# Transcriptomic biomarkers reveal jasmonic and salicylic acid state under field herbivory

**DOI:** 10.1101/2025.05.29.656841

**Authors:** Atsuki Tomita, Taro Maeda, Natsumi Mori-Moriyama, Yasuyuki Nomura, Yuko Kurita, Makoto Kashima, Shigeyuki Betsuyaku, Yasuhiro Sato, Atsushi J. Nagano

## Abstract

Plants coordinate defence against pathogens and herbivores through the crosstalk between salicylic acid (SA) and jasmonic acid (JA) signalling pathways. However, little is known about how plants integrate the phytohormone concentration profiles to shape defence states in complex field environments. Here we show that systematically varying combinatorial SA and JA concentrations in *Arabidopsis thaliana* reveals 43 distinct combination-specific transcriptional signatures beyond SA–JA antagonism. Leveraging these data, we developed and validated machine-learning-based transcriptomic biomarkers that enabled independent quantification of SA and JA response states. By applying these biomarkers to field-grown *A. thaliana*, we revealed that SA/JA response states were dynamically shaped by abundant and co-occurrence of herbivores on individual plants. Our study provides a strategy to decode hormonal responses to biotic environments from field transcriptomes, bridging the gap between molecular signalling and ecological dynamics.

## Introduction

In natural environments, plants are challenged by the community of diverse attackers such as herbivores and pathogens. This complex biotic environment necessitates dynamic integration and processing of stress signals to optimize defence strategies. Interactions between the salicylic acid (SA) and jasmonic acid (JA) signalling pathways have long been characterized by reciprocal antagonism^1,2^: SA promotes resistance to biotrophic pathogens^3^, whereas JA mediates defence against insects and necrotrophic pathogens^4^, partly through glucosinolate (GSL)-based anti-herbivore metabolites in Brassicaceae^5,6^. The SA-JA crosstalk, often described as reciprocal antagonism, allows plants to prioritise defence against attackers but can also compromise resistance to others^1,7^. Genetic and pharmacological studies have also shown the varying states of SA and JA, such that SA–JA crosstalk can turn into antagonistic, synergistic or additive relationships depending on phytohormone dose and timing^8,9^. However, we still lack a high-resolution quantification of the SA–JA interaction landscape that is necessary to understand plant responses to diverse attackers.

The plant SA/JA response to diverse biotic stresses is inherently context-dependent and highly dynamic^2,10^,. The states of SA and JA varies not only among species and genotypes but also within a single plant–herbivore interaction process through different life stages. In *Arabidopsis thaliana*, for example, oviposition by *Pieris* rapae induces SA-dependent defences^12^, whereas feeding by the hatched larvae subsequently triggers JA-dependent responses^13^. SA and JA signalling are not only host-regulated but are also actively manipulated by pathogens, herbivores, and associated organisms^14,15^. This dynamic defence states create a feedback loop that alters herbivore oviposition and host choice^15^. Furthermore, in natural settings, the simultaneous or sequential attack by diverse herbivore communities^10,11^ drives a far greater diversification of SA/JA balances and downstream defence outputs than single-attacker experiments can simulate^16^. Therefore, developing robust method to quantify the SA/JA signalling levels is pivotal for bridging the gap between laboratory and field conditions. To this end, it is essential to incorporate multiple-gene markers, which are themselves subject to SA-JA crosstalk.

To quantify plant responses to the diverse SA/JA balances, a systematic transcriptomic analysis using a wide range of SA and JA concentration combinations is essential^8,9^. Such an approach is crucial for comprehensively mapping the antagonistic, synergistic, and dose-dependent interactions between these pathways. However, it is challenging to systematically analyse the various combinations of SA and JA concentrations because large-scale RNA-seq experiments are required. To overcome this issue, many researchers have so far developed high-throughput RNA-Seq library preparation methods^17^. For instance, our previous study developed a Low-cost and eASY RNA-Seq method named Lasy-Seq^18^. This method has been applied to the systematic measurement of transcriptomic responses to multiple environmental inputs ^19^ but not to hormonal treatments and biotic stresses yet.

Here we show transcriptomic profiles of responses to 64 combinations of SA and JA concentrations in *A. thaliana* seedlings and derive transcriptomic biomarkers that quantify SA and JA response states. We applied these biomarkers to field transcriptomes from 2,381 *A. thaliana* individuals growing in distinct field in Otsu, Japan, and Zurich, Switzerland. By integrating these estimated response states with herbivore census data, we aimed to explore how variation in SA/JA signalling is closely linked to the specific assemblages and co-occurrence patterns of major herbivores on individual plants.

## Results

### Expression topography of SA/JA responses revealed by high-throughput RNA-seq

To evaluate how SA/JA concentration altered the interaction between the SA and JA signalling pathways, we performed transcriptome profiling of *A. thaliana* seedlings treated with 64 different SA/JA concentrations combinations for six hours (Fig. 1a,b; Supplementary Table 1). We employed an assay system that allows the simultaneous processing of 96 samples using 96-well plates (Fig. 1a,b), yielding data of 370 samples with 12,963 expressed genes (Supplementary Fig. 1). PCA analysis indicated that the first principal component explained 19.3% of variance and coordinated with the SA treatment concentrations (Fig. 1c; Supplementary Fig. 2a), while the second principal component explained 6.7% of variance and coordinated with the JA treatment concentrations (Fig. 1d; Supplementary Fig. 2b). These variations were unlikely attributable to experimental bias as no apparent differences were observed among the 96 well plates (Supplementary Fig. 2c–f). The number of DEGs between control and phytohormone-treated samples increased in a dose-dependent manner for SA and JA (Supplementary Fig. 3a–c). For both SA and JA, there was a larger number of up-regulated genes than down-regulated genes in most conditions from the 0 mM to 0.33 mM treatments (Supplementary Fig. 3d).

**Figure 1.**
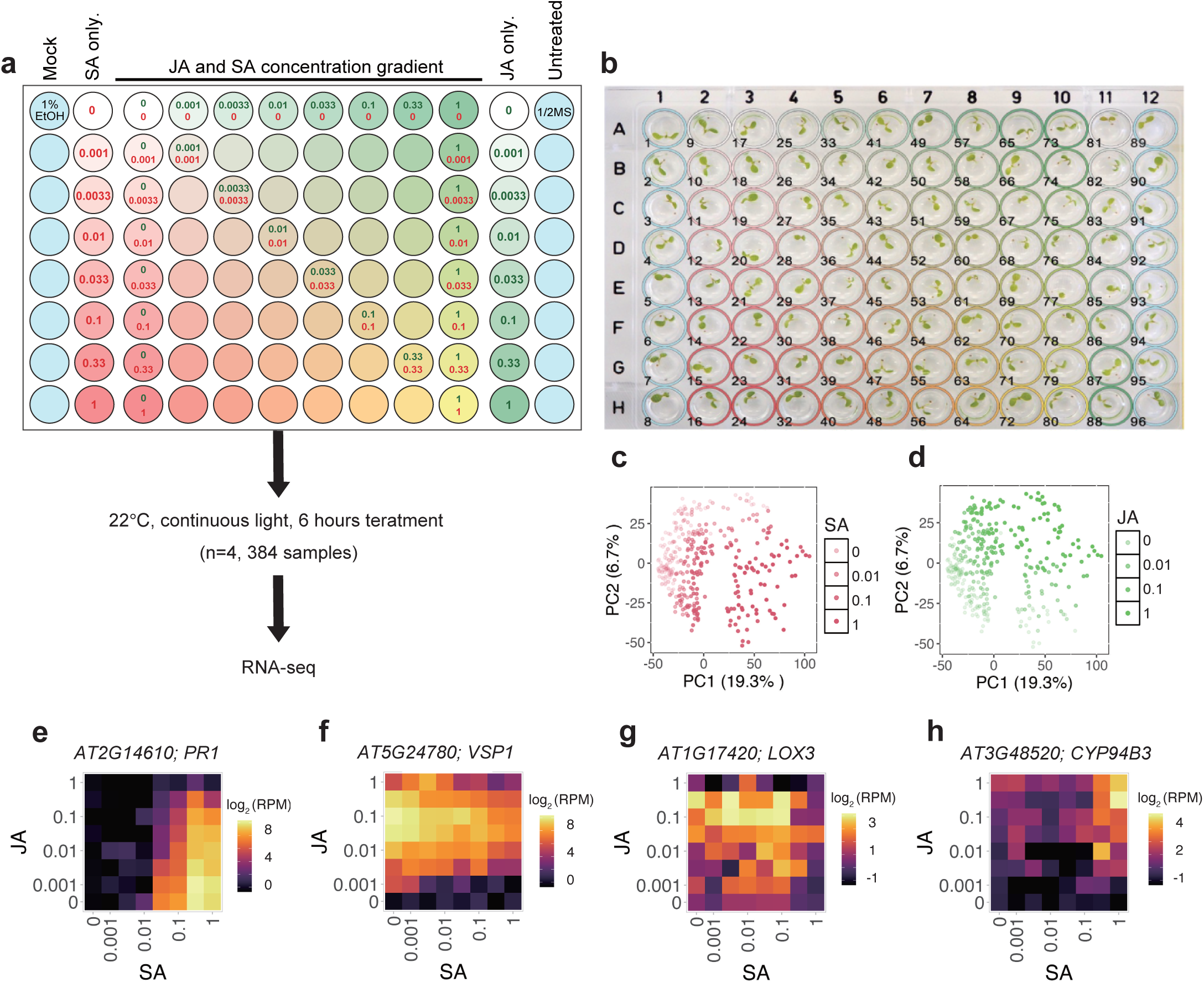
Treatments of 64 combinations of SA/JA concentrations (a) SA/JA treatment for *A. thaliana* seedling using 96-well plates. Each sample was treated with SA and JA at 64 conditions that were systematically combined among 8 concentrations: 0 mM, 0.001 mM, 0.0033 mM, 0.01 mM, 0.033 mM, 0.1 mM, 0.33 mM, and 1 mM. (b) A 96-well plate with seedling just prior to the SA/JA treatment. (c. d) Principal component analysis of samples. Each point represents samples. The color of each point indicates the log-transformed SA concentration (c), JA concentration (d). (e-h) Expression topography of genes related SA/JA responses. The horizontal axis of the heatmap corresponds to the concentration of SA, and the vertical axis to the concentration of JA. The color of each cell indicates the average log_2_(RPM + 1) for each concentration condition, n = 3∼11. *PR1* is a marker gene for SA responses. (e). *VSP1* is a marker gene for JA responses. (f). *LOX3* is encoding enzymes involved in JA synthesis (g). *CYP94B3* is involved in the downregulation of the JA response (h).

The systematic data then enabled us to depict a gene expression pattern in response to SA/JA concentration, which was designated as an expression topography hereafter (Fig. 1e–h). We evaluated the expression topography of well-known marker genes involved in the SA/JA response in addition to two genes associated with JA metabolism, such as *PR1*, *VSP1*, *LOX3* and *CYP94B3*. The SA response marker gene, *PR1*, exhibited an expression threshold at 0.033 mM SA with an optimal concentration of 0.33 mM, expression of which was completely suppressed under 1 mM JA (Fig. 1e; Supplementary Fig. 4a). The JA response marker gene, *VSP1*, exhibited an expression threshold at 0.001 mM JA and an optimum at 0.1 mM, expression of which was reduced under SA conditions (Fig. 1f; Supplementary Fig. 4b). These results are consistent with the qualitative understandings of SA/JA responses and SA-JA antagonism in previous studies^8,20^. Regarding JA metabolism, we detected expression changes of two key genes *LOX3* and *CYP94B3* in response to the SA/JA concentration. The expression of *LOX3* increased in a JA-concentration-dependent manner up to 0.33 mM JA (Fig. 1g; Supplementary Fig. 4c). This increase is consistent with previous evidence for autoregulation, since LOX3 encodes the enzyme for the first step of JA biosynthesis and accordingly LOX3-initiated pathway is known to exhibit a positive feedback loop^21^. Interestingly, *LOX3* expression was suppressed under 1 mM SA/JA treatment, suggesting that the positive feedback loop of *LOX3* is terminated once SA/JA levels reach substantial accumulation. In contrast, the expression of *CYP94B3*, a key regulator in the suppression of the JA signalling cascade, showed a slight increasing trend in a JA concentration-dependent manner (Fig. 1h; Supplementary Fig. 4d). Since CYP94B3 oxidizes the bioactive form of jasmonoyl-L-isoleucine, this oxidation by CYP94B3 regulates JA signalling levels and could affect SA/JA signalling levels ^22^. Notably, *CYP94B3* expression was maximally induced under high SA conditions, showing a 4.6-fold increase between 0 mM and 1 mM SA in the presence of 1 mM JA. These results indicate that these genes fine-tune the SA/JA crosstalk depending on the specific combination of concentrations.

### Diverse functional responses to the SA/JA concentrations in the expression topography

To characterize the diversity of expression topographies, we focused on 3,001 SA/JA-responsive genes (Supplementary Fig. 1) and performed affinity propagation clustering. This classification revealed 43 distinct gene clusters characterized by specific SA/JA concentration-dependent regulation patterns (Fig. 2a) and hierarchically aggregated into broad groups A–H to capture higher-order trends (Supplementary Fig. 5). These clusters contained 23 to 299 genes that shared highly coherent expression topographies (Fig. 2b). Cluster A-02 expression showed JA-dependent activation and suppression by SA. Conversely, cluster C-01 exhibited SA-dependent activation and JA-dependent suppression. To identify putative cis-regulatory elements among the expression topographies, we performed motif enrichment analysis on the upstream sequences of genes in each cluster (Fig. 2c; Supplementary Table 2). Clusters A-02 and E-03, which peaked at intermediate concentrations of both SA and JA, were strongly enriched for a GT3a type trihelix motif, suggesting that GT element-binding factors preferentially tune gene expression under moderate combinatorial phytohormone concentrations. By contrast, the SA dominated cluster C-01 was characterized by pronounced enrichment of TGA-class bZIP motifs, consistent with canonical NPR1-TGA-dependent SA signalling. Additional clusters were marked by motifs for other stress-associated transcription factors (e.g. other members of TGA, WRKY, ERF and MYB). Collectively, these results suggest that distinct *cis*-regulatory architectures provide the basis for complex SA/JA concentration-dependent outputs.

**Figure 2.**
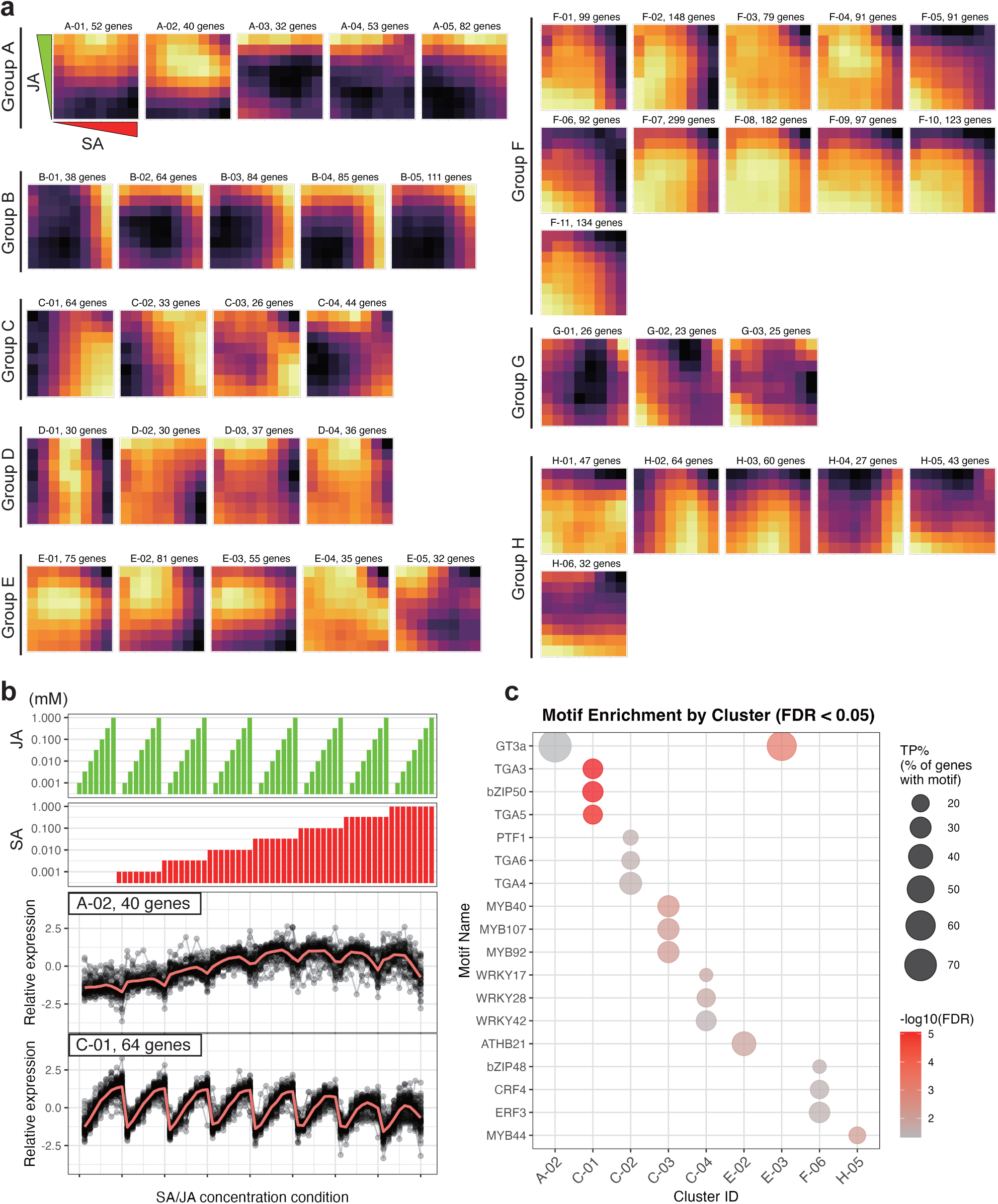
Forty-three gene clusters of expression topography represent various SA/JA responses (a) Average expression topography of each cluster. The horizontal axis is the concentration of SA, and the vertical axis is the concentration of JA. The colors represent the relative expression values of the genes in each cluster, with lighter colors indicating higher expression levels. (b) Upper 2 plots indicate concentration of SA/JA. Lower 2 plots show the relative expression levels of the cluster A-02 and cluster C-01. Expression profiles of 40 genes in A-02 and 64 genes in C-01 are indicated as gray points and lines. Mean expression is indicated as red lines. (c) Transcription factor binding motifs enriched in gene clusters. Bubbles represent motifs significantly enriched (FDR < 0.05, enrichment ratio ≥ 2, TP, > 10,) in each cluster. Bubble size indicates the percentage of target sequences containing the motif, and bubble color represents the statistical significance (-log10(FDR)). The top 3 most significant motifs per cluster are shown. A full list of enriched motifs was indicated in Supplemental Table 2.

To examine association between expression topographies and biological functions, we also conducted a GO enrichment analysis on 43 clusters. The analysis showed that 18 clusters were significantly enriched with 1 to 103 GO terms (adjusted *P* < 0.05, Fig. 3a; Supplementary Table 3). Genes in cluster A-02 were enriched in GO terms “response to jasmonic acid” (GO:0009753, adjusted *P* = 4.09 × 10^-4^). This cluster contained many JA-responsive genes reported in previous studies, such as *VSP2*, *JAZ8*, *JAZ10* and *JOX3* ^23^. Cluster C-01 showed the expression induced by SA and suppressed by JA. The *PR1* gene was contained in this cluster. Multiple GO terms were enriched in cluster C-01, including “response to bacterium” (GO:0009617, adjusted *P* = 1.18 × 10^-7^) and “response to salicylic acid” (GO:0009751, adjusted *P* = 3.55 × 10^-6^). When both SA and JA were treated in high concentrations, genes in cluster B-01 were up-regulated and genes in clusters F-07 and F-08 were down-regulated. In cluster B-01, GO terms such as “protein folding” (GO:0006457, adjusted *P* = 2.68 × 10^-5^) and “response to reactive oxygen species” (GO:0000302, adjusted *P* = 1.61 × 10^-3^) were enriched. The cluster B-01 mainly comprised genes encoding heat shock proteins such as HSP17.4 and HSP17.6. Clusters F-07 and F-08 were enriched in GO terms such as “ribosome biogenesis” (GO: 0042254, adjusted *P* = 7.71× 10^-30^ and 5.31 × 10^-4^) and “translation” (GO: 0006412, adjusted *P* = 1.52 × 10^-75^ and 3.28 × 10^-2^). GO terms associated with “photosynthesis” (GO:0015979, adjusted *P* = 1.57 × 10^-4^ and 4.22 × 10^-3^) were enriched in clusters F-05 and F-06. Genes in the clusters F-05 and F-06 were down-regulated in a dose-dependent and additive manner by SA/JA co-treatment (Fig. 2a). These results were consistent with previous studies, which demonstrated that co-treatment with SA and JA reduced the expression of photosynthesis-related genes ^23^.

**Figure 3.**
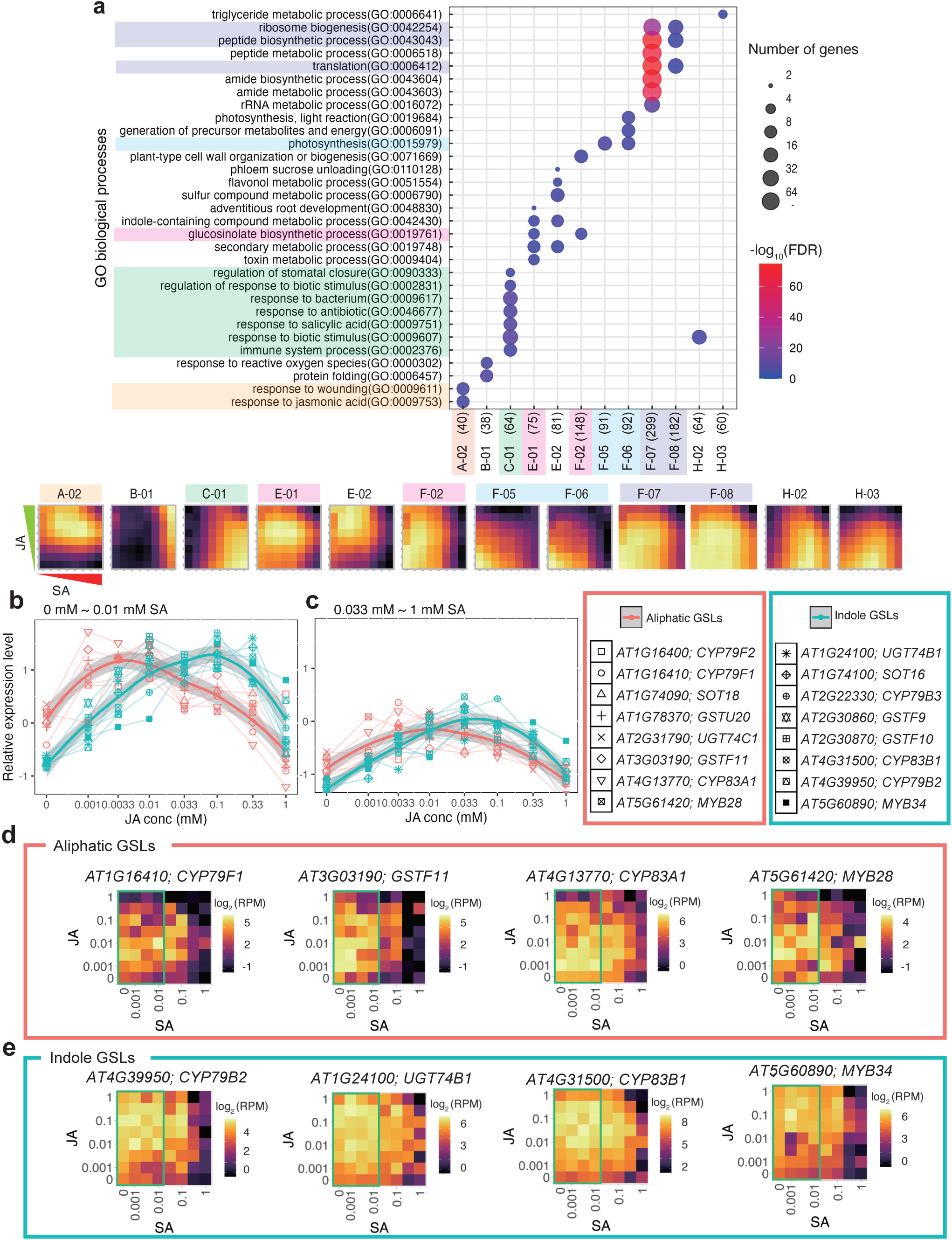
Genes with similar expression topographies were involved in same biological prosses. (a) GO enrichment in the gene clusters. The number of genes contained in each cluster is indicated in parentheses. The size of the circles indicates the number of genes. The color scale indicates the adjusted P-value (FDR) in the GO enrichment analysis. A full list of enriched GO terms was indicated in Supplemental Table 3. (b, c) Relative expression levels of glucosinolate (GSL) synthesis genes. Mean expressions for 0 mM to 0.01 mM SA conditions (b) or 0.033 mM to 1 mM SA conditions (c) are plotted. The horizontal axis indicates JA concentration (mM), and the vertical axis indicates relative expression level. Aliphatic GSL synthesis genes are shown in red. Indole GSL synthesis genes are shown in blue. Bold lines indicate LOESS curves drawn for each glucosinolate type. (d) Expression topographies of aliphatic GSL synthesis genes. (e) Expression topographies of indole GSL synthesis genes. Data used in the panel (b) are indicated by green boxes.

We found that two subtypes of GSLs, such as indole and aliphatic GSL, were induced by different concentrations of JA. In cluster E-01, the GO term “glucosinolate biosynthetic process” (GO:0019761, adjusted *P* = 5.32 × 10^-3^) was significantly enriched (Fig. 3a). Interestingly, “glucosinolate biosynthetic process” was also significantly enriched in cluster F-02 (adjusted *P* = 8.22 × 10^-3^). The cluster E-01 contained genes related to indole glucosinolate biosynthesis, *e.g. SUR1, CYP83B1, CYP79B3, MAM*. In contrast, the cluster F-02 contained genes related to aliphatic GSL biosynthesis, *e.g. MAM3, CYP79F2, BCAT4, SOT18*. The expression peak of indole GSL biosynthesis genes was found at lower concentration of JA than that of aliphatic GSL (Fig. 3b). In low SA concentrations (0 mM to 0.01 mM), the expression peaks of indole GSL biosynthesis genes were found around 0.1 mM JA, whereas those of aliphatic GSL biosynthesis genes were found around 0.003 mM JA (Fig. 3b). The expression peaks of indole GSL biosynthesis genes were found at approximately 30-fold higher JA concentrations than that of aliphatic GSL-related genes. This response was specific to low SA concentrations (Fig. 3c, Supplementary Fig. 6). Taken together, the ∼30-fold difference in JA sensitivity between these pathways, combined with their suppression by SA, suggests a defence strategy finely tuned by the quantitative balance of SA and JA.

### Multiple genes are required to account for the SA/JA response

Our comprehensive dataset allowed us to systematically evaluate how the expression of a single gene correlated with treated concentrations of SA and JA (Fig. 4a). Specifically, we focused on *PR1* and *VSP1* as they have long been used as major expression markers of SA and JA response^23^. Correlation coefficients between *PR1* and treatment concentrations of SA/JA were 0.63 and -0.30, respectively (Fig. 4a), expression of which had the 29th highest correlation with SA concentration. In contrast, *VSP1* had the 4,695th highest correlation with JA concentration. Correlation coefficients between *VSP1* and SA/JA treatment concentrations were -0.20 and 0.07, respectively (Fig. 4a). The genome-wide pattern of correlations indicates that correlations with SA and JA are associated with each other (Fig. 4a). We also focused on the other genes with high correlations. For example, *HRG2* exhibited the highest positive correlation with SA treatment concentration (*r* = 0.74), while that with JA was 0.14 (Fig. 4b). Meanwhile, UL4Z exhibited the highest negative correlation with SA, showing correlations of -0.70 and -0.22 with SA and JA, respectively (Fig. 4c). In contrast, the gene with the highest positive correlation with JA treatment concentration was *AT2G47950*. The correlation with JA was 0.85 and that with SA was 0.16 (Fig. 4d). The gene with the highest negative correlation with JA was *SAUR6*, with a correlation of -0.68 with JA and -0.44 with SA (Fig. 4e). These four genes responded to SA/JA at a specific threshold level (Fig. 4b–e), indicating that the expression level of a single gene alone cannot be markers to predict quantitative phytohormone levels These results showed that multiple genes responded to SA/JA treatments in a complex manner, thereby leading us to develop a quantitative response marker using multiple genes.

**Figure 4.**
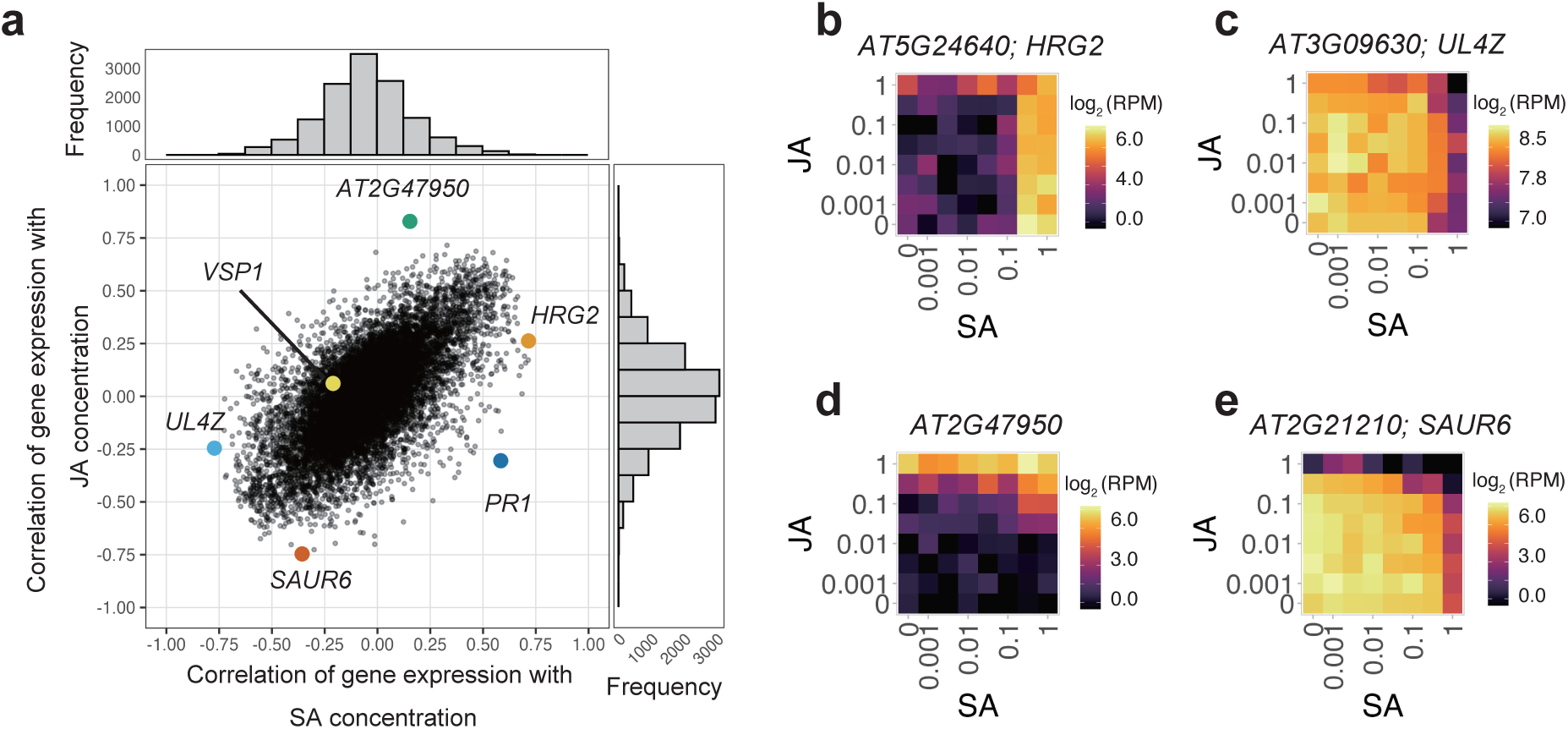
Dose-responsive gene expression and Transcriptomic biomarker of SA/JA response state (a,b) Histogram and scatter plot of correlation coefficients between gene expression levels and SA/JA concentrations (a), Genes with the highest or lowest correlation coefficient are indicated as color points. (b–e) Expression topographies of genes with the highest positive and negative correlation. b, H2O2 response gene *HRG2*, showing the highest positive correlation with SA. (c) Ribosomal protein gene *UL4Z*, showing the highest negative correlation with SA. (d) Uncharacterized protein gene *AT2G47950*, showing the highest positive correlation with JA. (e) Auxin-inducible gene *SAUR6*, showing the highest negative correlation with JA.

### Development of a transcriptomic biomarker of SA/JA response state

To develop a transcriptomic biomarker capable of estimating the quantitative state of SA/JA responses, we employed machine learning-based regressions on the expression levels of multiple genes. Specifically, we used an ensemble-learning approach with L1-regularized linear regression (least absolute shrinkage and selection operator, LASSO) to construct predictive models from the combinatorial SA/JA transcriptome dataset (Fig. 5a; Supplementary Fig. 7; see Methods for training/test data details). The resulting biomarkers demonstrated high predictive accuracy in an independent test dataset, yielding correlation coefficients of 0.87 for SA and 0.95 for JA between predicted and actual treatment concentrations (Fig. 5d,e). Notably, these values exceeded those of the best-correlated single genes (Fig. 4), although some underestimation was observed in the high-concentration range. These biomarkers also exhibited high specificity, such that the predicted SA and JA concentrations were weakly correlated with each other (*r* = 0.18; Fig. 5f) and vice versa (*r* = 0.38; Fig. 5g). To infer the functional basis of these markers, we determined the contribution of individual genes by calculating their selection frequency within the ensemble model (Supplementary Fig. 8). The distributions of selection frequencies indicated that limited number of genes were frequently used in the marker, but identified predominant contributors (selection frequency > 0.95) for the JA marker including canonical signaling components such as *JAZ1*, *JAZ2*, *JAZ3*, *JAZ5*, *JAZ6*, and *LOX2*. Similarly, predominant contributors for the SA marker included known SA-responsive genes, including *ZAR1*, *WRKY18*, *WRKY70*, *ROXY18*, *ECS1* and *ARR4* (Supplementary Table 4). Thus, our multi-gene transcriptomic markers provide a prospective predictor for the SA/JA response states.

**Figure 5.**
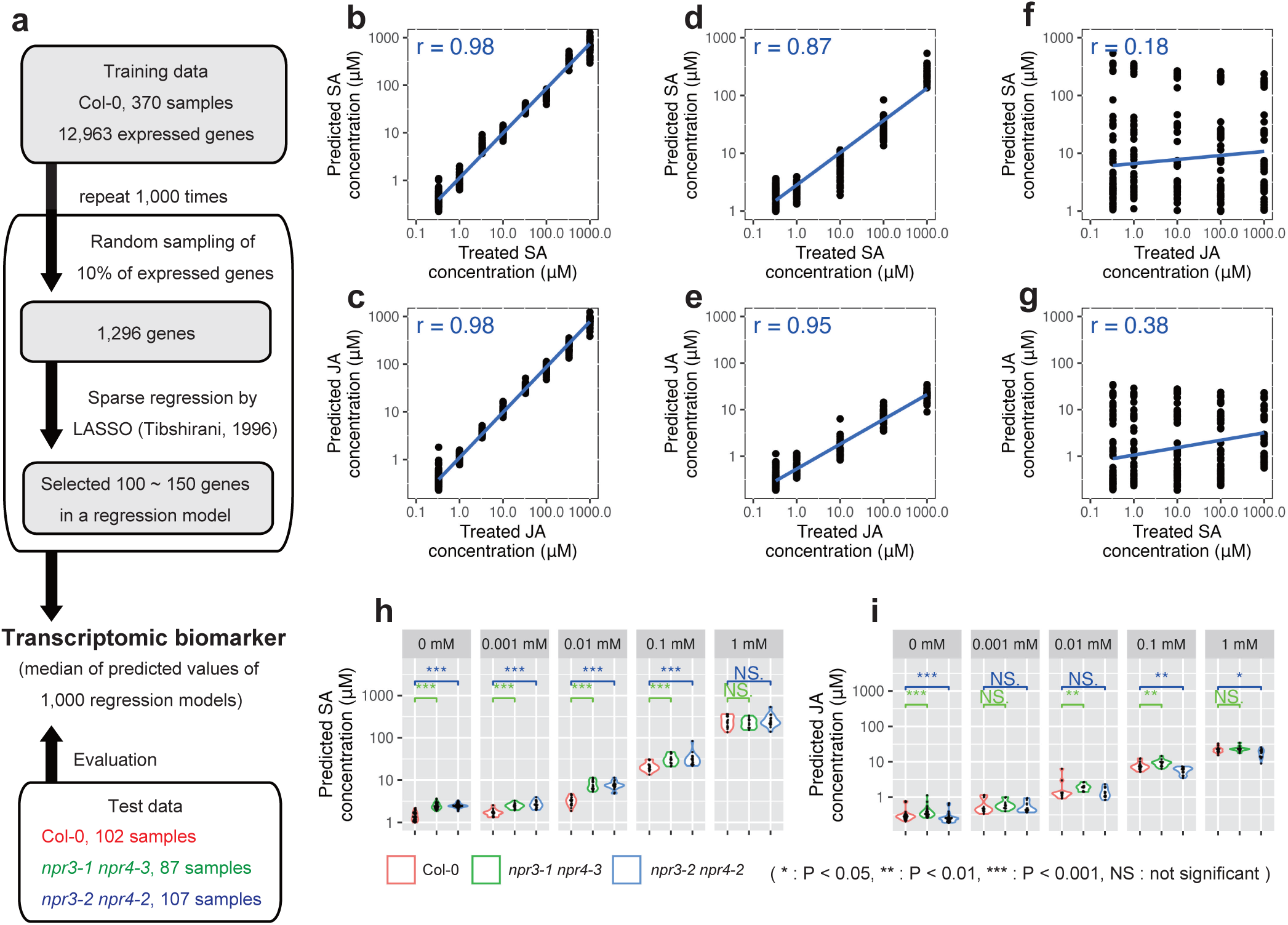
Transcriptomic biomarker of SA/JA response state (a) Scheme of development of a transcriptomic biomarker of SA/JA response state from RNA-seq data. See Materials and Methods for further details. (b,c) Predictions of the transcriptomic biomarkers for training data. Predicting of the marker for SA response state and treated SA concentration (b). Predicting of the marker for JA response state and treated JA concentration (c). (d-g) Predicting of the transcriptomic biomarkers for test data of Col-0. Prediction of the SA marker and treated SA concentration for test data (d). Prediction of the JA marker and treated JA concentration for test data (e). Prediction of the SA marker and treated JA concentration for test data (f). Prediction of the JA marker and treated SA concentration for test data (g). The horizontal axis is the predicted SA/JA concentration by the transcriptomic biomarker, and the vertical axis is the treated SA/JA concentration. The result of liner regression is shown in the blue line. (h, i) Predictions of the SA/JA marker for npr3/4 double mutants. Data obtained from 25 concentration conditions were grouped by the treated concentrations of SA (h) and JA (i). The vertical axis indicates the predicted concentrations by the transcriptomic biomarker. Results of Mann-Whitney’s U test were indicated as *: P < 0.05, **: P < 0.01, ***: P < 0.001 and NS: not significant.

To demonstrate the utility of our biomarkers for quantitative phenotyping, we applied them to investigate SA/JA response states in *npr3/4* double mutants (npr3-1npr4-3 and npr3-2npr4-2), which are deficient in SA perception^24,25^. We profiled the transcriptomes of these mutants under 25 combinatorial SA/JA conditions (total 194 samples; Supplementary Fig. 7) and estimated their response states using our transcriptomic biomarkers (Fig. 5h,i). These mutants showed significantly higher predicted SA concentration than that in Col-0 for all SA treatment conditions except 1 mM SA (Fig. 5h), indicating that the mutants seemed to have a higher sensitivity to SA than Col-0. This result suggested that NPR3/4 suppressed the SA signalling pathway in conditions with lower than 1mM SA, which was consistent with previous studies showing that NPR3 and NPR4 acted as transcriptional co-repressors of SA-responsive genes^24,25^. For JA responses, the mutants showed significant differences from Col-0 under several conditions, although these effects were not consistent among mutant alleles (Fig. 5i). Collectively, these results demonstrate that our transcriptomic biomarkers provide a robust, quantitative framework for decoding complex phytohormone response states, and suggest their applicability to phenotyping of diverse mutants.

### Application of biomarkers to transcriptome dataset under field herbivory

Lastly, we asked whether our biomarkers were able to capture JA and SA responses to herbivores in complex field environment. To address this question, we applied our biomarkers to field data including 2,381 *A. thaliana* individuals and their insect communities (1,195 samples in Otsu, and 1,186 samples in Zurich). SA and JA responses were uncorrelated in Zurich, whereas a significant positive correlation was detected in Otsu (Fig. 6a). We then tested whether these responses were associated with insect communities, which considerably differed between the two sites^26^ (Fig. 6b). The vast majority of herbivores in Zurich were caterpillars of diamondback moths (‘Px’) and flea beetles (‘Ps’ and ‘Pa’) that could jump between plants^27^, and those in Otsu were small white butterflies *Pieris rapae* (‘Pr’) and Px. While the variability of JA and SA responses was not directly explained by insect species richness (Supplementary Fig. 9), we detected specific responses to abundant herbivores (Fig. 6b). Specifically, in Zurich, the individual number of Px and wingless *Brevicoryne brassicae* (‘Bb’) were positively correlated with the JA response and negatively correlated with the SA response, consistent with a canonical SA-JA crosstalk pattern^1,2^. Conversely, Pa and Ps were positively associated with the SA response and negatively with the JA response. In Otsu, both eggs (‘Pr_e’) and larvae (Pr_l’) of small white butterflies correlated positively with SA and JA responses. These results suggest that herbivore communities may differentiate JA and SA responses through the species-specific influences between the two sites.

**Figure 6.**
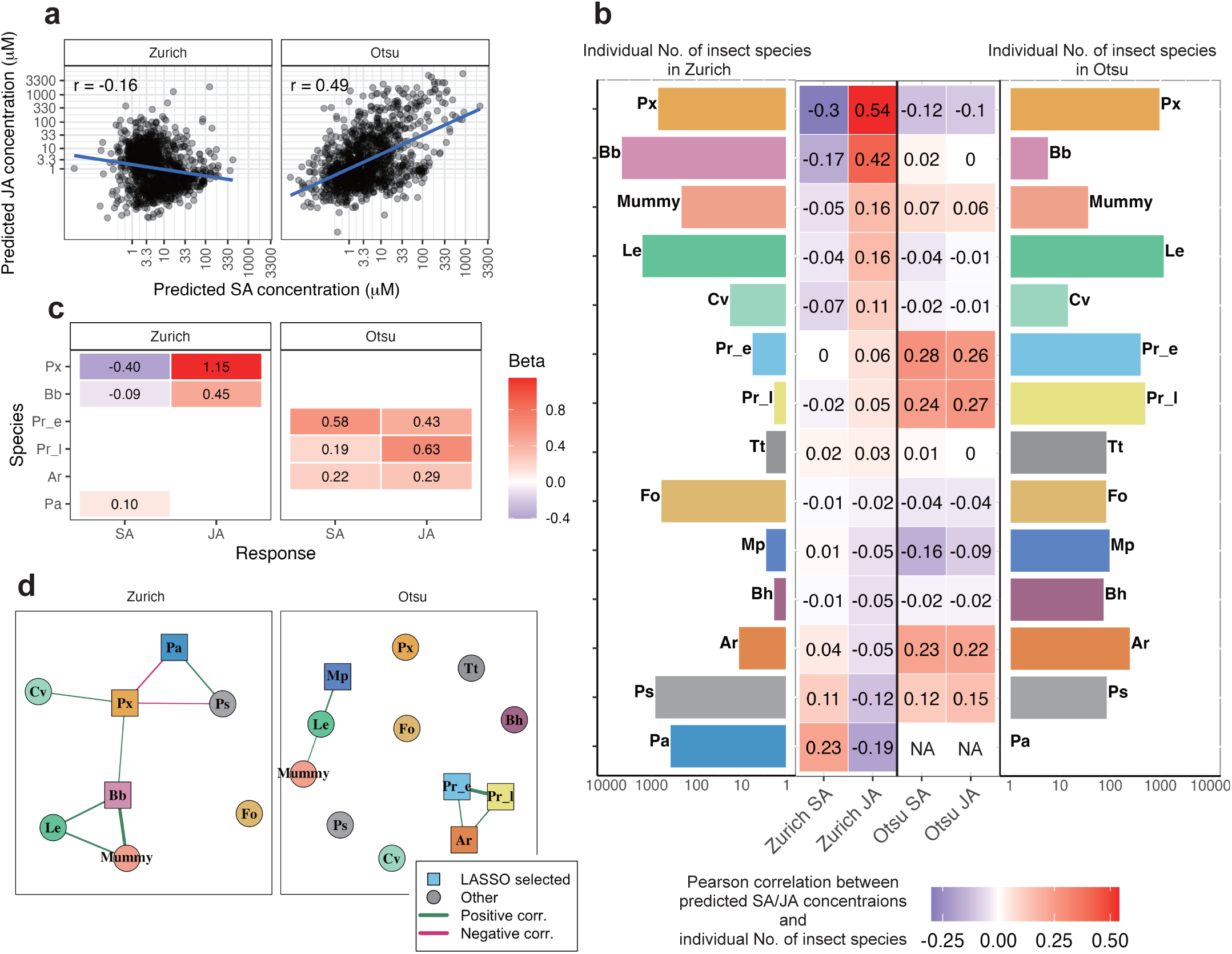
SA/JA response states of field transcriptomes and their association with herbivore communities. (a) Relationship between predicted SA and JA concentrations in Zurich and Otsu populations. Blue lines and r indicate linear regression fits and Pearson correlation coefficients, respectively. (b) Heatmap of Pearson correlation coefficients between predicted SA/JA concentrations and insect abundance. Side bars indicate total insect counts per population. Colors represent correlation strength (red: positive; blue: negative). (c) LASSO regression coefficients estimating insect effects on SA/JA responses. Only species with non-zero coefficients are shown. Colors represent the sign and magnitude of coefficients (red: positive; blue: negative). (d) Insect co-occurrence networks based on significant correlations (|r| > 0.15, p < 0.05). Nodes represent insect species. Square: LASSO-selected predictors. Circle: others. Edges represent correlations (green: positive/co-occurrence; pink: negative/segregation), with width proportional to correlation strength (|r|).

To further narrow down the herbivore-specific responses, we performed LASSO regression of herbivore abundance on the JA and SA responses (Fig. 6c). In Zurich, Px showed the strongest effects on both hormonal responses (β = 1.15 for JA, β = -0.40 for SA). Bb was also selected as a predictor of the JA response (β = 0.45). This result is consistent with the general pattern that chewing herbivores induce JA signaling ^28^. Pa exhibited the only positive coefficient for the SA response (β = 0.10). In Otsu, Pr_e and Pr_l emerged as the strongest predictors of the SA (β = 0.58) and JA (β = 0.63) responses, respectively, indicating life-stage-specific signatures within the same species. These patterns are consistent with SA induction by oviposition and JA induction by larval feeding^12,13^.

To corroborate these findings, we analysed herbivore co-occurrence networks on individual plants (Fig. 6d). Species selected by LASSO occupied central positions within their respective networks. In Zurich, Pa and Px were significantly negatively associated (r = -0.25; Supplementary Fig. 10a), whereas the Otsu network was cohesive with a strong co-occurrence between Pr_e and Pr_l (r = 0.59; Supplementary Fig. 10b). Taken together, these results indicate that the presence of major herbivores, coupled with their co-occurrence or exclusion patterns, shapes site-specific JA and SA responses.

## Discussion

Our systematic approach combining combinatorial phytohormone treatments with large-scale transcriptomics provides quantitative evidence for SA/JA responses. For example, our analysis revealed that genes encoding the biosynthesis of antimicrobial indole GSLs peaked at high JA concentrations, whereas those mediating aliphatic GSLs, which can deter a wide variety of herbivores^29^, peaked at lower JA levels (Fig. 3b). Given that high JA accumulation is characteristic of wounded sites^30^ while lower levels are typical of systemic tissues^31^, this differential regulation reflects an effective strategy whereby plants deploy localized antimicrobial defences at wound sites while priming systemic tissues against subsequent herbivory. As demonstrated by the concentration-specific regulation of GSL biosynthesis genes, our experimental system successfully captured fine-tuned transcriptomic changes triggered in response to different SA/JA concentrations. These findings provide testable hypotheses based on large-scale transcriptome data.

We successfully quantified the response of whole seedlings to a wide range of SA/JA concentrations and combinations, but this might obscure tissue-specific and cell-type-specific responses could be obscured. Our measurement of total RNA amounts of shoot and root indicated that approximately 70% of total RNA of seedling was derived from shoot and 30% from root (Supplementary Fig. 12a). Responses of root-specific highly expressed genes were successfully measured (Supplementary Fig. 12b). These findings suggest that tissue-specific gene expressions can be detected unless they exhibit opposite patterns and are thus offset among tissues. While capturing specific responses in minor cell types remains challenging, highly expressed genes in specialized cells (e.g., *TGG2* in myrosin cells^32^; Supplementary Fig. 12c) may still be detectable. Considering the rapid advances of single-cell RNA-Seq technologies, future integration of systematic SA/JA treatment with single-cell RNA-Seq analysis will be a promising direction to fully resolve these cell-type-specific responses.

The transcriptomic biomarker that we developed in this study offer a robust framework for predicting SA/JA response states across diverse contexts. Advances in methodology have enabled sensitive and accurate quantification of phytohormones^33^. However, these measured levels do not always correspond to response states^33^. The complementary use of our biomarkers with phytohormone quantification will help elucidate the mechanisms underlying the discrepancies. Furthermore, our biomarkers will be applicable to large-scale transcriptome data in emerging research areas, such as single-cell and field transcriptomics. A study combining single-nucleus RNA-Seq and spatial transcriptomics to analyse plant immune responses has been reported recently^34^. Applying the transcriptomic biomarkers to such datasets would reveal SA/JA response dynamics during plant immune responses at a single-cell resolution.

In field transcriptomics, massive transcriptome data acquired under natural conditions are modelled with environmental data to reveal the responses of organisms in real-world environments^35^. While laboratory studies have frequently reported apparent trade-offs between SA and JA response^2,8,9,36^, our field transcriptomics data reveal a more complex reality shaped by community composition. We show that the balance between SA and JA differs between the two field sites, exhibited significant differences in the insect community composition as shown by our previous fieldwork^26^. Specifically, in the Otsu site, the Pr was abundant, egg deposition and larval feeding of which are known to respectively induce SA and JA-mediated defence responses^12,13^. This simultaneous induction may account for why SA and JA are positively correlated in the Otsu site. Contrarily, Pa and Px were abundant in the Zurich site. Pa can jump between neighbouring plants while chewing leaves^26,27^, which may be associated with the plant’s SA response states. Importantly, integrating transcriptomic biomarkers with herbivore data provided testable hypotheses about plant-herbivore relationships. For instance, we detected the positive correlation between Pa abundance and SA responses, along with its segregation from Px. This suggests the selective colonization of highly mobile Pa onto plants that have already induced an SA response. Collectively, contrasting SA/JA balance between the two field sites might be accounted by herbivore specificity and mobility, which can be tested by excluding insect herbivores from the field.

In conclusion, our findings highlight the utility of combining transcriptomics and herbivore community data to understand phytohormonal responses to complex biotic stress in complex field environments. Previous field transcriptomic studies have primarily focused on analyses using meteorological data^19,37,38^. While these efforts have successfully elucidated plant responses to abiotic environmental factors, the responses to biotic factors remain largely unexplored. By integrating the transcriptome biomarkers developed in this study with field transcriptomics, it will deepen our understanding of the interplay between plant defence responses and meteorological factors in real-world environments.

## Materials and methods

### Plant materials and SA/JA treatment

We used *A. thaliana* (Col-0: CS70000, *npr3-1npr4-3*: CS72351 and *npr3-2npr4-2*: CS72352) to analyse the responses to SA & JA concentrations. The seeds of these lines were obtained from the Arabidopsis Biological Resource Center. Methyl jasmonate (135-14411, FUJIFILM Wako, Osaka, Japan) and sodium salicylate (S3007, Merck, Darmstadt, Germany) were dissolved to 1 M in ethanol or water, respectively. 100× SA or 100× JA stock solutions were then diluted from 1 M stock to 0 mM, 0.1 mM, 0.33 mM, 1 mM, 3.3 mM, 10 mM, 33 mM, and 100 mM in ethanol or water, respectively. Sixty-four variations of 50× SA/JA mixed stock solutions were prepared by combining eight concentrations of 100× SA and 100× JA, and mixing equal amounts of each. These 64 stock solutions for SA/JA treated samples, 50% EtOH in 1/2 MS medium for mock samples, and only 1/2 MS medium for untreated samples were transferred to a 96-well plate, and stored at −20°C until use. A single sterilized seed was sown in each well of 96-well plates and grown in 50 μL of liquid 1/2 MS medium. After 3 days of treatment at 4°C in the dark, the plants were cultivated at 20 °C for 5 days under continuous light. SA/JA treatments were performed by adding 2 μL of 50× SA/JA mixed stock solutions and 48 μL of liquid 1/2 MS medium to each well containing 5-day-old seedlings. This allowed the 5-day-old seedlings to be incubated for 6 h under 64 variations of SA and JA concentration conditions (Fig. 1a). To monitor the broad spectrum of early defence responses, we treated with SA/JA for 6 h^39^. Mock and untreated samples were treated for same time as above using 1% ethanol in 1/2 MS medium and only 1/2 MS medium, respectively. Four replicates of the plant were treated by each SA & JA condition, respectively. We removed the medium and added 100 μL of direct-RT buffer ^40^ and a zirconia bead (YTZ-4, AS-ONE, Japan) to each well. Samples were ground using the TissueLyser II (QIAGEN, MD, USA) by 2 rounds for 1.5 minutes of 2.5 Hz. The homogenates were centrifuged at 3,000 rpm for 10 min, and 50 μL of the supernatants were stored as lysates at -20 °C. For test data of the transcriptomic biomarkers of SA/JA response state, we performed the same procedure using Col-0, *npr3-1npr4-3* and *npr3-2npr4-2* under twenty-five SA & JA concentration conditions made by combining five concentrations: 0 mM, 0.001 mM, 0.01 mM, 0.1 mM and 1 mM (Supplementary Fig. 7).

### RNA-Seq experiment and basic data analysis

The 5 μL of lysate was used by RNA-Seq library preparation with the Lasy-seq version 1.1 method ^18^. The library was sequenced by HiSeqX (Illumina, San Diego, CA) with the paired-end 150+150bp mode. Trimomatic version 0.3.3 ^41^ was used for trimming read data, and bowtie version 1.1.2 ^42^ and RSEM version 1.3.0 ^43^ were used for expression quantification. For the following analysis, we used log_2_(RPM+1) values calculated by the same procedure of the previous study^18^. The expression topography of all SA/JA responsive genes is shown in Supplementary Fig. 13.

We excluded samples with fewer than 10^5.5^ total reads from the analysis (Supplementary Fig. 1b). We defined genes with more than 1.5 average log_2_ (RPM+1) as the expressed genes (Supplementary 1c). The consistency of the dataset across different 96-well plates was verified by PCA (Supplementary Fig. 2c-f). We calculated the standard deviation (SD) of the SA/JA-treated and control samples using the following procedure. To eliminate the effect of differences in sample numbers, 63 samples were randomly selected from each of the SA/JA-added samples and control samples. We calculated standard deviations among the 63 SA/JA-added samples as SD_treated_ and standard deviations among the 63 control samples as SD_control_. In the expressed genes, SA/JA responsive genes (3,001 genes in 370 samples) were selected based on the ratio of the SD_treated_ to SD_control_ (Supplementary Fig. 1d).

We detected differentially expressed genes between the control and each concentration treatment using a likelihood ratio test of the edgeR package ^44^. 0 mM SA and 0 mM JA, mock and untreated conditions were used as control samples. The FDR was controlled using Benjamini and Hochberg’s method^45^.

### Expression Topography

The average expression levels were calculated for each condition. The average expression levels of 8 × 8 conditions data for each gene were designated as an expression topography. To extract key characteristics of the expression topography, smoothing by a 3 × 3 moving average filter was applied to the expression data. Z-scores of the smoothed expression data were calculated using the scale function of R. Using the apcluster version 1.4.8 package ^46^, clustering was performed by the affinity propagation method based on Z-scores of the smoothed expression data of SA/JA-responsive genes. Pearson’s correlation coefficient (*r*) was calculated using the Z-scores averaged over each cluster of affinity propagation. Groups of clusters were determined by hierarchical clustering. The correlation coefficients were as the distance between clusters, and the optimal number of groups was estimated by the gap statistic.

### Gene Ontology analysis

Gene Ontology annotations were obtained from The Arabidopsis Information Resource (TAIR) on June 29, 2021. Fisher’s exact test was used to test the enrichment of GO terms. The multiple testing correction was performed with a FDR^45^. For these analyses, R 3.6.0 and 4.2.0 were used.

### Transcription Factor Binding Motif Enrichment Analysis

Transcription factor binding motif enrichment analysis was performed using the Simple Enrichment Analysis (SEA) tool from the MEME Suite version 5.5.0^47^. Promoter sequences (1,000 bp upstream of the transcription start site) were extracted for each gene identified through affinity propagation clustering. A background sequence set was constructed from all promoter sequences of expressed genes. The analysis utilized the Arabidopsis DAP-seq motif database^48^. SEA was run with default settings, and multiple testing was controlled using the FDR^45^. To ensure robustness, we focused on motifs satisfying stringent criteria: false discovery rate (FDR) < 0.05, enrichment ratio ≥ 2, and presence in >10% of target sequences (Supplemental Table 2).

### Transcriptomic biomarker of SA/JA response state

To create transcriptomic biomarkers correlated specifically with SA or JA, we employed the method using ensemble learning and L1-regularized linear regression. Expression data of the 12,963 expressed genes from 370 samples in 64 concentration conditions were used as training data. The 1,296 genes, 10% of the expressed genes, were randomly selected and used as input variables for LASSO^49^. The hyperparameter λ of LASSO was determined by cross-validation. The selection of subset of genes and LASSO were repeated 1,000 times. The median of the predicted values of the 1,000 regression models was used as the transcriptomic biomarker. Gene selection frequency was defined as the proportion of trials in which the gene had non-zero coefficient out of trials in which the gene was used as input variable (Supplementary Fig. 8; Supplemental Table 4). The test data of regression models were transcriptome data at 25 conditions of SA and JA combining 5 concentrations: 0, 0.001, 0.01, 0.1, and 1 mM.

### Field experiment under naturally occurring herbivores

In this study, we reused field data deposited by Sato et al. (2024) who observed naturally occurring herbivores on 199 *A. thaliana* accessions in the field. Briefly, 199 accessions were planted nearly 600 plants (= 199 accessions x 3 blocks) at outdoor gardens located in Zurich, Switzerland (47°23’N, 8°33’E; alt. ca. 500 m) and Otsu, Japan (35°06’N, 134°56’E; alt. ca. 200 m). Plants were initially cultivated under short-day conditions and then transferred to the field sites during early summer in 2017 and 2018. The 200 potted plants (199 accessions and an additional Col-0) were randomly placed within a 260 cm x 64 cm block in a checkered manner. Insect herbivores on individual plants were counted every two to three days throughout the growing season. The original phenotypic data utilized in this study are available in the Zenodo repository (doi:10.5281/zenodo.7945318).

### RNA-Seq experiment of Field samples

Leaf samples for transcriptomic analysis were harvested three weeks post-transplantation, corresponding to the endpoint of the herbivore observations. To minimize circadian variations in gene expression, all samples were collected within a two-hour window cantered around solar noon at each site. Harvested leaves were immediately immersed in an RNA preservation buffer containing 5.3 M (NH4)2SO4, 20 mM EDTA, and 25 mM trisodium citrate dihydrate (pH 5.2). Samples were incubated at 4℃ overnight and subsequently stored at -80℃ until processing. Total RNA was extracted using the Maxwell 16 Lev Plant RNA Kit (Promega Japan) according to the manufacturer’s protocol. RNA-seq libraries were prepared using the Lasy-Seq version 1.1^18^. These libraries were sequenced by Illumina HiSeq 2500 and HiSeq X platforms to generate 50-bp and 150-bp reads, from the 3’ end, respectively. Output fastq files were processed and mapped in the same way as described above. We excluded samples with fewer than 10^5.5^ total reads from the analysis. We defined genes with more than 1.5 average log_2_(RPM+1) as the expressed genes. Expressed genes in all three independent datasets (training, validation, and field datasets from Zurich and Otsu) were retained for biomarkers. Following the procedure for the transcriptomic biomarker developed in this study, the SA/JA response states were estimated for field-grown *A. thaliana* plants.

### Prediction of SA/JA response states from herbivore abundances

To quantify which herbivore species were most strongly associated with SA/JA response states, we tested whether the predicted SA/JA scores could be explained by herbivore community composition. To minimize the influence of strongly tissue- or stage-specific genes and to enrich for core components of SA/JA signalling, we used as biomarkers only expressed genes (log_2_(RPM + 1) > 1.5) in both the seedling training dataset and the field leaf dataset. We constructed four models in total, corresponding to each combination of field site (Zurich or Otsu) and response variable (SA or JA). In each model, the response variable was the predicted SA or JA response state, and the predictors were the abundances of 14 herbivore species recorded on individual plants. We fitted LASSO^49^ regression models using the glmnet package in R (family = “gaussian”, alpha = 1). The regularization parameter λ was selected by cross-validation using the default procedures implemented in glmnet.

### Analysis of co-occurrence patterns among herbivore species

log_2_-transformed count data were used to assess the association of herbivore species abundances on individual plants. We calculated Pearson’s correlation coefficients between all species pairs. Pearson’s correlation was tested using cor.test in R (Supplementary Fig. 10). Species pairs showing significant associations (|r| > 0.15 and p < 0.05) were retained and used to construct the abundance-based co-occurrence network (Fig. 6d). The network was visualized using the Fruchterman–Reingold algorithm implemented in the igraph package in R, such that species with stronger associations (|r|) were positioned closer to each other. Herbivore species that were extremely rare (maximum co-occurrence frequency < 0.01 across all pairs; Supplementary Fig. 11) were excluded from the network visualizations.

## Data availability

The RNA-Seq data of combinatorial treatments with SA and JA were submitted to the NCBI Sequence Read Archive repository under the BioProject PRJNA1232845. The RNA-Seq data from the field herbivory experiments were submitted to the NCBI Sequence Read Archive repository under the BioProject PRJNA1055060, PRJNA1055104, PRJNA1055317, PRJNA1055424, PRJNA1055734, PRJNA1055736, PRJNA1056126, and PRJNA1055939. R scripts and data used in the analysis (count data, etc.) are available via the figshare repository (https://doi.org/10.6084/m9.figshare.29323187). (These data are currently under embargo and will be made publicly available upon acceptance of the manuscript.)

## Acknowledgements

We thank Dynacom Co., Ltd. (Chiba, Japan) for technical assistance with the analysis of RNA-Seq data.

## Author contributions

AJN and SB conceived the study. YN, YK and MK developed the assay system. AT, YS and NMM conducted the RNA-Seq experiments. AT, TM, YS and AJN analysed the data. AT, YS and AJN wrote the manuscript with input from all co-authors.

