## Supplemental Figures S1-S12 for "Transcriptomic biomarkers reveal jasmonic and salicylic acid state under field herbivory"

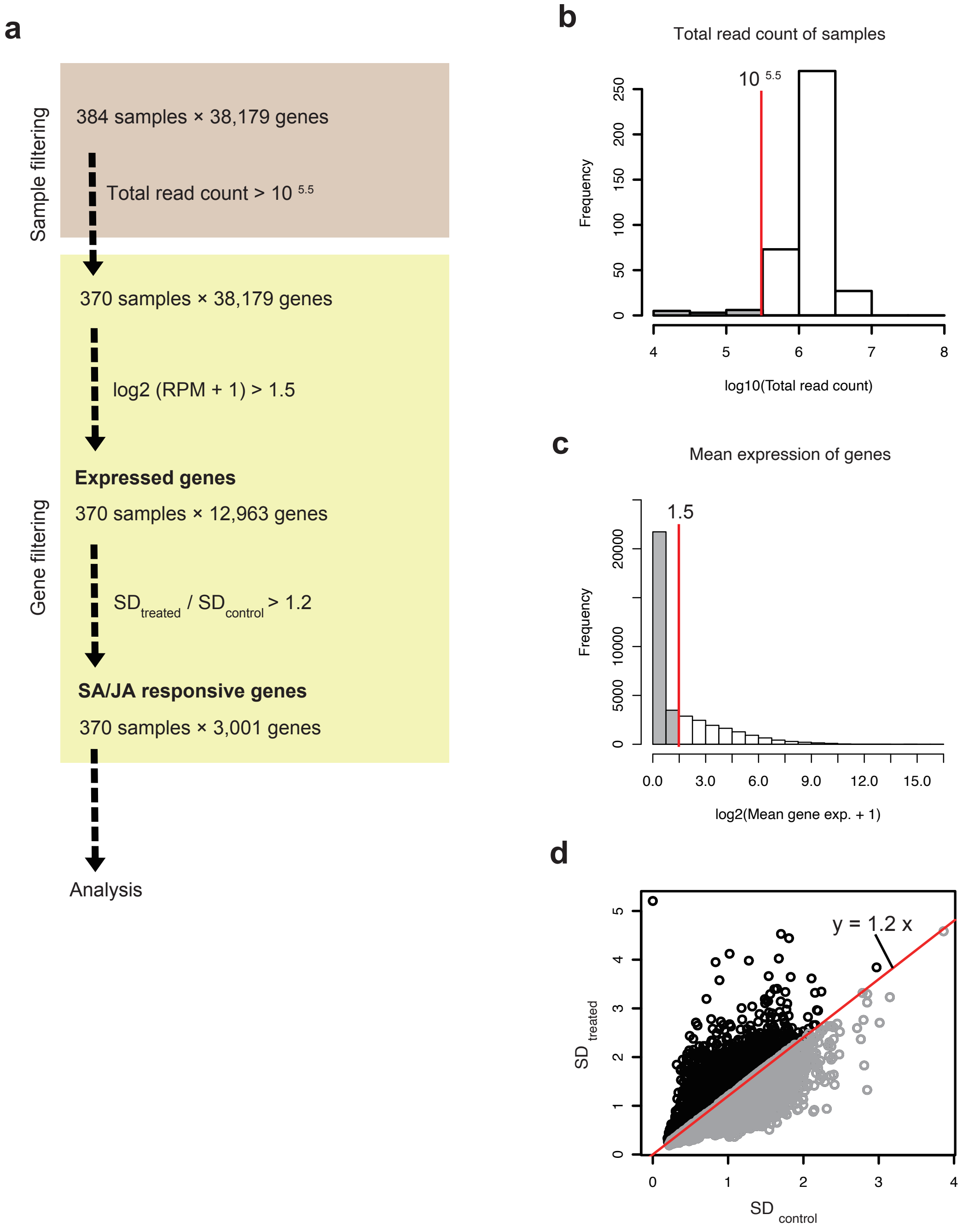


Supplementary Fig.1 Sample and gene filtering of RNA-seq data.

(a) Schematic flow of sample and gene filtering. (b) Histogram of total read counts for samples. (c) Histogram of average expression levels. Genes whose expressions were greater than 1.5 were defined as expressed genes. (d) Scatter plot of SDtreated and SDcontrol. Genes whose ratio of SDtreated to SDcontrol were greater than 1.2 were defined as SA/JA responsive genes. Red lines indicate thresholds.

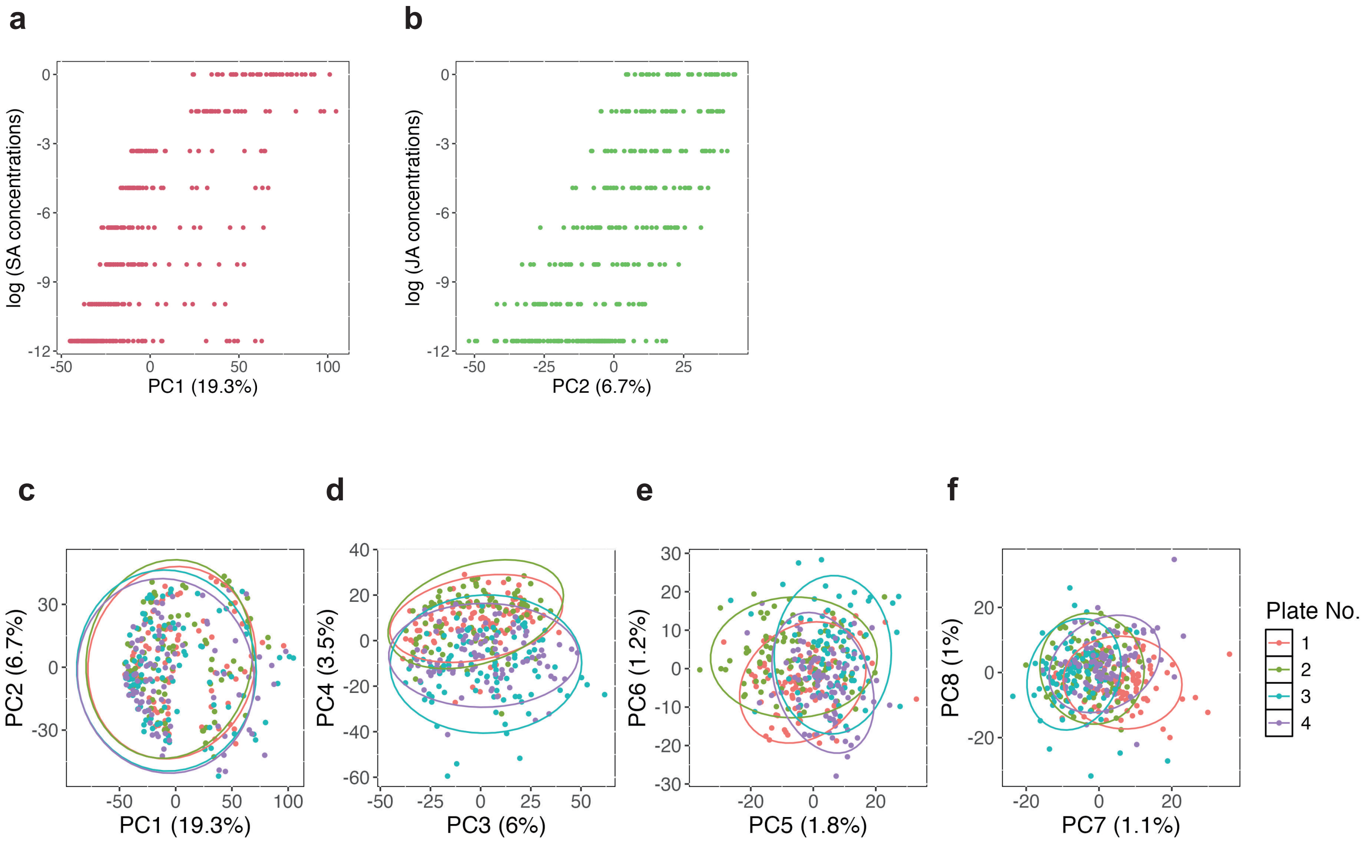


Supplementary Fig.2 Principal-component analysis of transcriptome samples.
(a) Scores of PC1 (19.3 % variance) plotted against log-transformed SA concentrations. The PC1 primarily captures SA dose-dependent effects. (b) Scores of PC2 (6.7 % variance) plotted against log-transformed JA concentrations. The PC2 primarily captures JA dose-dependent effects. (c–f) Pairwise combinations of the subsequent PCs (PC1–PC8) colored by 96-well plate number (Plate 1, red; Plate 2, green; Plate 3, blue; Plate 4, purple). The overlap among ellipses shows that the Top 8 PCs were not captured by plate-to-plate variation.

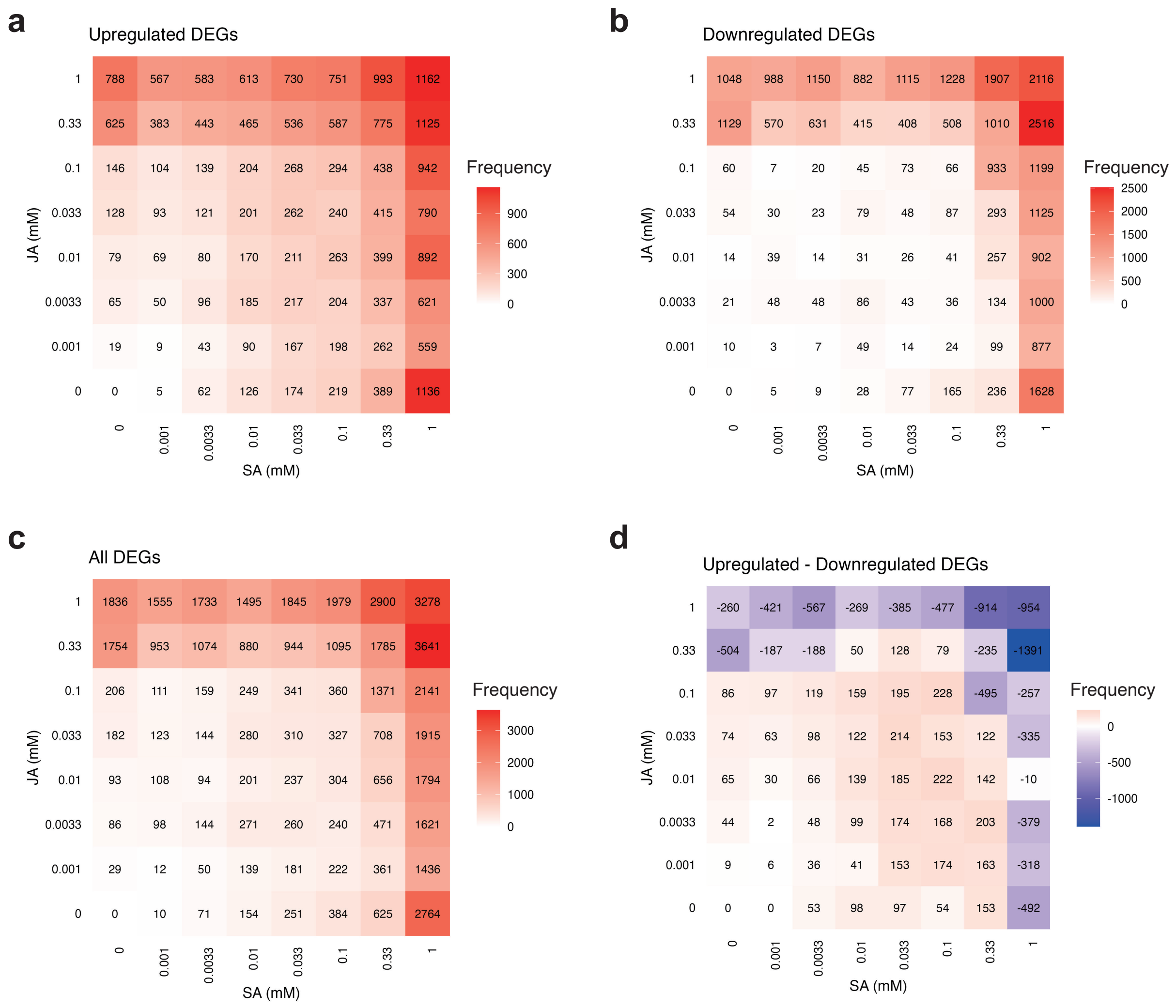


Supplementary Fig.3 Number of differentially expressed genes by SA/JA treatment.

The number of differentially expressed genes between the control group and other 63 SA/JA treatment conditions (FDR = 0.05). (a) Number of genes that were up-regulated by SA/JA treatments. (b) Number of genes that were down-regulated by SA/JA treatments. (c) Number of genes that were differentially expressed by SA/JA treatments. (d) Difference between the number of up-regulated genes and the number down-regulated by SA/JA treatments.

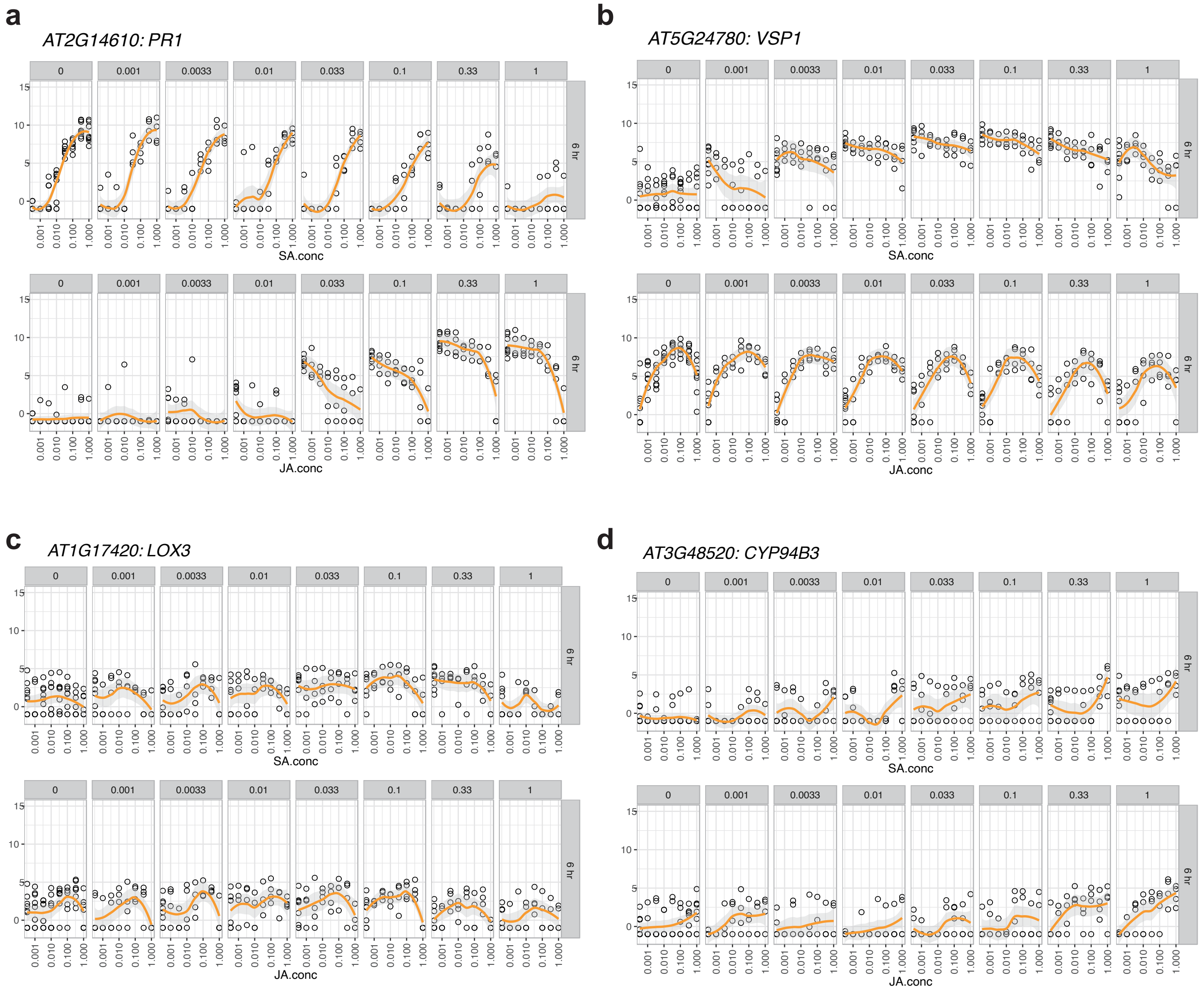


Supplementary Fig.4 Gene expression under combined SA/JA treatment.

(a–d) Scatter plot of gene expression levels under SA/JA treatment conditions (mM). Black circles represent $\log_2$ (RPM + 1) expression values for each individual RNA-seq sample. Upper panels of x-axis are SA concentration and JA fixed at the value indicated above each panel. Lower panels of x-axis are JA concentration and SA fixed at the value indicated above each panel. The orange line indicates LOESS curves, and the light-grey band shows the 95 % confidence interval.

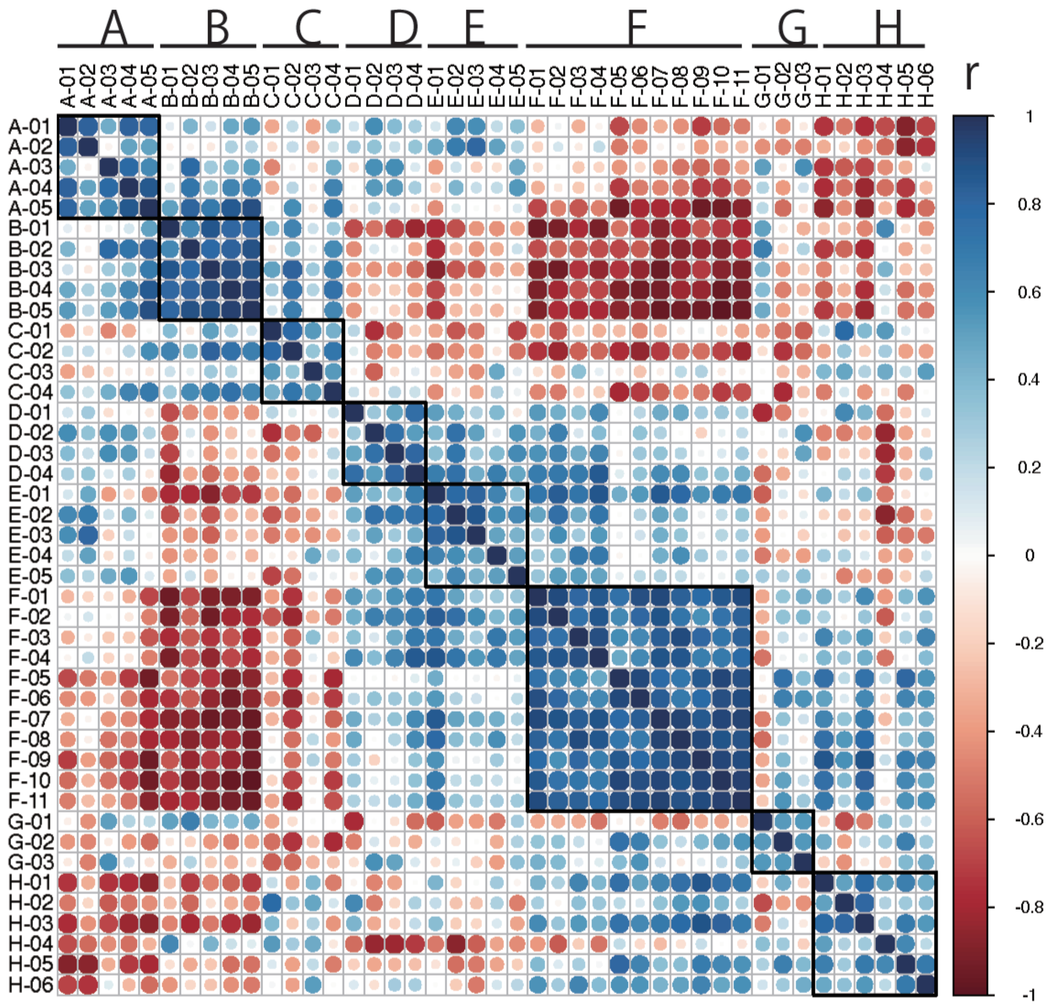


Supplementary Fig. 5 Matrix of Pearson's correlation between the means of each cluster.
Clusters sorted into eight groups by hierarchical clustering. Colour indicates Pearson's correlation.

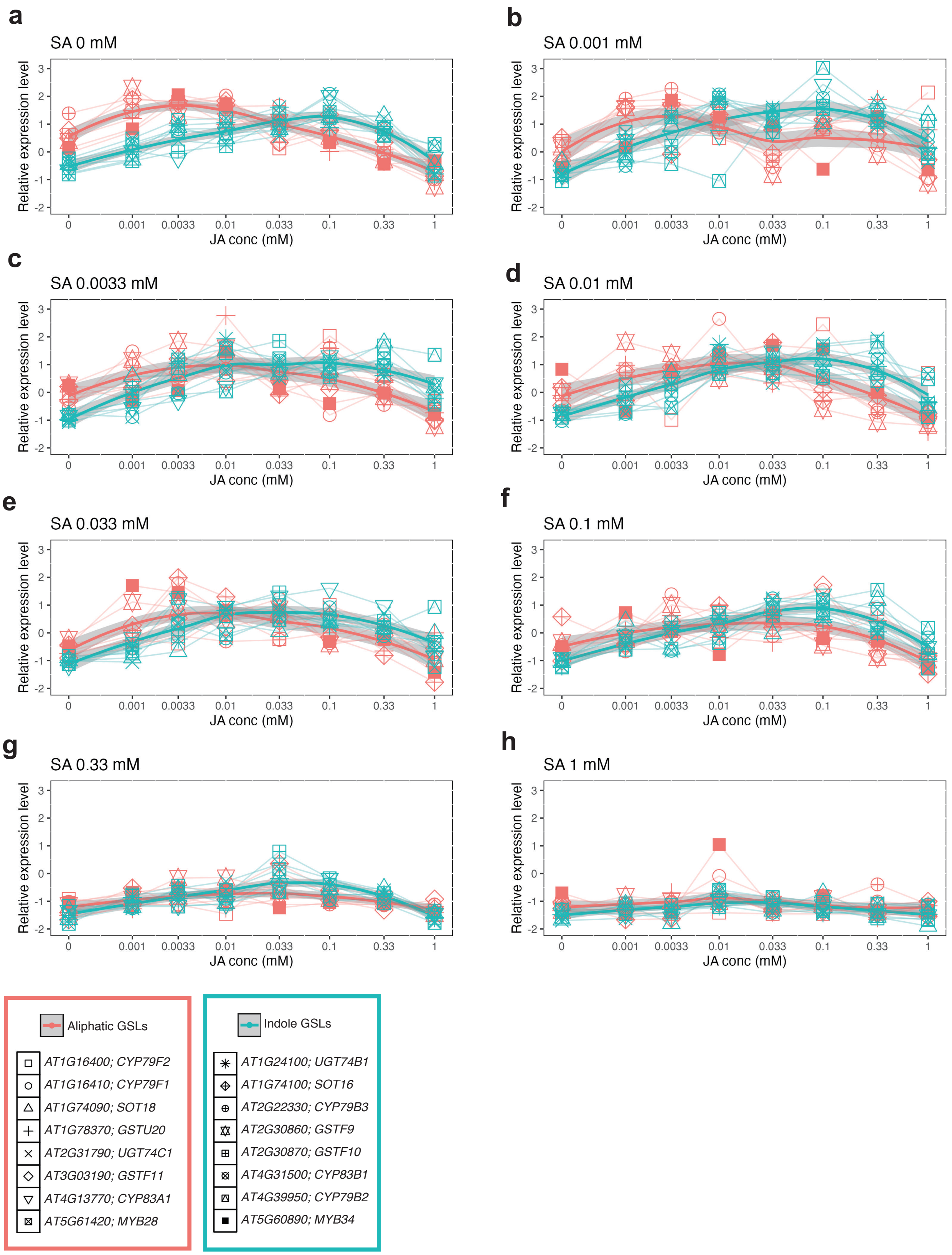


Supplementary Fig. 6 Relative expression levels of GSL synthesis genes.
(a–h) Relative expression levels of GSL synthesis genes at SA concentrations for 0 mM (a), 0.001 mM (b), 0.0033 mM (c), 0.01 mM (d), 0.033 mM (e), 0.1 mM (f), 0.33 mM (g), 1 mM (h). The horizontal axis indicates JA concentration (mM), and the vertical axis indicates relative expression level. Aliphatic GSL synthesis genes are shown in red. Indole GSL synthesis genes are shown in blue. Bold lines indicate LOESS curves drawn for each glucosinolate type.

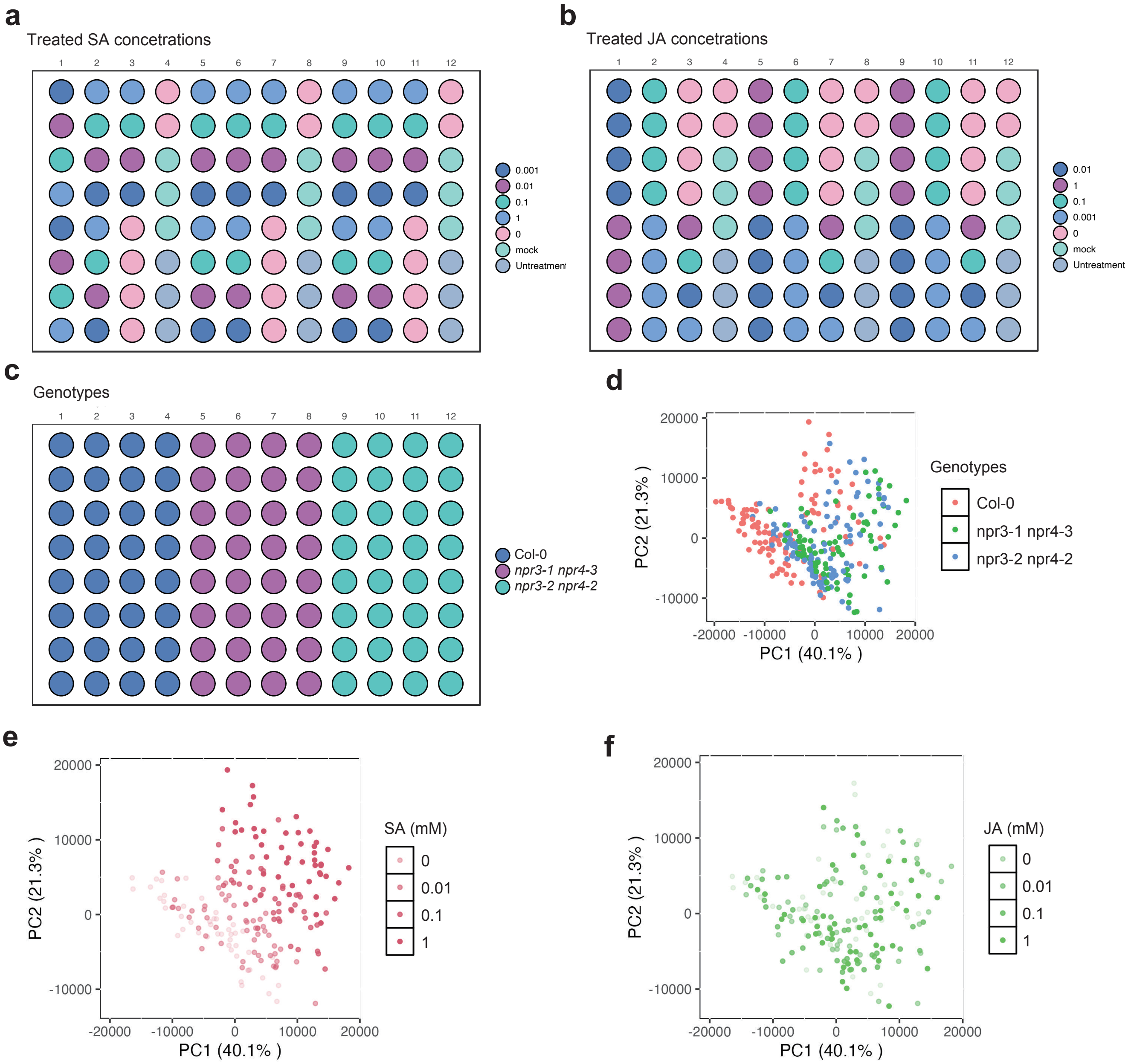


Supplementary Fig. 7 Summary of test data of transcriptomic biomarkers for SA/JA response states.

Layouts of SA treatment (a), JA treatment (b), and genotype (c) for test data. Seedlings were treated with SA and JA at 25 conditions that were made by systematically combining 5 concentrations of SA and JA: 0 mM, 0.001 mM, 0.01 mM, 0.1 mM, and 1 mM. (d–f) Principal component analysis of samples of test data. The color of each point indicates the genotypes (d), log-transformed SA concentrations (e) and JA concentrations (f).

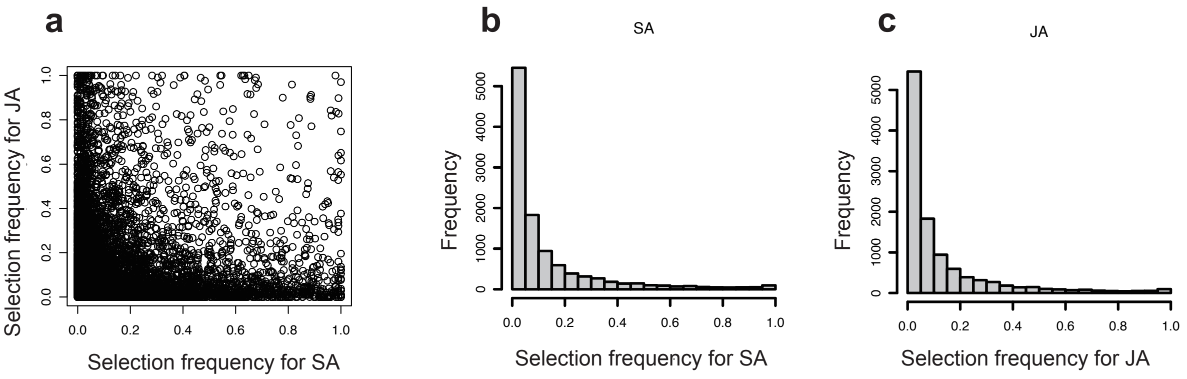


Supplementary Fig. 8 Selection frequency of genes in the transcriptomic biomarker.
(a) Scatter plot of selection frequency in the SA marker and JA marker. (b,c) Histogram of selection frequency in the SA marker (b) and JA marker (c).

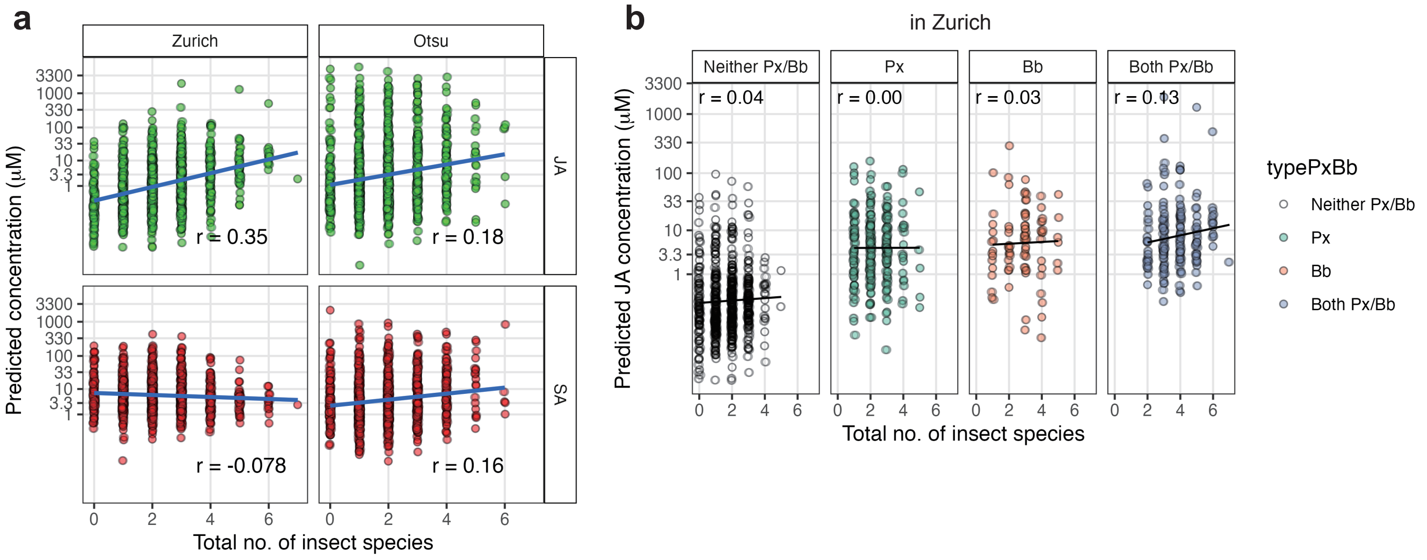


Supplementary Fig. 9 JA/SA responses was not directly explained by insect species richness.

In Zurich, the observed correlation between insect species richness and the JA response states was driven predominantly by a small number of strong JA-inducing species. These results suggest that the SA/JA responses are governed chiefly by the presence and functional traits of specific herbivores, rather than by overall community diversity.

(a) Relationship between insect species richness and predicted SA/JA reponse states across locations. X-axis: total number of insect species per plant; Y-axis: predicted SA/JA concentration (µM). Each point represents an individual plant. Blue lines show linear regression fits. r indicates Pearson correlation coefficient. Left column: Zurich (n = 1186); right column: Otsu (n = 1195).

(b) The relationship between total insect species richness and predicted JA concentration was analyzed separately for four categories based on the presence or absence of Plutella xylostella (Px) and Brevicoryne brassicae (Bb). Data shown are from Zurich samples. Each panel shows the scatter plot with regression line and Pearson correlation coefficient (r). Categories: Neither Px/Bb (neither species present), Px (only P. xylostella present), Bb (only B. brassicae present), and Both Px/Bb (both species present). Points represent individual plant samples.

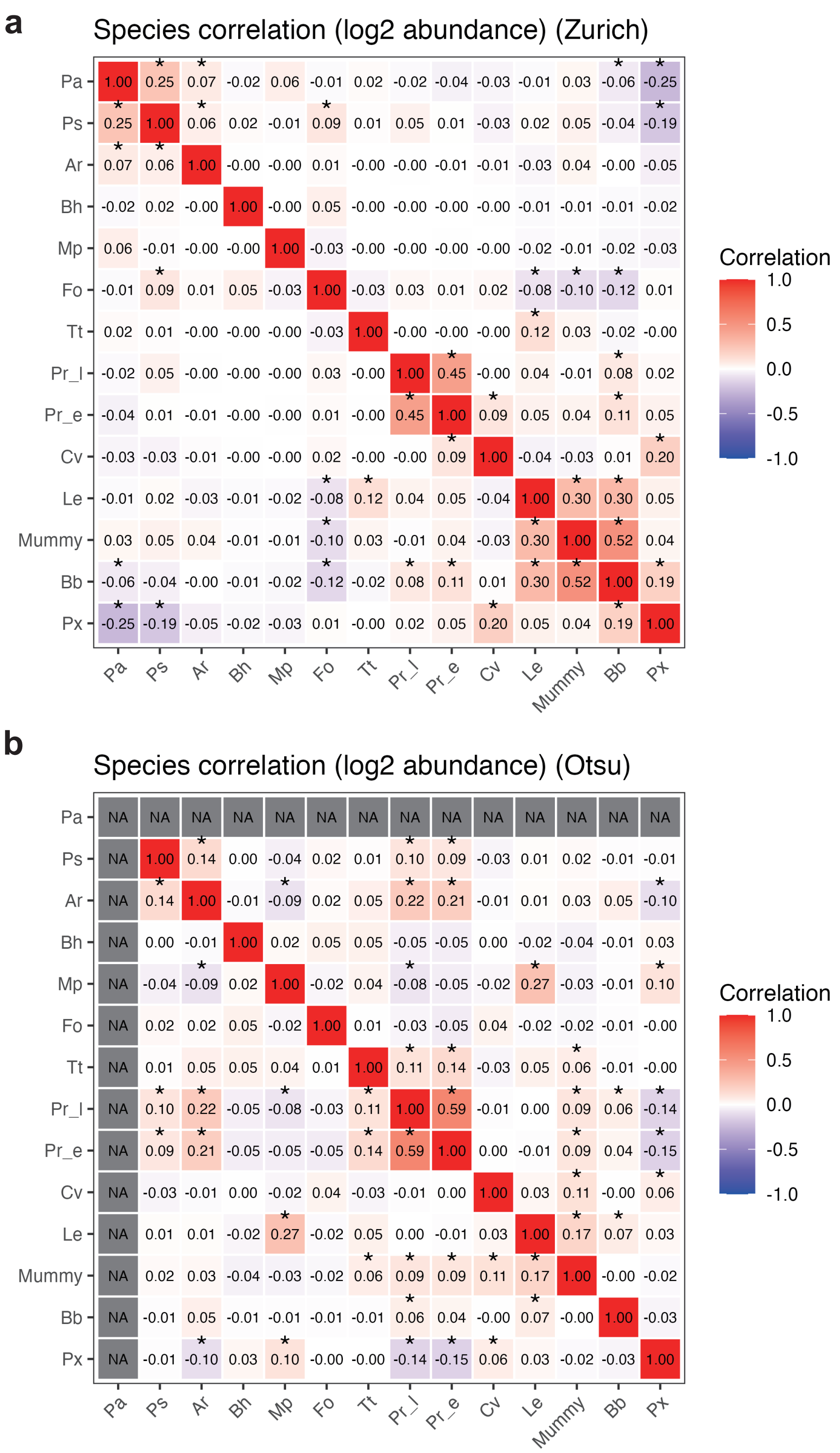


Supplementary Fig. 10 Comparison of insect herbivore site-specific co-occurrence patterns.

Correlation coefficient matrix in Zurich(a) and Otsu (c). Heatmap shows Pearson correlation coefficients (r) between $\log_2 x + 1$ -transformed No. of individual insect species for each species pair. Color gradient from blue (negative correlation) through white (no correlation) to red (positive correlation) represents association strength. Black asterisks denote statistical significance ($p < 0.05$). Positive correlations suggest co-occurrence tendencies, while negative correlations indicate competitive exclusion or habitat segregation. Across sites, wingless Lipaphis erysimi (‘Le’) and parasitoid-induced aphid mummies (‘Mummy’) consistently co-occurred (Zurich: r = 0.30; Otsu: r = 0.17), as expected for a host–parasitoid relationship, thereby providing an internal validation of the inferred co-occurrence patterns.

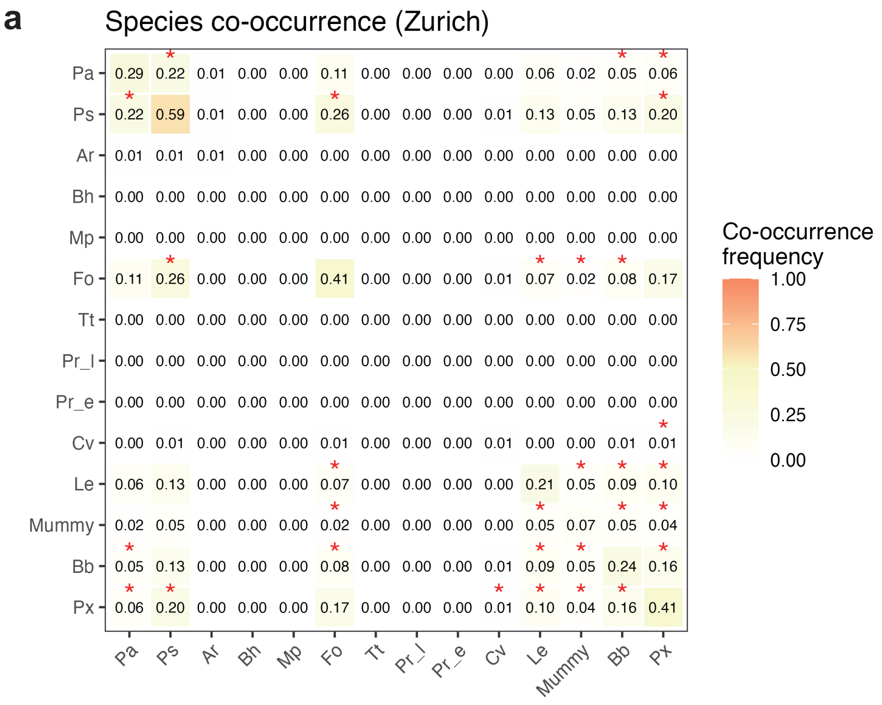


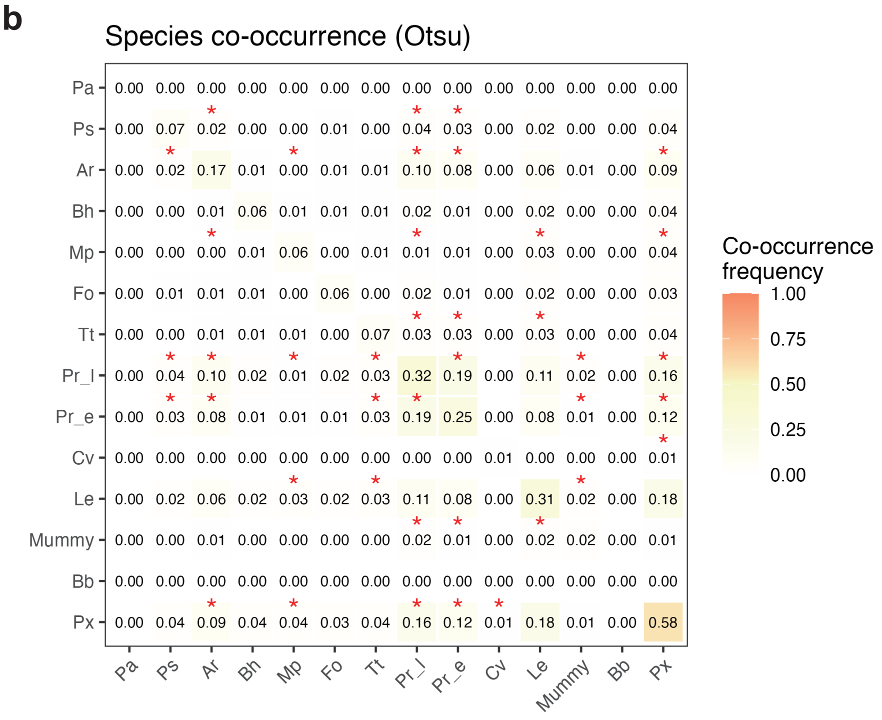


Supplementary Fig. 11 Comparison of insect herbivore site-specific co-occurrence frequency.

Co-occurrence frequency matrix in Zurich (b) and Otsu (d). Heatmap shows the frequency of co-occurrence for each species pair on individual plants. Values represent the proportion of samples (0-1) where both species were present simultaneously. Diagonal values indicate species prevalence. Color gradient from white (low) to orange (high) represents co-occurrence strength. Numbers indicate frequencies to two decimal places. Red asterisks denote significant associations ($\chi^2$ test, $p < 0.05$). Zurich: n = 1186 plants; Otsu: n = 1195 plants.

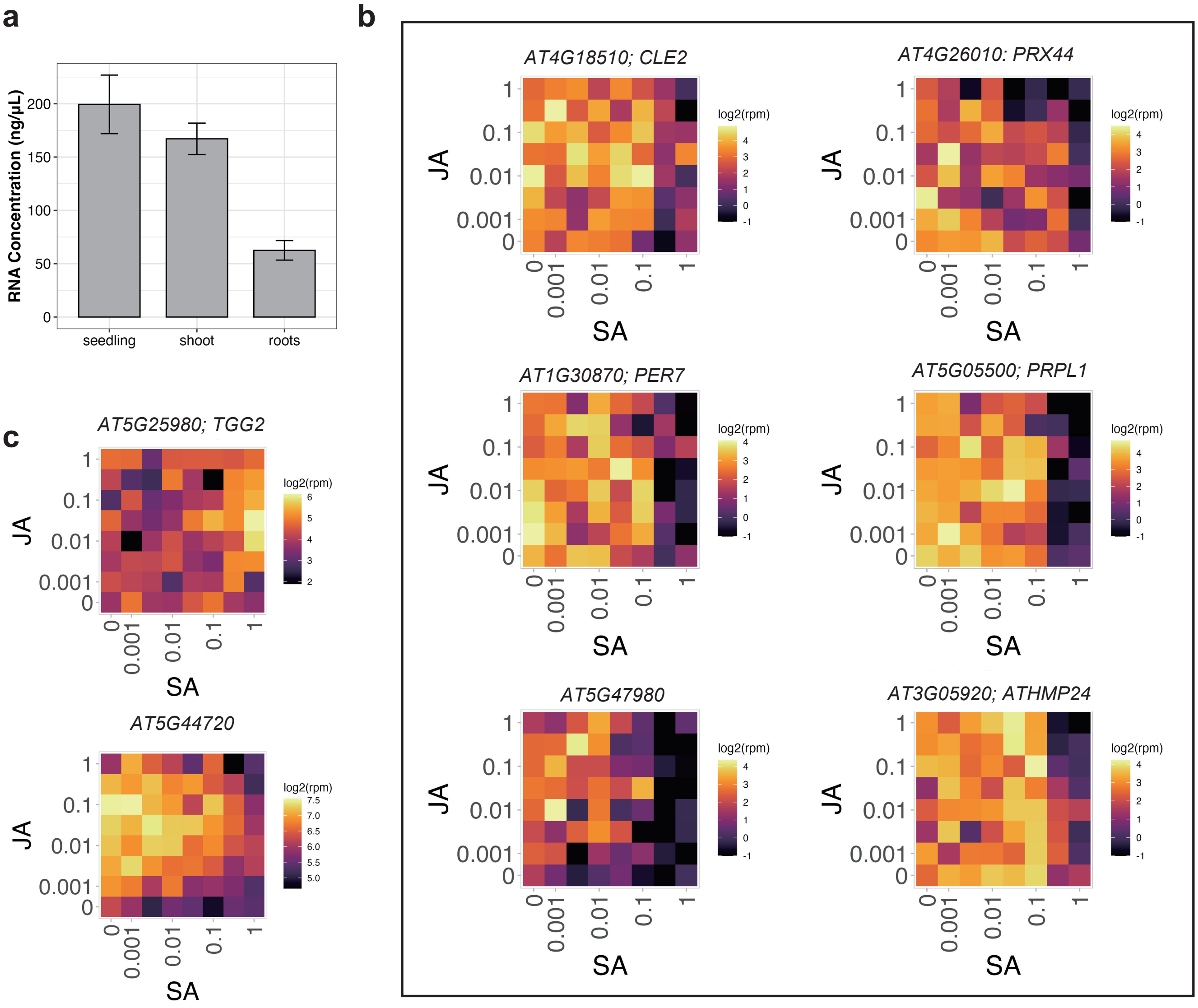


Supplementary Fig. 12 Gene expressions of root-specific genes were detectable.

(a) The total RNA contents of seedlings, shoots and roots. Error bars represent the standard deviation (n = 12). (b) Expression topographies of genes expressed specifically and highly in roots (A Christ et al., 2013). The horizontal axis of the heatmap corresponds to the concentration of SA, and the vertical axis to the concentration of JA. The color of each cel indicates the average log2(RPM + 1) for each concentration condition, n = 3~11.
