## Supplemental Figures S13 for "Transcriptomic biomarkers reveal jasmonic and salicylic acid state under field herbivory"

# AT1G02990.1 : NA

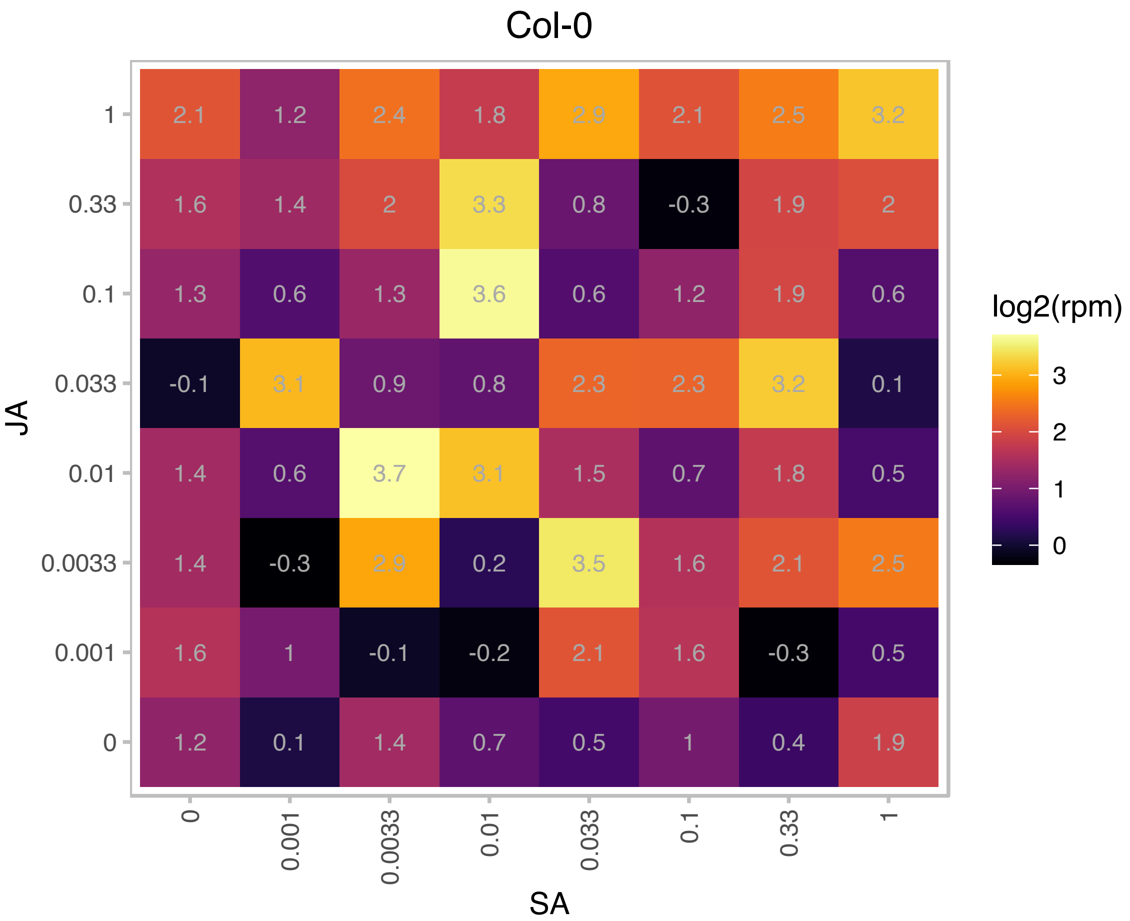


| | |
|---|---|
| Gene | AT1G02990 |
| is.expressed_genes | TRUE |
| is.responsive_genes | TRUE |
| is.apcluster | A-01 |

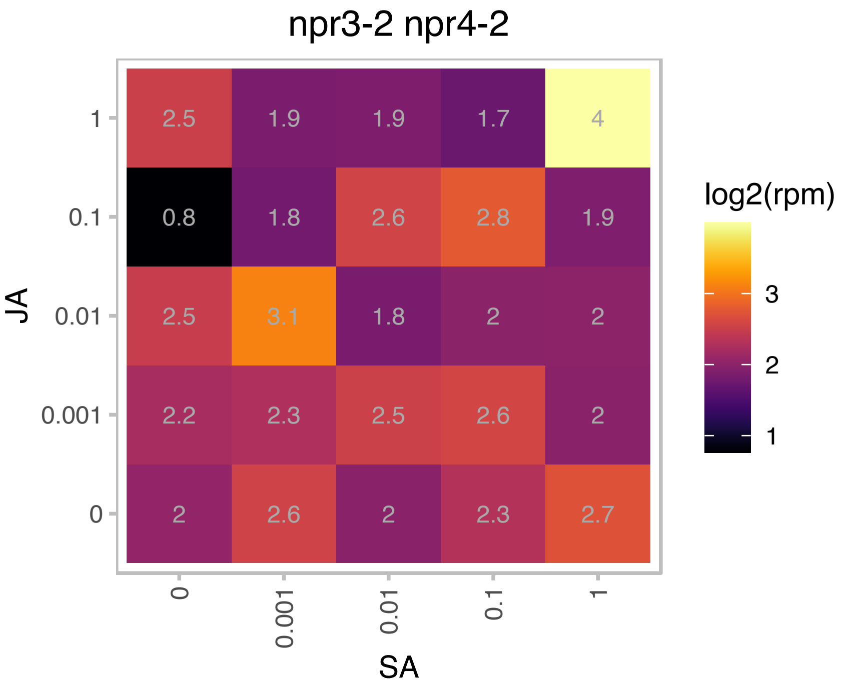


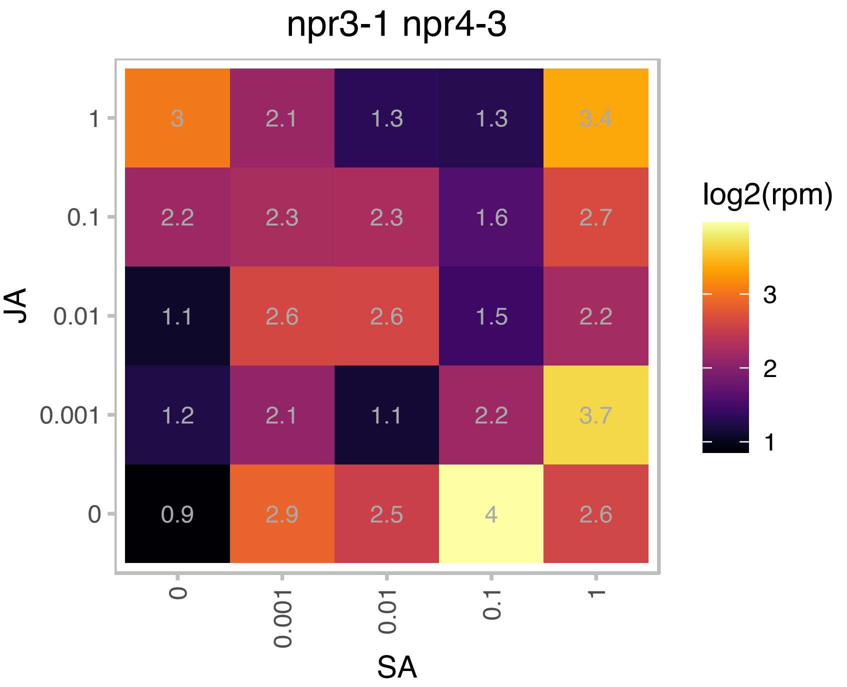

### AT1G05620.1 : NSH2, URH2

uridine-ribohydrolase 2

Encodes a cytosolic inosine nucleoside hydrolase. Transcript levels for this gene are elevated in older leaves suggesting that it may play a role in purine catabolism during senescence.

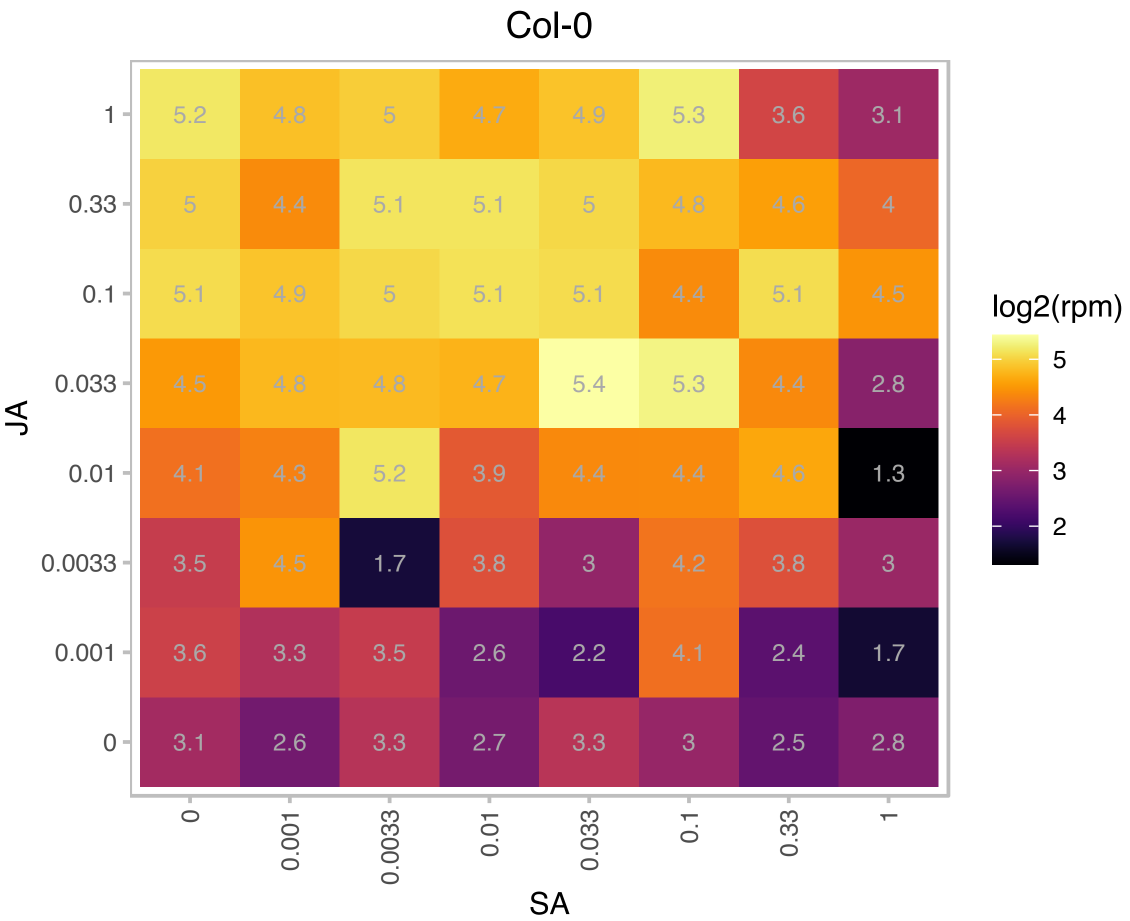


| | |
|---|---|
| Gene | AT1G05620 |
| is.expressed_genes | TRUE |
| is.responsive_genes | TRUE |
| is.apcluster | A-01 |

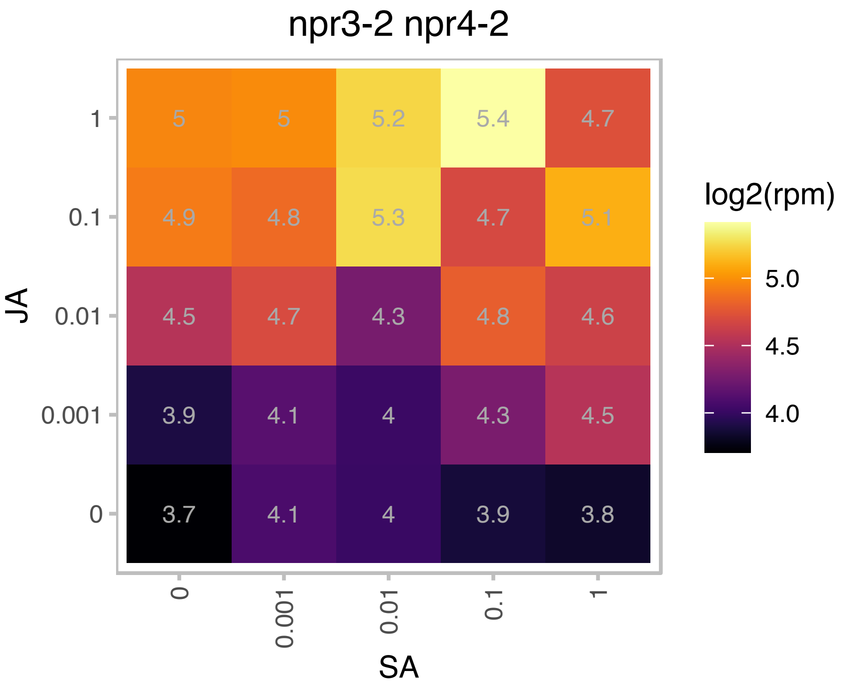


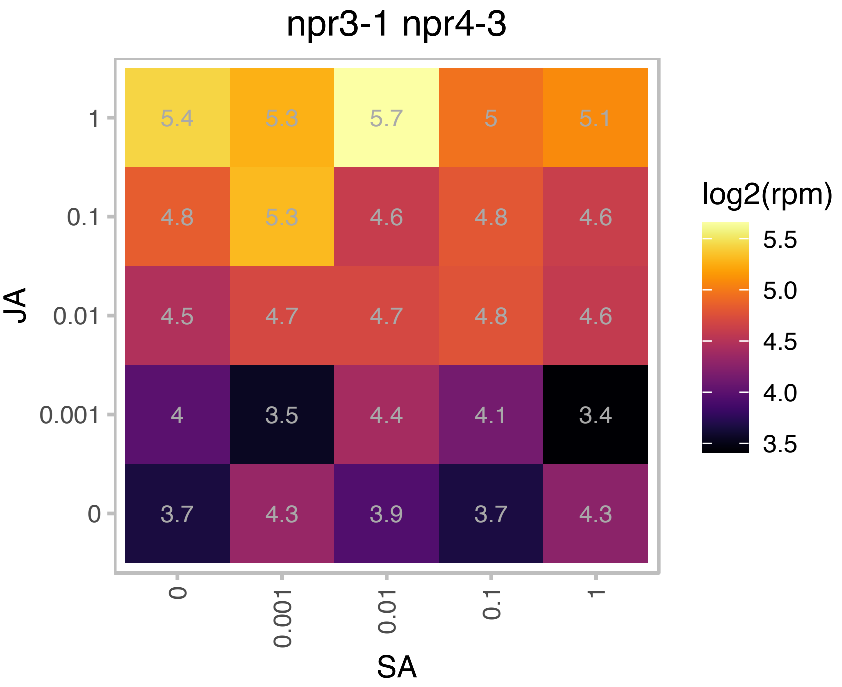

# AT1G07750.1 : NA

RmlC-like cupins superfamily protein

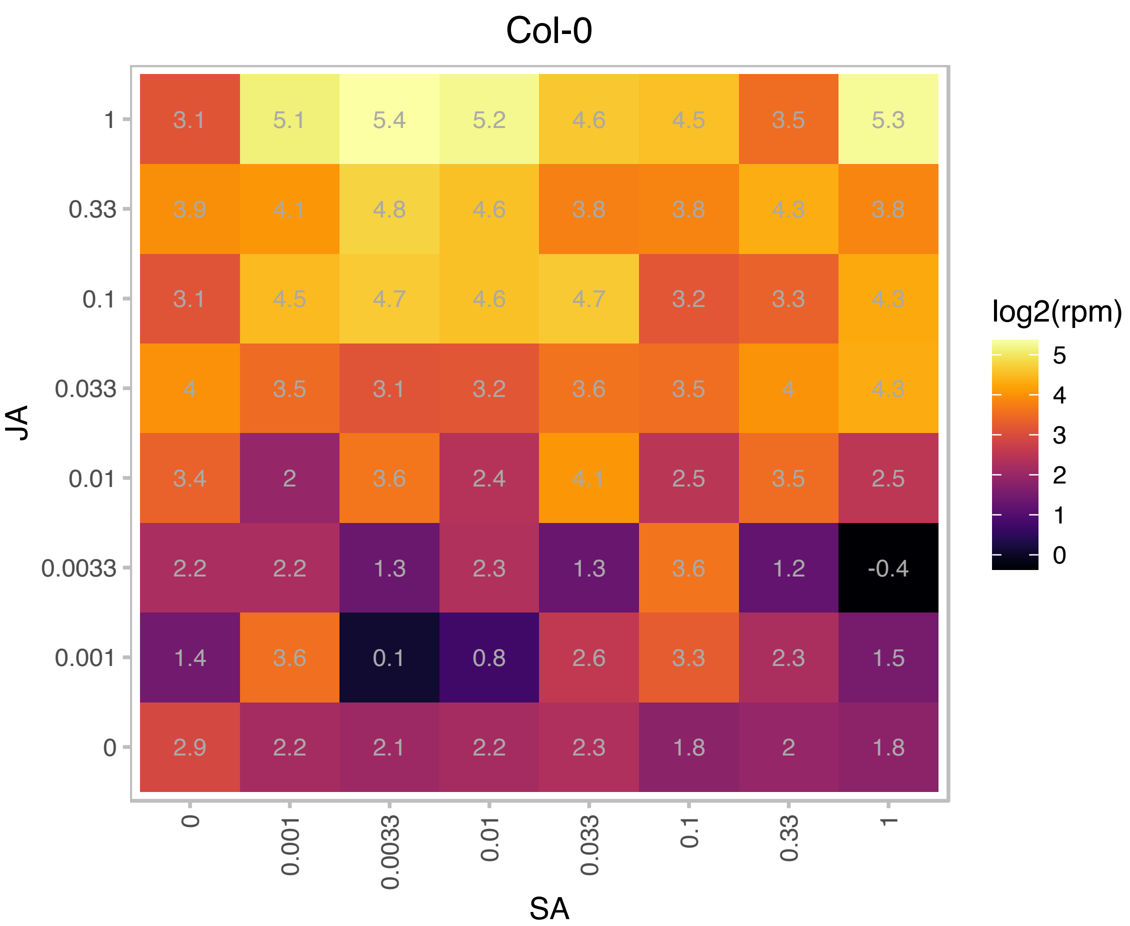


| | |
|---|---|
| Gene | AT1G07750 |
| is.expressed_genes | TRUE |
| is.responsive_genes | TRUE |
| is.apcluster | A-01 |

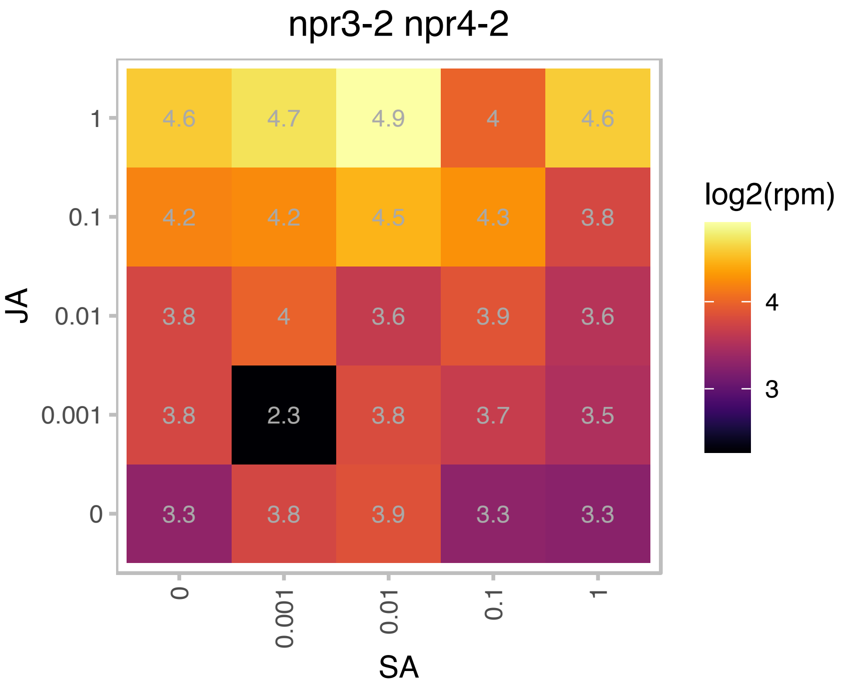


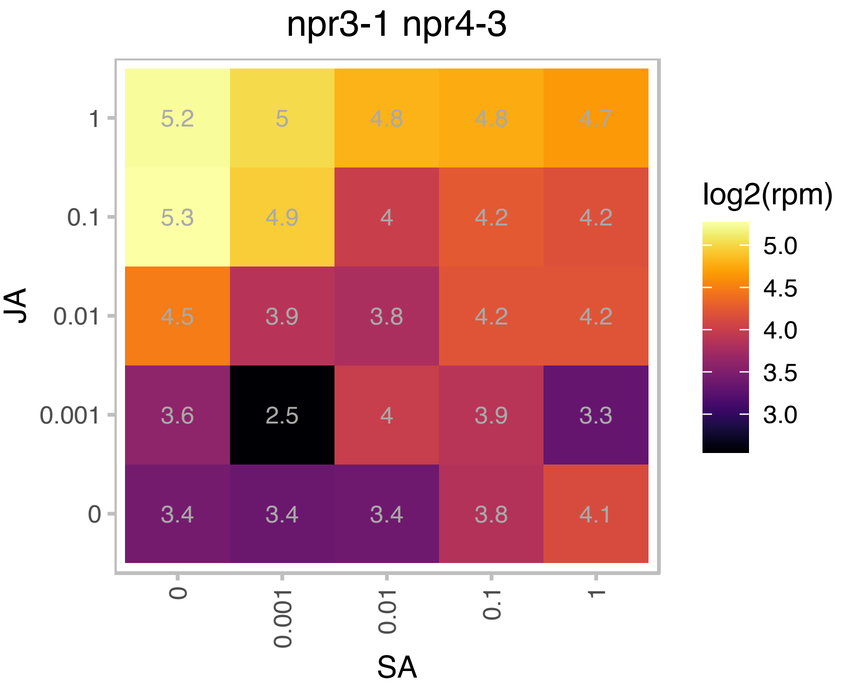

### AT1G14120.1 : AtDAO2

2-oxoglutarate (2OG) and Fe(II)-dependent oxygenase superfamily protein
DAO2 is an IAA oxidase expressed in root caps. it is a member of a family of dioxygenase and 2OG Fe(II) oxygenase domain and DAO domain containing proteins. It is expressed specifically in root cap cells and does not appear to be the major IAA oxidase in planta.

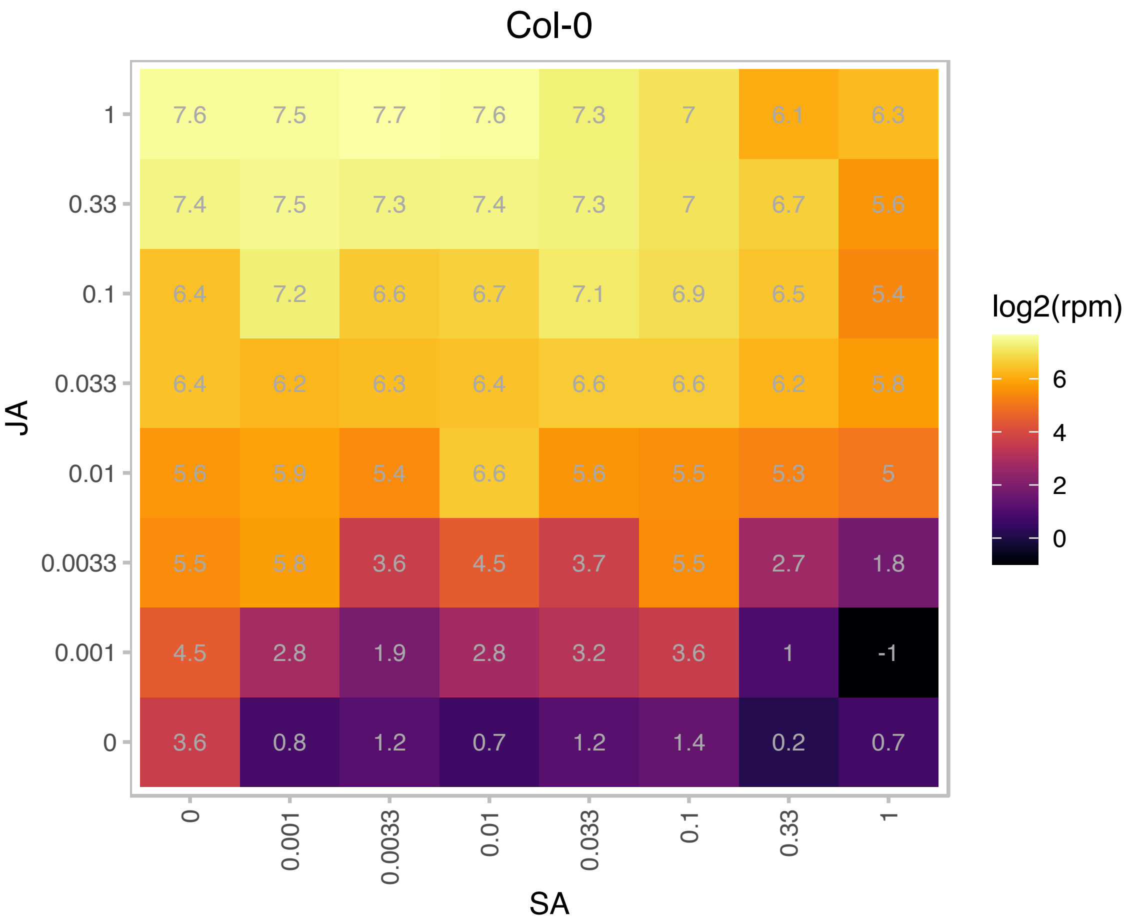


| | |
|---|---|
| Gene | AT1G14120 |
| is.expressed_genes | TRUE |
| is.responsive_genes | TRUE |
| is.apcluster | A-01 |

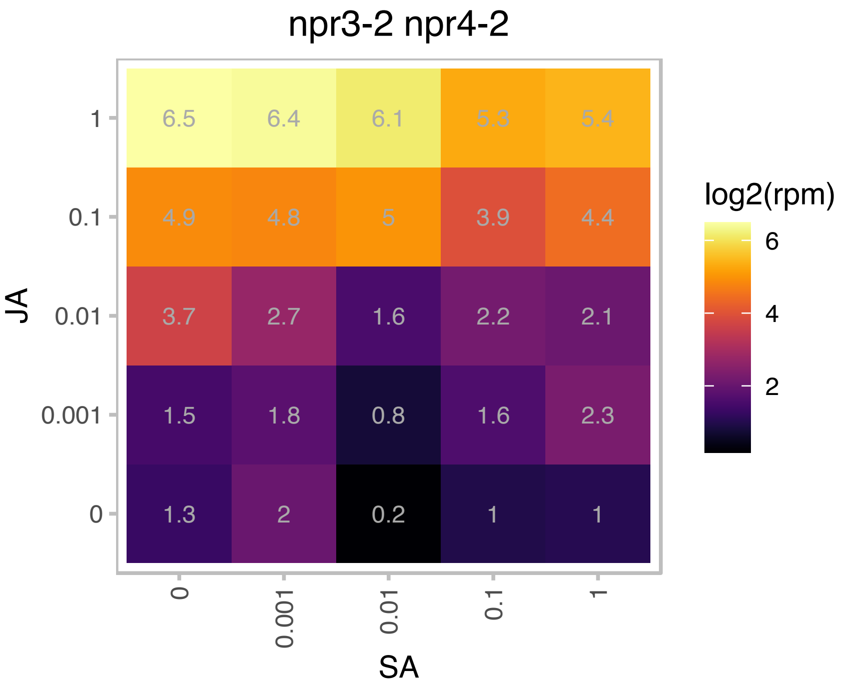


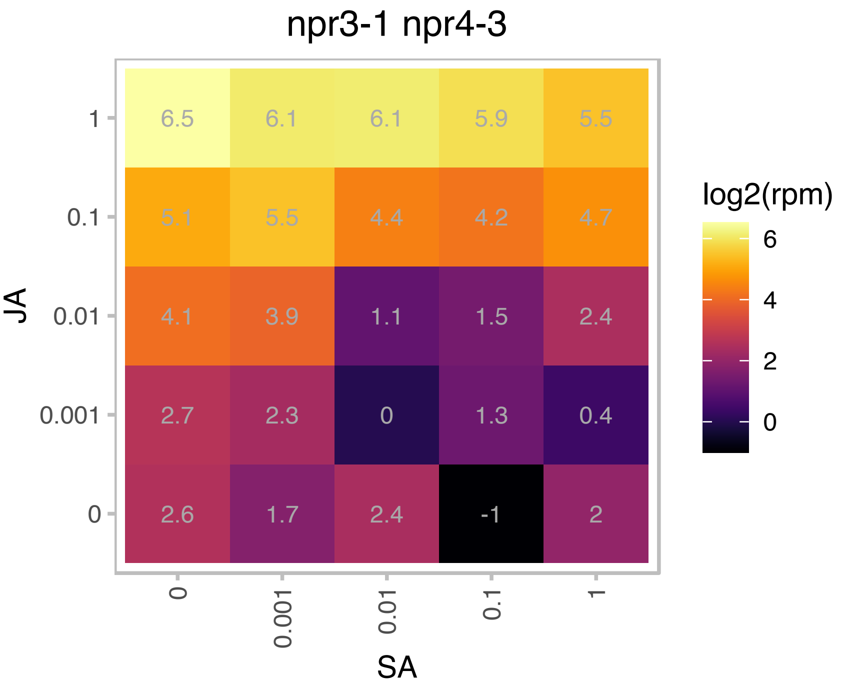

### AT1G18150.1 : ATMPK8, MPK8

Protein kinase superfamily protein

Encodes mitogen-activated protein kinase 8 (MPK8).  MPK8 connects protein phosphorylation, Ca2+, and ROS in the wound-signaling pathway.

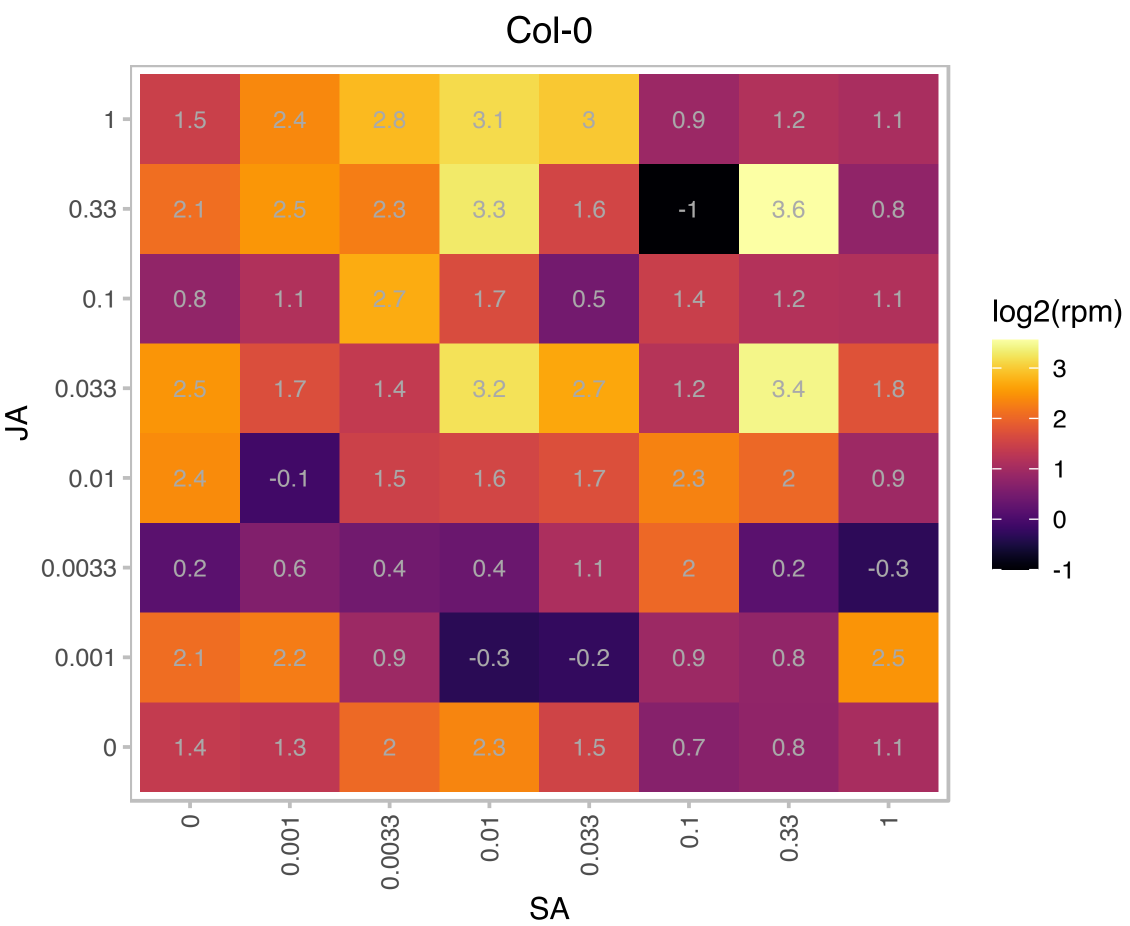


| | |
|---|---|
| Gene | AT1G18150 |
| is.expressed_genes | TRUE |
| is.responsive_genes | TRUE |
| is.apcluster | A-01 |

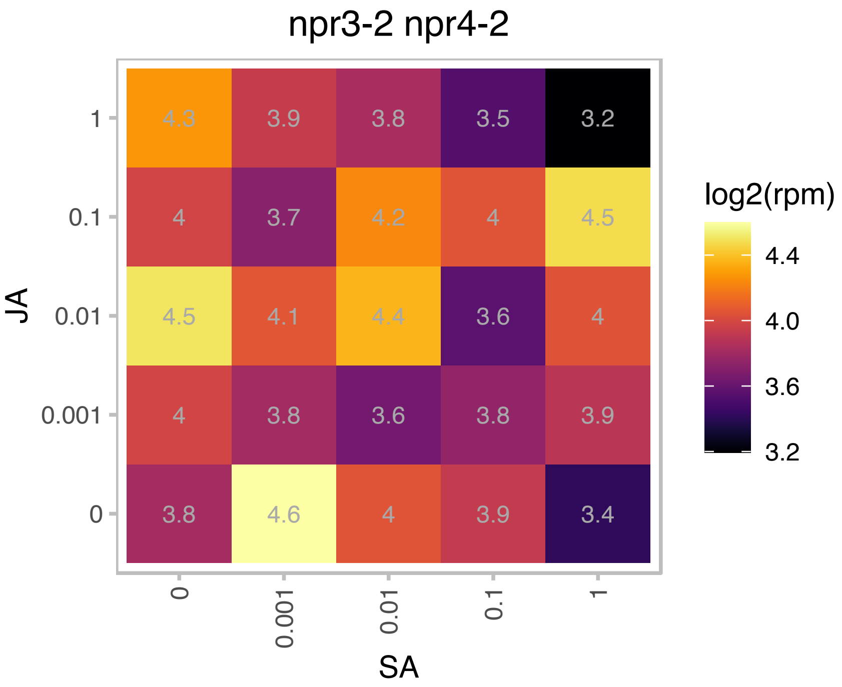


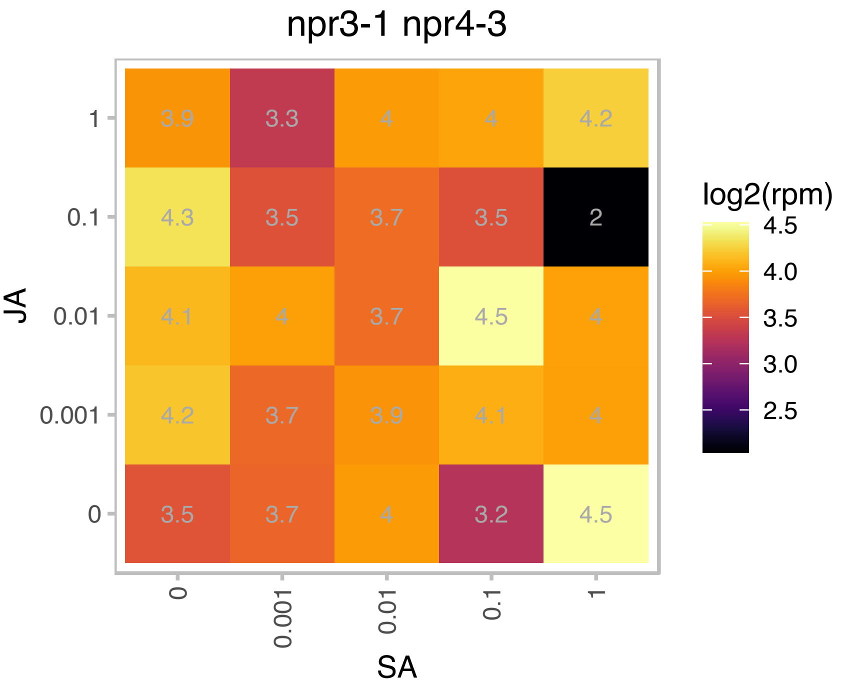

# AT1G23950.1 : NA

Protein of unknown function (DUF626)

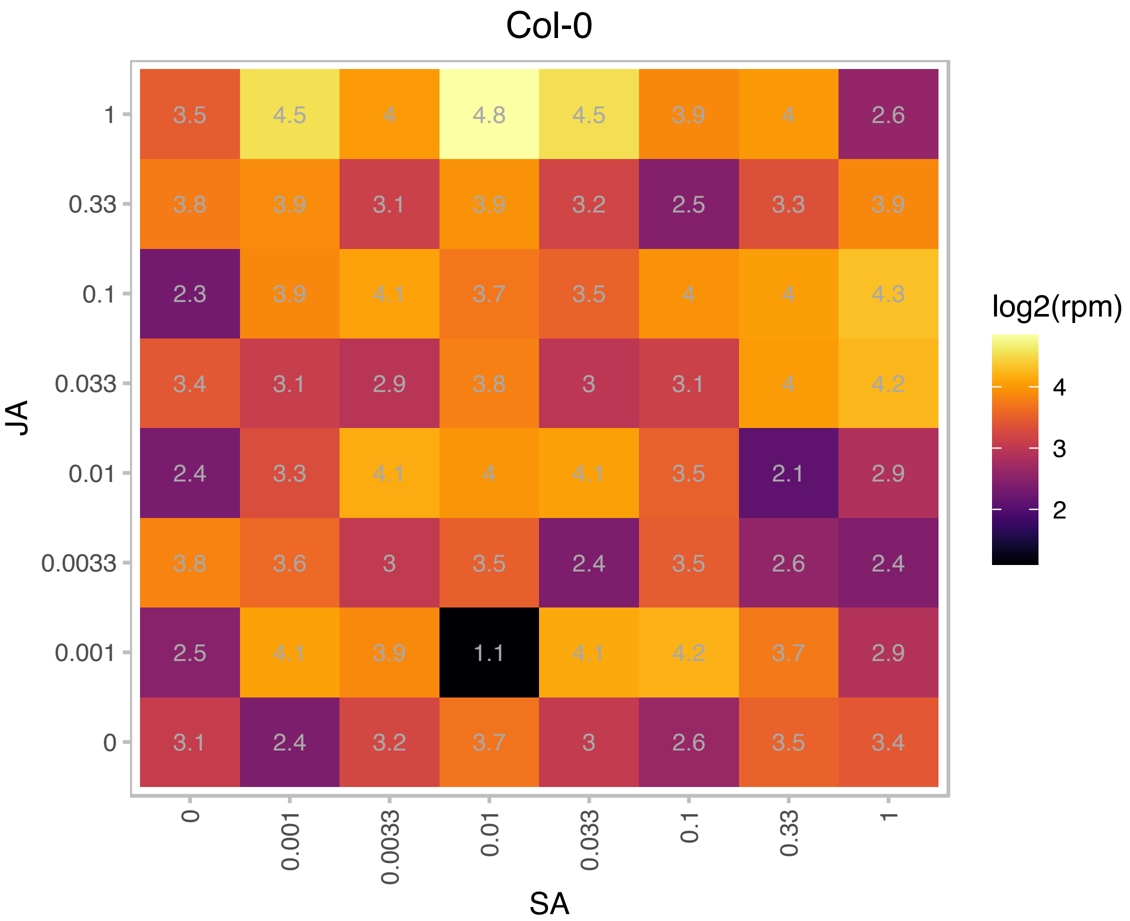


| | |
|---|---|
| Gene | AT1G23950 |
| is.expressed_genes | TRUE |
| is.responsive_genes | TRUE |
| is.apcluster | A-01 |

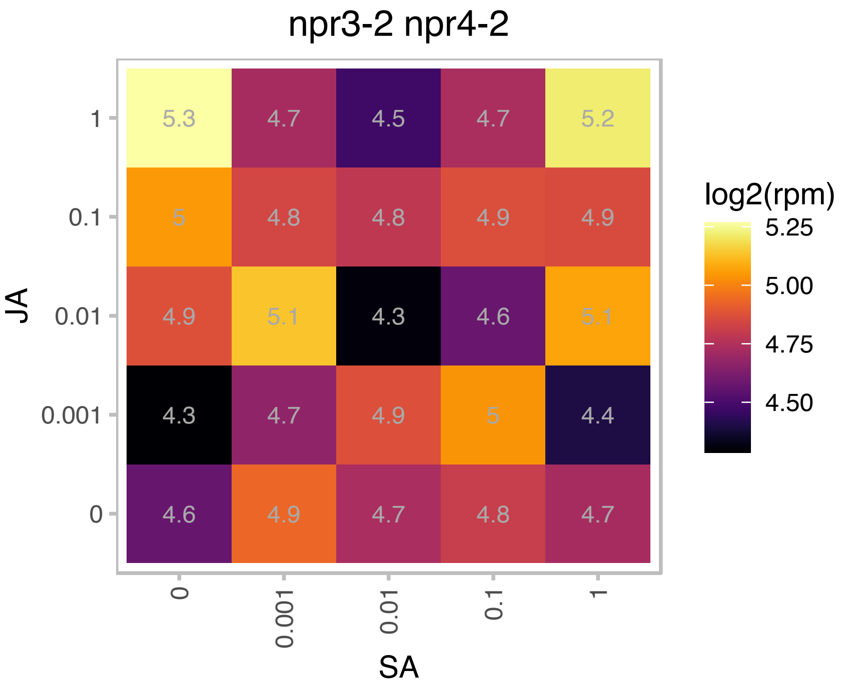


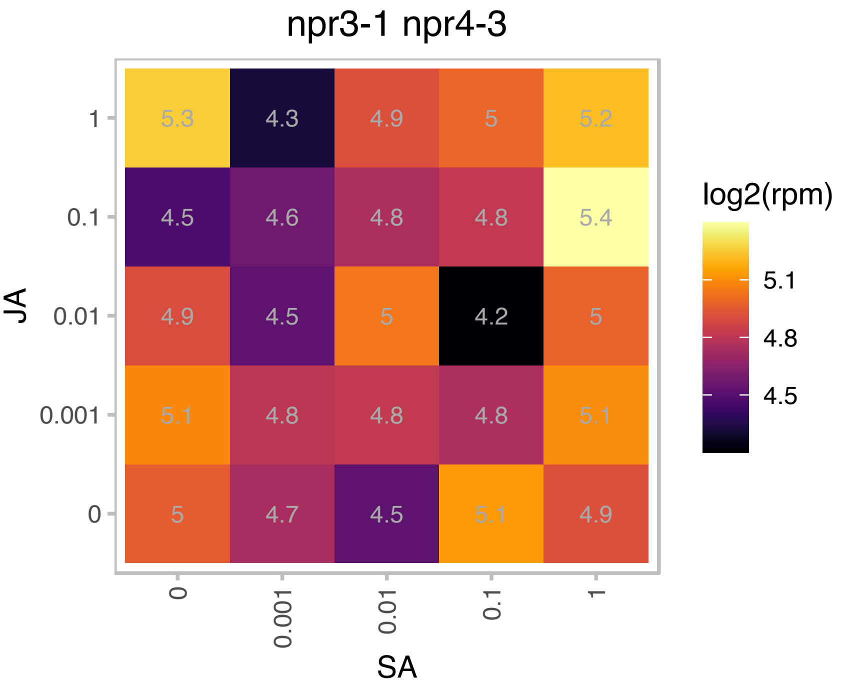

### AT1G30120.1 : PDH-E1 BETA

pyruvate dehydrogenase E1 beta

Encodes a putative plastid pyruvate dehydrogenase E1 beta subunit that is distinct from the mitochondrial pyruvate dehydrogenase E1 beta subunit.

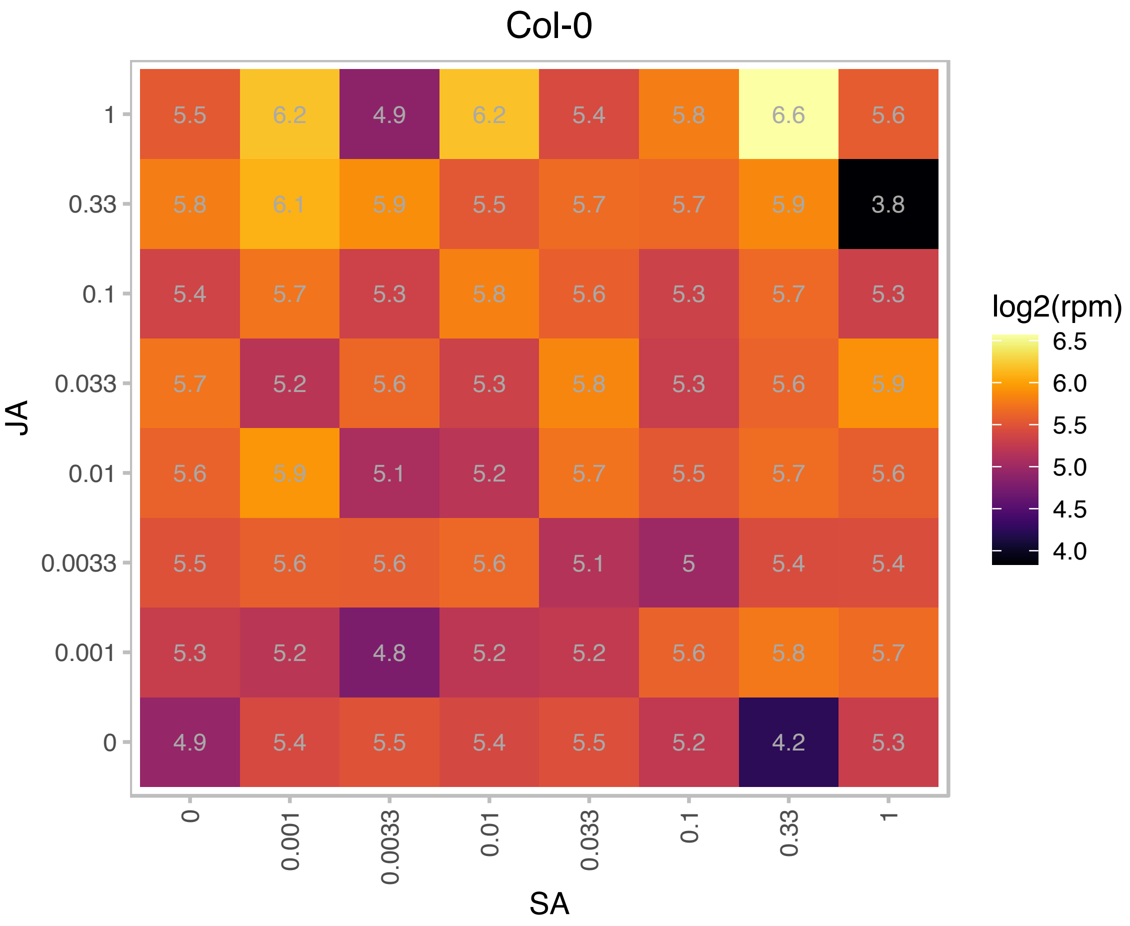


| | |
|---|---|
| Gene | AT1G30120 |
| is.expressed_genes | TRUE |
| is.responsive_genes | TRUE |
| is.apcluster | A-01 |

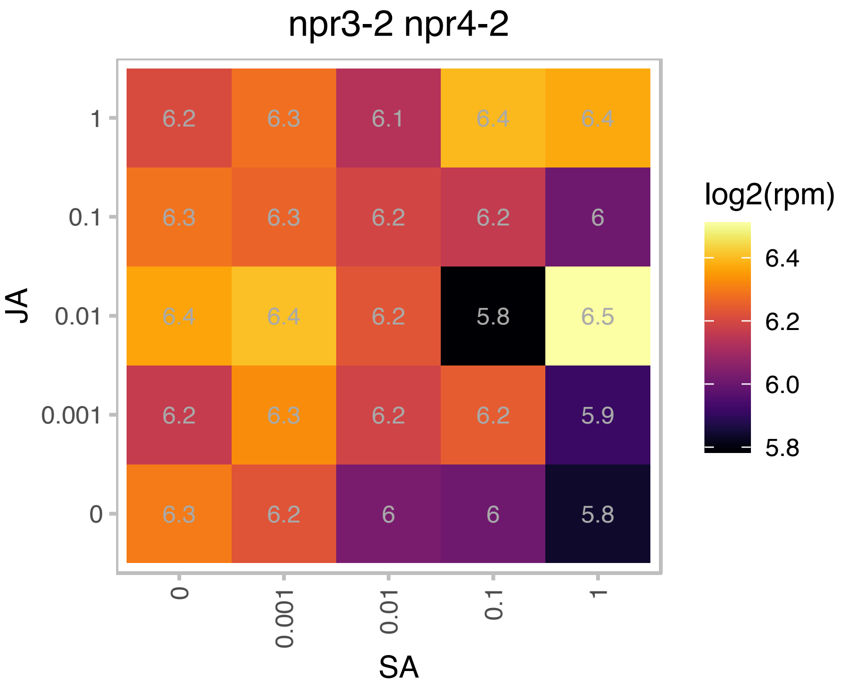


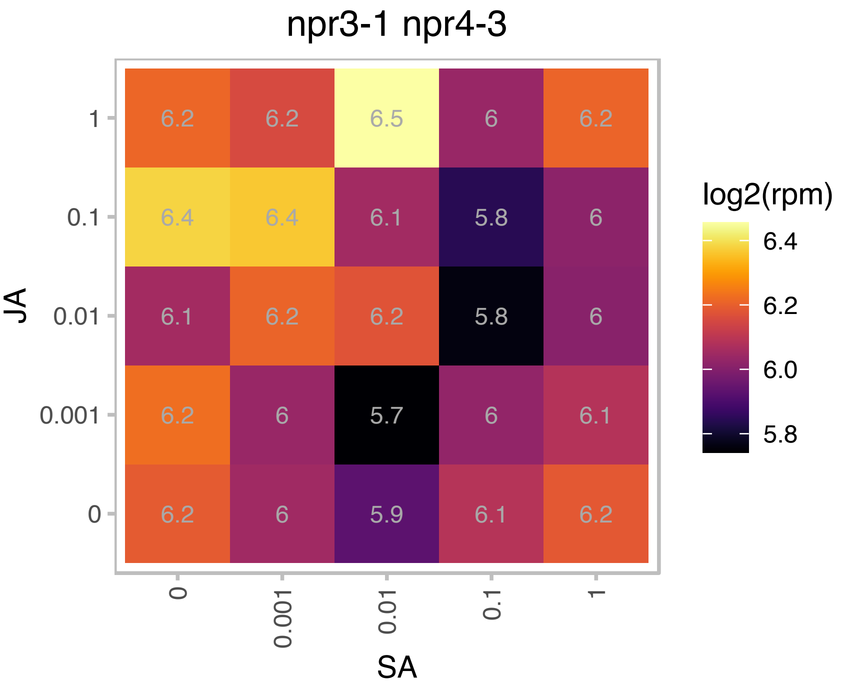

### AT1G44350.1 : ILL6

IAA-leucine resistant (ILR)-like gene 6

encodes a protein similar to IAA amino acid conjugate hydrolase.

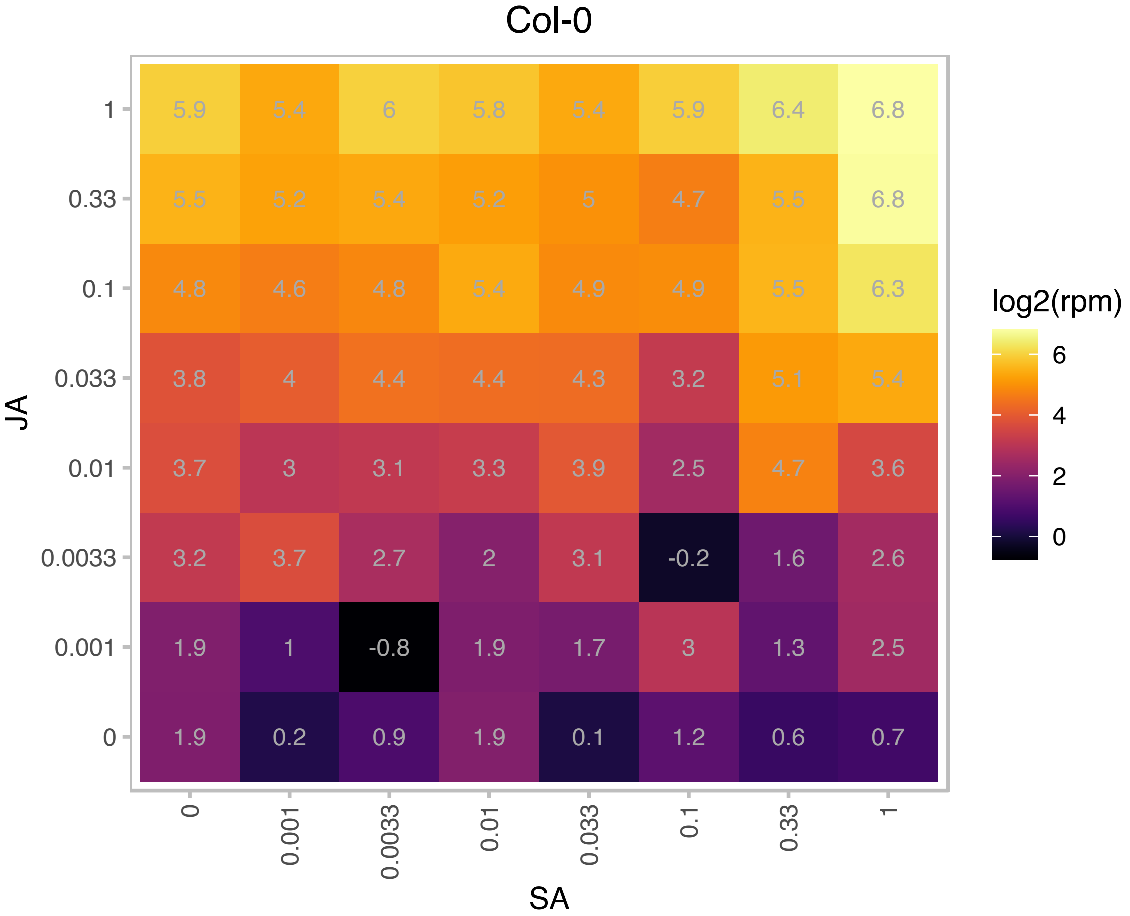


| | |
|---|---|
| Gene | AT1G44350 |
| is.expressed_genes | TRUE |
| is.responsive_genes | TRUE |
| is.apcluster | A-01 |

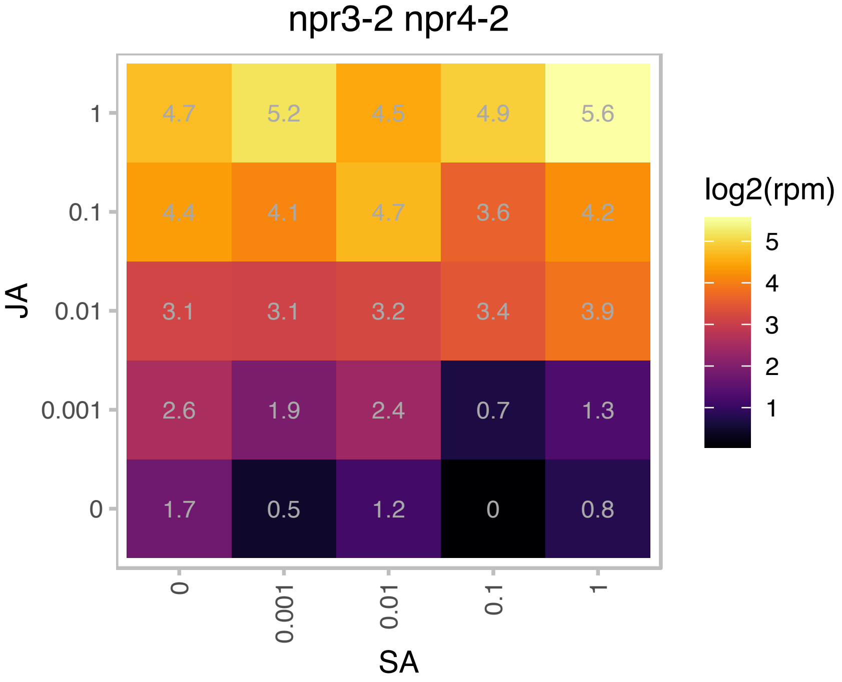


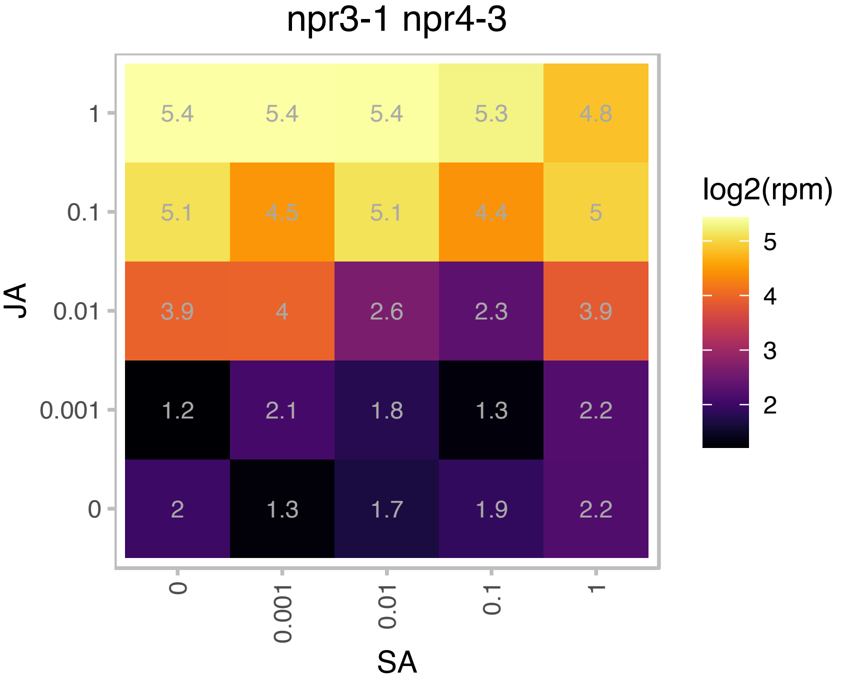

### AT1G50670.1 : OTU2

OTU-like cysteine protease family protein

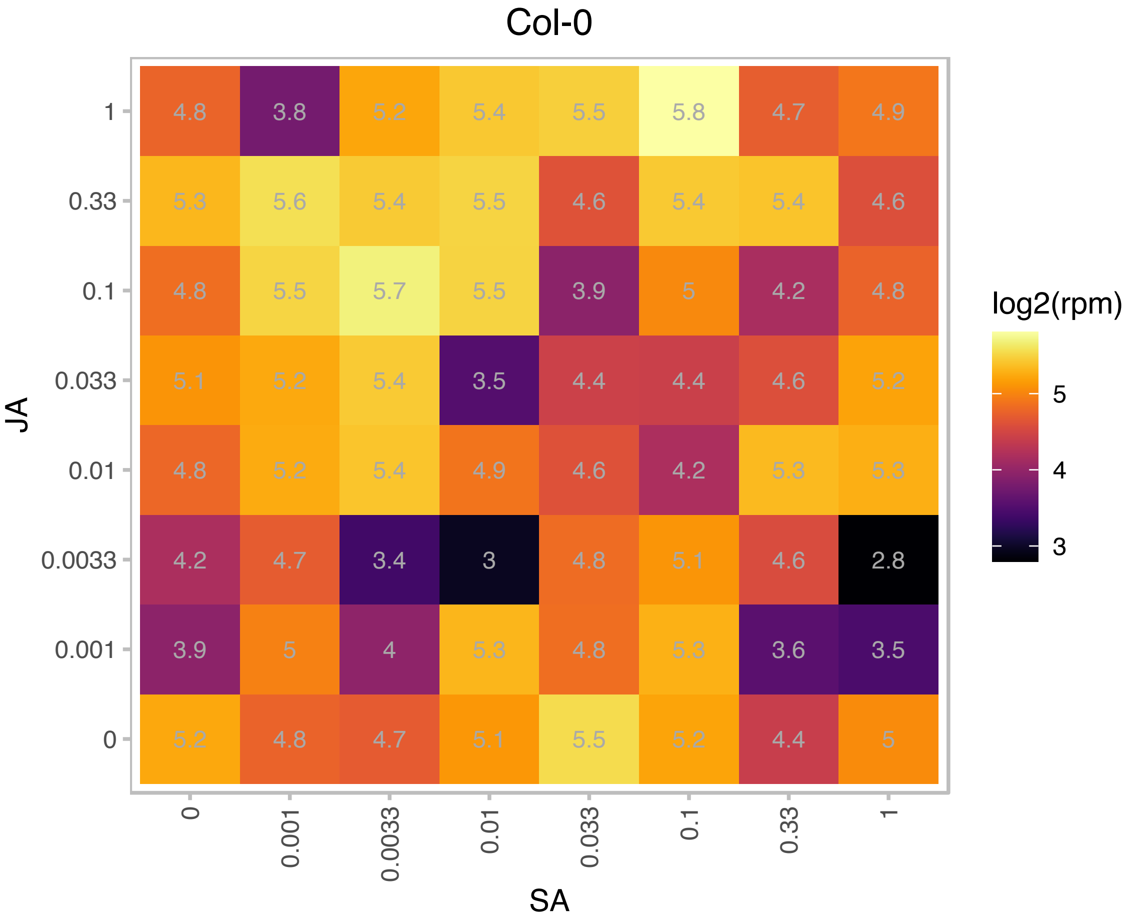


| | |
|---|---|
| Gene | AT1G50670 |
| is.expressed_genes | TRUE |
| is.responsive_genes | TRUE |
| is.apcluster | A-01 |

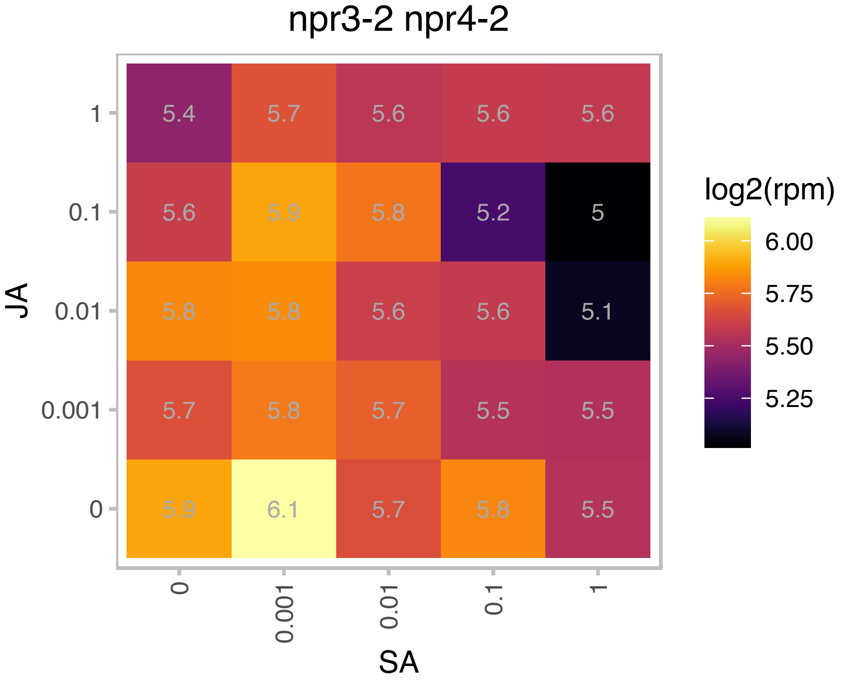


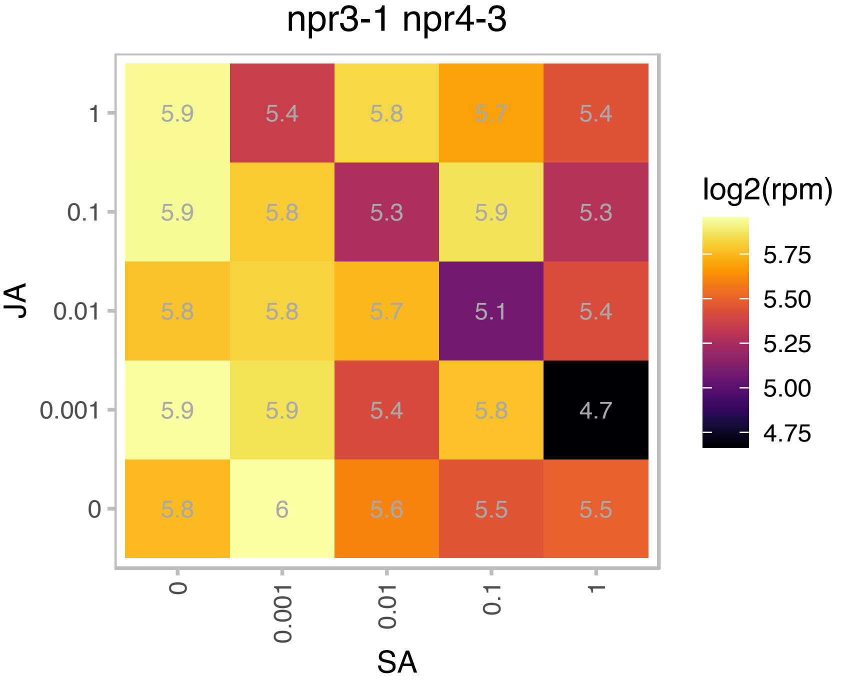

### AT1G52890.1 : ANAC019, ANAC19, NAC019

NAC domain containing protein 19

encodes a NAC transcription factor whose expression is induced by drought, high salt, and abscisic acid. This gene binds to ERD1 promoter in vitro.

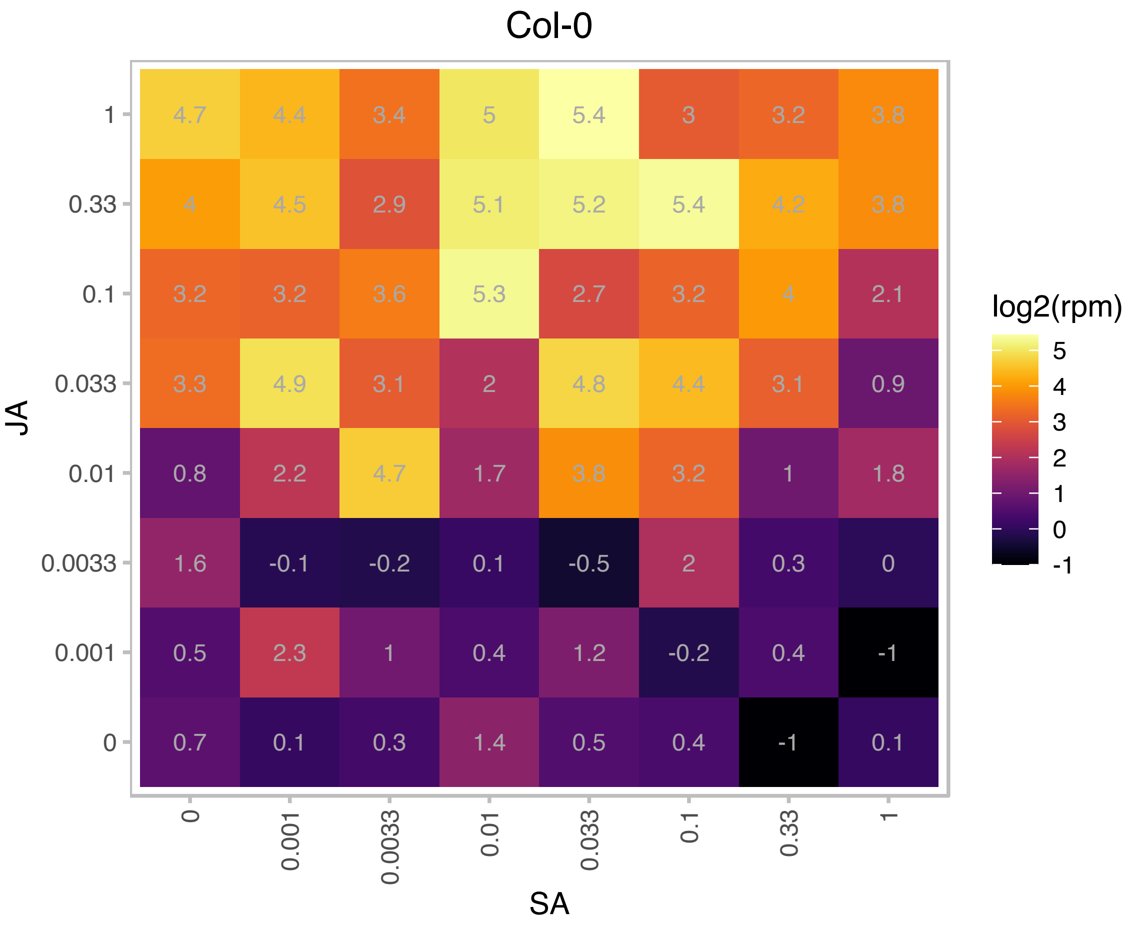


| | |
|---|---|
| Gene | AT1G52890 |
| is.expressed_genes | TRUE |
| is.responsive_genes | TRUE |
| is.apcluster | A-01 |

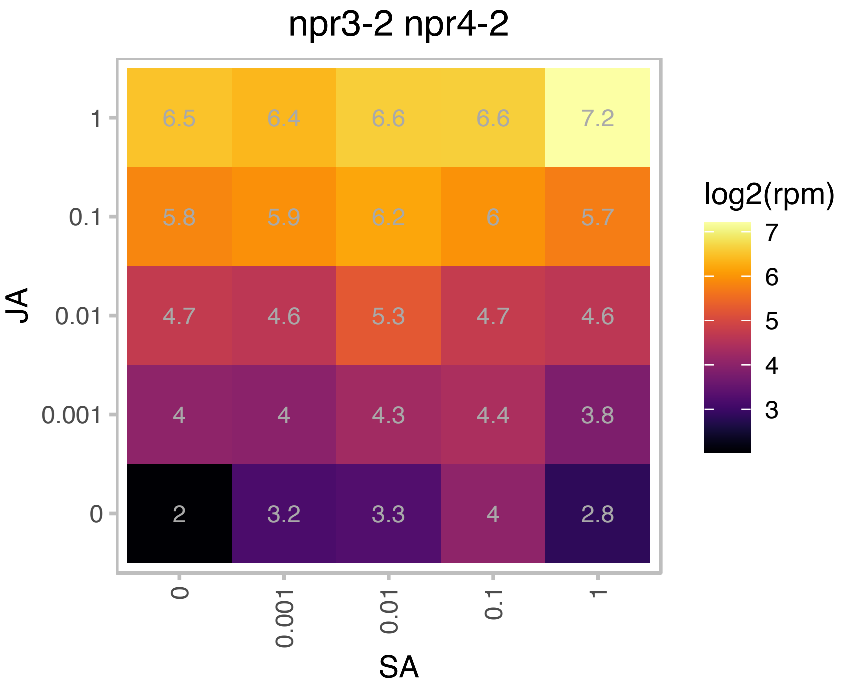


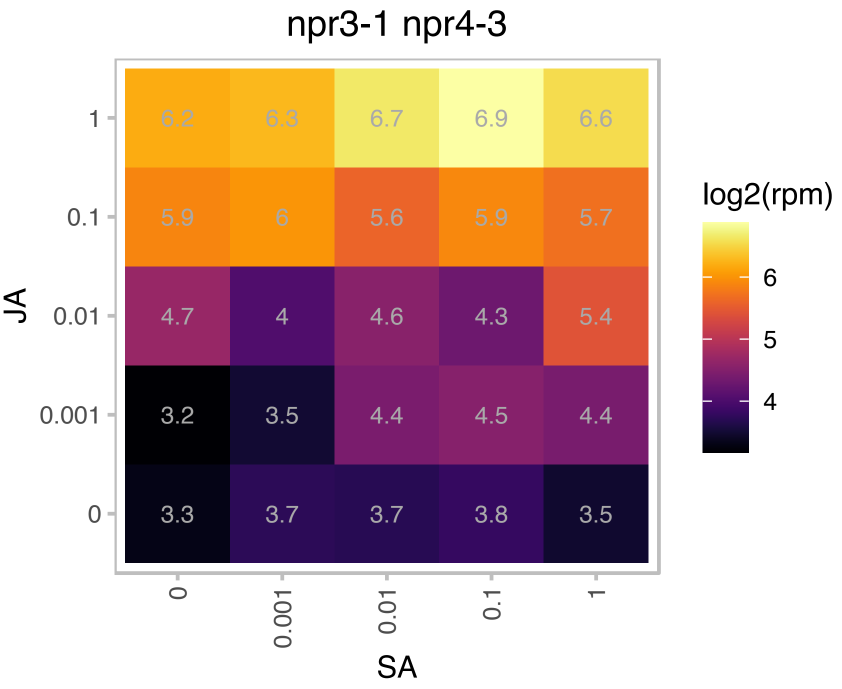

### AT1G67070.1 : DIN9, PMI2

Mannose-6-phosphate isomerase, type I

Encodes a protein with phosphomannose isomerase activity that is involved in synthesis of ascorbic acid. Expression is induced after 24 hours of dark treatment, in senescing leaves and treatment with exogenous photosynthesis inhibitor. Induction of gene expression was suppressed in excised leaves supplied with sugar. The authors suggest that the gene's expression pattern is responding to the level of sugar in the cell.

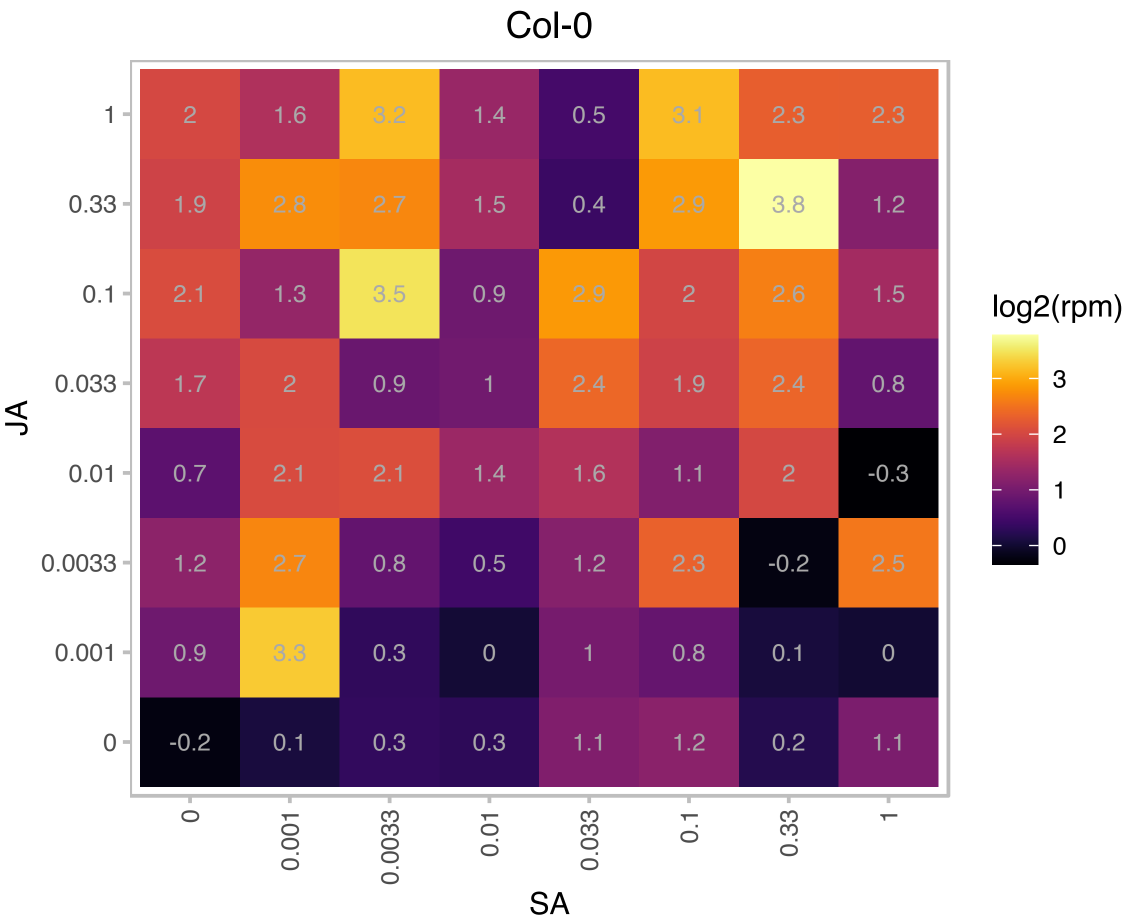


| | |
|---|---|
| Gene | AT1G67070 |
| is.expressed_genes | TRUE |
| is.responsive_genes | TRUE |
| is.apcluster | A-01 |

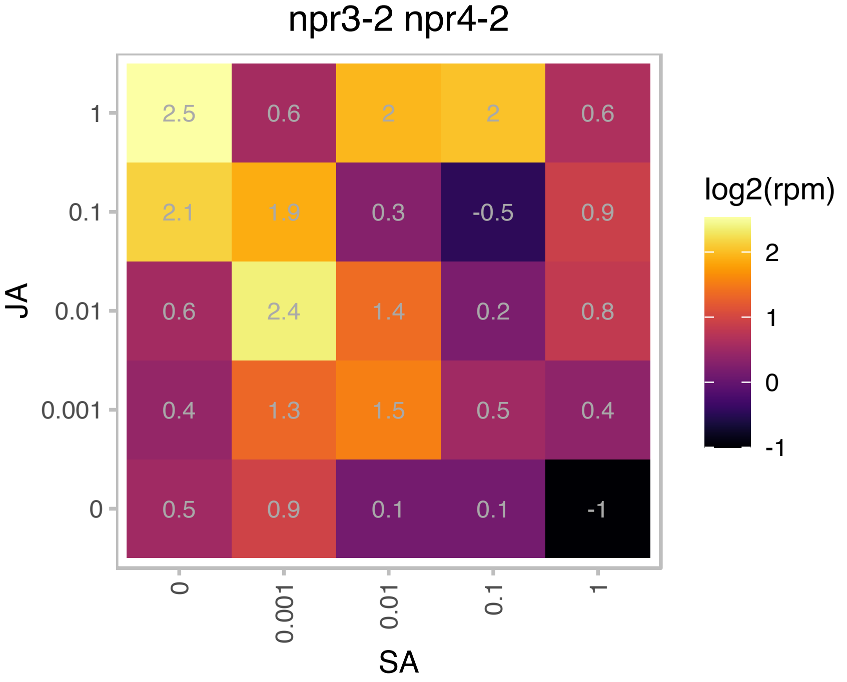


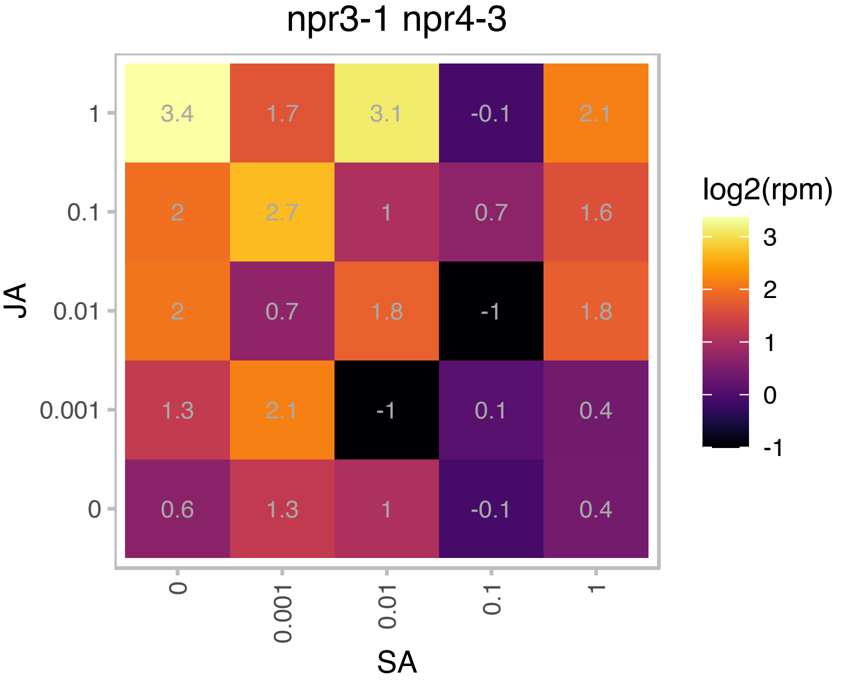

### AT1G69010.1 : BIM2

BES1-interacting Myc-like protein 2

Encodes BES1-INTERACTING MYC-LIKE 2 (BIM2), a PAR1 (PHYTOCHROME RAPIDLY REGULATED 1)-interacti
positively modulates the shade avoidance syndrome in Arabidopsis seedlings.

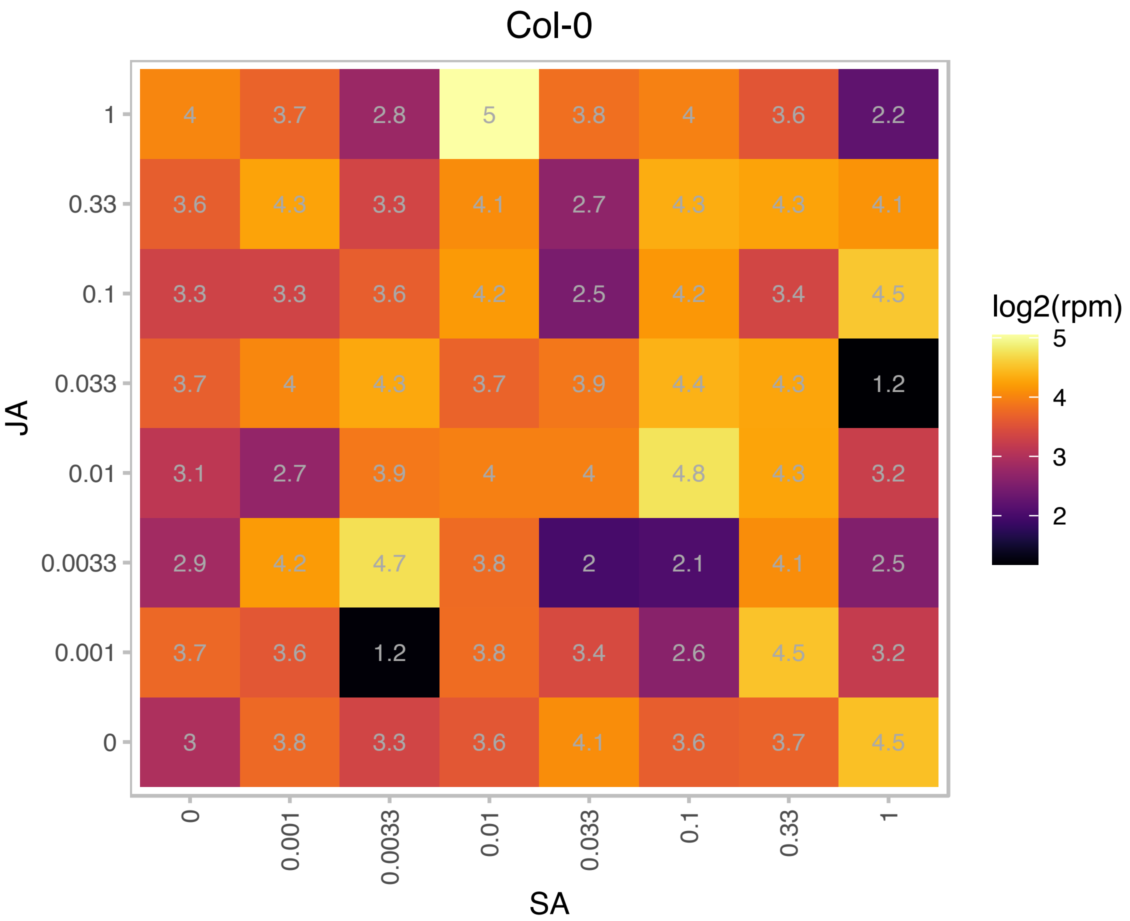


| | |
|---|---|
| Gene | AT1G69010 |
| is.expressed_genes | TRUE |
| is.responsive_genes | TRUE |
| is.apcluster | A-01 |

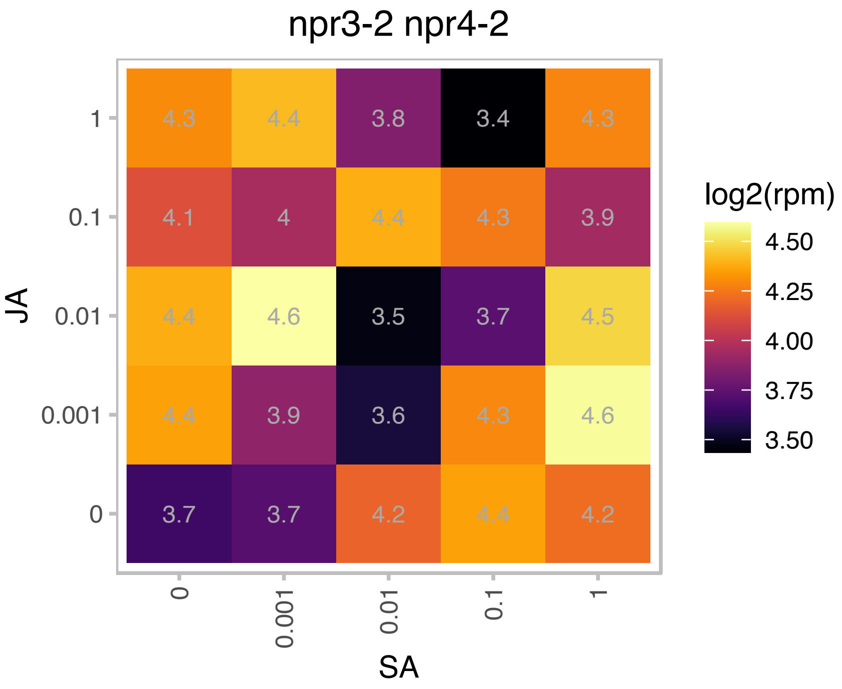


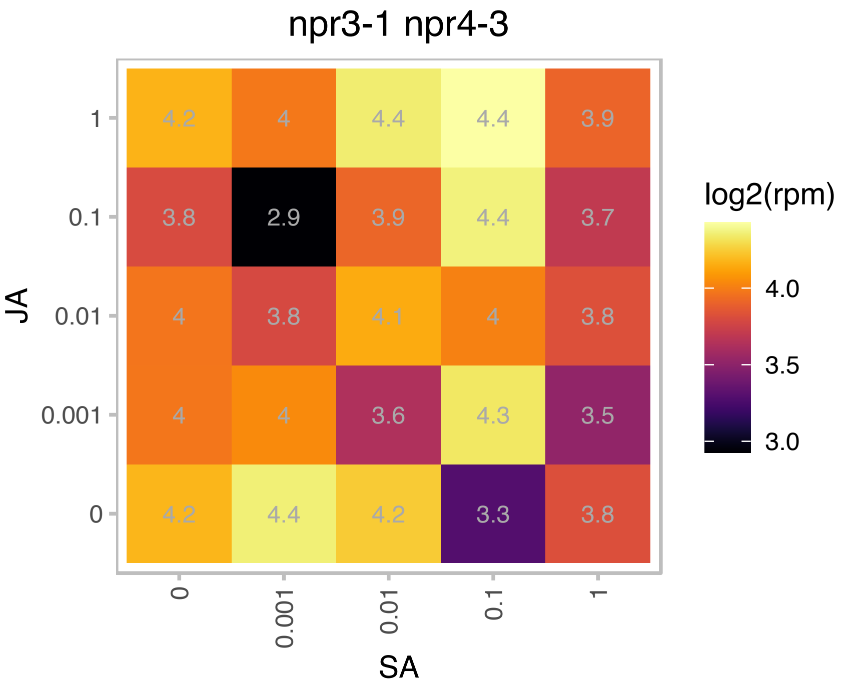

# AT1G69030.1 : NA

BSD domain-containing protein

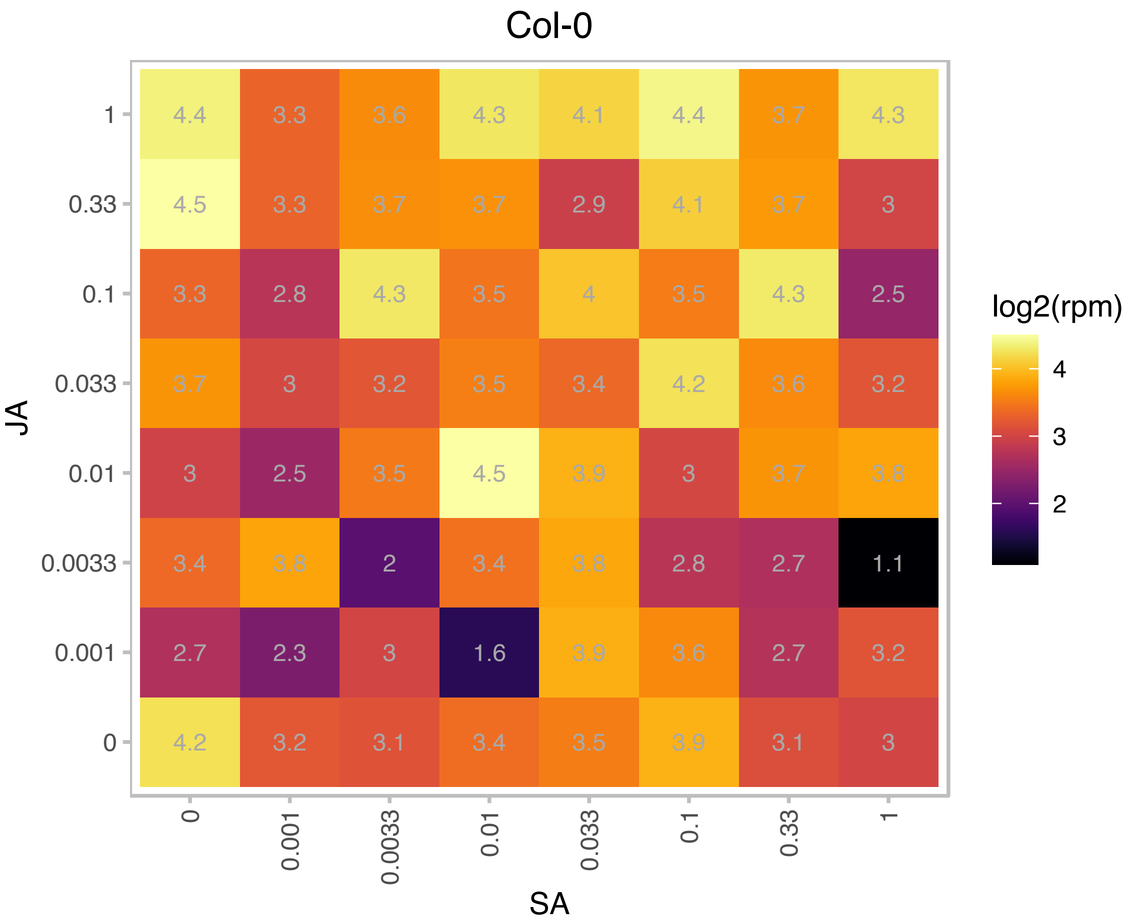


| | |
|---|---|
| Gene | AT1G69030 |
| is.expressed_genes | TRUE |
| is.responsive_genes | TRUE |
| is.apcluster | A-01 |

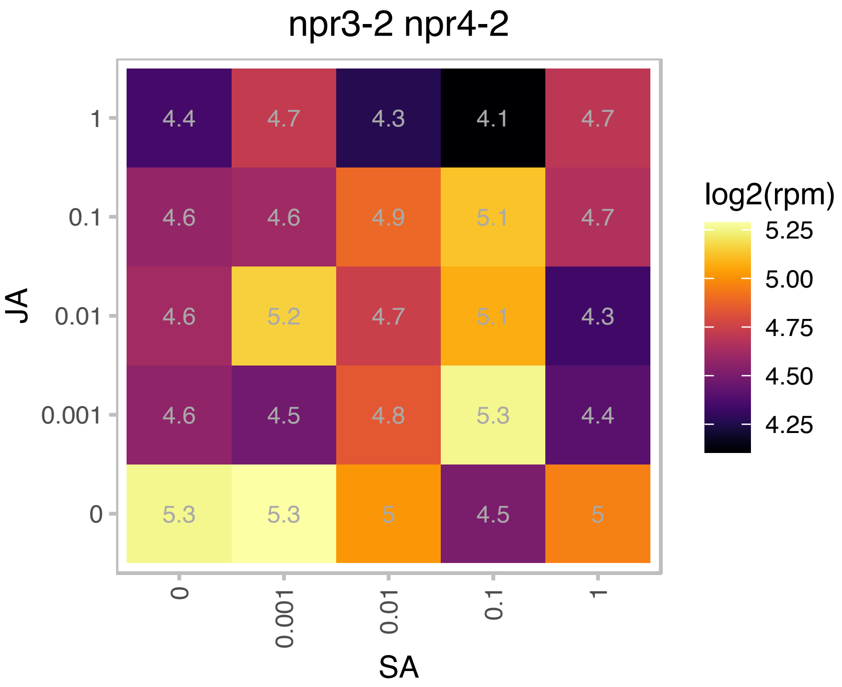


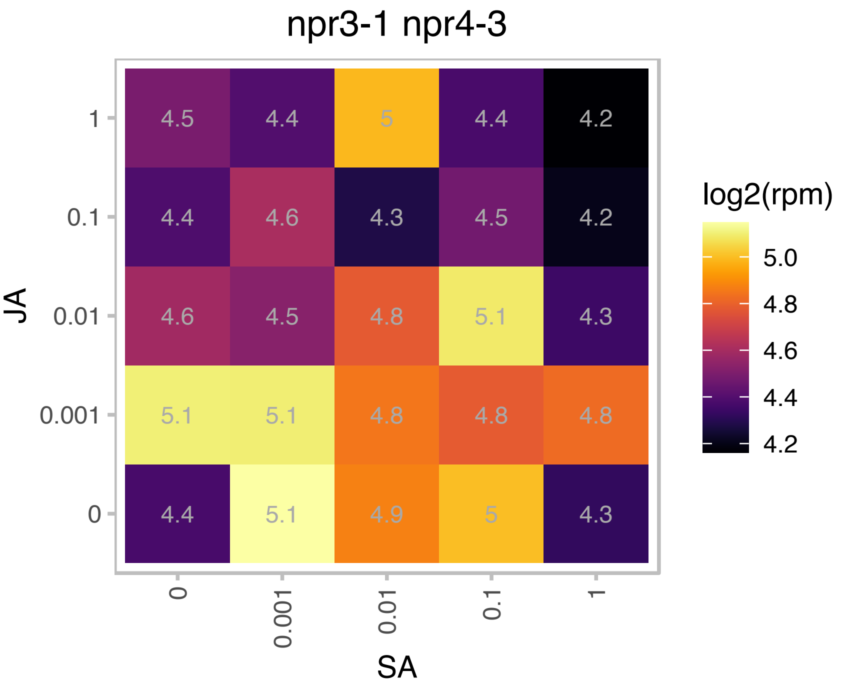

### AT1G72450.1 : JAZ6, TIFY11B

jasmonate-zim-domain protein 6
JAZ6 transcript levels rise in response to a jasmonate stimulus and a GFP:JAZ6 fusion protein localizes to the nucleus. Application of jasmonate methyl ester to Arabidopsis roots reduces the levels of a JAZ6:GUS fusion protein, presumably by stimulating ubiquitin-proteasome-mediated degradation.

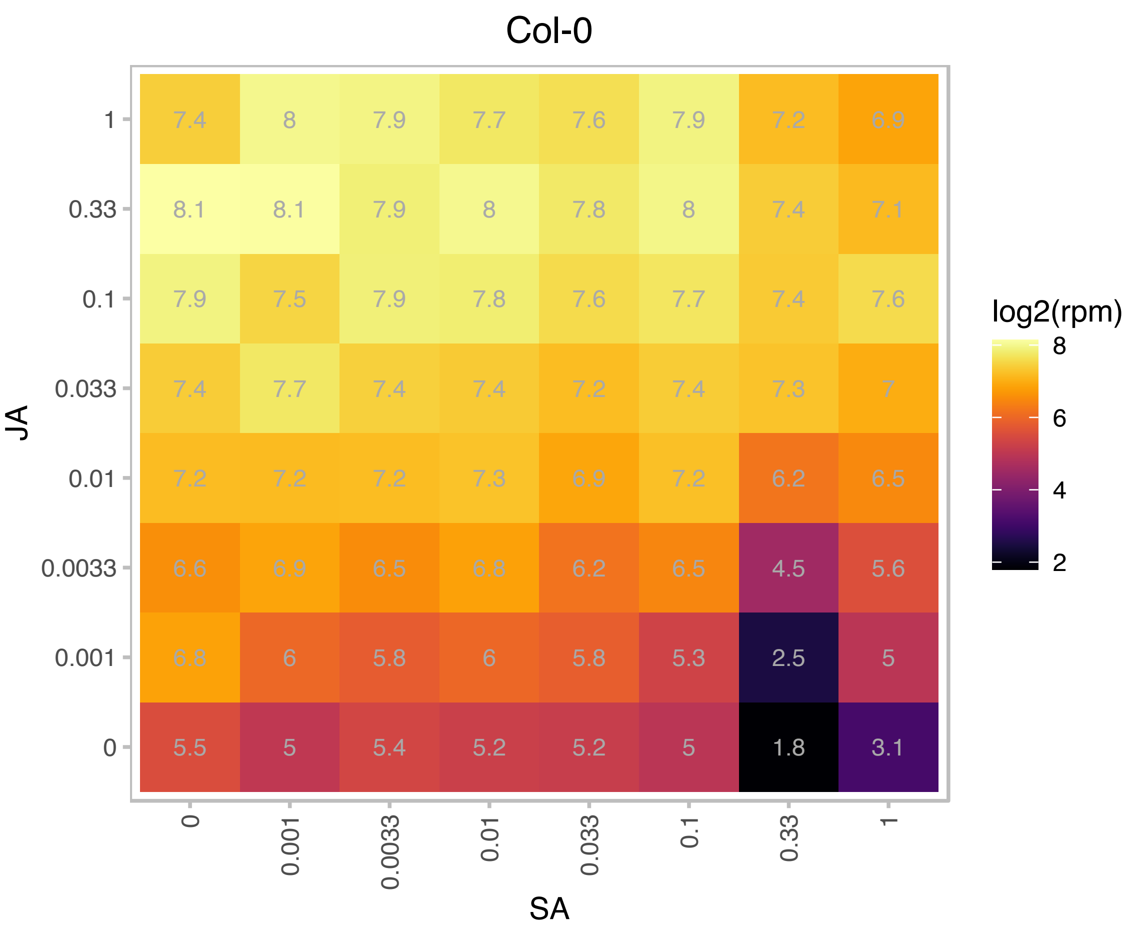


| | |
|---|---|
| Gene | AT1G72450 |
| is.expressed_genes | TRUE |
| is.responsive_genes | TRUE |
| is.apcluster | A-01 |

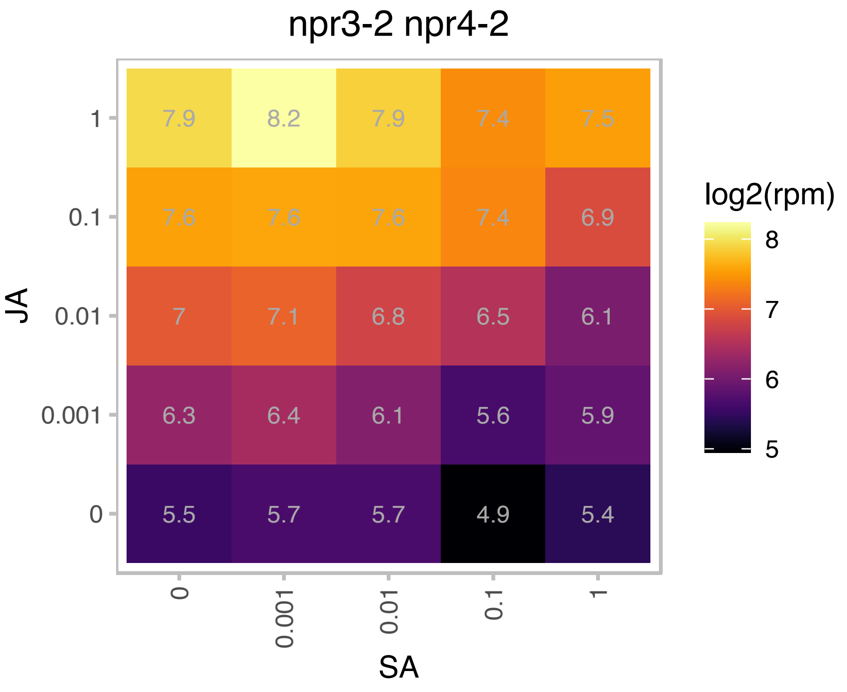


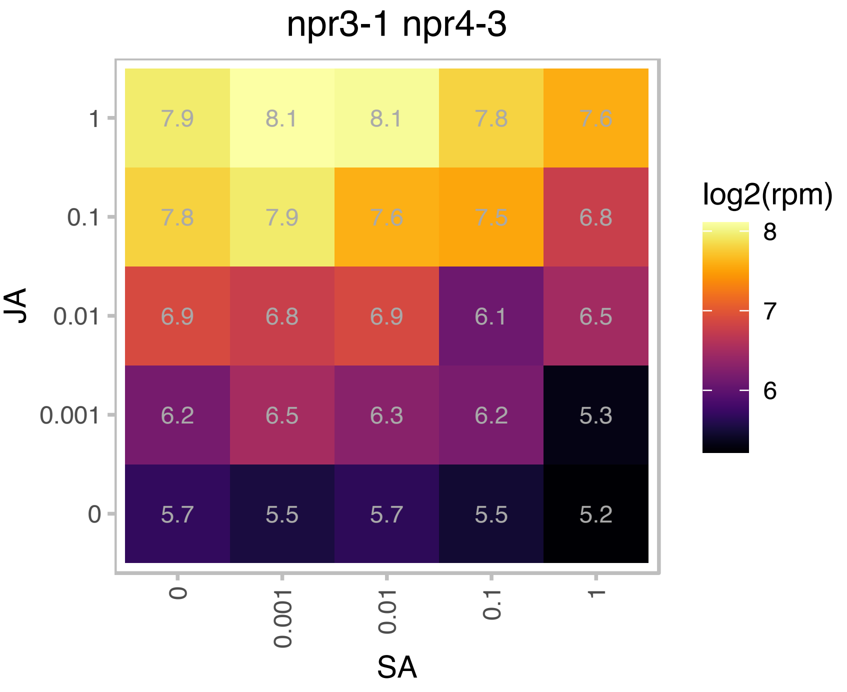

### AT1G73080.1 : ATPEPR1, PEPR1

PEP1 receptor 1

Encodes a leucine-rich repeat receptor kinase. Functions as a receptor for AtPep1 to amplify innate immunity response to pathogen attacks. The mRNA is cell-to-cell mobile.

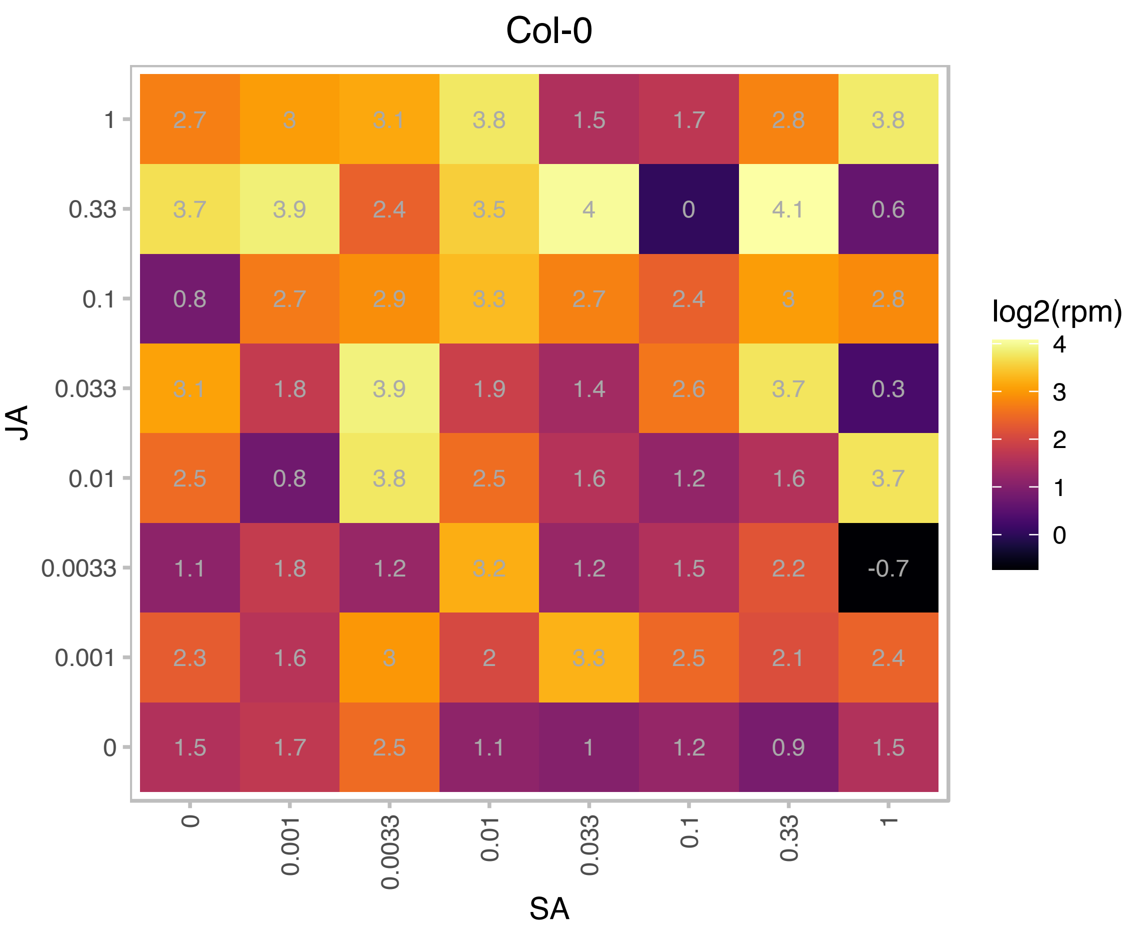


| | |
|---|---|
| Gene | AT1G73080 |
| is.expressed_genes | TRUE |
| is.responsive_genes | TRUE |
| is.apcluster | A-01 |

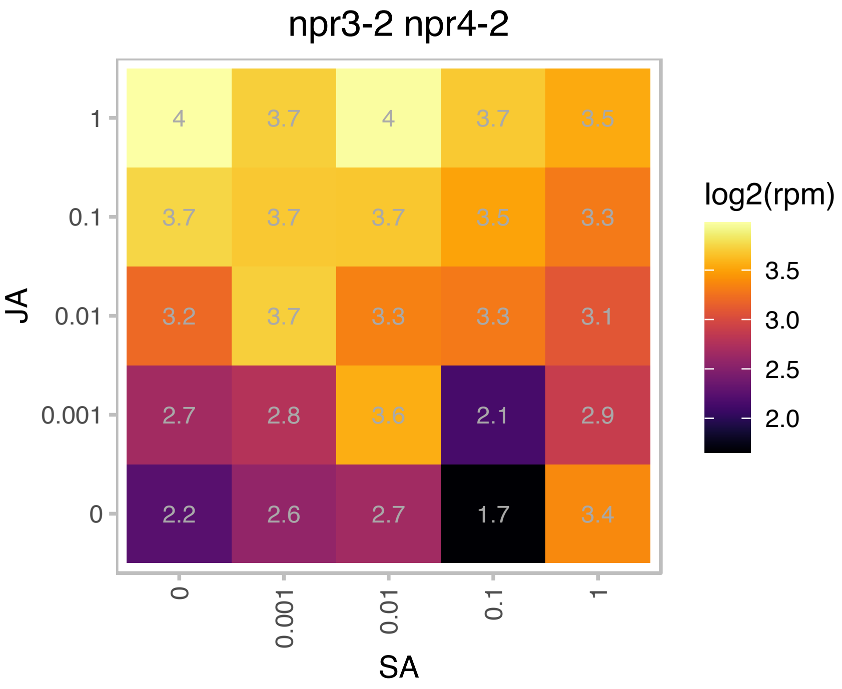


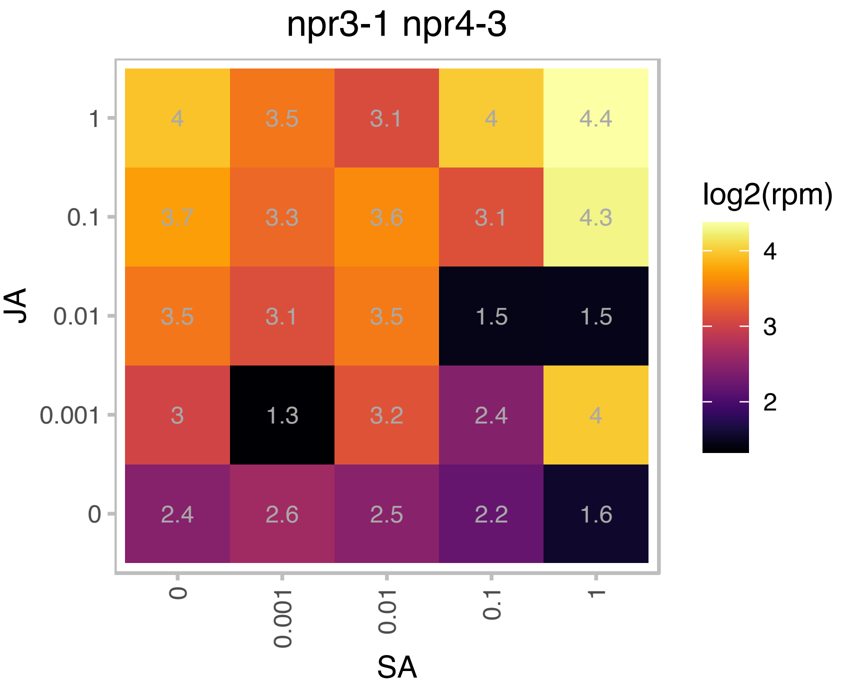

### AT1G76640.1 : CML39

Calcium-binding EF-hand family protein

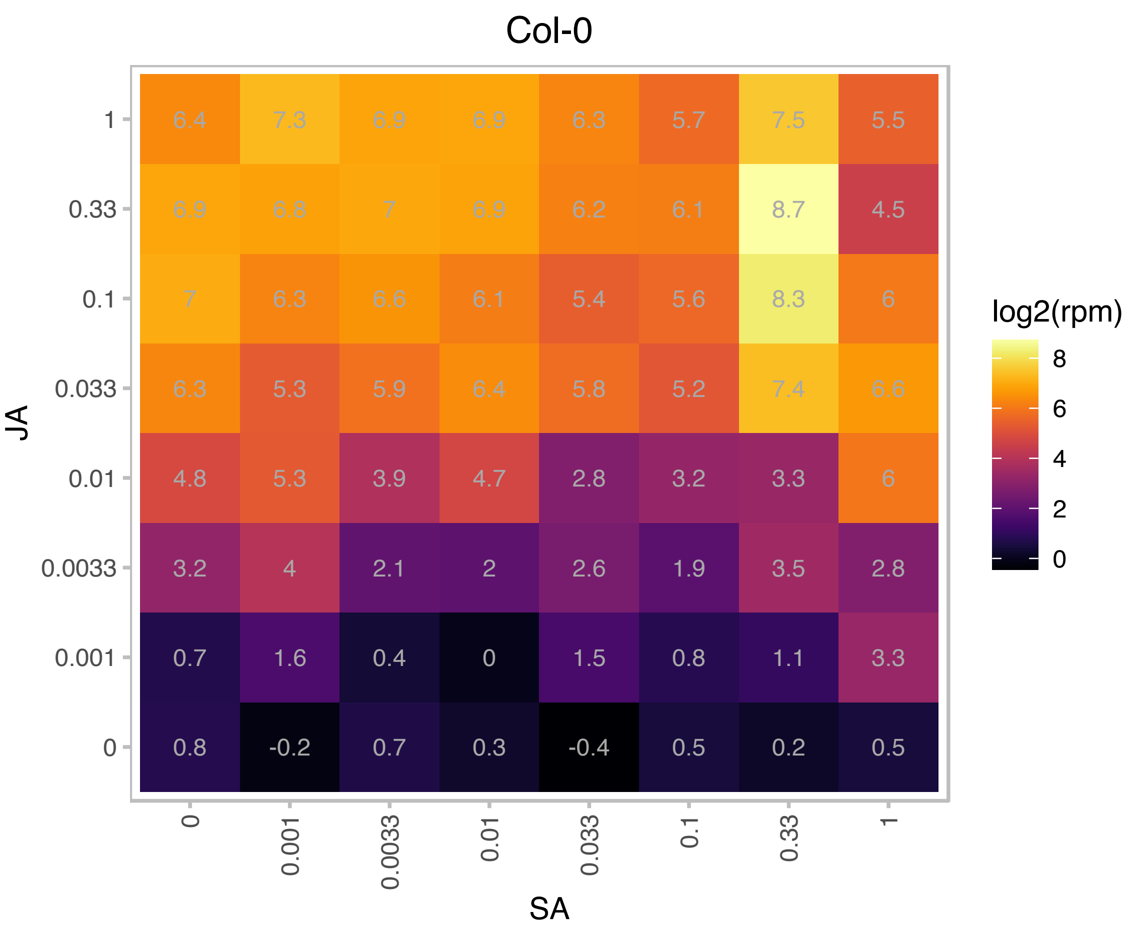


| | |
|---|---|
| Gene | AT1G76640 |
| is.expressed_genes | TRUE |
| is.responsive_genes | TRUE |
| is.apcluster | A-01 |

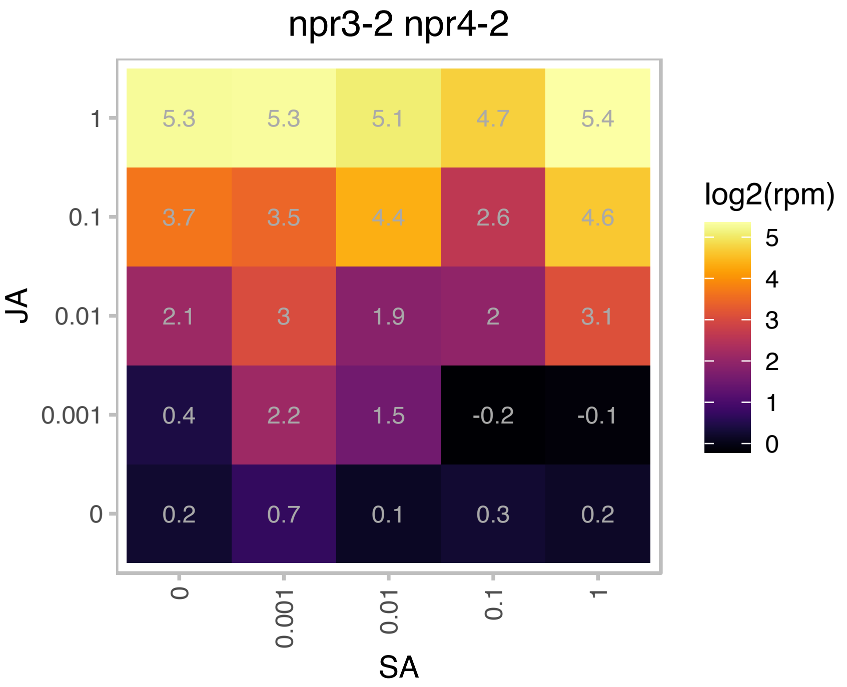


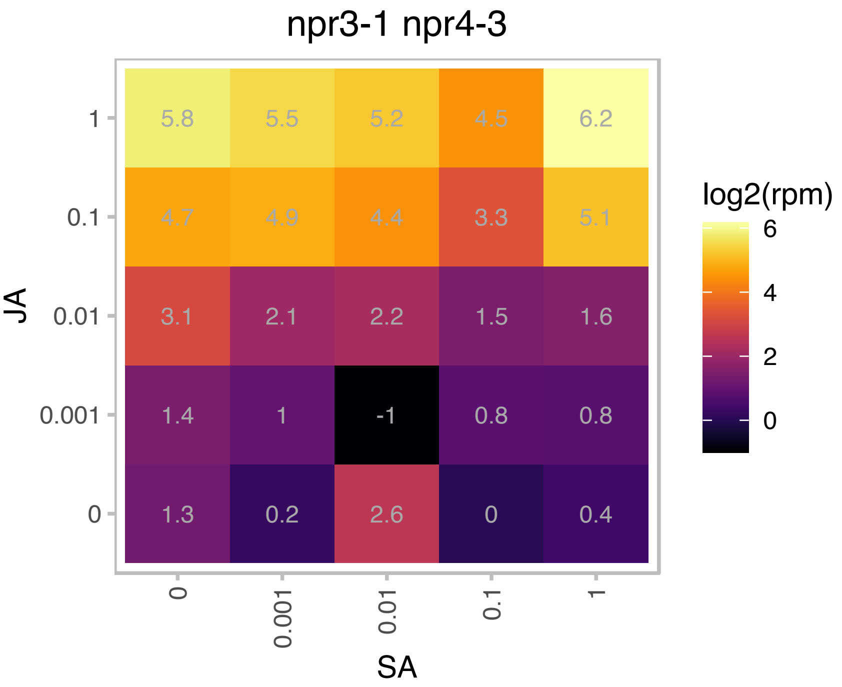

# AT1G78210.1 : NA

alpha/beta-Hydrolases superfamily protein

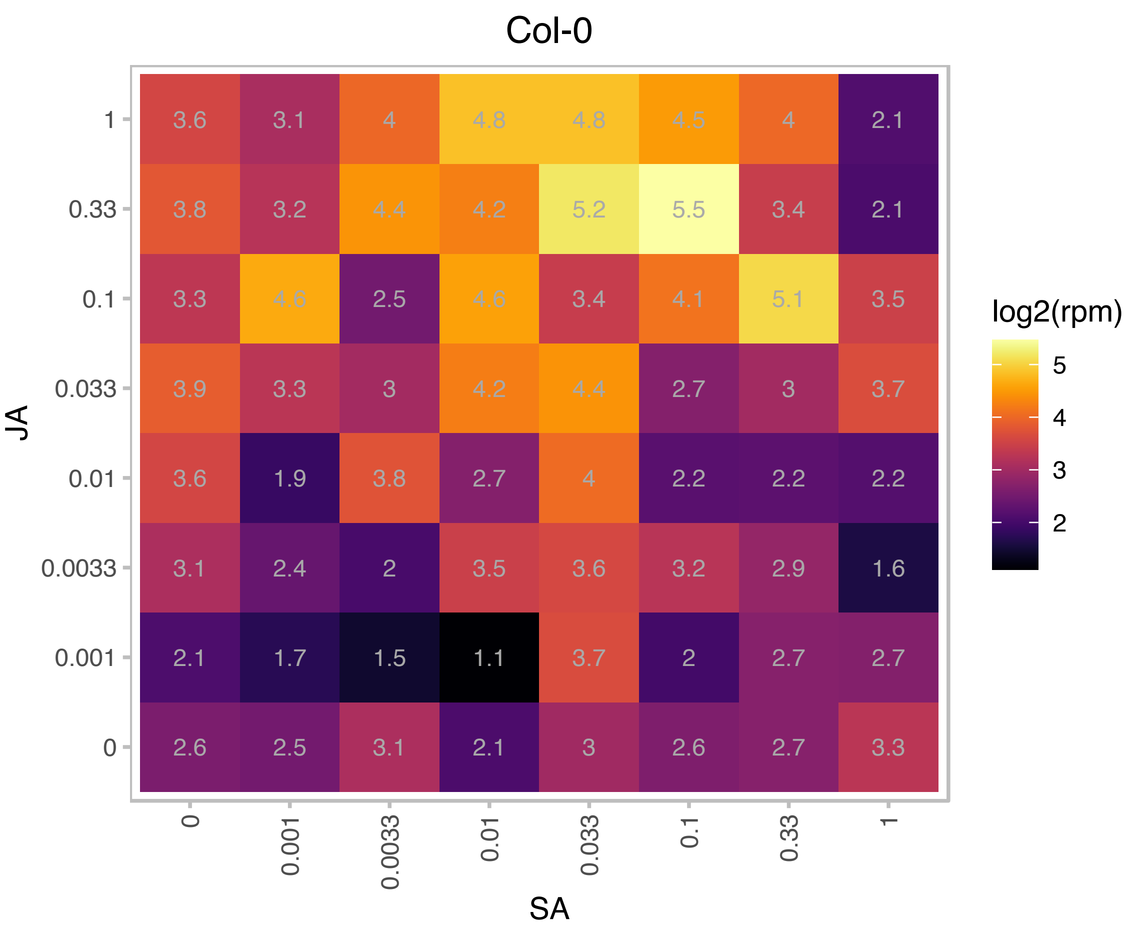


| | |
|---|---|
| Gene | AT1G78210 |
| is.expressed_genes | TRUE |
| is.responsive_genes | TRUE |
| is.apcluster | A-01 |

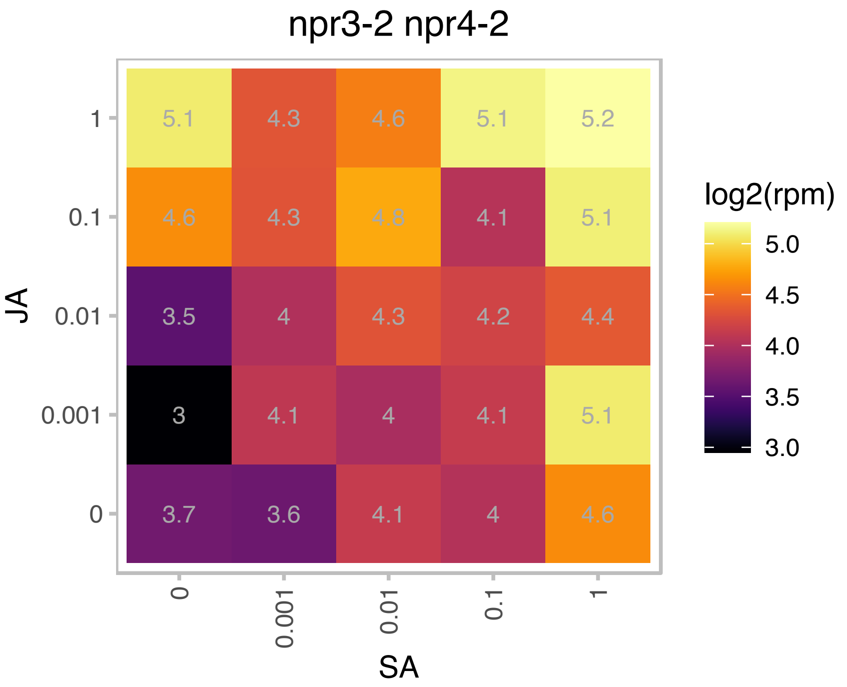


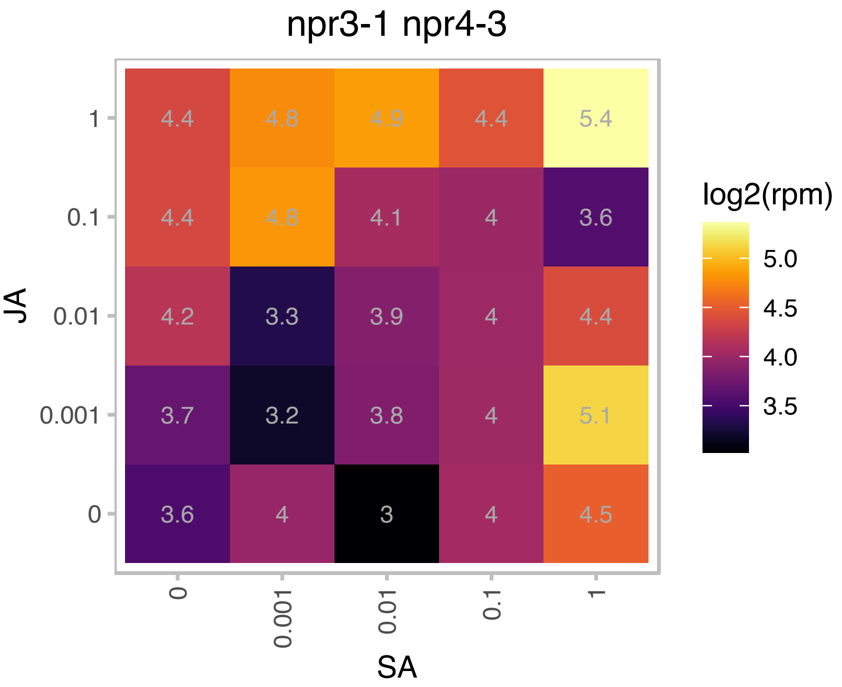

# AT2G07783.1 : NA

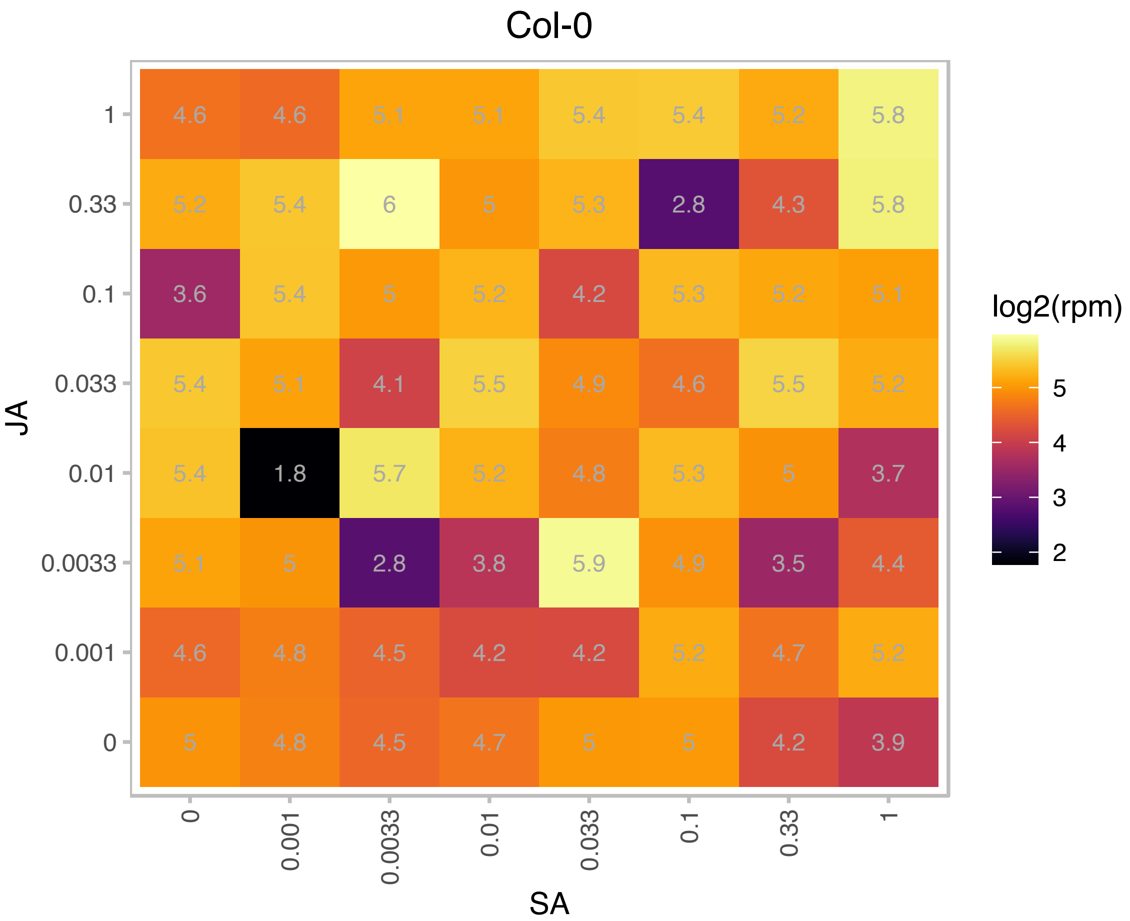


| | |
|---|---|
| Gene | AT2G07783 |
| is.expressed_genes | TRUE |
| is.responsive_genes | TRUE |
| is.apcluster | A-01 |

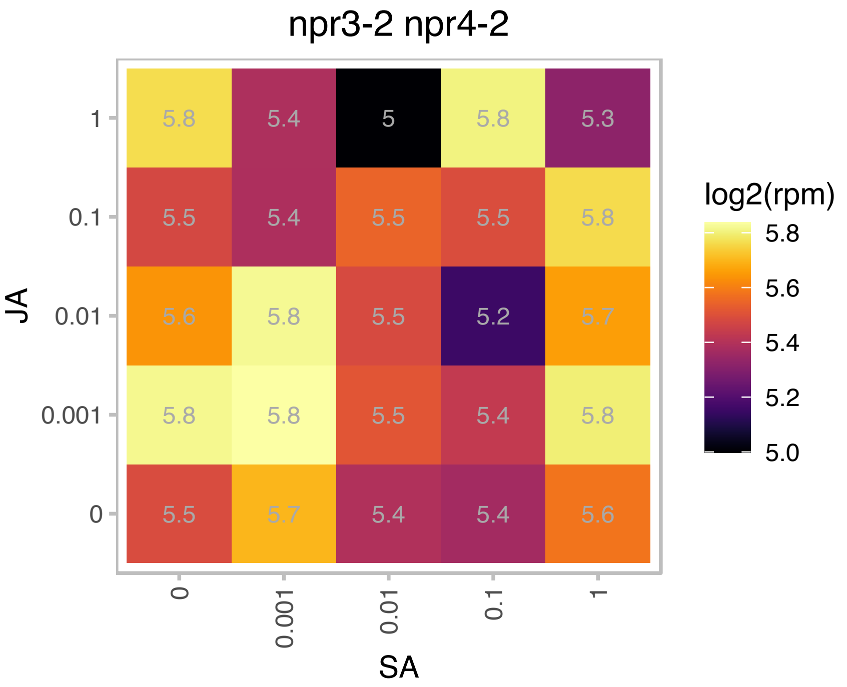


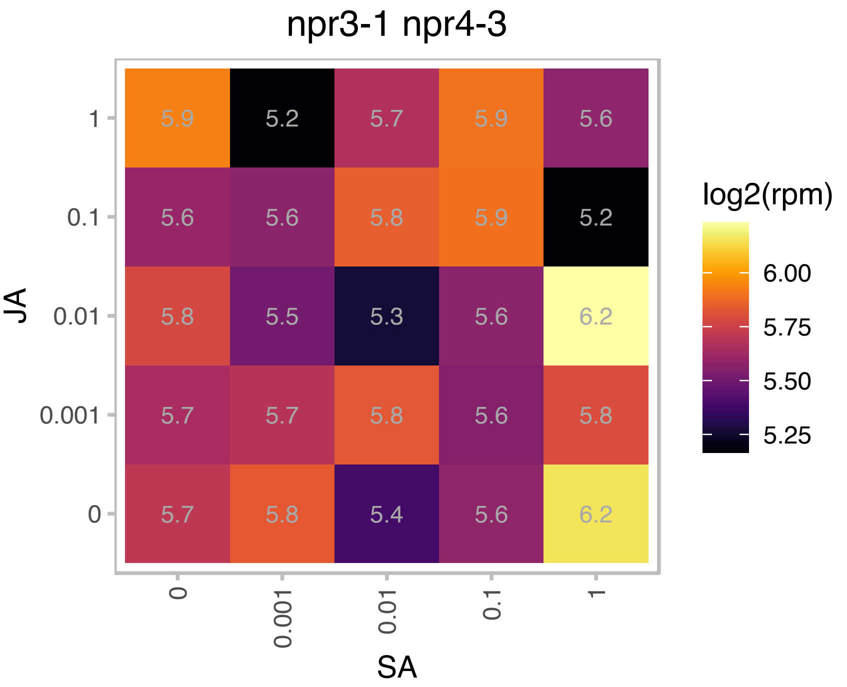

### AT2G26590.1 : RPN13

regulatory particle non-ATPase 13

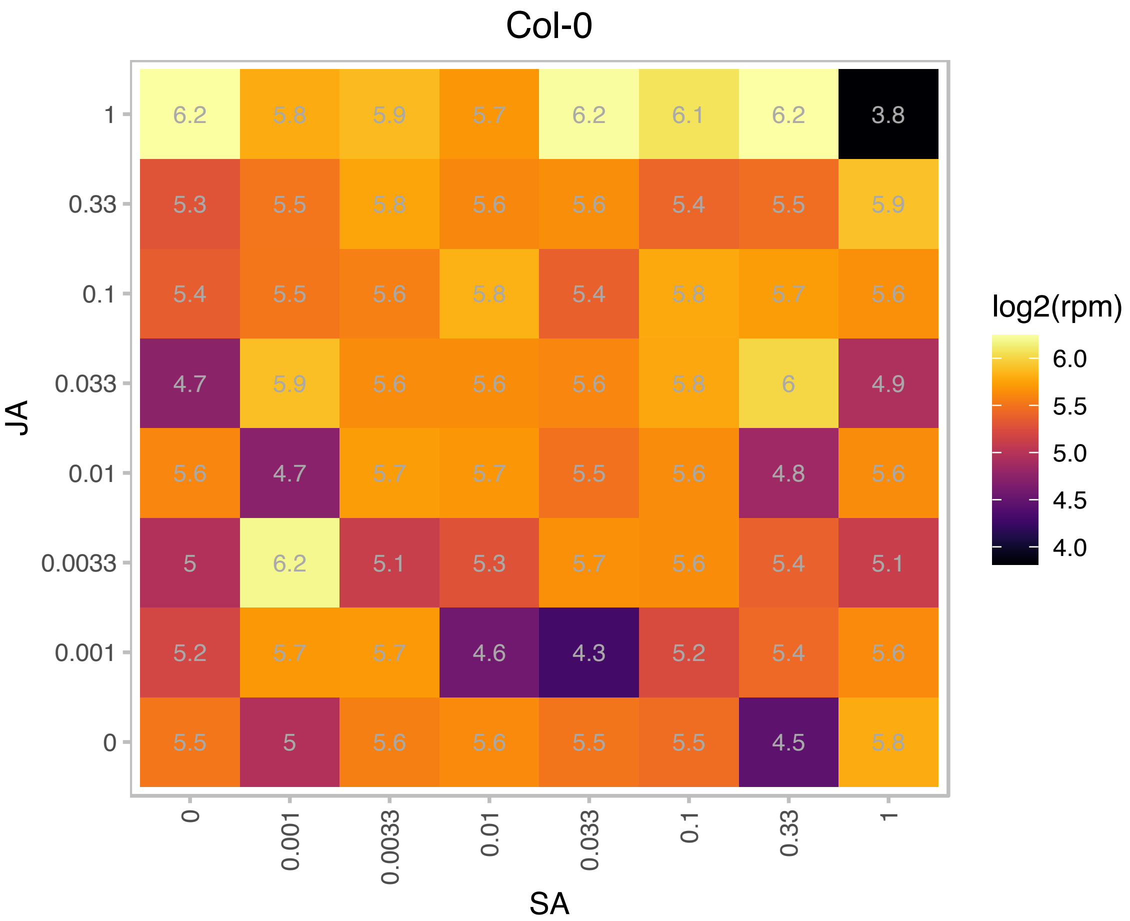


| | |
|---|---|
| Gene | AT2G26590 |
| is.expressed_genes | TRUE |
| is.responsive_genes | TRUE |
| is.apcluster | A-01 |

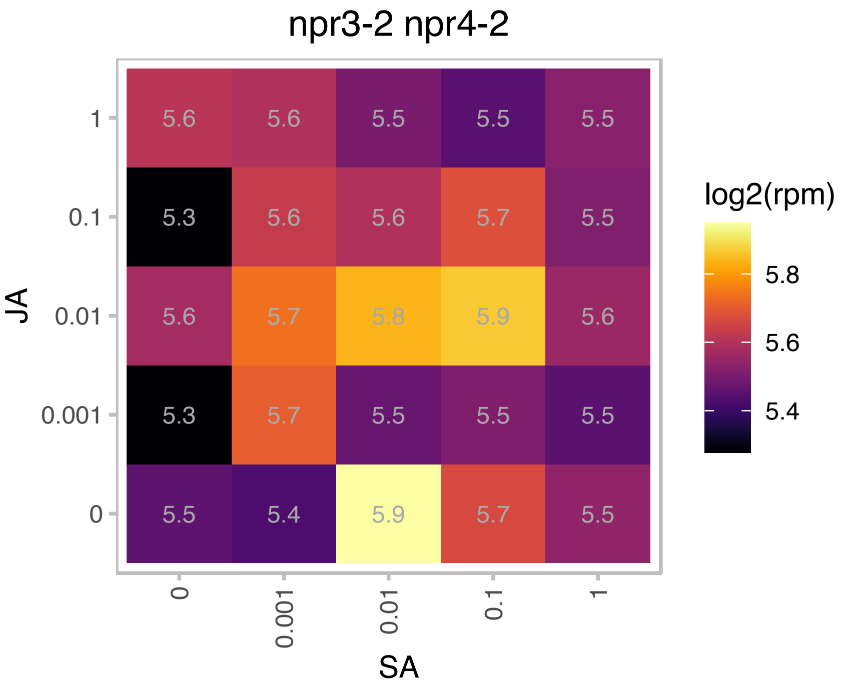


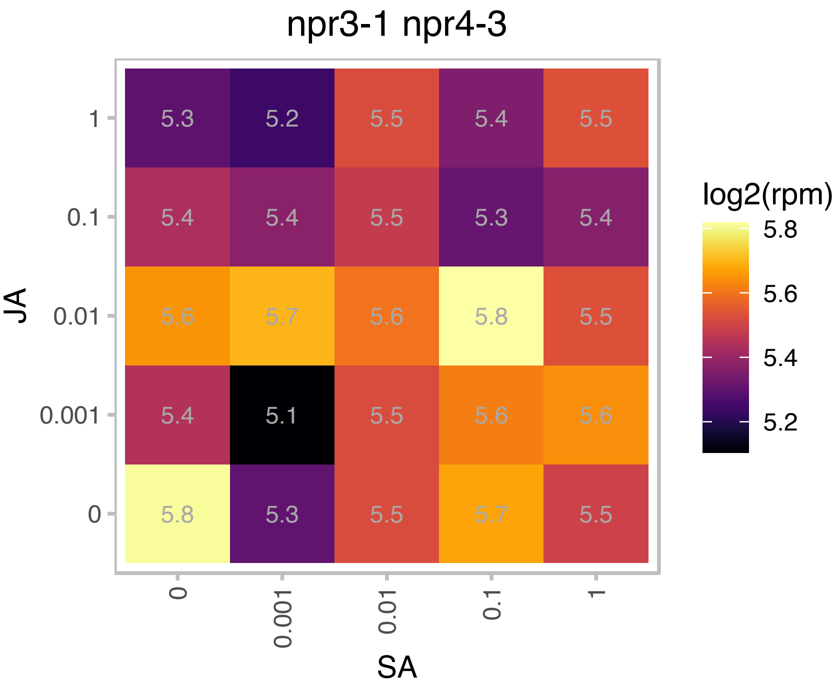

### AT2G29440.1 : ATGSTU6, GST24, GSTU6

glutathione S-transferase tau 6

Encodes glutathione transferase belonging to the tau class of GSTs. Naming convention according to Wagner et al. (2002).

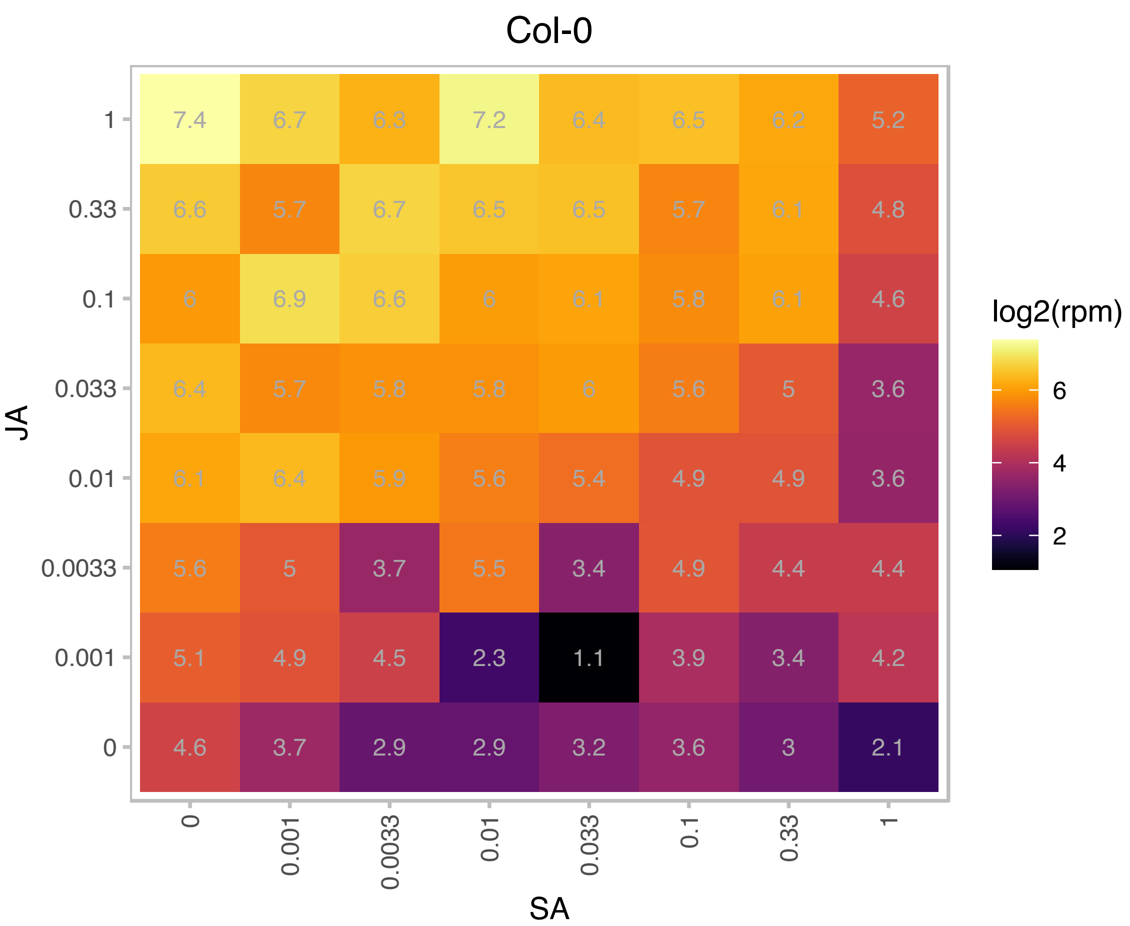


| | |
|---|---|
| Gene | AT2G29440 |
| is.expressed_genes | TRUE |
| is.responsive_genes | TRUE |
| is.apcluster | A-01 |

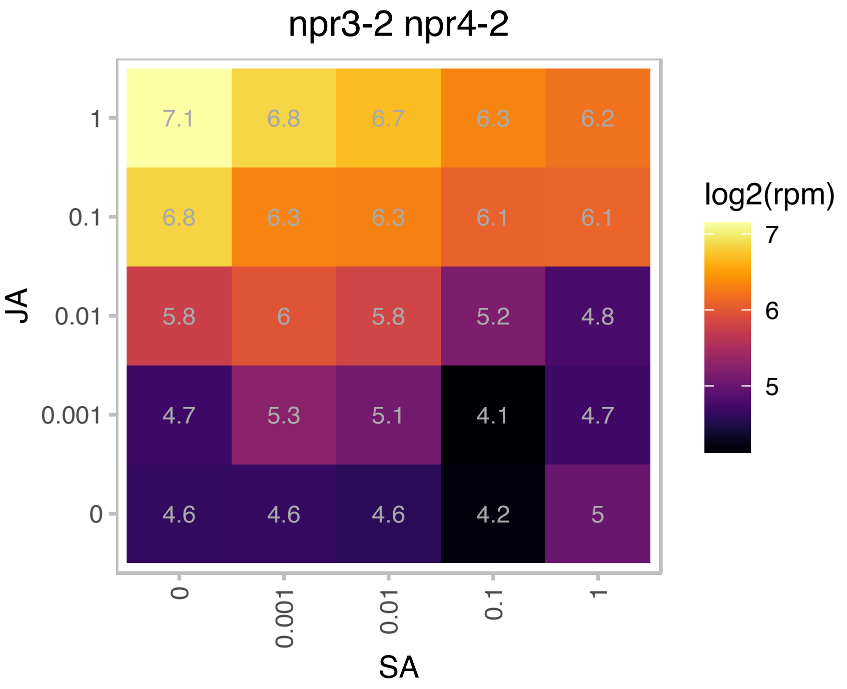


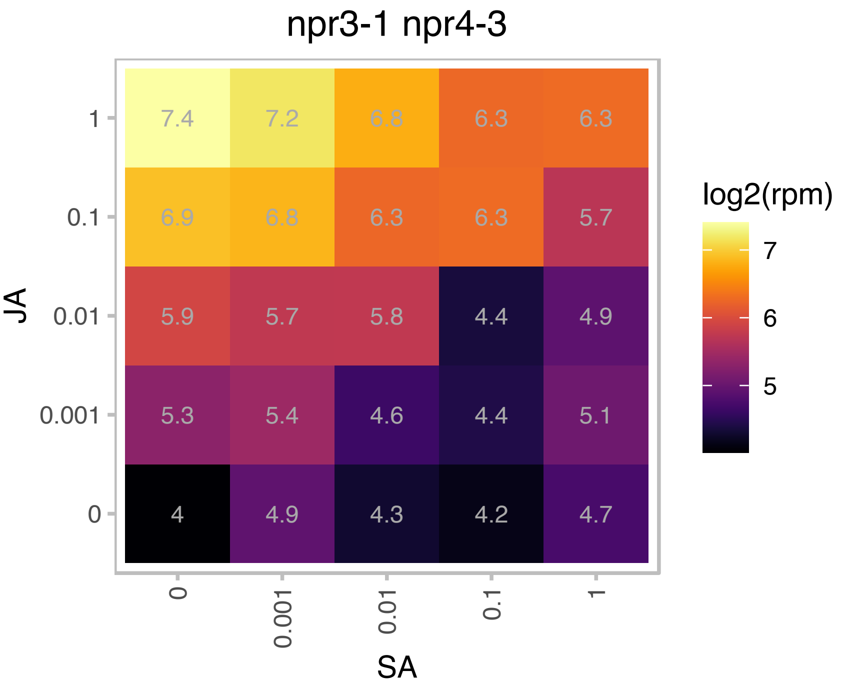

# AT2G32600.1 : NA

hydroxyproline-rich glycoprotein family protein

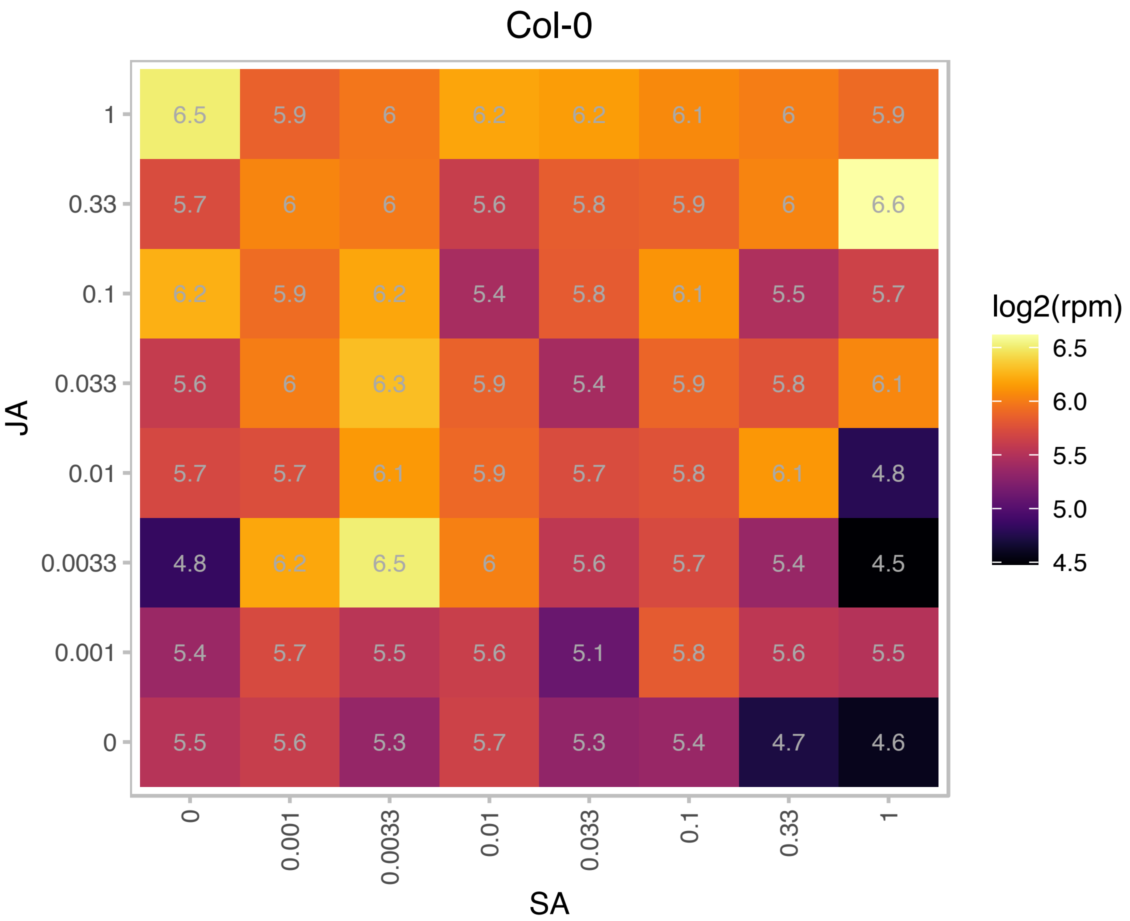


| | |
|---|---|
| Gene | AT2G32600 |
| is.expressed_genes | TRUE |
| is.responsive_genes | TRUE |
| is.apcluster | A-01 |

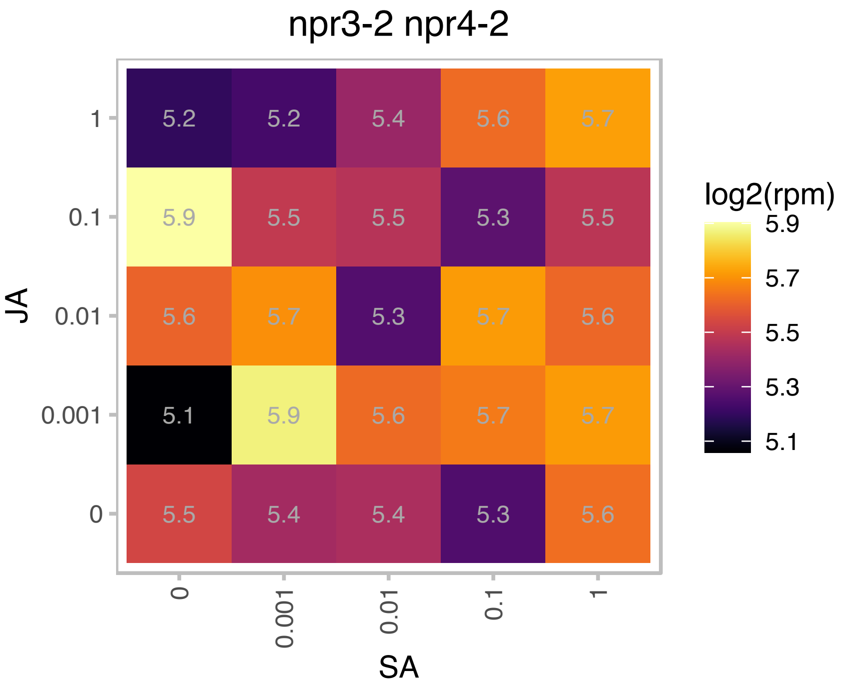


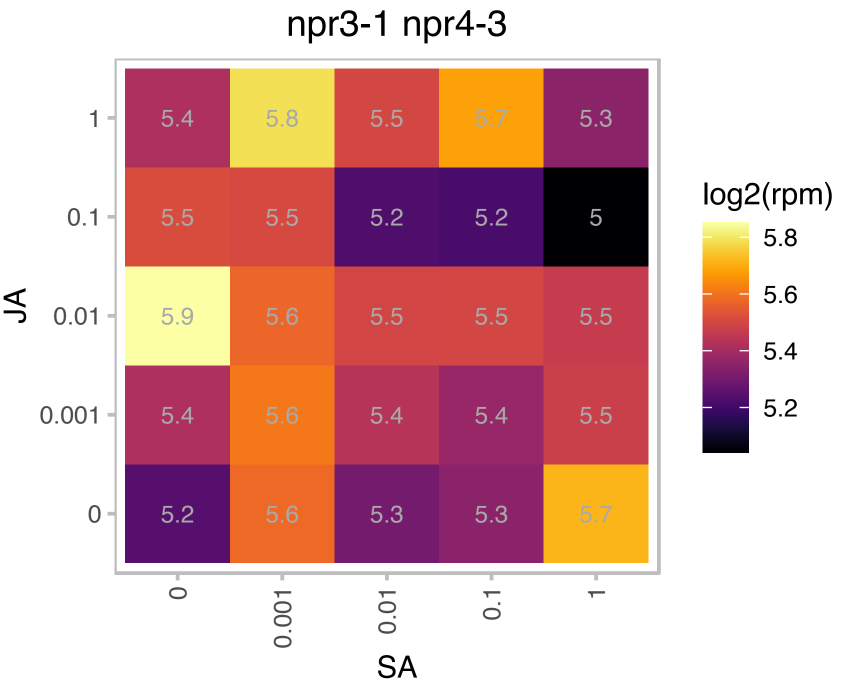

# AT2G34160.1 : NA

Alba DNA/RNA-binding protein

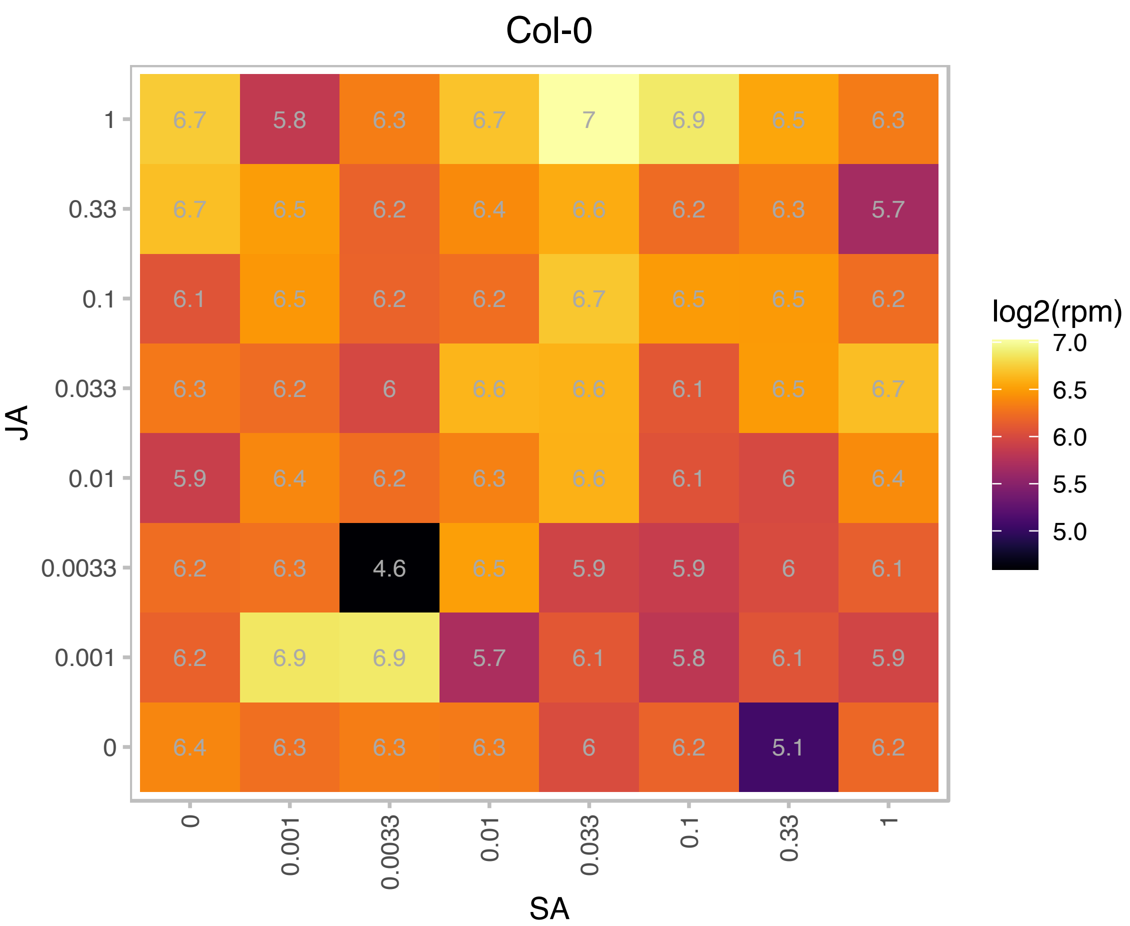


| | |
|---|---|
| Gene | AT2G34160 |
| is.expressed_genes | TRUE |
| is.responsive_genes | TRUE |
| is.apcluster | A-01 |

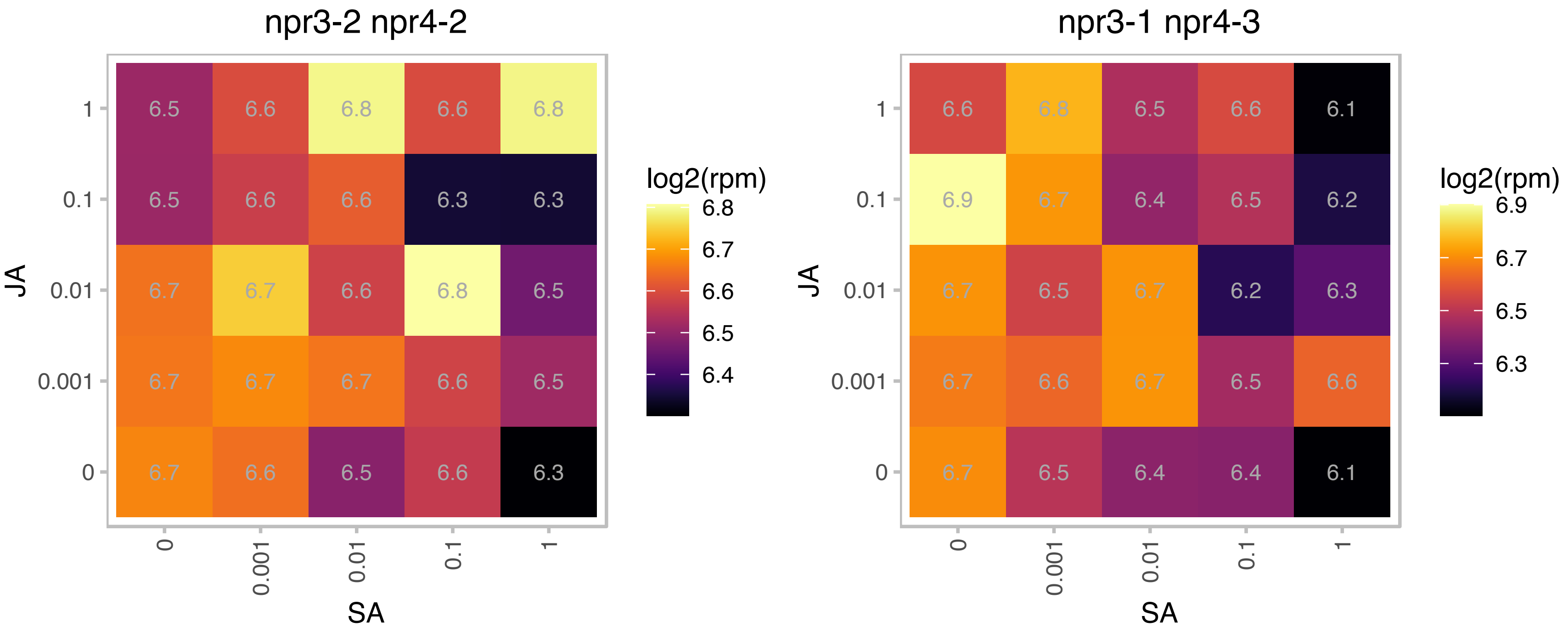

### AT2G38040.1 : CAC3

acetyl Co-enzyme a carboxylase carboxyltransferase alpha subunit

encodes the carboxyltransferase alpha subunit of acetyl-CoA carboxylase, involved in de novo fatty acid biosynthesis

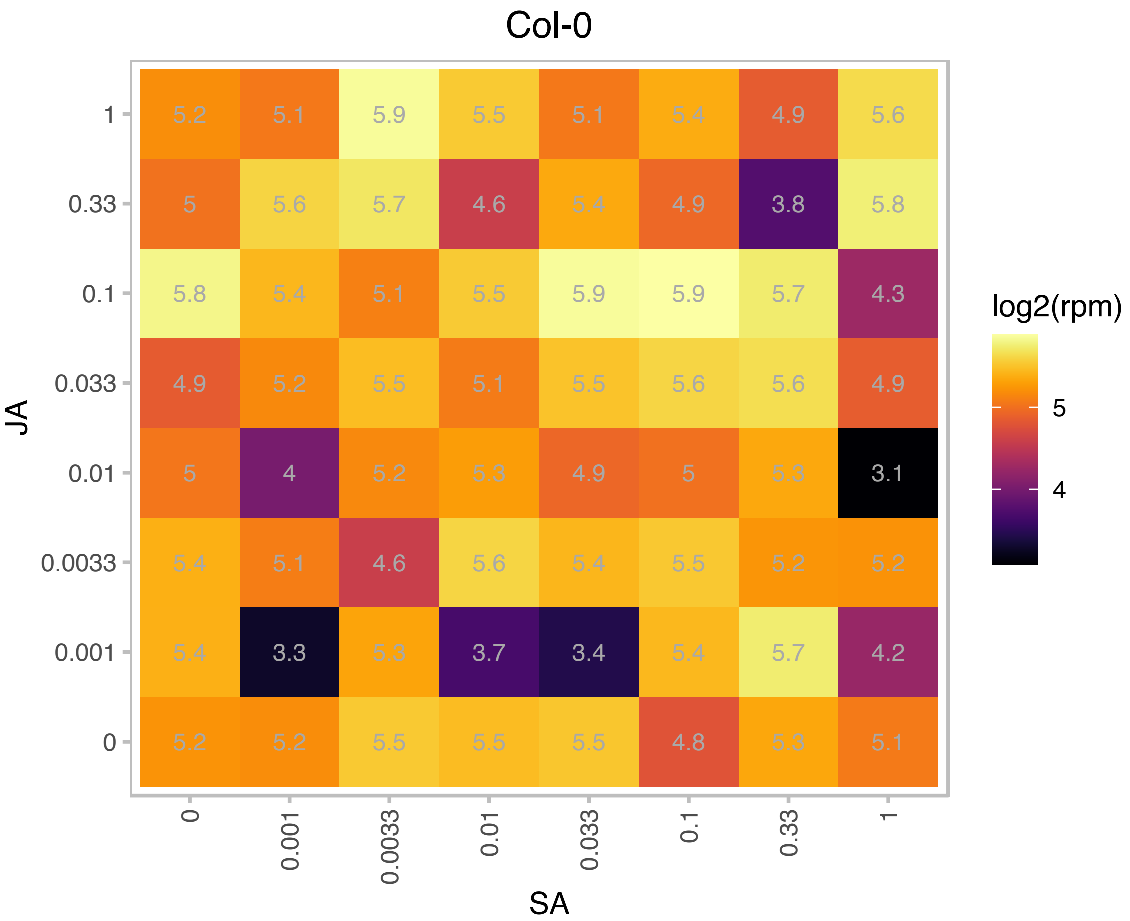


| | |
|---|---|
| Gene | AT2G38040 |
| is.expressed_genes | TRUE |
| is.responsive_genes | TRUE |
| is.apcluster | A-01 |

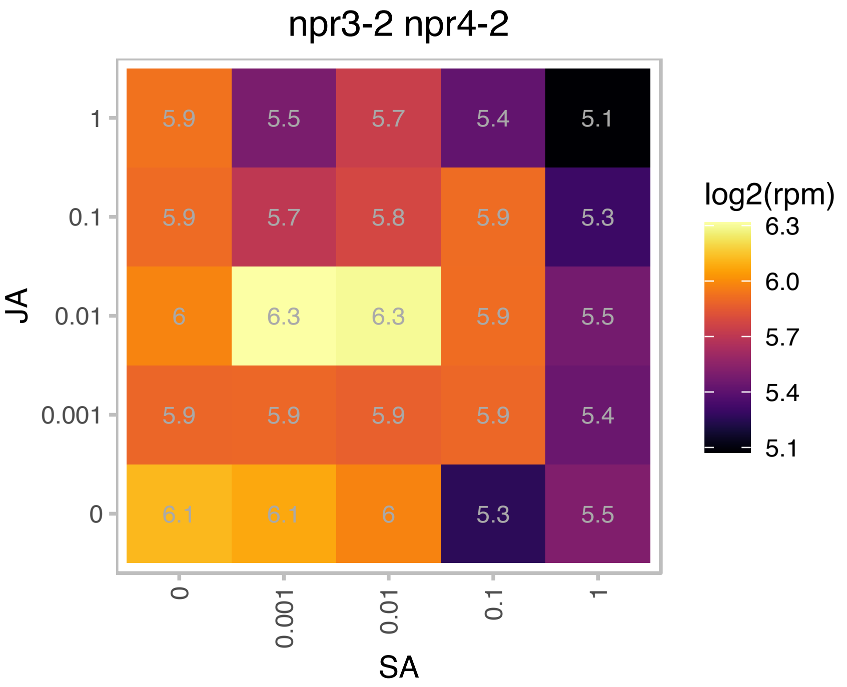


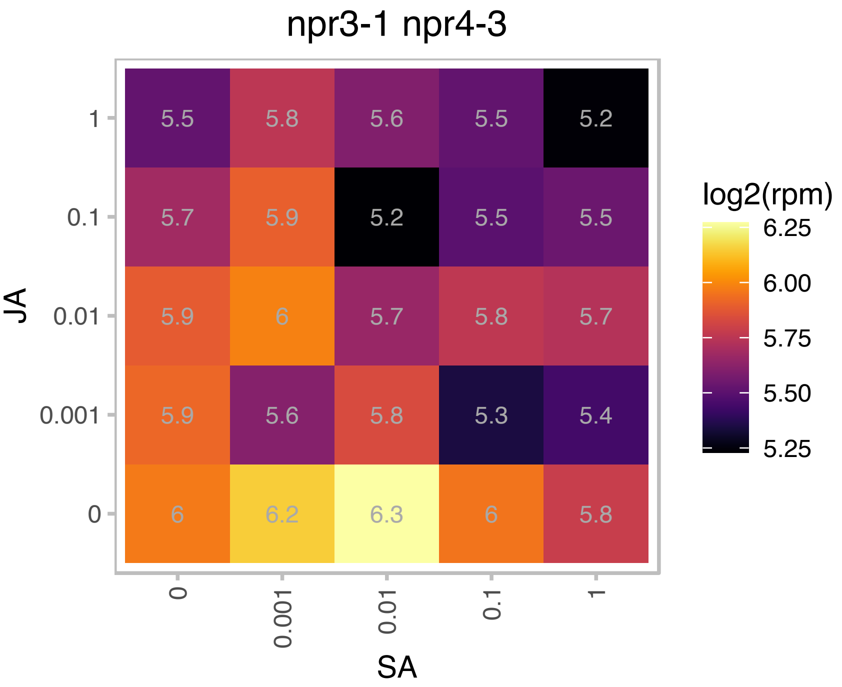

### AT2G38240.1 : JAO4, JOX4

2-oxoglutarate (2OG) and Fe(II)-dependent oxygenase superfamily protein

One of 4 paralogs encoding a 2-oxoglutarate/Fe(II)-dependent oxygenases that hydroxylates JA to 12-OH-JA.

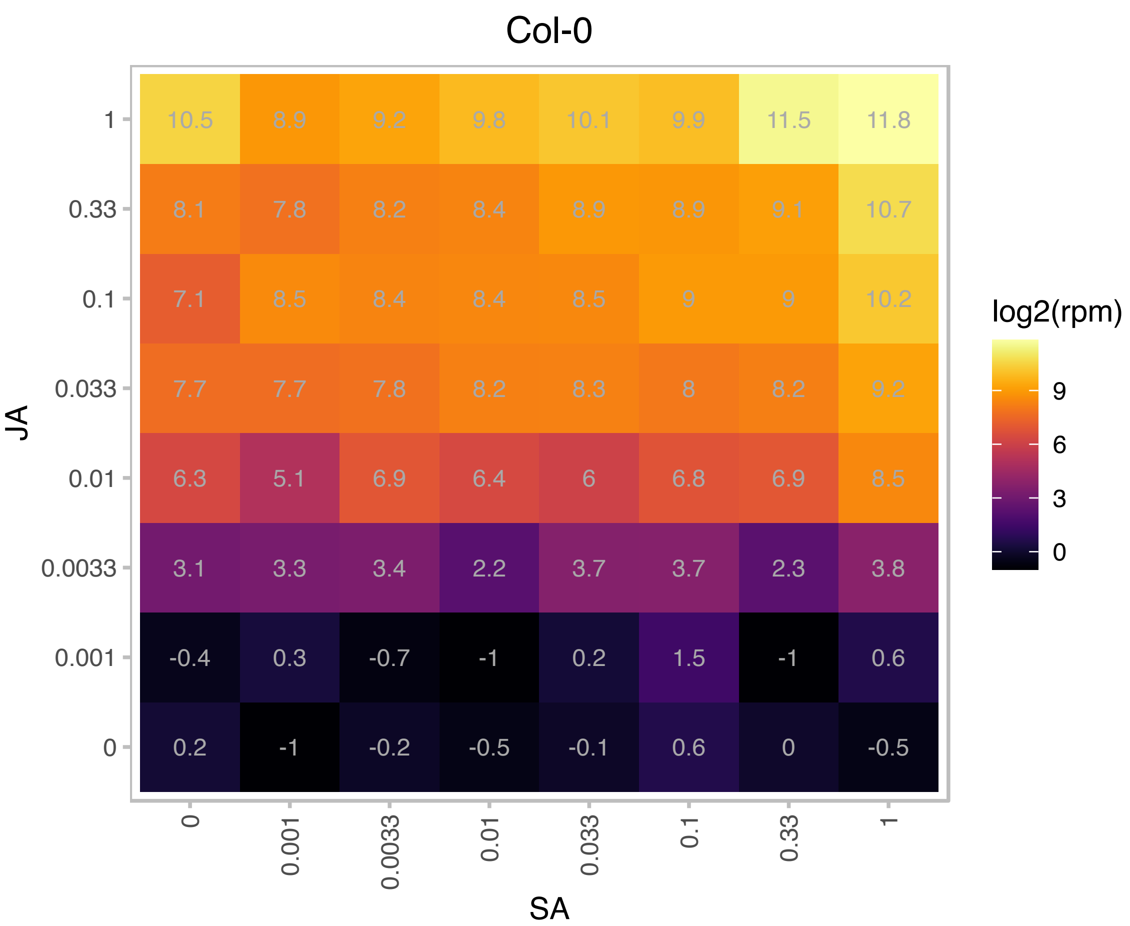


| | |
|---|---|
| Gene | AT2G38240 |
| is.expressed_genes | TRUE |
| is.responsive_genes | TRUE |
| is.apcluster | A-01 |

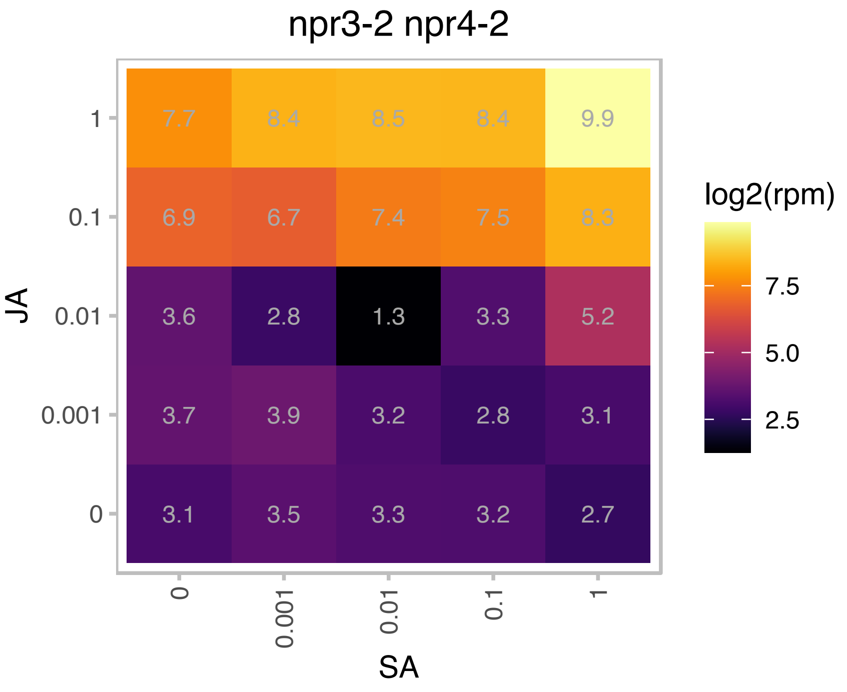


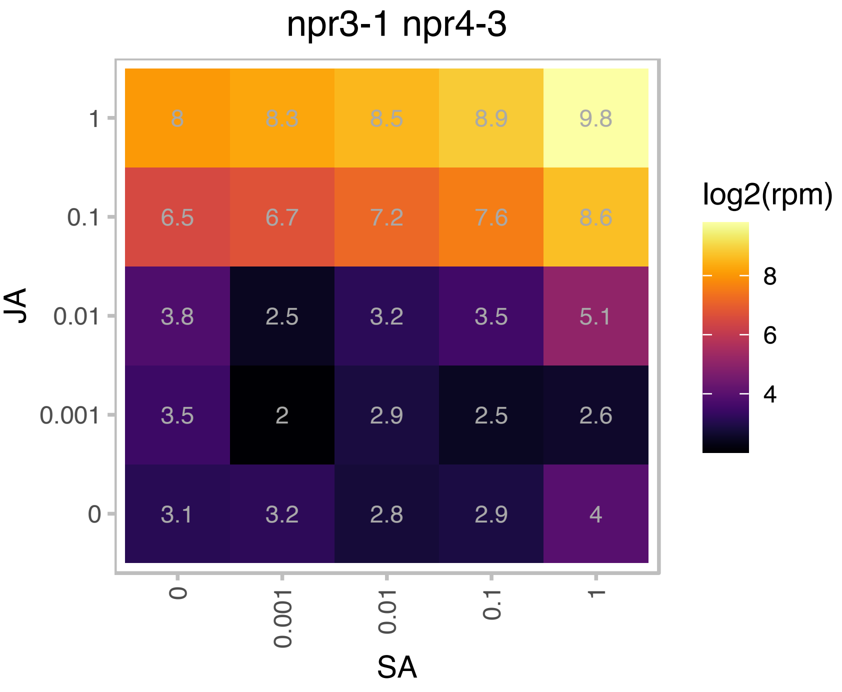

### AT3G02600.1 : ATLPP3, LPP3

lipid phosphate phosphatase 3

Encodes phosphatidic acid phosphatase. Expressed during germination.

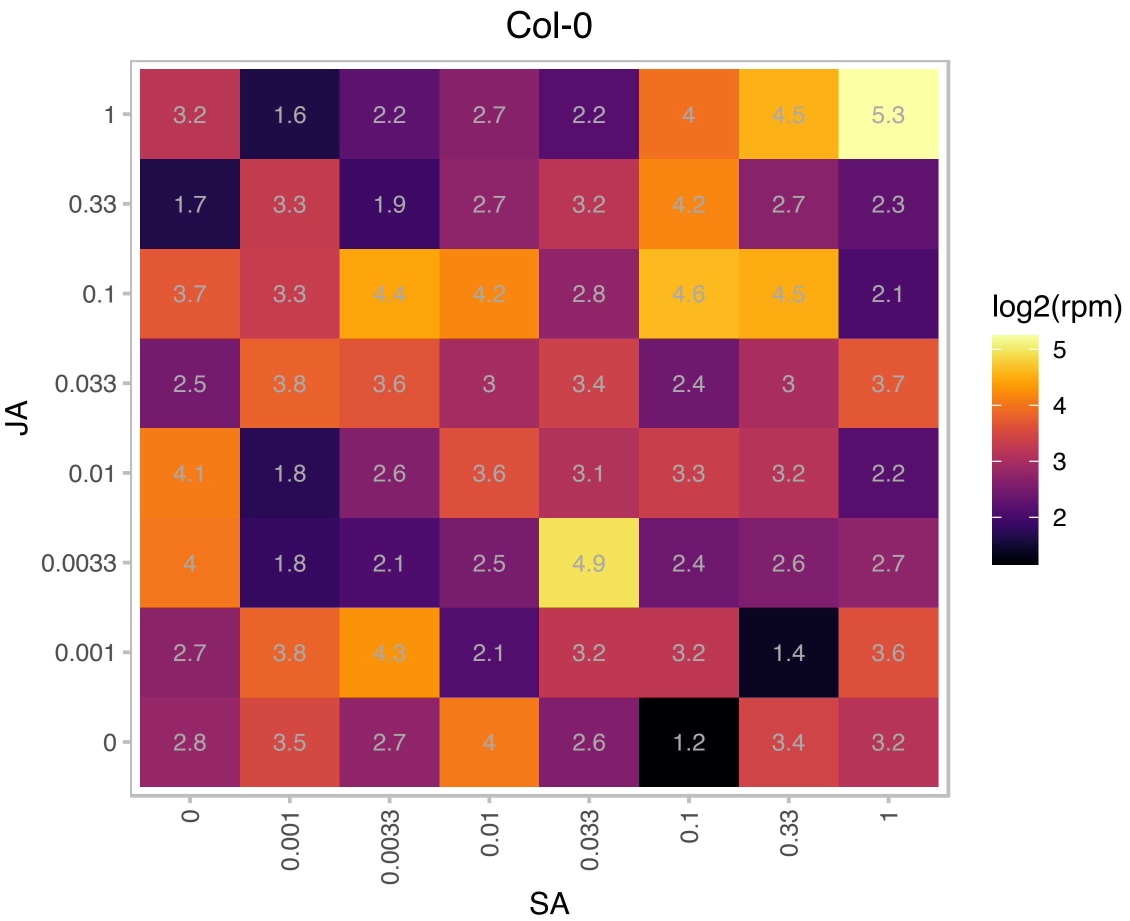


| | |
|---|---|
| Gene | AT3G02600 |
| is.expressed_genes | TRUE |
| is.responsive_genes | TRUE |
| is.apcluster | A-01 |

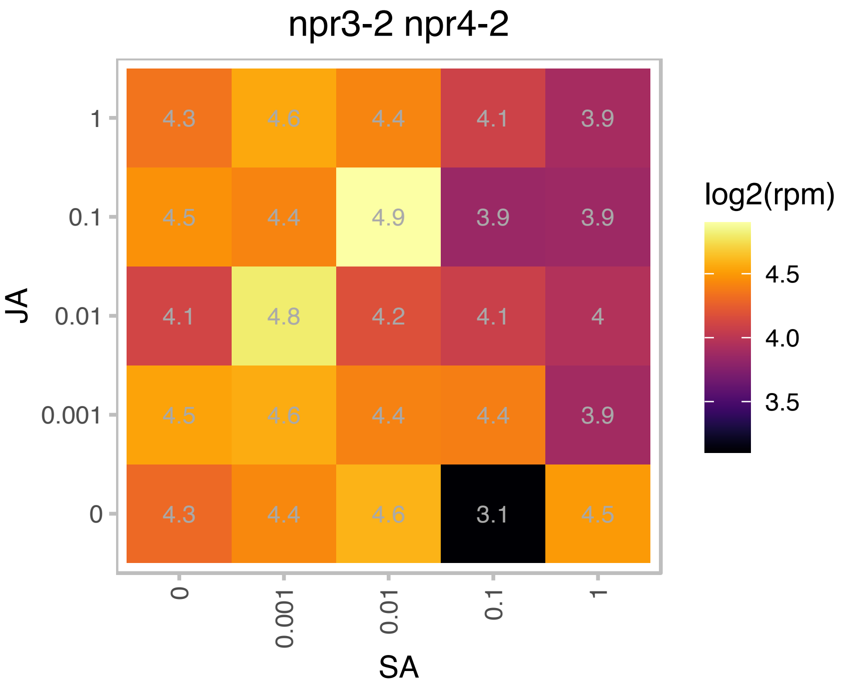


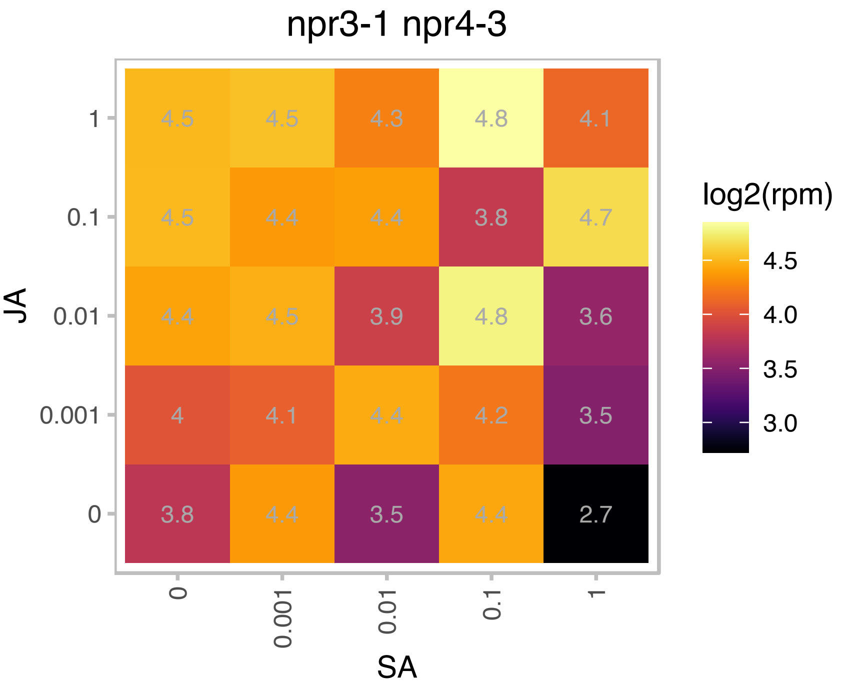

### AT3G08720.1 : ATPK19, ATPK2, ATS6K2, S6K2

serine/threonine protein kinase 2

Encodes a ribosomal-protein S6 kinase. Gene expression is induced by cold and salt (NaCl). Activation of AtS6k is regulated by 1-naphthylacetic acid and kinetin, at least in part, via a lipid kinase-dependent pathway. Phosphorylates specifically mammalian and plant S6 at 25 degrees C but not at 37 degrees C. Involved in translational up-regulation of ribosomal proteins.

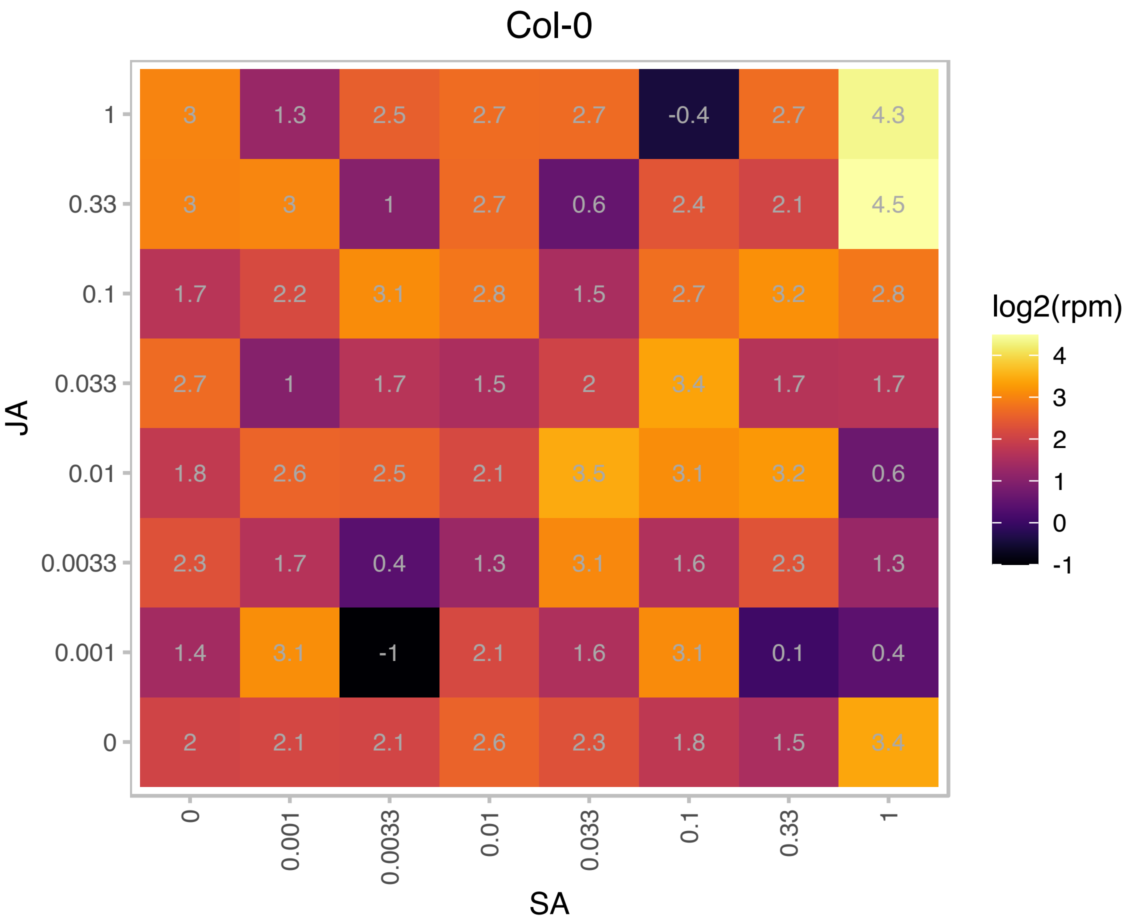


| | |
|---|---|
| Gene | AT3G08720 |
| is.expressed_genes | TRUE |
| is.responsive_genes | TRUE |
| is.apcluster | A-01 |

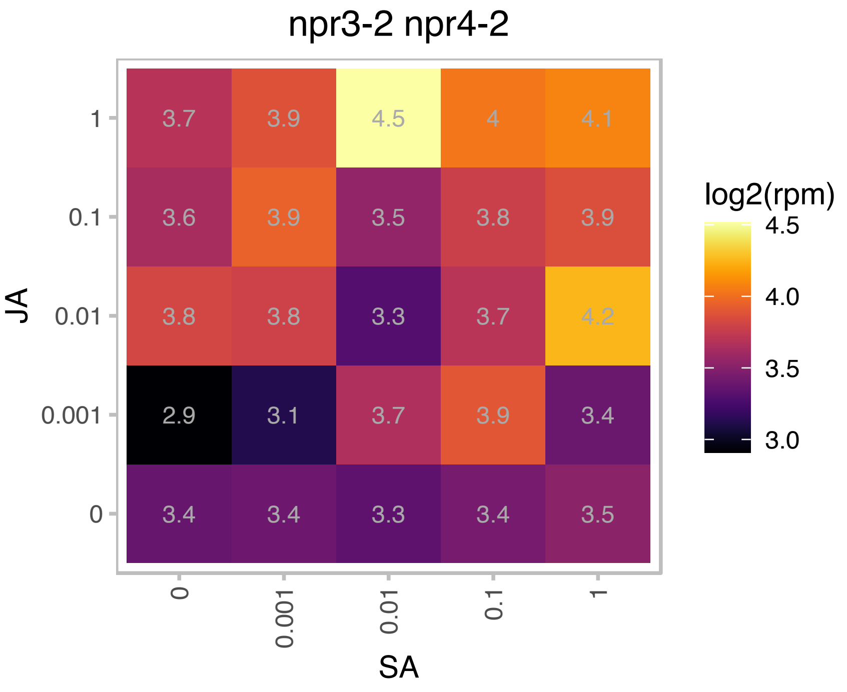


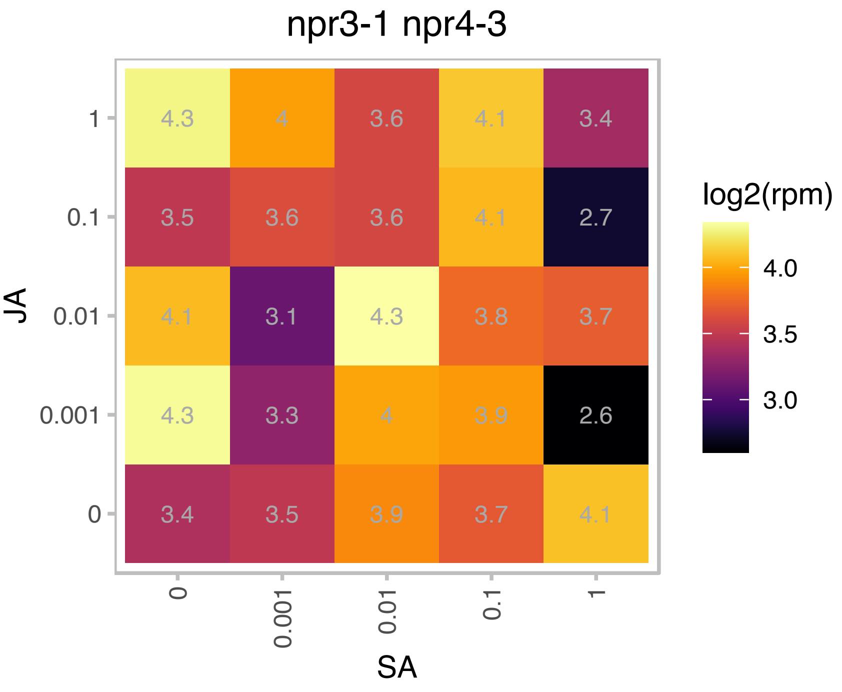

### AT3G11130.1 : AtCHC1, CHC1

Clathrin, heavy chain
CHC1 heavy chain subunit of clathrin. Involved in vesicle mediated trafficking. Mutants show reduced rates of endocytosis and defects clathrin mediated exocytosis. Mutants also have increased dehydration tolerance which may be related to the overall slower stomatal aperture dynamics. Overall growth is affected.

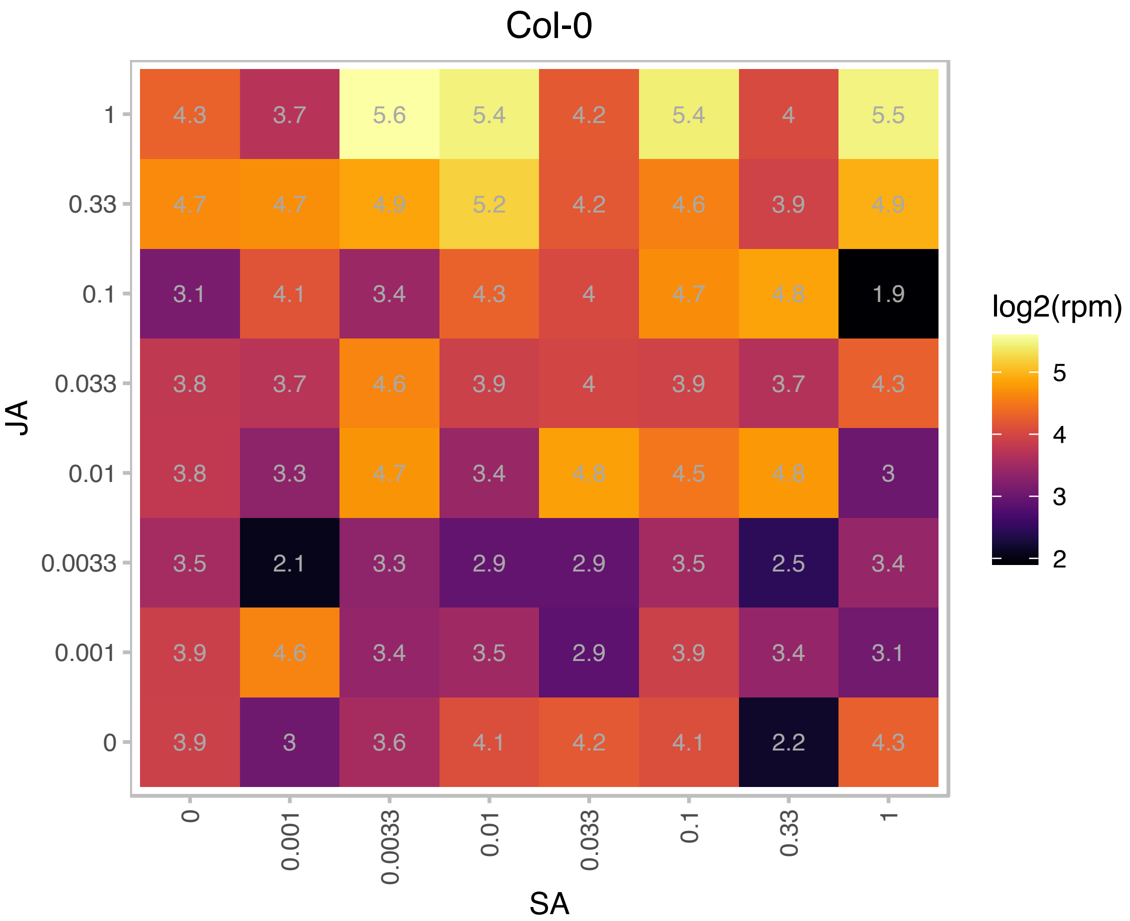


| | |
|---|---|
| Gene | AT3G11130 |
| is.expressed_genes | TRUE |
| is.responsive_genes | TRUE |
| is.apcluster | A-01 |

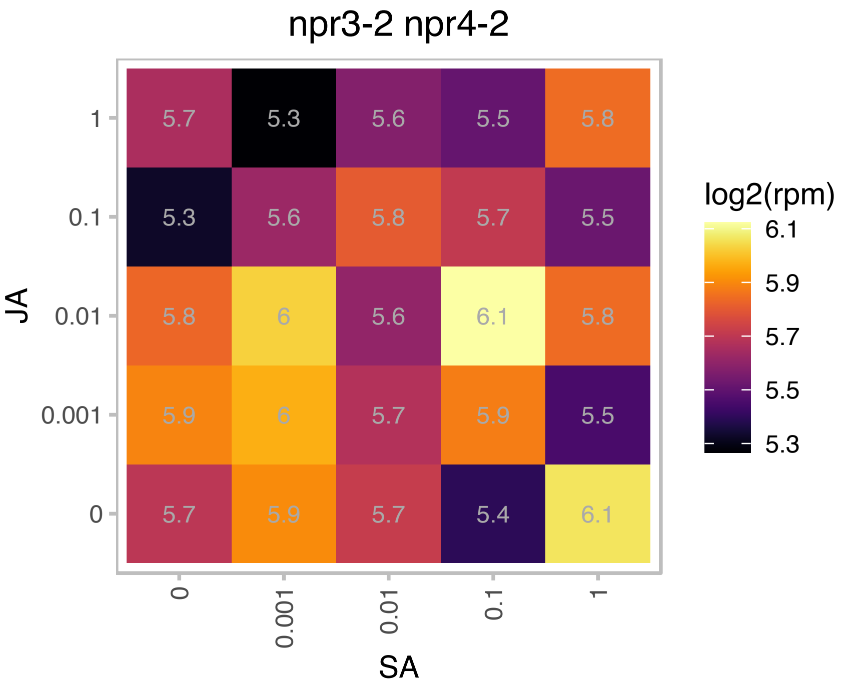


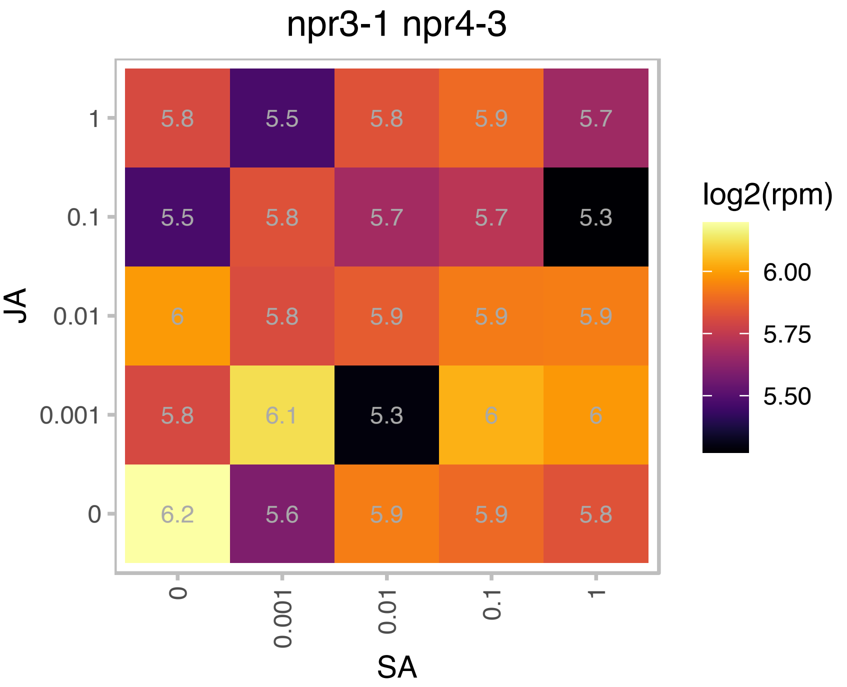

### AT3G23990.1 : HSP60, HSP60-3B

heat shock protein 60

mitochondrial chaperonin HSP. assist in rapid assembly of the oligomeric protein structures in the mitochondria.

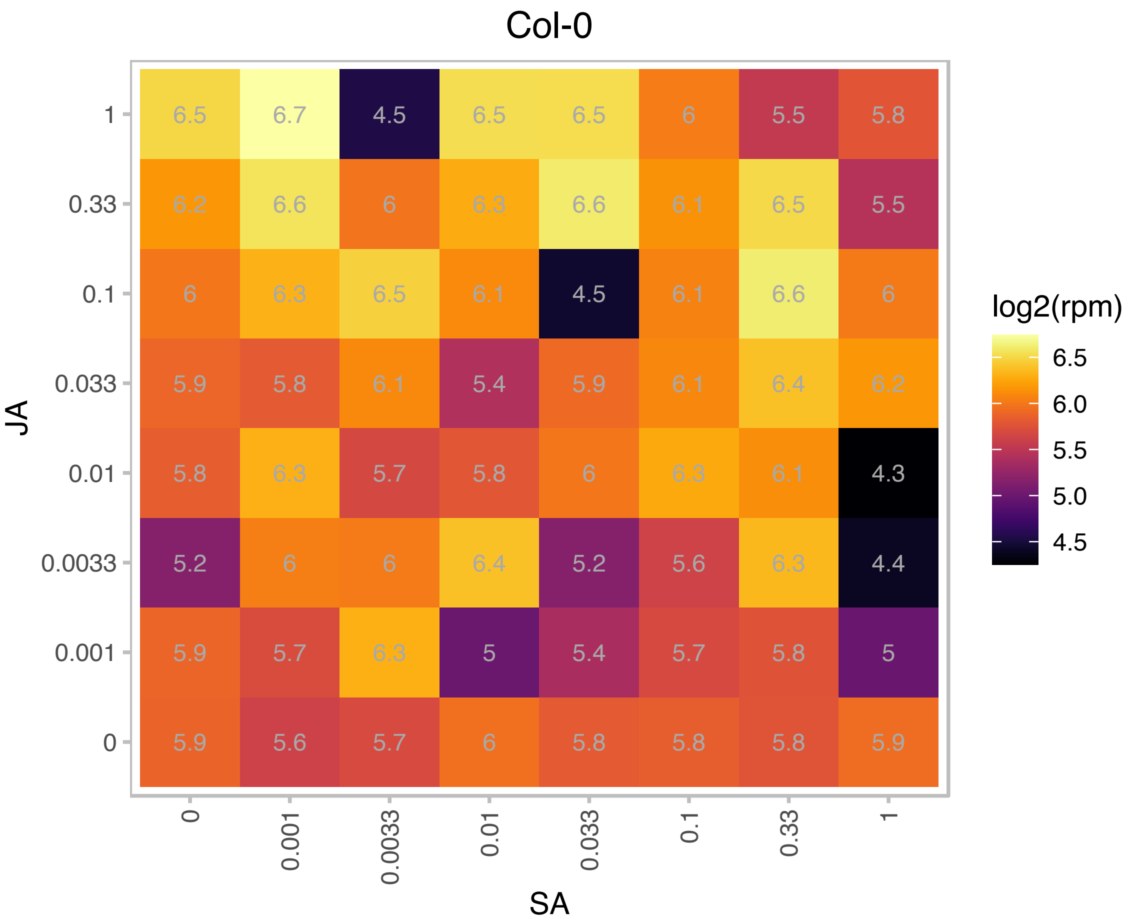


| | |
|---|---|
| Gene | AT3G23990 |
| is.expressed_genes | TRUE |
| is.responsive_genes | TRUE |
| is.apcluster | A-01 |

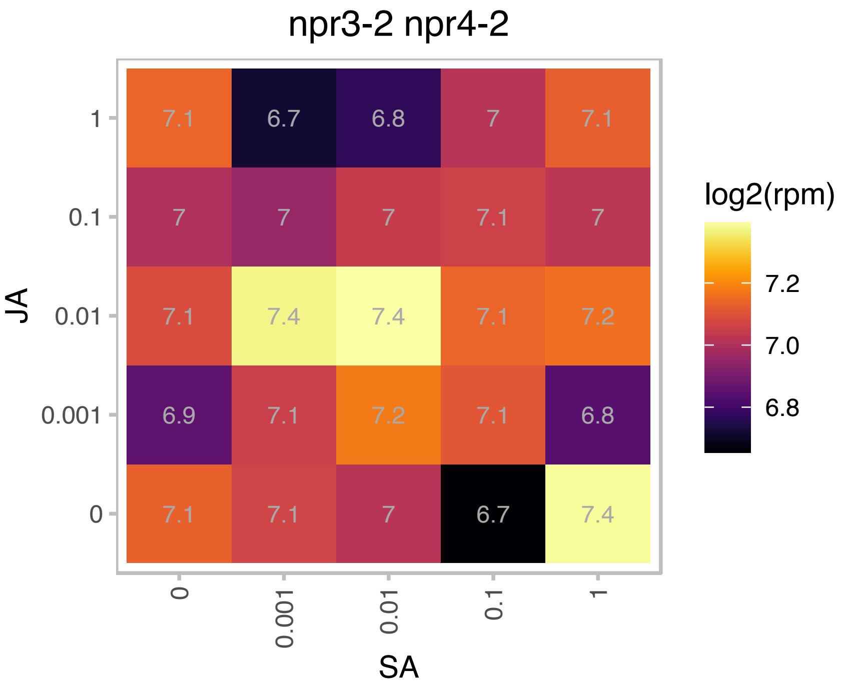


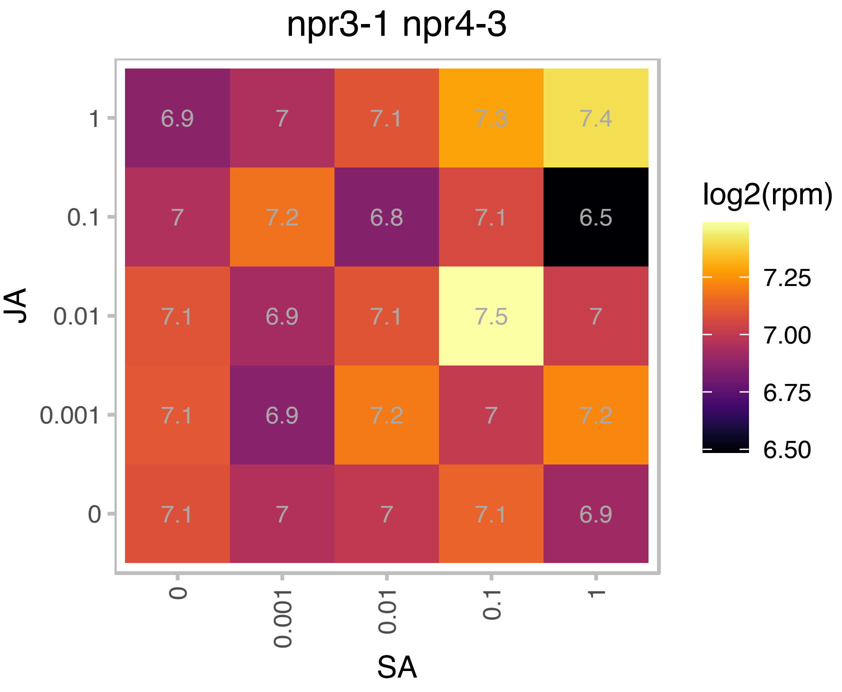

# AT3G26450.1 : NA

Polyketide cyclase/dehydrase and lipid transport superfamily protein

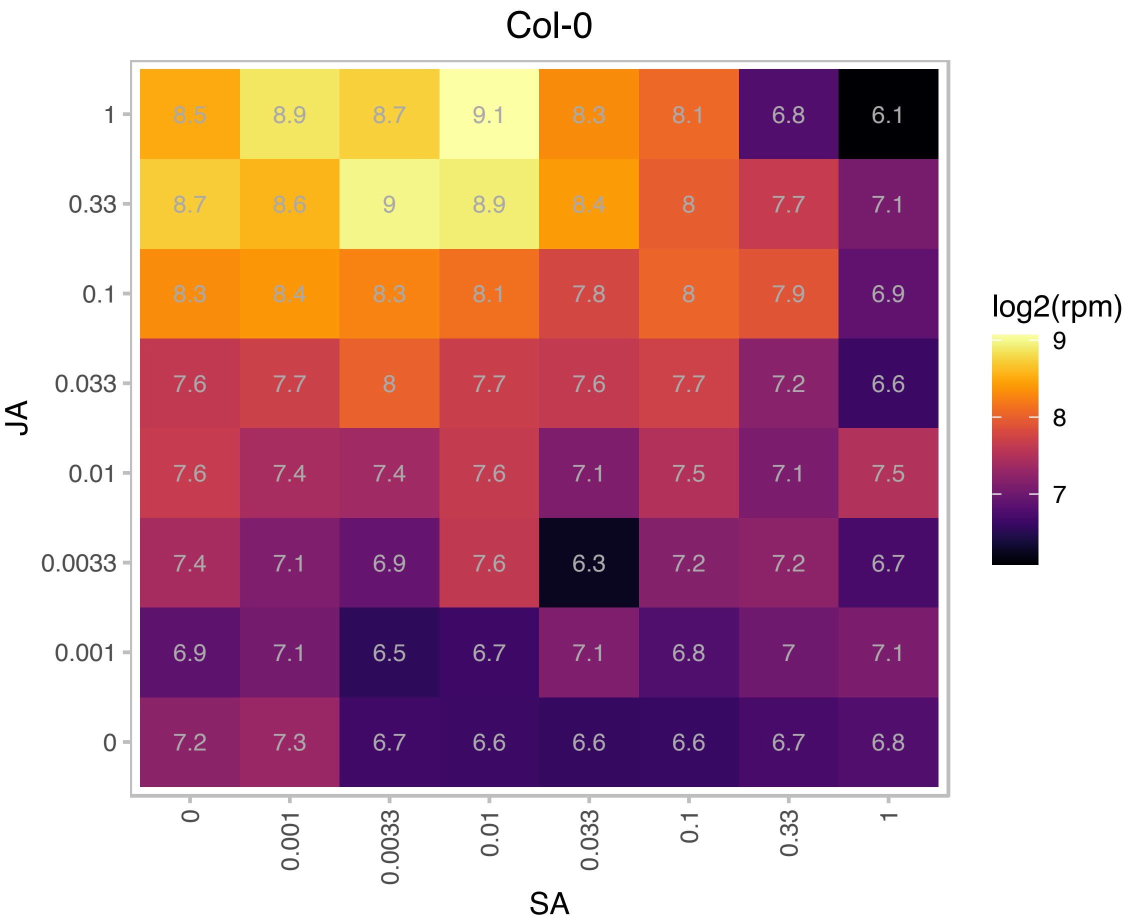


| | |
|---|---|
| Gene | AT3G26450 |
| is.expressed_genes | TRUE |
| is.responsive_genes | TRUE |
| is.apcluster | A-01 |

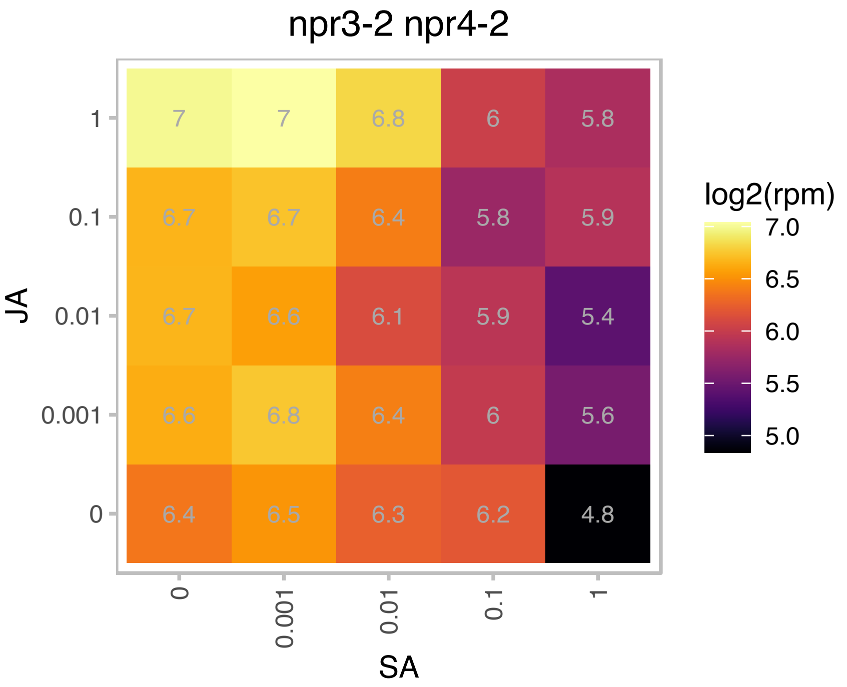


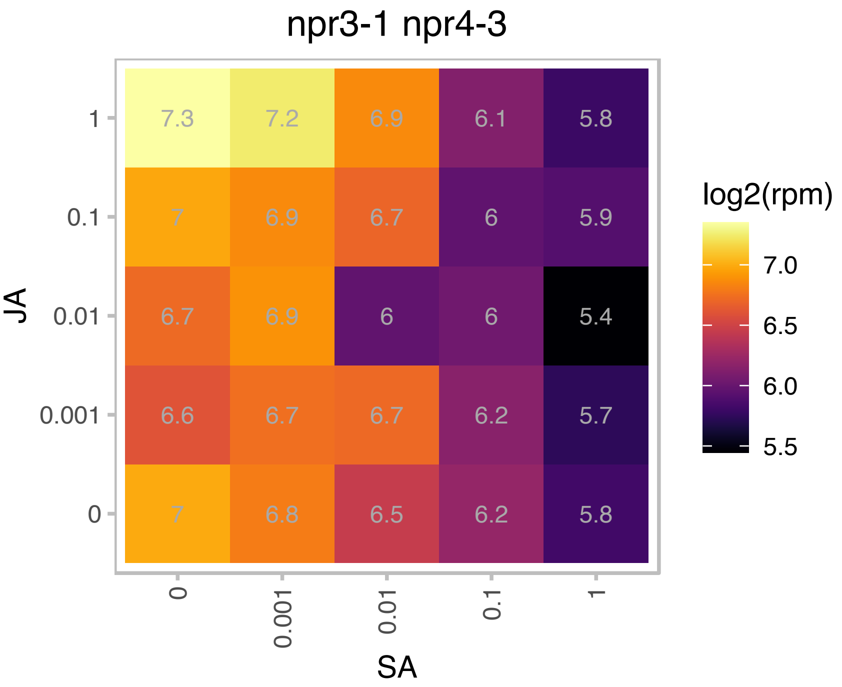

# AT3G43590.1 : NA

zinc knuckle (CCHC-type) family protein

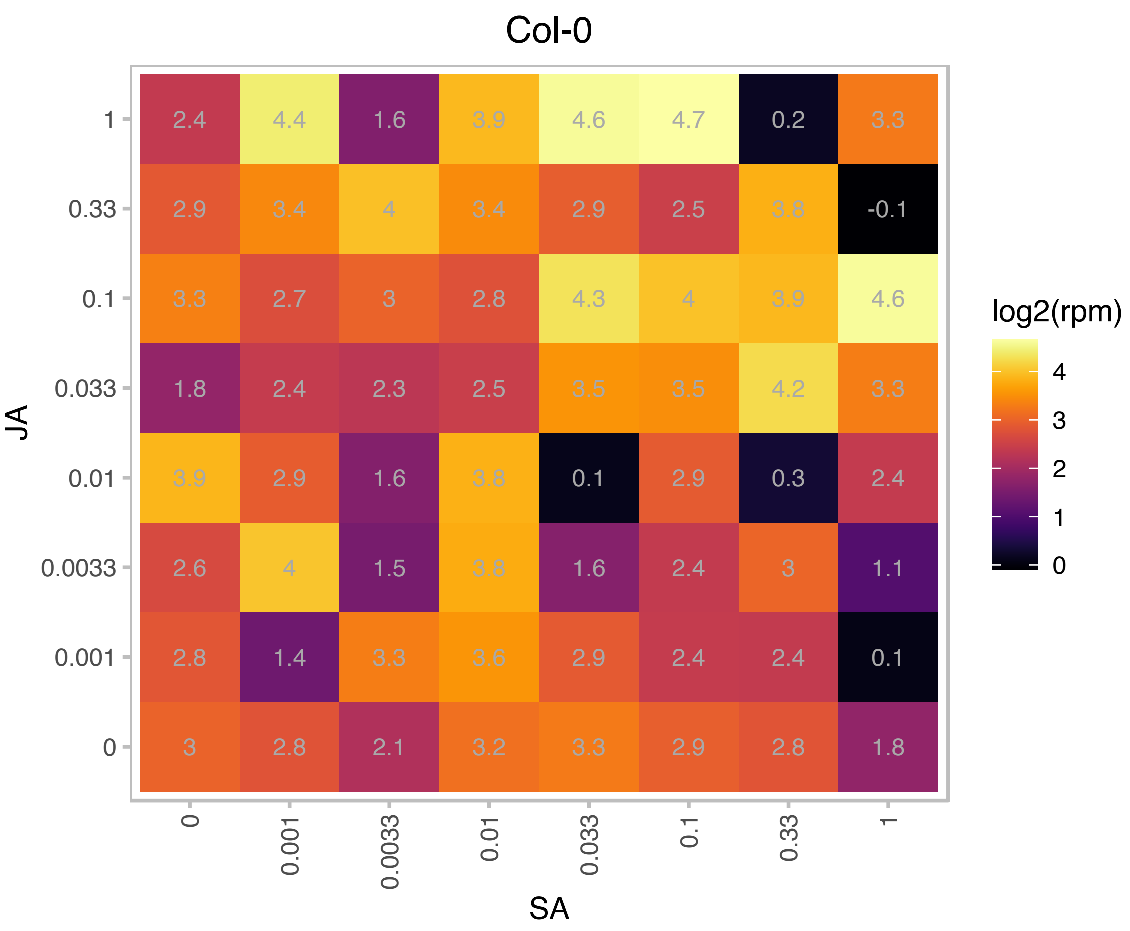


| | |
|---|---|
| Gene | AT3G43590 |
| is.expressed_genes | TRUE |
| is.responsive_genes | TRUE |
| is.apcluster | A-01 |

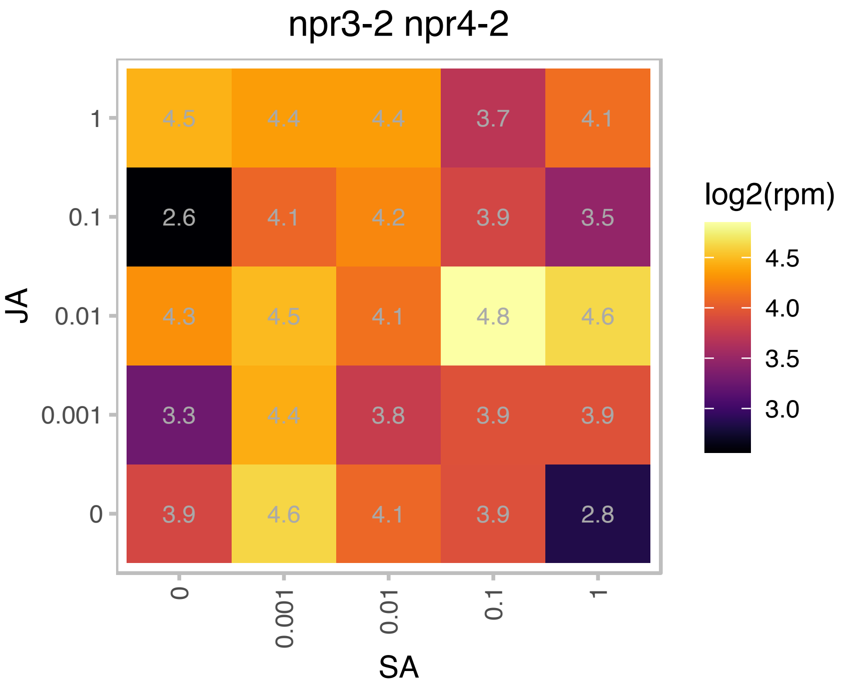


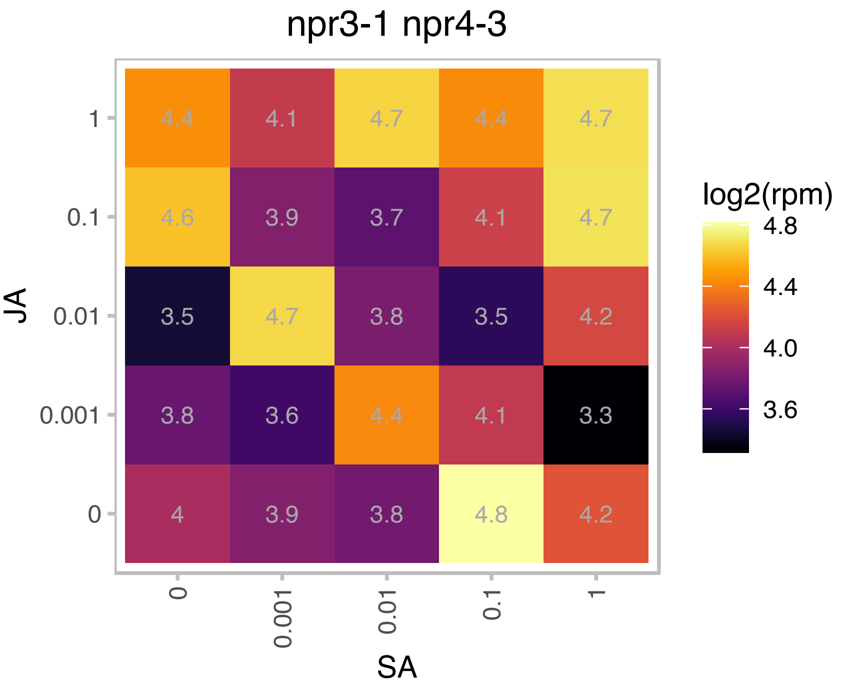

### AT3G47340.1 : ASN1, AT-ASN1, DIN6

glutamine-dependent asparagine synthase 1

encodes a glutamine-dependent asparagine synthetase, the predicted ASN1 peptide contains a purF-type glutamine-binding domain, and is expressed predominantly in shoot tissues, where light has a negative effect on its mRNA accumulation. Expression is induced within 3 hours of dark treatment, in senescing leaves and treatment with exogenous photosynthesis inhibitor. Induction of gene expression was suppressed in excised leaves supplied with sugar. The authors suggest that the gene's expression pattern is responding to the level of sugar in the cell.

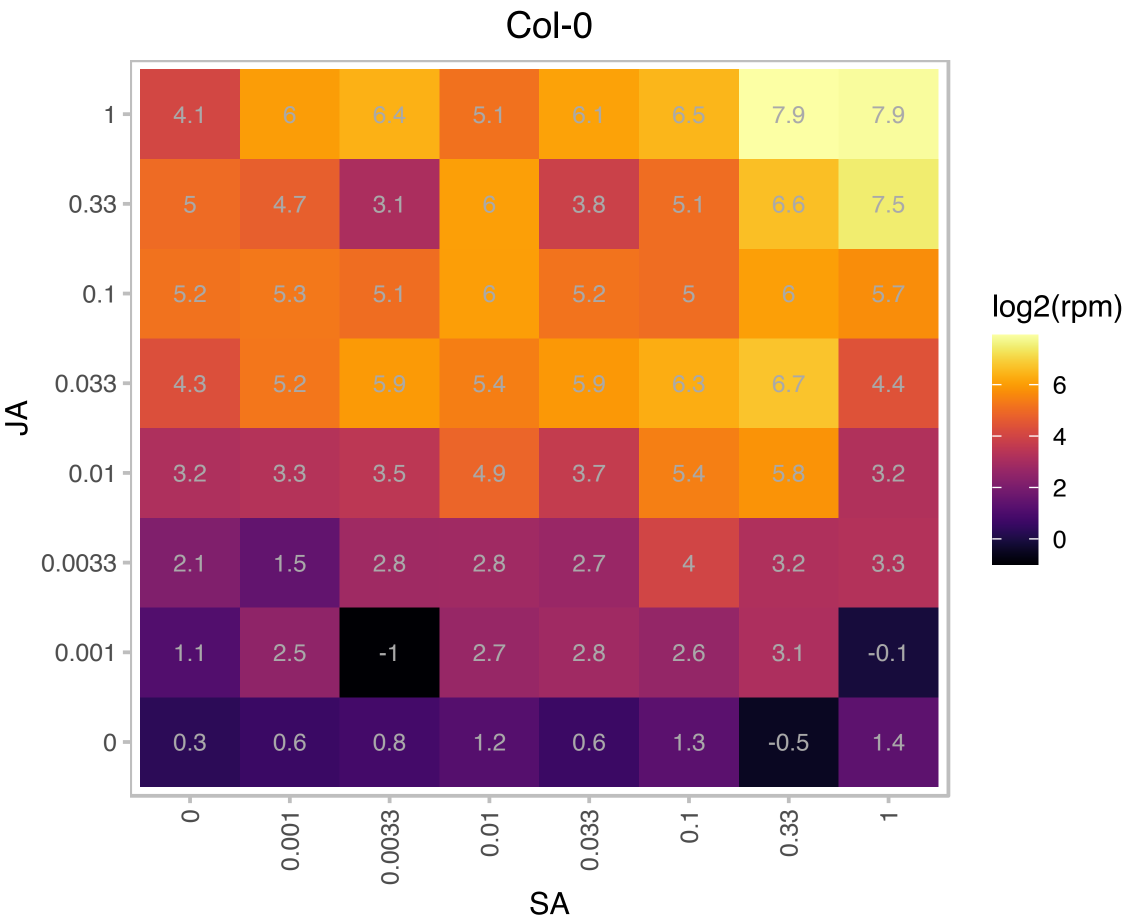


| | |
|---|---|
| Gene | AT3G47340 |
| is.expressed_genes | TRUE |
| is.responsive_genes | TRUE |
| is.apcluster | A-01 |

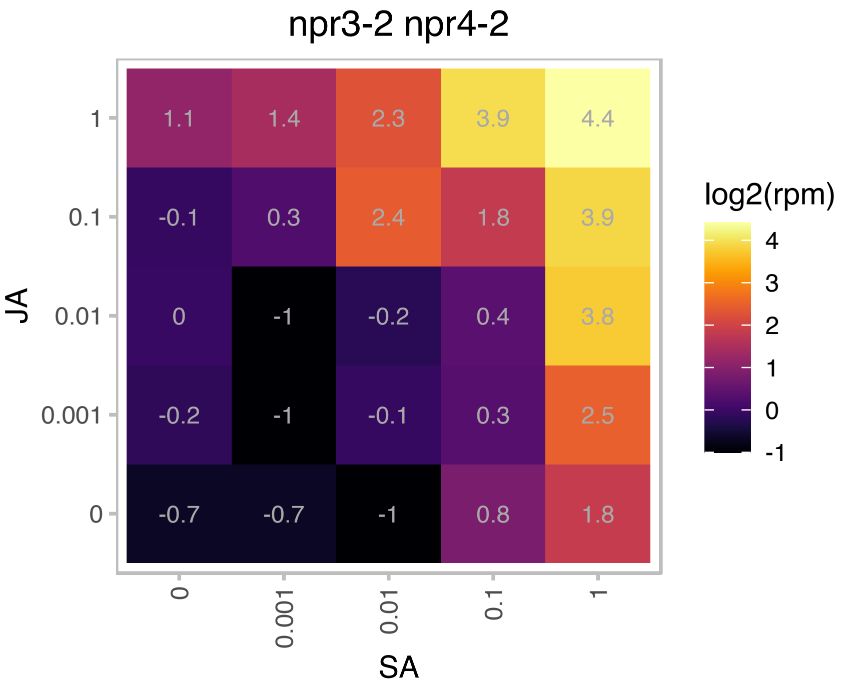


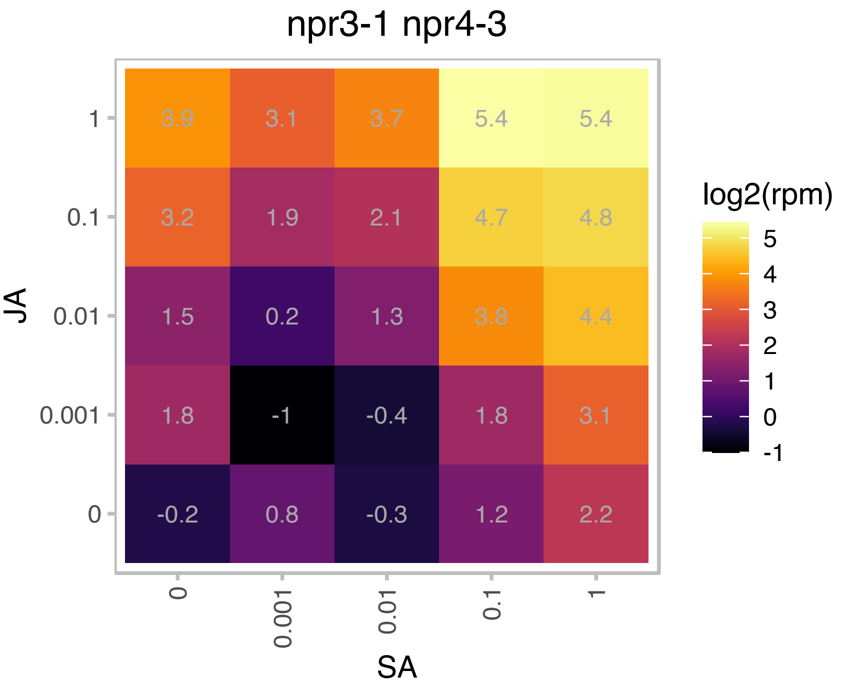

### AT3G55620.1 : eIF6A, emb1624

Translation initiation factor IF6

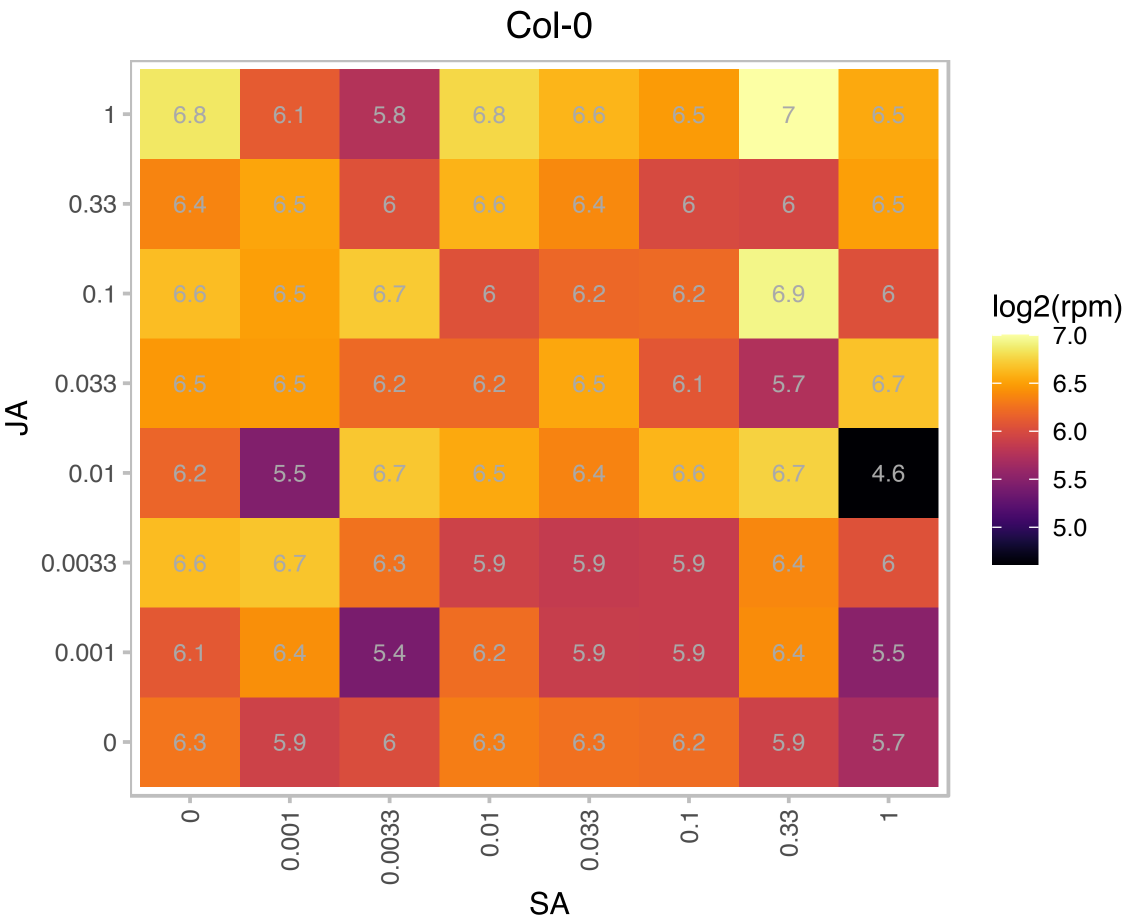


| | |
|---|---|
| Gene | AT3G55620 |
| is.expressed_genes | TRUE |
| is.responsive_genes | TRUE |
| is.apcluster | A-01 |

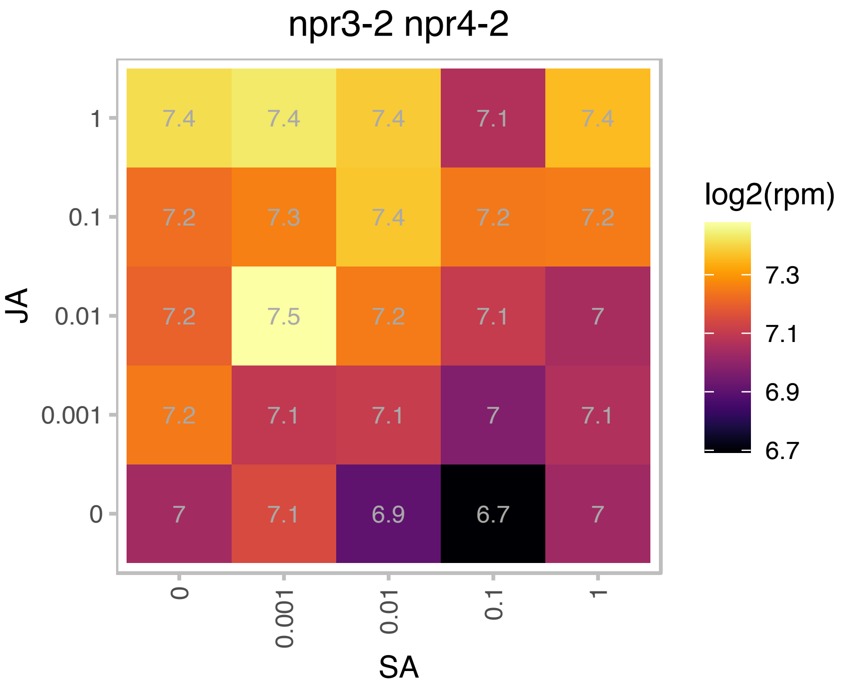


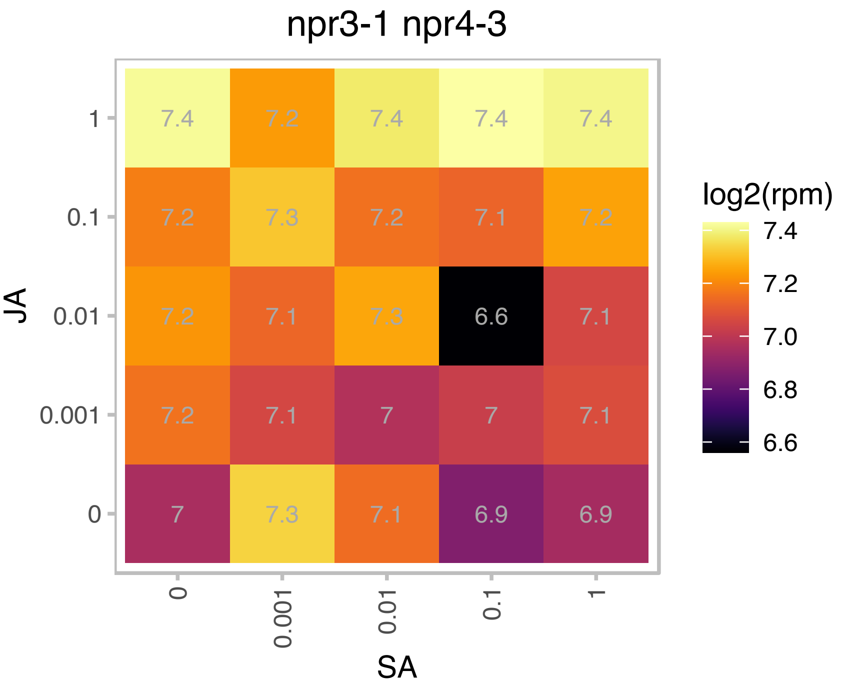

### AT3G55970.1 : ATJRG21, JAO3, JOX3, JRG21

jasmonate-regulated gene 21

One of 4 paralogs encoding a 2-oxoglutarate/Fe(II)-dependent oxygenases that hydroxylates JA to 12-OH-JA.

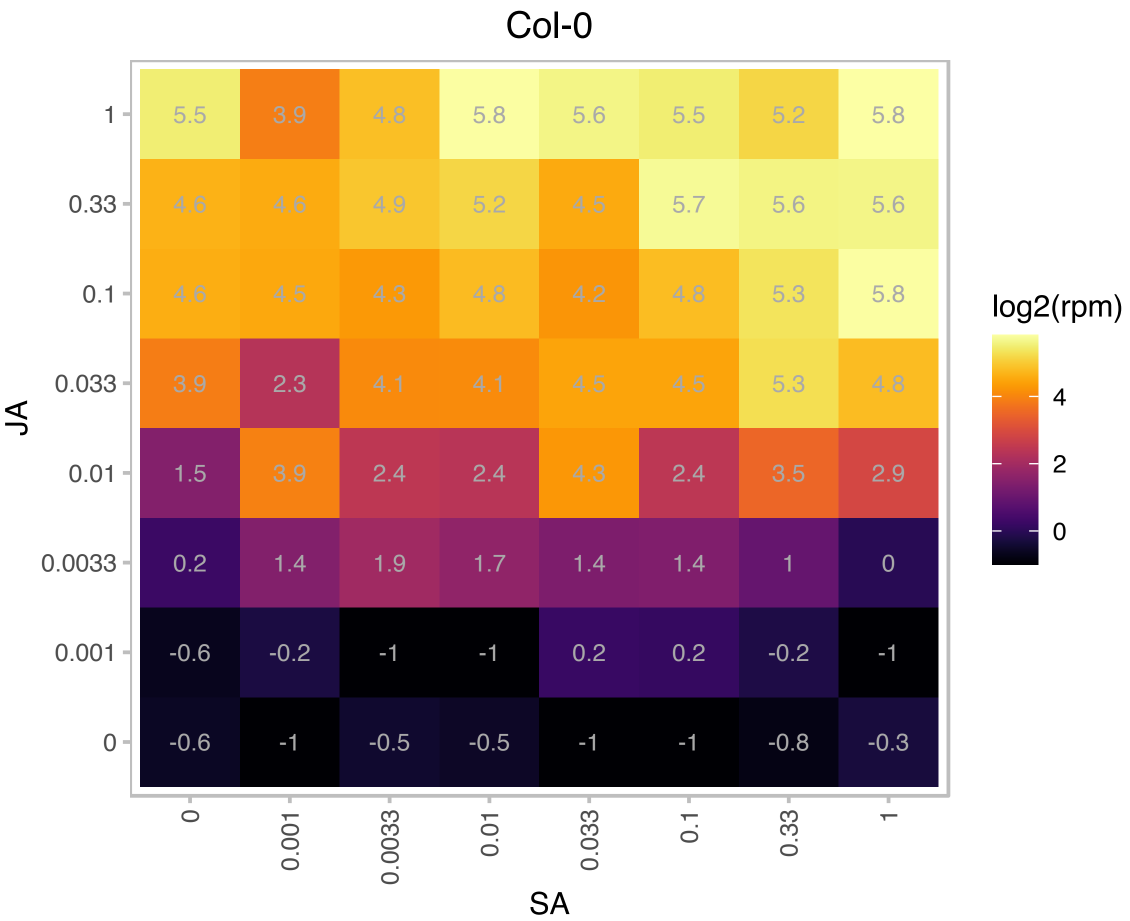


| | |
|---|---|
| Gene | AT3G55970 |
| is.expressed_genes | TRUE |
| is.responsive_genes | TRUE |
| is.apcluster | A-01 |

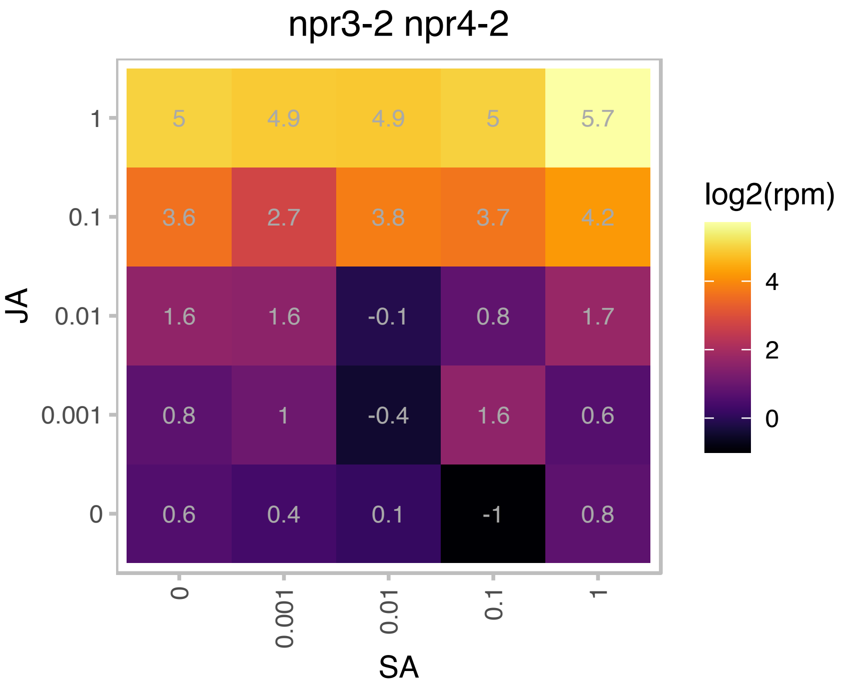


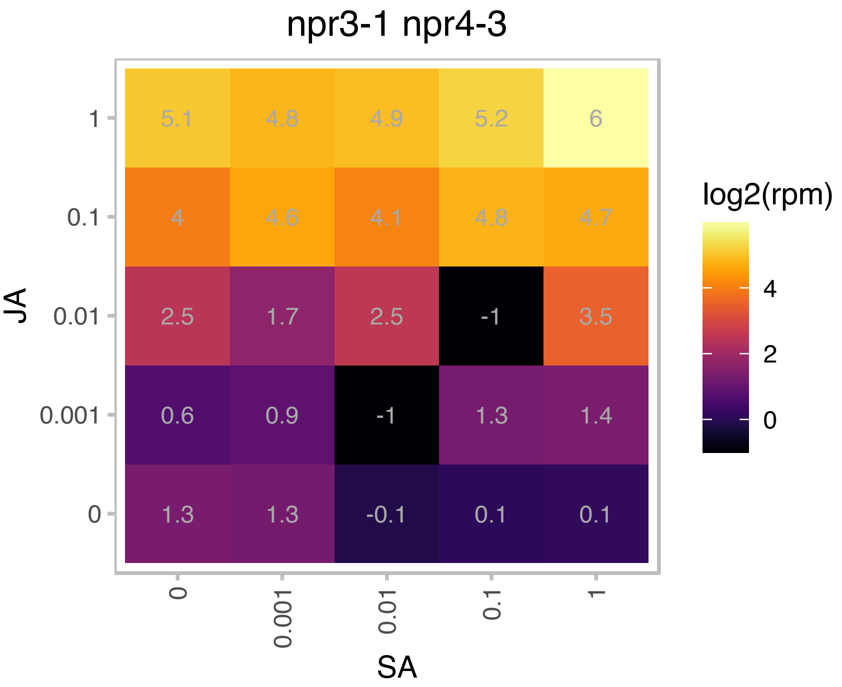

# AT3G57320.1 : NA

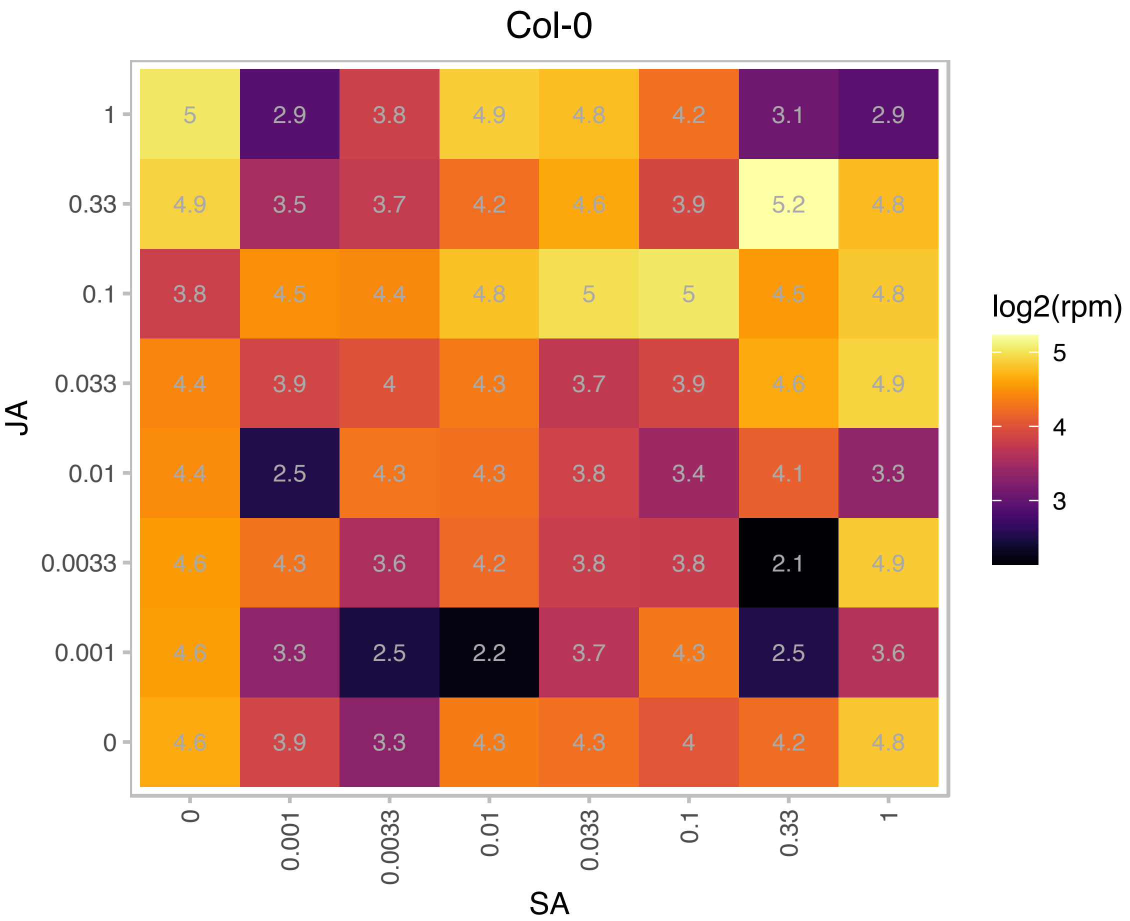


| | |
|---|---|
| Gene | AT3G57320 |
| is.expressed_genes | TRUE |
| is.responsive_genes | TRUE |
| is.apcluster | A-01 |

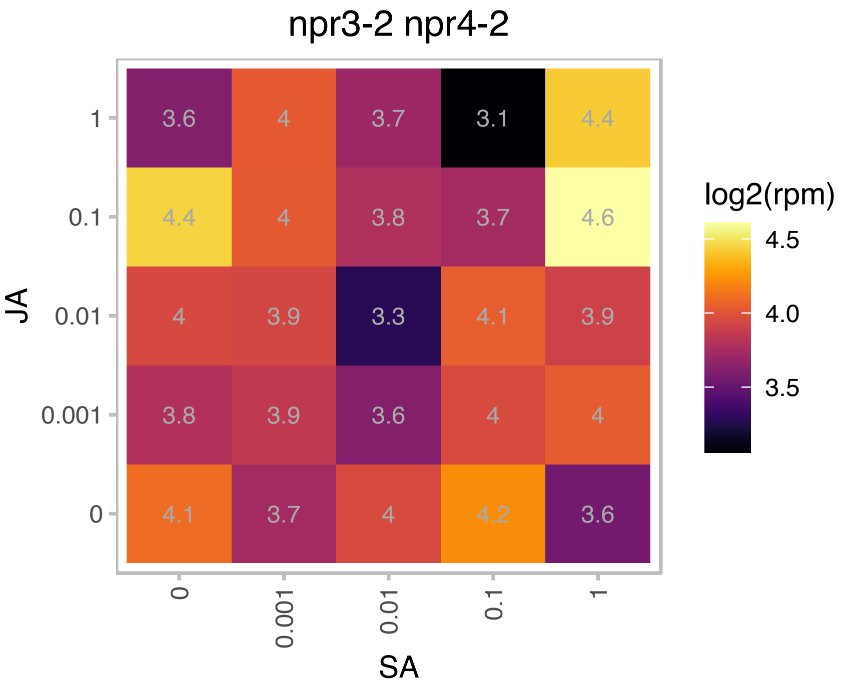


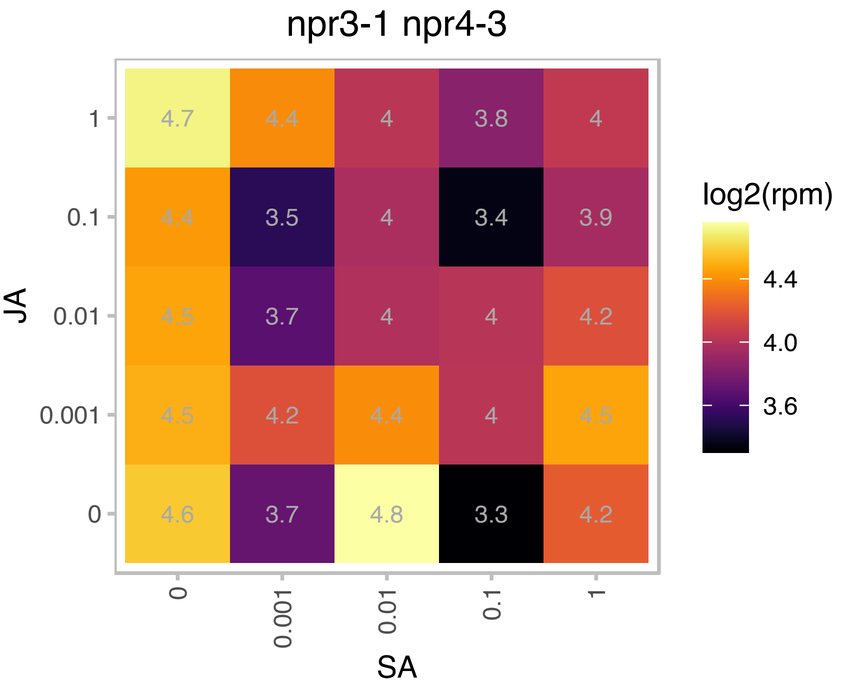

### AT4G05530.1 : IBR1, SDRA

indole-3-butyric acid response 1

Encodes a peroxisomal member of the short-chain dehydrogenase/reductase (SDR) family of enzymes. Loss of IBR function causes increased resistance to indole-3-butyric acid without affecting plant responses to IAA, NAA, and 2,4-D. This enzyme may be responsible for catalyzing a dehydrogenation step in the beta-oxidation-like conversion of IBA to IAA. The mRNA is cell-to-cell mobile.

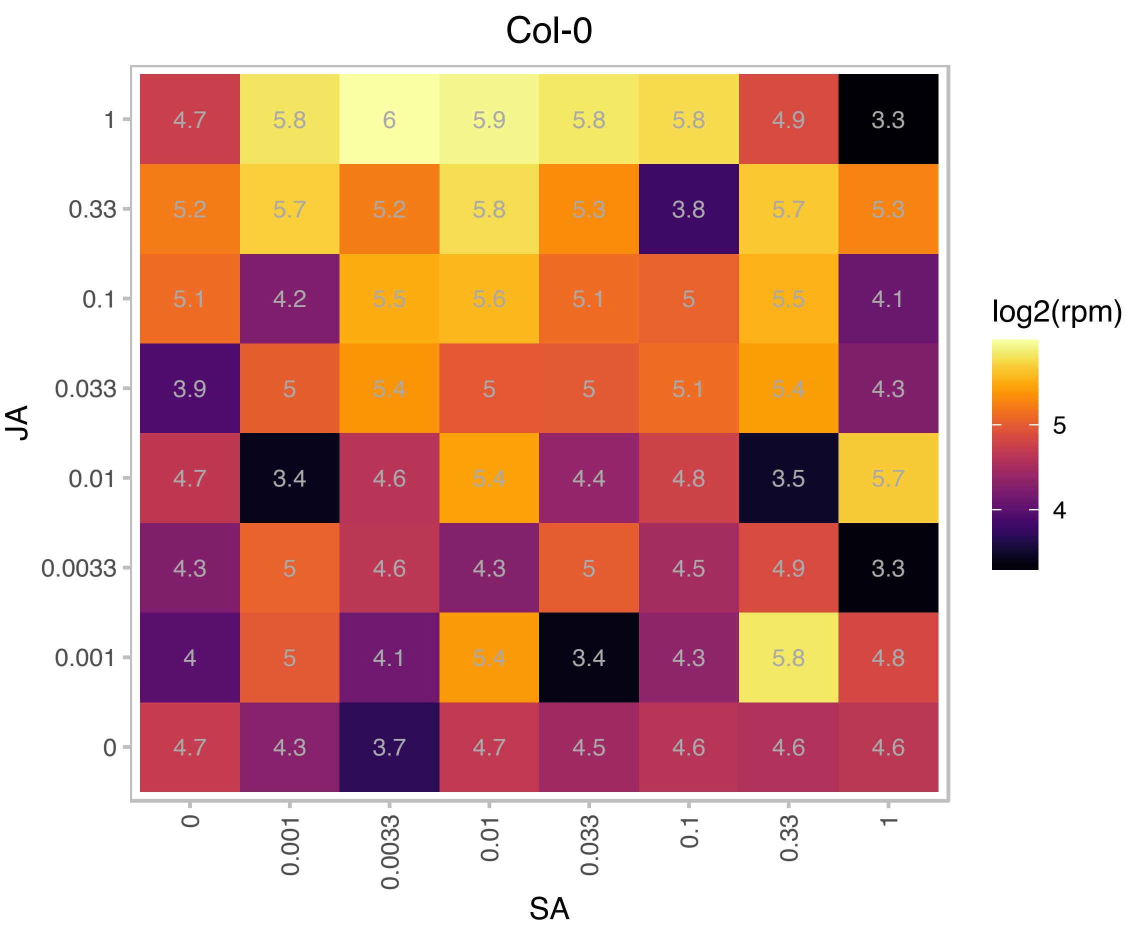


| | |
|---|---|
| Gene | AT4G05530 |
| is.expressed_genes | TRUE |
| is.responsive_genes | TRUE |
| is.apcluster | A-01 |

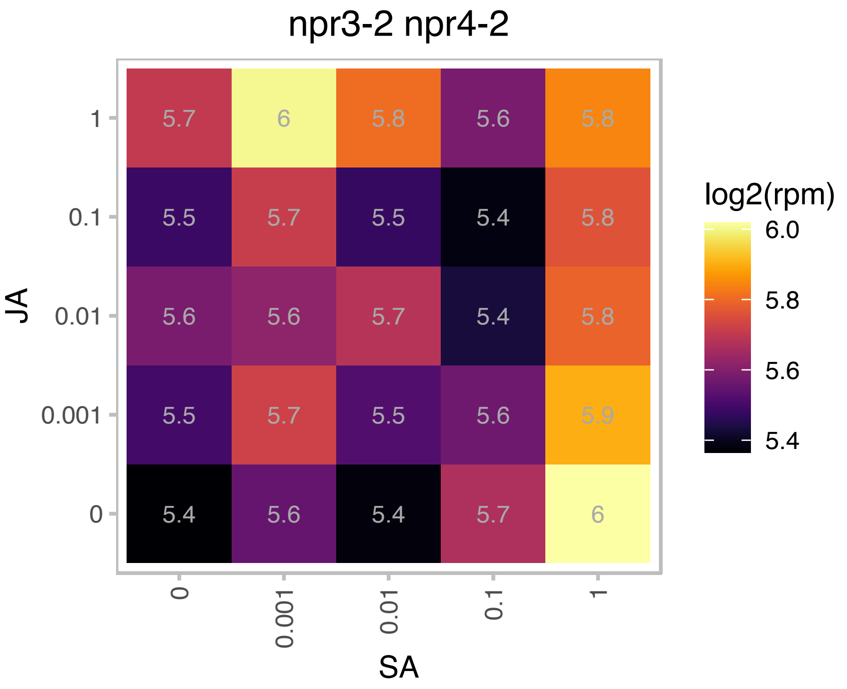


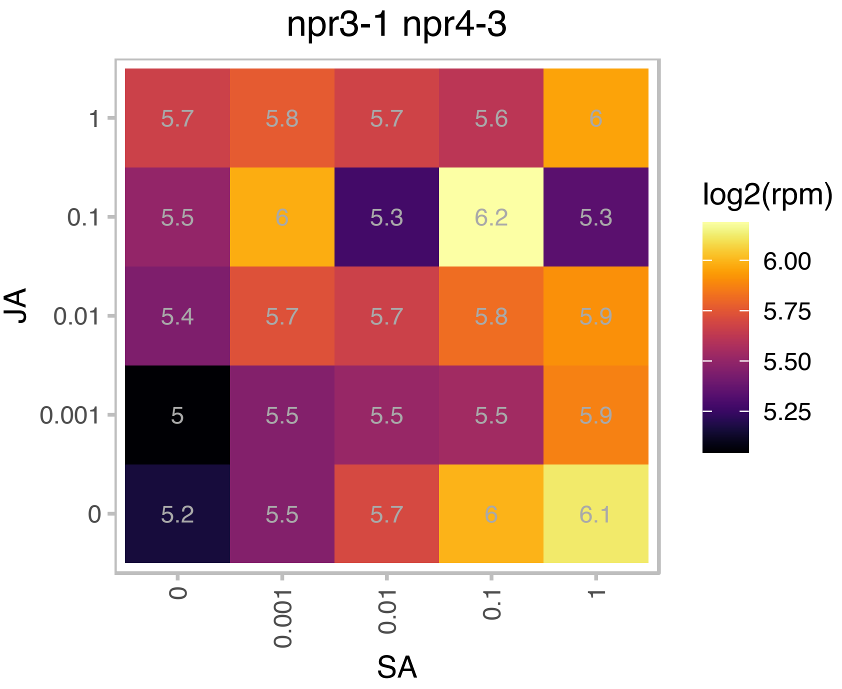

# AT4G32610.1 : NA

copper ion binding

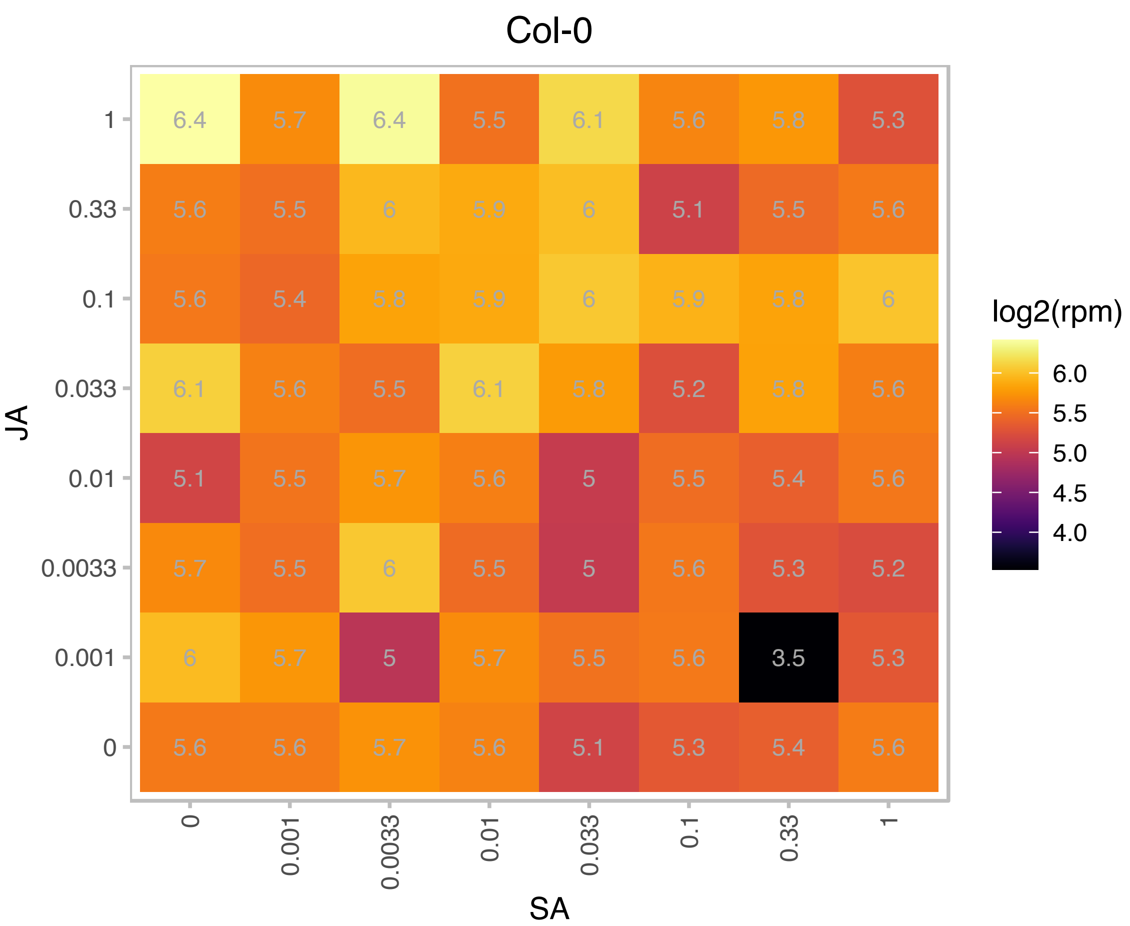


| | |
|---|---|
| Gene | AT4G32610 |
| is.expressed_genes | TRUE |
| is.responsive_genes | TRUE |
| is.apcluster | A-01 |

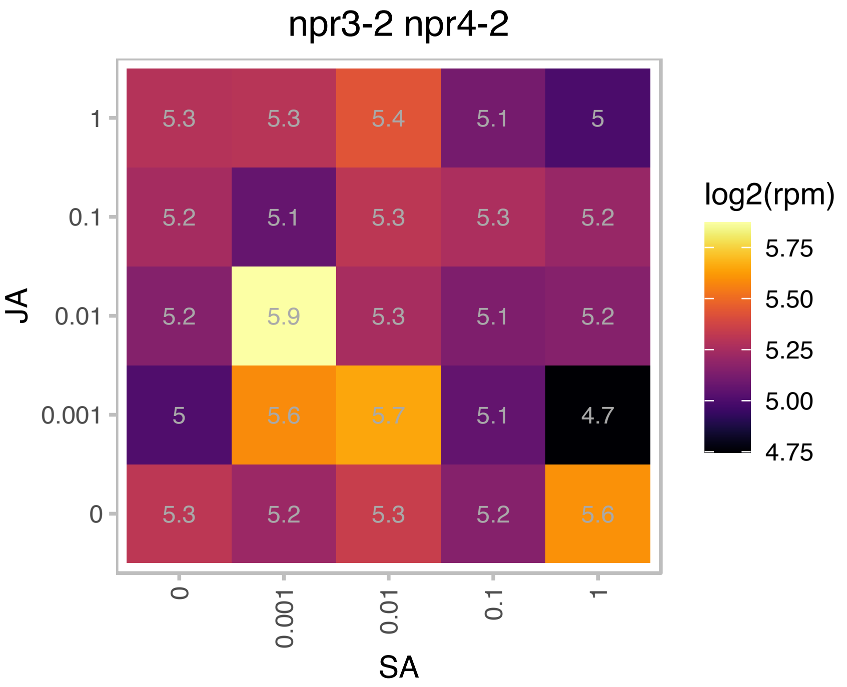


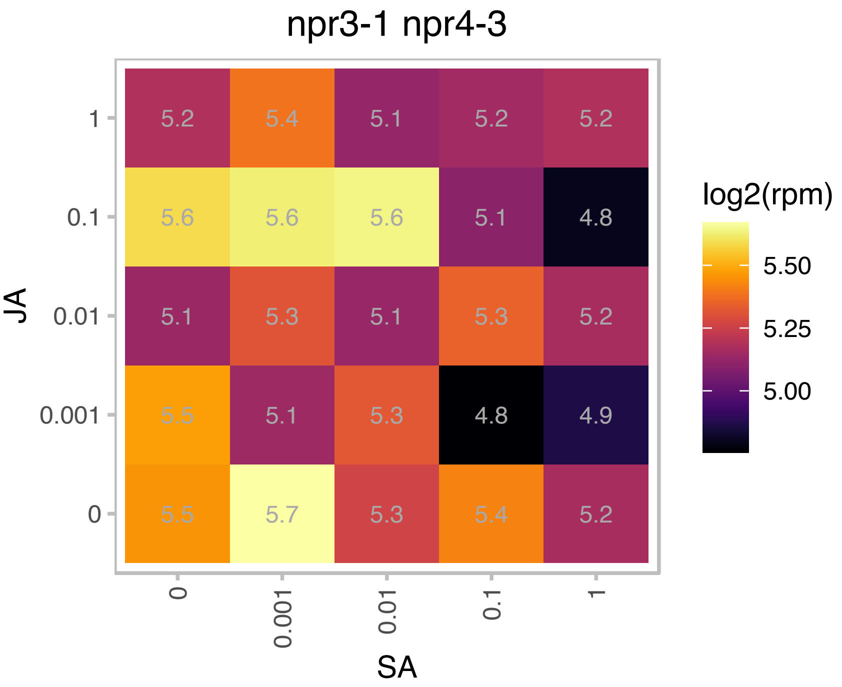

### AT4G34110.1 : ATPAB2, PAB2, PABP2

poly(A) binding protein 2

Putative poly-A binding protein. Member of a gene family .Expressed in stele and root meristem and post-fertilization ovules.Member of the class II family of PABP proteins. The mRNA is cell-to-cell mobile.

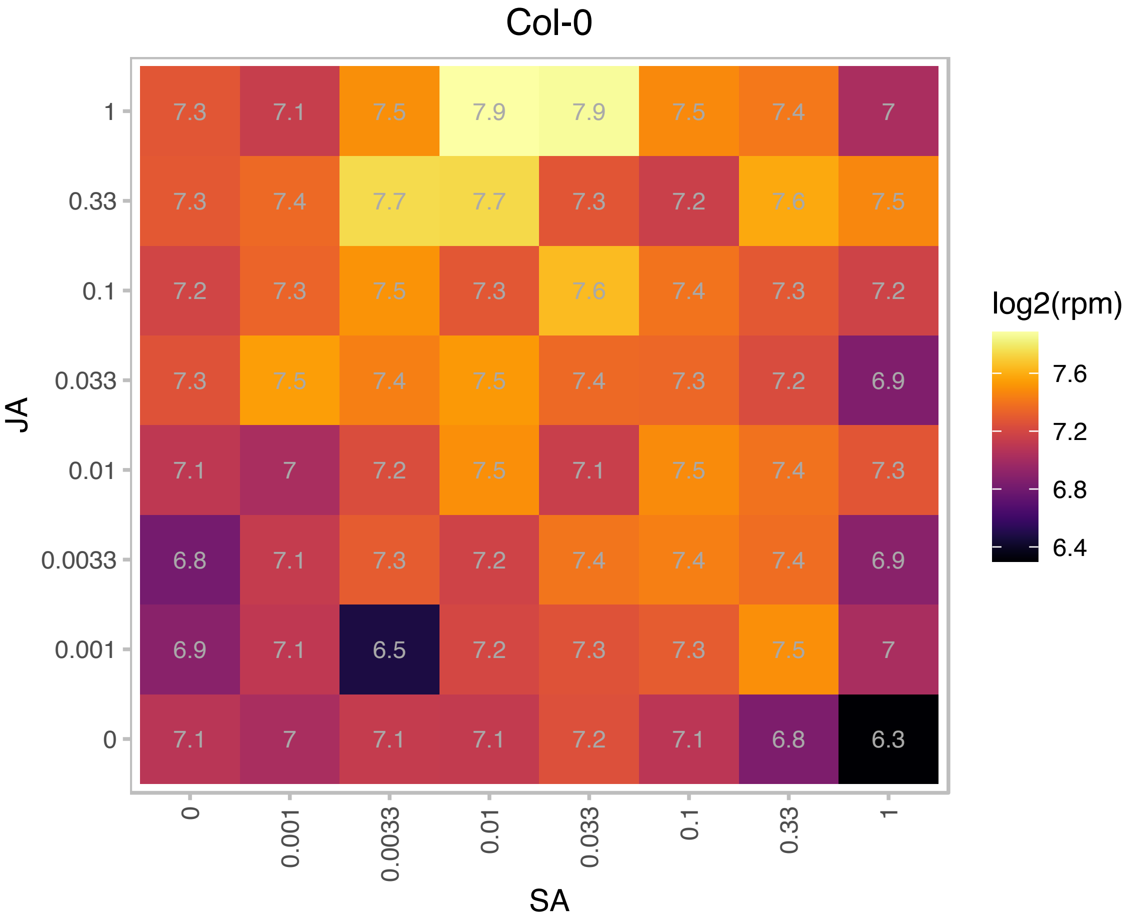


| | |
|---|---|
| Gene | AT4G34110 |
| is.expressed_genes | TRUE |
| is.responsive_genes | TRUE |
| is.apcluster | A-01 |

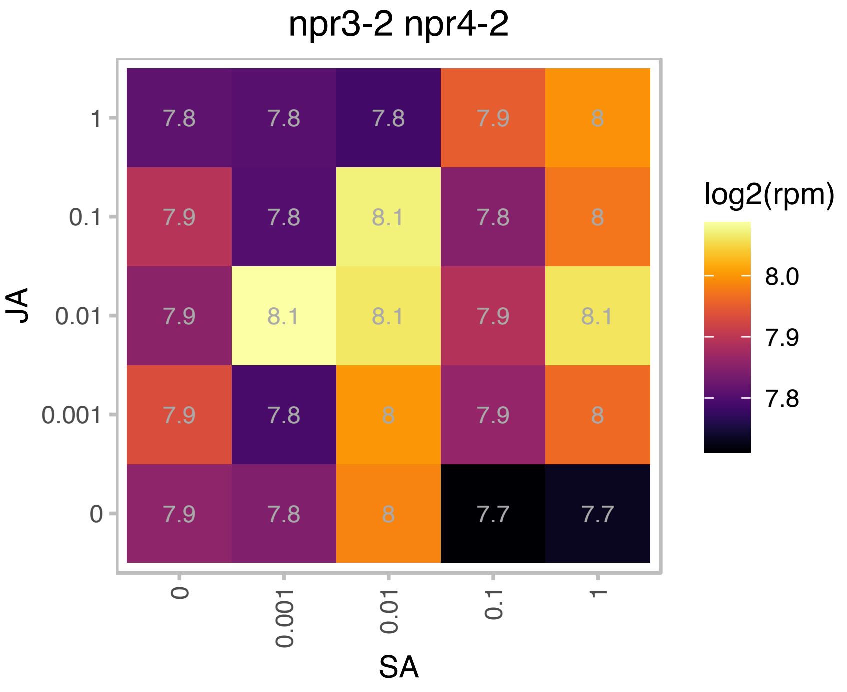


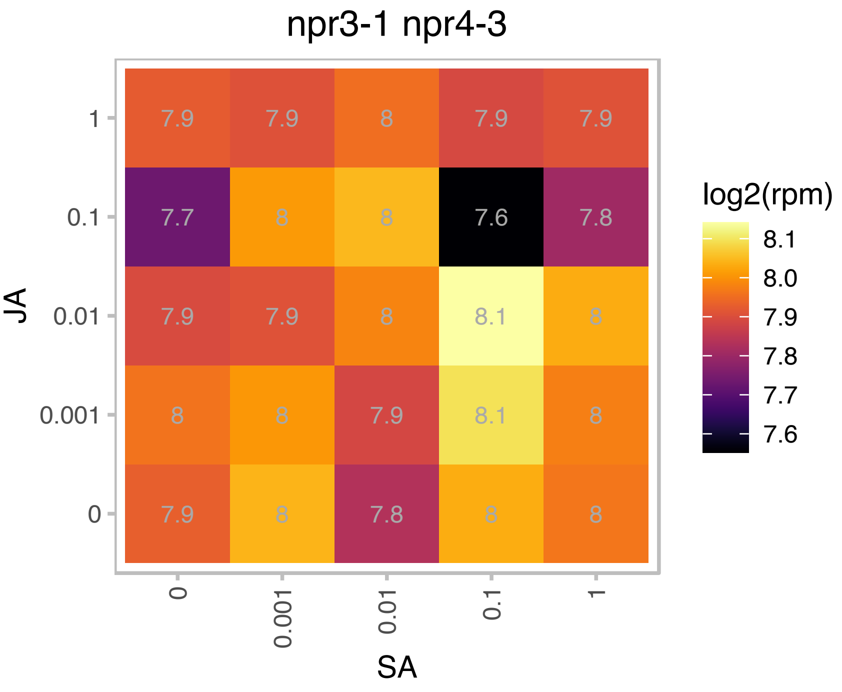

### AT4G35100.1 : PIP2;7, PIP3, PIP3A, SIMIP

plasma membrane intrinsic protein 3

a member of the plasma membrane intrinsic protein PIP. functions as aquaporin. Salt-stress-inducible MIP

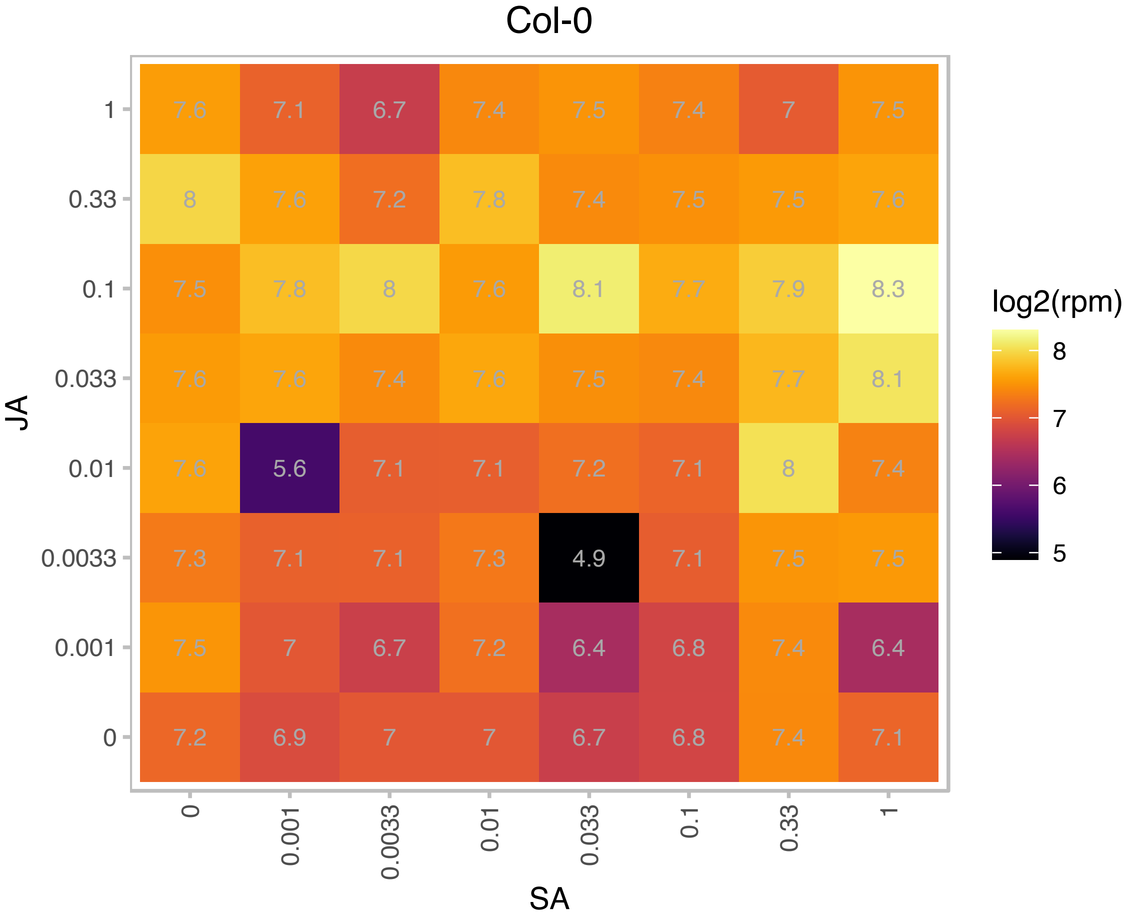


| | |
|---|---|
| Gene | AT4G35100 |
| is.expressed_genes | TRUE |
| is.responsive_genes | TRUE |
| is.apcluster | A-01 |

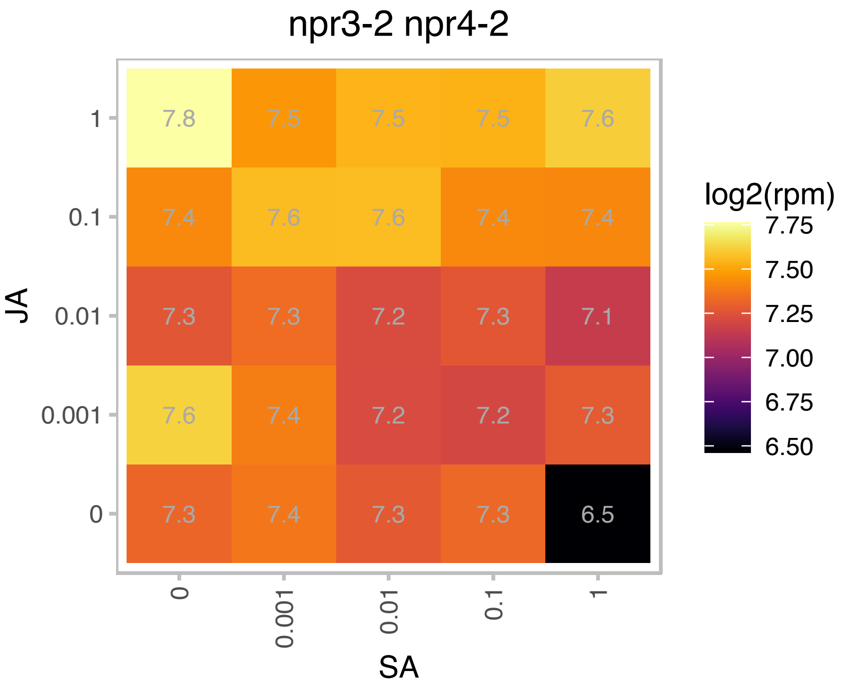


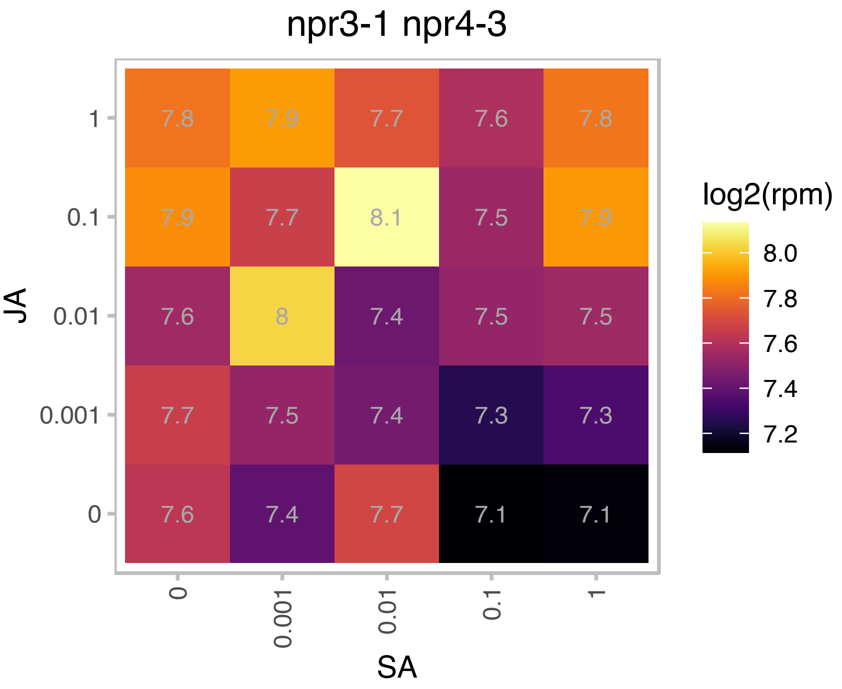

### AT4G36400.1 : D2HGDH

FAD-linked oxidases family protein

Encodes a (D)-2-hydroxyglutarate dehydrogenase.

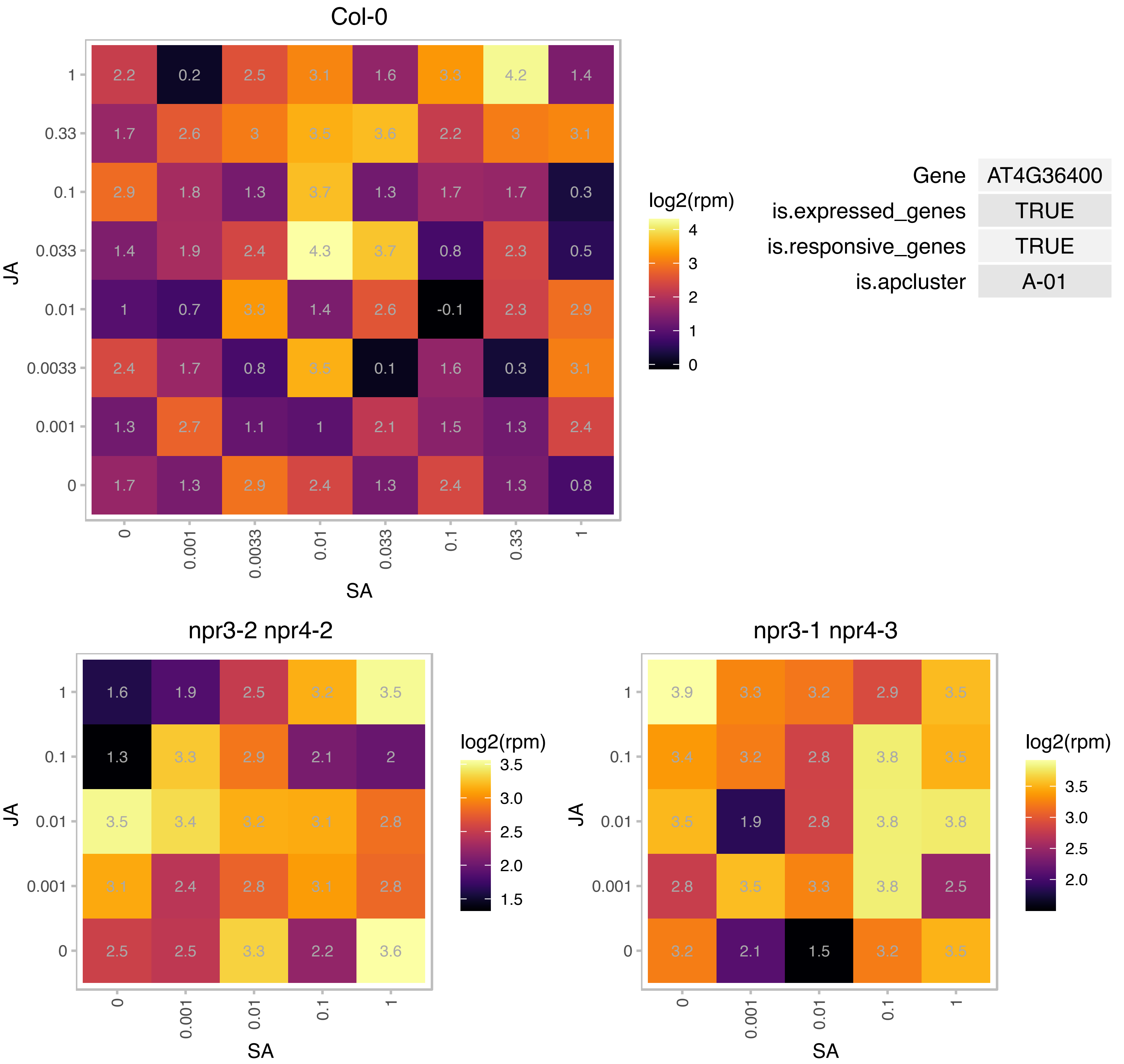


| | |
|---|---|
| Gene | AT4G36400 |
| is.expressed_genes | TRUE |
| is.responsive_genes | TRUE |
| is.apcluster | A-01 |

# AT4G40065.1 : NA

other RNA

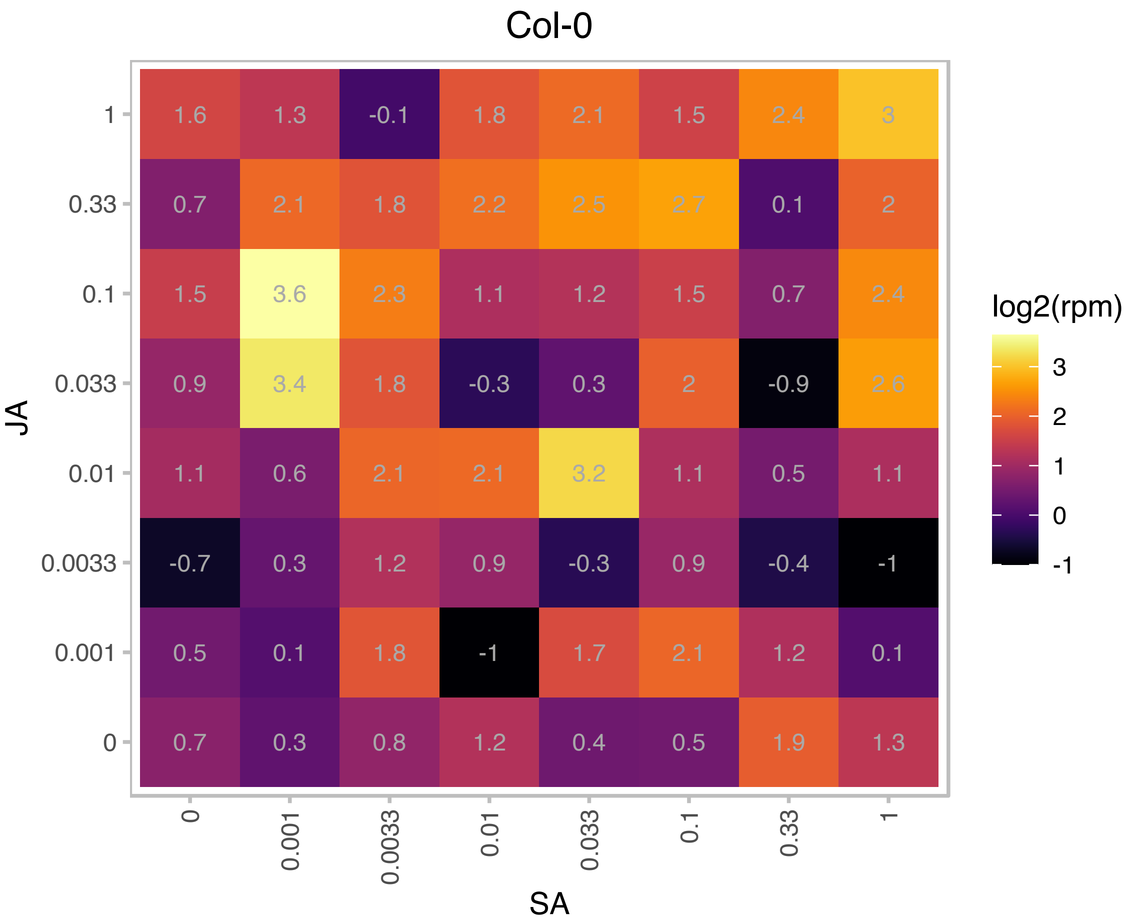


| | |
|---|---|
| Gene | AT4G40065 |
| is.expressed_genes | TRUE |
| is.responsive_genes | TRUE |
| is.apcluster | A-01 |

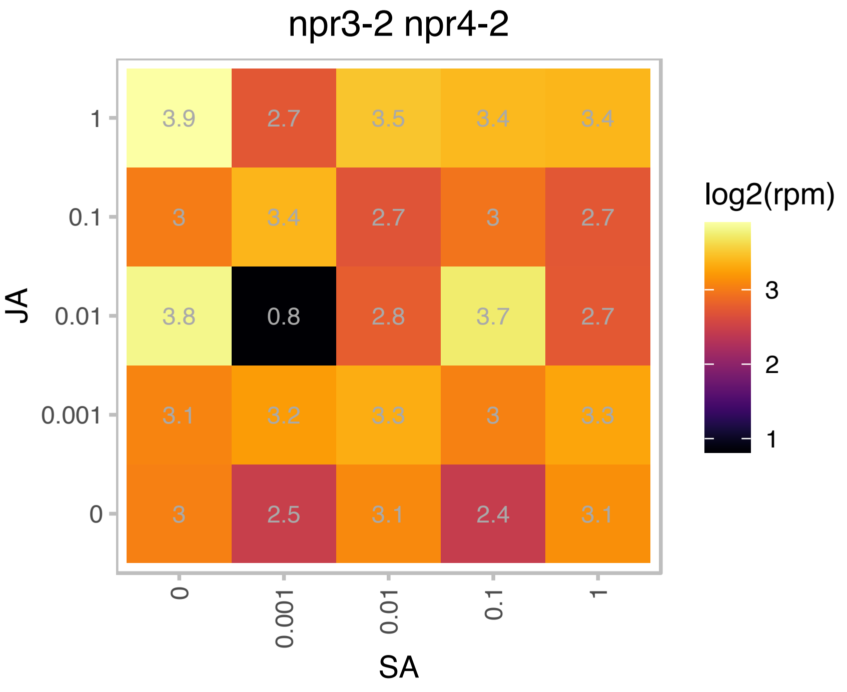


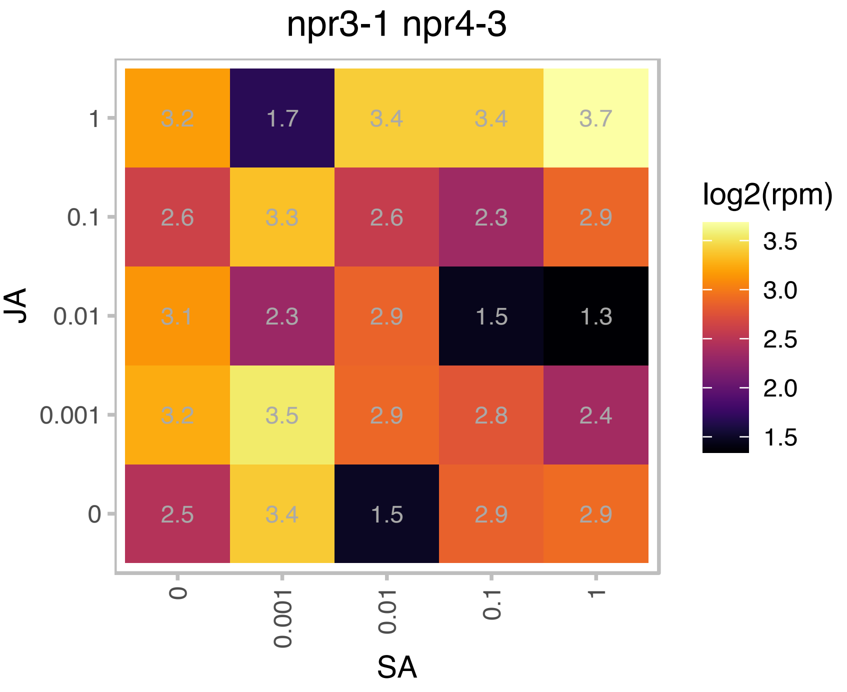

### AT5G01400.1 : ESP4

HEAT repeat-containing protein

Encodes a Symplekin/Pta1 homologue which would have the potential to interact with either ESP1 or AtCstF64.

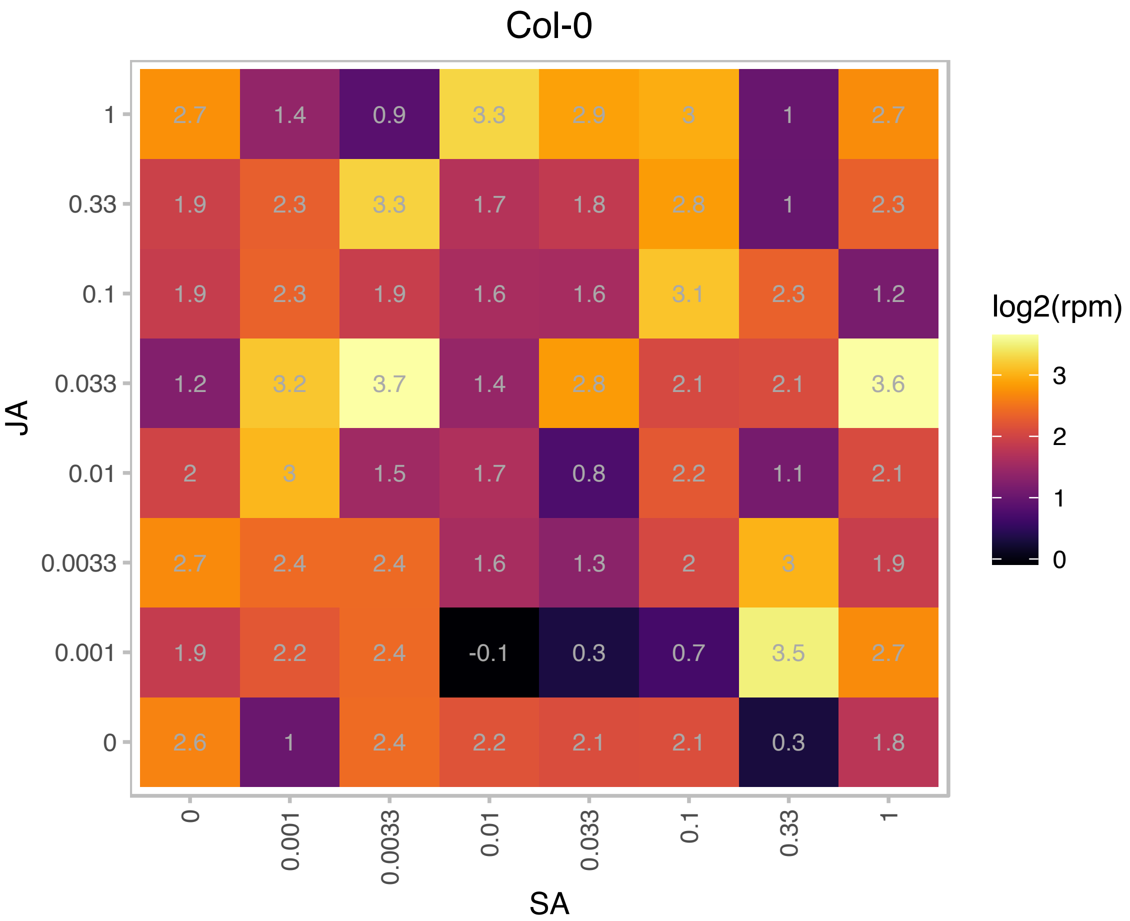


| | |
|---|---|
| Gene | AT5G01400 |
| is.expressed_genes | TRUE |
| is.responsive_genes | TRUE |
| is.apcluster | A-01 |

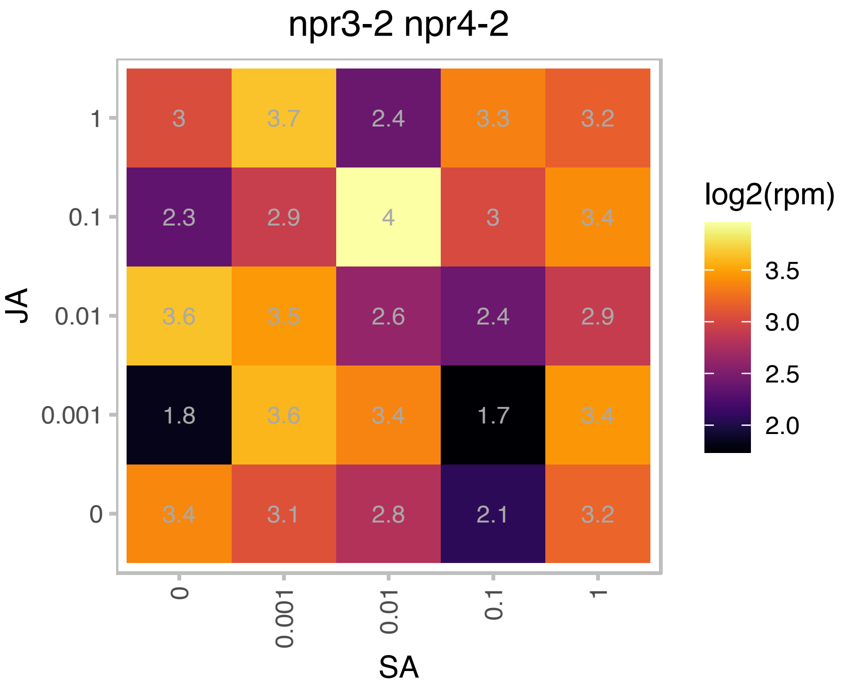


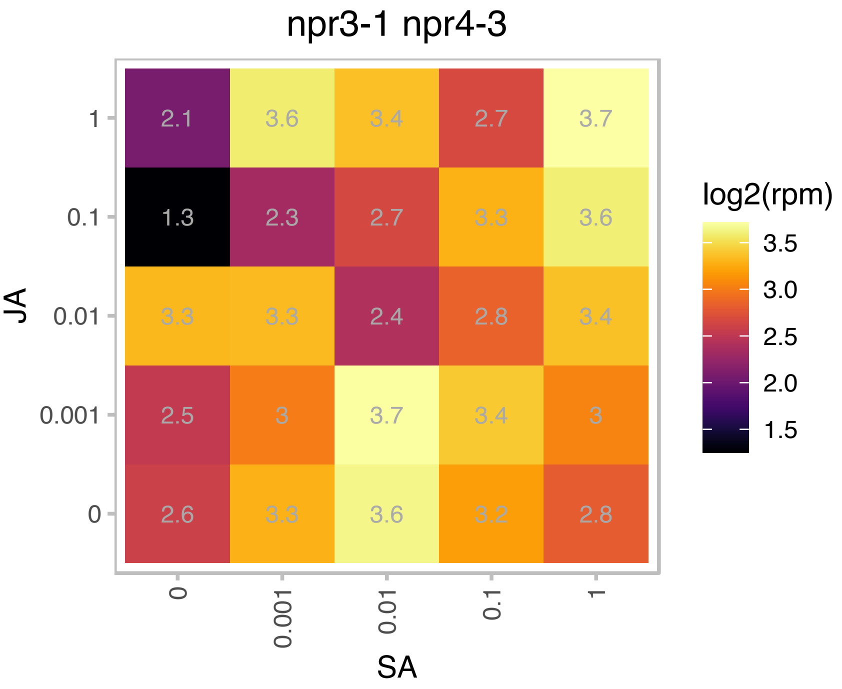

# AT5G07890.1 : NA

myosin heavy chain-related

Col-0

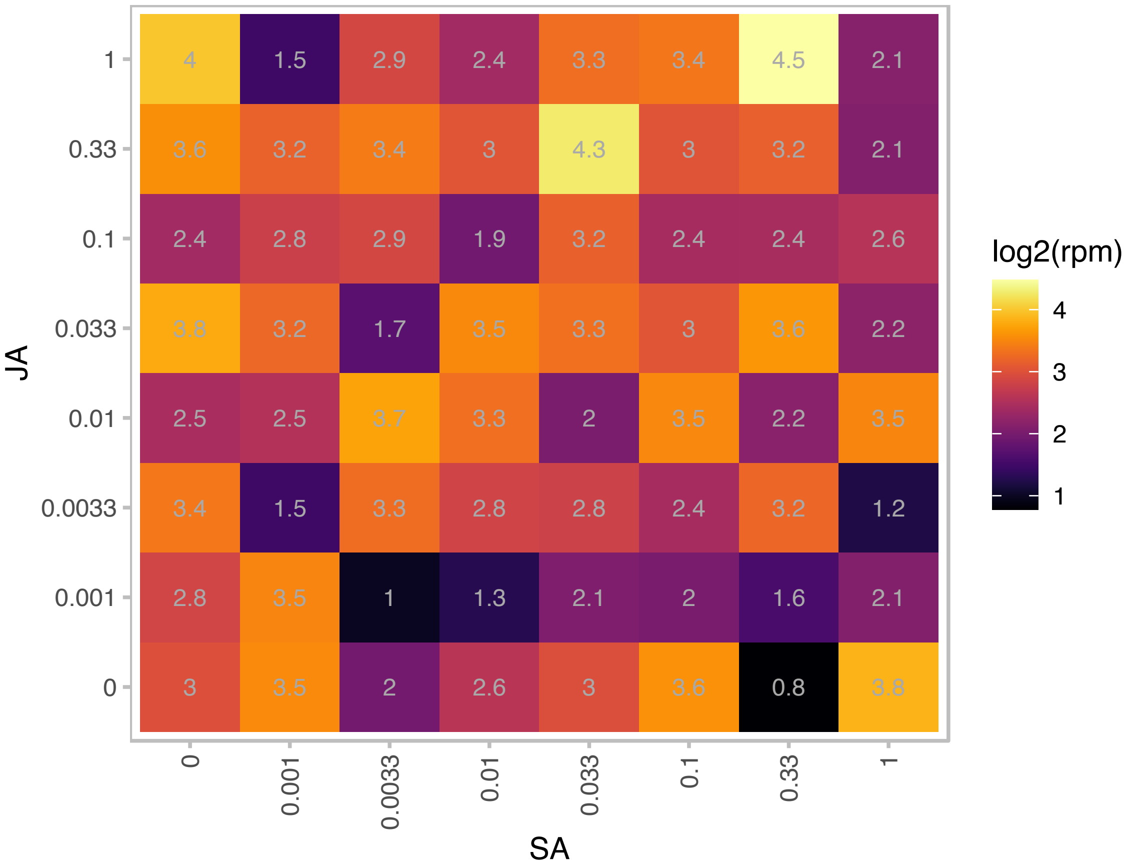


| | |
|---|---|
| Gene | AT5G07890 |
| is.expressed_genes | TRUE |
| is.responsive_genes | TRUE |
| is.apcluster | A-01 |

npr3-2 npr4-2

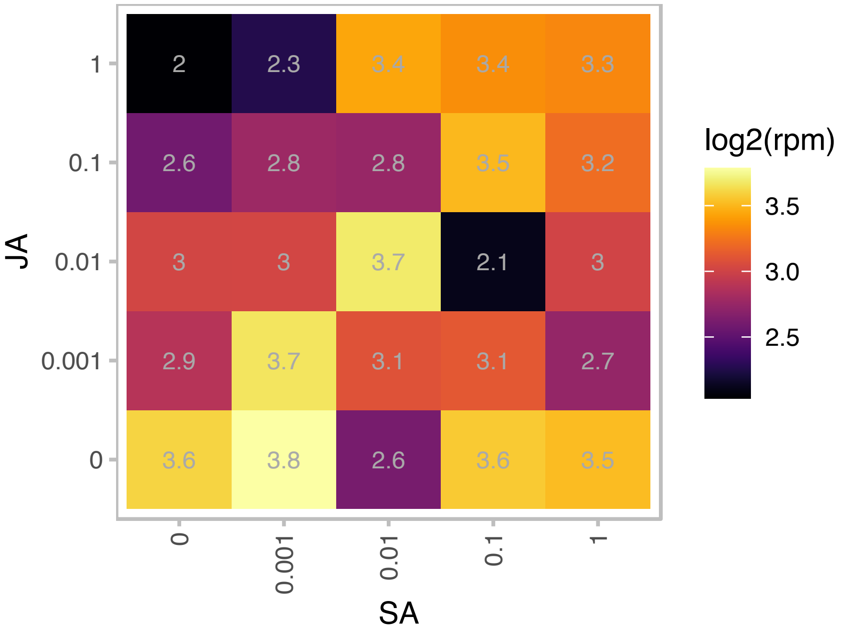


npr3-1 npr4-3

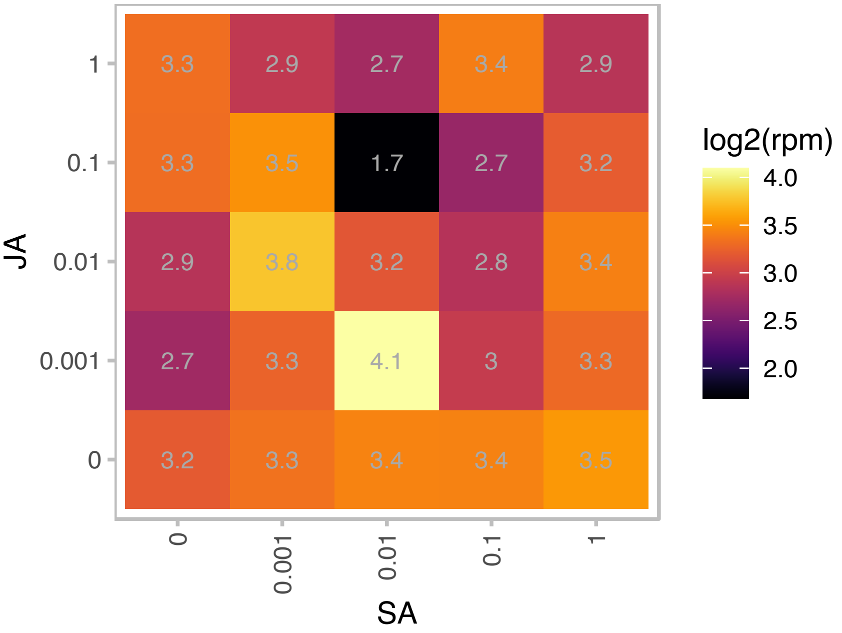

# AT5G10625.1 : NA

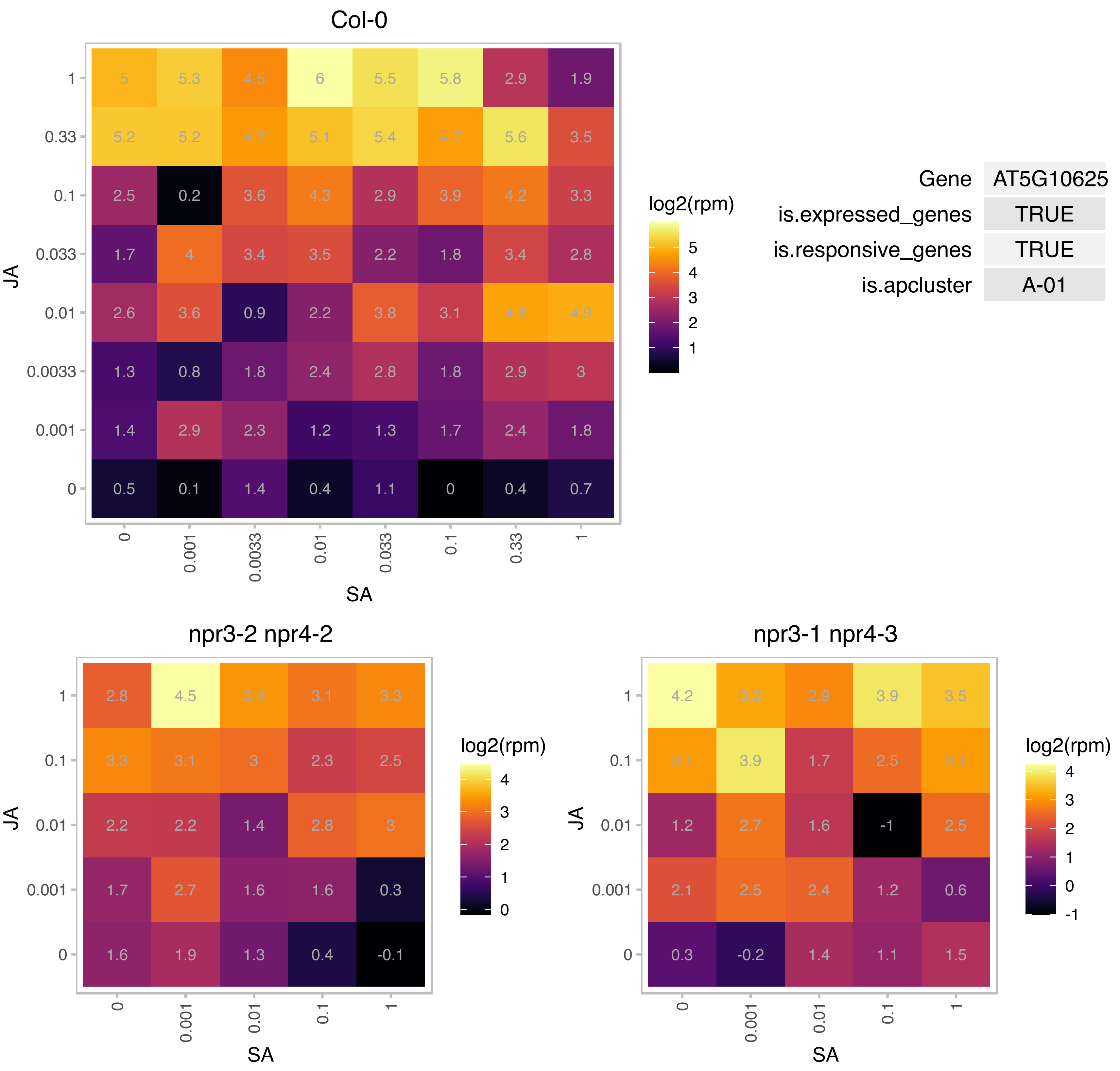

### AT5G12480.1 : CPK7

calmodulin-domain protein kinase 7

calmodulin-domain protein kinase CDPK isoform 7 (CPK7)

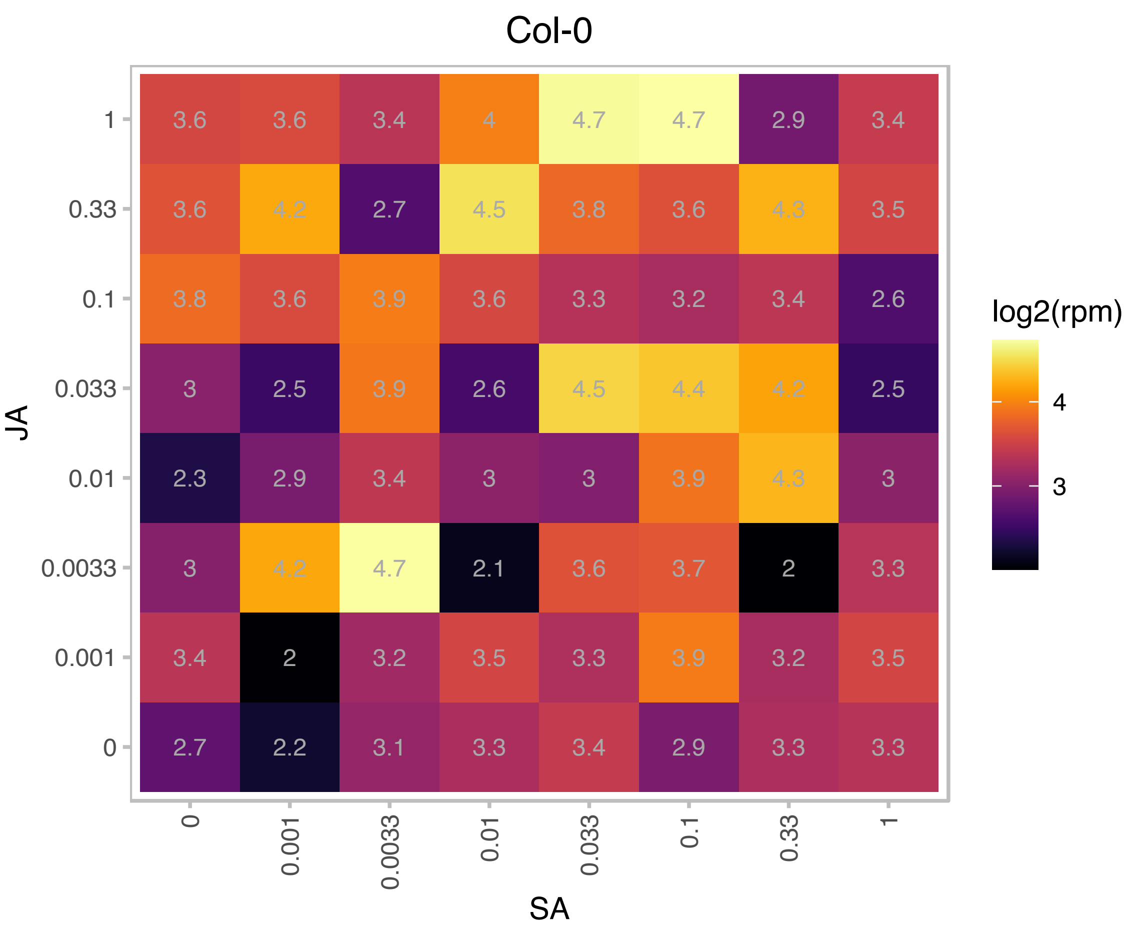


| | |
|---|---|
| Gene | AT5G12480 |
| is.expressed_genes | TRUE |
| is.responsive_genes | TRUE |
| is.apcluster | A-01 |

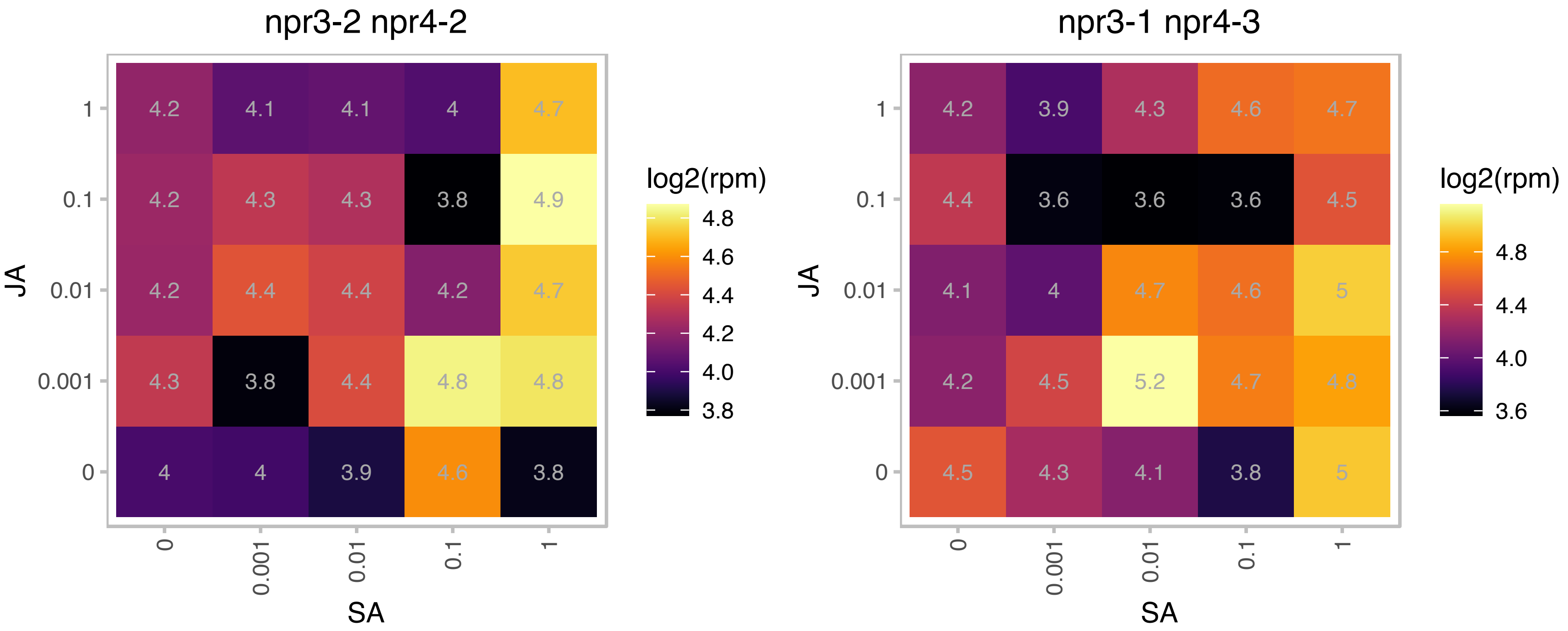

# AT5G23590.1 : NA

DNAJ heat shock N-terminal domain-containing protein

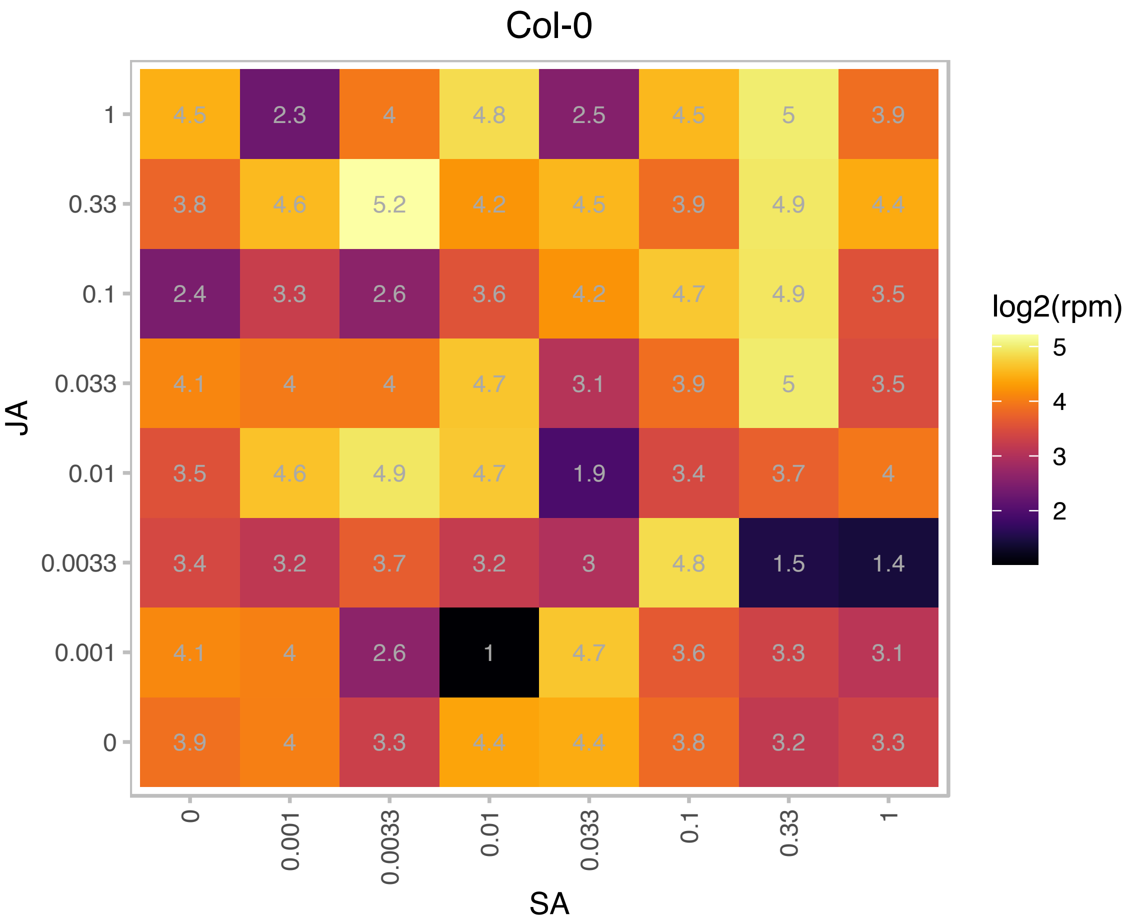


| | |
|---|---|
| Gene | AT5G23590 |
| is.expressed_genes | TRUE |
| is.responsive_genes | TRUE |
| is.apcluster | A-01 |

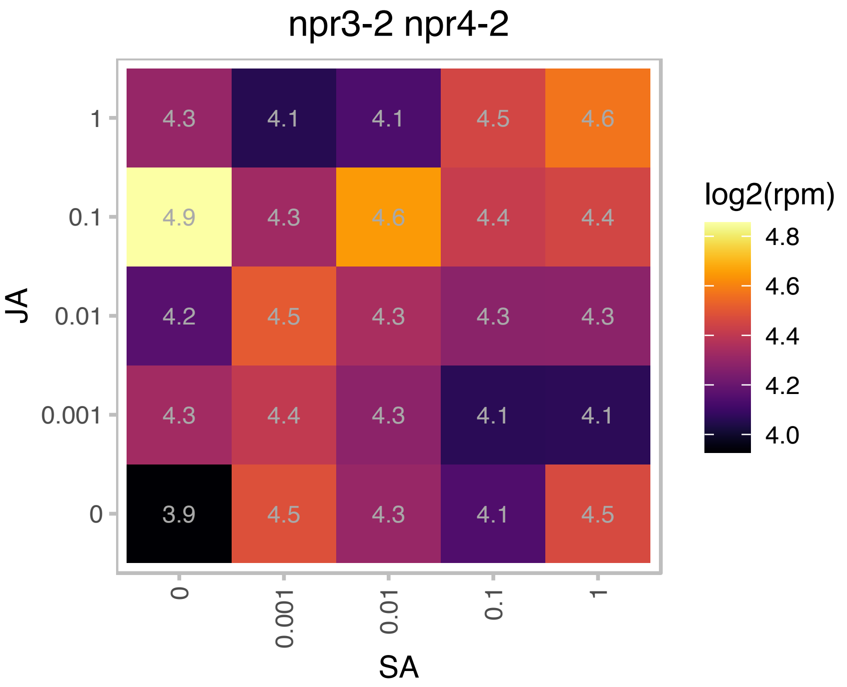


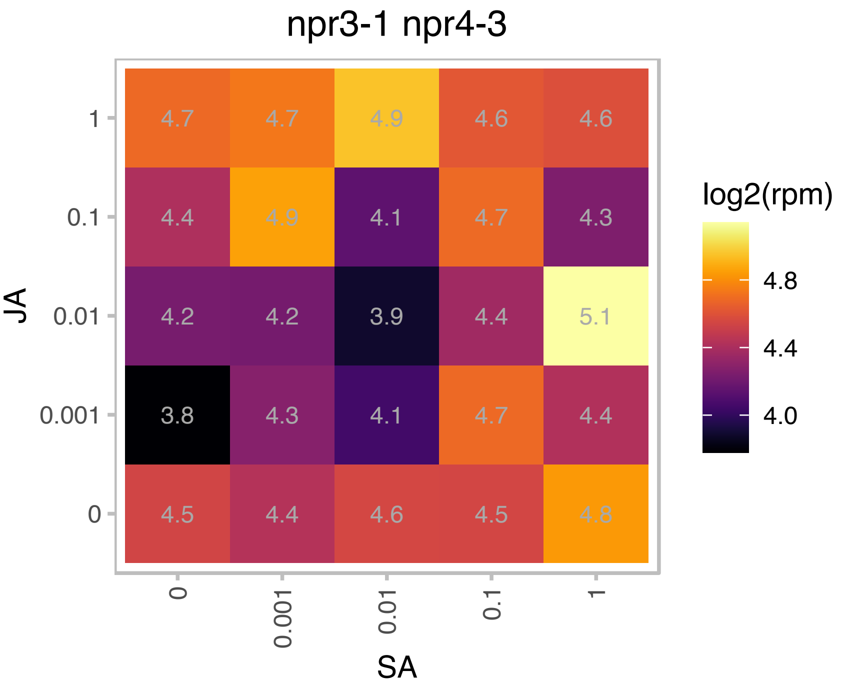

### AT5G26220.1 : AtGGCT2;1, GGCT2;1

ChaC-like family protein

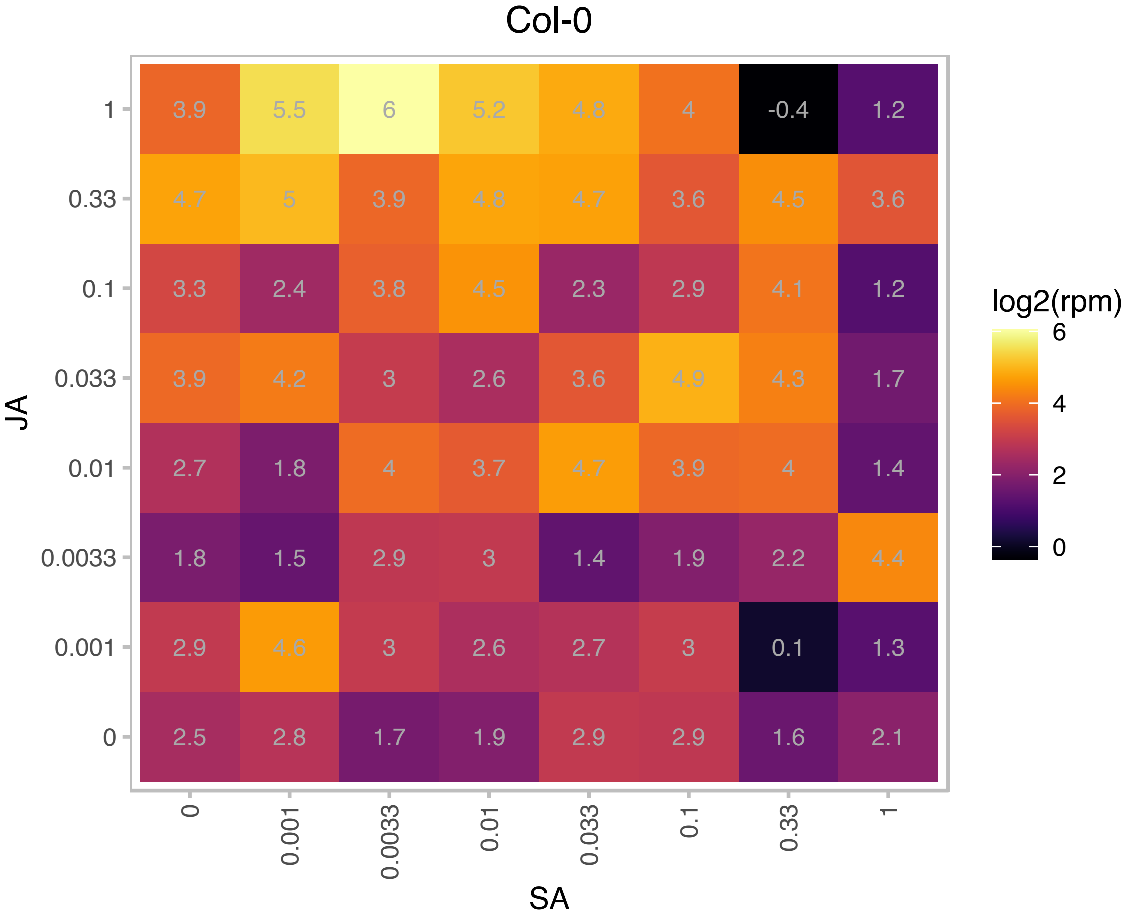


| | |
|---|---|
| Gene | AT5G26220 |
| is.expressed_genes | TRUE |
| is.responsive_genes | TRUE |
| is.apcluster | A-01 |

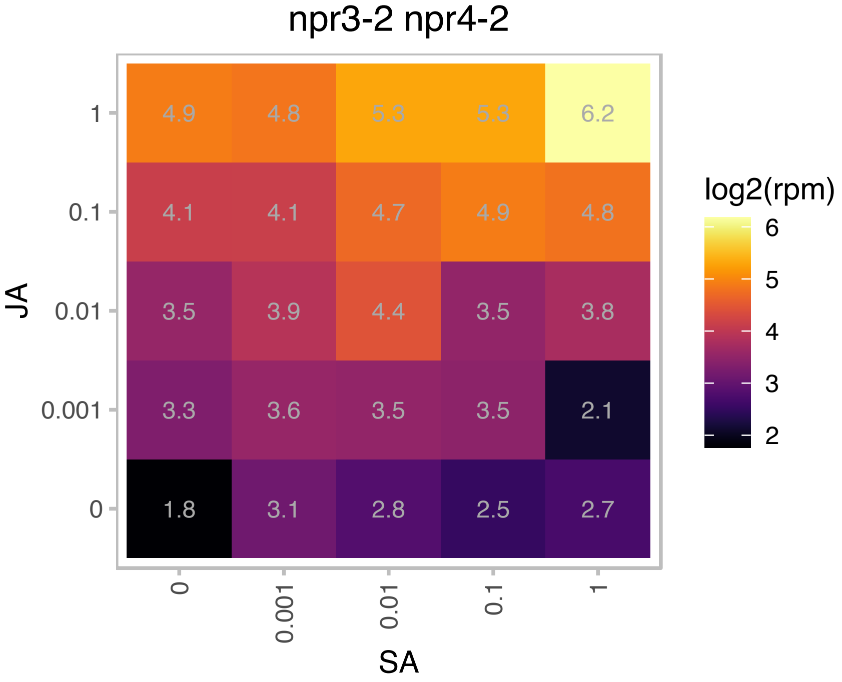


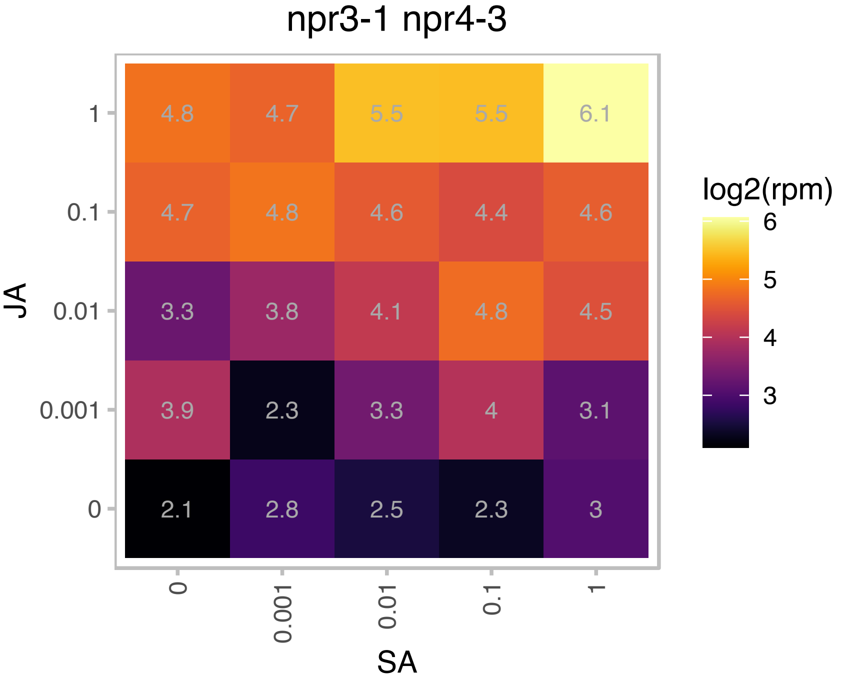

### AT5G38710.1 : PDH2

Methylenetetrahydrofolate reductase family protein

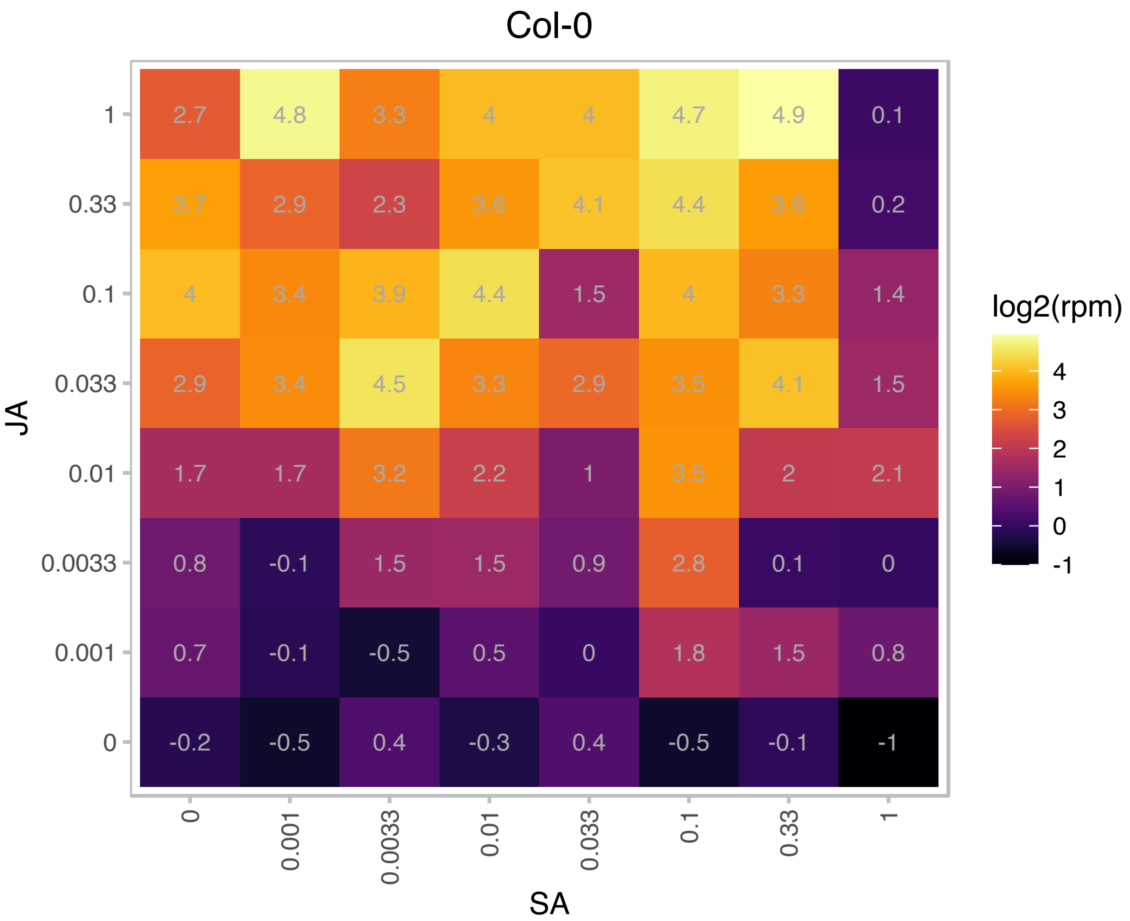


| Gene | AT5G38710 |
|---|---|
| is.expressed_genes | TRUE |
| is.responsive_genes | TRUE |
| is.apcluster | A-01 |

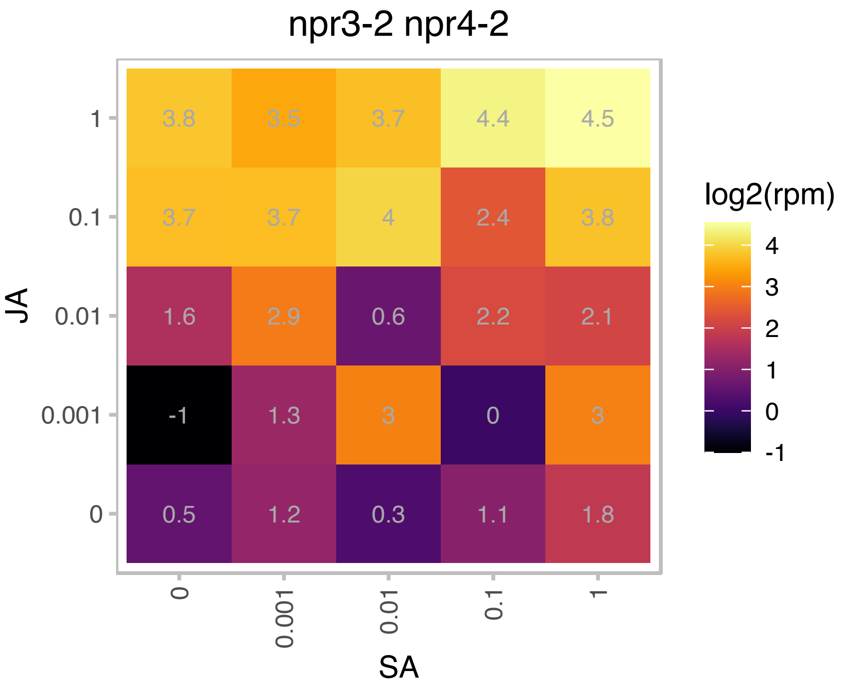


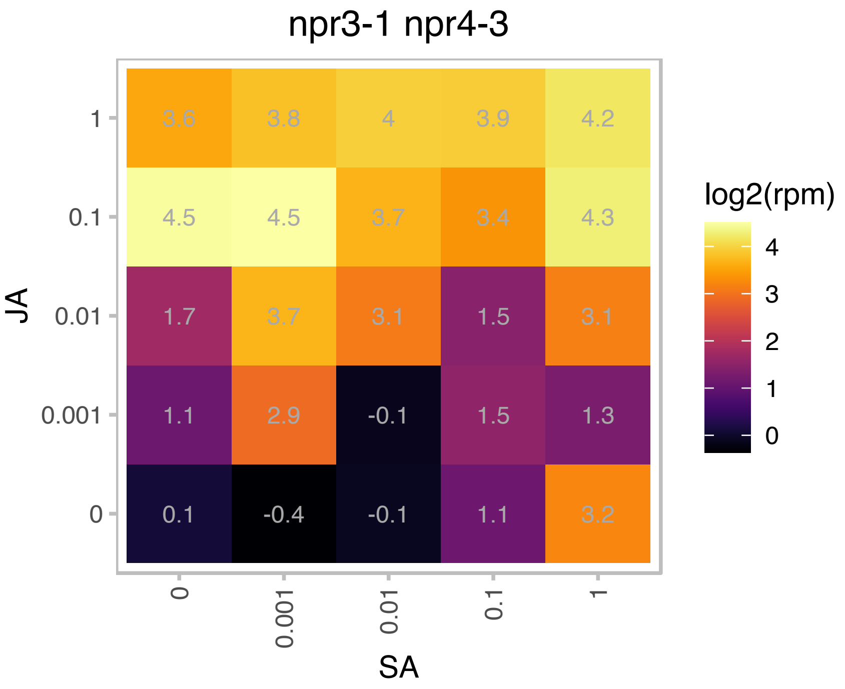

### AT5G40340.1 : PDP3

Tudor/PWWP/MBT superfamily protein

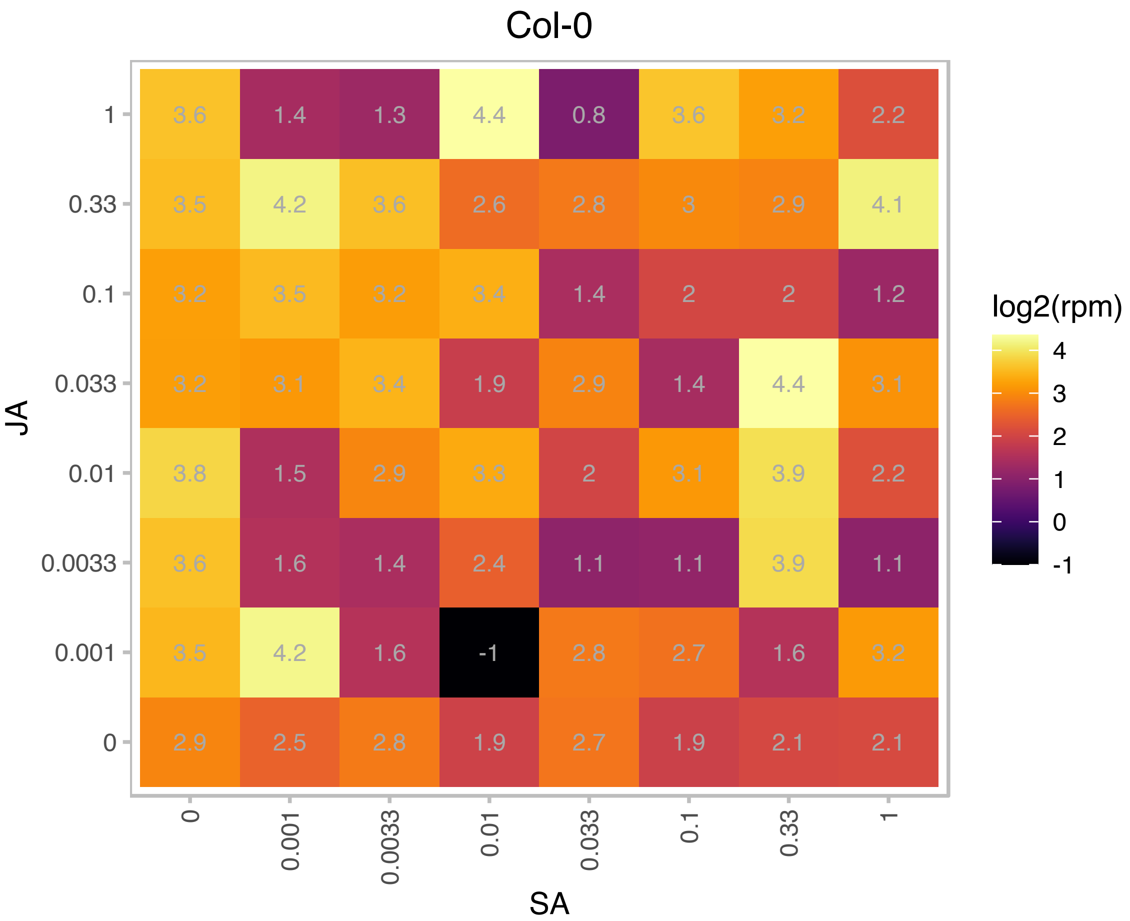


| | |
|---|---|
| Gene | AT5G40340 |
| is.expressed_genes | TRUE |
| is.responsive_genes | TRUE |
| is.apcluster | A-01 |

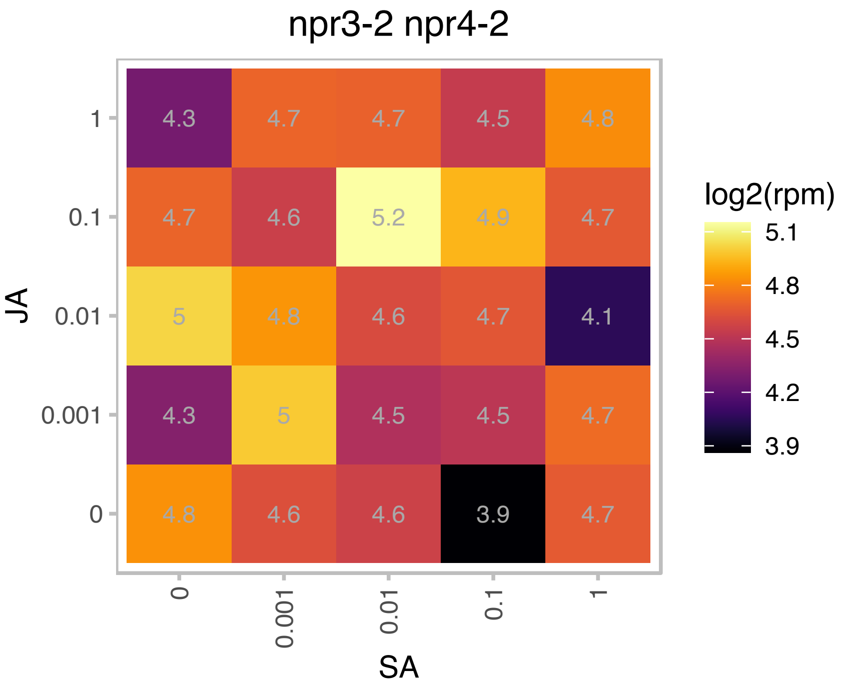


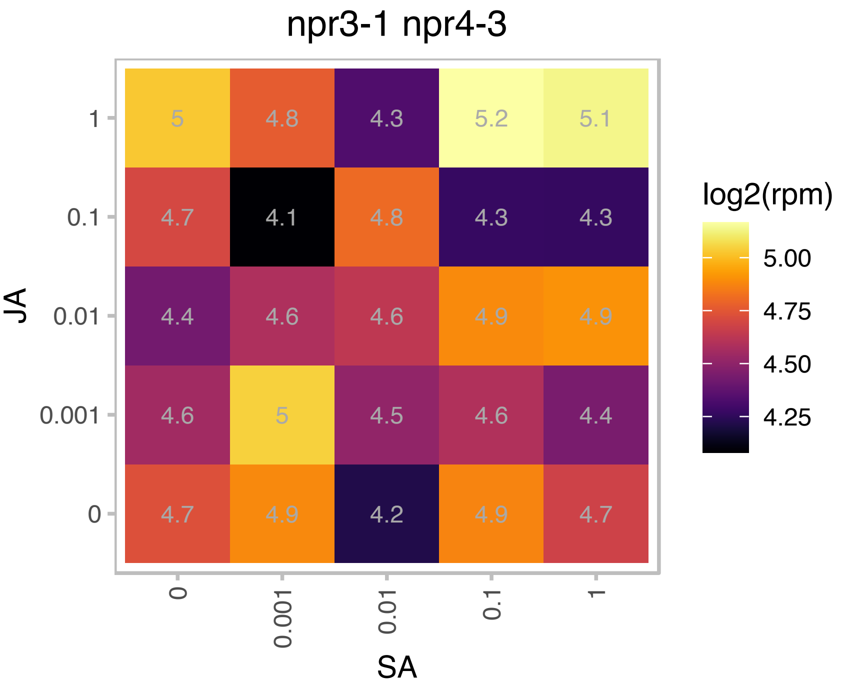

### AT5G42870.1 : ATPAH2, PAH2

phosphatidic acid phosphohydrolase 2

The PAH2 gene encodes a phosphatidate phosphohydrolase. Mutant analysis revealed its involvement in galactolipid synthesis pathway, and the membrane lipid remodeling. The pah1pah2 double-mutant showed enhance Al-susceptibility under low-P conditions, but there was no significant differences in Al tolerance between pah1pah2 and wild type when they were grown in a solution containing 35 μM Pi.

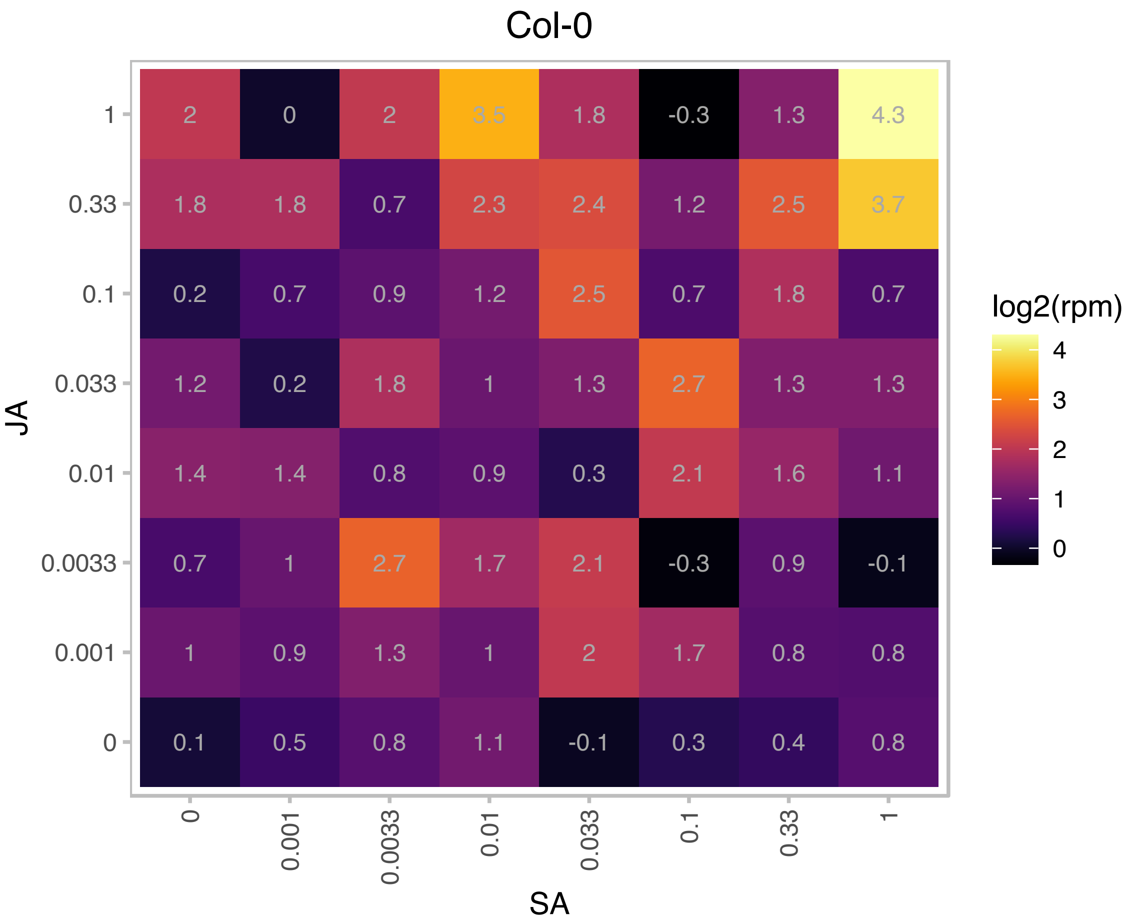


| | |
|---|---|
| Gene | AT5G42870 |
| is.expressed_genes | TRUE |
| is.responsive_genes | TRUE |
| is.apcluster | A-01 |

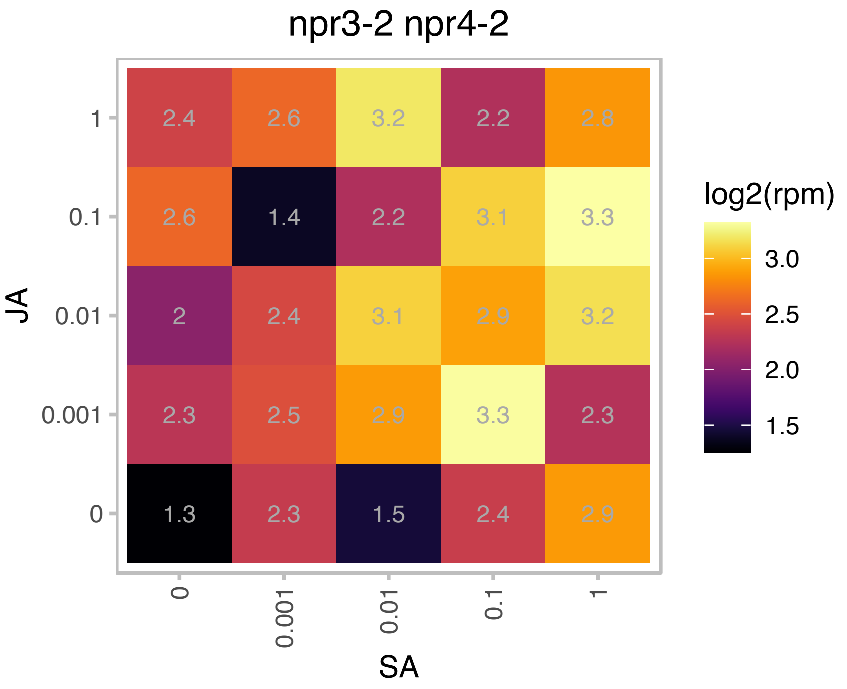


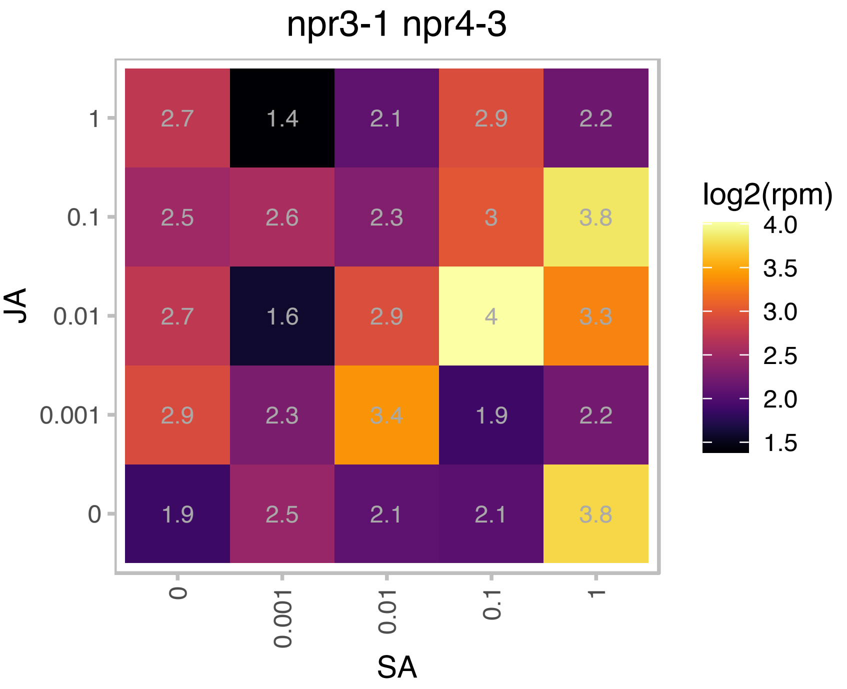

### AT5G63370.1 : CDKG1

Protein kinase superfamily protein

CDKG1 interacts with the splicing factor RSZ33 to regulate proper splicing of Cals5 Pre-mRNA.

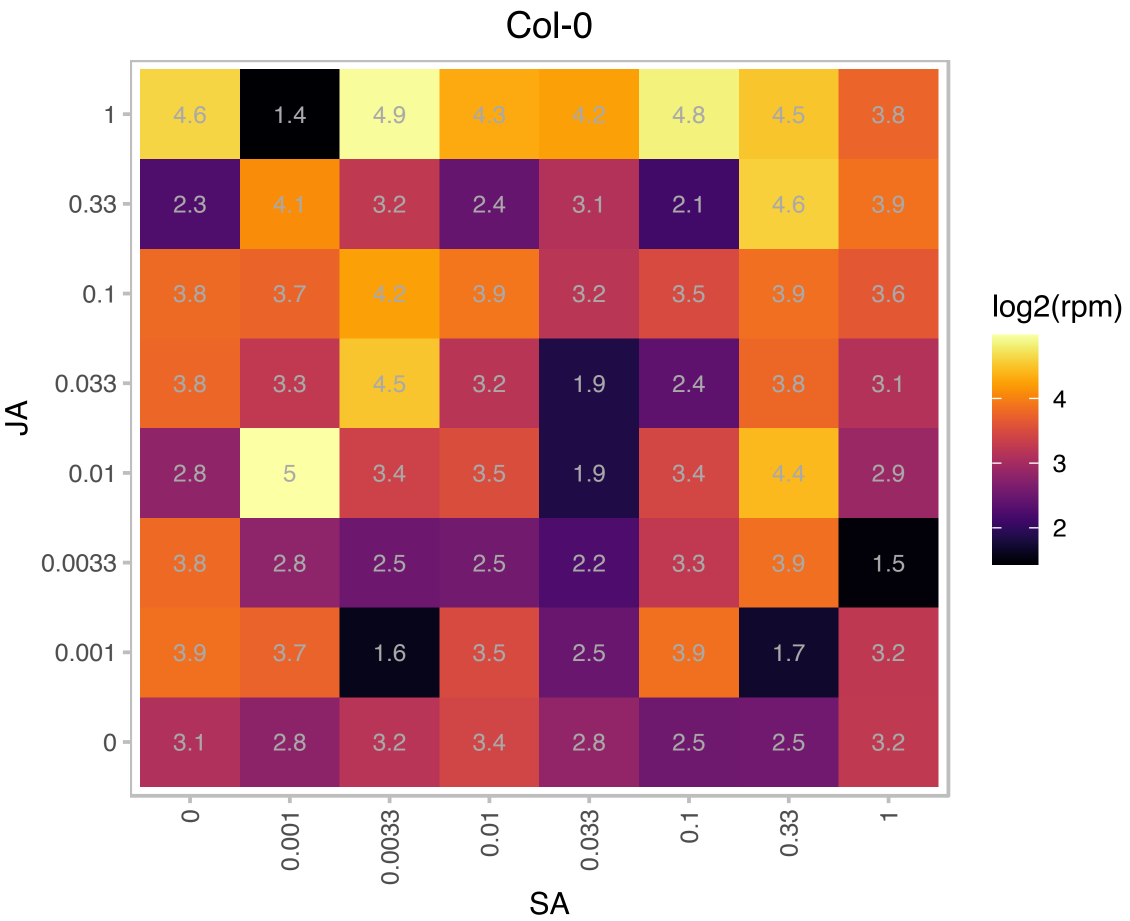


| | |
|---|---|
| Gene | AT5G63370 |
| is.expressed_genes | TRUE |
| is.responsive_genes | TRUE |
| is.apcluster | A-01 |

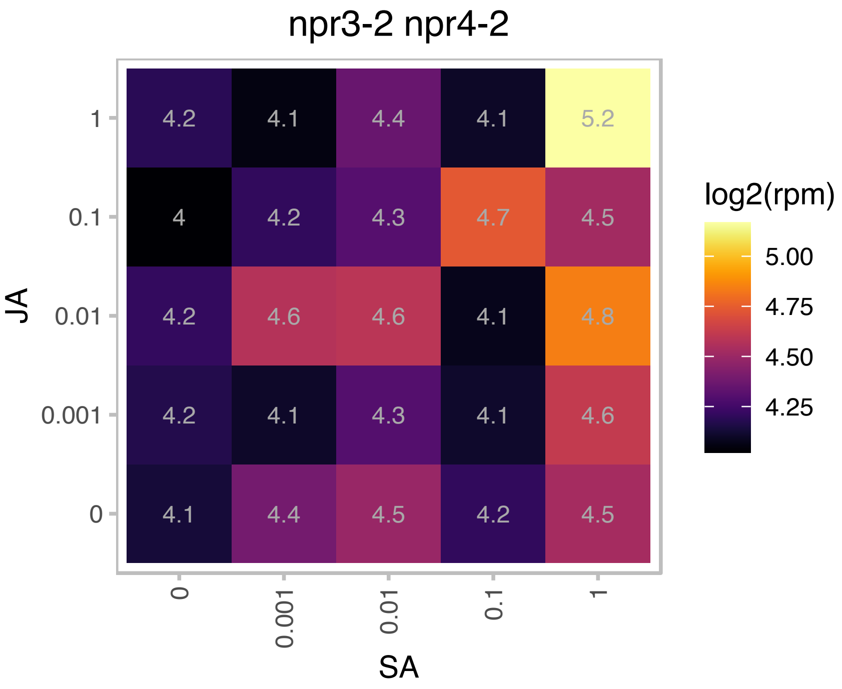


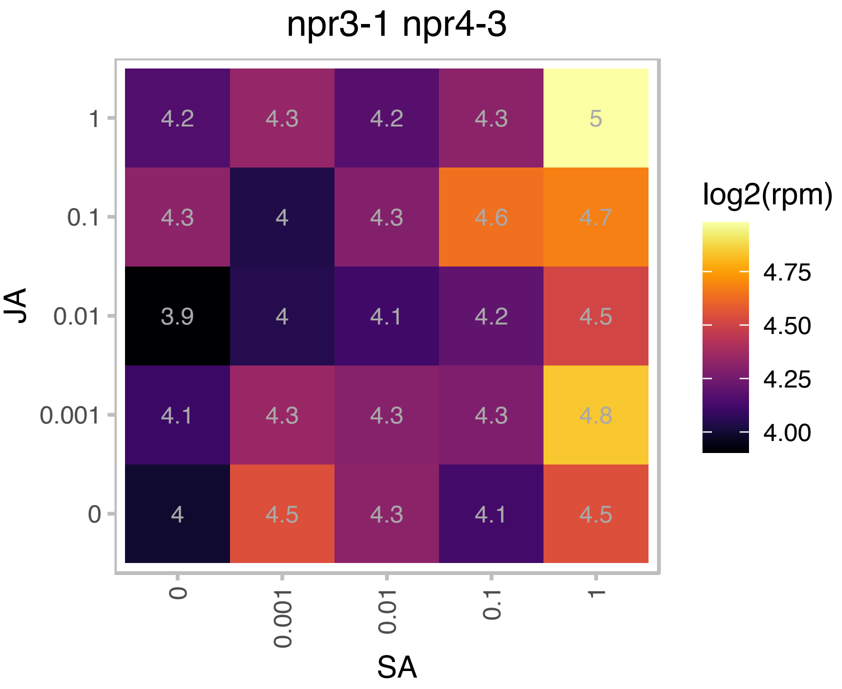

### AT5G65280.1 : GCL1

GCR2-like 1

Encodes a protein with reported similarity to GCR2 a putative G protein coupled receptor thought to be an ABA receptor. Loss of function mutations in GCL1 show no ABA response defects based on assays of seed germination and seedling development.GCL1 also has similarity to LANCL1 and LANCL2, human homologs of bacterial lanthionine synthetase.

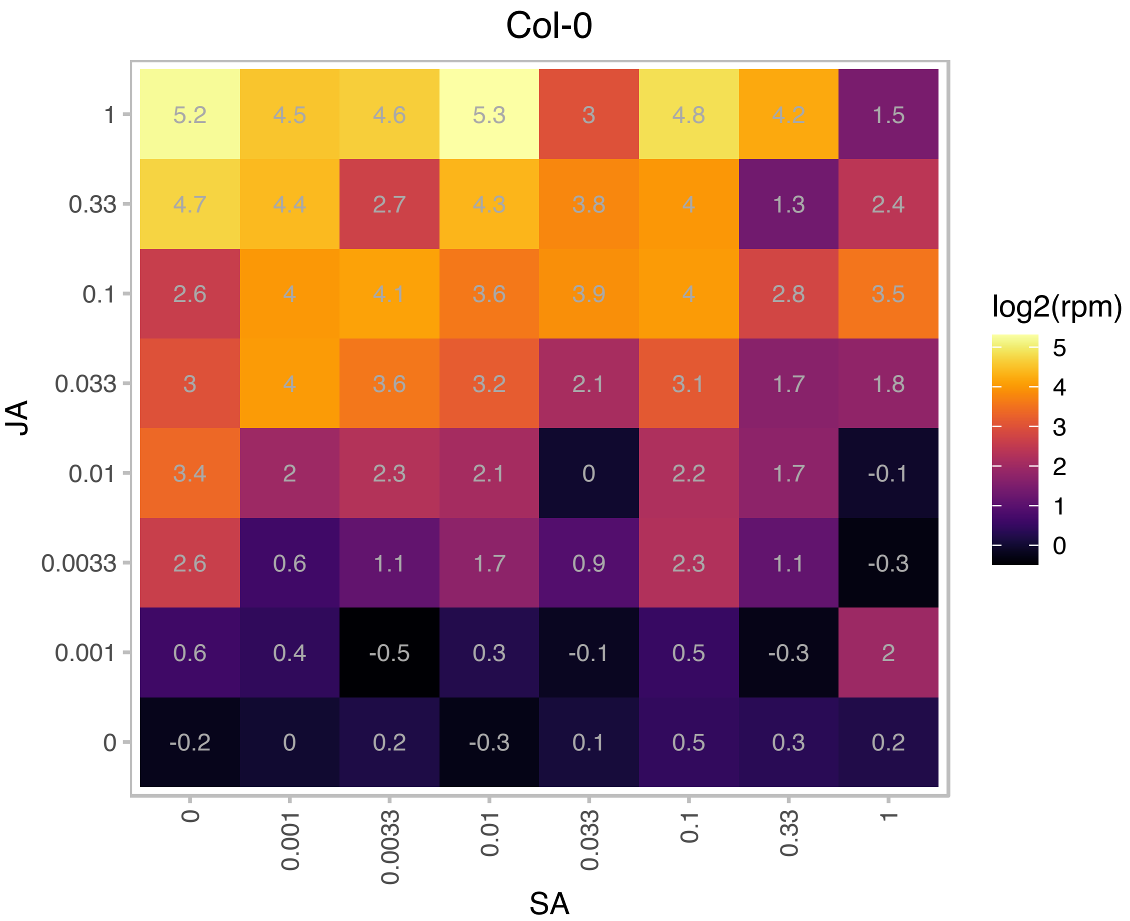


| | |
|---|---|
| Gene | AT5G65280 |
| is.expressed_genes | TRUE |
| is.responsive_genes | TRUE |
| is.apcluster | A-01 |

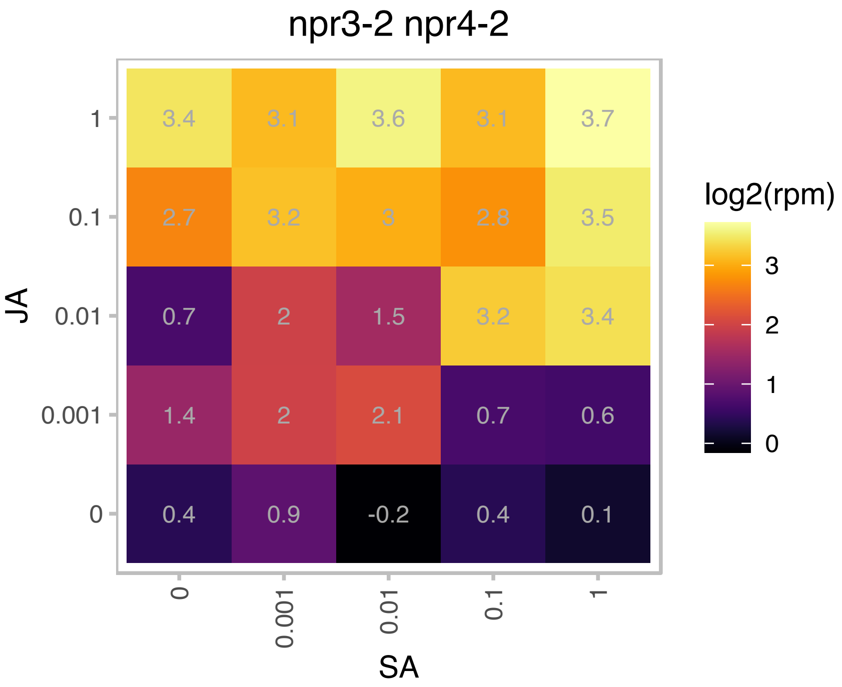


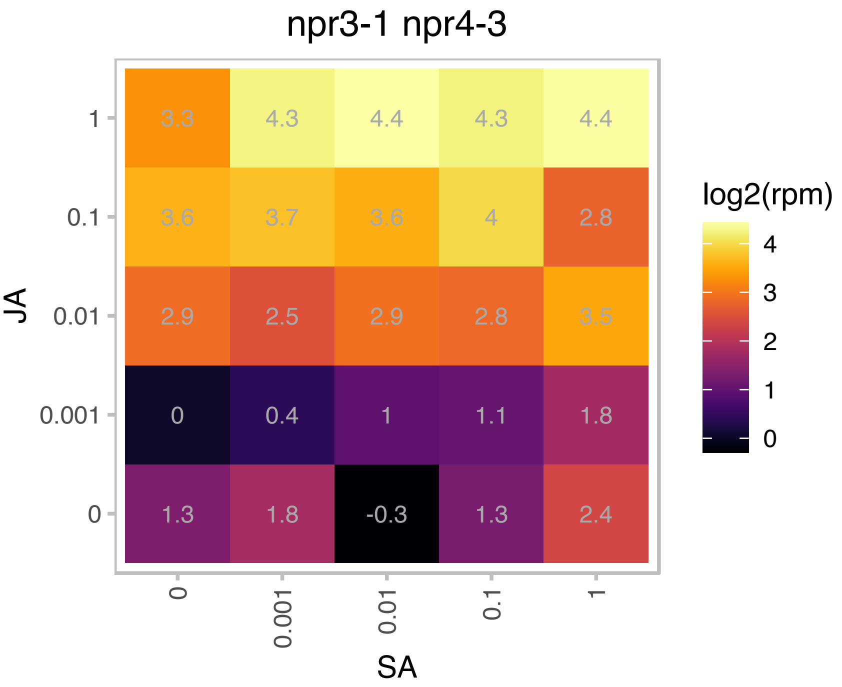

### AT5G67080.1 : MAPKKK19, MKK19

mitogen-activated protein kinase kinase kinase 19

member of MEKK subfamily

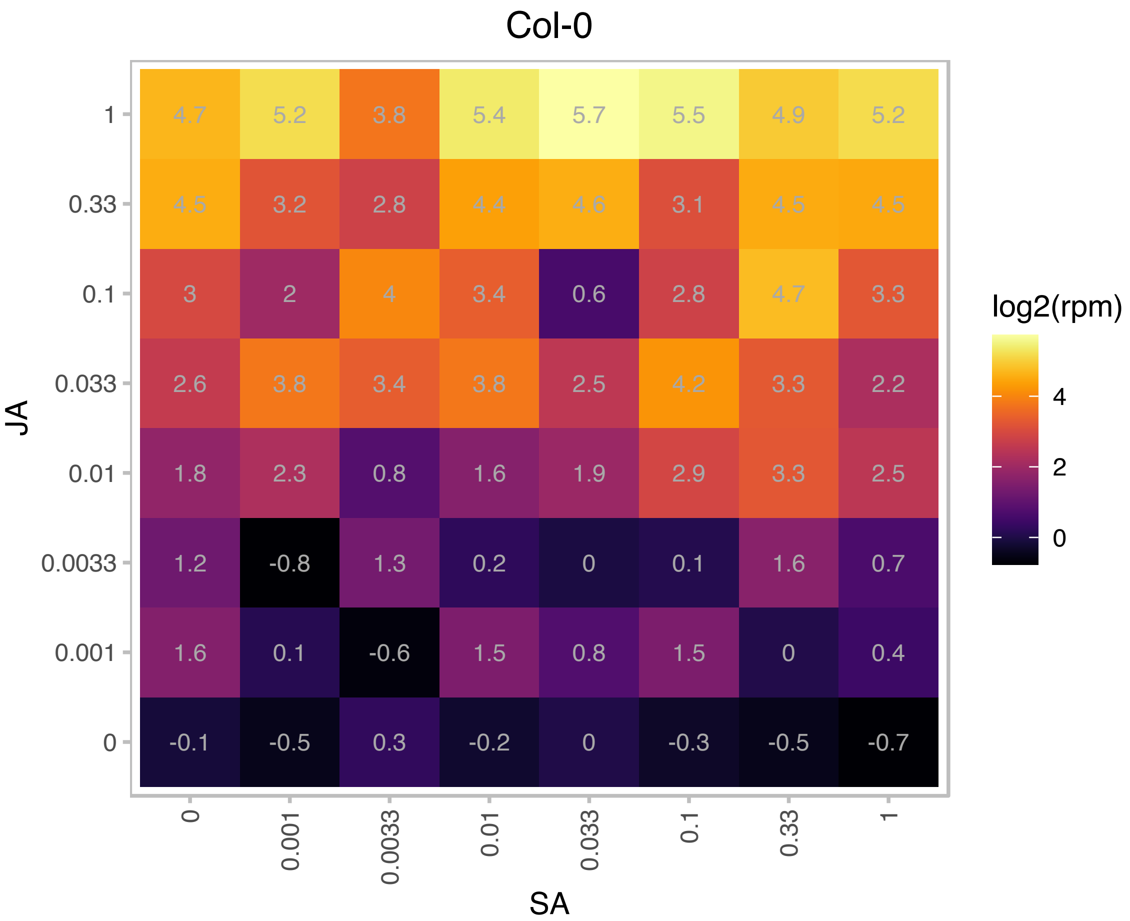


| | |
|---|---|
| Gene | AT5G67080 |
| is.expressed_genes | TRUE |
| is.responsive_genes | TRUE |
| is.apcluster | A-01 |

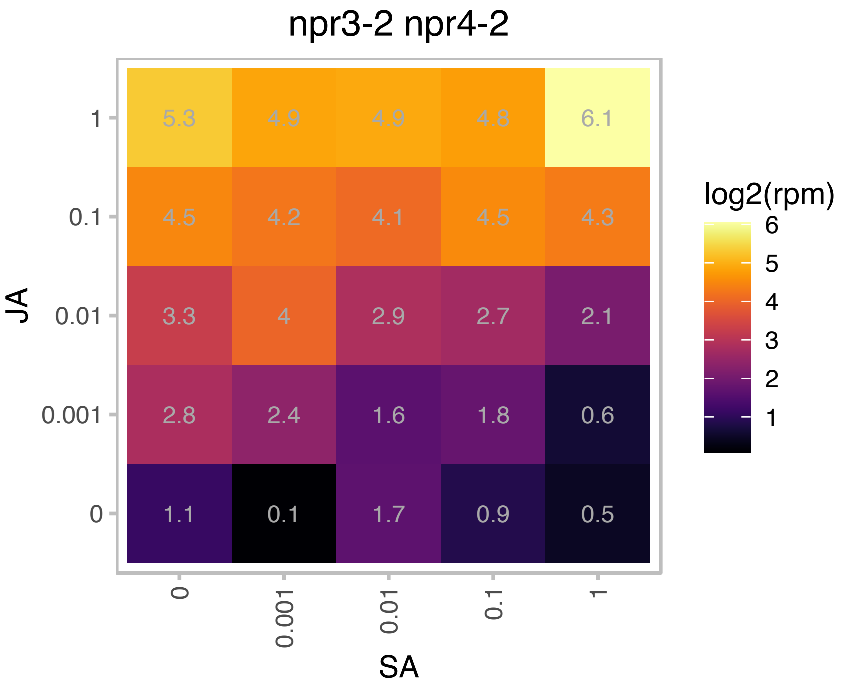


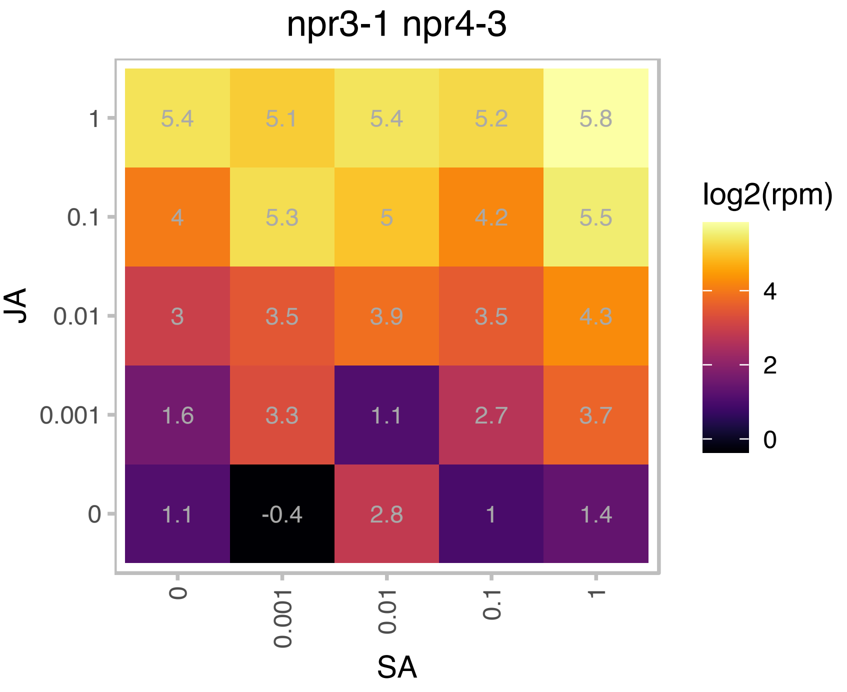

# AT1G06630.1 : NA

F-box/RNI-like superfamily protein

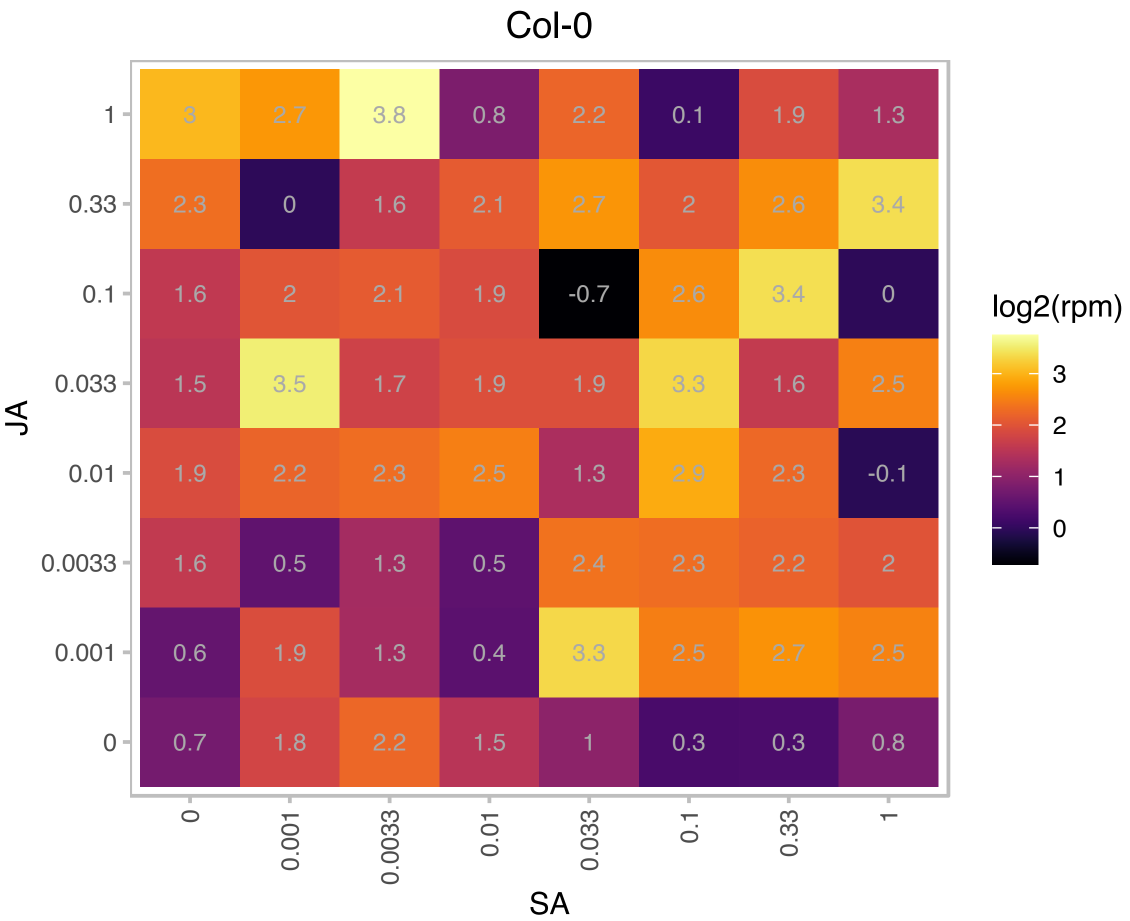


| | |
|---|---|
| Gene | AT1G06630 |
| is.expressed_genes | TRUE |
| is.responsive_genes | TRUE |
| is.apcluster | A-02 |

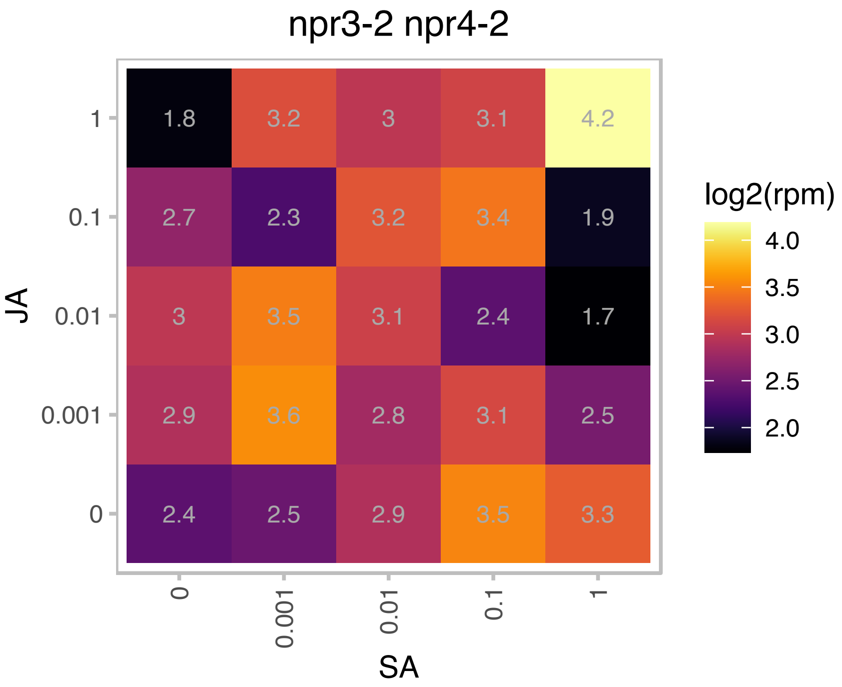


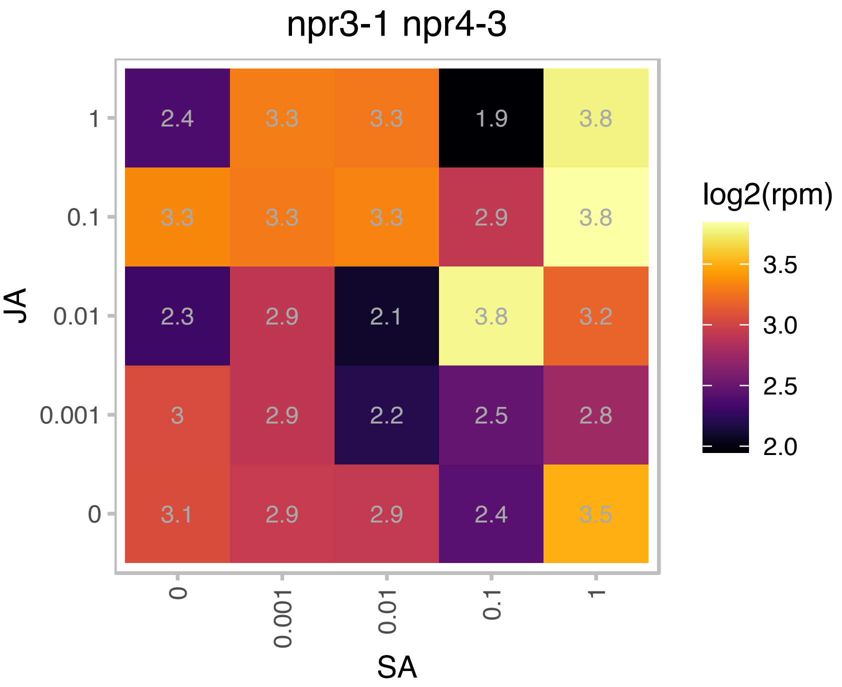

# AT1G20310.1 : NA

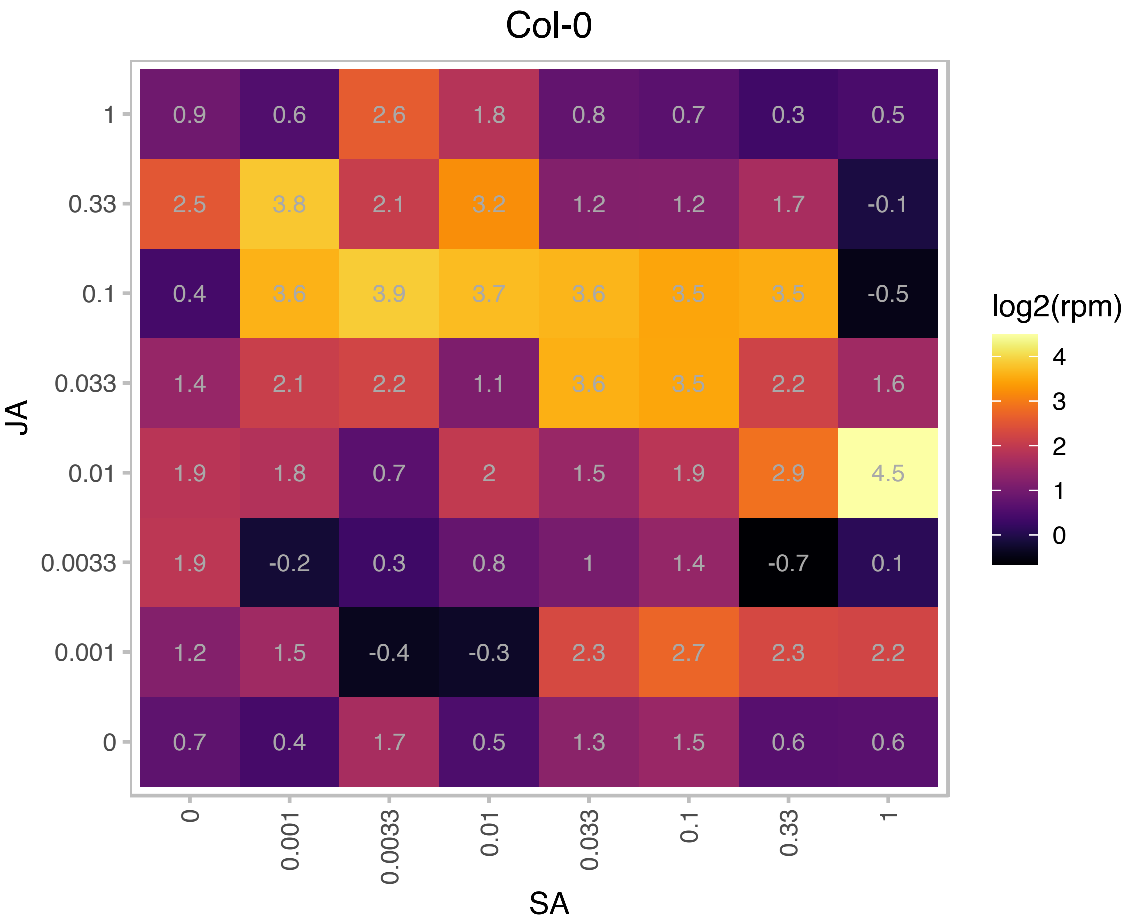


| | |
|---|---|
| Gene | AT1G20310 |
| is.expressed_genes | TRUE |
| is.responsive_genes | TRUE |
| is.apcluster | A-02 |

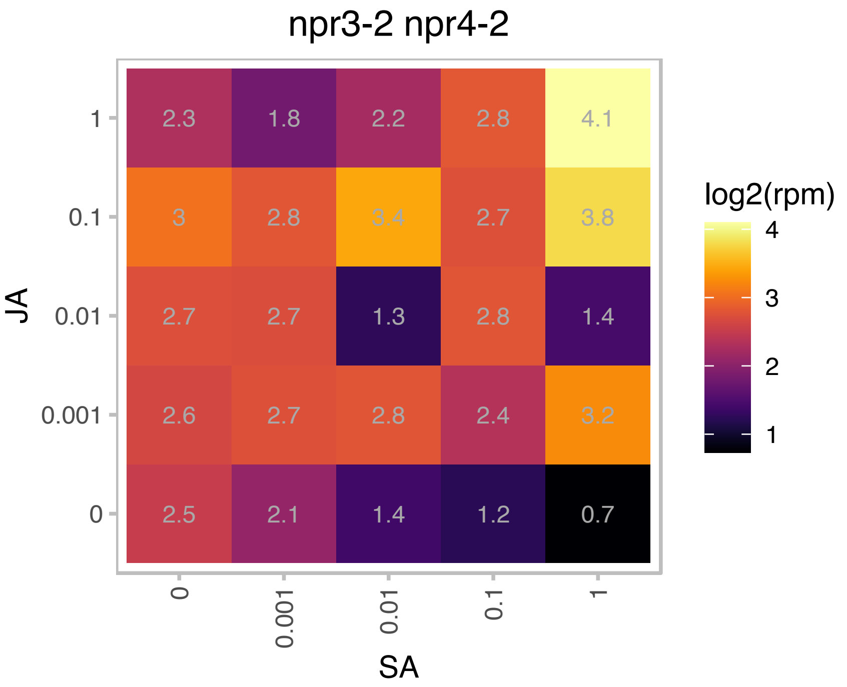


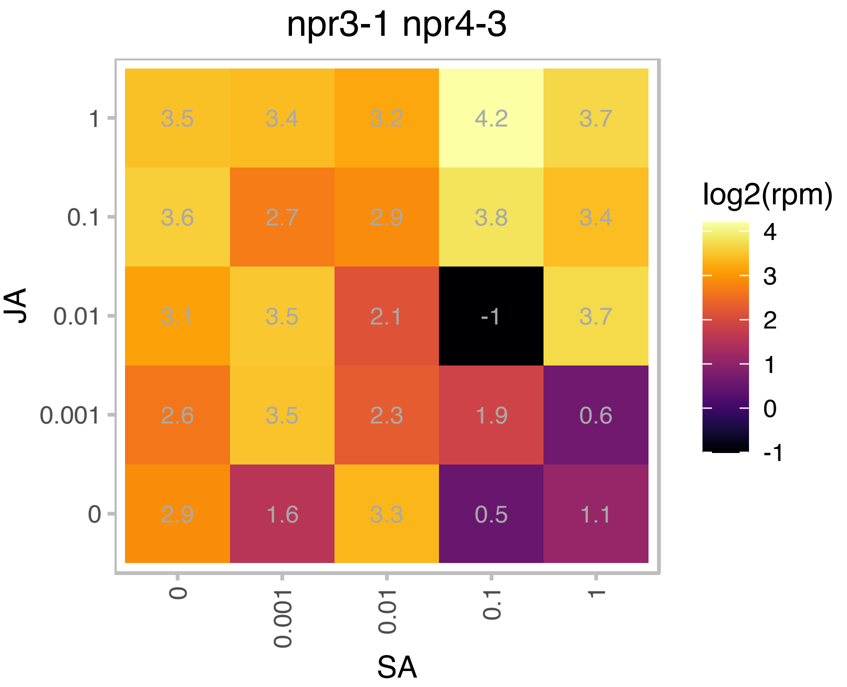

### AT1G30135.1 : JAZ8, TIFY5A

jasmonate-zim-domain protein 8

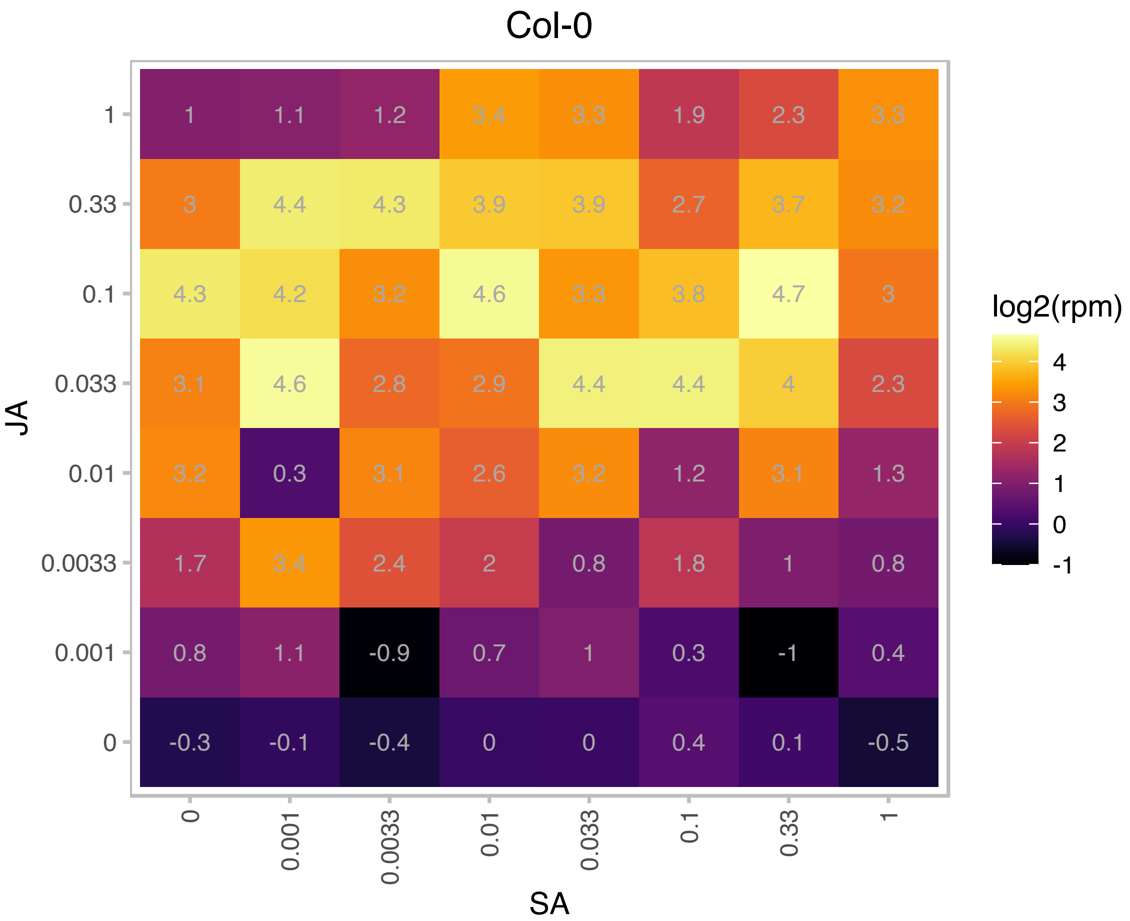


| | |
|---|---|
| Gene | AT1G30135 |
| is.expressed_genes | TRUE |
| is.responsive_genes | TRUE |
| is.apcluster | A-02 |

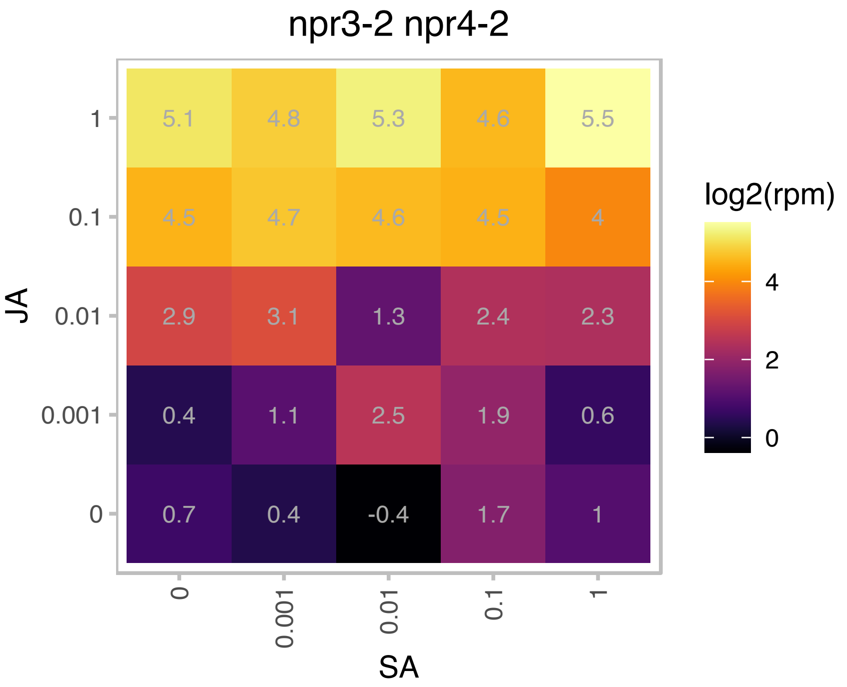


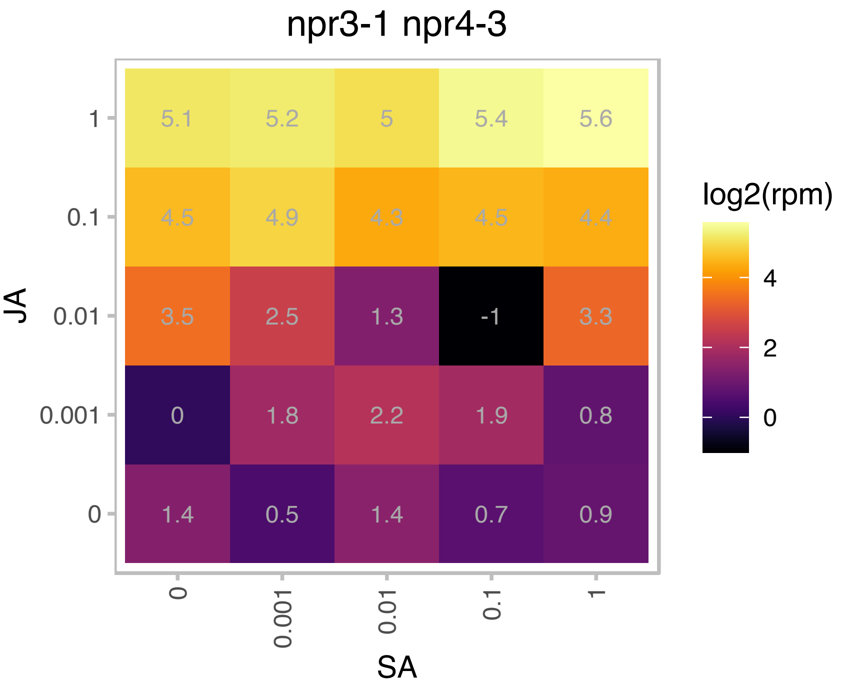

# AT1G54020.1 : NA

GDSL-like Lipase/Acylhydrolase superfamily protein

GDSL-motif esterase/acyltransferase/lipase. Enzyme group with broad substrate specificity that may catalyze acyltransfer or hydrolase reactions with lipid and non-lipid substrates.

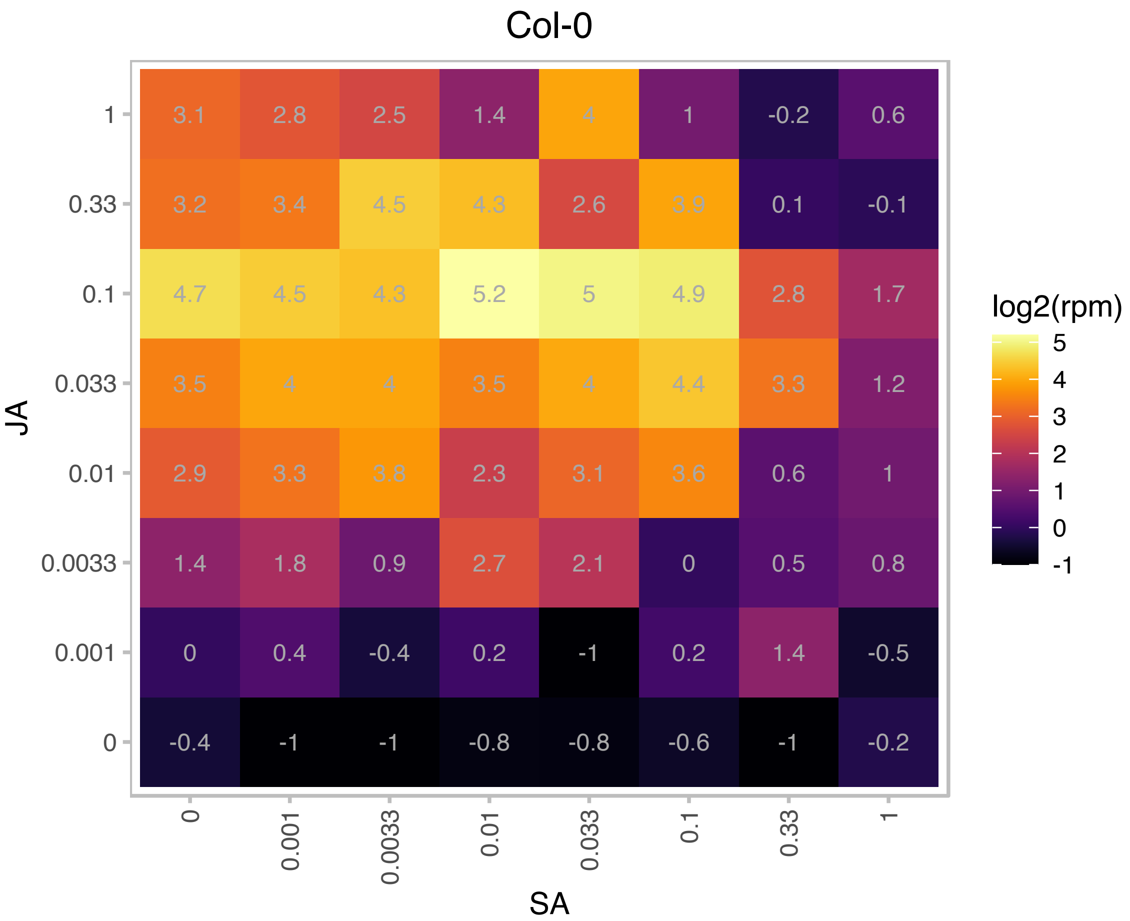


| | |
|---|---|
| Gene | AT1G54020 |
| is.expressed_genes | TRUE |
| is.responsive_genes | TRUE |
| is.apcluster | A-02 |

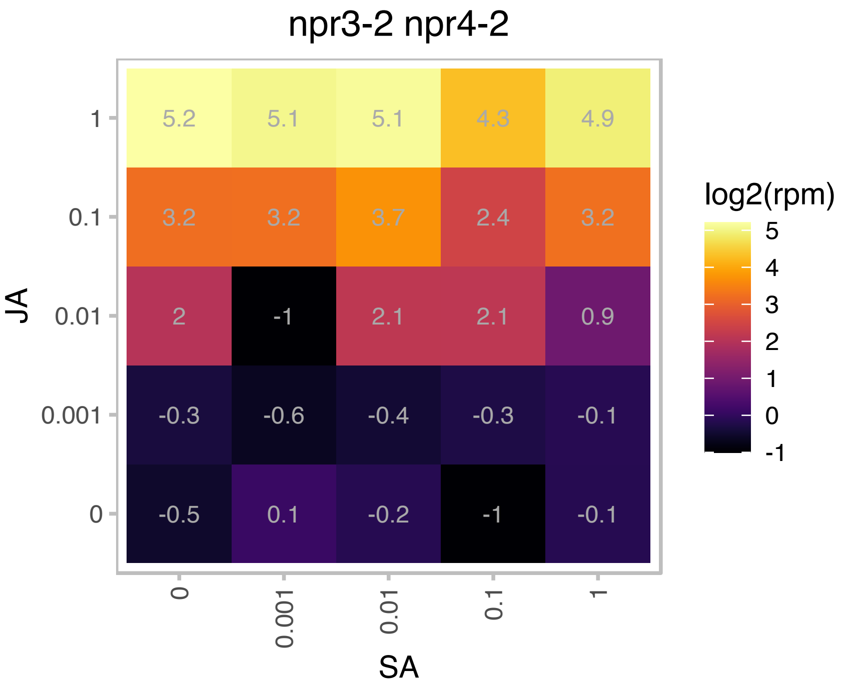


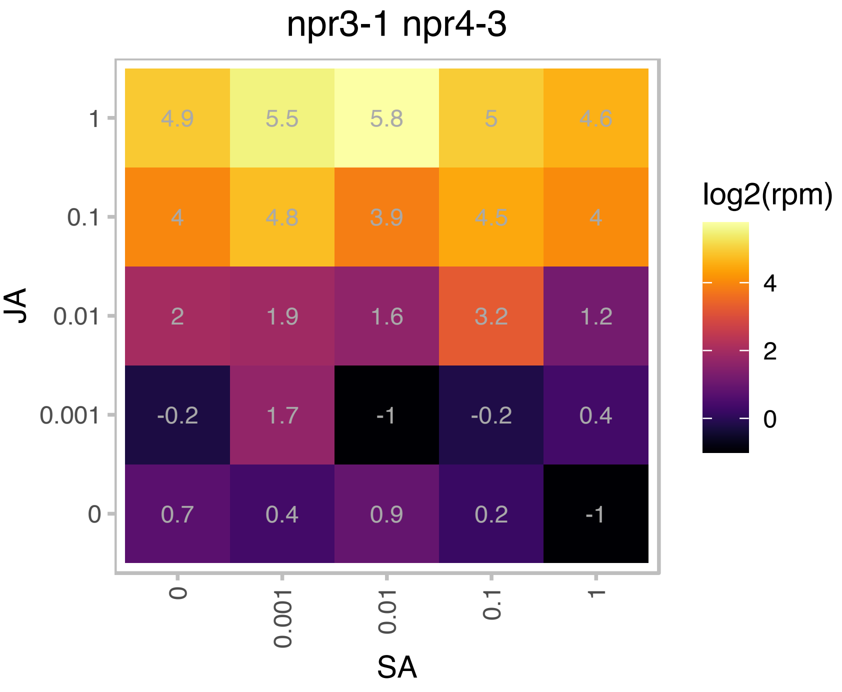

# AT1G66100.1 : NA

Plant thionin

Predicted to encode a PR (pathogenesis-related) protein.  Belongs to the plant thionin (PR-13) family with the following members: At1g66100, At5g36910, At1g72260, At2g15010, At1g12663, At1g12660.

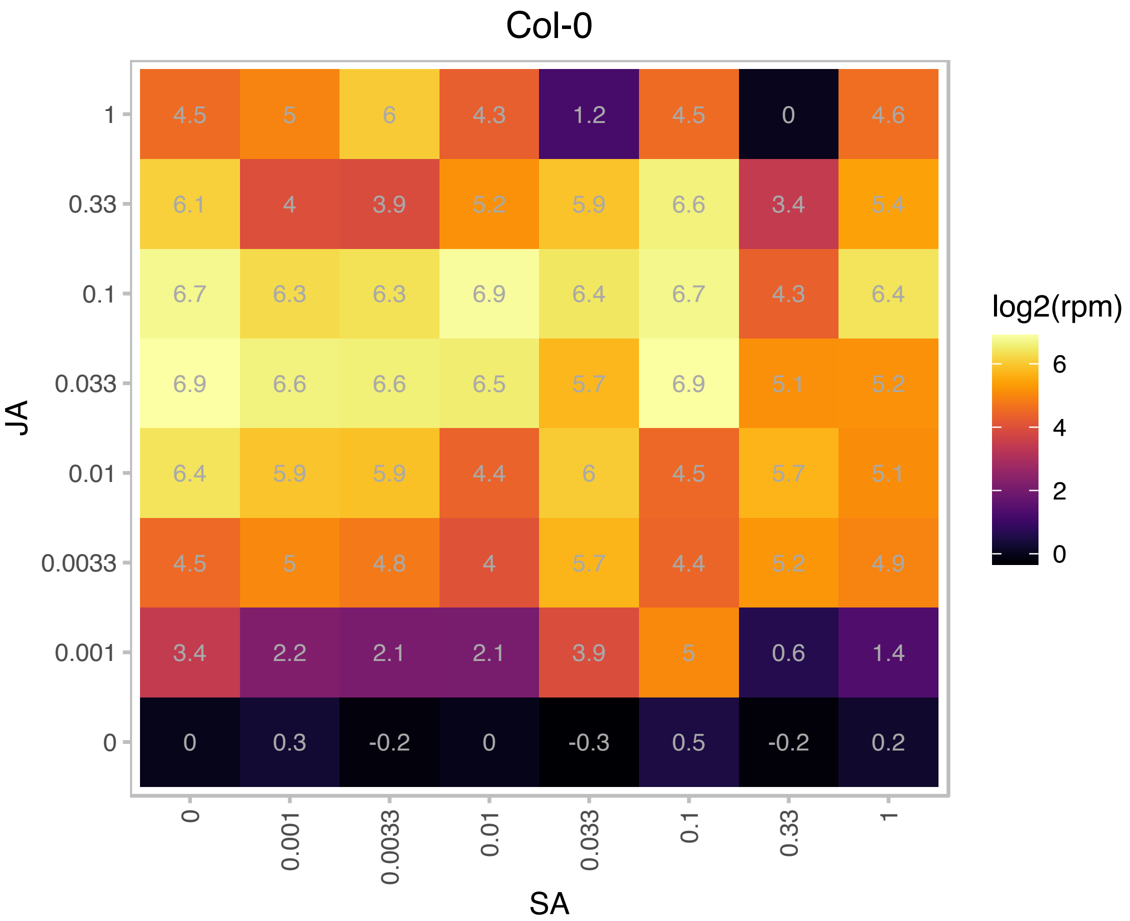


| | |
|---|---|
| Gene | AT1G66100 |
| is.expressed_genes | TRUE |
| is.responsive_genes | TRUE |
| is.apcluster | A-02 |

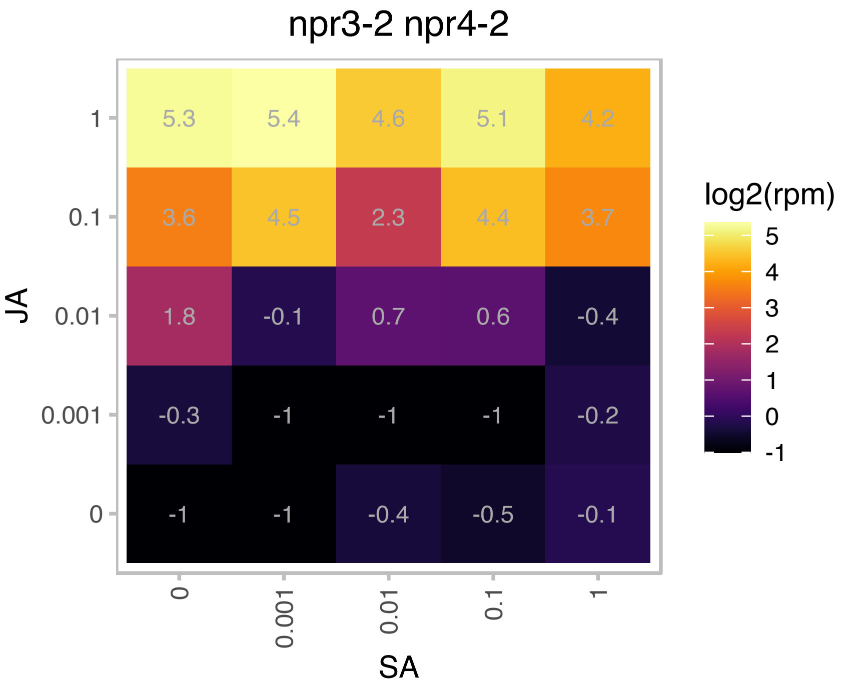


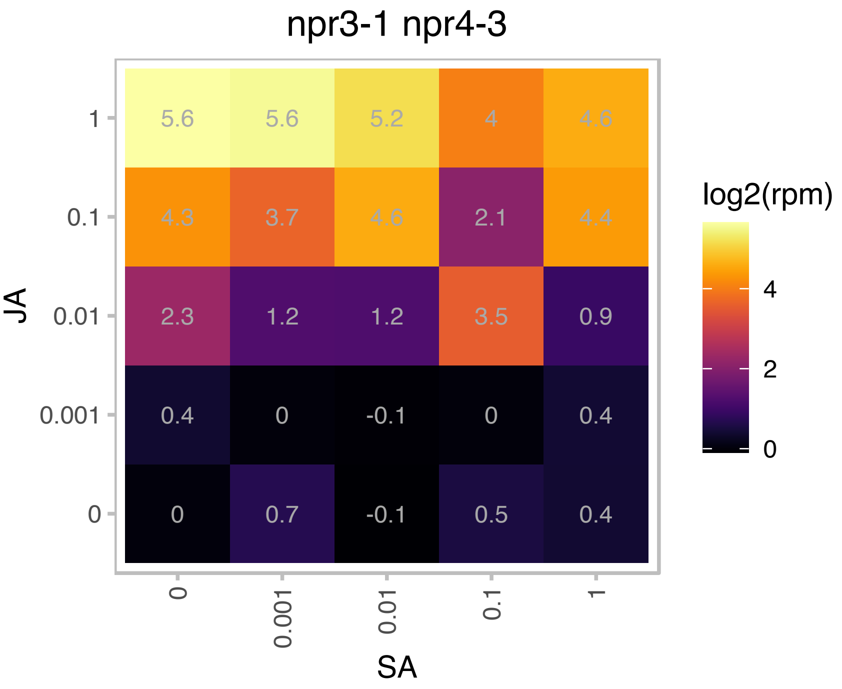

# AT2G15760.1 : NA

Protein of unknown function (DUF1645)

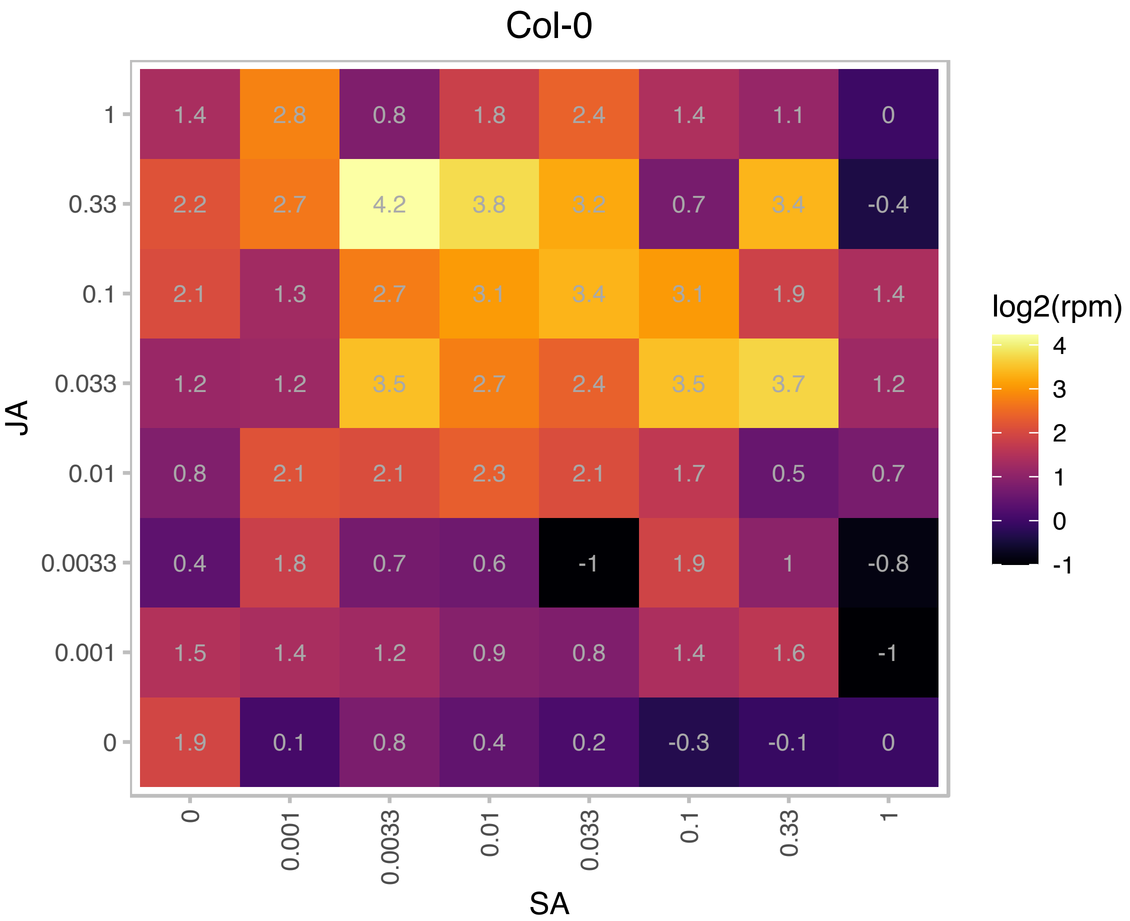


| | |
|---|---|
| Gene | AT2G15760 |
| is.expressed_genes | TRUE |
| is.responsive_genes | TRUE |
| is.apcluster | A-02 |

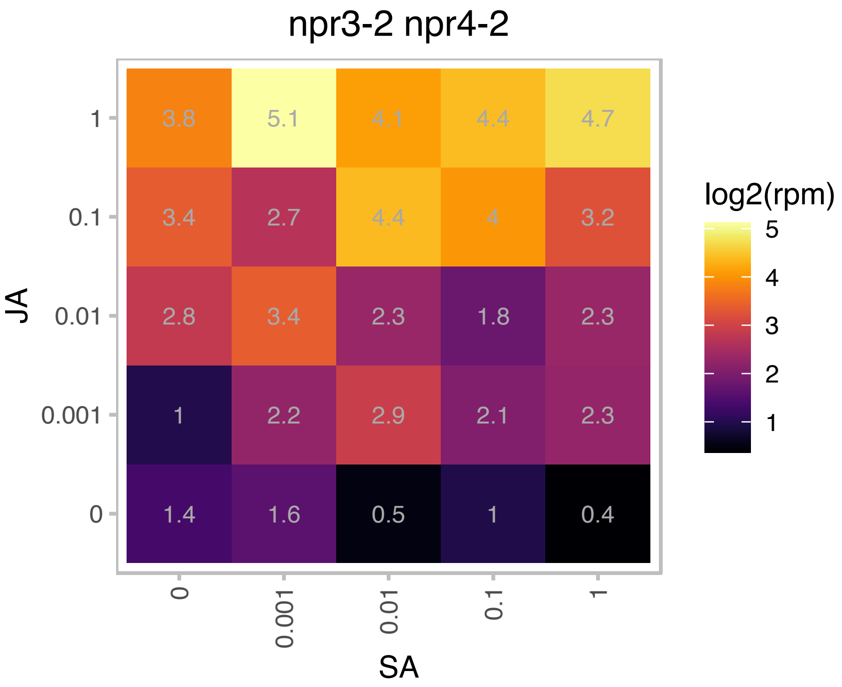


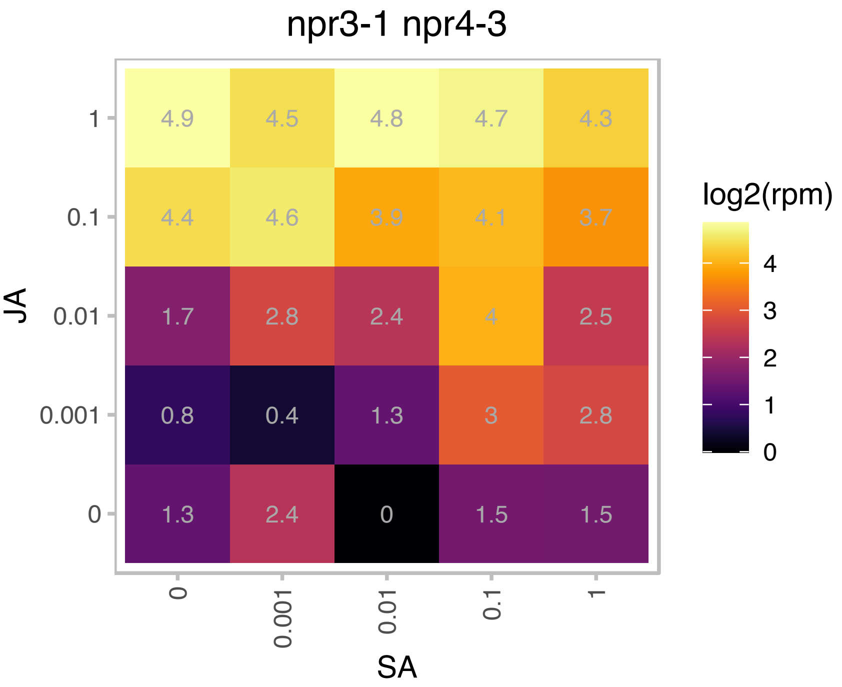

### AT2G24850.1 : TAT, TAT3

tyrosine aminotransferase 3

Encodes a tyrosine aminotransferase that is responsive to treatment with jasmonic acid.

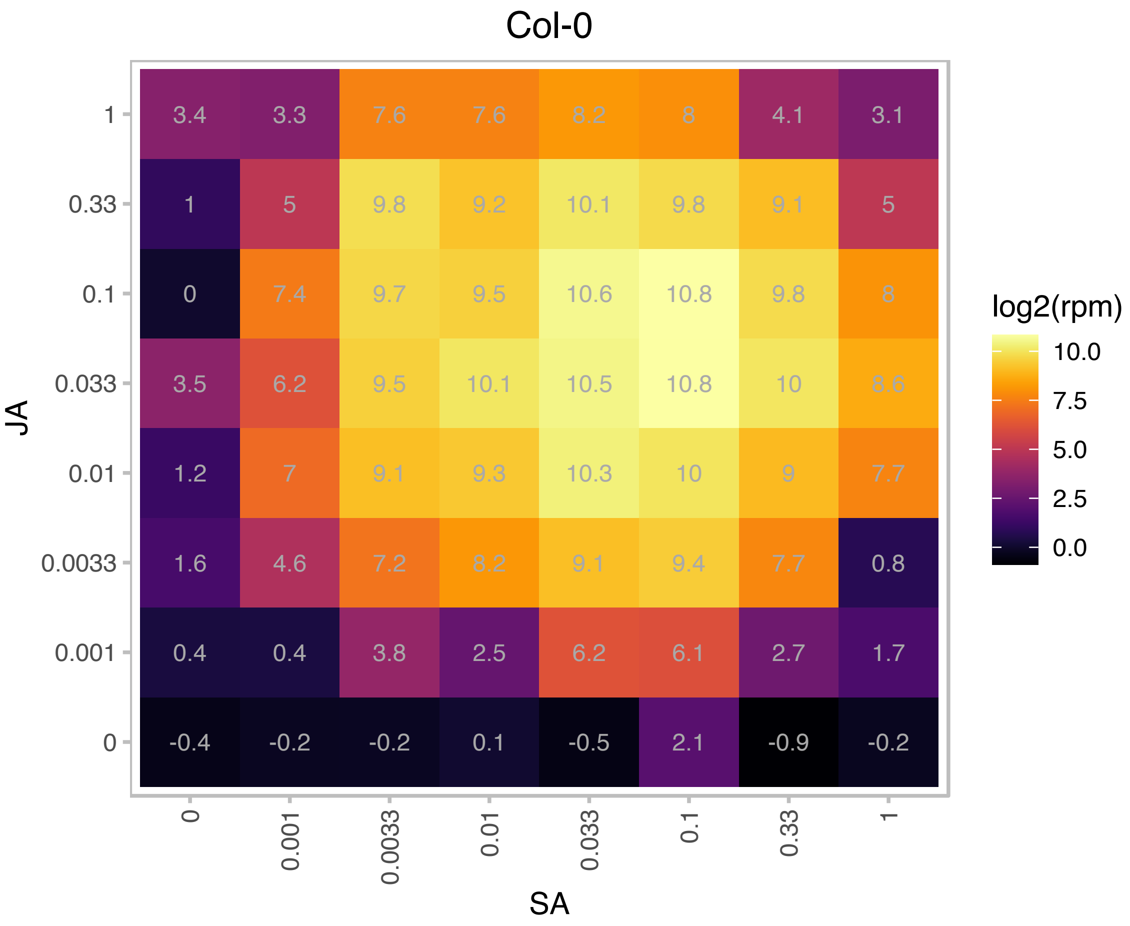


| | |
|---|---|
| Gene | AT2G24850 |
| is.expressed_genes | TRUE |
| is.responsive_genes | TRUE |
| is.apcluster | A-02 |

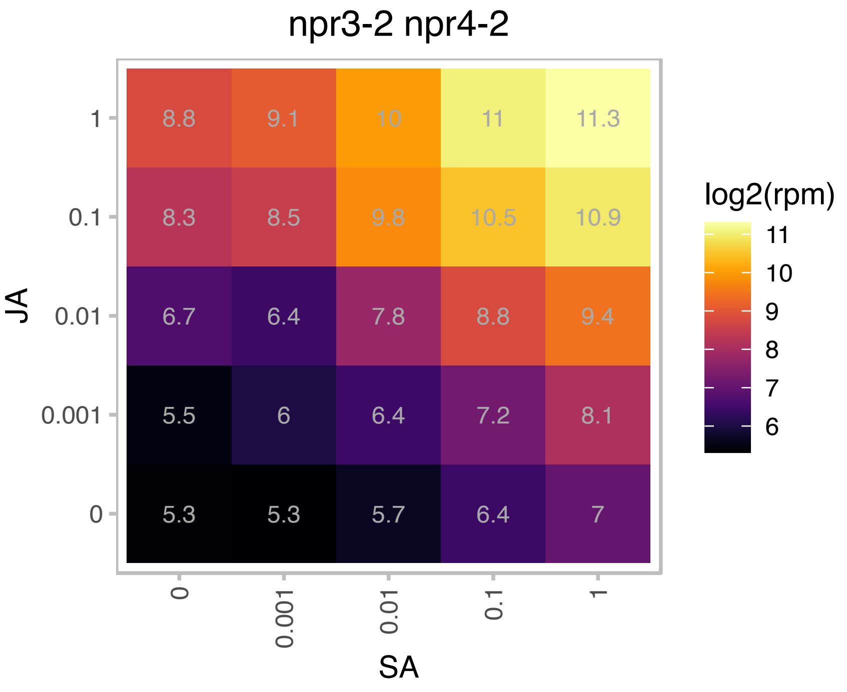


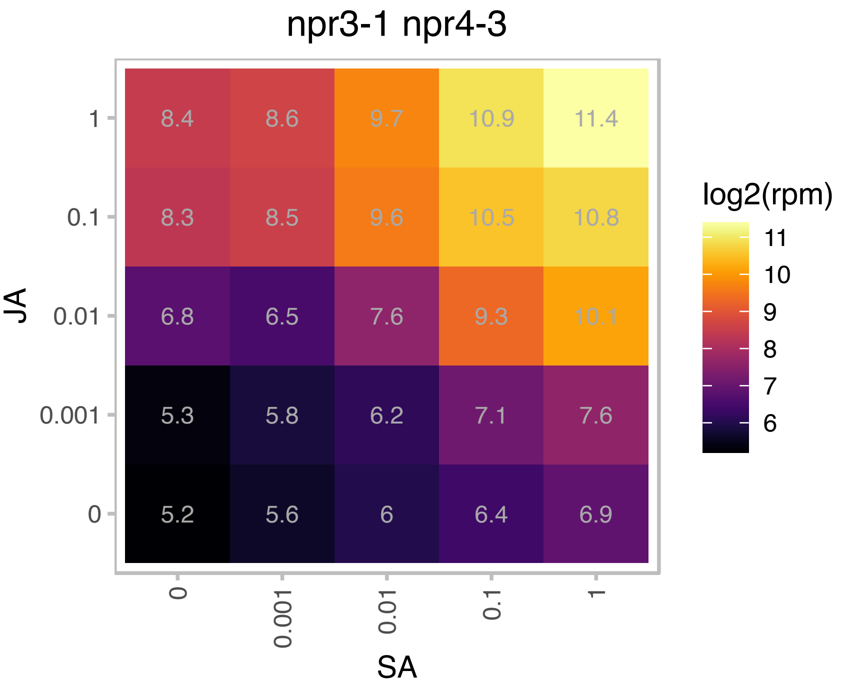

### AT2G27690.1 : CYP94C1

cytochrome P450, family 94, subfamily C, polypeptide 1
Encodes a CYP94C1. Has highest omega-hydroxylase activity with 9,10-epoxystearic acid, while also metabolized lauric acid (C12:0) and C18 unsaturated fatty acids. Gene expression is induced in response to wounding and jasmonic acid treatment.

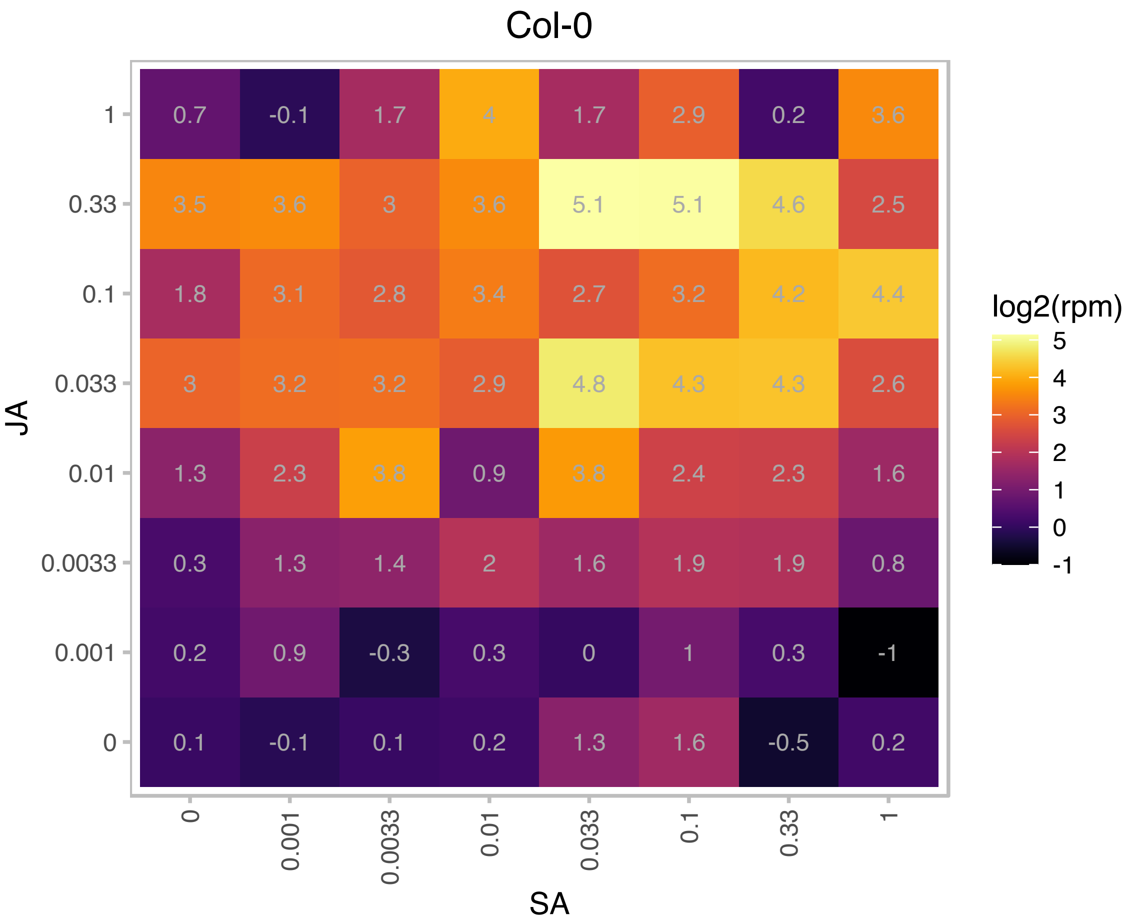


| | |
|---|---|
| Gene | AT2G27690 |
| is.expressed_genes | TRUE |
| is.responsive_genes | TRUE |
| is.apcluster | A-02 |

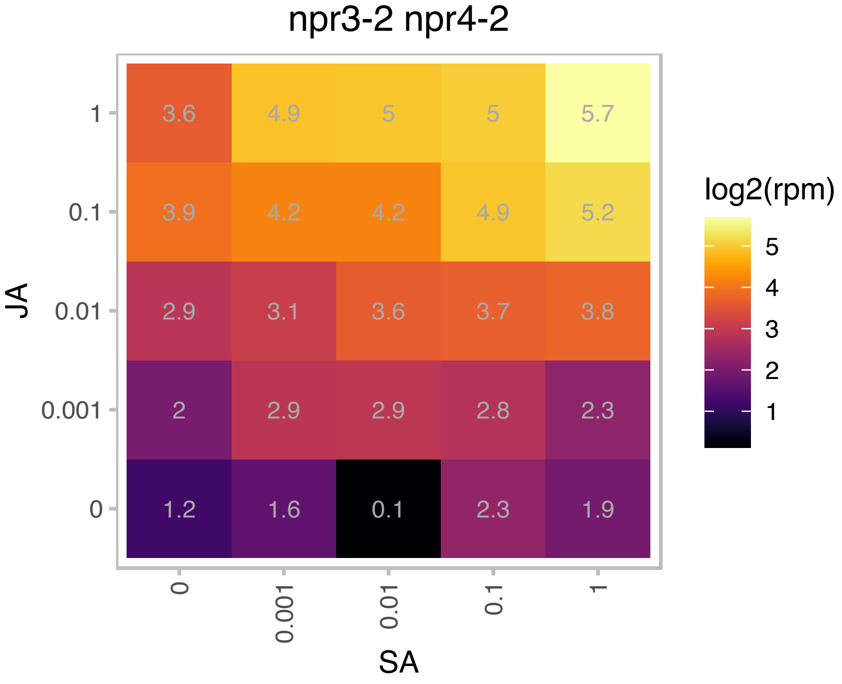


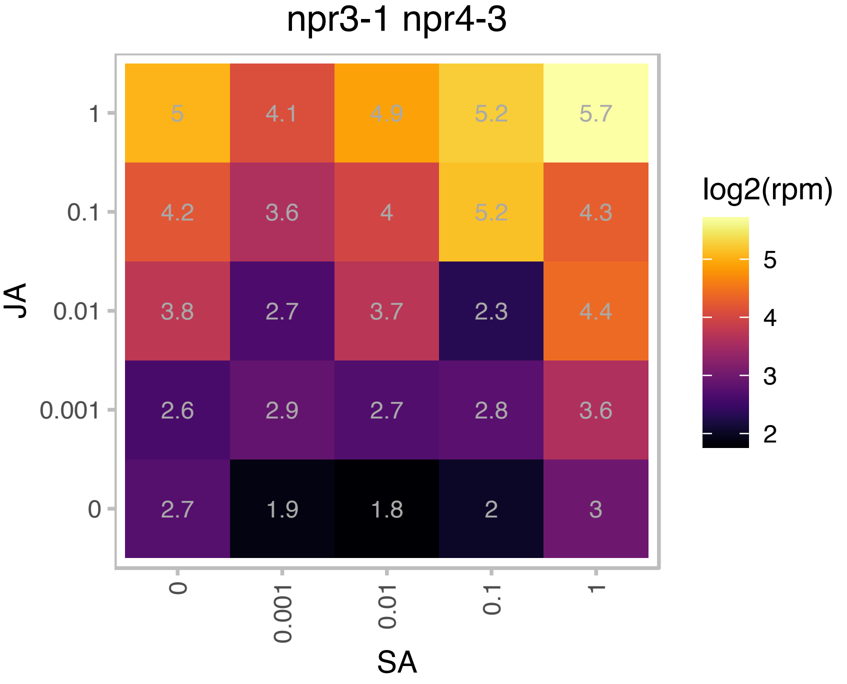

# AT2G37580.1 : NA

RING/U-box superfamily protein

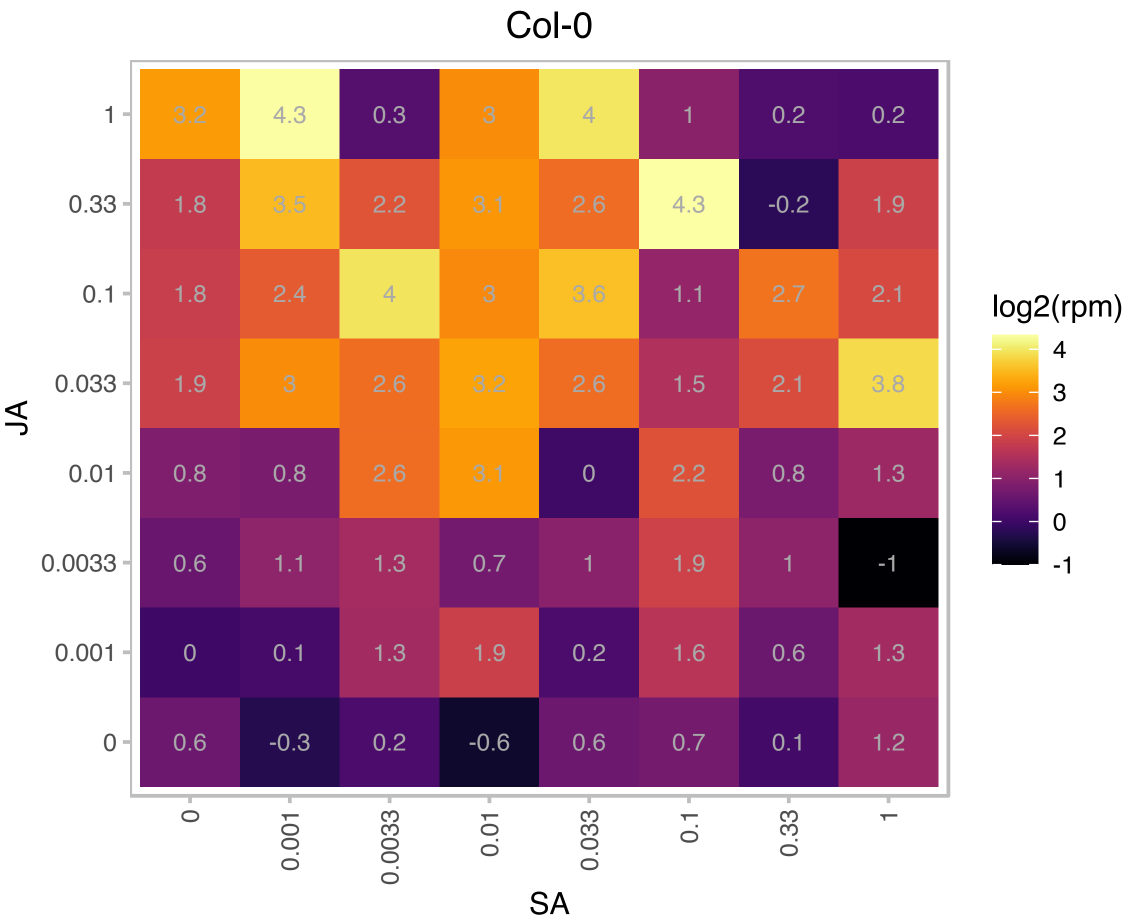


| | |
|---|---|
| Gene | AT2G37580 |
| is.expressed_genes | TRUE |
| is.responsive_genes | TRUE |
| is.apcluster | A-02 |

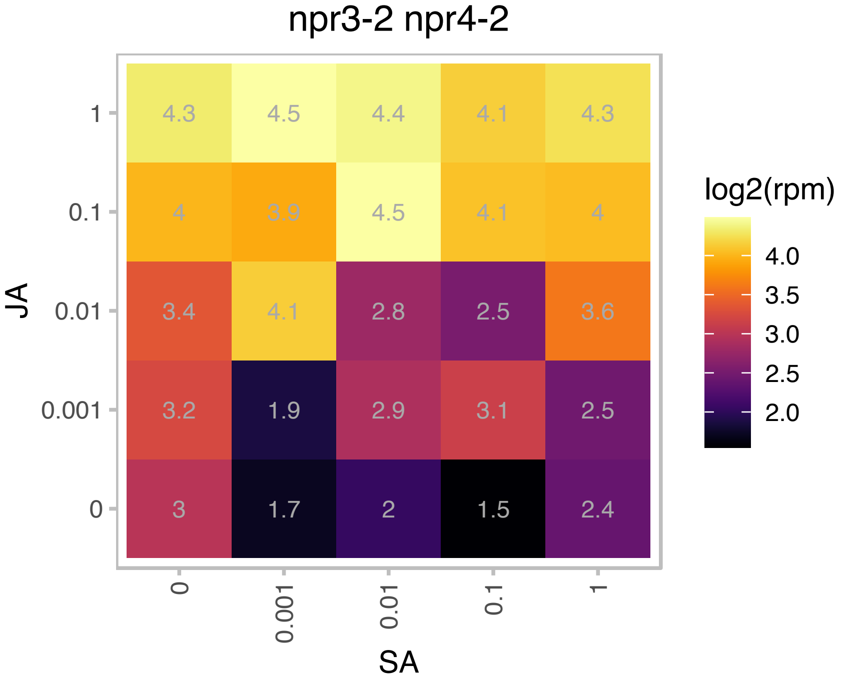


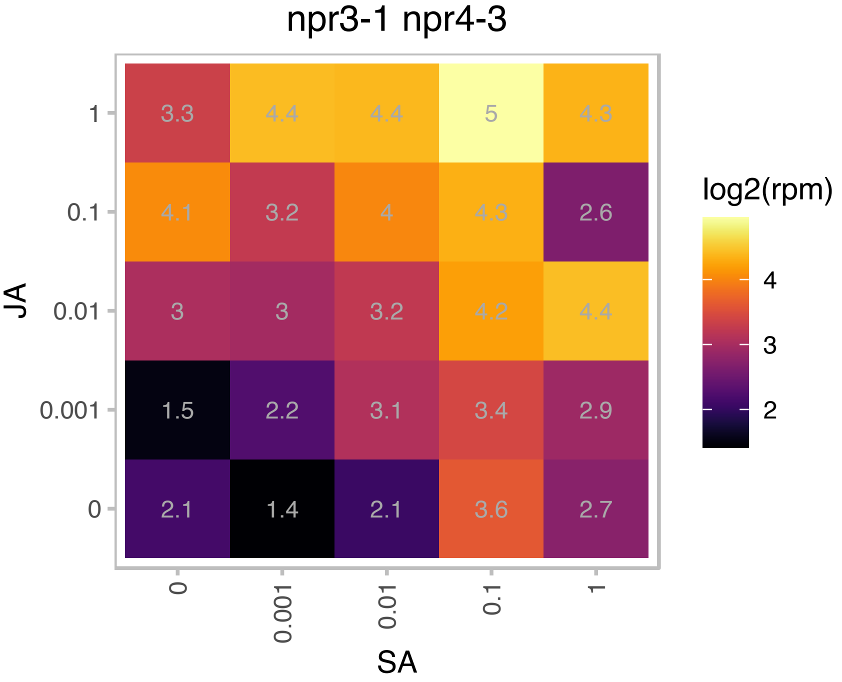

### AT2G38760.1 : ANN3, ANNAT3, AtANN3

annexin 3

Annexins are calcium binding proteins that are localized in the cytoplasm. When cytosolic Ca2+ increases, they relocate to the plasma membrane. The mRNA is cell-to-cell mobile.

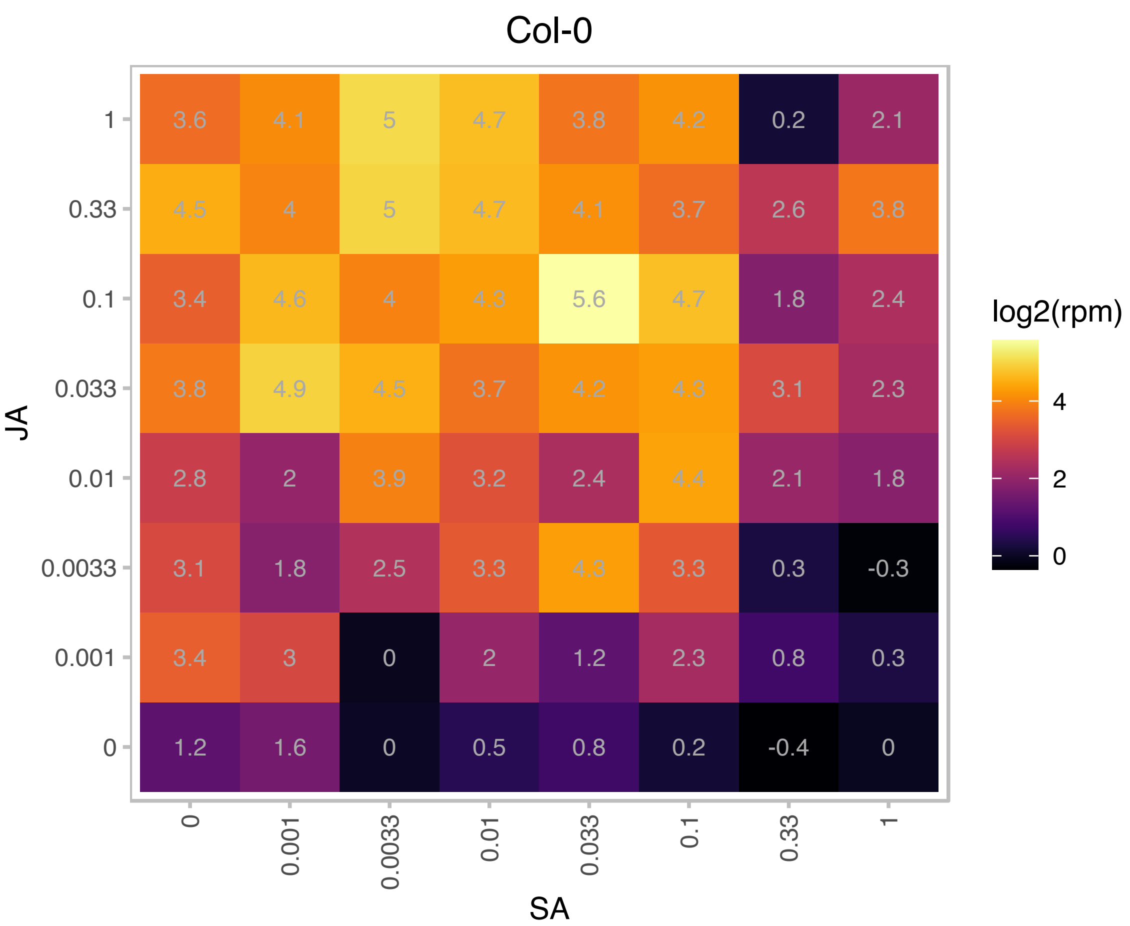


| | |
|---|---|
| Gene | AT2G38760 |
| is.expressed_genes | TRUE |
| is.responsive_genes | TRUE |
| is.apcluster | A-02 |

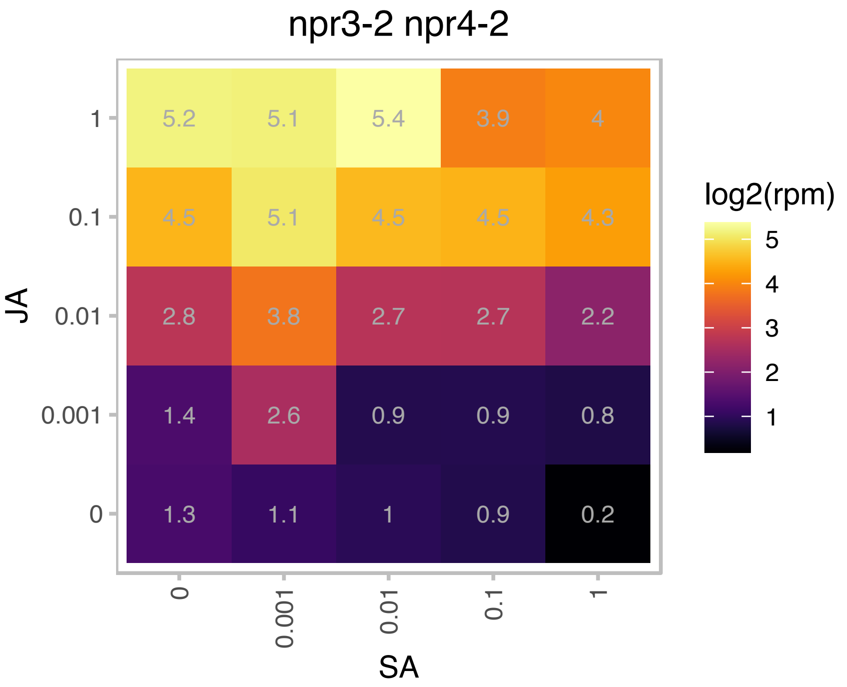


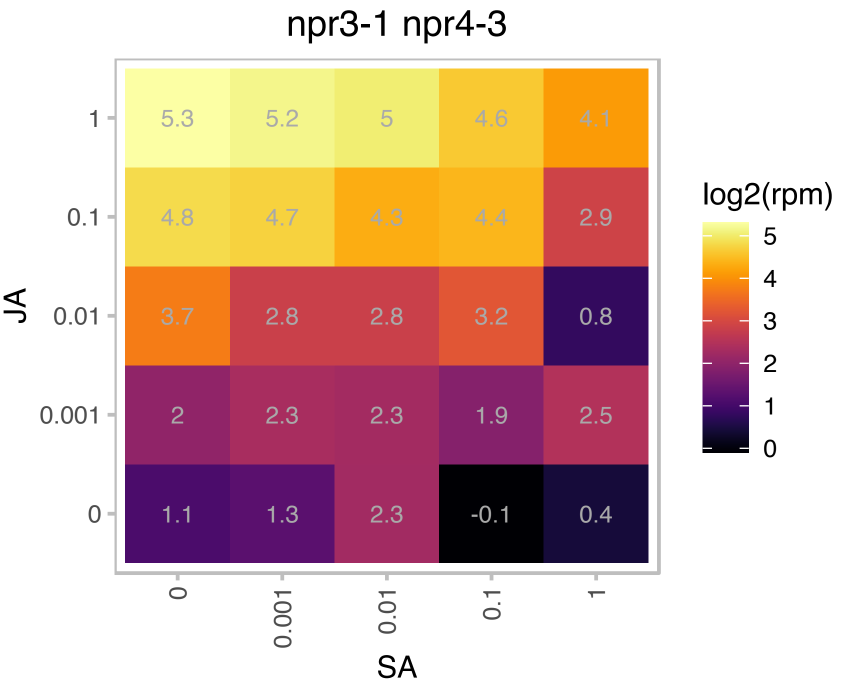

### AT2G39420.1 : MAGL8

alpha/beta-Hydrolases superfamily protein

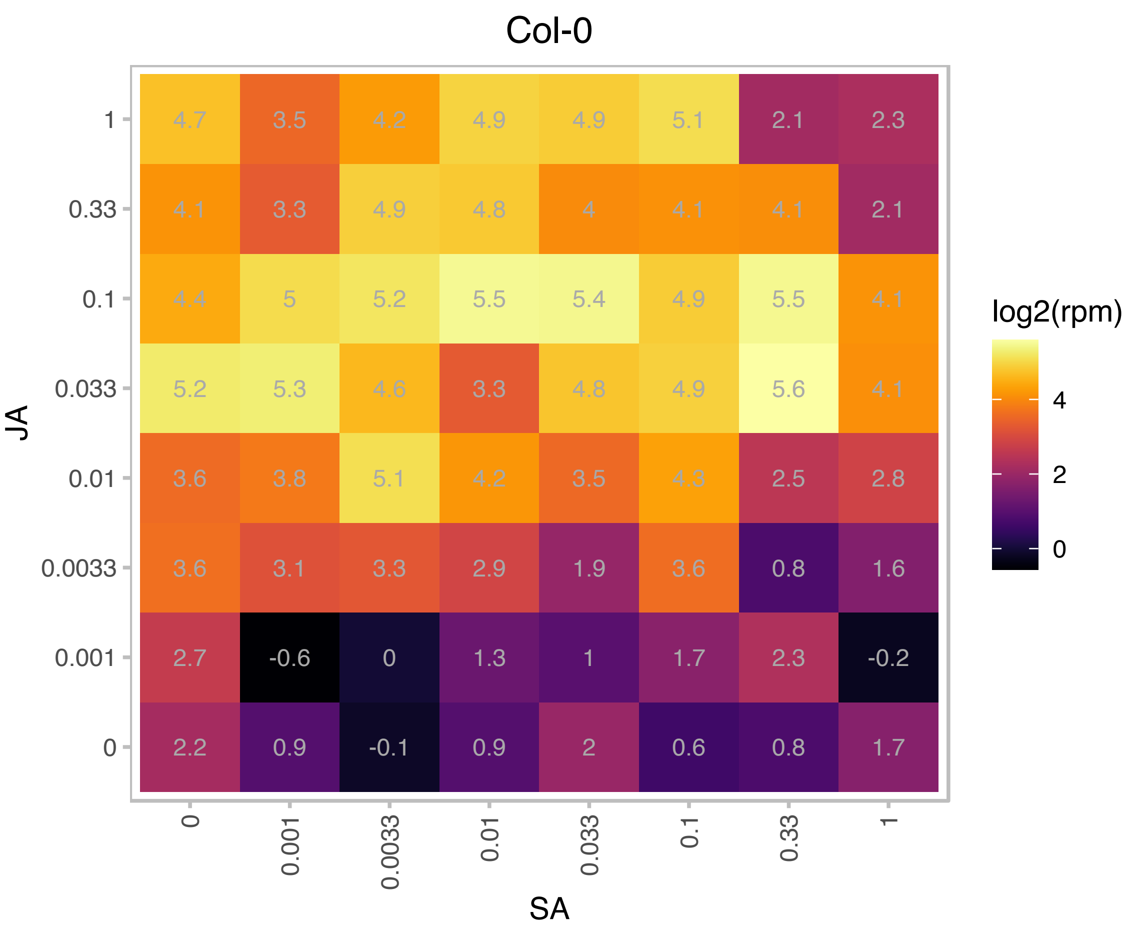


| | |
|---|---|
| Gene | AT2G39420 |
| is.expressed_genes | TRUE |
| is.responsive_genes | TRUE |
| is.apcluster | A-02 |

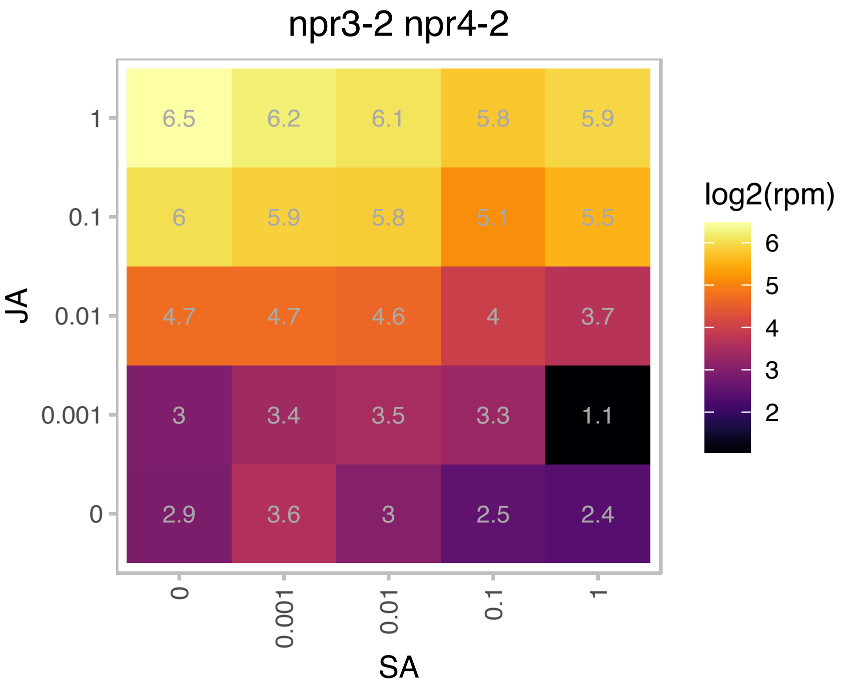


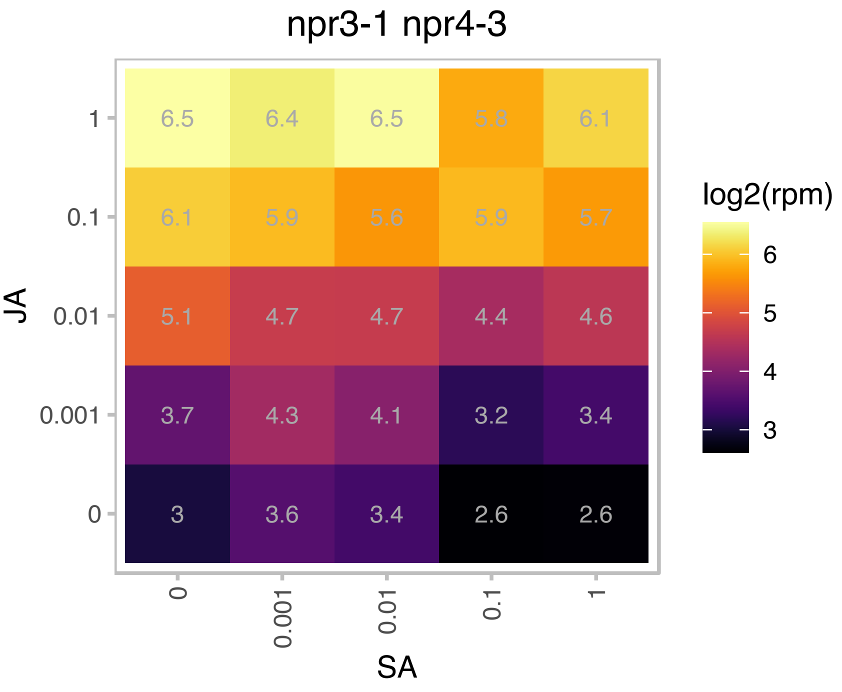

# AT2G40200.1 : NA

basic helix-loop-helix (bHLH) DNA-binding superfamily protein

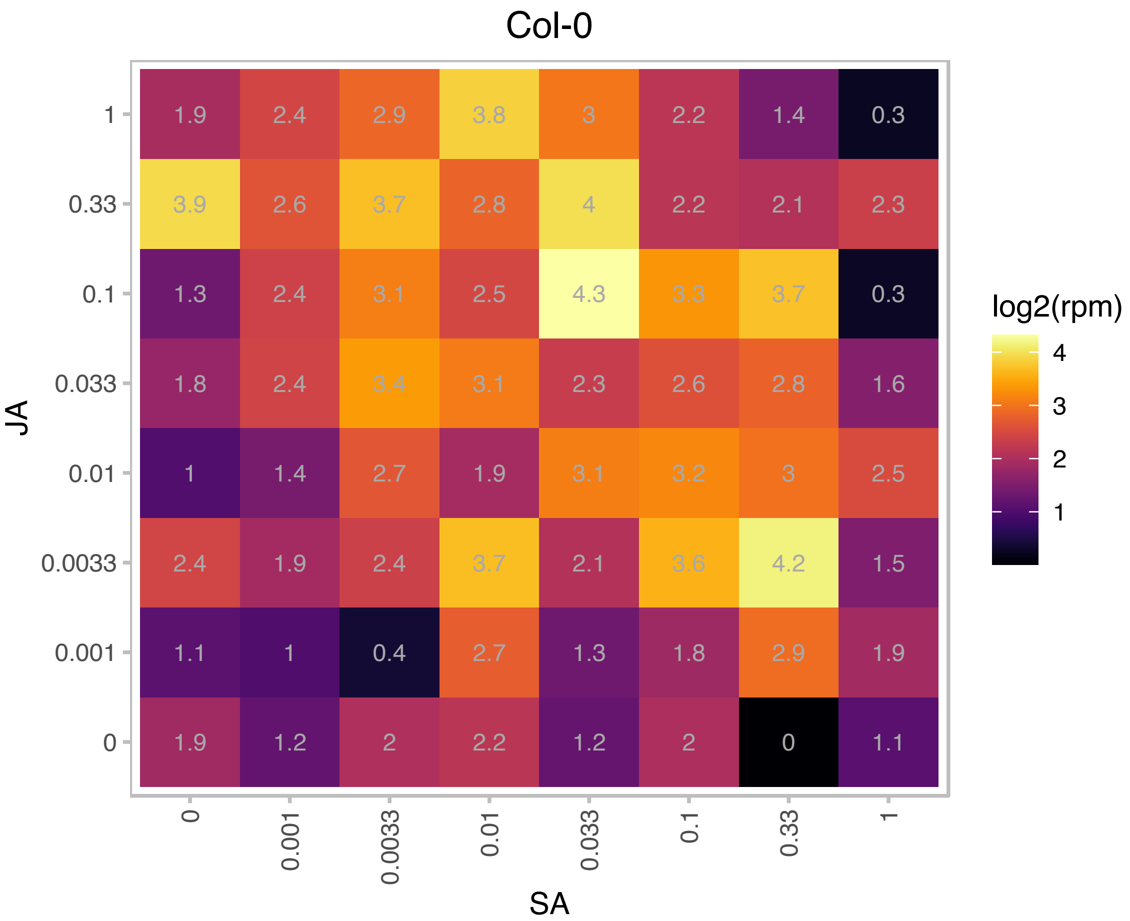


| Gene | AT2G40200 |
|---|---|
| is.expressed_genes | TRUE |
| is.responsive_genes | TRUE |
| is.apcluster | A-02 |

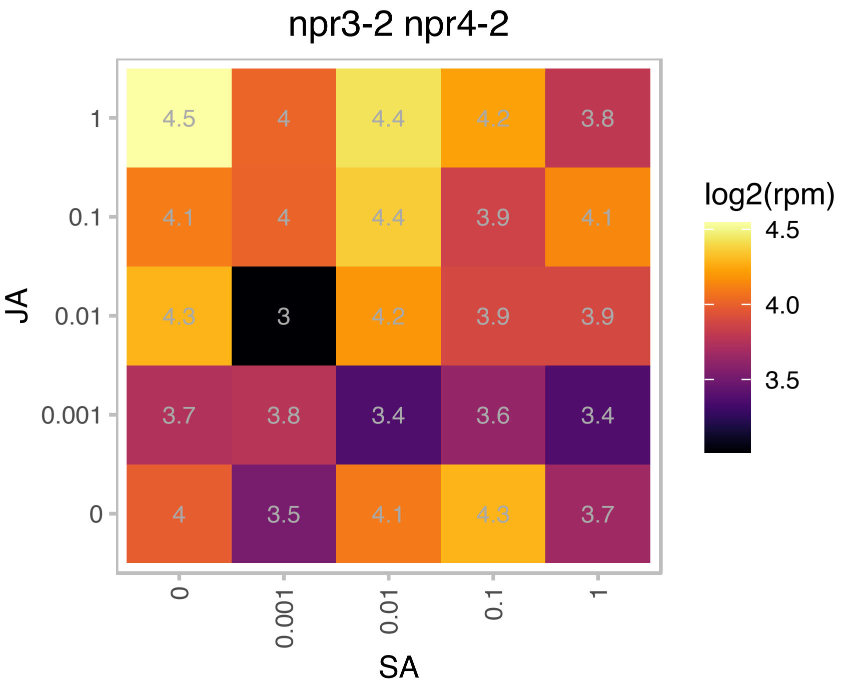


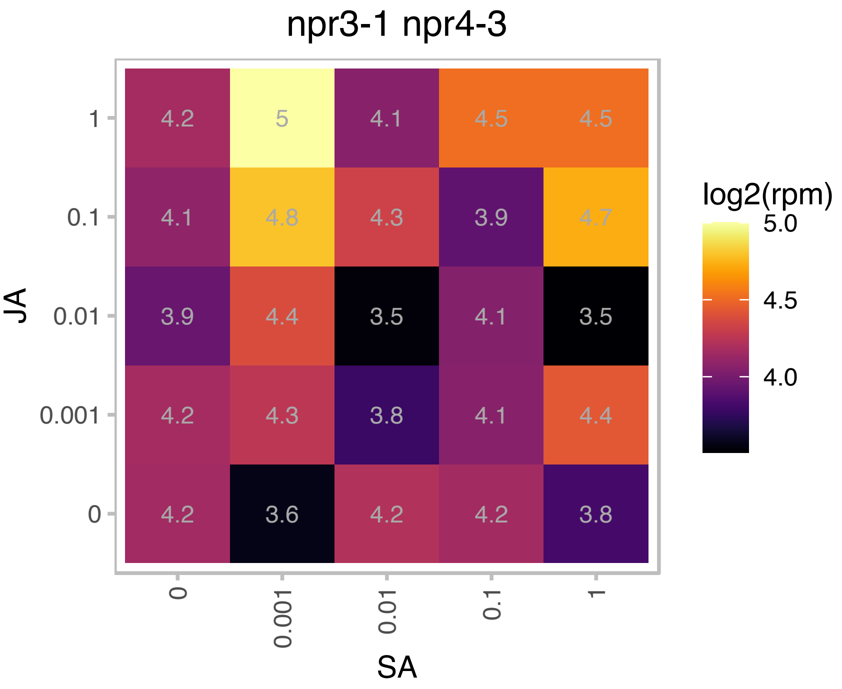

### AT3G10260.1 : RTNLB8

Reticulon family protein

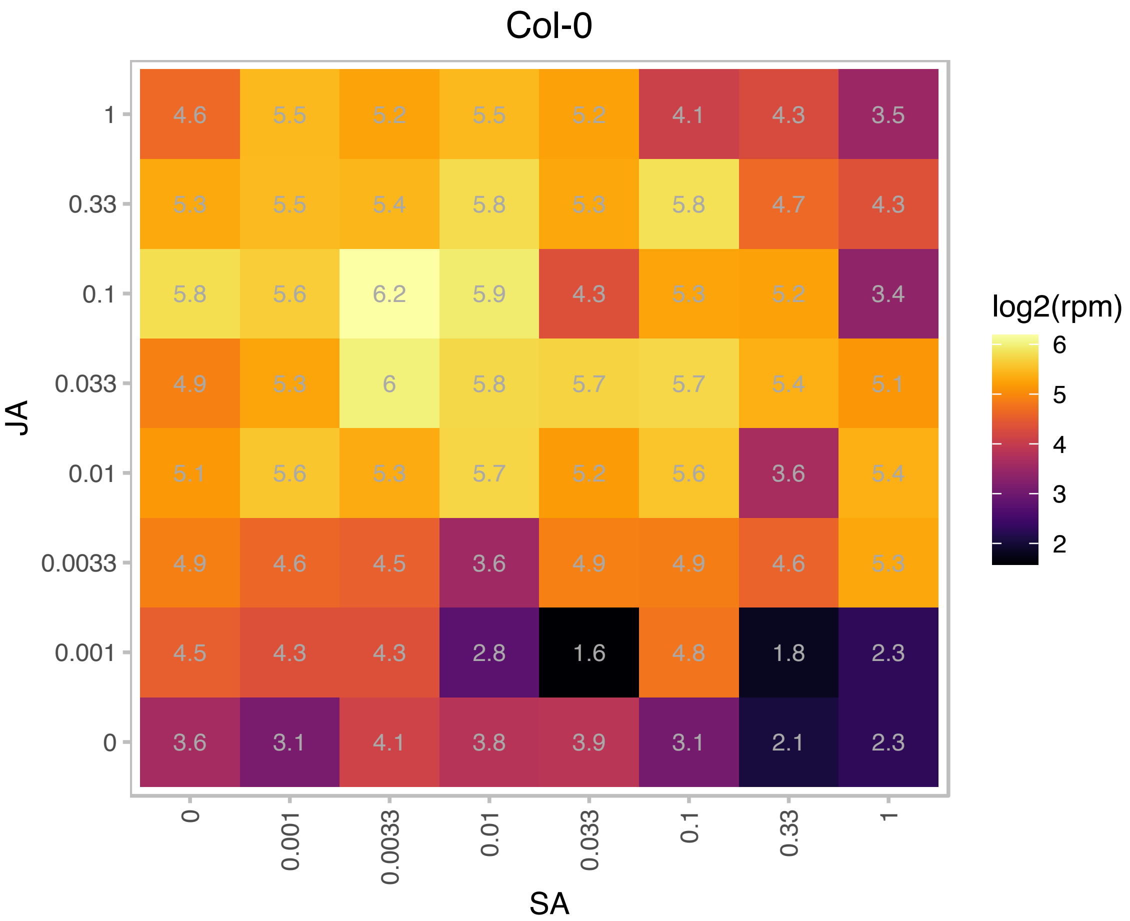


| | |
|---|---|
| Gene | AT3G10260 |
| is.expressed_genes | TRUE |
| is.responsive_genes | TRUE |
| is.apcluster | A-02 |

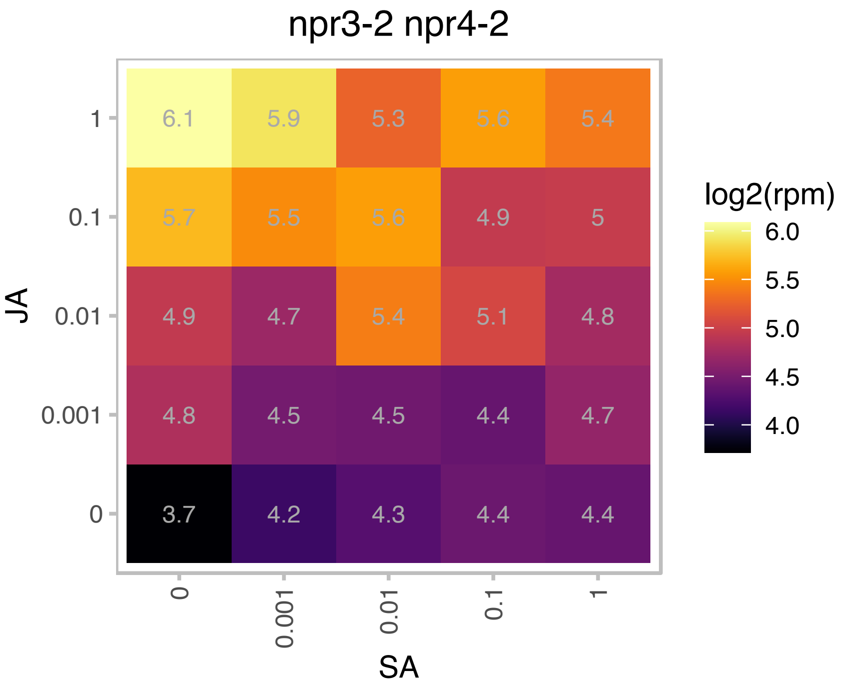


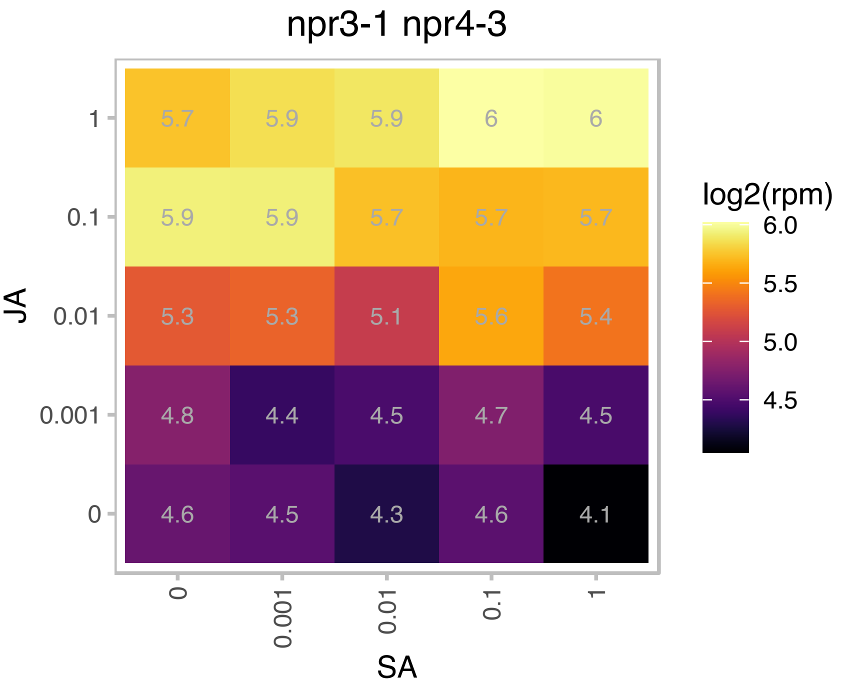

# AT3G14260.1 : NA

Protein of unknown function (DUF567)

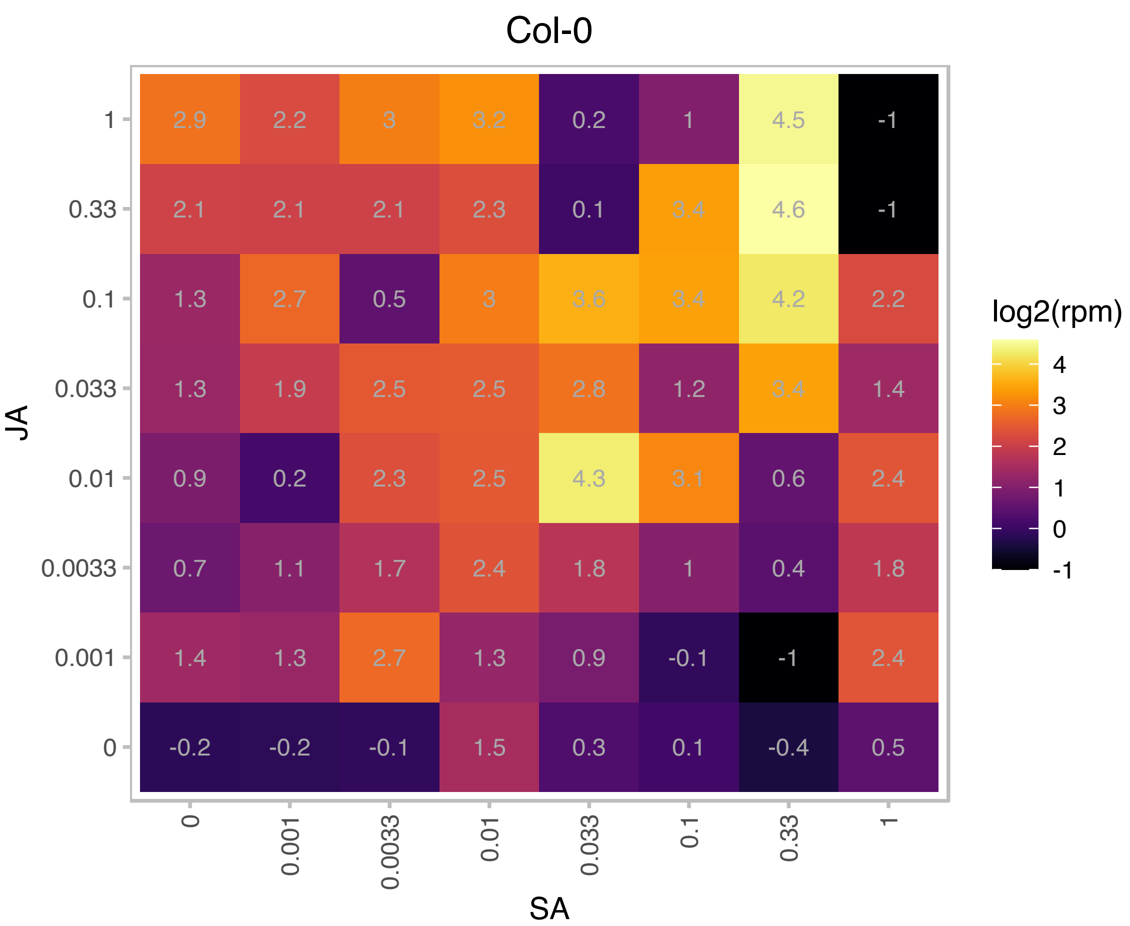


| | |
|---|---|
| Gene | AT3G14260 |
| is.expressed_genes | TRUE |
| is.responsive_genes | TRUE |
| is.apcluster | A-02 |

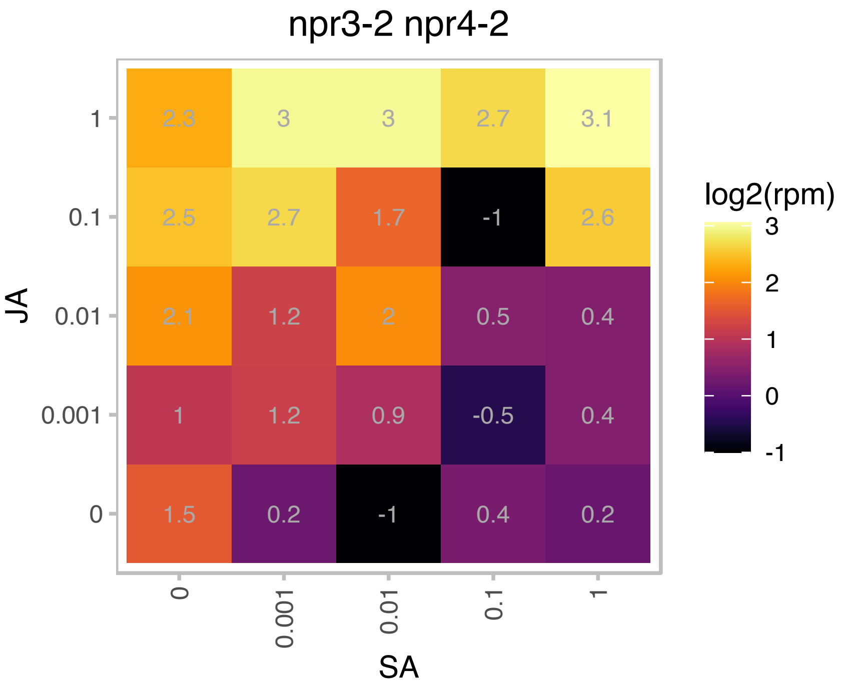


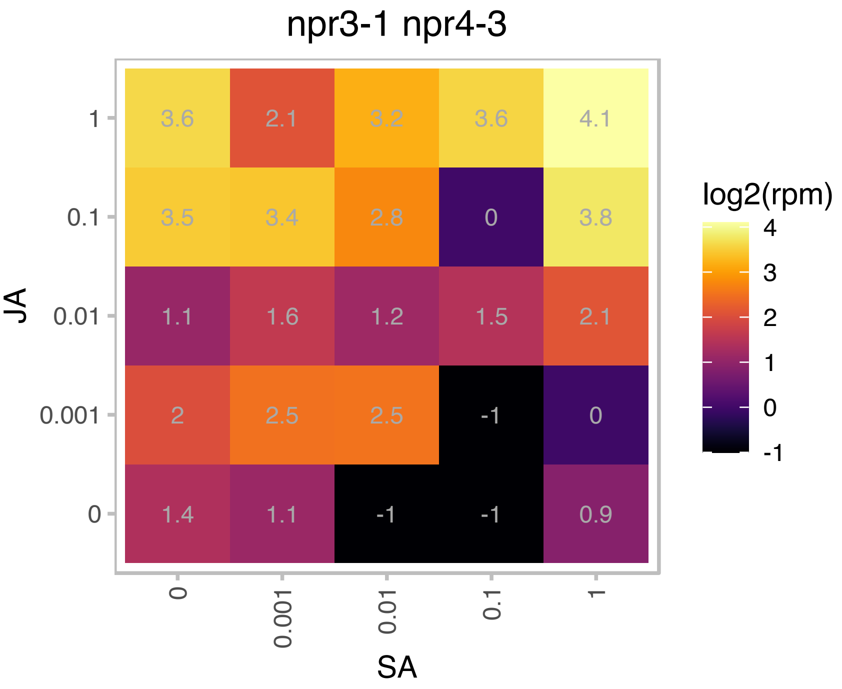

### AT3G16640.1 : AtTCTP1, TCTP1

Encodes a protein homologous to translationally controlled tumor protein (TCTP) from Drosophila. In flies, TCTP functions guanine nucleotide exchange factor in the TOR signaling pathway. TCTP is expressed throughout the plant with highest levels seen in meristematic regions of the shoot and root. Loss of function alleles are not transmitted through the male gametophyte due to defects in pollen tube growth. Hypomorphs, generated through RNAi, are dwarf and have smaller cells. These plants also have defects in lateral and primary root growth as well as root hair growth. The phenotypes are similar to TOR mutants suggesting that TCTP functions [illegible]

translationally controlled tumor protein

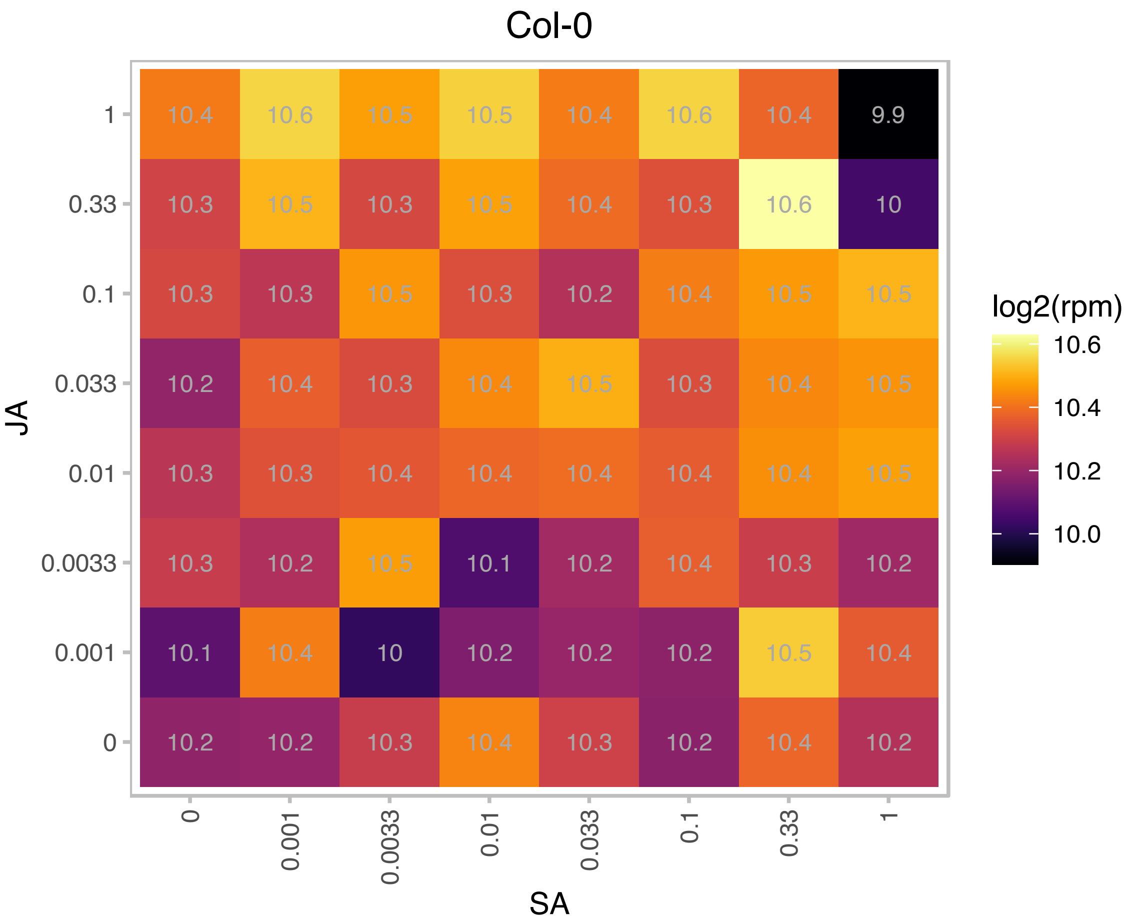


| | |
|---|---|
| Gene | AT3G16640 |
| is.expressed_genes | TRUE |
| is.responsive_genes | TRUE |
| is.apcluster | A-02 |

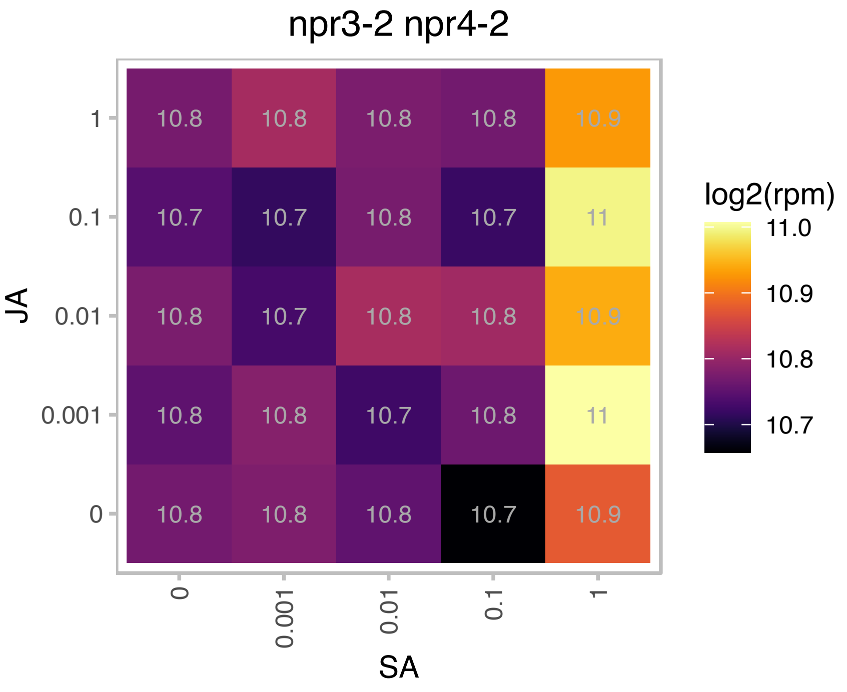


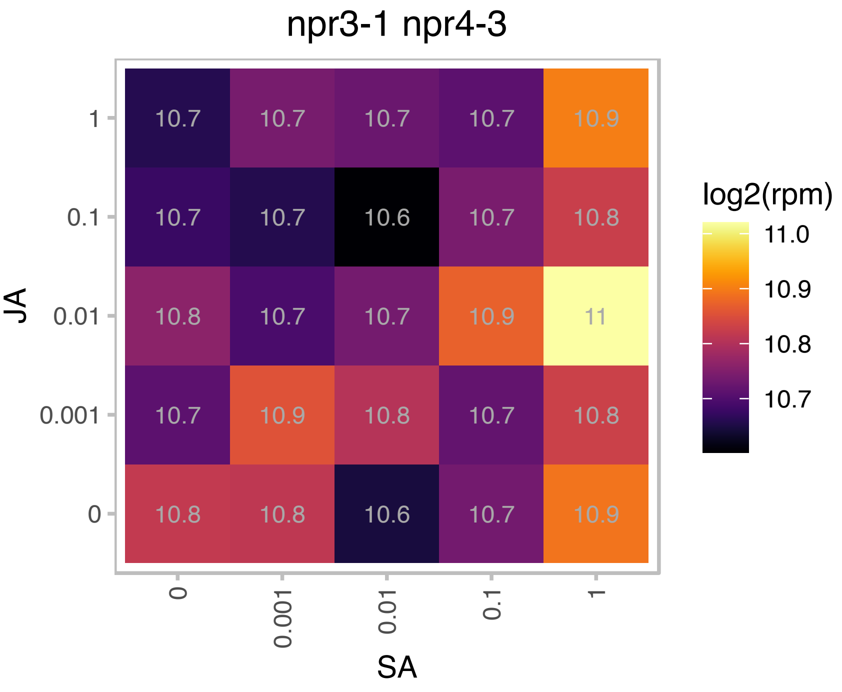

### AT3G16690.1 : AtSWEET16, SWEET16

Nodulin MtN3 family protein

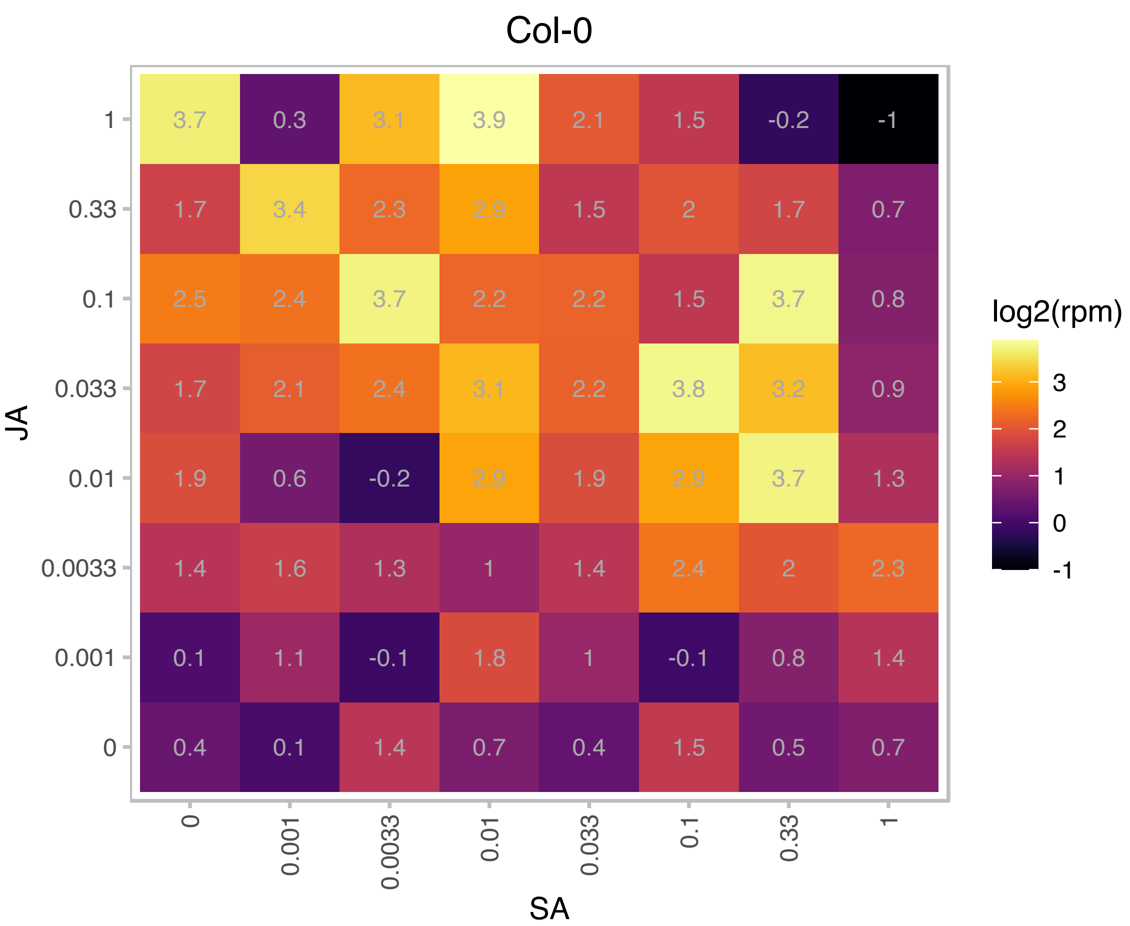


| | |
|---|---|
| Gene | AT3G16690 |
| is.expressed_genes | TRUE |
| is.responsive_genes | TRUE |
| is.apcluster | A-02 |

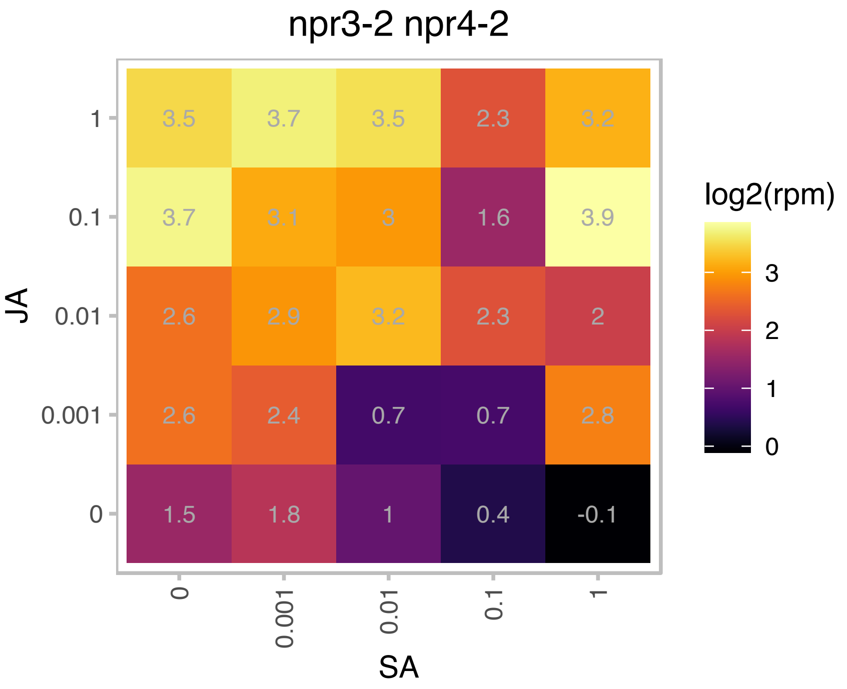


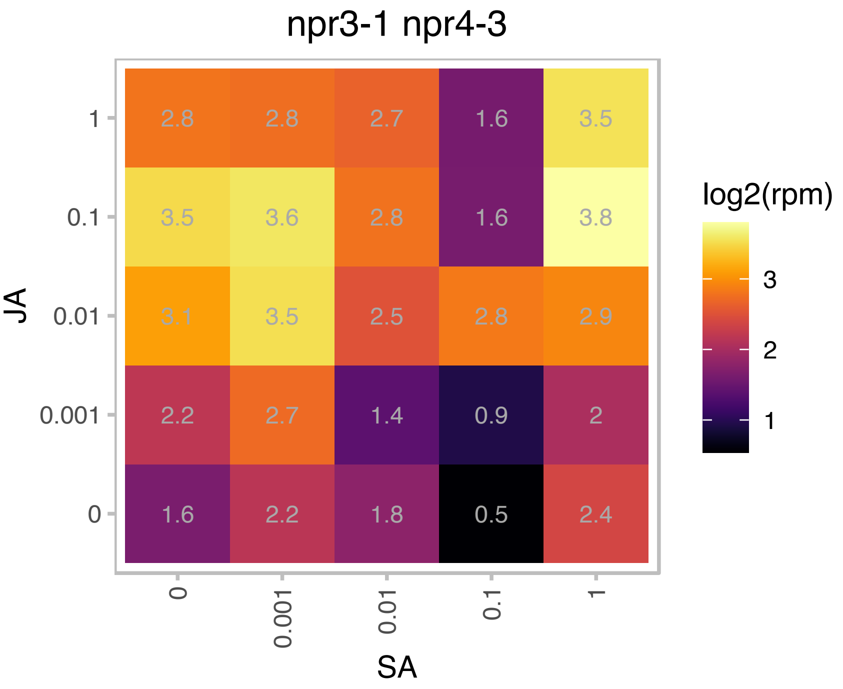

### AT3G26840.1 : PES2

Esterase/lipase/thioesterase family protein
Encodes a protein with phytyl ester synthesis and diacylglycerol acyltransferase activities that is involved in the deposition of free phytol and free fatty acids in the form of phytyl esters in chloroplasts, a process involved in maintaining the integrity of the photosynthetic membrane during abiotic stress and senescence.

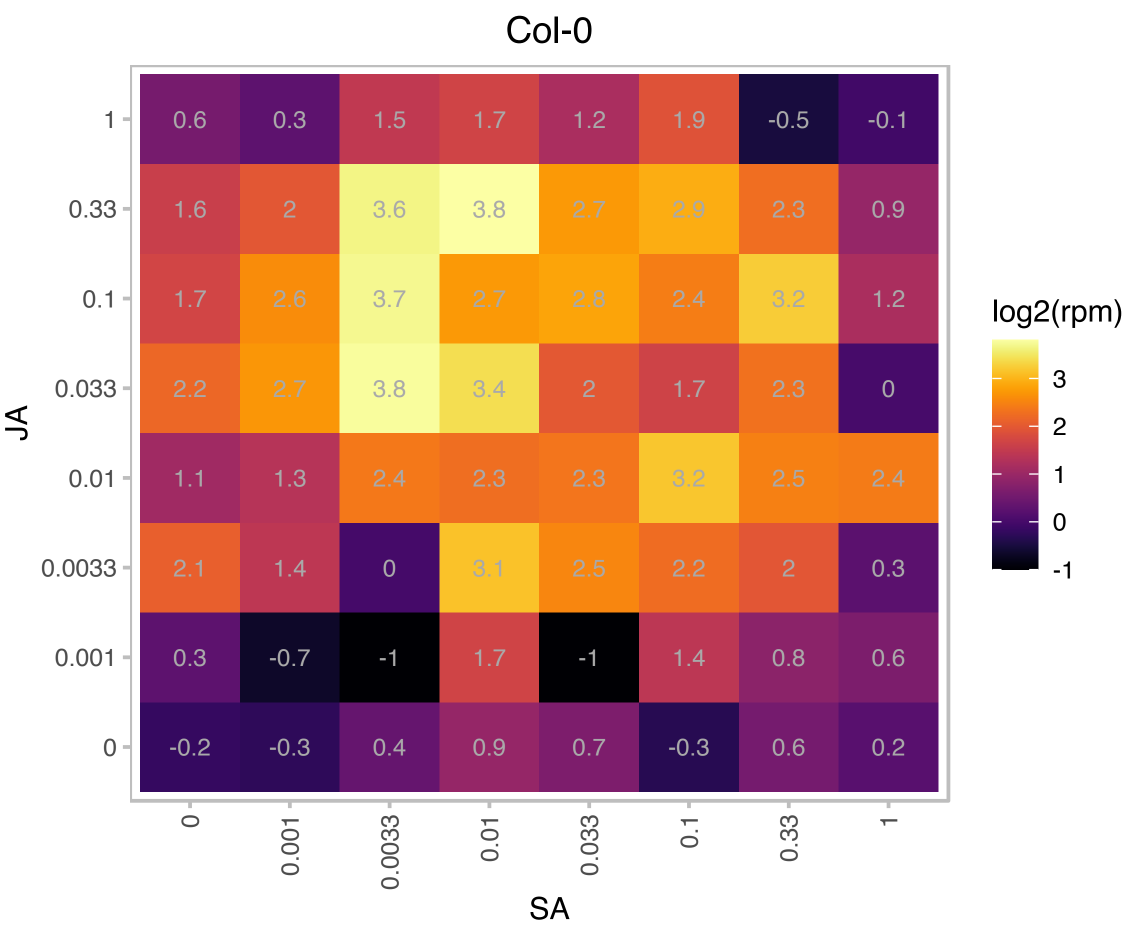


| | |
|---|---|
| Gene | AT3G26840 |
| is.expressed_genes | TRUE |
| is.responsive_genes | TRUE |
| is.apcluster | A-02 |

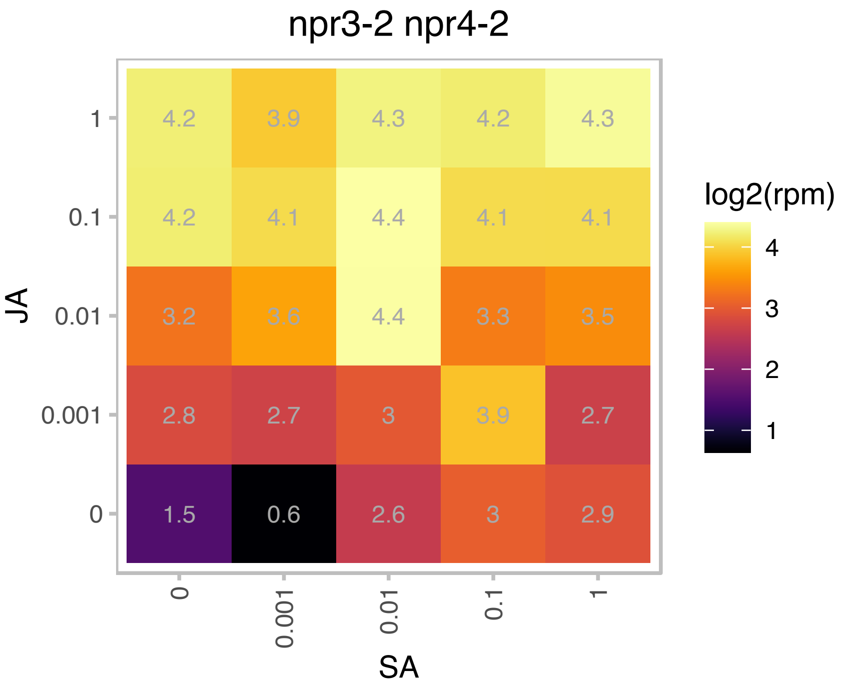


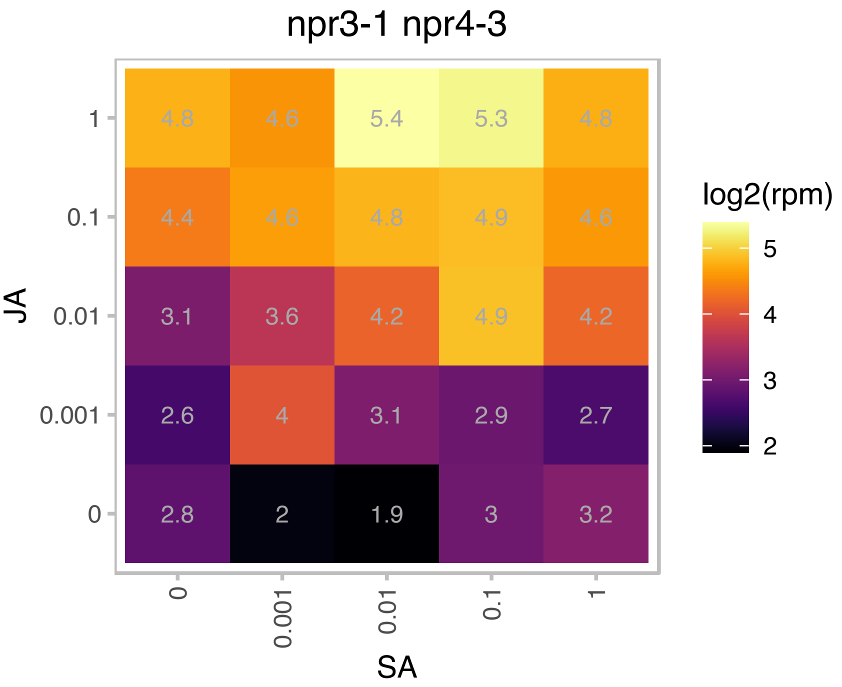

### AT3G27310.1 : PUX1

plant UBX domain-containing protein 1

encodes a protein that contains a UBX domain and regulates AtCDC48 by inhibiting its ATPase activity and by promoting the disassembly of the active hexamer. Phenotypic analysis of pux1 plants revealed that the loss of PUX1 accelerated the growth of various plant organs including roots and inflorescence shoots. AtCDC48 and SYP31 colocalize at the division plane during cytokinesis and to interact in vitro and in vivo.

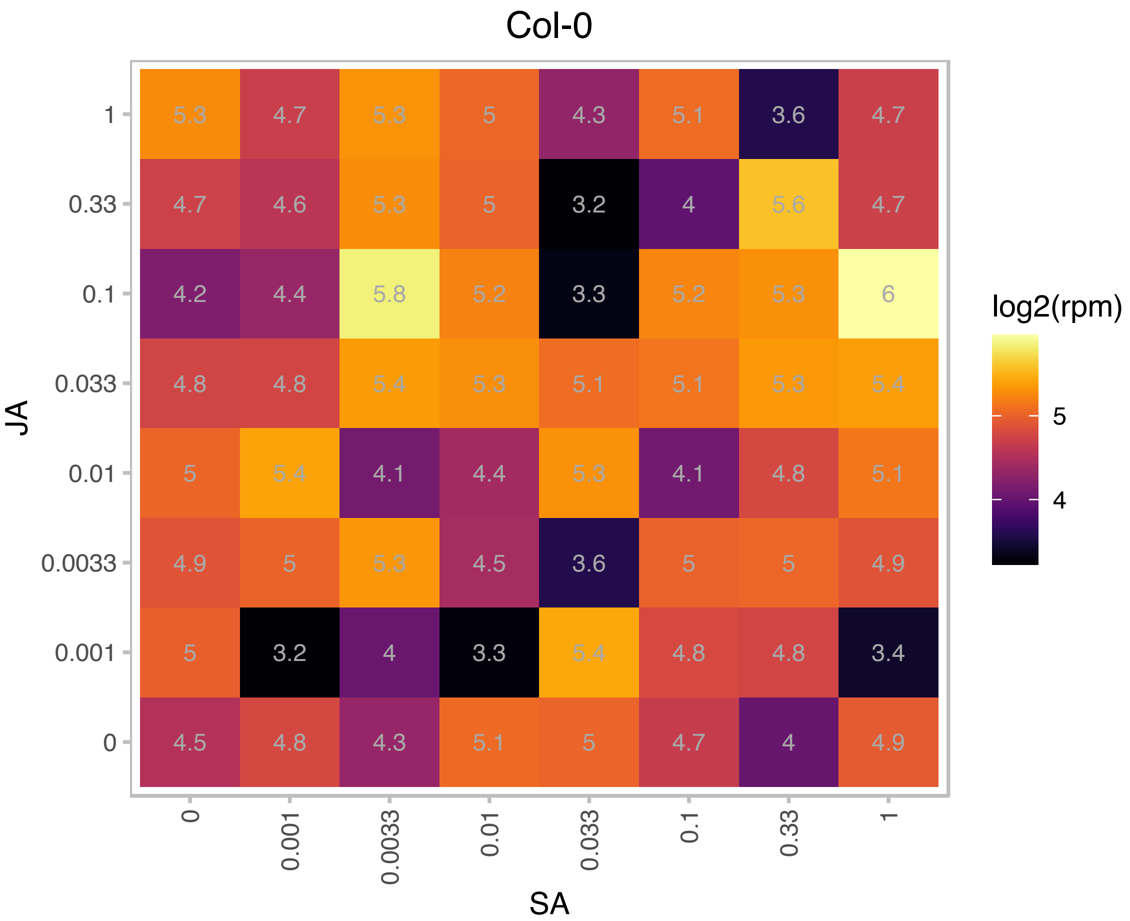


| | |
|---|---|
| Gene | AT3G27310 |
| is.expressed_genes | TRUE |
| is.responsive_genes | TRUE |
| is.apcluster | A-02 |

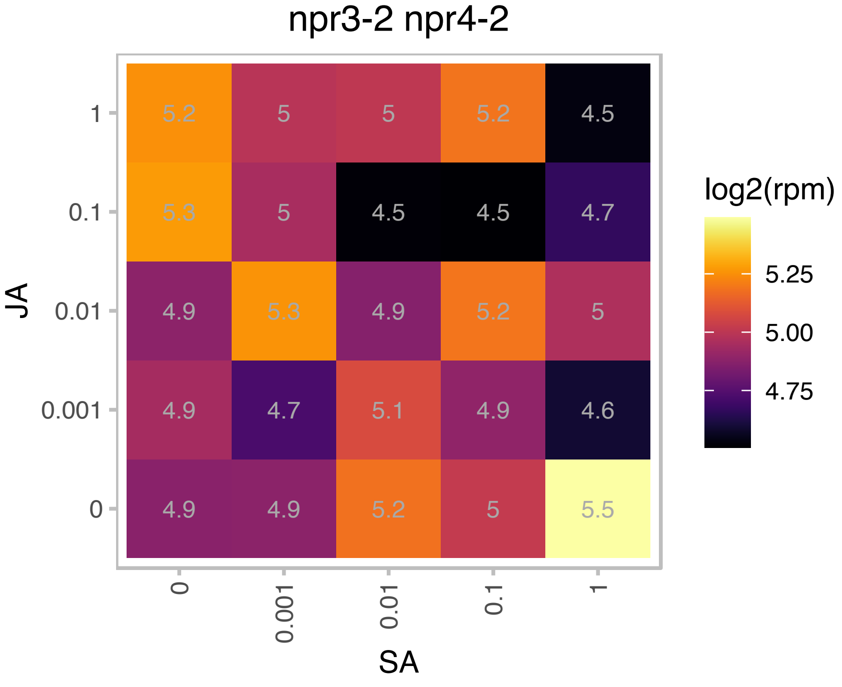


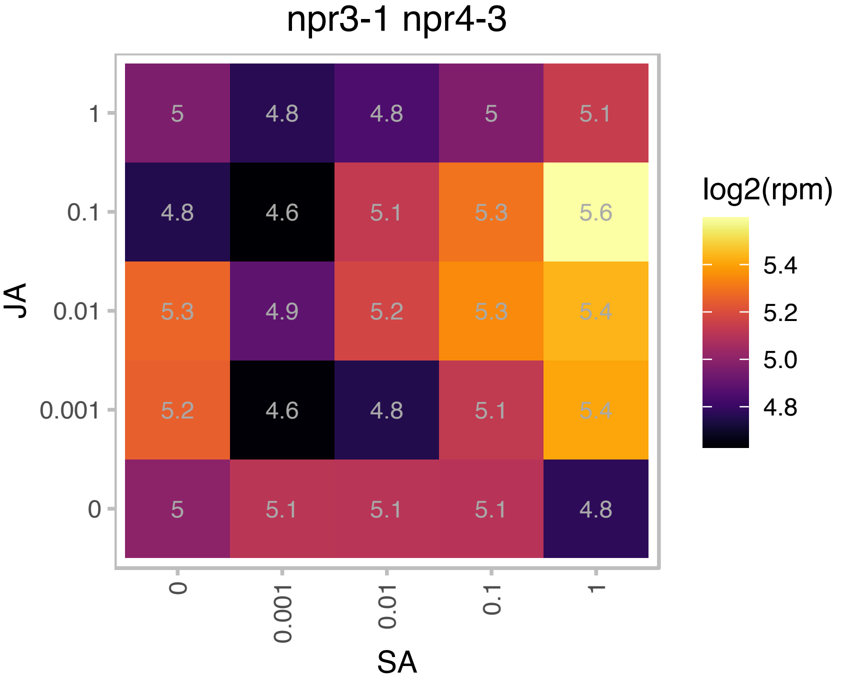

### AT3G44860.1 : FAMT

farnesoic acid carboxyl-O-methyltransferase

Encodes a farnesoic acid carboxyl-O-methyltransferase. The mRNA is cell-to-cell mobile.

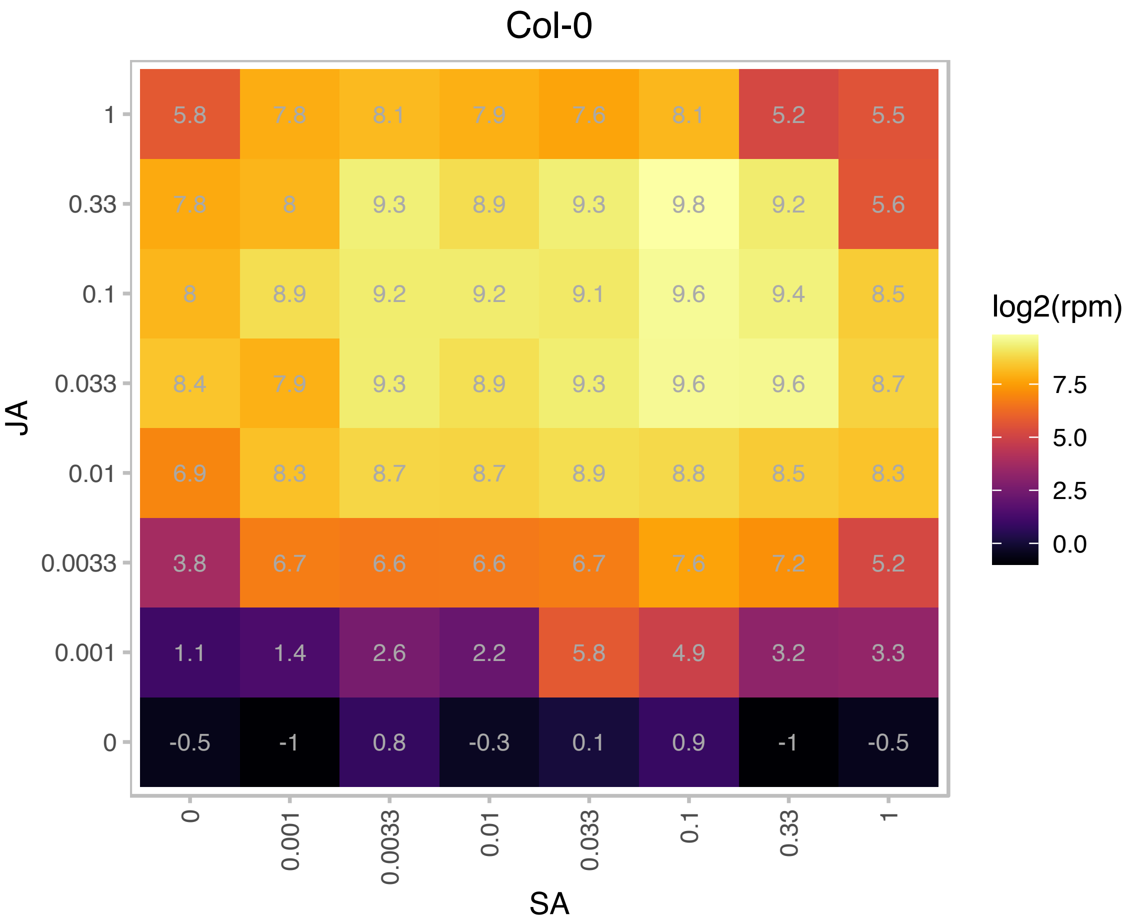


| | |
|---|---|
| Gene | AT3G44860 |
| is.expressed_genes | TRUE |
| is.responsive_genes | TRUE |
| is.apcluster | A-02 |

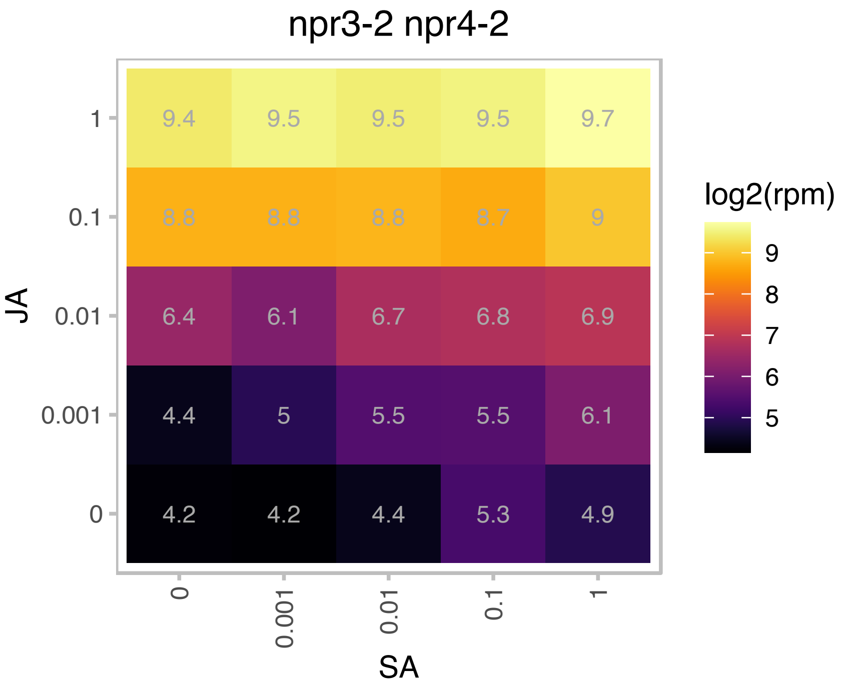


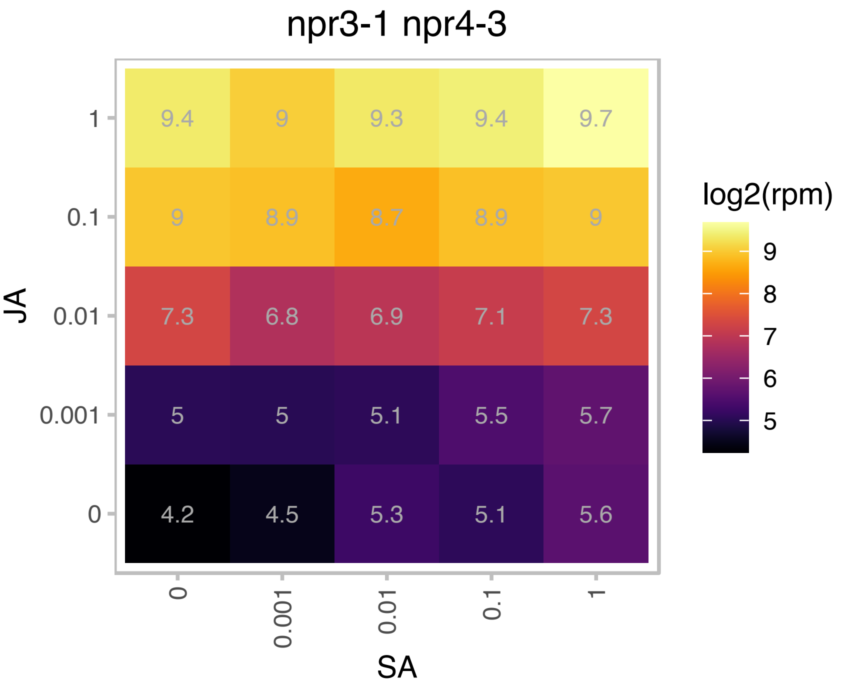

# AT3G51450.1 : NA

Calcium-dependent phosphotriesterase superfamily protein

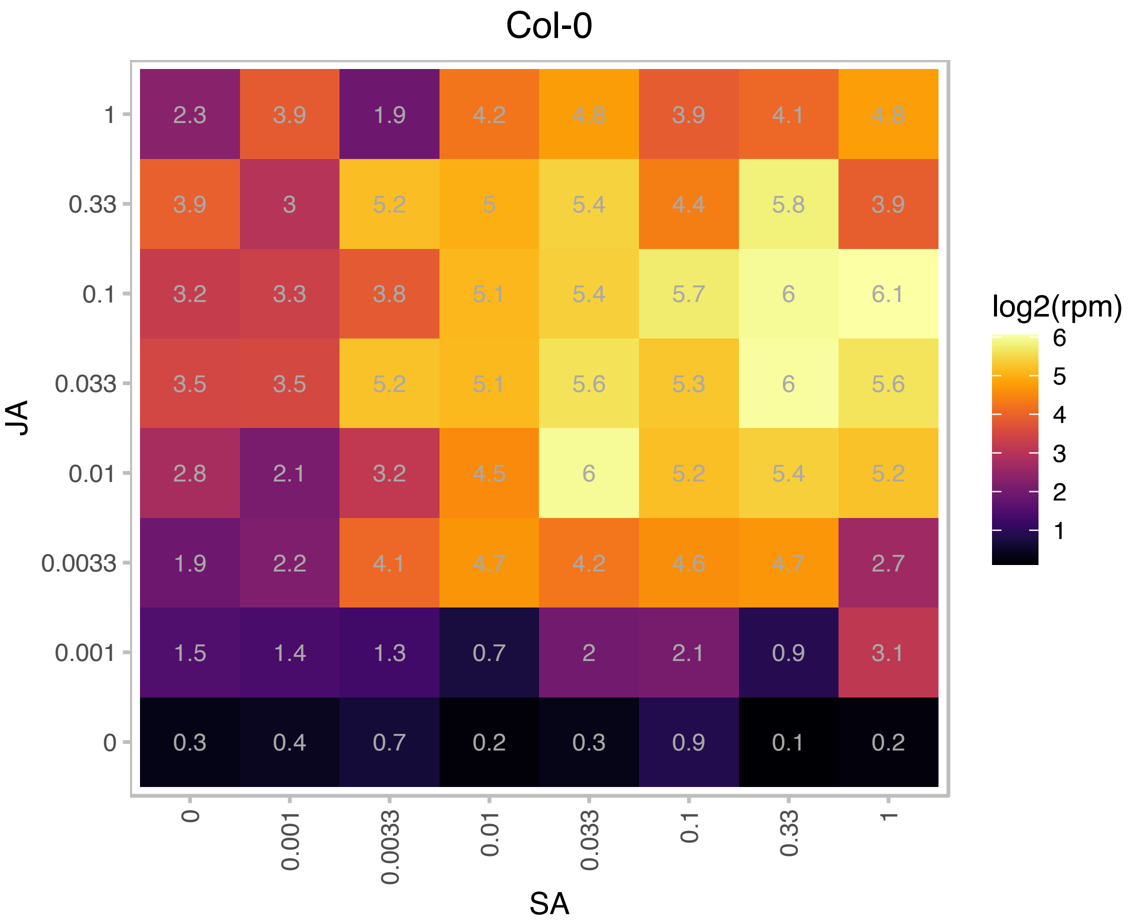


| Gene | AT3G51450 |
|---|---|
| is.expressed_genes | TRUE |
| is.responsive_genes | TRUE |
| is.apcluster | A-02 |

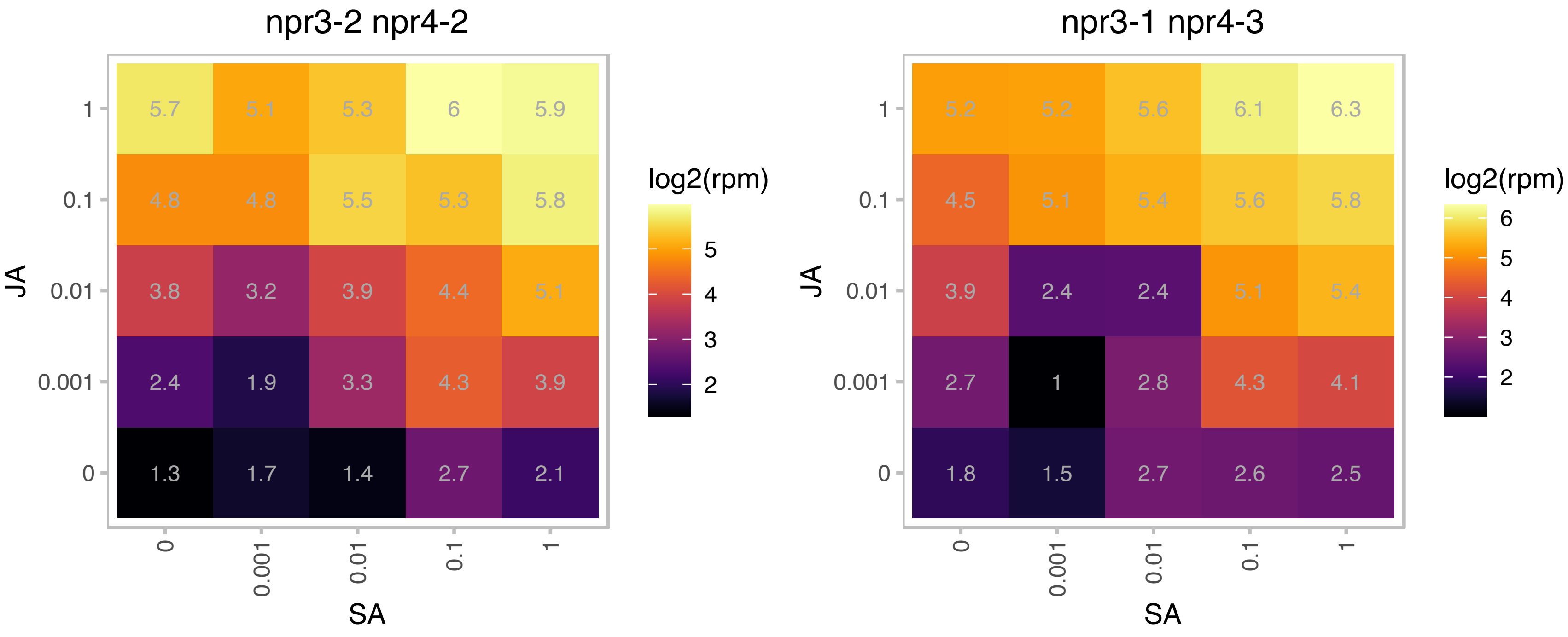

# AT4G01895.1 : NA

systemic acquired resistance (SAR) regulator protein NIMIN-1-related

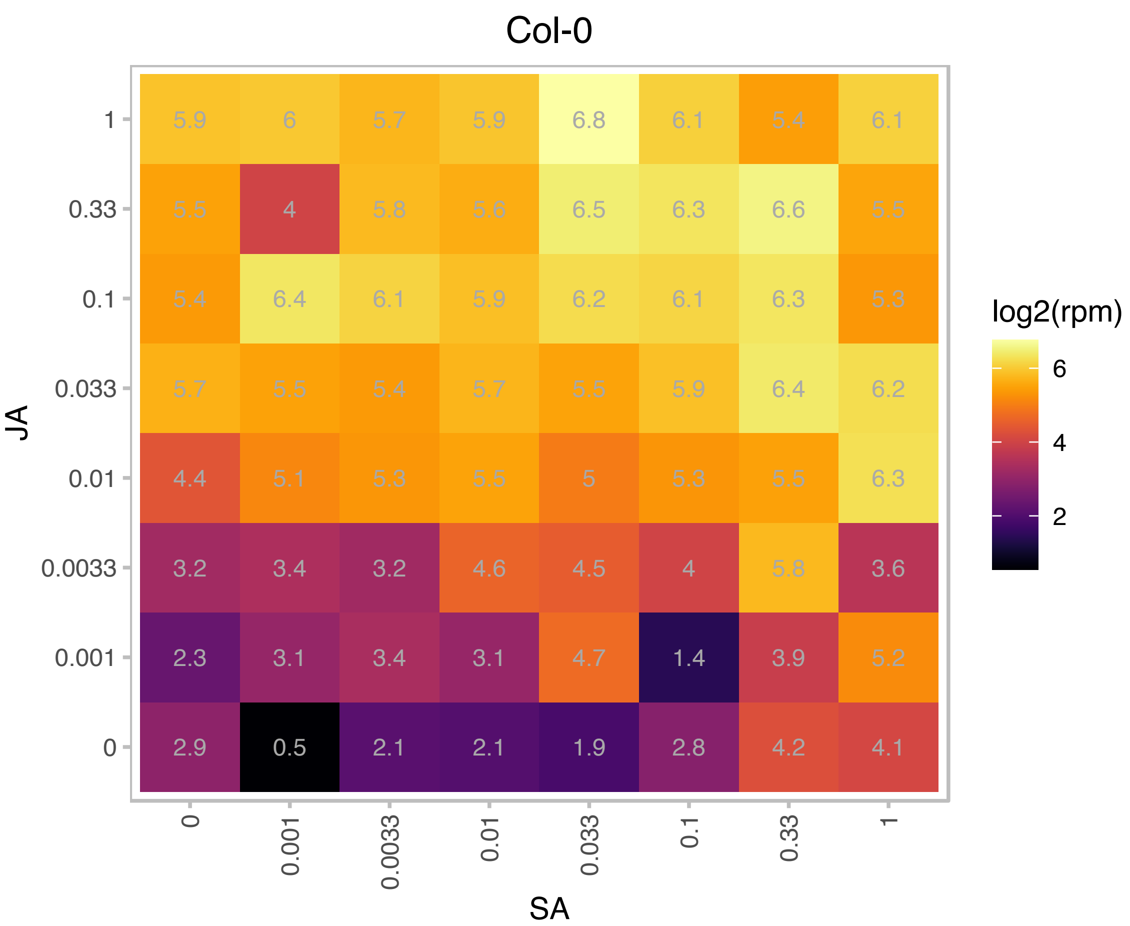


| | |
|---|---|
| Gene | AT4G01895 |
| is.expressed_genes | TRUE |
| is.responsive_genes | TRUE |
| is.apcluster | A-02 |

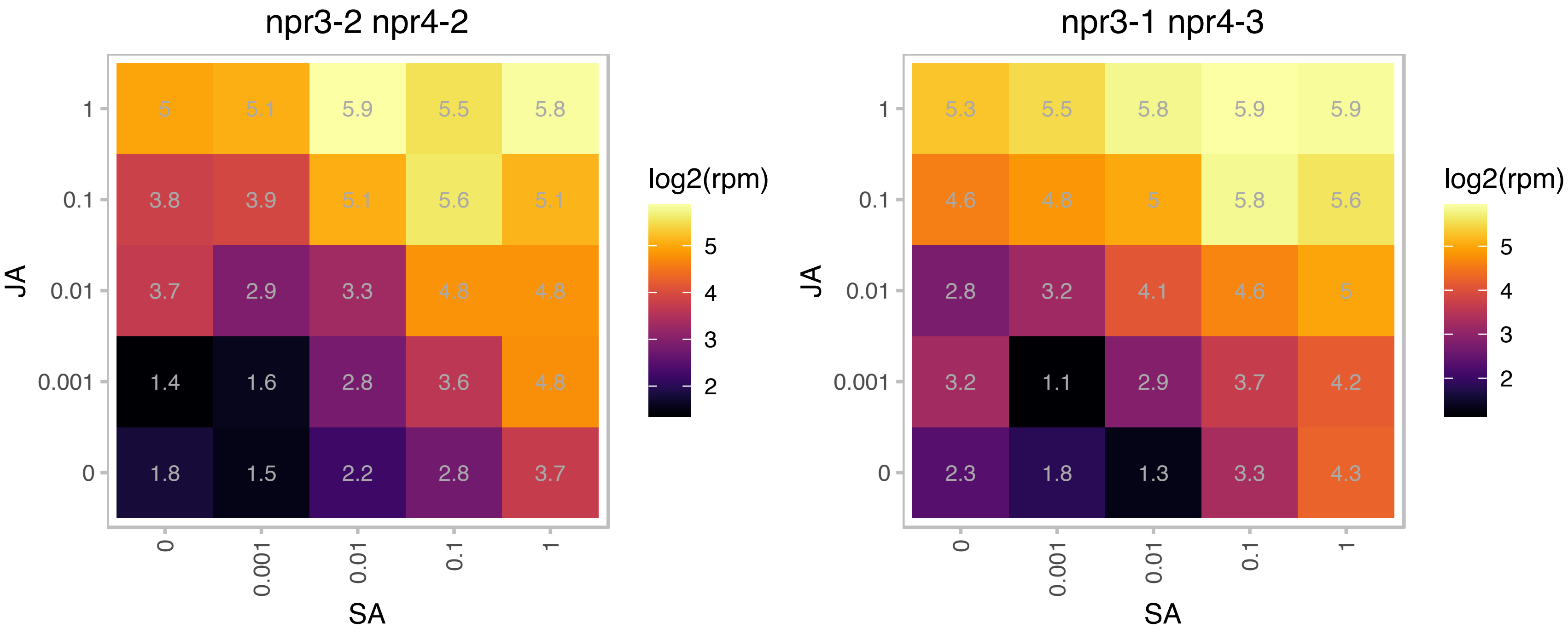

### AT4G04840.1 : ATMSRB6, MSRB6

methionine sulfoxide reductase B6

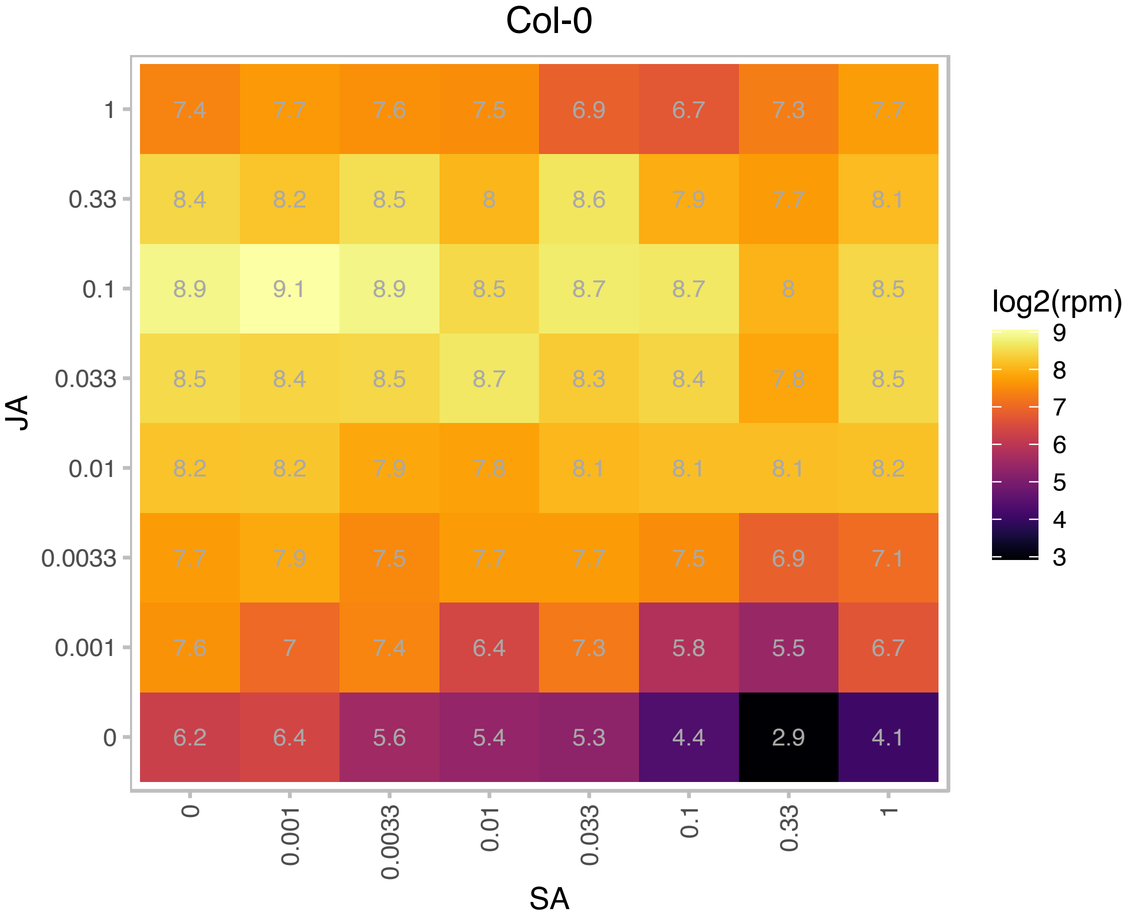


| | |
|---|---|
| Gene | AT4G04840 |
| is.expressed_genes | TRUE |
| is.responsive_genes | TRUE |
| is.apcluster | A-02 |

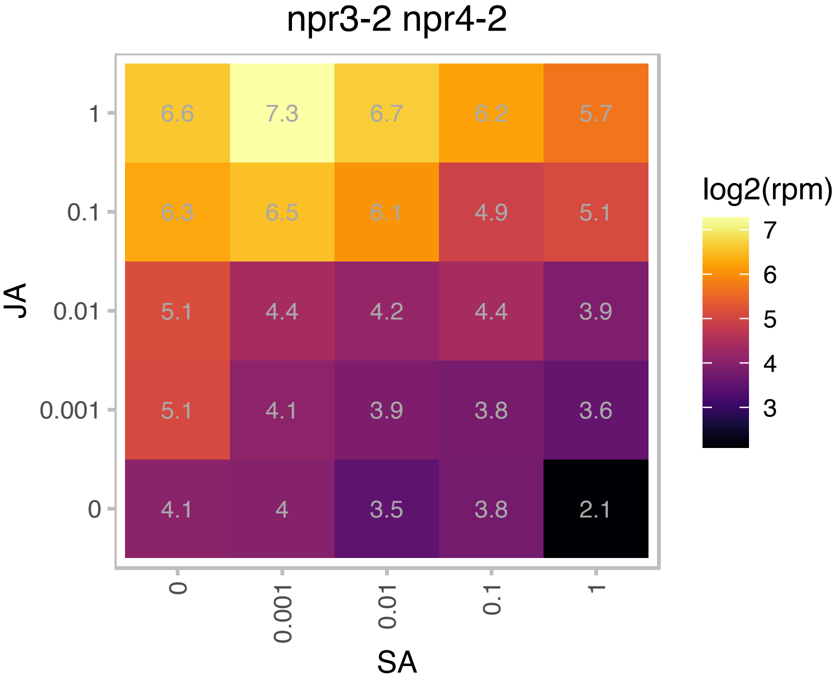


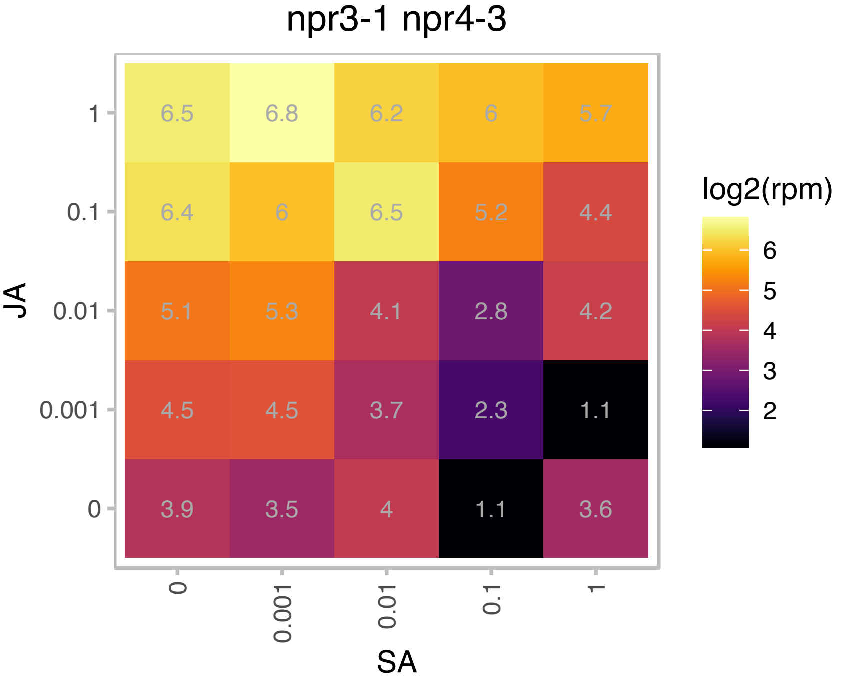

# AT4G09340.1 : NA

SPla/RYanodine receptor (SPRY) domain-containing protein

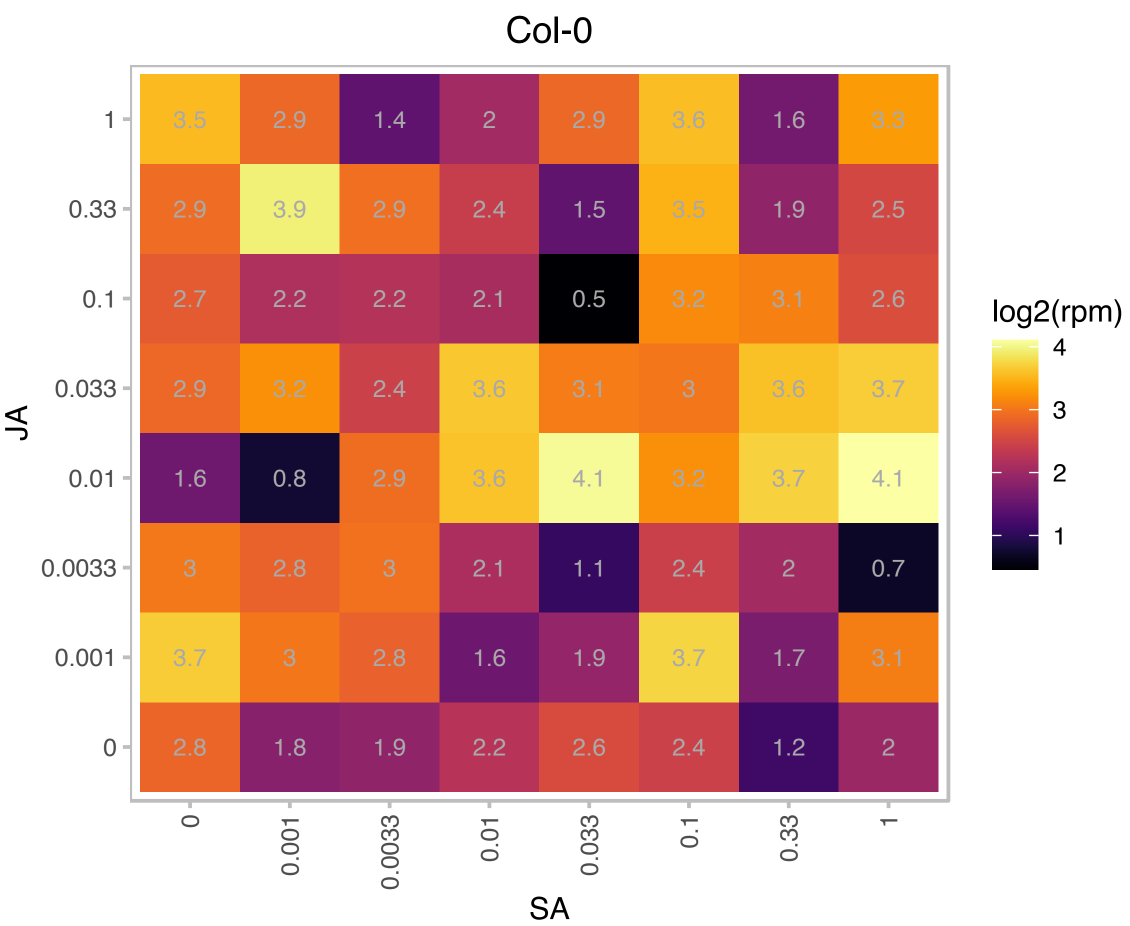


| | |
|---|---|
| Gene | AT4G09340 |
| is.expressed_genes | TRUE |
| is.responsive_genes | TRUE |
| is.apcluster | A-02 |

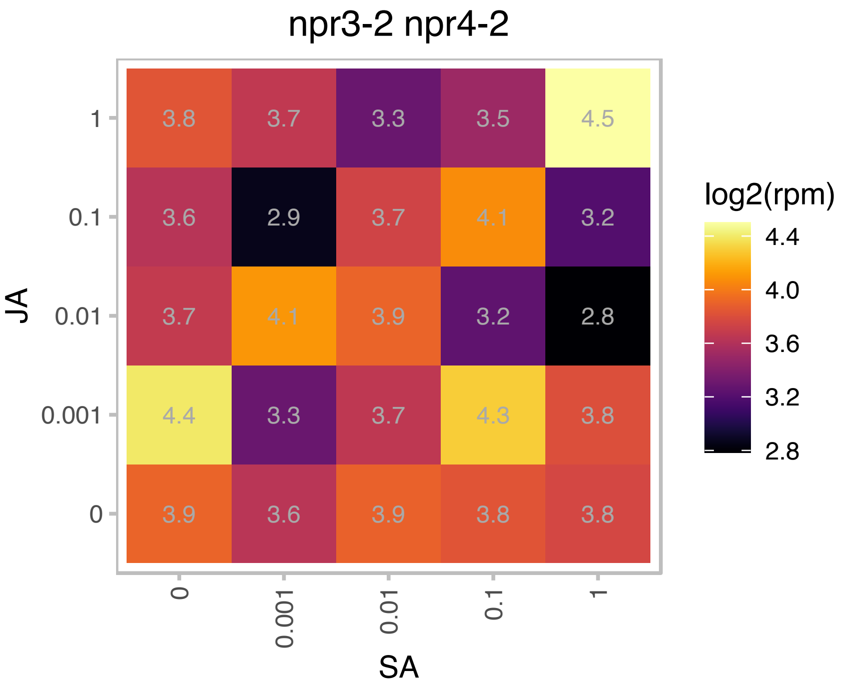


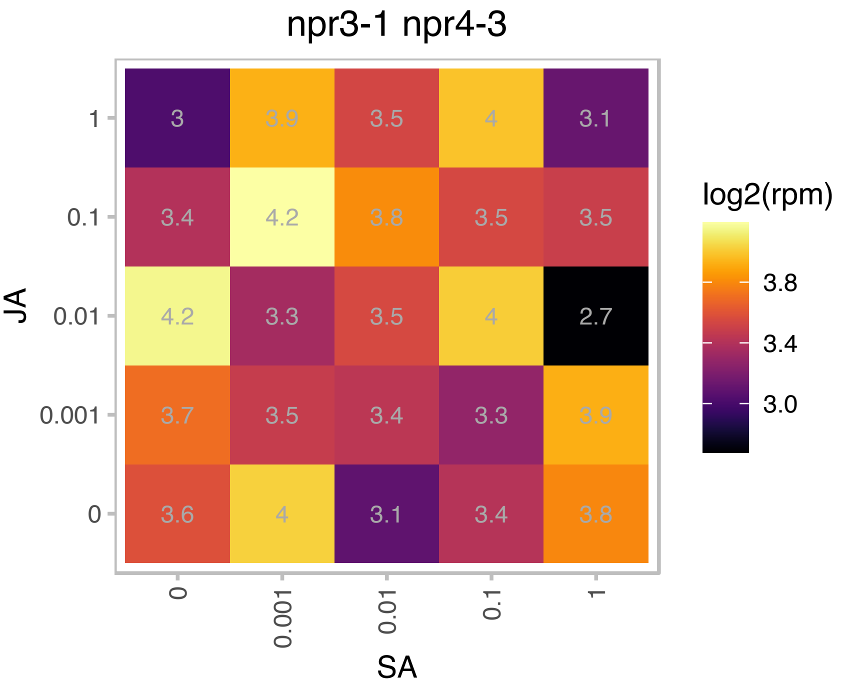

# AT4G19520.1 : NA

disease resistance protein (TIR-NBS-LRR class) family

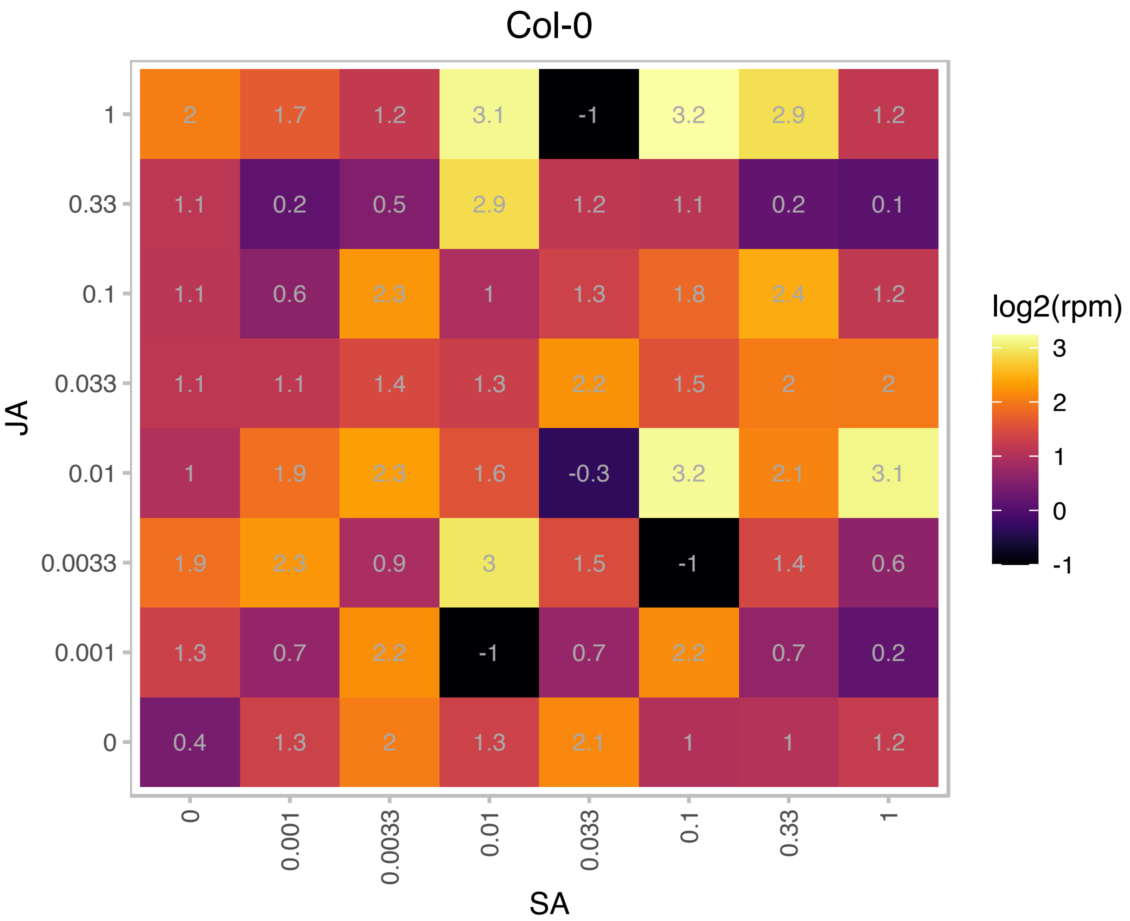


| | |
|---|---|
| Gene | AT4G19520 |
| is.expressed_genes | TRUE |
| is.responsive_genes | TRUE |
| is.apcluster | A-02 |

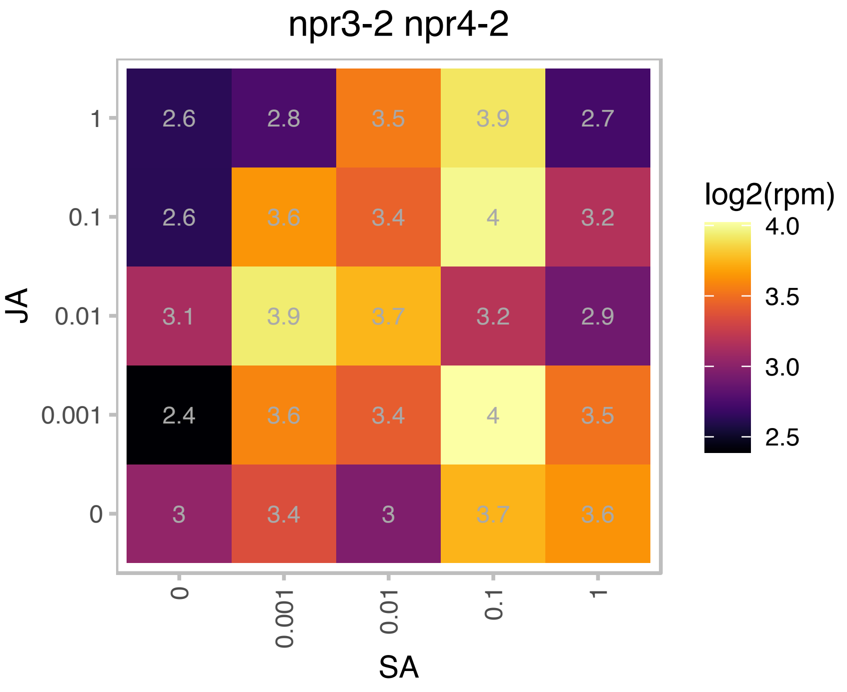


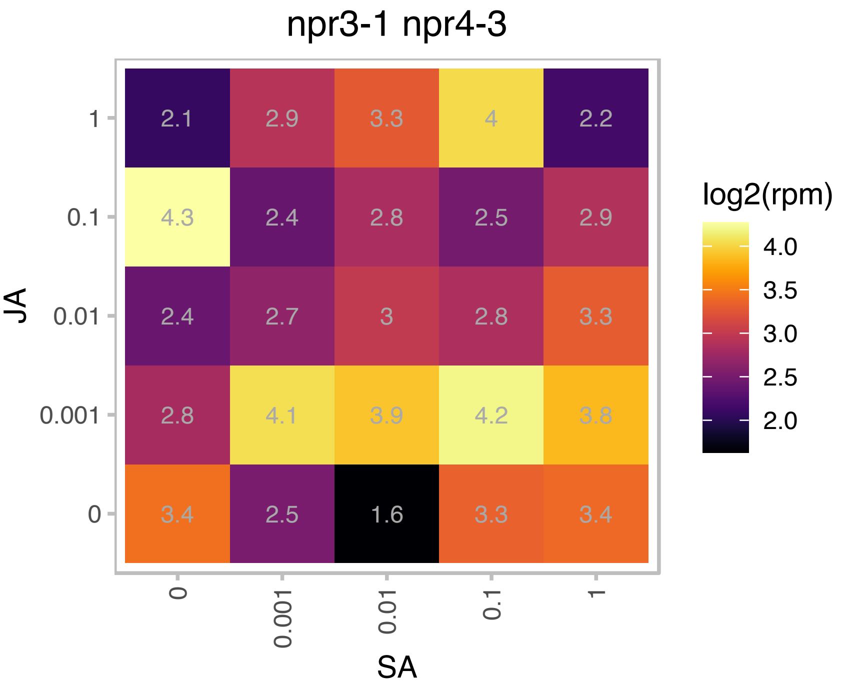

### AT4G21840.1 : ATMSRB8, MSRB8

methionine sulfoxide reductase B8

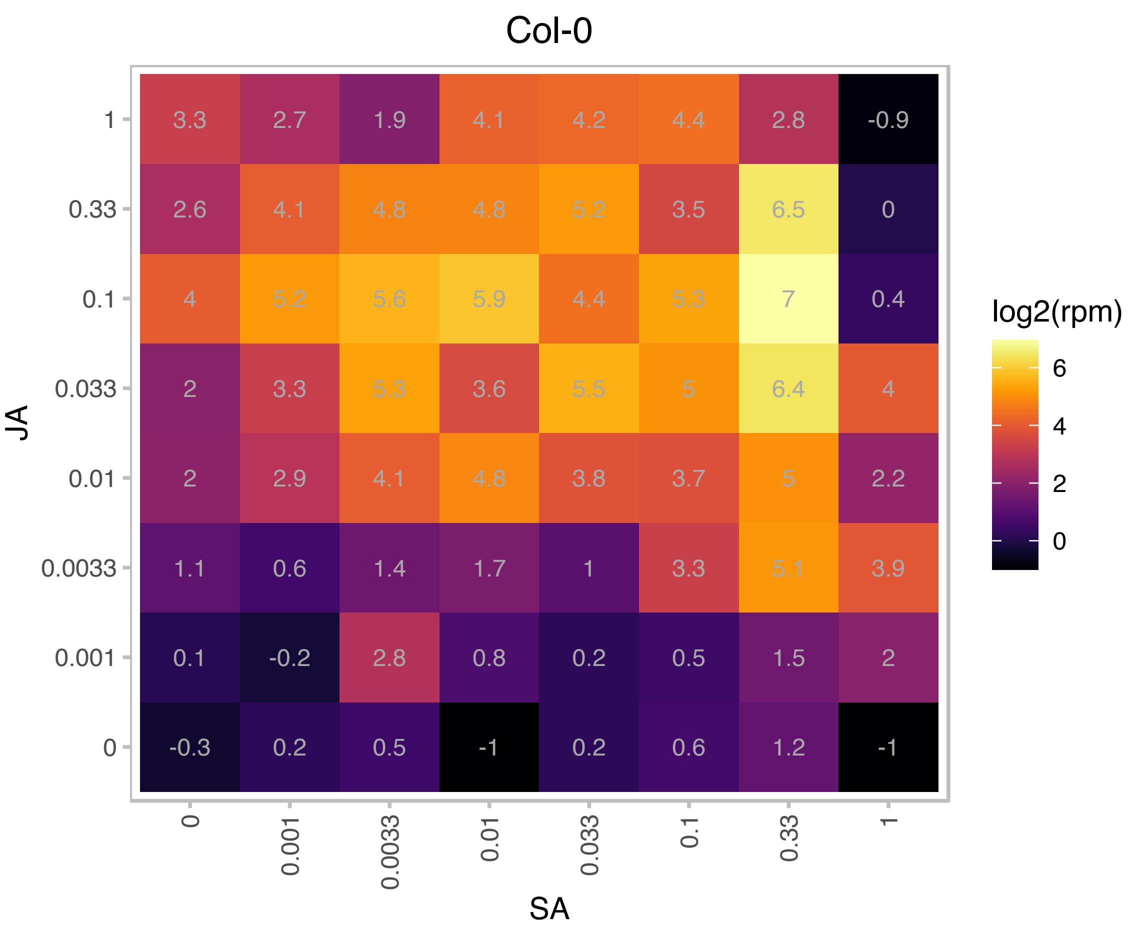


| | |
|---|---|
| Gene | AT4G21840 |
| is.expressed_genes | TRUE |
| is.responsive_genes | TRUE |
| is.apcluster | A-02 |

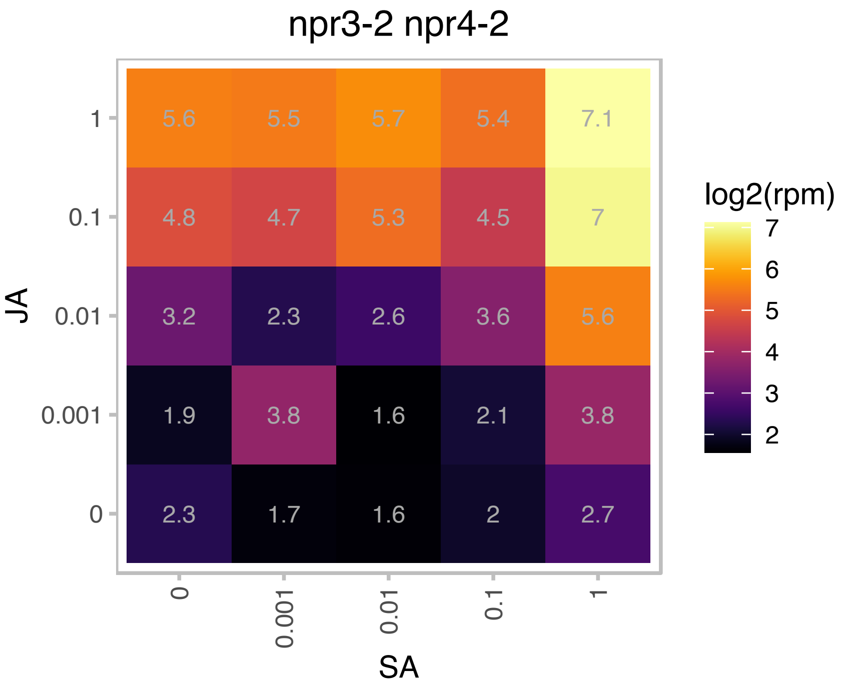


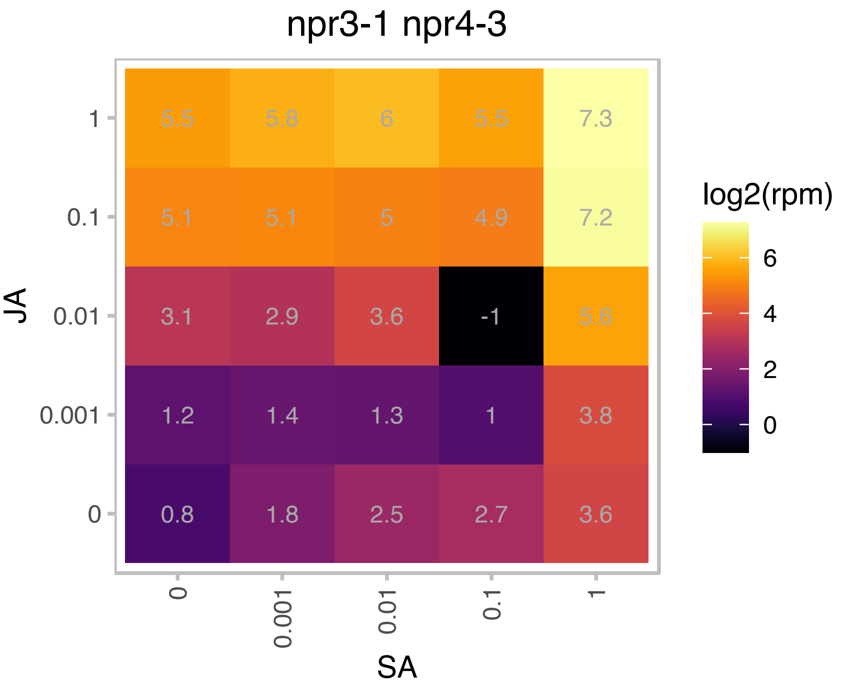

# AT4G21920.1 : NA

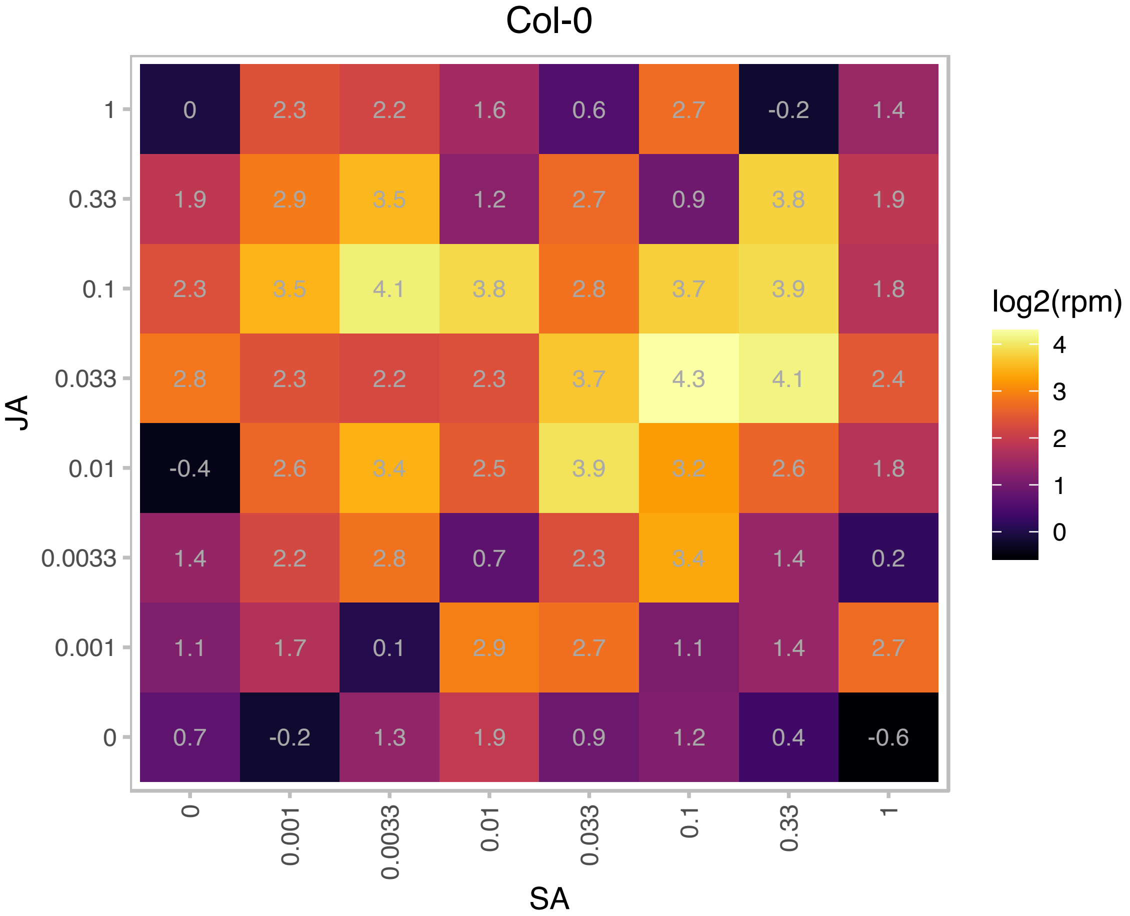


| | |
|---|---|
| Gene | AT4G21920 |
| is.expressed_genes | TRUE |
| is.responsive_genes | TRUE |
| is.apcluster | A-02 |

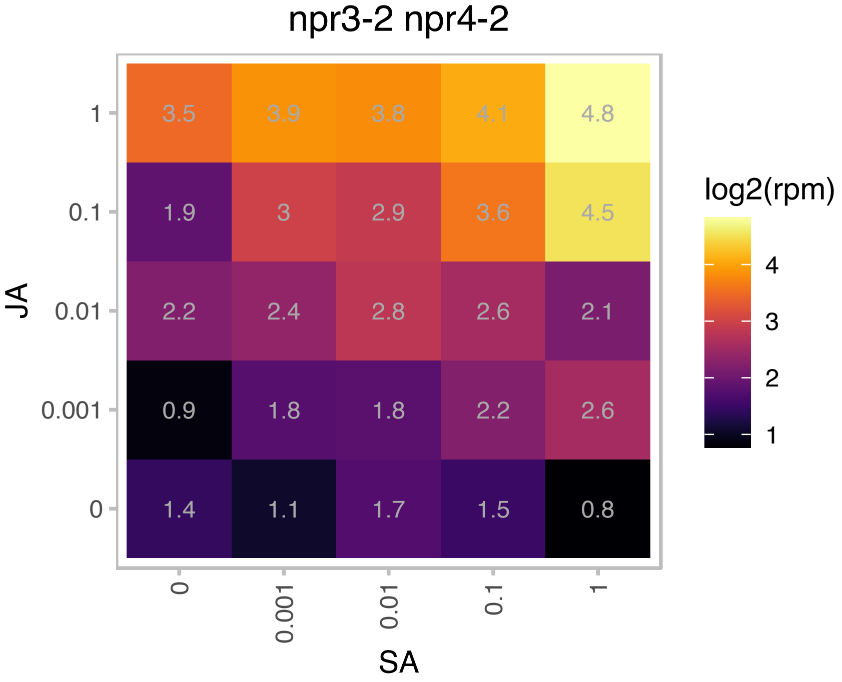


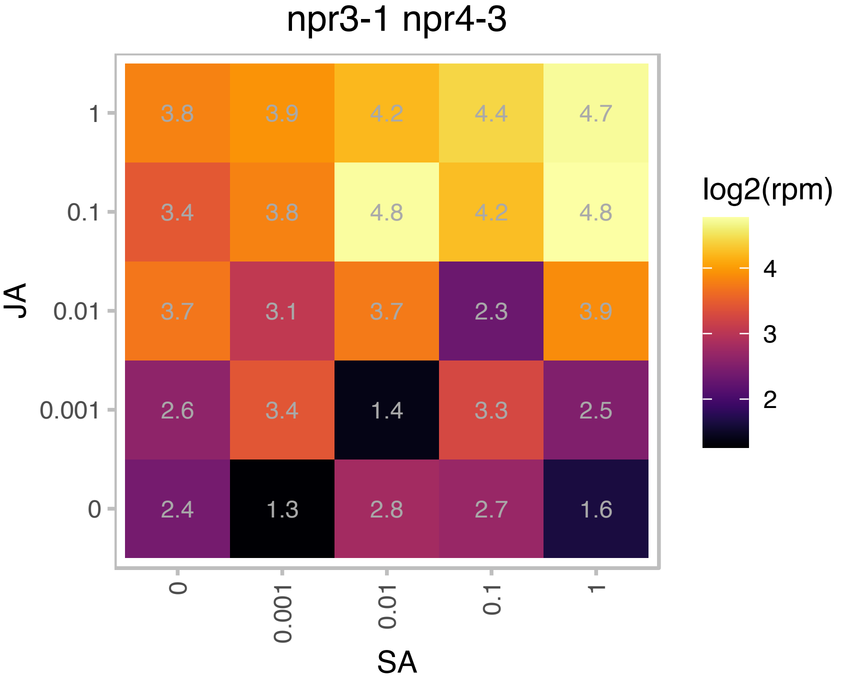

# AT4G23885.1 : NA

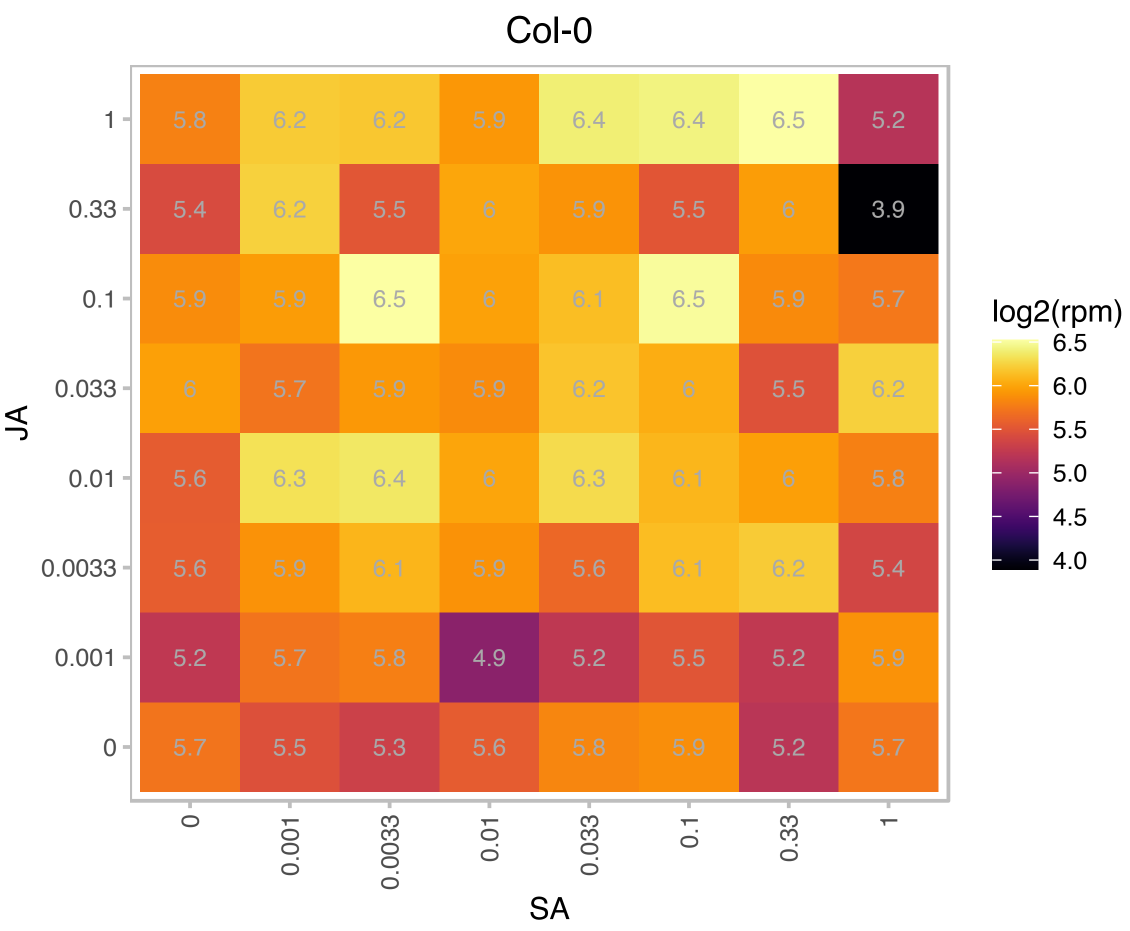


| | |
|---|---|
| Gene | AT4G23885 |
| is.expressed_genes | TRUE |
| is.responsive_genes | TRUE |
| is.apcluster | A-02 |

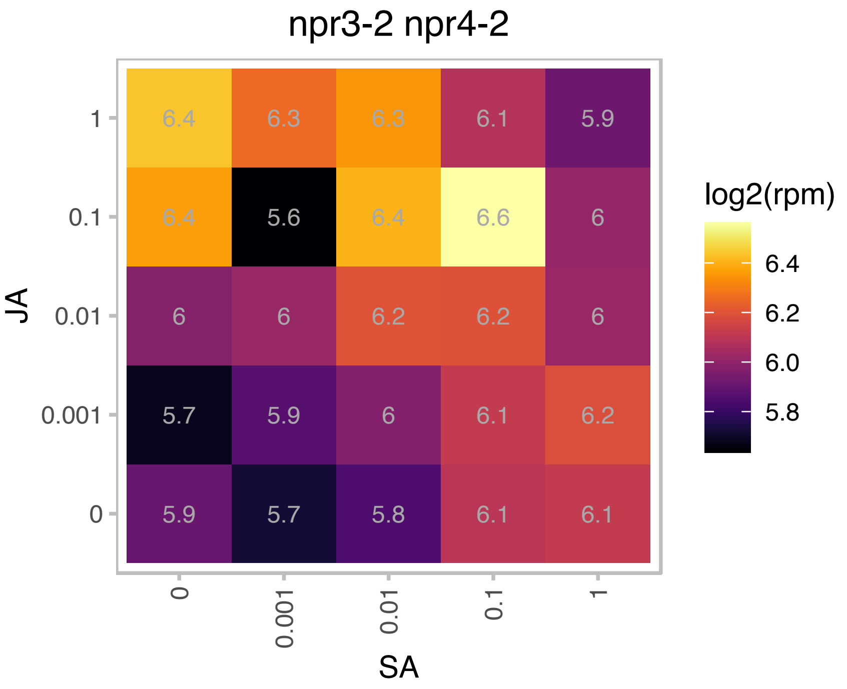


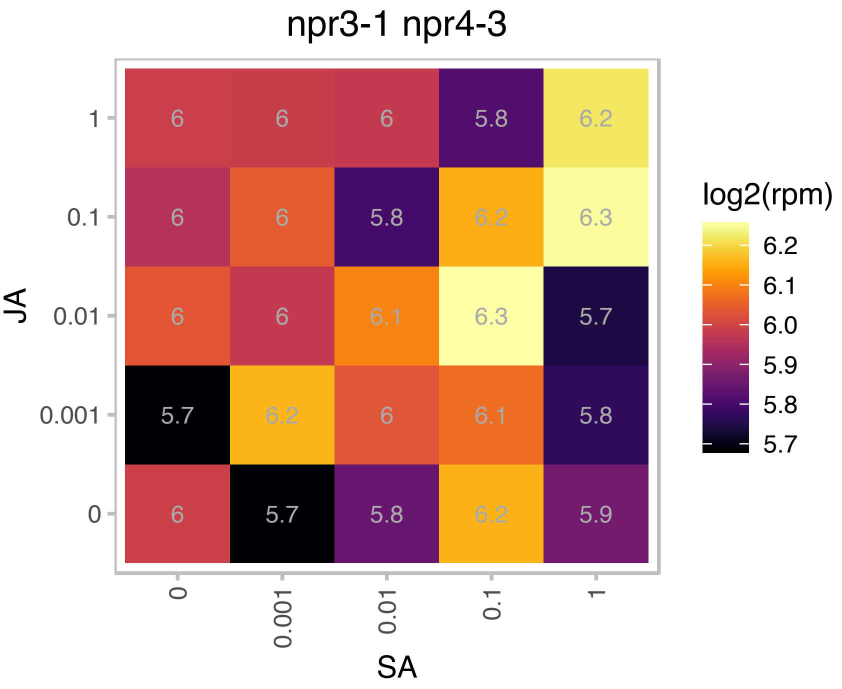

### AT4G39030.1 : EDS5, SCORD3, SID1

MATE efflux family protein
Encodes an orphan multidrug and toxin extrusion transporter. Essential component of salicylic acid-dependent signaling for disease resistance. Member of the MATE-transporter family. Expression induced by salicylic acid. Mutants are salicylic acid-deficient.

Col-0

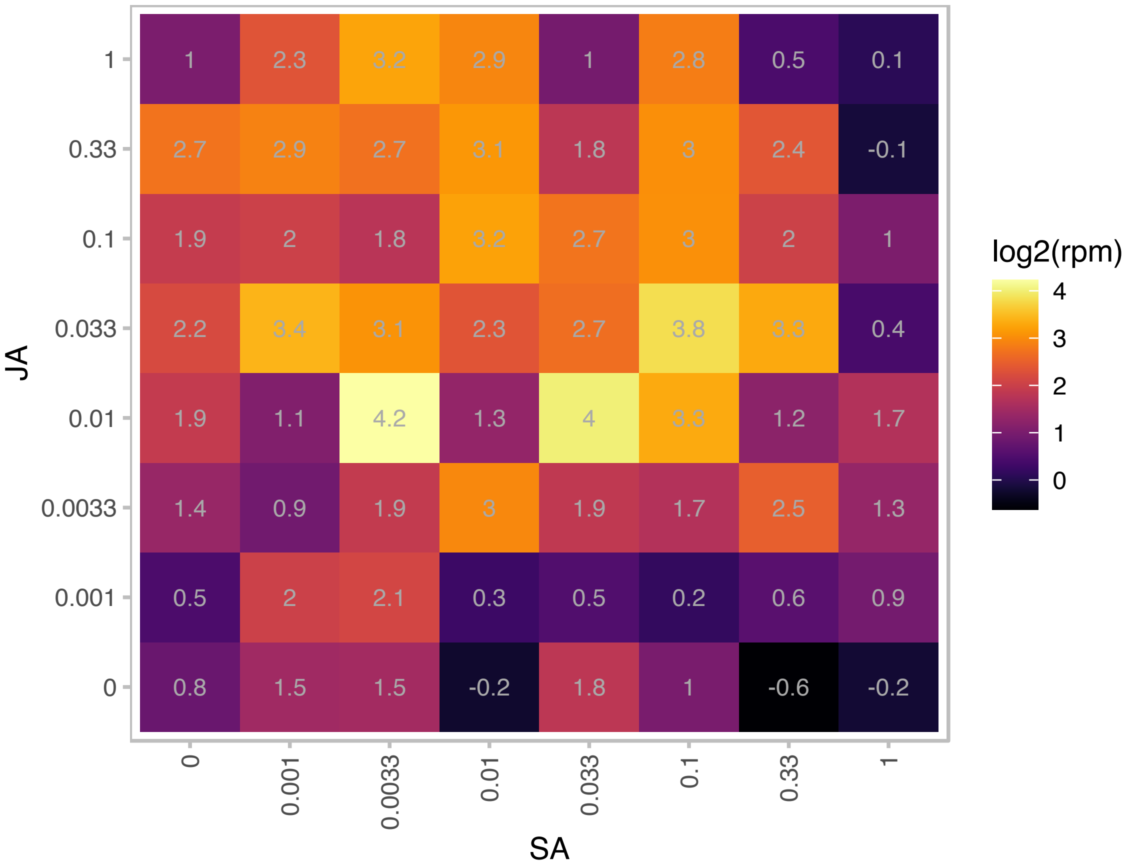


| | |
|---|---|
| Gene | AT4G39030 |
| is.expressed_genes | TRUE |
| is.responsive_genes | TRUE |
| is.apcluster | A-02 |

npr3-2 npr4-2

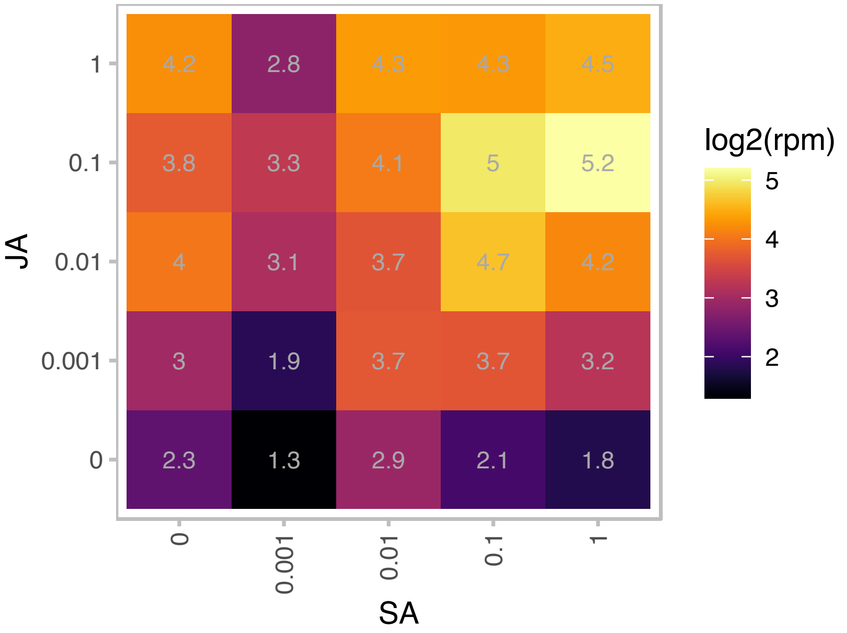


npr3-1 npr4-3

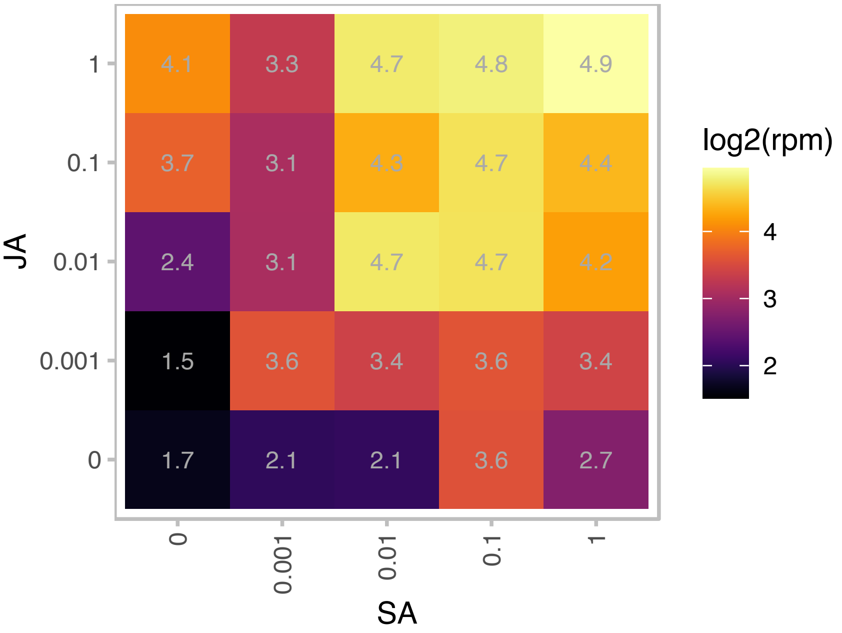

# AT5G04420.1 : NA

Galactose oxidase/kelch repeat superfamily protein

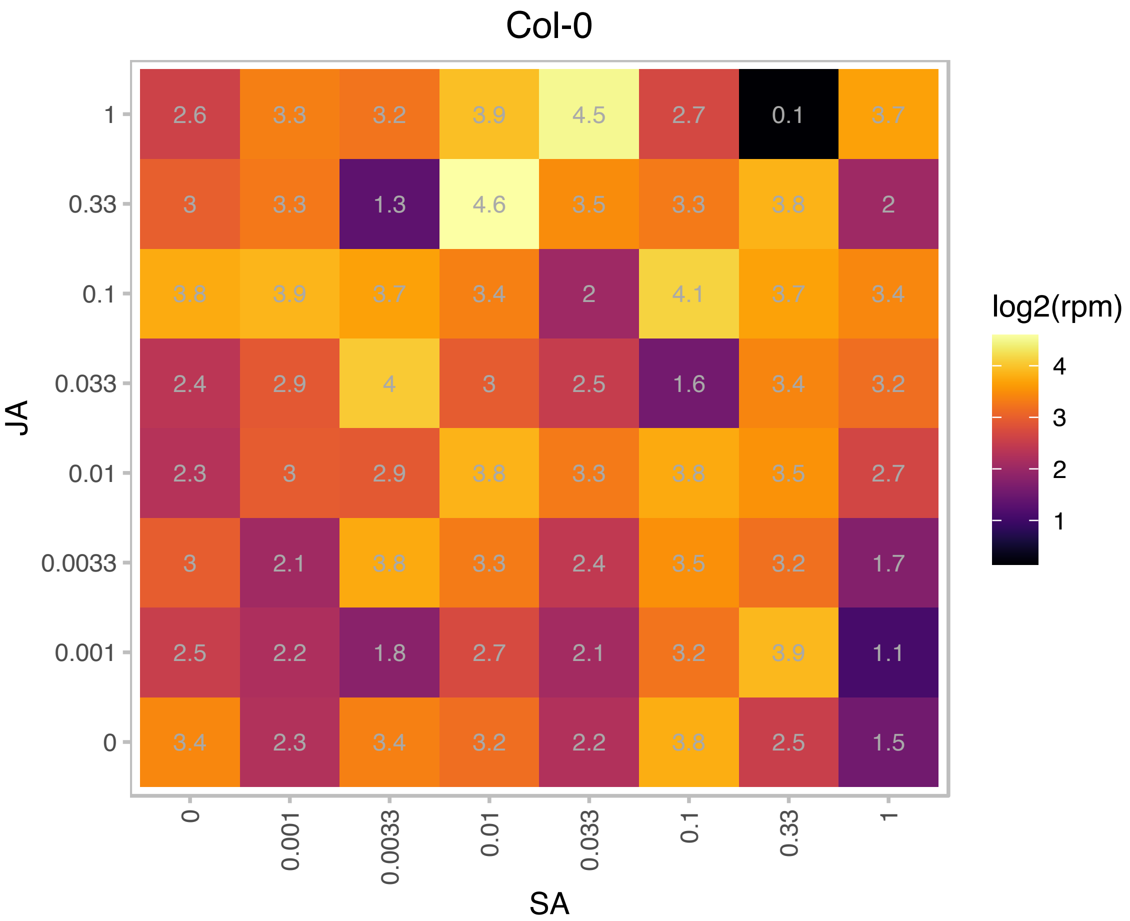


| | |
|---|---|
| Gene | AT5G04420 |
| is.expressed_genes | TRUE |
| is.responsive_genes | TRUE |
| is.apcluster | A-02 |

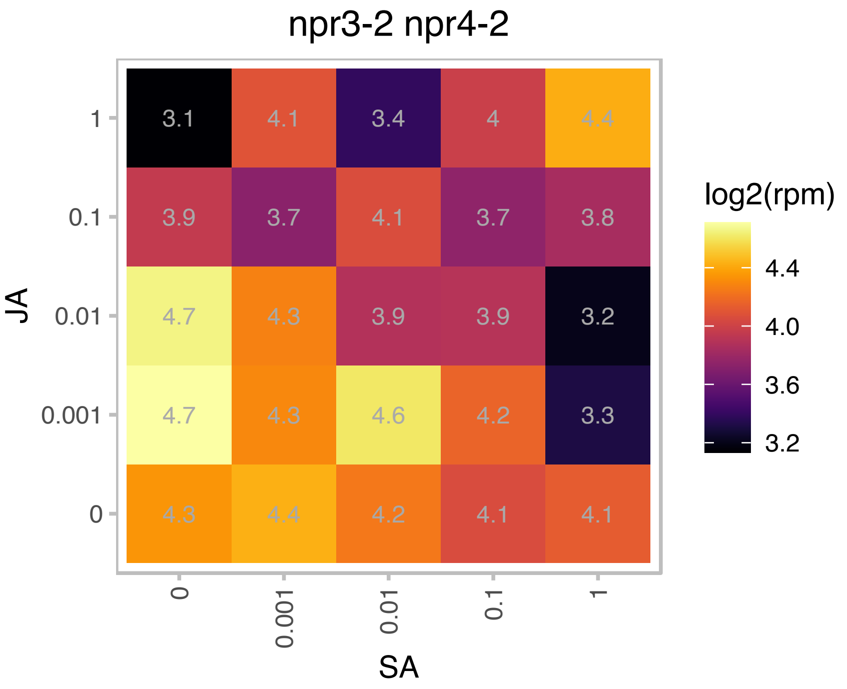


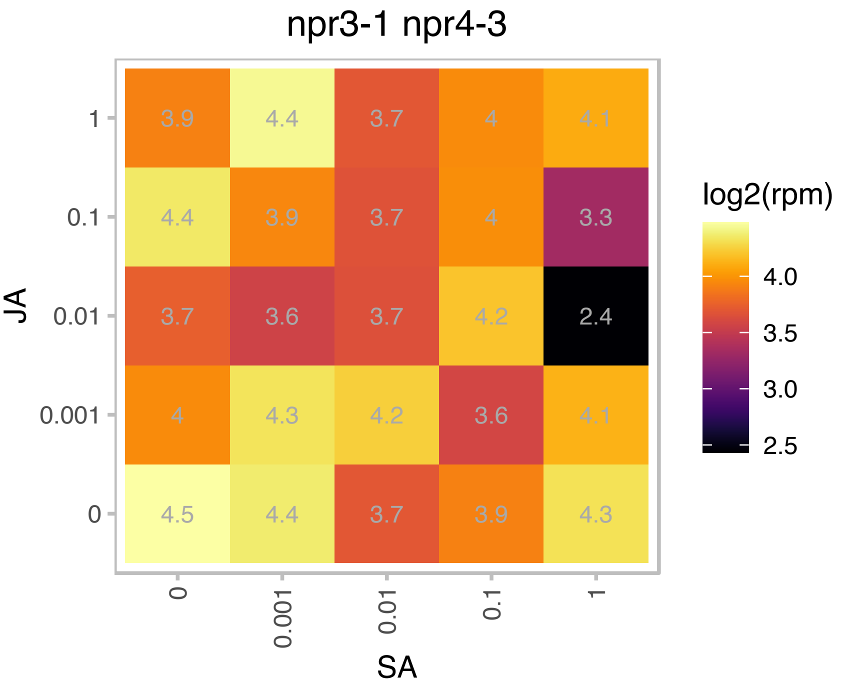

### AT5G05540.1 : SDN2

small RNA degrading nuclease 2

Col-0

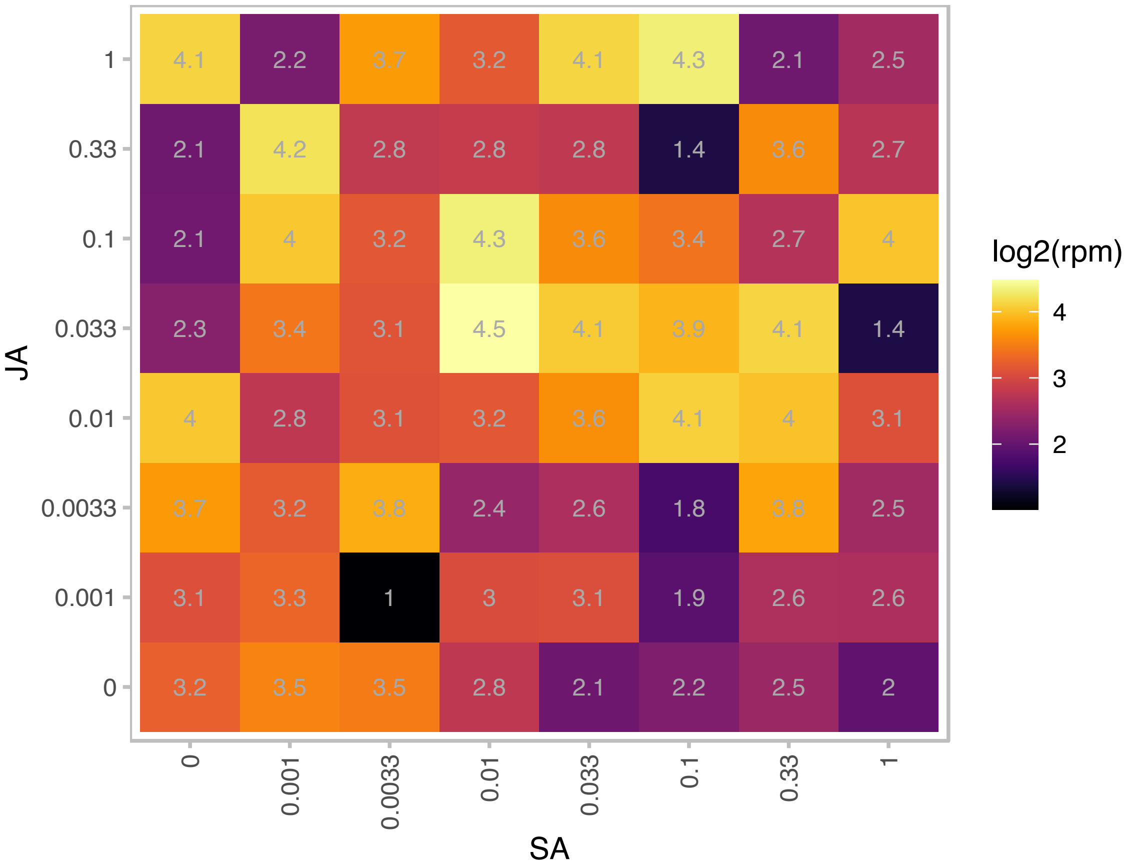


| | |
|---|---|
| Gene | AT5G05540 |
| is.expressed_genes | TRUE |
| is.responsive_genes | TRUE |
| is.apcluster | A-02 |

npr3-2 npr4-2

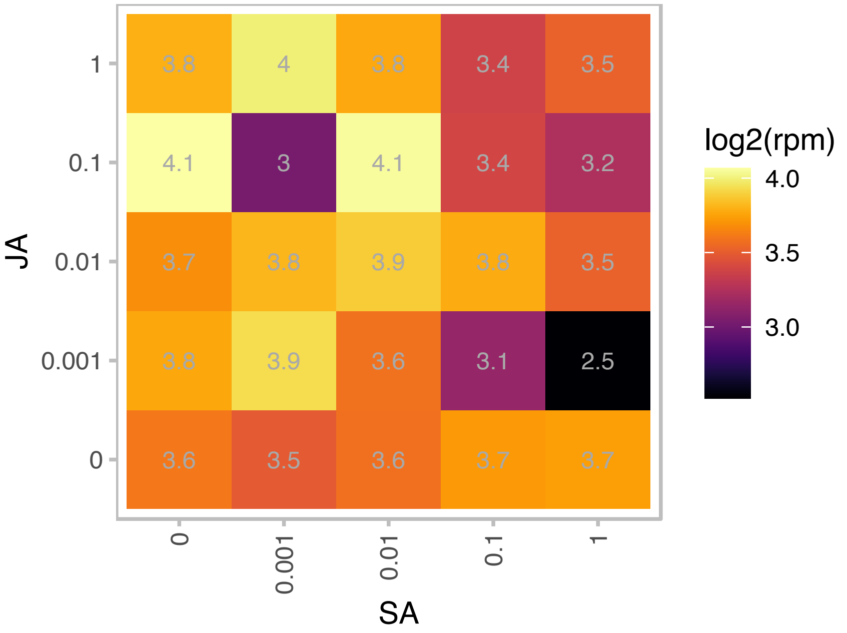


npr3-1 npr4-3

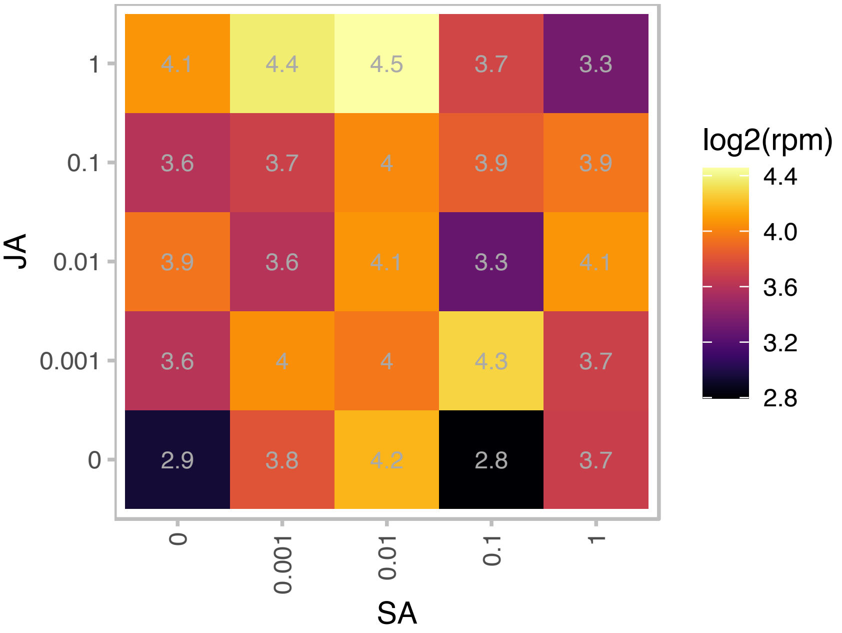

### AT5G07010.1 : ATST2A, ST2A

sulfotransferase 2A
Encodes a sulfotransferase that acts specifically on 11- and 12-hydroxyjasmonic acid. Transcript levels for this enzyme are increased by treatments with jasmonic acid (JA), 12-hydroxyJA, JA-isoleucine, and 12-oxyphytodienoic acid (a JA precursor).

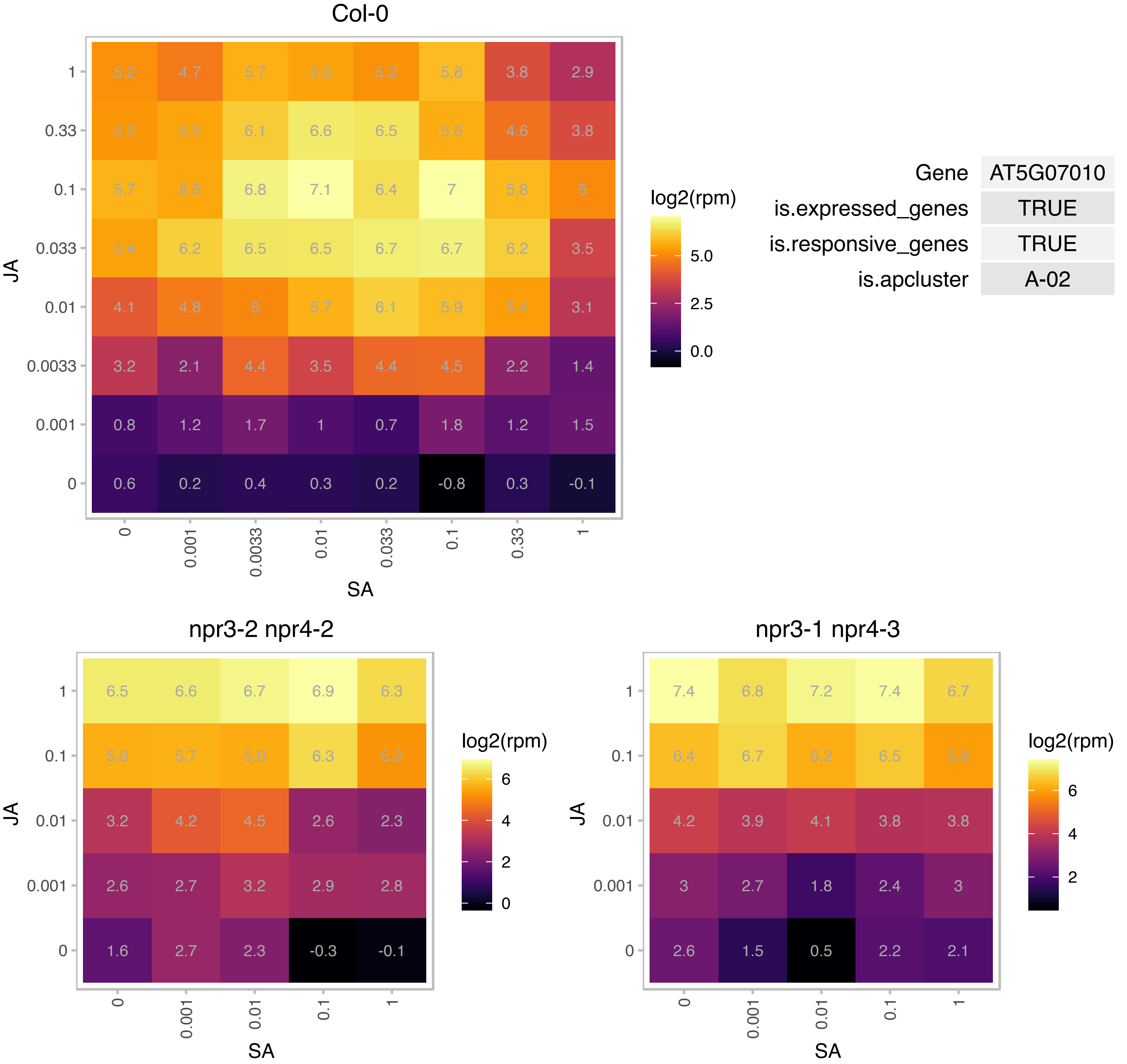


| | |
|---|---|
| Gene | AT5G07010 |
| is.expressed_genes | TRUE |
| is.responsive_genes | TRUE |
| is.apcluster | A-02 |

# AT5G12340.1 : NA

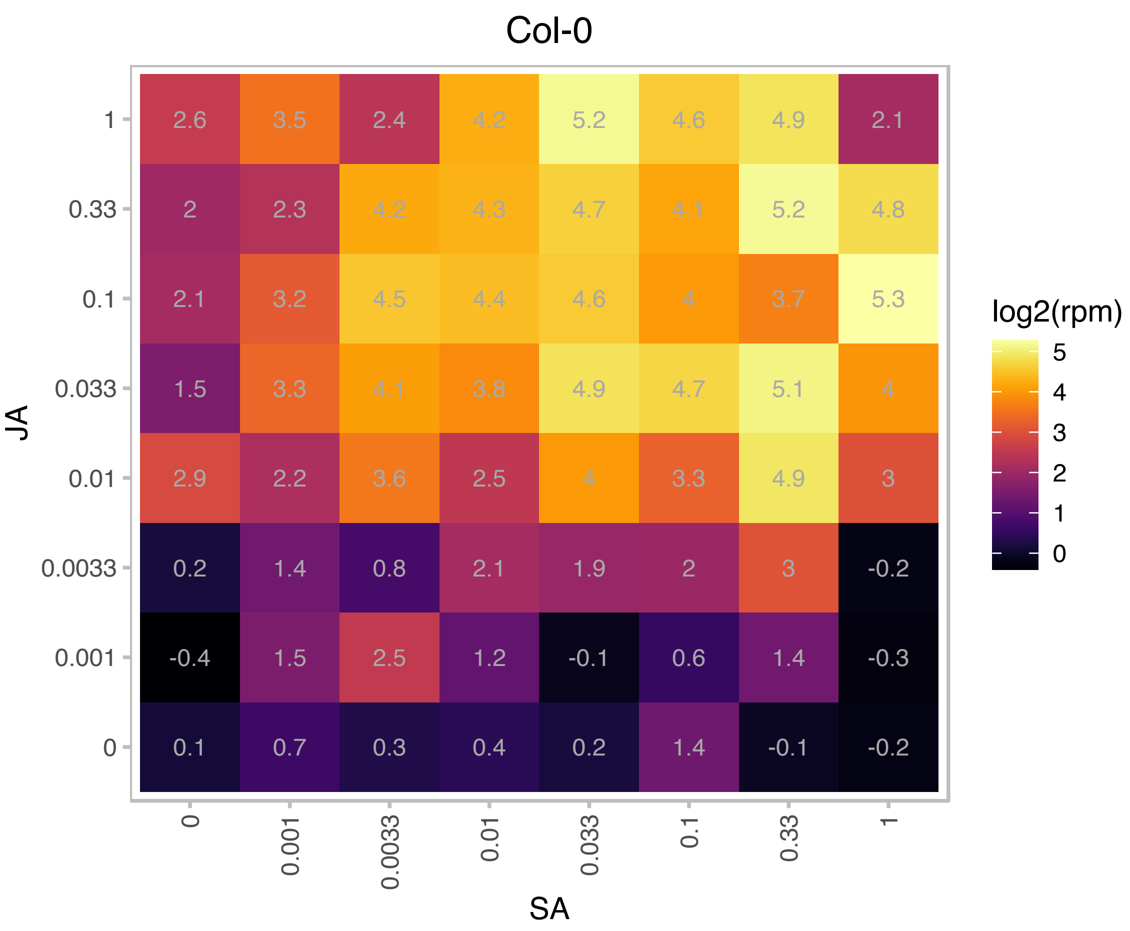


| | |
|---|---|
| Gene | AT5G12340 |
| is.expressed_genes | TRUE |
| is.responsive_genes | TRUE |
| is.apcluster | A-02 |

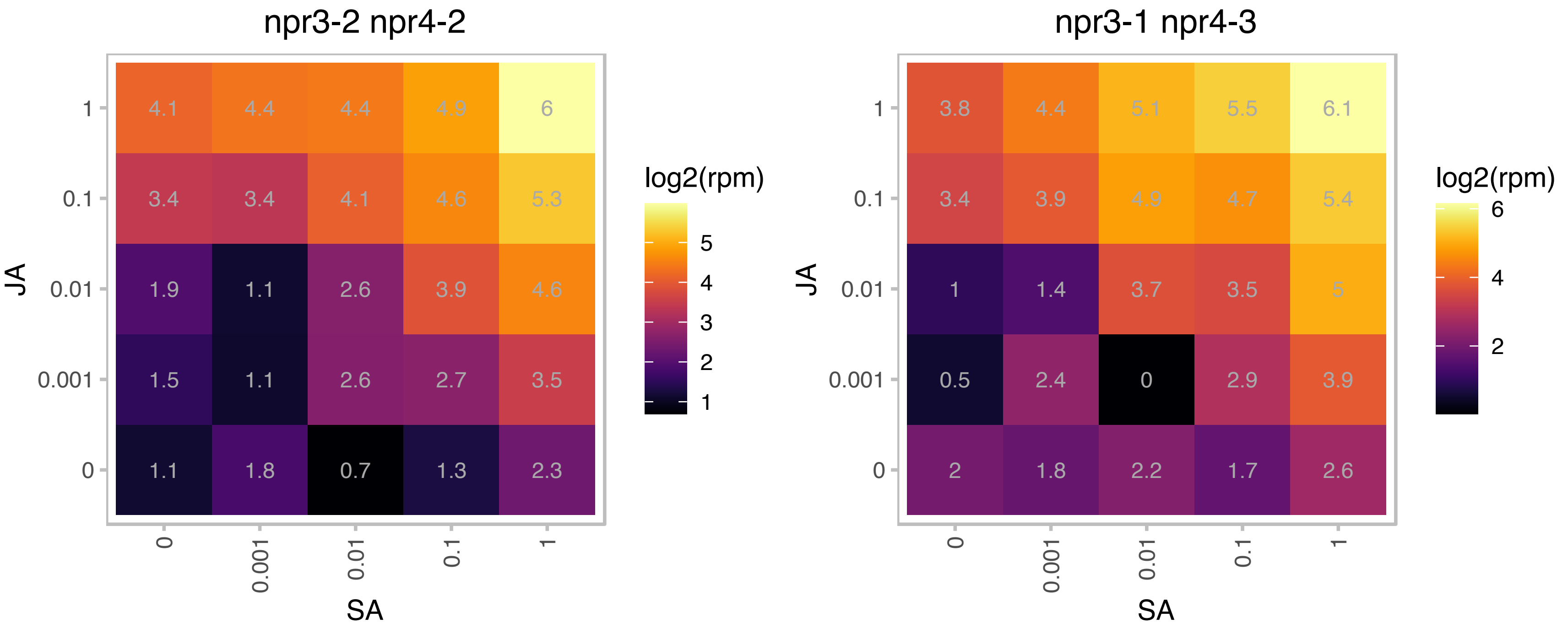

### AT5G13220.1 : JAS1, JAZ10, TIFY9

jasmonate-zim-domain protein 10

Plants overexpressing At5g13220.3, but not At5g13220.1 showed enhanced insensitivity to MeJa.

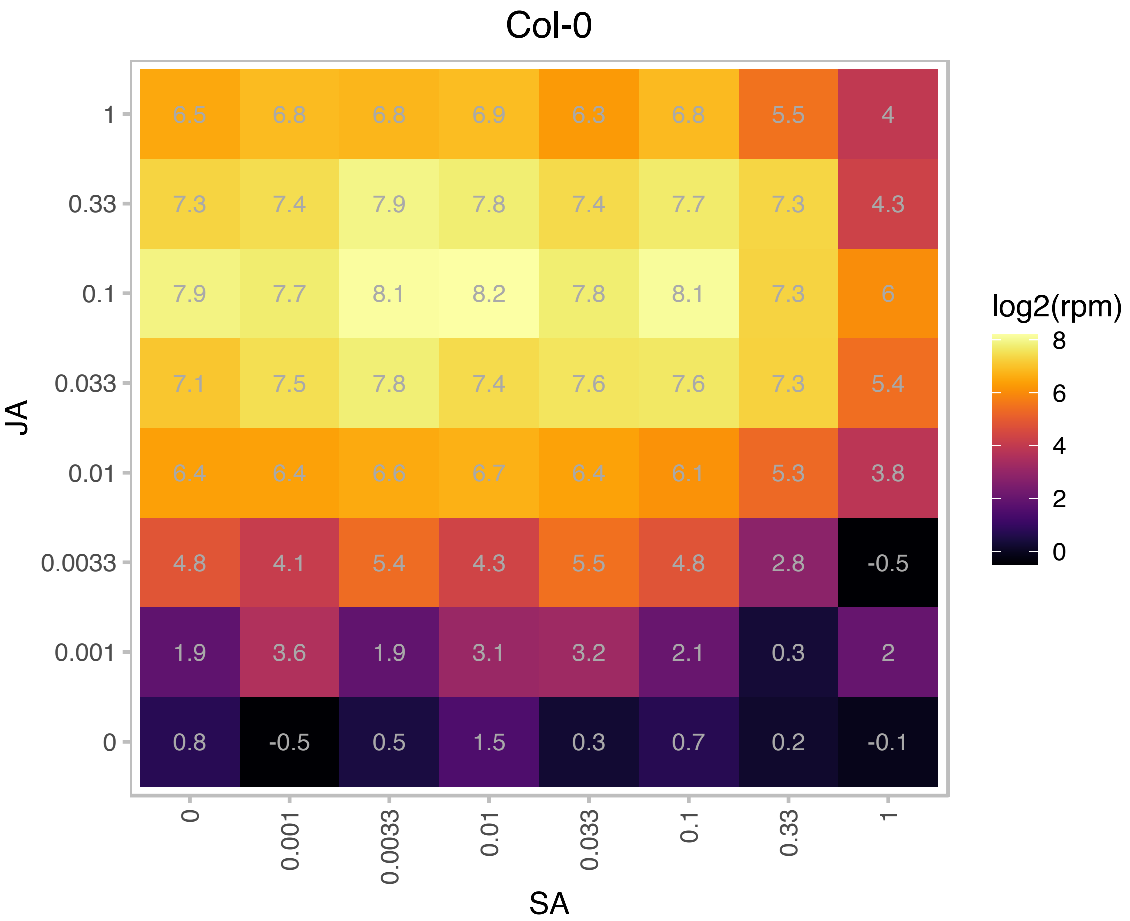


| | |
|---|---|
| Gene | AT5G13220 |
| is.expressed_genes | TRUE |
| is.responsive_genes | TRUE |
| is.apcluster | A-02 |

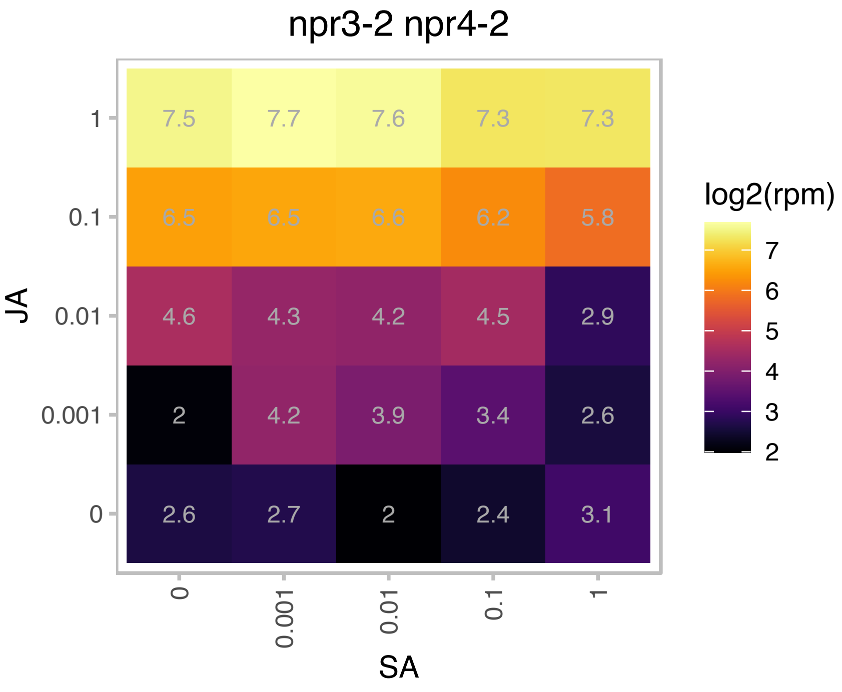


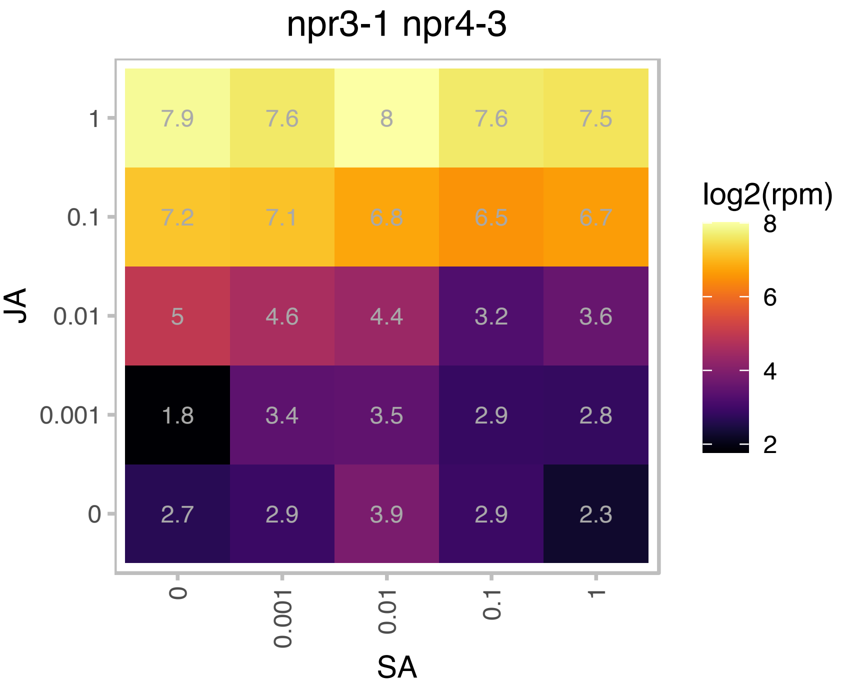

### AT5G14180.1 : MPL1

Myzus persicae-induced lipase 1

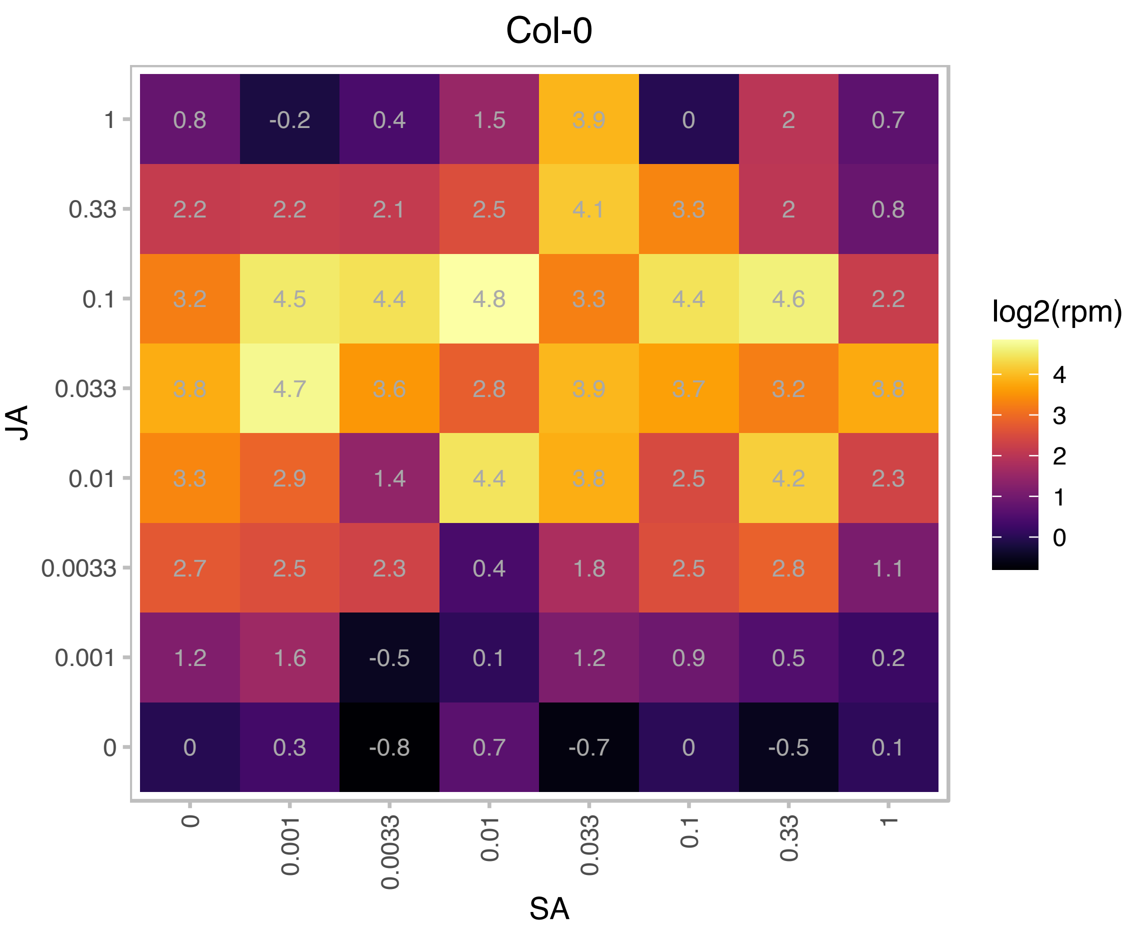


| | |
|---|---|
| Gene | AT5G14180 |
| is.expressed_genes | TRUE |
| is.responsive_genes | TRUE |
| is.apcluster | A-02 |

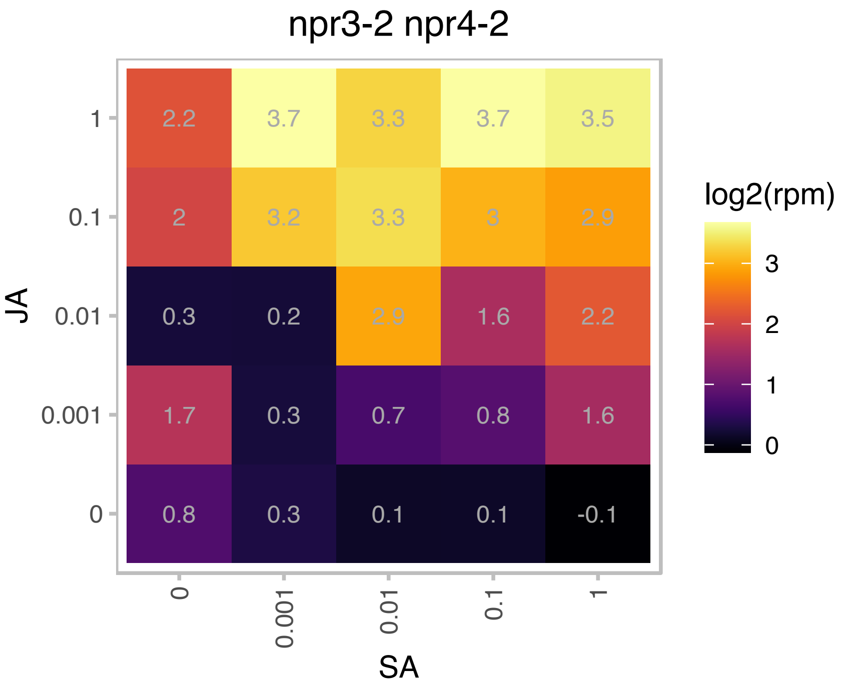


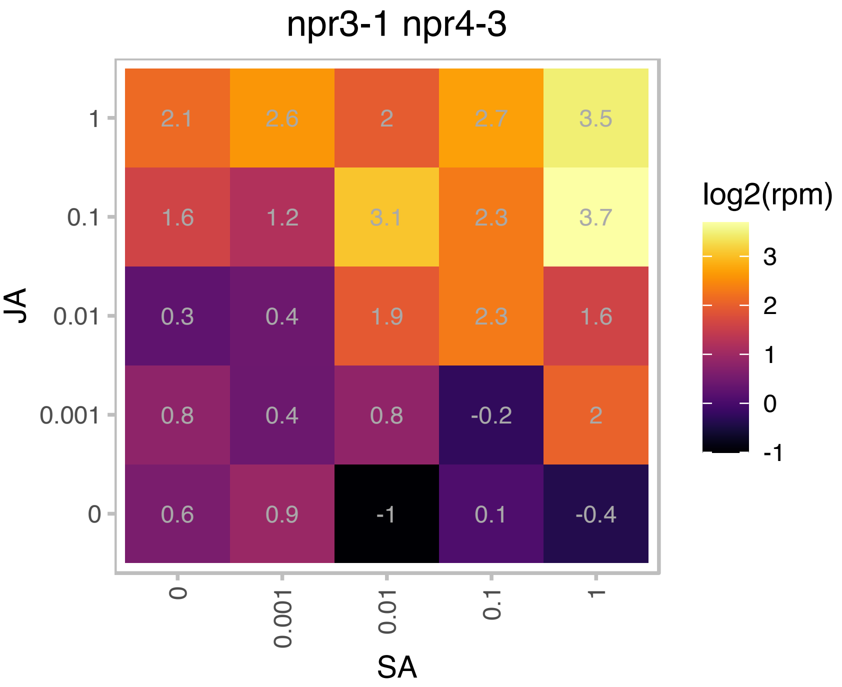

### AT5G18860.1 : NSH3

inosine-uridine preferring nucleoside hydrolase family protein

Encodes a purine nucleoside hydrolase active in the apoplast. It might play a role in salvaging extracellular ATP. NSH3 transcript levels rise in response to jasmonic acid and wounding.

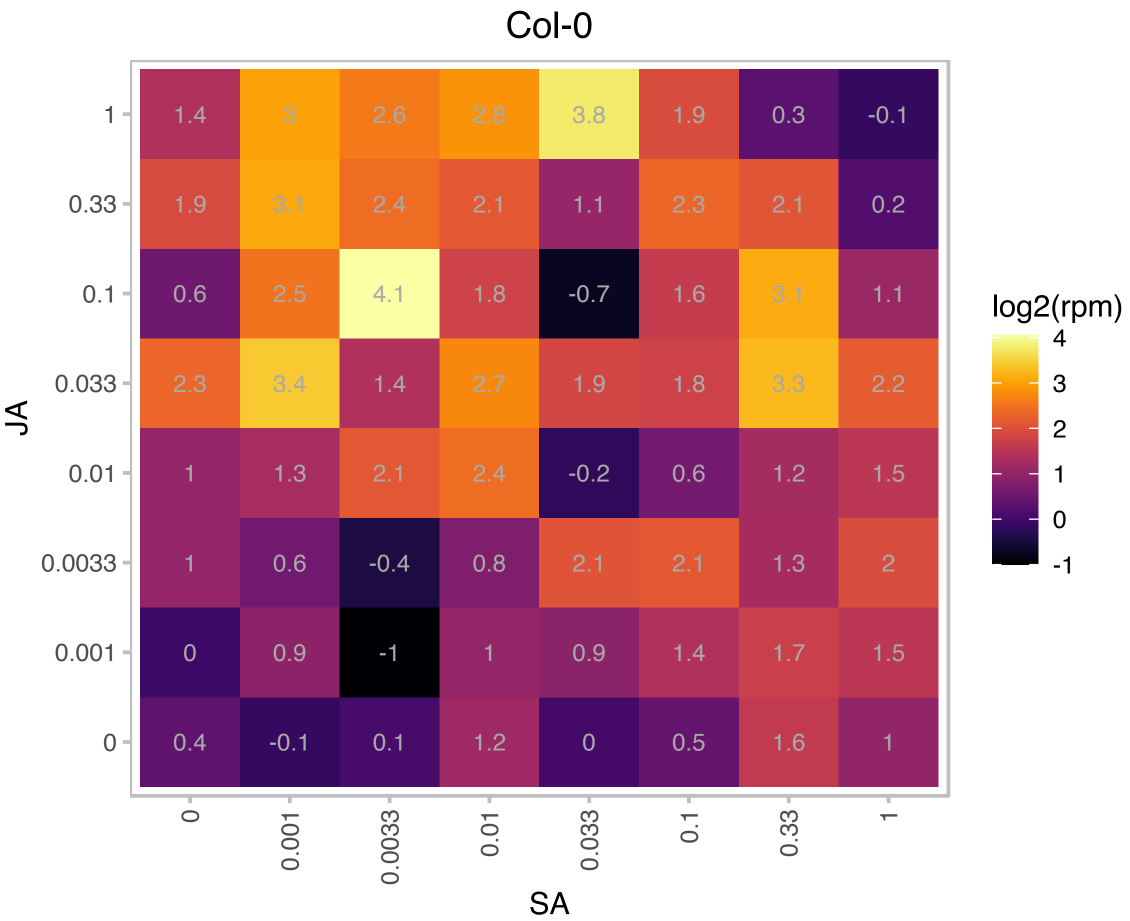


| | |
|---|---|
| Gene | AT5G18860 |
| is.expressed_genes | TRUE |
| is.responsive_genes | TRUE |
| is.apcluster | A-02 |

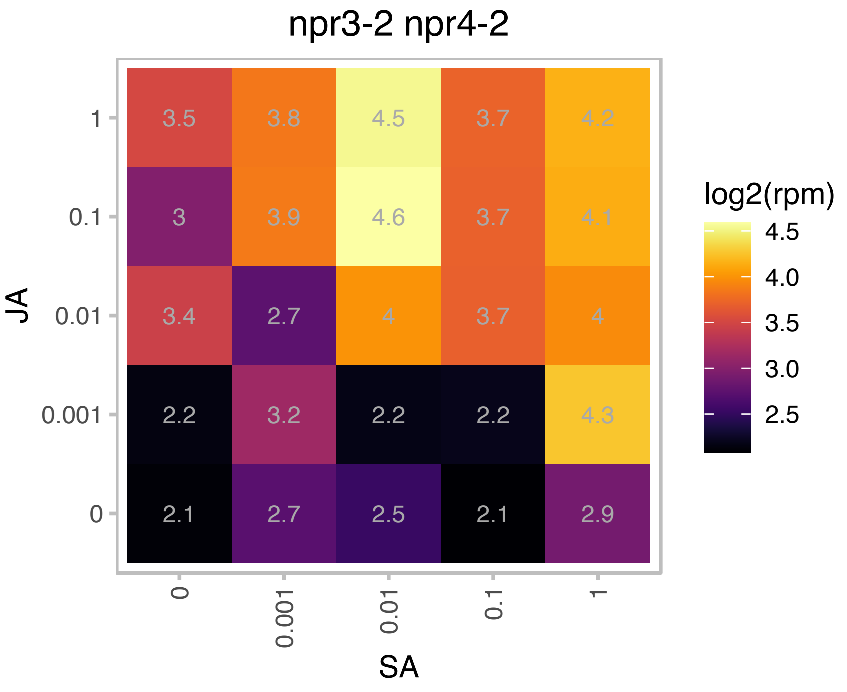


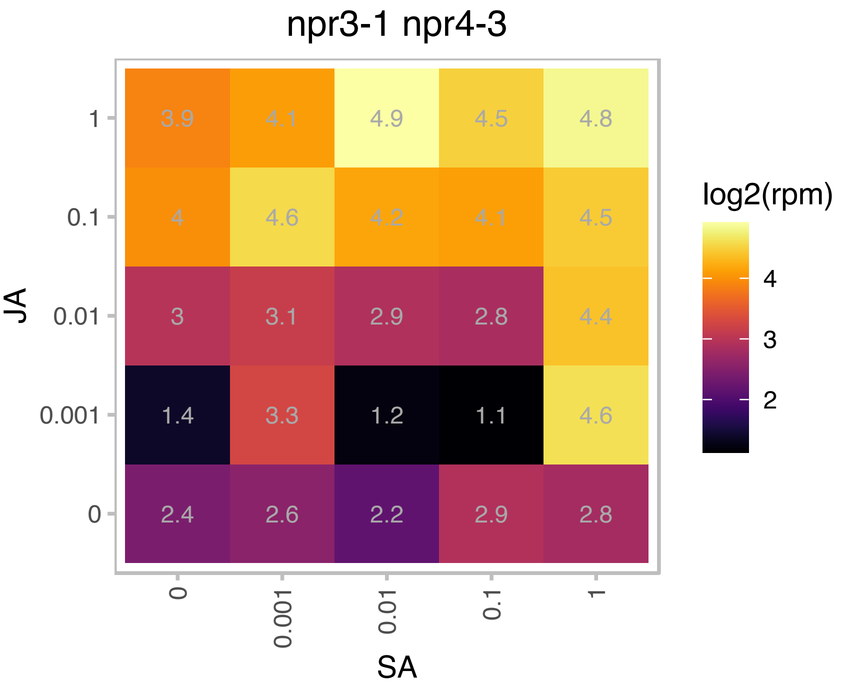

# AT5G19110.1 : NA

Eukaryotic aspartyl protease family protein

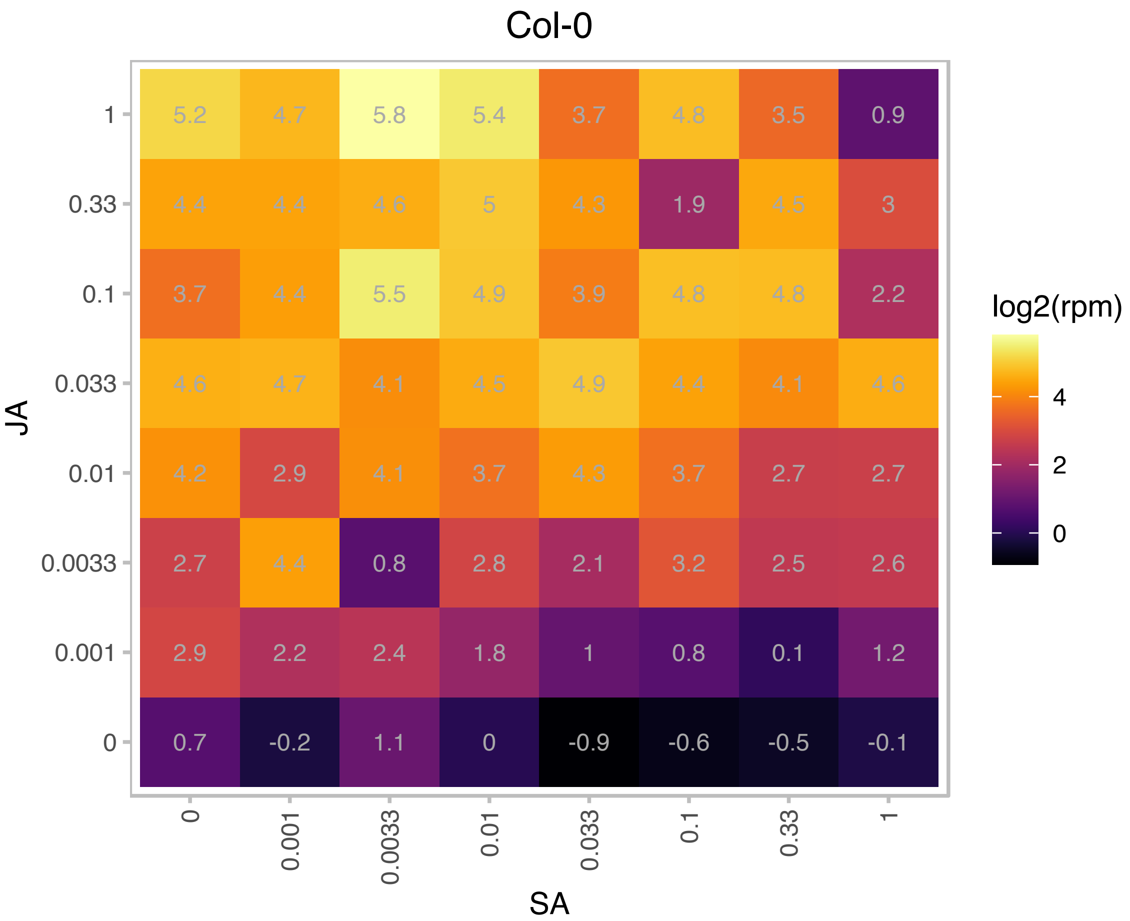


| | |
|---|---|
| Gene | AT5G19110 |
| is.expressed_genes | TRUE |
| is.responsive_genes | TRUE |
| is.apcluster | A-02 |

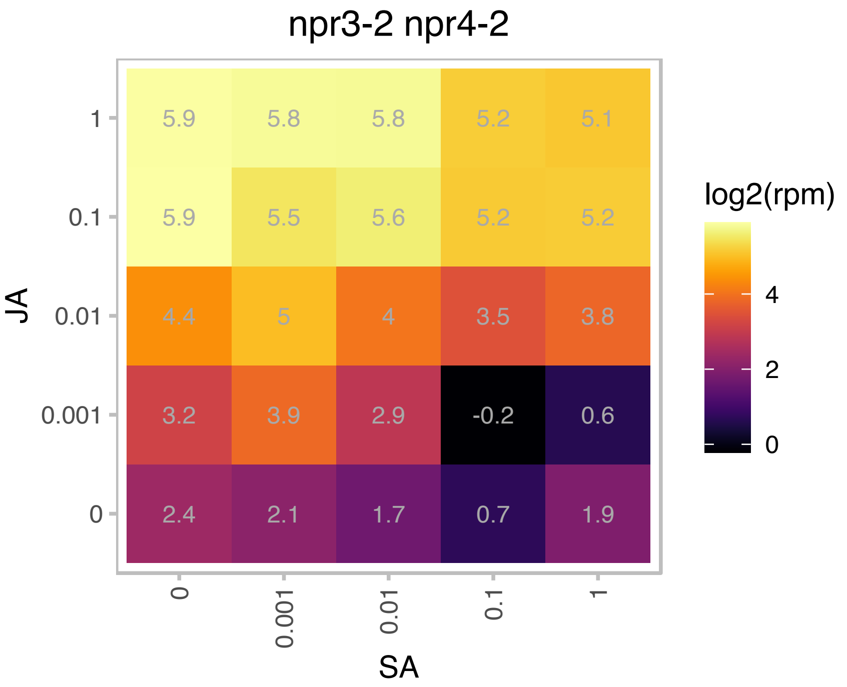


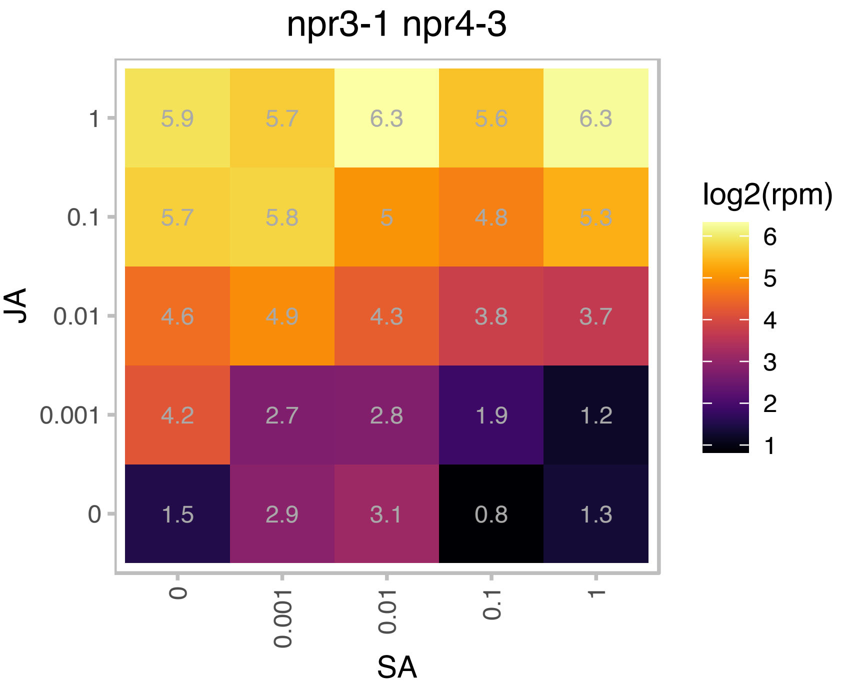

### AT5G24770.1 : ATVSP2, VSP2

vegetative storage protein 2
Has acid phosphatase activity dependent on the presence of divalent cations (Mg2+, Co2+, Zn2+, Mn2+) and anti-insect activity. Insects fed with the protein show a retarded development. Induced in response to abscisic acid, jasmonic acid, salt, water deficiency and wounding.

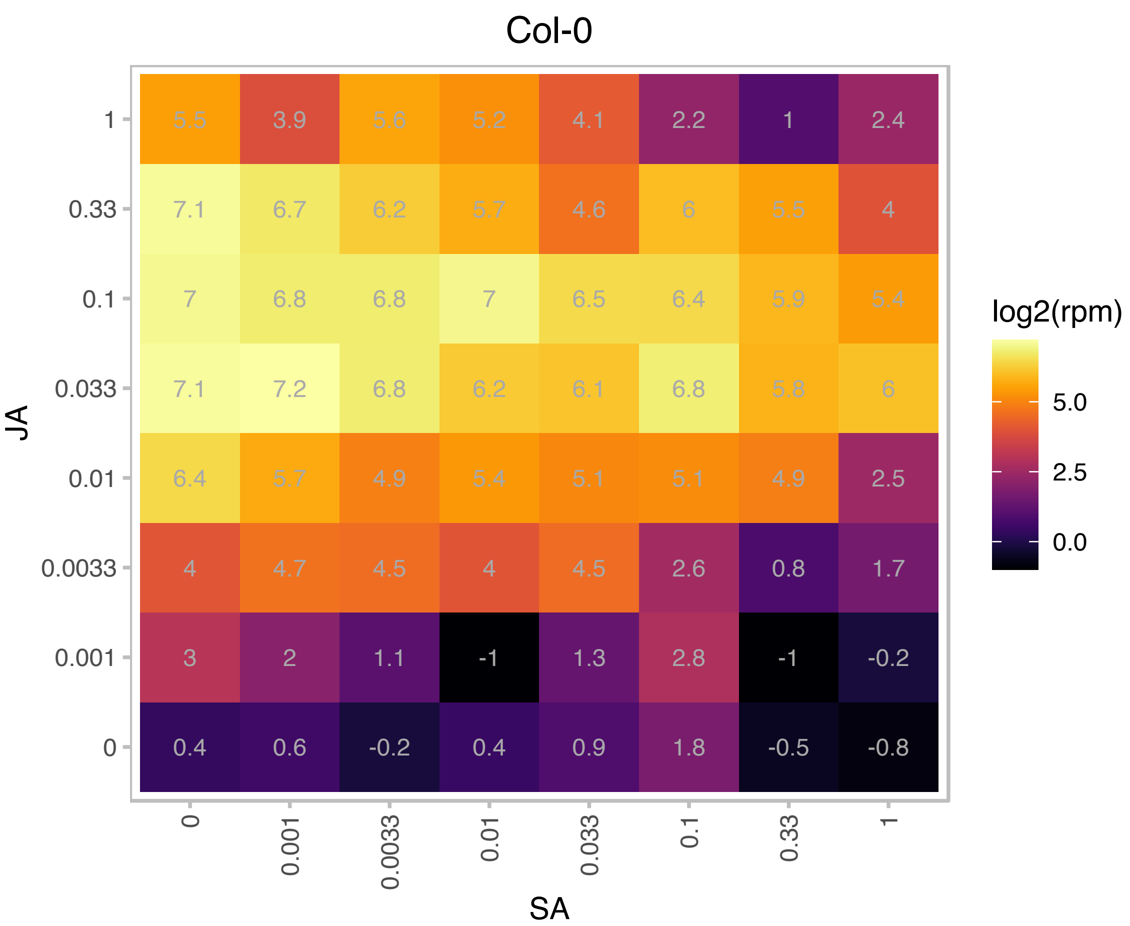


| | |
|---|---|
| Gene | AT5G24770 |
| is.expressed_genes | TRUE |
| is.responsive_genes | TRUE |
| is.apcluster | A-02 |

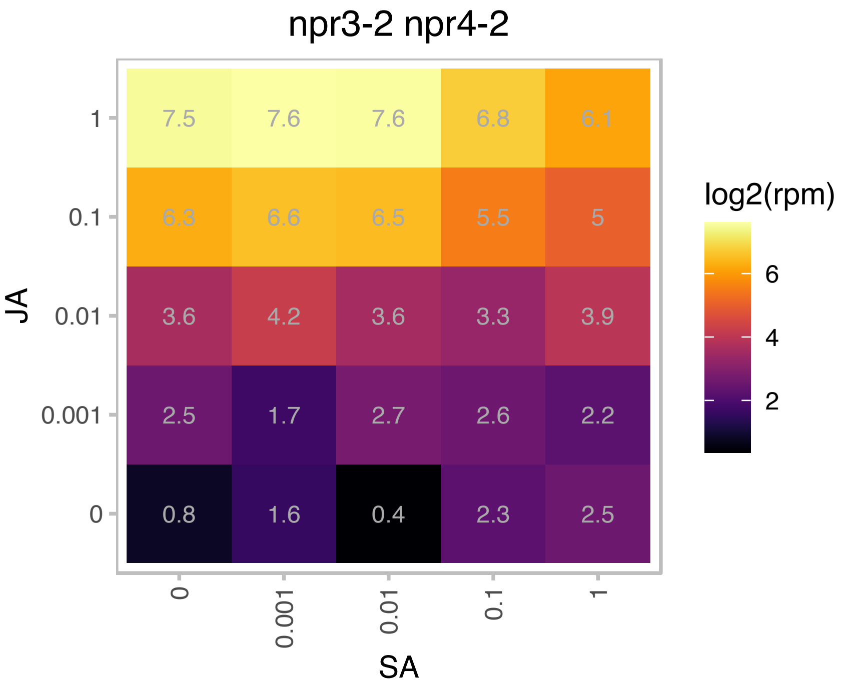


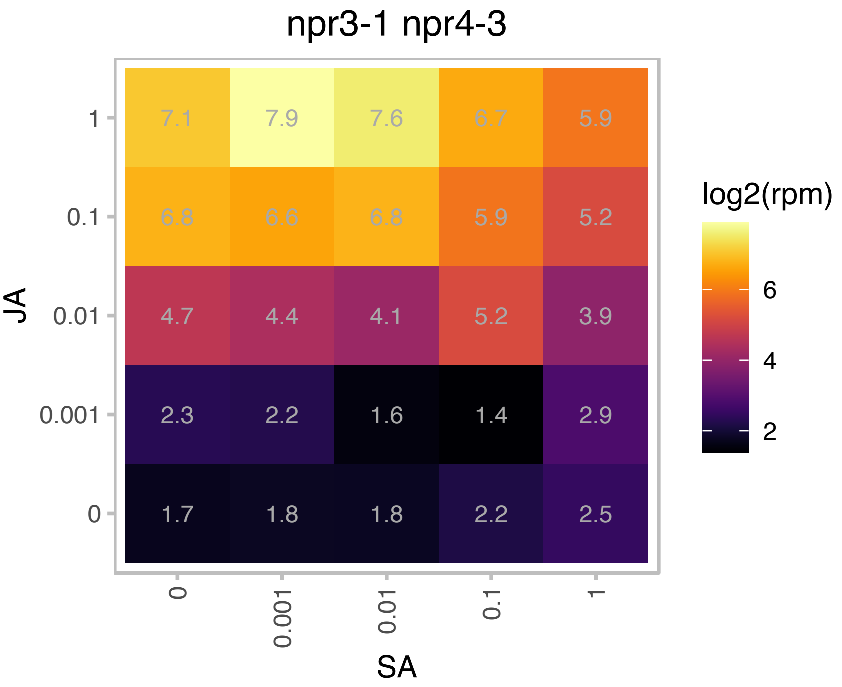

# AT5G43570.1 : NA

Serine protease inhibitor, potato inhibitor I-type family protein
Predicted to encode a PR (pathogenesis-related) peptide that belongs to the PR-6 proteinase inhibitor family. Six putative PR-6-type protein encoding genes are found in Arabidopsis: At2g38900, At2g38870, At5g43570, At5g43580, At3g50020 and At3g46860.

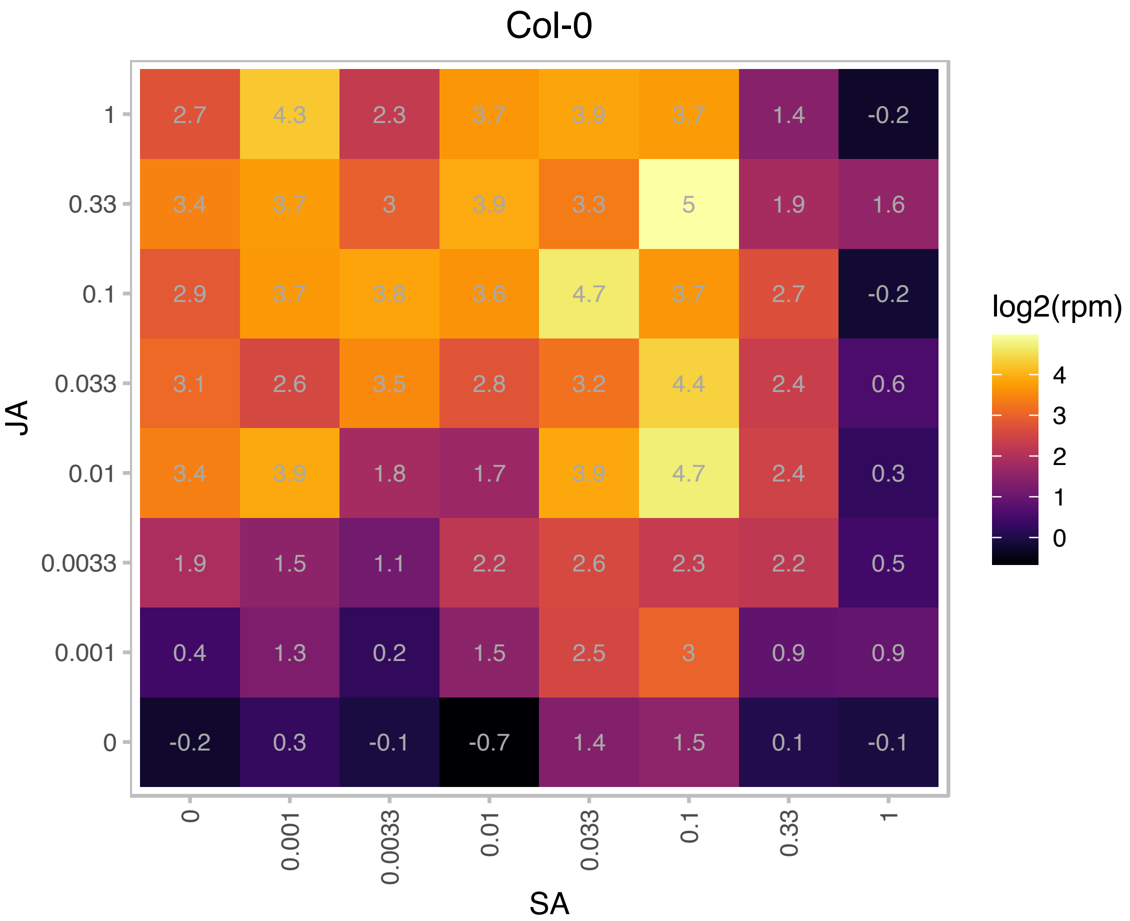


| | |
|---|---|
| Gene | AT5G43570 |
| is.expressed_genes | TRUE |
| is.responsive_genes | TRUE |
| is.apcluster | A-02 |

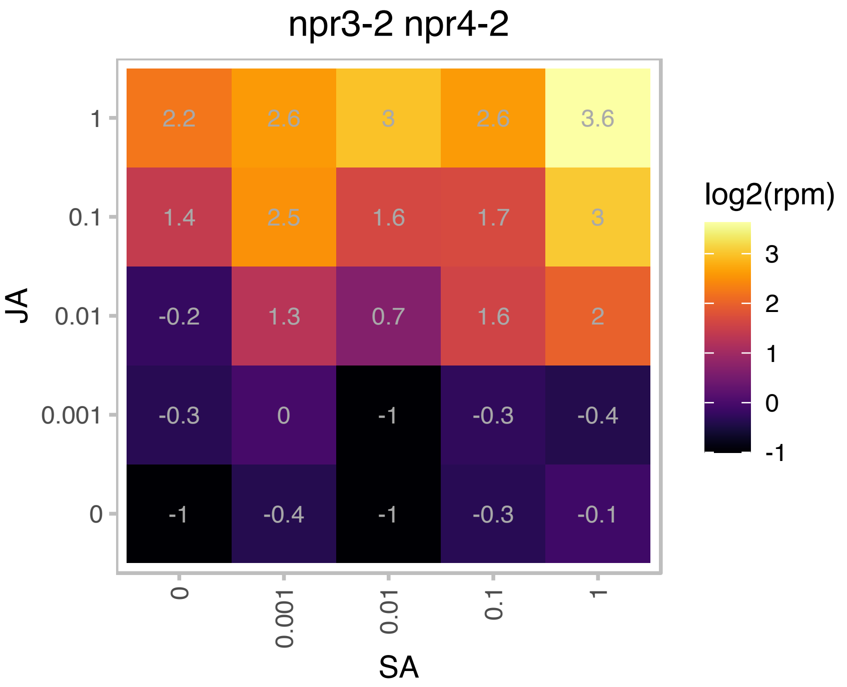


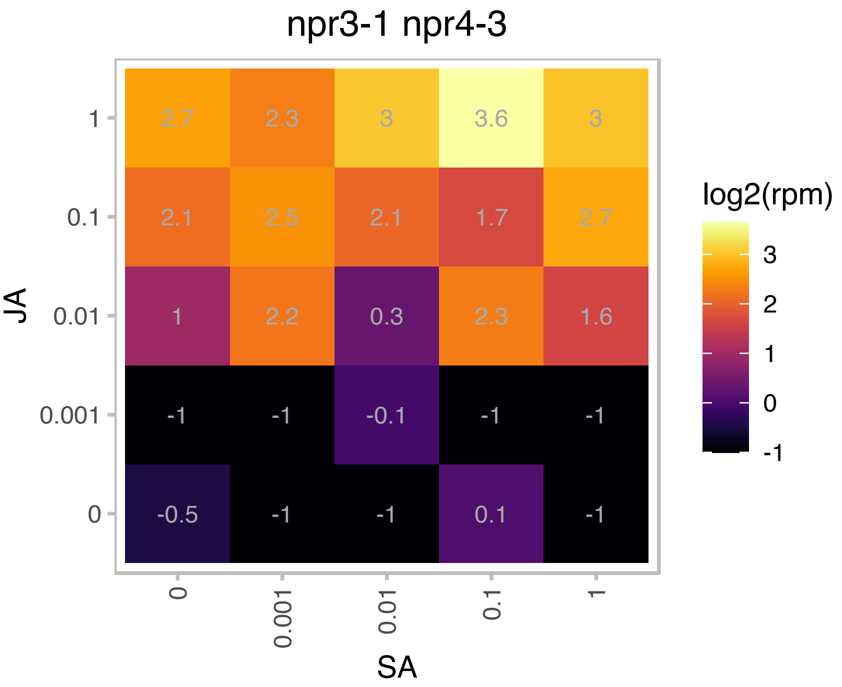

# AT5G59490.1 : NA

Haloacid dehalogenase-like hydrolase (HAD) superfamily protein

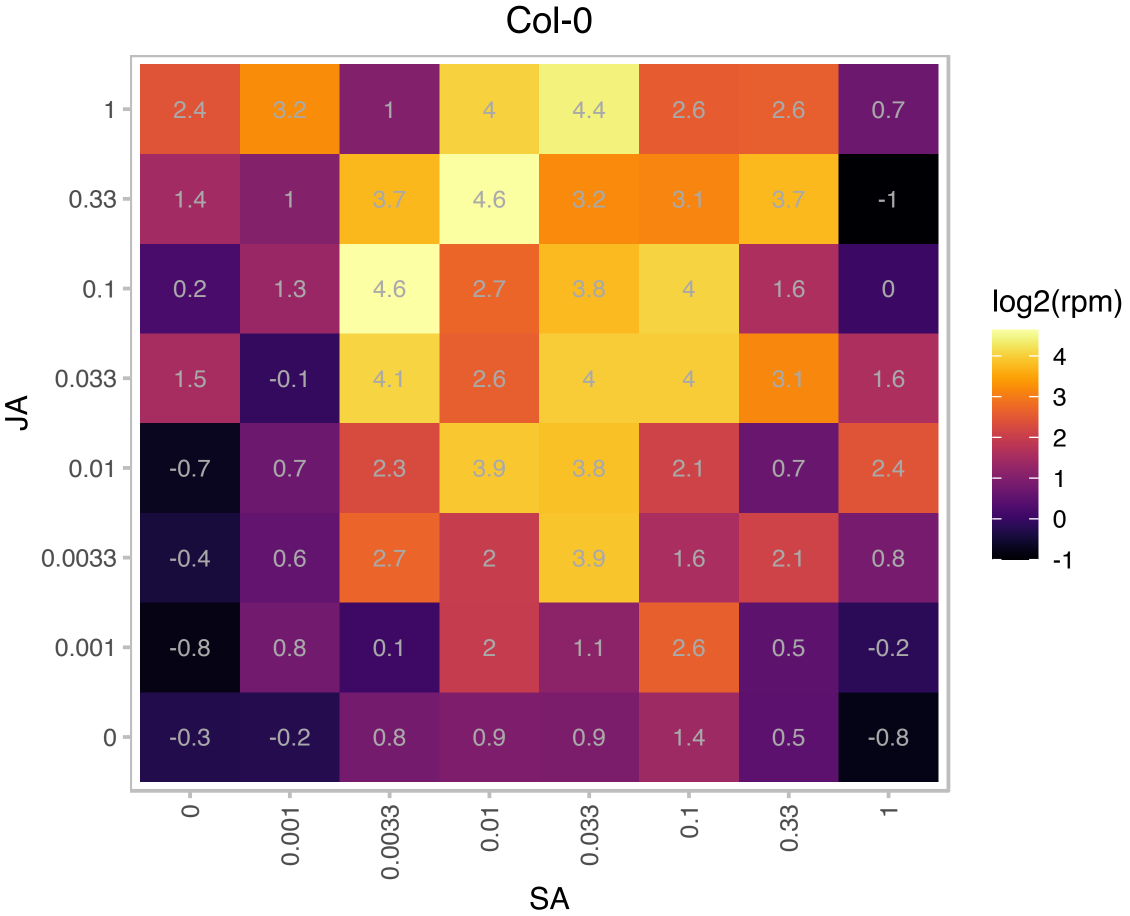


| | |
|---|---|
| Gene | AT5G59490 |
| is.expressed_genes | TRUE |
| is.responsive_genes | TRUE |
| is.apcluster | A-02 |

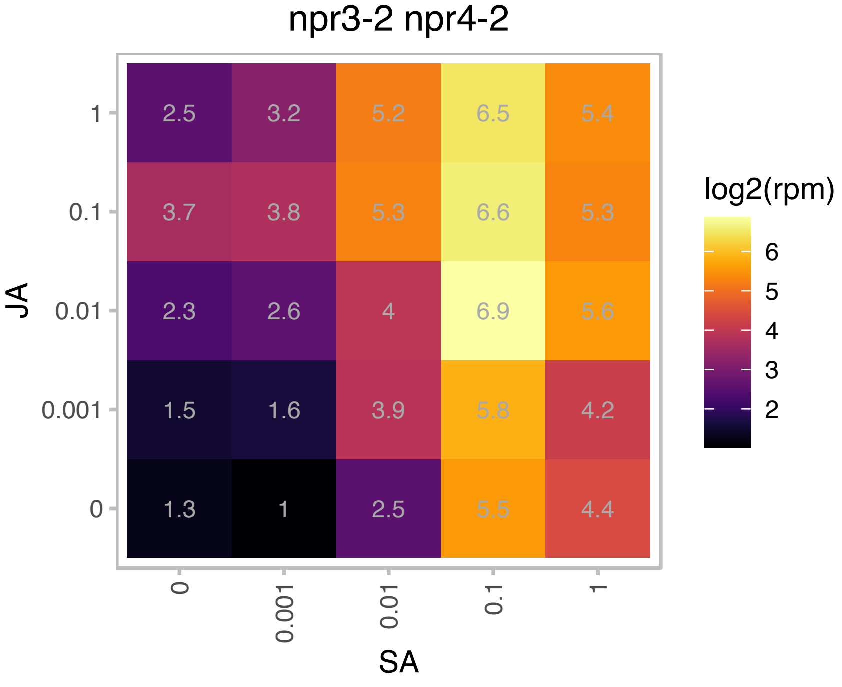


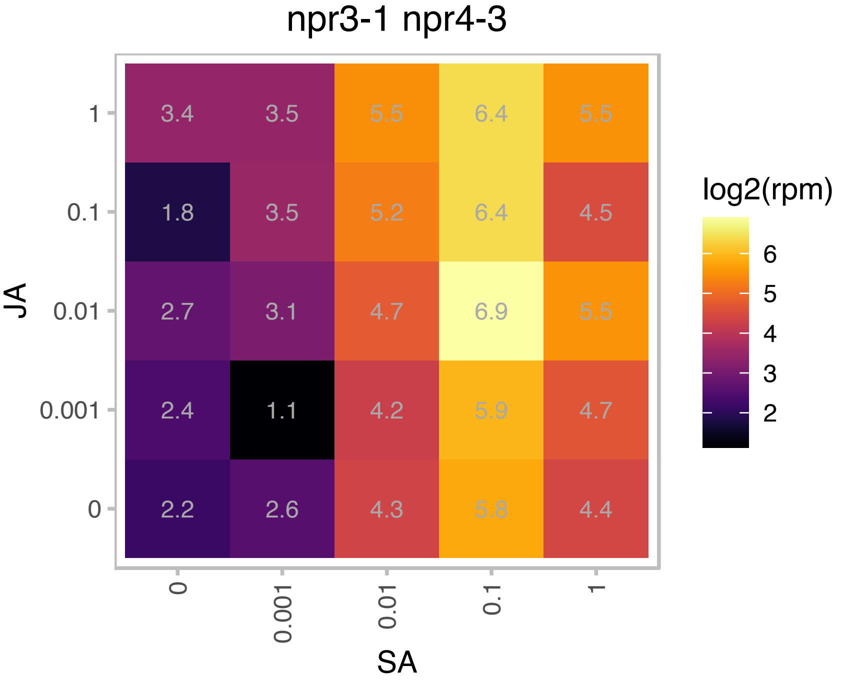

### ATCG00350.1 : PSAA

Photosystem I, PsaA/PsaB protein

Encodes psaA protein comprising the reaction center for photosystem I along with psaB protein; hydrophobic protein encoded by the chloroplast genome.

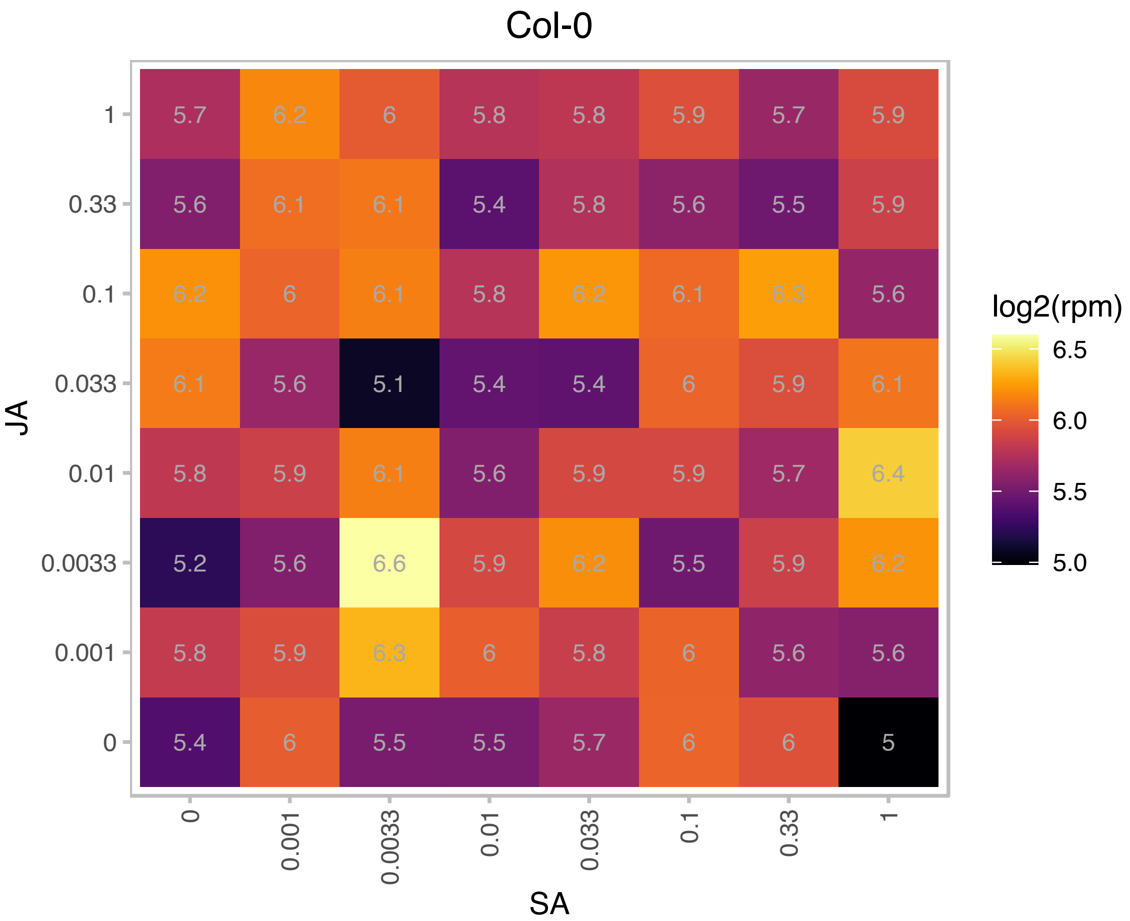


| | |
|---|---|
| Gene | ATCG00350 |
| is.expressed_genes | TRUE |
| is.responsive_genes | TRUE |
| is.apcluster | A-02 |

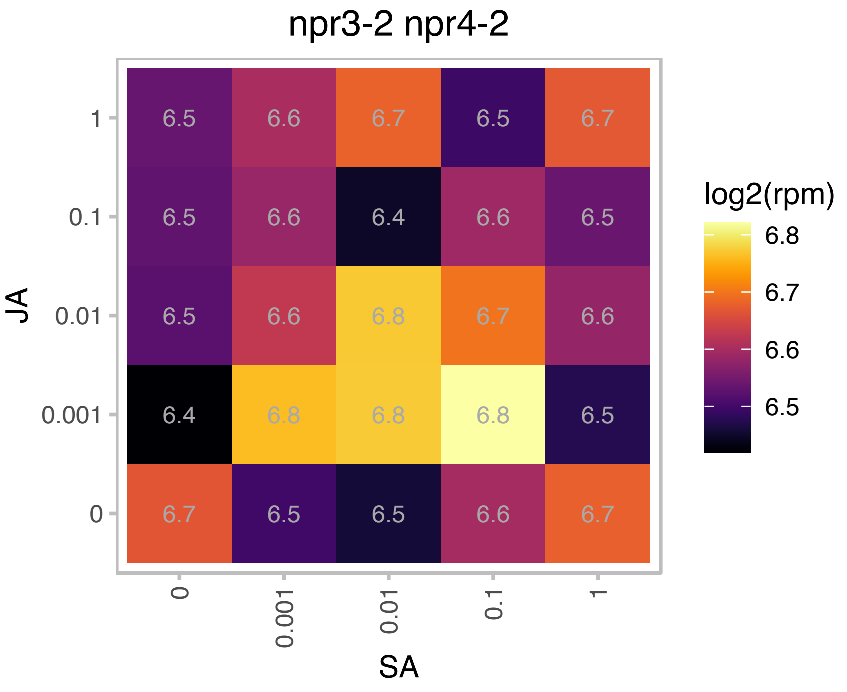


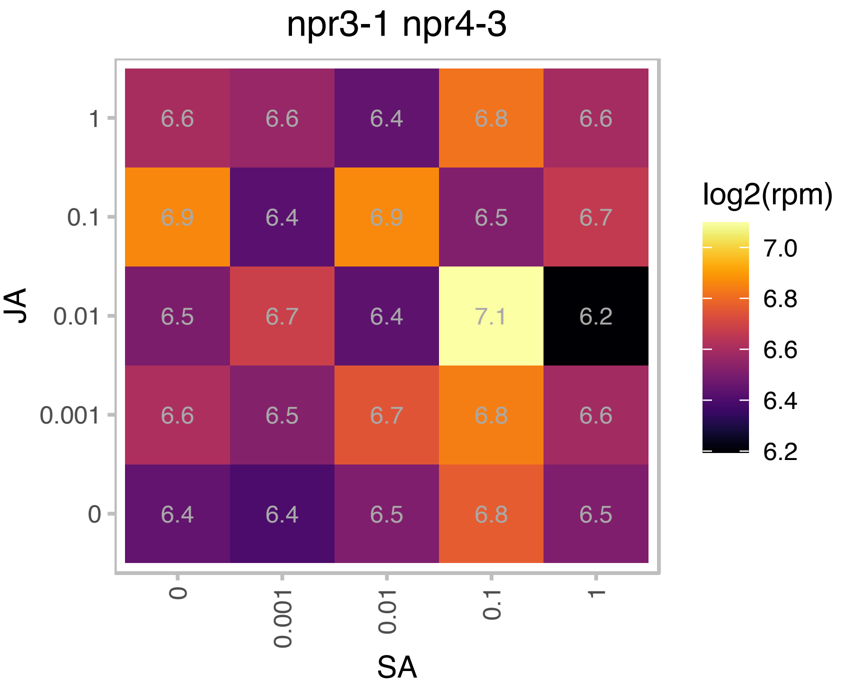

# AT1G07980.1 : NF-YC10

nuclear factor Y, subunit C10

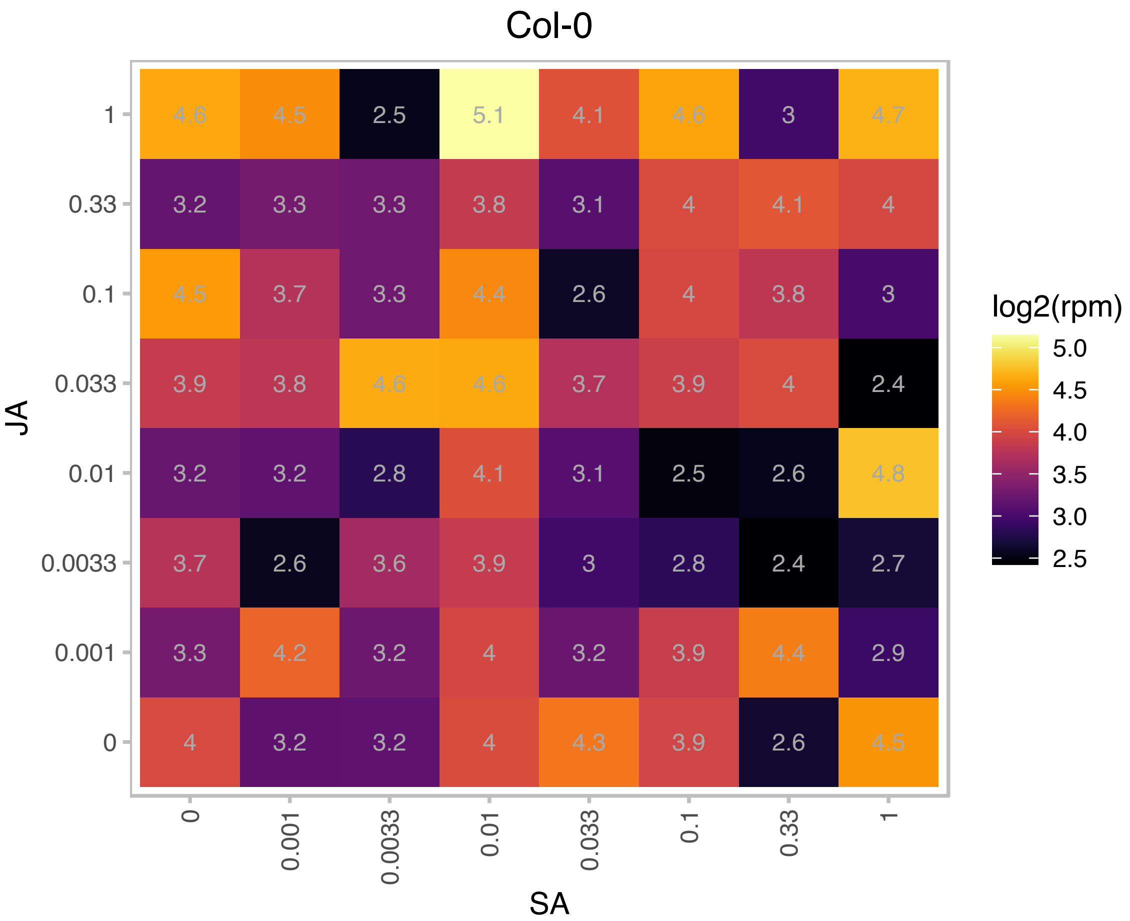


| | |
|---|---|
| Gene | AT1G07980 |
| is.expressed_genes | TRUE |
| is.responsive_genes | TRUE |
| is.apcluster | A-03 |

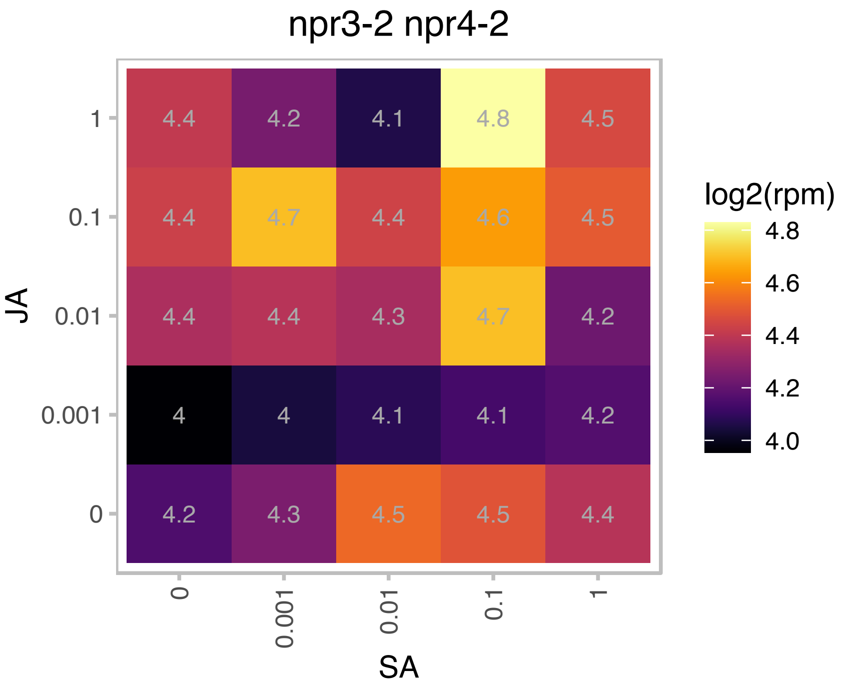


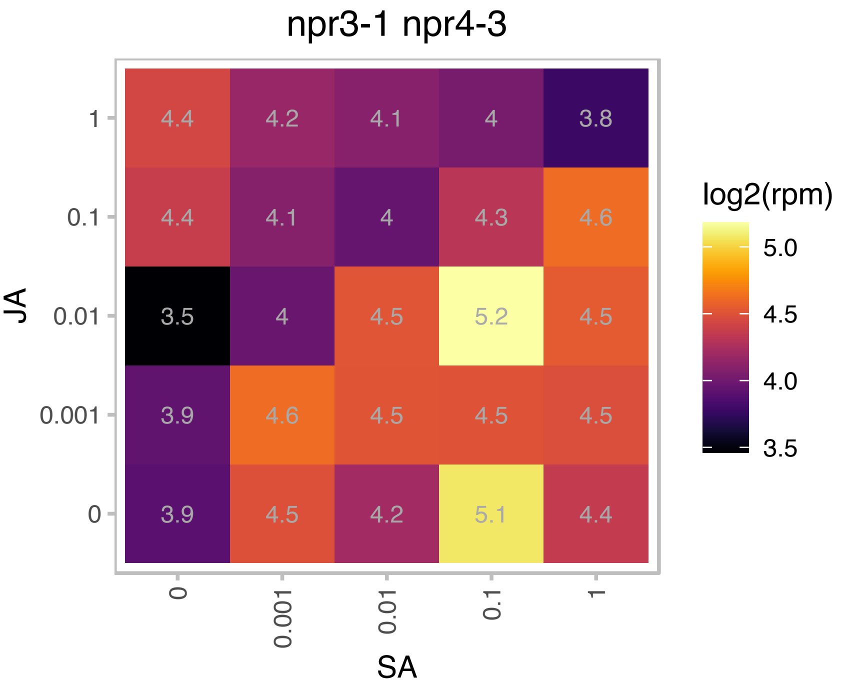

# AT1G17360.1 : NA

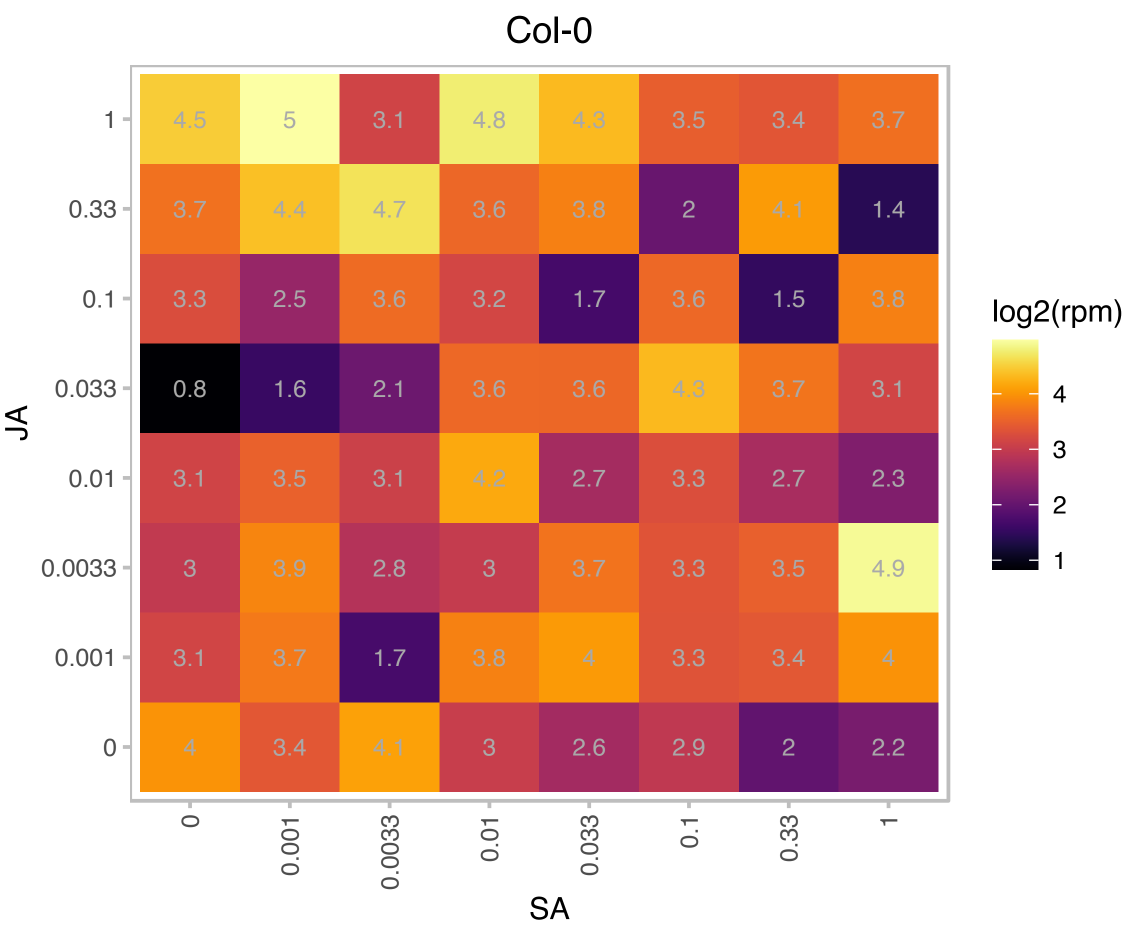


| | |
|---|---|
| Gene | AT1G17360 |
| is.expressed_genes | TRUE |
| is.responsive_genes | TRUE |
| is.apcluster | A-03 |

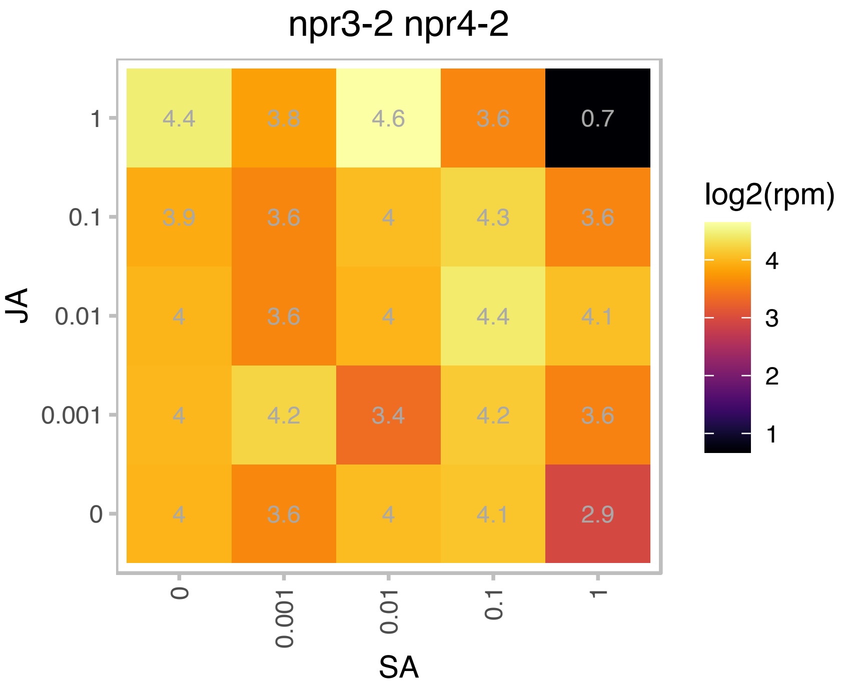


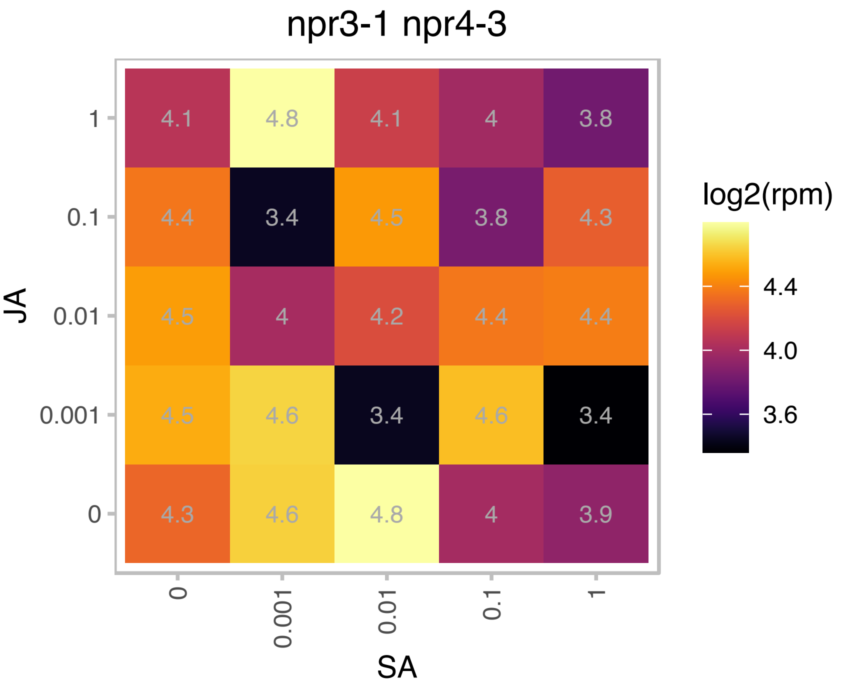

### AT1G21640.1 : ATNADK2, NADK2

NAD kinase 2

Encodes a protein with NAD kinase activity. The protein was also shown to bind calmodulin.

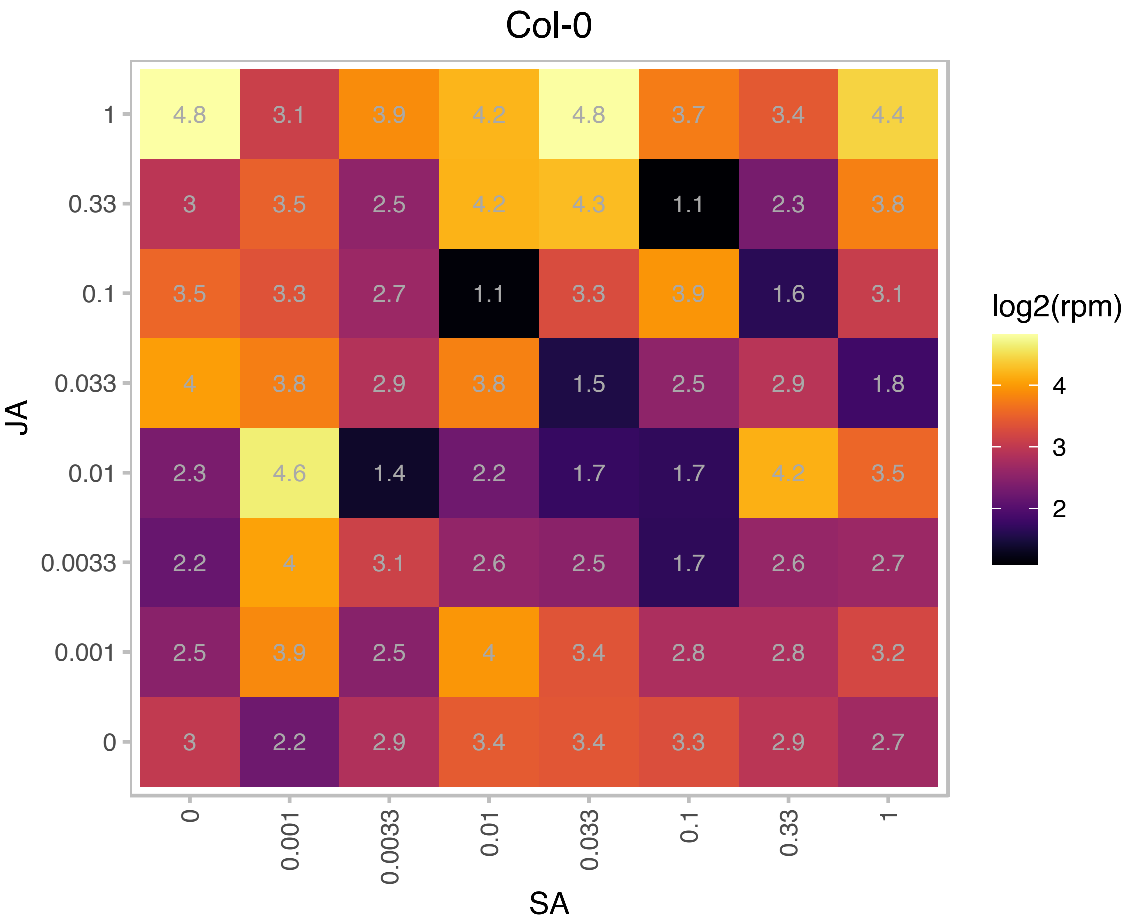


| | |
|---|---|
| Gene | AT1G21640 |
| is.expressed_genes | TRUE |
| is.responsive_genes | TRUE |
| is.apcluster | A-03 |

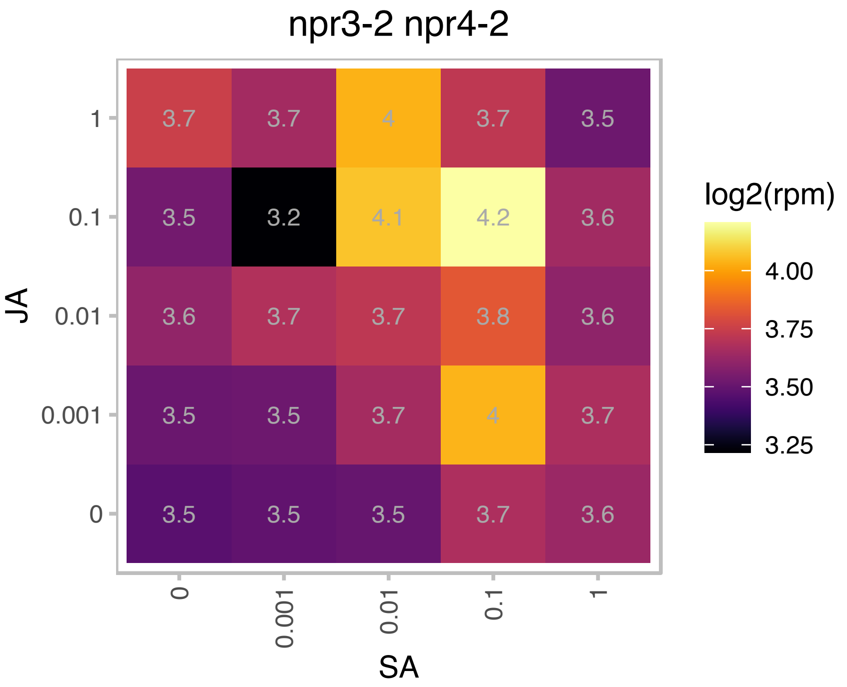


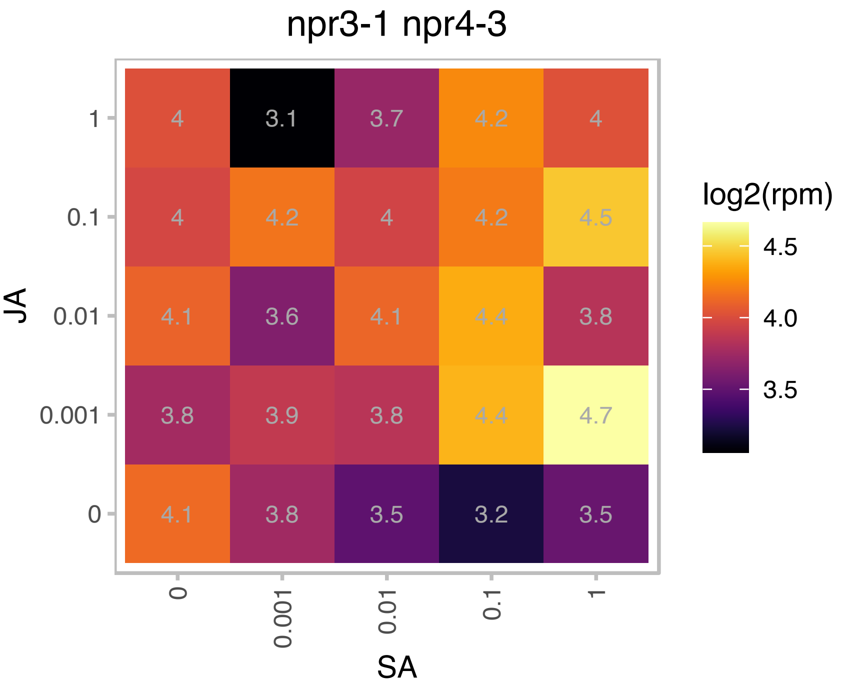

# AT1G30440.1 : NA

Phototropic-responsive NPH3 family protein

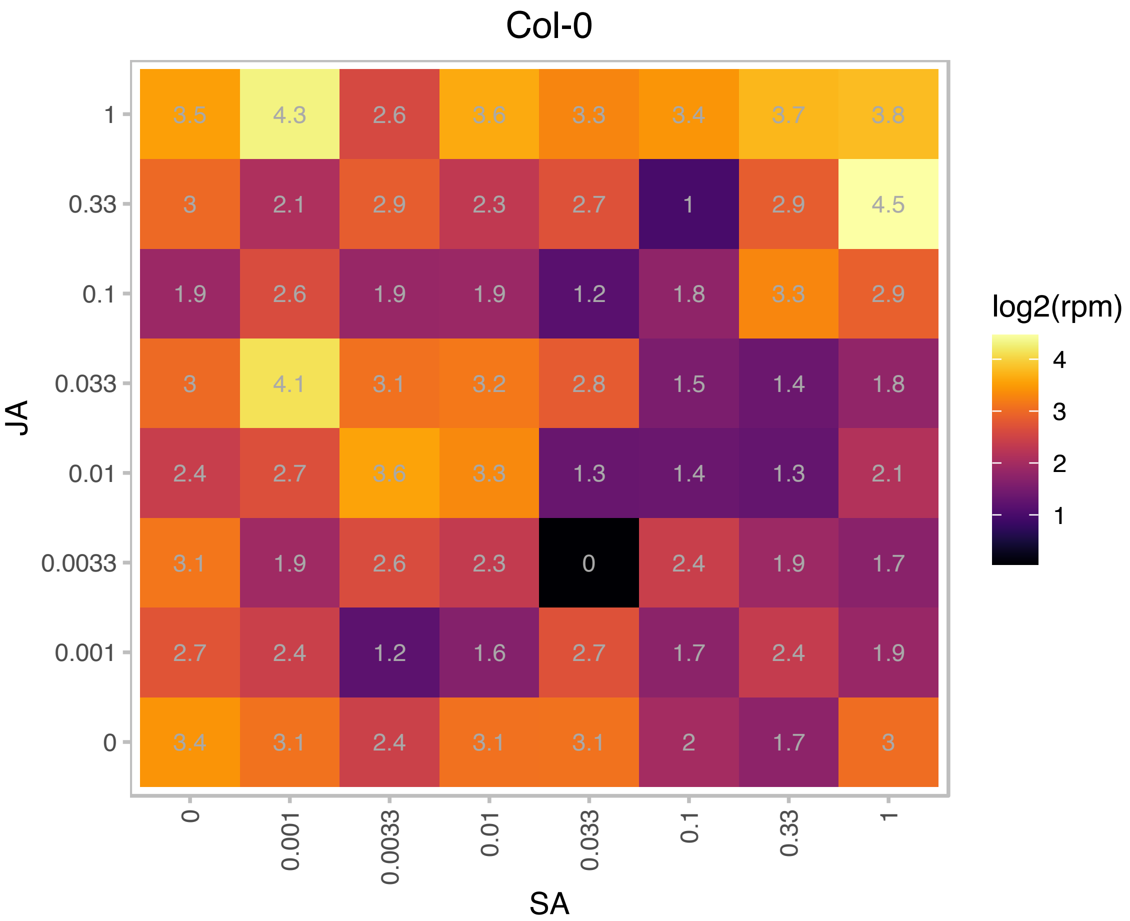


| Gene | AT1G30440 |
|---|---|
| is.expressed_genes | TRUE |
| is.responsive_genes | TRUE |
| is.apcluster | A-03 |

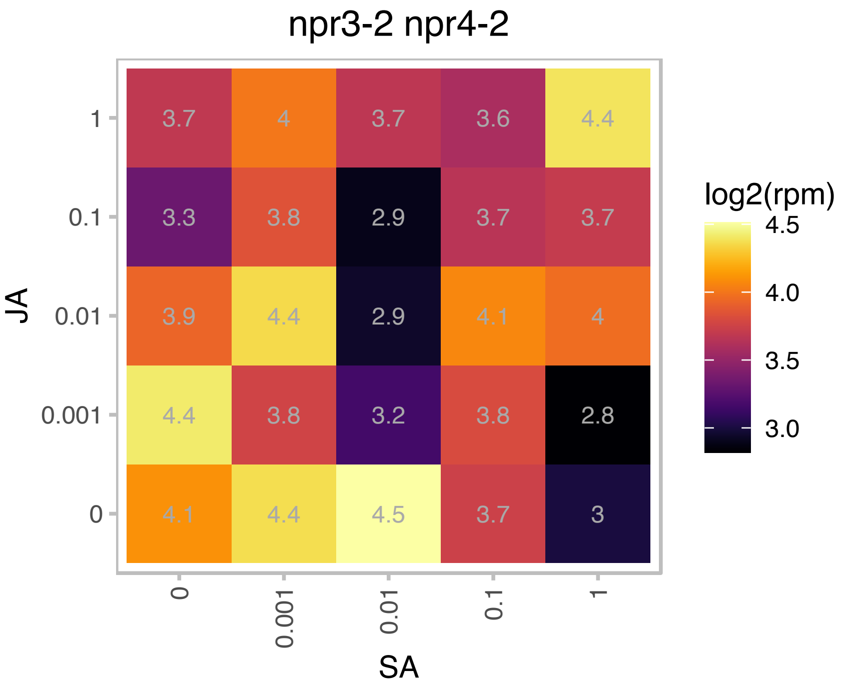


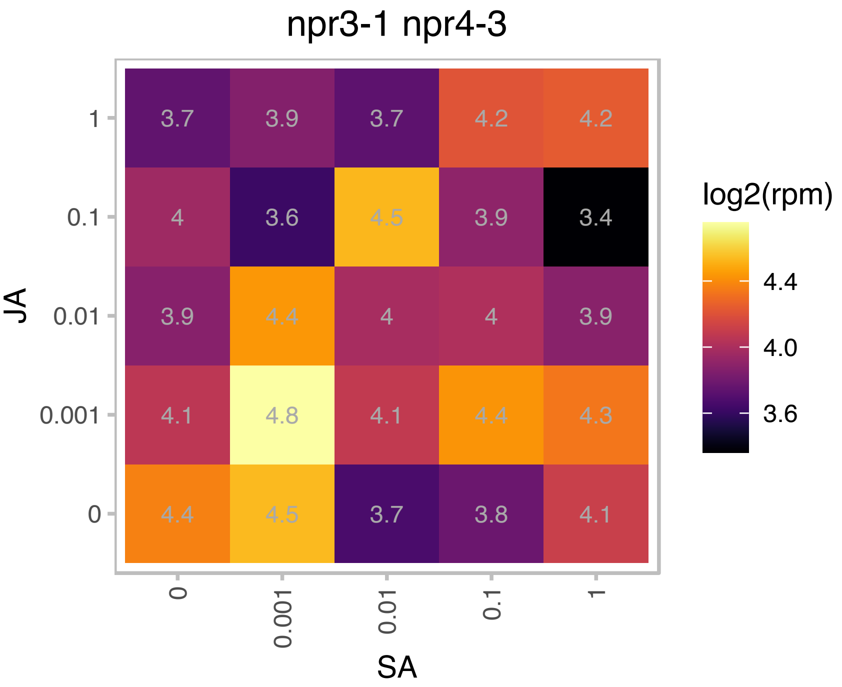

### AT1G62380.1 : ACO2, ATACO2

ACC oxidase 2

Encodes a protein similar to 1-aminocyclopropane-1-carboxylic oxidase (ACC oxidase). Expression of the AtACO2 transcripts is affected by ethylene.

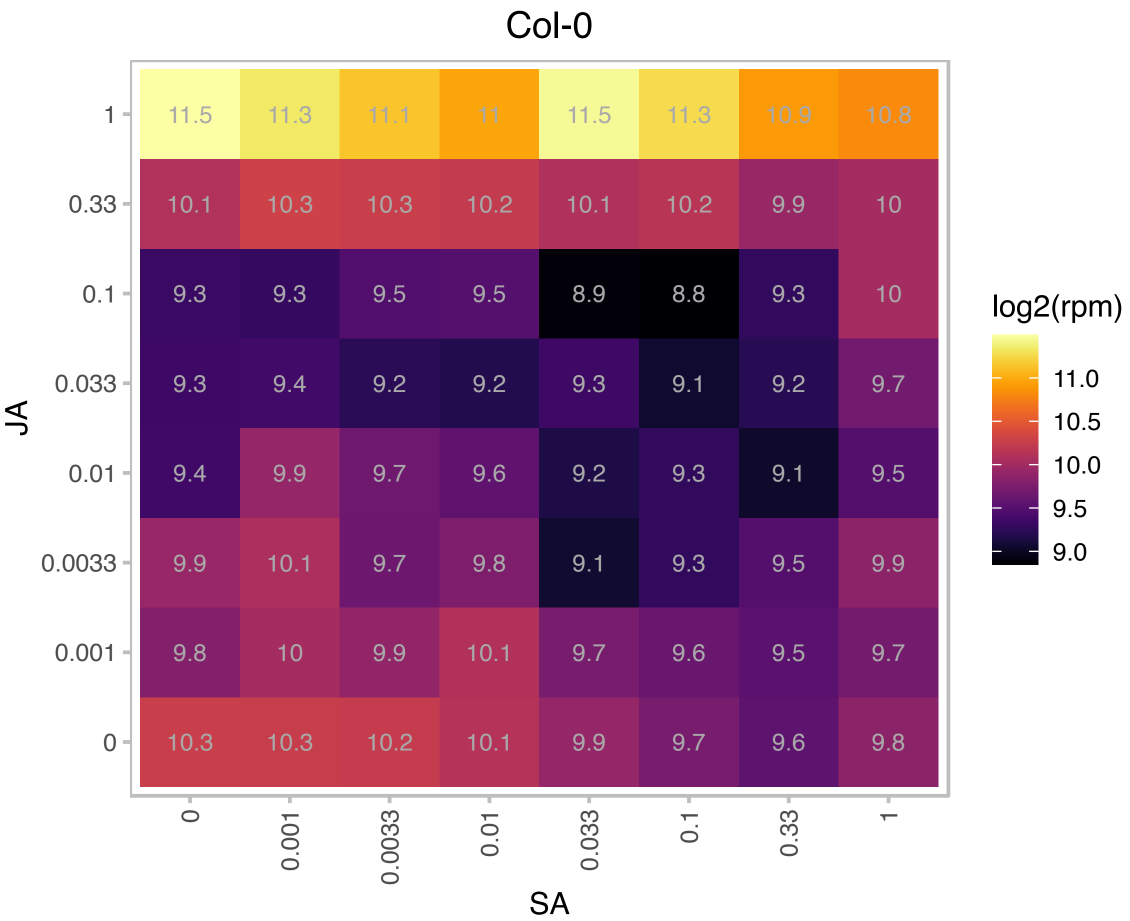


| | |
|---|---|
| Gene | AT1G62380 |
| is.expressed_genes | TRUE |
| is.responsive_genes | TRUE |
| is.apcluster | A-03 |

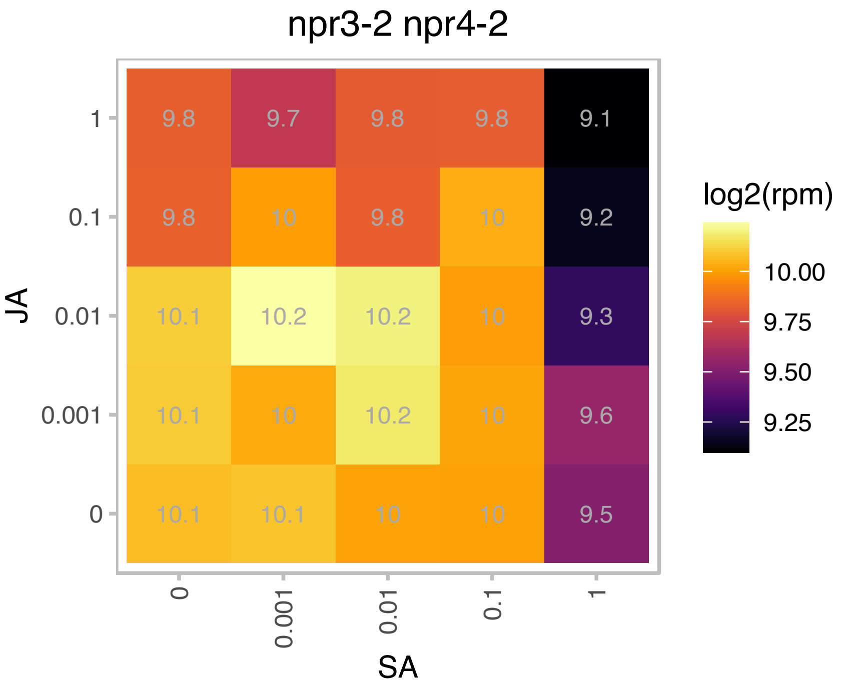


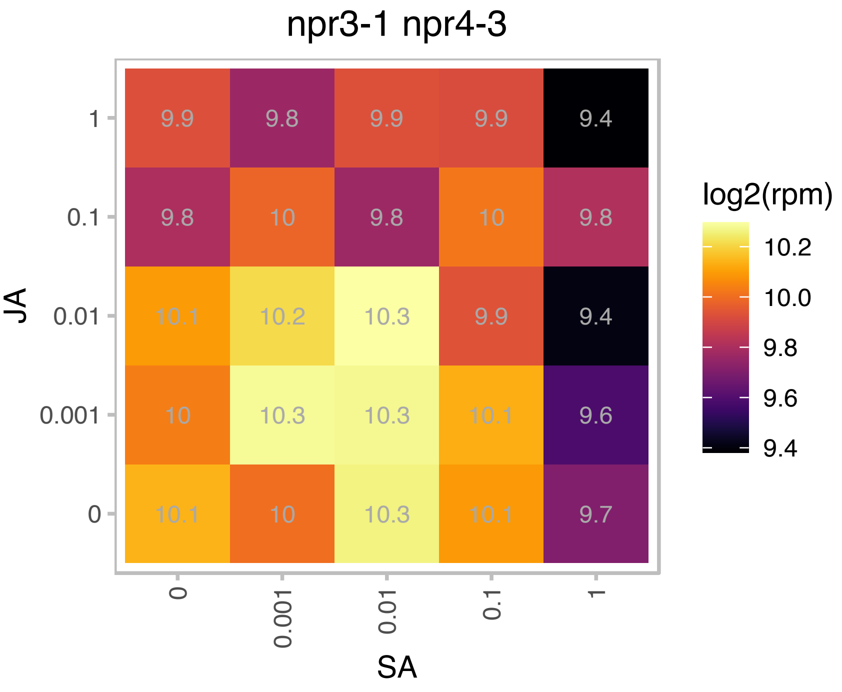

### AT1G65650.1 : UCH2

Peptidase C12, ubiquitin carboxyl-terminal hydrolase 1

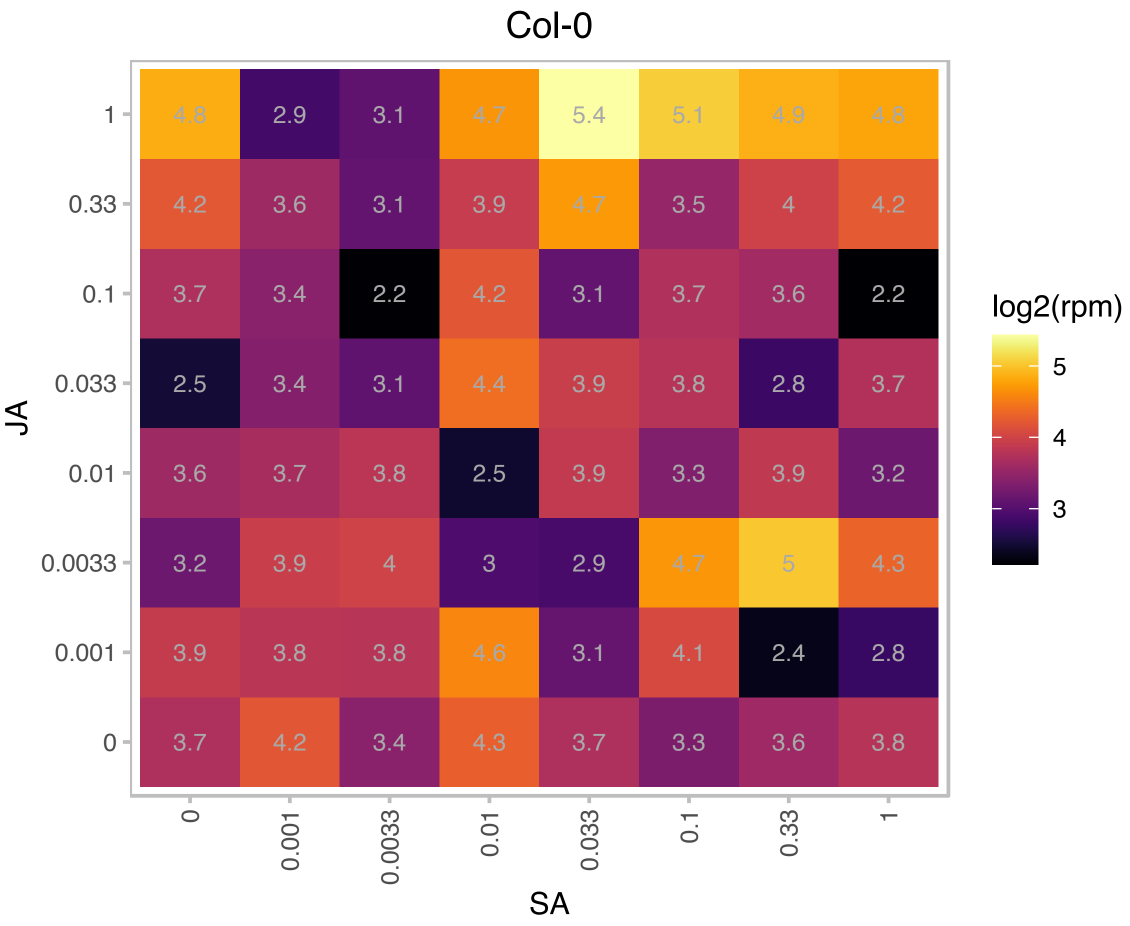


| | |
|---|---|
| Gene | AT1G65650 |
| is.expressed_genes | TRUE |
| is.responsive_genes | TRUE |
| is.apcluster | A-03 |

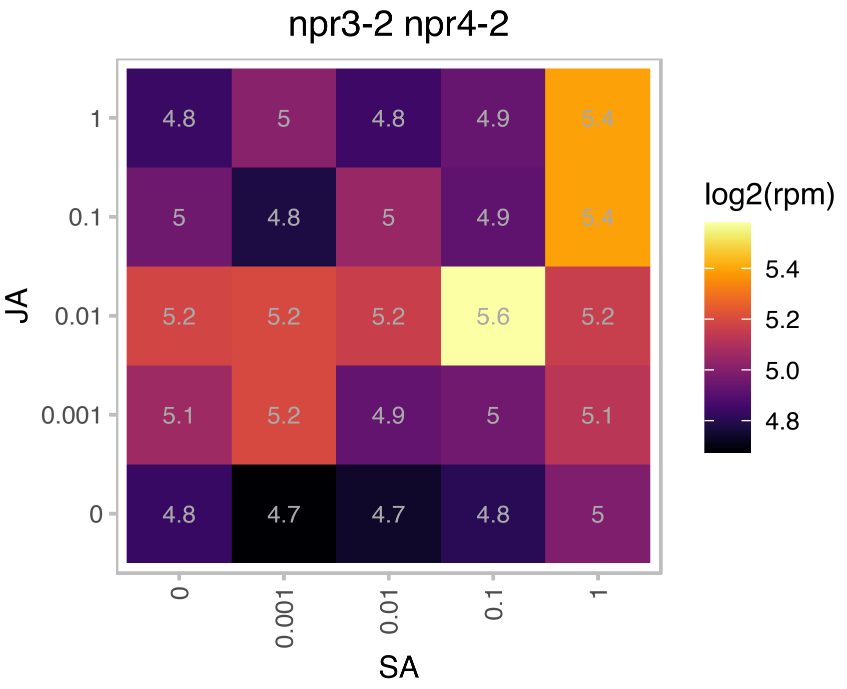


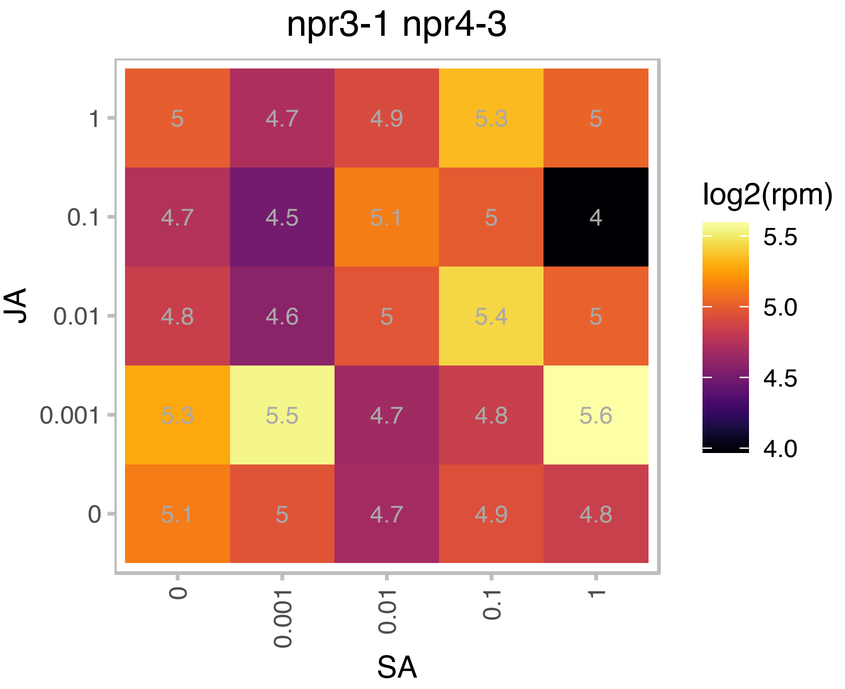

# AT2G02470.1 : AL6

alfin-like 6
Encodes a member of the Alfin1-like family of nuclear-localized PHD (plant homeodomain) domain containing proteins that acts as a novel upstream regulator of root hair formation during Pi starvation. All AL proteins except AL3 bind to di- or trimethylated histone H3 (H3K4me3/2). Members of this family include: AT5G05610 (AL1), AT3G11200 (AL2), AT3G42790 (AL3), AT5G26210 (AL4), AT5G20510 (AL5), AT2G02470 (AL6), AT1G14510

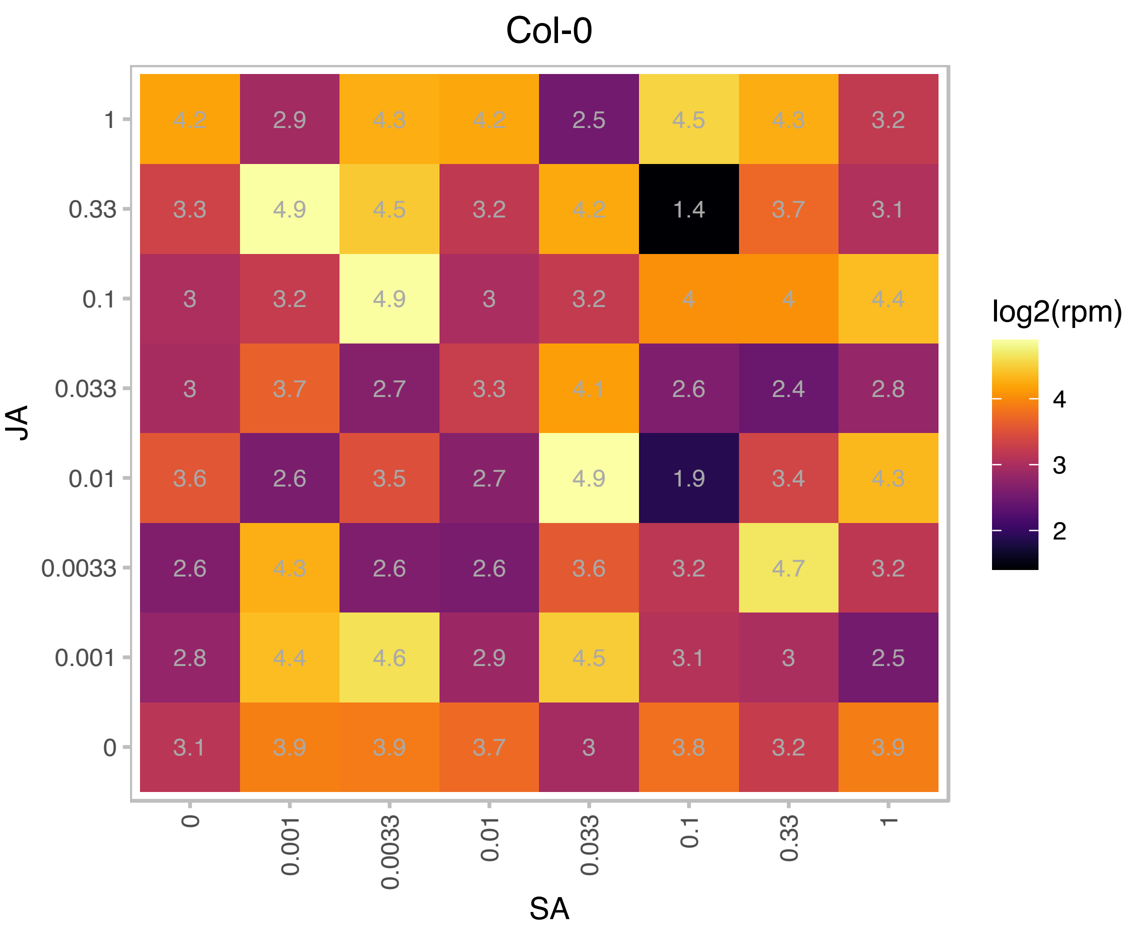


| | |
|---|---|
| Gene | AT2G02470 |
| is.expressed_genes | TRUE |
| is.responsive_genes | TRUE |
| is.apcluster | A-03 |

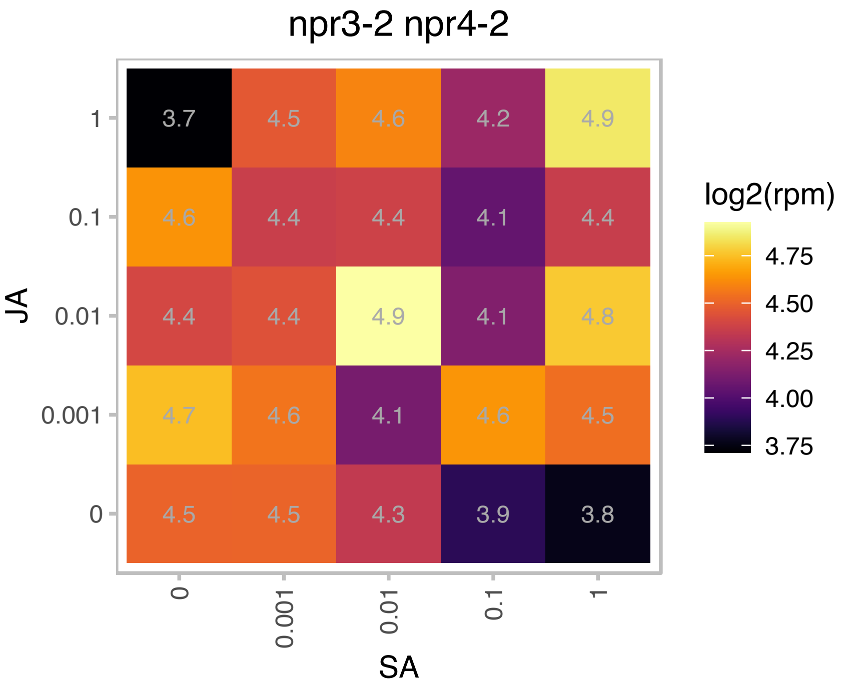


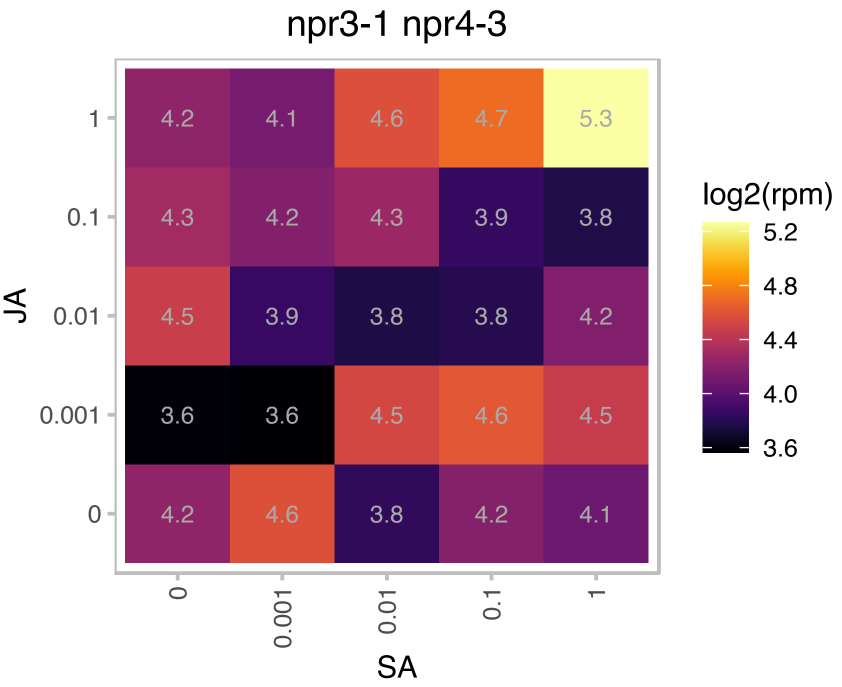

### AT2G15290.1 : AtTic21, CIA5, PIC1, TIC21

translocon at inner membrane of chloroplasts 21
Encodes a protein located in the chloroplast inner envelope. The study of mutant defective in the gene product suggests that the protein is involved in the translocation of protein across the envelope membrane into the chloroplast stroma.

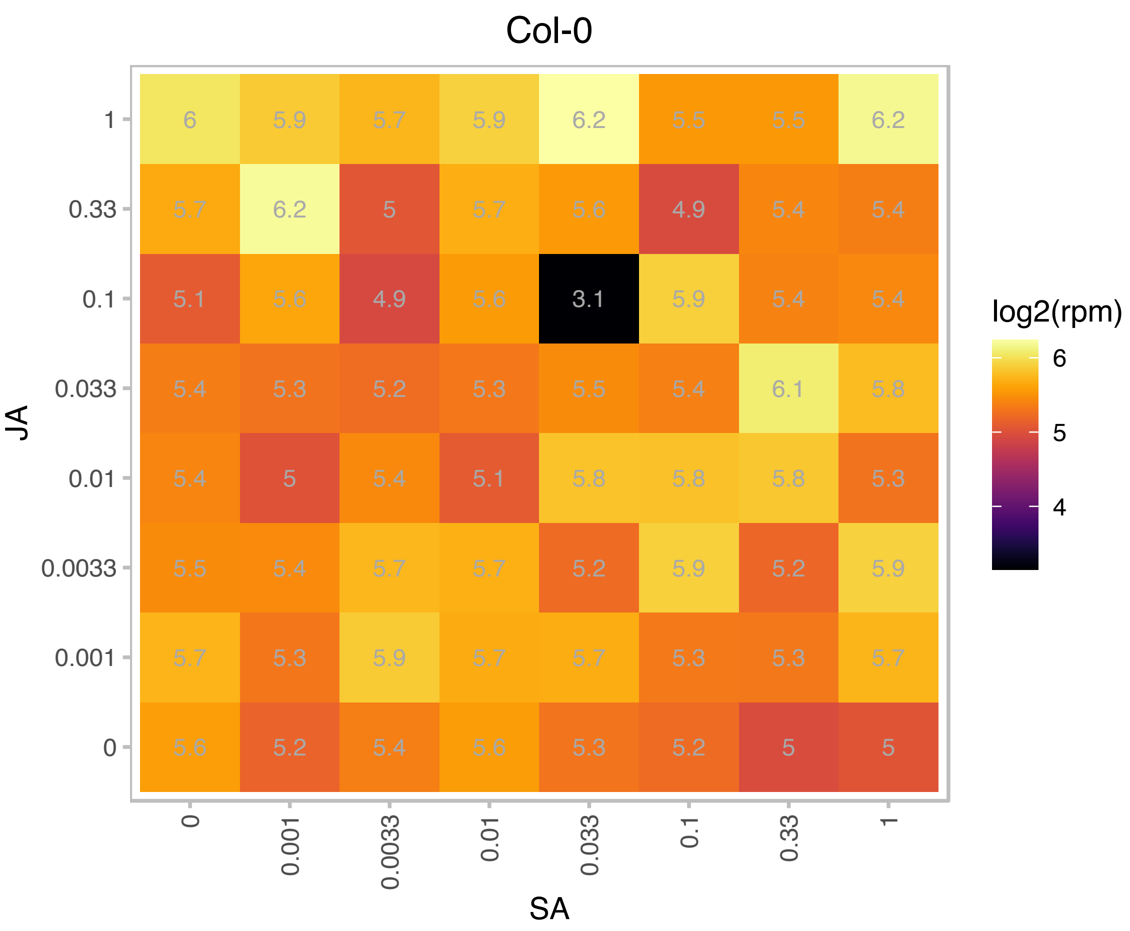


| | |
|---|---|
| Gene | AT2G15290 |
| is.expressed_genes | TRUE |
| is.responsive_genes | TRUE |
| is.apcluster | A-03 |

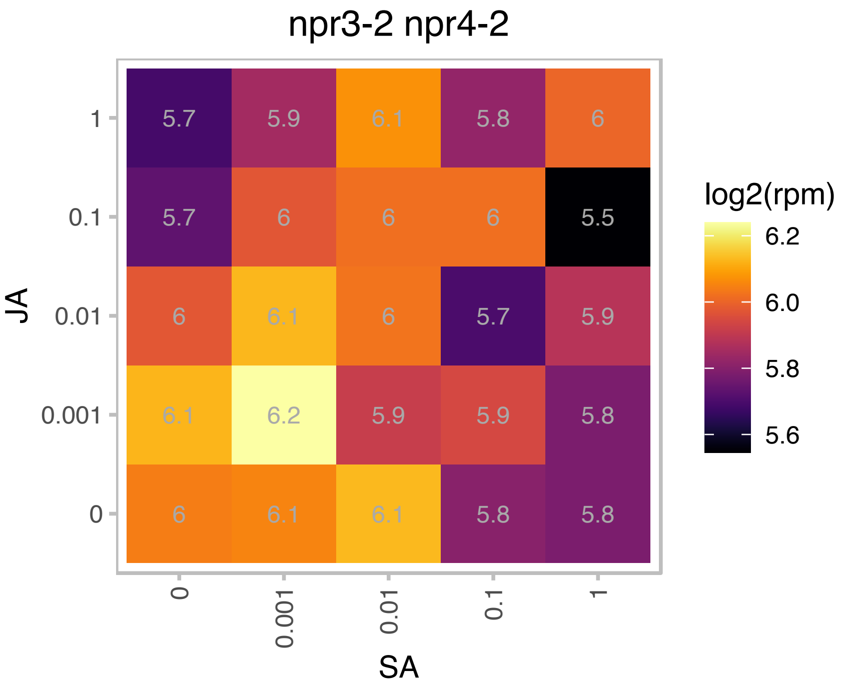


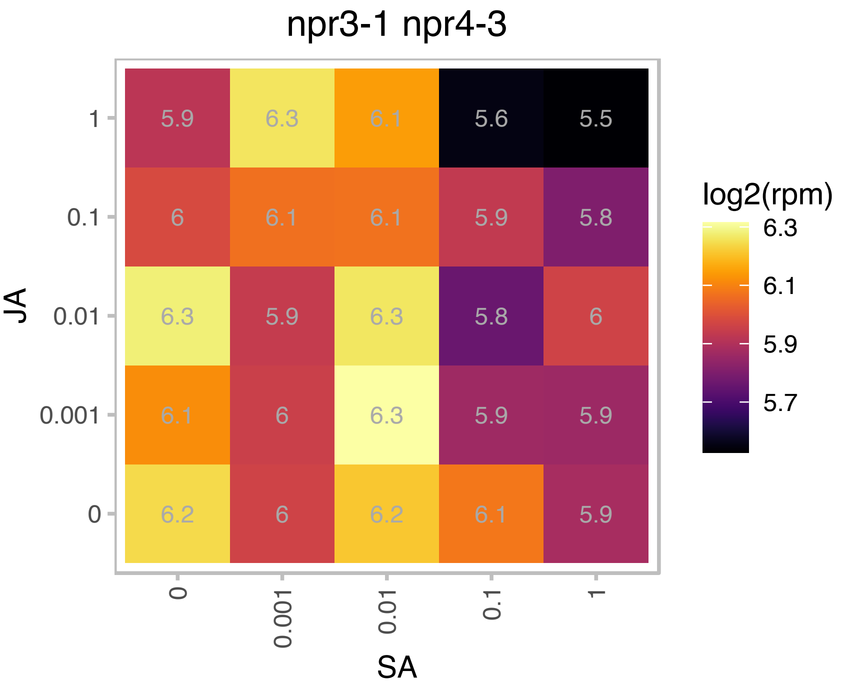

### AT2G17840.1 : ERD7

Senescence/dehydration-associated protein-related

Identified as drought-inducible gene by differential hybridization. Upregulated by high light, drought, cold and salt stress determined by microarray analysis.

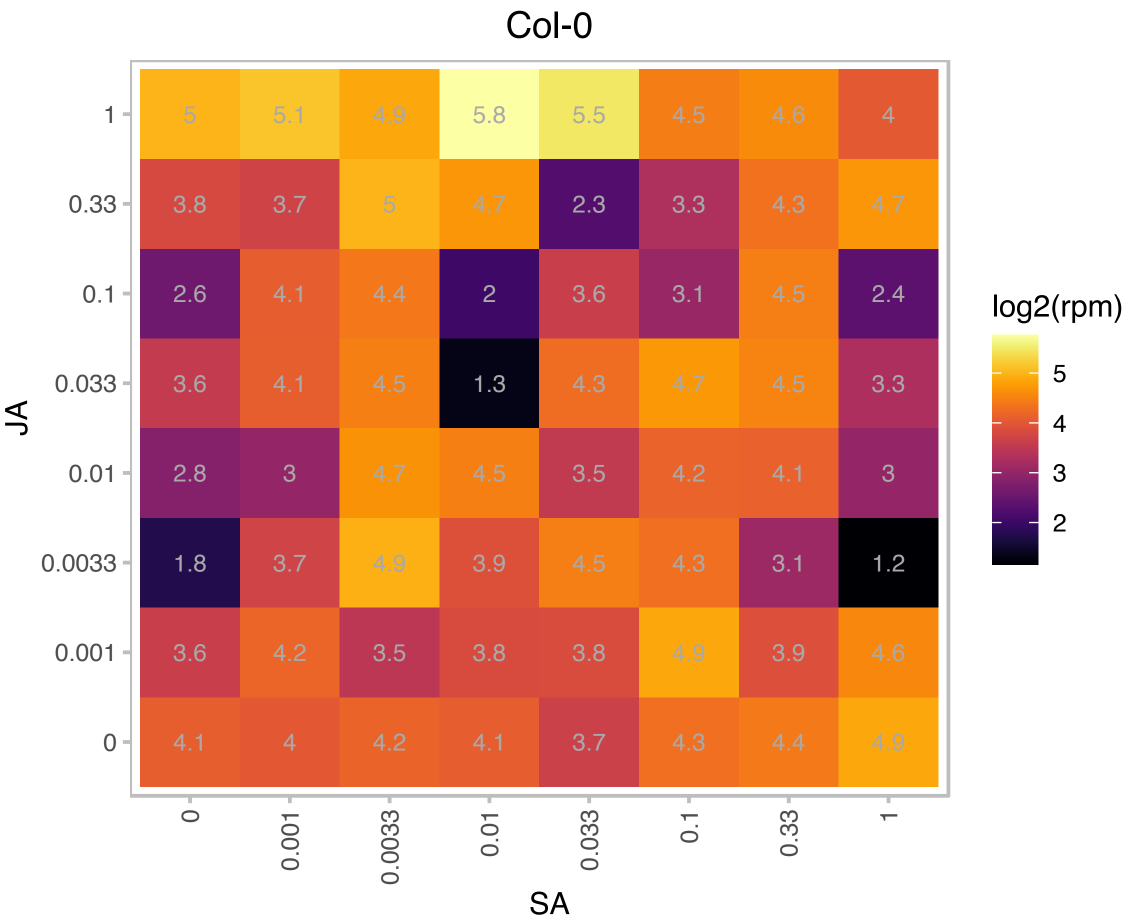


| | |
|---|---|
| Gene | AT2G17840 |
| is.expressed_genes | TRUE |
| is.responsive_genes | TRUE |
| is.apcluster | A-03 |

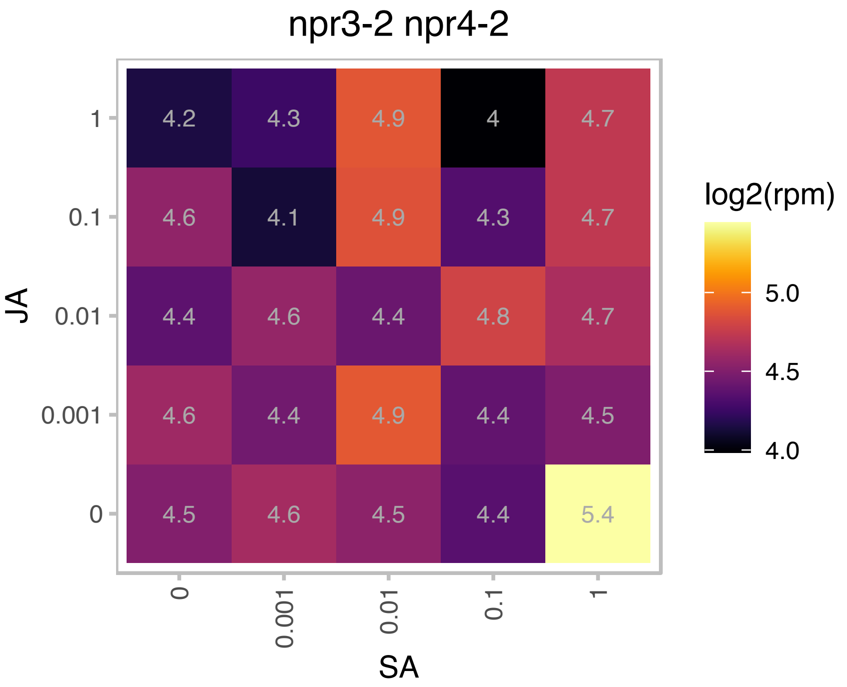


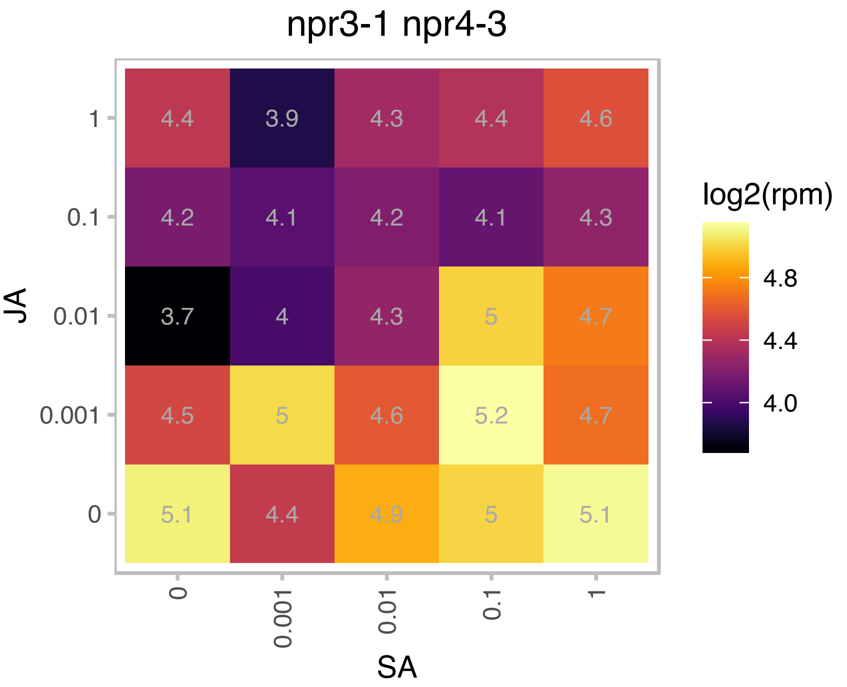

# AT2G22120.1 : NA

RING/FYVE/PHD zinc finger superfamily protein

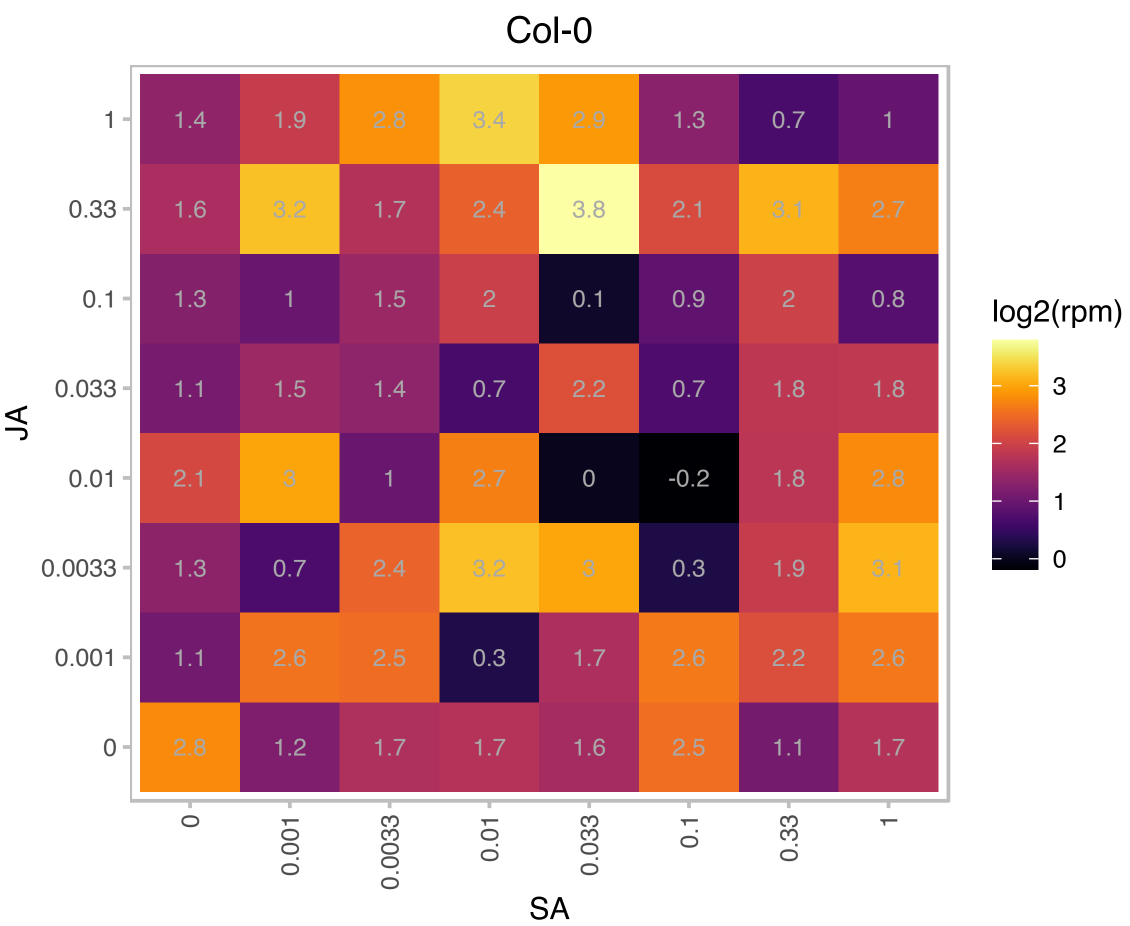


| | |
|---|---|
| Gene | AT2G22120 |
| is.expressed_genes | TRUE |
| is.responsive_genes | TRUE |
| is.apcluster | A-03 |

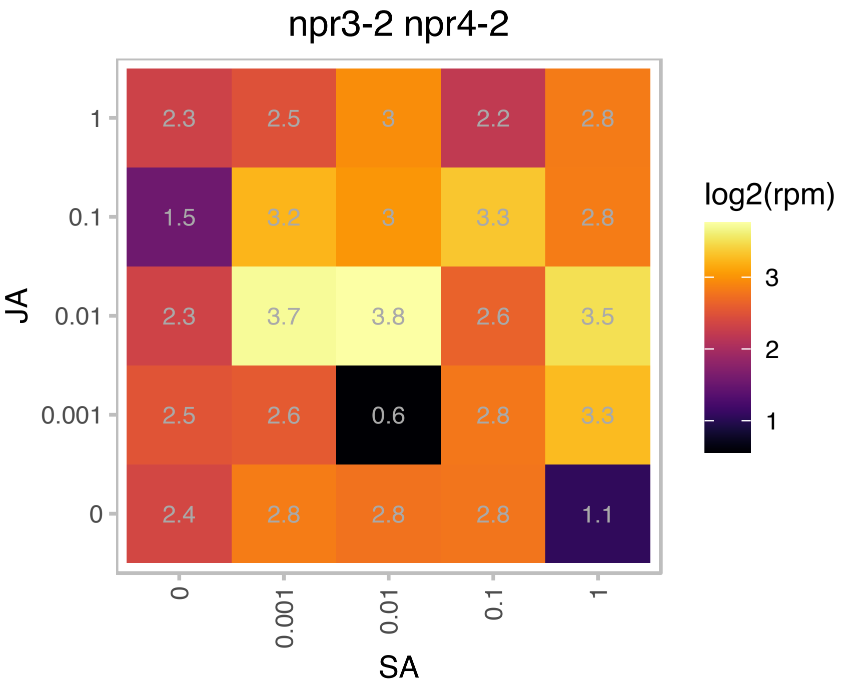


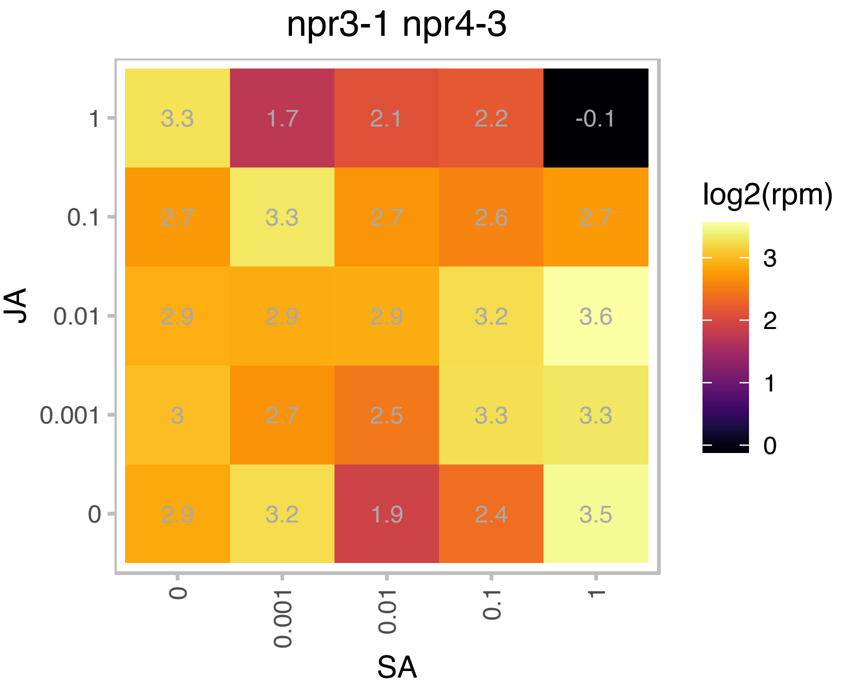

### AT2G40100.1 : LHCB4.3

light harvesting complex photosystem II

Lhcb4:3 protein (Lhcb4.3, light harvesting complex of photosystem II The mRNA is cell-to-cell mobile.

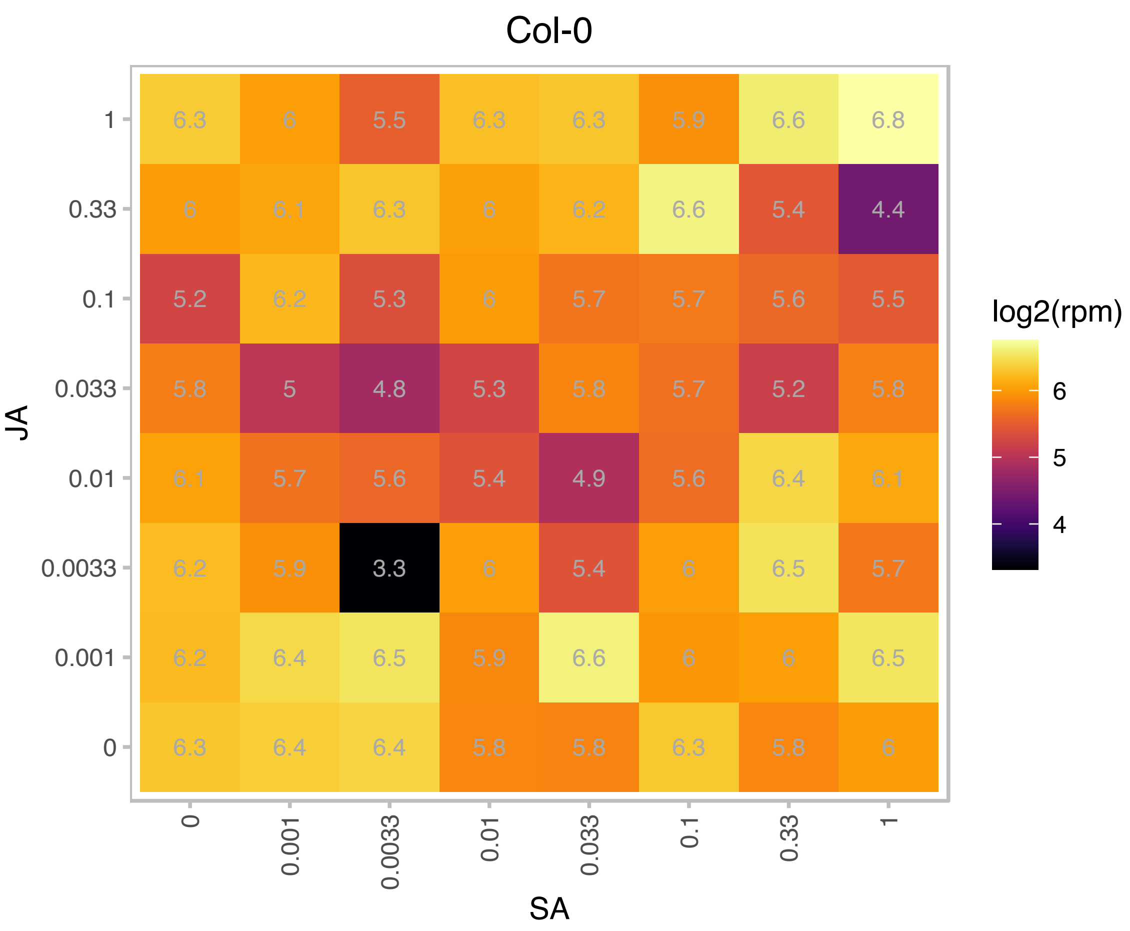


| | |
|---|---|
| Gene | AT2G40100 |
| is.expressed_genes | TRUE |
| is.responsive_genes | TRUE |
| is.apcluster | A-03 |

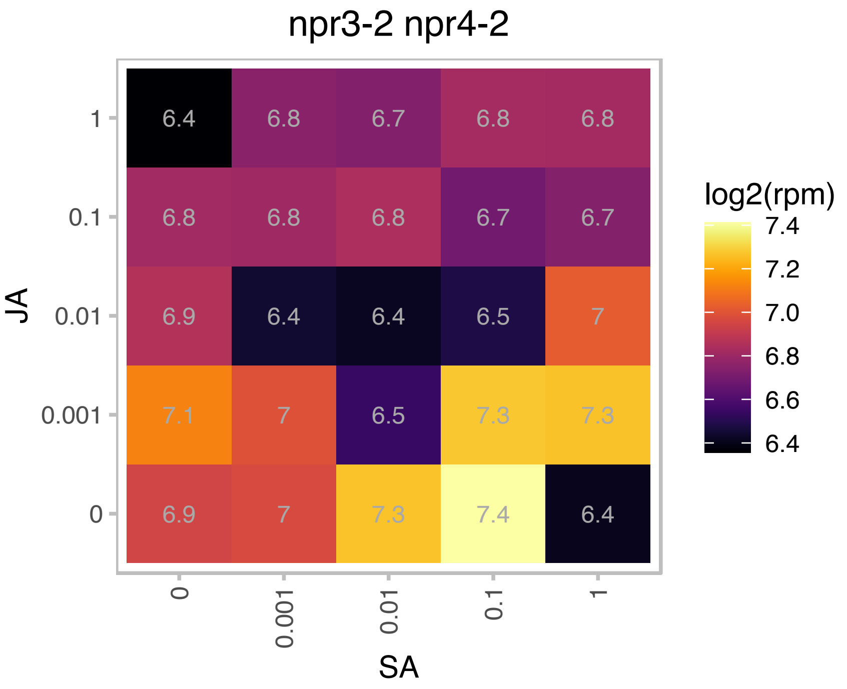


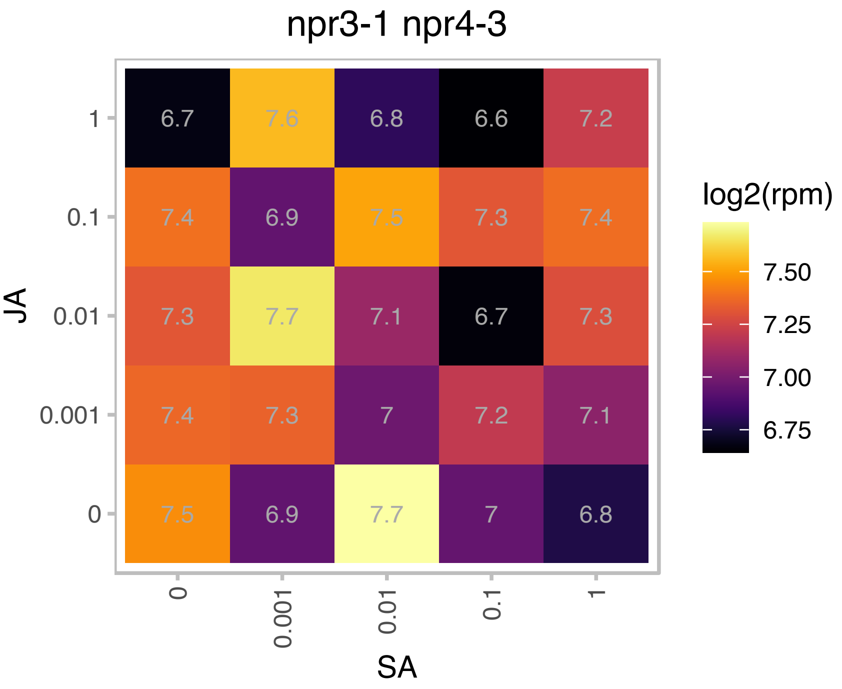

# AT3G03880.1 : NA

Protein of unknown function (DUF1639)

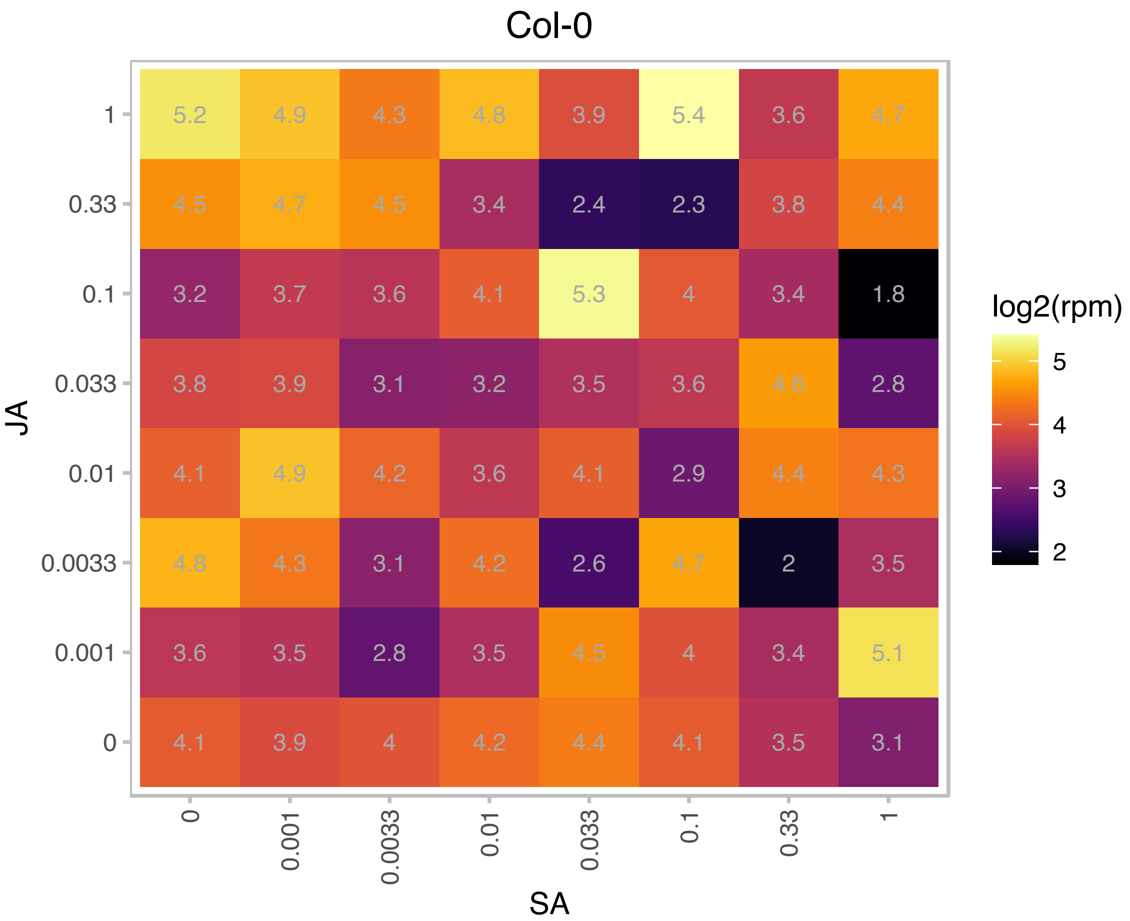


| | |
|---|---|
| Gene | AT3G03880 |
| is.expressed_genes | TRUE |
| is.responsive_genes | TRUE |
| is.apcluster | A-03 |

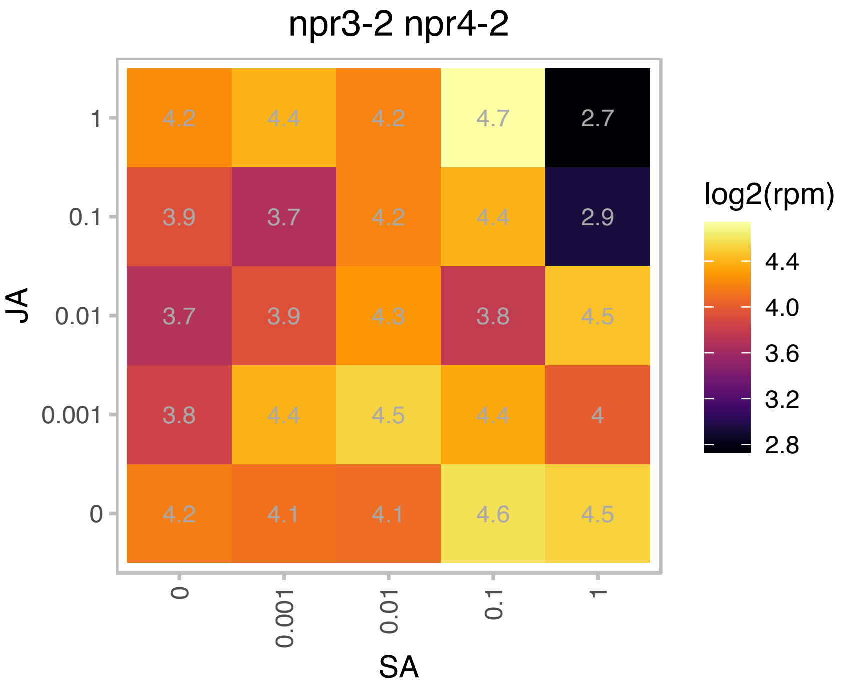


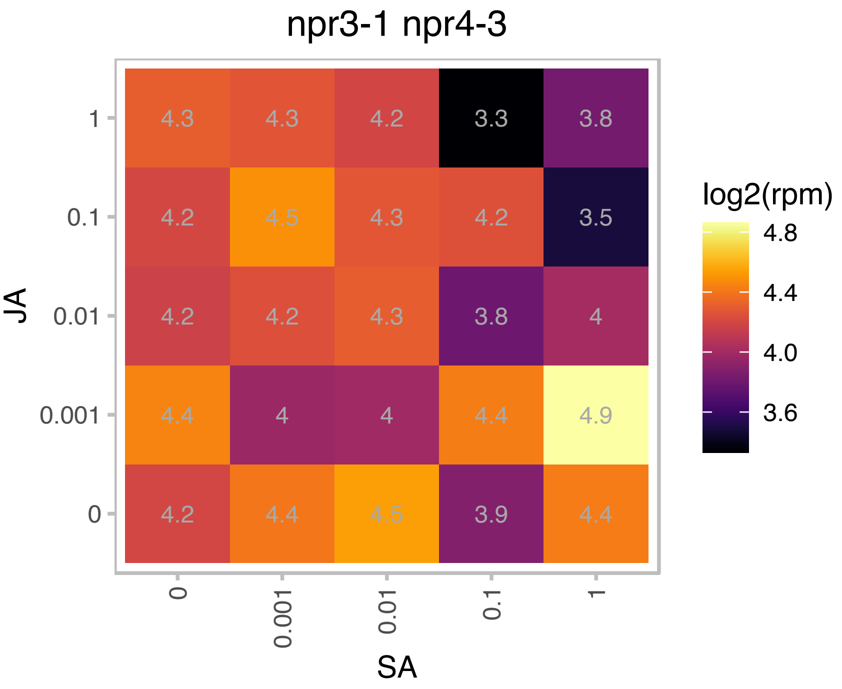

### AT3G22840.1 : ELIP, ELIP1

Chlorophyll A-B binding family protein

Encodes an early light-inducible protein.

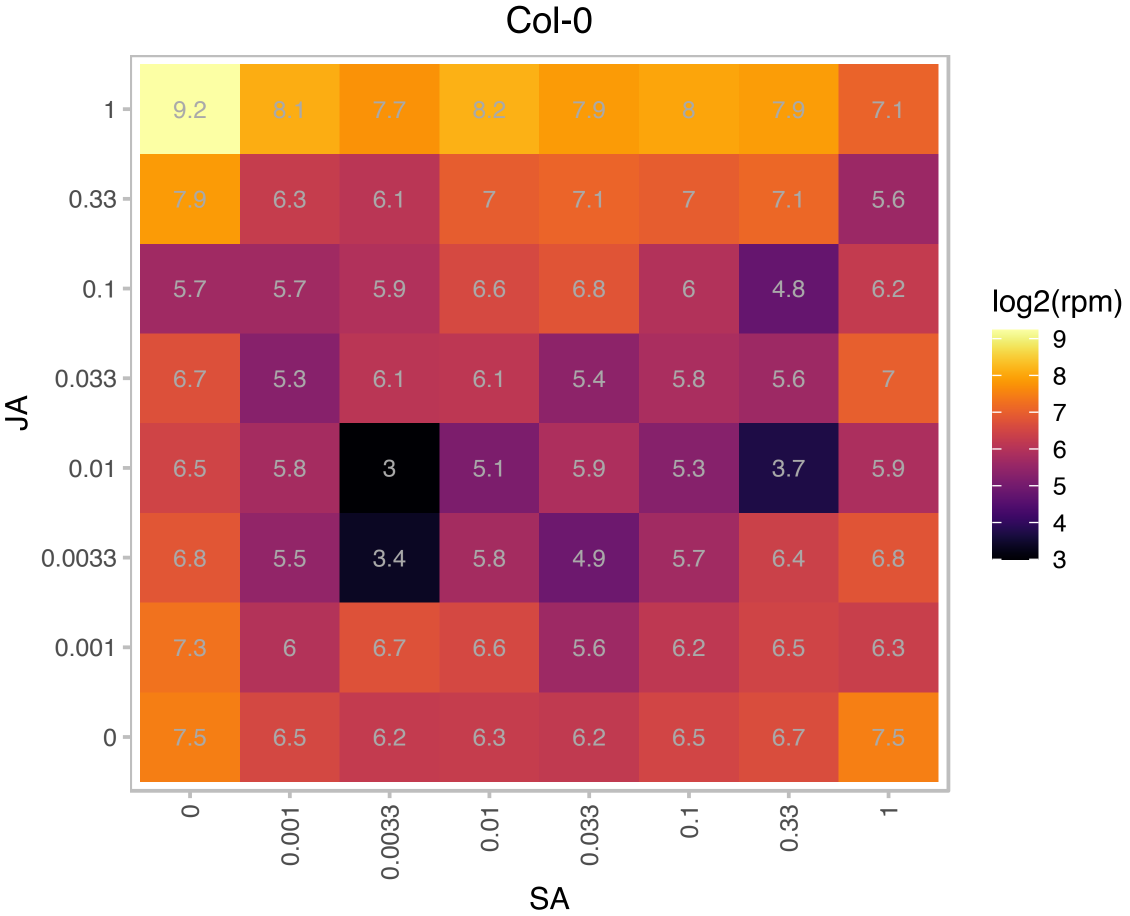


| | |
|---|---|
| Gene | AT3G22840 |
| is.expressed_genes | TRUE |
| is.responsive_genes | TRUE |
| is.apcluster | A-03 |

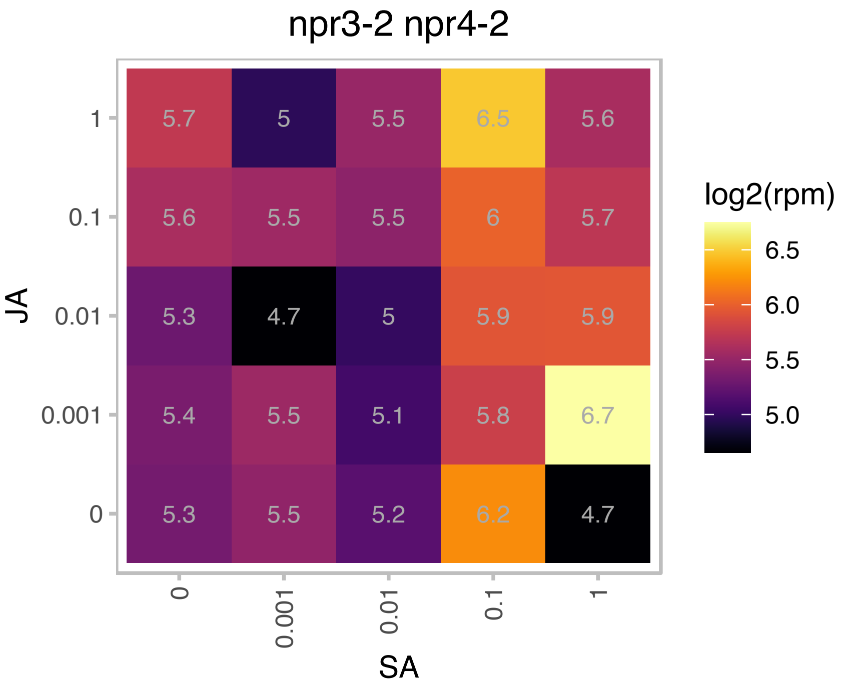


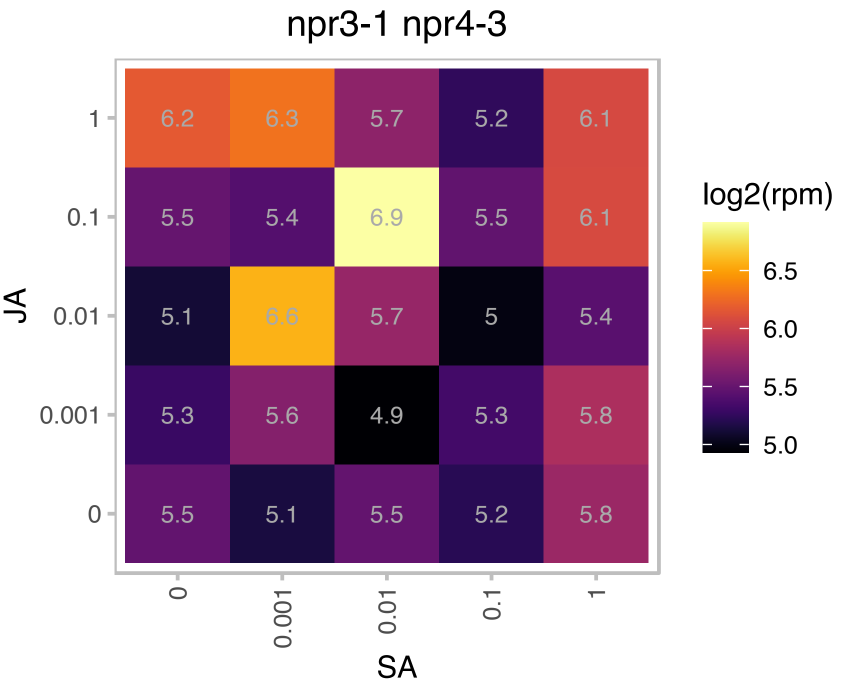

### AT3G27820.1 : ATMDAR4, MDAR4

monodehydroascorbate reductase 4

Encodes a peroxisome membrane-bound monodehydroascorbate reductase, involved in the ascorbate-glutathione cycle which removes toxic H2O2

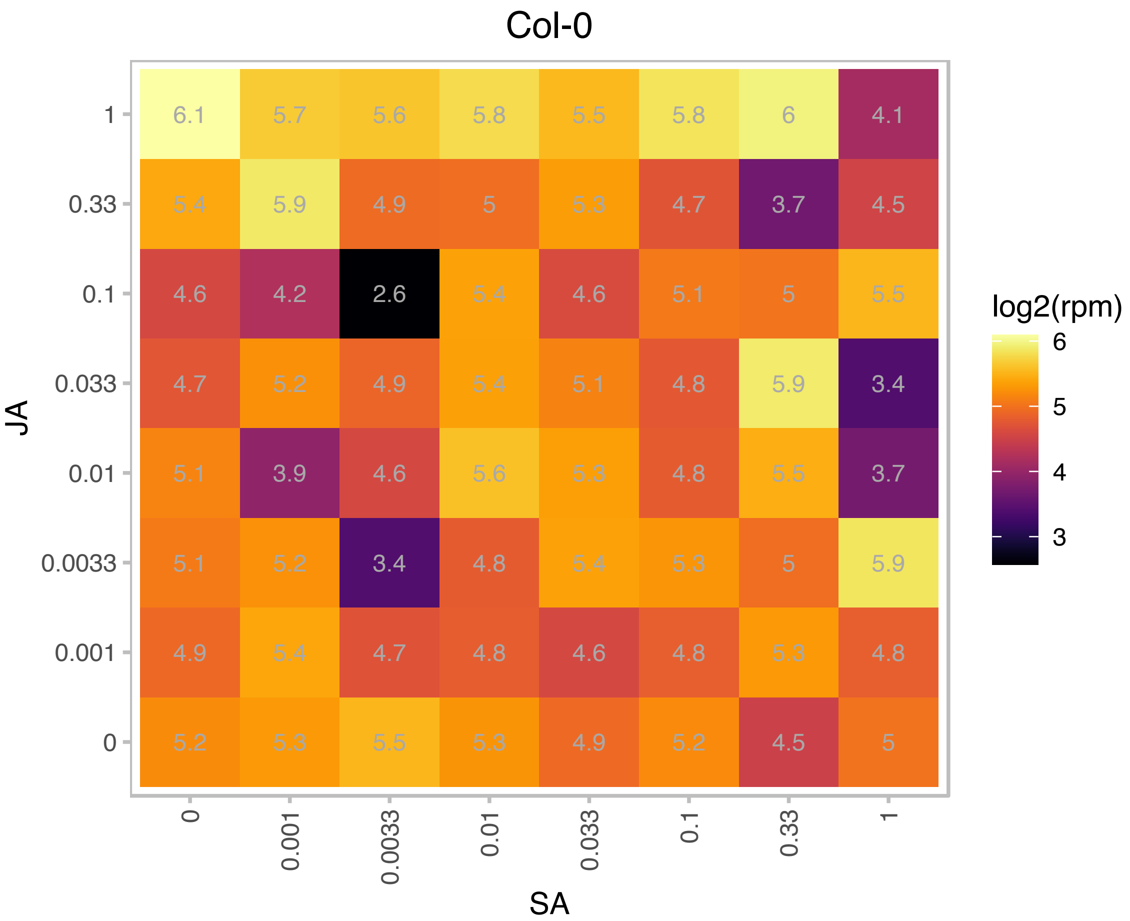


| | |
|---|---|
| Gene | AT3G27820 |
| is.expressed_genes | TRUE |
| is.responsive_genes | TRUE |
| is.apcluster | A-03 |

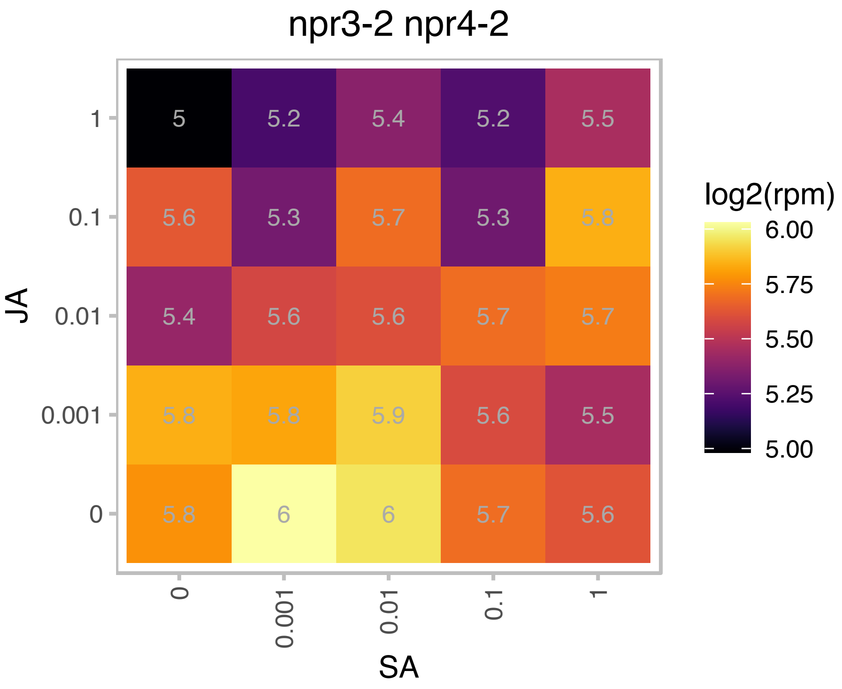


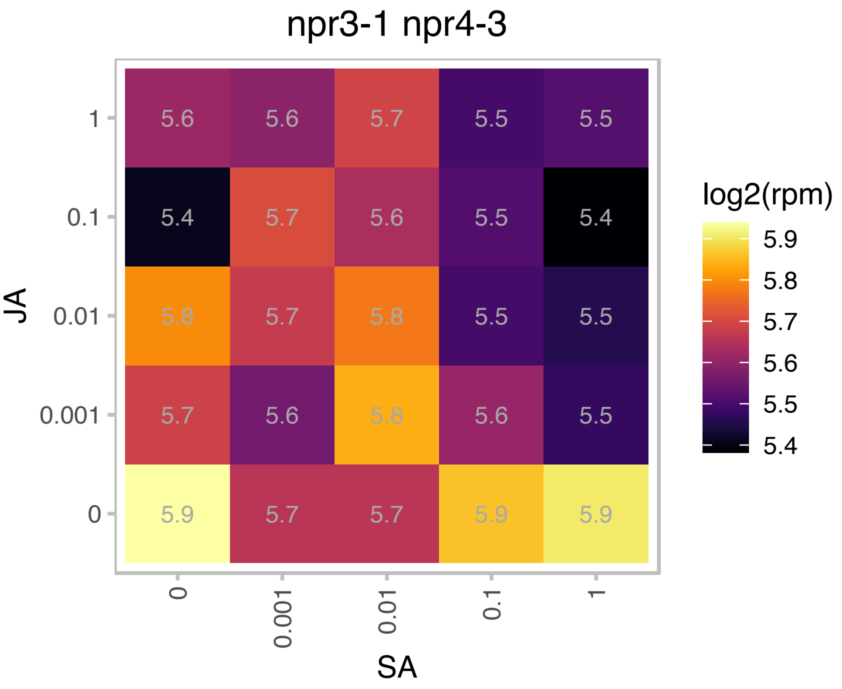

# AT3G52710.1 : NA

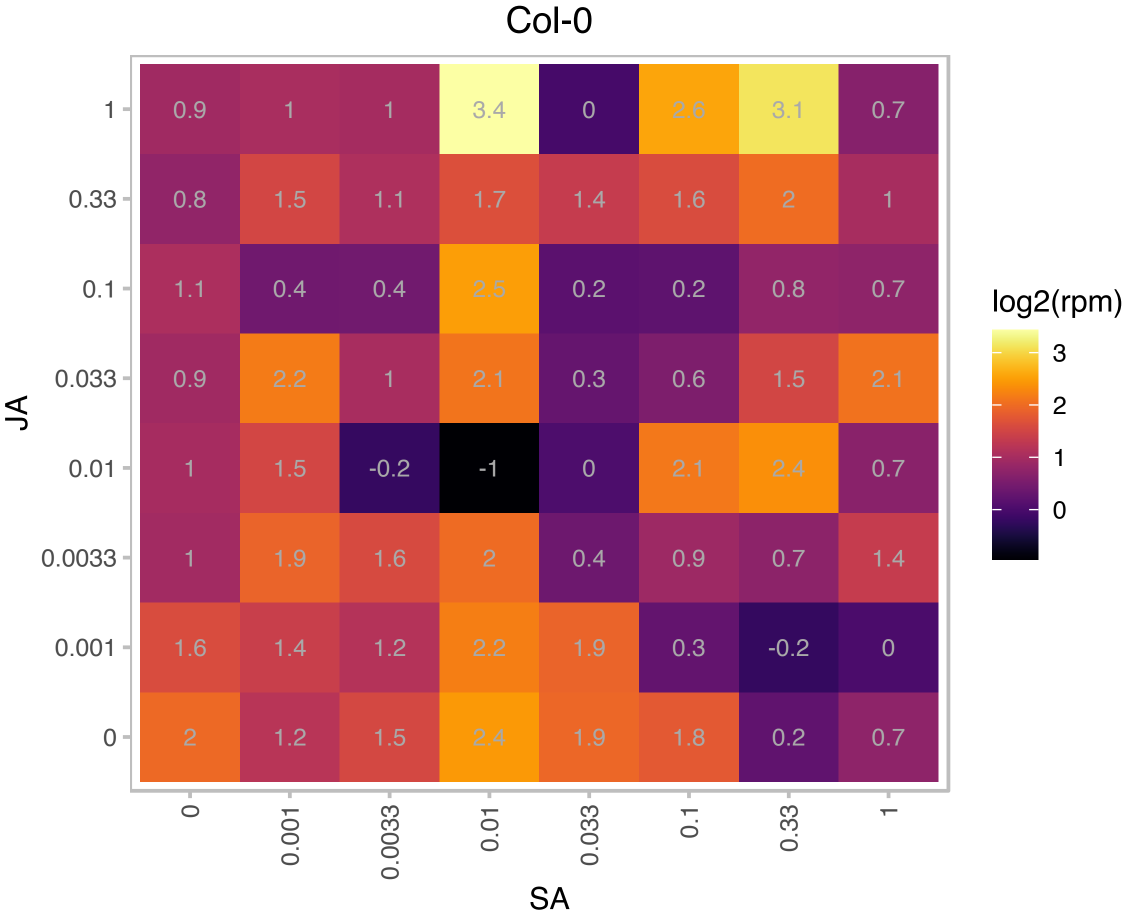


| | |
|---|---|
| Gene | AT3G52710 |
| is.expressed_genes | TRUE |
| is.responsive_genes | TRUE |
| is.apcluster | A-03 |

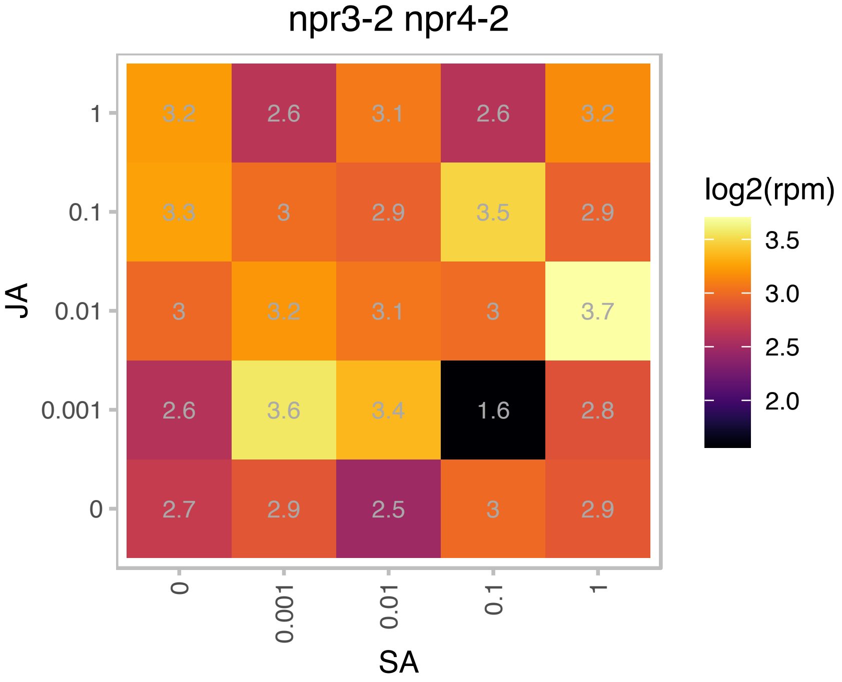


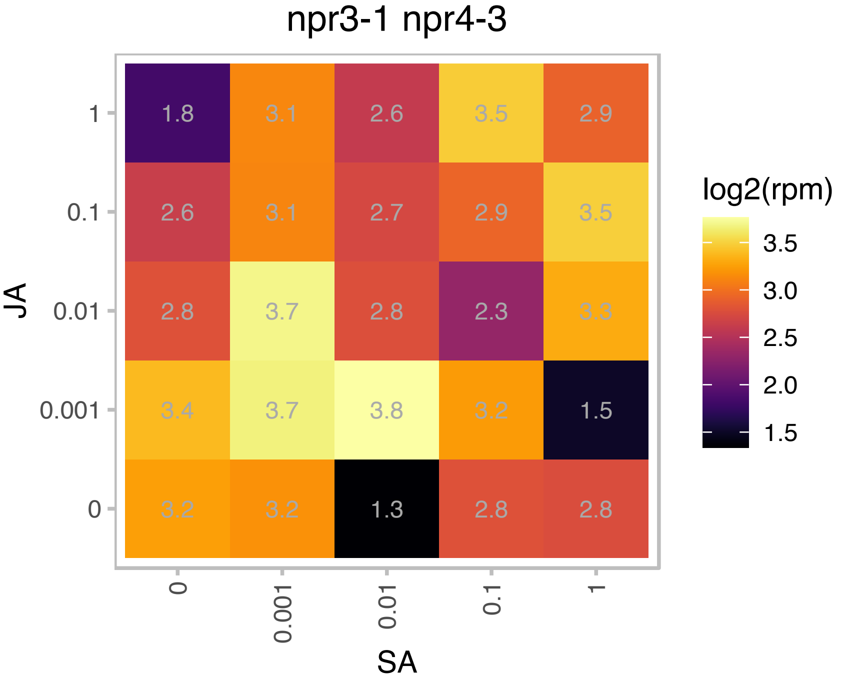

### AT3G62150.1 : ABCB21, PGP21

P-glycoprotein 21

Encodes a facultative transporter controlling auxin concentrations in plant cells.

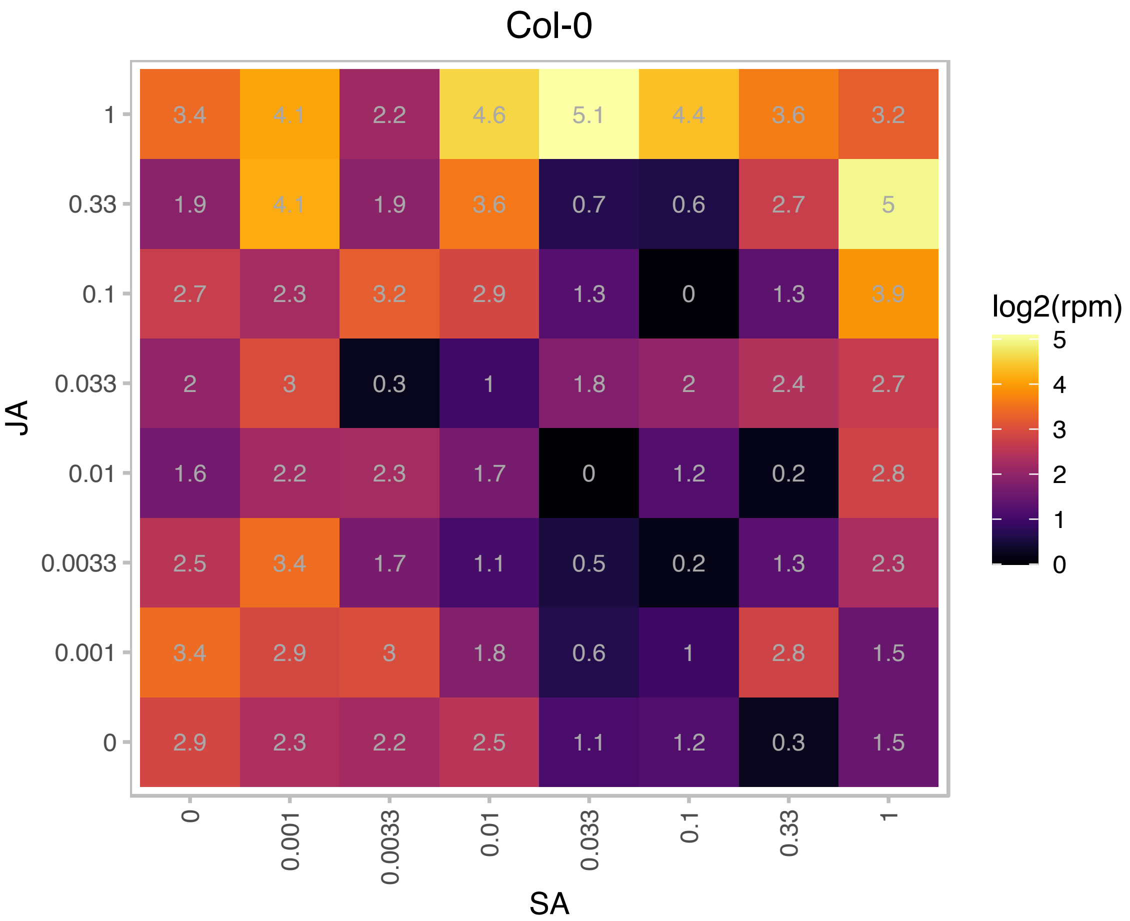


| | |
|---|---|
| Gene | AT3G62150 |
| is.expressed_genes | TRUE |
| is.responsive_genes | TRUE |
| is.apcluster | A-03 |

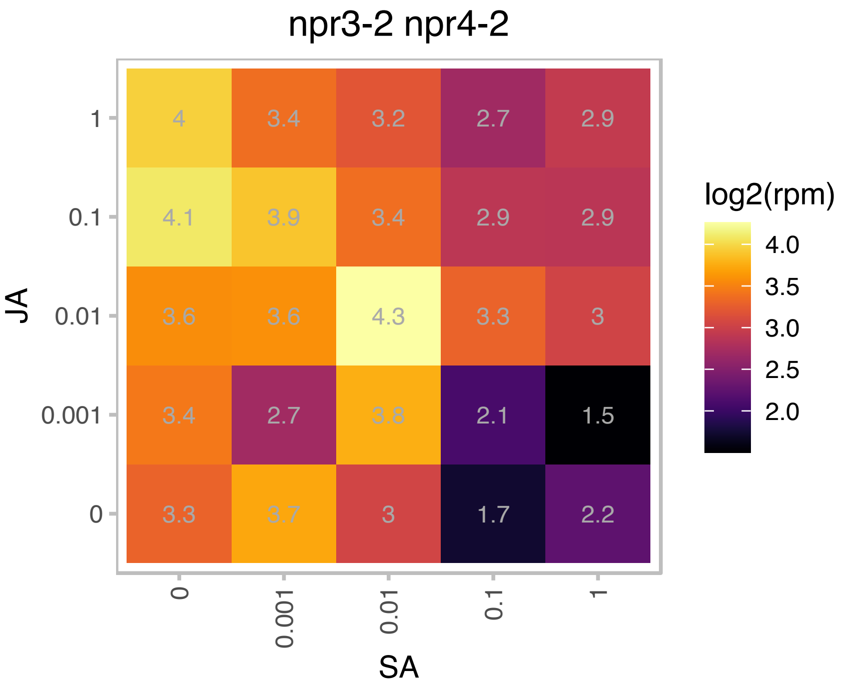


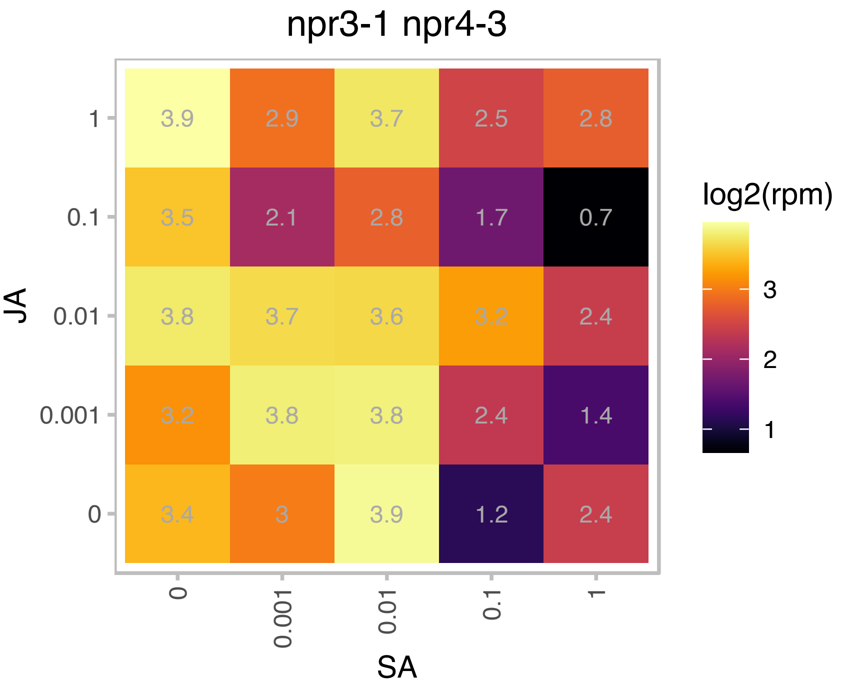

### AT4G02530.1 : MPH2

chloroplast thylakoid lumen protein

MPH2 is a green lineage-specific thylakoid lumen protein required for photosynthetic acclimation of PSII to stressful light conditions (PMID:28874535).

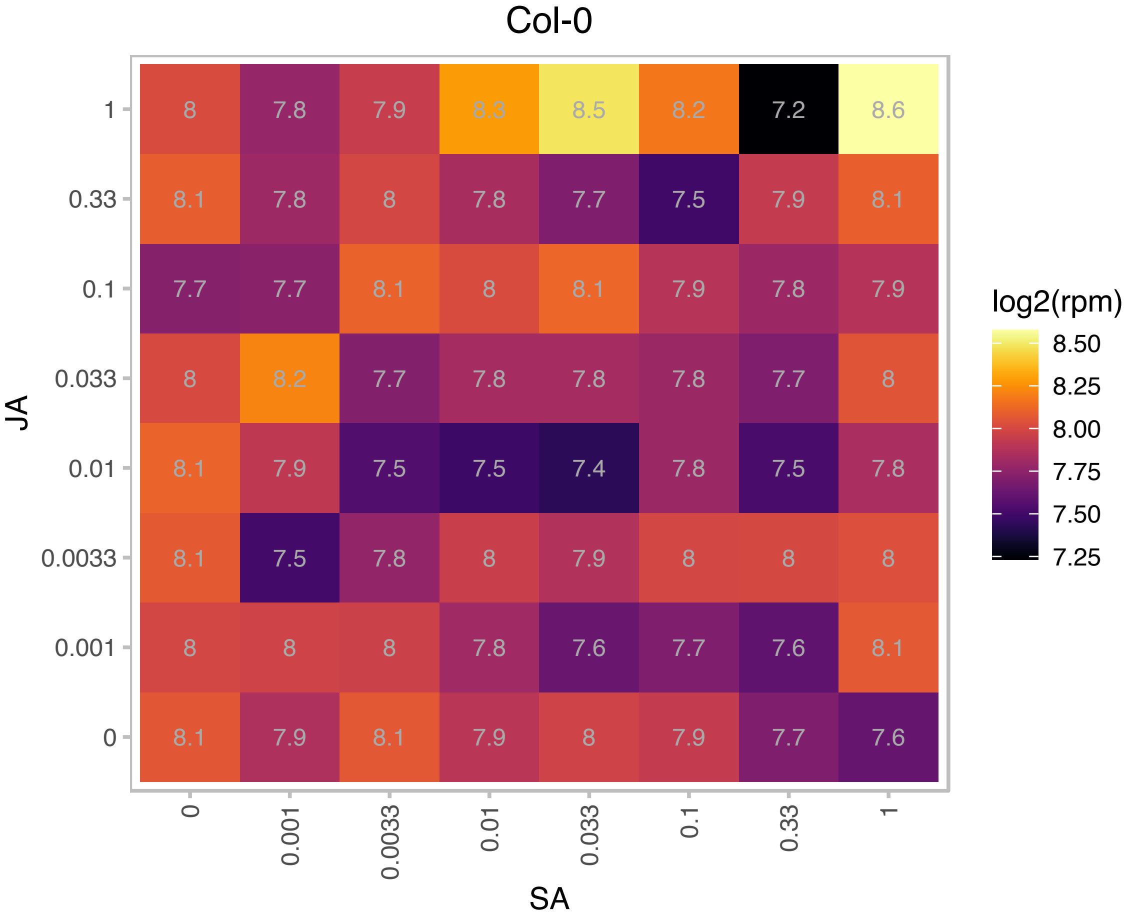


| | |
|---|---|
| Gene | AT4G02530 |
| is.expressed_genes | TRUE |
| is.responsive_genes | TRUE |
| is.apcluster | A-03 |

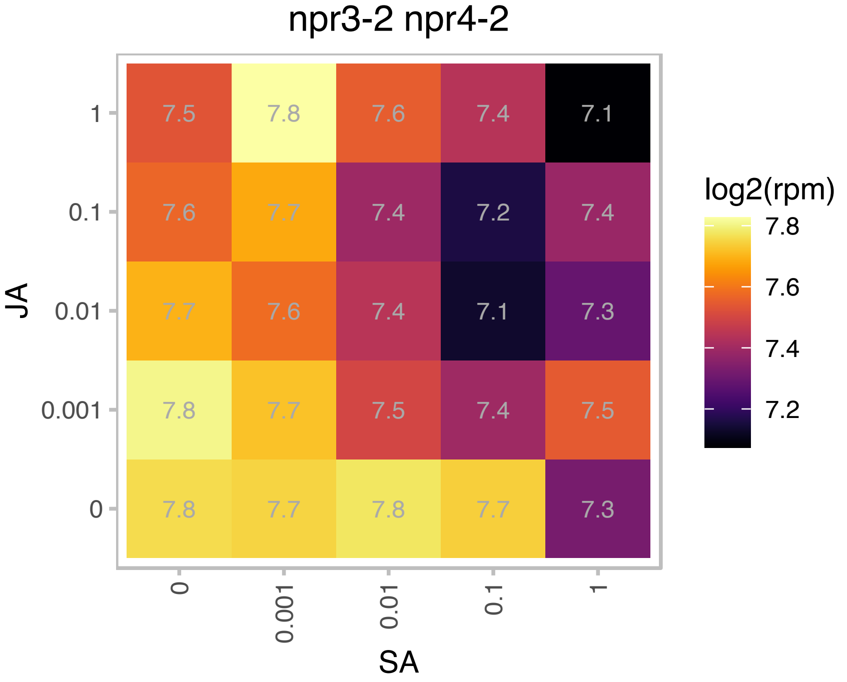


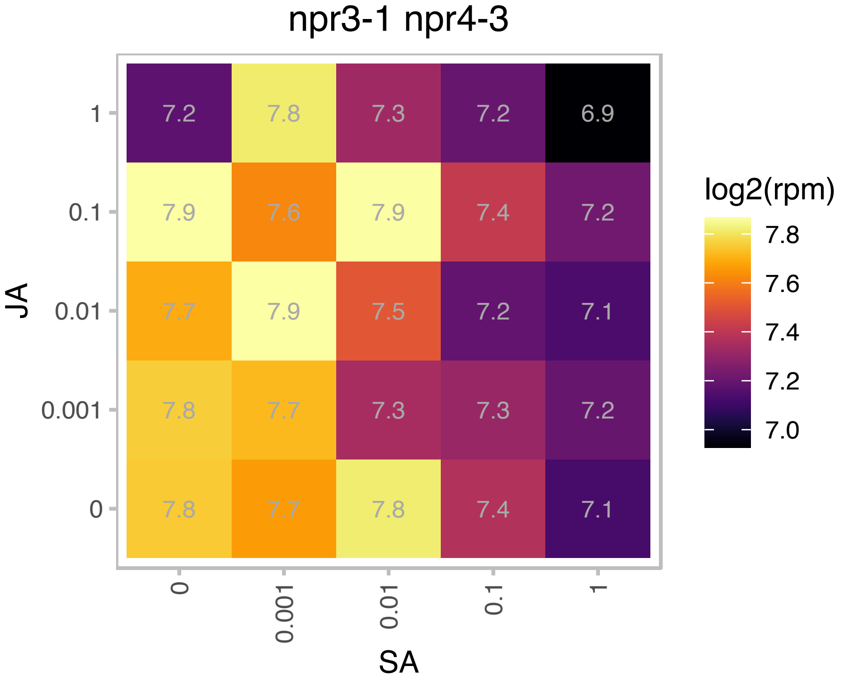

# AT4G02580.1 : NA

NADH-ubiquinone oxidoreductase 24 kDa subunit, putative

Col-0

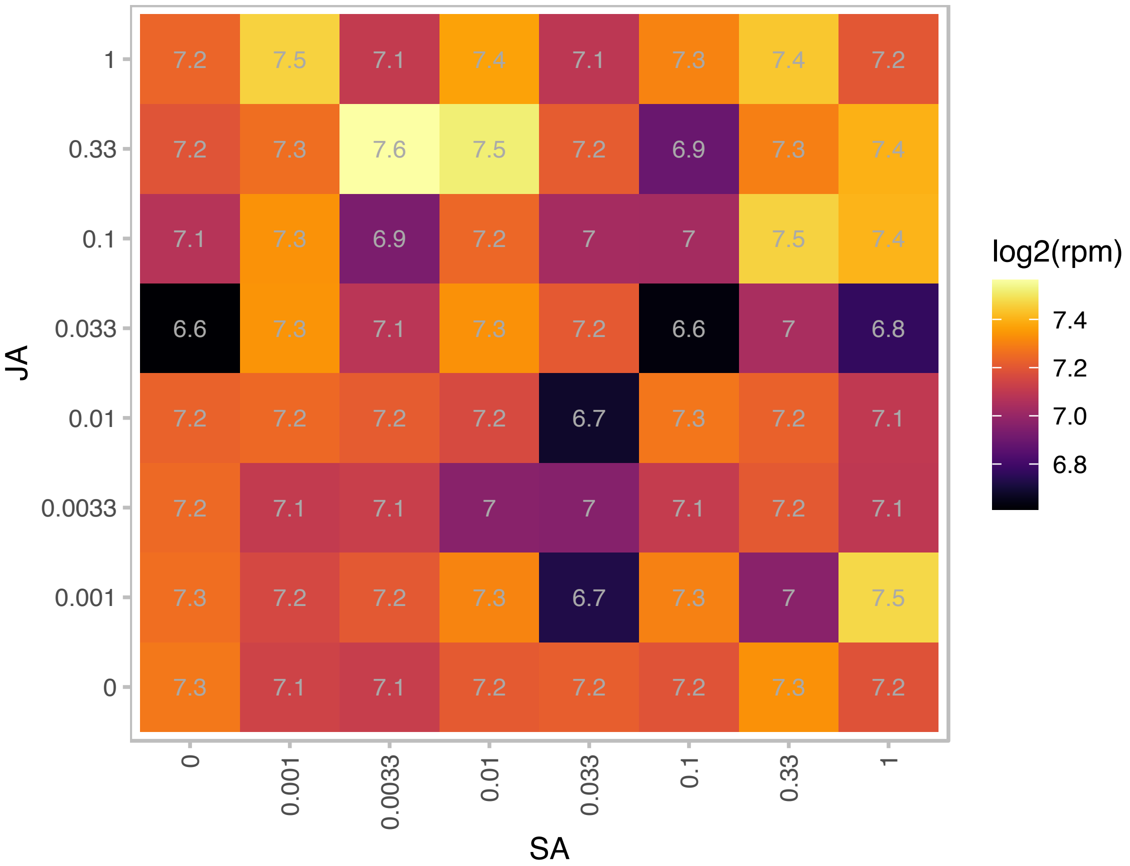


| | |
|---|---|
| Gene | AT4G02580 |
| is.expressed_genes | TRUE |
| is.responsive_genes | TRUE |
| is.apcluster | A-03 |

npr3-2 npr4-2

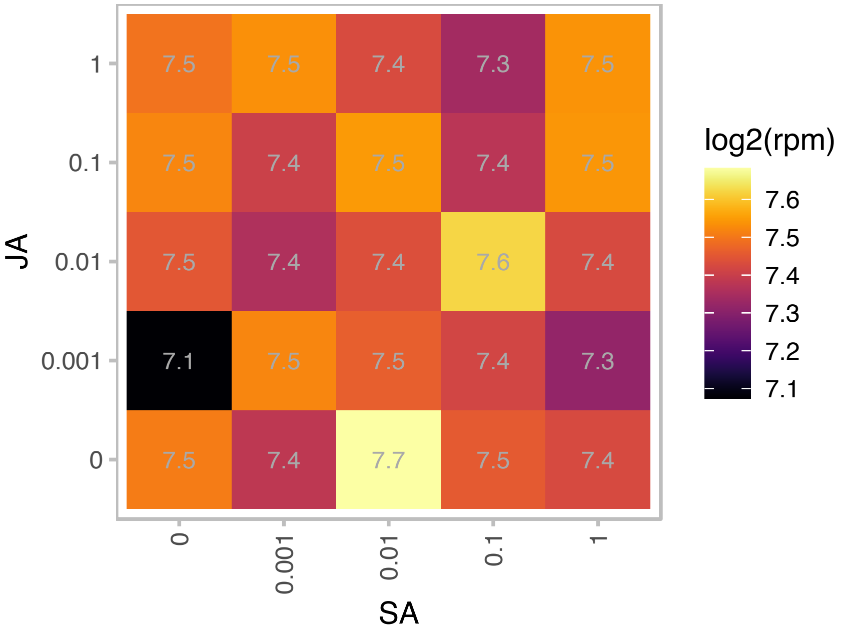


npr3-1 npr4-3

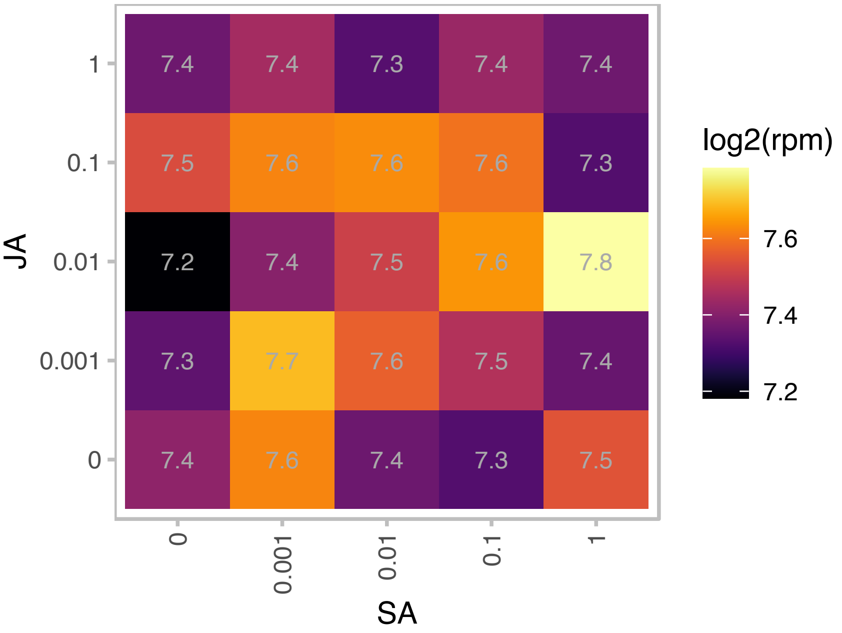

### AT4G20360.1 : ATRAB8D, ATRABE1B, RABE1b

RAB GTPase homolog E1B

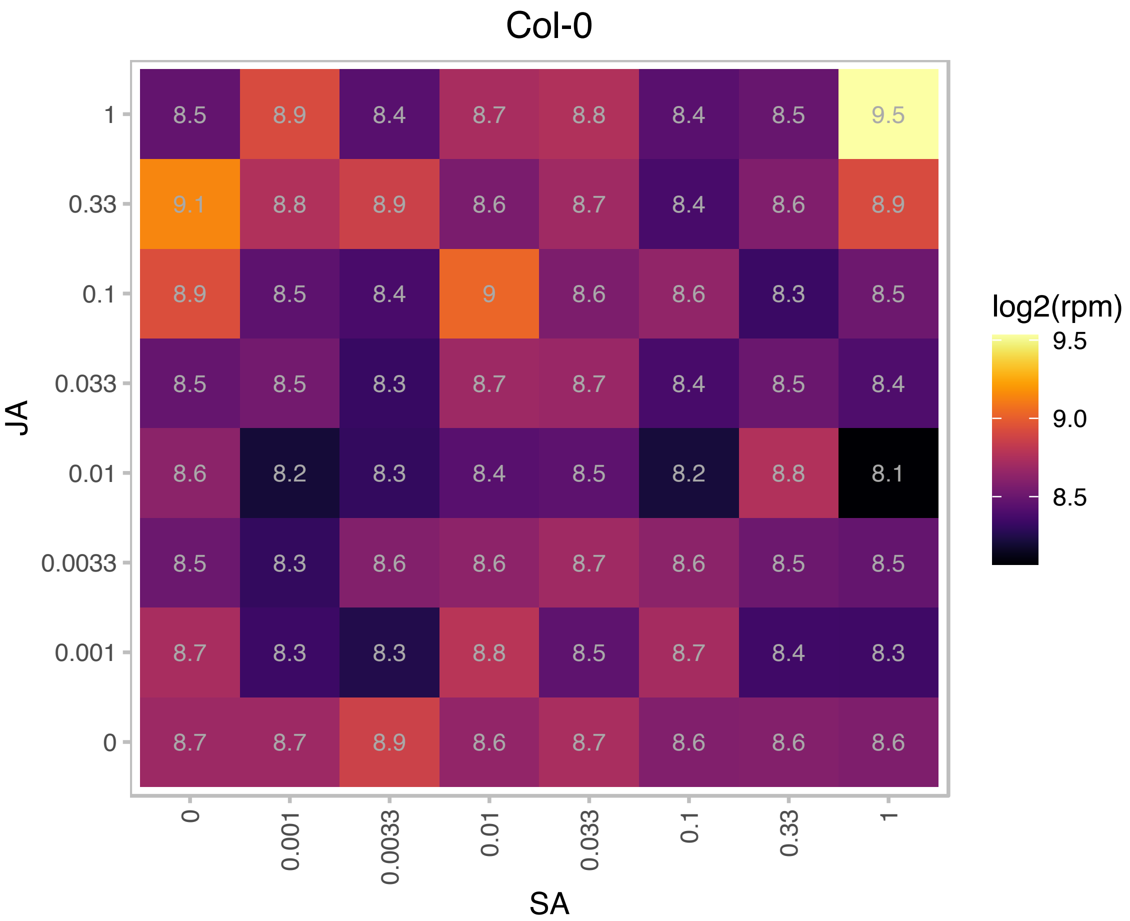


| | |
|---|---|
| Gene | AT4G20360 |
| is.expressed_genes | TRUE |
| is.responsive_genes | TRUE |
| is.apcluster | A-03 |

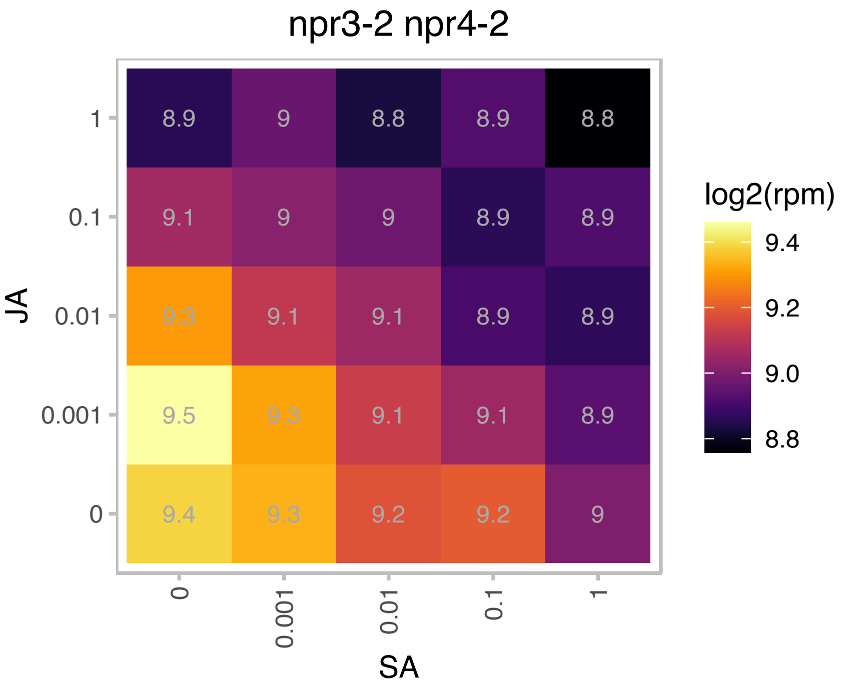


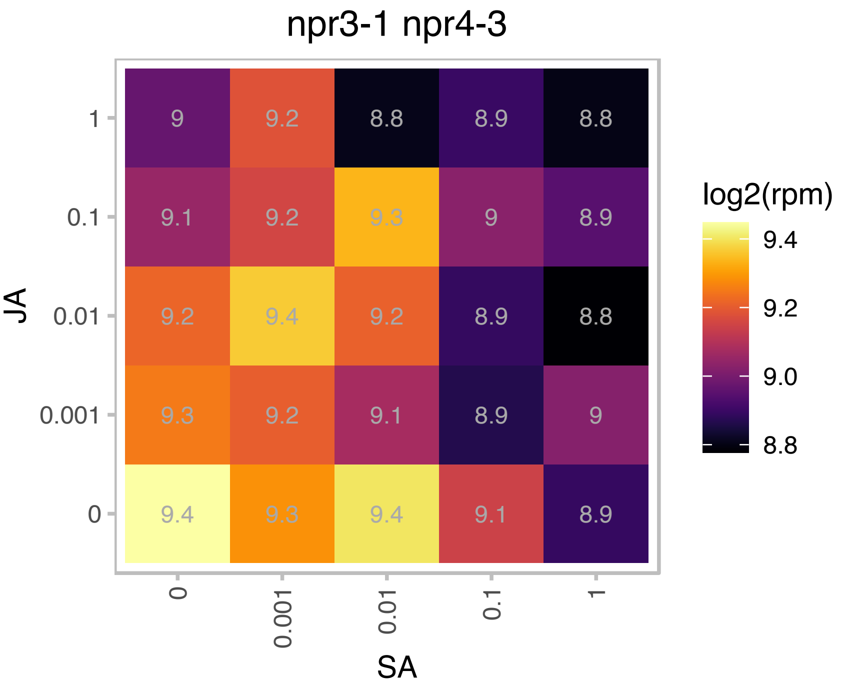

# AT4G22320.1 : NA

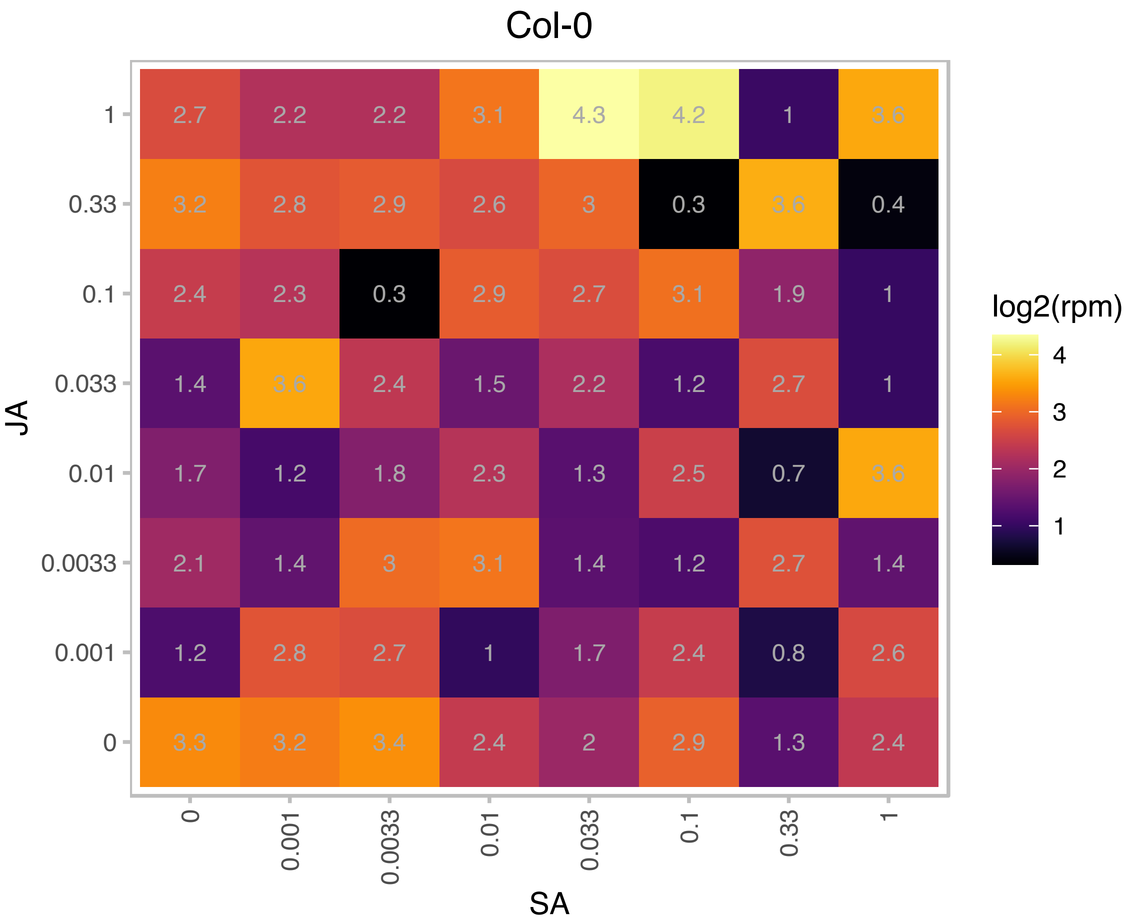


| | |
|---|---|
| Gene | AT4G22320 |
| is.expressed_genes | TRUE |
| is.responsive_genes | TRUE |
| is.apcluster | A-03 |

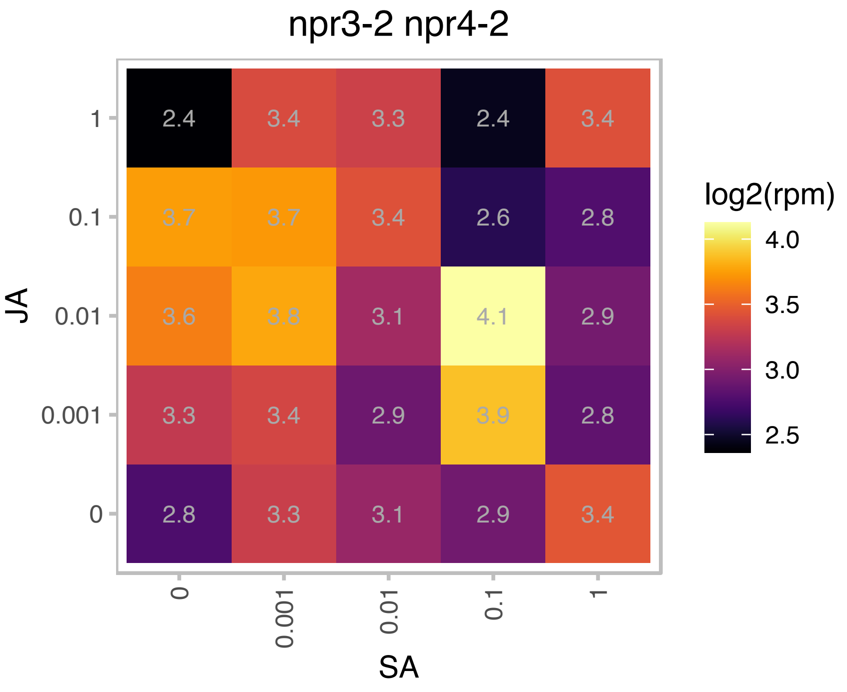


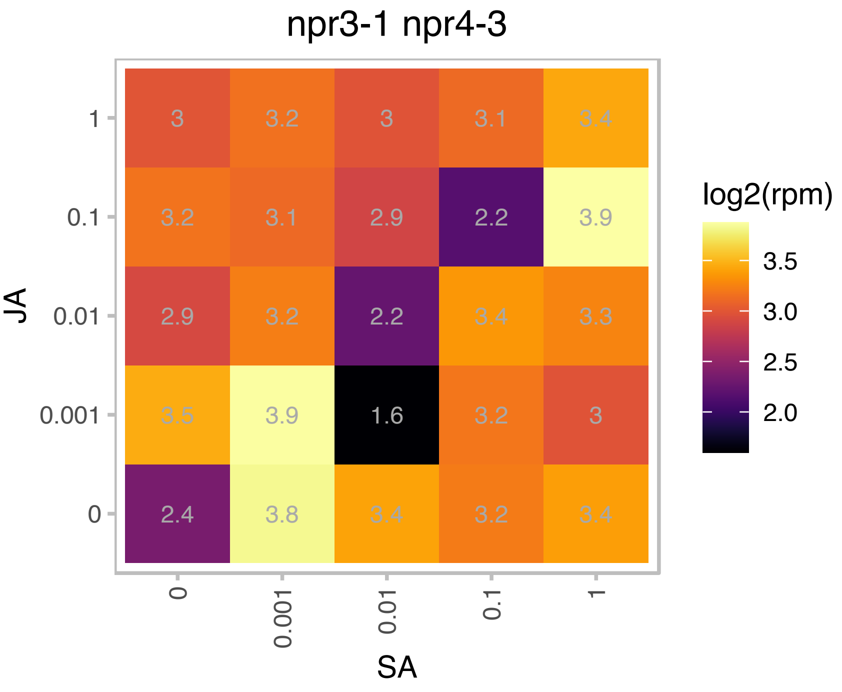

### AT4G36690.1 : ATU2AF65A

U2 snRNP auxilliary factor, large subunit, splicing factor

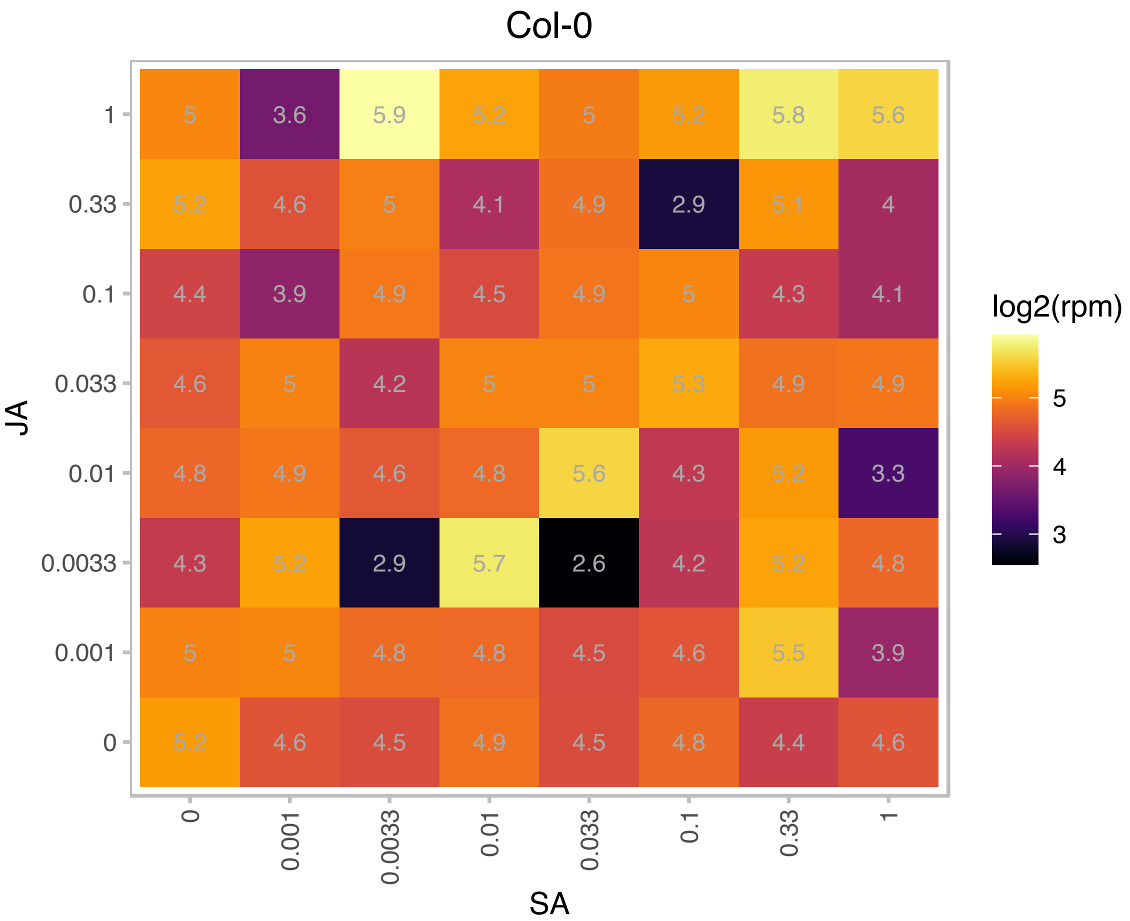


| | |
|---|---|
| Gene | AT4G36690 |
| is.expressed_genes | TRUE |
| is.responsive_genes | TRUE |
| is.apcluster | A-03 |

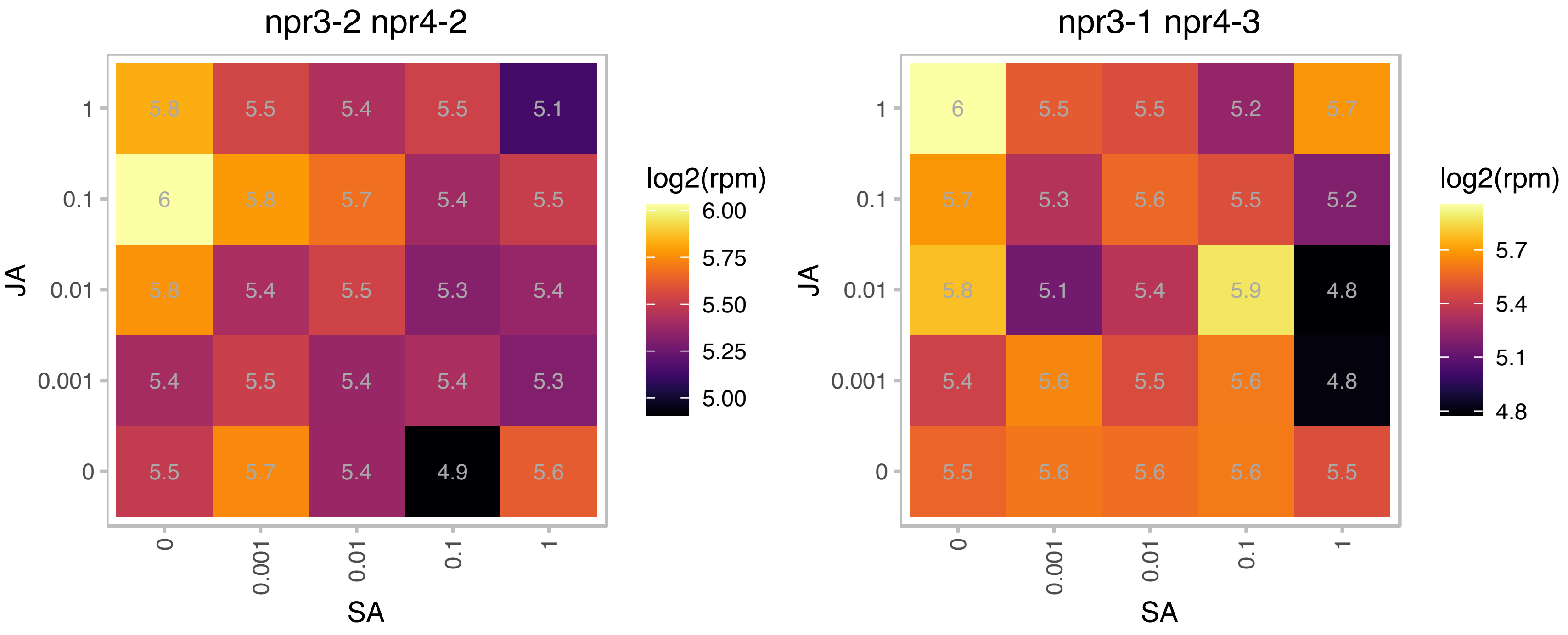

### AT5G05080.1 : ATUBC22, UBC22

ubiquitin-conjugating enzyme 22

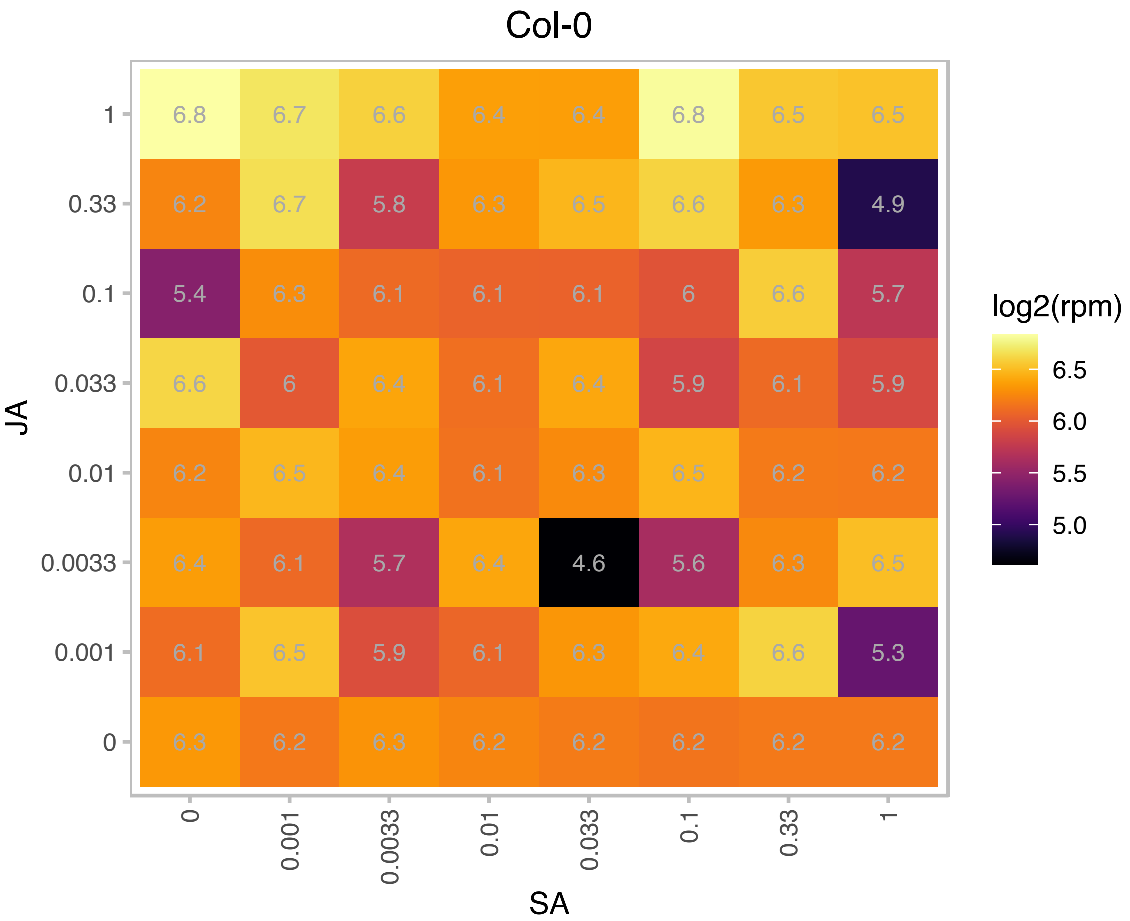


| | |
|---|---|
| Gene | AT5G05080 |
| is.expressed_genes | TRUE |
| is.responsive_genes | TRUE |
| is.apcluster | A-03 |

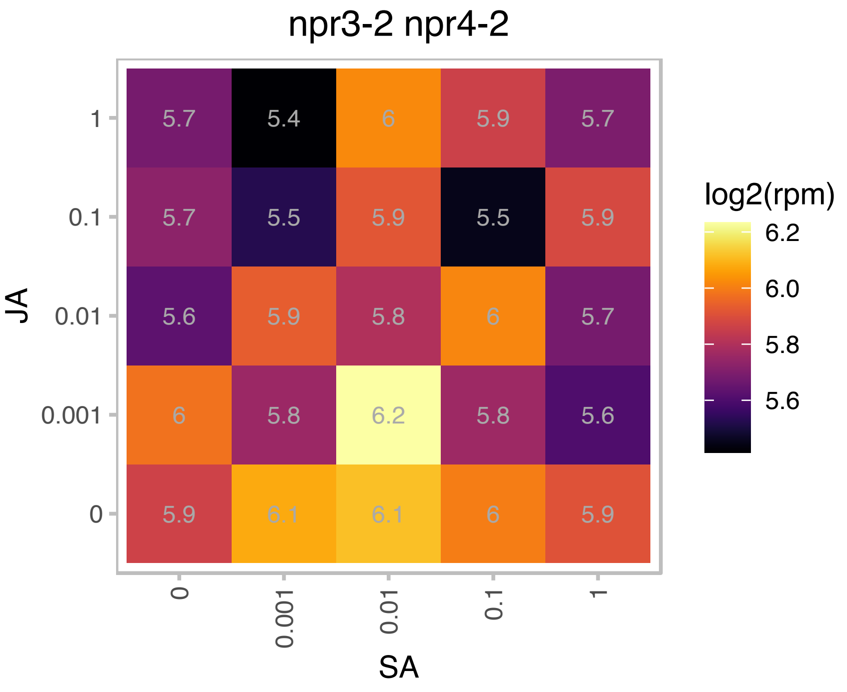


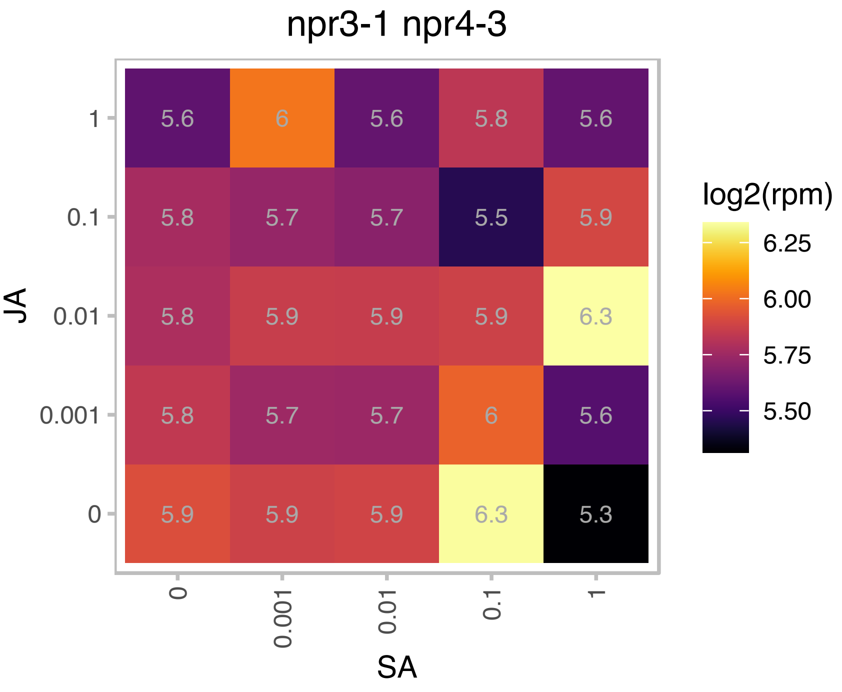

# AT5G08650.1 : NA

Small GTP-binding protein

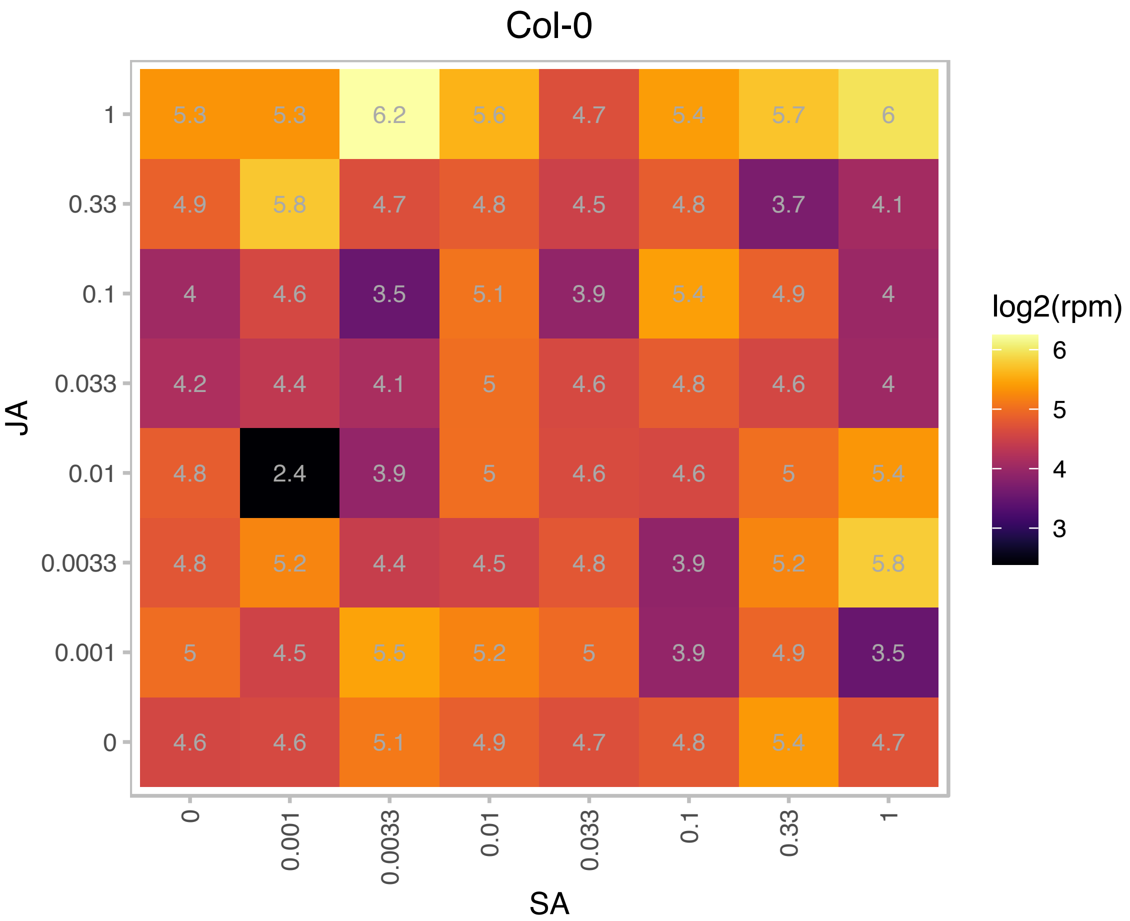


| | |
|---|---|
| Gene | AT5G08650 |
| is.expressed_genes | TRUE |
| is.responsive_genes | TRUE |
| is.apcluster | A-03 |

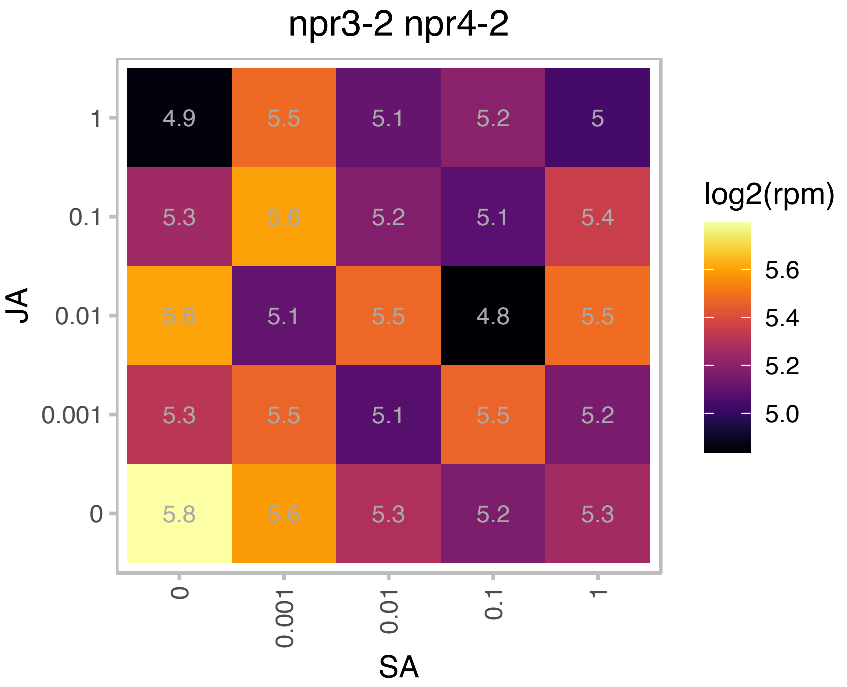


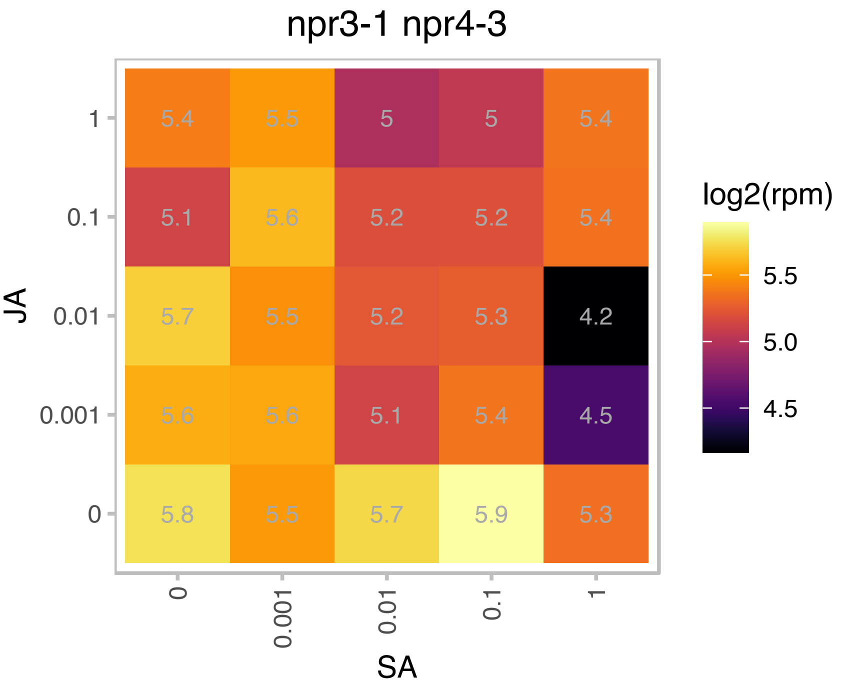

### AT5G09230.1 : AtSRT2, SRT2

sirtuin 2

Encodes SRT2, a member of the SIR2 (sirtuin) family HDAC (histone deacetylase) (SRT1/AT5g55760, SRT2/AT5G09230).

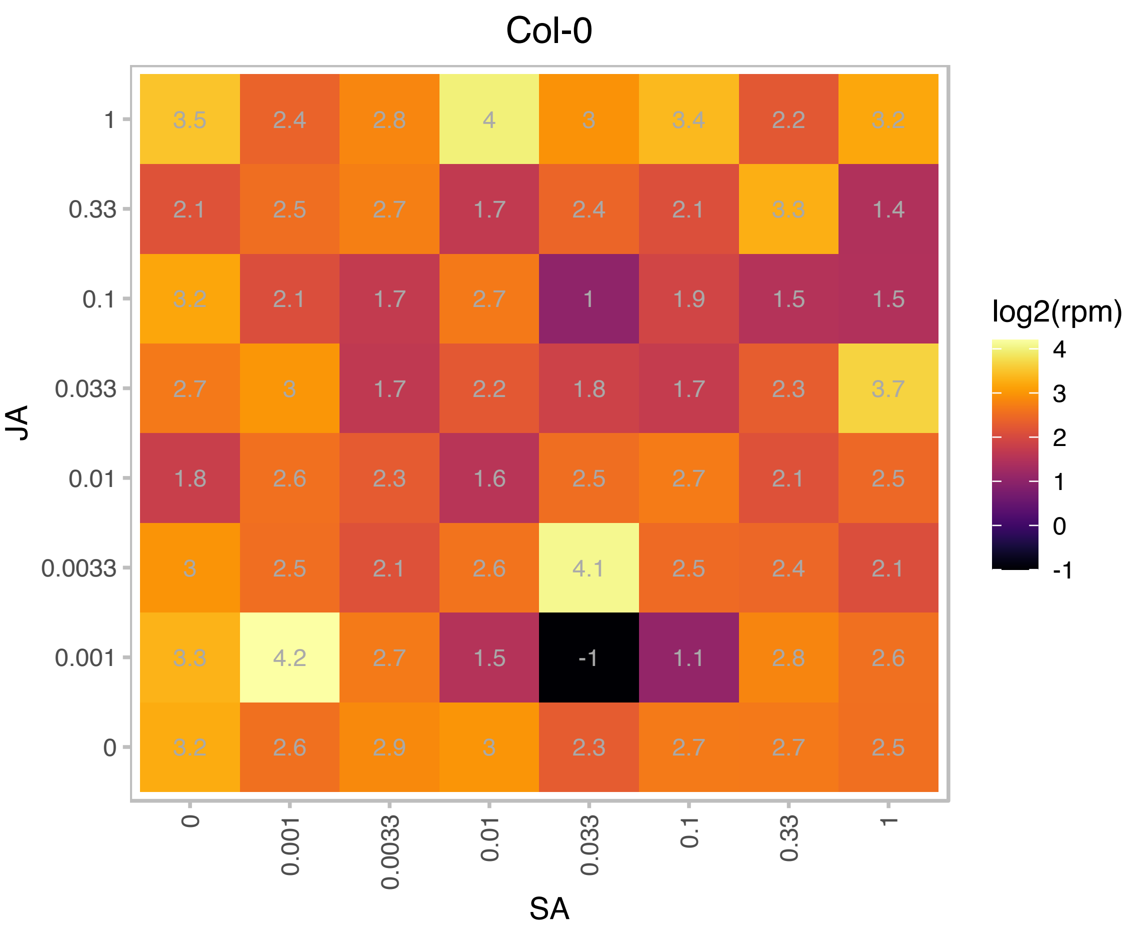


| | |
|---|---|
| Gene | AT5G09230 |
| is.expressed_genes | TRUE |
| is.responsive_genes | TRUE |
| is.apcluster | A-03 |

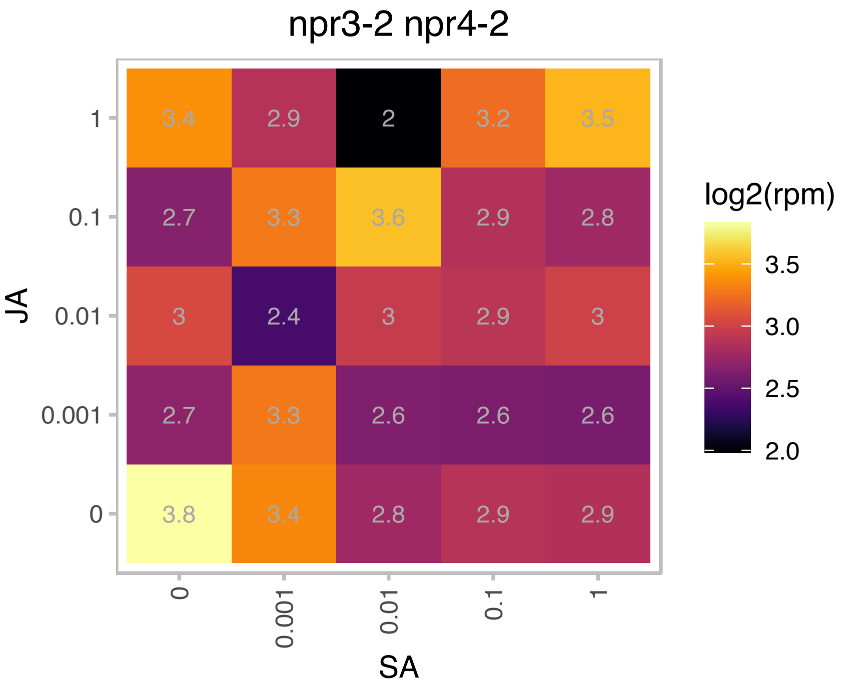


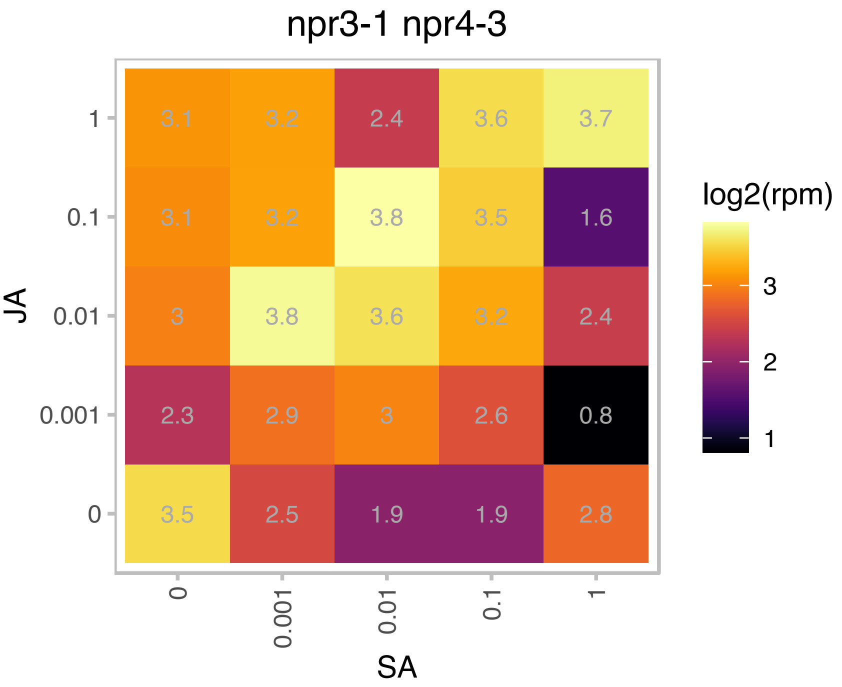

# AT5G11680.1 : NA

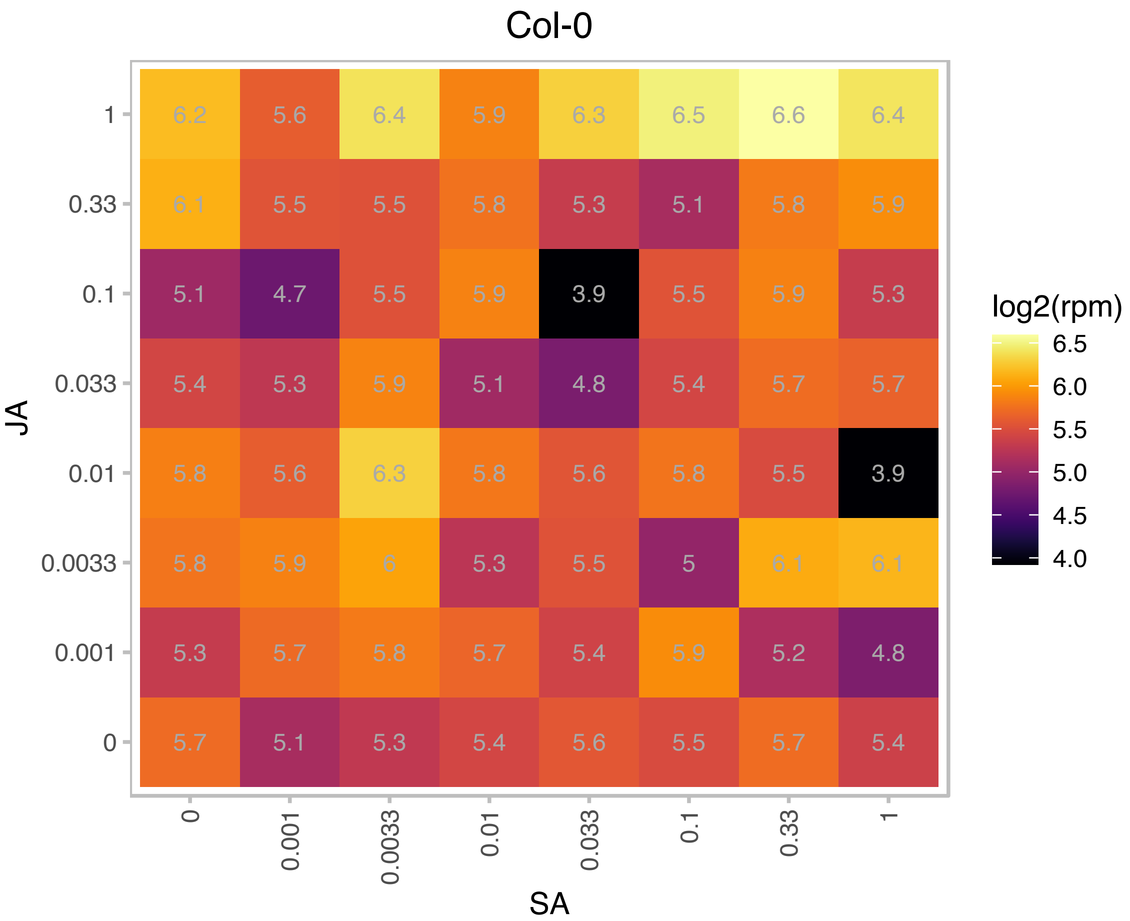


| | |
|---|---|
| Gene | AT5G11680 |
| is.expressed_genes | TRUE |
| is.responsive_genes | TRUE |
| is.apcluster | A-03 |

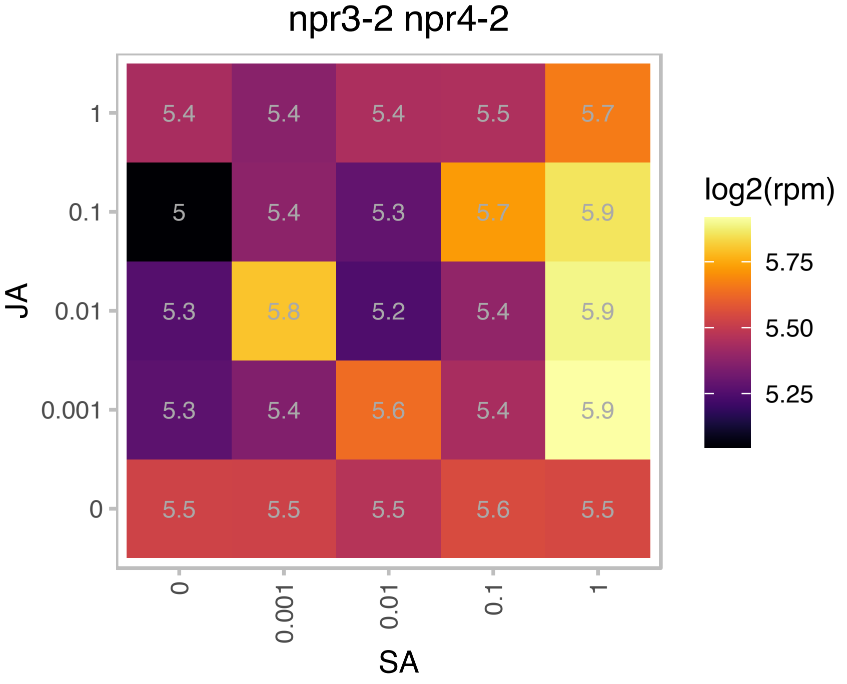


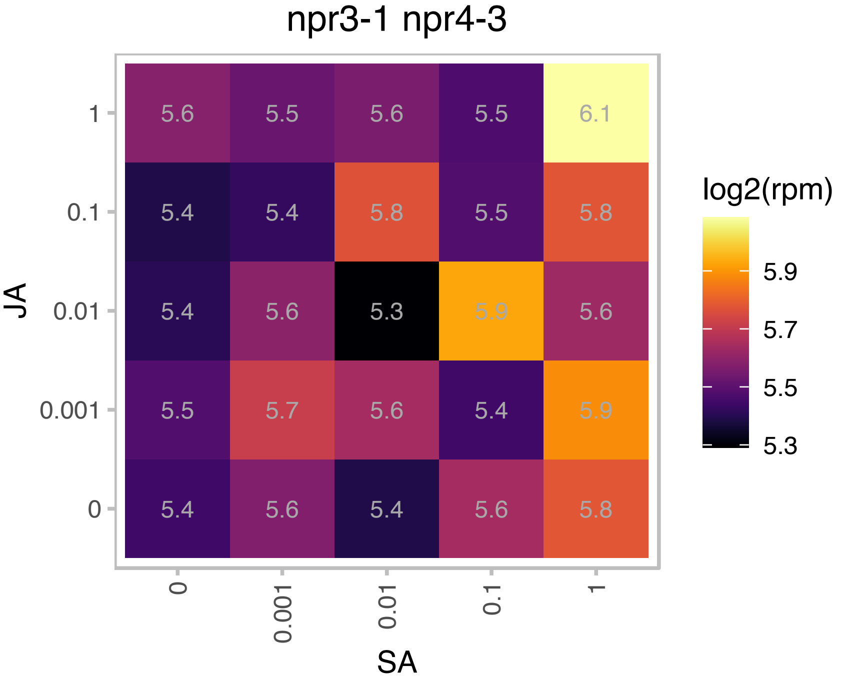

# AT5G13970.1 : NA

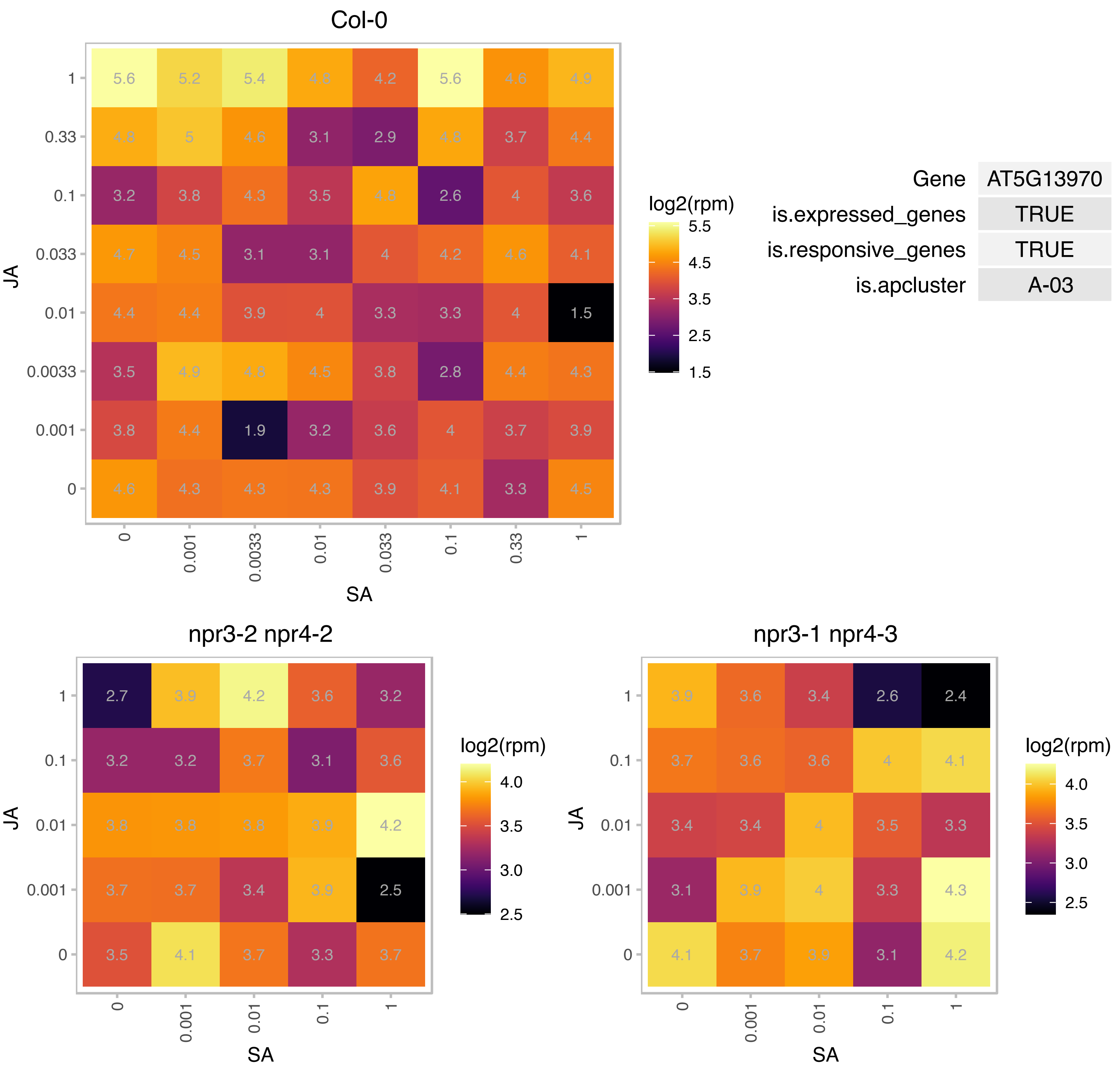

### AT5G15330.1 : ATSPX4, SPX4

SPX domain gene 4

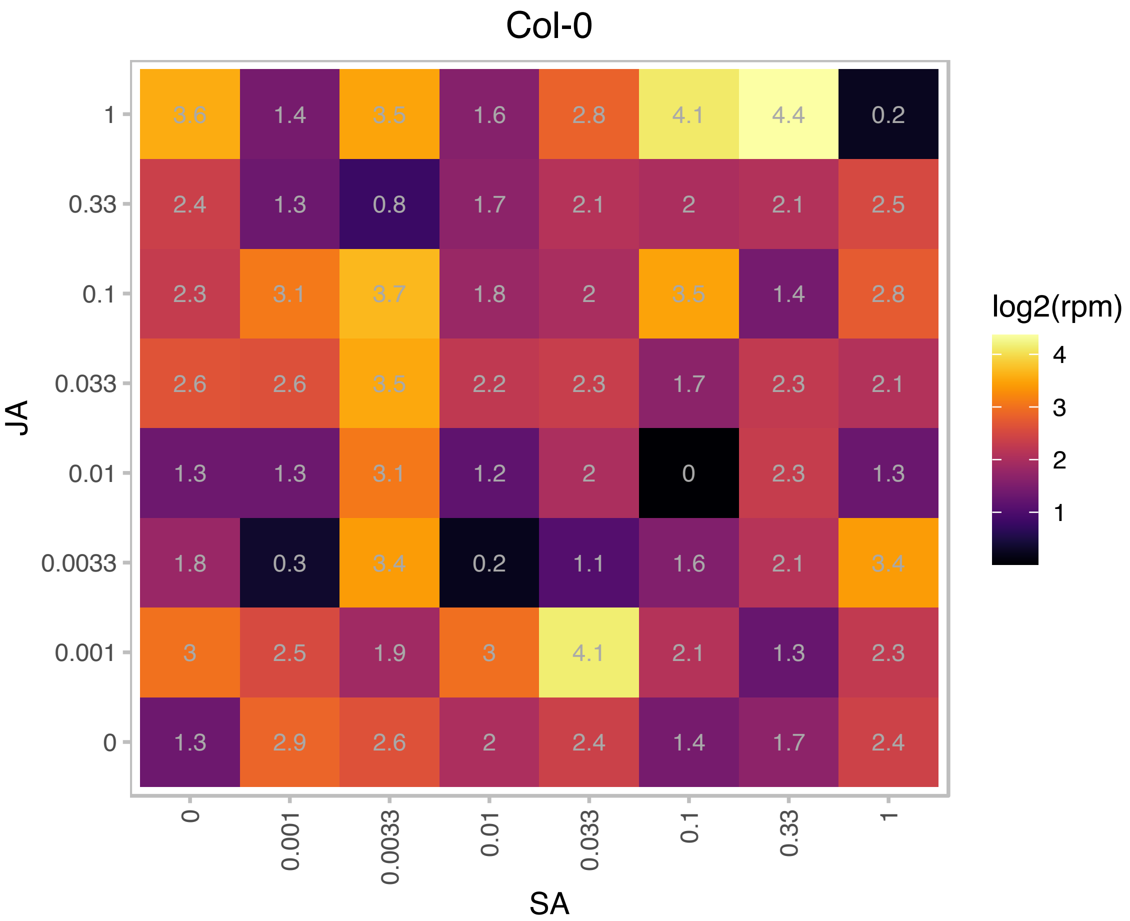


| | |
|---|---|
| Gene | AT5G15330 |
| is.expressed_genes | TRUE |
| is.responsive_genes | TRUE |
| is.apcluster | A-03 |

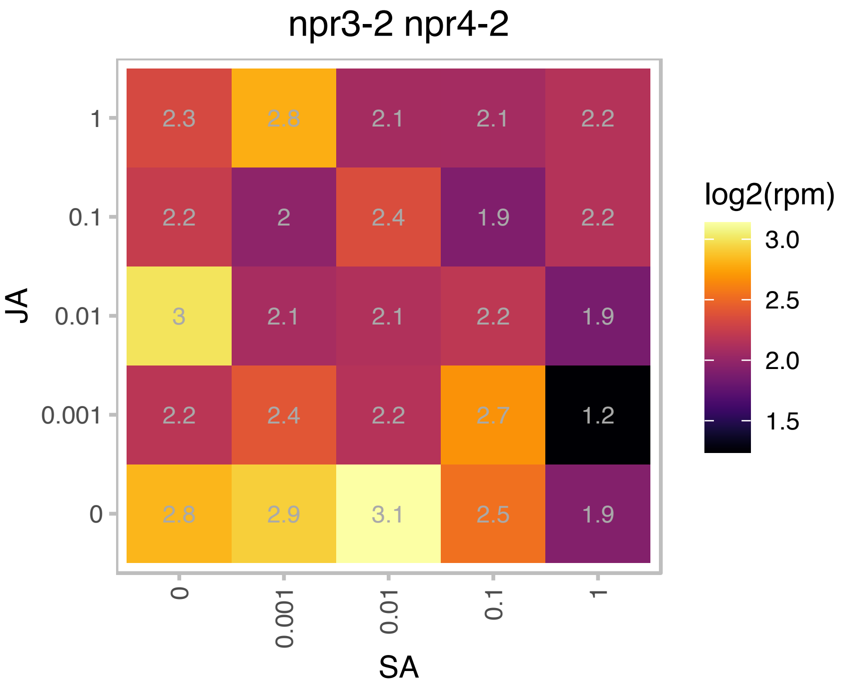


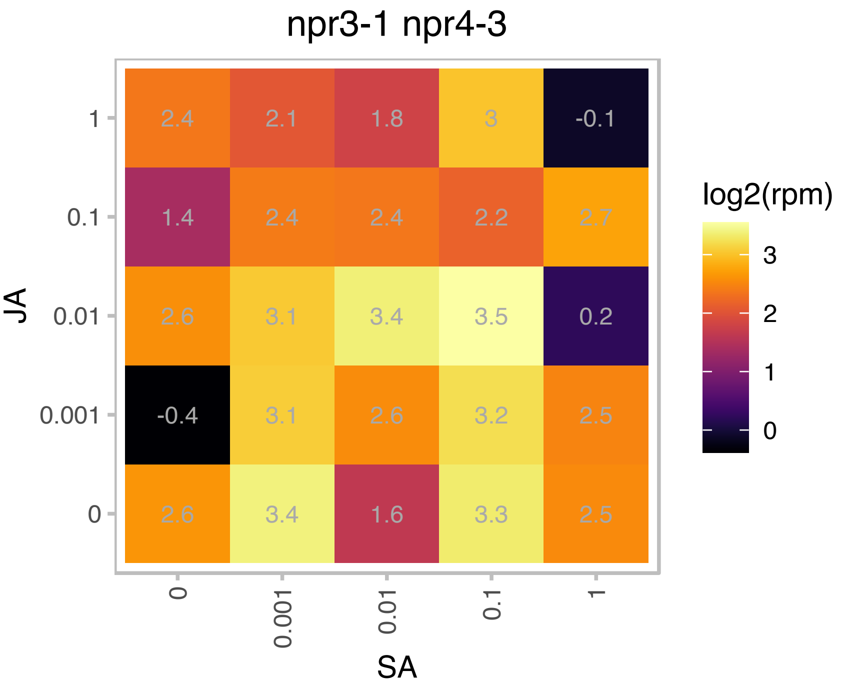

# AT5G15640.1 : NA

Mitochondrial substrate carrier family protein

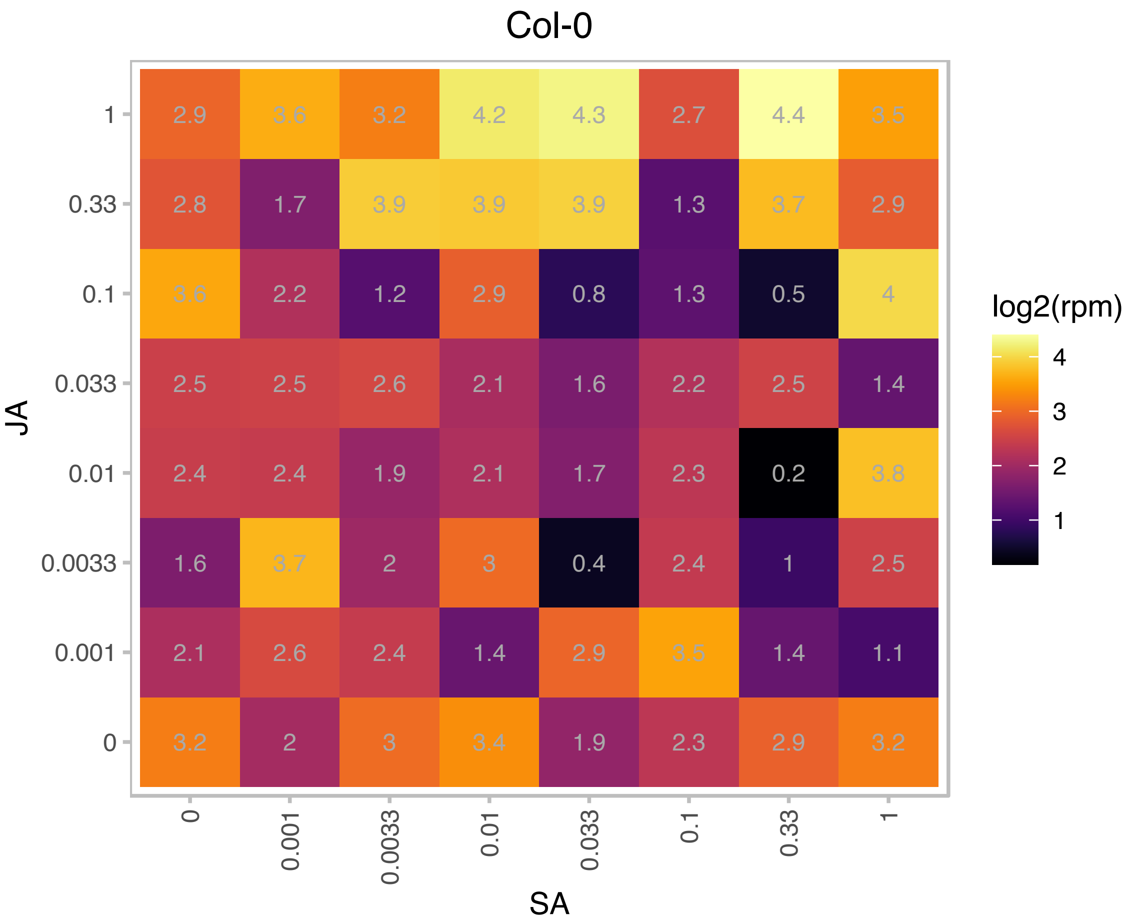


| | |
|---|---|
| Gene | AT5G15640 |
| is.expressed_genes | TRUE |
| is.responsive_genes | TRUE |
| is.apcluster | A-03 |

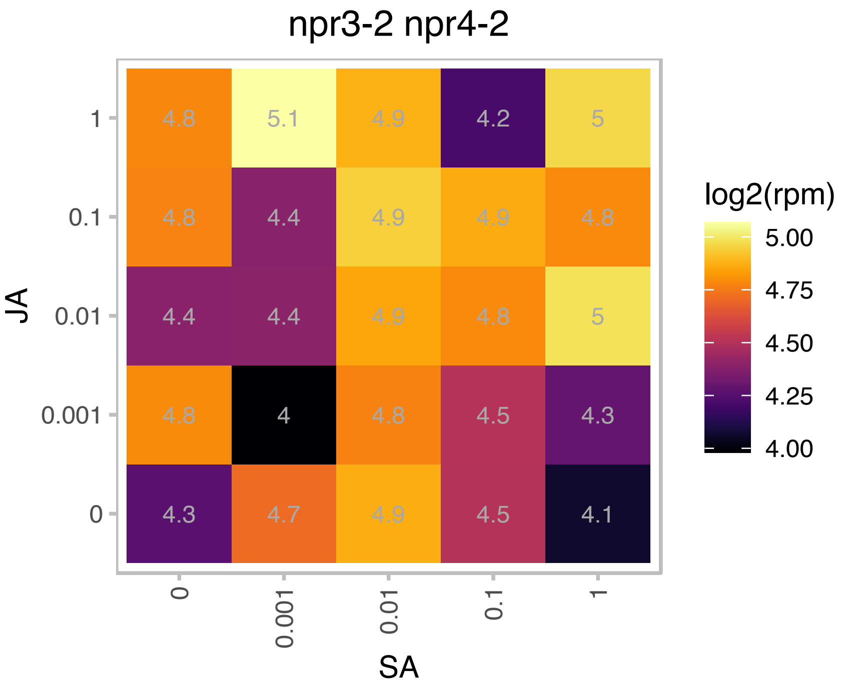


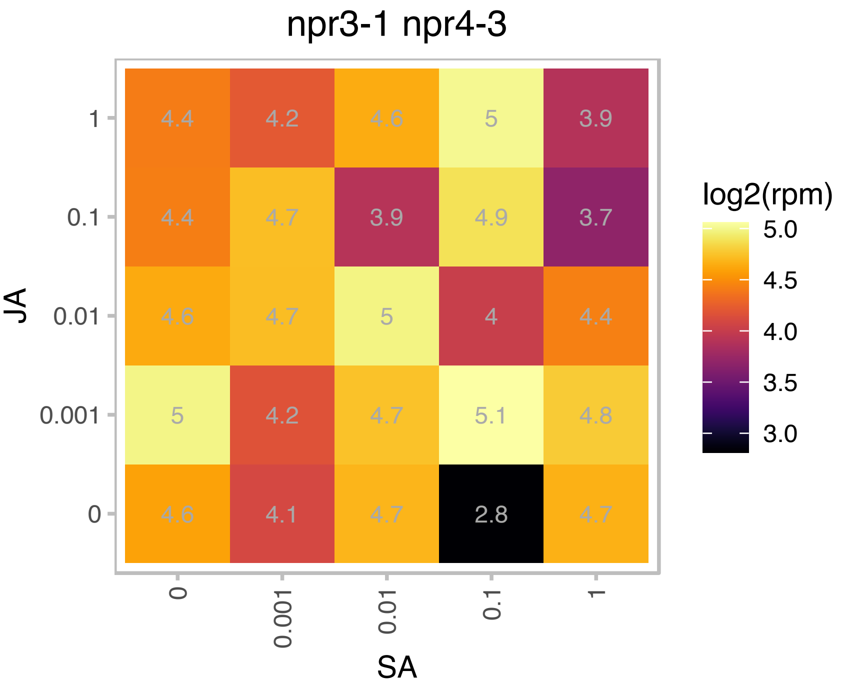

### AT5G22800.1 : EMB1030, EMB263, EMB86

Alanyl-tRNA synthetase, class IIc

A locus involved in embryogenesis. Mutations in this locus result in embryo lethality.

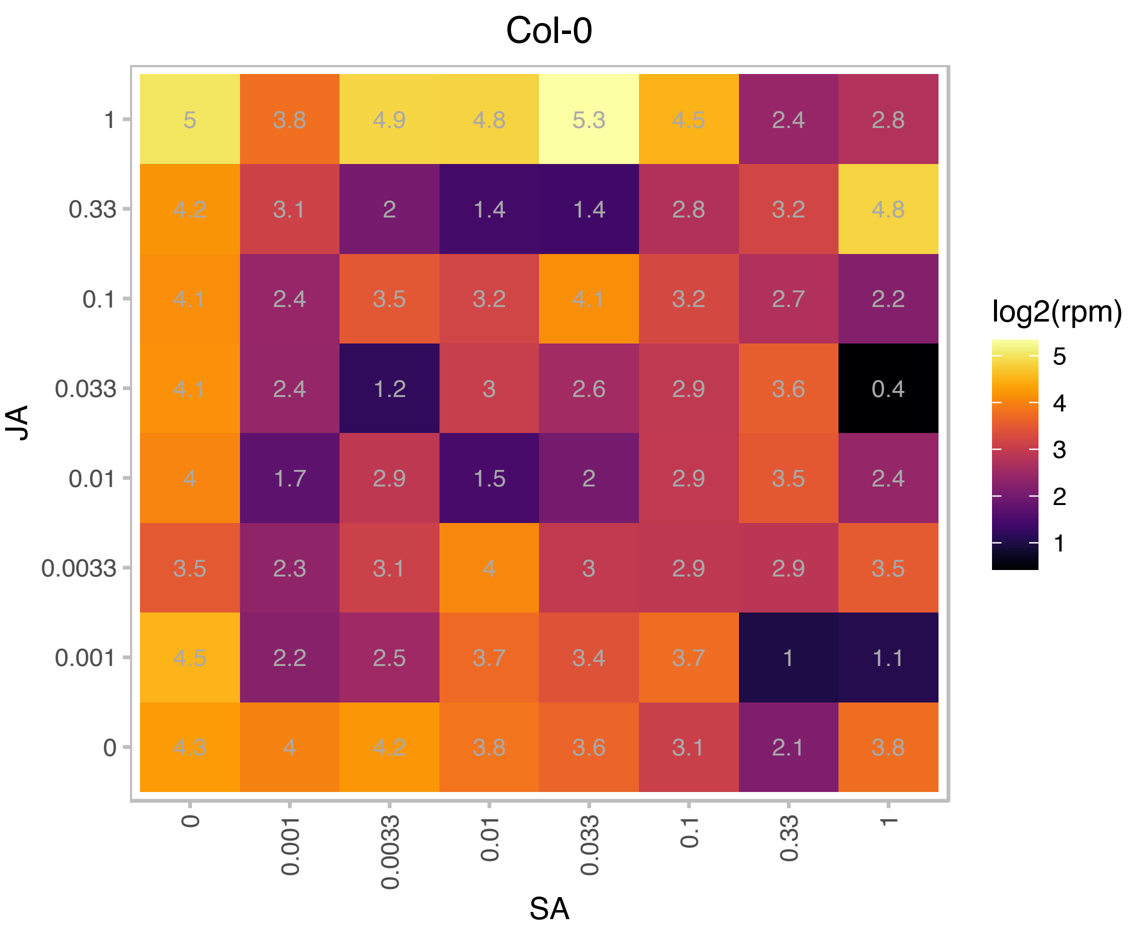


| | |
|---|---|
| Gene | AT5G22800 |
| is.expressed_genes | TRUE |
| is.responsive_genes | TRUE |
| is.apcluster | A-03 |

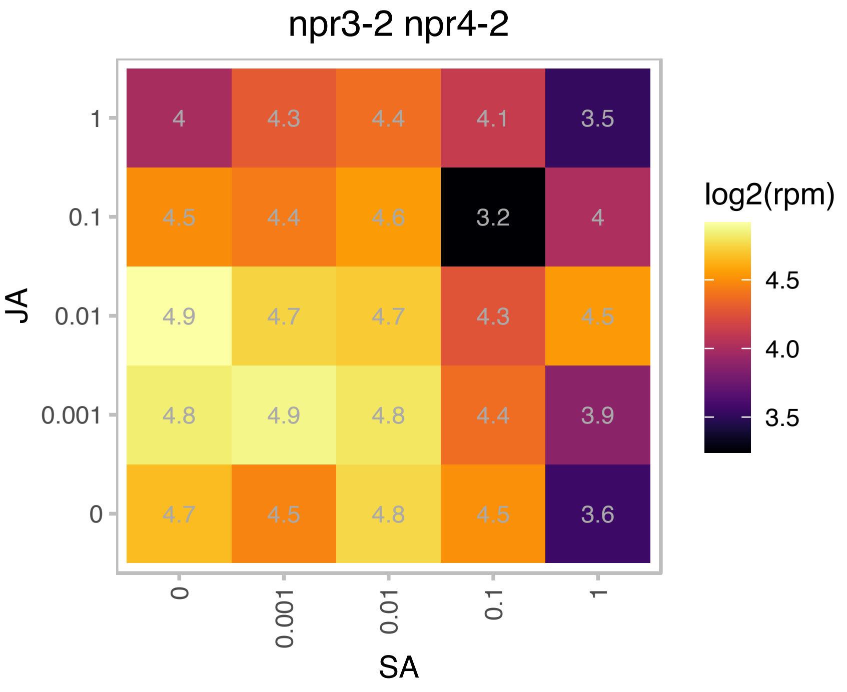


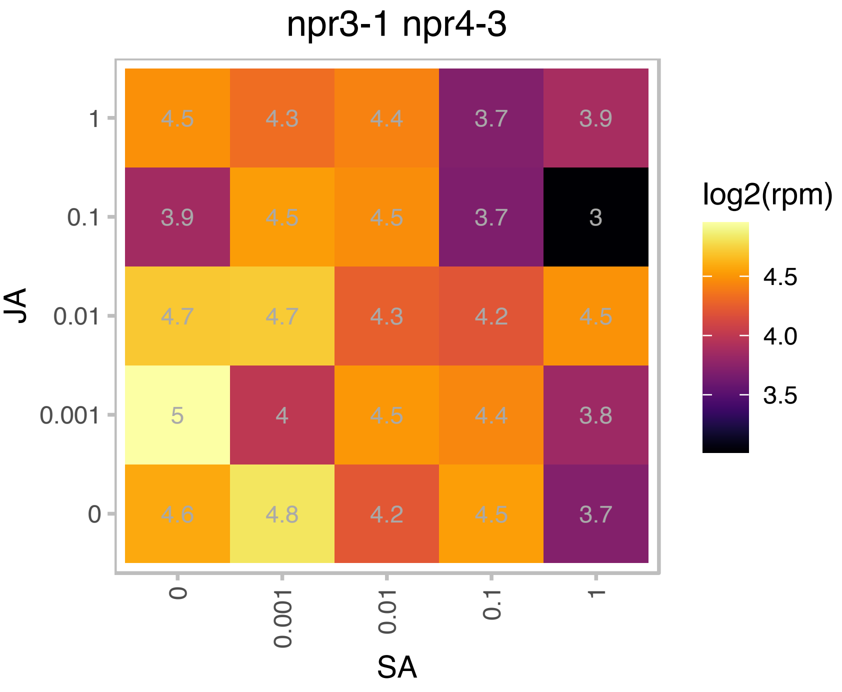

### AT5G40160.1 : EMB139, EMB506

Ankyrin repeat family protein

Encodes ankyrin repeat protein EMB506. Mutations in this locus result in embryo lethality.

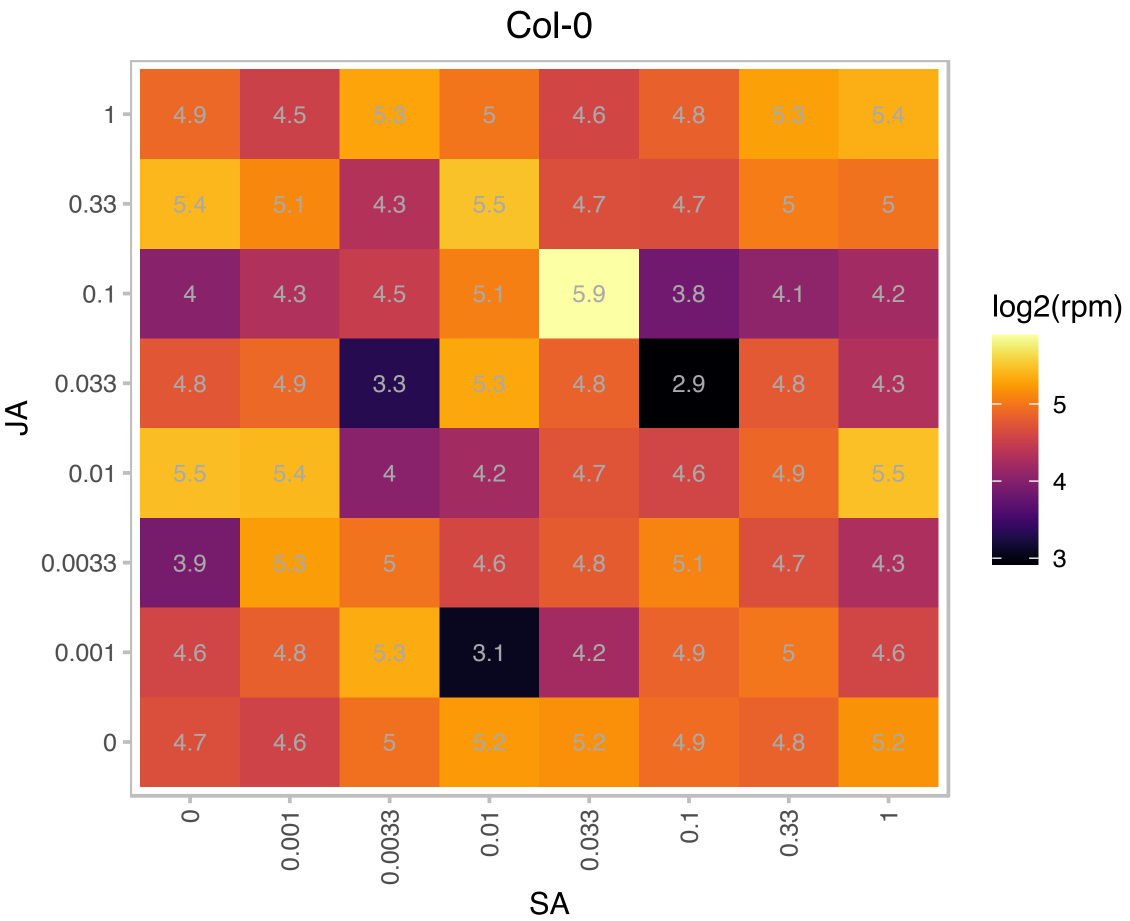


| | |
|---|---|
| Gene | AT5G40160 |
| is.expressed_genes | TRUE |
| is.responsive_genes | TRUE |
| is.apcluster | A-03 |

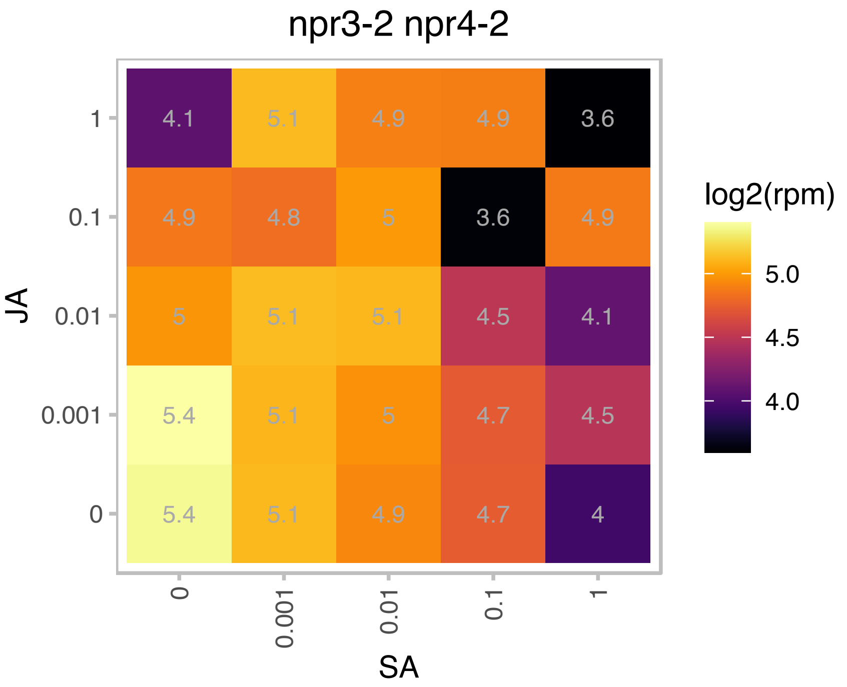


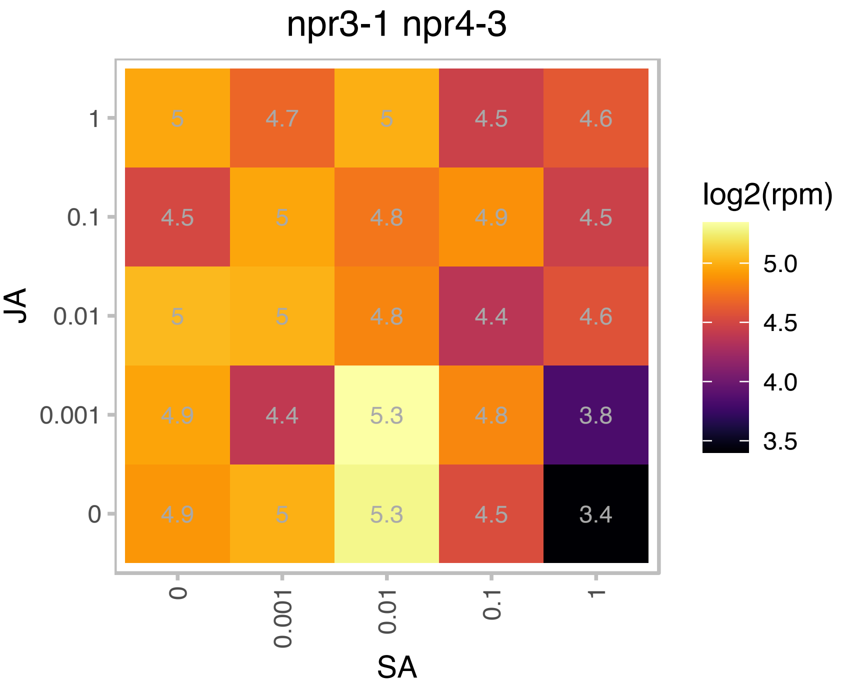

# AT5G45060.1 : NA

Disease resistance protein (TIR-NBS-LRR class) family

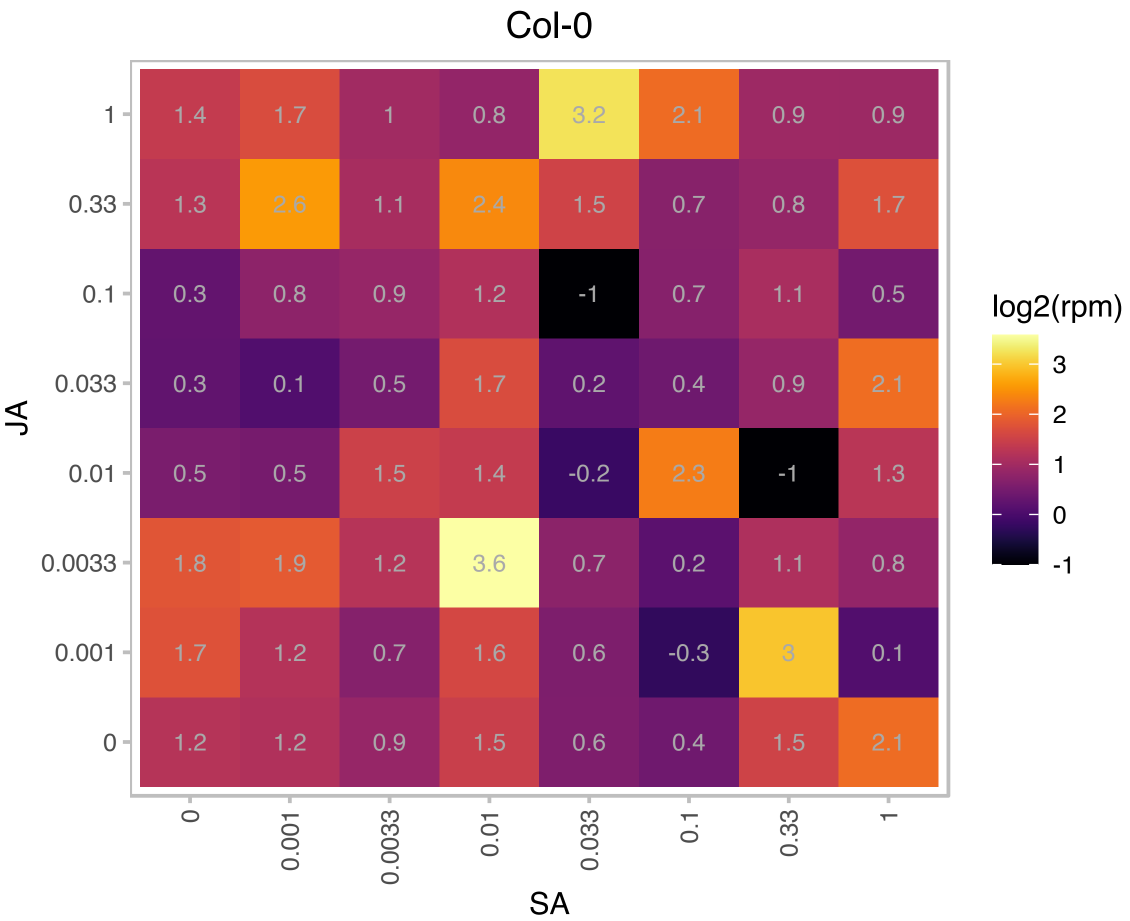


| | |
|---|---|
| Gene | AT5G45060 |
| is.expressed_genes | TRUE |
| is.responsive_genes | TRUE |
| is.apcluster | A-03 |

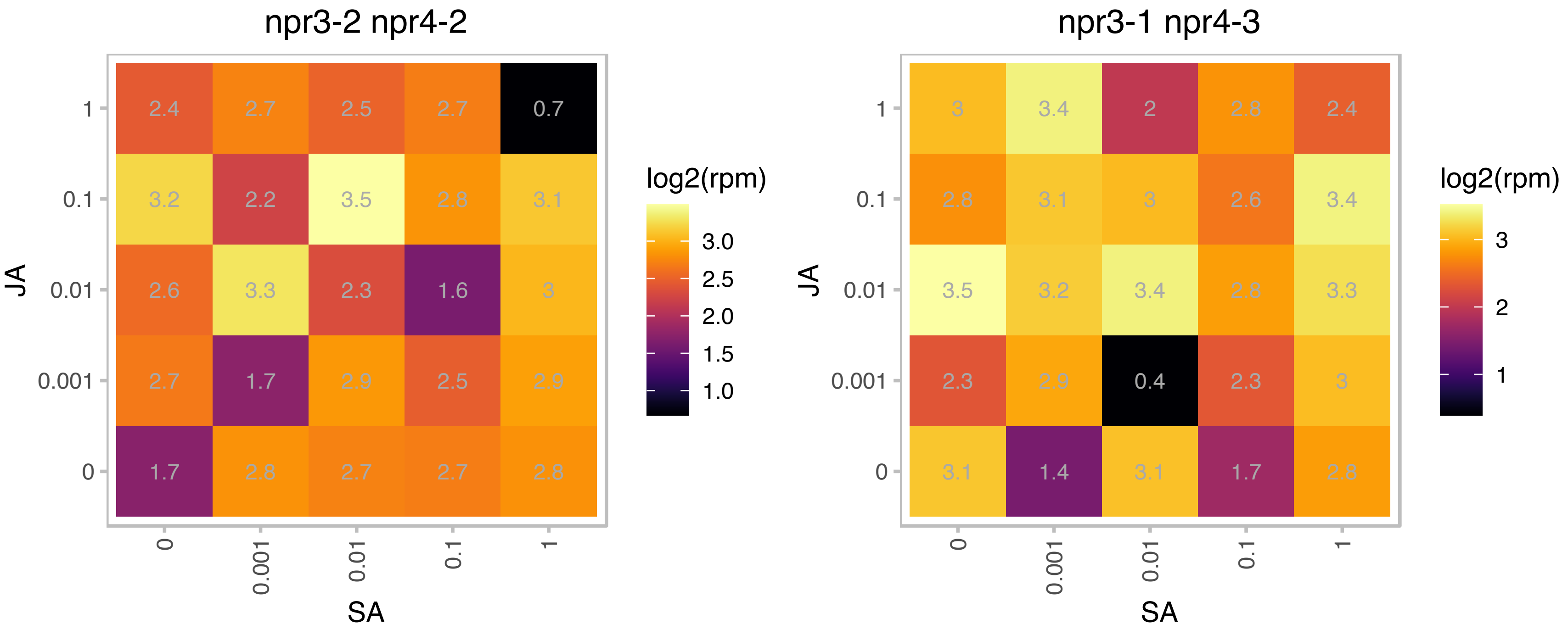

### AT5G62790.1 : DXR, PDE129

1-Deoxy-d-xylulose 5-phosphate reductoisomerase (DXR) catalyzes the first committed step of the 2-C-methyl-d-erythritol 4-phosphate pathway for isoprenoid biosynthesis. In Arabidopsis, DXR is encoded by a single-copy gene. Arabidopsis DXR is targeted to plastids and localizes into chloroplasts of leaf cells. DXR knockout or strongly silenced lines have a seedling lethal, albino phenotype. Transgenic, partially silenced lines expressing 35S:DXR have a variegated phenotype.

1-deoxy-D-xylulose 5-phosphate reductoisomerase

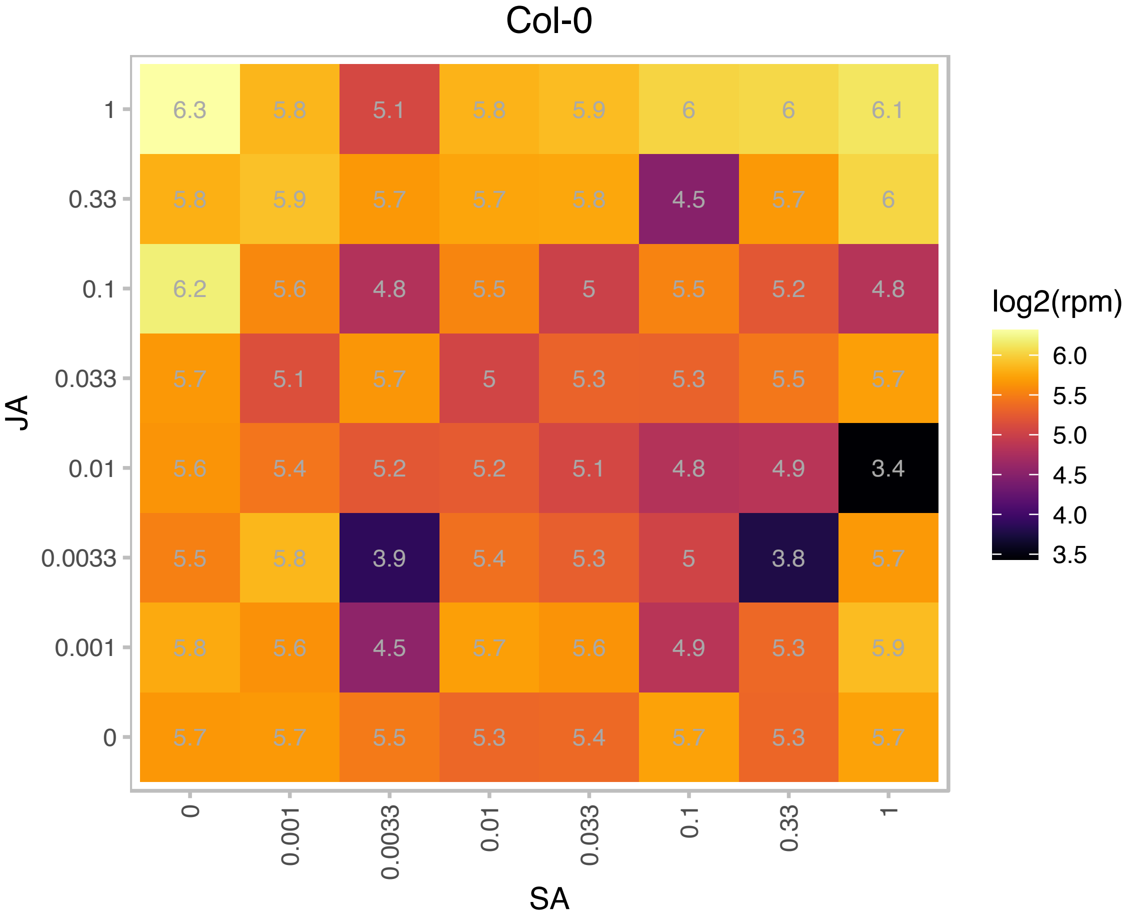


| | |
|---|---|
| Gene | AT5G62790 |
| is.expressed_genes | TRUE |
| is.responsive_genes | TRUE |
| is.apcluster | A-03 |

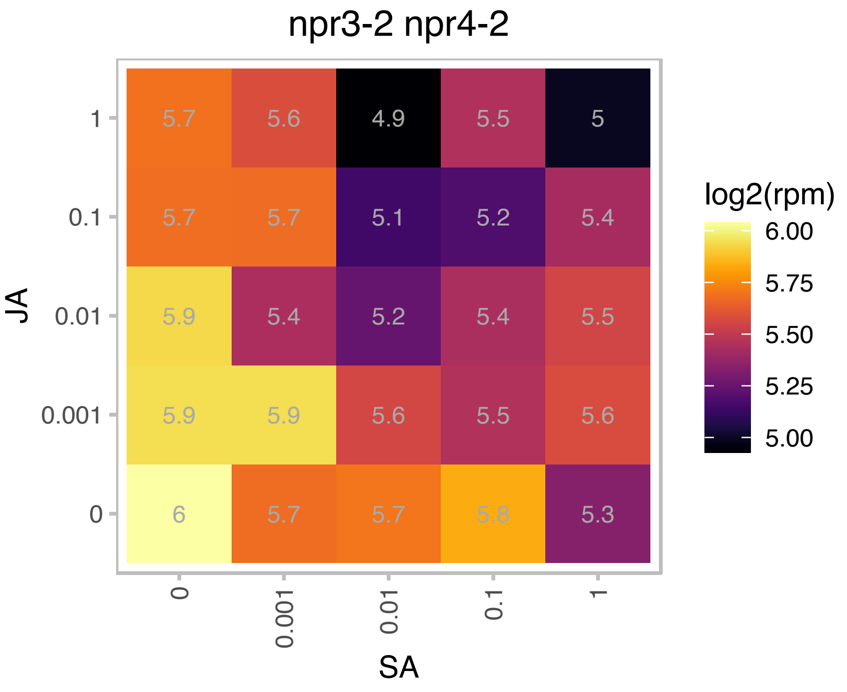


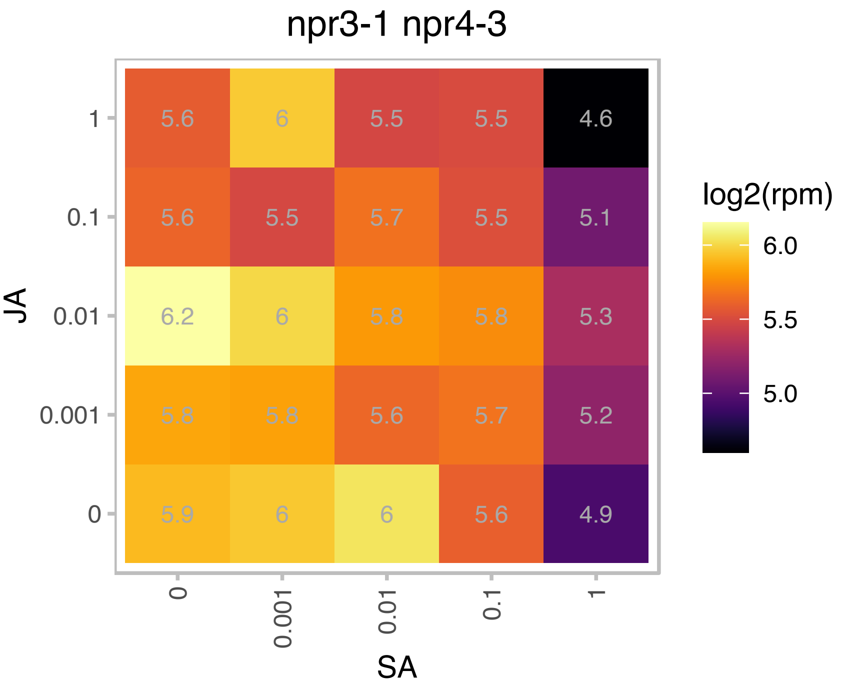

### AT1G17130.1 : CWC16b

Family of unknown function (DUF572)

DUF572 domain protein involved in alternative splicing.

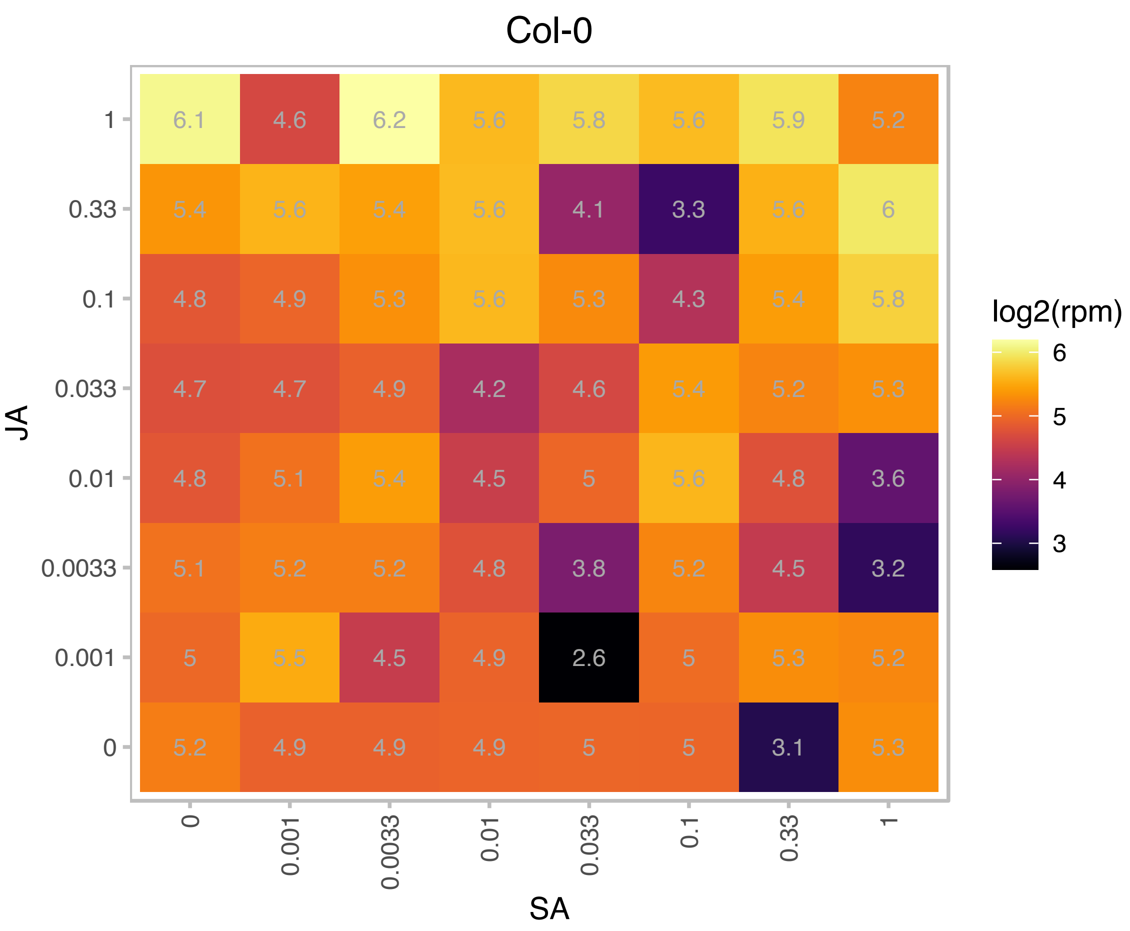


| | |
|---|---|
| Gene | AT1G17130 |
| is.expressed_genes | TRUE |
| is.responsive_genes | TRUE |
| is.apcluster | A-04 |

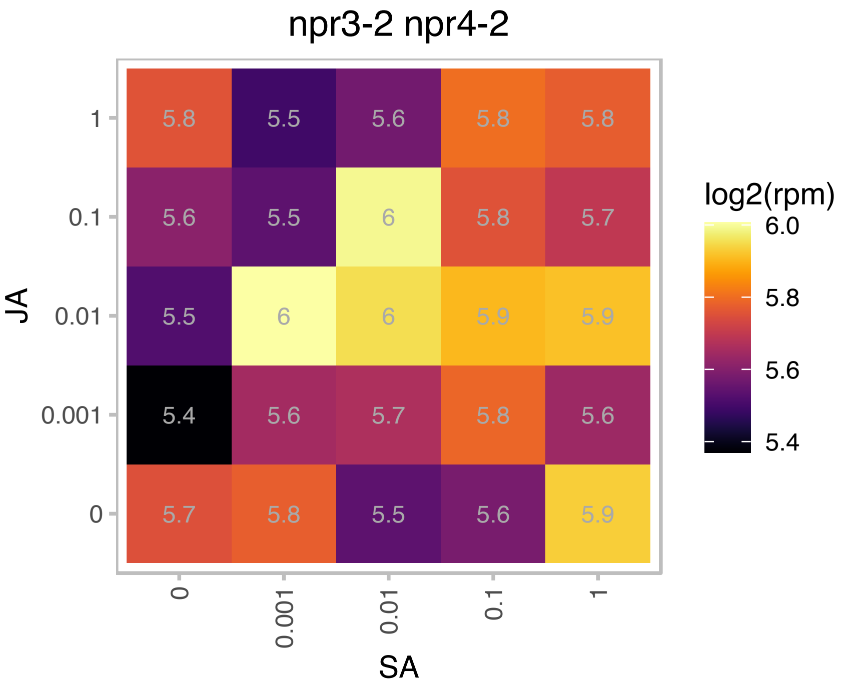


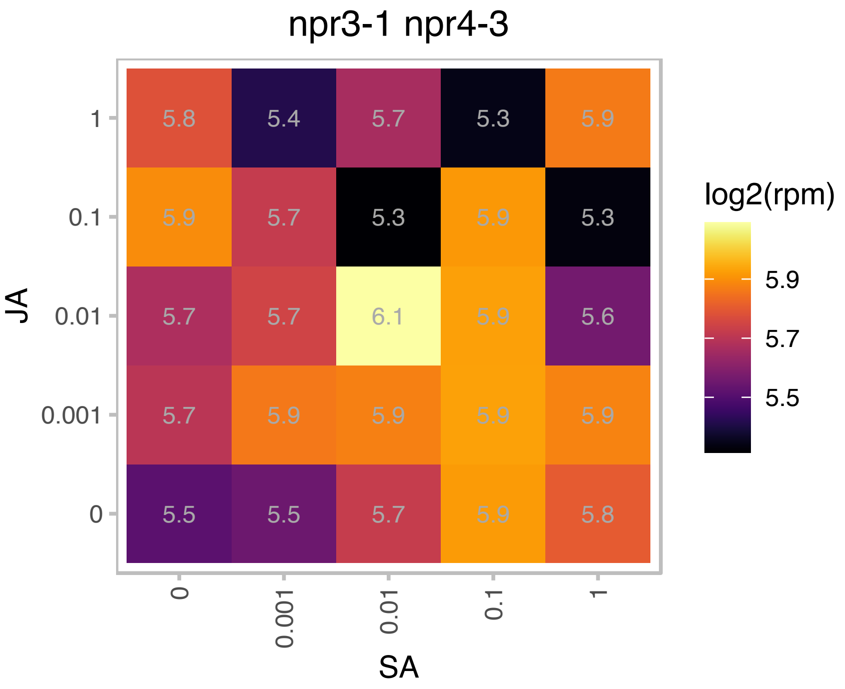

# AT1G18070.1 : NA

Translation elongation factor EF1A/initiation factor IF2gamma family protein

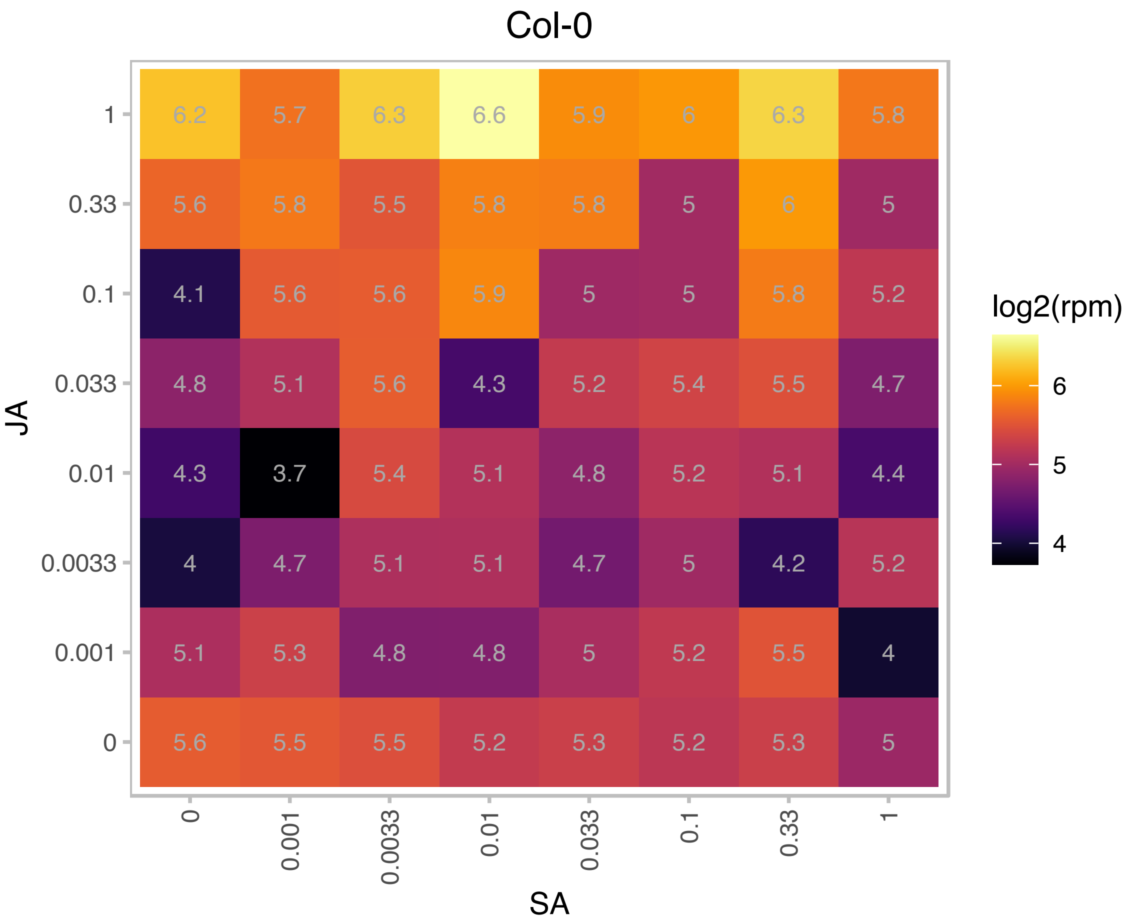


| | |
|---|---|
| Gene | AT1G18070 |
| is.expressed_genes | TRUE |
| is.responsive_genes | TRUE |
| is.apcluster | A-04 |

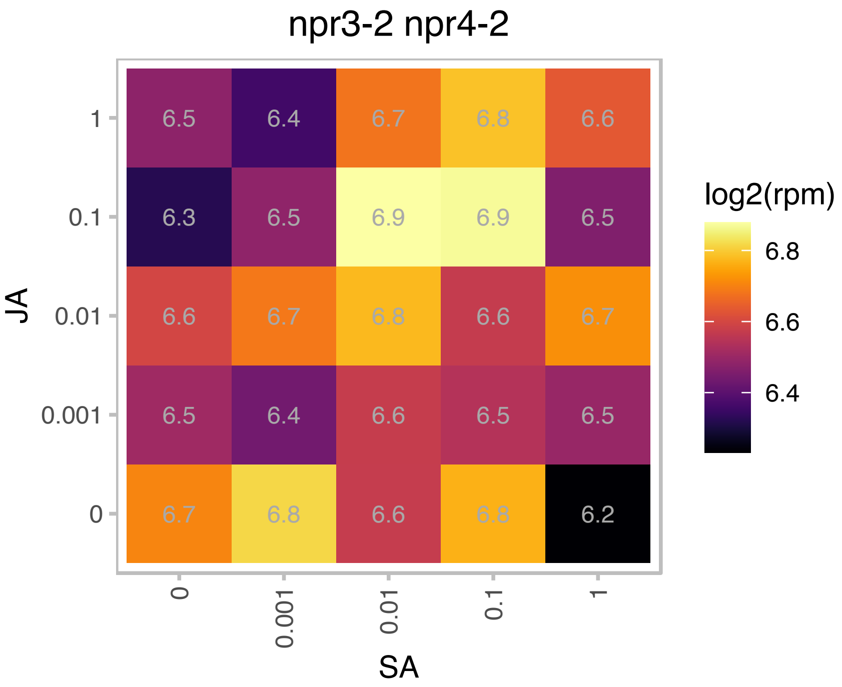


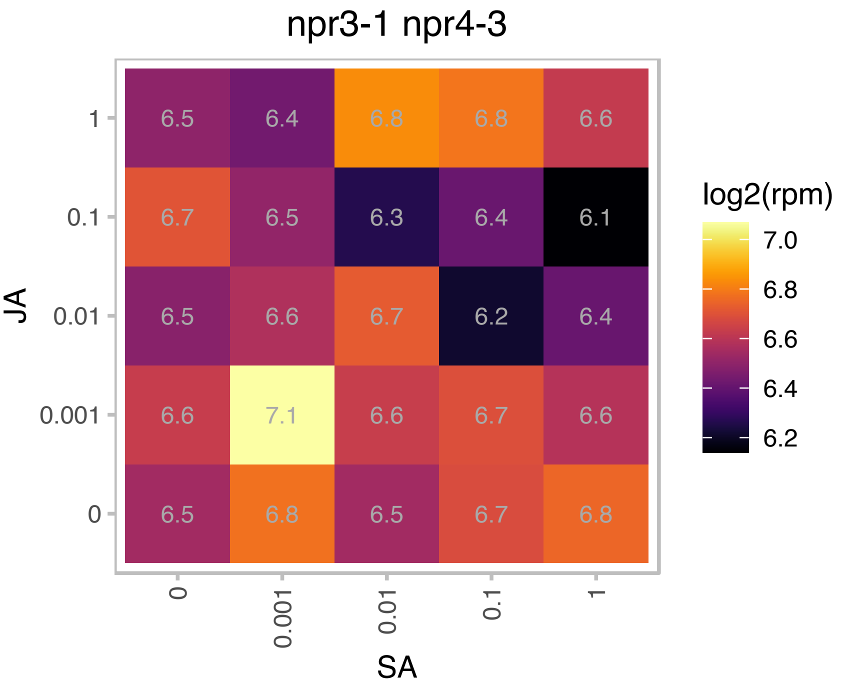

### AT1G23190.1 : PGM3

Phosphoglucomutase/phosphomannomutase family protein

Encodes a cytosolic phosphoglucomutase (PGM). Two Arabidopsis PGM proteins (AT1G70730/PGM2 and AT1G23190/PGM3) have high sequence similarities and redundant functions. Mature plants possessing a single cPGM allele had a major reduction in cPGM activity. Whereas pgm2 and pgm3 single mutants are undistinguishable from the wild type, loss of both PGM2 and PGM3 severely impairs male and female gametophyte development. The mRNA is cell-to-cell mobile.

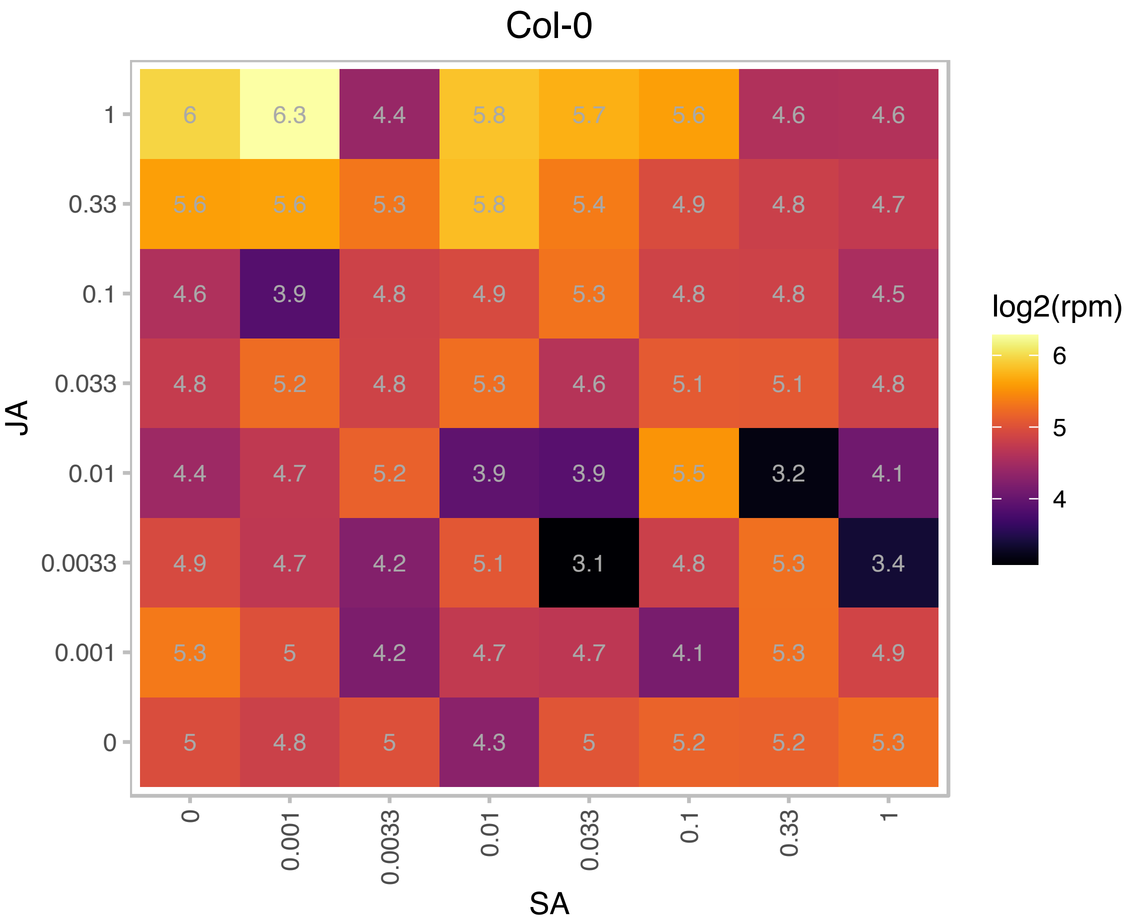


| | |
|---|---|
| Gene | AT1G23190 |
| is.expressed_genes | TRUE |
| is.responsive_genes | TRUE |
| is.apcluster | A-04 |

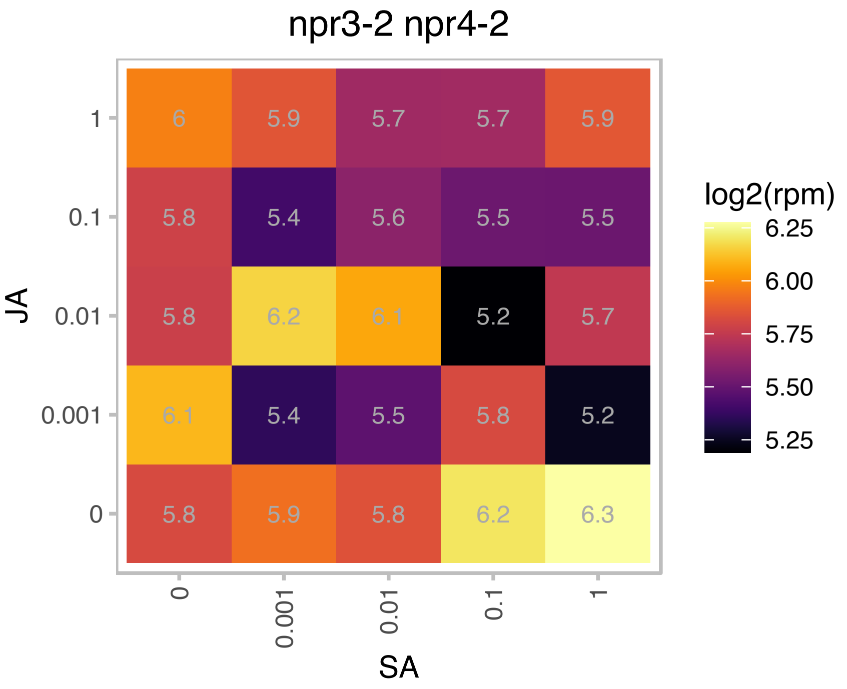


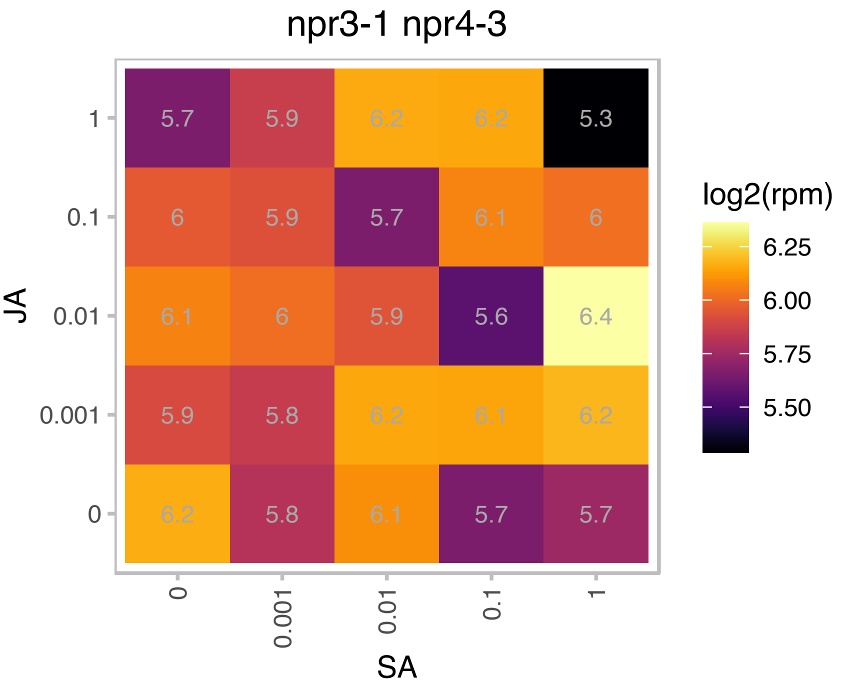

# AT1G32160.1 : NA

Protein of unknown function (DUF760)

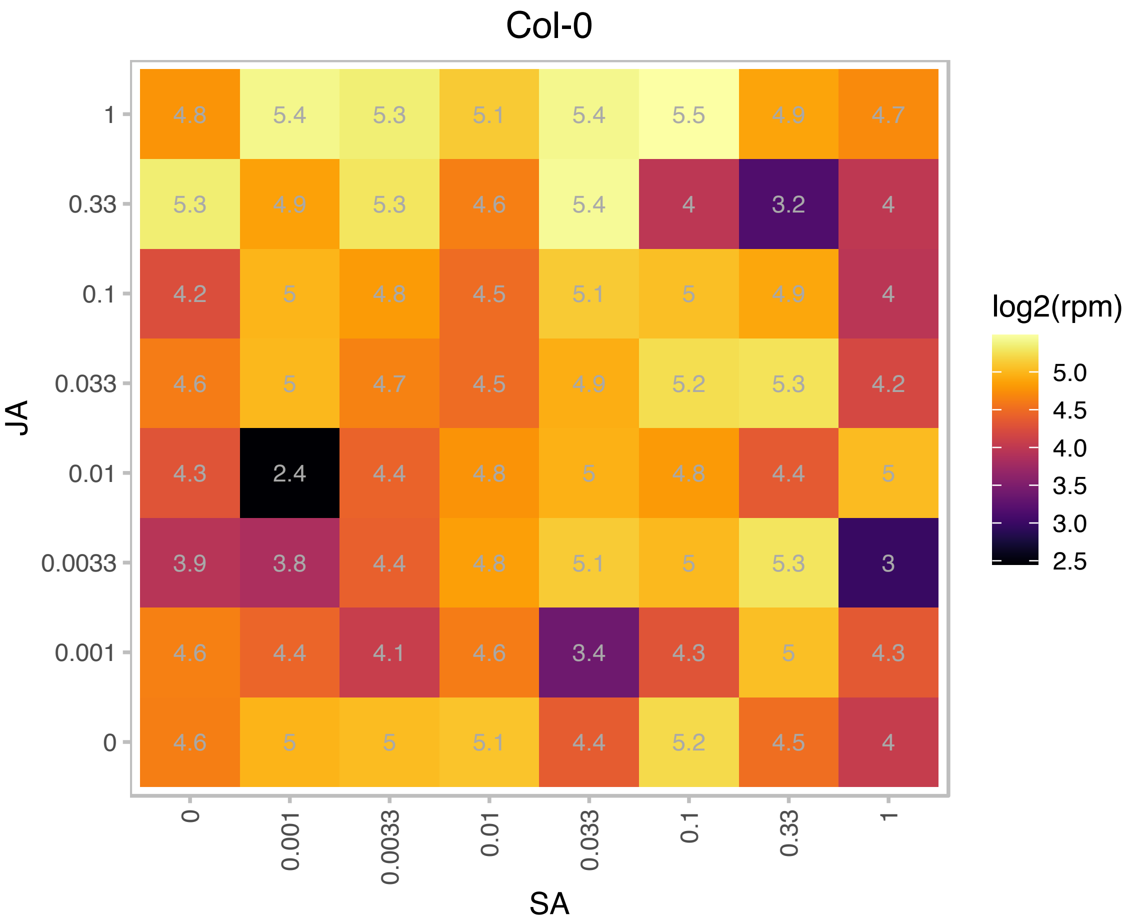


| | |
|---|---|
| Gene | AT1G32160 |
| is.expressed_genes | TRUE |
| is.responsive_genes | TRUE |
| is.apcluster | A-04 |

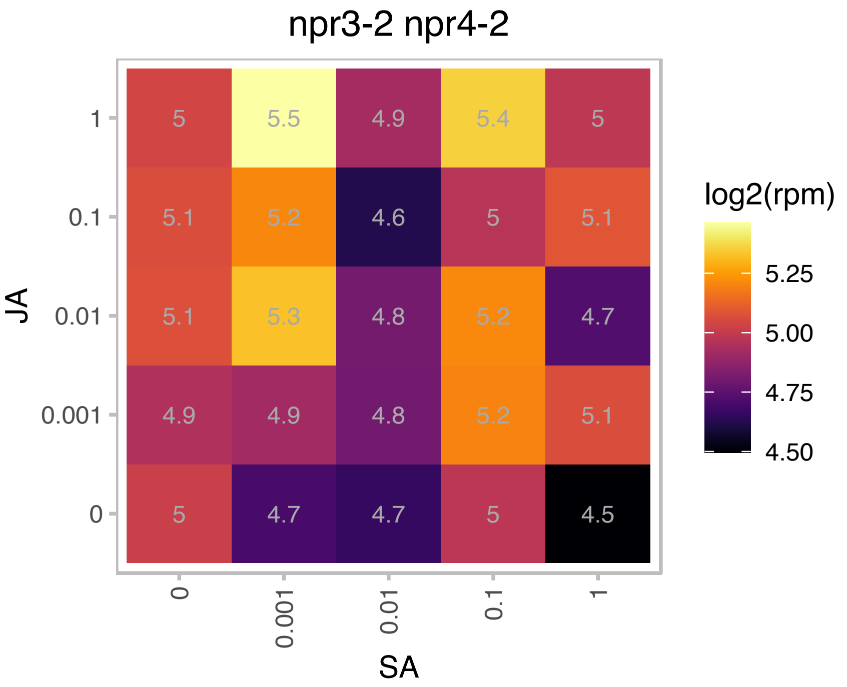


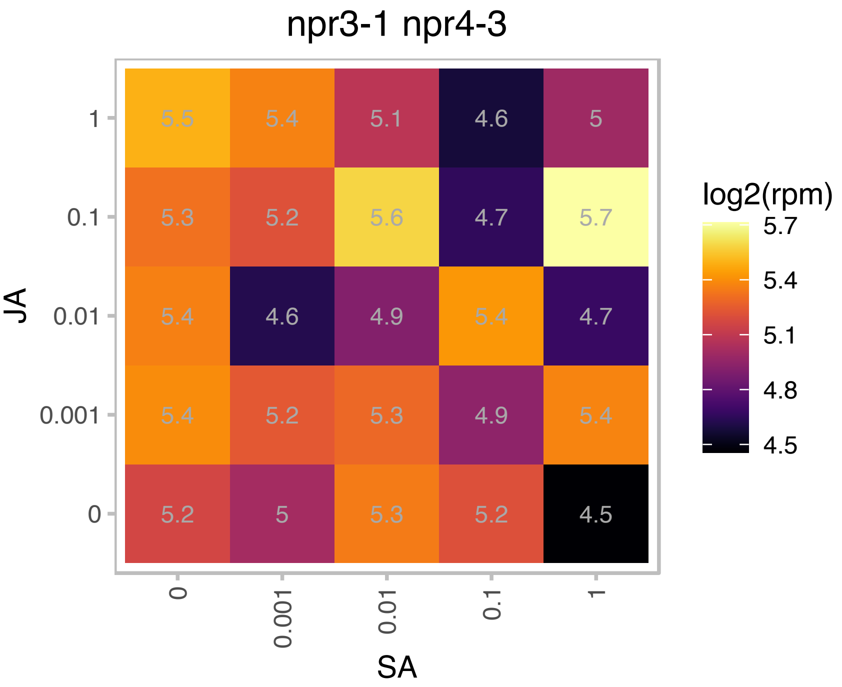

### AT1G36160.1 : ACC1, AT-ACC1, EMB22, GK, GSD1, PAS3

acetyl-CoA carboxylase 1

Encodes acetyl-CoA carboxylase. Mutant displays uncoordinated cell divisions which are enhanced by cytokinins. Mutant also has aberrant organization of the apical region in the embryo and abnormal root and shoot development and is deficient in freezing tolerance after cold acclimation. Essential for very long chain fatty acid elongation. The mRNA is cell-to-cell mobile.

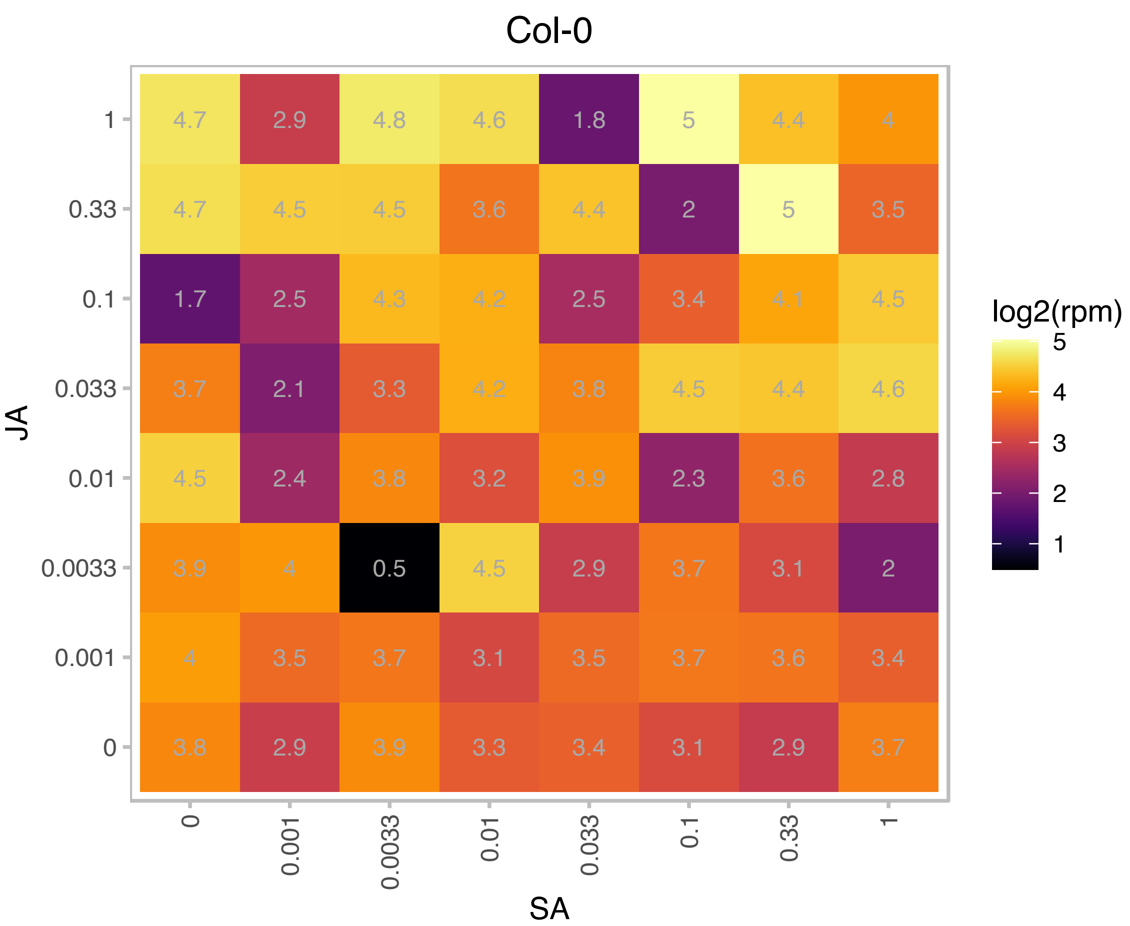


| | |
|---|---|
| Gene | AT1G36160 |
| is.expressed_genes | TRUE |
| is.responsive_genes | TRUE |
| is.apcluster | A-04 |

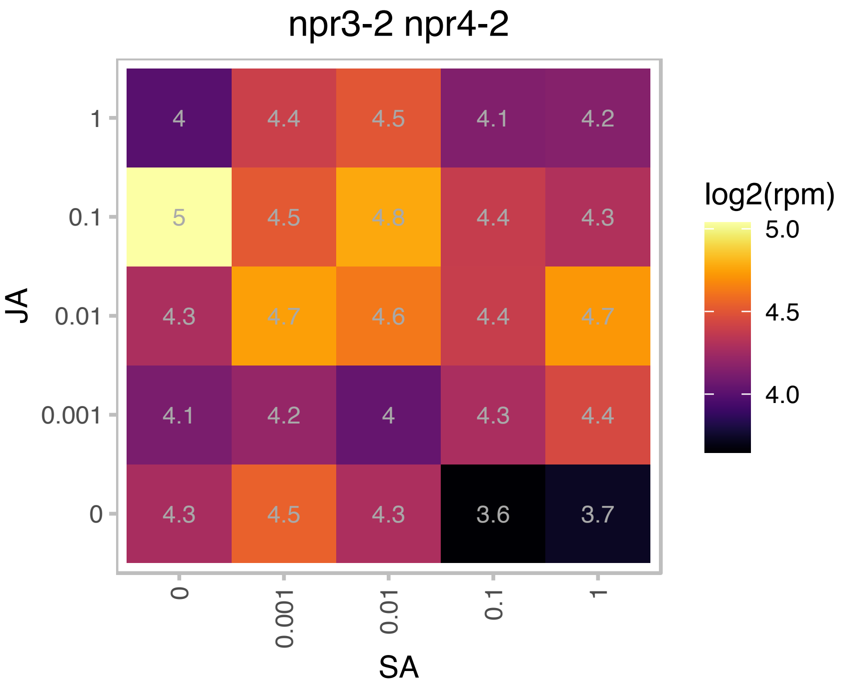


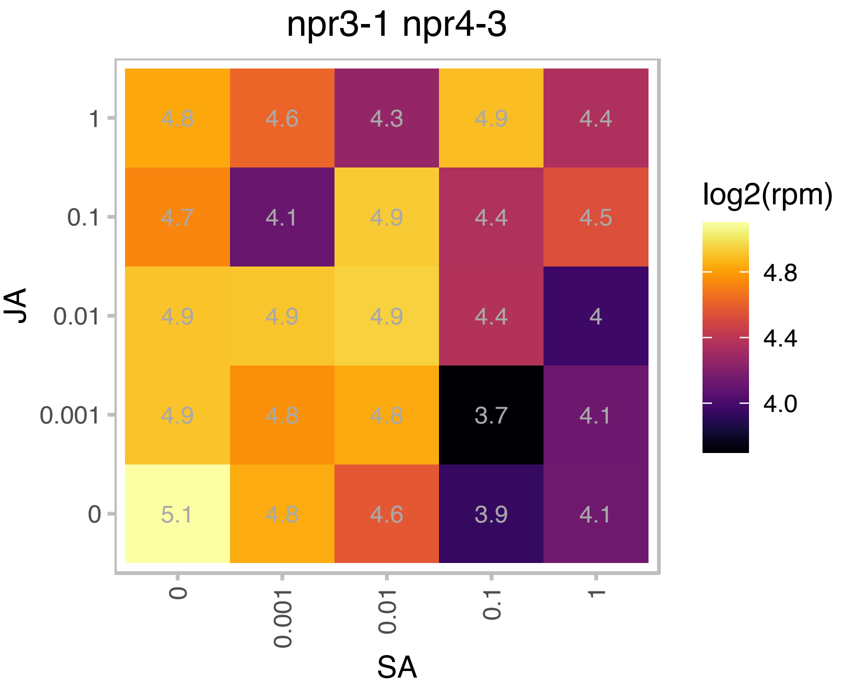

### AT1G51600.1 : GATA28, TIFY2A, ZML2

ZIM-LIKE 2

member of a novel family of plant-specific GATA-type transcription factors.

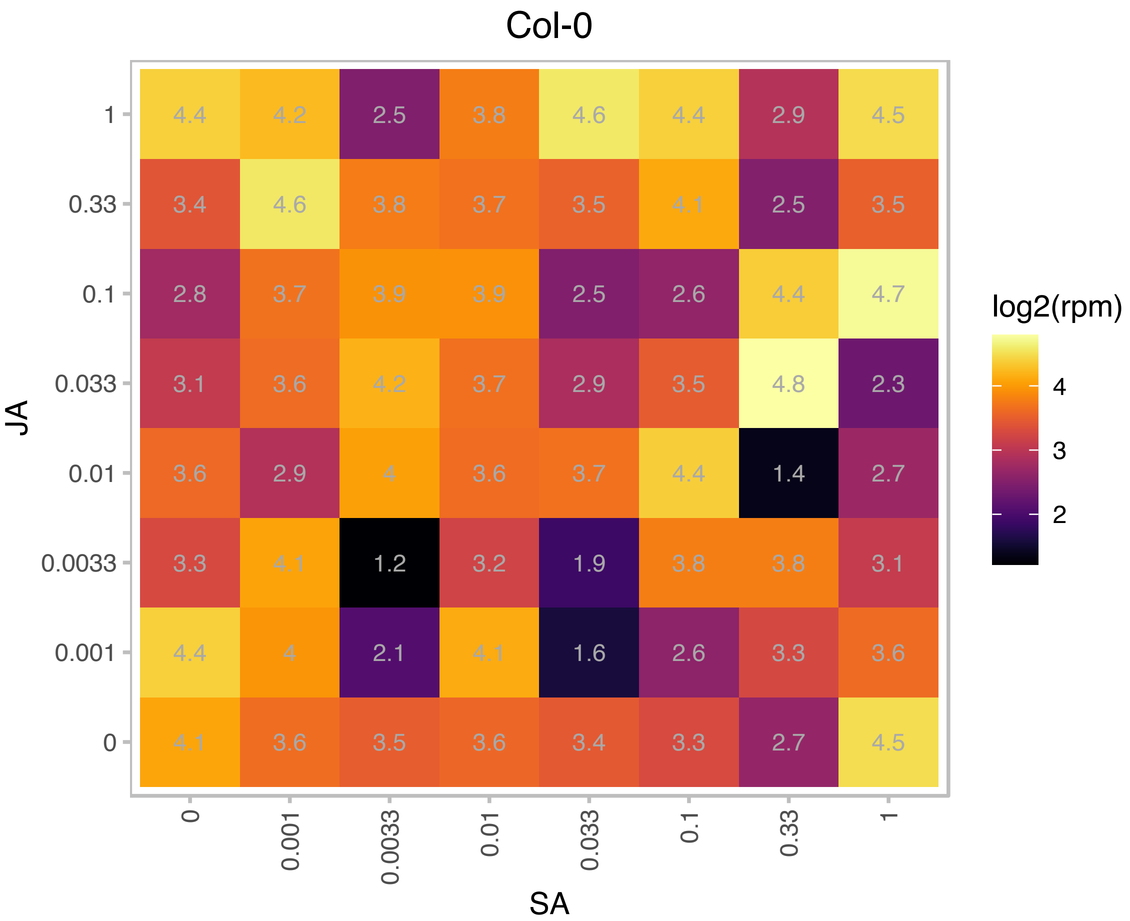


| | |
|---|---|
| Gene | AT1G51600 |
| is.expressed_genes | TRUE |
| is.responsive_genes | TRUE |
| is.apcluster | A-04 |

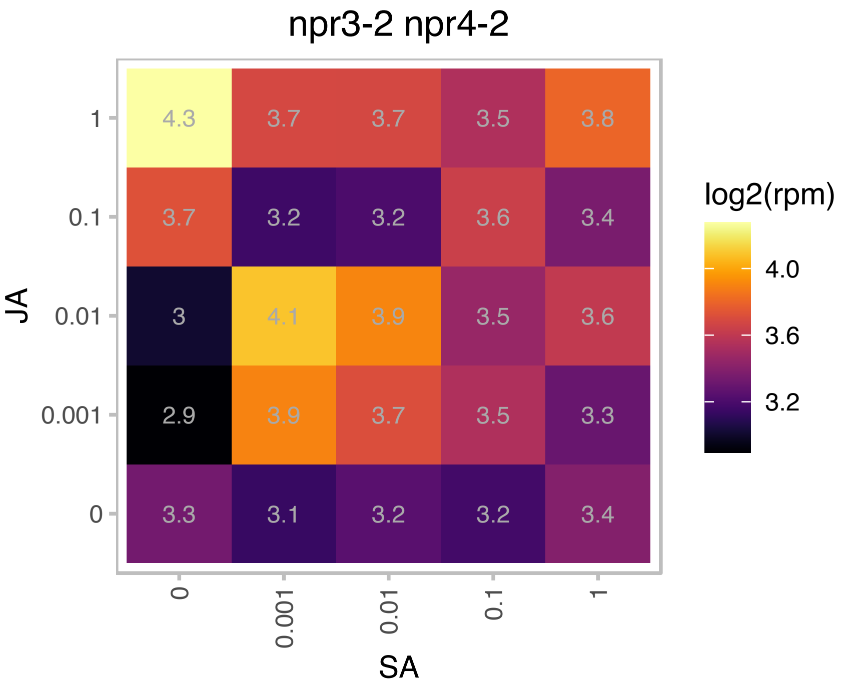


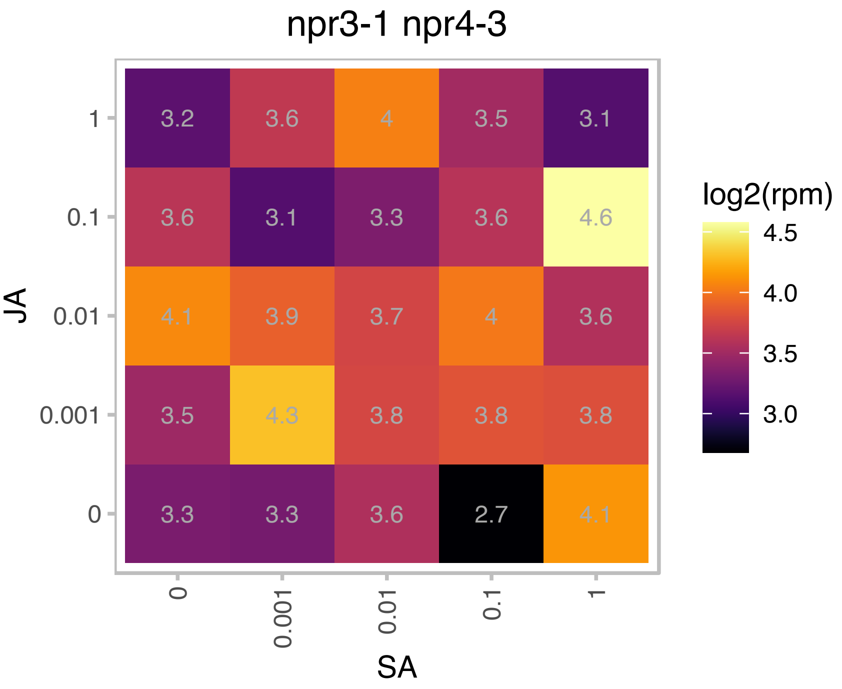

### AT1G53210.1 : AtNCL, NCL

sodium/calcium exchanger family protein / calcium-binding EF hand family protein

Encodes a Na+/Ca 2+ exchanger-like protein that participates in the maintenance of Ca 2+ homeostasis. The mRNA is cell-to-cell mobile.

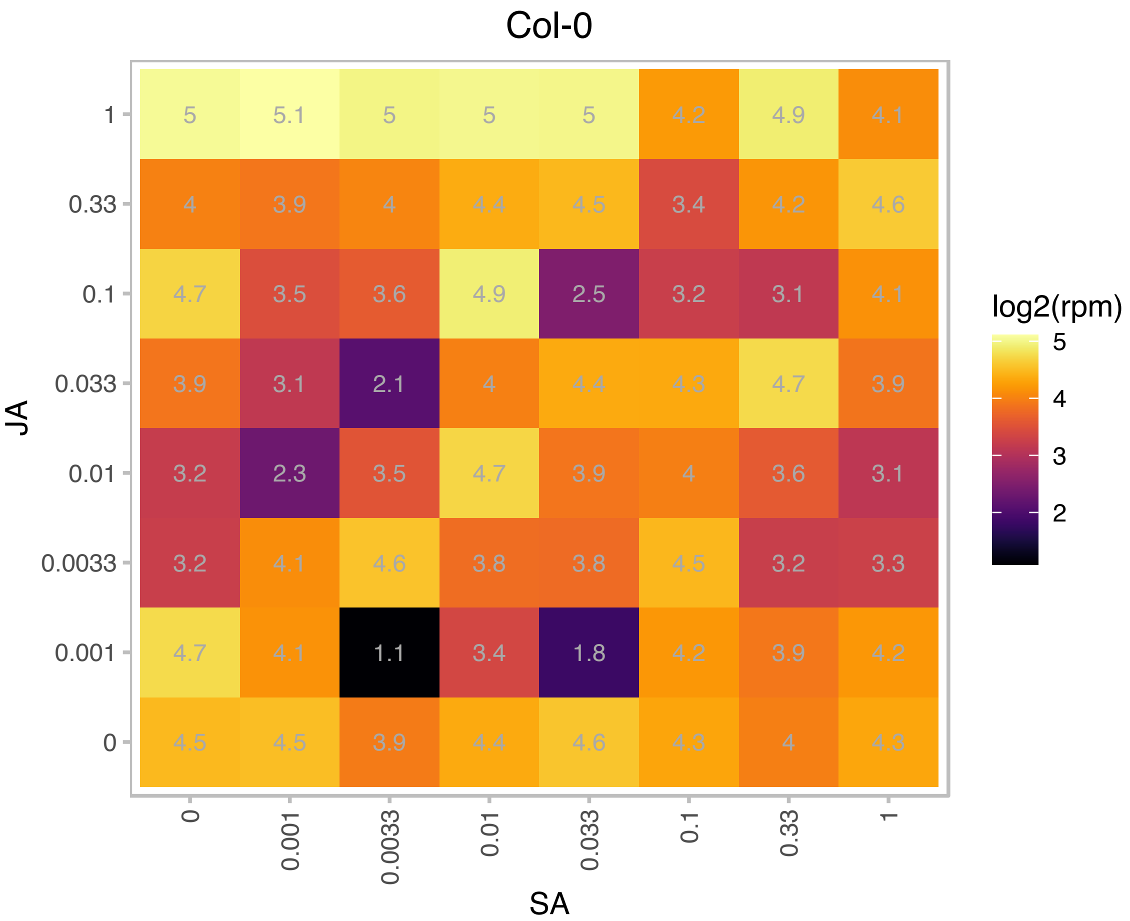


| | |
|---|---|
| Gene | AT1G53210 |
| is.expressed_genes | TRUE |
| is.responsive_genes | TRUE |
| is.apcluster | A-04 |

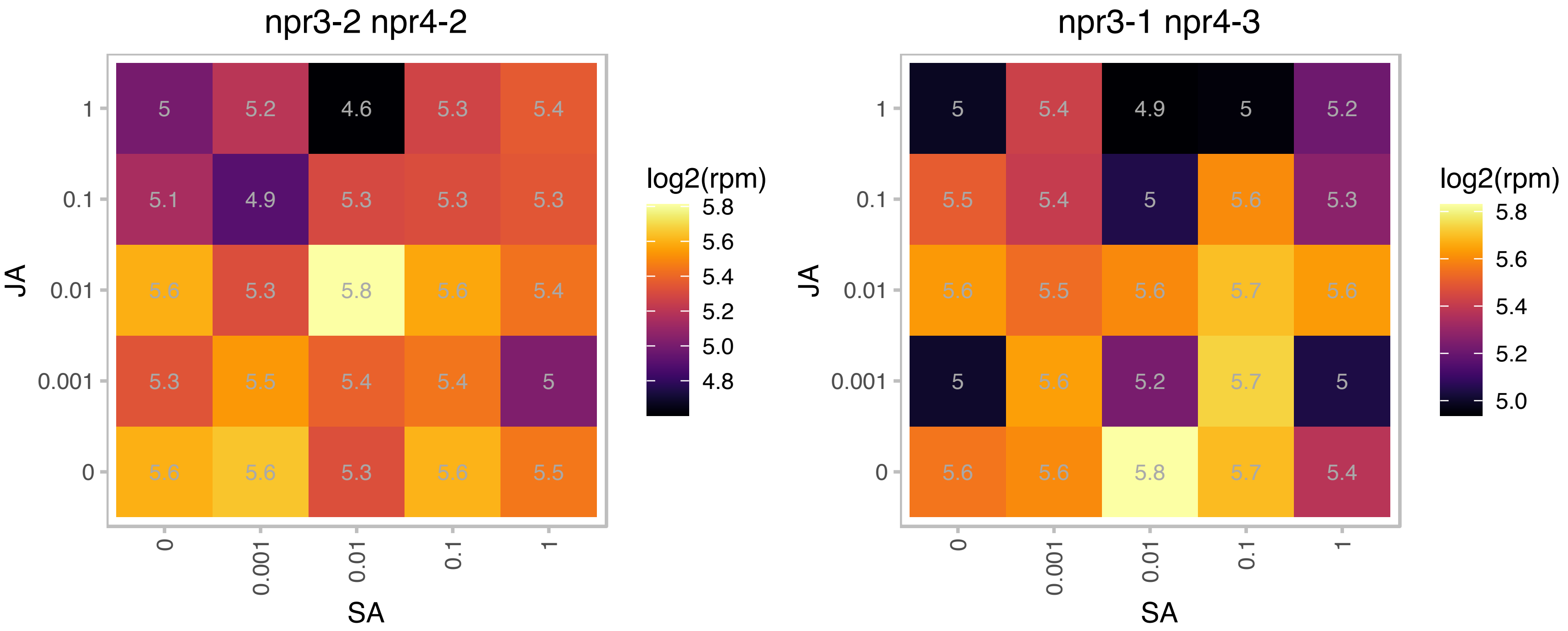

### AT1G70320.1 : UPL2

ubiquitin-protein ligase 2

encodes a ubiquitin-protein ligase-like protein containing a HECT domain. There are six other HECT-domain UPLs in Arabidopsis.

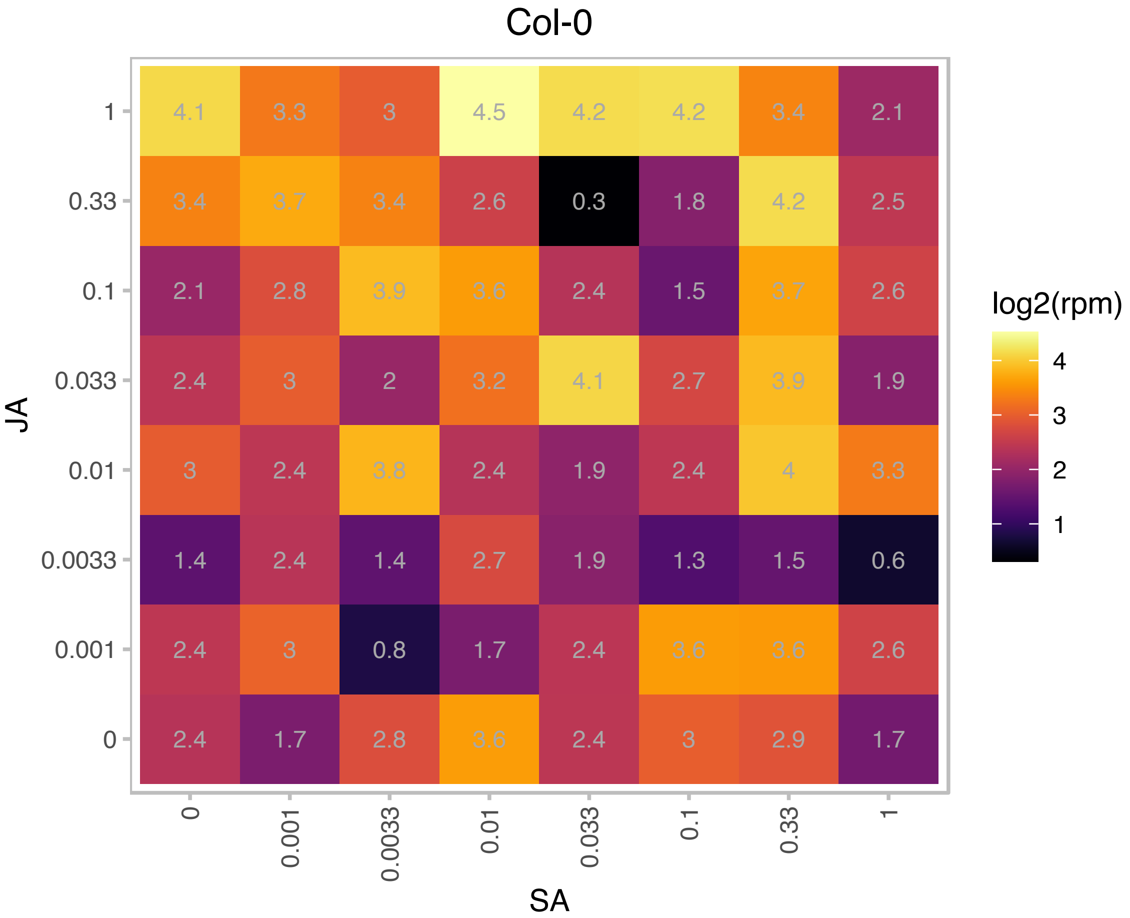


| | |
|---|---|
| Gene | AT1G70320 |
| is.expressed_genes | TRUE |
| is.responsive_genes | TRUE |
| is.apcluster | A-04 |

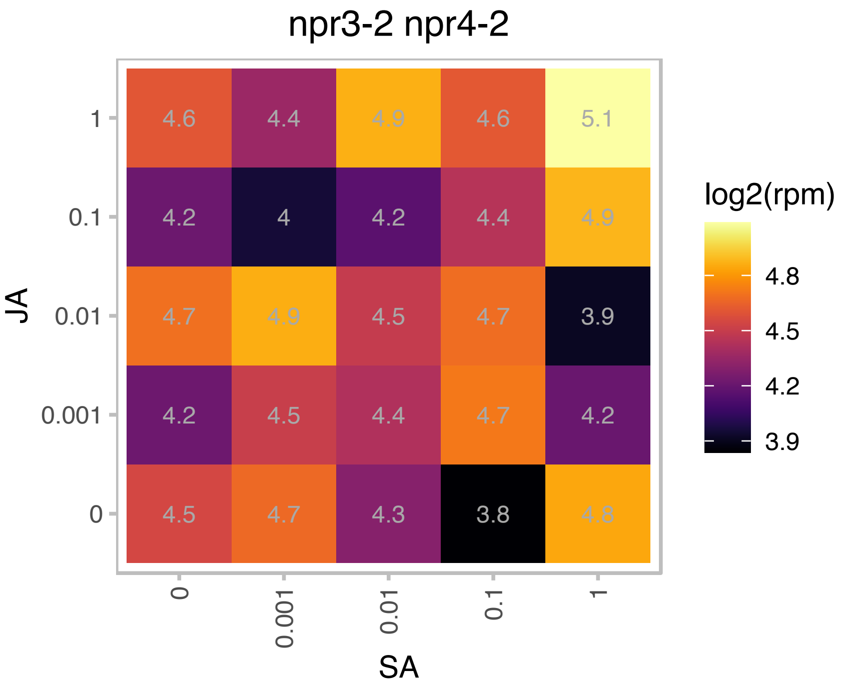


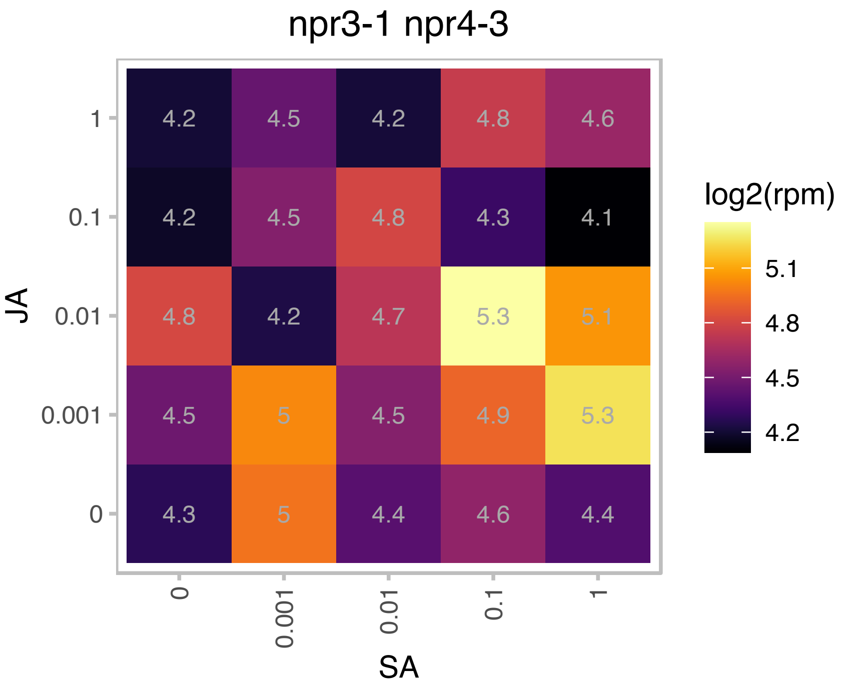

### AT1G78680.1 : ATGGH2, GGH2

gamma-glutamyl hydrolase 2

The Arabidopsis protein AtGGH2 is a gamma-glutamyl hydrolase acting specifically on monoglutamates. The enzyme is involved in the tetrahydrofolate metabolism and located to the vacuole.

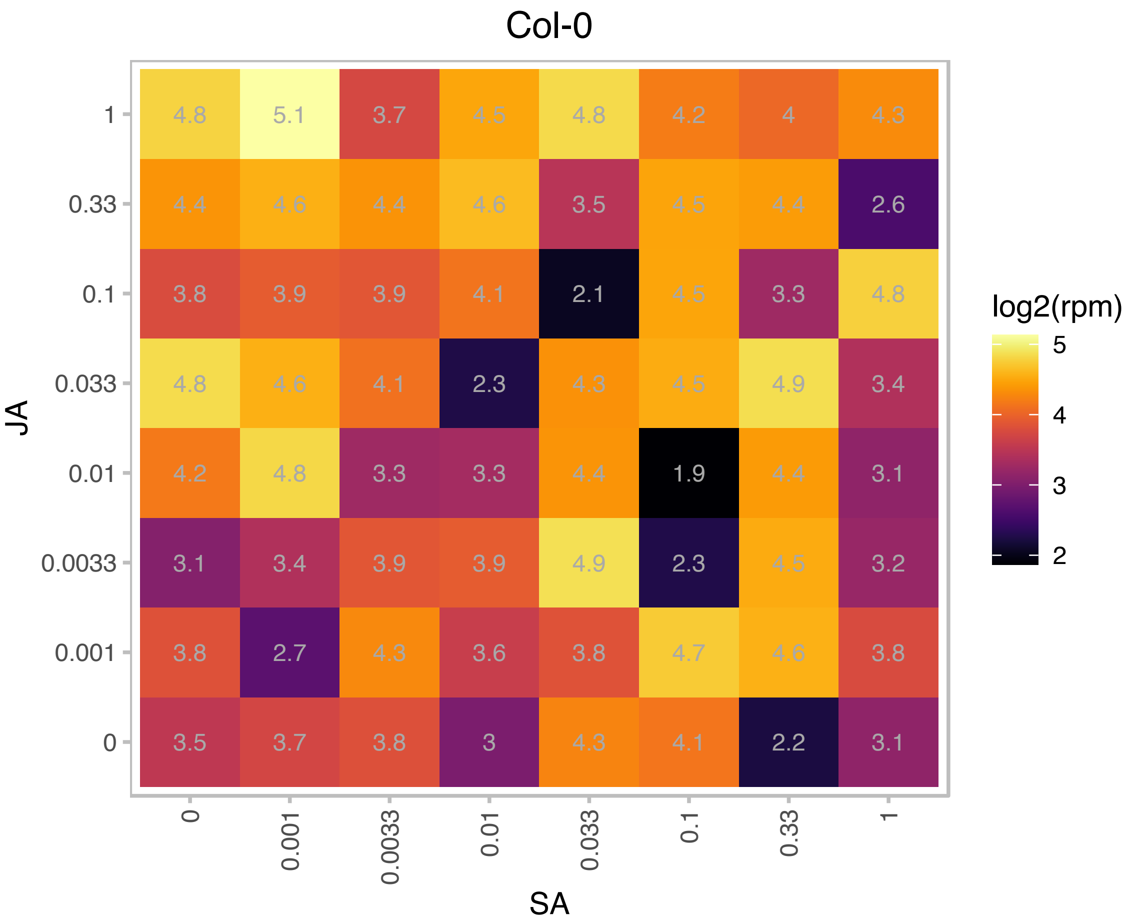


| | |
|---|---|
| Gene | AT1G78680 |
| is.expressed_genes | TRUE |
| is.responsive_genes | TRUE |
| is.apcluster | A-04 |

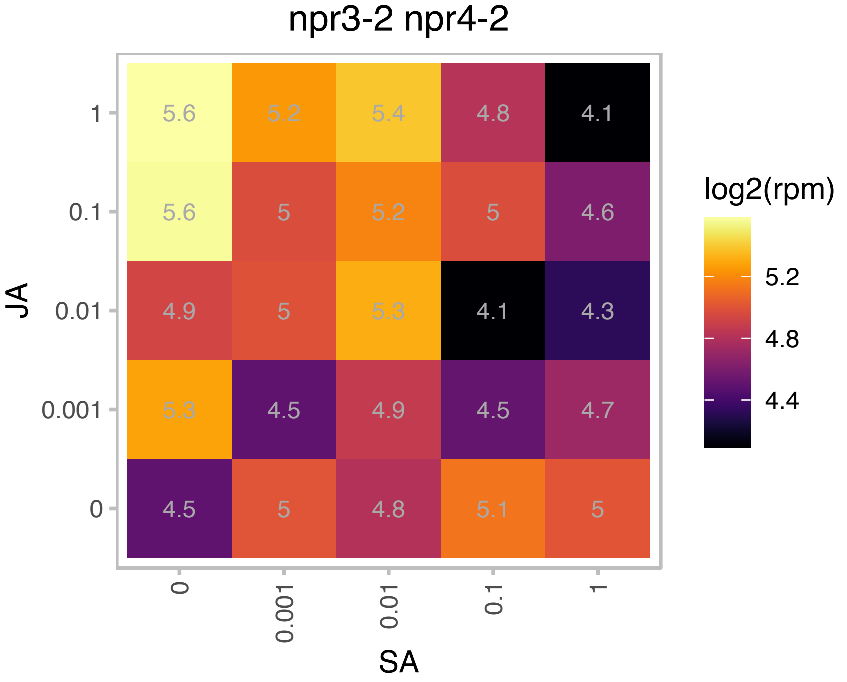


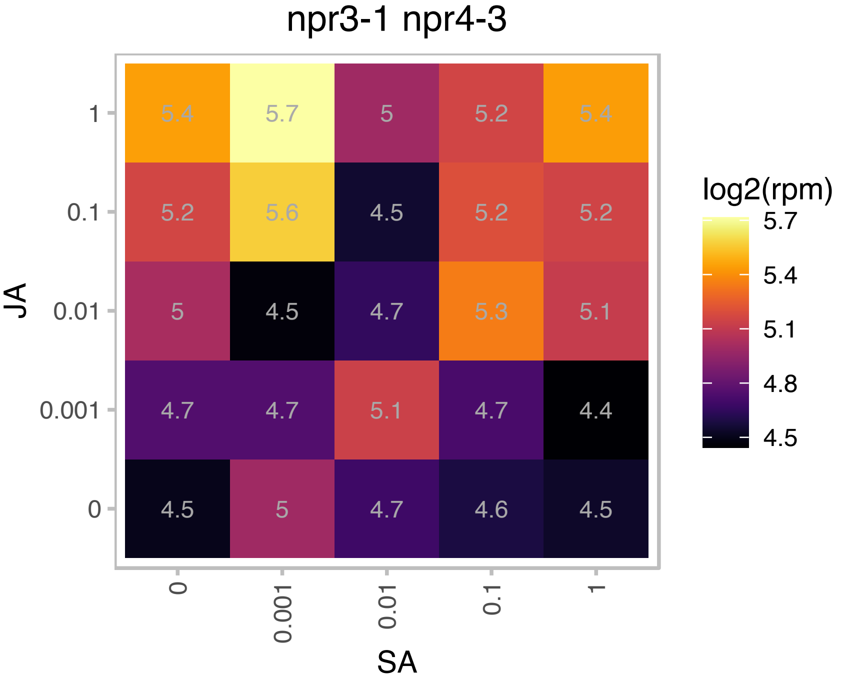

### AT1G79230.1 : ATMST1, ATRDH1, MST1, ST1, STR1

mercaptopyruvate sulfurtransferase 1

encodes a sulfurtransferase/rhodaneses, which belongs to a group of enzymes widely distributed in all three phyla that catalyze the transfer of sulfur from a donor to a thiophilic acceptor substrate. The protein and transcript levels are NOT affected by senescence or exogenous cyanide, suggesting that sulfurtransferases are involved in cyanide detoxification.

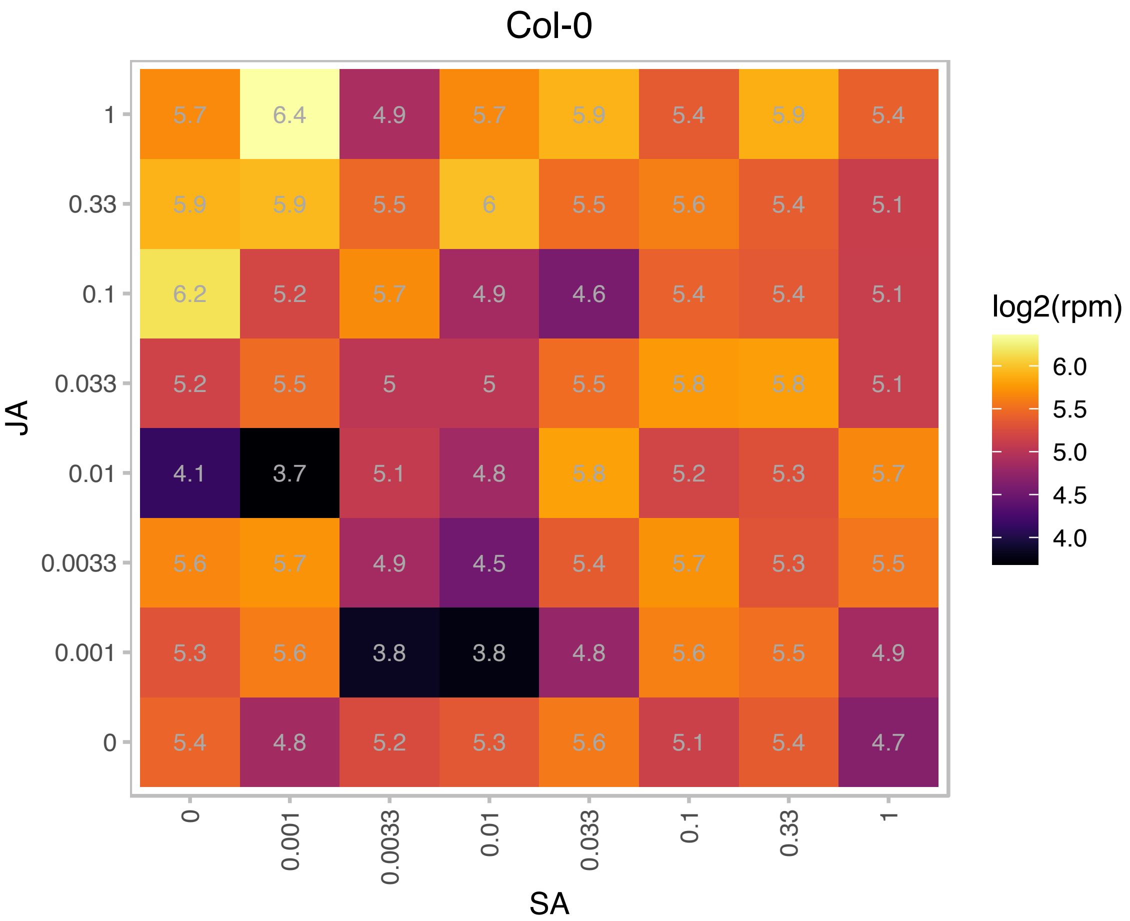


| | |
|---|---|
| Gene | AT1G79230 |
| is.expressed_genes | TRUE |
| is.responsive_genes | TRUE |
| is.apcluster | A-04 |

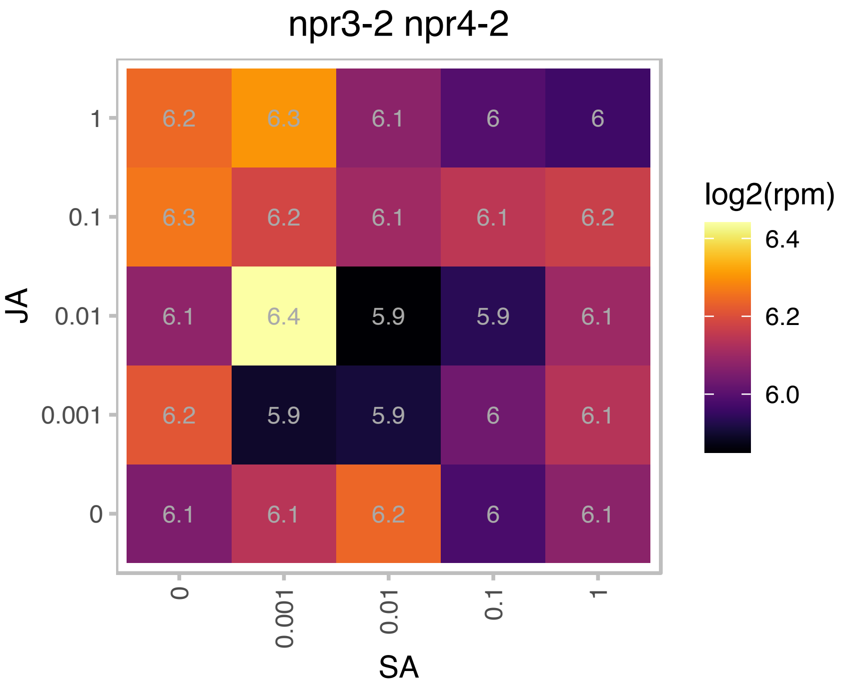


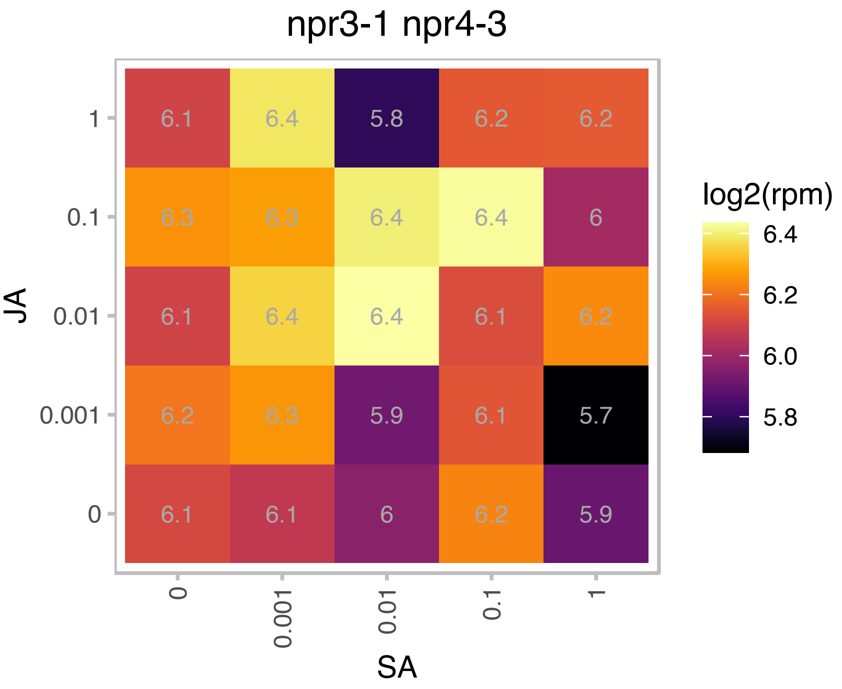

# AT1G80940.1 : NA

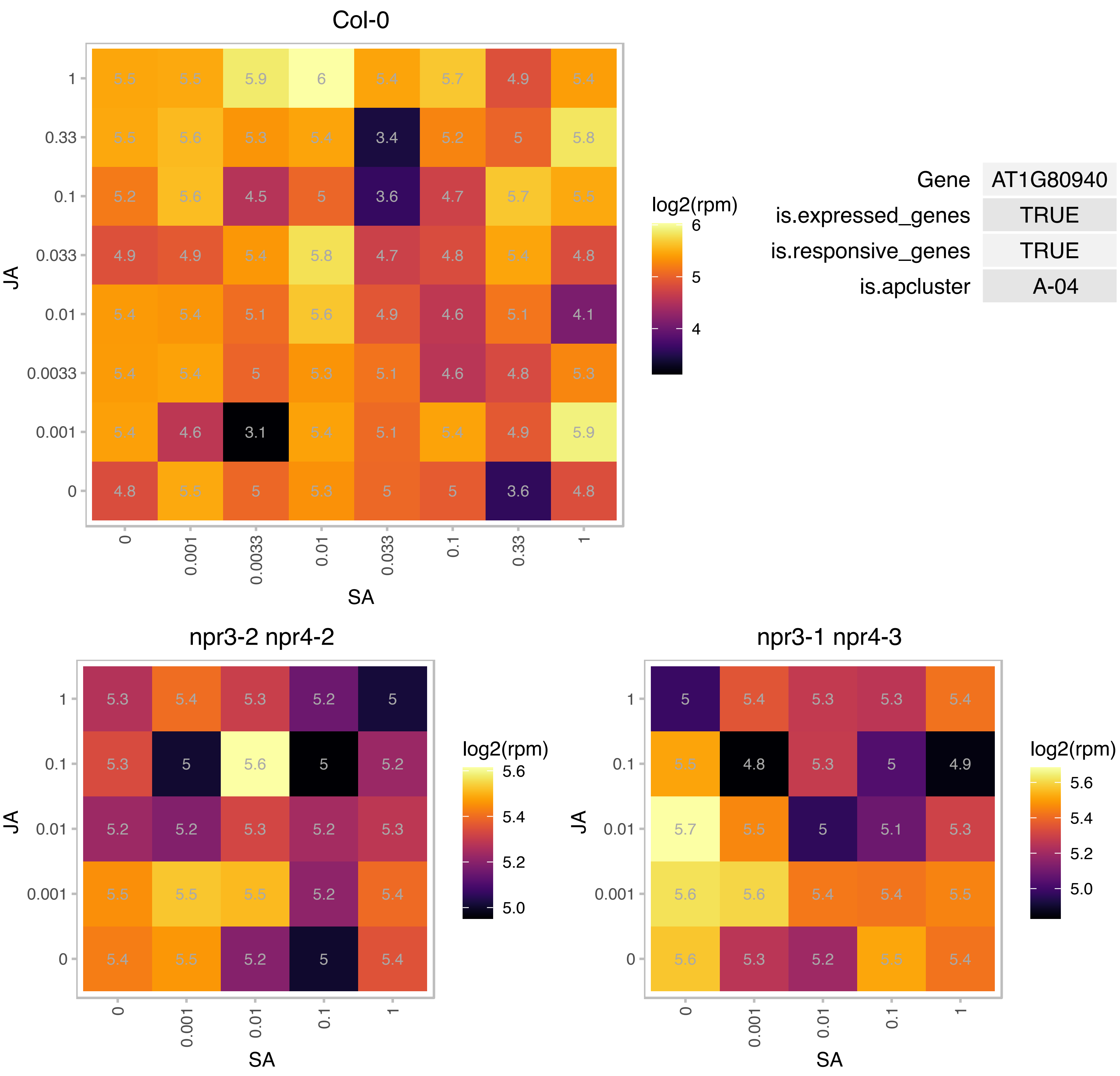

### AT2G17390.1 : AKR2B

ankyrin repeat-containing 2B
Highly homologous to AKR2A. Involved in chloroplast biogenesis. Double mutants of AKR2A and AKR2B have yello
leaves, significantly reduced chloroplast proteins, and no thylakoid membranes. The mRNA is cell-to-cell mobile.

Col-0

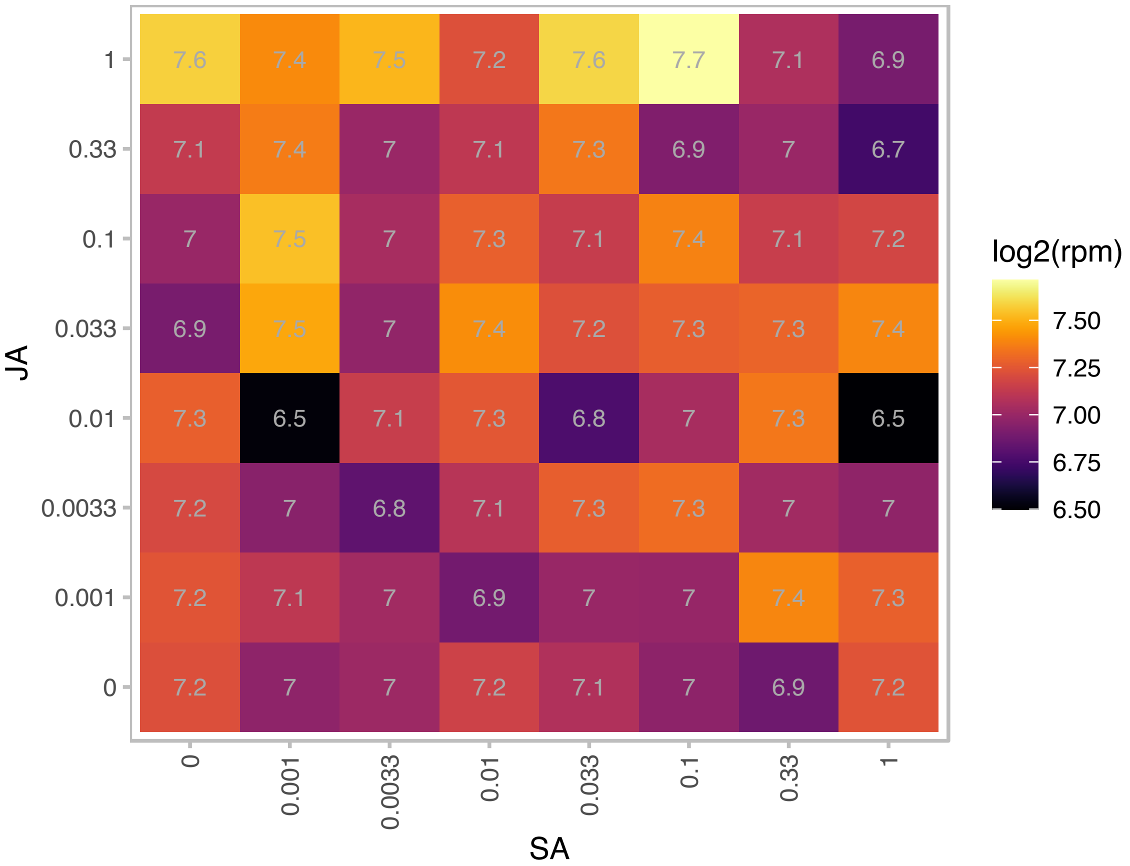


| | |
|---|---|
| Gene | AT2G17390 |
| is.expressed_genes | TRUE |
| is.responsive_genes | TRUE |
| is.apcluster | A-04 |

npr3-2 npr4-2

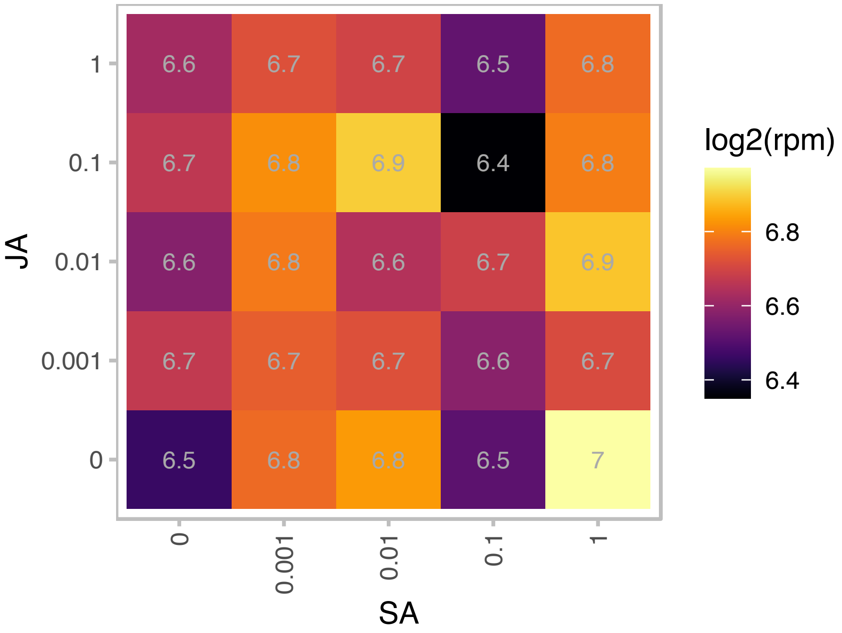


npr3-1 npr4-3

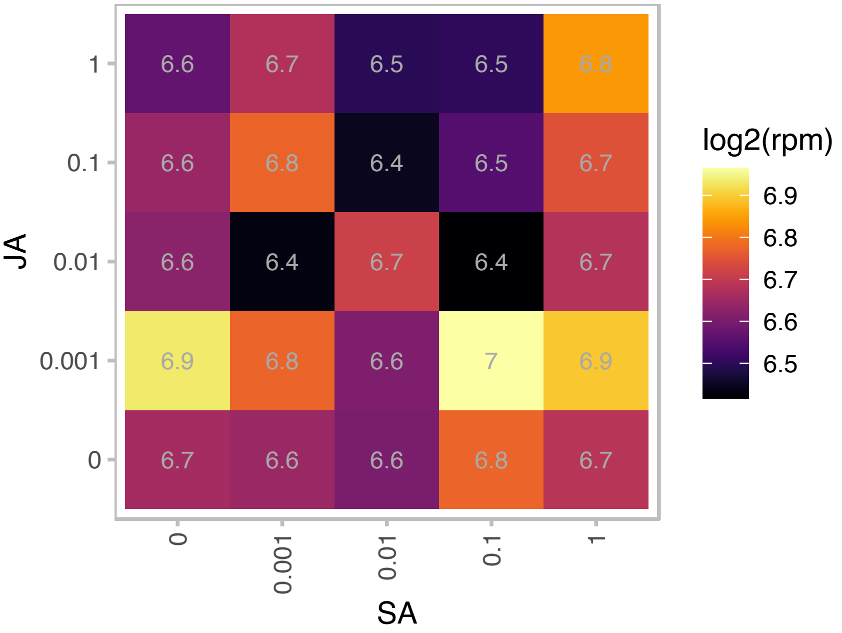

### AT2G20580.1 : ATRPN1A, RPN1A

26S proteasome regulatory subunit S2 1A

encoding the RPN subunits of the 26S proteasome The mRNA is cell-to-cell mobile.

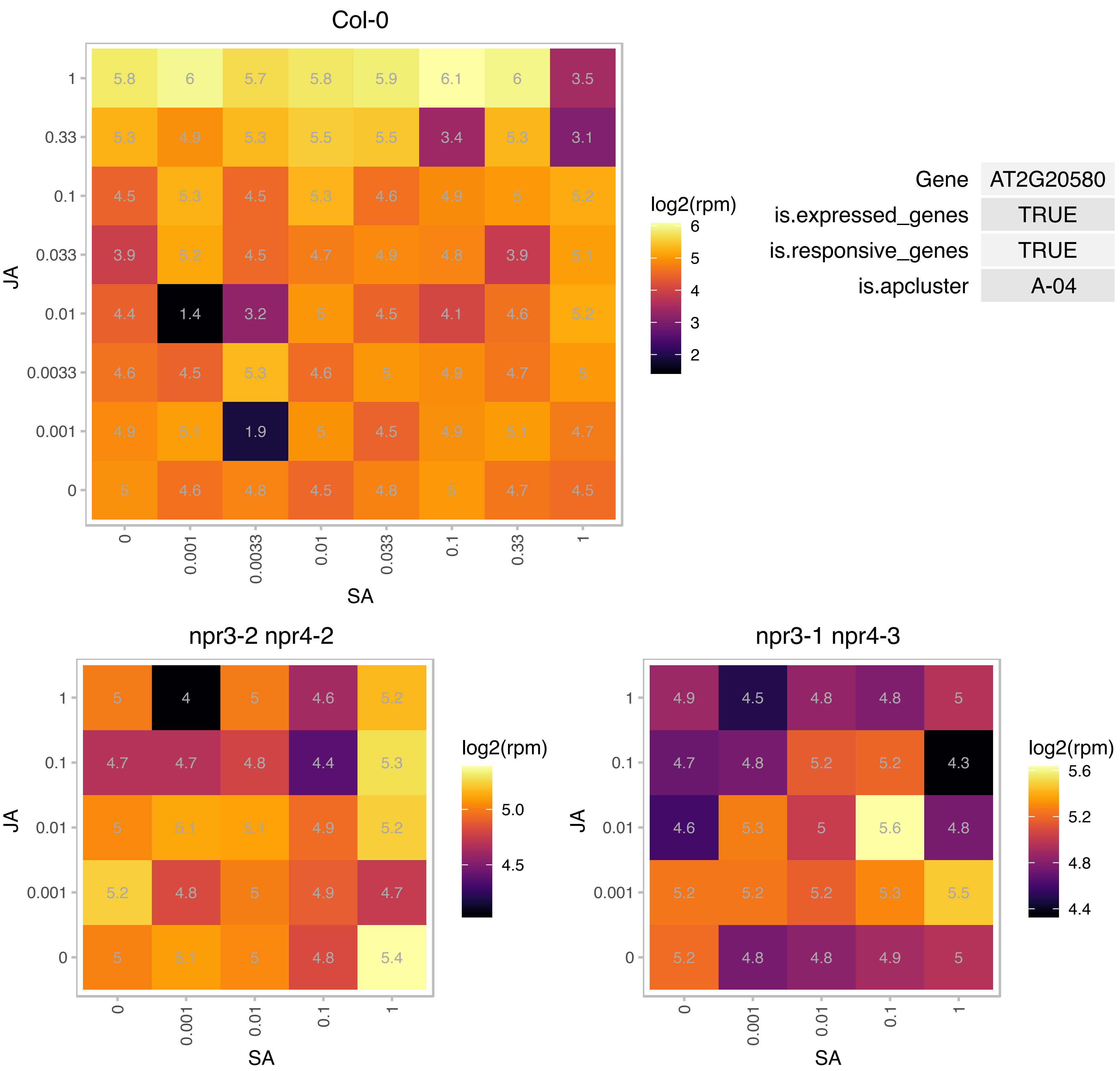


| | |
|---|---|
| Gene | AT2G20580 |
| is.expressed_genes | TRUE |
| is.responsive_genes | TRUE |
| is.apcluster | A-04 |

### AT2G21270.1 : UFD1

ubiquitin fusion degradation 1

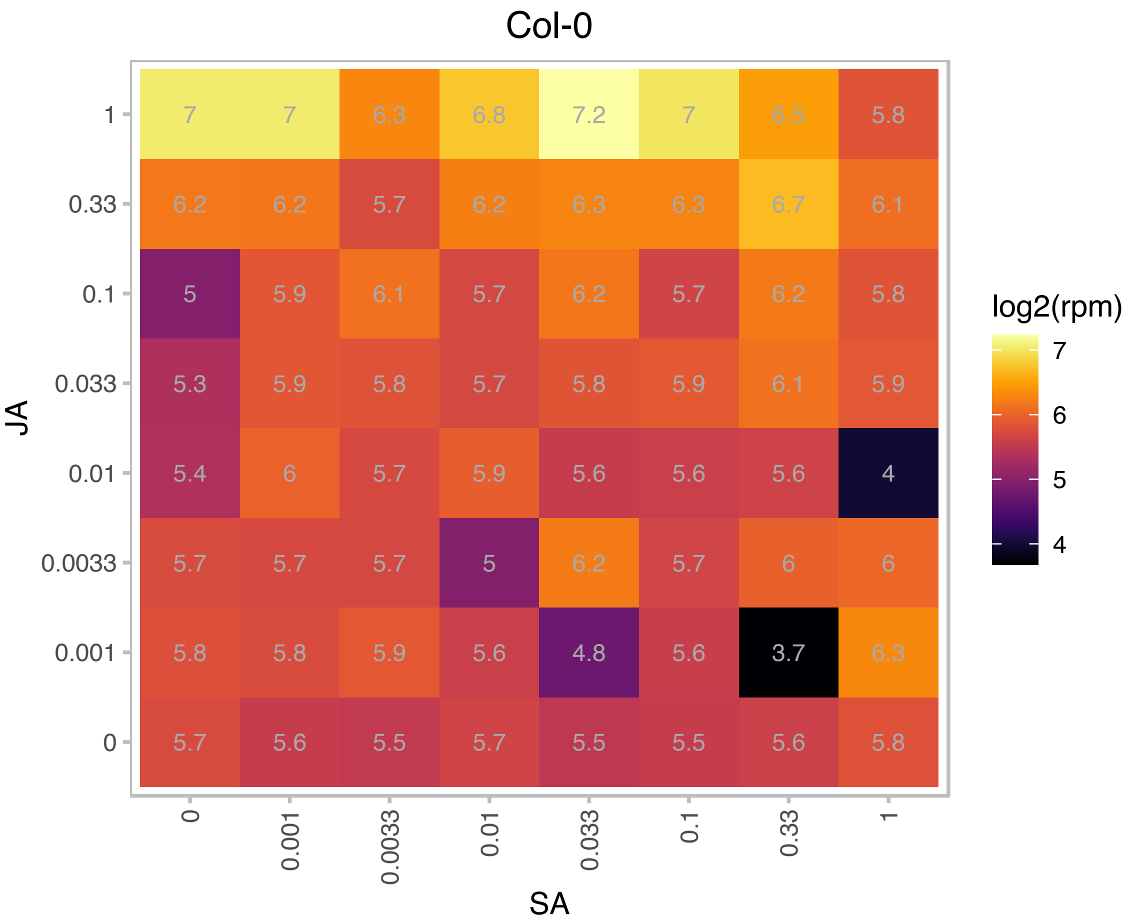


| | |
|---|---|
| Gene | AT2G21270 |
| is.expressed_genes | TRUE |
| is.responsive_genes | TRUE |
| is.apcluster | A-04 |

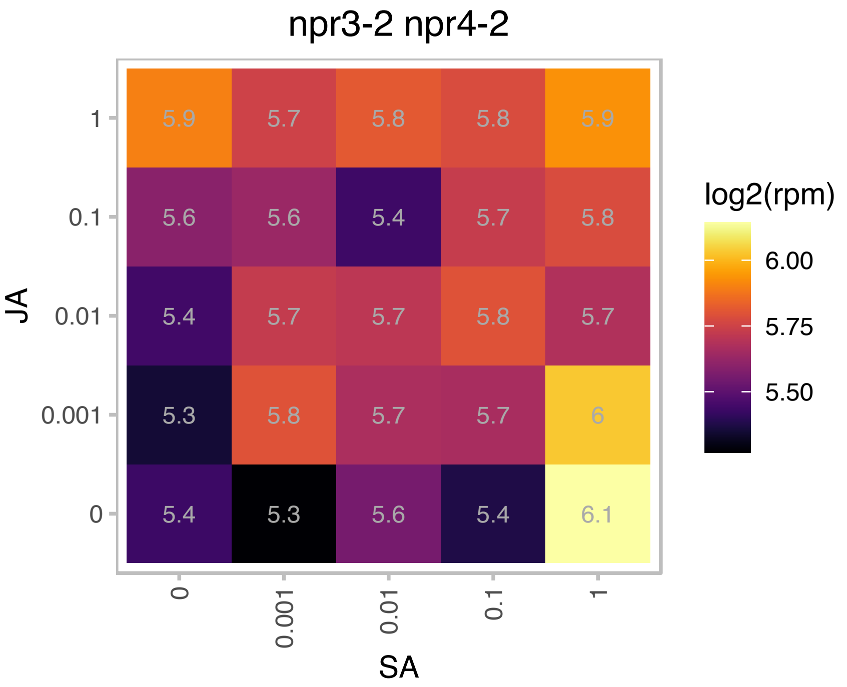


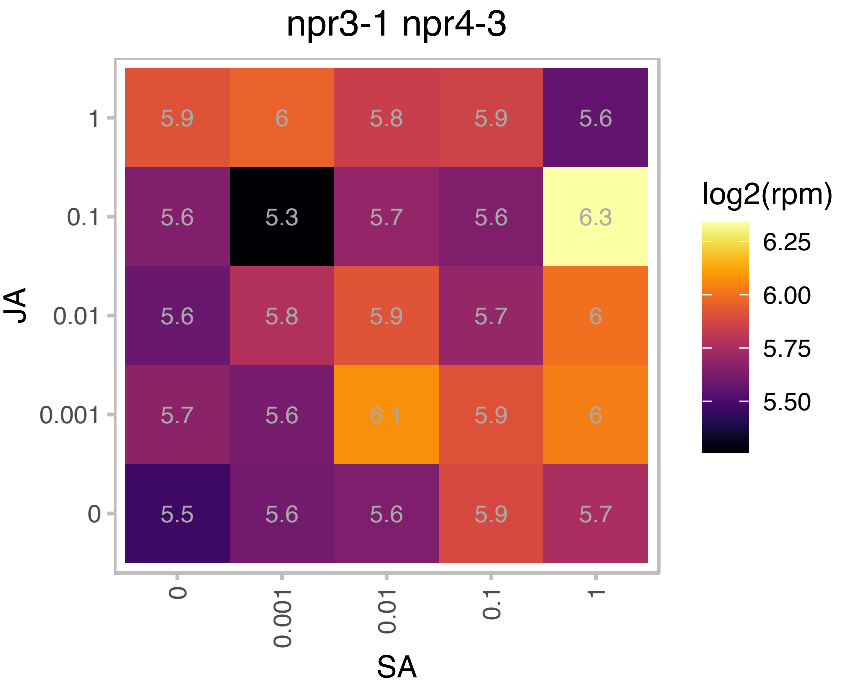

### AT2G22990.1 : SCPL8, SNG1

sinapoylglucose 1

sinapoylglucose:malate sinapoyltransferase. Catalyzes the formation of sinapoylmalate from sinapoylglucose. Mutants accumulate excess sinapoylglucose.

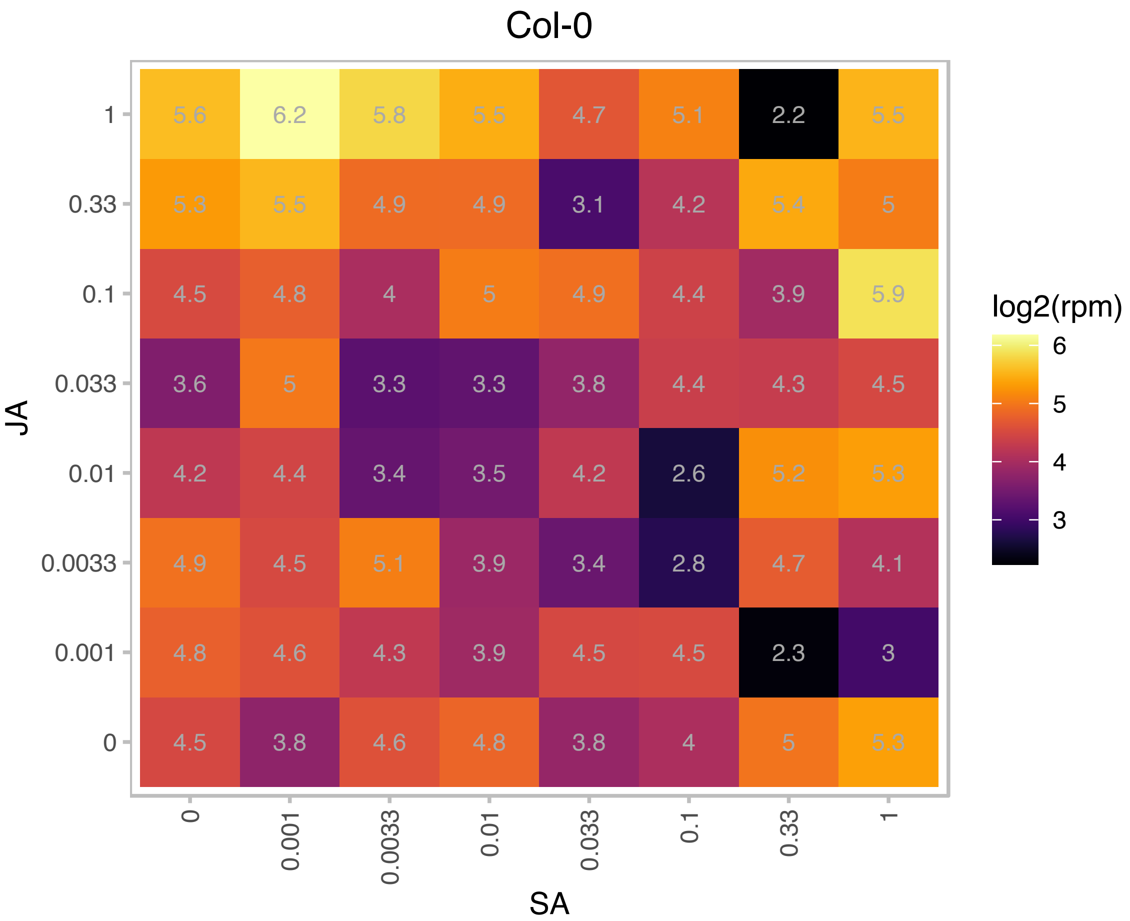


| | |
|---|---|
| Gene | AT2G22990 |
| is.expressed_genes | TRUE |
| is.responsive_genes | TRUE |
| is.apcluster | A-04 |

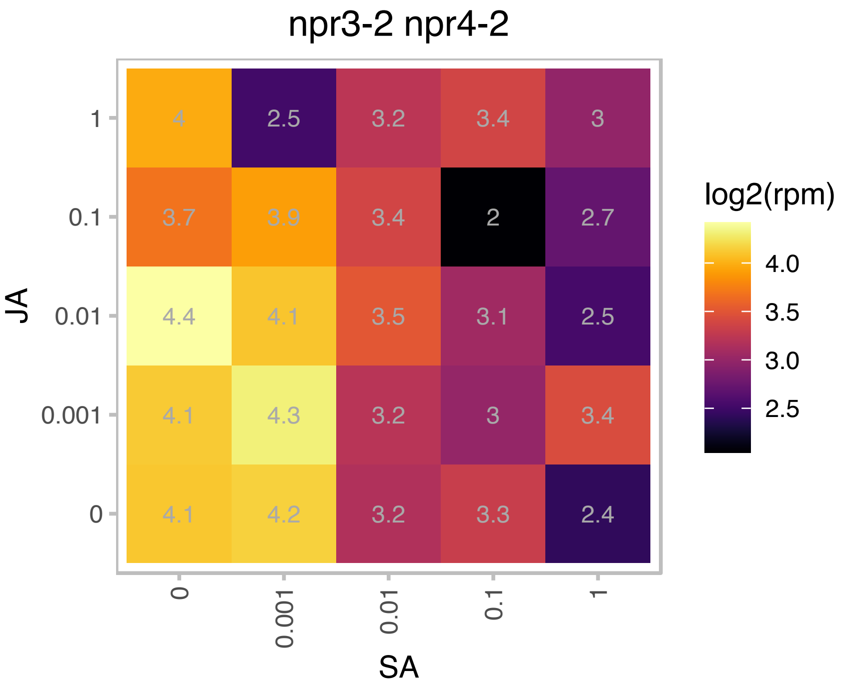


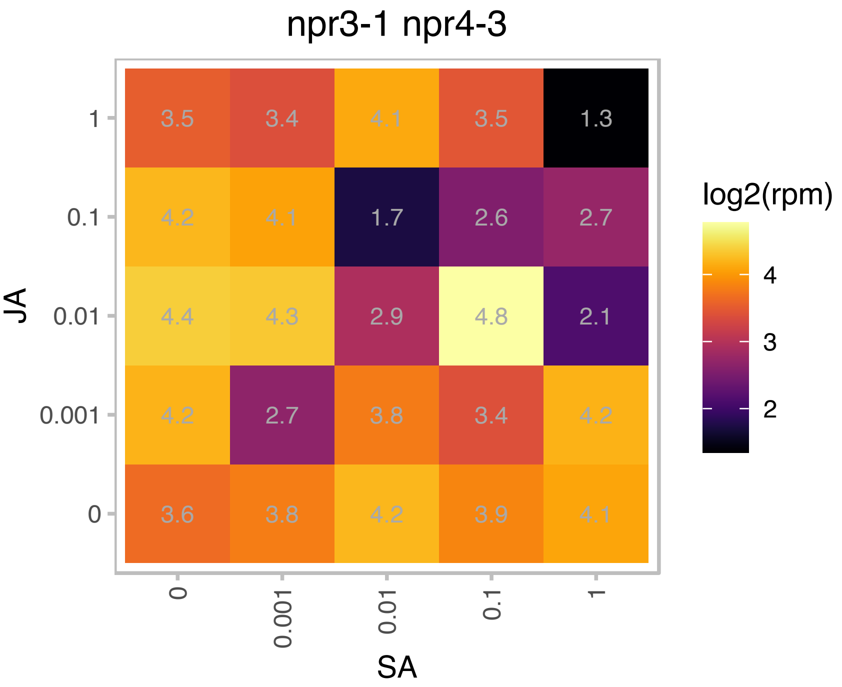

### AT2G41520.1 : DJC62, TPR15

Heat shock protein DnaJ with tetratricopeptide repeat

Encodes one of the 36 carboxylate clamp (CC)-tetratricopeptide repeat (TPR) proteins (Prasad 2010, Pubmed ID: 20856808) with potential to interact with Hsp90/Hsp70 as co-chaperones.

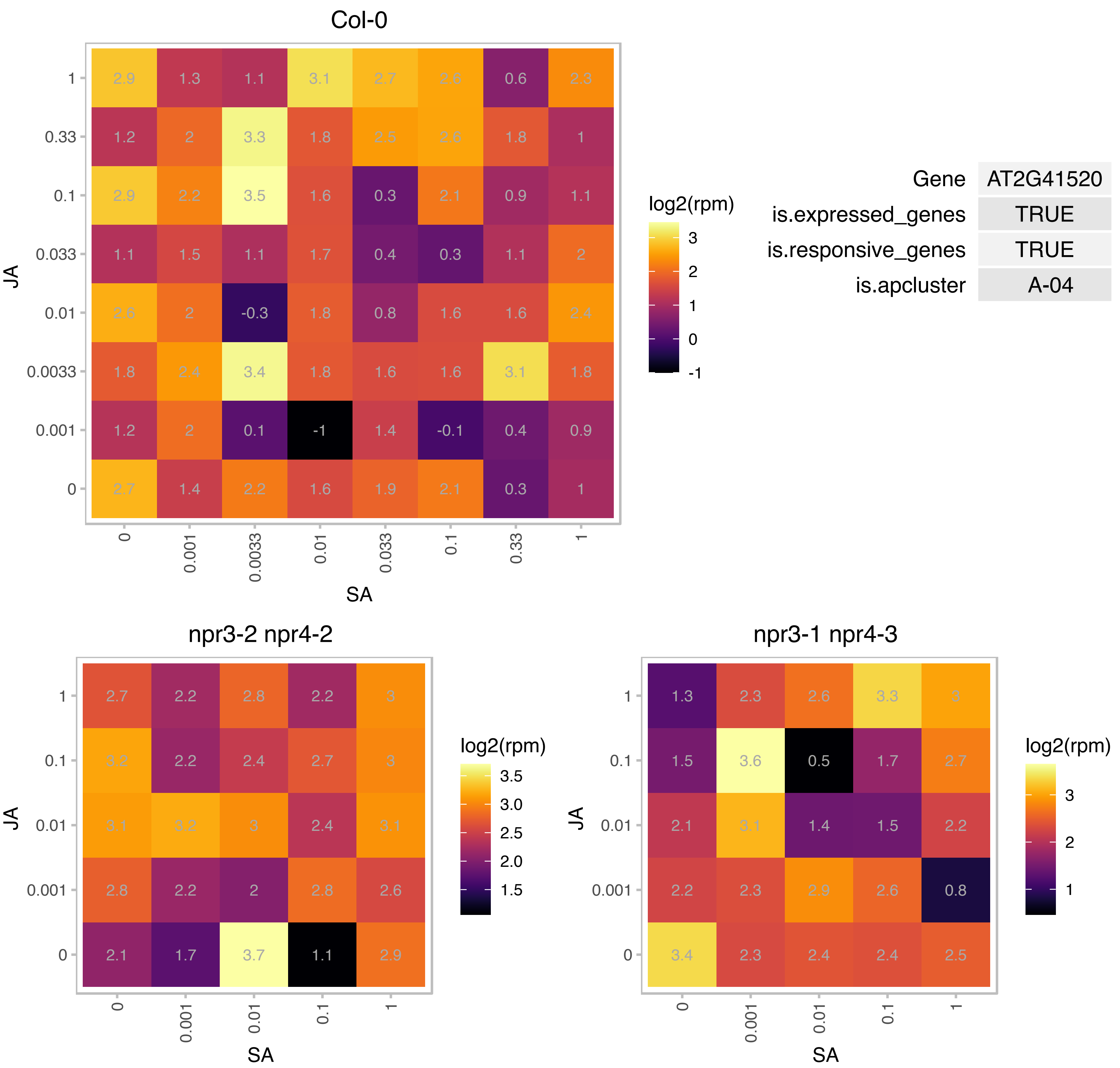


| | |
|---|---|
| Gene | AT2G41520 |
| is.expressed_genes | TRUE |
| is.responsive_genes | TRUE |
| is.apcluster | A-04 |

# AT2G44670.1 : NA

Protein of unknown function (DUF581)

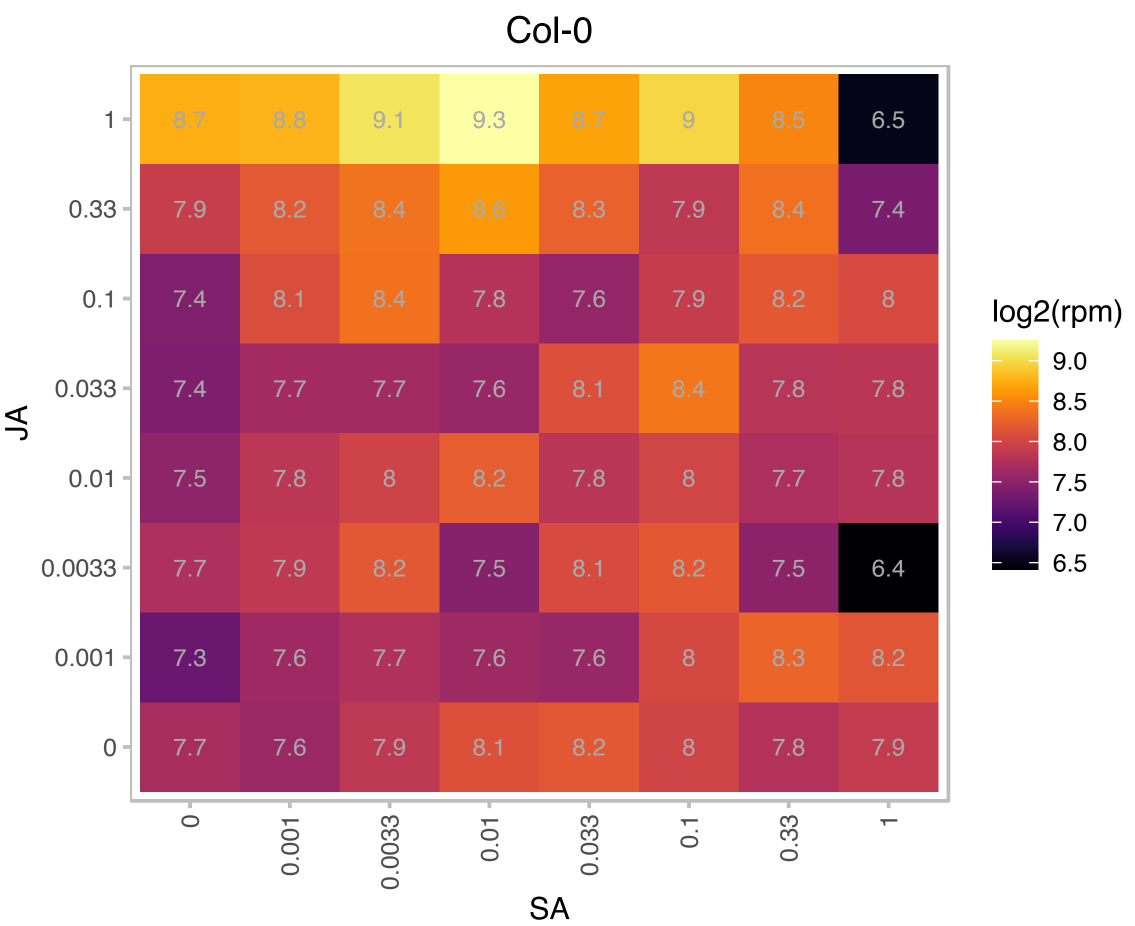


| | |
|---|---|
| Gene | AT2G44670 |
| is.expressed_genes | TRUE |
| is.responsive_genes | TRUE |
| is.apcluster | A-04 |

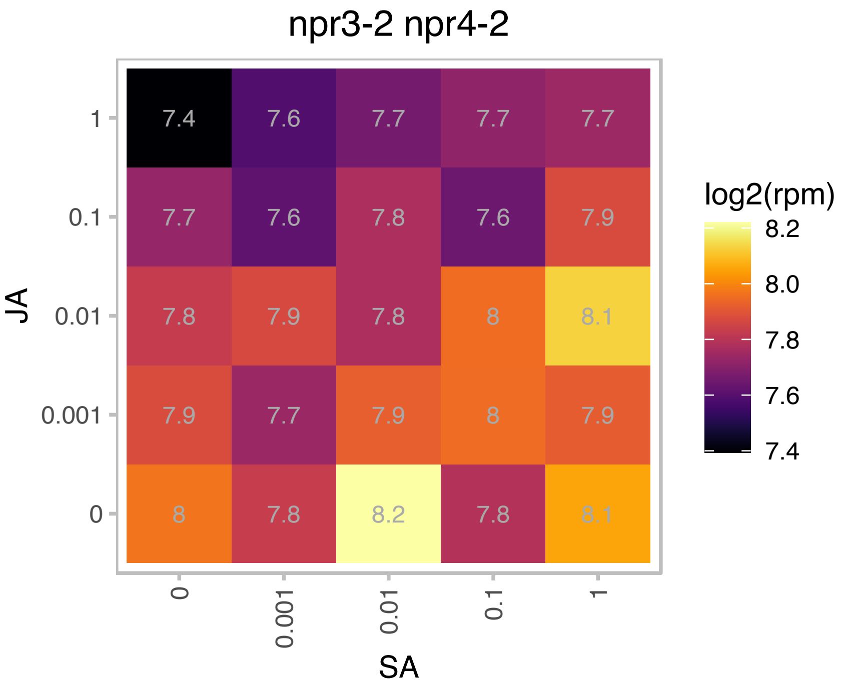


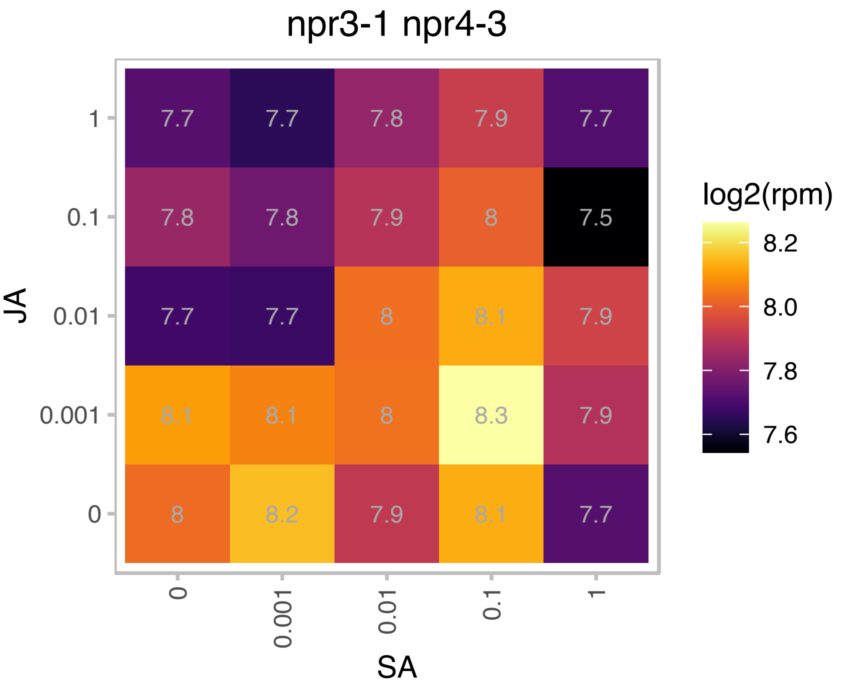

### AT3G03340.1 : UNE6

LUC7 related protein

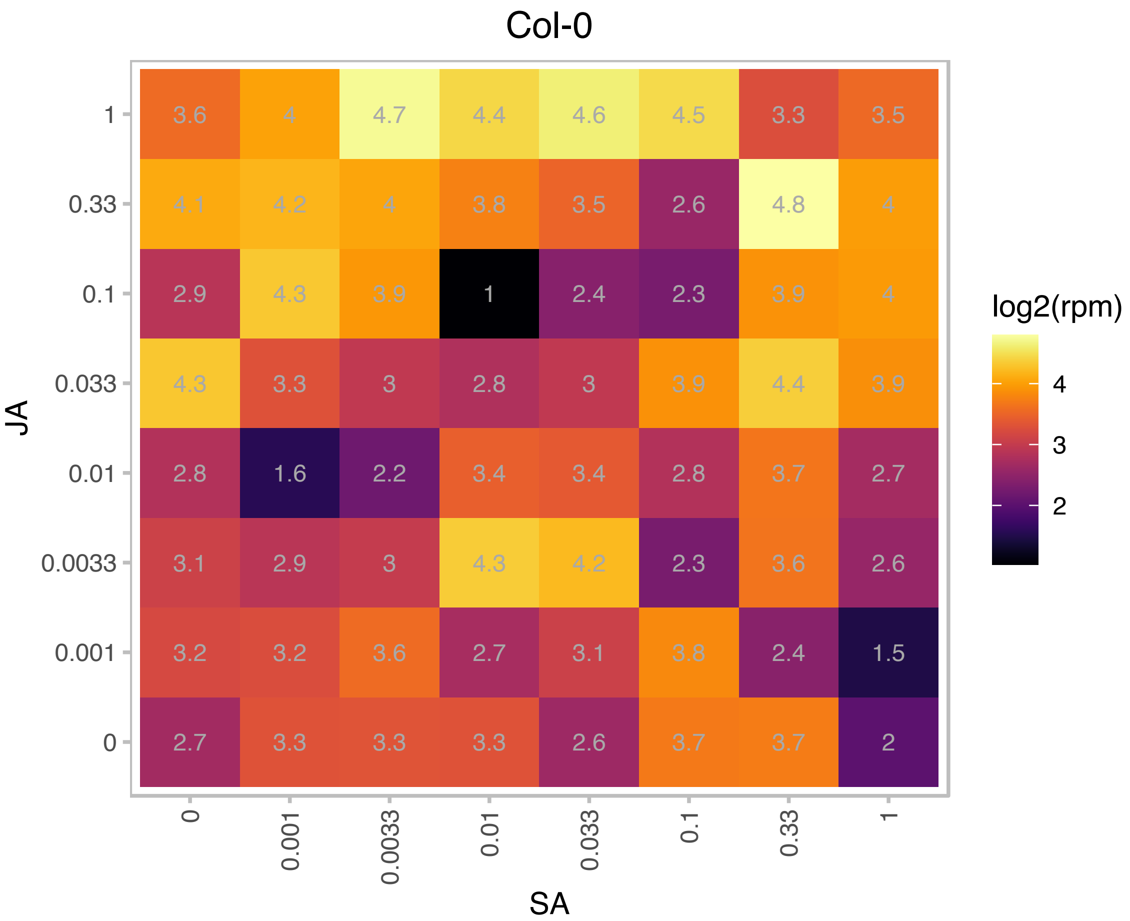


| | |
|---|---|
| Gene | AT3G03340 |
| is.expressed_genes | TRUE |
| is.responsive_genes | TRUE |
| is.apcluster | A-04 |

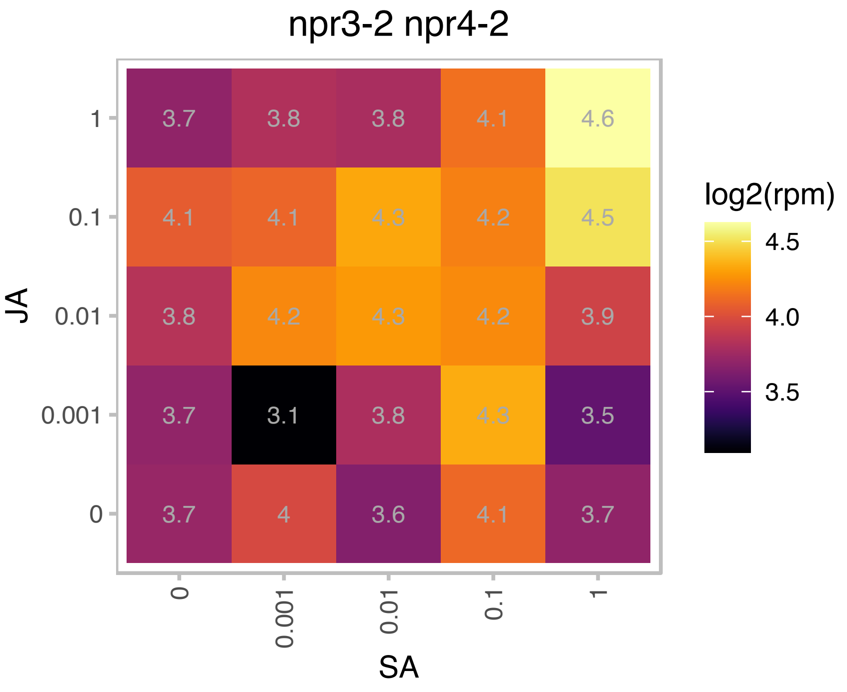


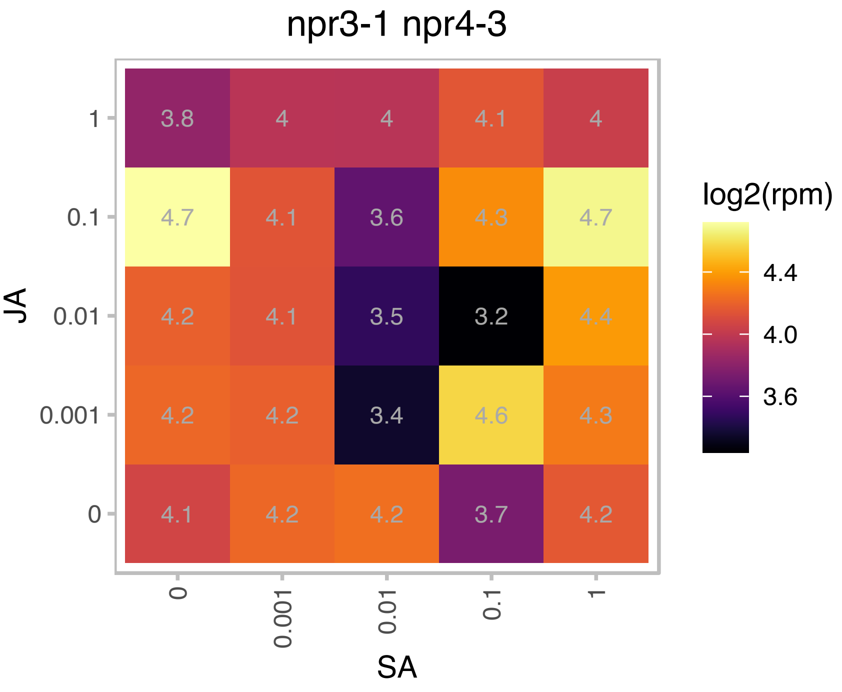

### AT3G05010.1 : Cand2

Protein of unknown function, transmembrane-40

Encodes a candidate G-protein Coupled Receptor that is involved in the regulation of root growth by bacterial N-acyl-homoserine lactones (AHLs) and plays a role in mediating interactions between plants and microbes.

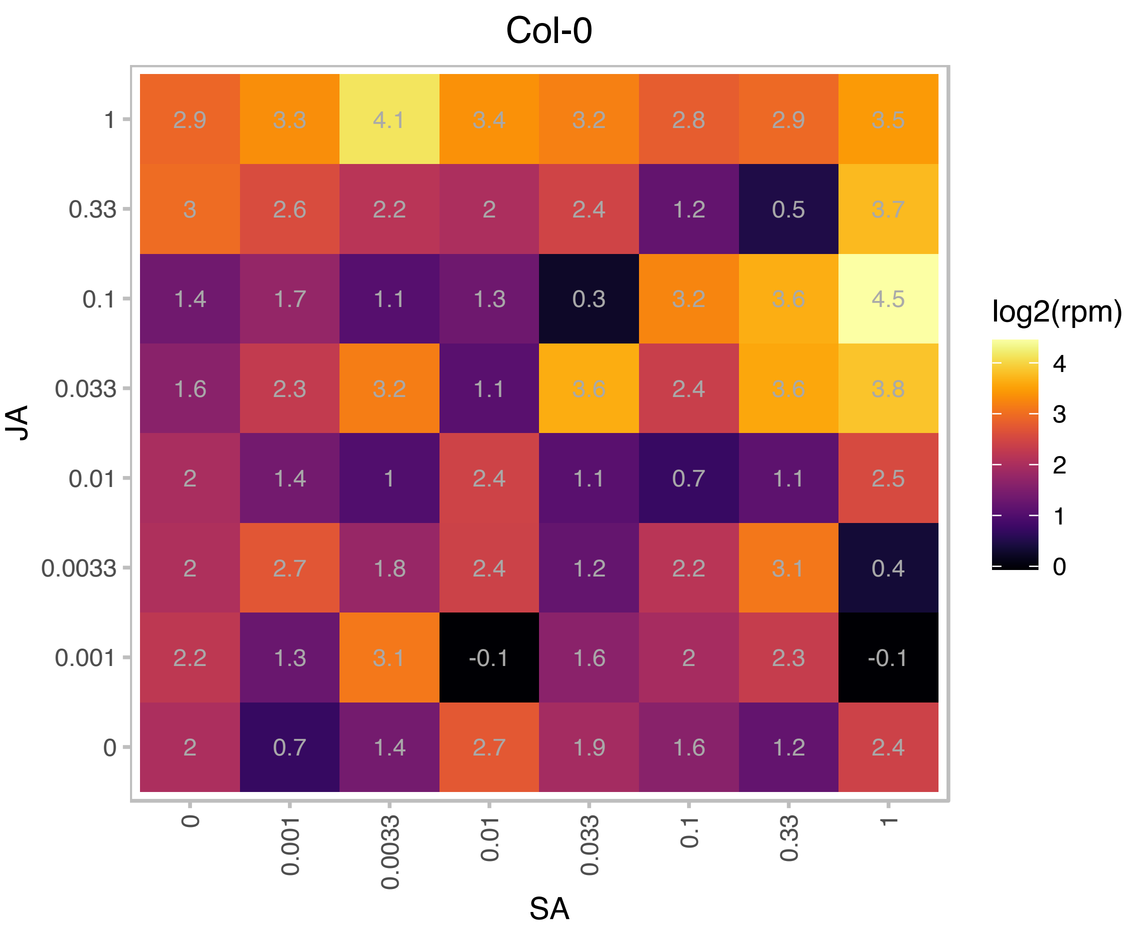


| | |
|---|---|
| Gene | AT3G05010 |
| is.expressed_genes | TRUE |
| is.responsive_genes | TRUE |
| is.apcluster | A-04 |

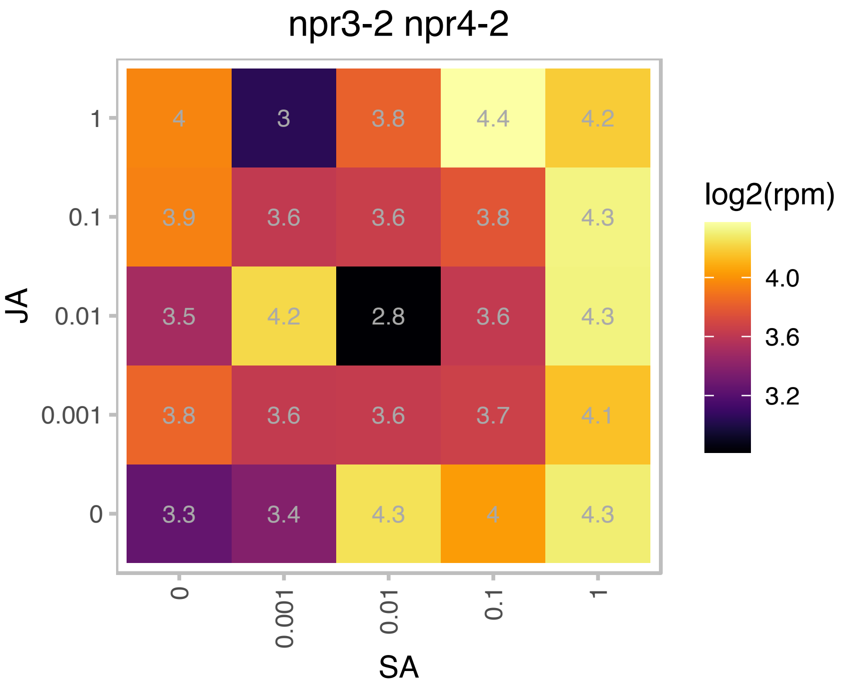


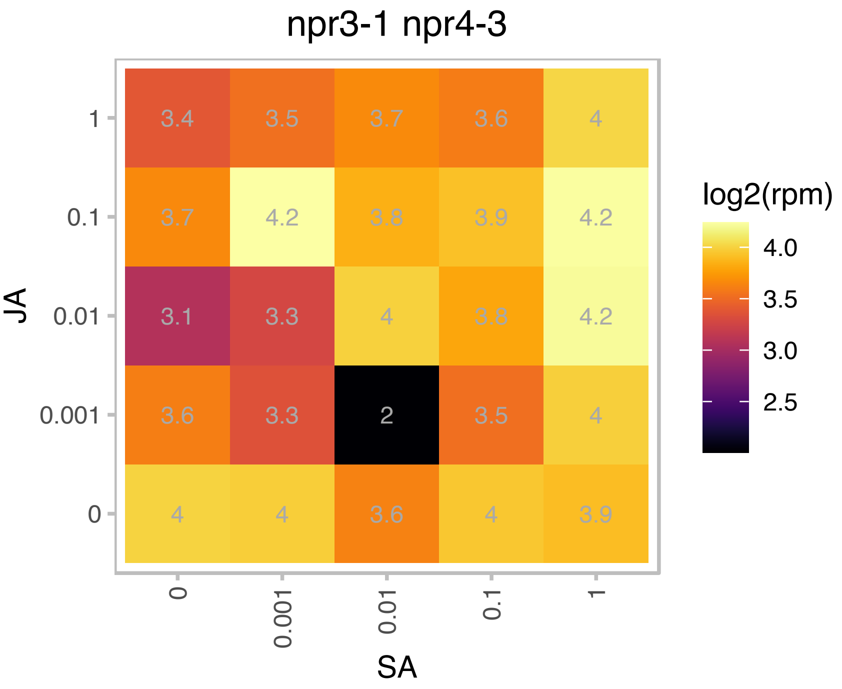

# AT3G05070.1 : NA

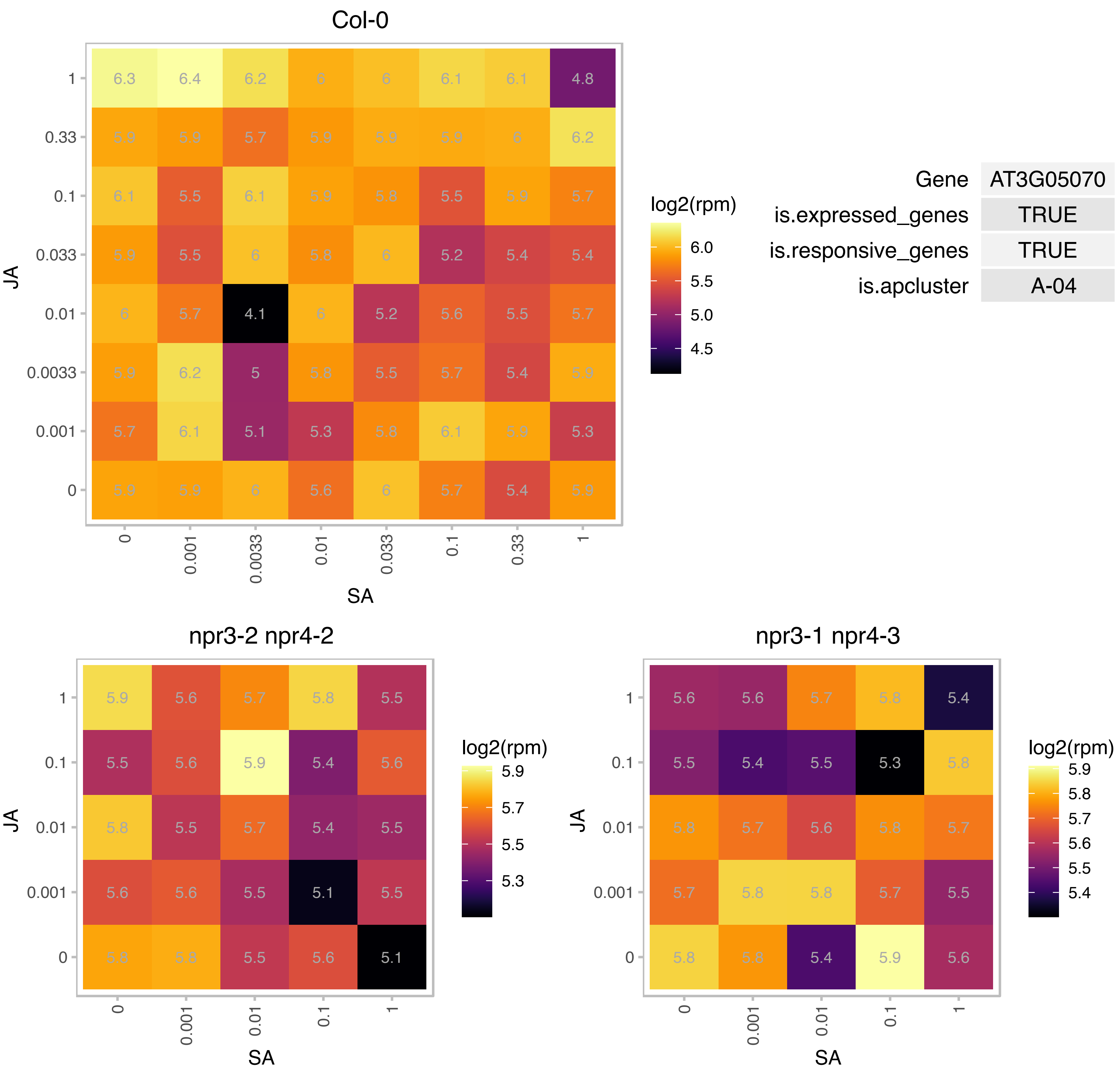

# AT3G06410.1 : NA

Zinc finger C-x8-C-x5-C-x3-H type family protein

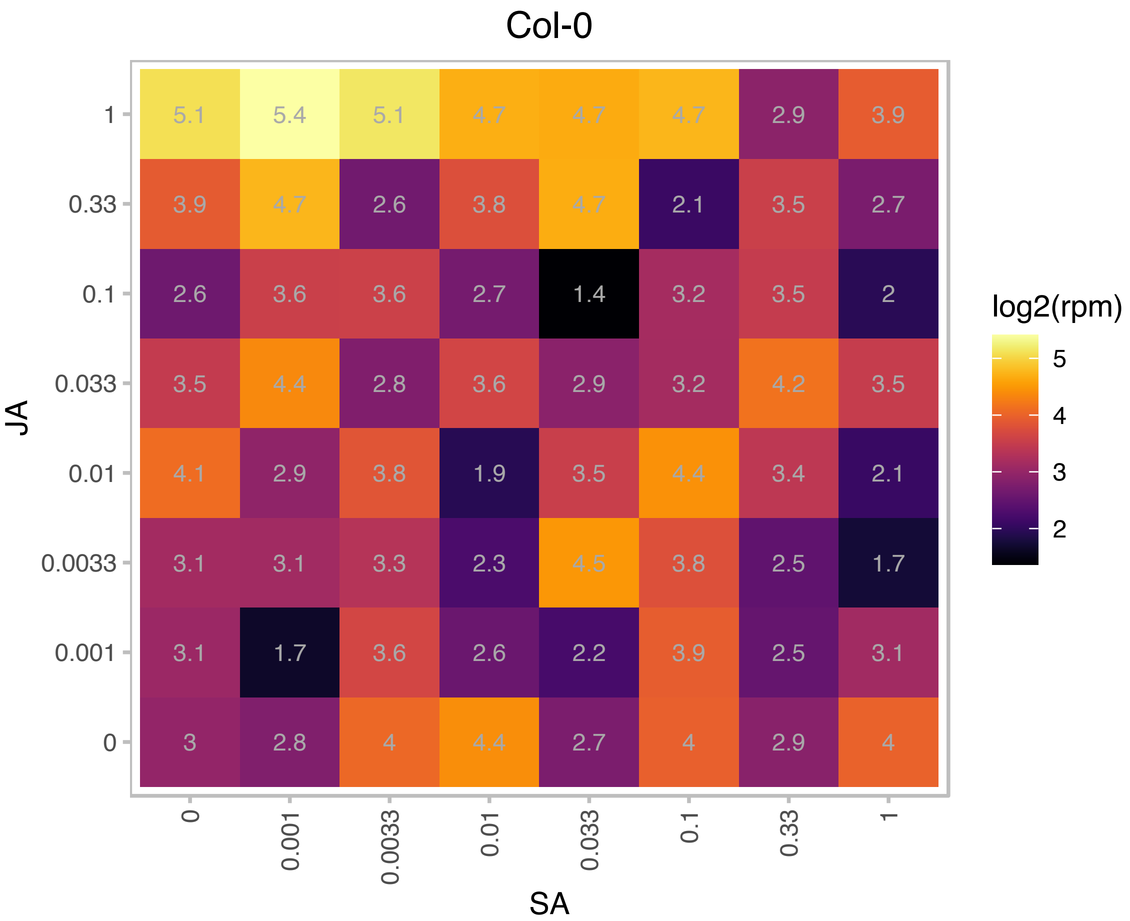


| | |
|---|---|
| Gene | AT3G06410 |
| is.expressed_genes | TRUE |
| is.responsive_genes | TRUE |
| is.apcluster | A-04 |

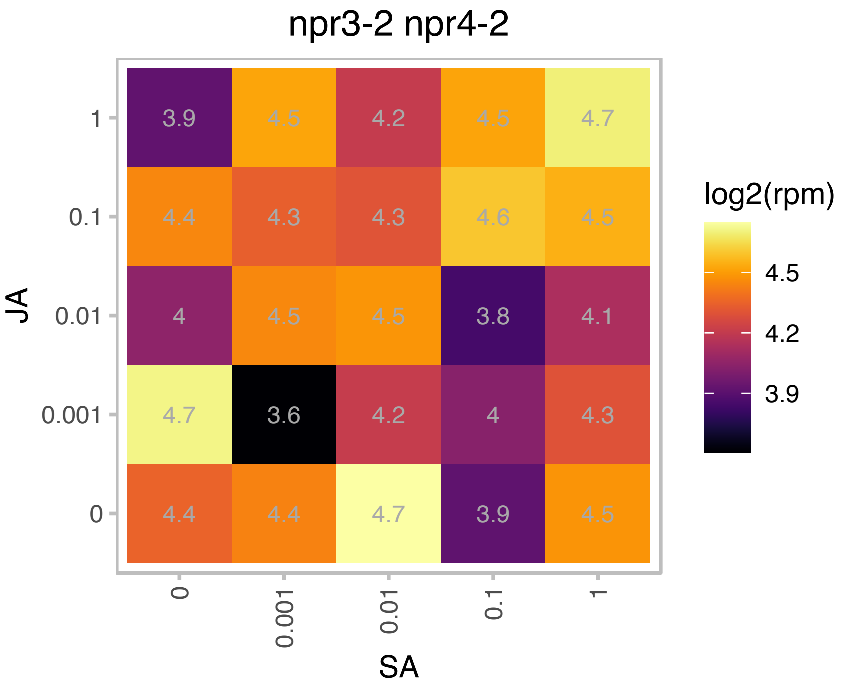


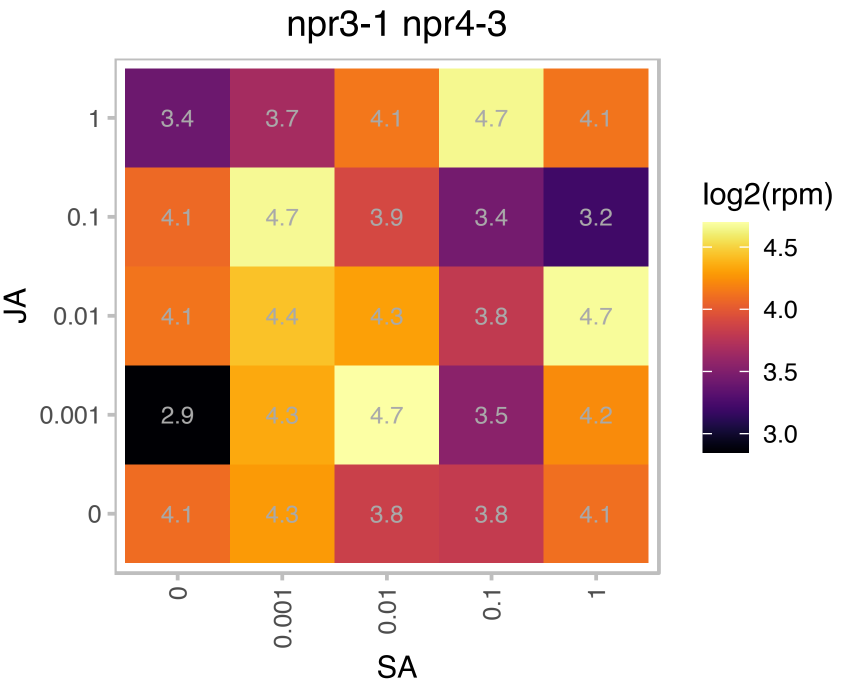

### AT3G14067.1 : SASP

Subtilase family protein
Encodes a protein with similarity to serine protease, subtilisin, that is upregulated during senescence and expressed in the arial portions of the plant.Loss of function mutations have increased branch number but normal silique length and seed set and therefore have increased fertility.

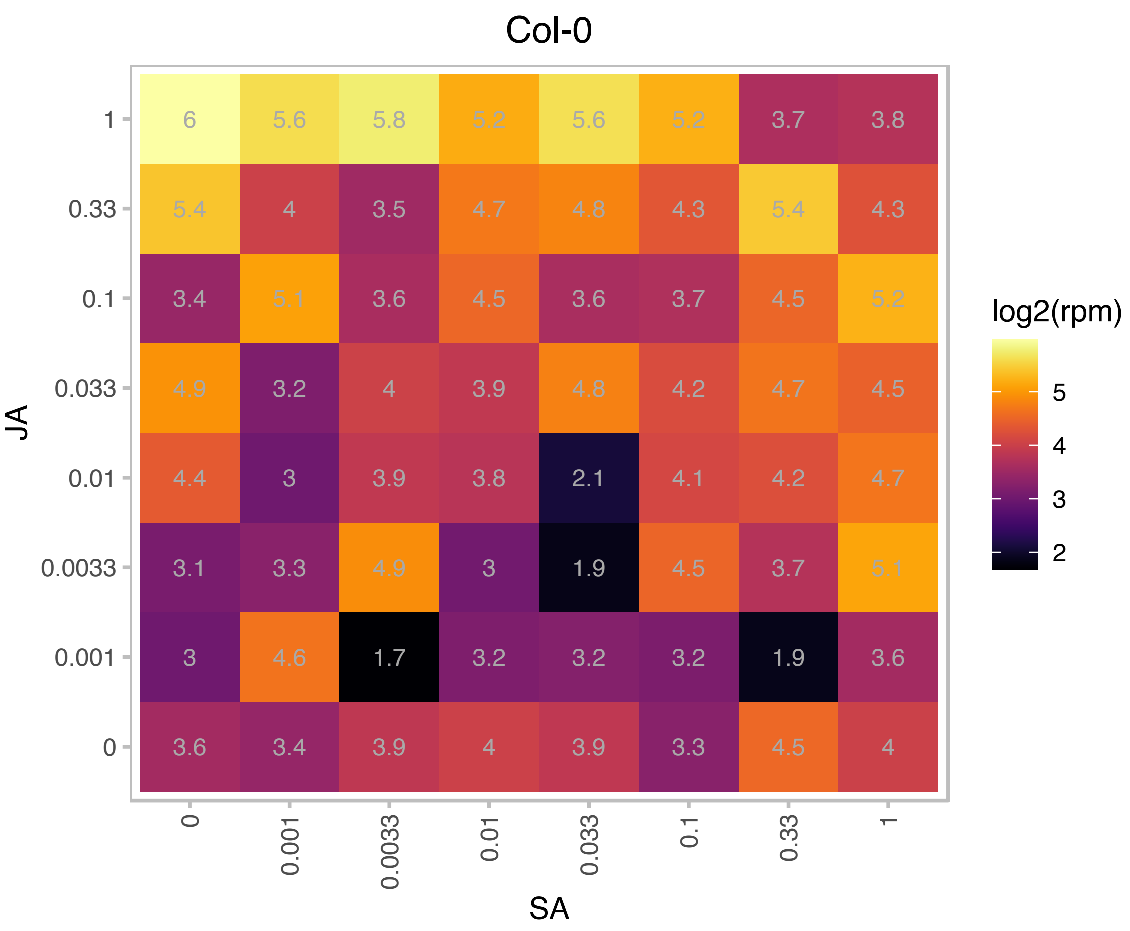


| | |
|---|---|
| Gene | AT3G14067 |
| is.expressed_genes | TRUE |
| is.responsive_genes | TRUE |
| is.apcluster | A-04 |

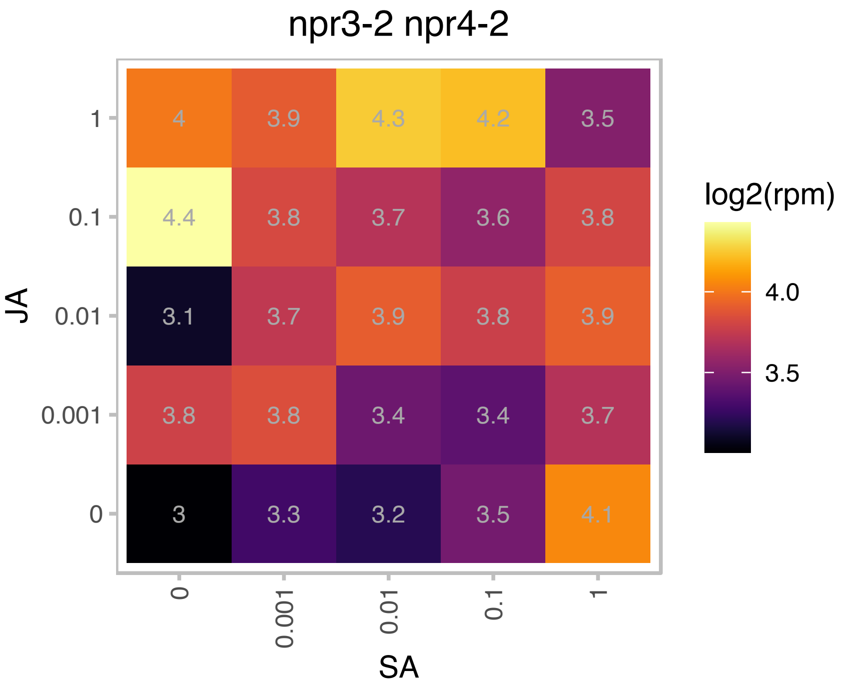


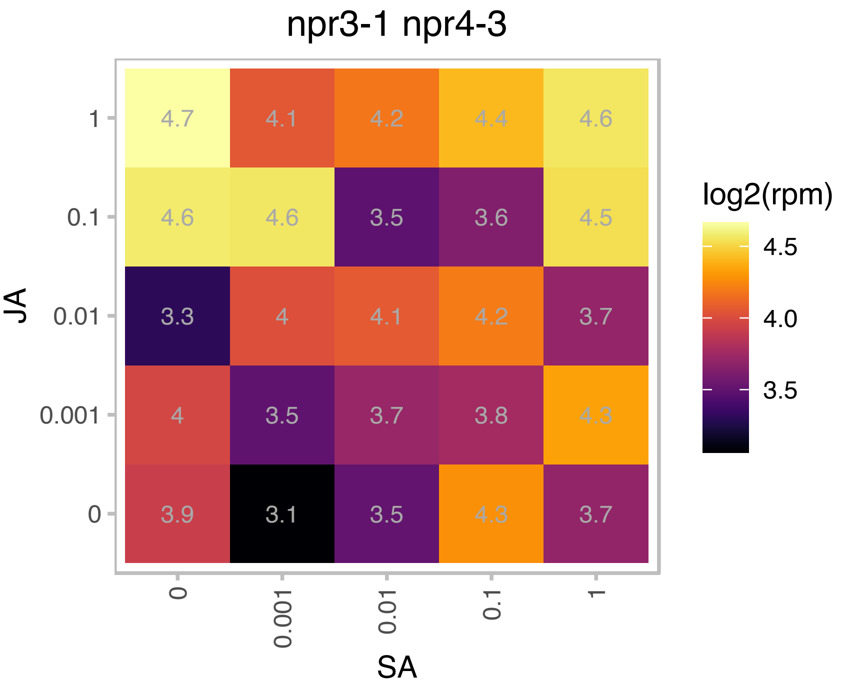

# AT3G15110.1 : NA

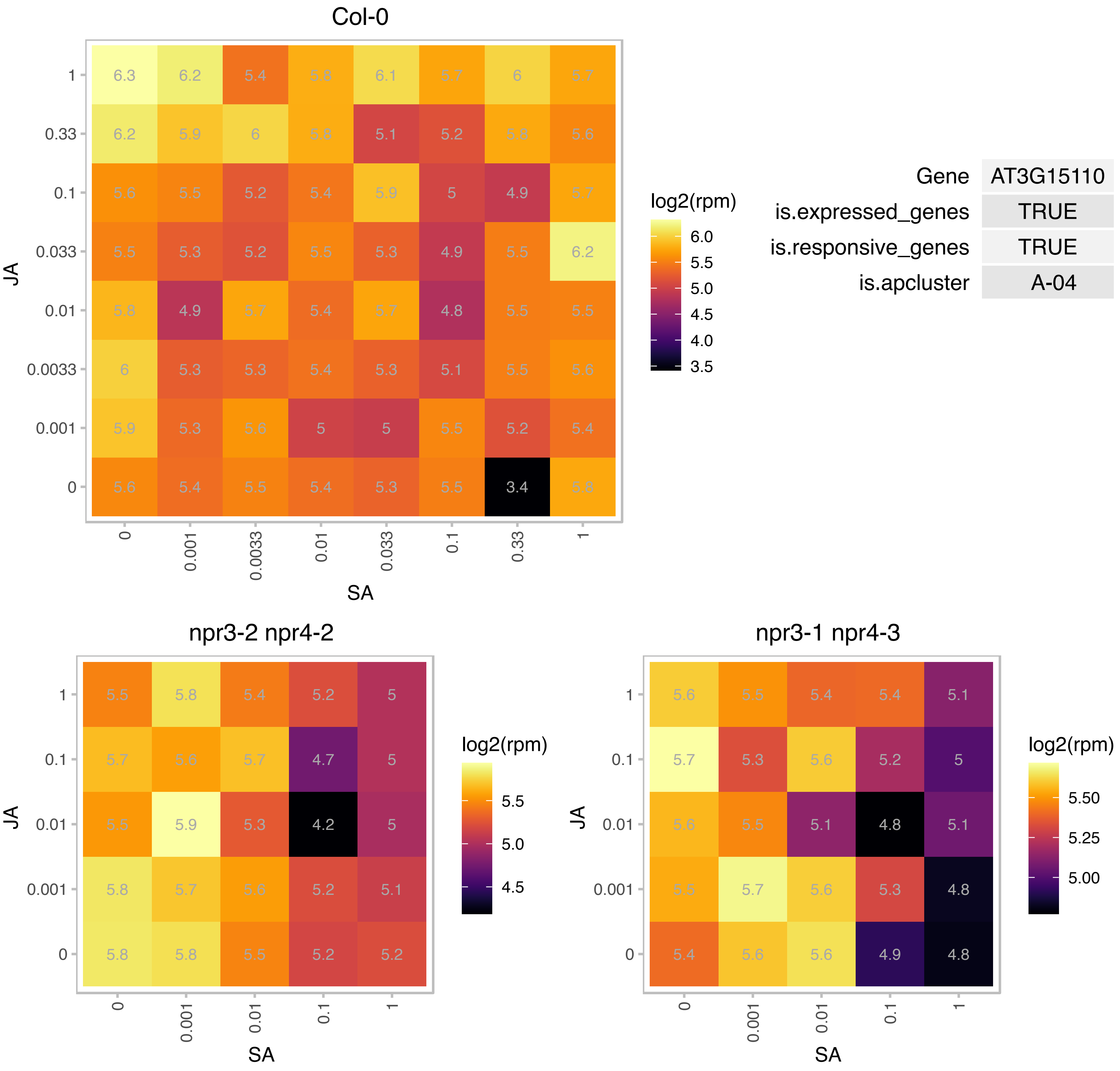


| | |
|---|---|
| Gene | AT3G15110 |
| is.expressed_genes | TRUE |
| is.responsive_genes | TRUE |
| is.apcluster | A-04 |

### AT3G19170.1 : ATPREP1, ATZNMP, PREP1

presequence protease 1

Zinc metalloprotease pitrilysin subfamily A. Signal peptide degrading enzyme targeted to mitochondria and chloroplasts. Expressed only in siliques and flowers

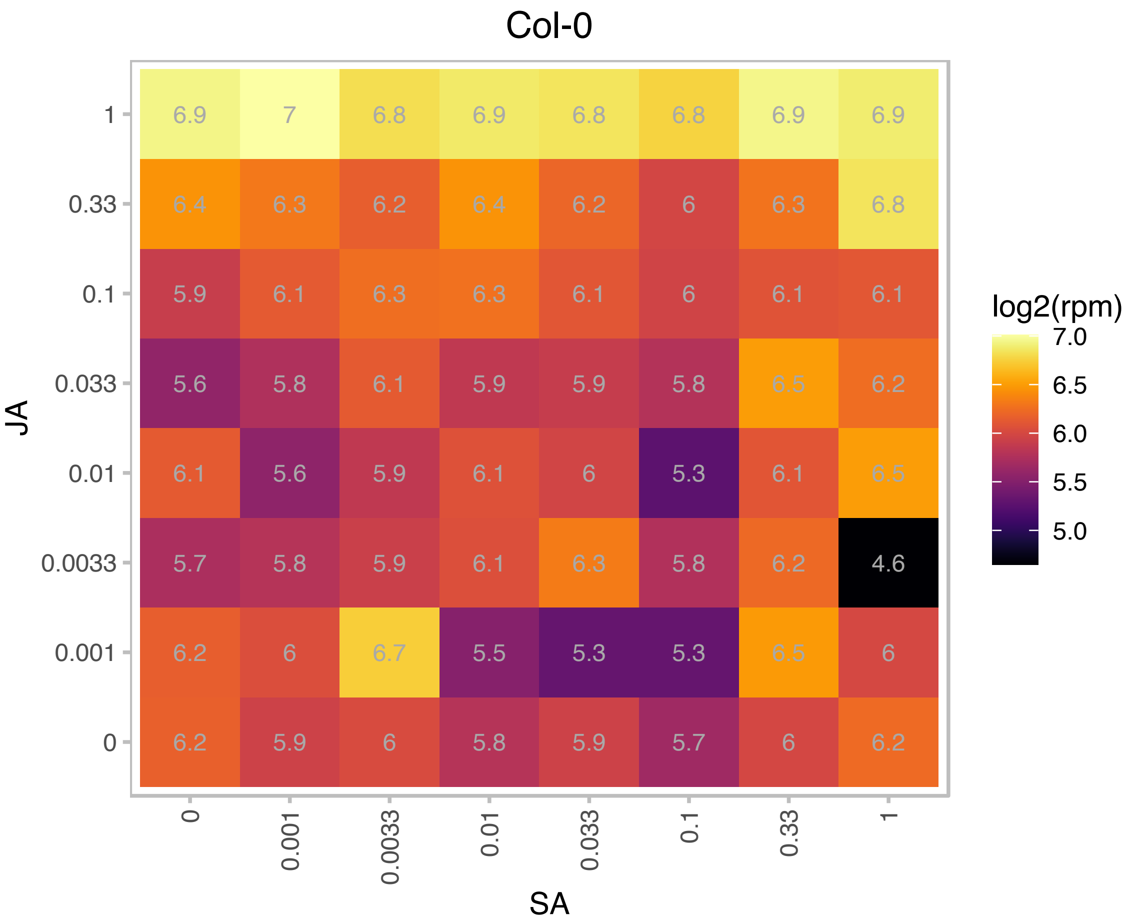


| | |
|---|---|
| Gene | AT3G19170 |
| is.expressed_genes | TRUE |
| is.responsive_genes | TRUE |
| is.apcluster | A-04 |

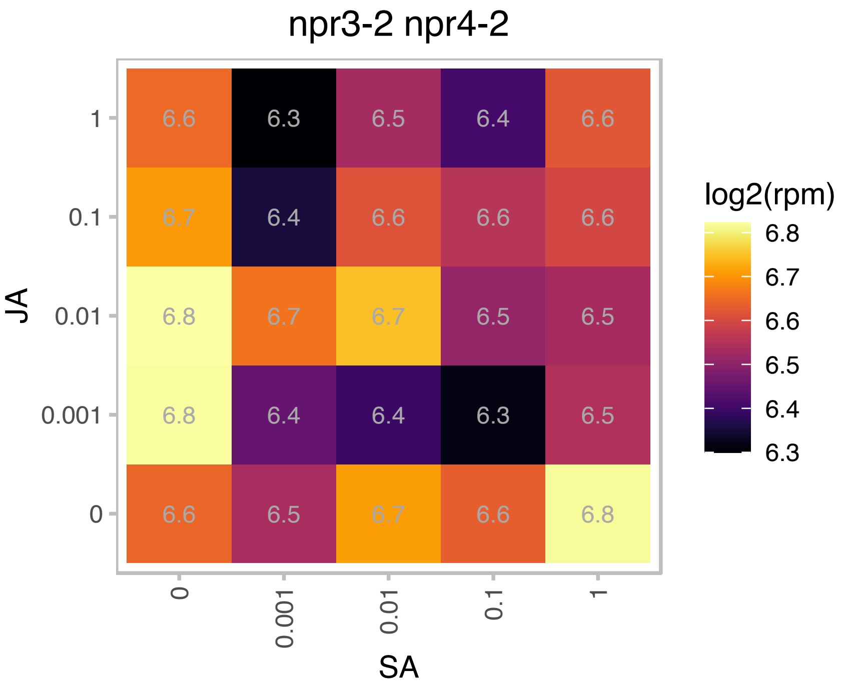


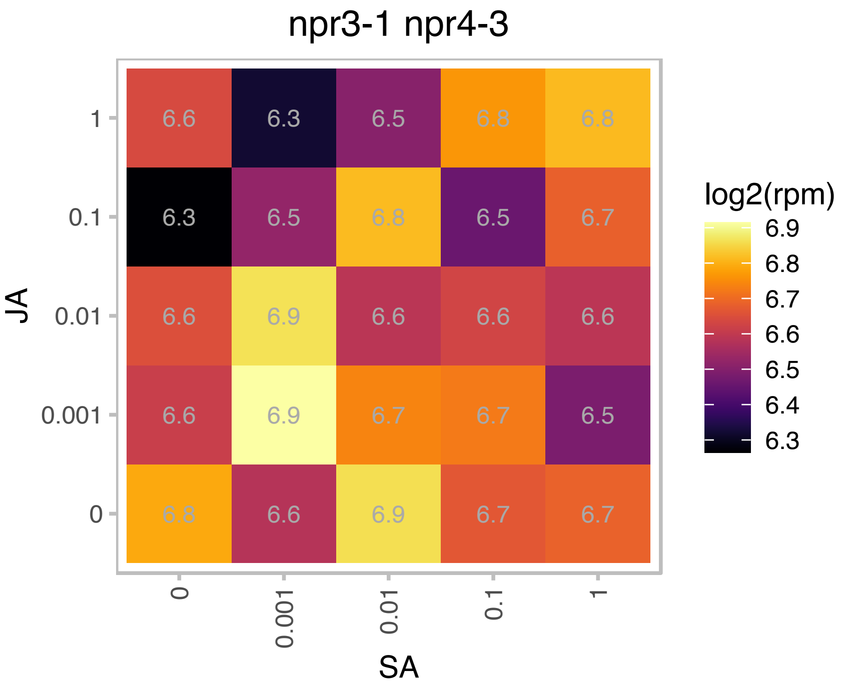

# AT3G24820.1 : NA

BSD domain-containing protein

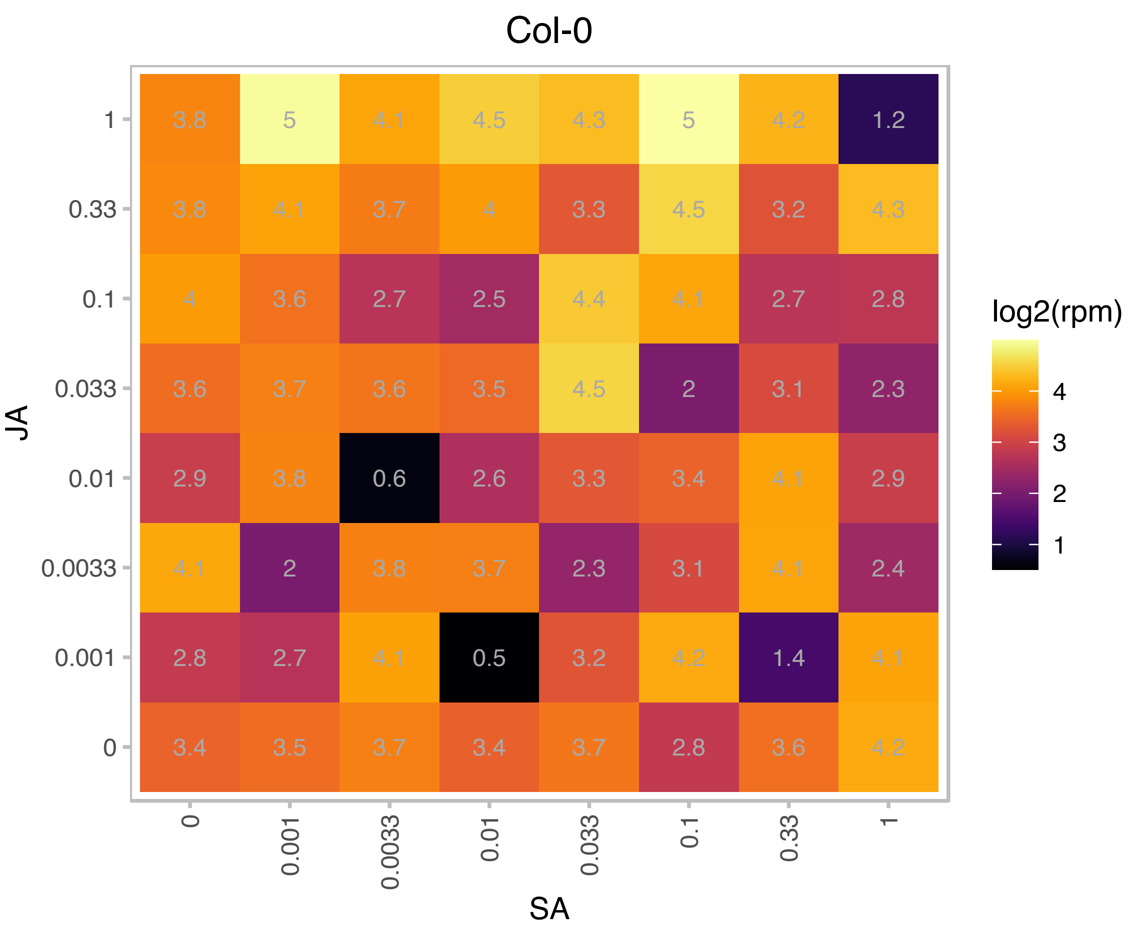


| | |
|---|---|
| Gene | AT3G24820 |
| is.expressed_genes | TRUE |
| is.responsive_genes | TRUE |
| is.apcluster | A-04 |

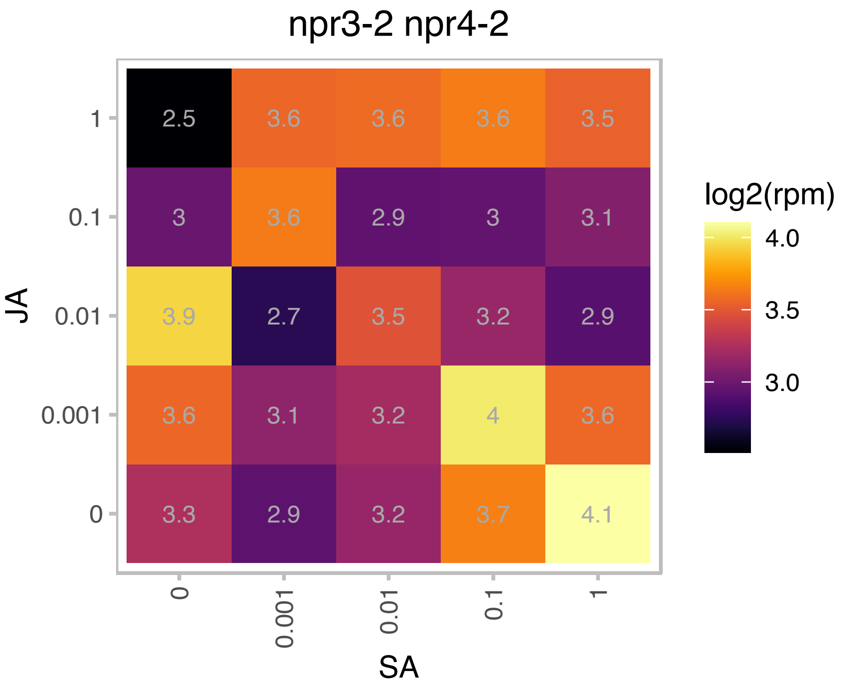


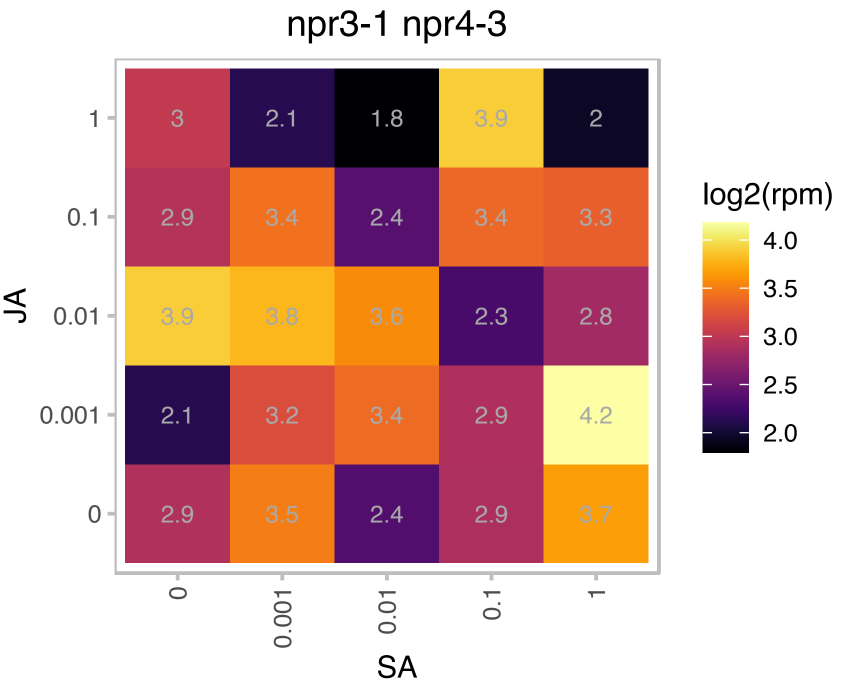

### AT3G45010.1 : scpl48

serine carboxypeptidase-like 48

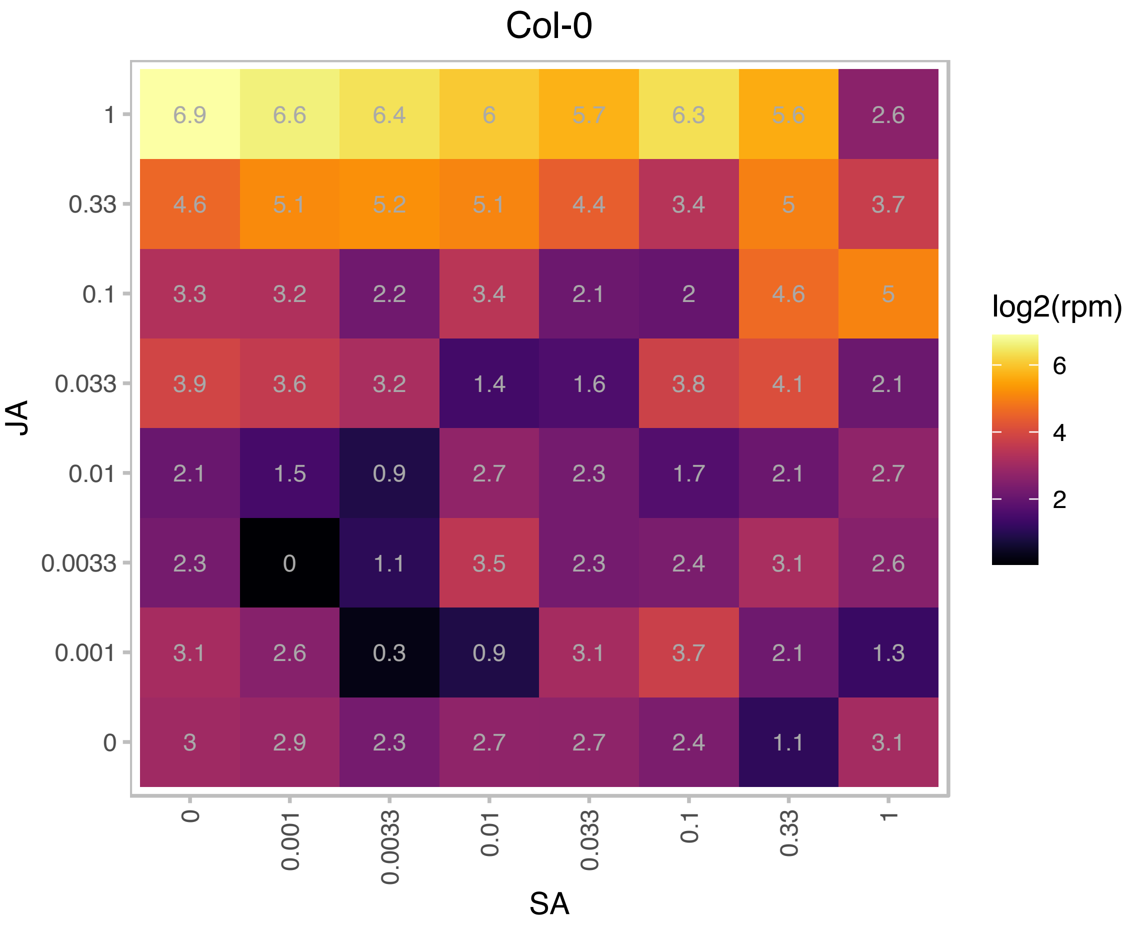


| | |
|---|---|
| Gene | AT3G45010 |
| is.expressed_genes | TRUE |
| is.responsive_genes | TRUE |
| is.apcluster | A-04 |

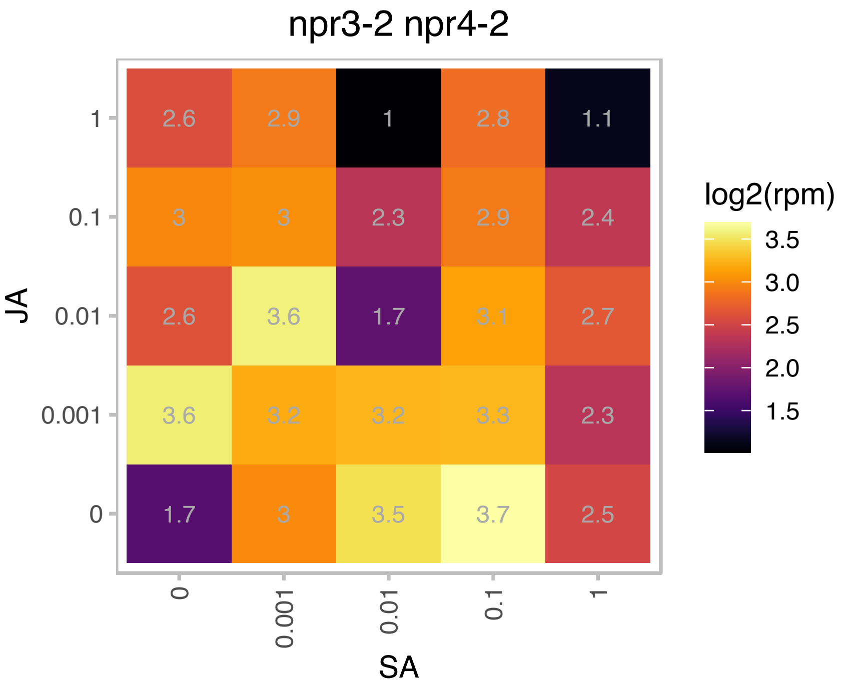


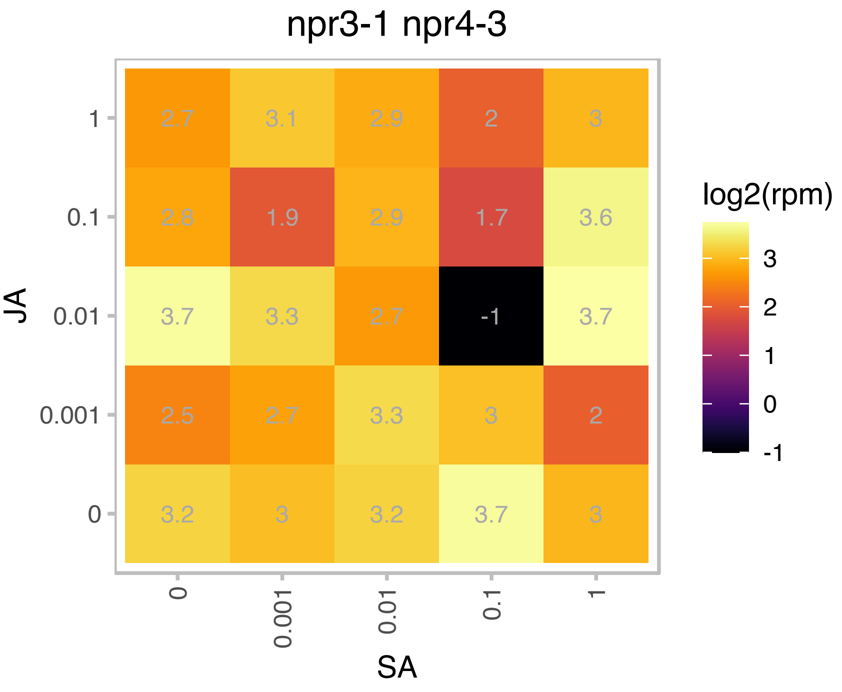

### AT3G46670.1 : UGT76E11

UDP-glucosyl transferase 76E11

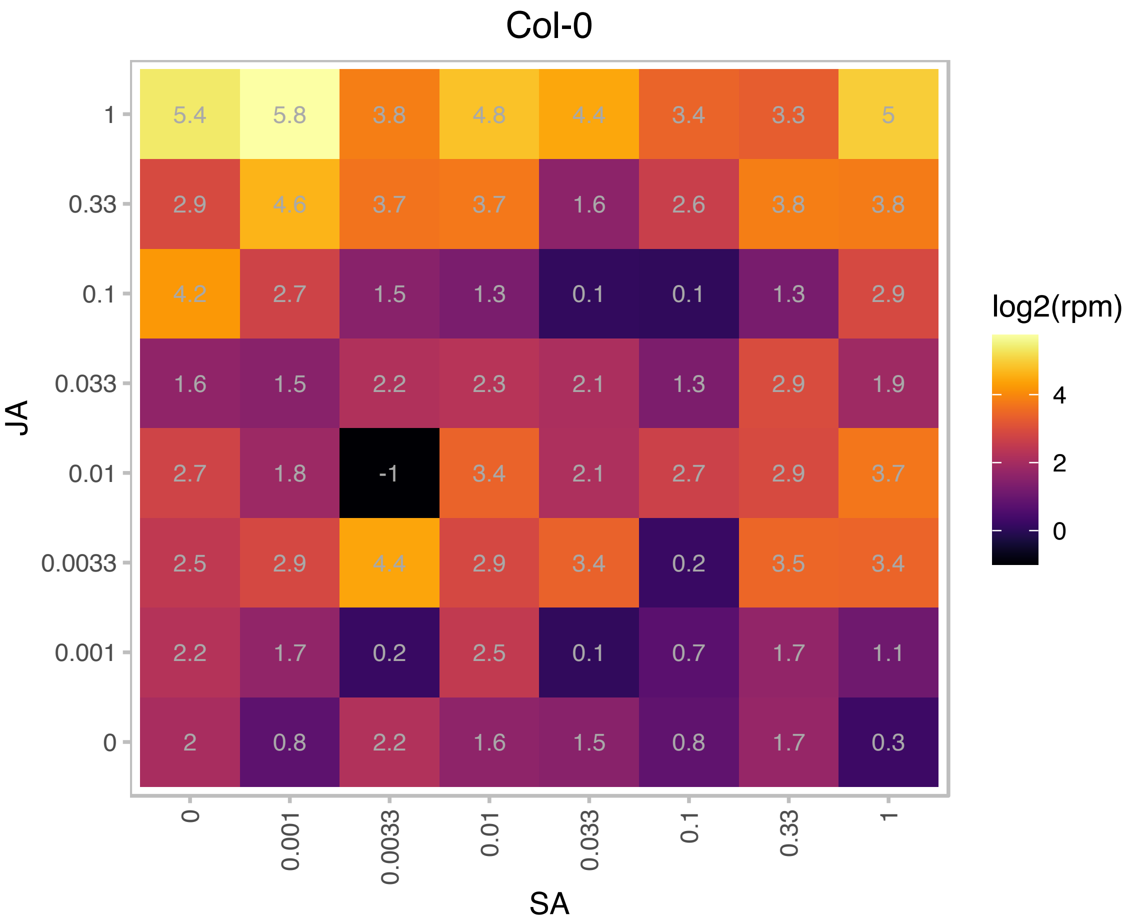


| | |
|---|---|
| Gene | AT3G46670 |
| is.expressed_genes | TRUE |
| is.responsive_genes | TRUE |
| is.apcluster | A-04 |

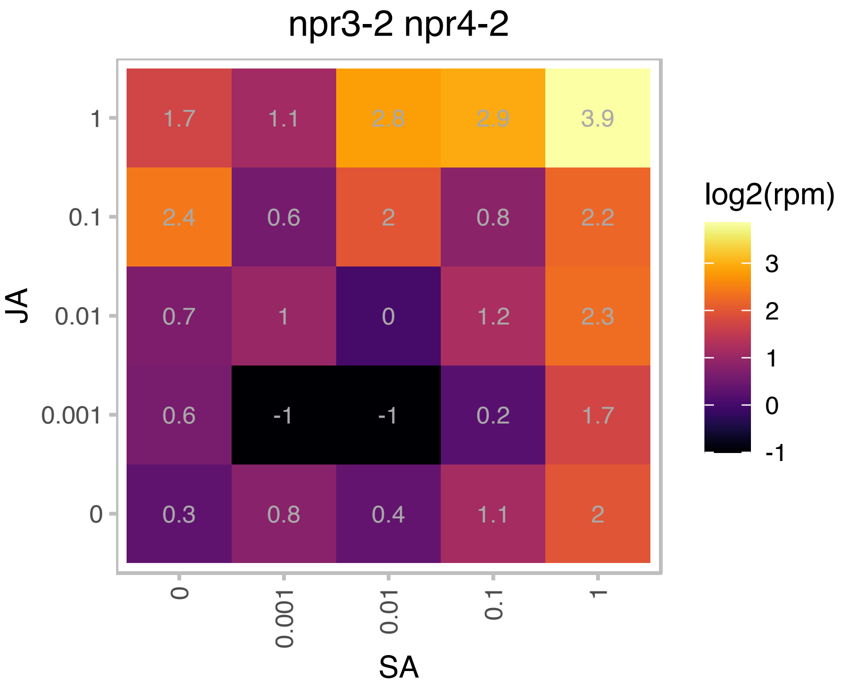


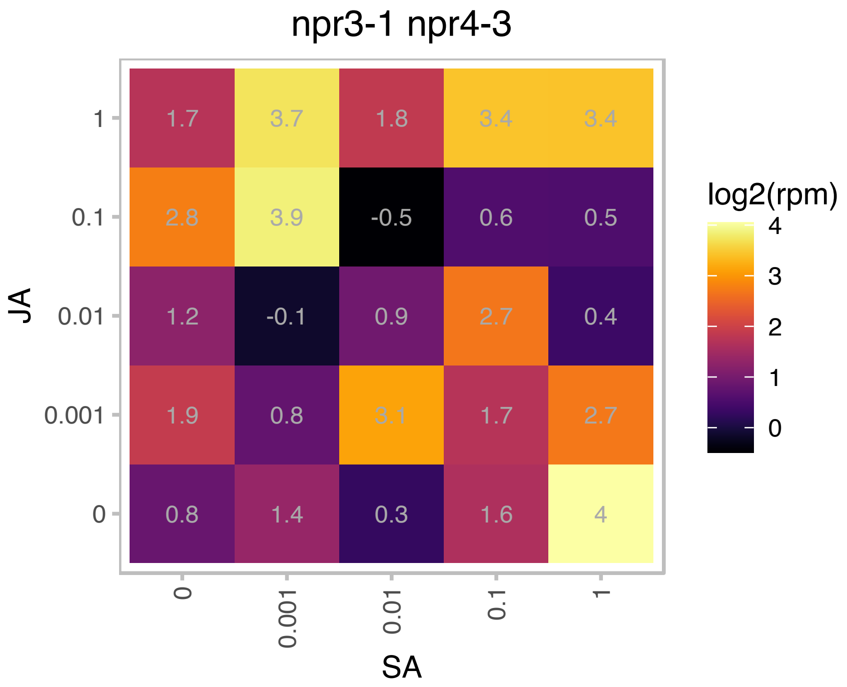

# AT3G50590.1 : NA

Transducin/WD40 repeat-like superfamily protein

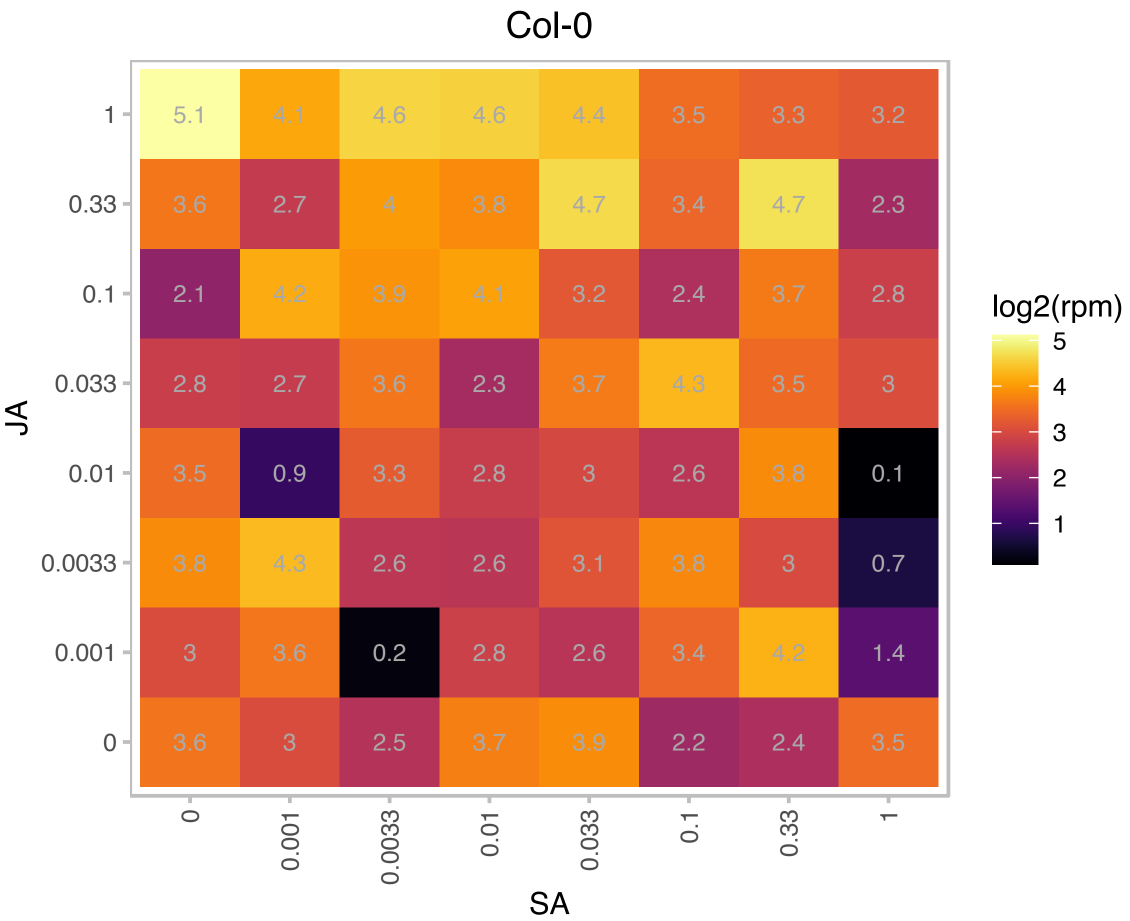


| | |
|---|---|
| Gene | AT3G50590 |
| is.expressed_genes | TRUE |
| is.responsive_genes | TRUE |
| is.apcluster | A-04 |

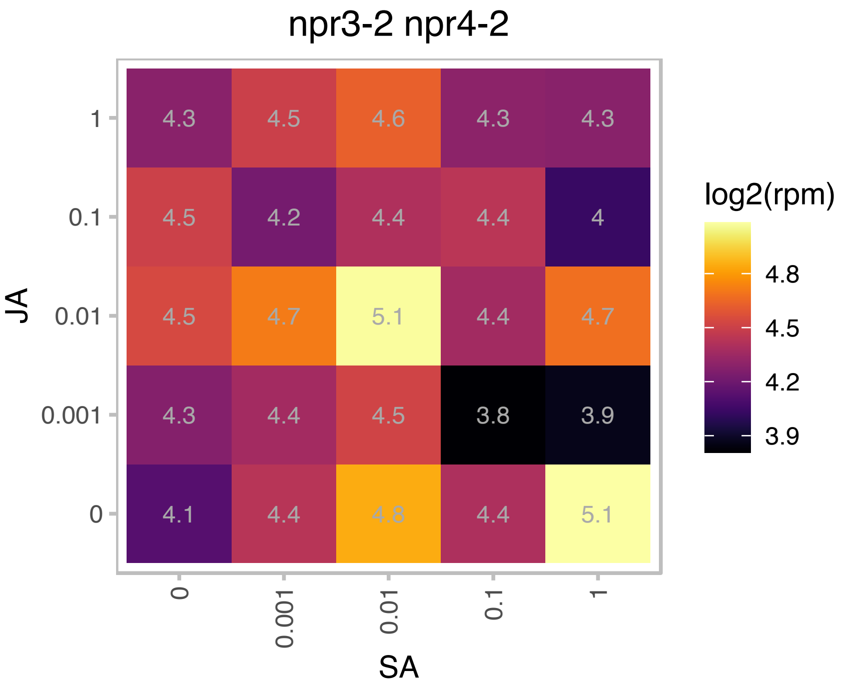


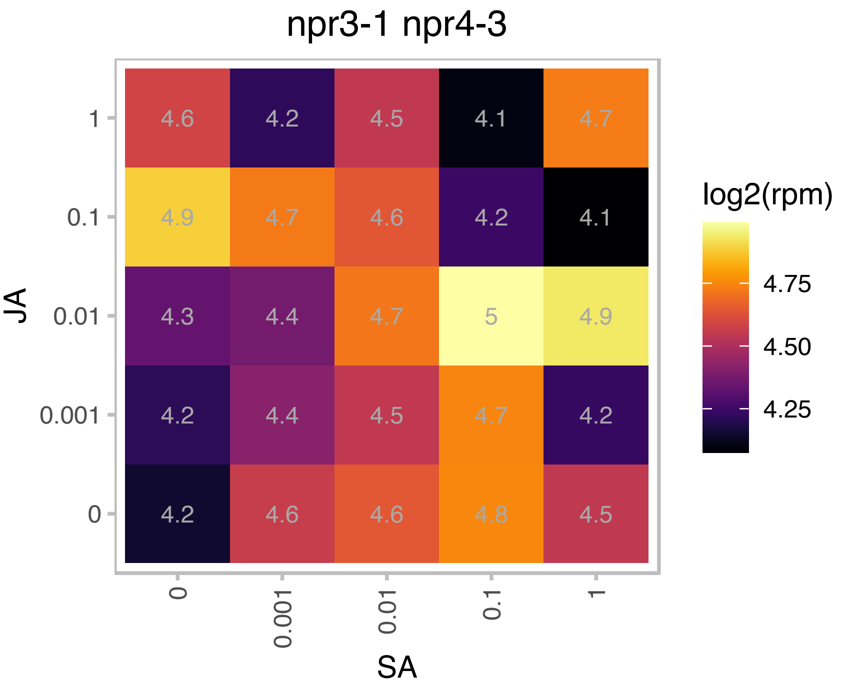

### AT3G53970.1 : PTRE1

proteasome inhibitor-related
PTRE1 was identified as homologous to human PI31. It has a conserved proline-rich domain at the C-terminus and a highly conserved FP (Fbxo7/PI31) dimerization domain at the N-terminus as well as some novel, conserved domains found only in plants. It regulates auxin signaling possibly via its proteosome suppressing activity.

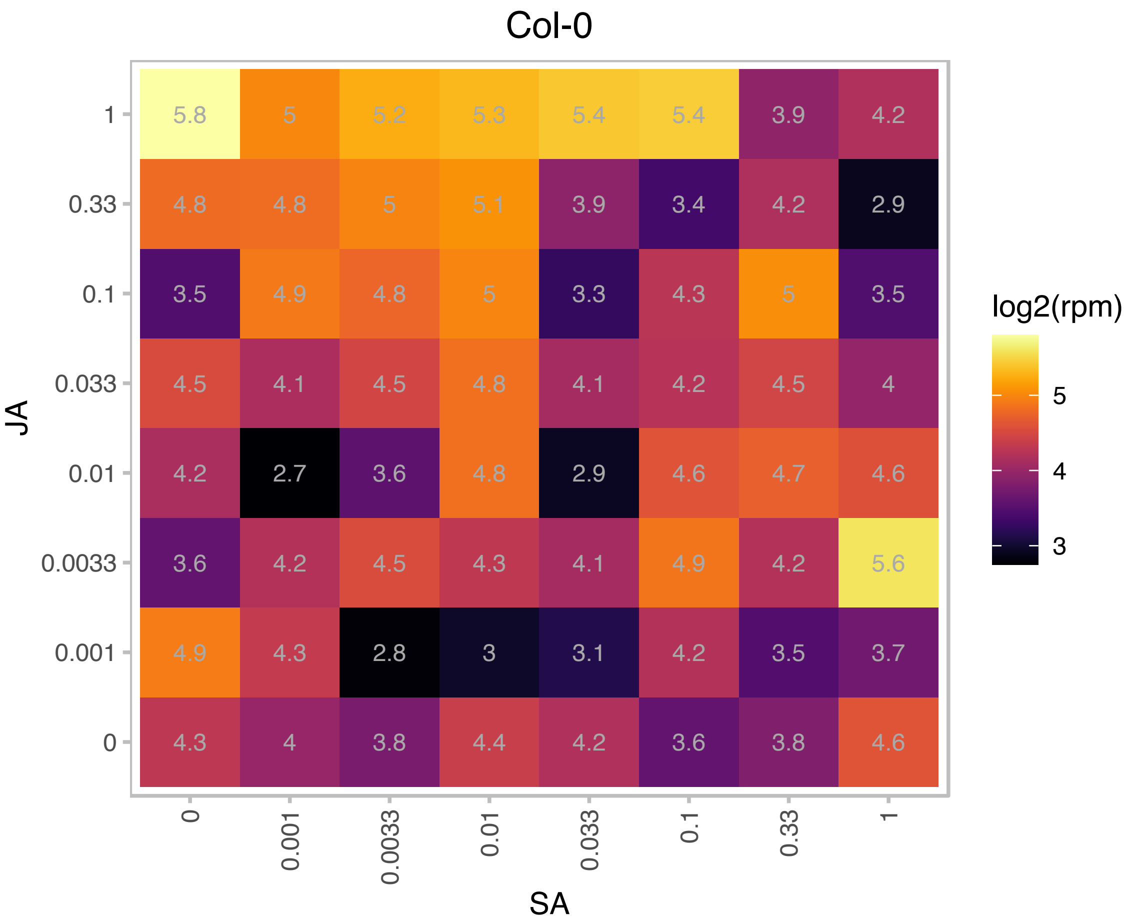


| | |
|---|---|
| Gene | AT3G53970 |
| is.expressed_genes | TRUE |
| is.responsive_genes | TRUE |
| is.apcluster | A-04 |

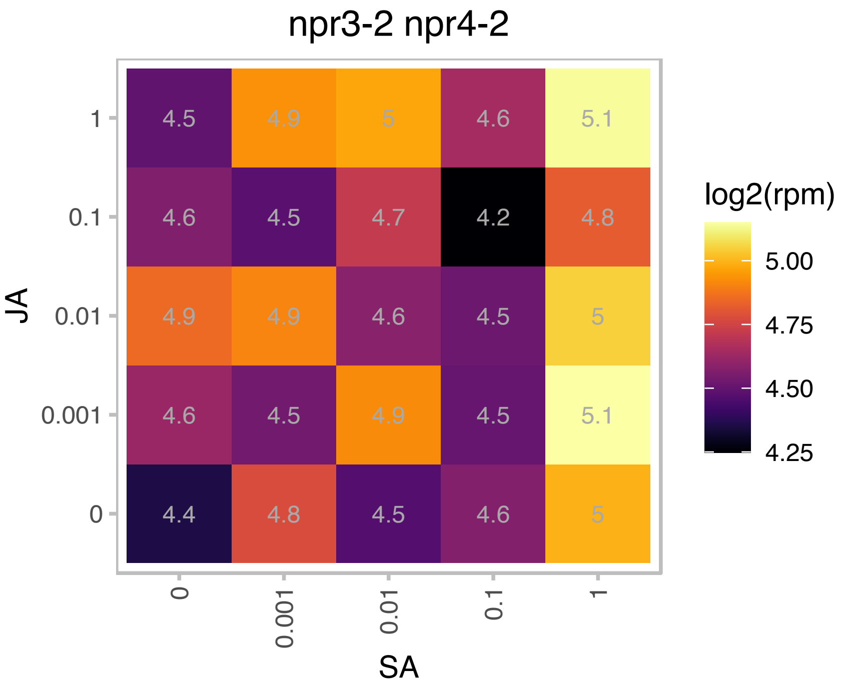


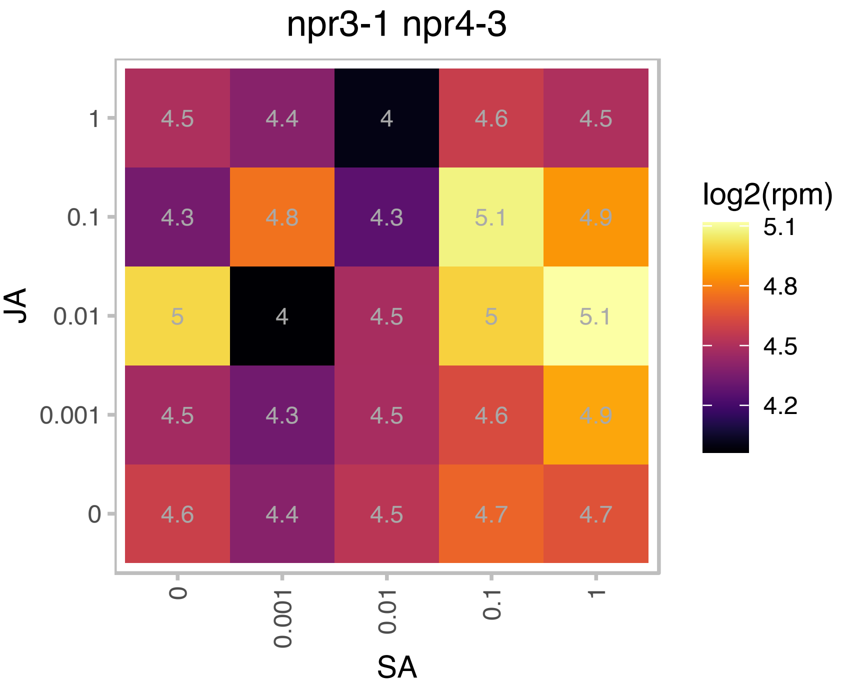

### AT3G55530.1 : AtSDIR1, SDIR1

RING/U-box superfamily protein
Encodes an intracellular membrane localized protein with E3 ligase activity, found in the ER with the C-terminus facing the cytoplasm. It is involved in regulation of ABA signaling. Loss of function alleles show decreased sensitivity to ABA. Overexpression results in increased sensitivity to ABA.

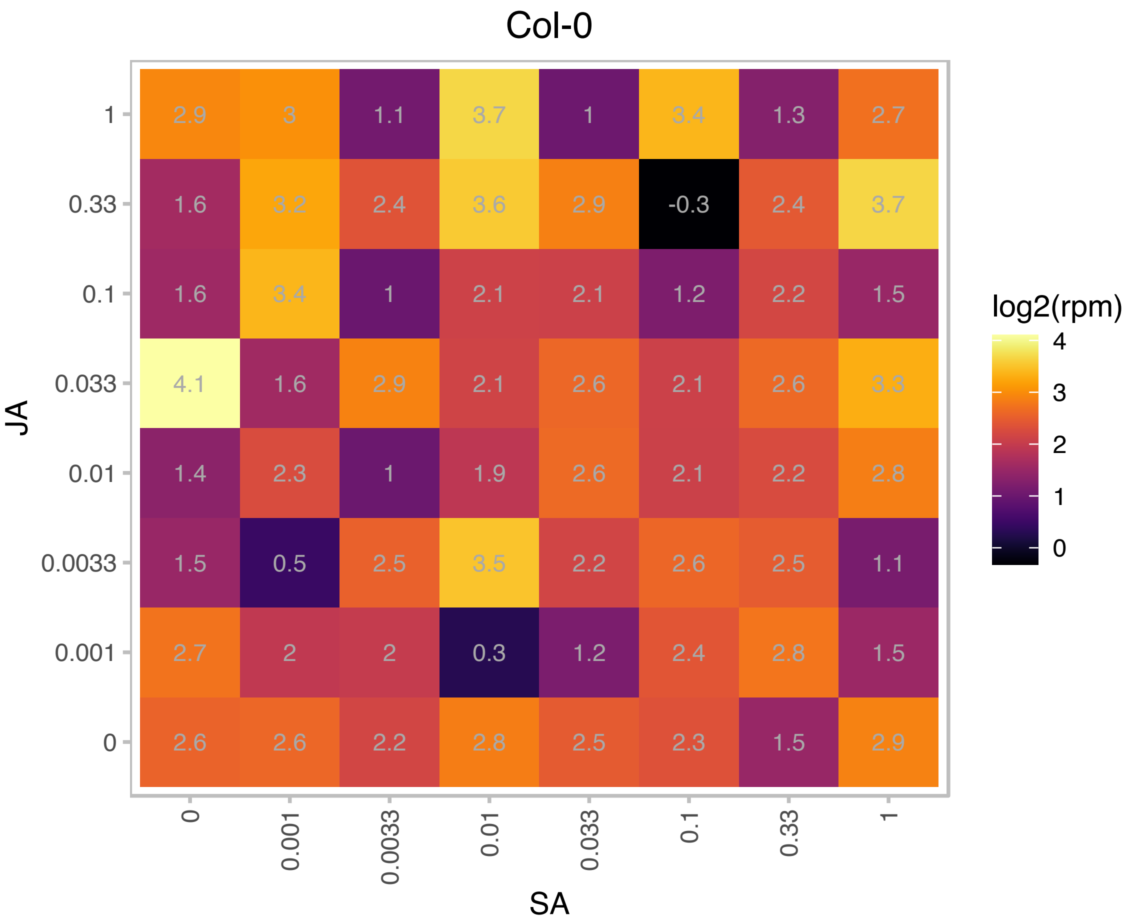


| | |
|---|---|
| Gene | AT3G55530 |
| is.expressed_genes | TRUE |
| is.responsive_genes | TRUE |
| is.apcluster | A-04 |

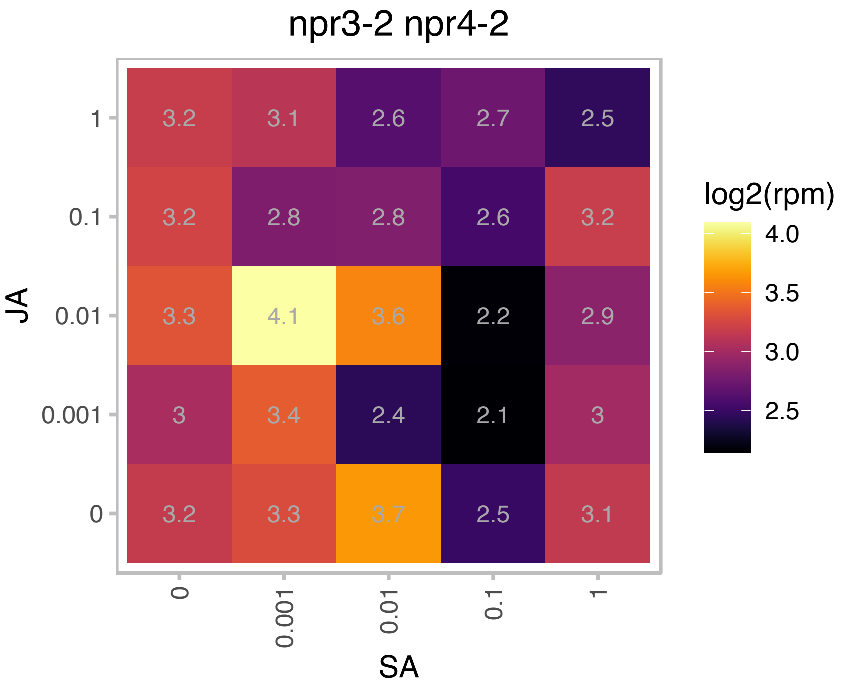


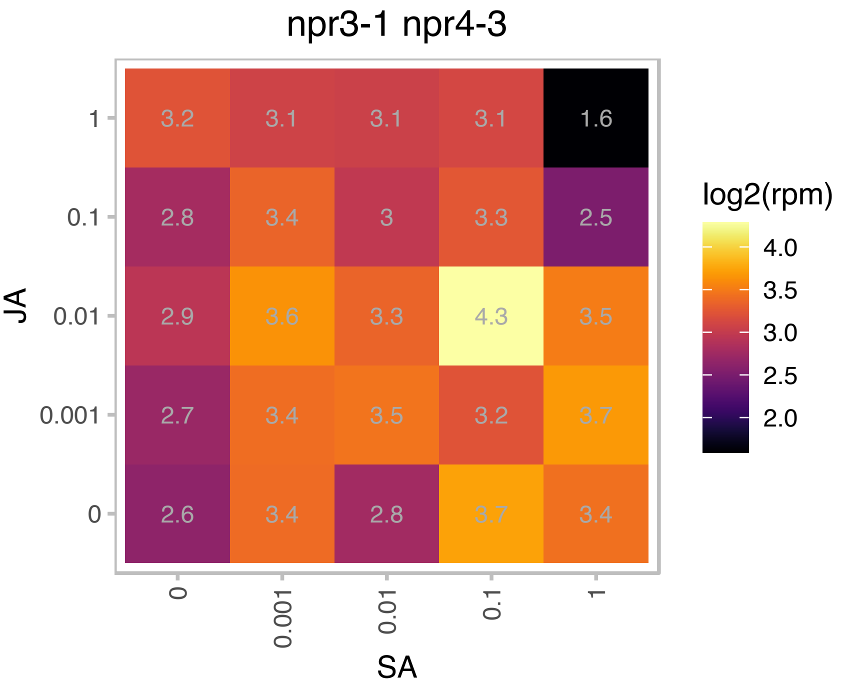

### AT3G59990.1 : MAP2B

methionine aminopeptidase 2B

Encodes a MAP2 like methionine aminopeptidase

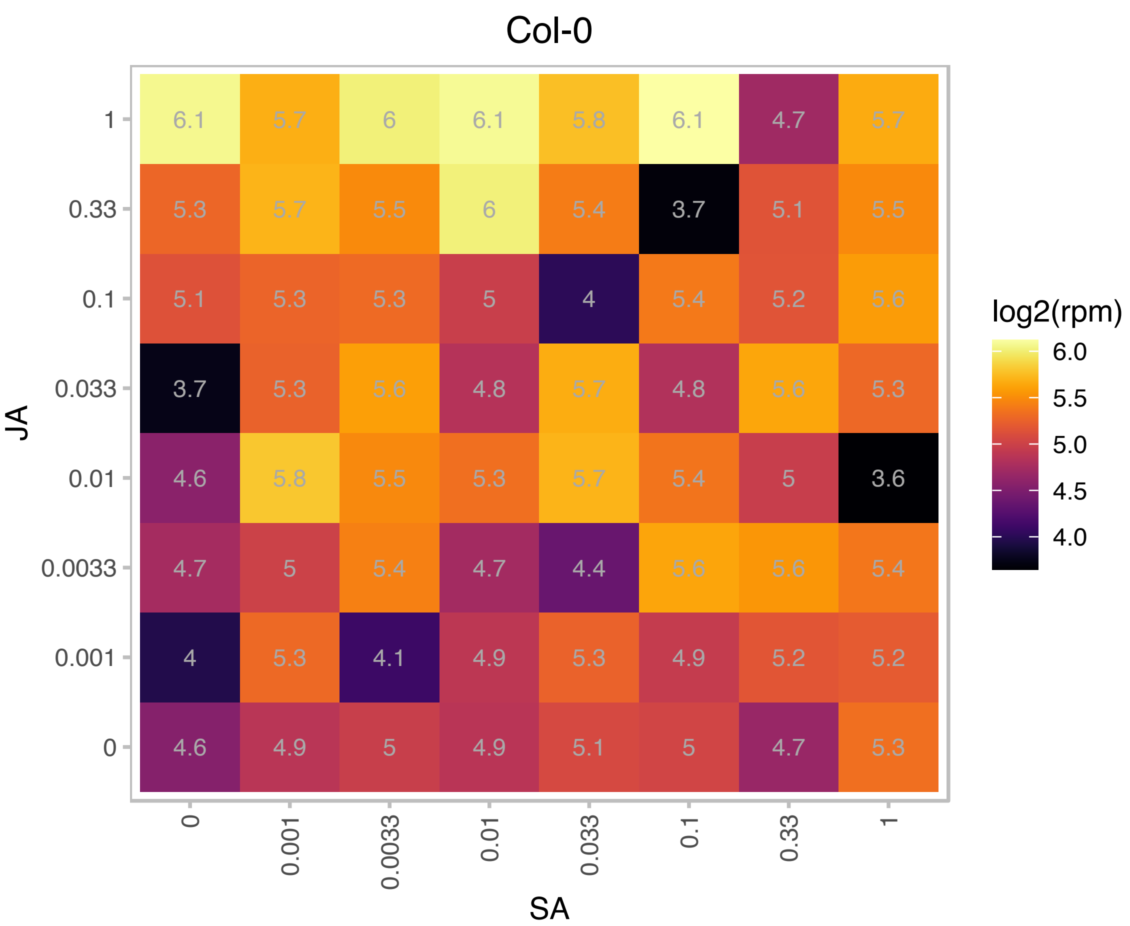


| | |
|---|---|
| Gene | AT3G59990 |
| is.expressed_genes | TRUE |
| is.responsive_genes | TRUE |
| is.apcluster | A-04 |

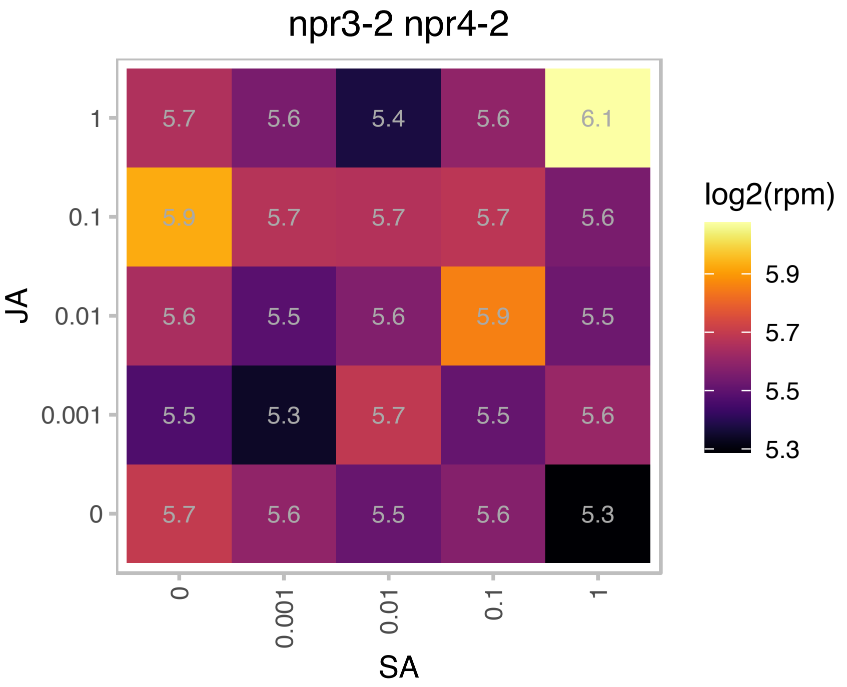


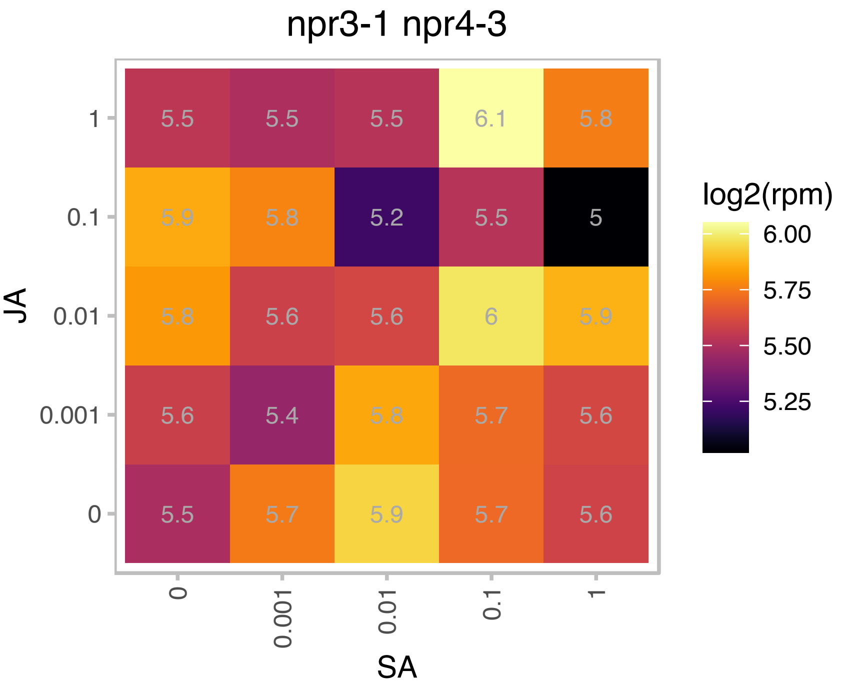

### AT4G11560.1 : AIPP3

bromo-adjacent homology (BAH) domain-containing protein

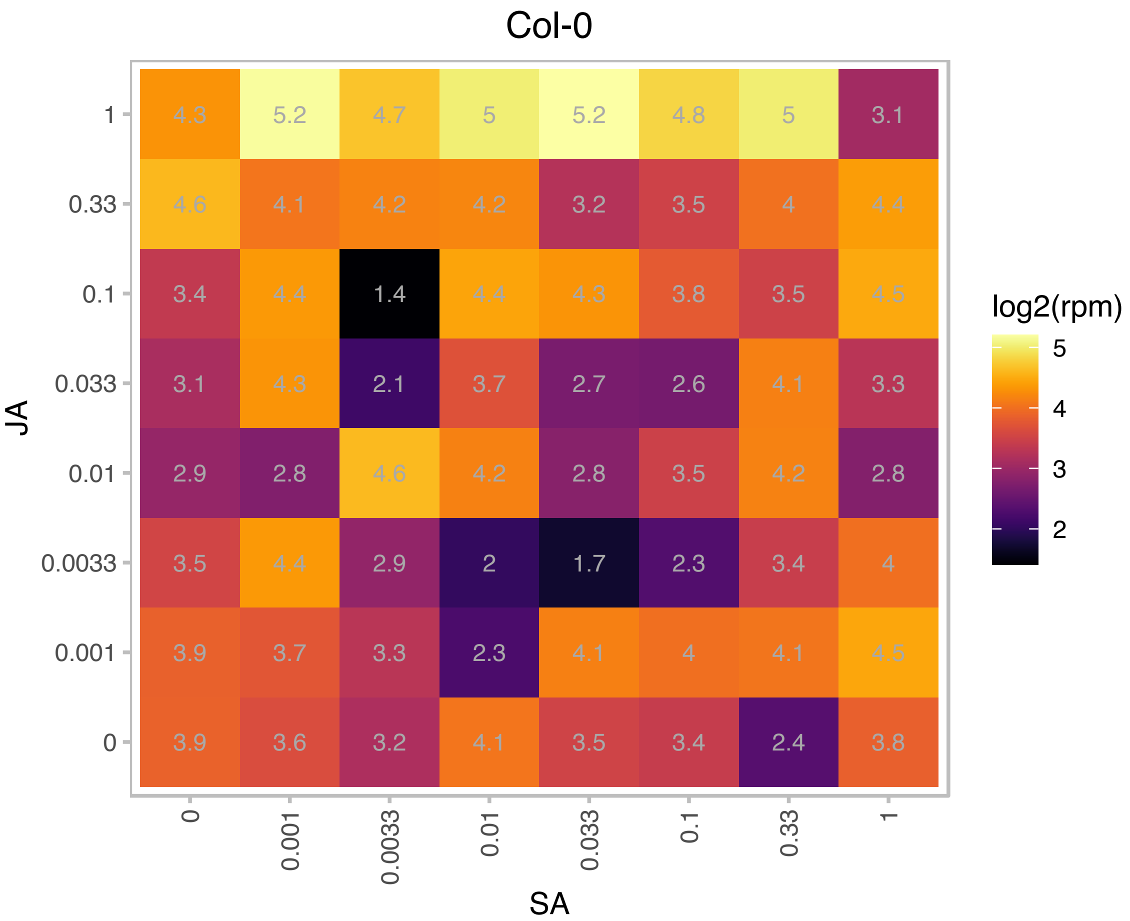


| | |
|---|---|
| Gene | AT4G11560 |
| is.expressed_genes | TRUE |
| is.responsive_genes | TRUE |
| is.apcluster | A-04 |

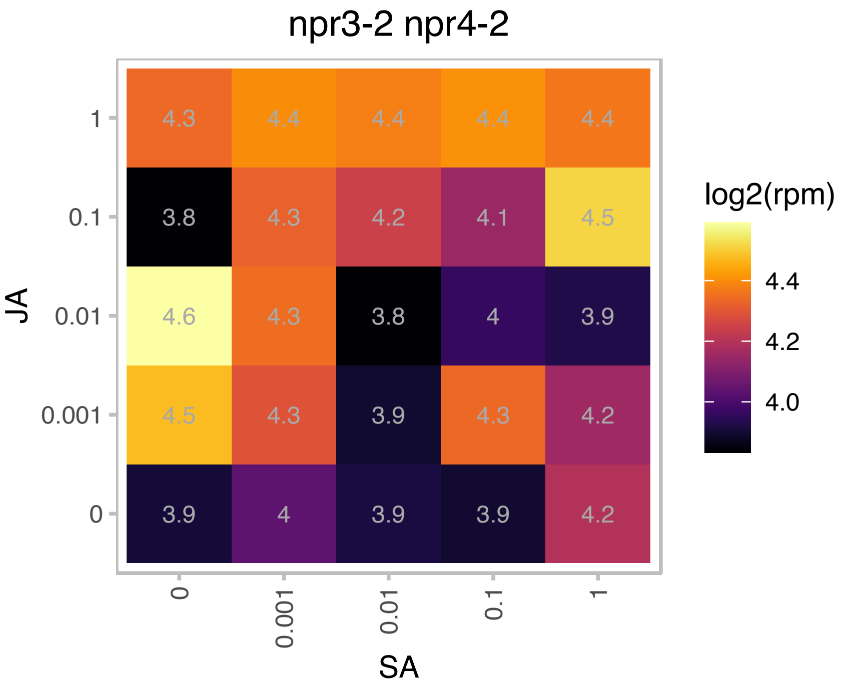


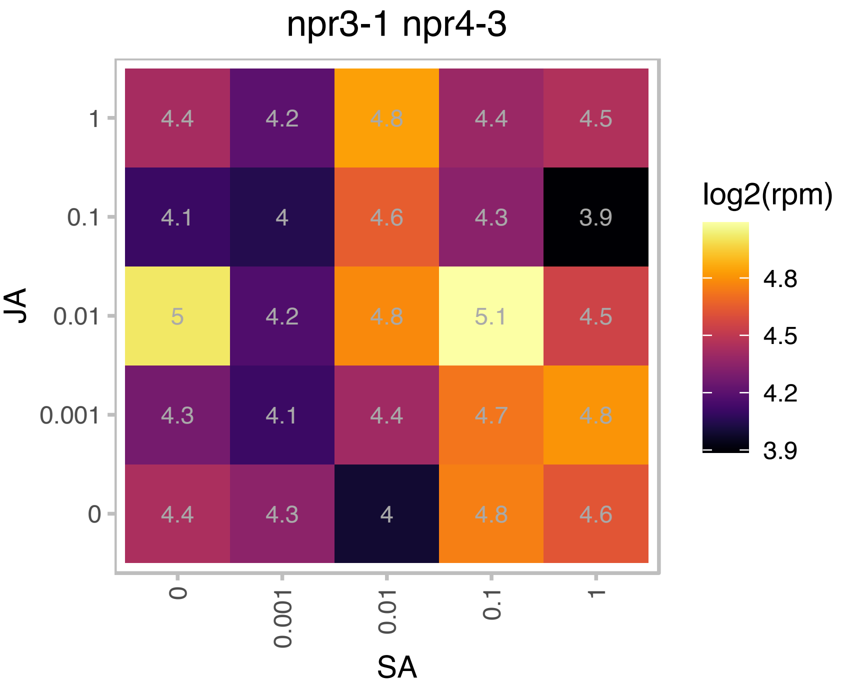

### AT4G17510.1 : UCH3

ubiquitin C-terminal hydrolase 3

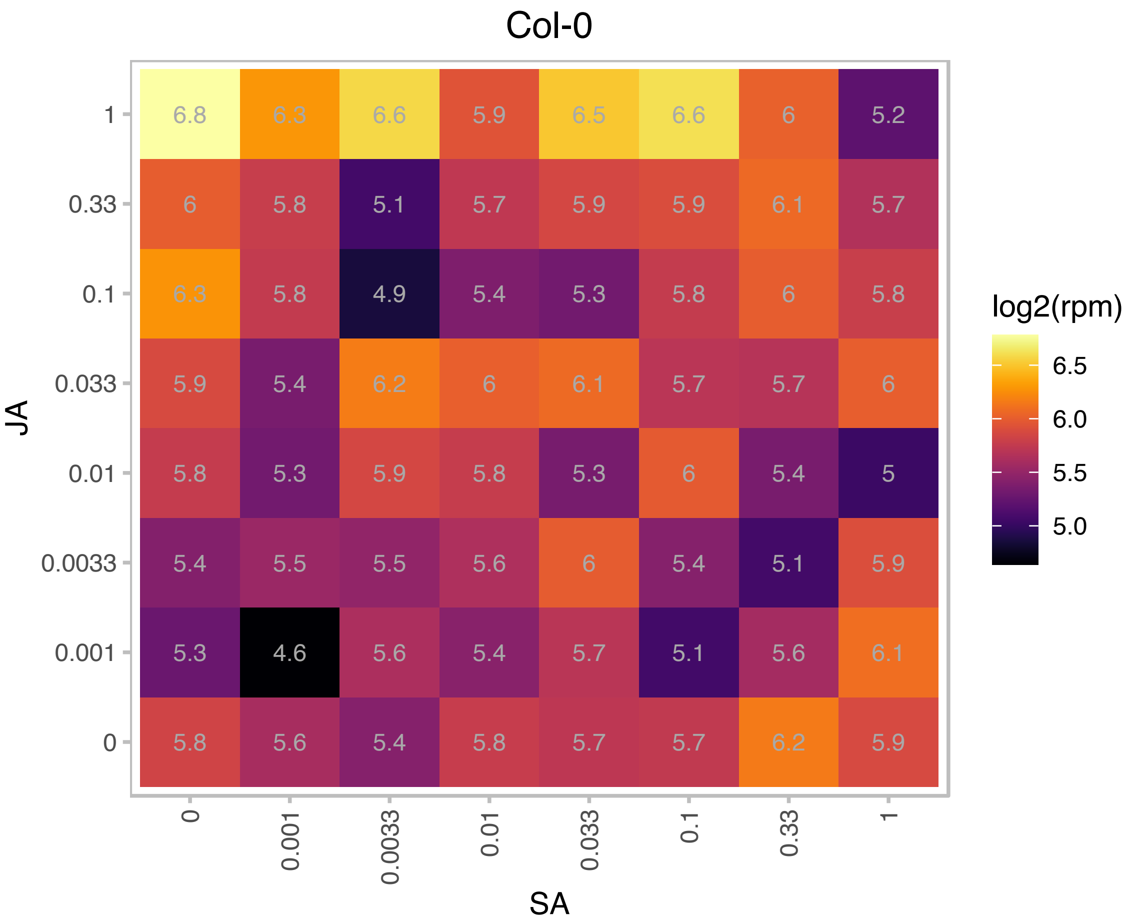


| | |
|---|---|
| Gene | AT4G17510 |
| is.expressed_genes | TRUE |
| is.responsive_genes | TRUE |
| is.apcluster | A-04 |

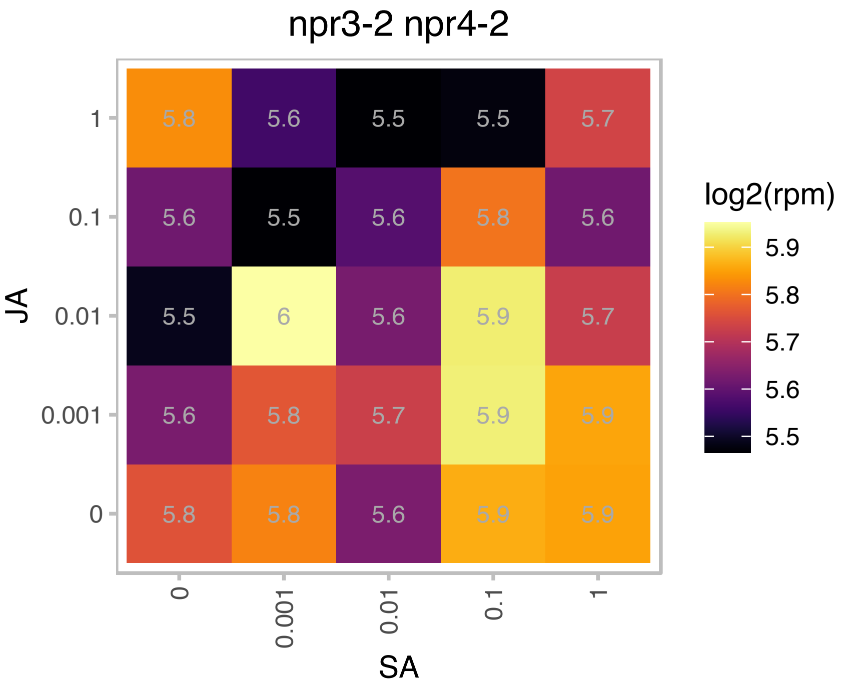


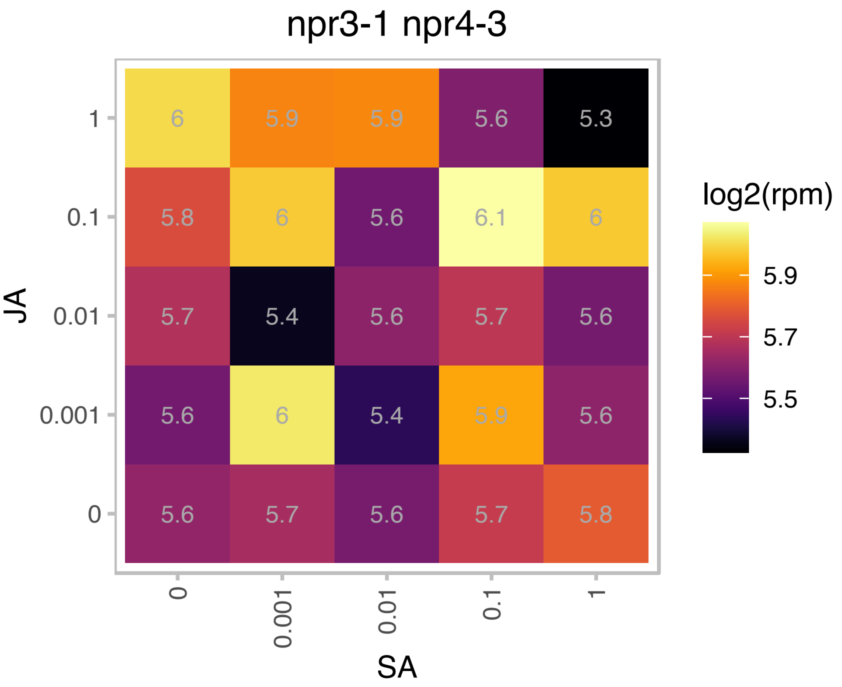

### AT4G20850.1 : TPP2

tripeptidyl peptidase ii
Tripeptidyl Peptidase II. Ser protease that assembles into a large oligomeric complex containing two proteins of 153 and 142 kD that are derived from a single TPP2 gene, with the smaller version missing part of the C-terminal end. Not essential, based on the lack of phenotype of a T-DNA disruption mutant.

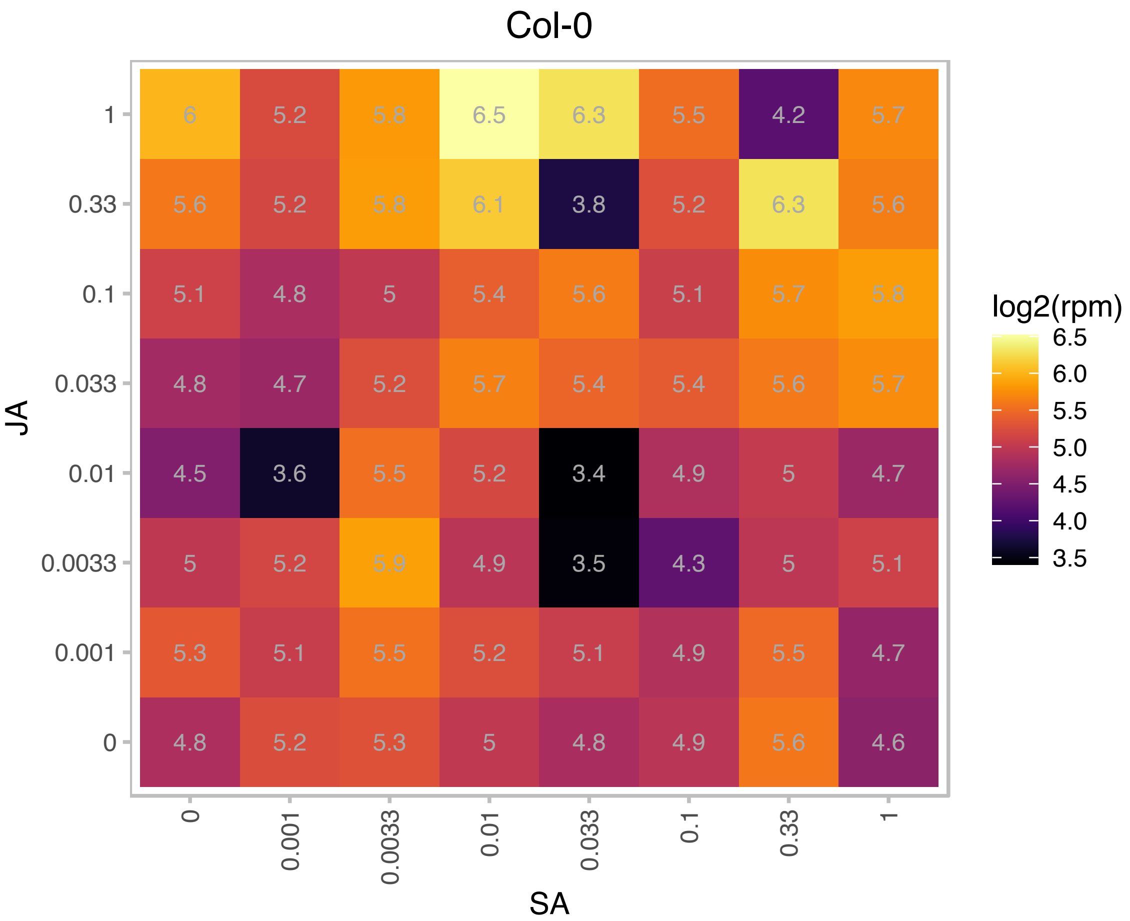


| | |
|---|---|
| Gene | AT4G20850 |
| is.expressed_genes | TRUE |
| is.responsive_genes | TRUE |
| is.apcluster | A-04 |

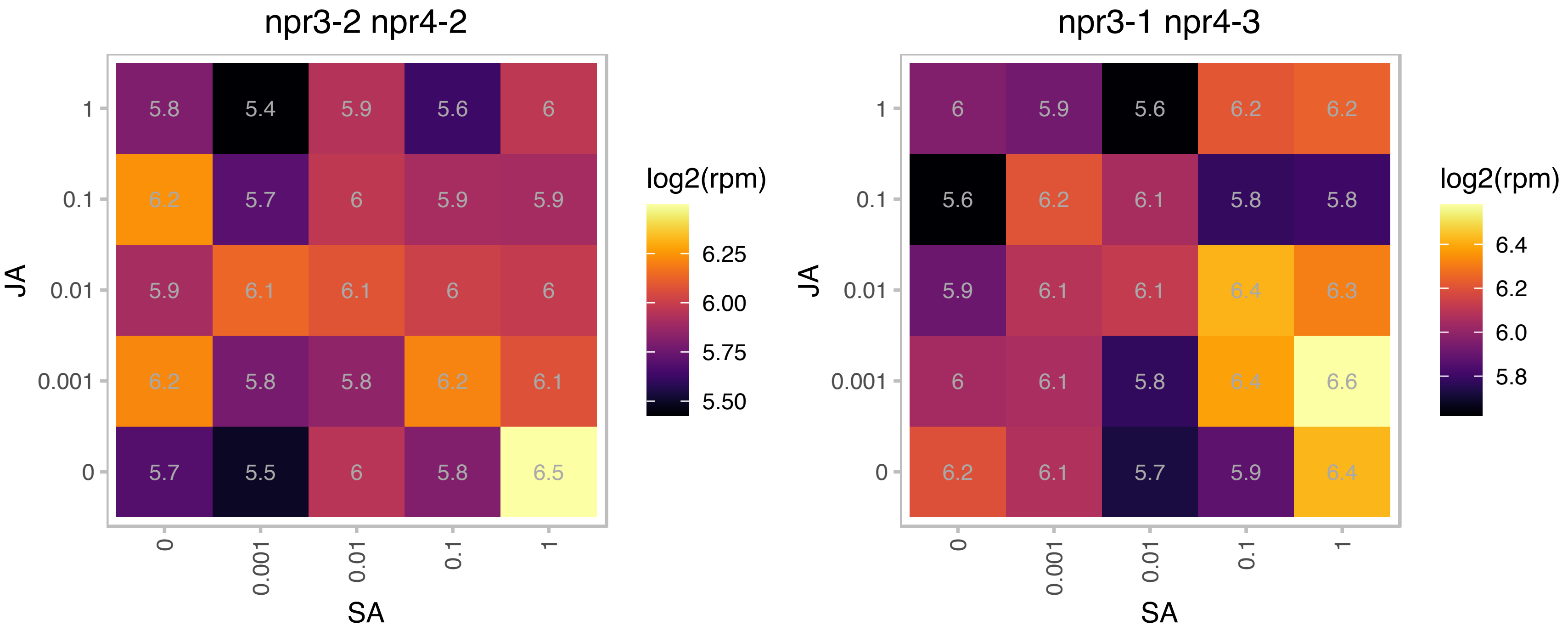

### AT4G33090.1 : APM1, ATAPM1

aminopeptidase M1

encodes an aminopeptidase, a ortholog of mouse microsomal AP (EC 3.4.11.2).

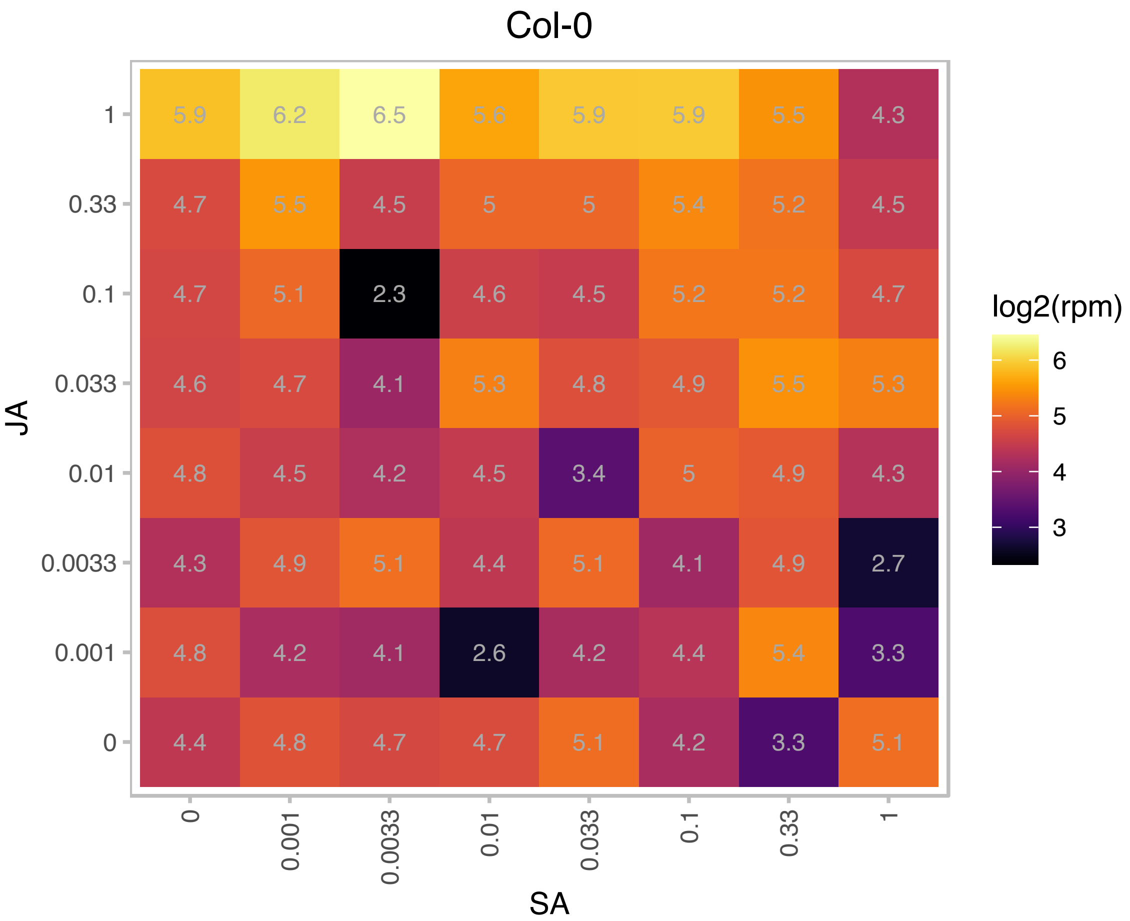


| | |
|---|---|
| Gene | AT4G33090 |
| is.expressed_genes | TRUE |
| is.responsive_genes | TRUE |
| is.apcluster | A-04 |

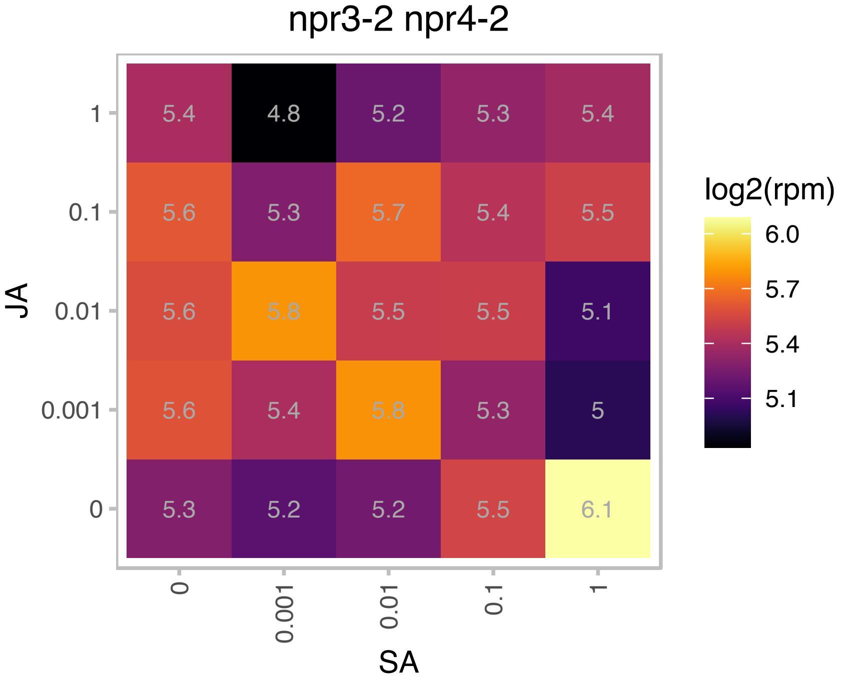


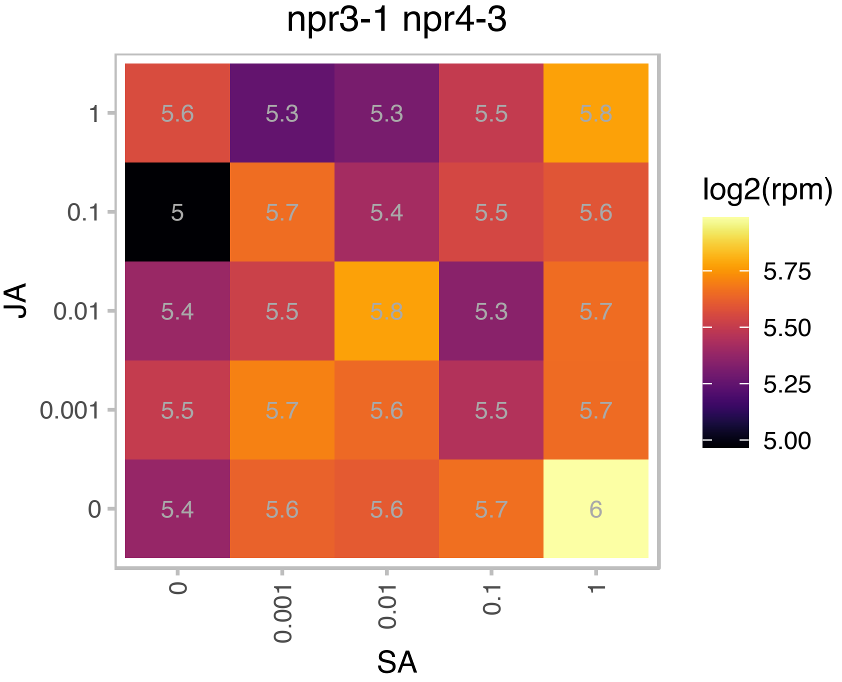

# AT4G35510.1 : NA

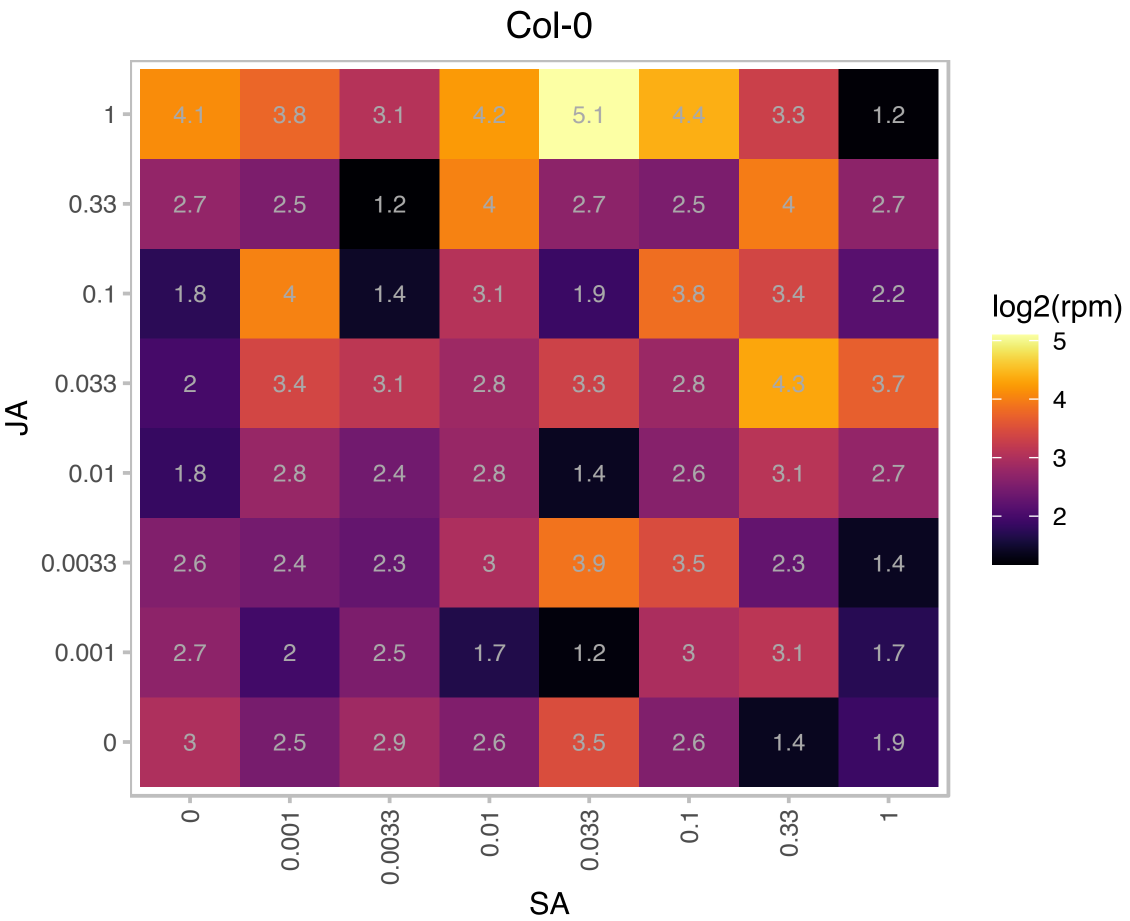


| | |
|---|---|
| Gene | AT4G35510 |
| is.expressed_genes | TRUE |
| is.responsive_genes | TRUE |
| is.apcluster | A-04 |

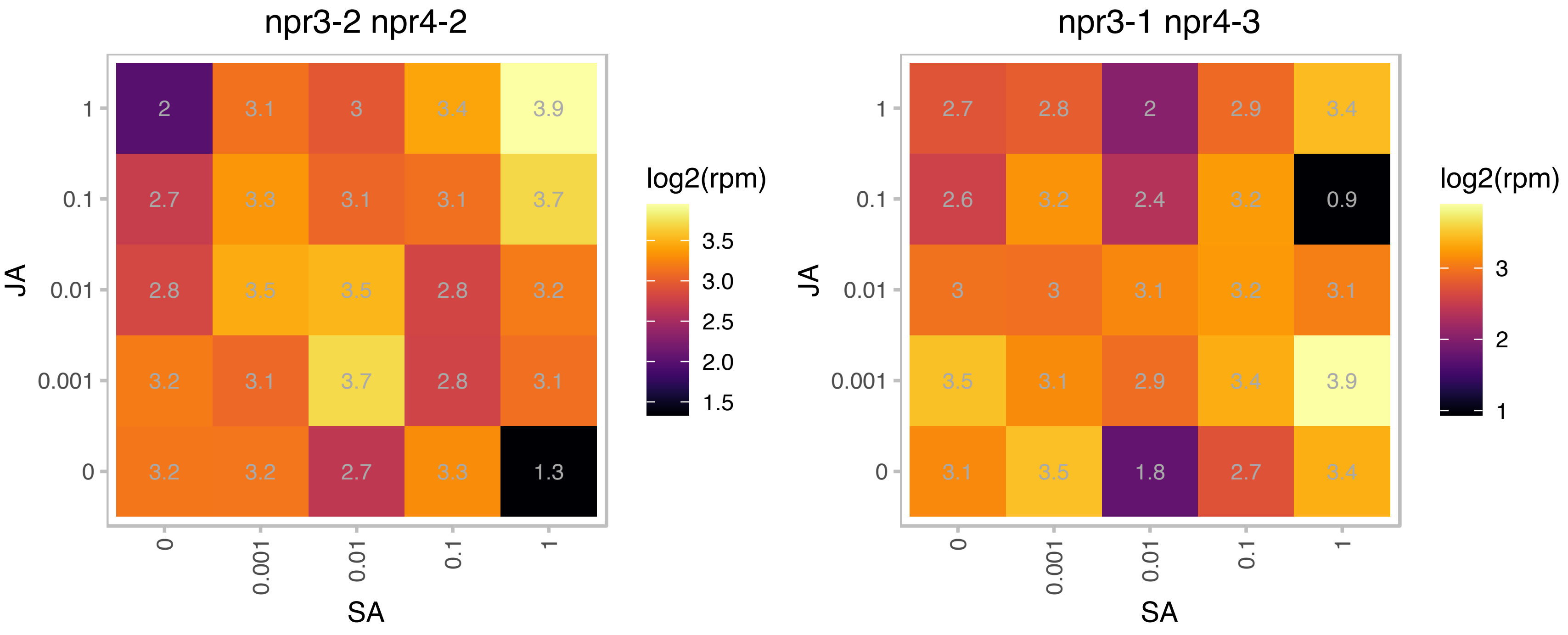

### AT5G03690.1 : AtFBA4, FBA4

Aldolase superfamily protein

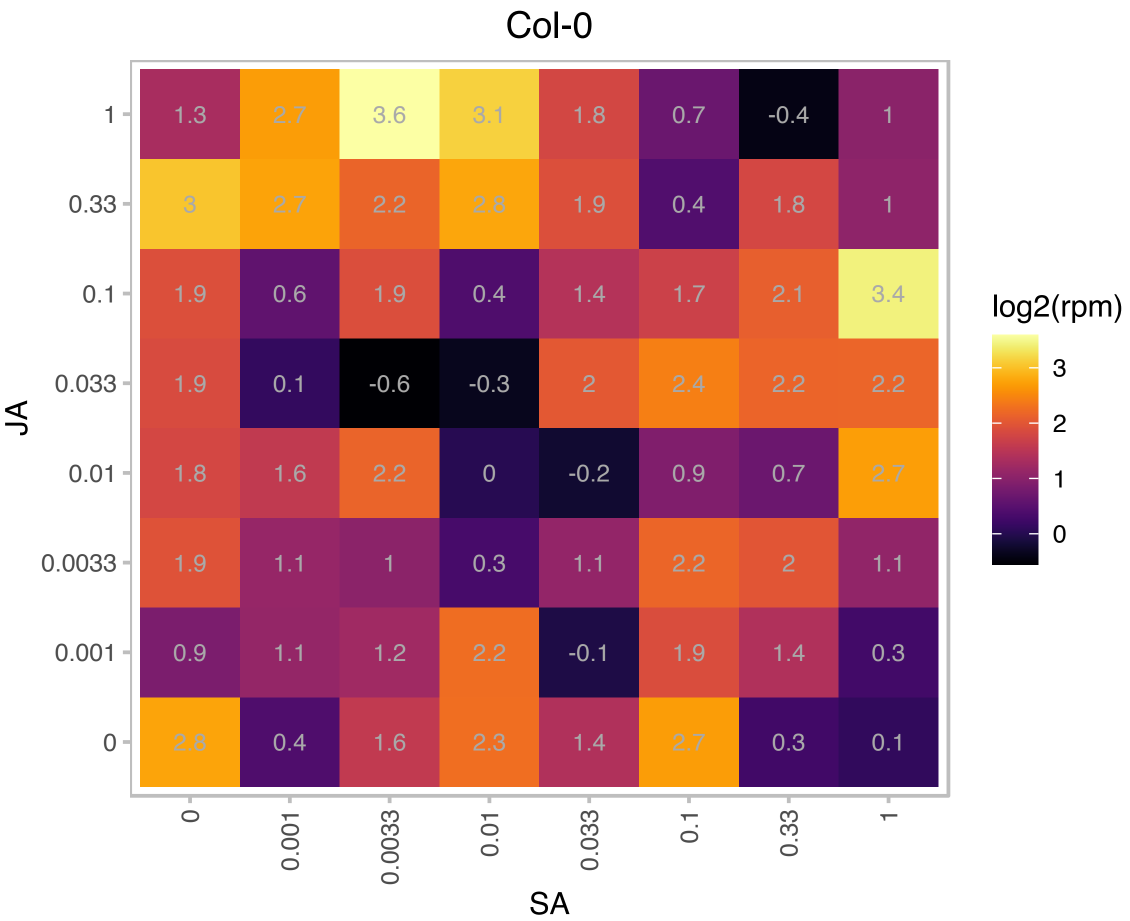


| | |
|---|---|
| Gene | AT5G03690 |
| is.expressed_genes | TRUE |
| is.responsive_genes | TRUE |
| is.apcluster | A-04 |

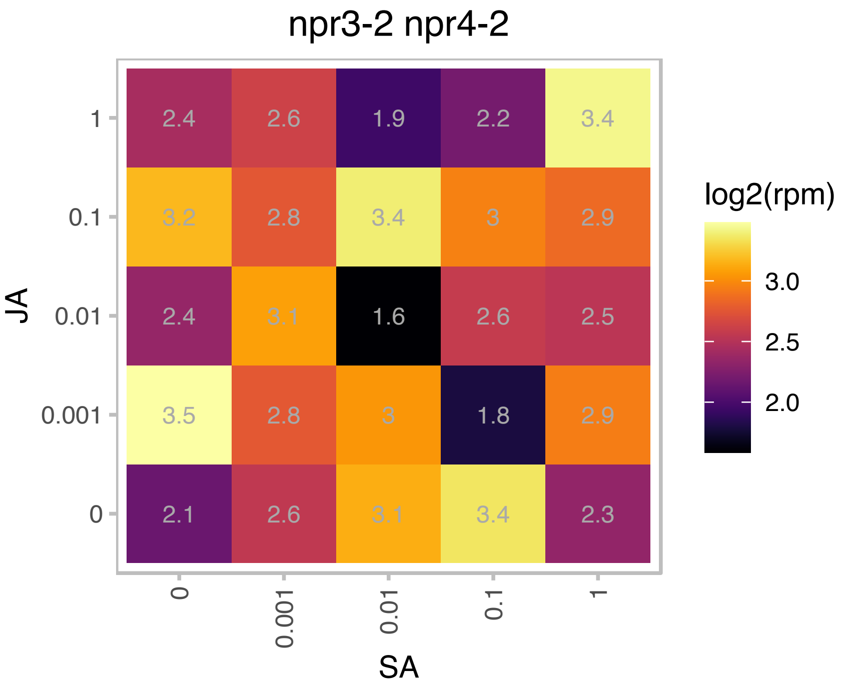


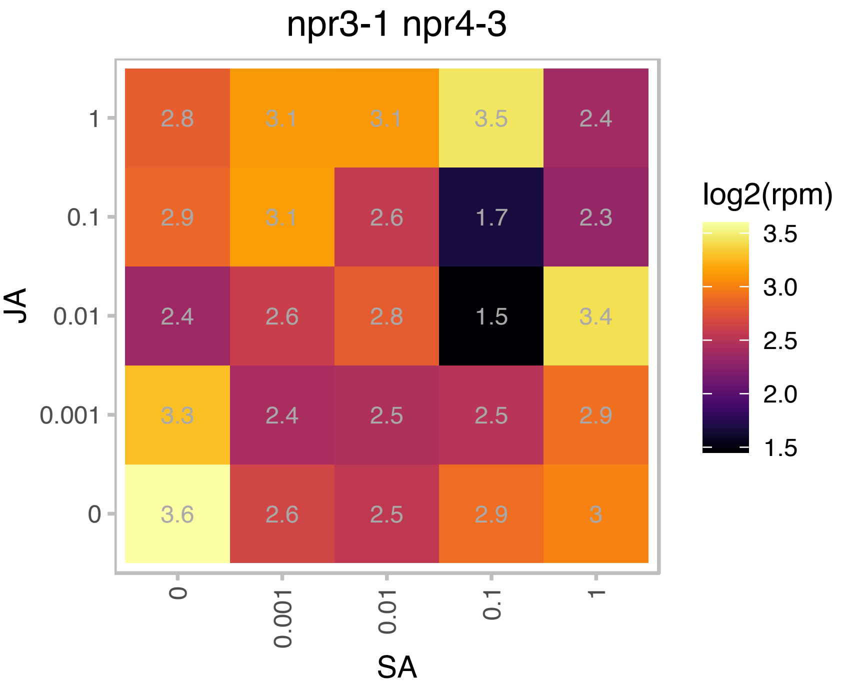

### AT5G09530.1 : PELPK1, PRP10

hydroxyproline-rich glycoprotein family protein

The gene encodes a unique protein which contains 36 repeats of a unique pentapeptide (Pro-Glu-LeuIleIVal-Pro-Lys). It has been shown tobe involved in growth and development.

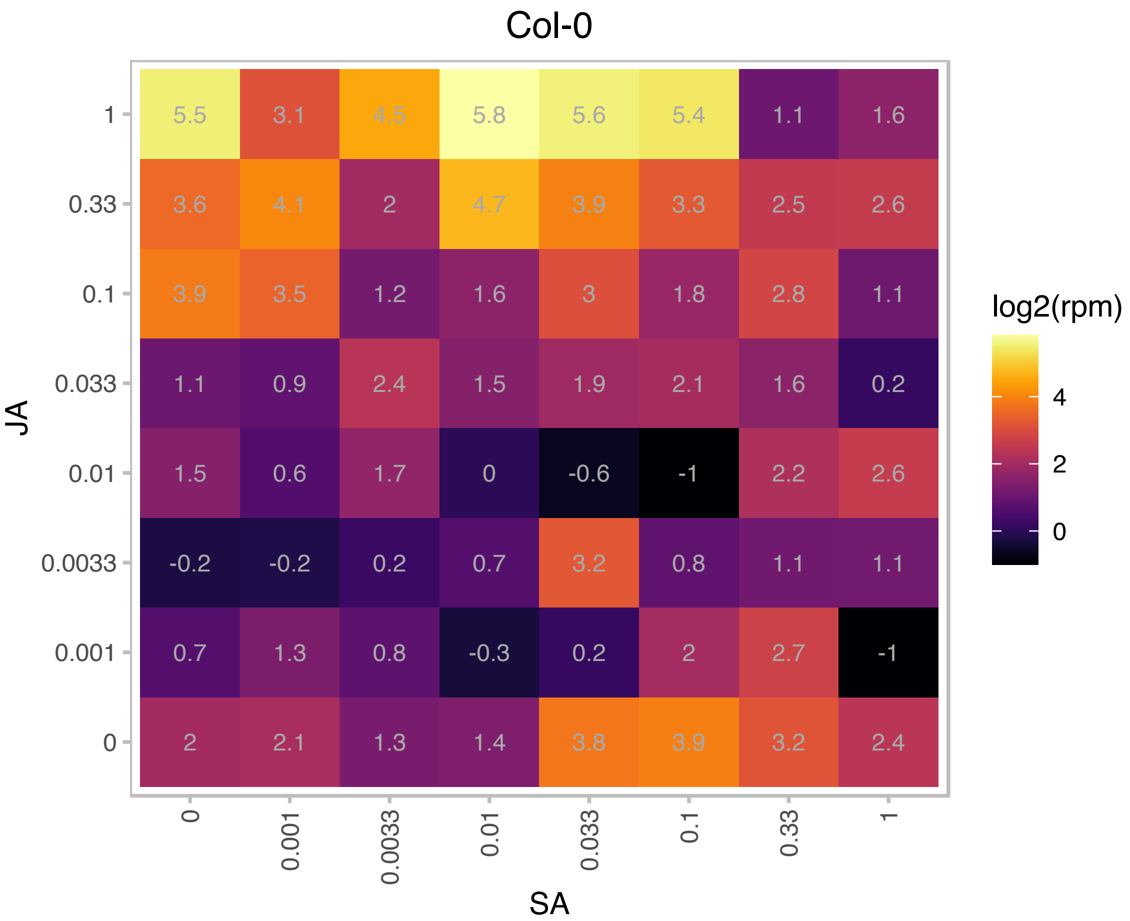


| | |
|---|---|
| Gene | AT5G09530 |
| is.expressed_genes | TRUE |
| is.responsive_genes | TRUE |
| is.apcluster | A-04 |

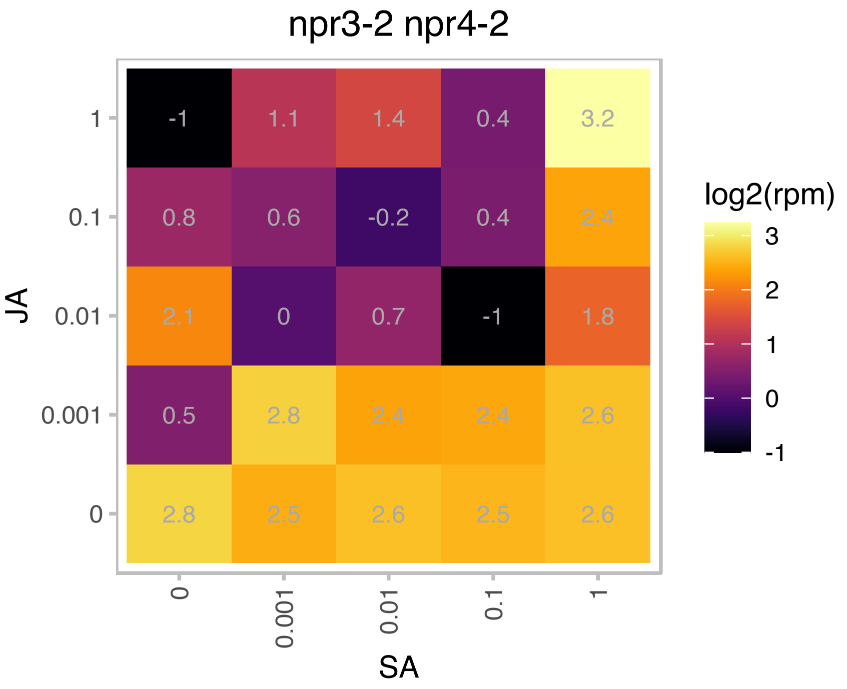


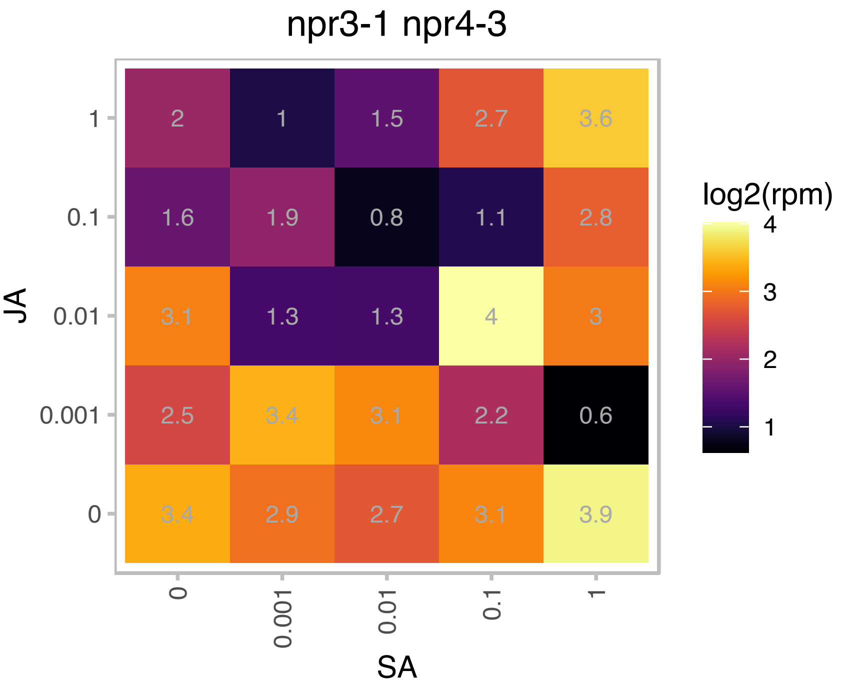

# AT5G16730.1 : NA

Plant protein of unknown function (DUF827)

Encodes a microtubule-associated protein. The mRNA is cell-to-cell mobile.

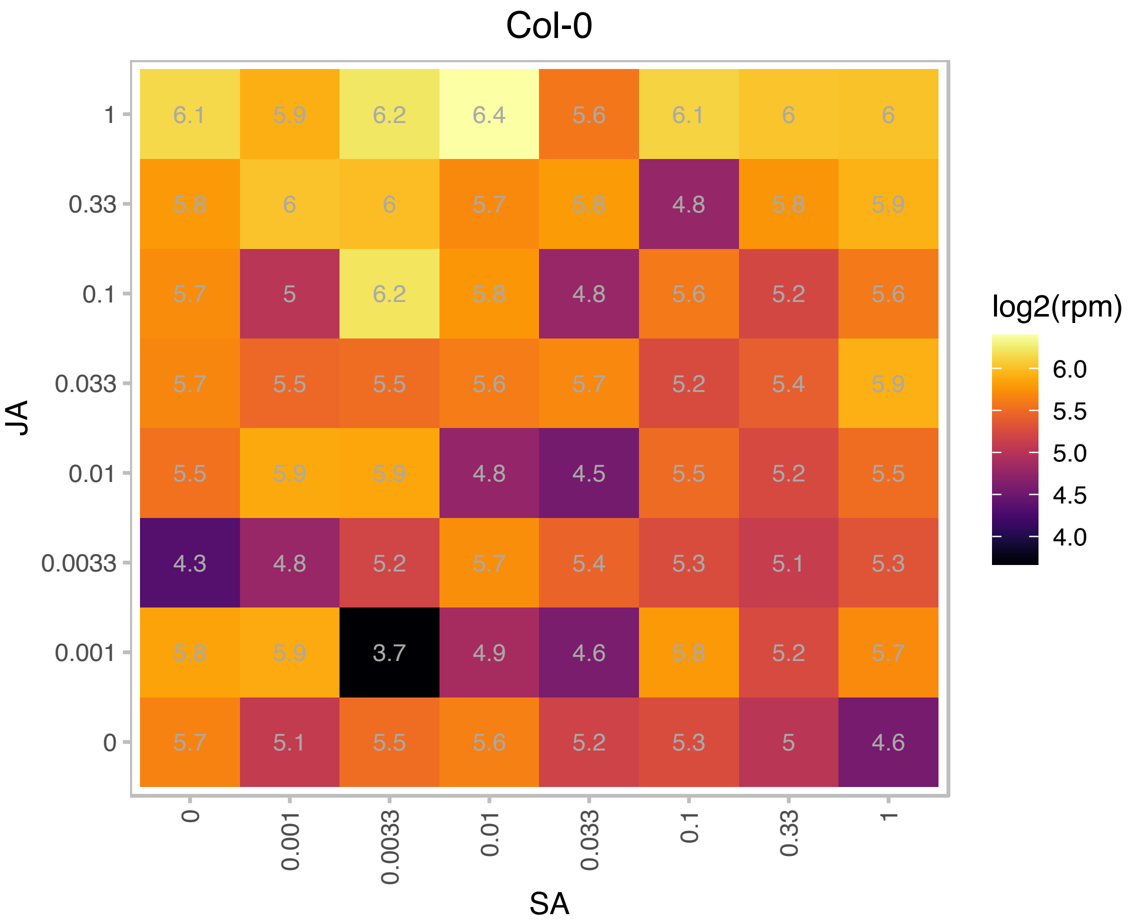


| | |
|---|---|
| Gene | AT5G16730 |
| is.expressed_genes | TRUE |
| is.responsive_genes | TRUE |
| is.apcluster | A-04 |

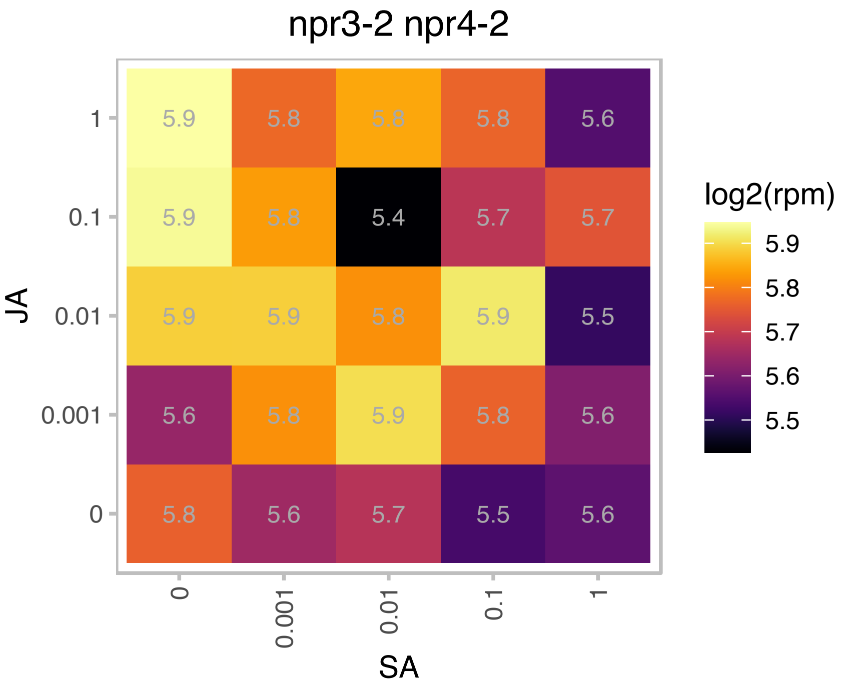


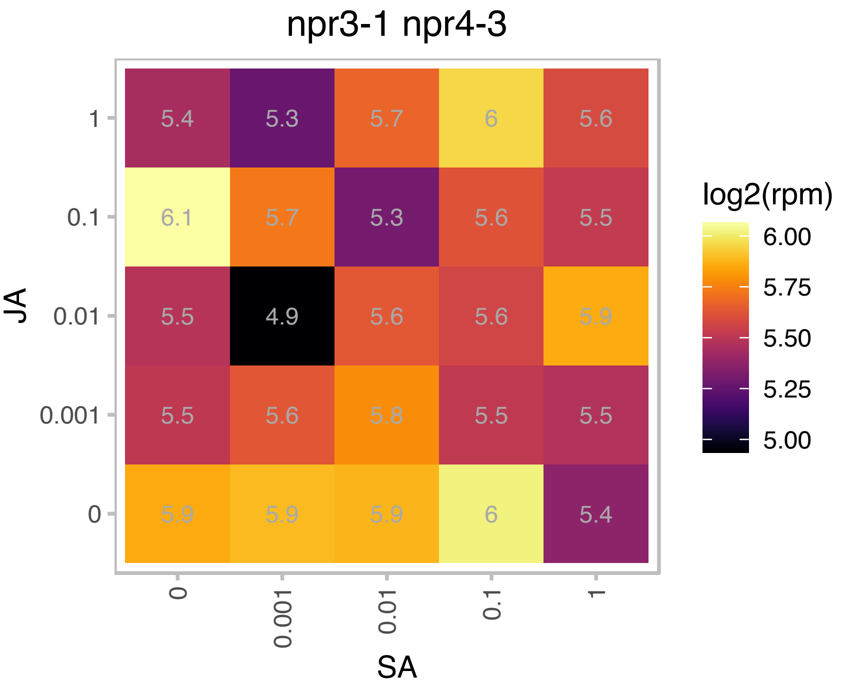

# AT5G21105.1 : NA

Plant L-ascorbate oxidase

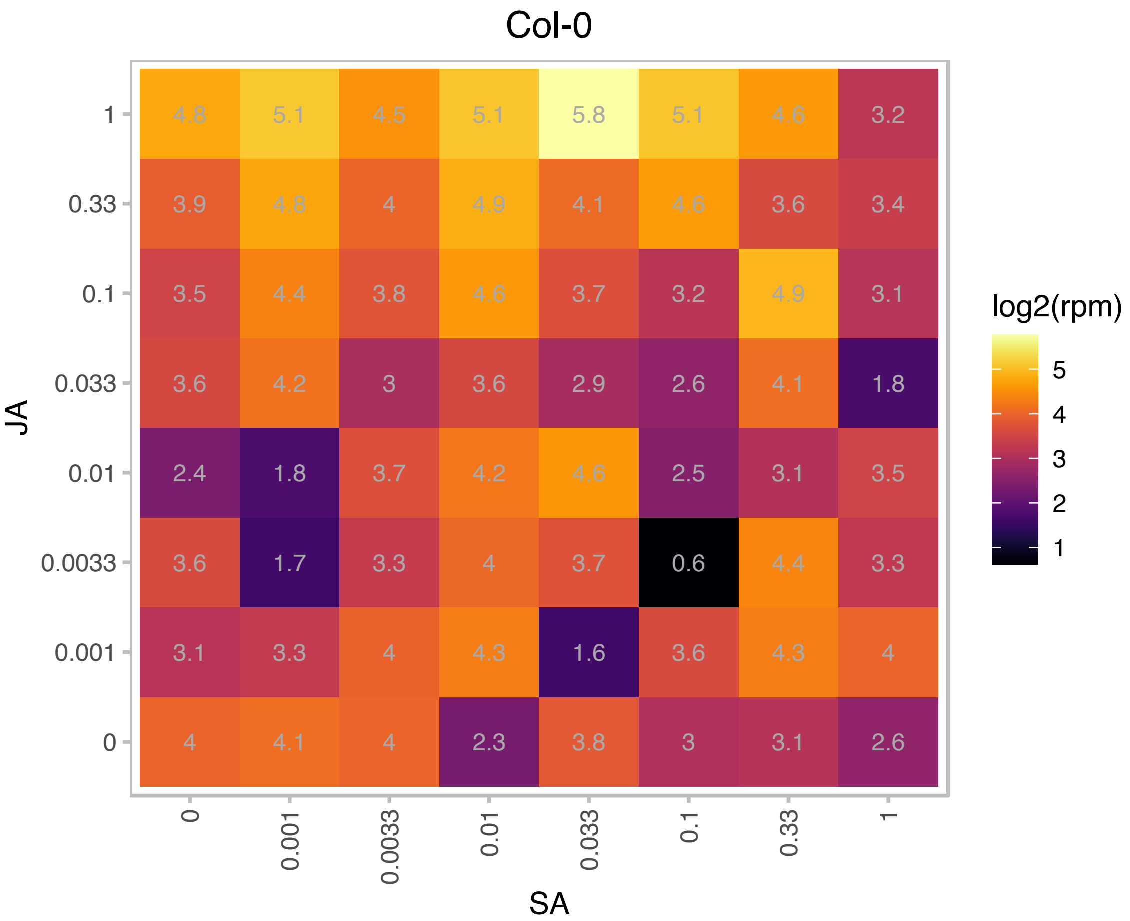


| | |
|---|---|
| Gene | AT5G21105 |
| is.expressed_genes | TRUE |
| is.responsive_genes | TRUE |
| is.apcluster | A-04 |

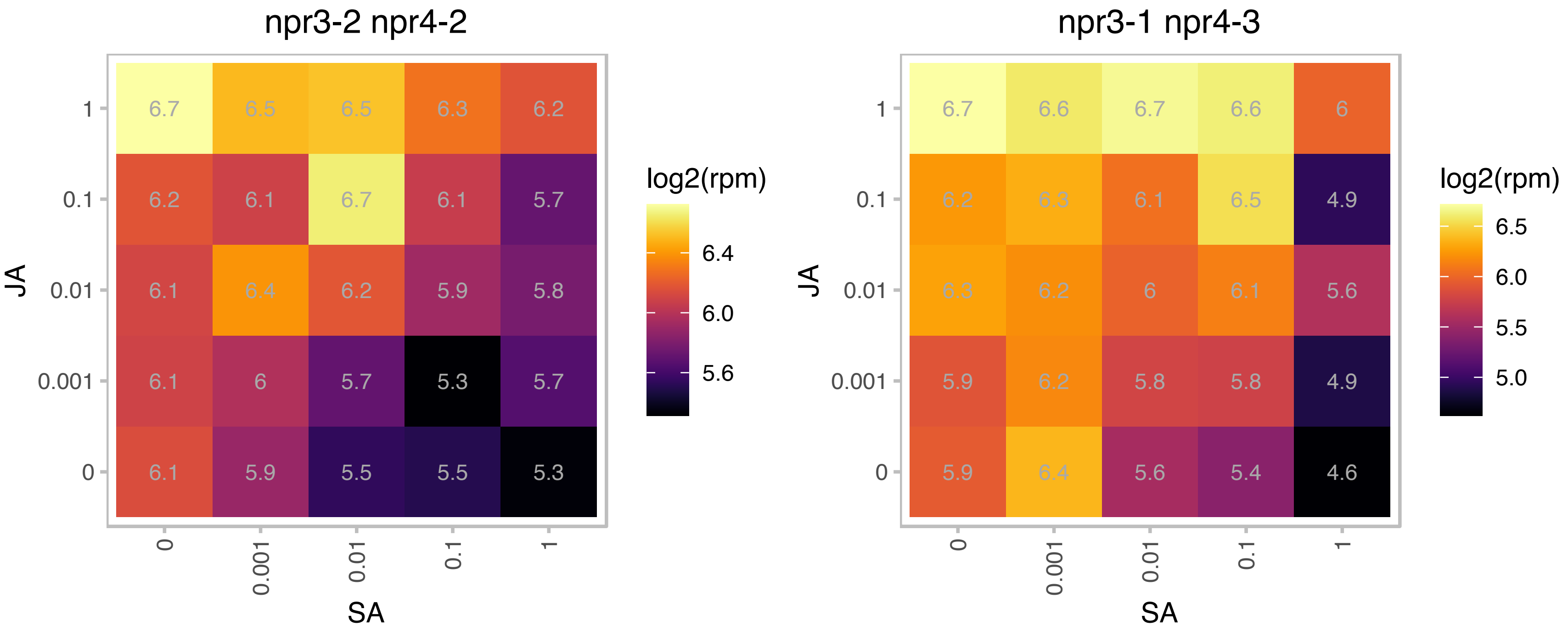

### AT5G24360.1 : ATIRE1-1, AtIRE1b, IRE1-1, IRE1B

inositol requiring 1-1

IRE1A and IRE1B catalyze bZIP60 mRNA splicing, producing the active bZIP60 transcription factor.

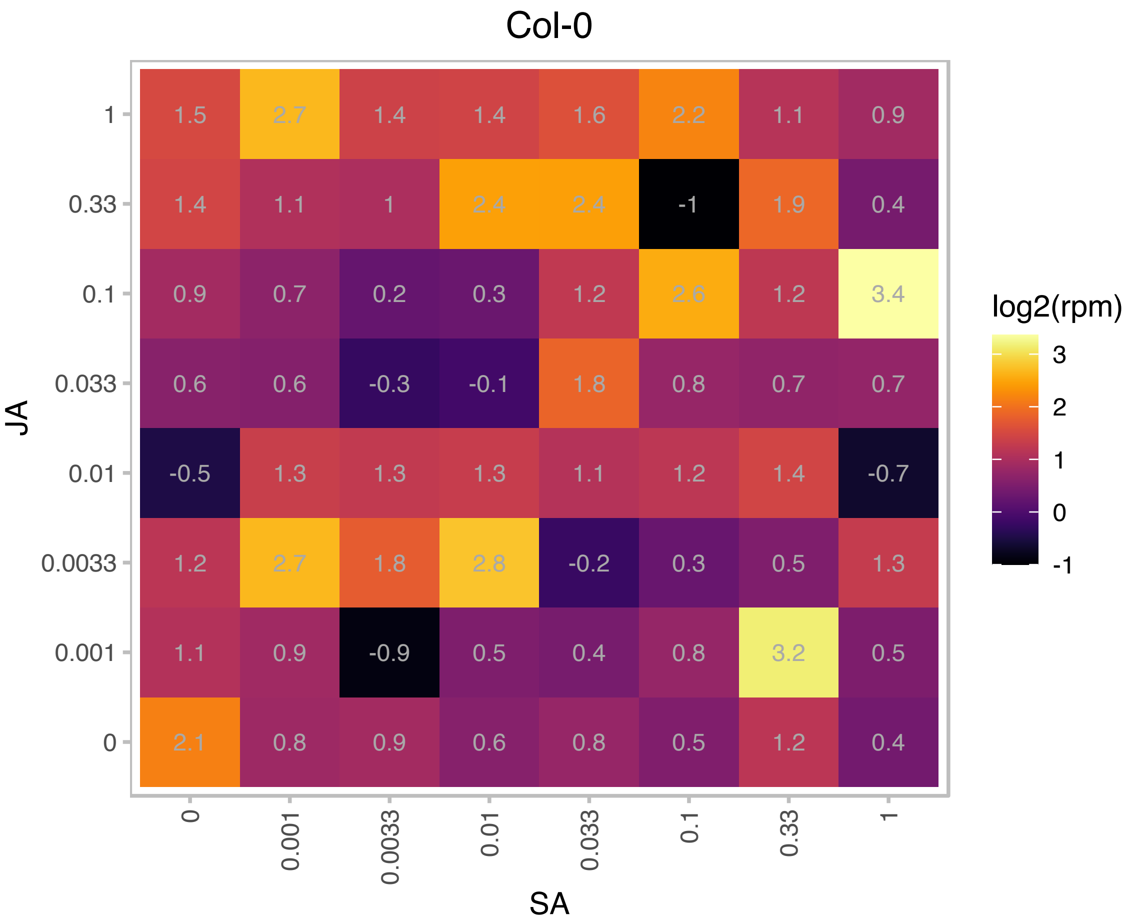


| | |
|---|---|
| Gene | AT5G24360 |
| is.expressed_genes | TRUE |
| is.responsive_genes | TRUE |
| is.apcluster | A-04 |

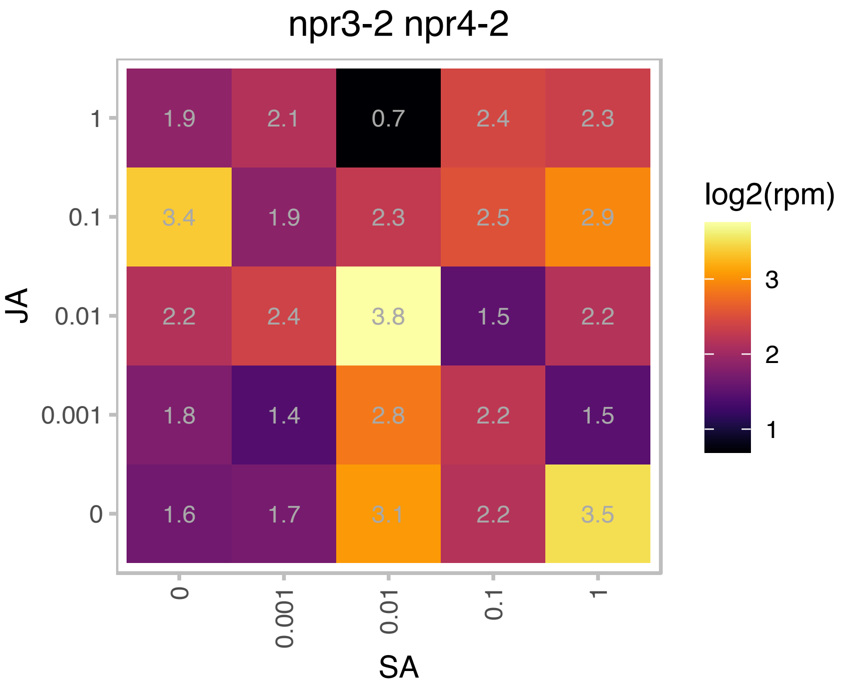


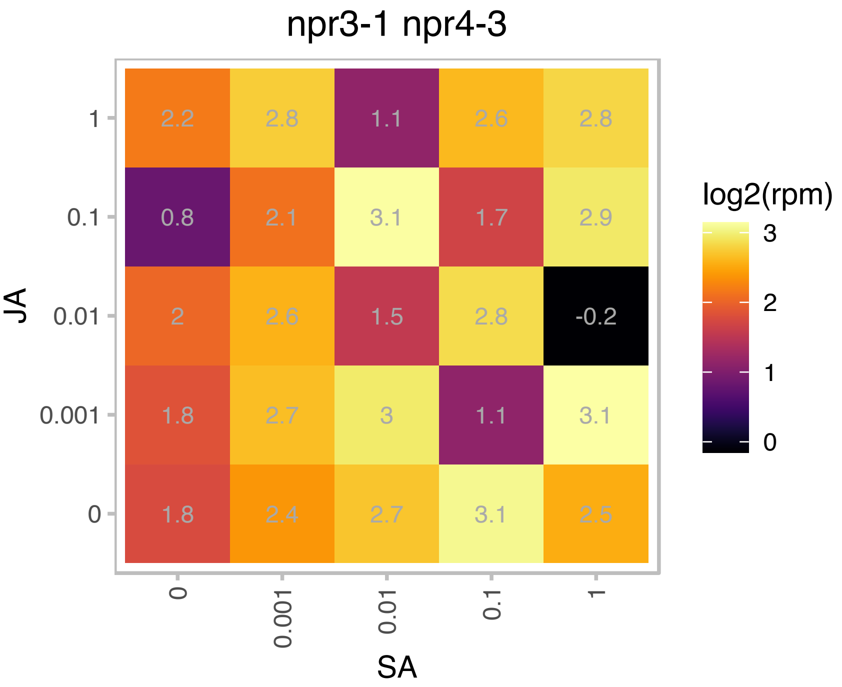

### AT5G24380.1 : ATYSL2, YSL2

YELLOW STRIPE like 2

closest Arabidopsis homolog of Zea maize metal-phytosiderophore/metal-nicotianamine transporter ZmYS1

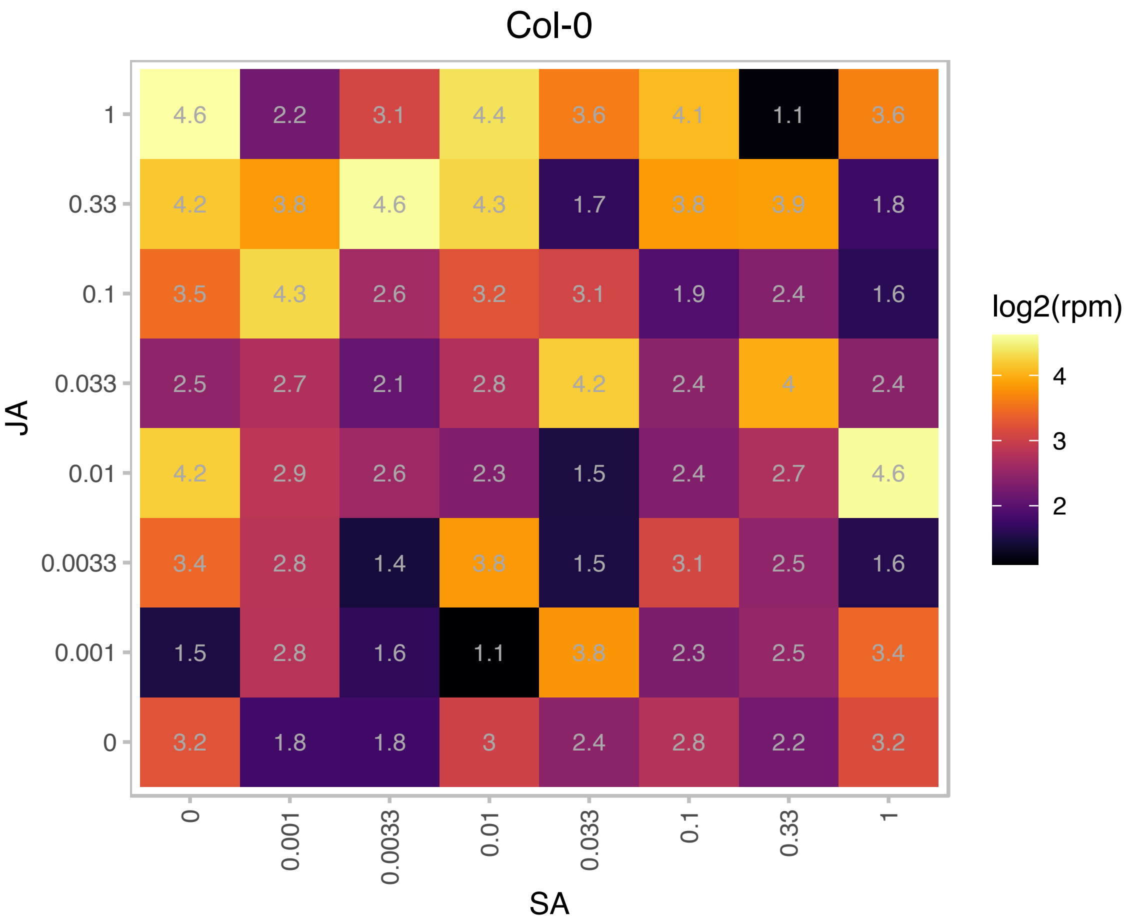


| | |
|---|---|
| Gene | AT5G24380 |
| is.expressed_genes | TRUE |
| is.responsive_genes | TRUE |
| is.apcluster | A-04 |

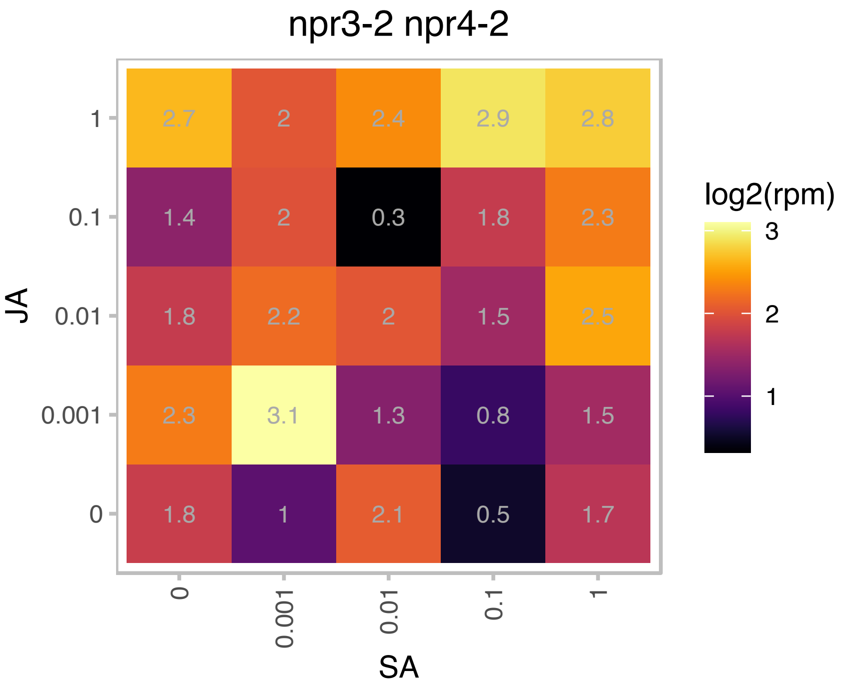


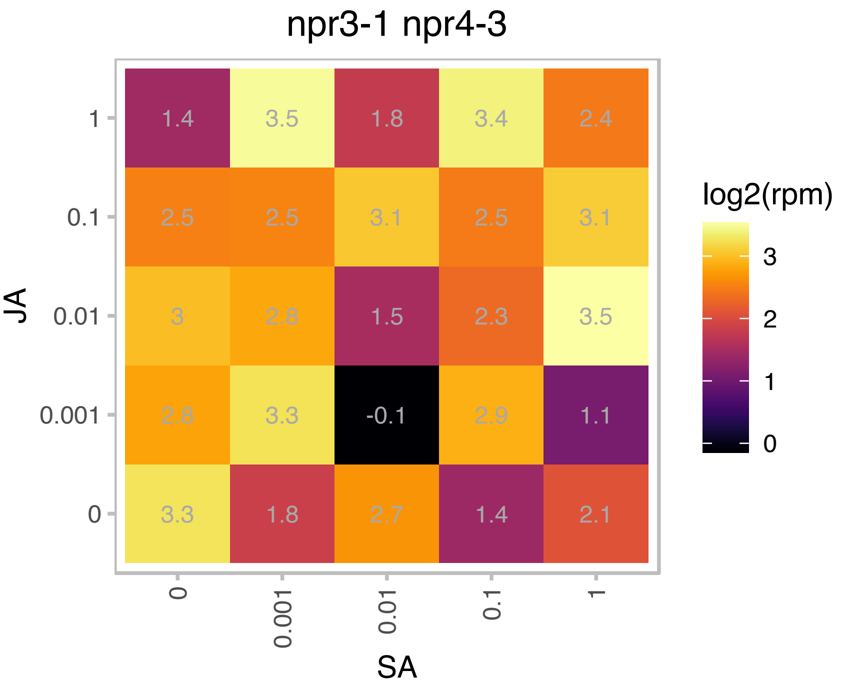

# AT5G38840.1 : NA

SMAD/FHA domain-containing protein

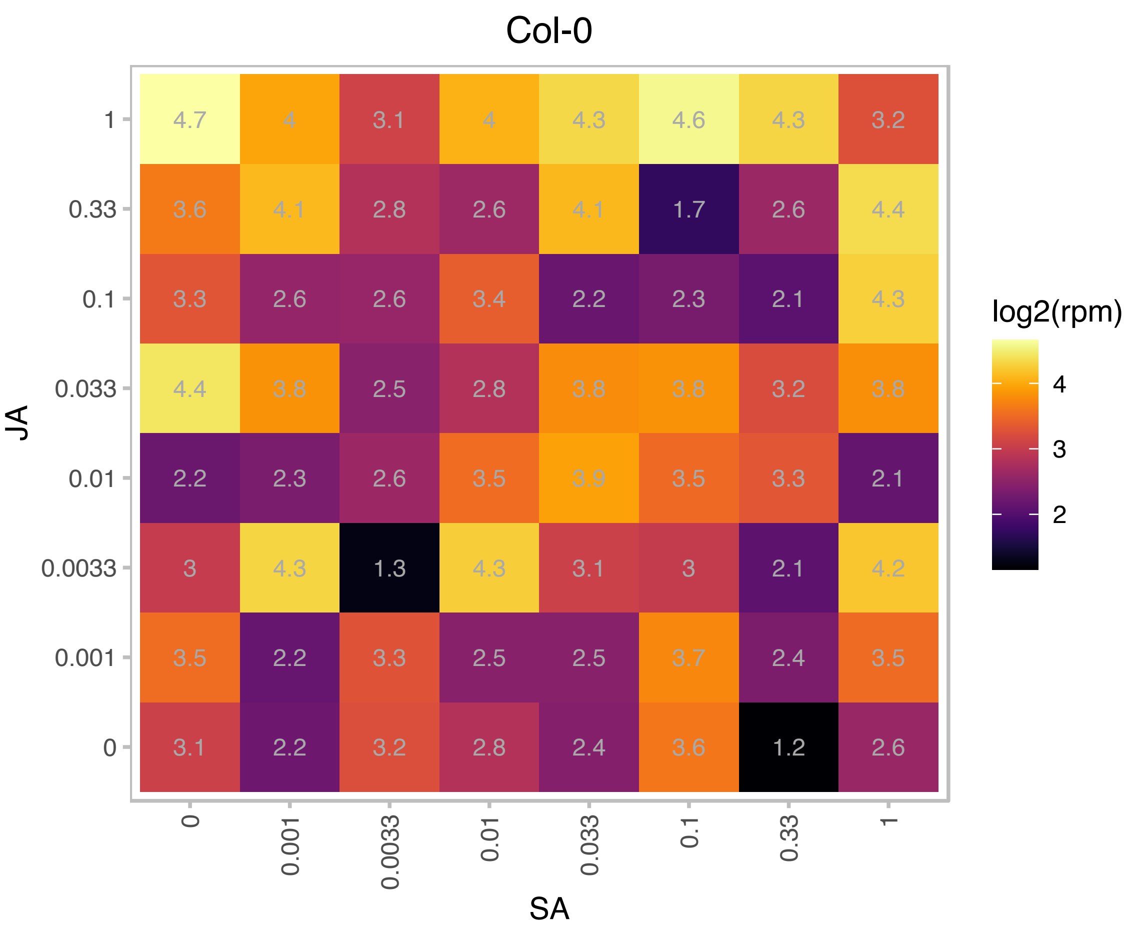


| | |
|---|---|
| Gene | AT5G38840 |
| is.expressed_genes | TRUE |
| is.responsive_genes | TRUE |
| is.apcluster | A-04 |

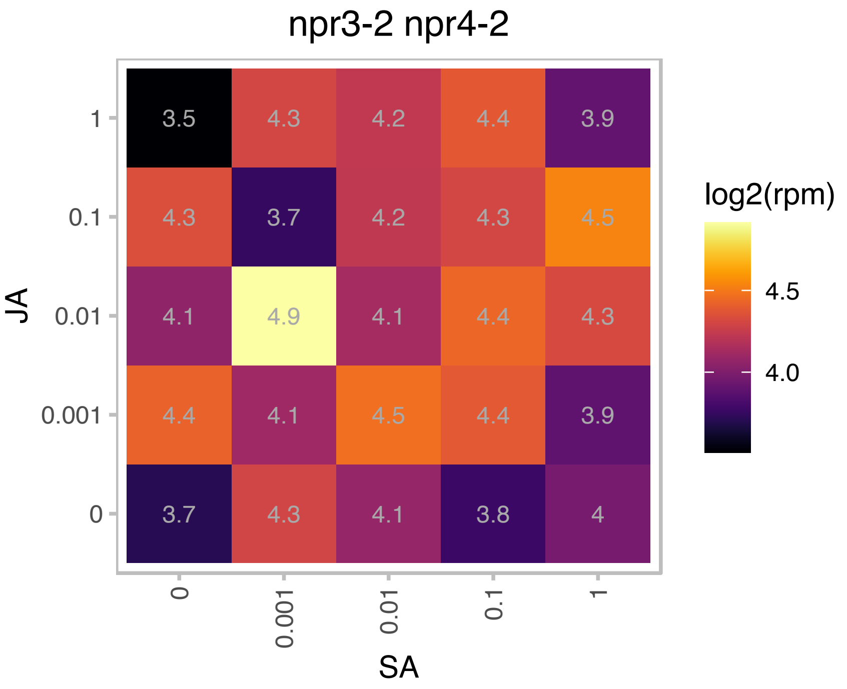


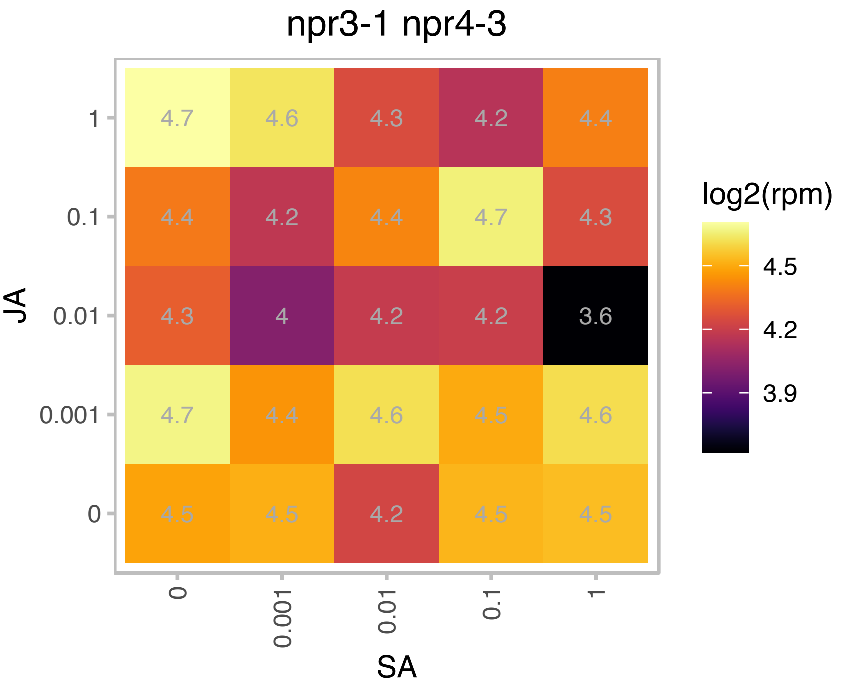

# AT5G40190.1 : NA

RNA ligase/cyclic nucleotide phosphodiesterase family protein

Identified in a screen for calmodulin-binding proteins obtained from an auxin treated cDNA library.

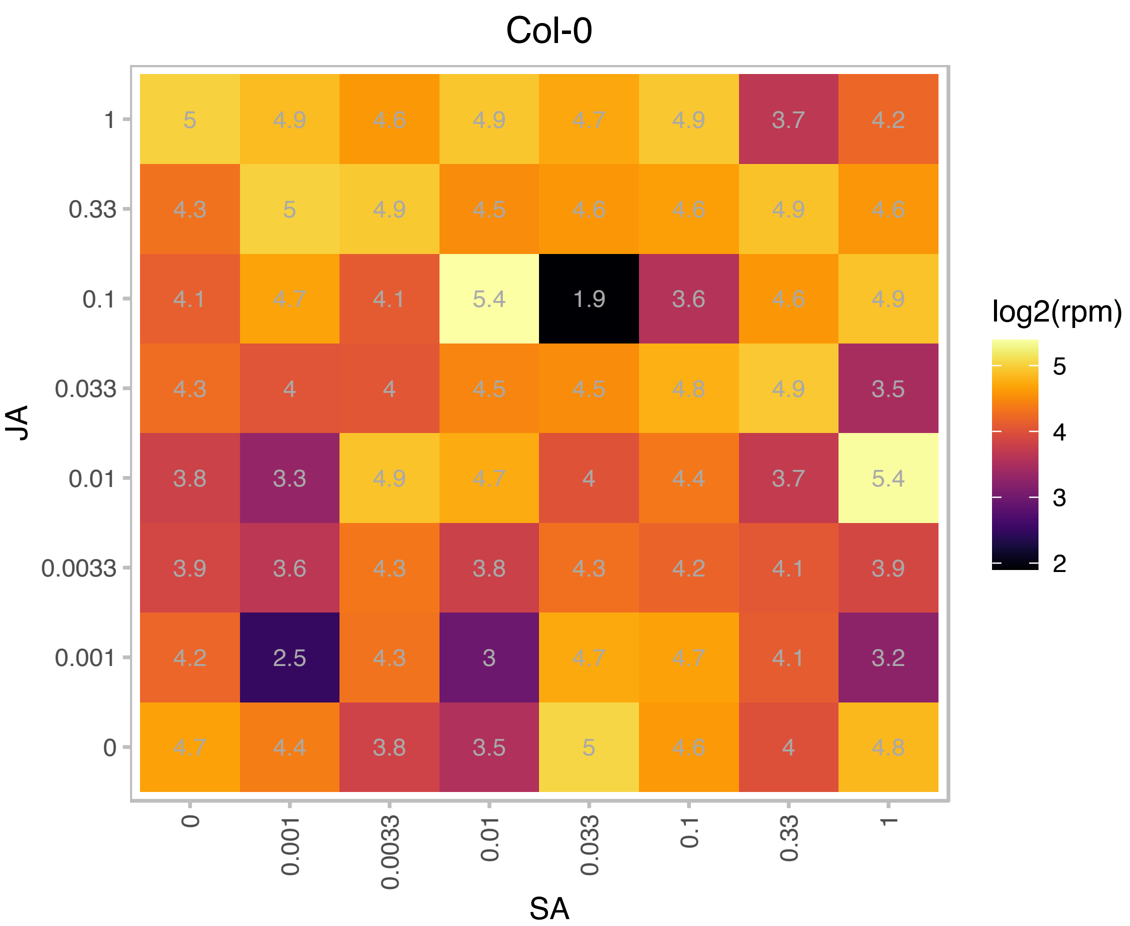


| | |
|---|---|
| Gene | AT5G40190 |
| is.expressed_genes | TRUE |
| is.responsive_genes | TRUE |
| is.apcluster | A-04 |

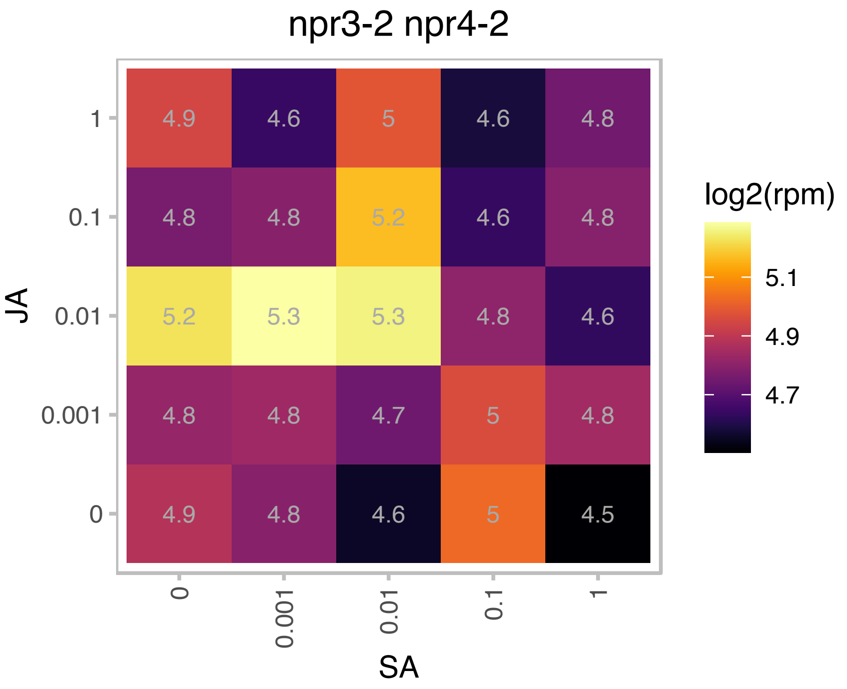


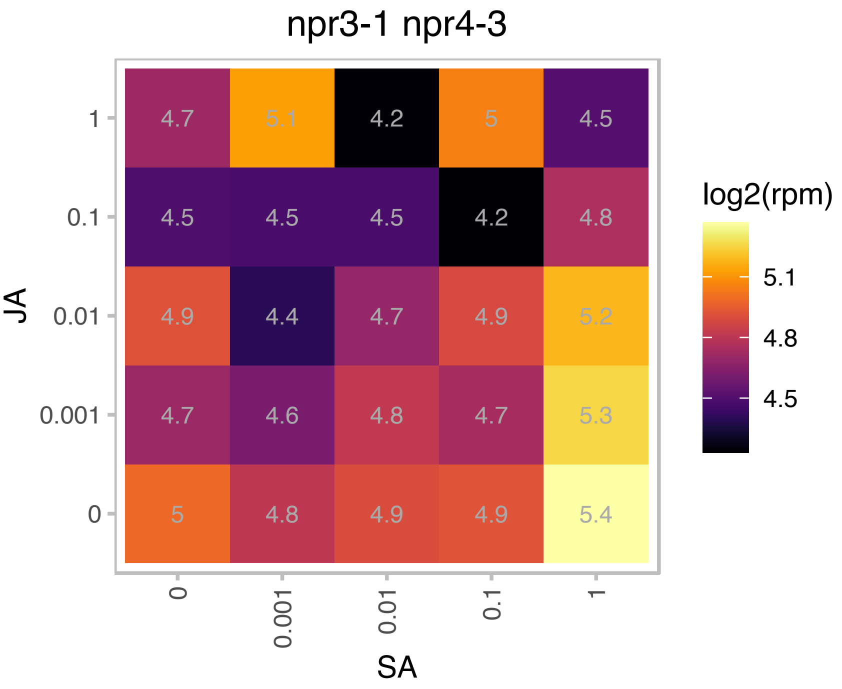

### AT5G40580.1 : PBB2

20S proteasome beta subunit PBB2

Encodes 20S proteasome beta subunit PBB2 (PBB2).

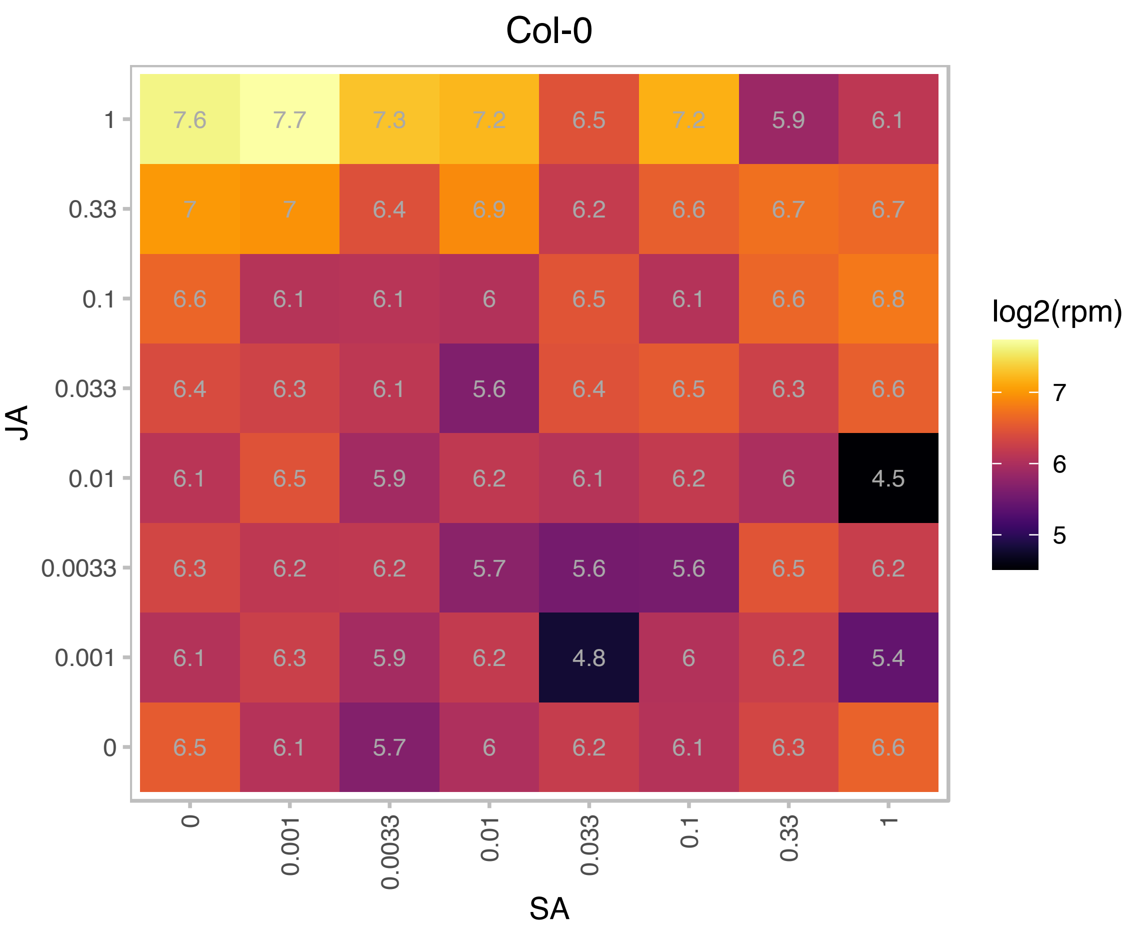


| | |
|---|---|
| Gene | AT5G40580 |
| is.expressed_genes | TRUE |
| is.responsive_genes | TRUE |
| is.apcluster | A-04 |

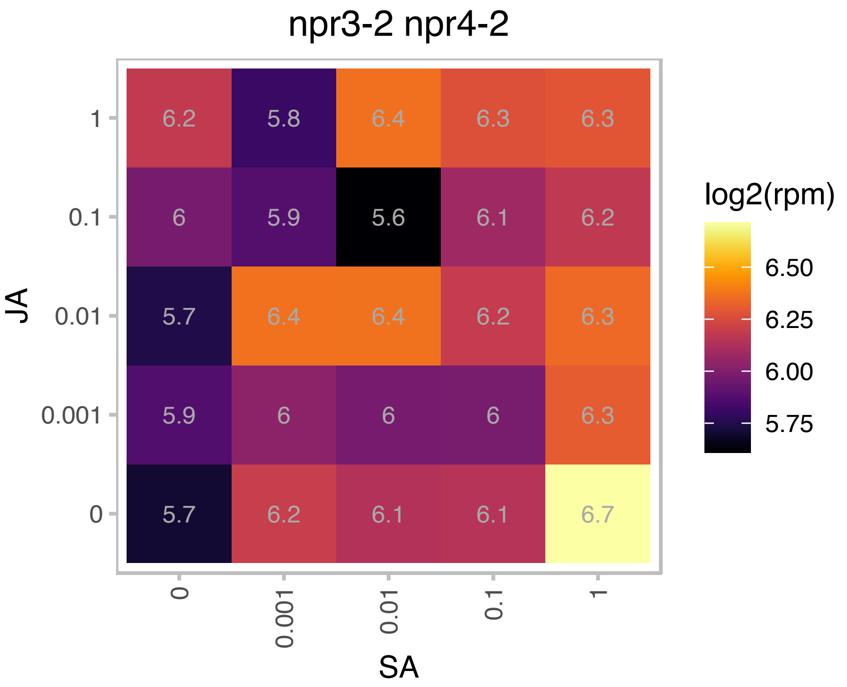


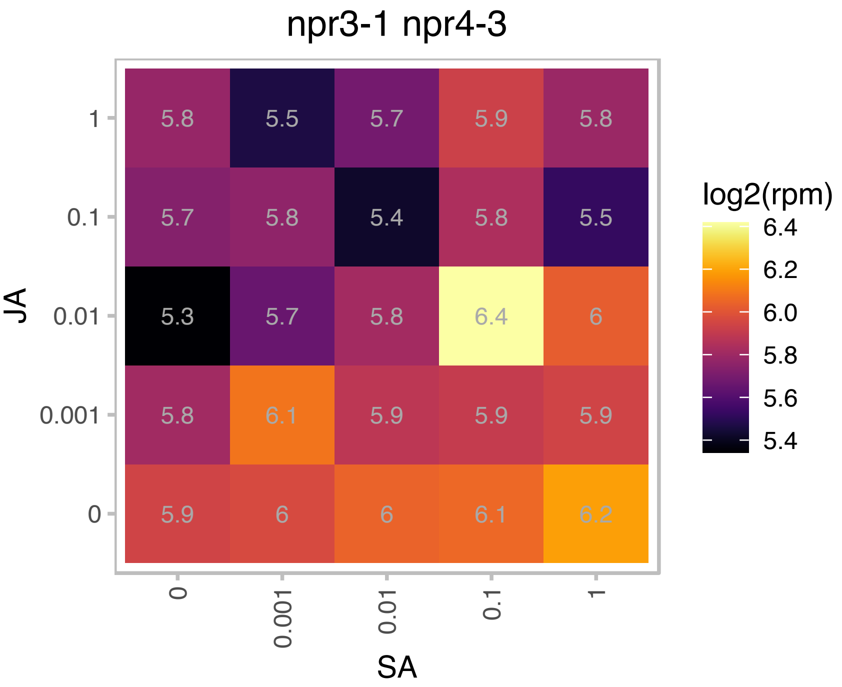

### AT5G43280.1 : ATDCI1, DCI1, DCI1

delta(3,5),delta(2,4)-dienoyl-CoA isomerase 1

Encodes the peroxisomal delta 3,5-delta2,4-dienoyl-CoA isomerase, a enzyme involved in degradation of unsaturated fatty acids. Gene expression is induced upon seed germination.

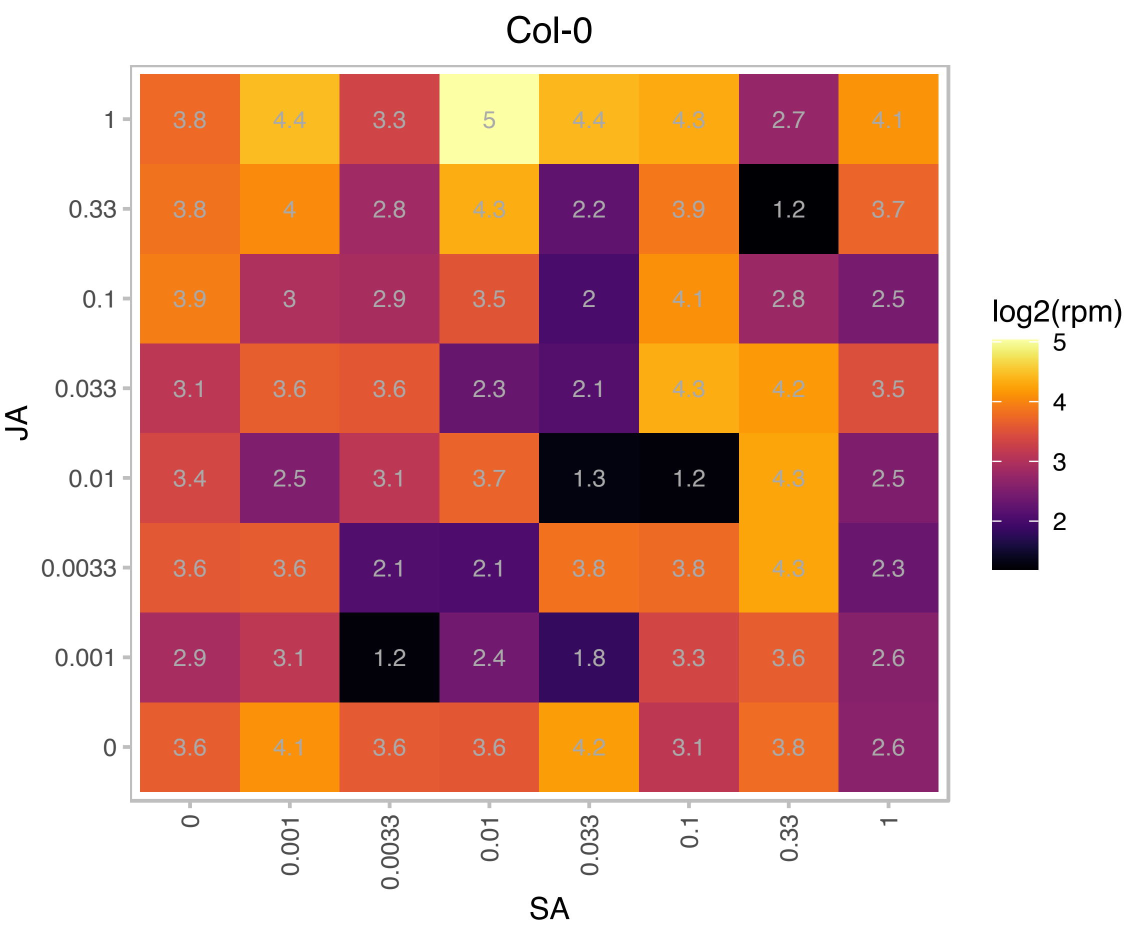


| | |
|---|---|
| Gene | AT5G43280 |
| is.expressed_genes | TRUE |
| is.responsive_genes | TRUE |
| is.apcluster | A-04 |

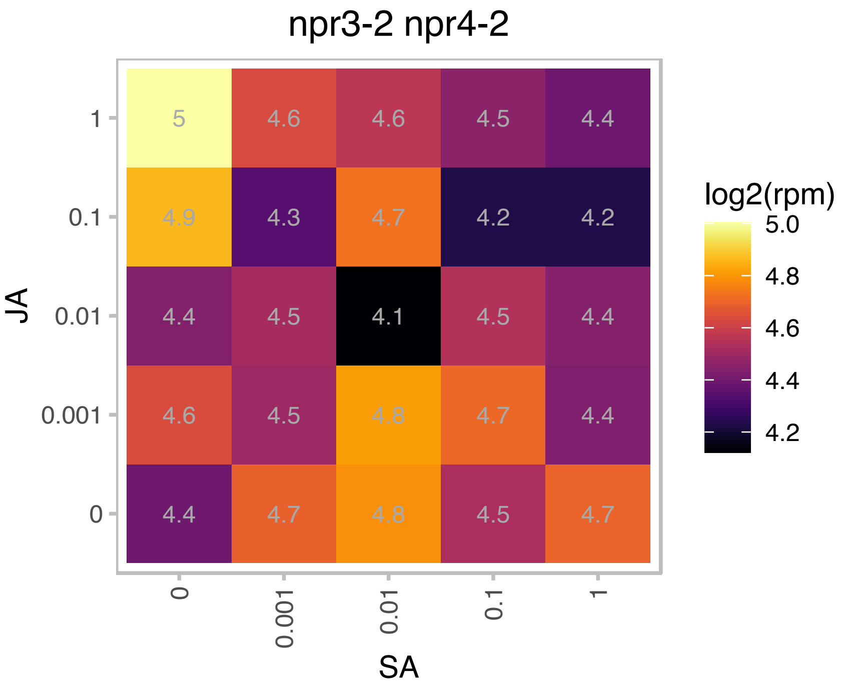


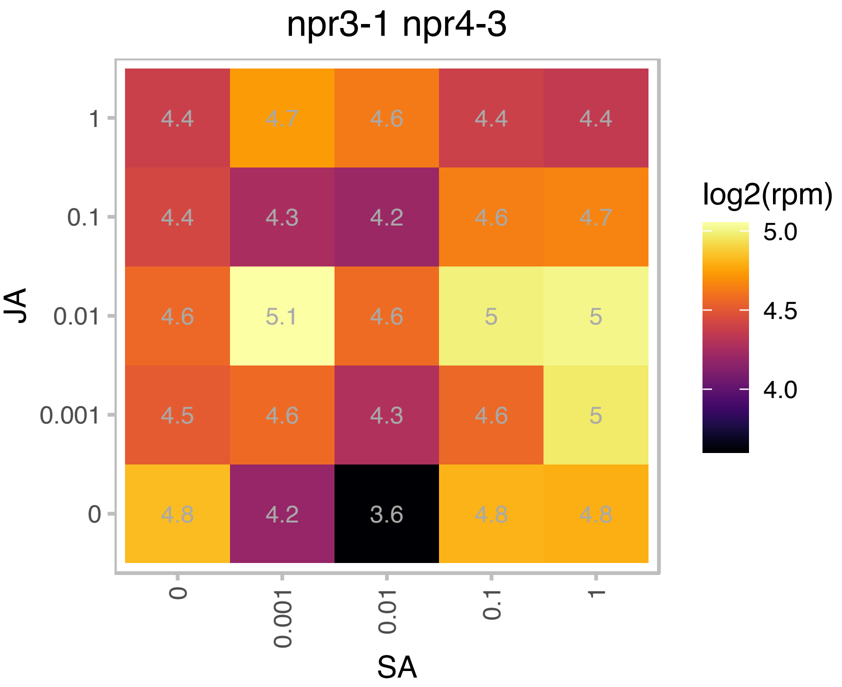

### AT5G47690.1 : AtPDS5A, PDS5, PDS5A

binding

One of 5 PO76/PDS5 cohesion cofactor orthologs of Arabidopsis.

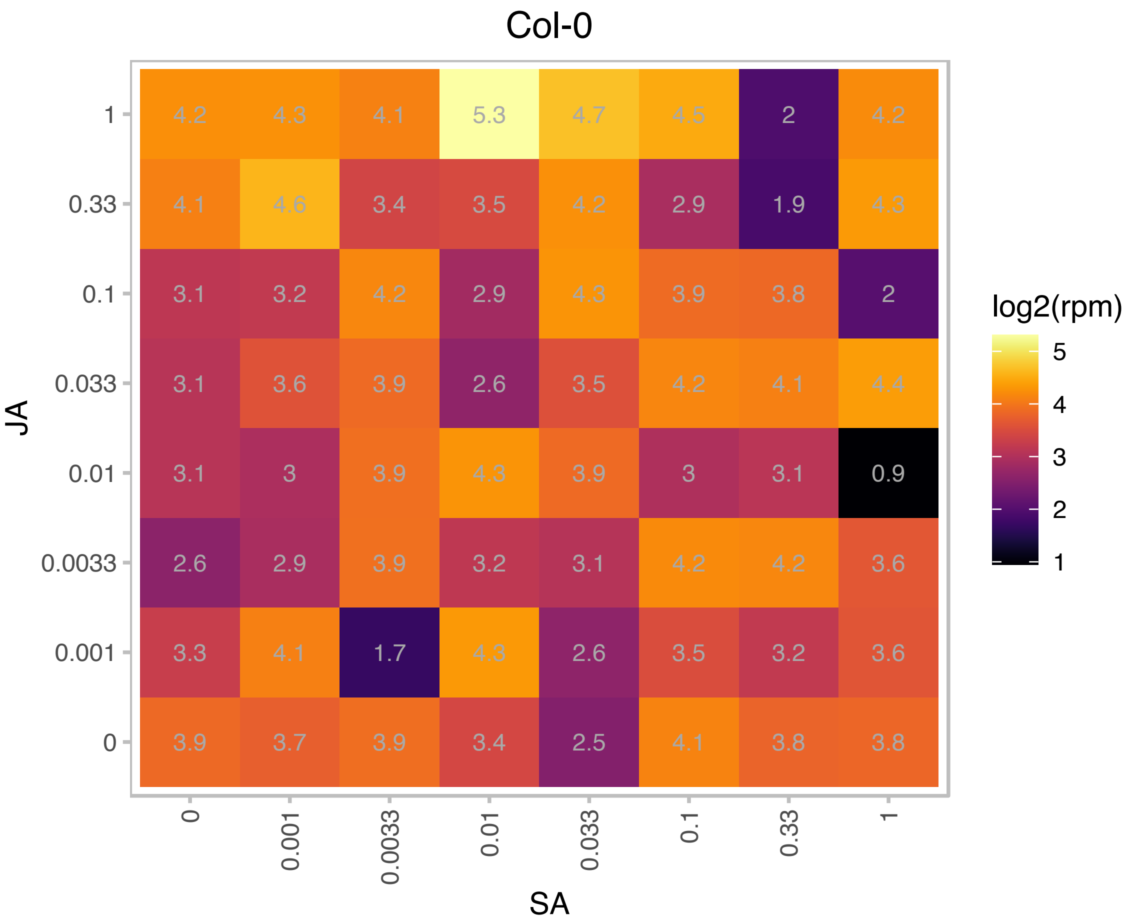


| | |
|---|---|
| Gene | AT5G47690 |
| is.expressed_genes | TRUE |
| is.responsive_genes | TRUE |
| is.apcluster | A-04 |

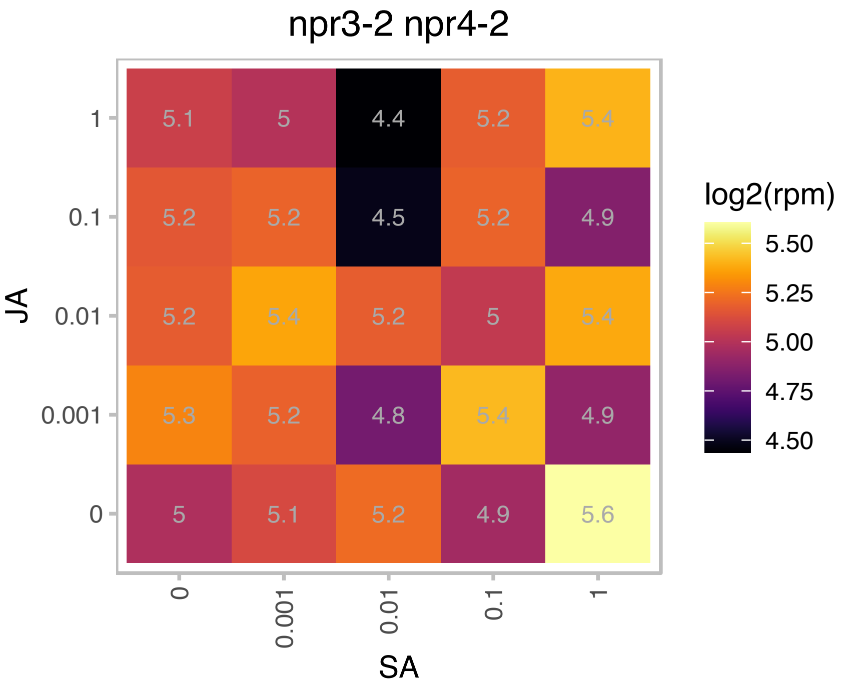


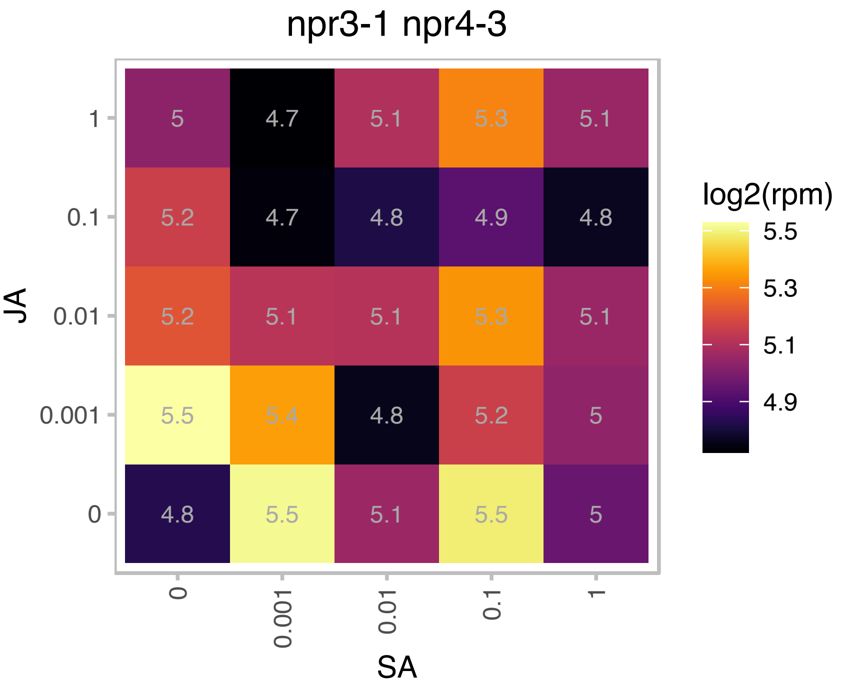

### AT5G50850.1 : MAB1

Transketolase family protein

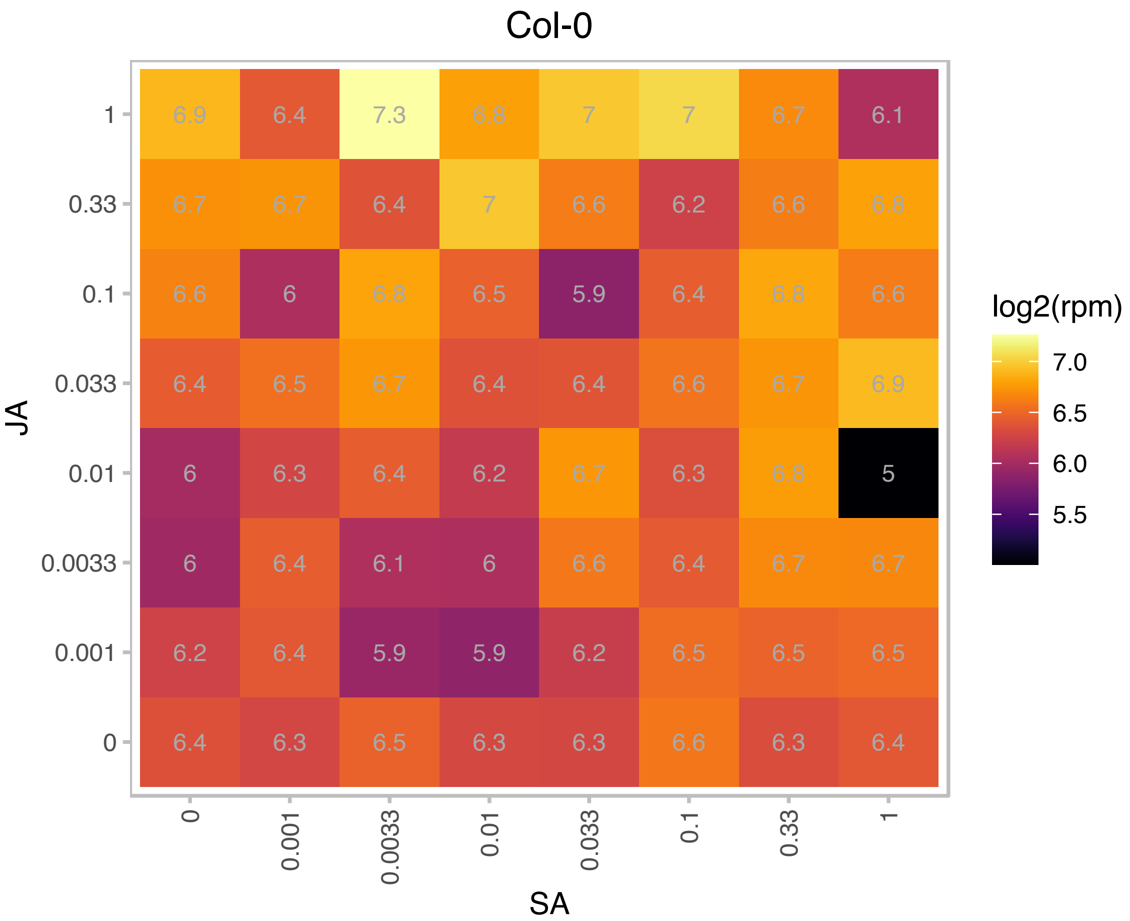


| | |
|---|---|
| Gene | AT5G50850 |
| is.expressed_genes | TRUE |
| is.responsive_genes | TRUE |
| is.apcluster | A-04 |

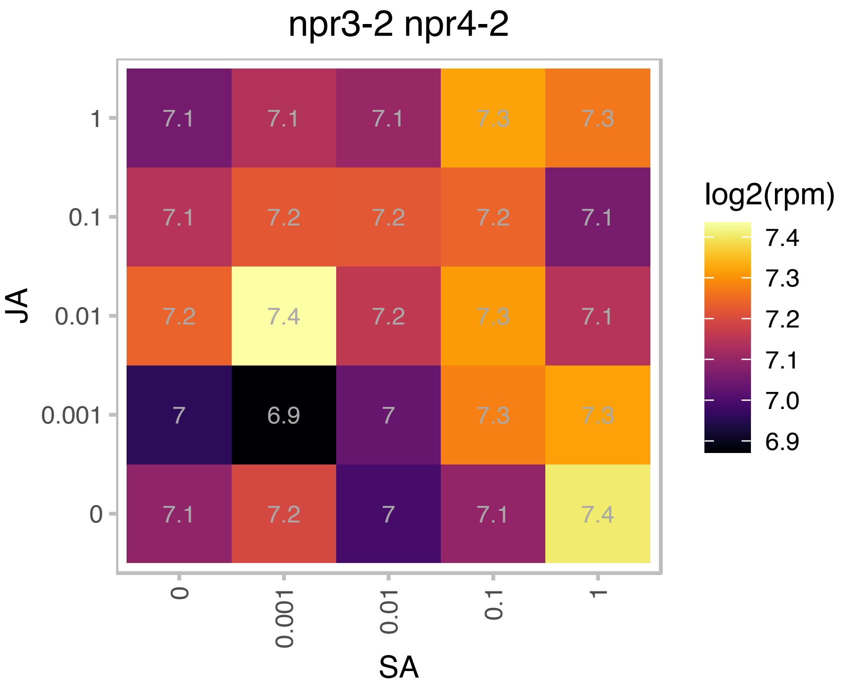


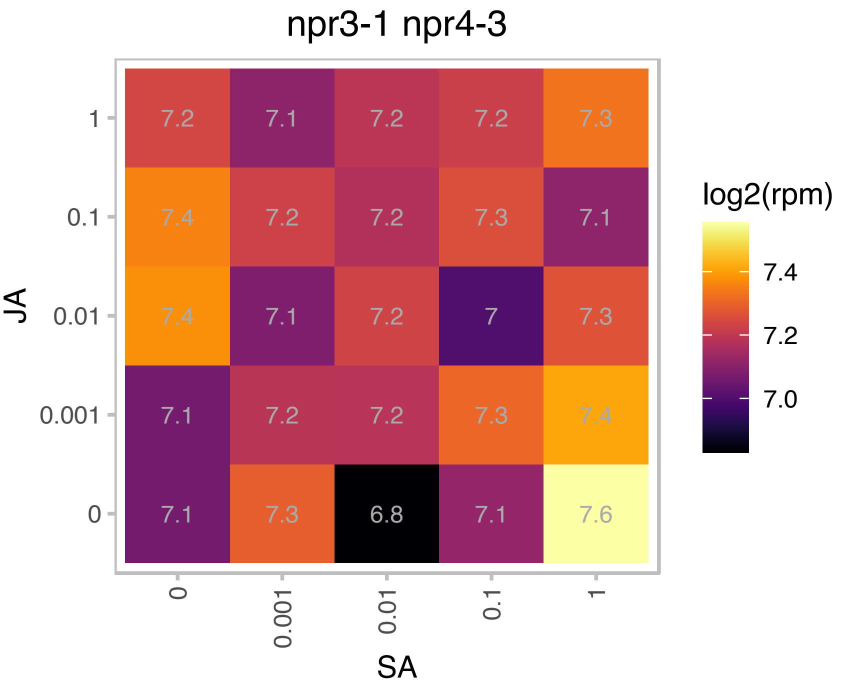

### AT5G53850.1 : DEP1

haloacid dehalogenase-like hydrolase family protein

Encodes a trifunctional dehydratase / enolase / phosphatase involved in the methionine salvage.

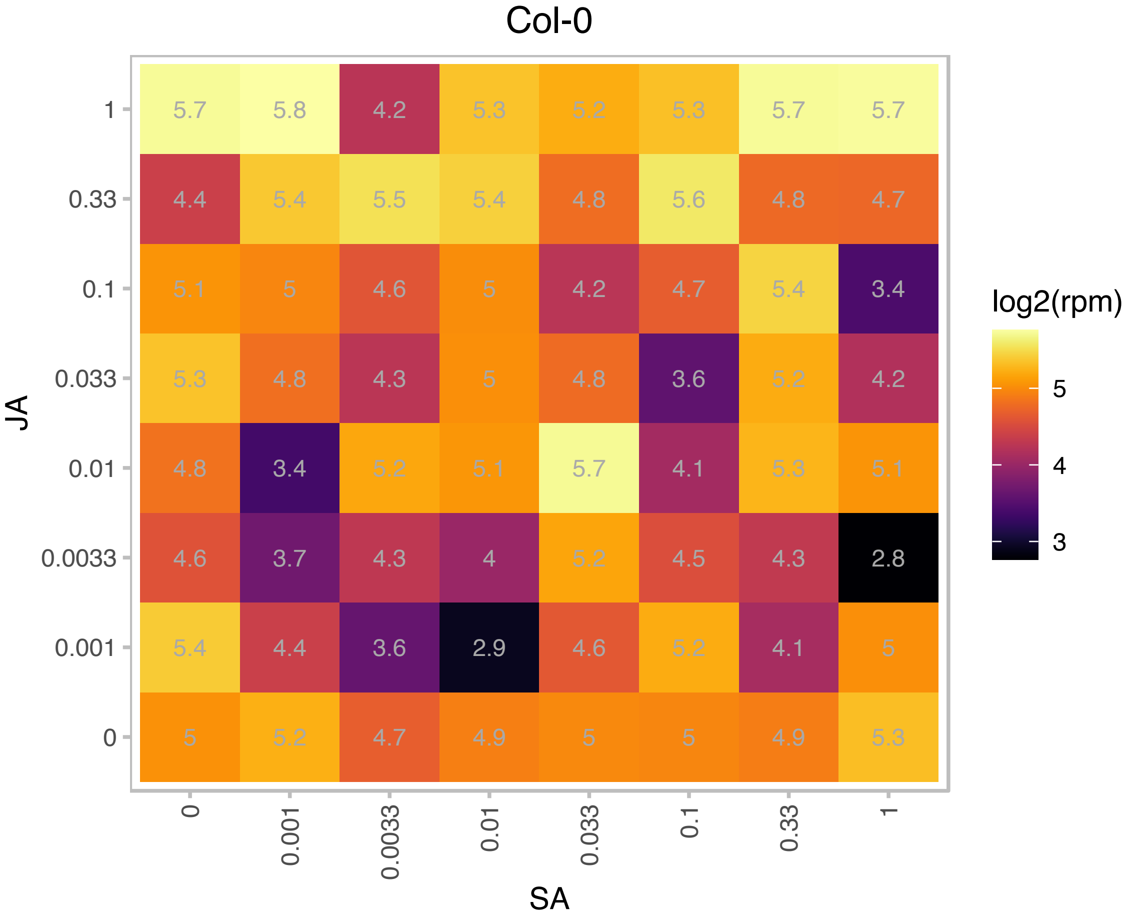


| | |
|---|---|
| Gene | AT5G53850 |
| is.expressed_genes | TRUE |
| is.responsive_genes | TRUE |
| is.apcluster | A-04 |

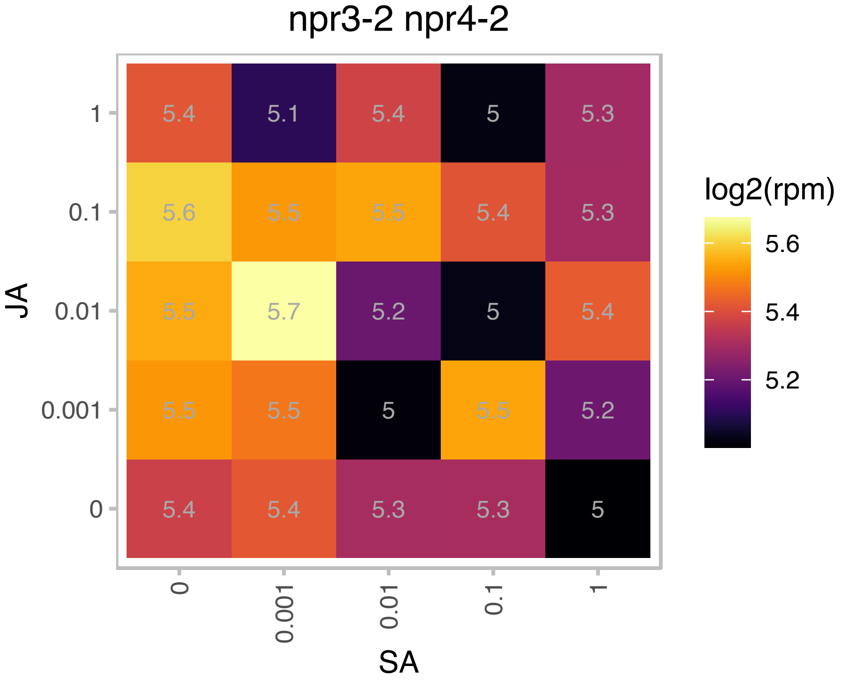


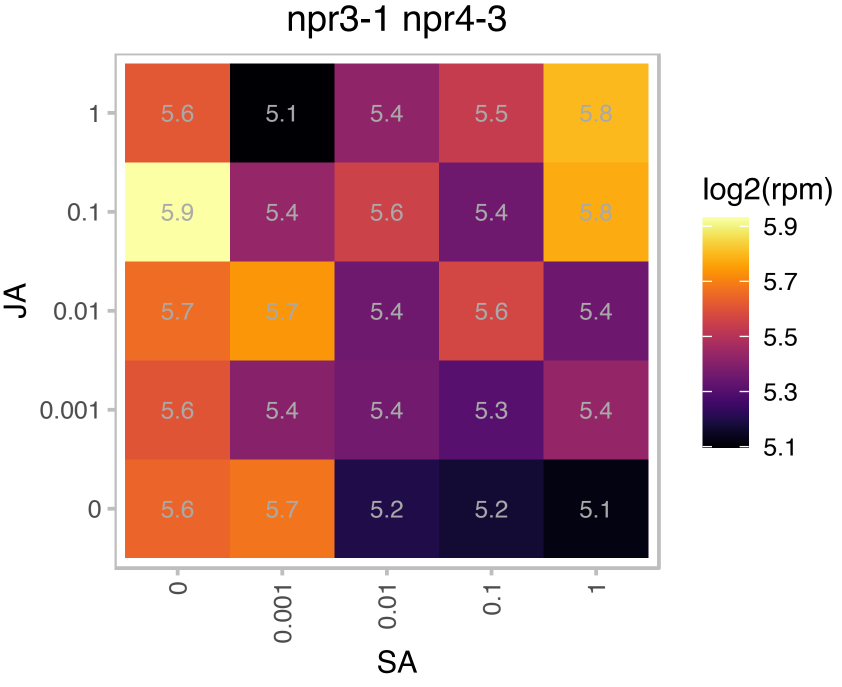

### AT5G56010.1 : AtHsp90-3, AtHsp90.3, HSP81-3, Hsp81.3

heat shock protein 81-3
A member of heat shock protein 90 (HSP90) gene family. Expressed in all tissues and abundant in root apical meristem, pollen and tapetum. Expression is NOT heat-induced but induced by IAA and NaCl. Overexpression reduced tolerance to heat and conferred higher tolerance to calcium. The mRNA is cell-to-cell mobile.

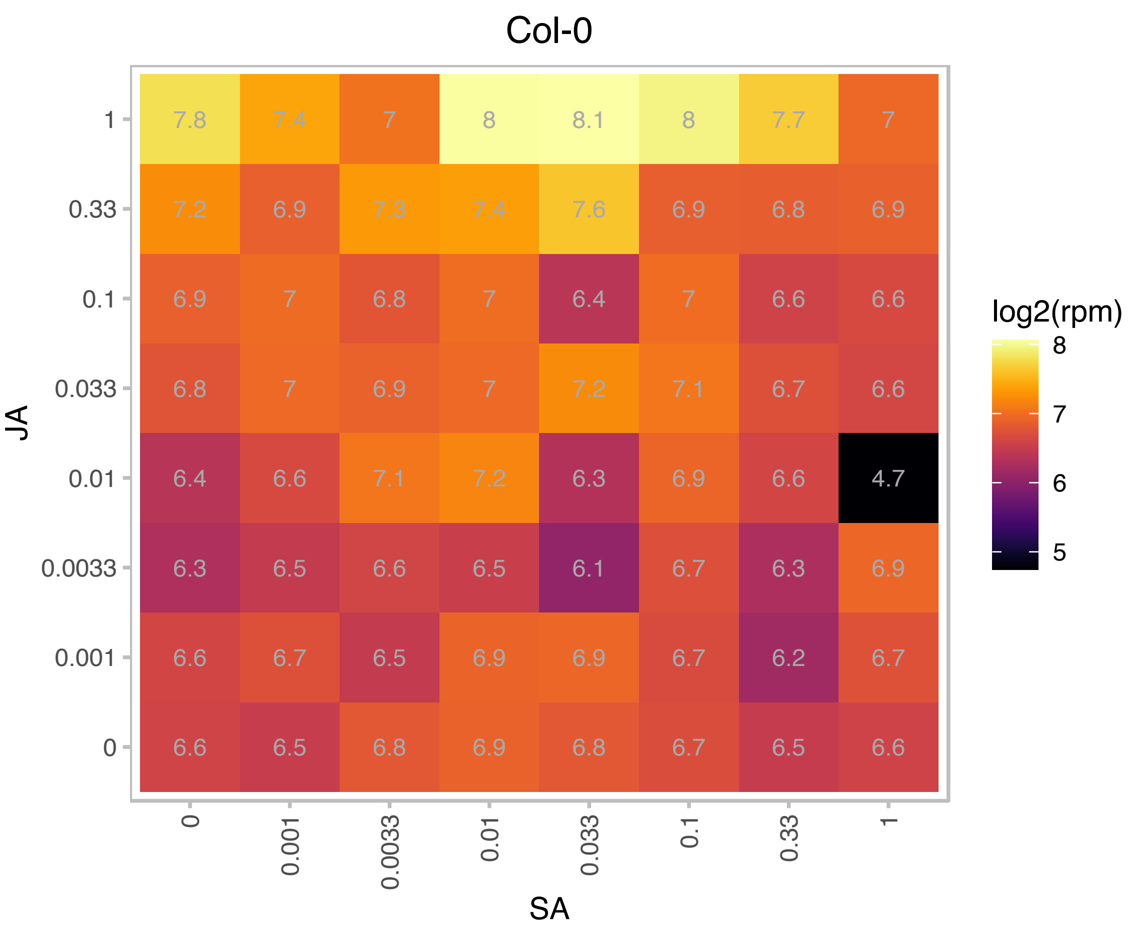


| | |
|---|---|
| Gene | AT5G56010 |
| is.expressed_genes | TRUE |
| is.responsive_genes | TRUE |
| is.apcluster | A-04 |

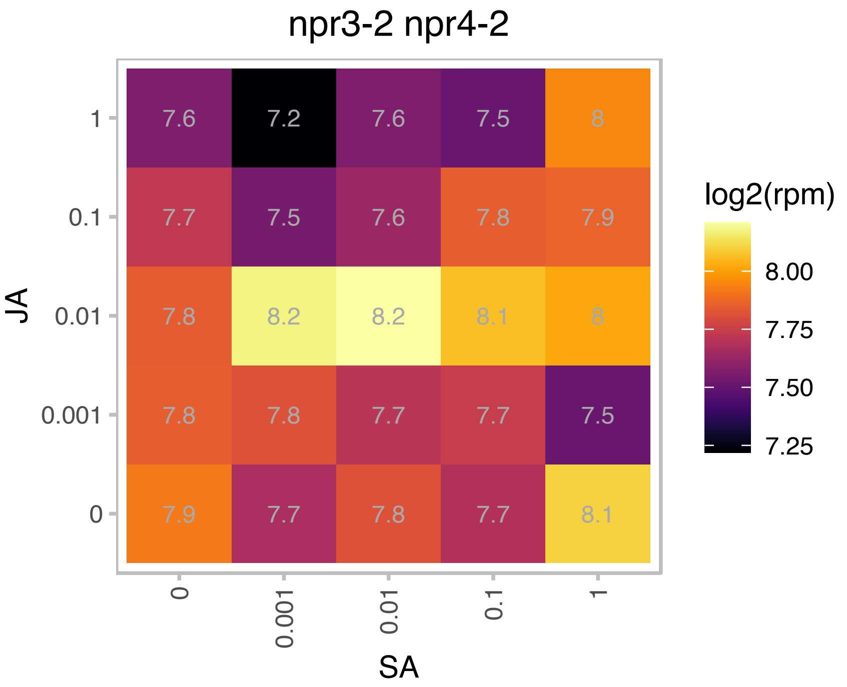


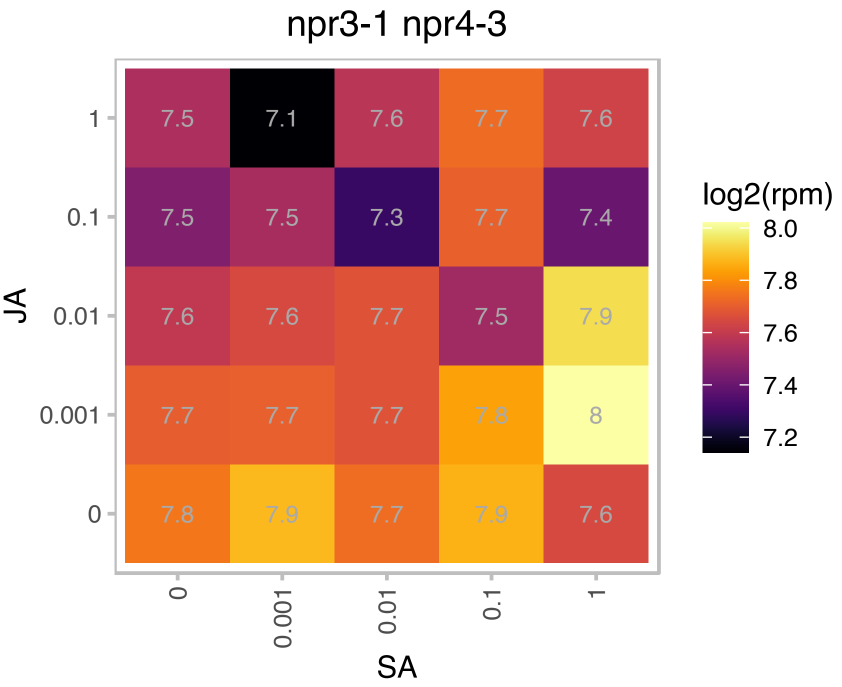

# AT5G56190.1 : NA

Transducin/WD40 repeat-like superfamily protein

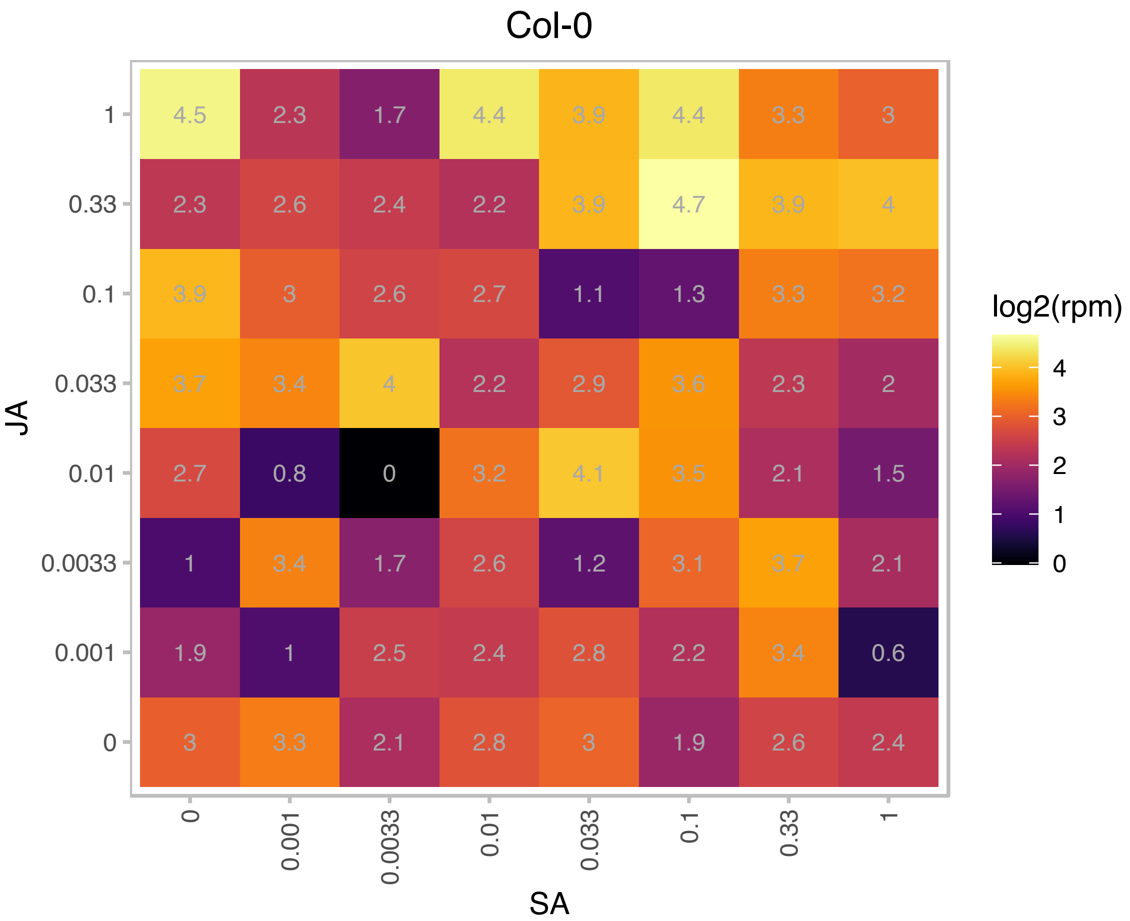


| | |
|---|---|
| Gene | AT5G56190 |
| is.expressed_genes | TRUE |
| is.responsive_genes | TRUE |
| is.apcluster | A-04 |

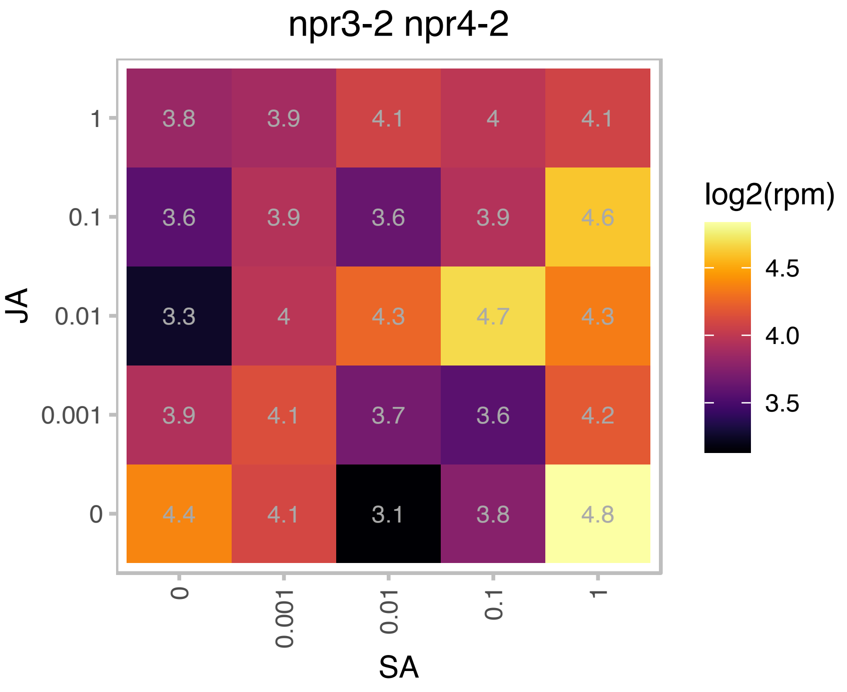


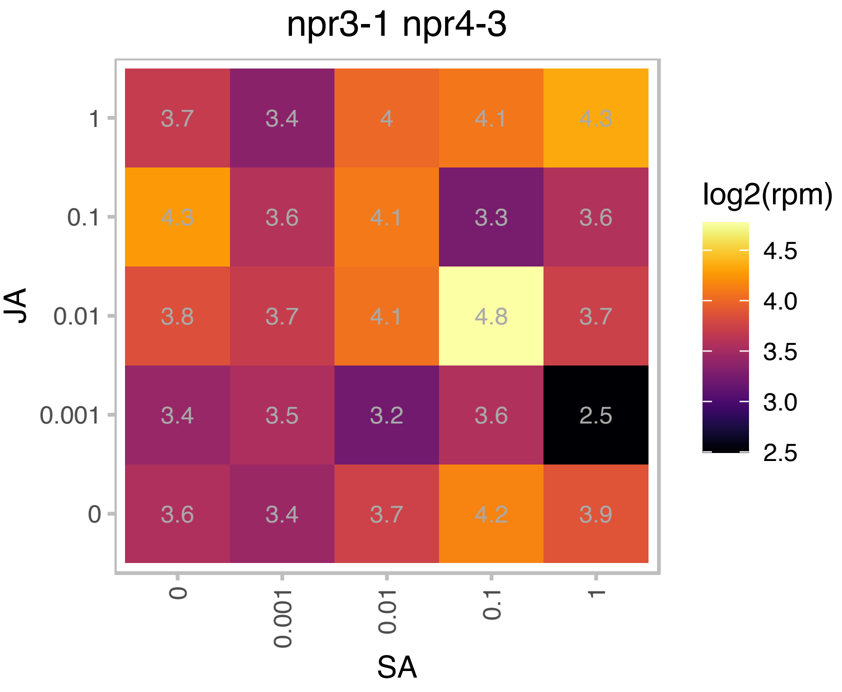

### AT5G57990.1 : UBP23

ubiquitin-specific protease 23

Encodes a ubiquitin-specific protease.

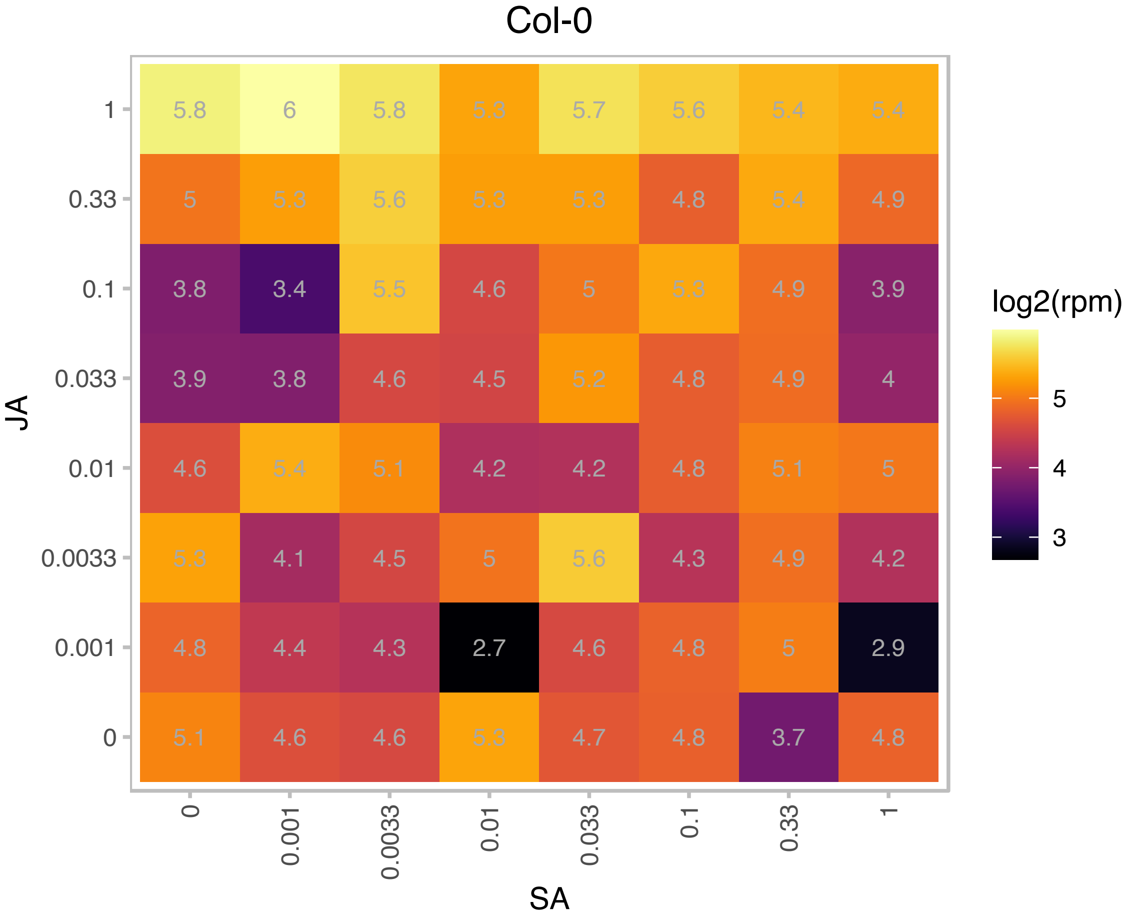


| | |
|---|---|
| Gene | AT5G57990 |
| is.expressed_genes | TRUE |
| is.responsive_genes | TRUE |
| is.apcluster | A-04 |

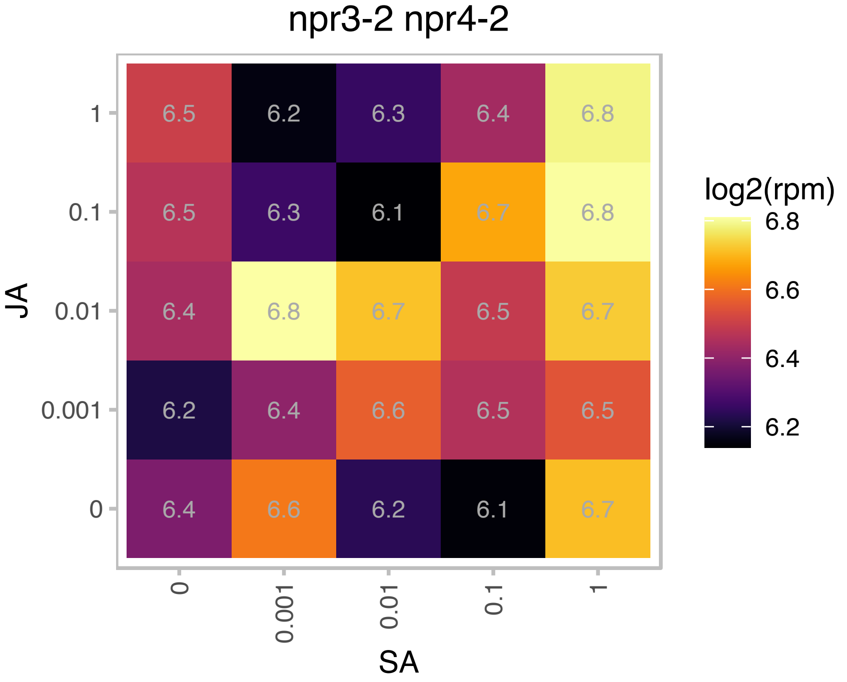


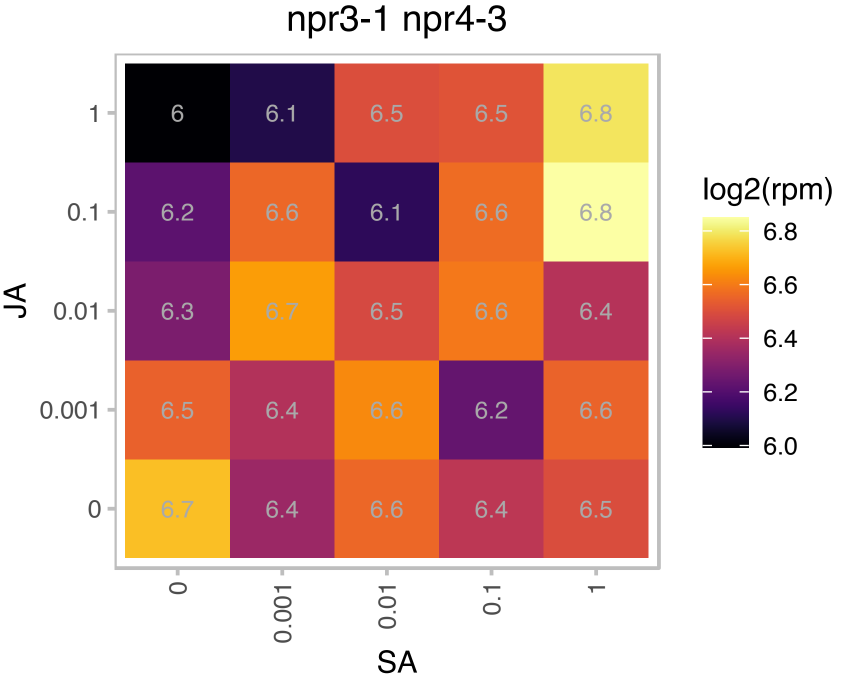

# AT5G65495.1 : NA

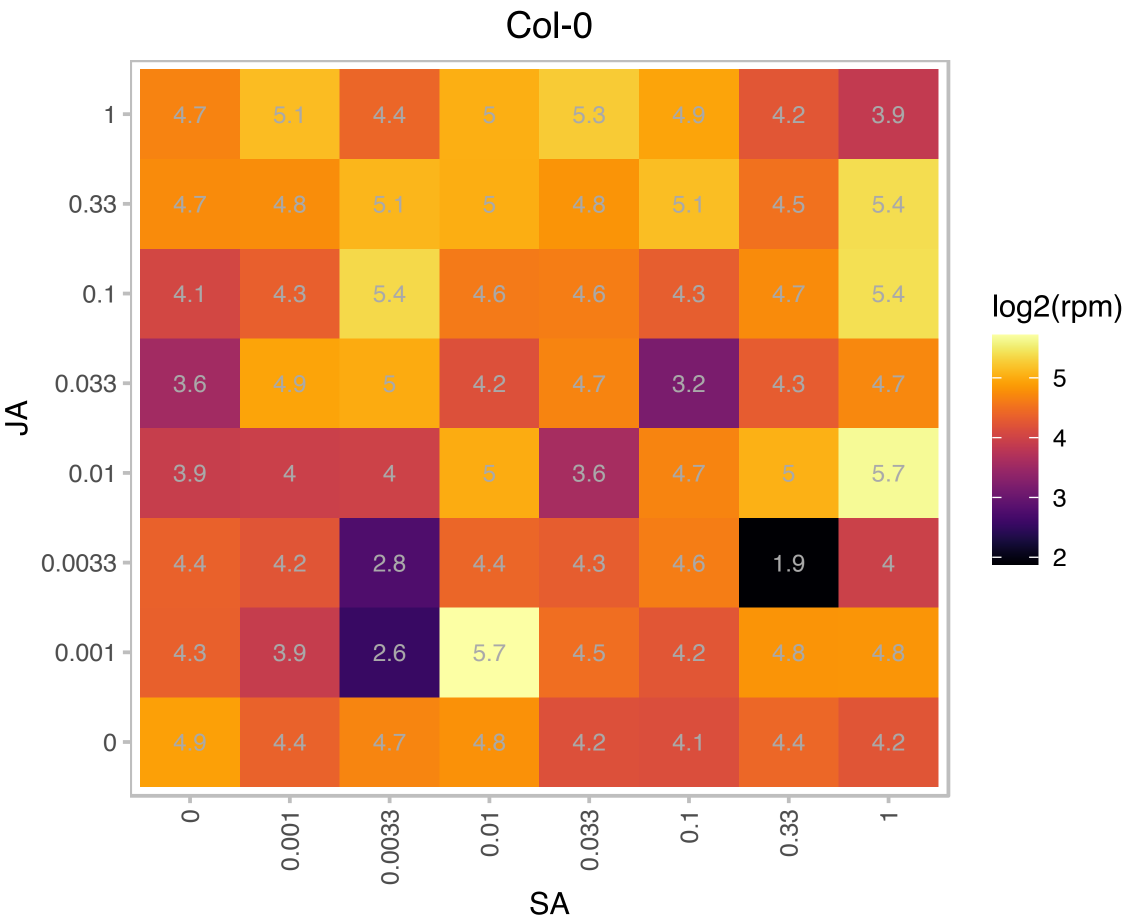


| | |
|---|---|
| Gene | AT5G65495 |
| is.expressed_genes | TRUE |
| is.responsive_genes | TRUE |
| is.apcluster | A-04 |

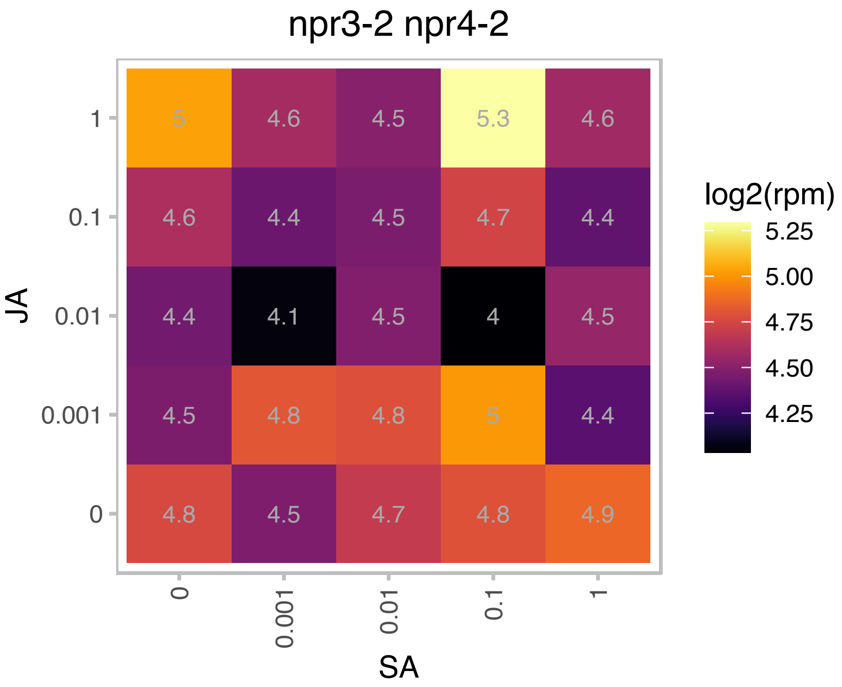


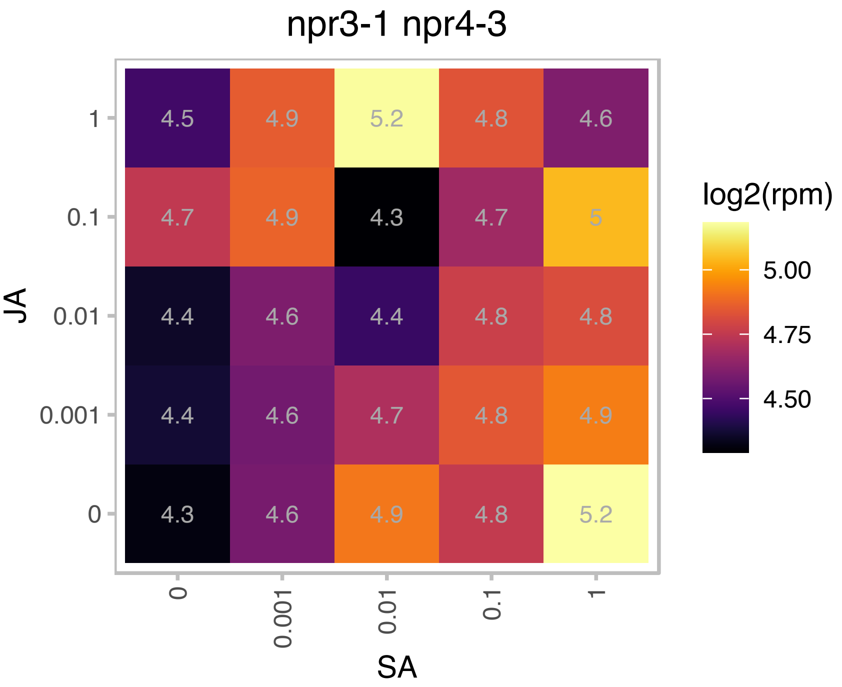

# AT1G03350.1 : NA

BSD domain-containing protein

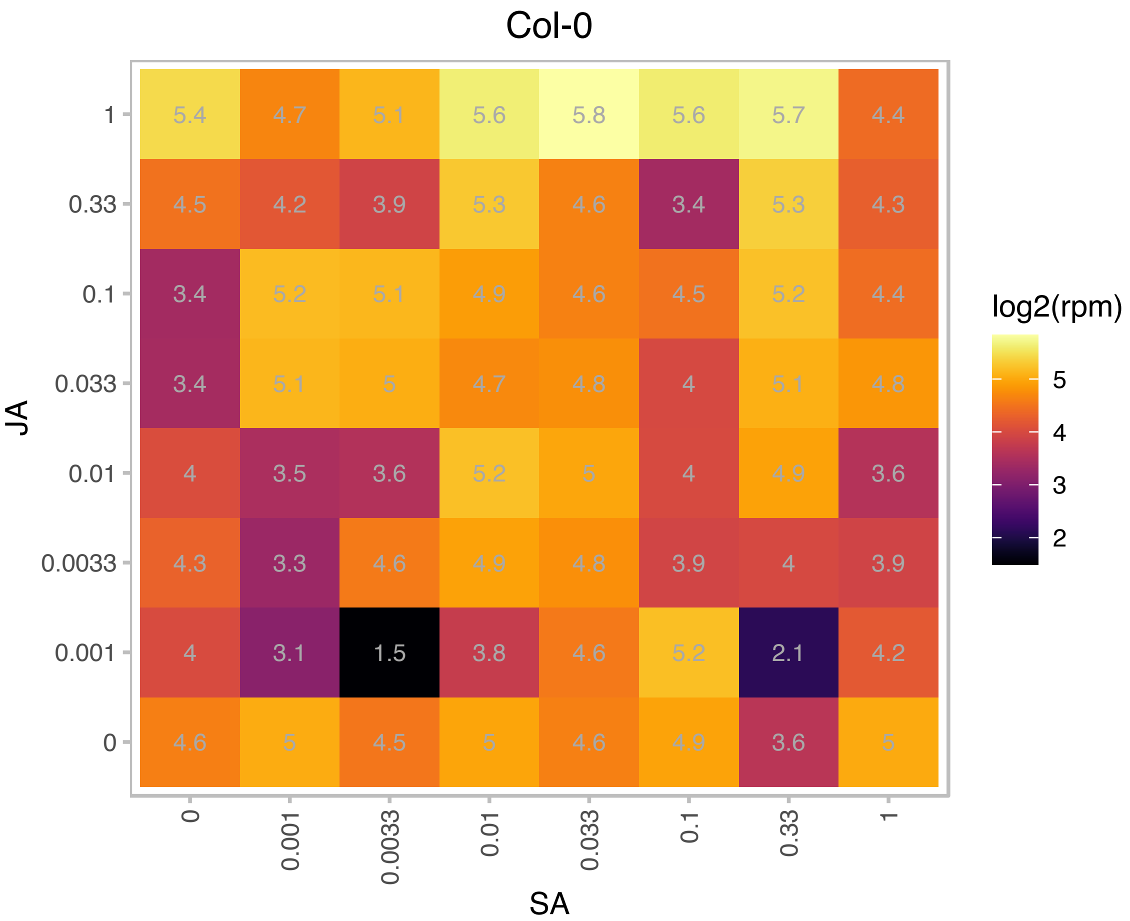


| | |
|---|---|
| Gene | AT1G03350 |
| is.expressed_genes | TRUE |
| is.responsive_genes | TRUE |
| is.apcluster | A-05 |

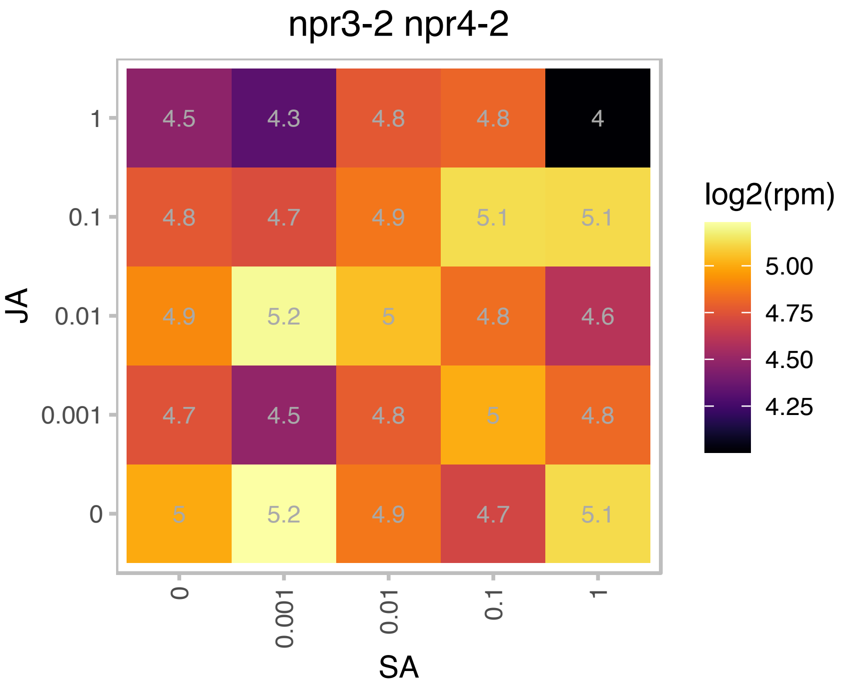


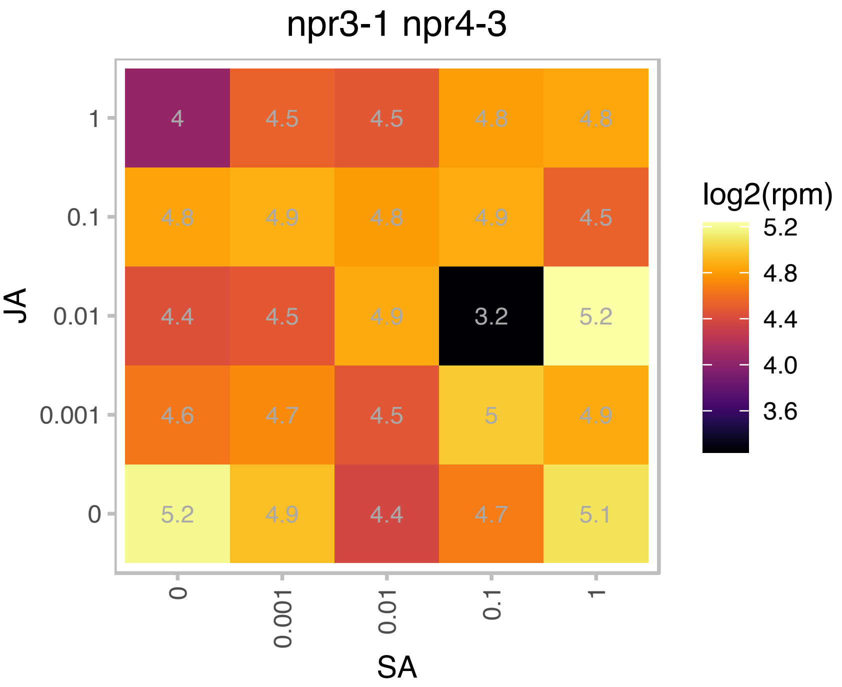

### AT1G03970.1 : GBF4

G-box binding factor 4

encodes a basic leucine zipper G-box binding factor that can bind to G-box motifs only as heterodimers with GBF2 or GBF3. A single amino acid change can confer G-box binding as homodimers.

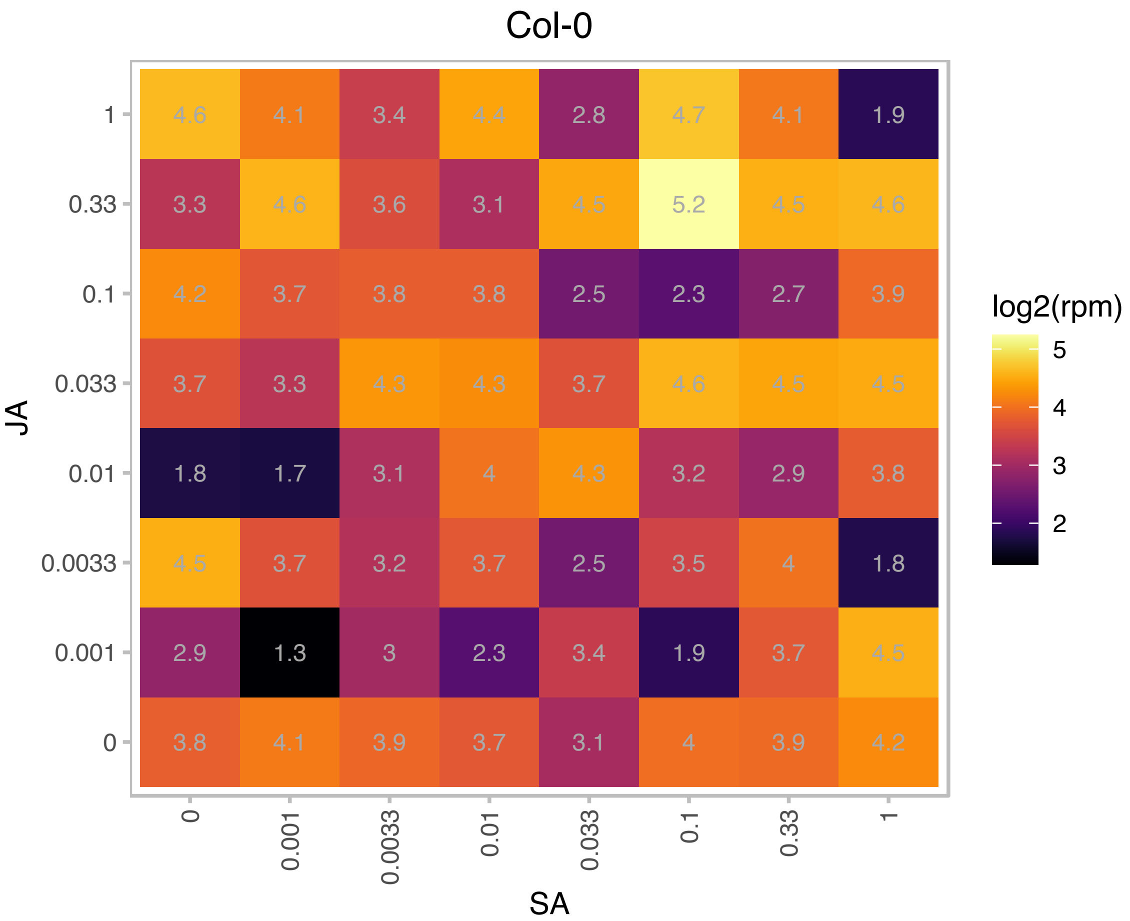


| | |
|---|---|
| Gene | AT1G03970 |
| is.expressed_genes | TRUE |
| is.responsive_genes | TRUE |
| is.apcluster | A-05 |

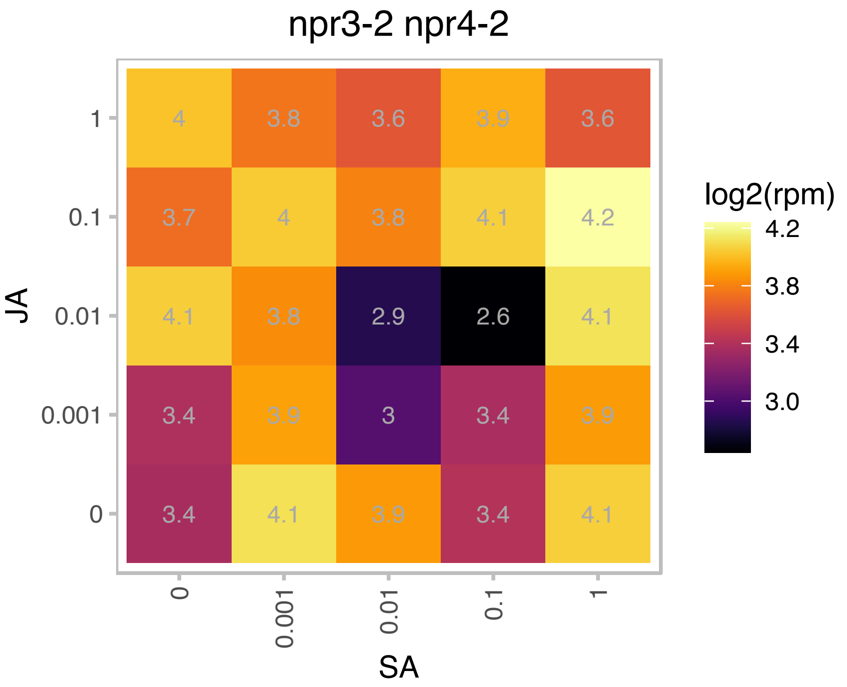


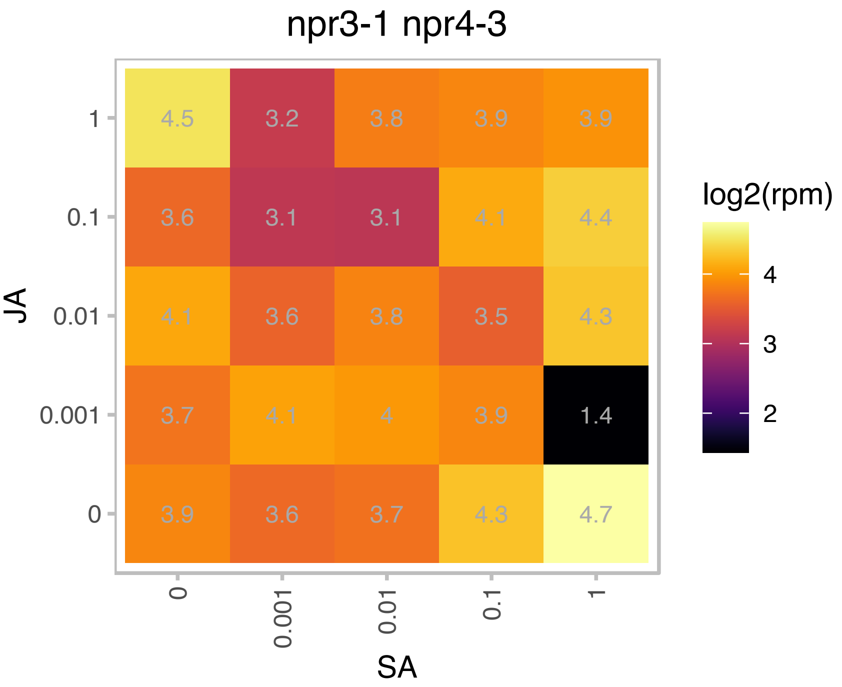

# AT1G07350.1 : SR45a

RNA-binding (RRM/RBD/RNP motifs) family protein

Encodes a serine/arginine rich-like protein, SR45a. Involved in the regulation of stress-responsive alternative splicing.

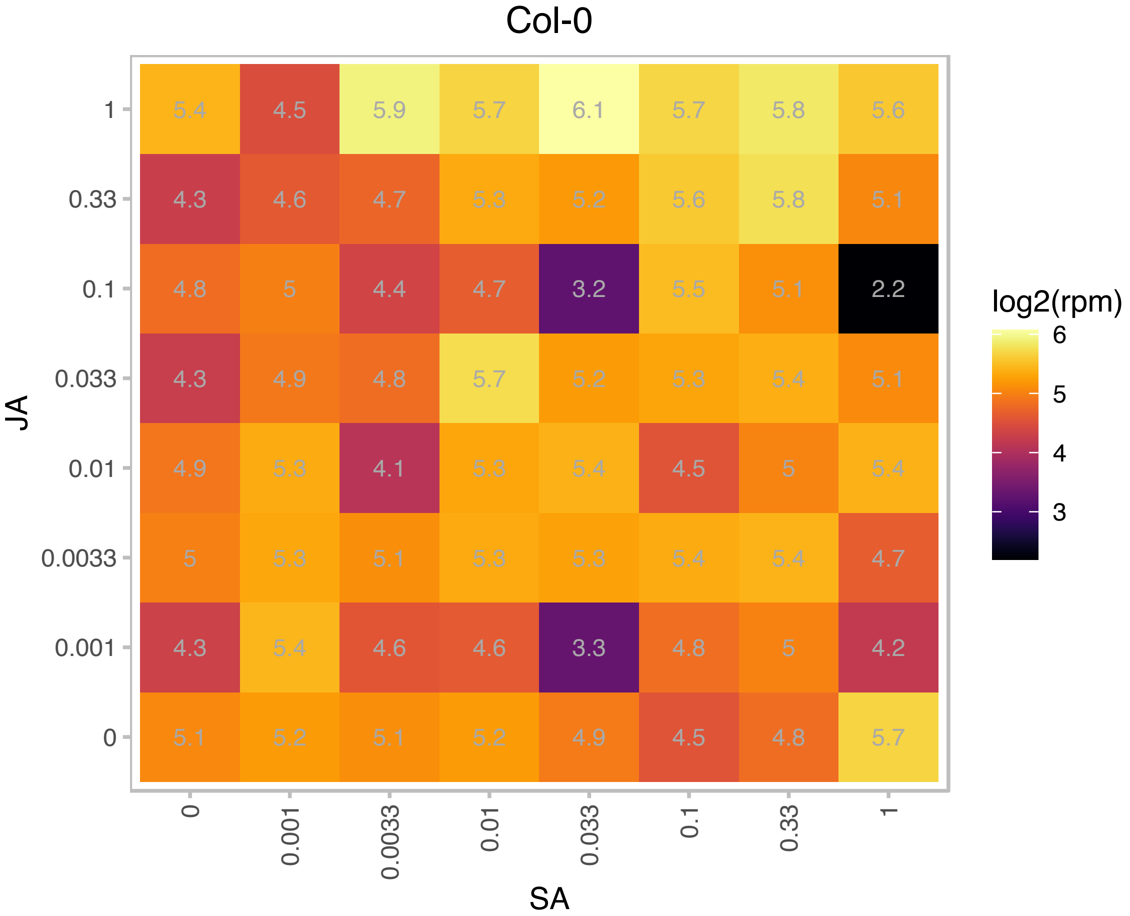


| | |
|---|---|
| Gene | AT1G07350 |
| is.expressed_genes | TRUE |
| is.responsive_genes | TRUE |
| is.apcluster | A-05 |

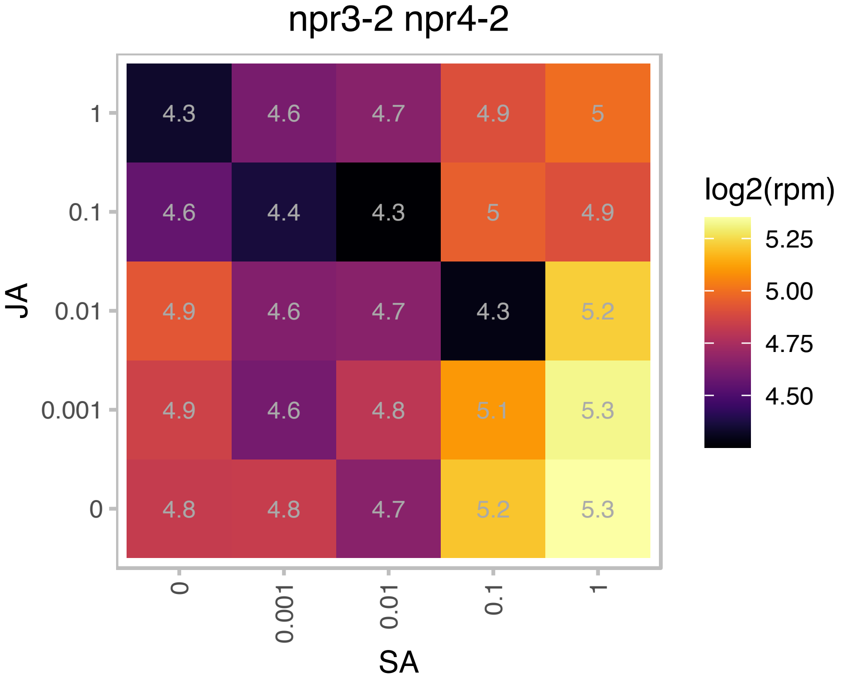


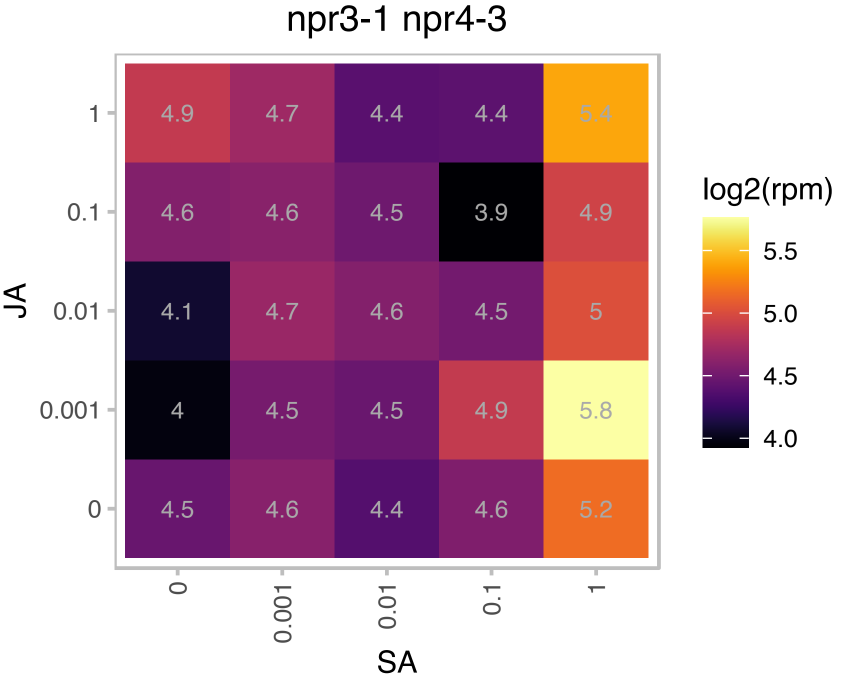

### AT1G07670.1 : ATECA4, ECA4

endomembrane-type CA-ATPase 4

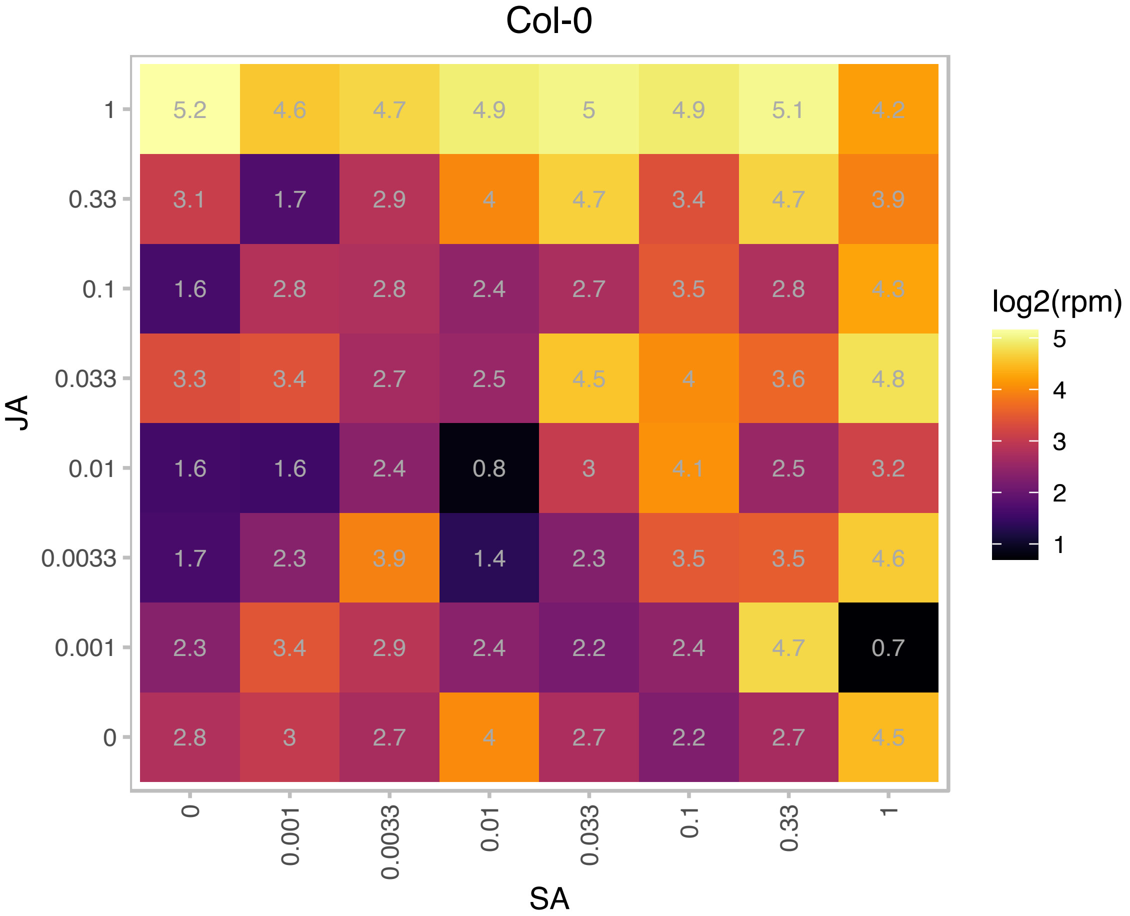


| | |
|---|---|
| Gene | AT1G07670 |
| is.expressed_genes | TRUE |
| is.responsive_genes | TRUE |
| is.apcluster | A-05 |

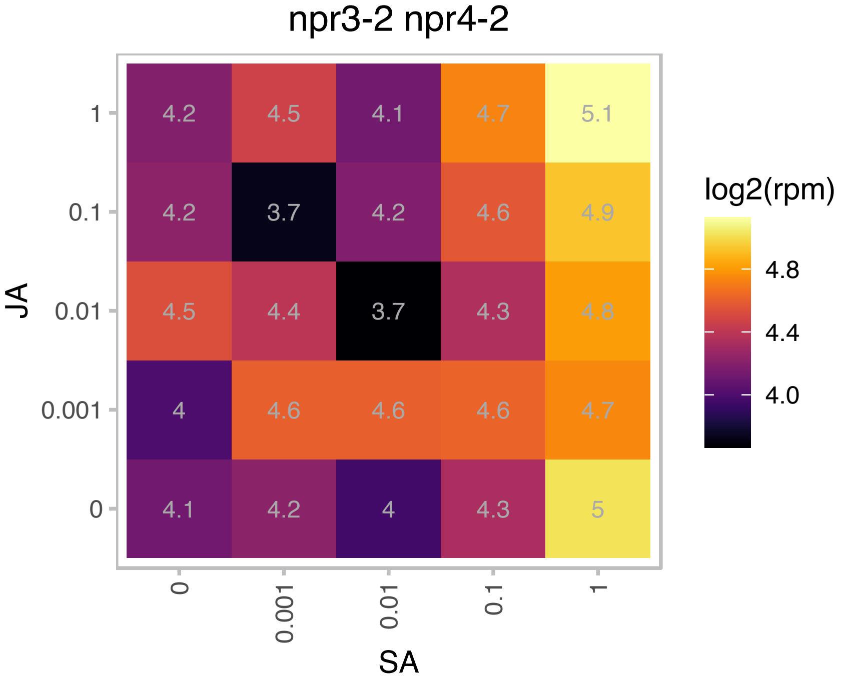


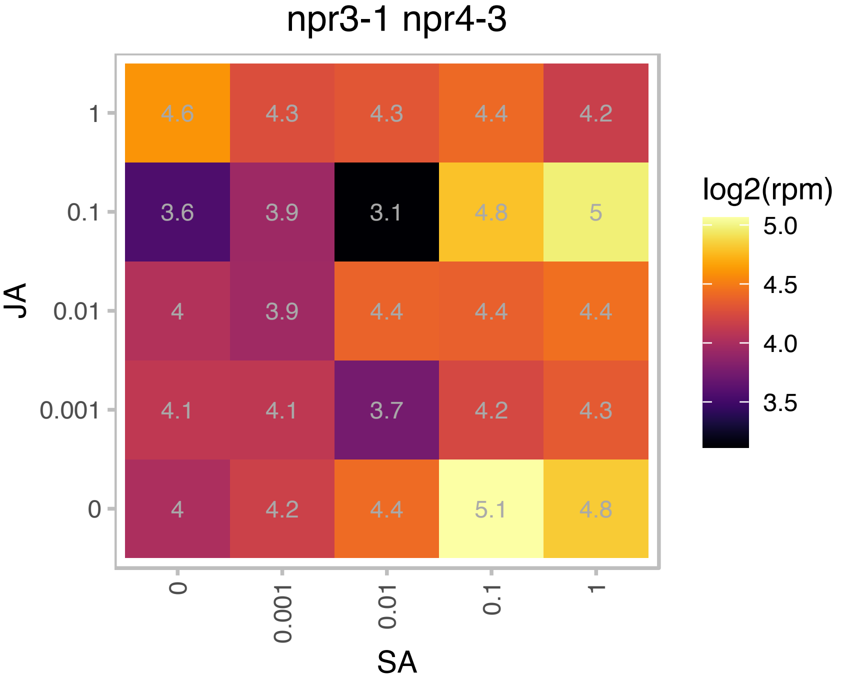

### AT1G07810.1 : ACA3, ATECA1, ECA1

ER-type Ca2+-ATPase 1

Encodes an ER-type Ca2+-pumping ATPase. The mRNA is cell-to-cell mobile.

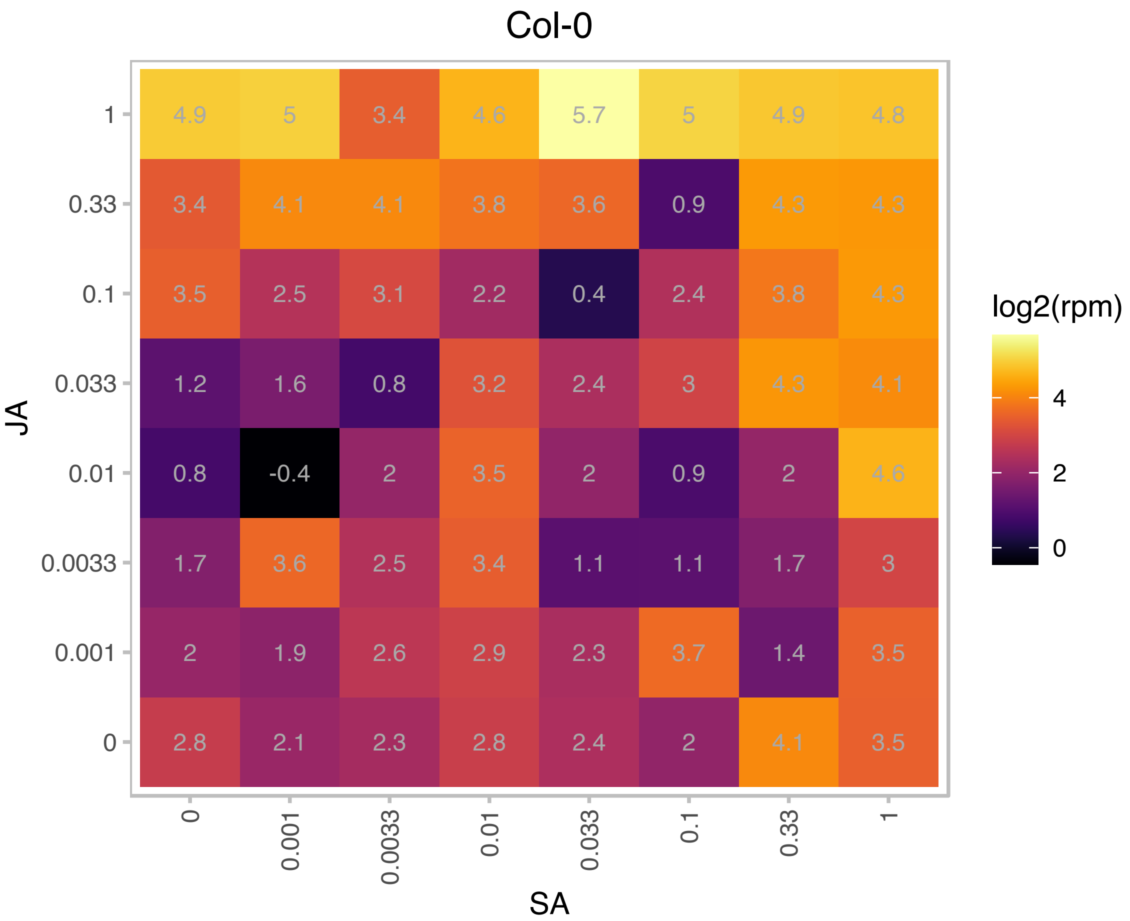


| | |
|---|---|
| Gene | AT1G07810 |
| is.expressed_genes | TRUE |
| is.responsive_genes | TRUE |
| is.apcluster | A-05 |

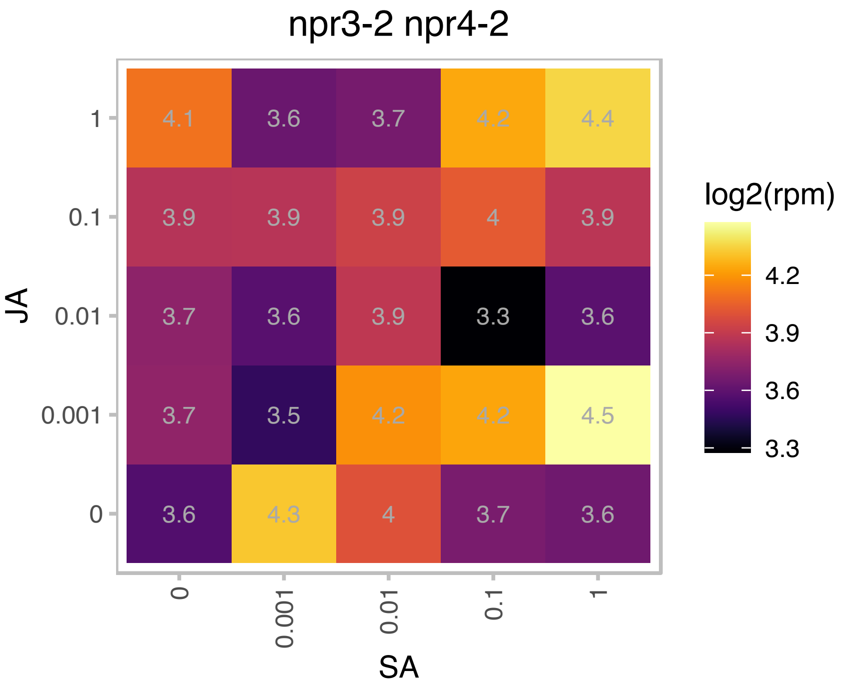


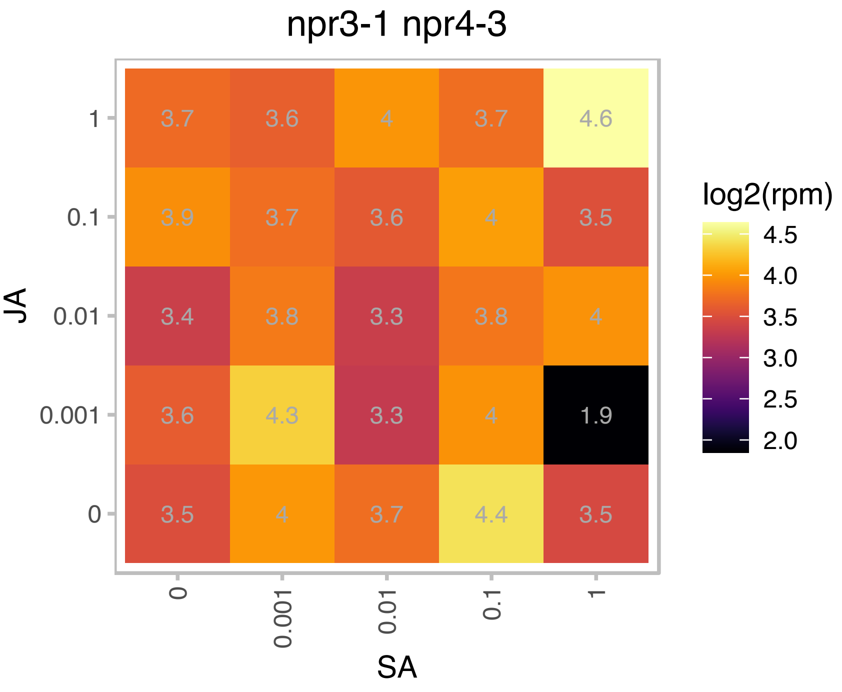

# AT1G09130.1 : NA

ATP-dependent caseinolytic (Clp) protease/crotonase family protein

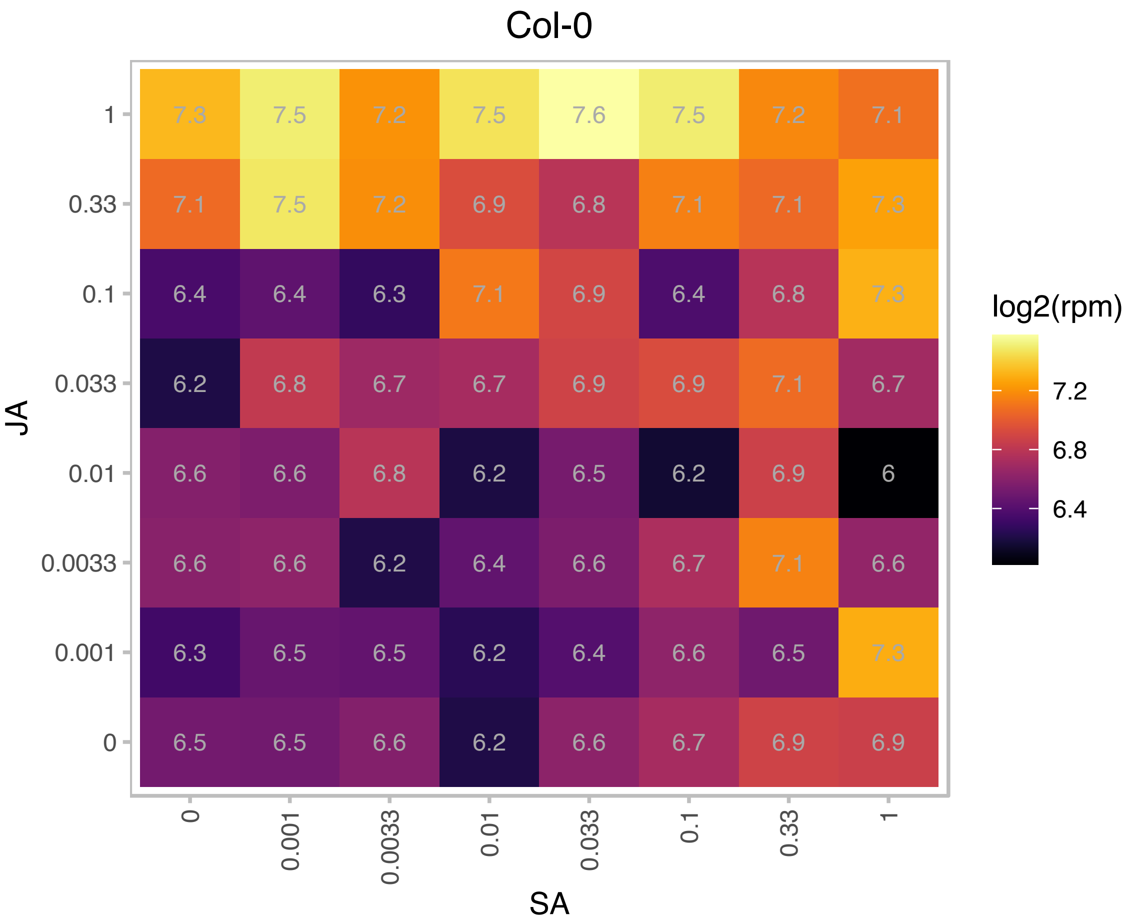


| | |
|---|---|
| Gene | AT1G09130 |
| is.expressed_genes | TRUE |
| is.responsive_genes | TRUE |
| is.apcluster | A-05 |

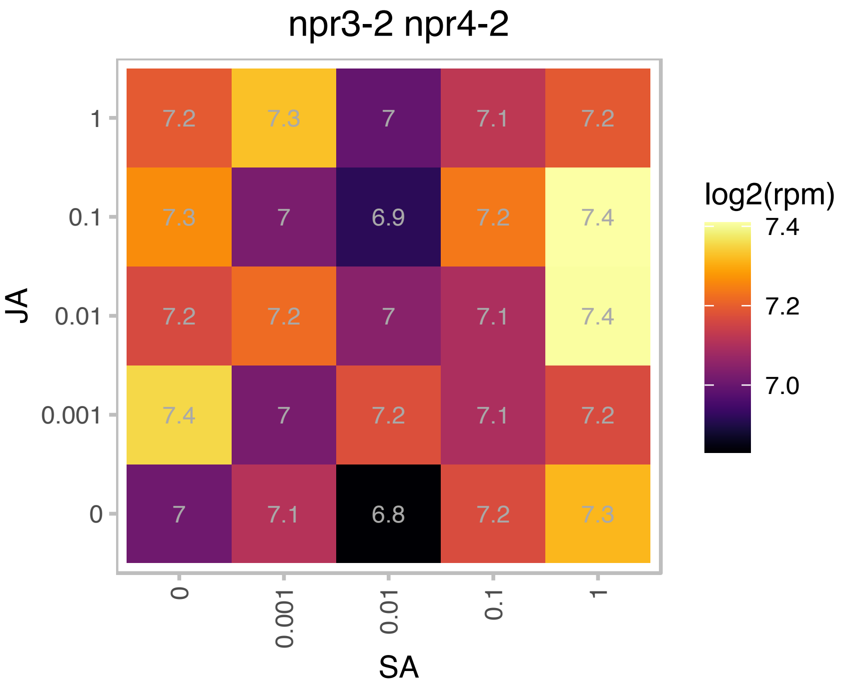


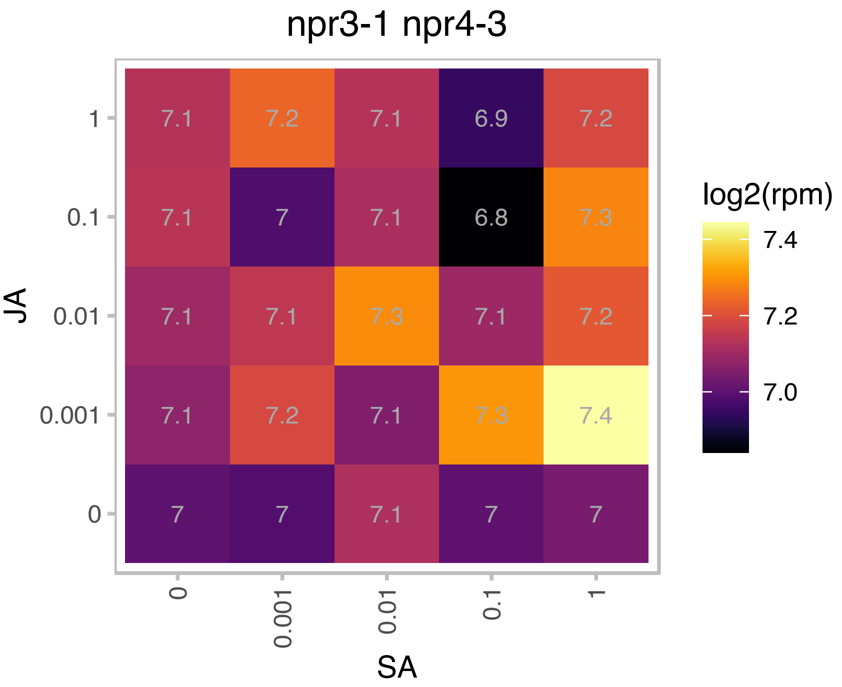

# AT1G09870.1 : NA

histidine acid phosphatase family protein

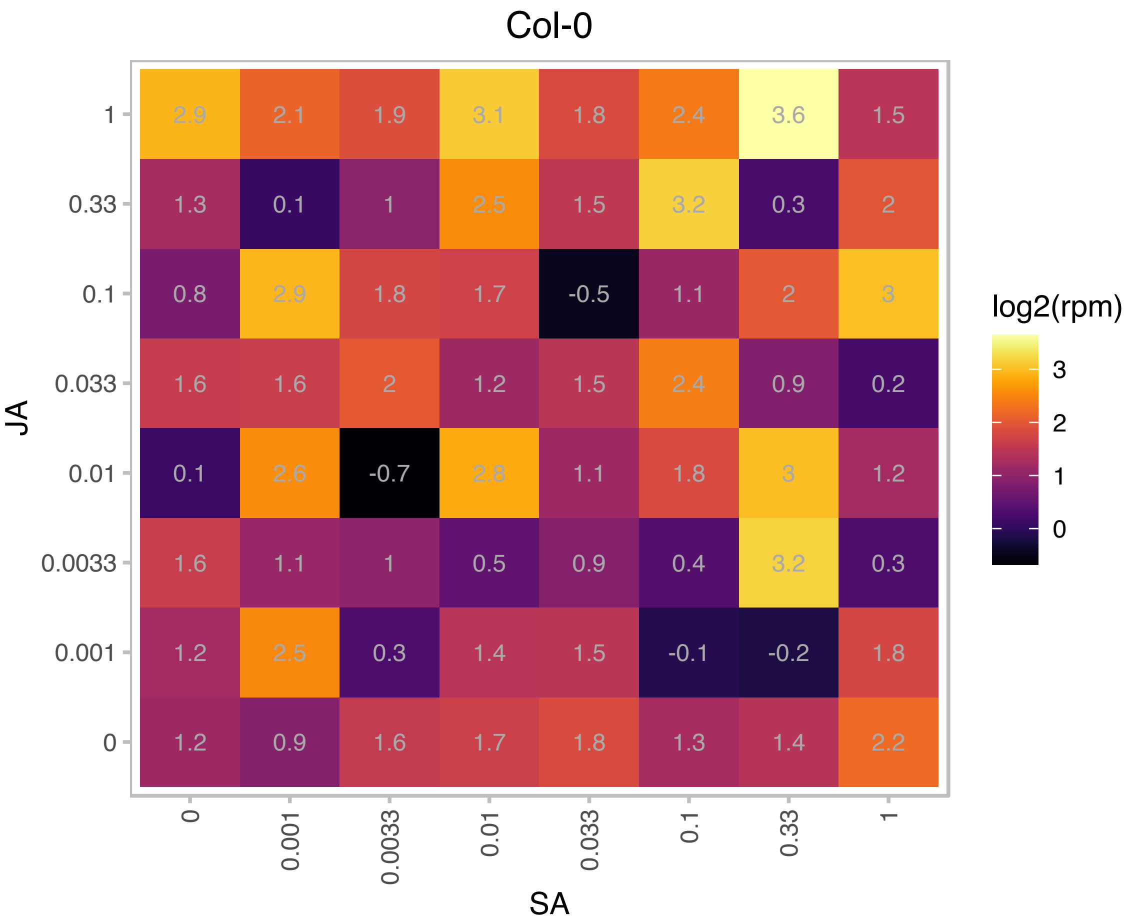


| | |
|---|---|
| Gene | AT1G09870 |
| is.expressed_genes | TRUE |
| is.responsive_genes | TRUE |
| is.apcluster | A-05 |

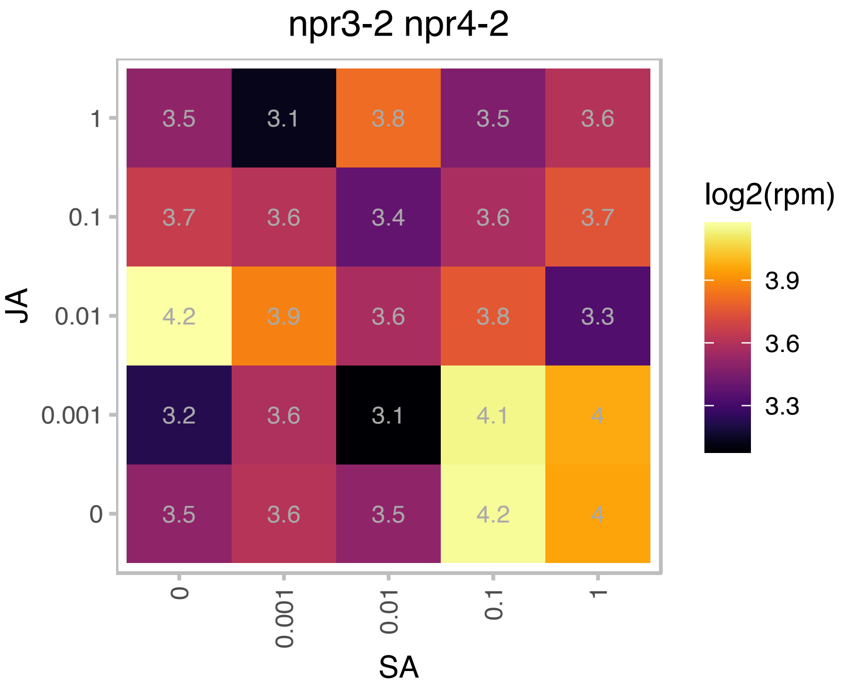


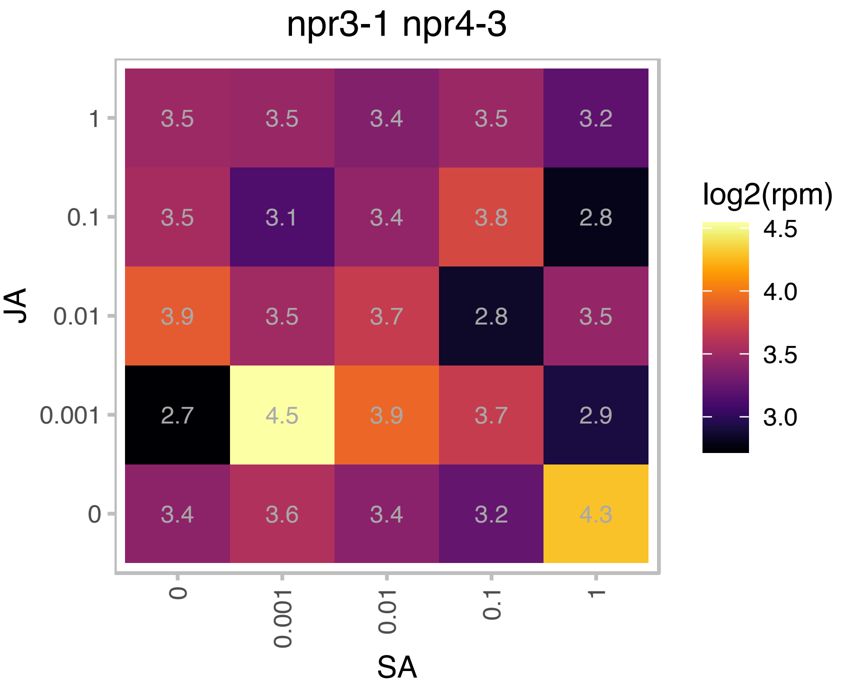

# AT1G10580.1 : NA

Transducin/WD40 repeat-like superfamily protein

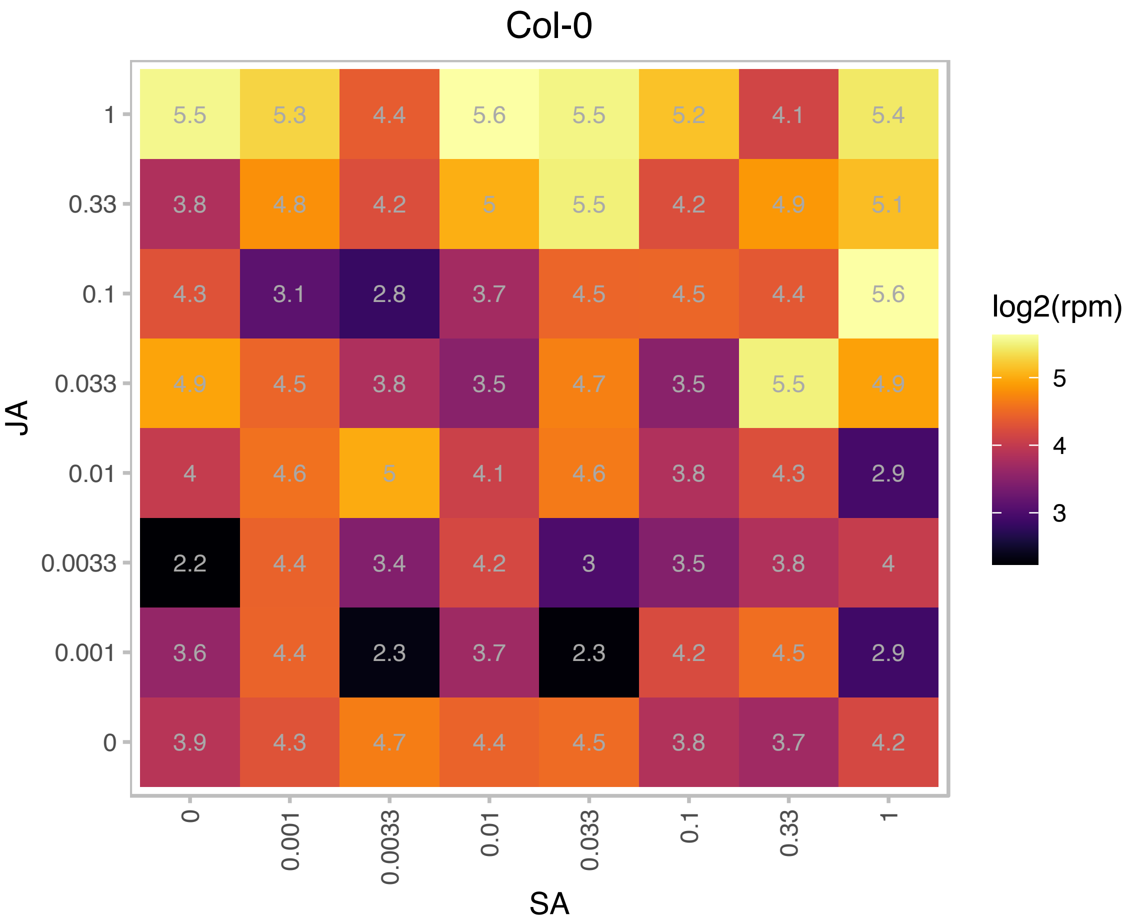


| | |
|---|---|
| Gene | AT1G10580 |
| is.expressed_genes | TRUE |
| is.responsive_genes | TRUE |
| is.apcluster | A-05 |

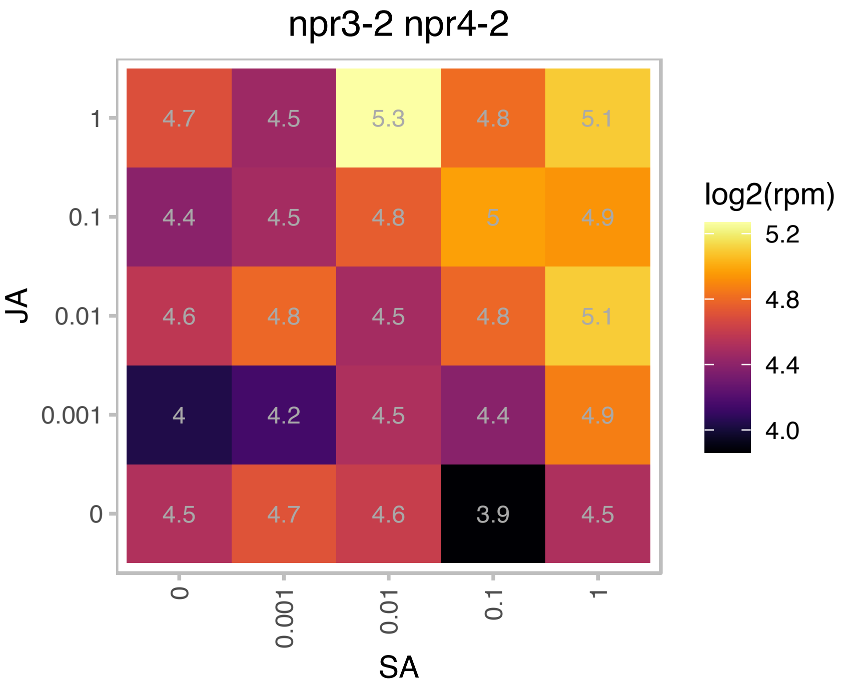


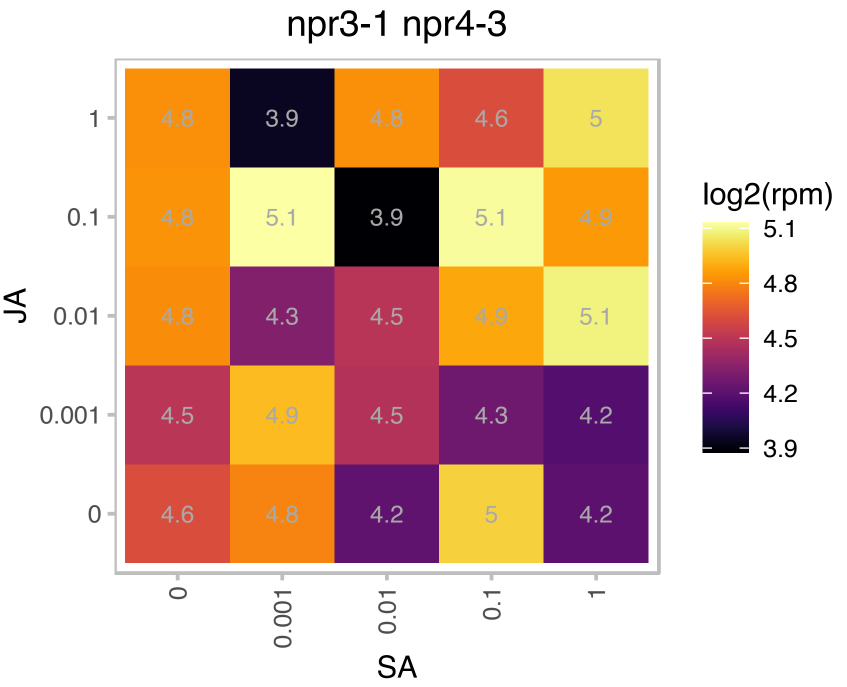

# AT1G10585.1 : NA

basic helix-loop-helix (bHLH) DNA-binding superfamily protein

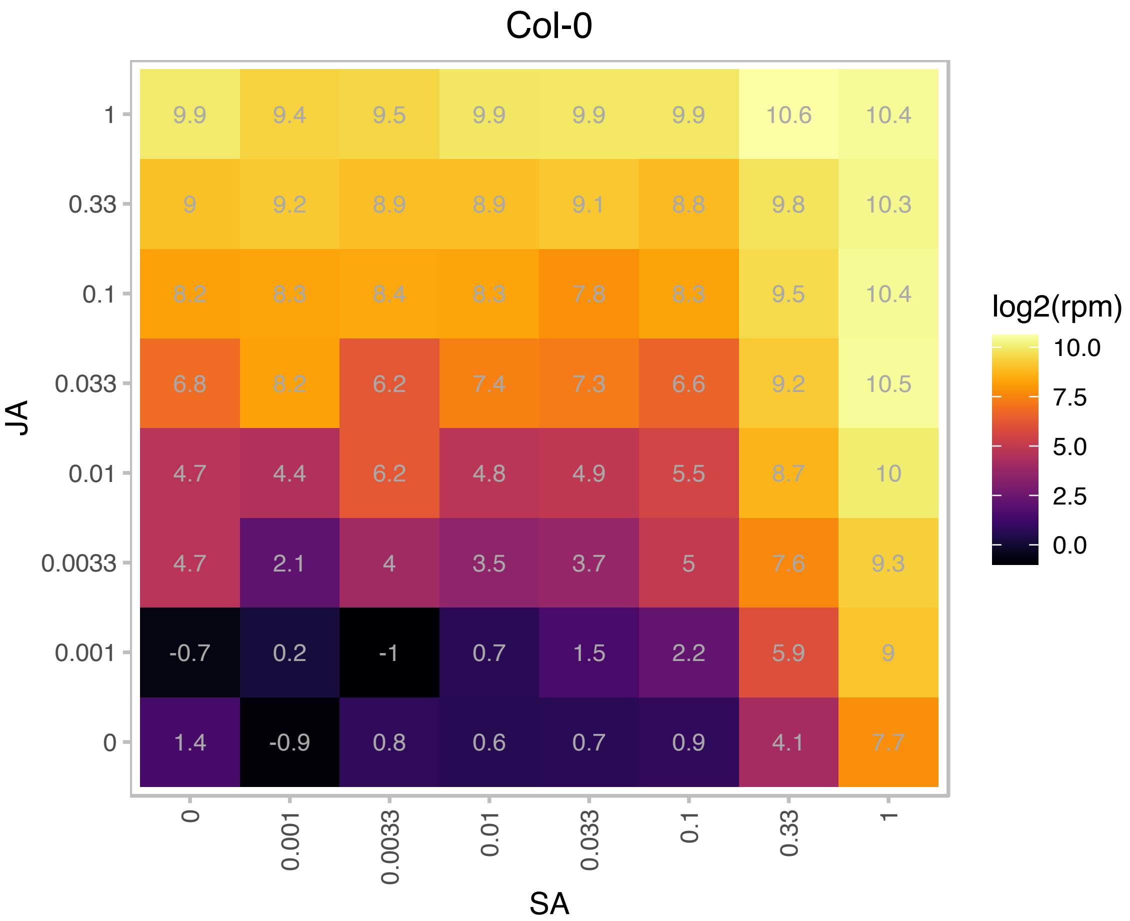


| | |
|---|---|
| Gene | AT1G10585 |
| is.expressed_genes | TRUE |
| is.responsive_genes | TRUE |
| is.apcluster | A-05 |

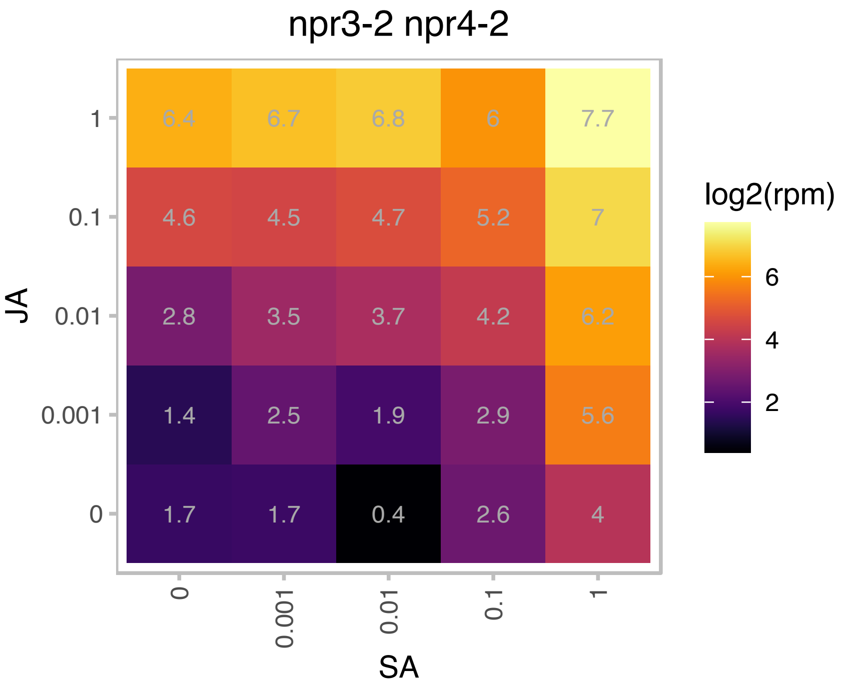


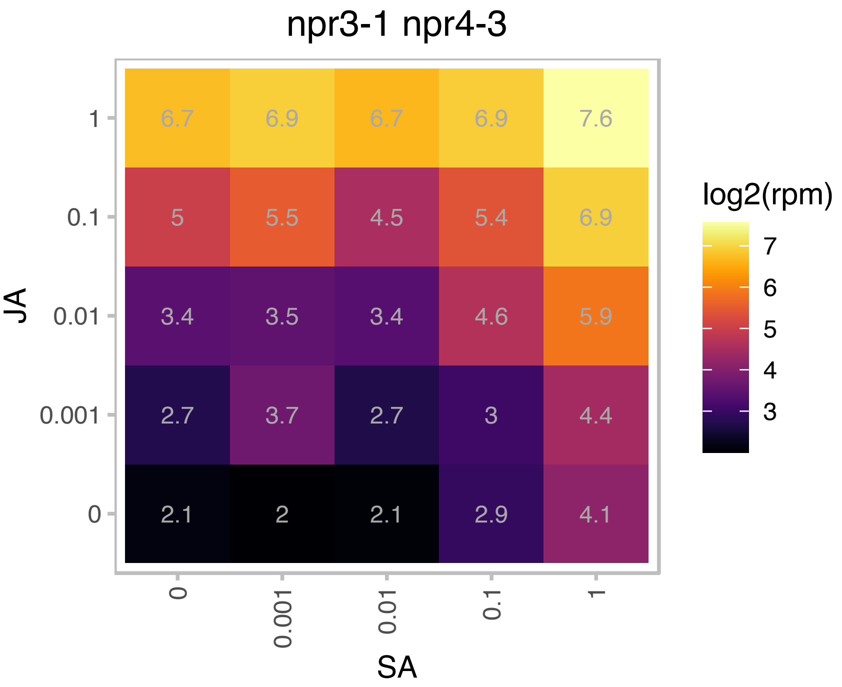

### AT1G14370.1 : APK2A, Kin1, PBL2

protein kinase 2A

Encodes protein kinase APK2a. Protein is N-myristoylated.

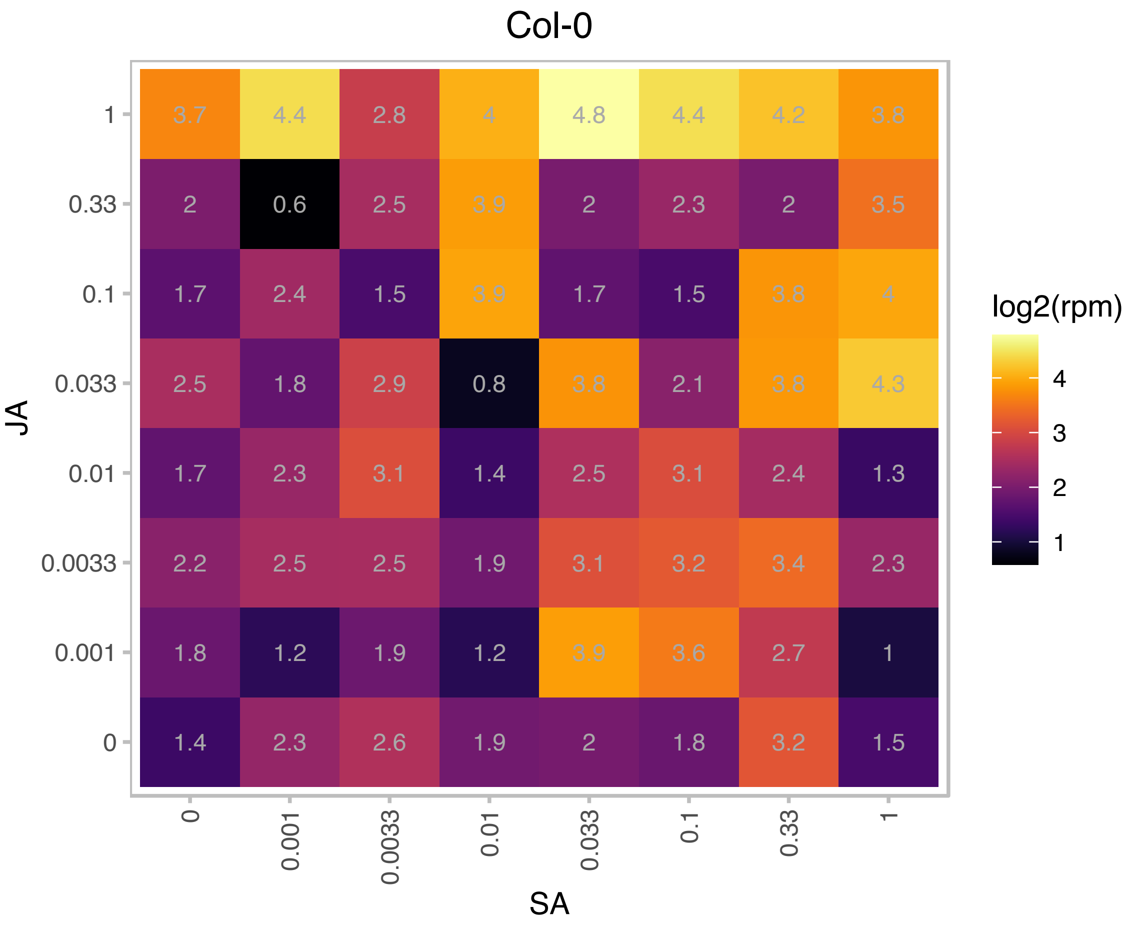


| | |
|---|---|
| Gene | AT1G14370 |
| is.expressed_genes | TRUE |
| is.responsive_genes | TRUE |
| is.apcluster | A-05 |

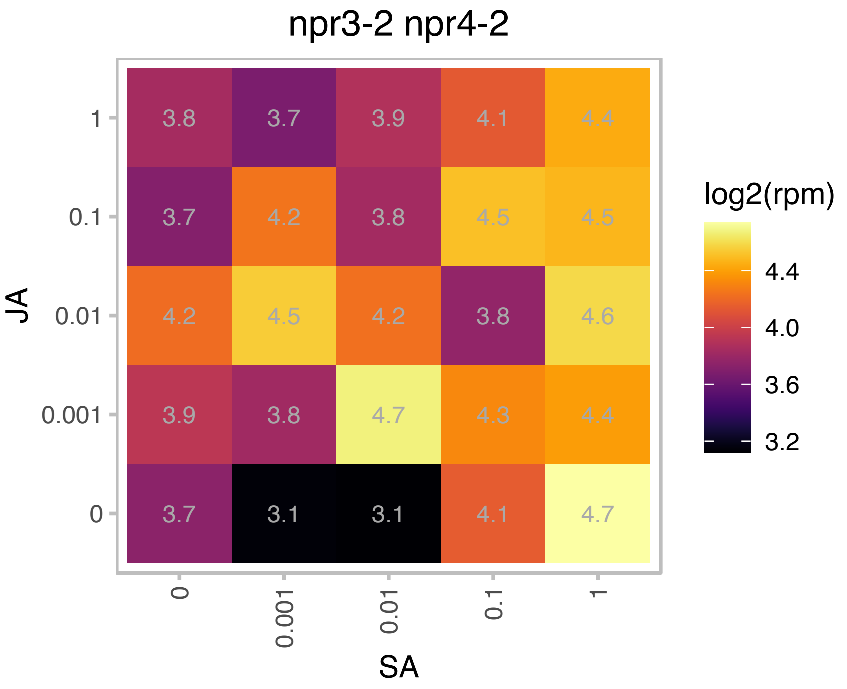


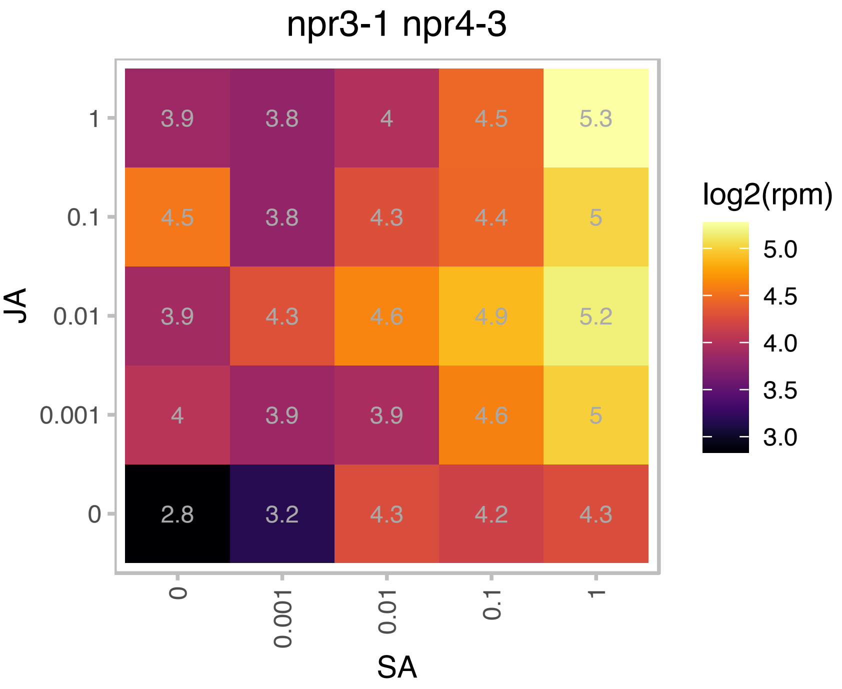

### AT1G20575.1 : DPMS1

Nucleotide-diphospho-sugar transferases superfamily protein

Encodes the catalytic core of the dolichol phosphate mannase synthase (DPMS) complex. It is not active on its own but requires the presence of DPMS2 and DPMS3 for full activity. It is localized in the ER and mediates isoprenyl-linked glycan biogenesis, inﬂuences development, stress response, and ammonium hypersensitivity.

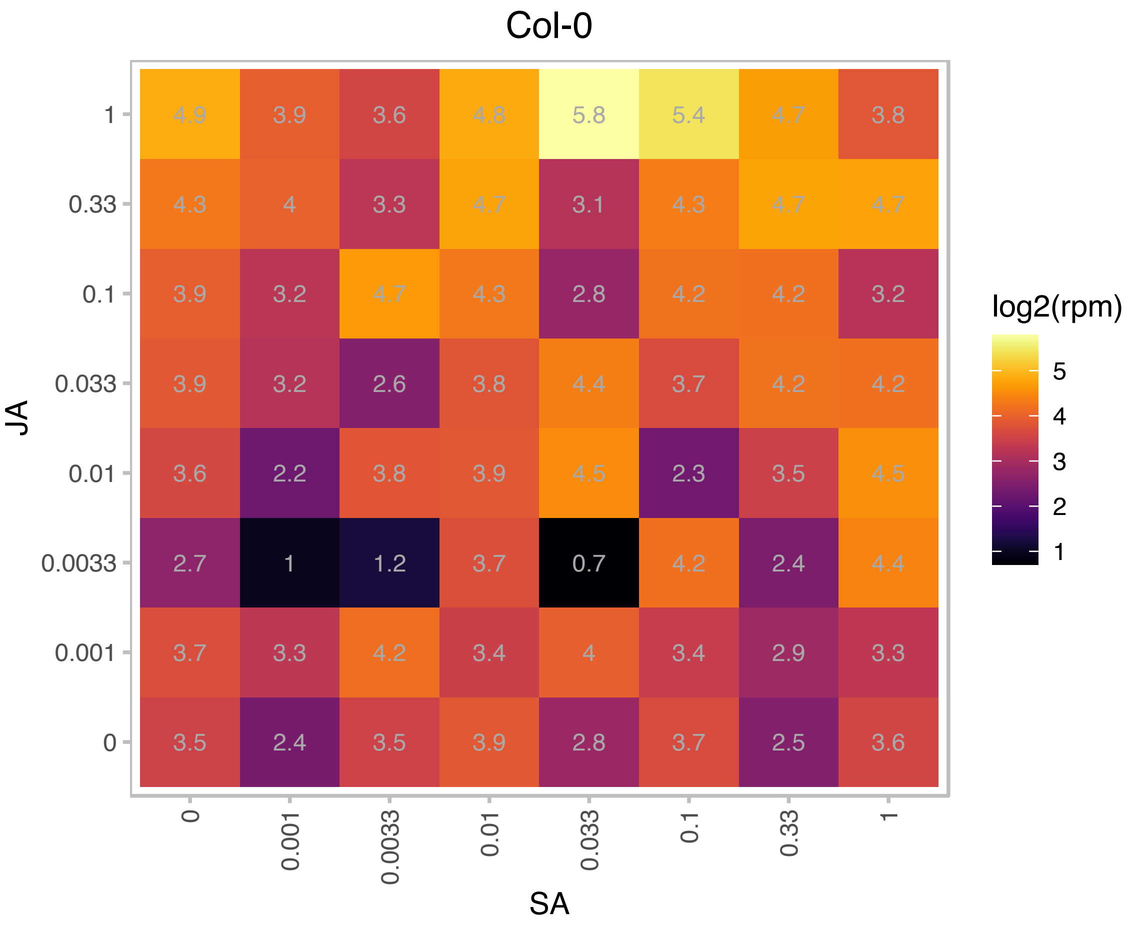


| | |
|---|---|
| Gene | AT1G20575 |
| is.expressed_genes | TRUE |
| is.responsive_genes | TRUE |
| is.apcluster | A-05 |

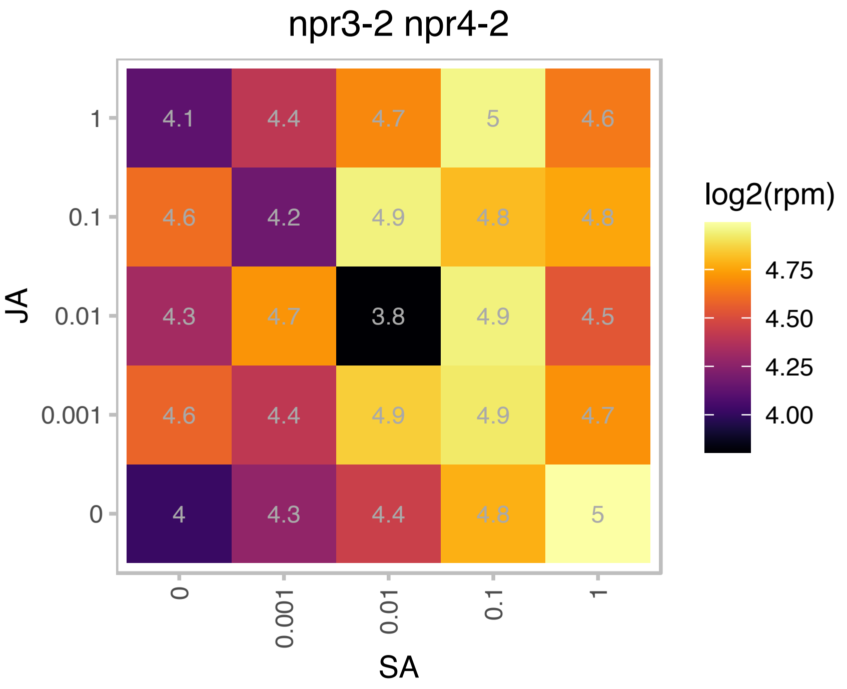


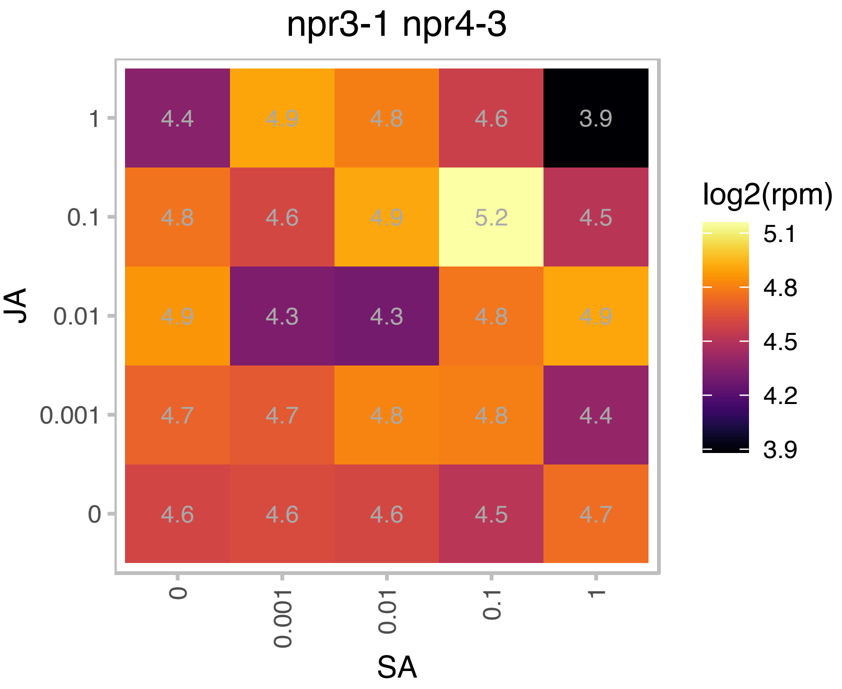

# AT1G21350.1 : NA

Thioredoxin superfamily protein

| | |
|---|---|
| Gene | AT1G21350 |
| is.expressed_genes | TRUE |
| is.responsive_genes | TRUE |
| is.apcluster | A-05 |

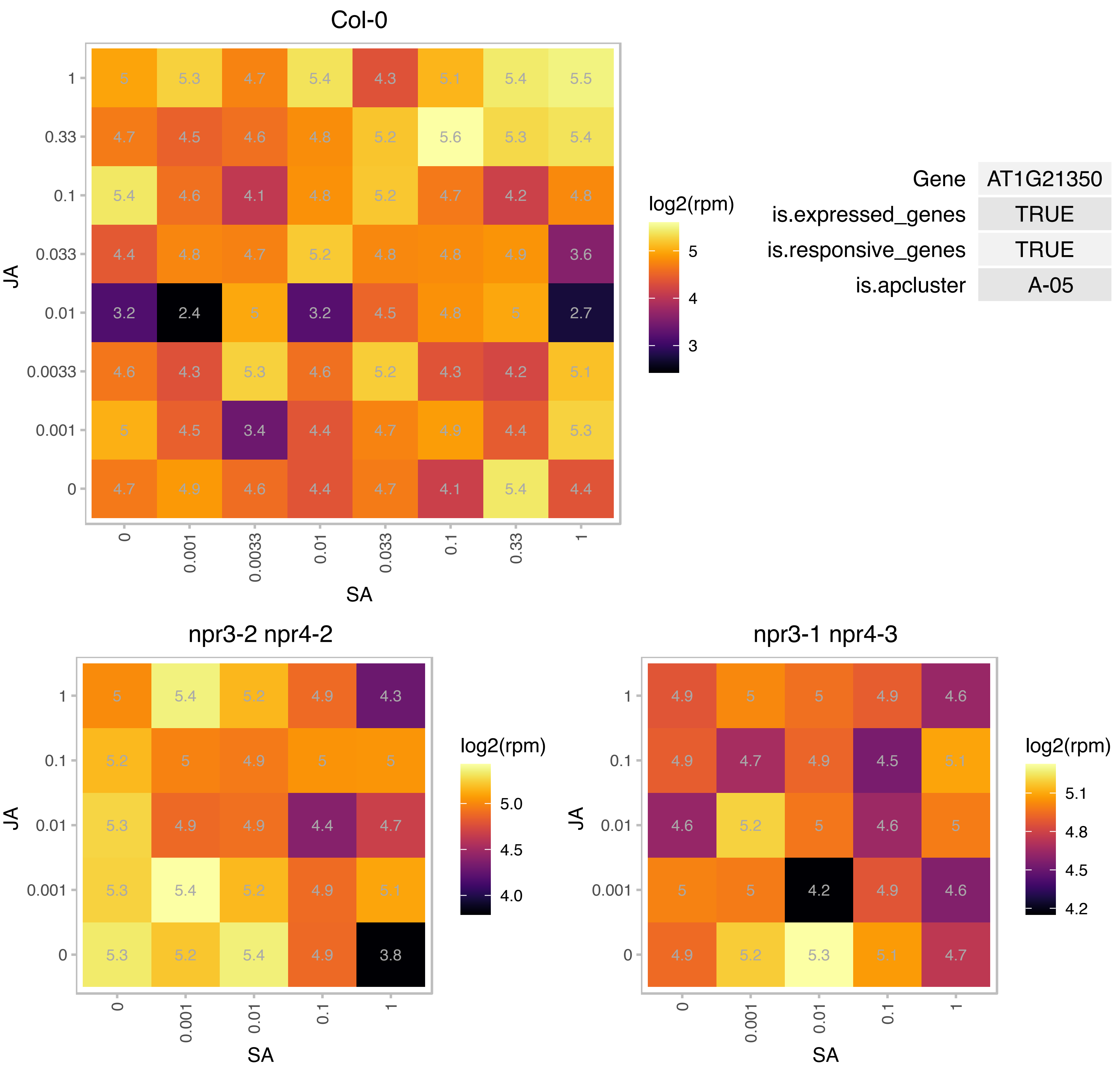

### AT1G24460.1 : TNO1

Encodes a novel protein that co-immunoprecipitates with SYP41. It is involved in vacuolar trafficking and salt tolerance, potentially via a role in vesicle fusion and in maintaining TGN structure or identity. Mutants display delayed formation of the Brefeldin A (BFA) compartment in cotyledons upon application of BFA.

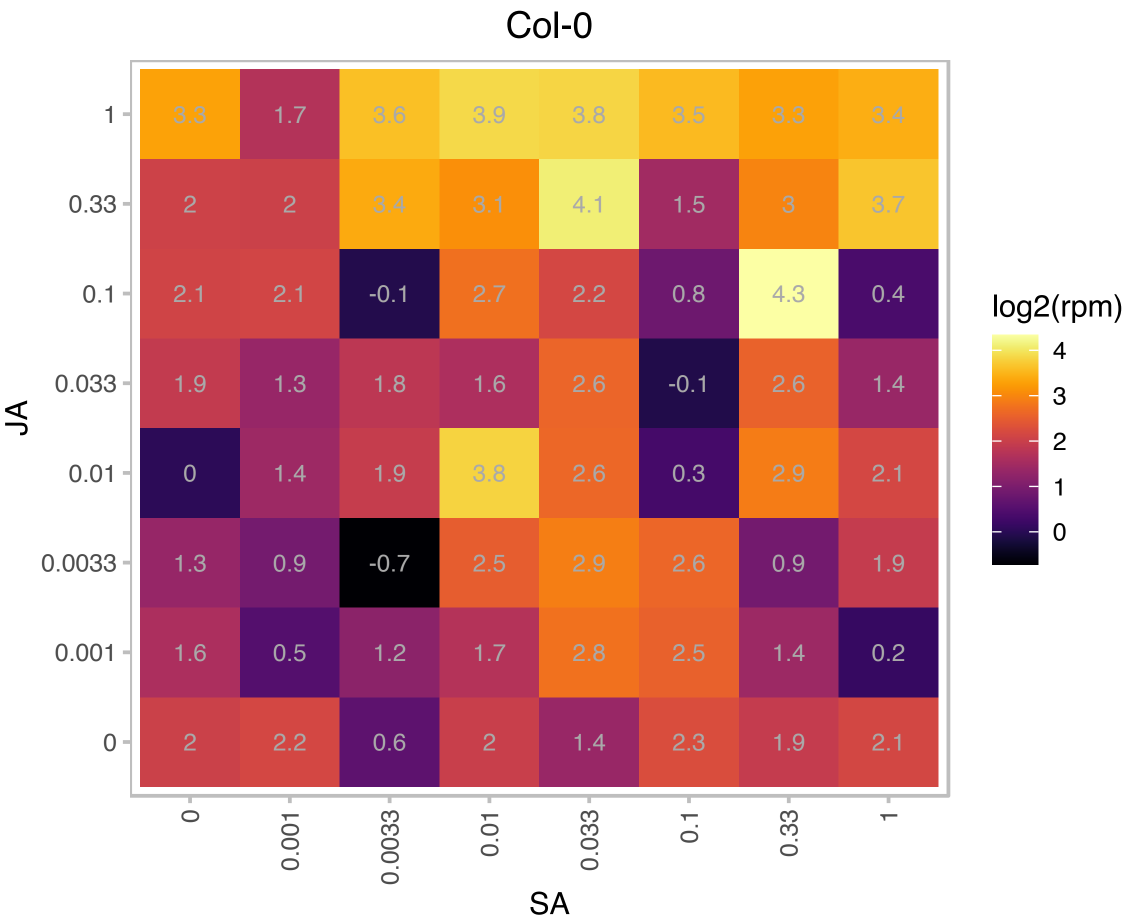


| | |
|---|---|
| Gene | AT1G24460 |
| is.expressed_genes | TRUE |
| is.responsive_genes | TRUE |
| is.apcluster | A-05 |

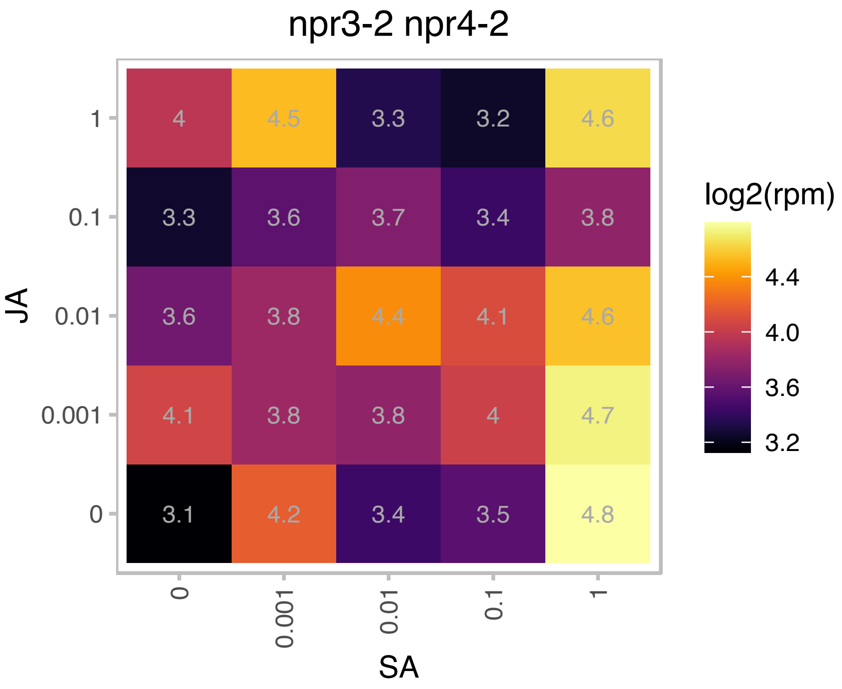


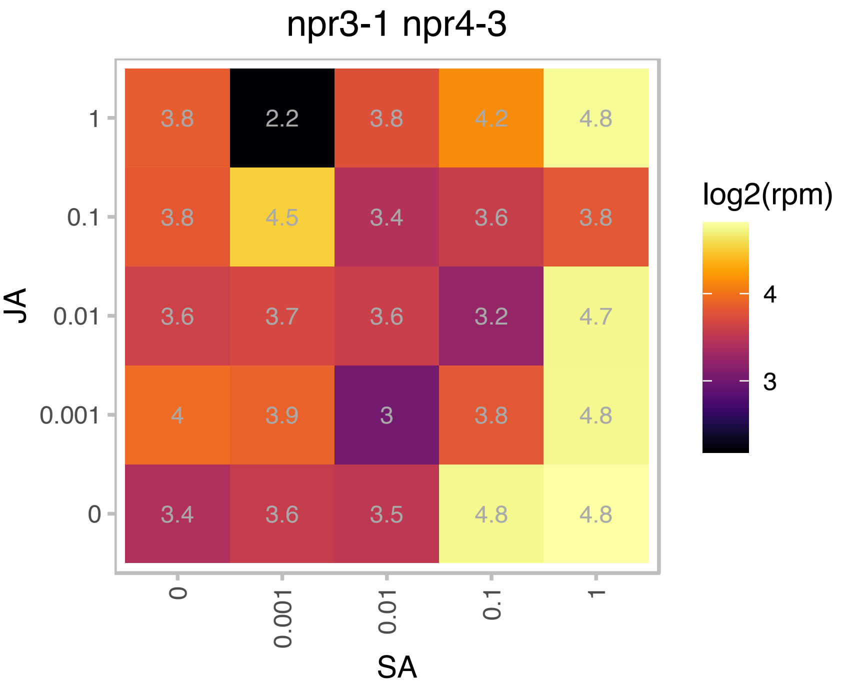

### AT1G27730.1 : STZ, ZAT10

Salt tolerance zinc finger

Related to Cys2/His2-type zinc-finger proteins found in higher plants. Compensated for a subset of calcineurin deficiency in yeast. Salt tolerance produced by ZAT10 appeared to be partially dependent on ENA1/PMR2, a P-type ATPase required for Li+ and Na+ efflux in yeast. The protein is localized to the nucleus, acts as a transcriptional repressor and is responsive to chitin oligomers. Also involved in response to photooxidative stress.

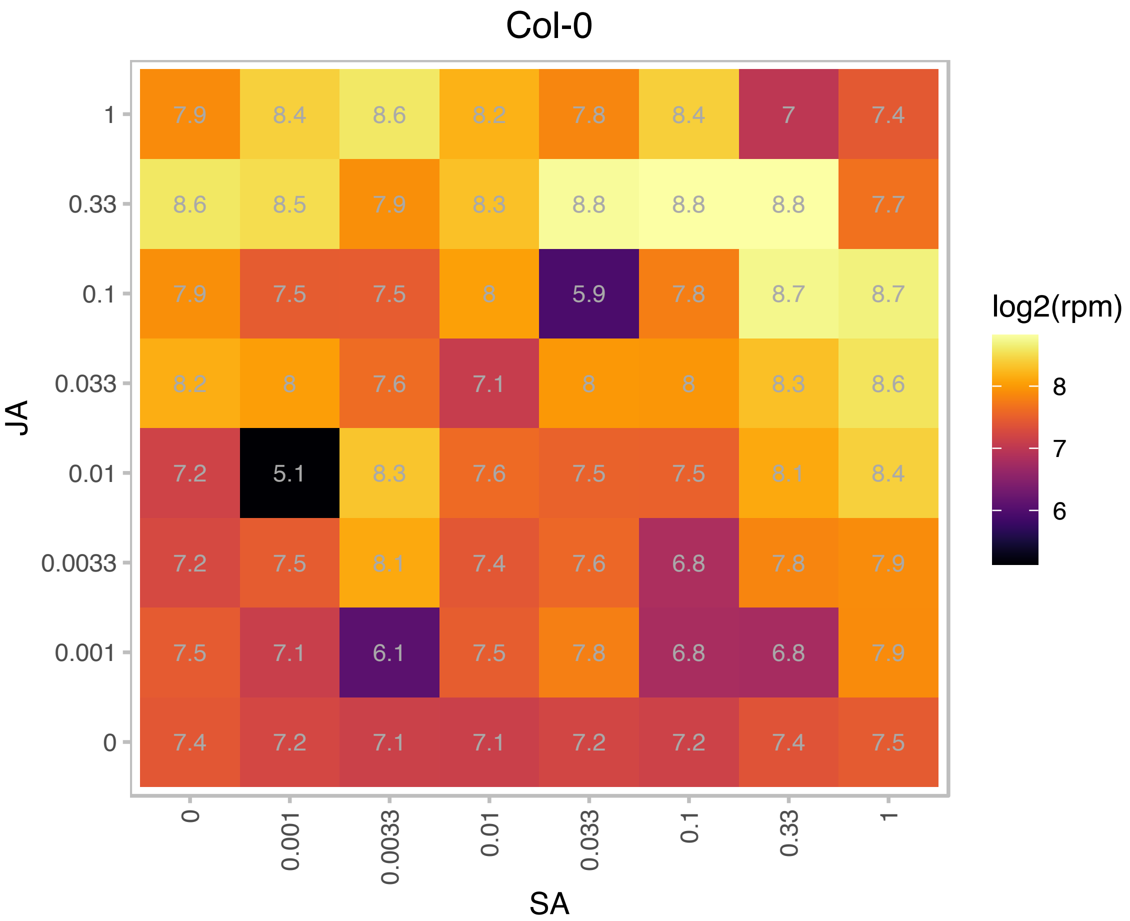


| | |
|---|---|
| Gene | AT1G27730 |
| is.expressed_genes | TRUE |
| is.responsive_genes | TRUE |
| is.apcluster | A-05 |

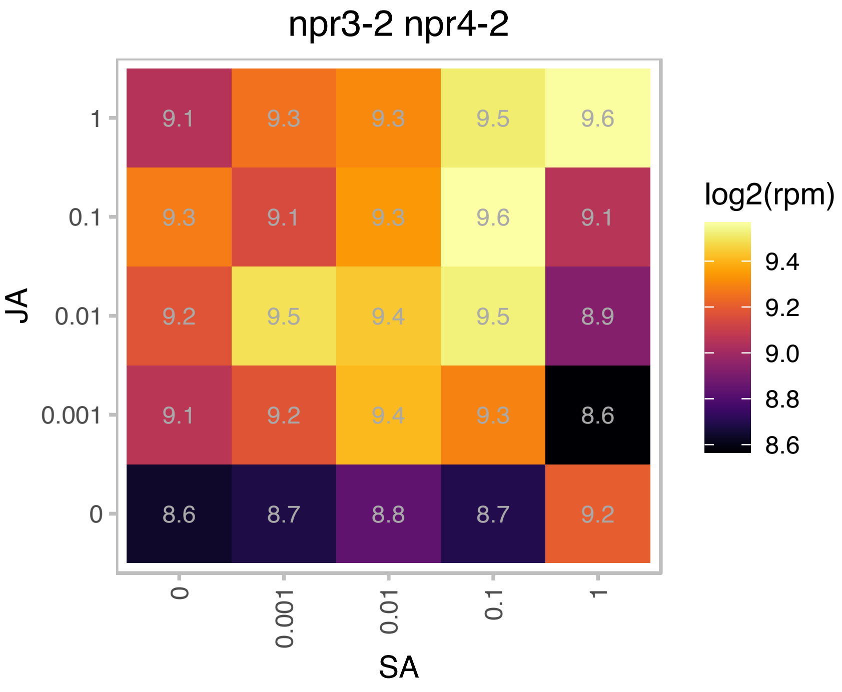


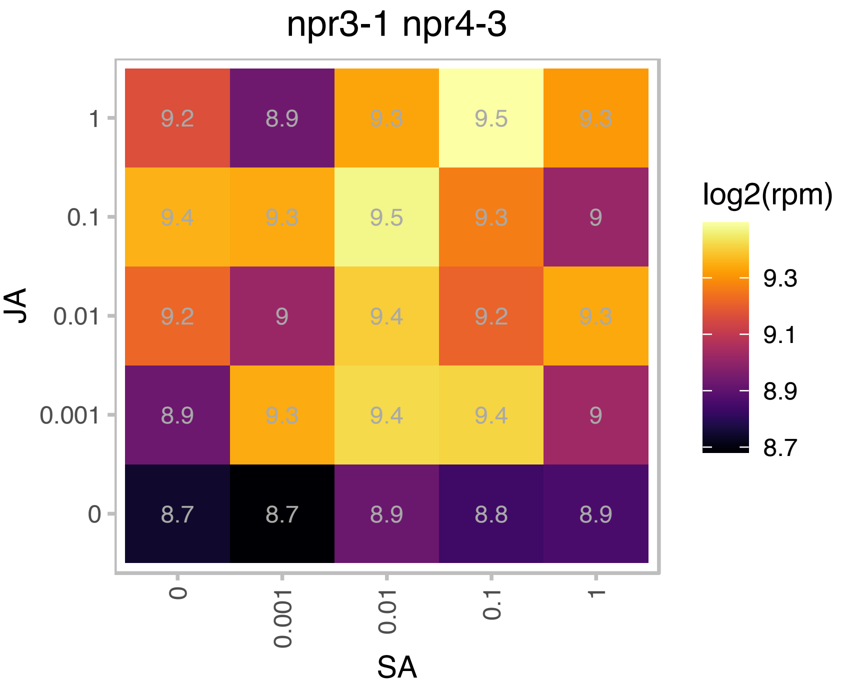

### AT1G27970.1 : NTF2B

nuclear transport factor 2B

Encodes an ortholog of yeast NTF2, a nuclear envelop transport protein that functions as the nuclear import receptor for RanGDP, an essential player in nucleocytoplasmic transport. The mRNA is cell-to-cell mobile.

| | |
|---|---|
| Gene | AT1G27970 |
| is.expressed_genes | TRUE |
| is.responsive_genes | TRUE |
| is.apcluster | A-05 |

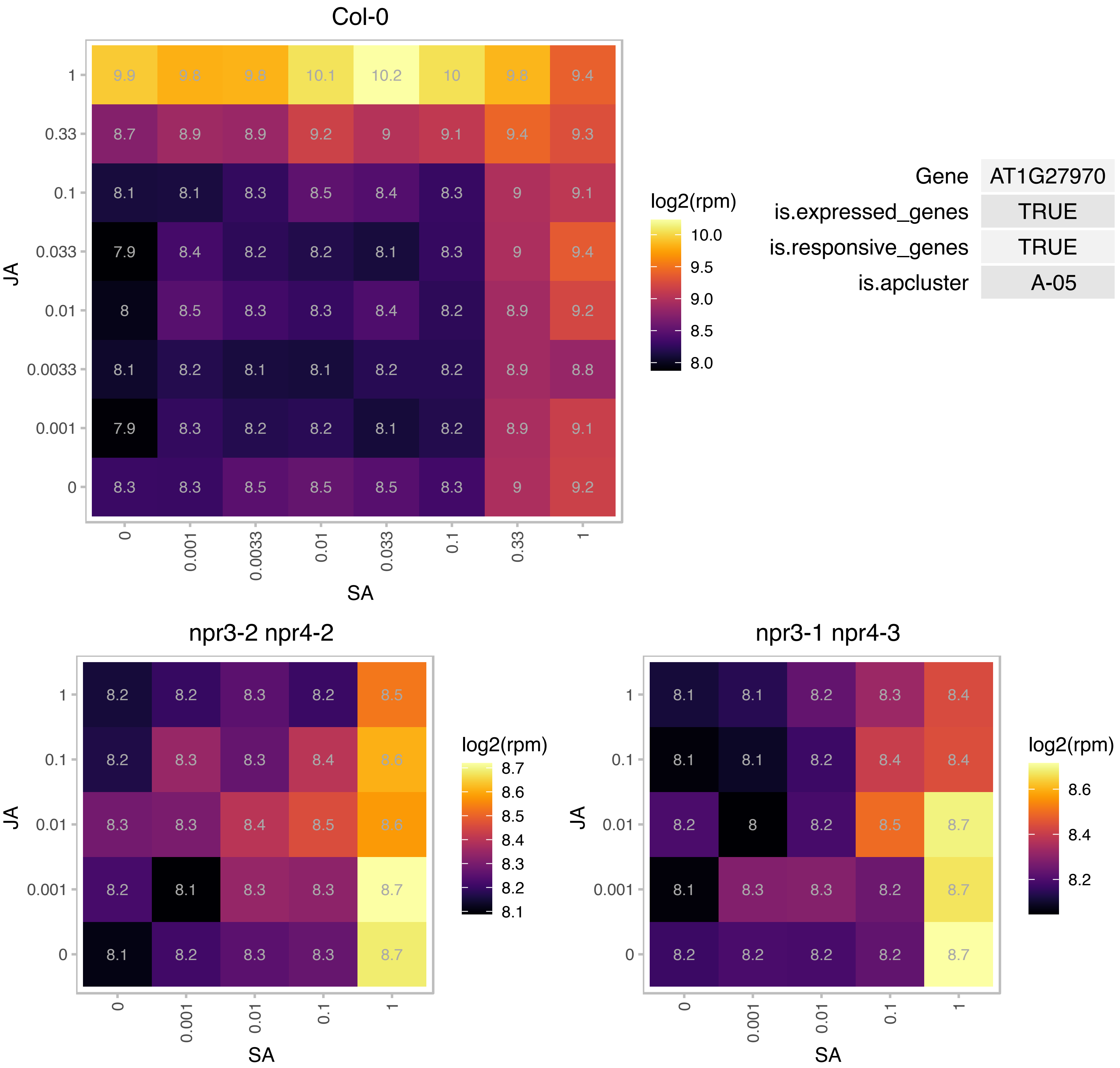

### AT1G35910.1 : AtTPPD, TPPD

Haloacid dehalogenase-like hydrolase (HAD) superfamily protein

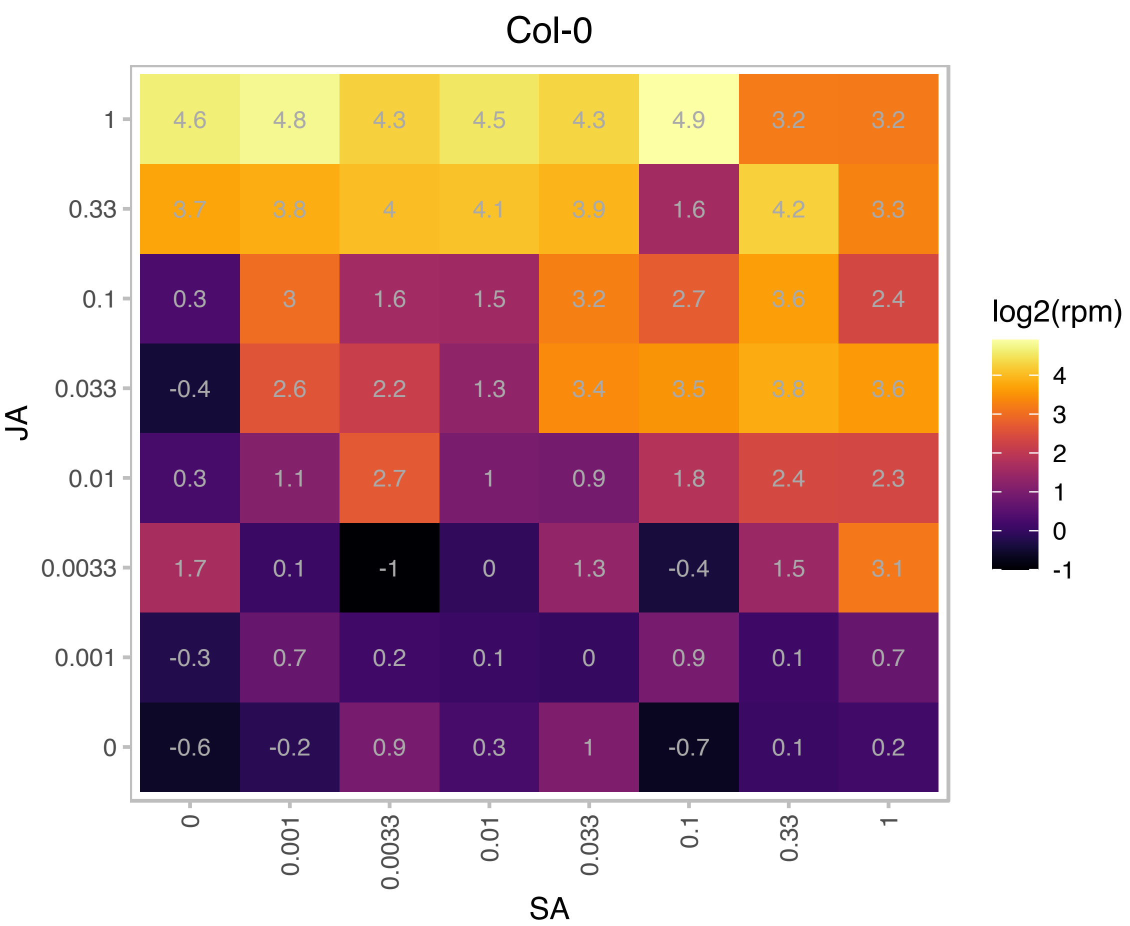


| | |
|---|---|
| Gene | AT1G35910 |
| is.expressed_genes | TRUE |
| is.responsive_genes | TRUE |
| is.apcluster | A-05 |

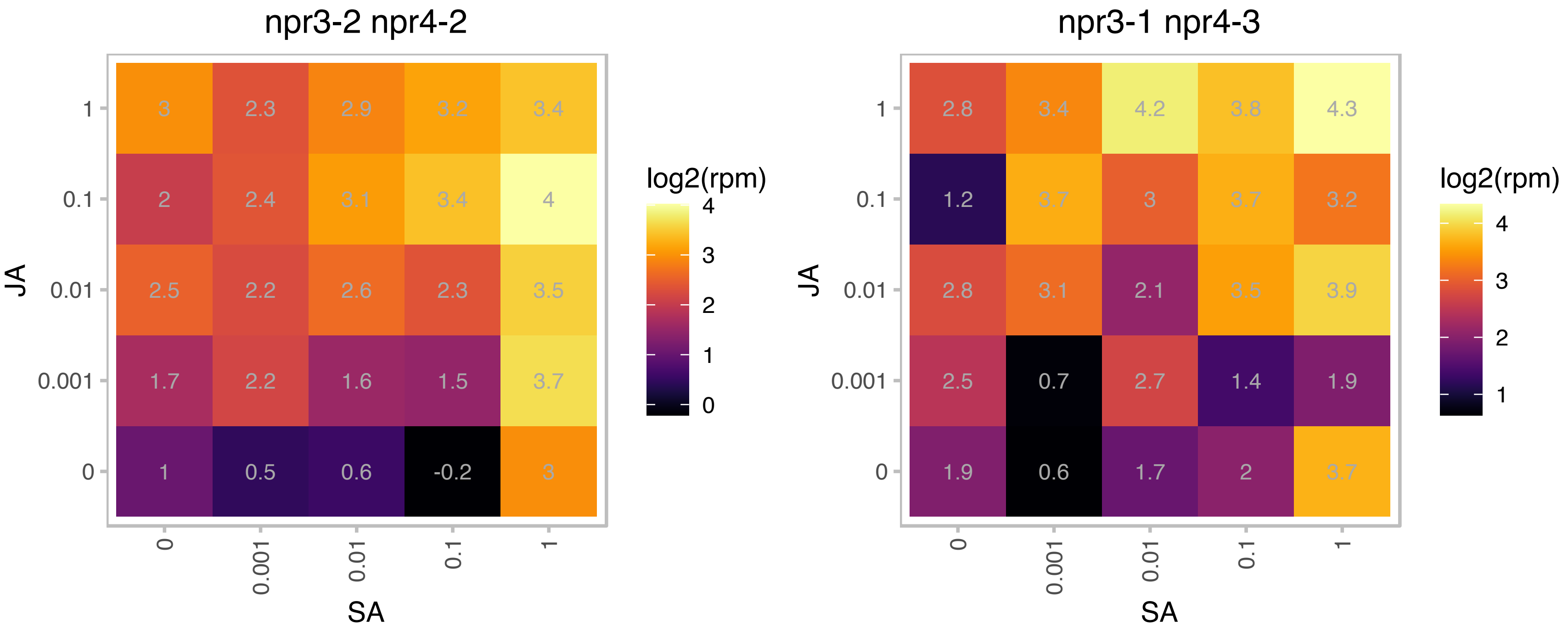

# AT1G45000.1 : NA

AAA-type ATPase family protein

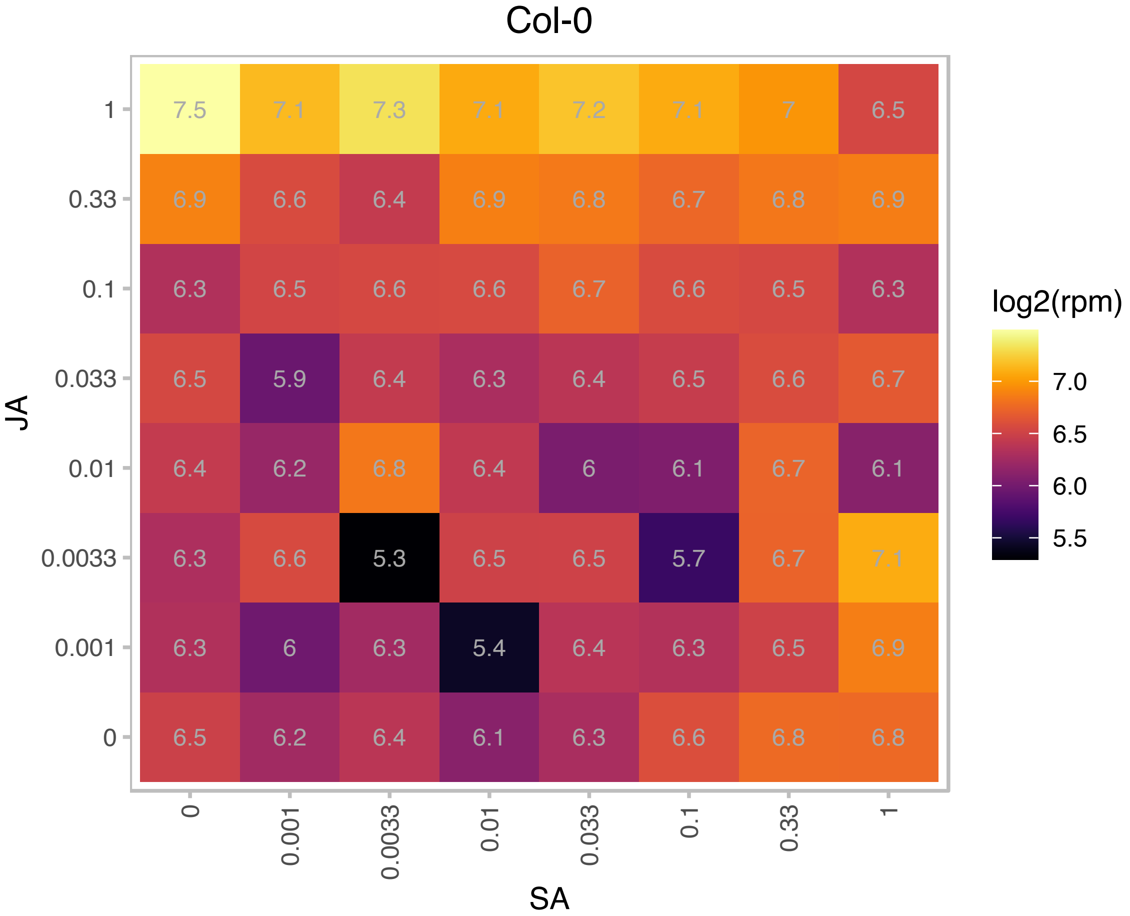


| | |
|---|---|
| Gene | AT1G45000 |
| is.expressed_genes | TRUE |
| is.responsive_genes | TRUE |
| is.apcluster | A-05 |

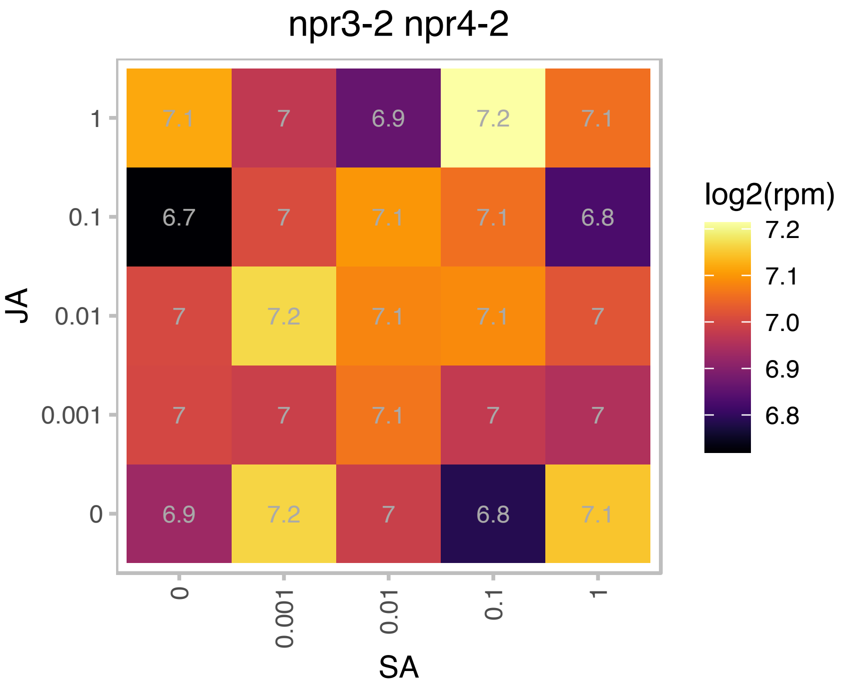


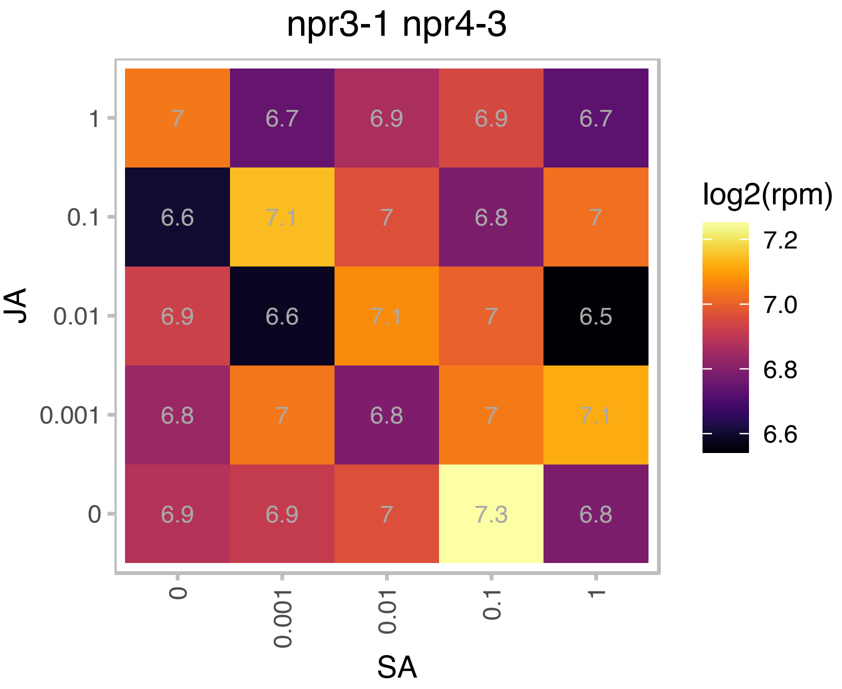

# AT1G48900.1 : NA

Signal recognition particle, SRP54 subunit protein

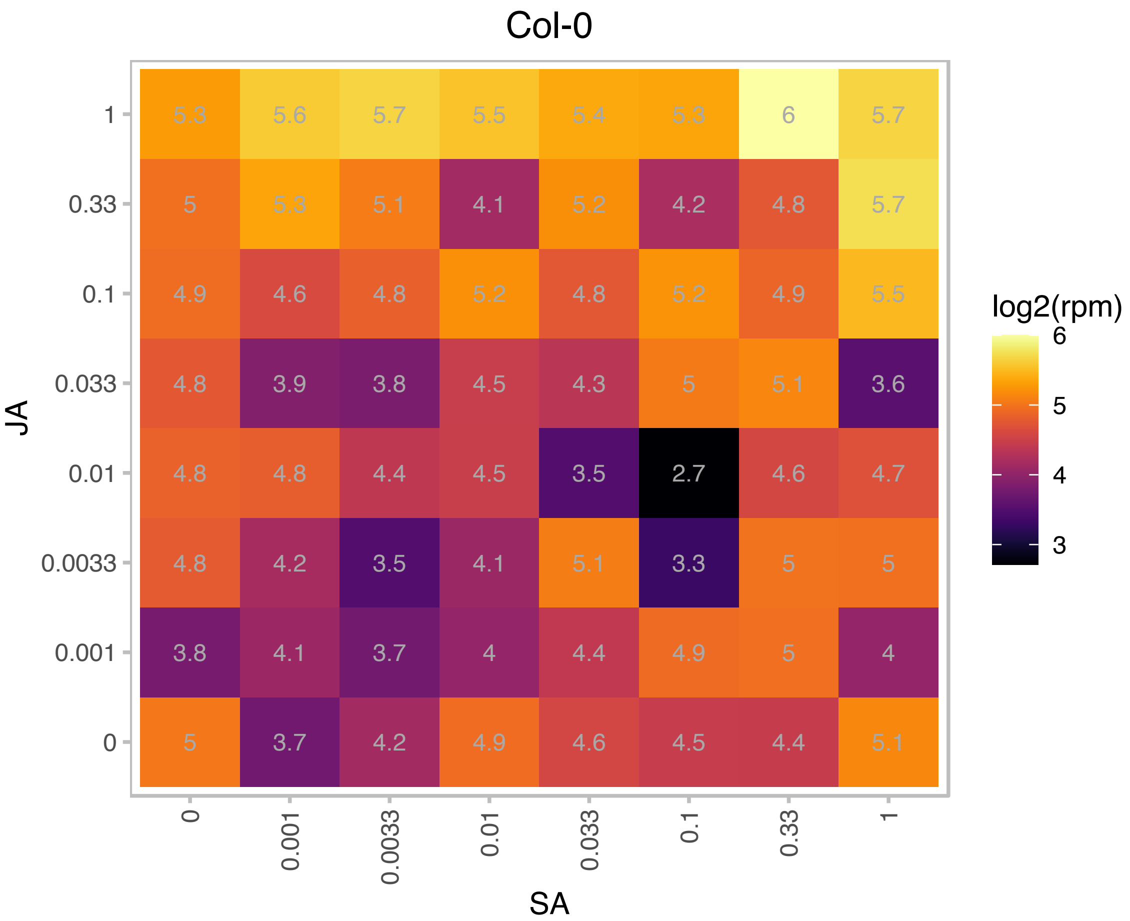


| | |
|---|---|
| Gene | AT1G48900 |
| is.expressed_genes | TRUE |
| is.responsive_genes | TRUE |
| is.apcluster | A-05 |

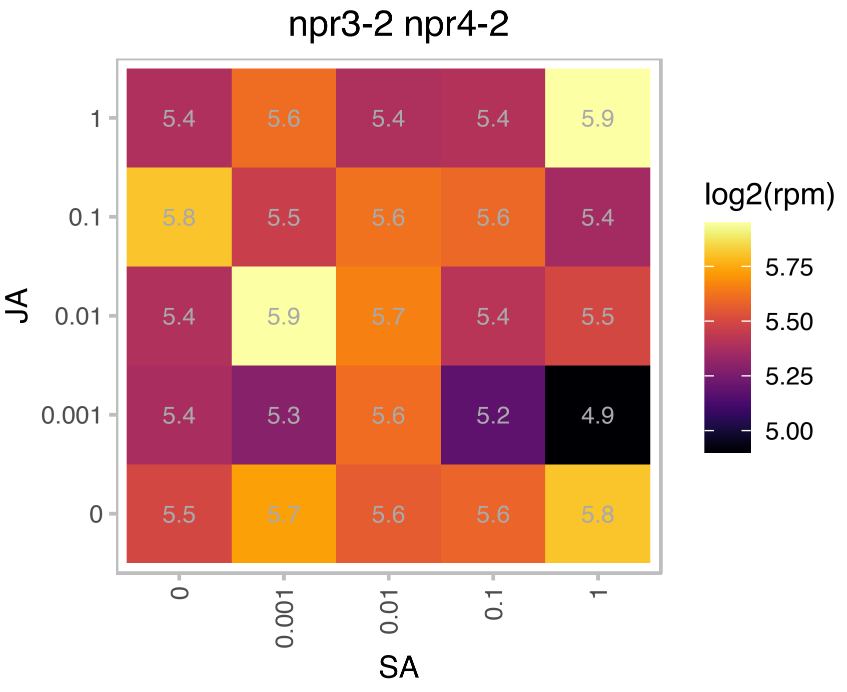


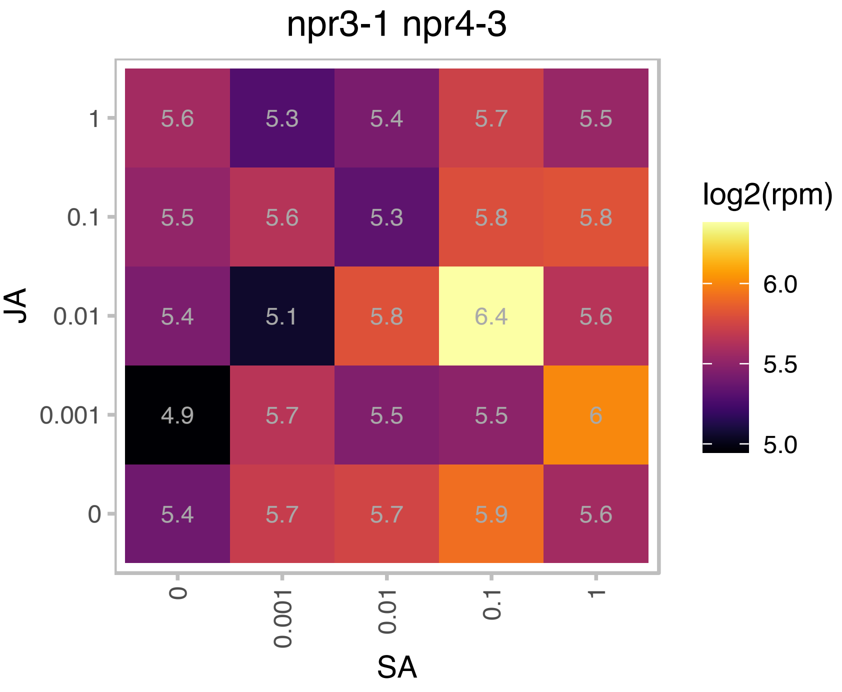

### AT1G49300.1 : ATRAB7, ATRABG3E, RABG3E

RAB GTPase homolog G3E

encodes a small GTPase involved in membrane trafficking. Gene expression is induced by hydrogen peroxide and
lines. Lines overexpressing the gene are more tolerant to high salt and hyperosmotic conditions.

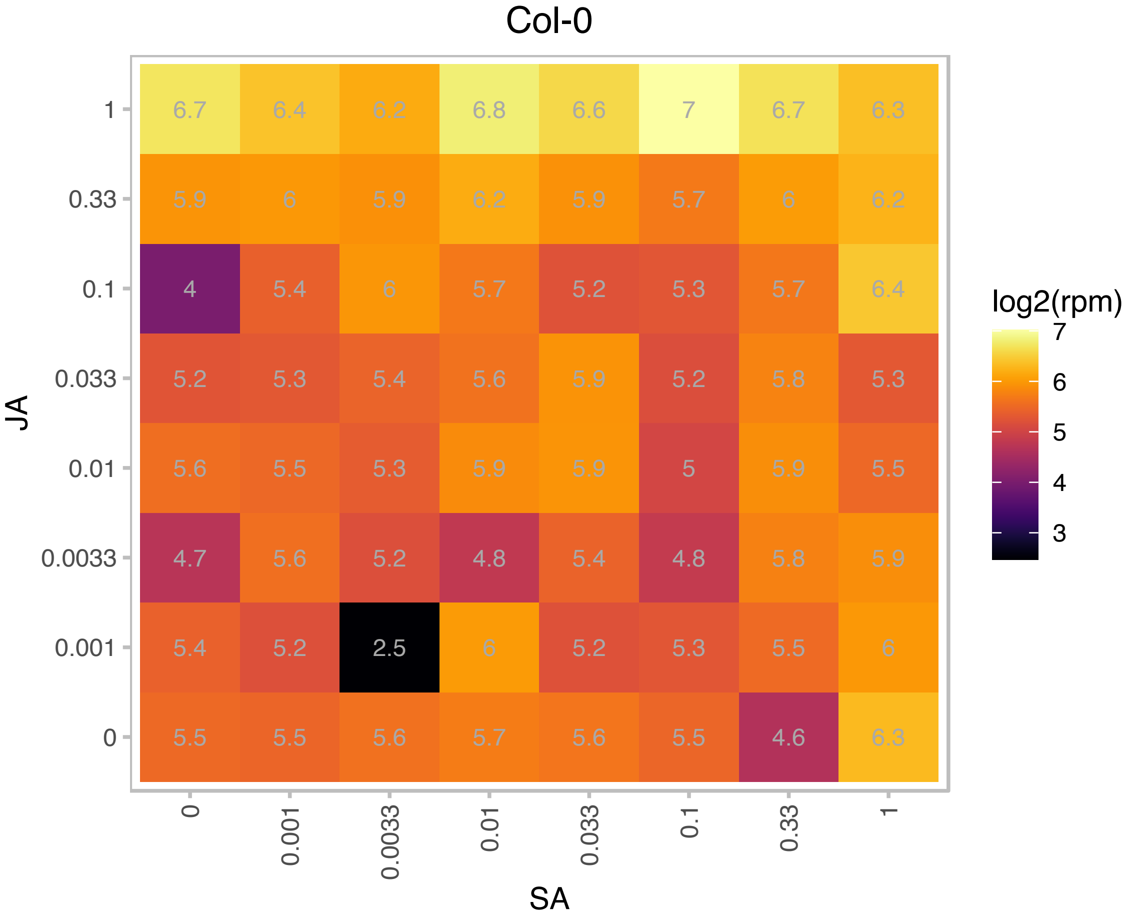


| | |
|---|---|
| Gene | AT1G49300 |
| is.expressed_genes | TRUE |
| is.responsive_genes | TRUE |
| is.apcluster | A-05 |

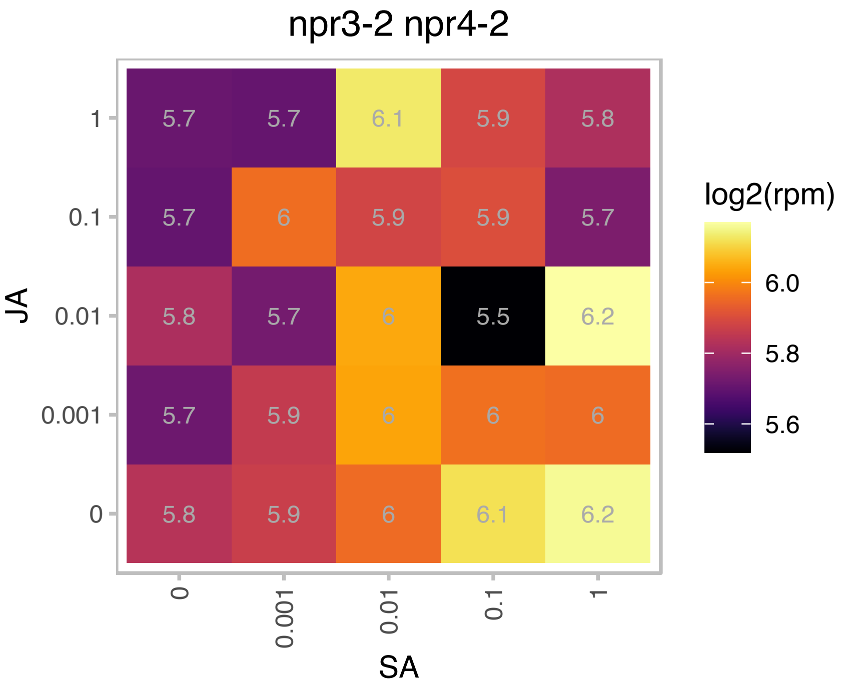


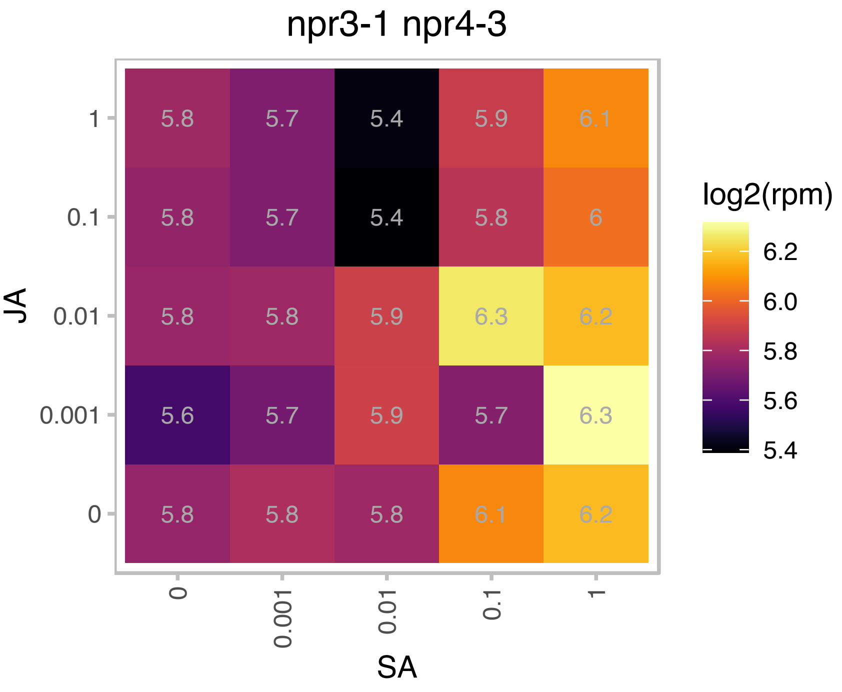

# AT1G51350.1 : NA

ARM repeat superfamily protein

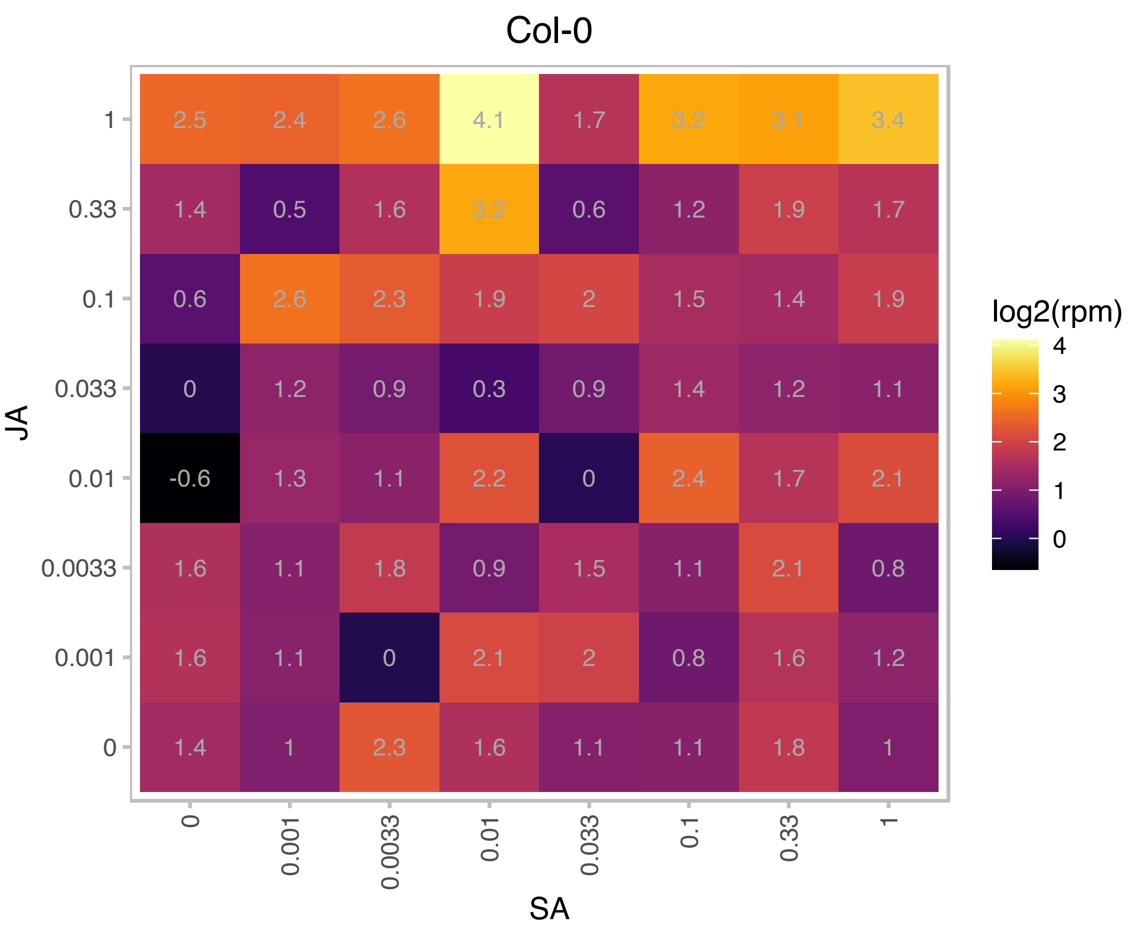


| | |
|---|---|
| Gene | AT1G51350 |
| is.expressed_genes | TRUE |
| is.responsive_genes | TRUE |
| is.apcluster | A-05 |

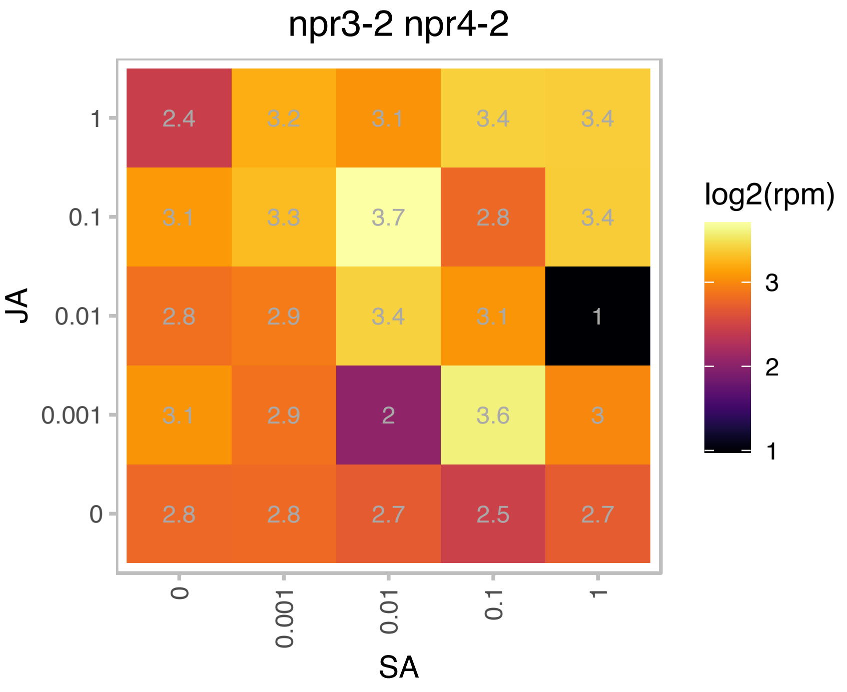


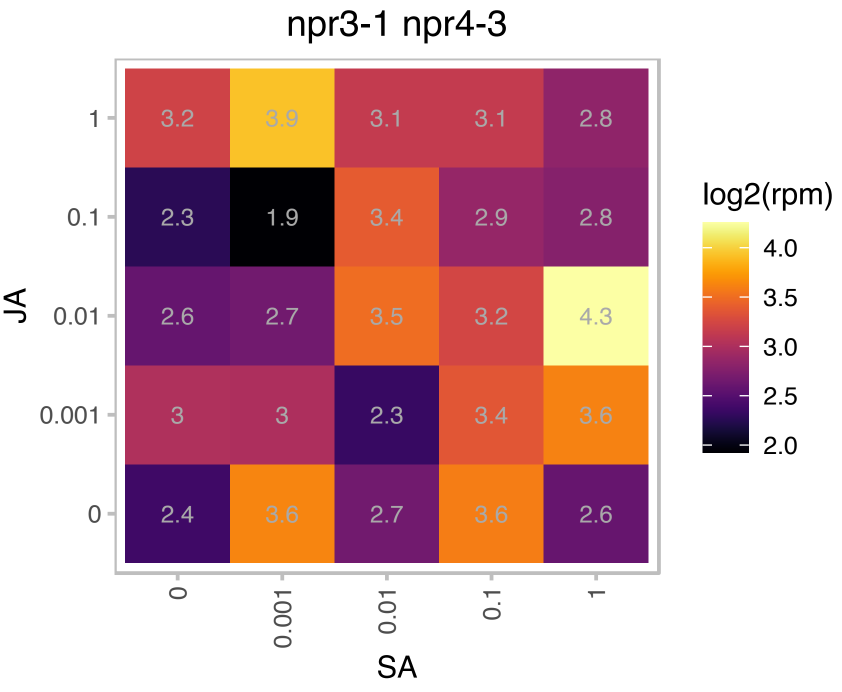

### AT1G61800.1 : ATGPT2, GPT2

glucose-6-phosphate/phosphate translocator 2

glucose6-Phosphate/phosphate transporter 2. Expression is upregulated in the shoot of cax1/cax3 mutant. The mRNA is cell-to-cell mobile.

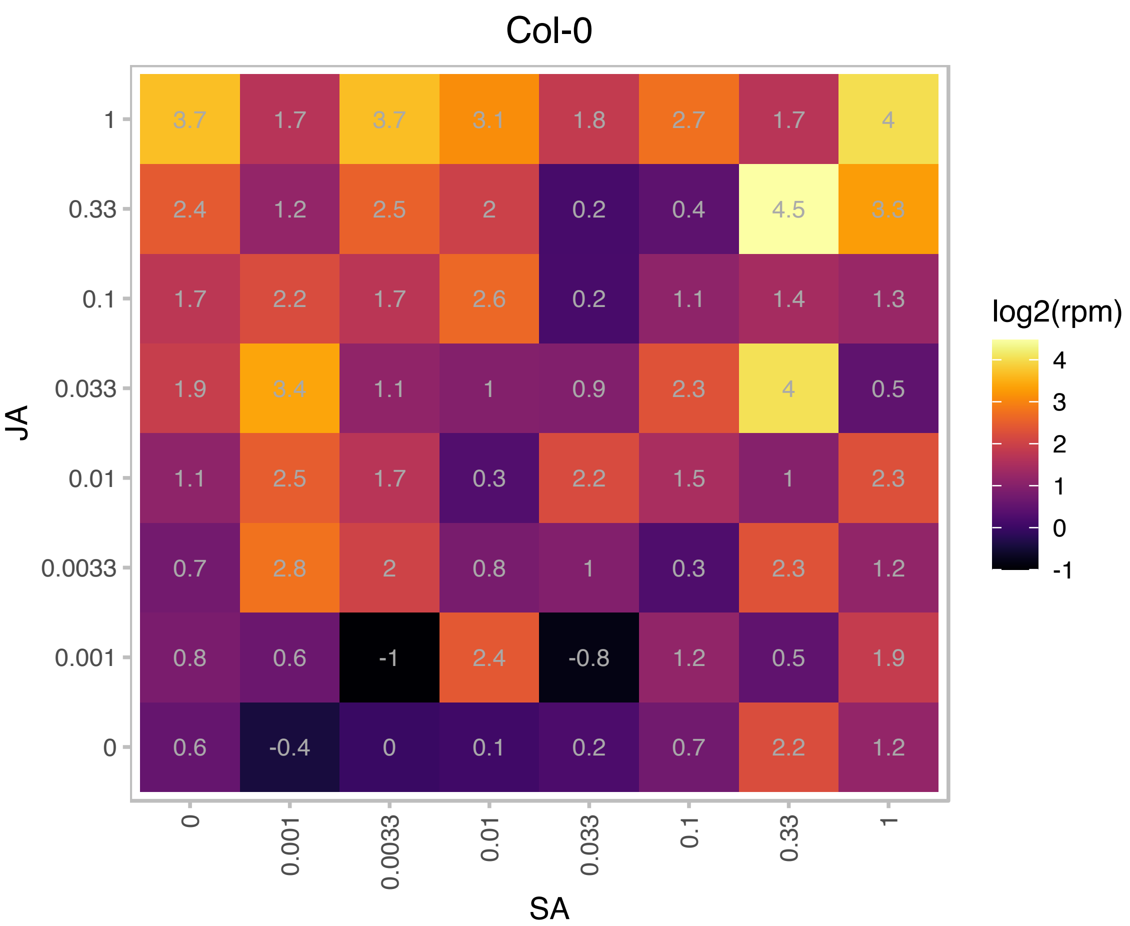


| | |
|---|---|
| Gene | AT1G61800 |
| is.expressed_genes | TRUE |
| is.responsive_genes | TRUE |
| is.apcluster | A-05 |

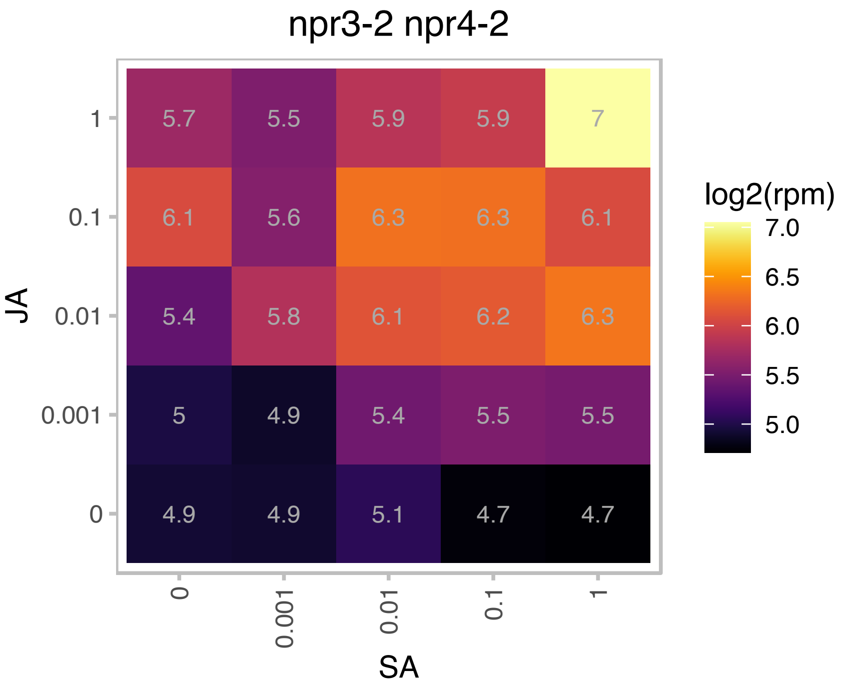


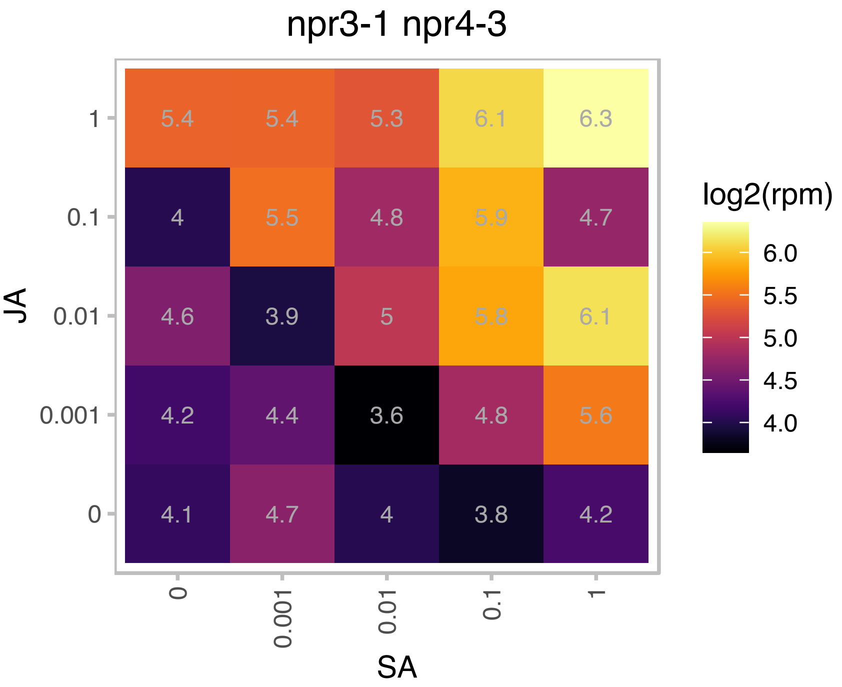

# AT1G62480.1 : NA

Vacuolar calcium-binding protein-related

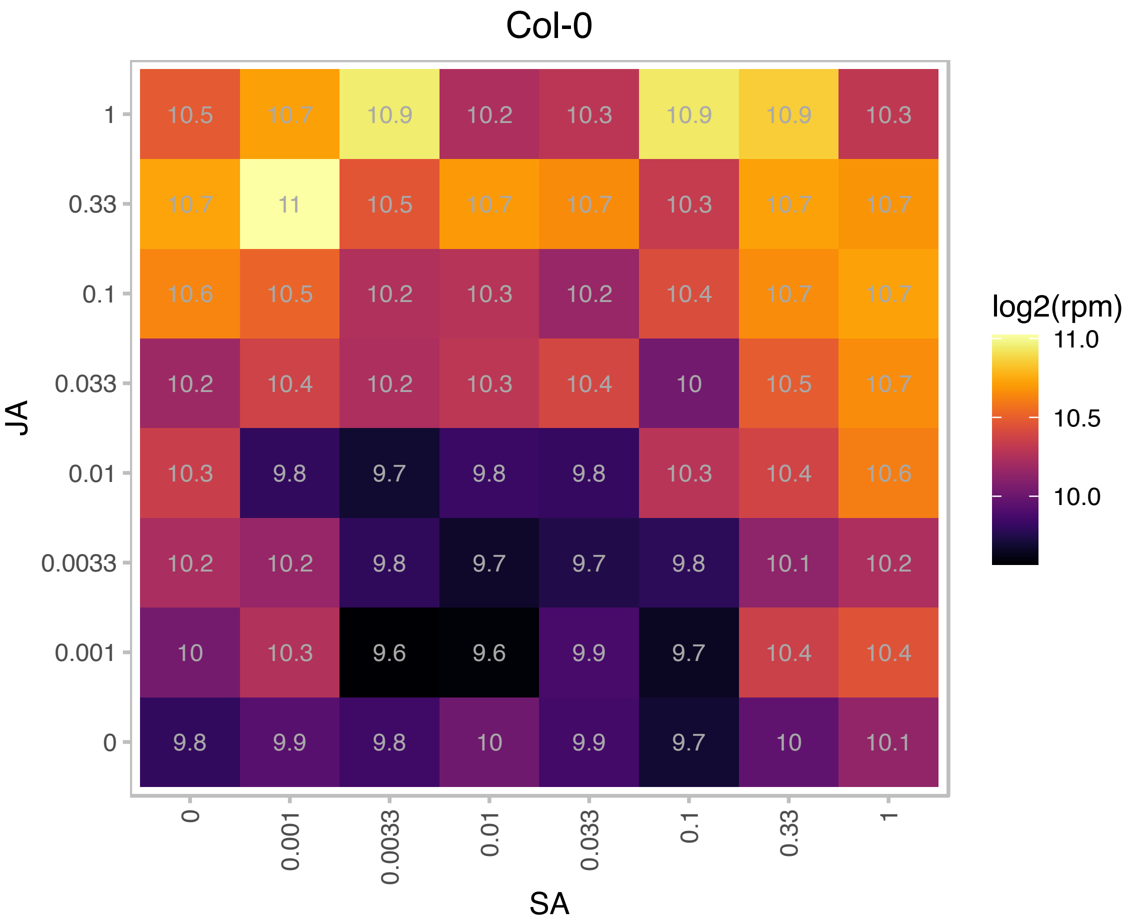


| | |
|---|---|
| Gene | AT1G62480 |
| is.expressed_genes | TRUE |
| is.responsive_genes | TRUE |
| is.apcluster | A-05 |

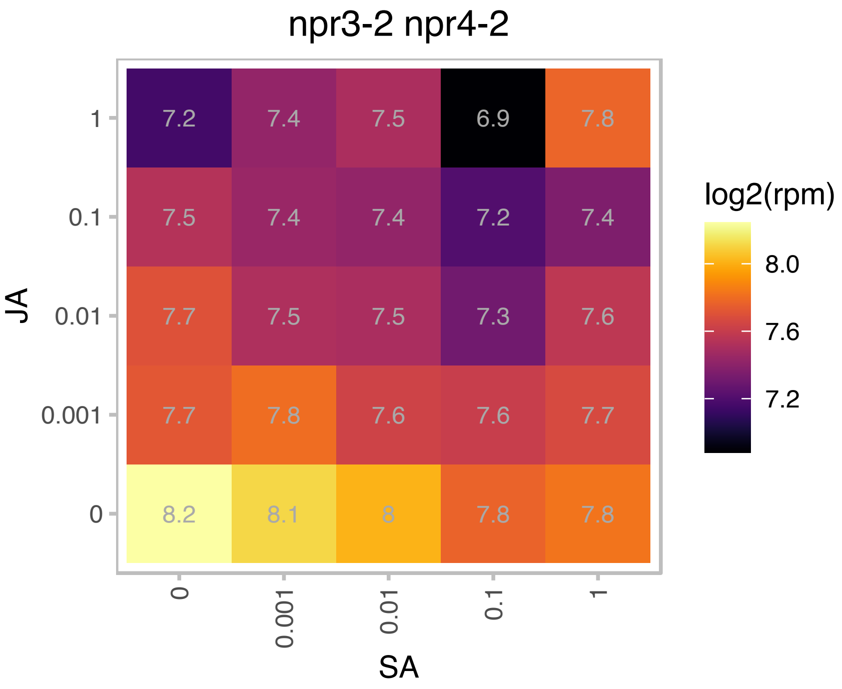


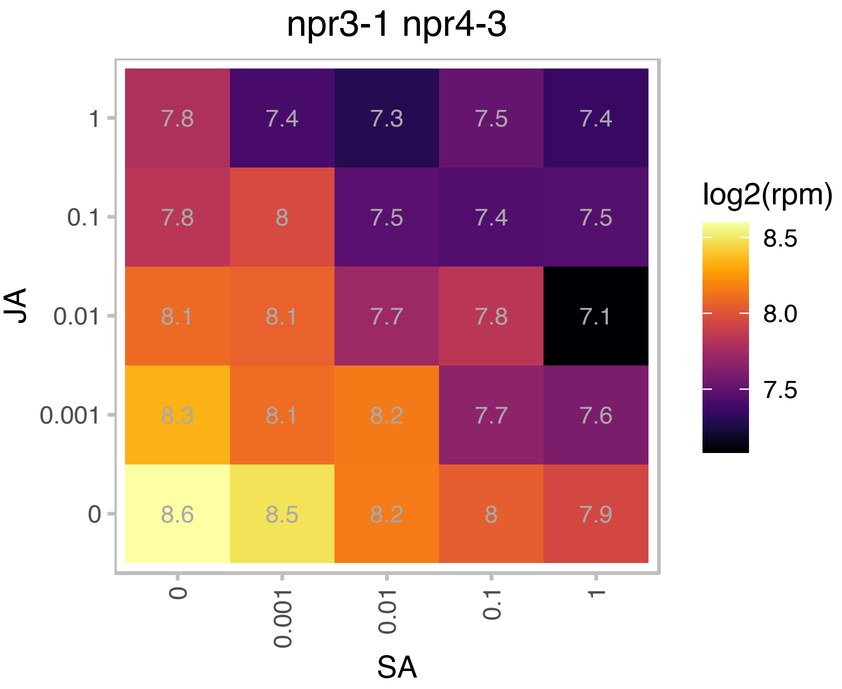

# AT1G66760.1 : NA

MATE efflux family protein

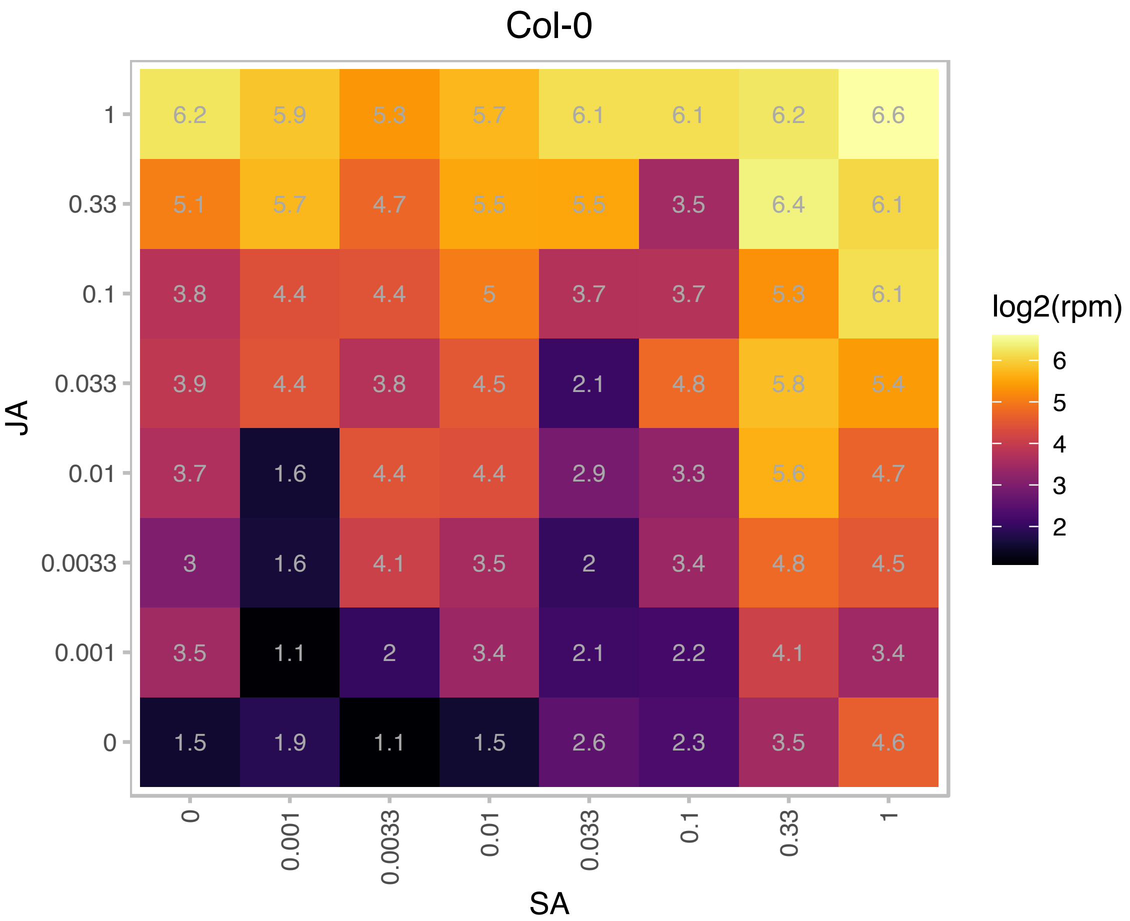


| | |
|---|---|
| Gene | AT1G66760 |
| is.expressed_genes | TRUE |
| is.responsive_genes | TRUE |
| is.apcluster | A-05 |

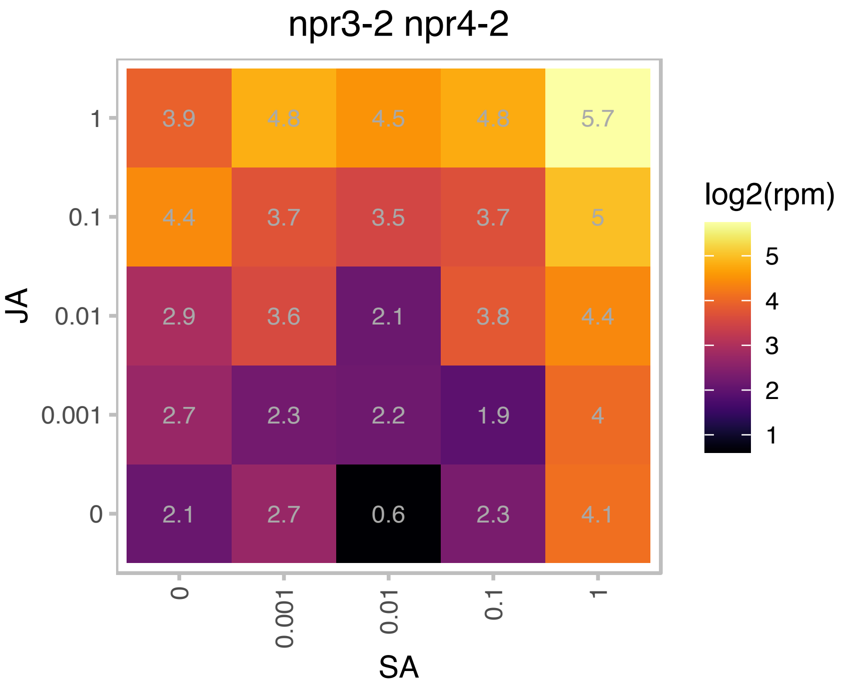


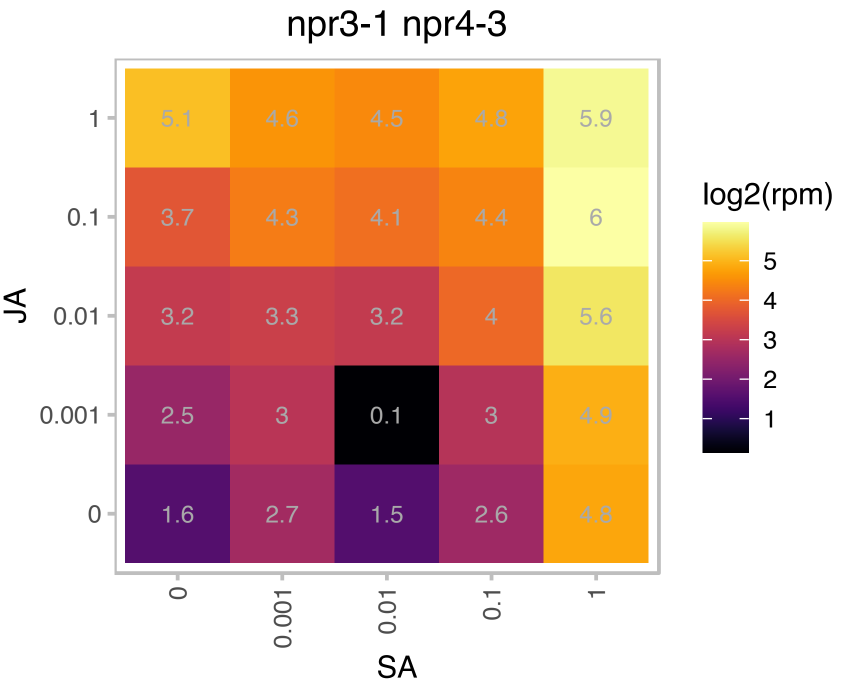

# AT1G68620.1 : NA

alpha/beta-Hydrolases superfamily protein

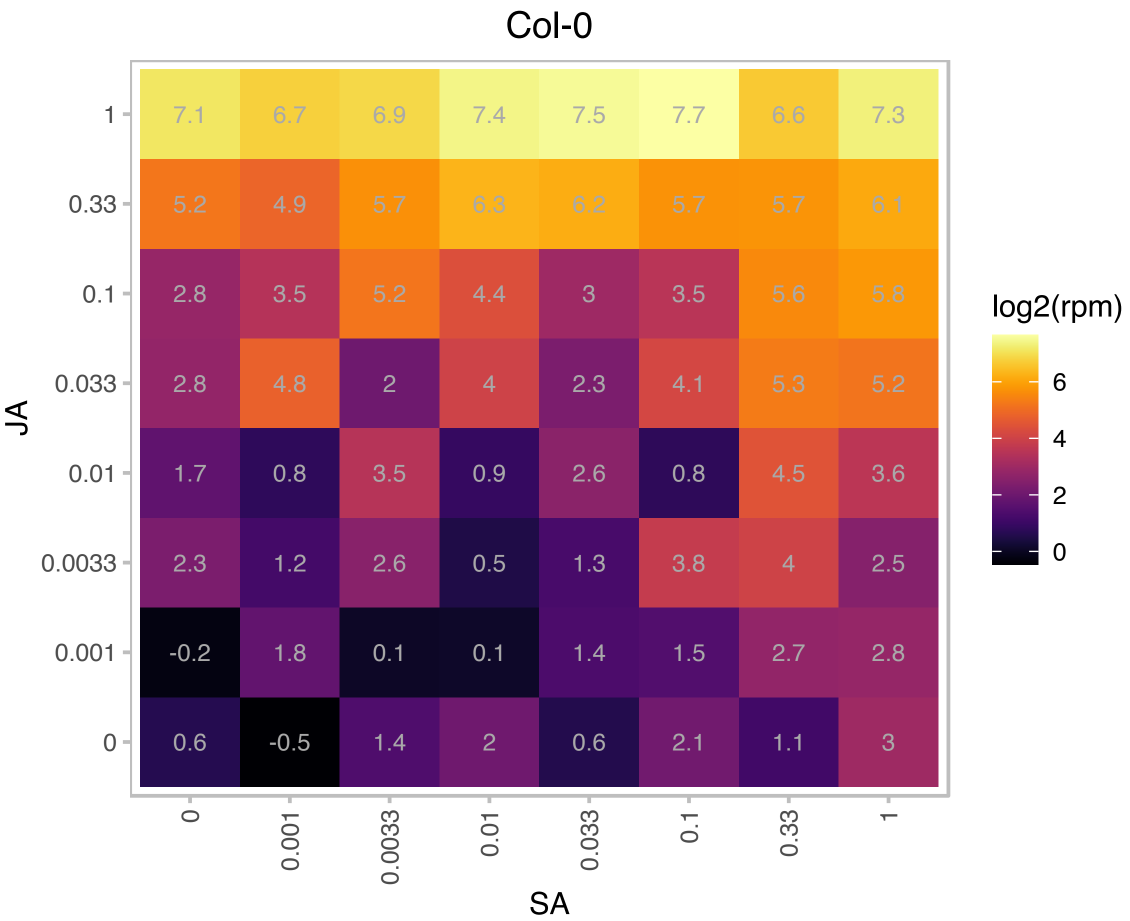


| | |
|---|---|
| Gene | AT1G68620 |
| is.expressed_genes | TRUE |
| is.responsive_genes | TRUE |
| is.apcluster | A-05 |

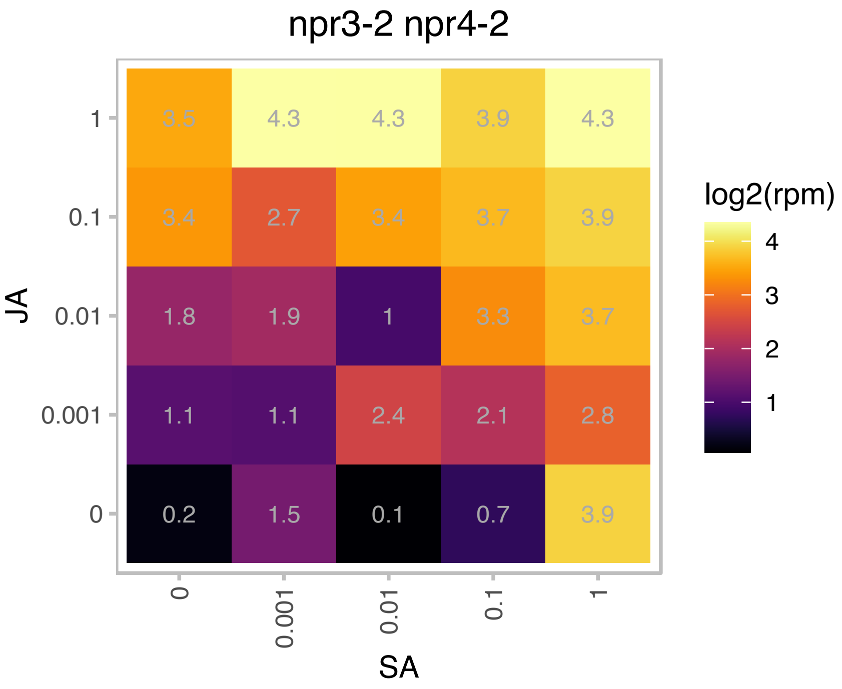


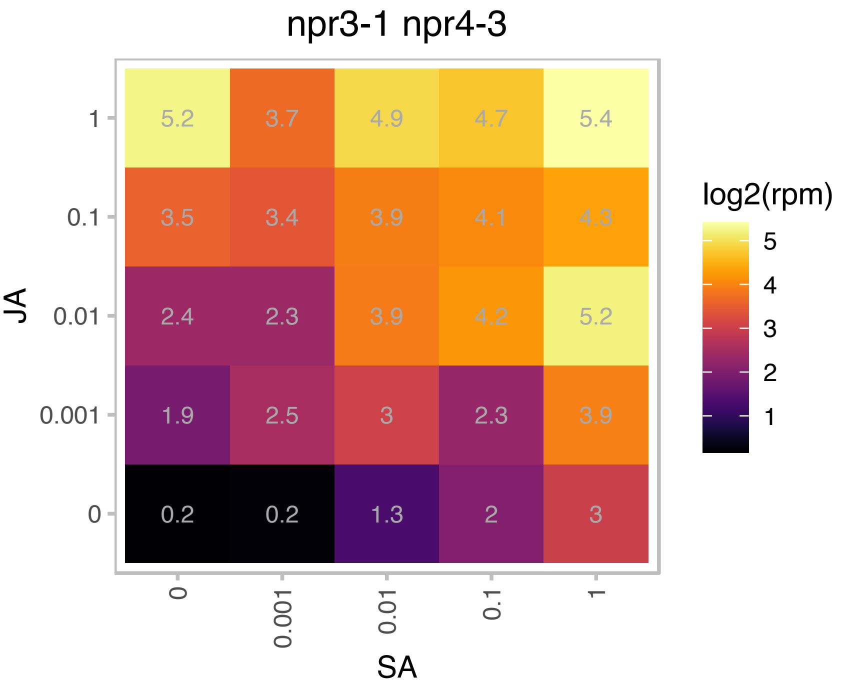

# AT1G75990.1 : NA

PAM domain (PCI/PINT associated module) protein

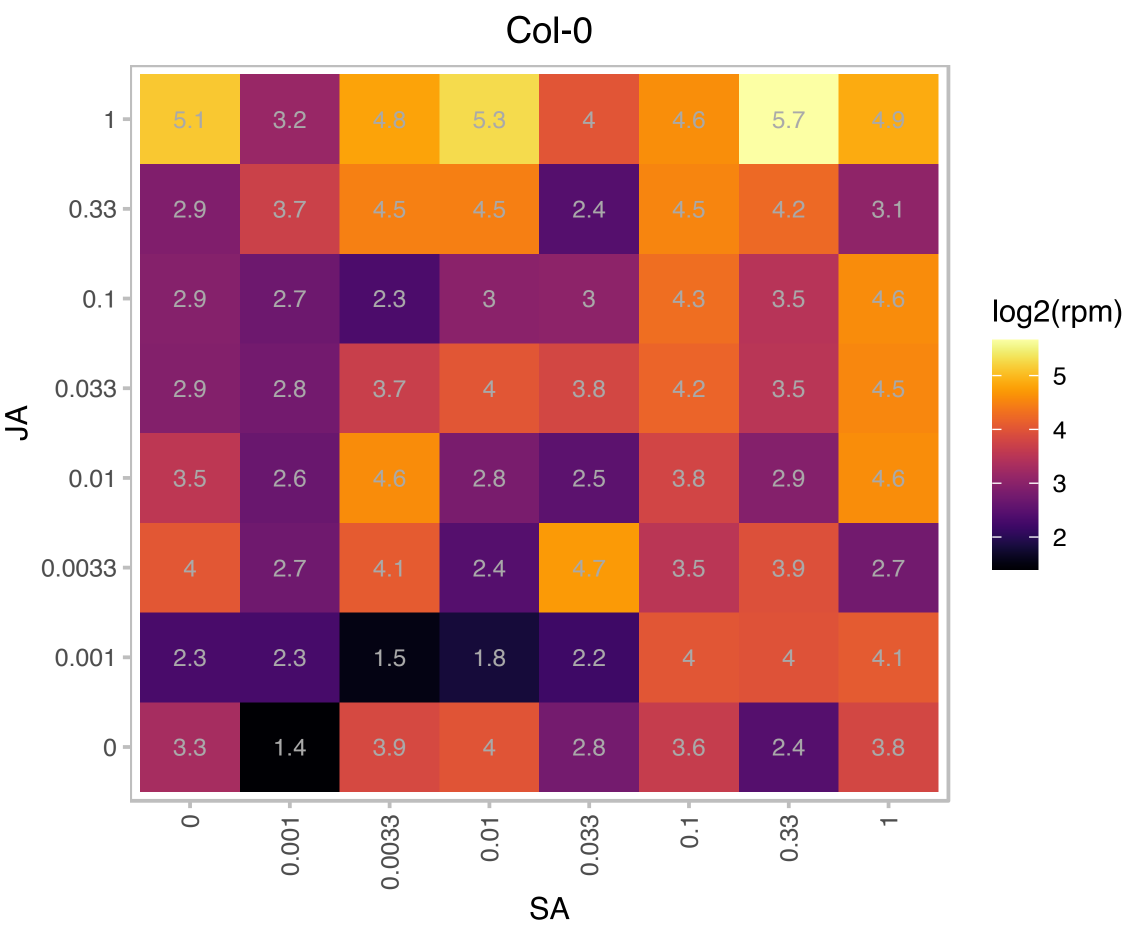


| | |
|---|---|
| Gene | AT1G75990 |
| is.expressed_genes | TRUE |
| is.responsive_genes | TRUE |
| is.apcluster | A-05 |

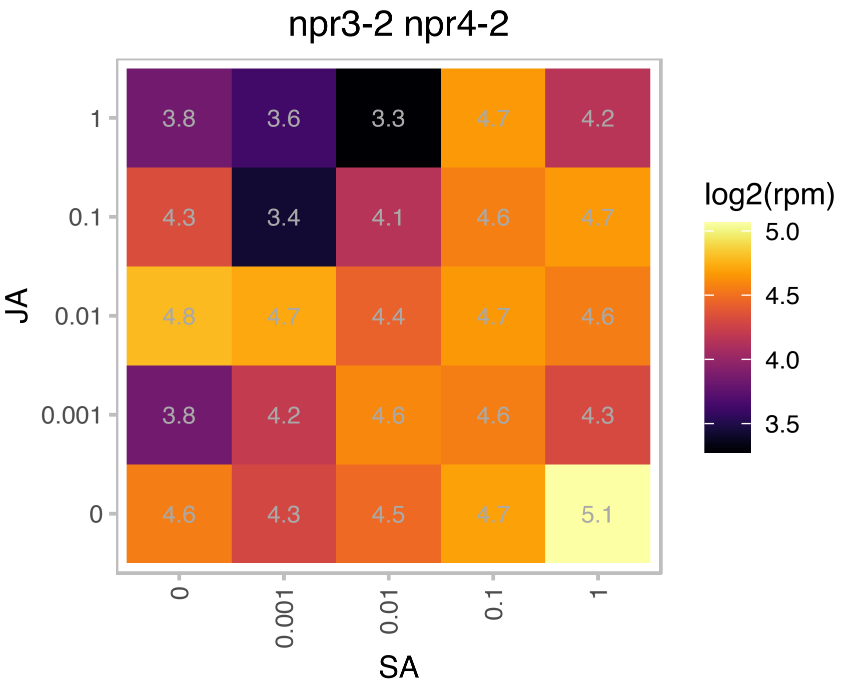


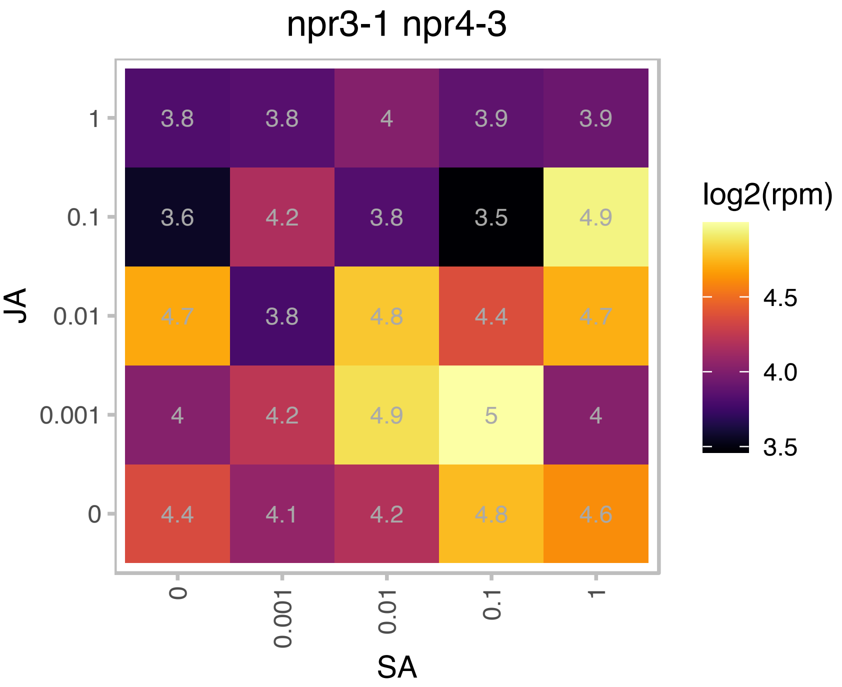

### AT1G79930.1 : AtHsp70-14, HSP91

heat shock protein 91

encodes high molecular weight heat shock protein 70 not a HSP90 homolog, mRNA is constitutively expressed but transiently induced after heat shock

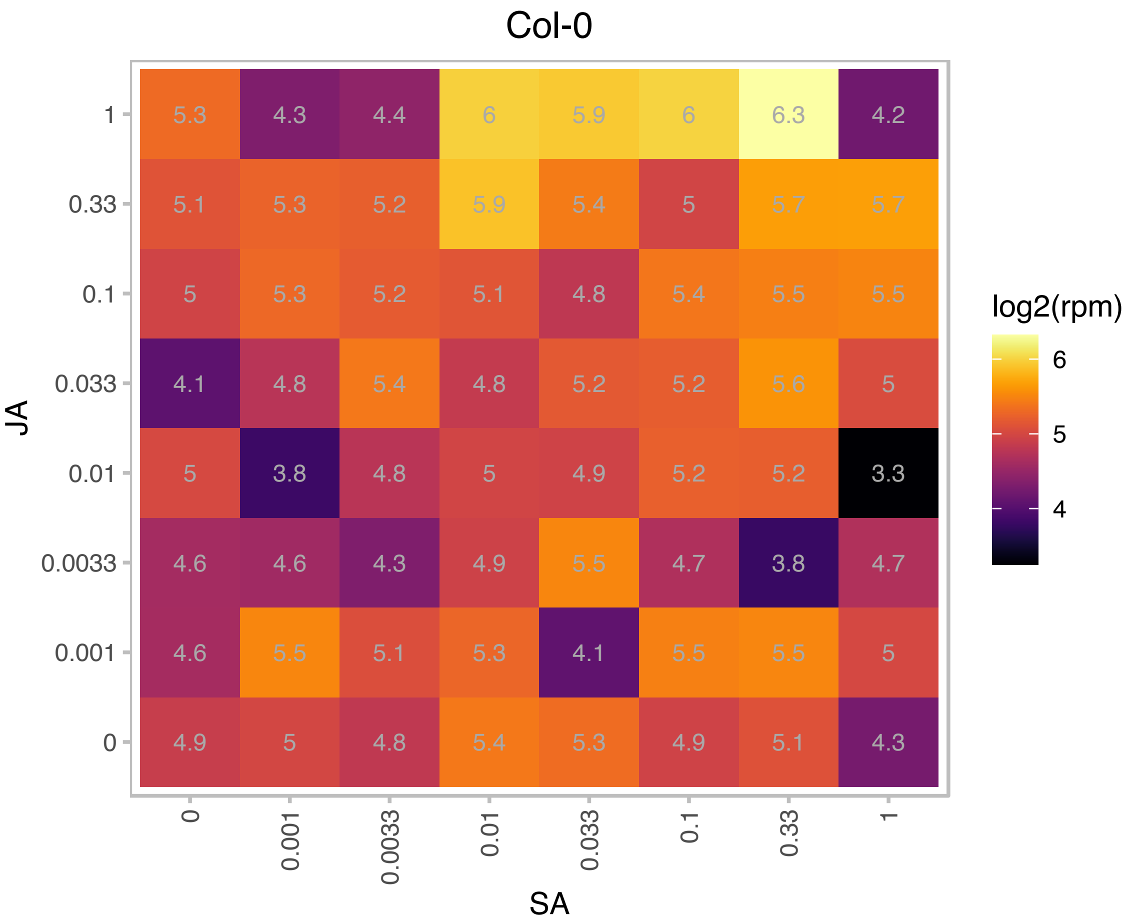


| | |
|---|---|
| Gene | AT1G79930 |
| is.expressed_genes | TRUE |
| is.responsive_genes | TRUE |
| is.apcluster | A-05 |

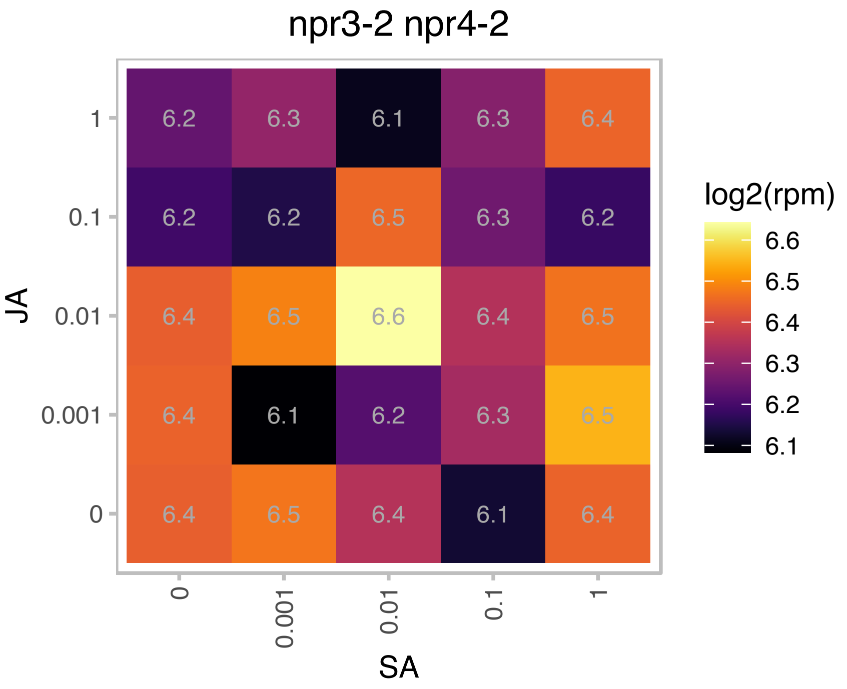


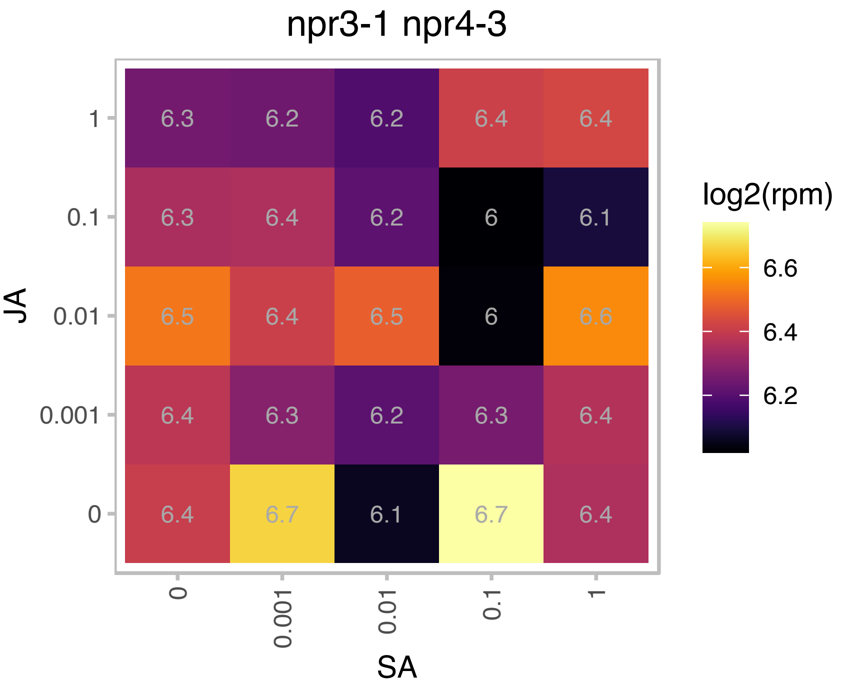

### AT1G80160.1 : GLYI7

Lactoylglutathione lyase / glyoxalase I family protein

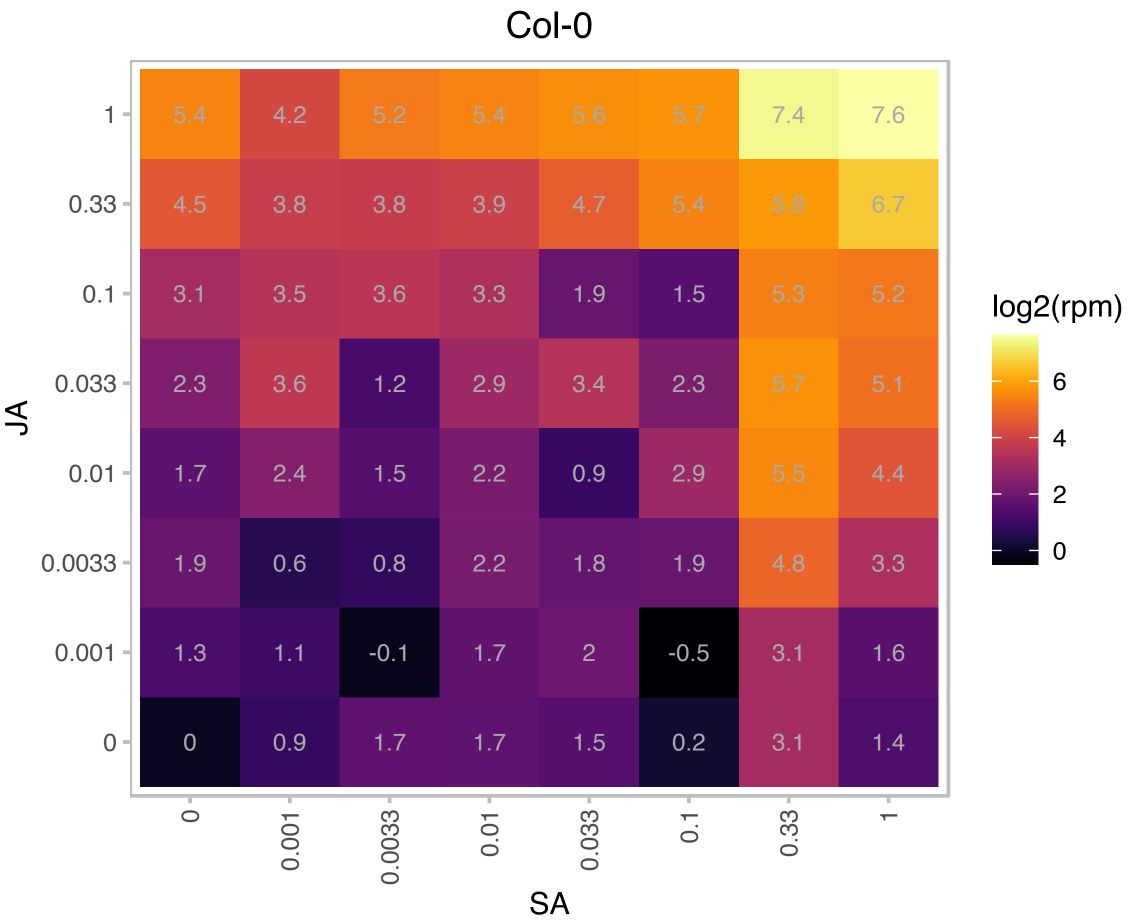


| | |
|---|---|
| Gene | AT1G80160 |
| is.expressed_genes | TRUE |
| is.responsive_genes | TRUE |
| is.apcluster | A-05 |

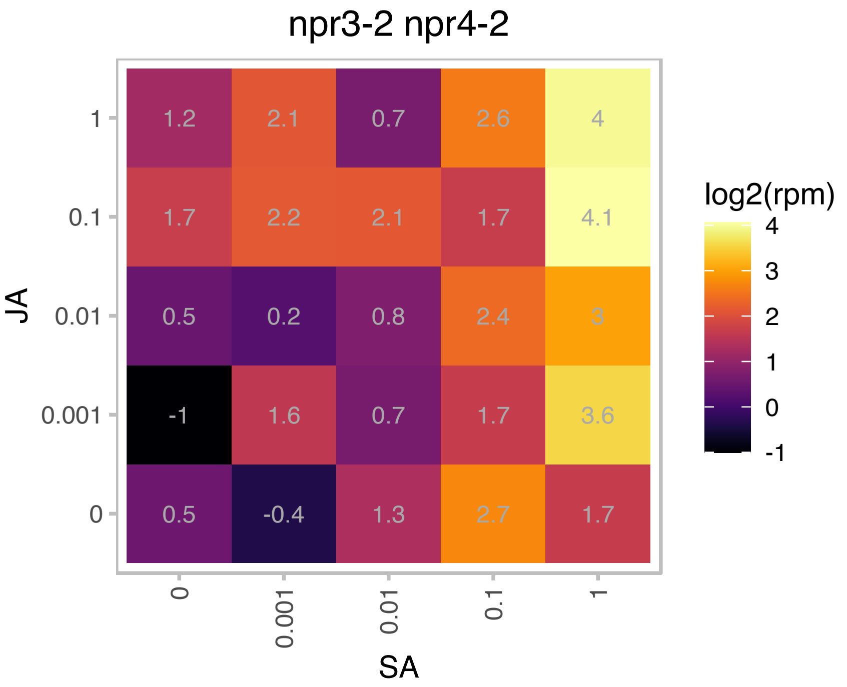


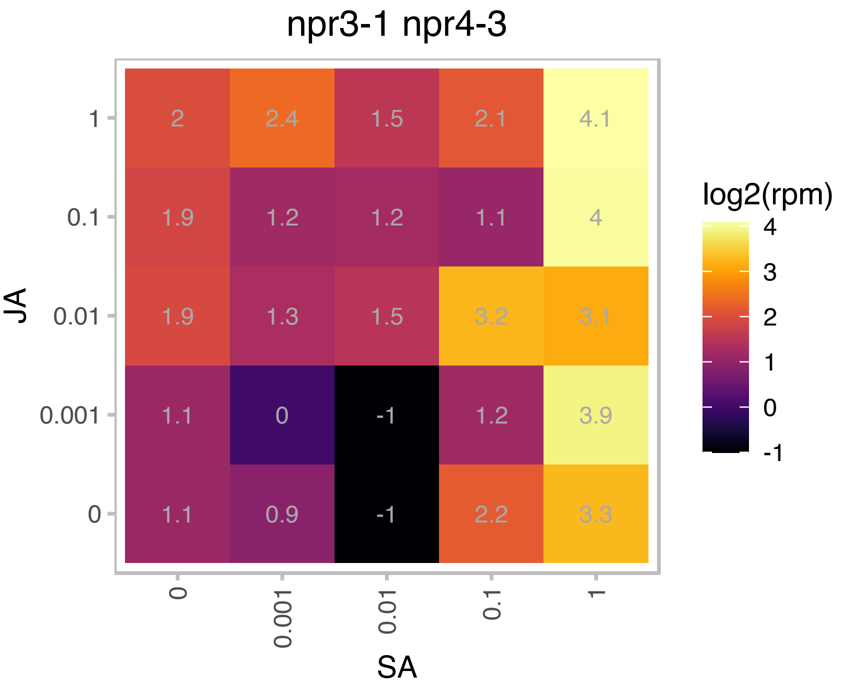

### AT2G16720.1 : ATMYB7, ATY49, MYB7

myb domain protein 7

Encodes a member of MYB3R- and R2R3- type MYB- encoding gene family that acts as a repressor of flavonol biosynthesis. AtMYB7 gene expression is induced by salt treatment.

Col-0

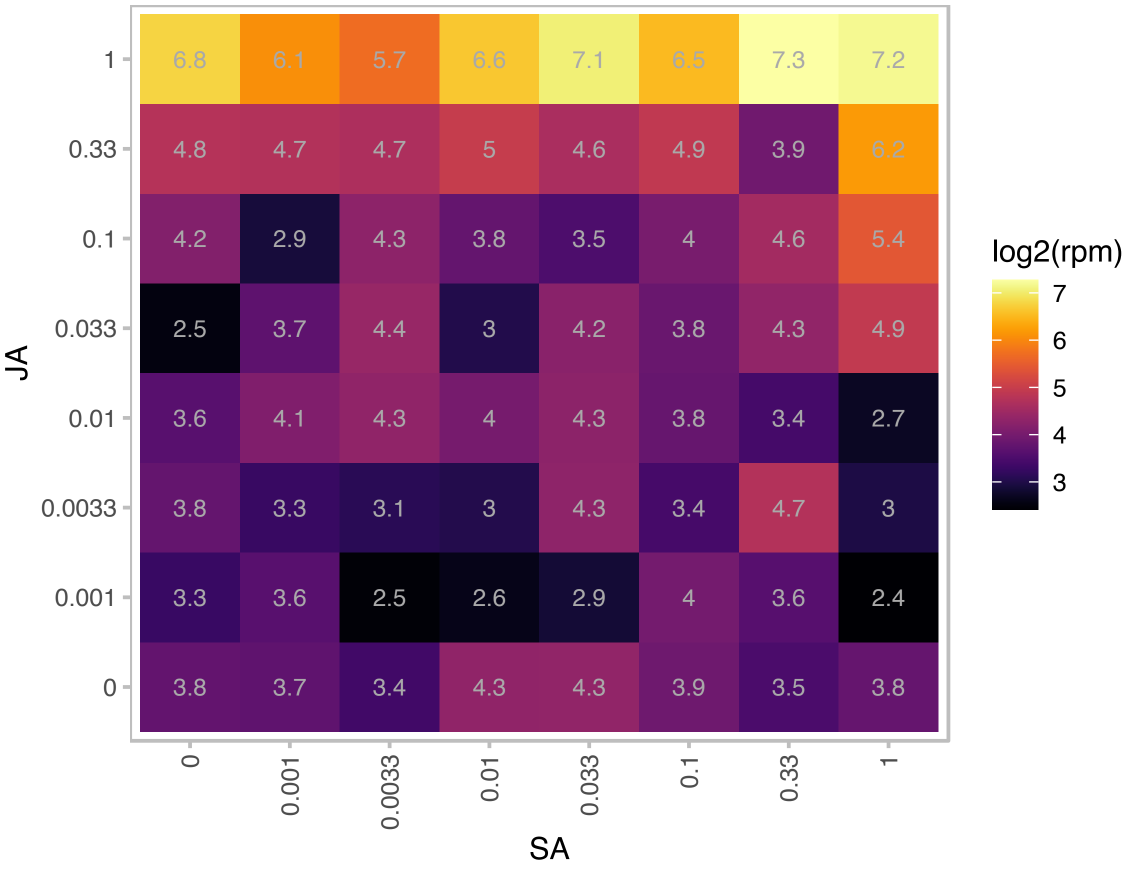


| | |
|---|---|
| Gene | AT2G16720 |
| is.expressed_genes | TRUE |
| is.responsive_genes | TRUE |
| is.apcluster | A-05 |

npr3-2 npr4-2

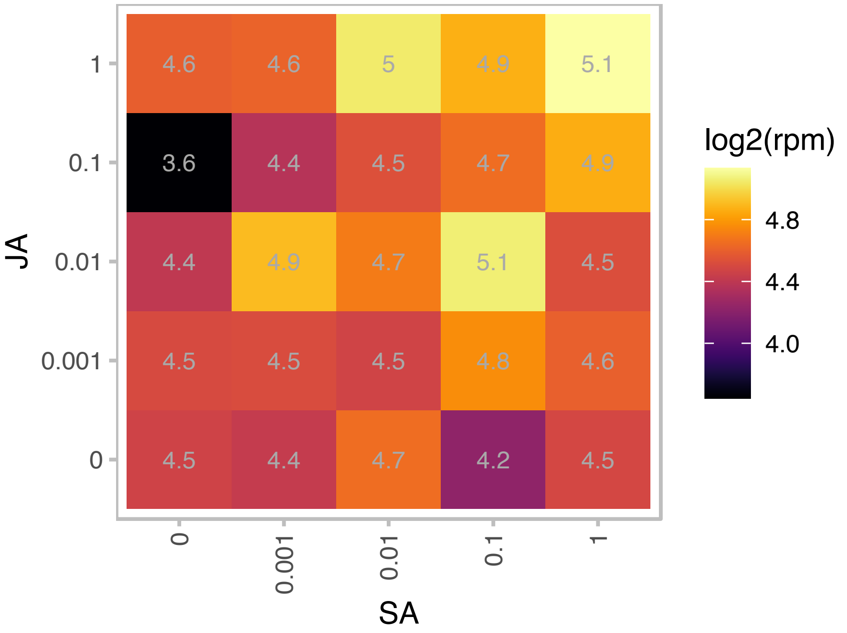


npr3-1 npr4-3

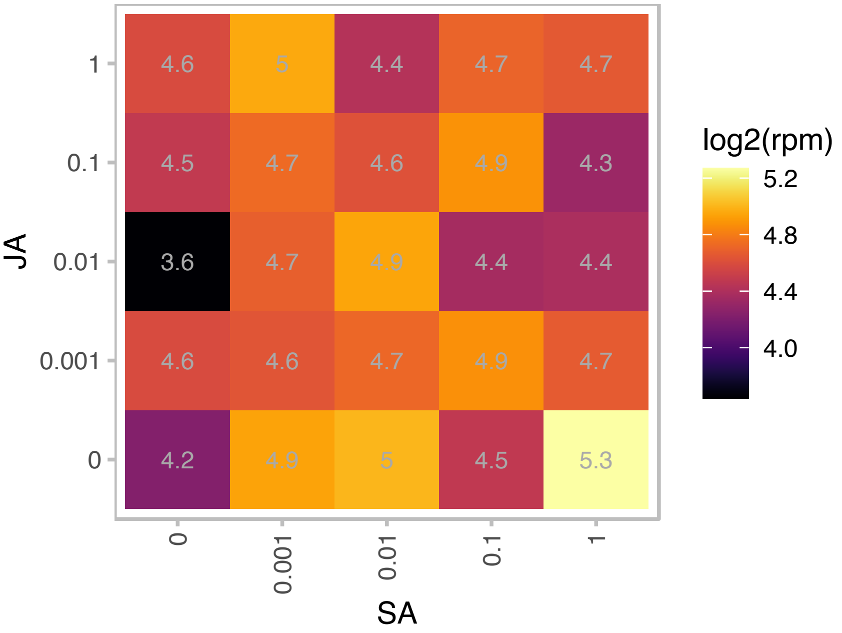

### AT2G17700.1 : STY8

ACT-like protein tyrosine kinase family protein

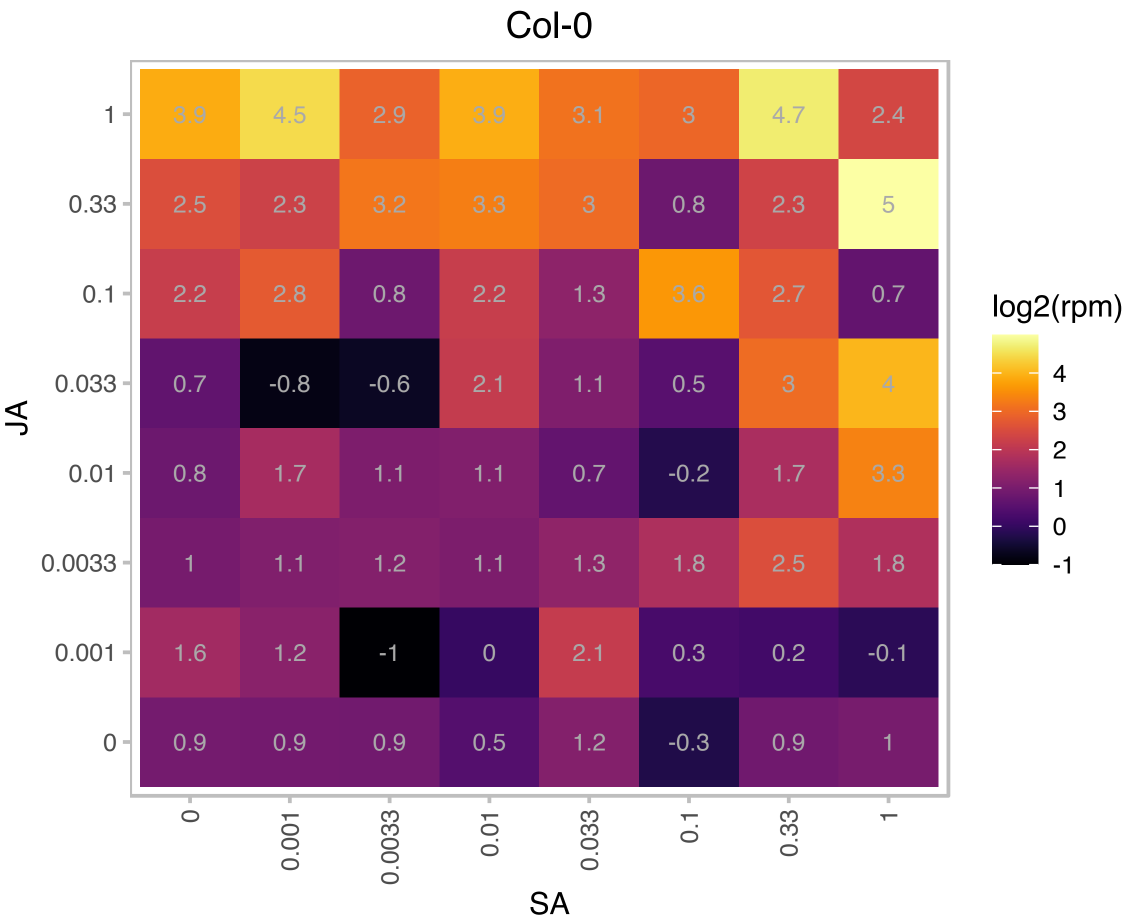


| | |
|---|---|
| Gene | AT2G17700 |
| is.expressed_genes | TRUE |
| is.responsive_genes | TRUE |
| is.apcluster | A-05 |

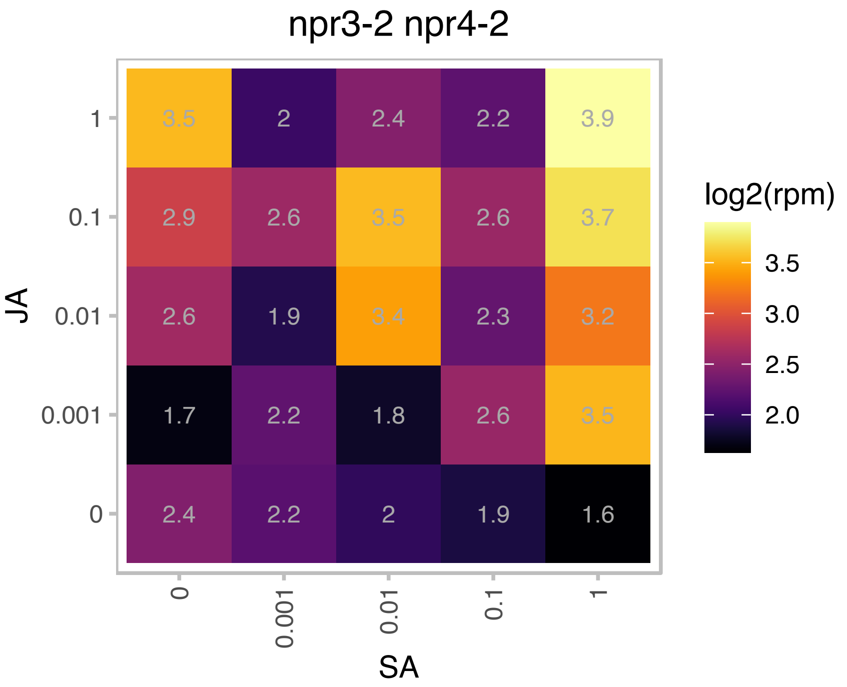


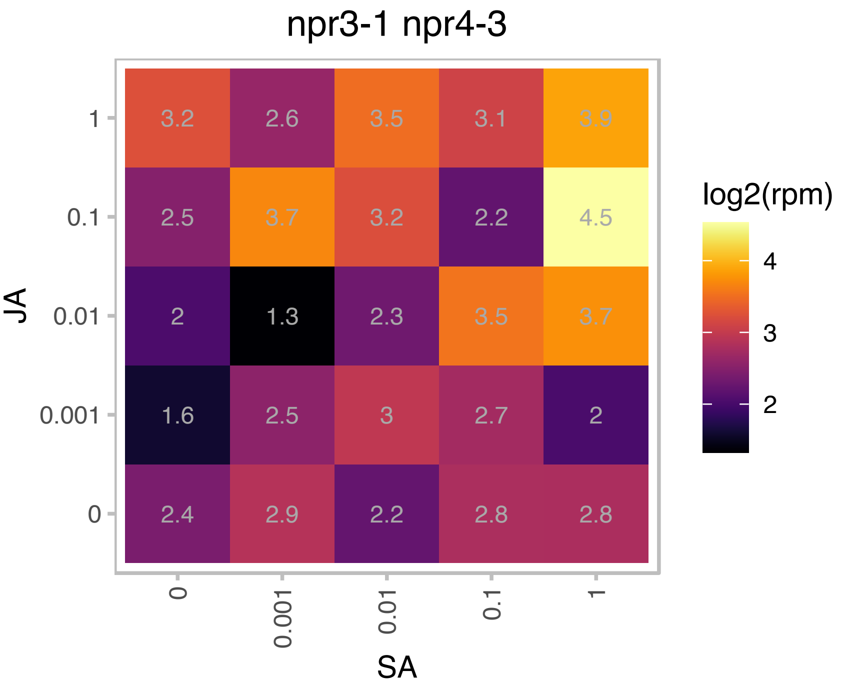

### AT2G19710.1 : ISTL5

Regulator of Vps4 activity in the MVB pathway protein

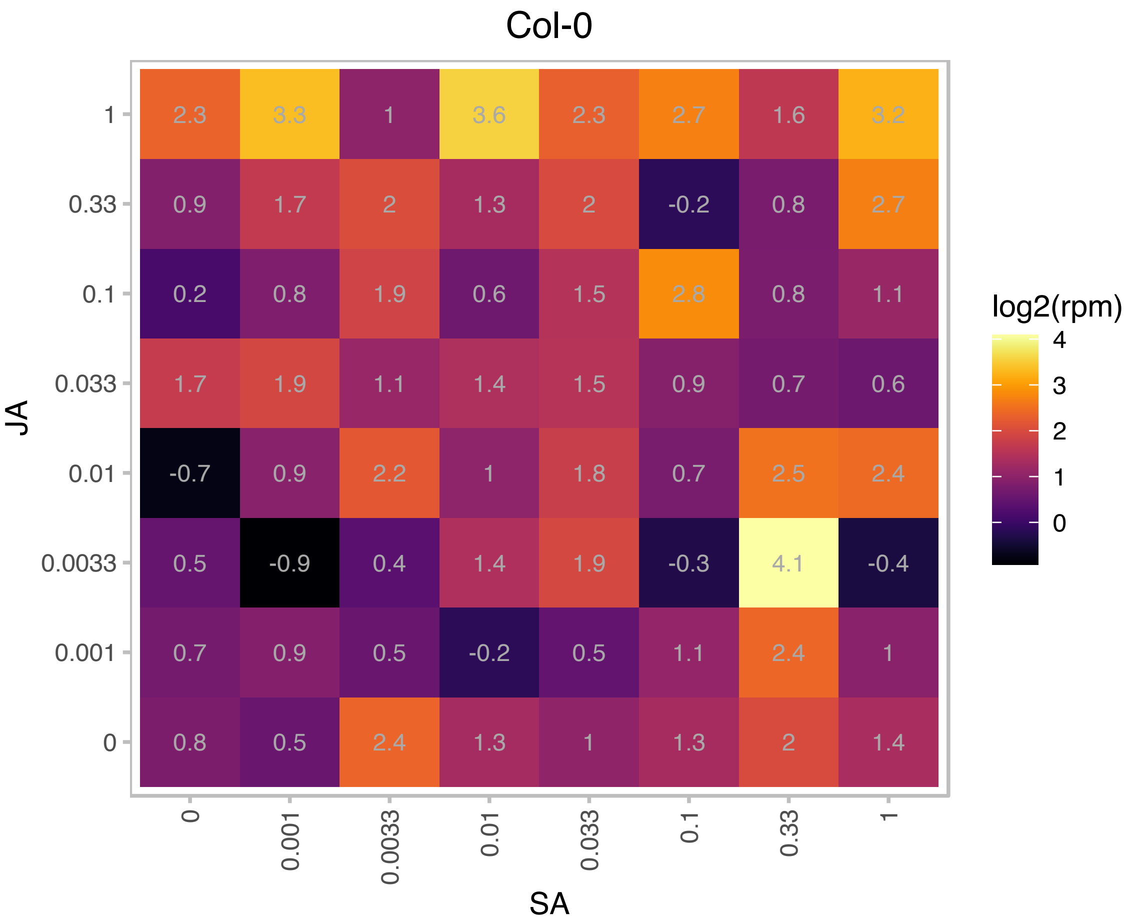


| | |
|---|---|
| Gene | AT2G19710 |
| is.expressed_genes | TRUE |
| is.responsive_genes | TRUE |
| is.apcluster | A-05 |

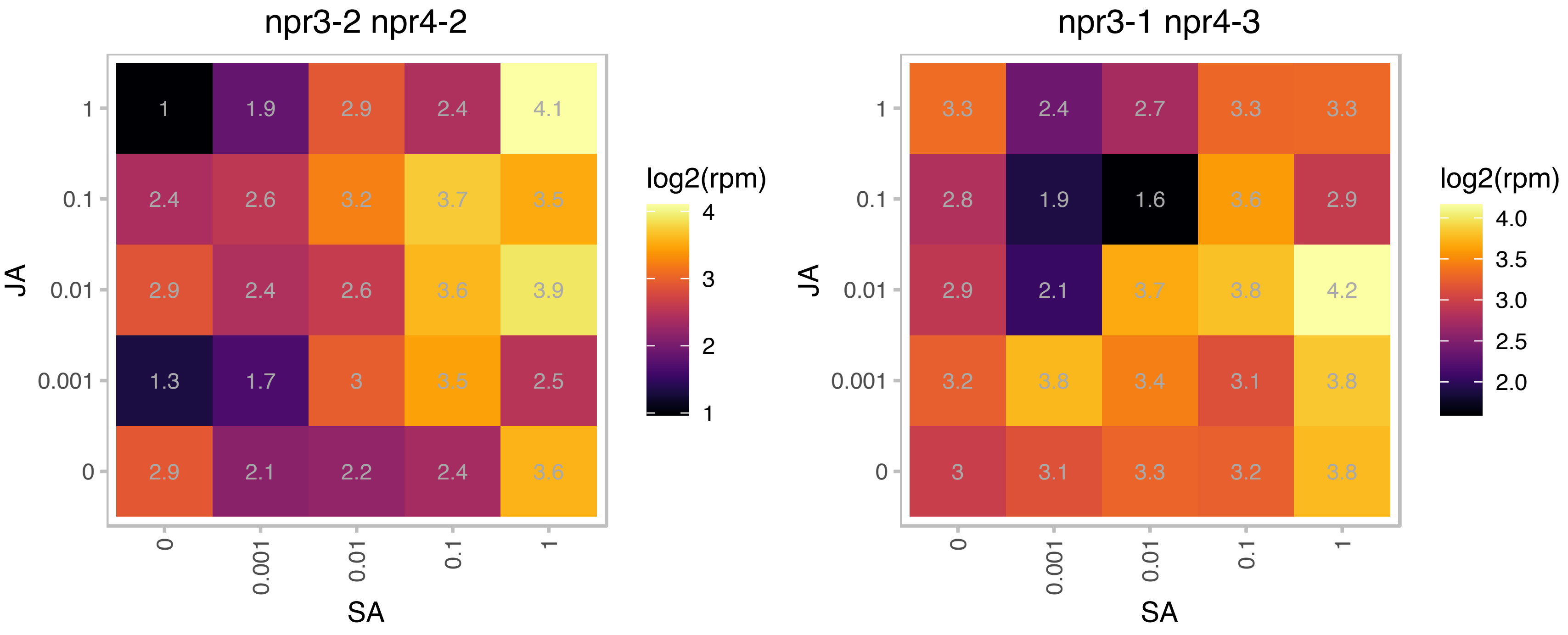

# AT2G23110.1 : NA

Late embryogenesis abundant protein, group 6

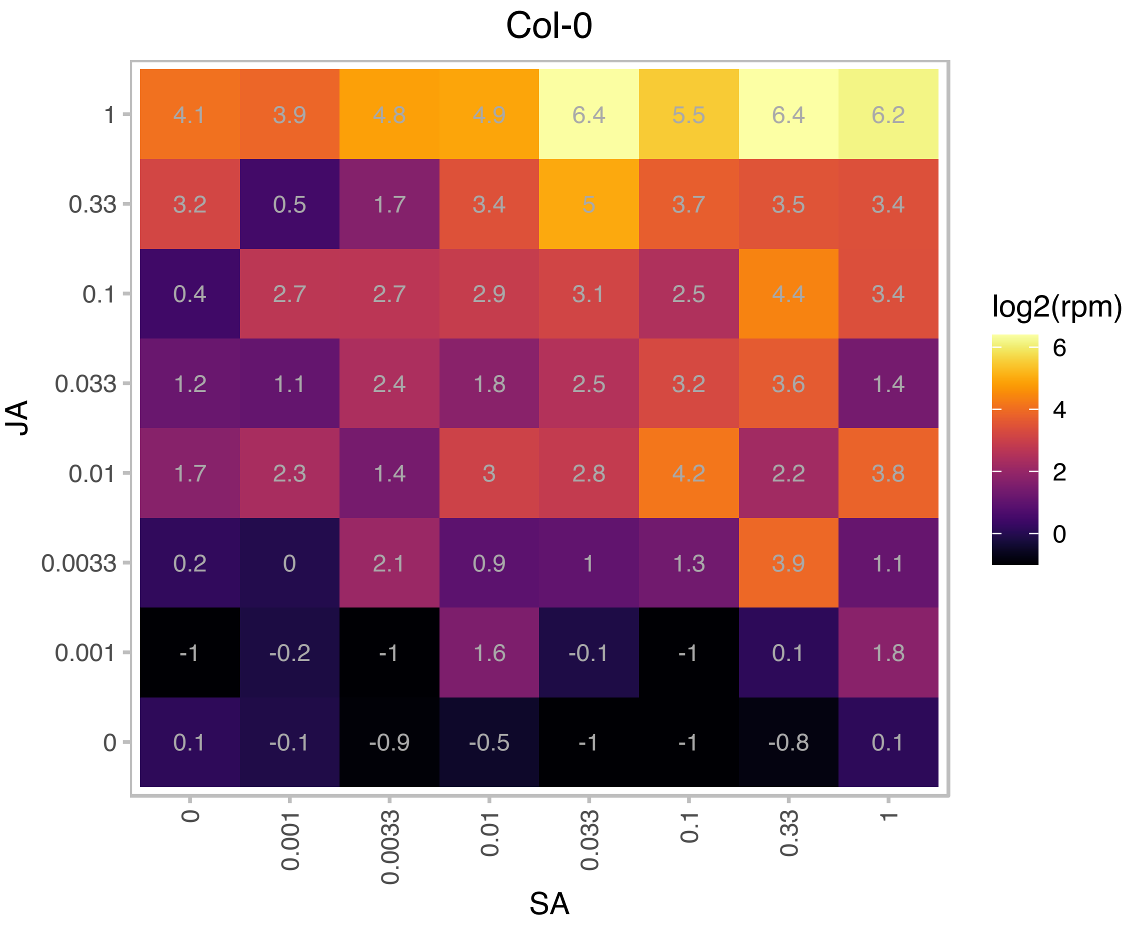


| | |
|---|---|
| Gene | AT2G23110 |
| is.expressed_genes | TRUE |
| is.responsive_genes | TRUE |
| is.apcluster | A-05 |

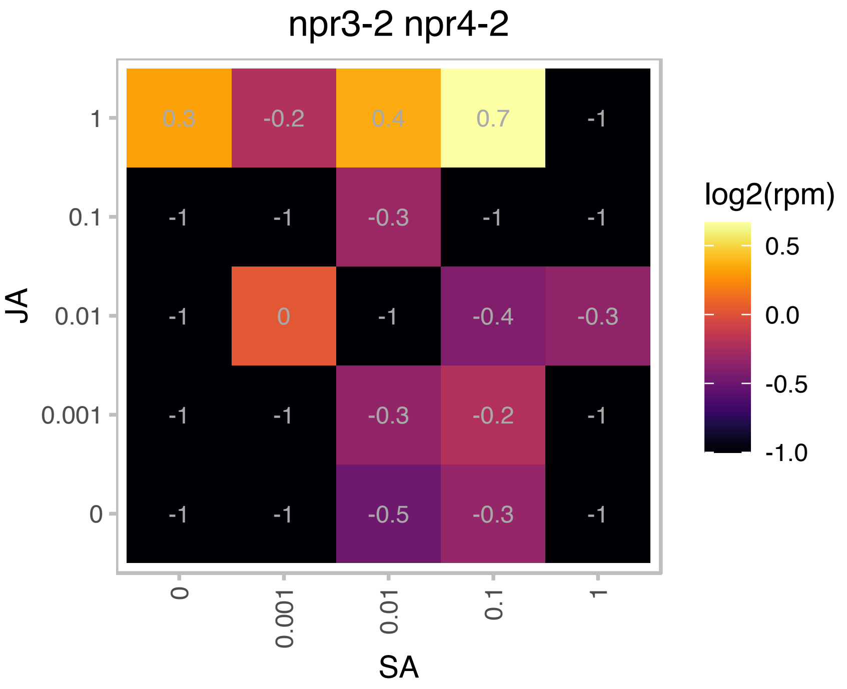


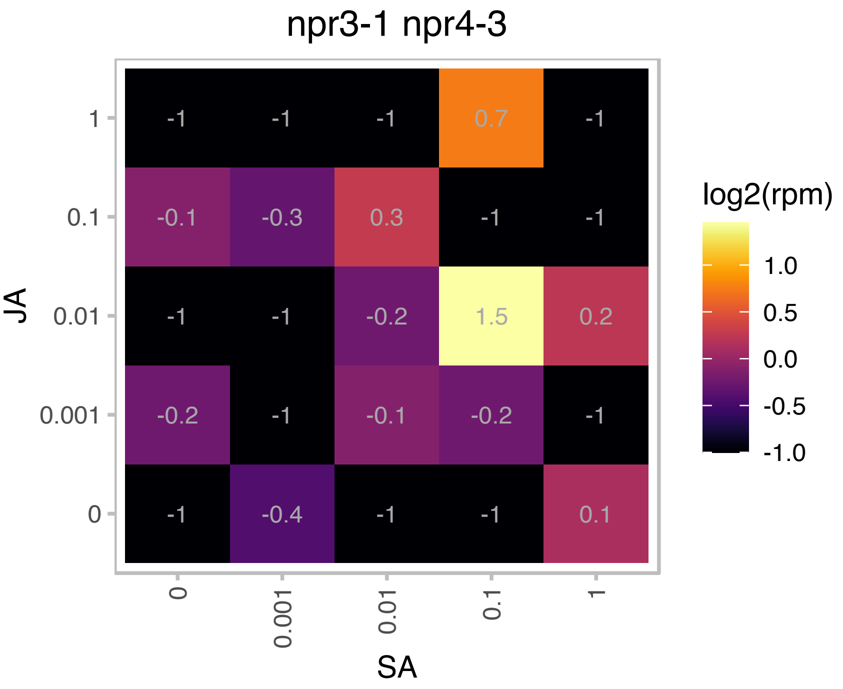

# AT2G25070.1 : NA

Protein phosphatase 2C family protein

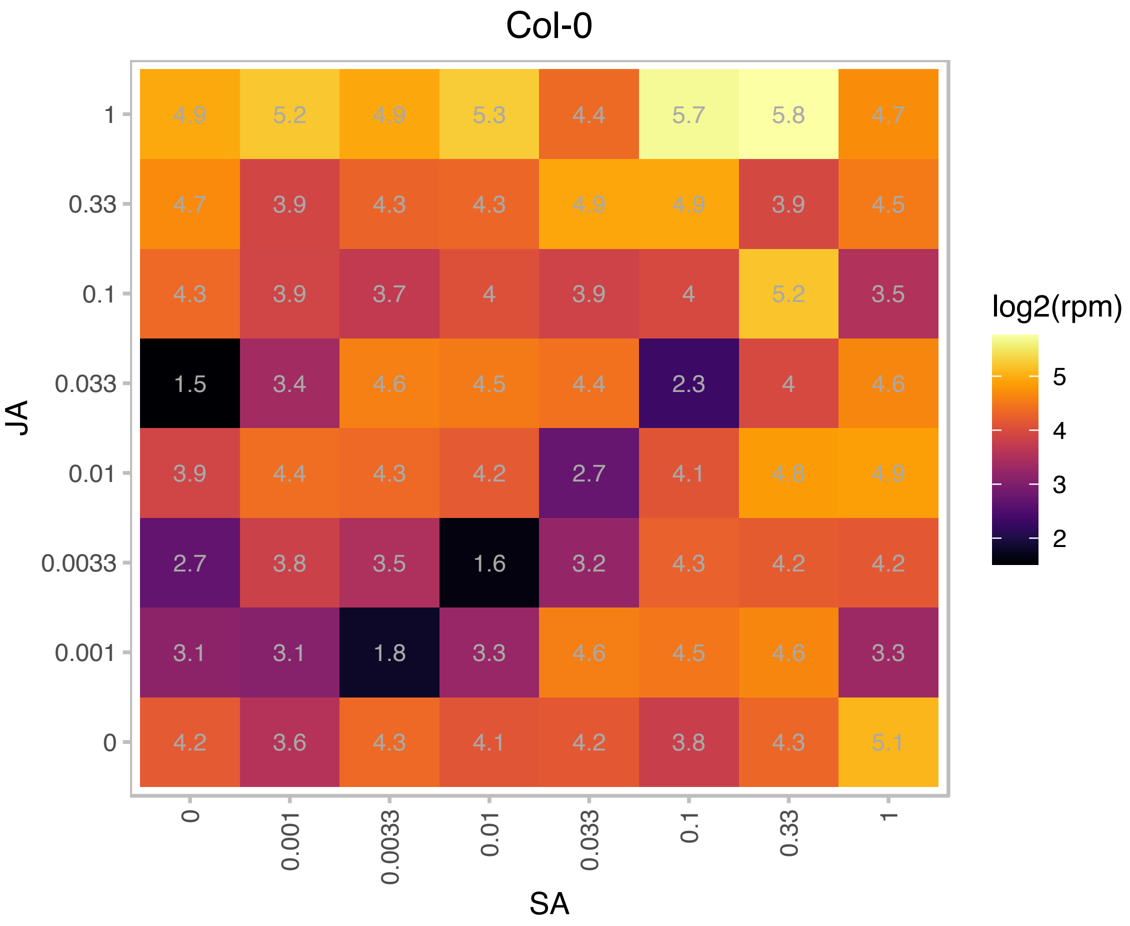


| | |
|---|---|
| Gene | AT2G25070 |
| is.expressed_genes | TRUE |
| is.responsive_genes | TRUE |
| is.apcluster | A-05 |

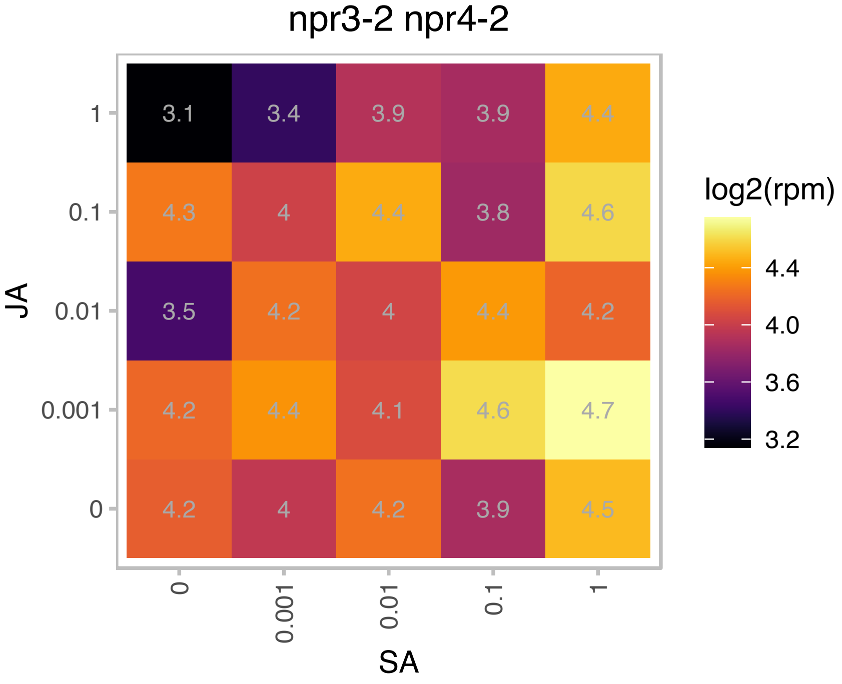


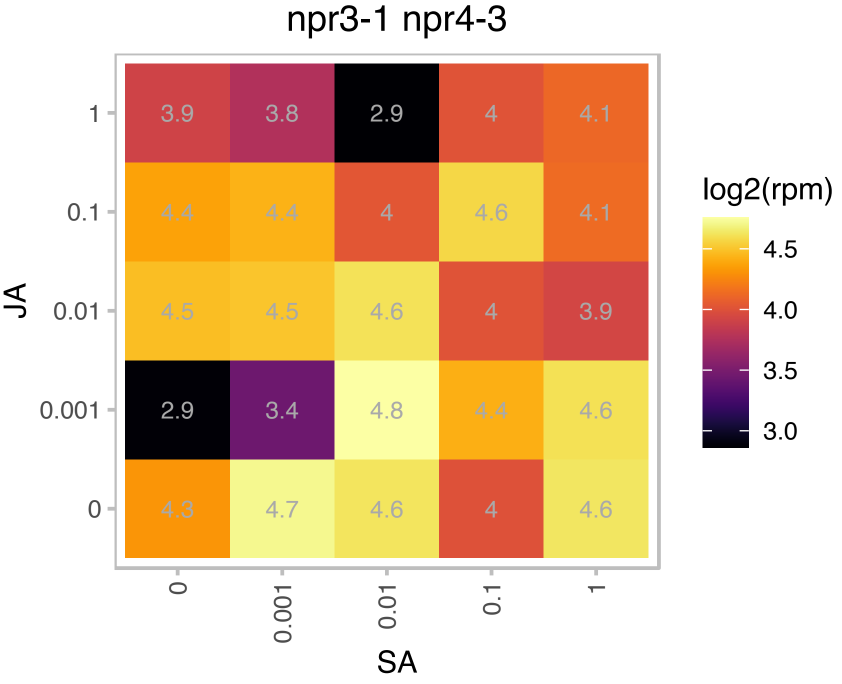

### AT2G26770.1 : SCAB1

plectin-related

Encodes a plant-specific actin binding protein SCAB1 (STOMATAL CLOSURE-RELATED ACTIN BINDING PROTEI
SCAB1 stabilizes actin filaments and regulates stomatal movement.

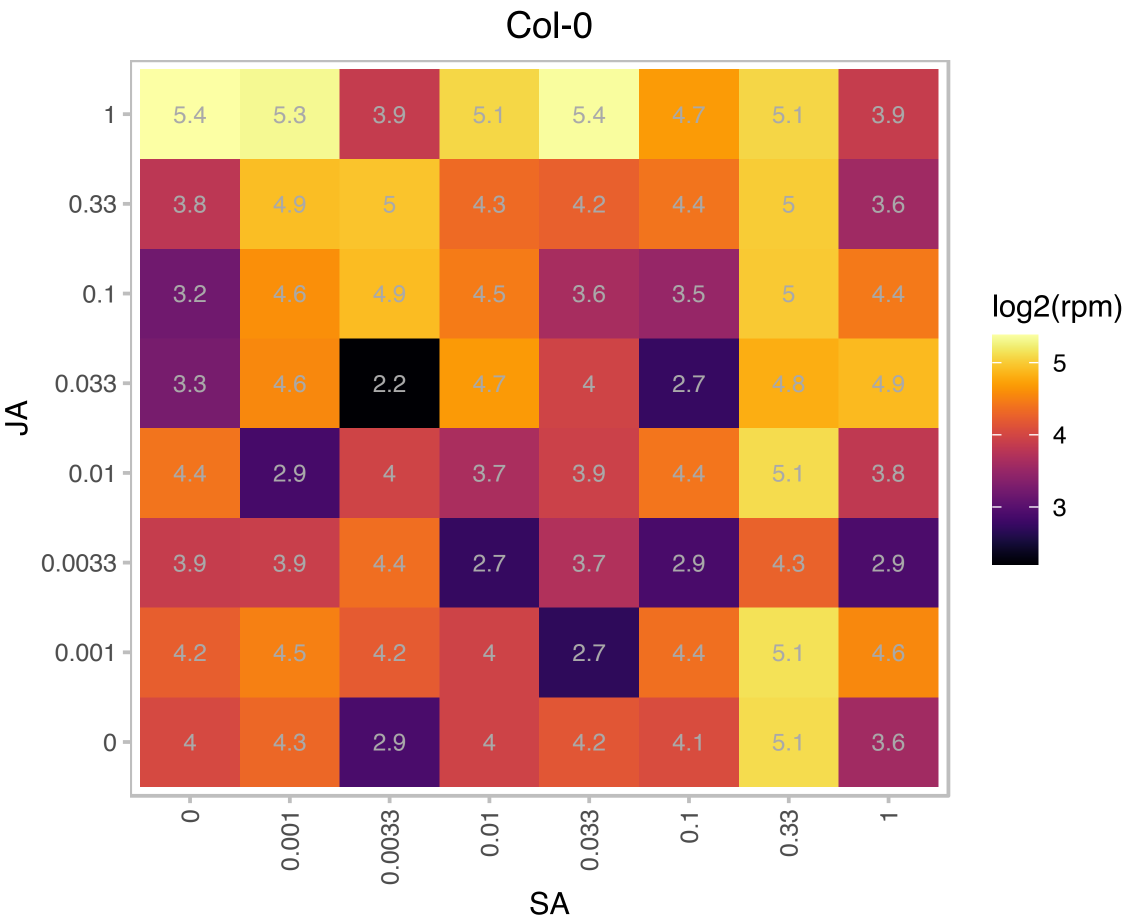


| | |
|---|---|
| Gene | AT2G26770 |
| is.expressed_genes | TRUE |
| is.responsive_genes | TRUE |
| is.apcluster | A-05 |

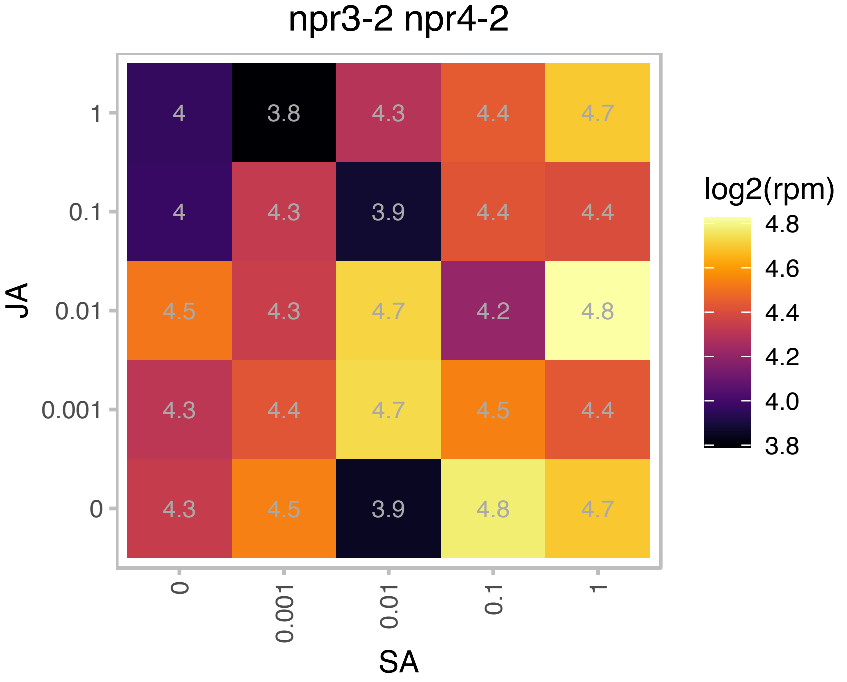


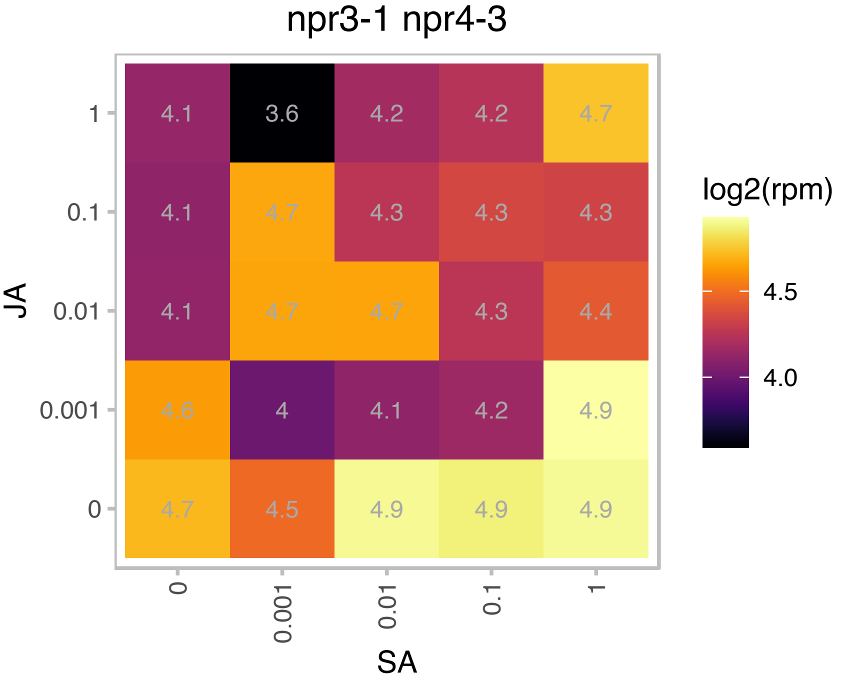

# AT2G27090.1 : NA

Protein of unknown function (DUF630 and DUF632)

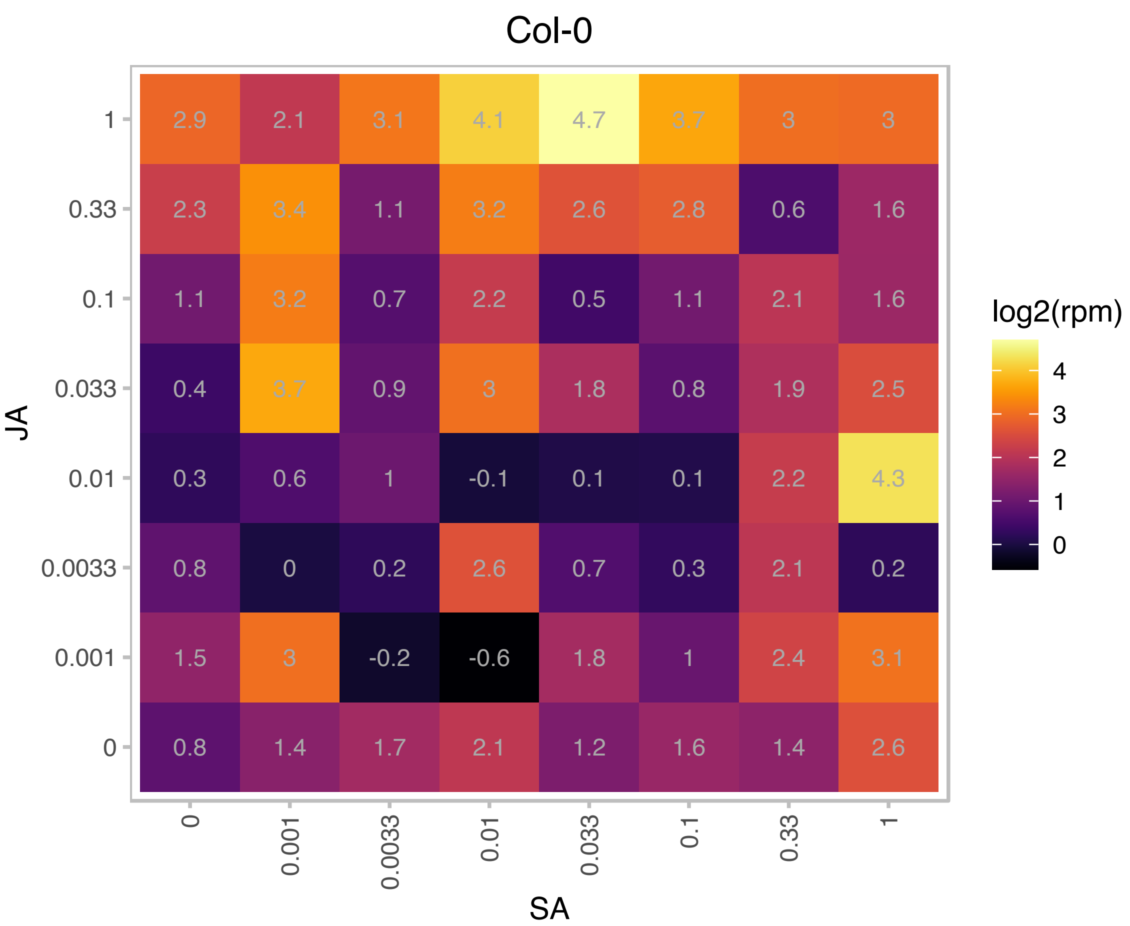


| | |
|---|---|
| Gene | AT2G27090 |
| is.expressed_genes | TRUE |
| is.responsive_genes | TRUE |
| is.apcluster | A-05 |

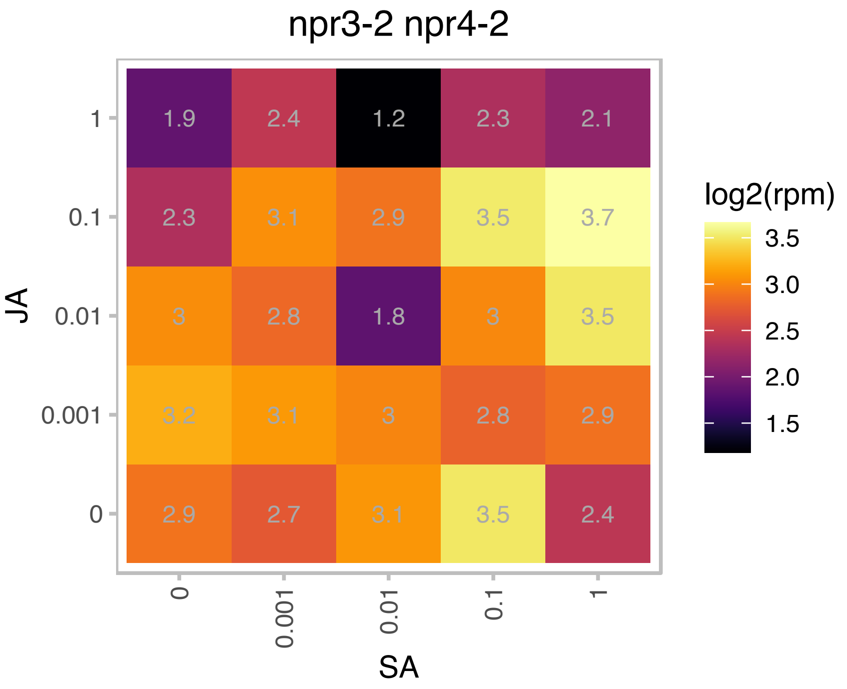


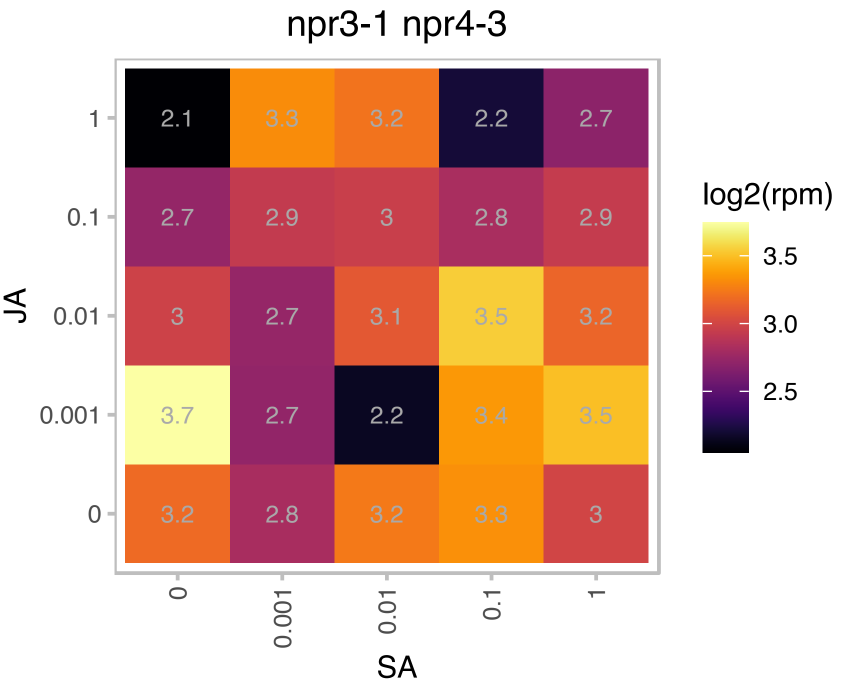

### AT2G29470.1 : ATGSTU3, GST21, GSTU3

glutathione S-transferase tau 3

Encodes glutathione transferase belonging to the tau class of GSTs. Naming convention according to Wagner et al. (2002).

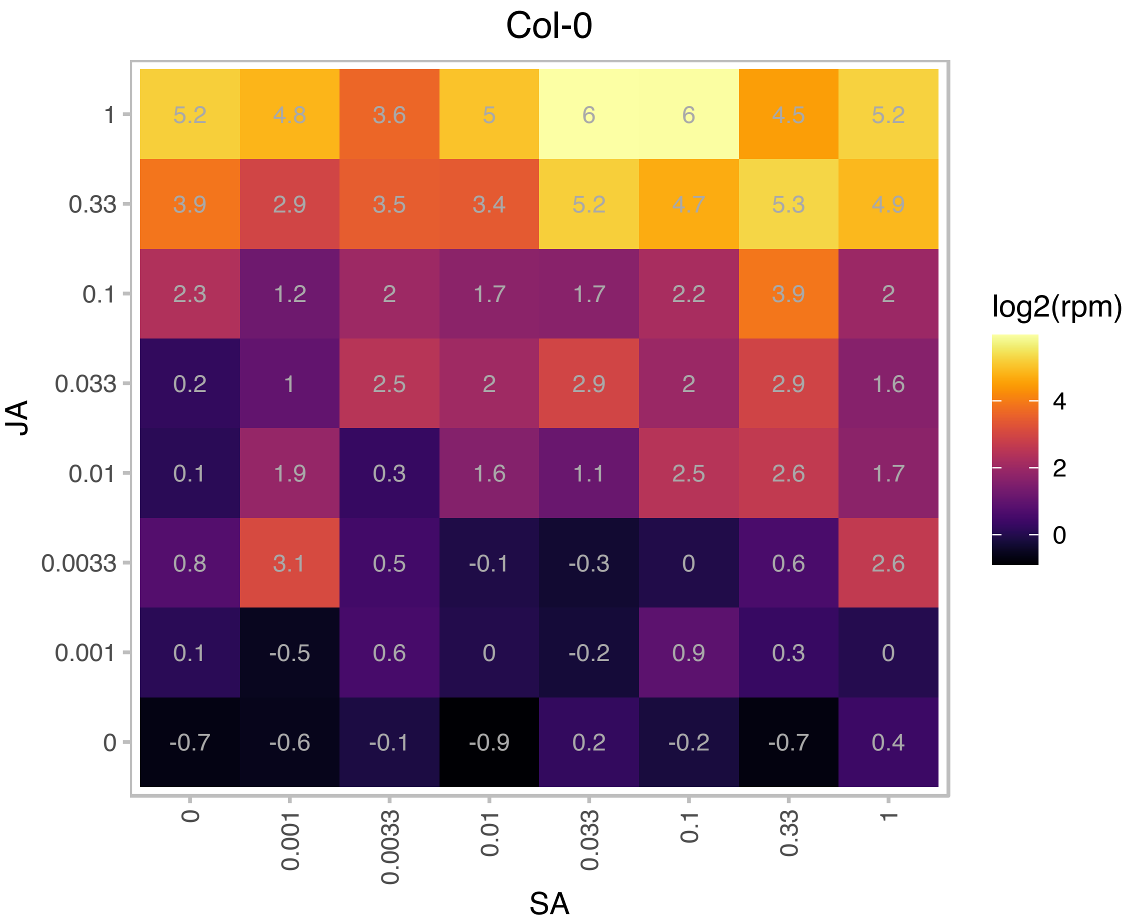


| | |
|---|---|
| Gene | AT2G29470 |
| is.expressed_genes | TRUE |
| is.responsive_genes | TRUE |
| is.apcluster | A-05 |

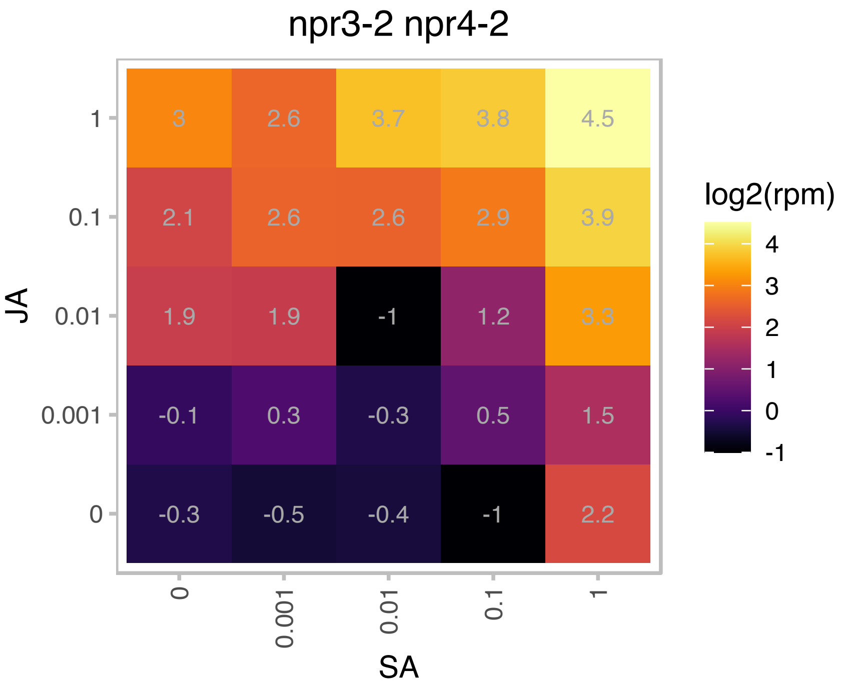


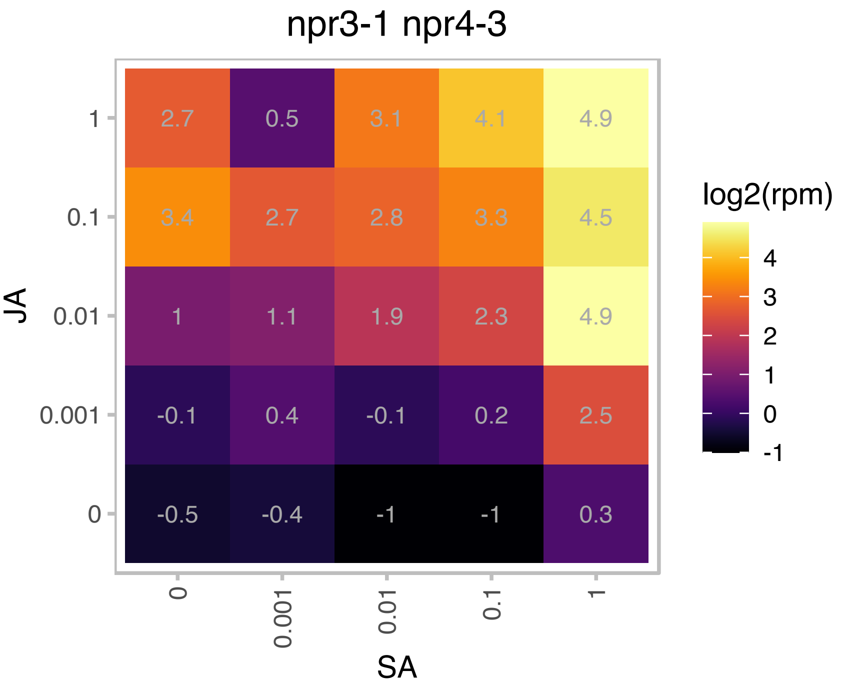

### AT2G42160.1 : BRIZ1

zinc finger (ubiquitin-hydrolase) domain-containing protein

Encodes a RING domain containing protein BRIZ1. BRIZ1 (At2g42160) and BRIZ2 (At2g26000) proteins form a heteromeric E3 ligase complex required for seed germination and post-germination growth.

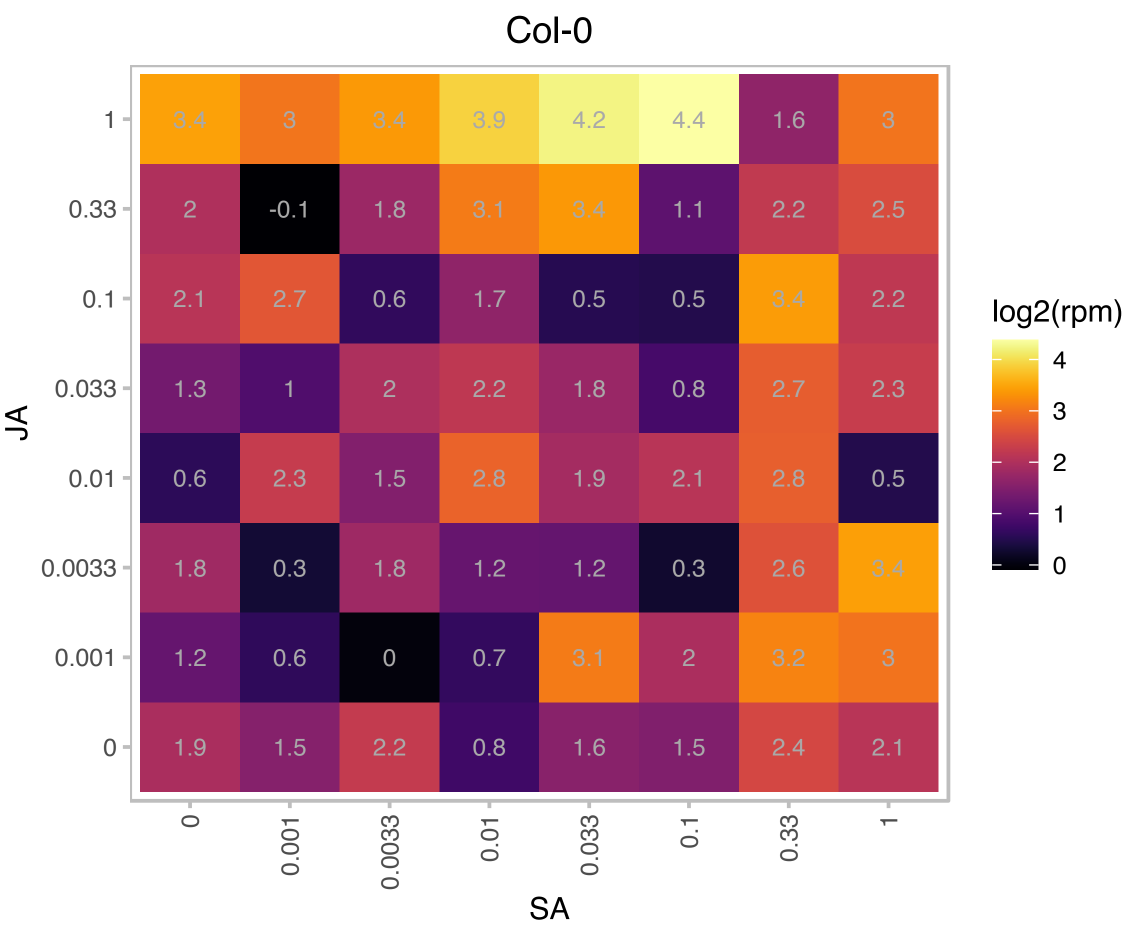


| | |
|---|---|
| Gene | AT2G42160 |
| is.expressed_genes | TRUE |
| is.responsive_genes | TRUE |
| is.apcluster | A-05 |

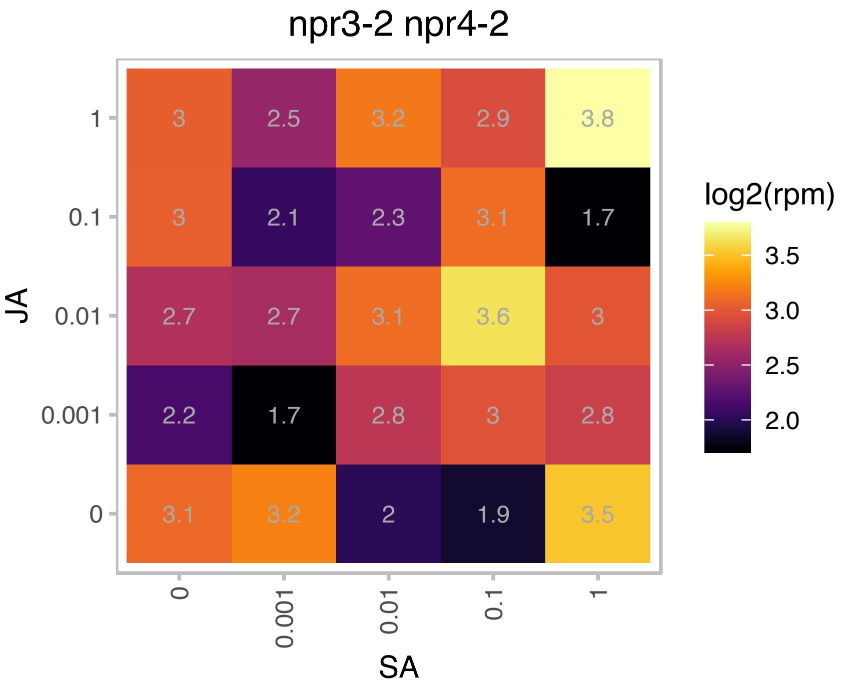


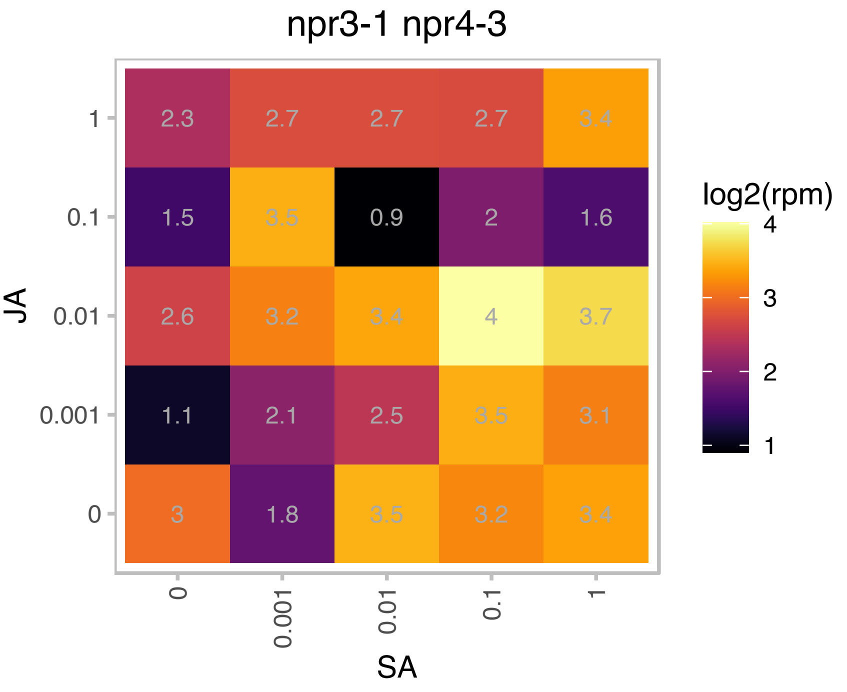

### AT2G44100.1 : AT-GDI1, ATGDI1, GDI1

guanosine nucleotide diphosphate dissociation inhibitor 1

GDP dissociation inhibitor involved in vesicular membrane traffic

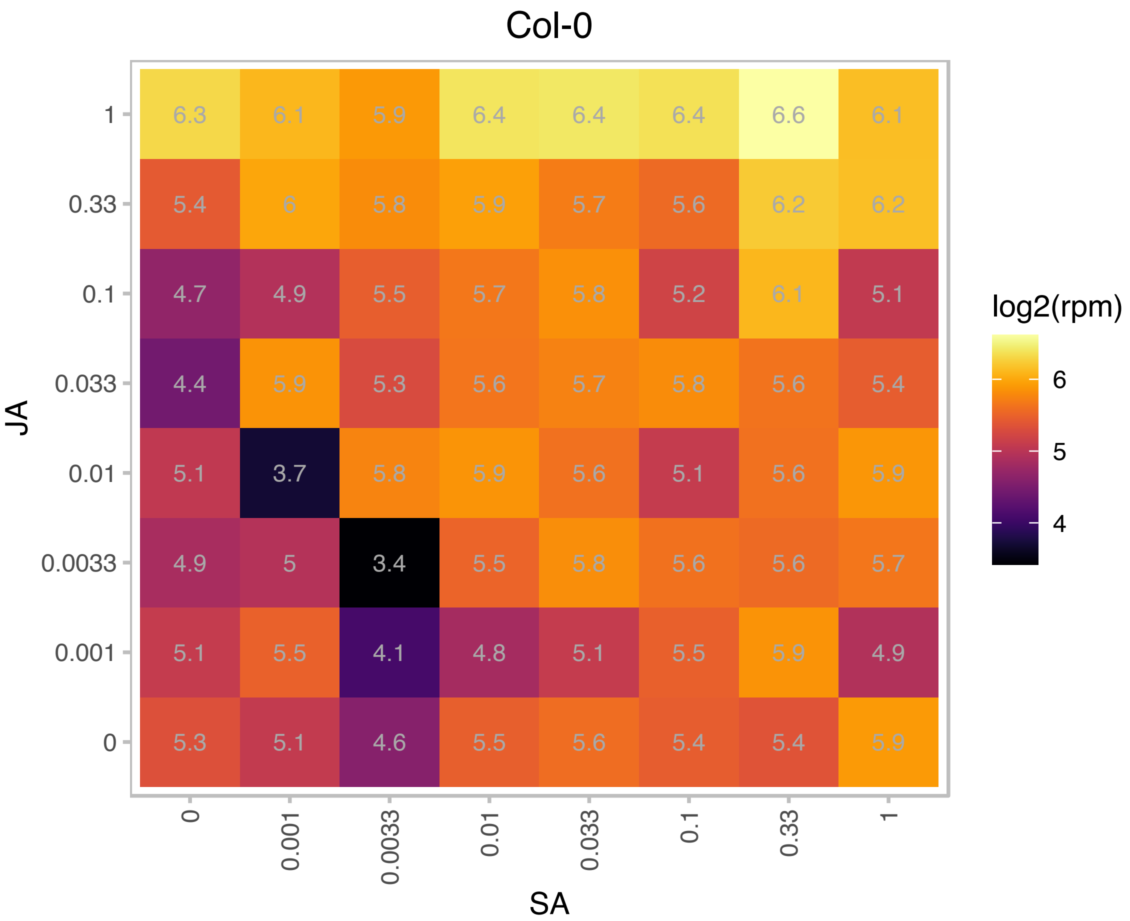


| | |
|---|---|
| Gene | AT2G44100 |
| is.expressed_genes | TRUE |
| is.responsive_genes | TRUE |
| is.apcluster | A-05 |

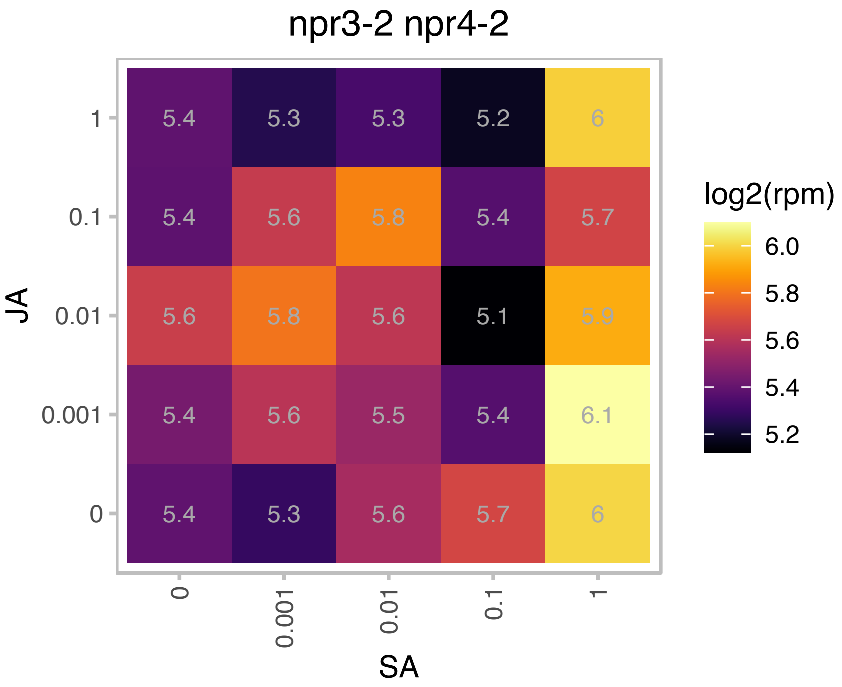


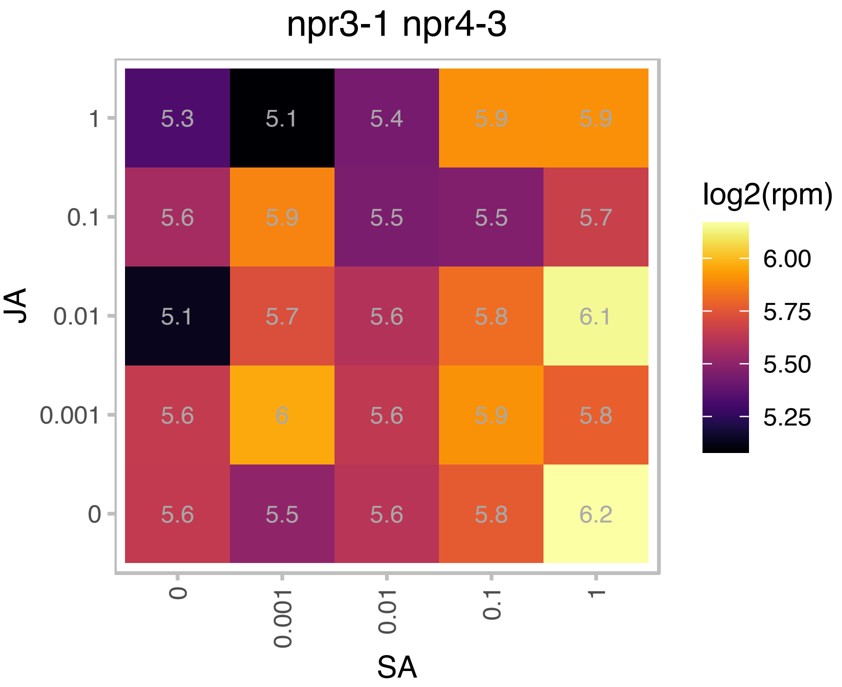

# AT2G47950.1 : NA

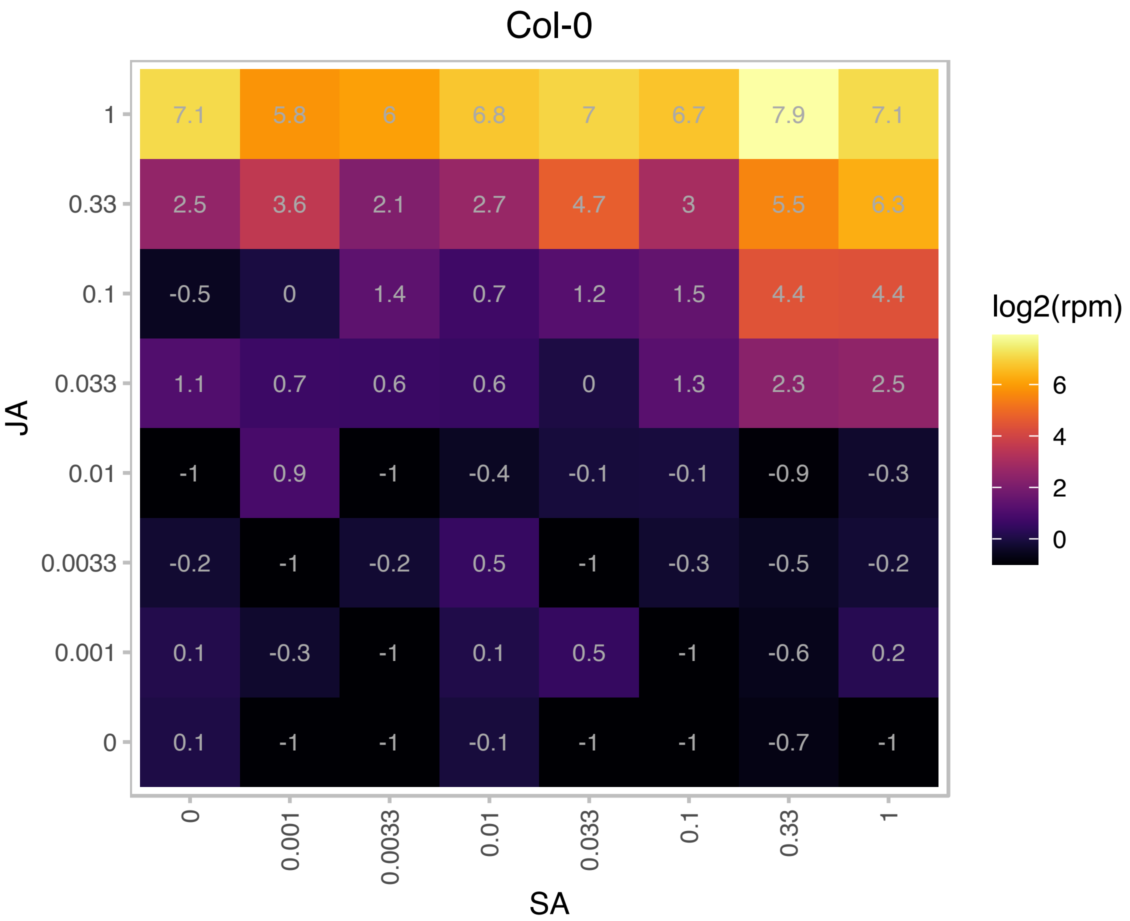


| | |
|---|---|
| Gene | AT2G47950 |
| is.expressed_genes | TRUE |
| is.responsive_genes | TRUE |
| is.apcluster | A-05 |

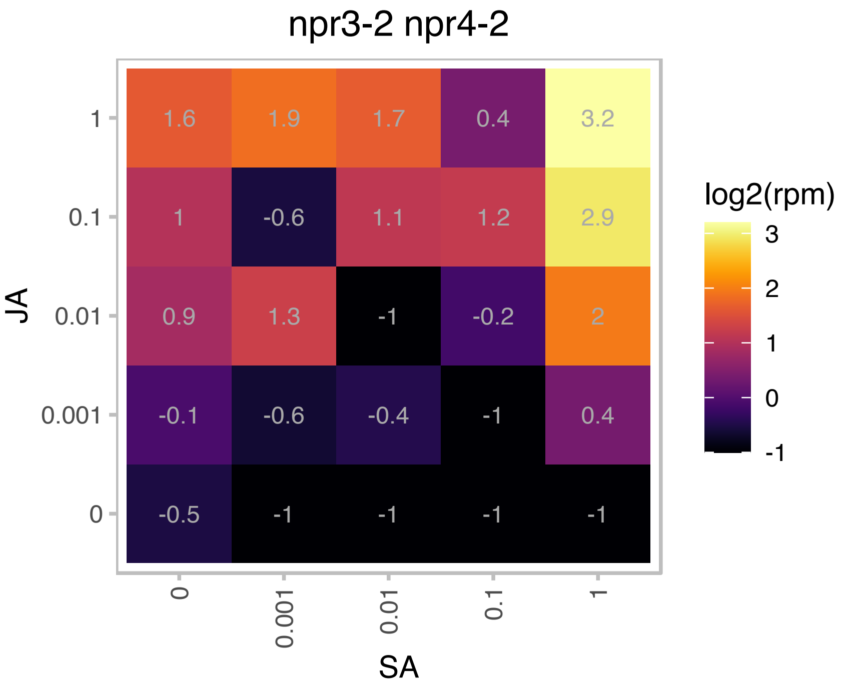


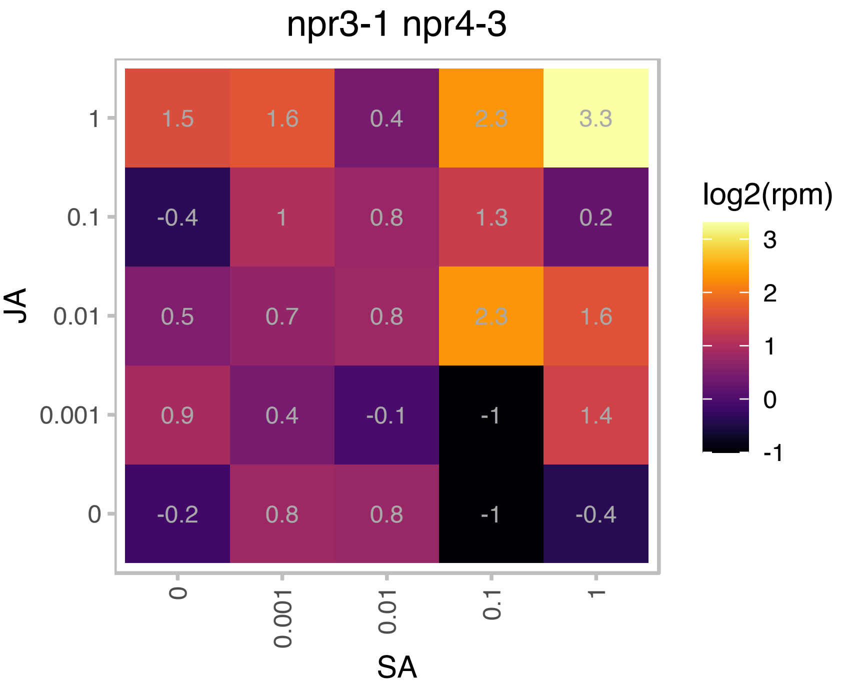

### AT3G02875.1 : ILR1

Peptidase M20/M25/M40 family protein

Hydrolyzes amino acid conjugates of the plant growth regulator indole-3-acetic acid (IAA), including IAA-Leu and IAA-Phe. Uses Mg and Co ions as cofactors.

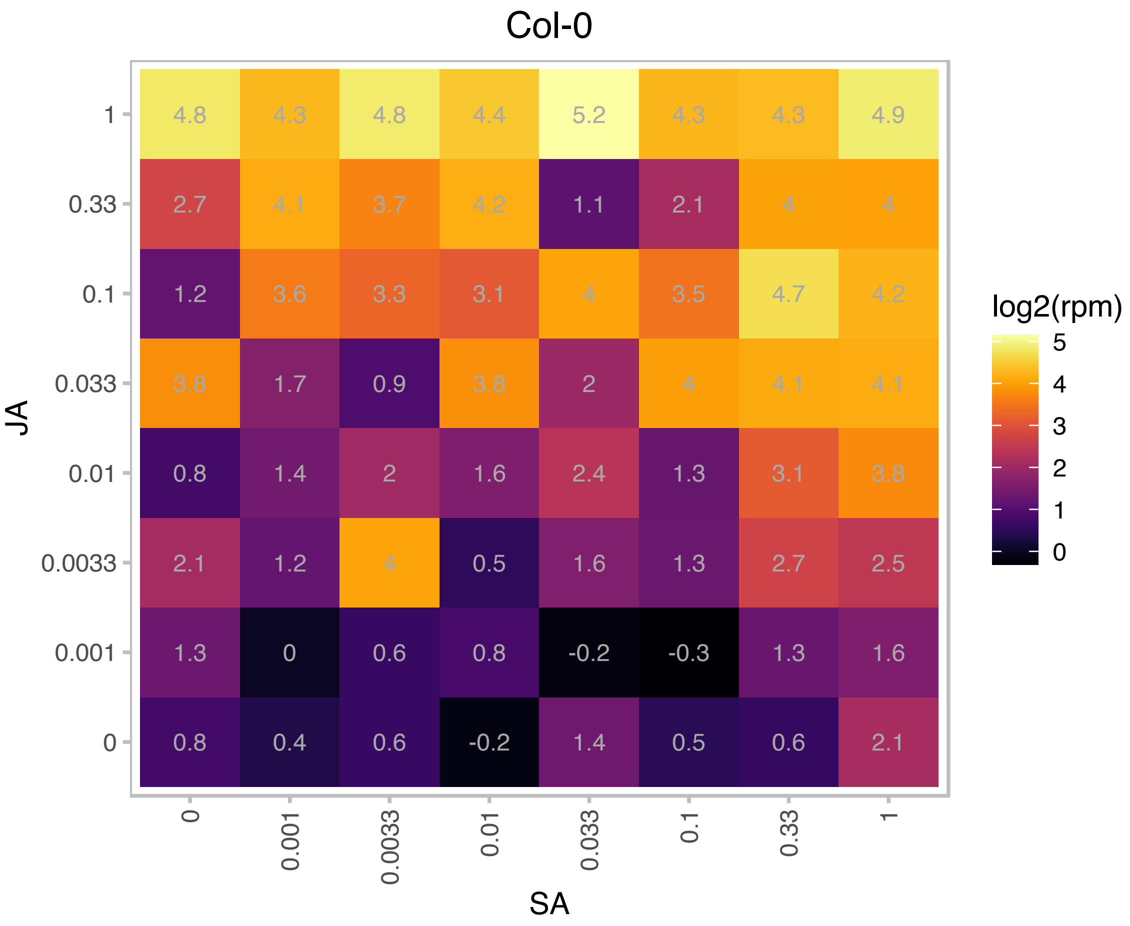


| | |
|---|---|
| Gene | AT3G02875 |
| is.expressed_genes | TRUE |
| is.responsive_genes | TRUE |
| is.apcluster | A-05 |

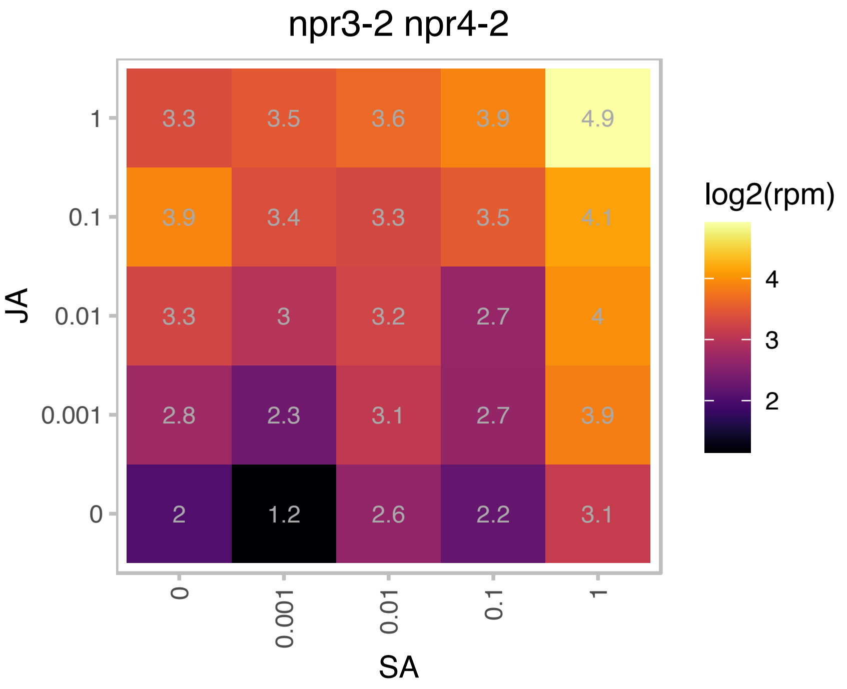


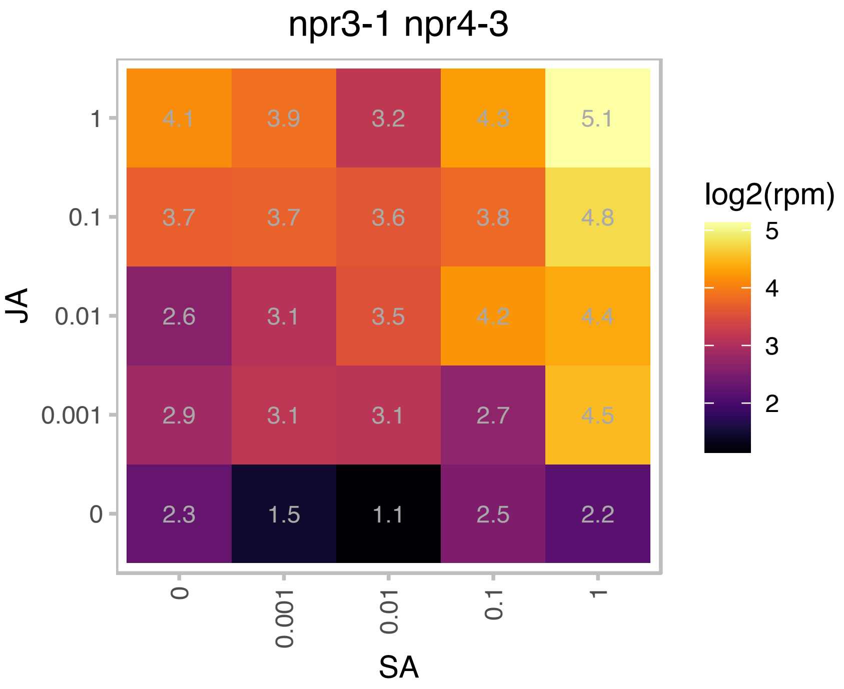

# AT3G05170.1 : NA

Phosphoglycerate mutase family protein

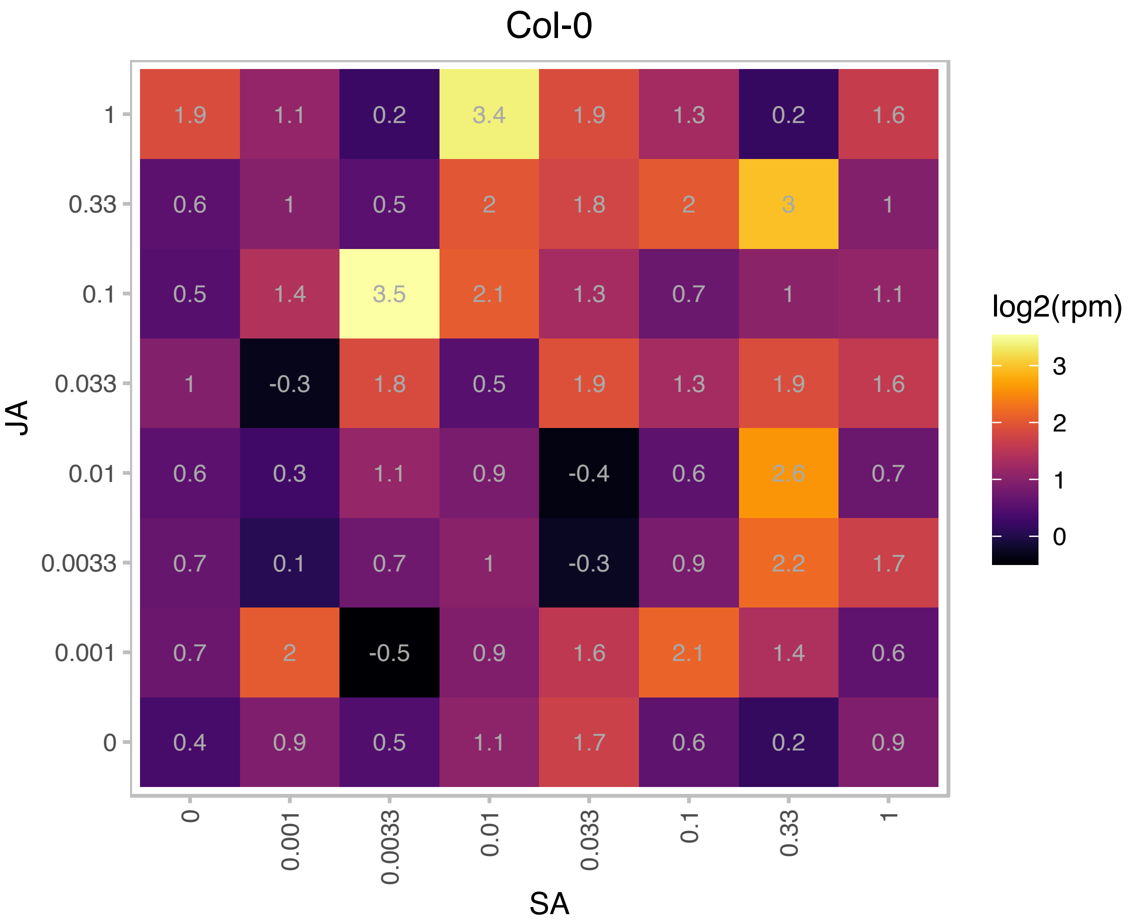


| | |
|---|---|
| Gene | AT3G05170 |
| is.expressed_genes | TRUE |
| is.responsive_genes | TRUE |
| is.apcluster | A-05 |

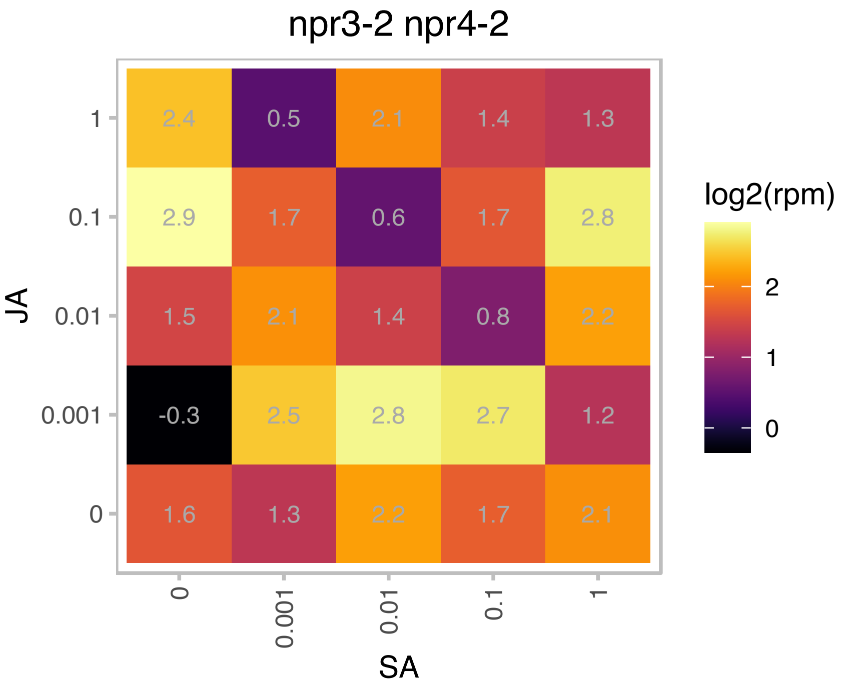


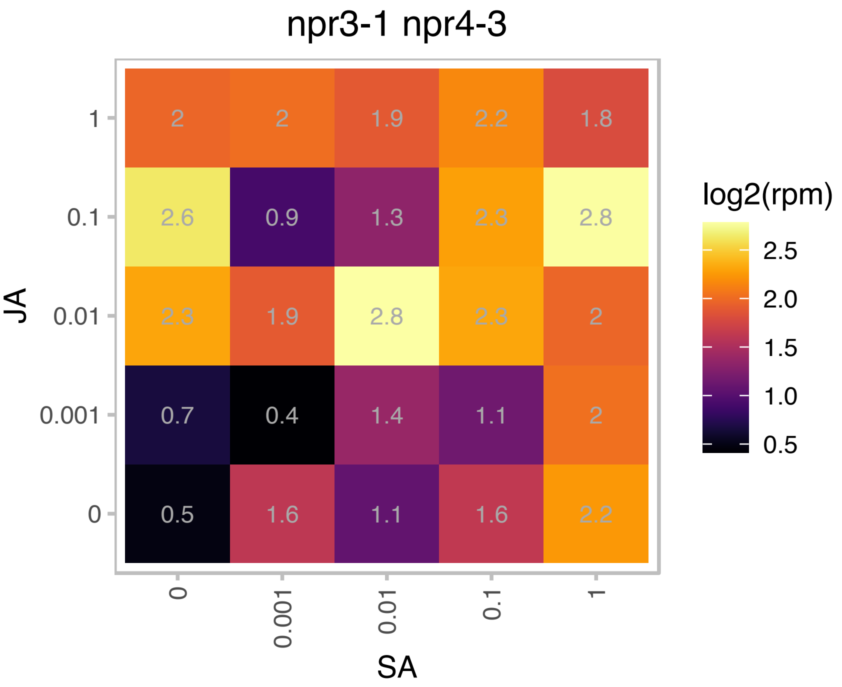

### AT3G06490.1 : AtMYB108, BOS1, MYB108

myb domain protein 108

putative transcription factor MYB108 (MYB108) mRNA,

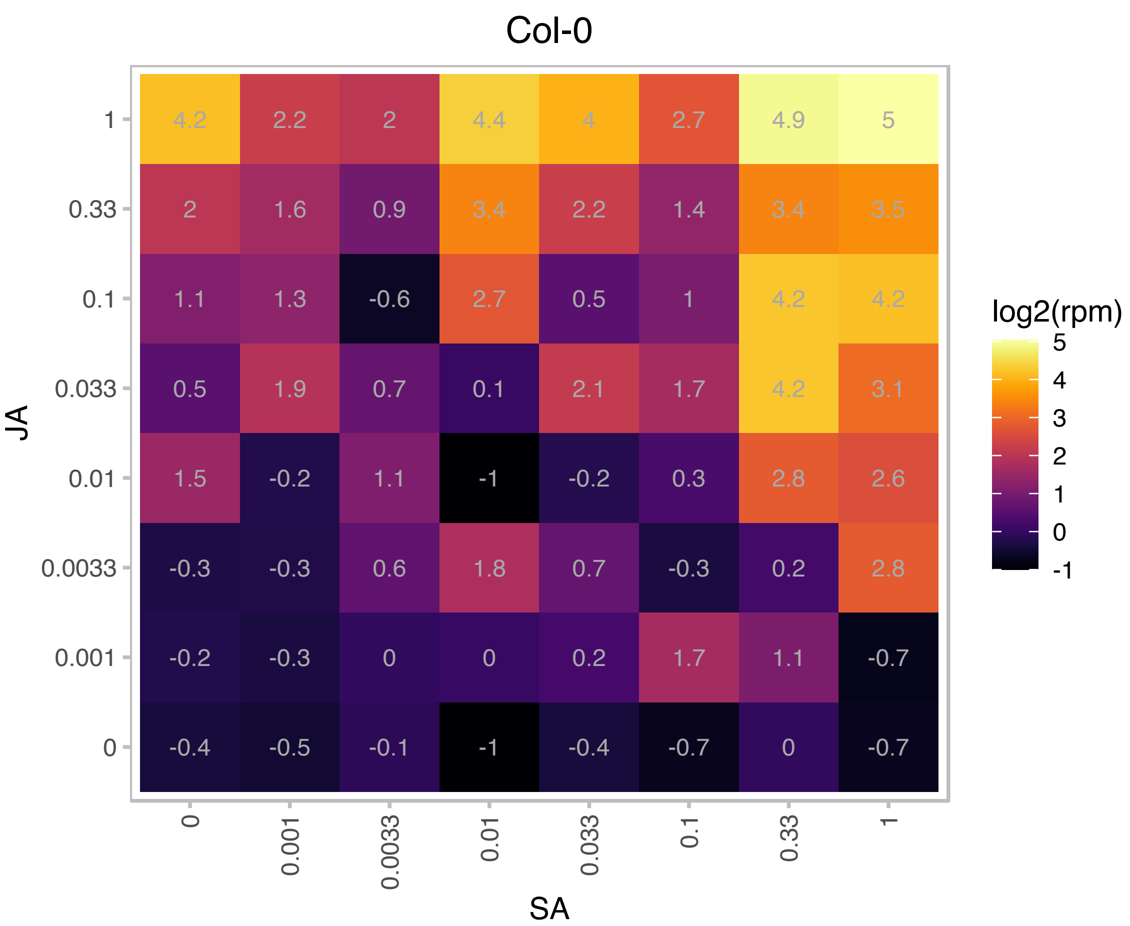


| | |
|---|---|
| Gene | AT3G06490 |
| is.expressed_genes | TRUE |
| is.responsive_genes | TRUE |
| is.apcluster | A-05 |

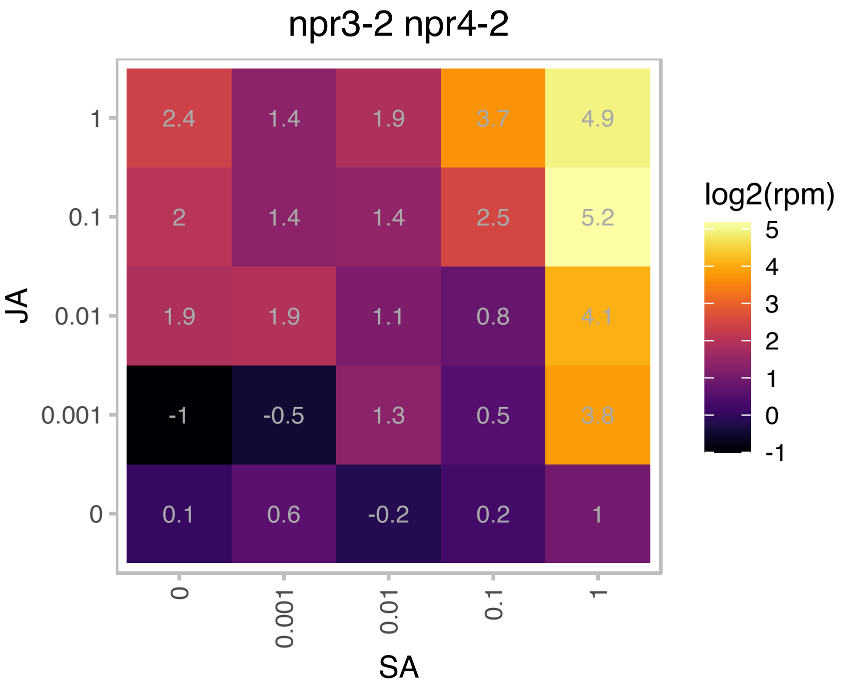


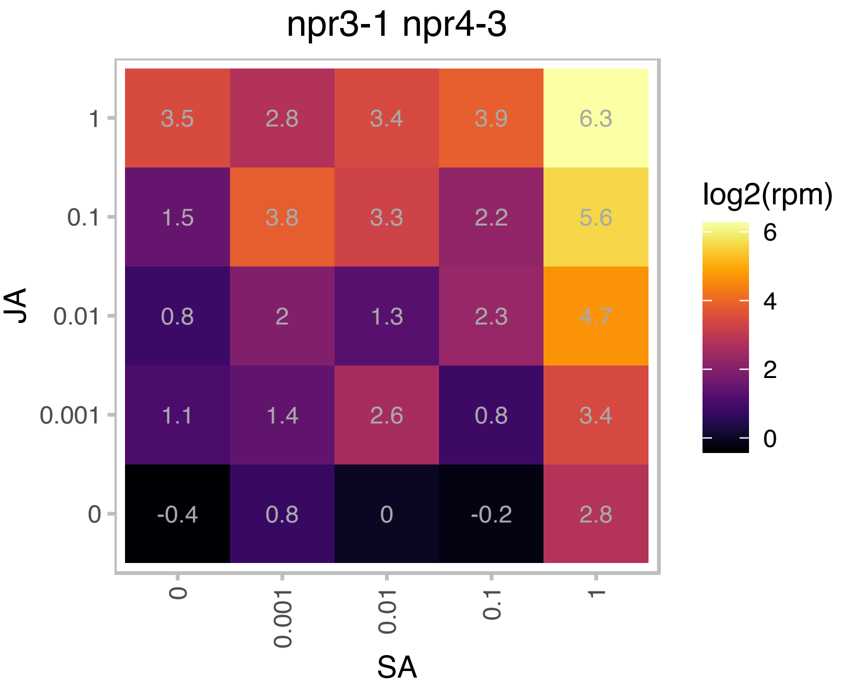

# AT3G14960.1 : NA

Galactosyltransferase family protein

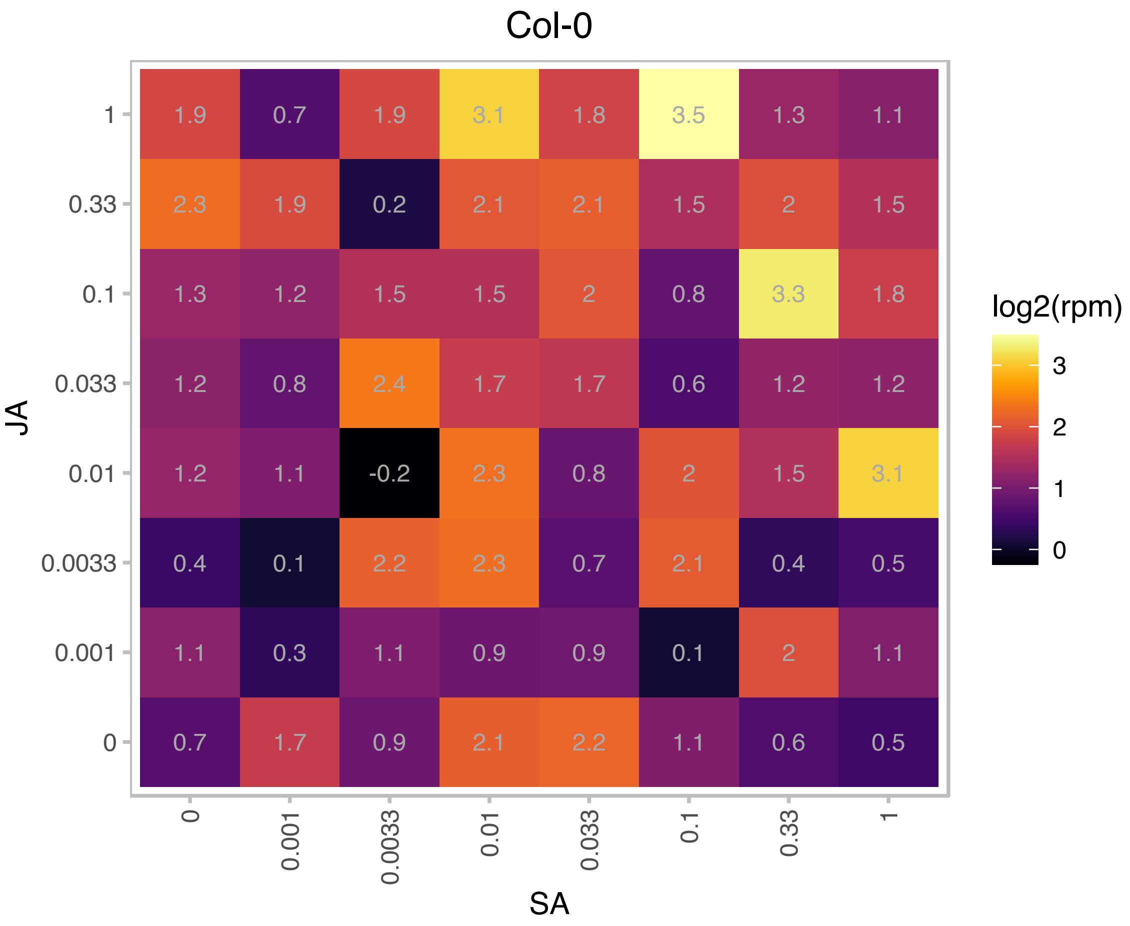


| Gene | AT3G14960 |
|---|---|
| is.expressed_genes | TRUE |
| is.responsive_genes | TRUE |
| is.apcluster | A-05 |

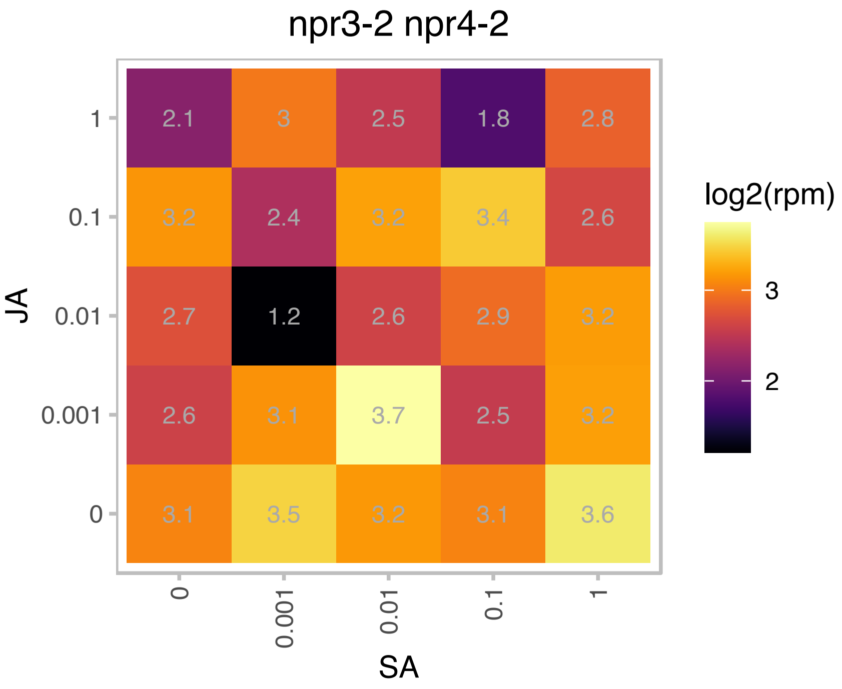


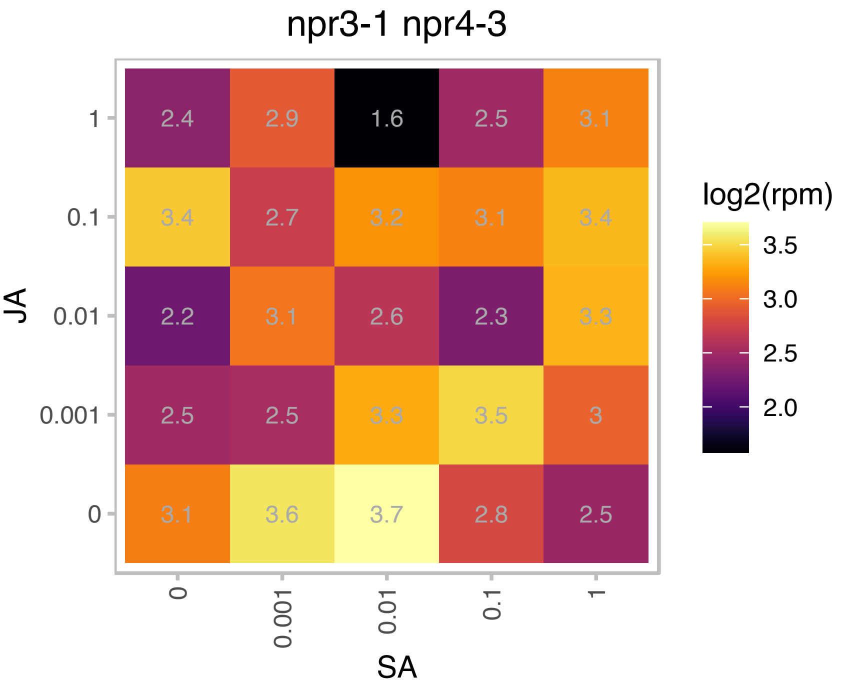

# AT3G15290.1 : NA

3-hydroxyacyl-CoA dehydrogenase family protein

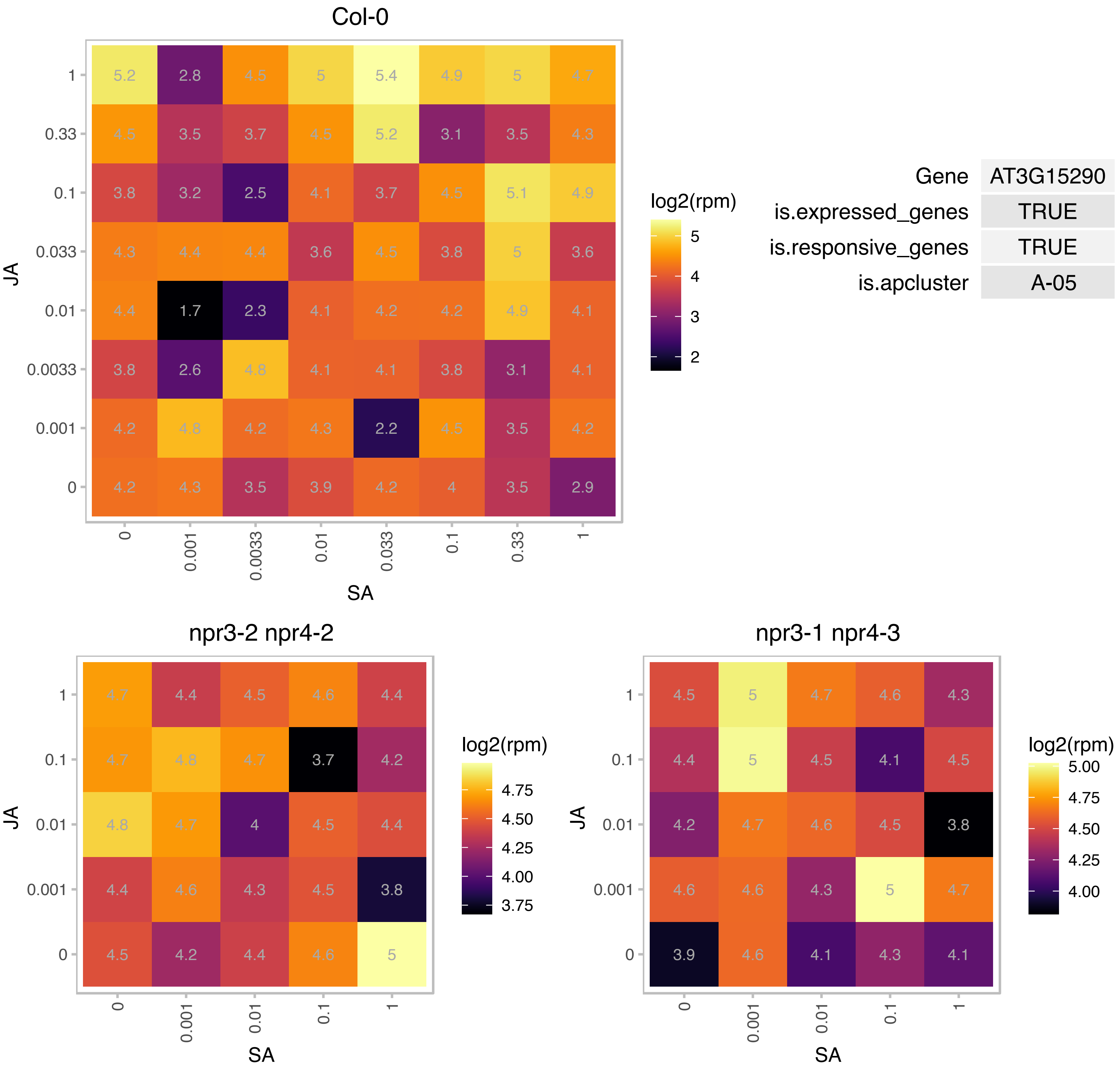


| | |
|---|---|
| Gene | AT3G15290 |
| is.expressed_genes | TRUE |
| is.responsive_genes | TRUE |
| is.apcluster | A-05 |

# AT3G21295.1 : NA

Tudor/PWWP/MBT superfamily protein

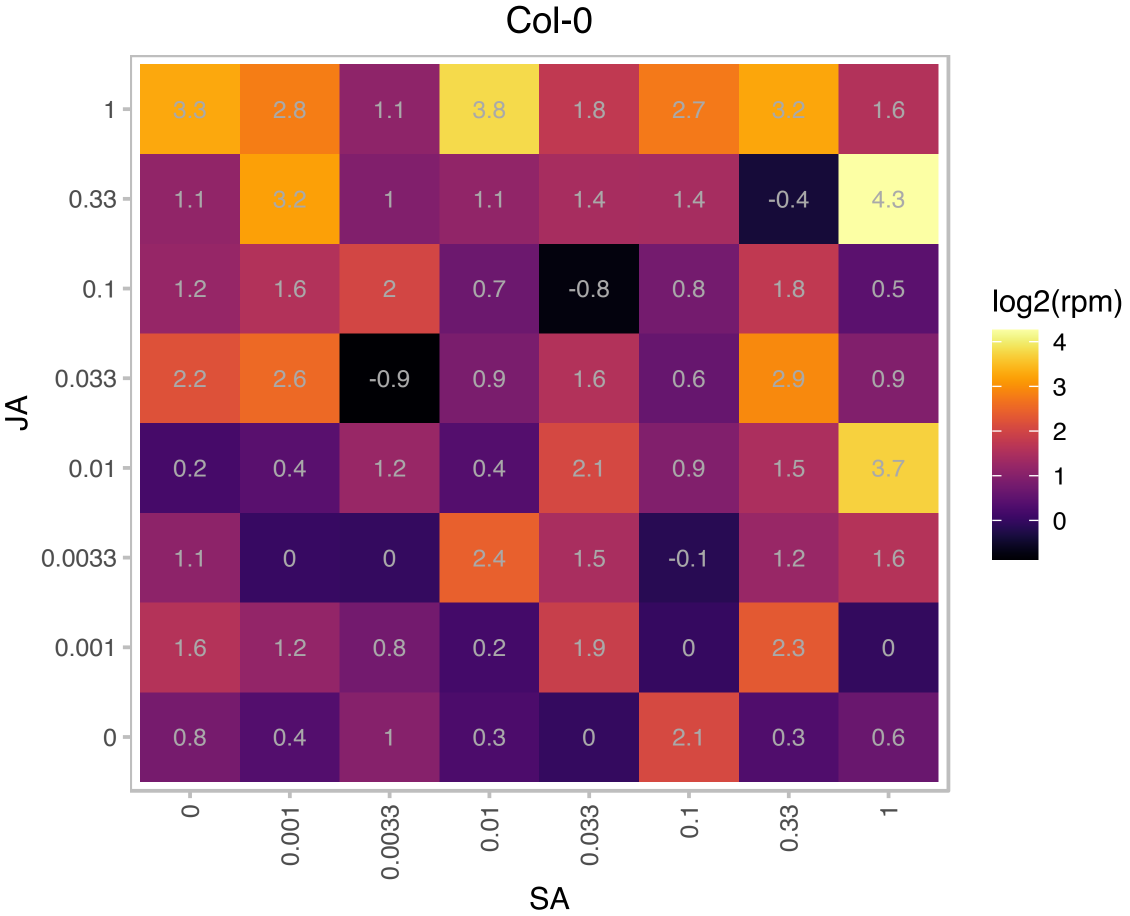


| | |
|---|---|
| Gene | AT3G21295 |
| is.expressed_genes | TRUE |
| is.responsive_genes | TRUE |
| is.apcluster | A-05 |

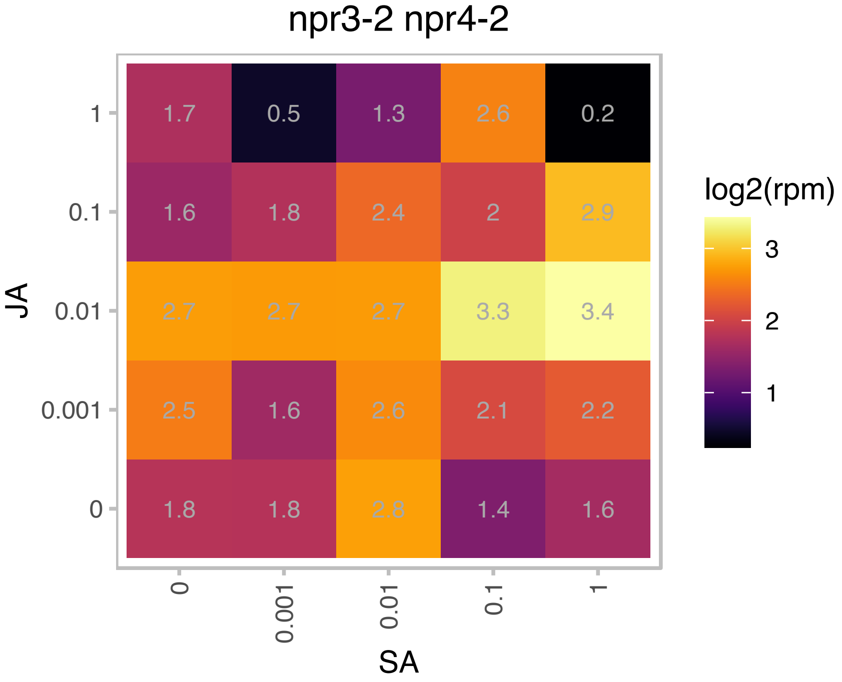


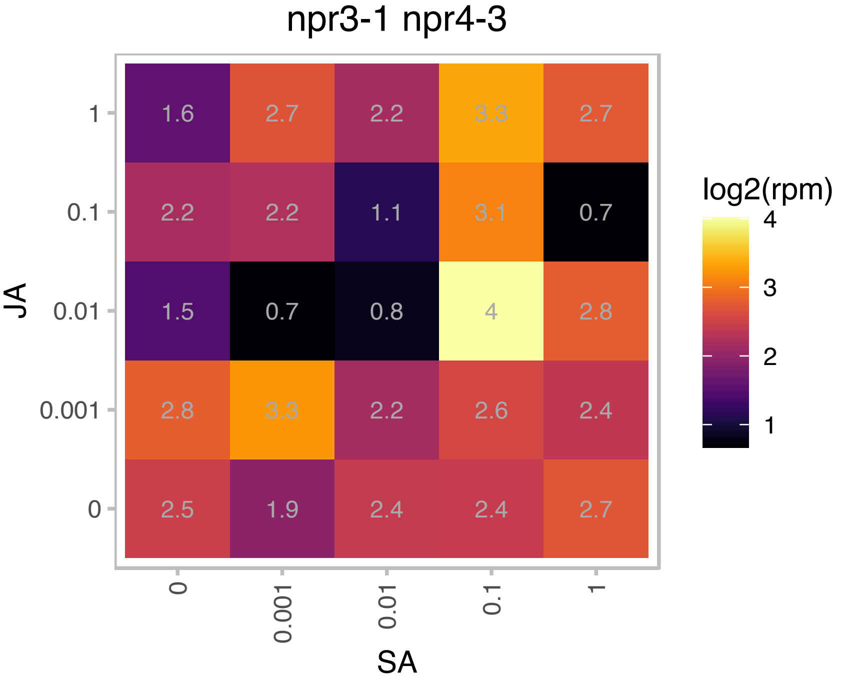

### AT3G22170.1 : CPD45, FHY3

far-red elongated hypocotyls 3

A component of the PHYA signaling network, mediates the FR-HIR response to far-red light in concert with FAR1.

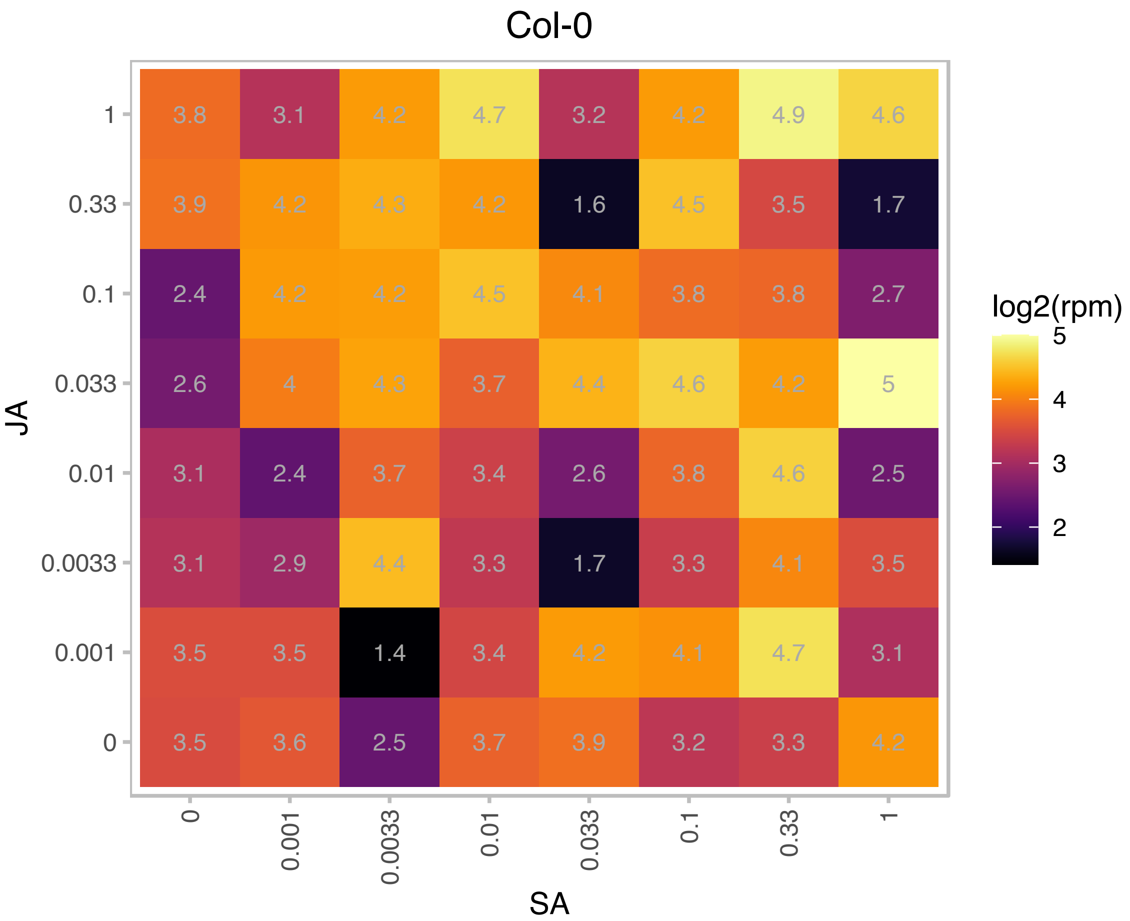


| | |
|---|---|
| Gene | AT3G22170 |
| is.expressed_genes | TRUE |
| is.responsive_genes | TRUE |
| is.apcluster | A-05 |

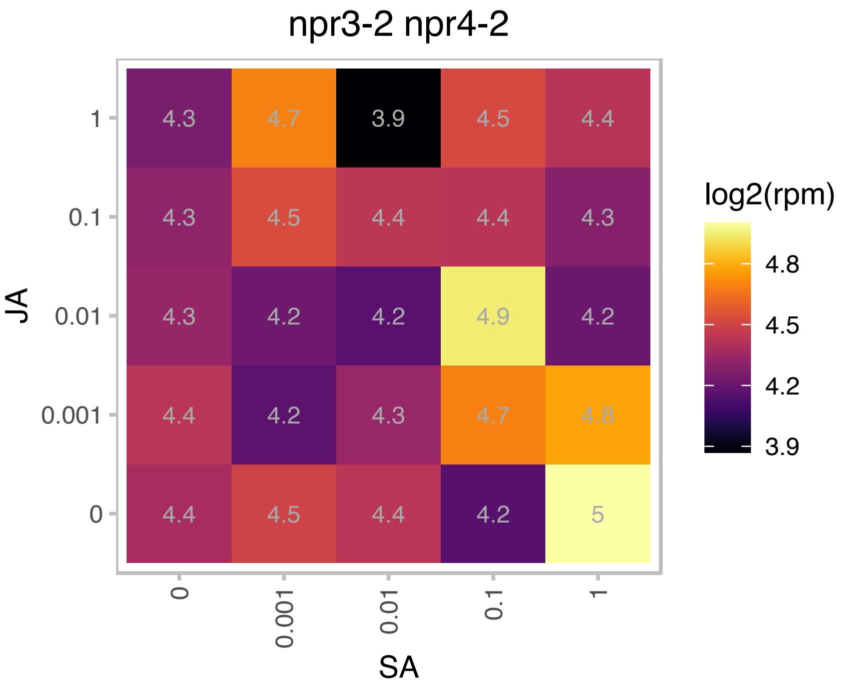


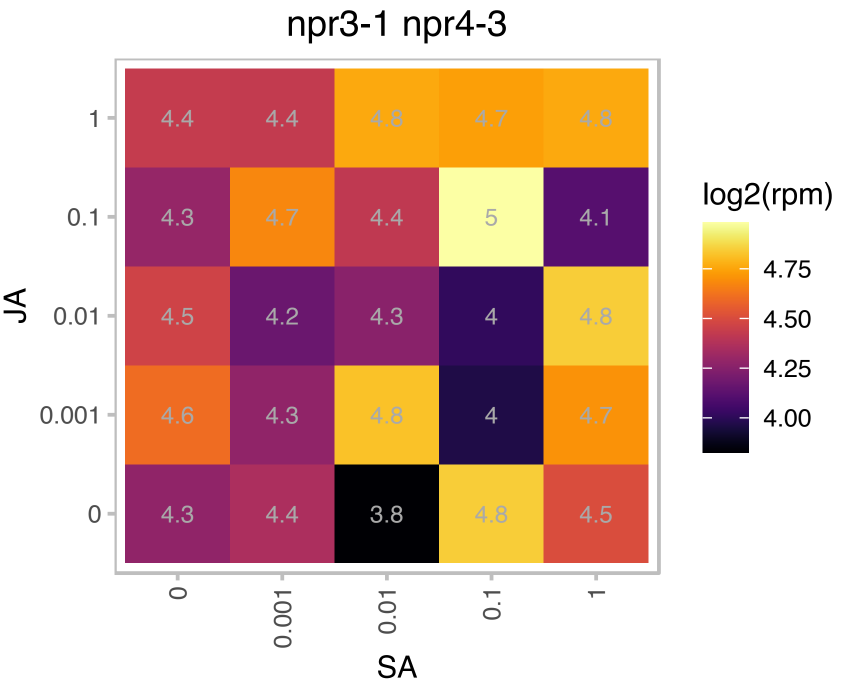

# AT3G22980.1 : NA

Ribosomal protein S5/Elongation factor G/III/V family protein

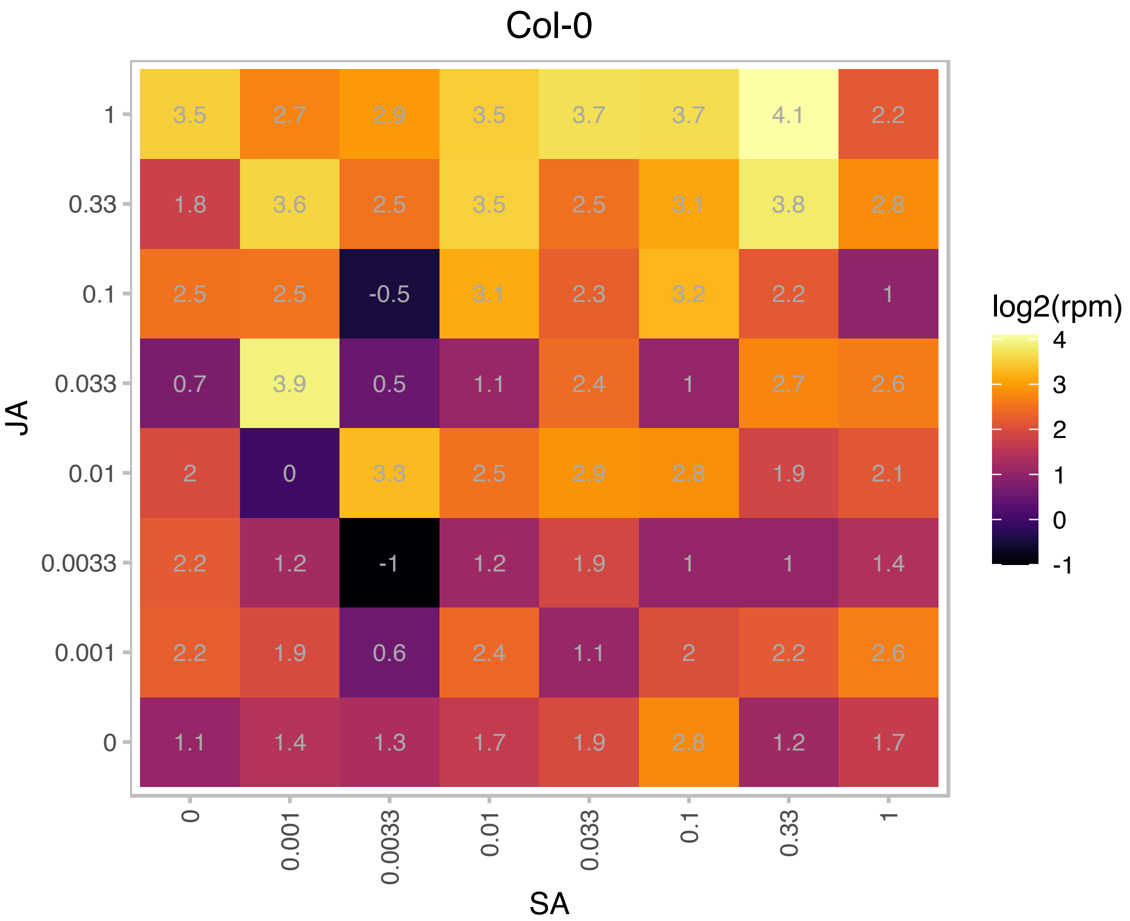


| | |
|---|---|
| Gene | AT3G22980 |
| is.expressed_genes | TRUE |
| is.responsive_genes | TRUE |
| is.apcluster | A-05 |

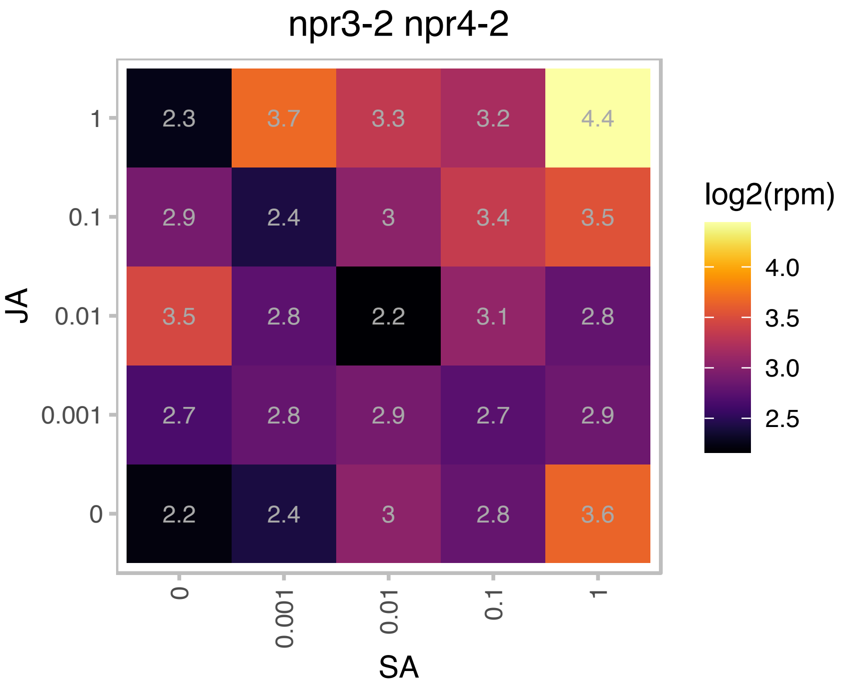


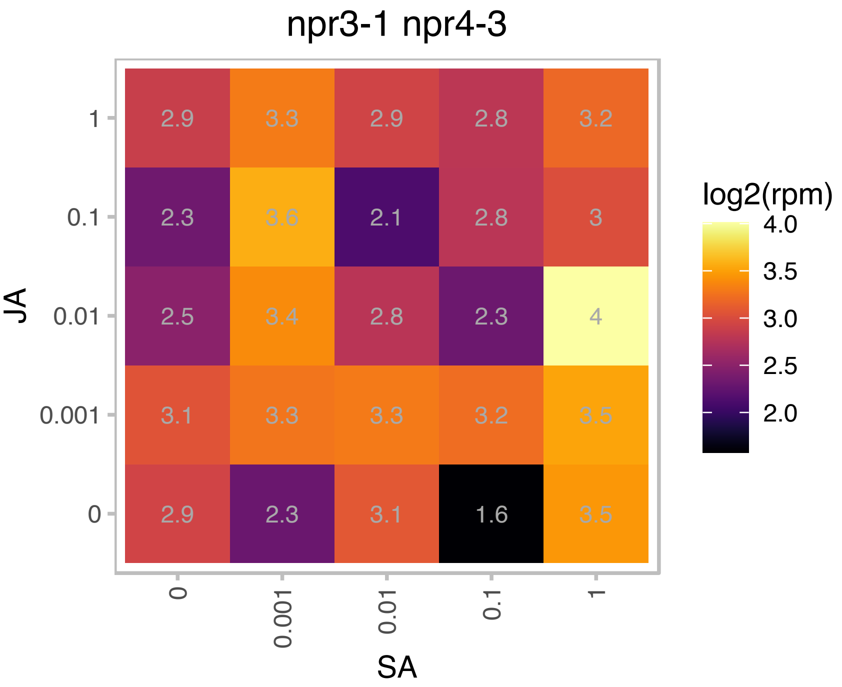

### AT3G25610.1 : ALA10

ATPase E1-E2 type family protein / haloacid dehalogenase-like hydrolase family protein

Encodes aminophospholipid ATPase10 (ALA10), a P4-type ATPase flippase that internalizes exogenous phospholipids across the plasma membrane.

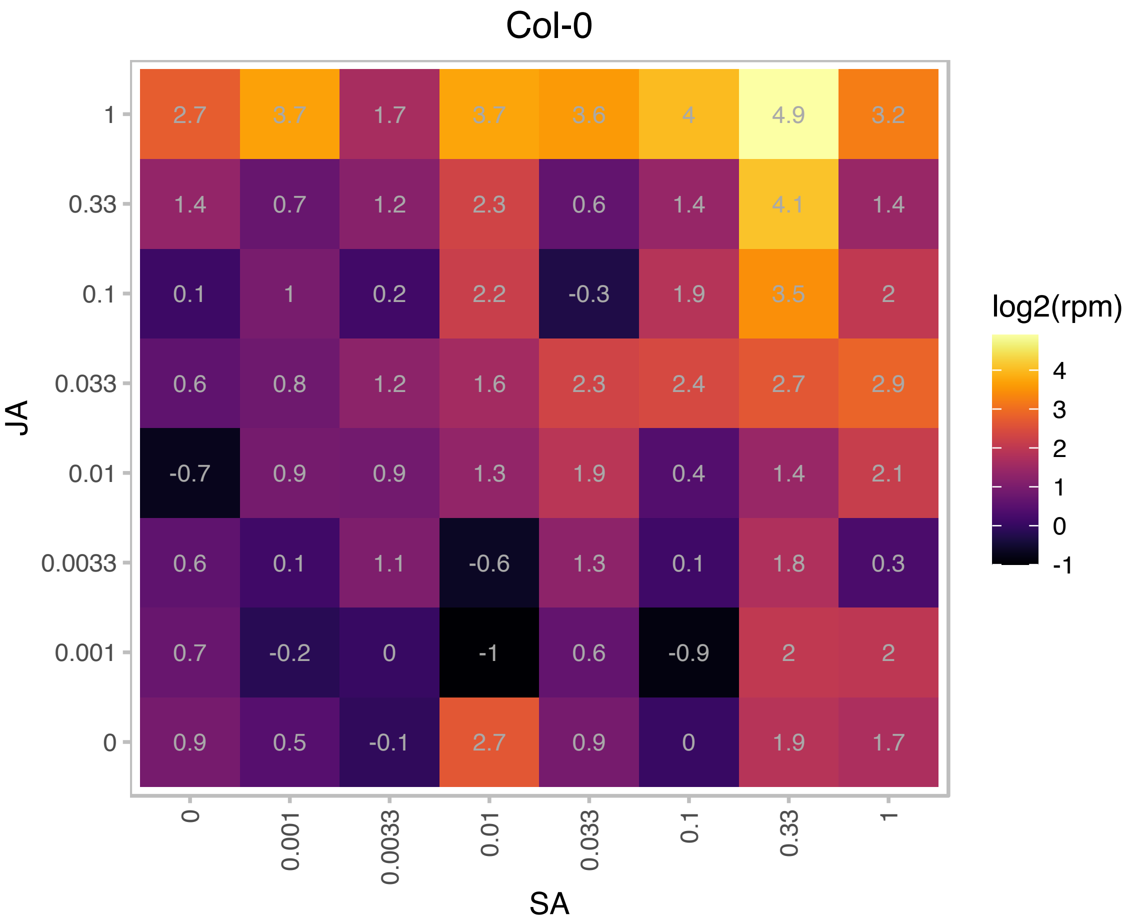


| | |
|---|---|
| Gene | AT3G25610 |
| is.expressed_genes | TRUE |
| is.responsive_genes | TRUE |
| is.apcluster | A-05 |

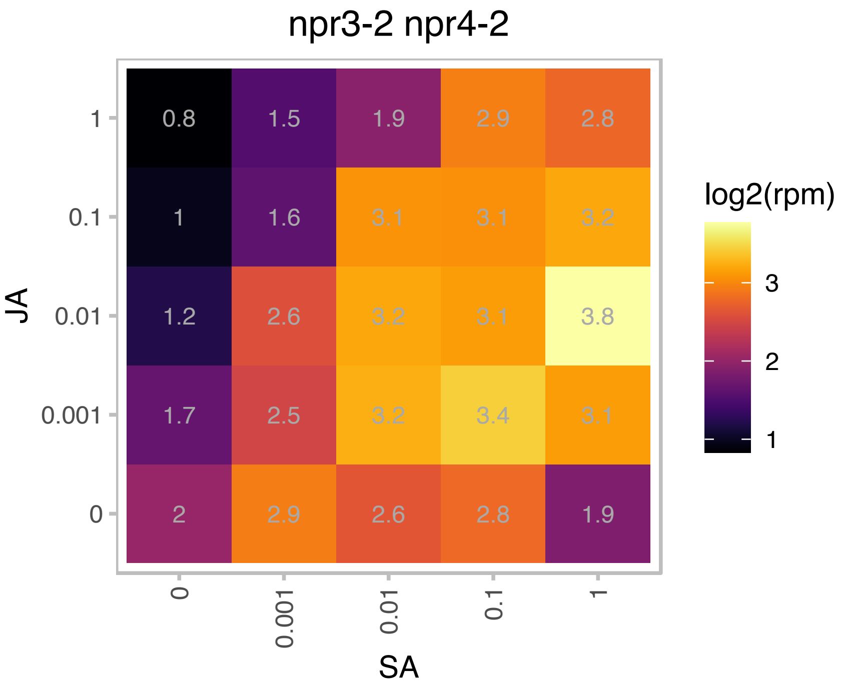


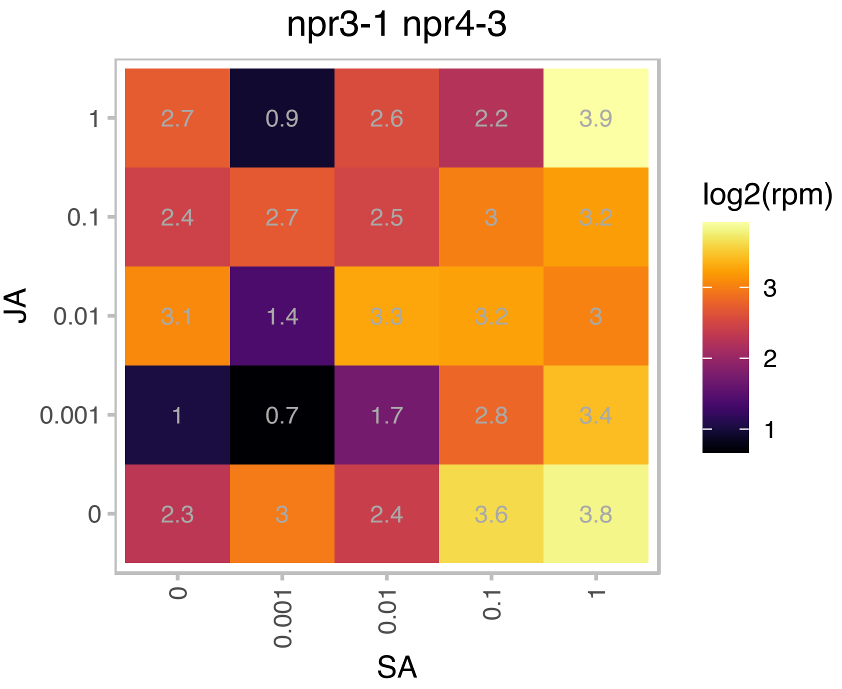

### AT3G26170.1 : CYP71B19

cytochrome P450, family 71, subfamily B, polypeptide 19

putative cytochrome P450

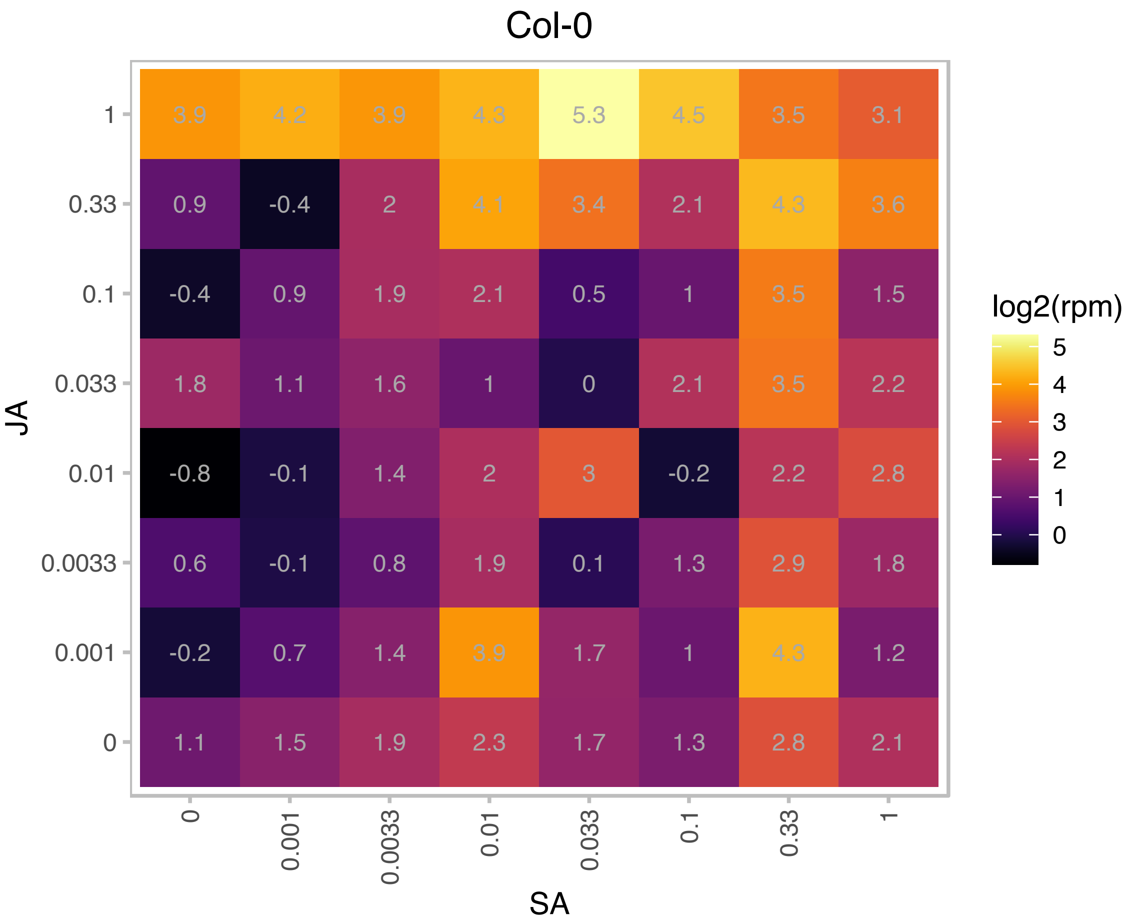


| Gene | AT3G26170 |
|---|---|
| is.expressed_genes | TRUE |
| is.responsive_genes | TRUE |
| is.apcluster | A-05 |

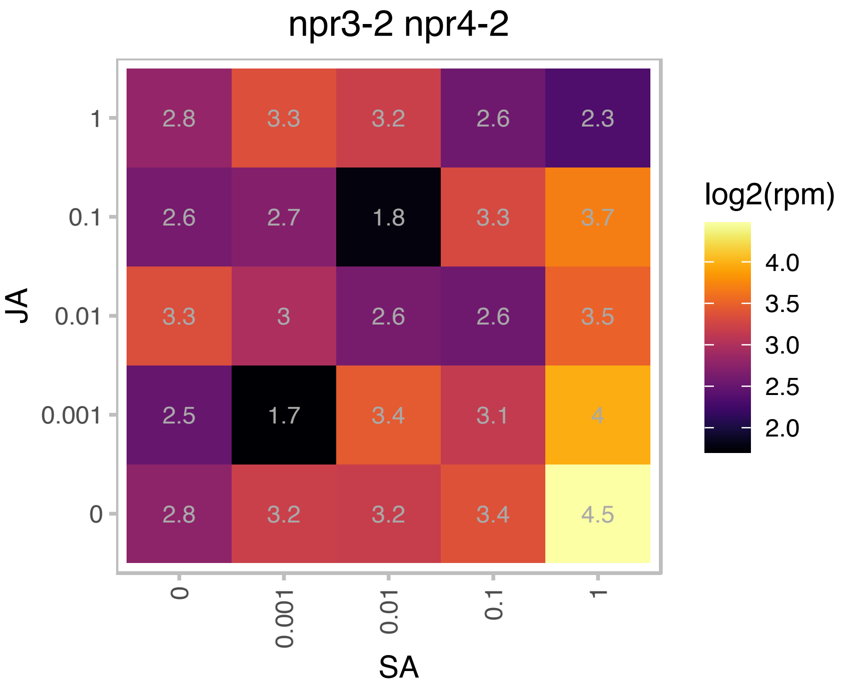


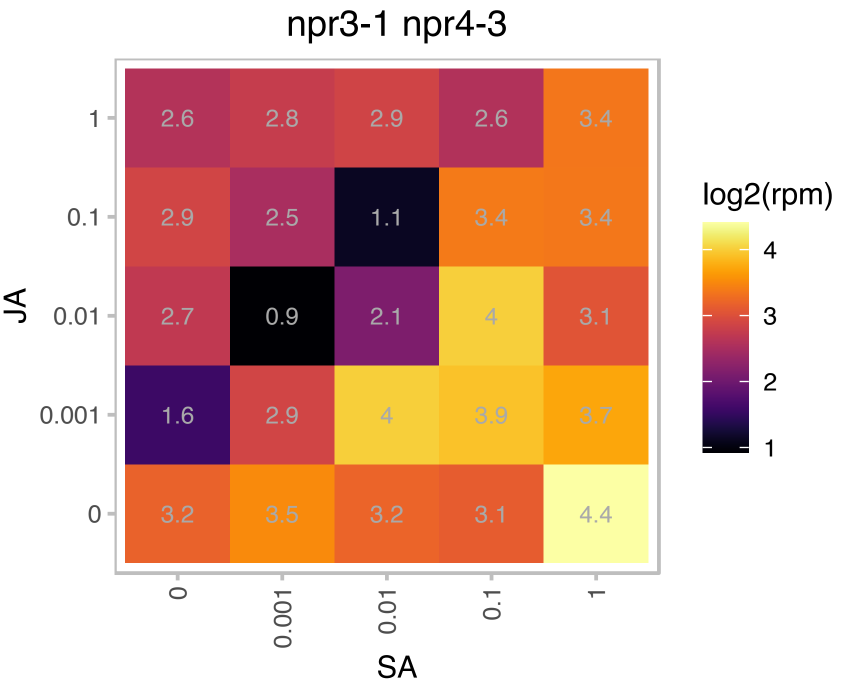

### AT3G46660.1 : UGT76E12

UDP-glucosyl transferase 76E12

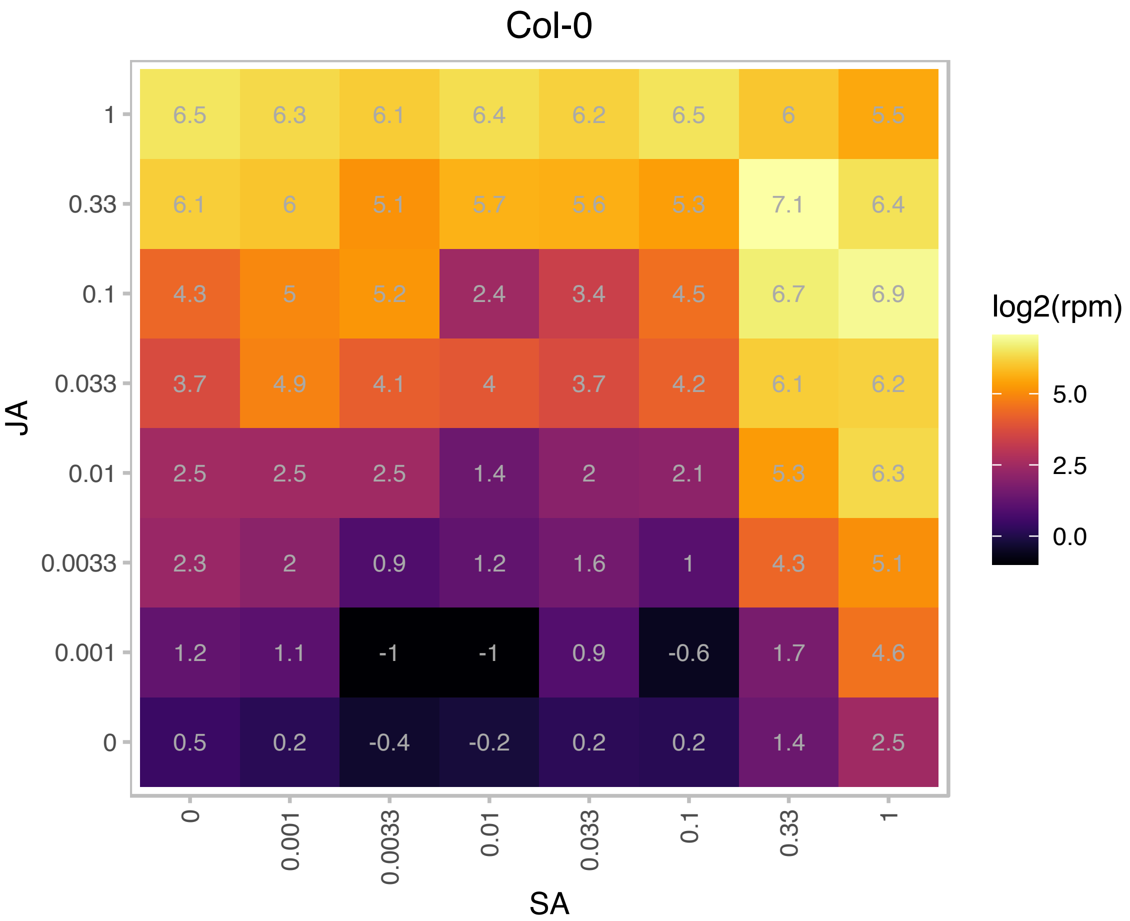


| | |
|---|---|
| Gene | AT3G46660 |
| is.expressed_genes | TRUE |
| is.responsive_genes | TRUE |
| is.apcluster | A-05 |

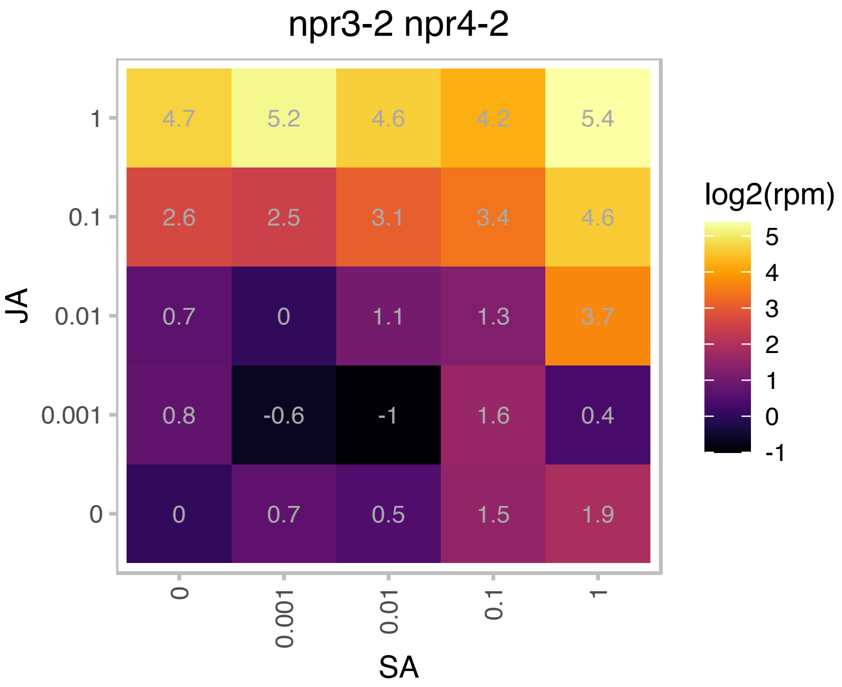


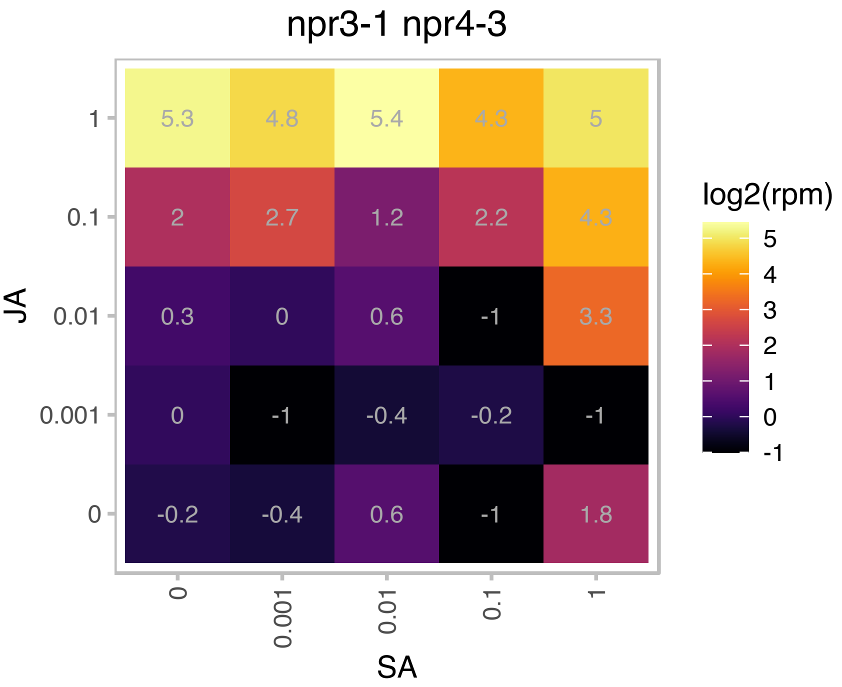

### AT3G48880.1 : SNIPER4

RNI-like superfamily protein
Encodes an F-box protein, SNIPER4, that regulates the turnover of MUSE13 and MUSE14, redundant TRAF protei
serving as adaptors in the SCFCRP1 complex to facilitate the turnover of nucleotide-binding domain and
leucine-rich repeats (NLR) immune receptors.

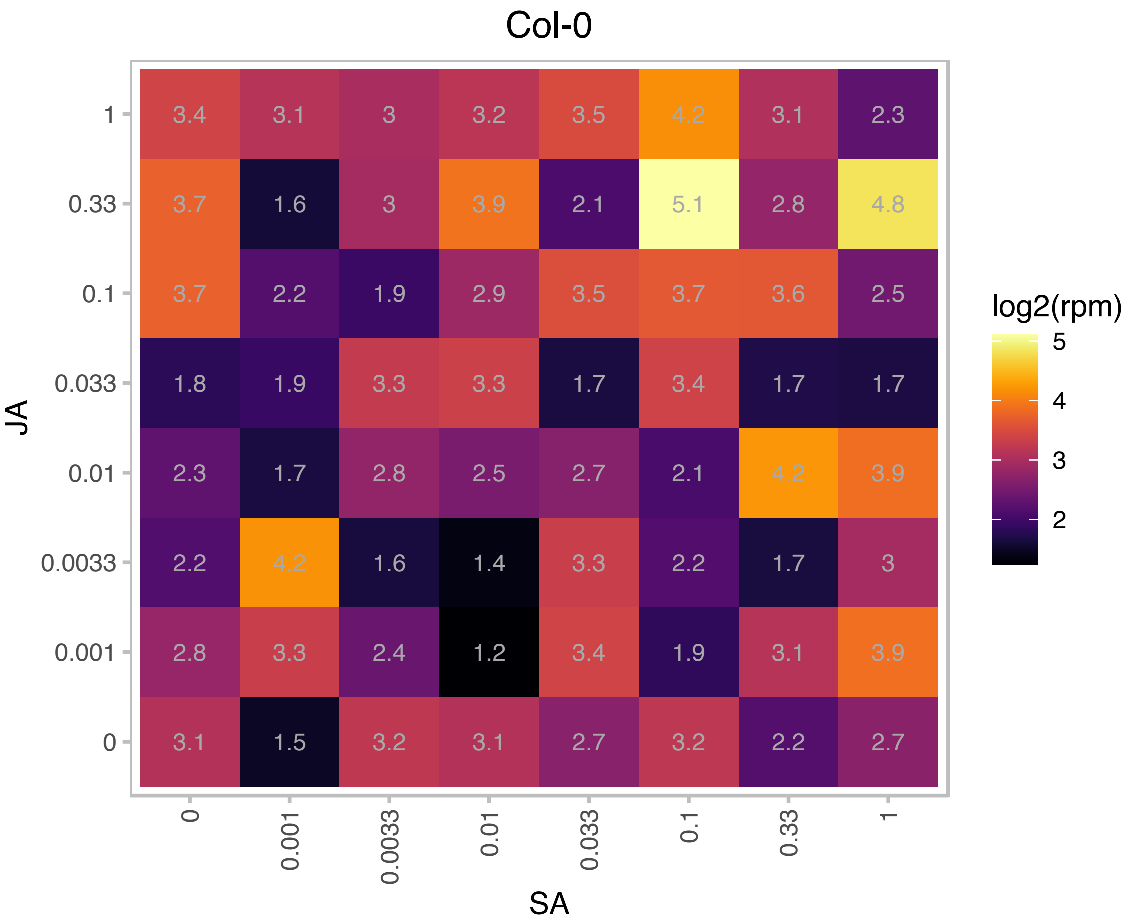


| | |
|---|---|
| Gene | AT3G48880 |
| is.expressed_genes | TRUE |
| is.responsive_genes | TRUE |
| is.apcluster | A-05 |

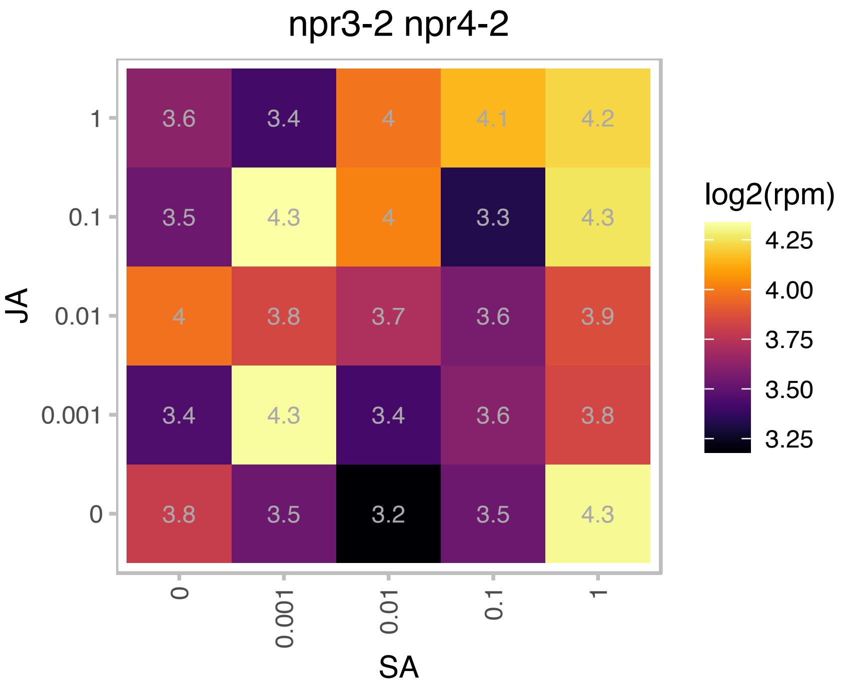


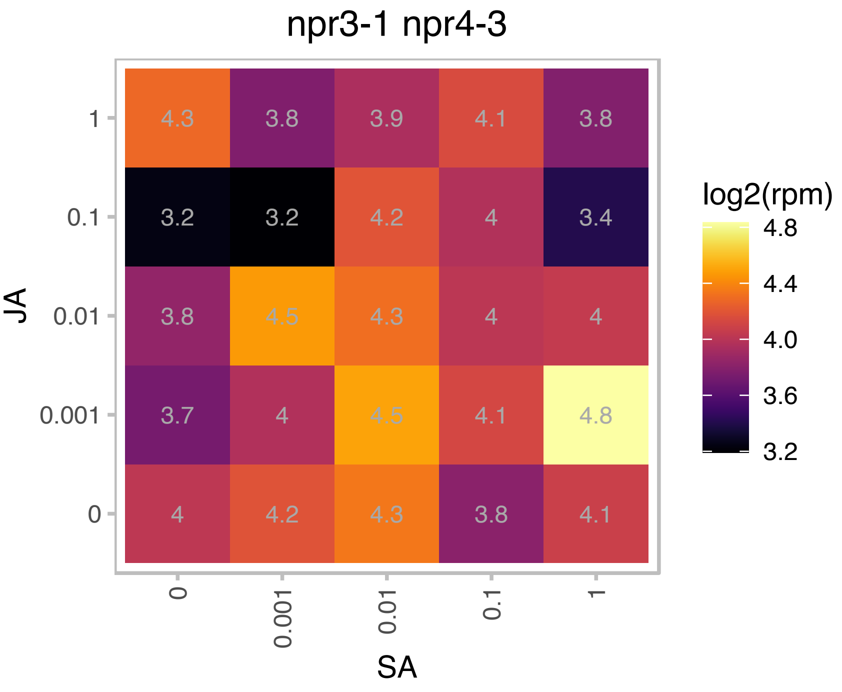

# AT3G51090.1 : NA

Protein of unknown function (DUF1640)

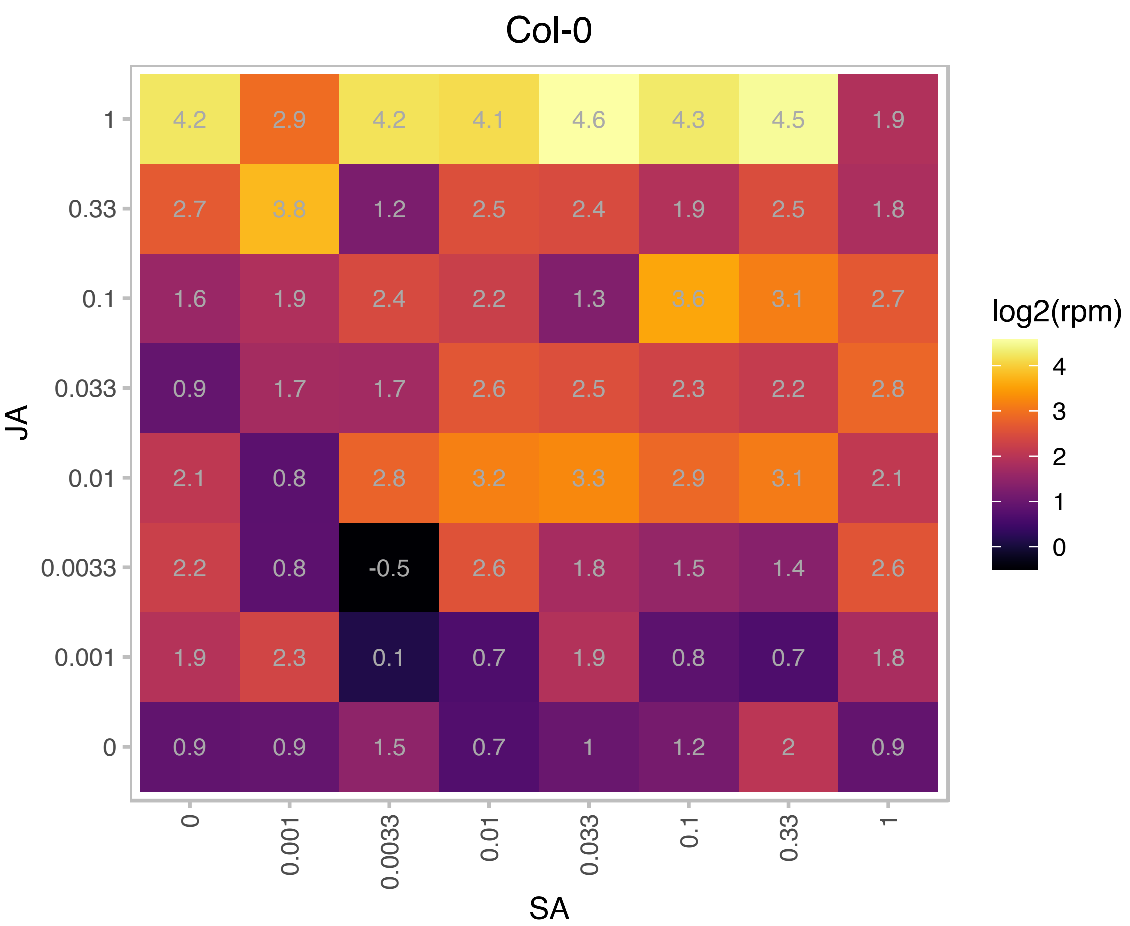


| | |
|---|---|
| Gene | AT3G51090 |
| is.expressed_genes | TRUE |
| is.responsive_genes | TRUE |
| is.apcluster | A-05 |

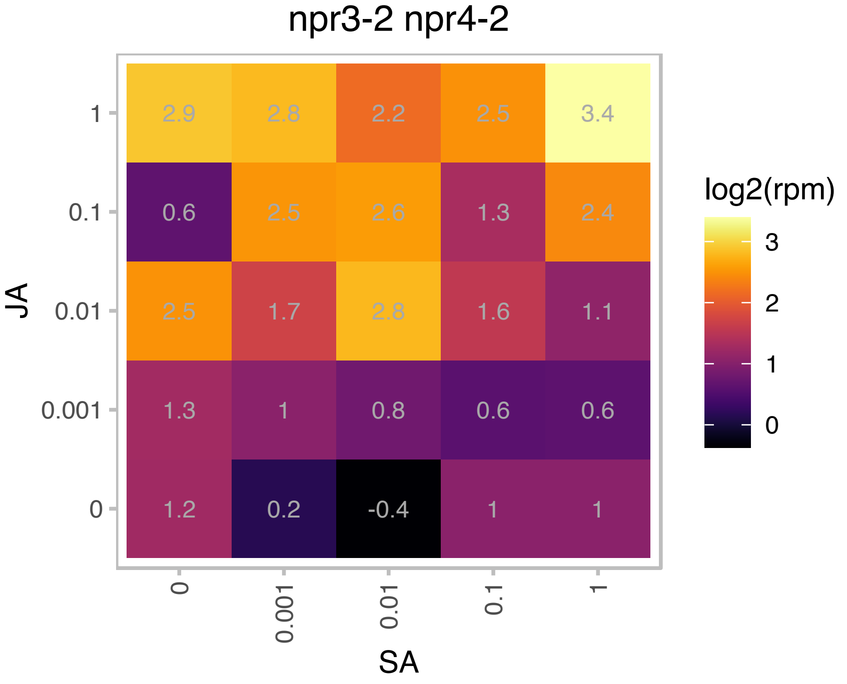


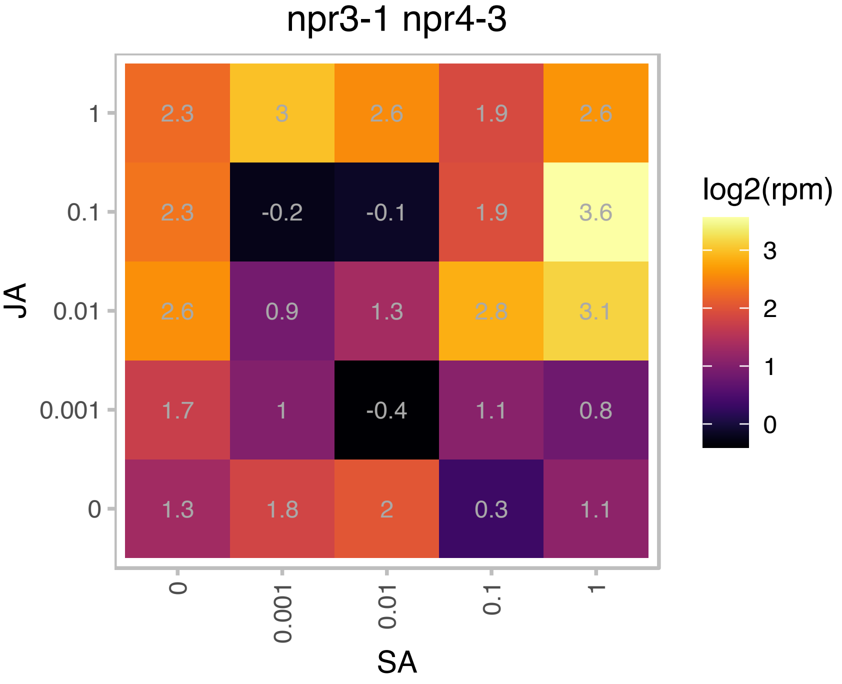

### AT3G51460.1 : RHD4

Phosphoinositide phosphatase family protein

Encodes RHD4 (ROOT HAIR DEFECTIVE4), a phosphatidylinositol-4-phosphate phosphatase required for root hair development. The mRNA is cell-to-cell mobile.

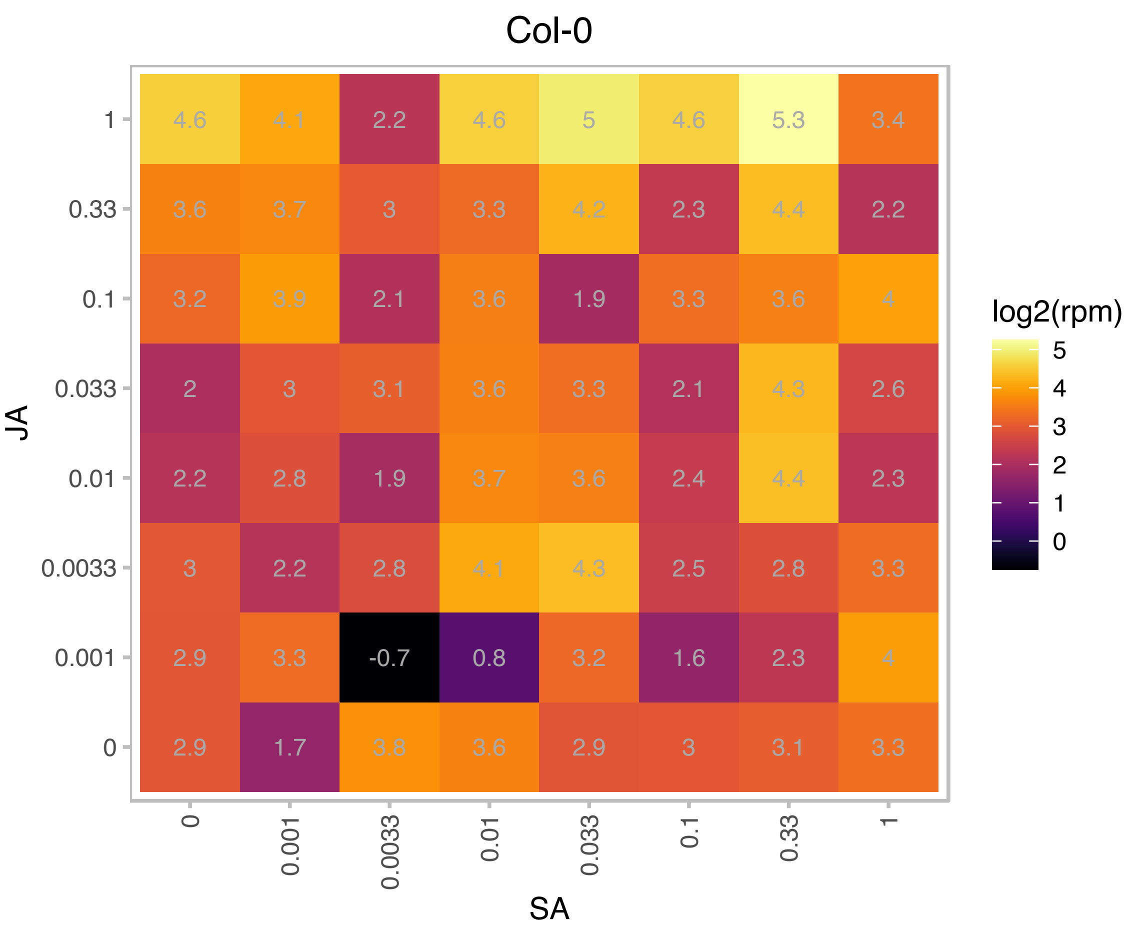


| | |
|---|---|
| Gene | AT3G51460 |
| is.expressed_genes | TRUE |
| is.responsive_genes | TRUE |
| is.apcluster | A-05 |

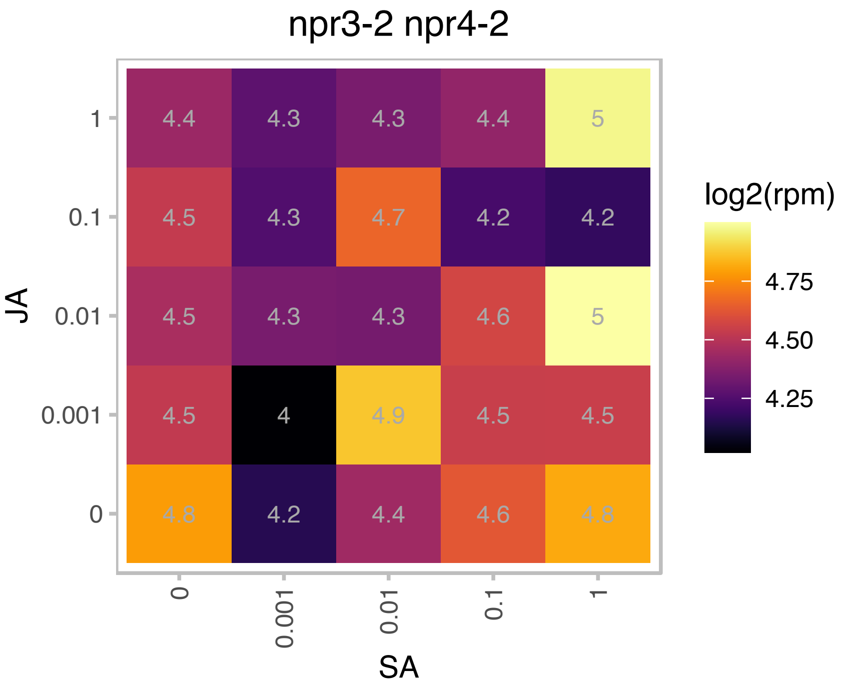


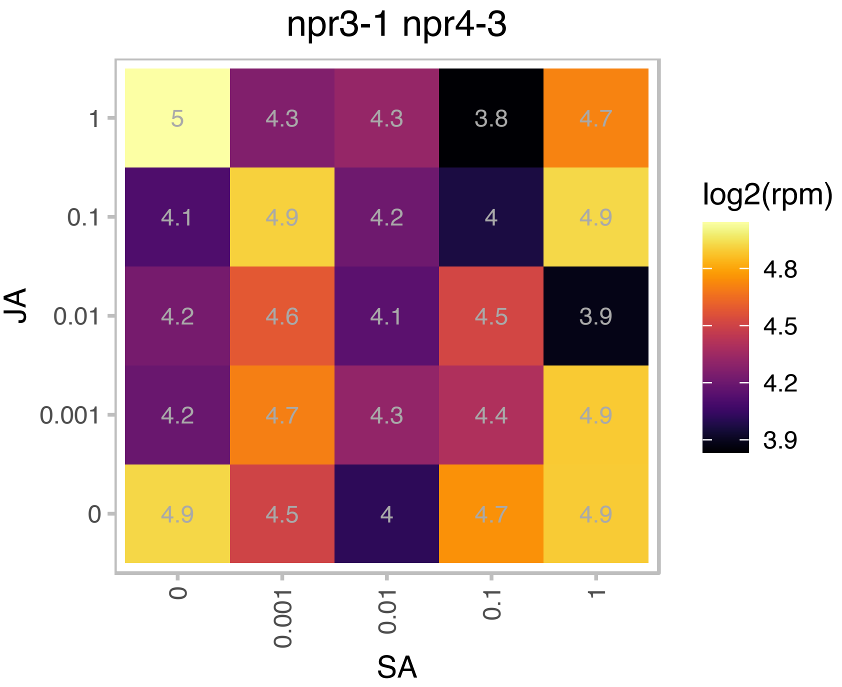

# AT3G51990.1 : NA

Protein kinase superfamily protein

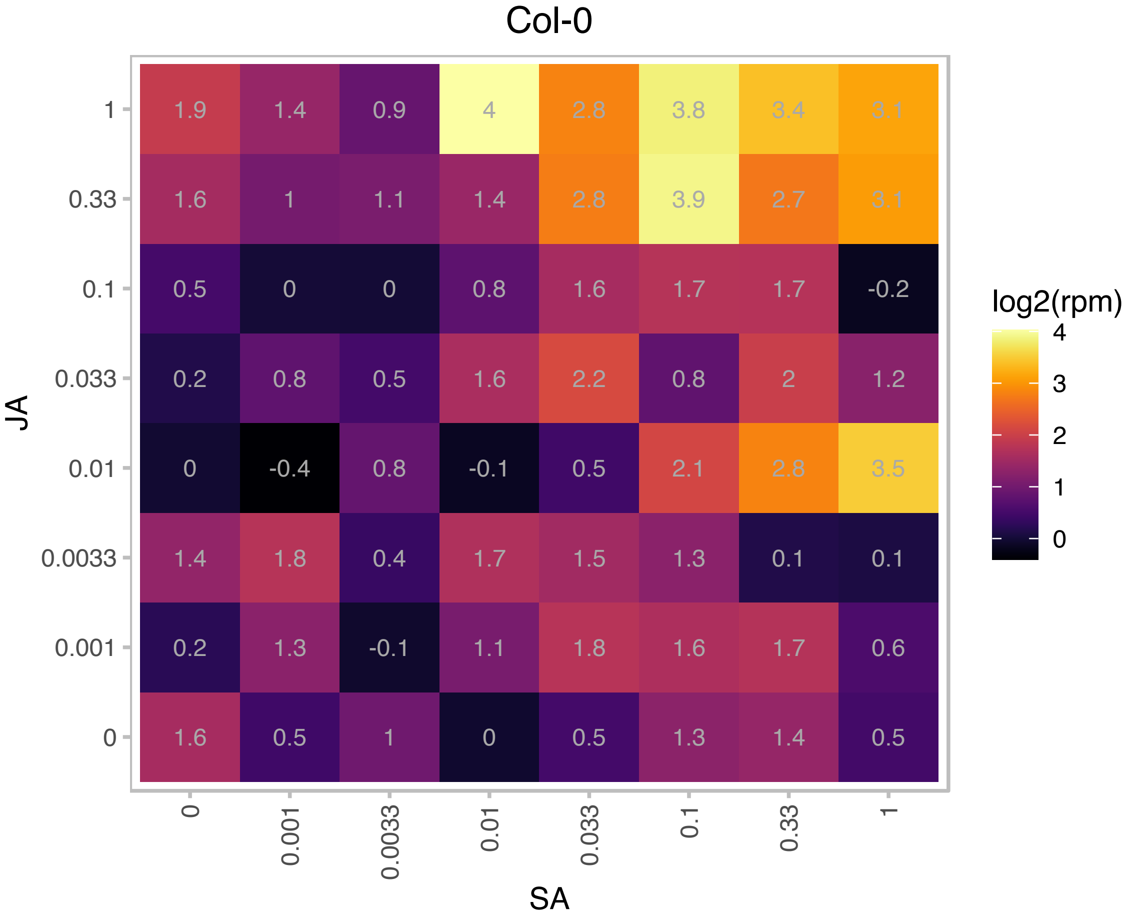


| | |
|---|---|
| Gene | AT3G51990 |
| is.expressed_genes | TRUE |
| is.responsive_genes | TRUE |
| is.apcluster | A-05 |

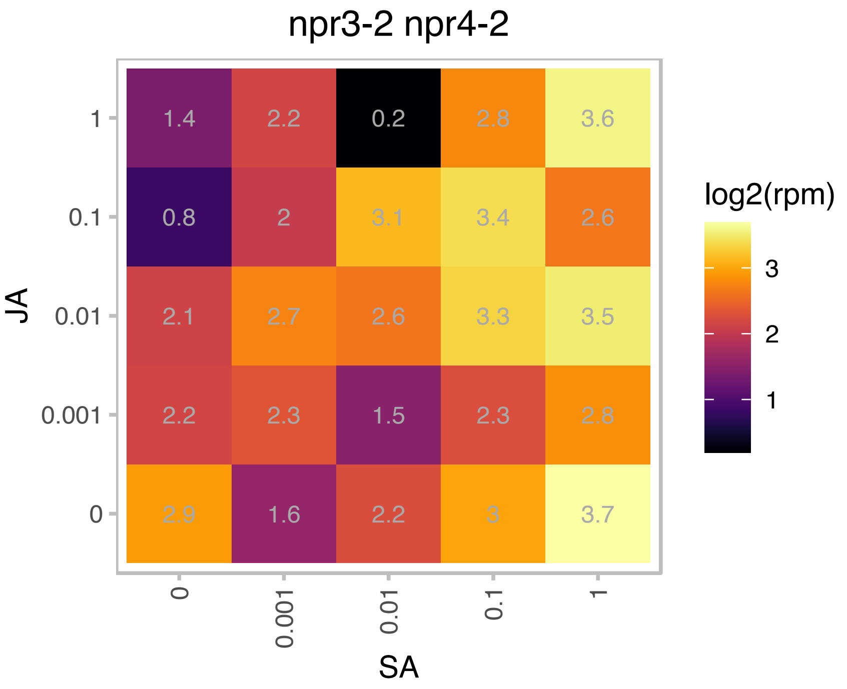


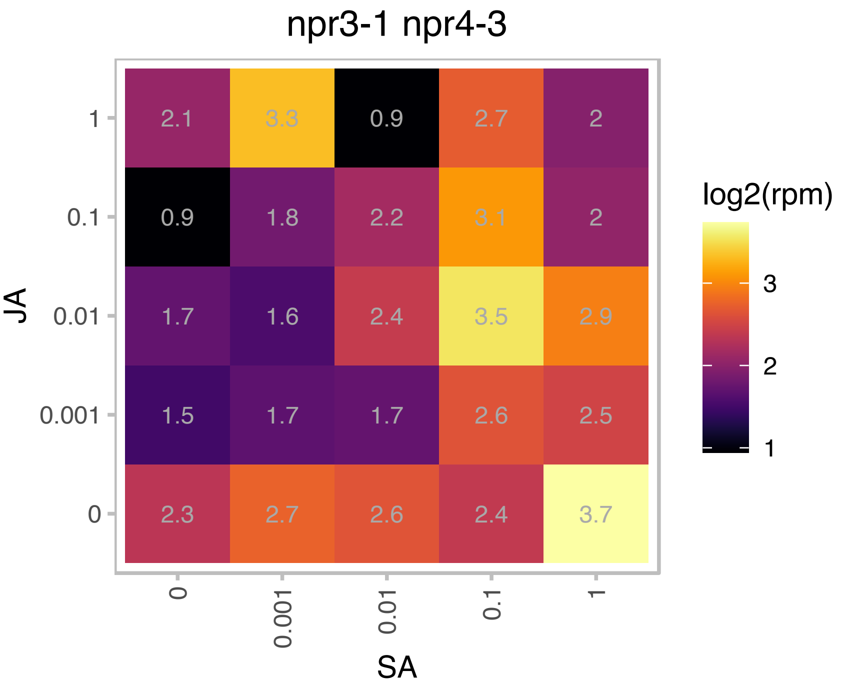

### AT3G55400.1 : OVA1

methionyl-tRNA synthetase / methionine--tRNA ligase / MetRS (cpMetRS)

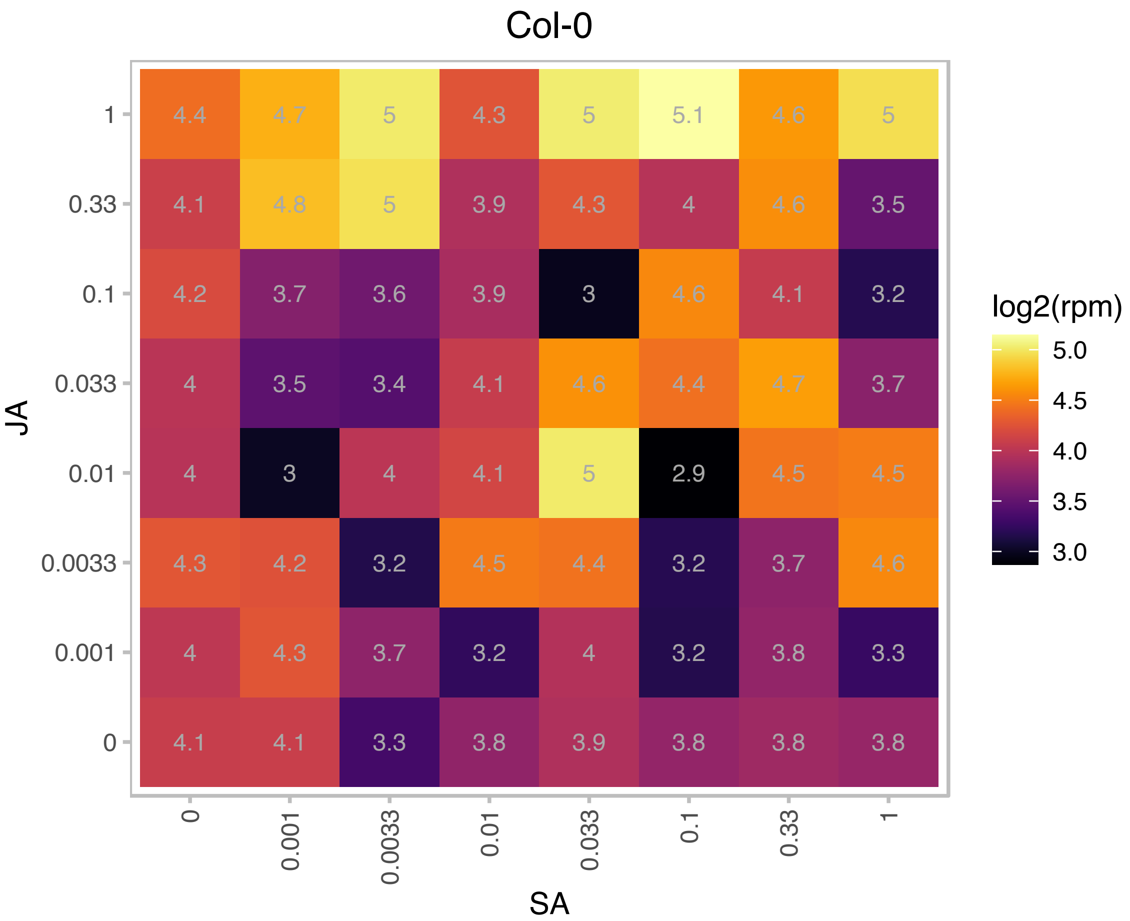


| | |
|---|---|
| Gene | AT3G55400 |
| is.expressed_genes | TRUE |
| is.responsive_genes | TRUE |
| is.apcluster | A-05 |

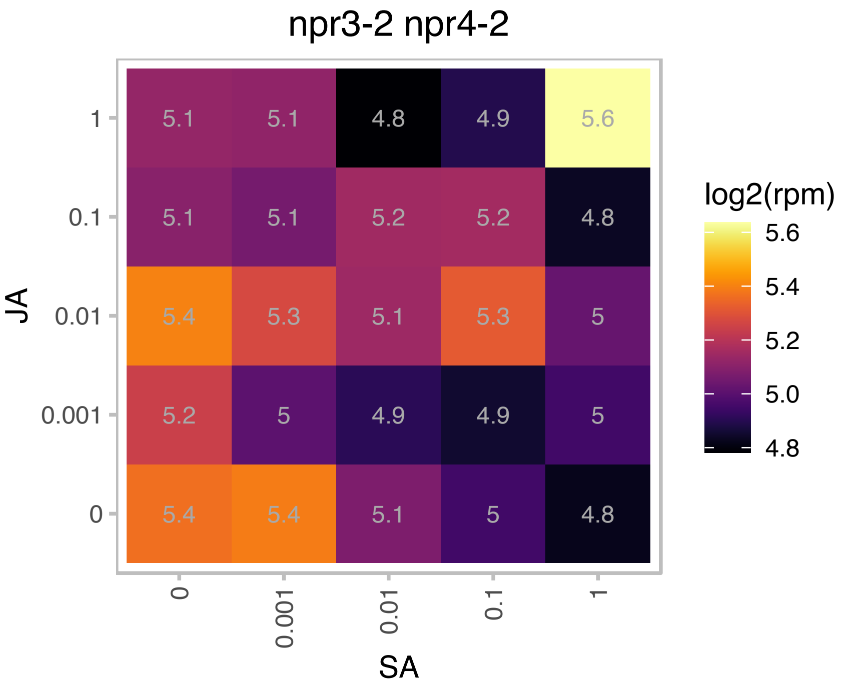


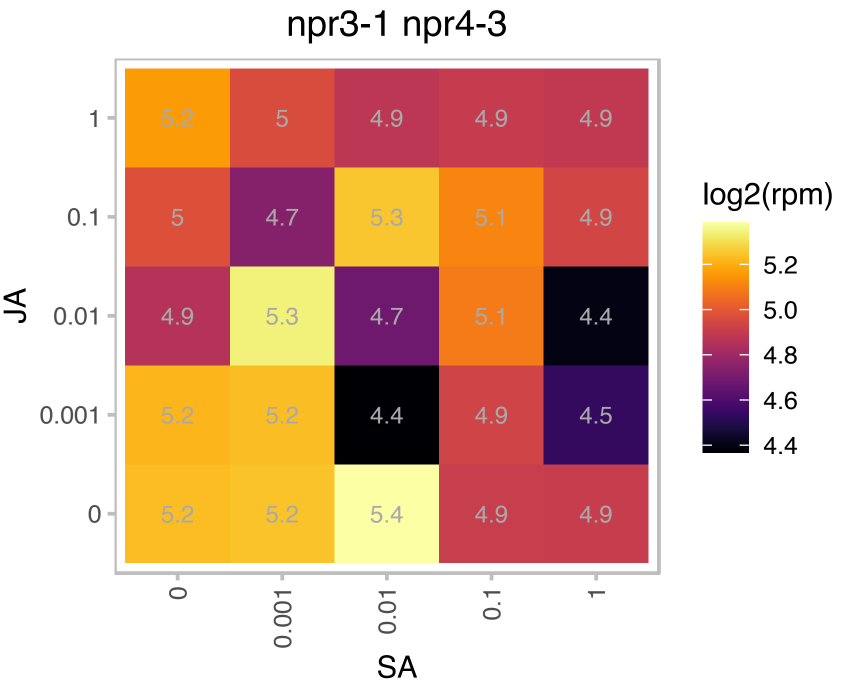

# AT3G57890.1 : NA

Tubulin binding cofactor C domain-containing protein

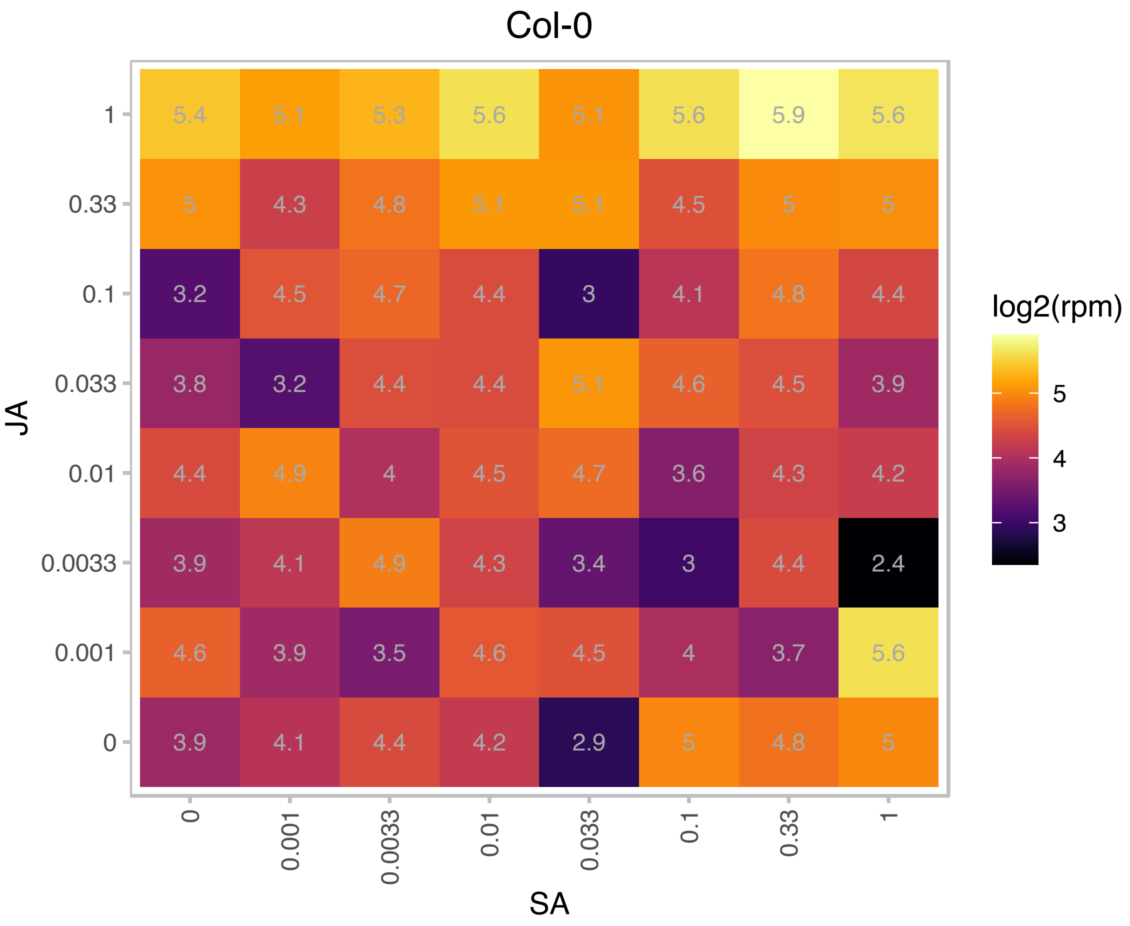


| | |
|---|---|
| Gene | AT3G57890 |
| is.expressed_genes | TRUE |
| is.responsive_genes | TRUE |
| is.apcluster | A-05 |

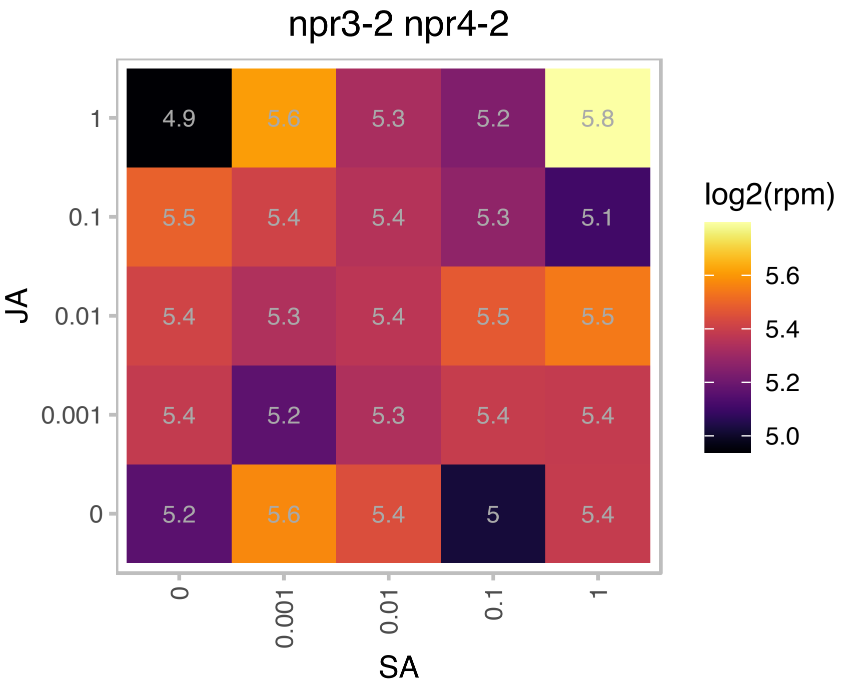


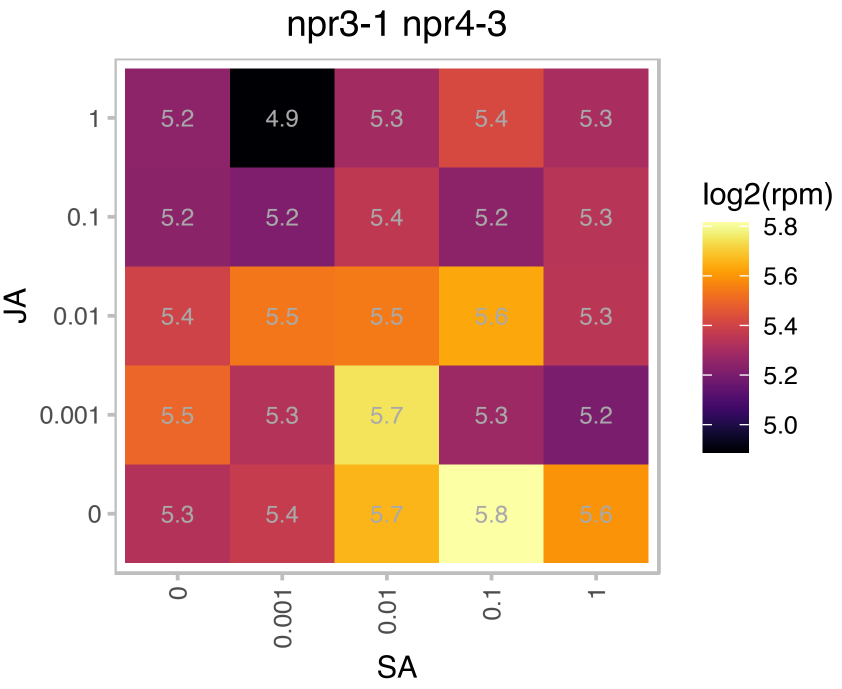

### AT3G63010.1 : ATGID1B, GID1B

alpha/beta-Hydrolases superfamily protein
Encodes a gibberellin (GA) receptor ortholog of the rice GA receptor gene (OsGID1). Has GA-binding activity, showing higher affinity to GA4. Interacts with DELLA proteins in vivo in the presence of GA4. The mRNA is cell-to-cell mobile.

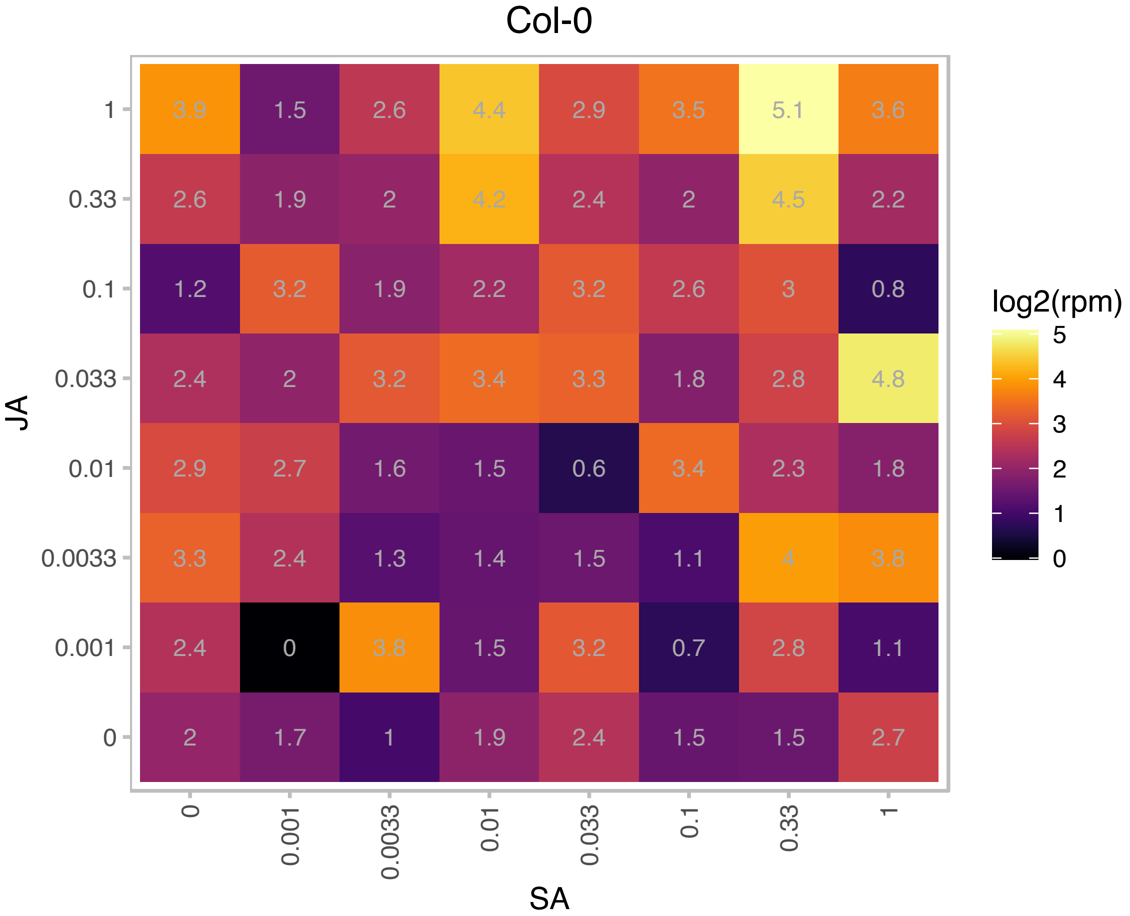


| | |
|---|---|
| Gene | AT3G63010 |
| is.expressed_genes | TRUE |
| is.responsive_genes | TRUE |
| is.apcluster | A-05 |

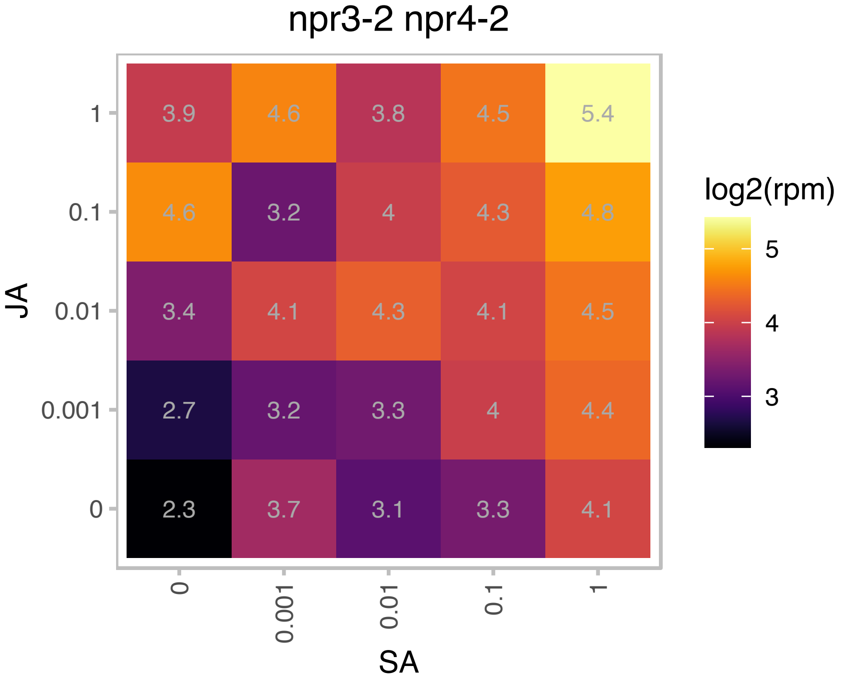


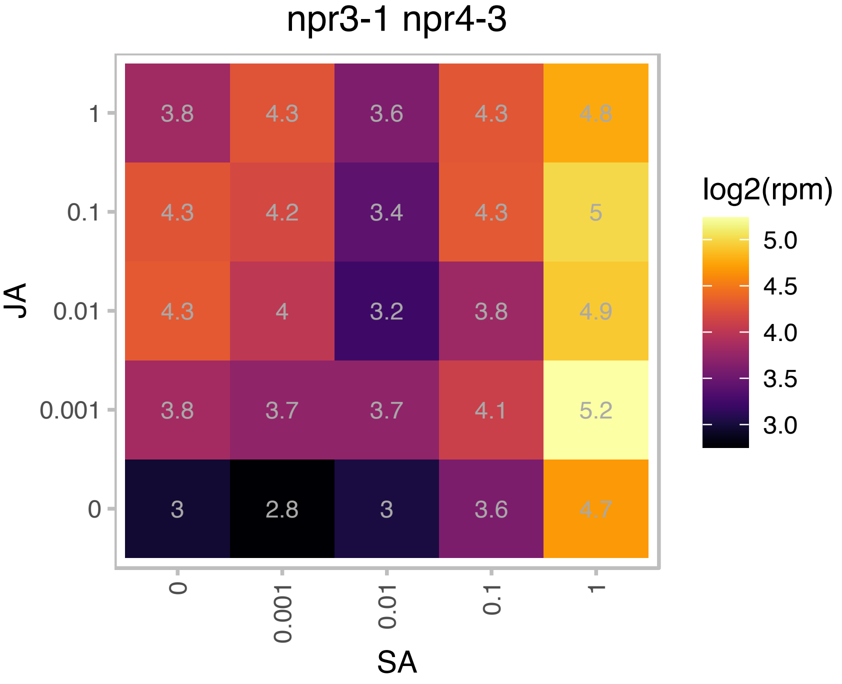

# AT4G02740.1 : NA

F-box/RNI-like superfamily protein

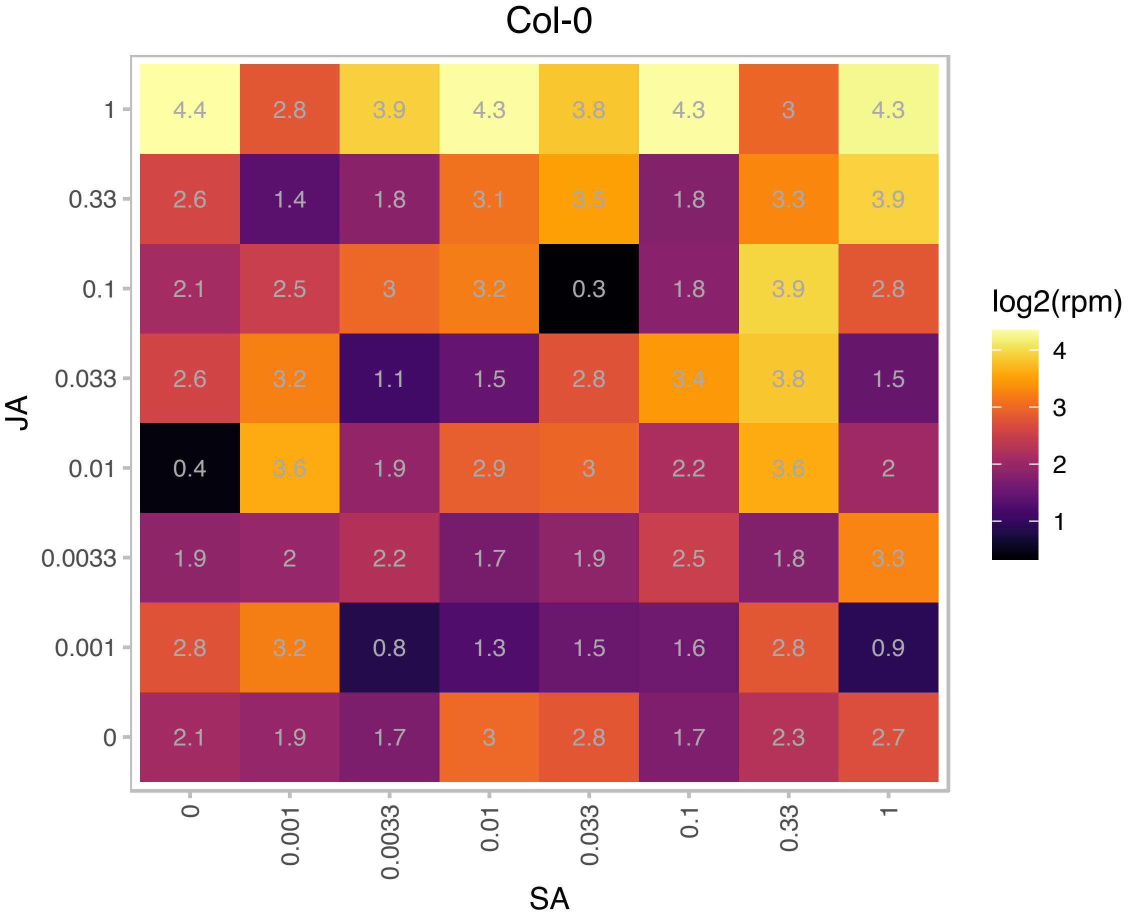


| | |
|---|---|
| Gene | AT4G02740 |
| is.expressed_genes | TRUE |
| is.responsive_genes | TRUE |
| is.apcluster | A-05 |

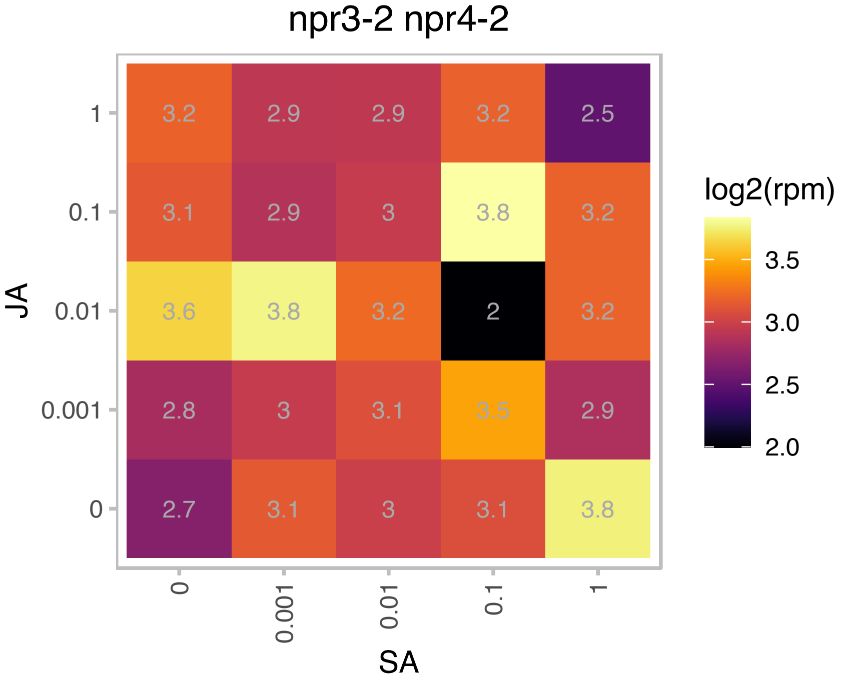


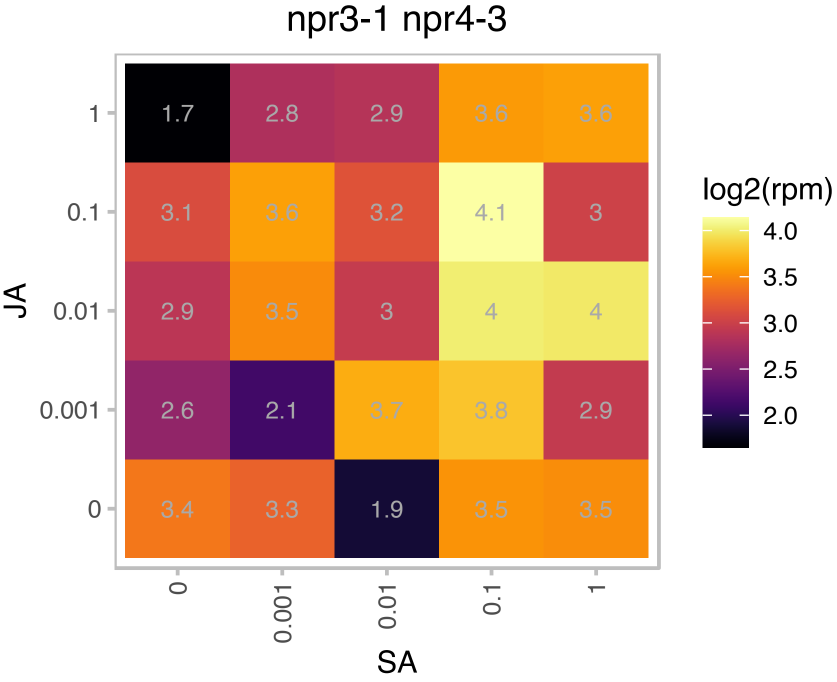

### AT4G04910.1 : NSF

AAA-type ATPase family protein

N-ethylmaleimide sensitive factor

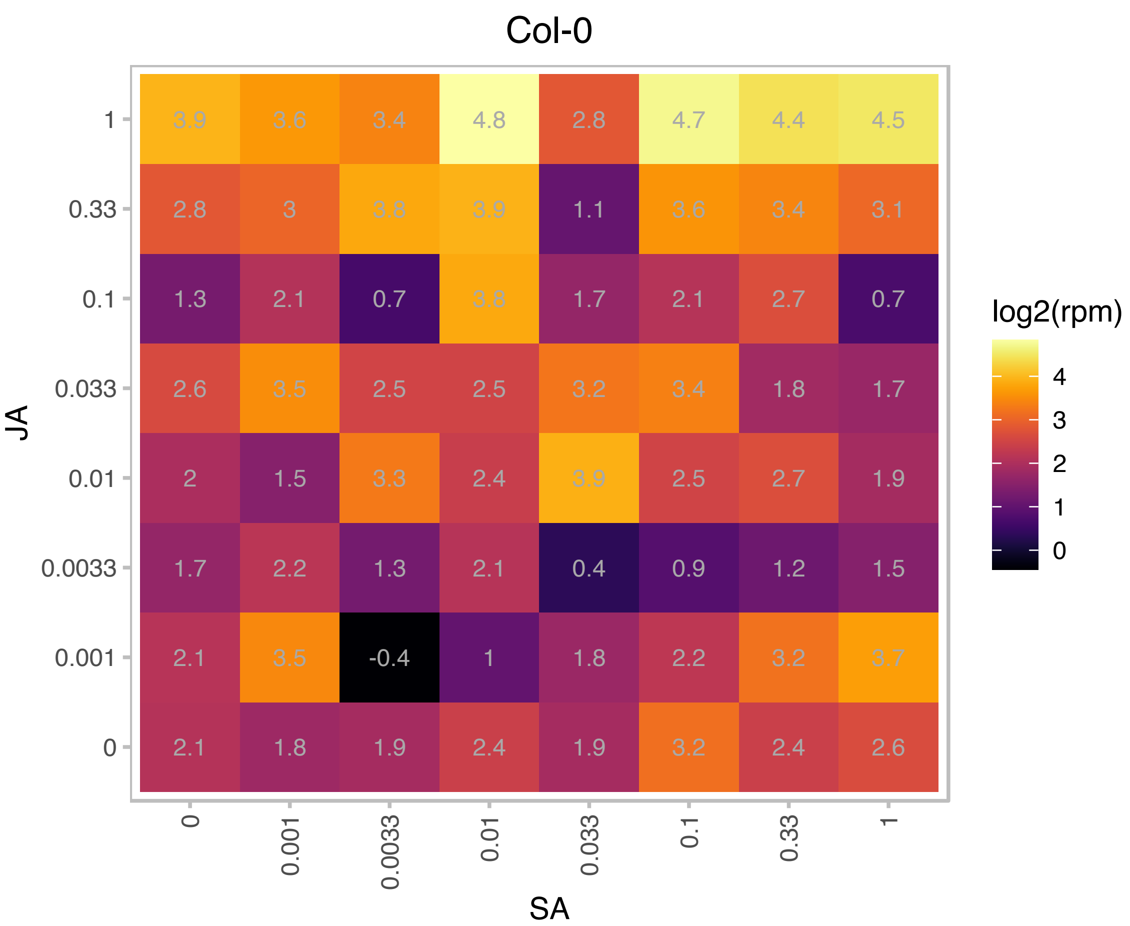


| | |
|---|---|
| Gene | AT4G04910 |
| is.expressed_genes | TRUE |
| is.responsive_genes | TRUE |
| is.apcluster | A-05 |

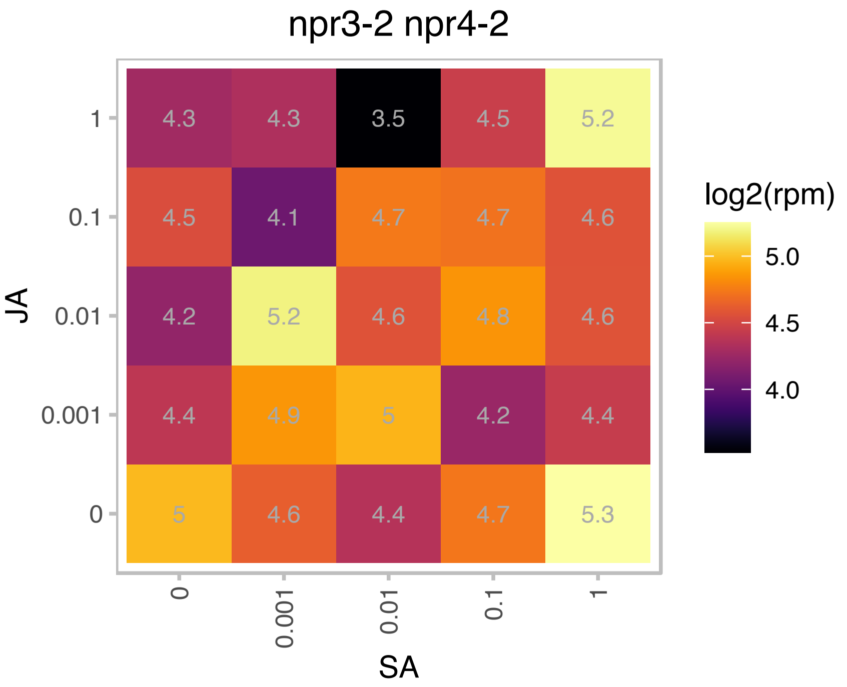


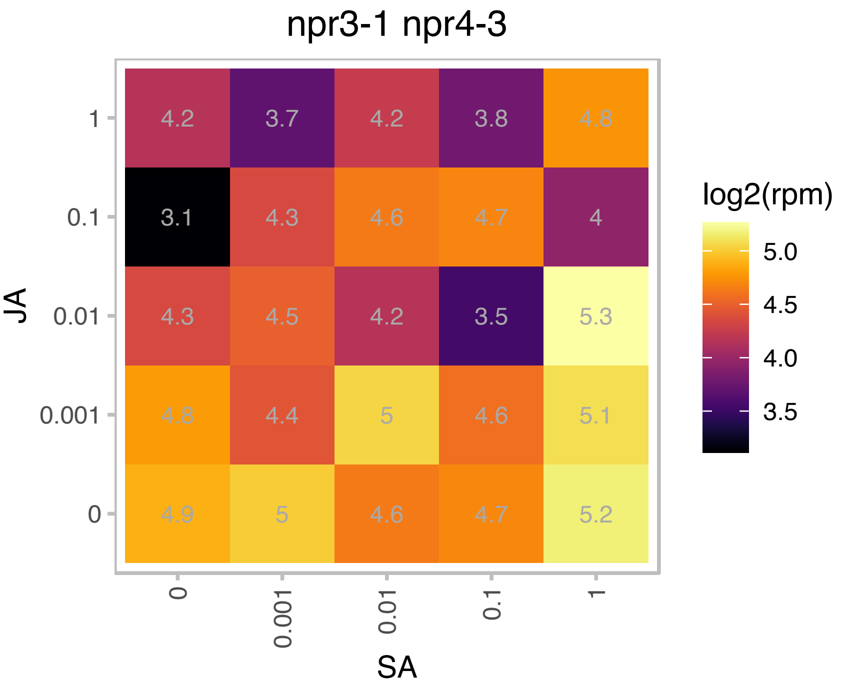

### AT4G11150.1 : emb2448, TUF, TUFF, VHA-E1

vacuolar ATP synthase subunit E1

Encodes a vacuolar H+-ATPase subunit E isoform 1 which is required for Golgi organization and vacuole function in embryogenesis. The mRNA is cell-to-cell mobile.

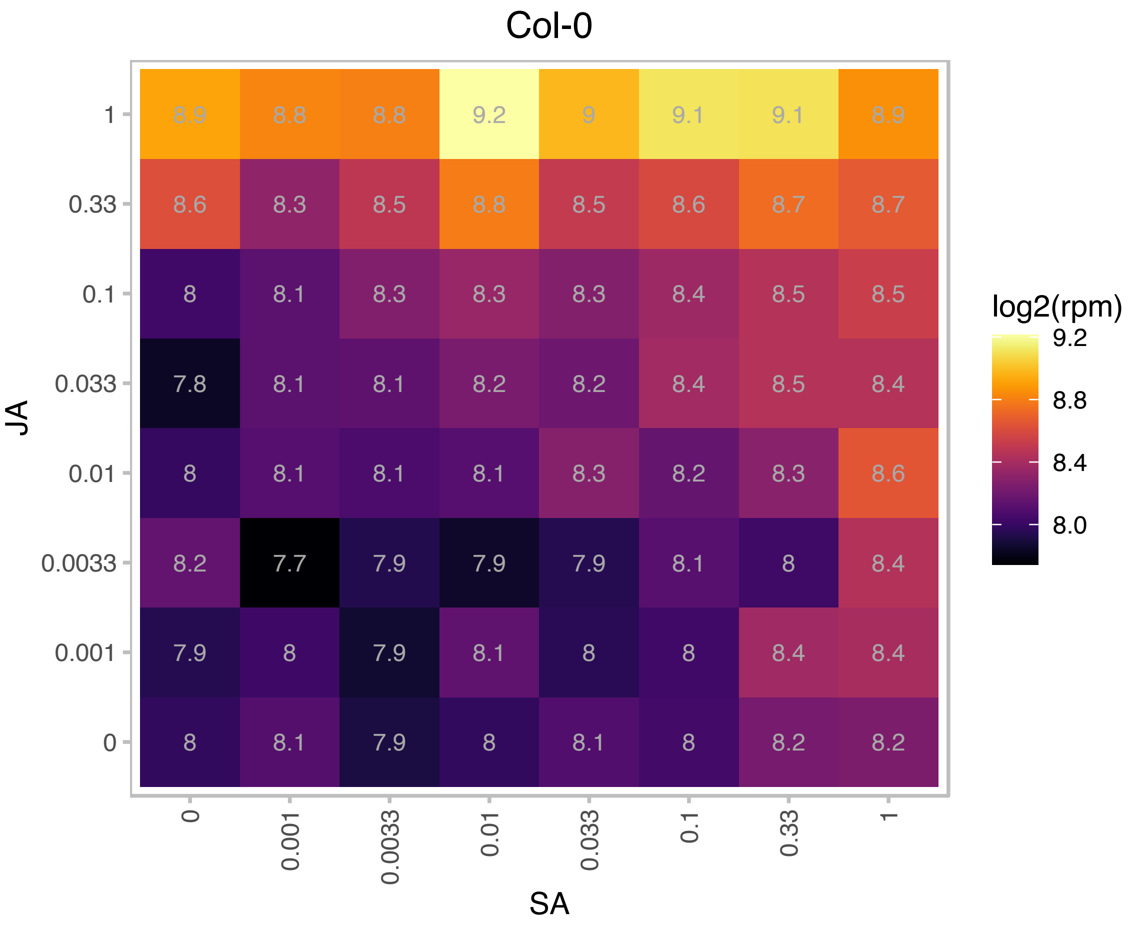


| | |
|---|---|
| Gene | AT4G11150 |
| is.expressed_genes | TRUE |
| is.responsive_genes | TRUE |
| is.apcluster | A-05 |

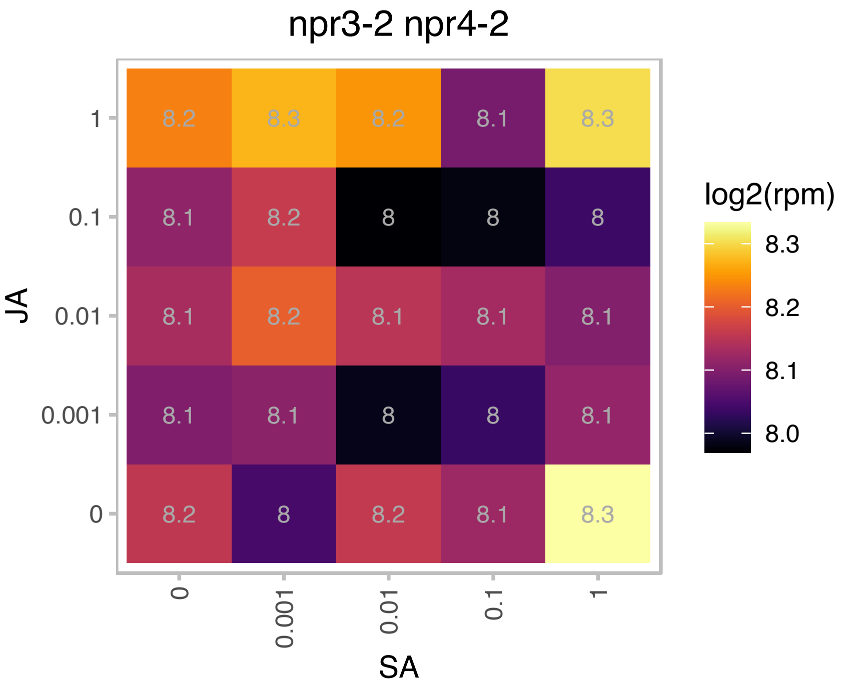


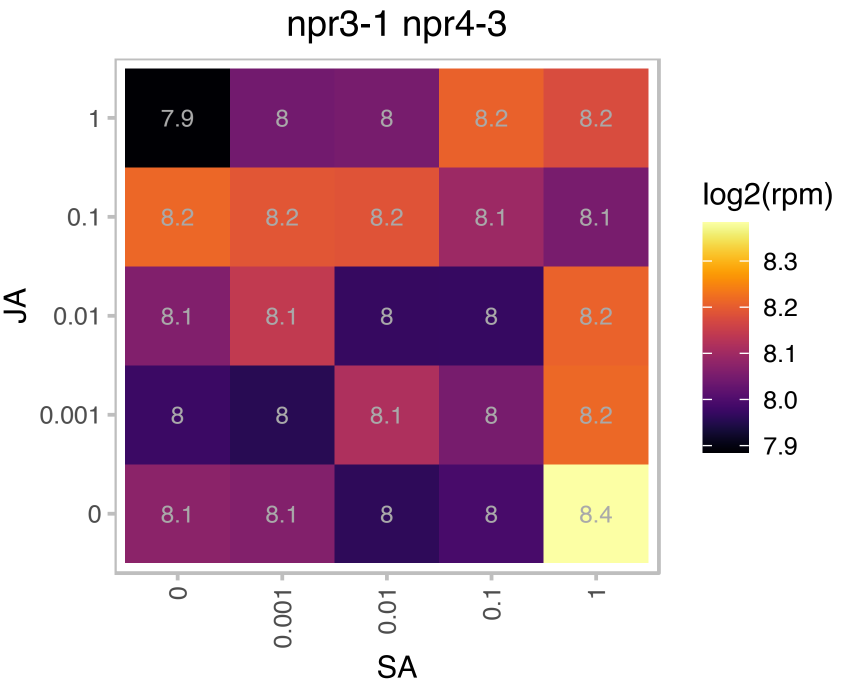

# AT4G14500.1 : NA

Polyketide cyclase/dehydrase and lipid transport superfamily protein

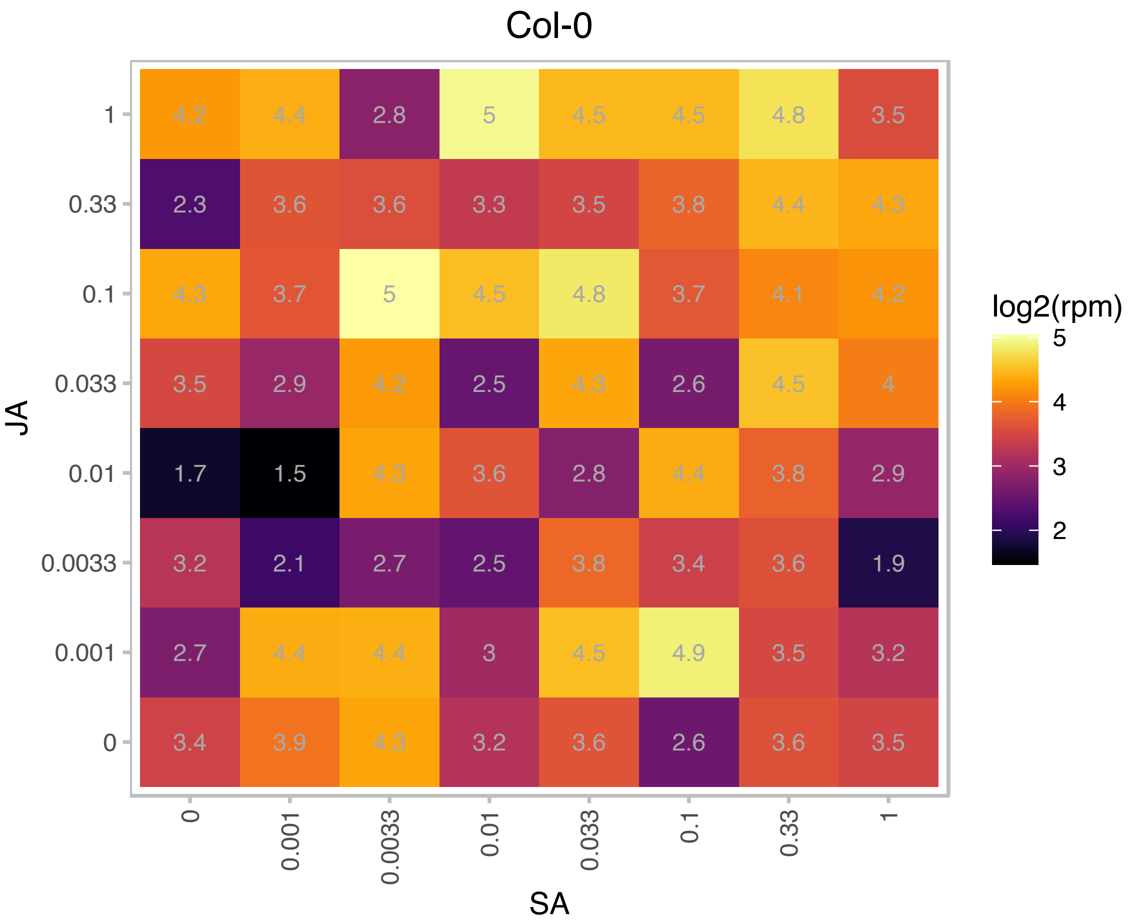


| | |
|---|---|
| Gene | AT4G14500 |
| is.expressed_genes | TRUE |
| is.responsive_genes | TRUE |
| is.apcluster | A-05 |

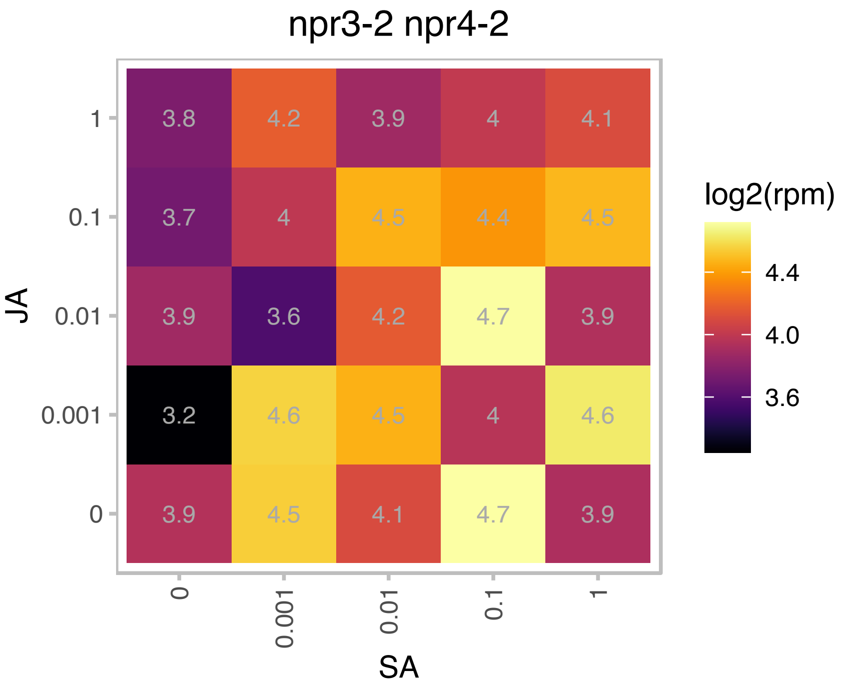


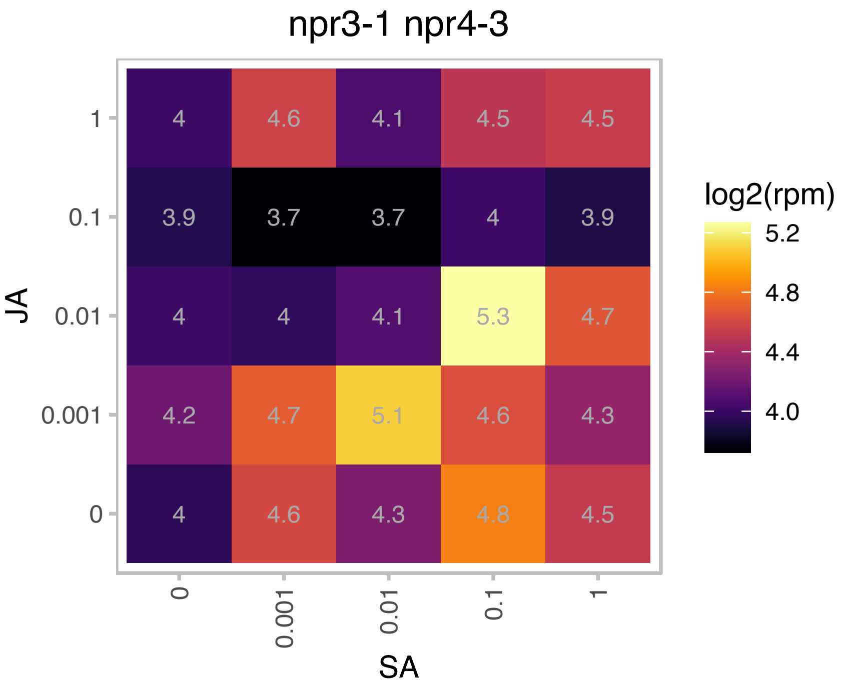

### AT4G21680.1 : AtNPF7.2, NPF7.2, NRT1.8

NITRATE TRANSPORTER 1.8

Encodes a nitrate transporter (NRT1.8). Functions in nitrate removal from the xylem sap. Mediates cadmium tolerance.

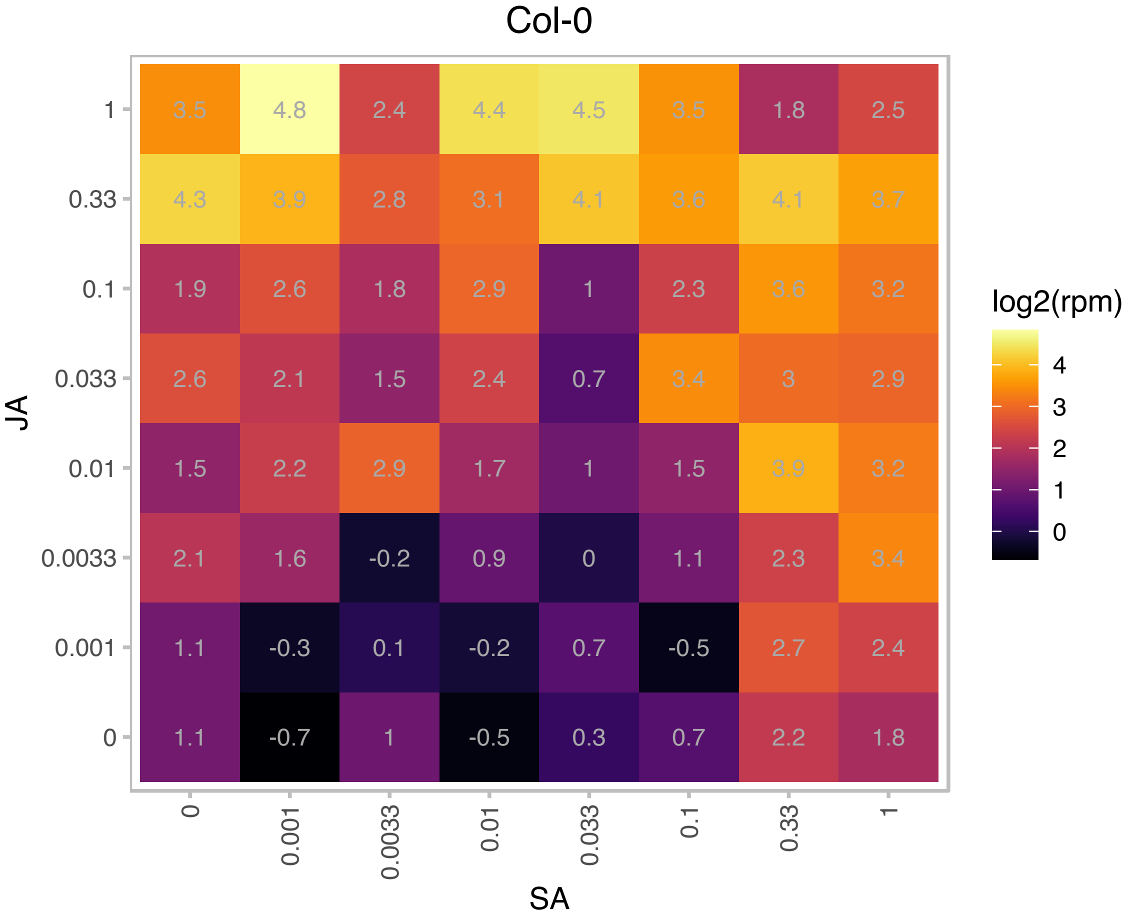


| | |
|---|---|
| Gene | AT4G21680 |
| is.expressed_genes | TRUE |
| is.responsive_genes | TRUE |
| is.apcluster | A-05 |

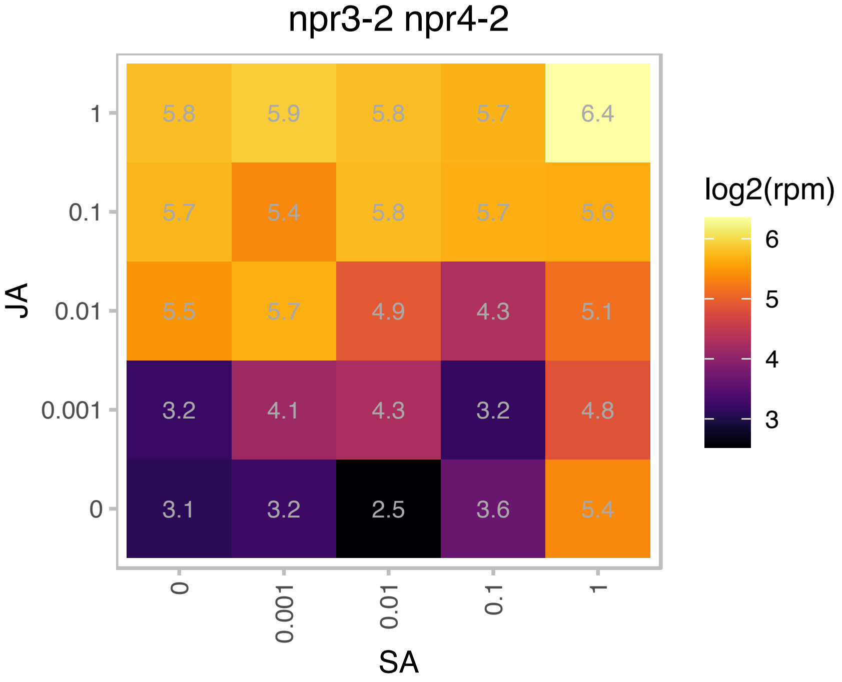


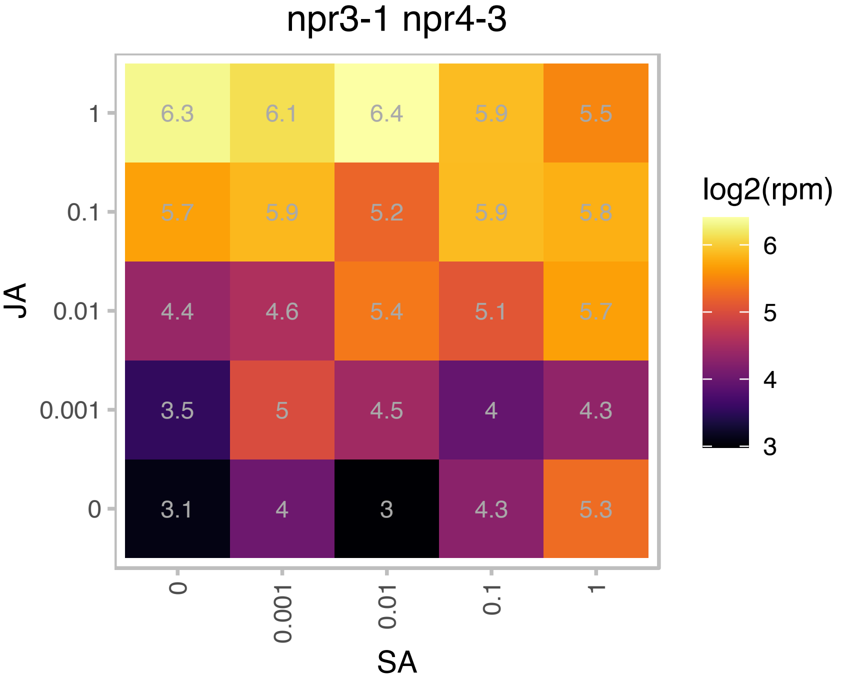

# AT4G24380.1 : NA

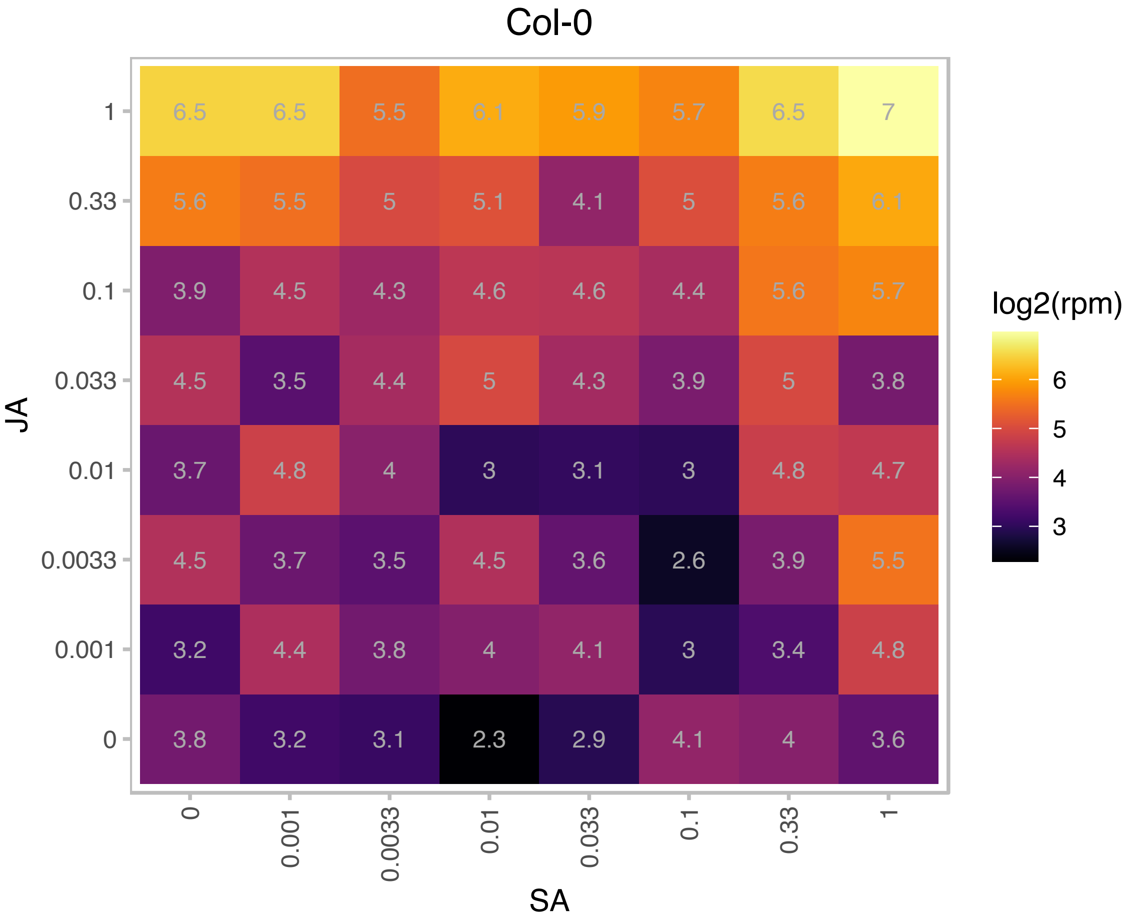


| | |
|---|---|
| Gene | AT4G24380 |
| is.expressed_genes | TRUE |
| is.responsive_genes | TRUE |
| is.apcluster | A-05 |

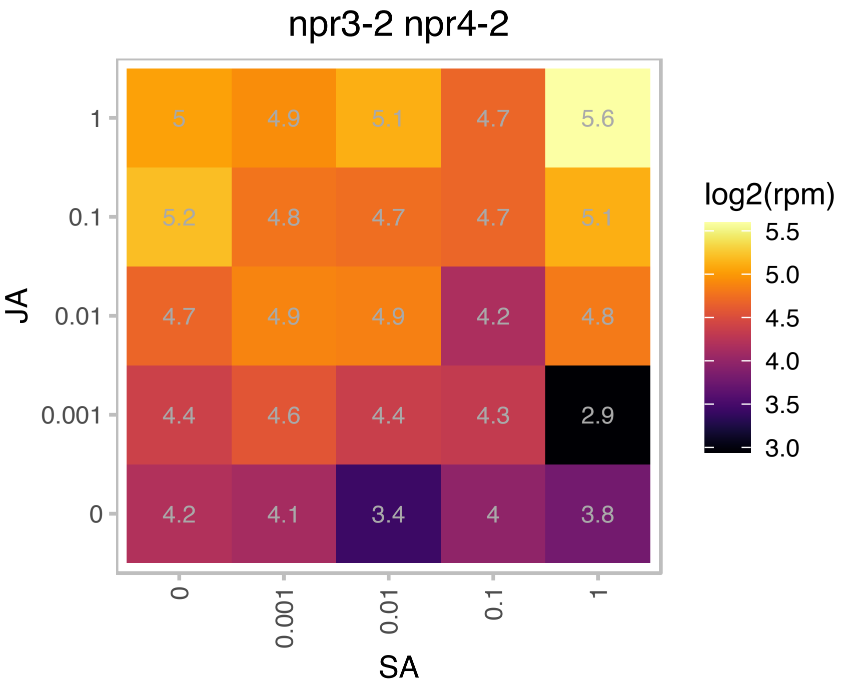


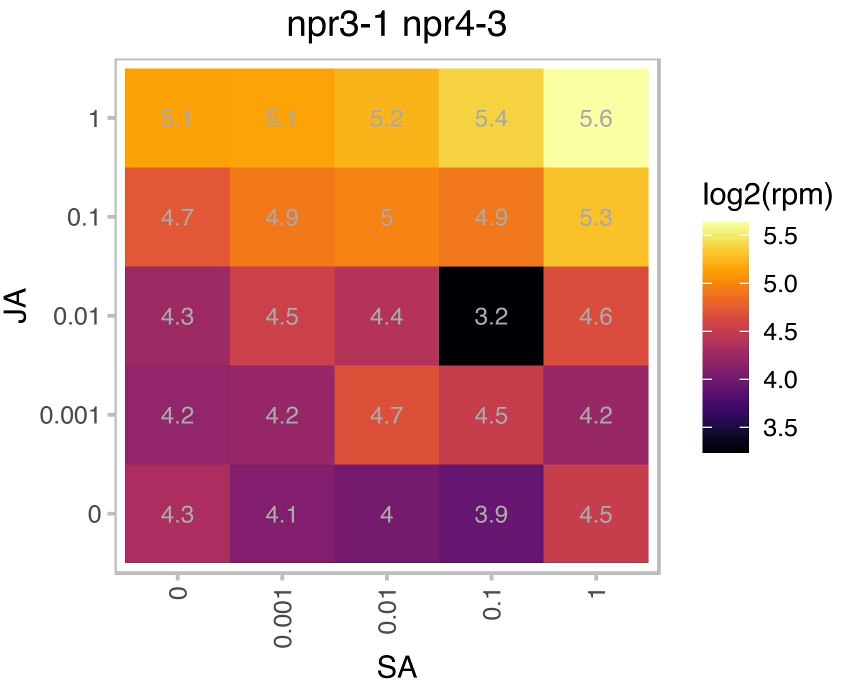

### AT4G27990.1 : ATYLMG1-2, YLMG1-2

YGGT family protein

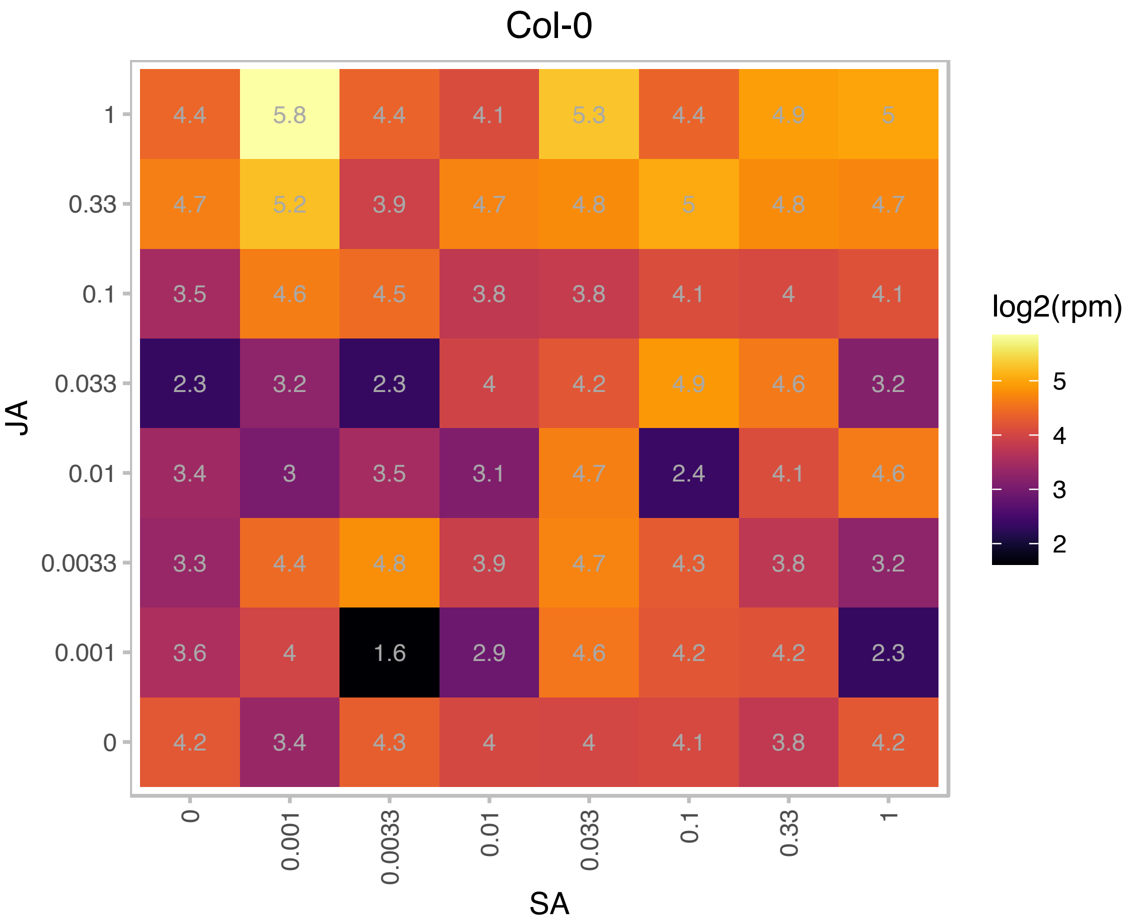


| | |
|---|---|
| Gene | AT4G27990 |
| is.expressed_genes | TRUE |
| is.responsive_genes | TRUE |
| is.apcluster | A-05 |

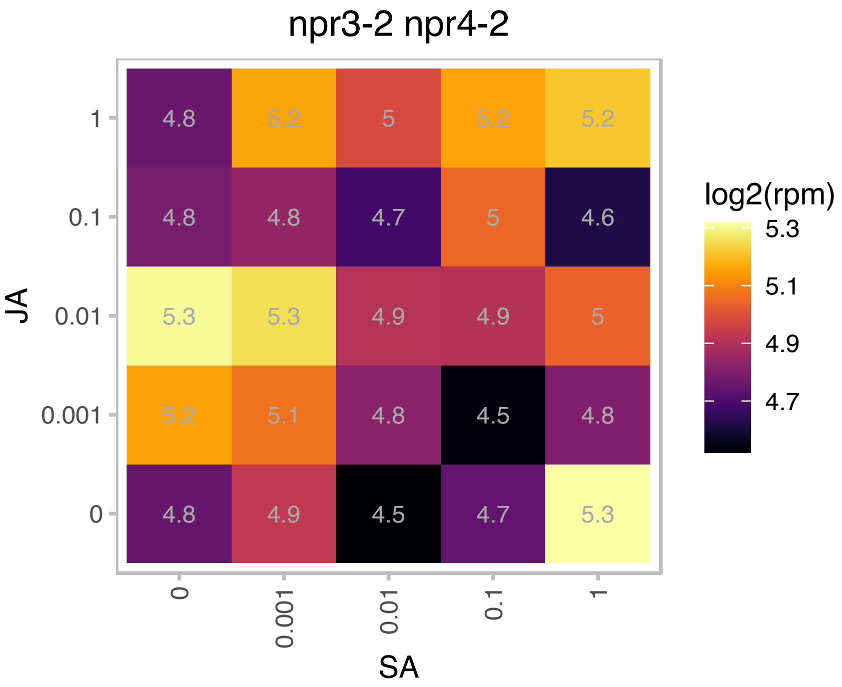


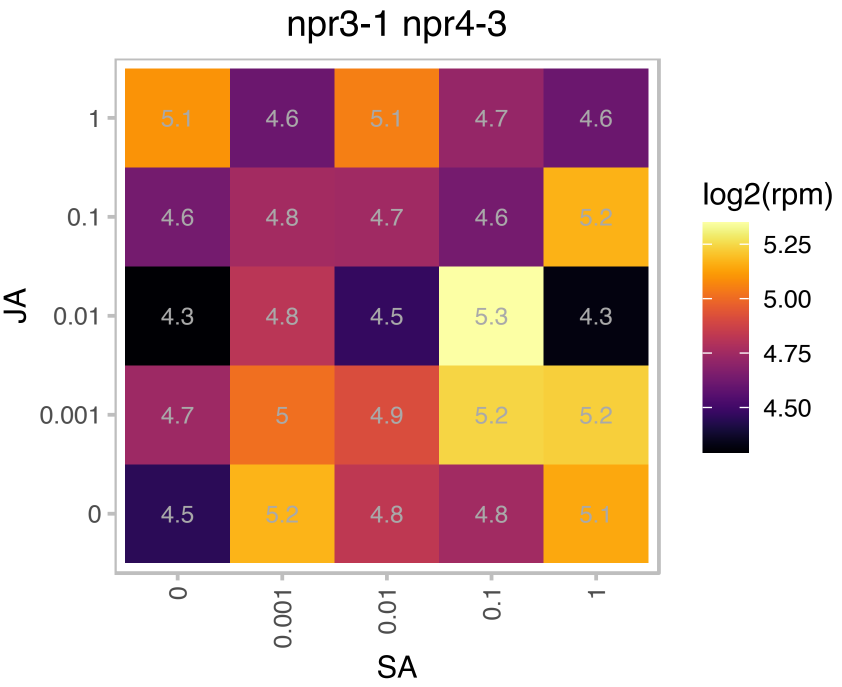

# AT4G31480.1 : NA

Coatomer, beta subunit

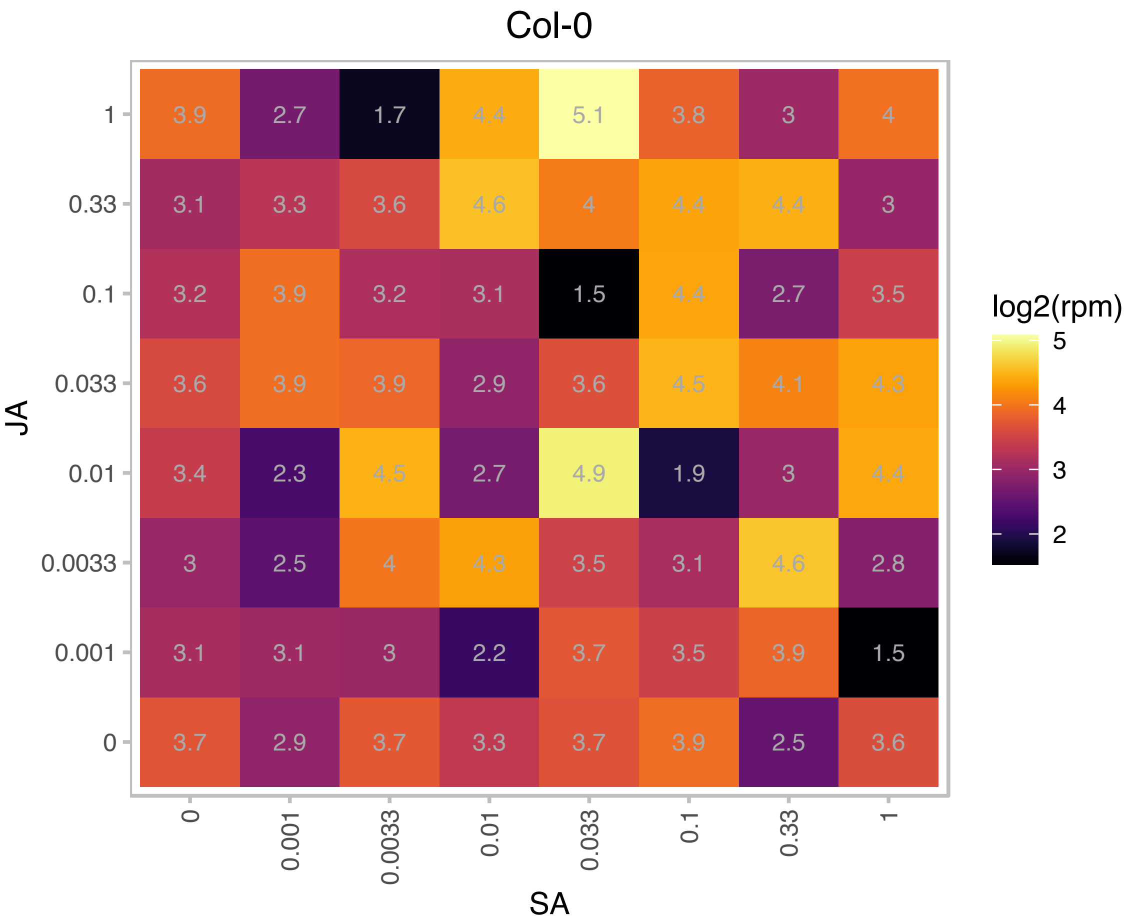


| | |
|---|---|
| Gene | AT4G31480 |
| is.expressed_genes | TRUE |
| is.responsive_genes | TRUE |
| is.apcluster | A-05 |

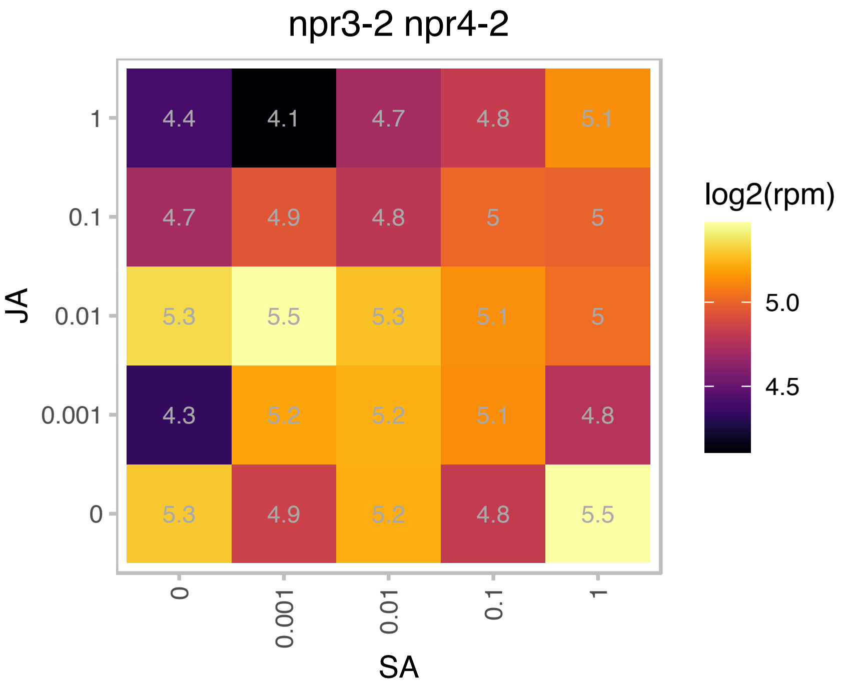


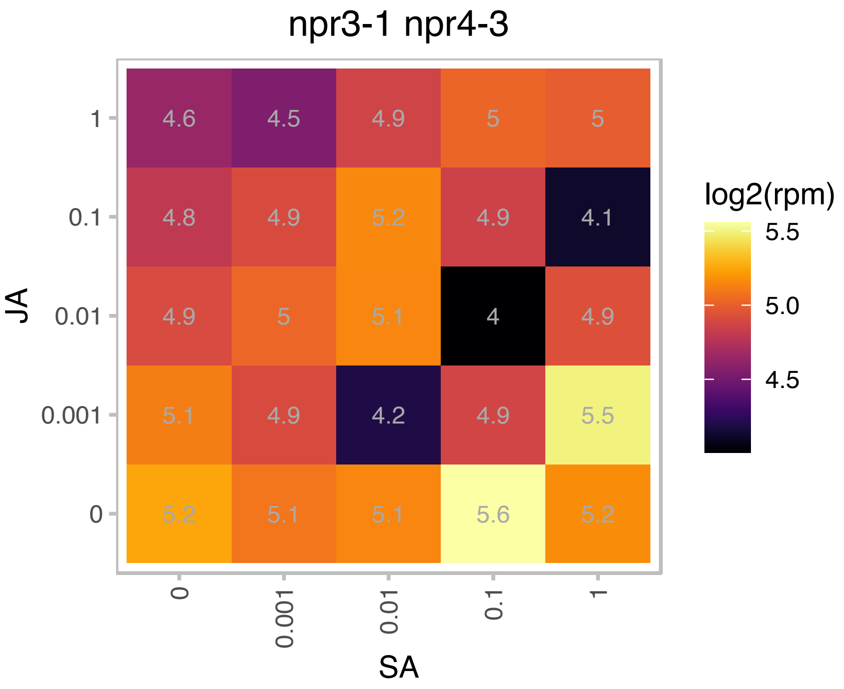

### AT4G33680.1 : AGD2

Pyridoxal phosphate (PLP)-dependent transferases superfamily protein
Encodes an L,L-diaminopimelate aminotransferase. Involved in disease resistance against Pseudomonas syringae. mutants have elevated SA levels, a low level of spontaneous cell death, callose deposition, and enlarged cells in leaves. genetically maps on chr 4 between L23H3 and nga1139.

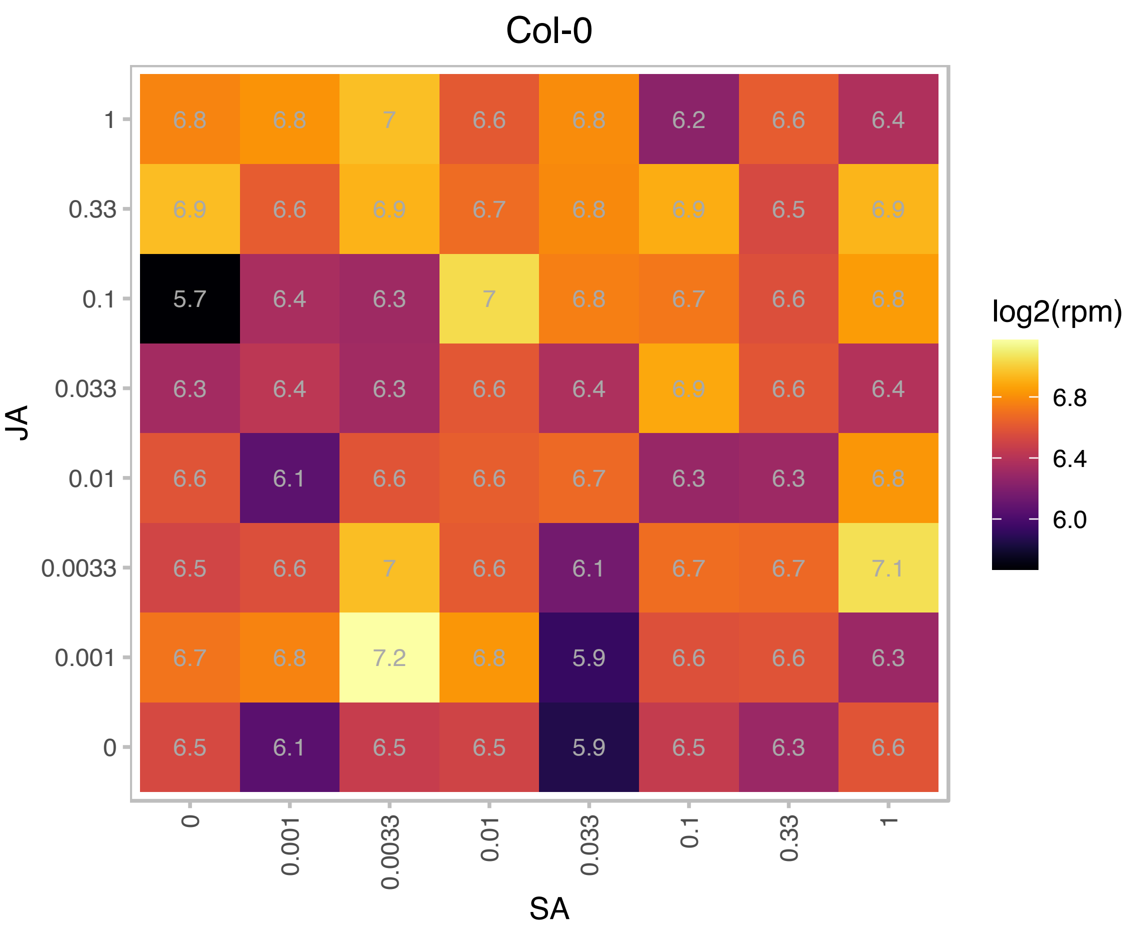


| | |
|---|---|
| Gene | AT4G33680 |
| is.expressed_genes | TRUE |
| is.responsive_genes | TRUE |
| is.apcluster | A-05 |

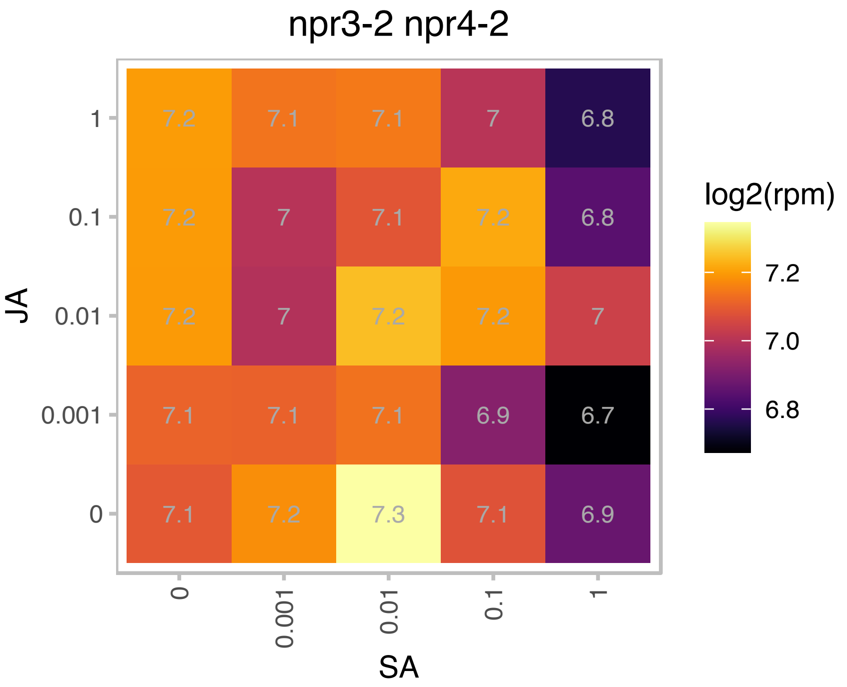


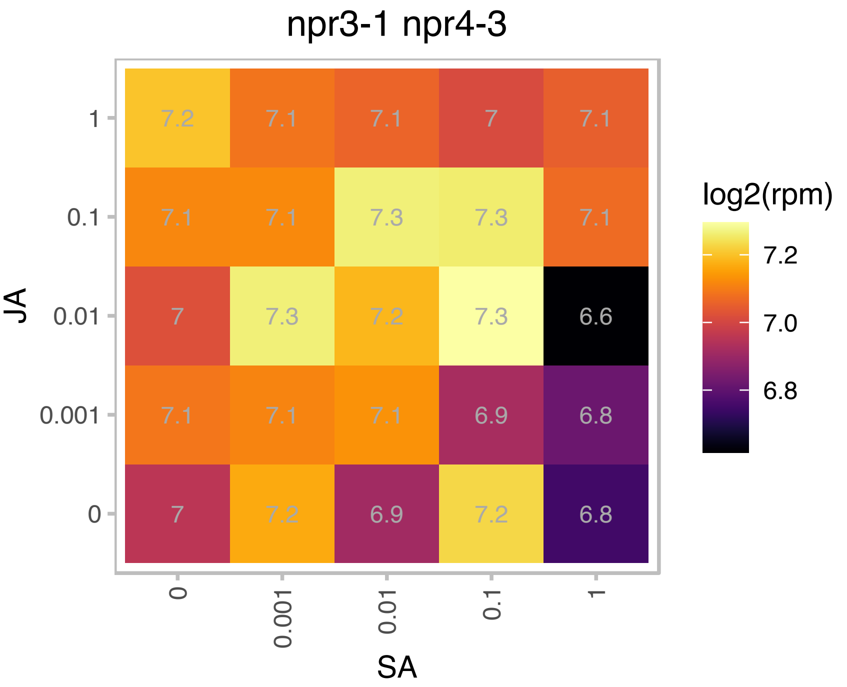

### AT4G36890.1 : IRX14

IRX14 was identified as MUCI64 in a reverse genetic screen for MUCILAGE-RELATED genes. IRX14/MUCI64 is a G
Nucleotide-diphospho-sugar transferases superfamily protein
protein essential for xylan elongation in seed coat mucilage. The xylan backbone maintains the attachment of mucilage to the seed surface and the distribution of cellulose. It was identified based on its gene expression co-variance with the IRX3 gene involved in secondary cell wall synthesis.  A biochemical assay using the irx14 mutant indicates that IRX14 might function in xylose chain elongation.

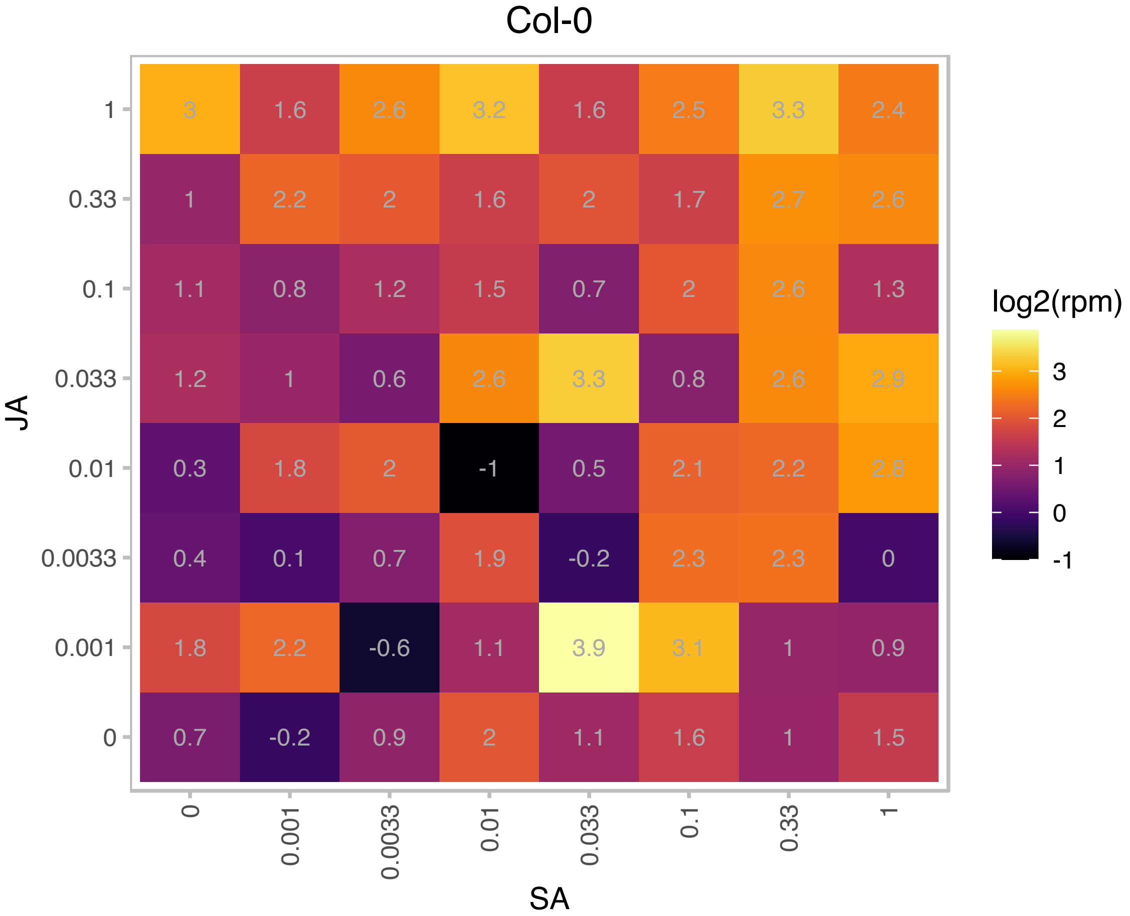


| | |
|---|---|
| Gene | AT4G36890 |
| is.expressed_genes | TRUE |
| is.responsive_genes | TRUE |
| is.apcluster | A-05 |

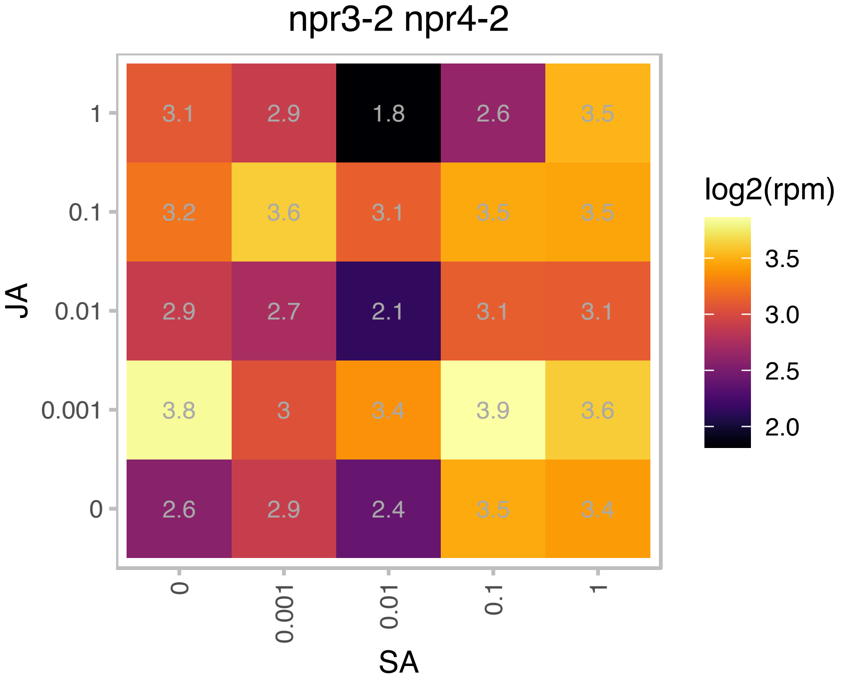


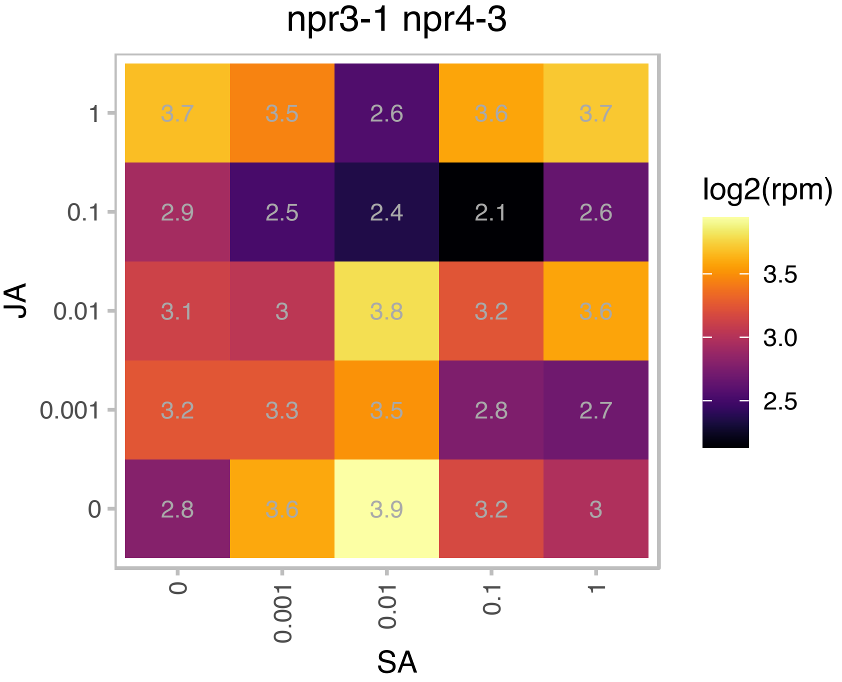

### AT5G03210.1 : AtDIP2, DIP2

Encodes a small polypeptide contributing to resistance to potyvirus.

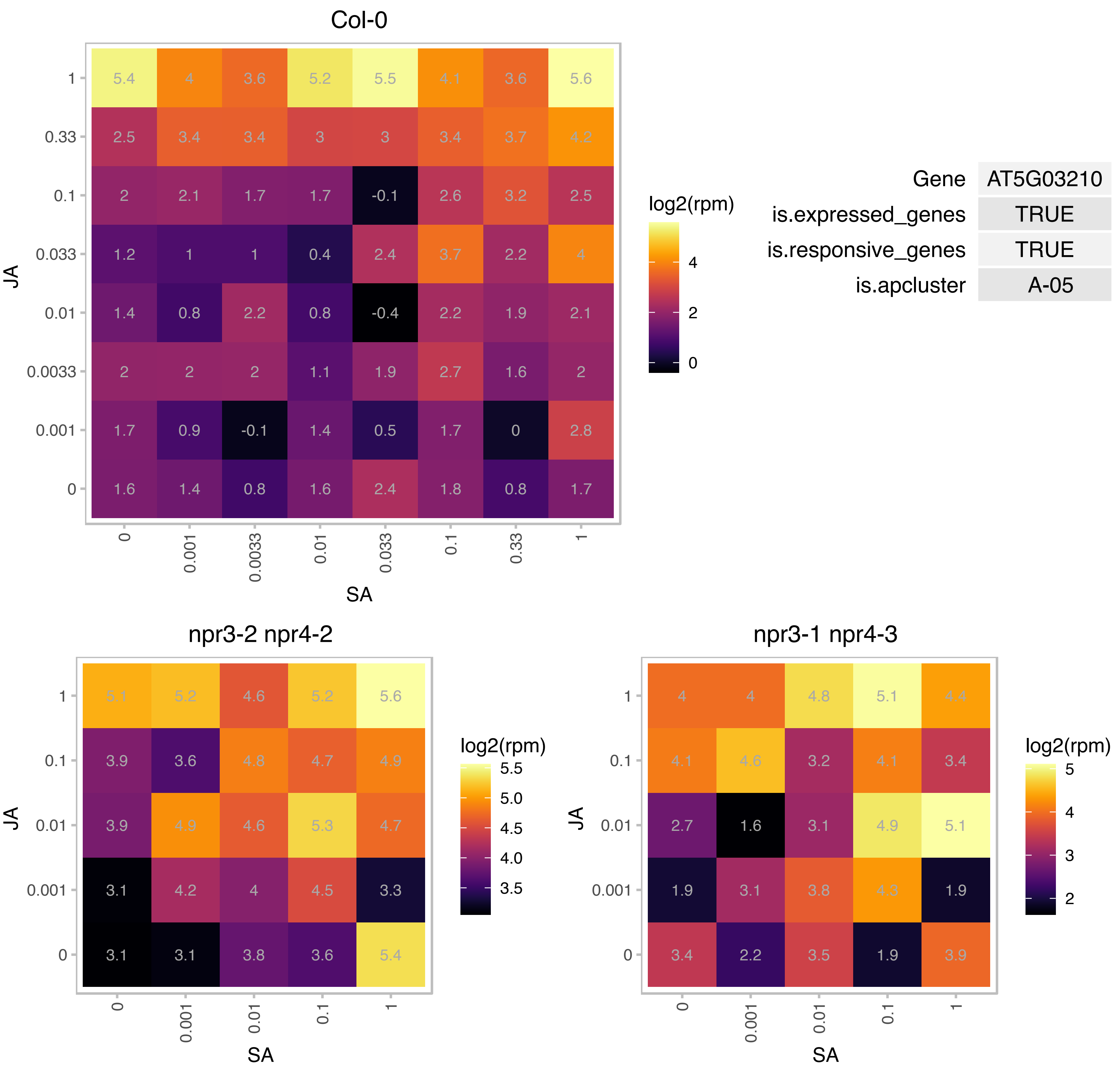


| | |
|---|---|
| Gene | AT5G03210 |
| is.expressed_genes | TRUE |
| is.responsive_genes | TRUE |
| is.apcluster | A-05 |

# AT5G07740.1 : NA

actin binding

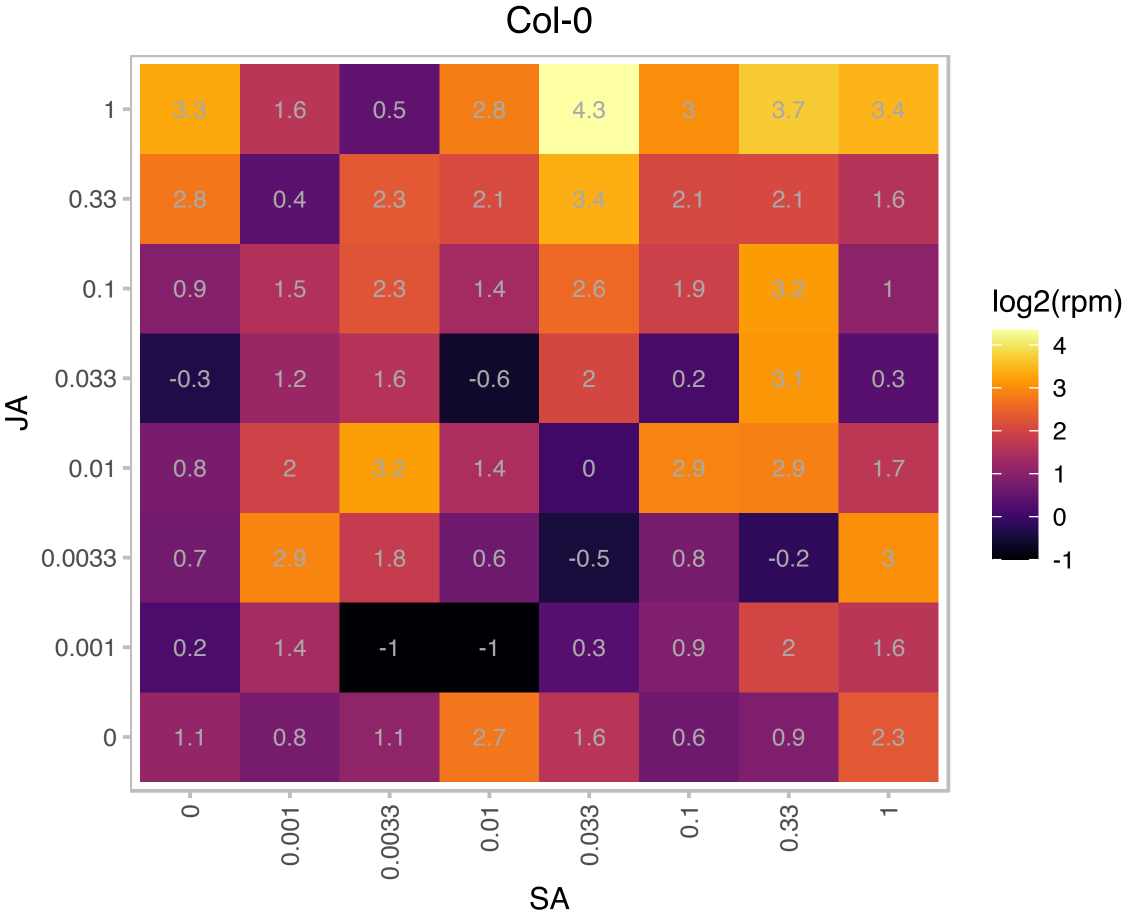


| | |
|---|---|
| Gene | AT5G07740 |
| is.expressed_genes | TRUE |
| is.responsive_genes | TRUE |
| is.apcluster | A-05 |

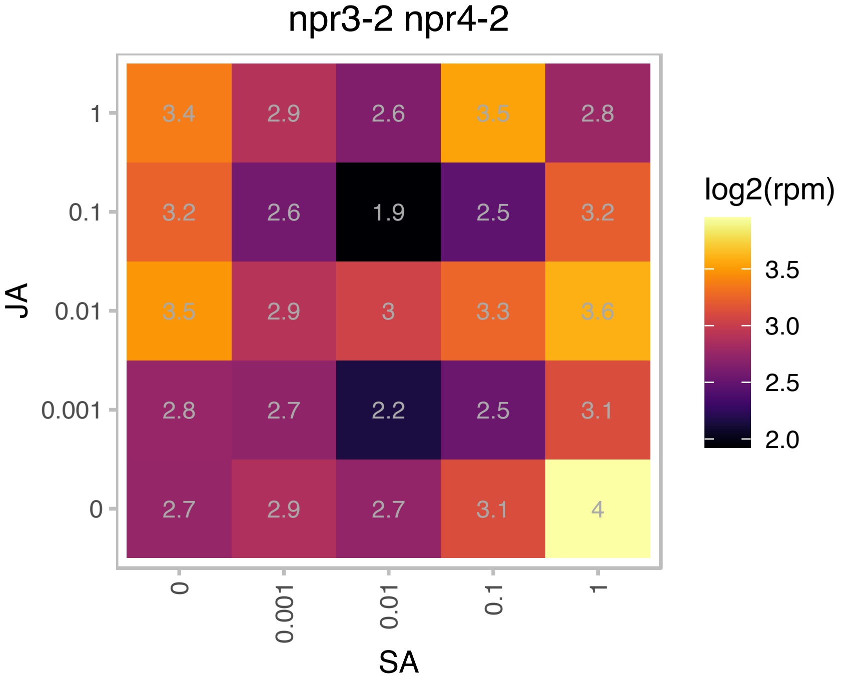


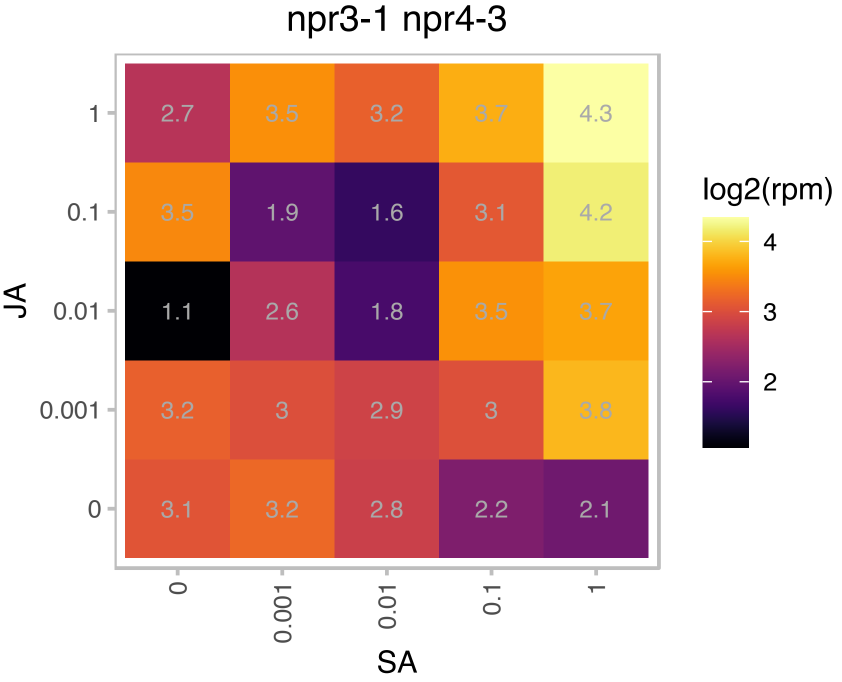

# AT5G12010.1 : NA

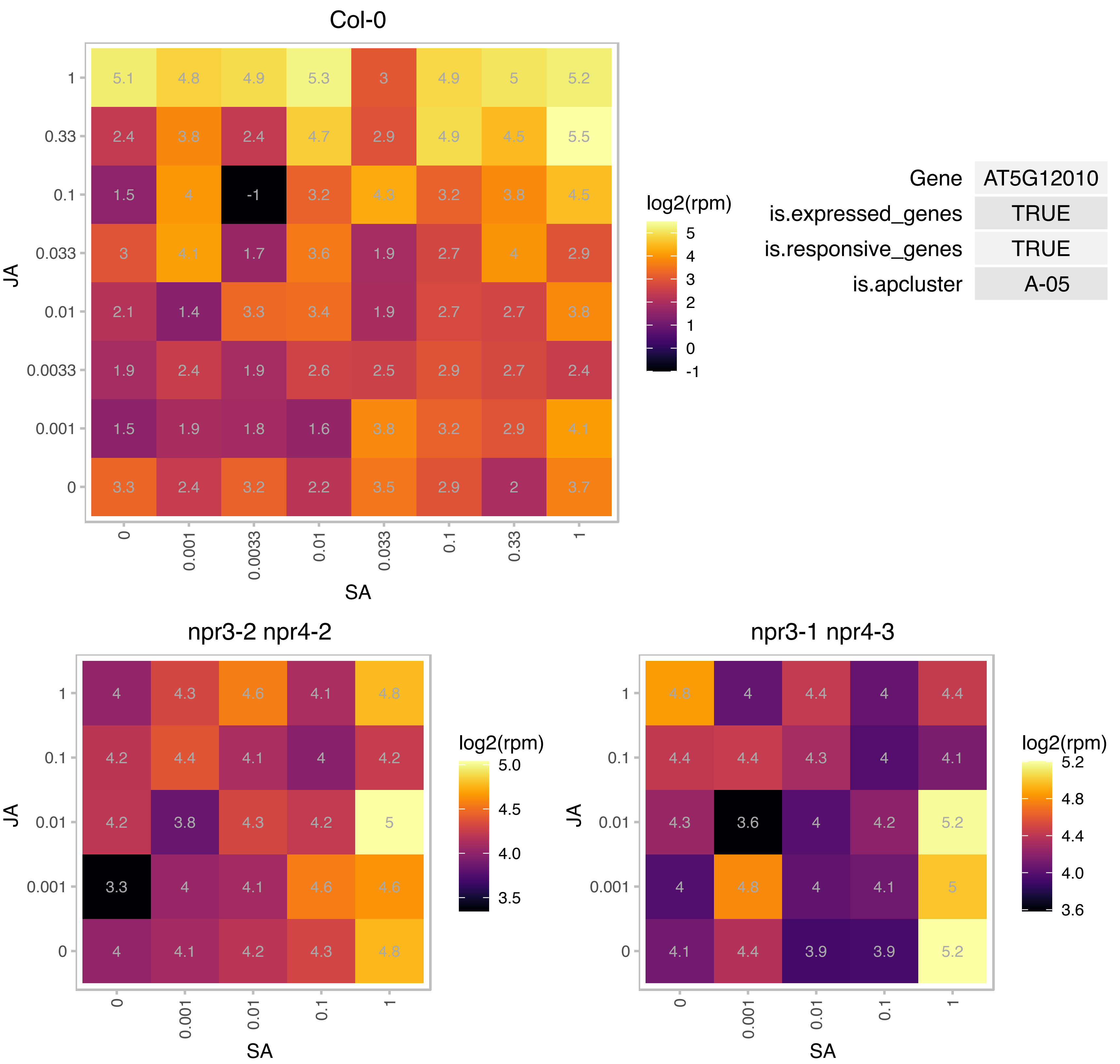


| | |
|---|---|
| Gene | AT5G12010 |
| is.expressed_genes | TRUE |
| is.responsive_genes | TRUE |
| is.apcluster | A-05 |

### AT5G13280.1 : AK, AK-LYS1, AK1

aspartate kinase 1
Asp kinase inhibited by Lys and S-adenosylmethionine. Contains regulatory domains that belong to the ACT domain family, which allow binding to a extreme variety of ligands. Can function as a monomer or as a dimer with acetohydroxyacid synthase (HSDH).

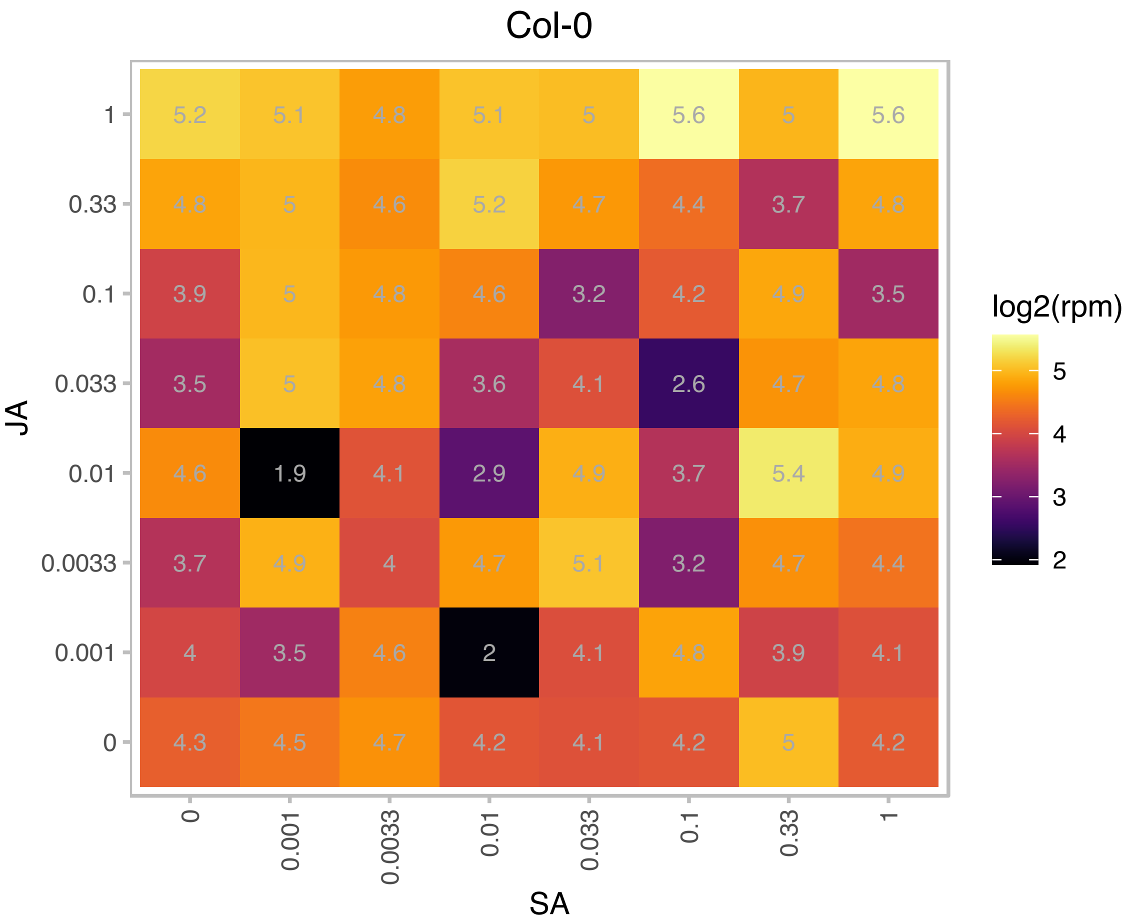


| | |
|---|---|
| Gene | AT5G13280 |
| is.expressed_genes | TRUE |
| is.responsive_genes | TRUE |
| is.apcluster | A-05 |

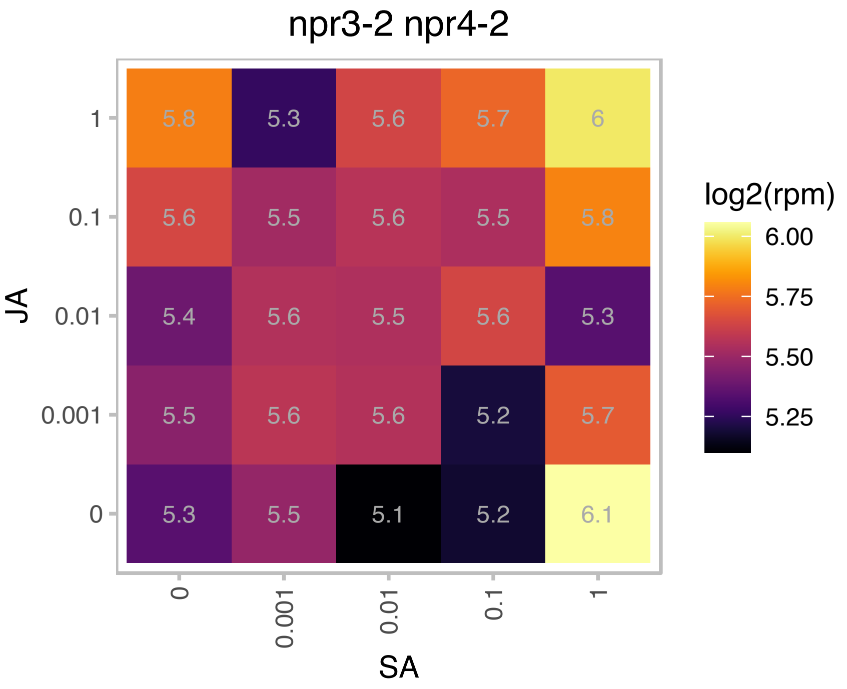


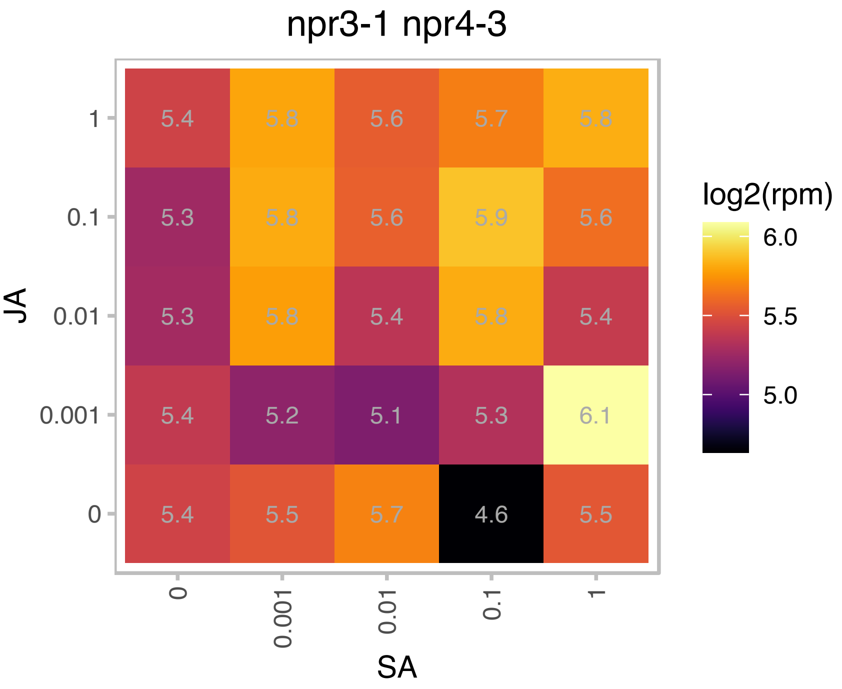

### AT5G17640.1 : ASG1

Protein of unknown function (DUF1005)

Expression of this gene is induced by abscisic acid and salt stress.

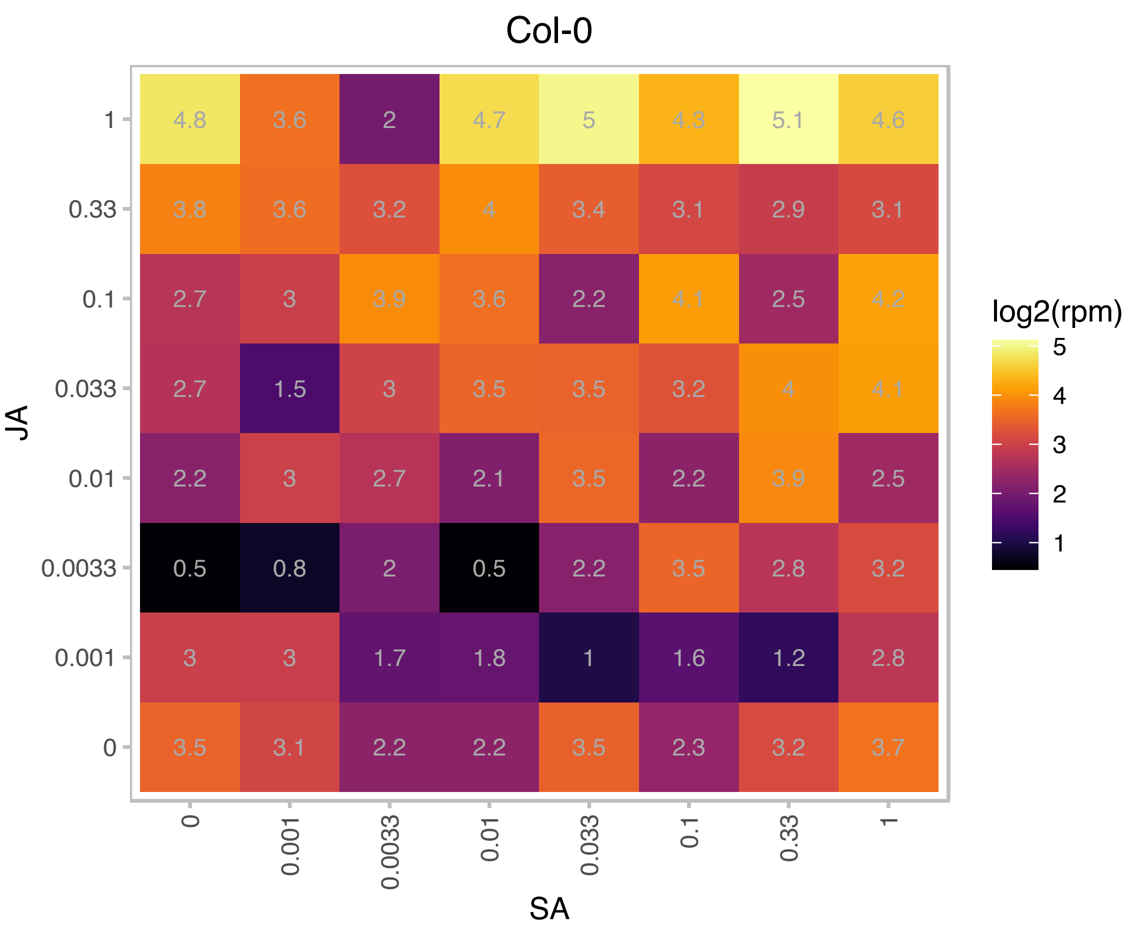


| | |
|---|---|
| Gene | AT5G17640 |
| is.expressed_genes | TRUE |
| is.responsive_genes | TRUE |
| is.apcluster | A-05 |

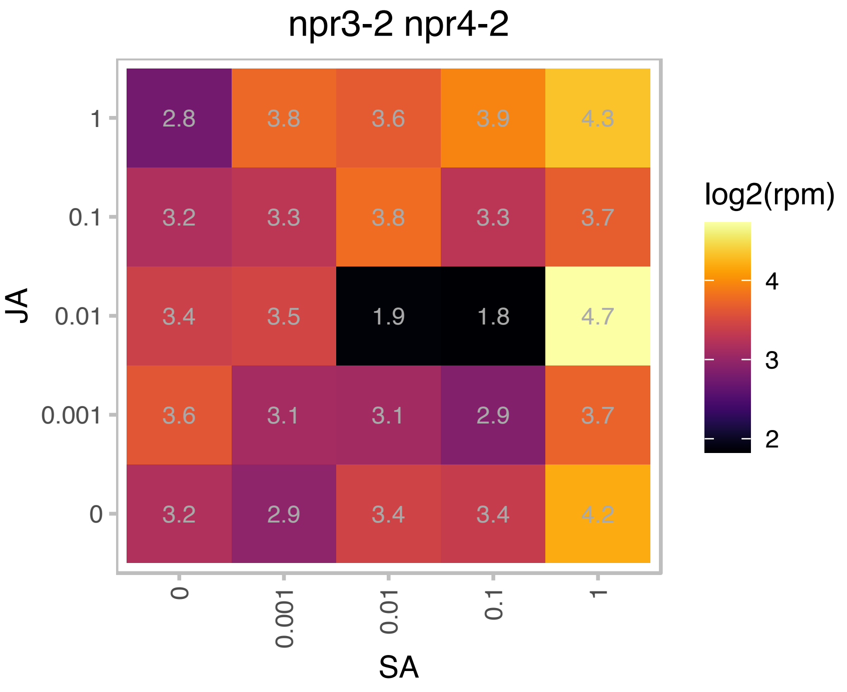


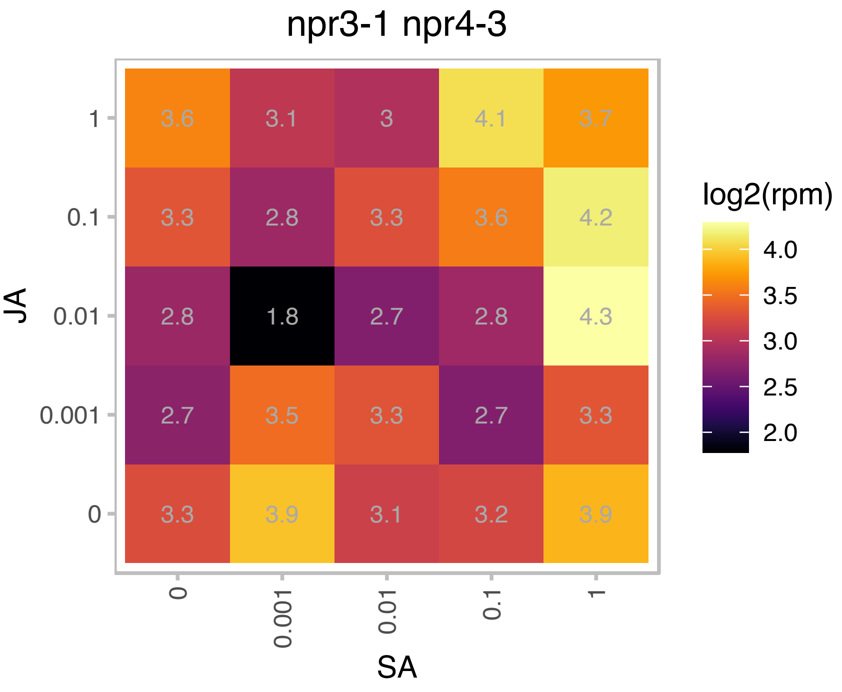

# AT5G21280.1 : NA

hydroxyproline-rich glycoprotein family protein

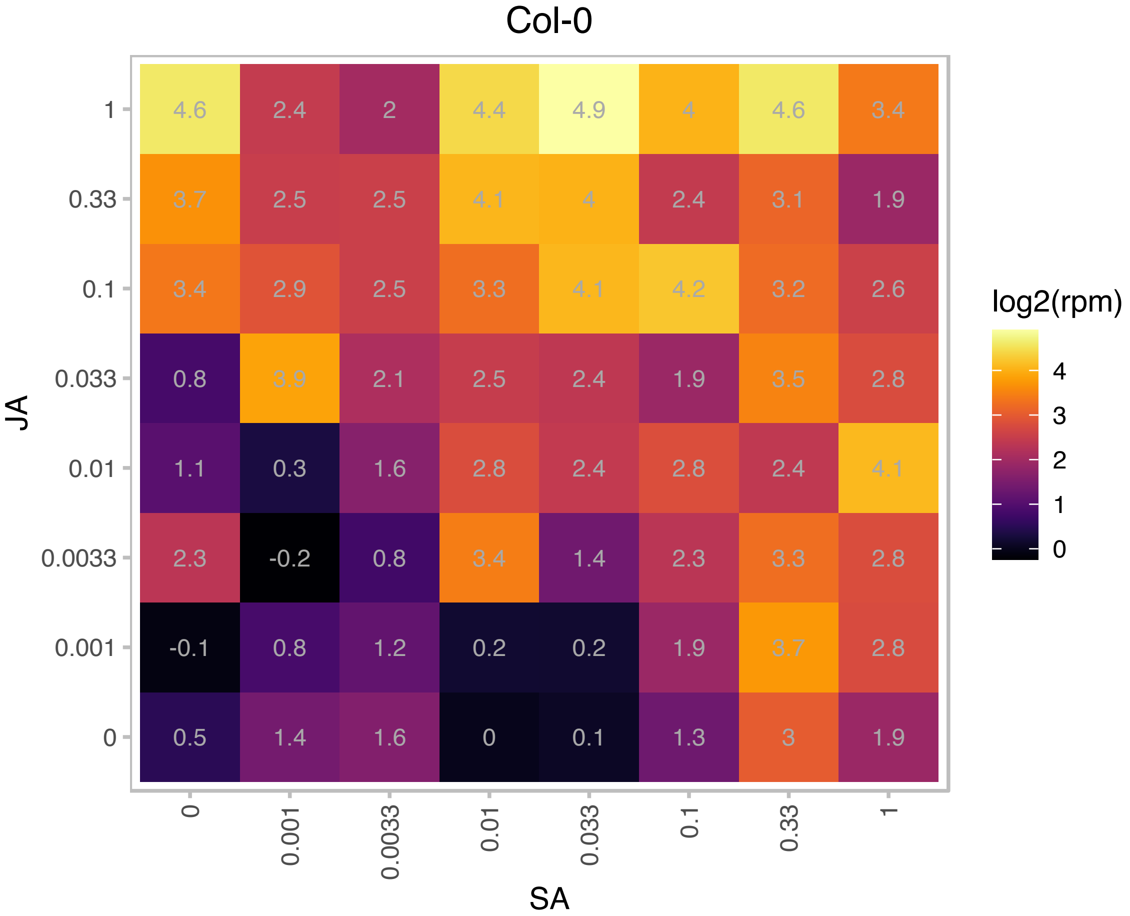


| | |
|---|---|
| Gene | AT5G21280 |
| is.expressed_genes | TRUE |
| is.responsive_genes | TRUE |
| is.apcluster | A-05 |

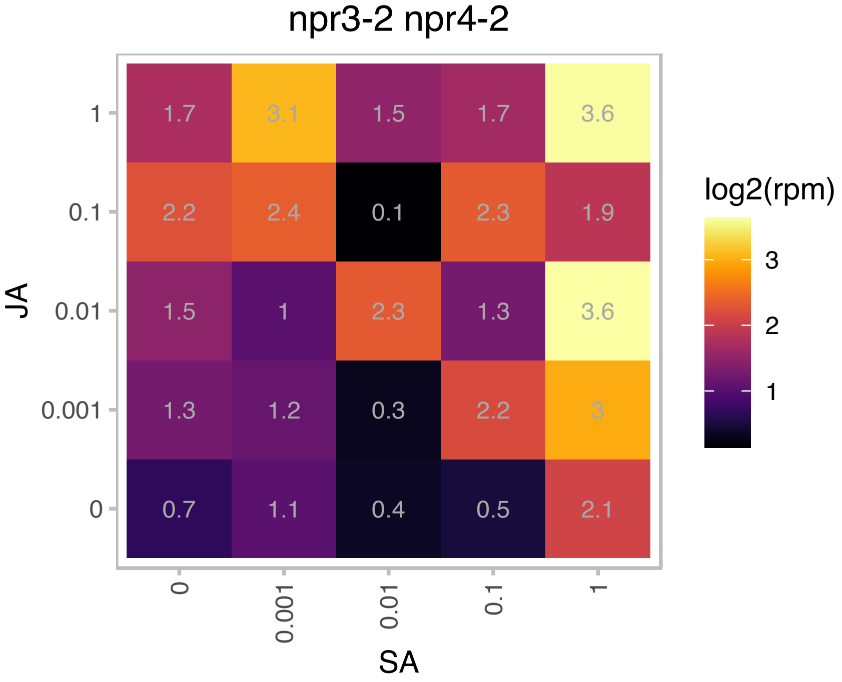


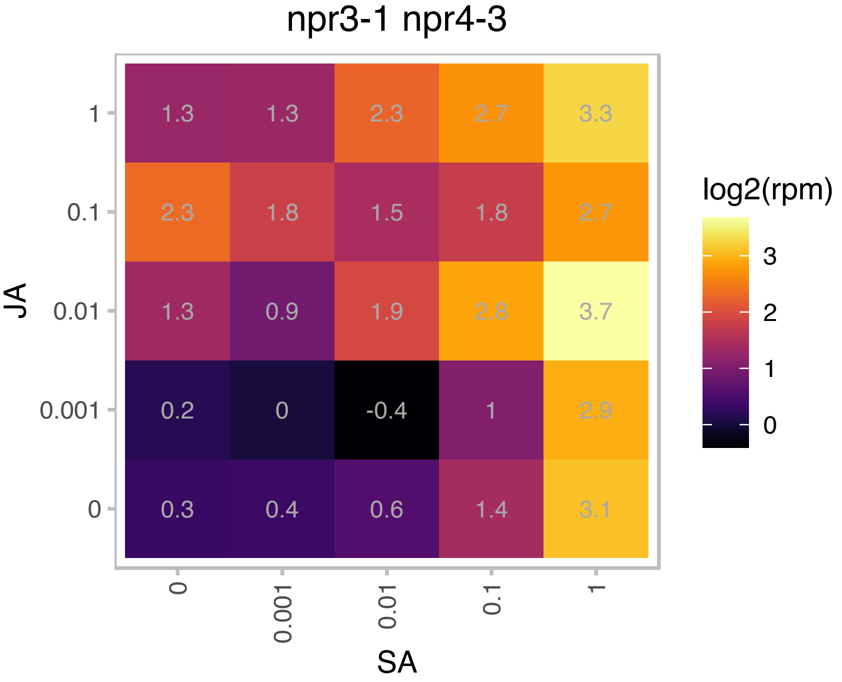

### AT5G27520.1 : AtPNC2, PNC2

peroxisomal adenine nucleotide carrier 2

encodes a peroxisomal adenine nucleotide transporter, involved in fatty acid beta-oxidation during early stage of postgerminative growth.

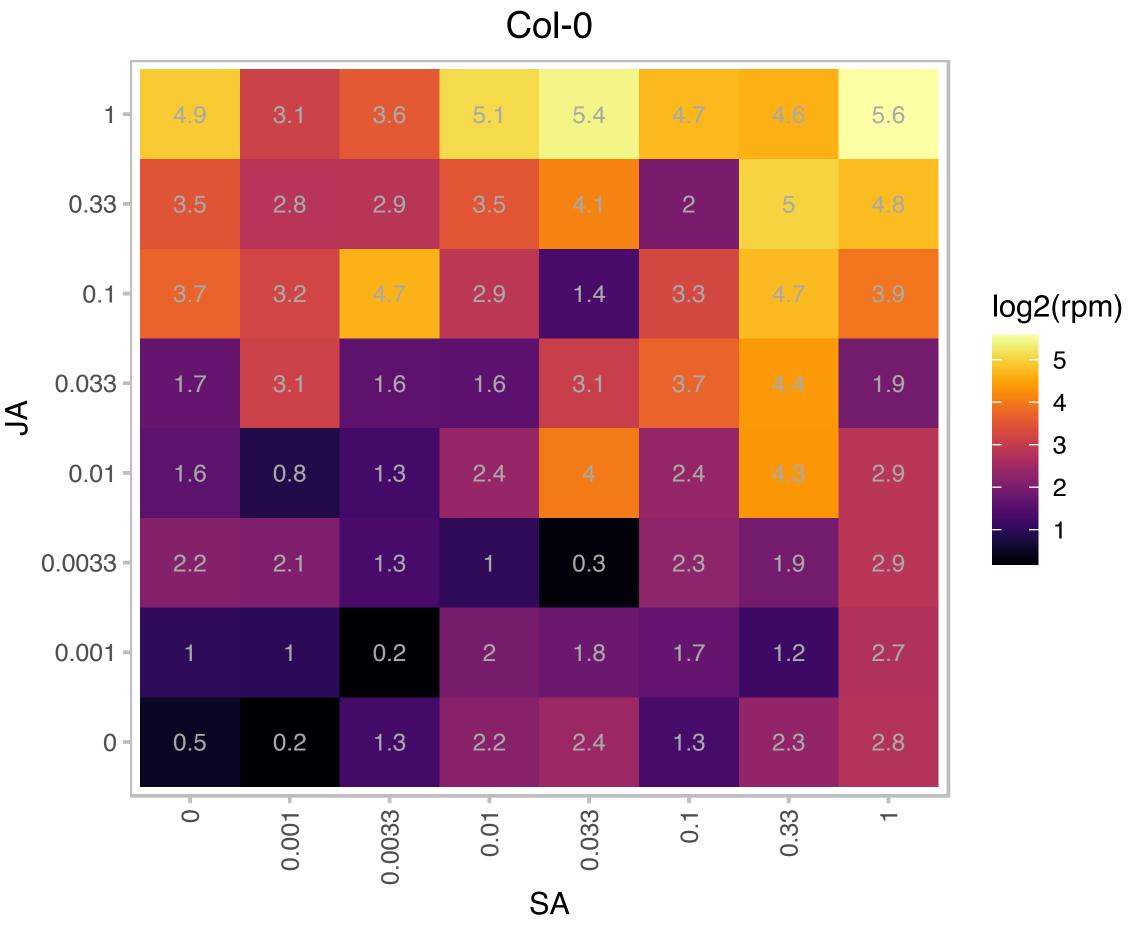


| | |
|---|---|
| Gene | AT5G27520 |
| is.expressed_genes | TRUE |
| is.responsive_genes | TRUE |
| is.apcluster | A-05 |

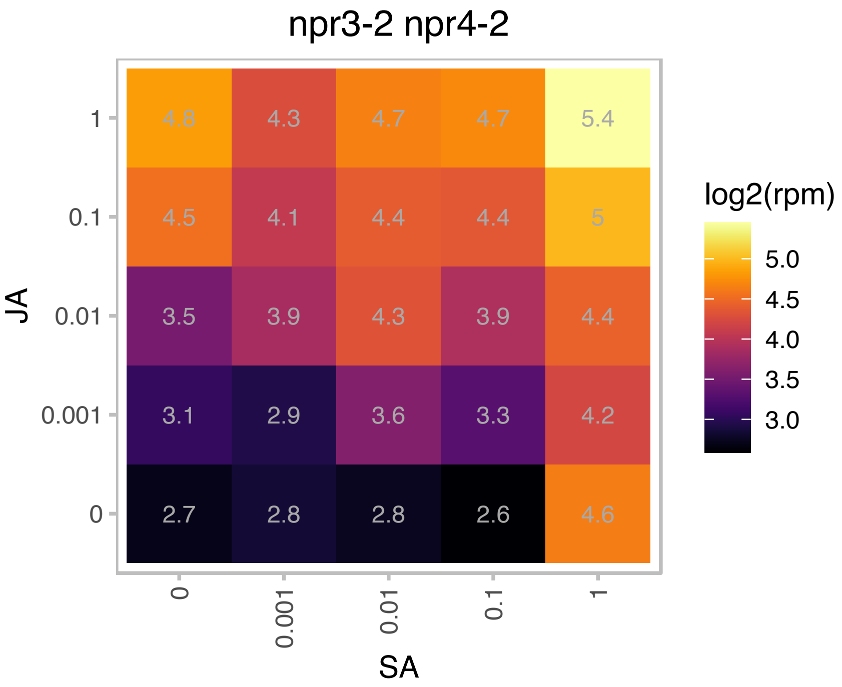


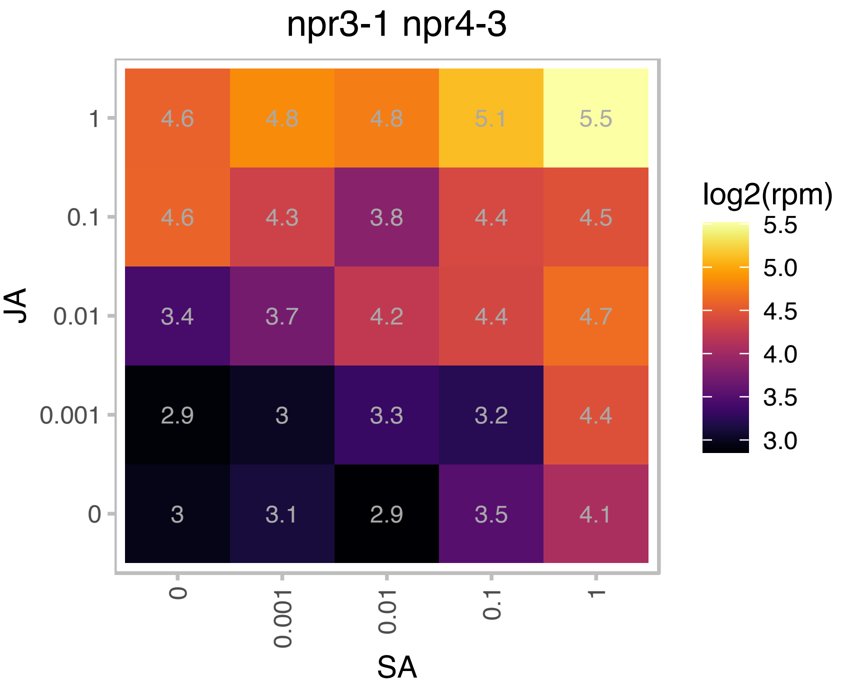

### AT5G27540.1 : emb2473, MIRO1

MIRO-related GTP-ase 1

Encodes a protein with similarity to GTPases that is localized to the mitochondrion. Involved in embryogenesis, pollen tube growth and required for mitochondrial development.

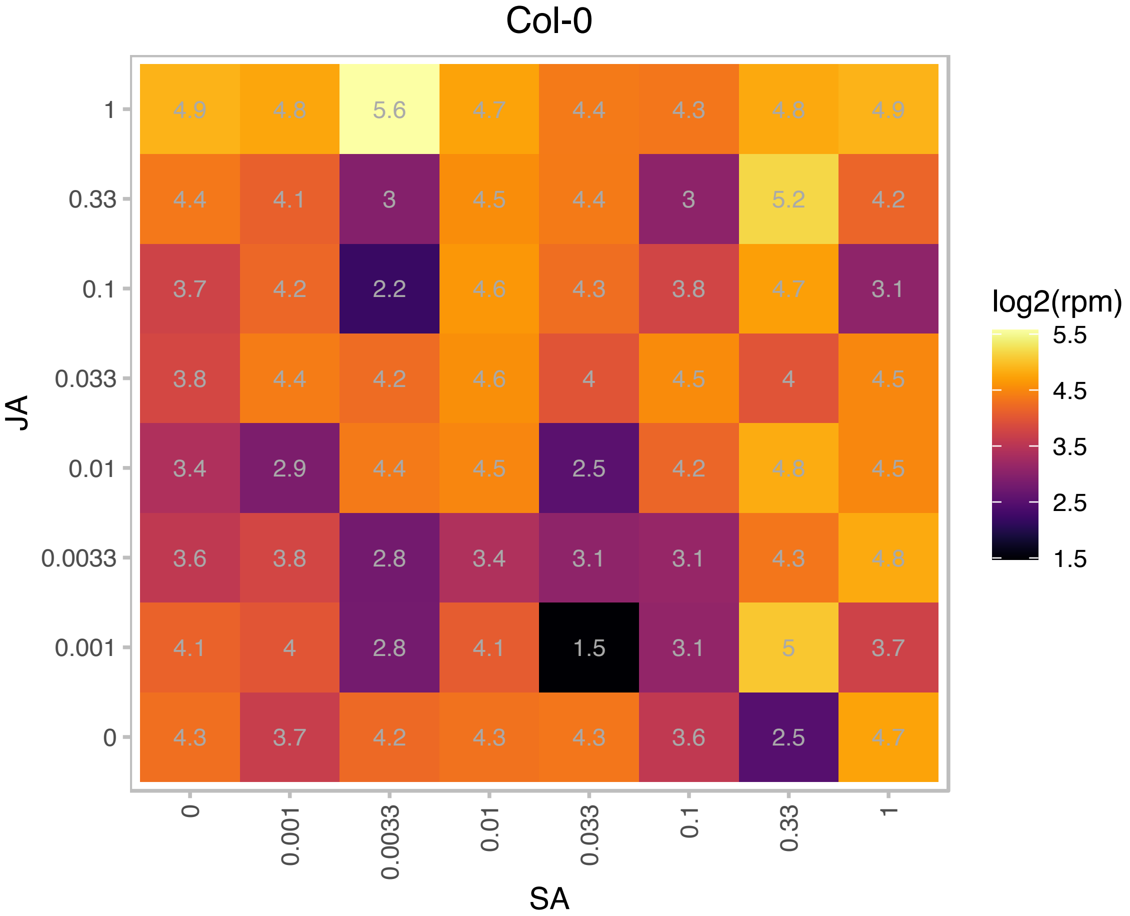


| | |
|---|---|
| Gene | AT5G27540 |
| is.expressed_genes | TRUE |
| is.responsive_genes | TRUE |
| is.apcluster | A-05 |

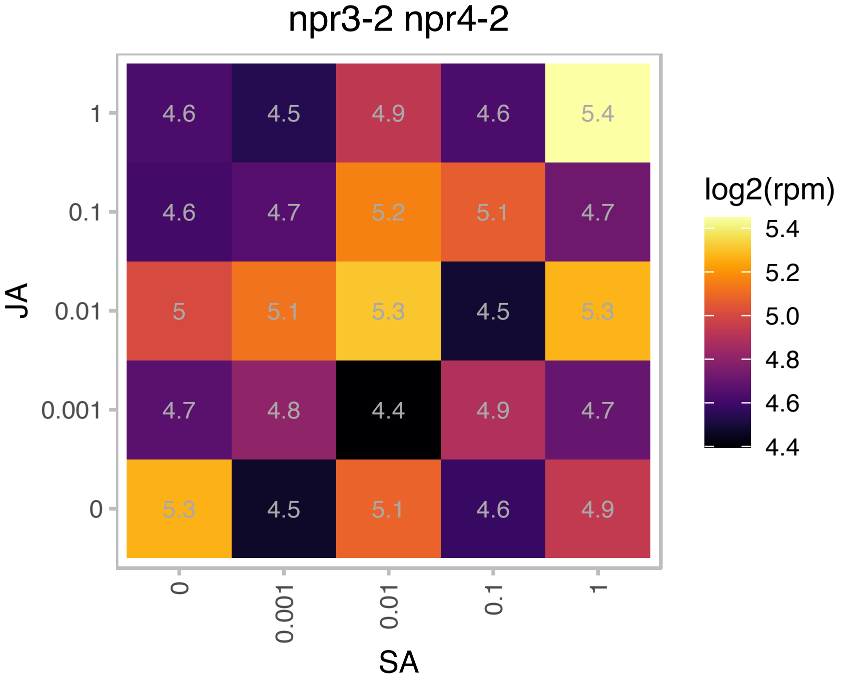


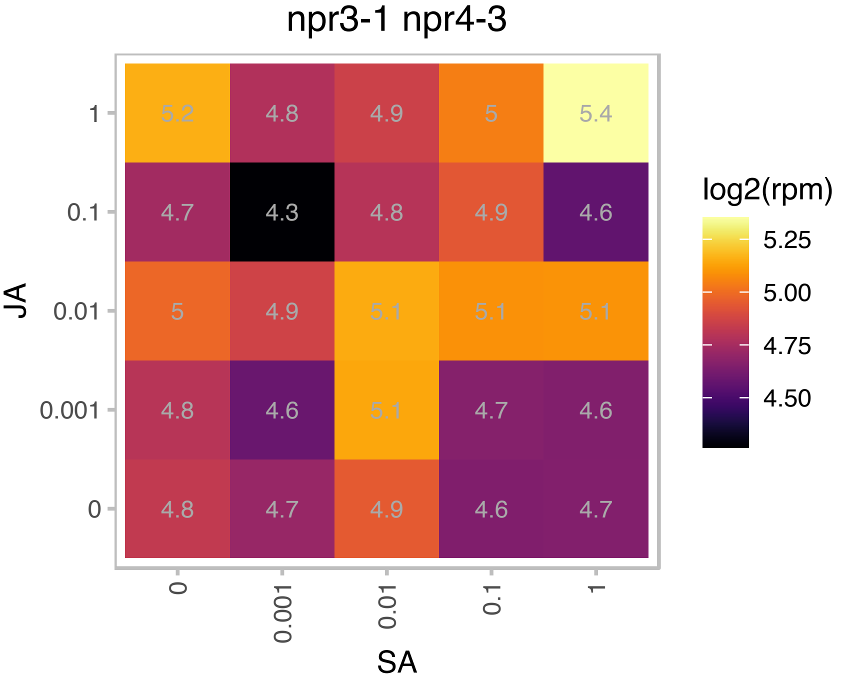

# AT5G35680.1 : NA

Nucleic acid-binding, OB-fold-like protein

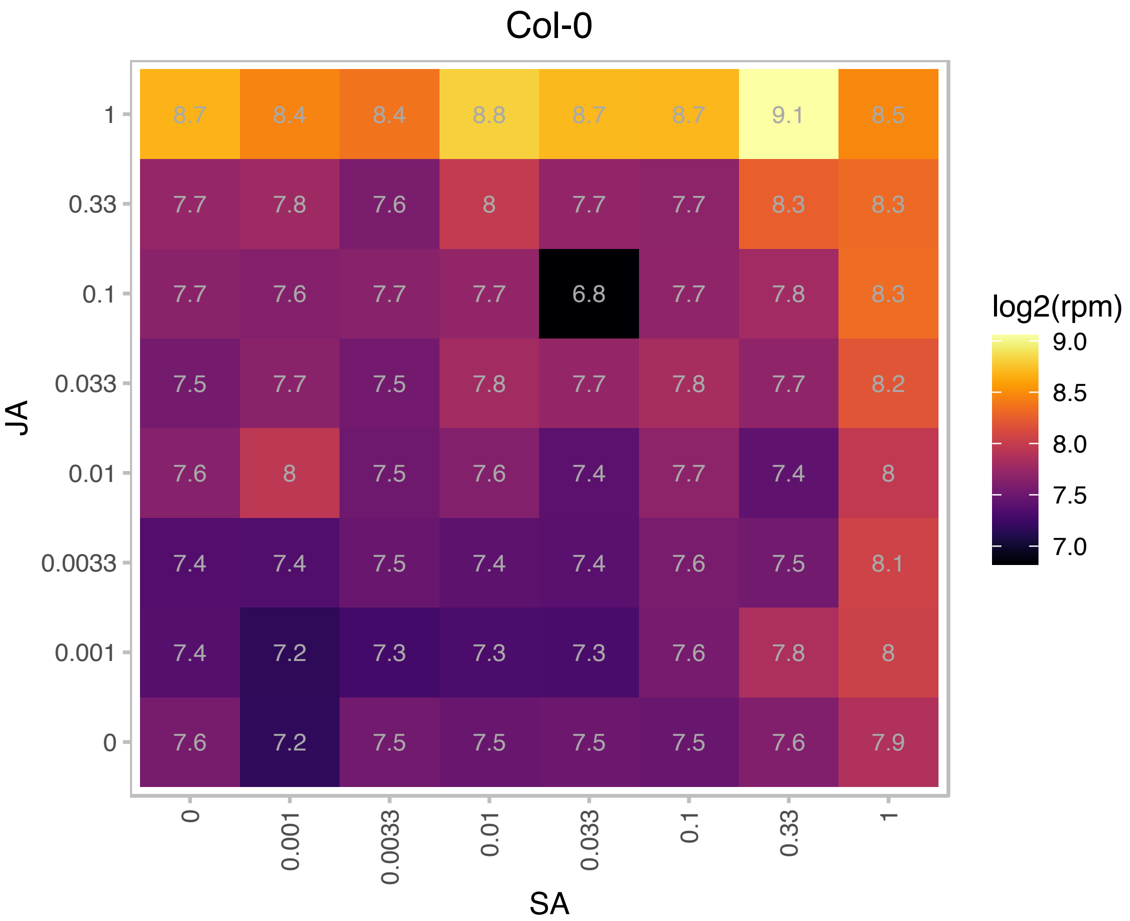


| | |
|---|---|
| Gene | AT5G35680 |
| is.expressed_genes | TRUE |
| is.responsive_genes | TRUE |
| is.apcluster | A-05 |

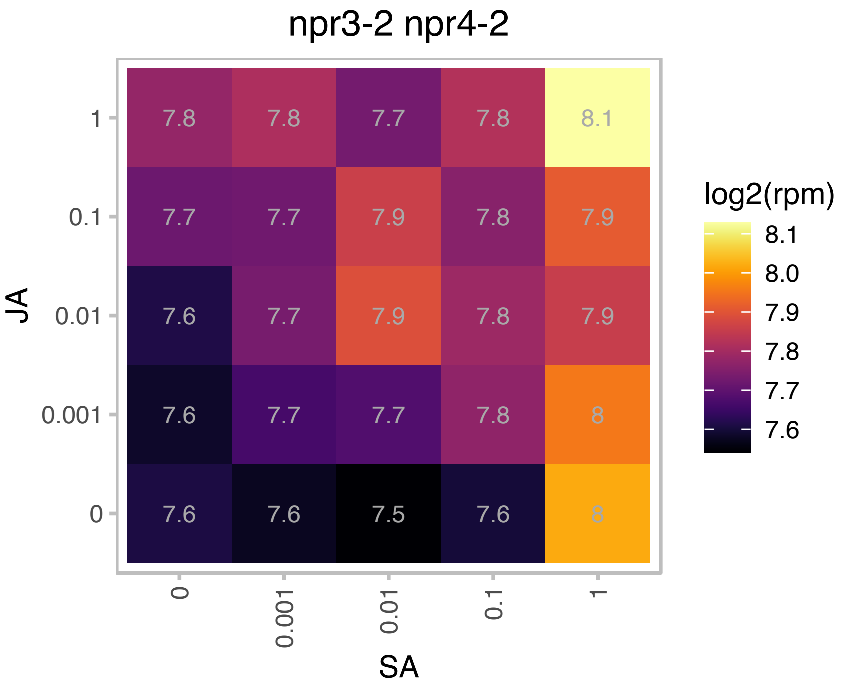


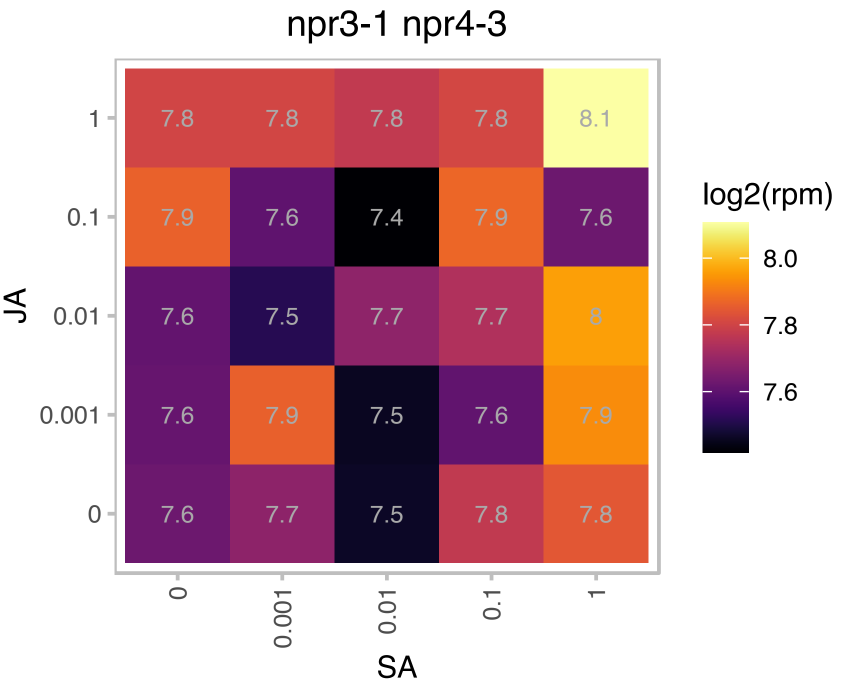

### AT5G37510.1 : CI76, EMB1467

NADH-ubiquinone dehydrogenase, mitochondrial, putative

Encodes a subunit of the 400 kDa subcomplex of the mitochondrial NADH dehydrogenase (complex I). The protein has been isolated in the male gametophyte. The mRNA is cell-to-cell mobile.

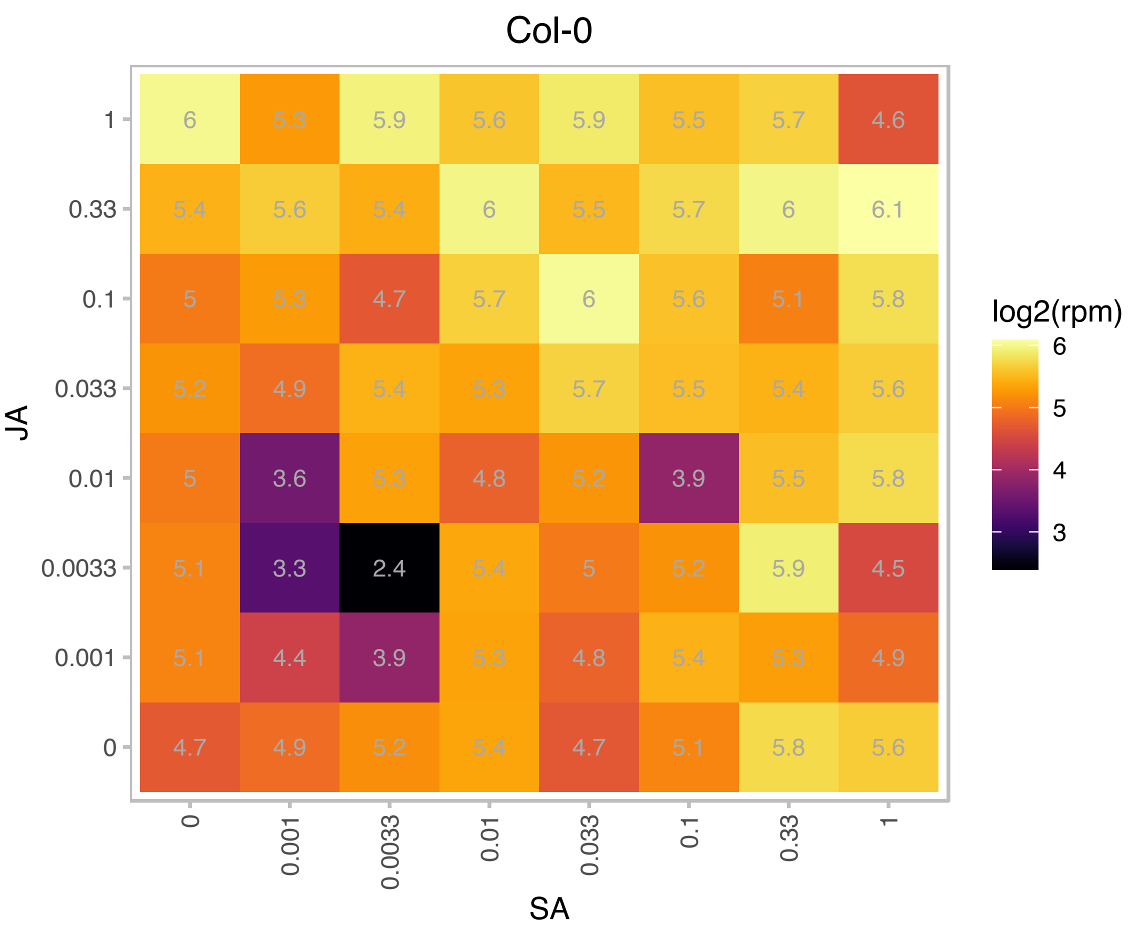


| | |
|---|---|
| Gene | AT5G37510 |
| is.expressed_genes | TRUE |
| is.responsive_genes | TRUE |
| is.apcluster | A-05 |

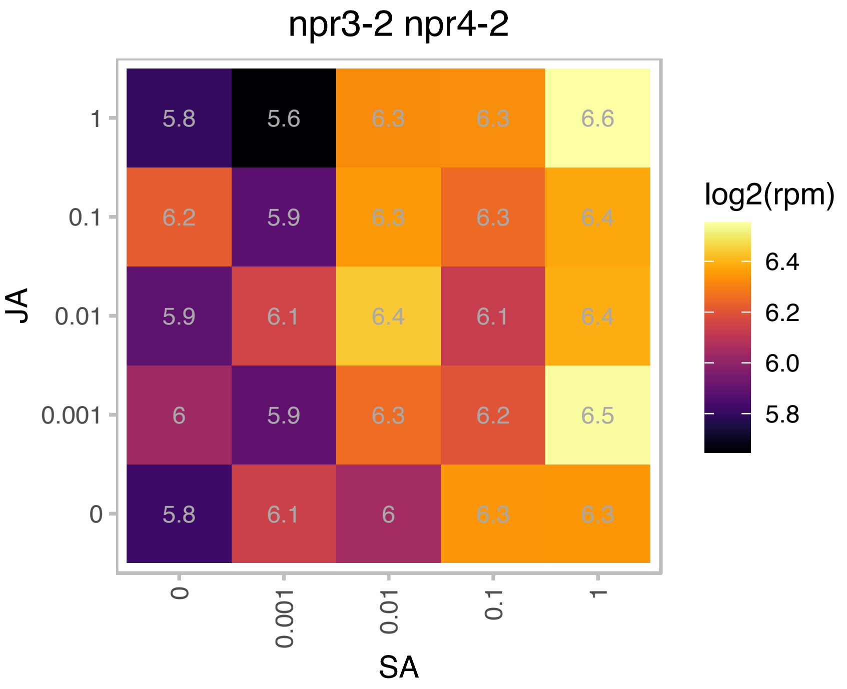


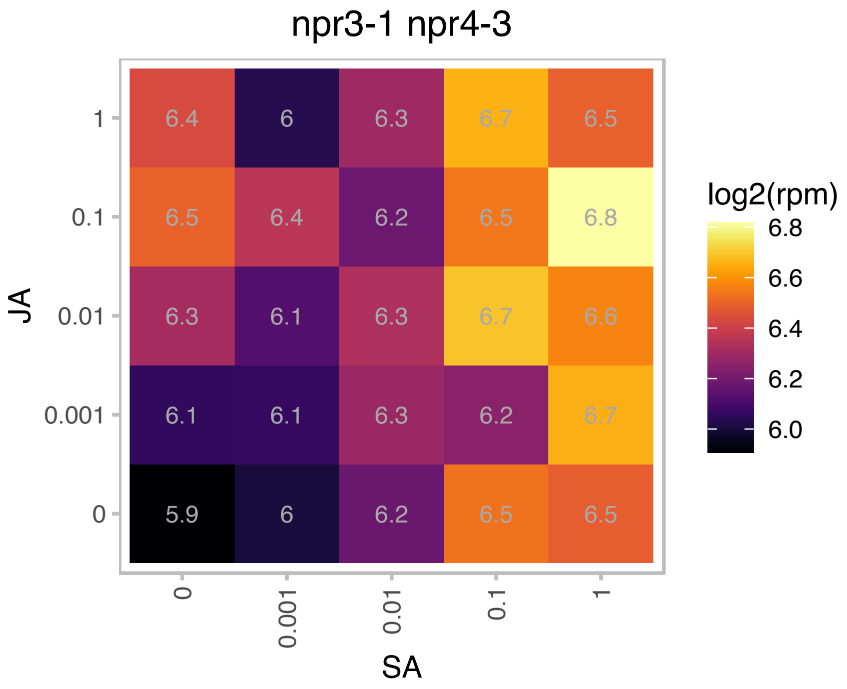

### AT5G43320.1 : CK1.8, ckl8

casein kinase I-like 8

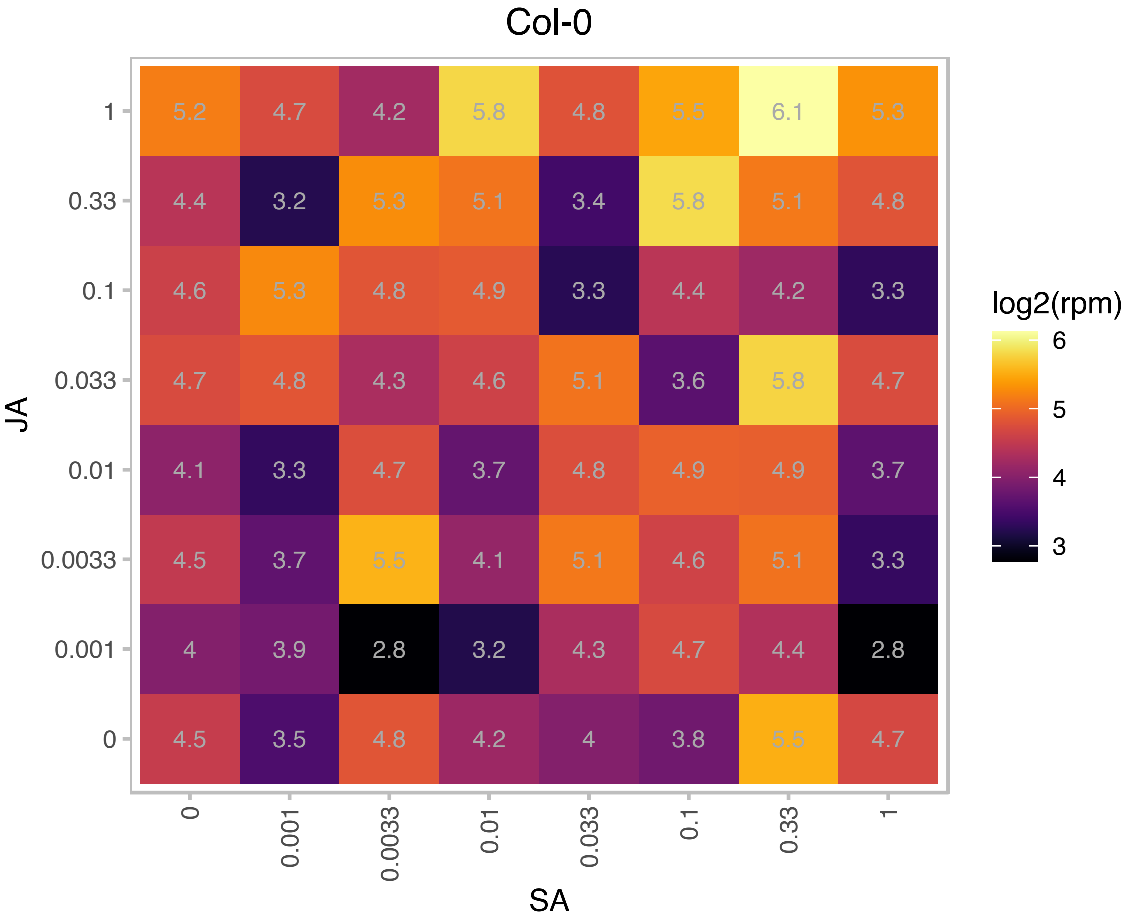


| | |
|---|---|
| Gene | AT5G43320 |
| is.expressed_genes | TRUE |
| is.responsive_genes | TRUE |
| is.apcluster | A-05 |

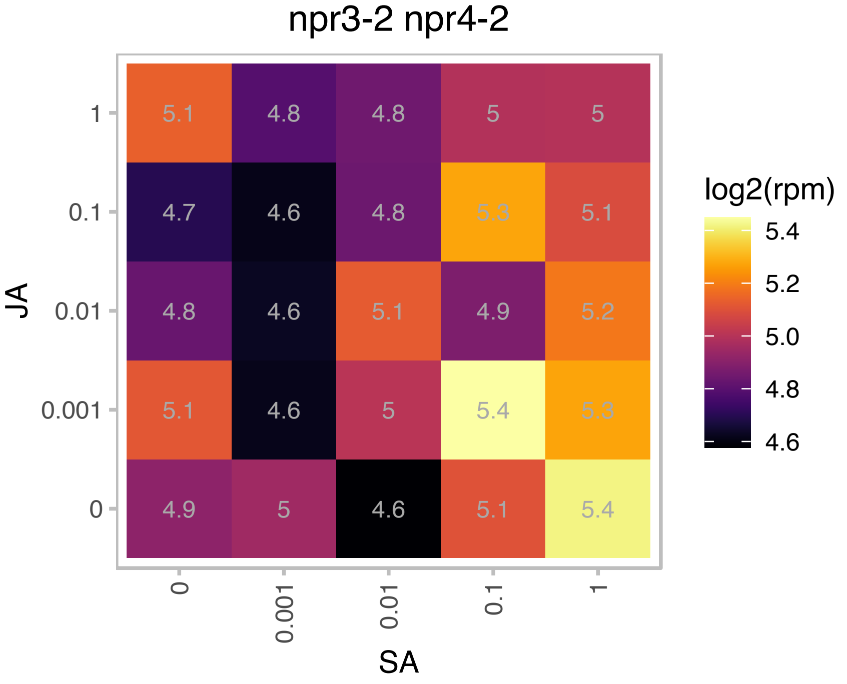


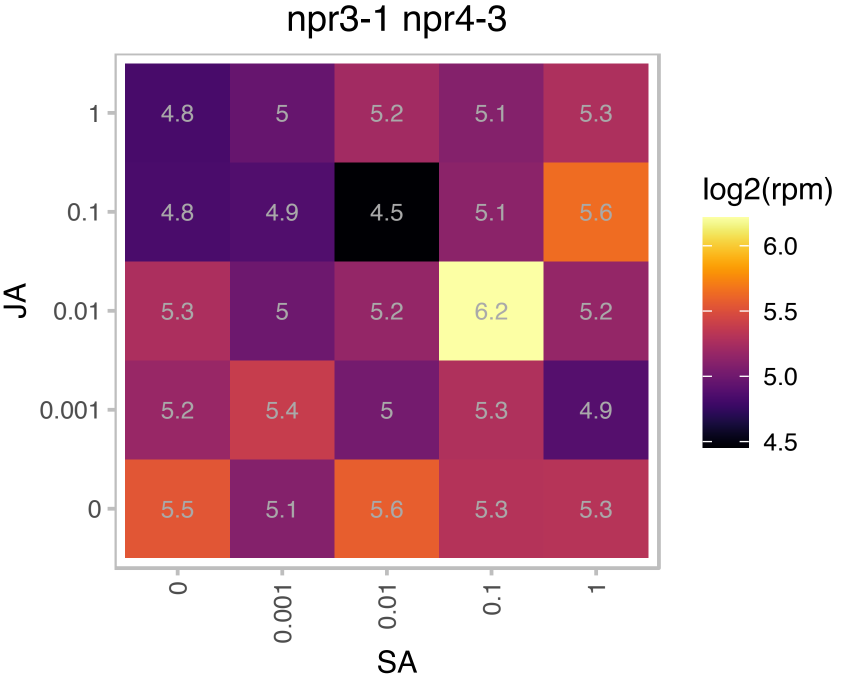

# AT5G44320.1 : NA

Eukaryotic translation initiation factor 3 subunit 7 (eIF-3)

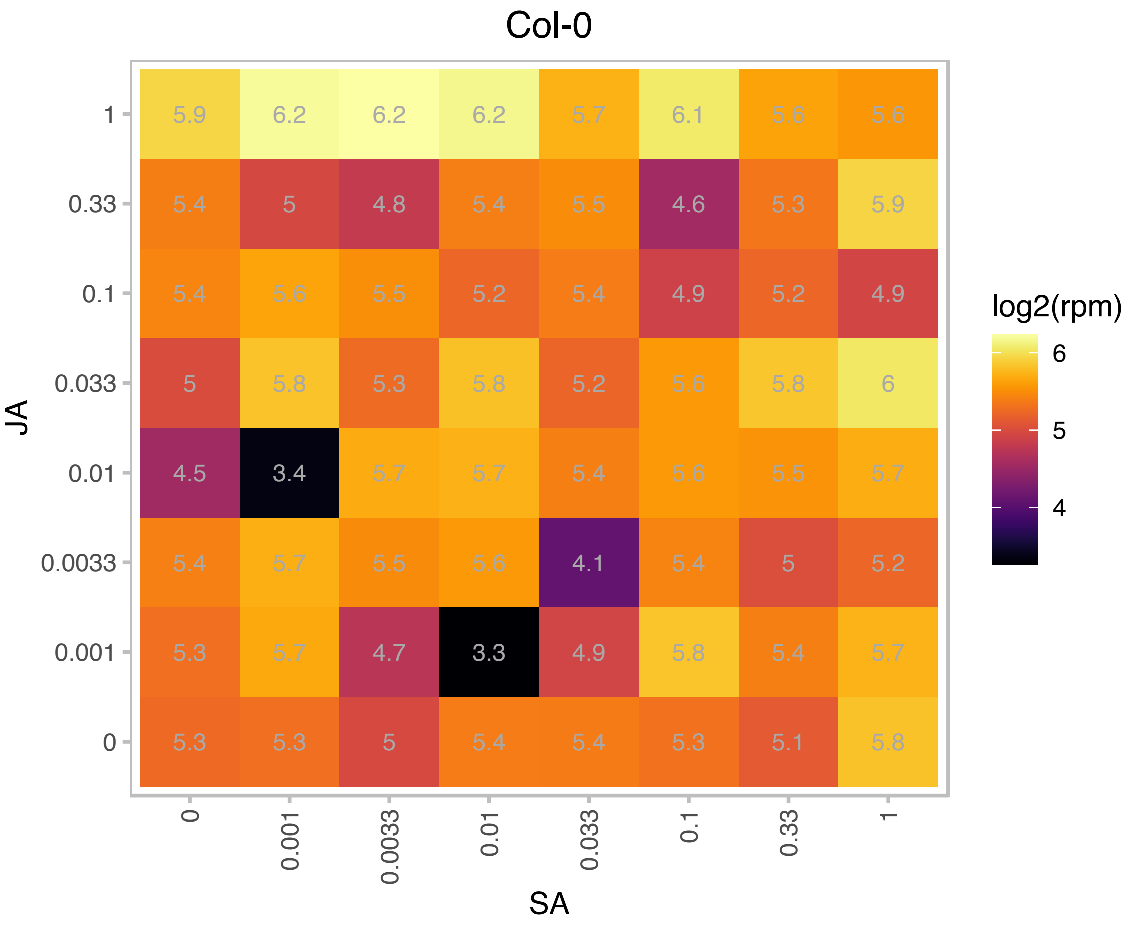


| | |
|---|---|
| Gene | AT5G44320 |
| is.expressed_genes | TRUE |
| is.responsive_genes | TRUE |
| is.apcluster | A-05 |

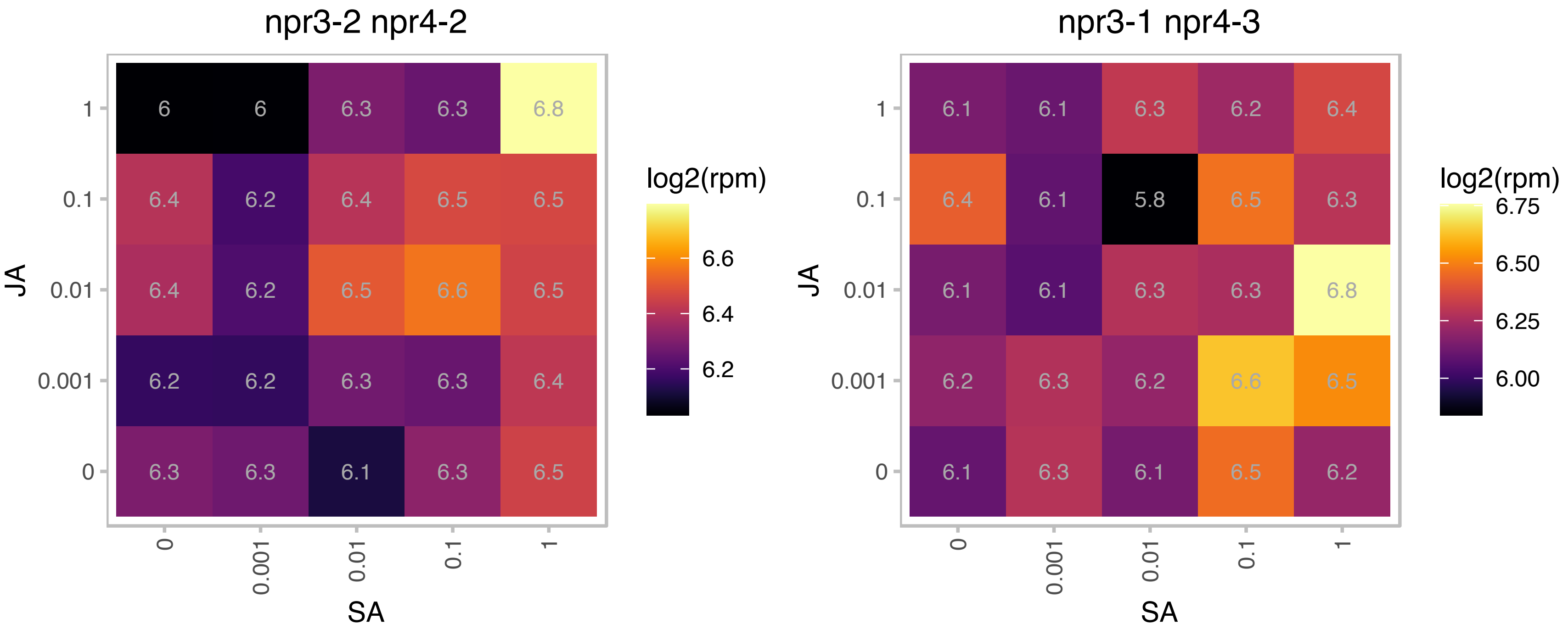

### AT5G44340.1 : TUB4

tubulin beta chain 4

beta tubulin gene

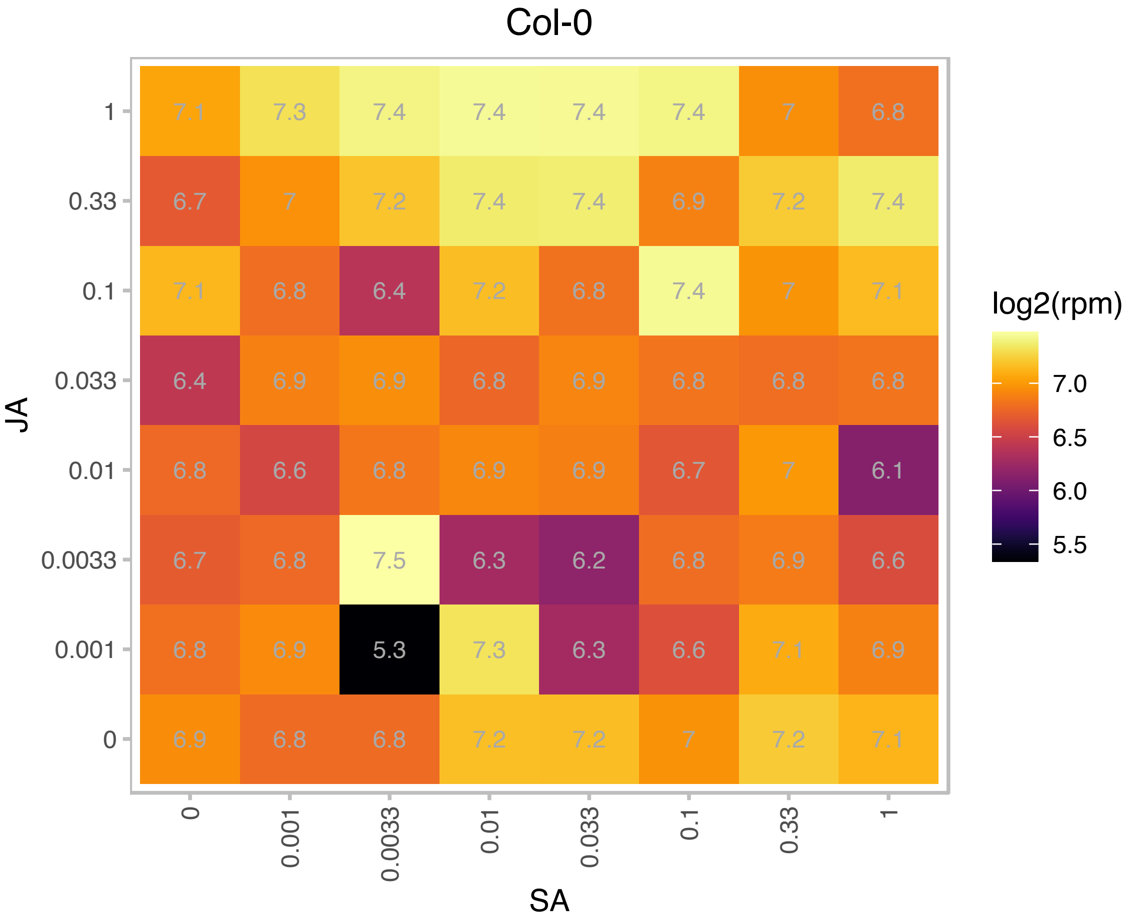


| | |
|---|---|
| Gene | AT5G44340 |
| is.expressed_genes | TRUE |
| is.responsive_genes | TRUE |
| is.apcluster | A-05 |

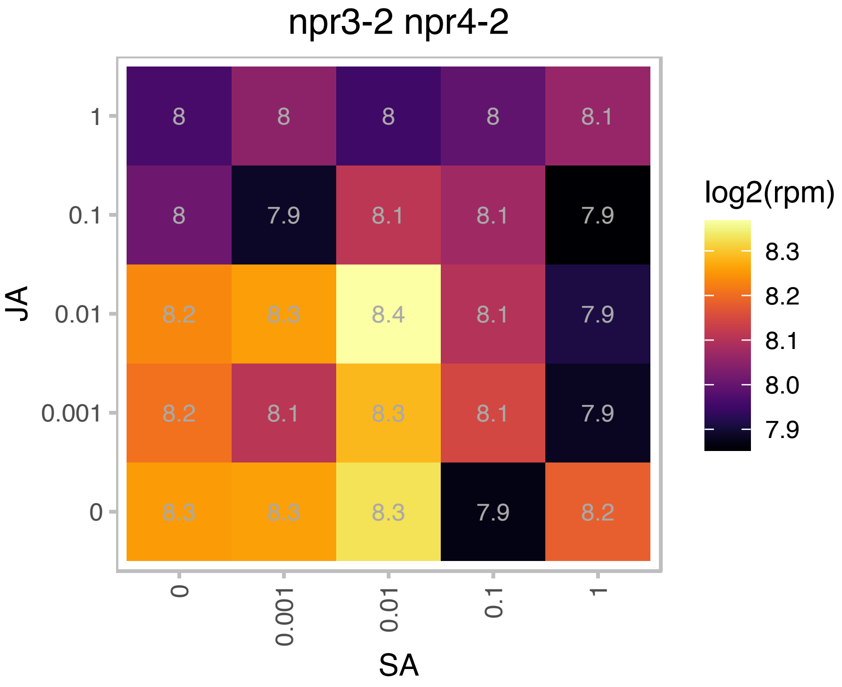


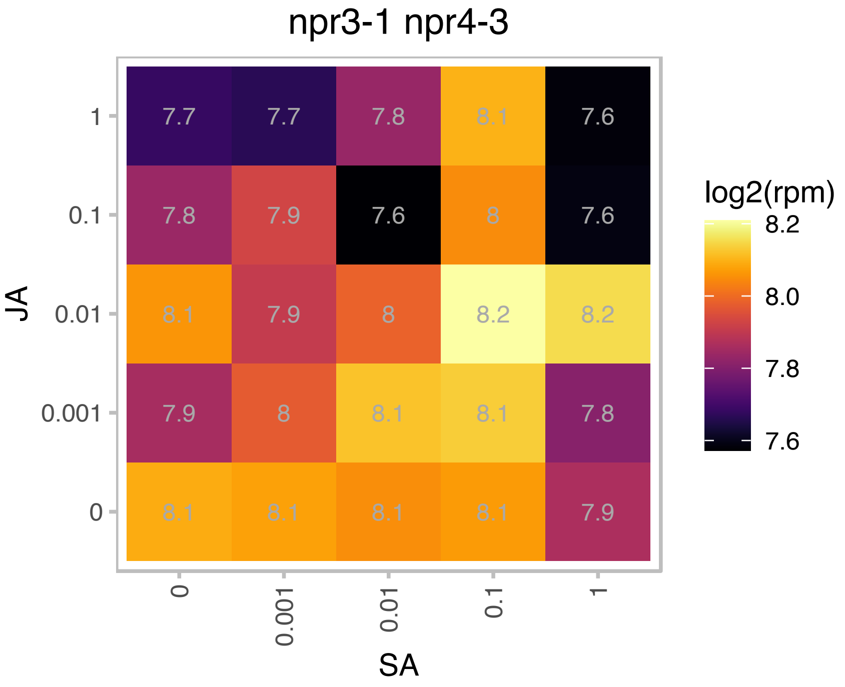

# AT5G59480.1 : NA

Haloacid dehalogenase-like hydrolase (HAD) superfamily protein

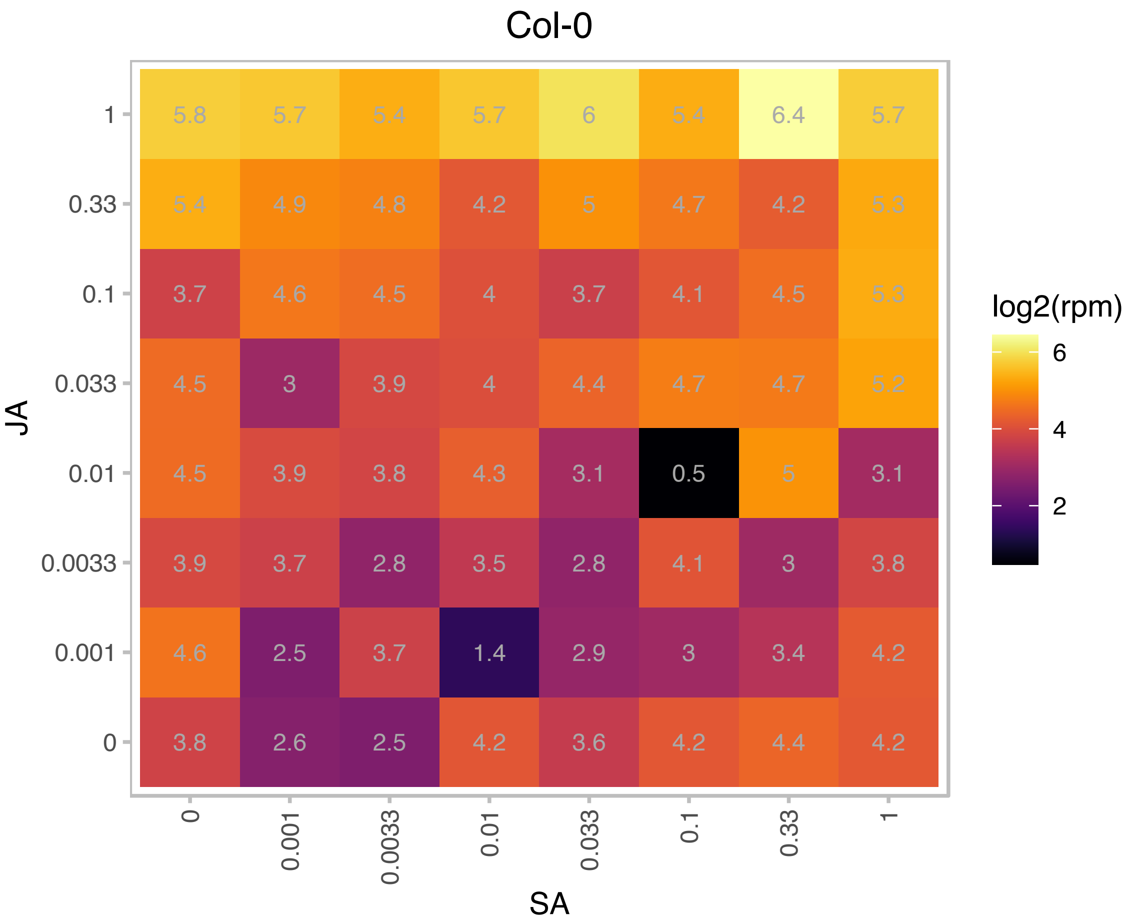


| | |
|---|---|
| Gene | AT5G59480 |
| is.expressed_genes | TRUE |
| is.responsive_genes | TRUE |
| is.apcluster | A-05 |

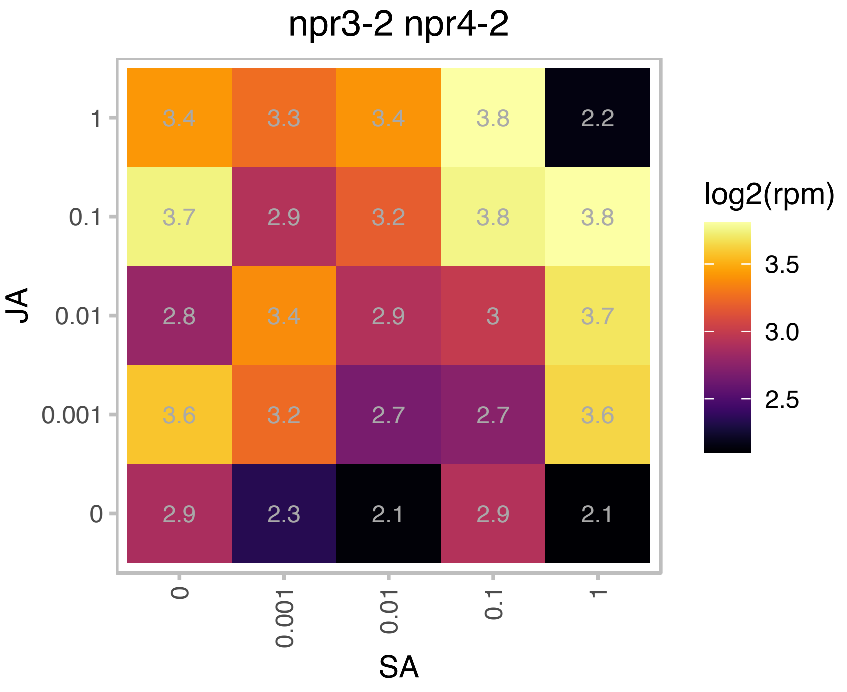


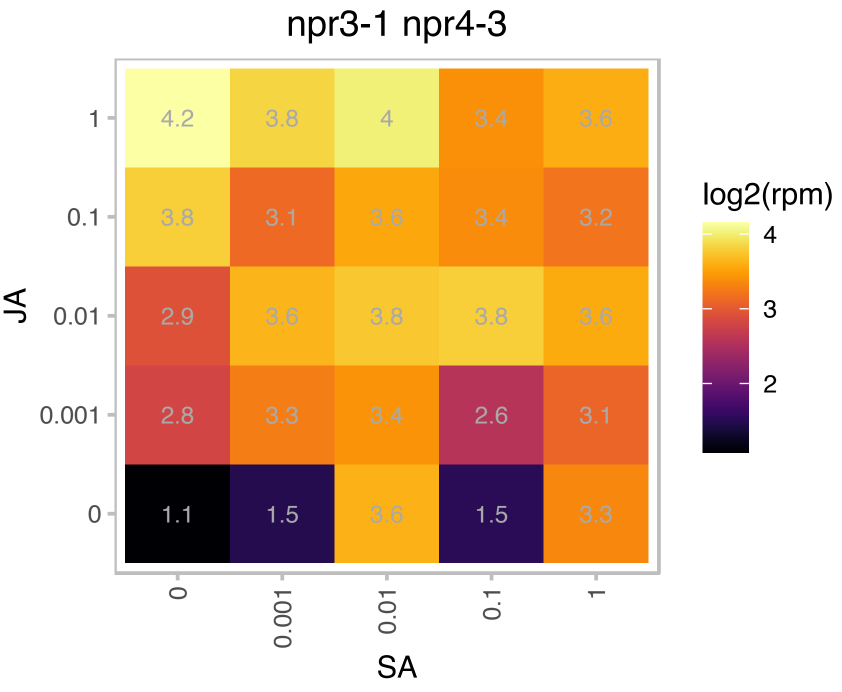

### AT5G63970.1 : RGLG3

Copine (Calcium-dependent phospholipid-binding protein) family

Encodes a ubiquitin ligase that is an essential upstream modulator of JA signaling in response to various stimuli.

| | |
|---|---|
| Gene | AT5G63970 |
| is.expressed_genes | TRUE |
| is.responsive_genes | TRUE |
| is.apcluster | A-05 |

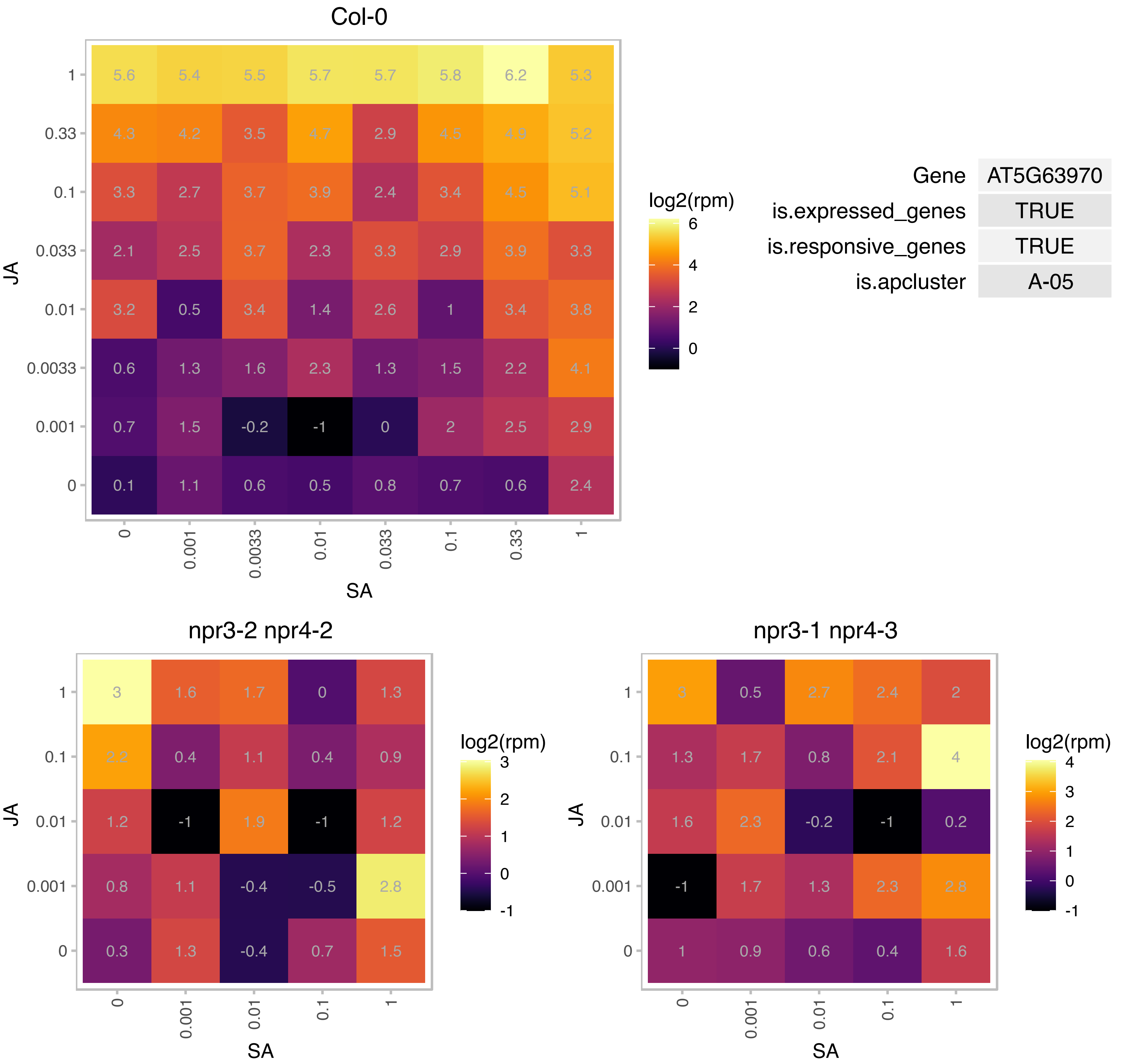

### AT5G65270.1 : AtRABA4a, RABA4a

RAB GTPase homolog A4A

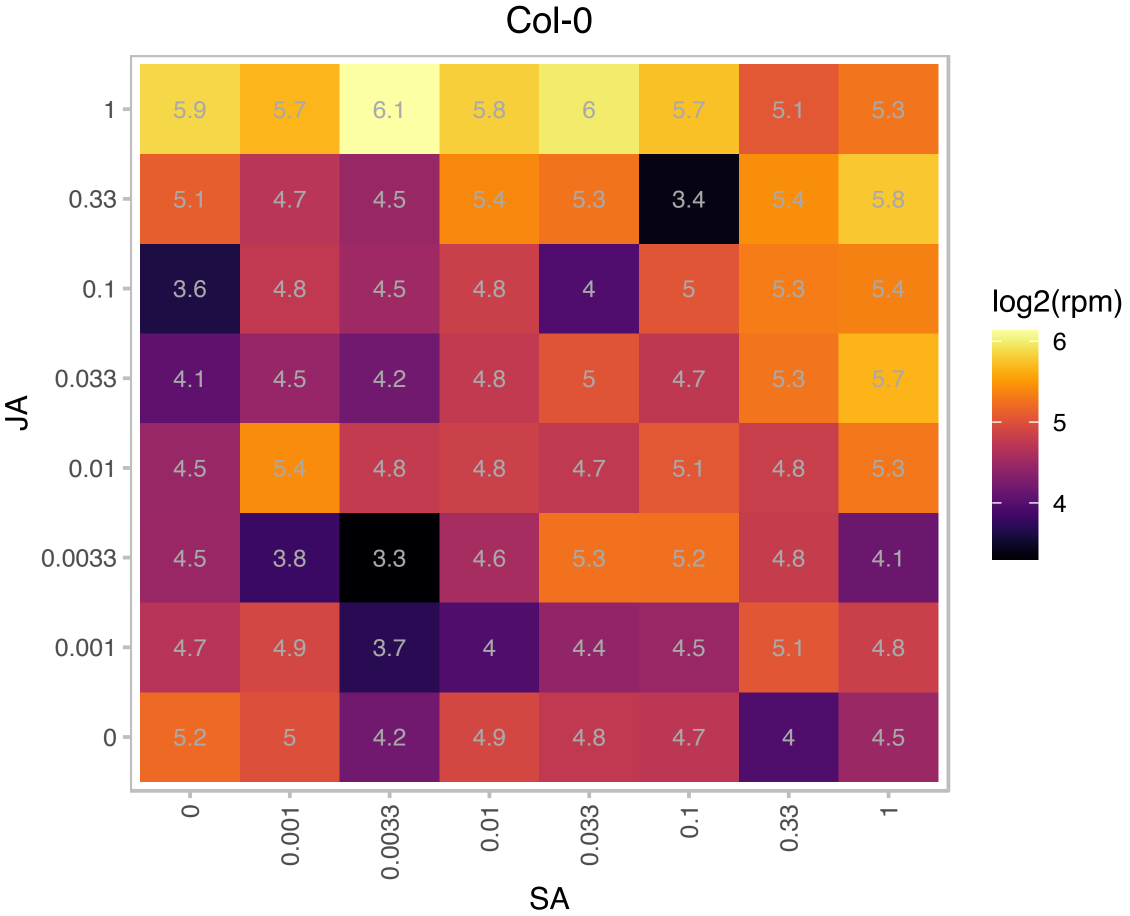


| | |
|---|---|
| Gene | AT5G65270 |
| is.expressed_genes | TRUE |
| is.responsive_genes | TRUE |
| is.apcluster | A-05 |

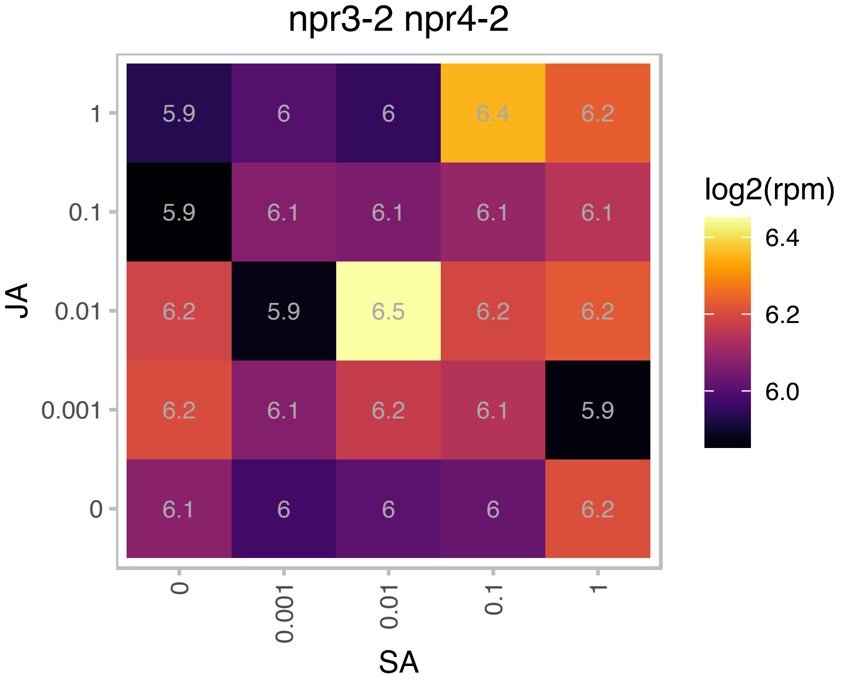


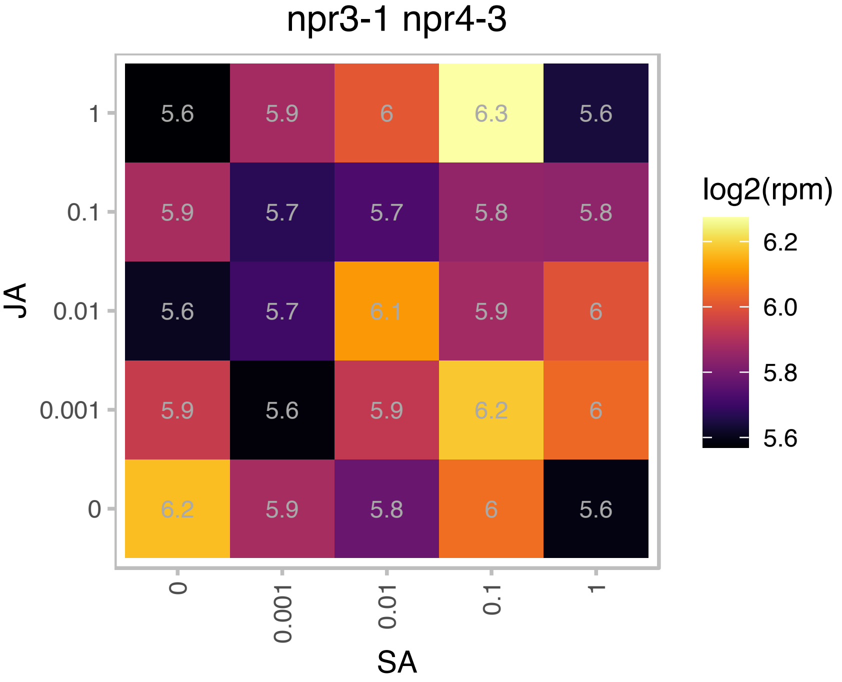

### AT1G06530.1 : PMD2

Tropomyosin-related

Encodes PEROXISOMAL AND MITOCHONDRIAL DIVISION FACTOR2. Involved in mitochondrial morphogenesis.

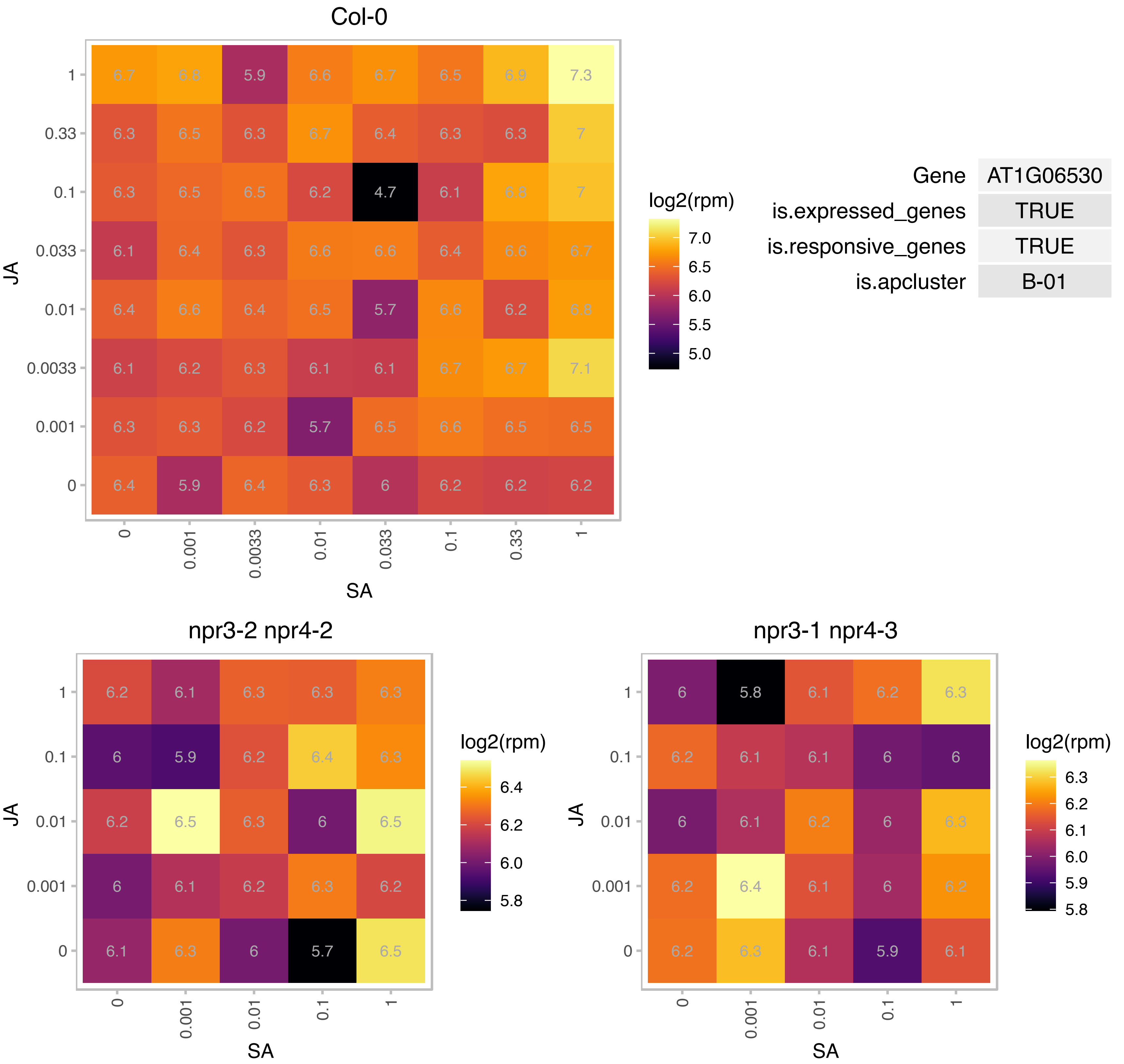


| Gene | AT1G06530 |
|---|---|
| is.expressed_genes | TRUE |
| is.responsive_genes | TRUE |
| is.apcluster | B-01 |

### AT1G08110.1 : AtGLYI2

lactoylglutathione lyase family protein / glyoxalase I family protein

Encodes a Zn2+ dependent glyoxylase.

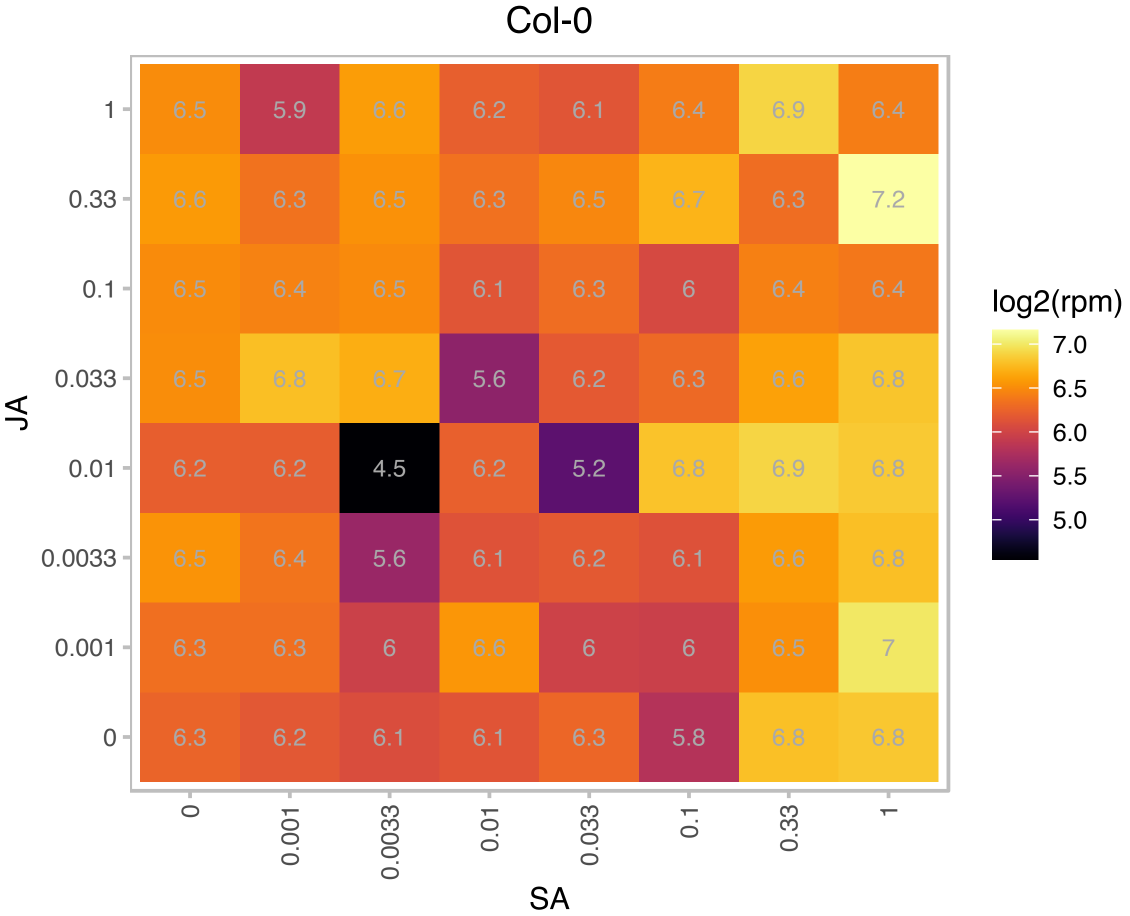


| | |
|---|---|
| Gene | AT1G08110 |
| is.expressed_genes | TRUE |
| is.responsive_genes | TRUE |
| is.apcluster | B-01 |

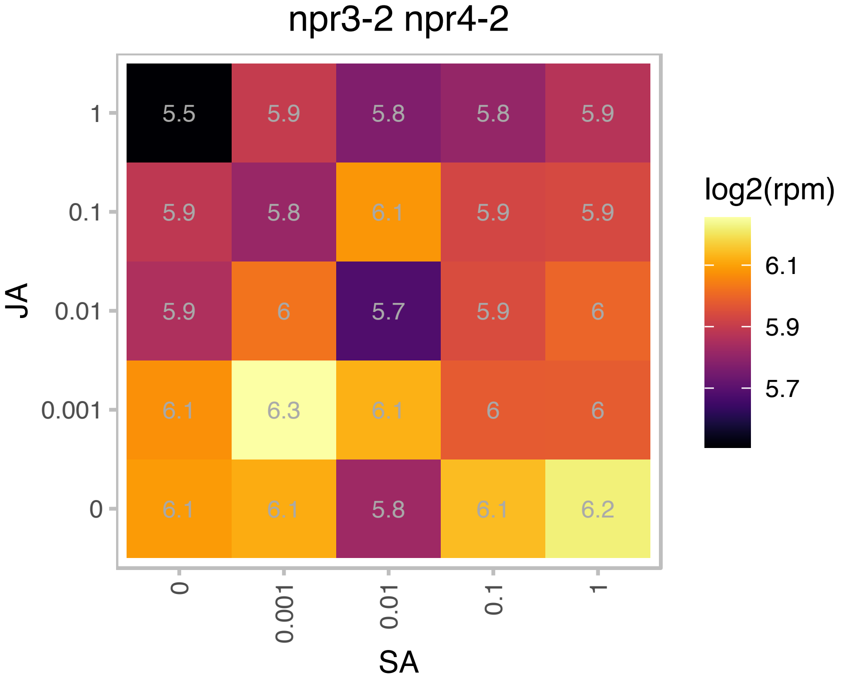


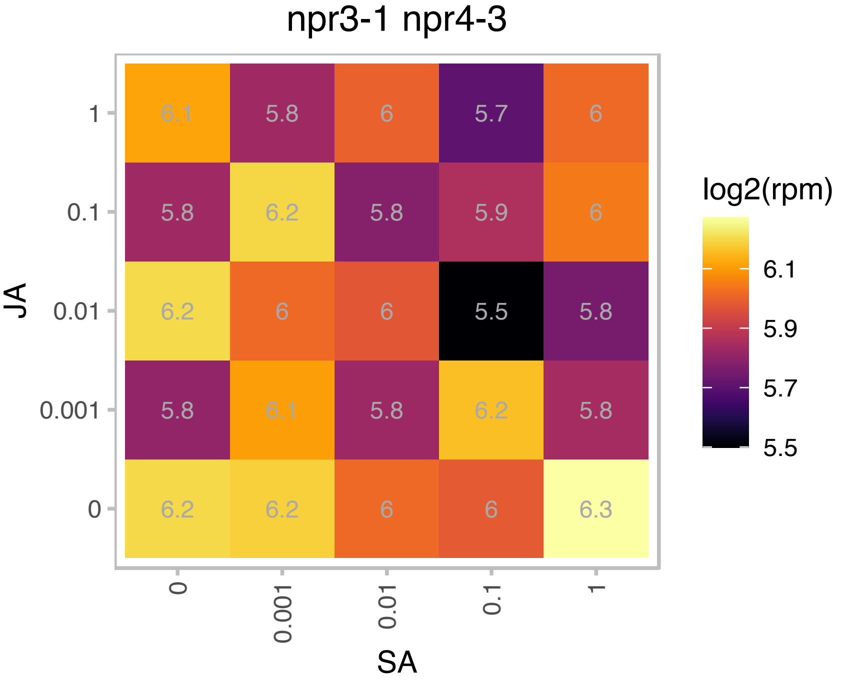

### AT1G16030.1 : Hsp70b

heat shock protein 70B

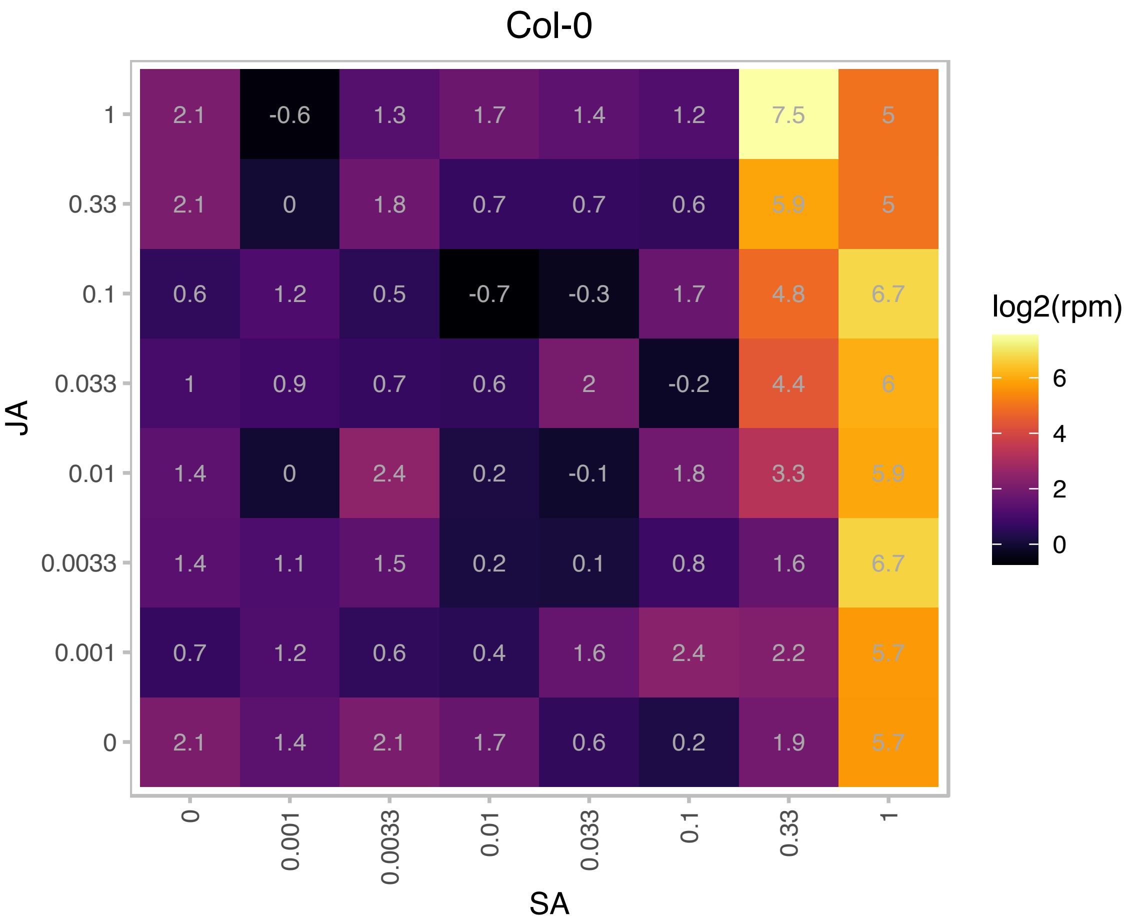


| | |
|---|---|
| Gene | AT1G16030 |
| is.expressed_genes | TRUE |
| is.responsive_genes | TRUE |
| is.apcluster | B-01 |

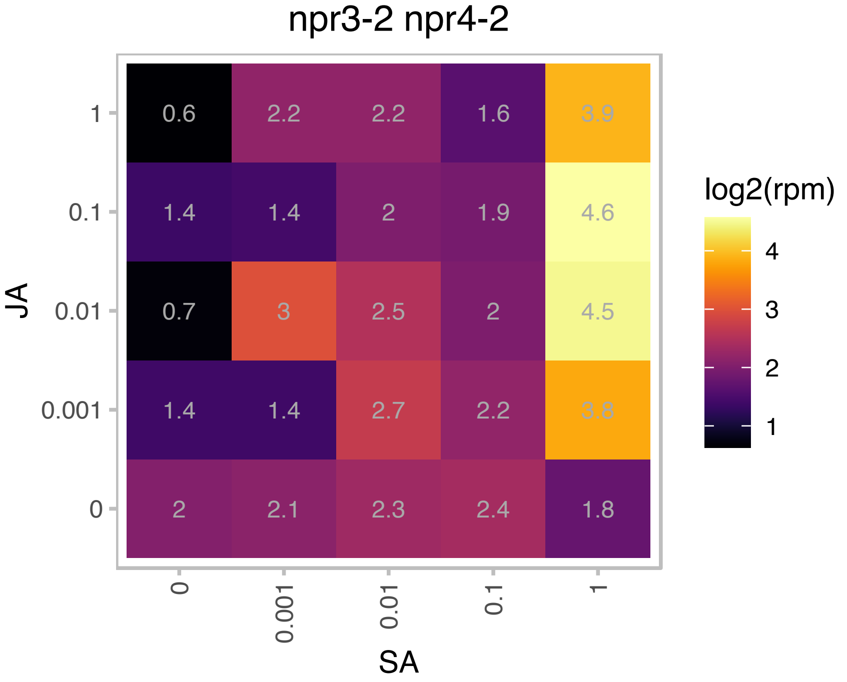


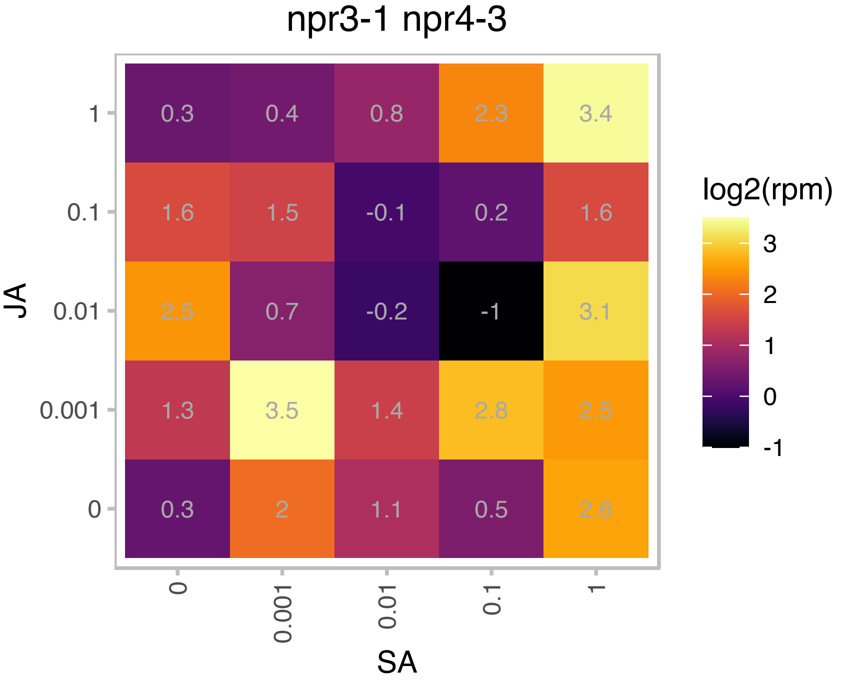

# AT1G28540.1 : NA

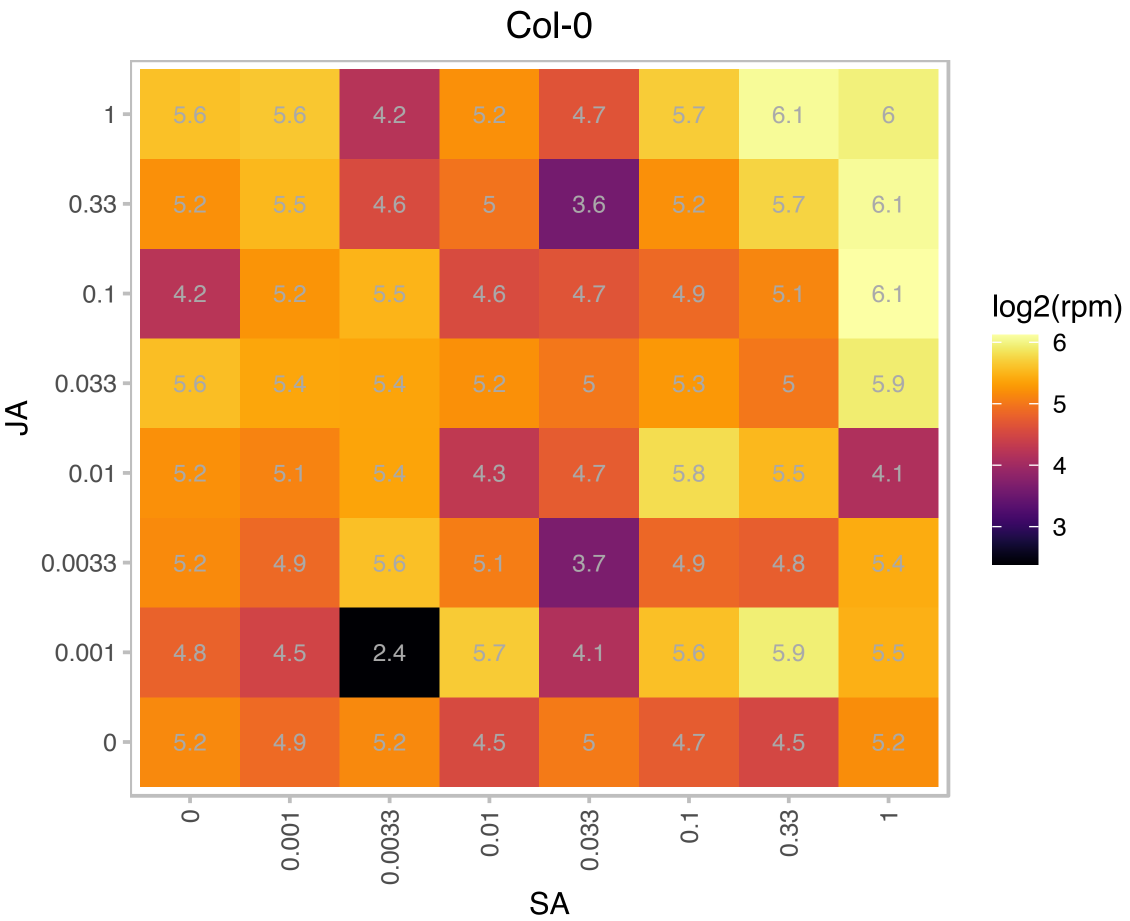


| | |
|---|---|
| Gene | AT1G28540 |
| is.expressed_genes | TRUE |
| is.responsive_genes | TRUE |
| is.apcluster | B-01 |

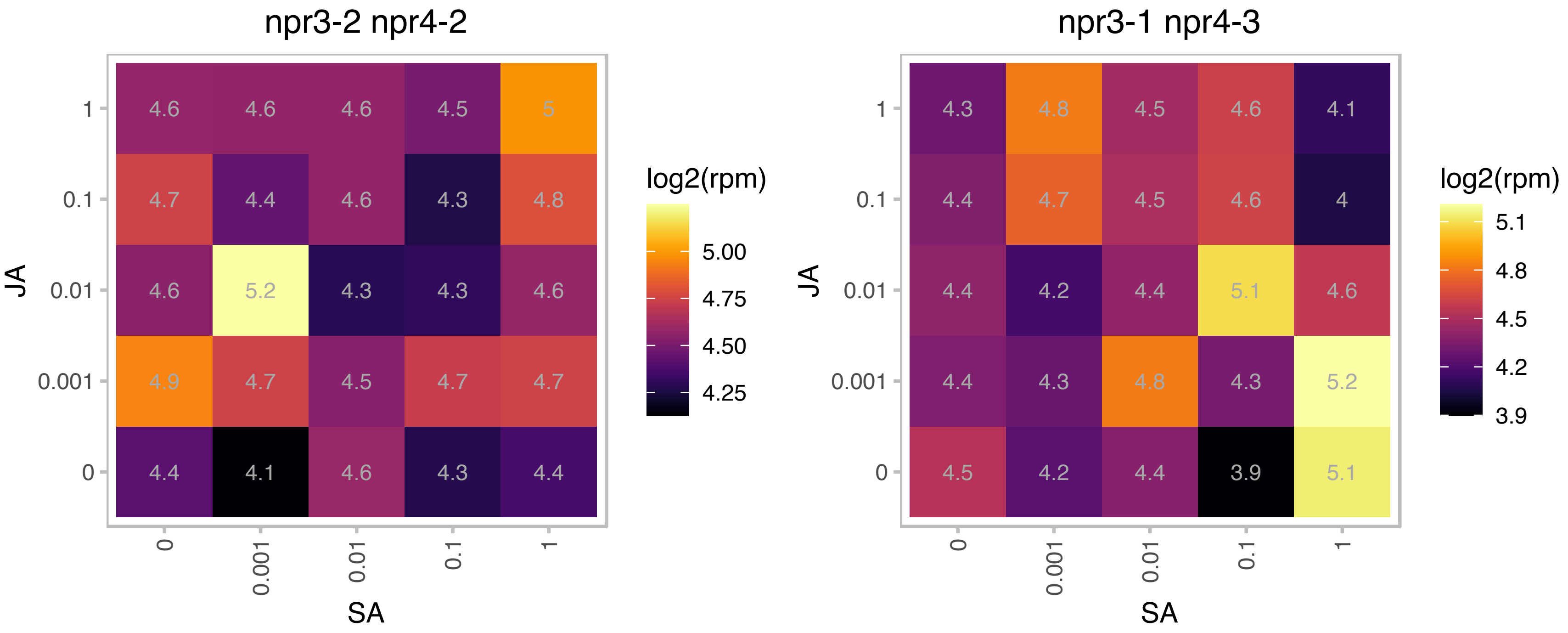

# AT1G44608.1 : NA

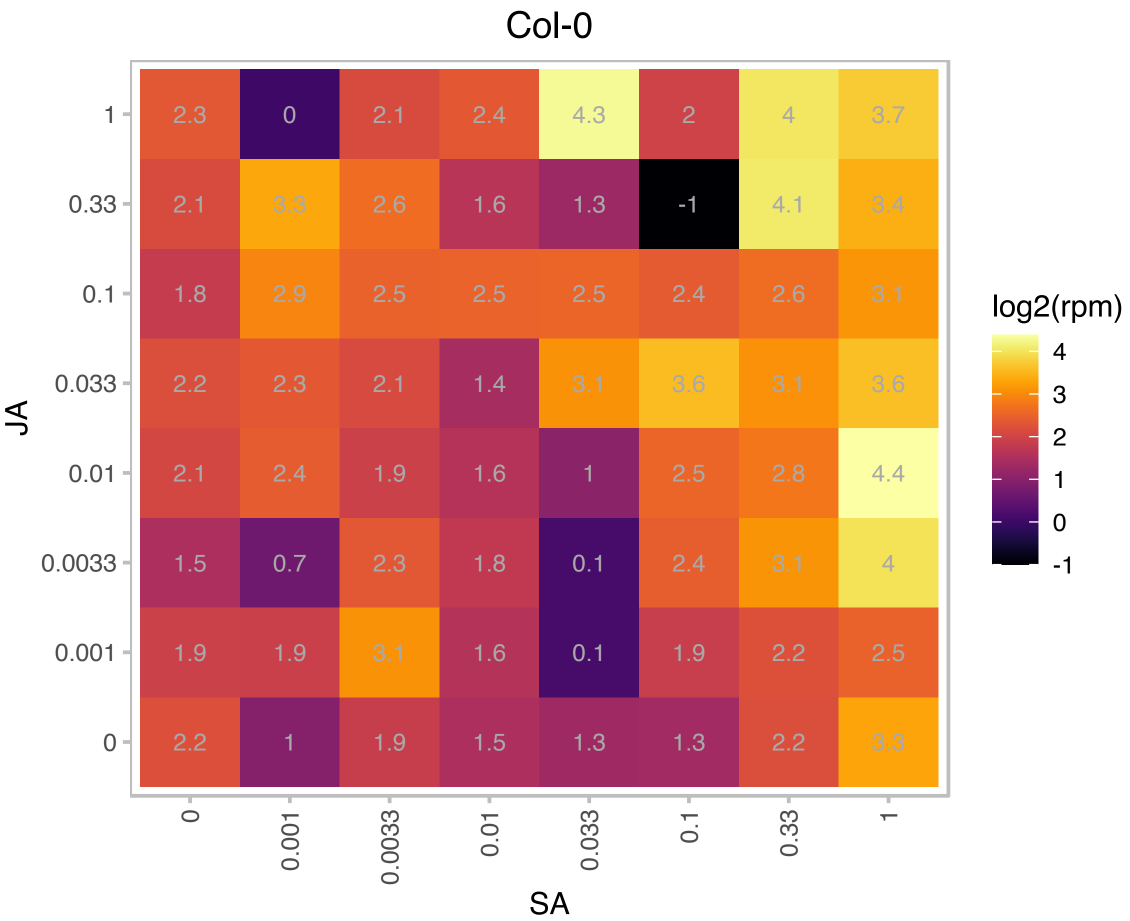


| | |
|---|---|
| Gene | AT1G44608 |
| is.expressed_genes | TRUE |
| is.responsive_genes | TRUE |
| is.apcluster | B-01 |

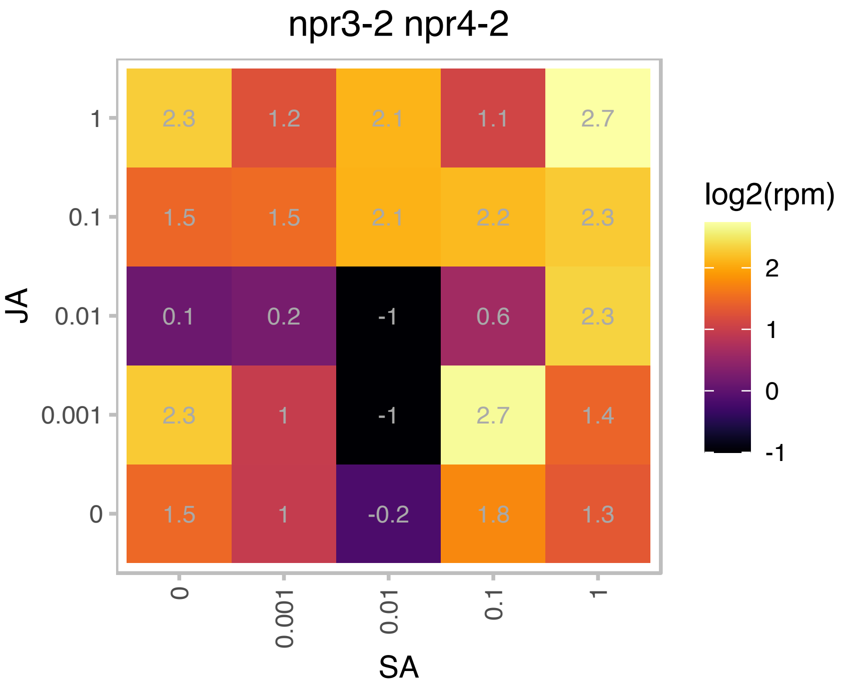


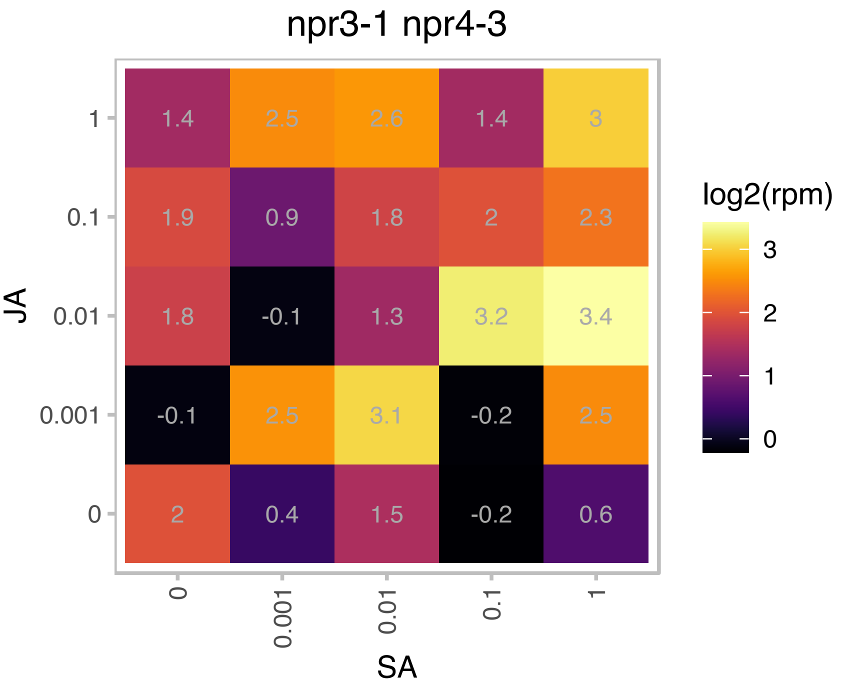

# AT1G44890.1 : NA

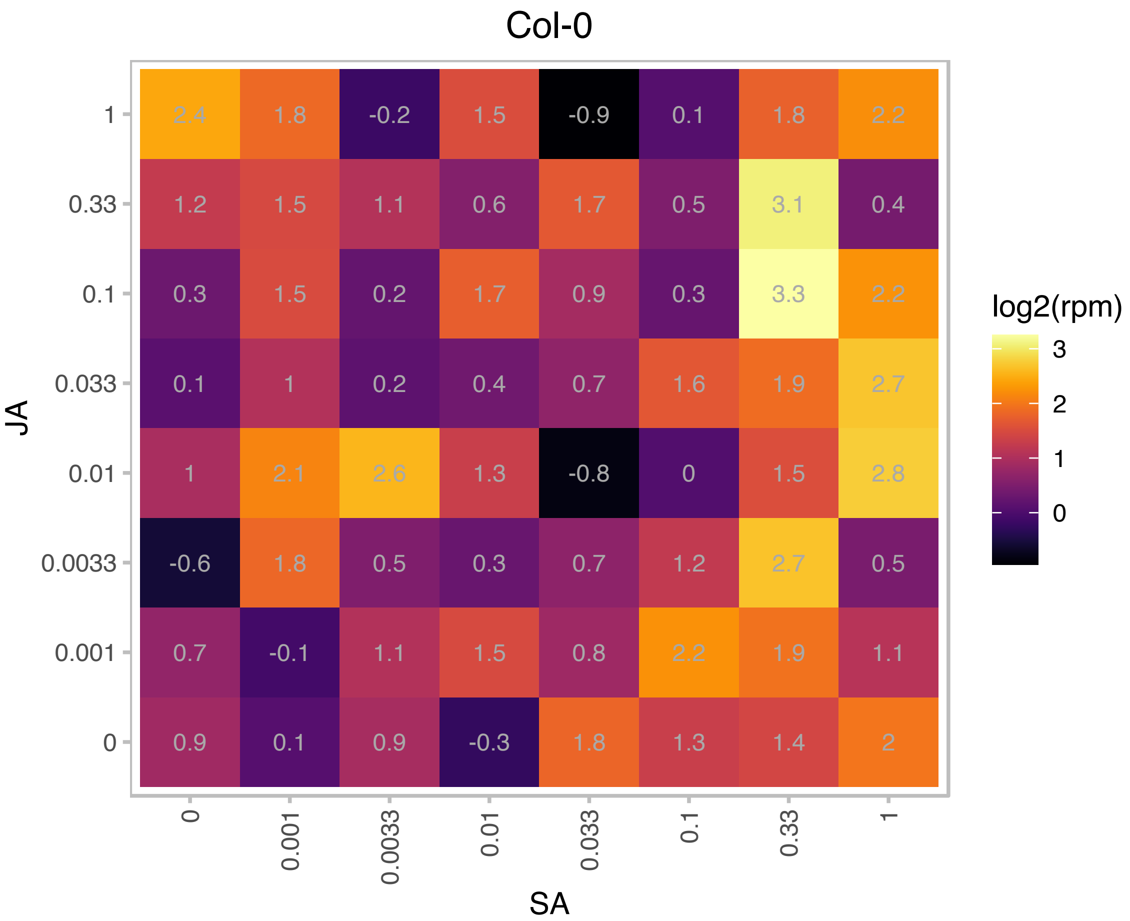


| | |
|---|---|
| Gene | AT1G44890 |
| is.expressed_genes | TRUE |
| is.responsive_genes | TRUE |
| is.apcluster | B-01 |

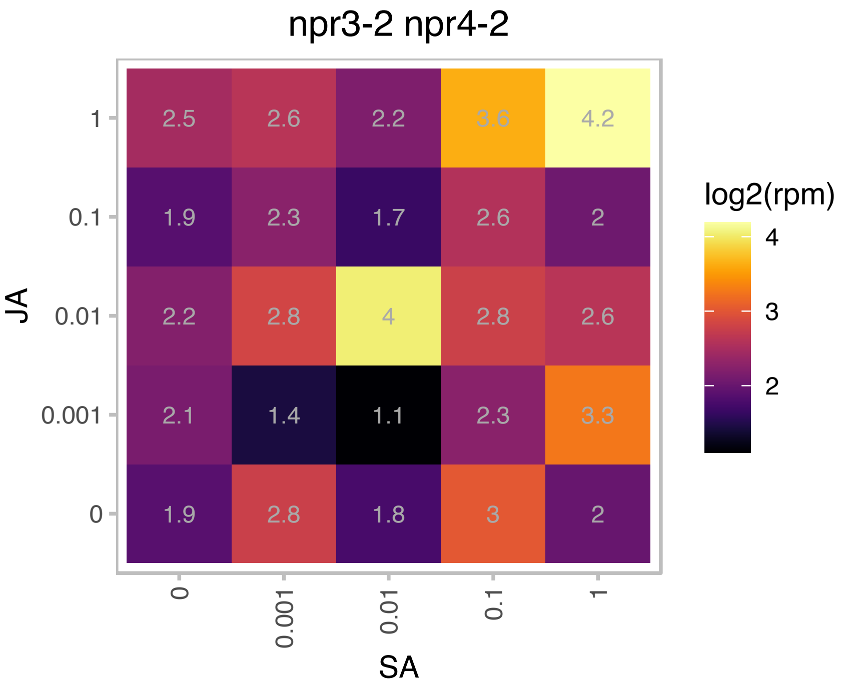


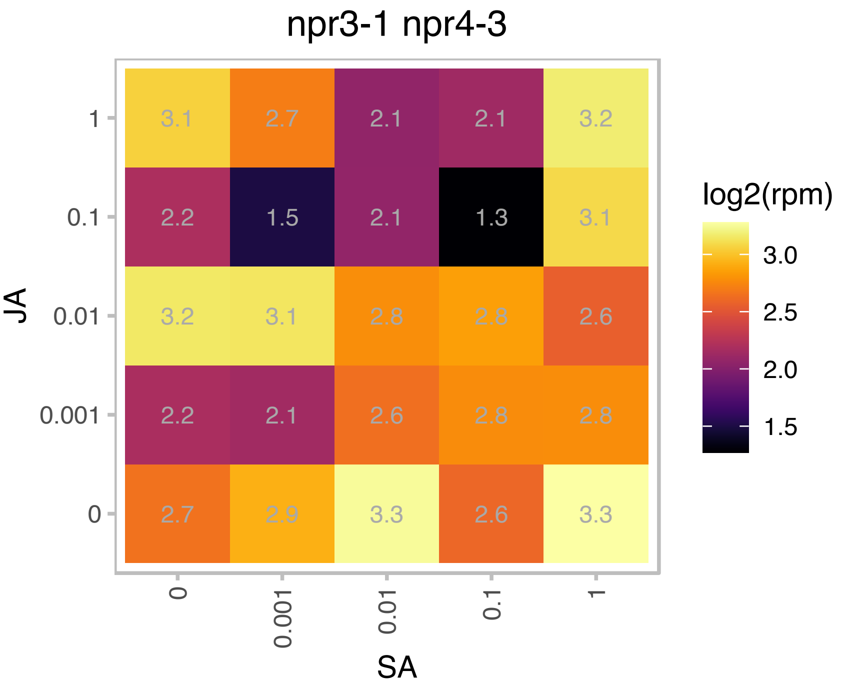

# AT1G45233.2 : -

-

-

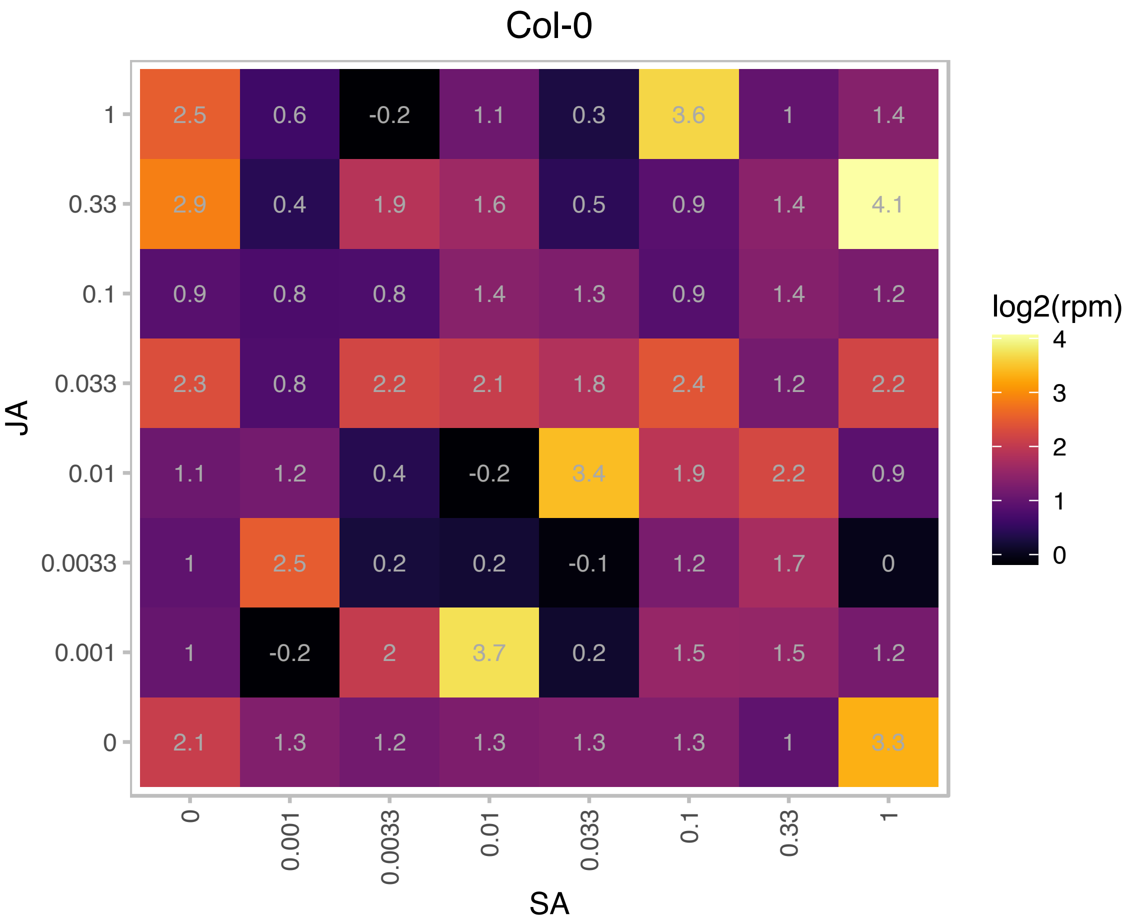


| | |
|---|---|
| Gene | AT1G45233 |
| is.expressed_genes | TRUE |
| is.responsive_genes | TRUE |
| is.apcluster | B-01 |

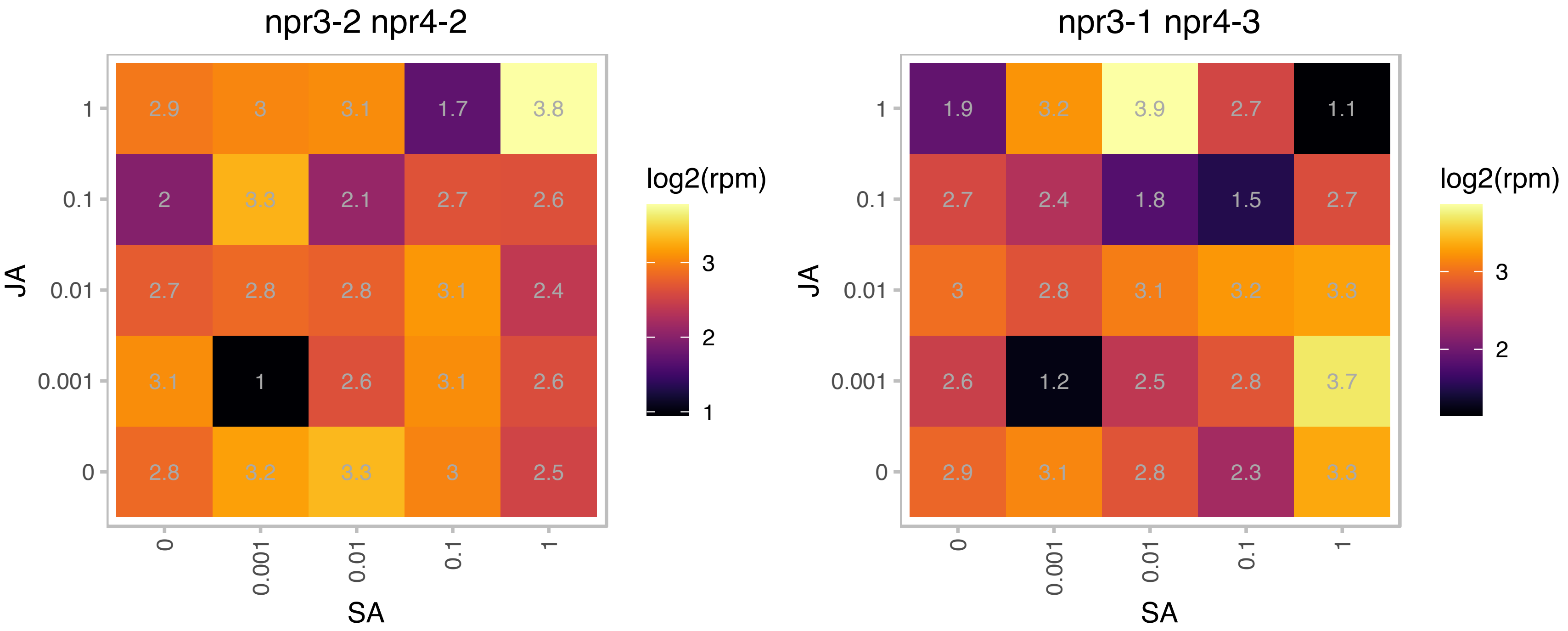

### AT1G52827.1 : ATCDT1, AthCYSTM2, CDT1

cadmium tolerance 1

Cysteine-rich protein that confers cadmium tolerance to yeast. Homologs are also found in rice, Brassica rapa and pine.

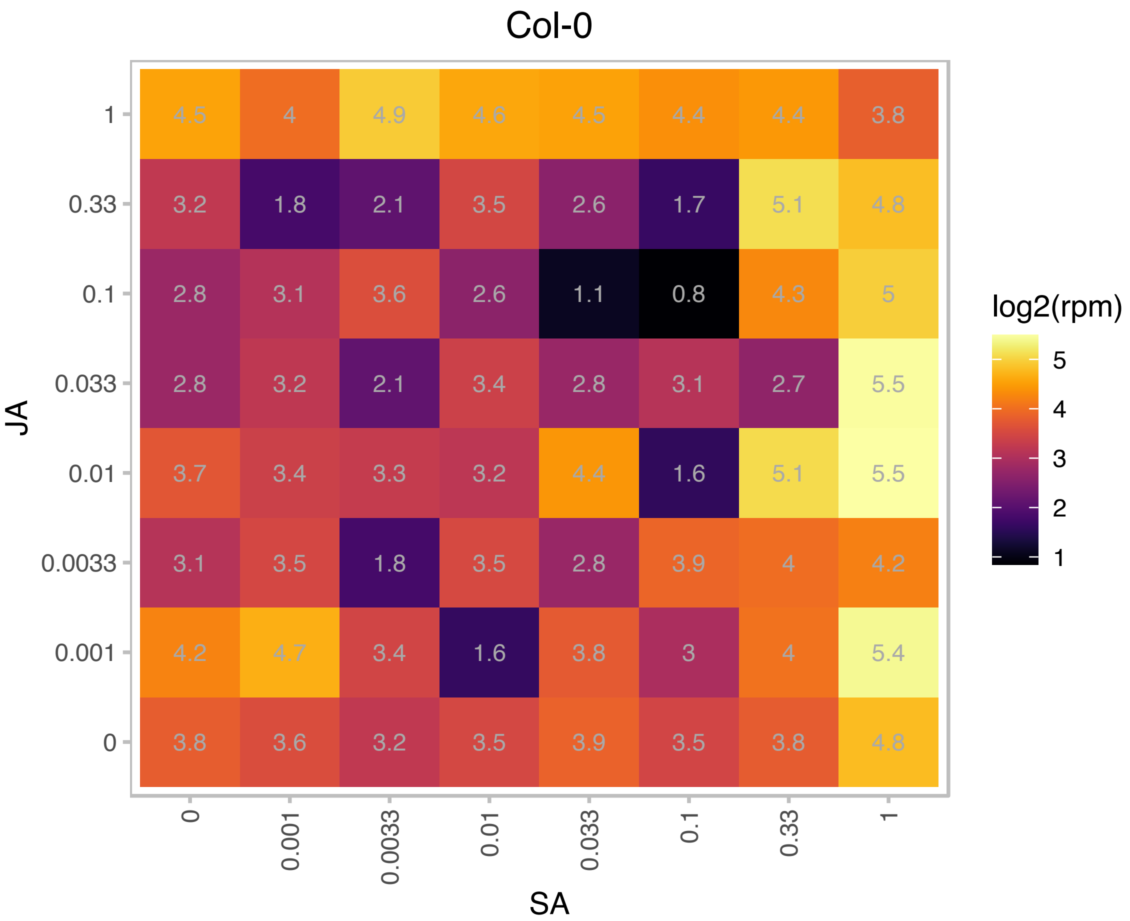


| | |
|---|---|
| Gene | AT1G52827 |
| is.expressed_genes | TRUE |
| is.responsive_genes | TRUE |
| is.apcluster | B-01 |

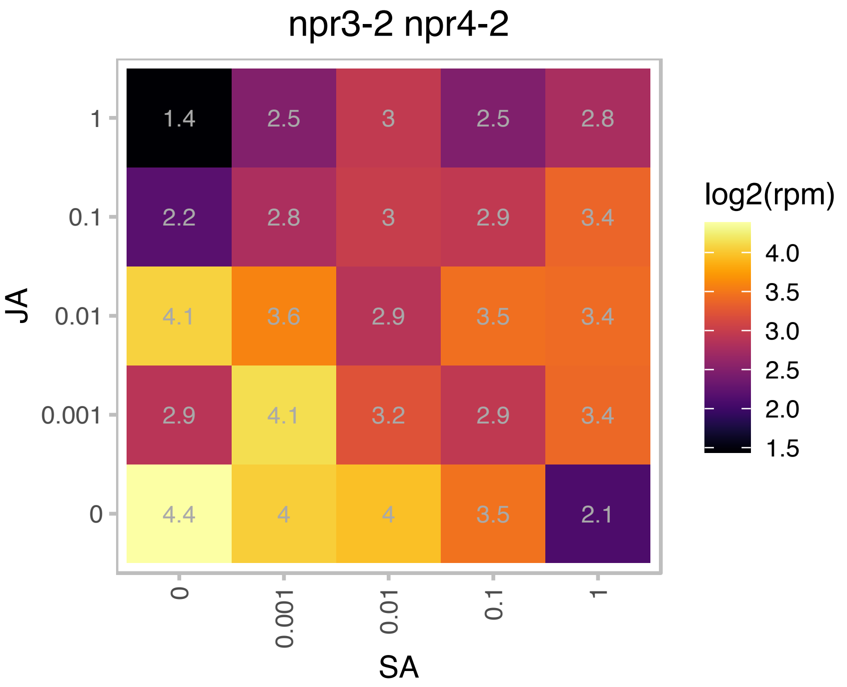


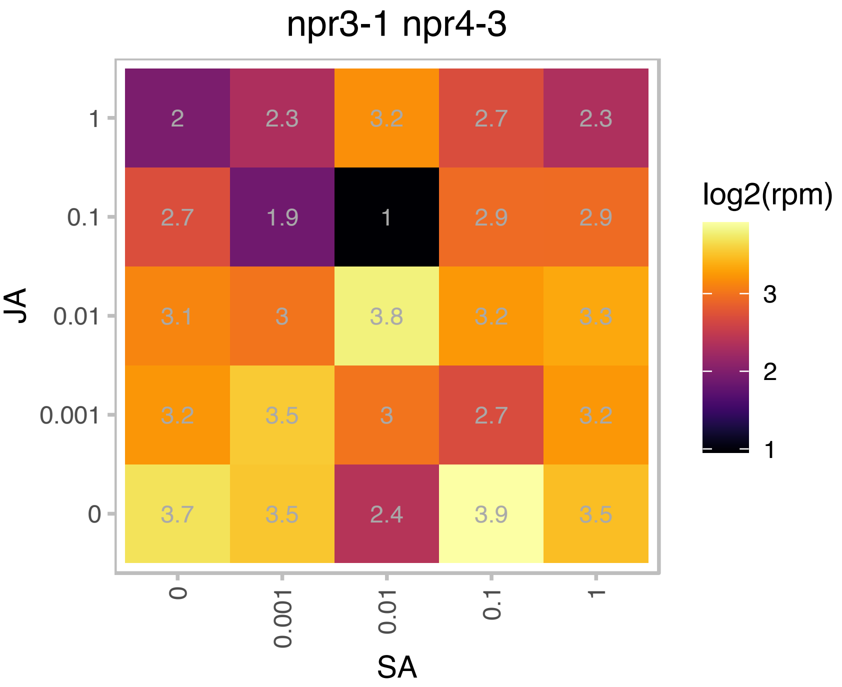

# AT1G53540.1 : NA

HSP20-like chaperones superfamily protein

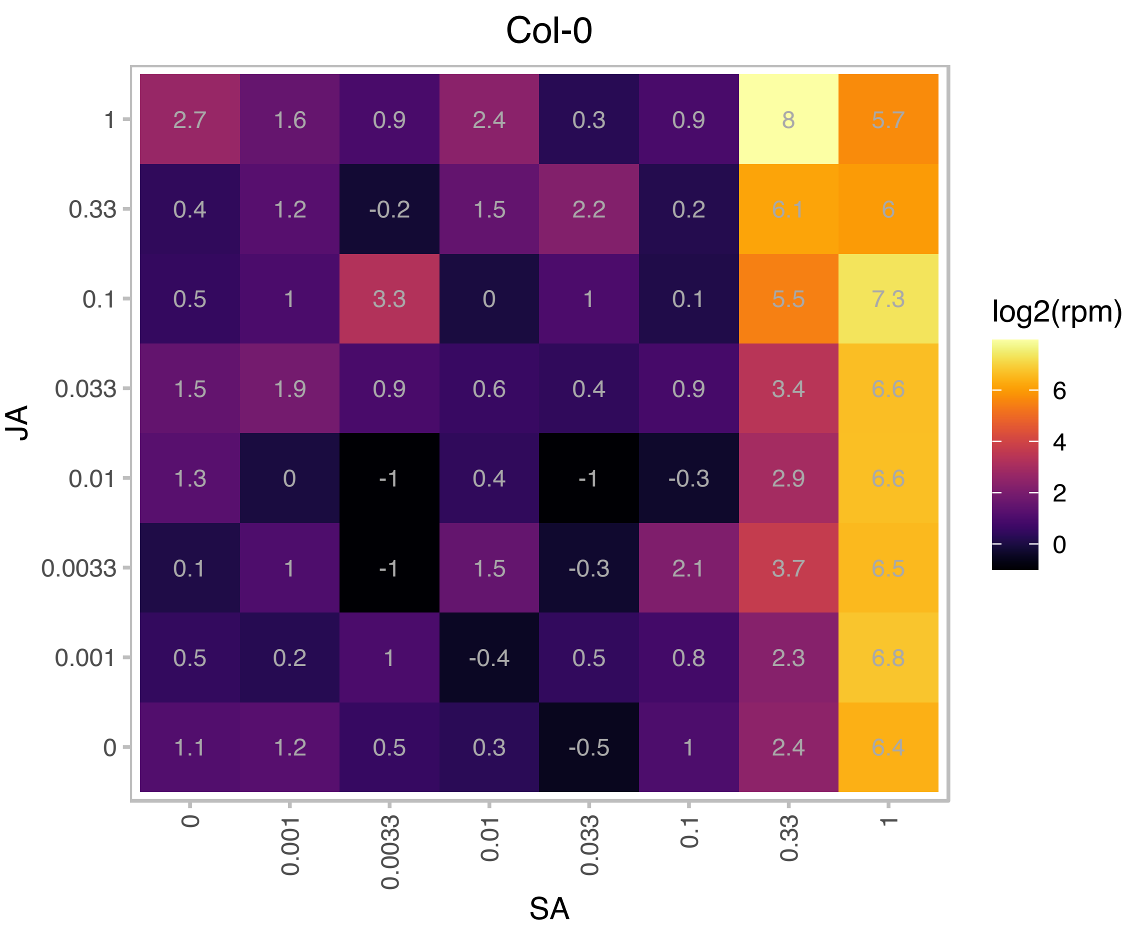


| | |
|---|---|
| Gene | AT1G53540 |
| is.expressed_genes | TRUE |
| is.responsive_genes | TRUE |
| is.apcluster | B-01 |

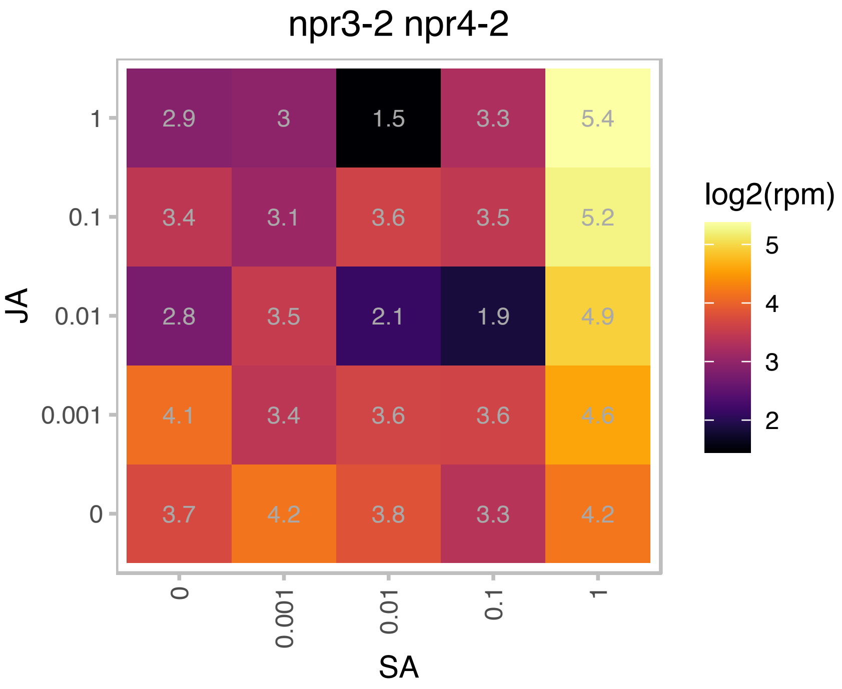


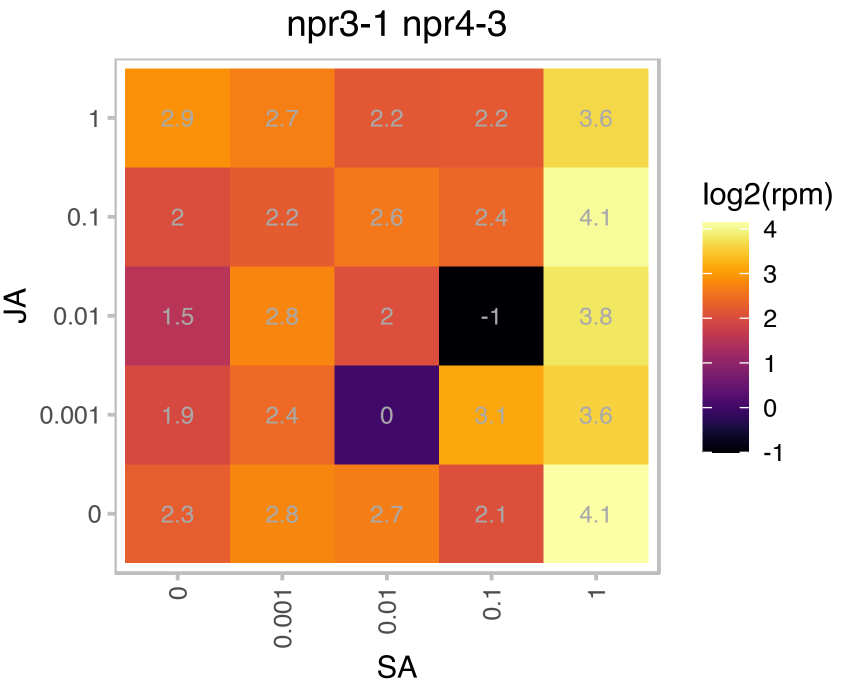

# AT1G54050.1 : NA

HSP20-like chaperones superfamily protein

| | |
|---|---|
| Gene | AT1G54050 |
| is.expressed_genes | TRUE |
| is.responsive_genes | TRUE |
| is.apcluster | B-01 |

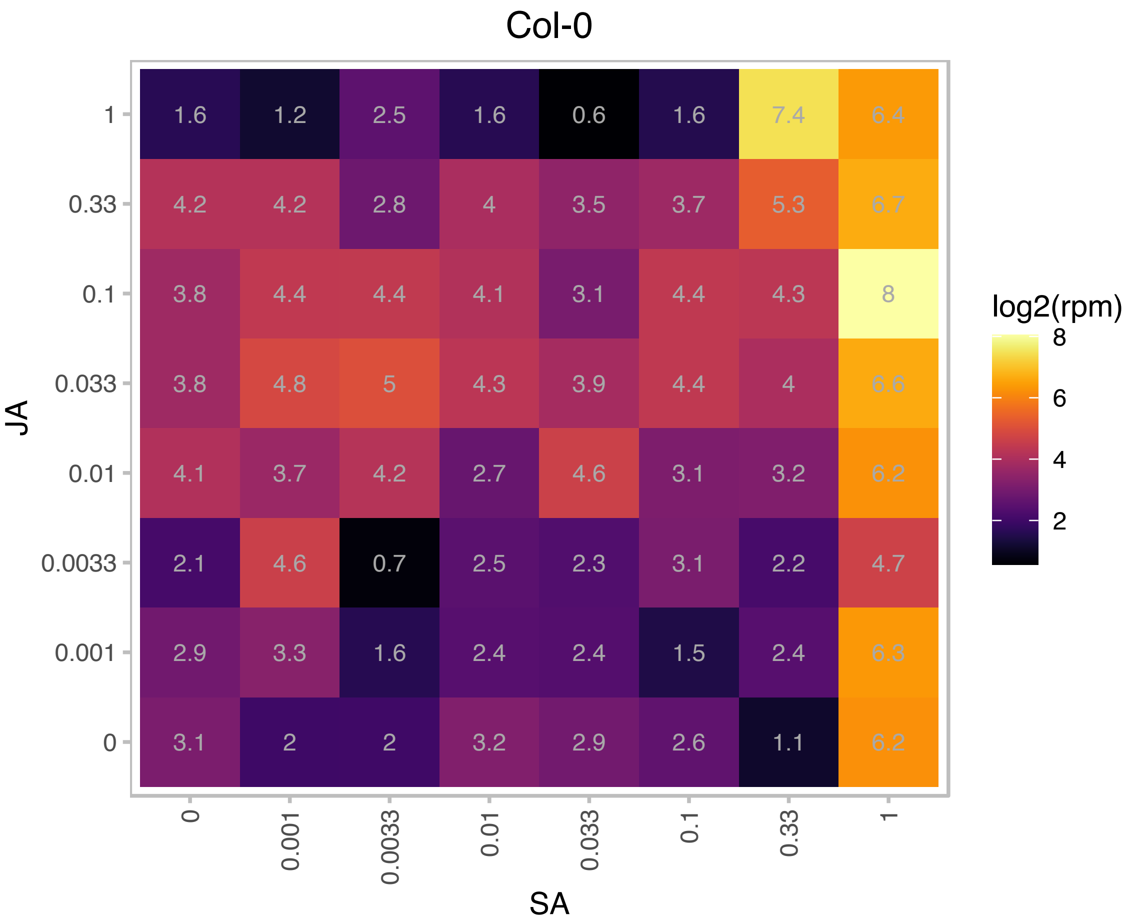


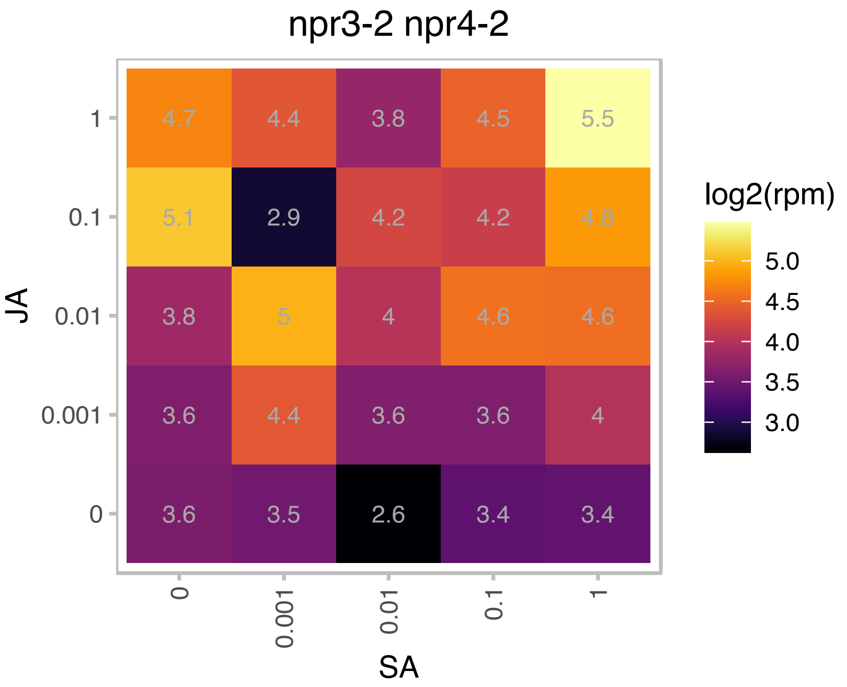


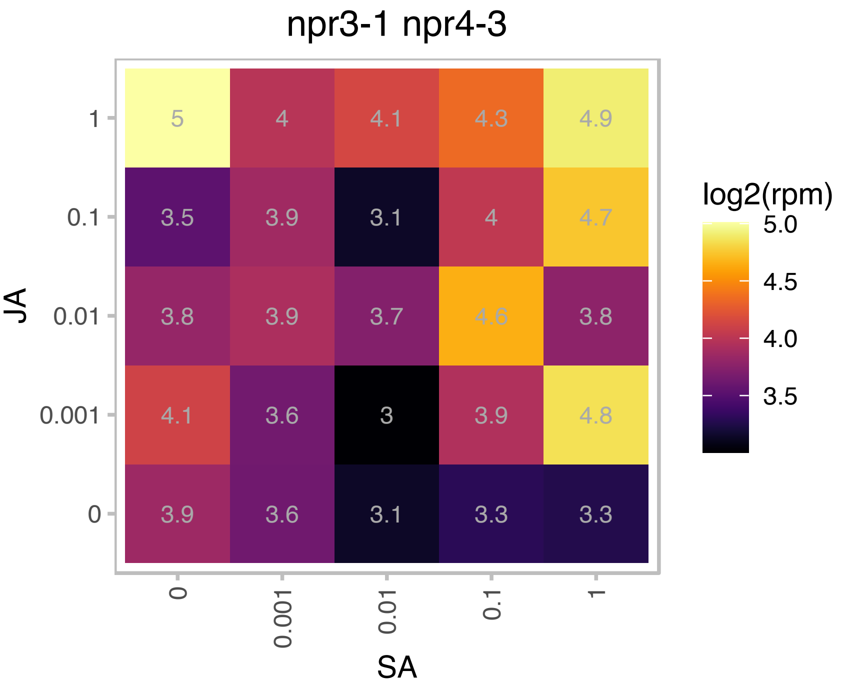

### AT1G56580.1 : SVB

Protein of unknown function, DUF538

Encodes SMALLER WITH VARIABLE BRANCHES (SVB), a protein with a conserved domain of unknown function (
The trichomes of the SVB mutants are smaller and exhibit branches of variable length and number.

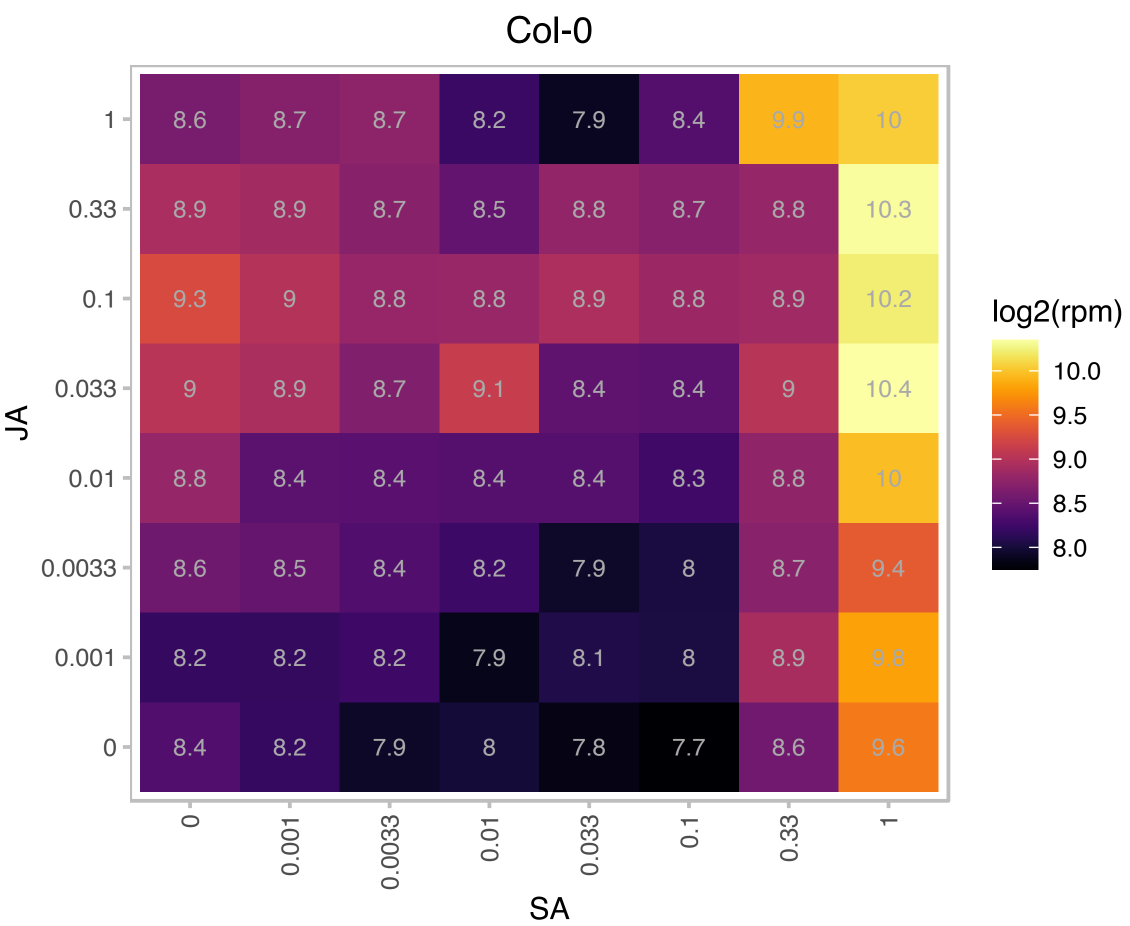


| | |
|---|---|
| Gene | AT1G56580 |
| is.expressed_genes | TRUE |
| is.responsive_genes | TRUE |
| is.apcluster | B-01 |

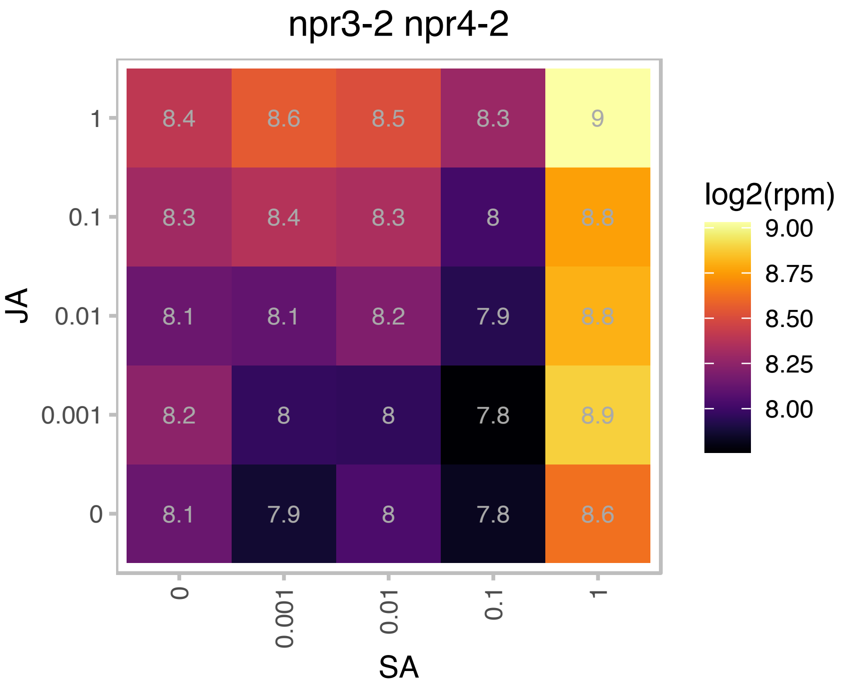


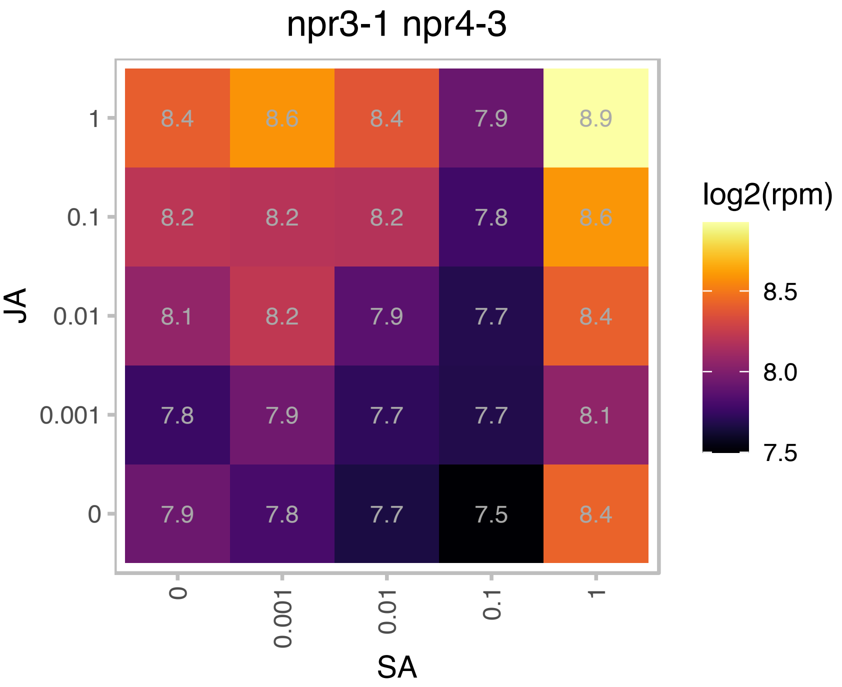

# AT1G67365.1 : NA

other RNA

Potential natural antisense gene, locus overlaps with AT1G67370

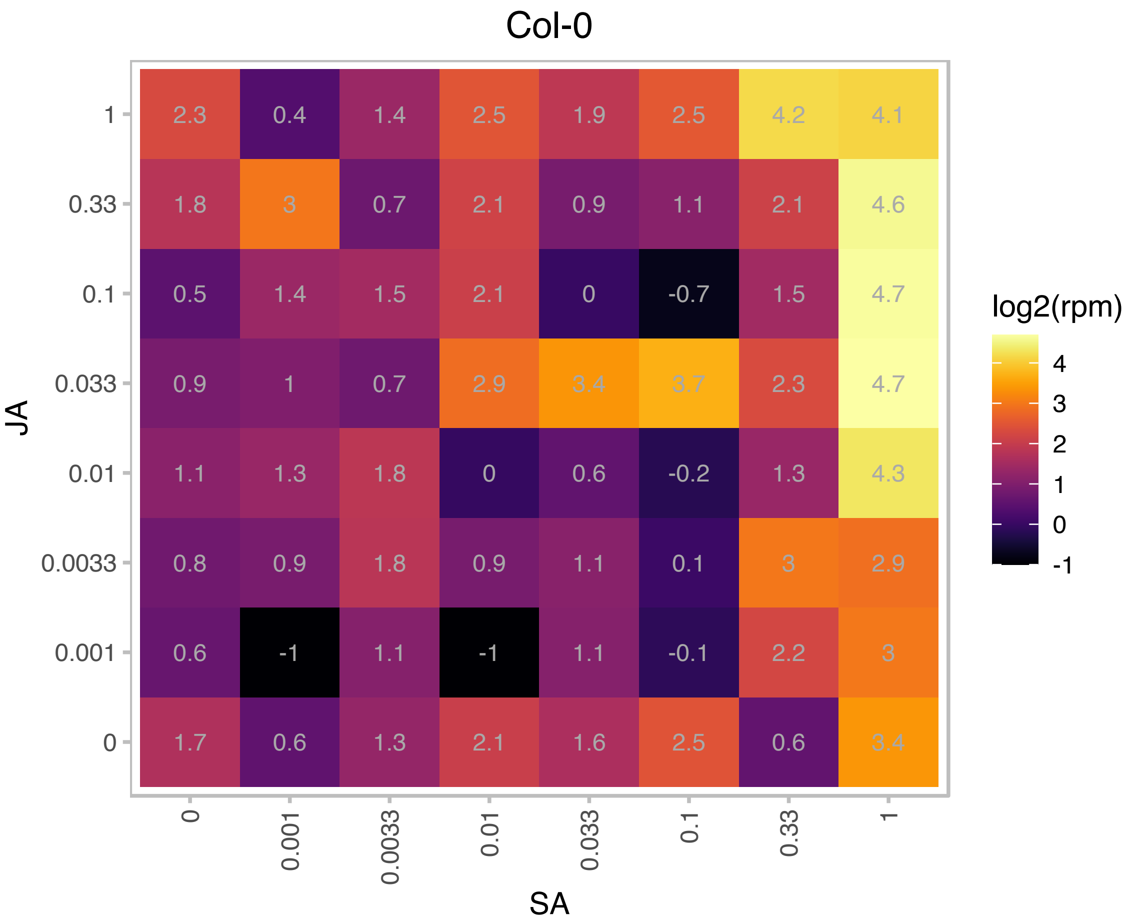


| | |
|---|---|
| Gene | AT1G67365 |
| is.expressed_genes | TRUE |
| is.responsive_genes | TRUE |
| is.apcluster | B-01 |

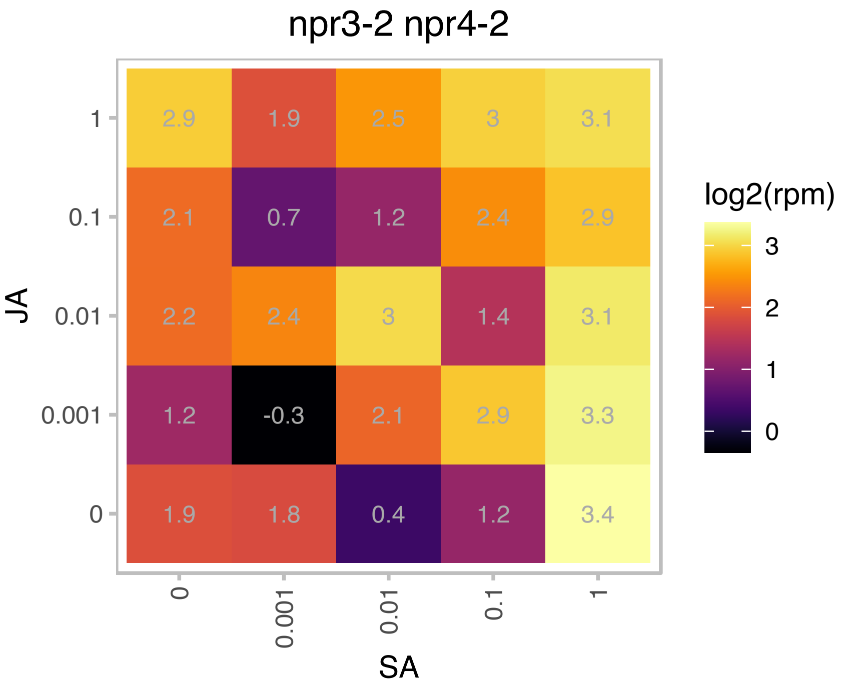


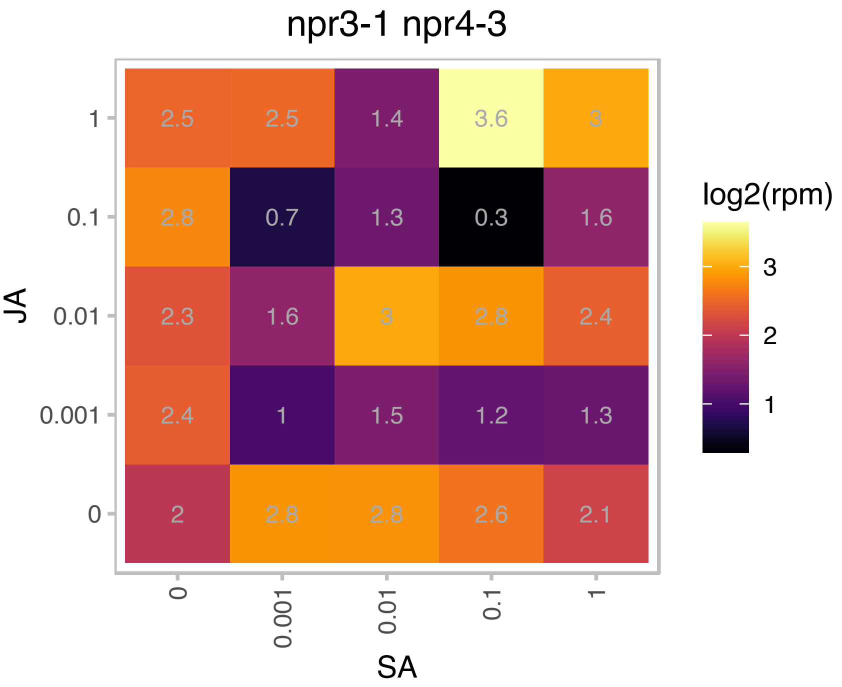

### AT1G68260.1 : ALT3

Thioesterase superfamily protein

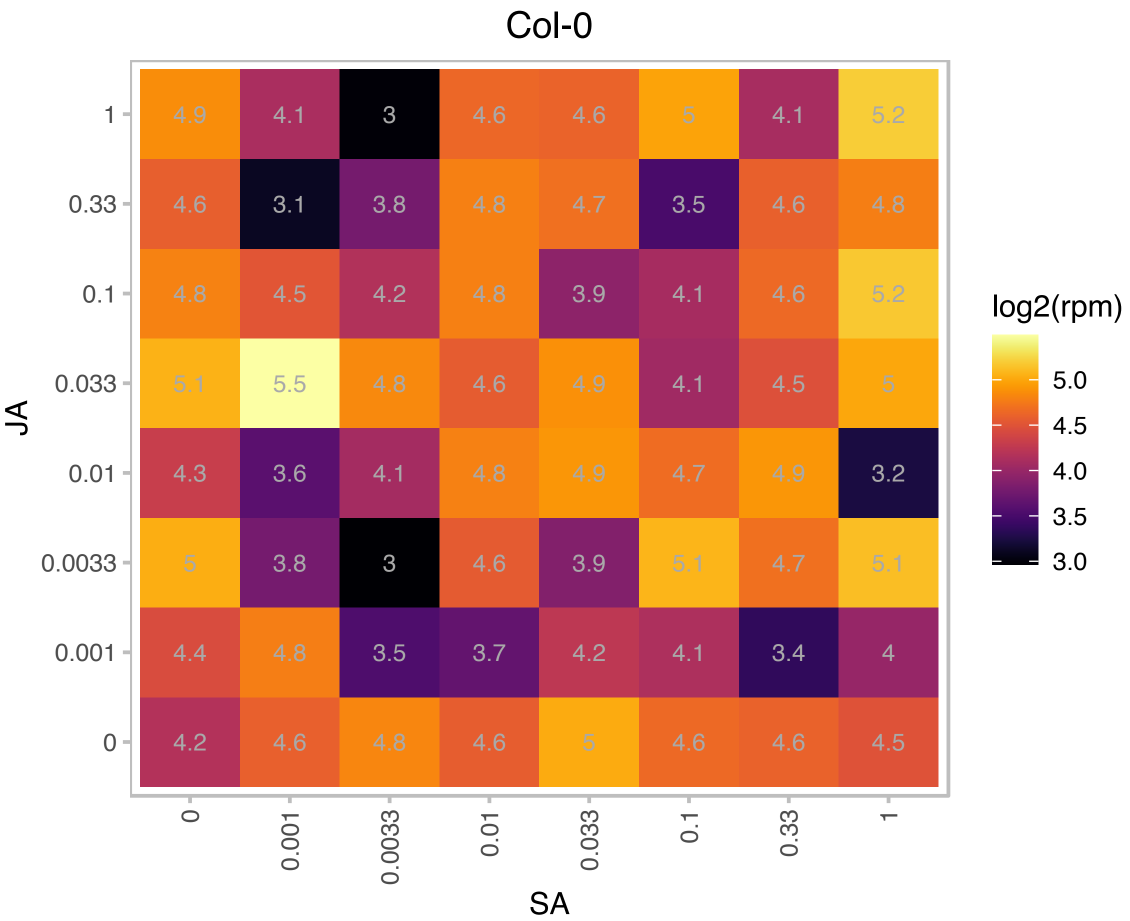


| | |
|---|---|
| Gene | AT1G68260 |
| is.expressed_genes | TRUE |
| is.responsive_genes | TRUE |
| is.apcluster | B-01 |

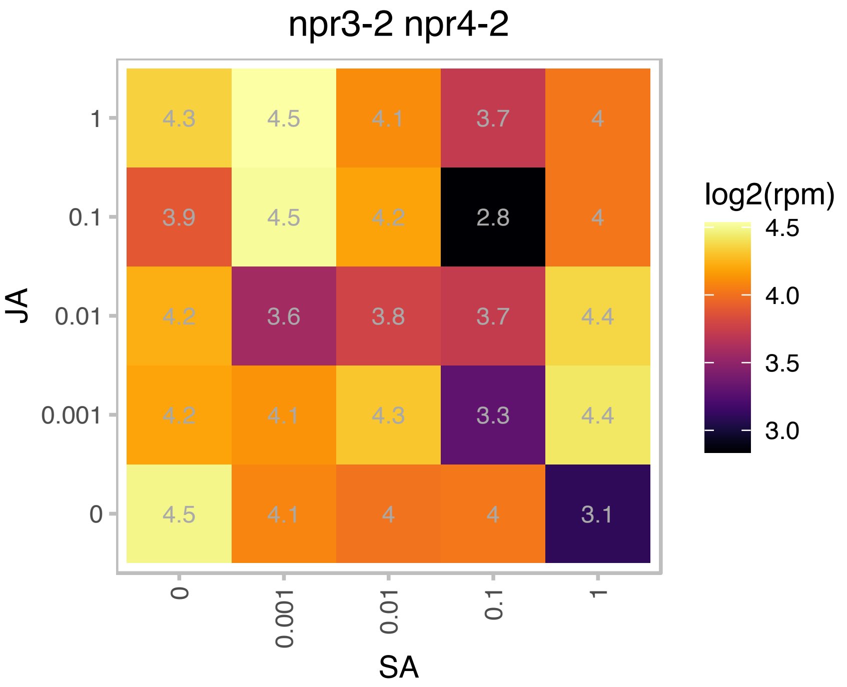


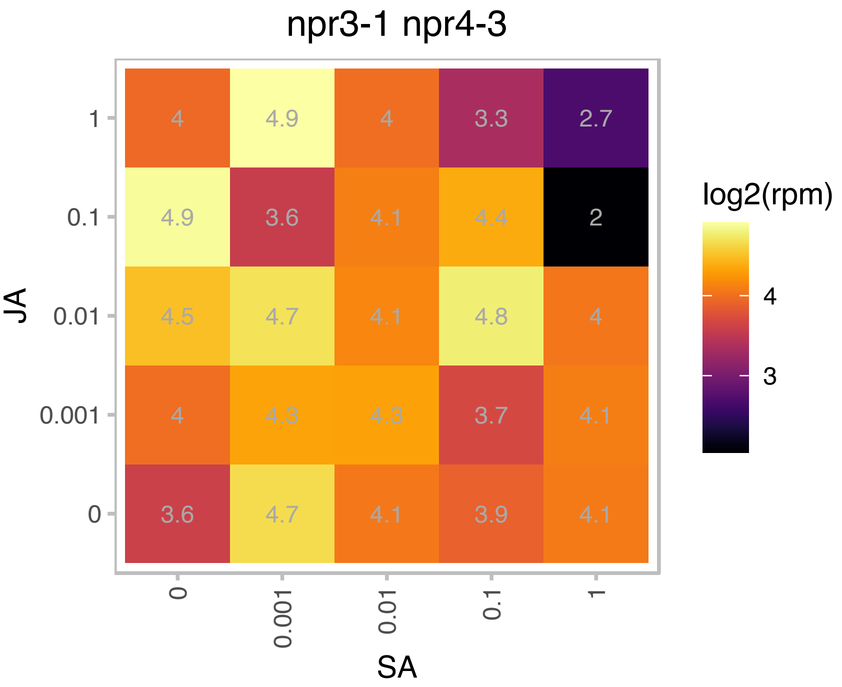

# AT1G79245.1 : NA

| | |
|---|---|
| Gene | AT1G79245 |
| is.expressed_genes | TRUE |
| is.responsive_genes | TRUE |
| is.apcluster | B-01 |

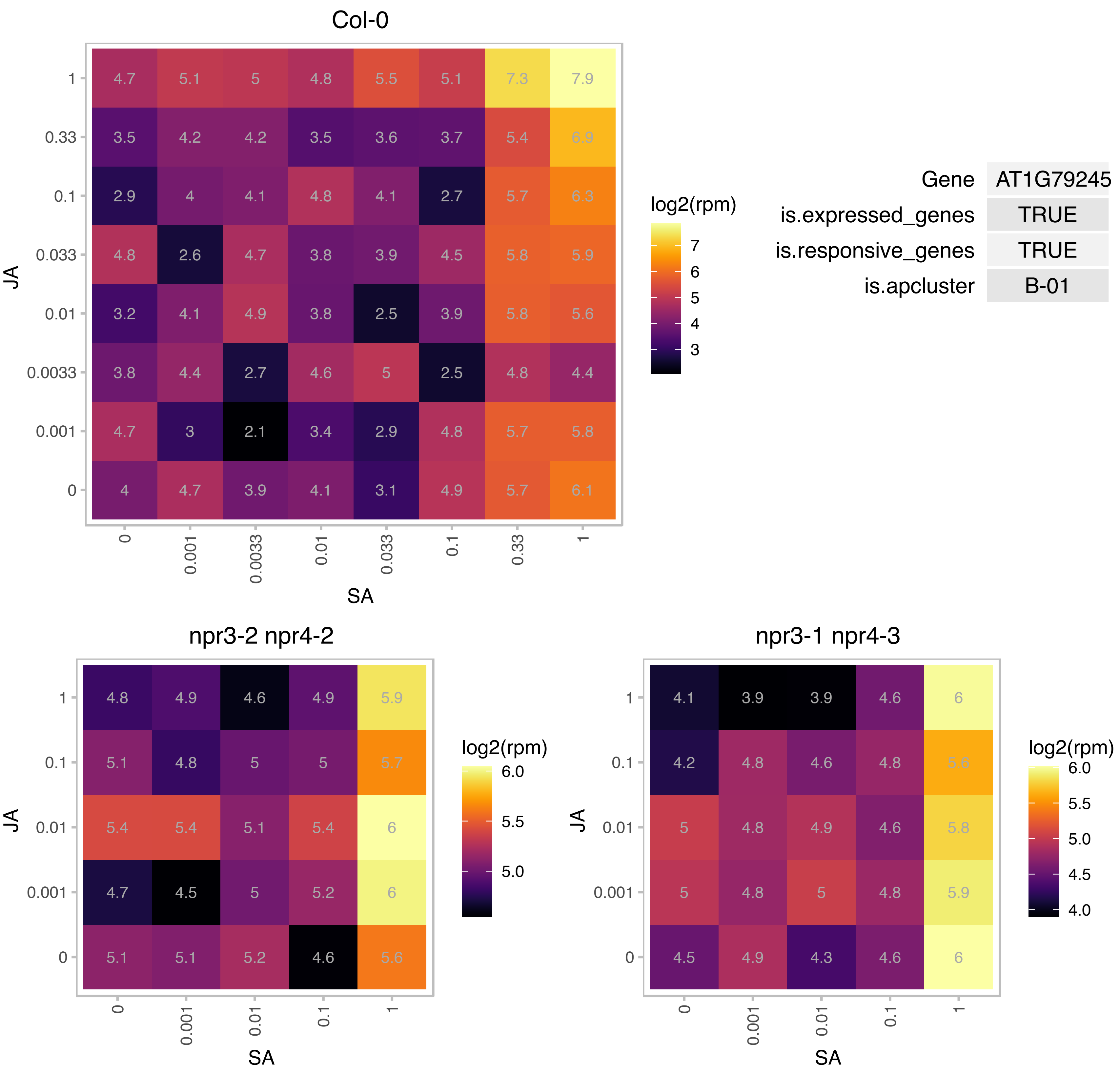

# AT2G05632.1 : NA

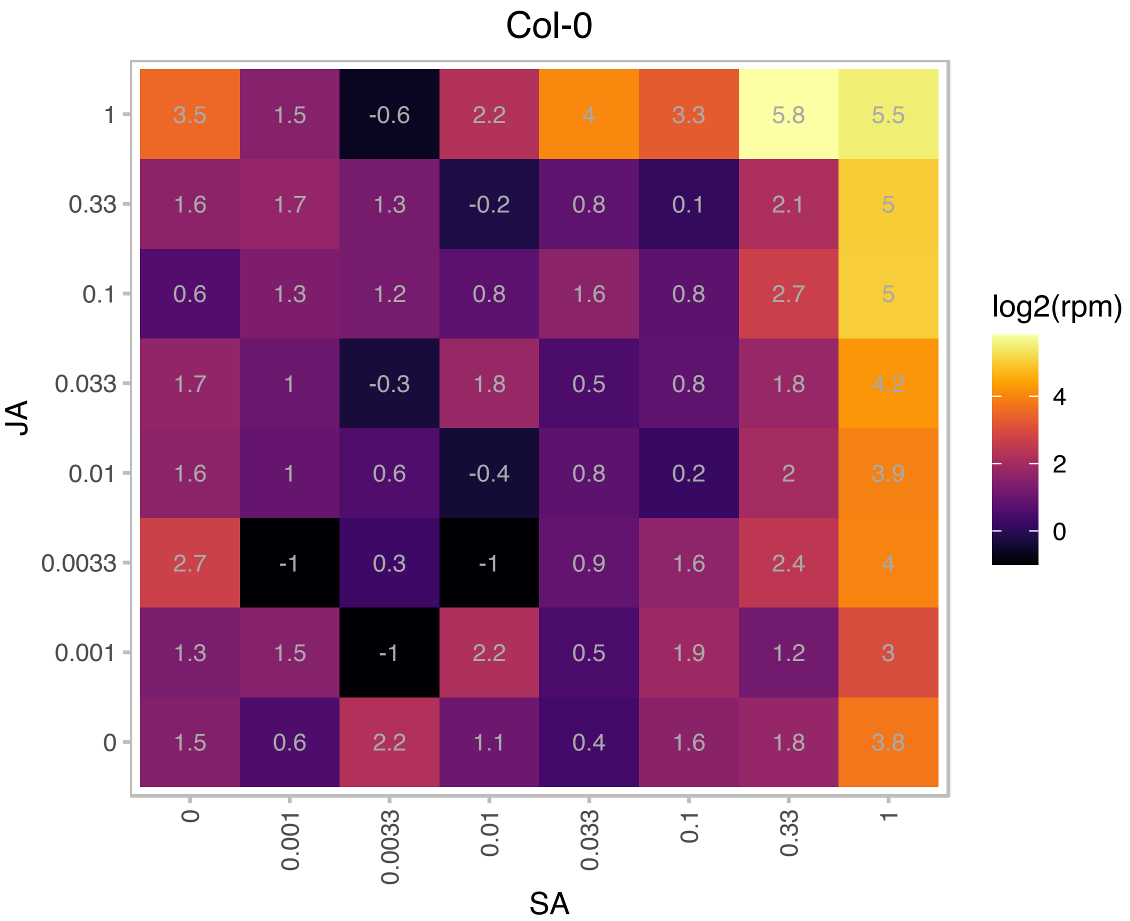


| | |
|---|---|
| Gene | AT2G05632 |
| is.expressed_genes | TRUE |
| is.responsive_genes | TRUE |
| is.apcluster | B-01 |

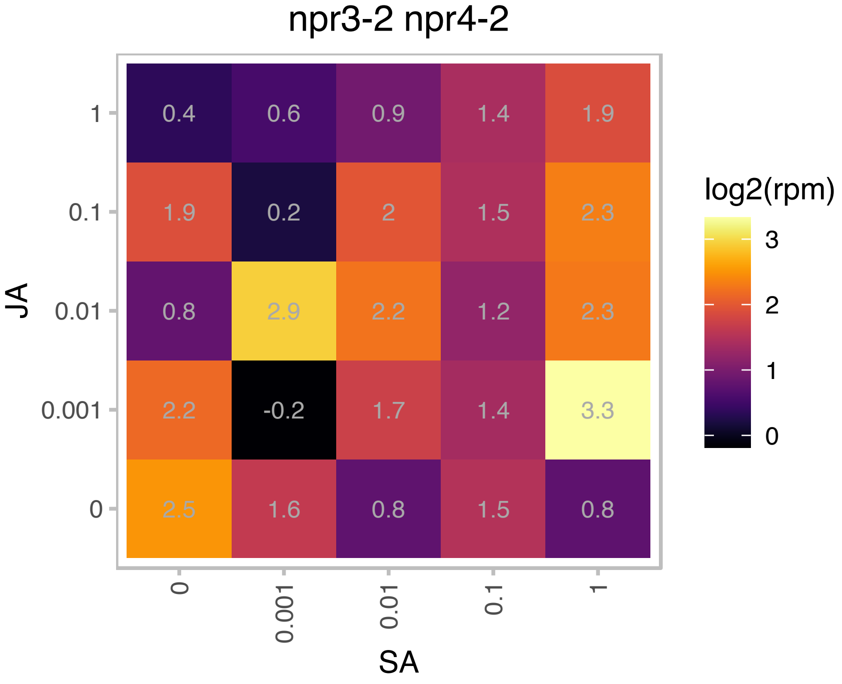


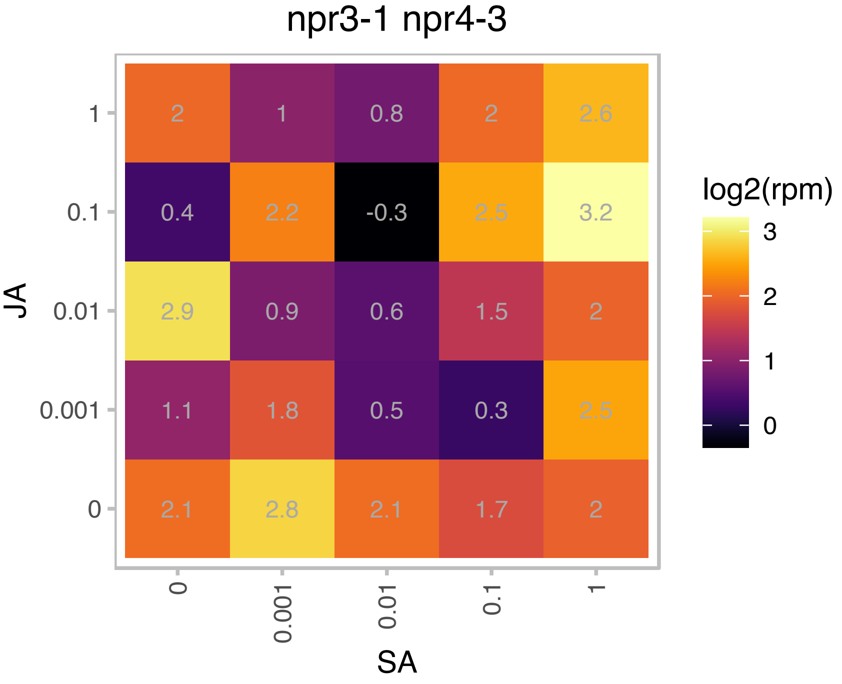

### AT2G20530.1 : ATPHB6, PHB6

prohibitin 6

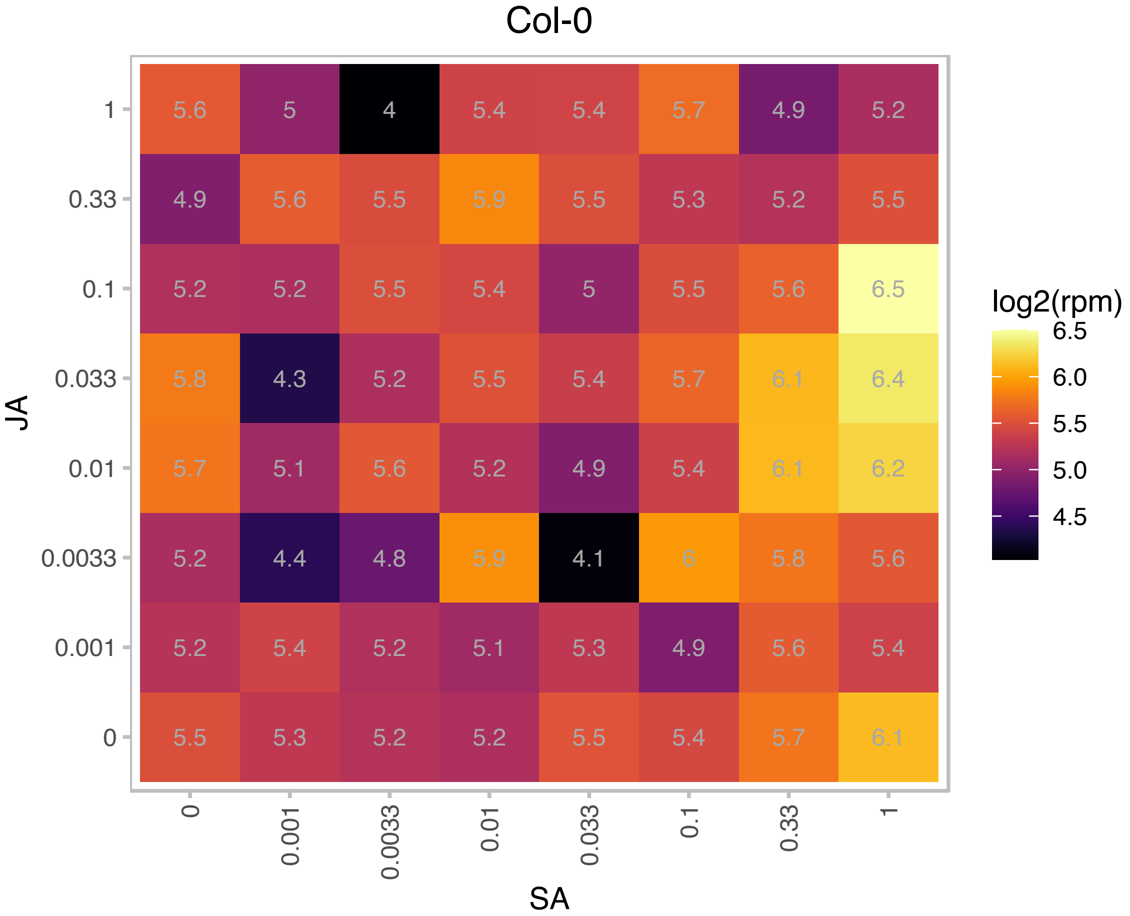


| | |
|---|---|
| Gene | AT2G20530 |
| is.expressed_genes | TRUE |
| is.responsive_genes | TRUE |
| is.apcluster | B-01 |

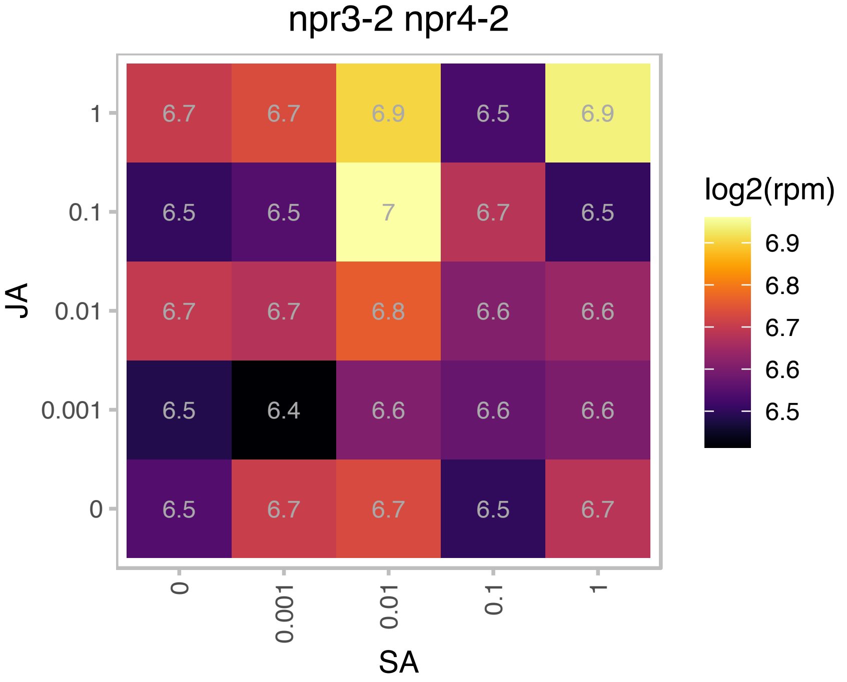


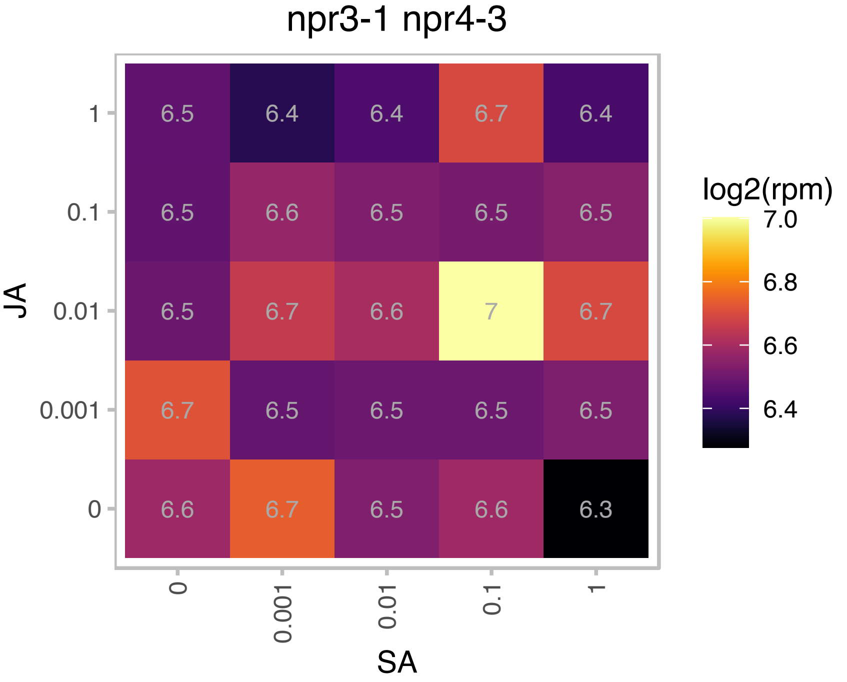

# AT2G21640.1 : NA

Encodes a protein of unknown function that is a marker for oxidative stress response.Expression in rosette leaves is activated by high concentration of boron.

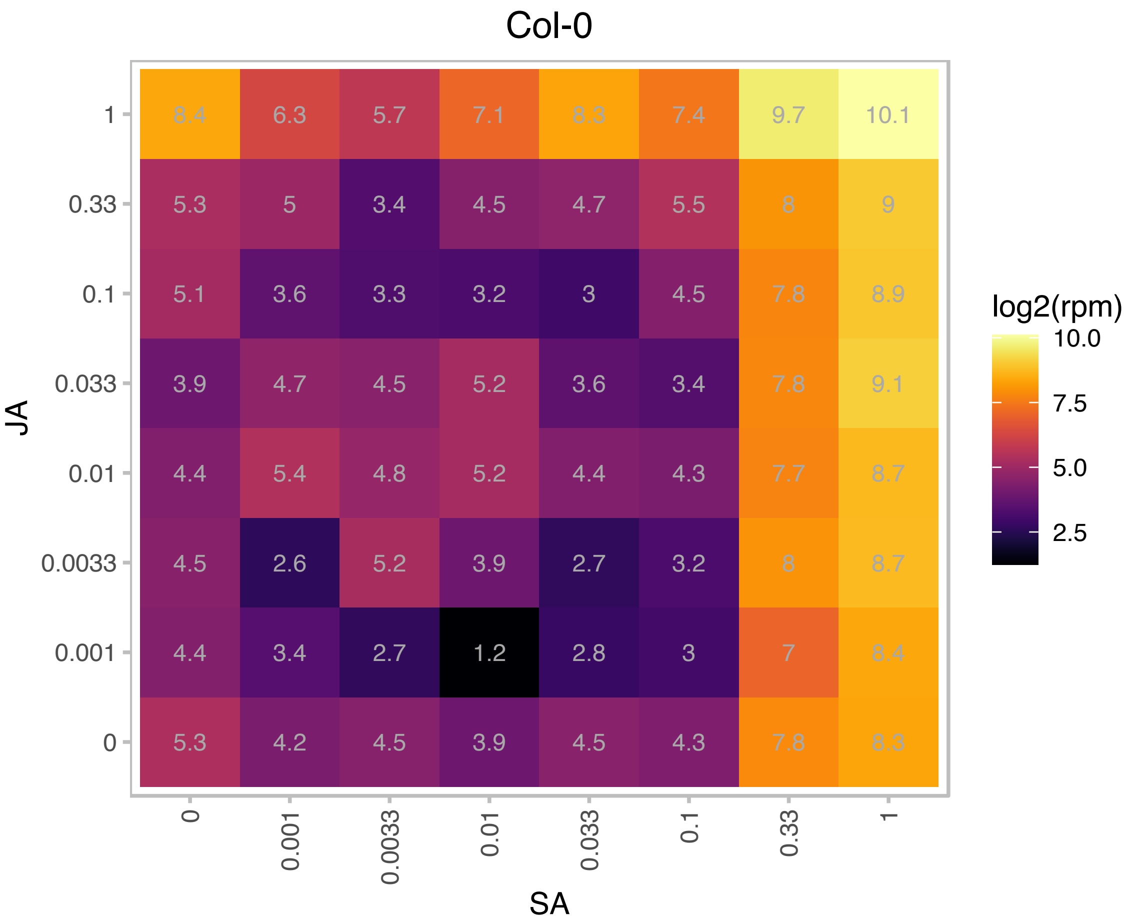


| | |
|---|---|
| Gene | AT2G21640 |
| is.expressed_genes | TRUE |
| is.responsive_genes | TRUE |
| is.apcluster | B-01 |

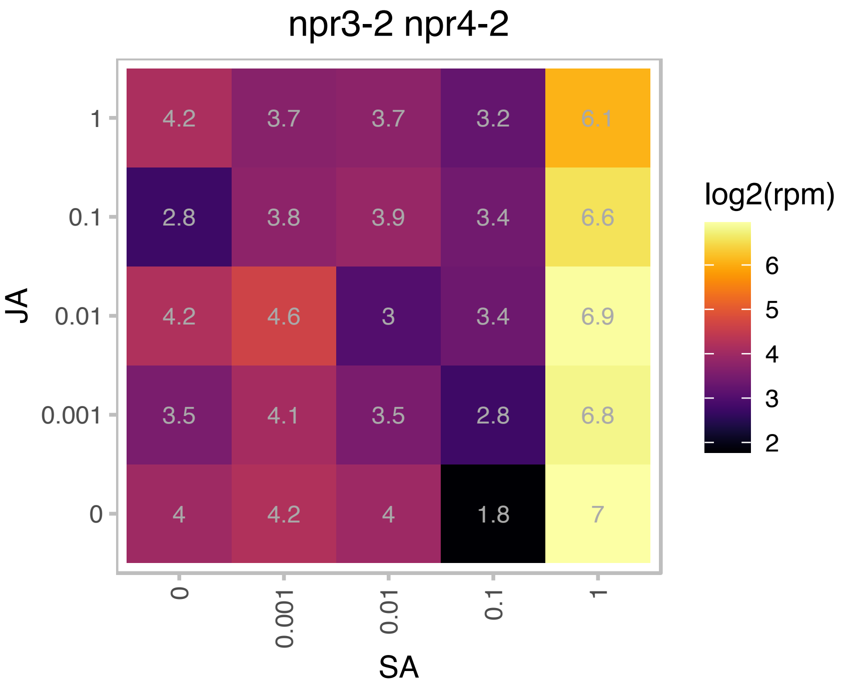


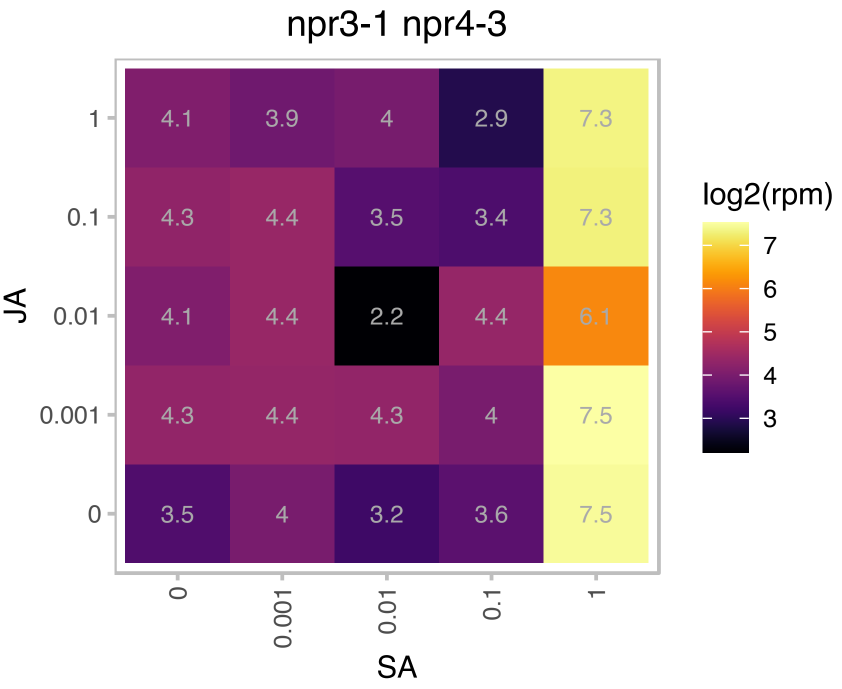

# AT2G23170.1 : GH3.3

Auxin-responsive GH3 family protein

encodes an IAA-amido synthase that conjugates Asp and other amino acids to auxin in vitro.

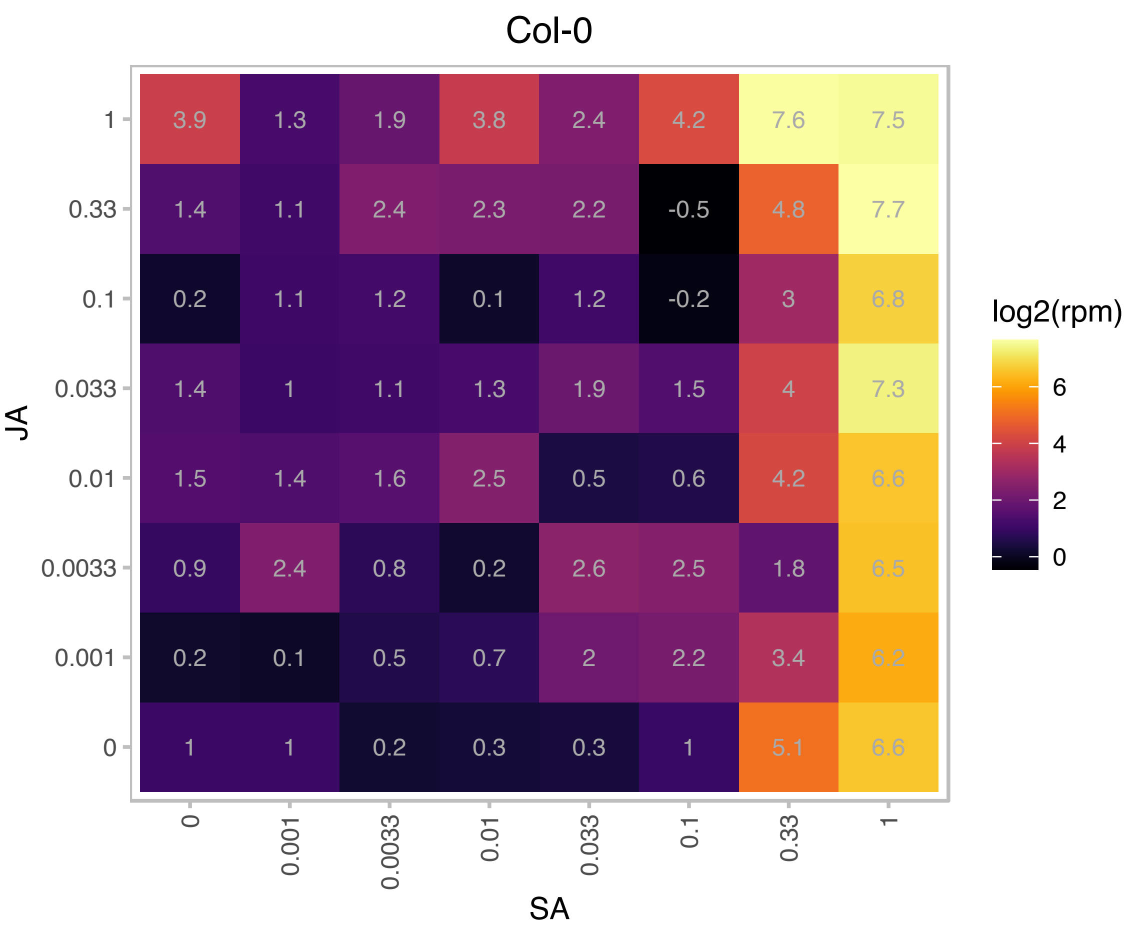


| | |
|---|---|
| Gene | AT2G23170 |
| is.expressed_genes | TRUE |
| is.responsive_genes | TRUE |
| is.apcluster | B-01 |

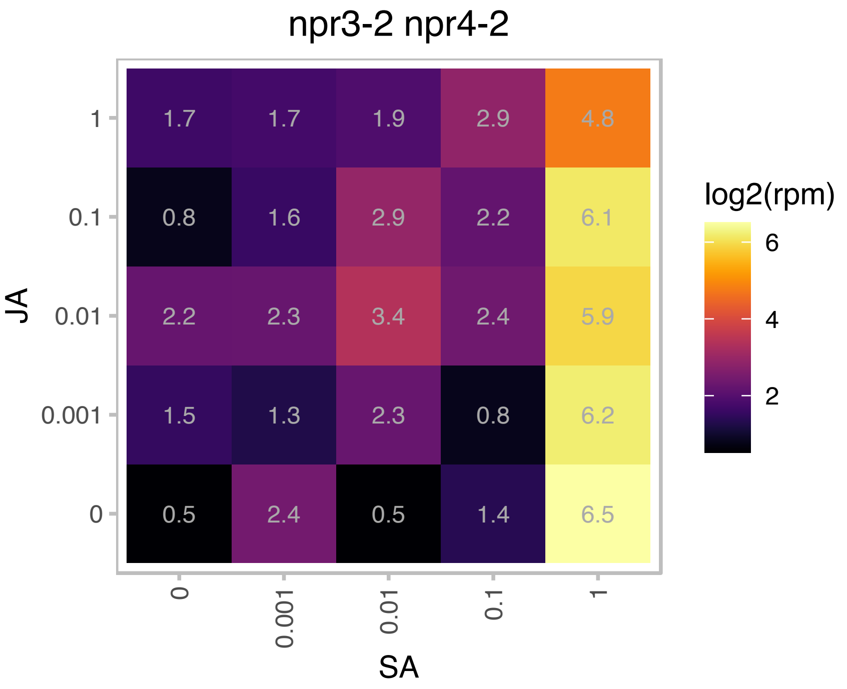


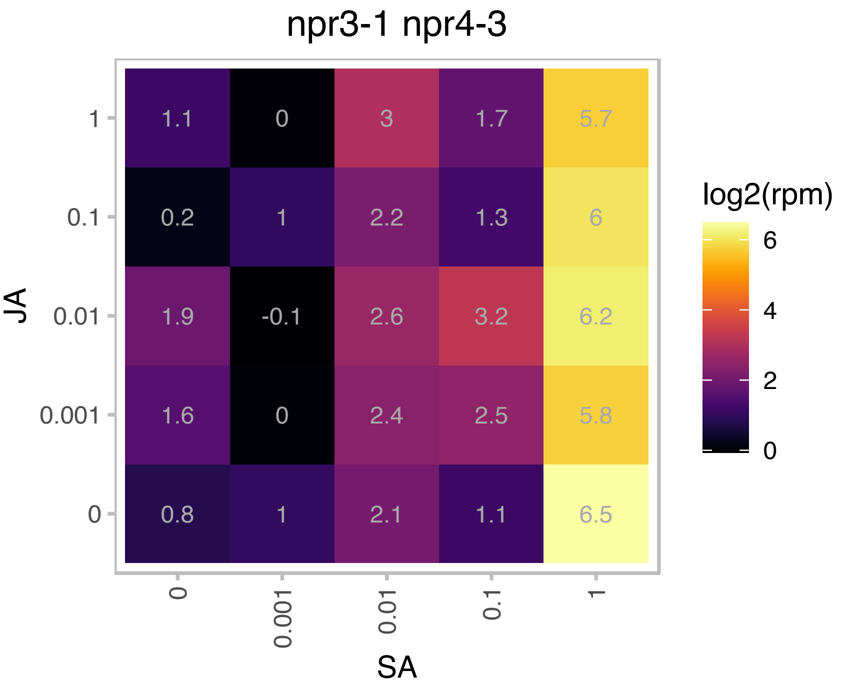

### AT2G30130.1 : ASL5, LBD12, PCK1

Lateral organ boundaries (LOB) domain family protein

Overexpression/activation tagged allele has epinastic leaves, reduced apical dominance and is sterile. Gene is similar to asymmetric leaves (AS)/lateral organ boundary (LOB) genes which repress KNOX gene expression.

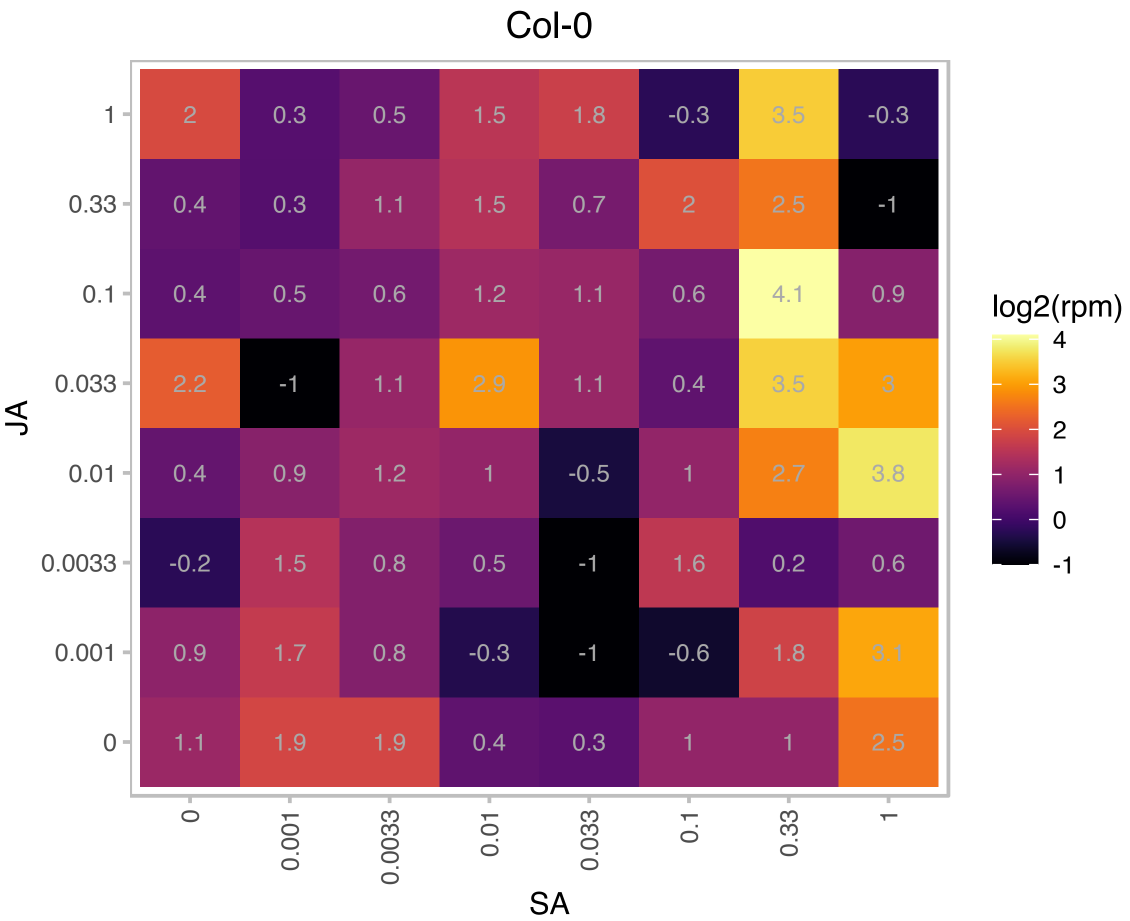


| | |
|---|---|
| Gene | AT2G30130 |
| is.expressed_genes | TRUE |
| is.responsive_genes | TRUE |
| is.apcluster | B-01 |

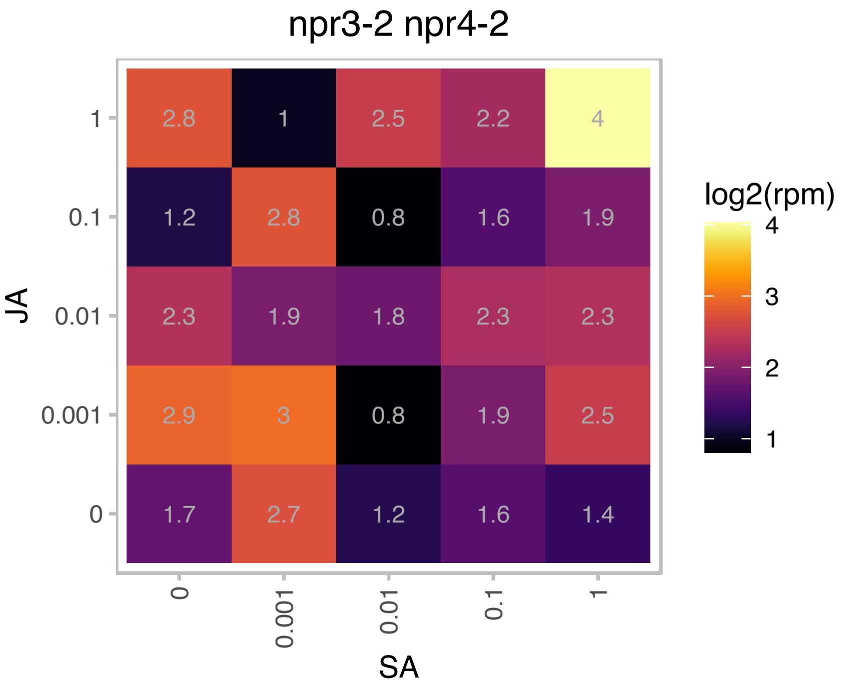


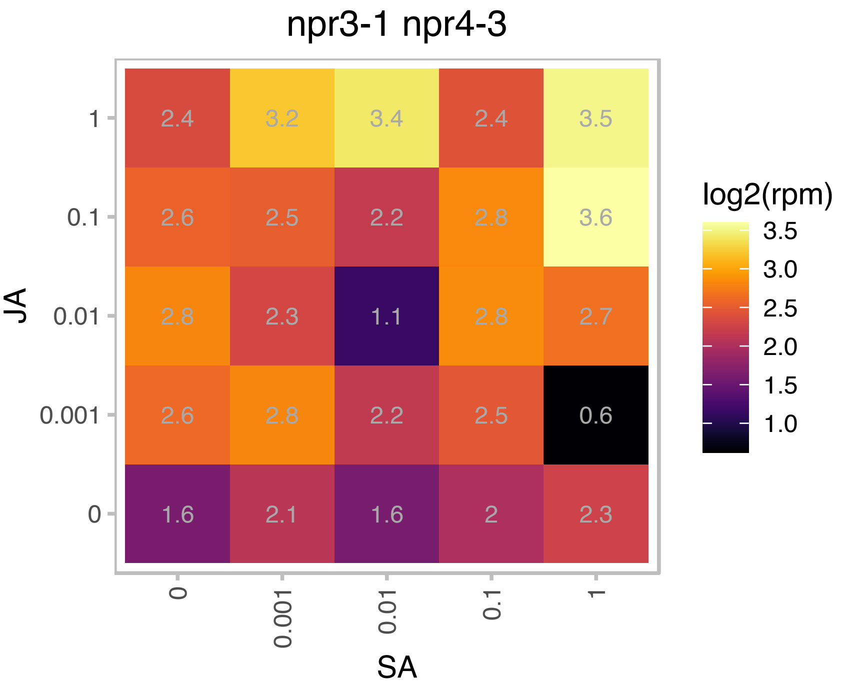

# AT2G34460.1 : NA

NAD(P)-binding Rossmann-fold superfamily protein

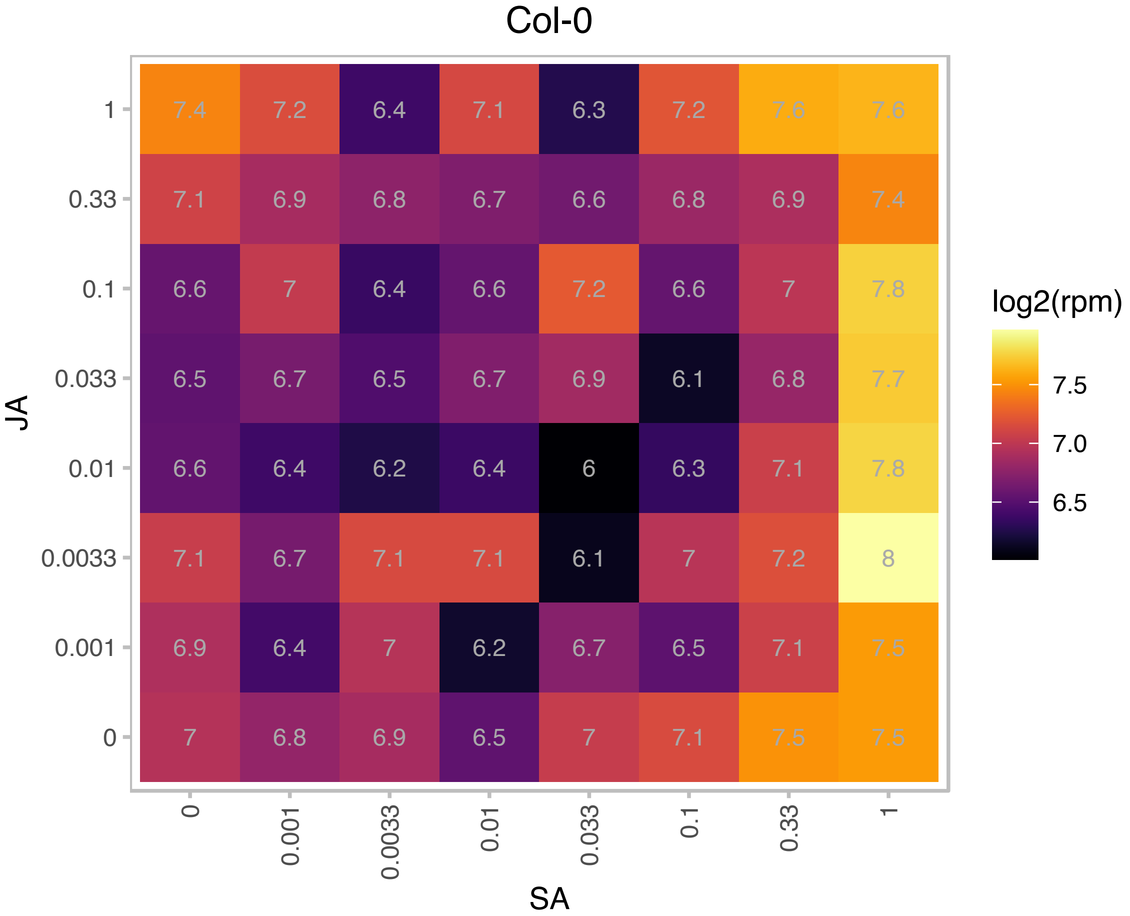


| Gene | AT2G34460 |
|---|---|
| is.expressed_genes | TRUE |
| is.responsive_genes | TRUE |
| is.apcluster | B-01 |

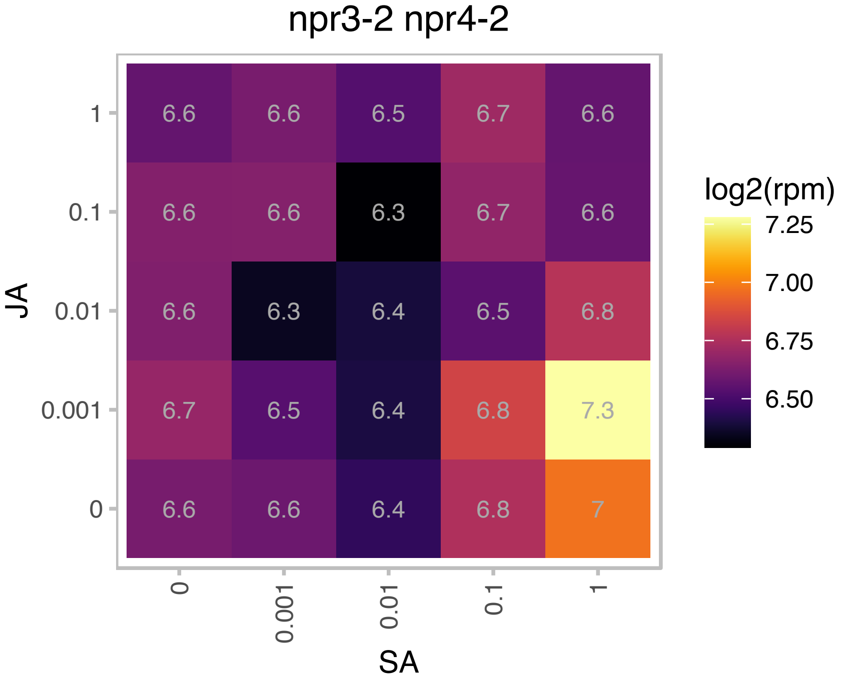


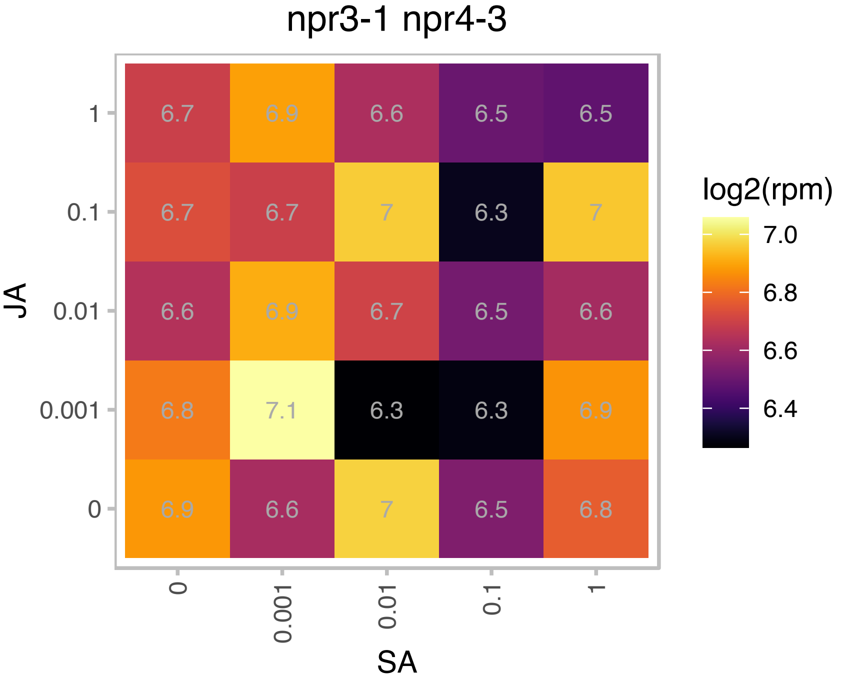

# AT2G37130.1 : NA

Peroxidase superfamily protein

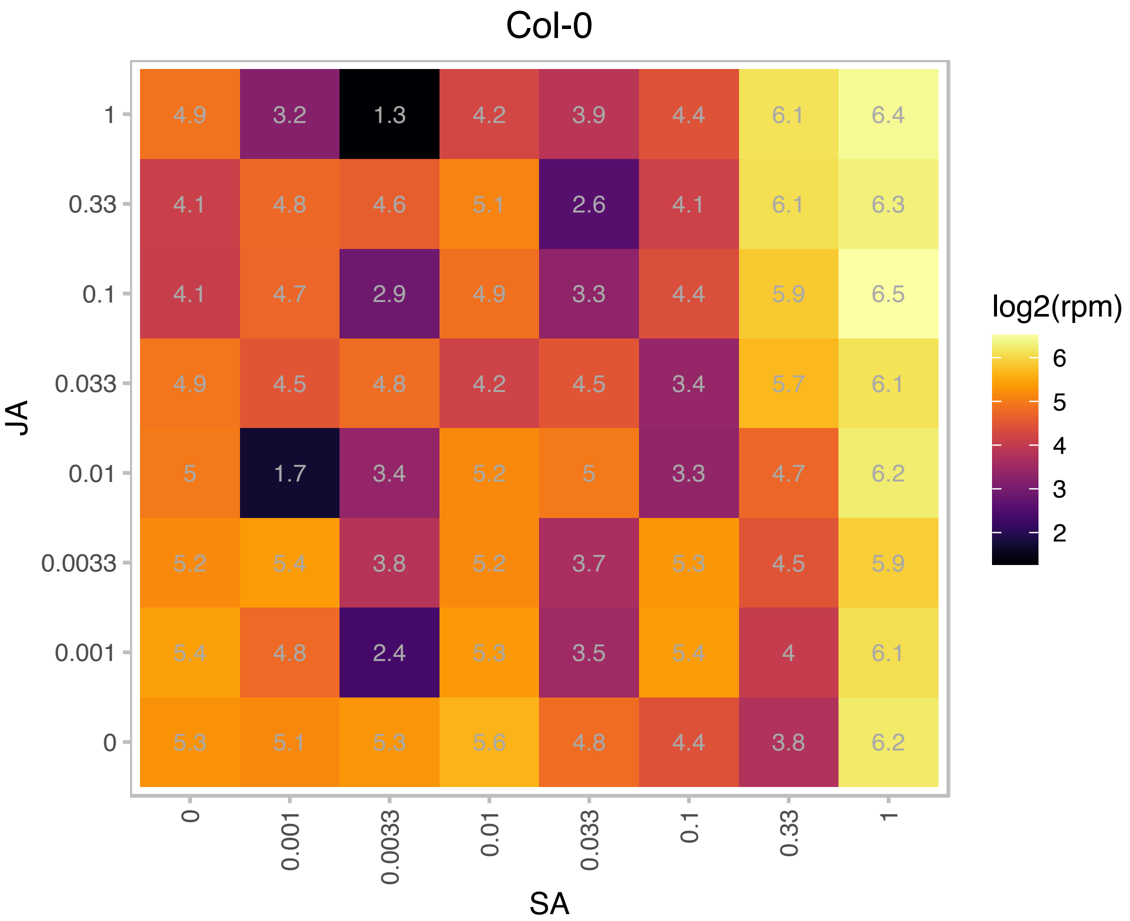


| | |
|---|---|
| Gene | AT2G37130 |
| is.expressed_genes | TRUE |
| is.responsive_genes | TRUE |
| is.apcluster | B-01 |

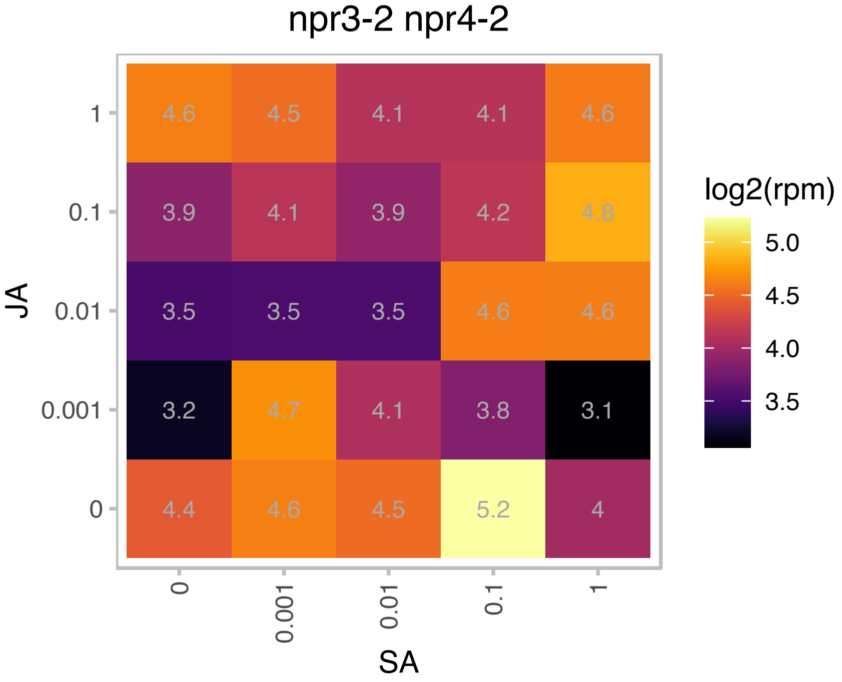


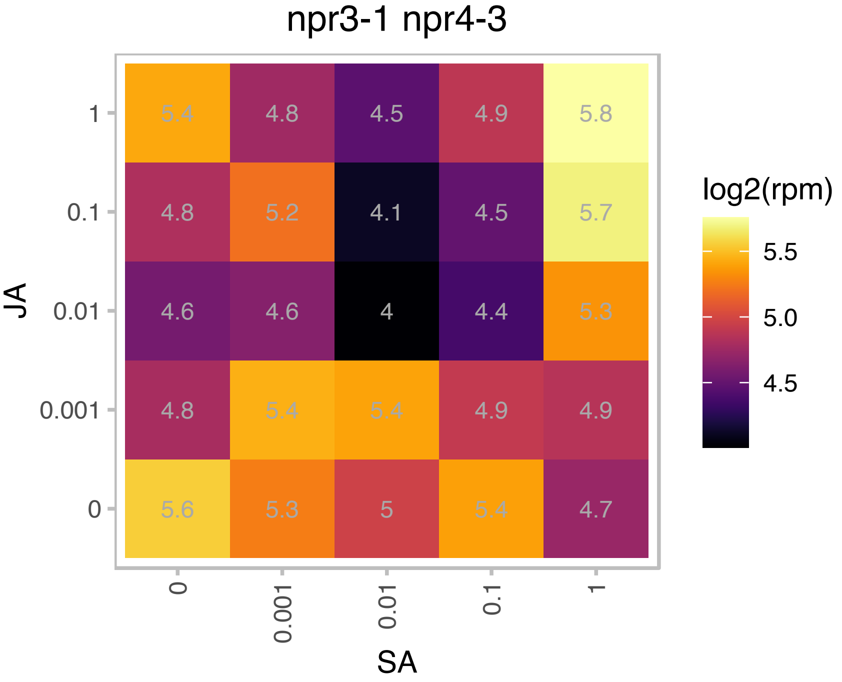

### AT3G04630.1 : WDL1

WVD2-like 1

Member of a small gene family which have a KLEEK domain which may be involved in protein- protein interactions. Over expression of WDL1 results in abnormal root development.

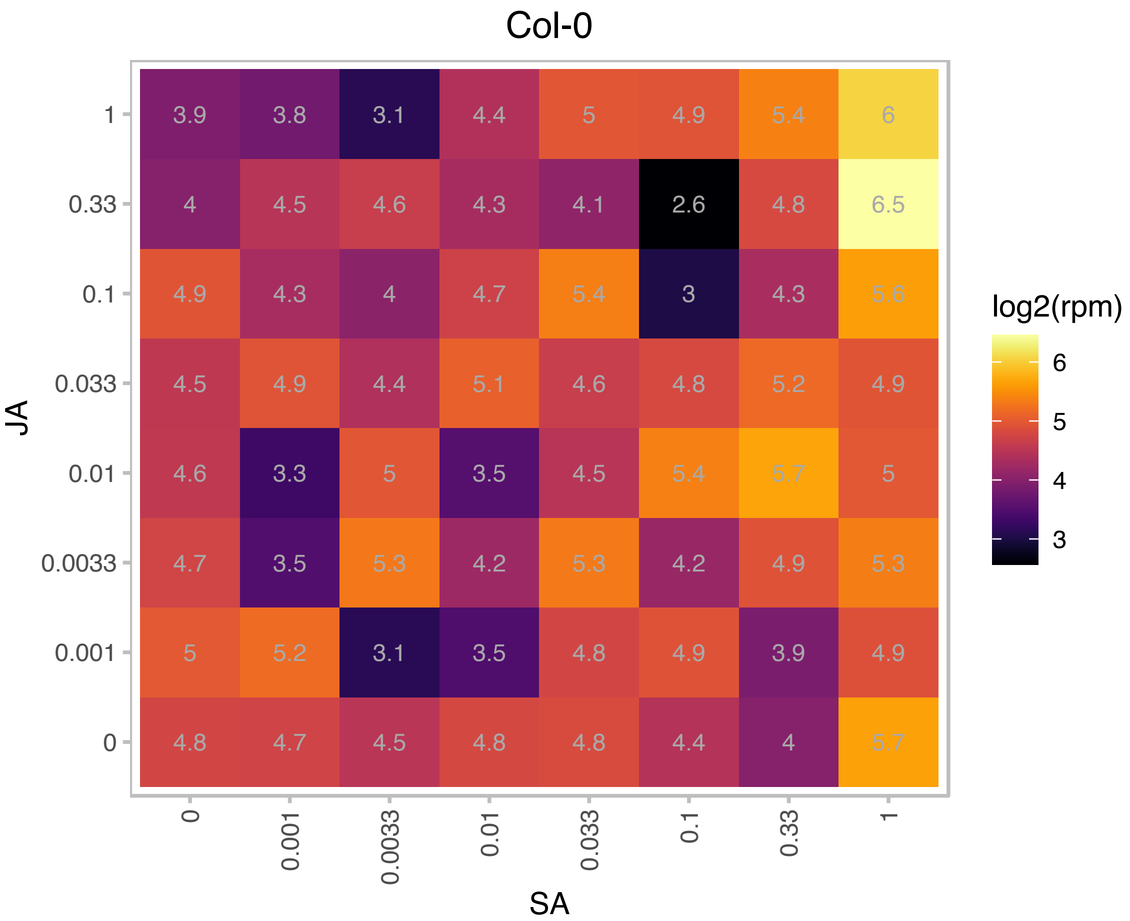


| | |
|---|---|
| Gene | AT3G04630 |
| is.expressed_genes | TRUE |
| is.responsive_genes | TRUE |
| is.apcluster | B-01 |

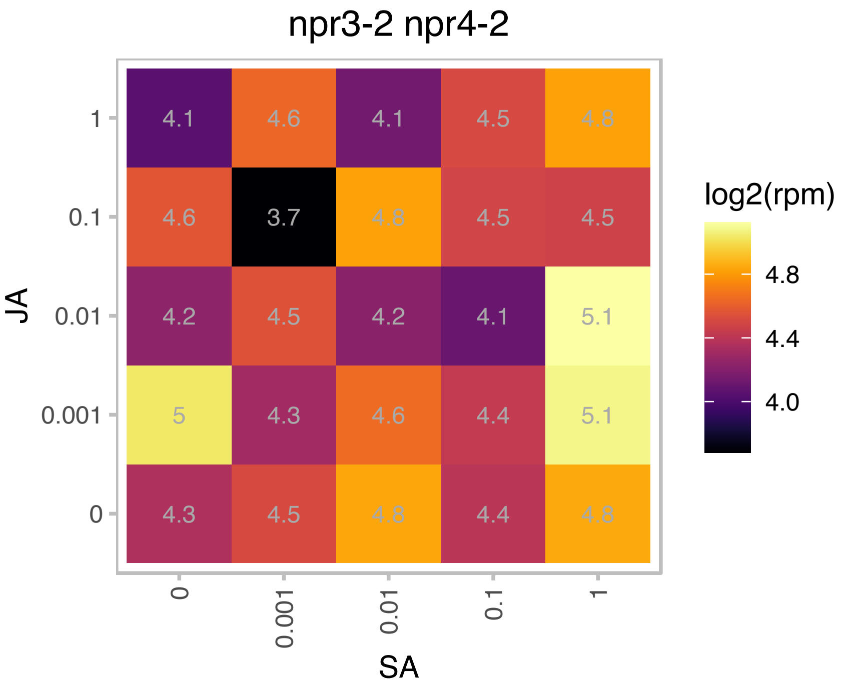


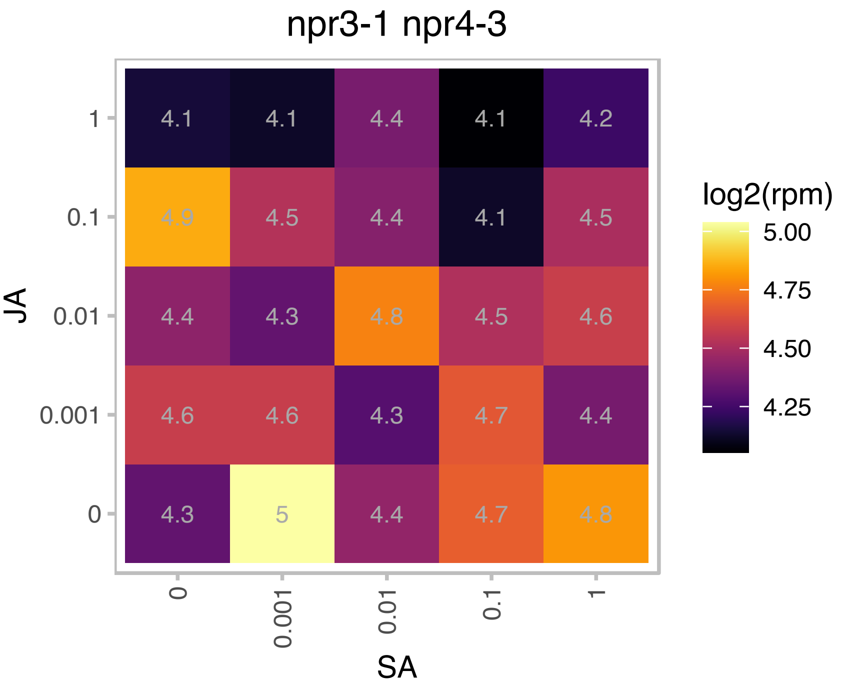

### AT3G16150.1 : ASPGB1

N-terminal nucleophile aminohydrolases (Ntn hydrolases) superfamily protein

Encodes an asparaginase that catalyzes the degradation of L-asparagine to L-aspartic acid and ammonia. The mRNA is cell-to-cell mobile.

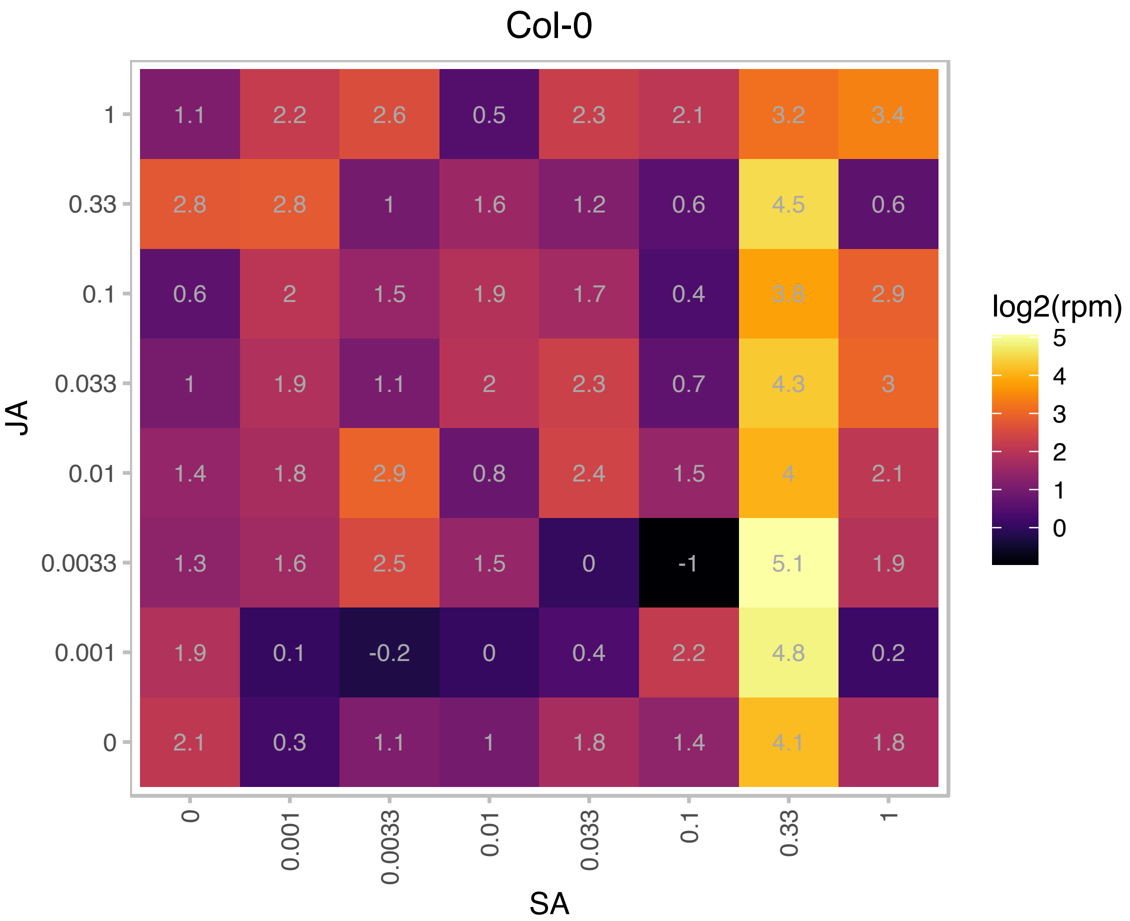


| | |
|---|---|
| Gene | AT3G16150 |
| is.expressed_genes | TRUE |
| is.responsive_genes | TRUE |
| is.apcluster | B-01 |

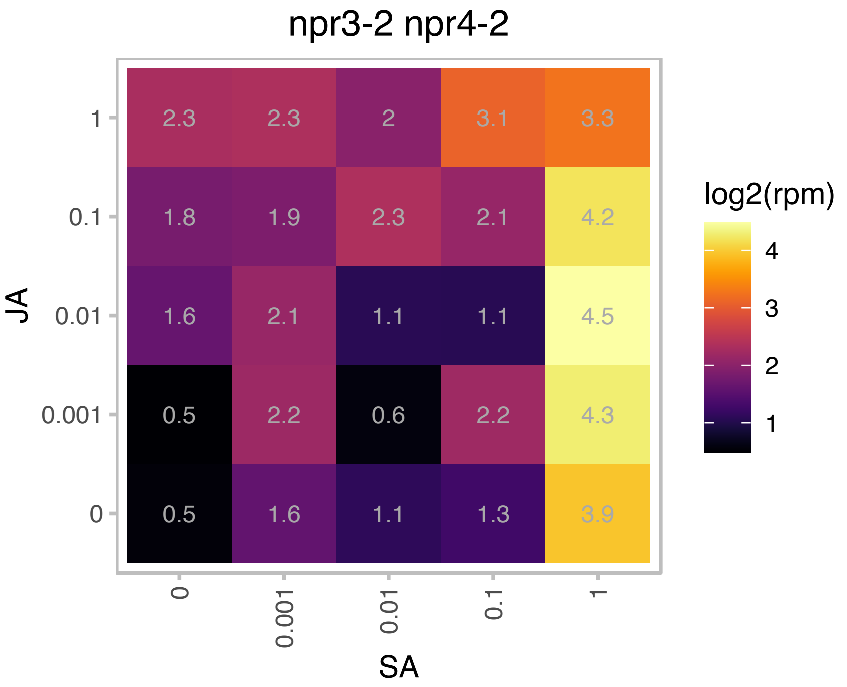


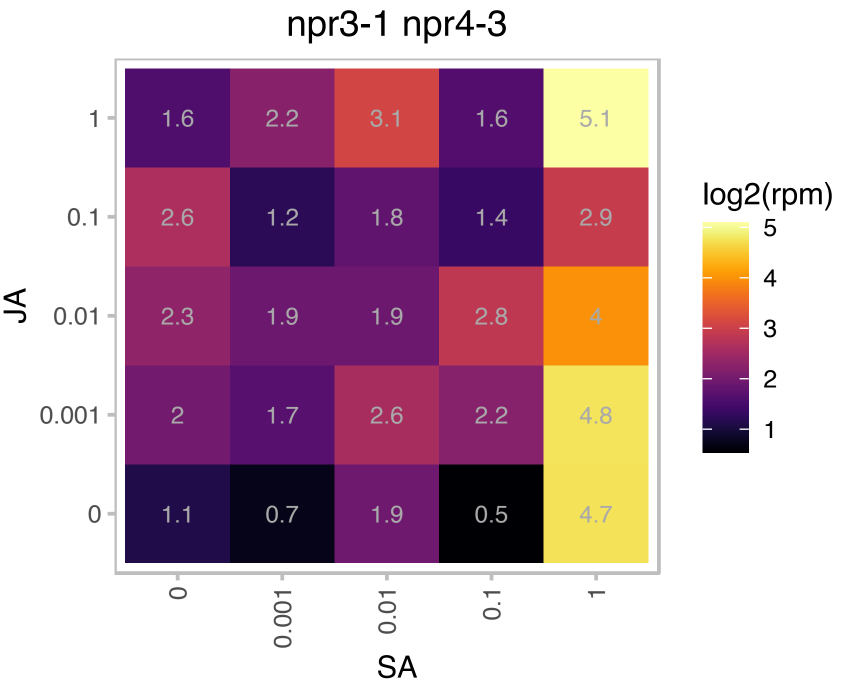

### AT3G18140.1 : LST8-1

Transducin/WD40 repeat-like superfamily protein

Encodes LST8-1, involved in regulating amino acid accumulation and the synthesis of myo-inositol and raffinose during plant adaptation to long days.

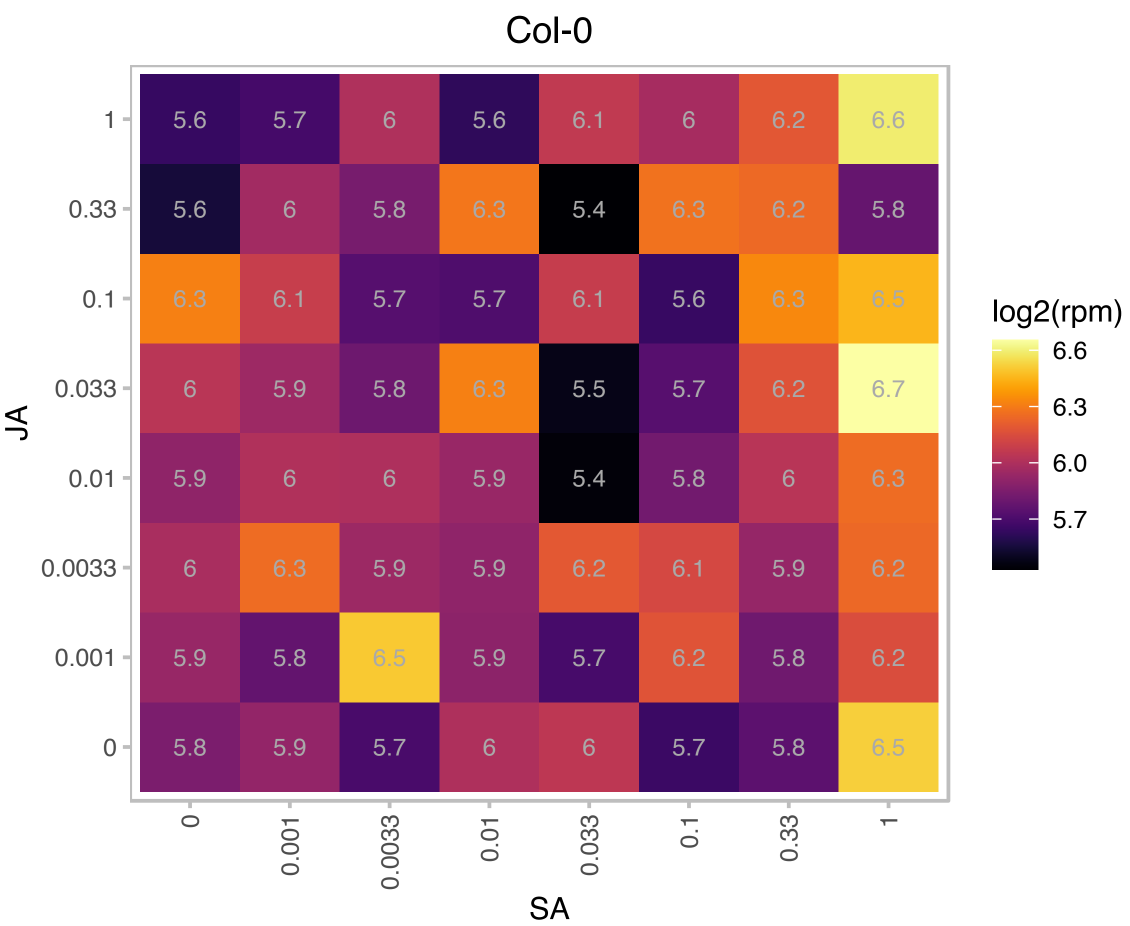


| | |
|---|---|
| Gene | AT3G18140 |
| is.expressed_genes | TRUE |
| is.responsive_genes | TRUE |
| is.apcluster | B-01 |

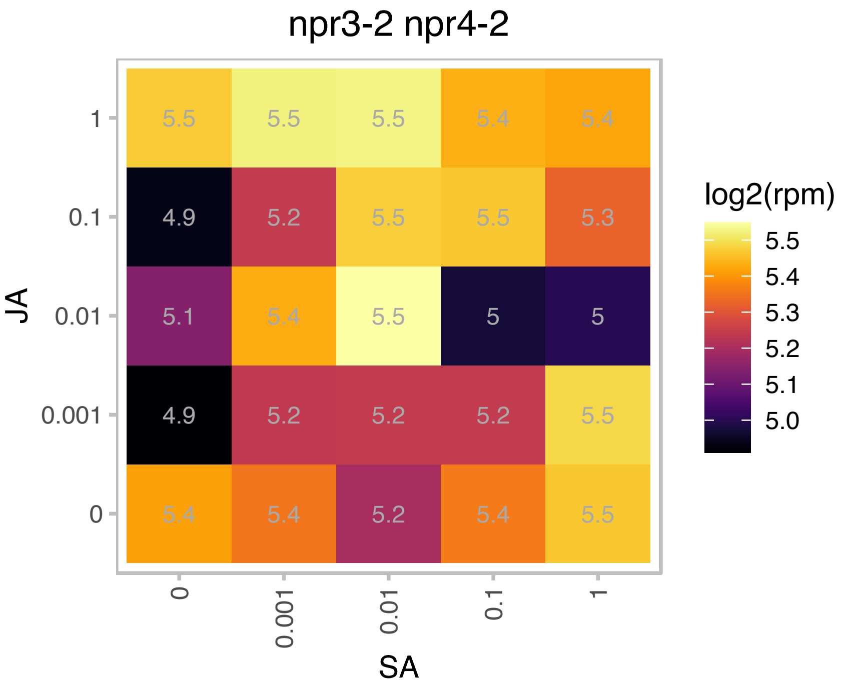


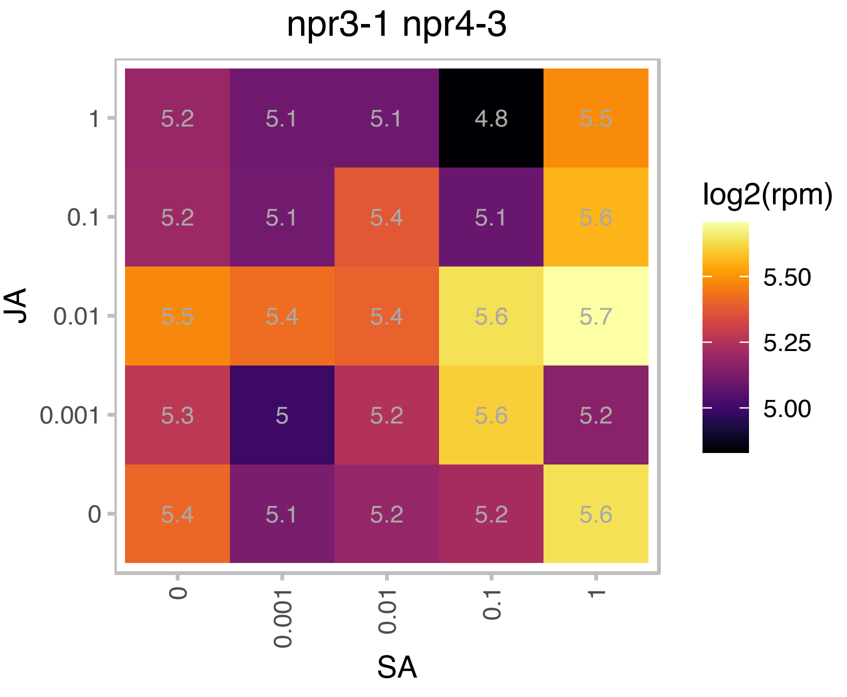

### AT3G46230.1 : ATHSP17.4, HSP17.4

heat shock protein 17.4

member of the class I small heat-shock protein (sHSP) family, which accounts for the majority of sHSPs in maturing seeds

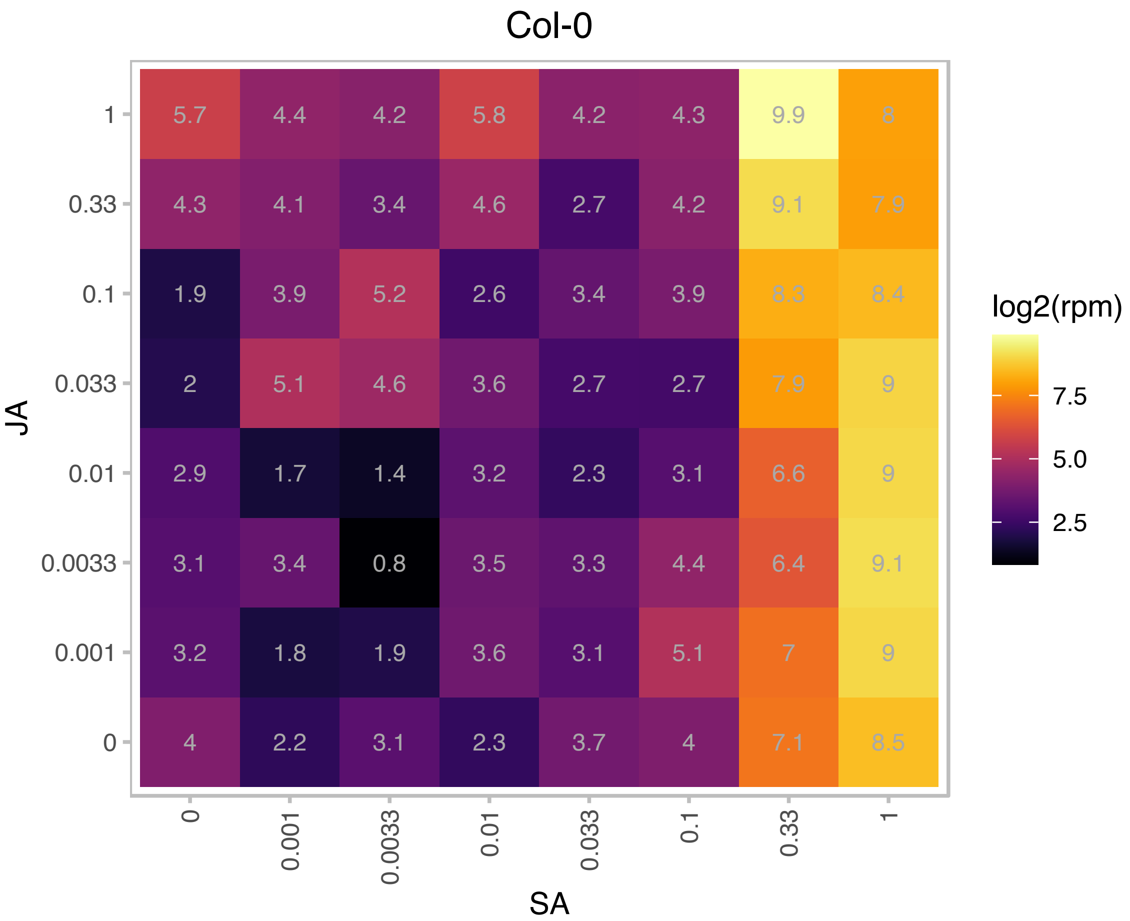


| | |
|---|---|
| Gene | AT3G46230 |
| is.expressed_genes | TRUE |
| is.responsive_genes | TRUE |
| is.apcluster | B-01 |

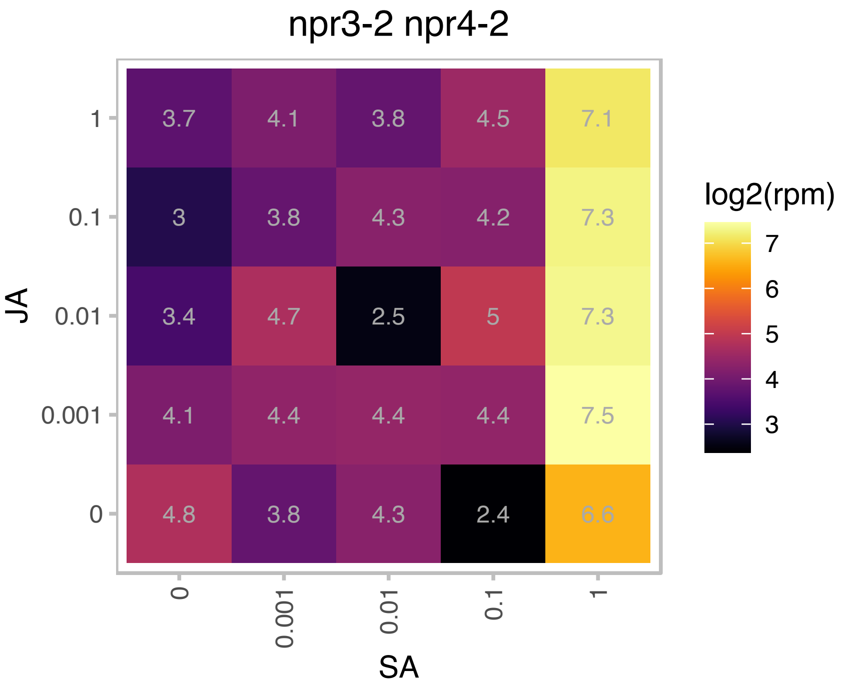


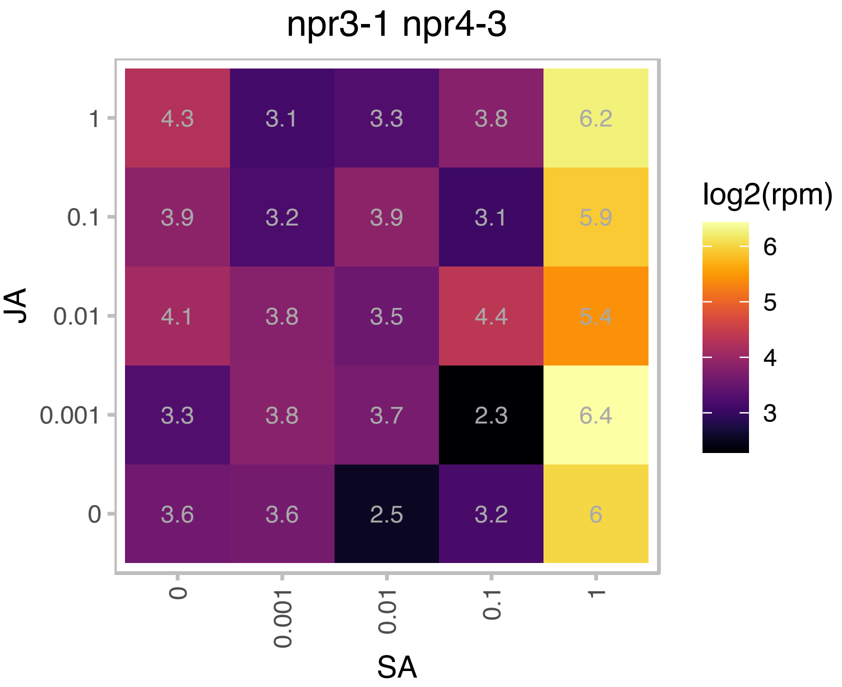

# AT3G48115.1 : NA

other RNA

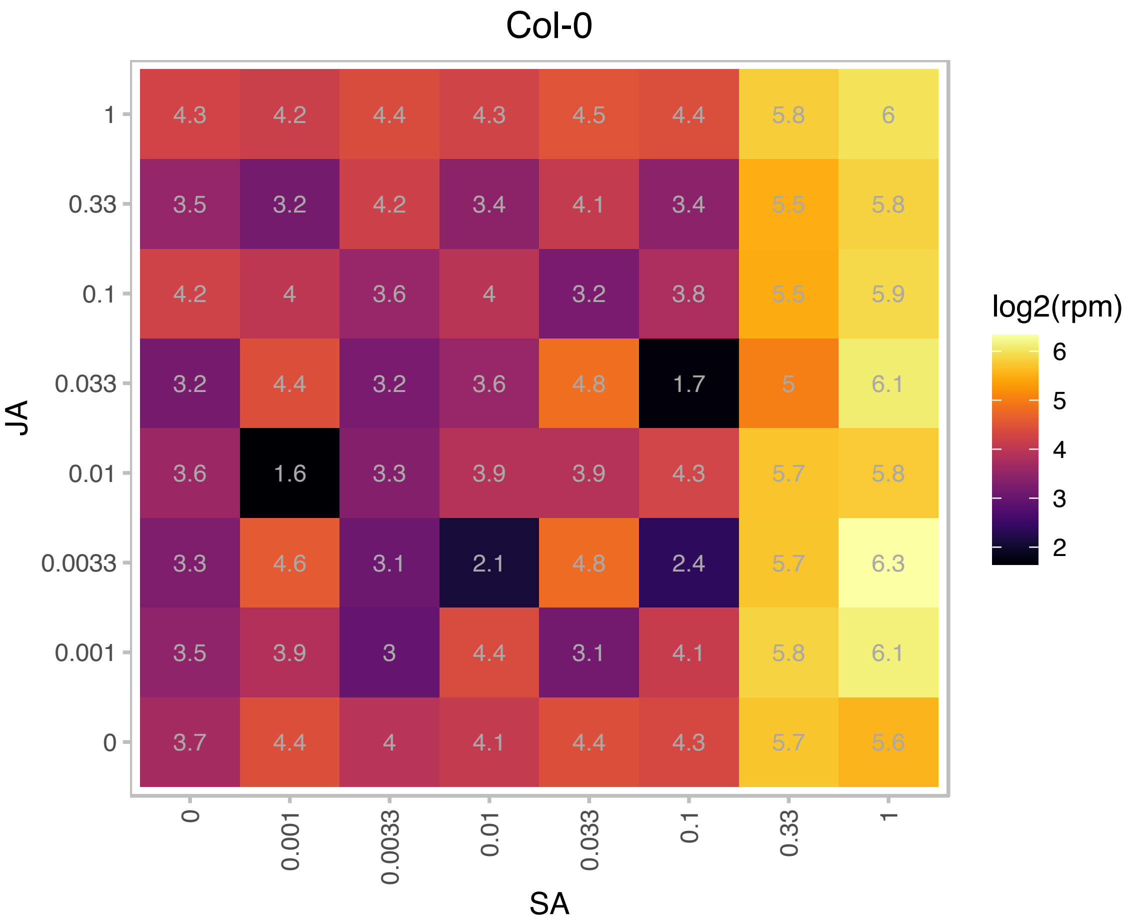


| | |
|---|---|
| Gene | AT3G48115 |
| is.expressed_genes | TRUE |
| is.responsive_genes | TRUE |
| is.apcluster | B-01 |

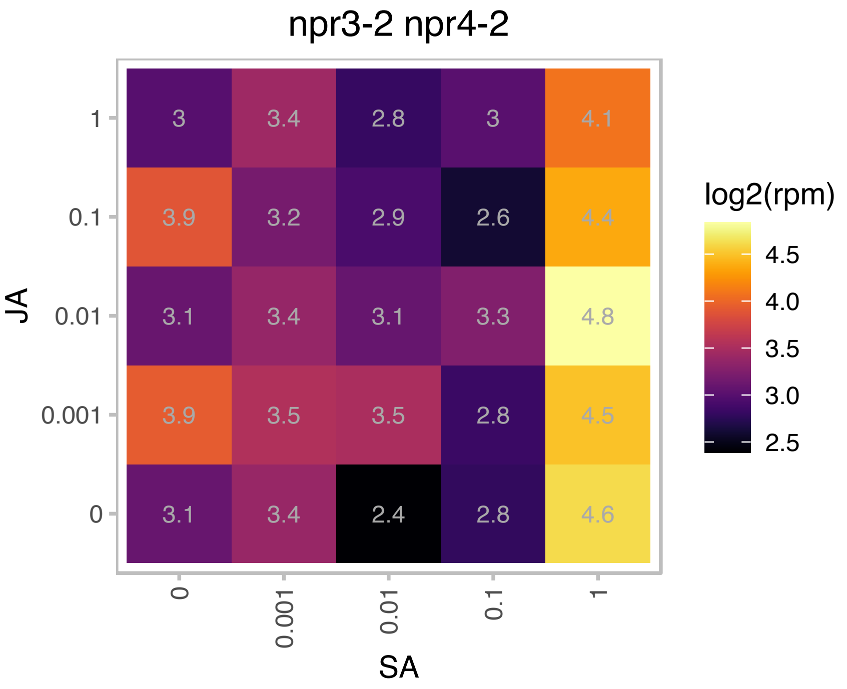


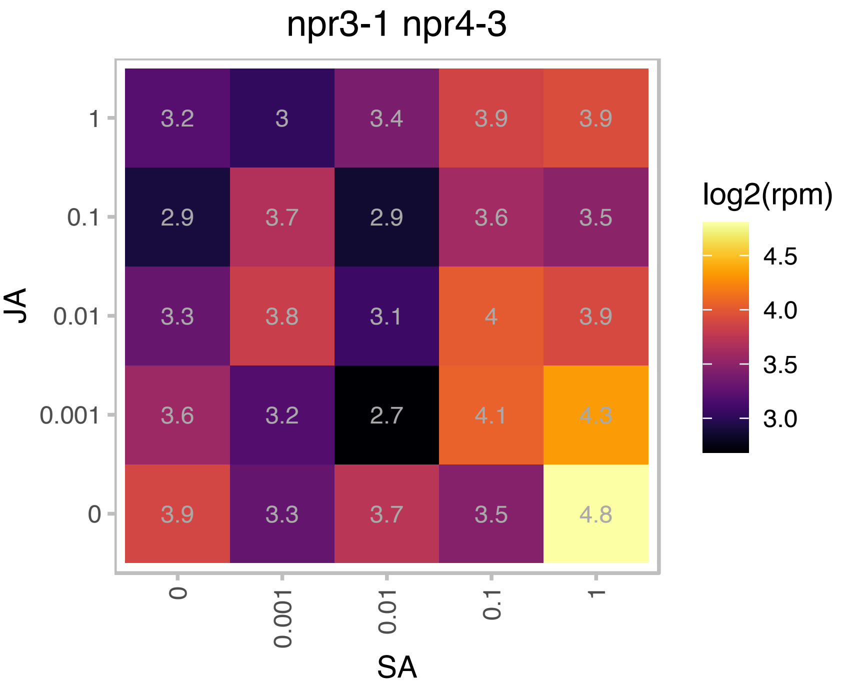

### AT3G62030.1 : CYP20-3, ROC4

rotamase CYP 4
nuclear-encoded chloroplast stromal cyclophilin CYP20-3 (also known as ROC4). Protein is tyrosine-phosphorylated and its phosphorylation state is modulated in response to ABA in Arabidopsis thaliana seeds.

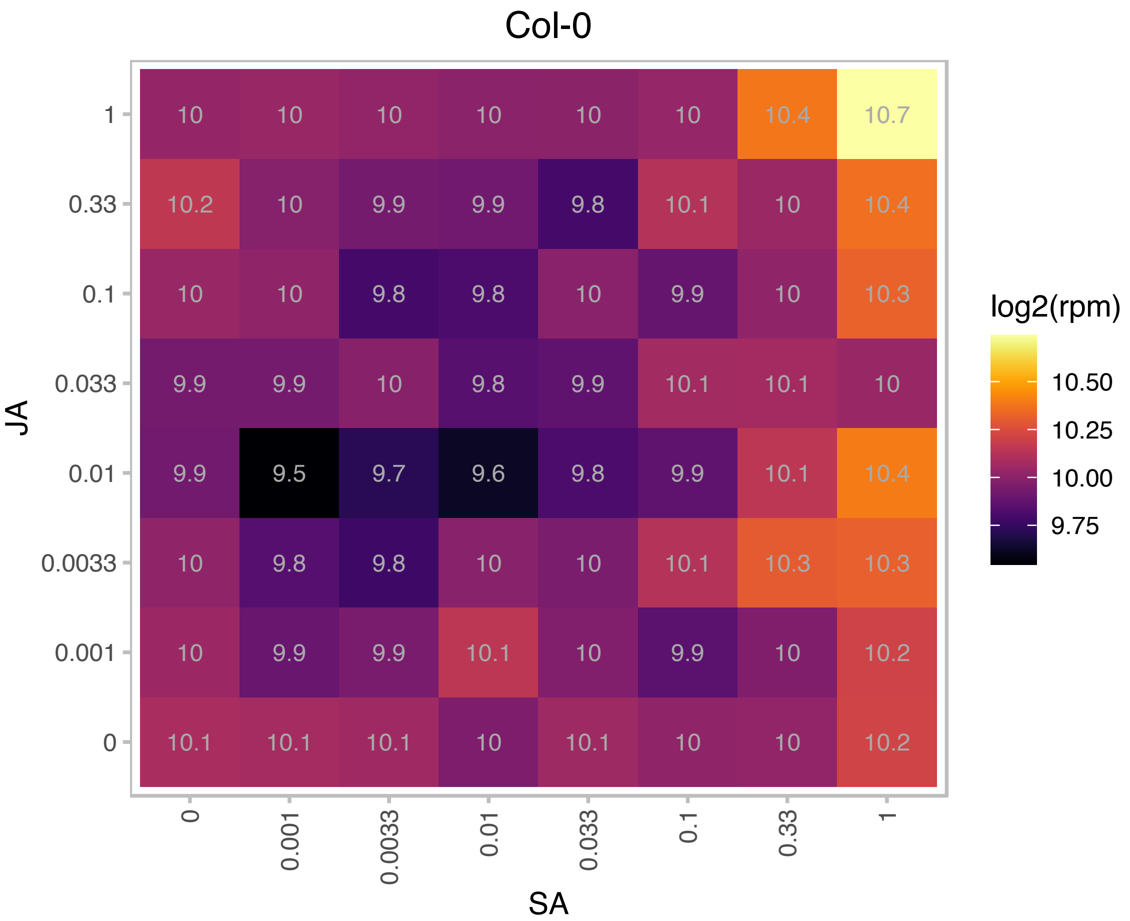


| | |
|---|---|
| Gene | AT3G62030 |
| is.expressed_genes | TRUE |
| is.responsive_genes | TRUE |
| is.apcluster | B-01 |

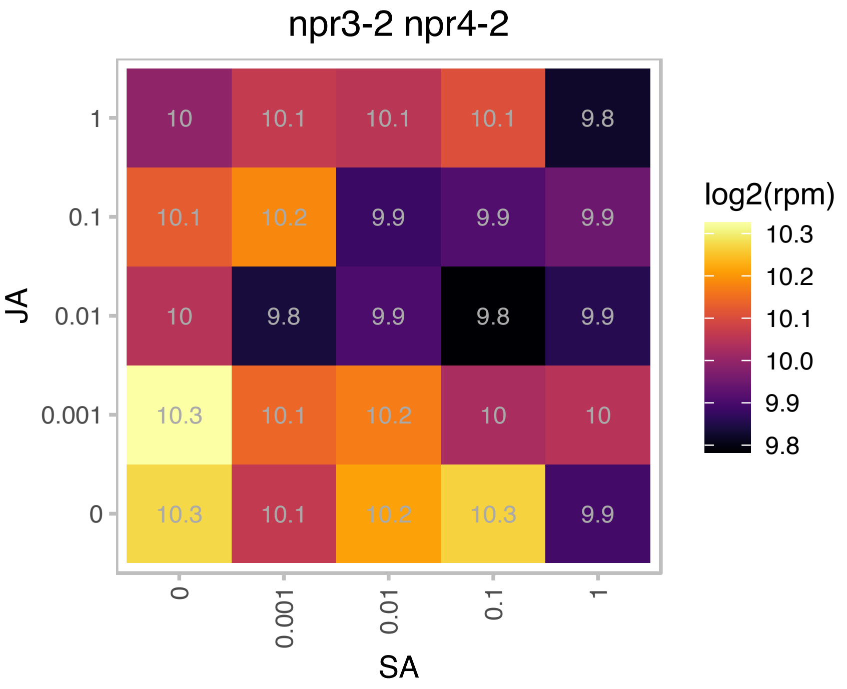


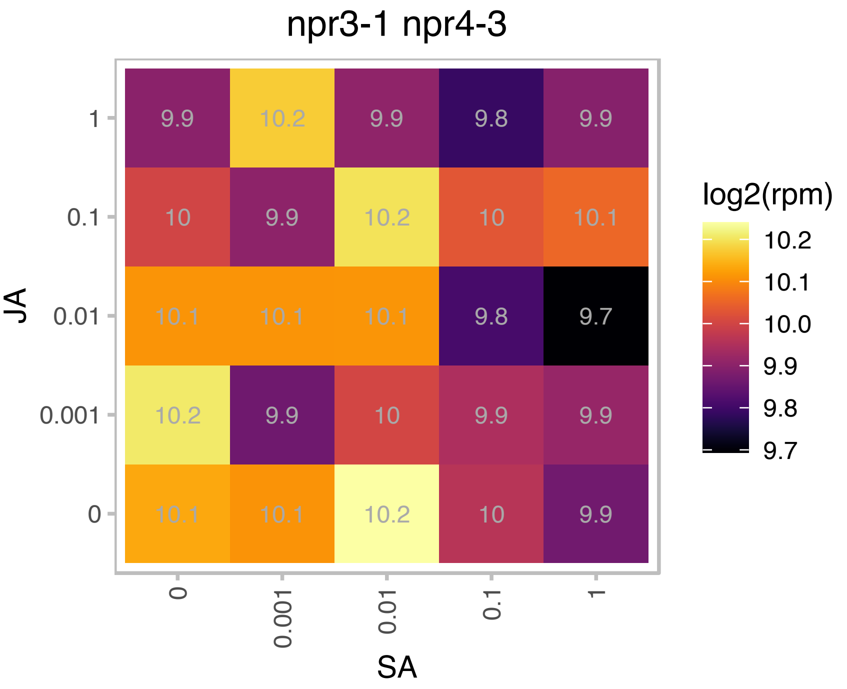

# AT4G08695.1 : NA

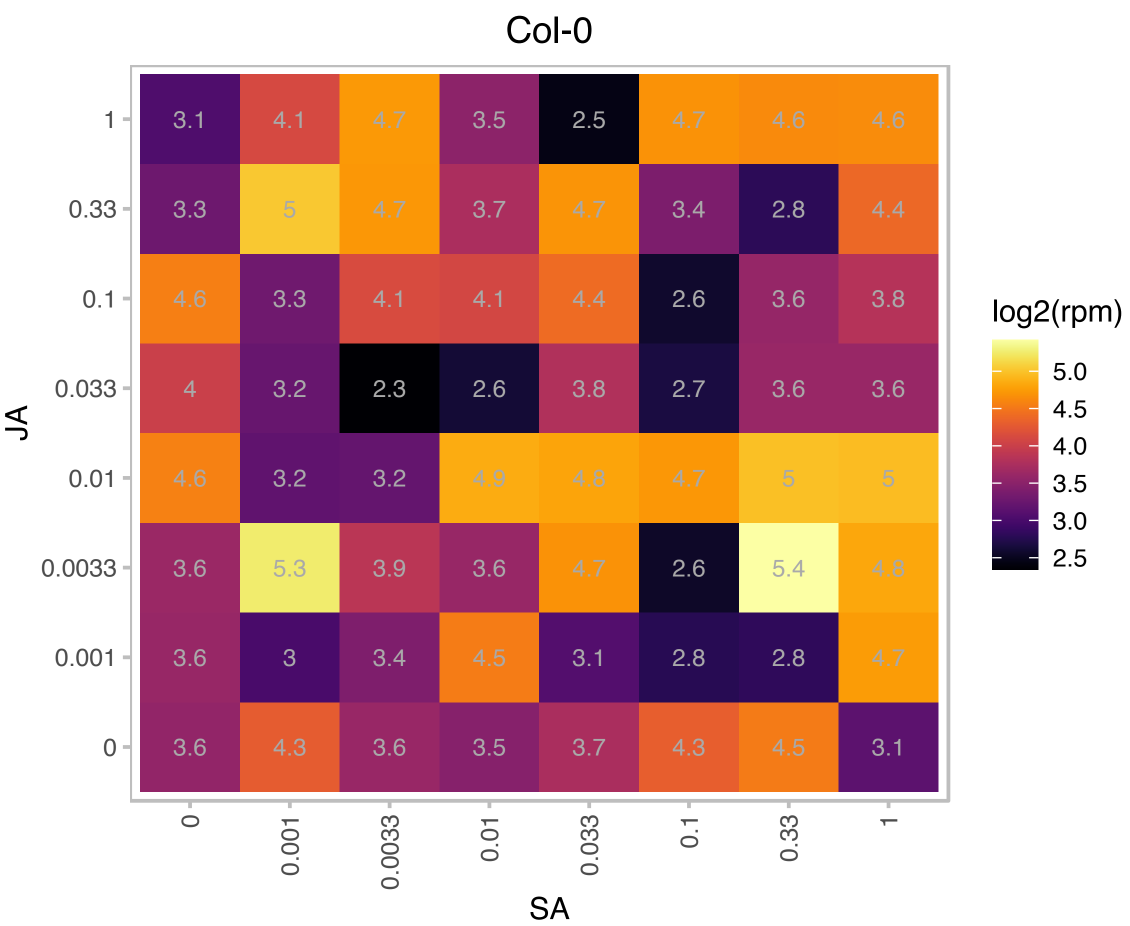


| | |
|---|---|
| Gene | AT4G08695 |
| is.expressed_genes | TRUE |
| is.responsive_genes | TRUE |
| is.apcluster | B-01 |

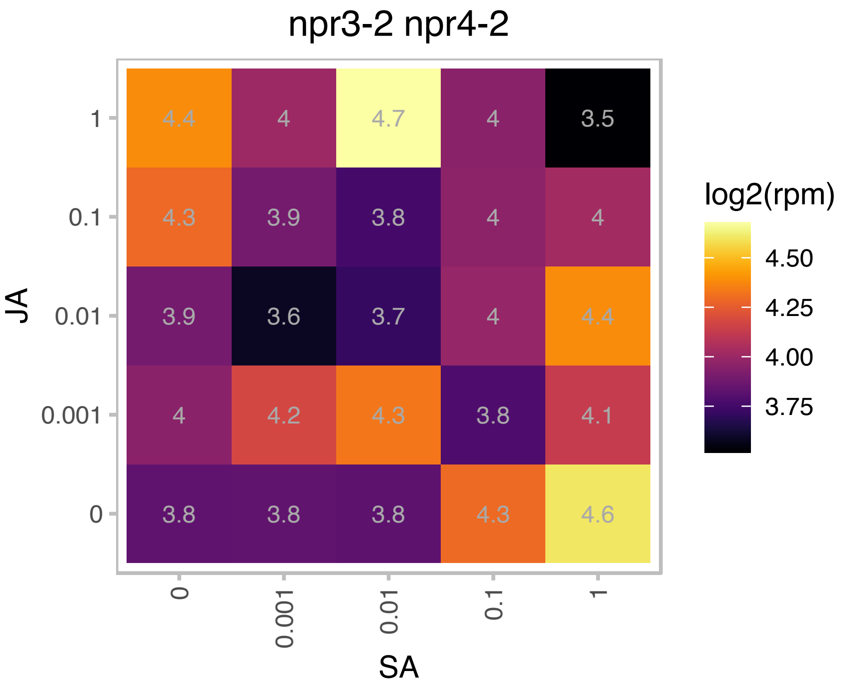


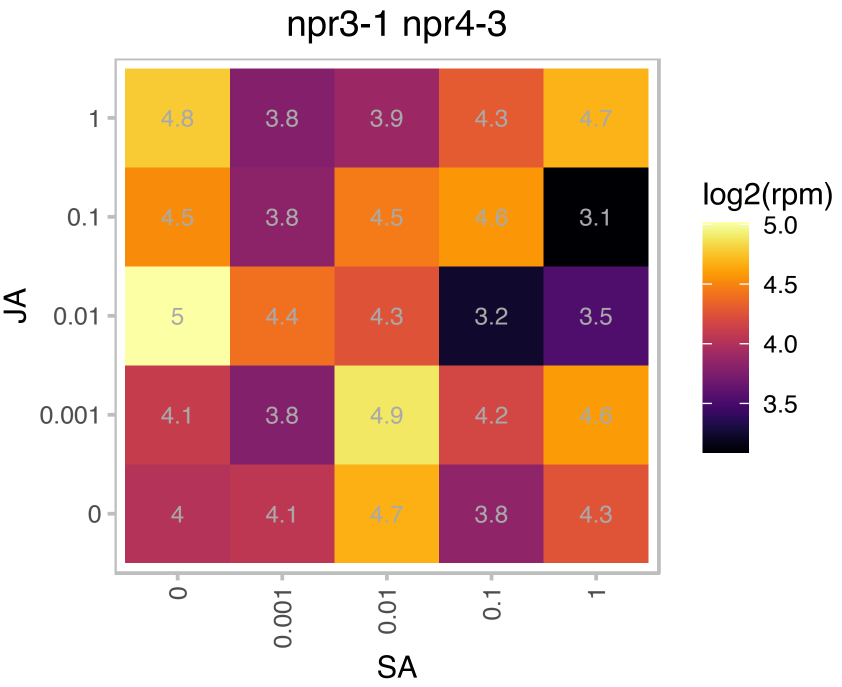

# AT4G26555.1 : NA

FKBP-like peptidyl-prolyl cis-trans isomerase family protein

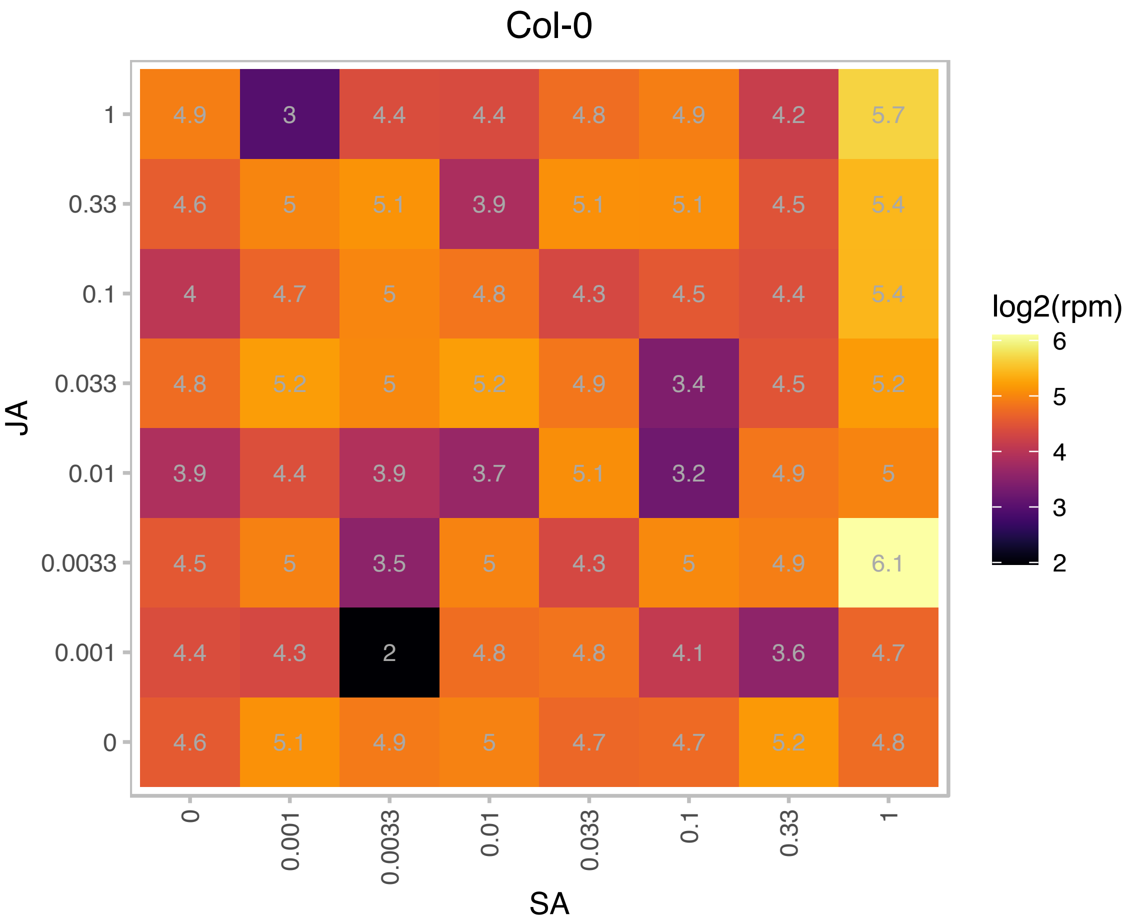


| | |
|---|---|
| Gene | AT4G26555 |
| is.expressed_genes | TRUE |
| is.responsive_genes | TRUE |
| is.apcluster | B-01 |

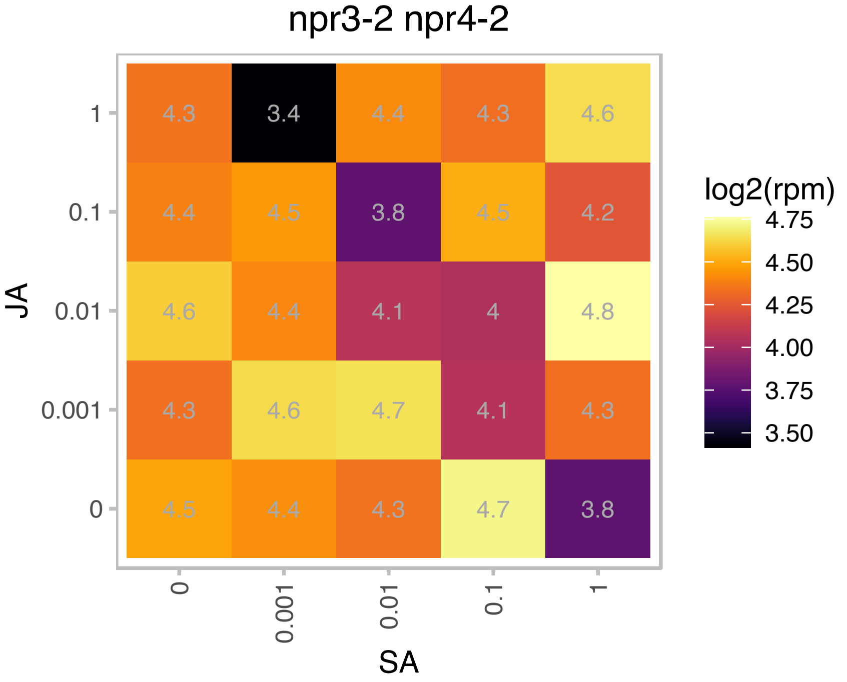


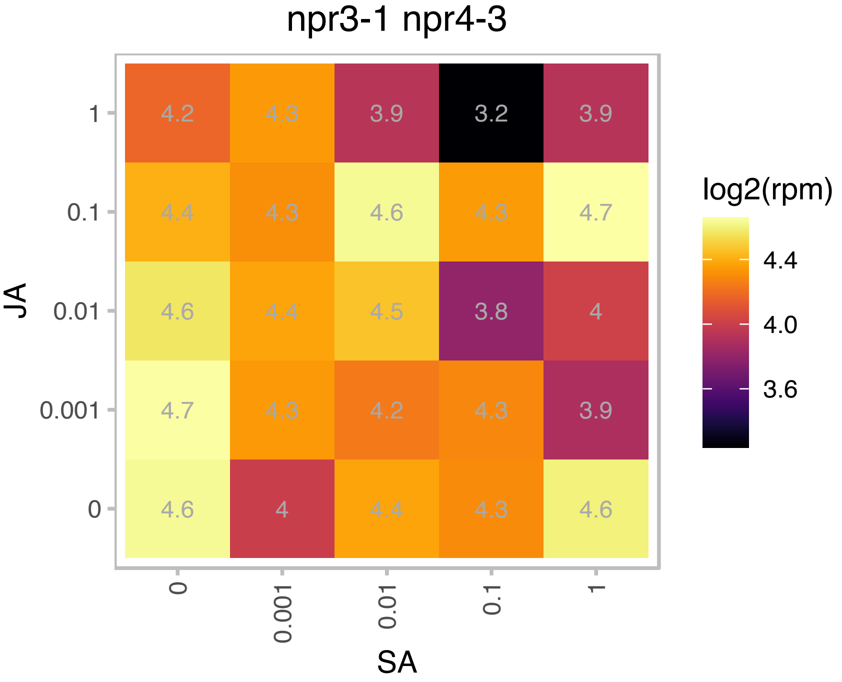

### AT4G34410.1 : ERF109, RRTF1

Encodes a member of the ERF (ethylene response factor) subfamily B-3 of ERF/AP2 transcription factor family. The protein contains one AP2 domain. There are 18 members in this subfamily including ATERF-1, ATERF-2, AND ATERF-5. Regulates programmed cell death (PCD) inhibitor genes. Involved in retarding programmed cell death under salt stress due to the regulation of processes participating in ROS inhibition. ERF-regulated transcripts belong to the tryptophan biosynthesis, tryptophan metabolism, and downstream plant hormone signal transduction pathways, where ERF109 potentially acts as a ?master switch? mediator of a cascade of consecutive events across the three pathways, promoting plant growth and re-adjustment to homeostasis due the

redox responsive transcription factor 1

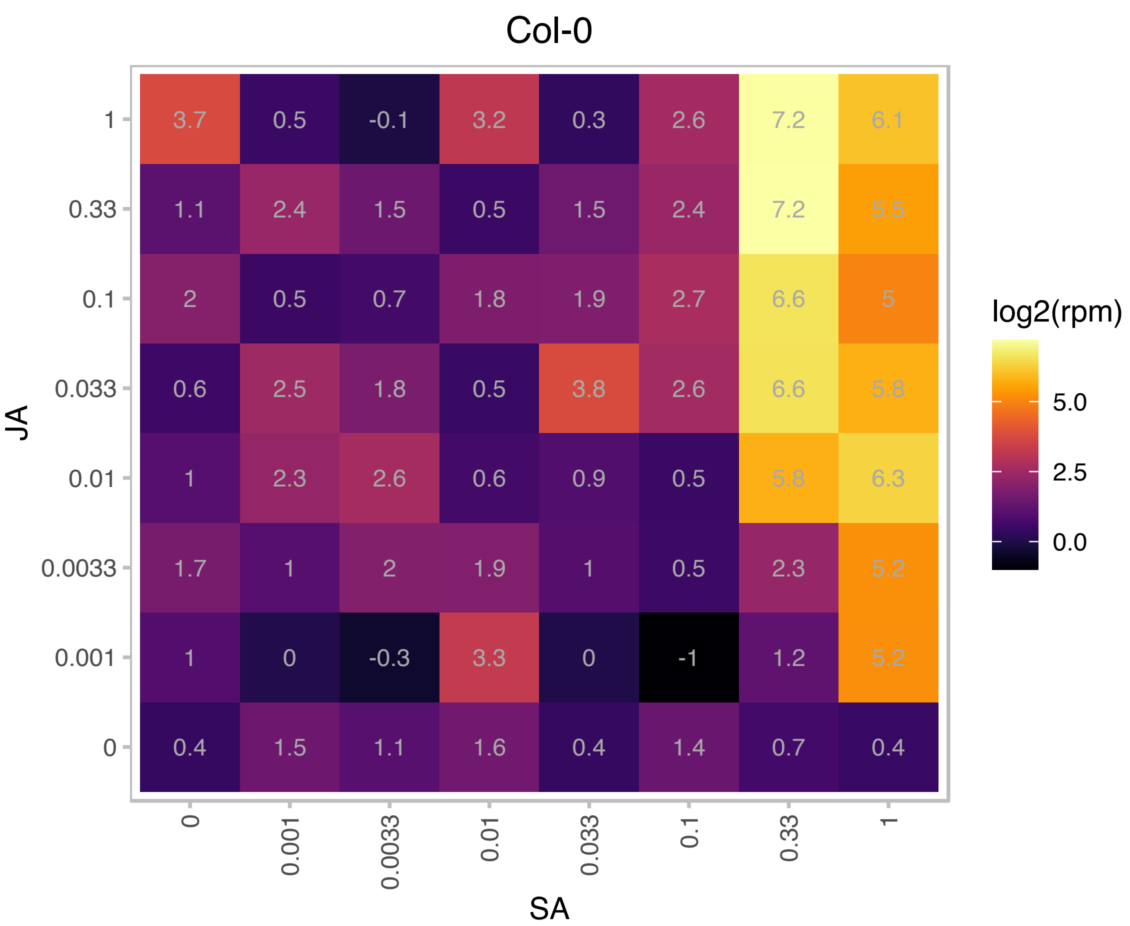


| | |
|---|---|
| Gene | AT4G34410 |
| is.expressed_genes | TRUE |
| is.responsive_genes | TRUE |
| is.apcluster | B-01 |

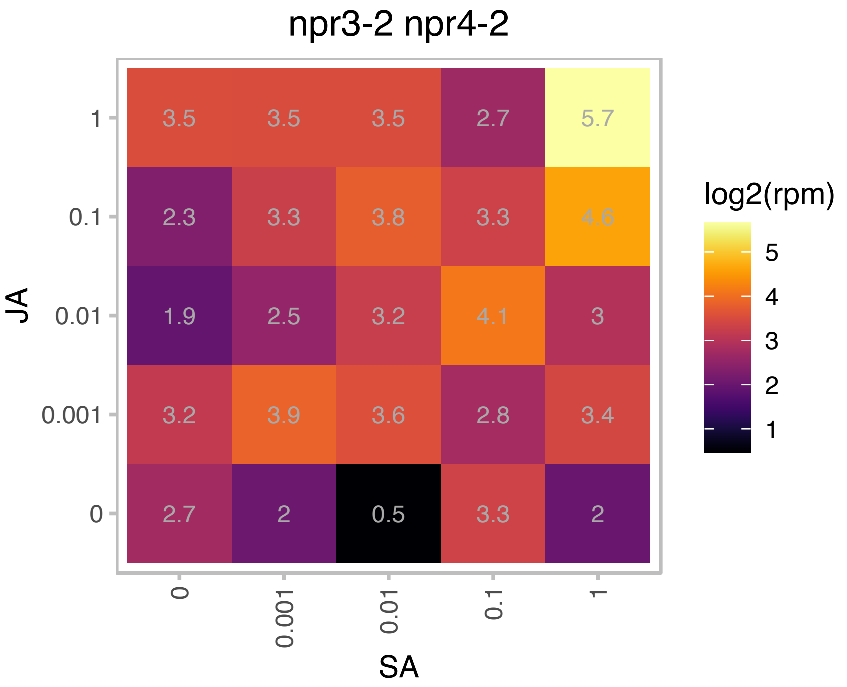


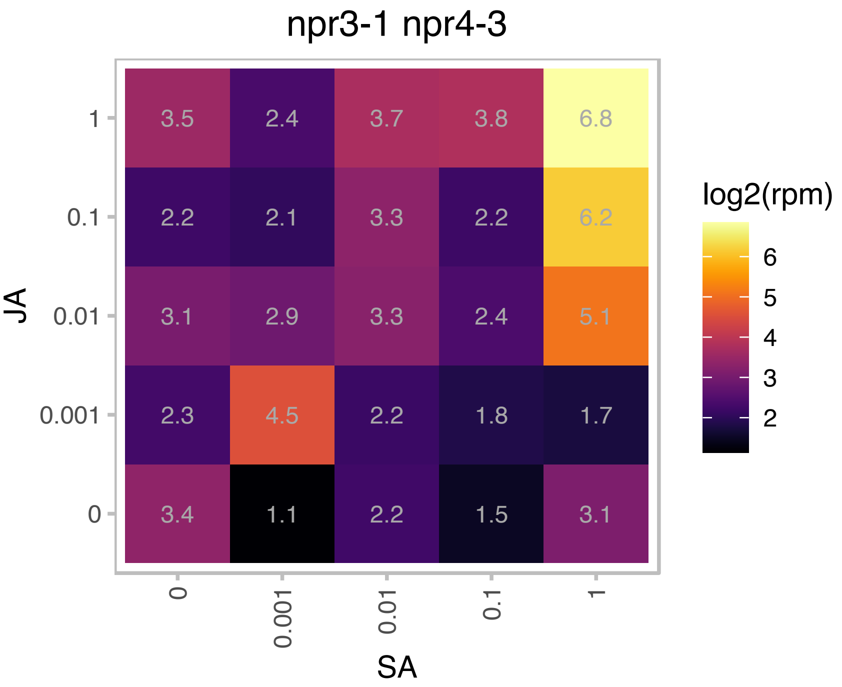

### AT5G12020.1 : HSP17.6II

17.6 kDa class II heat shock protein

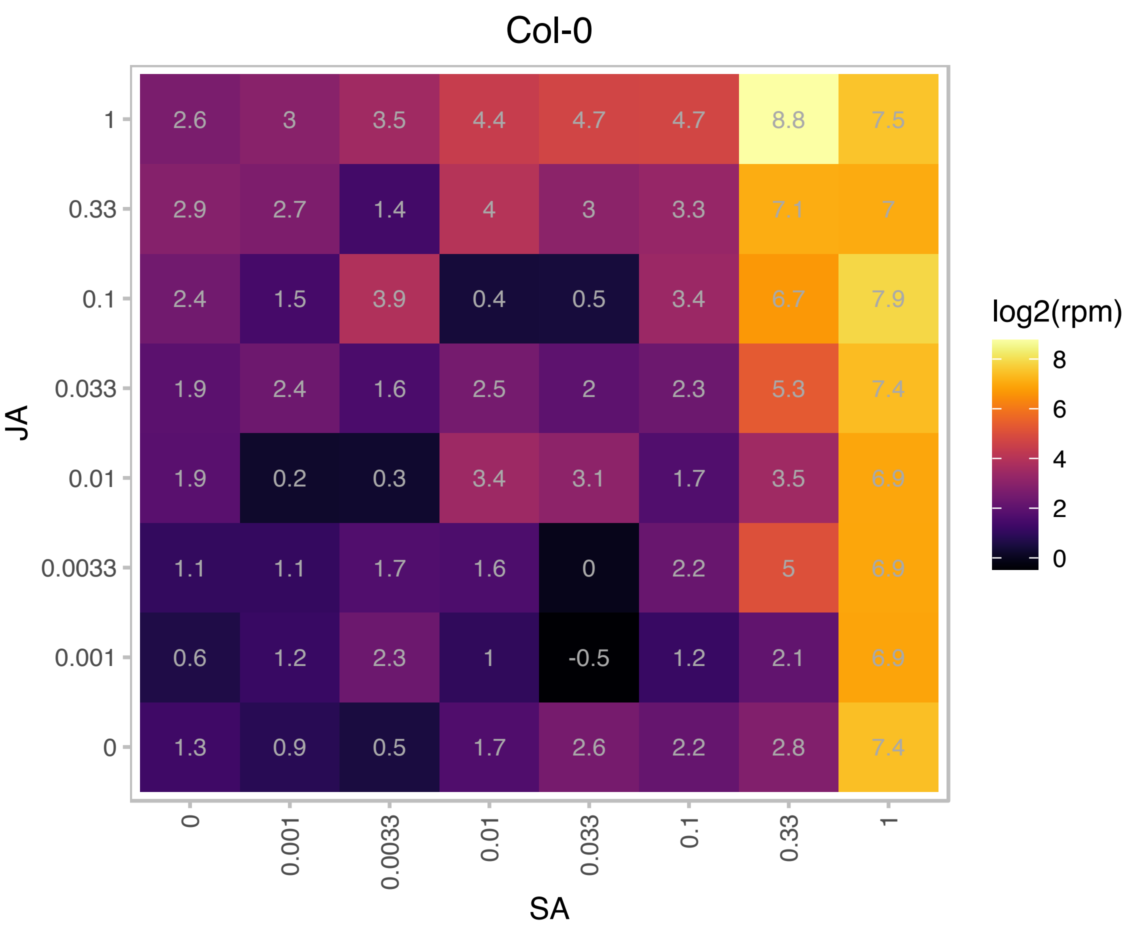


| | |
|---|---|
| Gene | AT5G12020 |
| is.expressed_genes | TRUE |
| is.responsive_genes | TRUE |
| is.apcluster | B-01 |

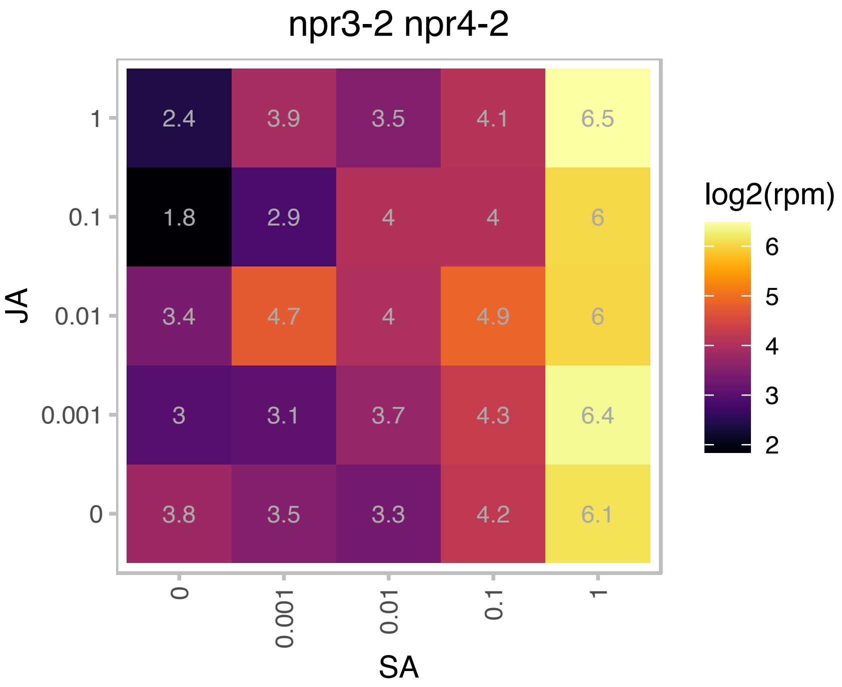


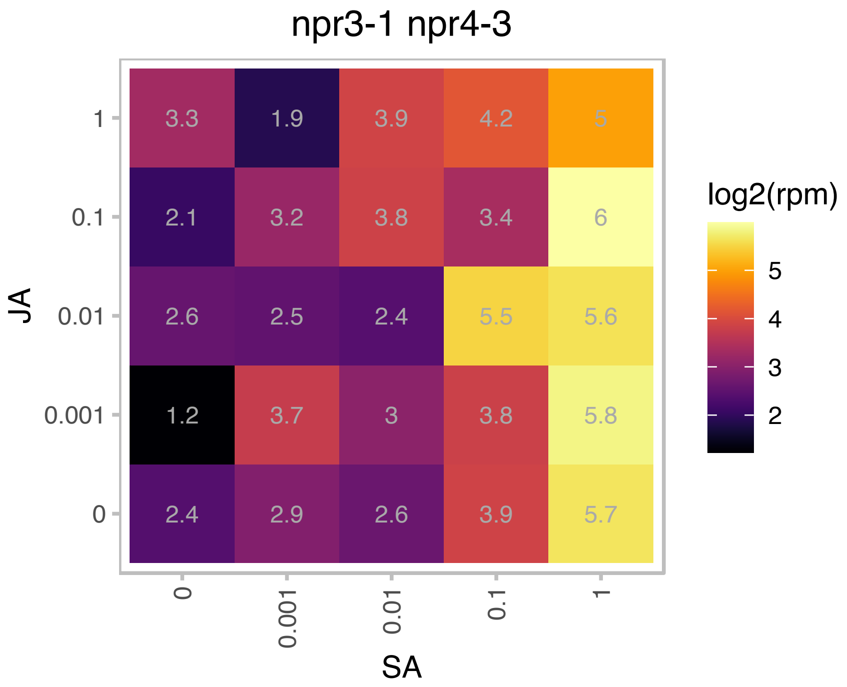

### AT5G12030.1 : AT-HSP17.6A, HSP17.6, HSP17.6A

heat shock protein 17.6A

Encodes a cytosolic small heat shock protein with chaperone activity that is induced by heat and osmotic stress and is also expressed late in seed development.

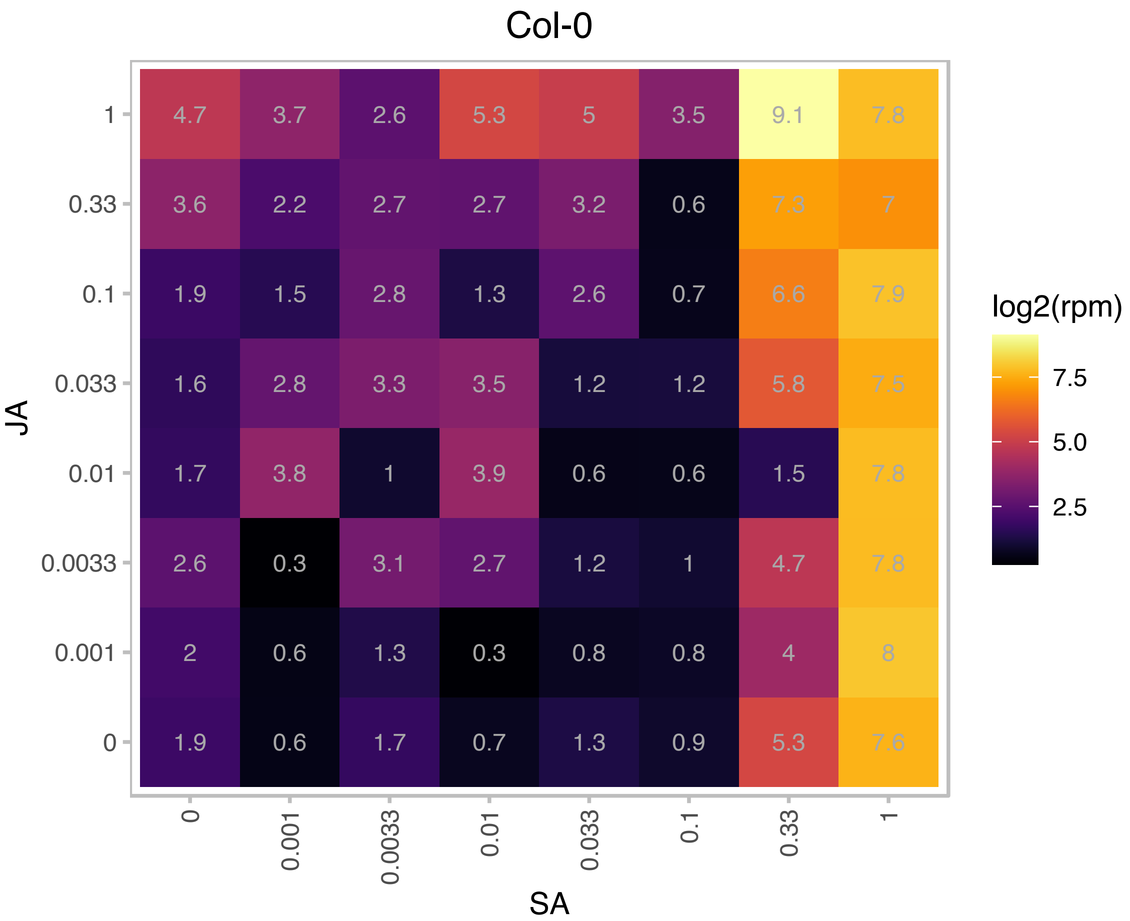


| | |
|---|---|
| Gene | AT5G12030 |
| is.expressed_genes | TRUE |
| is.responsive_genes | TRUE |
| is.apcluster | B-01 |

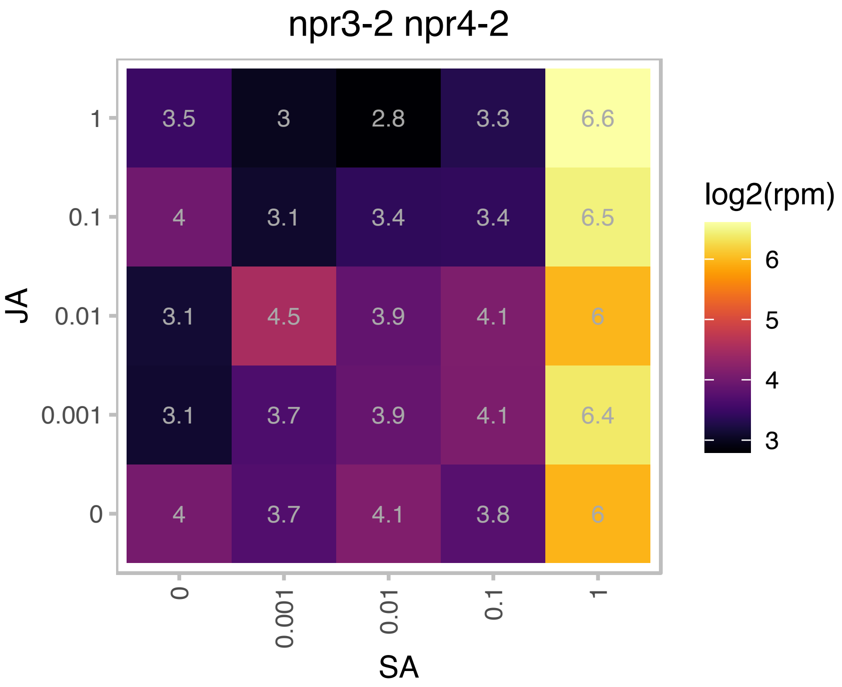


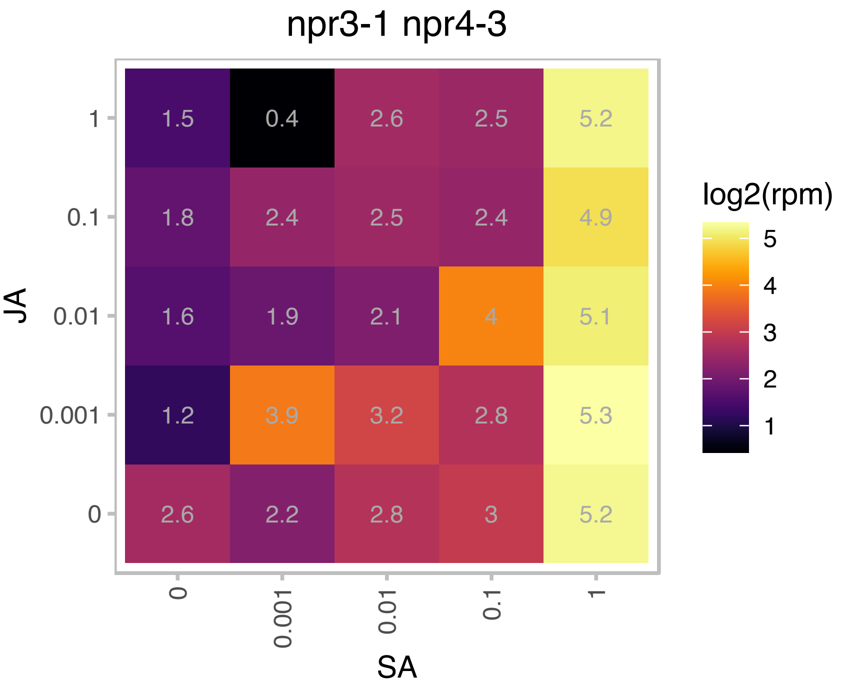

# AT5G24640.1 : NA

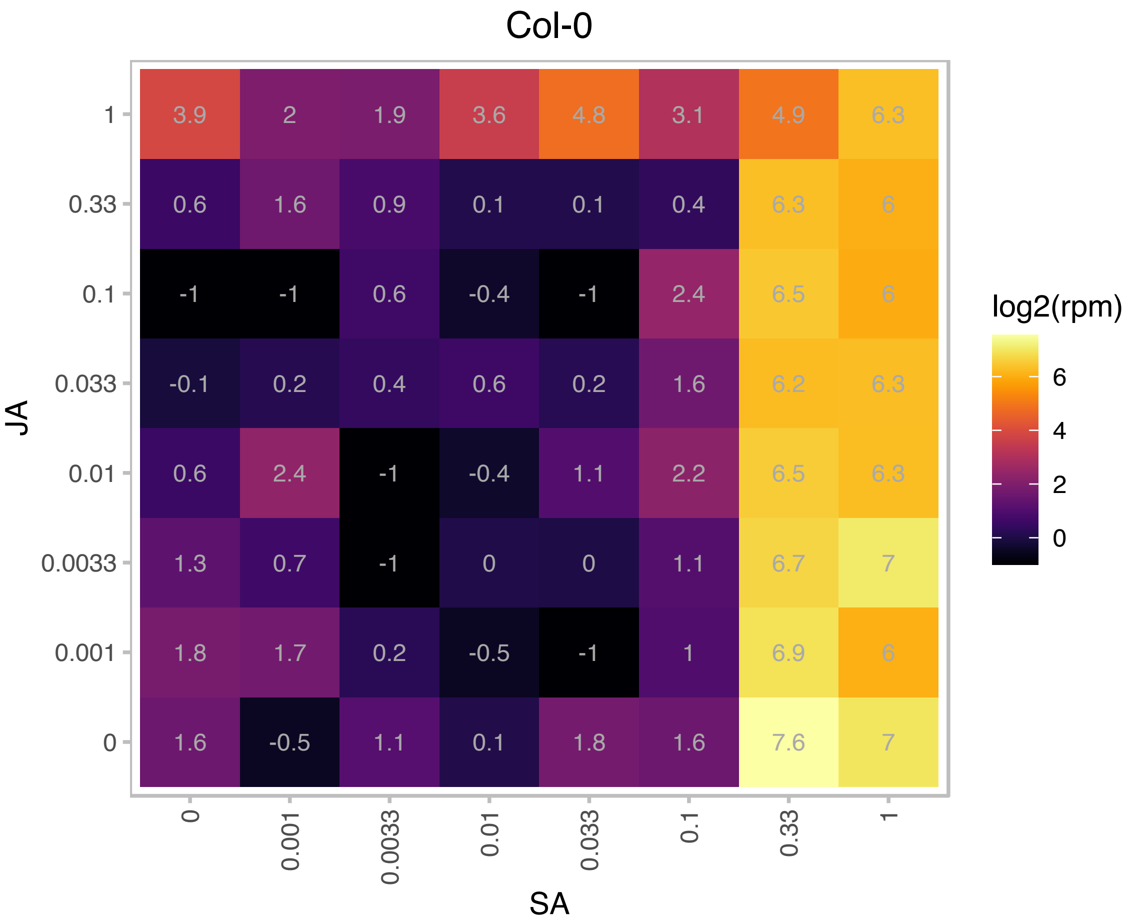


| | |
|---|---|
| Gene | AT5G24640 |
| is.expressed_genes | TRUE |
| is.responsive_genes | TRUE |
| is.apcluster | B-01 |

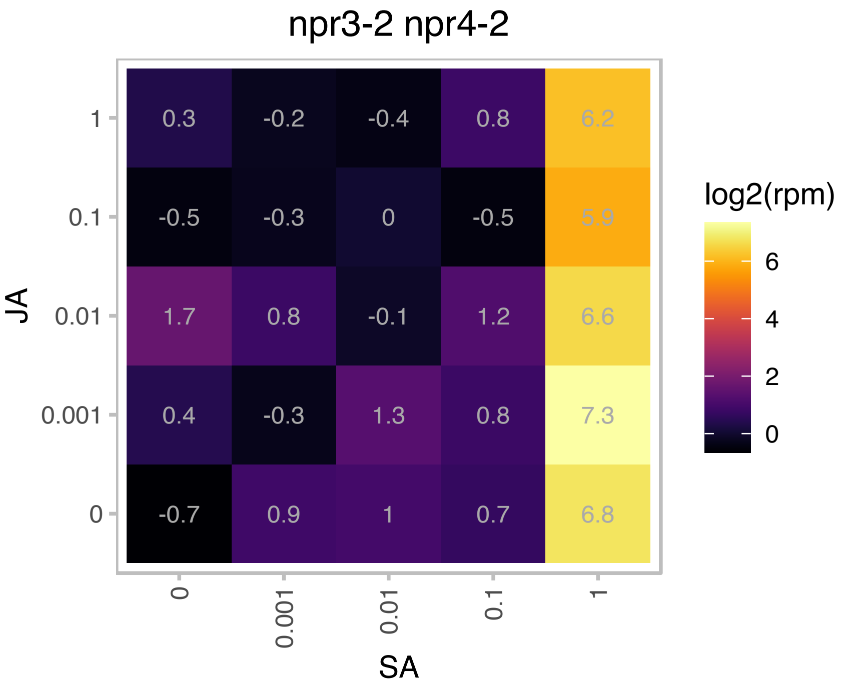


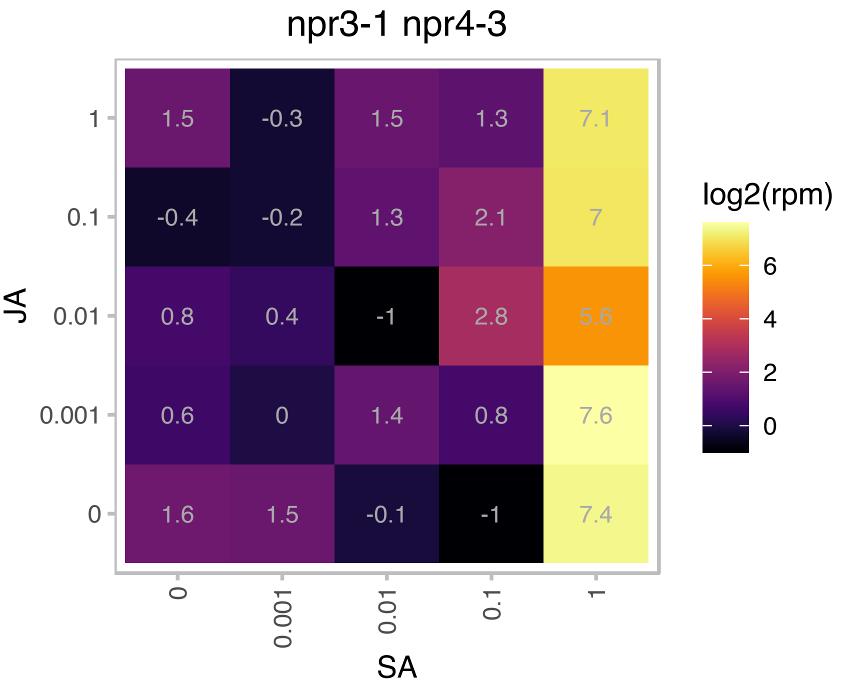

# AT5G52020.1 : NA

Integrase-type DNA-binding superfamily protein

encodes a member of the DREB subfamily A-4 of ERF/AP2 transcription factor family. The protein contains one AP2 domain. There are 17 members in this subfamily including TINY.

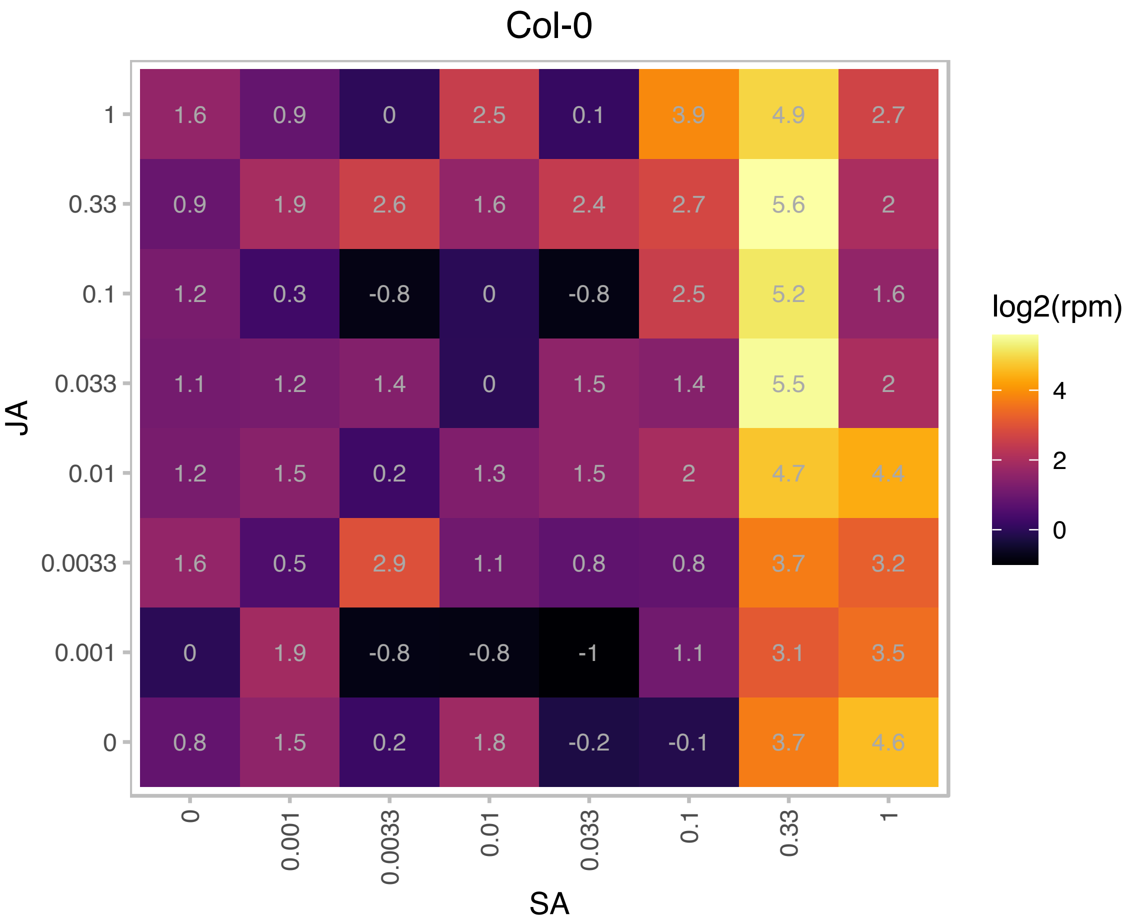


| | |
|---|---|
| Gene | AT5G52020 |
| is.expressed_genes | TRUE |
| is.responsive_genes | TRUE |
| is.apcluster | B-01 |

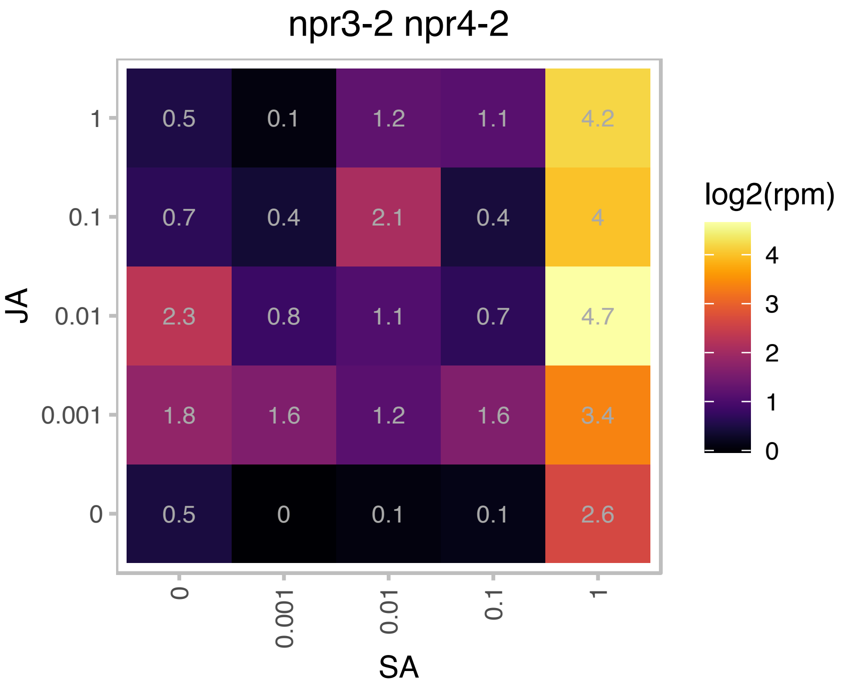


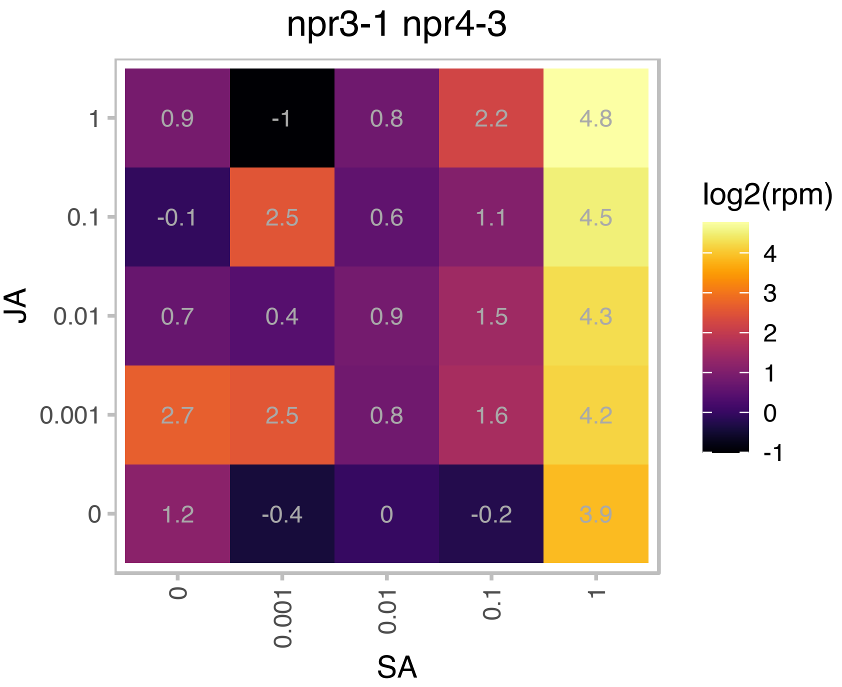

### AT5G53360.1 : SINAT5

TRAF-like superfamily protein

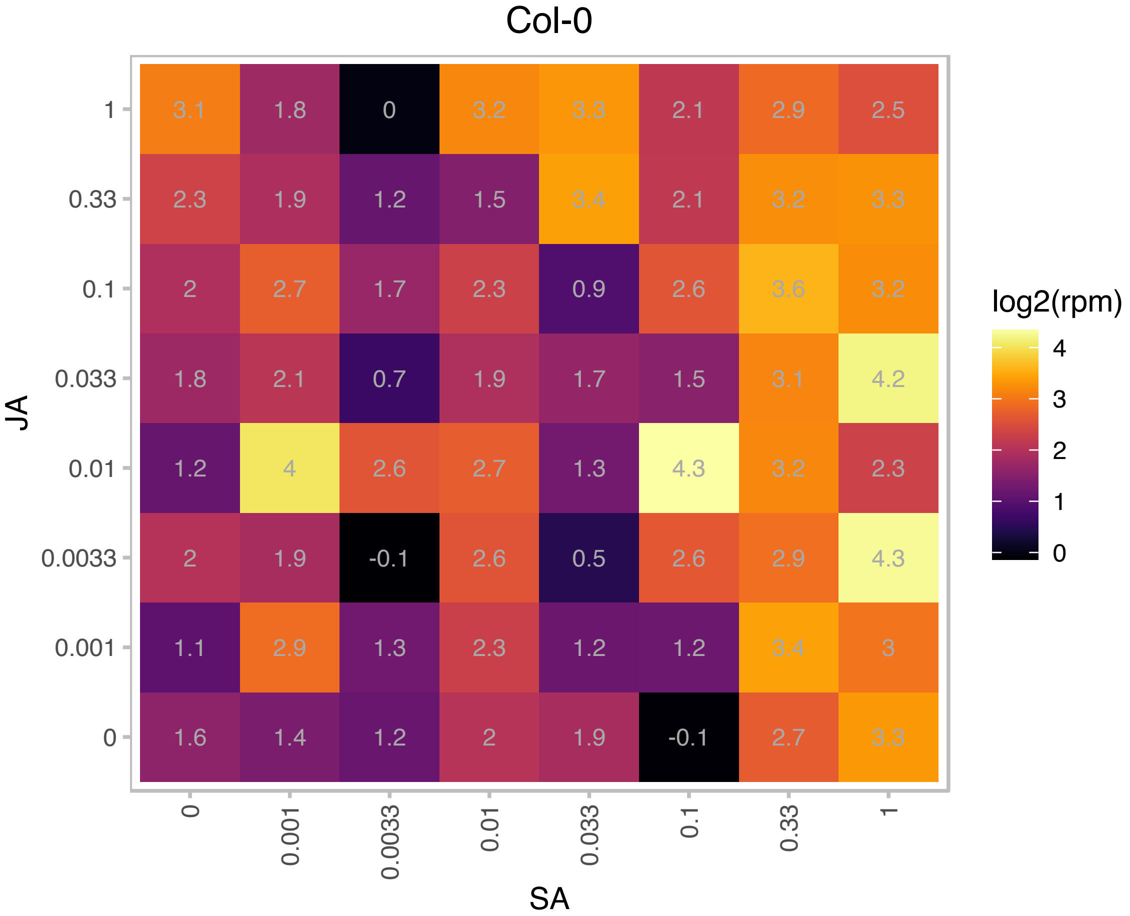


| | |
|---|---|
| Gene | AT5G53360 |
| is.expressed_genes | TRUE |
| is.responsive_genes | TRUE |
| is.apcluster | B-01 |

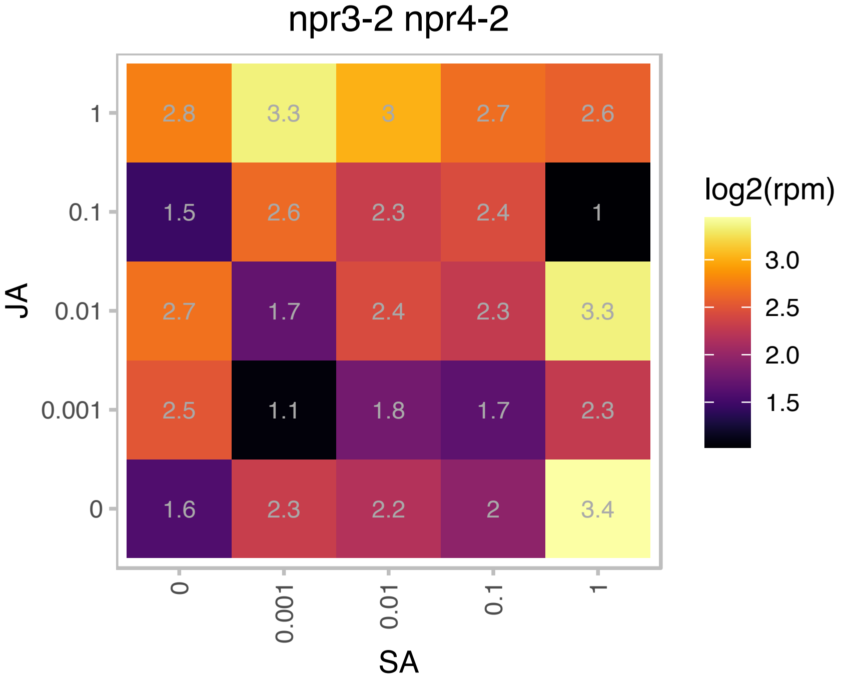


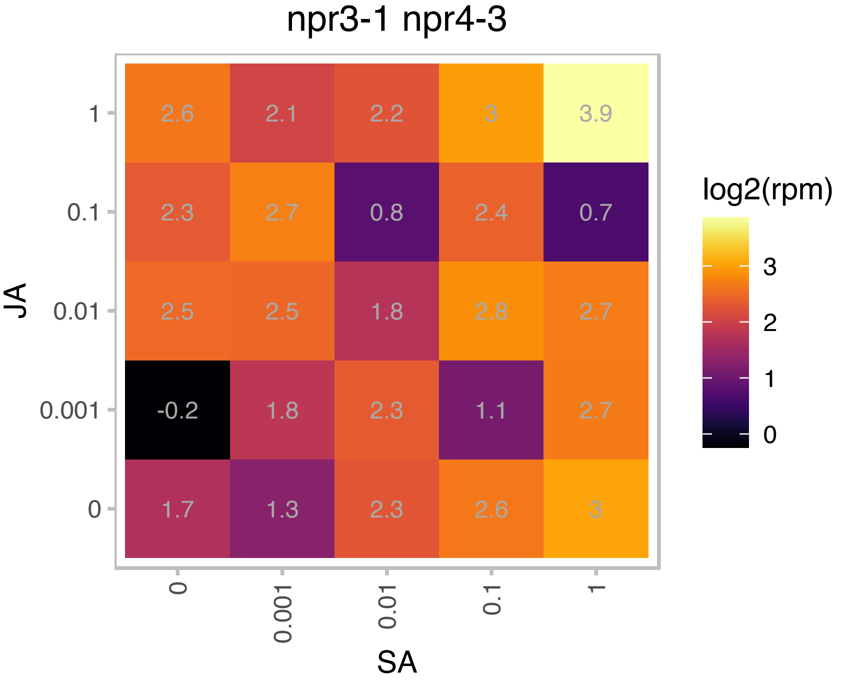

### AT5G54490.1 : PBP1

pinoid-binding protein 1

Encodes a PINOID (PID)-binding protein containing putative EF-hand calcium-binding motifs. The interaction is dependent on the presence of calcium. mRNA expression is up-regulated by auxin. Not a phosphorylation target of PID, likely acts upstream of PID to regulate the activity of this protein in response to changes in calcium levels.

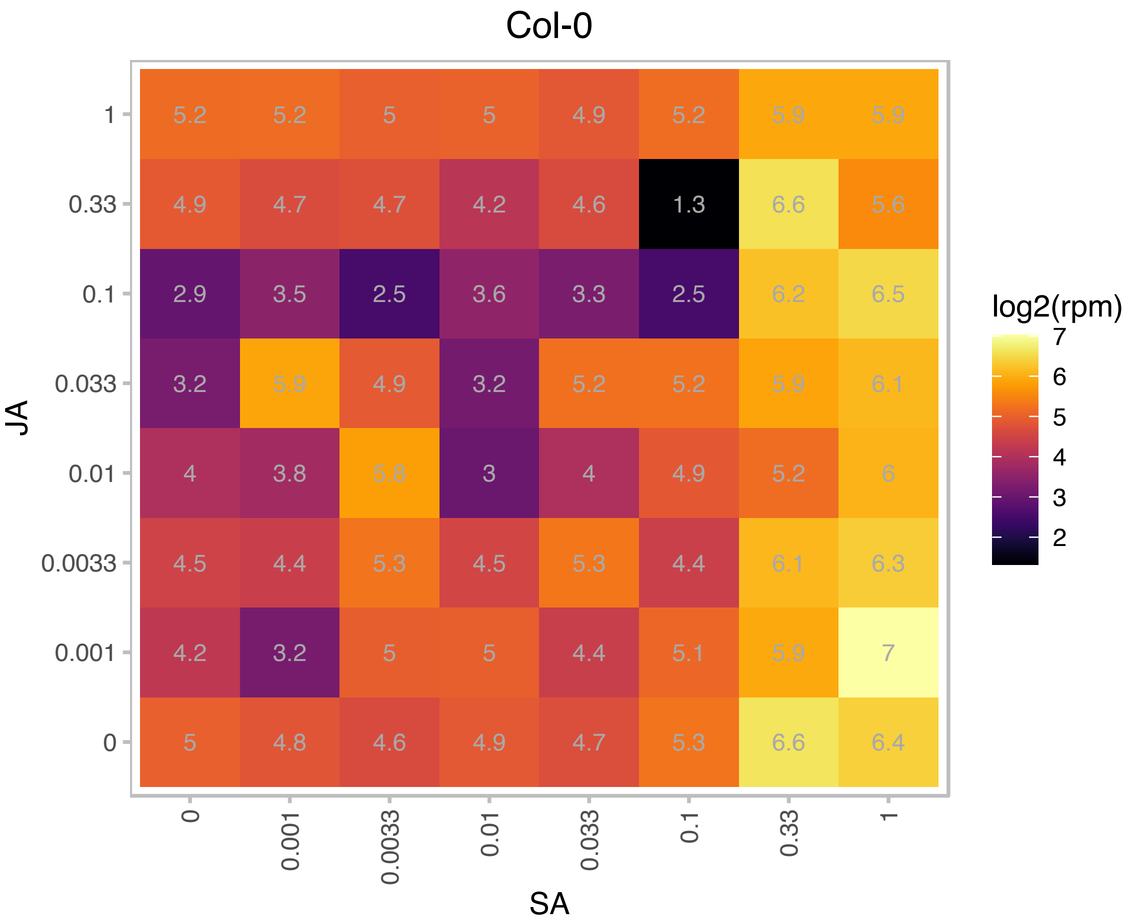


| | |
|---|---|
| Gene | AT5G54490 |
| is.expressed_genes | TRUE |
| is.responsive_genes | TRUE |
| is.apcluster | B-01 |

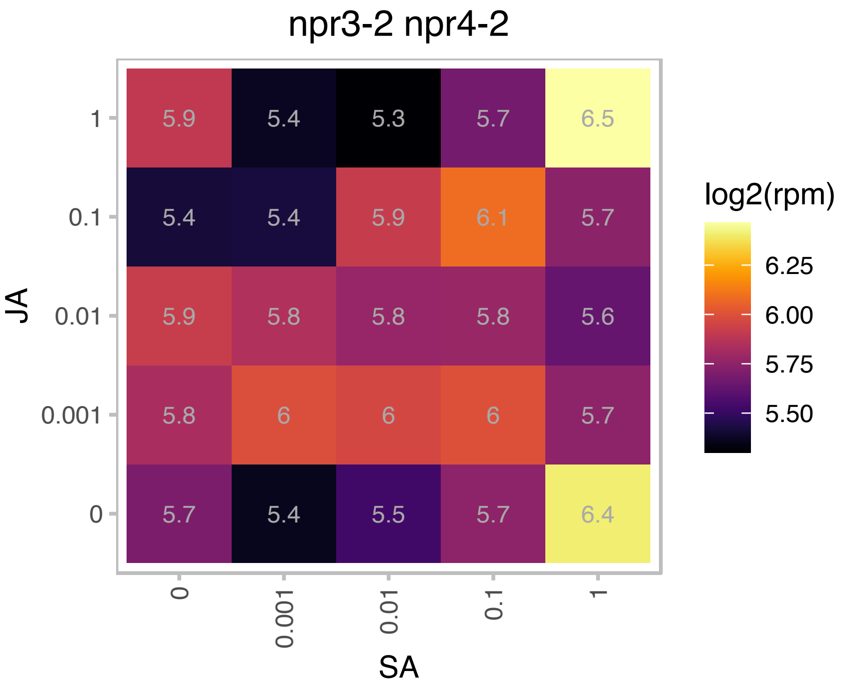


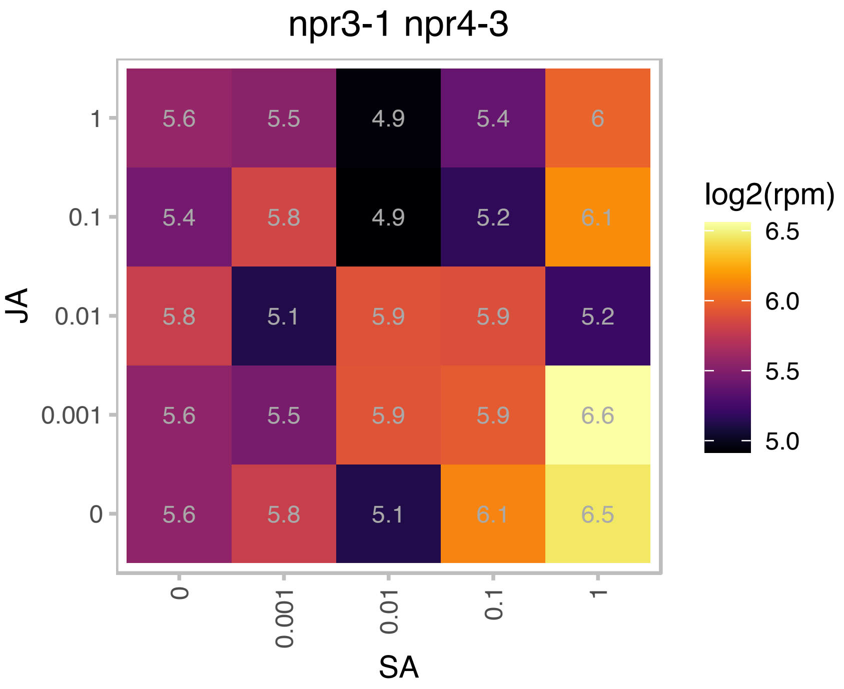

### AT5G59720.1 : HSP18.2

heat shock protein 18.2

encodes a low molecular weight heat shock protein that contains the heat shock element in the promoter region. Expression is induced in response to heat shock.

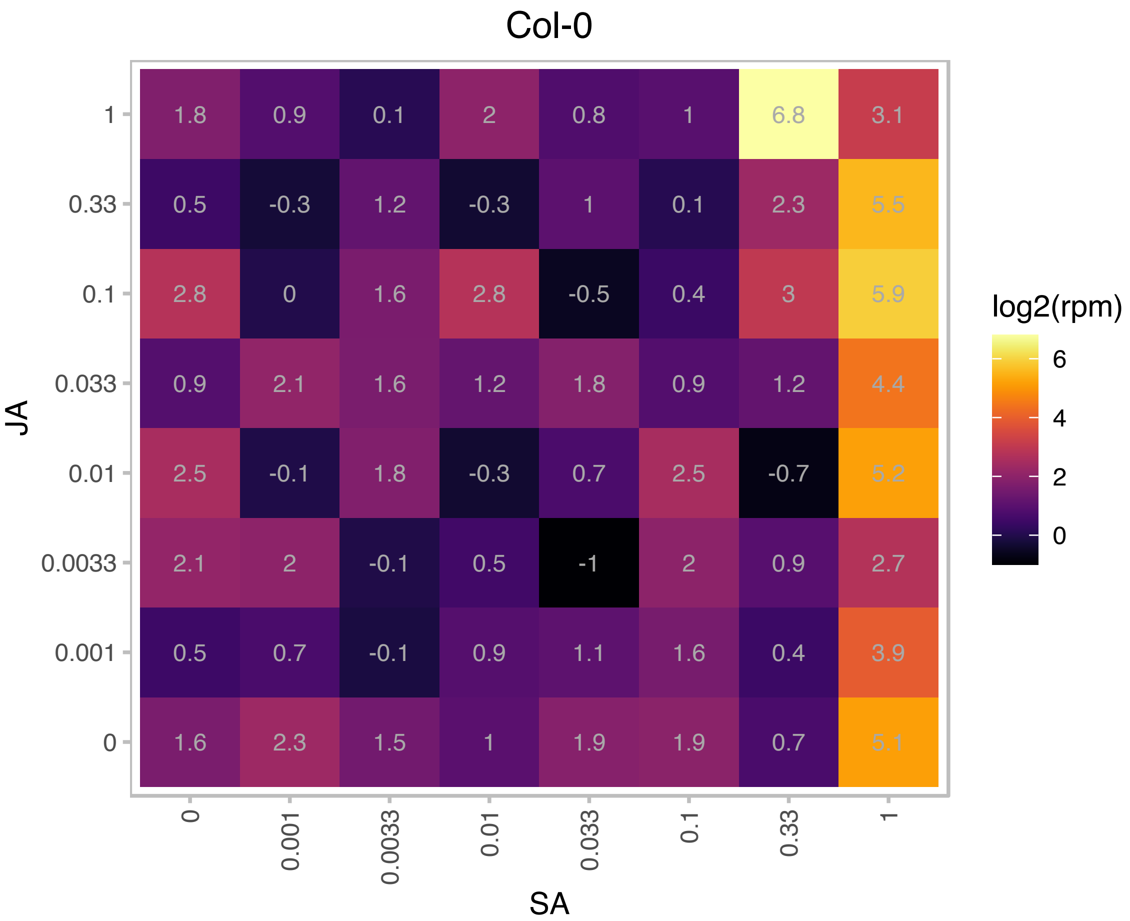


| | |
|---|---|
| Gene | AT5G59720 |
| is.expressed_genes | TRUE |
| is.responsive_genes | TRUE |
| is.apcluster | B-01 |

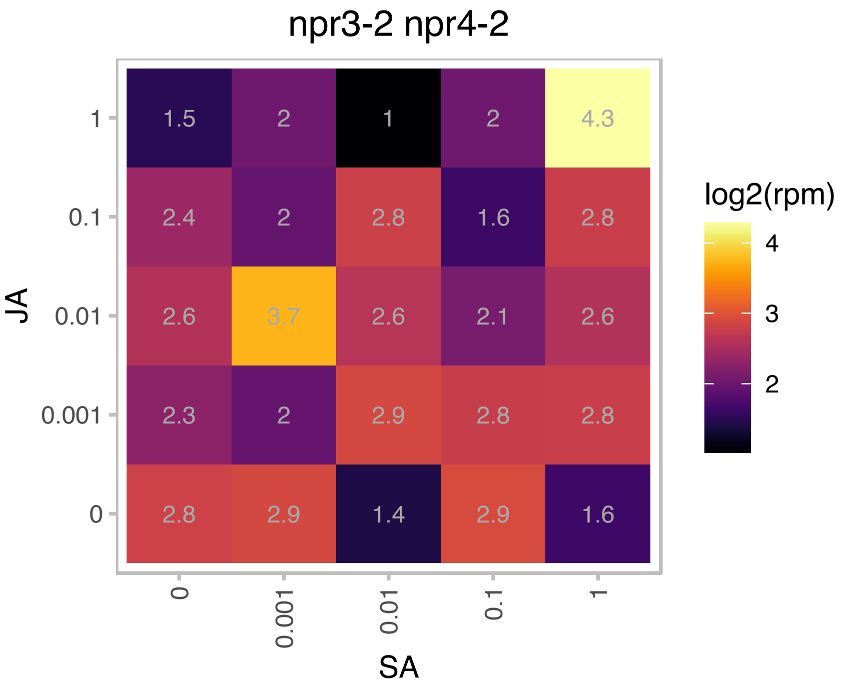


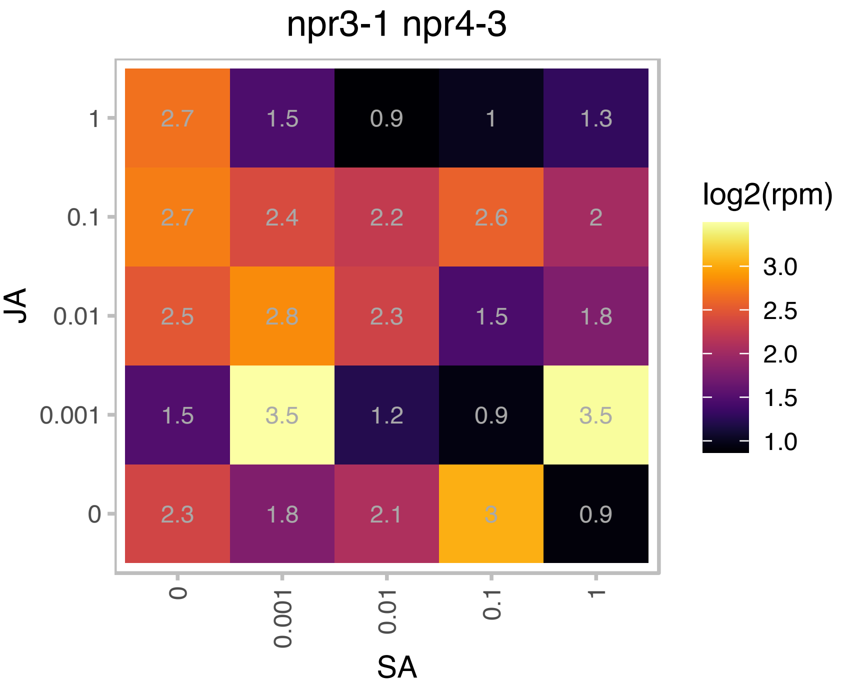

### ATCG00190.1 : RPOB

RNA polymerase subunit beta

Chloroplast DNA-dependent RNA polymerase B subunit. The transcription of this gene is regulated by a nuclear encoded RNA polymerase. This gene has been transferred to mitochondrial genome during crucifer evolution.

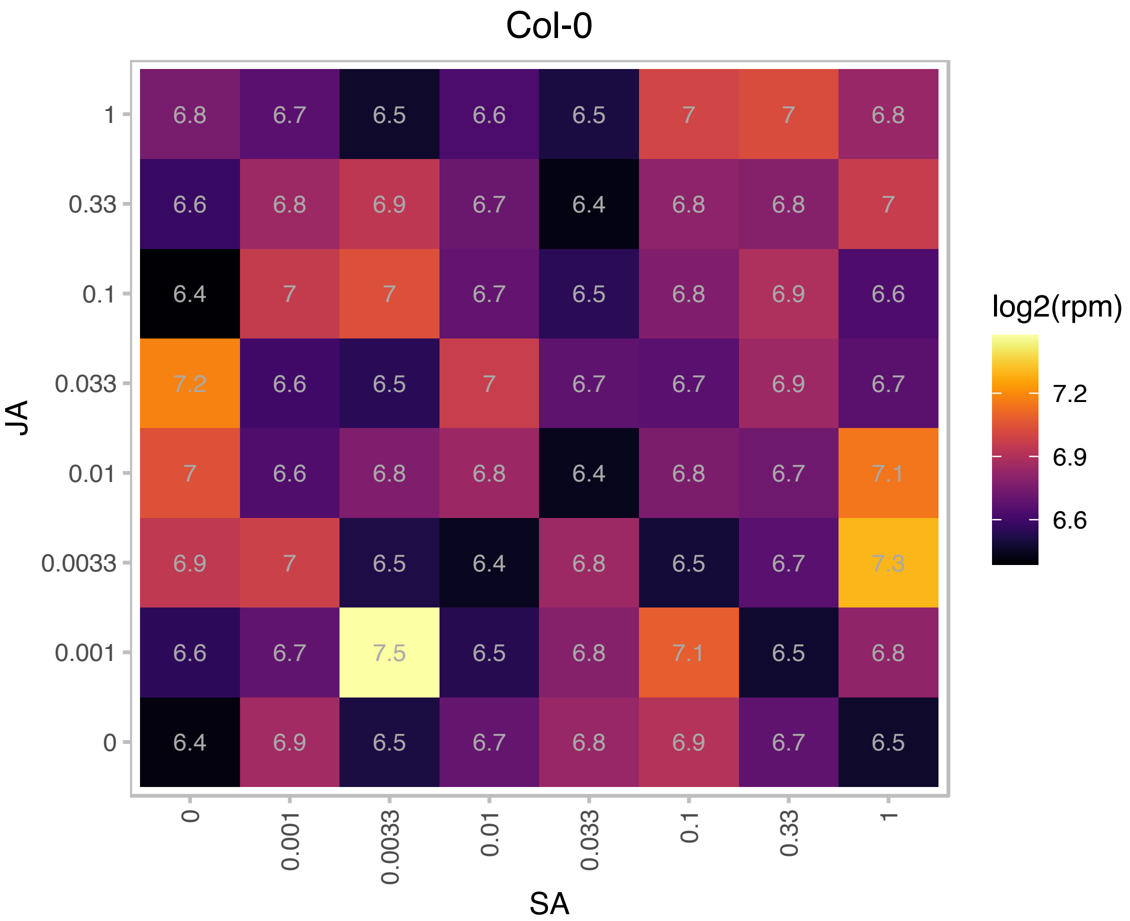


| | |
|---|---|
| Gene | ATCG00190 |
| is.expressed_genes | TRUE |
| is.responsive_genes | TRUE |
| is.apcluster | B-01 |

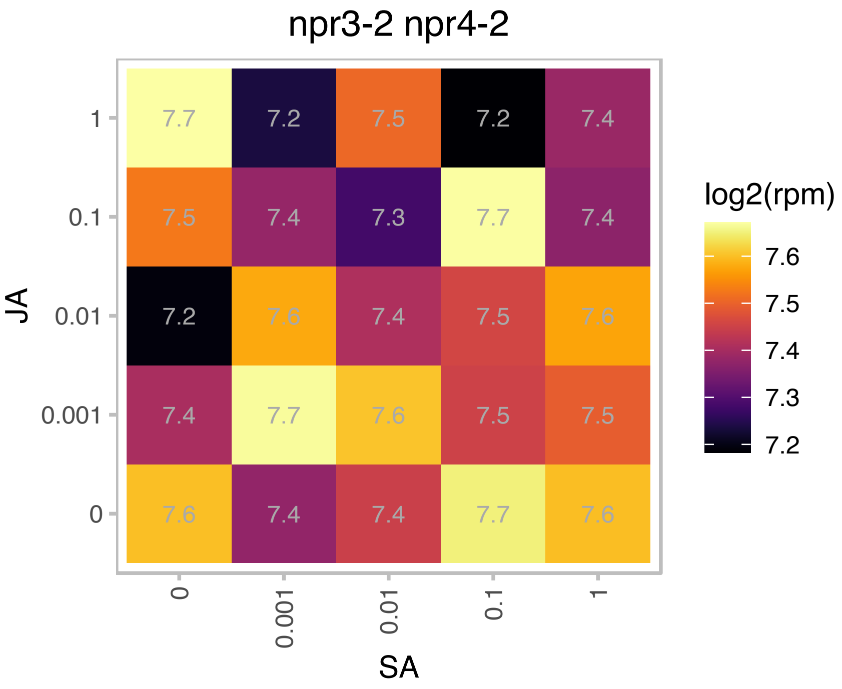


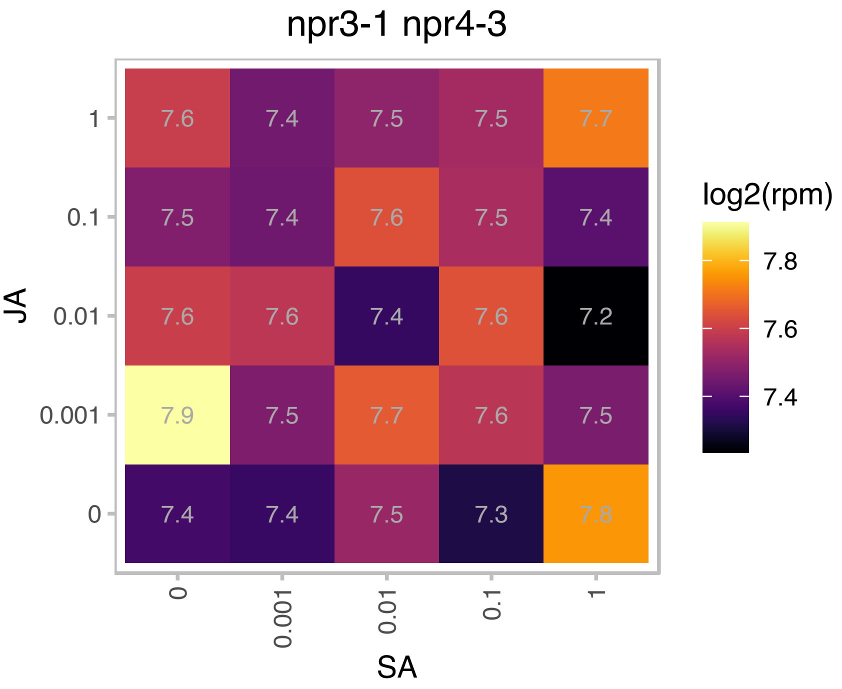

# AT1G04350.1 : NA

2-oxoglutarate (2OG) and Fe(II)-dependent oxygenase superfamily protein

encodes a protein whose sequence is similar to 2-oxoglutarate-dependent dioxygenase

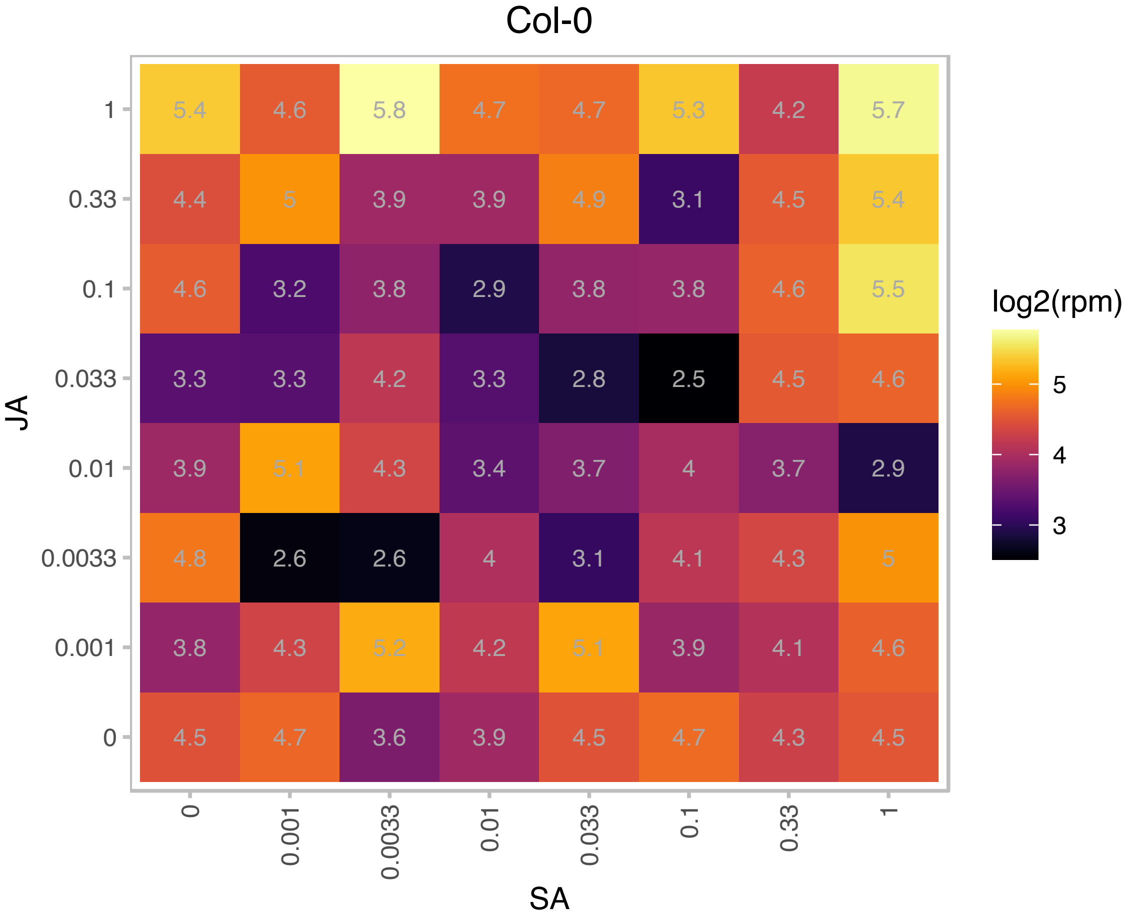


| | |
|---|---|
| Gene | AT1G04350 |
| is.expressed_genes | TRUE |
| is.responsive_genes | TRUE |
| is.apcluster | B-02 |

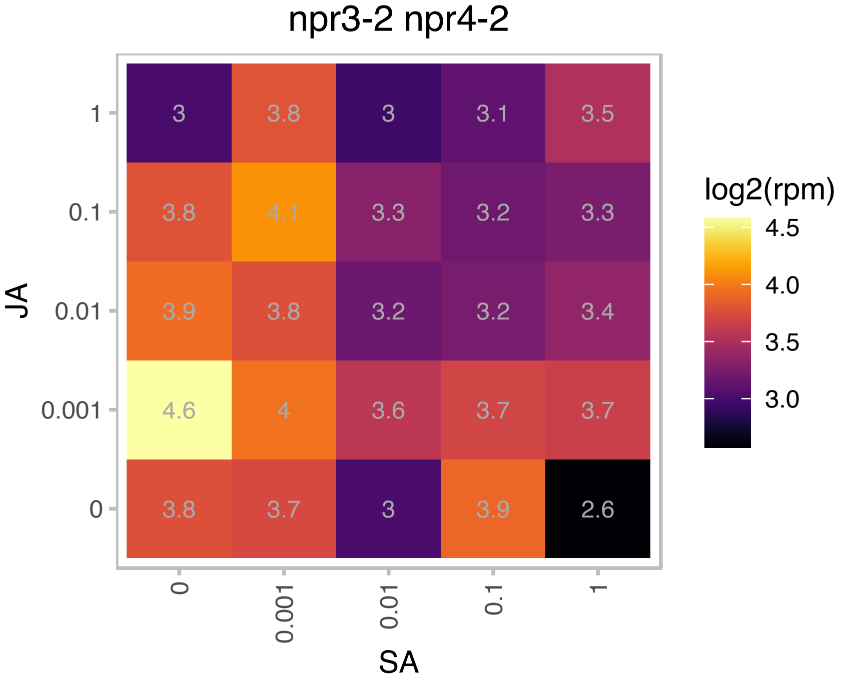


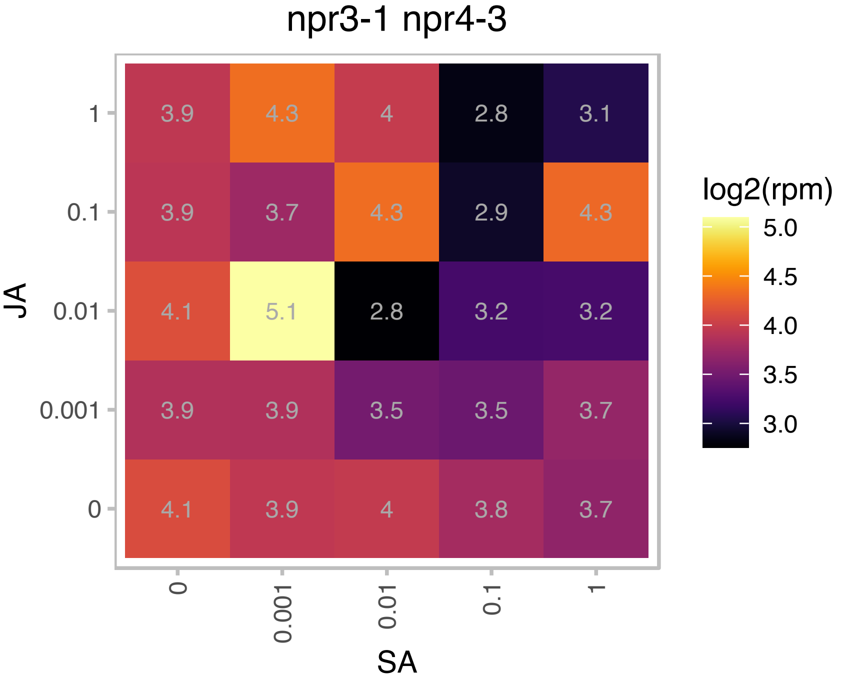

### AT1G05850.1 : ATCTL1, CTL1, ELP, ELP1, ERH2, HOT2, [illegible]

Chitinase family protein

Encodes an endo chitinase-like protein AtCTL1. Essential for tolerance to heat, salt and drought stresses. Also involved in root hair development, cell expansion and response to cytokinin. Allelic to erh2. 11 alleles described in Hauser (1995). Mutant is defective in acquired thermotolerance, appears semidwarf throughout its life cycle and has extra lateral branches. There are two EMS alleles. Expression of AtHSP101 is not affected in the mutants.

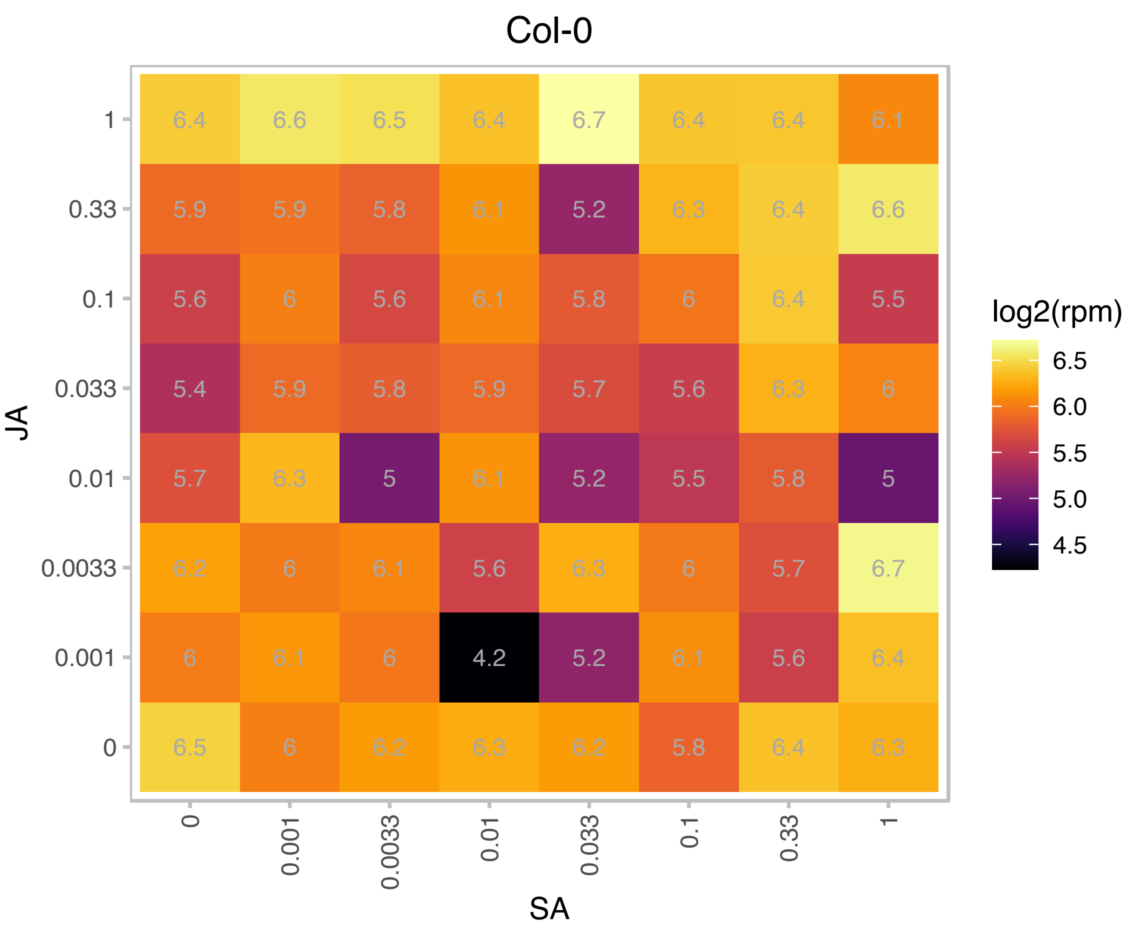


| | |
|---|---|
| Gene | AT1G05850 |
| is.expressed_genes | TRUE |
| is.responsive_genes | TRUE |
| is.apcluster | B-02 |

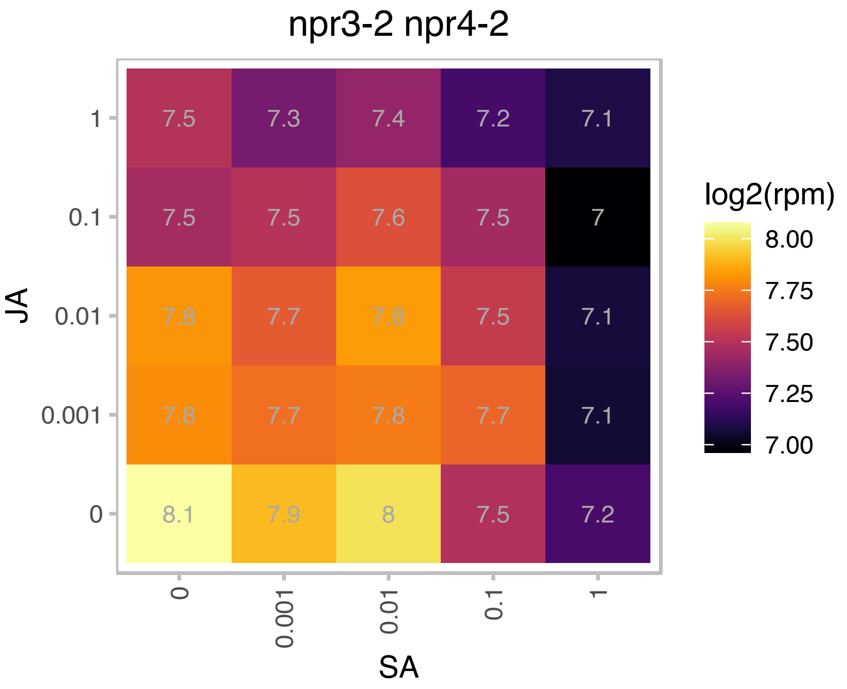


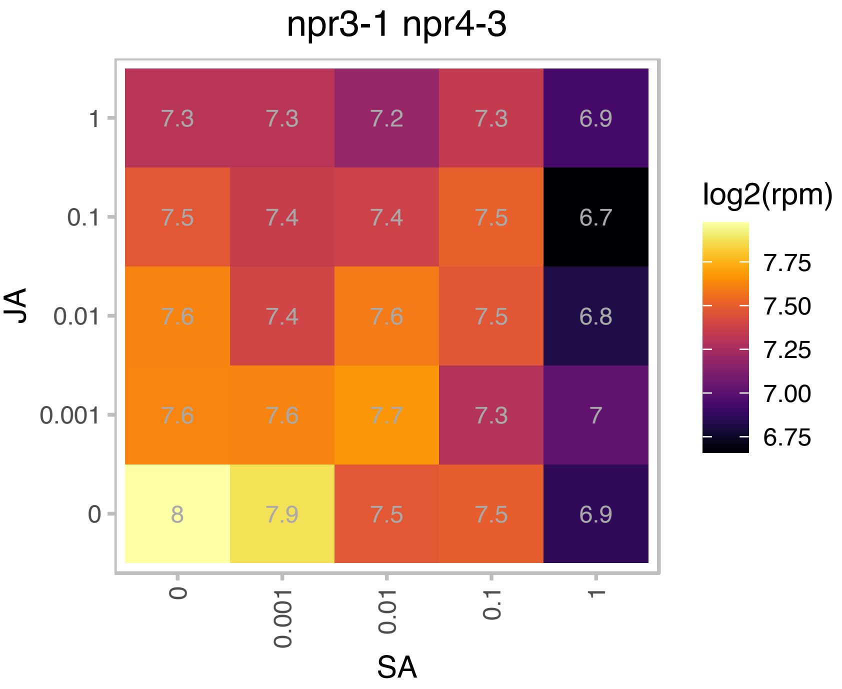

### AT1G10670.1 : ACLA-1

ATP-citrate lyase A-1

One of the three genes encoding subunit A of the trimeric protein ATP Citrate Lyase. Antisense ACLA-1 plants cause a reduction in cytosolic acetyl-CoA metabolism and have upregulation of stress-related genes and down-regulation of primary metabolism and growth genes, suggesting the mutation restricts normal growth and developmental processes and puts the plant into a state of stress.

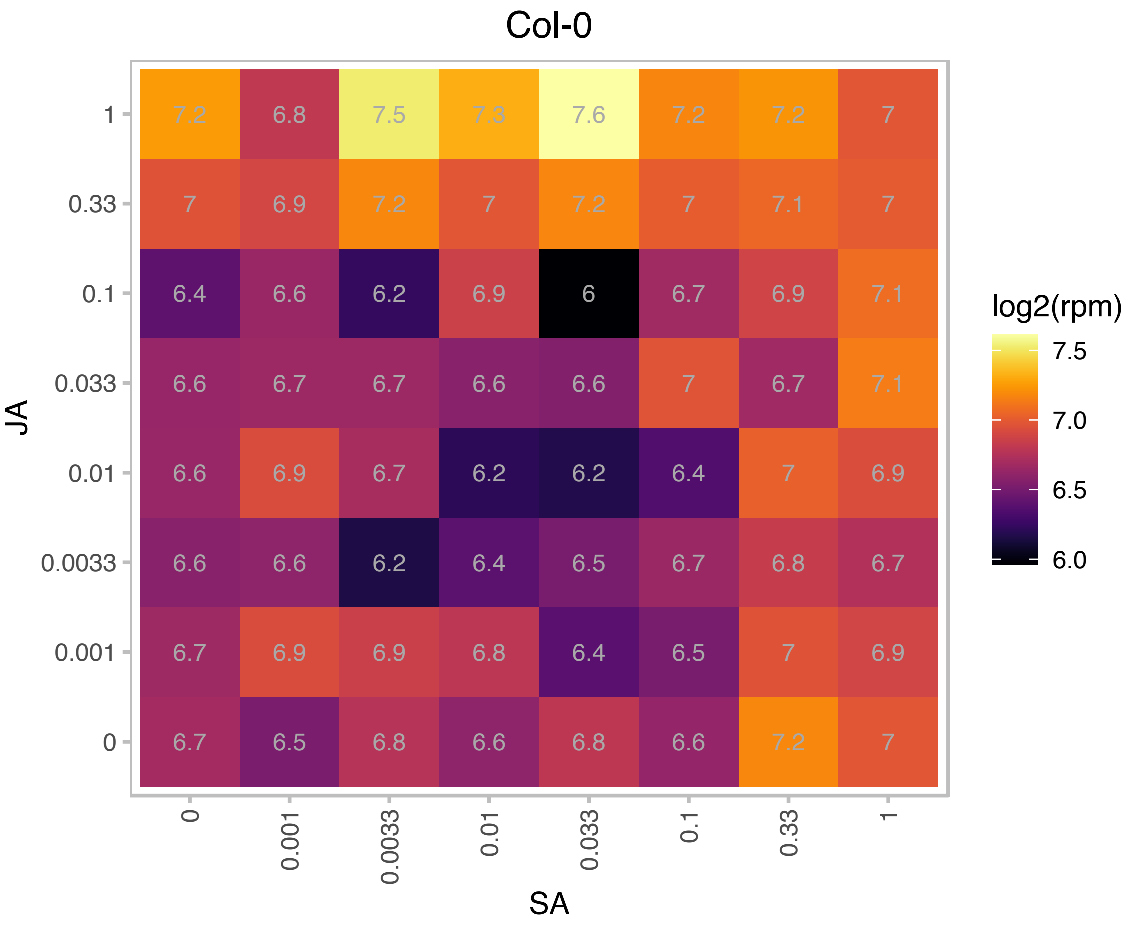


| | |
|---|---|
| Gene | AT1G10670 |
| is.expressed_genes | TRUE |
| is.responsive_genes | TRUE |
| is.apcluster | B-02 |

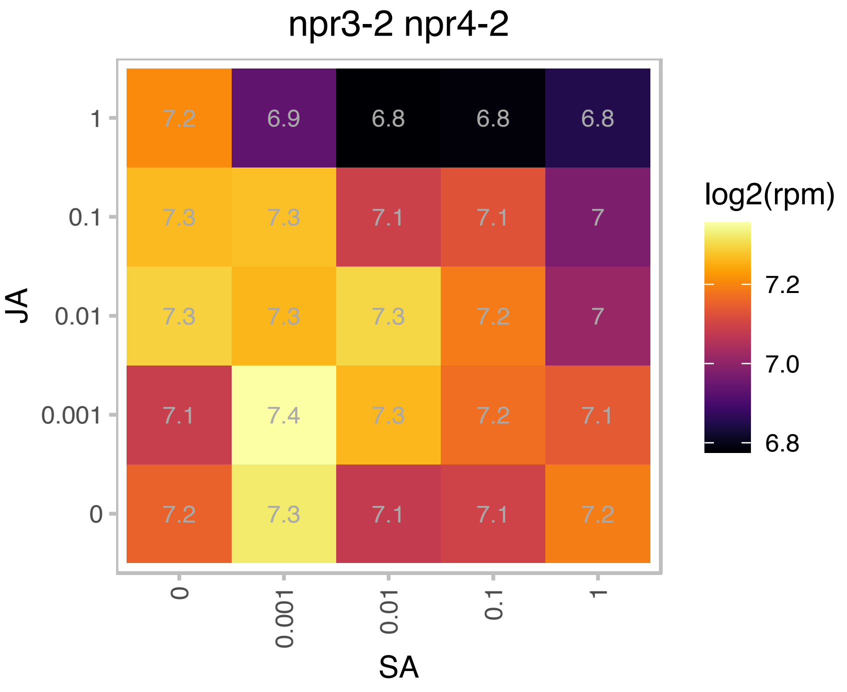


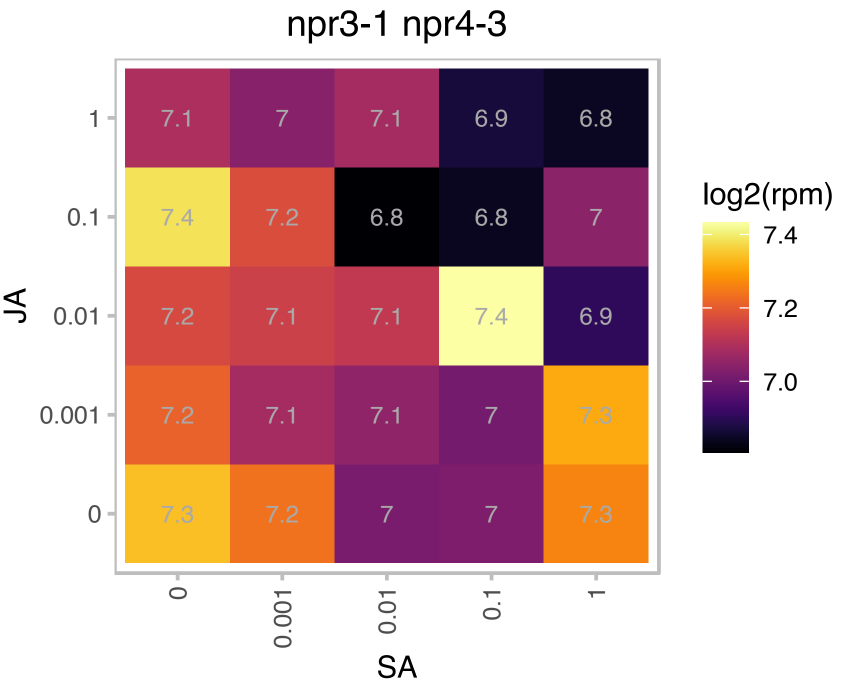

# AT1G13820.1 : NA

alpha/beta-Hydrolases superfamily protein

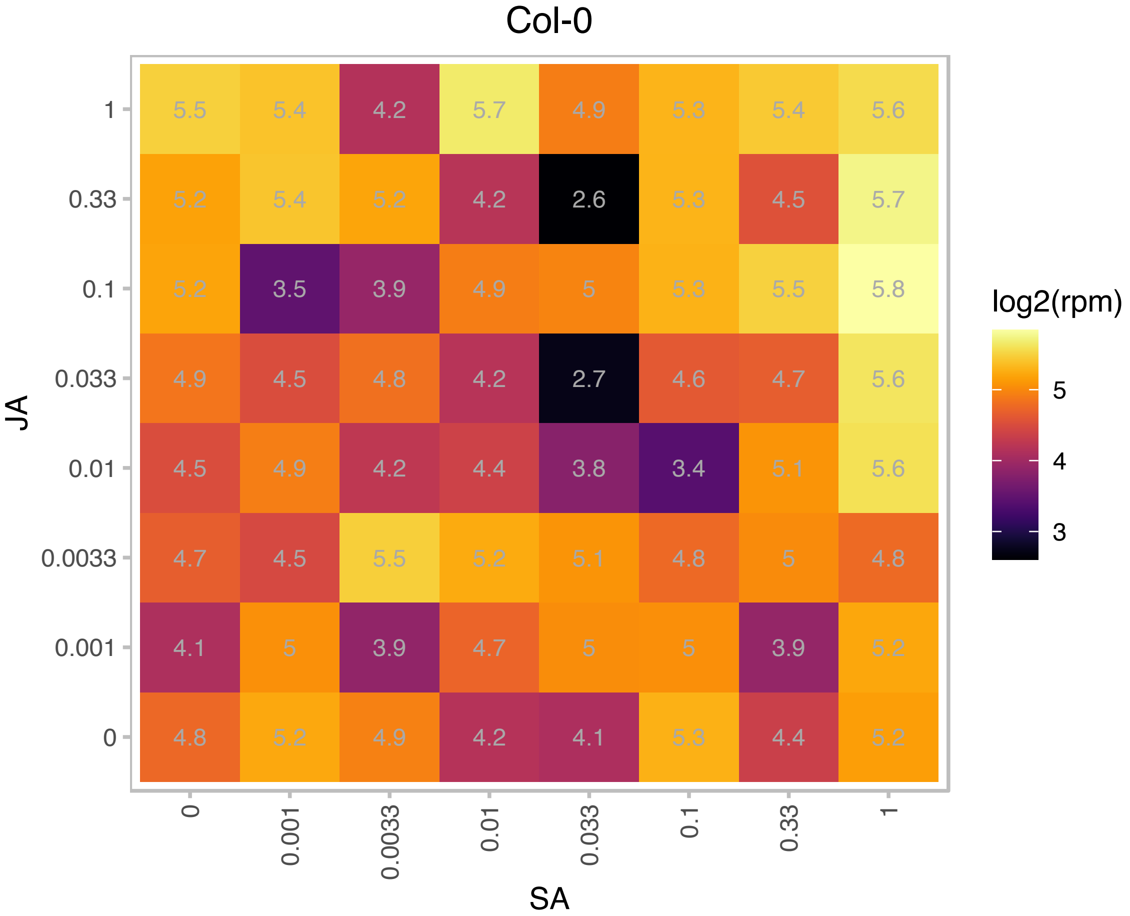


| | |
|---|---|
| Gene | AT1G13820 |
| is.expressed_genes | TRUE |
| is.responsive_genes | TRUE |
| is.apcluster | B-02 |

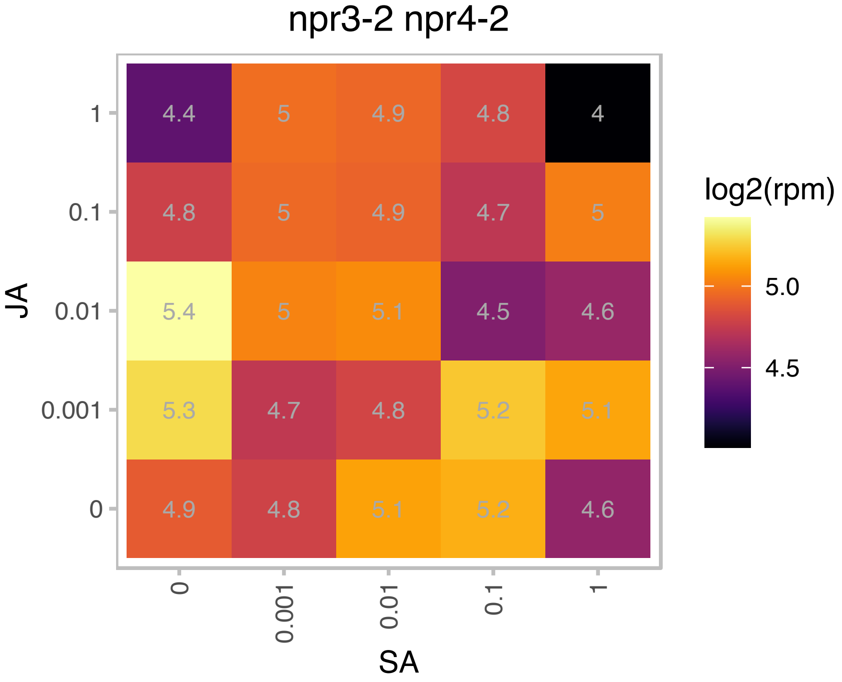


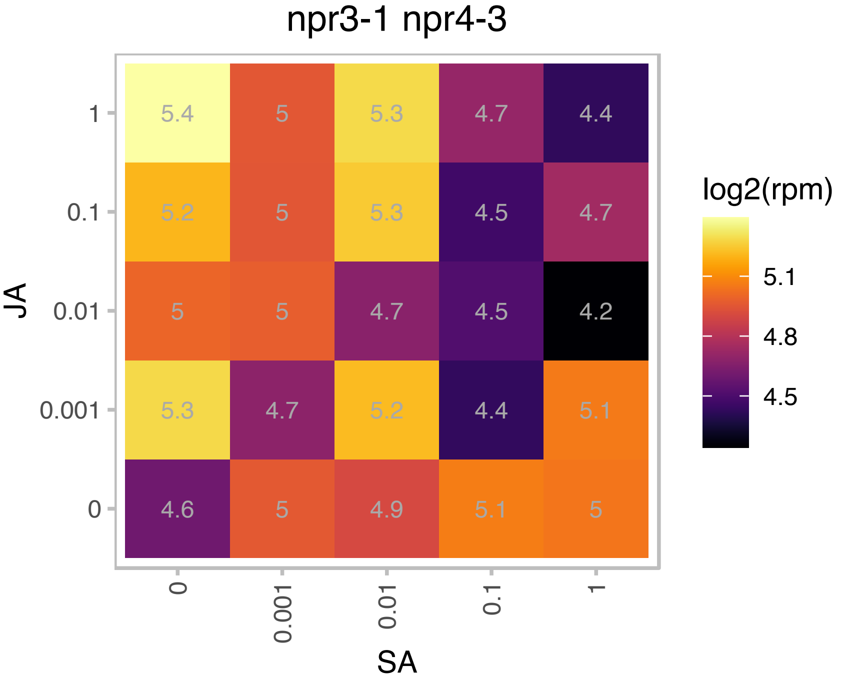

# AT1G19200.1 : NA

Protein of unknown function (DUF581)

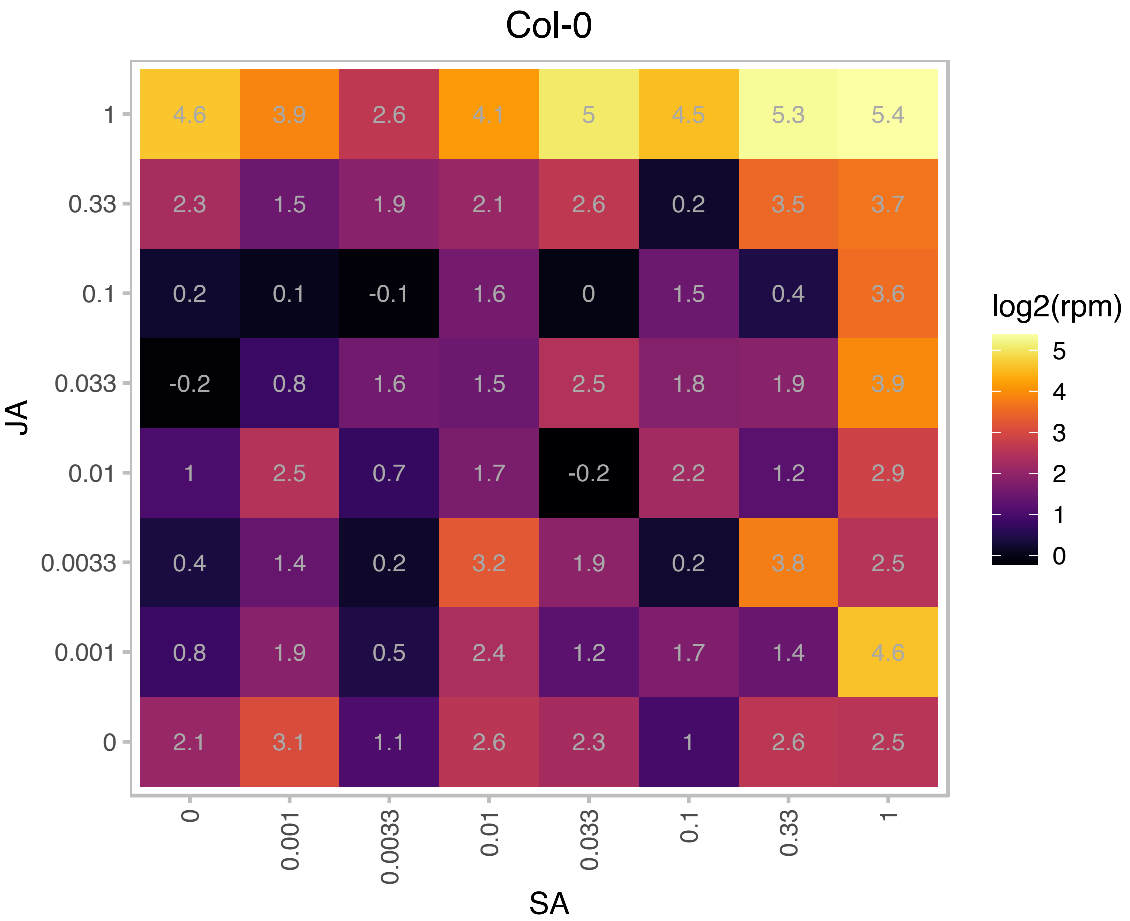


| | |
|---|---|
| Gene | AT1G19200 |
| is.expressed_genes | TRUE |
| is.responsive_genes | TRUE |
| is.apcluster | B-02 |

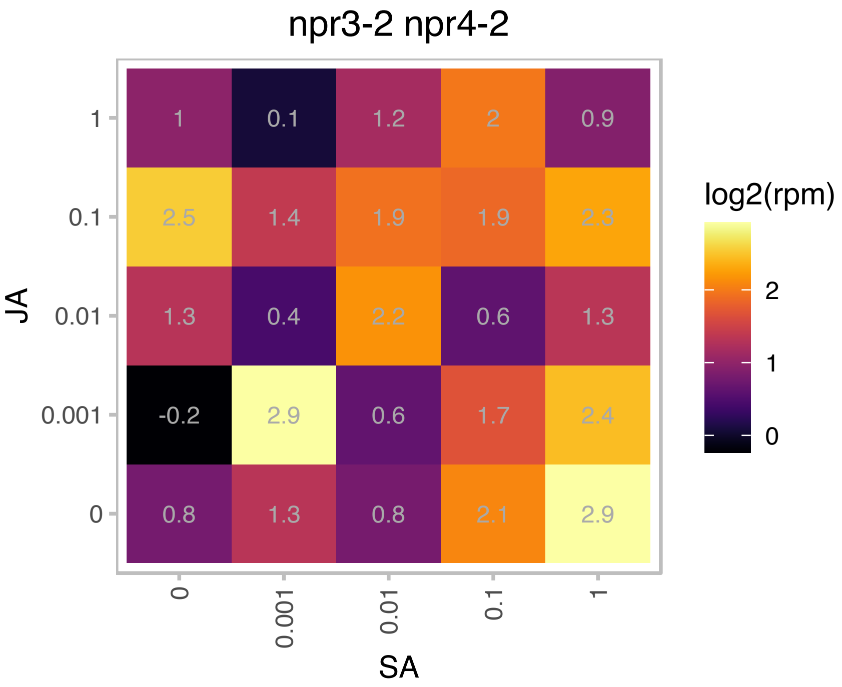


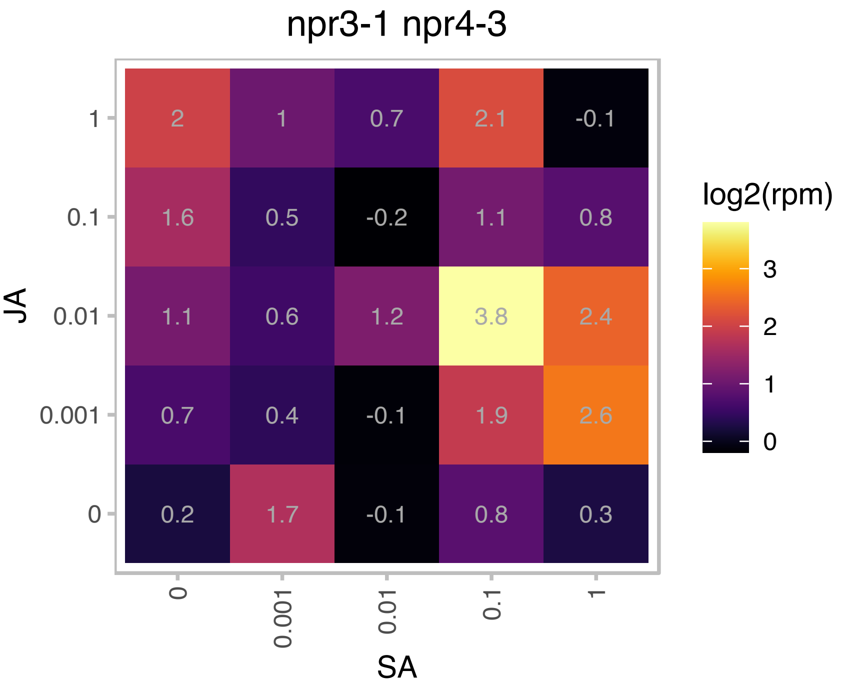

### AT1G20440.1 : AtCOR47, COR47, RD17

cold-regulated 47

Belongs to the dehydrin protein family, which contains highly conserved stretches of 7-17 residues that are repetitively scattered in their sequences, the K-, S-, Y- and lysine rich segments. Cold regulated gene, amino acid sequence homology with Group II LEA (late embryogenesis abundant) proteins. Also responds to osmotic stress, ABA, dehydration and inhibits e.coli growth while overexpressed. COR47 and RAB18 double overexpressor plants are cold tolerant.

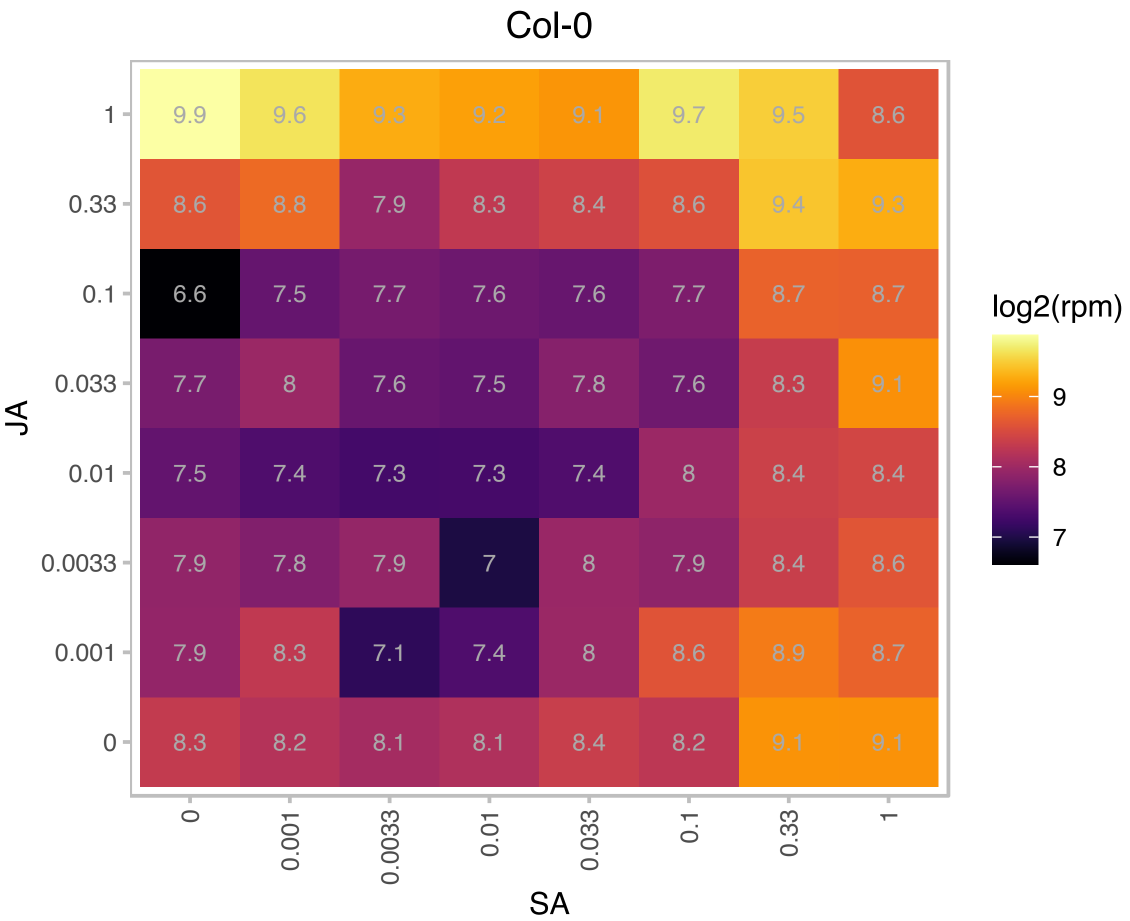


| | |
|---|---|
| Gene | AT1G20440 |
| is.expressed_genes | TRUE |
| is.responsive_genes | TRUE |
| is.apcluster | B-02 |

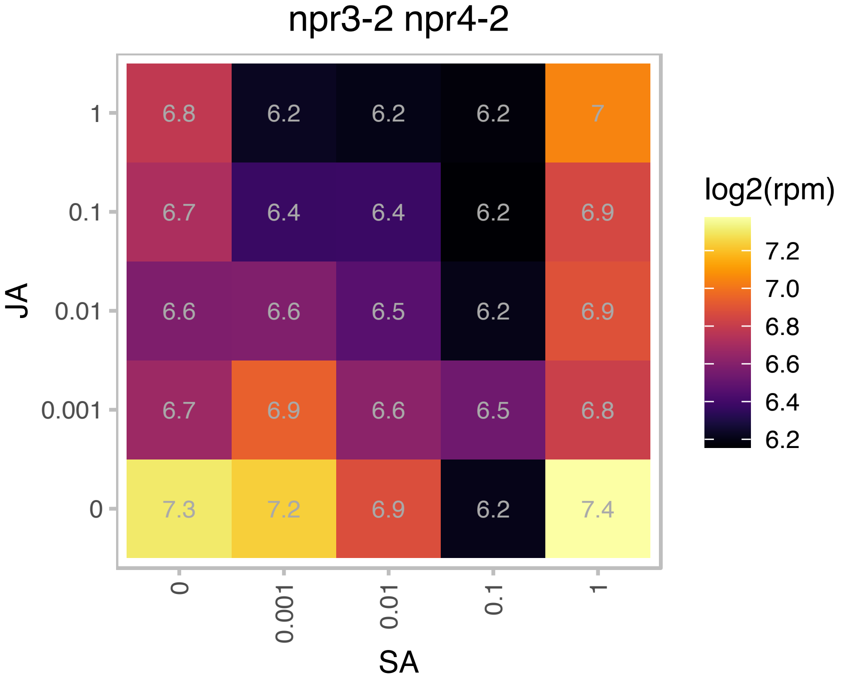


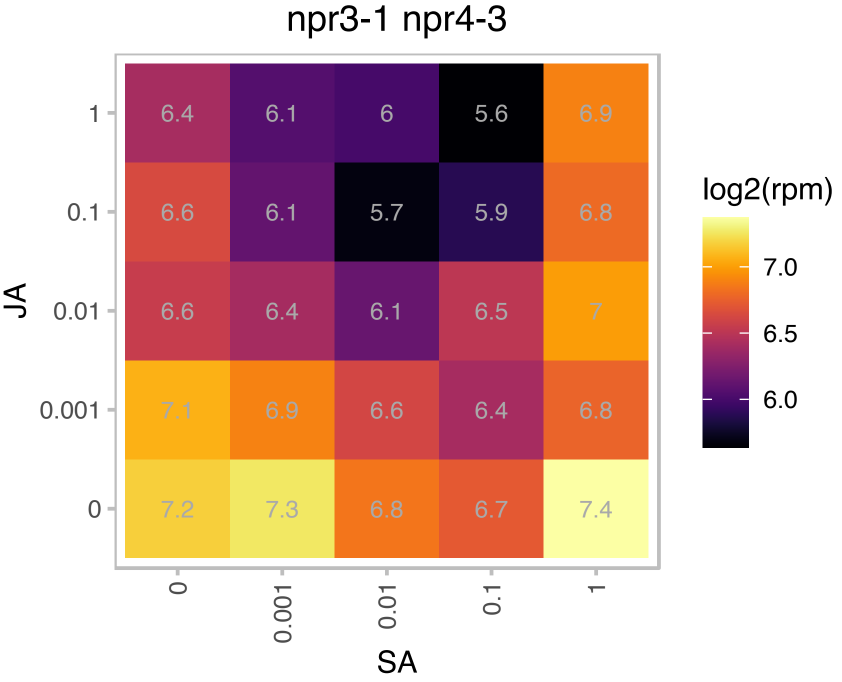

### AT1G23550.1 : SRO2

similar to RCD one 2

Encodes a protein with similarity to RCD1 but without the WWE domain. The protein does have a PARP signature upstream of the C-terminal protein interaction domain. The PARP signature may bind NAD+ and attach the ADP-ribose-moiety from NAD+ to the target molecule. Its presence suggests a role for the protein in ADP ribosylation. Its transcript level is up-regulated by tunicamycin (N-linked glycosylation inhibitor causing ER stress).

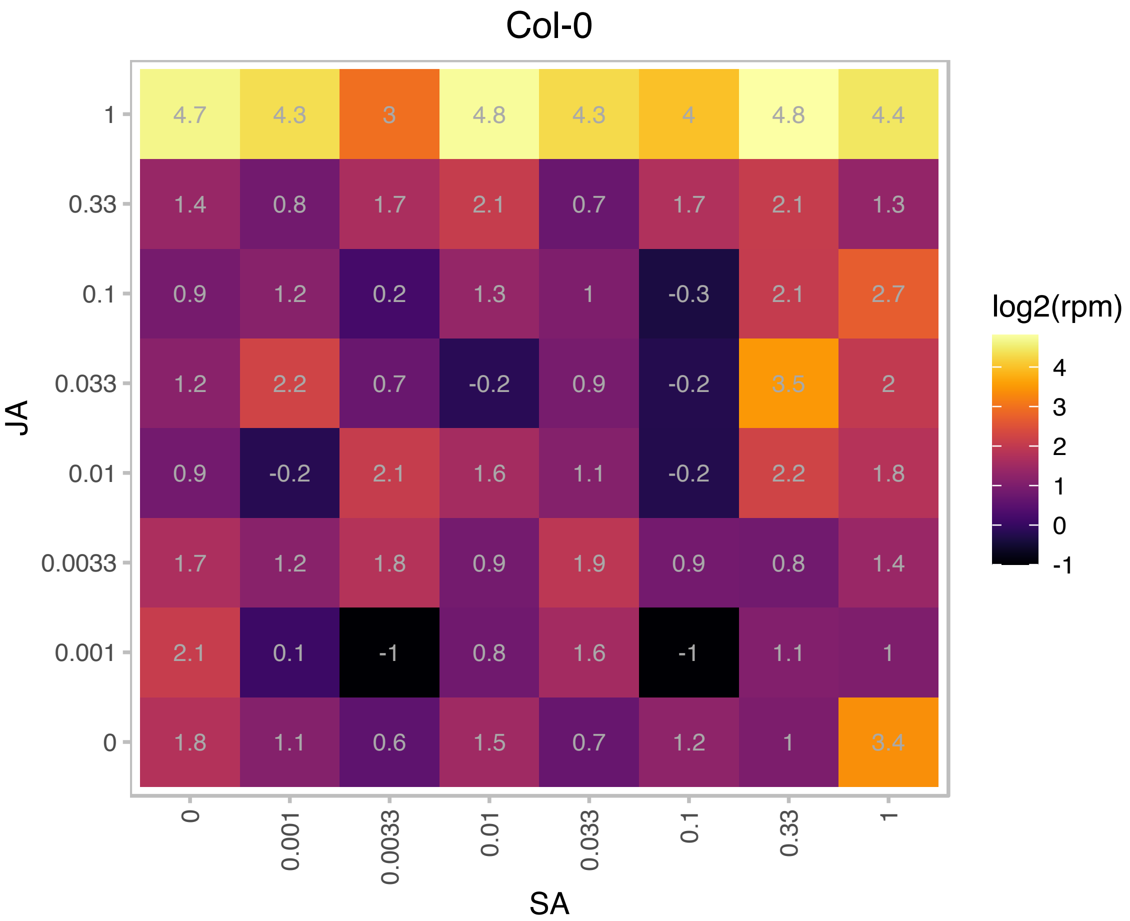


| | |
|---|---|
| Gene | AT1G23550 |
| is.expressed_genes | TRUE |
| is.responsive_genes | TRUE |
| is.apcluster | B-02 |

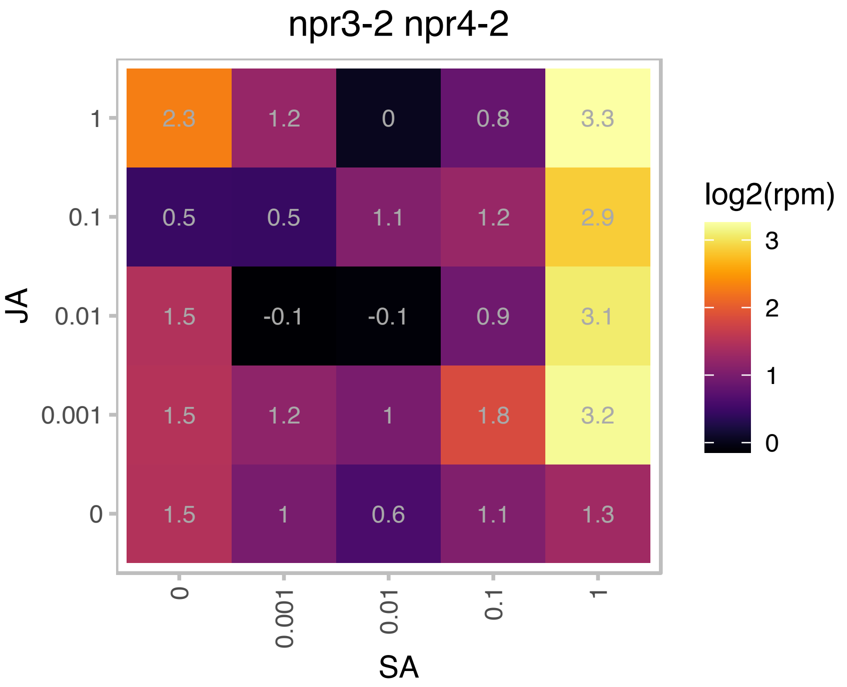


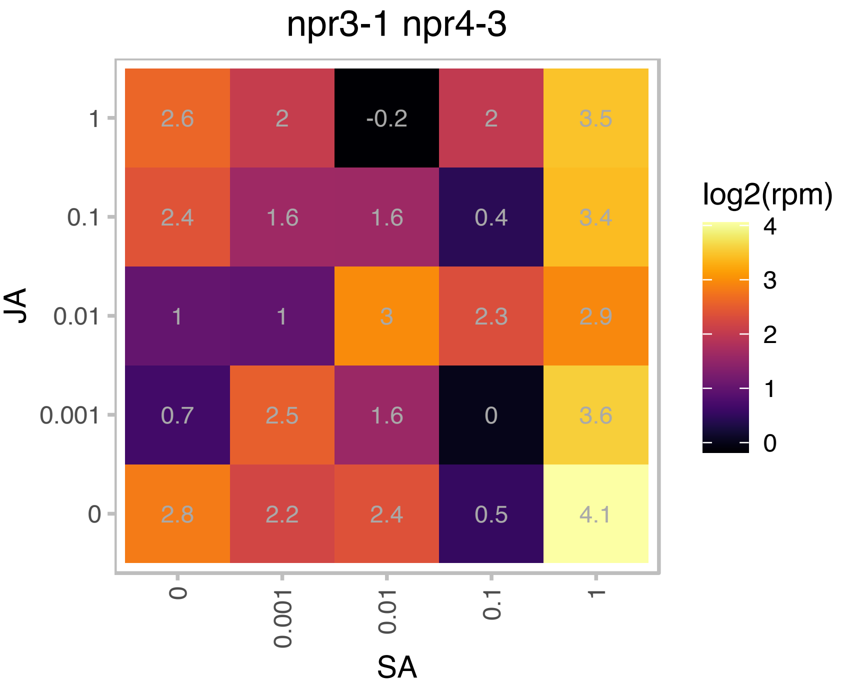

### AT1G35470.1 : RanBPM

SPla/RYanodine receptor (SPRY) domain-containing protein

Encodes a homologue of human RanBPM that belongs to the uncharacterized family of plant SPRY, LisH, CTLH an[illegible] CRA domain-containing proteins.

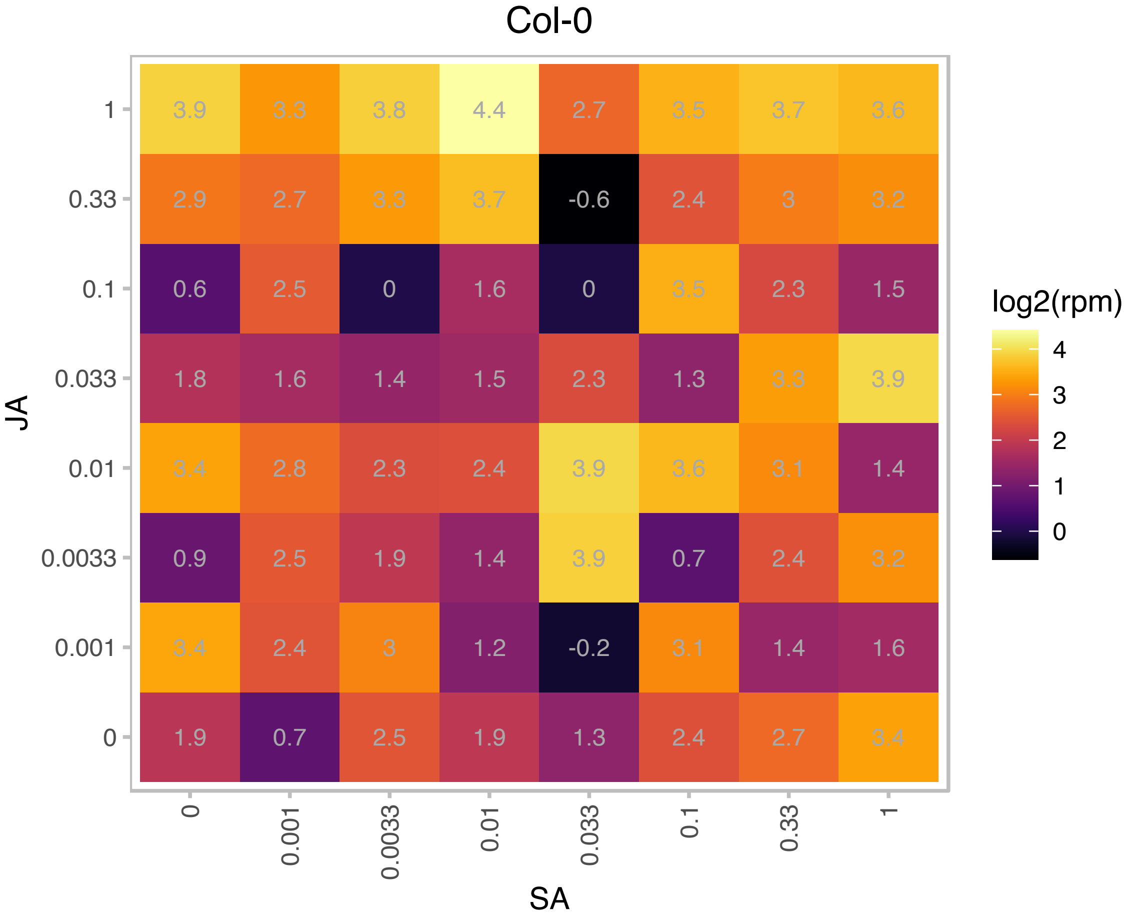


| | |
|---|---|
| Gene | AT1G35470 |
| is.expressed_genes | TRUE |
| is.responsive_genes | TRUE |
| is.apcluster | B-02 |

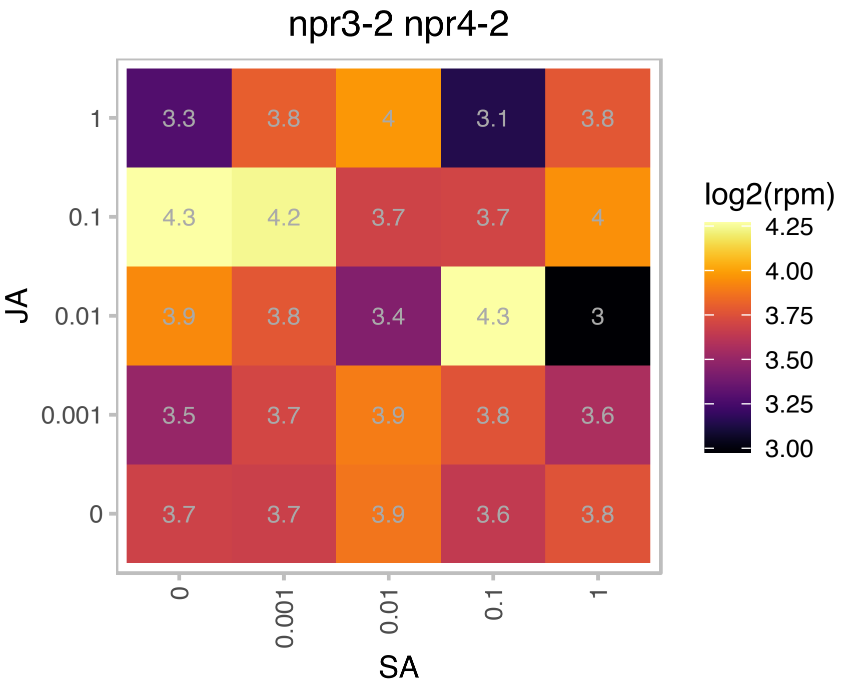


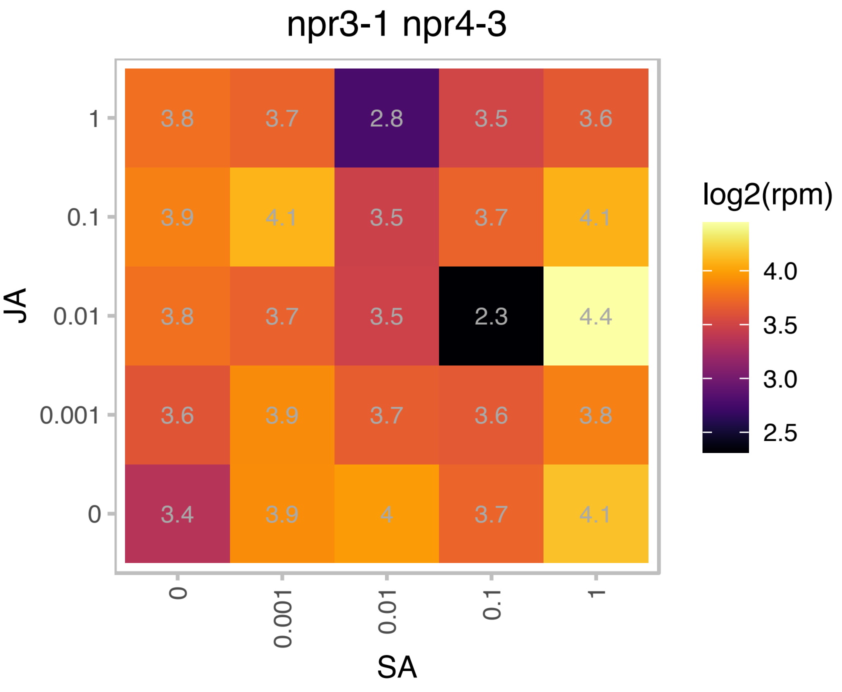

# AT1G43690.1 : NA

ubiquitin interaction motif-containing protein

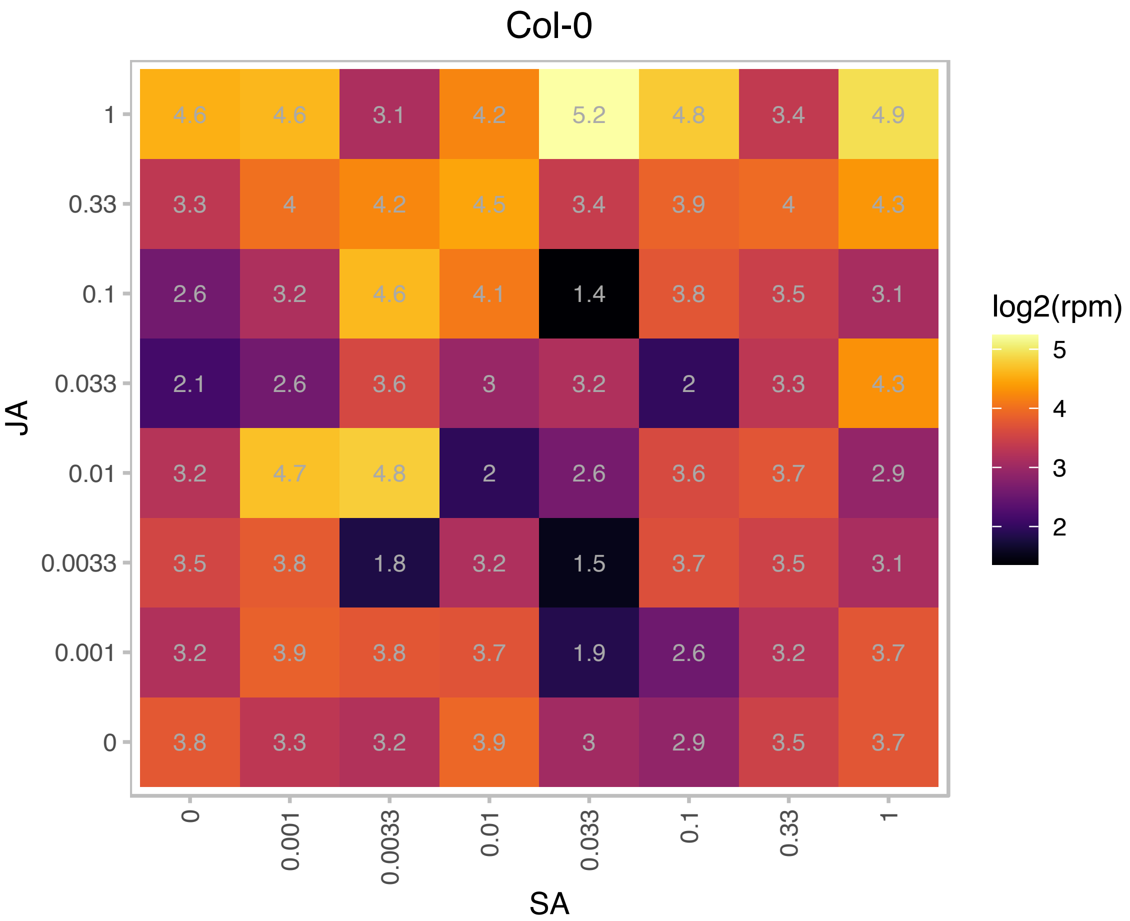


| | |
|---|---|
| Gene | AT1G43690 |
| is.expressed_genes | TRUE |
| is.responsive_genes | TRUE |
| is.apcluster | B-02 |

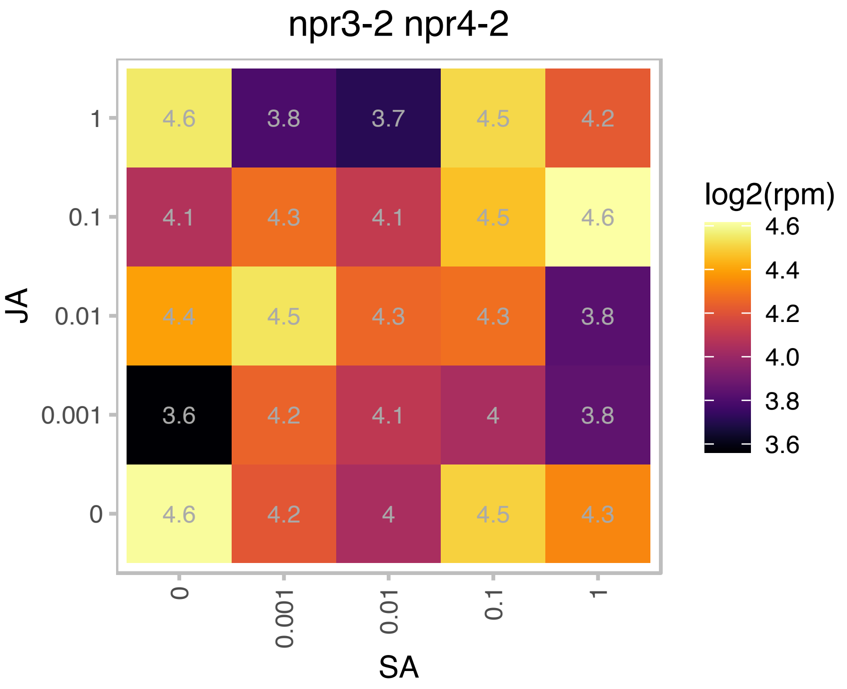


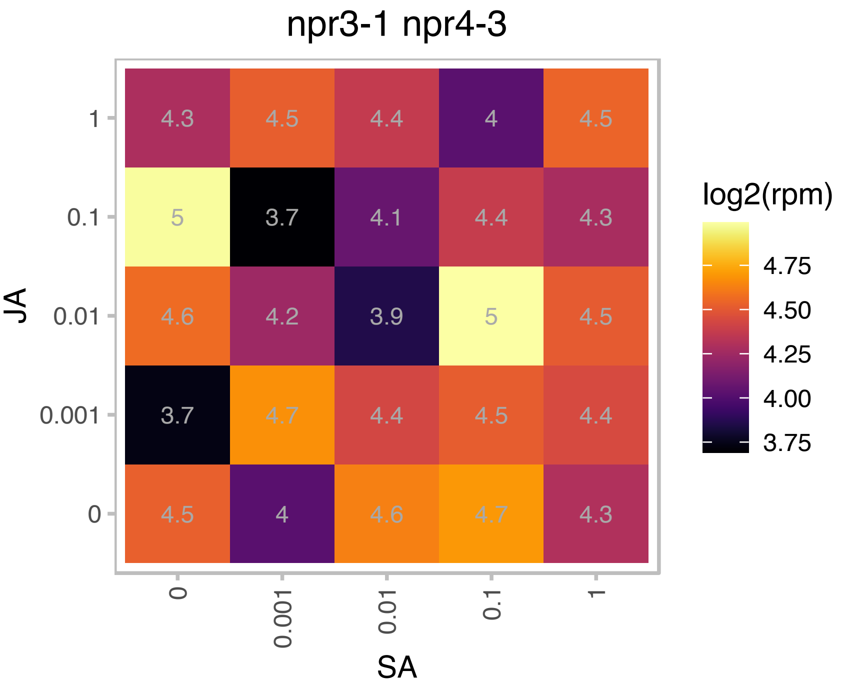

# AT1G45688.1 : CC1

CC1 is a plant specific gene that interacts with with the cellulose synthase complex and microtubules. It appears to play a role in localizing CESA to the membrane, microtuble dynamics , particularly during salt stress.

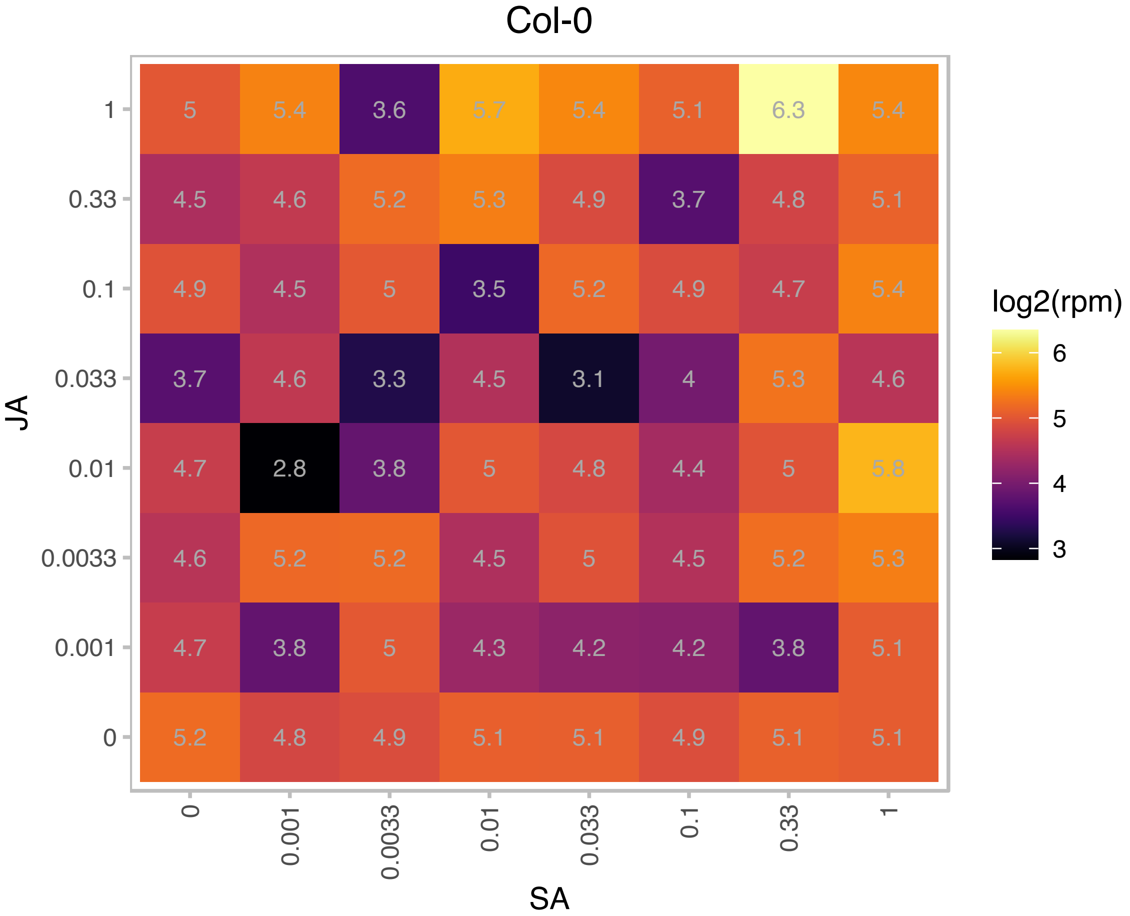


| | |
|---|---|
| Gene | AT1G45688 |
| is.expressed_genes | TRUE |
| is.responsive_genes | TRUE |
| is.apcluster | B-02 |

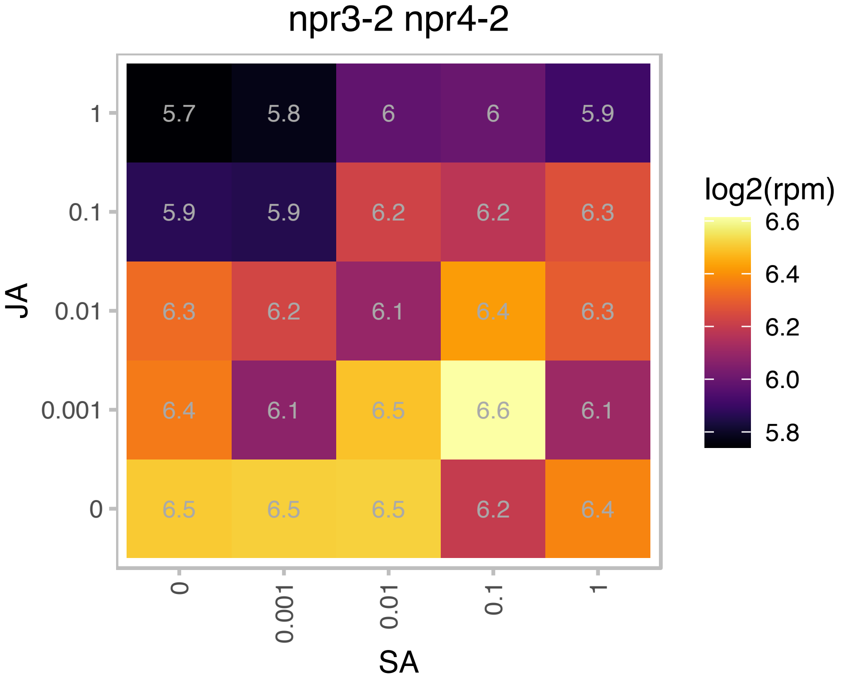


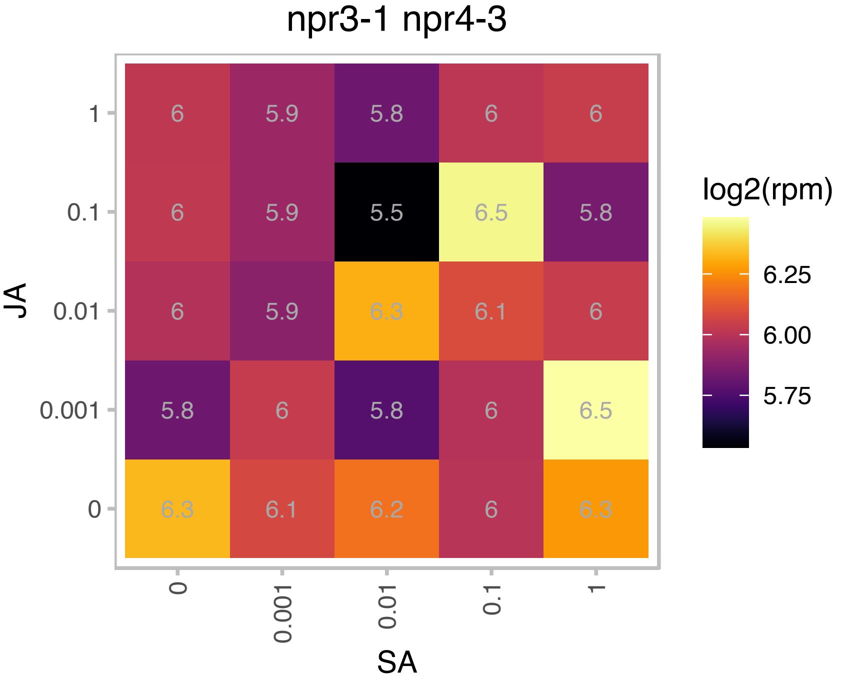

### AT1G47250.1 : PAF2

20S proteasome alpha subunit F2

Encodes 20S proteasome subunit PAF2 (PAF2).

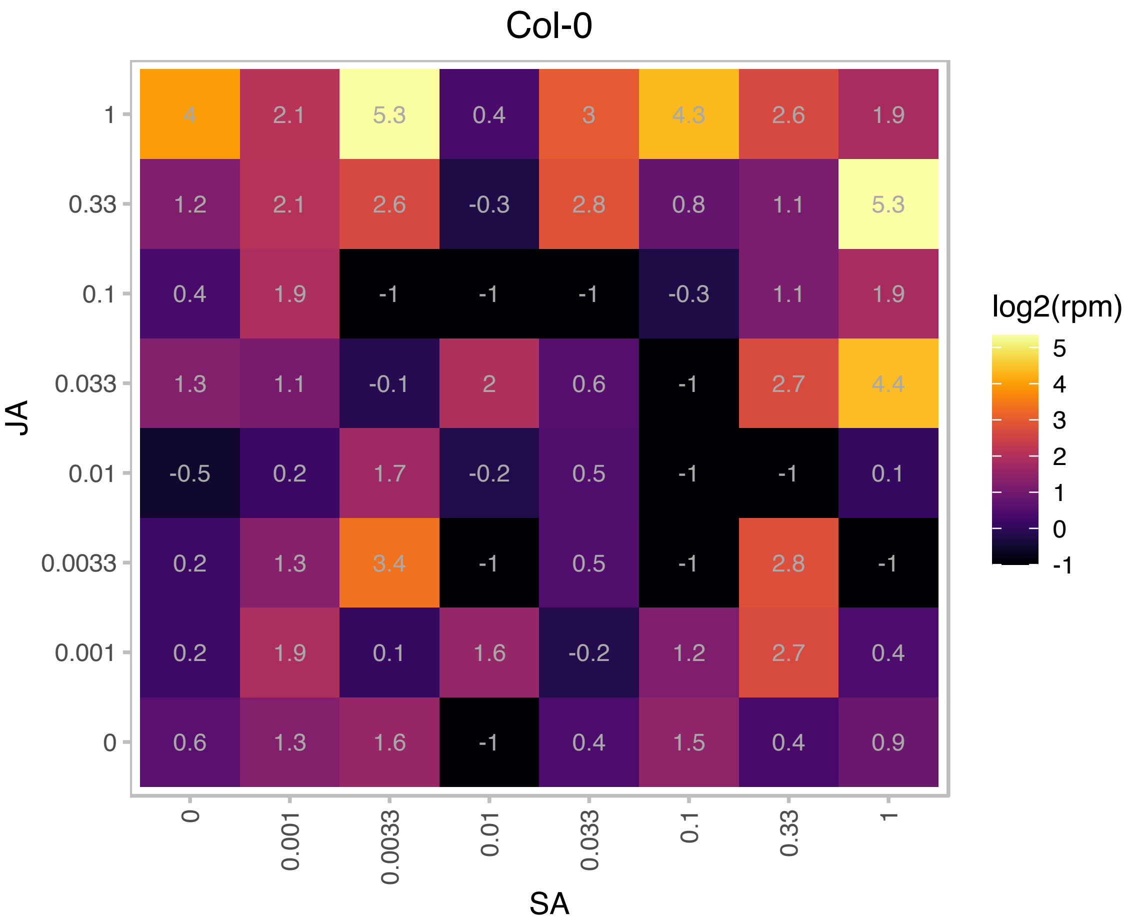


| | |
|---|---|
| Gene | AT1G47250 |
| is.expressed_genes | TRUE |
| is.responsive_genes | TRUE |
| is.apcluster | B-02 |

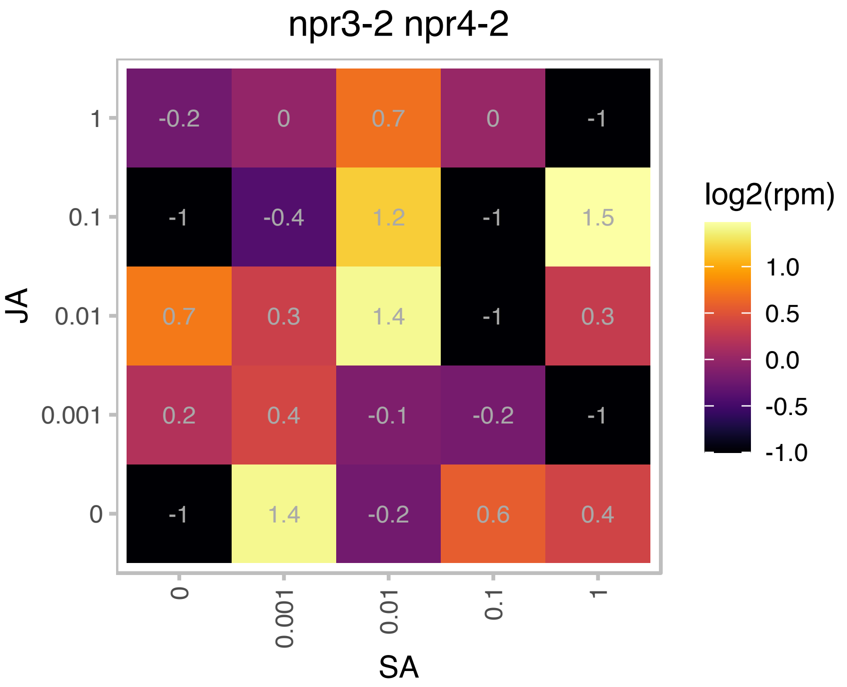


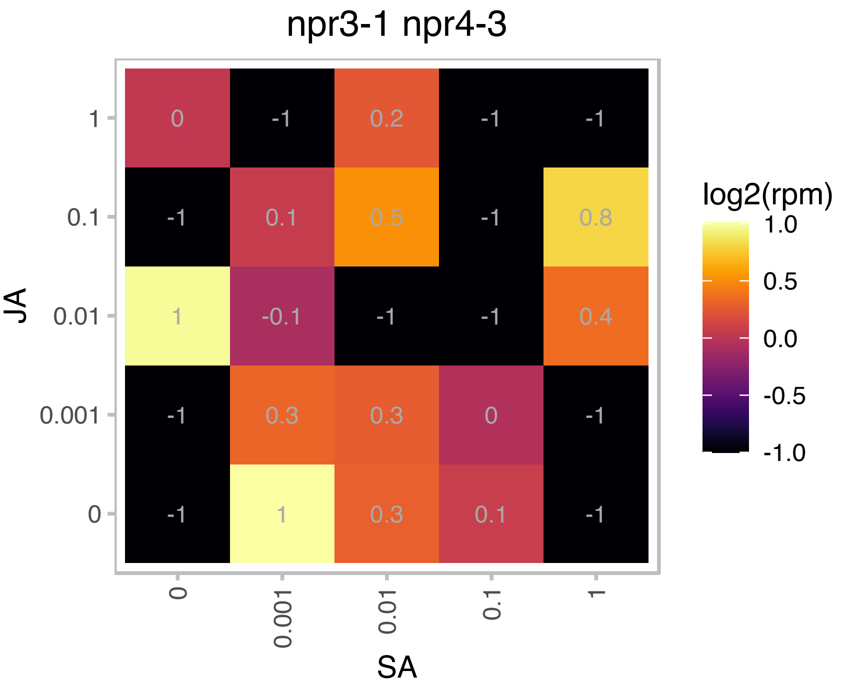

### AT1G48030.1 : mtLPD1

mitochondrial lipoamide dehydrogenase 1

Encodes a mitochondrial lipoamide dehydrogenase whose expression is induced by light.

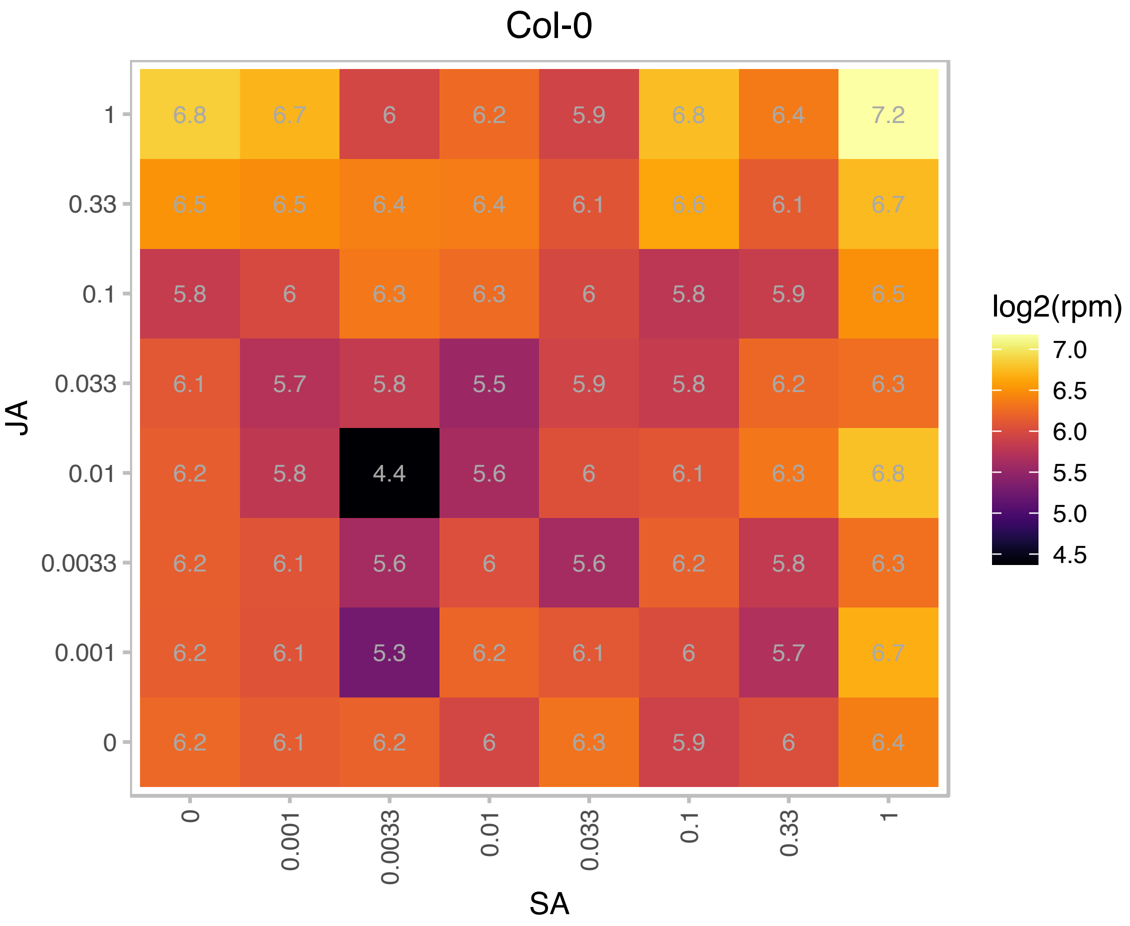


| | |
|---|---|
| Gene | AT1G48030 |
| is.expressed_genes | TRUE |
| is.responsive_genes | TRUE |
| is.apcluster | B-02 |

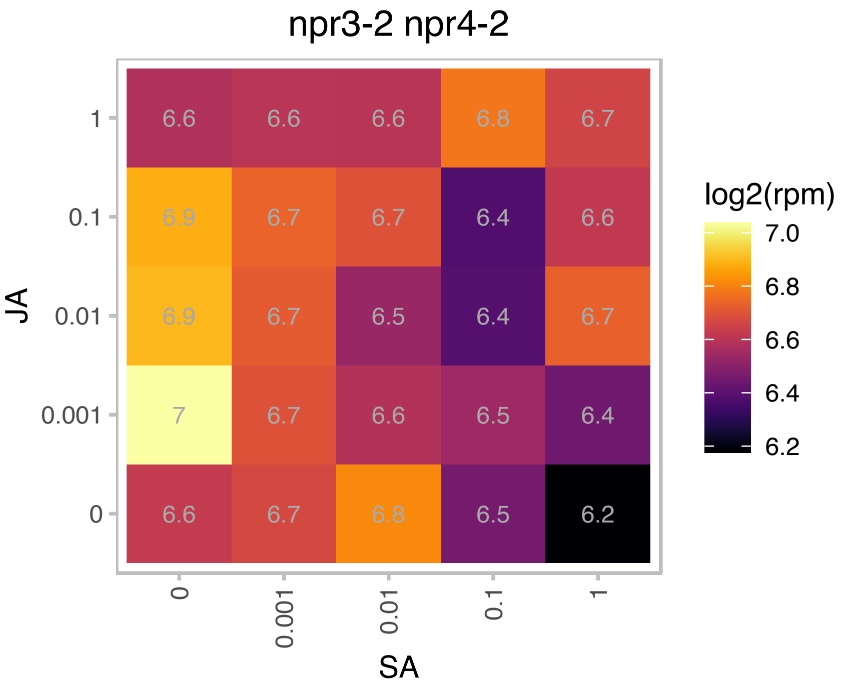


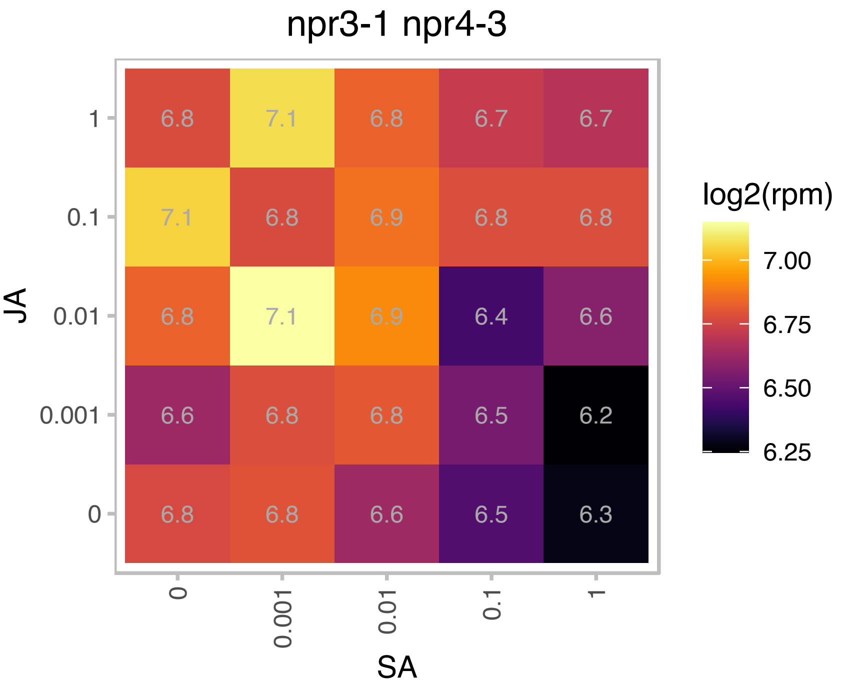

### AT1G49620.1 : AtKRP7, ICK5, ICN6, KRP7

Cyclin-dependent kinase inhibitor family protein
Kip-related protein (KRP) gene, encodes CDK (cyclin-dependent kinase) inhibitor (CKI), negative regulator of cell division. A member of seven KRP genes found in Arabidopsis thaliana. Differential expression patterns for distinct KRPs were revealed by in situ hybridization. Binds to D type cyclins and may inhibit cell cycle.

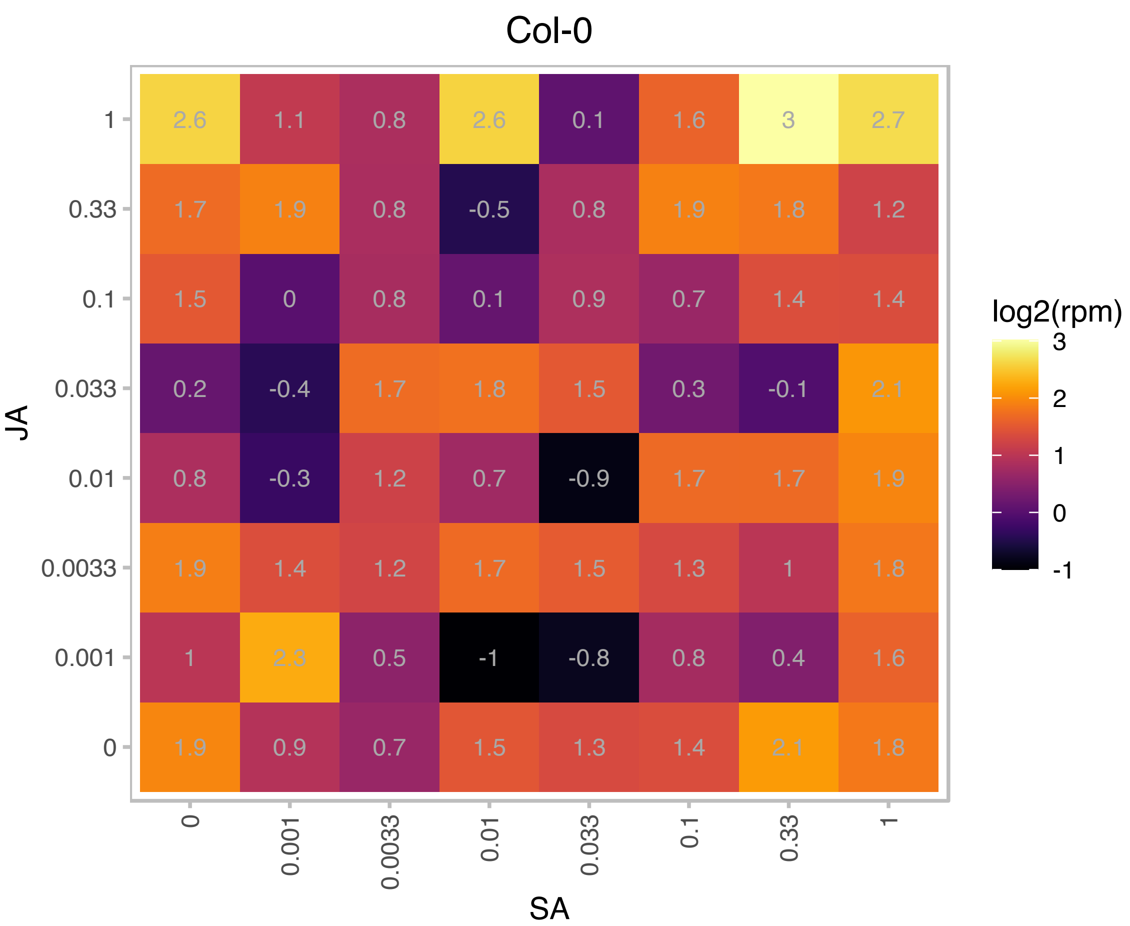


| | |
|---|---|
| Gene | AT1G49620 |
| is.expressed_genes | TRUE |
| is.responsive_genes | TRUE |
| is.apcluster | B-02 |

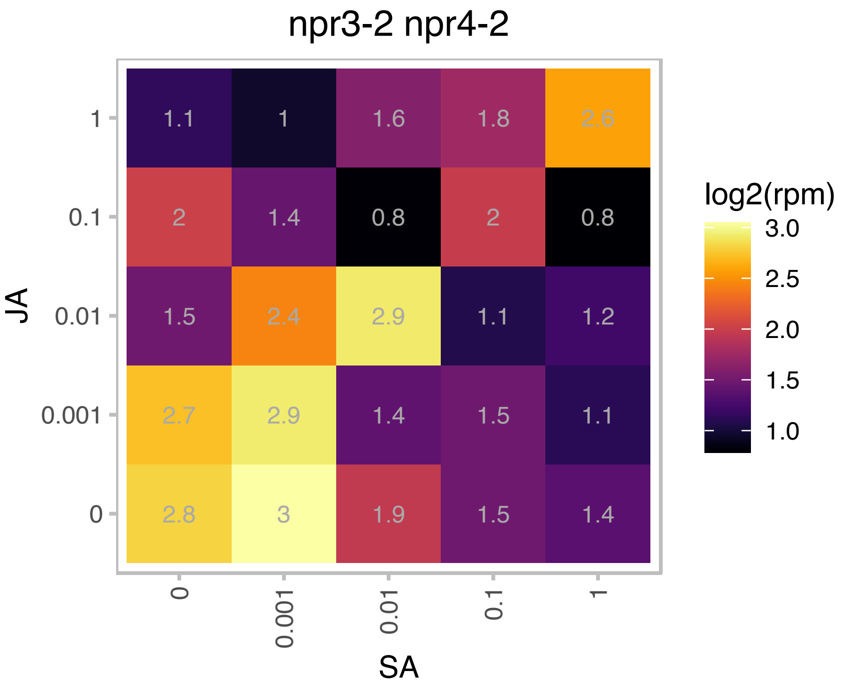


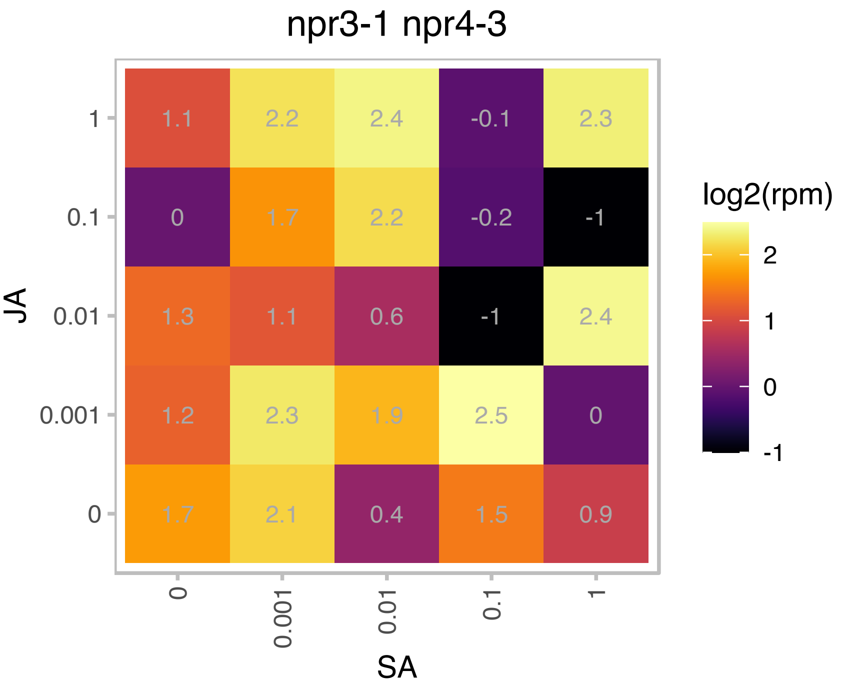

### AT1G56500.1 : SOQ1

haloacid dehalogenase-like hydrolase family protein

Encodes a thylakoid membrane protein with thioredoxin-like and beta-propeller domains located in the lumen and a haloacid-dehalogenase domain exposed to the chloroplast stroma. The protein's role may be to prevent formation of a slowly reversible form of antenna quenching, thereby maintaining the efficiency of light harvesting. The mRNA is cell-to-cell mobile.

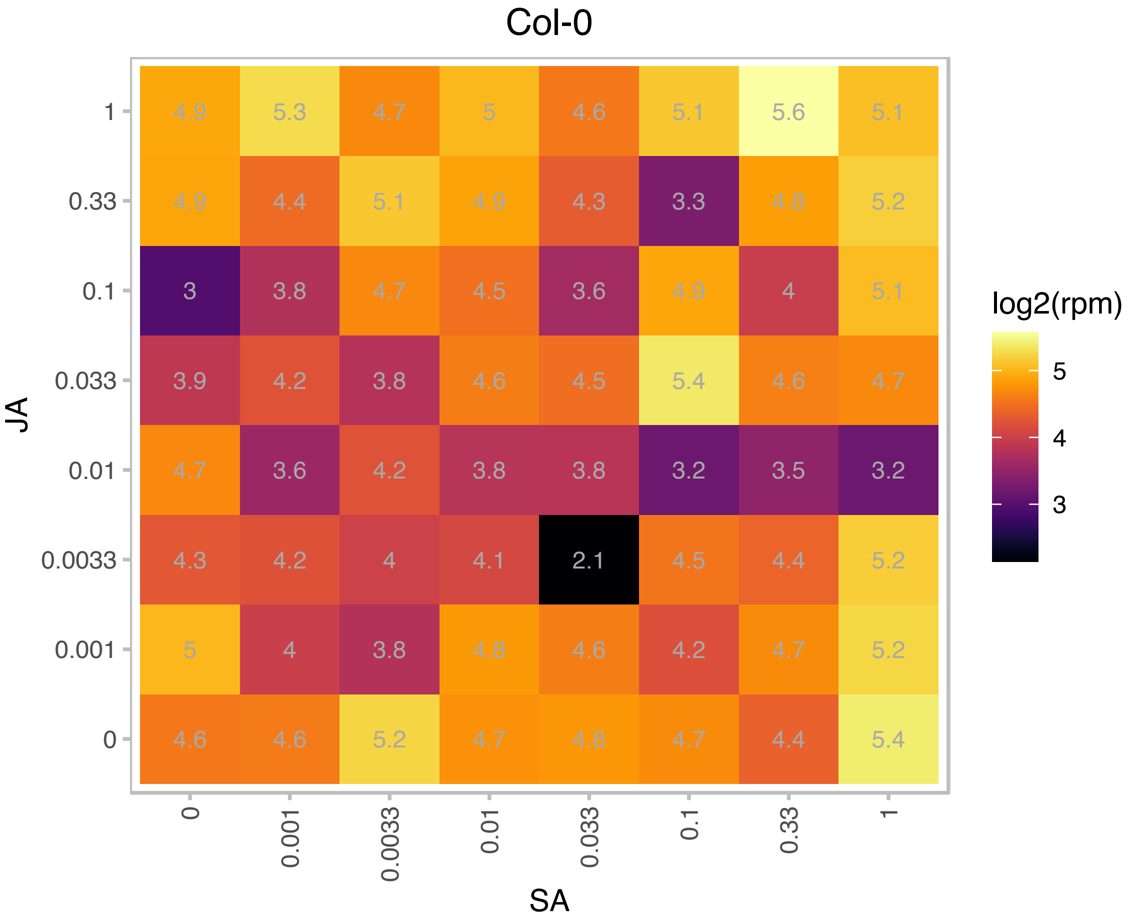


| | |
|---|---|
| Gene | AT1G56500 |
| is.expressed_genes | TRUE |
| is.responsive_genes | TRUE |
| is.apcluster | B-02 |

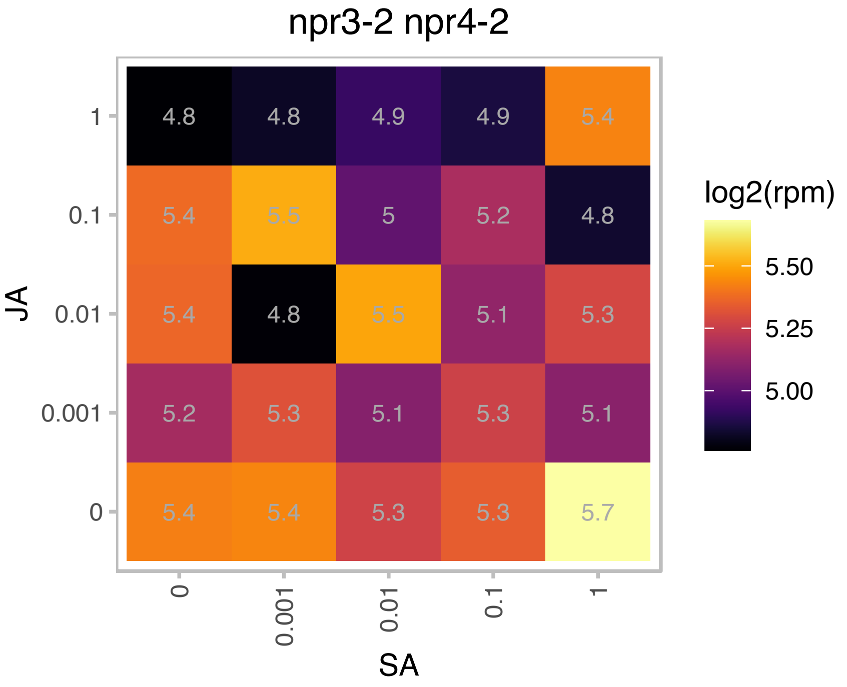


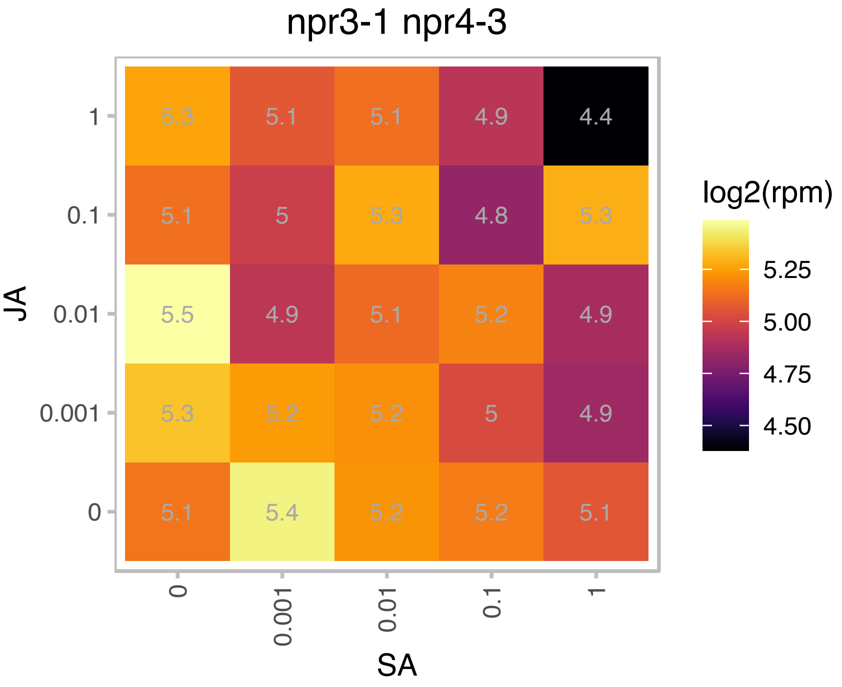

### AT1G69270.1 : AtRPK1, RPK1

receptor-like protein kinase 1

RPK1 is a leucine-rich receptor-like kinase located in the plasma membrane which is upregulated by abscisic acid, dehydration, high salt, low temperature, but not by other plant hormones. RPK1 knock-out and antisense plants show an ABA-insensitive phenotype. RPK1 plays a role in ABA-controlled cell proliferation and is a regulator of the ABA signal transduction pathway. Overexpression of the LRR domain has a dominant negative effect on RPK1. Mutations in RPK1 uncouple cotyledon anlagen and primordia by modulating epidermal cell shape and polarity.

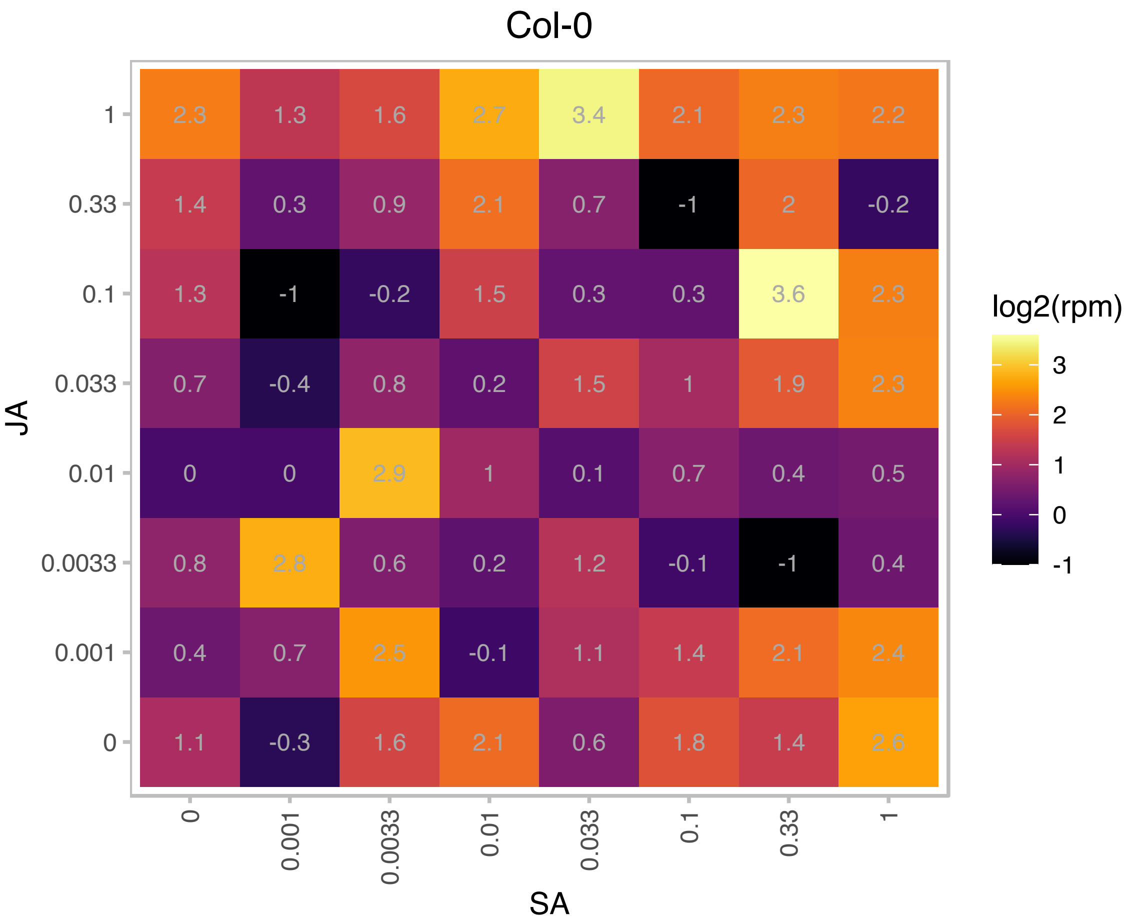


| | |
|---|---|
| Gene | AT1G69270 |
| is.expressed_genes | TRUE |
| is.responsive_genes | TRUE |
| is.apcluster | B-02 |

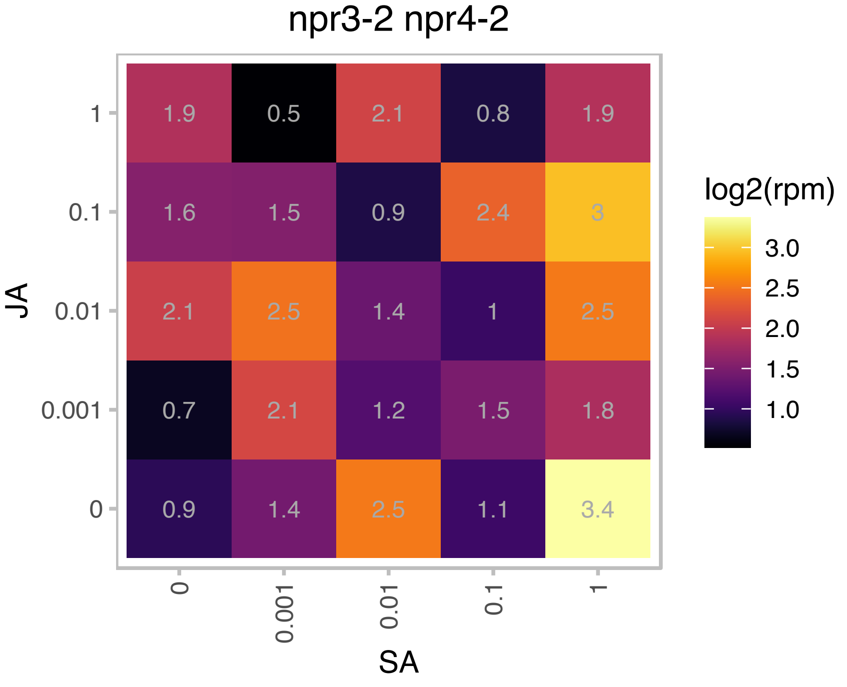


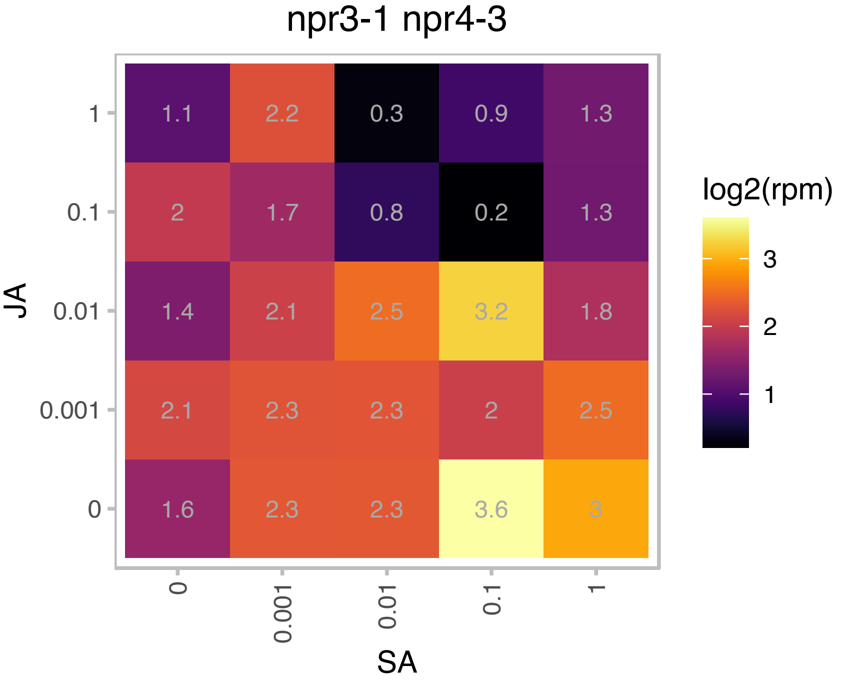

### AT1G72150.1 : PATL1

PATELLIN 1

novel cell-plate-associated protein that is related in sequence to proteins involved in membrane trafficking in other eukaryotes The mRNA is cell-to-cell mobile.

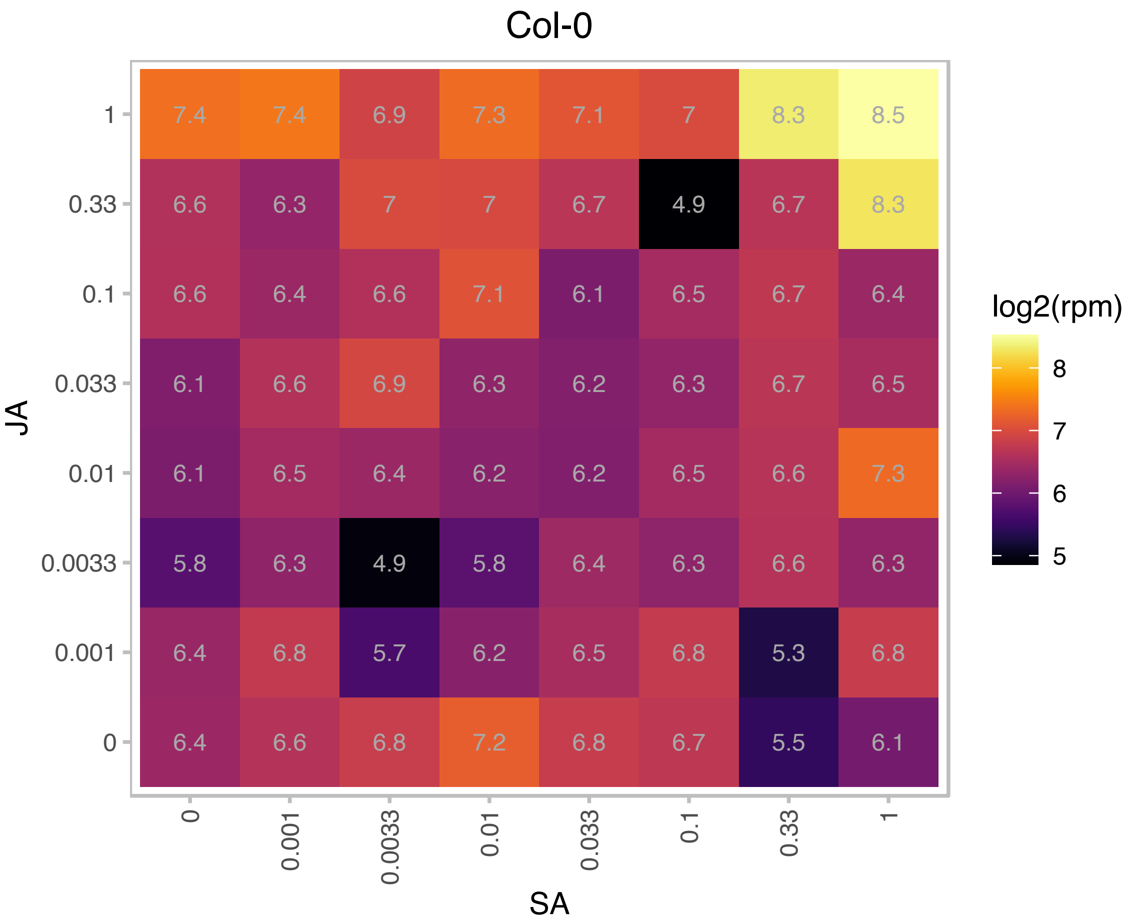


| | |
|---|---|
| Gene | AT1G72150 |
| is.expressed_genes | TRUE |
| is.responsive_genes | TRUE |
| is.apcluster | B-02 |

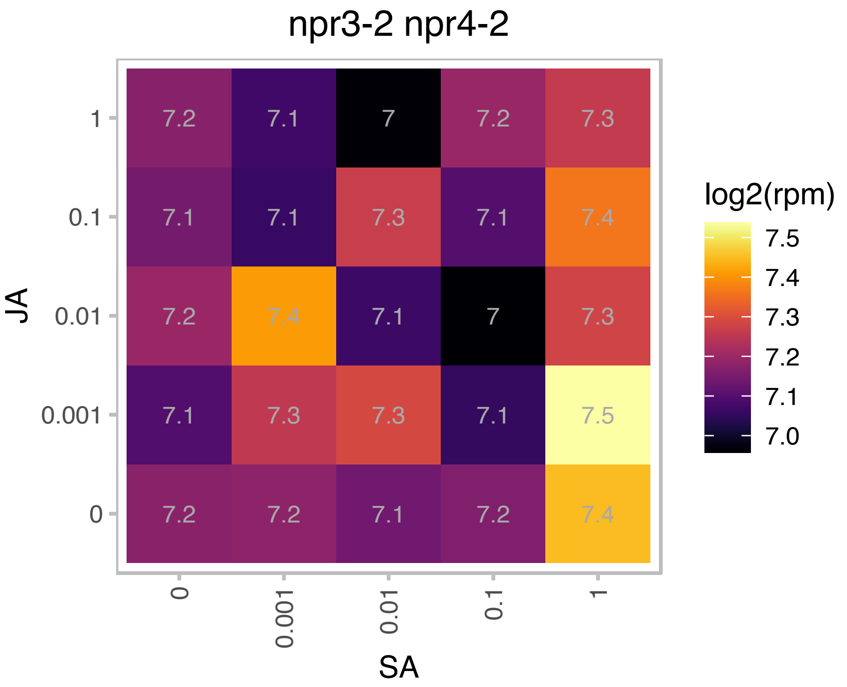


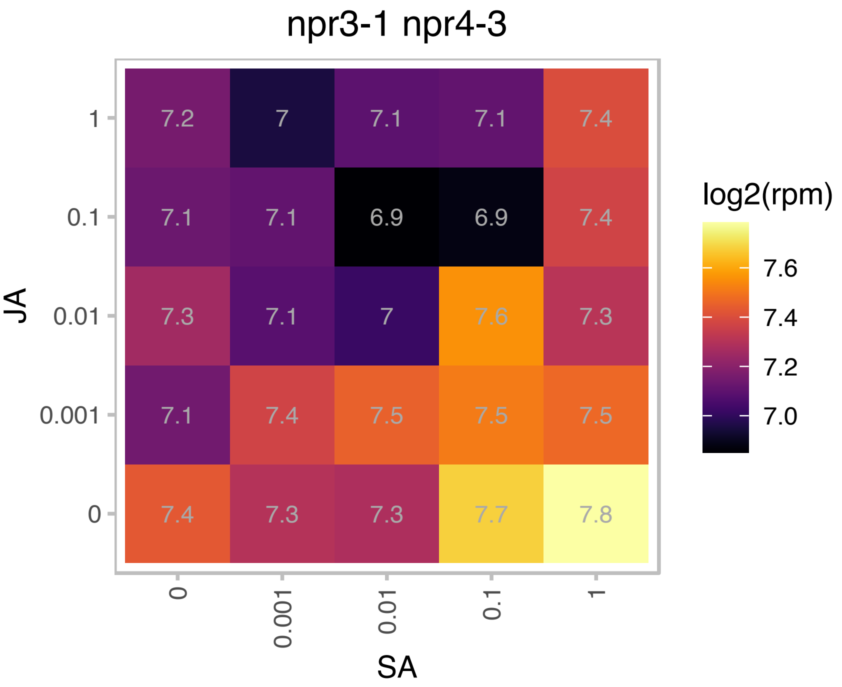

# AT1G73240.1 : NA

Col-0

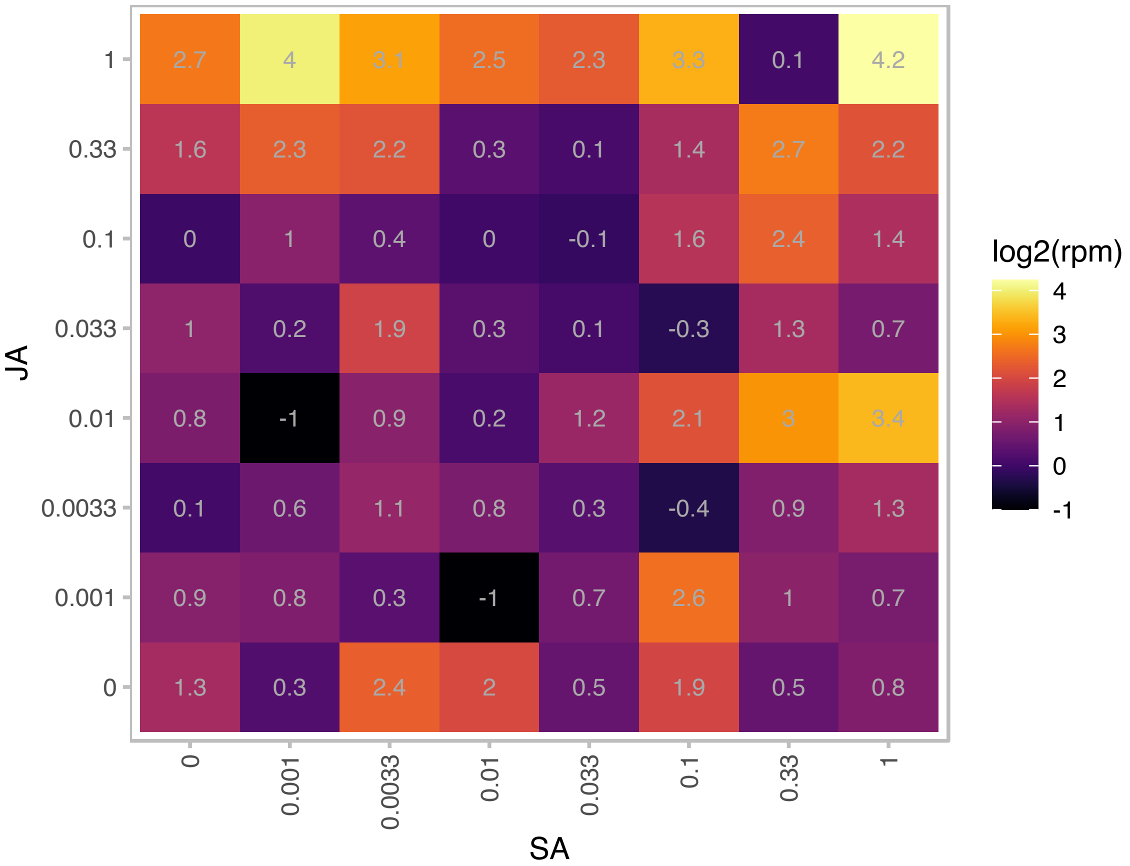


| | |
|---|---|
| Gene | AT1G73240 |
| is.expressed_genes | TRUE |
| is.responsive_genes | TRUE |
| is.apcluster | B-02 |

npr3-2 npr4-2

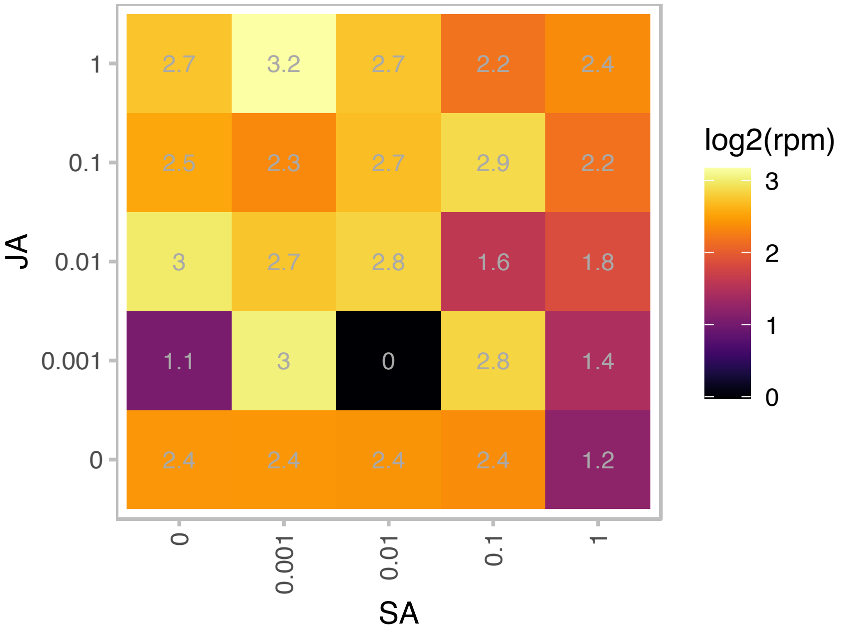


npr3-1 npr4-3

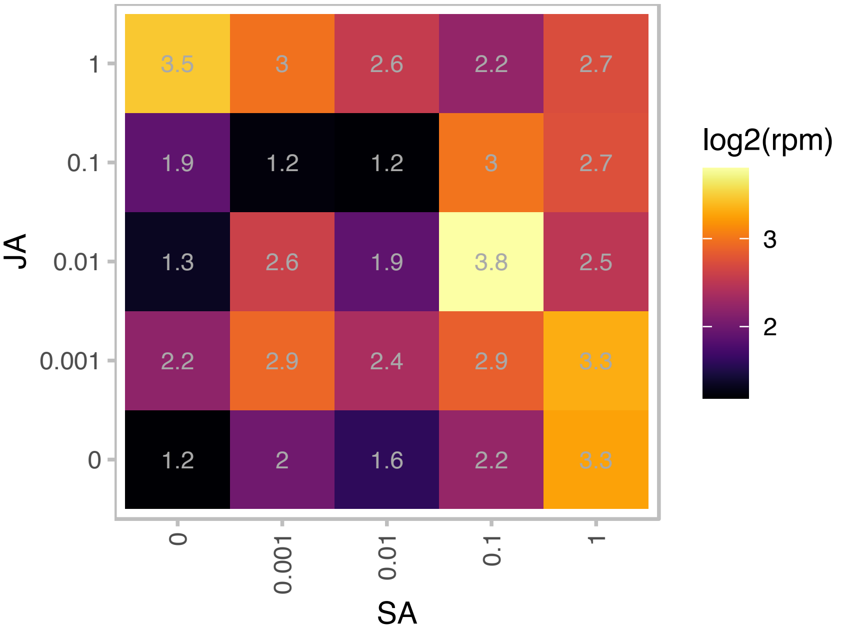

# AT1G74250.1 : NA

DNAJ heat shock N-terminal domain-containing protein

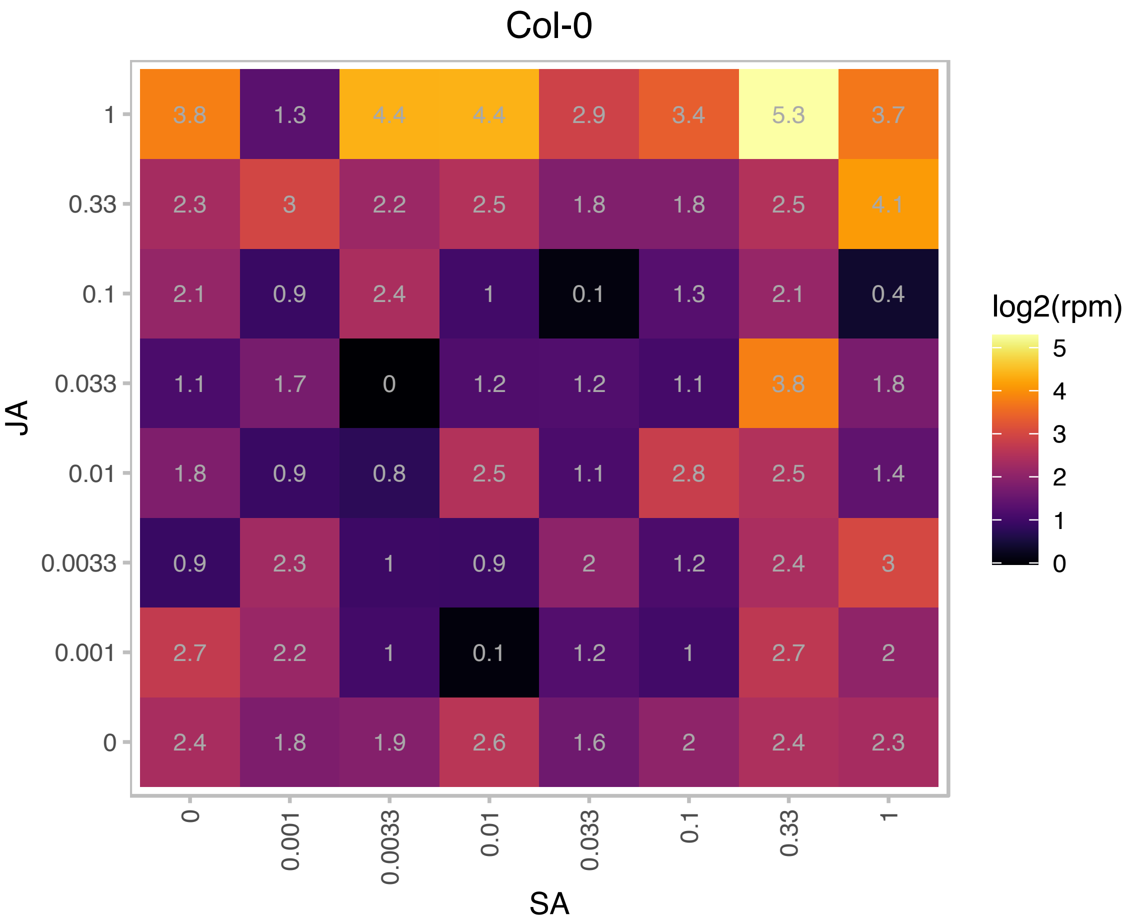


| | |
|---|---|
| Gene | AT1G74250 |
| is.expressed_genes | TRUE |
| is.responsive_genes | TRUE |
| is.apcluster | B-02 |

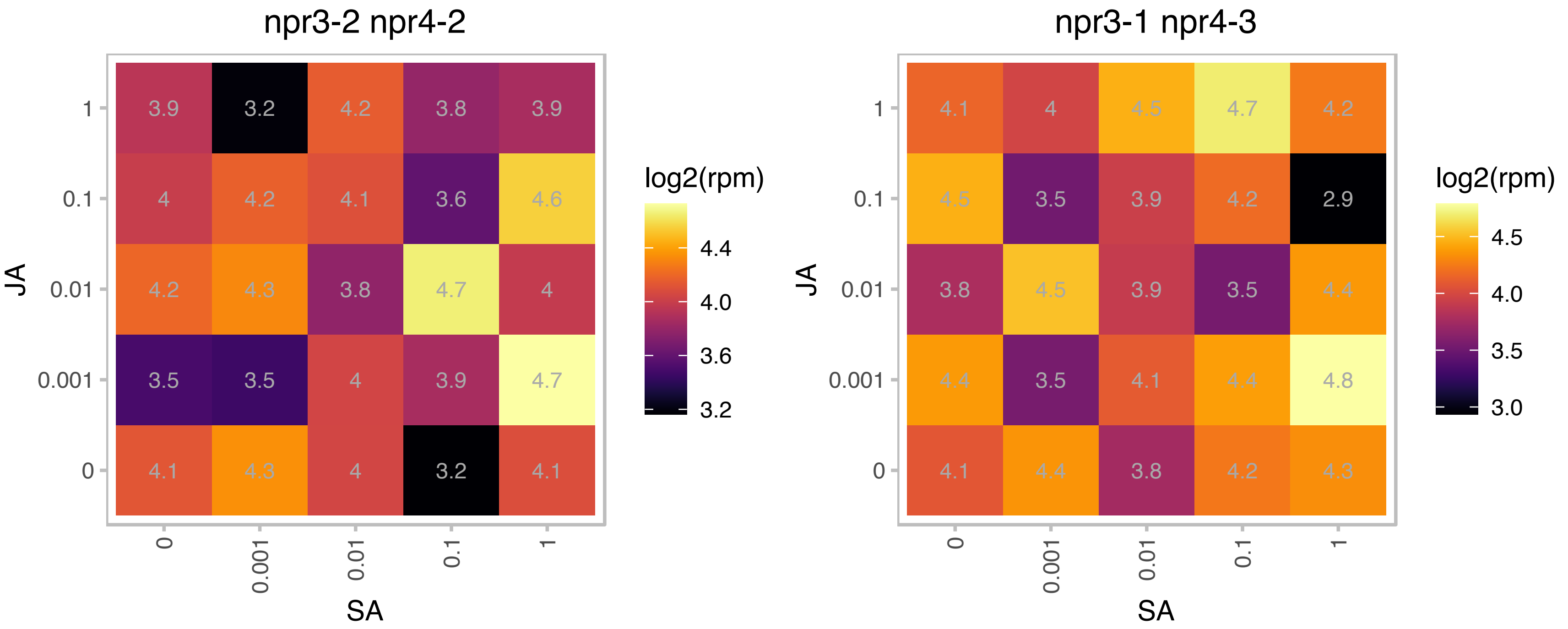

### AT1G75750.1 : GASA1

GAST1 protein homolog 1

GA-responsive GAST1 protein homolog regulated by BR and GA antagonistically.  Possibly involved in cell elongation based on expression data The mRNA is cell-to-cell mobile.

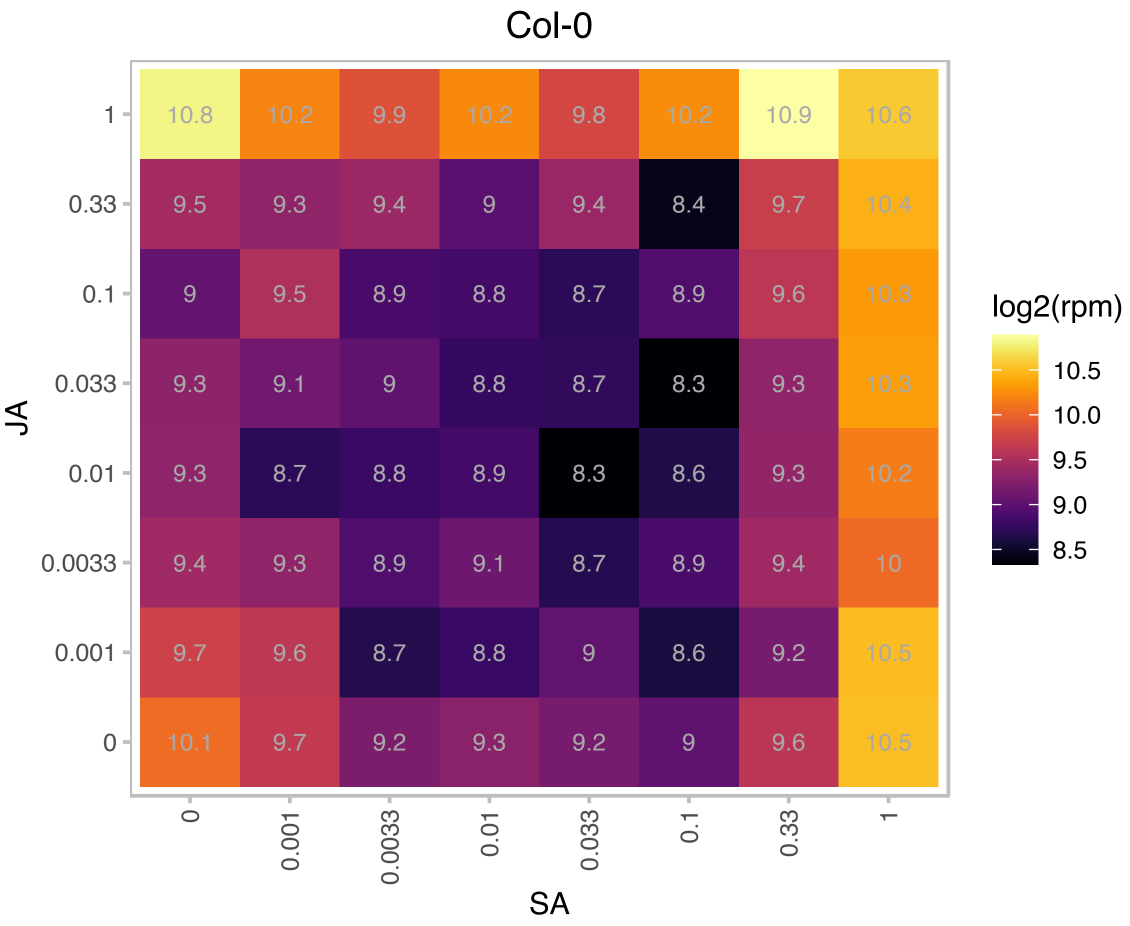


| | |
|---|---|
| Gene | AT1G75750 |
| is.expressed_genes | TRUE |
| is.responsive_genes | TRUE |
| is.apcluster | B-02 |

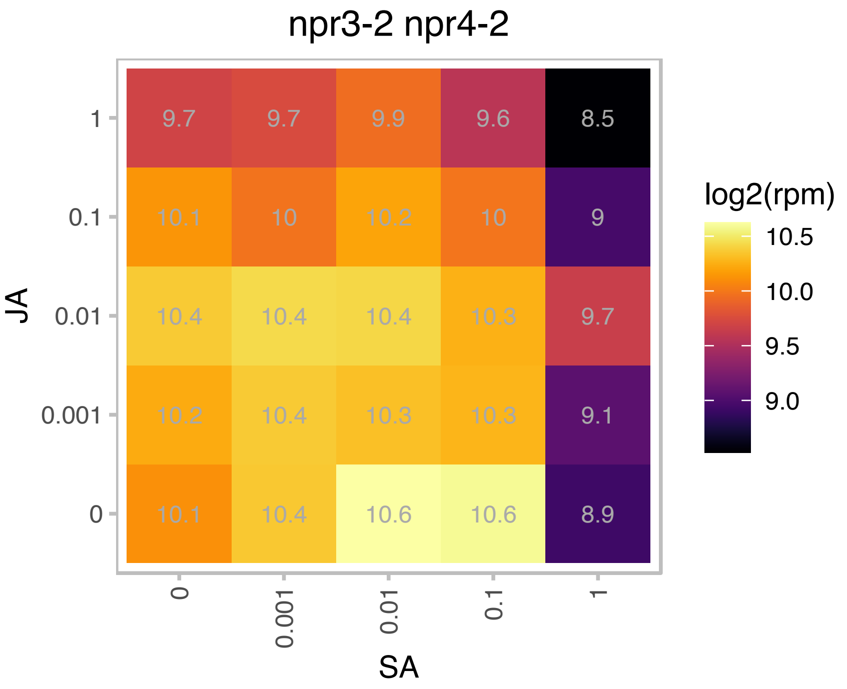


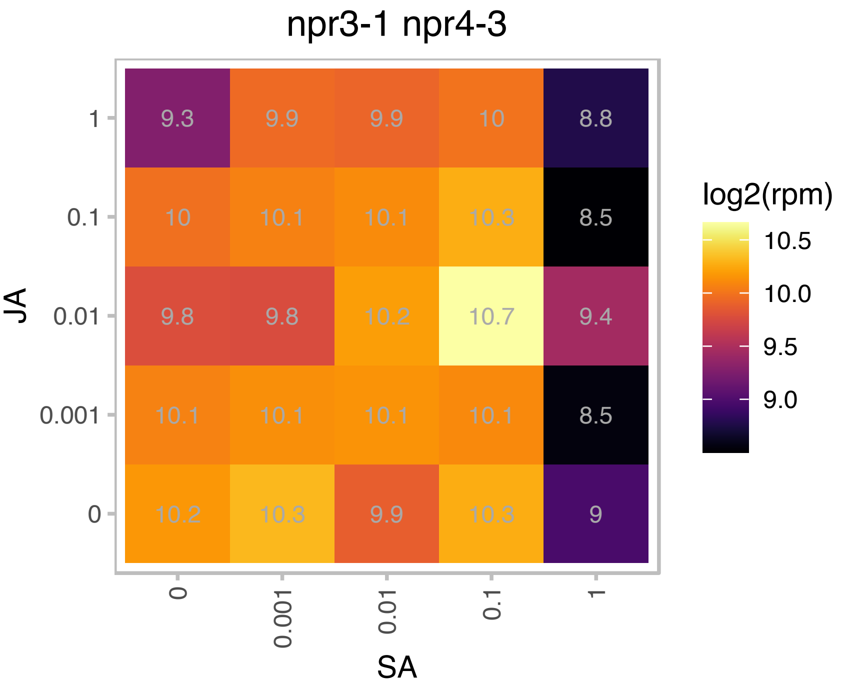

### AT1G78600.1 : BBX22, DBB3, LZF1, STH3

light-regulated zinc finger protein 1

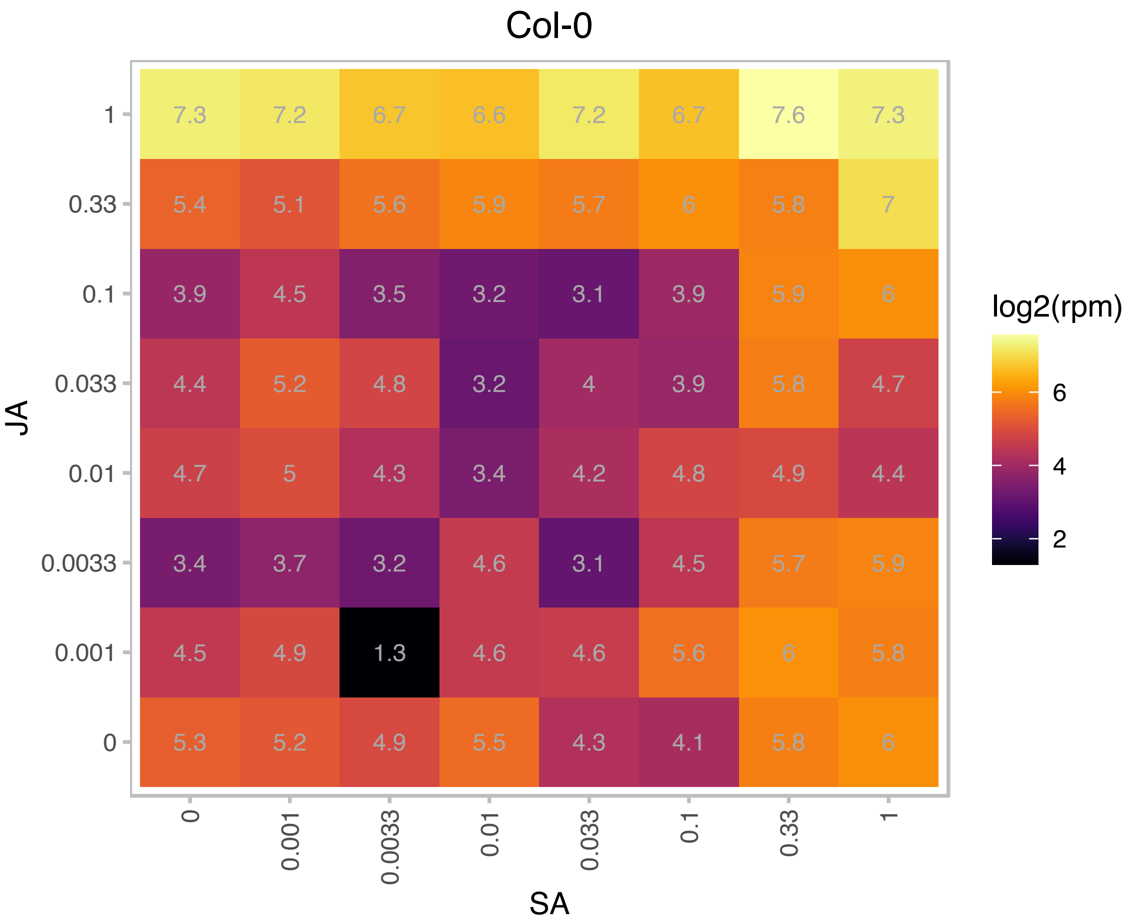


| | |
|---|---|
| Gene | AT1G78600 |
| is.expressed_genes | TRUE |
| is.responsive_genes | TRUE |
| is.apcluster | B-02 |

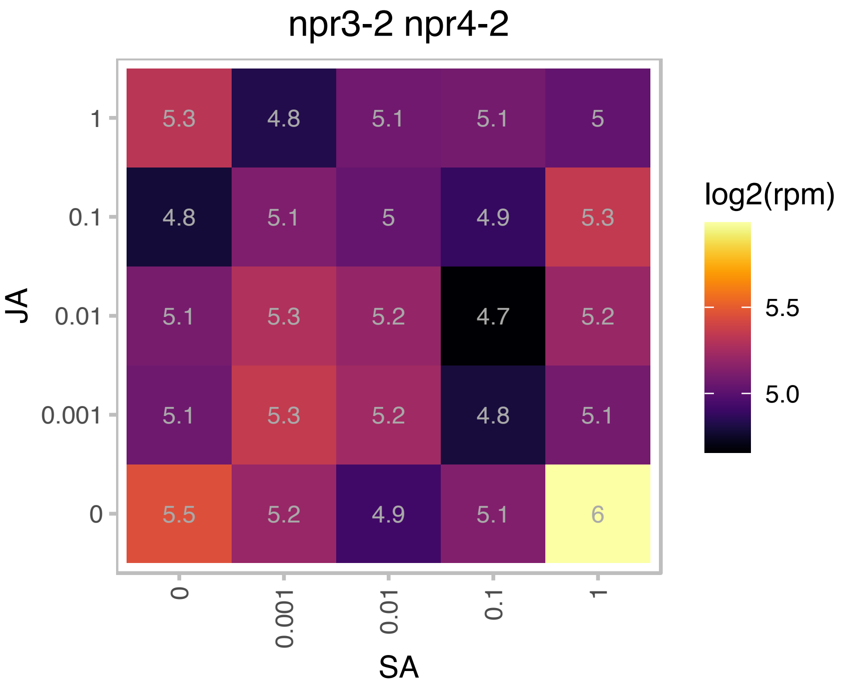


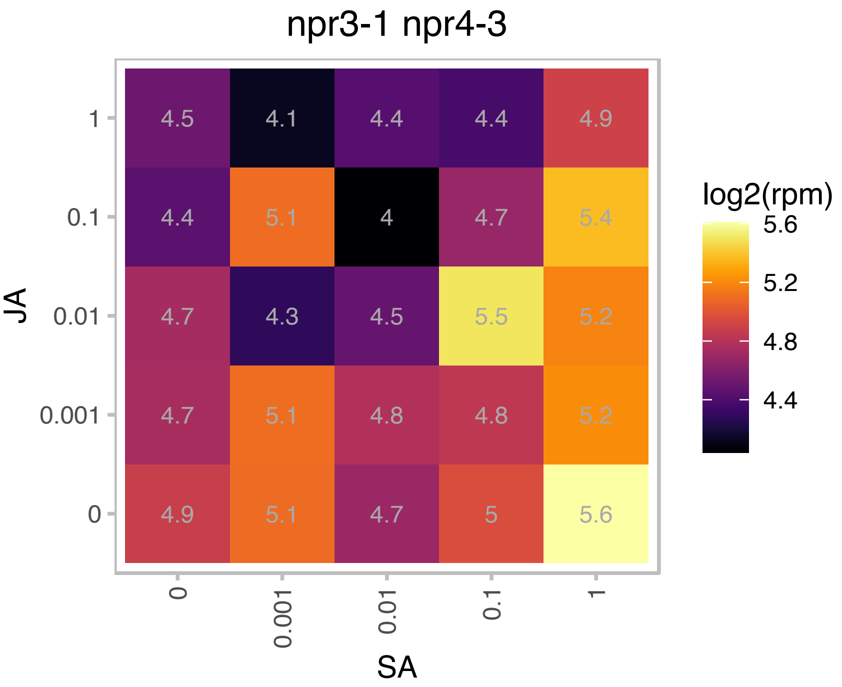

### AT1G79410.1 : AtOCT5, OCT5

organic cation/carnitine transporter5

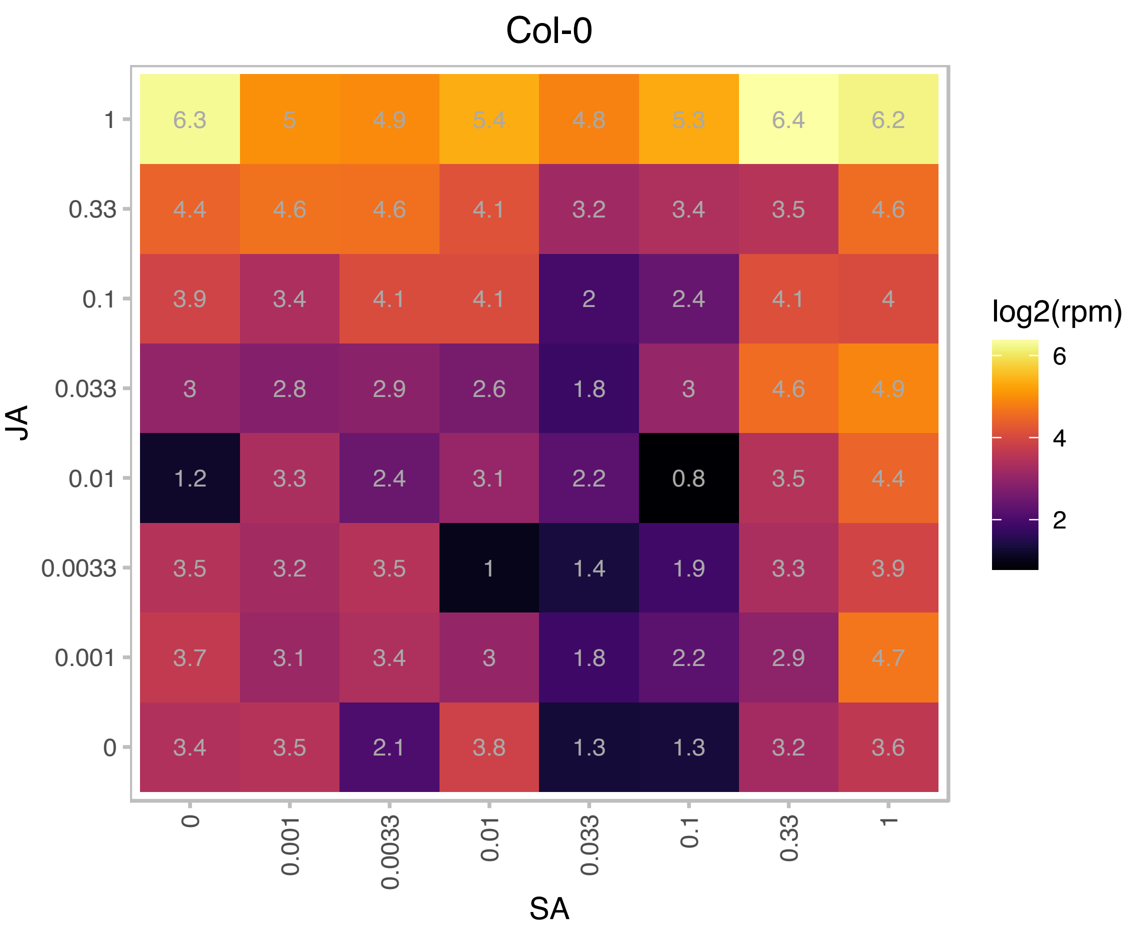


| | |
|---|---|
| Gene | AT1G79410 |
| is.expressed_genes | TRUE |
| is.responsive_genes | TRUE |
| is.apcluster | B-02 |

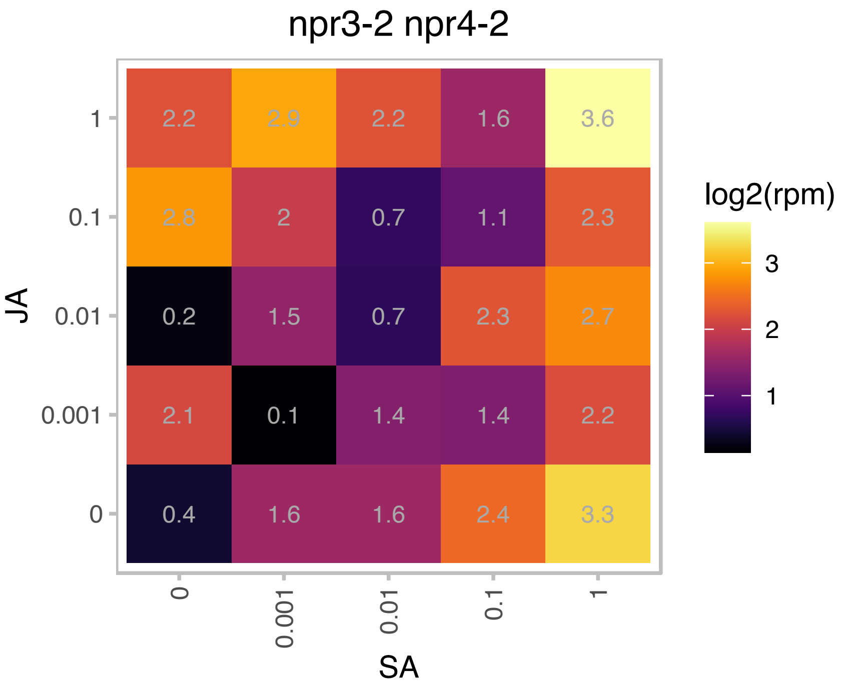


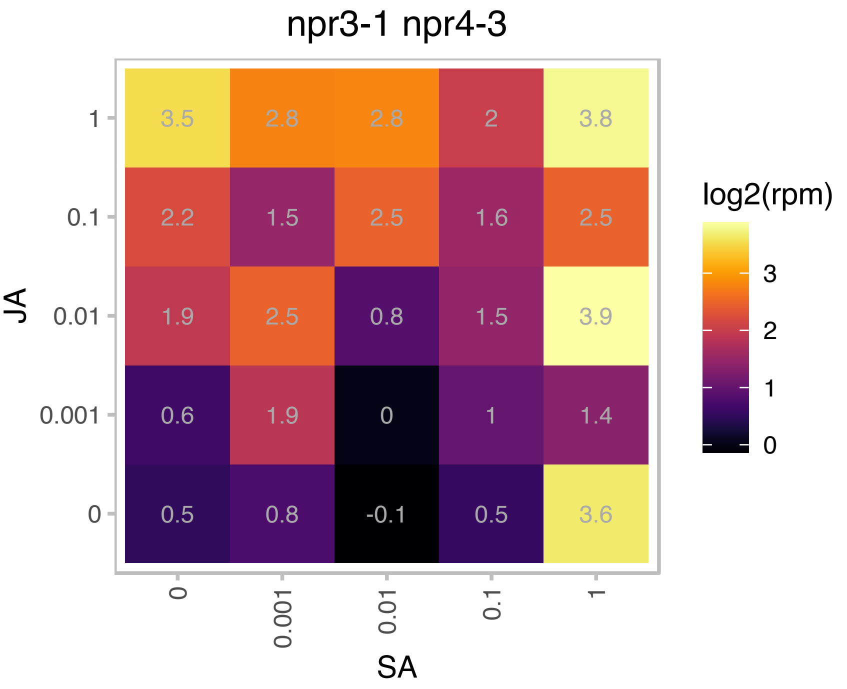

### AT1G80380.1 : GLYK

P-loop containing nucleoside triphosphate hydrolases superfamily protein

encodes a glycerate kinase which catalyzes the last step of photorespiration C2 cycle.

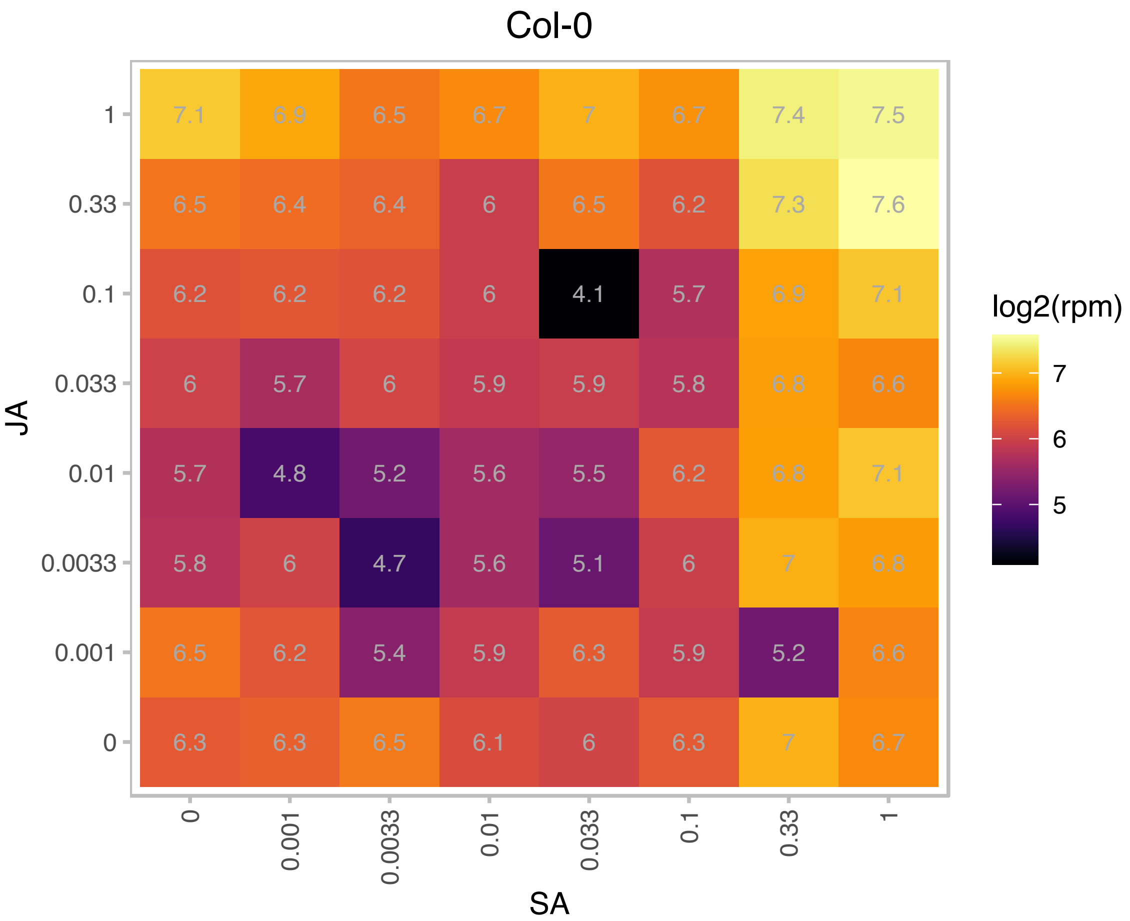


| | |
|---|---|
| Gene | AT1G80380 |
| is.expressed_genes | TRUE |
| is.responsive_genes | TRUE |
| is.apcluster | B-02 |

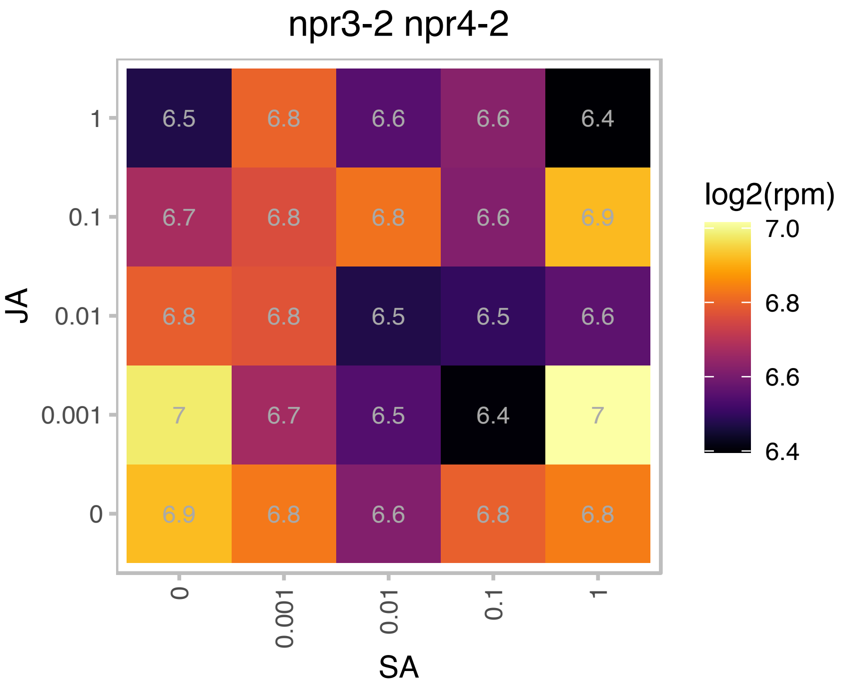


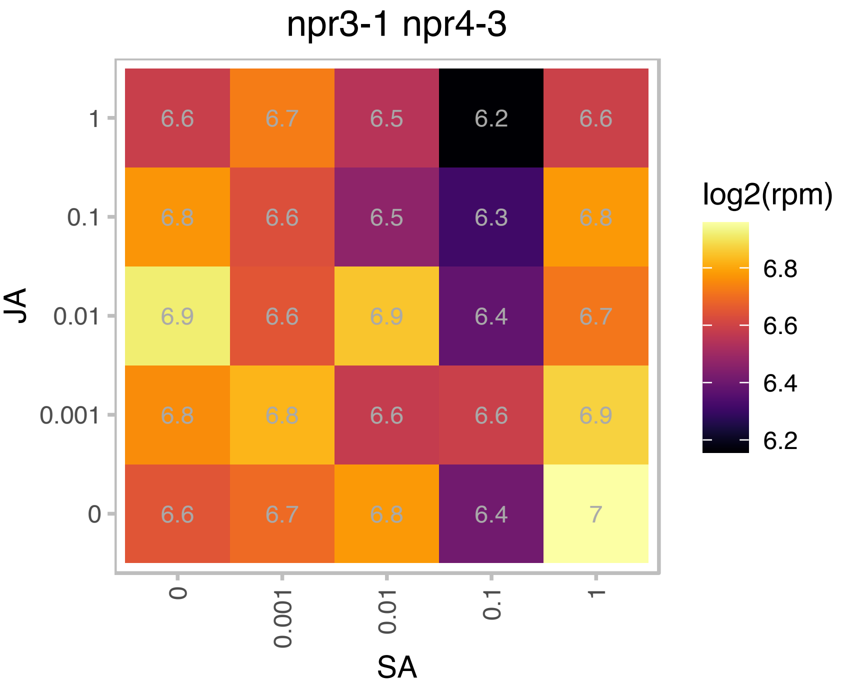

### AT2G01850.1 : ATXTH27, EXGT-A3, XTH27

endoxyloglucan transferase A3

EXGT-A3 has homology to xyloglucan endotransglucosylases/hydrolases (XTHs). Mutants in this gene show a lesion mimic phenotype associated with leaf maturation and a reduction in the number of tertiary veins. Individual tracheary elements in the mutants are shorter, but phloem transport activity is not severely affected. EXGT-A3 plays a role in xyloglucan degradation in the differentiating tracheary elements of rosette leaves. The mRNA is cell-to-cell mobile.

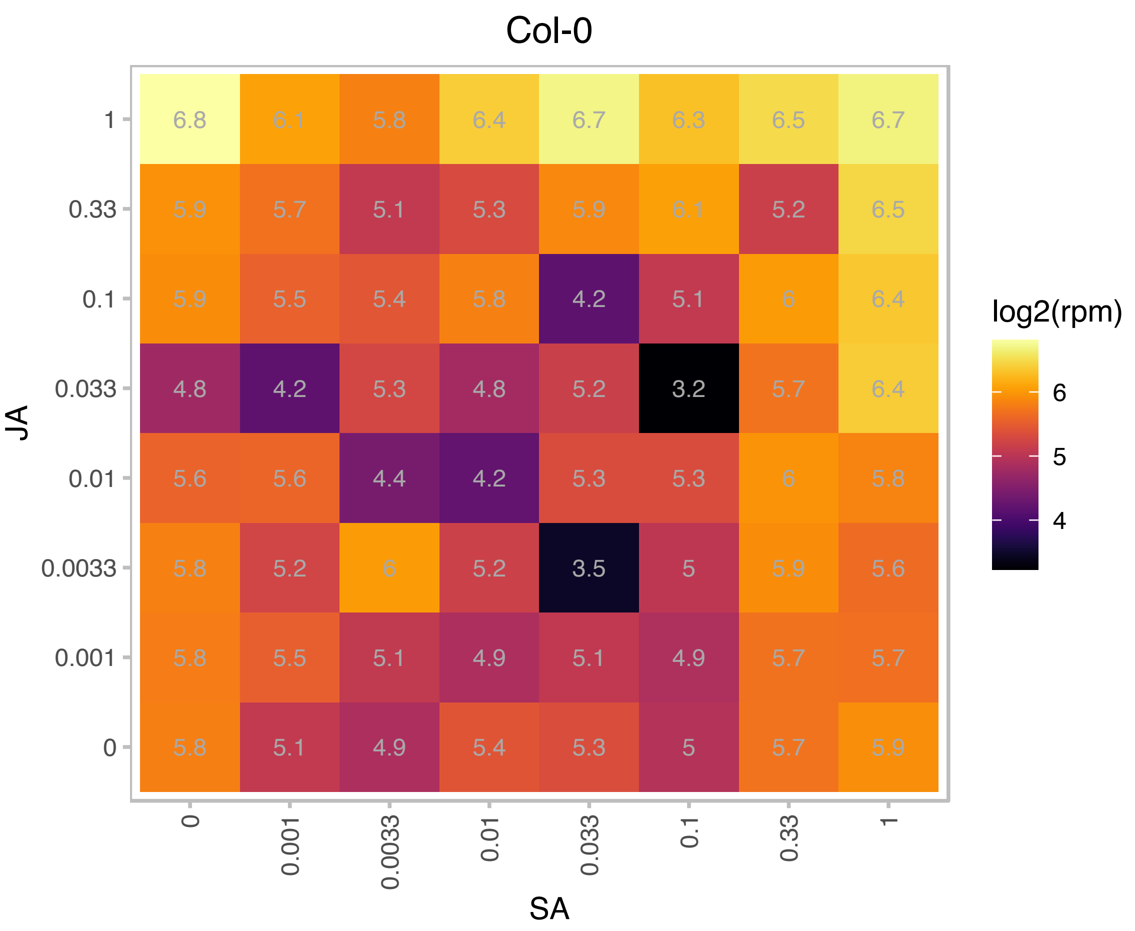


| | |
|---|---|
| Gene | AT2G01850 |
| is.expressed_genes | TRUE |
| is.responsive_genes | TRUE |
| is.apcluster | B-02 |

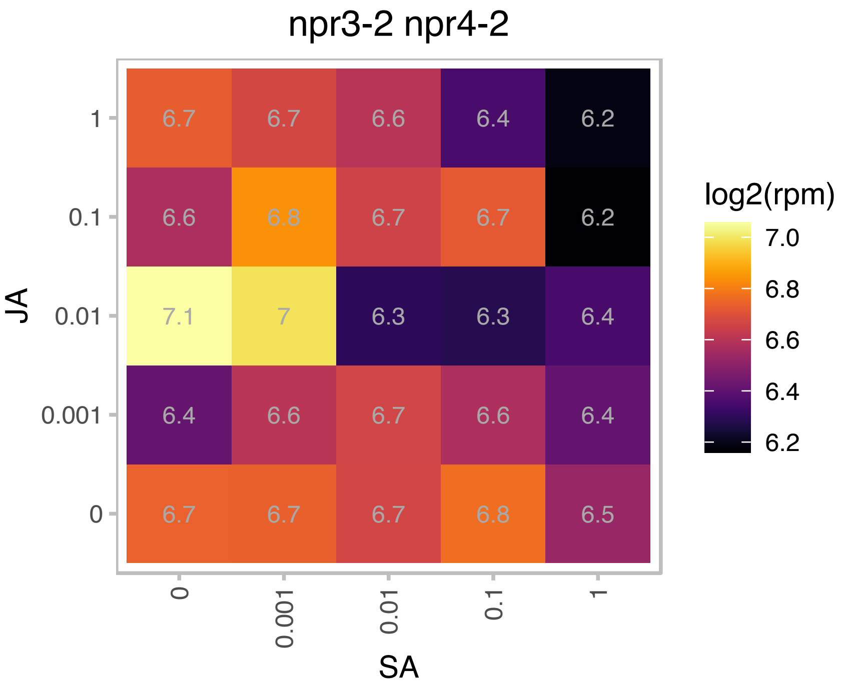


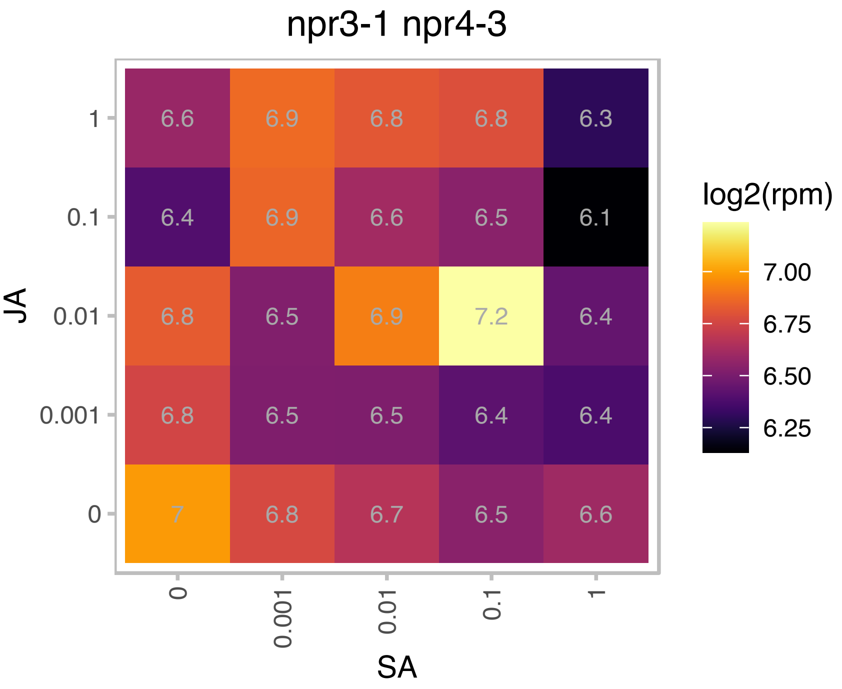

# AT2G04795.1 : NA

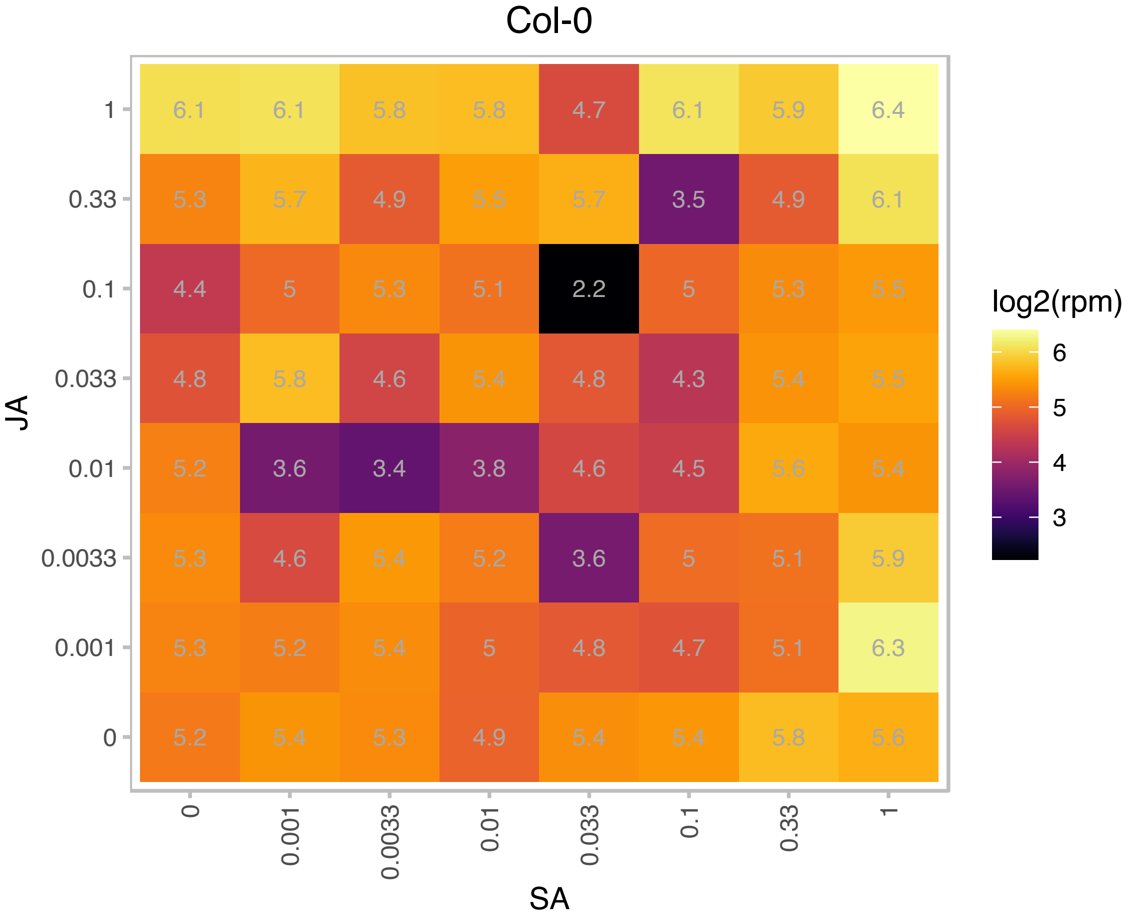


| | |
|---|---|
| Gene | AT2G04795 |
| is.expressed_genes | TRUE |
| is.responsive_genes | TRUE |
| is.apcluster | B-02 |

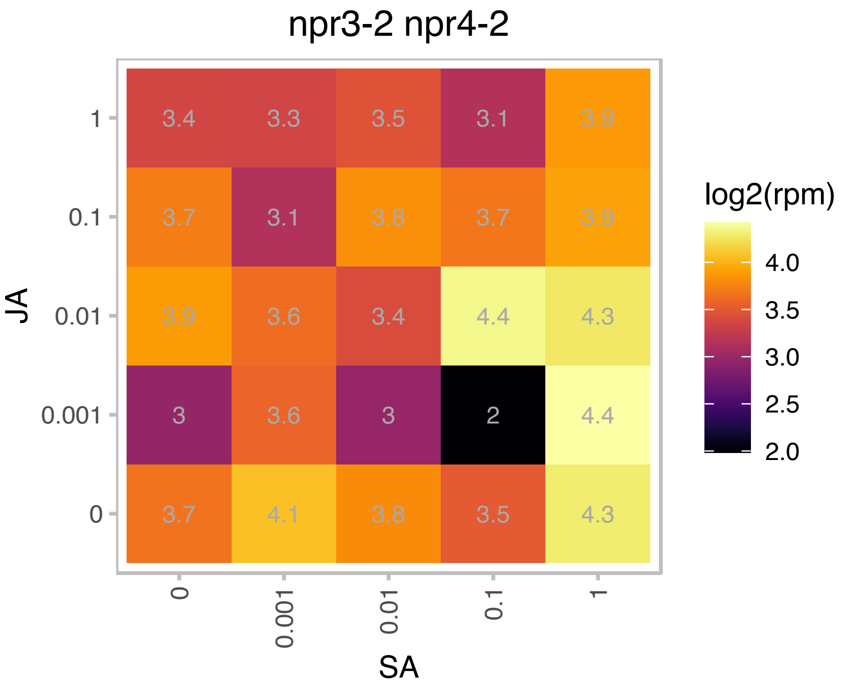


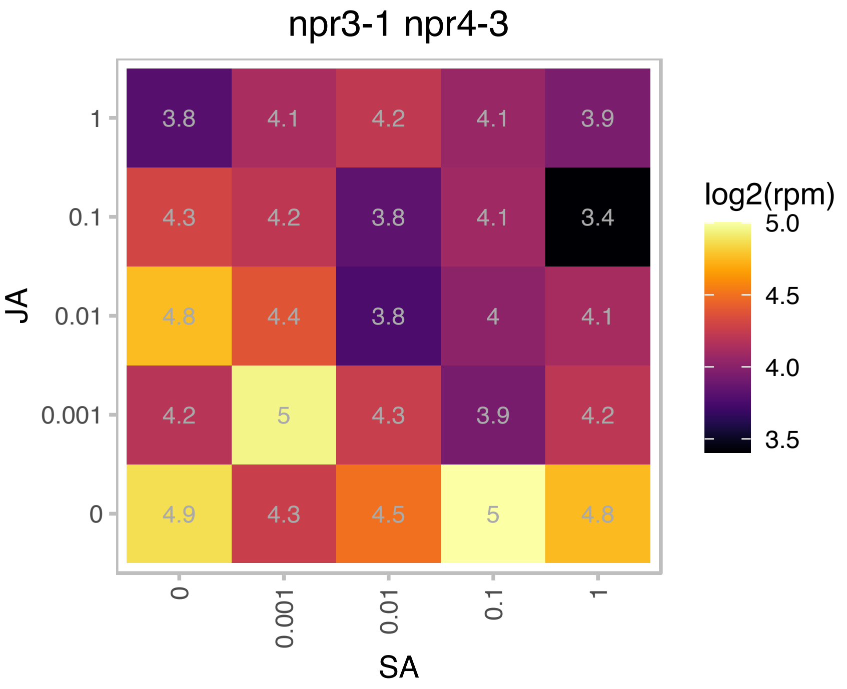

# AT2G20740.1 : NA

Tetraspanin family protein

| | |
|---|---|
| Gene | AT2G20740 |
| is.expressed_genes | TRUE |
| is.responsive_genes | TRUE |
| is.apcluster | B-02 |

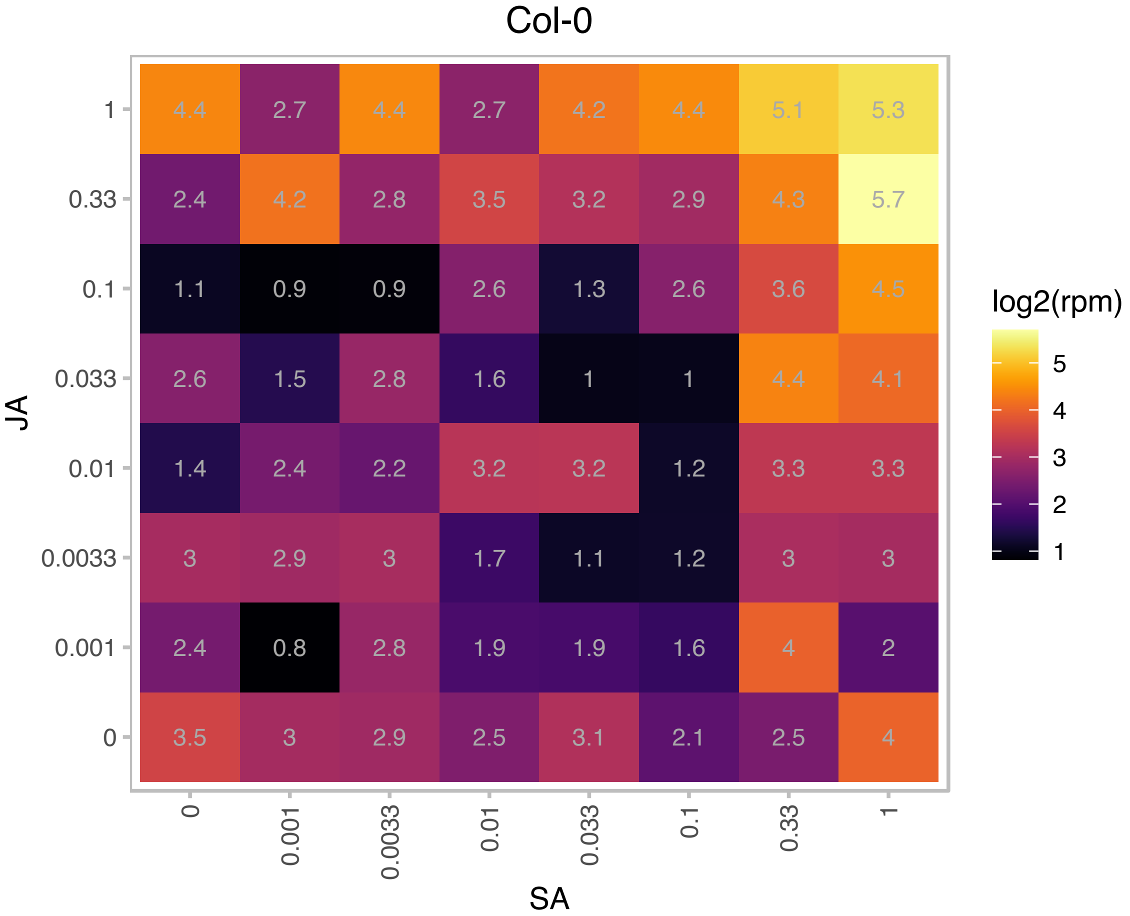


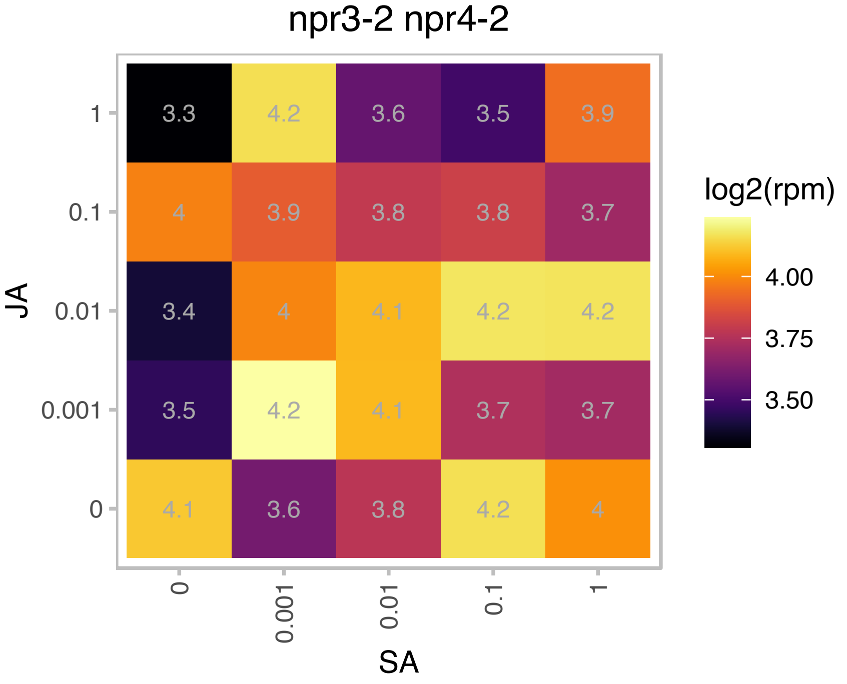


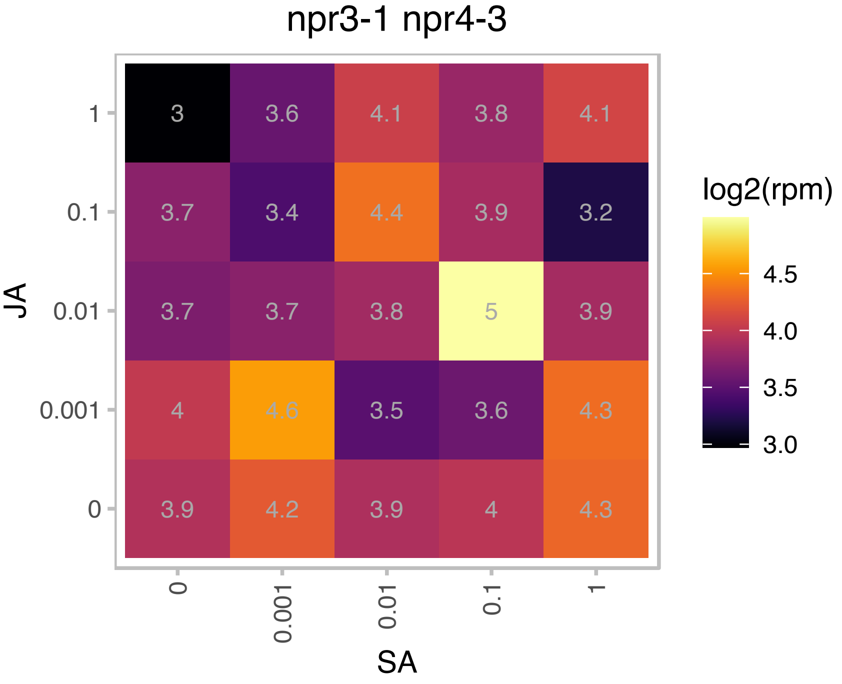

### AT2G25930.1 : ELF3, PYK20

hydroxyproline-rich glycoprotein family protein
Encodes a nuclear protein that is expressed rhythmically and interacts with phytochrome B to control plant development and flowering through a signal transduction pathway. Required component of the core circadian clock regardless of light conditions.

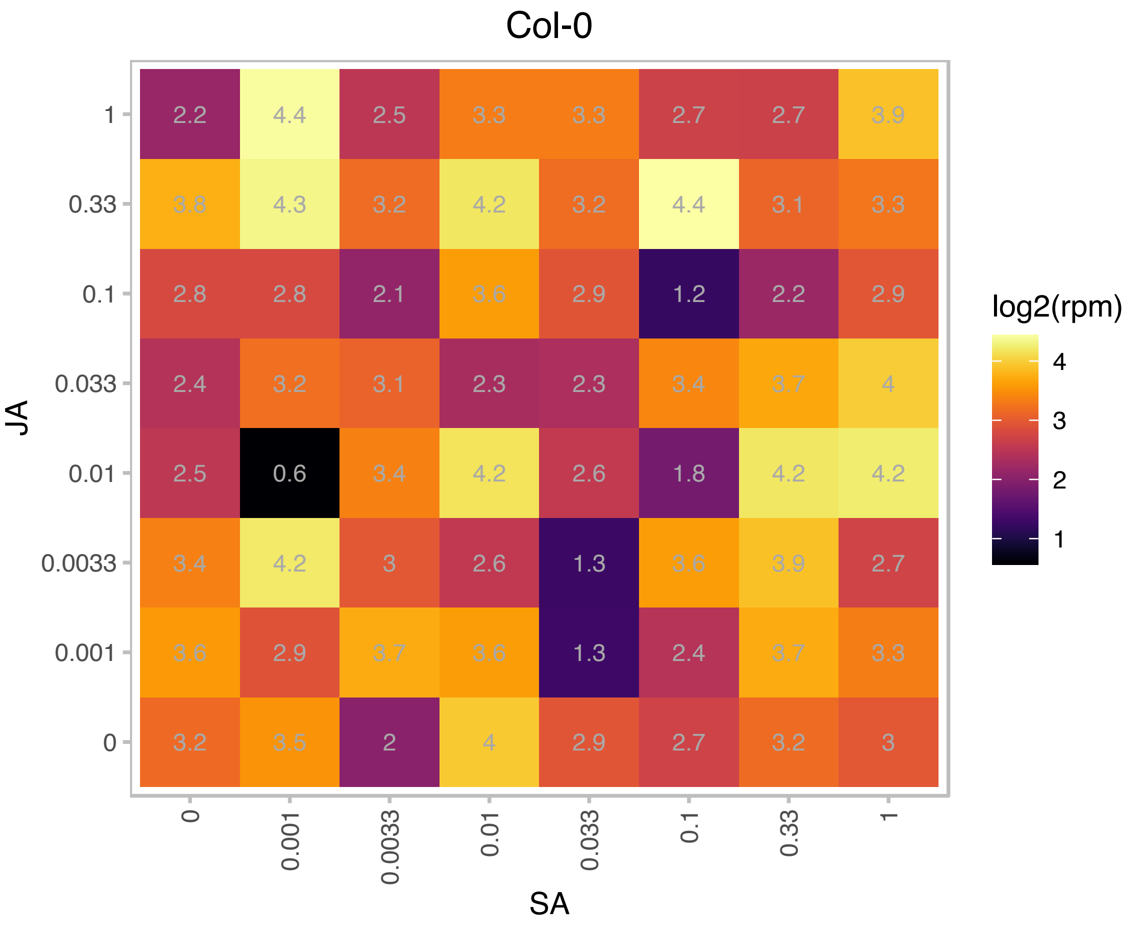


| | |
|---|---|
| Gene | AT2G25930 |
| is.expressed_genes | TRUE |
| is.responsive_genes | TRUE |
| is.apcluster | B-02 |

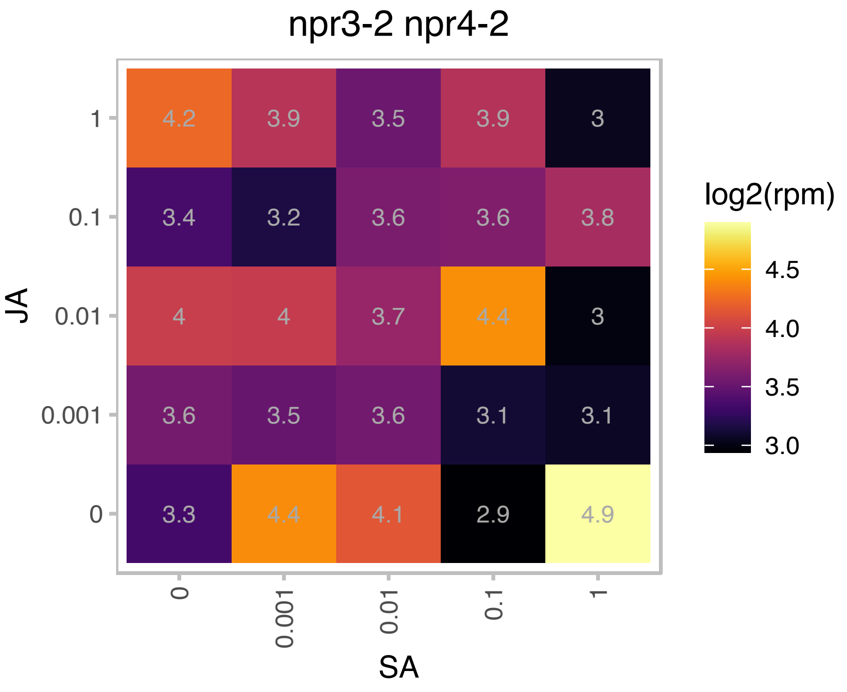


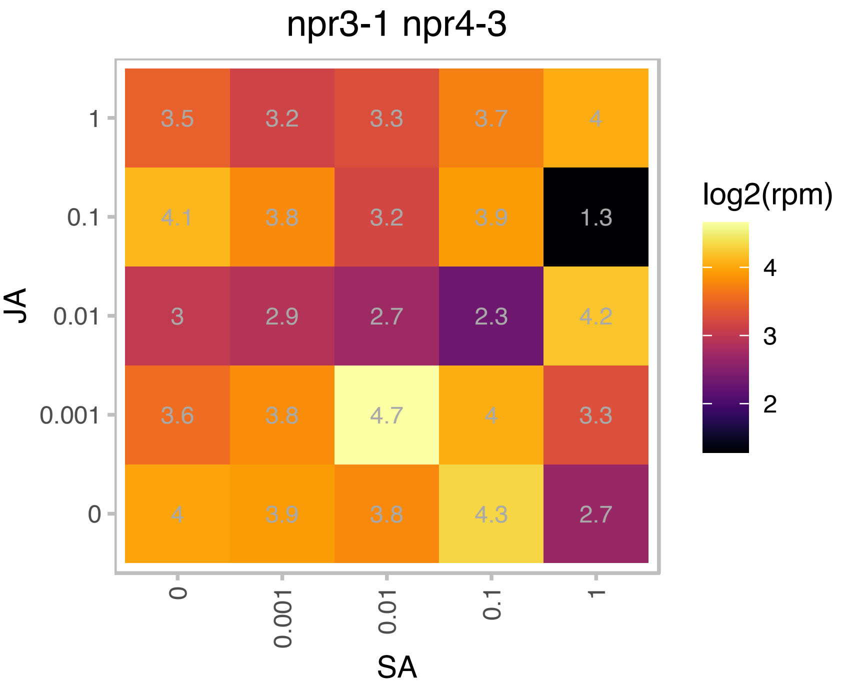

### AT2G31280.1 : CPUORF7, LHL2, LL2

conserved peptide upstream open reading frame 7

Encodes a LHW-like protein with 80% amino acid identity to LHW.

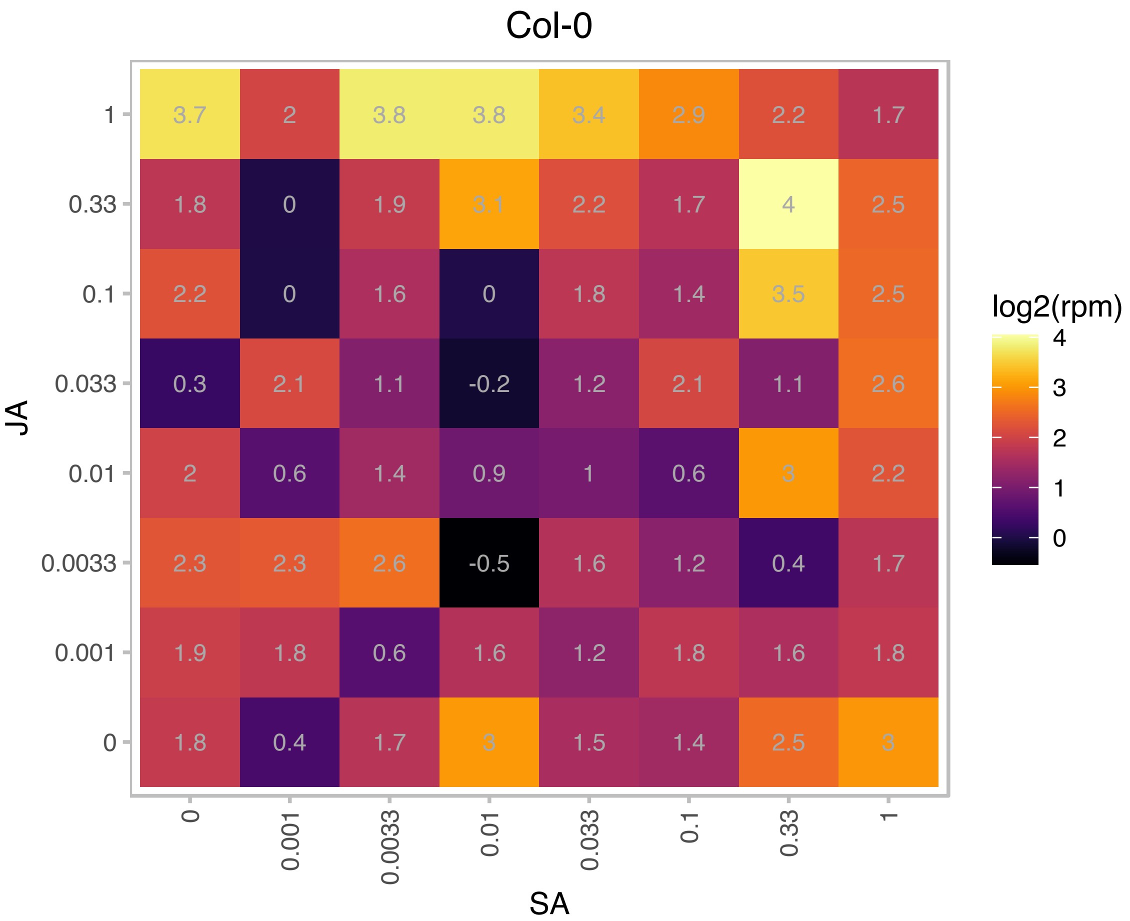


| | |
|---|---|
| Gene | AT2G31280 |
| is.expressed_genes | TRUE |
| is.responsive_genes | TRUE |
| is.apcluster | B-02 |

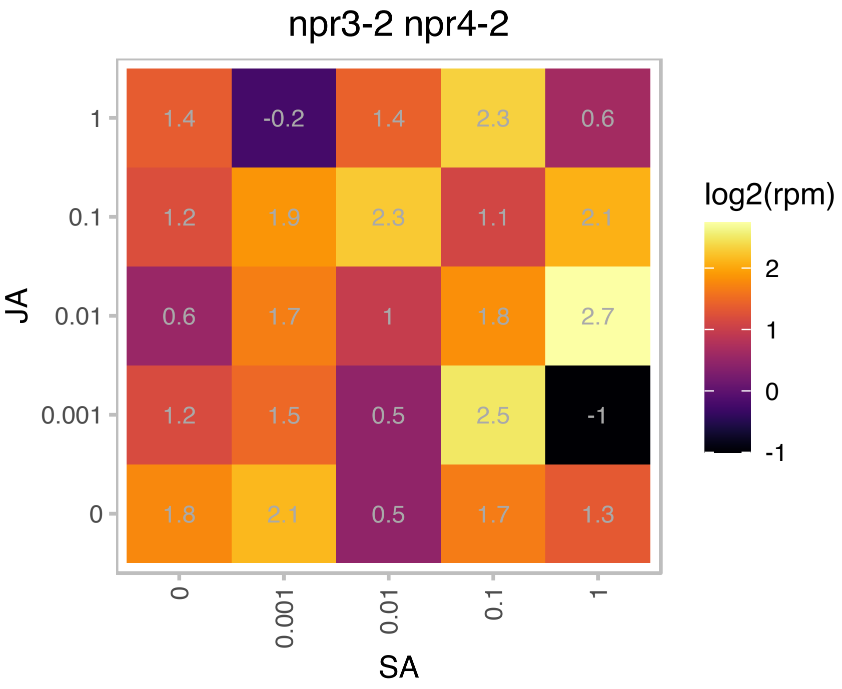


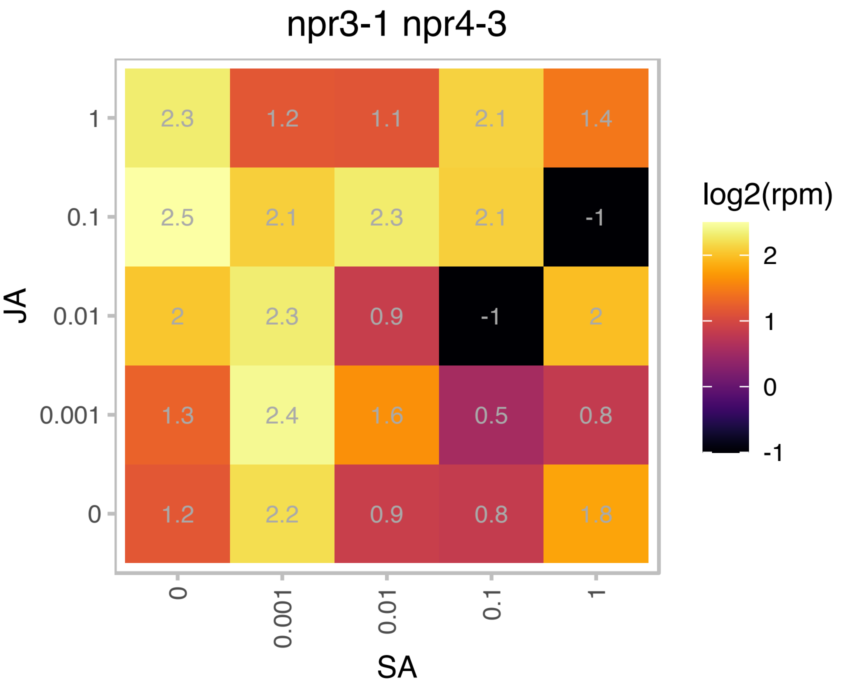

### AT2G32950.1 : ATCOP1, COP1, DET340, EMB168, FUS1

Transducin/WD40 repeat-like superfamily protein

Represses photomorphogenesis and induces skotomorphogenesis in the dark. Contains a ring finger zinc-binding motif, a coiled-coil domain, and several WD-40 repeats, similar to G-beta proteins. The C-terminus has homology to TAFII80, a subunit of the TFIID component of the RNA polymerase II of Drosophila. Nuclear localization in the dark and cytoplasmic in the light. The mRNA is cell-to-cell mobile.

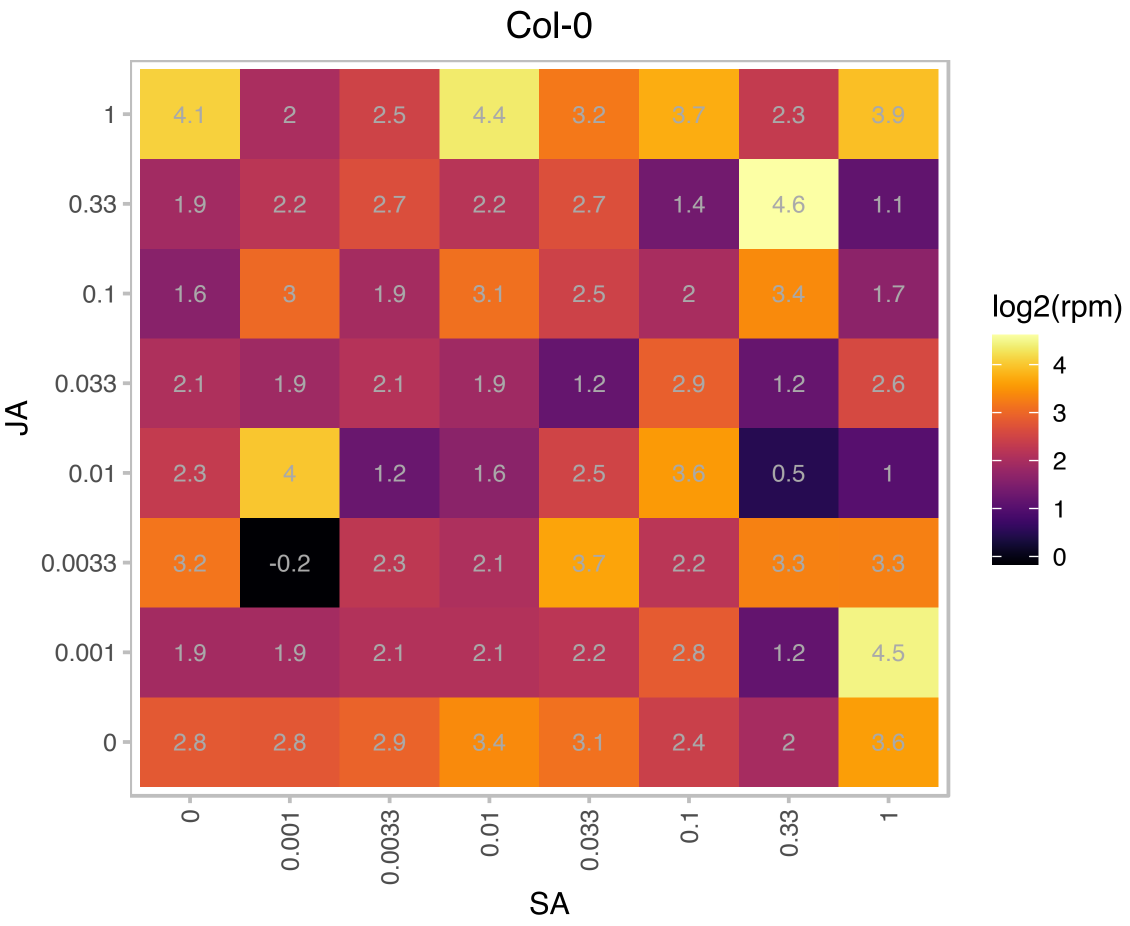


| | |
|---|---|
| Gene | AT2G32950 |
| is.expressed_genes | TRUE |
| is.responsive_genes | TRUE |
| is.apcluster | B-02 |

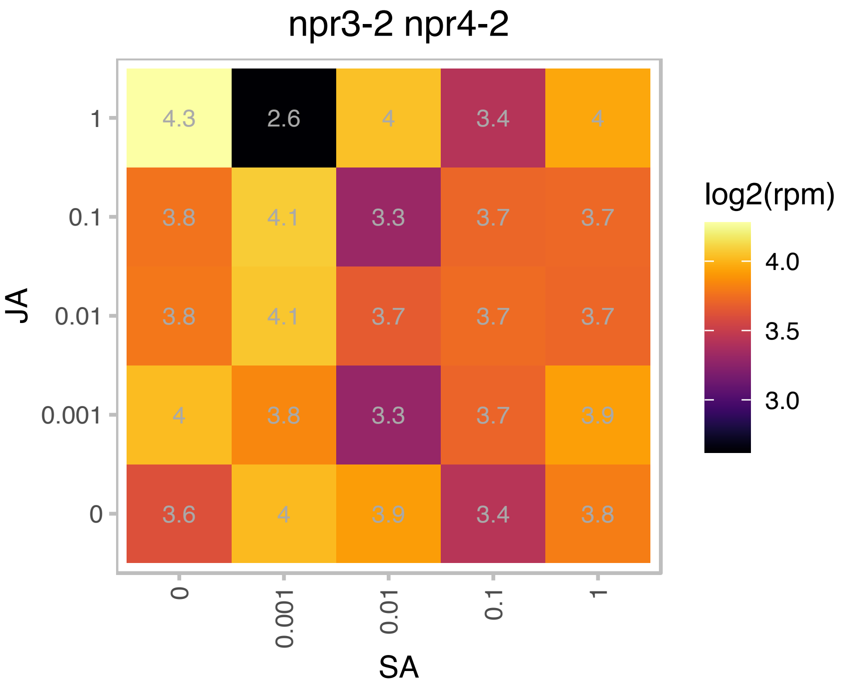


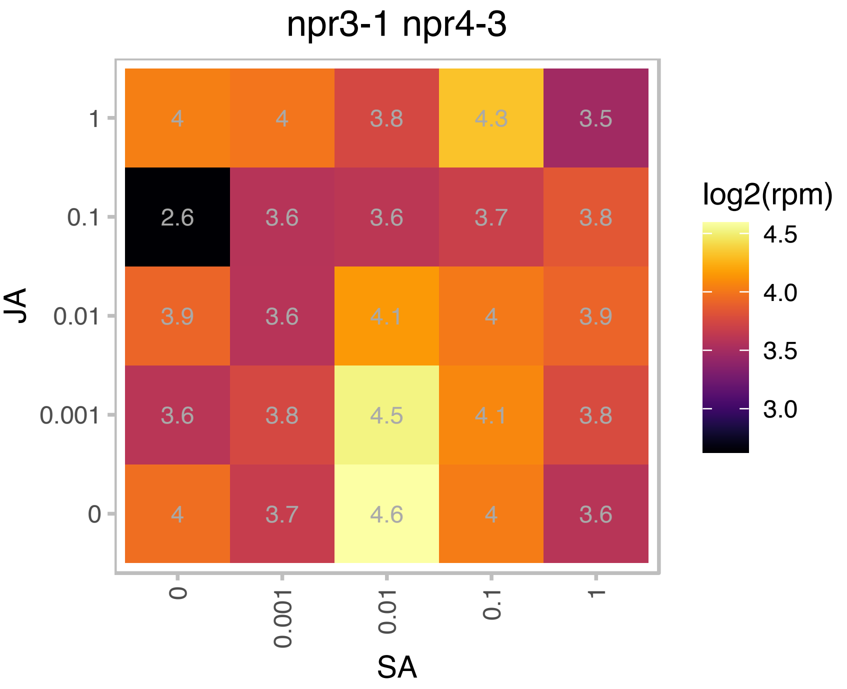

# AT2G35490.1 : NA

Plastid-lipid associated protein PAP / fibrillin family protein

| | |
|---|---|
| Gene | AT2G35490 |
| is.expressed_genes | TRUE |
| is.responsive_genes | TRUE |
| is.apcluster | B-02 |

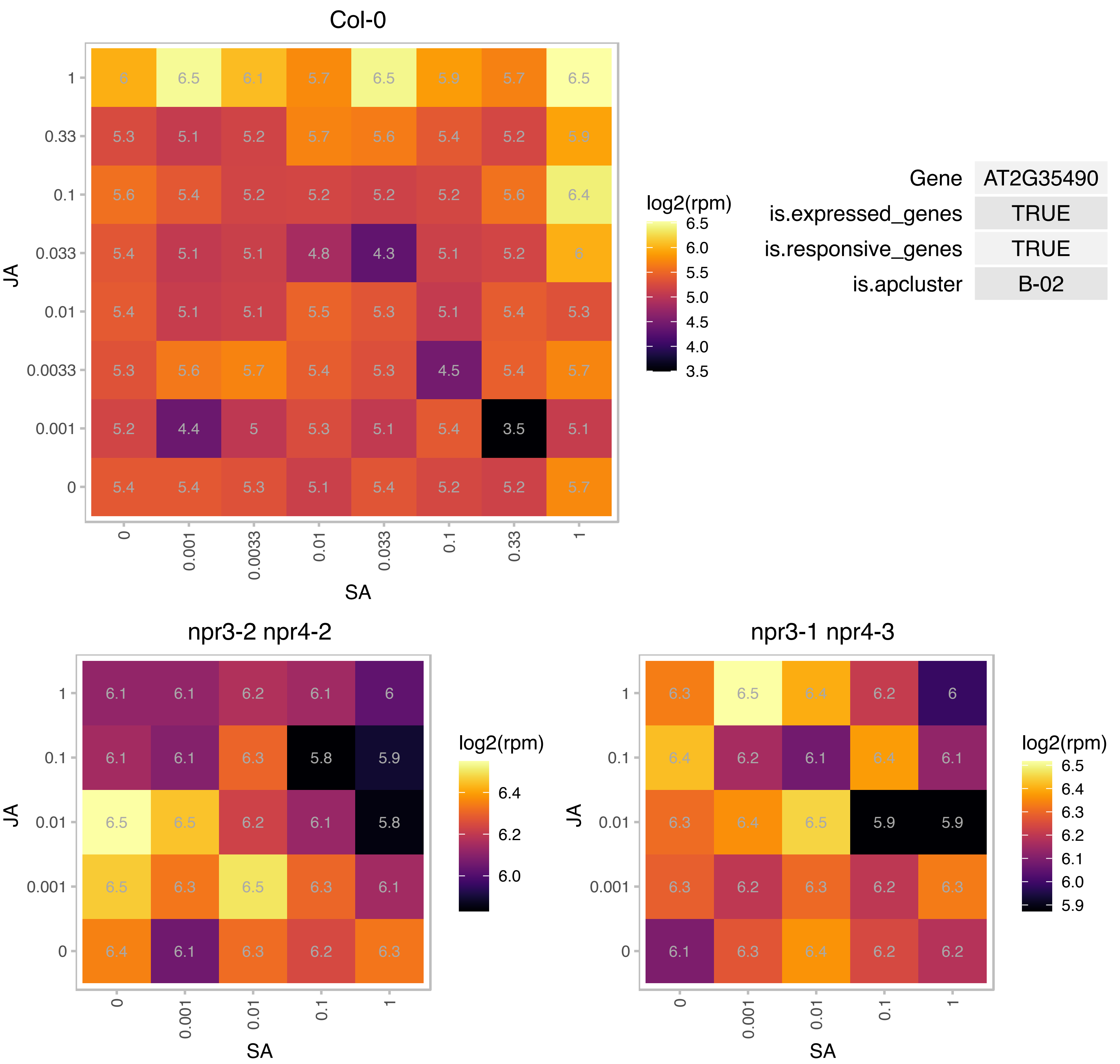

# AT2G47935.1 : NA

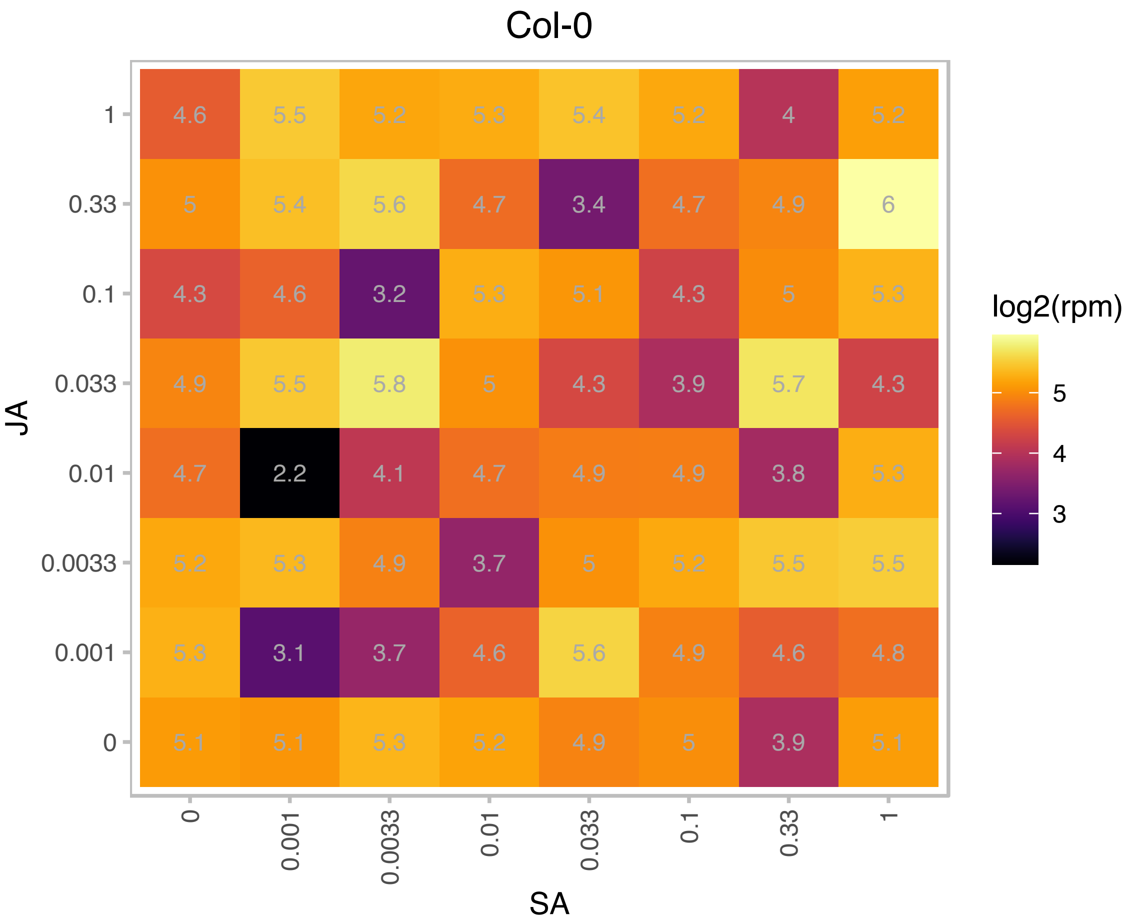


| | |
|---|---|
| Gene | AT2G47935 |
| is.expressed_genes | TRUE |
| is.responsive_genes | TRUE |
| is.apcluster | B-02 |

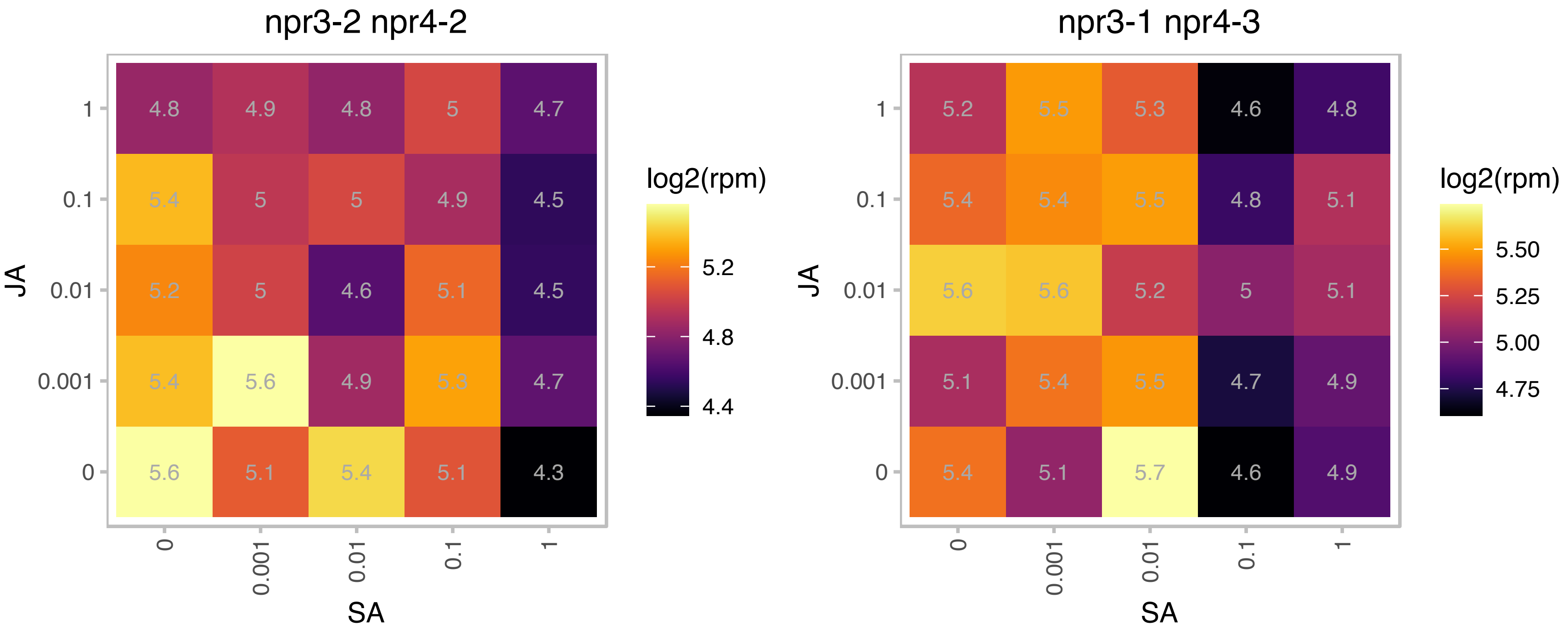

# AT3G05400.1 : NA

Major facilitator superfamily protein

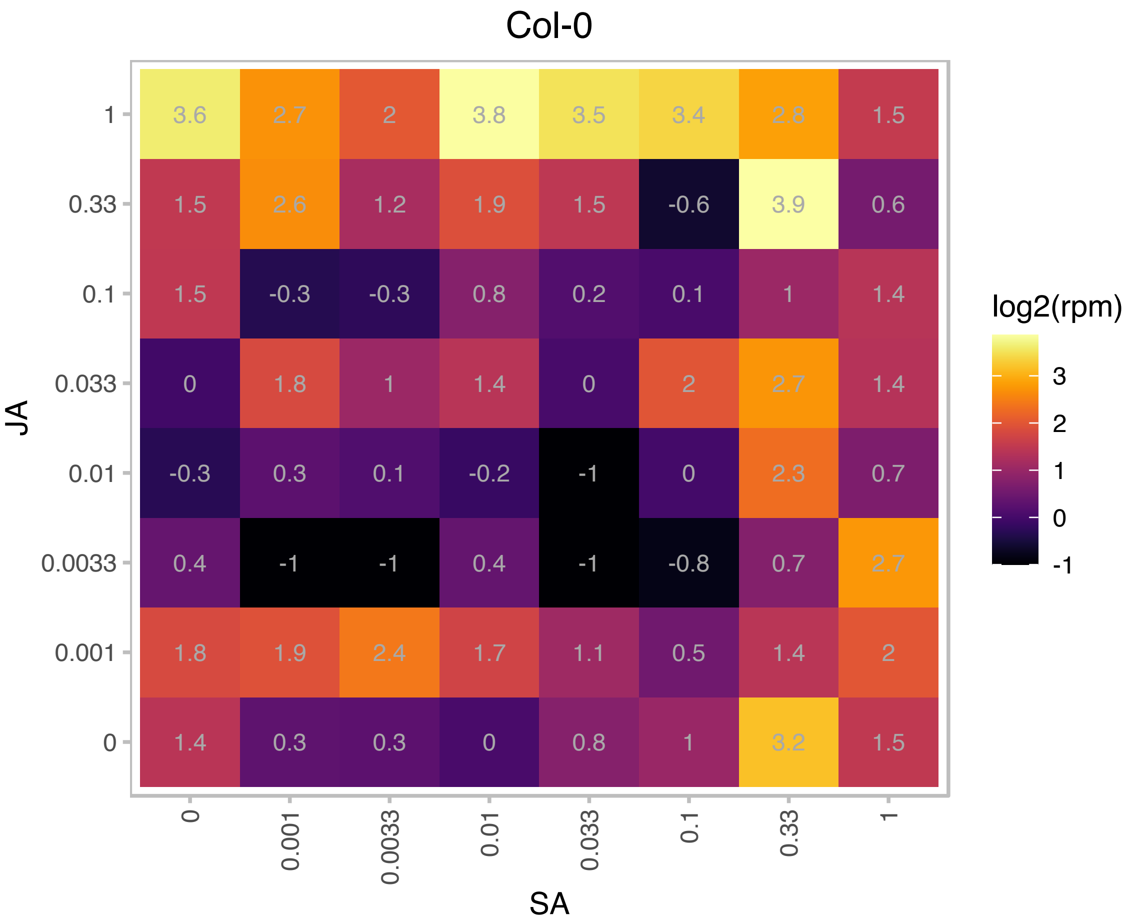


| | |
|---|---|
| Gene | AT3G05400 |
| is.expressed_genes | TRUE |
| is.responsive_genes | TRUE |
| is.apcluster | B-02 |

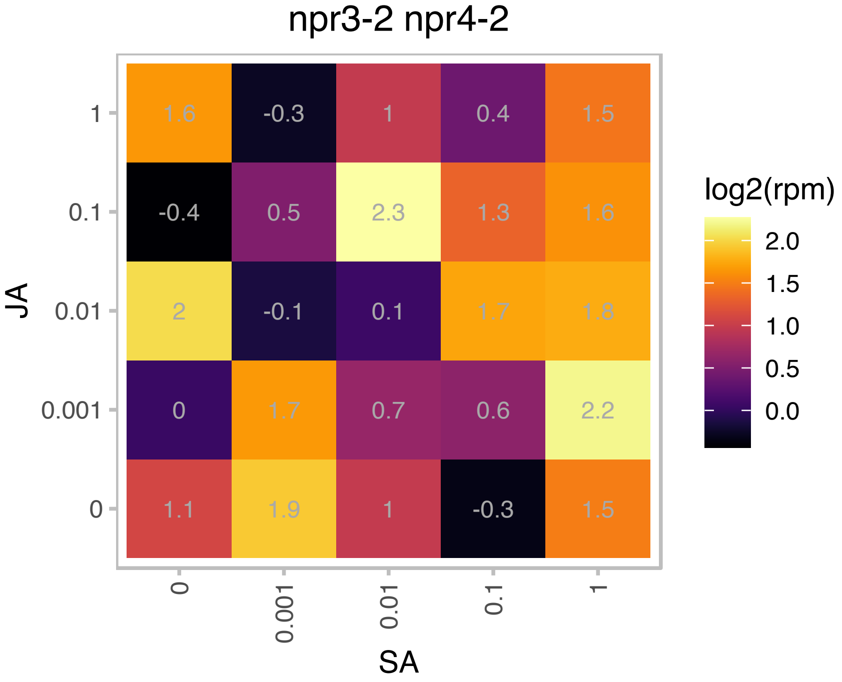


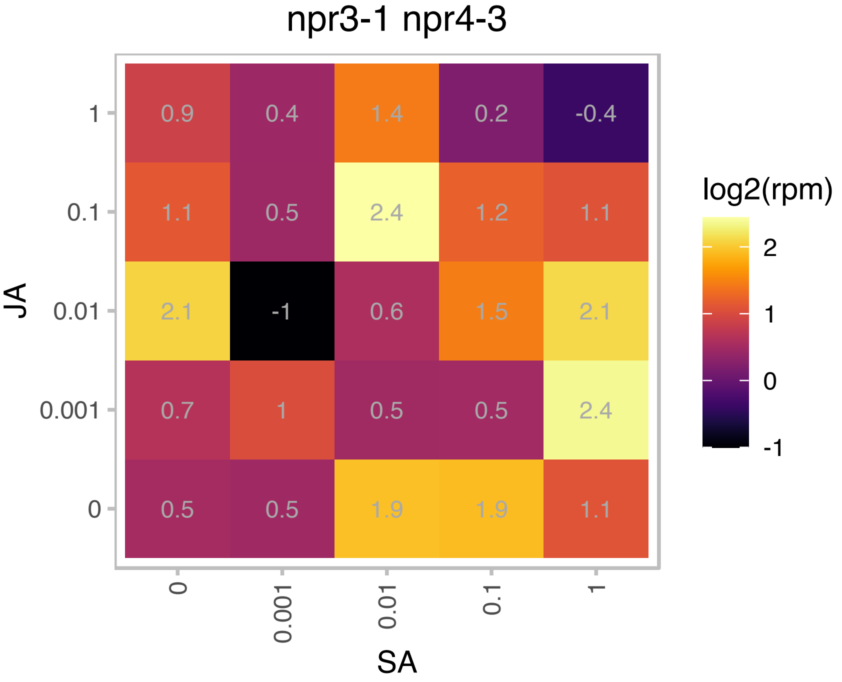

# AT3G07280.1 : NA

pseudogene of unknown protein

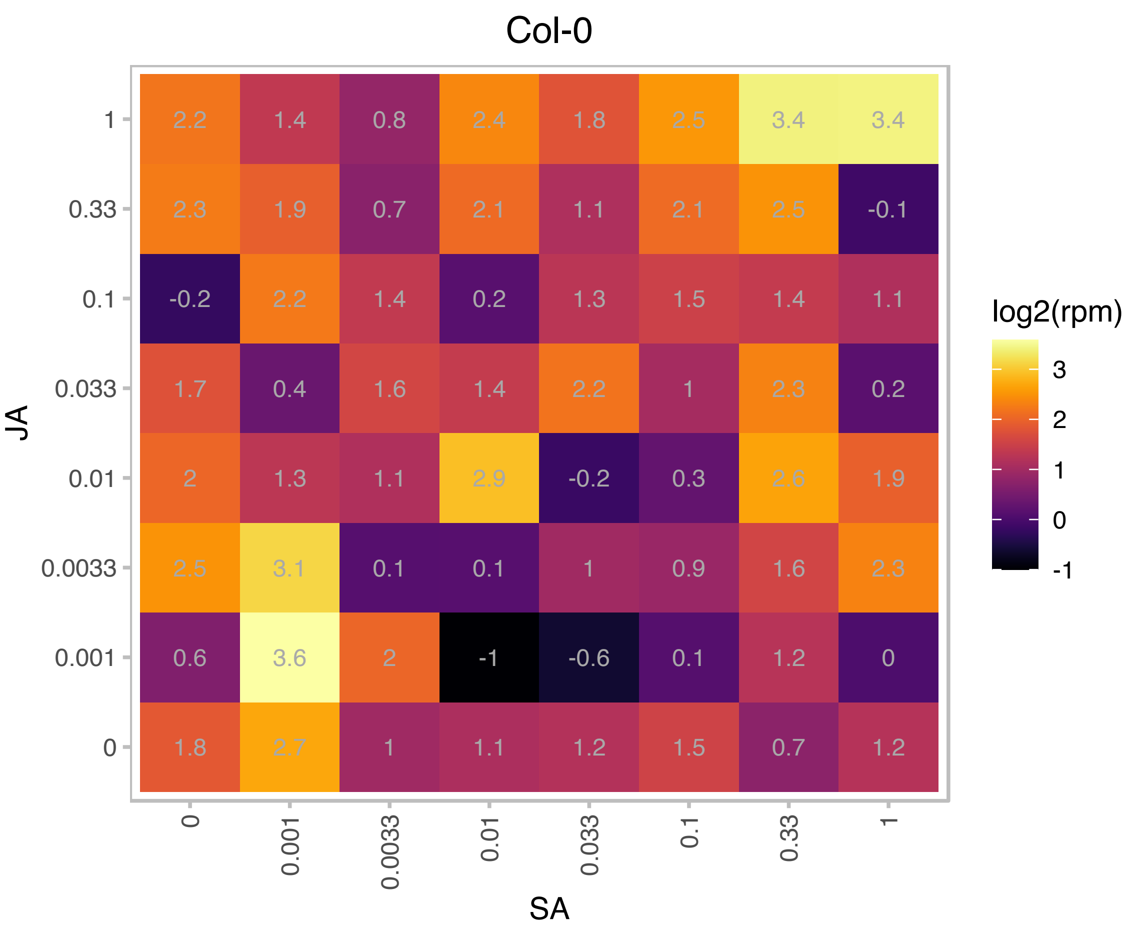


| | |
|---|---|
| Gene | AT3G07280 |
| is.expressed_genes | TRUE |
| is.responsive_genes | TRUE |
| is.apcluster | B-02 |

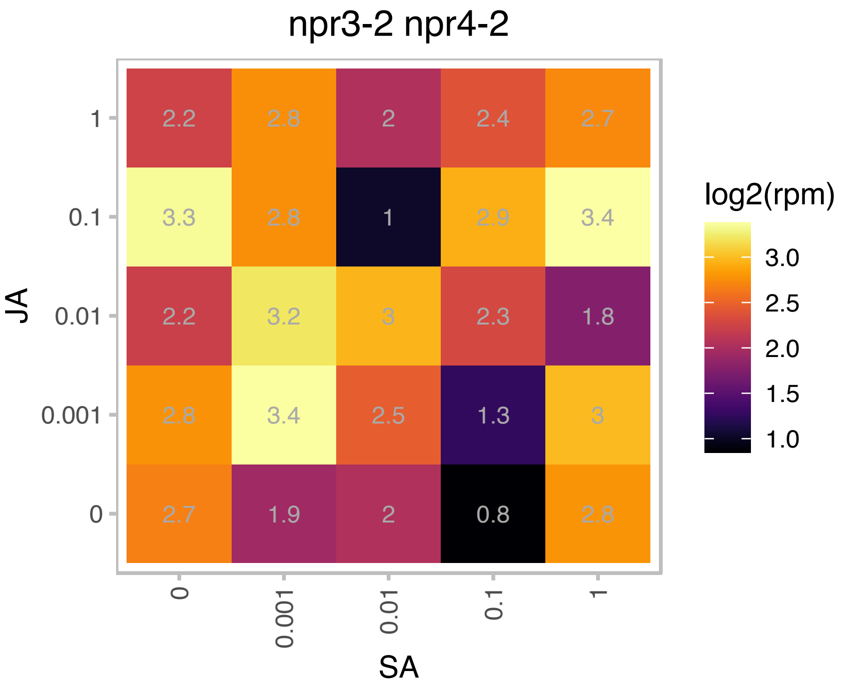


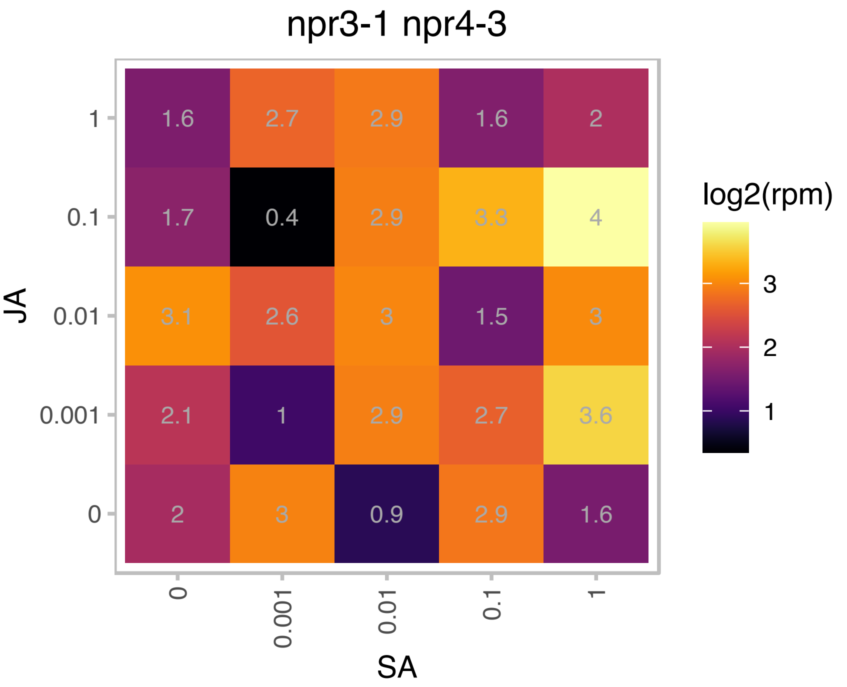

### AT3G09030.1 : EAP3

BTB/POZ domain-containing protein

EAP3 is a cytolsolic BTB/POZ-domain protein involved in trafficking of PEN3.

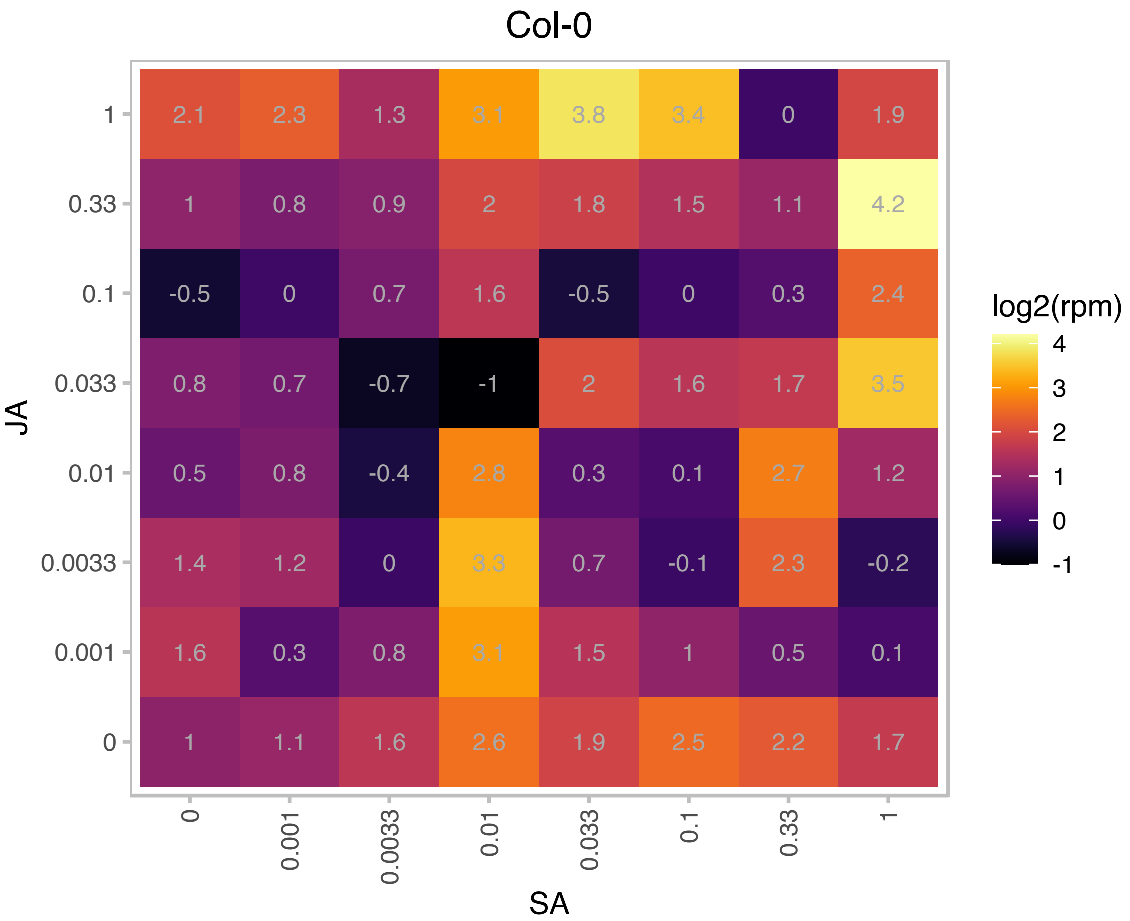


| | |
|---|---|
| Gene | AT3G09030 |
| is.expressed_genes | TRUE |
| is.responsive_genes | TRUE |
| is.apcluster | B-02 |

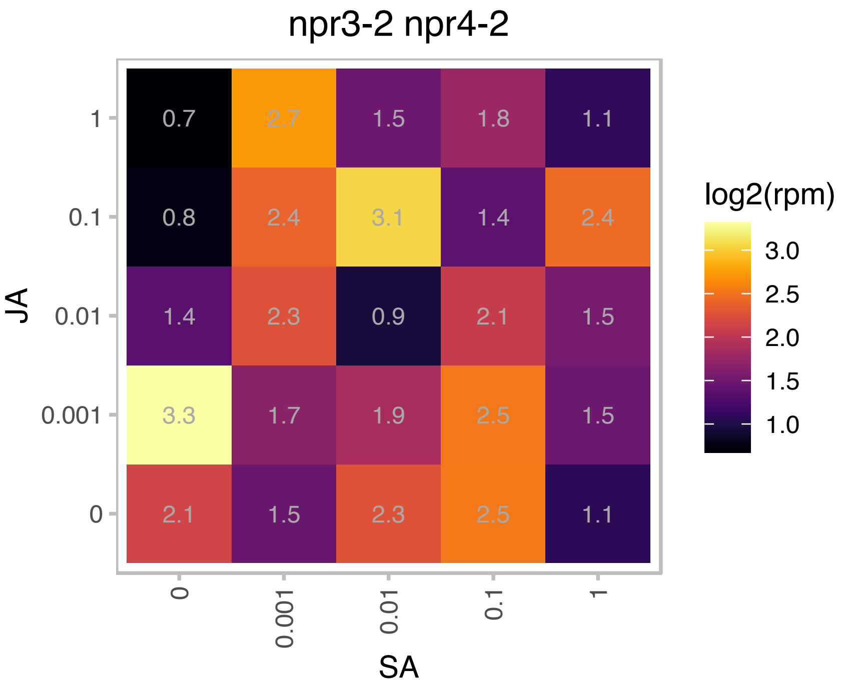


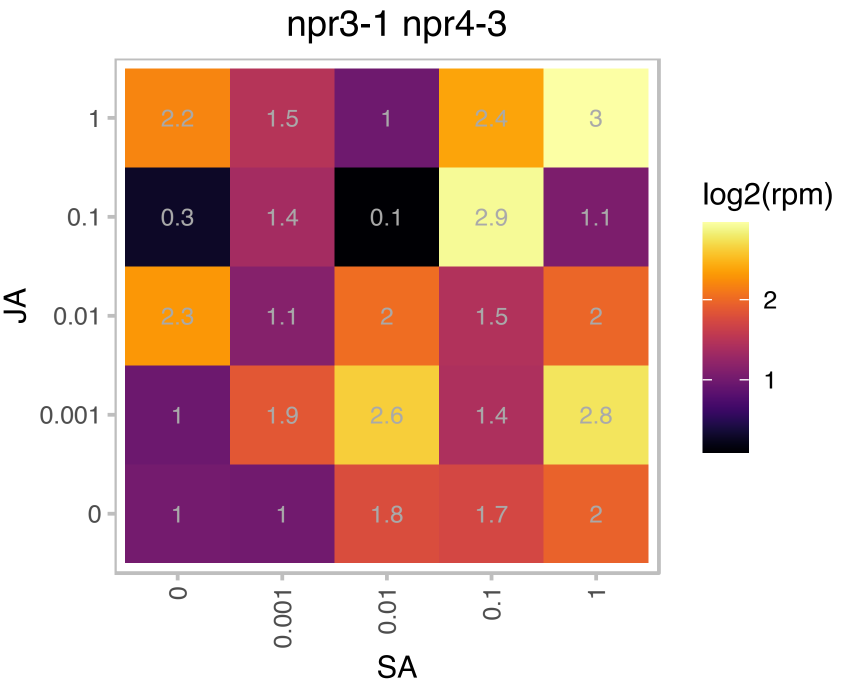

# AT3G15590.1 : NA

Tetratricopeptide repeat (TPR)-like superfamily protein

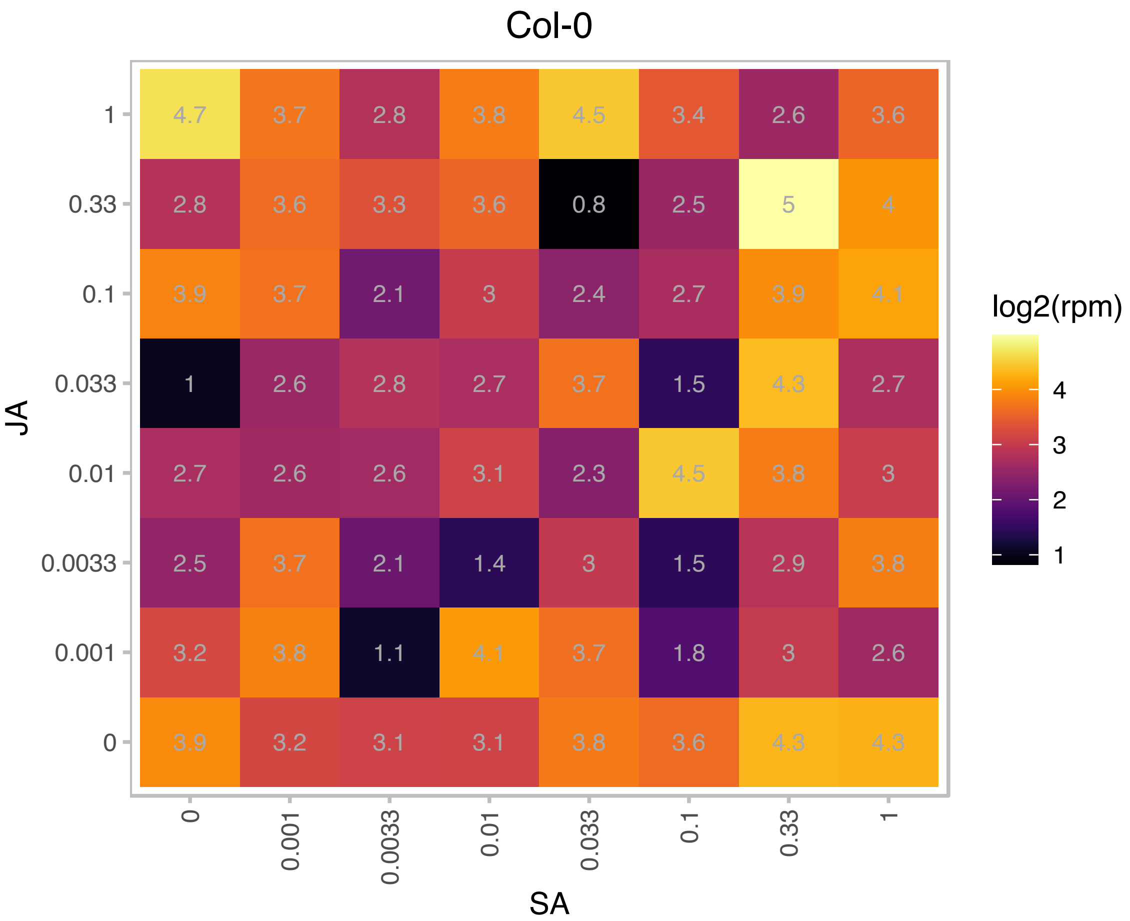


| | |
|---|---|
| Gene | AT3G15590 |
| is.expressed_genes | TRUE |
| is.responsive_genes | TRUE |
| is.apcluster | B-02 |

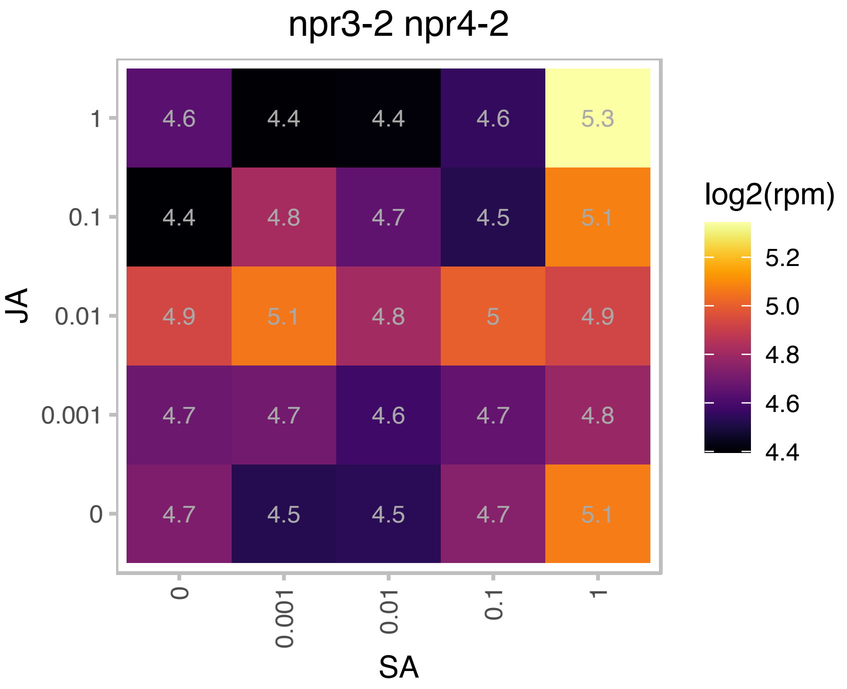


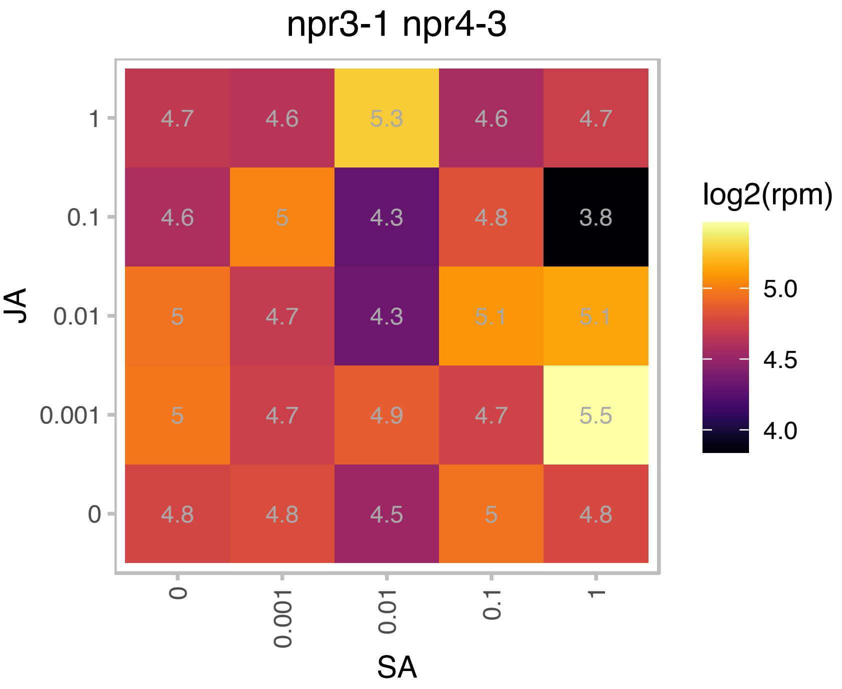

### AT3G26060.1 : ATPRX Q, PRXQ

Thioredoxin superfamily protein

encodes periredoxin Q which decomposes peroxides and plays a role in the protection of the photosynthetic apparatus

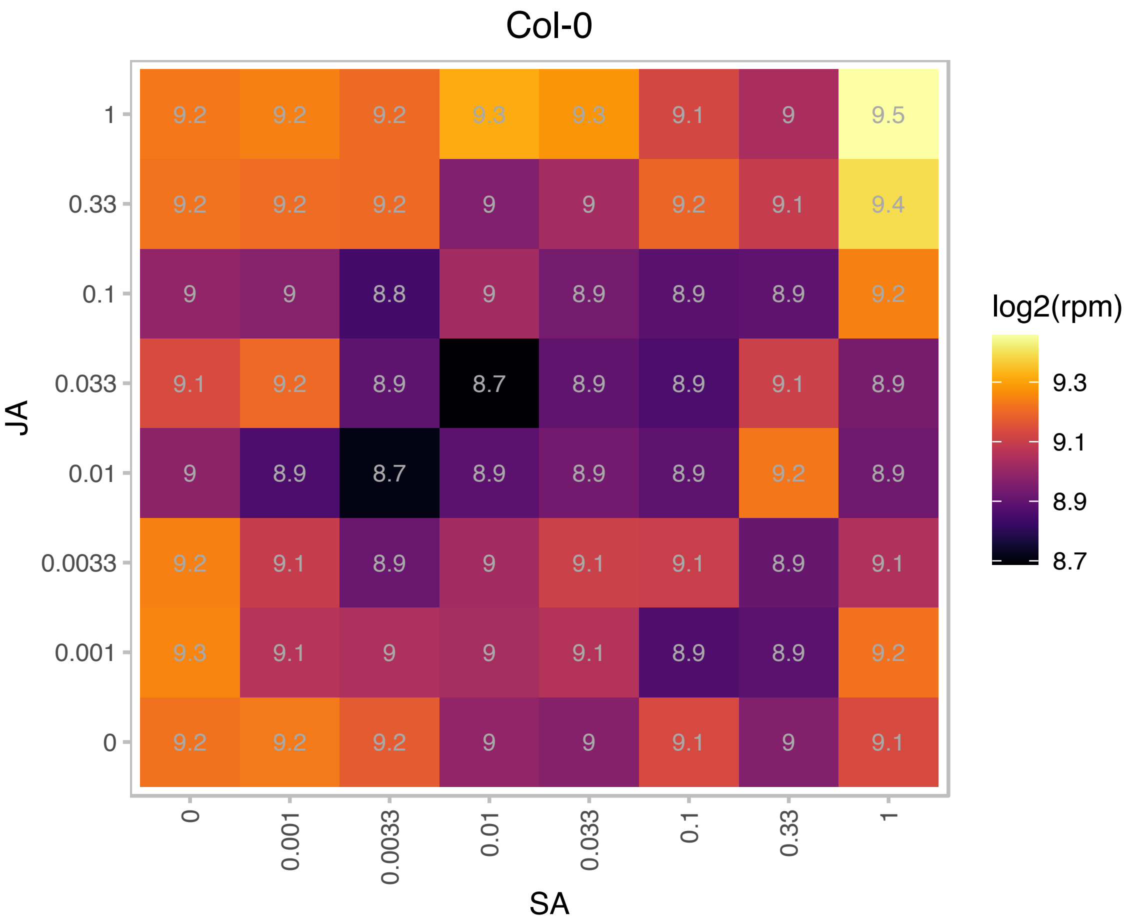


| | |
|---|---|
| Gene | AT3G26060 |
| is.expressed_genes | TRUE |
| is.responsive_genes | TRUE |
| is.apcluster | B-02 |

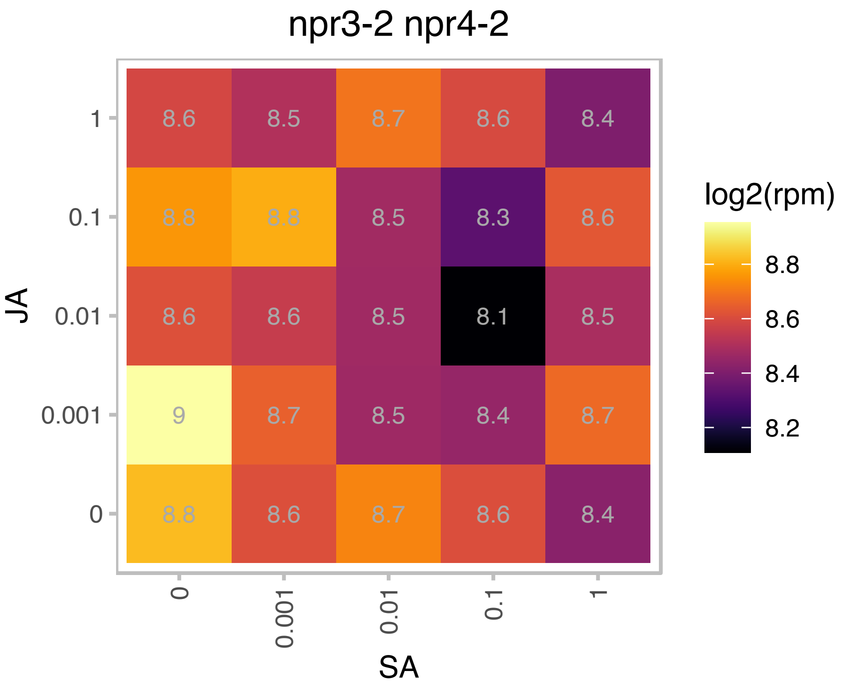


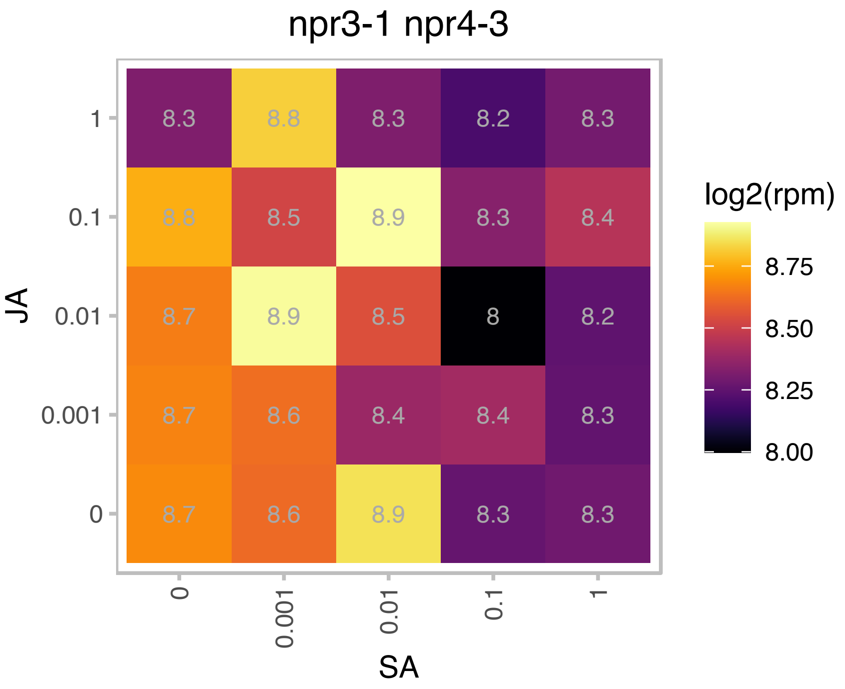

# AT3G27350.1 : NA

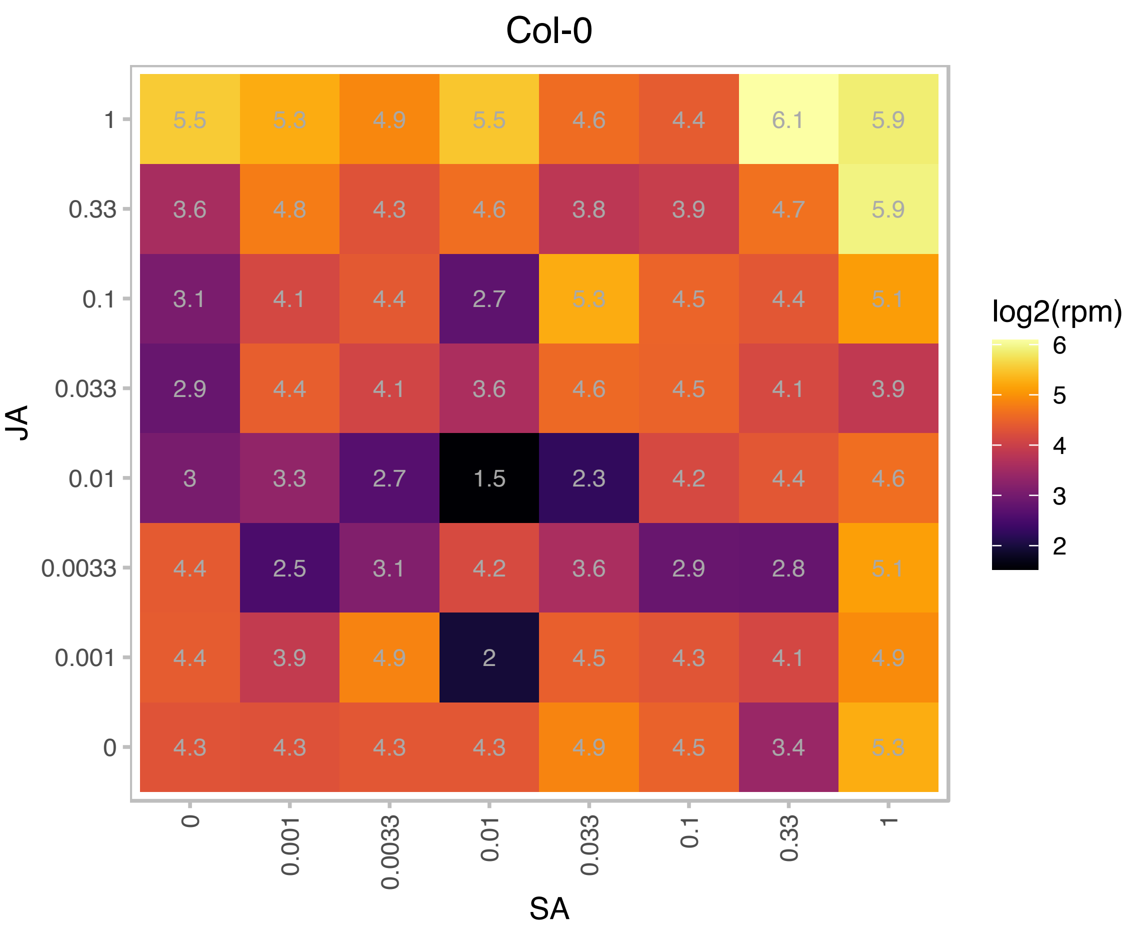


| | |
|---|---|
| Gene | AT3G27350 |
| is.expressed_genes | TRUE |
| is.responsive_genes | TRUE |
| is.apcluster | B-02 |

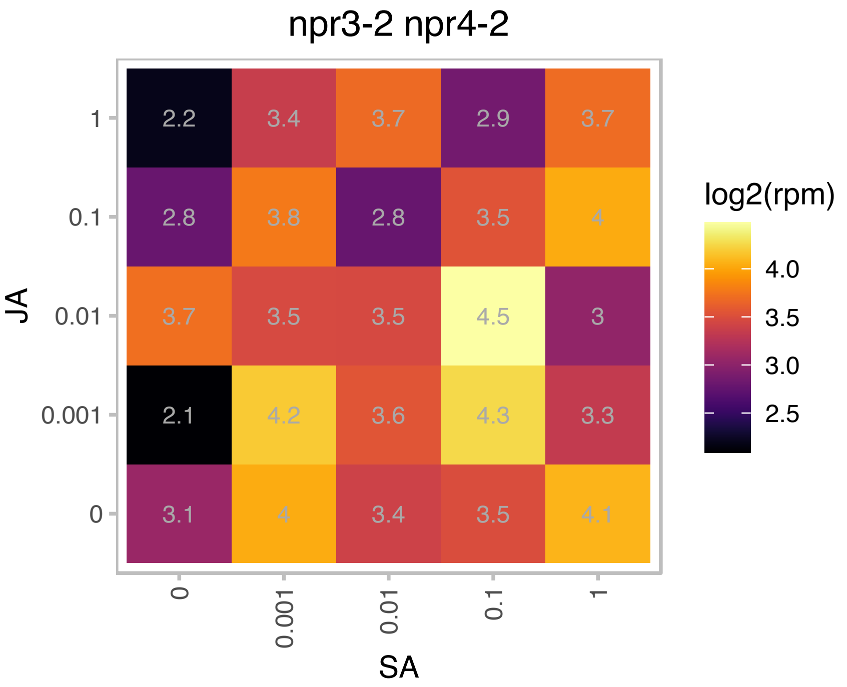


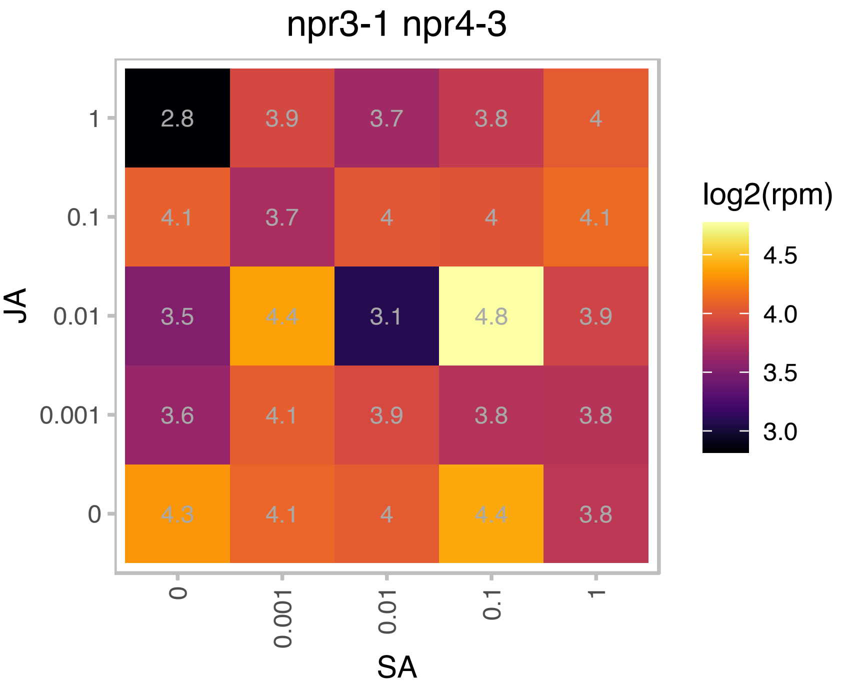

### AT3G54360.1 : NCA1

zinc ion binding

Encodes a catalase chaperon that is essential for catalase activity. Required for multiple stress responses.

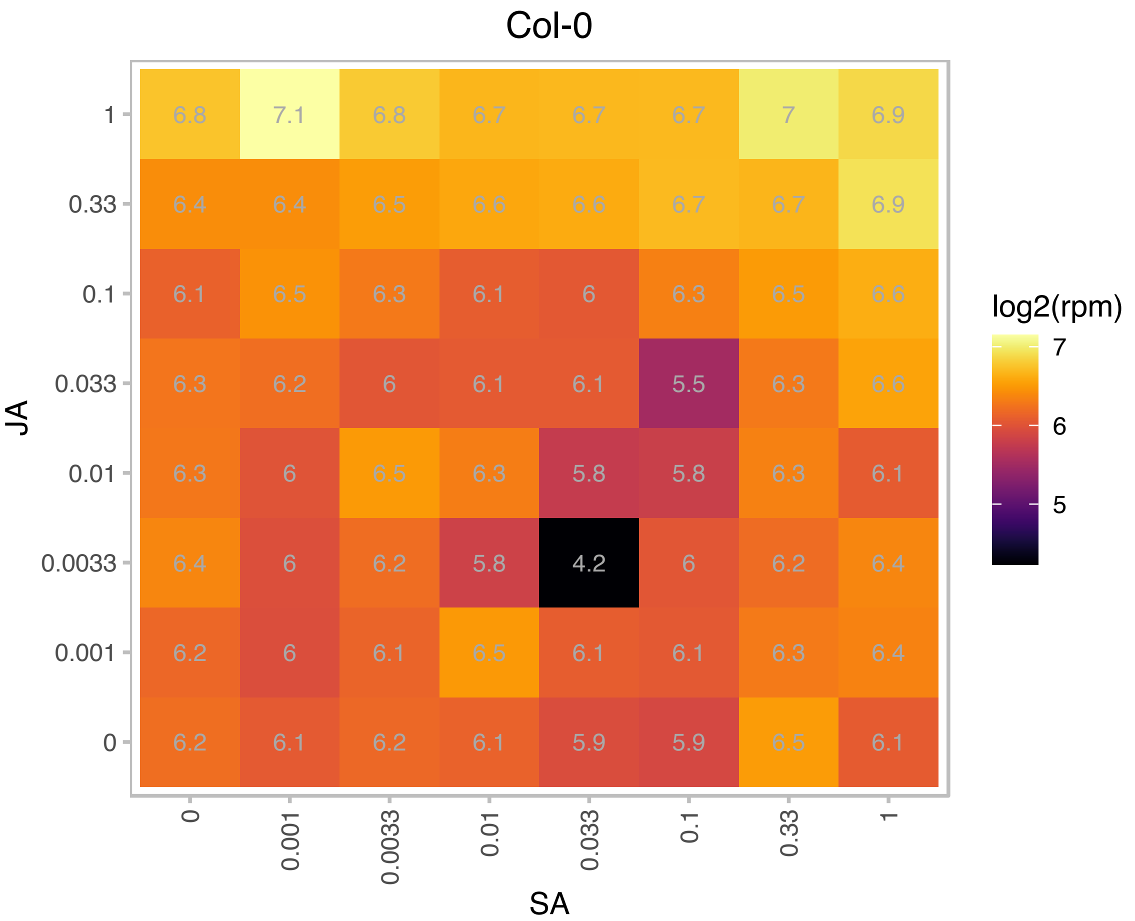


| | |
|---|---|
| Gene | AT3G54360 |
| is.expressed_genes | TRUE |
| is.responsive_genes | TRUE |
| is.apcluster | B-02 |

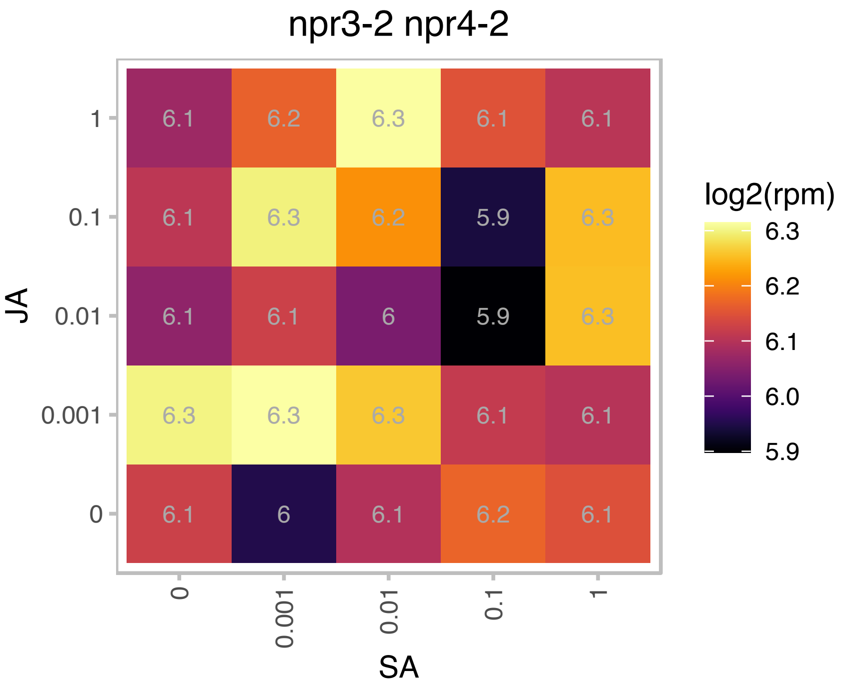


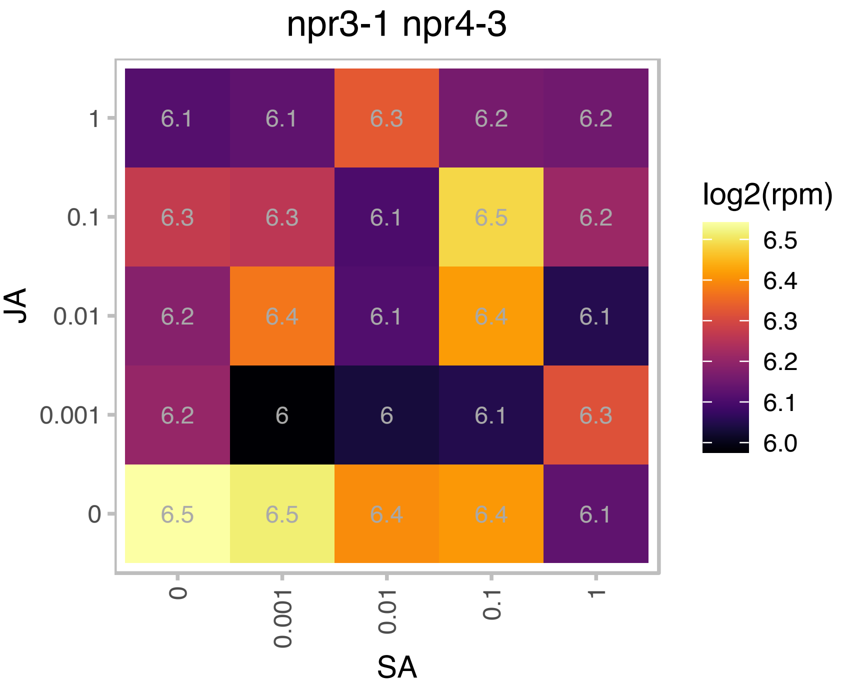

### AT3G58010.1 : PGL34

plastoglobulin 34kD

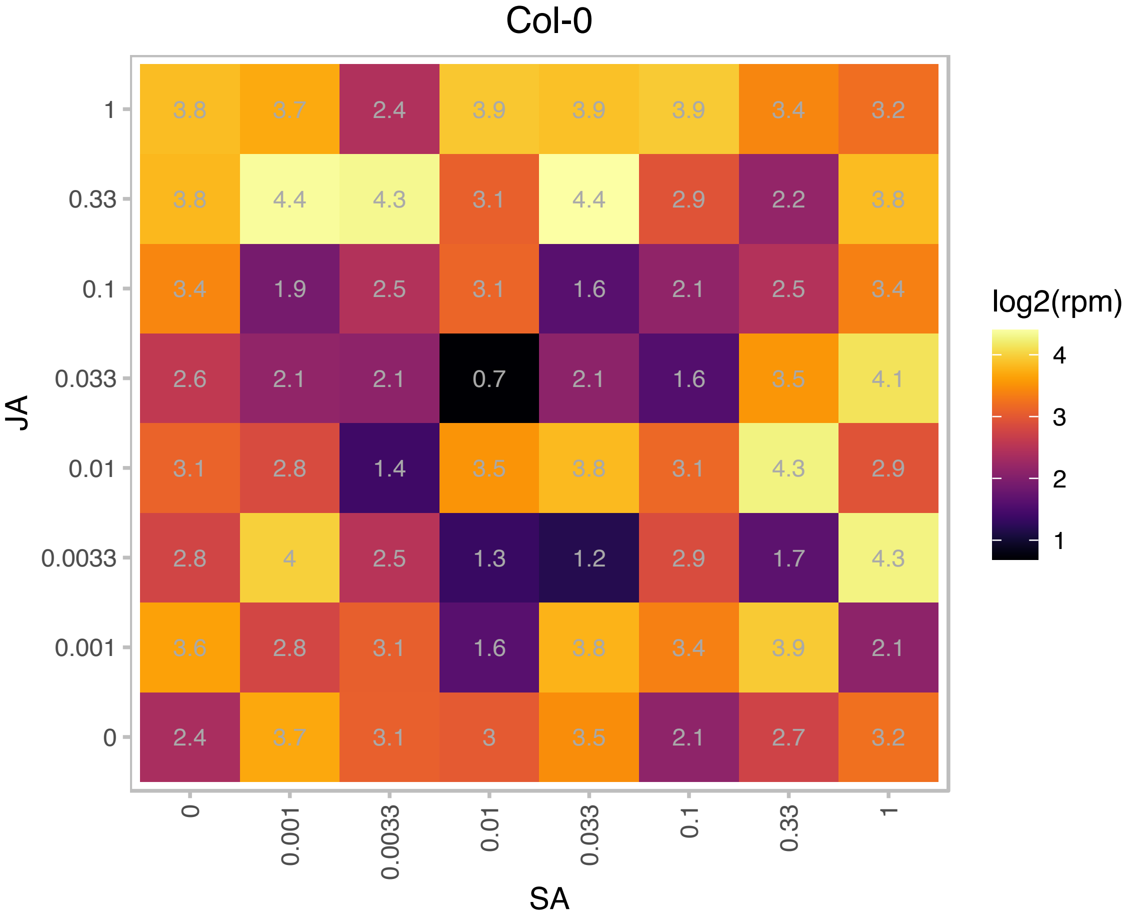


| | |
|---|---|
| Gene | AT3G58010 |
| is.expressed_genes | TRUE |
| is.responsive_genes | TRUE |
| is.apcluster | B-02 |

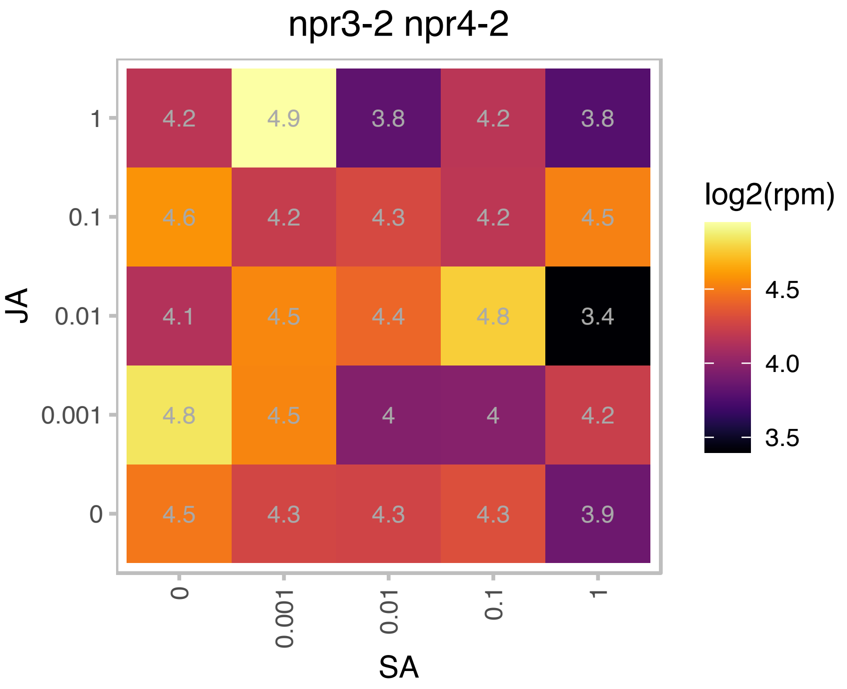


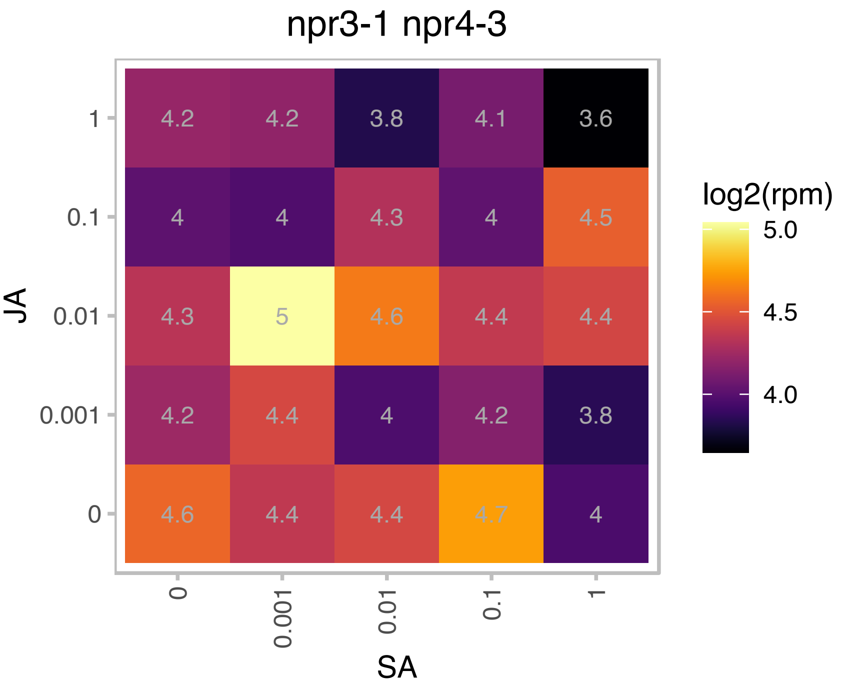

### AT3G60140.1 : BGLU30, DIN2, SRG2

Glycosyl hydrolase superfamily protein

Encodes a protein similar to beta-glucosidase and is a member of glycoside hydrolase family 1. Expression is induced after 24 hours of dark treatment, in senescing leaves and treatment with exogenous photosynthesis inhibitor. Induction of gene expression was suppressed in excised leaves supplied with sugar. The authors suggest that the gene's expression pattern is responding to the level of sugar in the cell. The mRNA is cell-to-cell mobile.

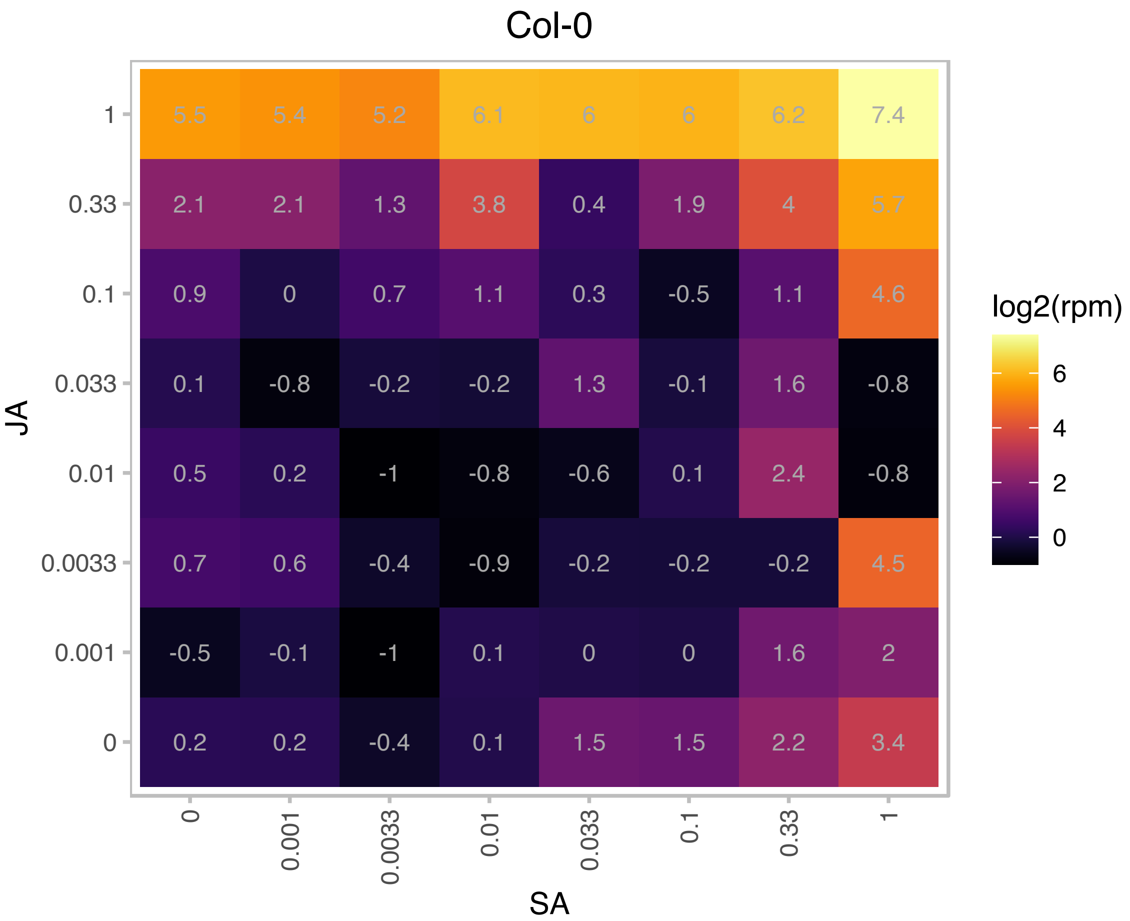


| | |
|---|---|
| Gene | AT3G60140 |
| is.expressed_genes | TRUE |
| is.responsive_genes | TRUE |
| is.apcluster | B-02 |

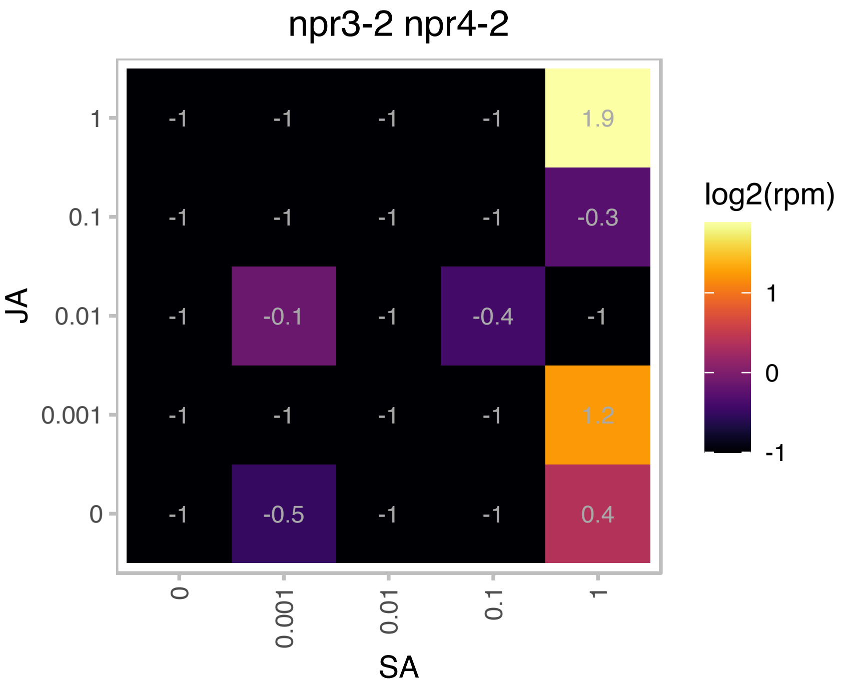


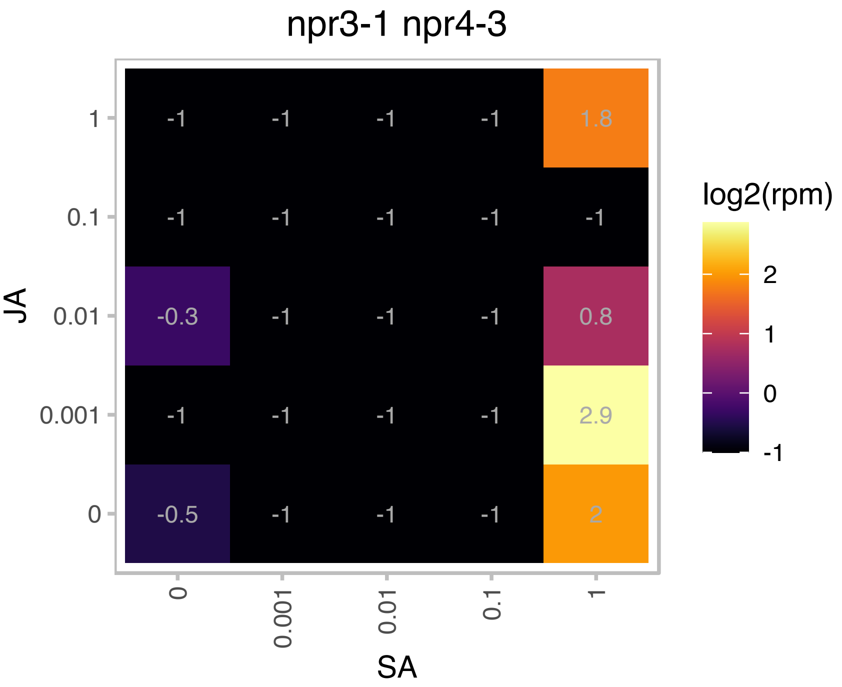

# AT3G61260.1 : NA

Remorin family protein

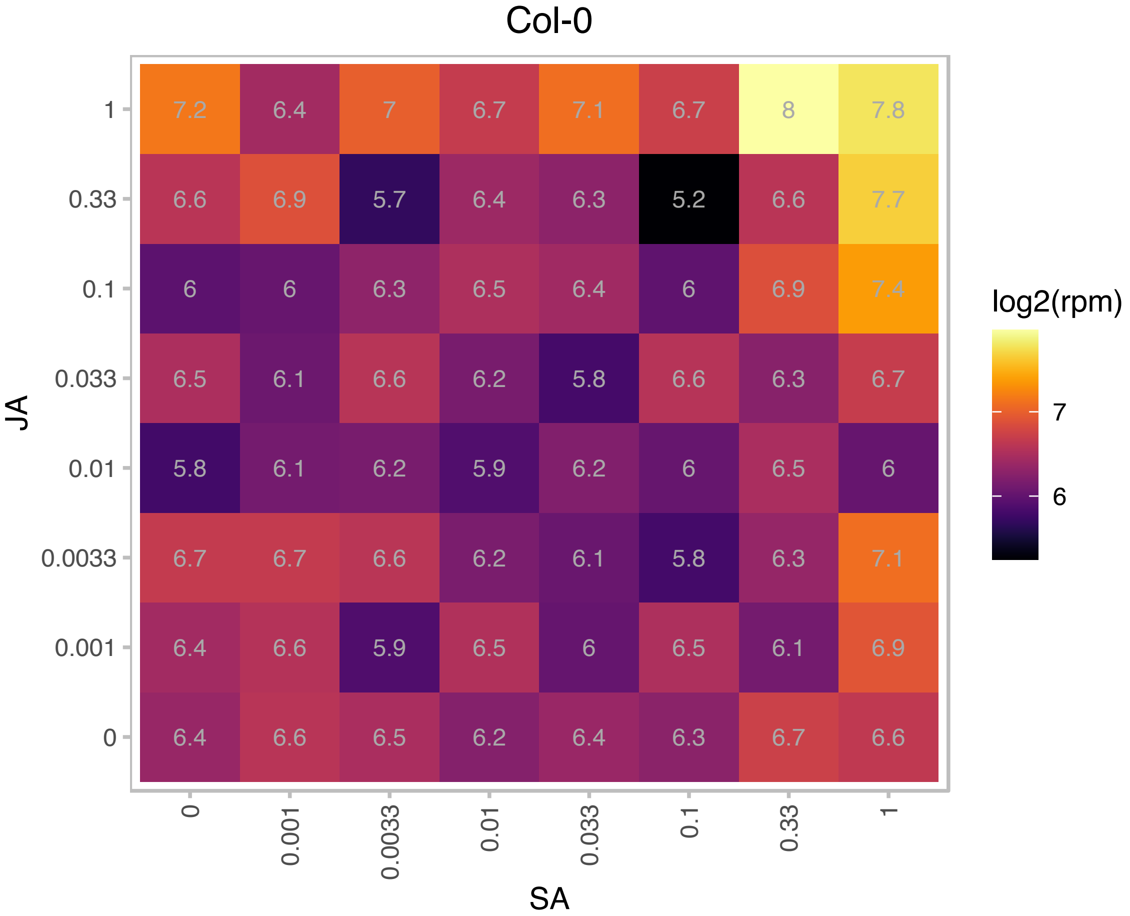


| | |
|---|---|
| Gene | AT3G61260 |
| is.expressed_genes | TRUE |
| is.responsive_genes | TRUE |
| is.apcluster | B-02 |

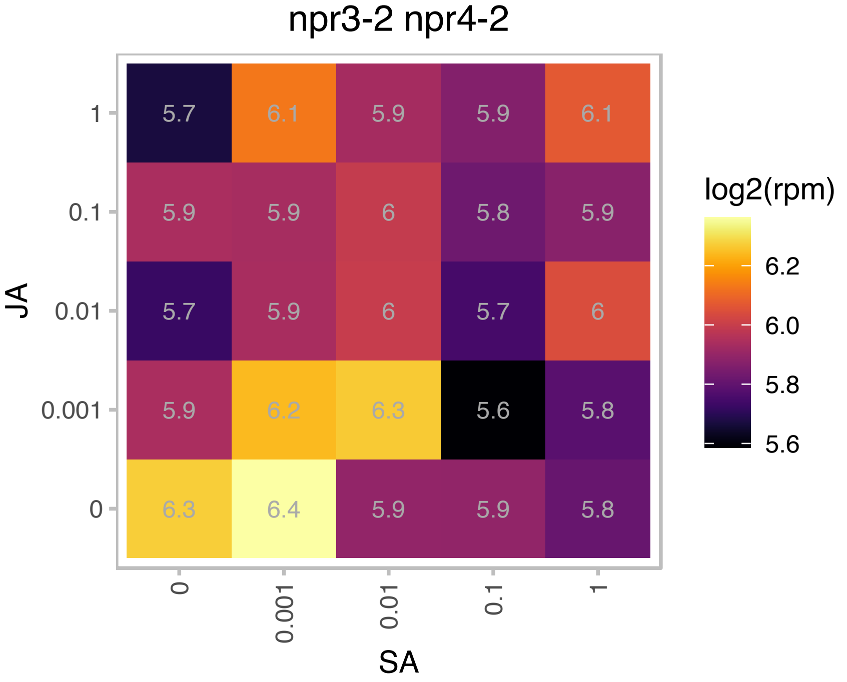


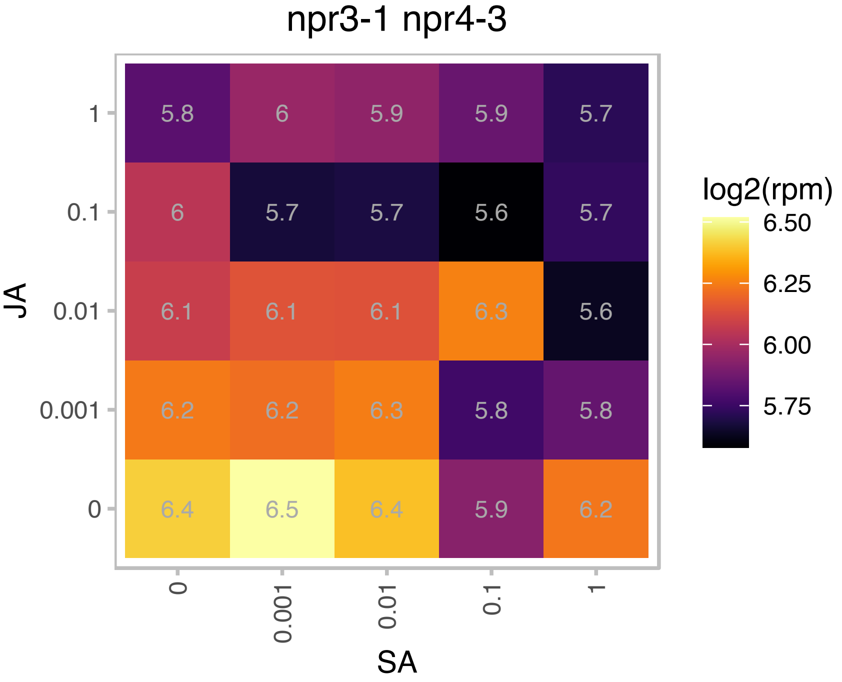

### AT4G12610.1 : ATRAP74, RAP74

transcription activators;DNA binding;RNA polymerase II transcription factors;catalytics;transcription initiation factors

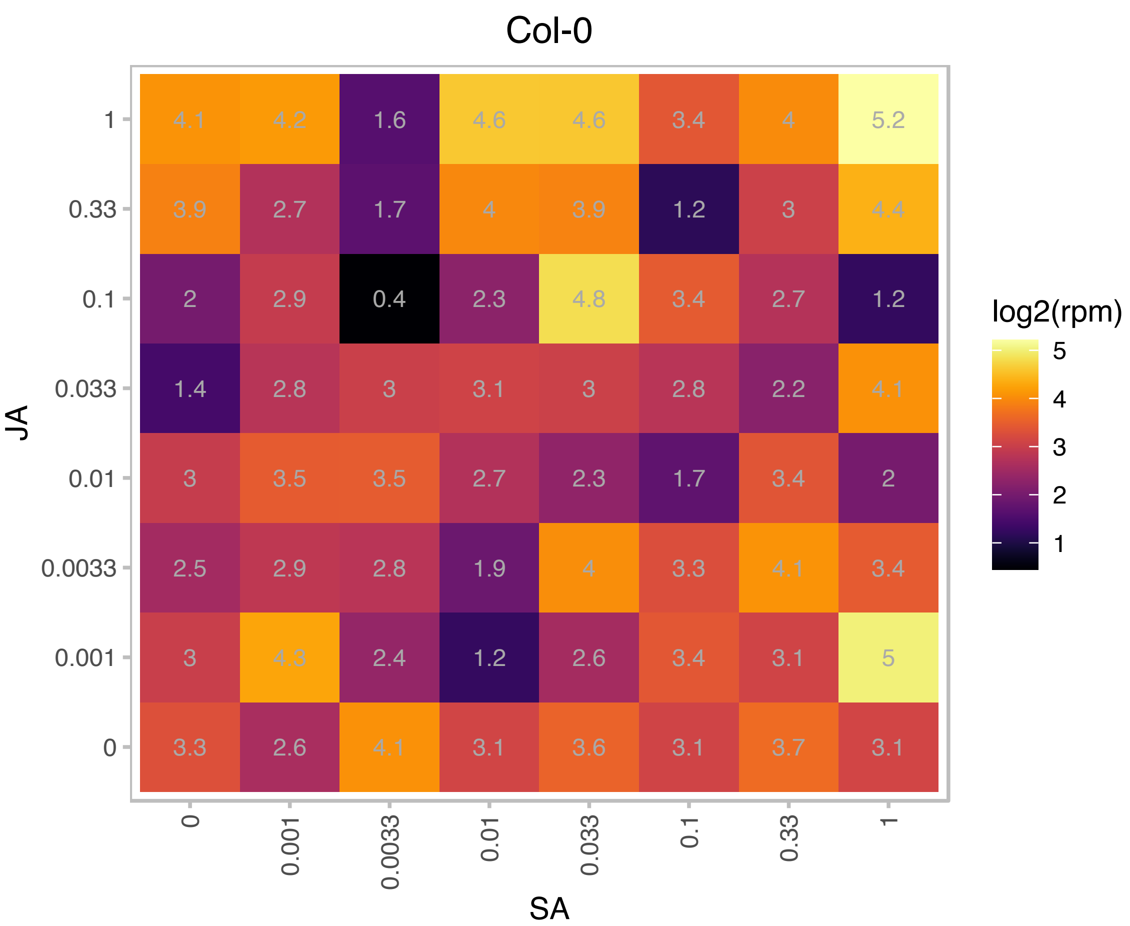


| | |
|---|---|
| Gene | AT4G12610 |
| is.expressed_genes | TRUE |
| is.responsive_genes | TRUE |
| is.apcluster | B-02 |

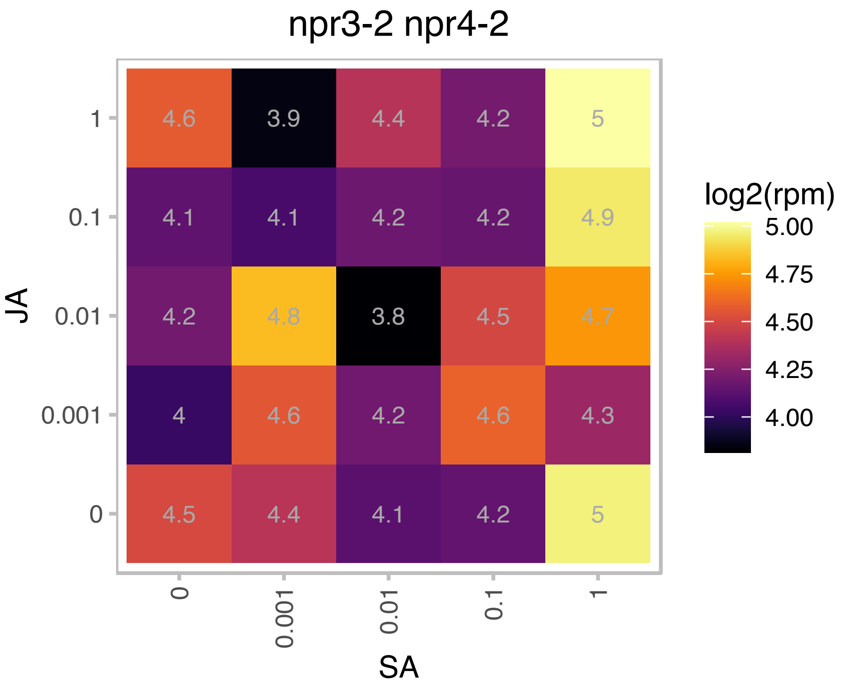


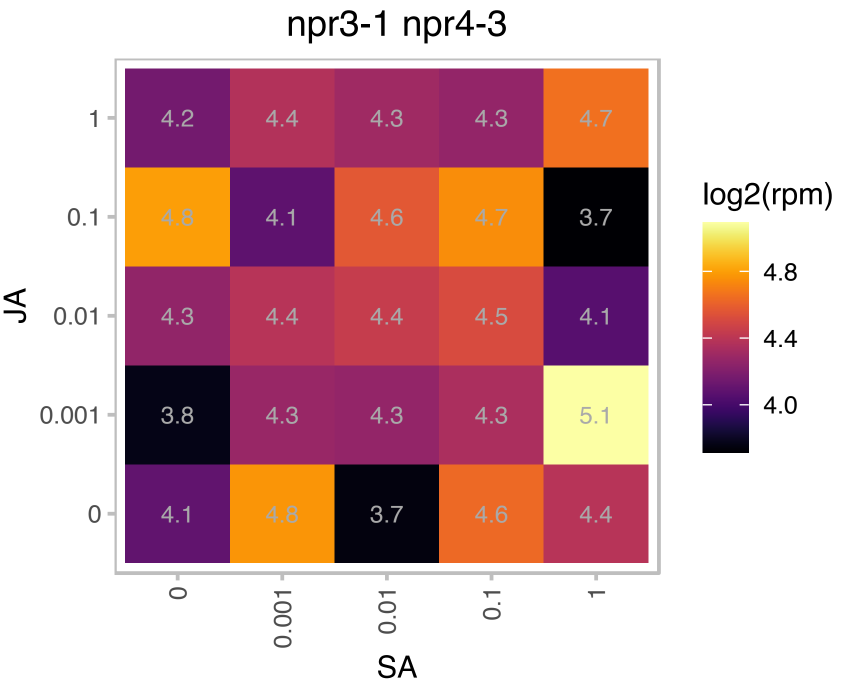

### AT4G20260.1 : ATPCAP1, MDP25, PCAP1

Encodes a Ca2+ and Cu2+ binding protein. N-terminal myristylation on glycine 2 appears to enable it to associate tightly with the plasma membrane. Recombinant PCaP1 interacts strongly with phosphatidylinositol 3,5-bisphosphate (PtdIns(3,5)P2) and PtdIns (3,4,5)P3, and weakly with PtdIns(3,5)P2 and PtdIns(4,5). It also interacts with calmodulin (CaM) in a calcium-dependent manner. CaM does not interfere with PCaP1 membrane localization but does weaken interactions between it and the PtdInsPs. PCaP1 has an apparent Kd of 10 uM for Cu2+ and can bind six ions per protein. Transcript levels for PCaP1 first fall and then rise following exposure to CuCl2. Mannitol, sorbitol, and the flg22 oligopeptide also increase expression levels. The mRN

plasma-membrane-associated cation-binding protein 1

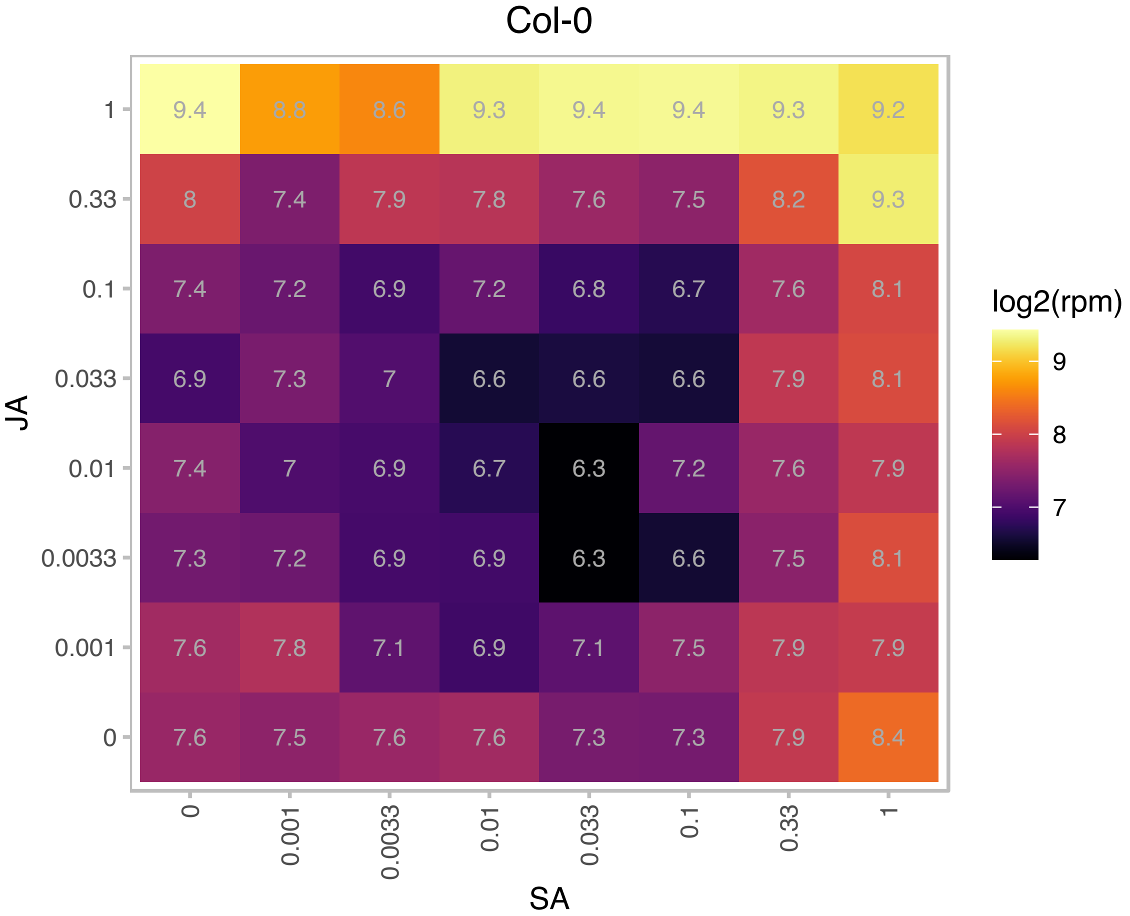


| | |
|---|---|
| Gene | AT4G20260 |
| is.expressed_genes | TRUE |
| is.responsive_genes | TRUE |
| is.apcluster | B-02 |

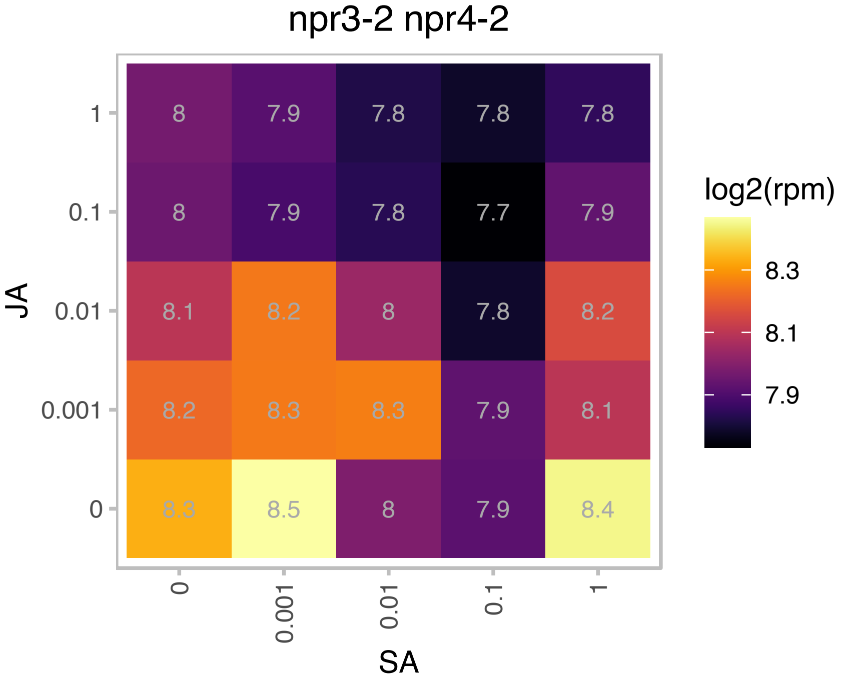


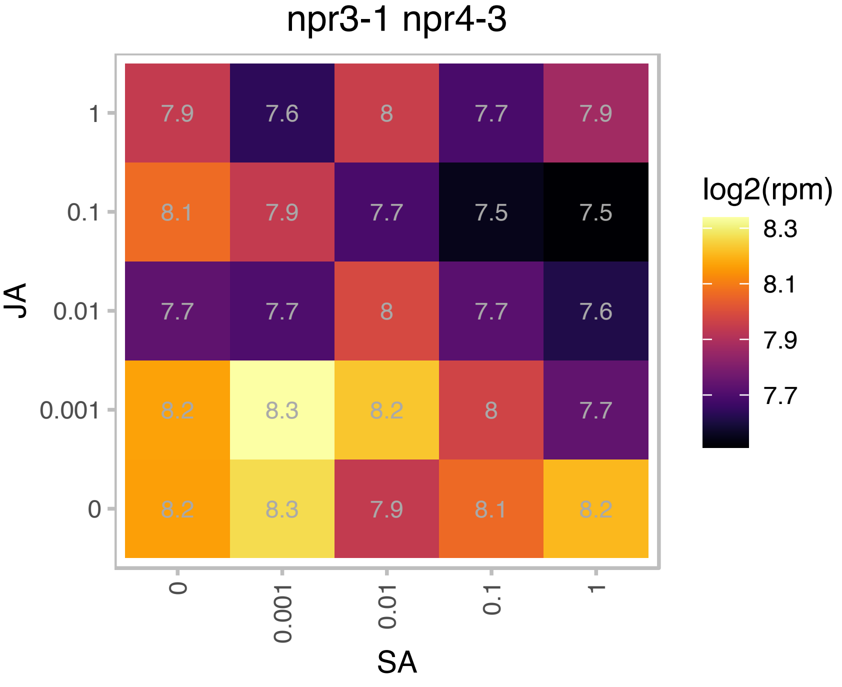

### AT4G21100.1 : DDB1B

damaged DNA binding protein 1B
One of two closely related genes similar to a damaged DNA binding protein originally described in mammals. May form a complex with DET1 to regulate photomorphogenesis. Loss of function mutations are lethal. The DDB1b protein binds with a number of DWD-containing proteins and may form part of a CUL4-based E3 ubiquitin ligase.

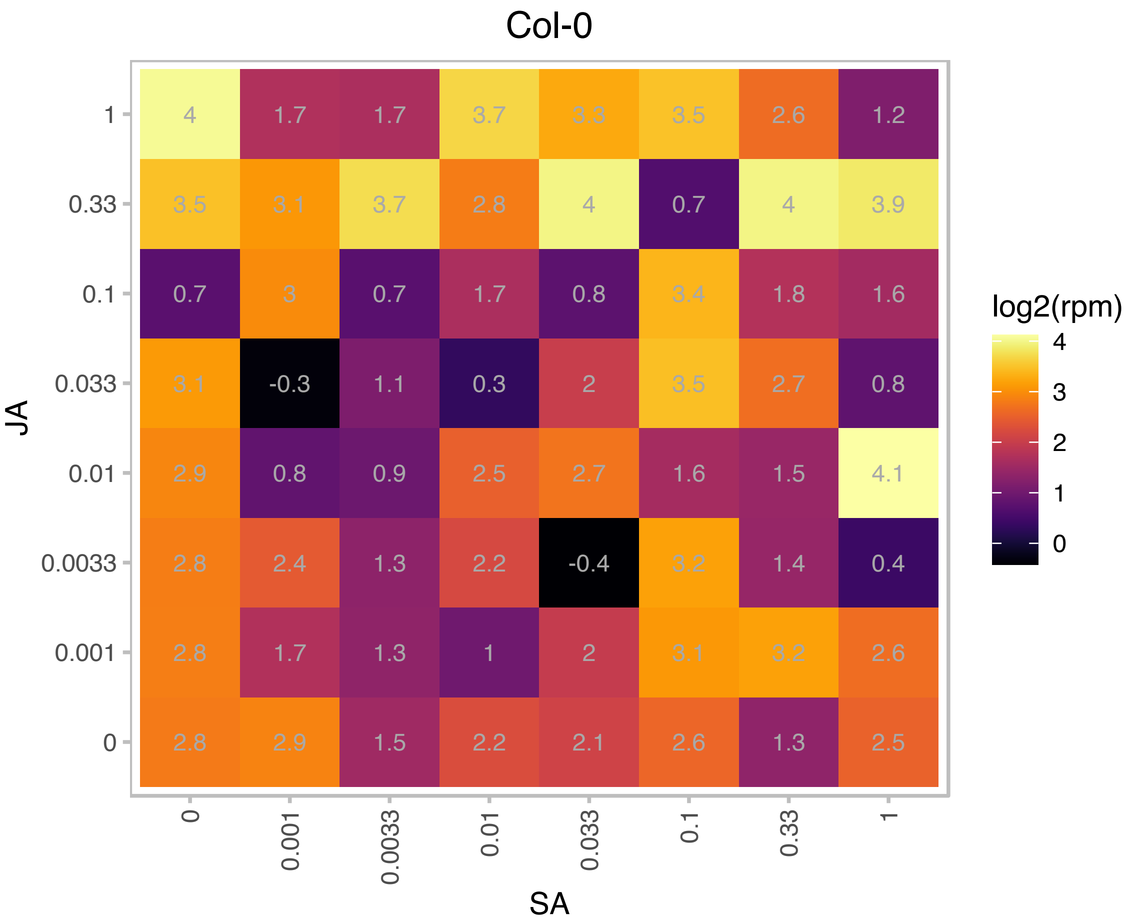


| | |
|---|---|
| Gene | AT4G21100 |
| is.expressed_genes | TRUE |
| is.responsive_genes | TRUE |
| is.apcluster | B-02 |

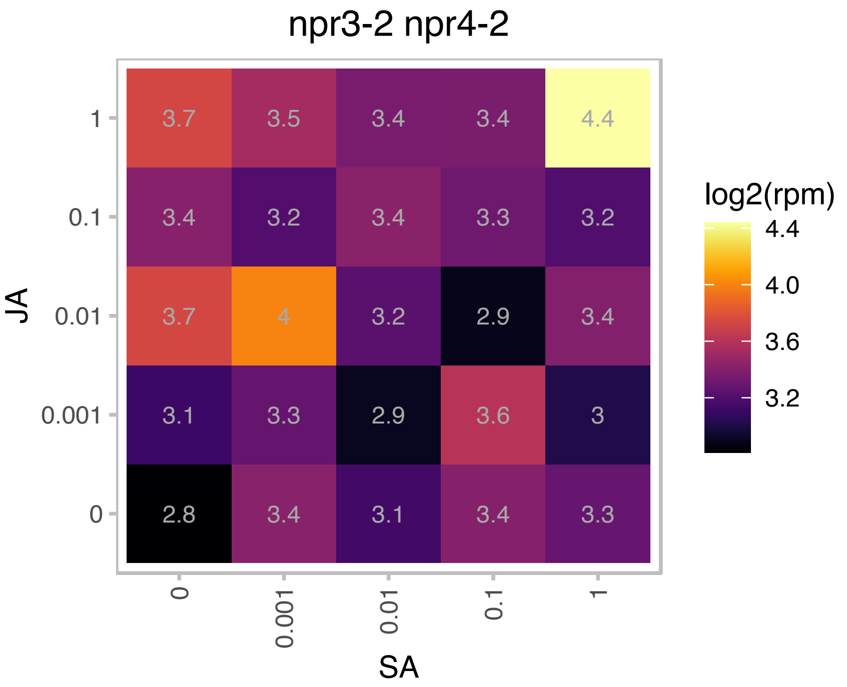


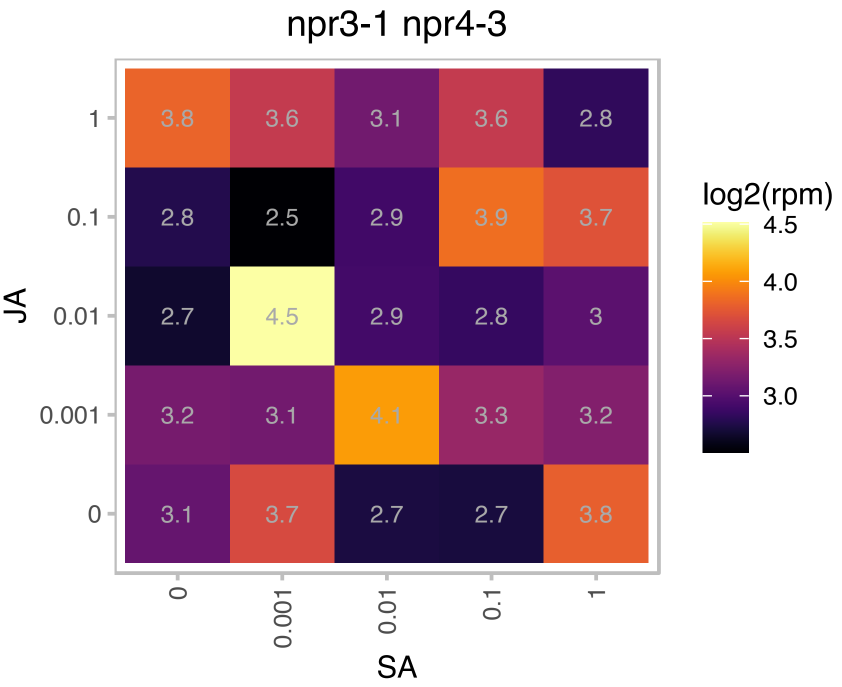

### AT4G25700.1 : B1, BCH1, BETA-OHASE 1, chy1

beta-hydroxylase 1

Converts beta-carotene to zeaxanthin via cryptoxanthin.

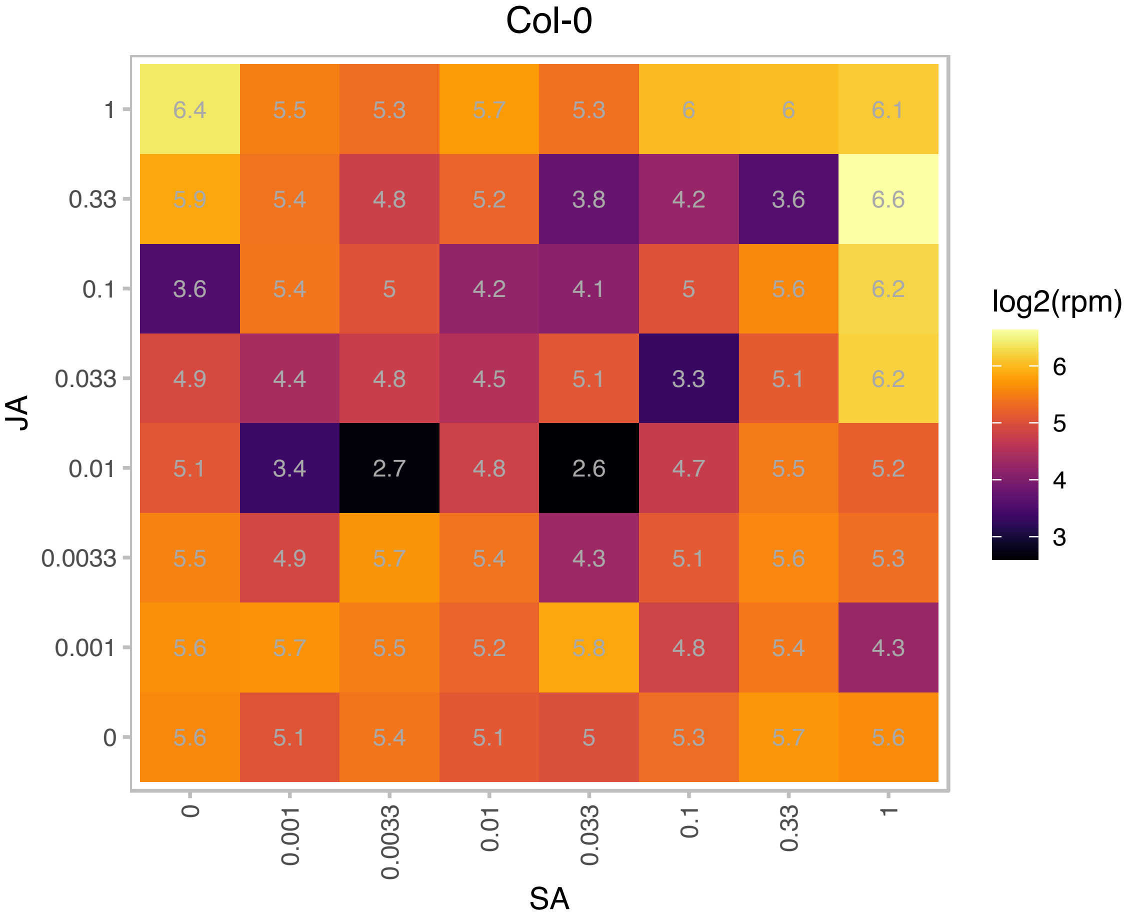


| | |
|---|---|
| Gene | AT4G25700 |
| is.expressed_genes | TRUE |
| is.responsive_genes | TRUE |
| is.apcluster | B-02 |

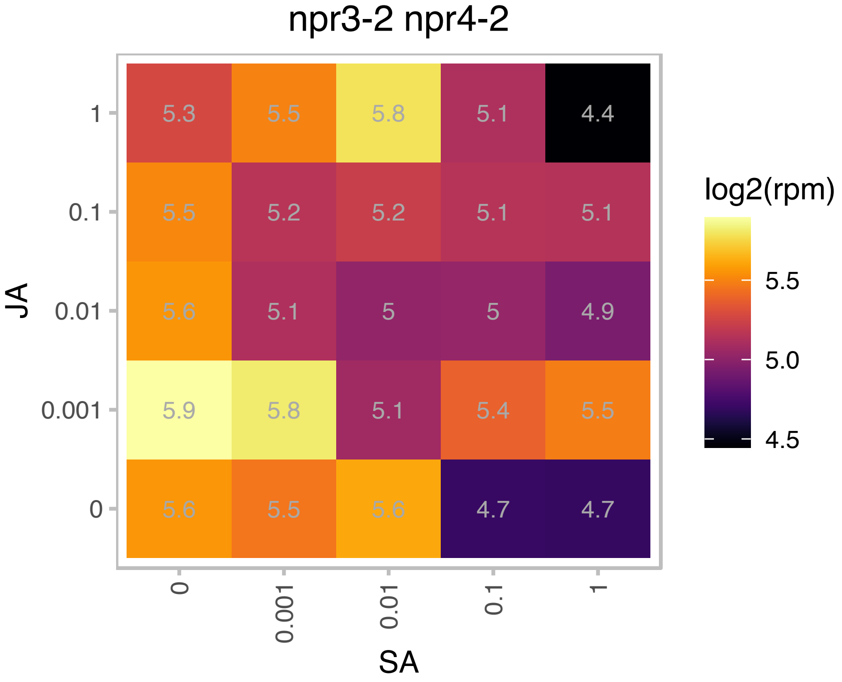


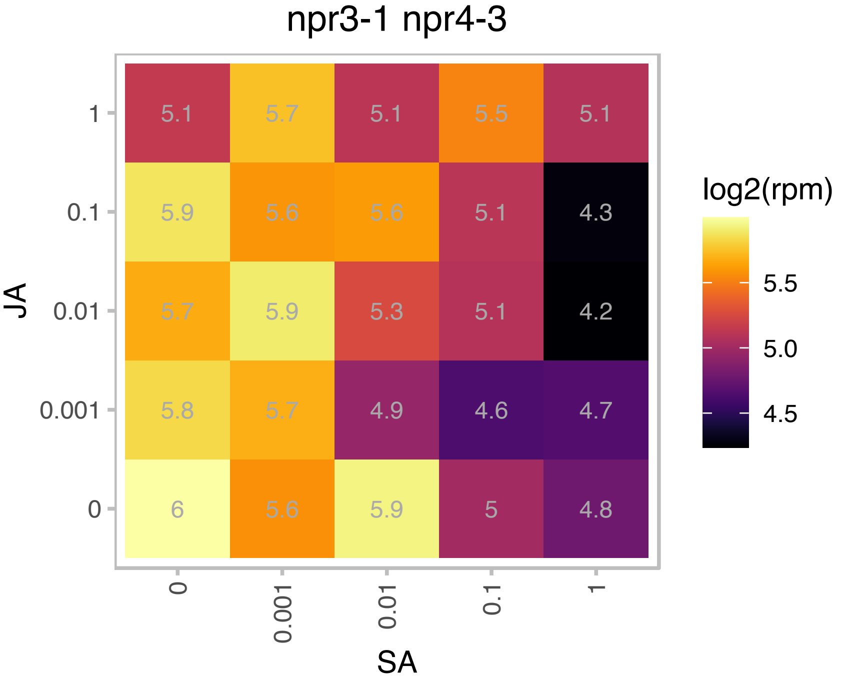

### AT4G25810.1 : XTH23, XTR6

xyloglucan endotransglycosylase 6

xyloglucan endotransglycosylase-related protein (XTR6)

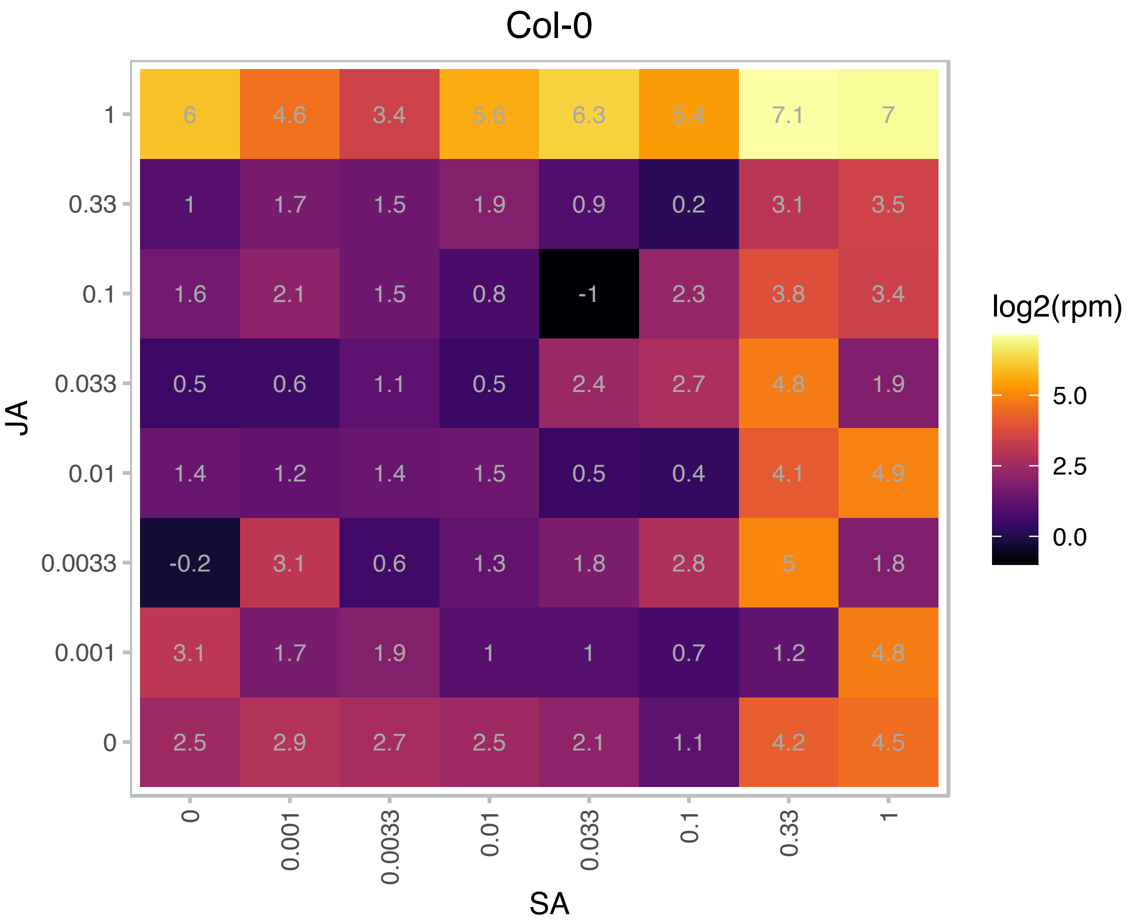


| | |
|---|---|
| Gene | AT4G25810 |
| is.expressed_genes | TRUE |
| is.responsive_genes | TRUE |
| is.apcluster | B-02 |

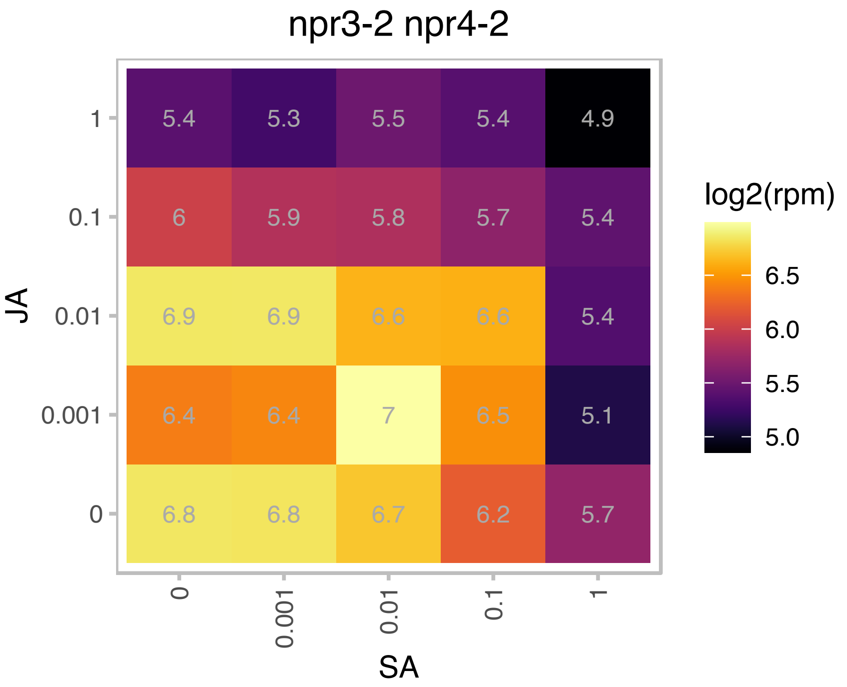


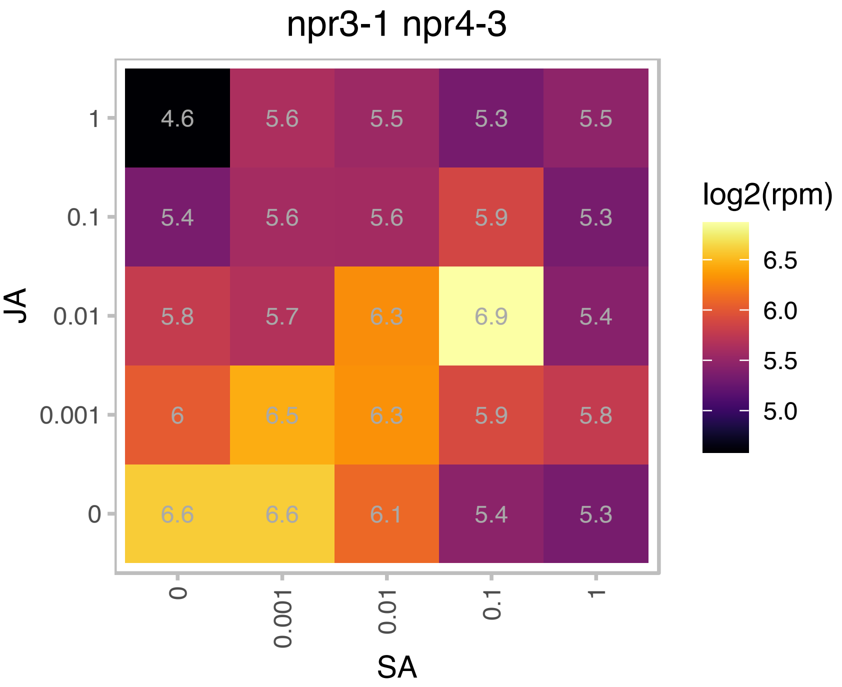

# AT4G33760.1 : NA

tRNA synthetase class II (D, K and N) family protein

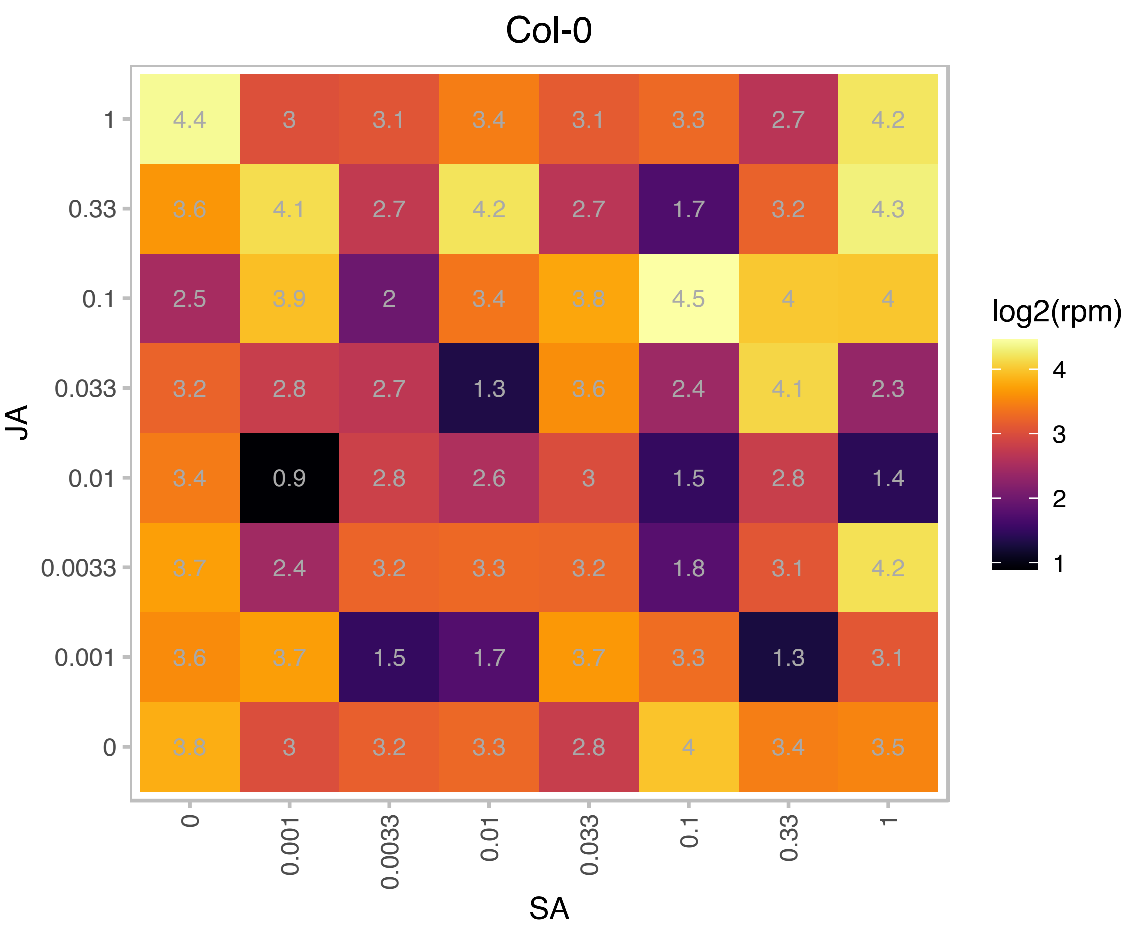


| | |
|---|---|
| Gene | AT4G33760 |
| is.expressed_genes | TRUE |
| is.responsive_genes | TRUE |
| is.apcluster | B-02 |

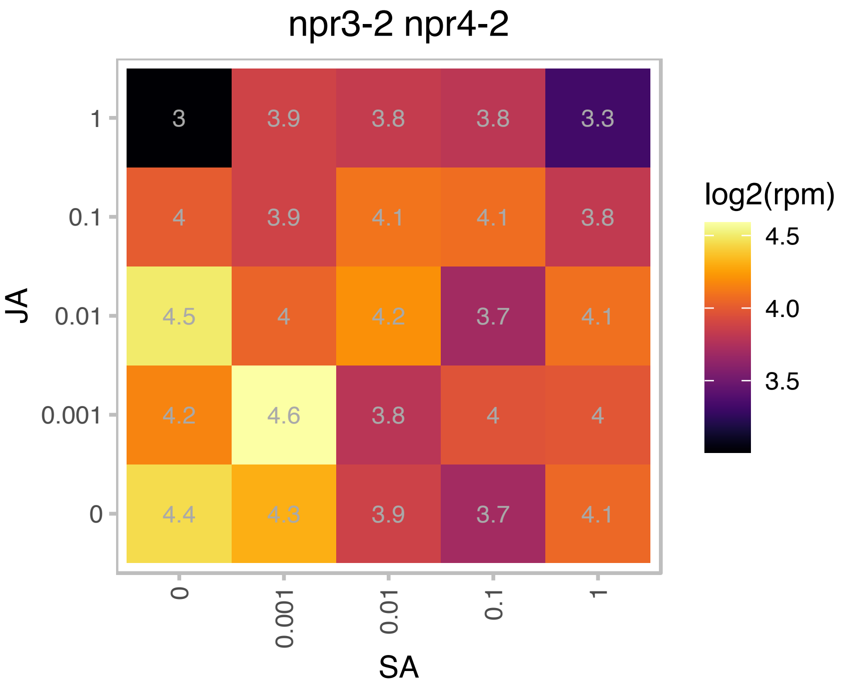


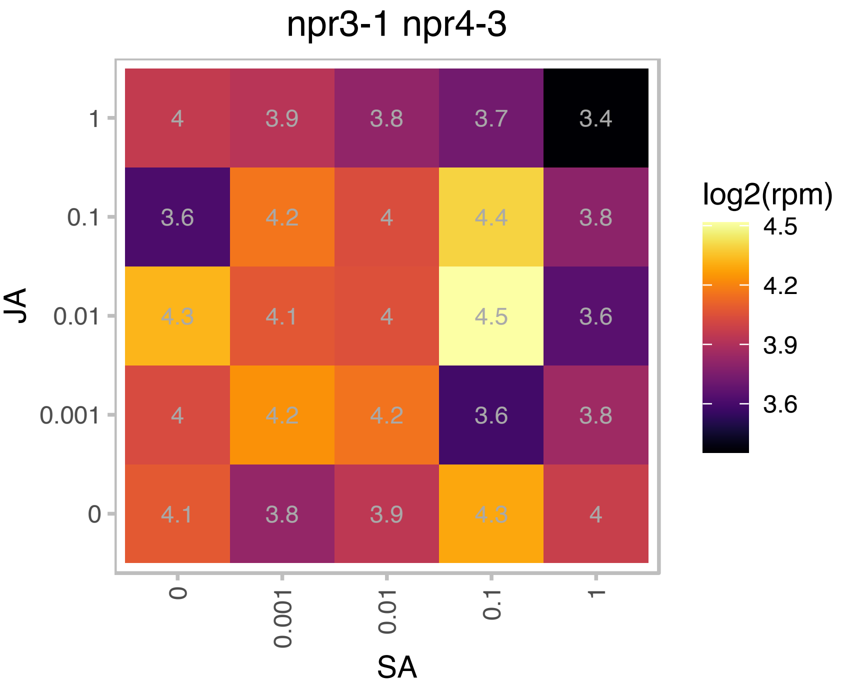

### AT4G37200.1 : HCF164

Thioredoxin superfamily protein
Encodes thioredoxin-like protein with disulfide reductase activity that is involved in the biogenesis of the plastid cytochrome b6f complex.  Protein is located in the thylakoid membrane with the C-terminal hydrophilic portion, containing the thioredoxin like domain, extending into the thylakoid lumen.

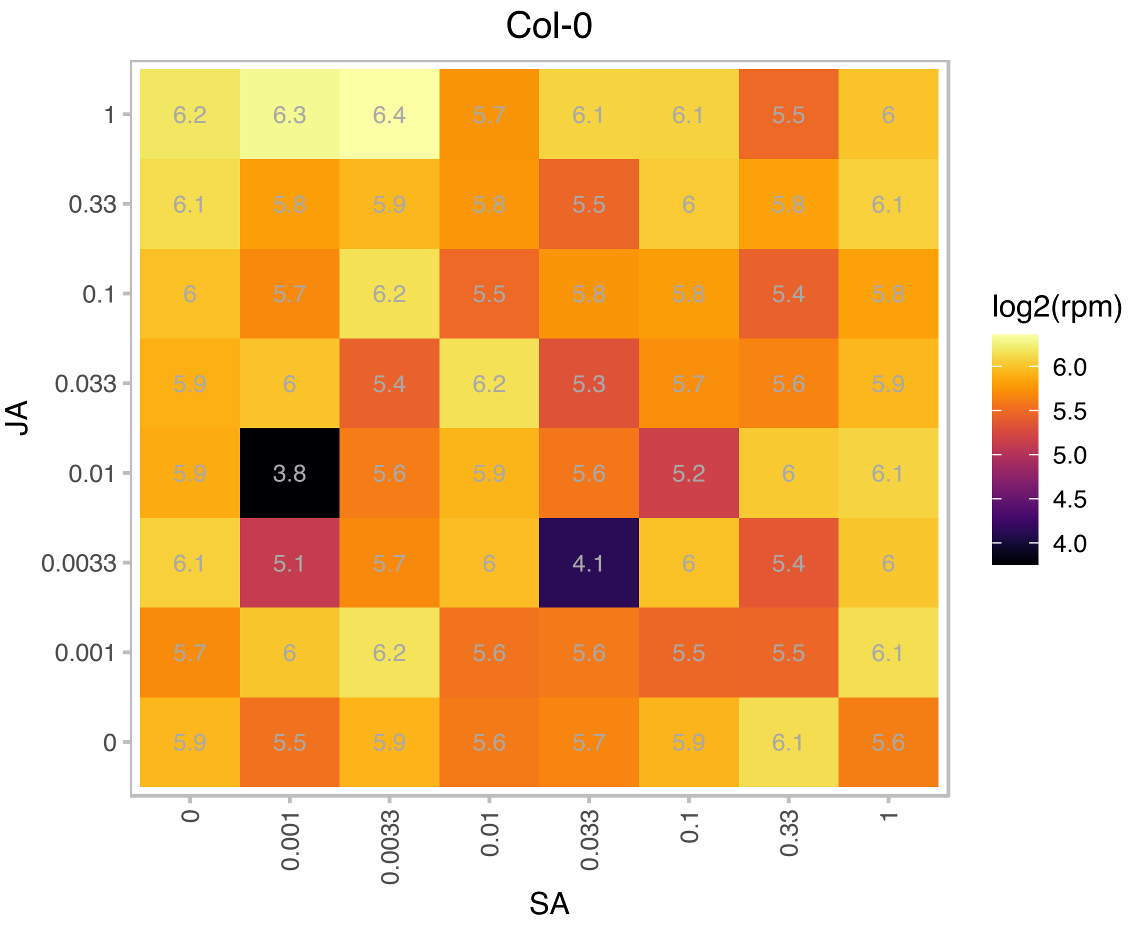


| | |
|---|---|
| Gene | AT4G37200 |
| is.expressed_genes | TRUE |
| is.responsive_genes | TRUE |
| is.apcluster | B-02 |

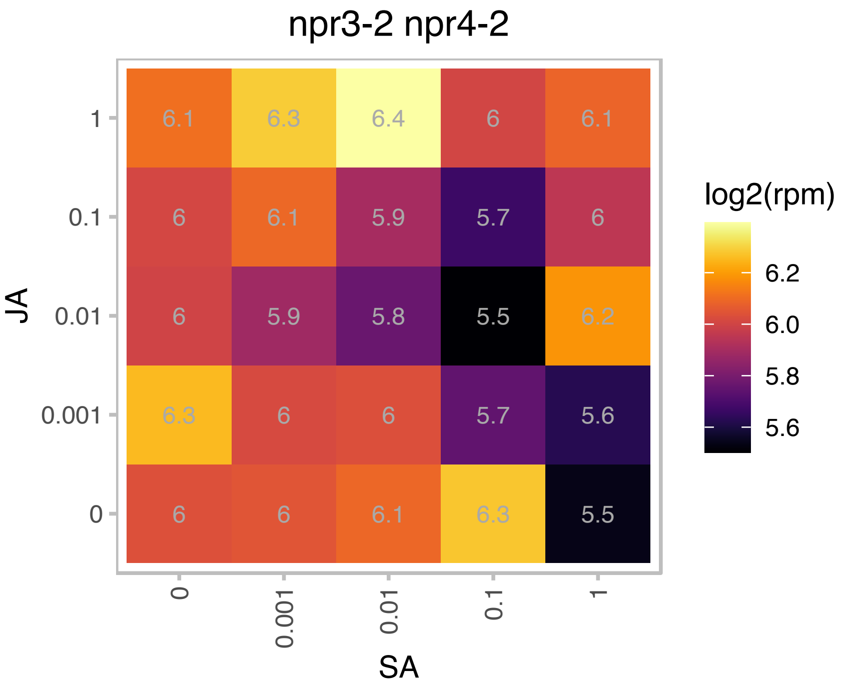


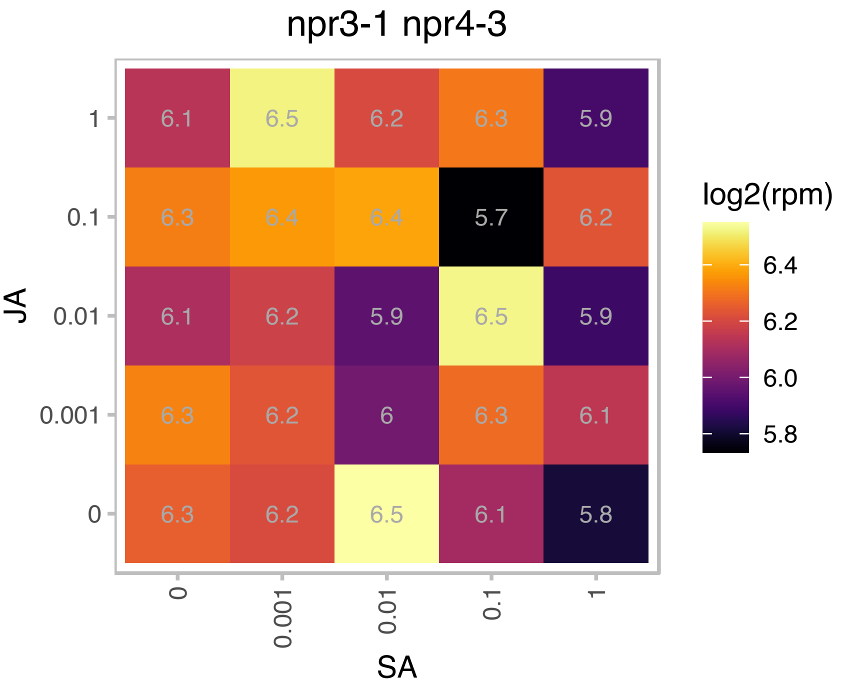

### AT4G38920.1 : ATVHA-C3, AVA-P3, VHA-C3

vacuolar-type H(+)-ATPase C3

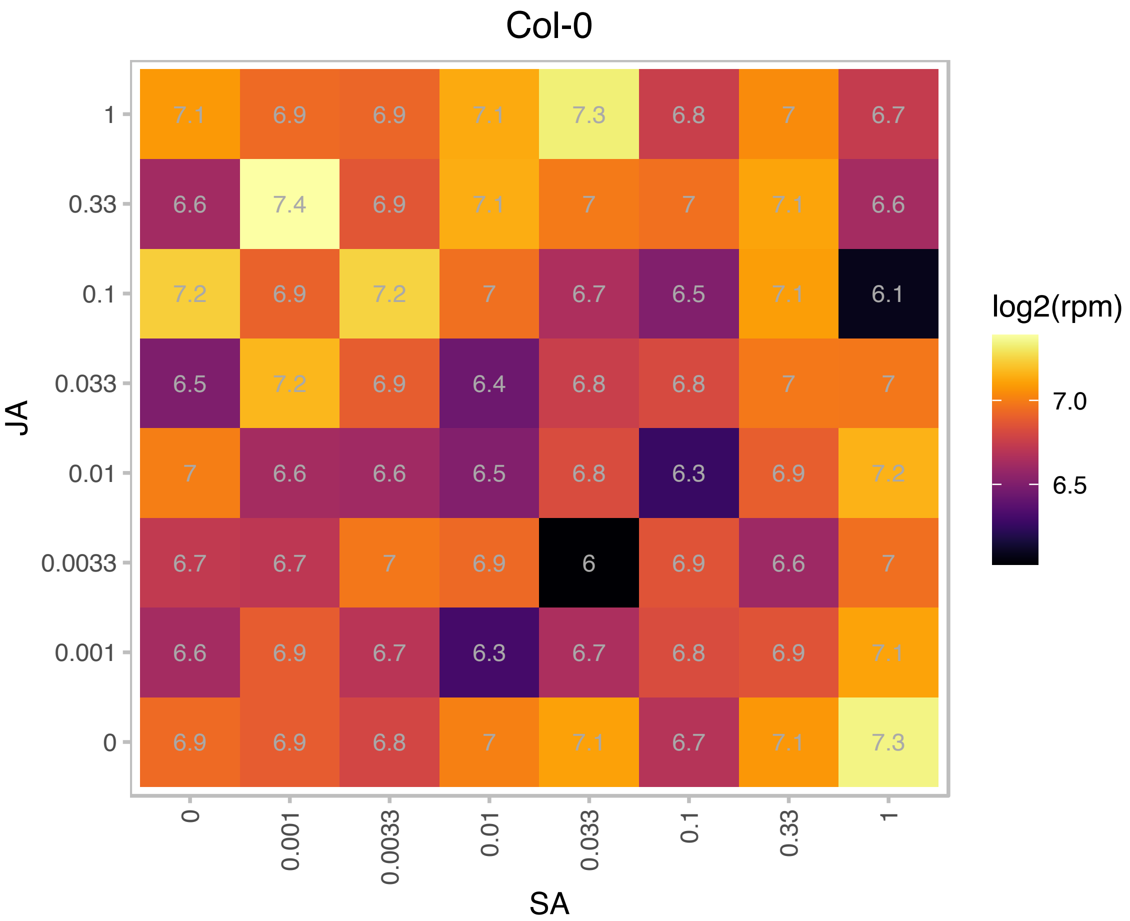


| | |
|---|---|
| Gene | AT4G38920 |
| is.expressed_genes | TRUE |
| is.responsive_genes | TRUE |
| is.apcluster | B-02 |

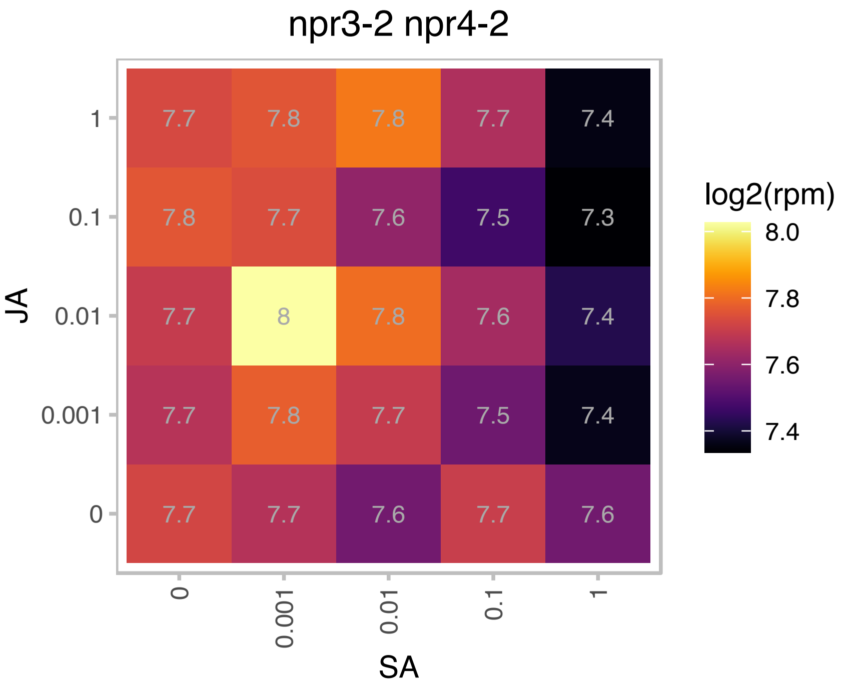


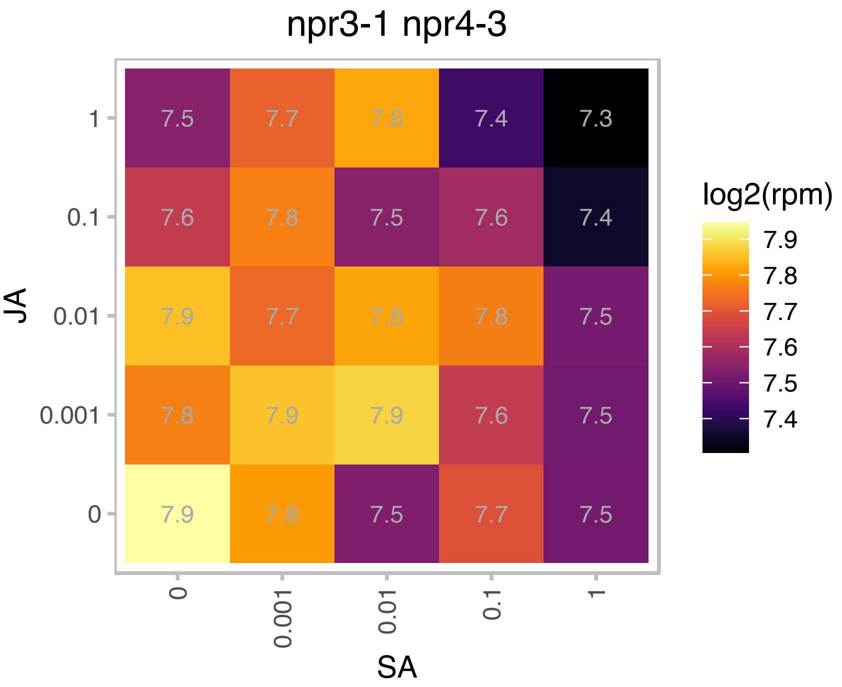

### AT4G39800.1 : ATIPS1, ATMIPS1, MI-1-P SYNTHASE, MI

myo-inositol-1-phosphate synthase 1
** Referred to as MIPS2 in Mitsuhashi et al 2008. myo-inositol-1-phosphate synthase isoform 1.Expressed in leaf, root and silique. Immunolocalization experiments with an antibody recognizing MIPS1, MIPS2, and MIPS3 showed endosperm localization.

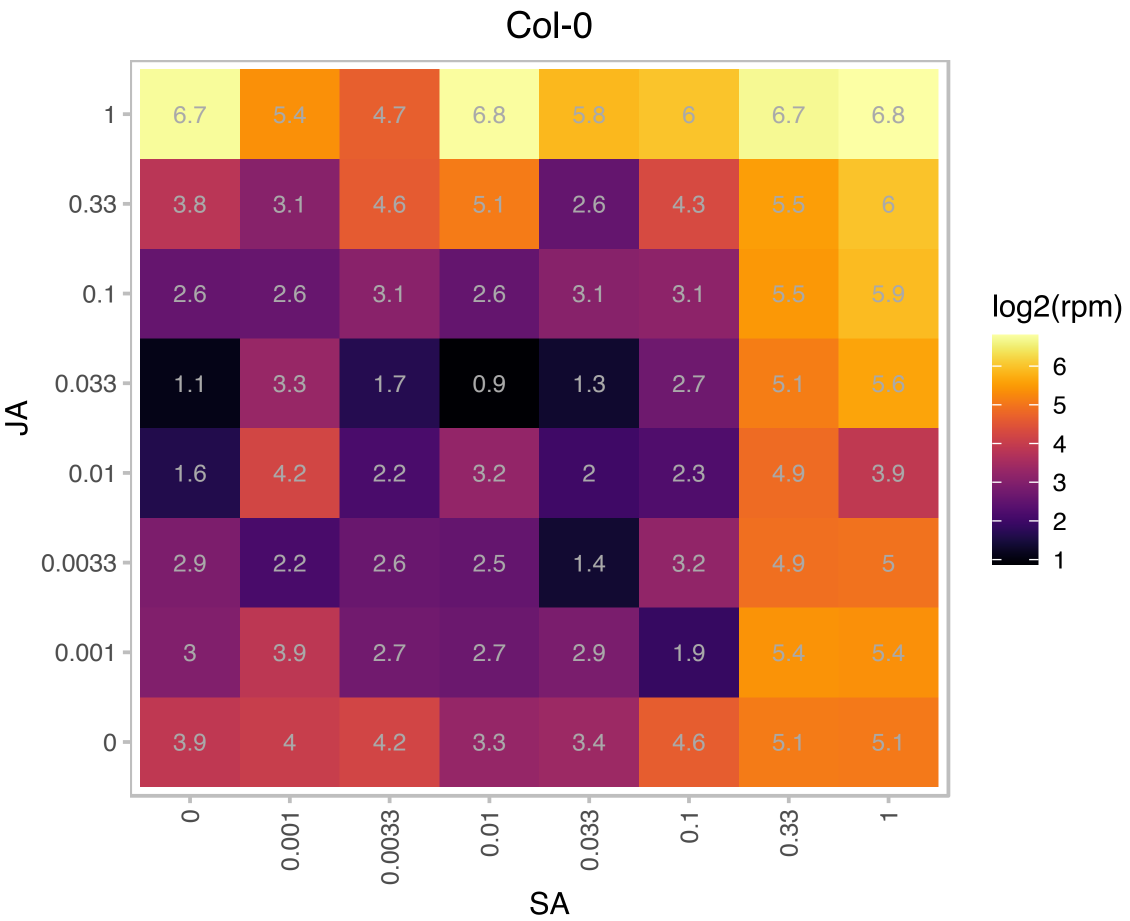


| | |
|---|---|
| Gene | AT4G39800 |
| is.expressed_genes | TRUE |
| is.responsive_genes | TRUE |
| is.apcluster | B-02 |

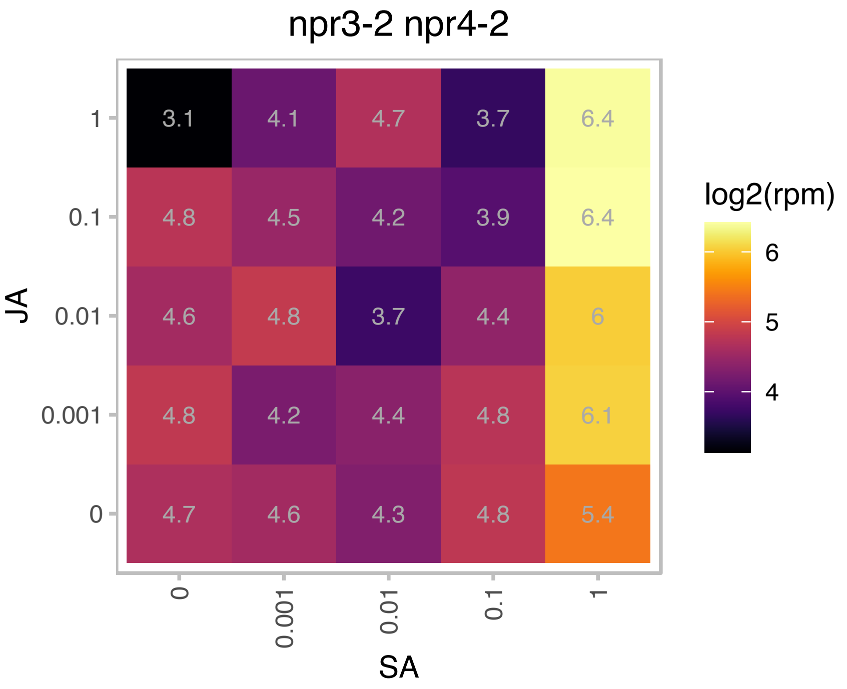


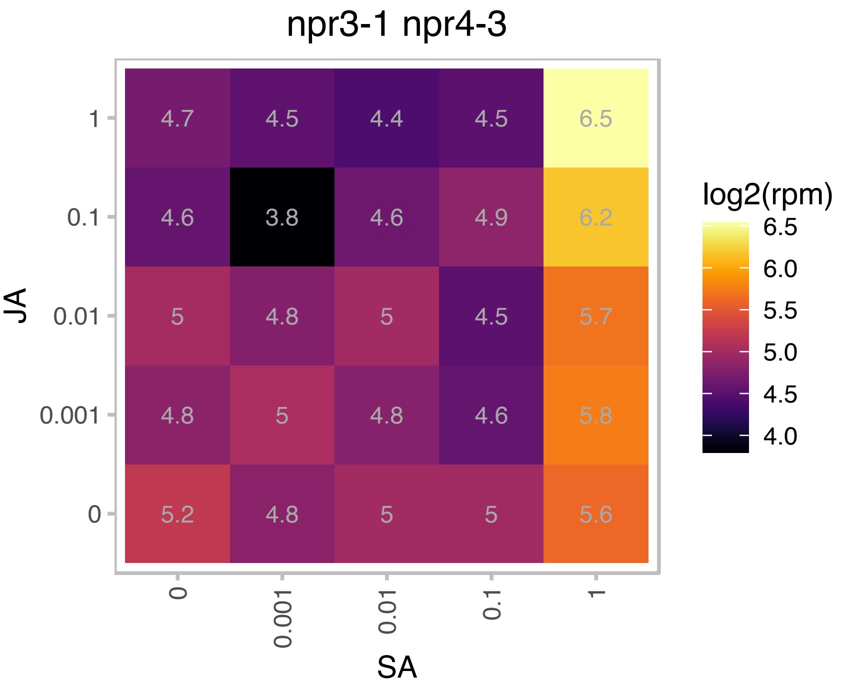

# AT5G03380.1 : NA

Heavy metal transport/detoxification superfamily protein

| | |
|---|---|
| Gene | AT5G03380 |
| is.expressed_genes | TRUE |
| is.responsive_genes | TRUE |
| is.apcluster | B-02 |

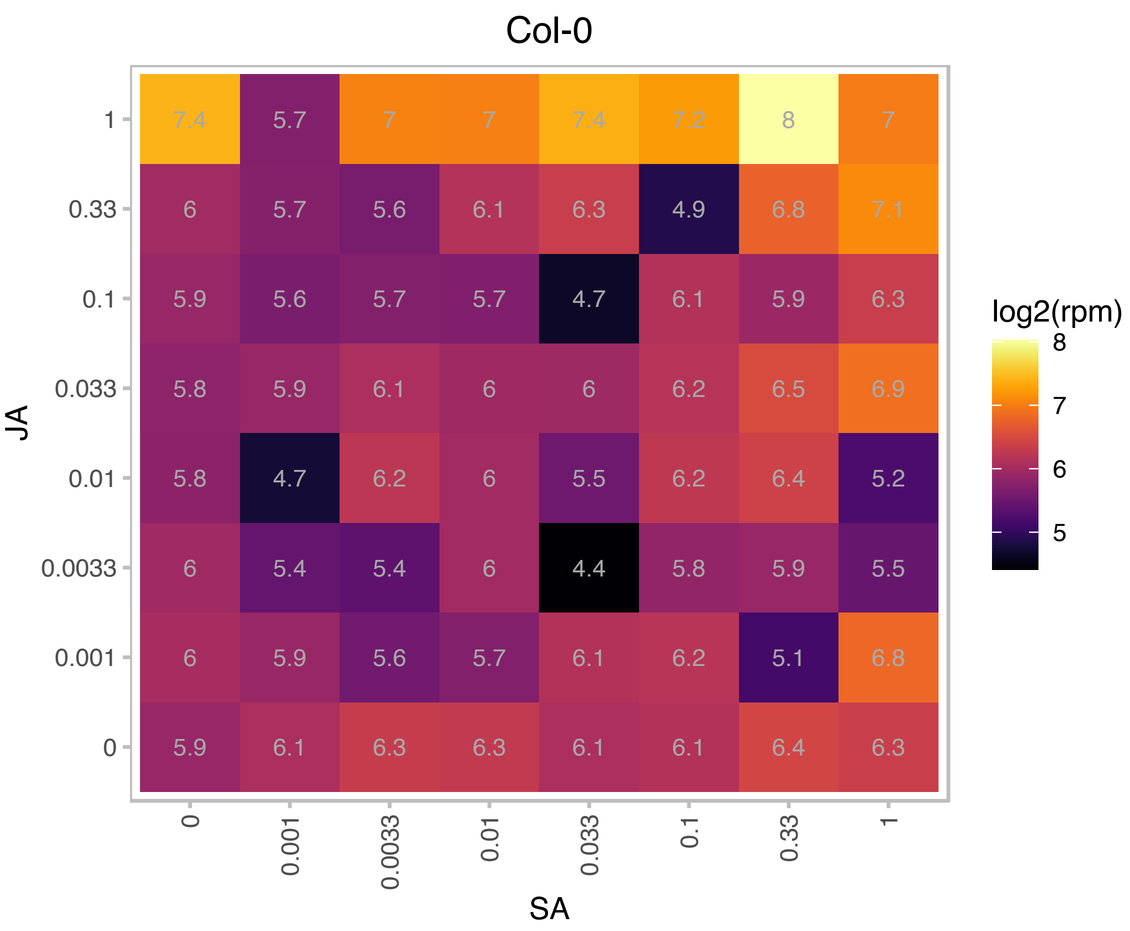


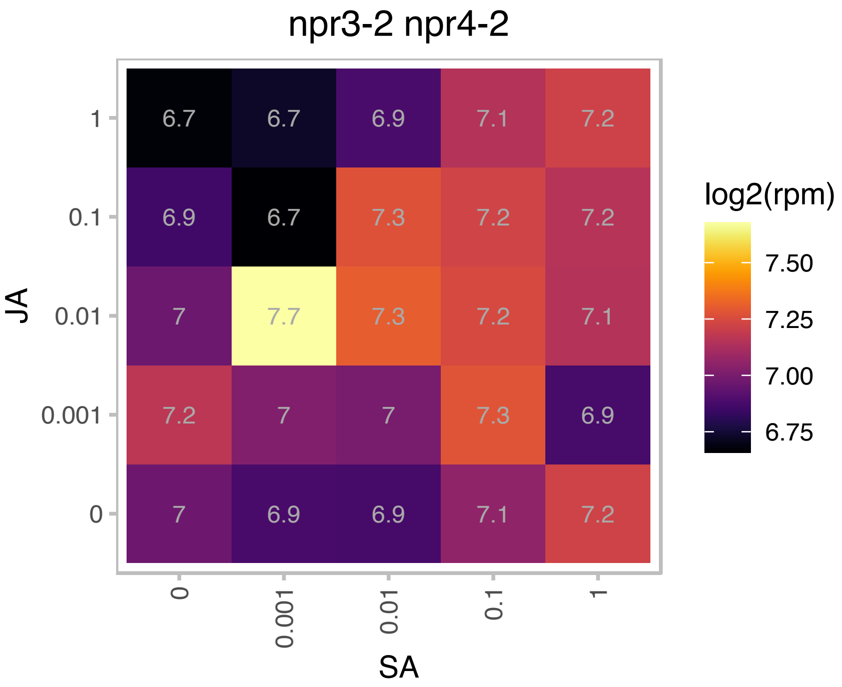


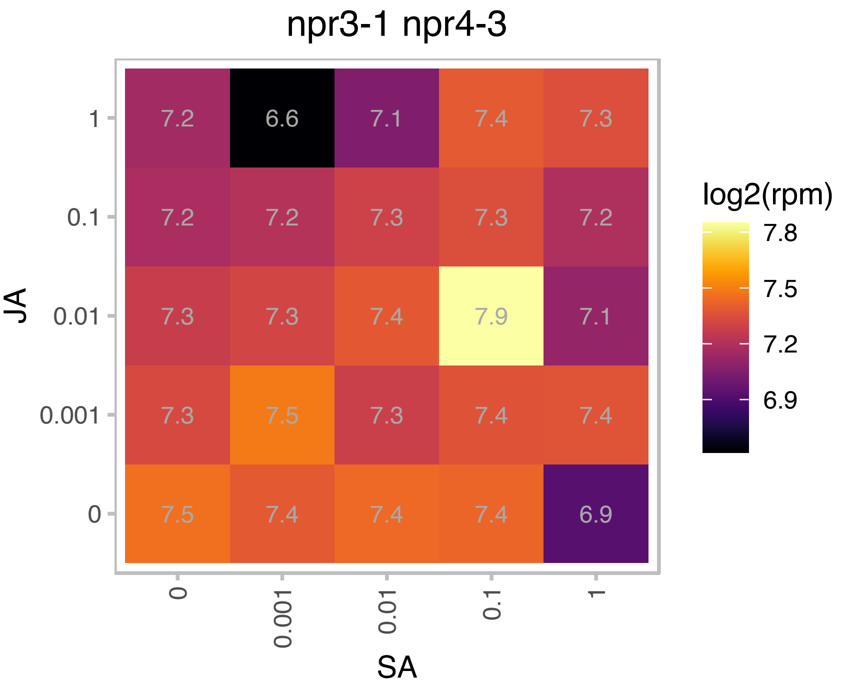

# AT5G06385.1 : NA

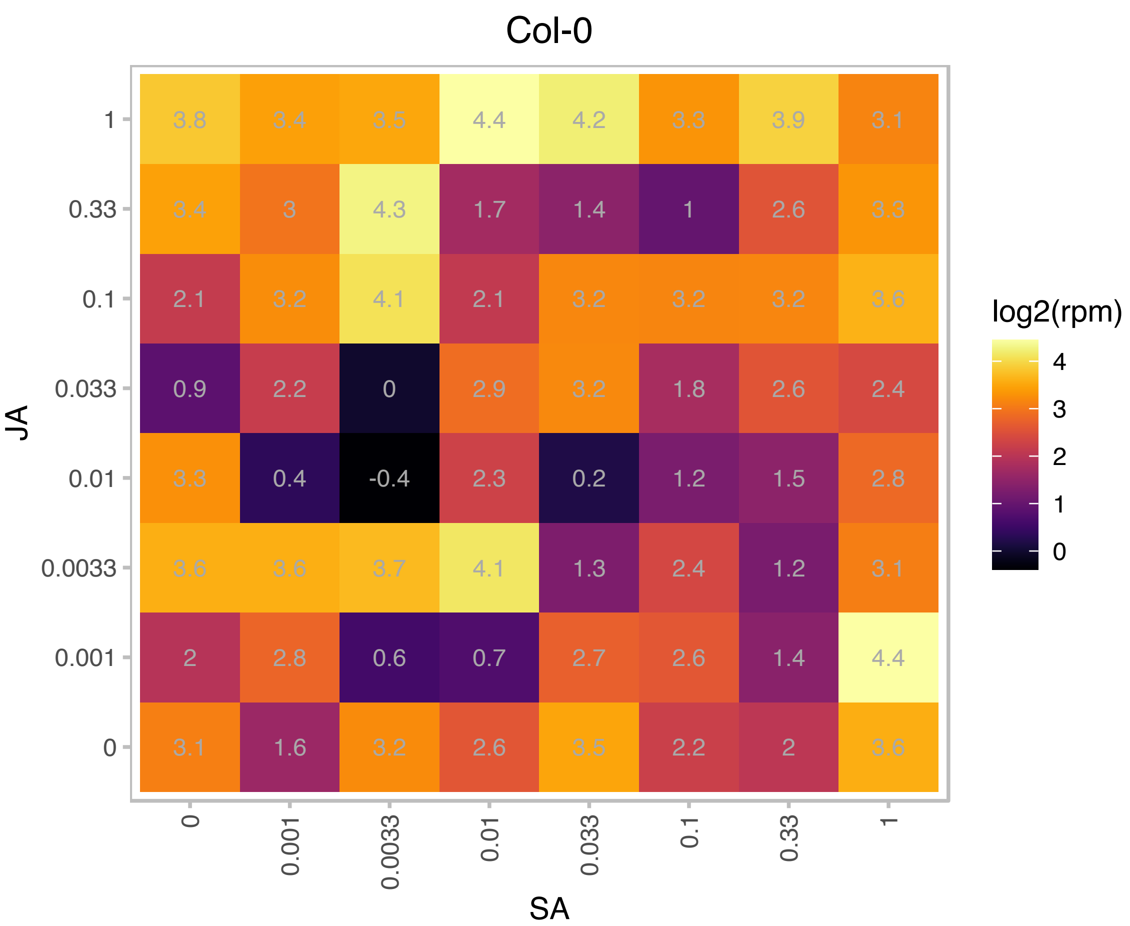


| | |
|---|---|
| Gene | AT5G06385 |
| is.expressed_genes | TRUE |
| is.responsive_genes | TRUE |
| is.apcluster | B-02 |

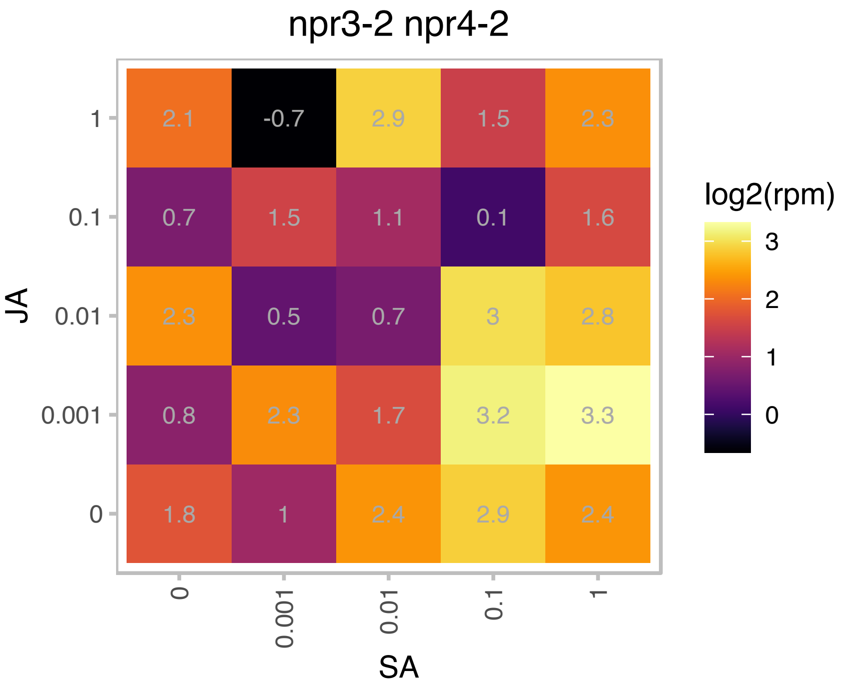


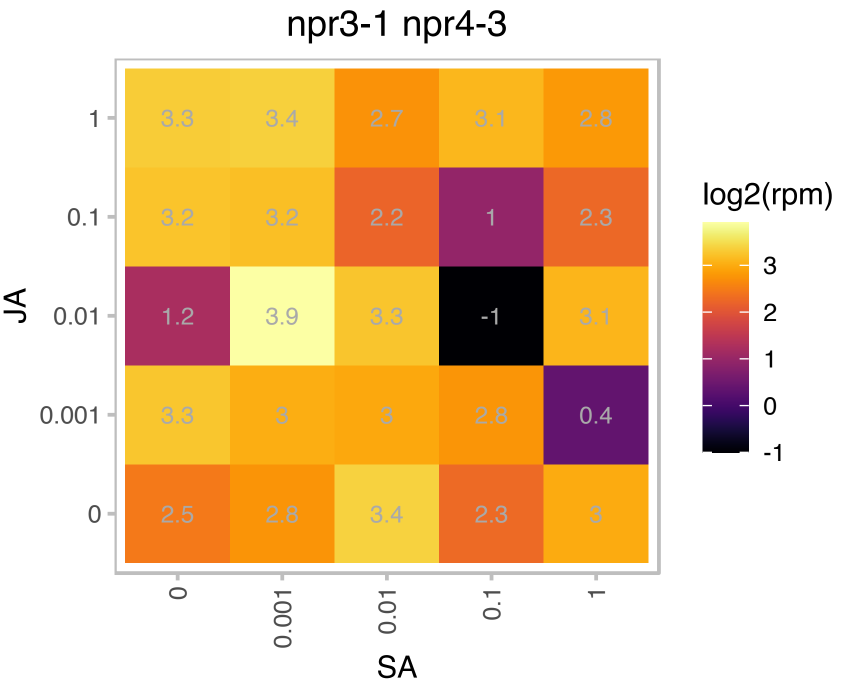

### AT5G13120.1 : ATCYP20-2, CYP20-2, Pnsl5

cyclophilin 20-2

Encodes a lumenal cyclophilin with peptidyl-prolyl isomerase activity that is associated with the NAD(P)H dehydrogenase complex in stromal regions of the thylakoid membrane. It is likely to be important for the accumulation of the hydrophobic domain of the NAD(P)H dehydrogenase complex. This complex is associated with PSI and is responsible for the reduction of plastoquinone.

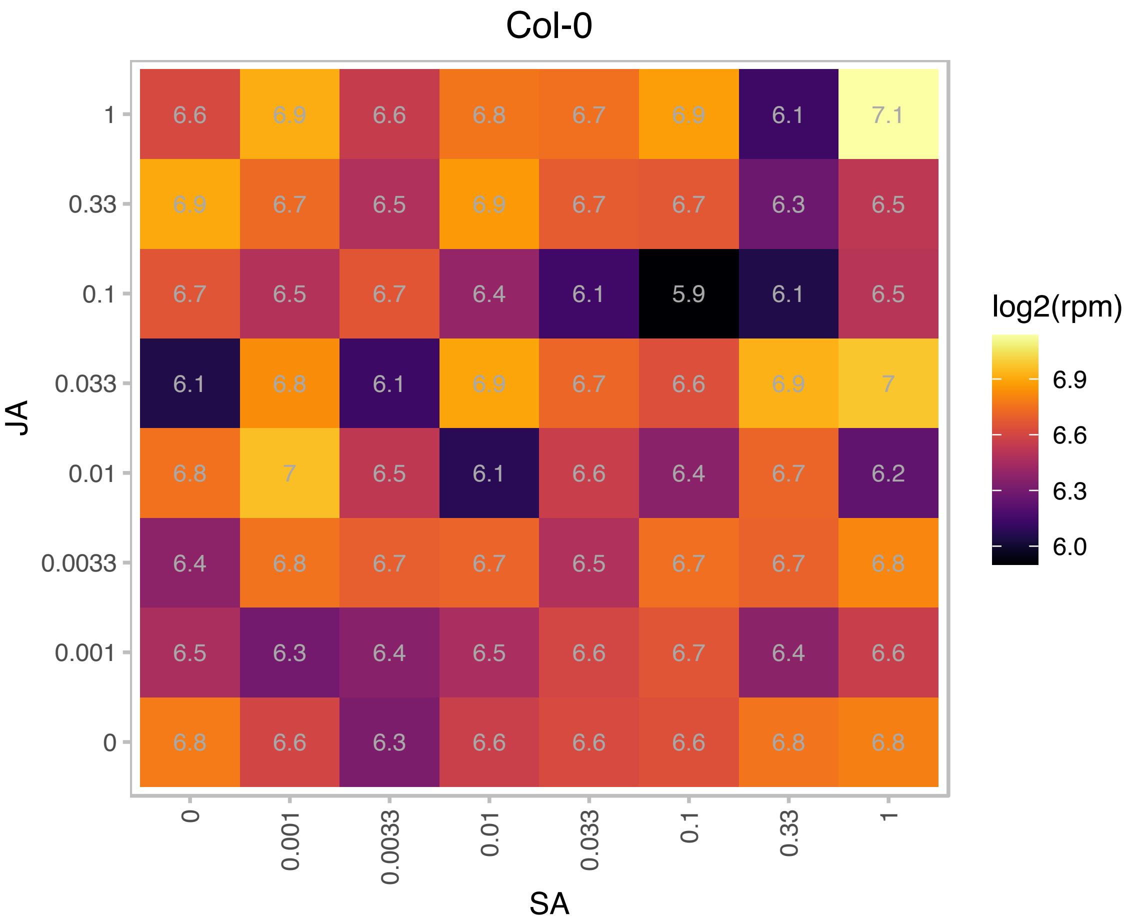


| | |
|---|---|
| Gene | AT5G13120 |
| is.expressed_genes | TRUE |
| is.responsive_genes | TRUE |
| is.apcluster | B-02 |

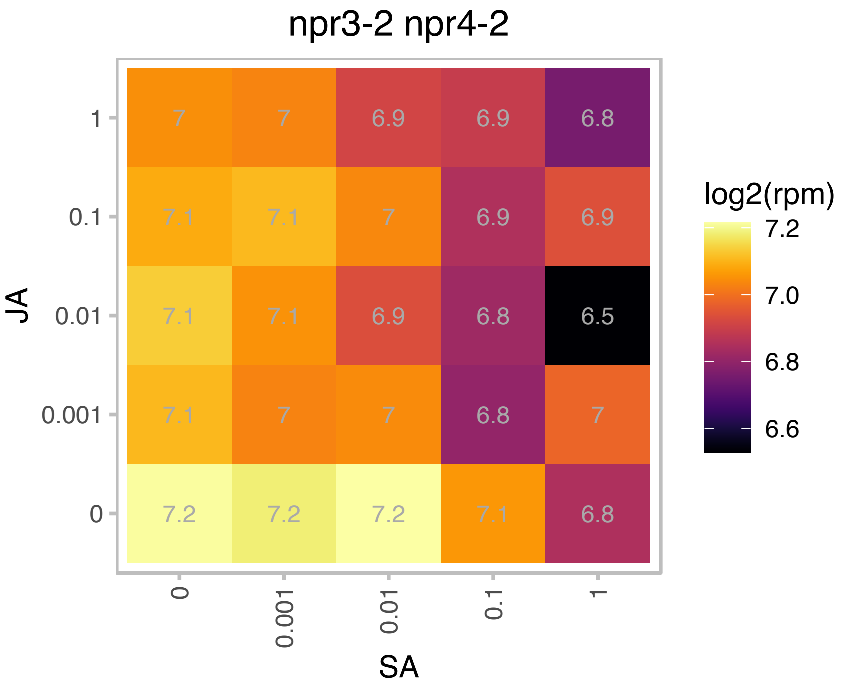


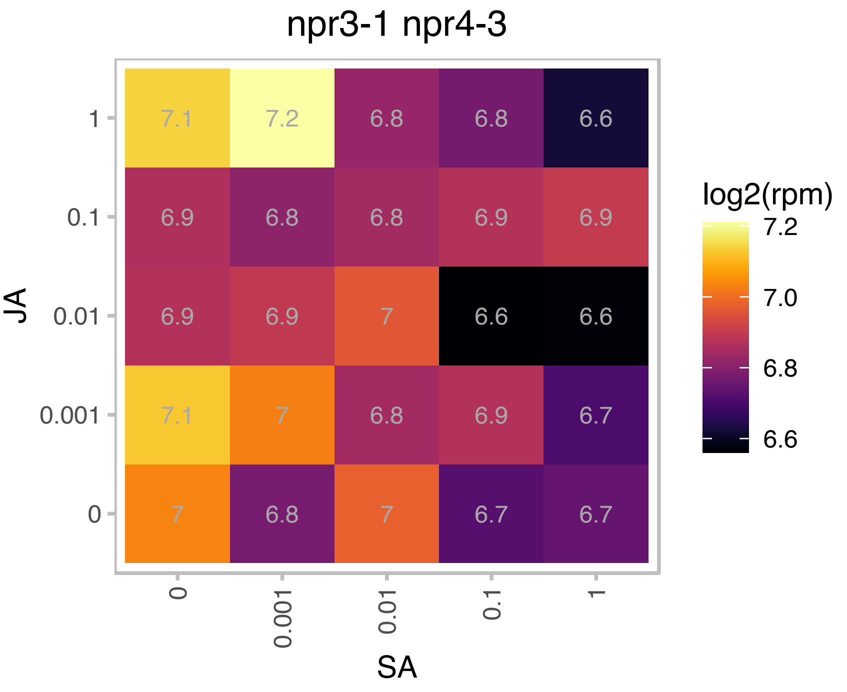

### AT5G24655.1 : LSU4

response to low sulfur 4

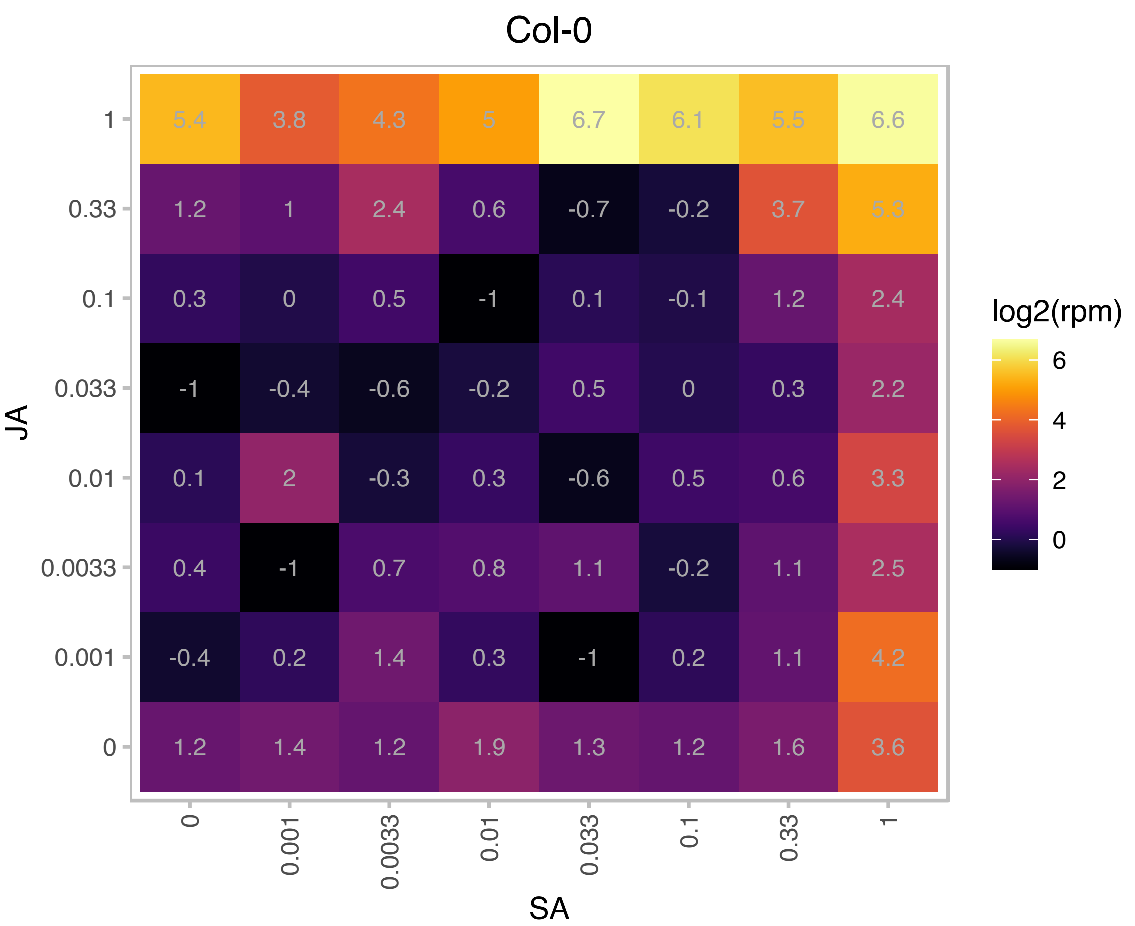


| | |
|---|---|
| Gene | AT5G24655 |
| is.expressed_genes | TRUE |
| is.responsive_genes | TRUE |
| is.apcluster | B-02 |

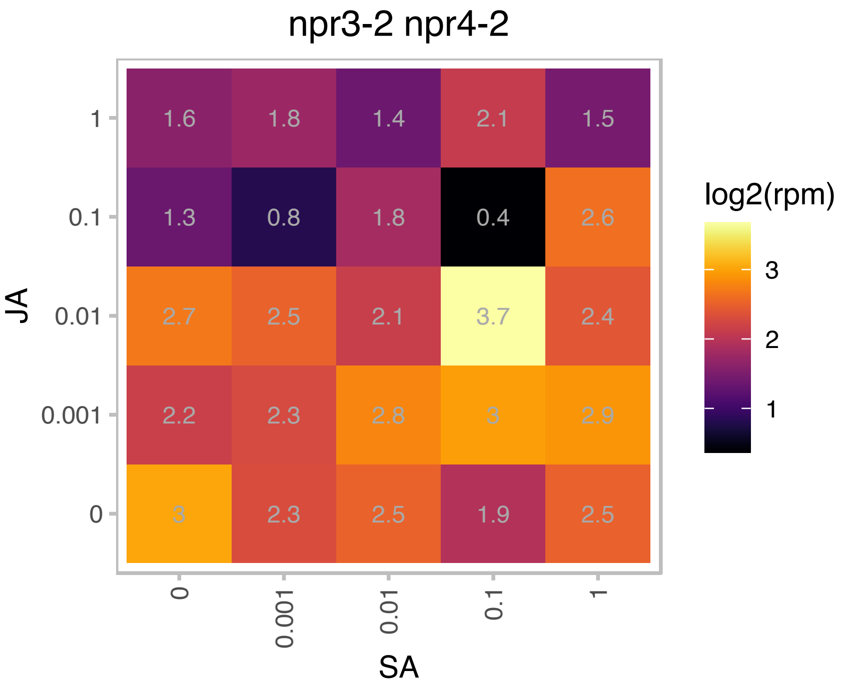


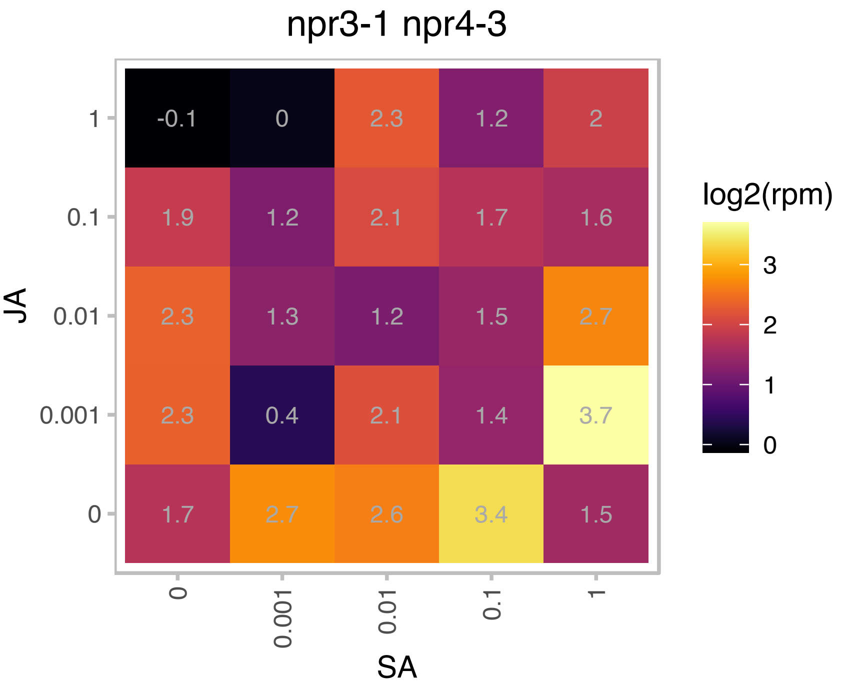

### AT5G25610.1 : ATRD22, RD22

BURP domain-containing protein

responsive to dehydration 22 (RD22) mediated by ABA

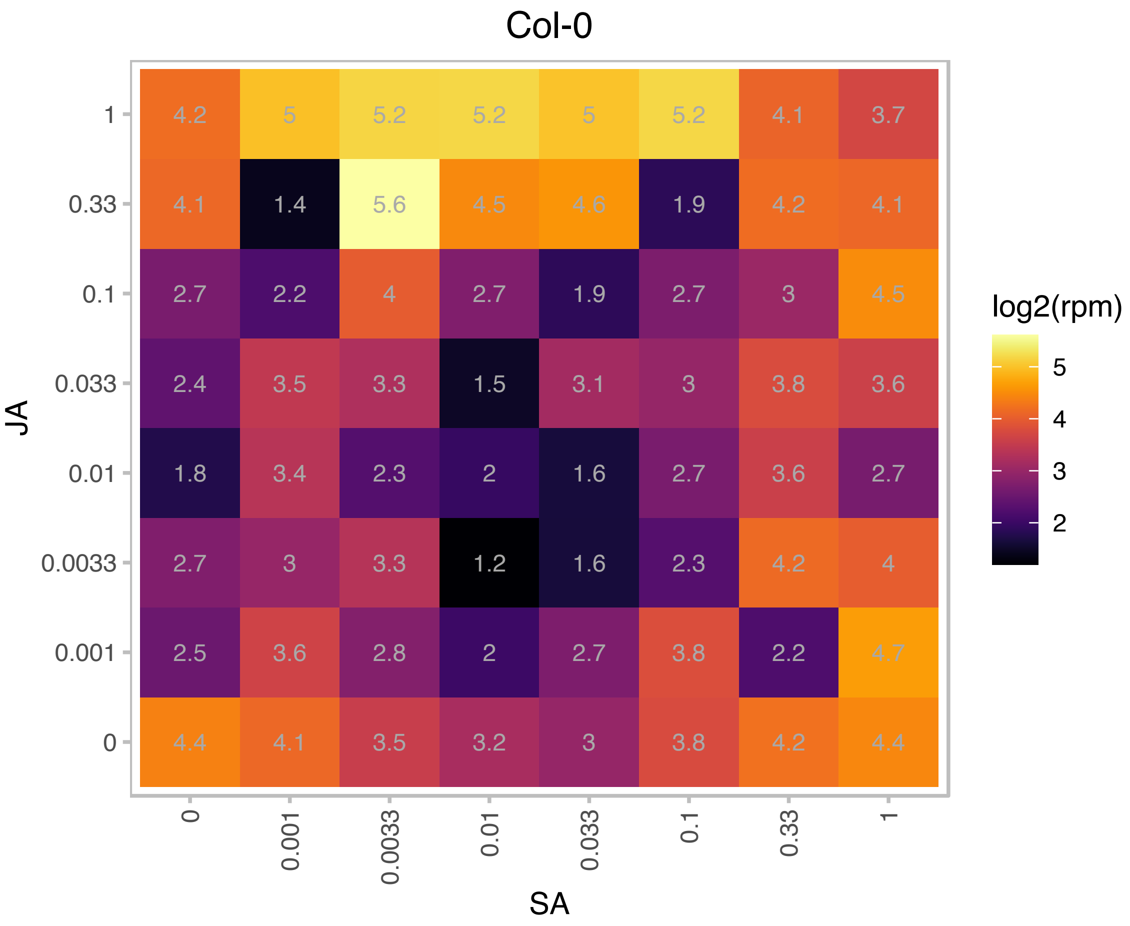


| | |
|---|---|
| Gene | AT5G25610 |
| is.expressed_genes | TRUE |
| is.responsive_genes | TRUE |
| is.apcluster | B-02 |

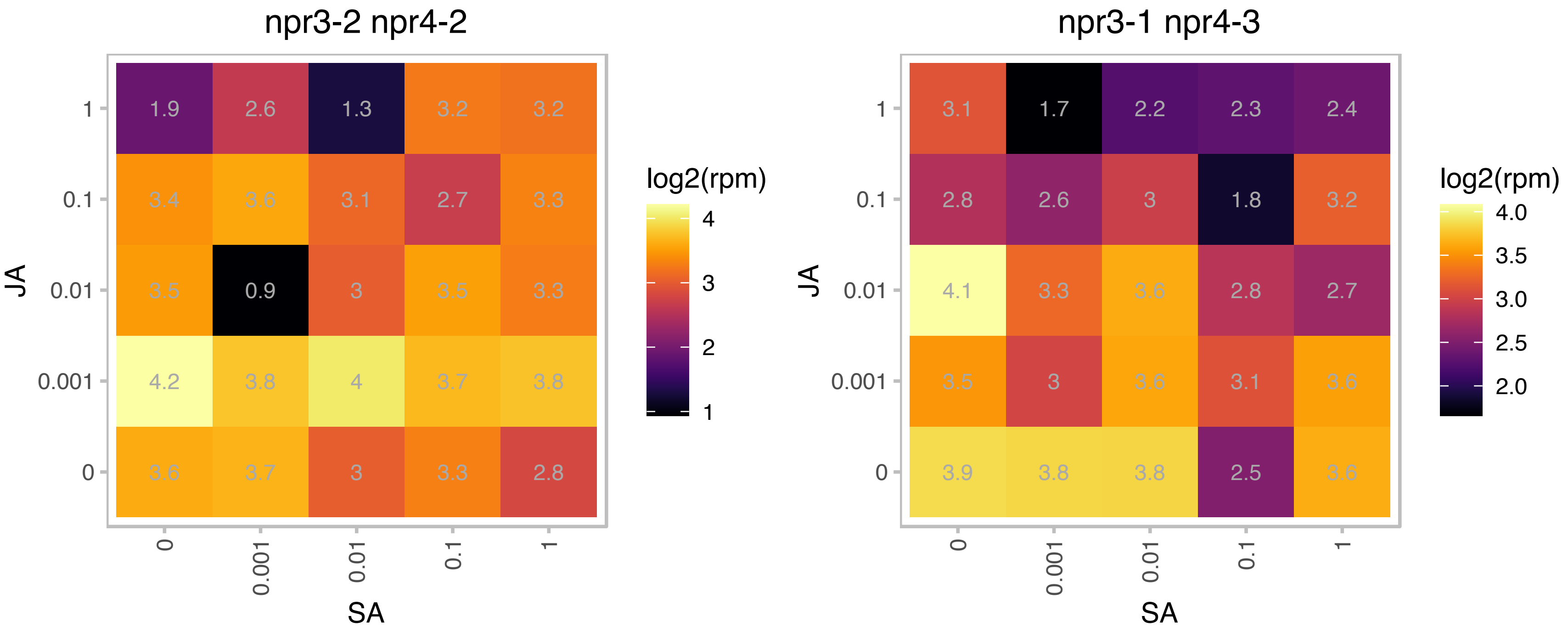

### AT5G27420.1 : ATL31, CNI1

carbon/nitrogen insensitive 1
Encodes CNI1 (Carbon/Nitrogen Insensitive1) (also named as ATL31), a RING type ubiquitin ligase that functions in the Carbon/Nitrogen response for growth phase transition in Arabidopsis seedlings. The mRNA is cell-to-cell mobile.

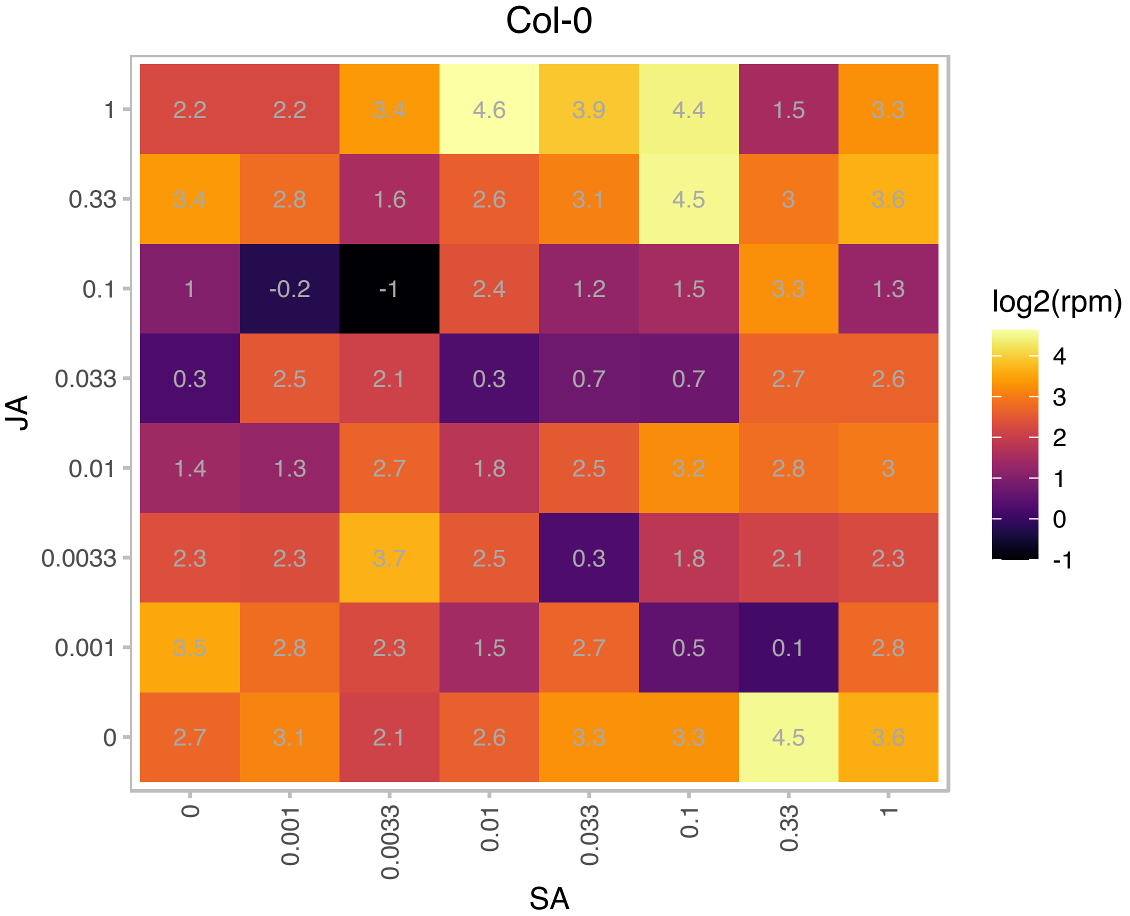


| | |
|---|---|
| Gene | AT5G27420 |
| is.expressed_genes | TRUE |
| is.responsive_genes | TRUE |
| is.apcluster | B-02 |

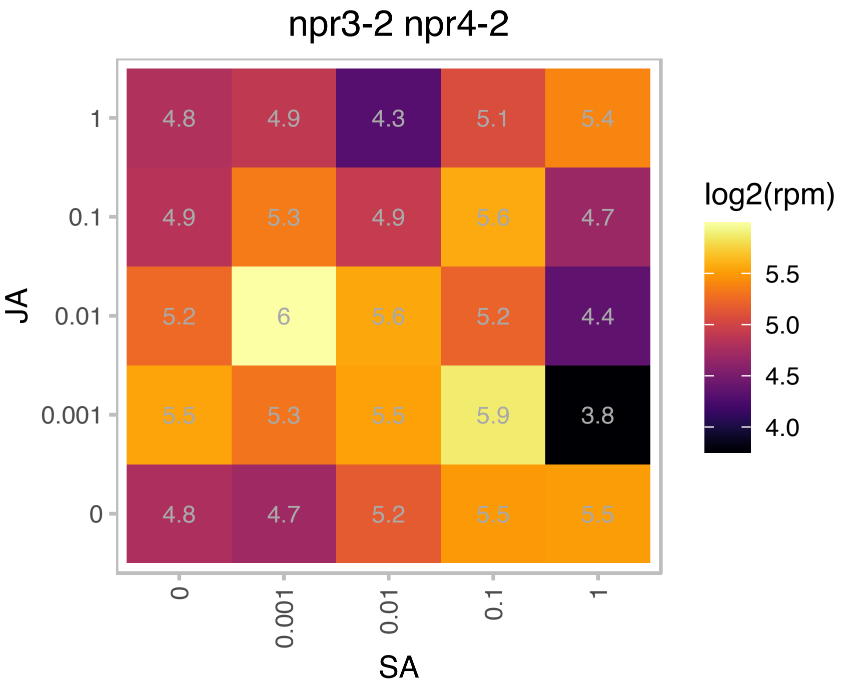


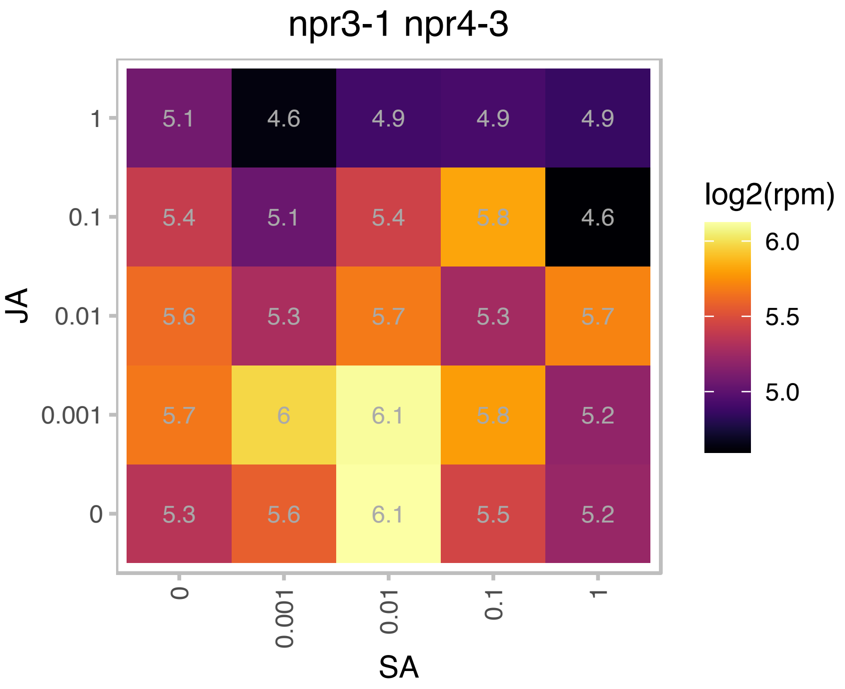

### AT5G27640.1 : ATEIF3B-1, ATTIF3B1, EIF3B, EIF3B-1, TIF

translation initiation factor 3B1

encodes a member of eukaryotic translation initiation factor 3B family.

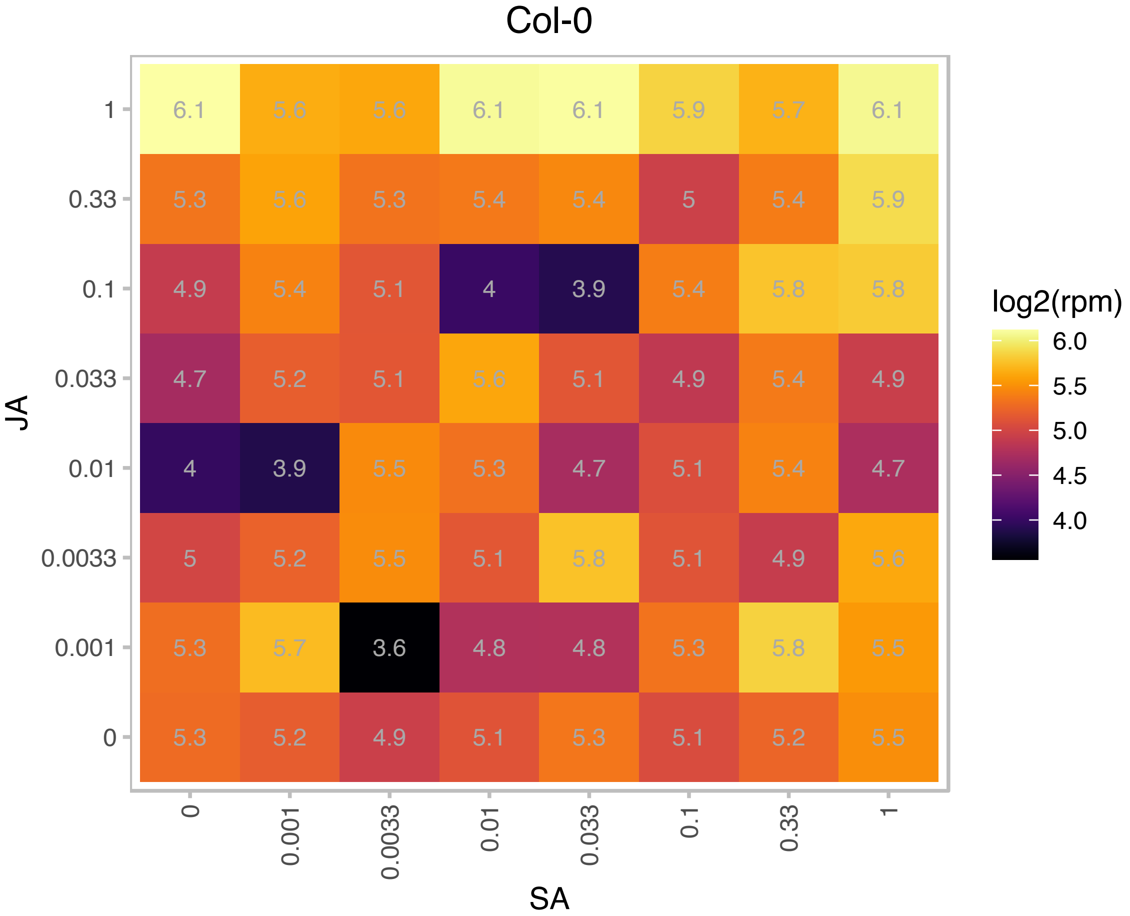


| | |
|---|---|
| Gene | AT5G27640 |
| is.expressed_genes | TRUE |
| is.responsive_genes | TRUE |
| is.apcluster | B-02 |

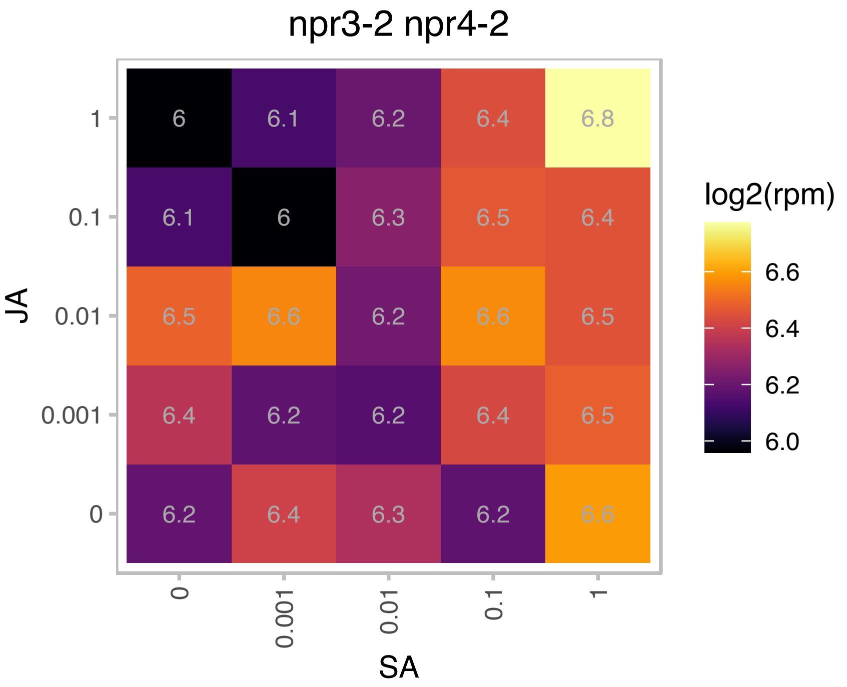


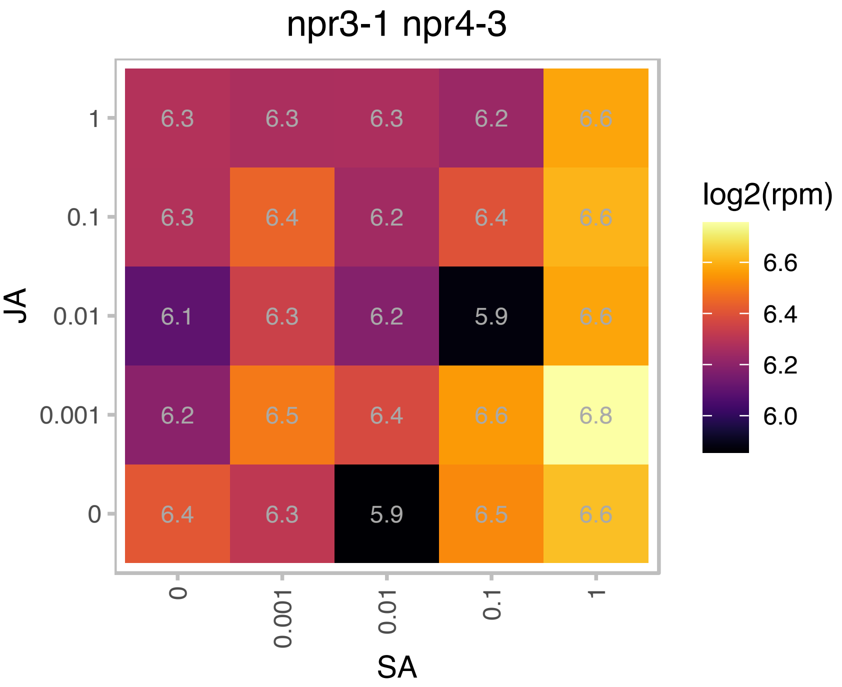

### AT5G41790.1 : CIP1

COP1-interactive protein 1
encodes a protein that physically interacts specifically with the putative coiled-coil region of COP1 in vitro. In hypocotyl and cotyledon protoplasts, it is associated to the cytoskeleton, but not in the root. expression is not regulated by light. The mRNA is cell-to-cell mobile.

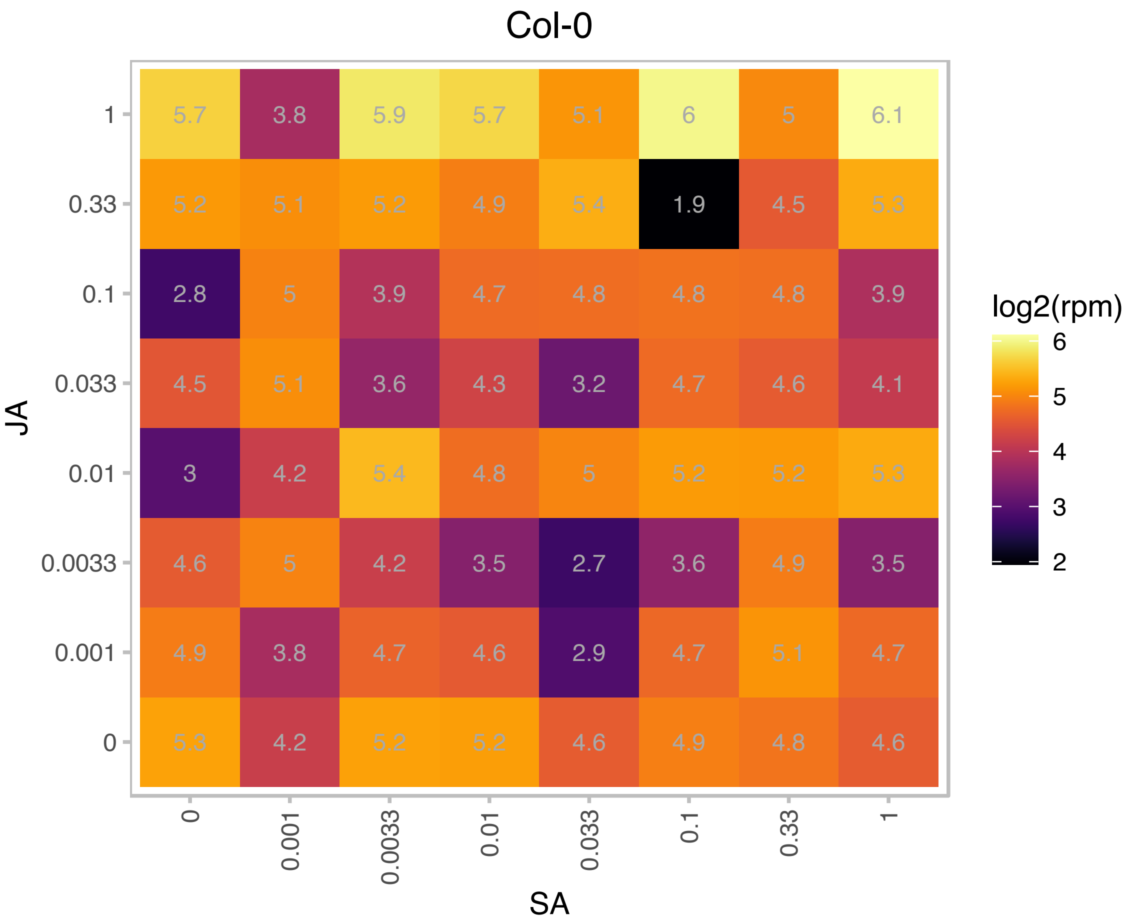


| | |
|---|---|
| Gene | AT5G41790 |
| is.expressed_genes | TRUE |
| is.responsive_genes | TRUE |
| is.apcluster | B-02 |

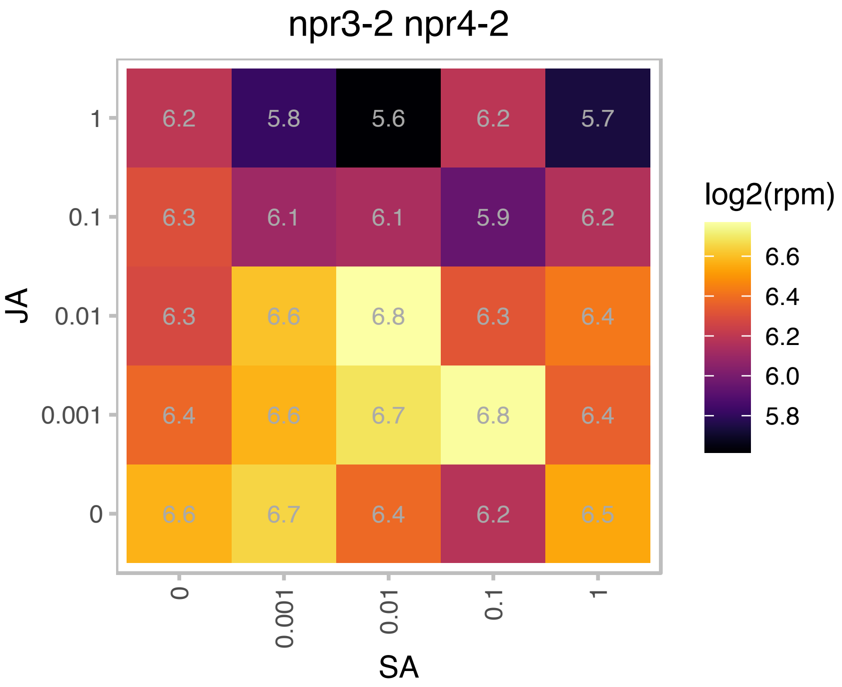


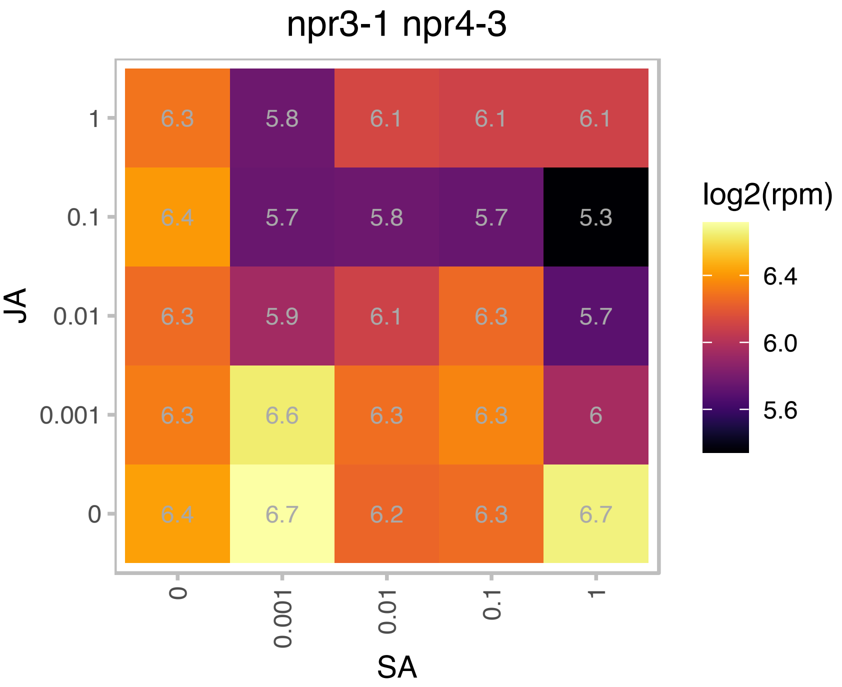

### AT5G42190.1 : ASK2, SKP1B

E3 ubiquitin ligase SCF complex subunit SKP1/ASK1 family protein

Similar to SKP1 in yeast and humans which are involved in mitotic cell cycle control and ubiquitin mediated proteolysis.

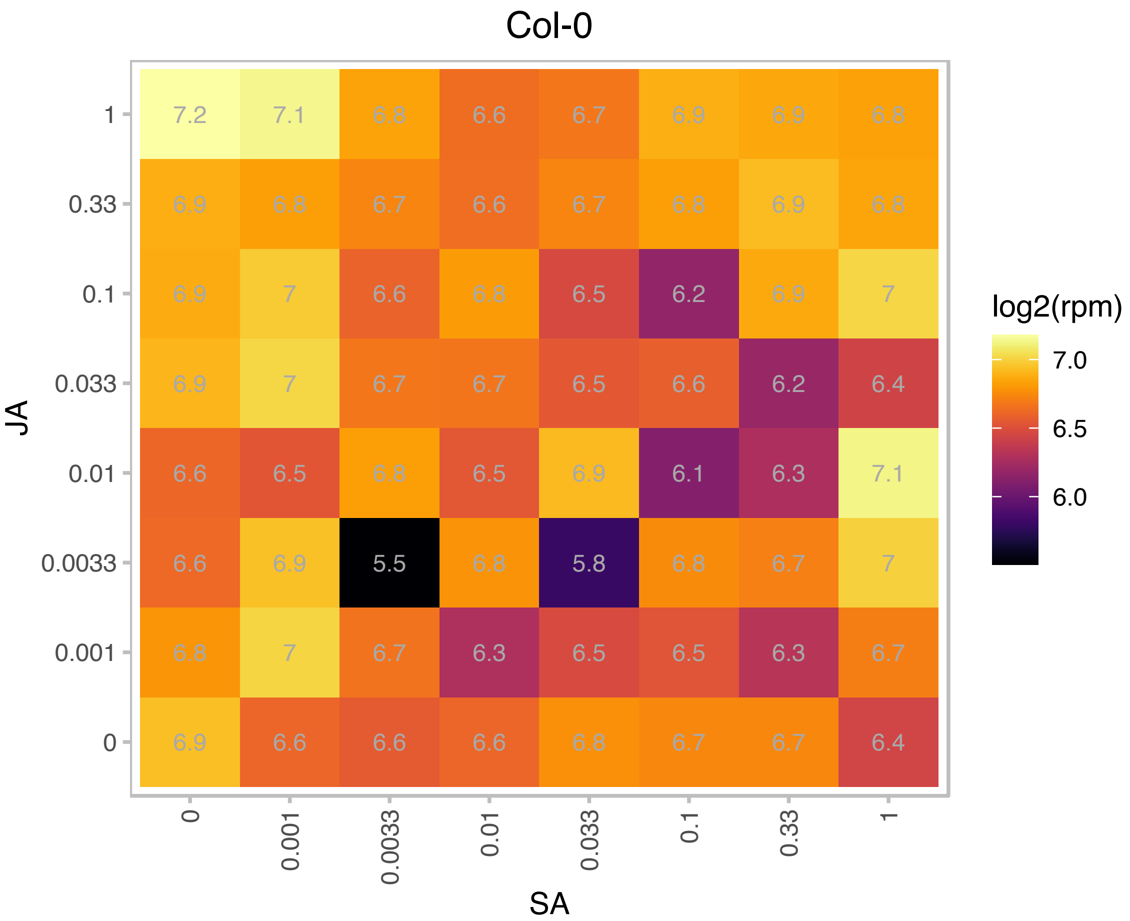


| | |
|---|---|
| Gene | AT5G42190 |
| is.expressed_genes | TRUE |
| is.responsive_genes | TRUE |
| is.apcluster | B-02 |

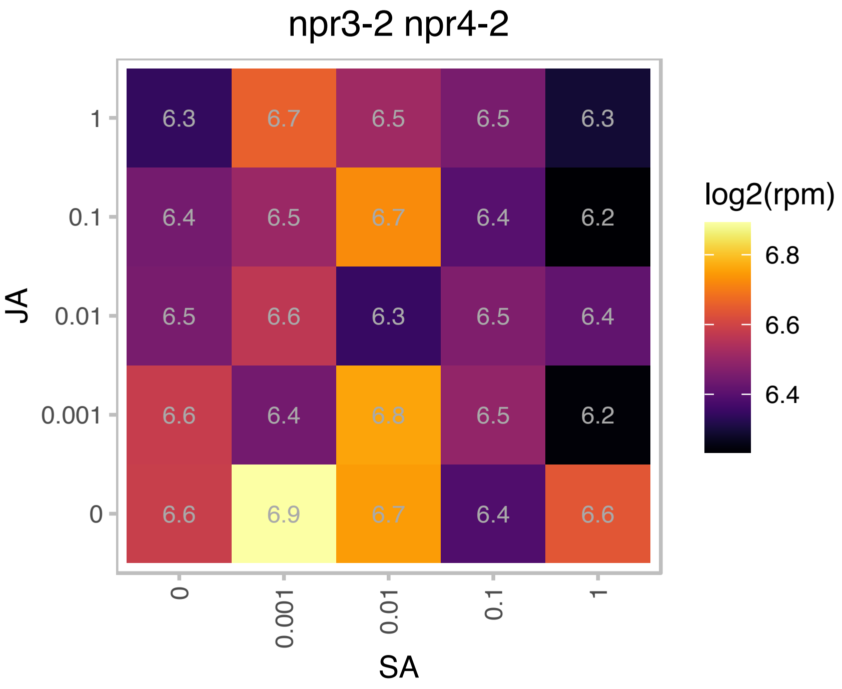


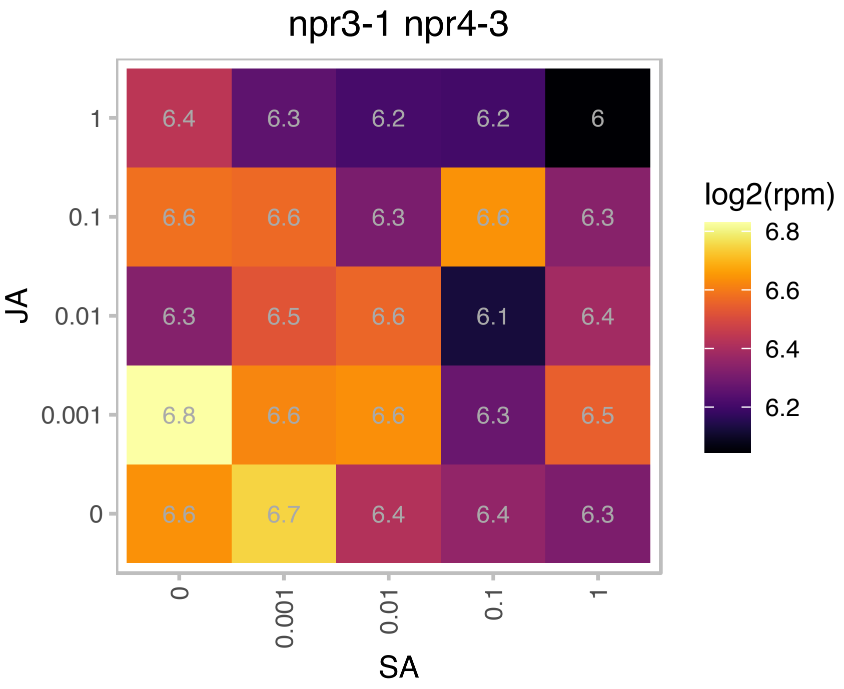

### AT5G51820.1 : ATPGMP, PGM, PGM1, STF1

phosphoglucomutase
Encodes a plastid isoform of the enzyme phosphoglucomutase involved in controlling photosynthetic carbon flow. Effective petiole movement against the direction of the gravity requires functional PGM activity that is required for full development of amyloplasts.

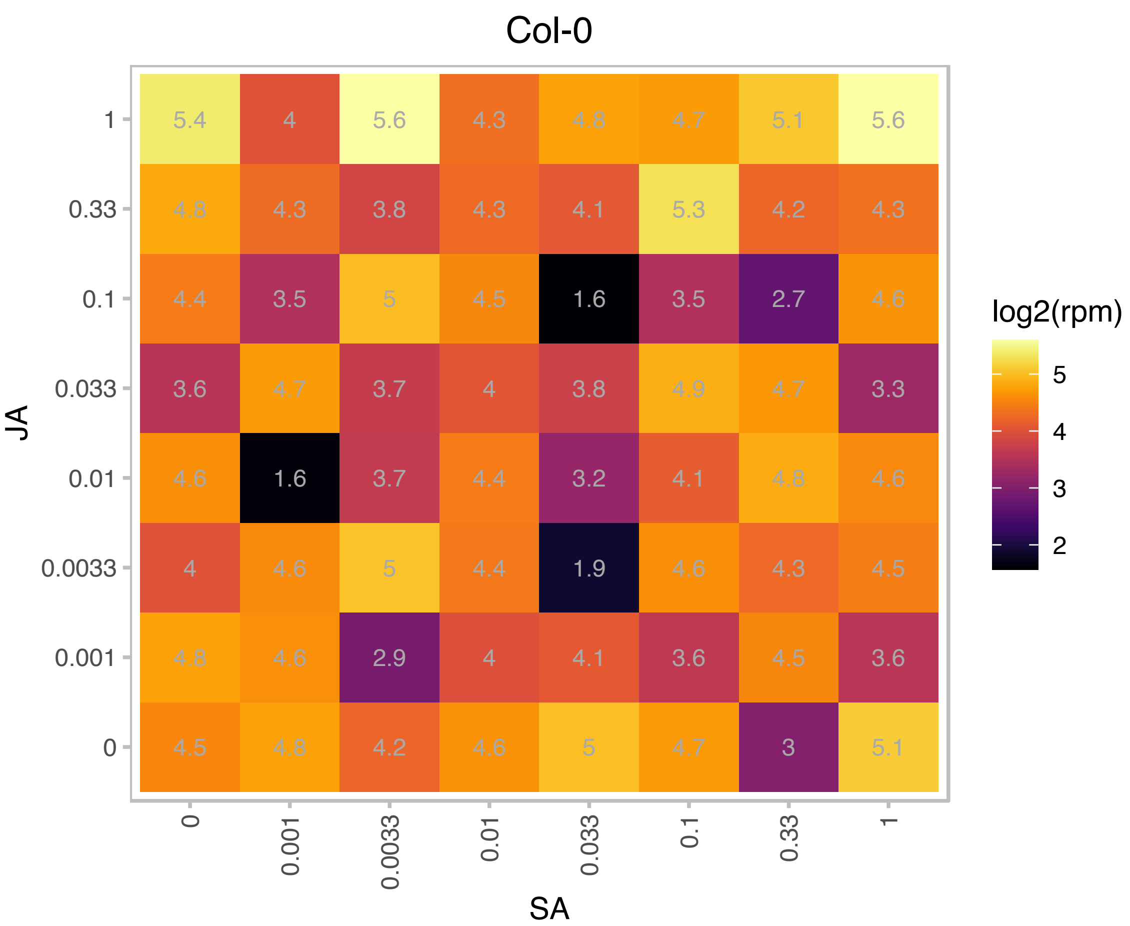


| | |
|---|---|
| Gene | AT5G51820 |
| is.expressed_genes | TRUE |
| is.responsive_genes | TRUE |
| is.apcluster | B-02 |

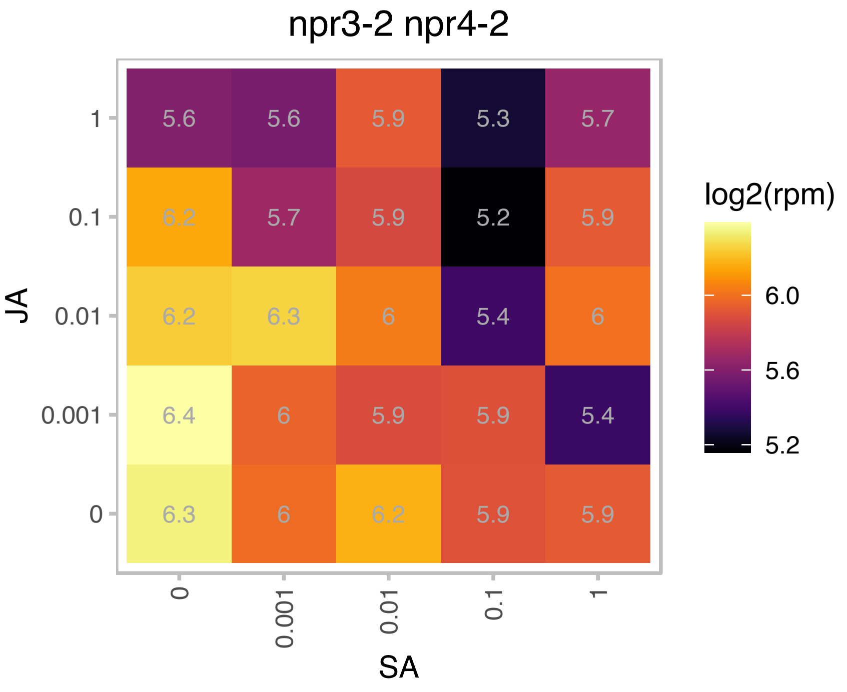


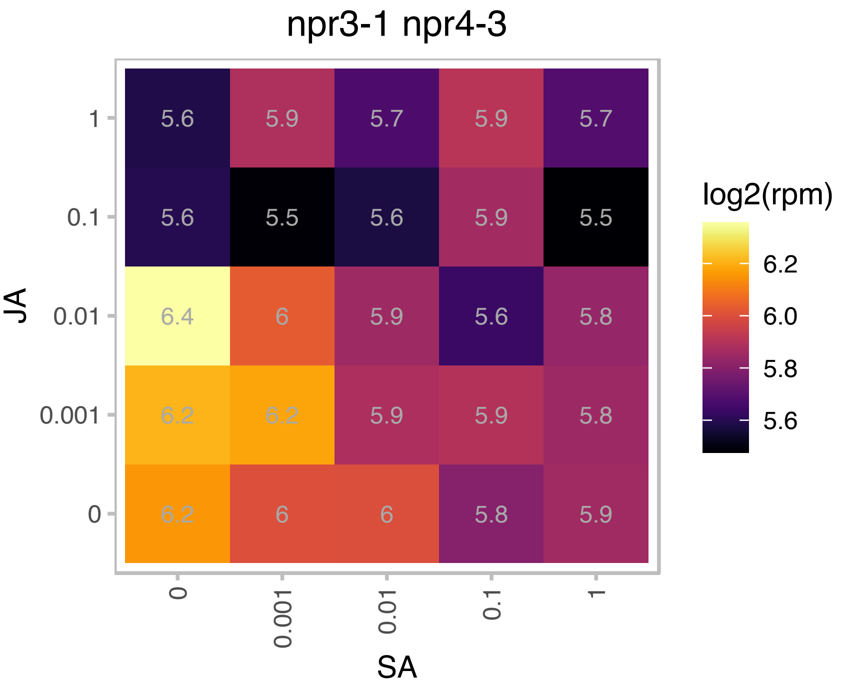

# AT5G54090.1 : NA

DNA mismatch repair protein MutS, type 2

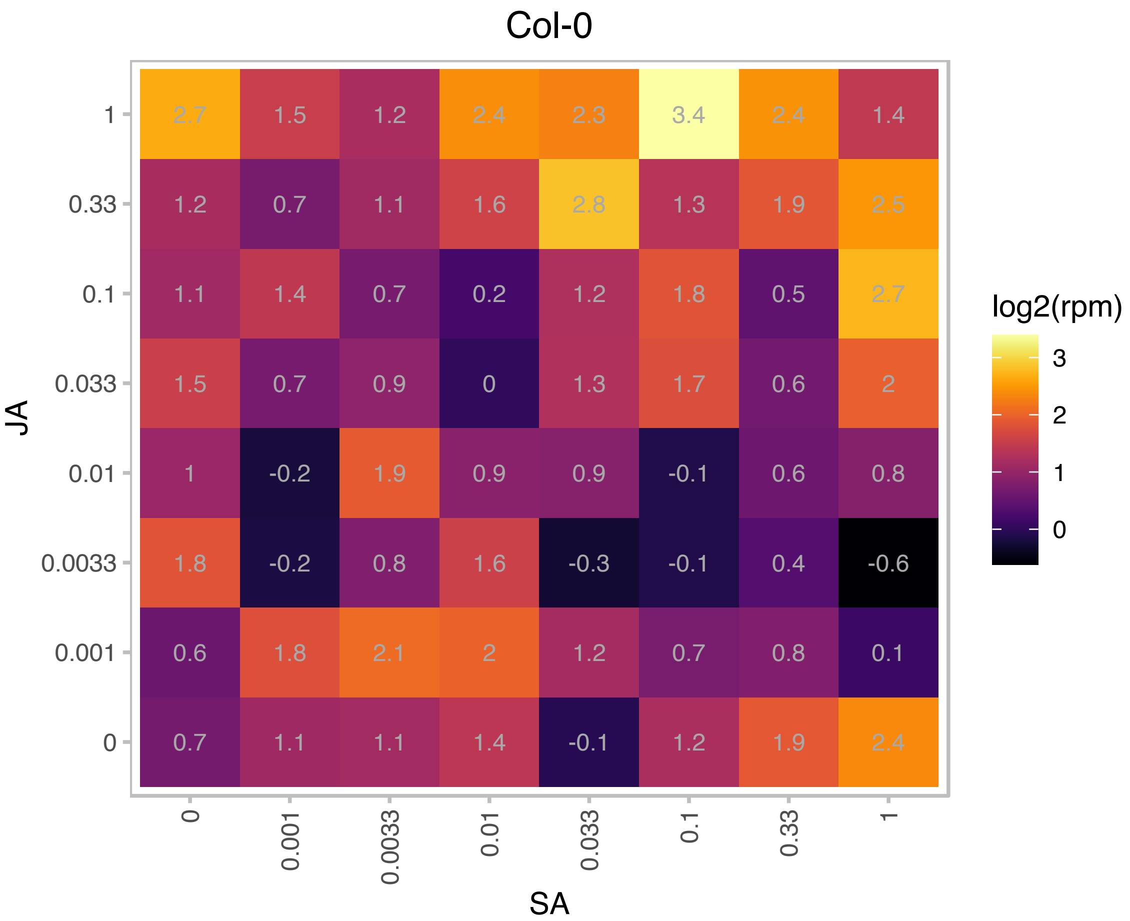


| | |
|---|---|
| Gene | AT5G54090 |
| is.expressed_genes | TRUE |
| is.responsive_genes | TRUE |
| is.apcluster | B-02 |

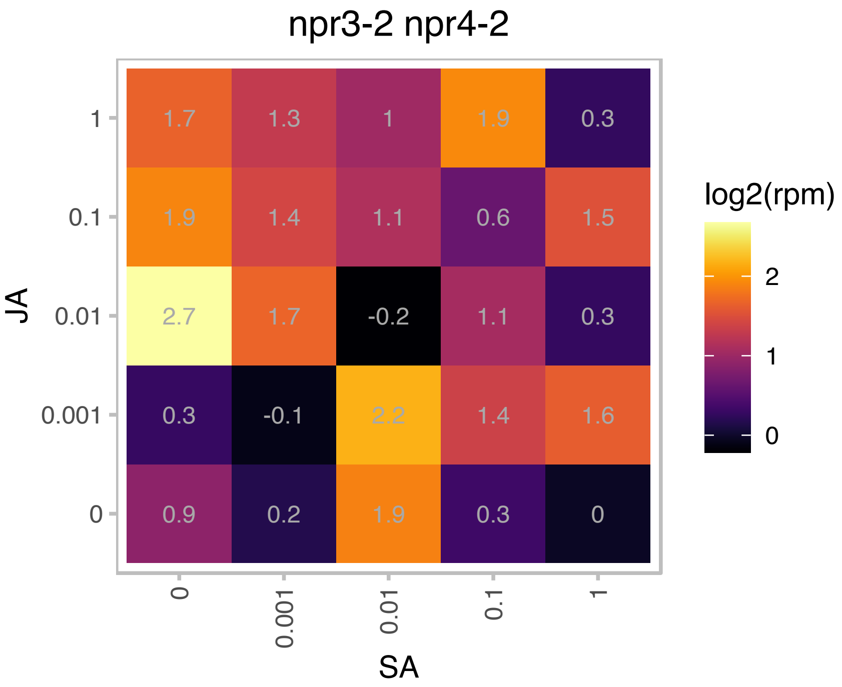


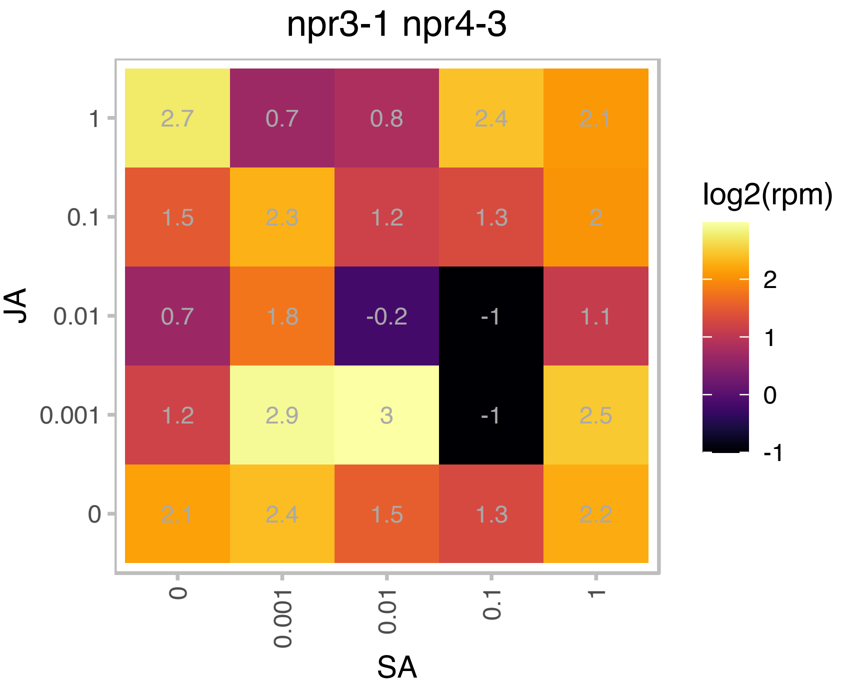

# AT5G55220.1 : NA

trigger factor type chaperone family protein

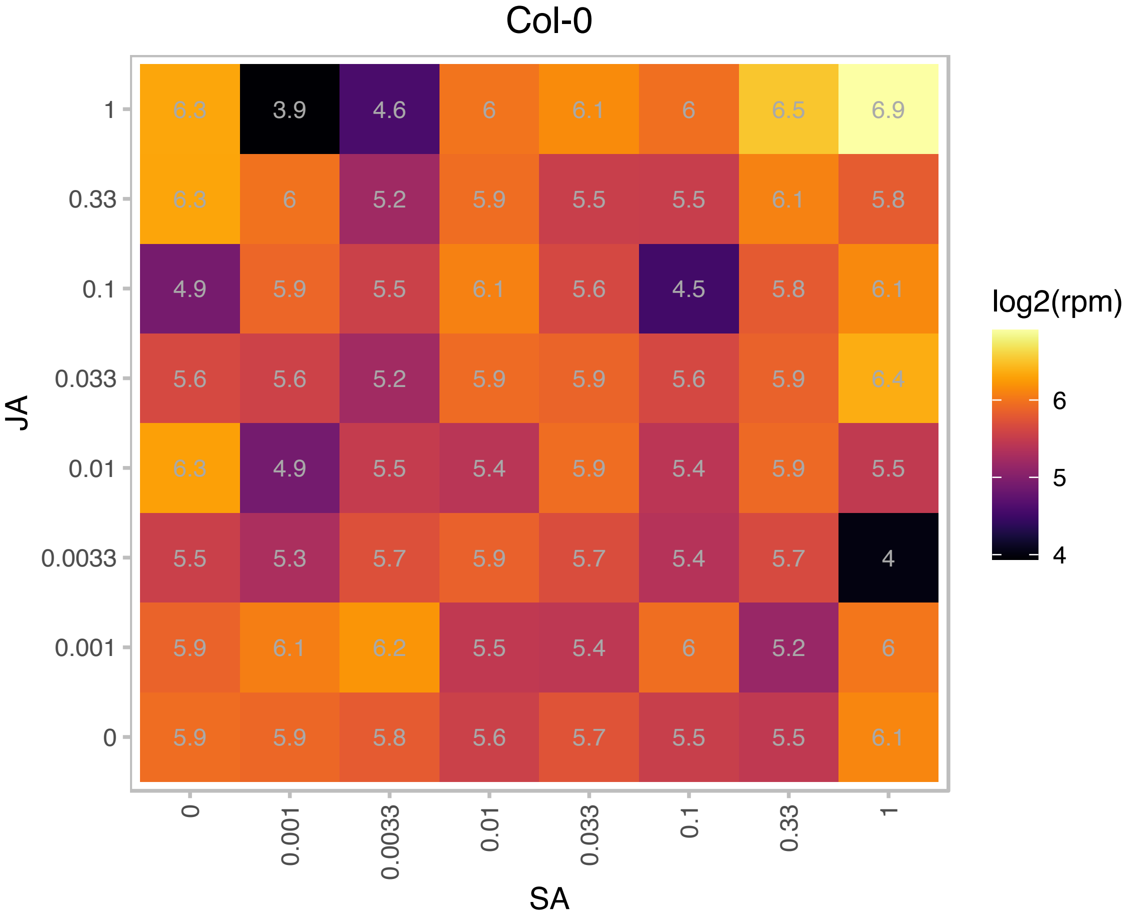


| | |
|---|---|
| Gene | AT5G55220 |
| is.expressed_genes | TRUE |
| is.responsive_genes | TRUE |
| is.apcluster | B-02 |

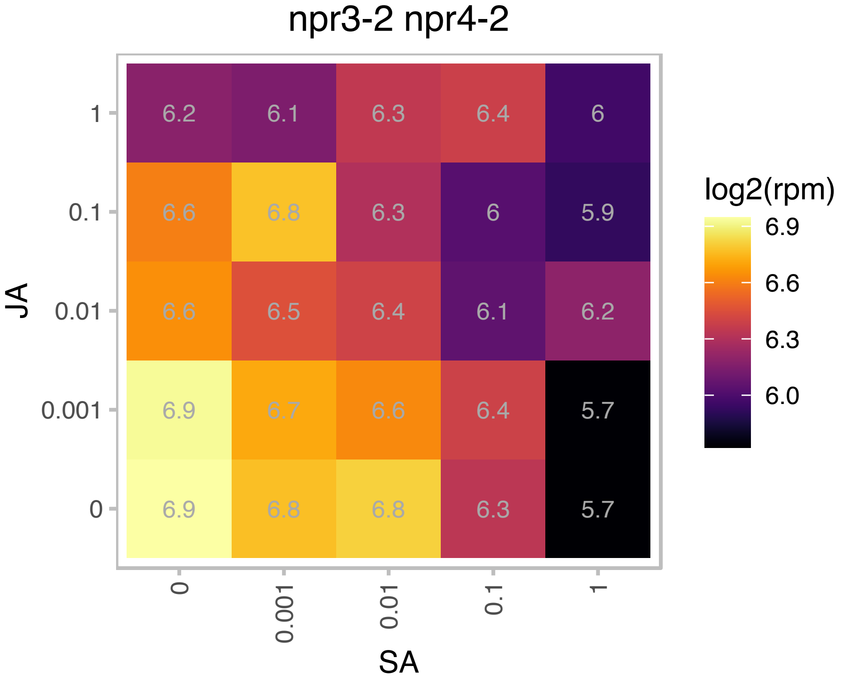


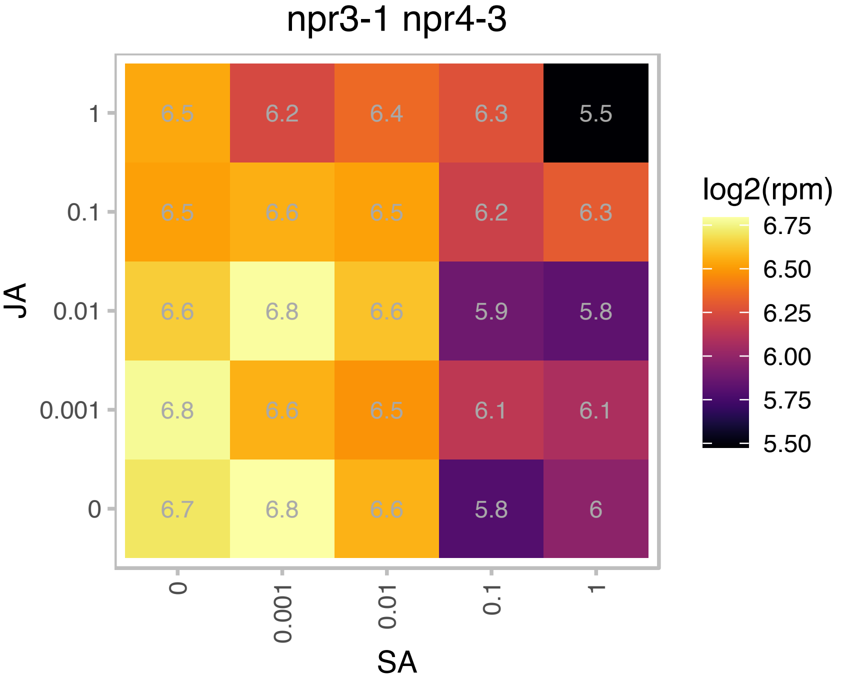

### AT5G60600.1 : CLB4, CSB3, GCPE, HDS, ISPG

Encodes a chloroplast-localized hydroxy-2-methyl-2-(E)-butenyl 4-diphosphate (HMBPP) synthase (HDS), catalyzes the formation of HMBPP from 2-C-methyl-D-erythrytol 2,4-cyclodiphosphate (MEcPP). The HDS enzyme controls the penultimate steps of the biosynthesis of IPP and dimethylallyl diphosphate (DMAPP) via the MEP pathway and may serve as a metabolic control point for SA-mediated disease resistance. In the light, the electrons required for the reaction catalyzed by HDS are directly provided by the electron flow from photosynthesis via ferredoxin. In the dark however, the enzyme requires an electron shuttle: [illegible]

4-hydroxy-3-methylbut-2-enyl diphosphate synthase

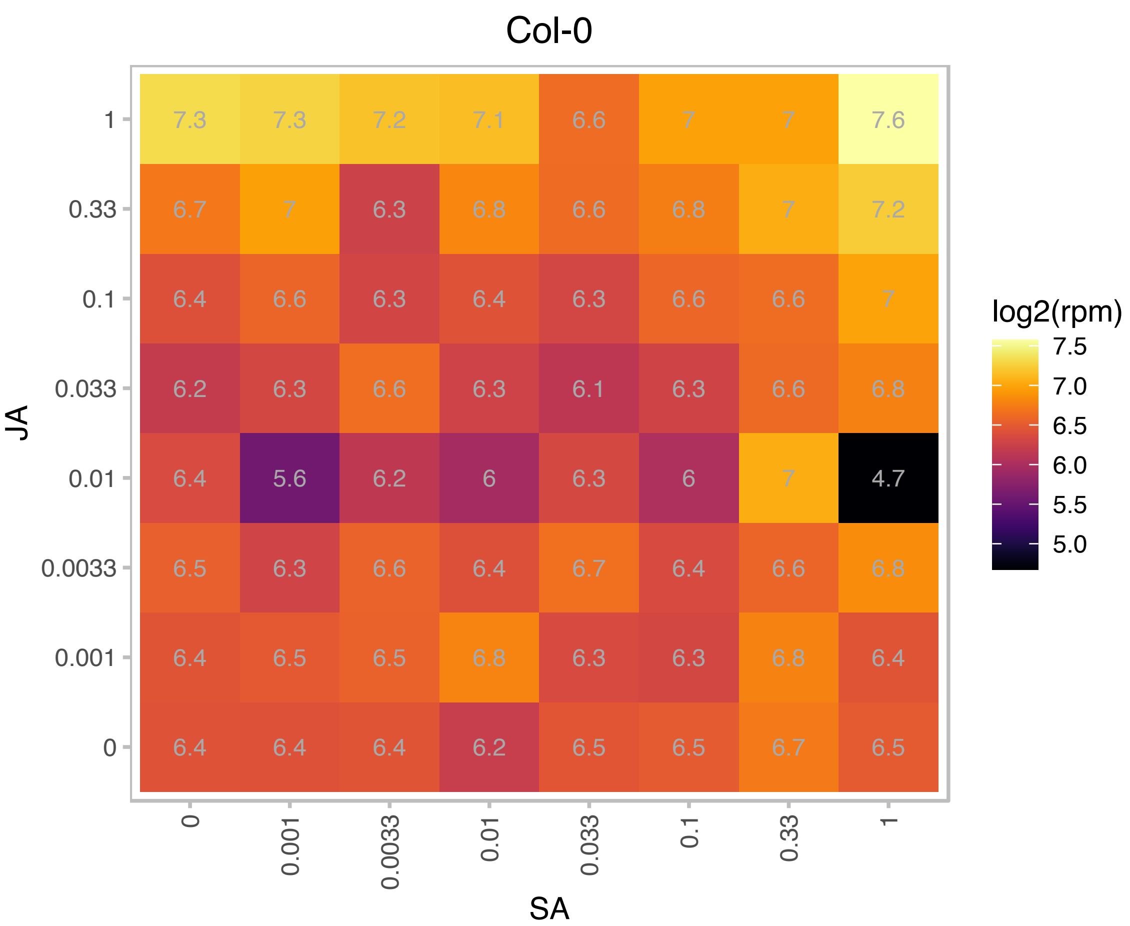


| | |
|---|---|
| Gene | AT5G60600 |
| is.expressed_genes | TRUE |
| is.responsive_genes | TRUE |
| is.apcluster | B-02 |

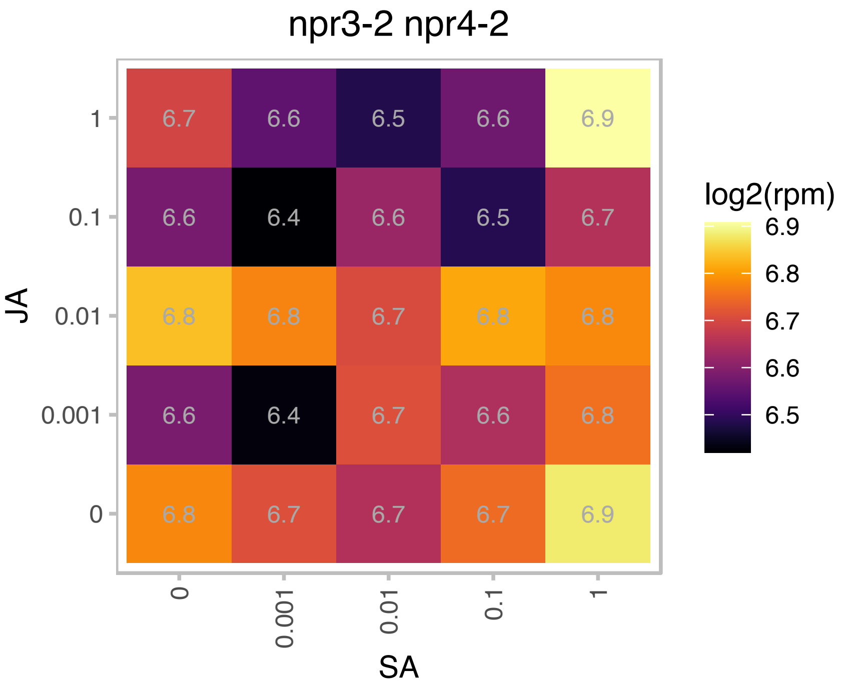


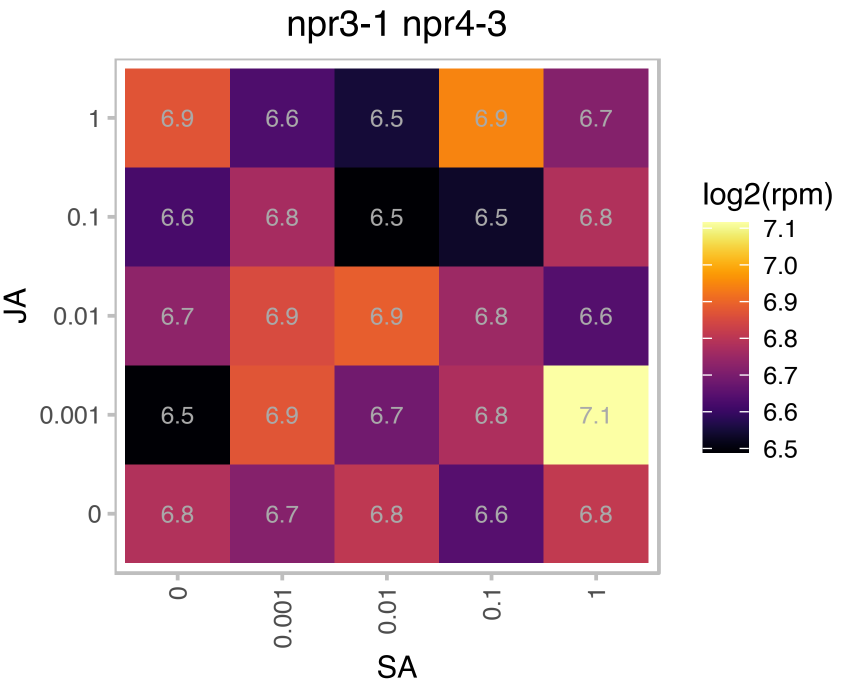

# AT5G65840.1 : NA

Thioredoxin superfamily protein

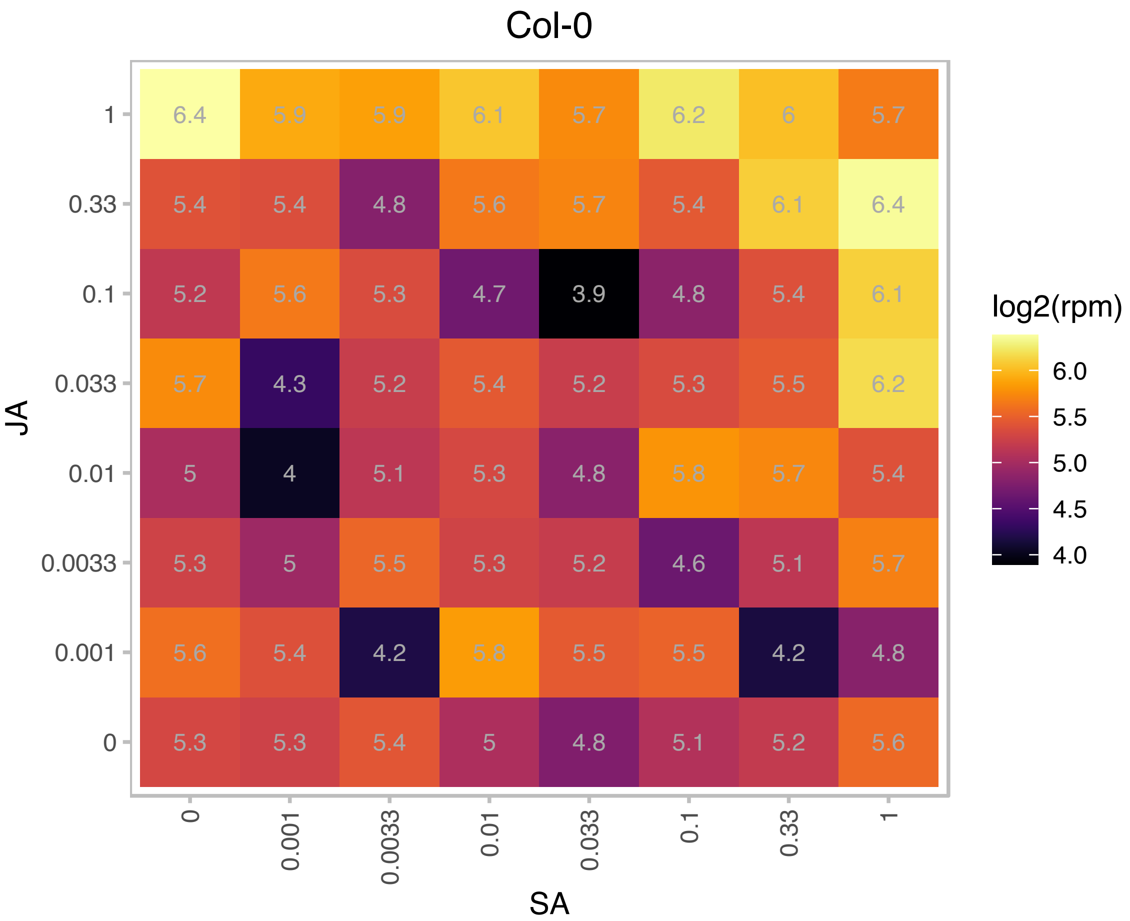


| | |
|---|---|
| Gene | AT5G65840 |
| is.expressed_genes | TRUE |
| is.responsive_genes | TRUE |
| is.apcluster | B-02 |

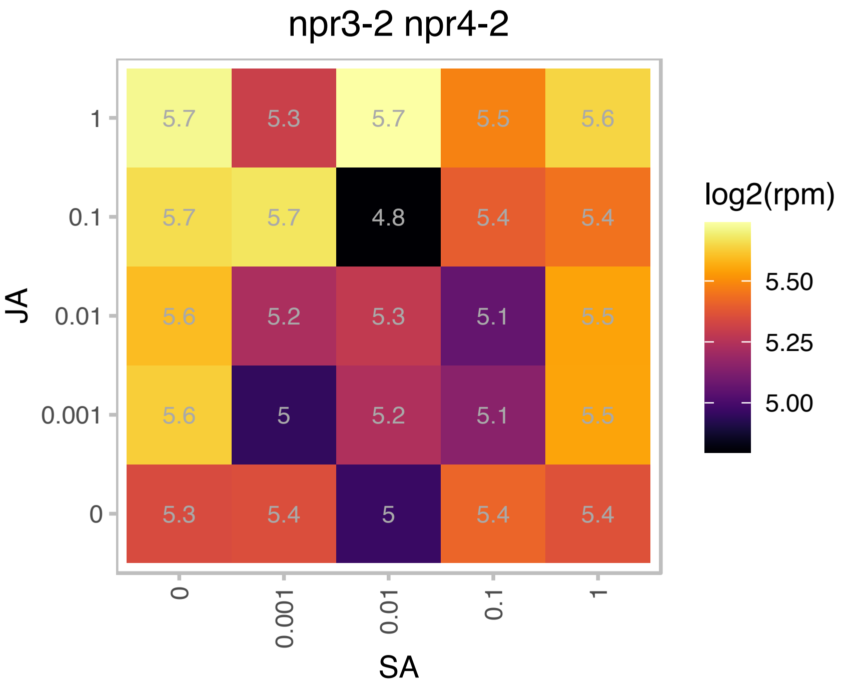


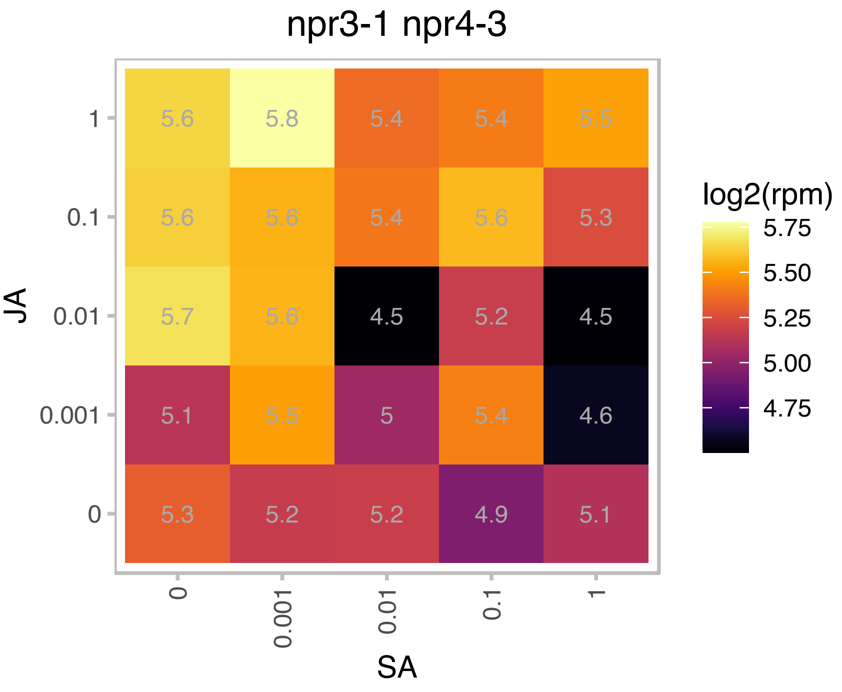

### AT5G66190.1 : ATLFNR1, FNR1, LFNR1

ferredoxin-NADP(+)-oxidoreductase 1

Encodes a leaf-type ferredoxin:NADP(H) oxidoreductase. It is present in both chloroplast stroma and thylakoid membranes but is more abundant in the thylakoid. The affinity of this enzyme for ferredoxin is slightly, but significantly, higher than AtLFNR2, an isoform of the same enzyme. AtLFNR1 forms a heterodimer with AtFNR2 and is also a prerequisite to attach AtFNR2 to the thylakoid membrane.

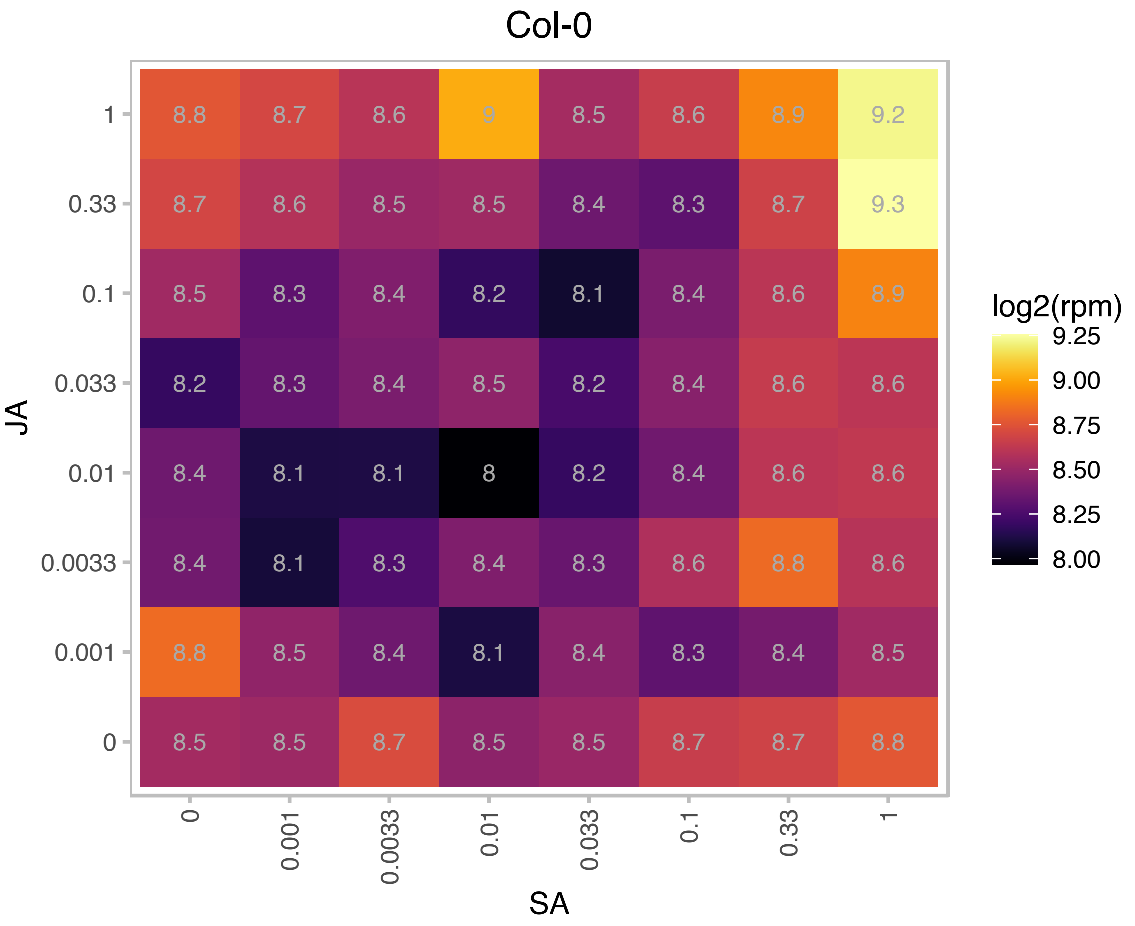


| | |
|---|---|
| Gene | AT5G66190 |
| is.expressed_genes | TRUE |
| is.responsive_genes | TRUE |
| is.apcluster | B-02 |

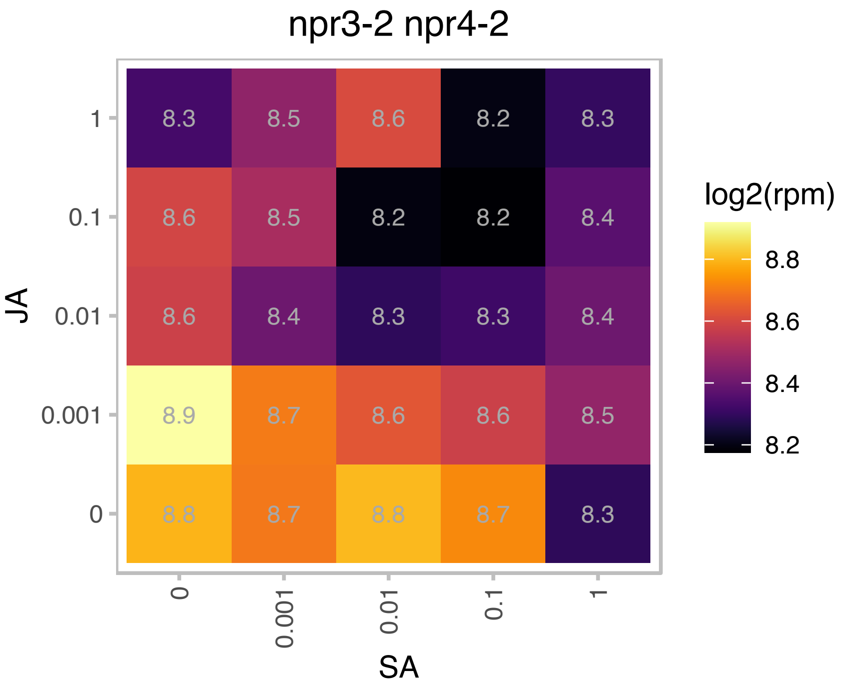


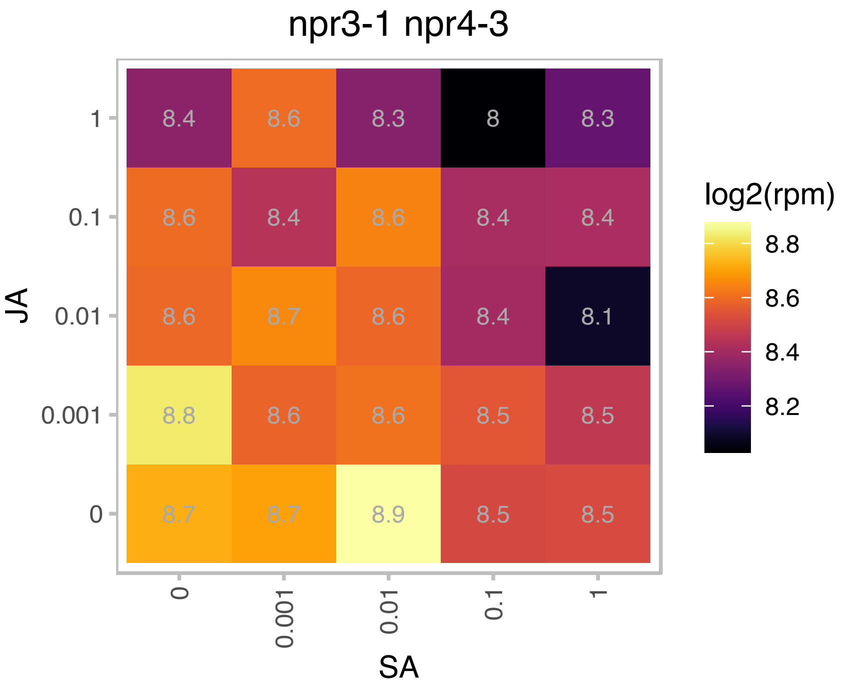

### AT1G02220.1 : ANAC003, NAC003

NAC domain containing protein 3

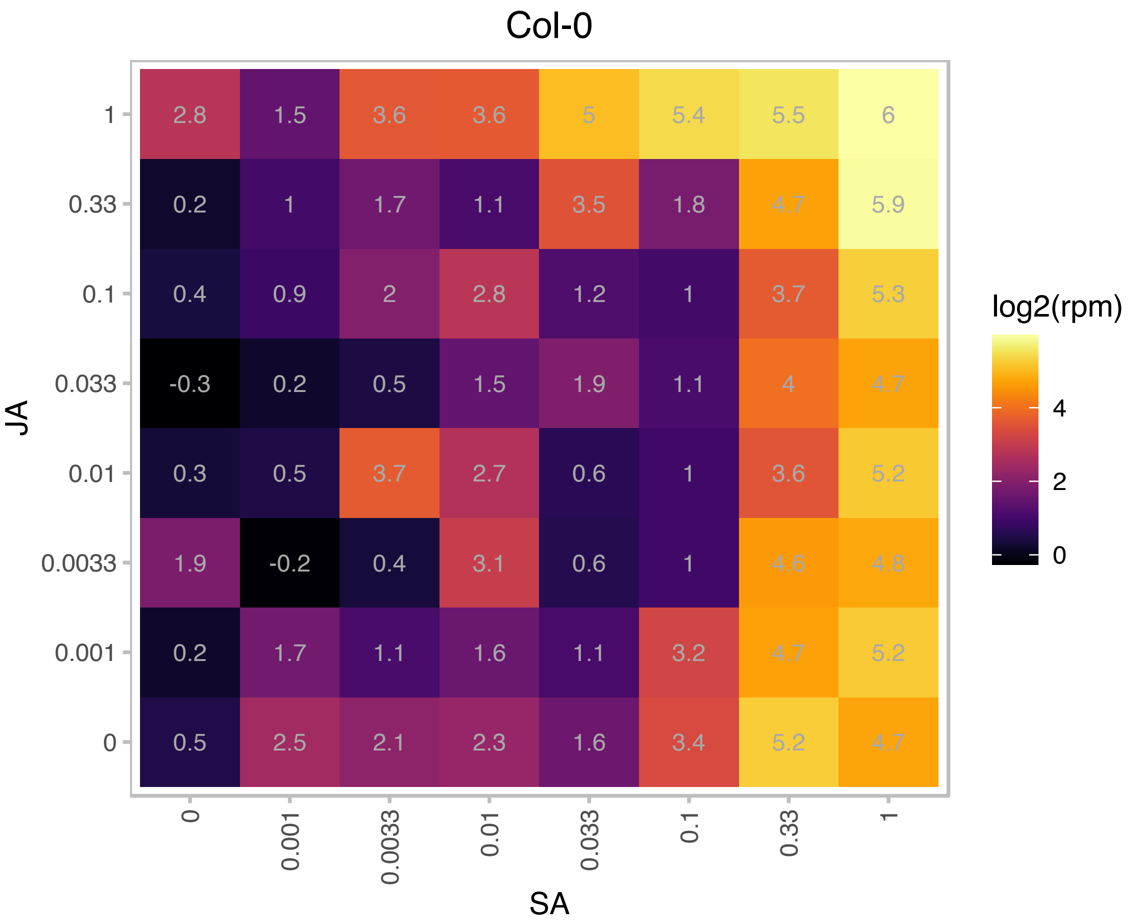


| | |
|---|---|
| Gene | AT1G02220 |
| is.expressed_genes | TRUE |
| is.responsive_genes | TRUE |
| is.apcluster | B-03 |

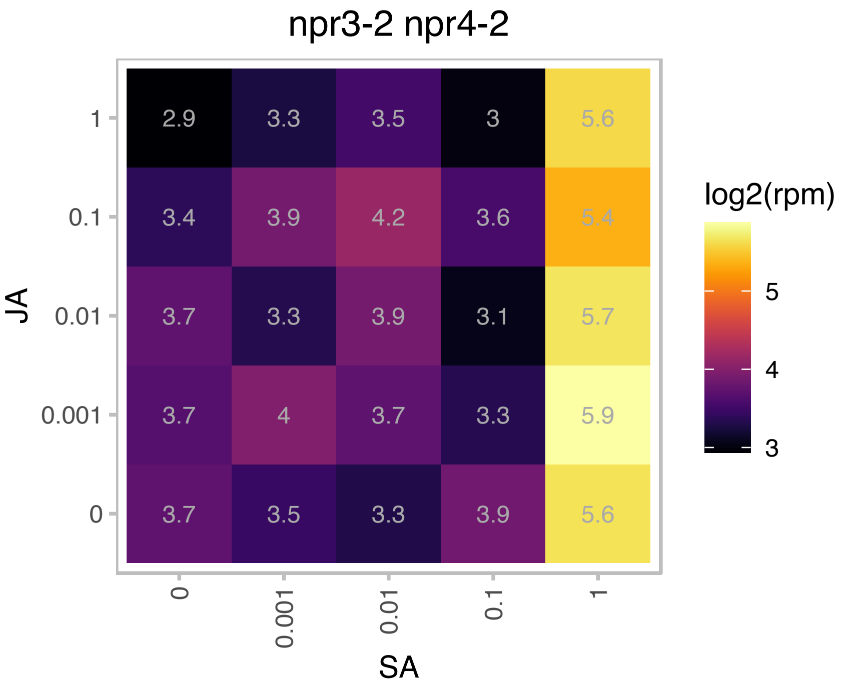


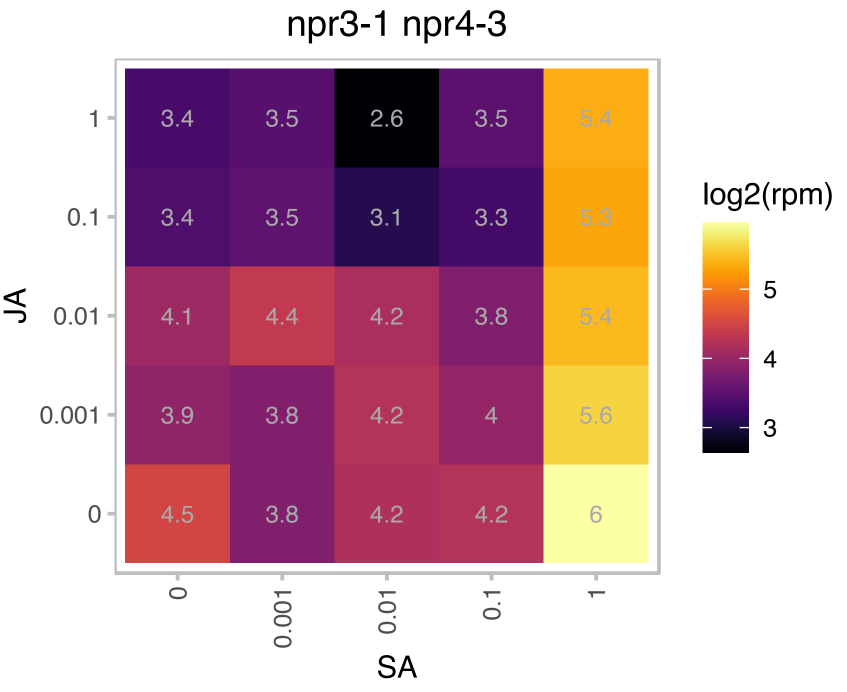

# AT1G05170.1 : NA

Galactosyltransferase family protein

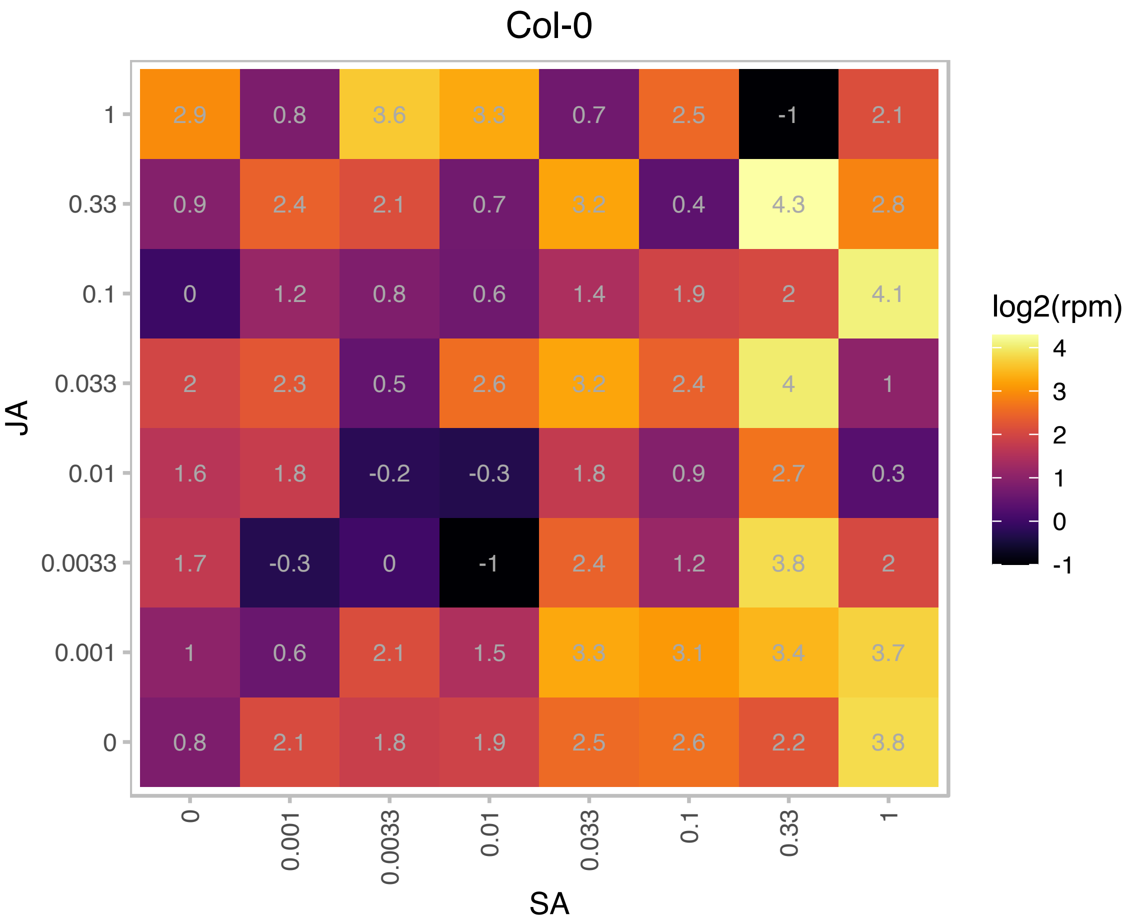


| | |
|---|---|
| Gene | AT1G05170 |
| is.expressed_genes | TRUE |
| is.responsive_genes | TRUE |
| is.apcluster | B-03 |

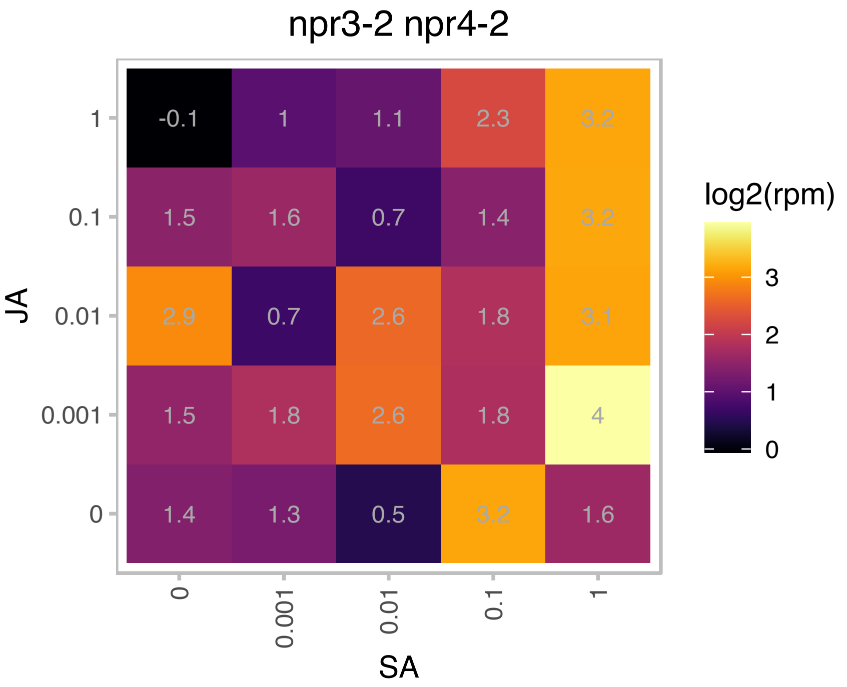


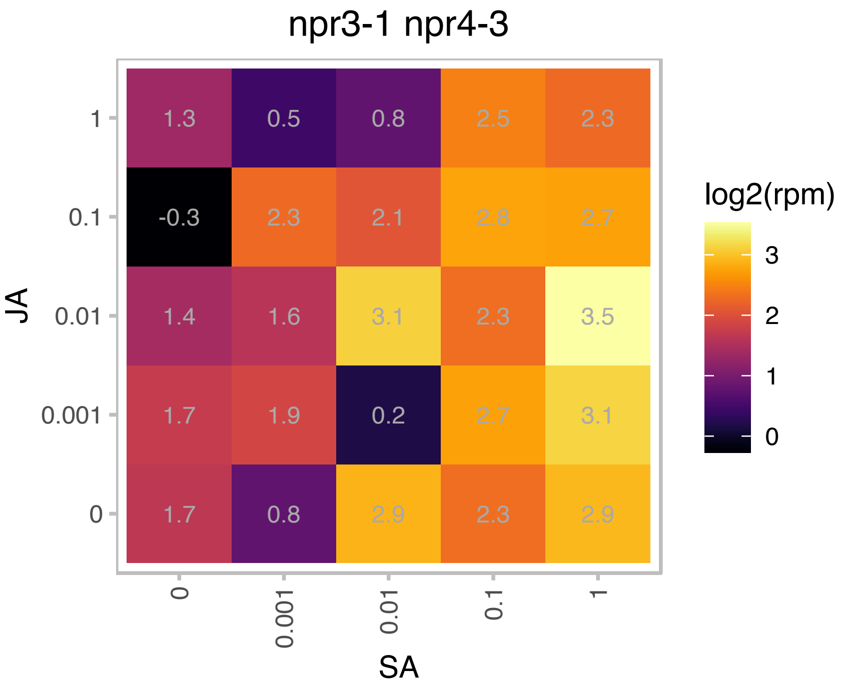

### AT1G07620.1 : ATOBGM, TRM32

GTP-binding protein Obg/CgtA

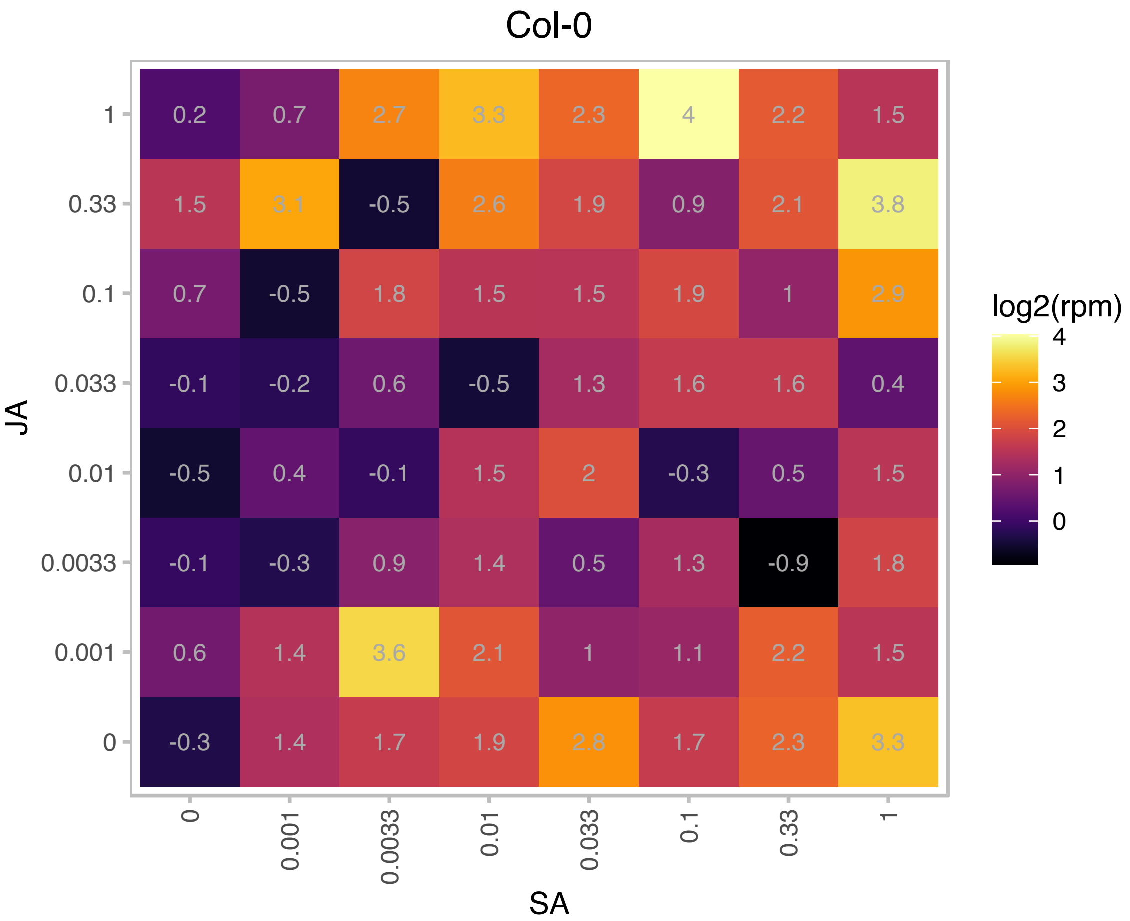


| | |
|---|---|
| Gene | AT1G07620 |
| is.expressed_genes | TRUE |
| is.responsive_genes | TRUE |
| is.apcluster | B-03 |

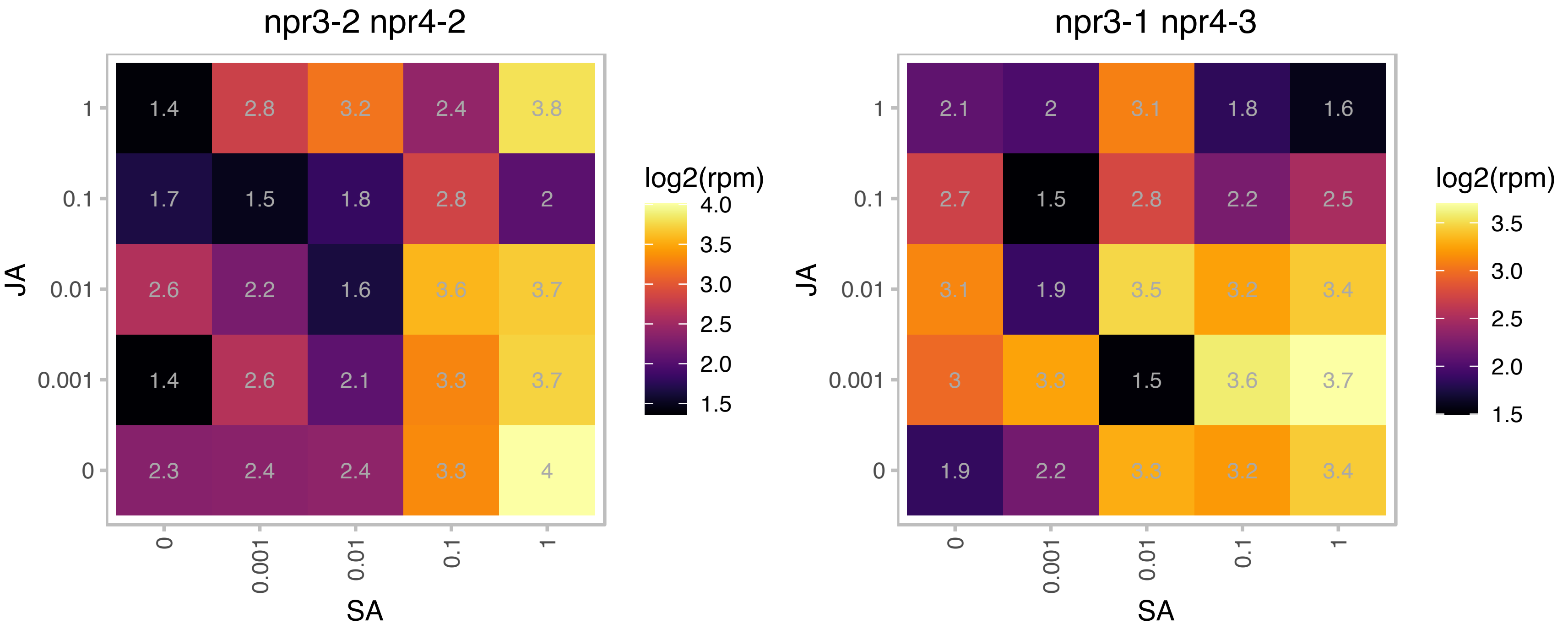

# AT1G08050.1 : NA

Zinc finger (C3HC4-type RING finger) family protein

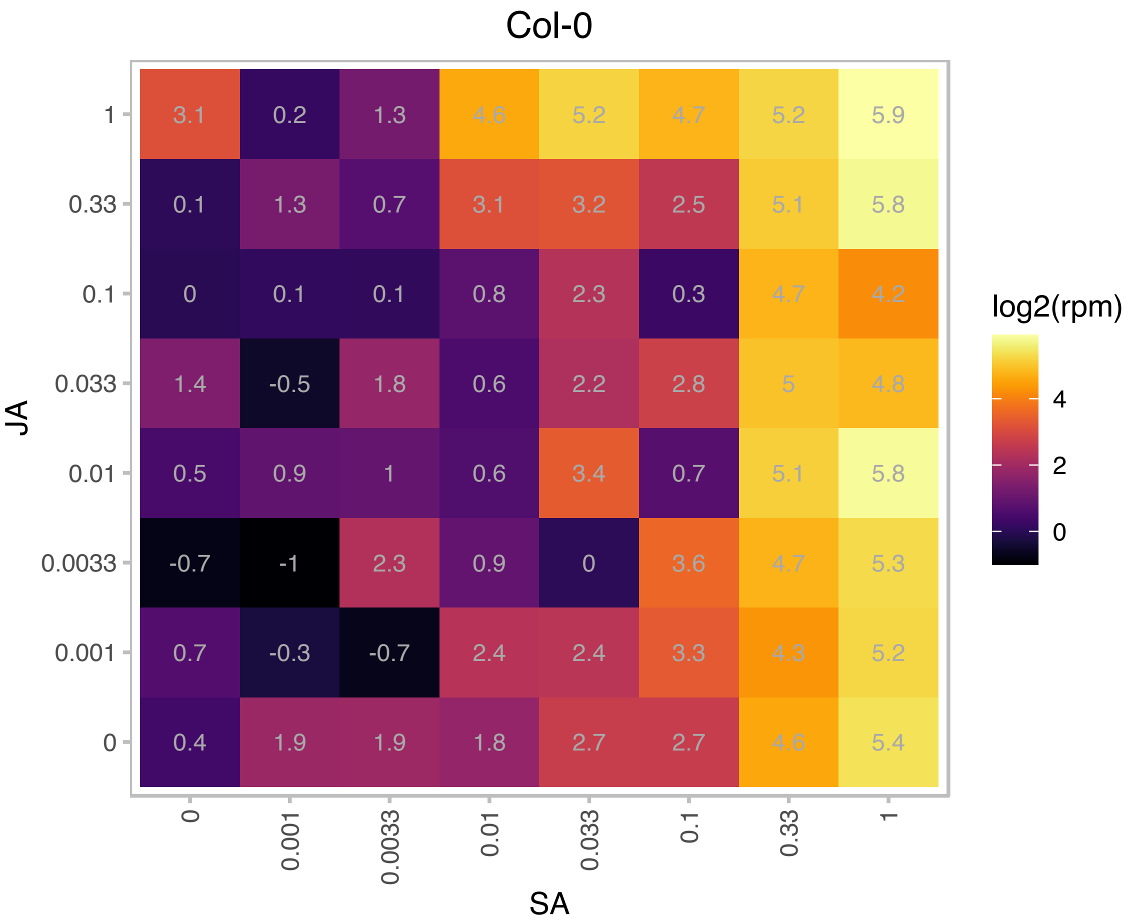


| | |
|---|---|
| Gene | AT1G08050 |
| is.expressed_genes | TRUE |
| is.responsive_genes | TRUE |
| is.apcluster | B-03 |

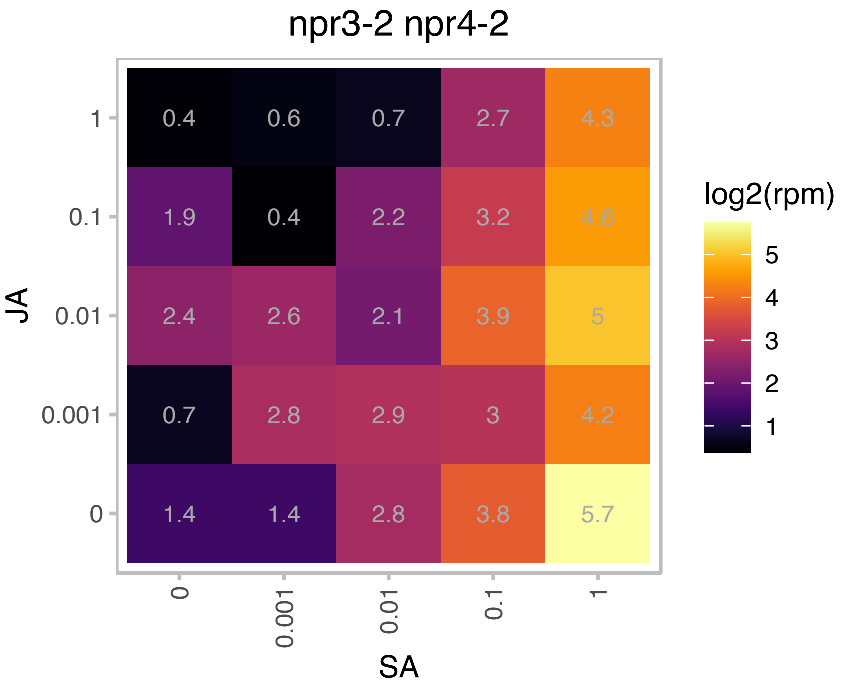


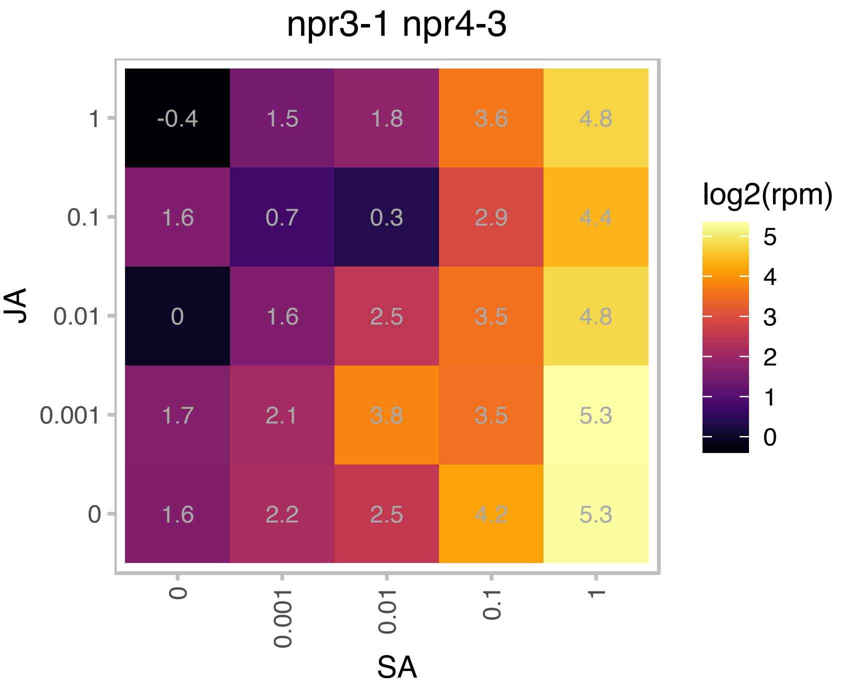

### AT1G16510.1 : SAUR41

SAUR-like auxin-responsive protein family.

Encodes a clade III SAUR gene with a distinctive expression pattern in root meristems. It is normally expressed in the quiescent center and cortex/endodermis initials and upon auxin stimulation, the expression is found in the endodermal layer. Overexpression studies suggest roles in cell expansion and auxin transport.

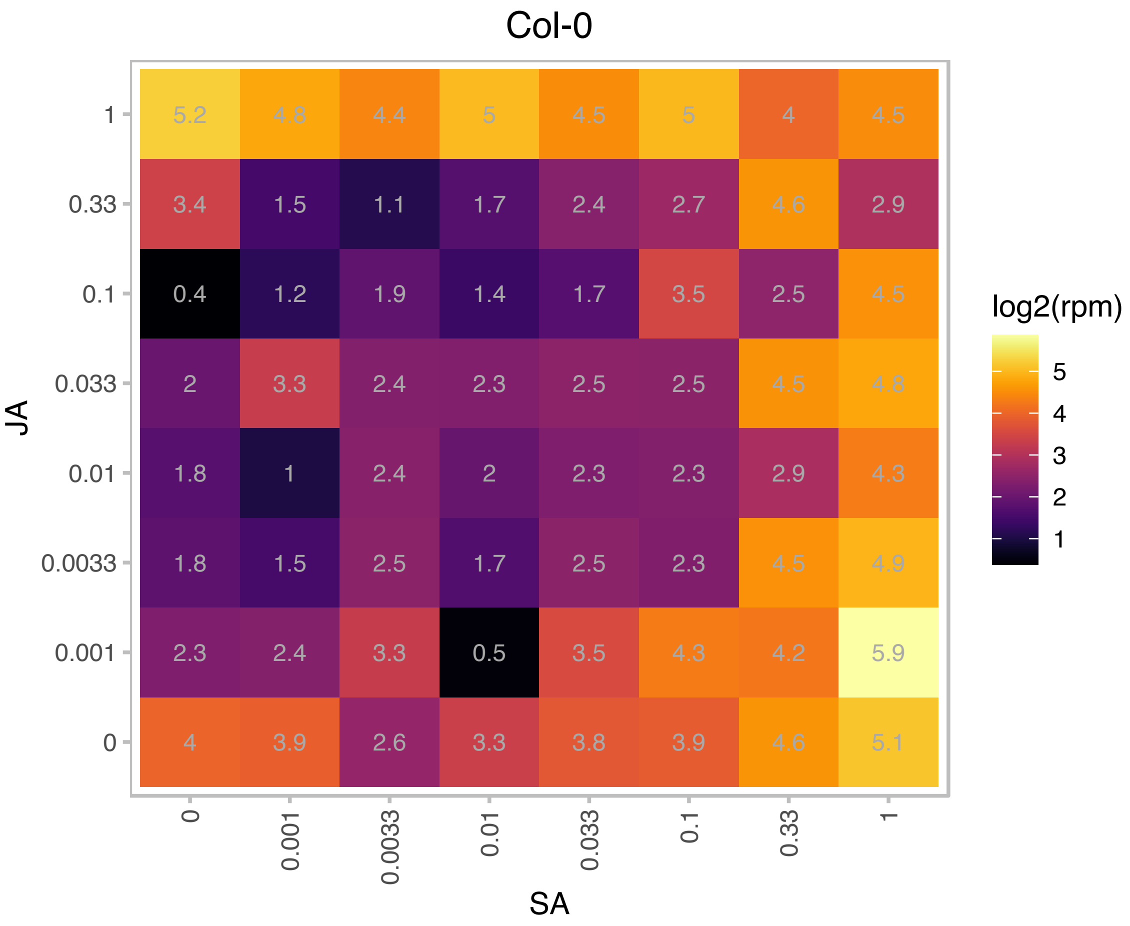


| | |
|---|---|
| Gene | AT1G16510 |
| is.expressed_genes | TRUE |
| is.responsive_genes | TRUE |
| is.apcluster | B-03 |

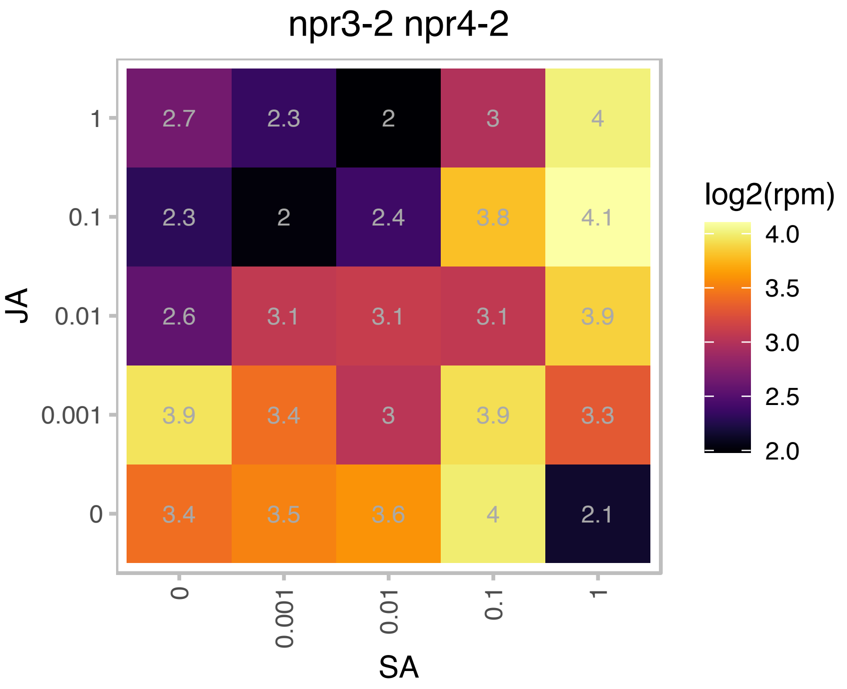


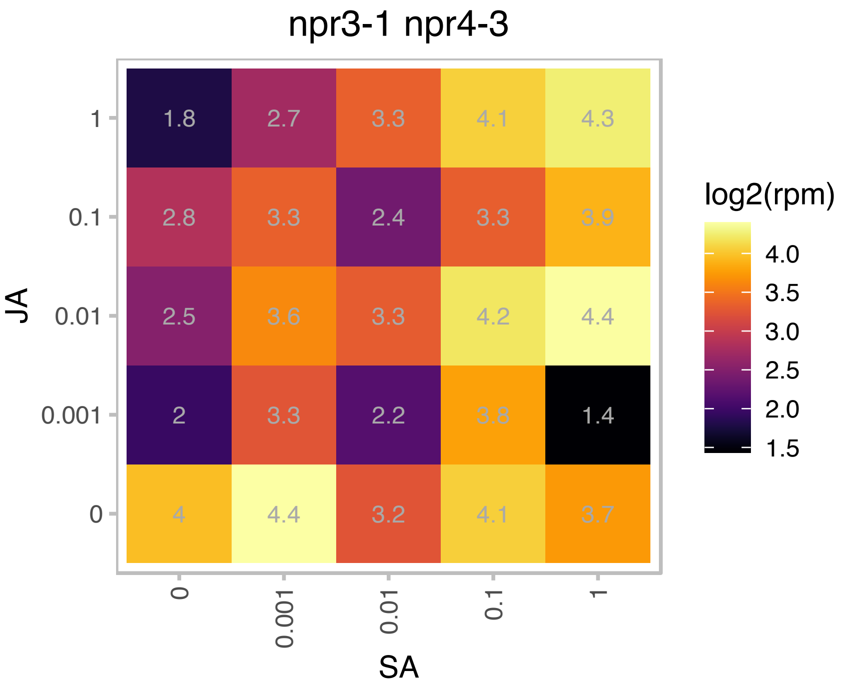

# AT1G18020.1 : NA

FMN-linked oxidoreductases superfamily protein

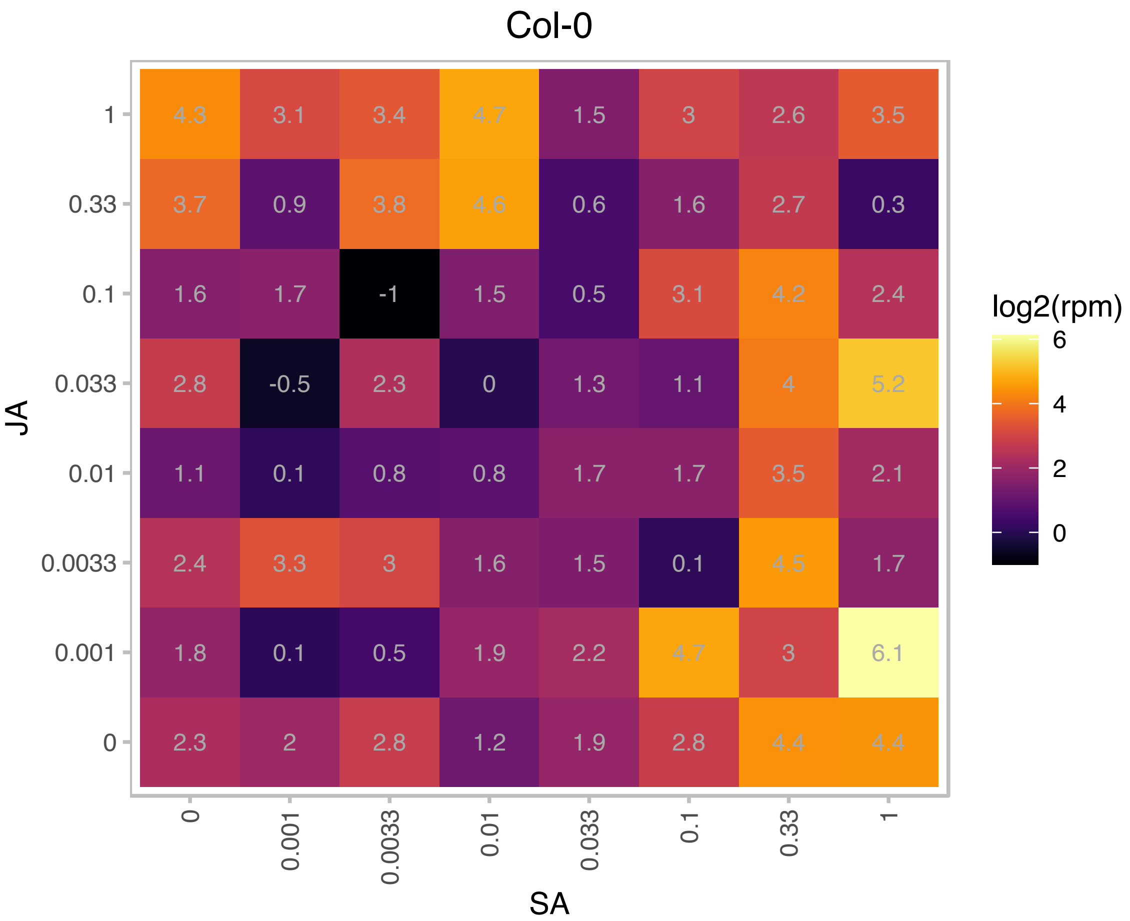


| | |
|---|---|
| Gene | AT1G18020 |
| is.expressed_genes | TRUE |
| is.responsive_genes | TRUE |
| is.apcluster | B-03 |

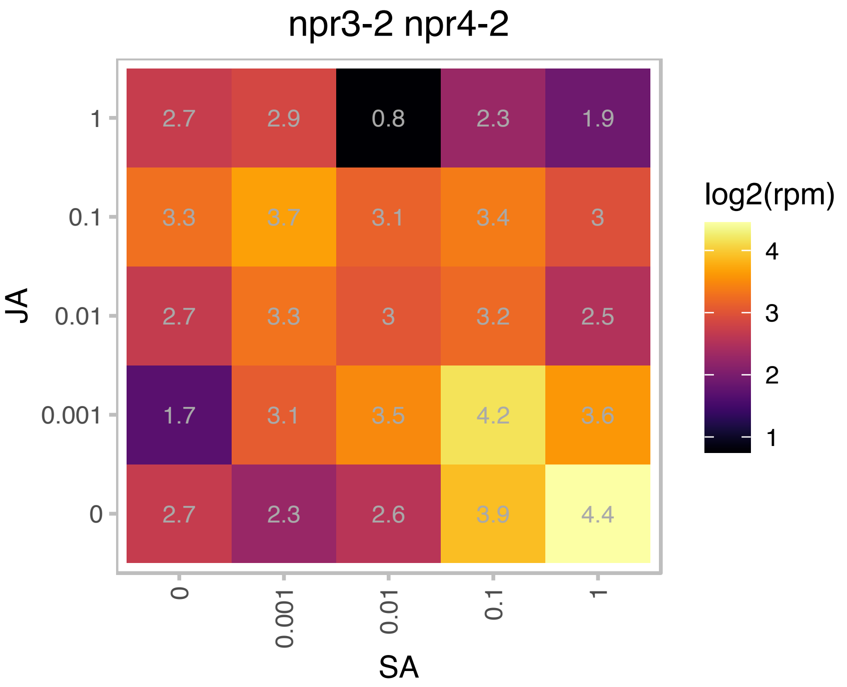


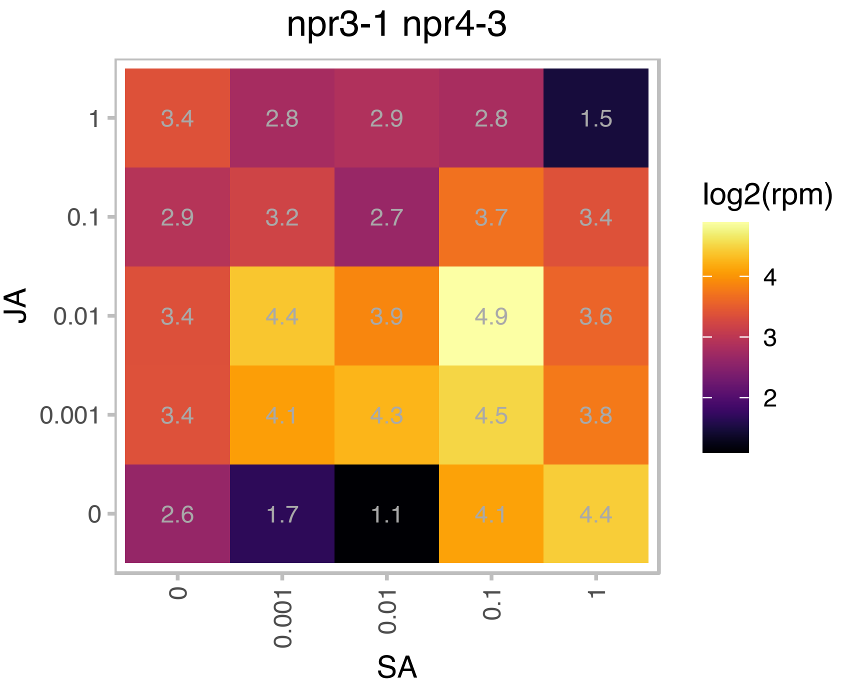

### AT1G21390.1 : emb2170

embryo defective 2170

#### Col-0

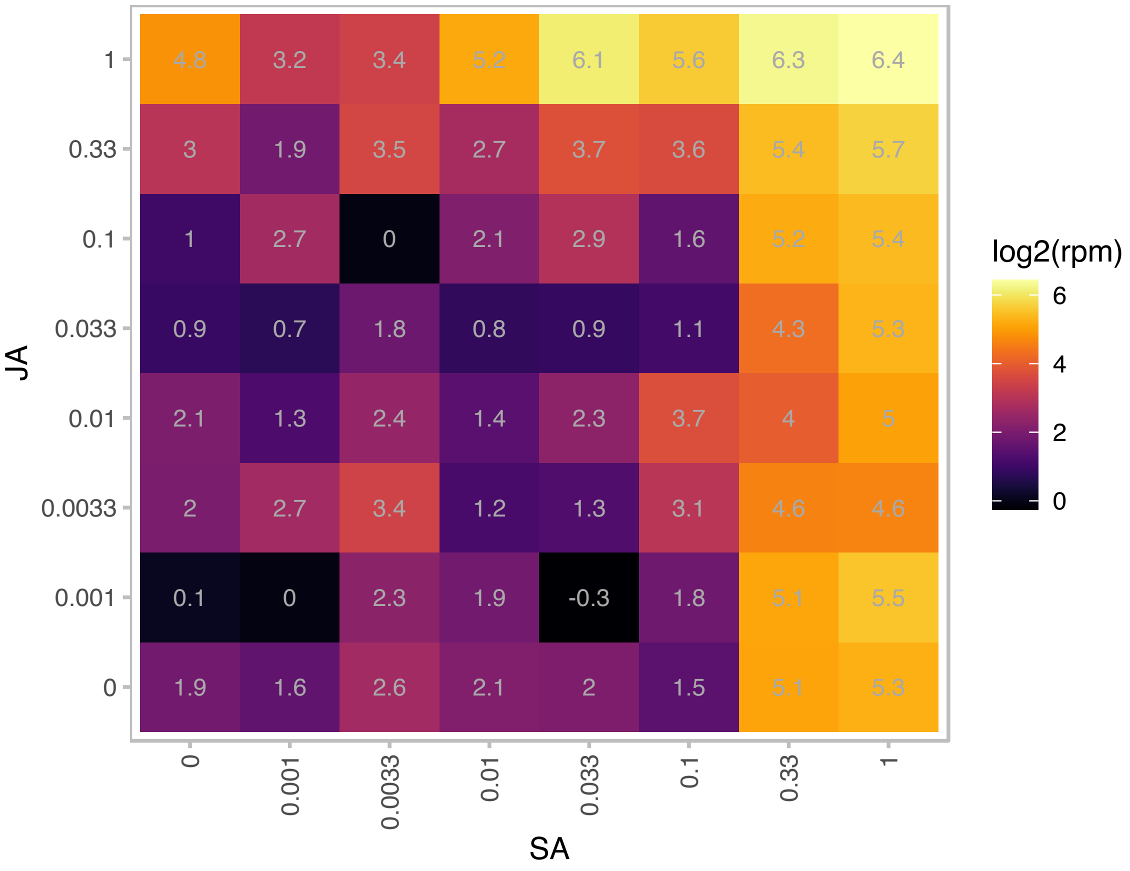


| | |
|---|---|
| Gene | AT1G21390 |
| is.expressed_genes | TRUE |
| is.responsive_genes | TRUE |
| is.apcluster | B-03 |

#### npr3-2 npr4-2

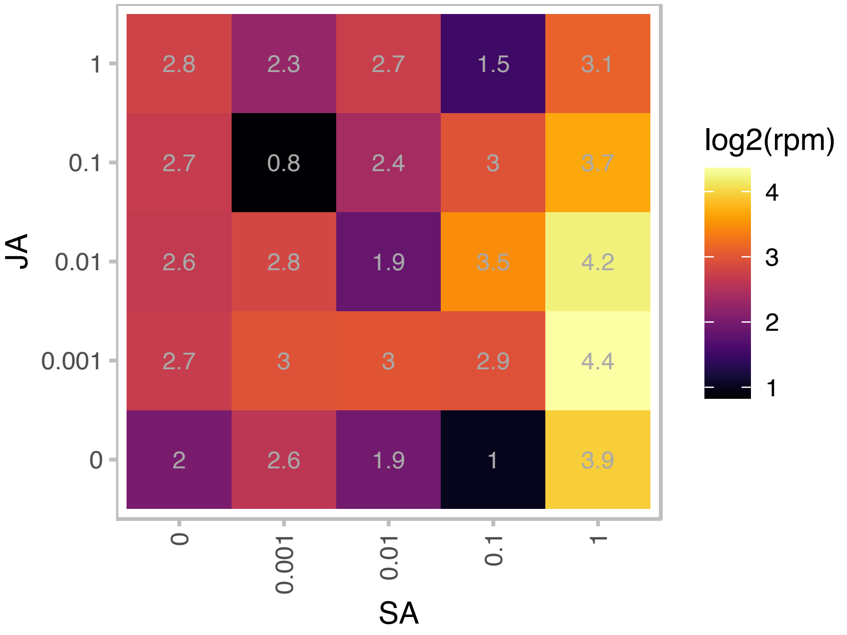


#### npr3-1 npr4-3

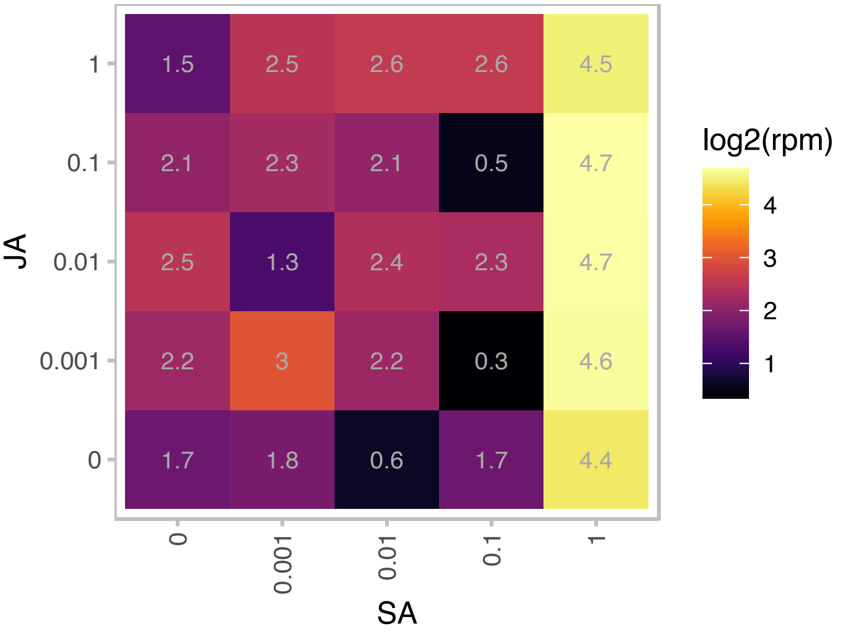

# AT1G22470.1 : NA

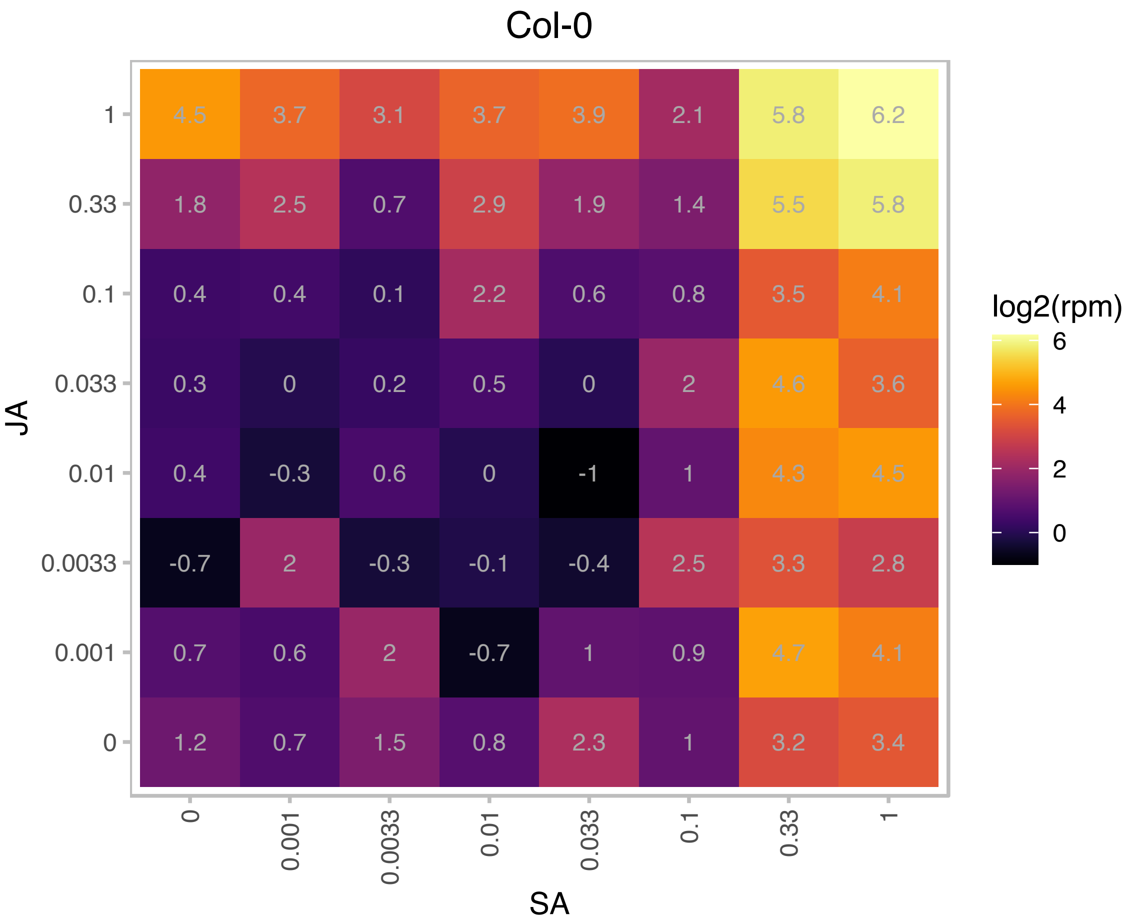


| | |
|---|---|
| Gene | AT1G22470 |
| is.expressed_genes | TRUE |
| is.responsive_genes | TRUE |
| is.apcluster | B-03 |

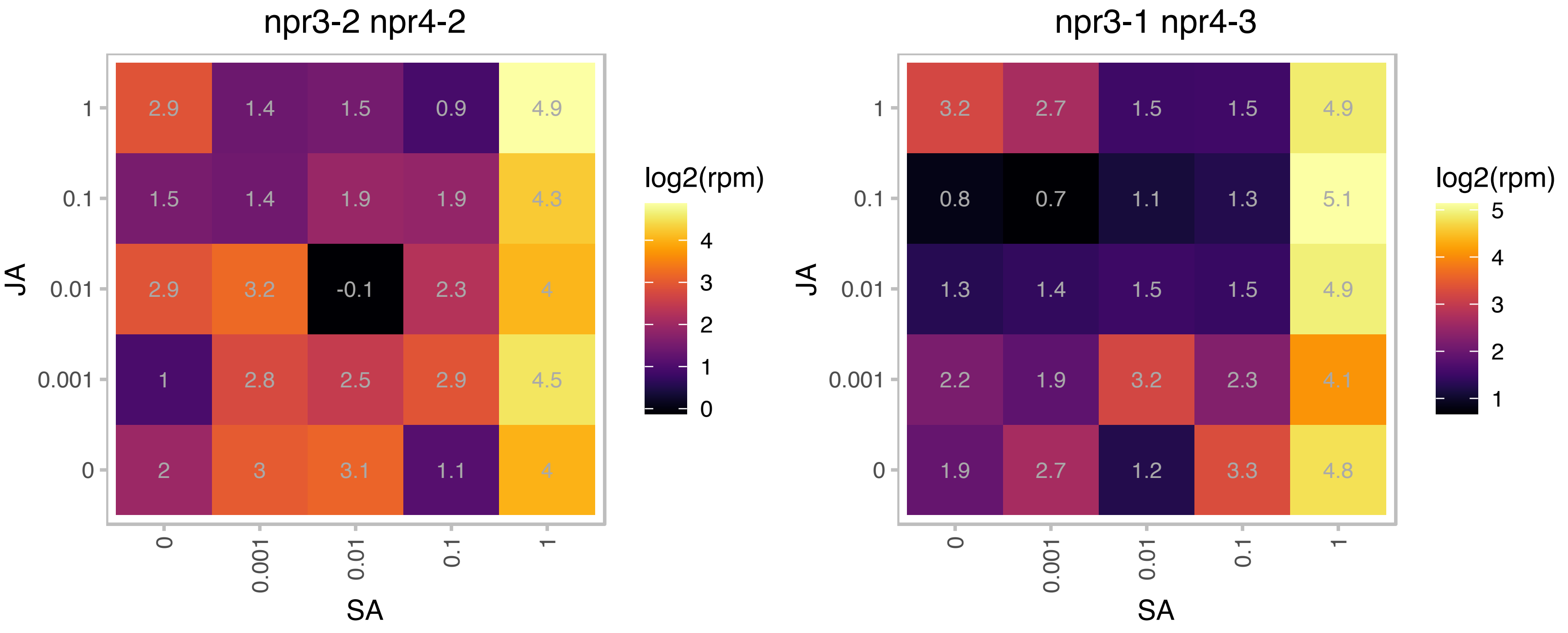

### AT1G32940.1 : AtSBT3.5, SBT3.5

Subtilase family protein

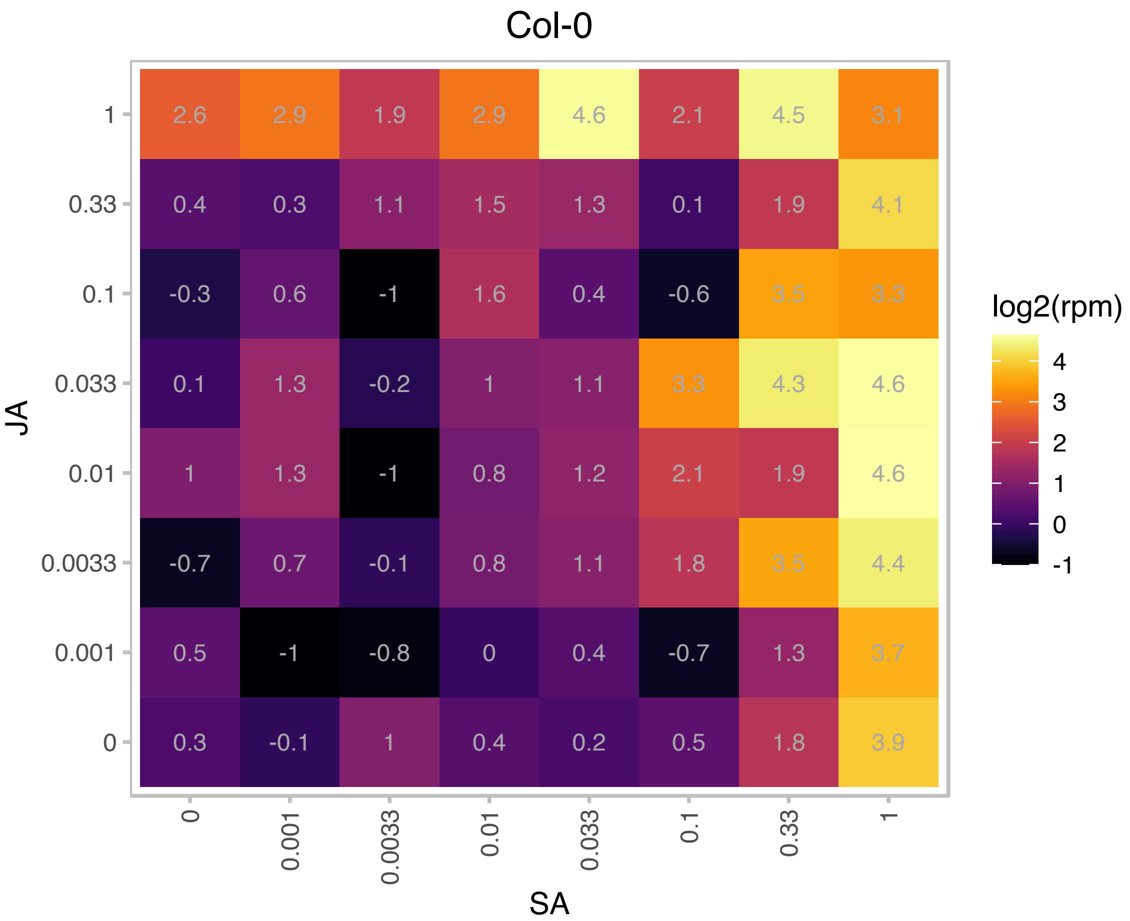


| | |
|---|---|
| Gene | AT1G32940 |
| is.expressed_genes | TRUE |
| is.responsive_genes | TRUE |
| is.apcluster | B-03 |

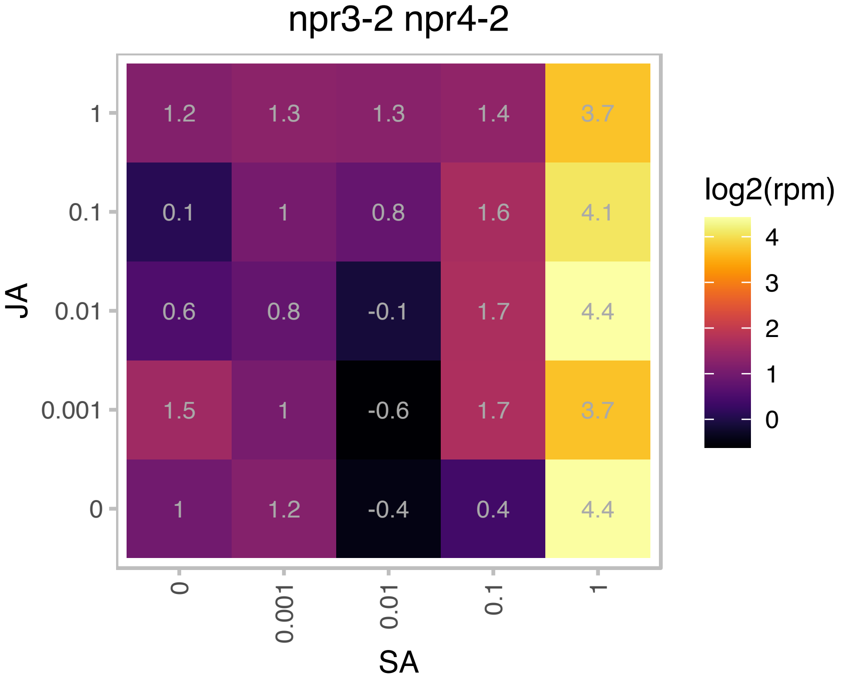


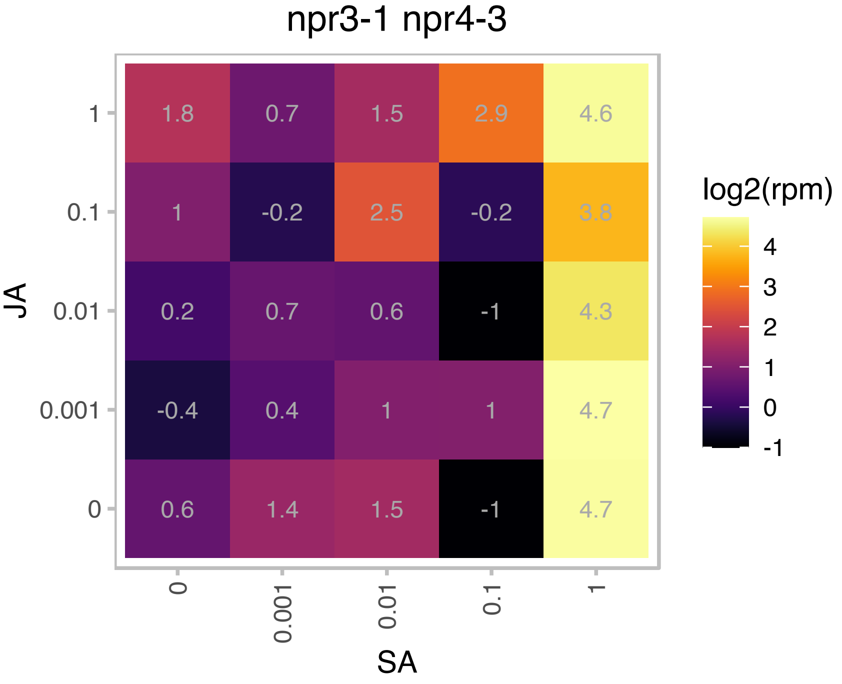

# AT1G35516.1 : NA

myb-like transcription factor family protein

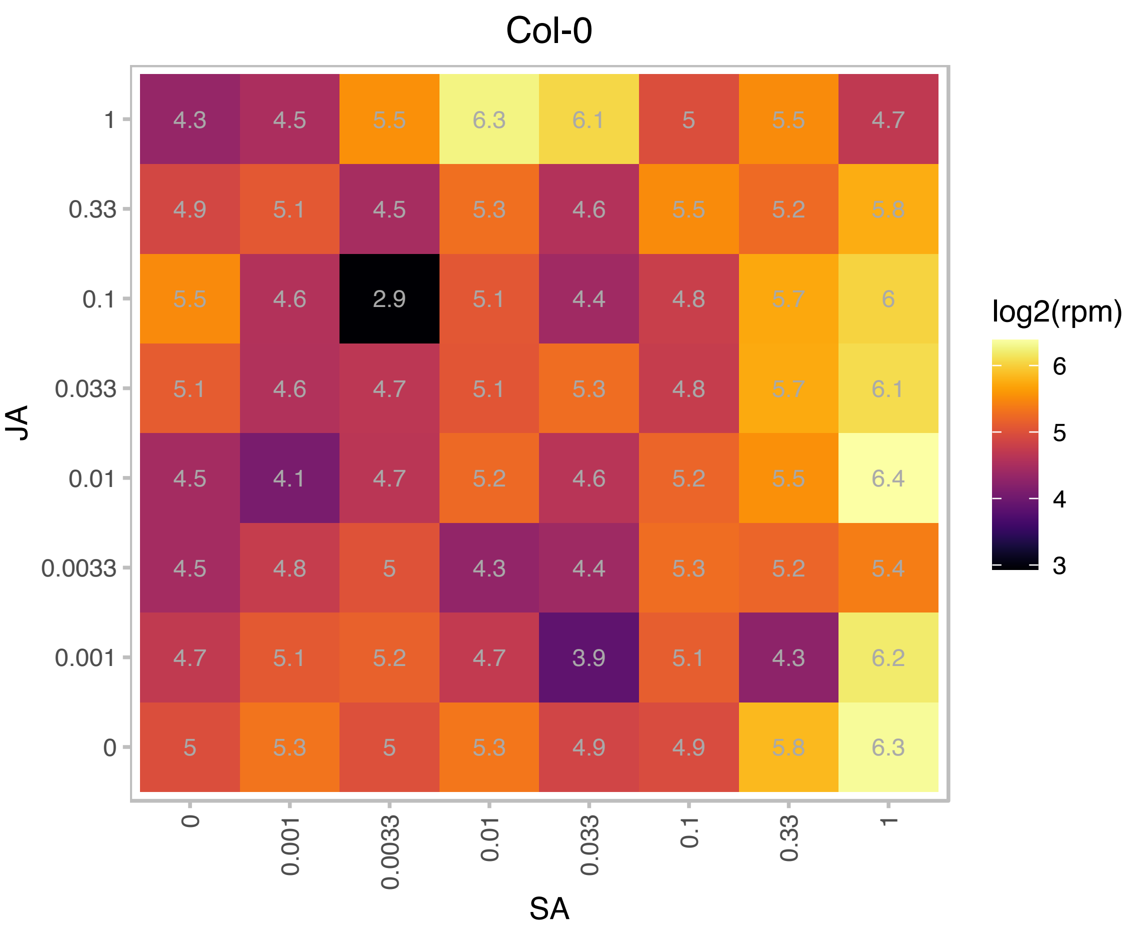


| | |
|---|---|
| Gene | AT1G35516 |
| is.expressed_genes | TRUE |
| is.responsive_genes | TRUE |
| is.apcluster | B-03 |

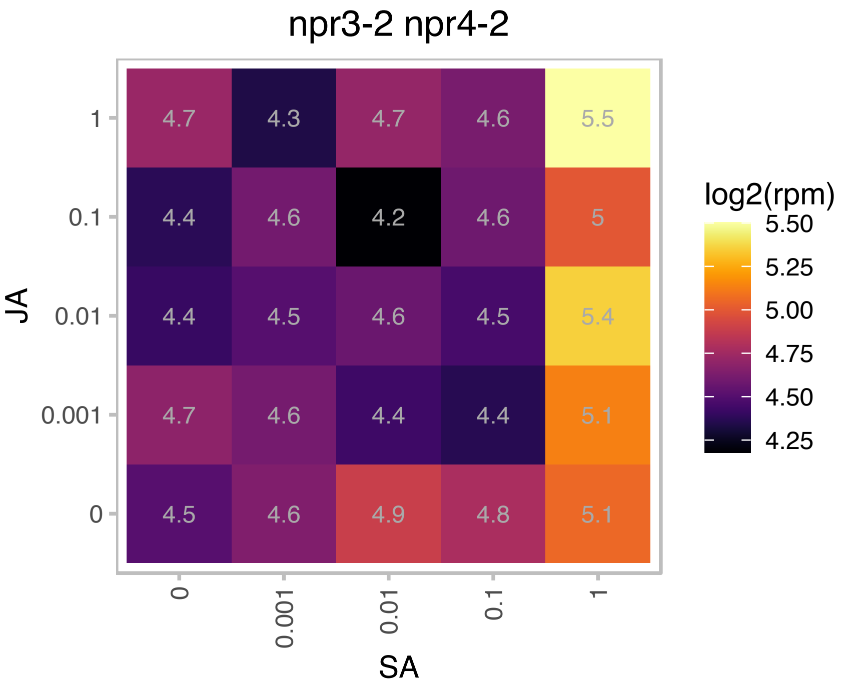


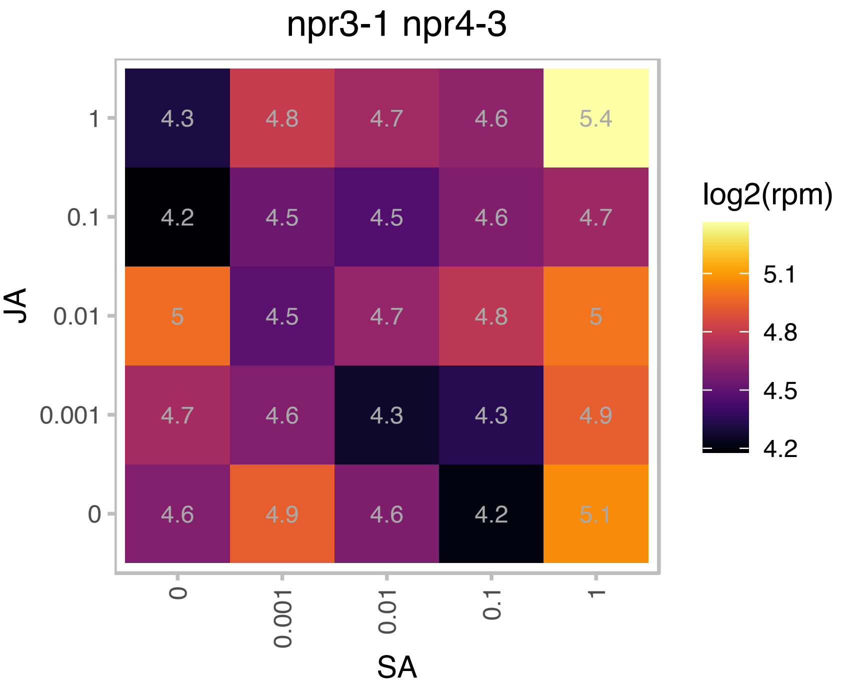

### AT1G43710.1 : ATSDC1, EMB1075, SDC1

Pyridoxal phosphate (PLP)-dependent transferases superfamily protein

Encodes a serine decarboxylase that is involved in ethanolamine metabolism and is crucial for plant growth.

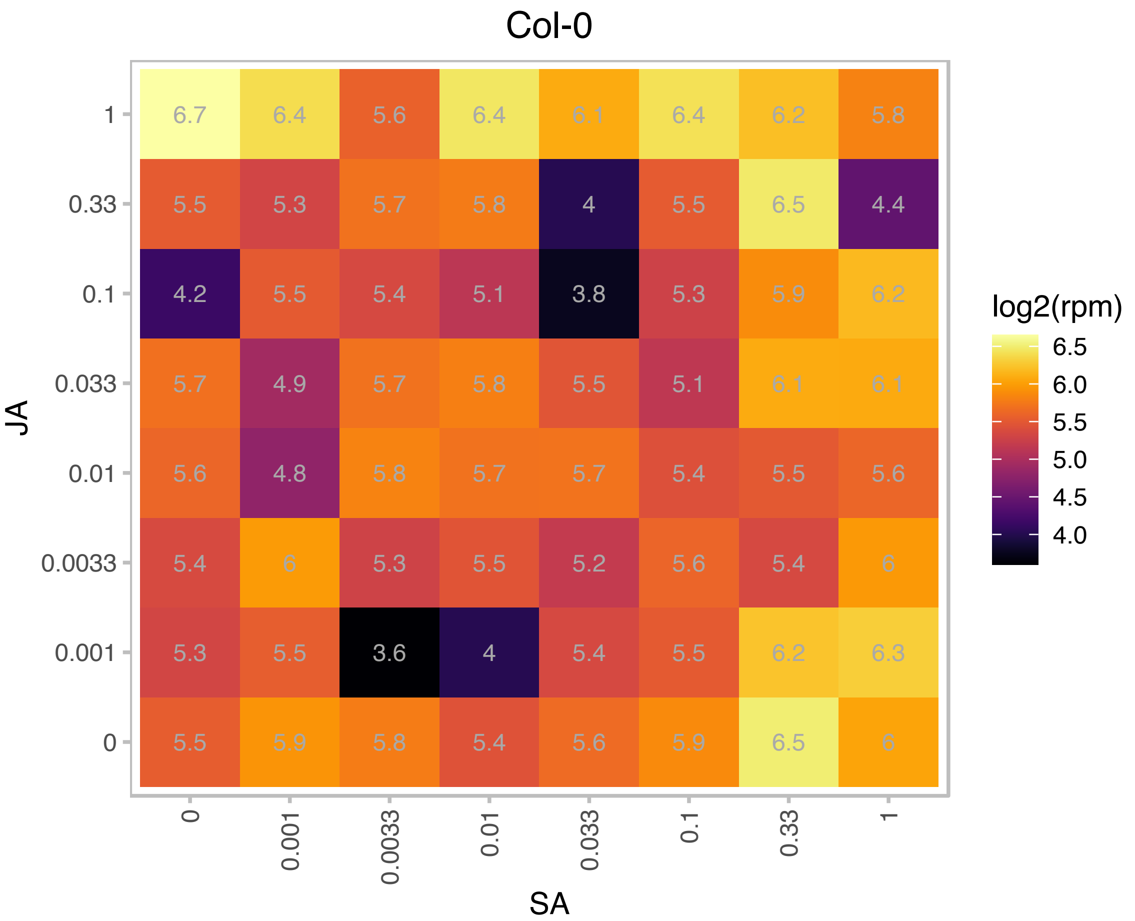


| | |
|---|---|
| Gene | AT1G43710 |
| is.expressed_genes | TRUE |
| is.responsive_genes | TRUE |
| is.apcluster | B-03 |

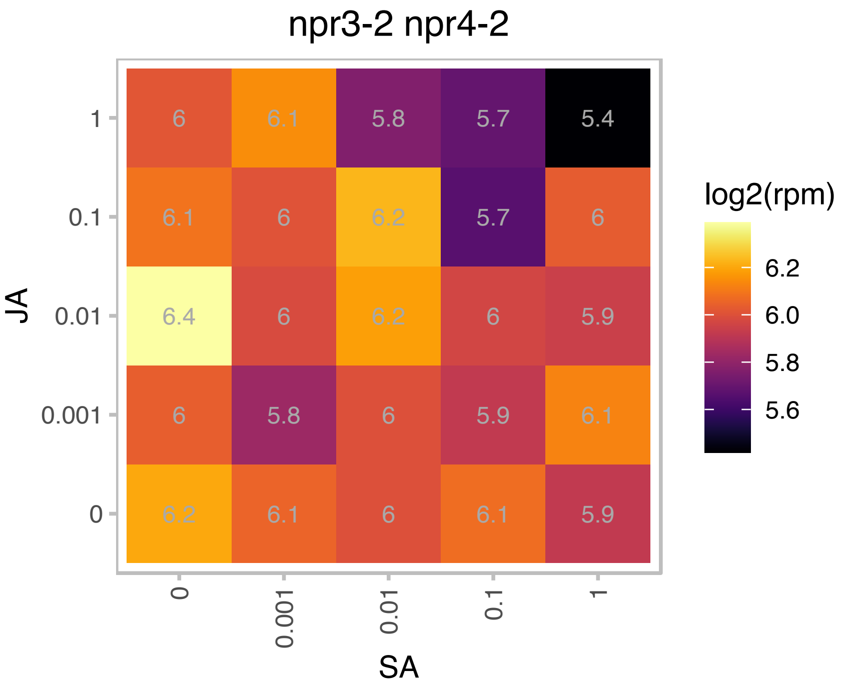


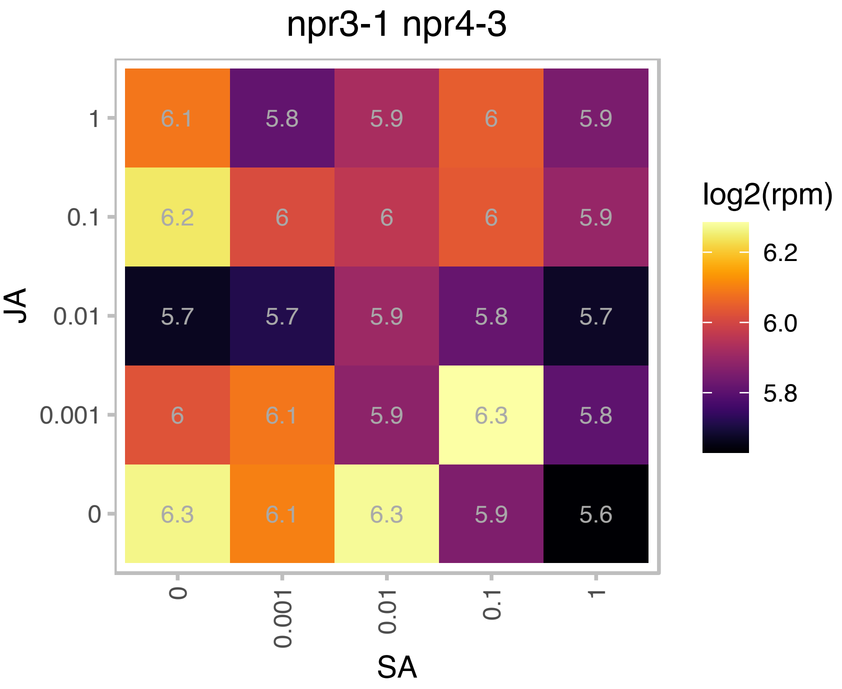

# AT1G52420.1 : NA

UDP-Glycosyltransferase superfamily protein

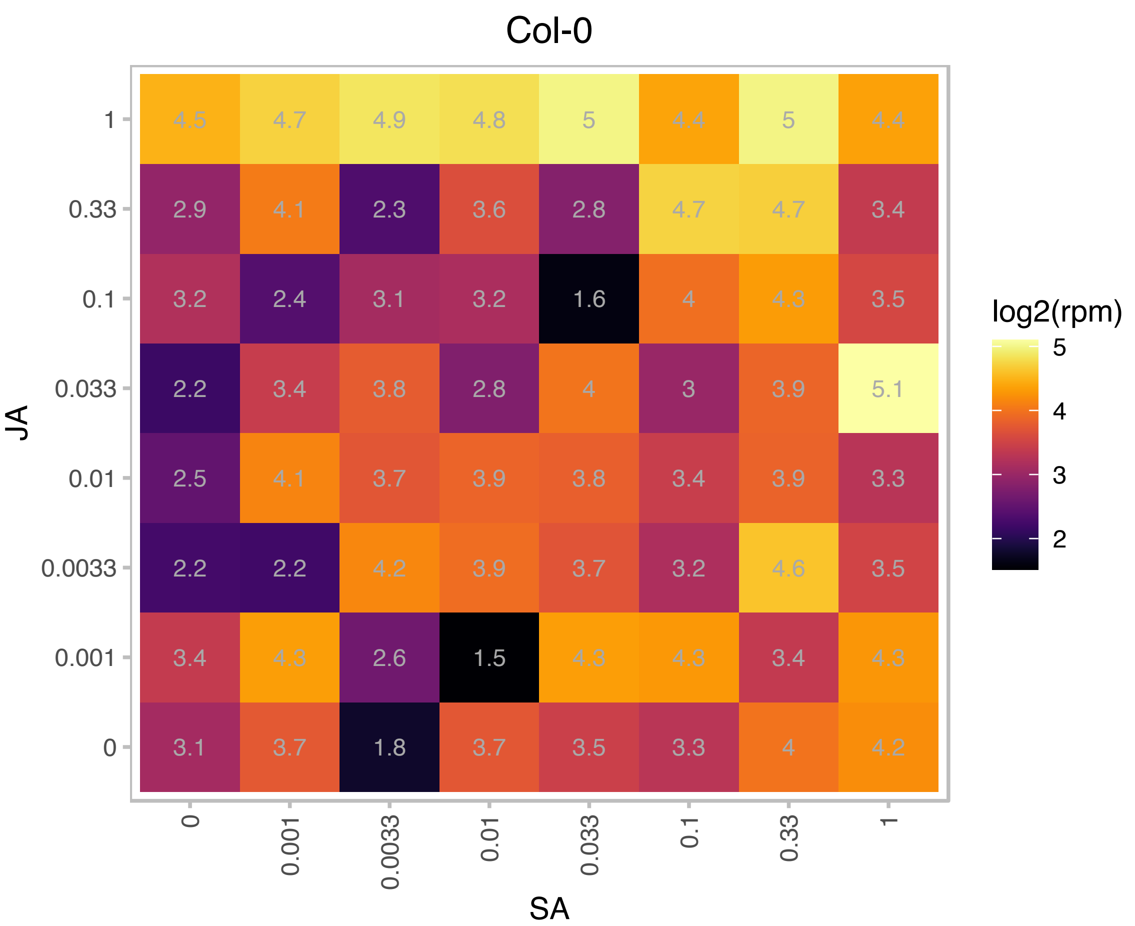


| Gene | AT1G52420 |
|---|---|
| is.expressed_genes | TRUE |
| is.responsive_genes | TRUE |
| is.apcluster | B-03 |

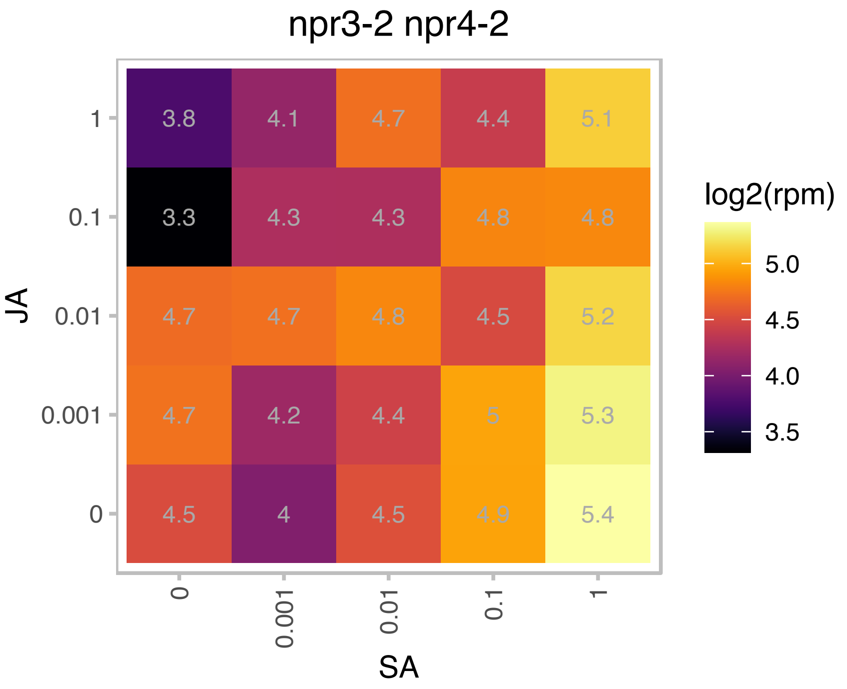


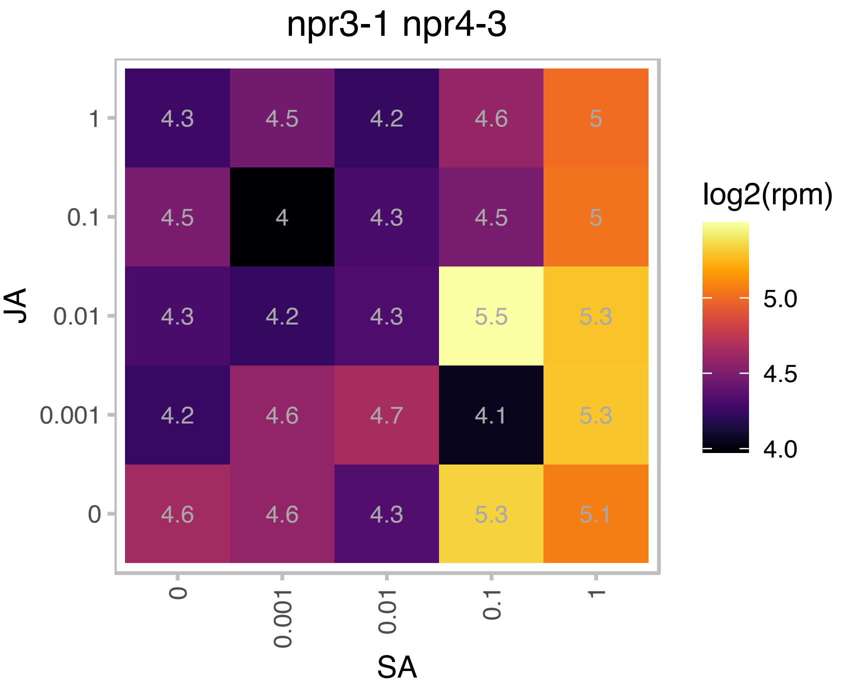

# AT1G54120.1 : NA

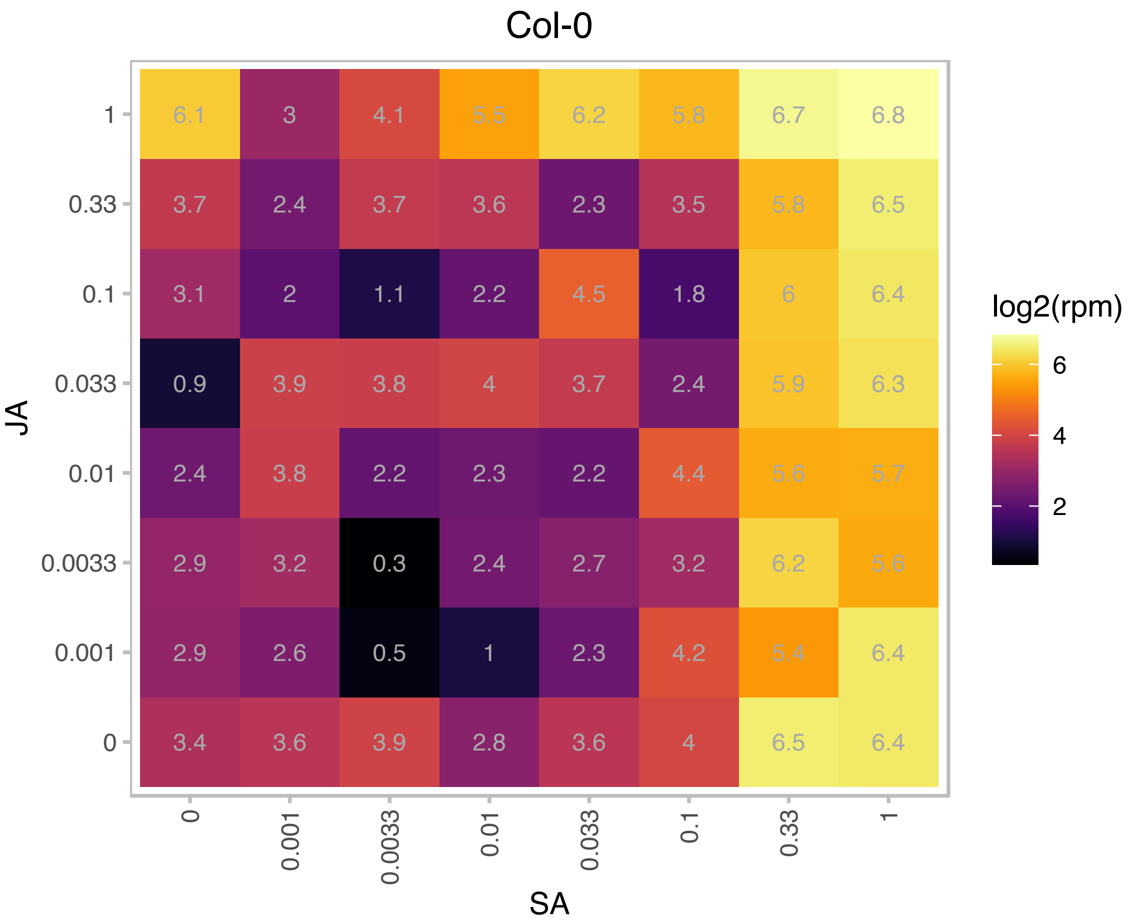


| | |
|---|---|
| Gene | AT1G54120 |
| is.expressed_genes | TRUE |
| is.responsive_genes | TRUE |
| is.apcluster | B-03 |

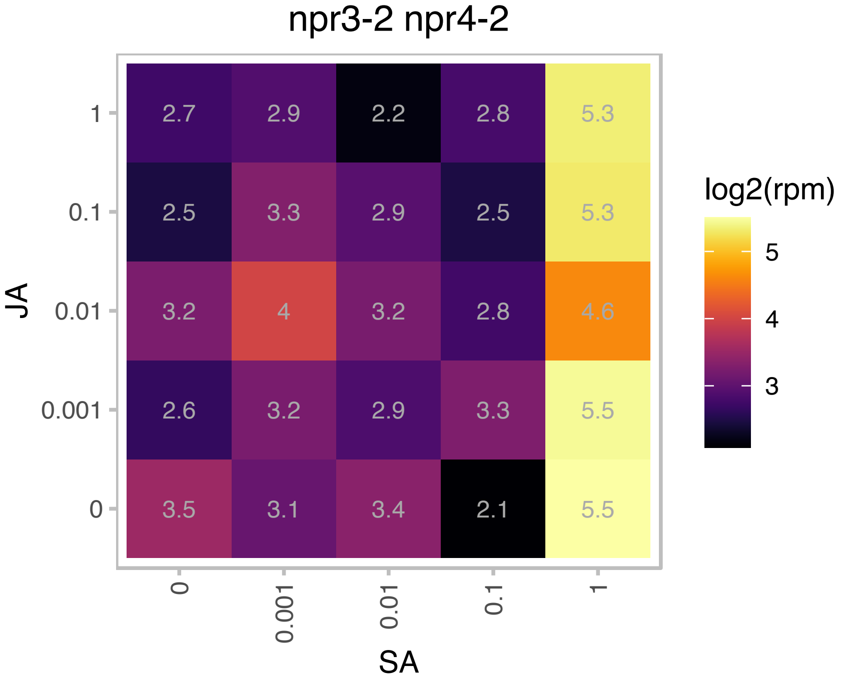


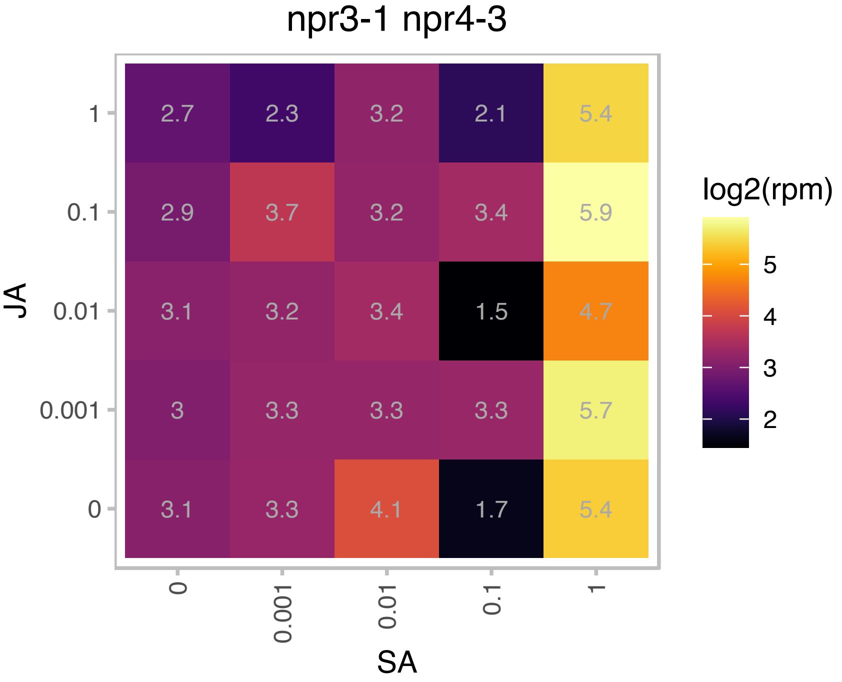

# AT1G55535.1 : NA

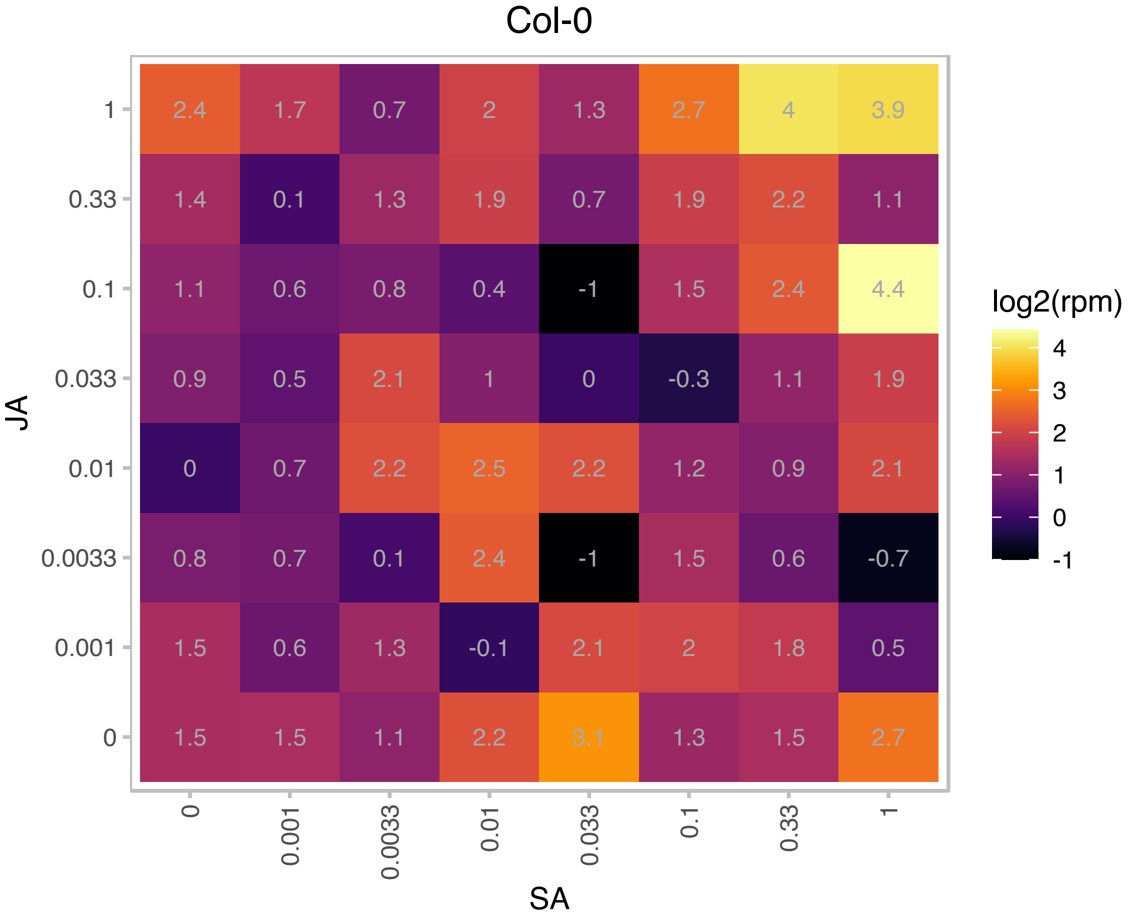


| | |
|---|---|
| Gene | AT1G55535 |
| is.expressed_genes | TRUE |
| is.responsive_genes | TRUE |
| is.apcluster | B-03 |

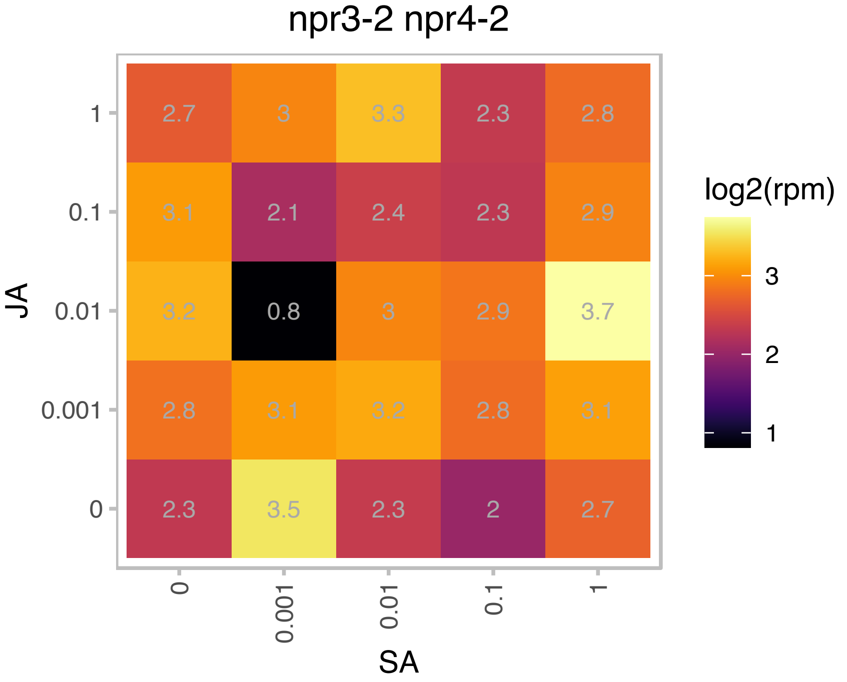


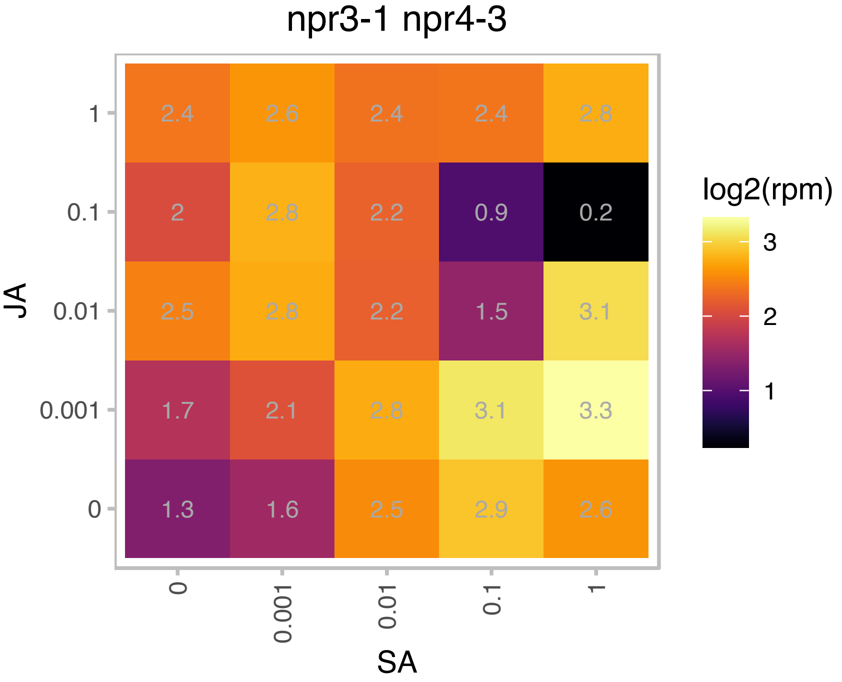

### AT1G59590.1 : ZCF37

ZCF37

ZCF37 mRNA, complete cds The mRNA is cell-to-cell mobile.

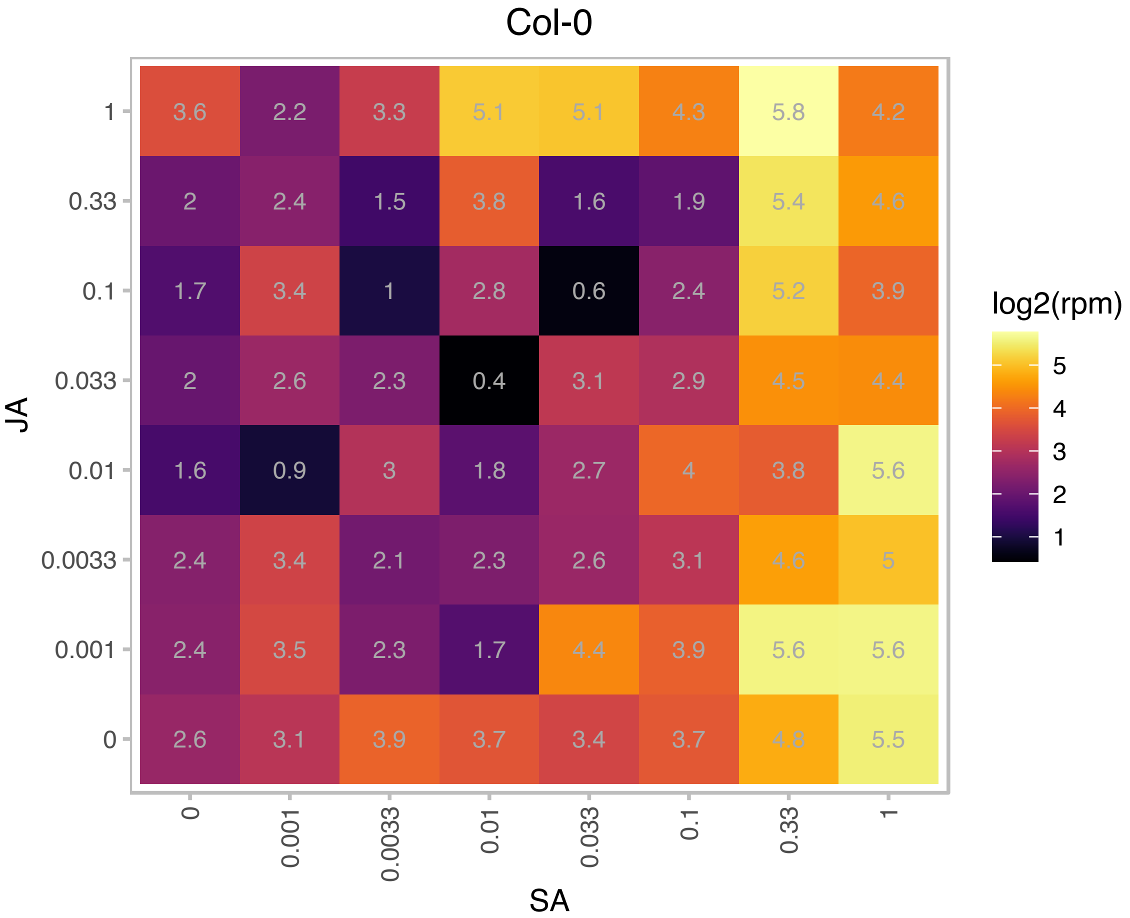


| | |
|---|---|
| Gene | AT1G59590 |
| is.expressed_genes | TRUE |
| is.responsive_genes | TRUE |
| is.apcluster | B-03 |

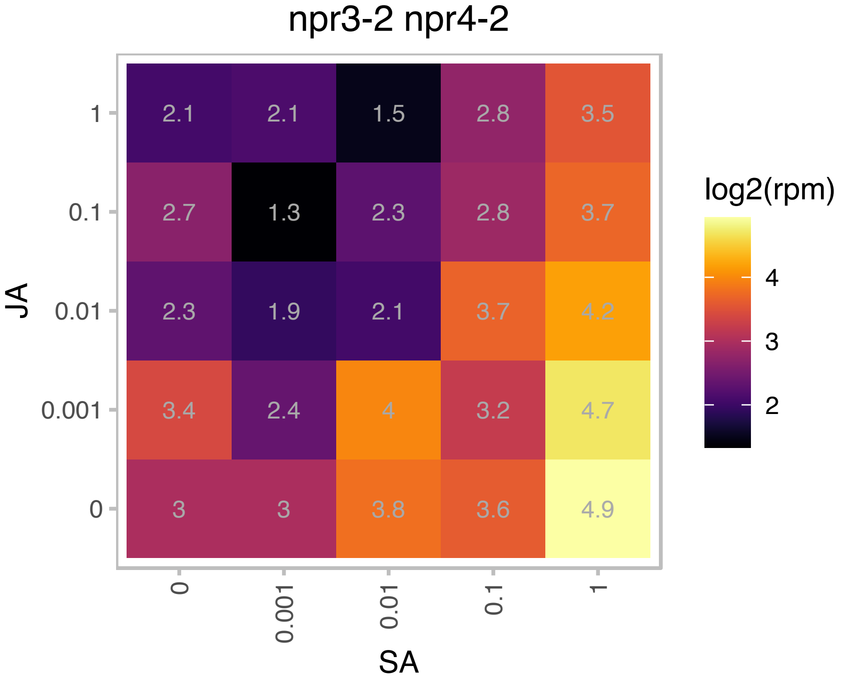


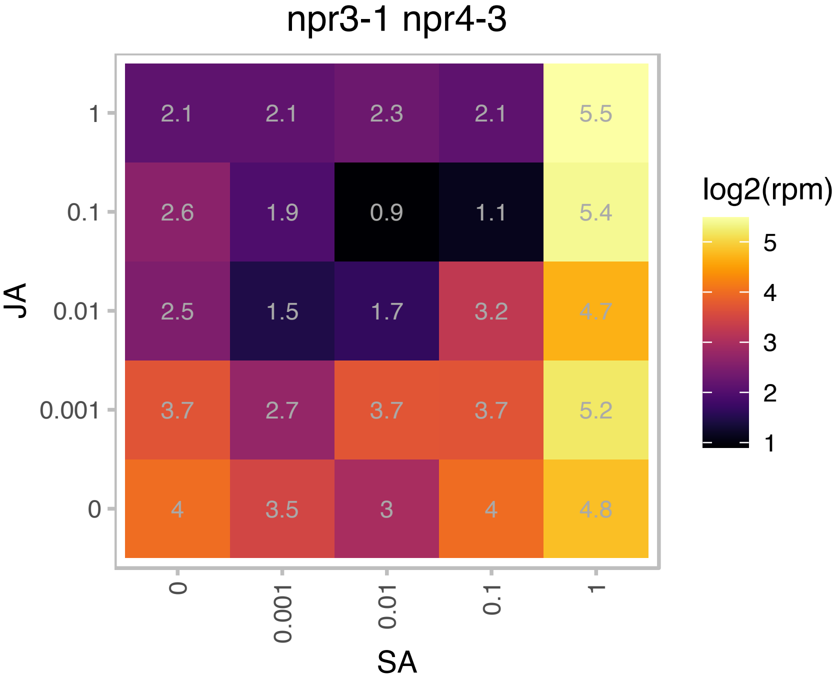

### AT1G59900.1 : AT-E1 ALPHA, E1 ALPHA

pyruvate dehydrogenase complex E1 alpha subunit

encodes the e1 alpha subunit of the pyruvate dehydrogenase complex (PDC) The mRNA is cell-to-cell mobile.

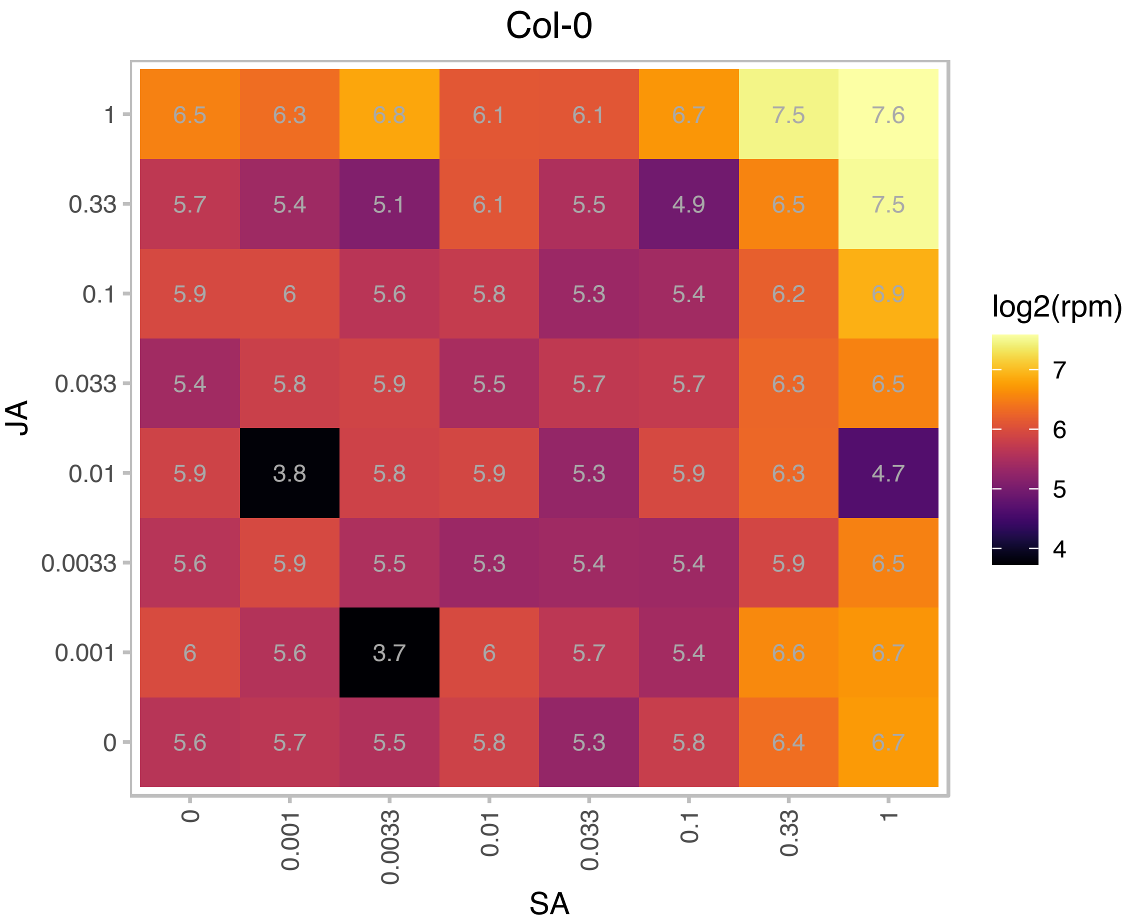


| | |
|---|---|
| Gene | AT1G59900 |
| is.expressed_genes | TRUE |
| is.responsive_genes | TRUE |
| is.apcluster | B-03 |

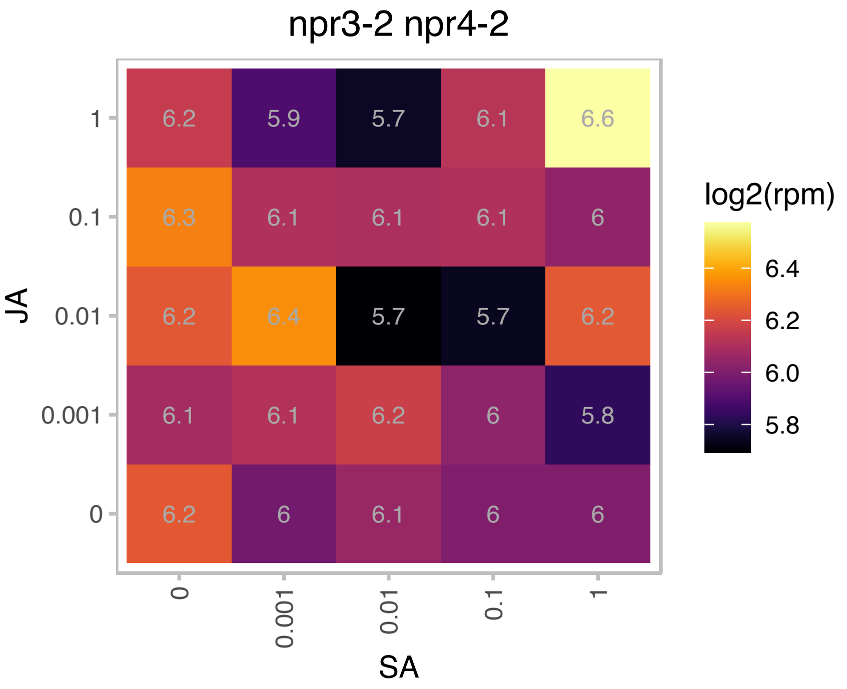


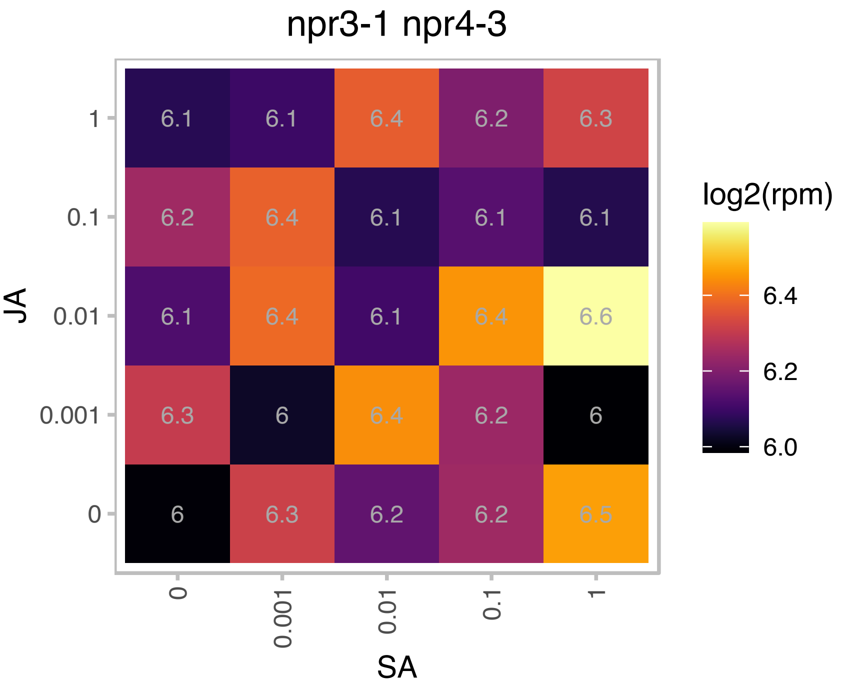

# AT1G65920.1 : NA

Regulator of chromosome condensation (RCC1) family with FYVE zinc finger domain

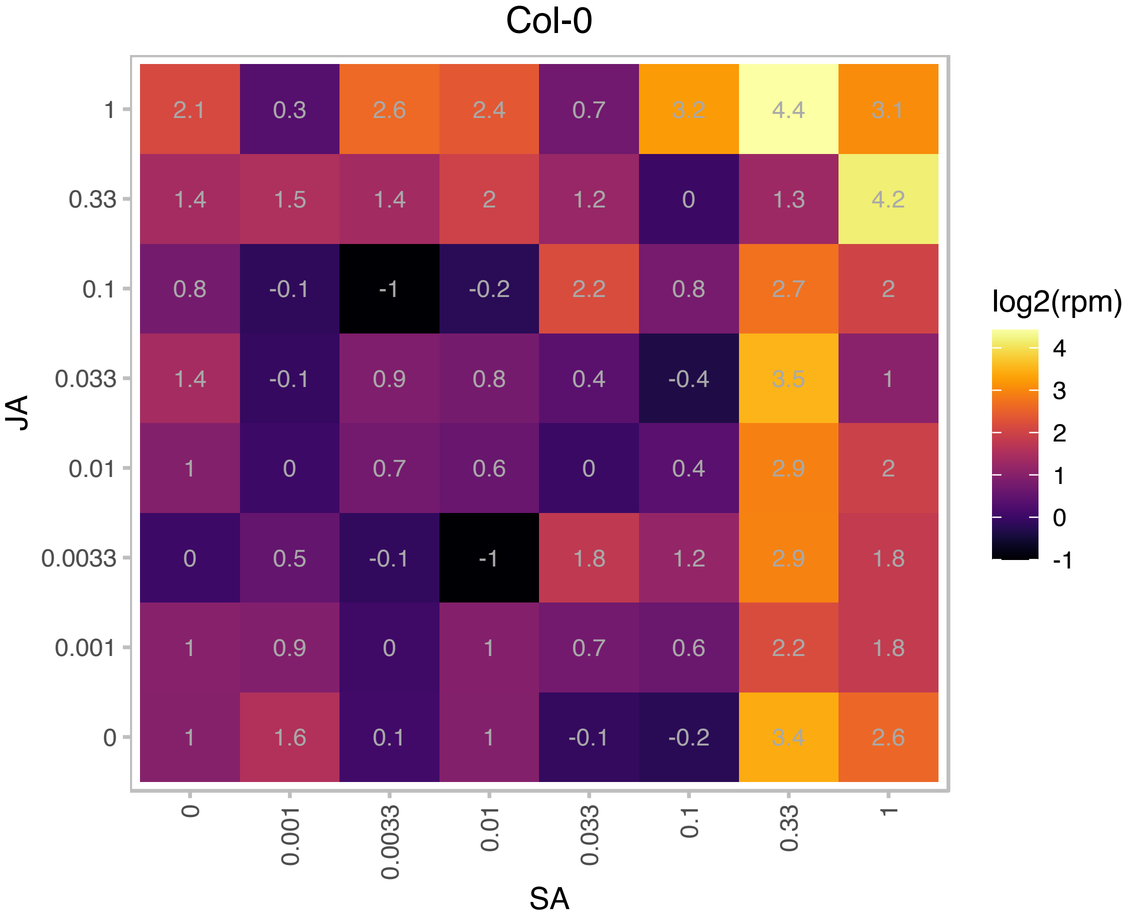


| | |
|---|---|
| Gene | AT1G65920 |
| is.expressed_genes | TRUE |
| is.responsive_genes | TRUE |
| is.apcluster | B-03 |

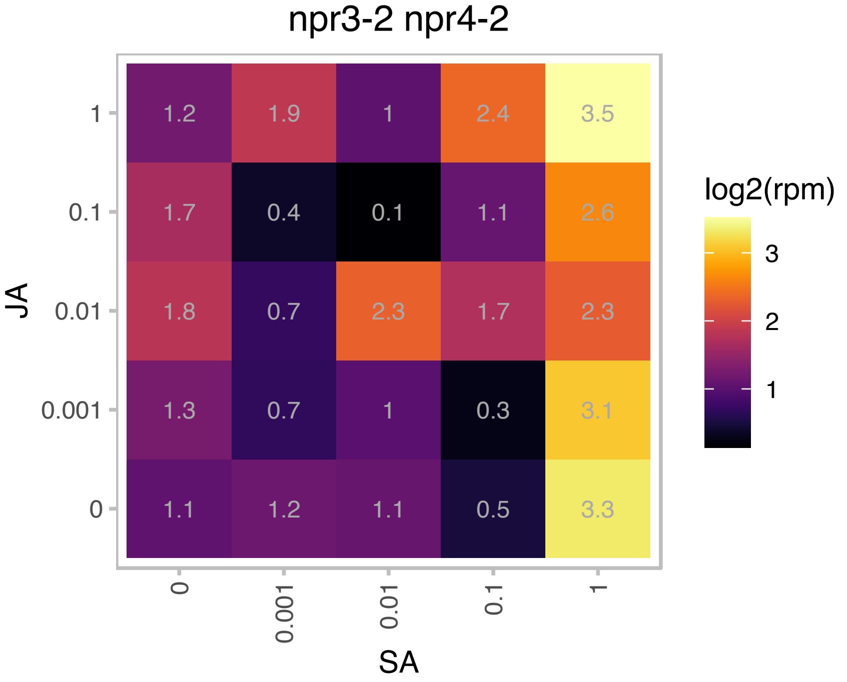


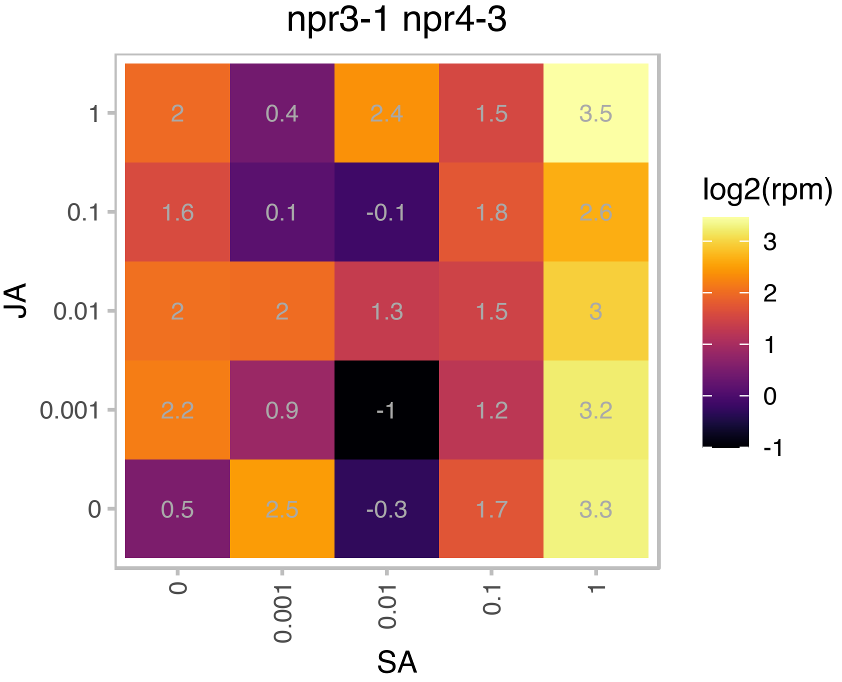

# AT1G68820.1 : NA

Transmembrane Fragile-X-F-associated protein

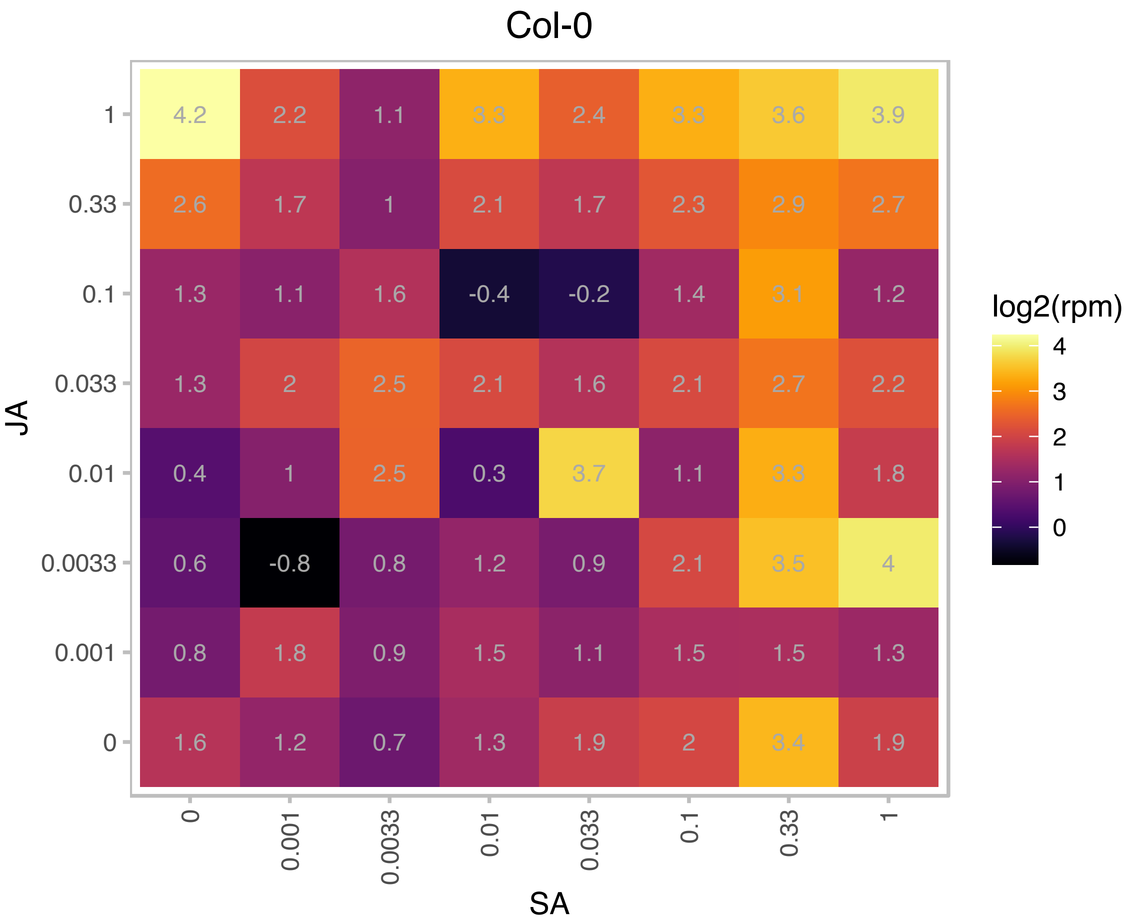


| | |
|---|---|
| Gene | AT1G68820 |
| is.expressed_genes | TRUE |
| is.responsive_genes | TRUE |
| is.apcluster | B-03 |

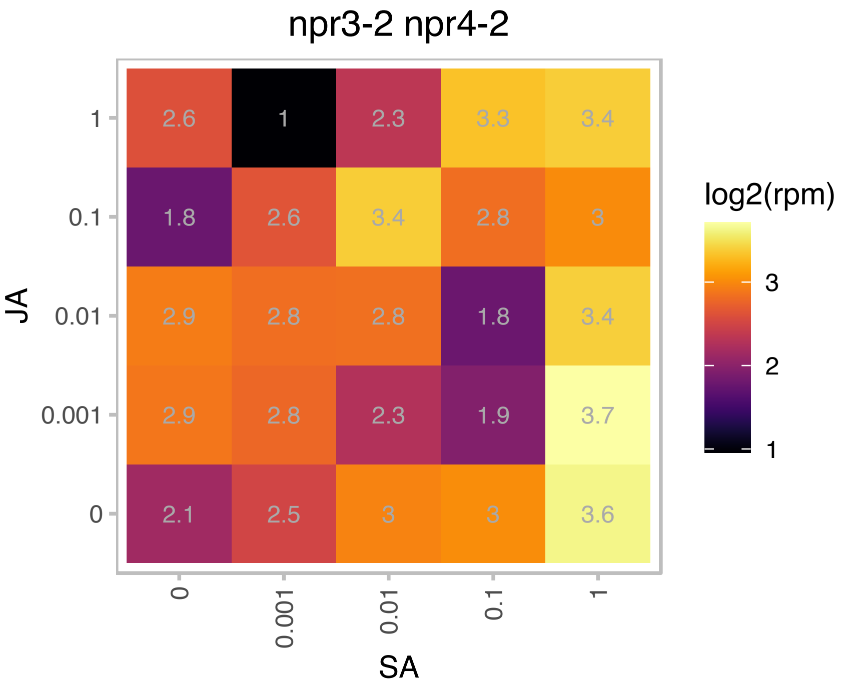


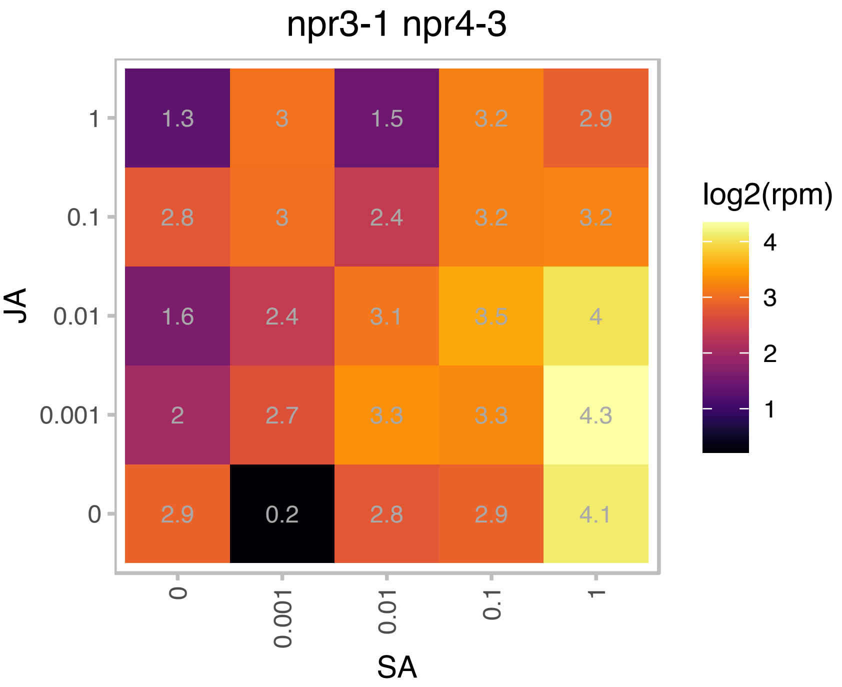

### AT1G68880.1 : AtbZIP, bZIP

basic leucine-zipper 8

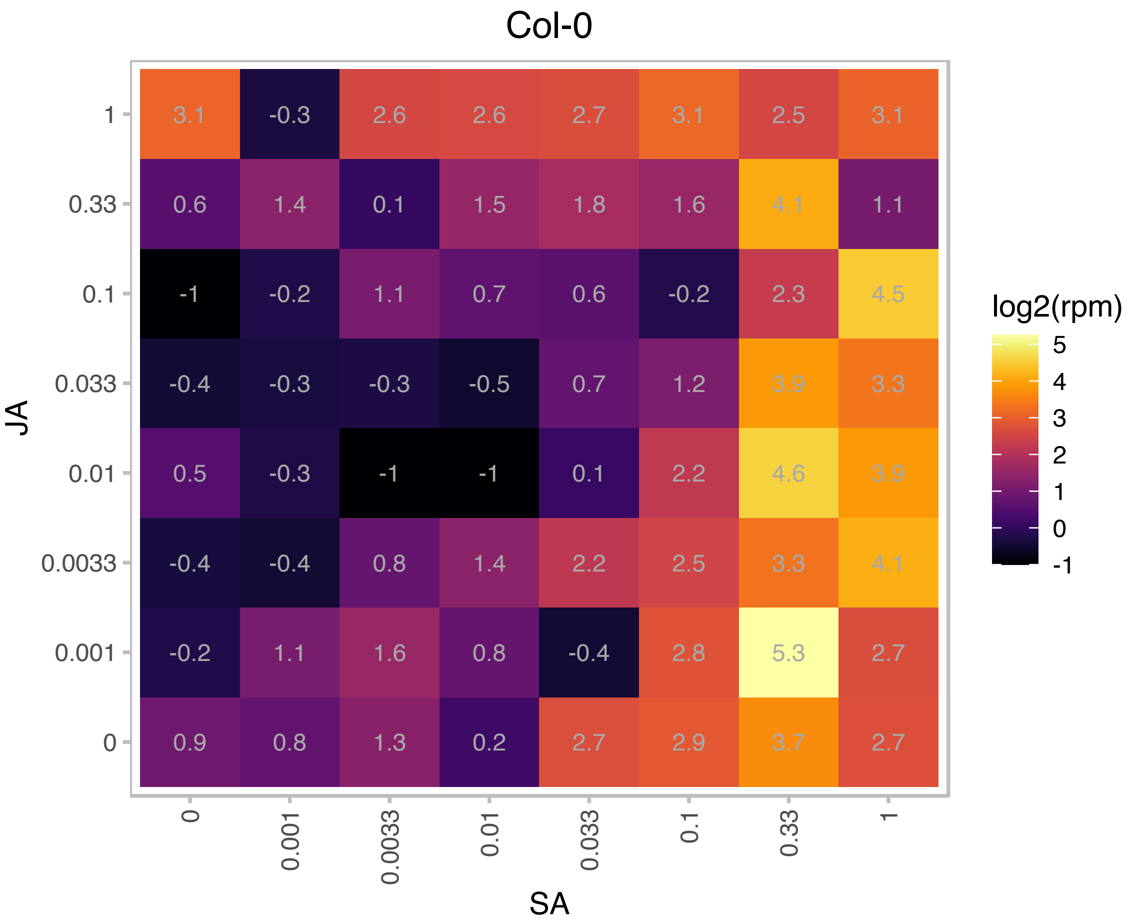


| | |
|---|---|
| Gene | AT1G68880 |
| is.expressed_genes | TRUE |
| is.responsive_genes | TRUE |
| is.apcluster | B-03 |

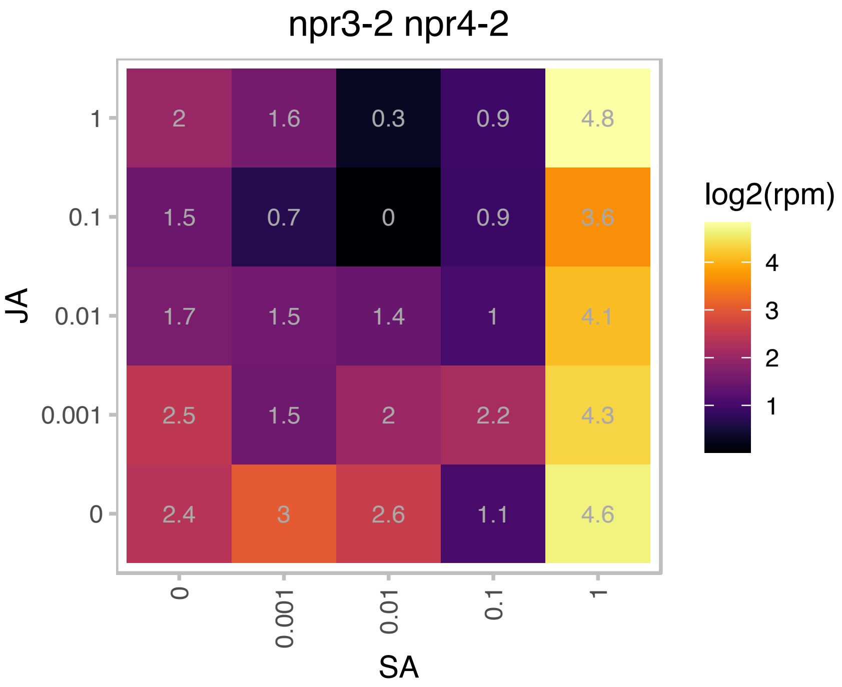


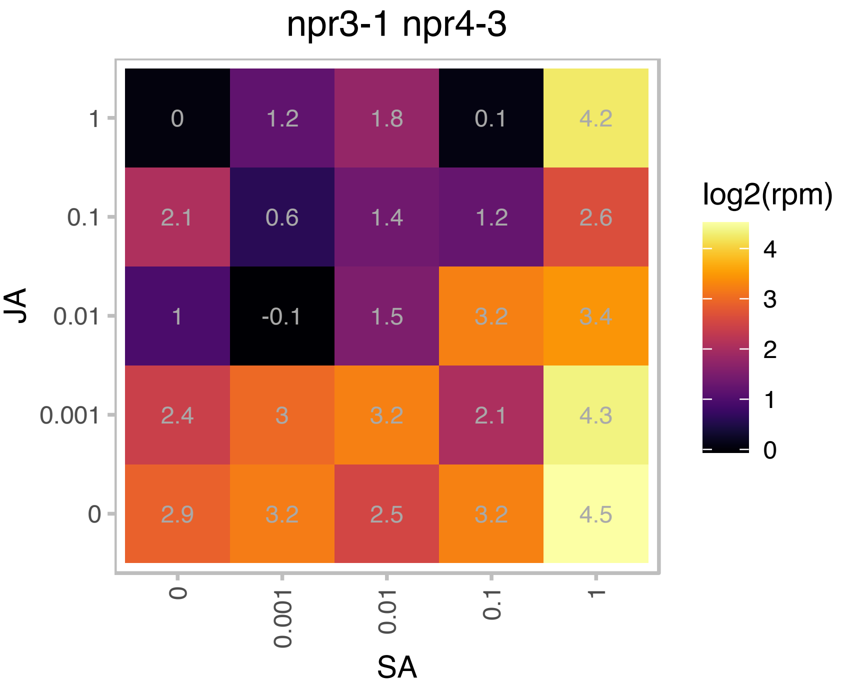

# AT1G70740.1 : NA

Protein kinase superfamily protein

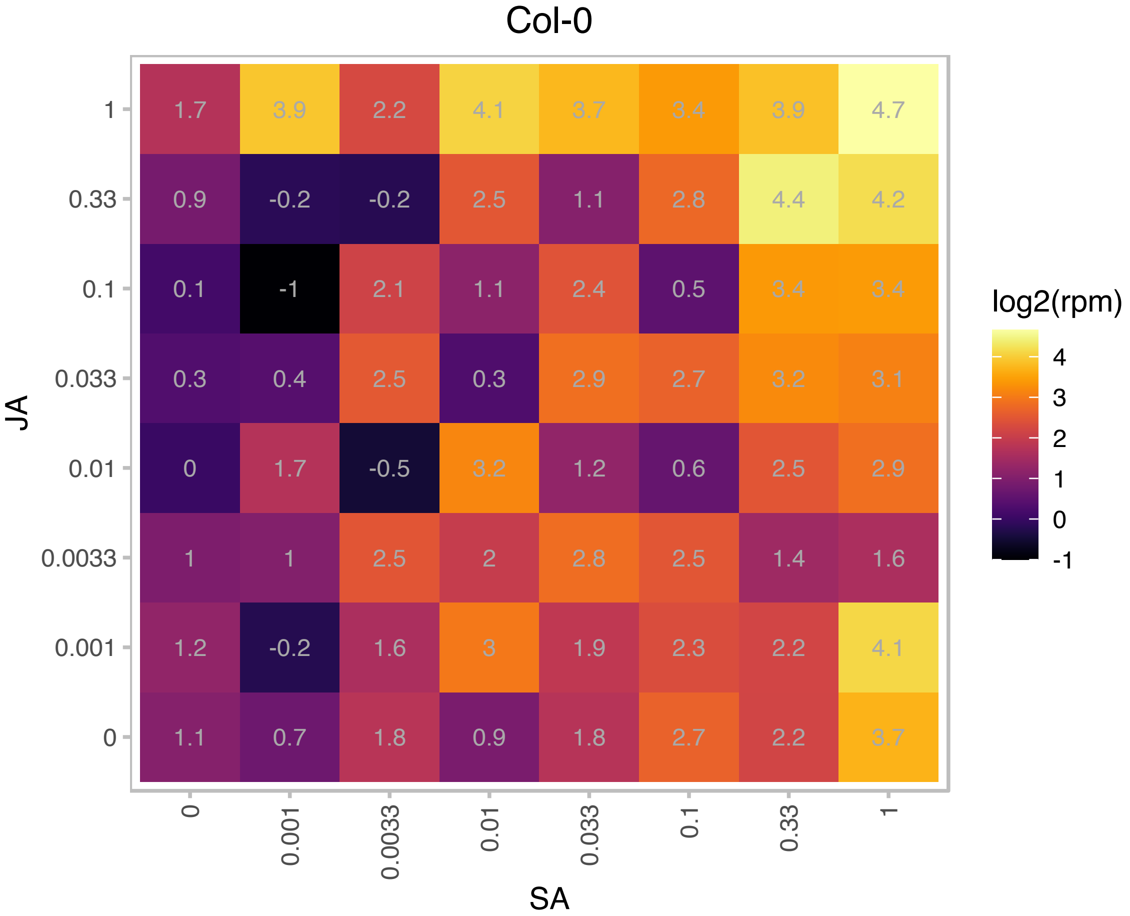


| | |
|---|---|
| Gene | AT1G70740 |
| is.expressed_genes | TRUE |
| is.responsive_genes | TRUE |
| is.apcluster | B-03 |

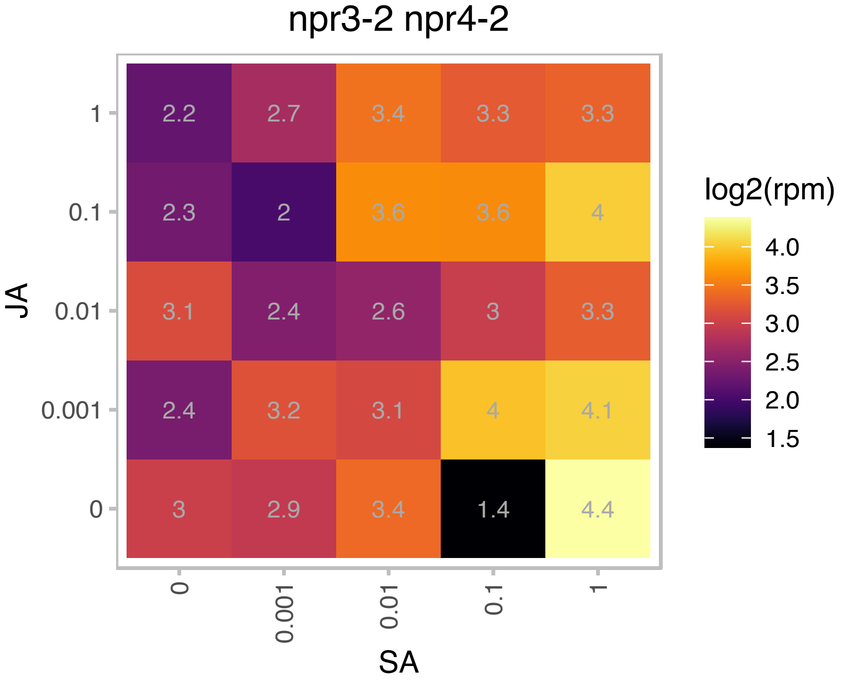


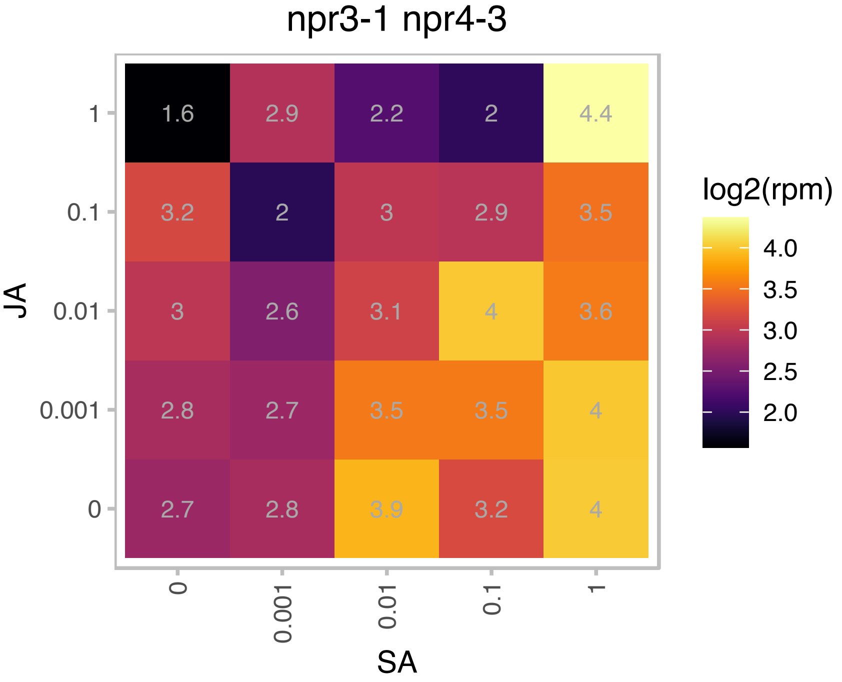

# AT1G76920.1 : NA

F-box family protein

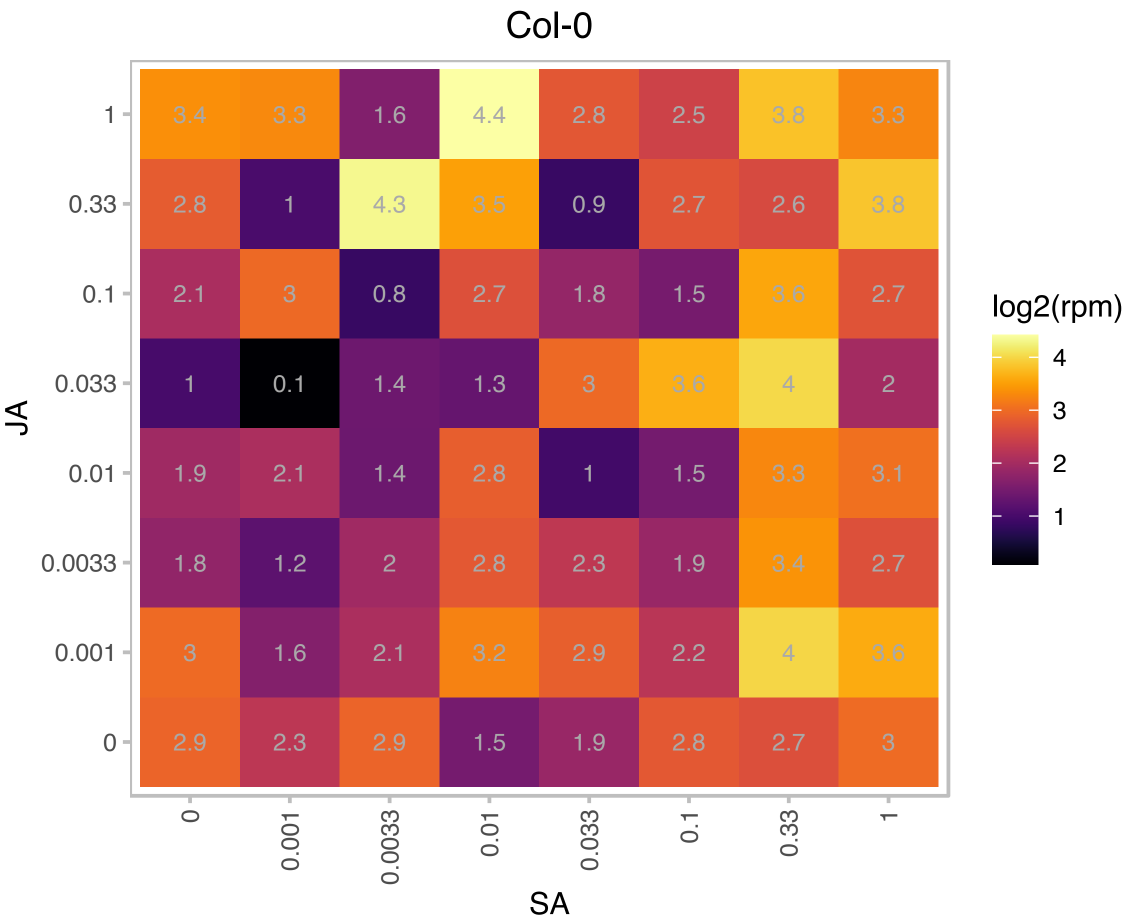


| | |
|---|---|
| Gene | AT1G76920 |
| is.expressed_genes | TRUE |
| is.responsive_genes | TRUE |
| is.apcluster | B-03 |

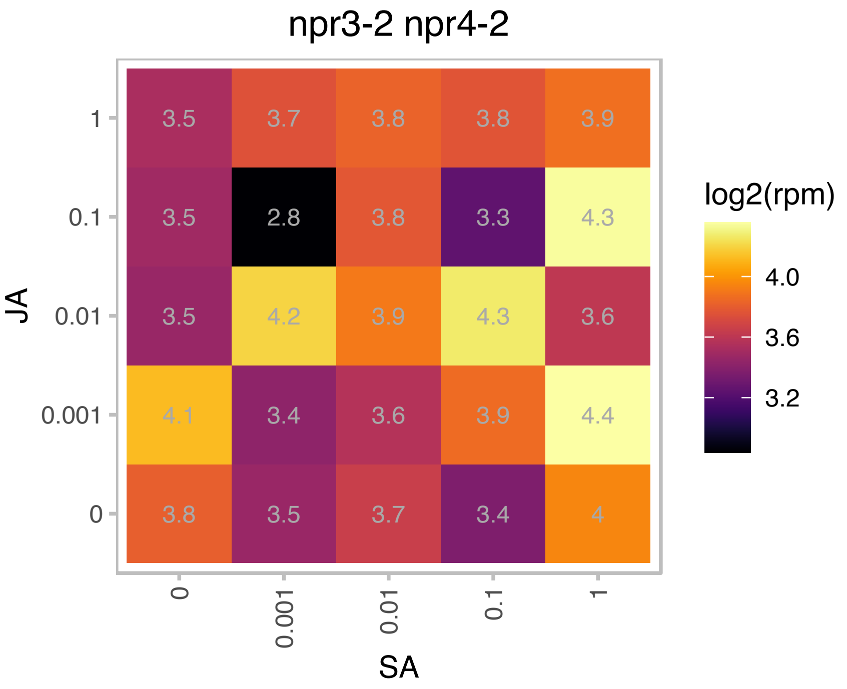


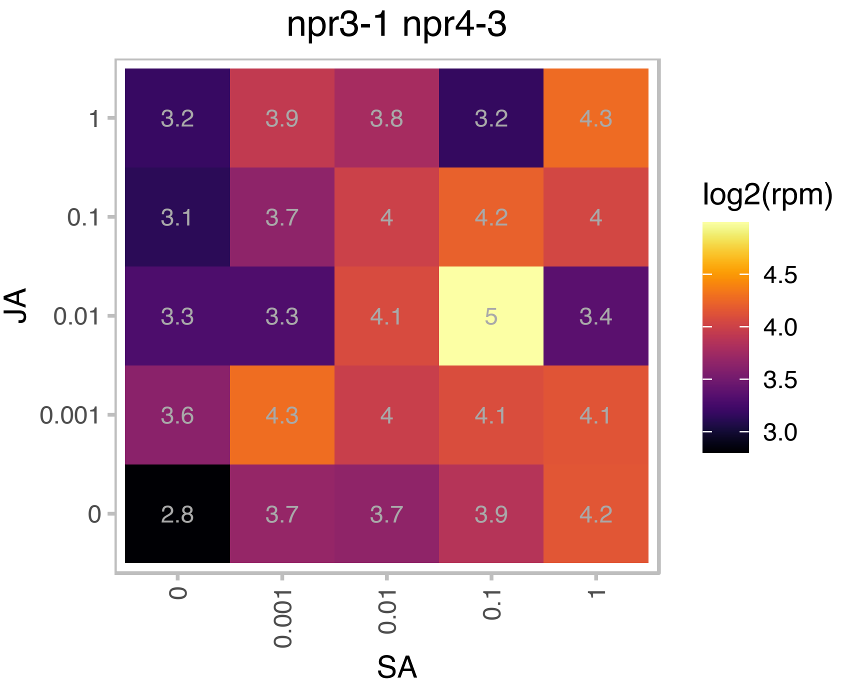

# AT1G76970.1 : NA

Target of Myb protein 1

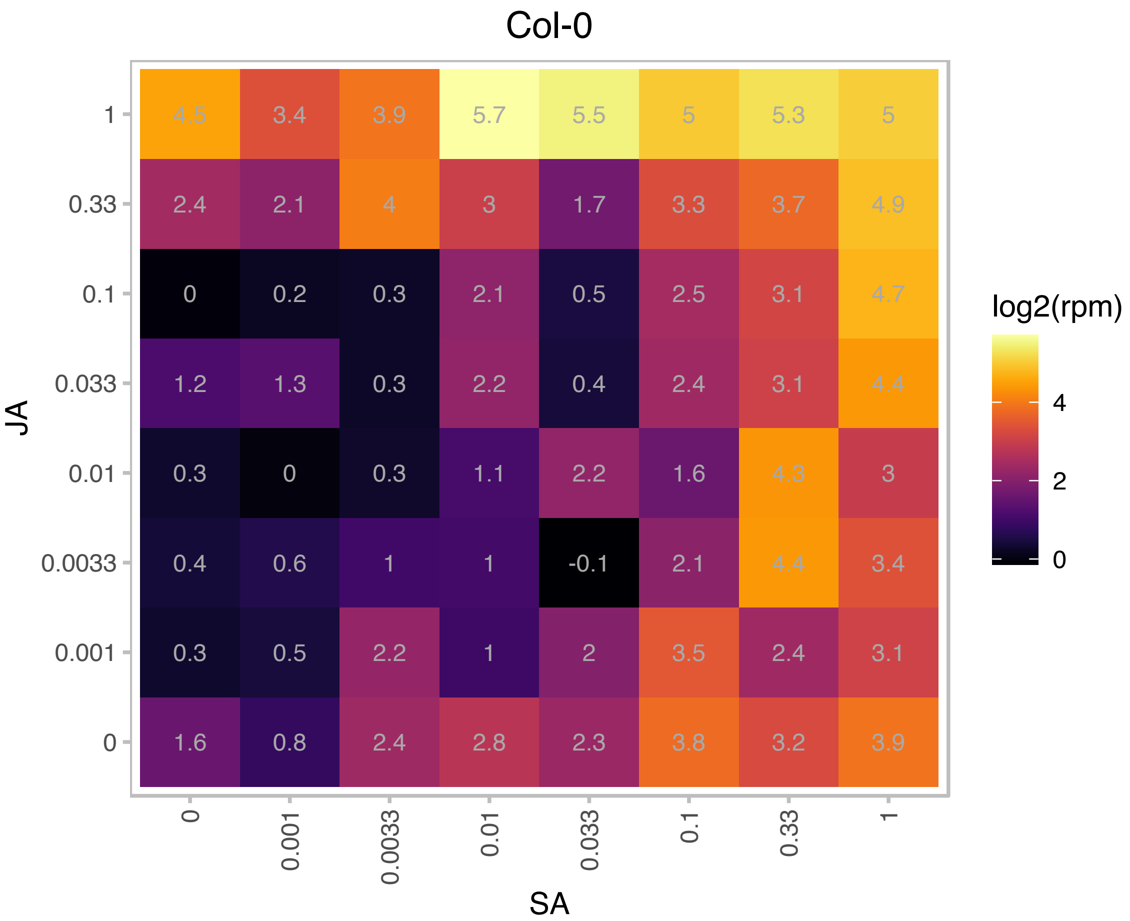


| | |
|---|---|
| Gene | AT1G76970 |
| is.expressed_genes | TRUE |
| is.responsive_genes | TRUE |
| is.apcluster | B-03 |

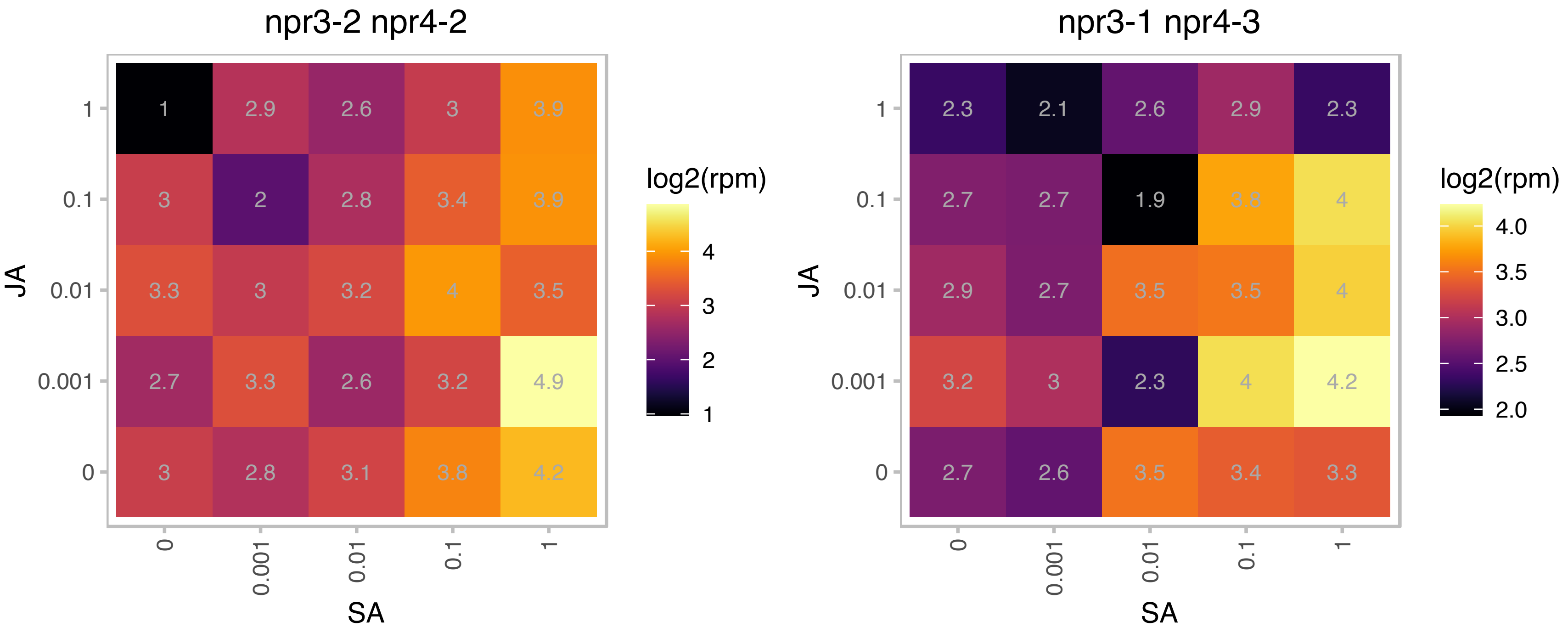

# AT1G78420.1 : DA2

RING/U-box superfamily protein

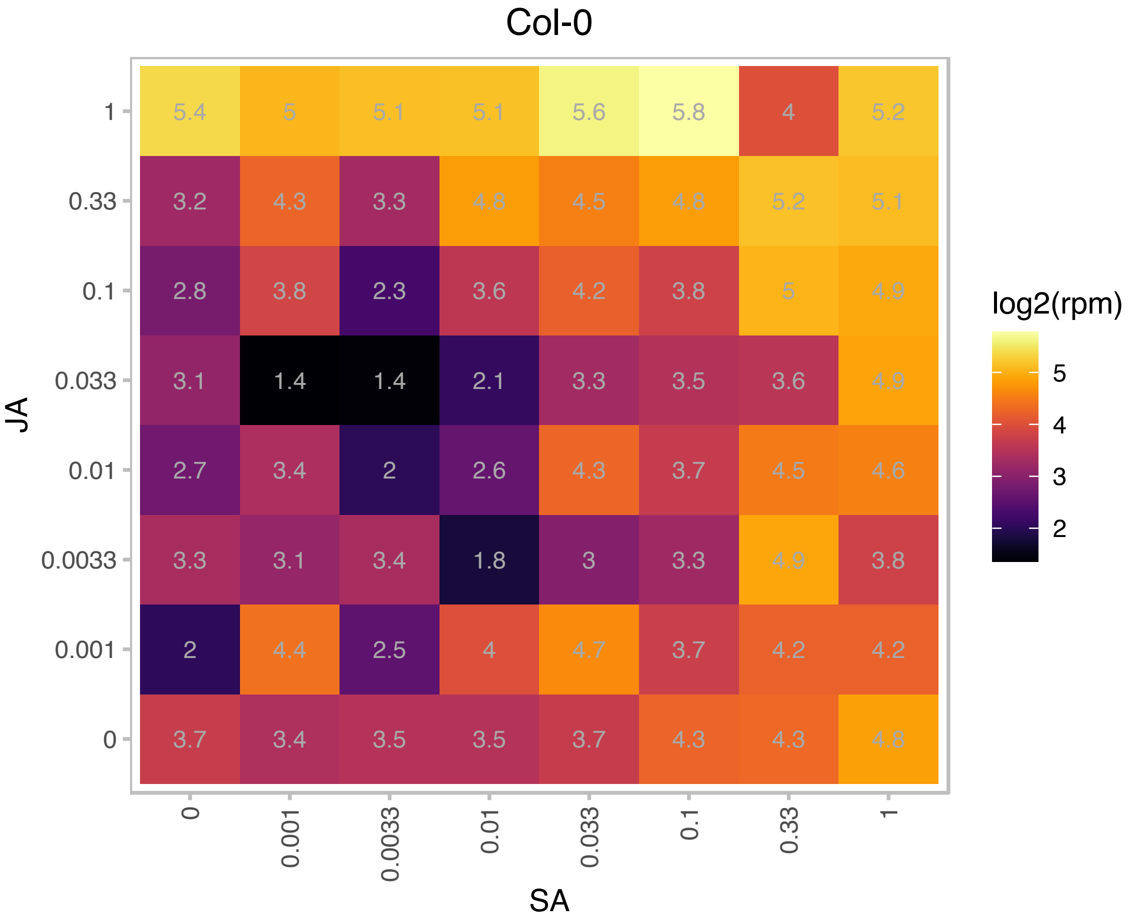


| | |
|---|---|
| Gene | AT1G78420 |
| is.expressed_genes | TRUE |
| is.responsive_genes | TRUE |
| is.apcluster | B-03 |

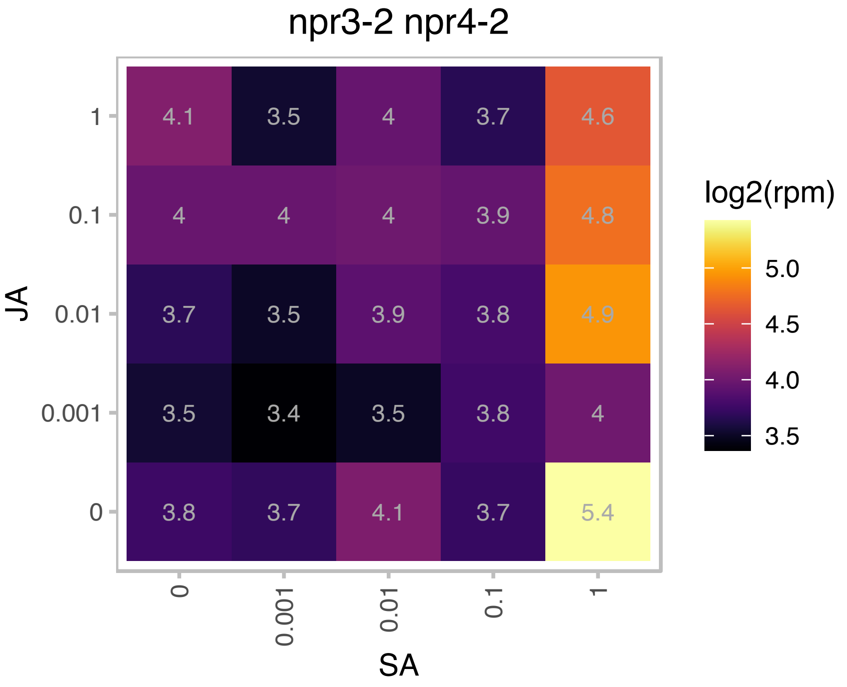


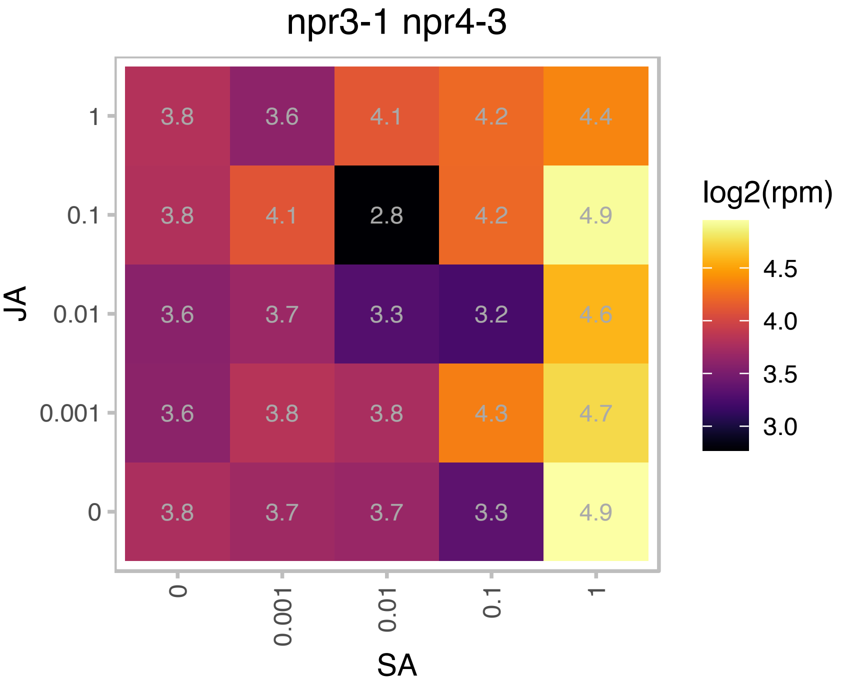

# AT2G01820.1 : NA

Leucine-rich repeat protein kinase family protein

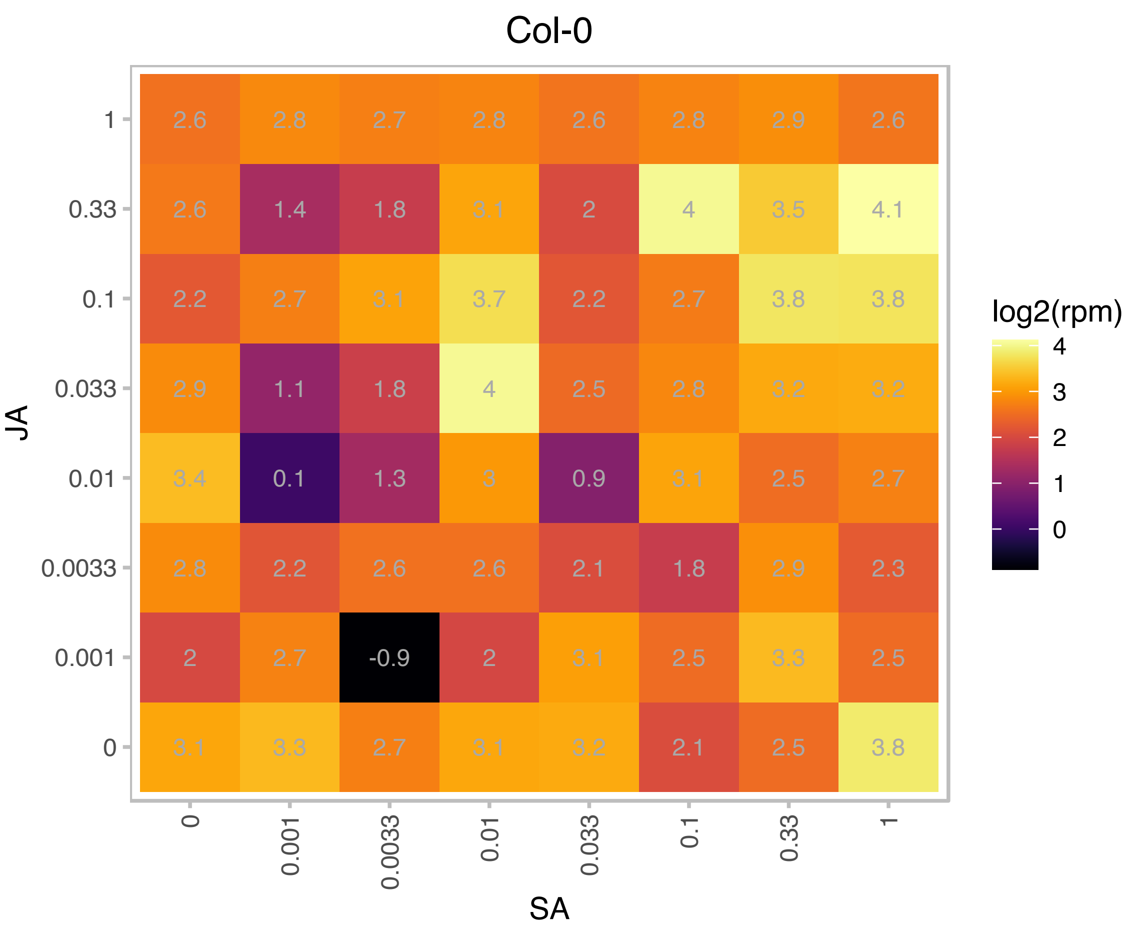


| | |
|---|---|
| Gene | AT2G01820 |
| is.expressed_genes | TRUE |
| is.responsive_genes | TRUE |
| is.apcluster | B-03 |

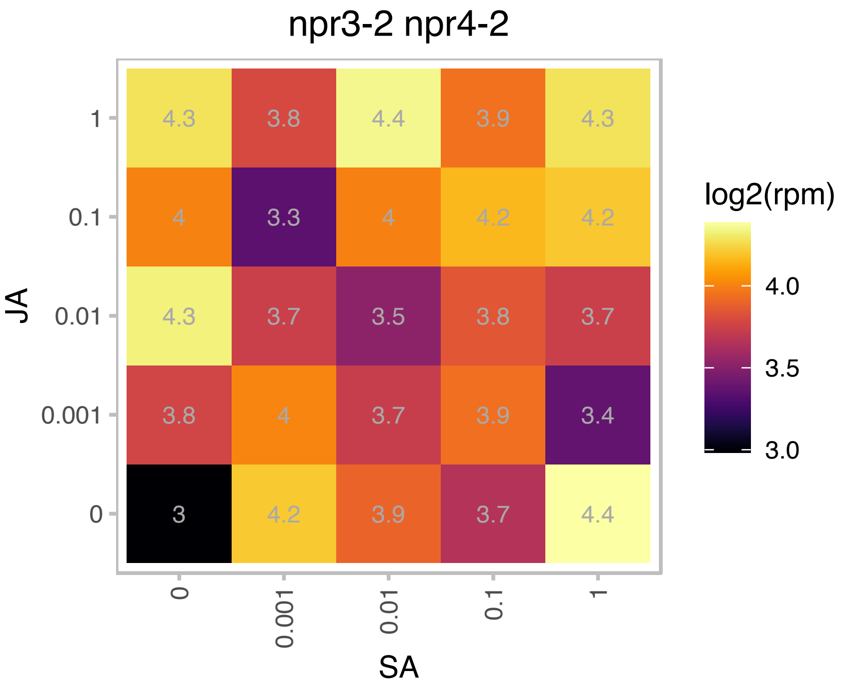


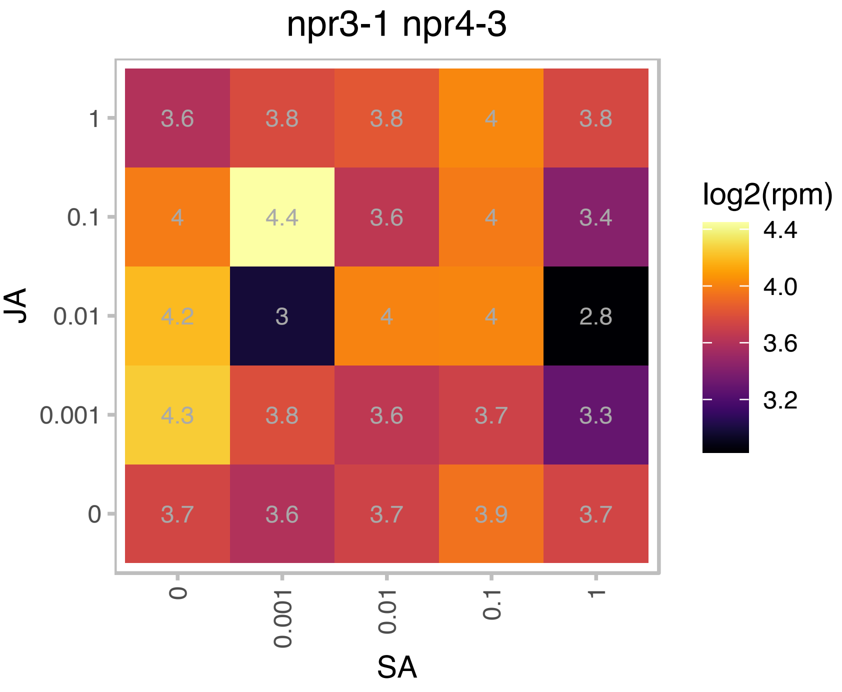

### AT2G01930.1 : ATBPC1, BBR, BPC1

basic pentacysteine1
BASIC PENTACYSTEINE1 (BPC1) is a regulator of the homeotic Arabidopsis thaliana gene SEEDSTICK (STK), wh
controls ovule identity. BPC1 induces conformational changes by cooperative binding to purine-rich elements present in the STK regulatory sequence. STK is upregulated in bpc1 mutant.

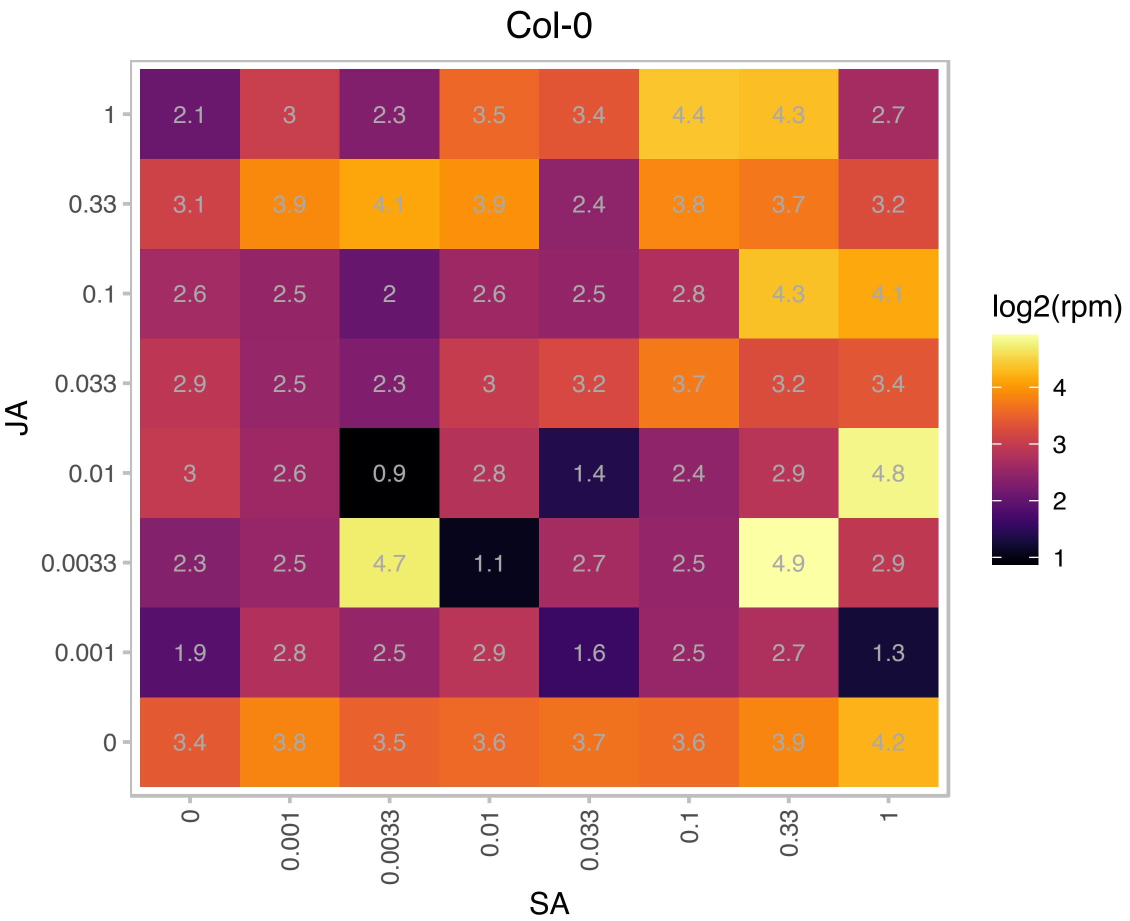


| | |
|---|---|
| Gene | AT2G01930 |
| is.expressed_genes | TRUE |
| is.responsive_genes | TRUE |
| is.apcluster | B-03 |

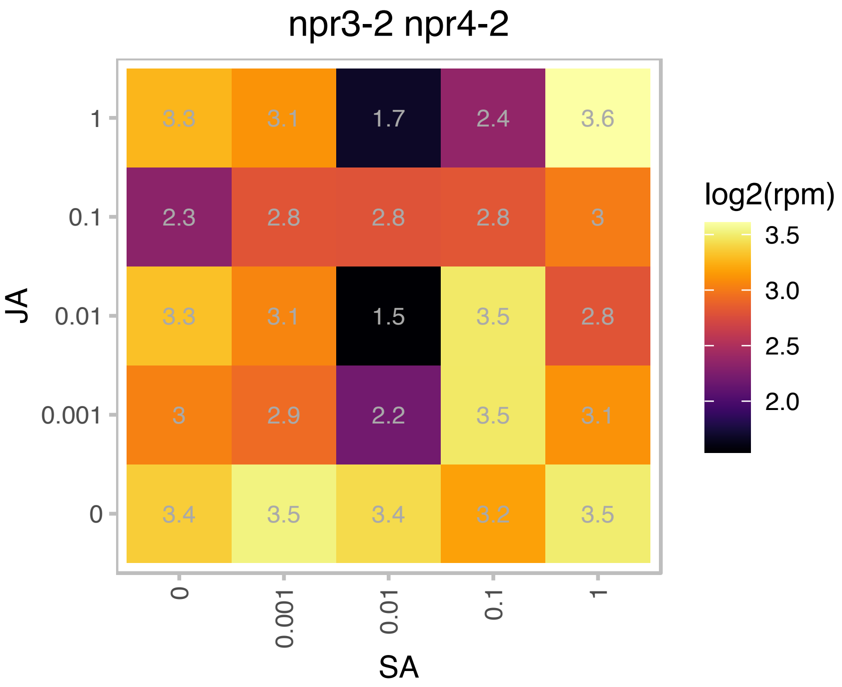


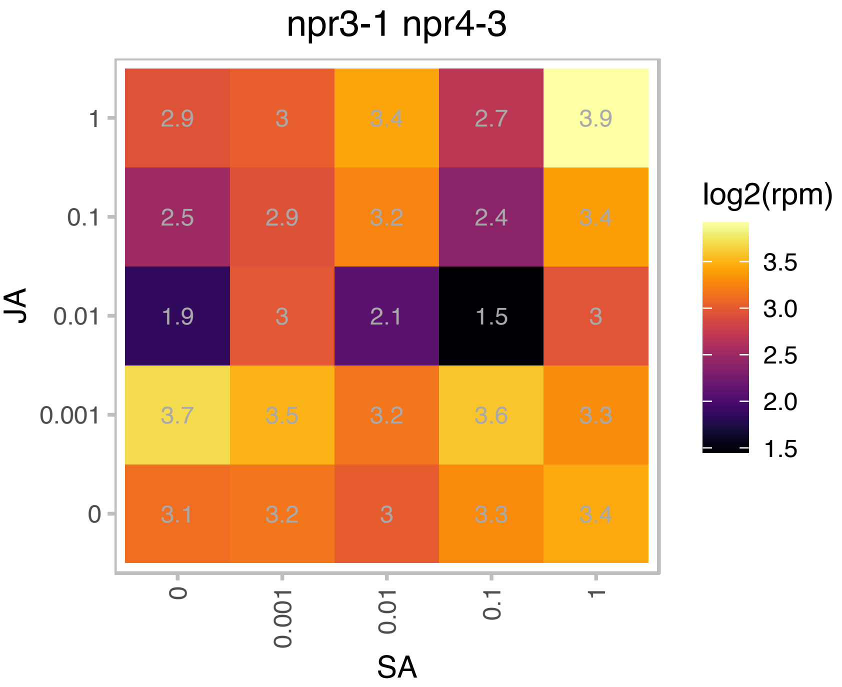

# AT2G04050.1 : NA

MATE efflux family protein

Col-0

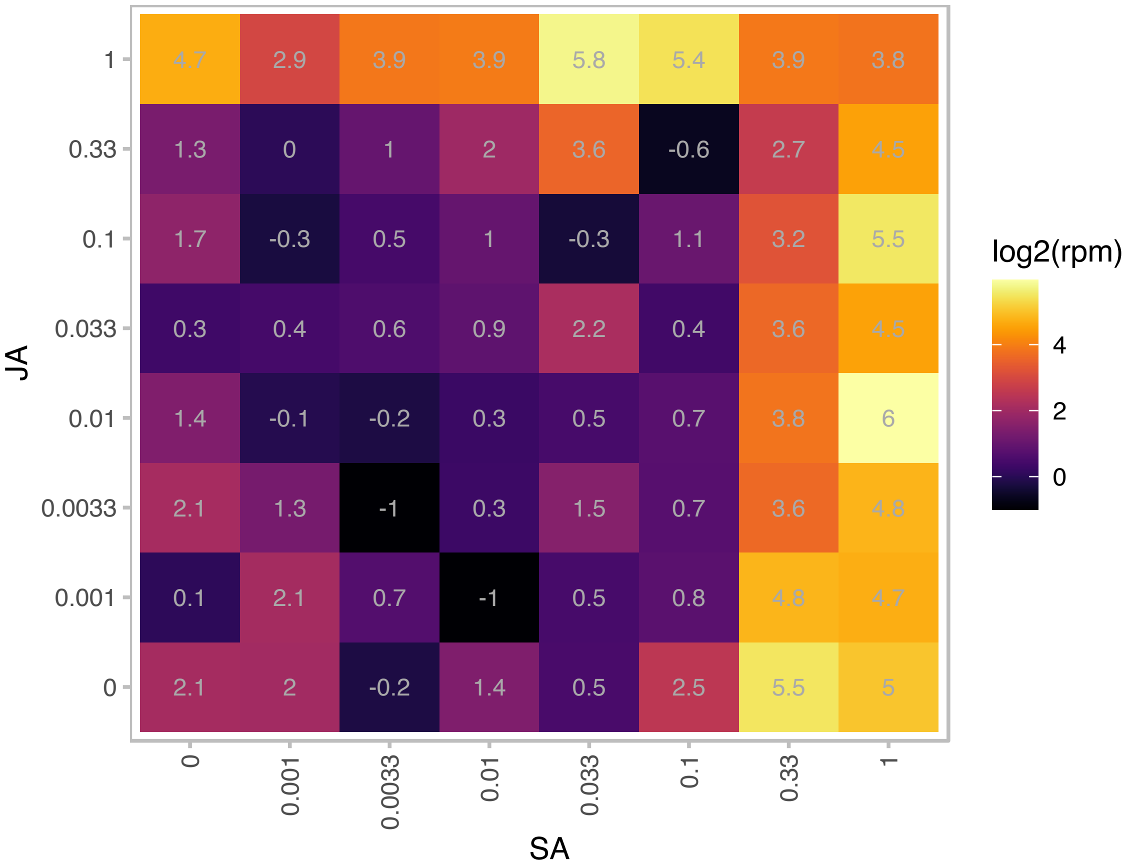


| | |
|---|---|
| Gene | AT2G04050 |
| is.expressed_genes | TRUE |
| is.responsive_genes | TRUE |
| is.apcluster | B-03 |

npr3-2 npr4-2

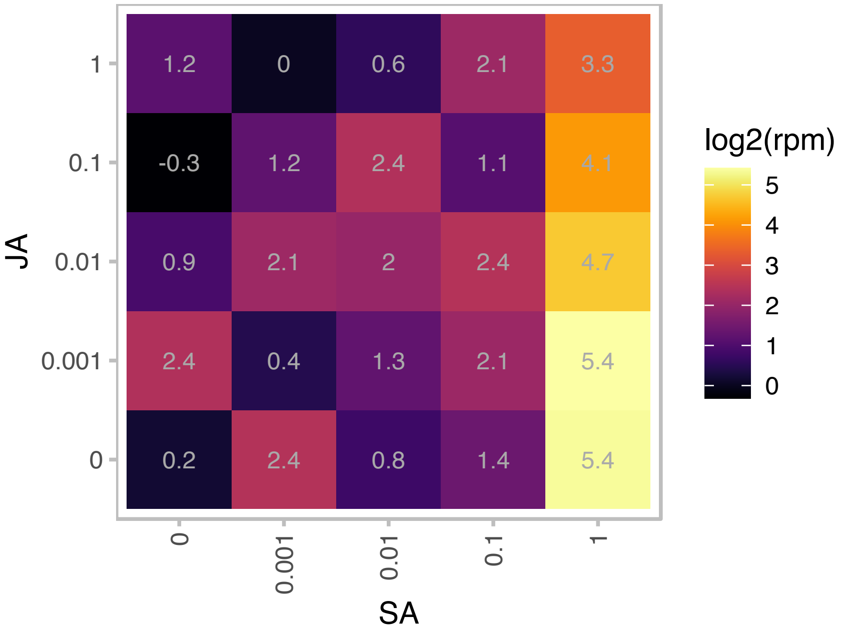


npr3-1 npr4-3

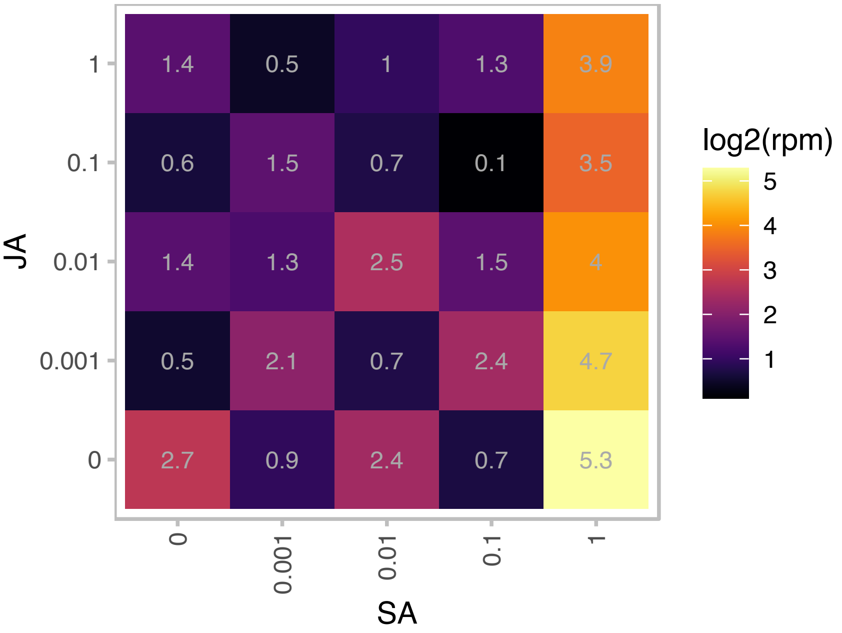

# AT2G14115.1 : NA

other RNA

Potential natural antisense gene, locus overlaps with AT2G14120

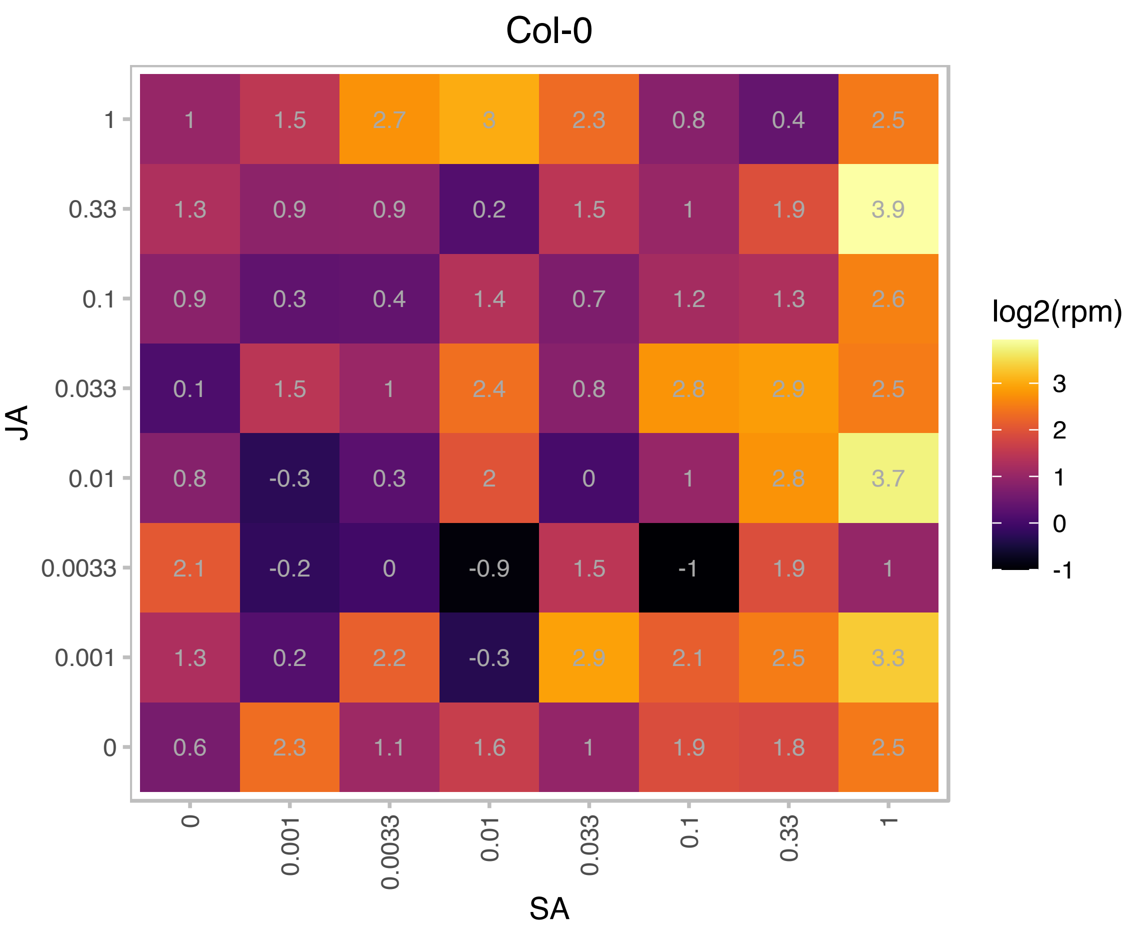


| | |
|---|---|
| Gene | AT2G14115 |
| is.expressed_genes | TRUE |
| is.responsive_genes | TRUE |
| is.apcluster | B-03 |

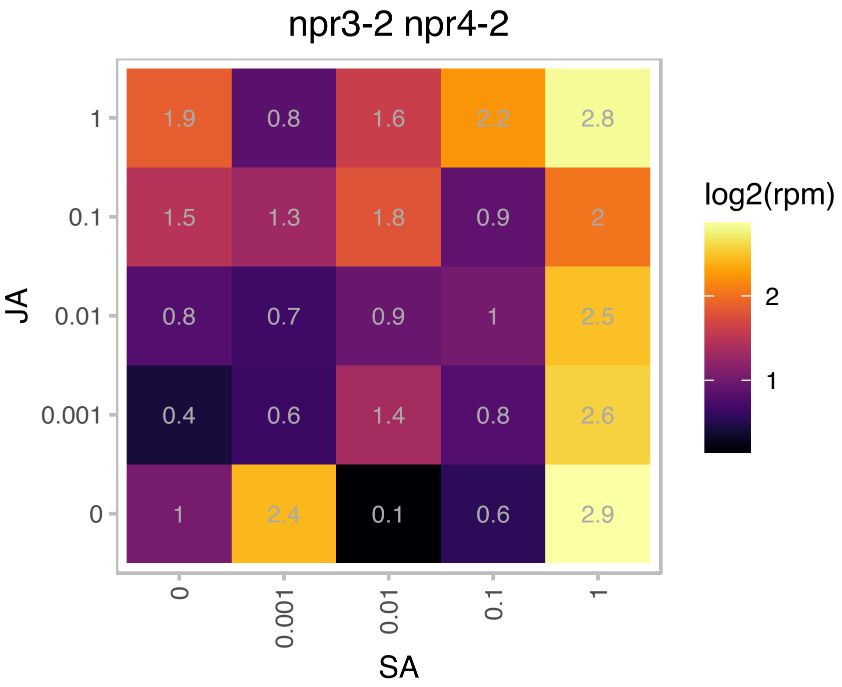


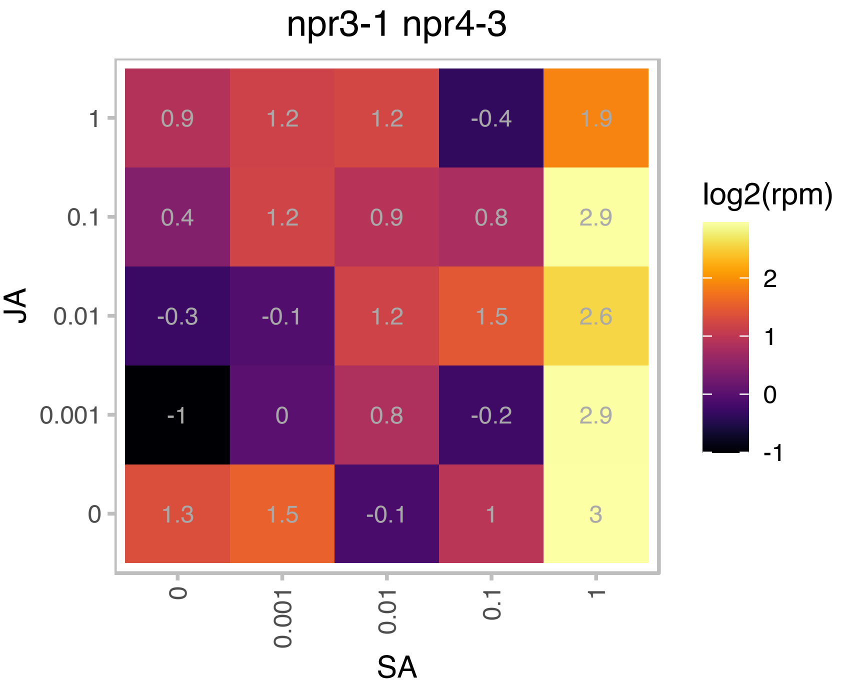

### AT2G19450.1 : ABX45, AS11, ATDGAT, AtDGAT1, DGAT1,

membrane bound O-acyl transferase (MBOAT) family protein
Encodes Acyl-CoA:diacylglycerol acyltransferase (DGAT) catalyzes the final step of the triacylglycerol synthesis pathway. An insertion mutation in the TAG1 gene results in altered lipid phenotype. Role in senescence and seed development. Its preferred substrate is linolenoyl-CoA (C18:3-CoA).

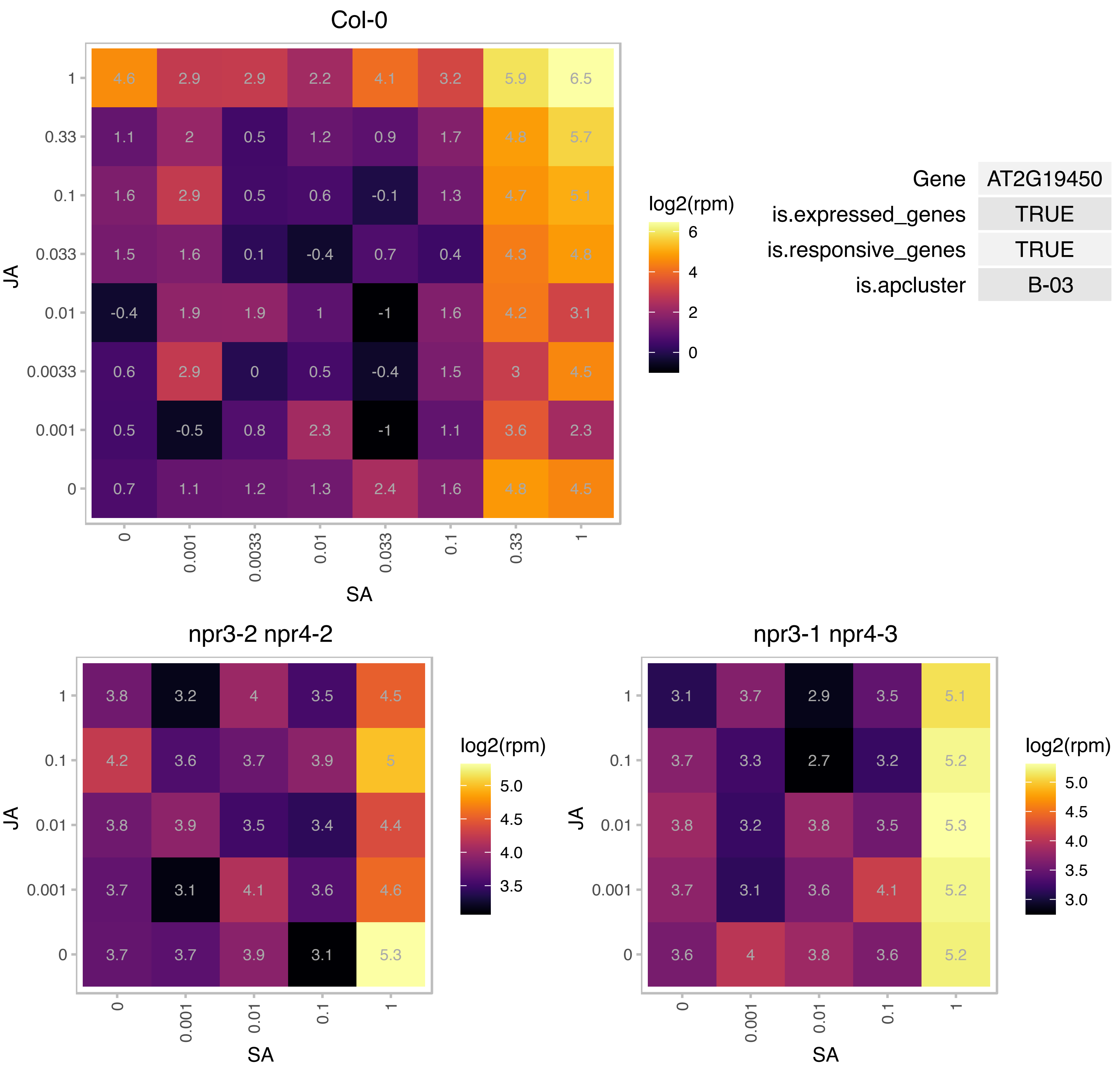

# AT2G19460.1 : NA

Protein of unknown function (DUF3511)

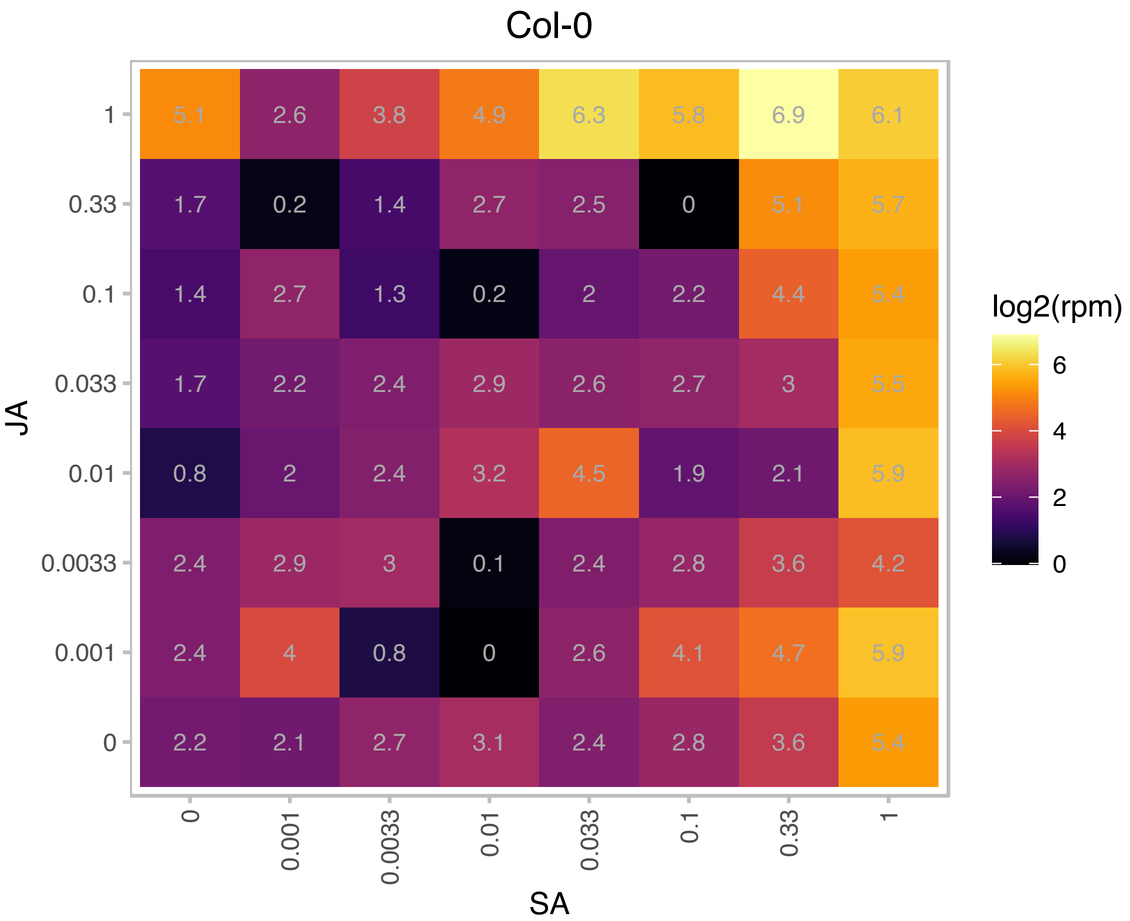


| | |
|---|---|
| Gene | AT2G19460 |
| is.expressed_genes | TRUE |
| is.responsive_genes | TRUE |
| is.apcluster | B-03 |

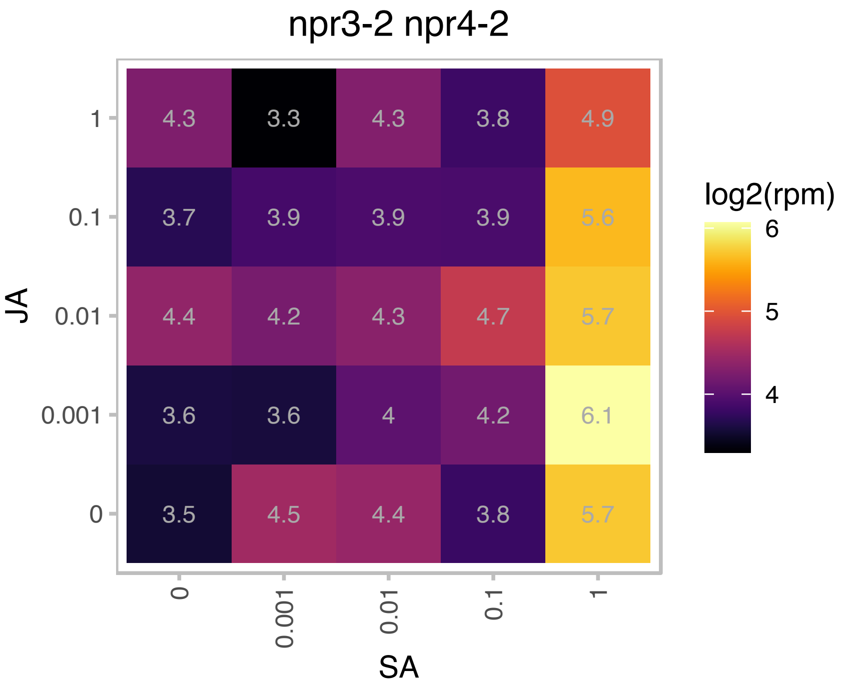


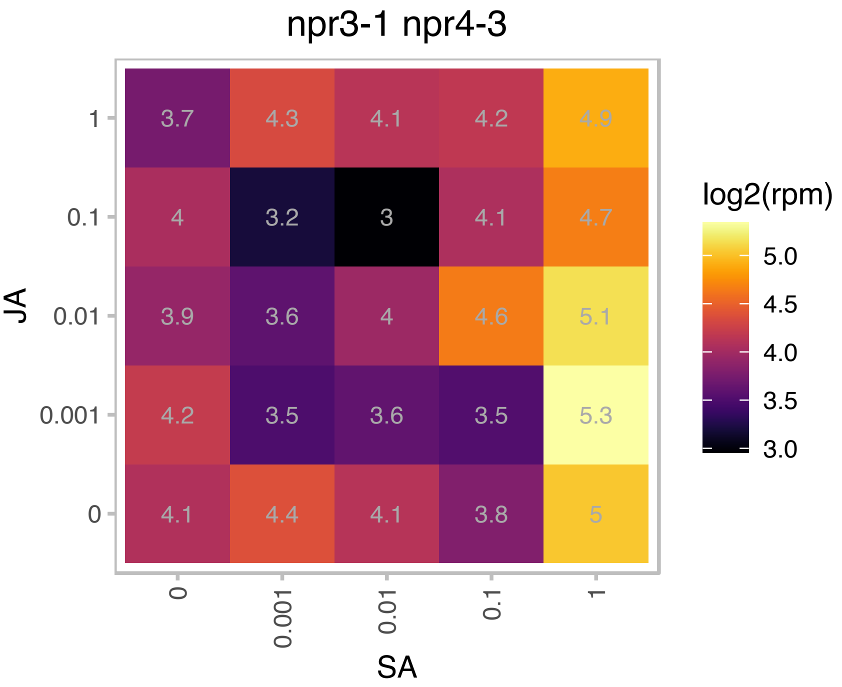

### AT2G20120.1 : COV1

Protein of unknown function (DUF502)

Encodes an integral membrane protein of unknown function, highly conserved between plants and bacteria; is likely to be involved in a mechanism that negatively regulates the differentiation of vascular tissue in the stem. Mutants display a dramatic increase in vascular tissue development in the stem in place of the interfascicular region that normally separates the vascular bundles.

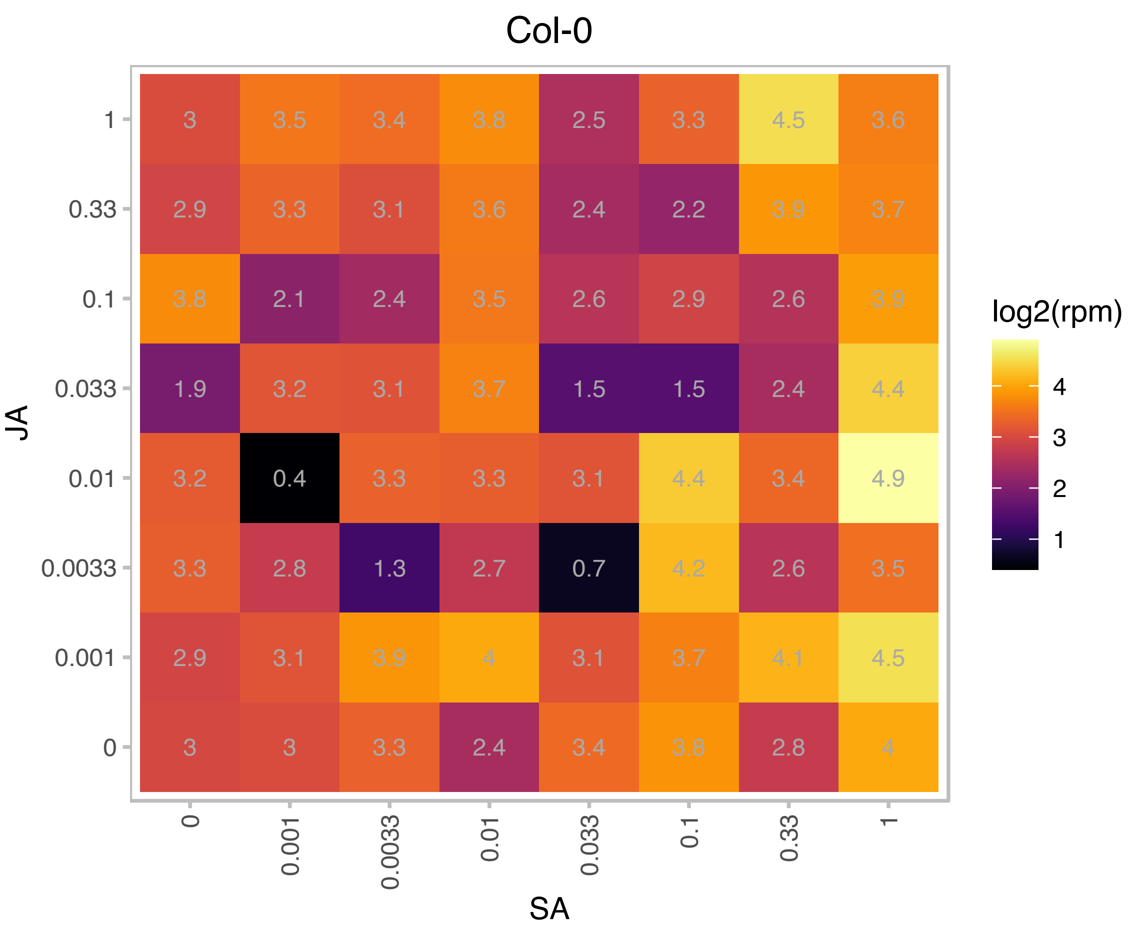


| | |
|---|---|
| Gene | AT2G20120 |
| is.expressed_genes | TRUE |
| is.responsive_genes | TRUE |
| is.apcluster | B-03 |

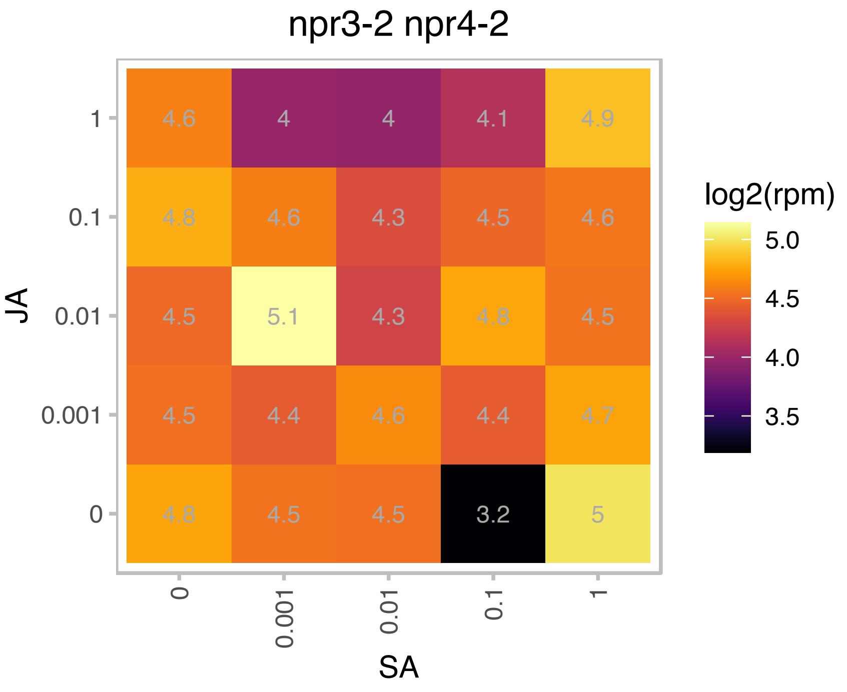


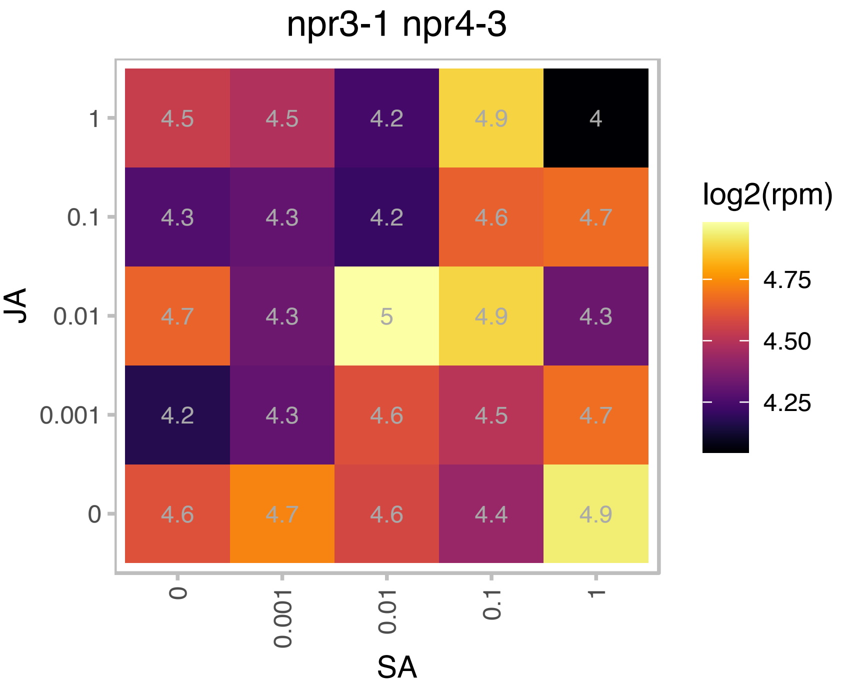

# AT2G25737.1 : NA

Sulfite exporter TauE/SafE family protein

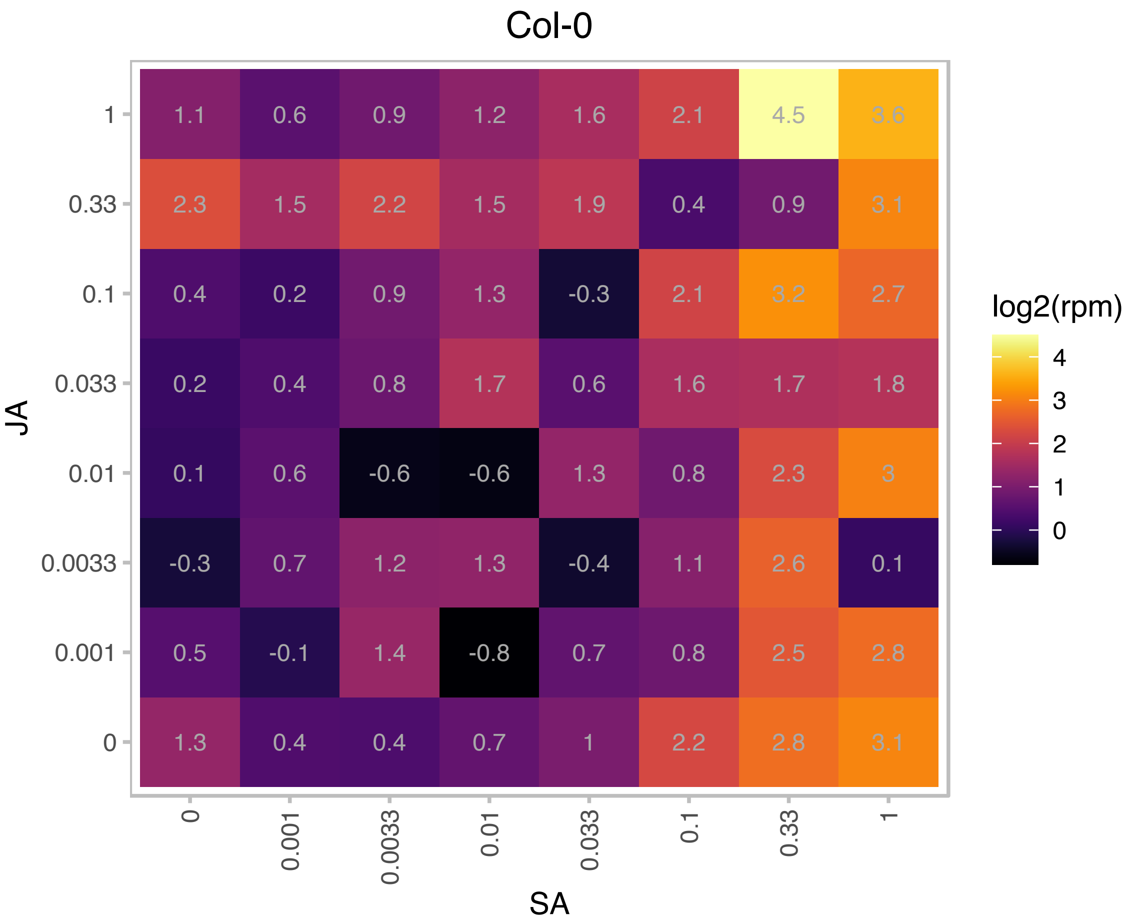


| | |
|---|---|
| Gene | AT2G25737 |
| is.expressed_genes | TRUE |
| is.responsive_genes | TRUE |
| is.apcluster | B-03 |

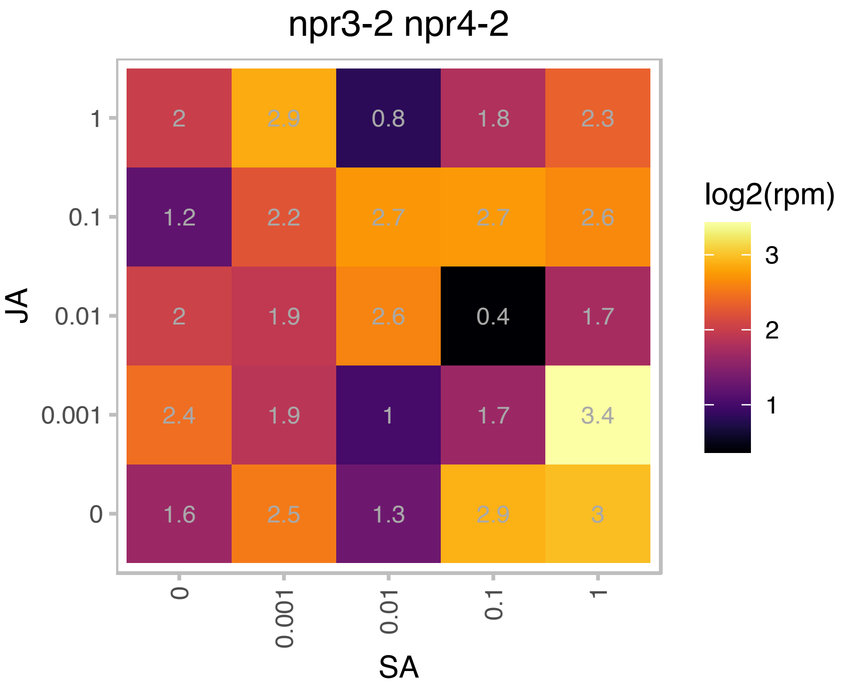


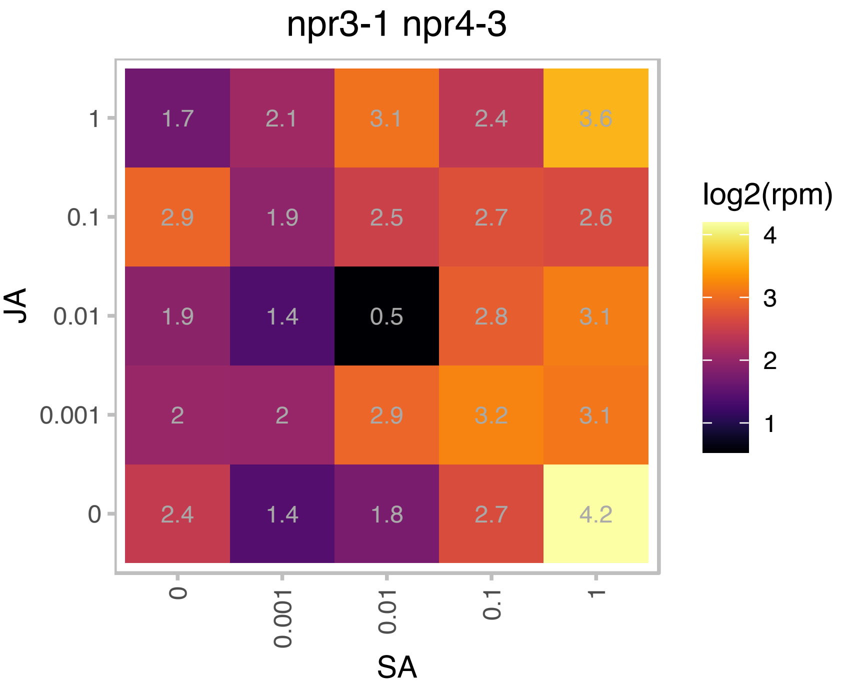

### AT2G29990.1 : NDA2

alternative NAD(P)H dehydrogenase 2

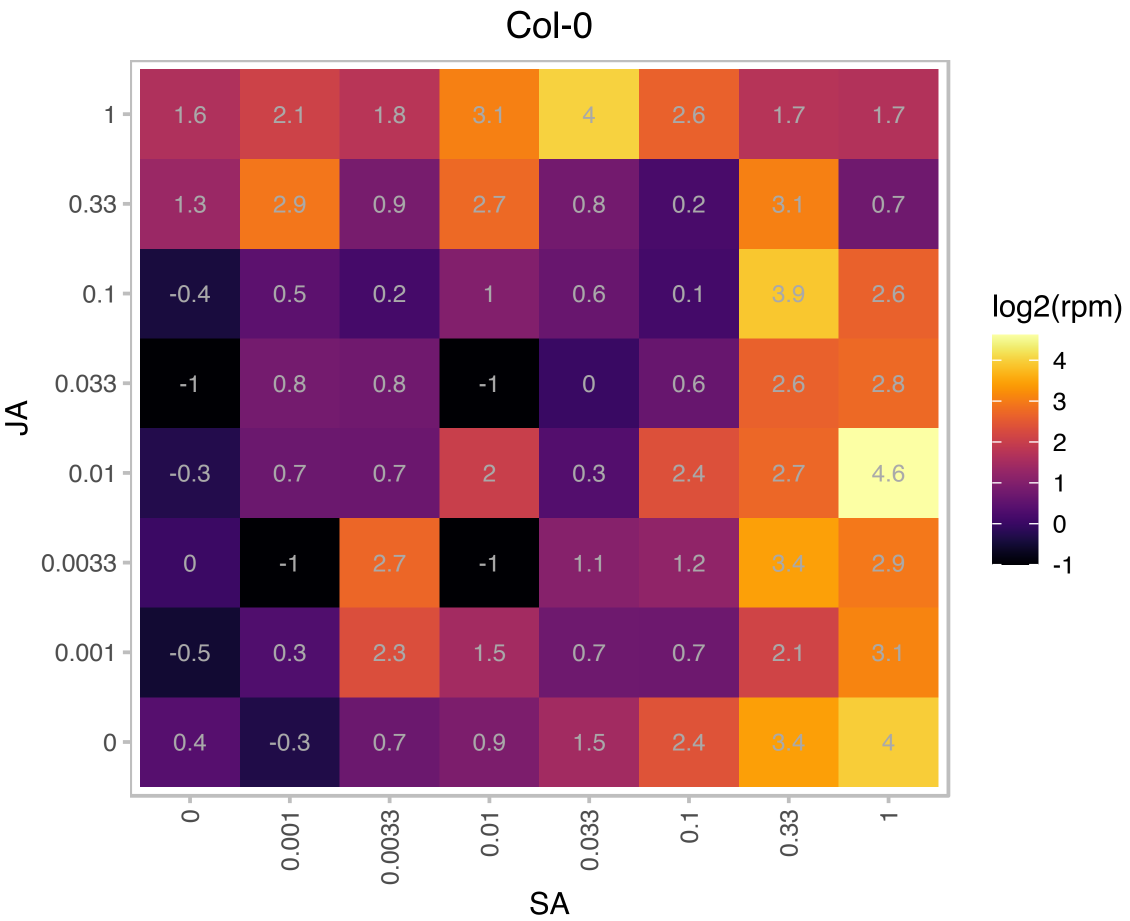


| | |
|---|---|
| Gene | AT2G29990 |
| is.expressed_genes | TRUE |
| is.responsive_genes | TRUE |
| is.apcluster | B-03 |

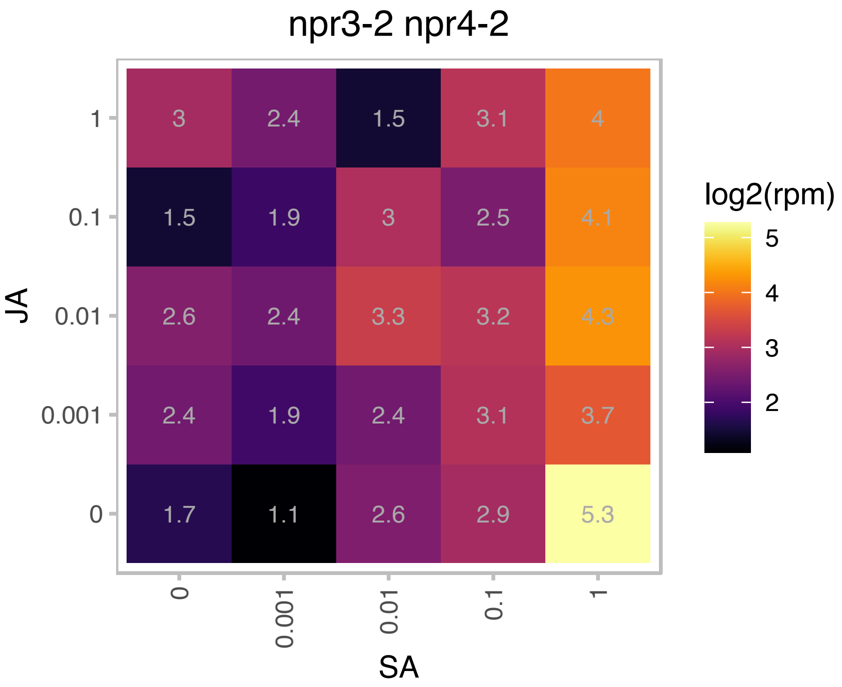


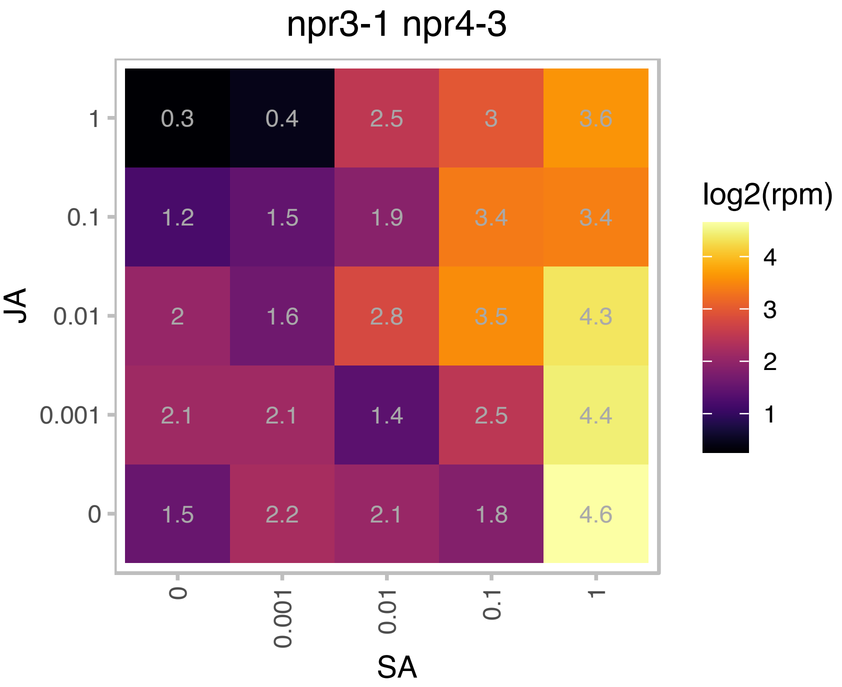

### AT2G31510.1 : ARI7, ATARI7

IBR domain-containing protein

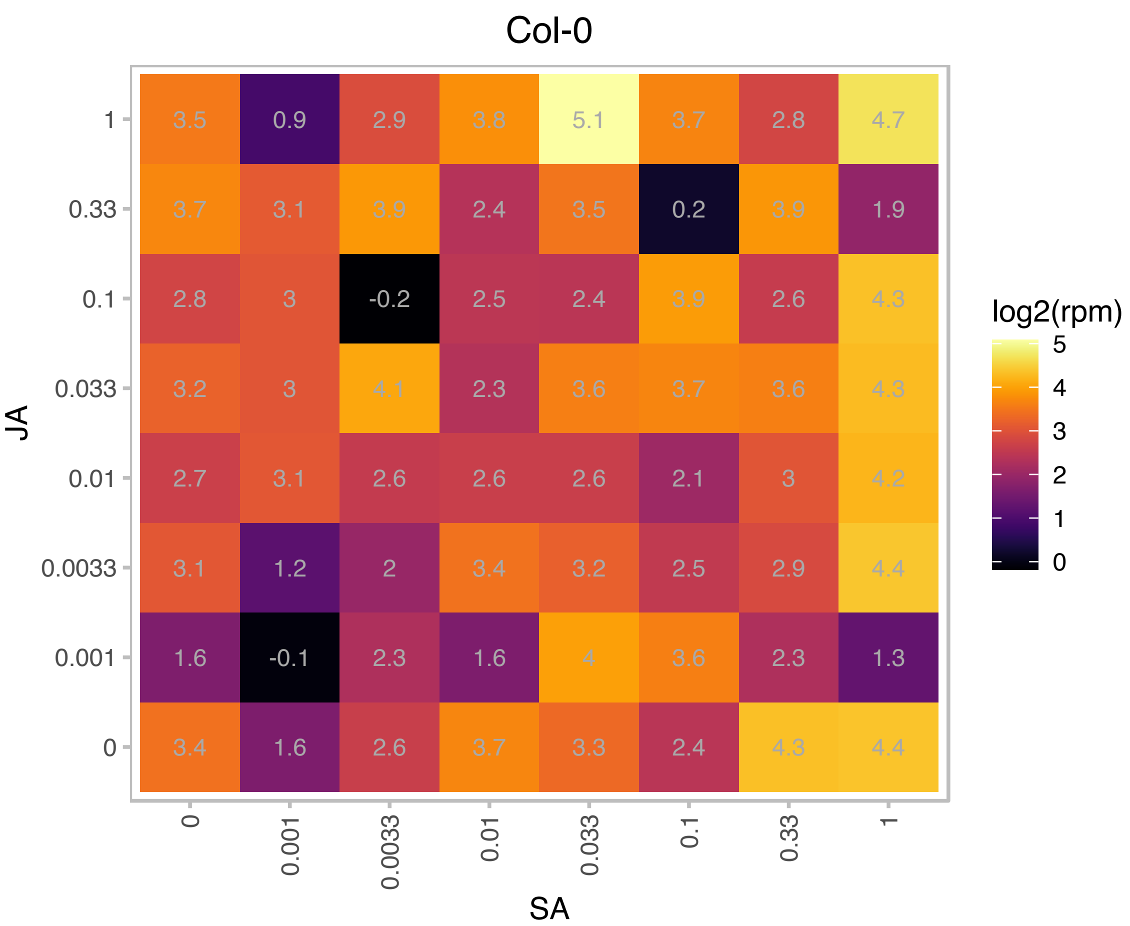


| | |
|---|---|
| Gene | AT2G31510 |
| is.expressed_genes | TRUE |
| is.responsive_genes | TRUE |
| is.apcluster | B-03 |

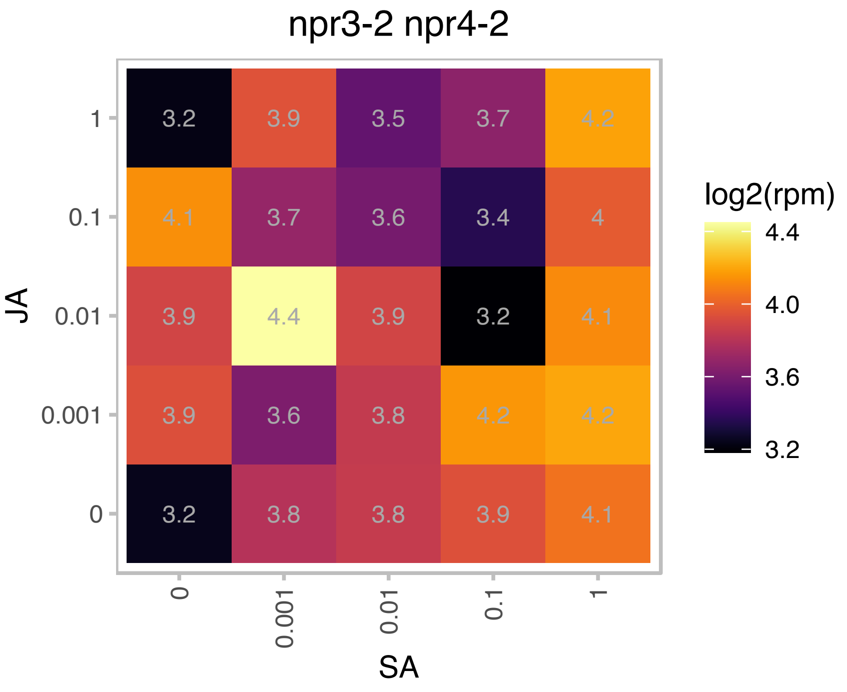


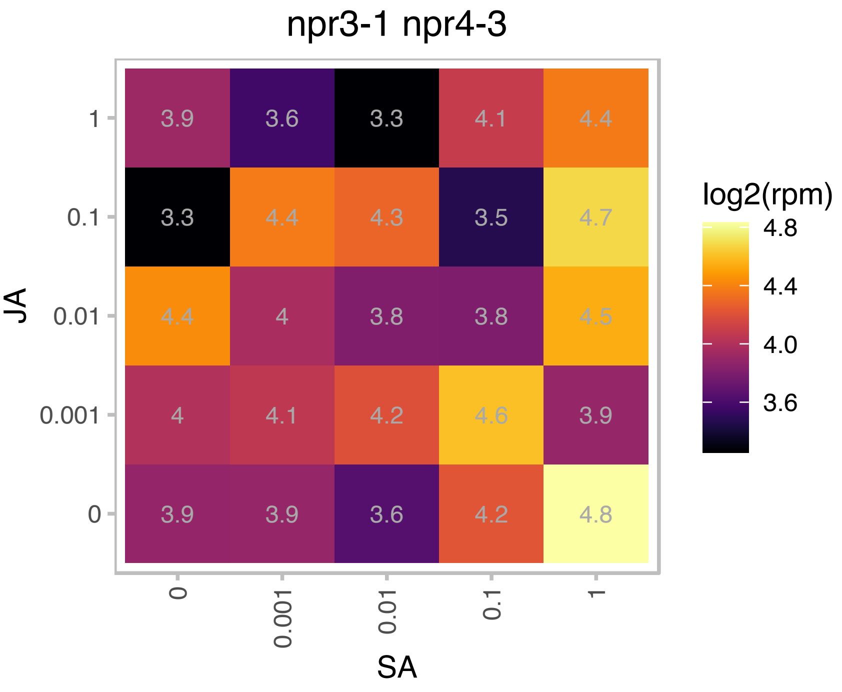

# AT2G37060.1 : NF-YB8

nuclear factor Y, subunit B8

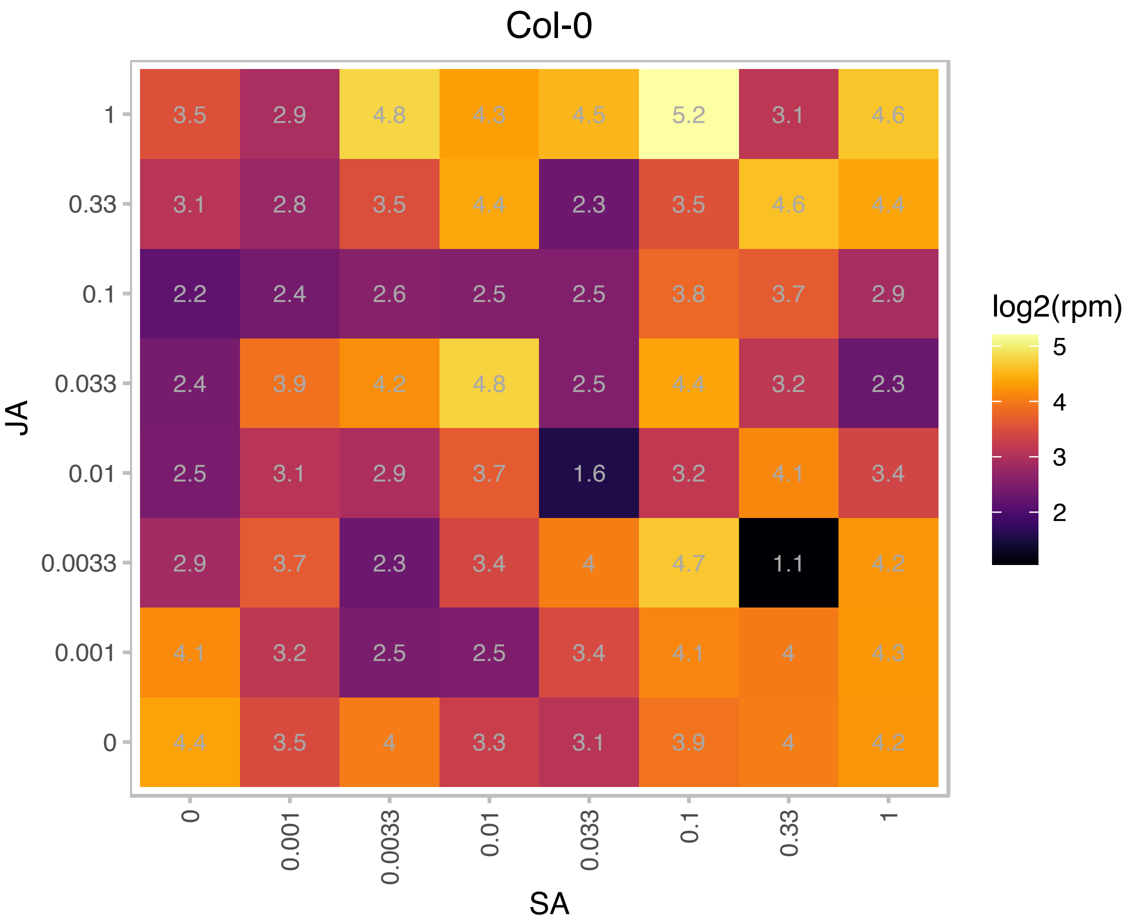


| | |
|---|---|
| Gene | AT2G37060 |
| is.expressed_genes | TRUE |
| is.responsive_genes | TRUE |
| is.apcluster | B-03 |

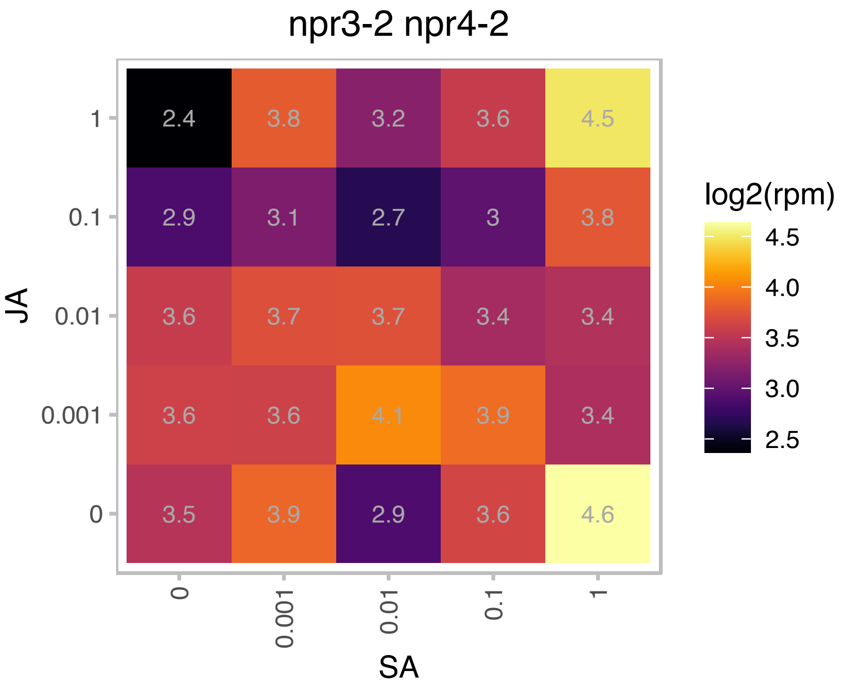


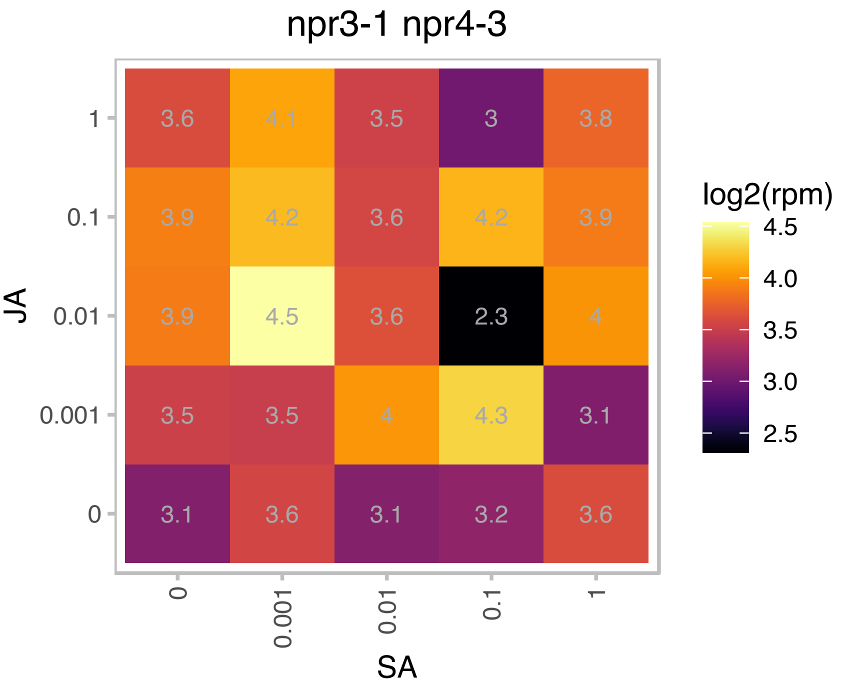

# AT2G37240.1 : NA

Thioredoxin superfamily protein

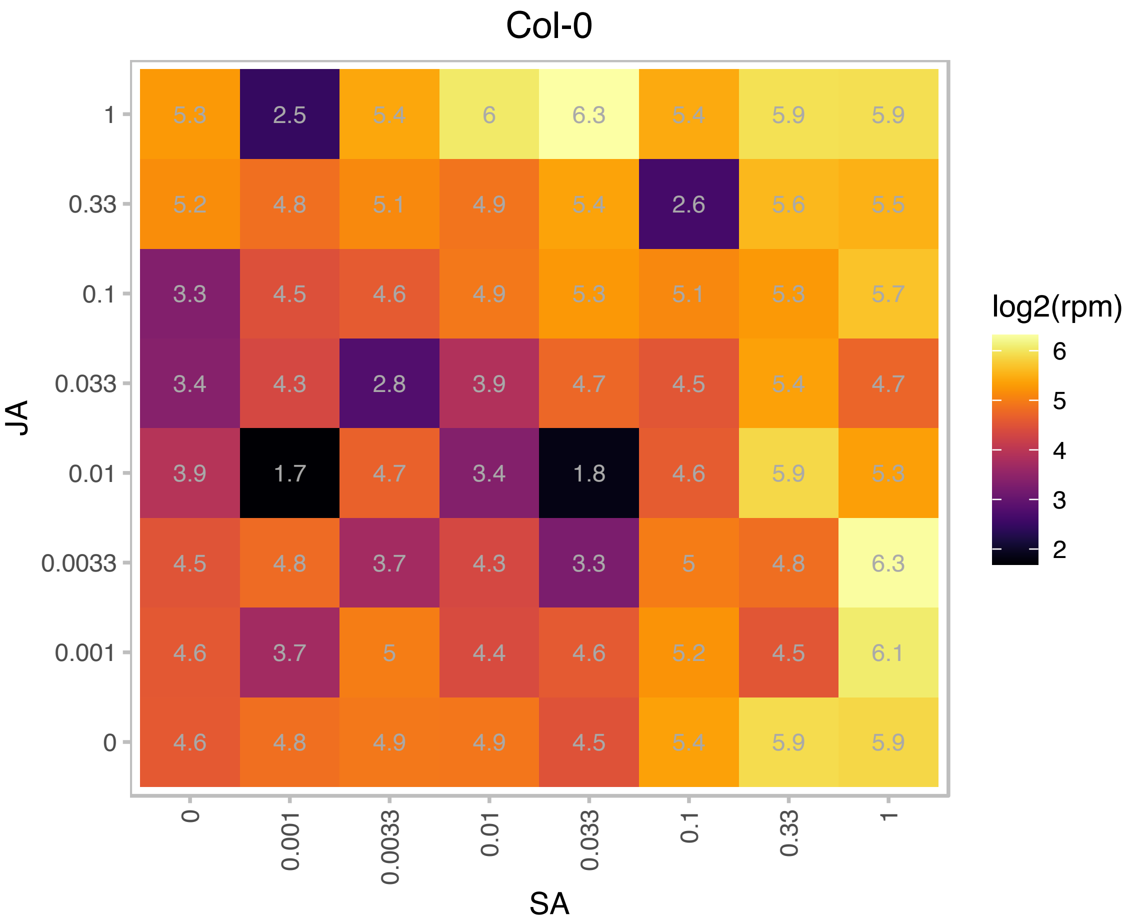

| | |
|---|---|
| Gene | AT2G37240 |
| is.expressed_genes | TRUE |
| is.responsive_genes | TRUE |
| is.apcluster | B-03 |

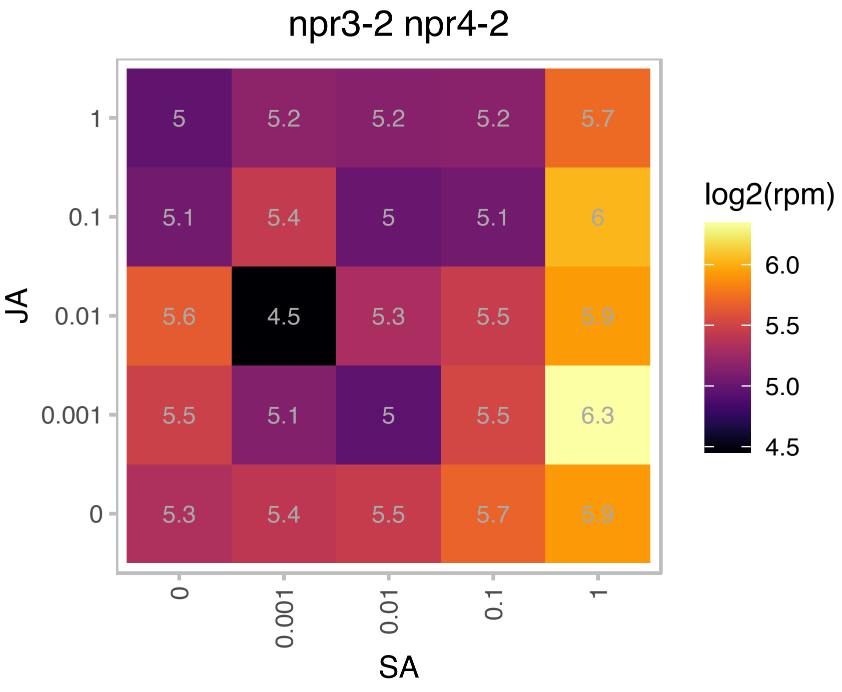

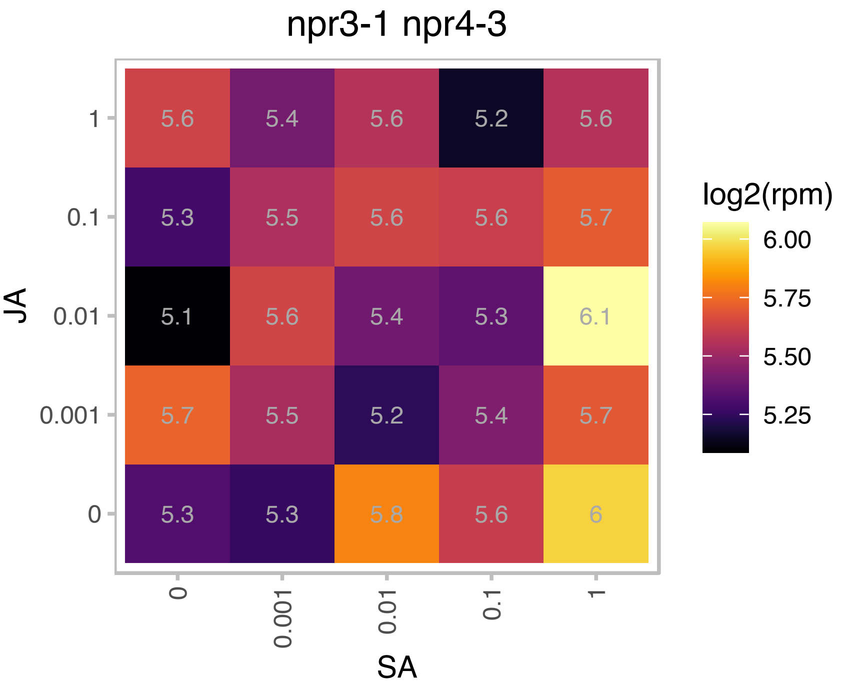

# AT2G38823.1 : NA

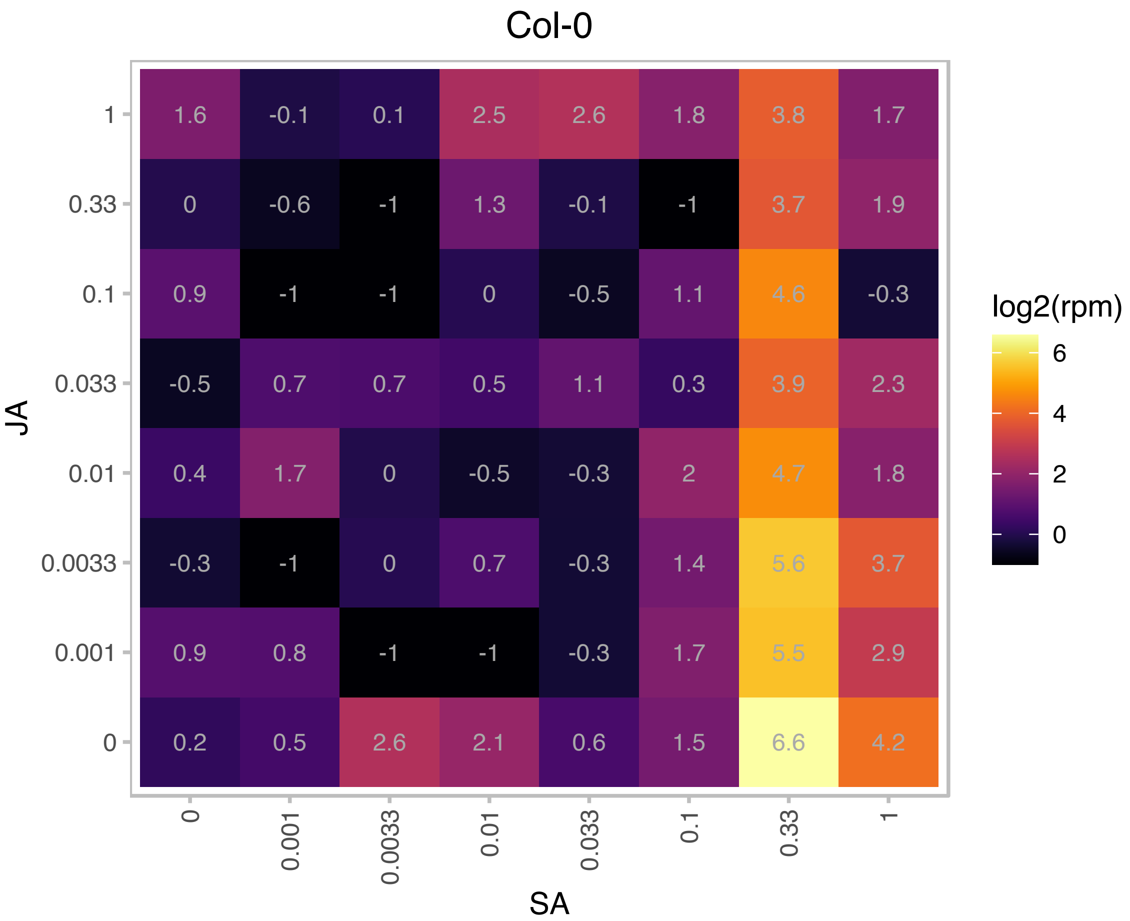


| | |
|---|---|
| Gene | AT2G38823 |
| is.expressed_genes | TRUE |
| is.responsive_genes | TRUE |
| is.apcluster | B-03 |

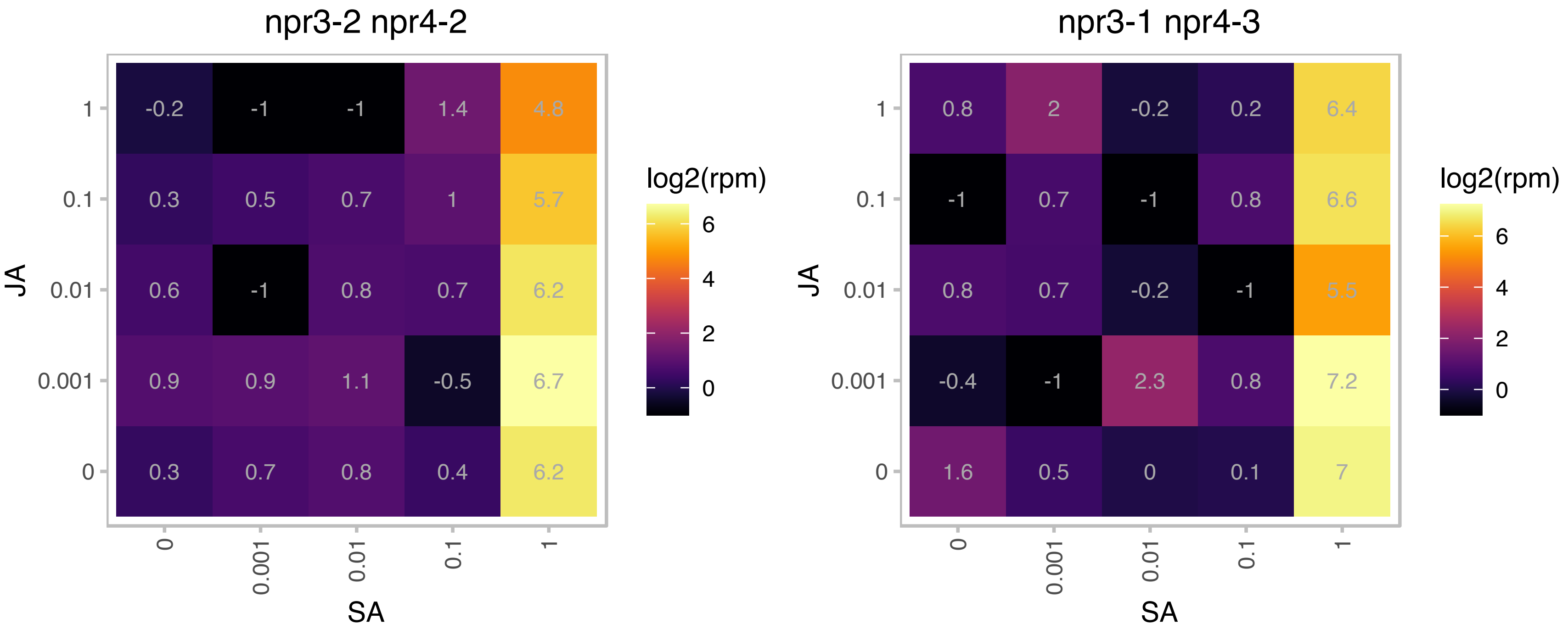

### AT2G40340.1 : AtERF48, DREB2C

Integrase-type DNA-binding superfamily protein
Encodes a member of the DREB subfamily A-2 of ERF/AP2 transcription factor family. The protein contains one AP2 domain. There are eight members in this subfamily including DREB2A AND DREB2B that are involved in response to drought.

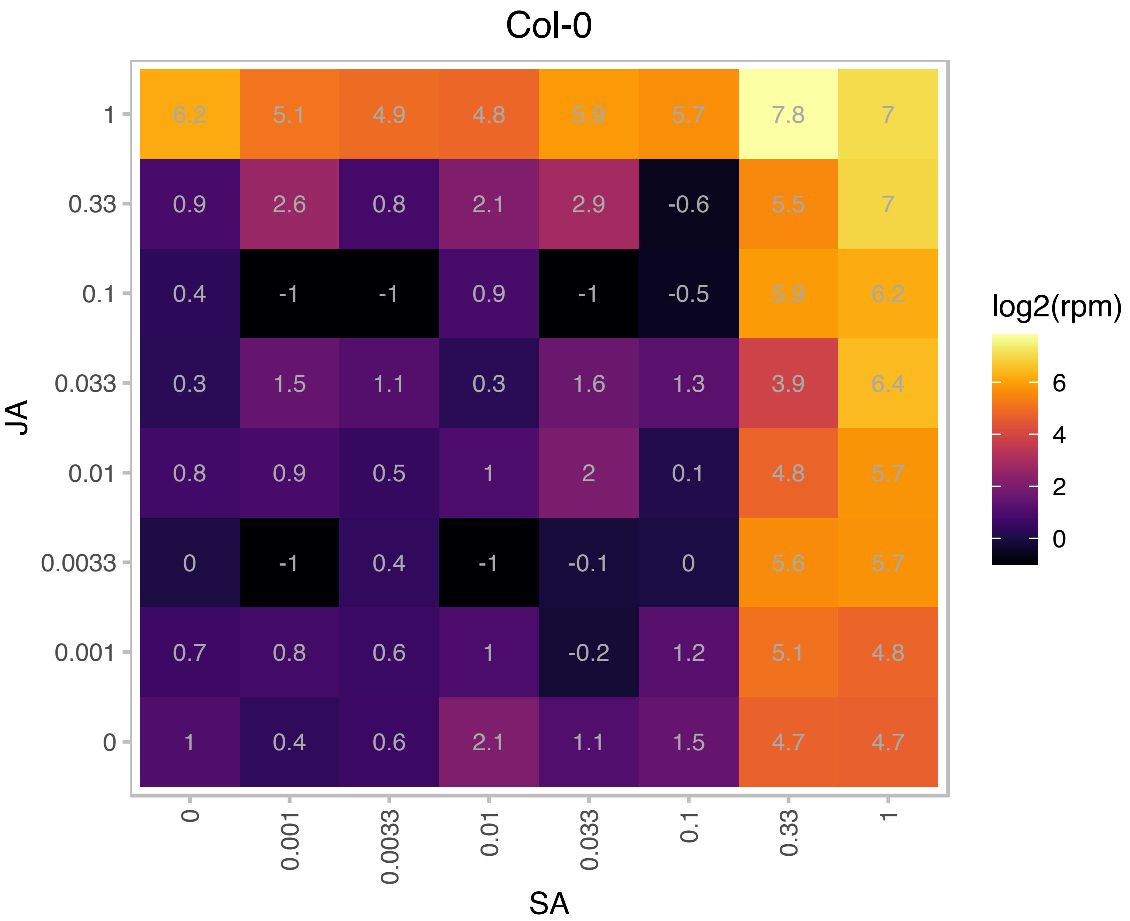


| | |
|---|---|
| Gene | AT2G40340 |
| is.expressed_genes | TRUE |
| is.responsive_genes | TRUE |
| is.apcluster | B-03 |

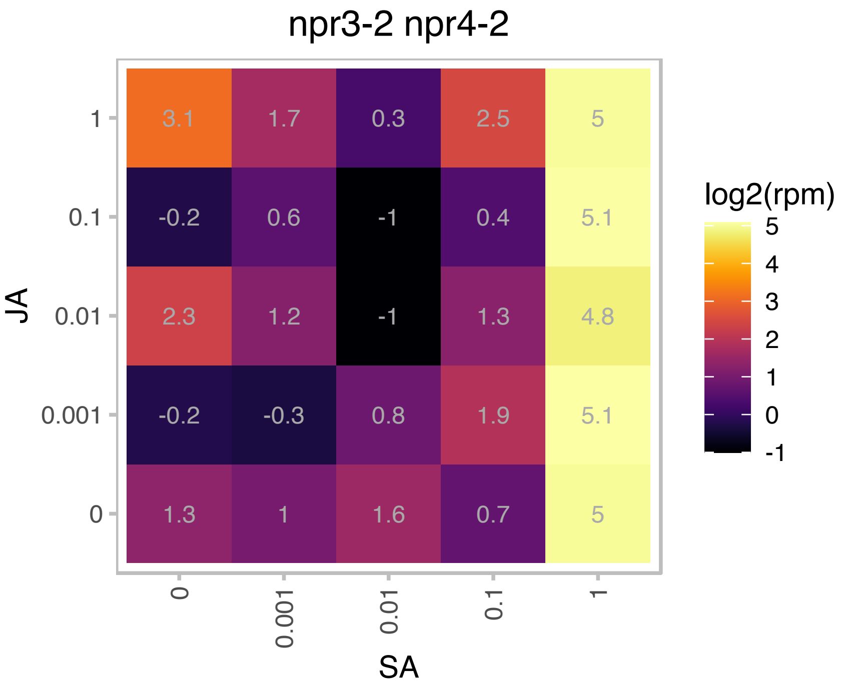


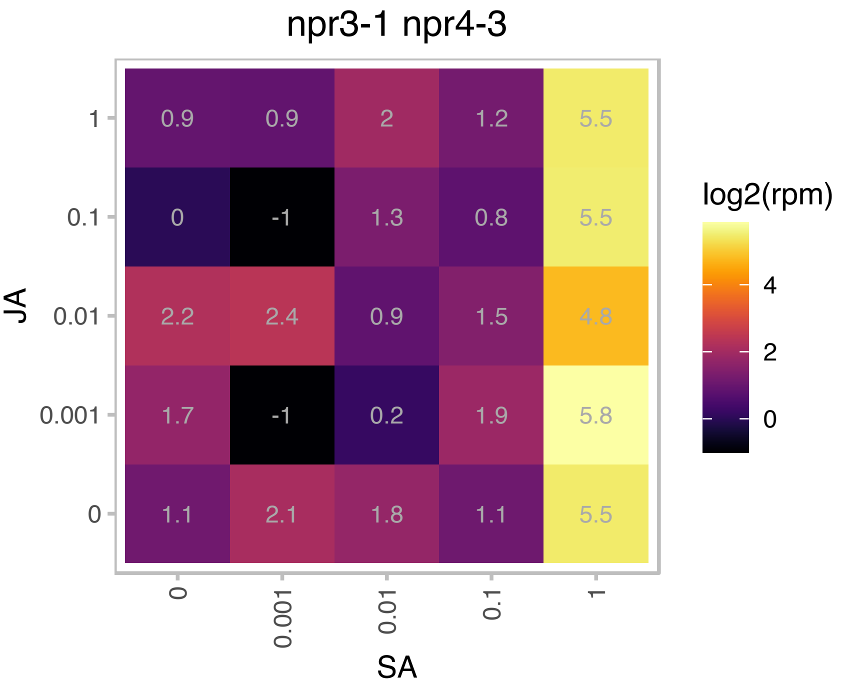

### AT2G40880.1 : ATCYSA, CYSA, FL3-27

cystatin A

Encodes a protein with cysteine proteinase inhibitor activity. Overexpression increases tolerance to abiotic stressors (i.e.salt,osmotic, cold stress). The mRNA is cell-to-cell mobile.

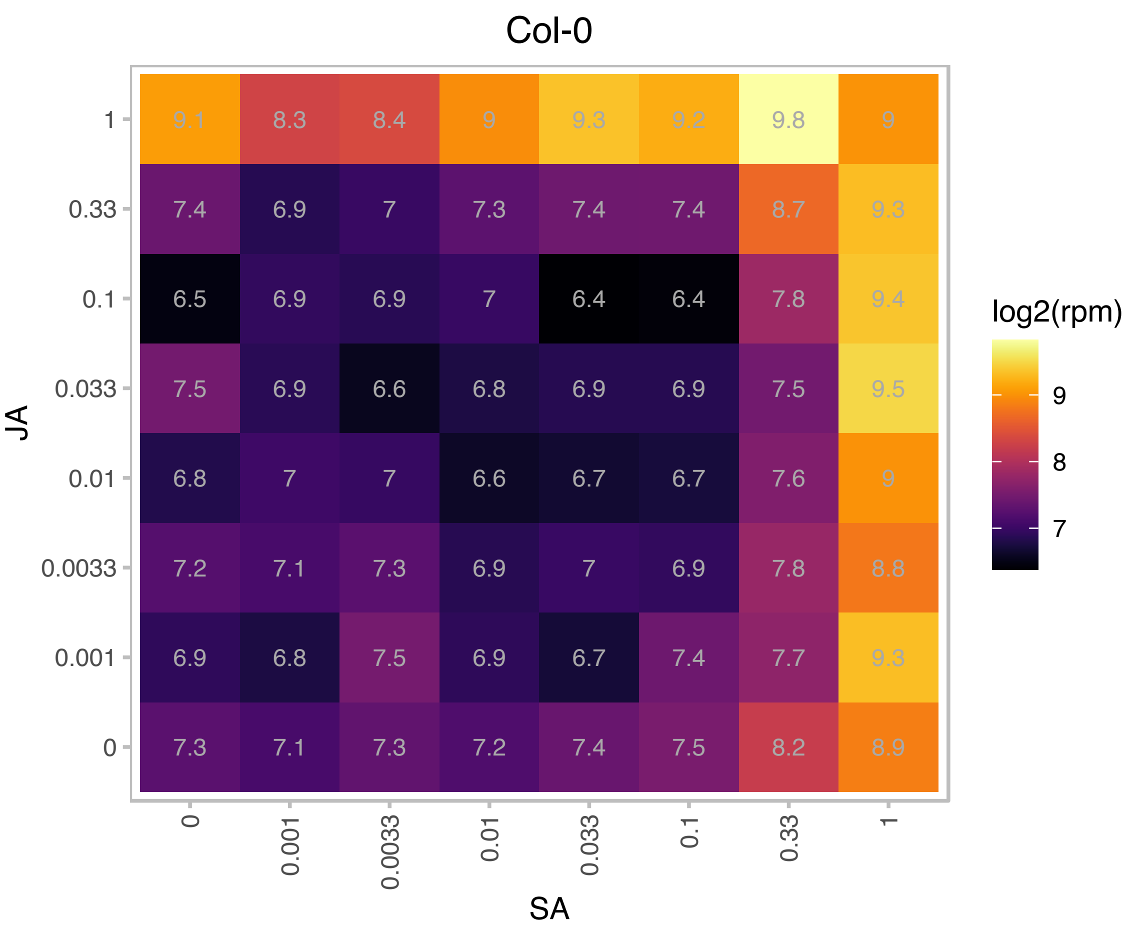


| | |
|---|---|
| Gene | AT2G40880 |
| is.expressed_genes | TRUE |
| is.responsive_genes | TRUE |
| is.apcluster | B-03 |

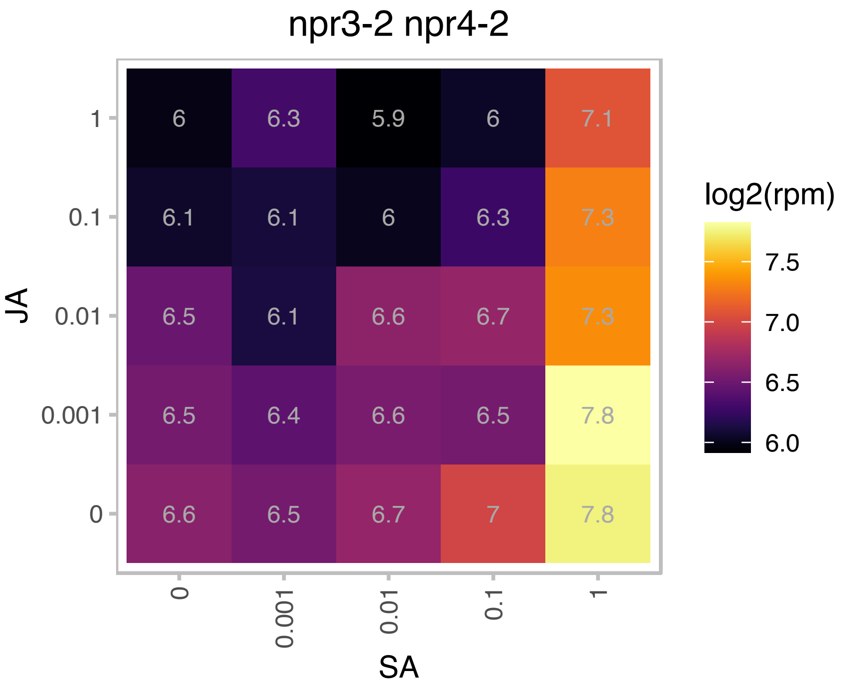


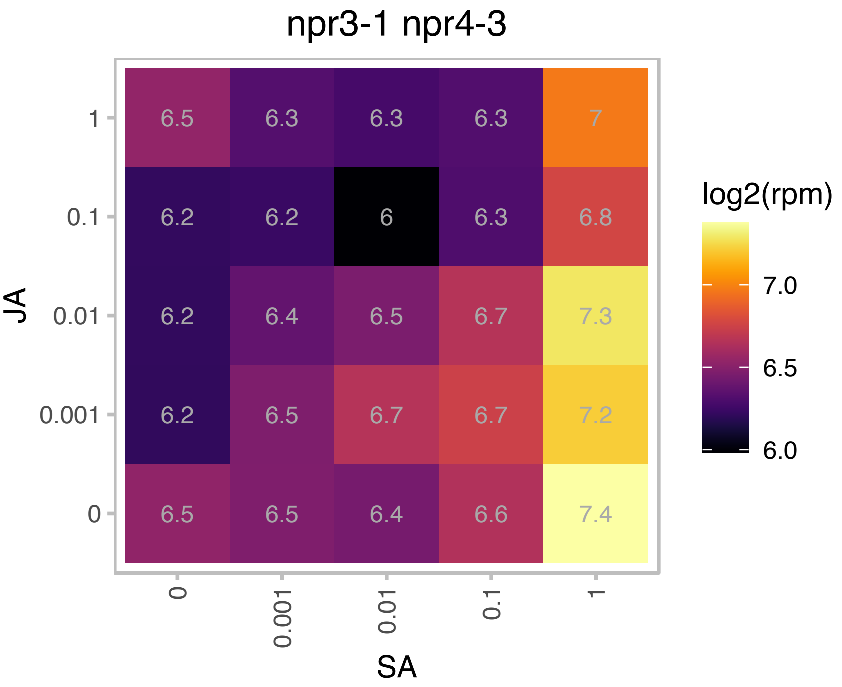

### AT2G41540.1 : GPDHC1

6-phosphogluconate dehydrogenase family protein
Encodes a protein with NAD-dependent glycerol-3-phosphate (G3P) dehydrogenase which was shown to compleme
an Escherichia coli strain: BB20-14, auxotrophic for glycerol/G3P due to a loss-of-function mutation in the
gpsA gene.

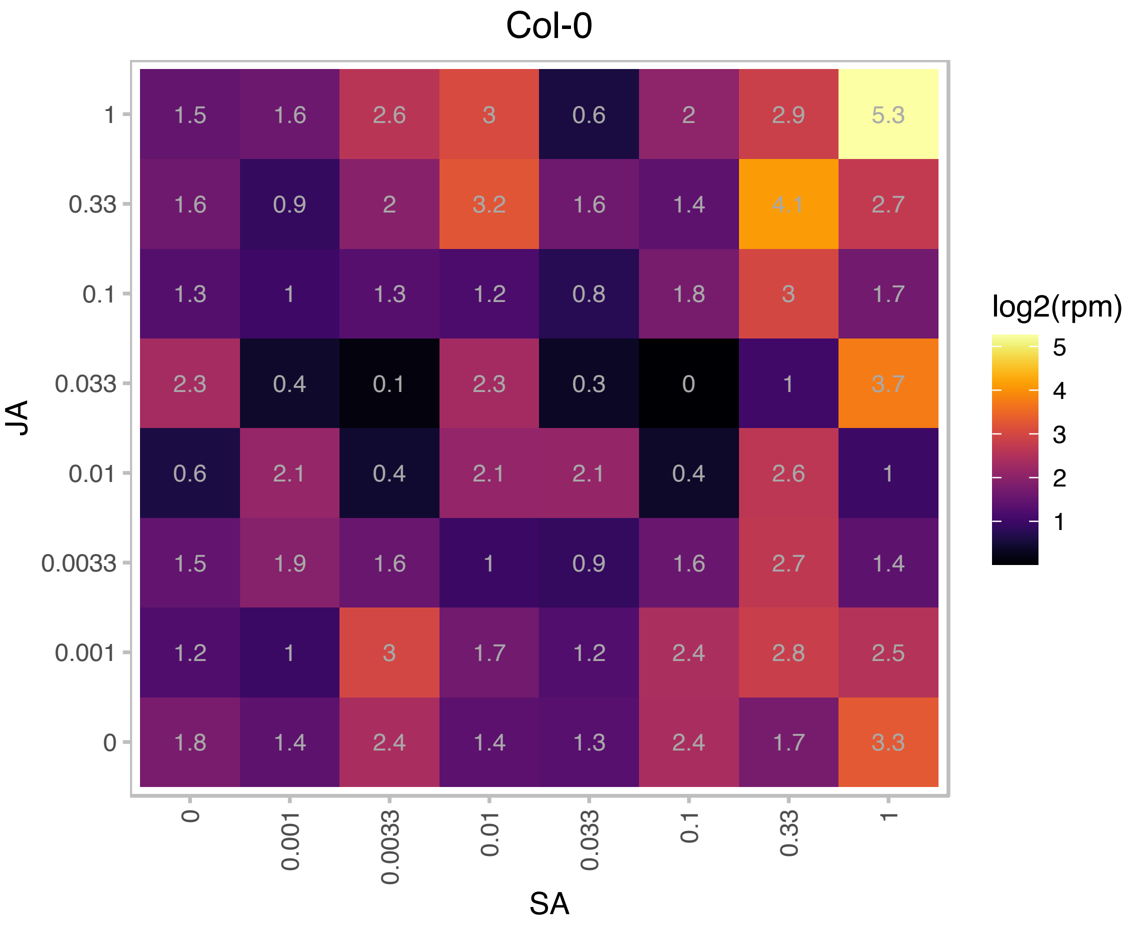


| | |
|---|---|
| Gene | AT2G41540 |
| is.expressed_genes | TRUE |
| is.responsive_genes | TRUE |
| is.apcluster | B-03 |

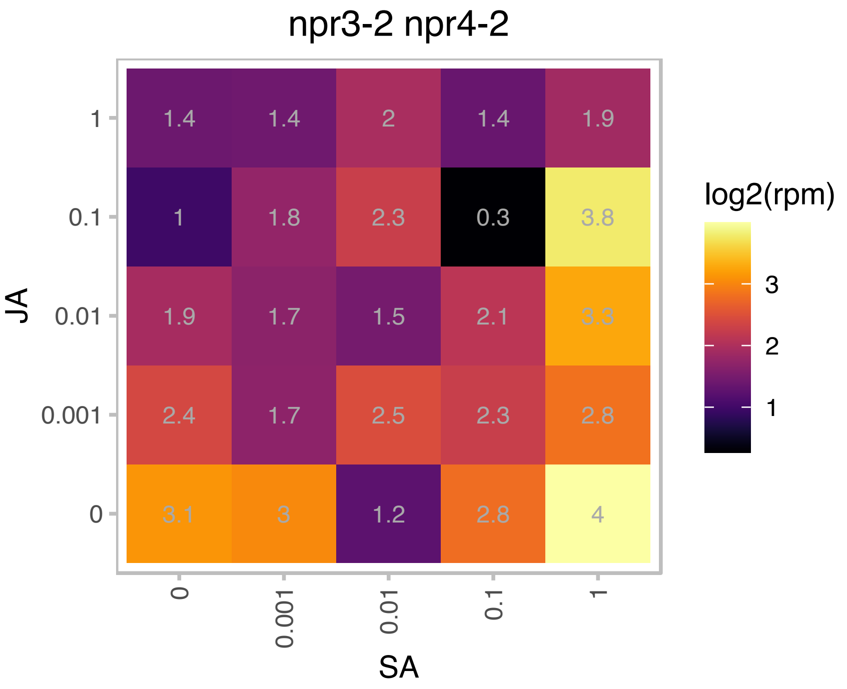


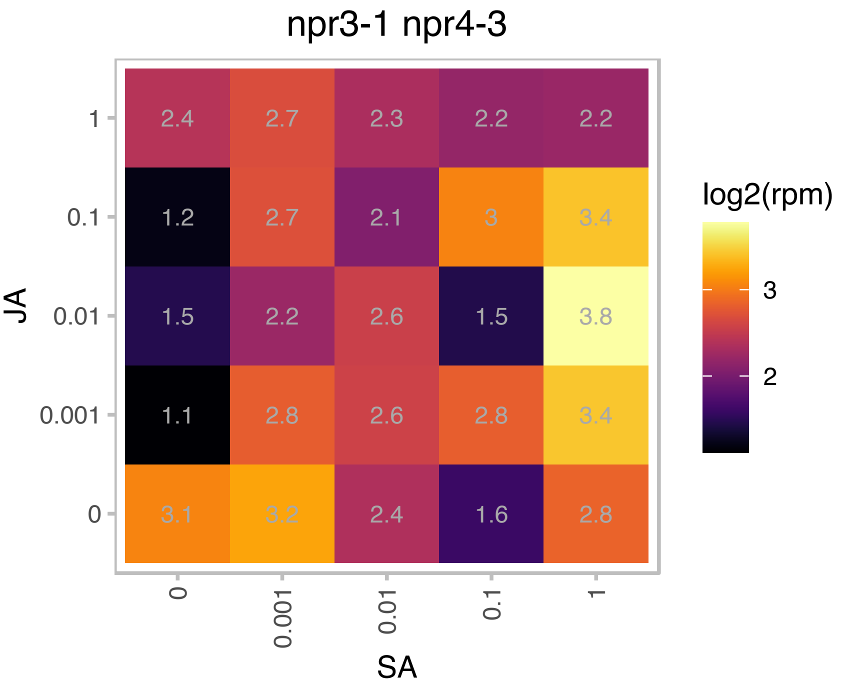

### AT2G47520.1 : AtERF71, ERF71, HRE2

Integrase-type DNA-binding superfamily protein
encodes a member of the ERF (ethylene response factor) subfamily B-2 of ERF/AP2 transcription factor family. The protein contains one AP2 domain. There are 5 members in this subfamily including RAP2.2 AND RAP2.12. It plays a role in hypoxia-induced root slanting.

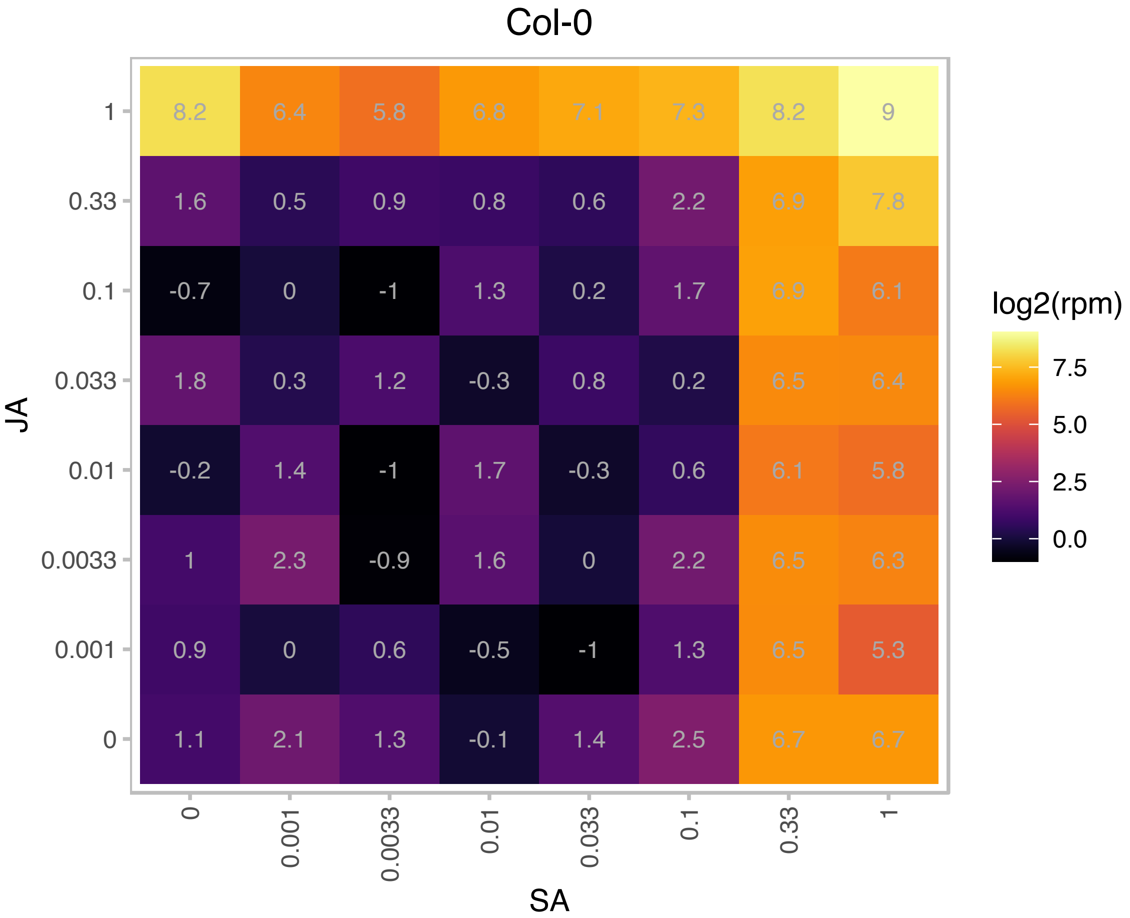


| | |
|---|---|
| Gene | AT2G47520 |
| is.expressed_genes | TRUE |
| is.responsive_genes | TRUE |
| is.apcluster | B-03 |

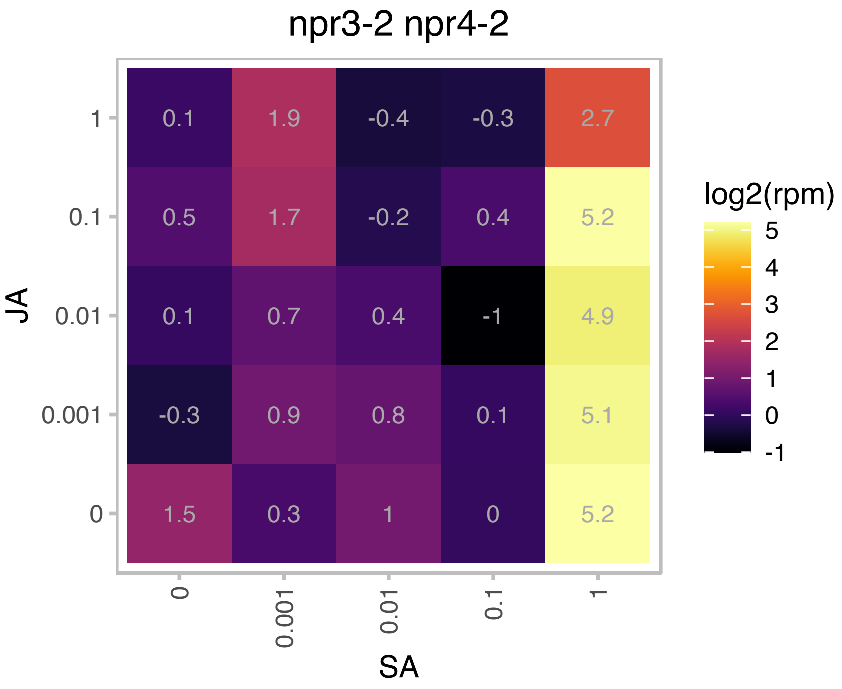


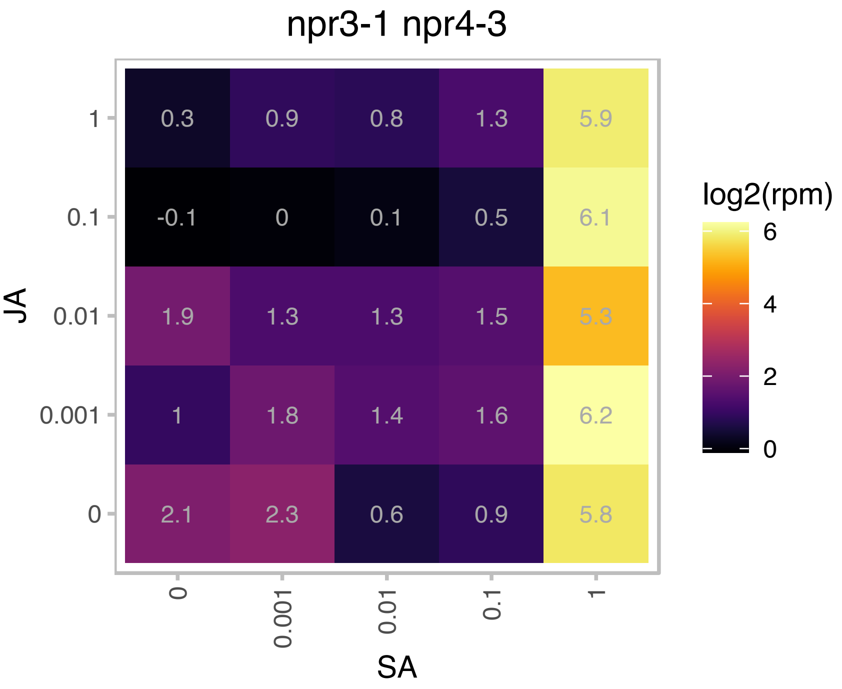

# AT2G47890.1 : NA

B-box type zinc finger protein with CCT domain

Col-0

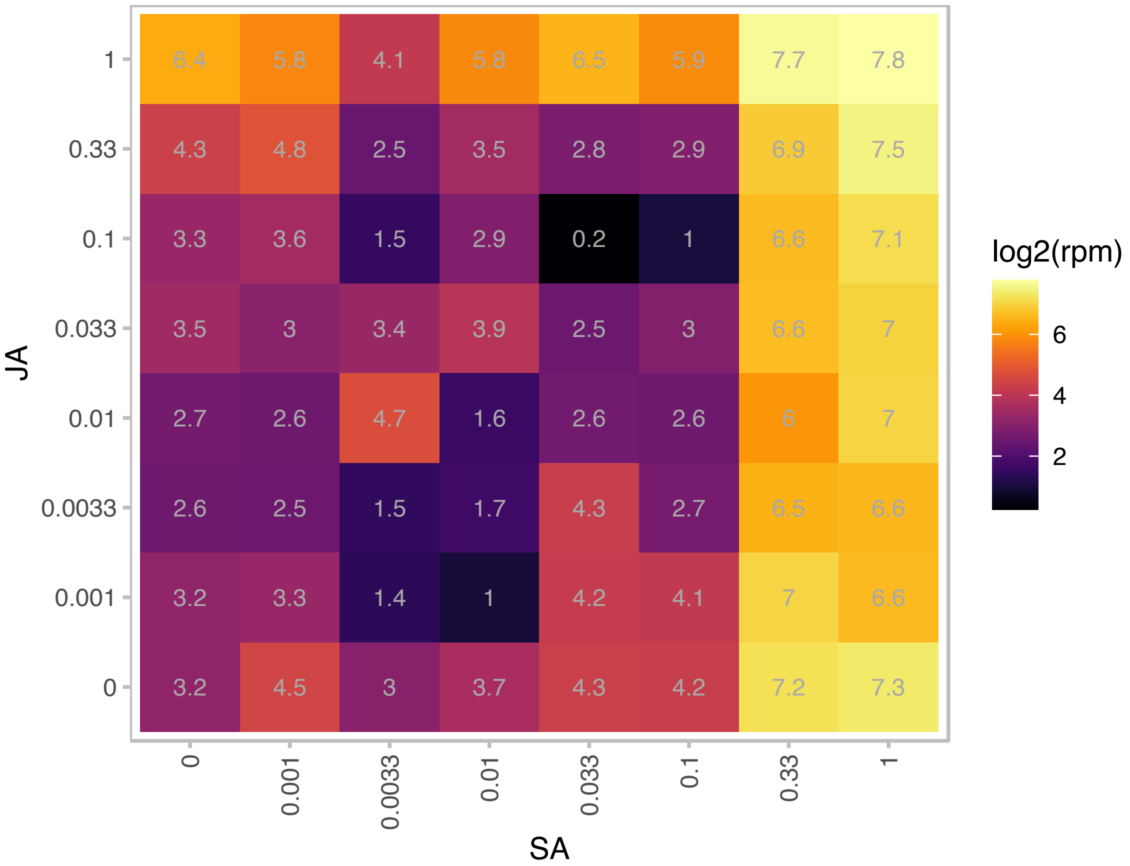


| | |
|---|---|
| Gene | AT2G47890 |
| is.expressed_genes | TRUE |
| is.responsive_genes | TRUE |
| is.apcluster | B-03 |

npr3-2 npr4-2

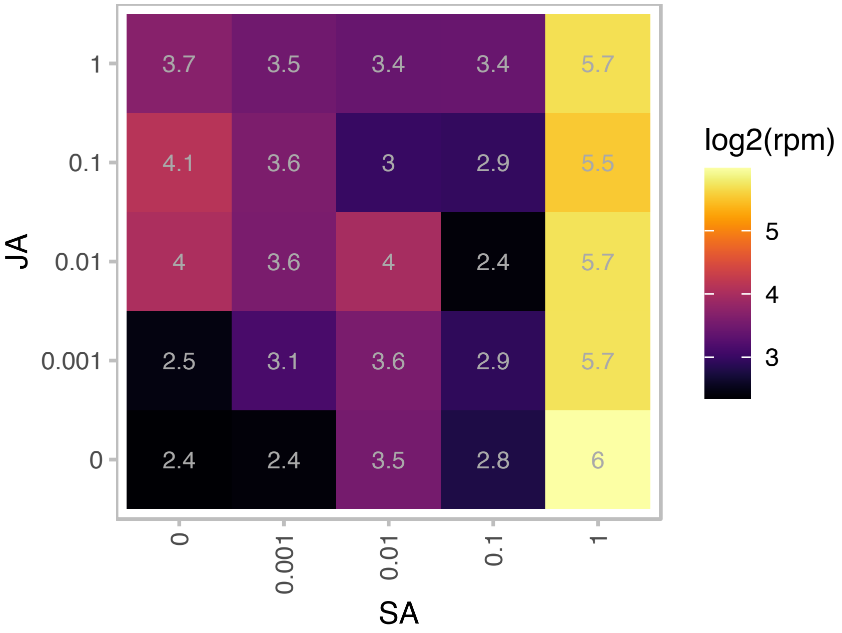


npr3-1 npr4-3

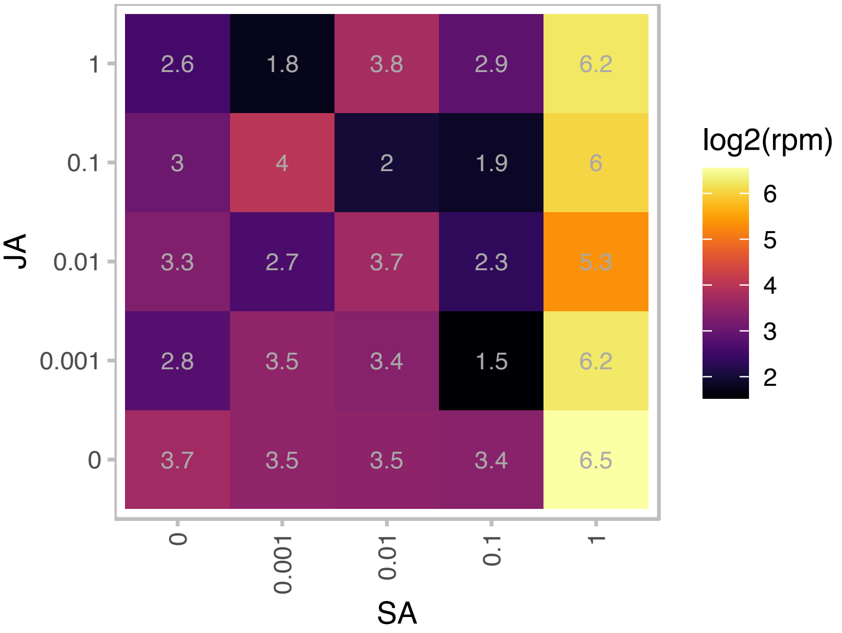

### AT3G03270.1 : HRU1

Adenine nucleotide alpha hydrolases-like superfamily protein

HRU1 is a hypoxia induced universal stress protein. It exists as two splice variants with AT3G03270.2 , which contains a putative dimerization domain, the predominant transcript found under anoxia. It is induced by RAP2.12. Subcellular localization is dynamic; under anoxia the localization of HRU1 shifts from cytoplasm to the plasma membrane.

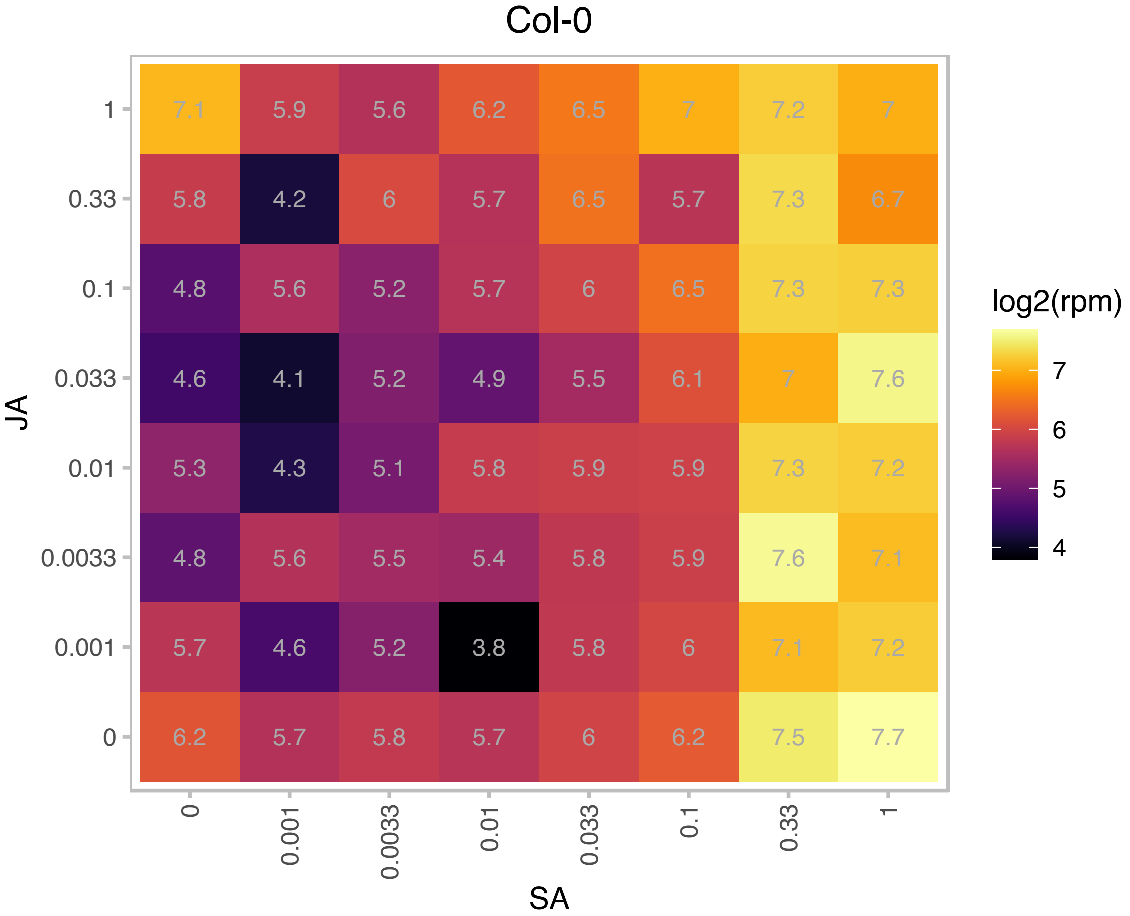


| | |
|---|---|
| Gene | AT3G03270 |
| is.expressed_genes | TRUE |
| is.responsive_genes | TRUE |
| is.apcluster | B-03 |

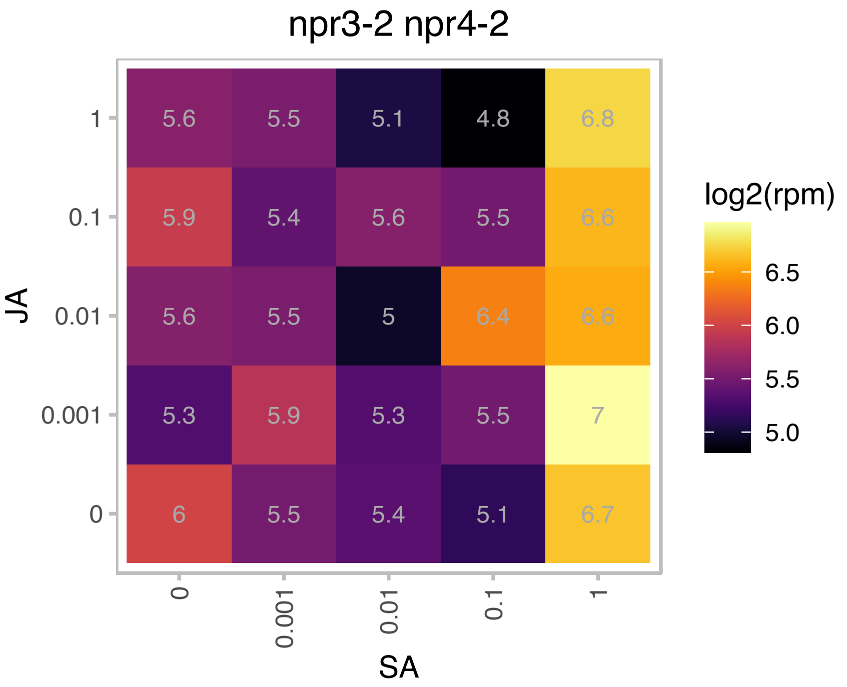


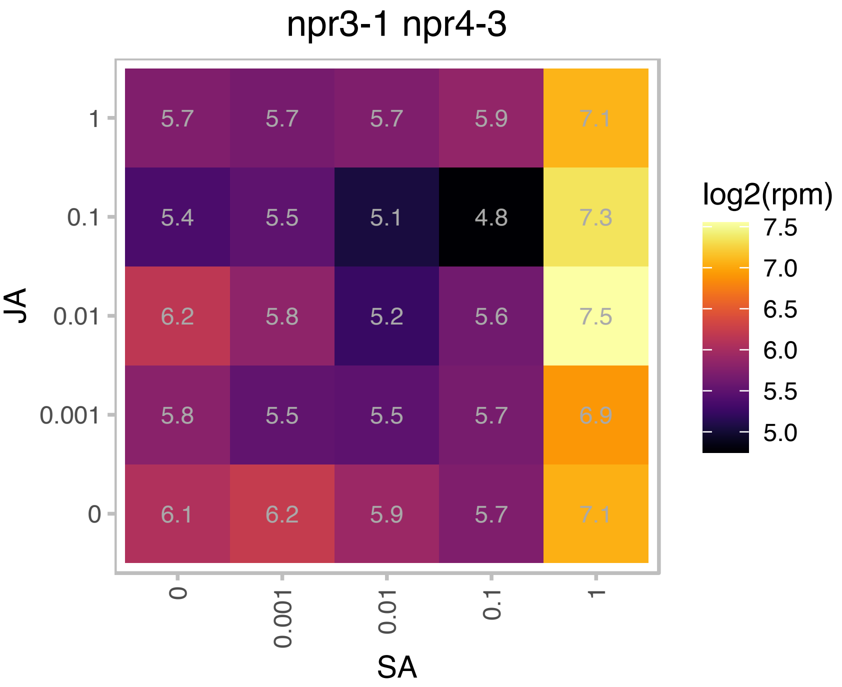

### AT3G03310.1 : ATLCAT3, LCAT3

lecithin:cholesterol acyltransferase 3

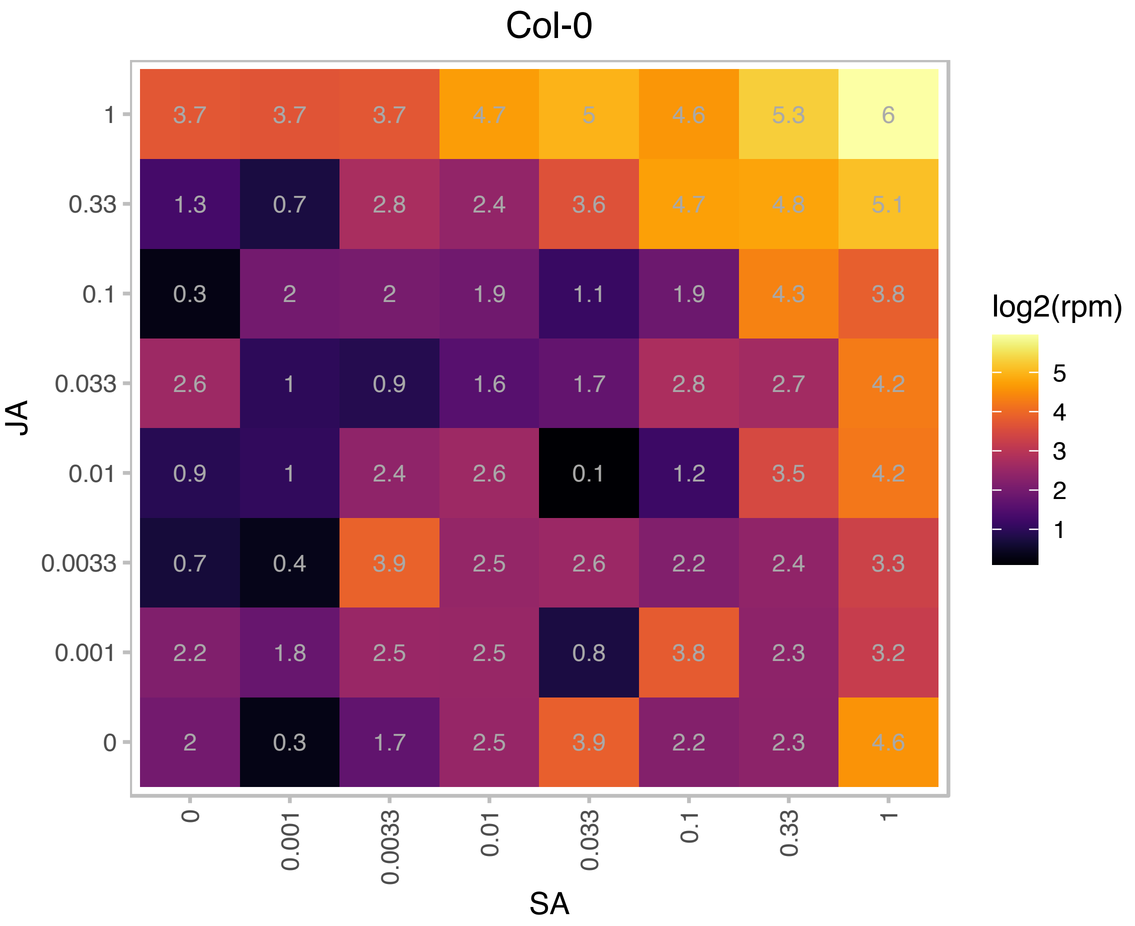


| | |
|---|---|
| Gene | AT3G03310 |
| is.expressed_genes | TRUE |
| is.responsive_genes | TRUE |
| is.apcluster | B-03 |

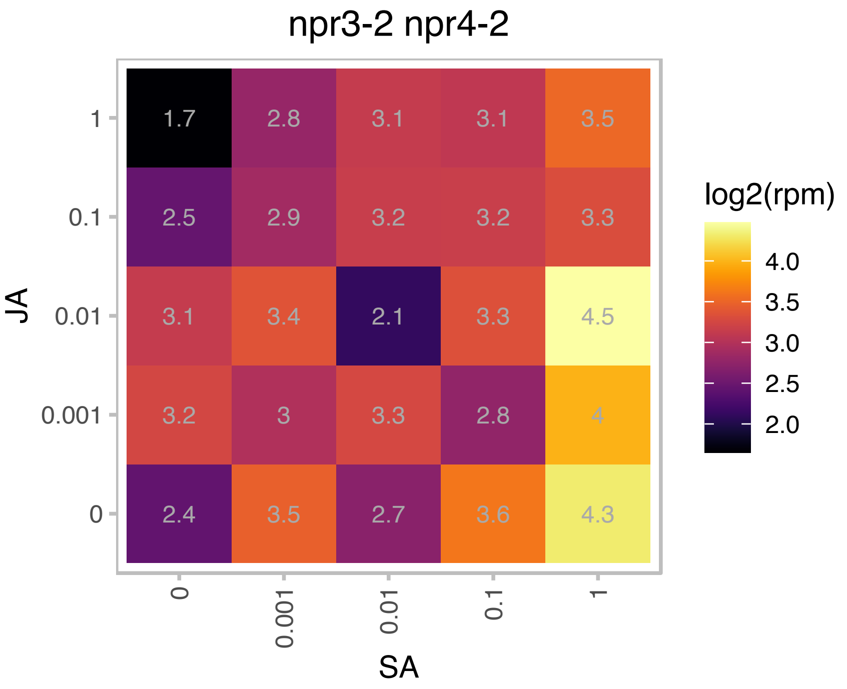


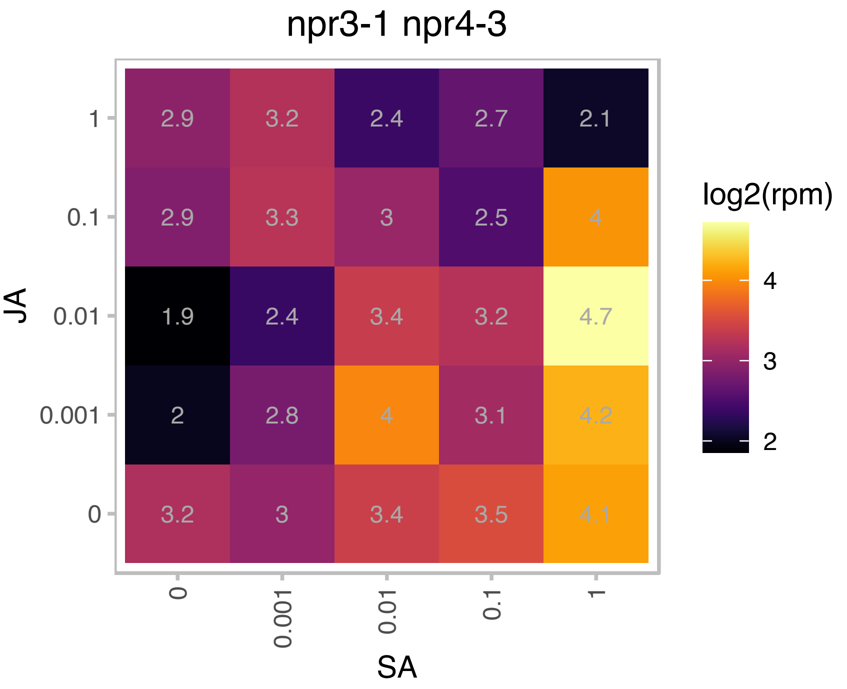

### AT3G04060.1 : anac046, NAC046

NAC domain containing protein 46
NAC046 is a member of the NAC domain containing family of transcription factors. It was identified in a screen for regulators of chlorophyll protein gene expression. Mutants in NAC046 have delayed senescence and increased CHL content suggesting a role in regulation of senescence and chlorophyll degradation.

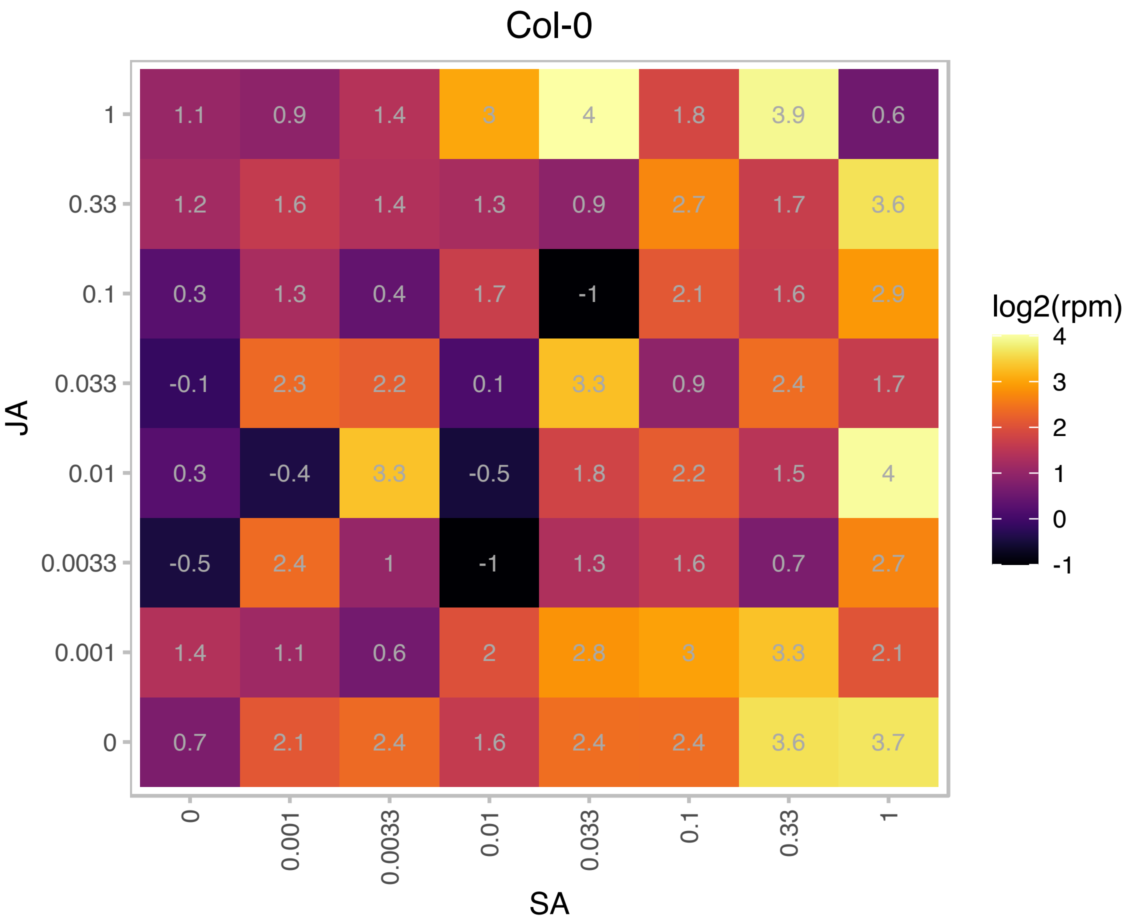


| | |
|---|---|
| Gene | AT3G04060 |
| is.expressed_genes | TRUE |
| is.responsive_genes | TRUE |
| is.apcluster | B-03 |

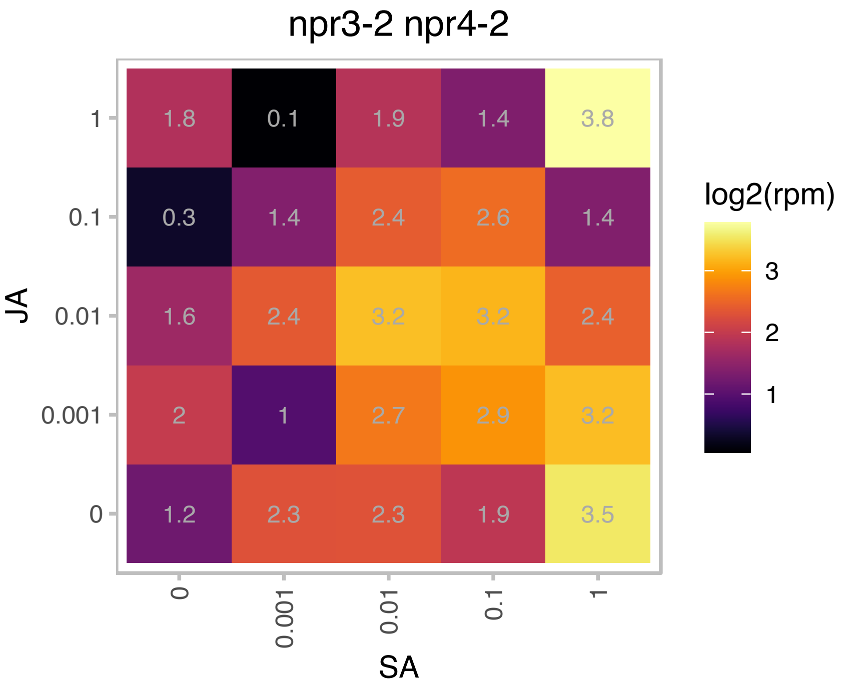


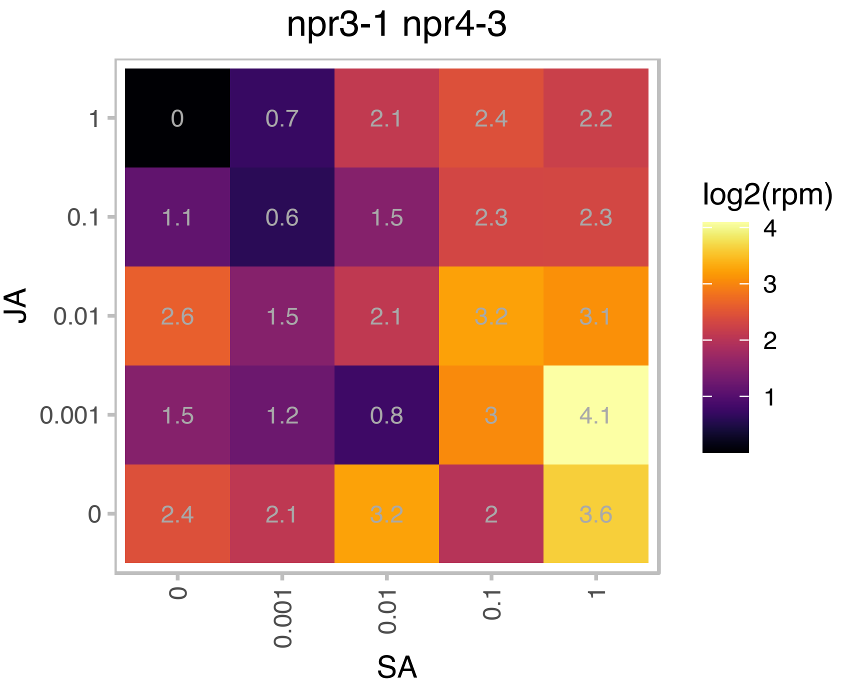

# AT3G04600.1 : NA

Nucleotidylyl transferase superfamily protein

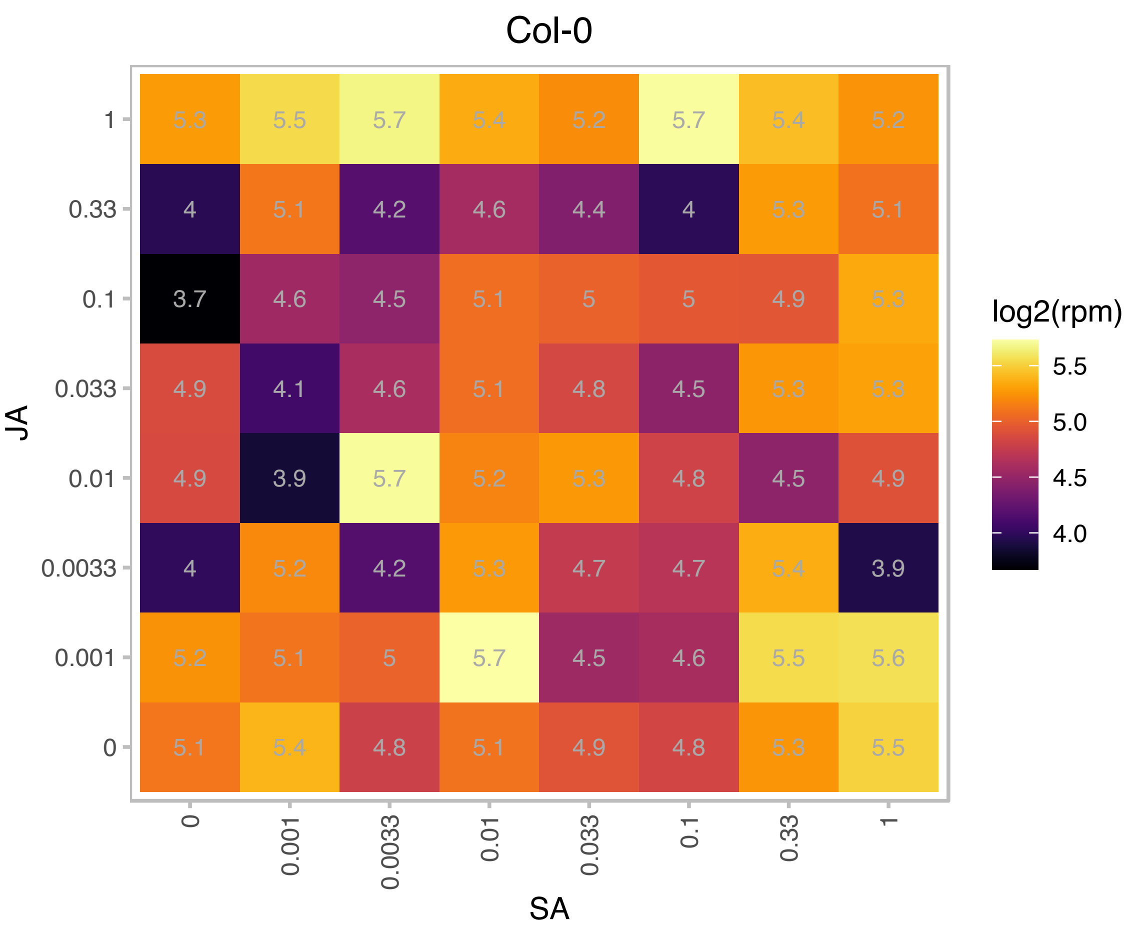


| | |
|---|---|
| Gene | AT3G04600 |
| is.expressed_genes | TRUE |
| is.responsive_genes | TRUE |
| is.apcluster | B-03 |

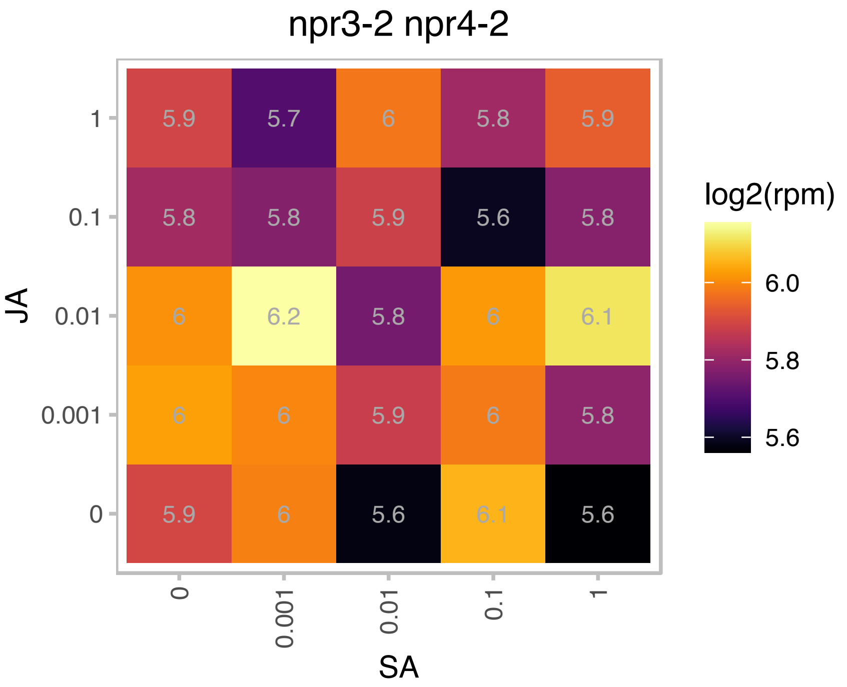


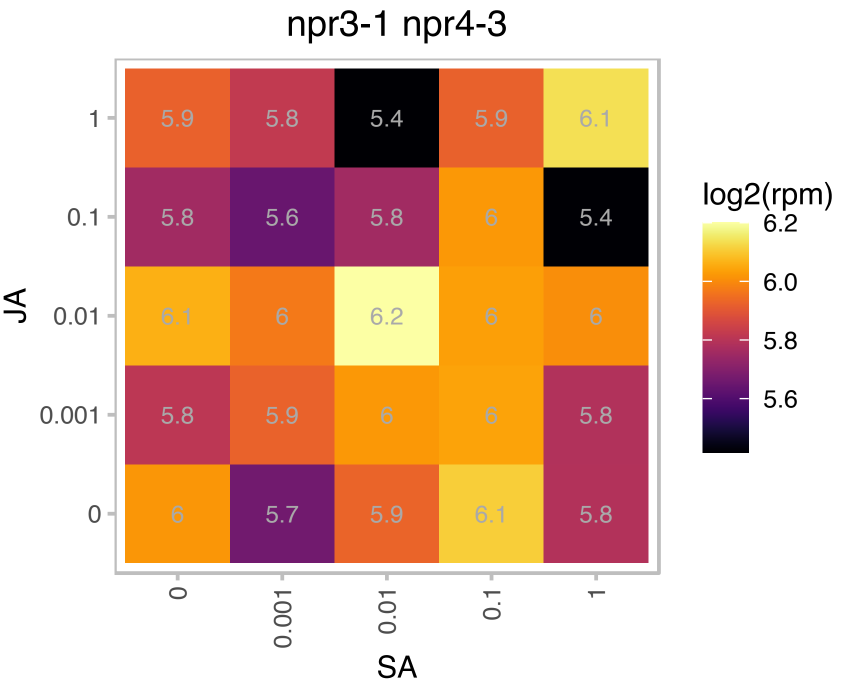

# AT3G05685.1 : NA

Cystatin/monellin superfamily protein

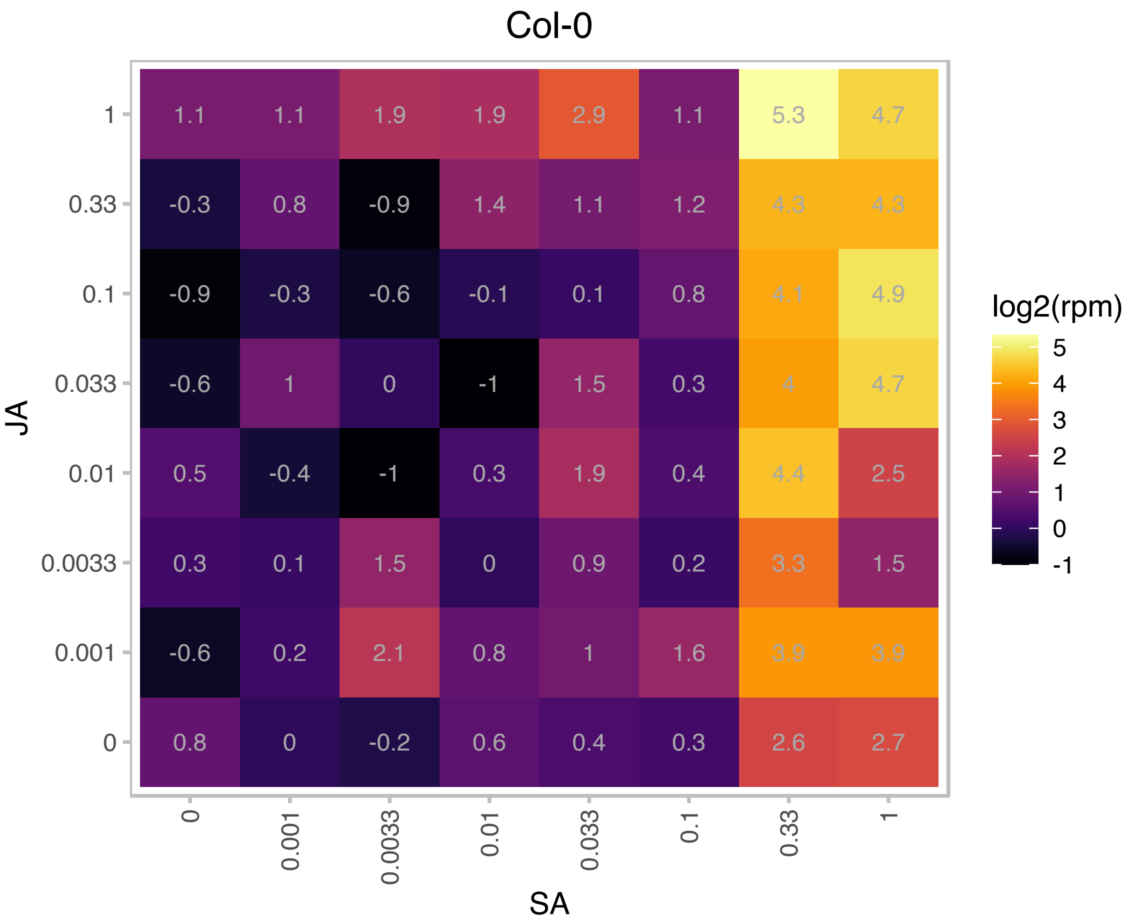


| | |
|---|---|
| Gene | AT3G05685 |
| is.expressed_genes | TRUE |
| is.responsive_genes | TRUE |
| is.apcluster | B-03 |

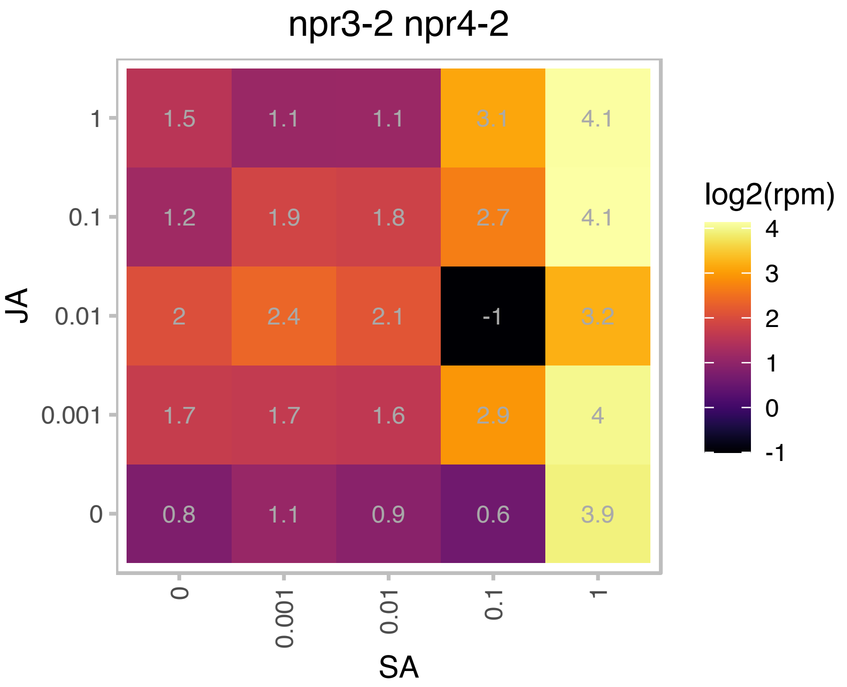


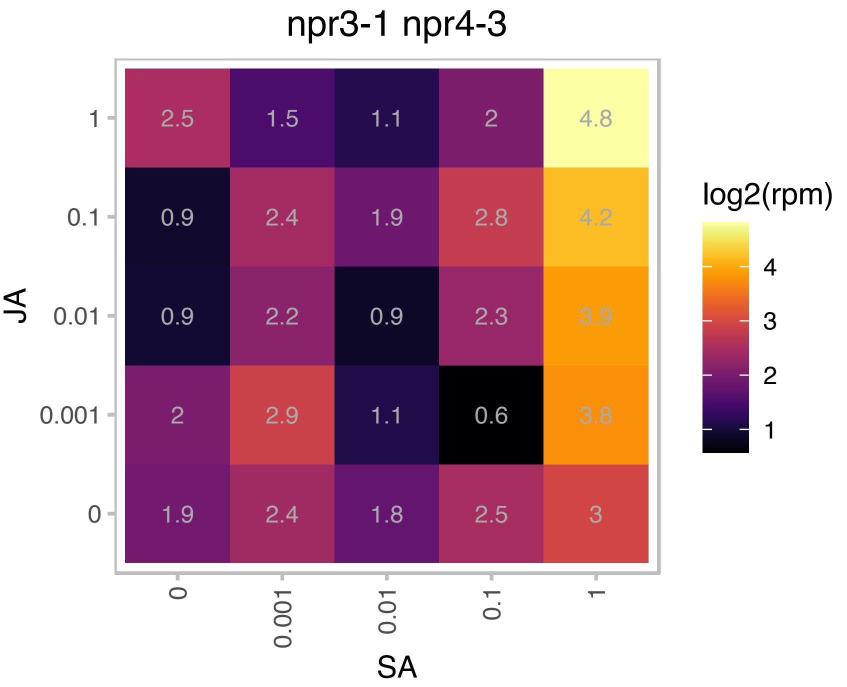

### AT3G08590.1 : iPGAM2

Phosphoglycerate mutase, 2,3-bisphosphoglycerate-independent

Encodes a 2,3-biphosphoglycerate-independent phosphoglycerate mutase that is involved in pollen development and stomatal movement.

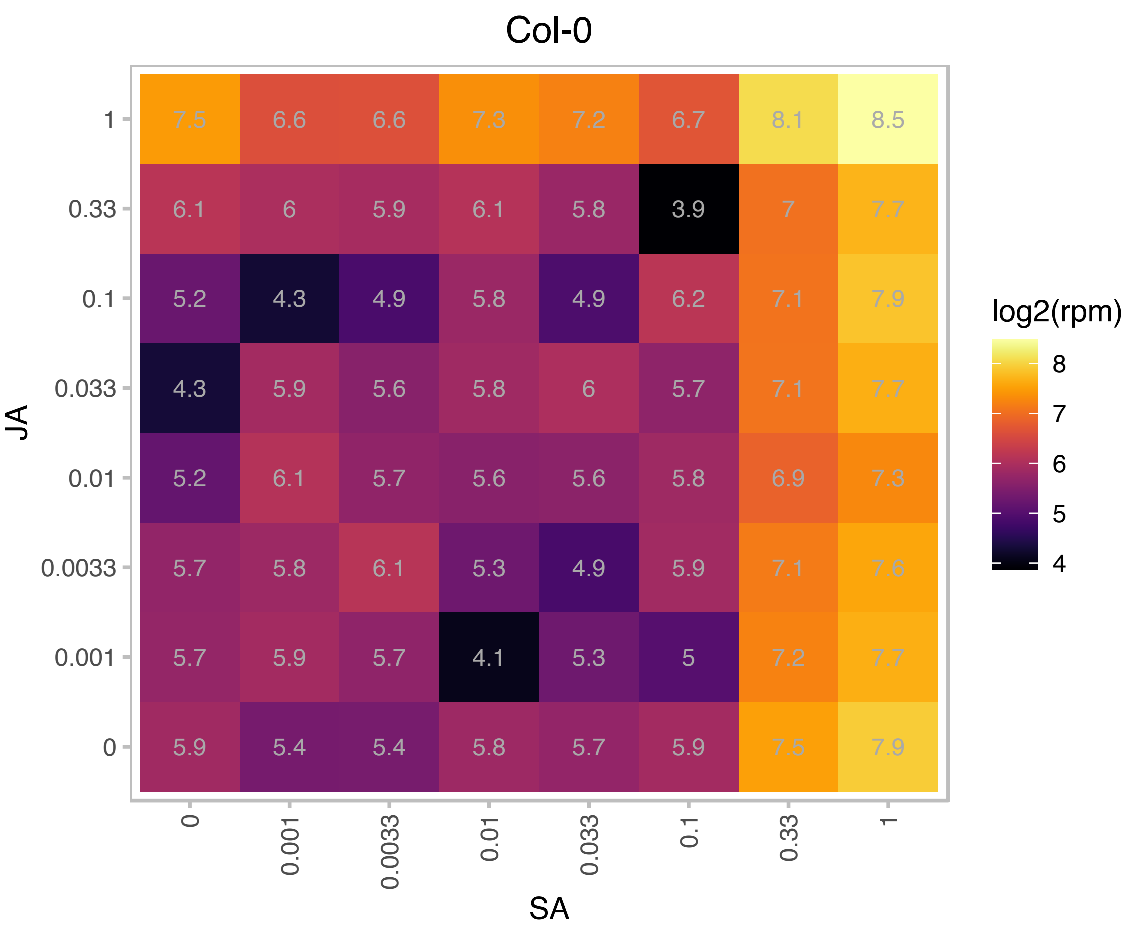


| | |
|---|---|
| Gene | AT3G08590 |
| is.expressed_genes | TRUE |
| is.responsive_genes | TRUE |
| is.apcluster | B-03 |

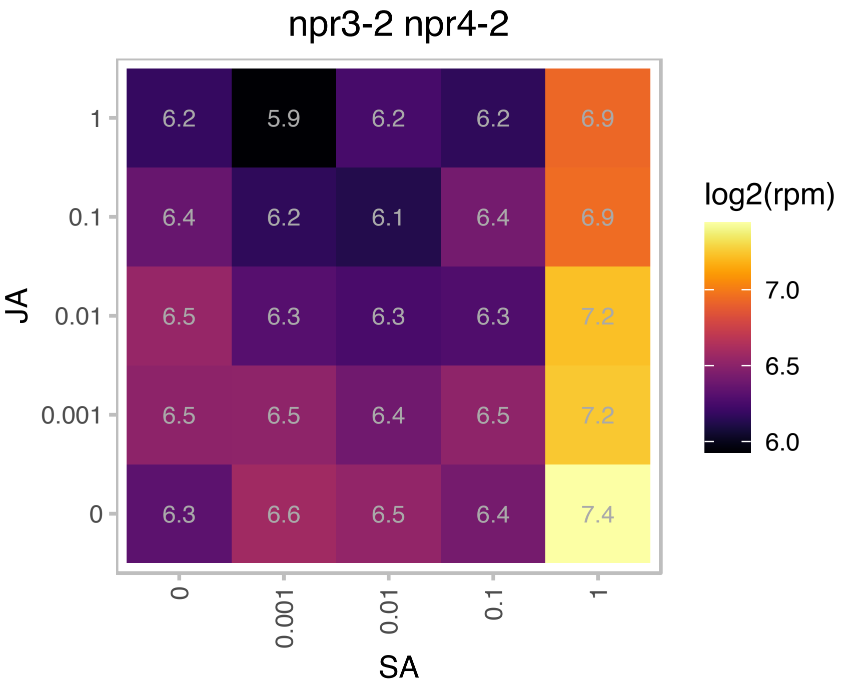


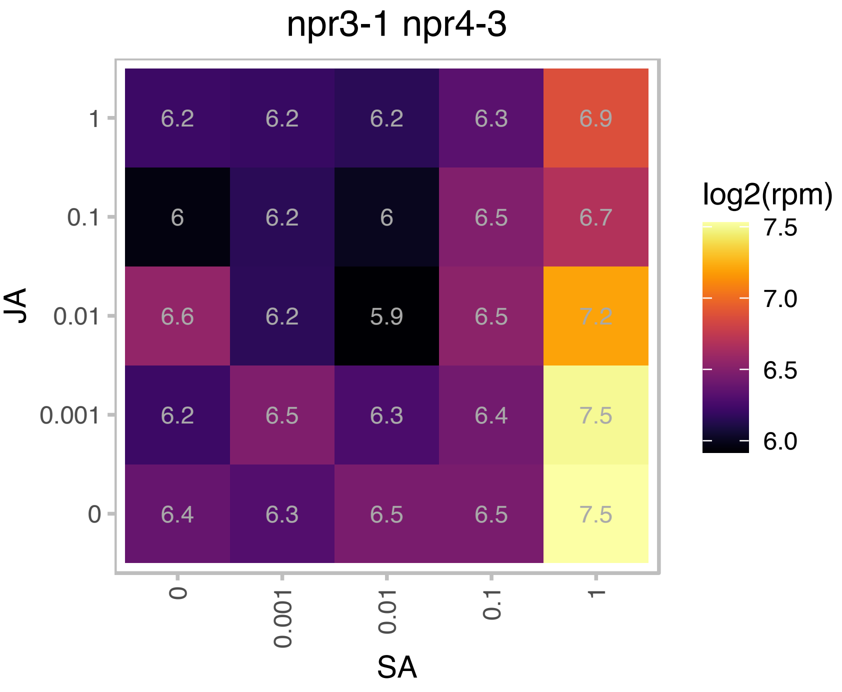

# AT3G09010.1 : NA

Protein kinase superfamily protein

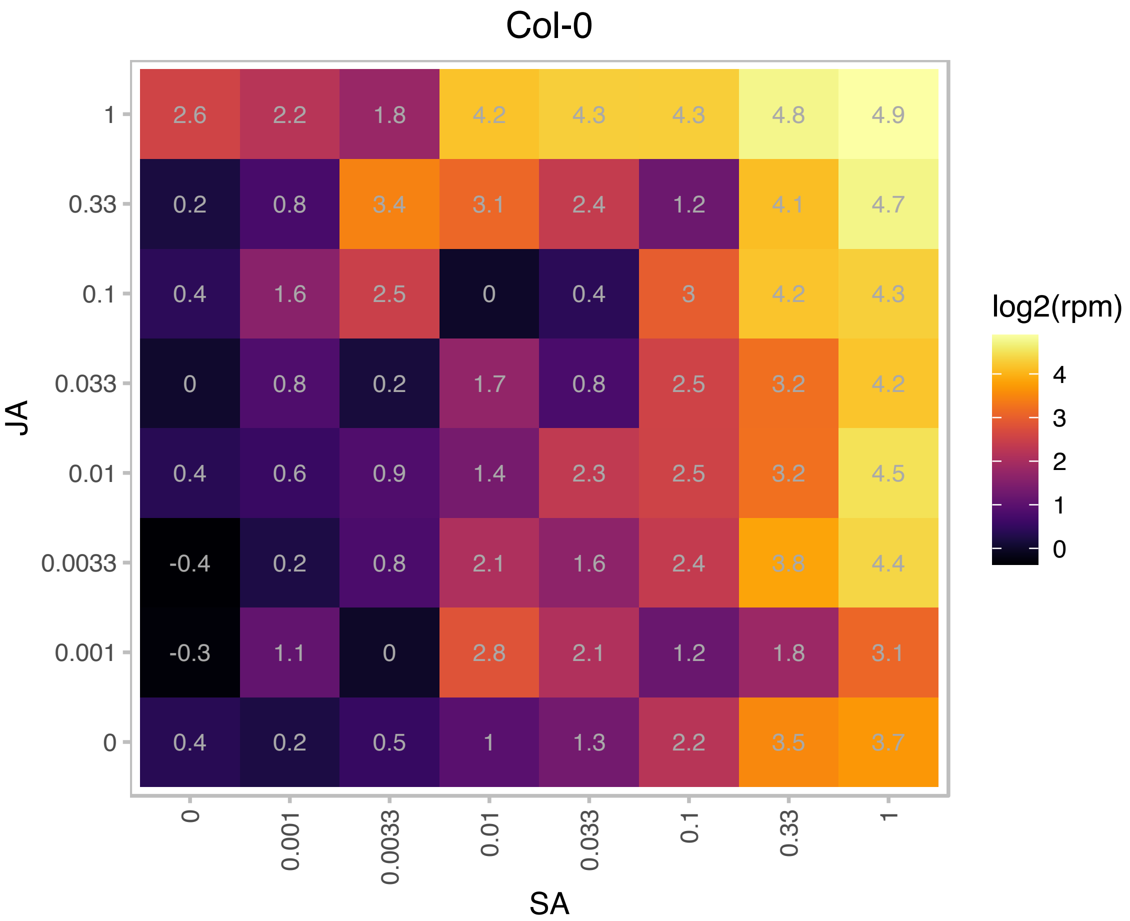


| | |
|---|---|
| Gene | AT3G09010 |
| is.expressed_genes | TRUE |
| is.responsive_genes | TRUE |
| is.apcluster | B-03 |

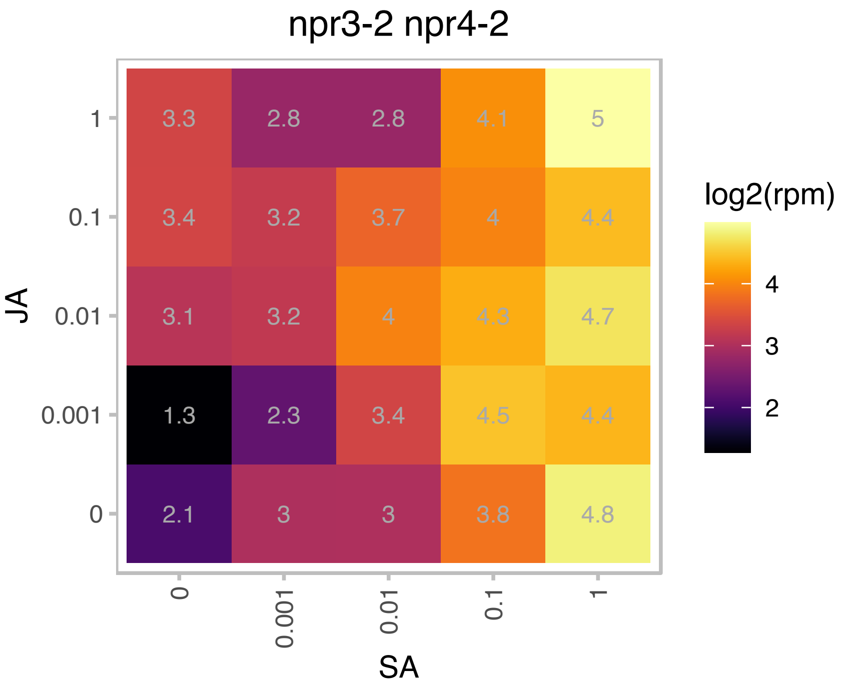


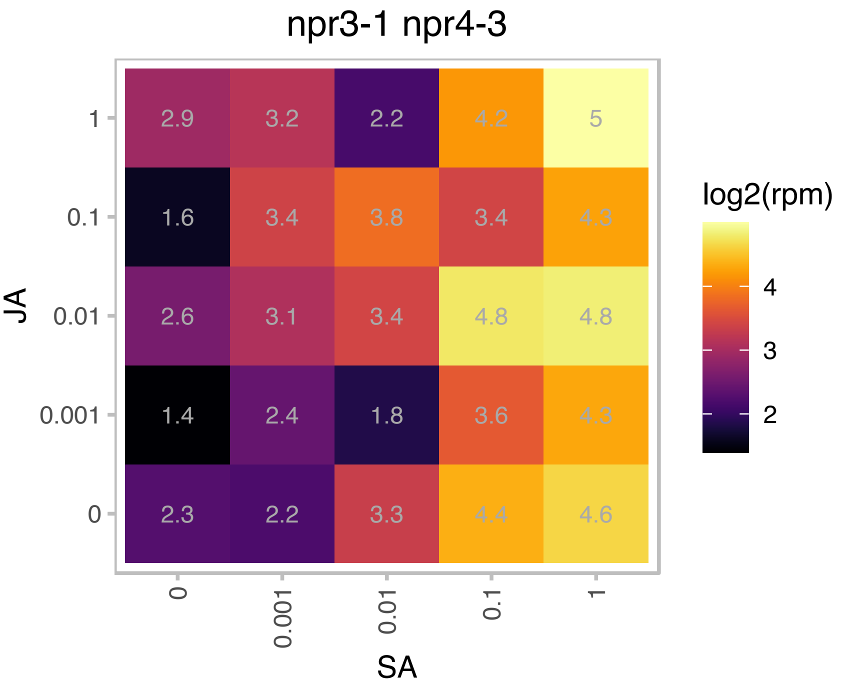

### AT3G09810.1 : IDH-VI

isocitrate dehydrogenase VI

Encodes a catalytic subunit of the mitochondrially-localized NAD+- dependent isocitrate dehydrogenase

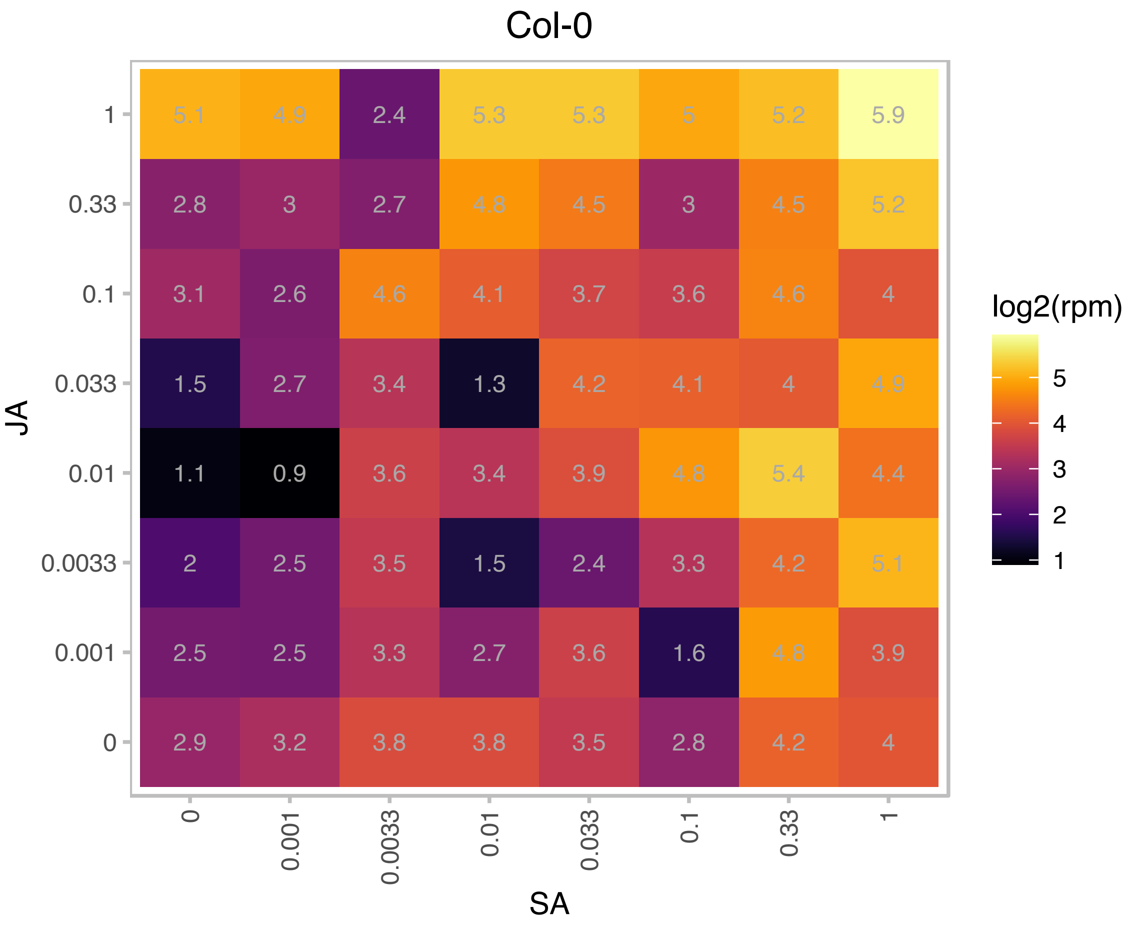


| | |
|---|---|
| Gene | AT3G09810 |
| is.expressed_genes | TRUE |
| is.responsive_genes | TRUE |
| is.apcluster | B-03 |

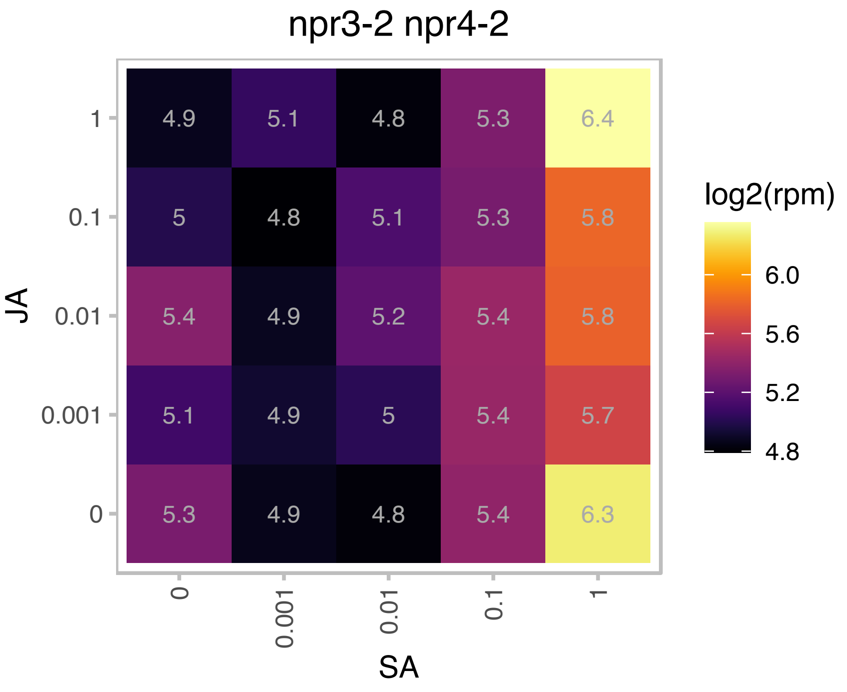


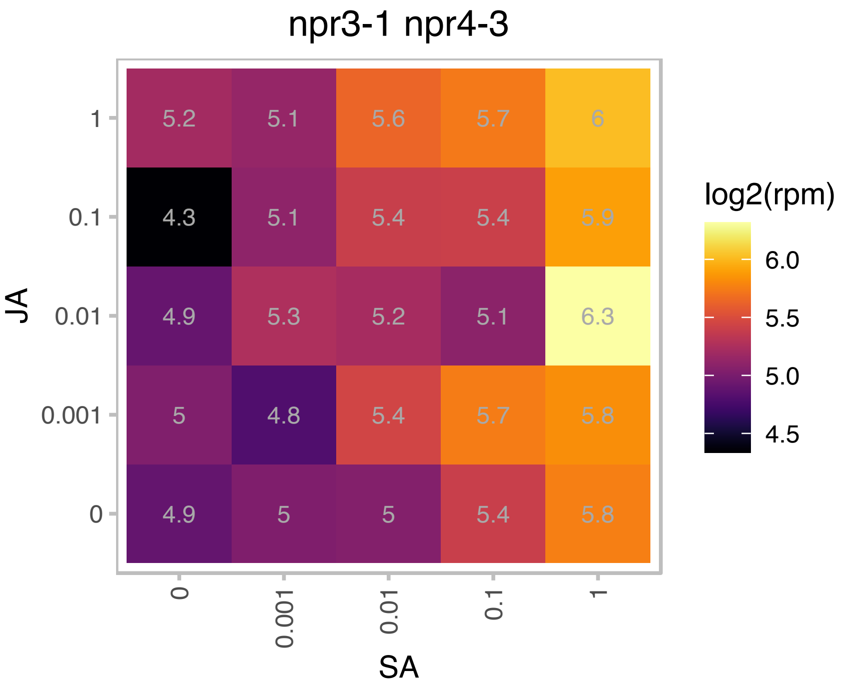

# AT3G14595.1 : NA

Ribosomal protein L18ae family

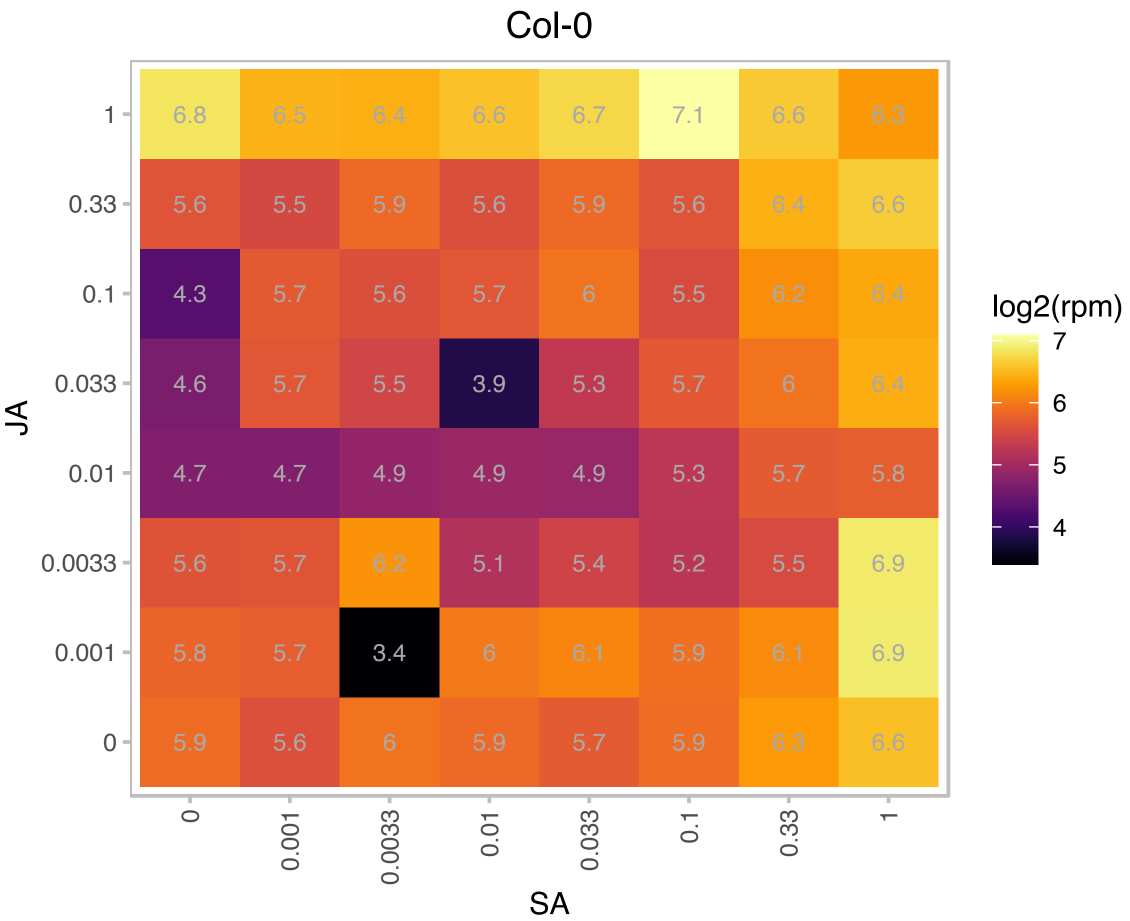


| | |
|---|---|
| Gene | AT3G14595 |
| is.expressed_genes | TRUE |
| is.responsive_genes | TRUE |
| is.apcluster | B-03 |

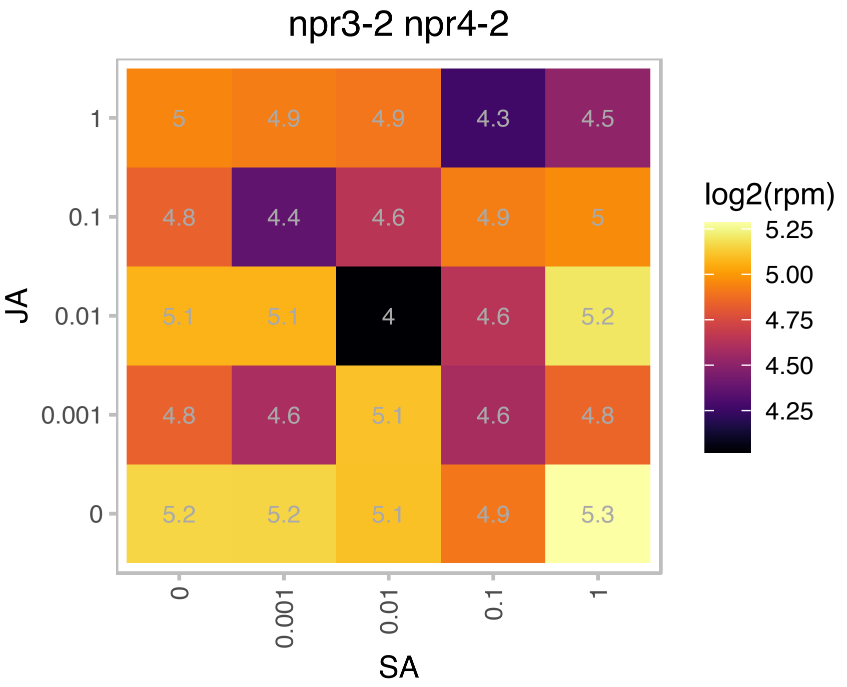


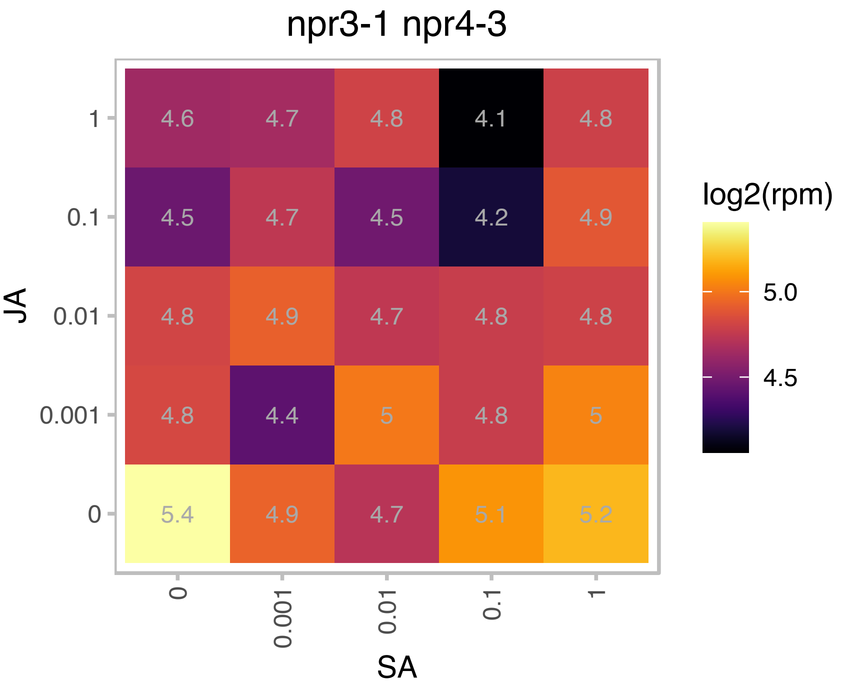

### AT3G15355.1 : PFU1, UBC25

ubiquitin-conjugating enzyme 25

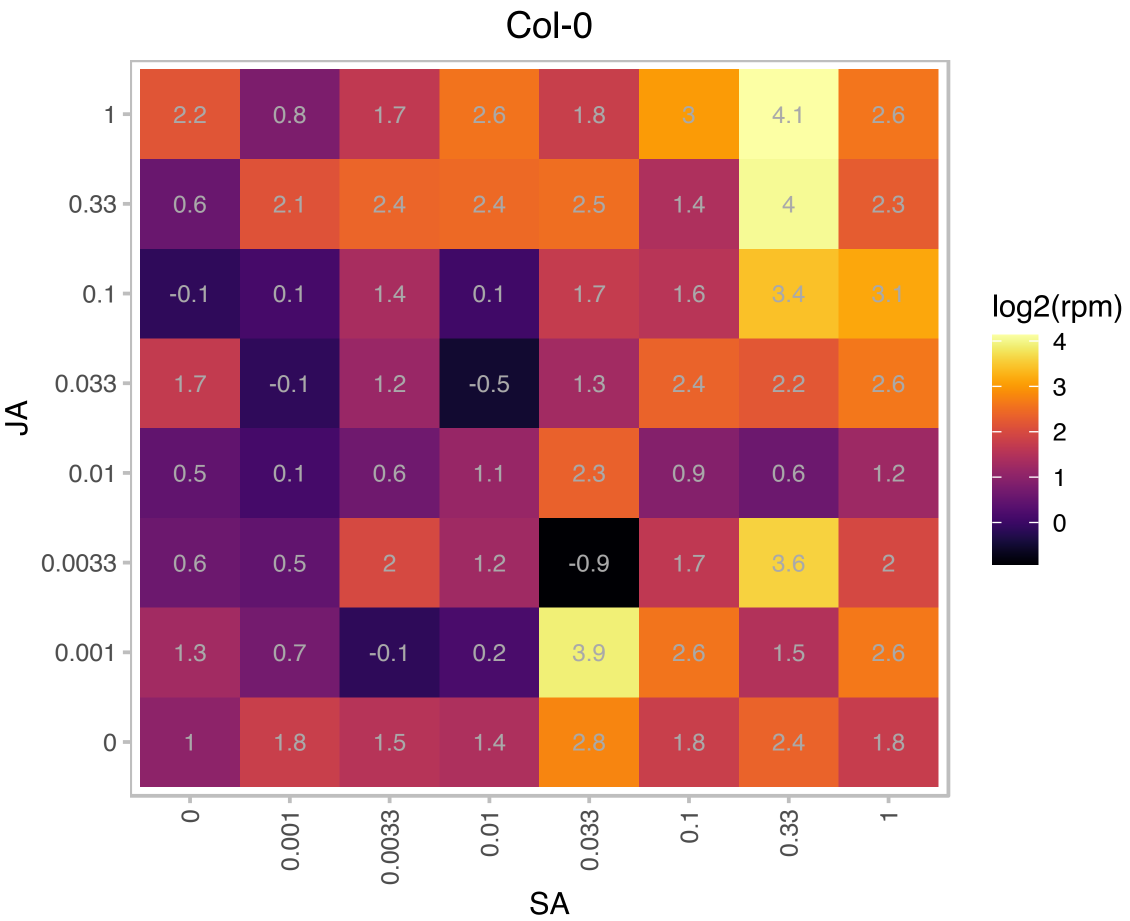


| | |
|---|---|
| Gene | AT3G15355 |
| is.expressed_genes | TRUE |
| is.responsive_genes | TRUE |
| is.apcluster | B-03 |

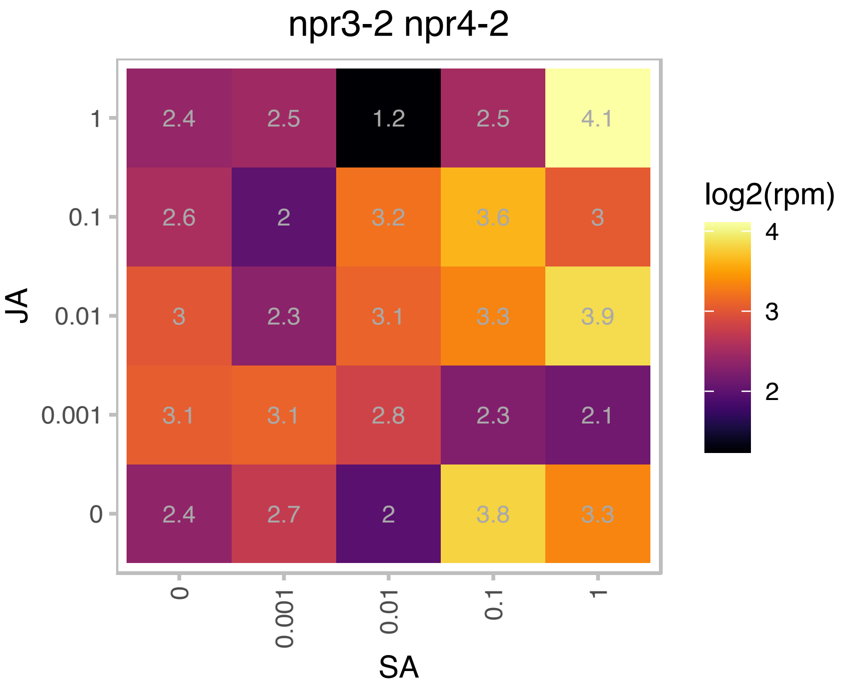


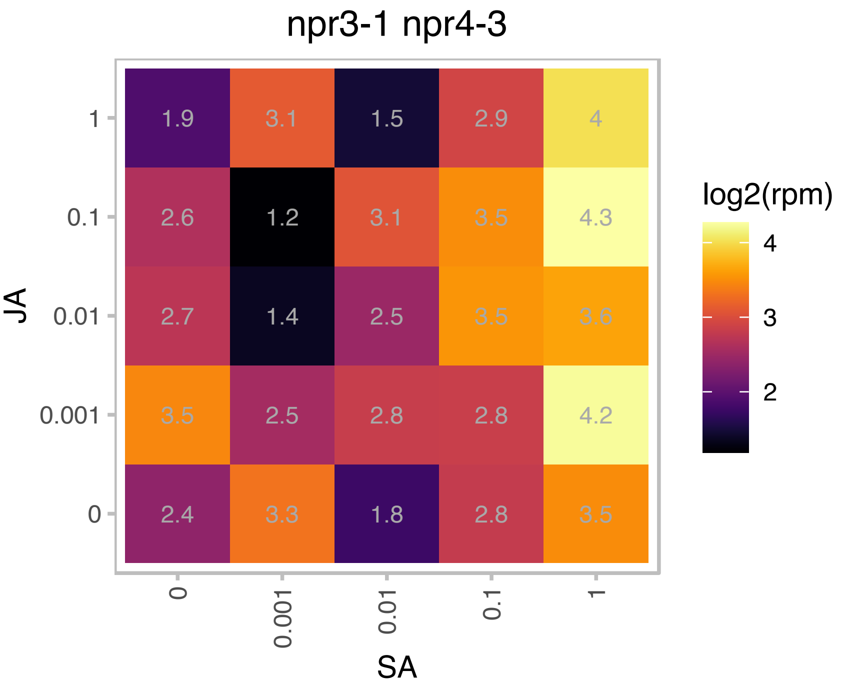

### AT3G22370.1 : AOX1A, ATAOX1A, AtHSR3, HSR3

alternative oxidase 1A

Encodes AOX1a, an isoform of alternative oxidase that is expressed in rosettes, flowers, and root. The alternative oxidase of plant mitochondria transfers electrons from the ubiquinone pool to oxygen without energy conservations. It is regulated through transcriptional control and by pyruvate. Plays a role in shoot acclimation to low temperature. Also is capable of ameliorating reactive oxygen species production when the cytochrome pathway is inhibited. AOX1a also functions as a marker for mitochondrial retrograde response. The mRNA is cell-to-cell mobile.

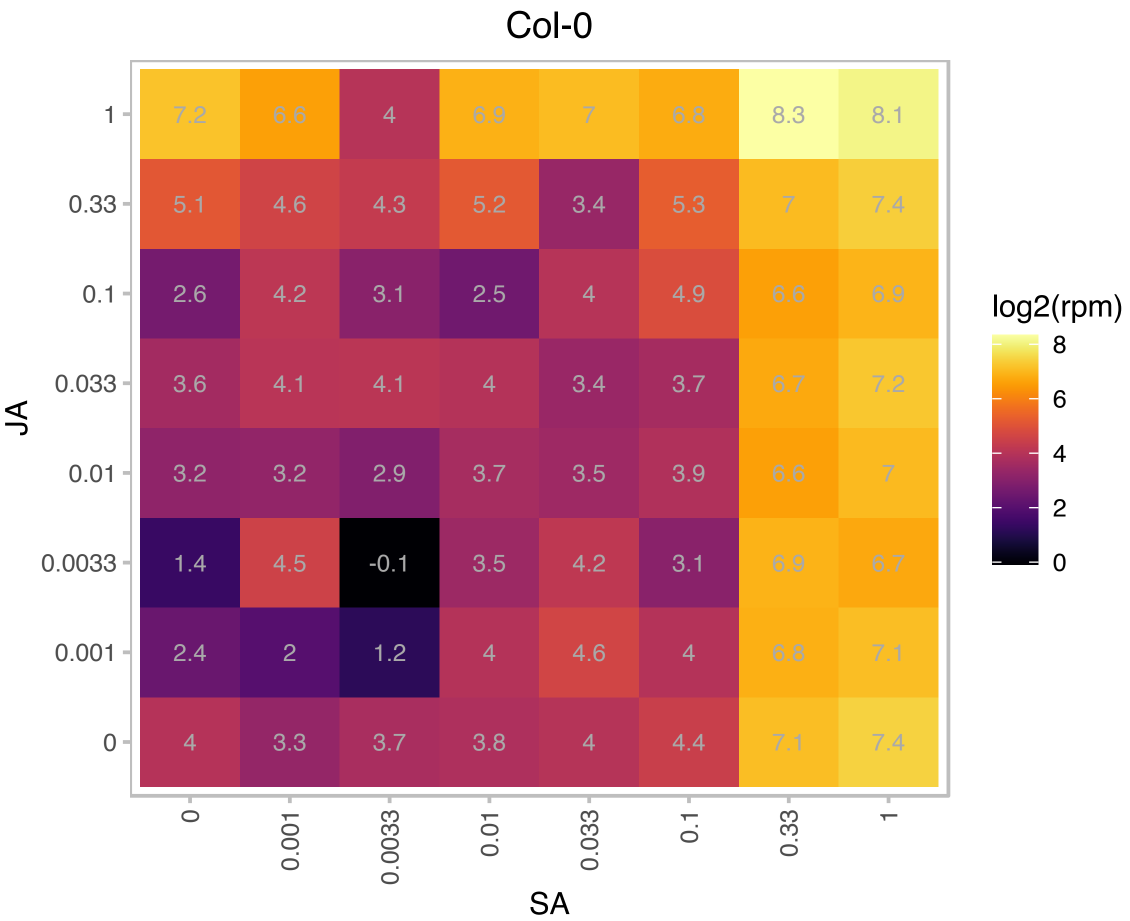


| Gene | AT3G22370 |
|---|---|
| is.expressed_genes | TRUE |
| is.responsive_genes | TRUE |
| is.apcluster | B-03 |

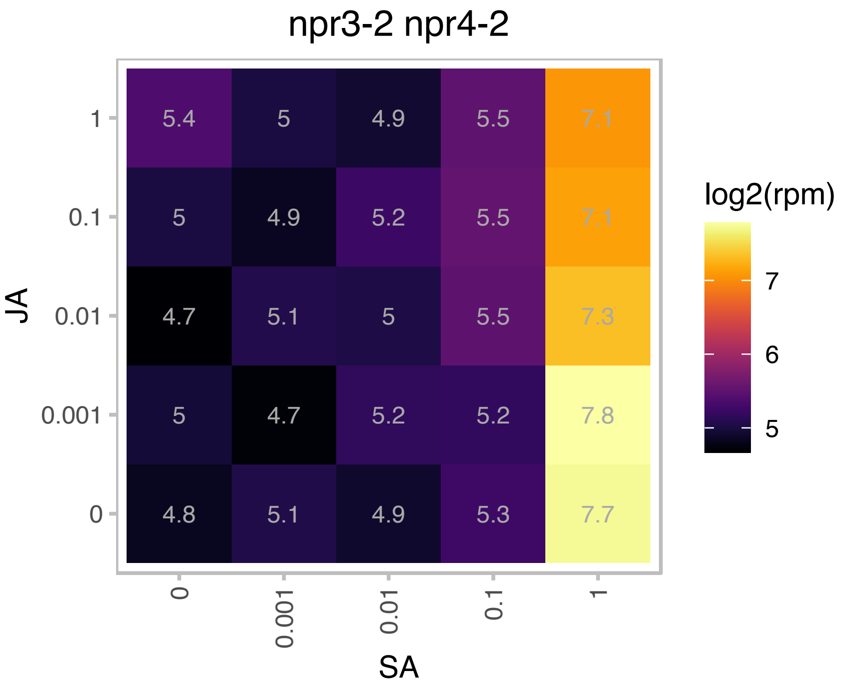


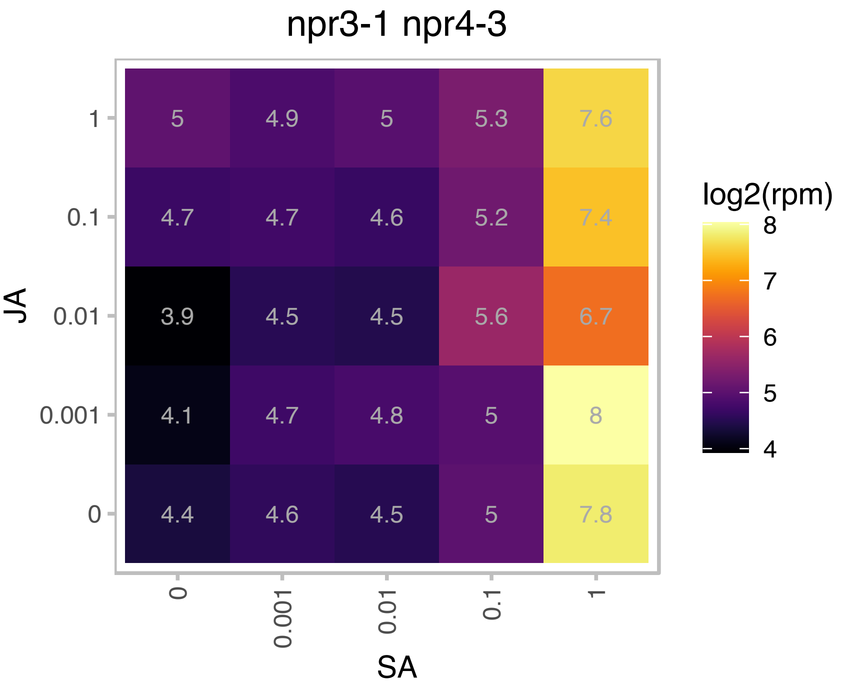

### AT3G24420.1 : DLK2

alpha/beta-Hydrolases superfamily protein

DLK2 is a divergent member of the DWARF14 family. It's expression is dependent on D14 and KAI2 but it does not appear to play a role in stringolactone metabolism.

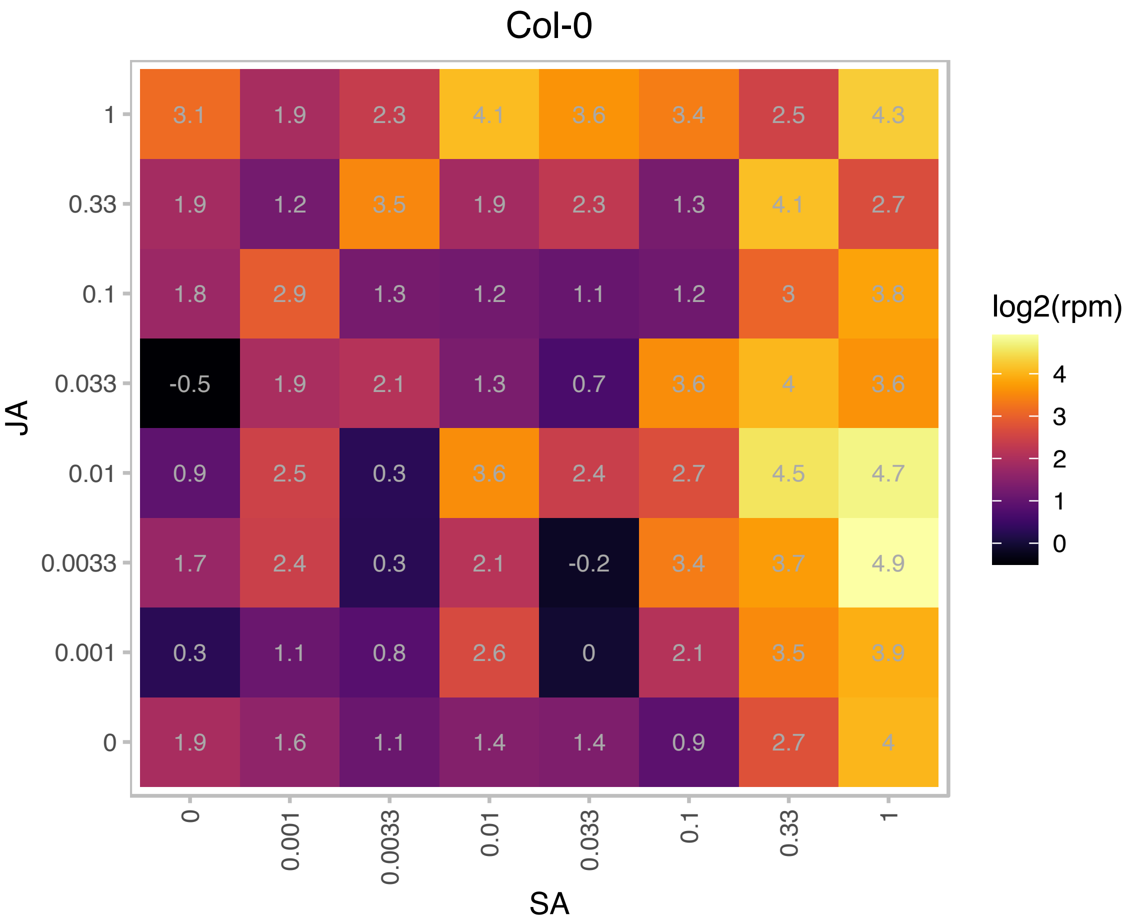


| | |
|---|---|
| Gene | AT3G24420 |
| is.expressed_genes | TRUE |
| is.responsive_genes | TRUE |
| is.apcluster | B-03 |

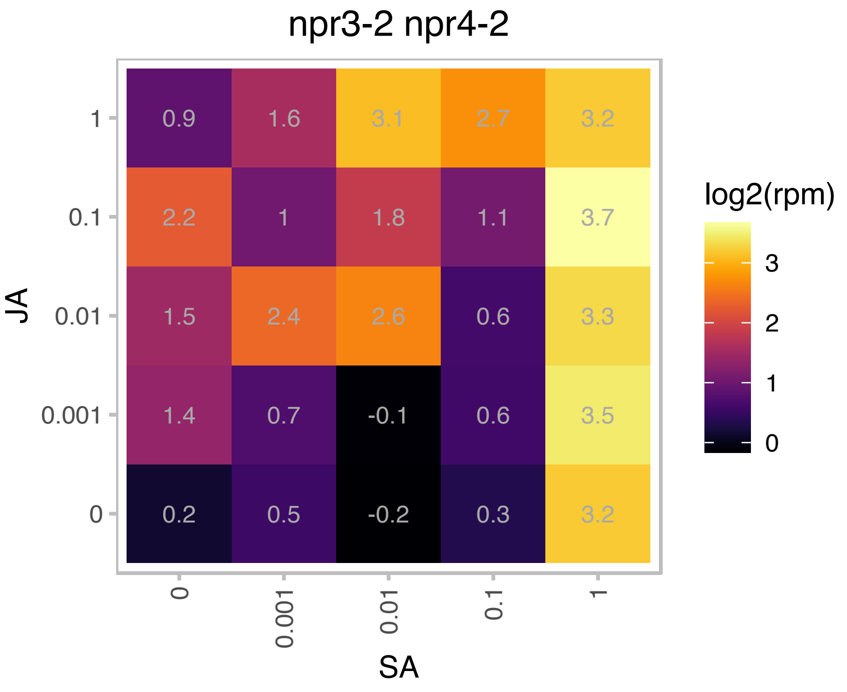


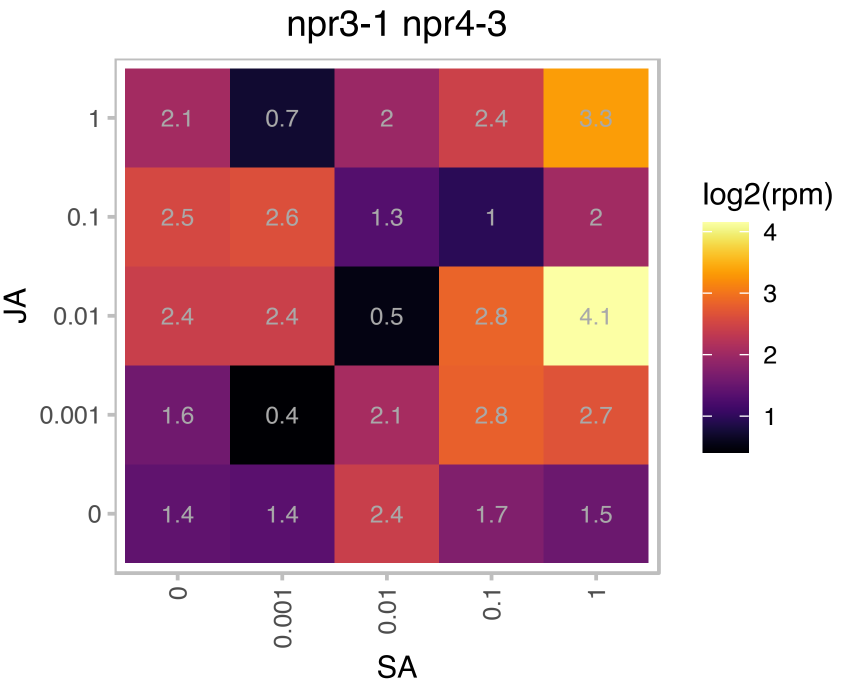

# AT3G48020.1 : NA

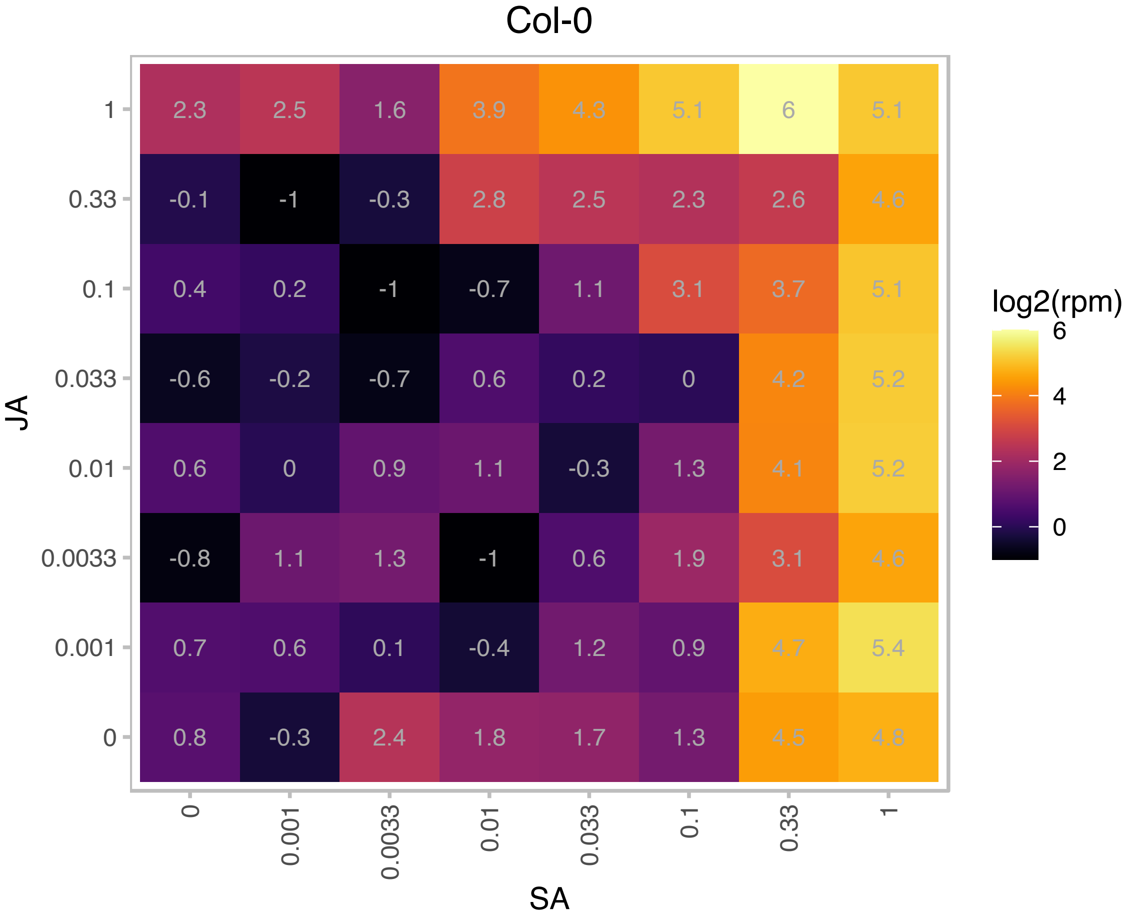


| | |
|---|---|
| Gene | AT3G48020 |
| is.expressed_genes | TRUE |
| is.responsive_genes | TRUE |
| is.apcluster | B-03 |

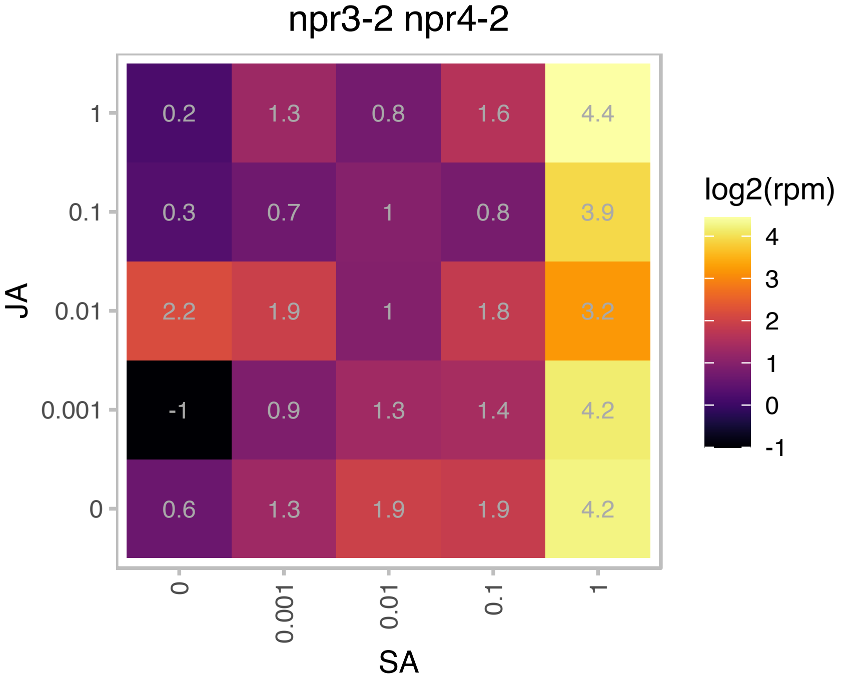


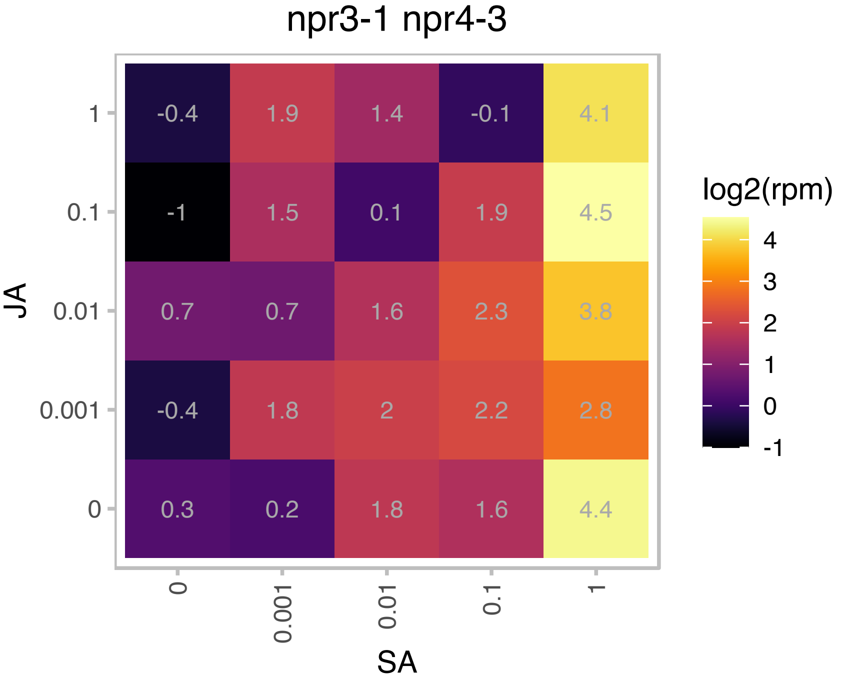

# AT3G48240.1 : NA

Octicosapeptide/Phox/Bem1p family protein

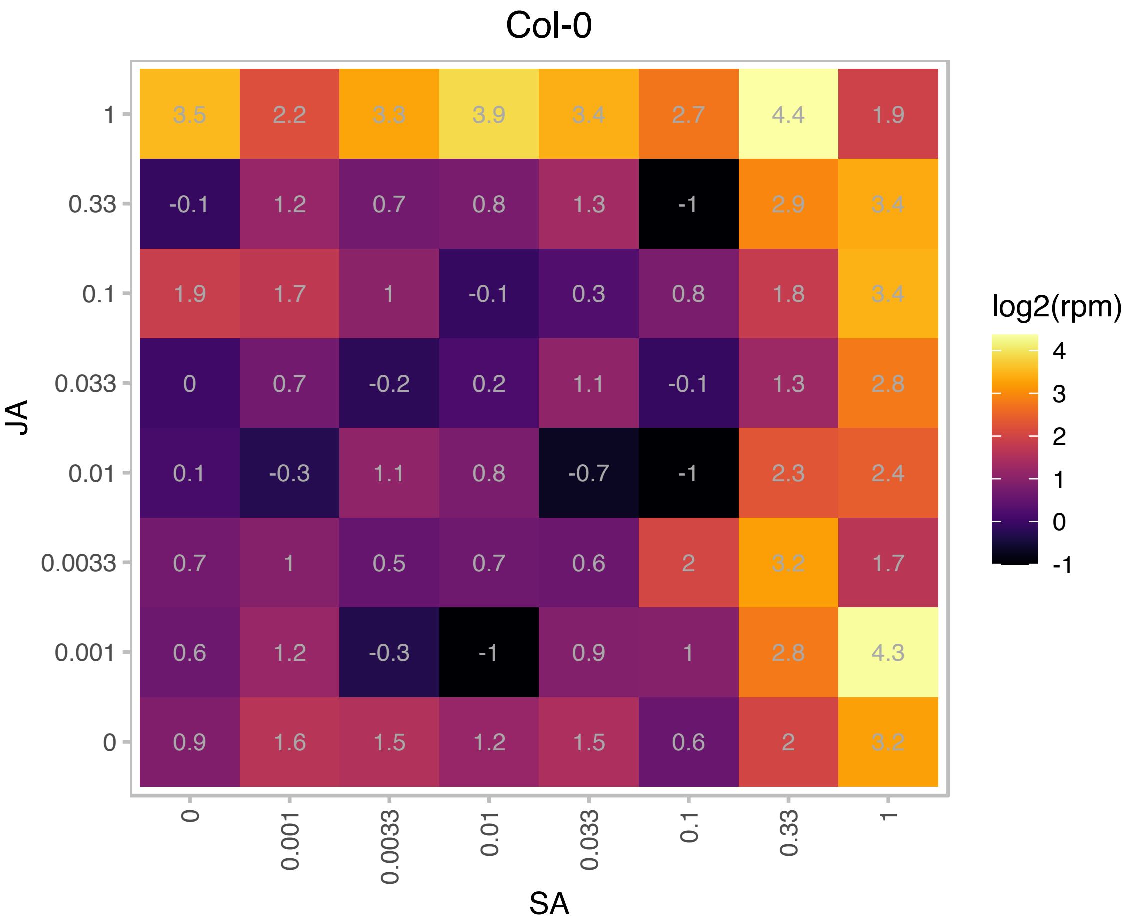


| | |
|---|---|
| Gene | AT3G48240 |
| is.expressed_genes | TRUE |
| is.responsive_genes | TRUE |
| is.apcluster | B-03 |

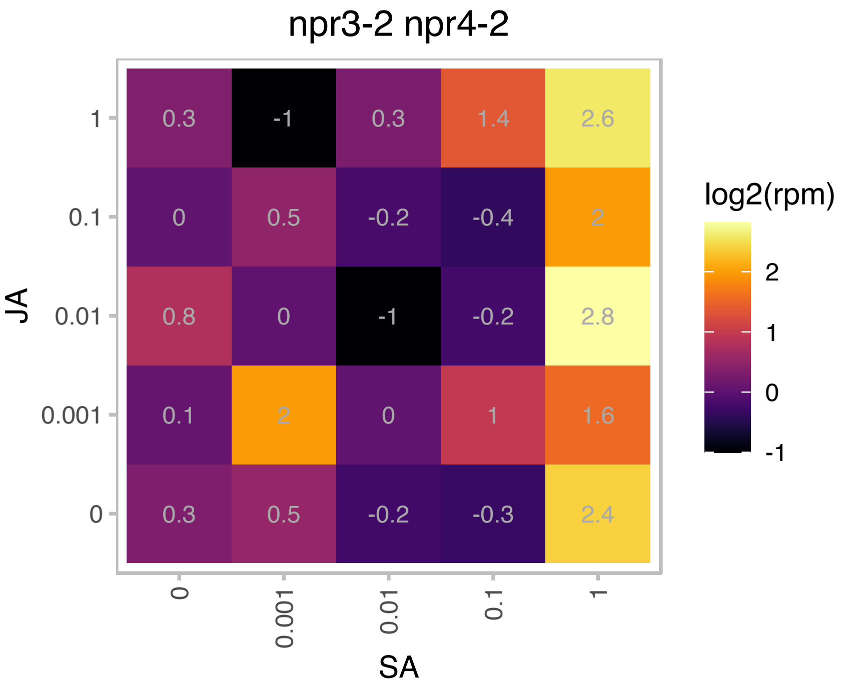


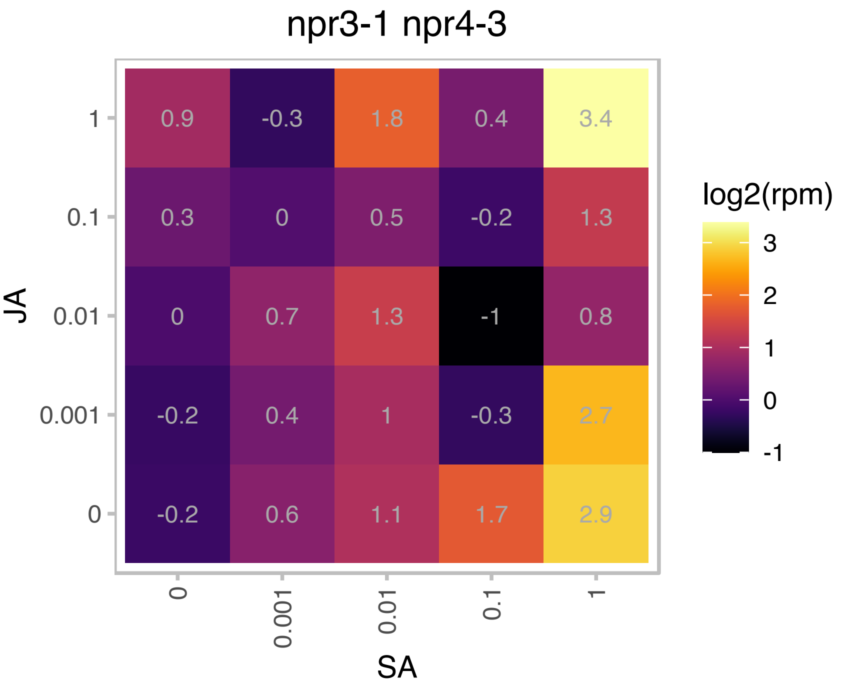

### AT3G54950.1 : PLA IIIA, PLP7, pPLAIIIbeta

patatin-like protein 6
Encodes pPLAIIIbeta, a member of the Group 3 patatin-related phospholipases. pPLAIIIbeta hydrolyzes phospholipids and galactolipids and additionally has acyl-CoA thioesterase activity. Alterations of pPLAIIIβ result in changes in lipid levels and composition.

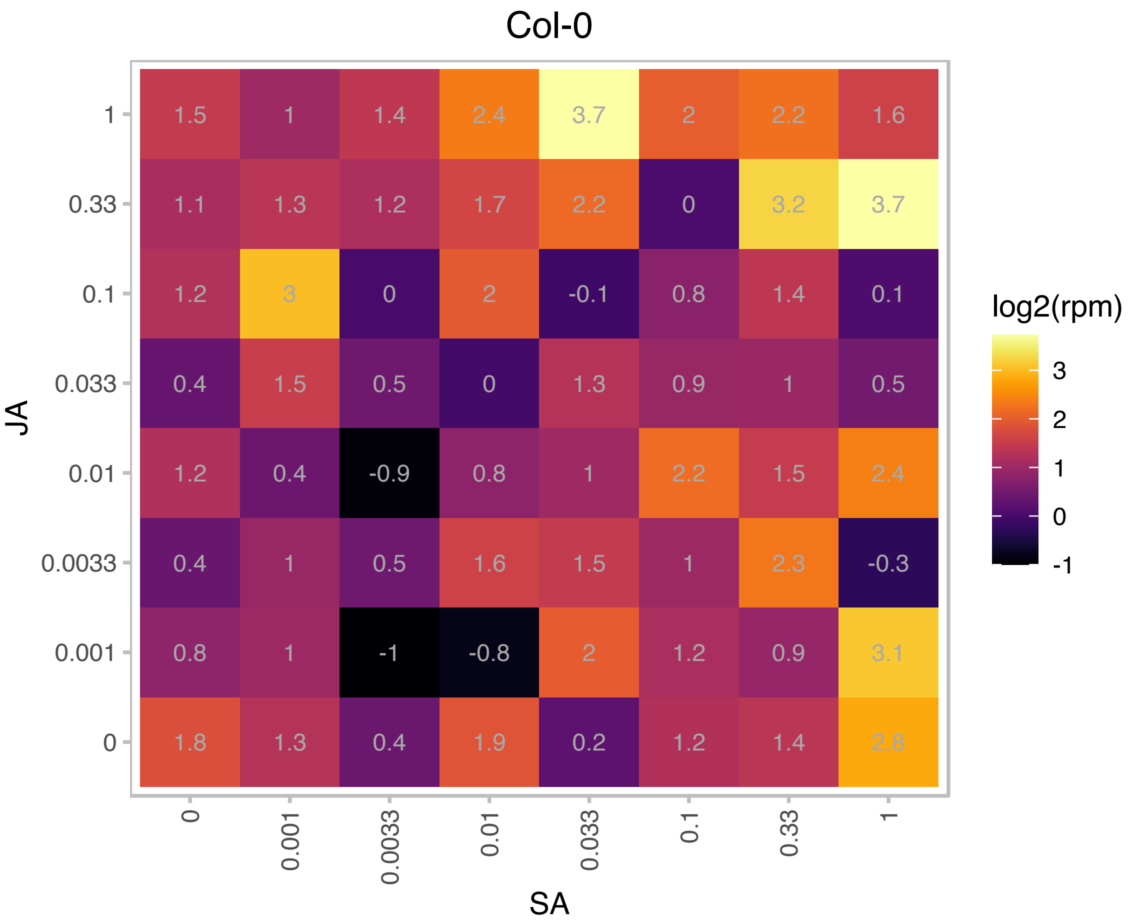


| | |
|---|---|
| Gene | AT3G54950 |
| is.expressed_genes | TRUE |
| is.responsive_genes | TRUE |
| is.apcluster | B-03 |

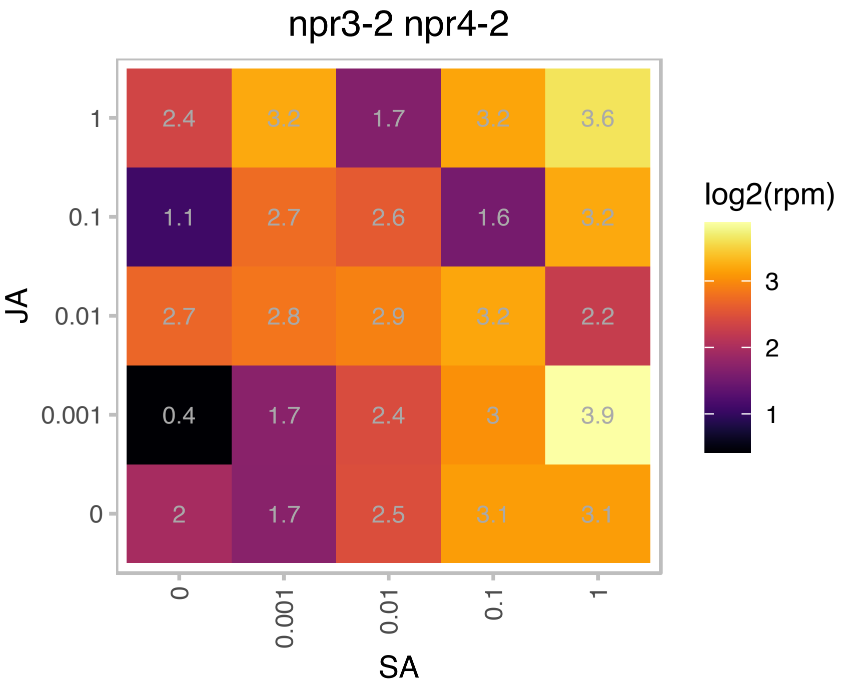


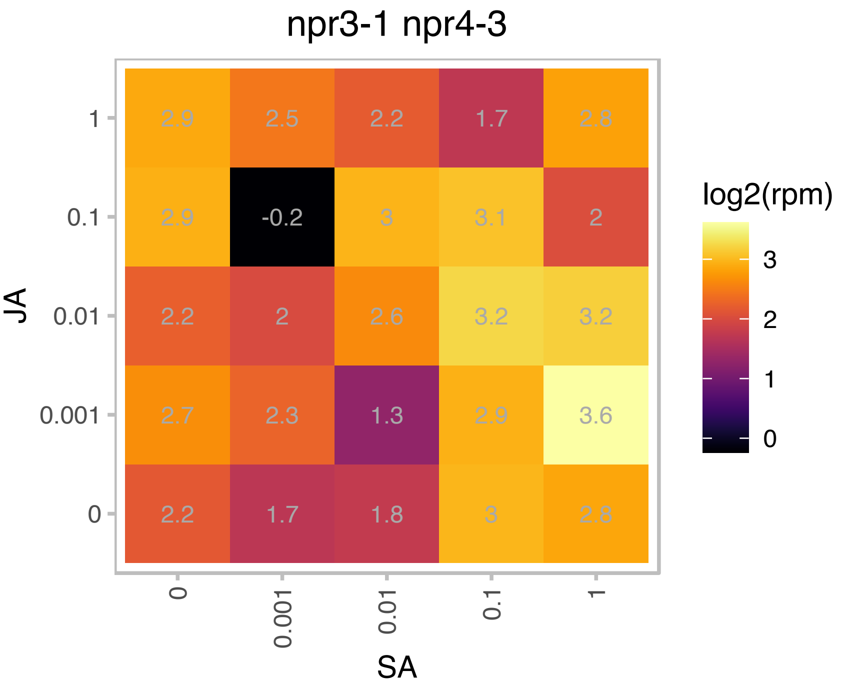

### AT3G60250.1 : CKB3

casein kinase II beta chain 3

Regulatory (beta) subunit of the protein kinase CK2. Involved in regulation of the circadian clock in Arabidopsis

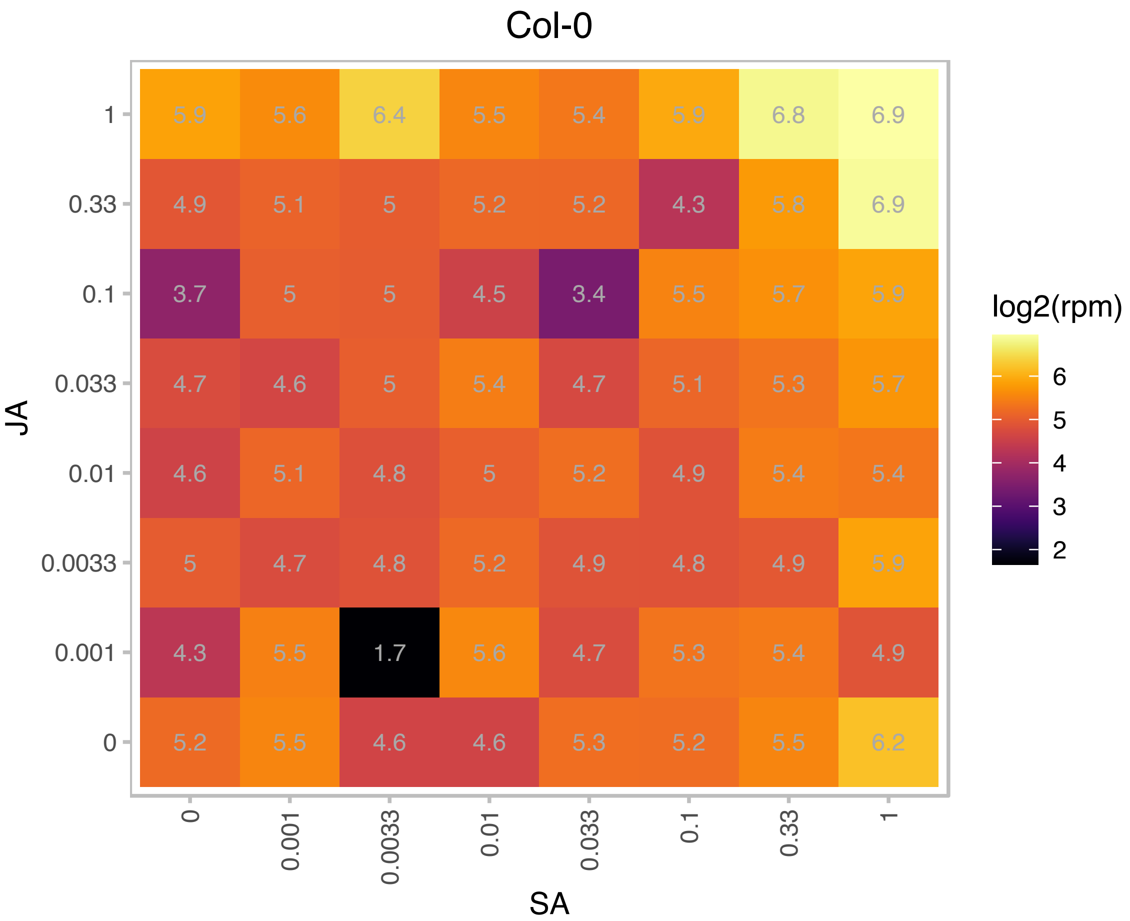


| | |
|---|---|
| Gene | AT3G60250 |
| is.expressed_genes | TRUE |
| is.responsive_genes | TRUE |
| is.apcluster | B-03 |

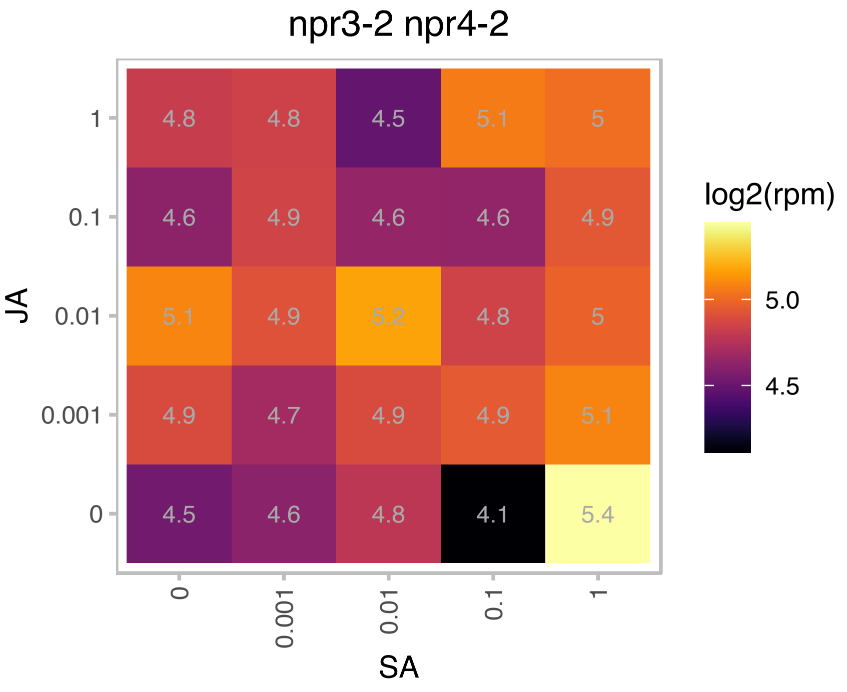


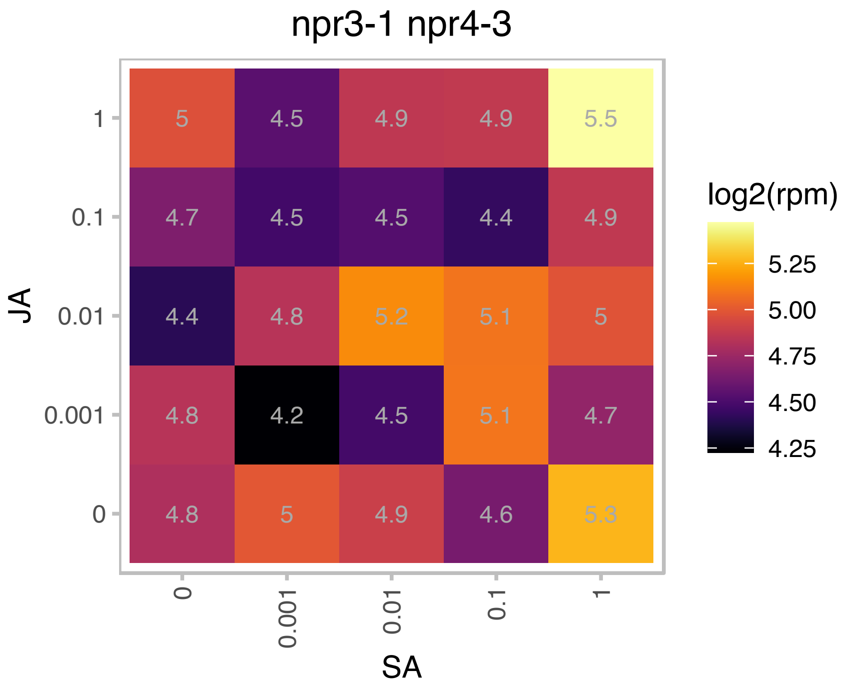

### AT3G61113.1 : URM12

Ubiquitin related modifier 1

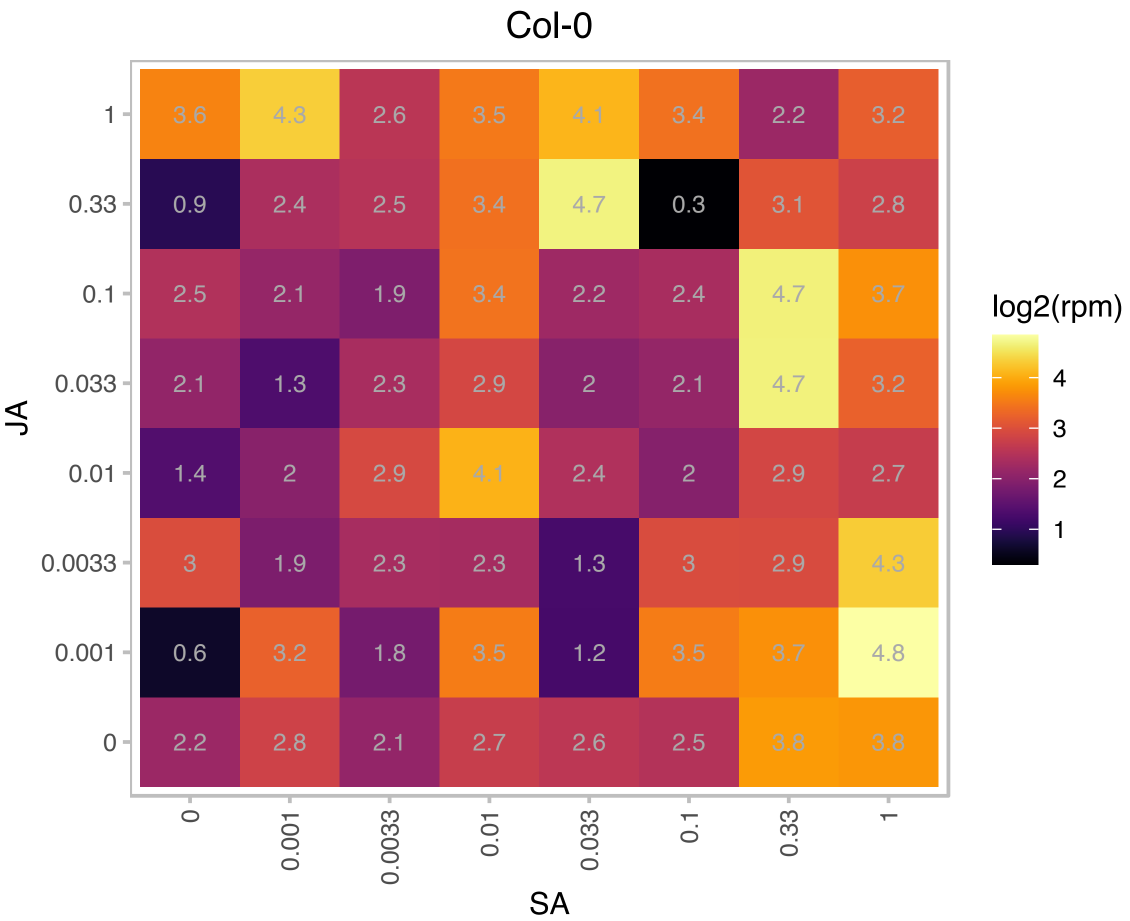


| | |
|---|---|
| Gene | AT3G61113 |
| is.expressed_genes | TRUE |
| is.responsive_genes | TRUE |
| is.apcluster | B-03 |

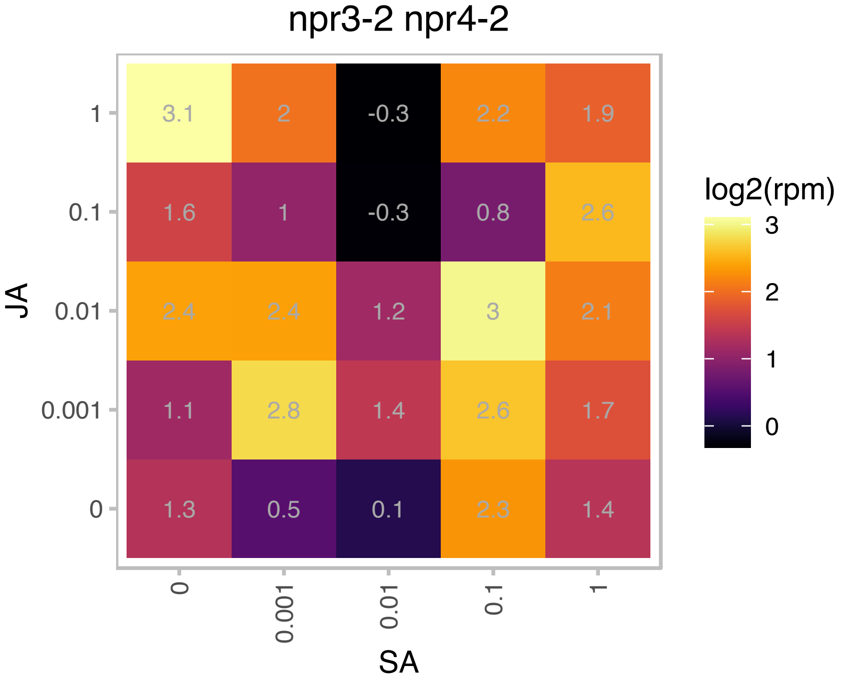


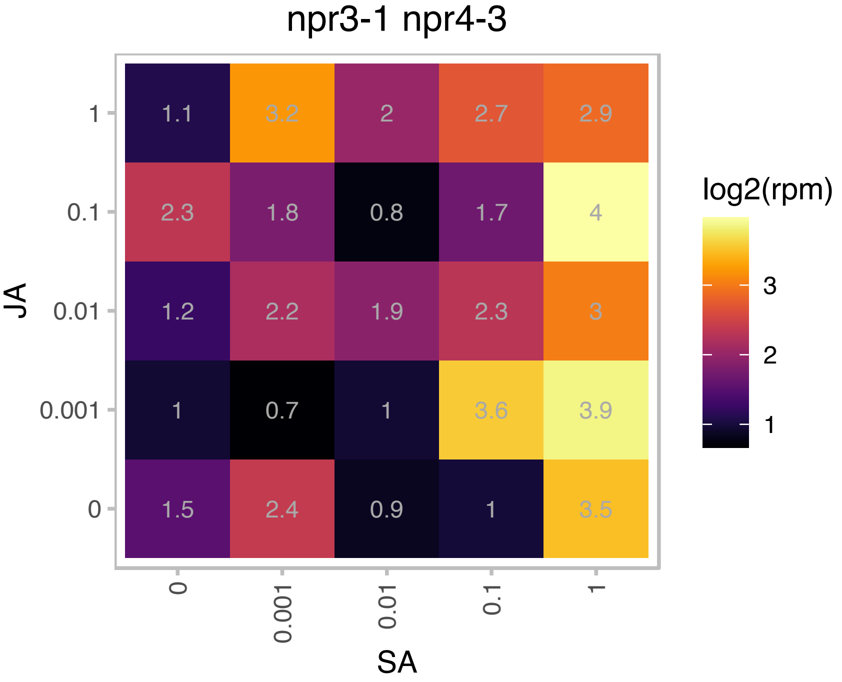

# AT4G11300.1 : NA

Protein of unknown function (DUF793)

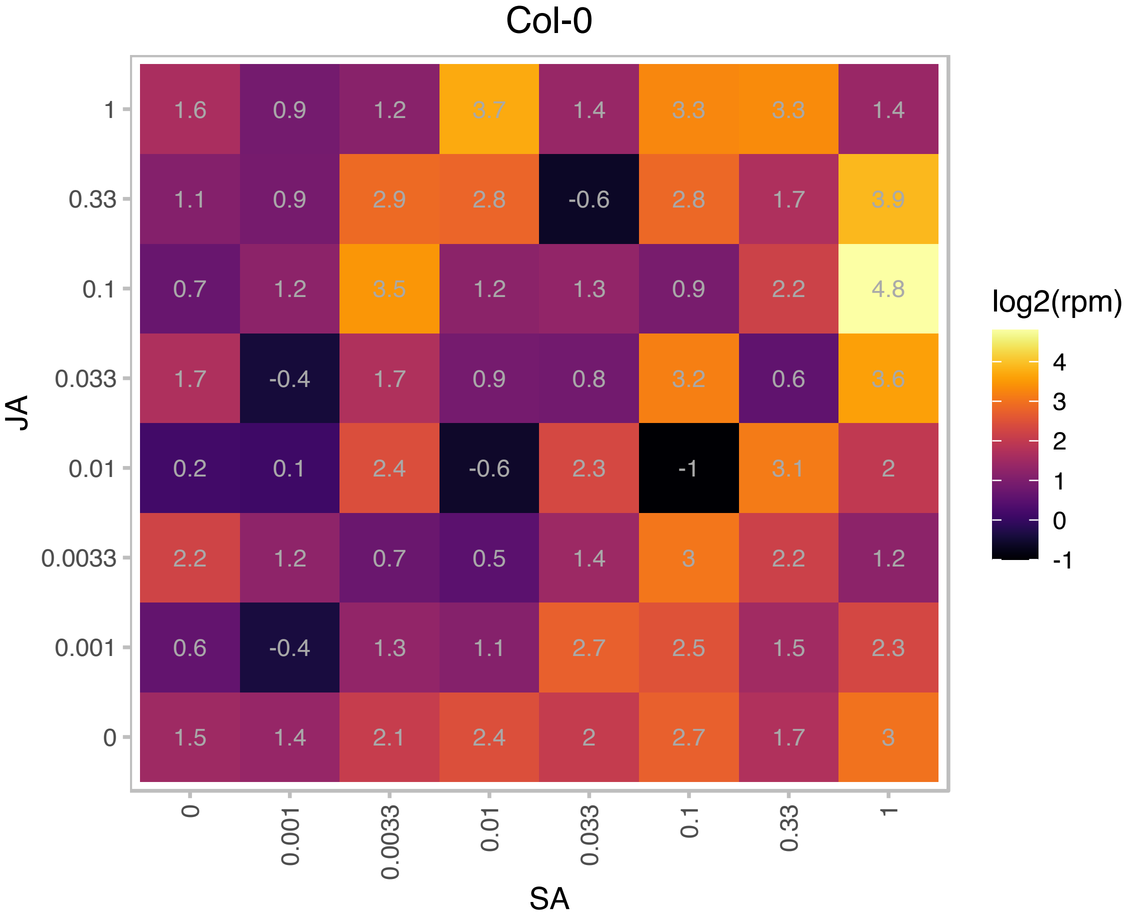


| | |
|---|---|
| Gene | AT4G11300 |
| is.expressed_genes | TRUE |
| is.responsive_genes | TRUE |
| is.apcluster | B-03 |

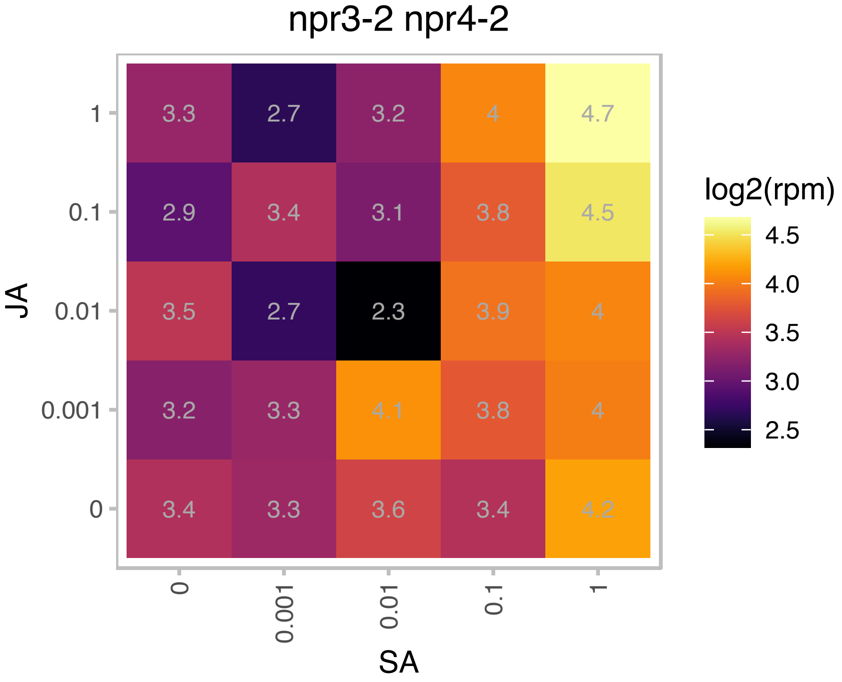


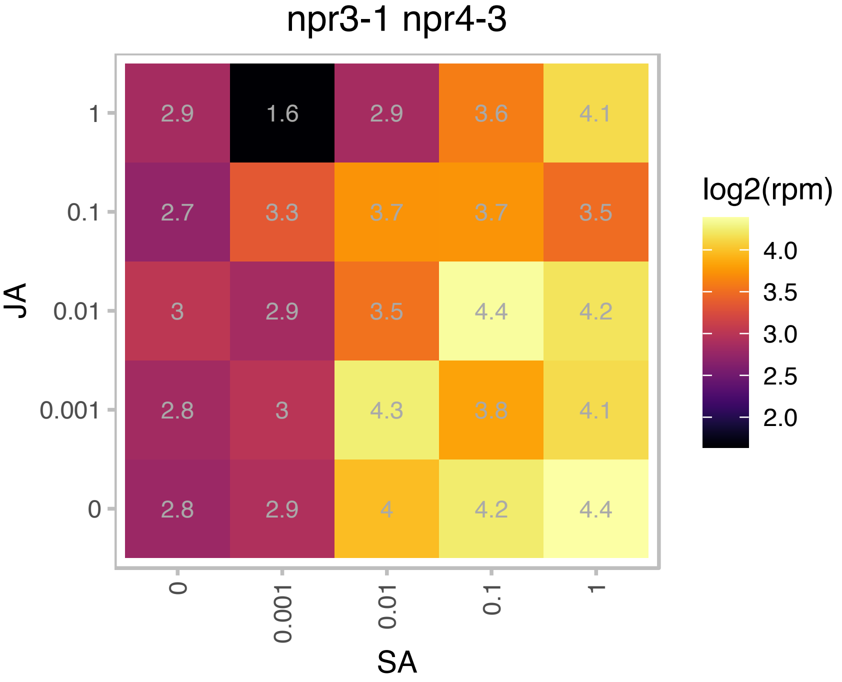

# AT4G14746.1 : NA

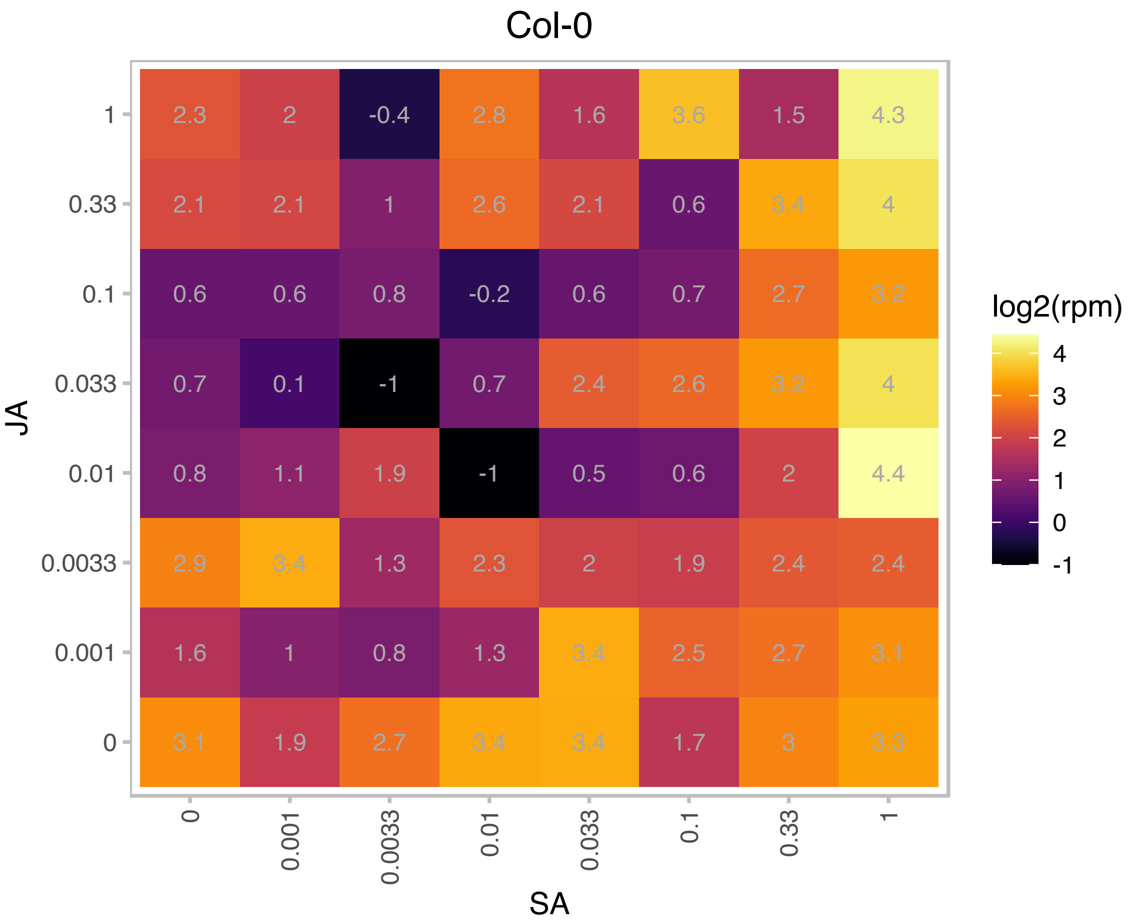


| | |
|---|---|
| Gene | AT4G14746 |
| is.expressed_genes | TRUE |
| is.responsive_genes | TRUE |
| is.apcluster | B-03 |

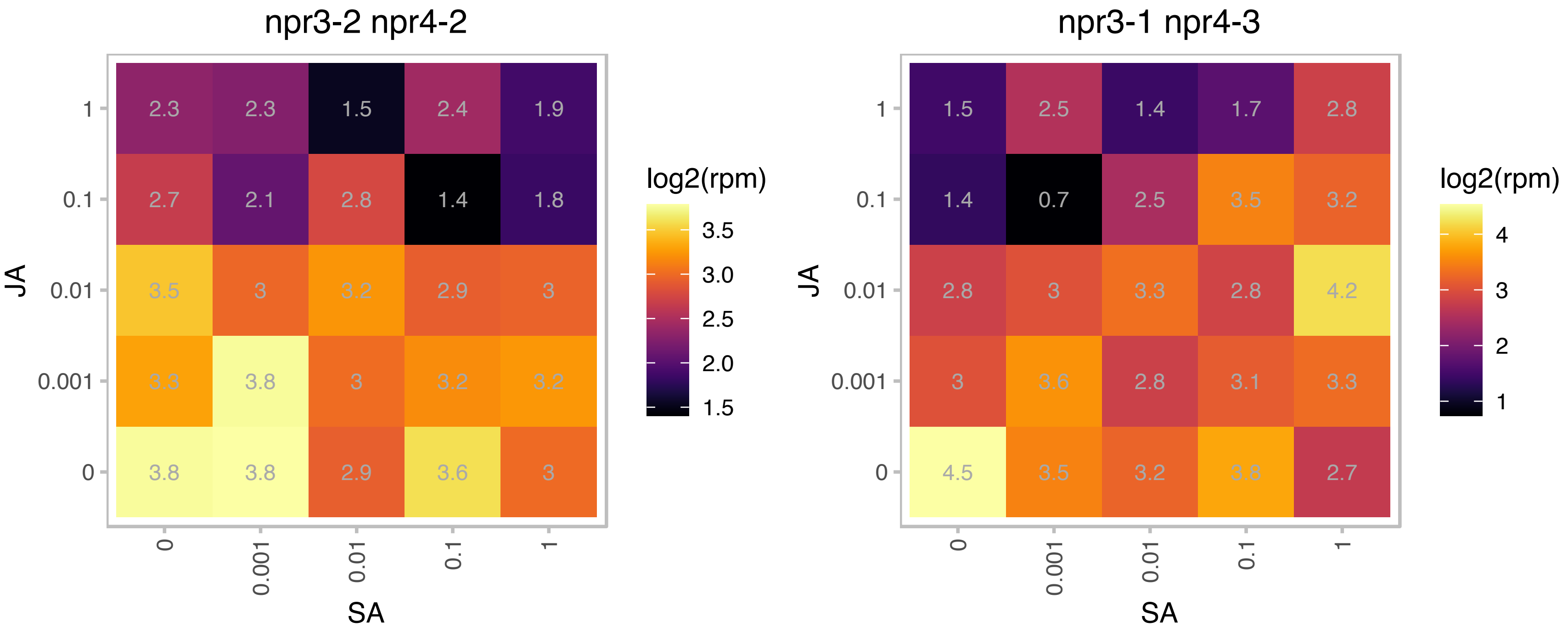

### AT4G17615.1 : ATCBL1, CBL1, SCABP5

calcineurin B-like protein 1

Member of AtCBL (Calcineurin B-like Calcium Sensor Proteins) family. Protein level is increased upon high salt, mannitol, and cold stresses. CBL1 interacts with CIPK23 and recruits the kinase to the plasma membrane where the substrate(s) of CIPK23 may reside. CBL1 localization is regulated by protein modification including myristolation and acylation.

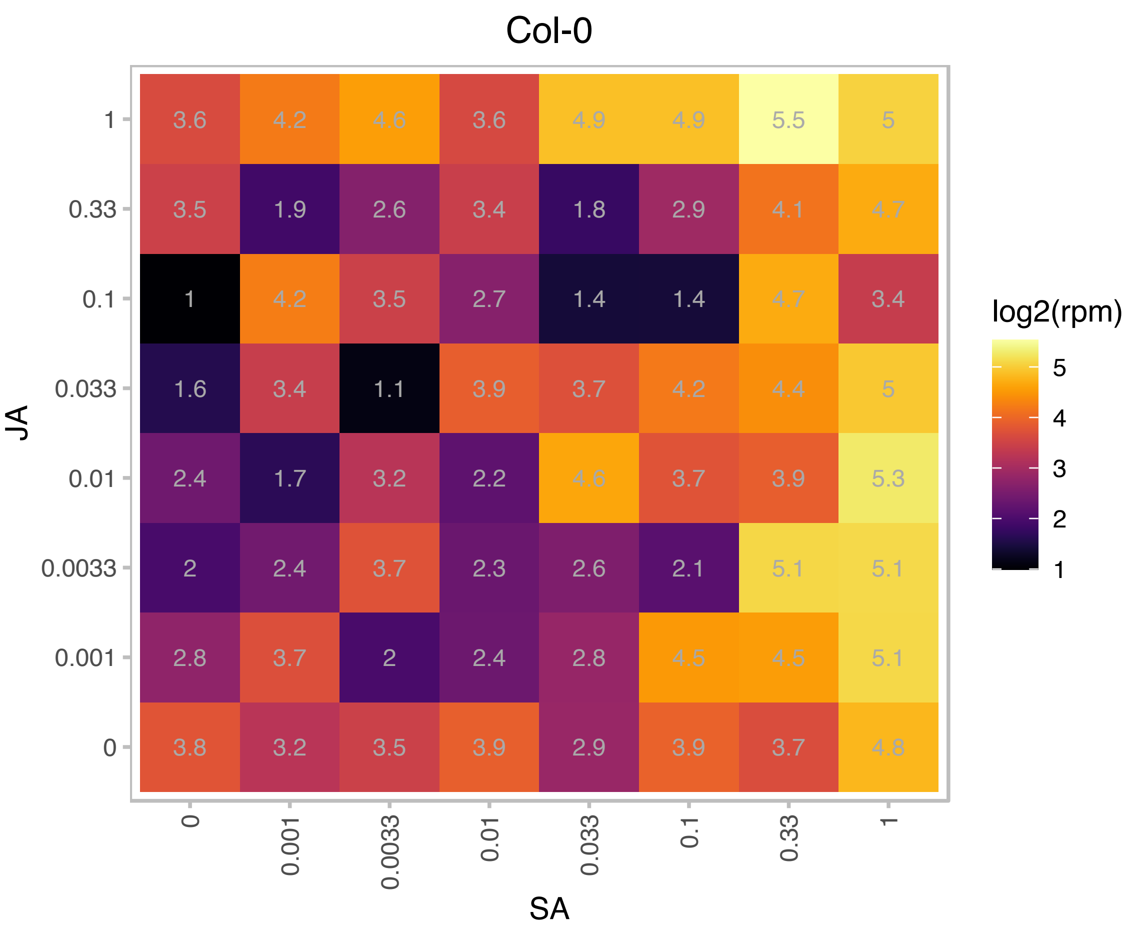


| | |
|---|---|
| Gene | AT4G17615 |
| is.expressed_genes | TRUE |
| is.responsive_genes | TRUE |
| is.apcluster | B-03 |

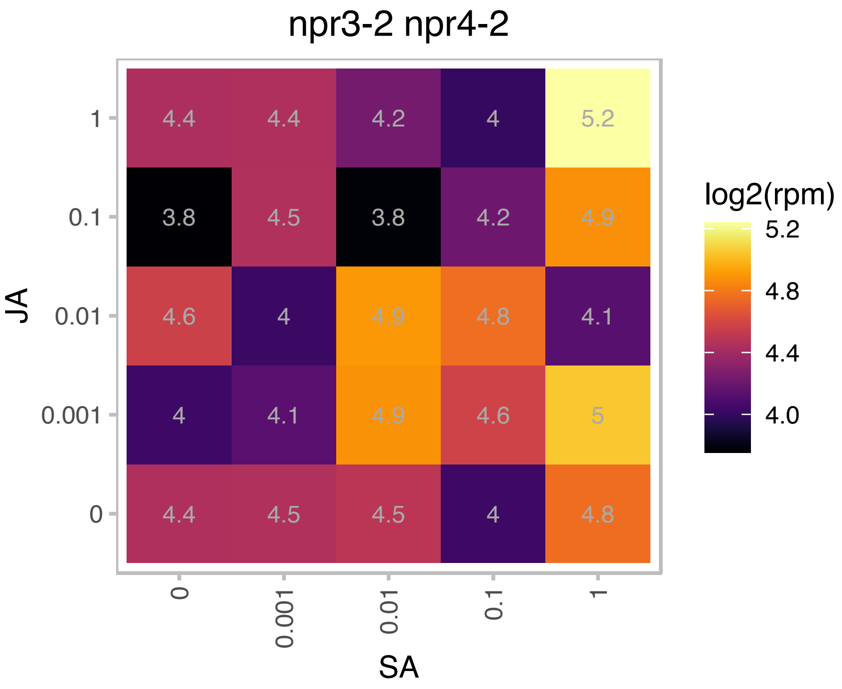


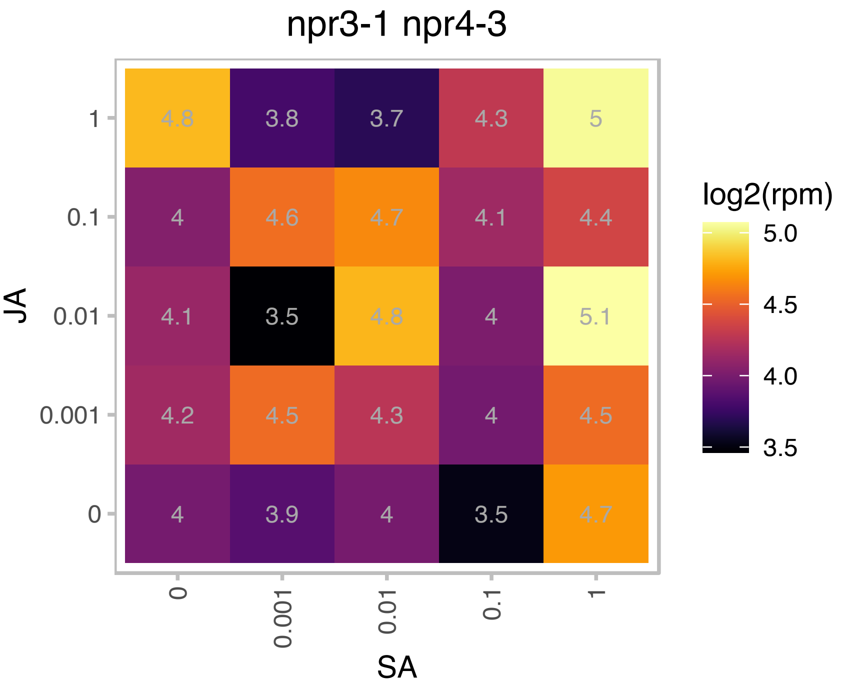

### AT4G23450.1 : AIRP1, AtAIRP1

RING/U-box superfamily protein

AtAIRP1 gene encodes a C3H2C3-type RING E3 Ub ligase. It has been shown to be a positive regulator in the Arabidopsis ABA-dependent drought response.

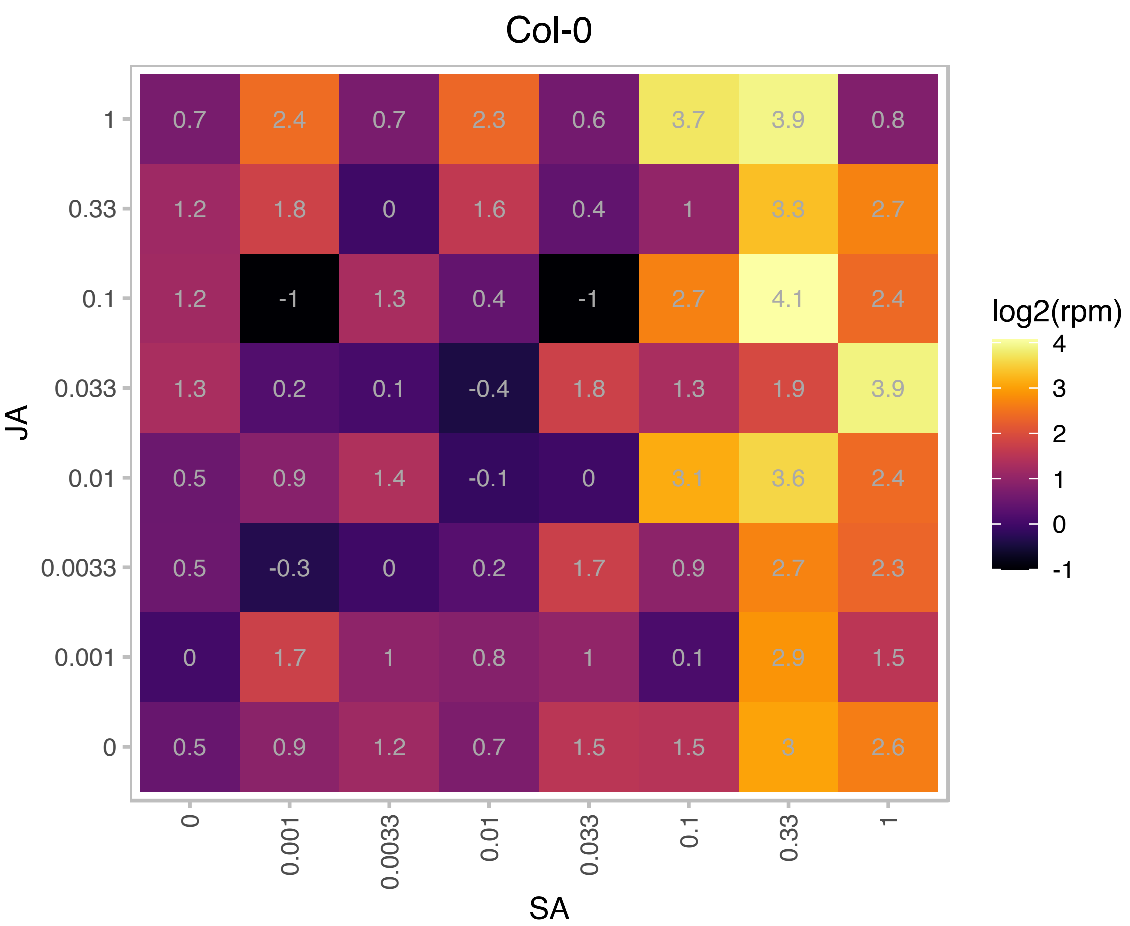


| JA \ SA | 0 | 0.001 | 0.0033 | 0.01 | 0.033 | 0.1 | 0.33 | 1 |
|---|---|---|---|---|---|---|---|---|
| 1 | 0.7 | 2.4 | 0.7 | 2.3 | 0.6 | 3.7 | 3.9 | 0.8 |
| 0.33 | 1.2 | 1.8 | 0 | 1.6 | 0.4 | 1 | 3.3 | 2.7 |
| 0.1 | 1.2 | -1 | 1.3 | 0.4 | -1 | 2.7 | 4.1 | 2.4 |
| 0.033 | 1.3 | 0.2 | 0.1 | -0.4 | 1.8 | 1.3 | 1.9 | 3.9 |
| 0.01 | 0.5 | 0.9 | 1.4 | -0.1 | 0 | 3.1 | 3.6 | 2.4 |
| 0.0033 | 0.5 | -0.3 | 0 | 0.2 | 1.7 | 0.9 | 2.7 | 2.3 |
| 0.001 | 0 | 1.7 | 1 | 0.8 | 1 | 0.1 | 2.9 | 1.5 |
| 0 | 0.5 | 0.9 | 1.2 | 0.7 | 1.5 | 1.5 | 3 | 2.6 |

| | |
|---|---|
| Gene | AT4G23450 |
| is.expressed_genes | TRUE |
| is.responsive_genes | TRUE |
| is.apcluster | B-03 |

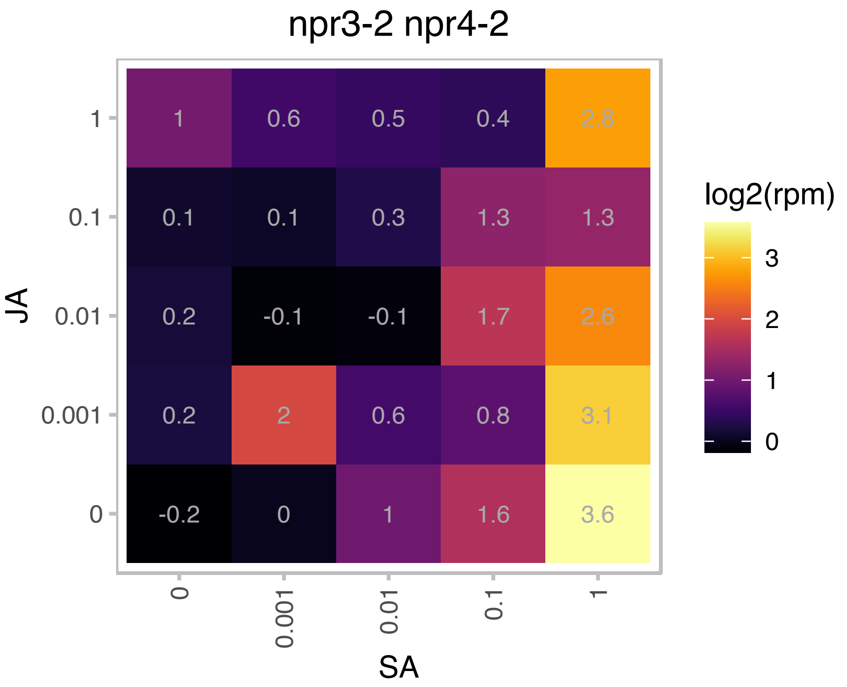


| JA \ SA | 0 | 0.001 | 0.01 | 0.1 | 1 |
|---|---|---|---|---|---|
| 1 | 1 | 0.6 | 0.5 | 0.4 | 2.8 |
| 0.1 | 0.1 | 0.1 | 0.3 | 1.3 | 1.3 |
| 0.01 | 0.2 | -0.1 | -0.1 | 1.7 | 2.6 |
| 0.001 | 0.2 | 2 | 0.6 | 0.8 | 3.1 |
| 0 | -0.2 | 0 | 1 | 1.6 | 3.6 |

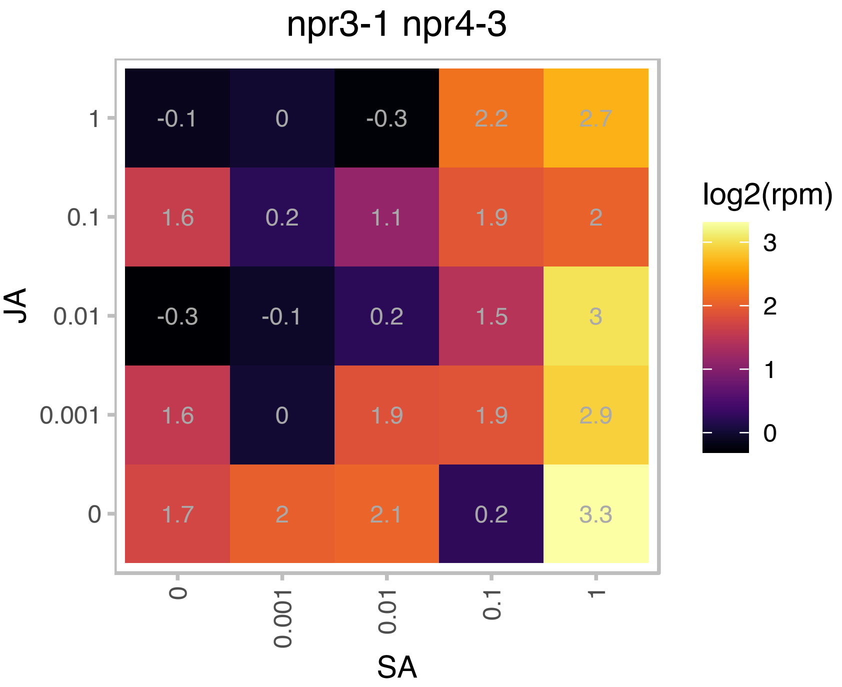


| JA \ SA | 0 | 0.001 | 0.01 | 0.1 | 1 |
|---|---|---|---|---|---|
| 1 | -0.1 | 0 | -0.3 | 2.2 | 2.7 |
| 0.1 | 1.6 | 0.2 | 1.1 | 1.9 | 2 |
| 0.01 | -0.3 | -0.1 | 0.2 | 1.5 | 3 |
| 0.001 | 1.6 | 0 | 1.9 | 1.9 | 2.9 |
| 0 | 1.7 | 2 | 2.1 | 0.2 | 3.3 |

### AT4G33040.1 : ROXY21

Thioredoxin superfamily protein

Encodes a member of the CC-type glutaredoxin (ROXY) family that has been shown to interact with the transcription factor TGA2.

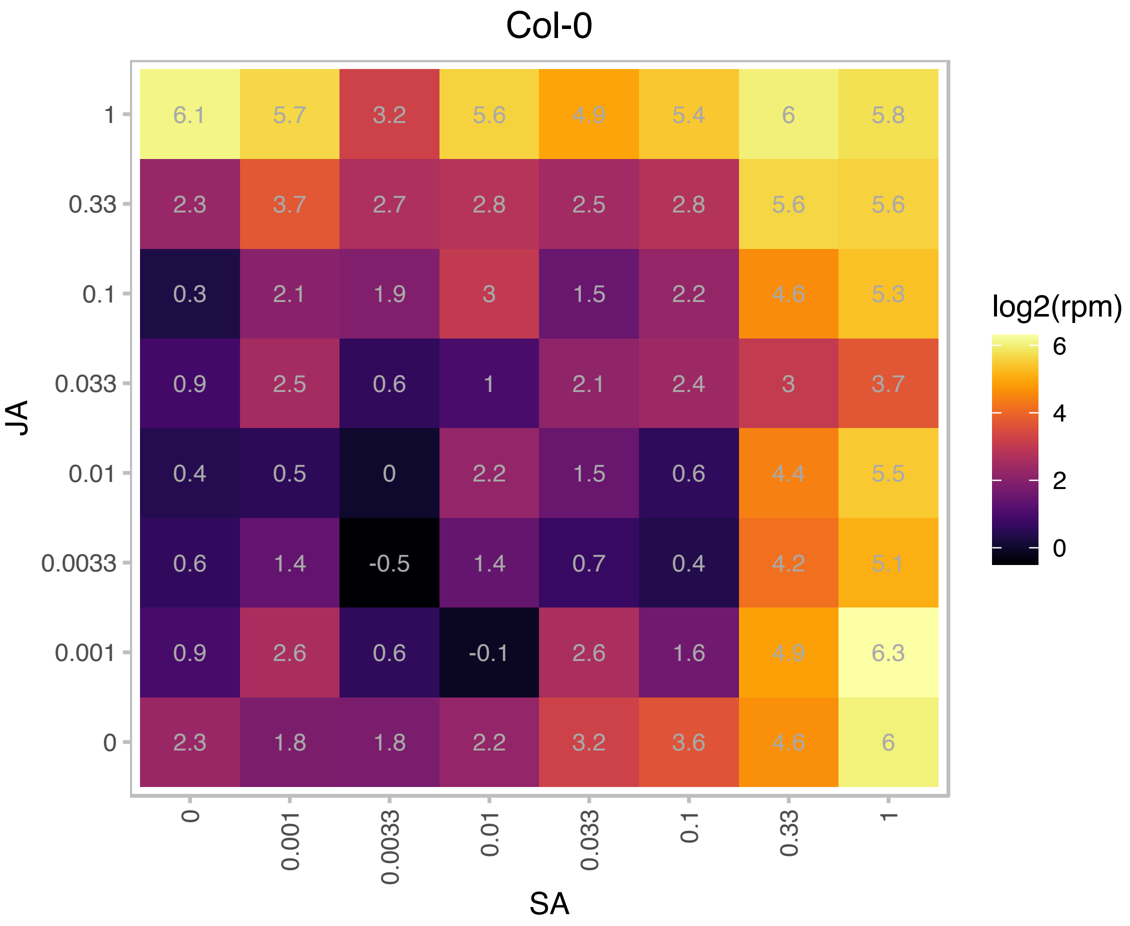


| | |
|---|---|
| Gene | AT4G33040 |
| is.expressed_genes | TRUE |
| is.responsive_genes | TRUE |
| is.apcluster | B-03 |

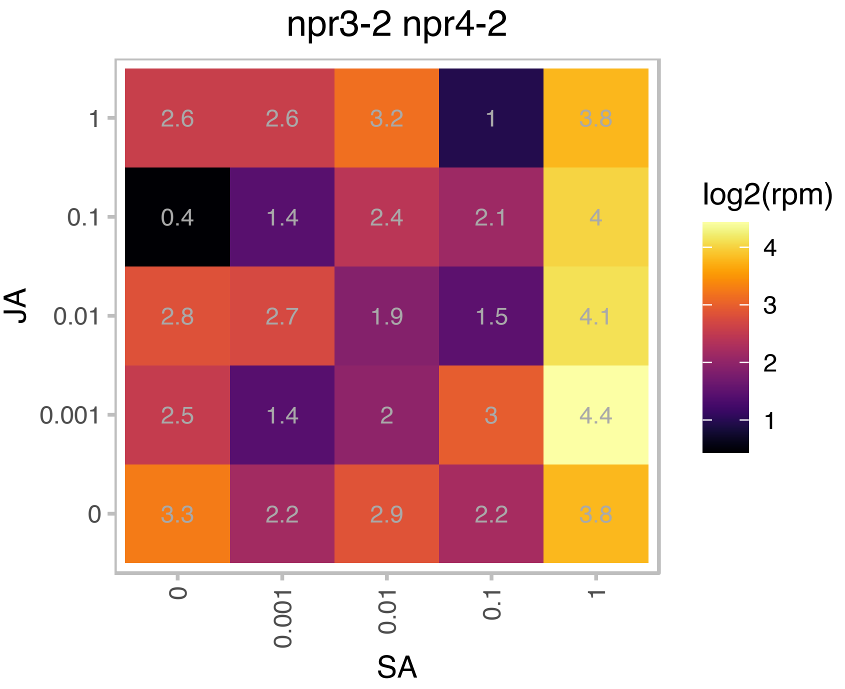


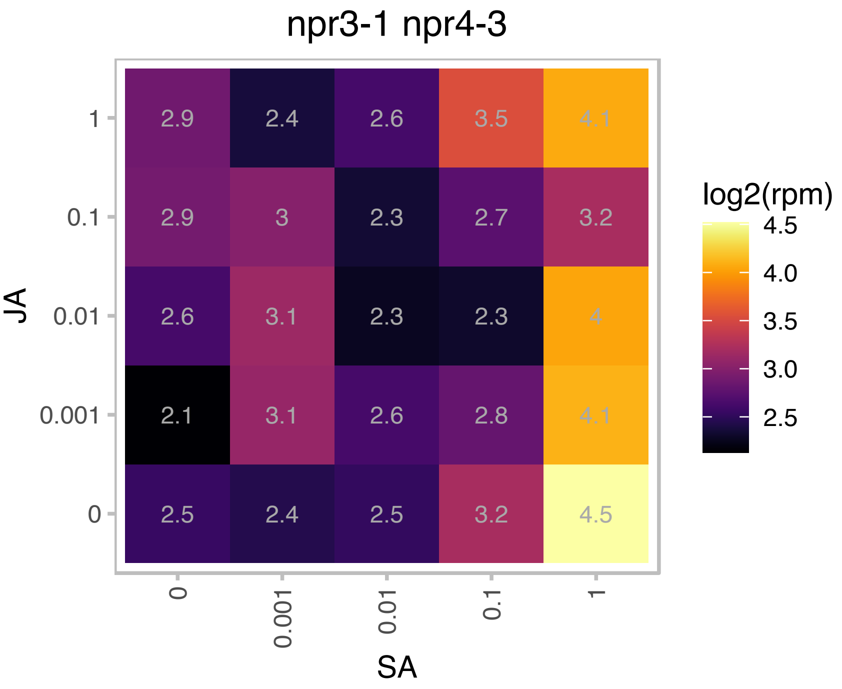

### AT4G37370.1 : CYP81D8

cytochrome P450, family 81, subfamily D, polypeptide 8

member of CYP81D

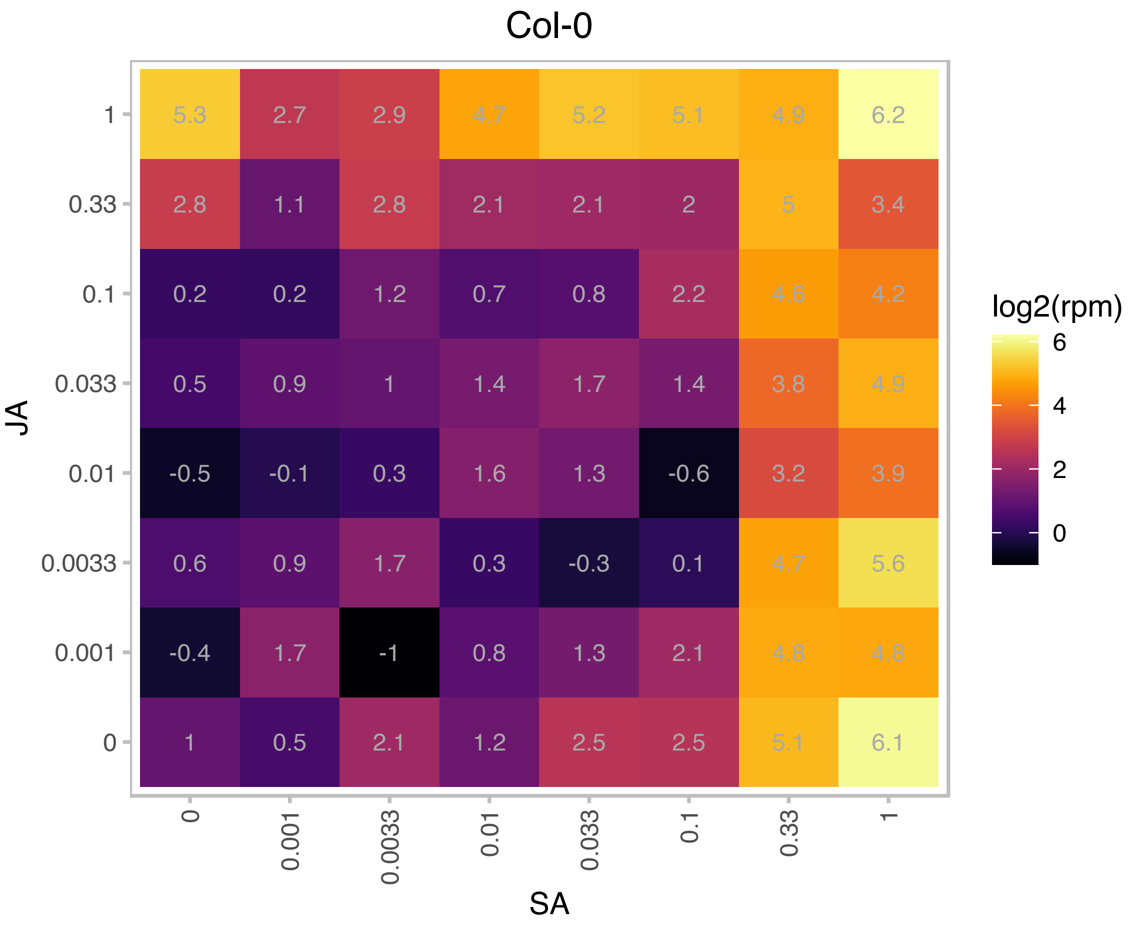


| | |
|---|---|
| Gene | AT4G37370 |
| is.expressed_genes | TRUE |
| is.responsive_genes | TRUE |
| is.apcluster | B-03 |

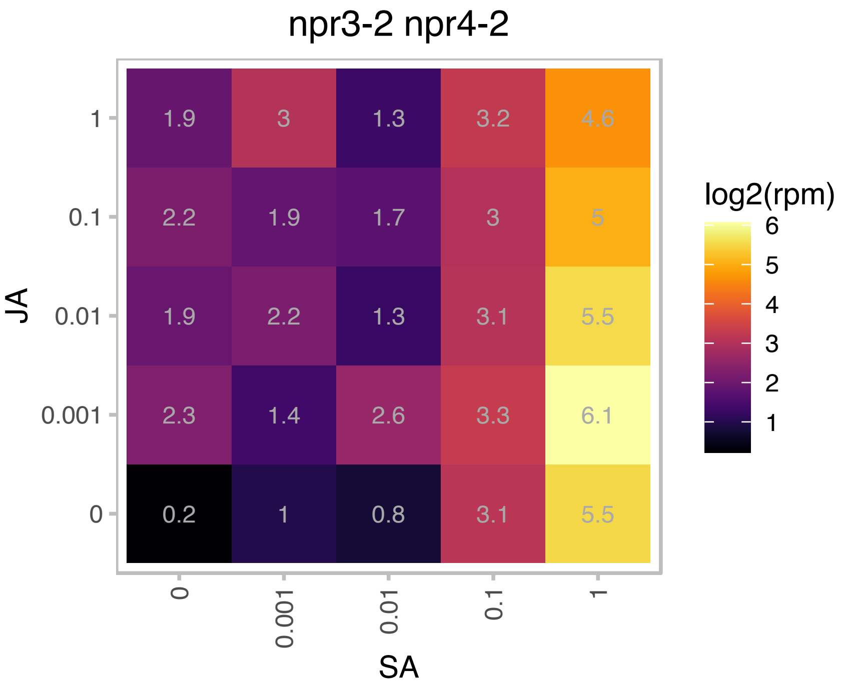


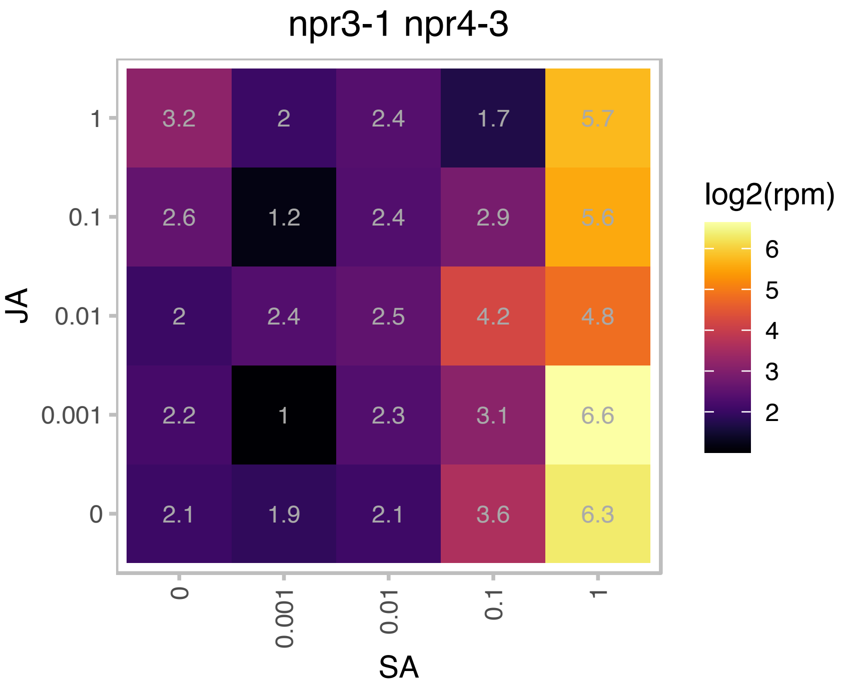

# AT5G03030.1 : NA

Chaperone DnaJ-domain superfamily protein

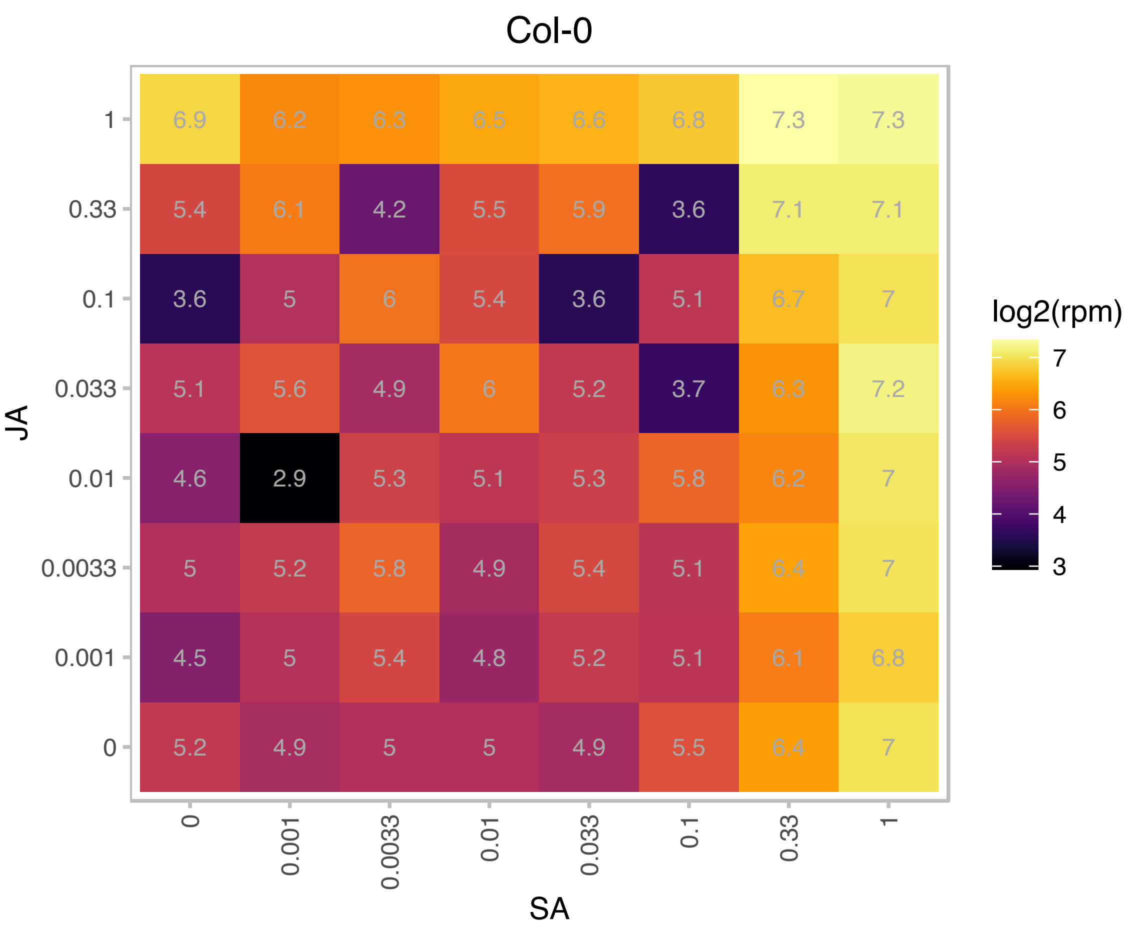


| | |
|---|---|
| Gene | AT5G03030 |
| is.expressed_genes | TRUE |
| is.responsive_genes | TRUE |
| is.apcluster | B-03 |

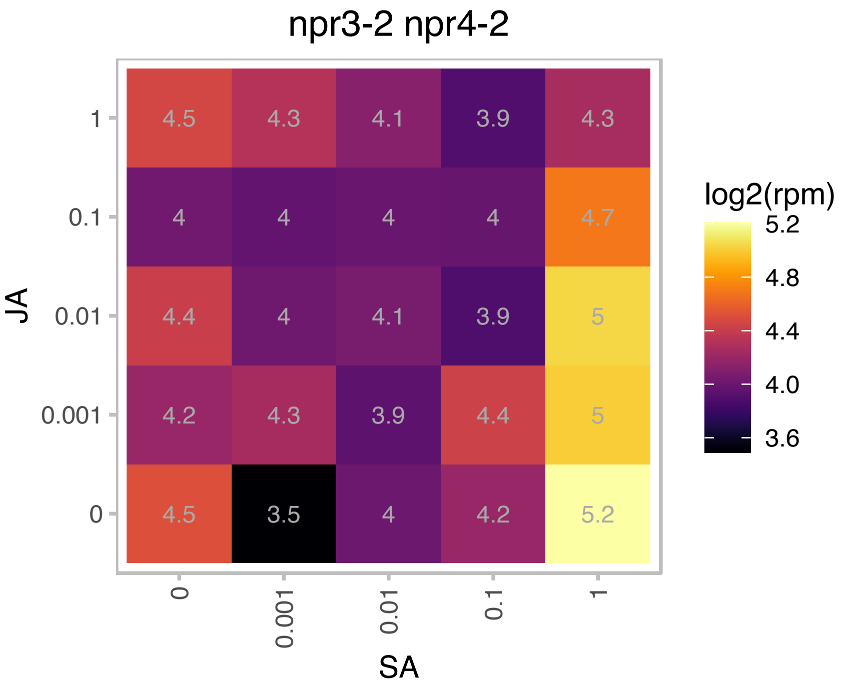


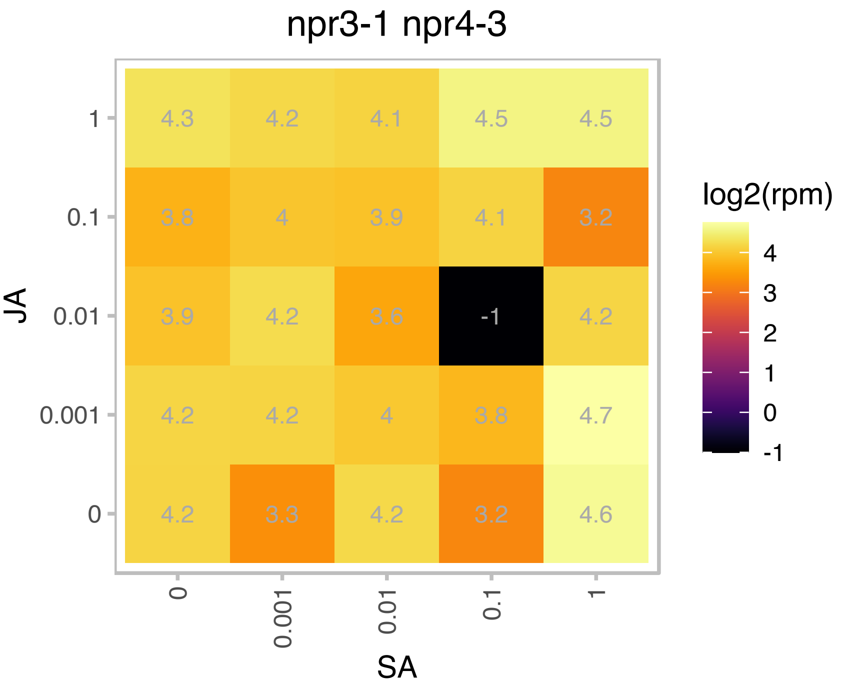

### AT5G03240.1 : UBQ3

polyubiquitin 3

encodes ubiquitin that is attached to proteins destined for degradation. UBQ3 is most homologous with UBQ4, and is expressed in higher levels in vegetative tissue but lower levels in flowers than UBQ4. UBQ3 encodes different number of ubiquitins in different ecotypes. UBQ3 transcript level is modulated by UV-B and light/dark treatments.

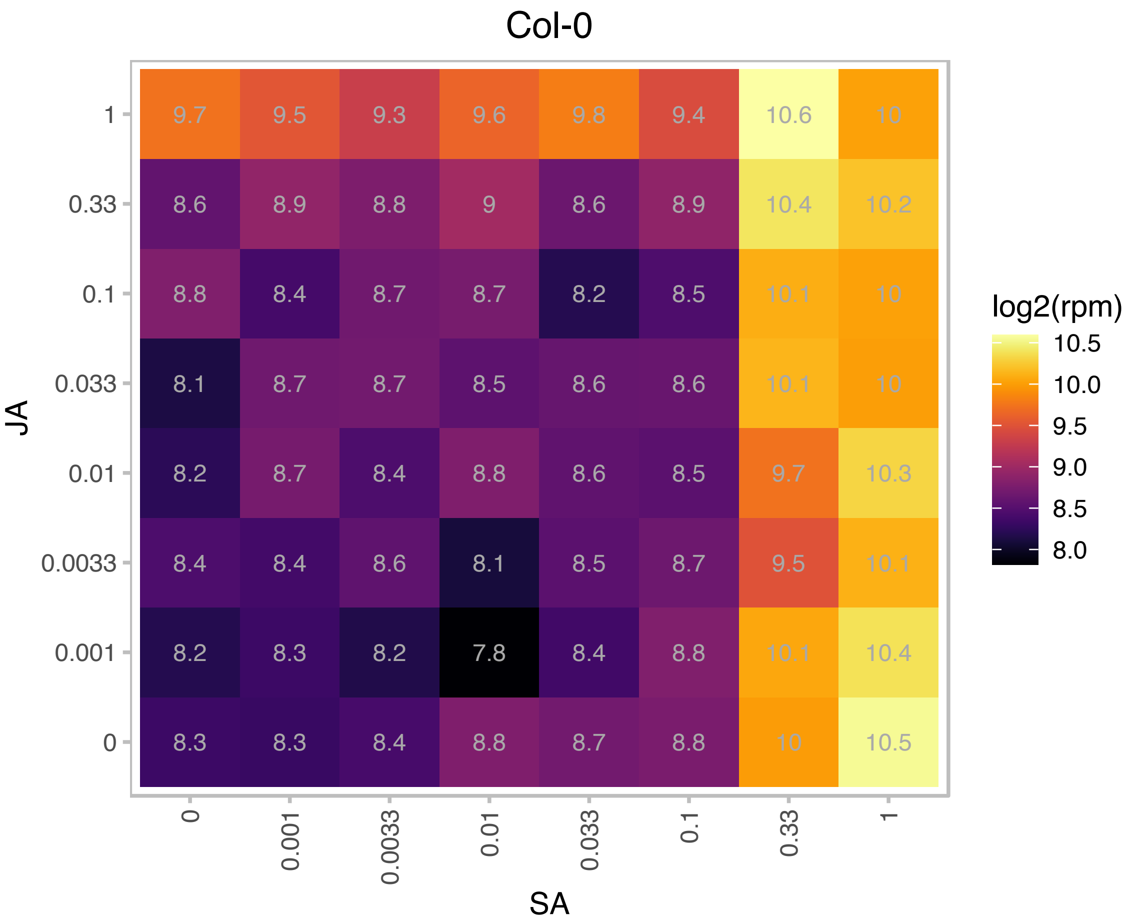


| | |
|---|---|
| Gene | AT5G03240 |
| is.expressed_genes | TRUE |
| is.responsive_genes | TRUE |
| is.apcluster | B-03 |

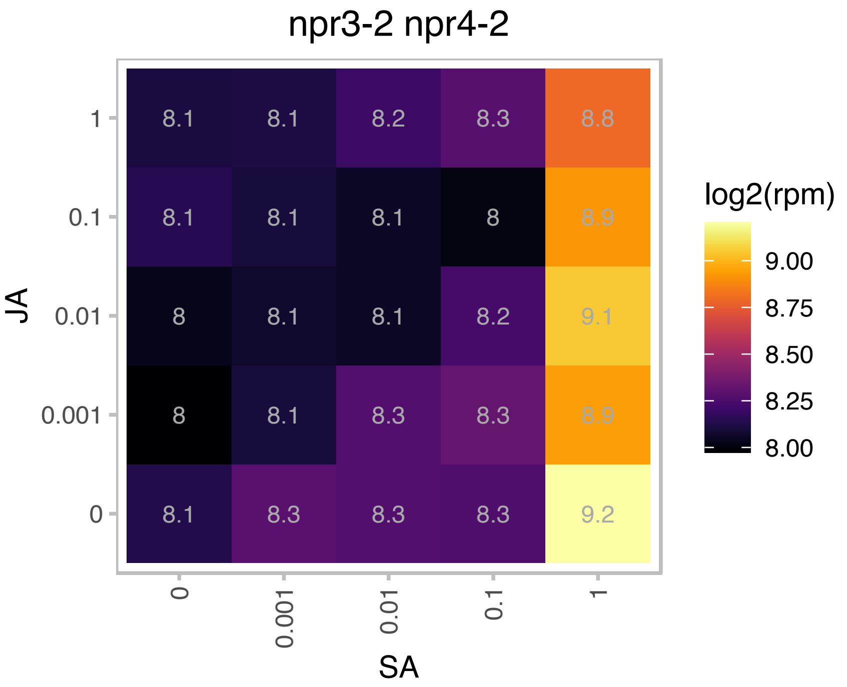


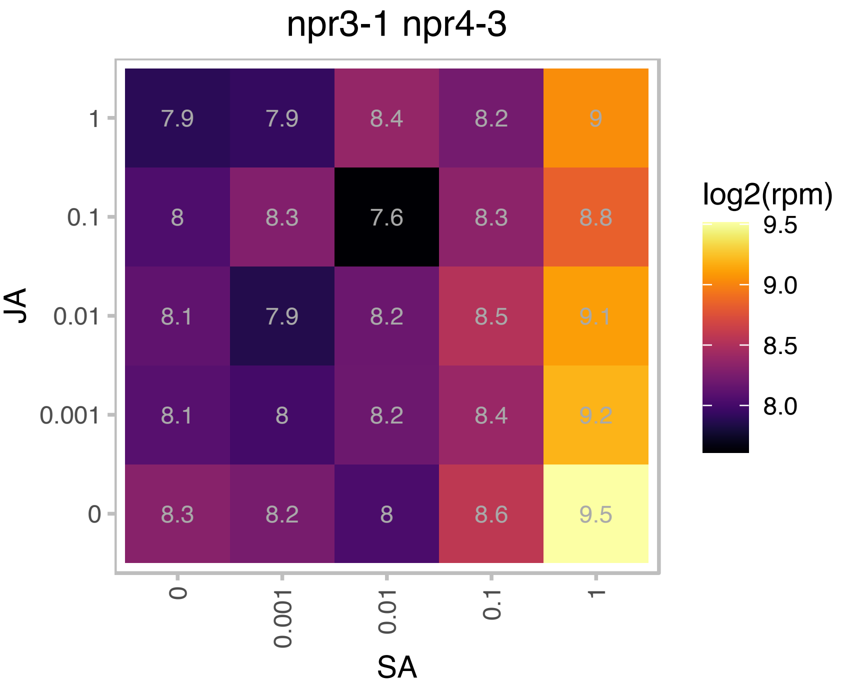

# AT5G07870.1 : NA

HXXXD-type acyl-transferase family protein

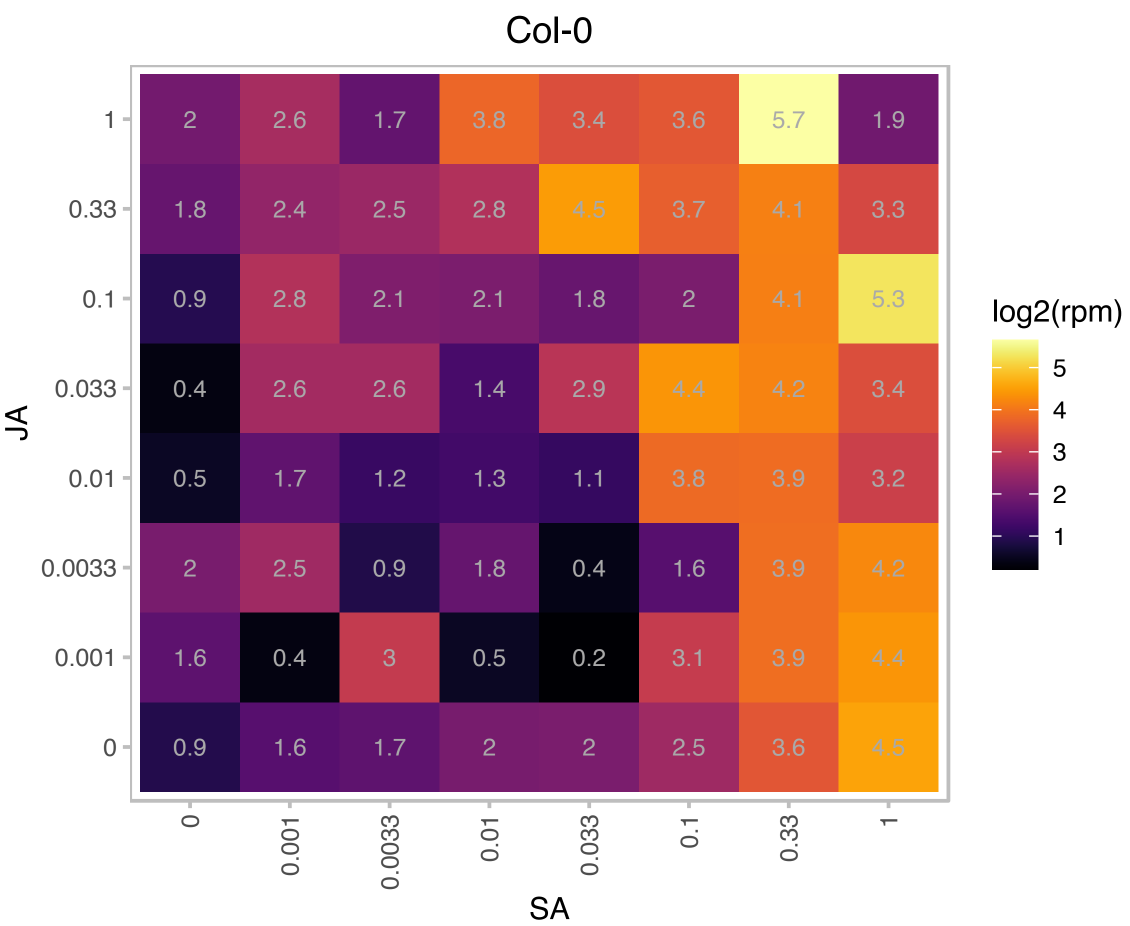


| | |
|---|---|
| Gene | AT5G07870 |
| is.expressed_genes | TRUE |
| is.responsive_genes | TRUE |
| is.apcluster | B-03 |

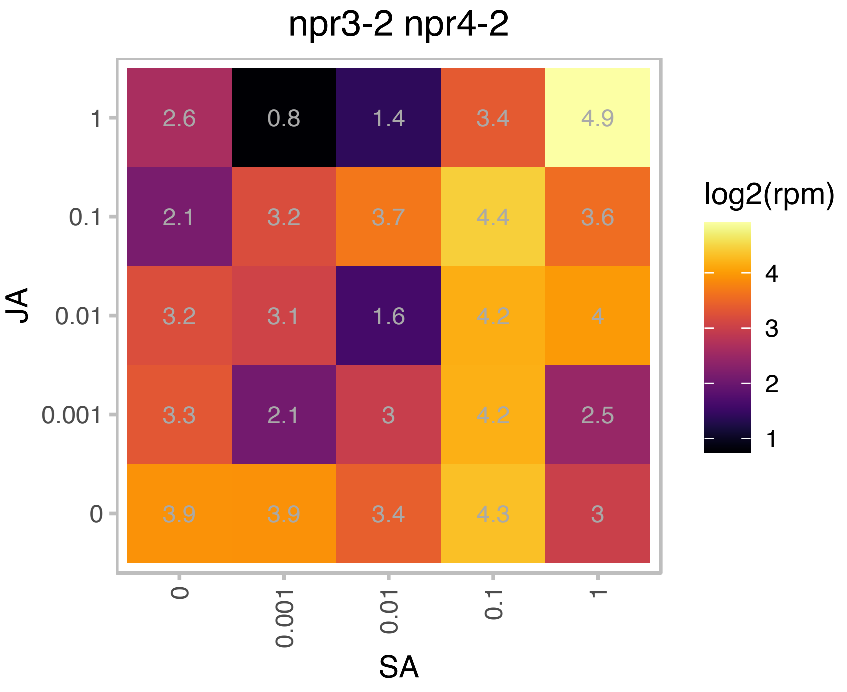


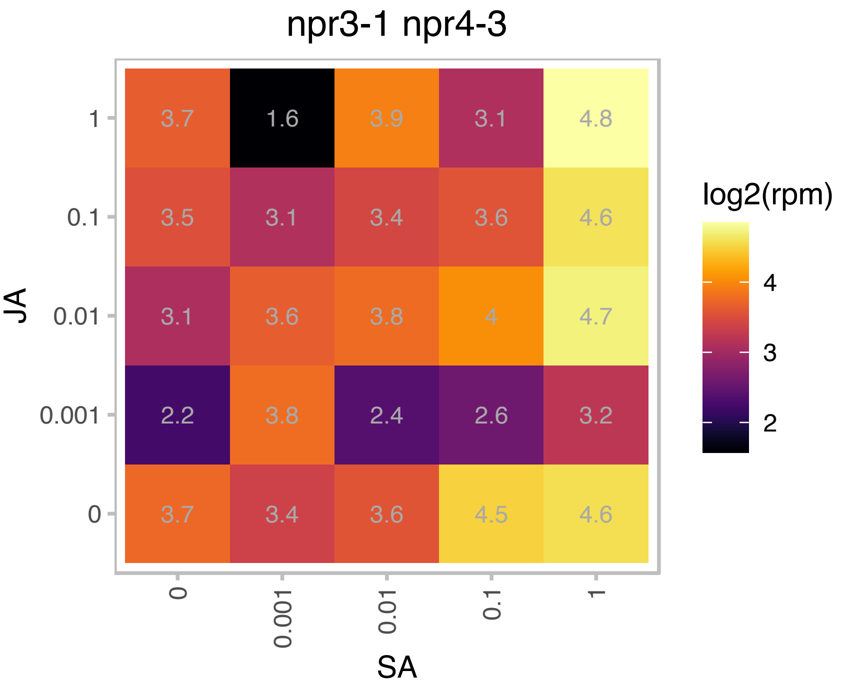

# AT5G10695.1 : NA

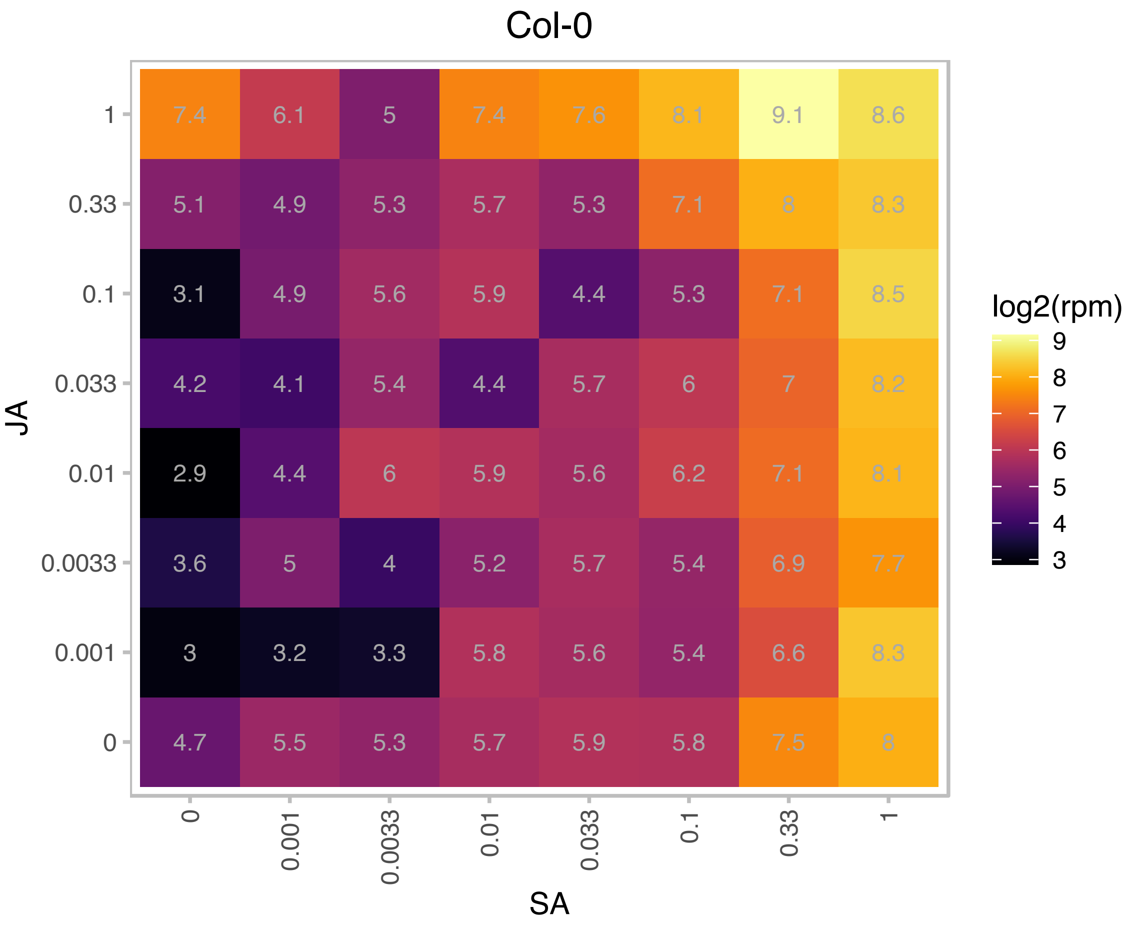


| | |
|---|---|
| Gene | AT5G10695 |
| is.expressed_genes | TRUE |
| is.responsive_genes | TRUE |
| is.apcluster | B-03 |

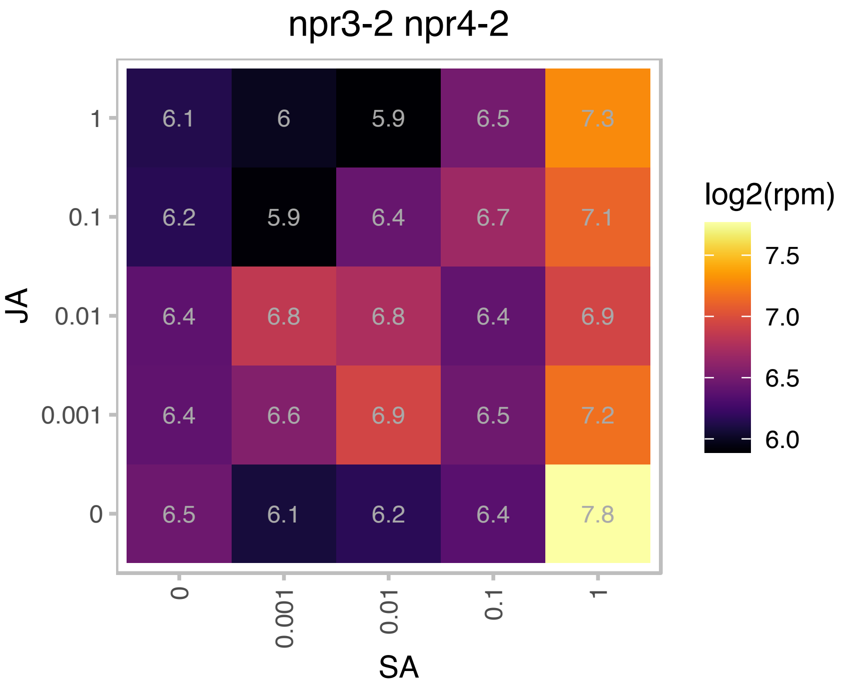


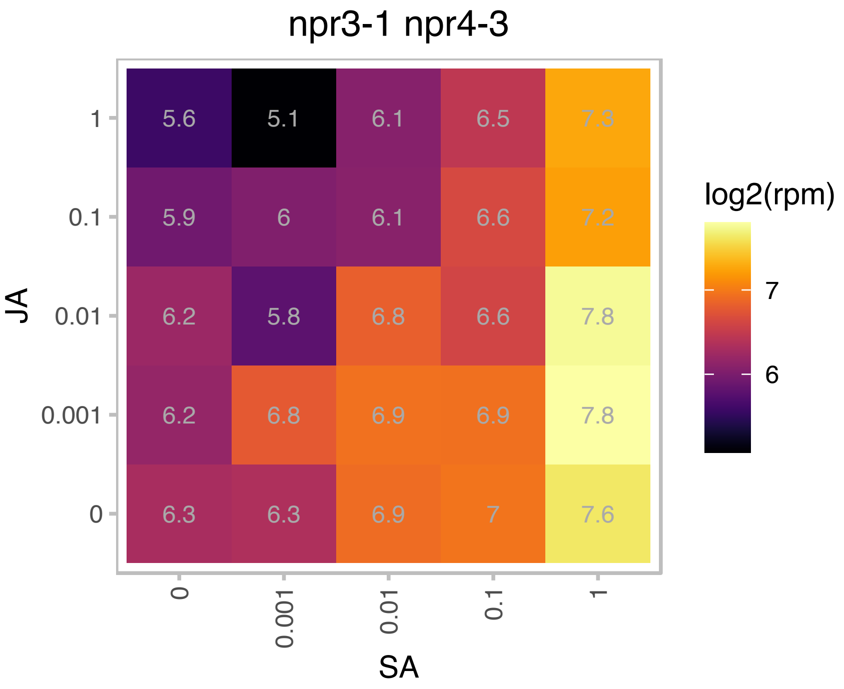

### AT5G18270.1 : ANAC087

Arabidopsis NAC domain containing protein 87

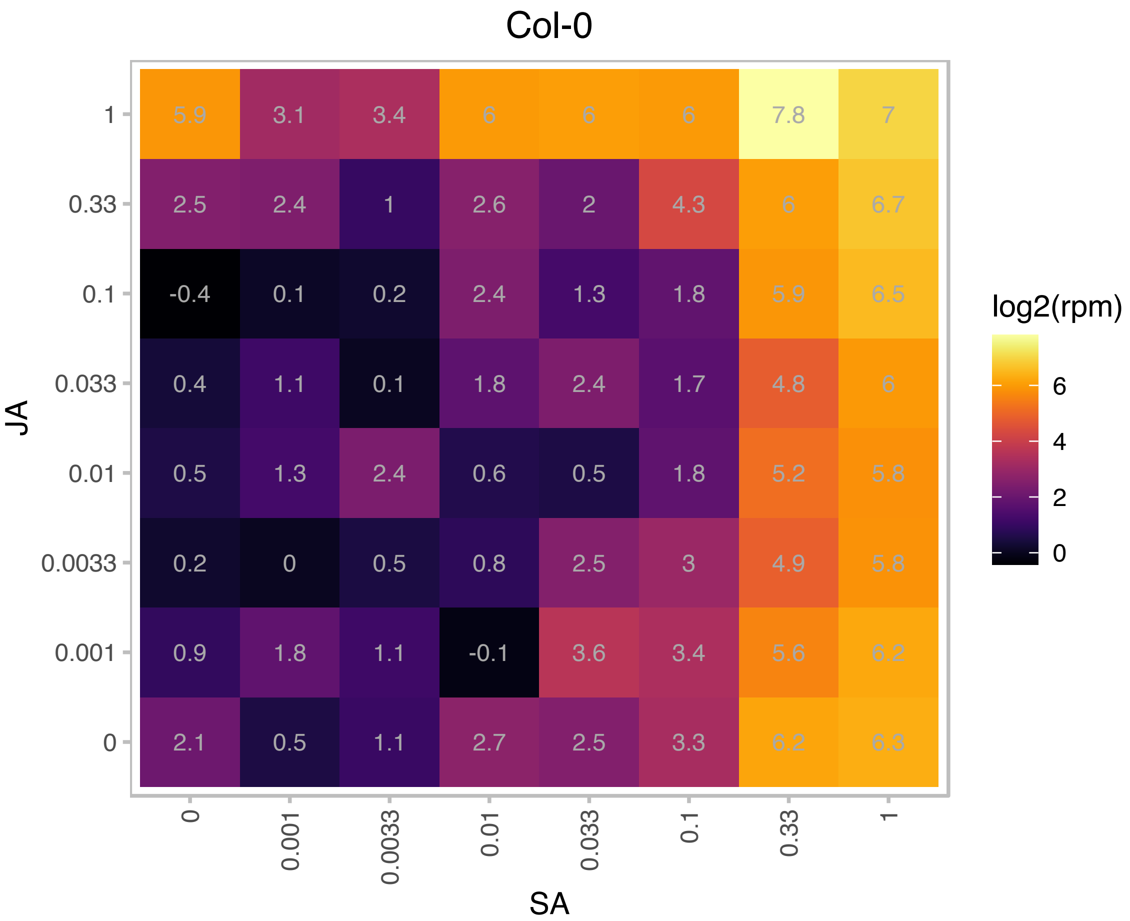


| | |
|---|---|
| Gene | AT5G18270 |
| is.expressed_genes | TRUE |
| is.responsive_genes | TRUE |
| is.apcluster | B-03 |

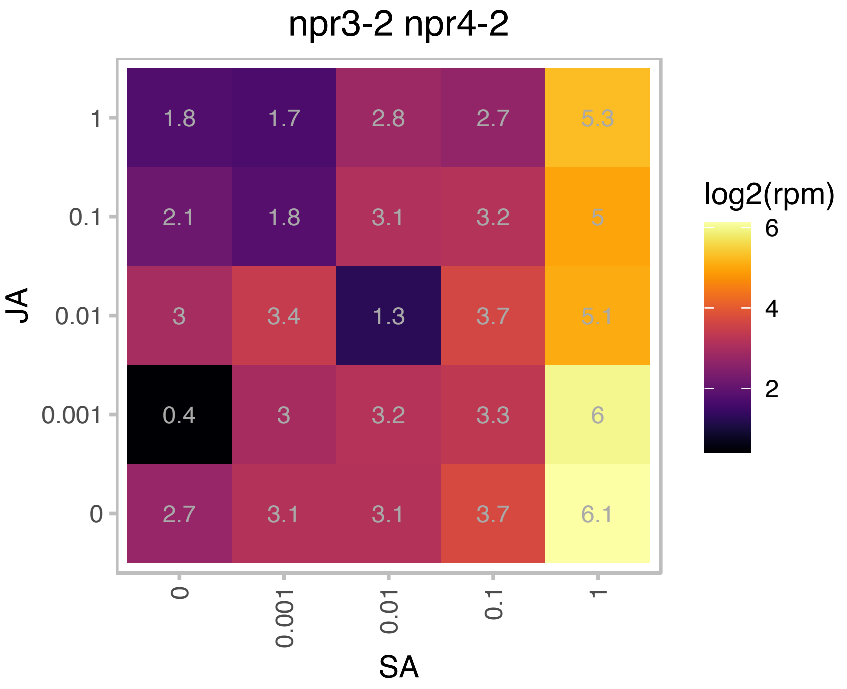


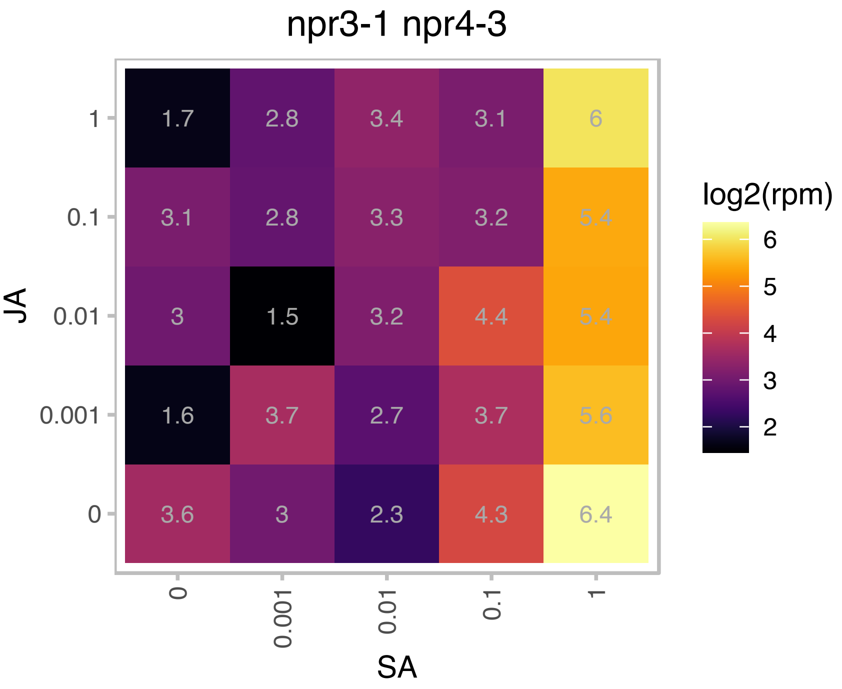

# AT5G24570.1 : NA

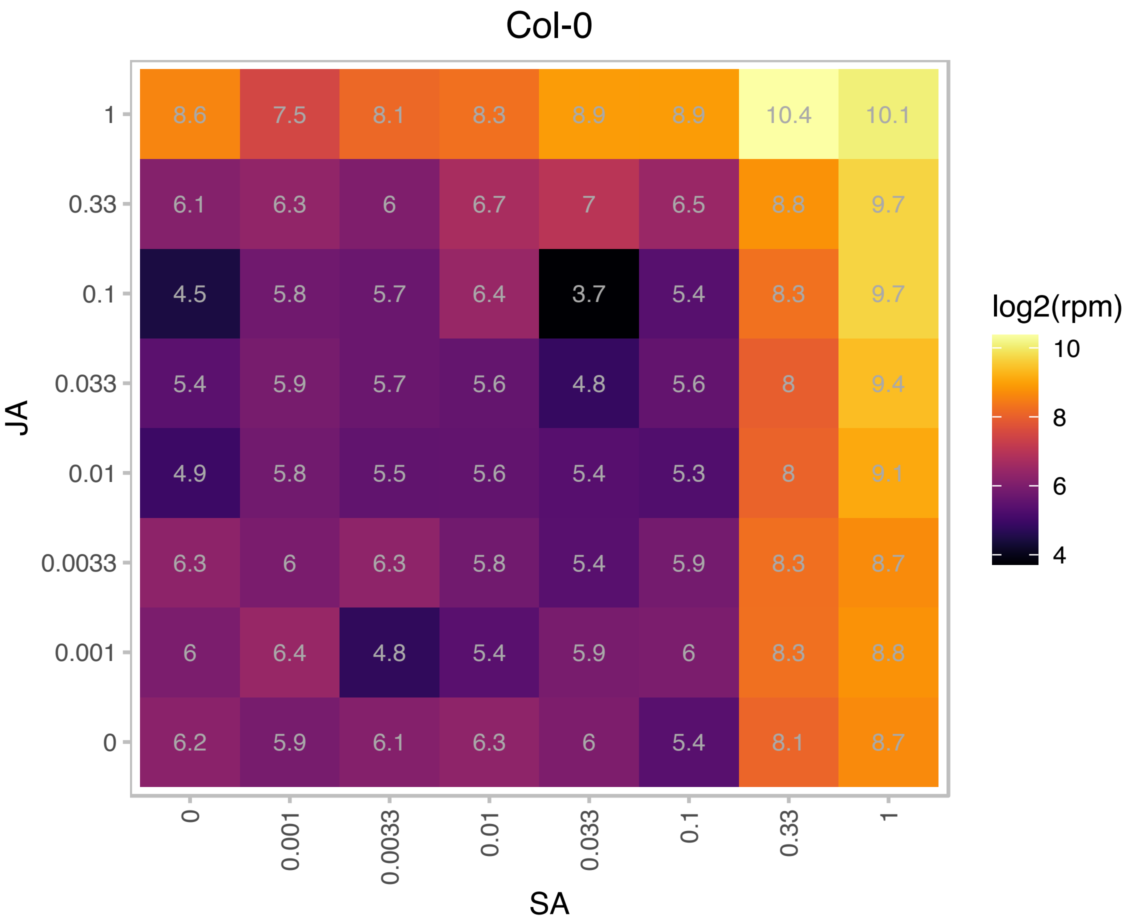


| | |
|---|---|
| Gene | AT5G24570 |
| is.expressed_genes | TRUE |
| is.responsive_genes | TRUE |
| is.apcluster | B-03 |

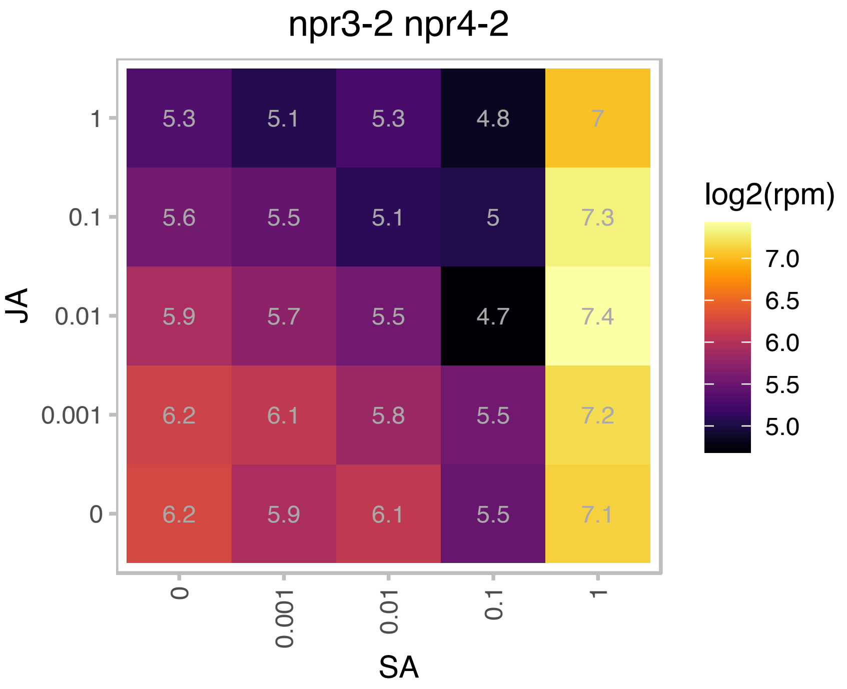


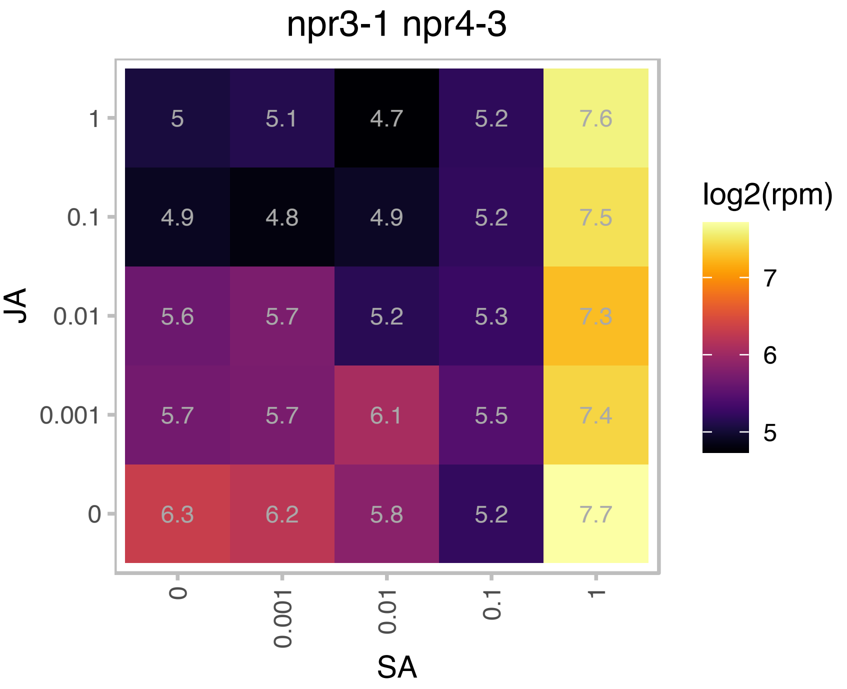

# AT5G37930.1 : NA

Protein with RING/U-box and TRAF-like domains

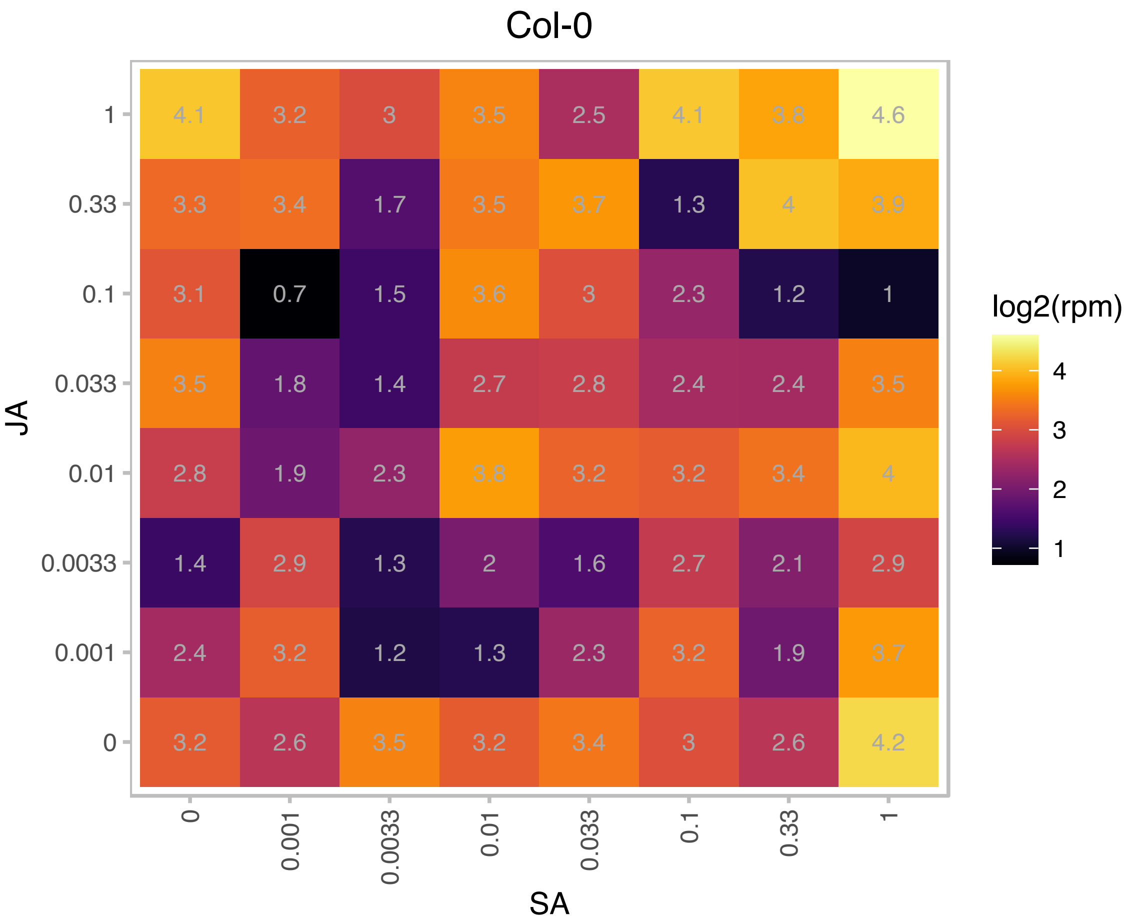


| | |
|---|---|
| Gene | AT5G37930 |
| is.expressed_genes | TRUE |
| is.responsive_genes | TRUE |
| is.apcluster | B-03 |

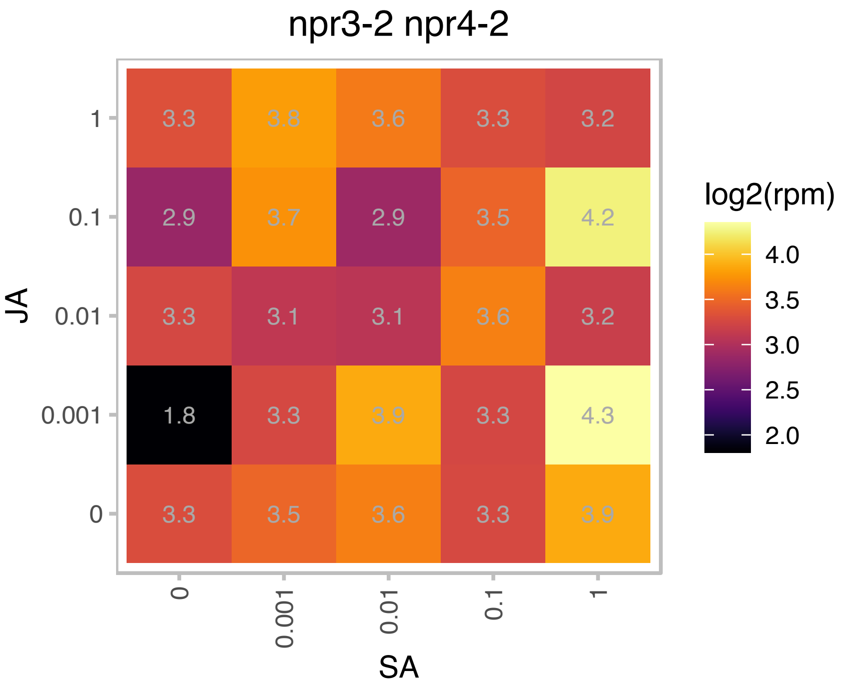


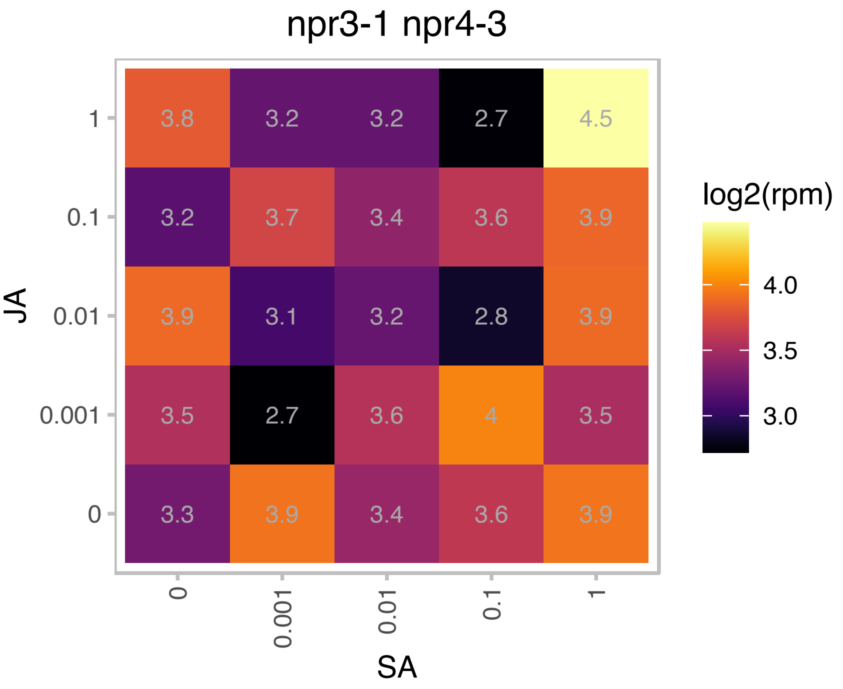

# AT5G38200.1 : NA

Class I glutamine amidotransferase-like superfamily protein

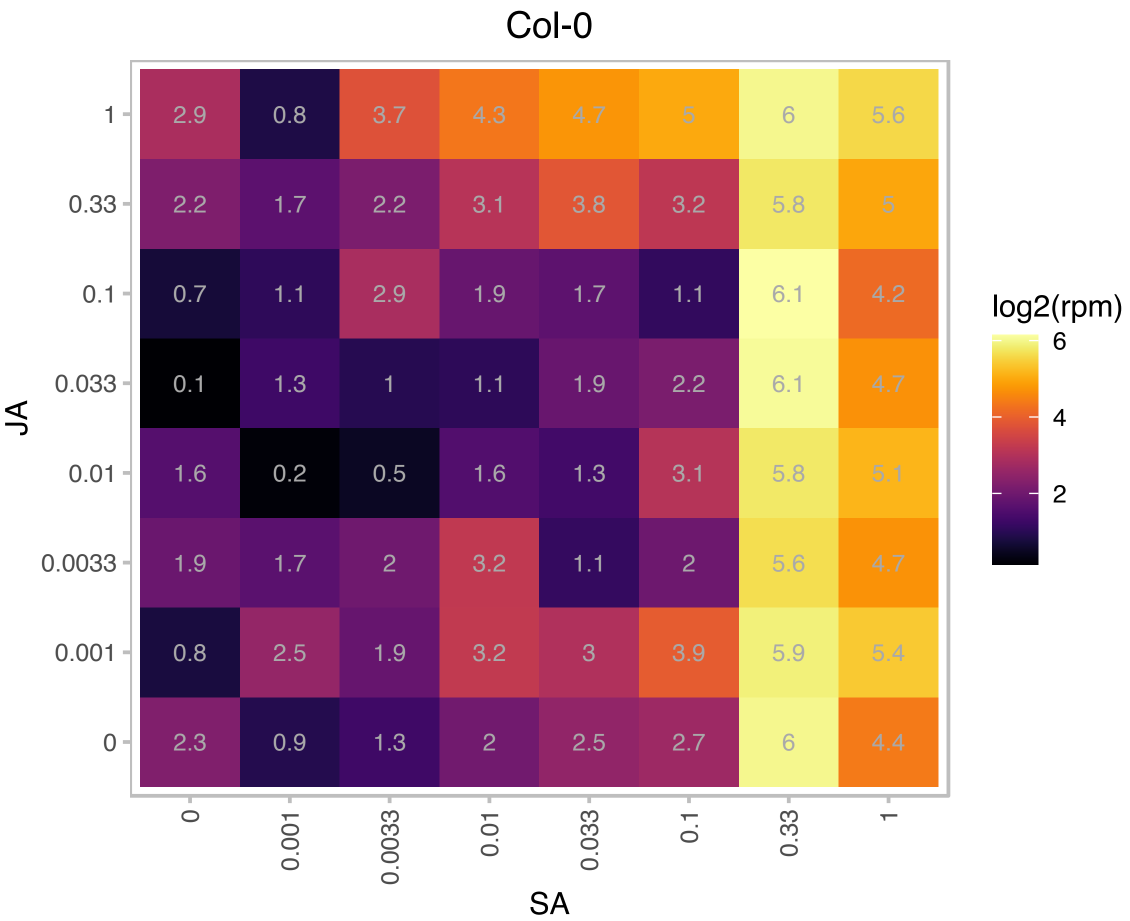


| | |
|---|---|
| Gene | AT5G38200 |
| is.expressed_genes | TRUE |
| is.responsive_genes | TRUE |
| is.apcluster | B-03 |

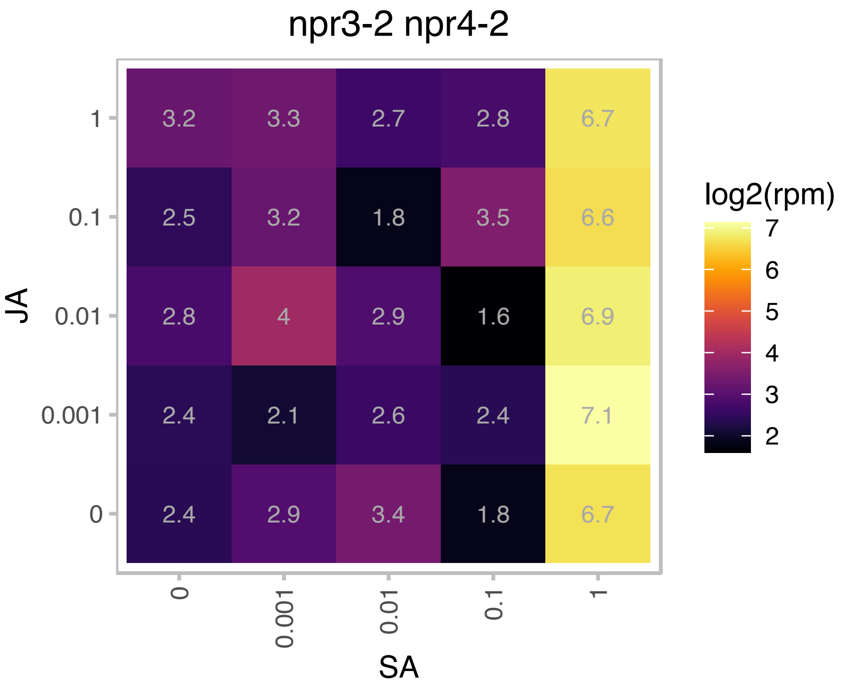


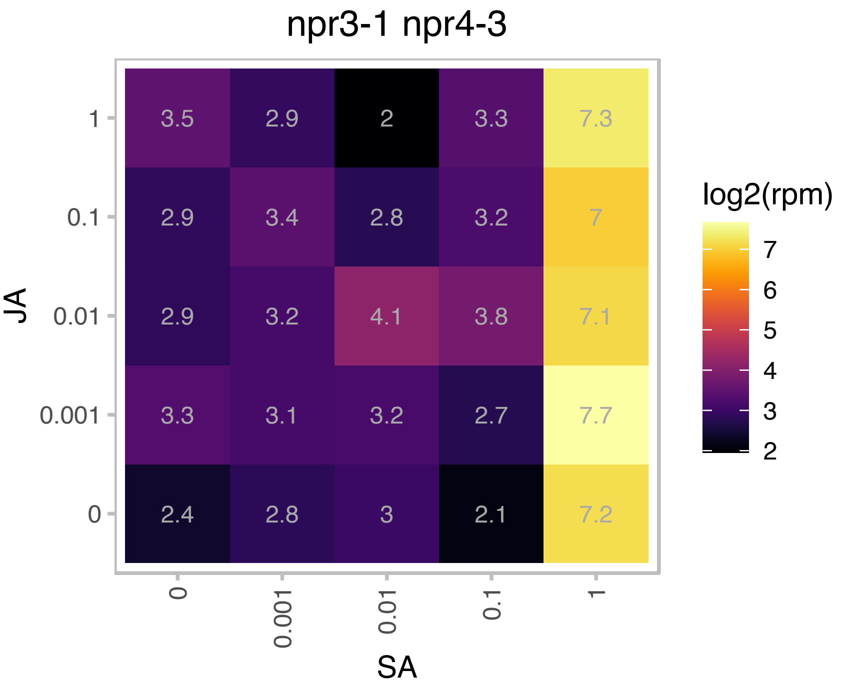

# AT5G40690.1 : NA

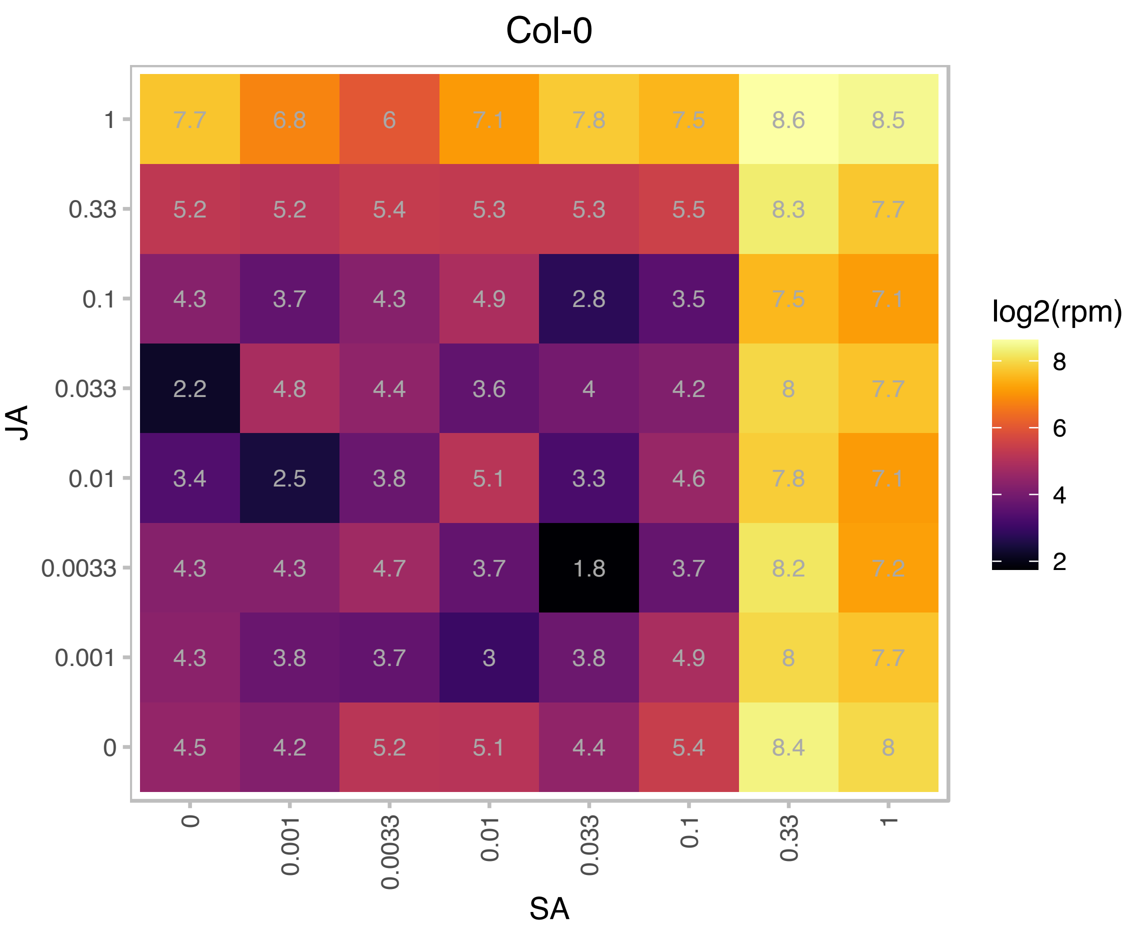


| | |
|---|---|
| Gene | AT5G40690 |
| is.expressed_genes | TRUE |
| is.responsive_genes | TRUE |
| is.apcluster | B-03 |

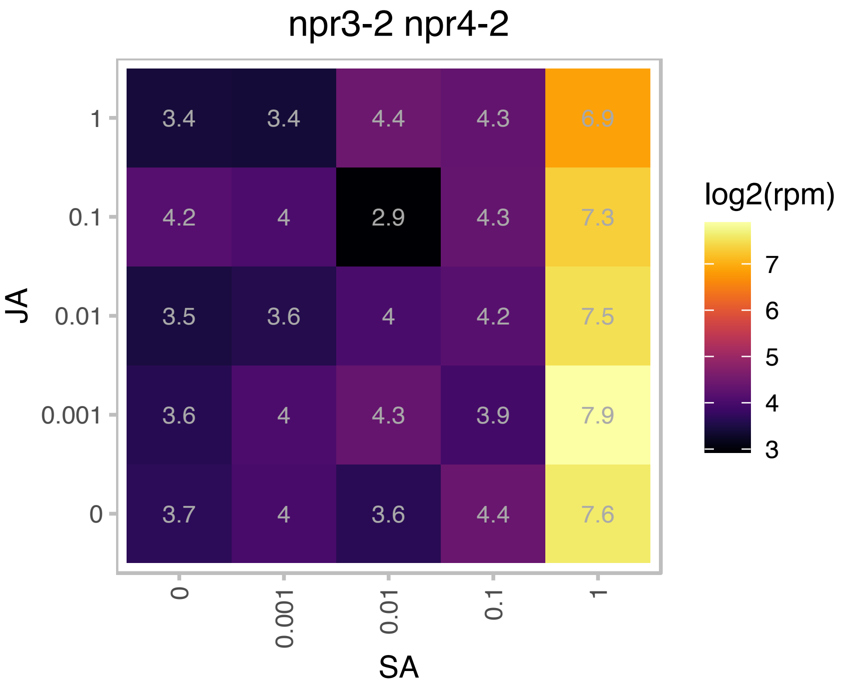


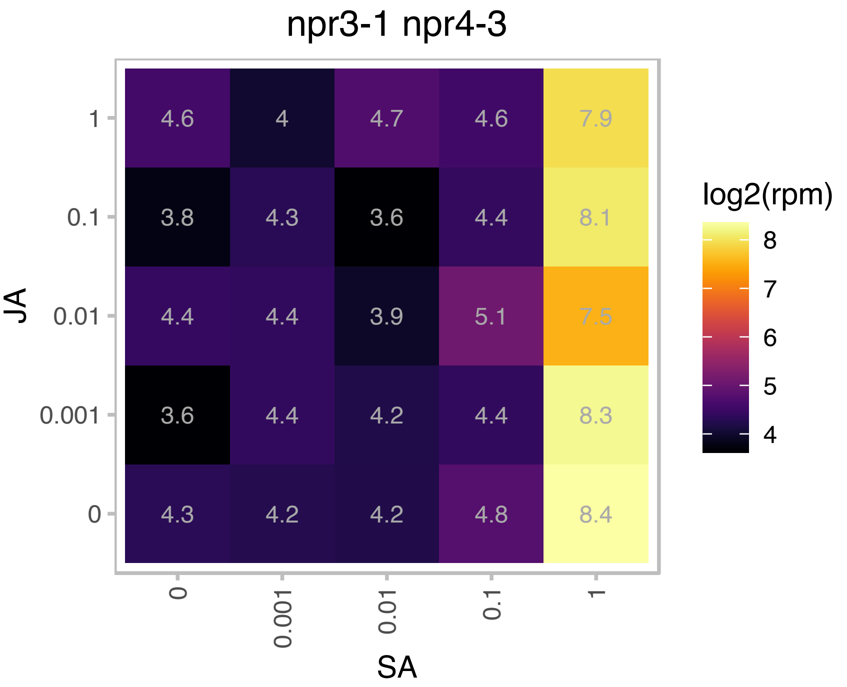

# AT5G40880.1 : NA

WD-40 repeat family protein / zfwd3 protein (ZFWD3)

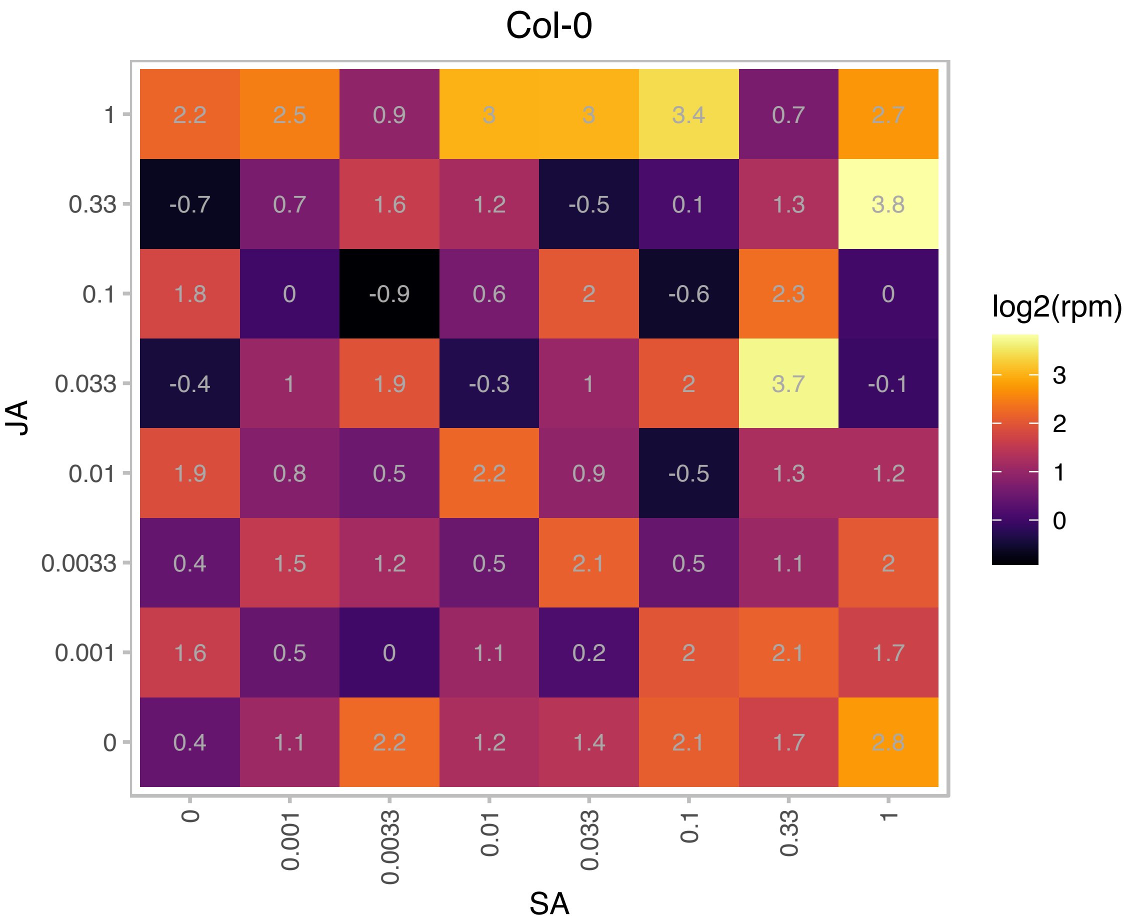


| | |
|---|---|
| Gene | AT5G40880 |
| is.expressed_genes | TRUE |
| is.responsive_genes | TRUE |
| is.apcluster | B-03 |

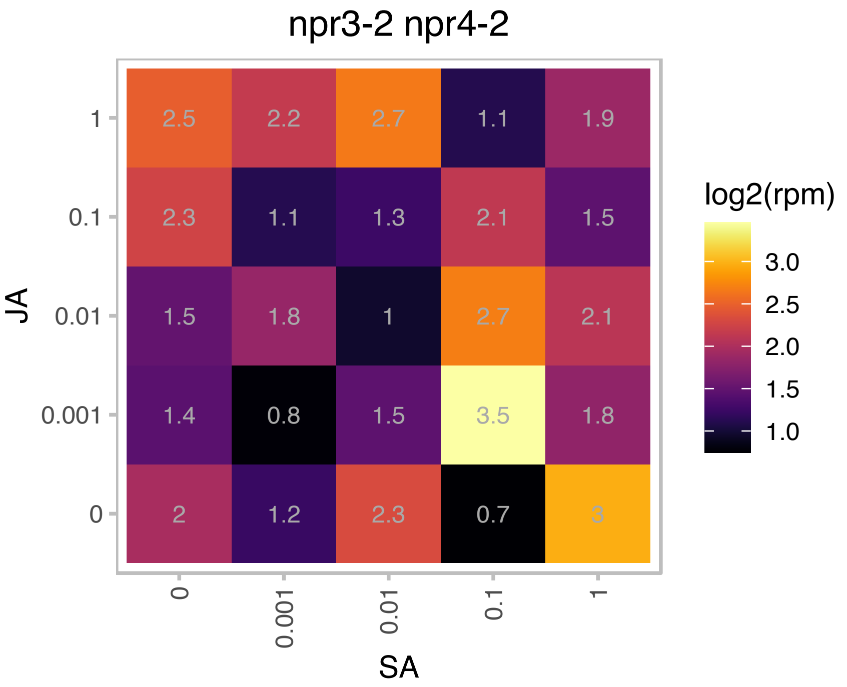


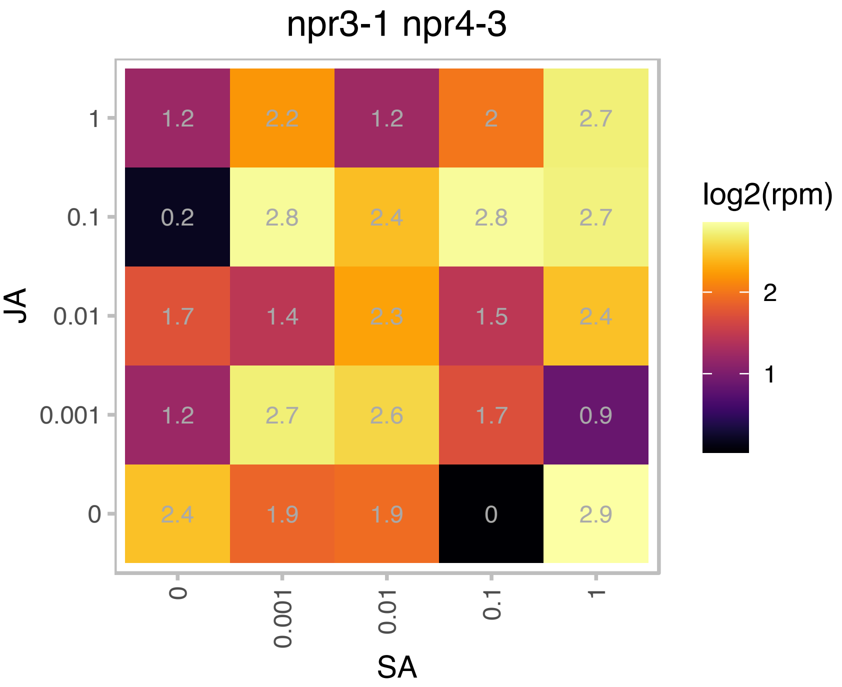

# AT5G47730.1 : NA

Sec14p-like phosphatidylinositol transfer family protein

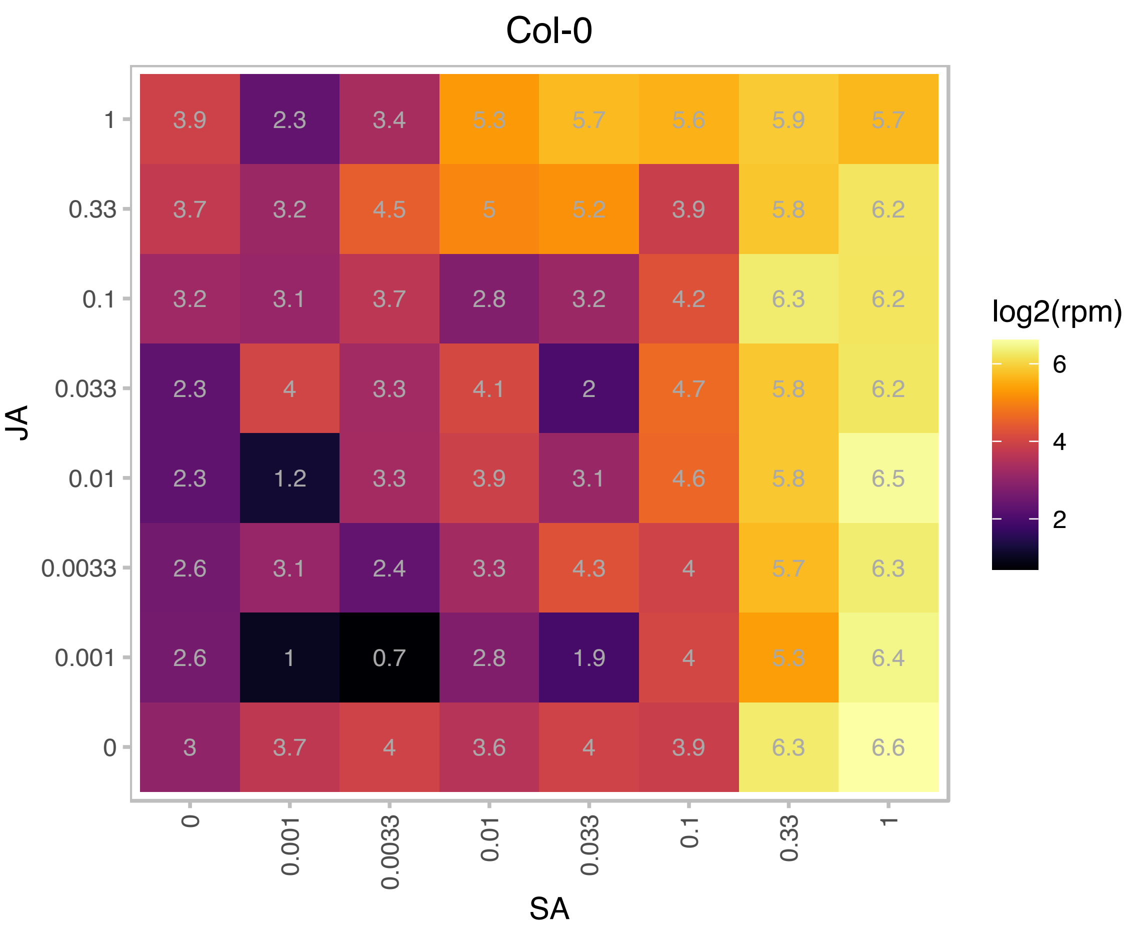


| | |
|---|---|
| Gene | AT5G47730 |
| is.expressed_genes | TRUE |
| is.responsive_genes | TRUE |
| is.apcluster | B-03 |

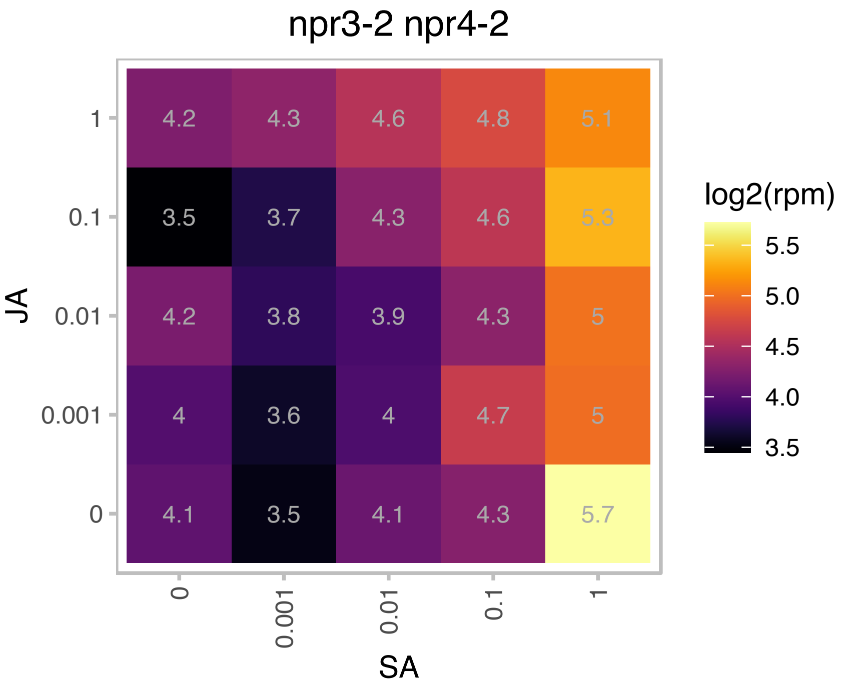


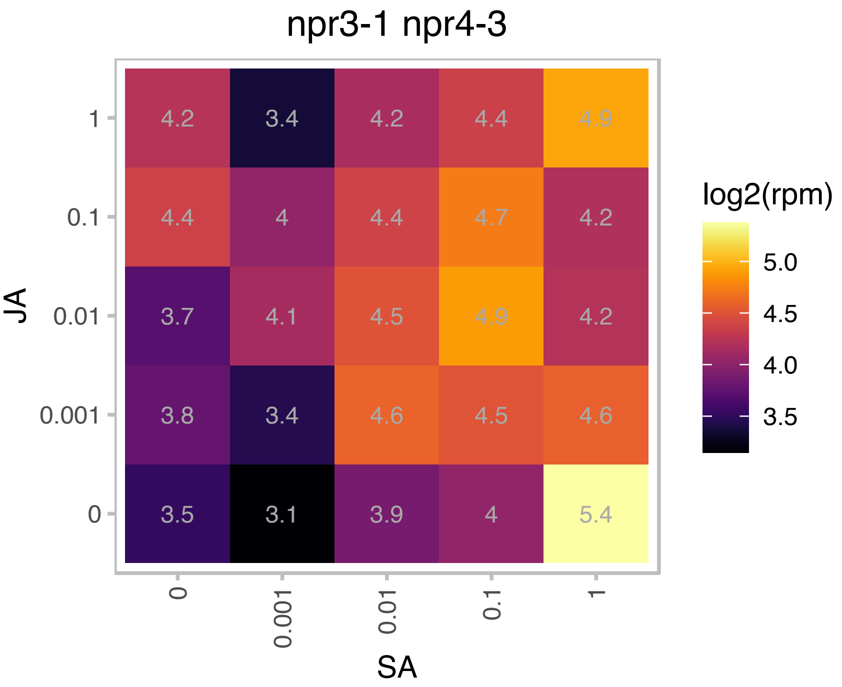

### AT5G47770.1 : FPS1

farnesyl diphosphate synthase 1

Encodes a protein with farnesyl diphosphate synthase activity.

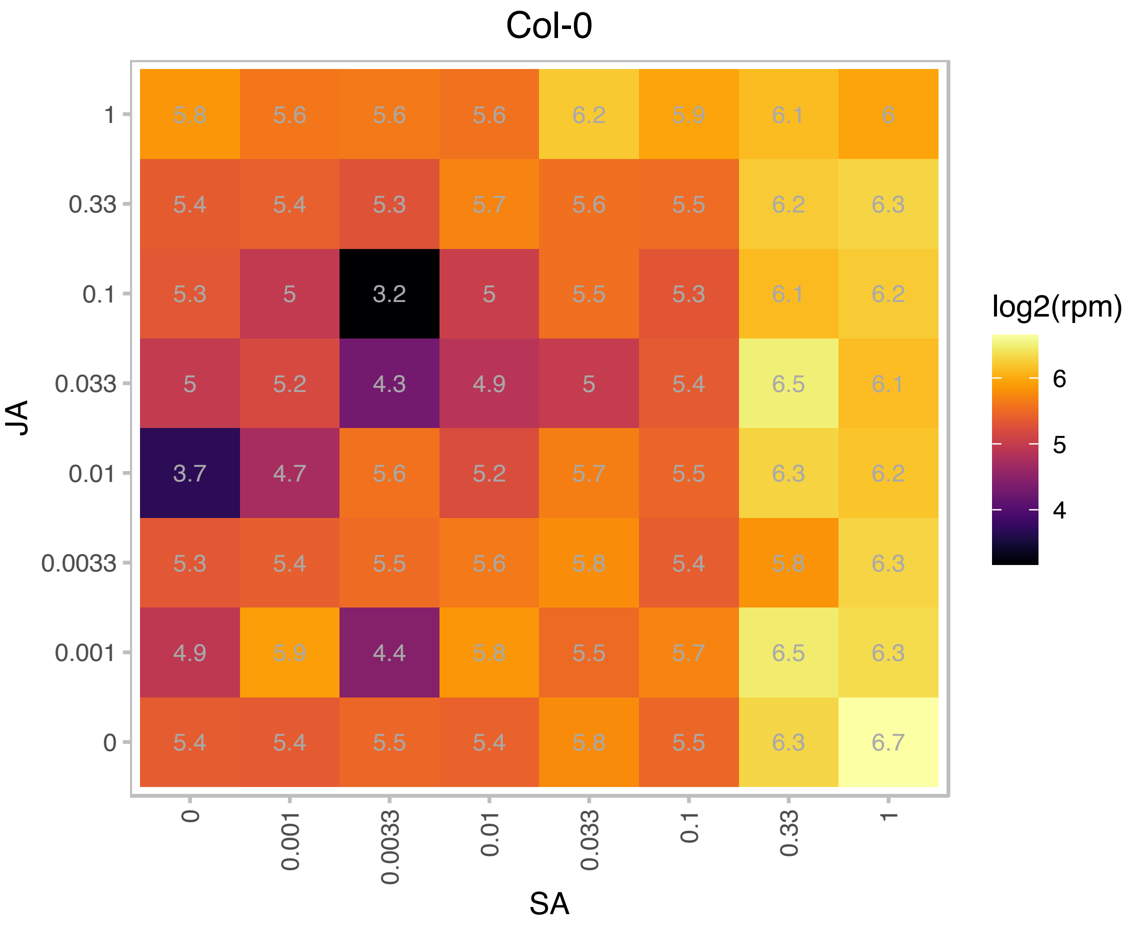


| | |
|---|---|
| Gene | AT5G47770 |
| is.expressed_genes | TRUE |
| is.responsive_genes | TRUE |
| is.apcluster | B-03 |

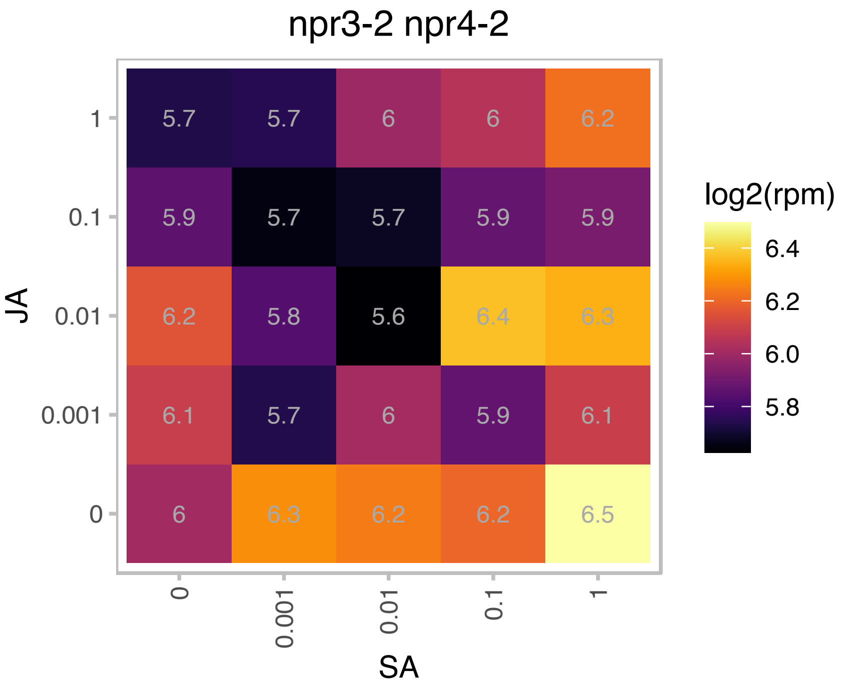


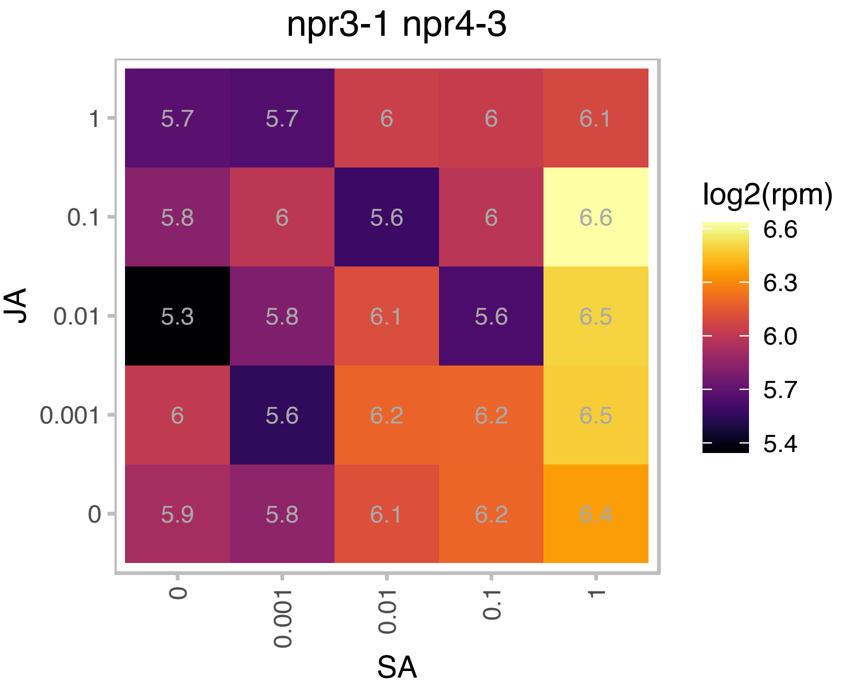

### AT5G48570.1 : ATFKBP65, FKBP65, ROF2

FKBP-type peptidyl-prolyl cis-trans isomerase family protein

Encodes one of the 36 carboxylate clamp (CC)-tetratricopeptide repeat (TPR) proteins (Prasad 2010, Pubmed ID: 20856808) with potential to interact with Hsp90/Hsp70 as co-chaperones.

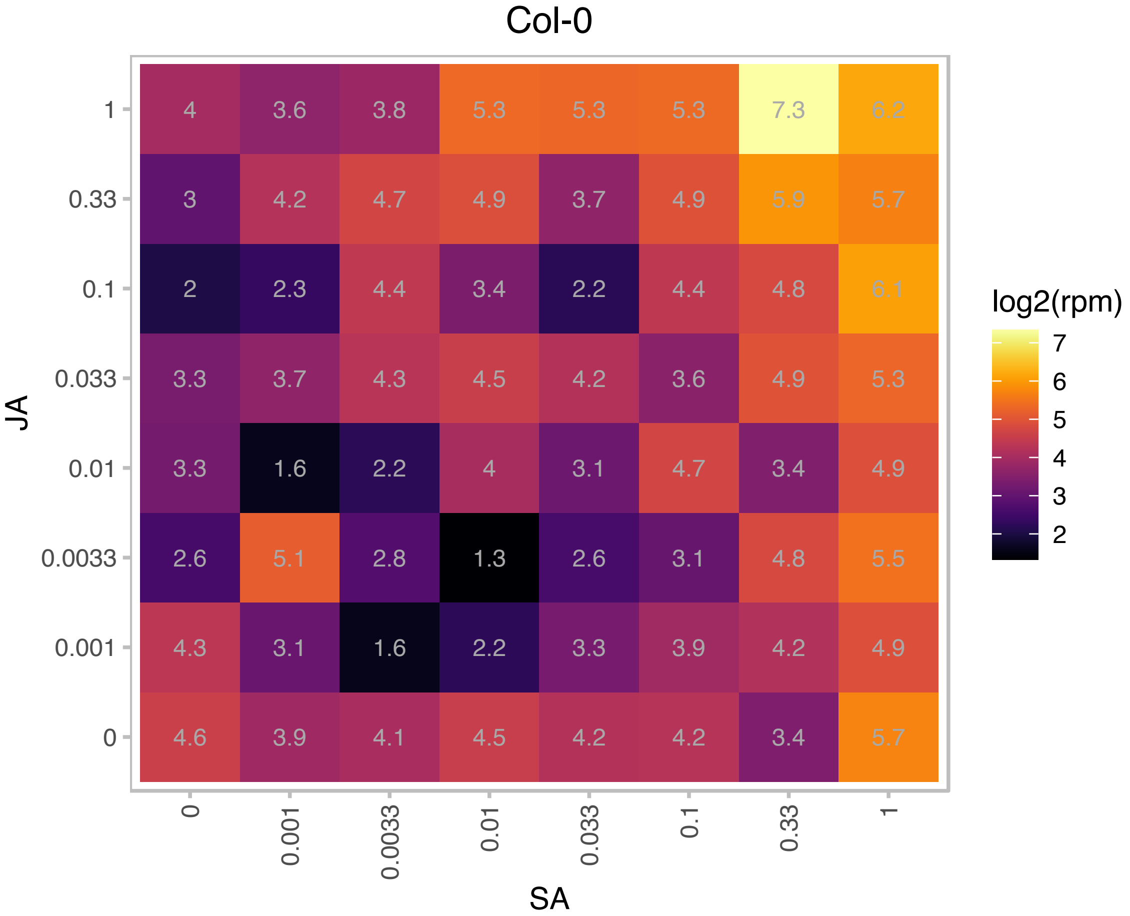


| | |
|---|---|
| Gene | AT5G48570 |
| is.expressed_genes | TRUE |
| is.responsive_genes | TRUE |
| is.apcluster | B-03 |

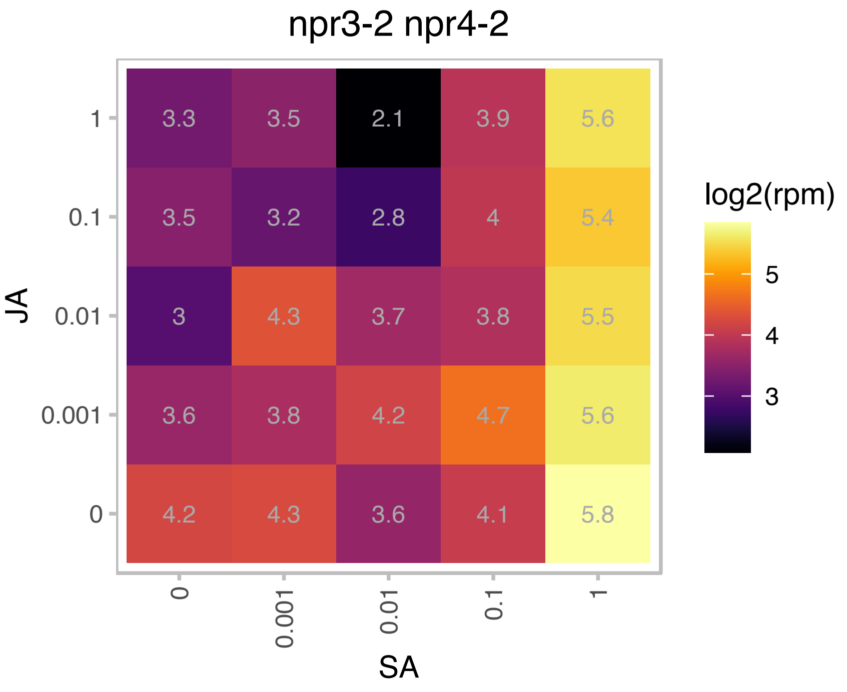


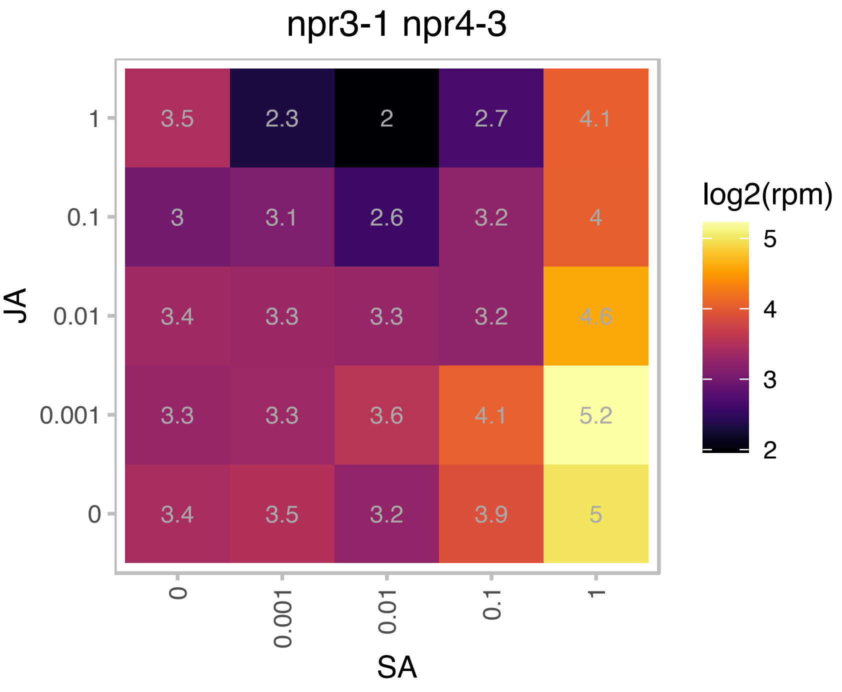

# AT5G49665.1 : NA

Zinc finger (C3HC4-type RING finger) family protein

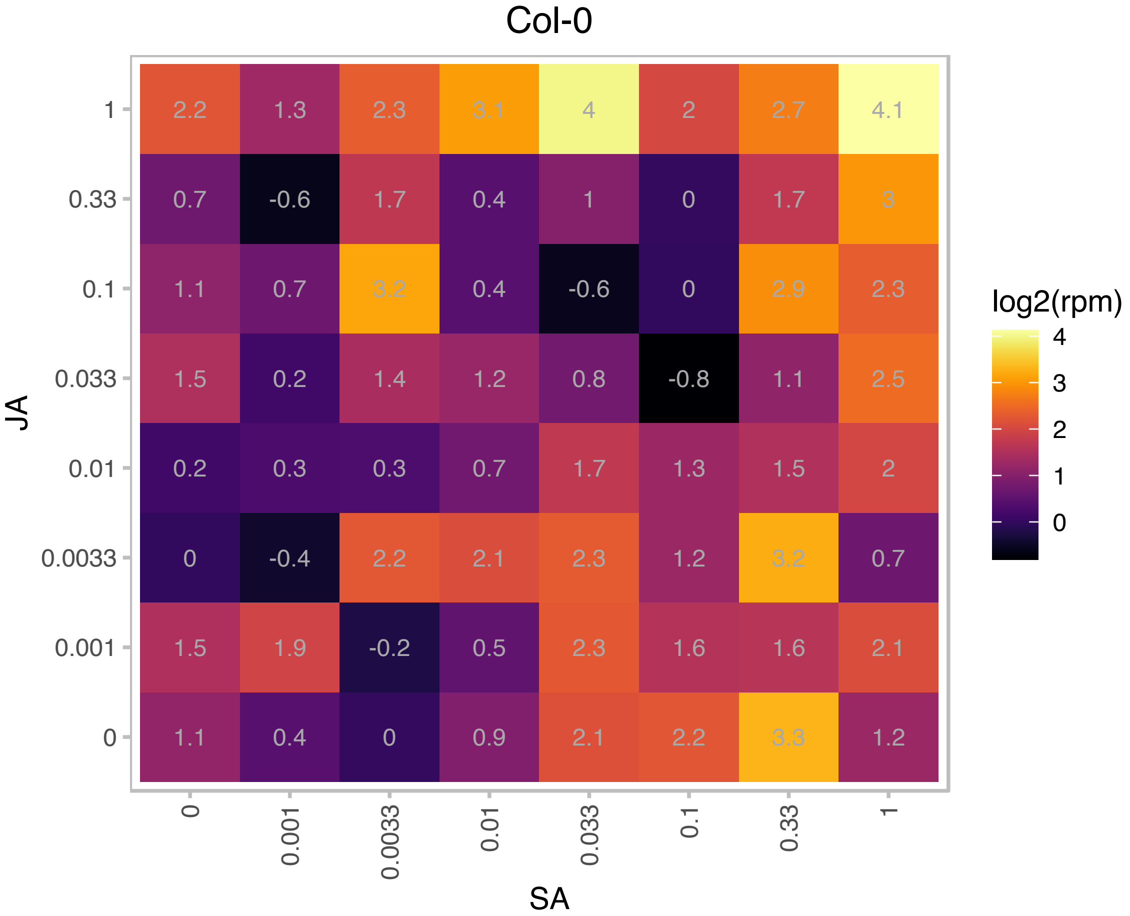


| | |
|---|---|
| Gene | AT5G49665 |
| is.expressed_genes | TRUE |
| is.responsive_genes | TRUE |
| is.apcluster | B-03 |

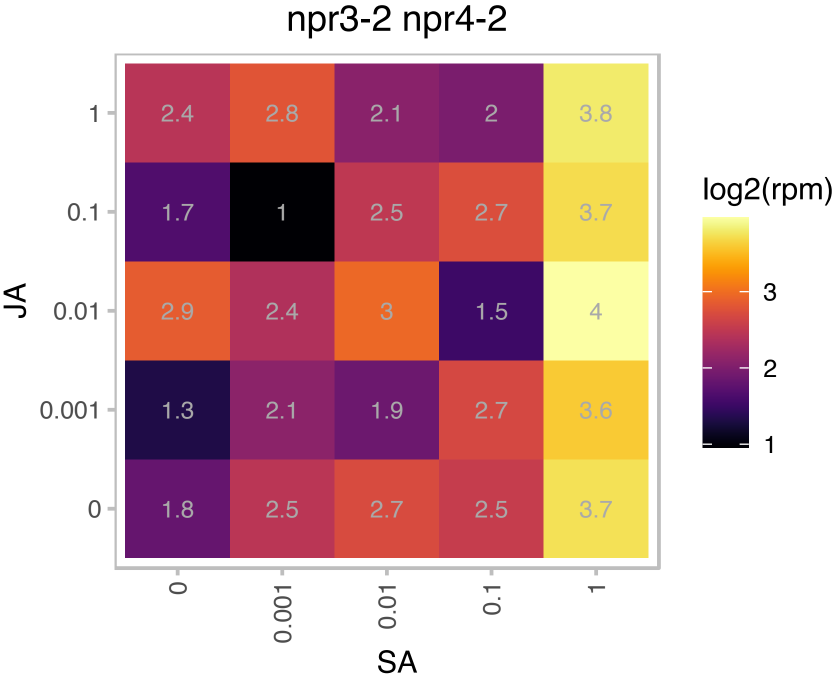


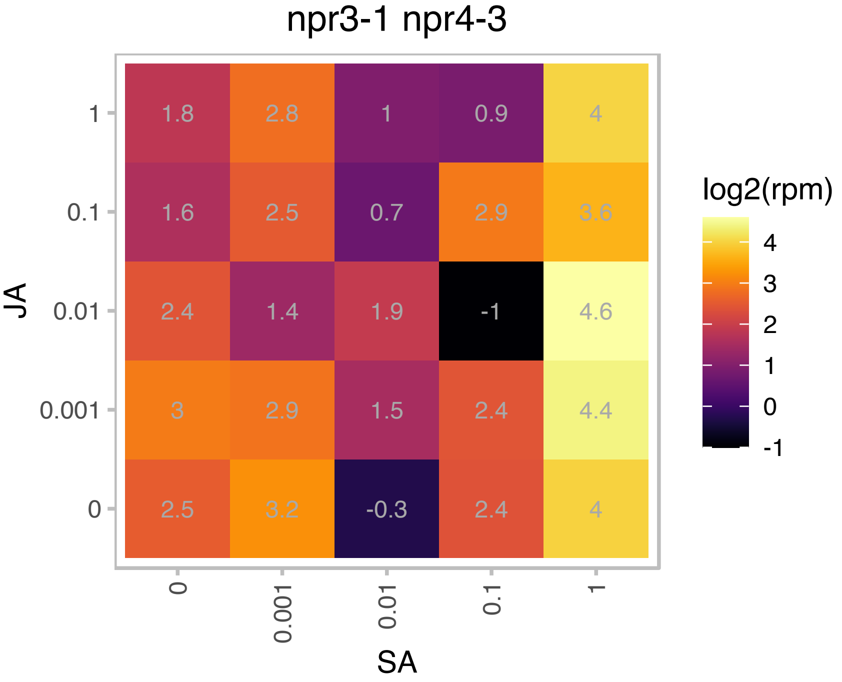

### AT5G53120.1 : ATSPDS3, SPDS3, SPMS

spermidine synthase 3

encodes a novel spermine synthase and is a paralog of previously characterized spermidine synthases, SPDS1 and SPDS2. SPDS3 forms heterodimers with SDPS2, which in turn forms heterodimers with SDPS1 in vivo. The ge[illegible] does not complement speDelta3 deficiency of spermidine synthase in yeast but DOES complement speDelta4 deficiency.

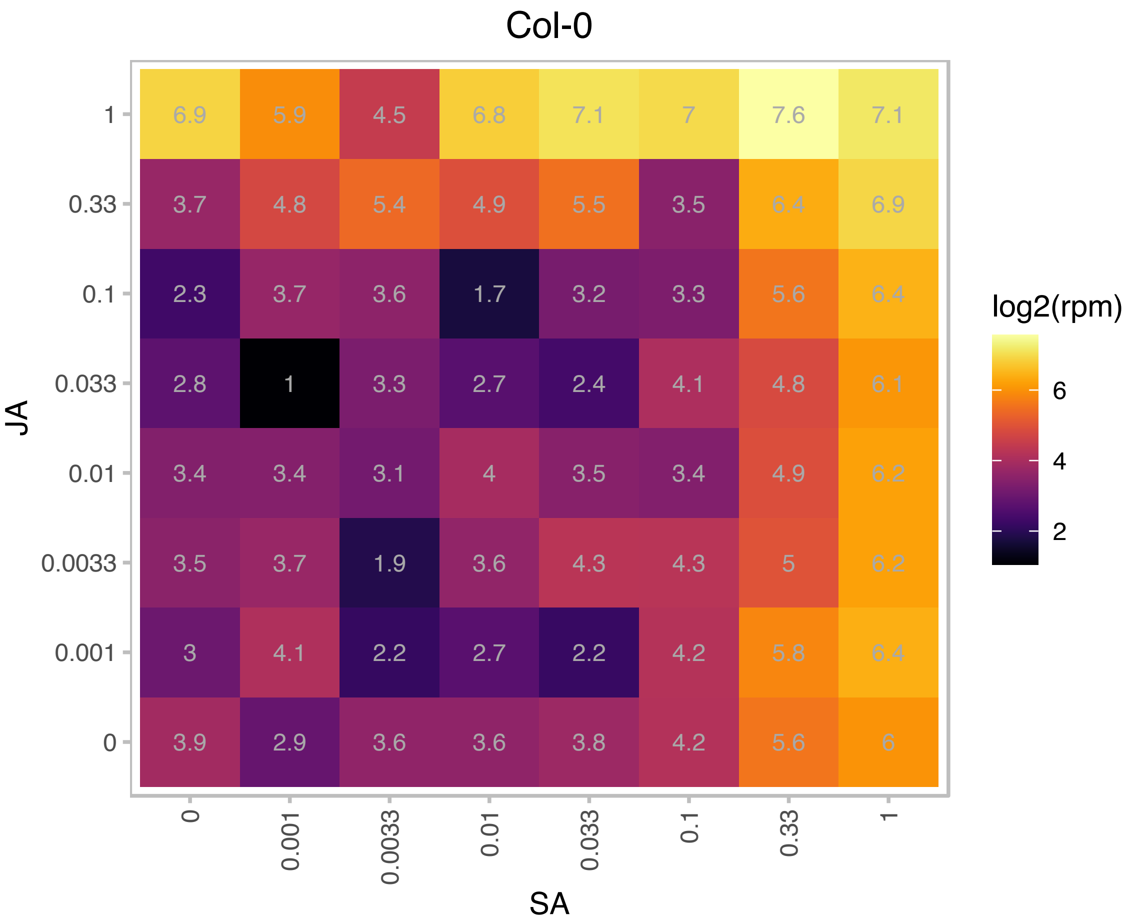


| | |
|---|---|
| Gene | AT5G53120 |
| is.expressed_genes | TRUE |
| is.responsive_genes | TRUE |
| is.apcluster | B-03 |

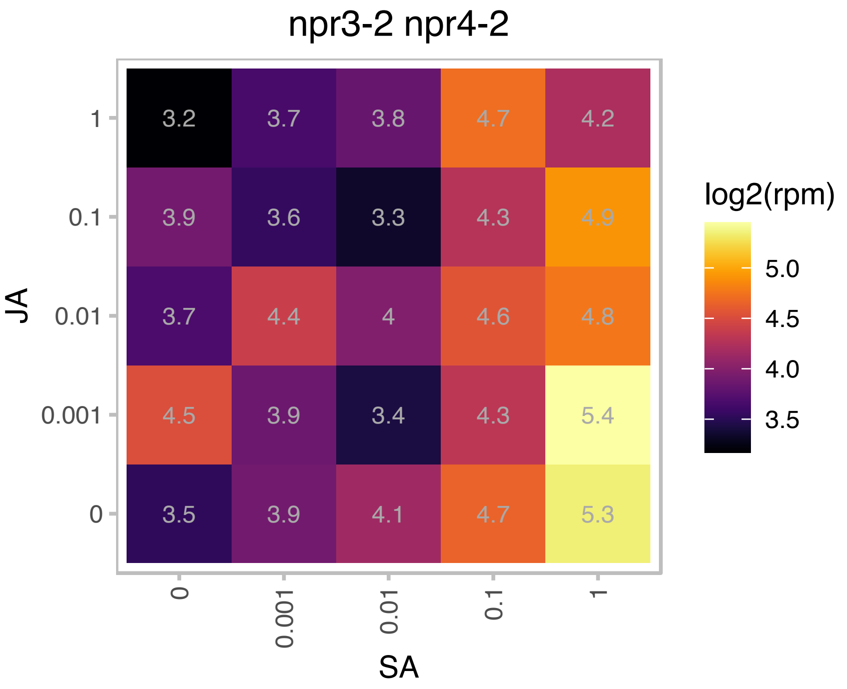


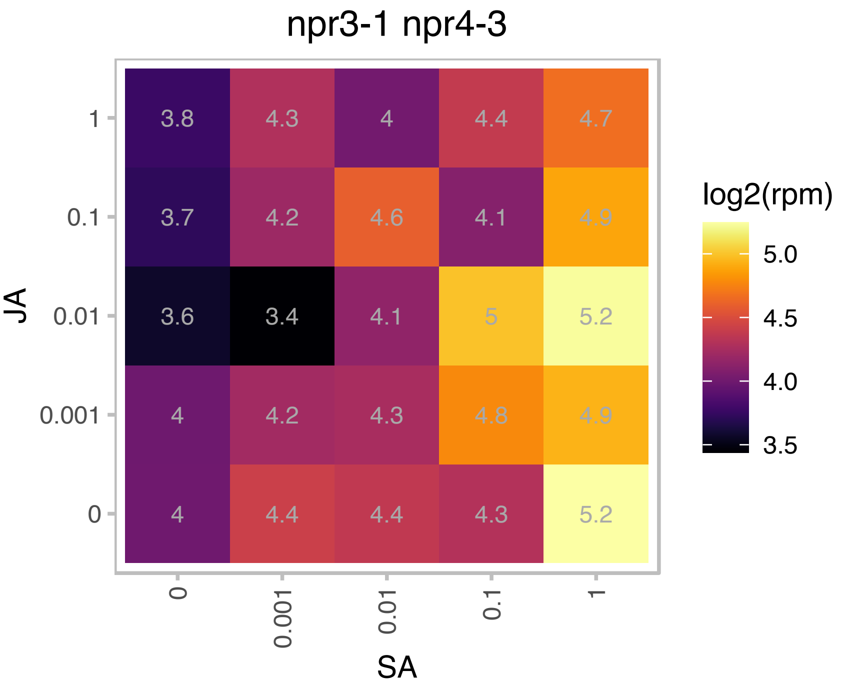

### AT5G56090.1 : COX15

cytochrome c oxidase 15

Encodes a homolog of COX15. Microarray analysis show a 3.2 fold increase in transcription after treatment with rotenone, an electron transport chain inhibitor.

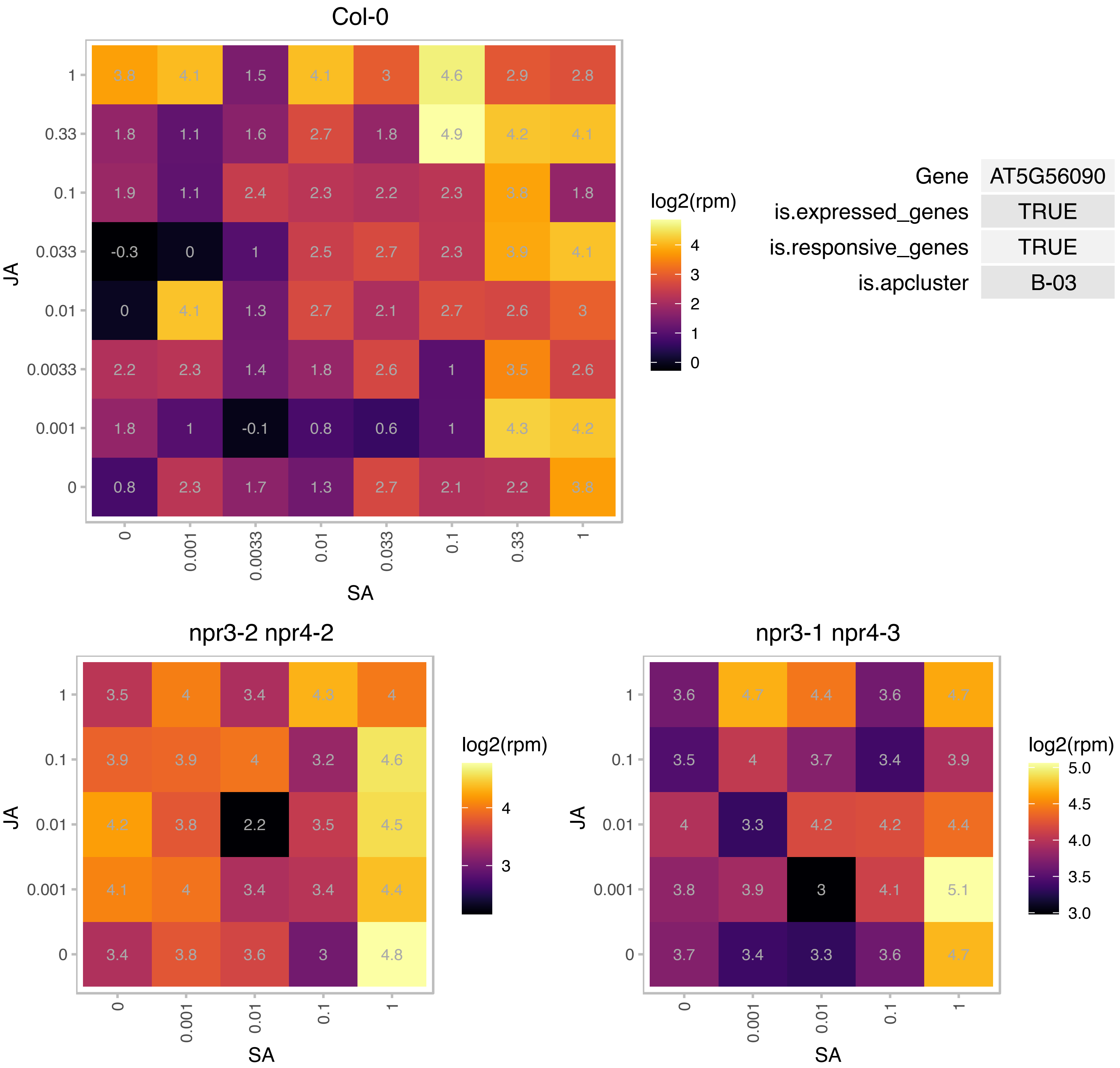


| | |
|---|---|
| Gene | AT5G56090 |
| is.expressed_genes | TRUE |
| is.responsive_genes | TRUE |
| is.apcluster | B-03 |

### AT5G59030.1 : COPT1

copper transporter 1

encodes a putative copper transport protein that contains copper-binding motif and functionally complements in copper-transport defective yeast strains

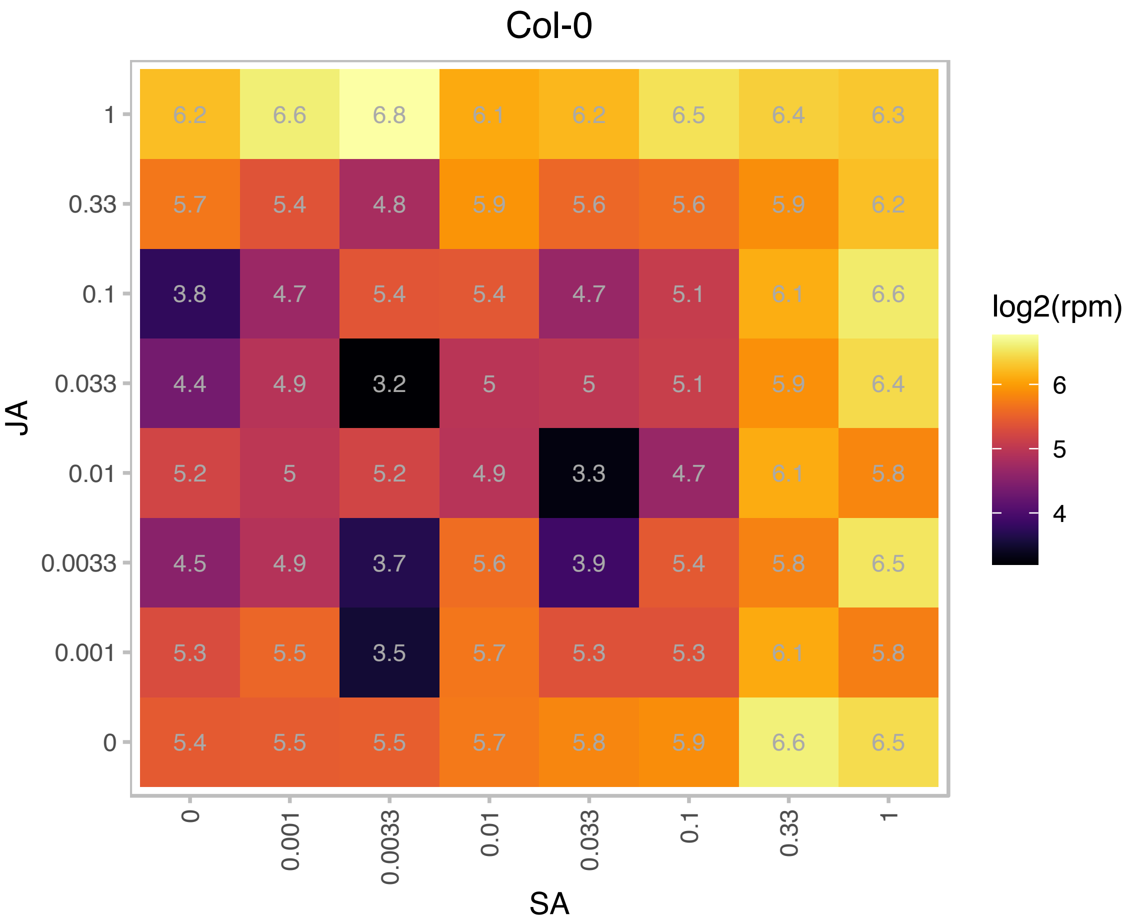


| | |
|---|---|
| Gene | AT5G59030 |
| is.expressed_genes | TRUE |
| is.responsive_genes | TRUE |
| is.apcluster | B-03 |

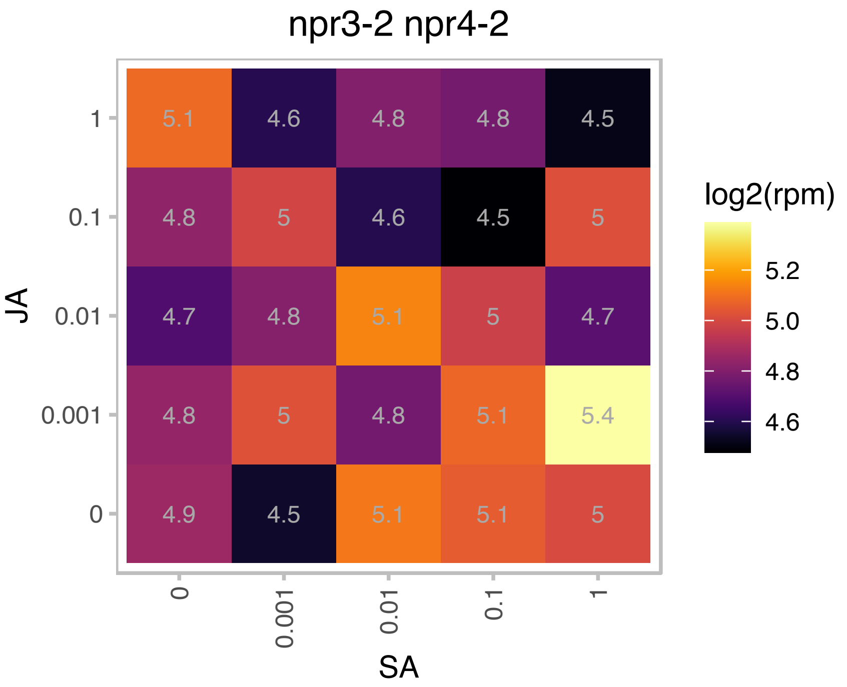


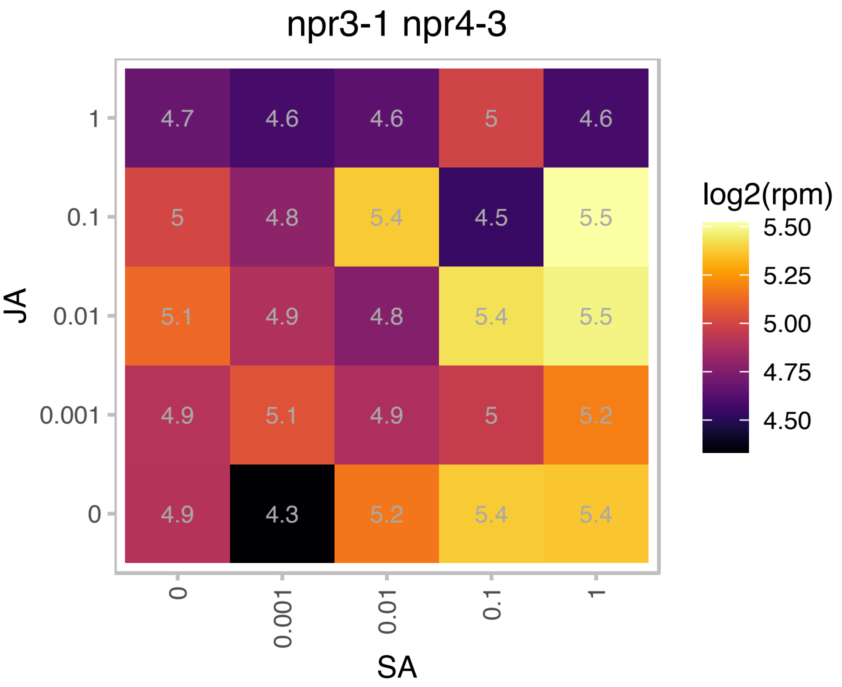

# AT5G62865.1 : NA

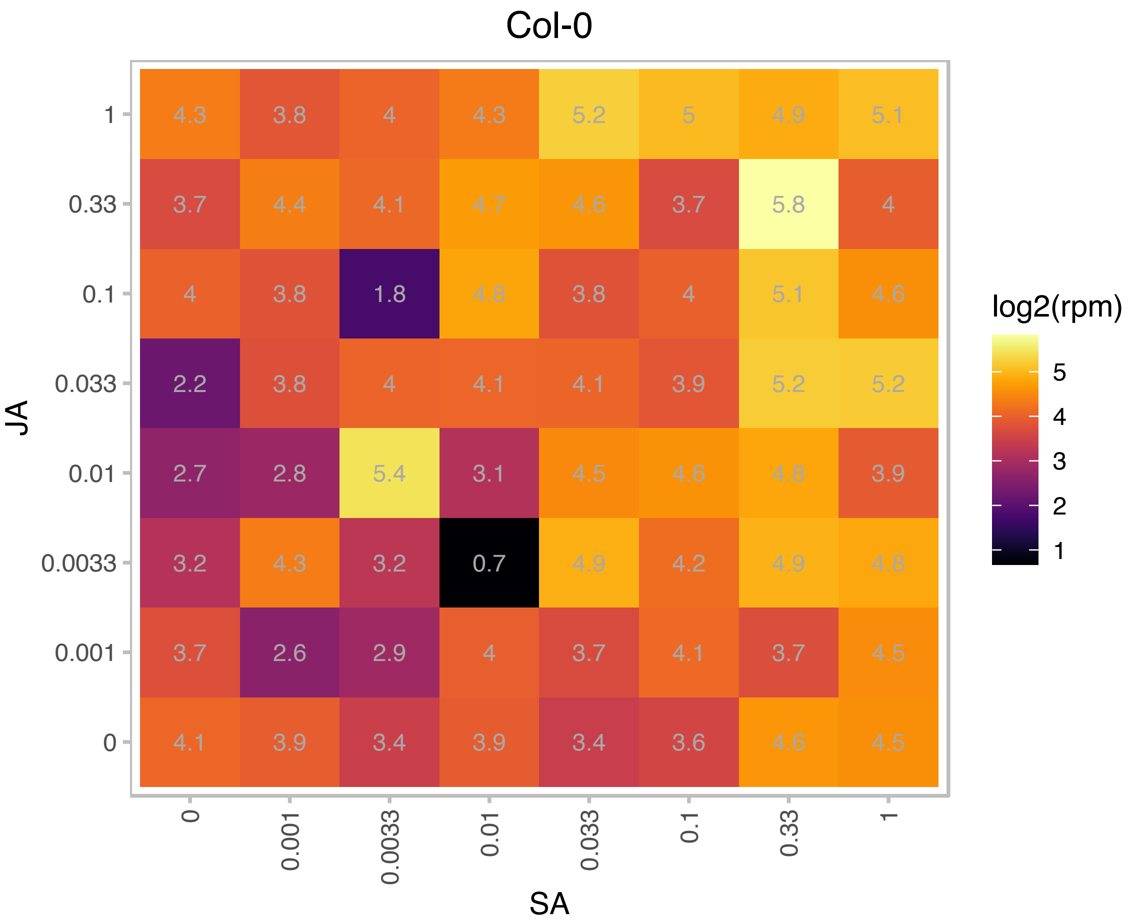


| | |
|---|---|
| Gene | AT5G62865 |
| is.expressed_genes | TRUE |
| is.responsive_genes | TRUE |
| is.apcluster | B-03 |

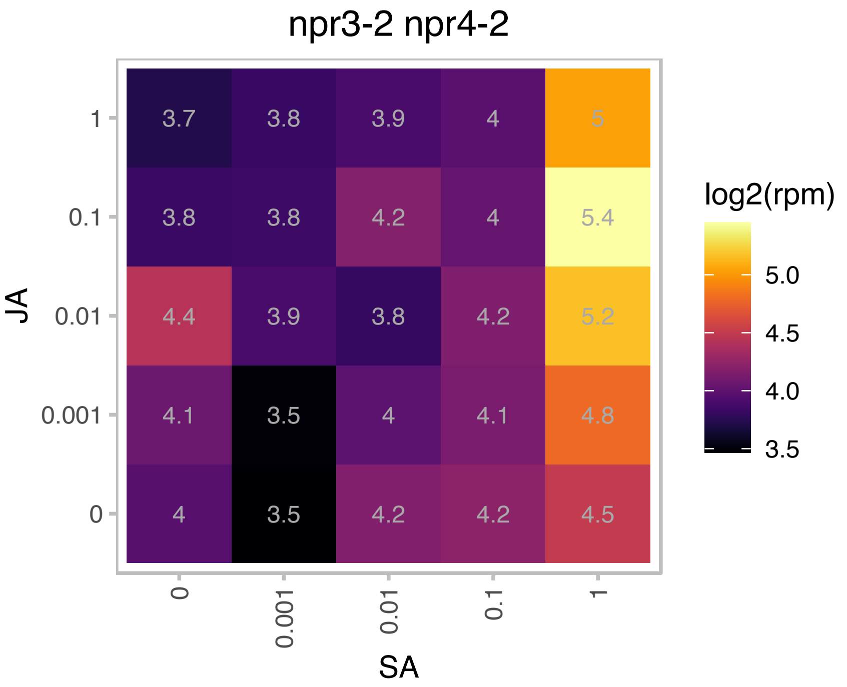


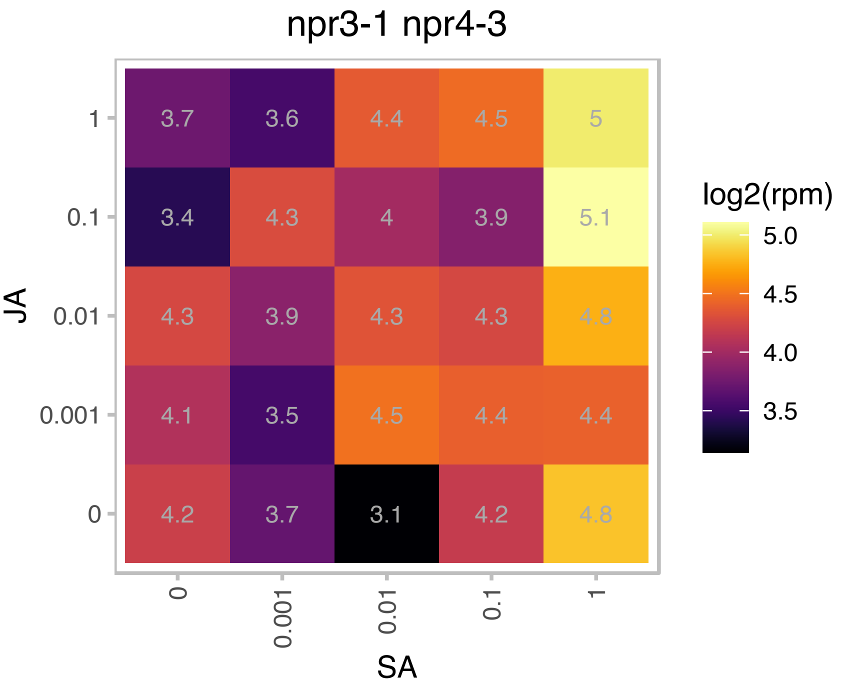

### AT5G64750.1 : ABR1

Integrase-type DNA-binding superfamily protein

Encodes a putative transcription factor containing an AP2 domain. Is a member of the ERF (ethylene response factor) subfamily B-4 of ERF/AP2 transcription factor family. Expressed in response to ABA, osmotic stress, sugar stress and drought. Mutants are hypersensitive to these stresses. May be involved in regulation of ABA mediated stress response. The mRNA is cell-to-cell mobile.

Col-0

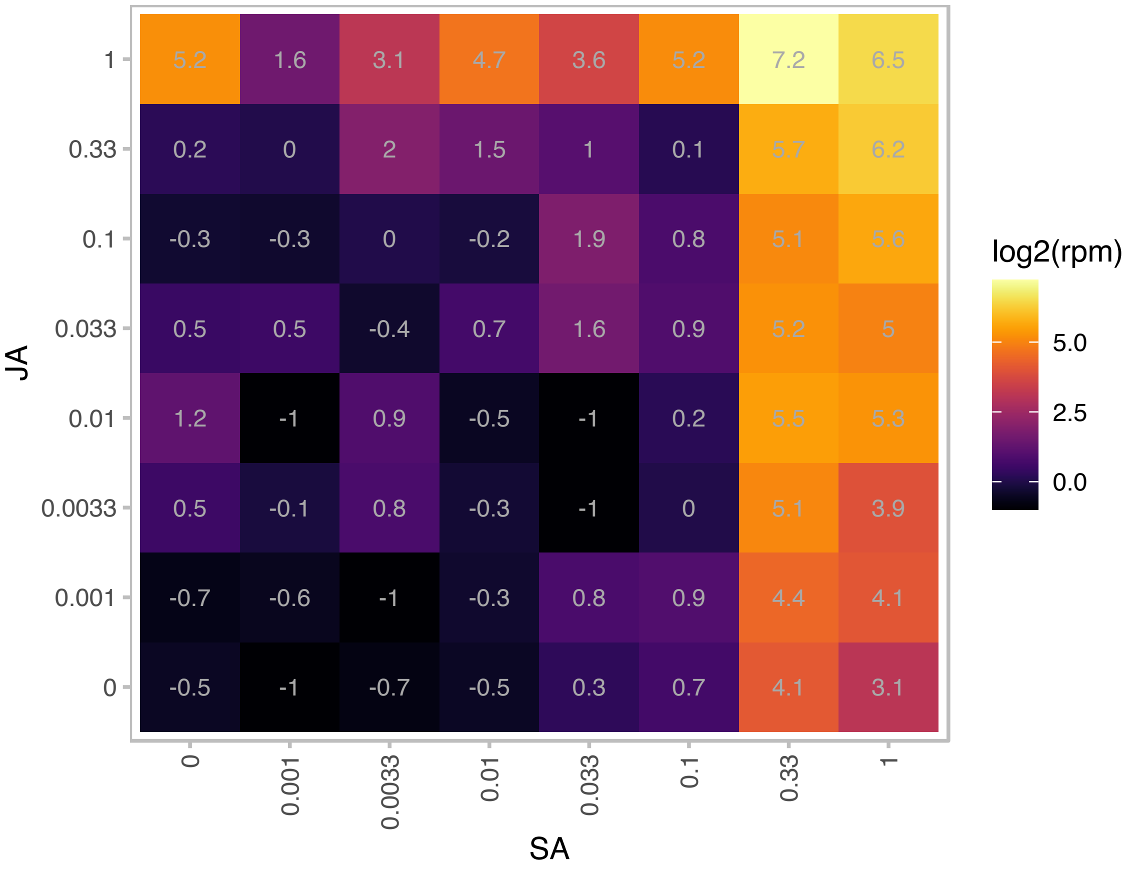


| | |
|---|---|
| Gene | AT5G64750 |
| is.expressed_genes | TRUE |
| is.responsive_genes | TRUE |
| is.apcluster | B-03 |

npr3-2 npr4-2

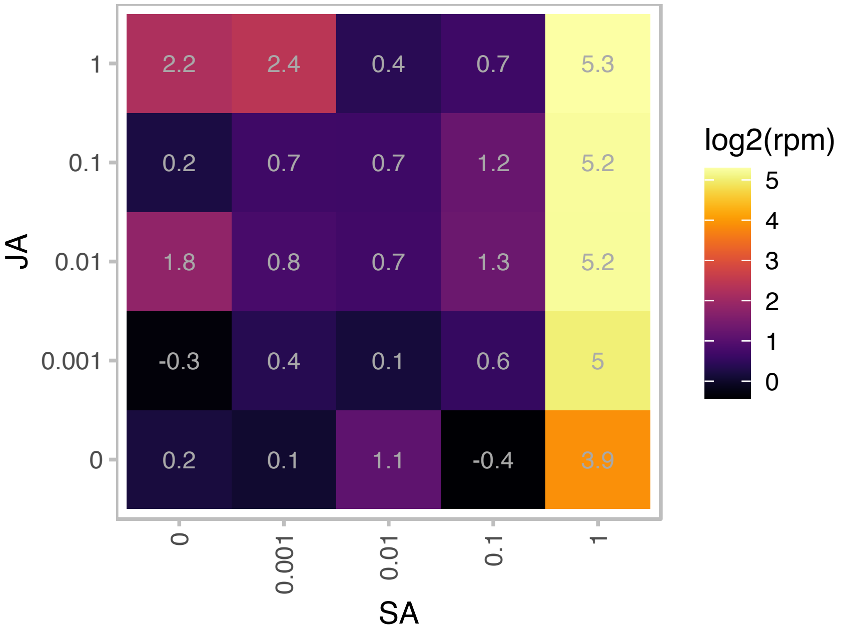


npr3-1 npr4-3

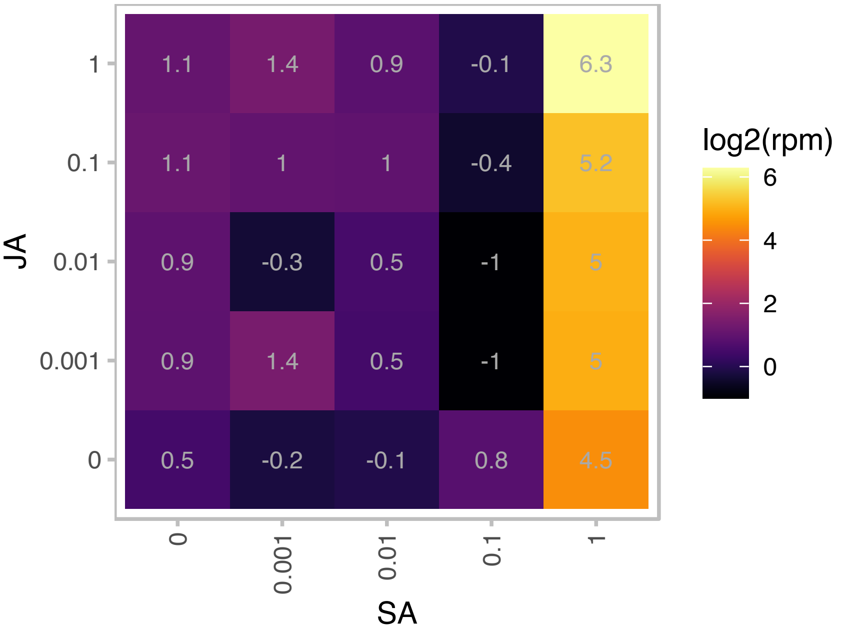

# AT5G66440.1 : NA

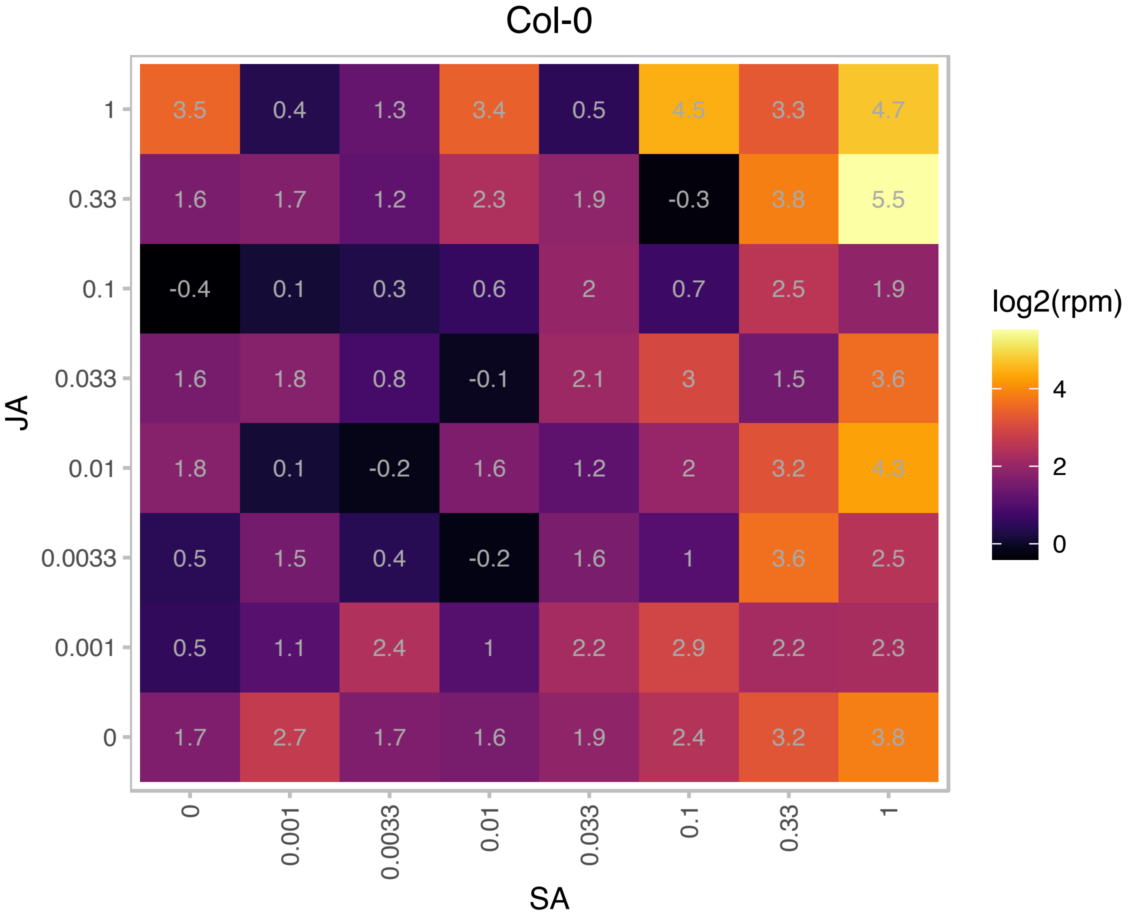


| | |
|---|---|
| Gene | AT5G66440 |
| is.expressed_genes | TRUE |
| is.responsive_genes | TRUE |
| is.apcluster | B-03 |

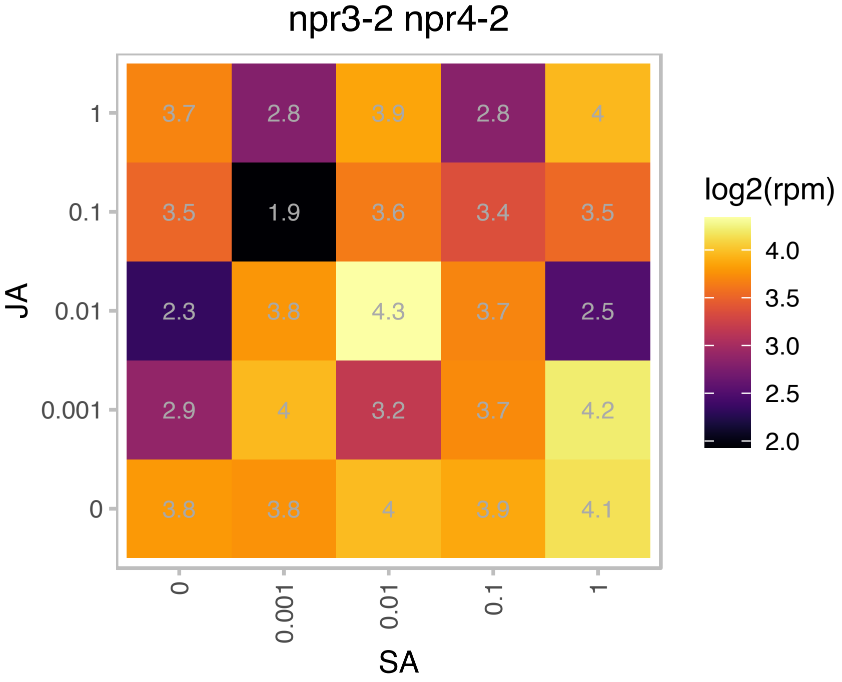


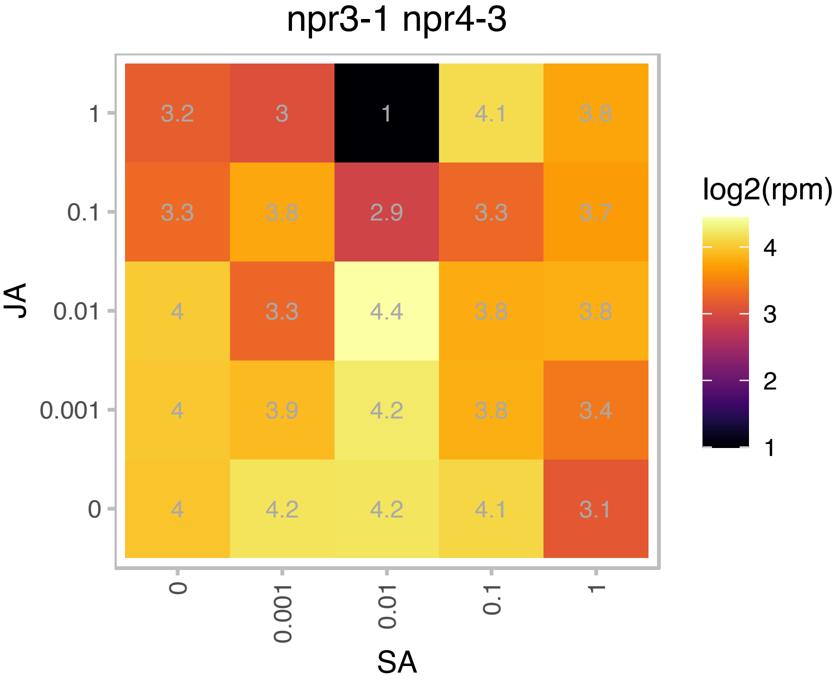

### AT1G03770.1 : ATRING1B, RING1B

RING 1B
Encodes a nuclear localized protein with similarity to animal polycomb repressive core complex1 (PRC1) core component RING.Appears to function redundantly with ATRING1a, a close paralog. Both interact physically with CLF and LHP1 and appear to function together to repress class I KNOX gene expression.

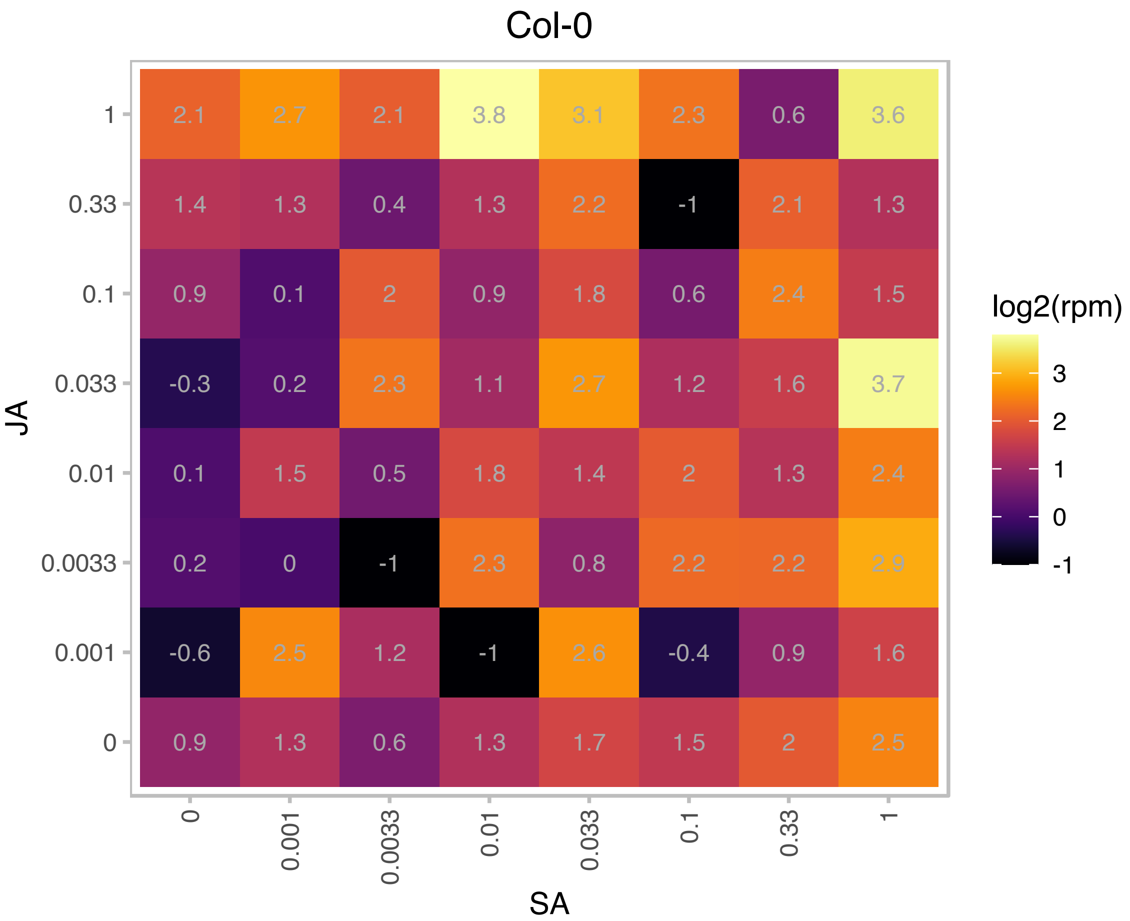


| | |
|---|---|
| Gene | AT1G03770 |
| is.expressed_genes | TRUE |
| is.responsive_genes | TRUE |
| is.apcluster | B-04 |

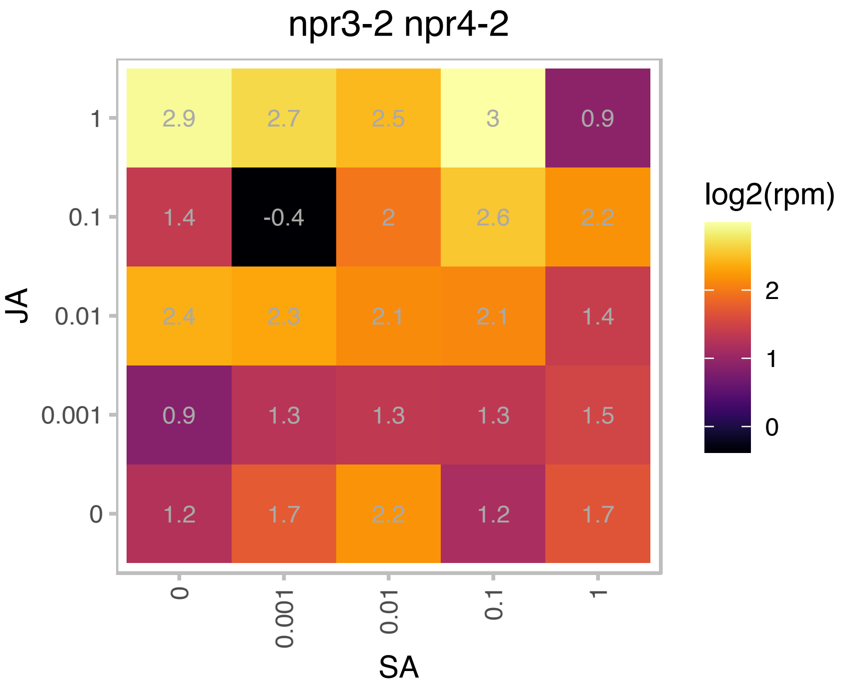


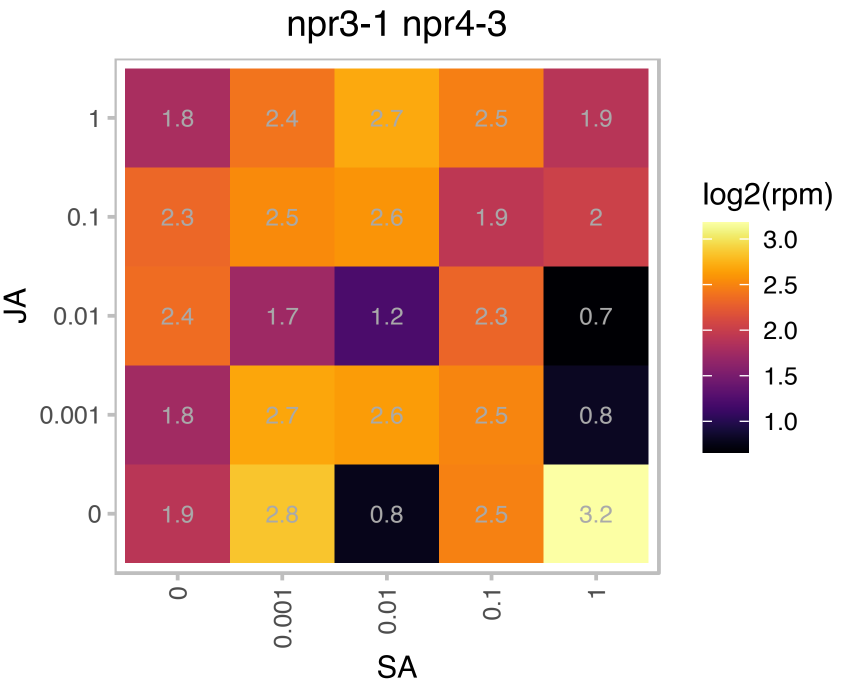

### AT1G03860.1 : ATPHB2, PHB2

prohibitin 2

prohibitin 2

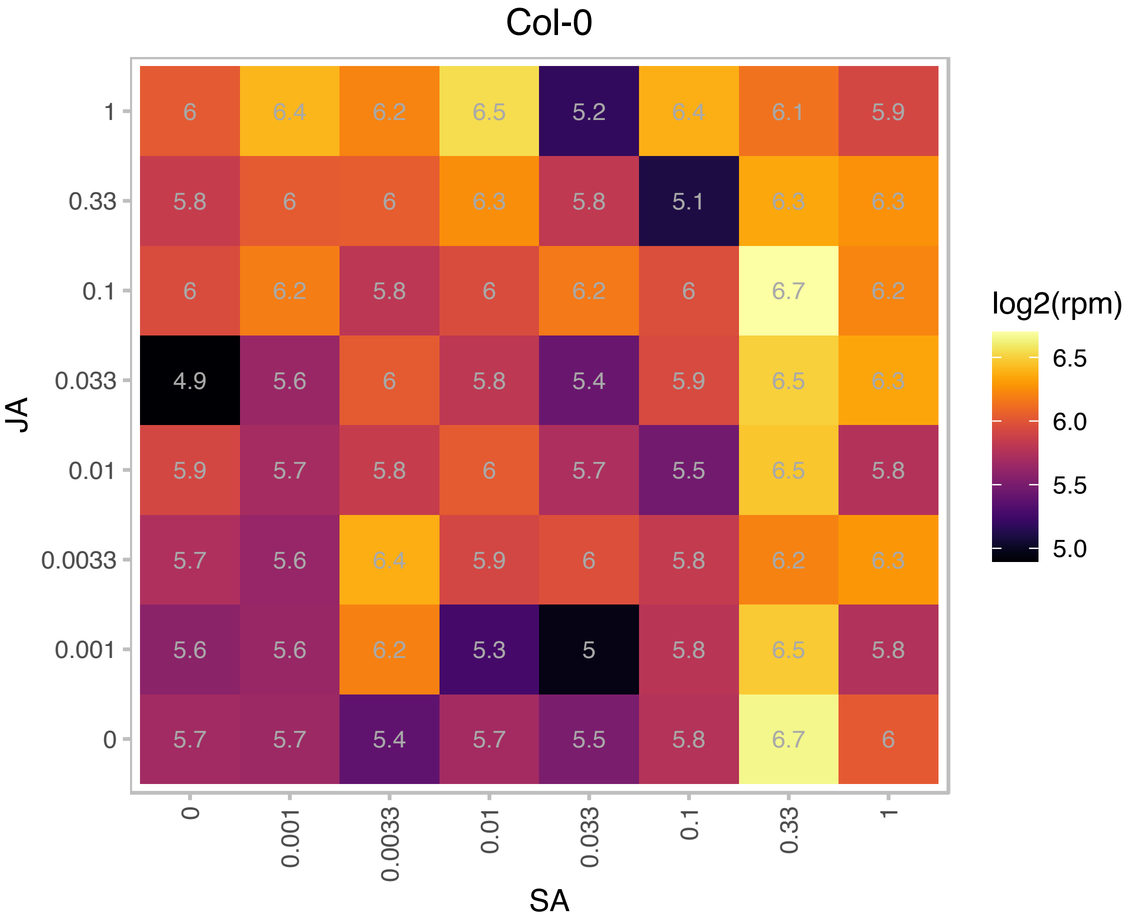


| | |
|---|---|
| Gene | AT1G03860 |
| is.expressed_genes | TRUE |
| is.responsive_genes | TRUE |
| is.apcluster | B-04 |

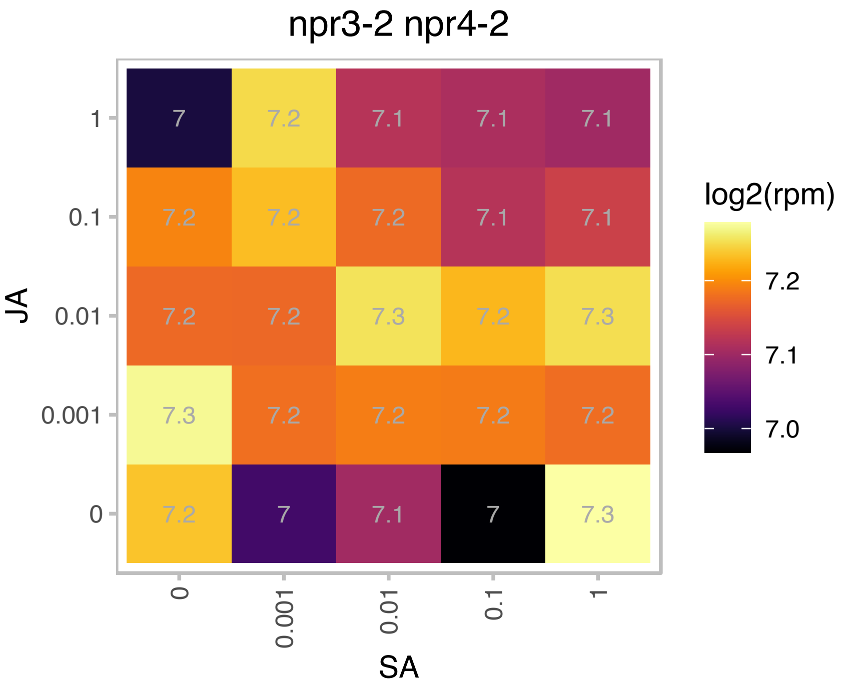


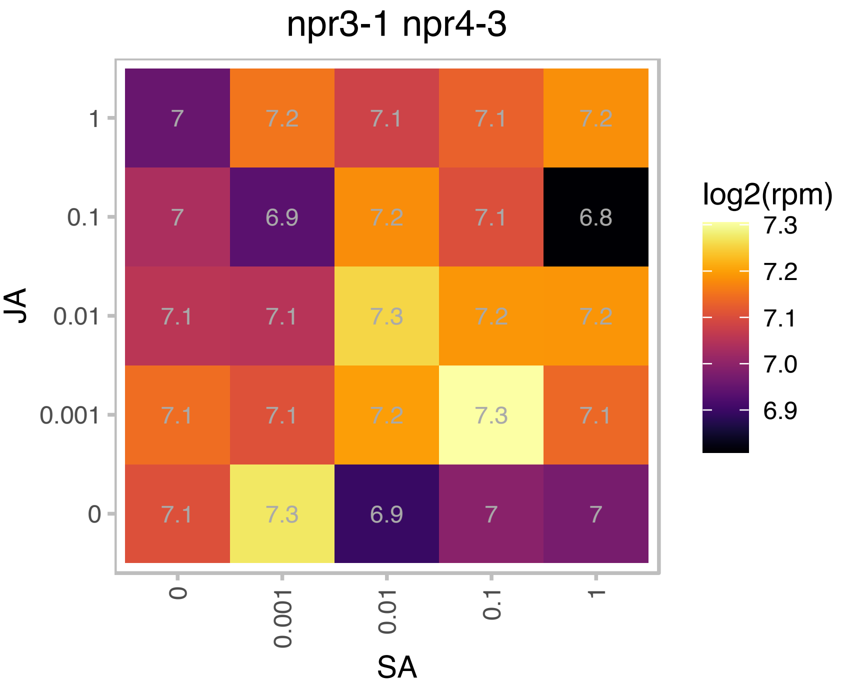

# AT1G04990.1 : NA

Zinc finger C-x8-C-x5-C-x3-H type family protein

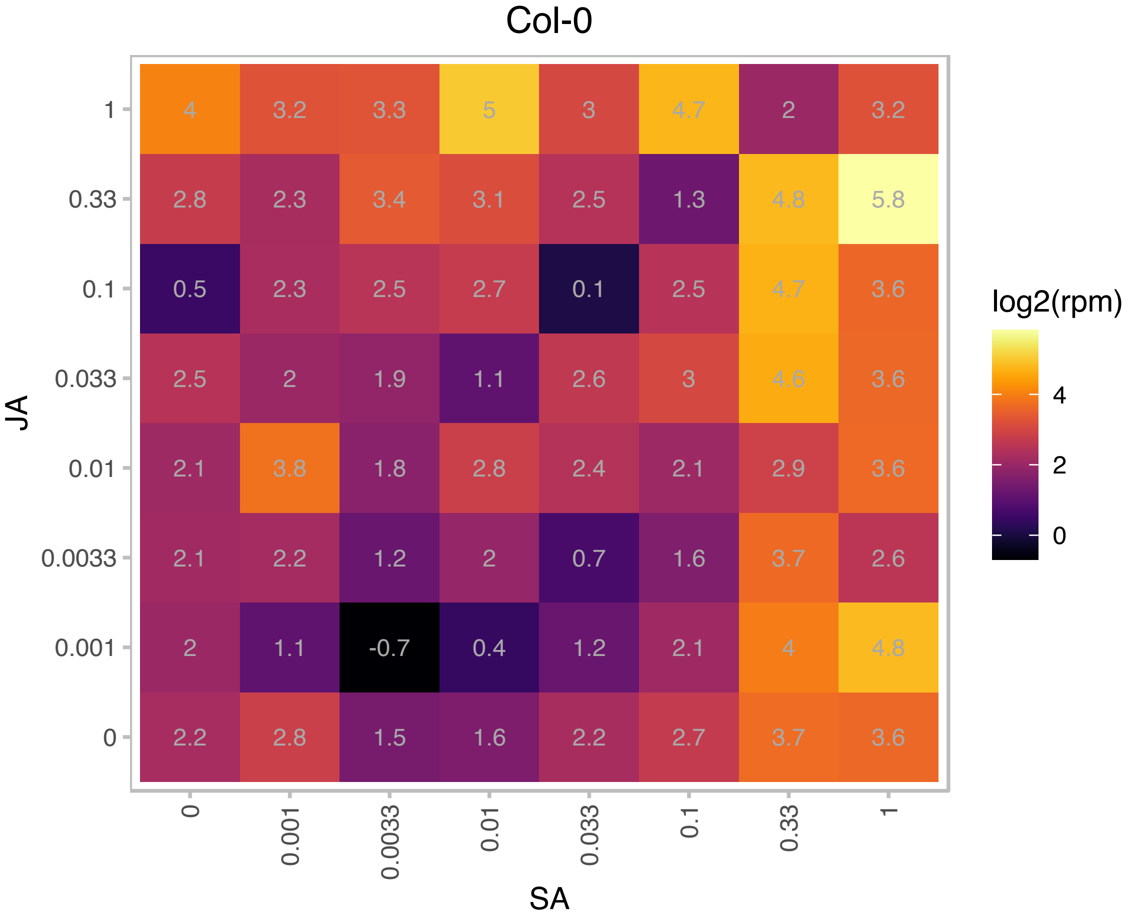


| | |
|---|---|
| Gene | AT1G04990 |
| is.expressed_genes | TRUE |
| is.responsive_genes | TRUE |
| is.apcluster | B-04 |

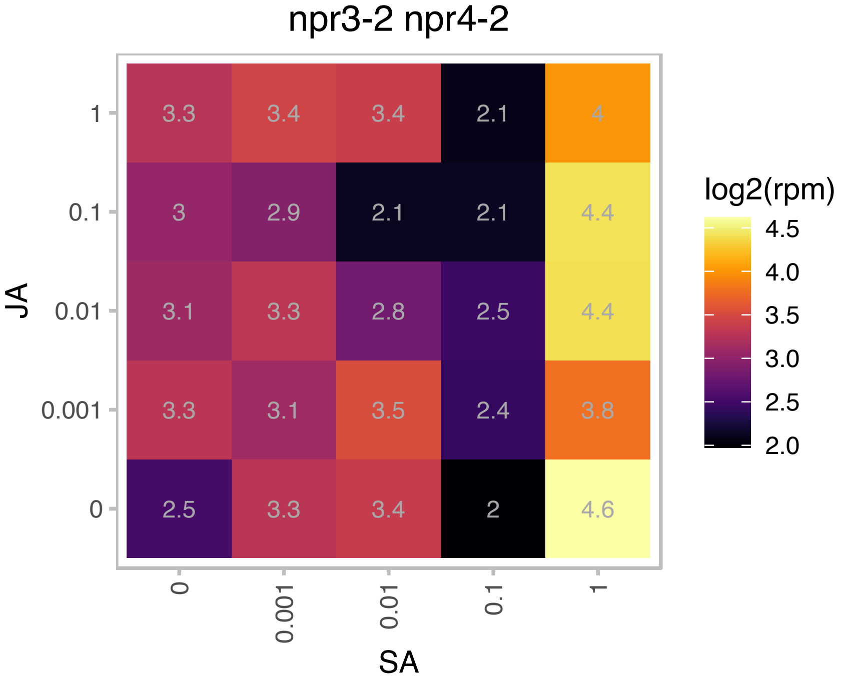


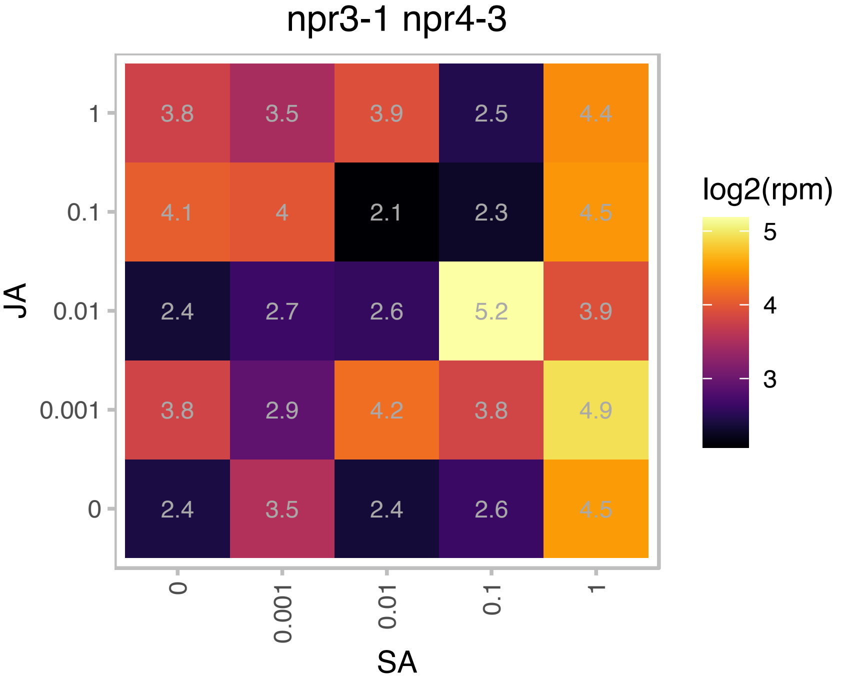

# AT1G05562.1 : NA

Potential natural antisense gene, locus overlaps with AT1G05560

#### Col-0

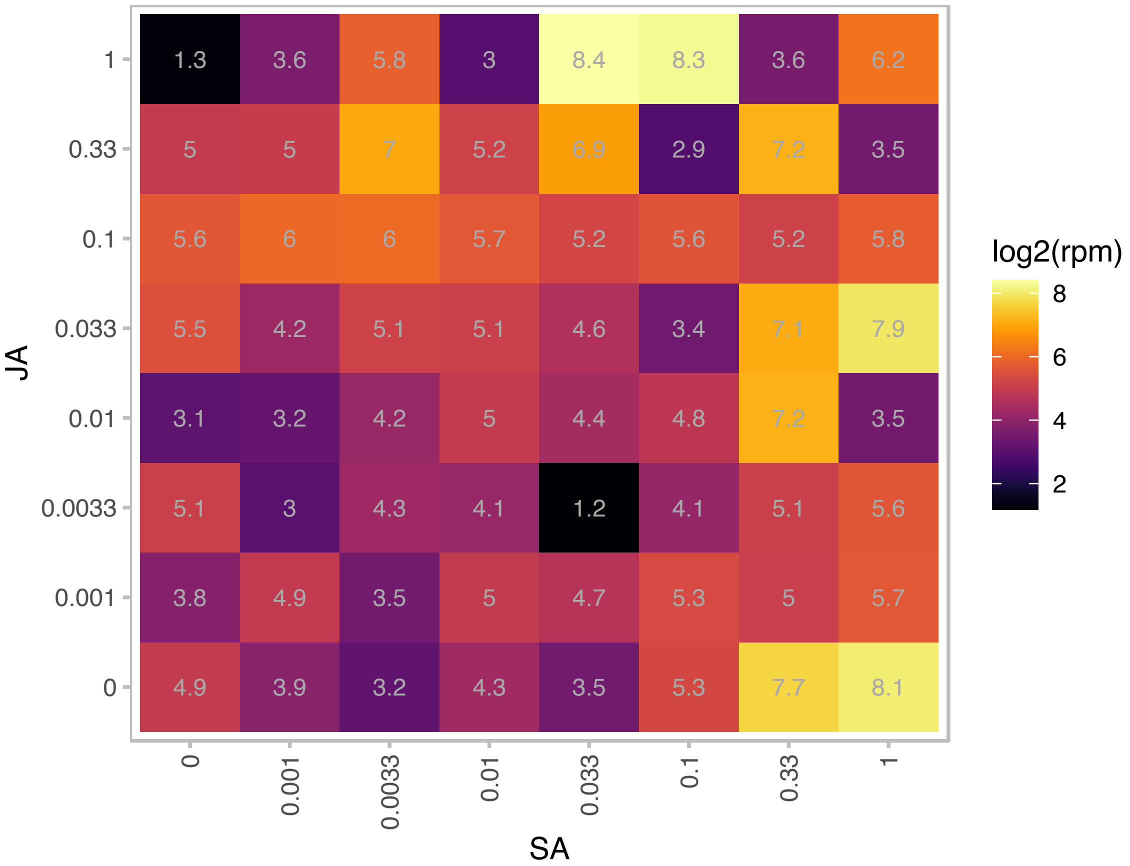


| | |
|---|---|
| Gene | AT1G05562 |
| is.expressed_genes | TRUE |
| is.responsive_genes | TRUE |
| is.apcluster | B-04 |

#### npr3-2 npr4-2

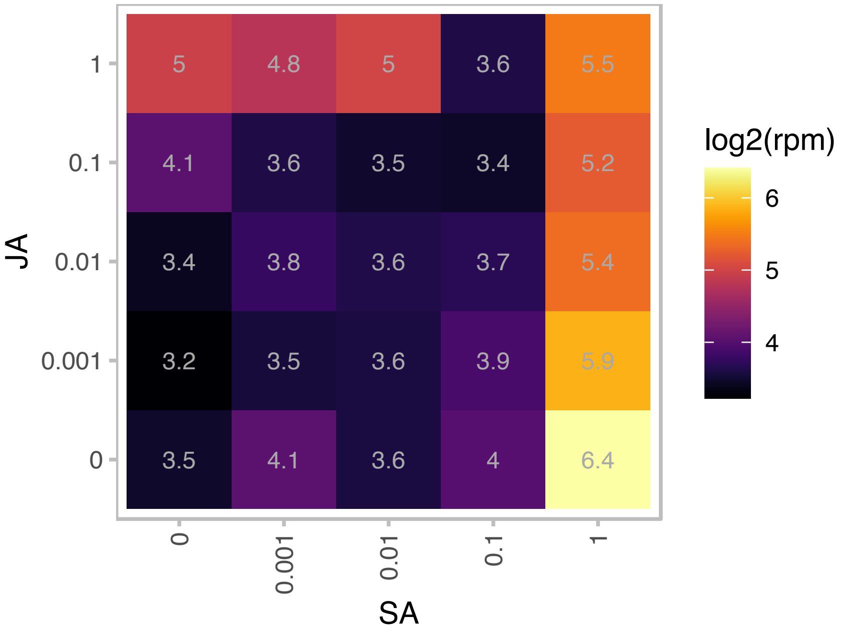


#### npr3-1 npr4-3

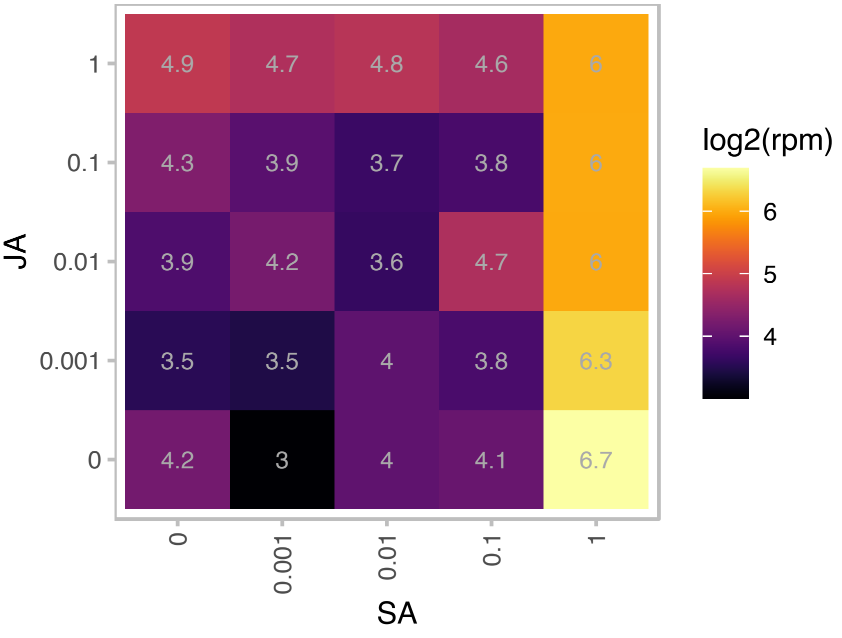

### AT1G05680.1 : UGT74E2

Uridine diphosphate glycosyltransferase 74E2

Encodes a UDP-glucosyltransferase, UGT74E2, that acts on IBA (indole-3-butyric acid) and affects auxin homeostasis. The transcript and protein levels of this enzyme are strongly induced by H2O2 and may allow integration of ROS (reactive oxygen species) and auxin signaling. This enzyme can also transfer glycosyl groups to several compounds related to the explosive TNT when this synthetic compound is taken up from the environment.

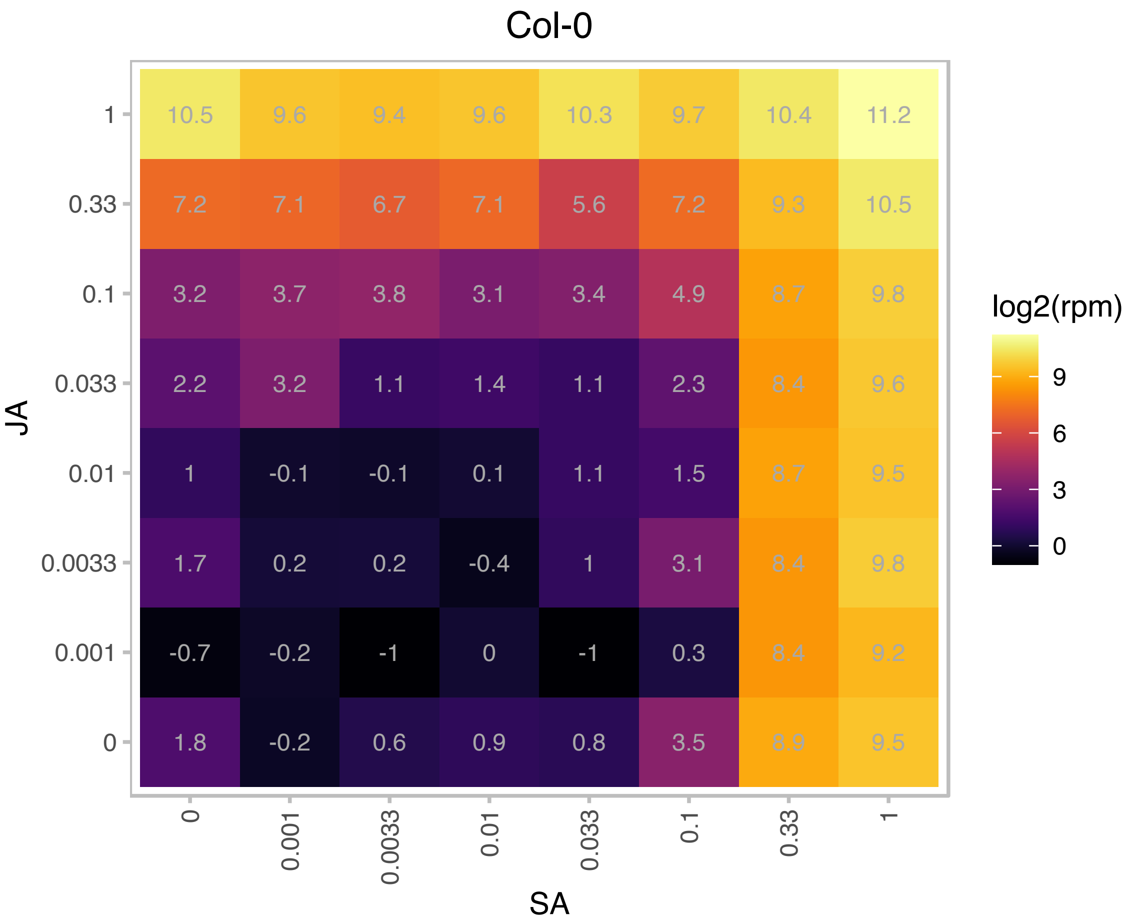


| | |
|---|---|
| Gene | AT1G05680 |
| is.expressed_genes | TRUE |
| is.responsive_genes | TRUE |
| is.apcluster | B-04 |

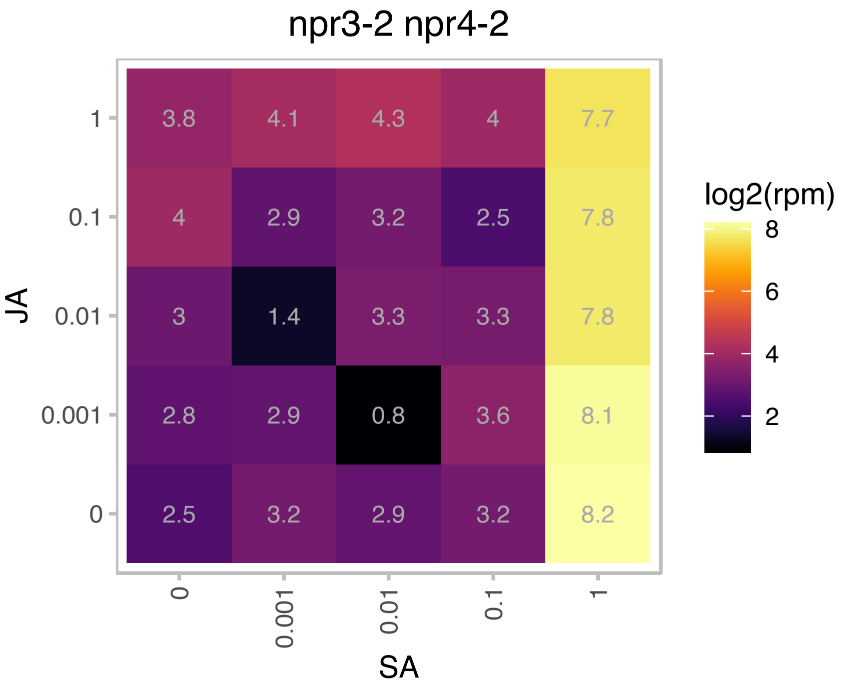


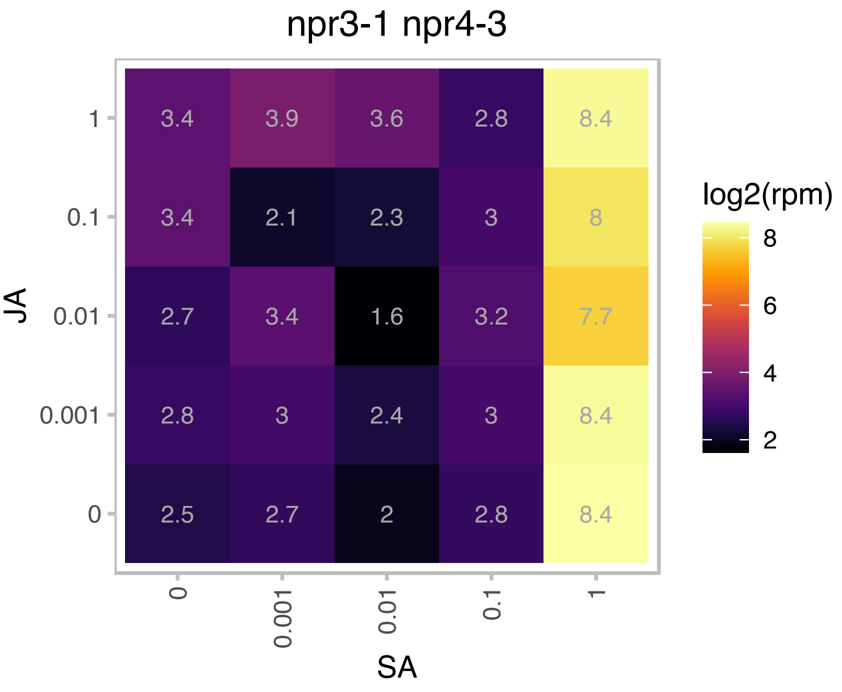

# AT1G09500.1 : NA

NAD(P)-binding Rossmann-fold superfamily protein

similar to Eucalyptus gunnii alcohol dehydrogenase of unknown physiological function (GI:1143445), Vigna unguiculata (gi:1854445), NOT a cinnamyl-alcohol dehydrogenase

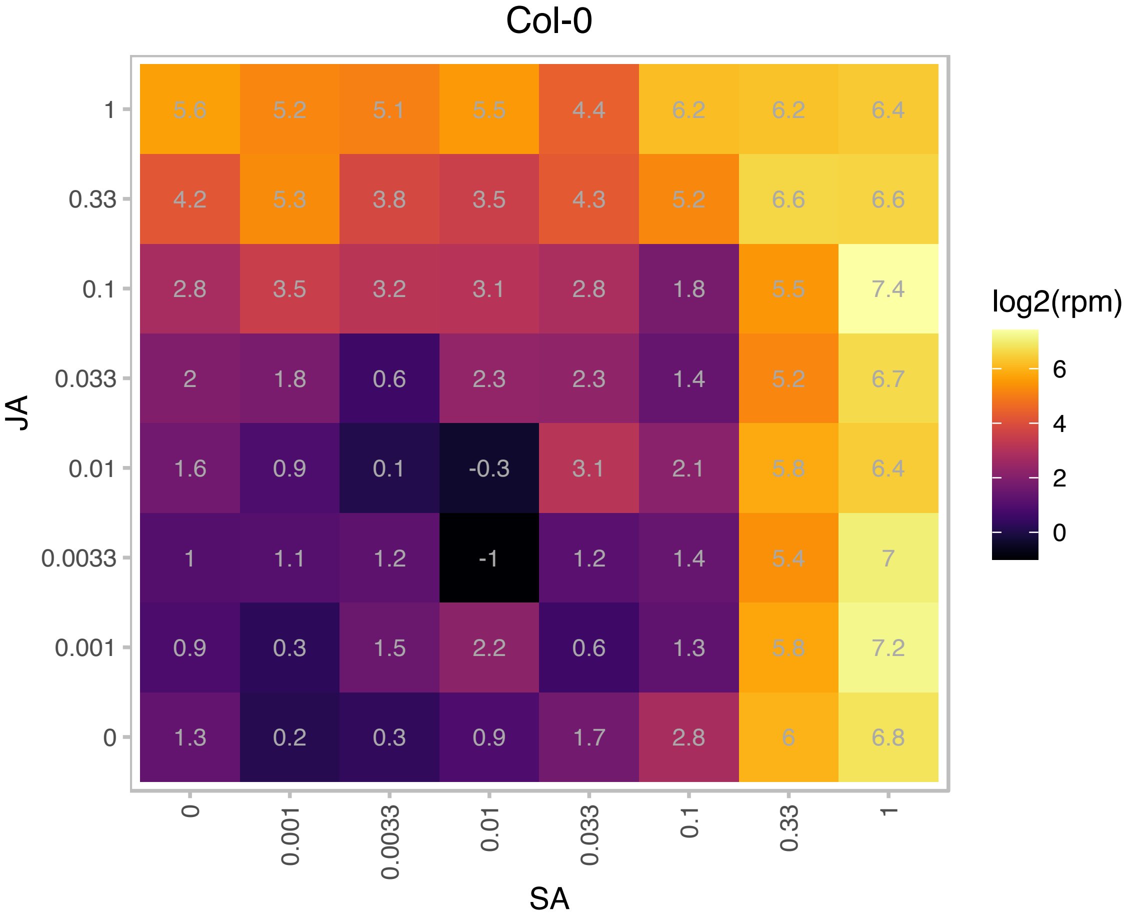


| | |
|---|---|
| Gene | AT1G09500 |
| is.expressed_genes | TRUE |
| is.responsive_genes | TRUE |
| is.apcluster | B-04 |

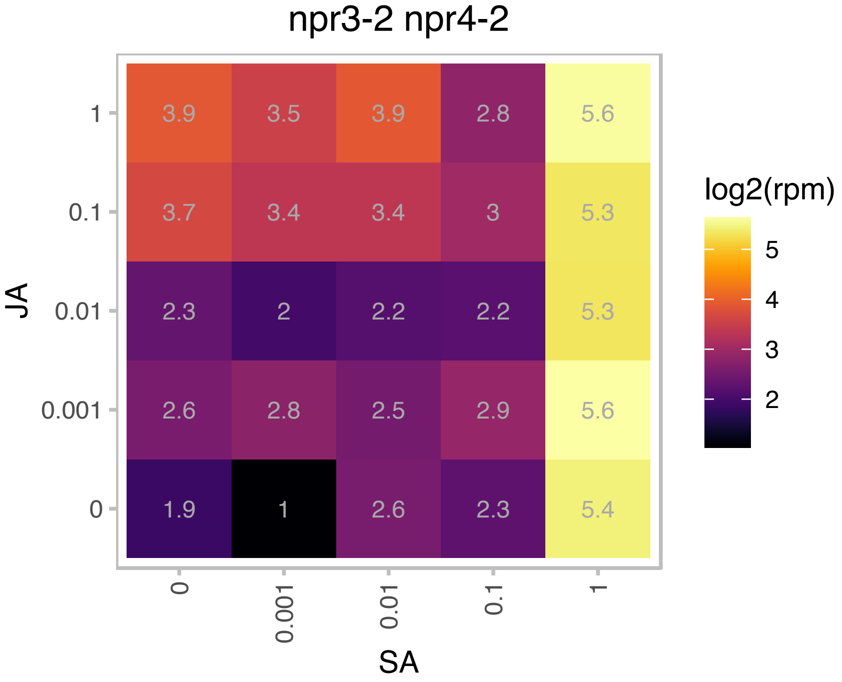


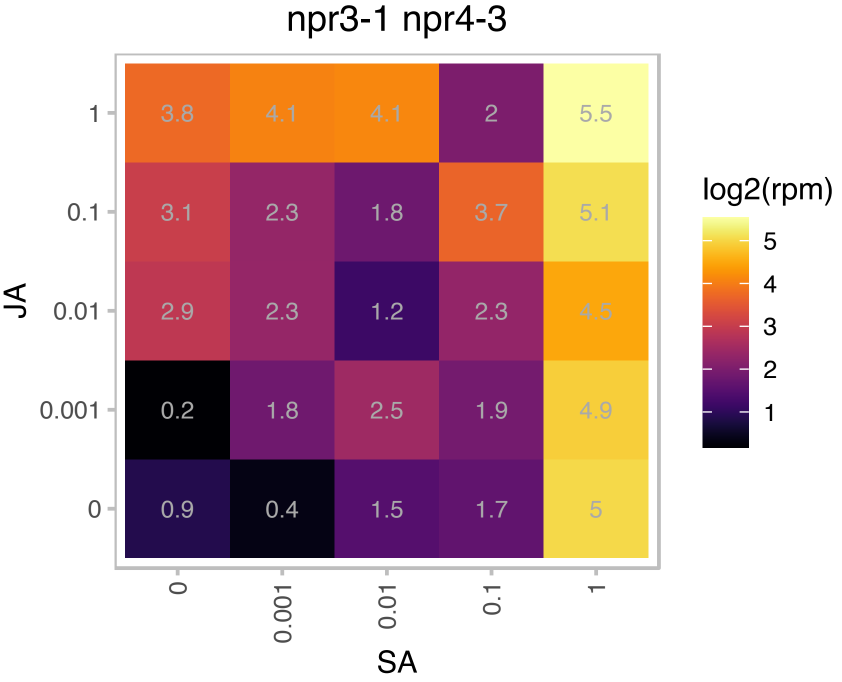

### AT1G12350.1 : ATCOAB, COAB

4-phospho-panto-thenoylcysteine synthetase

At1g12350 encodes phosphopantothenoylcysteine synthetase (phosphopantothenoylcysteine ligase). Its molecular function was shown to add cysteine forming (R)-4'-phospho-N-pantothenoylcysteine (PPC)

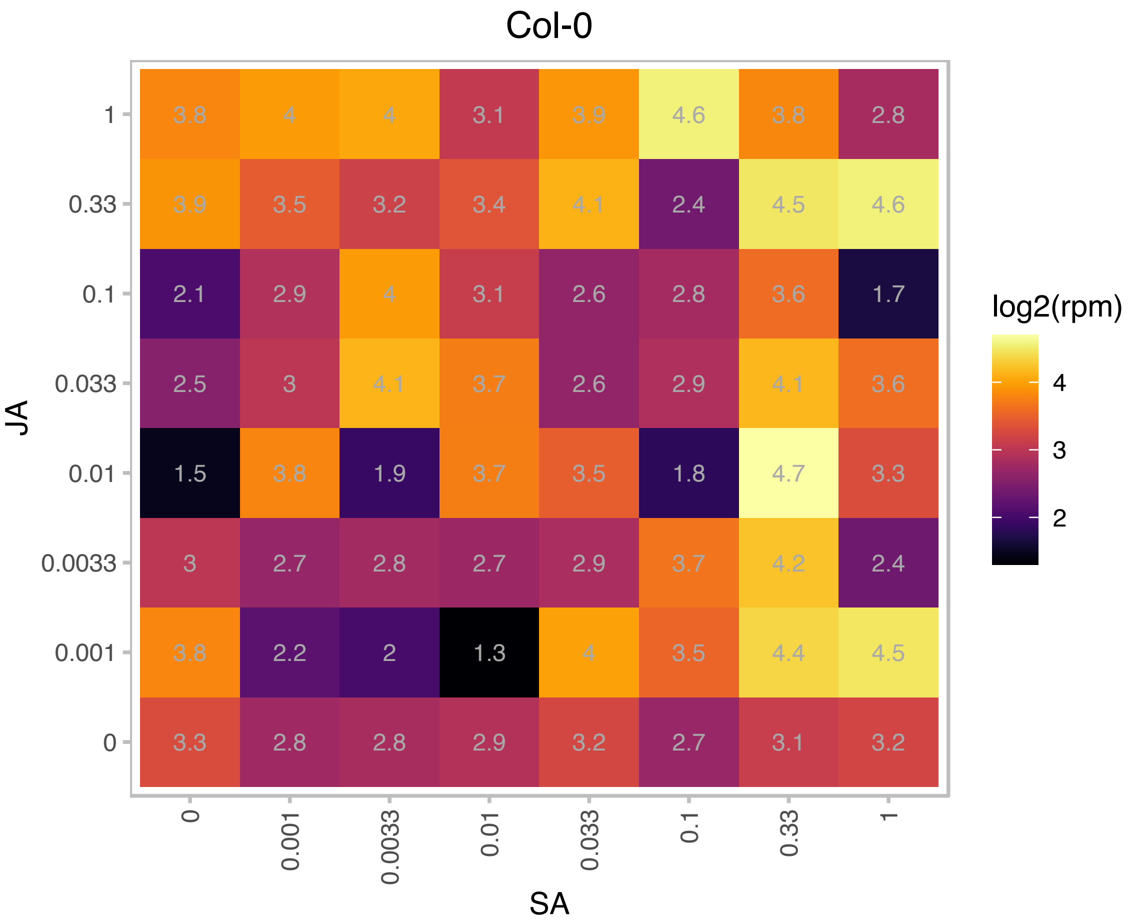


| | |
|---|---|
| Gene | AT1G12350 |
| is.expressed_genes | TRUE |
| is.responsive_genes | TRUE |
| is.apcluster | B-04 |

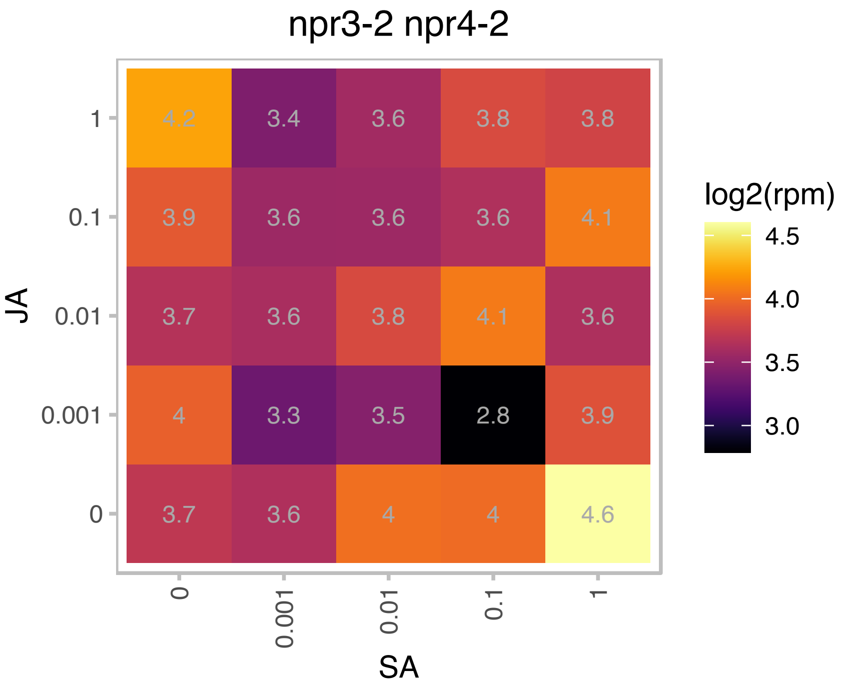


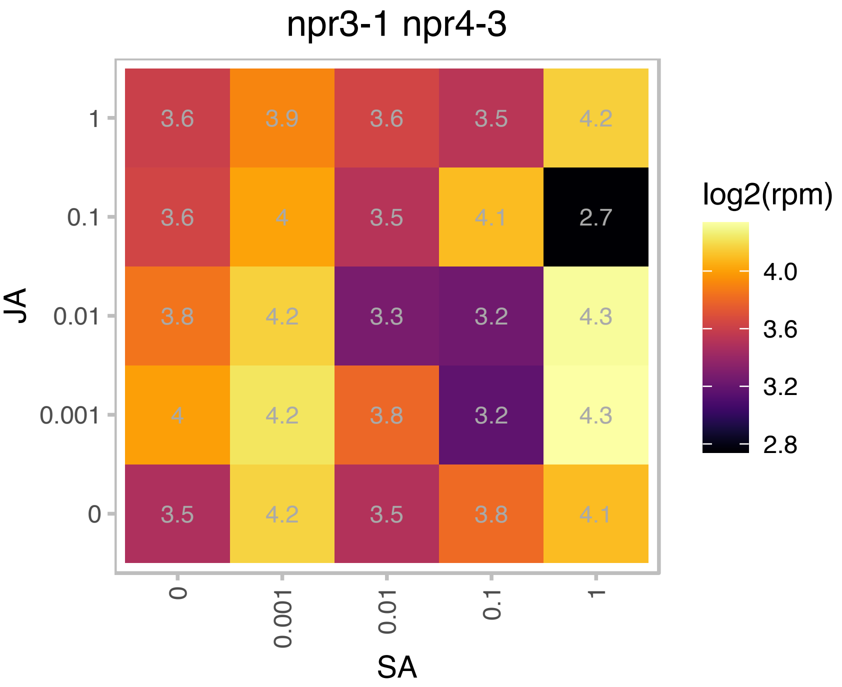

### AT1G13060.1 : PBE1

20S proteasome beta subunit E1

Encodes 20S proteasome beta subunit PBE1 (PBE1).

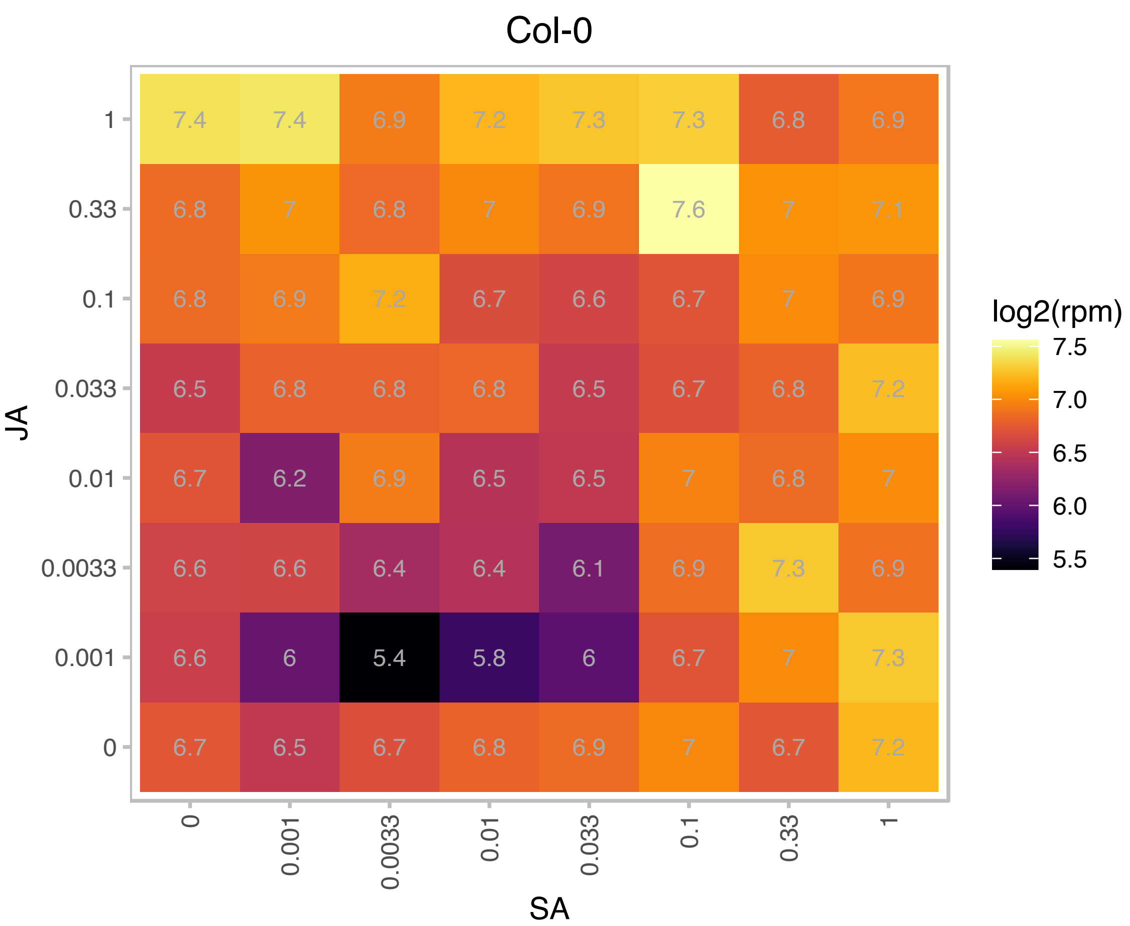


| | |
|---|---|
| Gene | AT1G13060 |
| is.expressed_genes | TRUE |
| is.responsive_genes | TRUE |
| is.apcluster | B-04 |

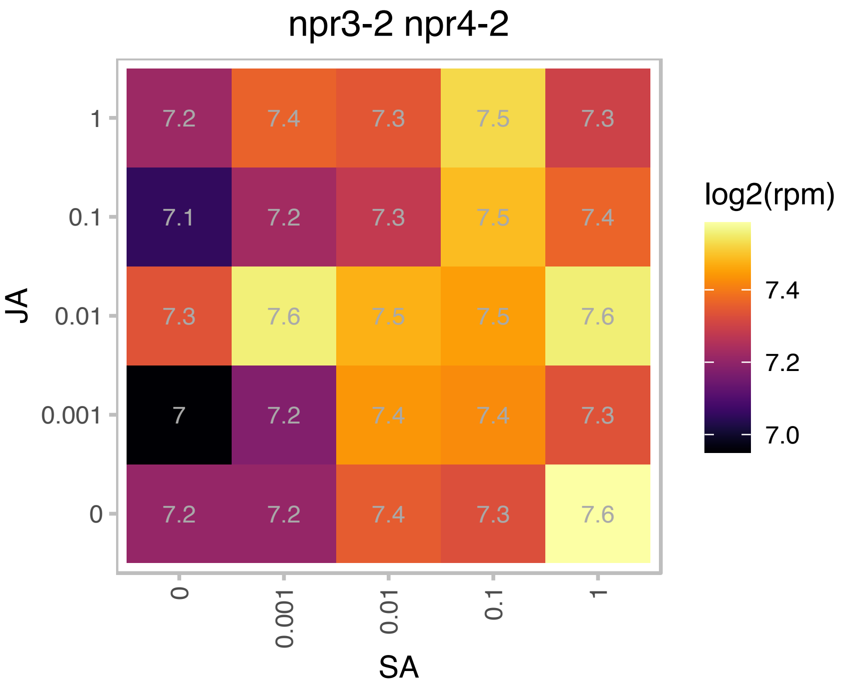


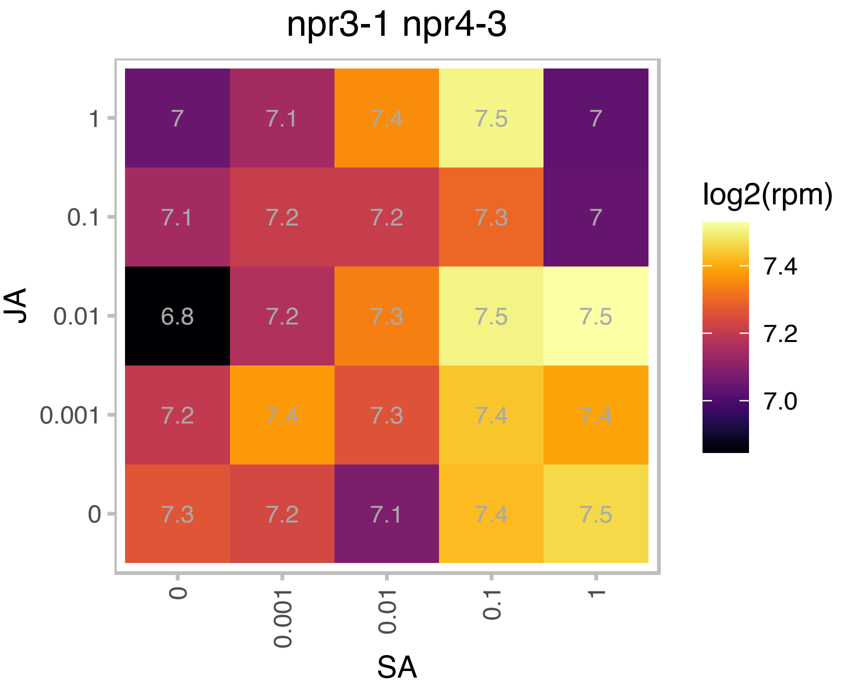

### AT1G17020.1 : ATSRG1, SRG1

senescence-related gene 1

Encodes a novel member of the Fe(II)/ascorbate oxidase gene family; senescence-related gene.

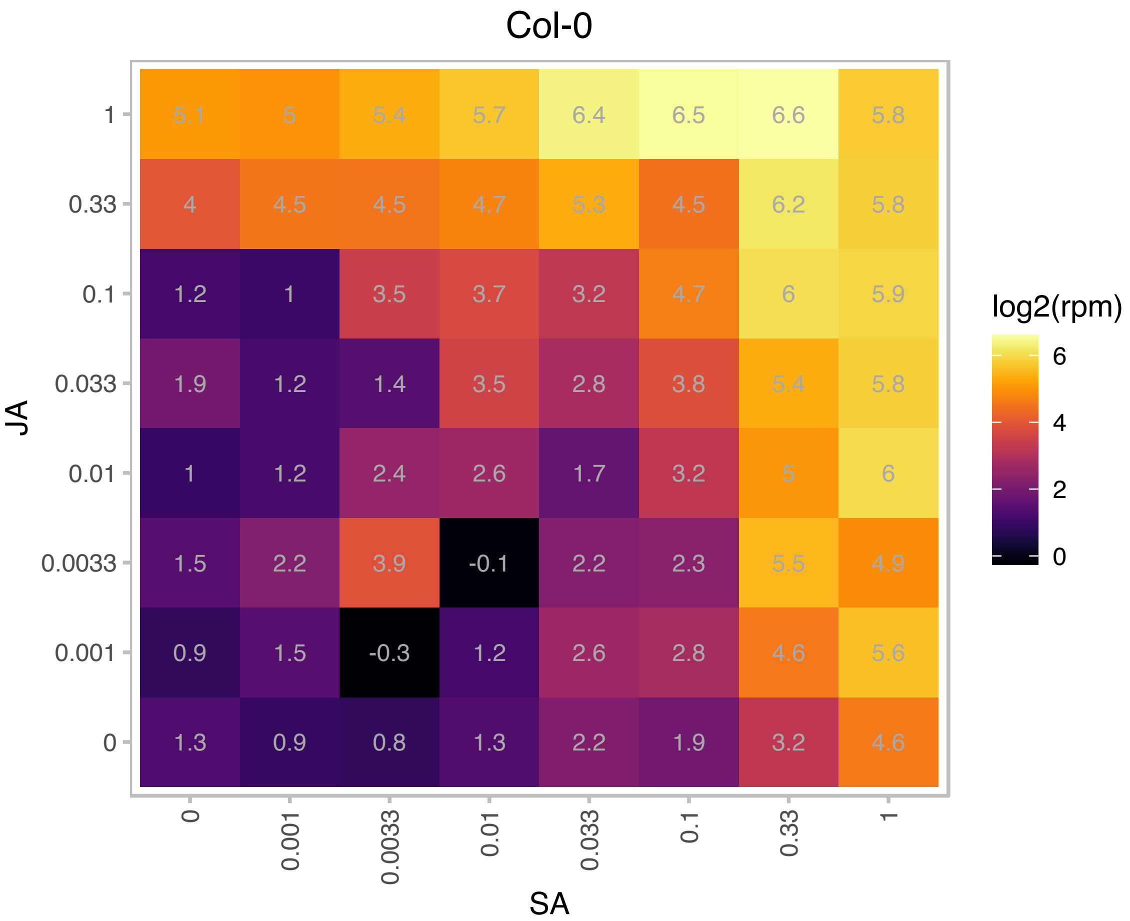


| | |
|---|---|
| Gene | AT1G17020 |
| is.expressed_genes | TRUE |
| is.responsive_genes | TRUE |
| is.apcluster | B-04 |

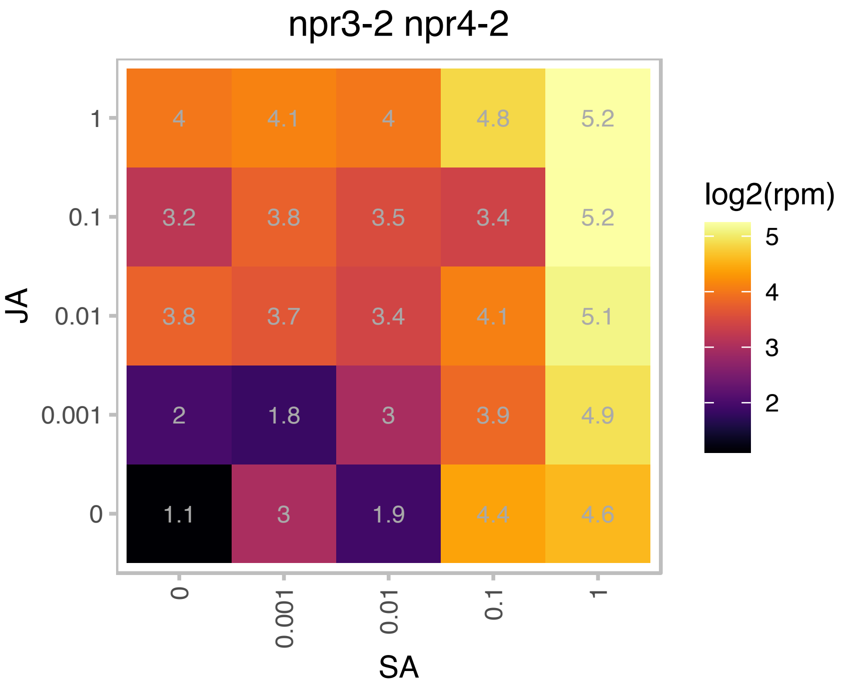


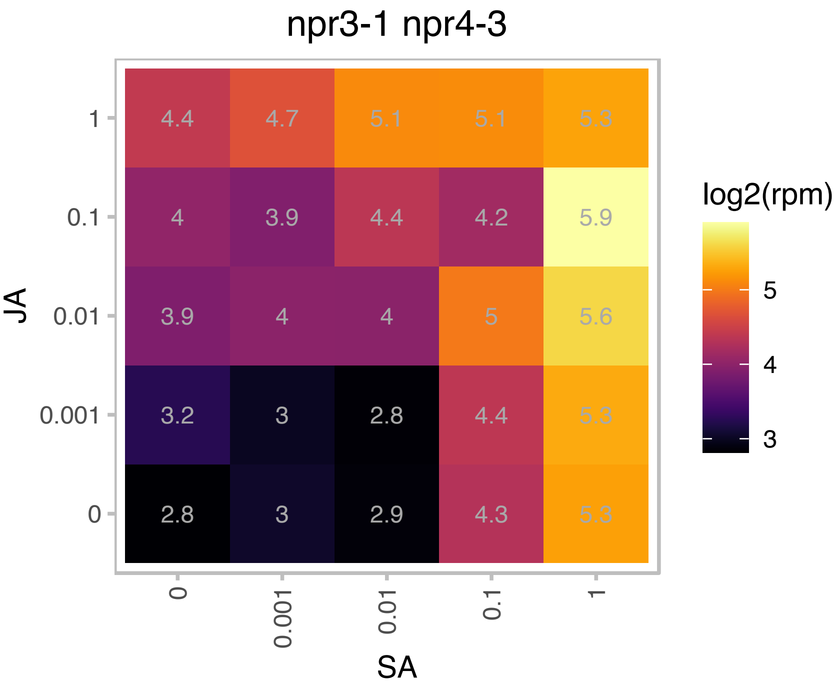

### AT1G17340.1 : SAC5

Phosphoinositide phosphatase family protein

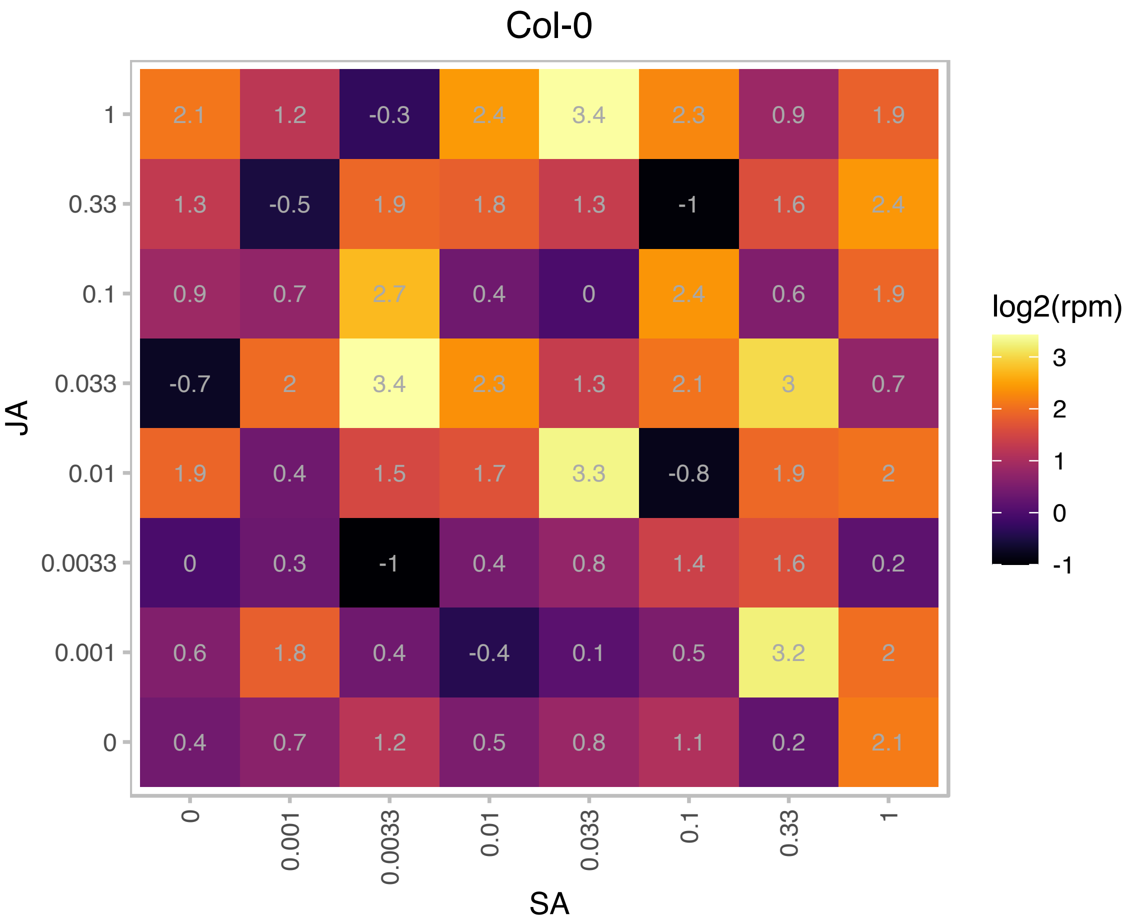


| | |
|---|---|
| Gene | AT1G17340 |
| is.expressed_genes | TRUE |
| is.responsive_genes | TRUE |
| is.apcluster | B-04 |

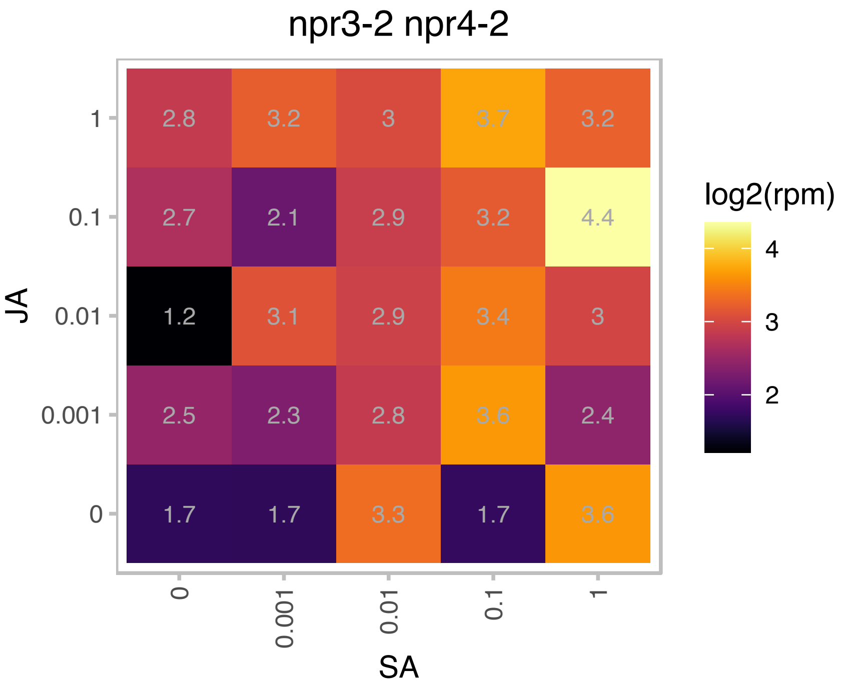


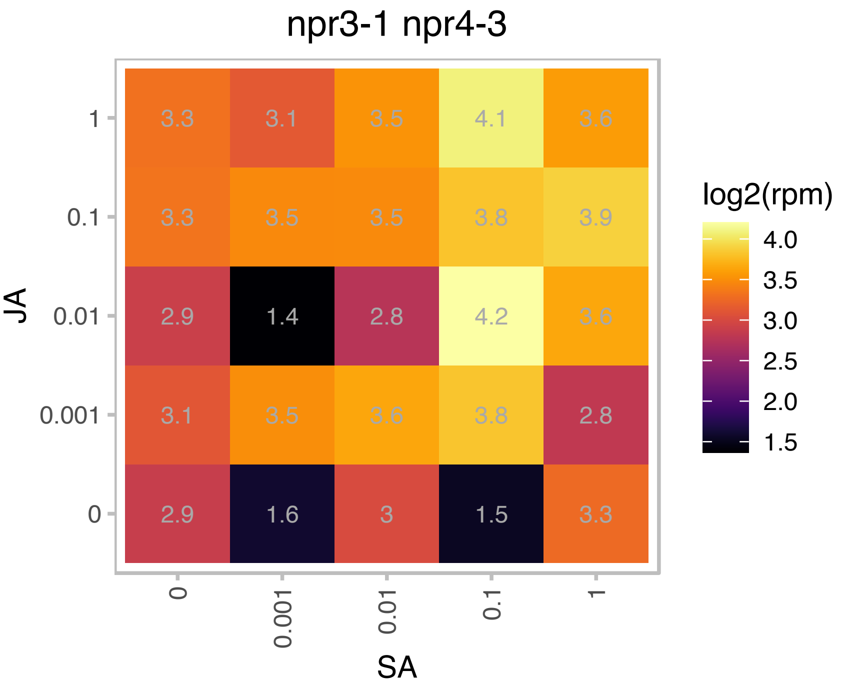

### AT1G17650.1 : AtGLYR2, GLYR2, GR2

glyoxylate reductase 2

Glyoxylate reductase located in chloroplasts.

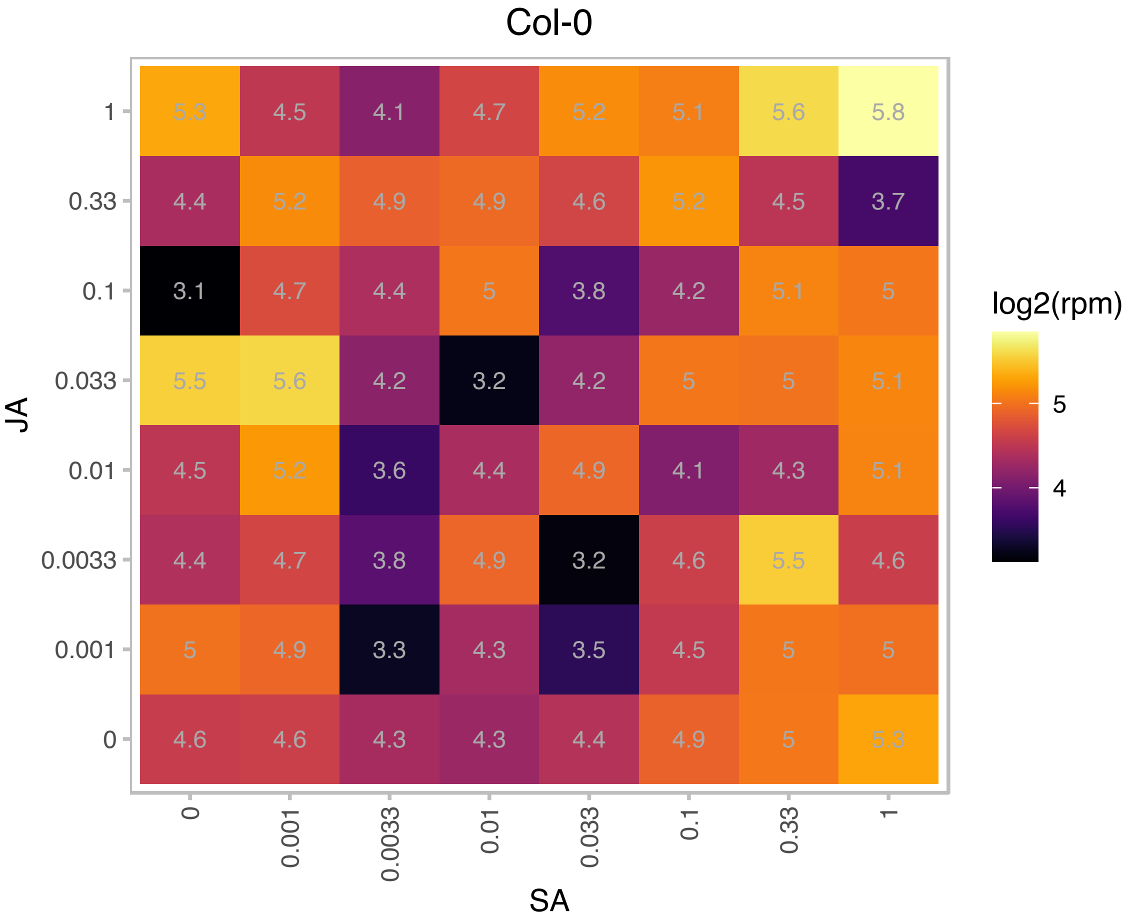


| | |
|---|---|
| Gene | AT1G17650 |
| is.expressed_genes | TRUE |
| is.responsive_genes | TRUE |
| is.apcluster | B-04 |

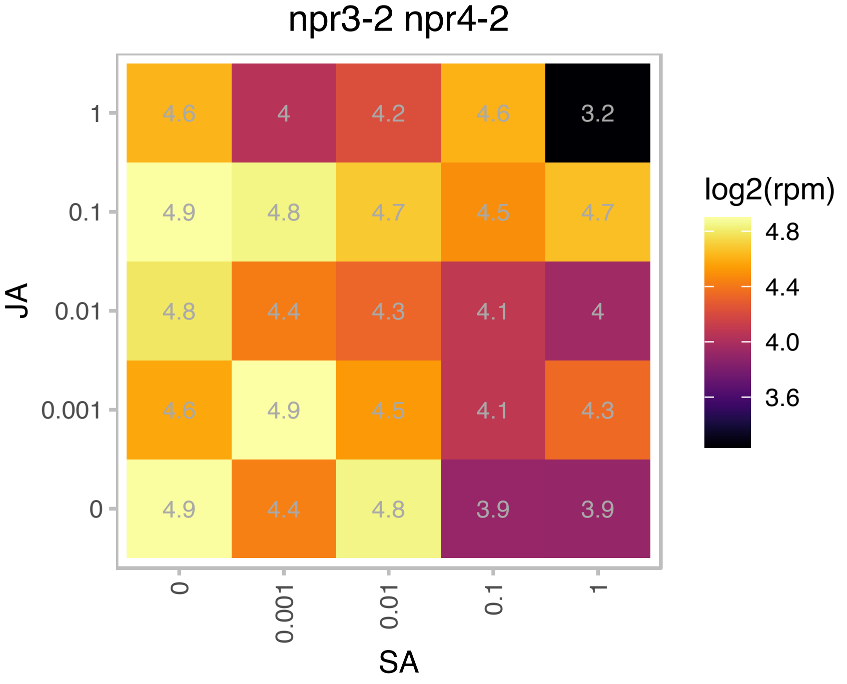


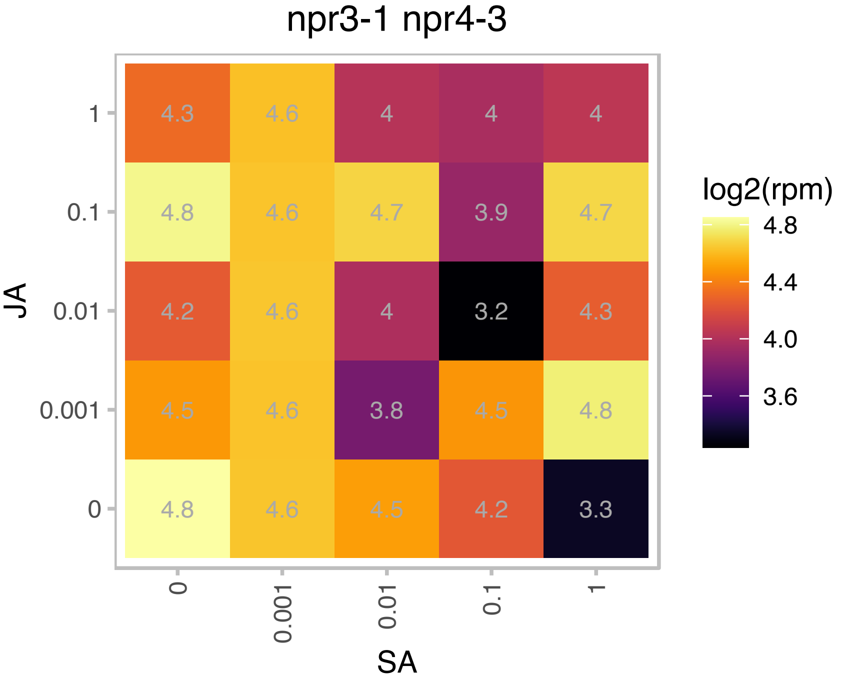

# AT1G29195.1 : NA

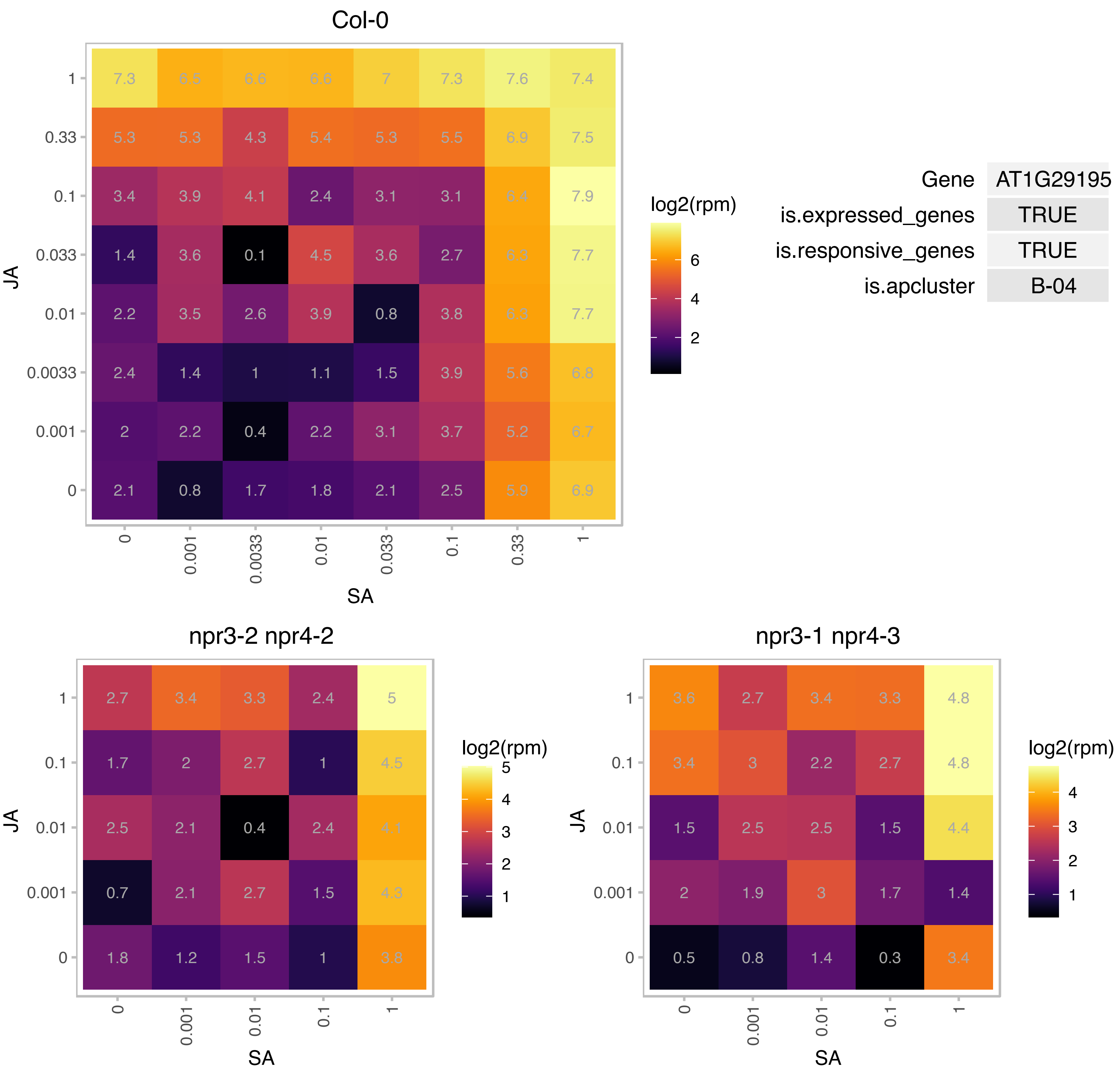

### AT1G29395.1 : COR413-TM1, COR413IM1, COR414-TM1

COLD REGULATED 314 INNER MEMBRANE 1

Integral membrane protein in the inner envelope of chloroplasts. Provide freezing tolerance. Expression is induced by short-term cold-treatment, water deprivation, and abscisic acid treatment.

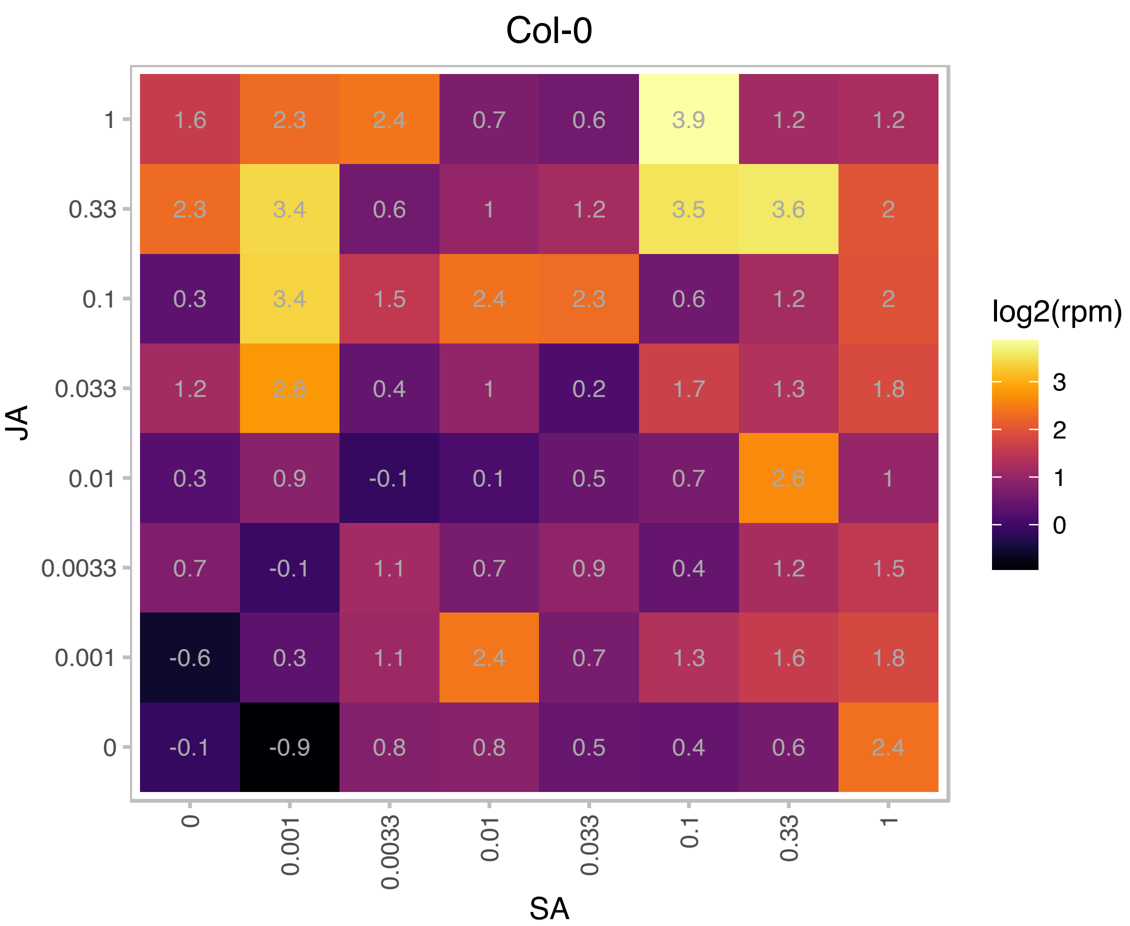


| | |
|---|---|
| Gene | AT1G29395 |
| is.expressed_genes | TRUE |
| is.responsive_genes | TRUE |
| is.apcluster | B-04 |

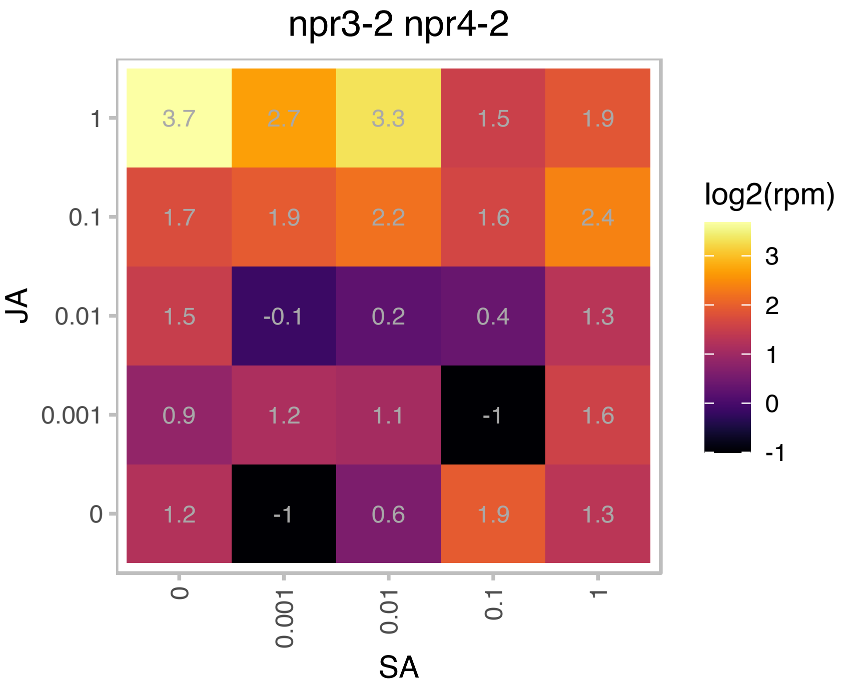


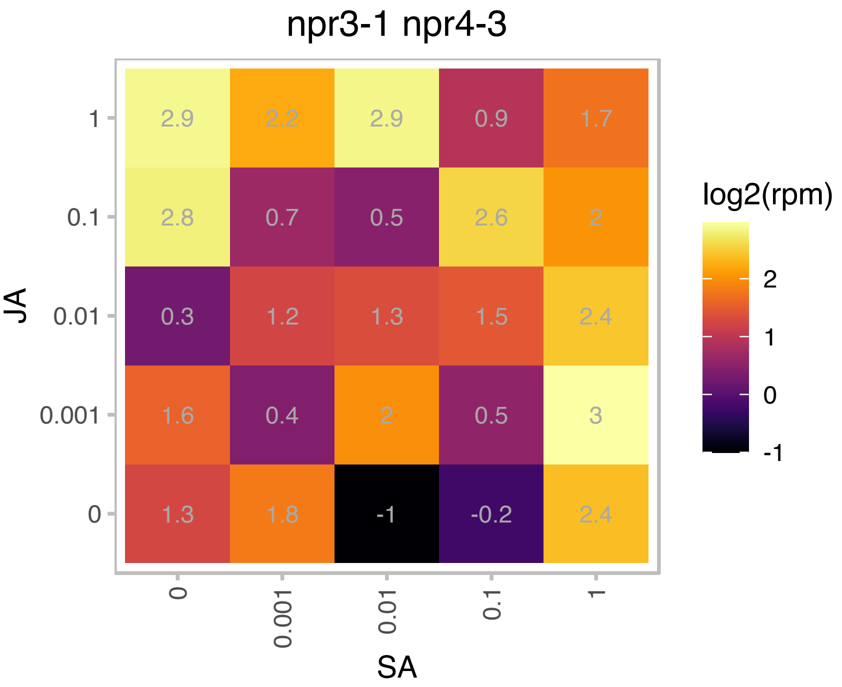

# AT1G47128.1 : RD21, RD21A

Granulin repeat cysteine protease family protein
cysteine proteinase precursor-like protein/ dehydration stress-responsive gene (RD21). Has been shown to have peptide ligase activity and protease activity in vitro. RD21 is involved in immunity to the necrotrophic fungal pathogen Botrytis cinerea.Activity detected in root, leaf, flower and cell culture.

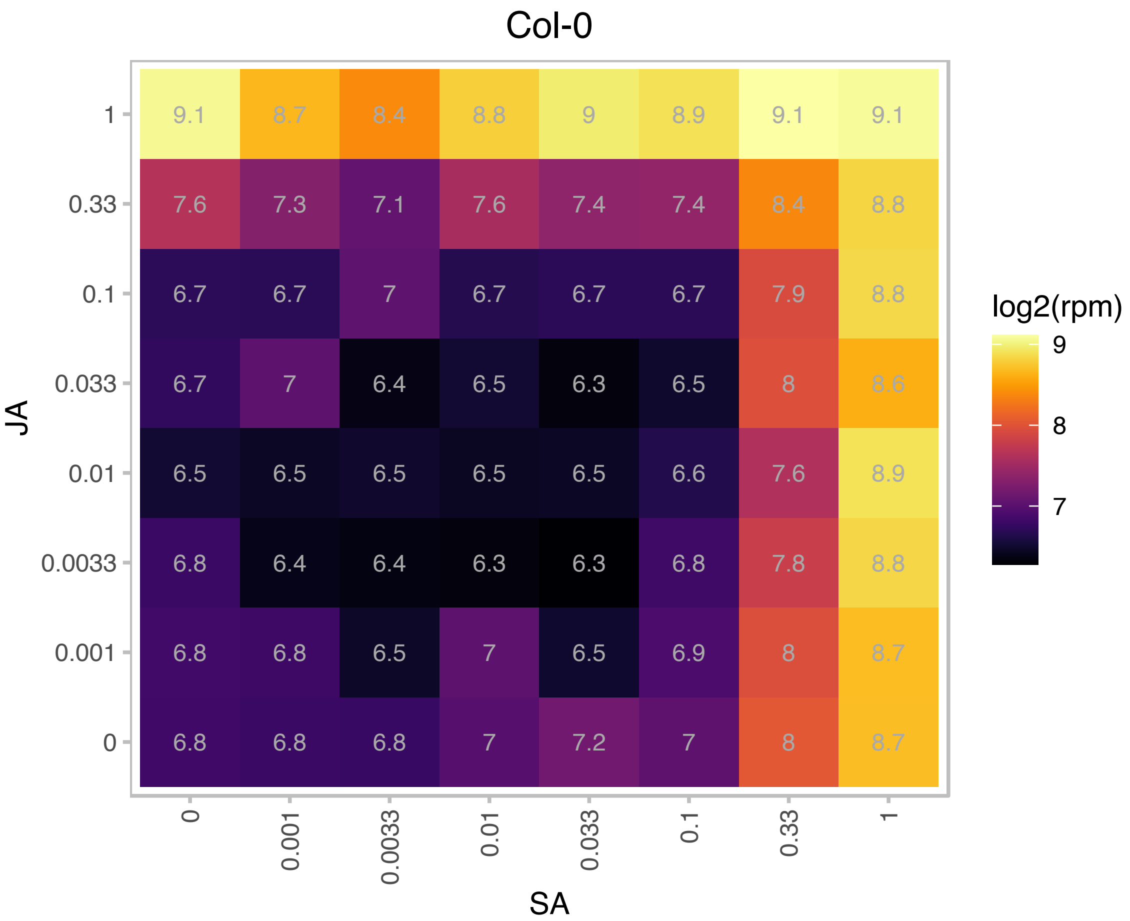


| | |
|---|---|
| Gene | AT1G47128 |
| is.expressed_genes | TRUE |
| is.responsive_genes | TRUE |
| is.apcluster | B-04 |

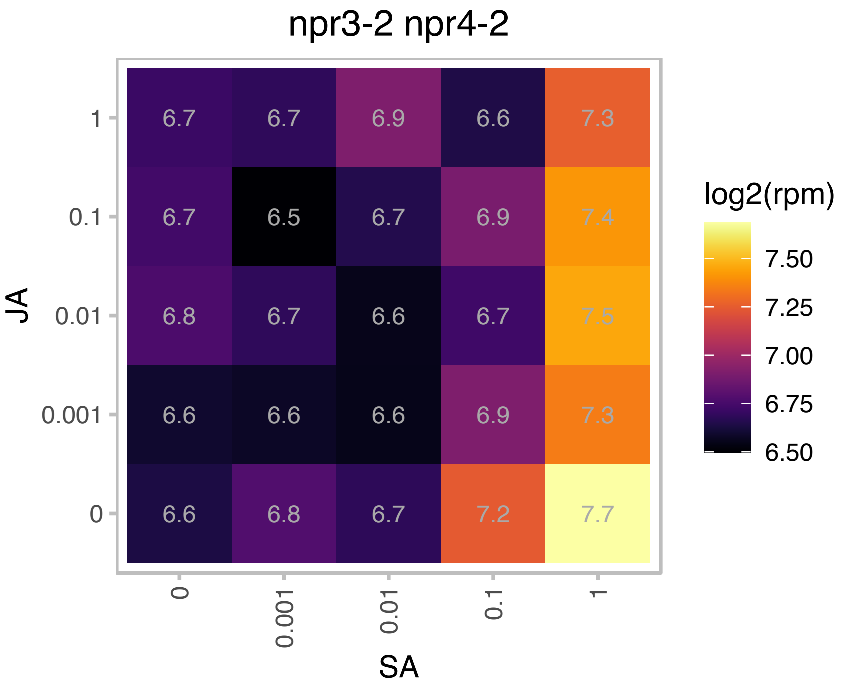


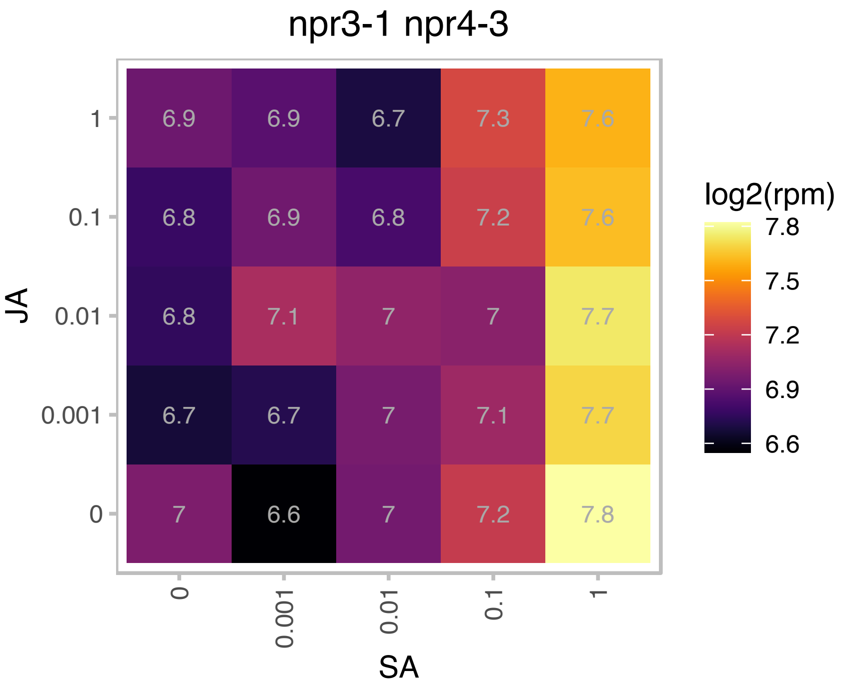

### AT1G51070.1 : bHLH115

basic helix-loop-helix (bHLH) DNA-binding superfamily protein

bHLH115 is a basic helix loop helix protein of the IVc subgroup that plays a role in iron homeostasis. It interacts with related family members and targets PYE and other genes involved in response to Fe.

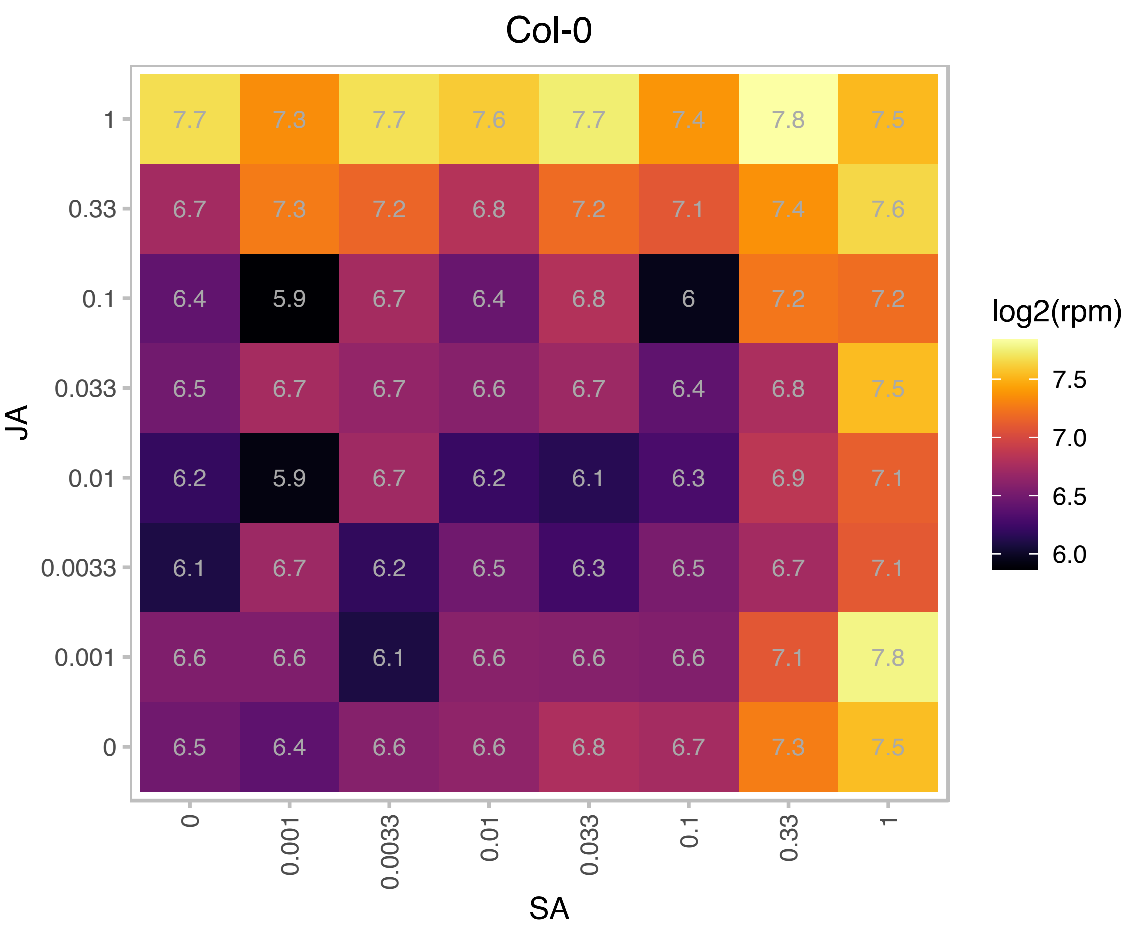


| | |
|---|---|
| Gene | AT1G51070 |
| is.expressed_genes | TRUE |
| is.responsive_genes | TRUE |
| is.apcluster | B-04 |

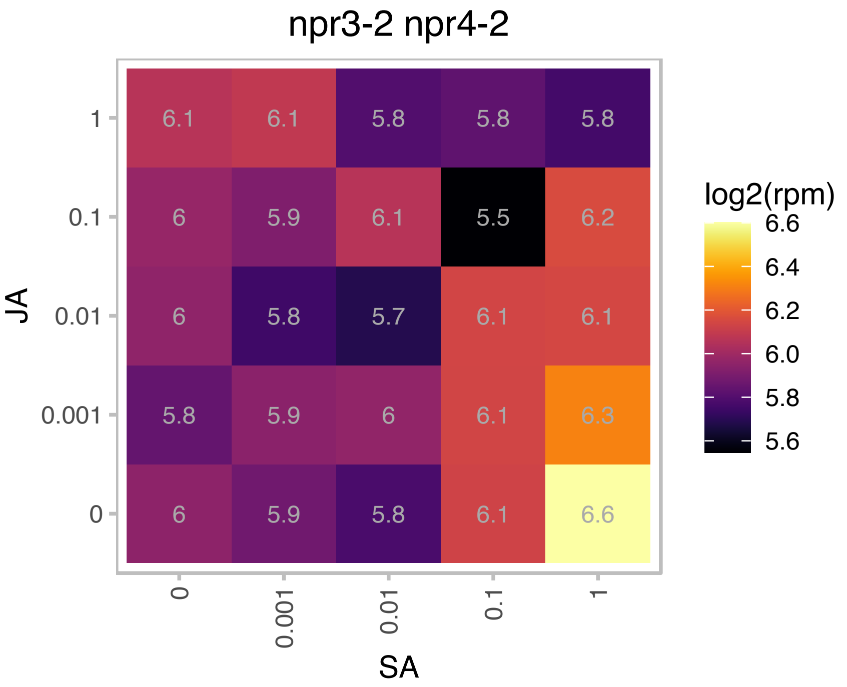


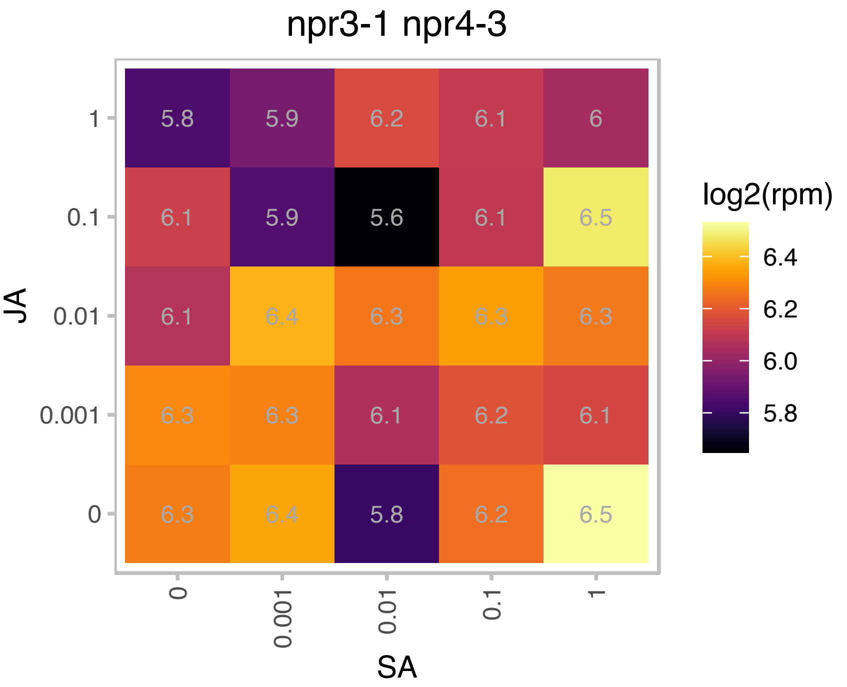

### AT1G60940.1 : SNRK2-10, SNRK2.10, SRK2B

SNF1-related protein kinase 2.10

encodes a member of SNF1-related protein kinases (SnRK2) whose activity is activated by ionic (salt) and non-ionic (mannitol) osmotic stress.

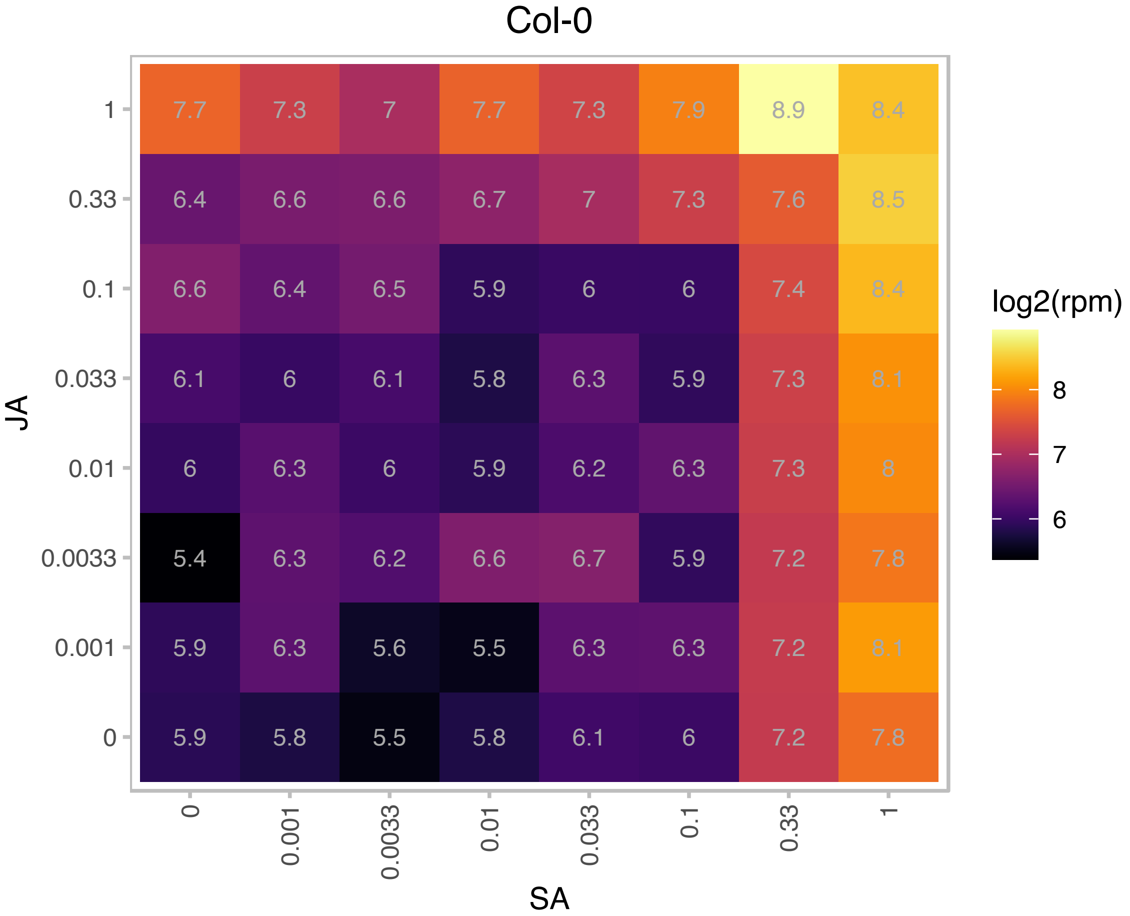


| | |
|---|---|
| Gene | AT1G60940 |
| is.expressed_genes | TRUE |
| is.responsive_genes | TRUE |
| is.apcluster | B-04 |

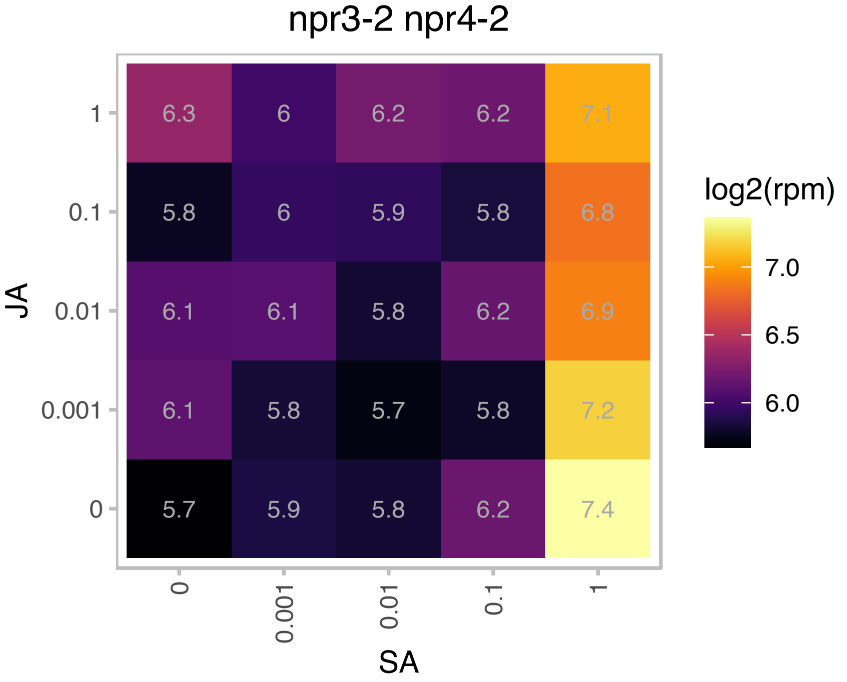


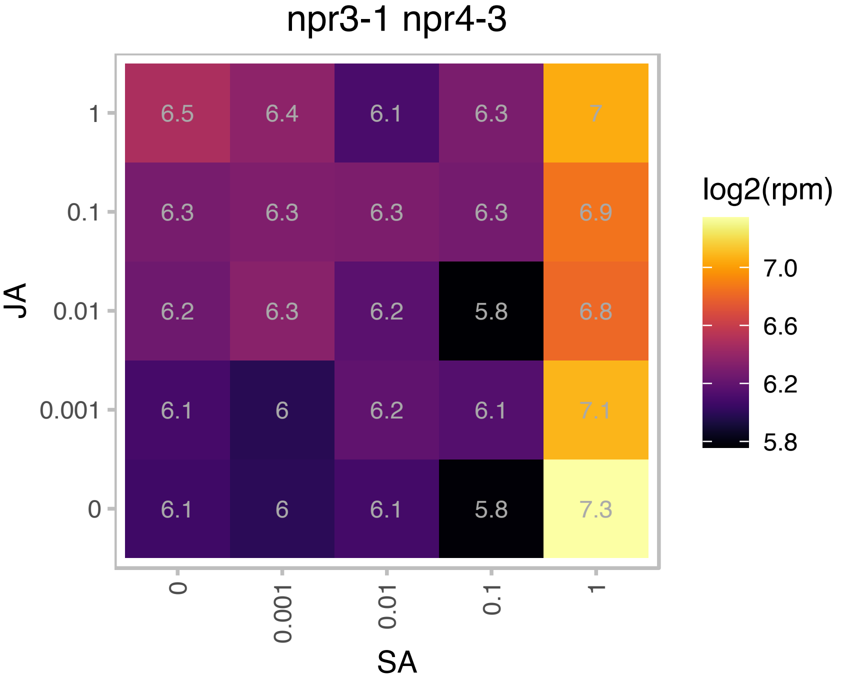

# AT1G61150.1 : NA

LisH and RanBPM domains containing protein

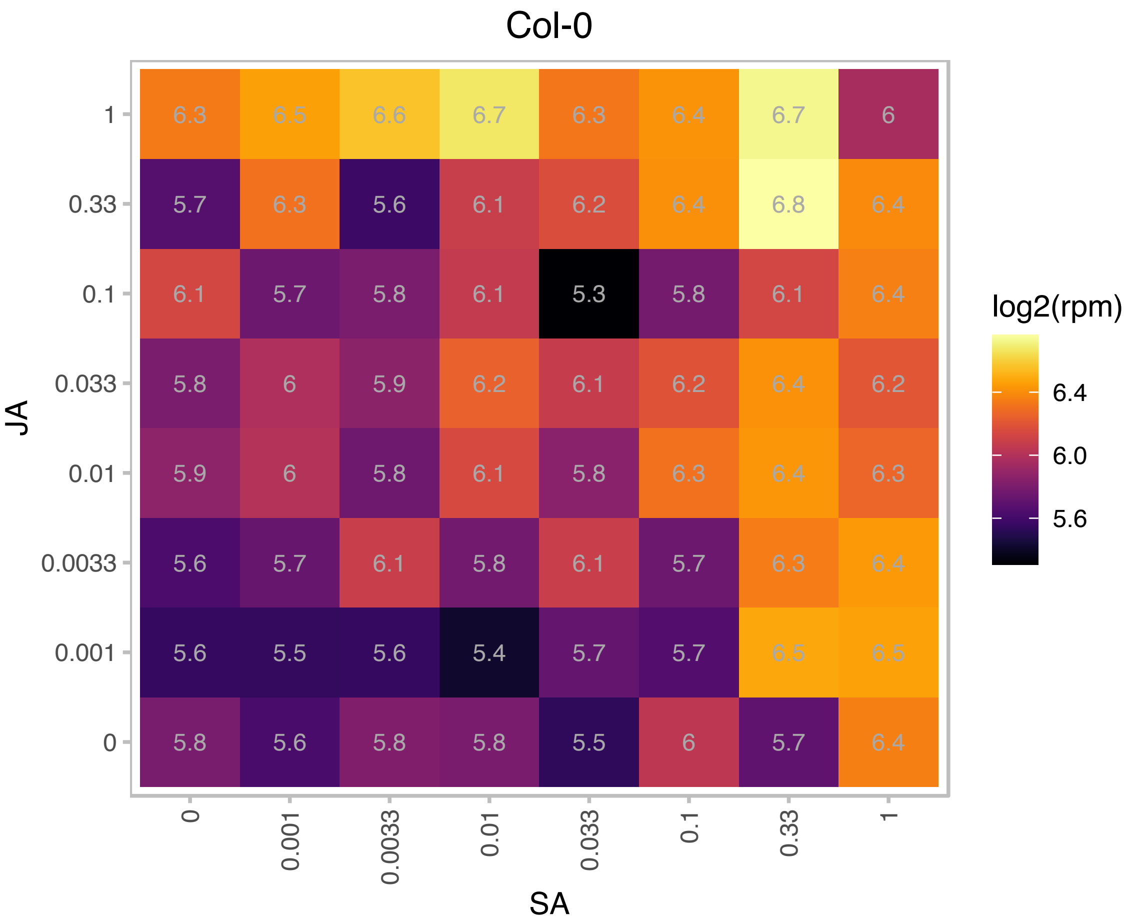


| | |
|---|---|
| Gene | AT1G61150 |
| is.expressed_genes | TRUE |
| is.responsive_genes | TRUE |
| is.apcluster | B-04 |

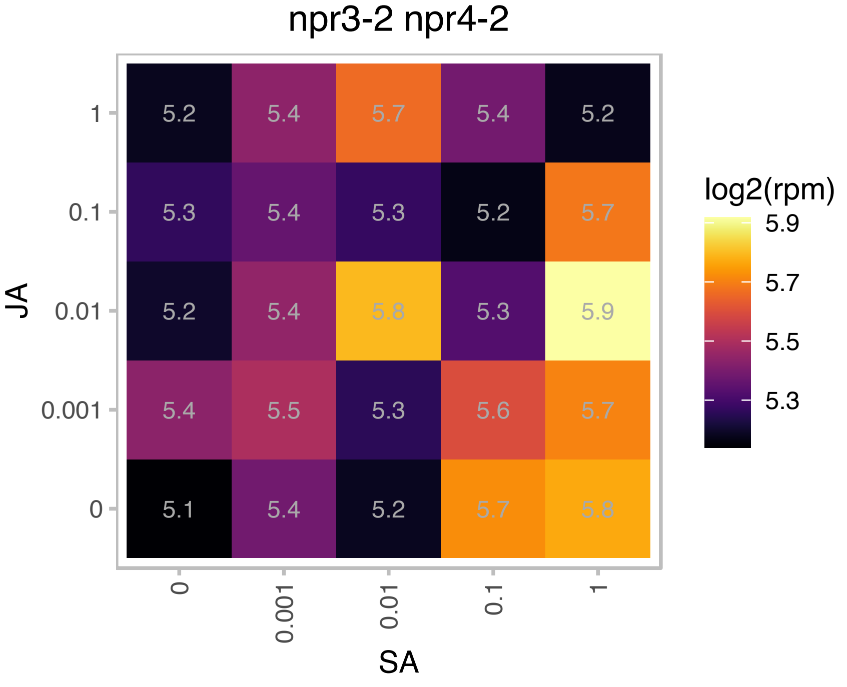


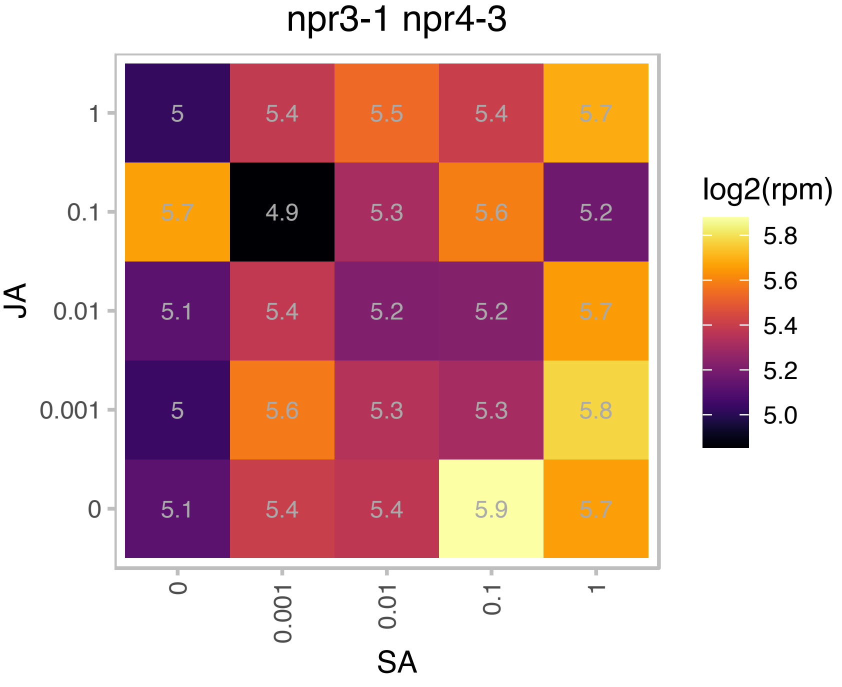

### AT1G62180.1 : APR2, APSR, ATAPR2, PRH, PRH43

5'adenylylphosphosulfate reductase 2

encodes a adenosine 5'-phosphosulfate reductase, involved in sulfate assimilation. Is a major effect locus for natural variation of shoot sulfate content in Arabidopsis.

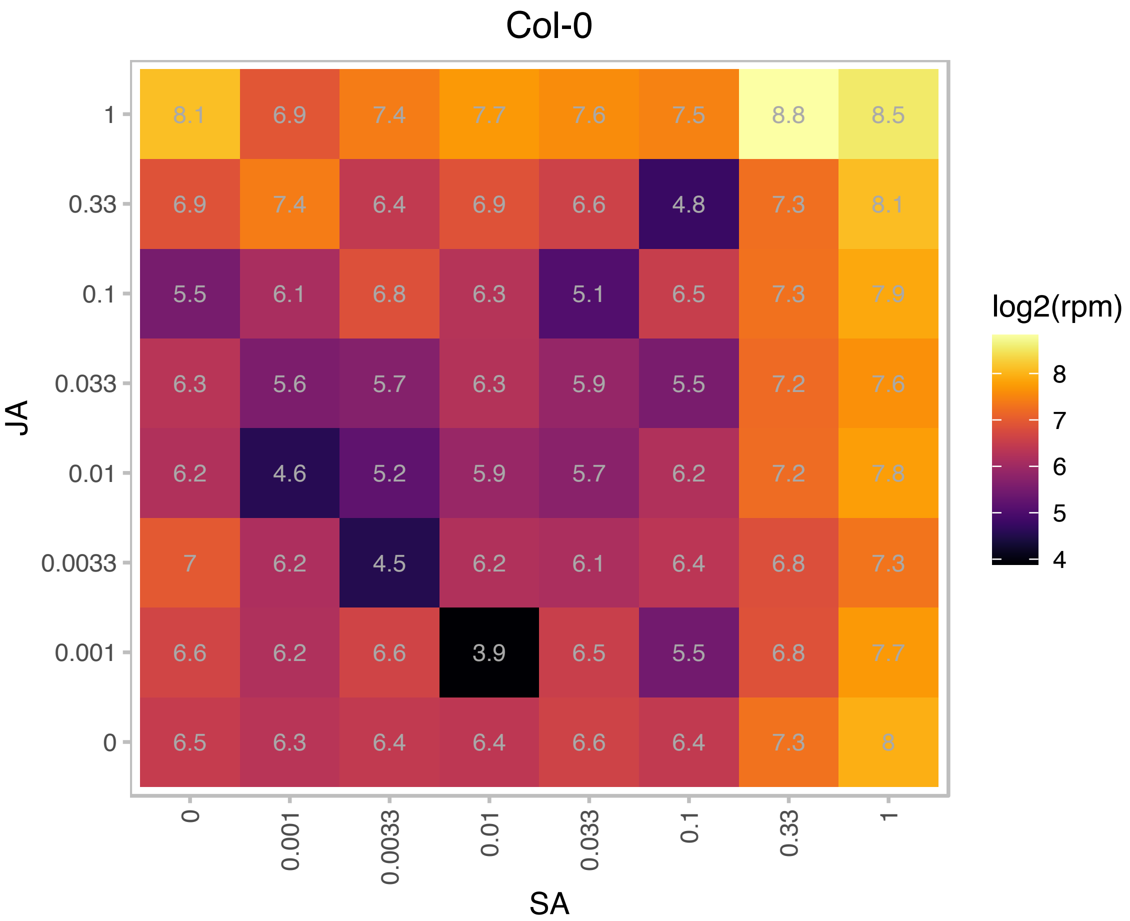


| | |
|---|---|
| Gene | AT1G62180 |
| is.expressed_genes | TRUE |
| is.responsive_genes | TRUE |
| is.apcluster | B-04 |

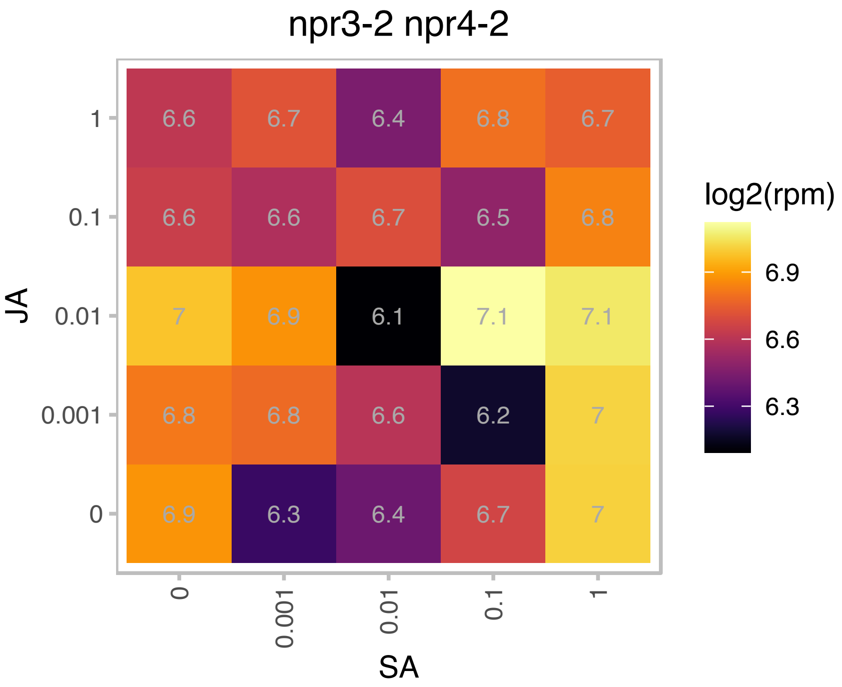


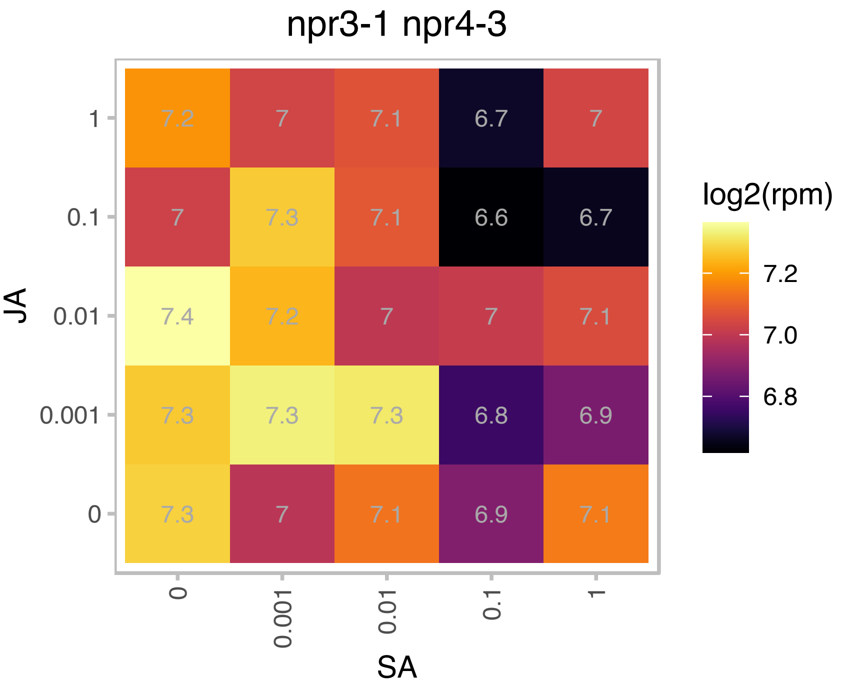

### AT1G62200.1 : AtNPF8.5, AtPTR6, NPF8.5, PTR6

Major facilitator superfamily protein

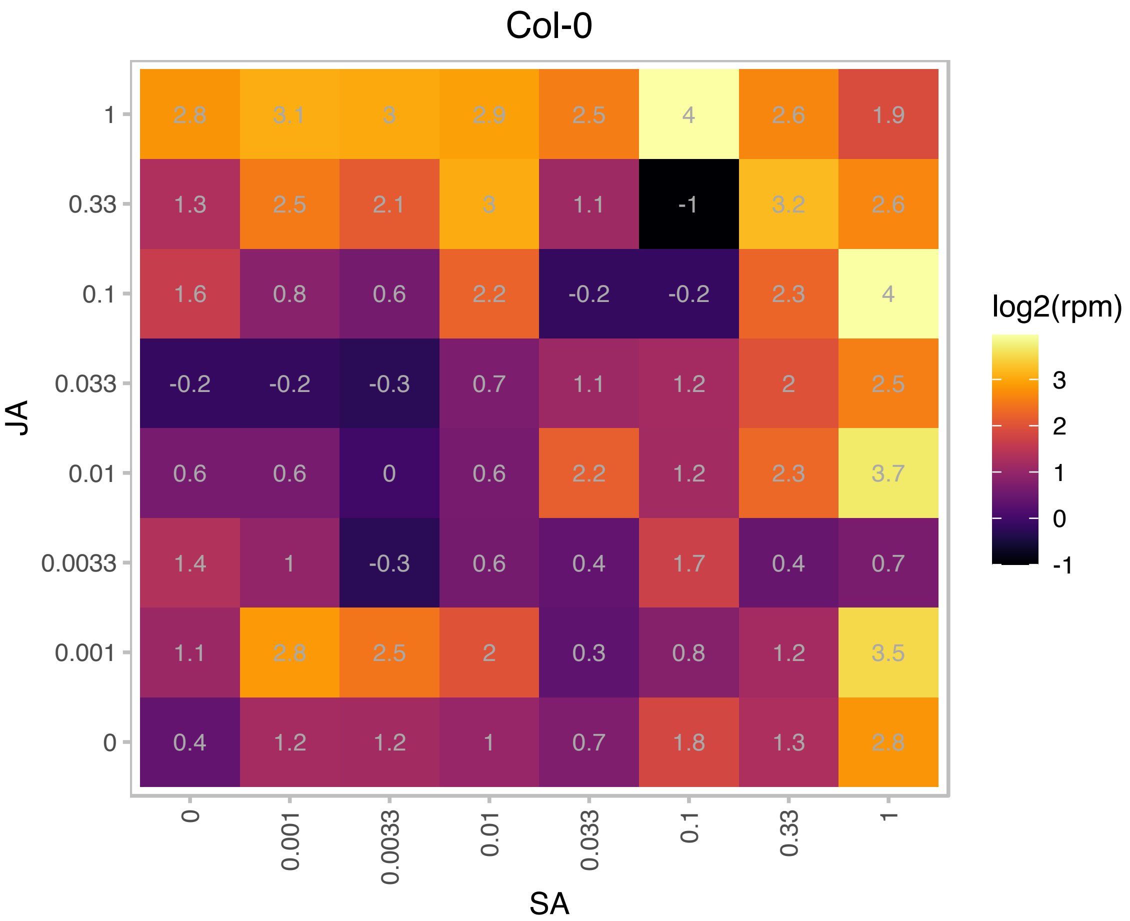


| | |
|---|---|
| Gene | AT1G62200 |
| is.expressed_genes | TRUE |
| is.responsive_genes | TRUE |
| is.apcluster | B-04 |

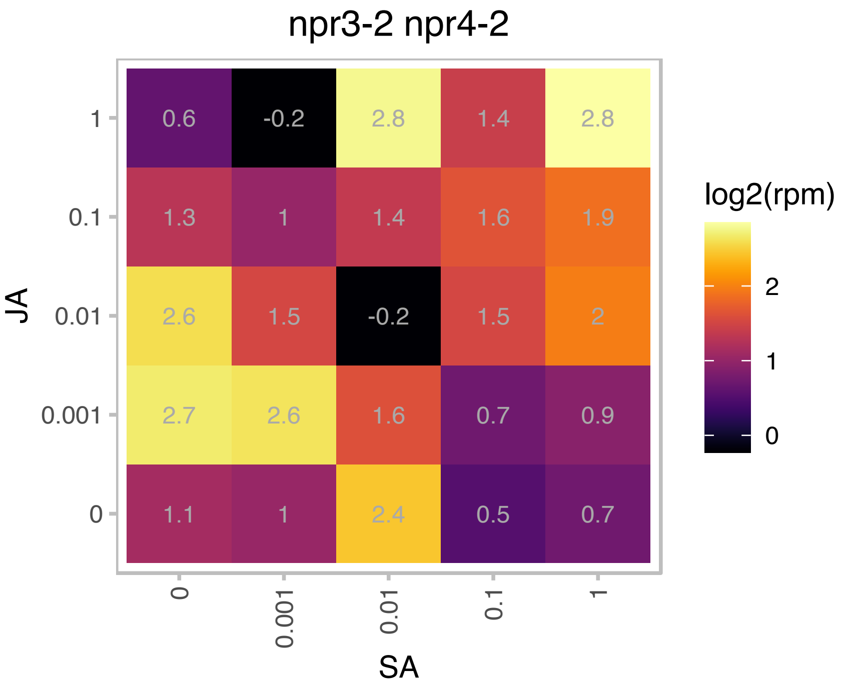


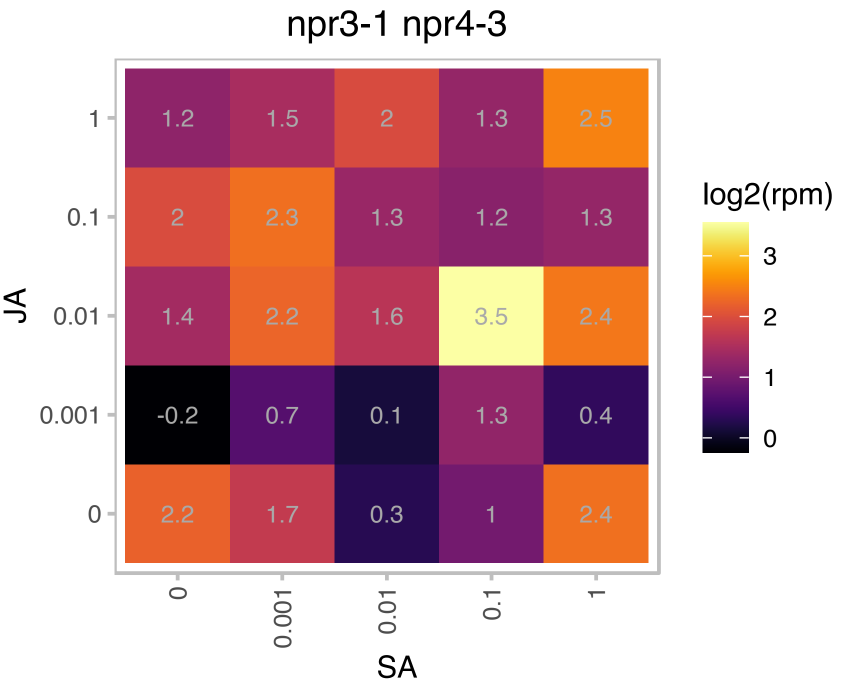

# AT1G63530.1 : NA

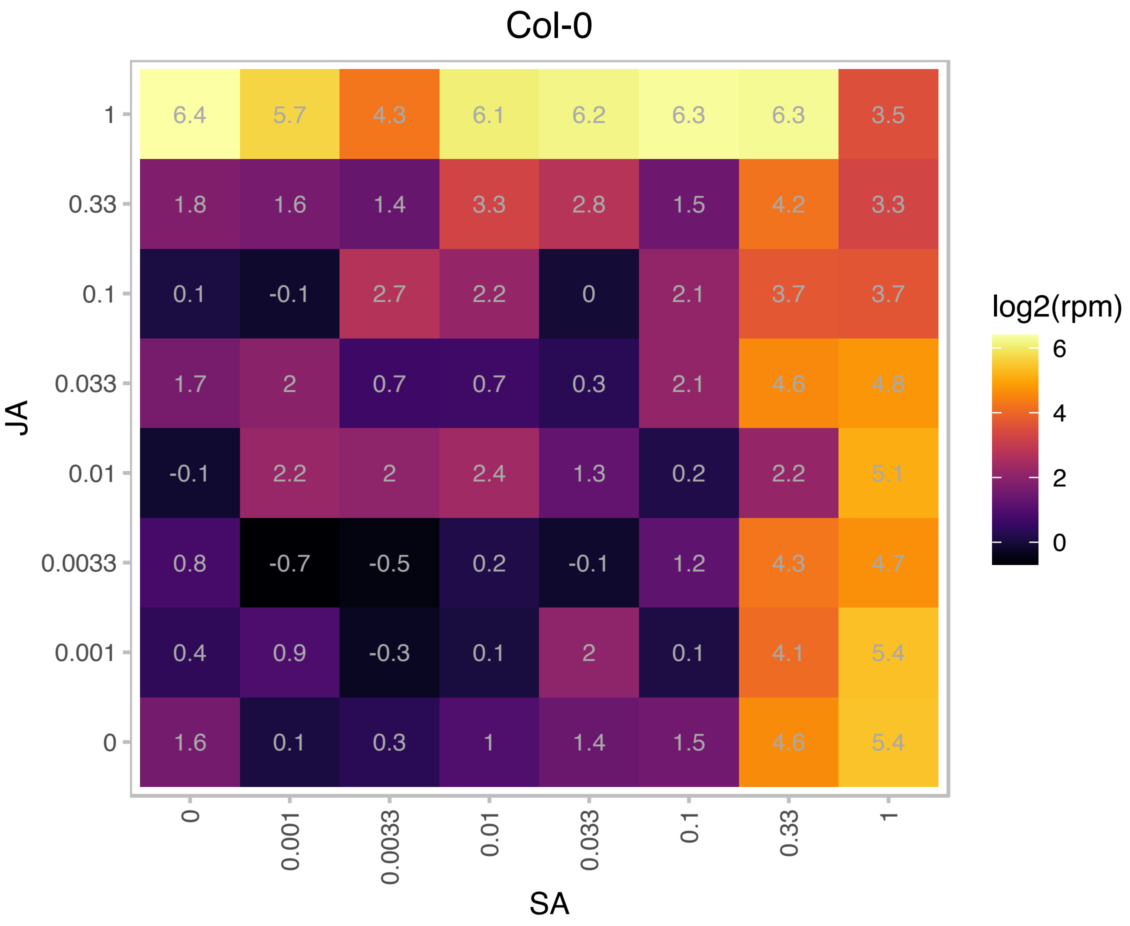


| | |
|---|---|
| Gene | AT1G63530 |
| is.expressed_genes | TRUE |
| is.responsive_genes | TRUE |
| is.apcluster | B-04 |

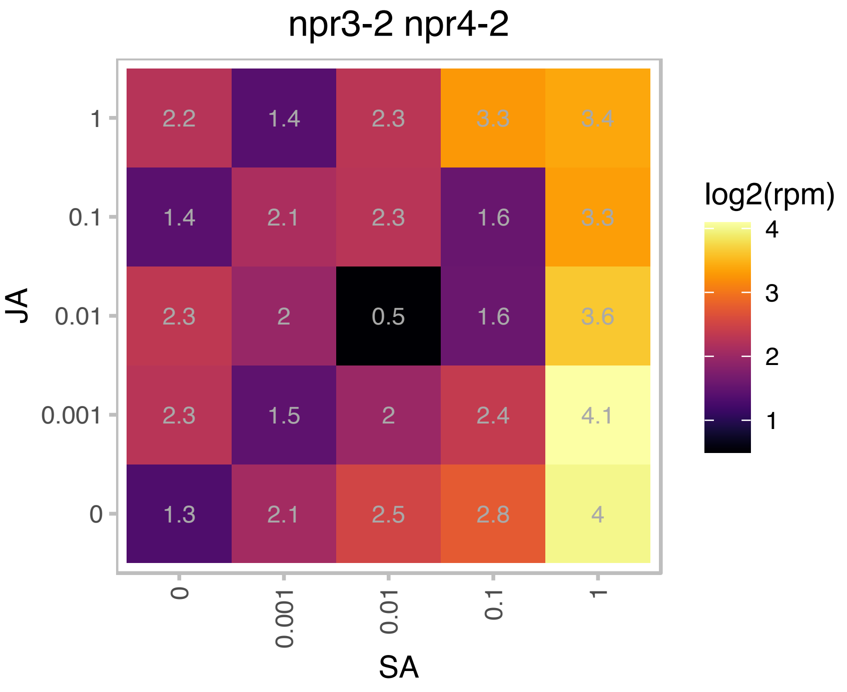


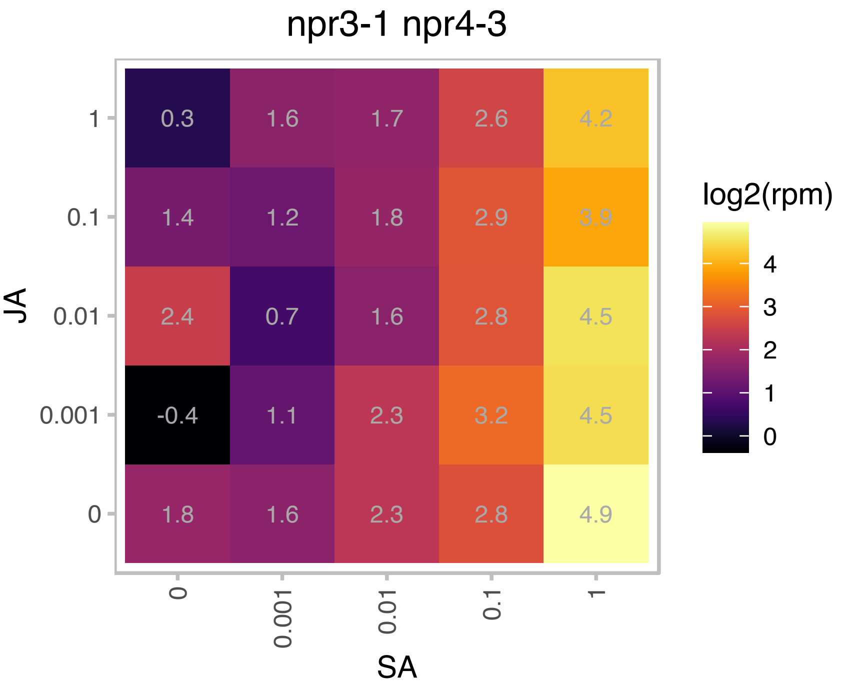

### AT1G63970.1 : ISPF, MECPS

isoprenoid F
Encodes a protein with 2C-methyl-D-erythritol 2,4-cyclodiphosphate synthase activity. The protein's activity was confirmed by heterologous expression of phenotypic complementation of the E. coli ispF mutant. Plants defective in this gene display an albino lethal phenotype.Homolog of E. coli IspF

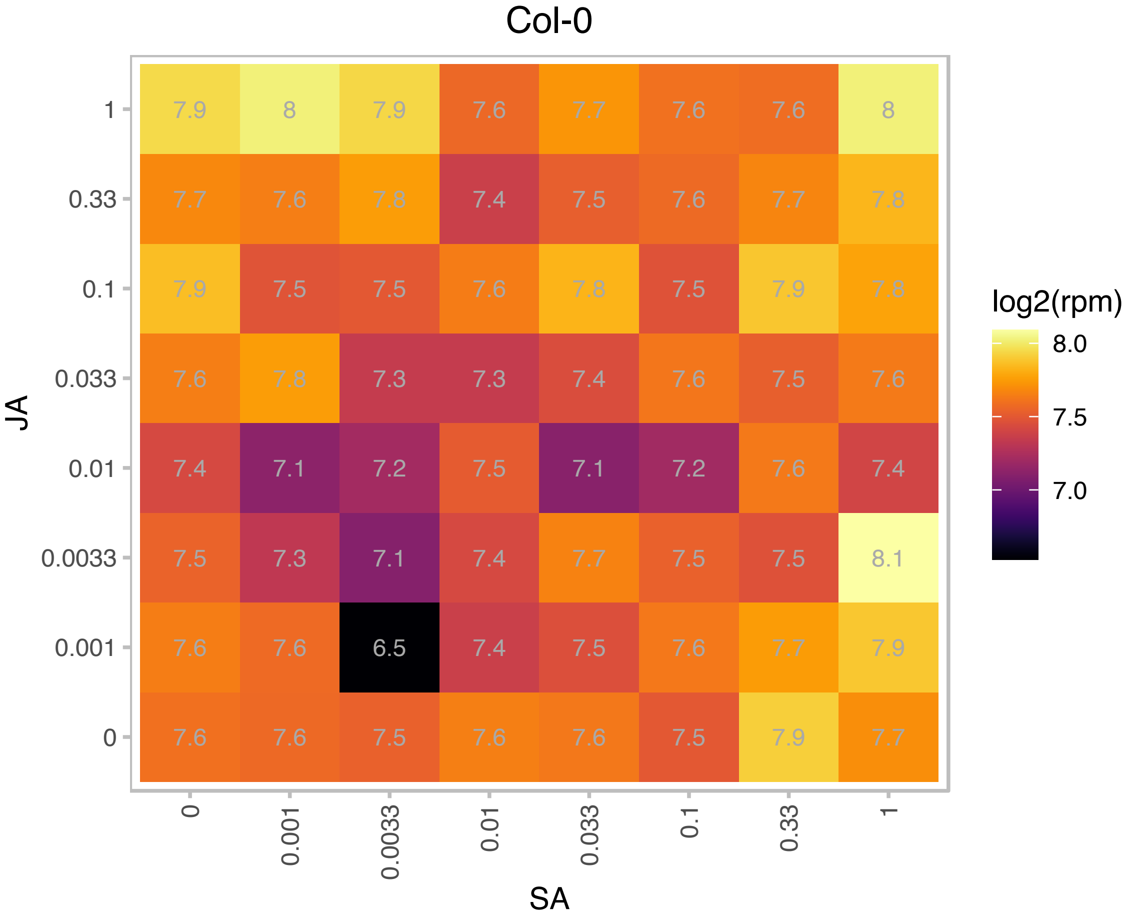


| | |
|---|---|
| Gene | AT1G63970 |
| is.expressed_genes | TRUE |
| is.responsive_genes | TRUE |
| is.apcluster | B-04 |

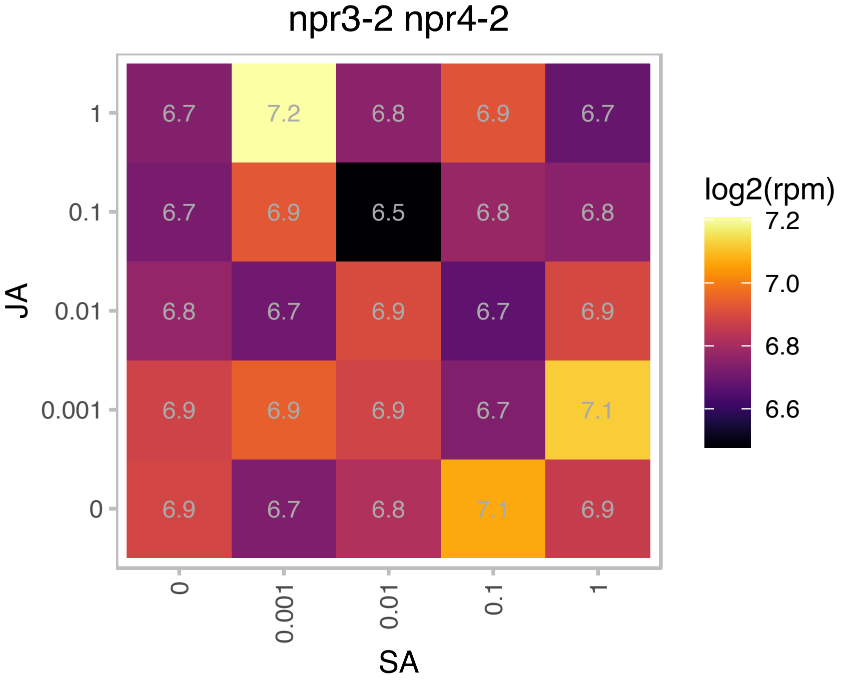


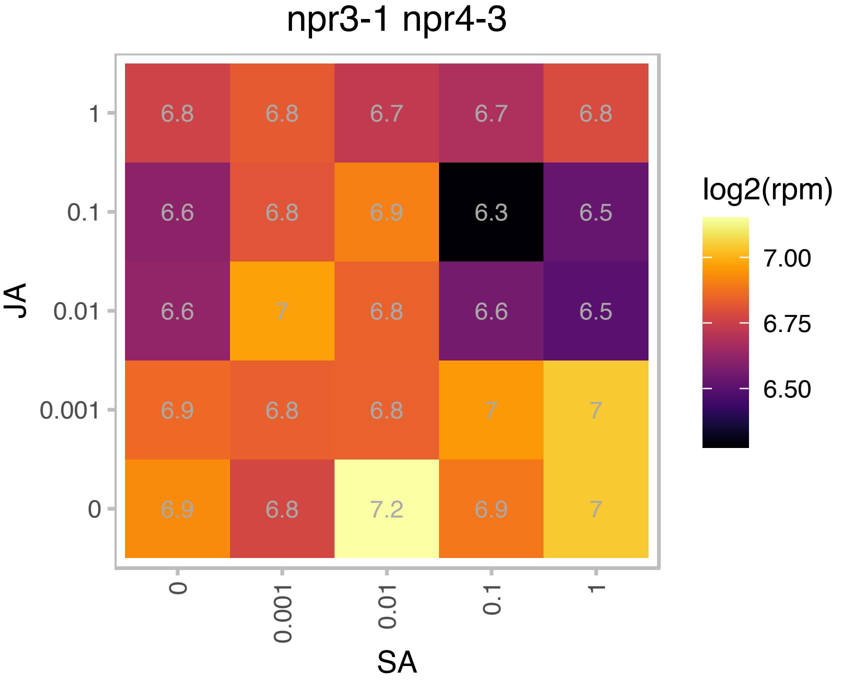

### AT1G64660.1 : ATMGL, MGL

methionine gamma-lyase
Encodes a functional methionine gamma-lyase, a cytosolic enzyme catalyzes the degradation of methionine into methanethiol, alpha-ketobutyrate and ammonia. The catabolism of excess methionine is important to methionine homeostasis. The mRNA is cell-to-cell mobile.

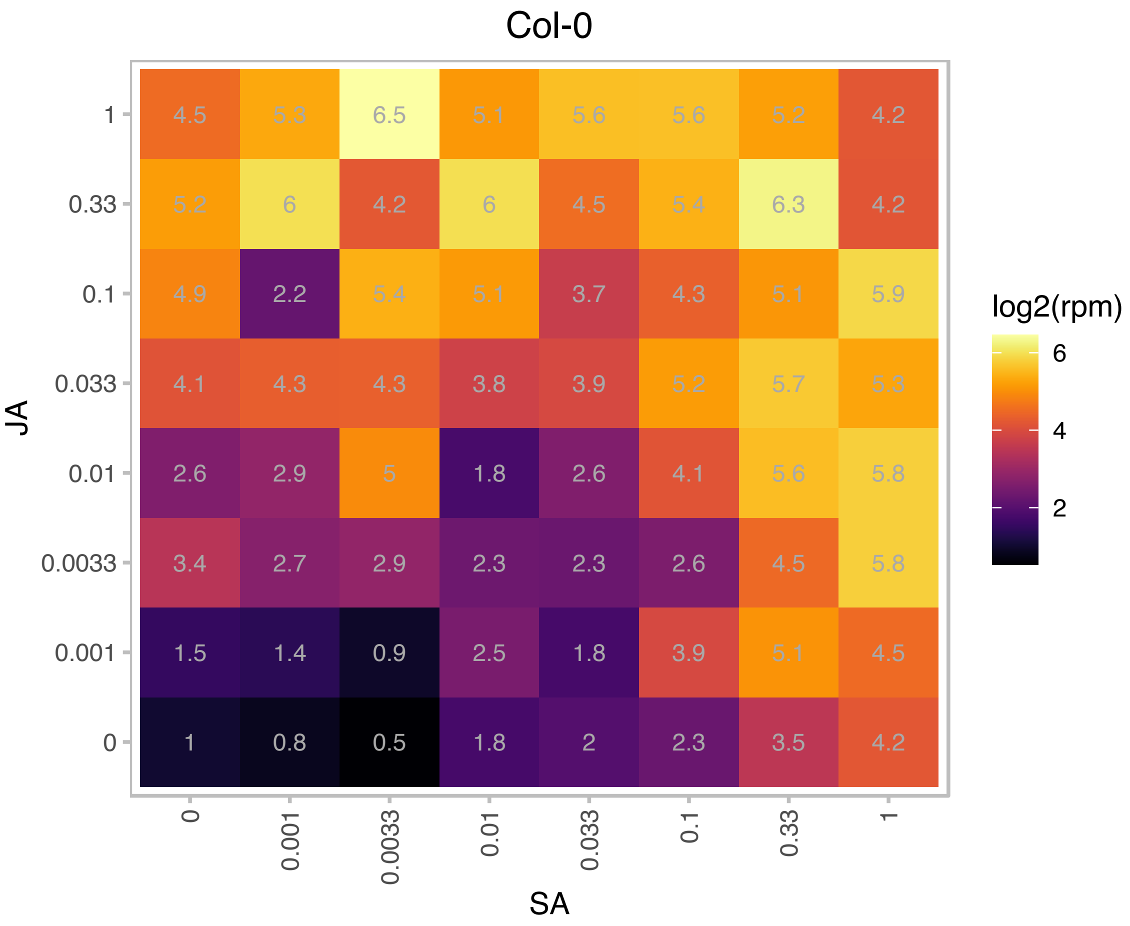

| | |
|---|---|
| Gene | AT1G64660 |
| is.expressed_genes | TRUE |
| is.responsive_genes | TRUE |
| is.apcluster | B-04 |

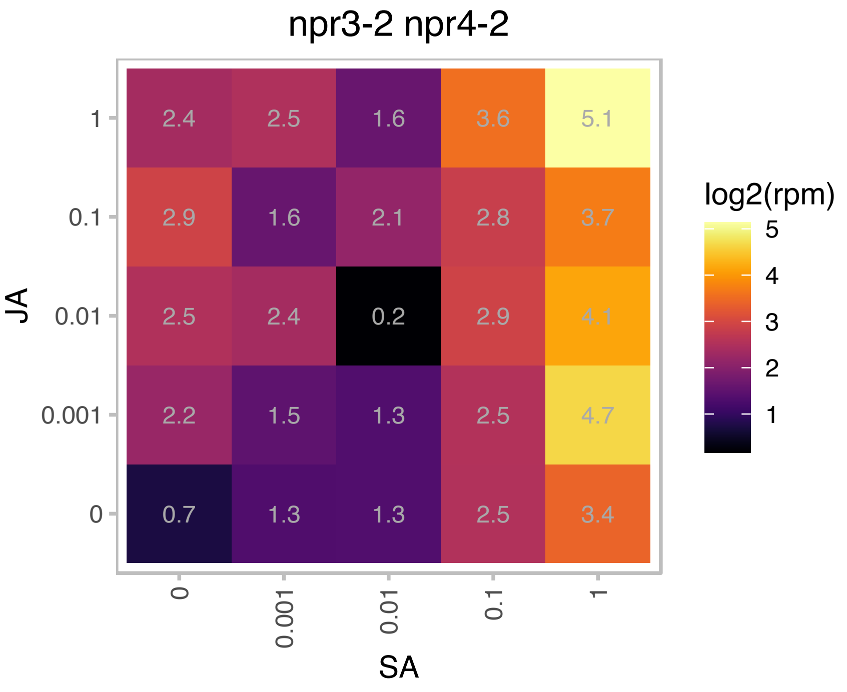

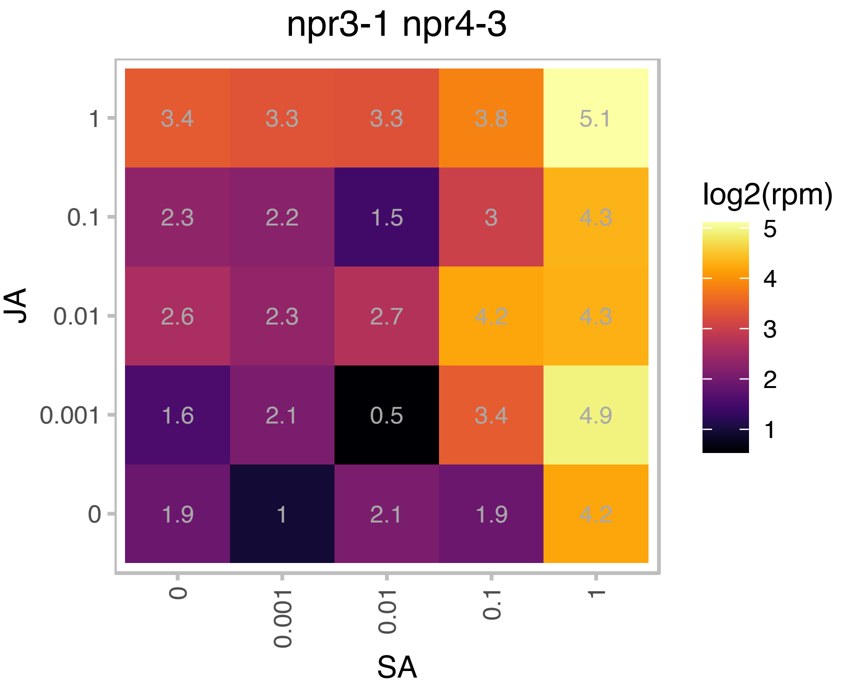

# AT1G69680.1 : NA

Mog1/PsbP/DUF1795-like photosystem II reaction center PsbP family protein

Col-0

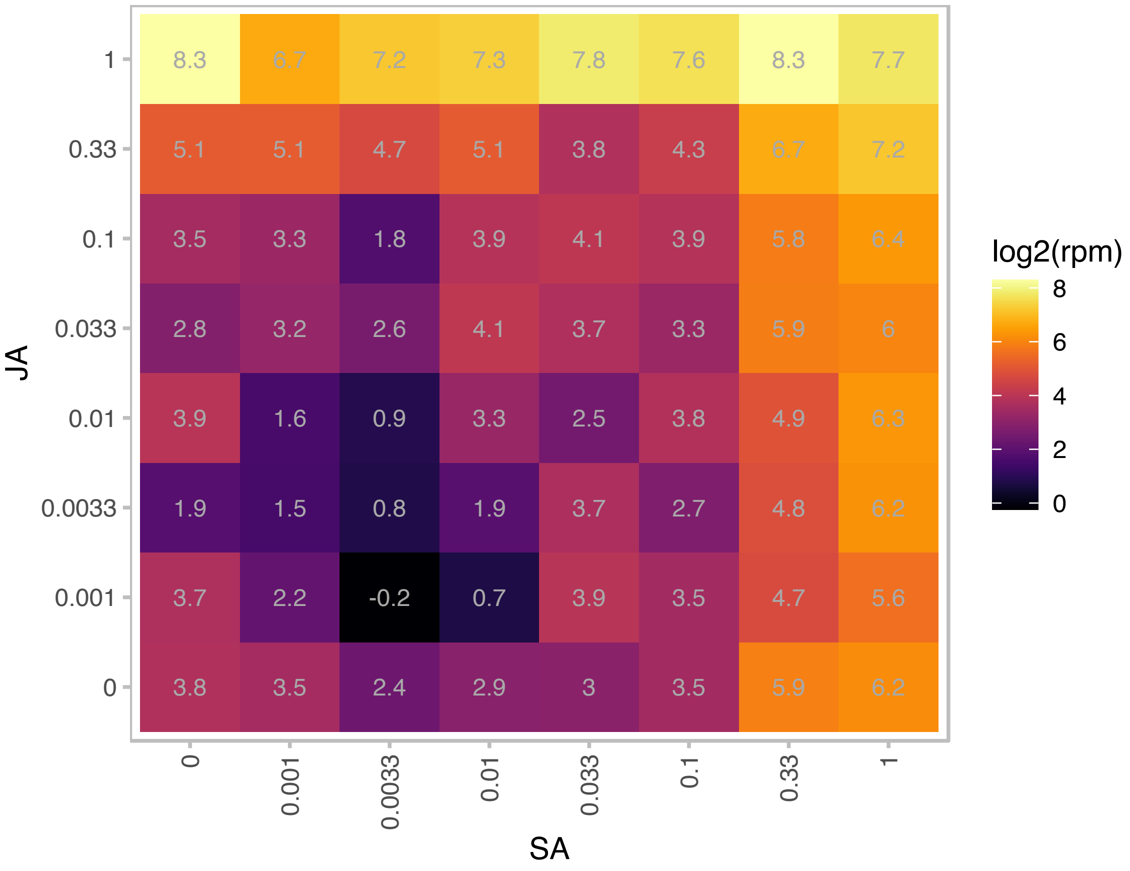


| | |
|---|---|
| Gene | AT1G69680 |
| is.expressed_genes | TRUE |
| is.responsive_genes | TRUE |
| is.apcluster | B-04 |

npr3-2 npr4-2

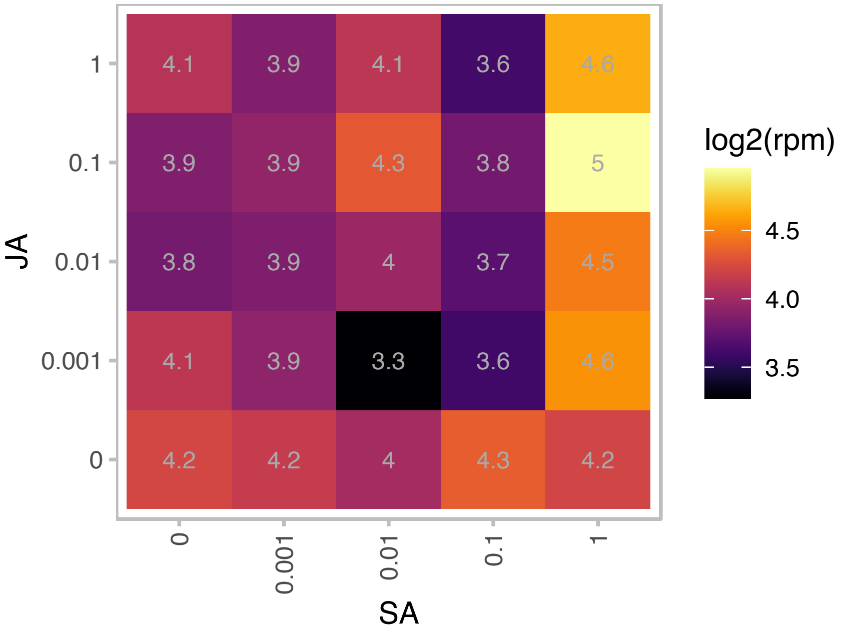


npr3-1 npr4-3

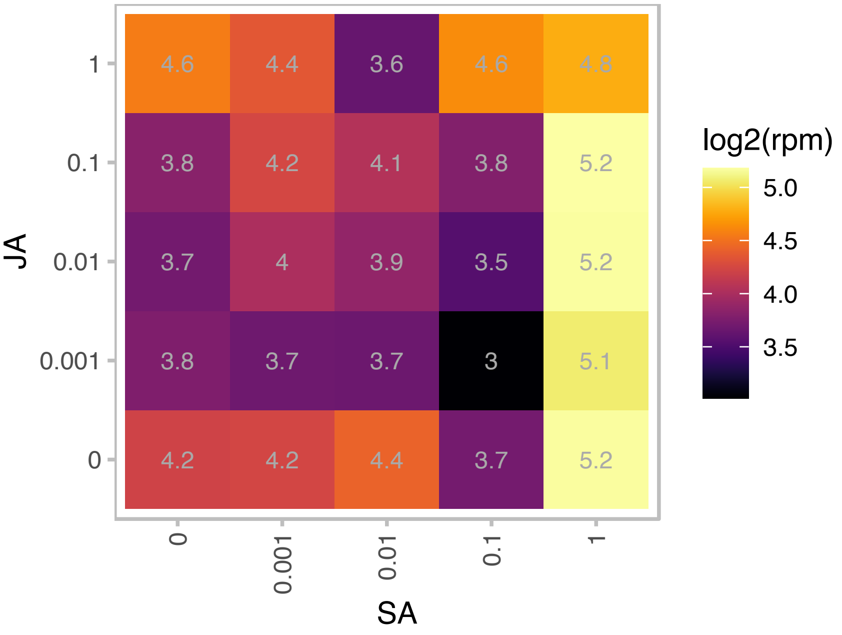

### AT1G69930.1 : ATGSTU11, GSTU11

glutathione S-transferase TAU 11

Encodes glutathione transferase belonging to the tau class of GSTs. Naming convention according to Wagner et al. (2002).

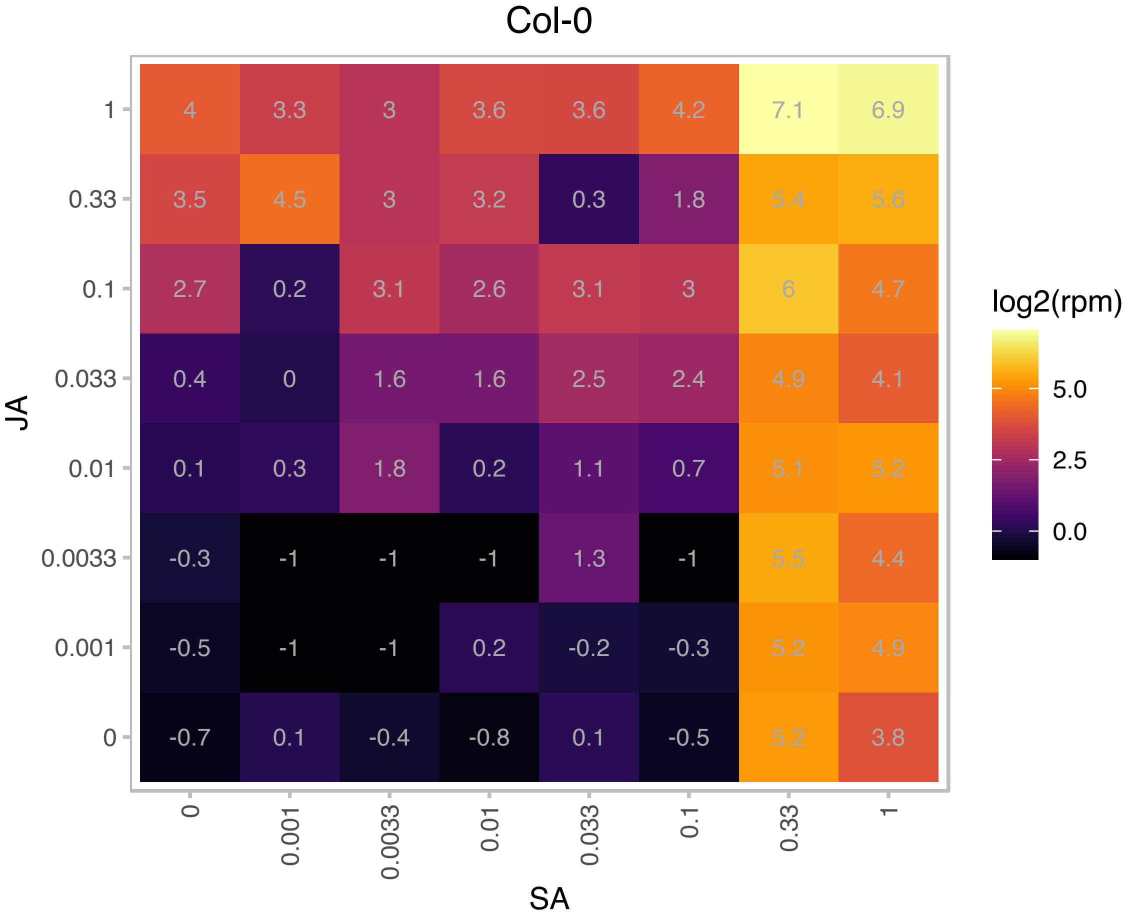


| | |
|---|---|
| Gene | AT1G69930 |
| is.expressed_genes | TRUE |
| is.responsive_genes | TRUE |
| is.apcluster | B-04 |

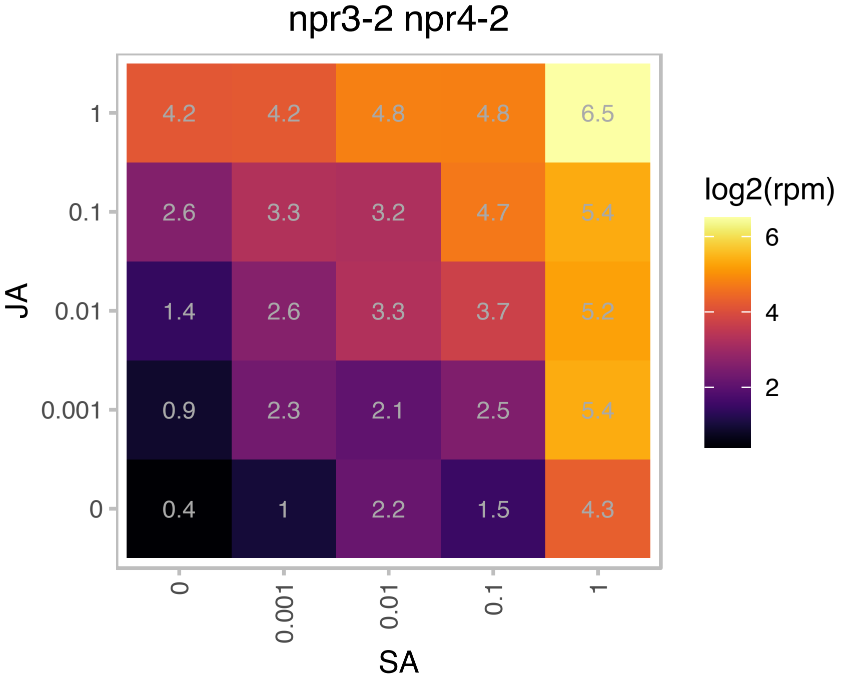


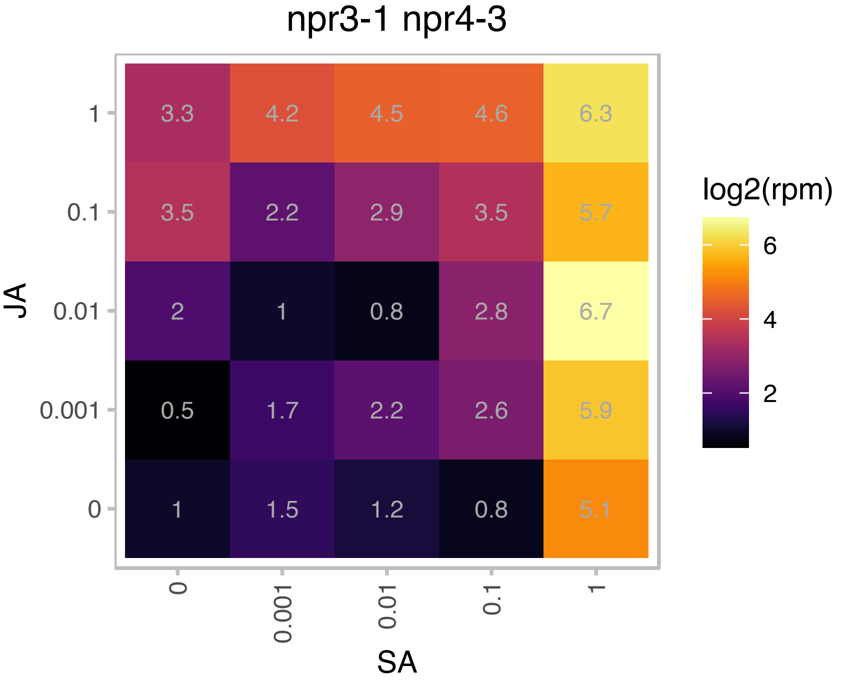

# AT1G71520.1 : NA

Integrase-type DNA-binding superfamily protein

encodes a member of the DREB subfamily A-5 of ERF/AP2 transcription factor family. The protein contains one AP2 domain. There are 16 members in this subfamily including RAP2.1, RAP2.9 and RAP2.10.

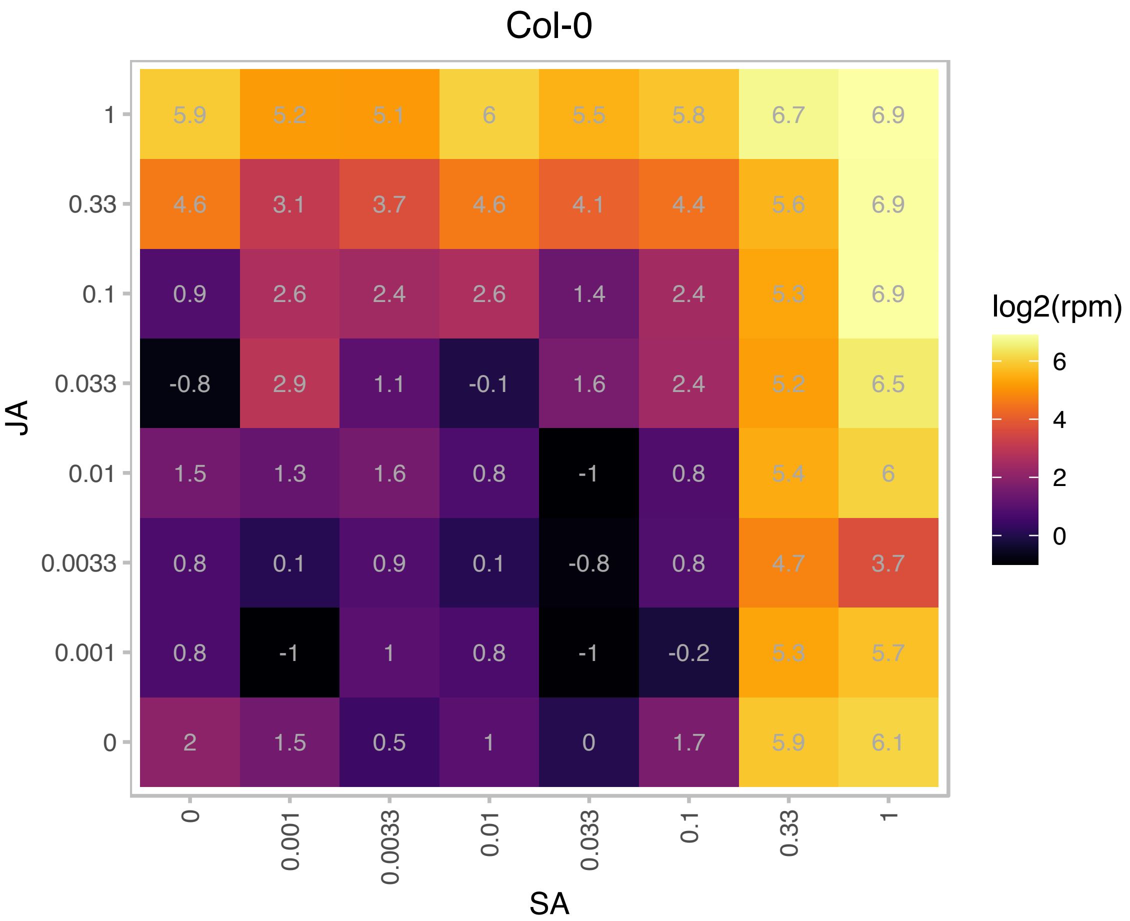


| | |
|---|---|
| Gene | AT1G71520 |
| is.expressed_genes | TRUE |
| is.responsive_genes | TRUE |
| is.apcluster | B-04 |

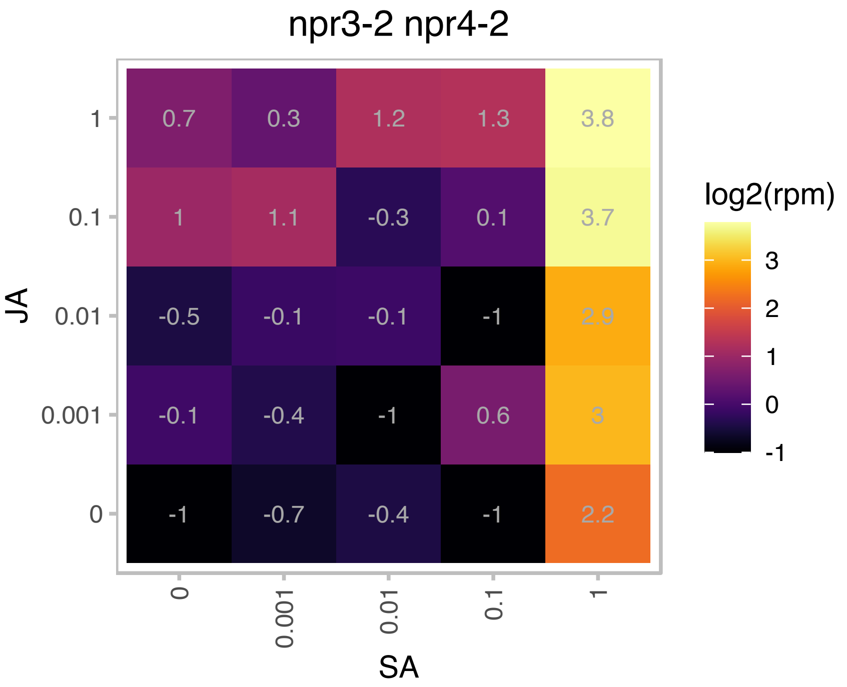


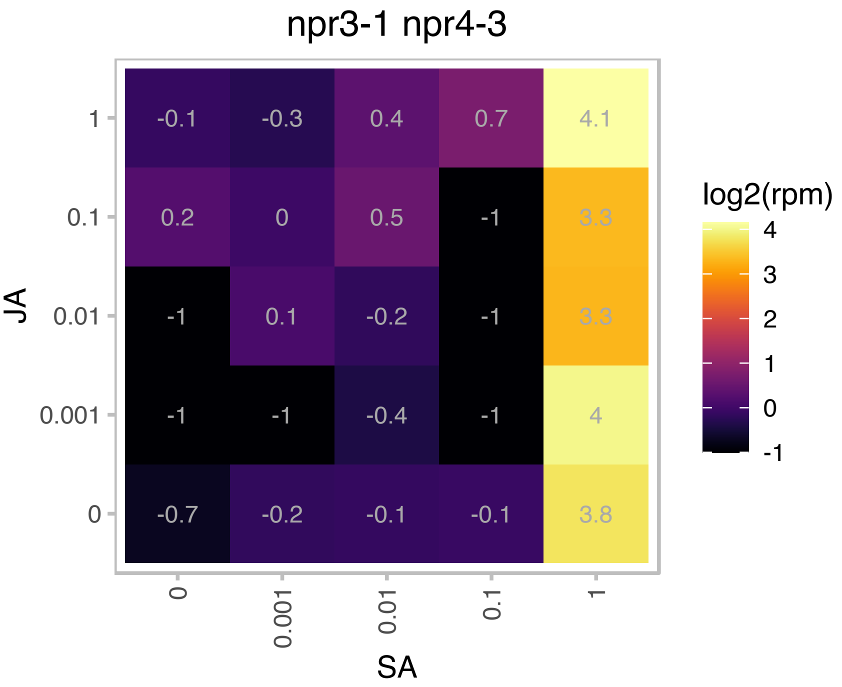

### AT1G72490.1 : AtNGR2, DRO1, LAZY4

DRO1 is a member of the IGT gene family and has a unknown function . It is expressed in roots and involved in leaf root architecture, specifically the orientation of lateral root angles. Involved in determining lateral root branch angle.

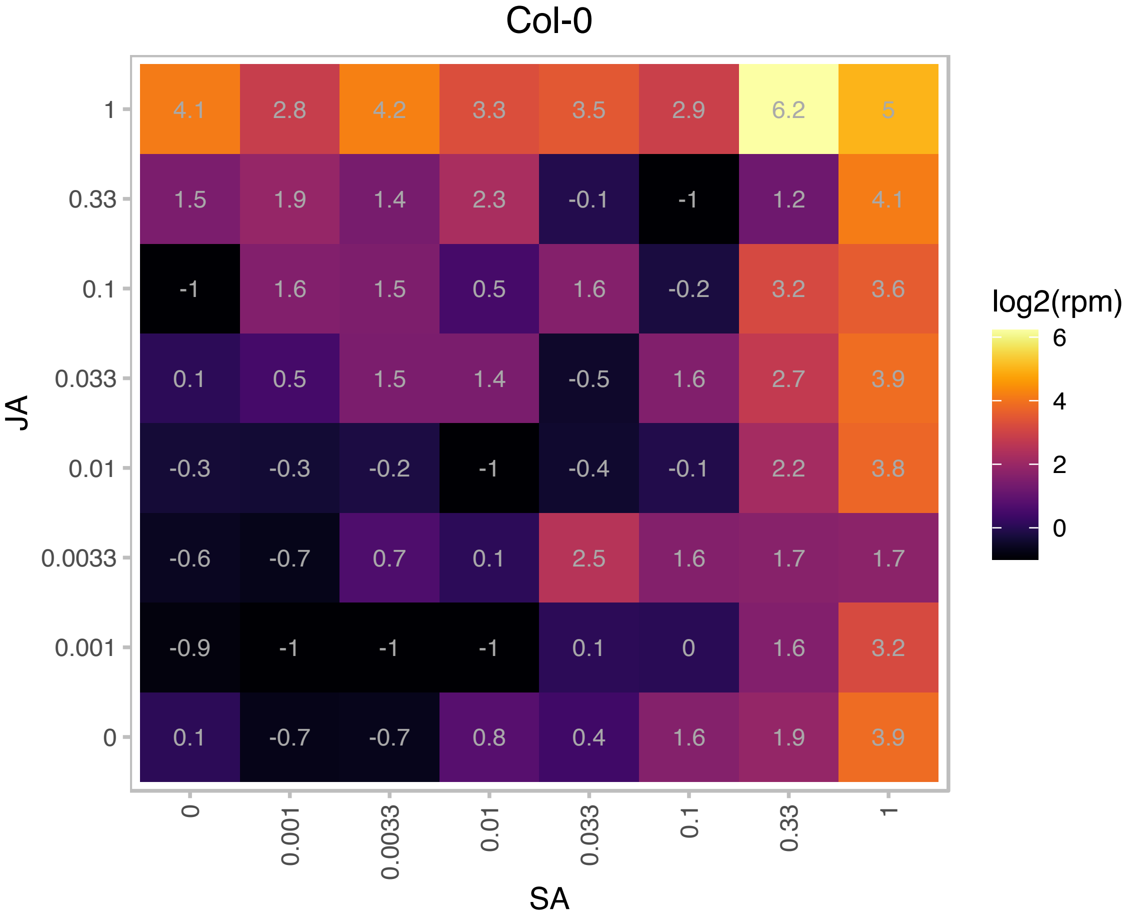


| | |
|---|---|
| Gene | AT1G72490 |
| is.expressed_genes | TRUE |
| is.responsive_genes | TRUE |
| is.apcluster | B-04 |

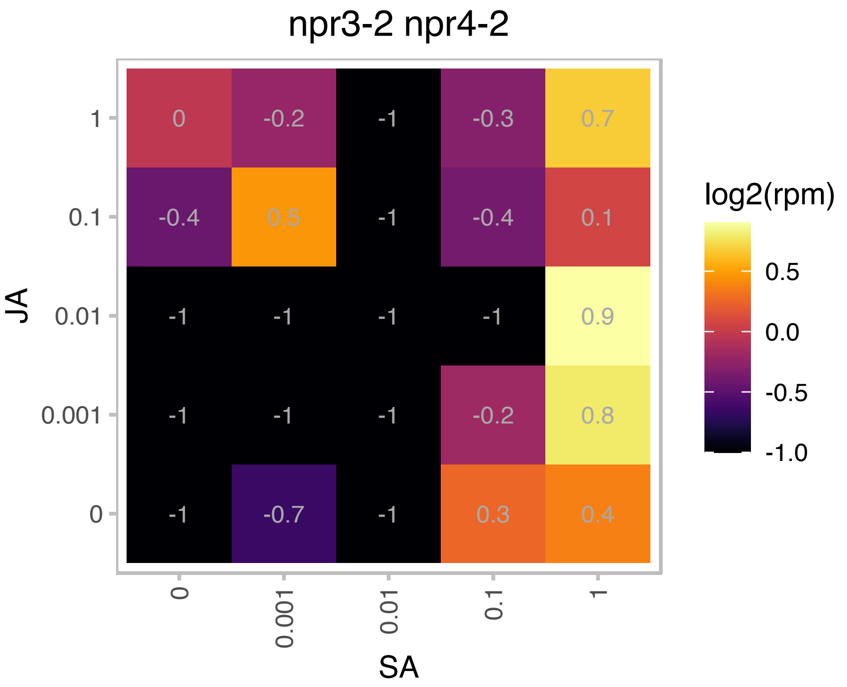


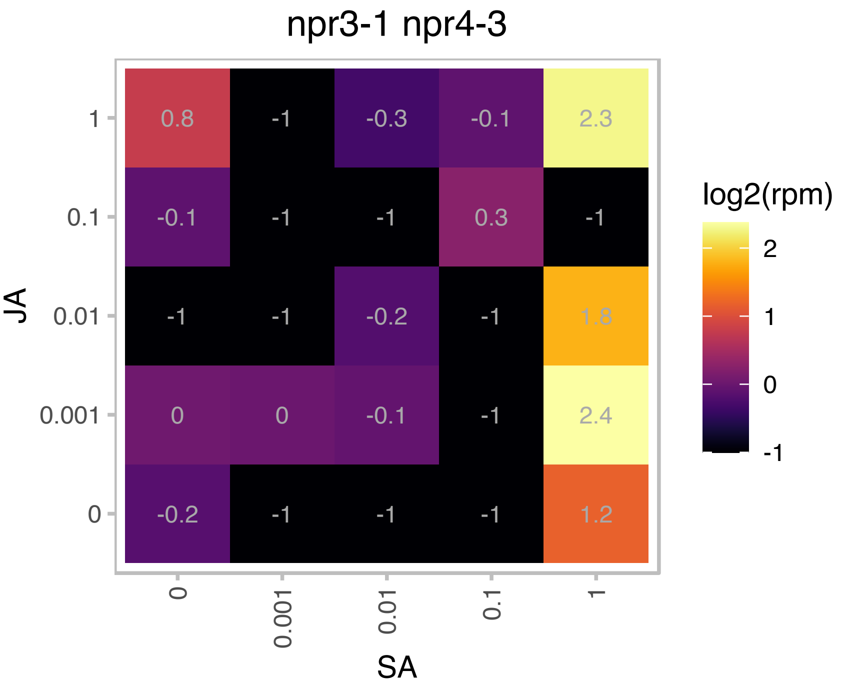

# AT1G75860.1 : NA

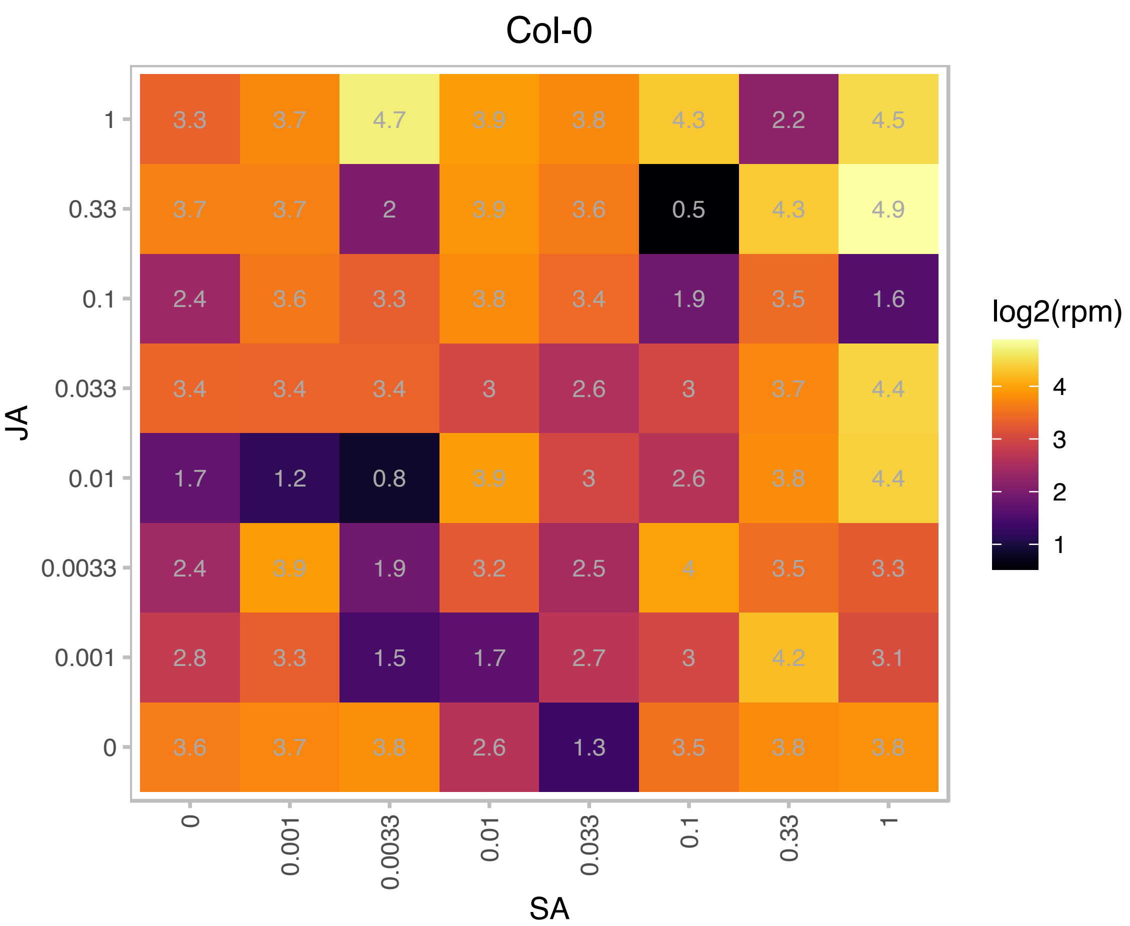


| | |
|---|---|
| Gene | AT1G75860 |
| is.expressed_genes | TRUE |
| is.responsive_genes | TRUE |
| is.apcluster | B-04 |

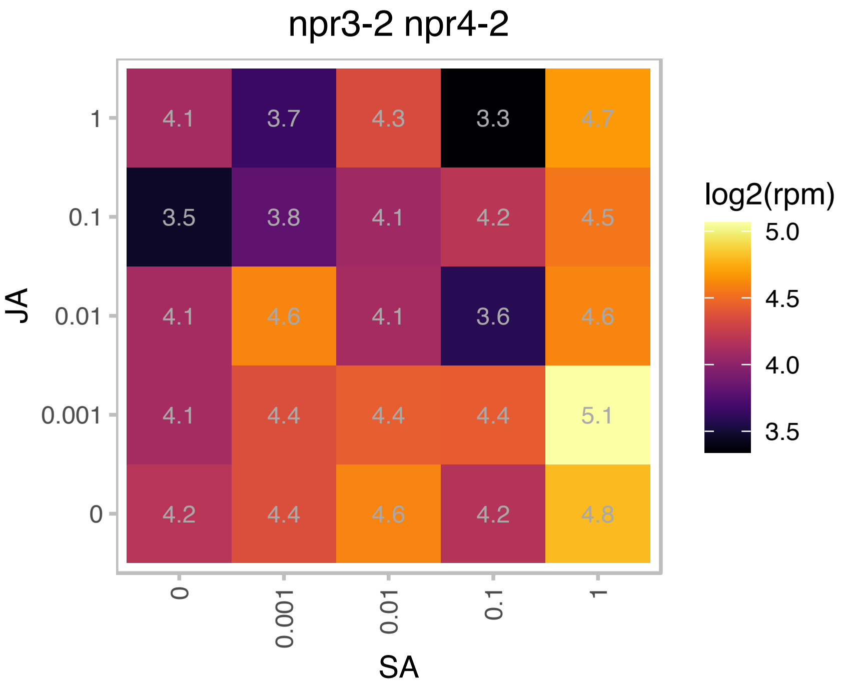


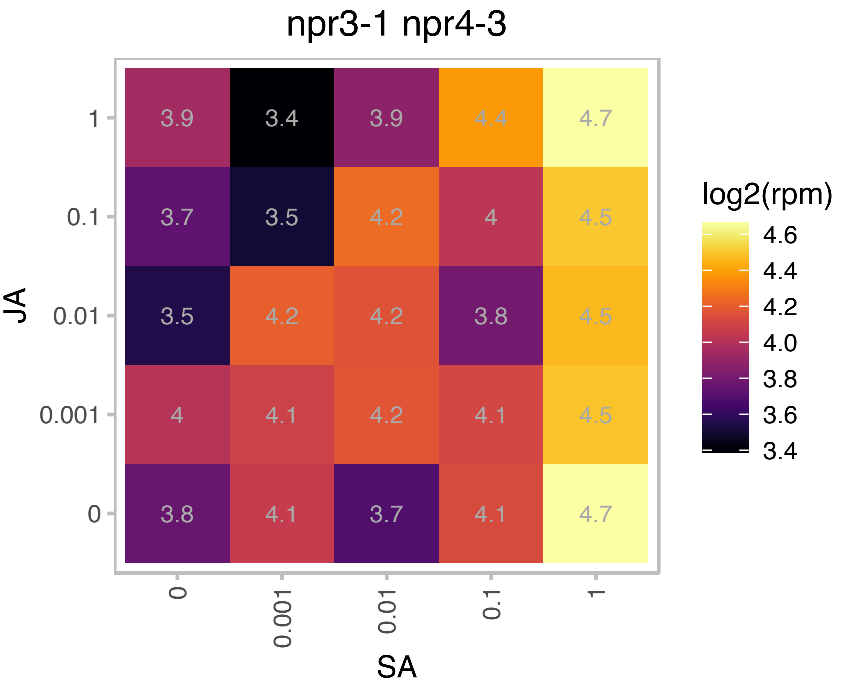

# AT1G76980.1 : NA

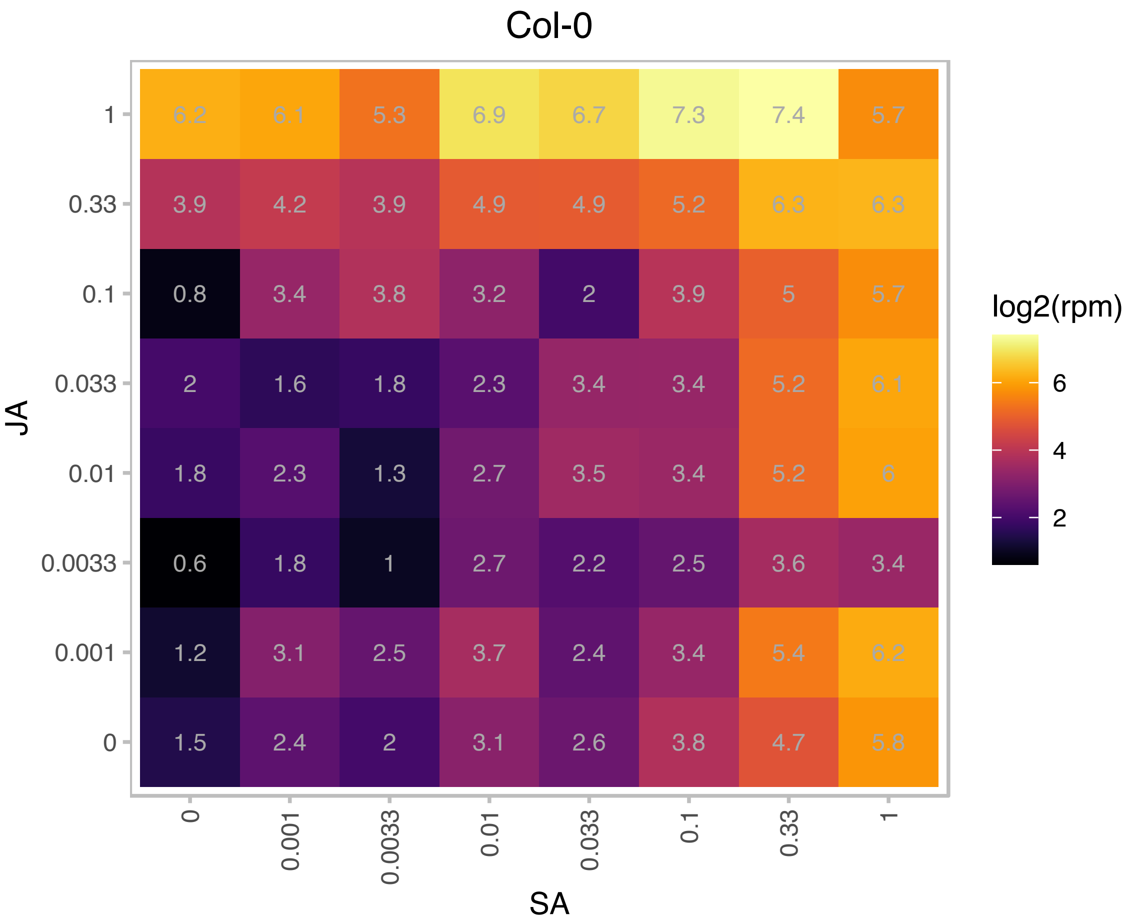


| | |
|---|---|
| Gene | AT1G76980 |
| is.expressed_genes | TRUE |
| is.responsive_genes | TRUE |
| is.apcluster | B-04 |

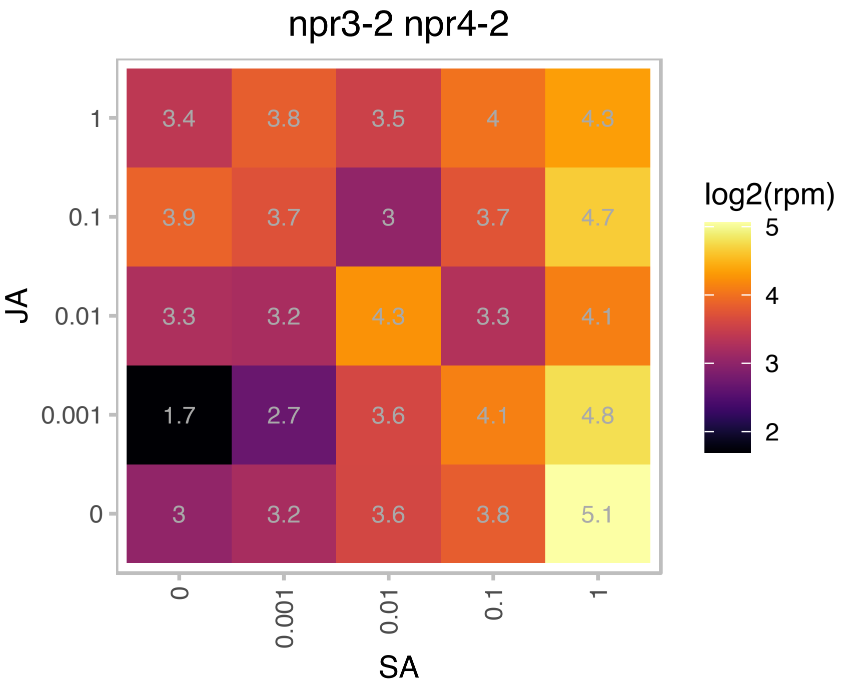


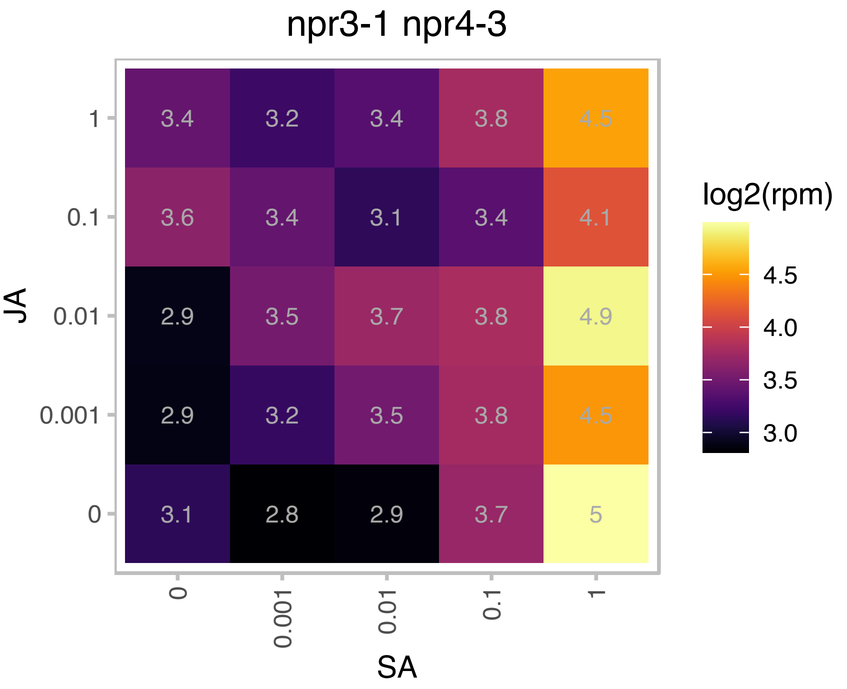

# AT2G07671.1 : NA

ATP synthase subunit C family protein

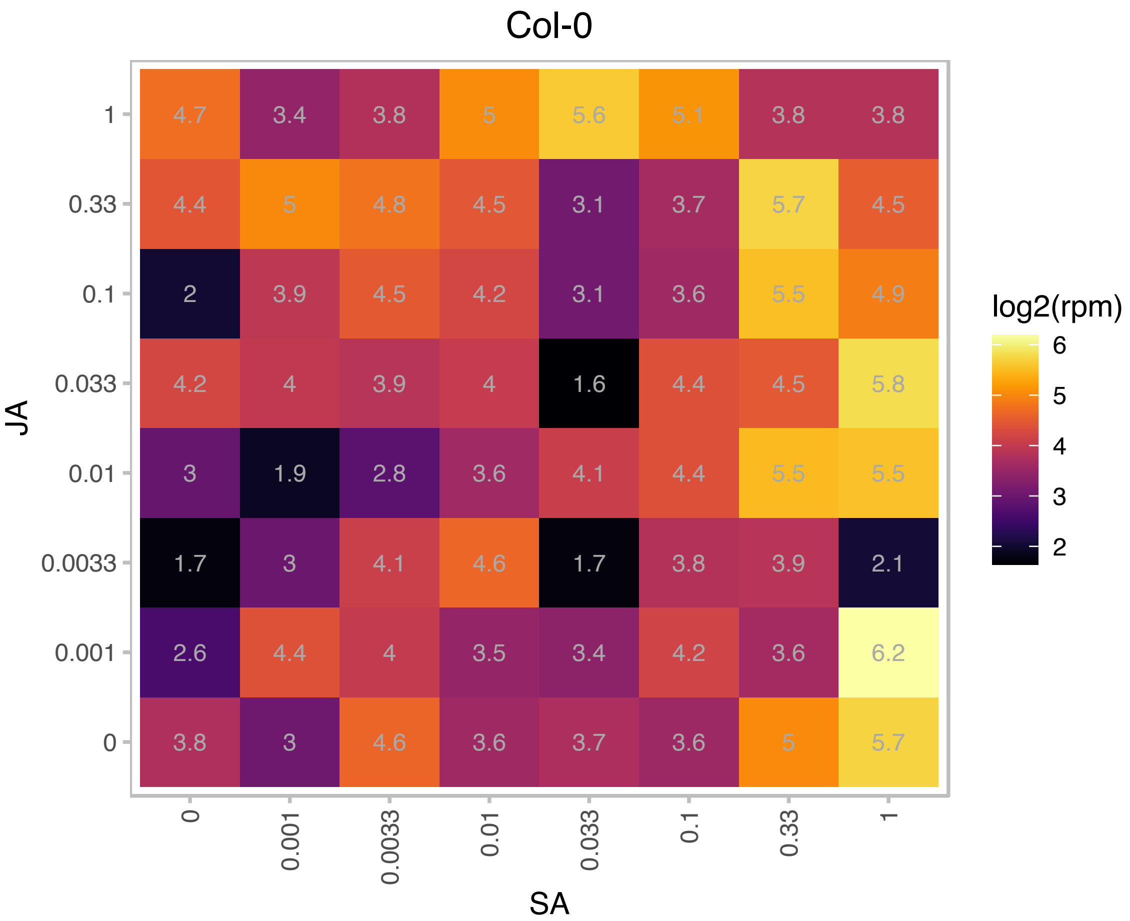


| | |
|---|---|
| Gene | AT2G07671 |
| is.expressed_genes | TRUE |
| is.responsive_genes | TRUE |
| is.apcluster | B-04 |

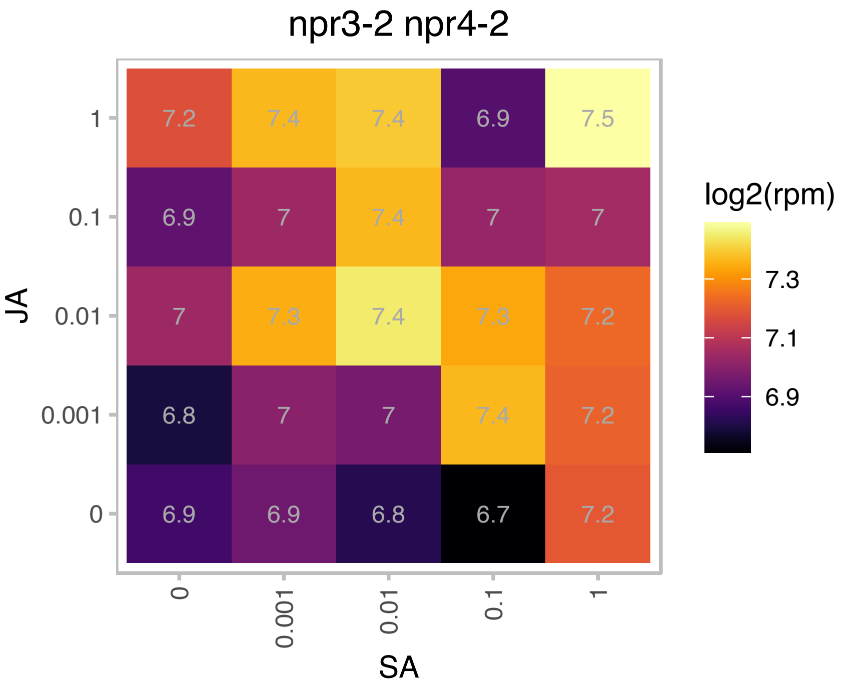


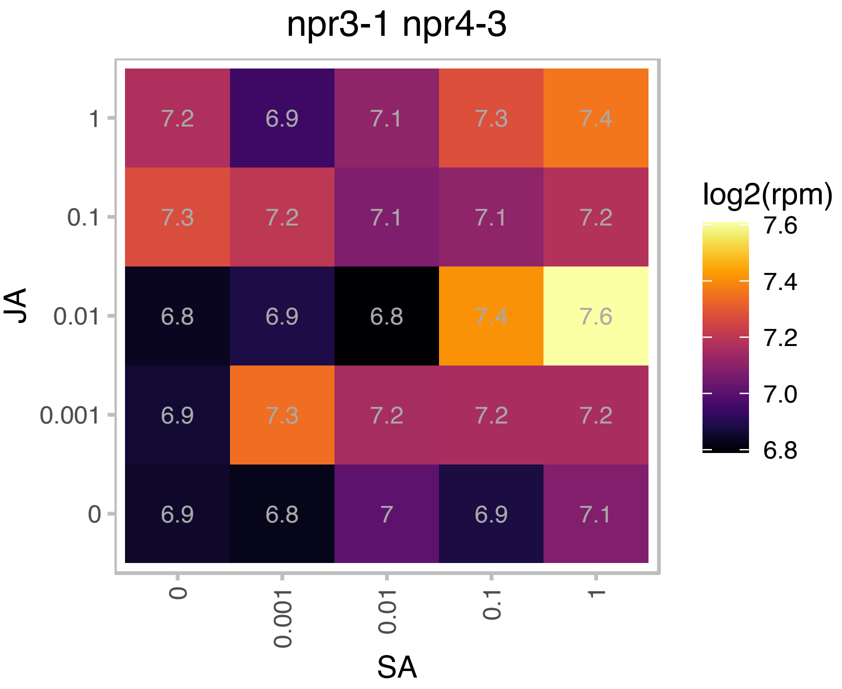

### AT2G15490.1 : UGT73B4

UDP-glycosyltransferase 73B4

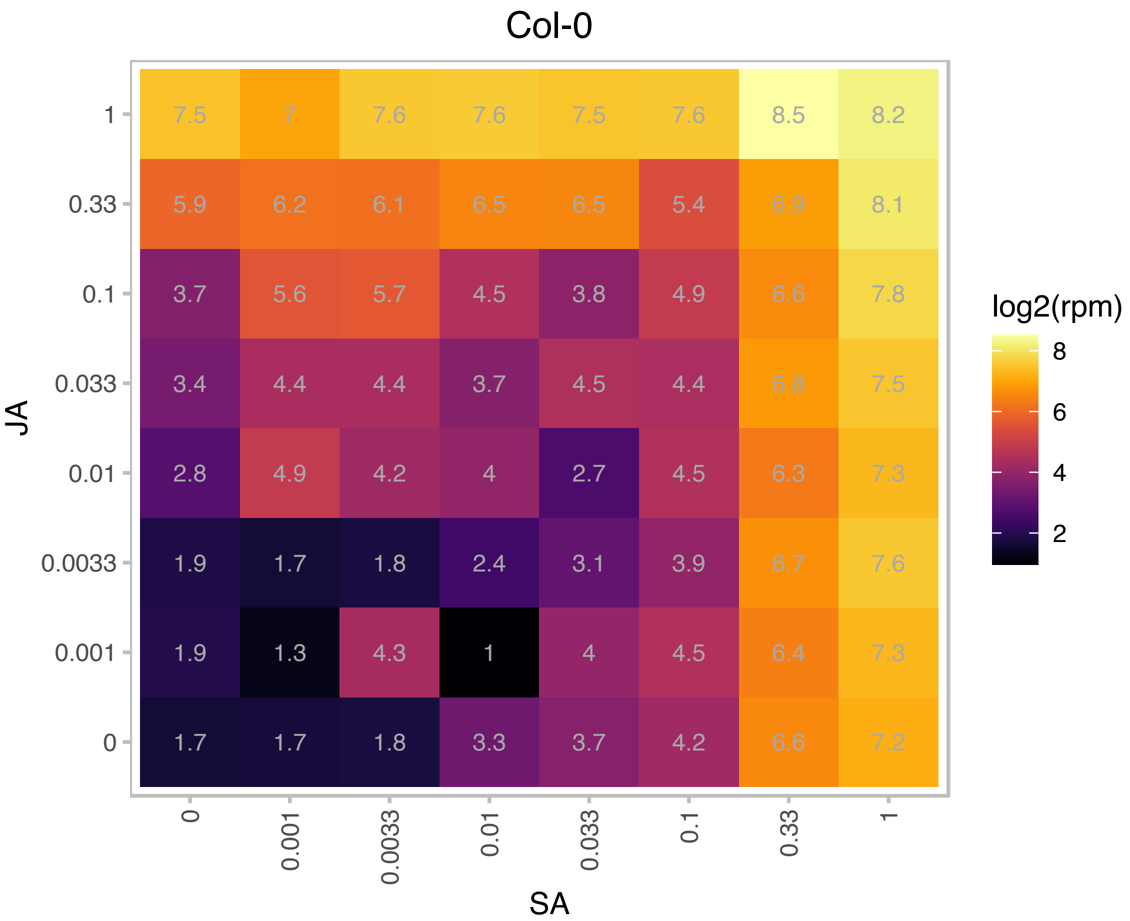


| | |
|---|---|
| Gene | AT2G15490 |
| is.expressed_genes | TRUE |
| is.responsive_genes | TRUE |
| is.apcluster | B-04 |

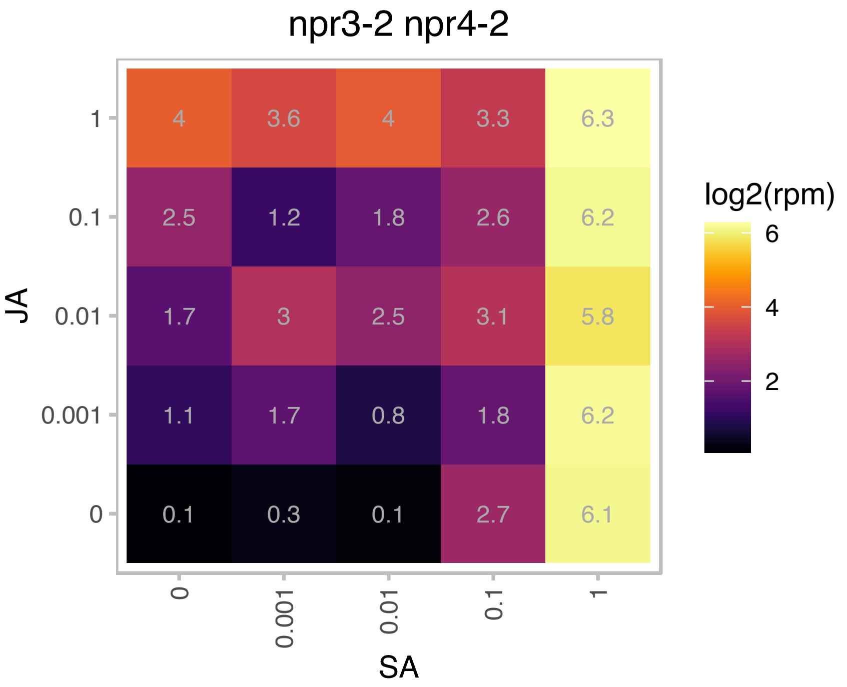


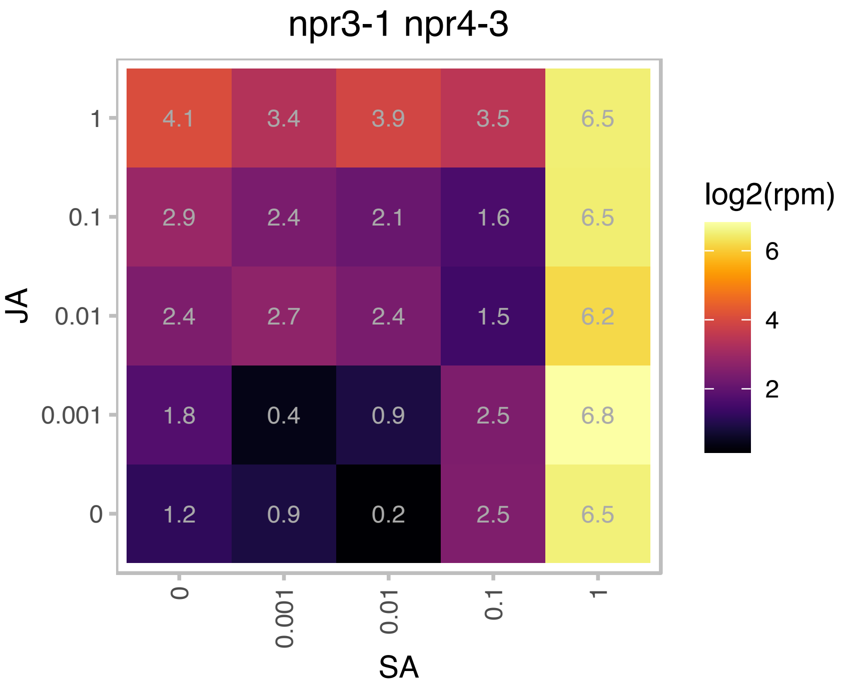

# AT2G18193.1 : NA

P-loop containing nucleoside triphosphate hydrolases superfamily protein

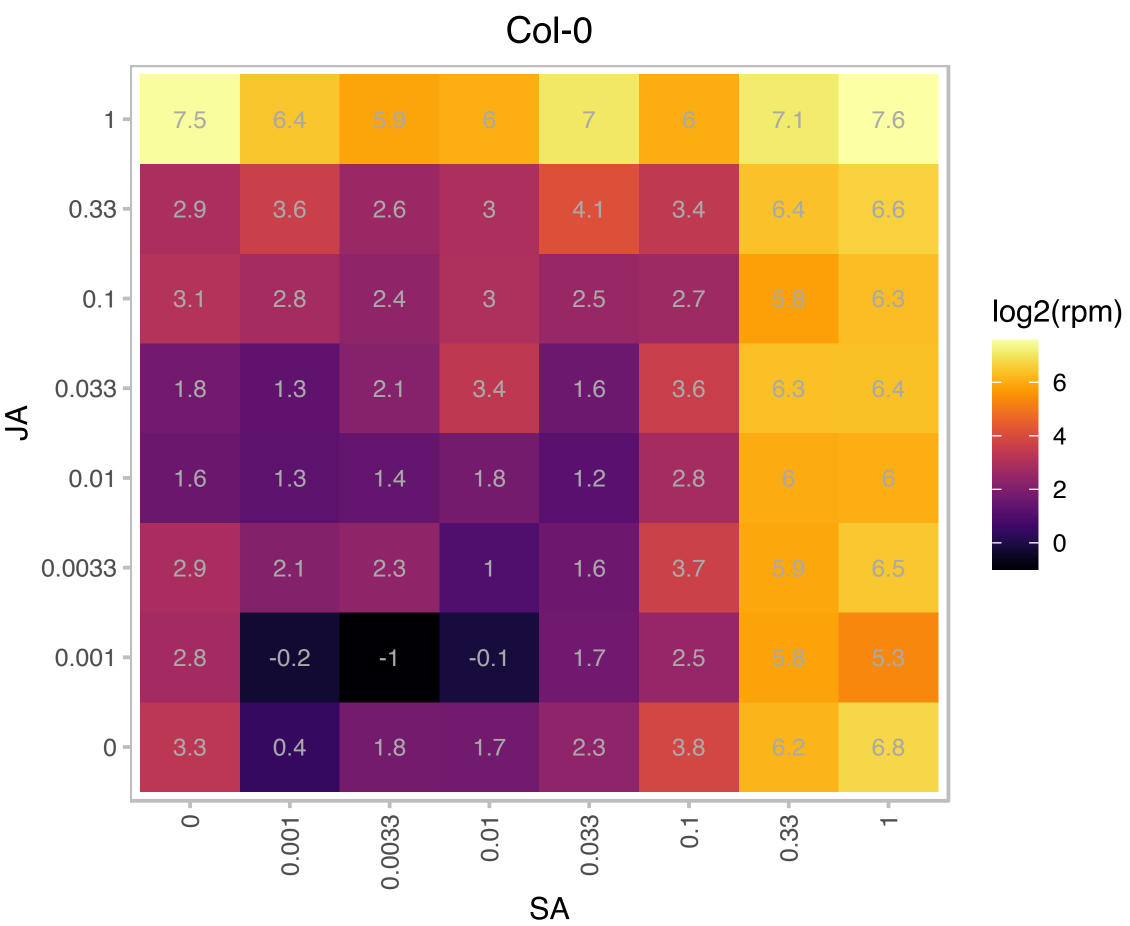


| | |
|---|---|
| Gene | AT2G18193 |
| is.expressed_genes | TRUE |
| is.responsive_genes | TRUE |
| is.apcluster | B-04 |

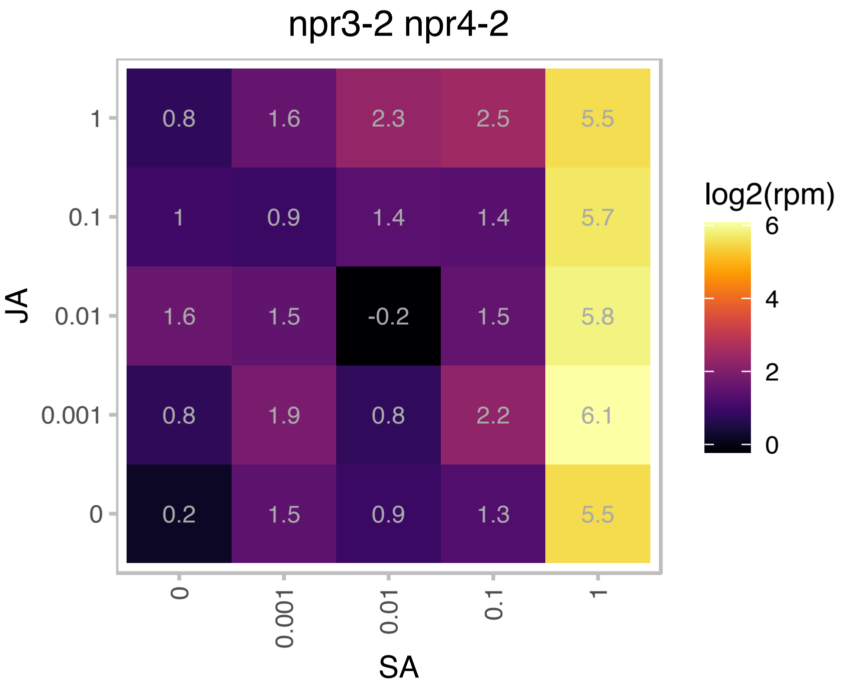


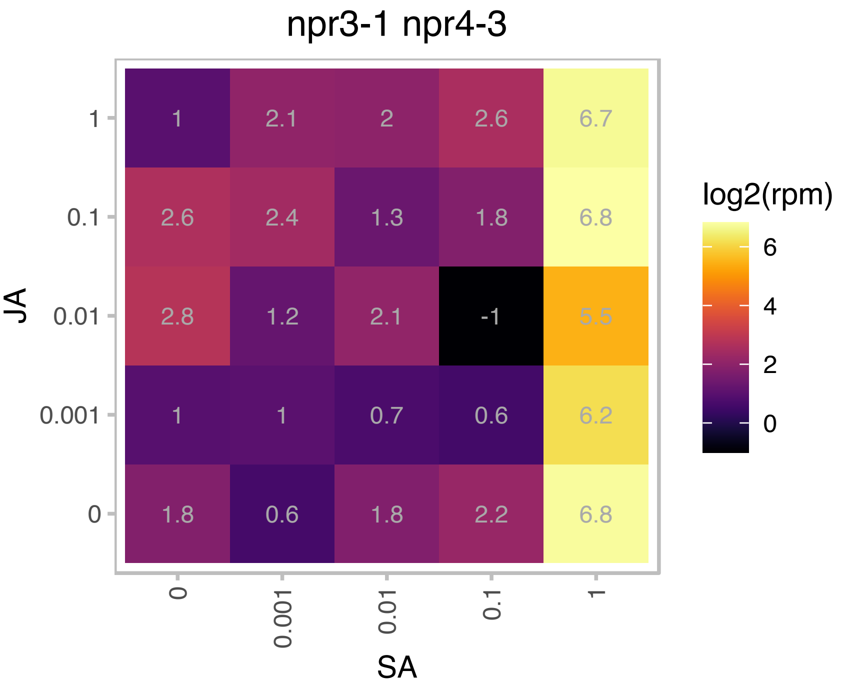

# AT2G20500.1 : NA

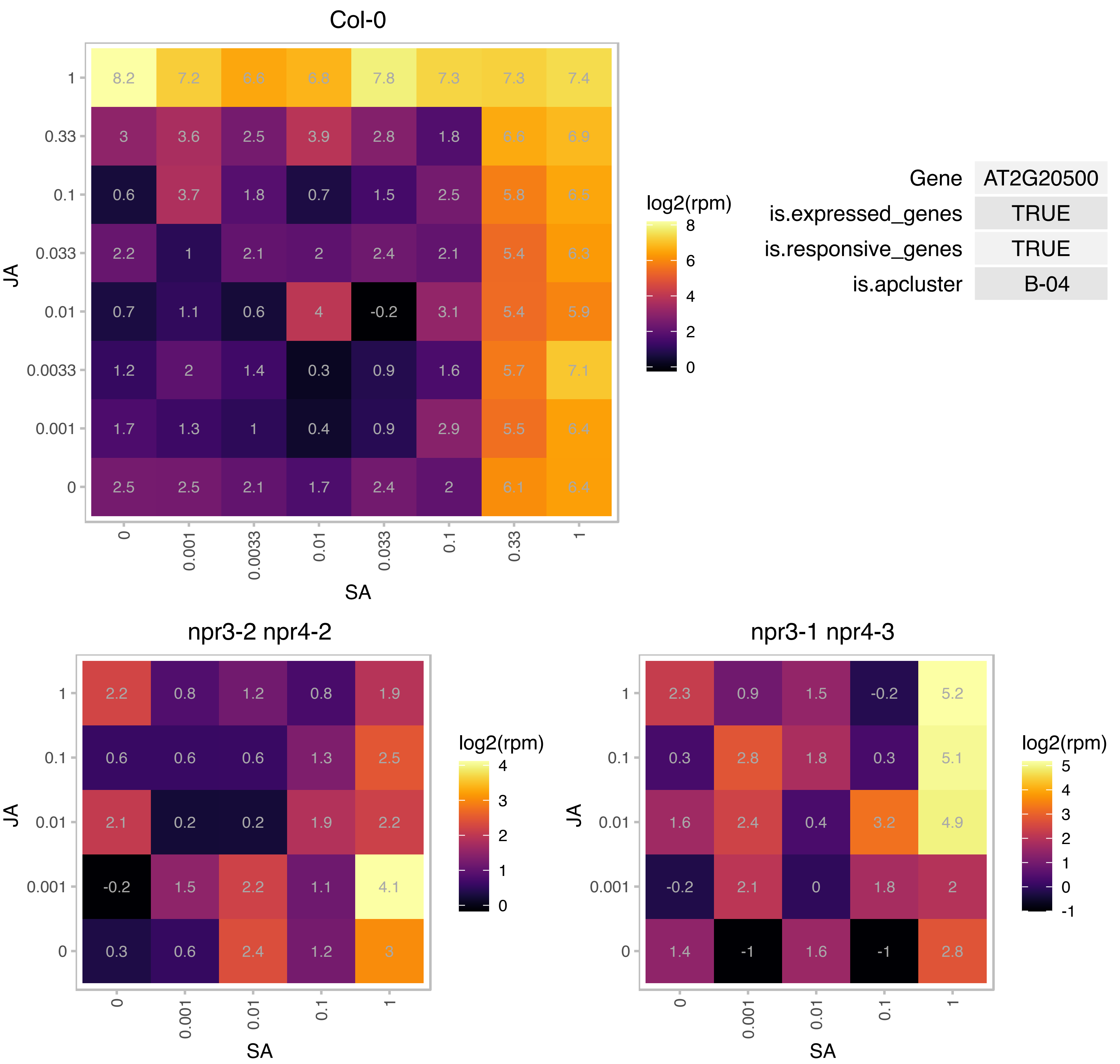


| | |
|---|---|
| Gene | AT2G20500 |
| is.expressed_genes | TRUE |
| is.responsive_genes | TRUE |
| is.apcluster | B-04 |

# AT2G25625.1 : CV

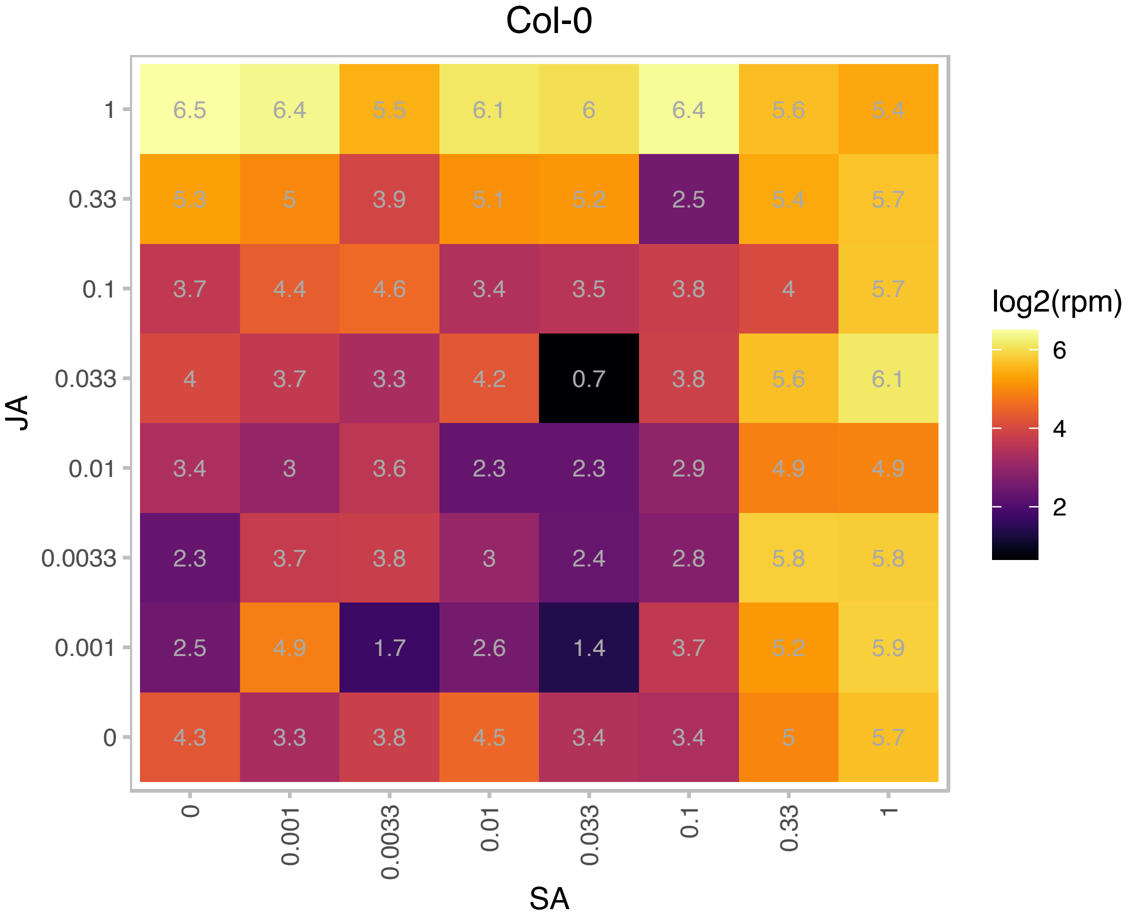


| | |
|---|---|
| Gene | AT2G25625 |
| is.expressed_genes | TRUE |
| is.responsive_genes | TRUE |
| is.apcluster | B-04 |

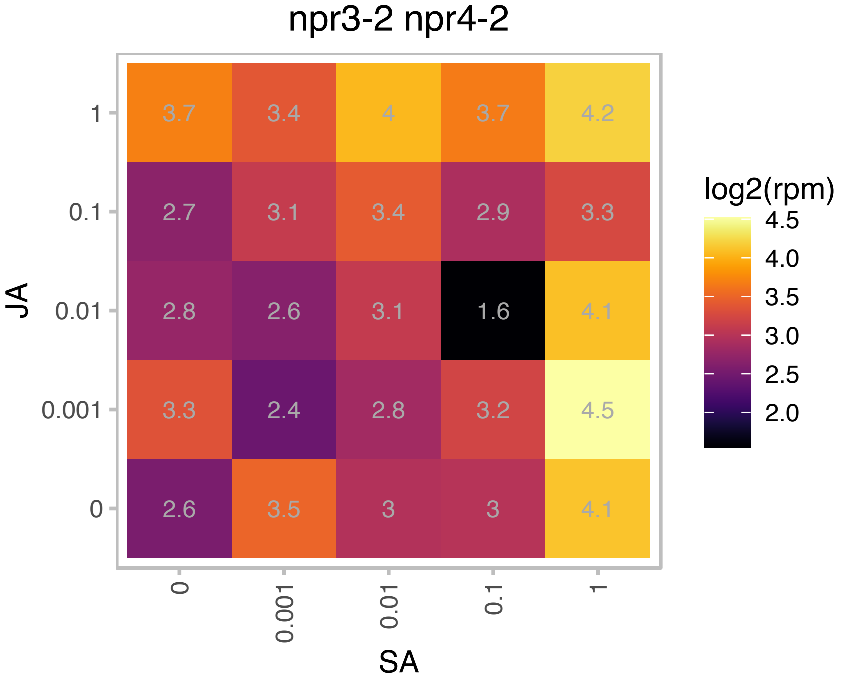


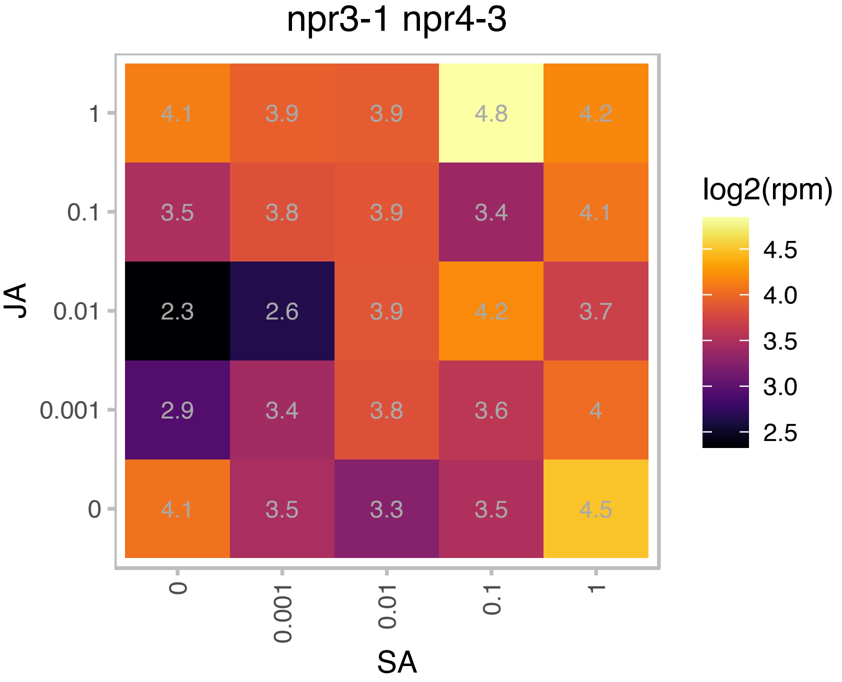

### AT2G29480.1 : ATGSTU2, GST20, GSTU2

glutathione S-transferase tau 2

Encodes glutathione transferase belonging to the tau class of GSTs. Naming convention according to Wagner et al. (2002).

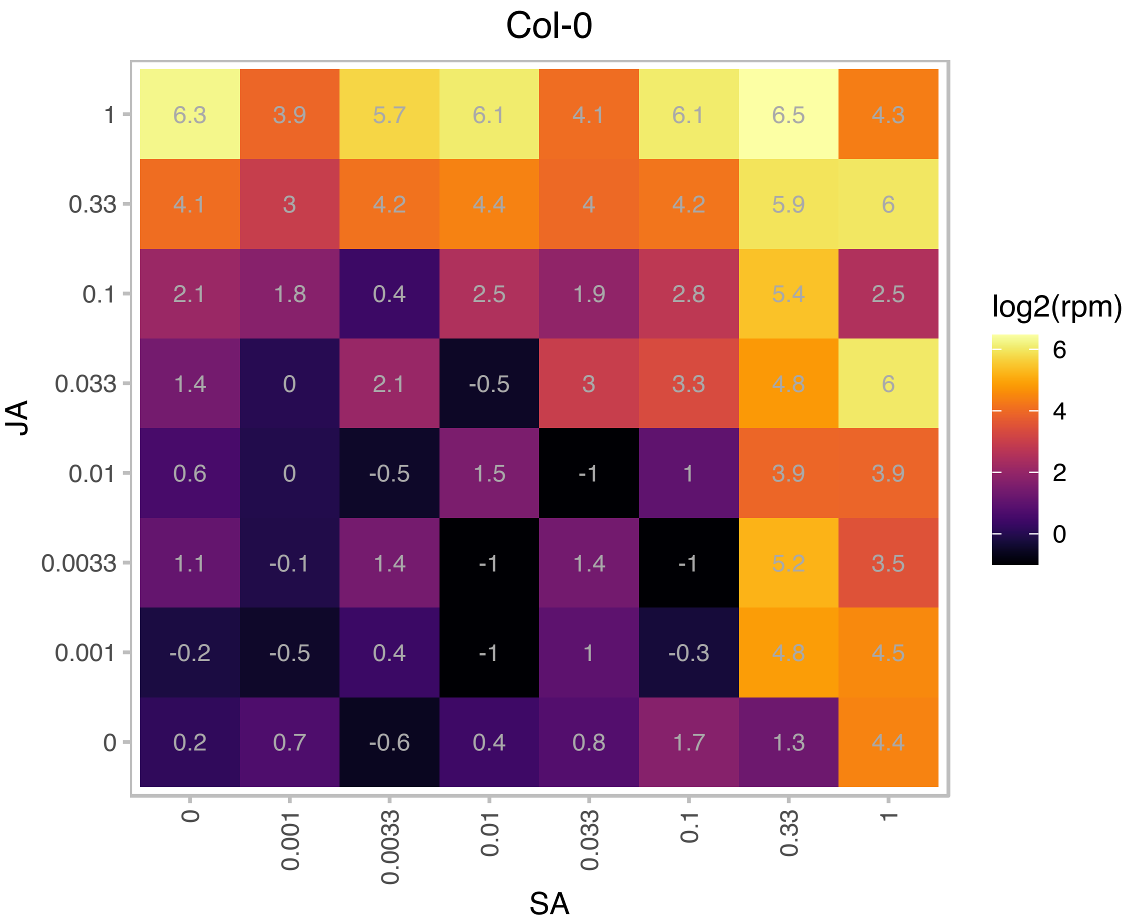


| | |
|---|---|
| Gene | AT2G29480 |
| is.expressed_genes | TRUE |
| is.responsive_genes | TRUE |
| is.apcluster | B-04 |

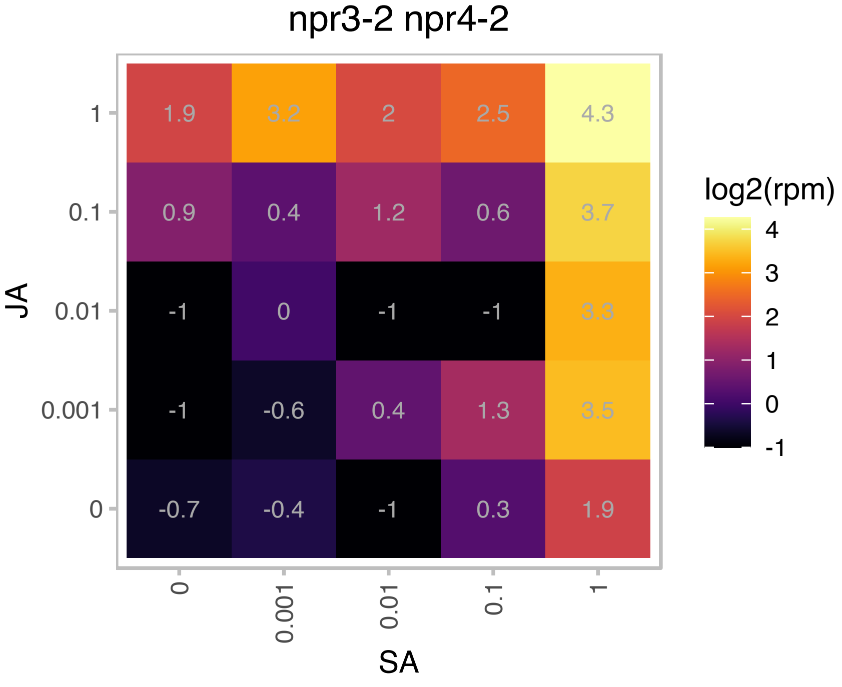


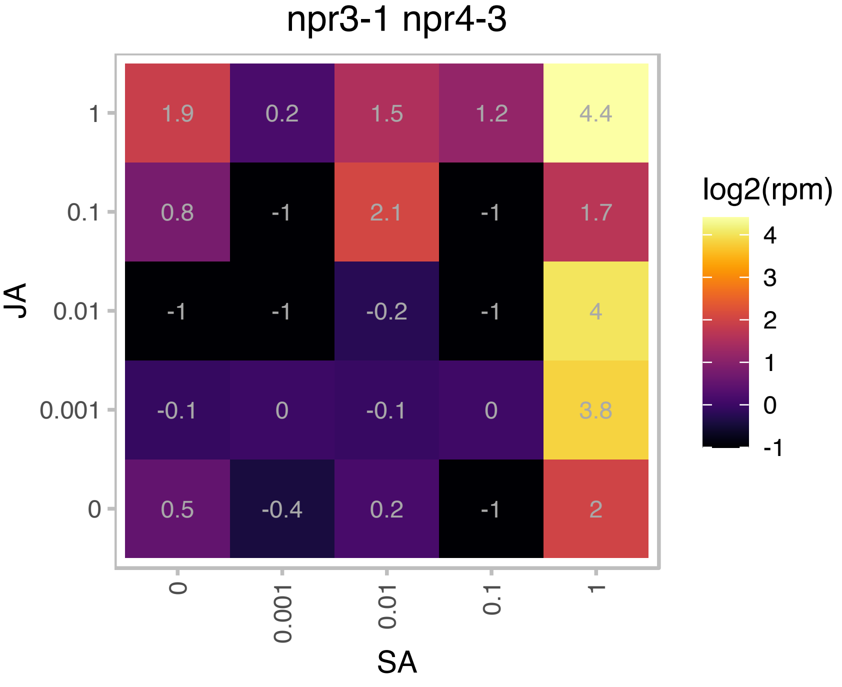

### AT2G30170.1 : PBCP

Protein phosphatase 2C family protein

Encodes a chloroplast PP2C phosphatase that is required for efficient dephosphorylation of PSII proteins and involved in light acclimation.

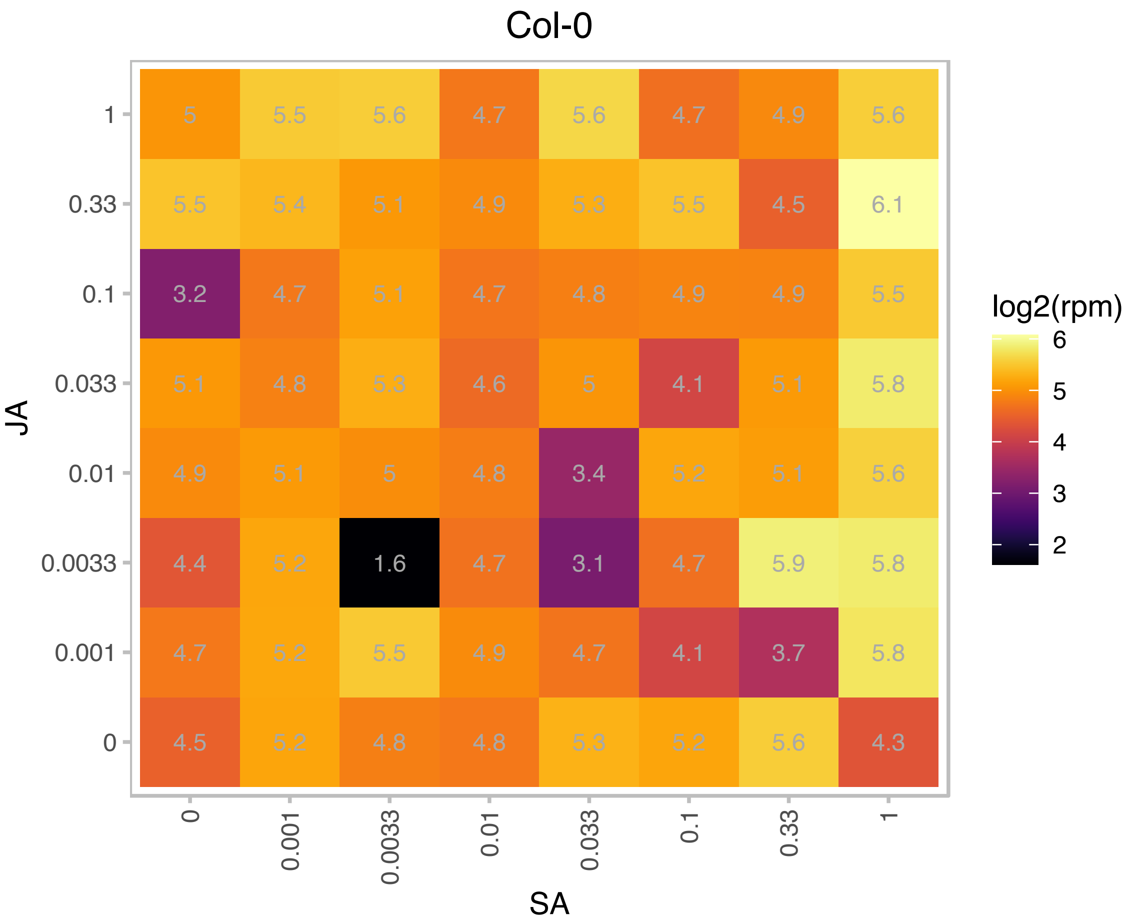


| | |
|---|---|
| Gene | AT2G30170 |
| is.expressed_genes | TRUE |
| is.responsive_genes | TRUE |
| is.apcluster | B-04 |

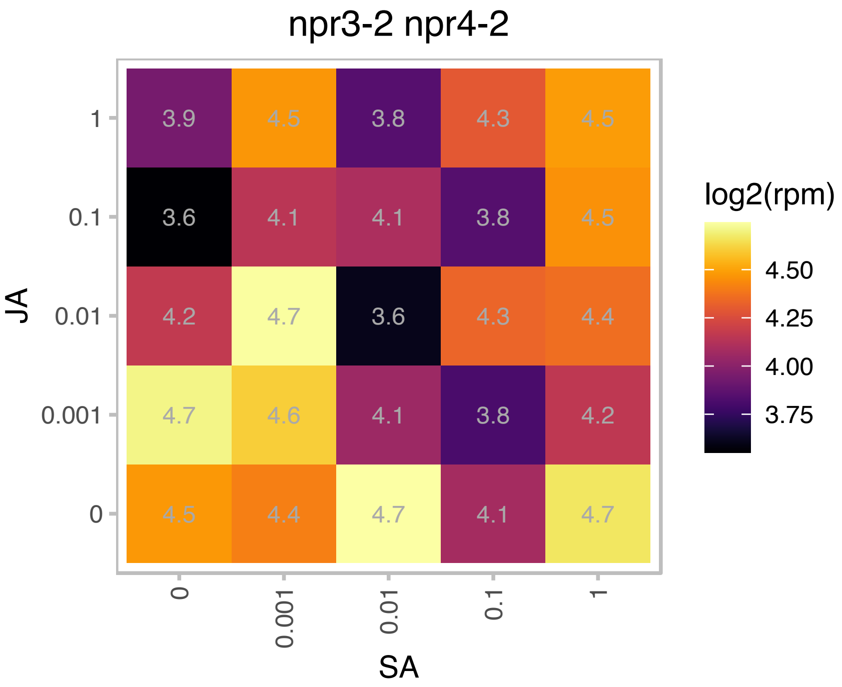


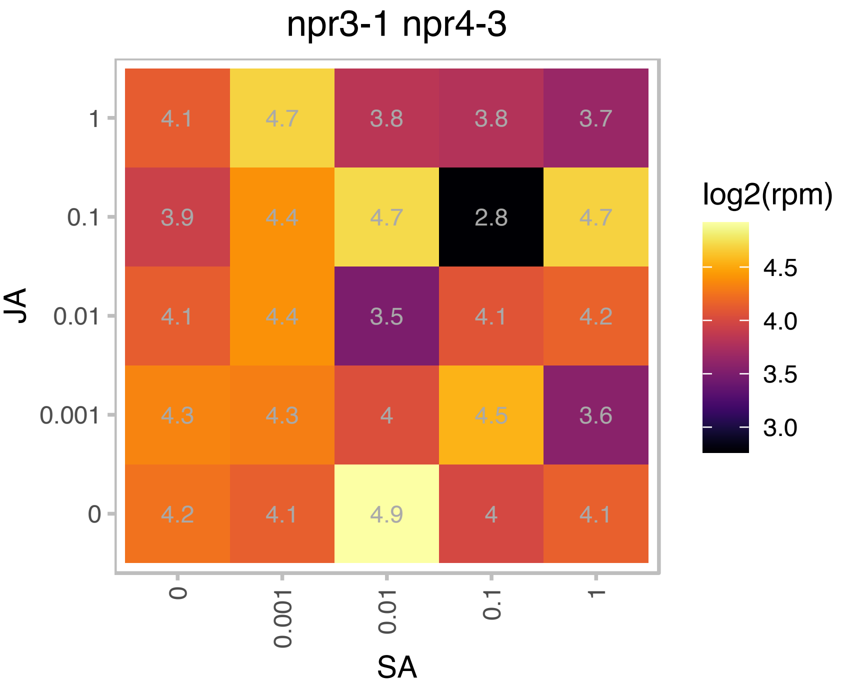

### AT2G31380.1 : BBX25, STH

salt tolerance homologue

a B-box zinc finger protein that interacts with COP1. contains a novel 11 amino acid motif at the C-terminus (also found at the N-terminus of HY5) that is involved in the COP1 interaction.

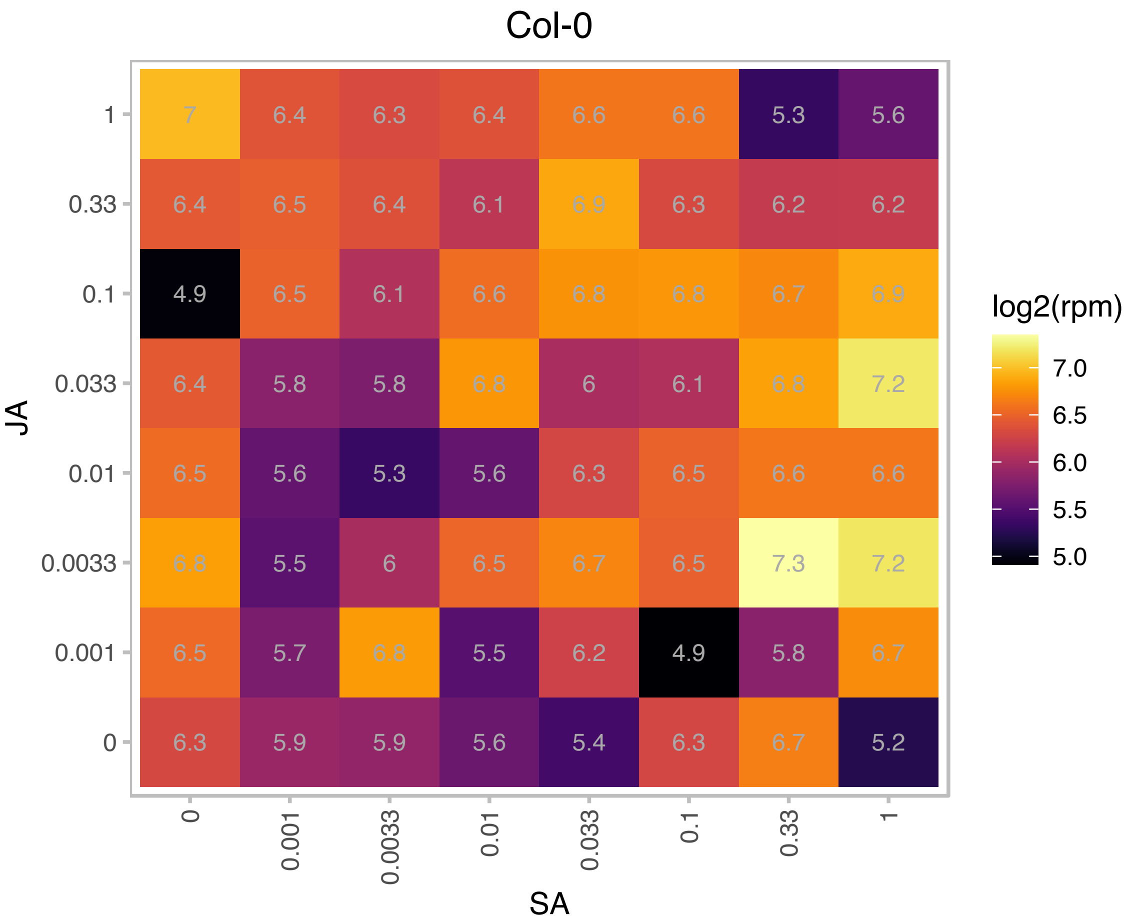


| Gene | AT2G31380 |
|---|---|
| is.expressed_genes | TRUE |
| is.responsive_genes | TRUE |
| is.apcluster | B-04 |

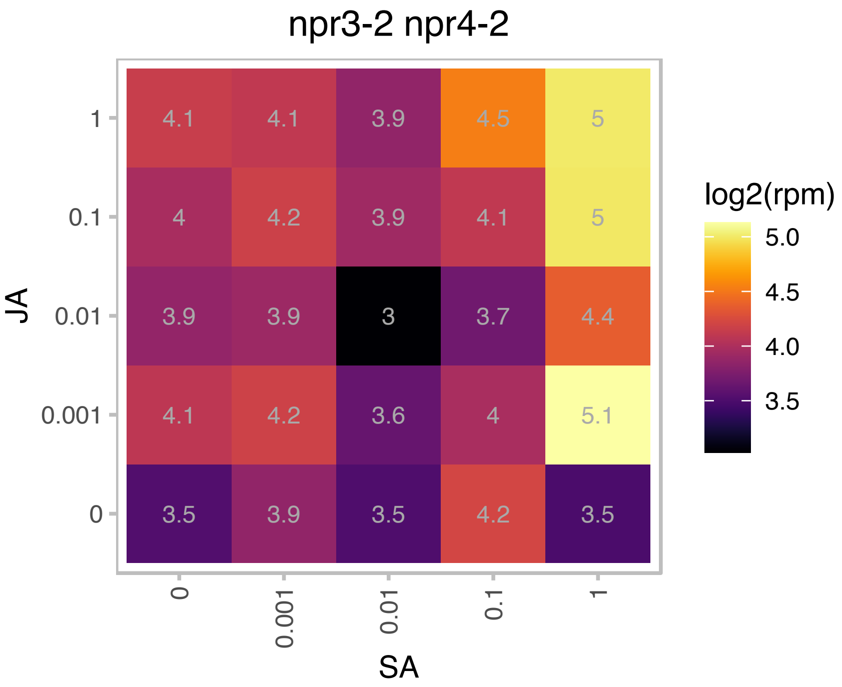


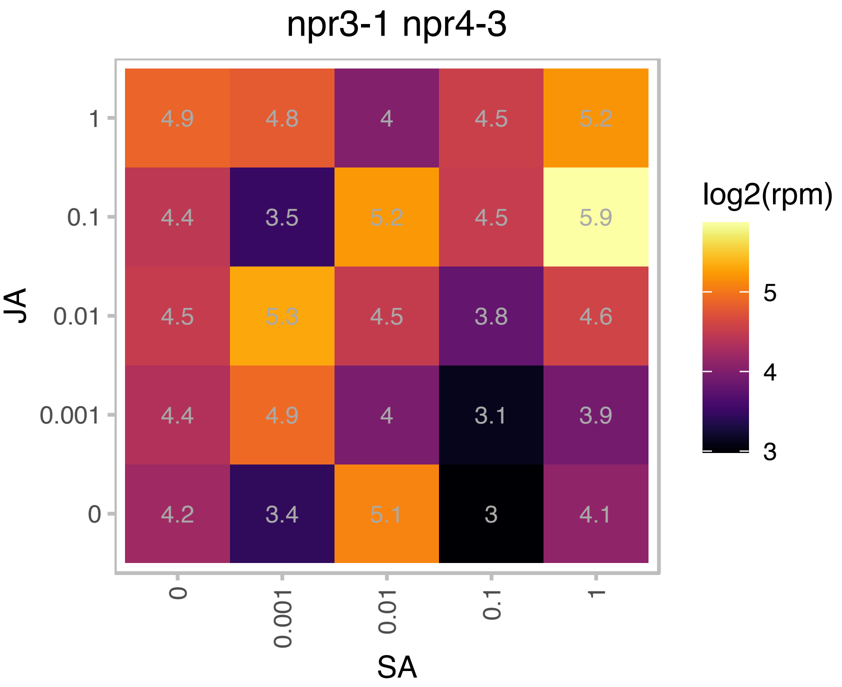

### AT2G31570.1 : ATGPX2, GPX2, GPXL2

glutathione peroxidase 2

glutathione peroxidase GPx

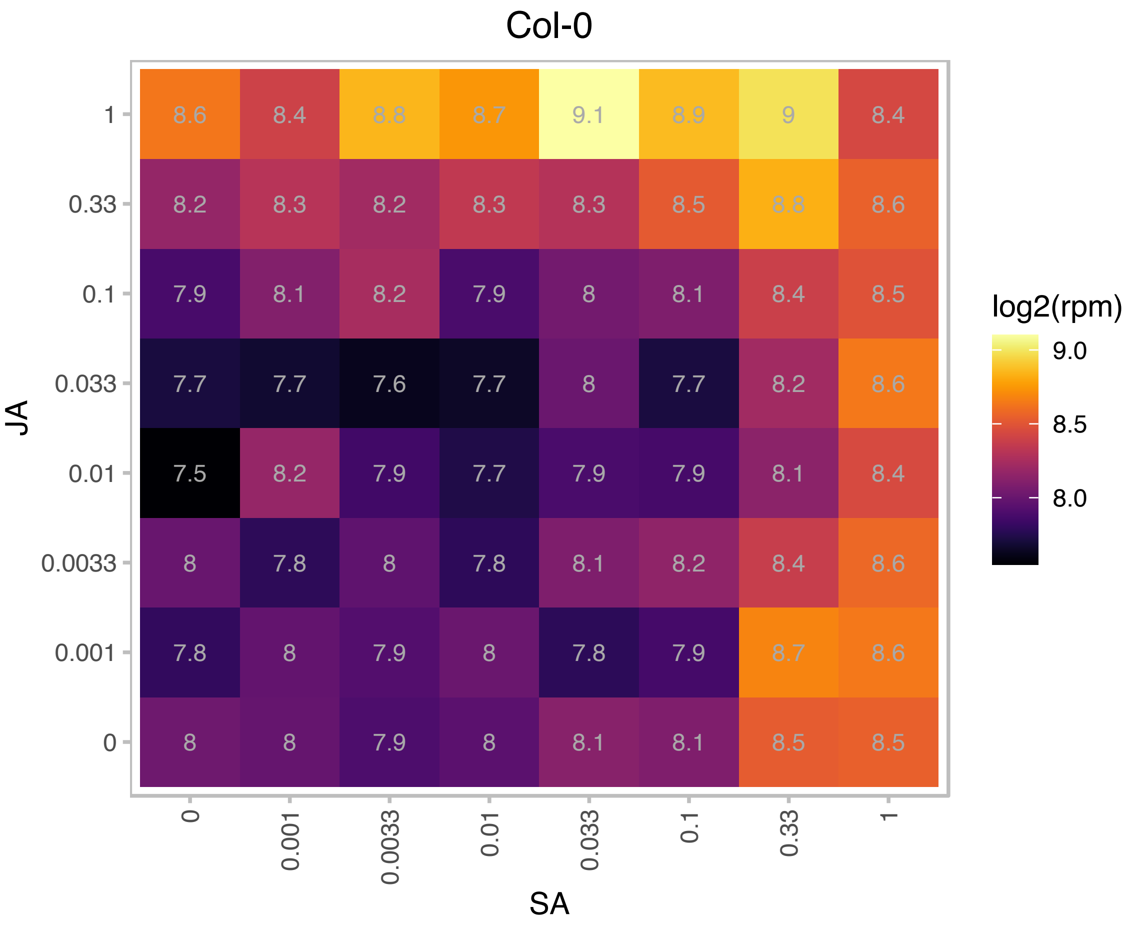


| | |
|---|---|
| Gene | AT2G31570 |
| is.expressed_genes | TRUE |
| is.responsive_genes | TRUE |
| is.apcluster | B-04 |

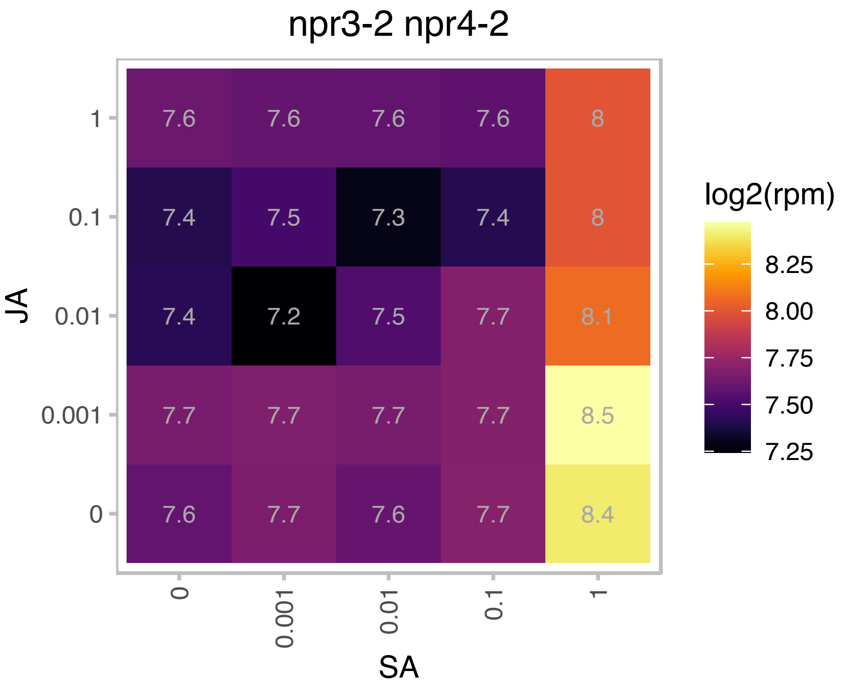


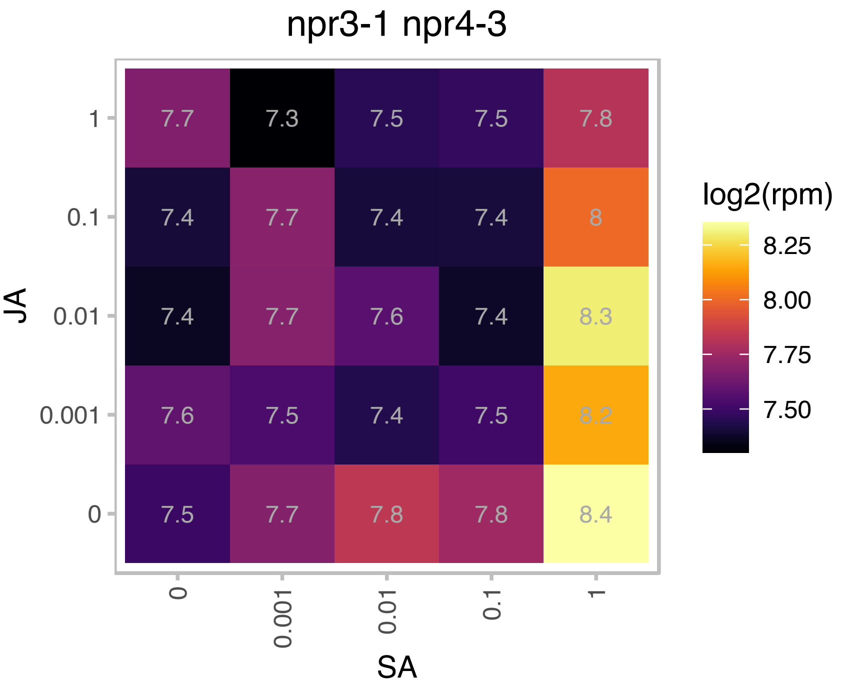

# AT2G36780.1 : NA

UDP-Glycosyltransferase superfamily protein

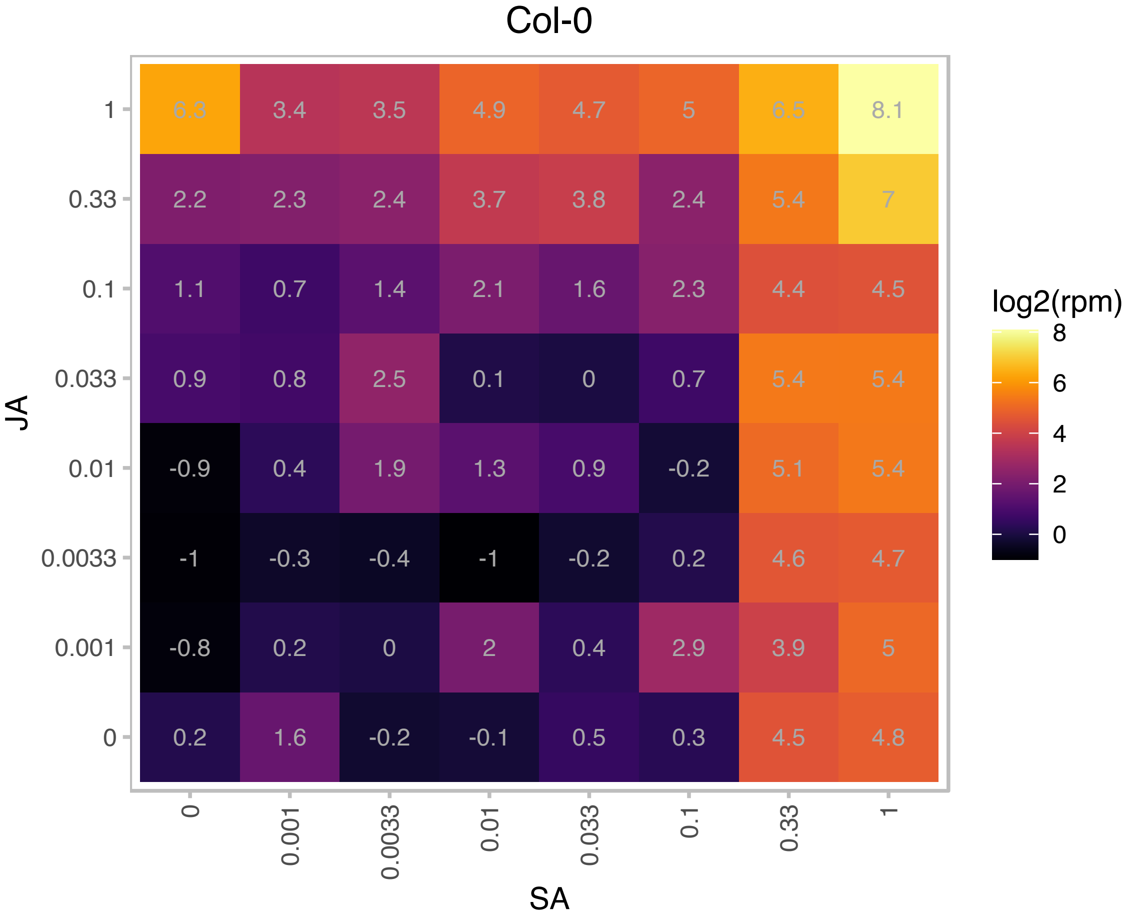


| | |
|---|---|
| Gene | AT2G36780 |
| is.expressed_genes | TRUE |
| is.responsive_genes | TRUE |
| is.apcluster | B-04 |

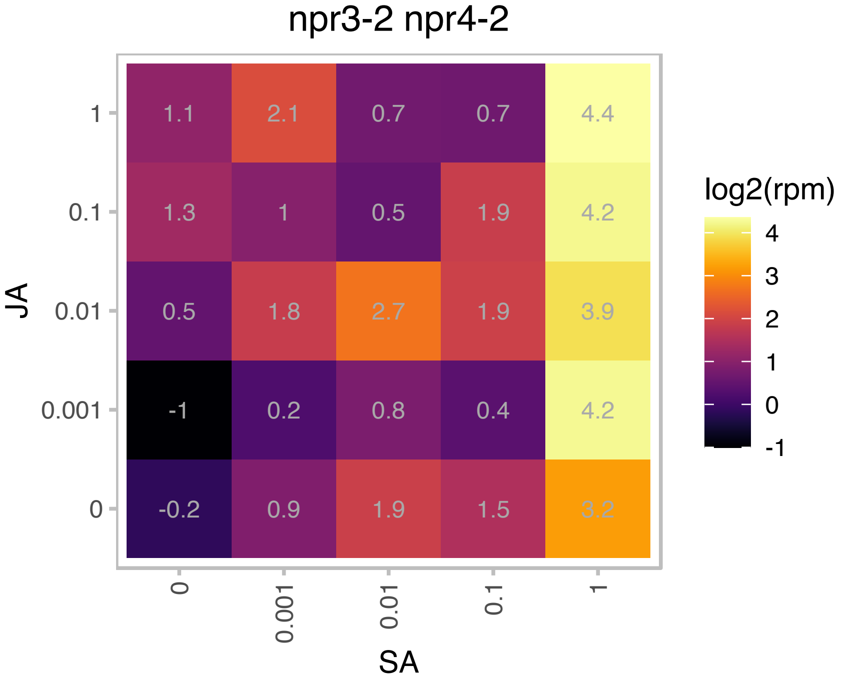


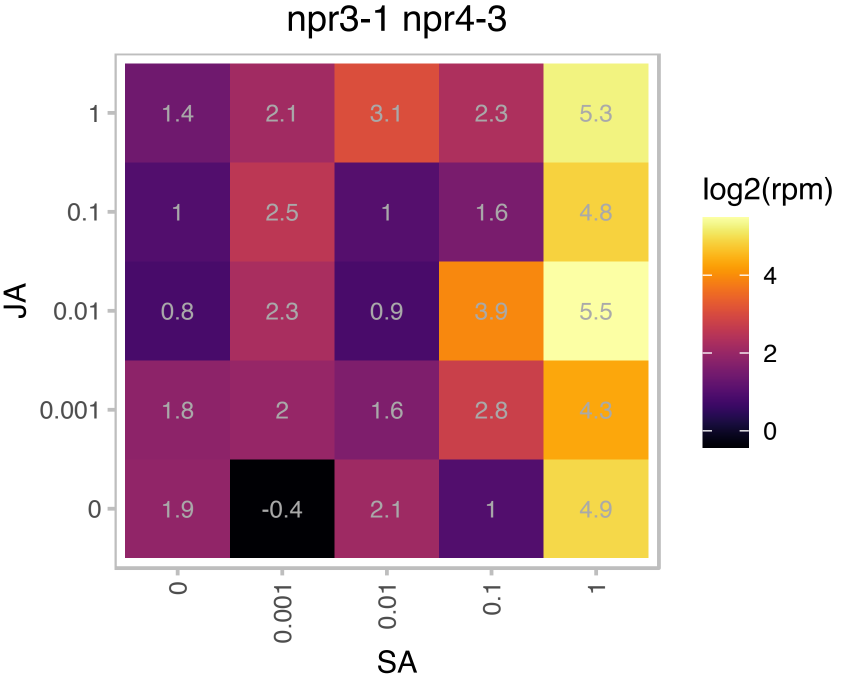

# AT2G38000.1 : NA

chaperone protein dnaJ-related

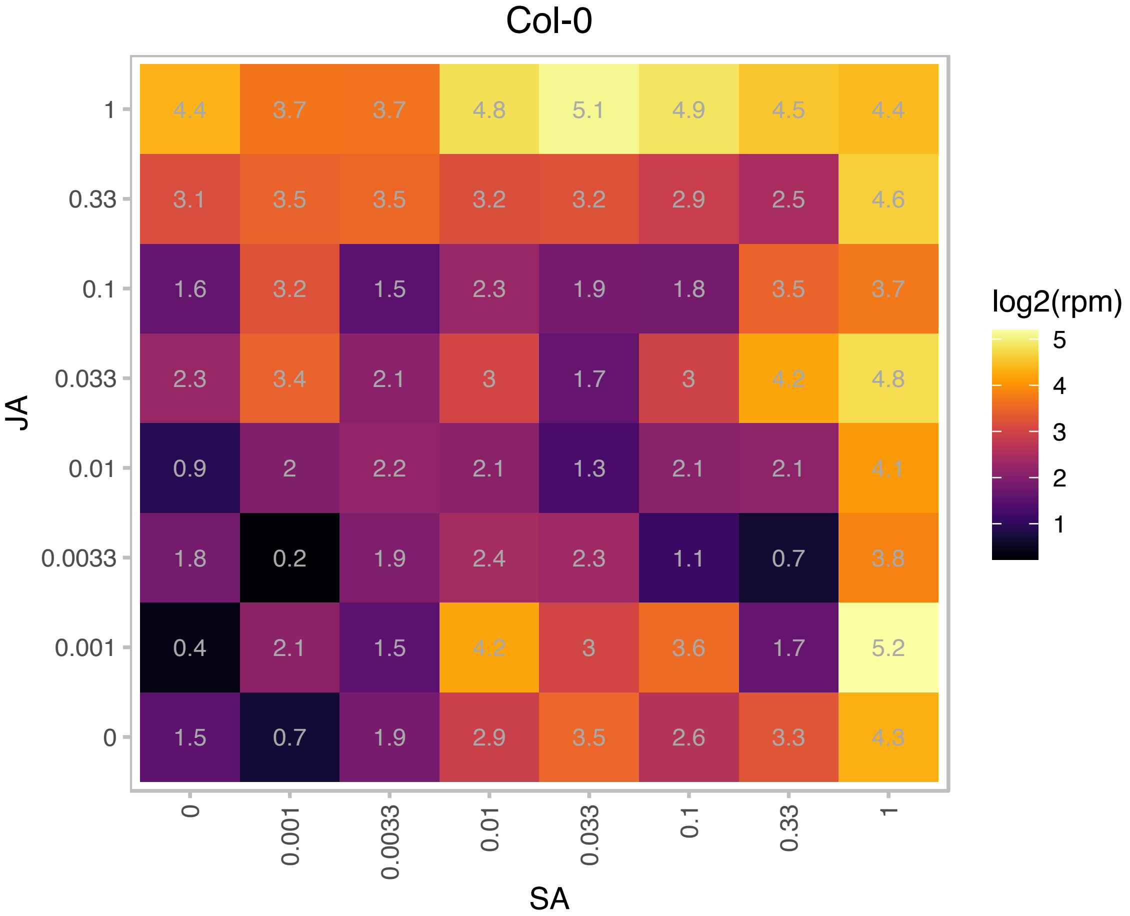


| | |
|---|---|
| Gene | AT2G38000 |
| is.expressed_genes | TRUE |
| is.responsive_genes | TRUE |
| is.apcluster | B-04 |

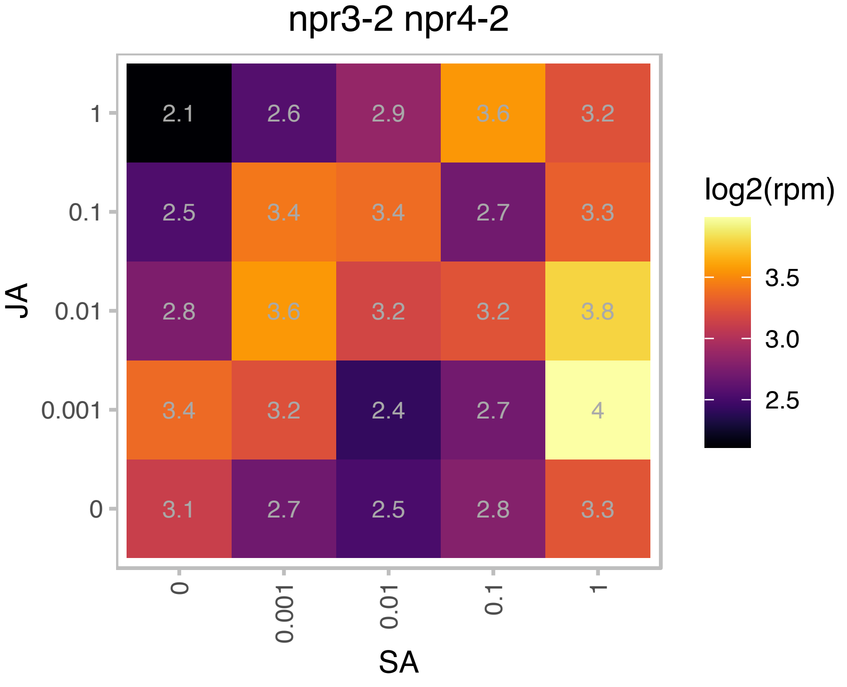


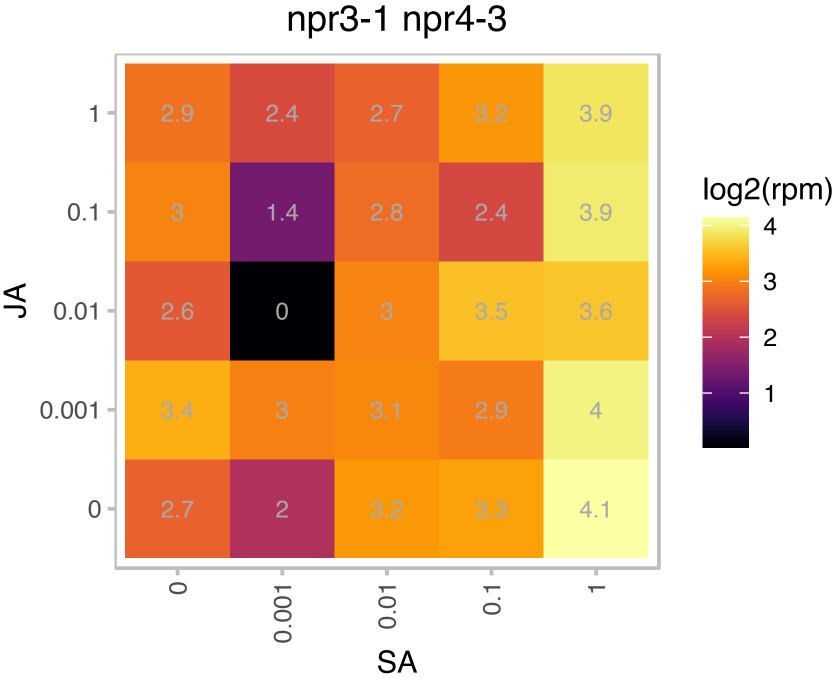

### AT2G44460.1 : BGLU28

beta glucosidase 28

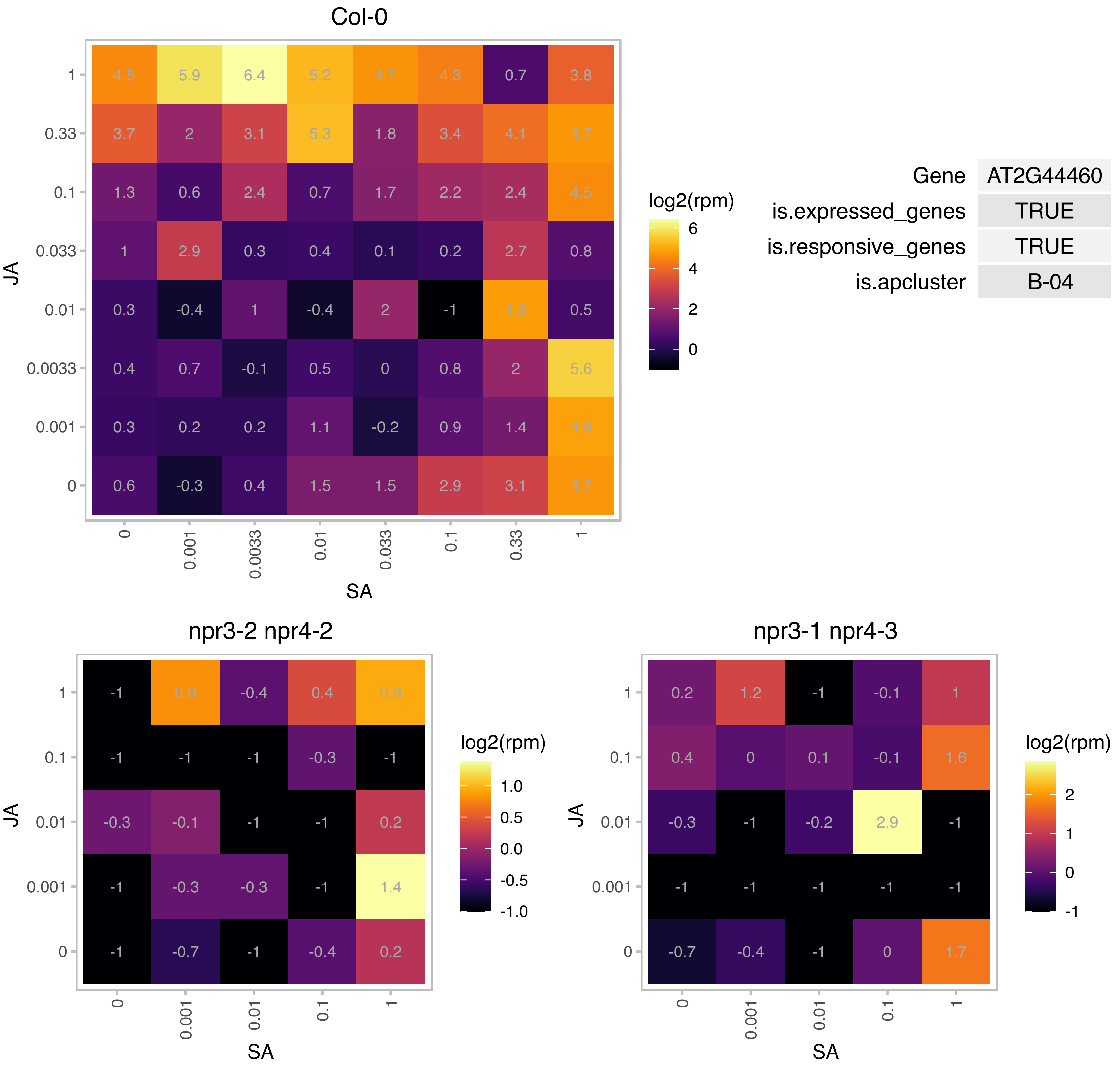


| | |
|---|---|
| Gene | AT2G44460 |
| is.expressed_genes | TRUE |
| is.responsive_genes | TRUE |
| is.apcluster | B-04 |

### AT2G46240.1 : ATBAG6, BAG6

BCL-2-associated athanogene 6

A member of Arabidopsis BAG (Bcl-2-associated athanogene) proteins, plant homologs of mammalian regulators of apoptosis. Expression of BAG6 in leaves was strongly induced by heat stress. Knockout mutants exhibited enhanced susceptibility to fungal pathogen Botrytis cinerea. Plant BAG proteins are multi-functional and remarkably similar to their animal counterparts, as they regulate apoptotic-like processes ranging from pathogen attack, to abiotic stress, to plant development. The mRNA is cell-to-cell mobile.

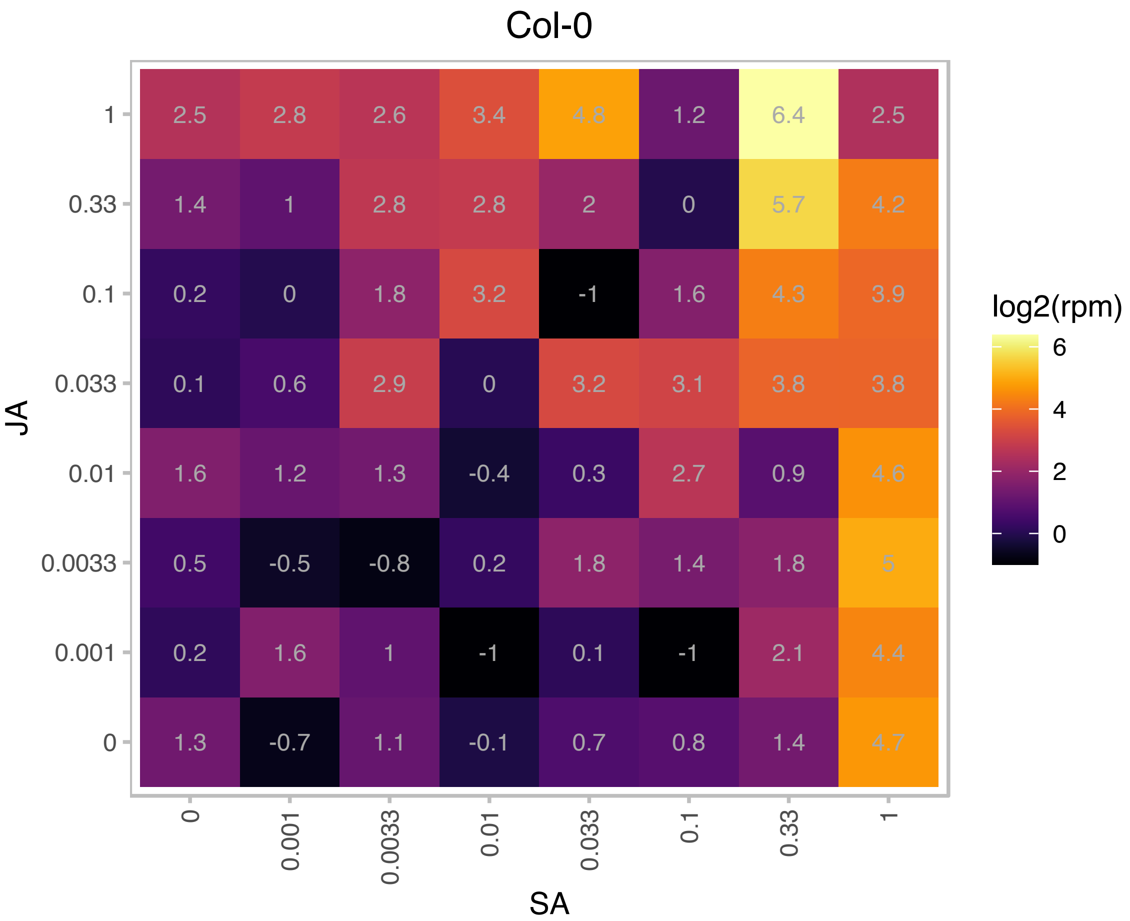


| | |
|---|---|
| Gene | AT2G46240 |
| is.expressed_genes | TRUE |
| is.responsive_genes | TRUE |
| is.apcluster | B-04 |

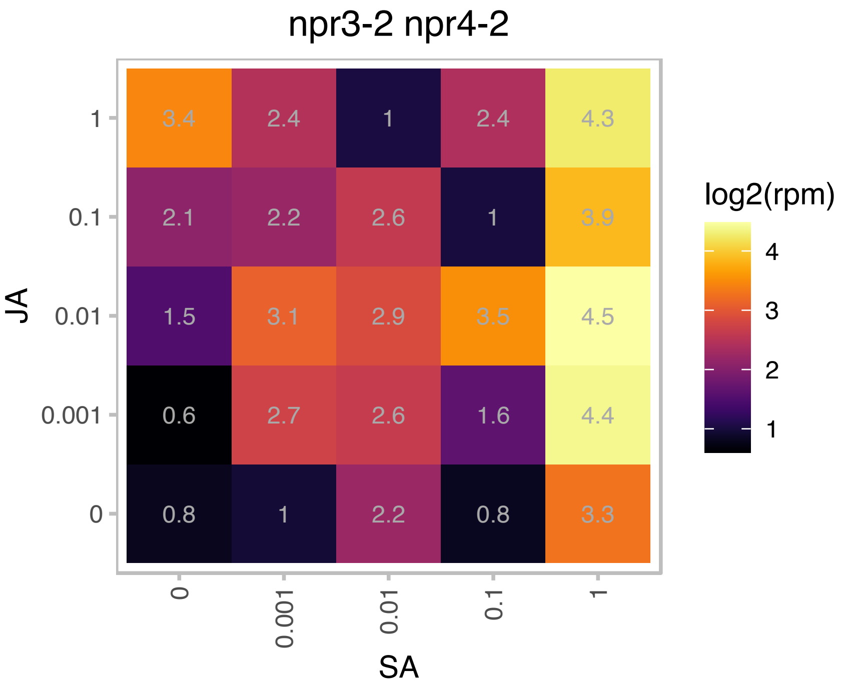


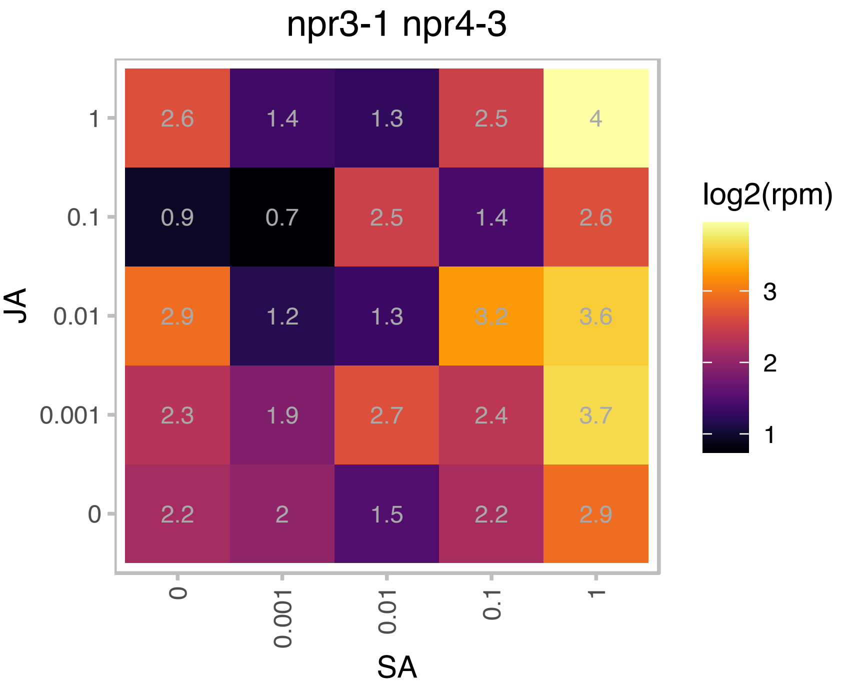

### AT2G47000.1 : ABCB4, AtABCB4, ATPGP4, MDR4, PGP4

ATP binding cassette subfamily B4
Encodes an auxin efflux transmembrane transporter that is a member of the multidrug resistance P-glycoprotein (MDR/PGP) subfamily of ABC transporters. Functions in the basipetal redirection of auxin from the root tip. Exhibits apolar plasma membrane localization in the root cap and polar localization in tissues above and is involved in root hair elongation.

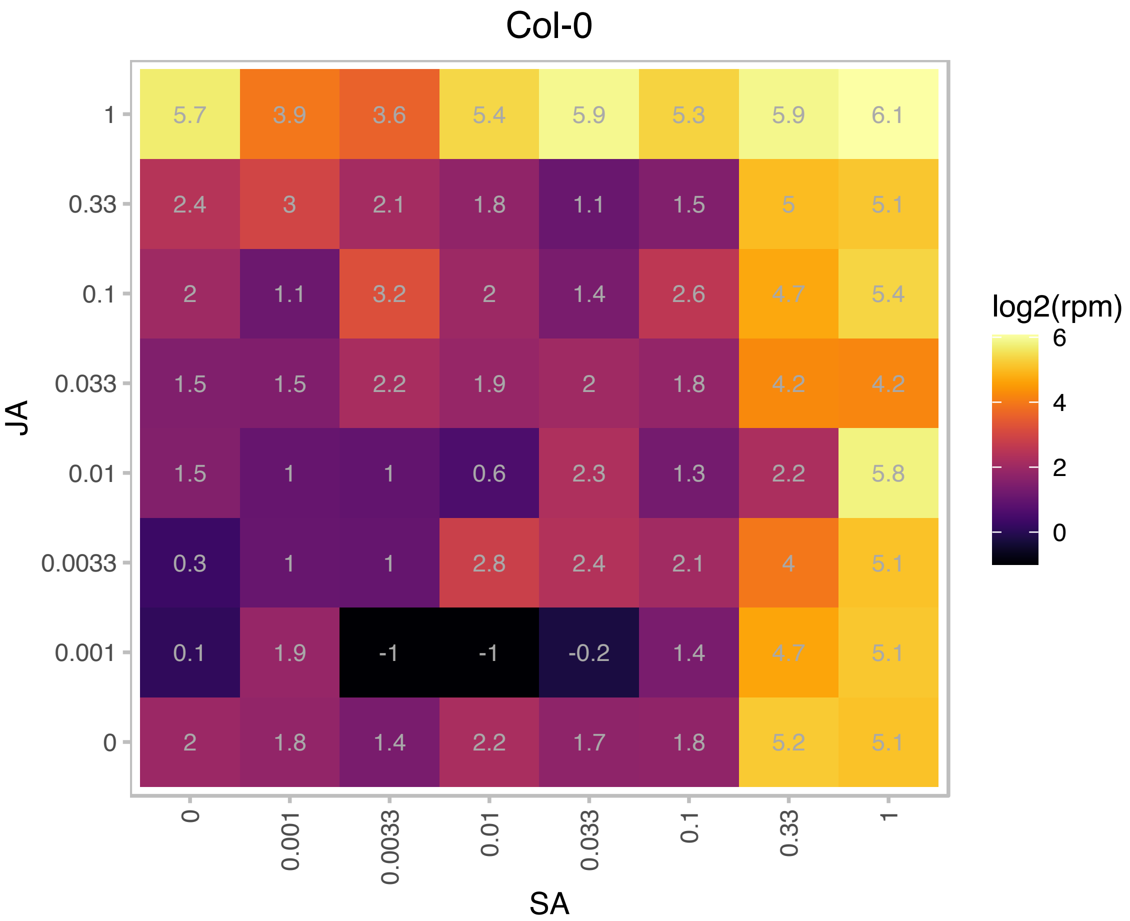


| | |
|---|---|
| Gene | AT2G47000 |
| is.expressed_genes | TRUE |
| is.responsive_genes | TRUE |
| is.apcluster | B-04 |

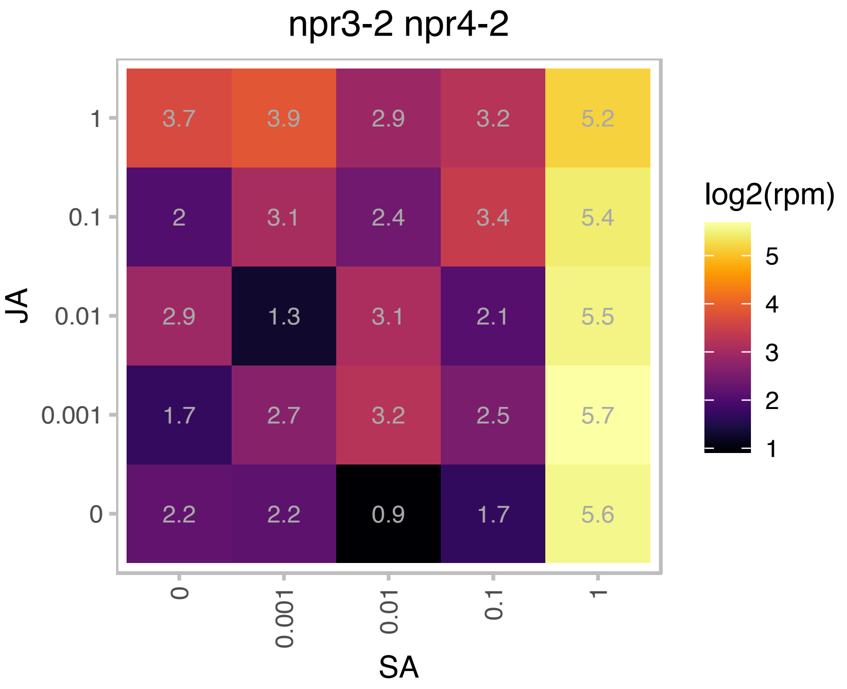


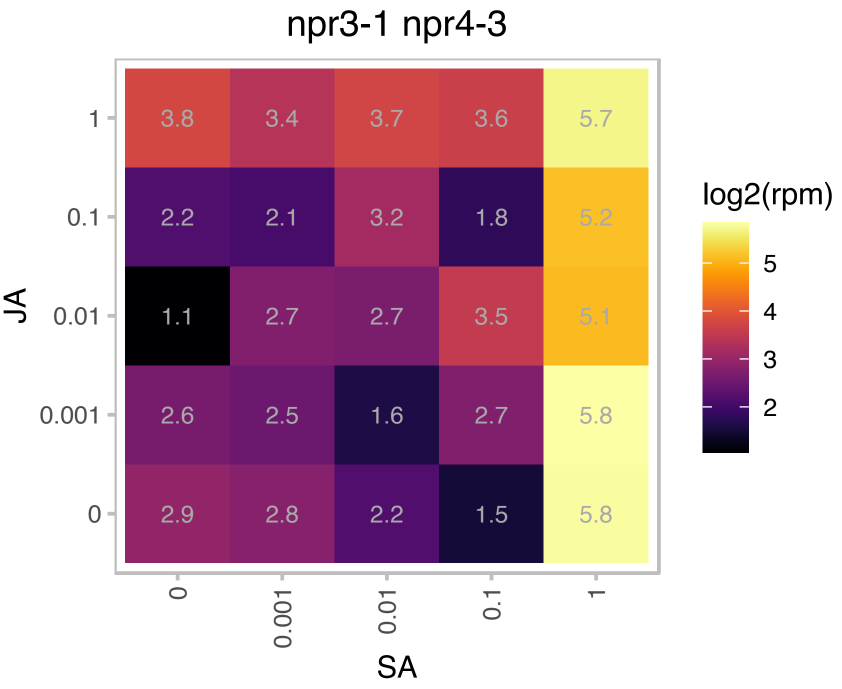

# AT3G03890.1 : NA

FMN binding

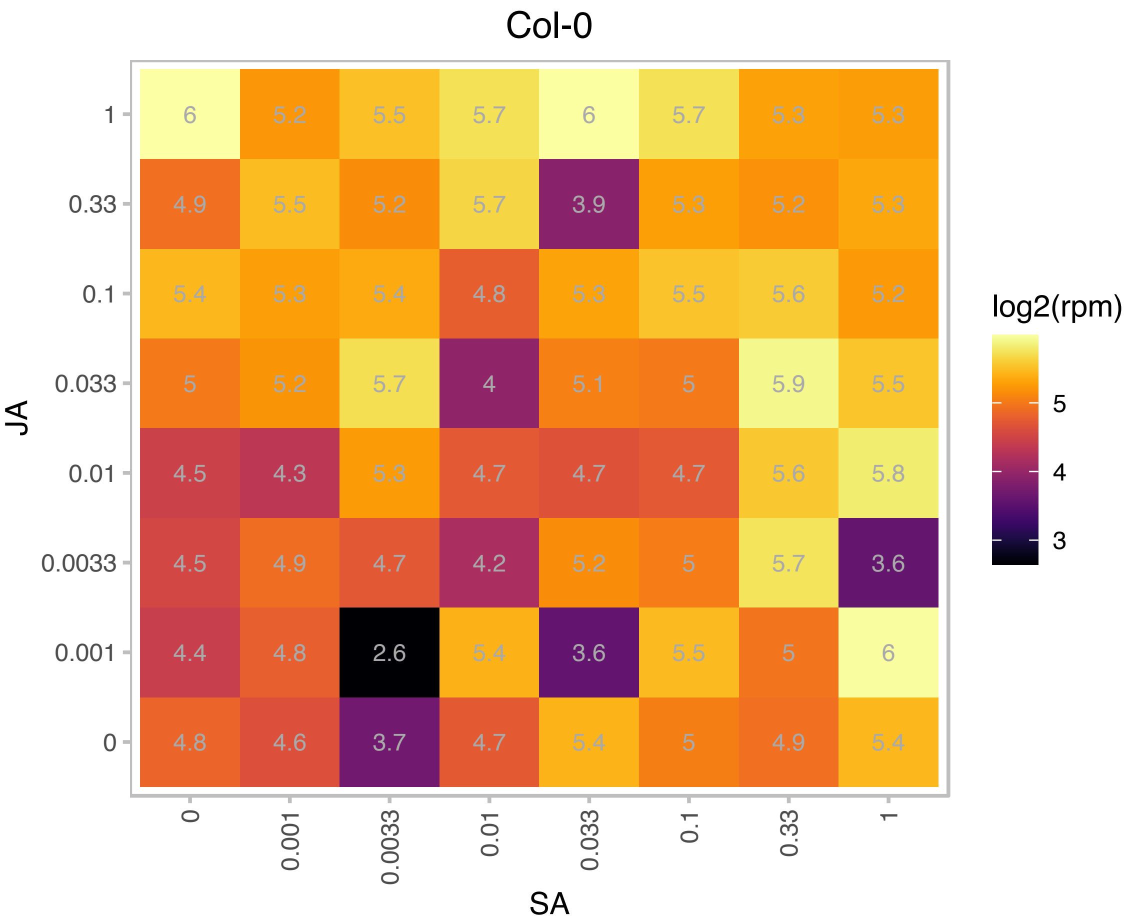


| | |
|---|---|
| Gene | AT3G03890 |
| is.expressed_genes | TRUE |
| is.responsive_genes | TRUE |
| is.apcluster | B-04 |

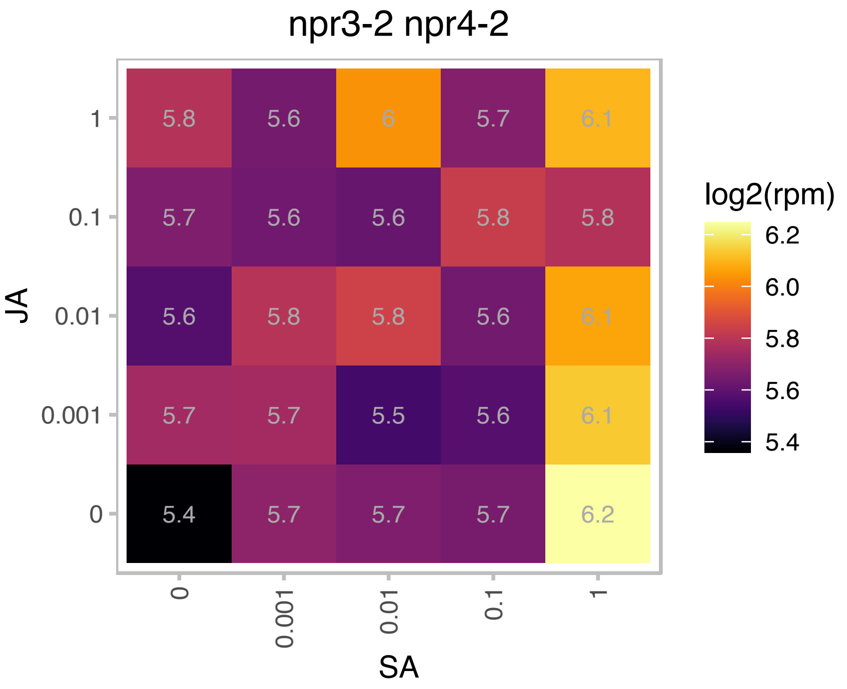


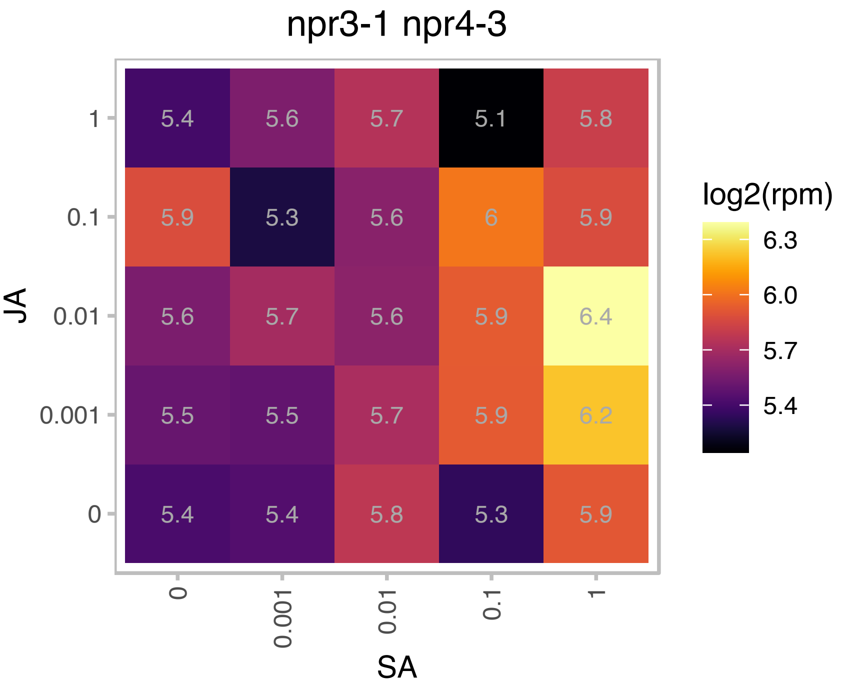

### AT3G12120.1 : AtFAD2, FAD2

fatty acid desaturase 2
Major enzyme responsible for the synthesis of 18:2 fatty acids in the endoplasmic reticulum. Contains His-rich motifs, which contribute to the interaction with the electron donor cytochrome b5. Mutations in this gene suppress the low temperature-induced phenotype of Arabidopsis tocopherol-deficient mutant vte2.

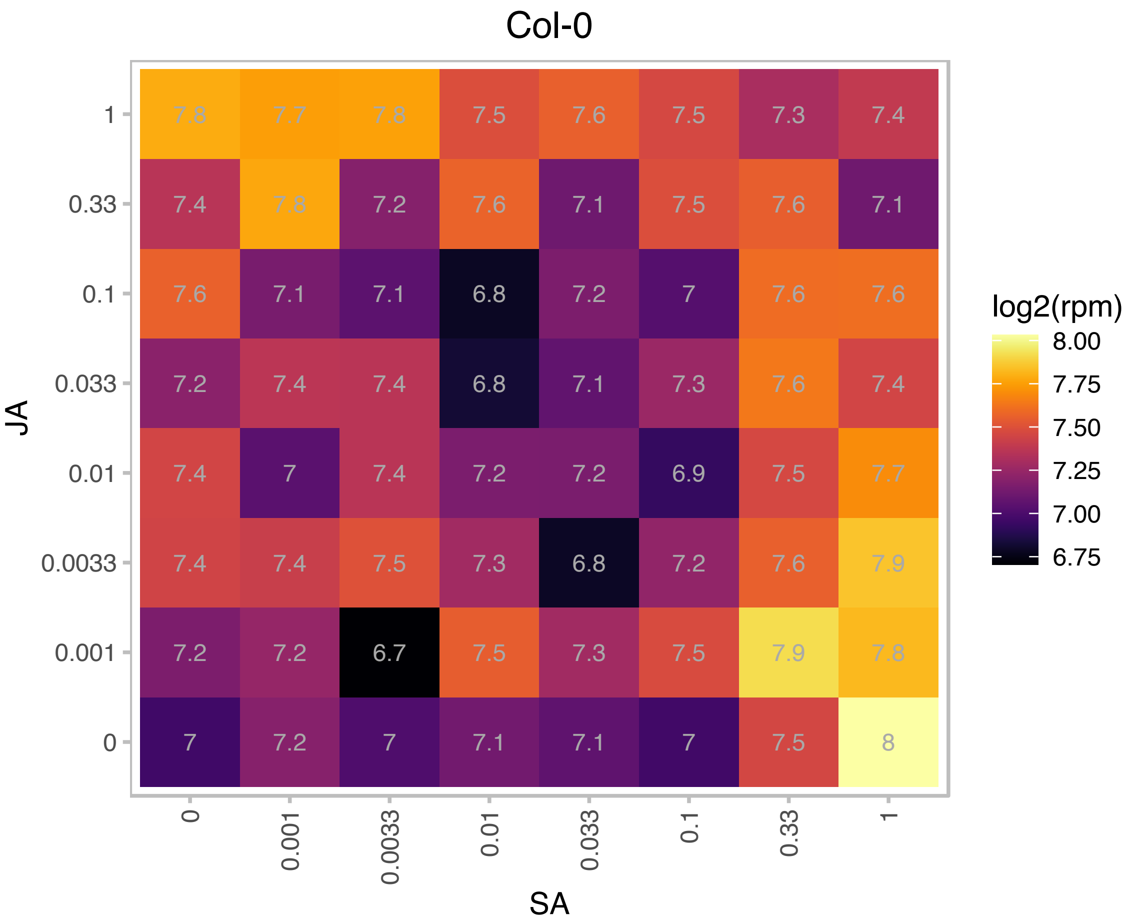


| | |
|---|---|
| Gene | AT3G12120 |
| is.expressed_genes | TRUE |
| is.responsive_genes | TRUE |
| is.apcluster | B-04 |

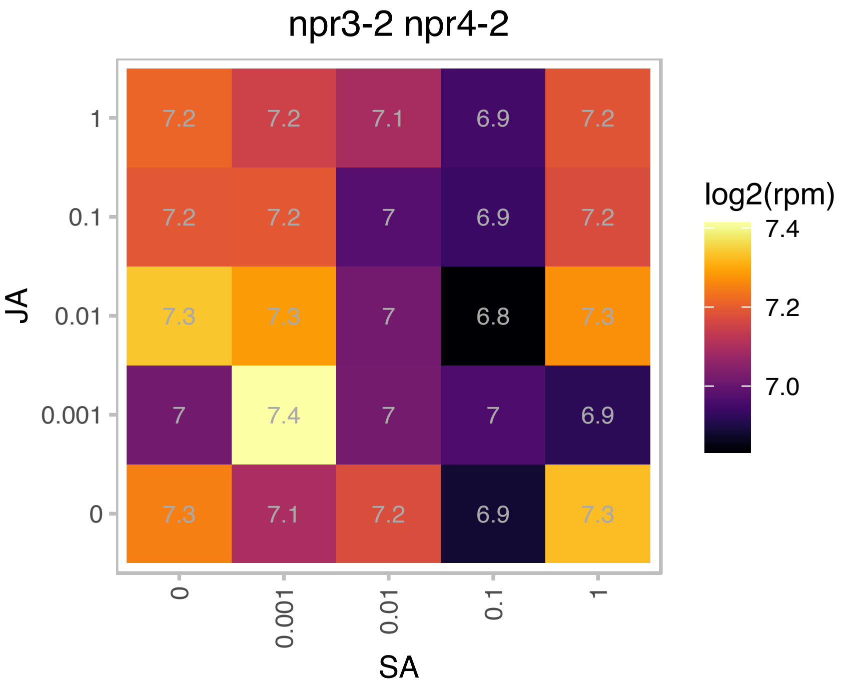


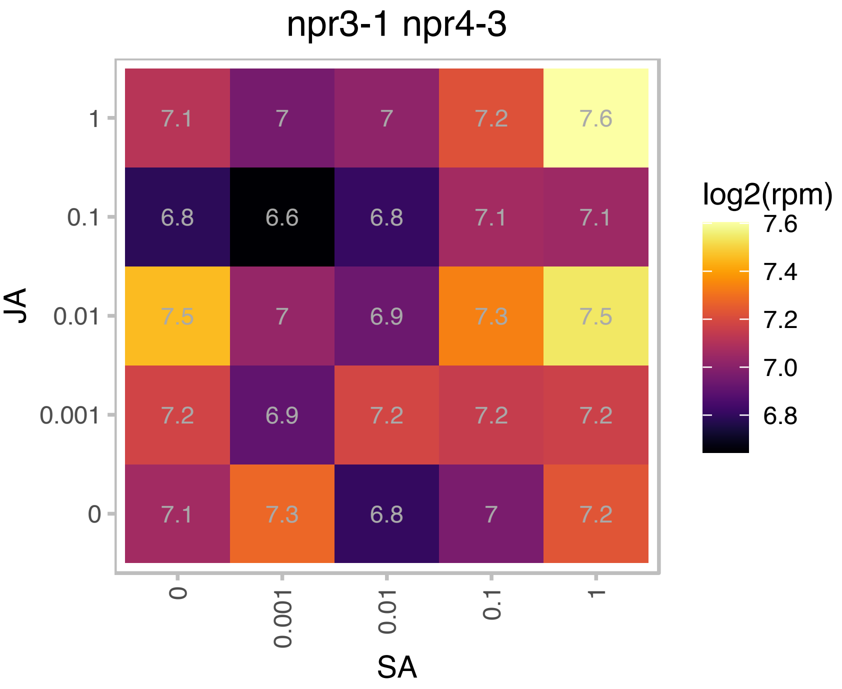

# AT3G12350.1 : NA

F-box family protein

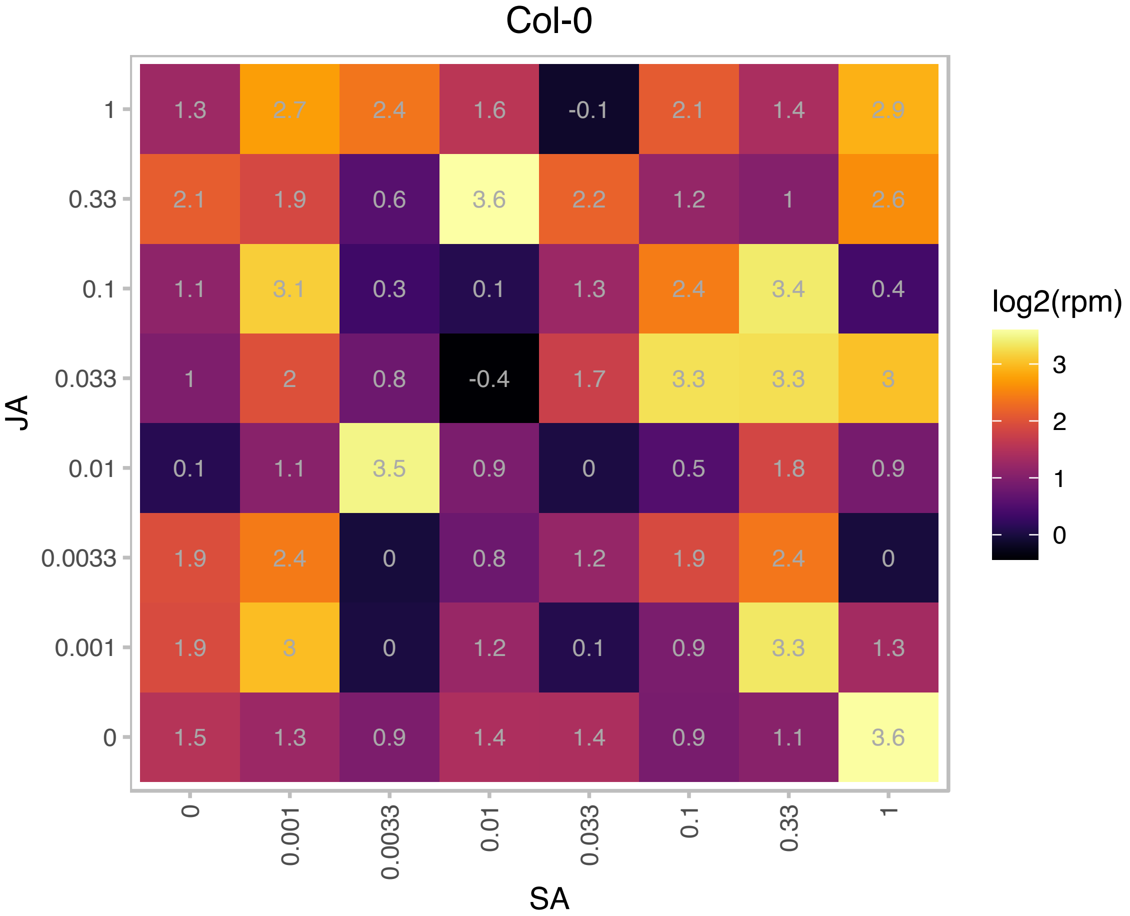


| | |
|---|---|
| Gene | AT3G12350 |
| is.expressed_genes | TRUE |
| is.responsive_genes | TRUE |
| is.apcluster | B-04 |

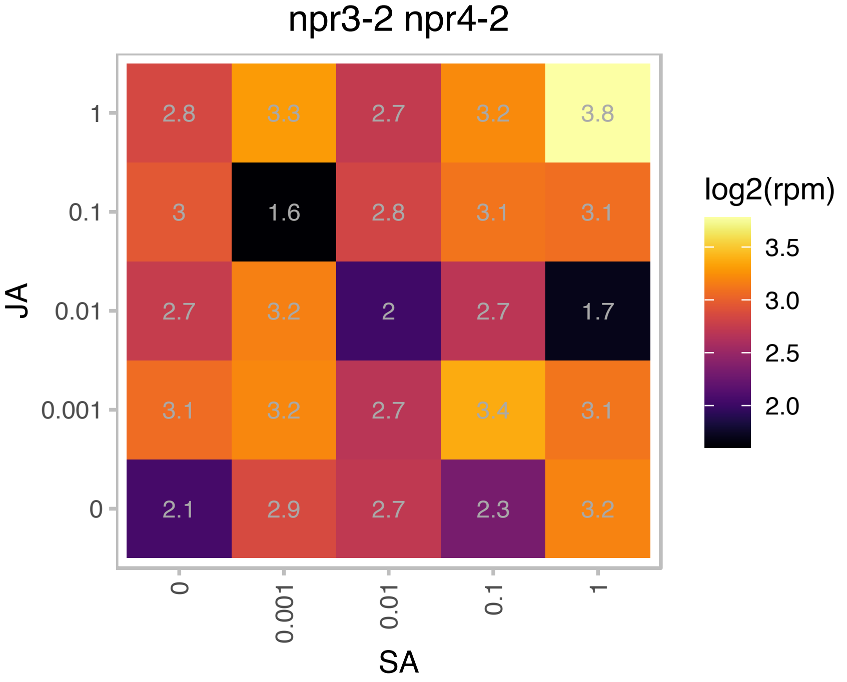


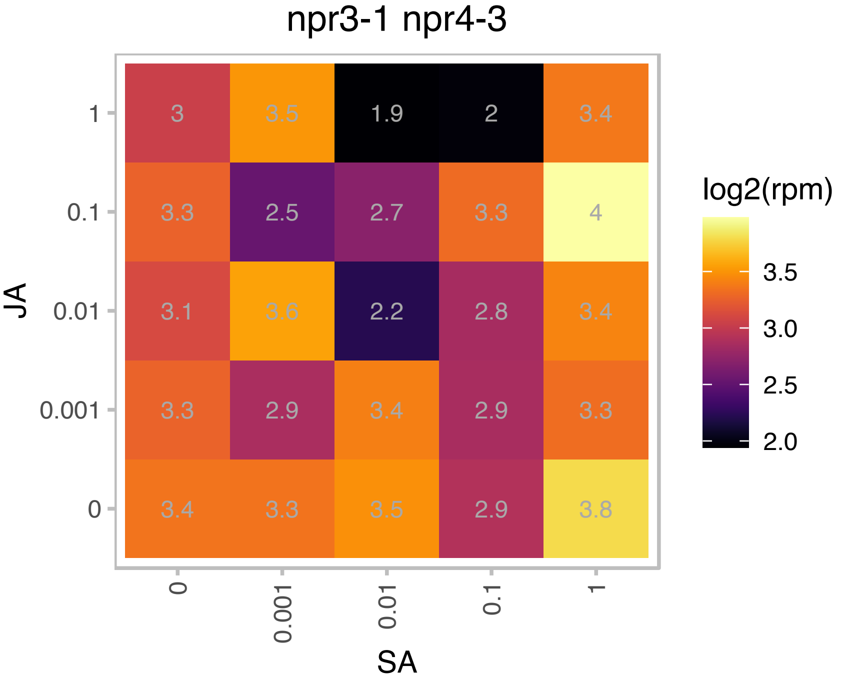

# AT3G14780.1 : NA

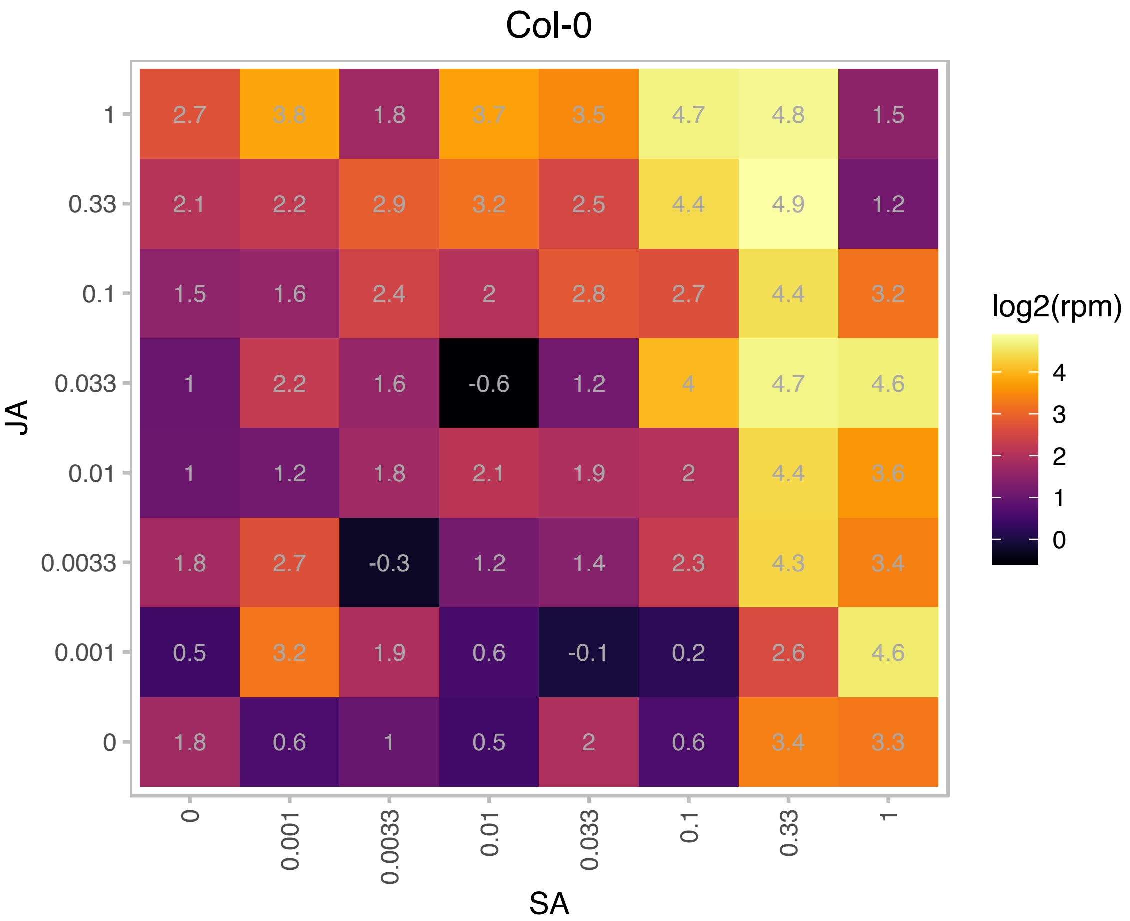


| | |
|---|---|
| Gene | AT3G14780 |
| is.expressed_genes | TRUE |
| is.responsive_genes | TRUE |
| is.apcluster | B-04 |

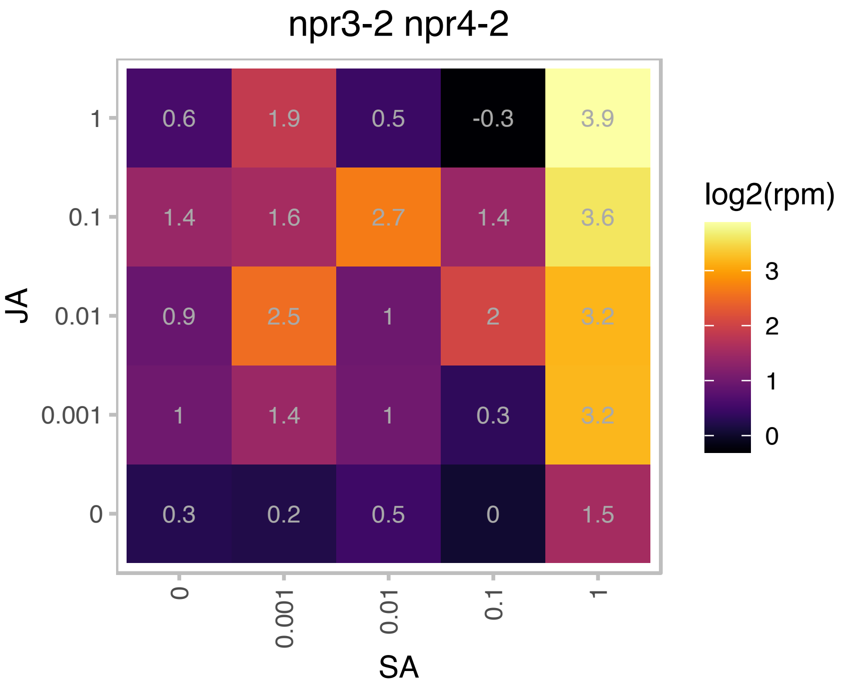


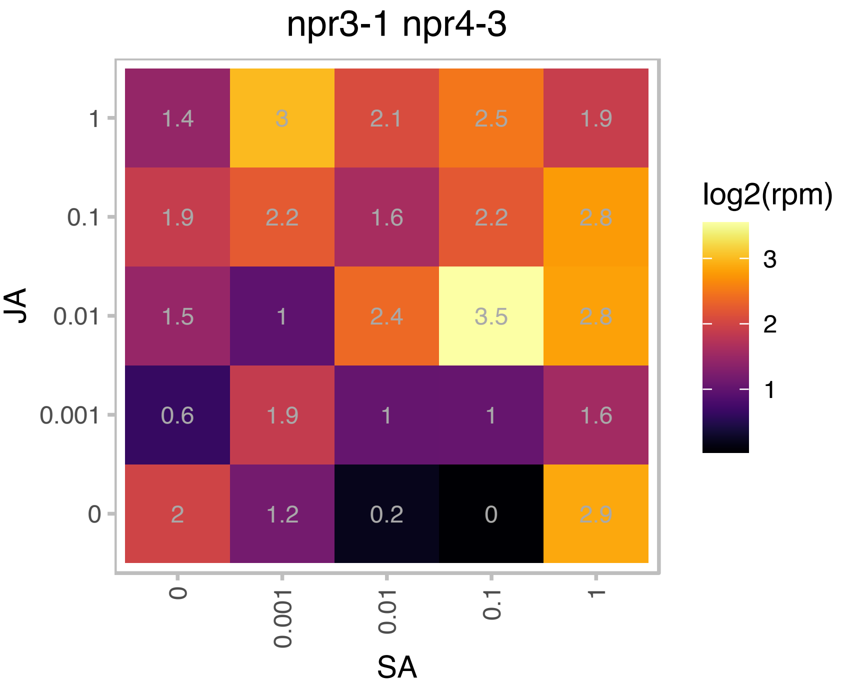

# AT3G16330.1 : NA

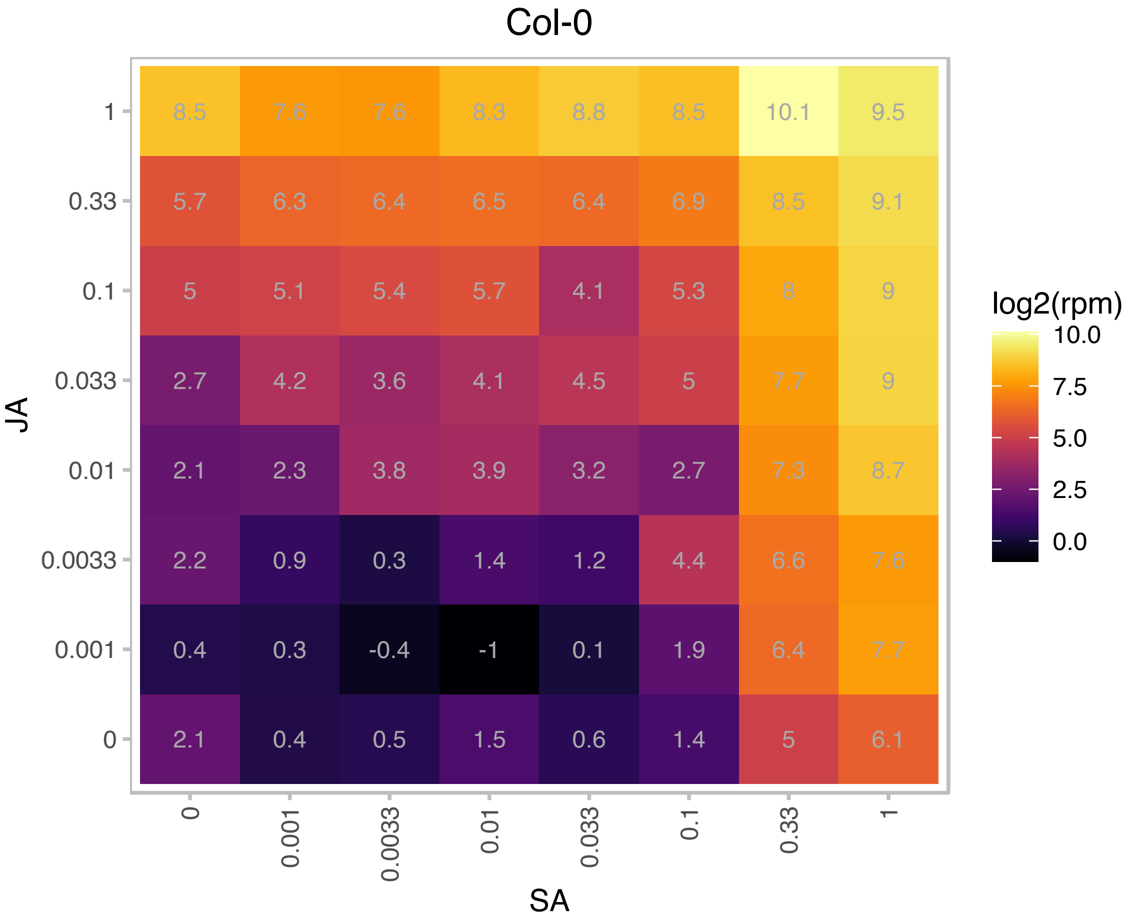


| | |
|---|---|
| Gene | AT3G16330 |
| is.expressed_genes | TRUE |
| is.responsive_genes | TRUE |
| is.apcluster | B-04 |

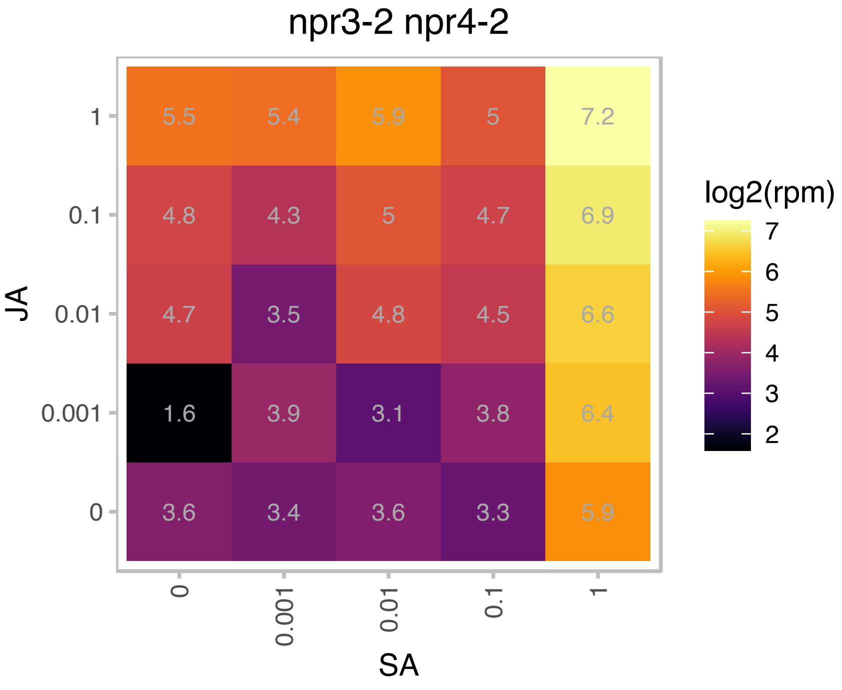


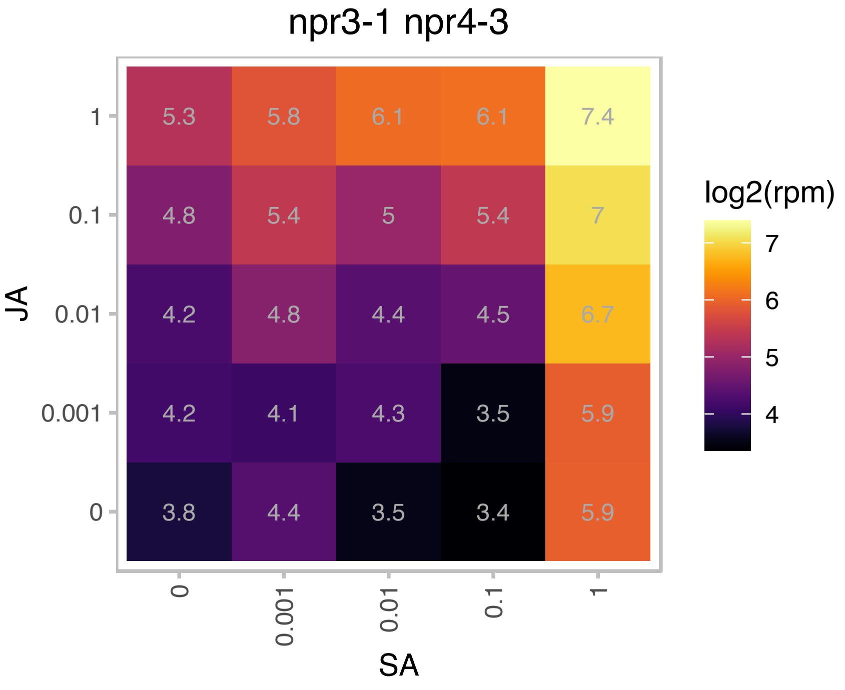

### AT3G16800.1 : EGR3

Protein phosphatase 2C family protein
EGR3 functions as a negative regulator of plant growth with prominent effect on plant growth during drought stress, EGR3 regulates microtubule organization and likely affects additional cytoskeleton and trafficking processes along the plasma membrane.

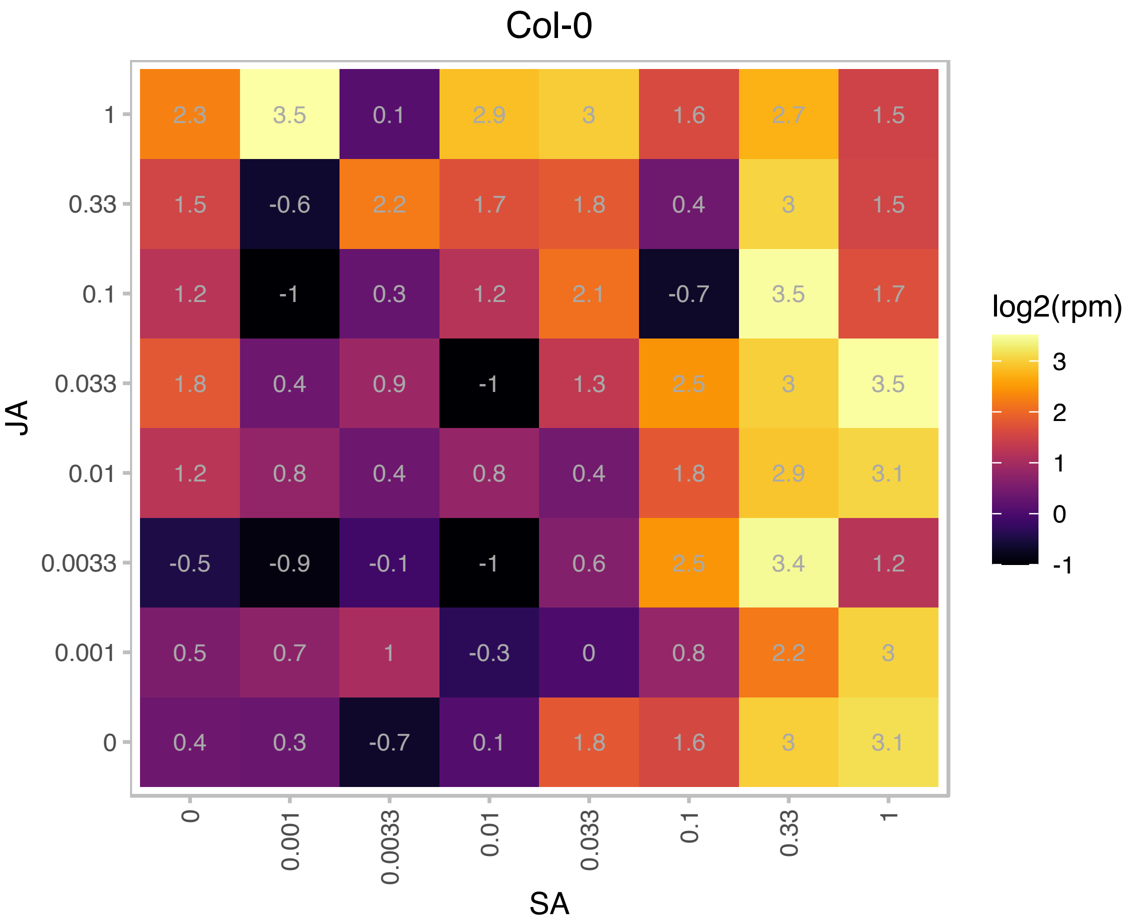


| | |
|---|---|
| Gene | AT3G16800 |
| is.expressed_genes | TRUE |
| is.responsive_genes | TRUE |
| is.apcluster | B-04 |

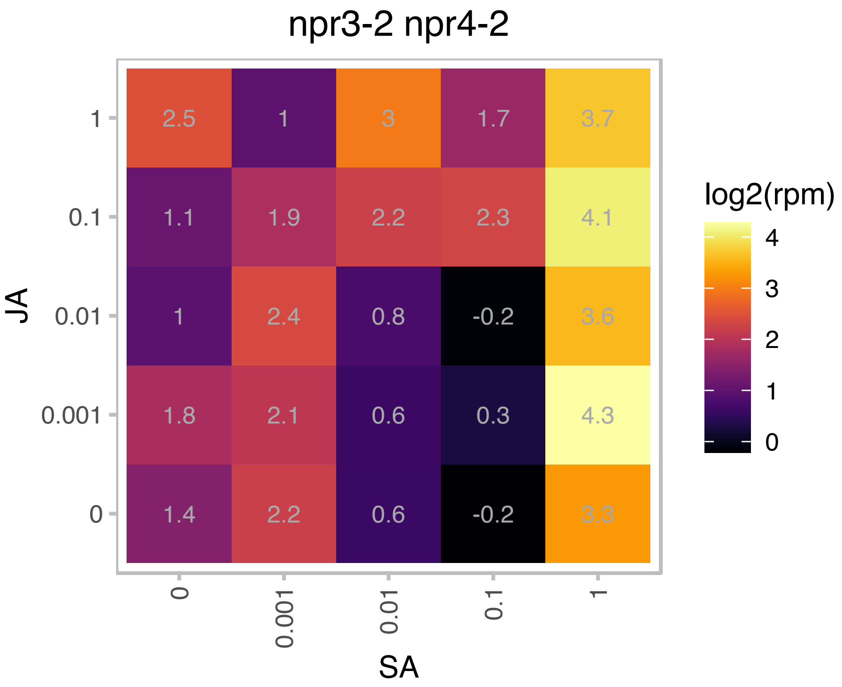


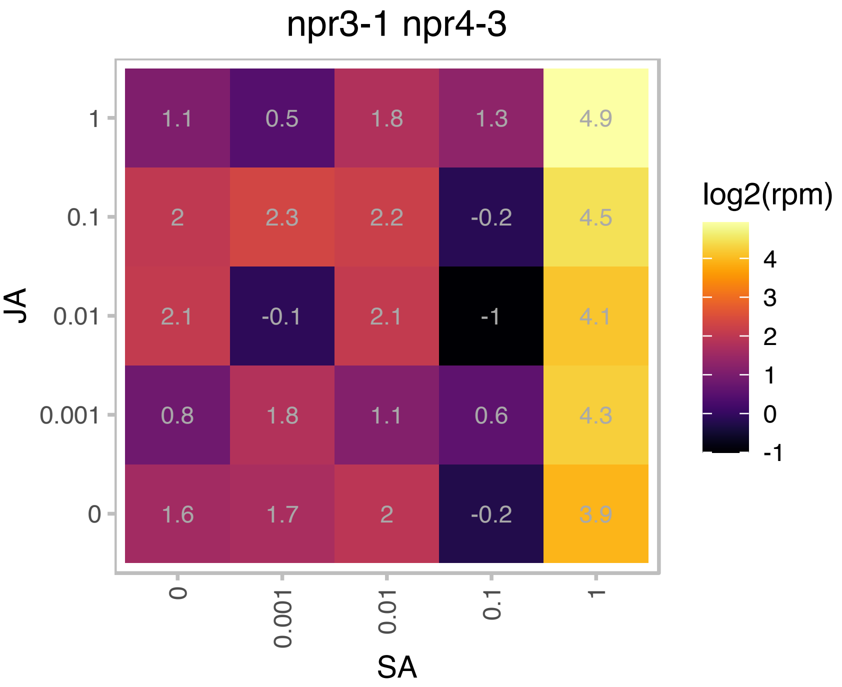

# AT3G19910.1 : NA

RING/U-box superfamily protein

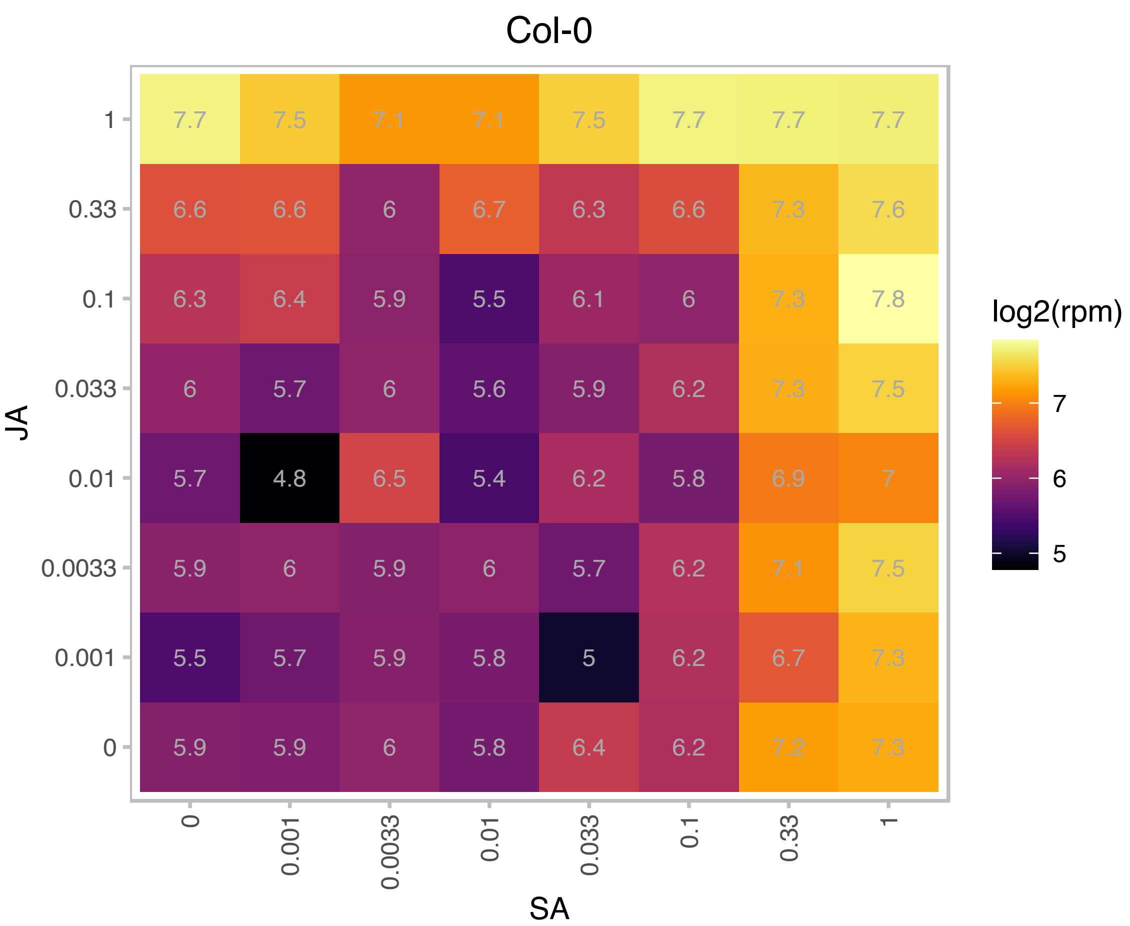


| | |
|---|---|
| Gene | AT3G19910 |
| is.expressed_genes | TRUE |
| is.responsive_genes | TRUE |
| is.apcluster | B-04 |

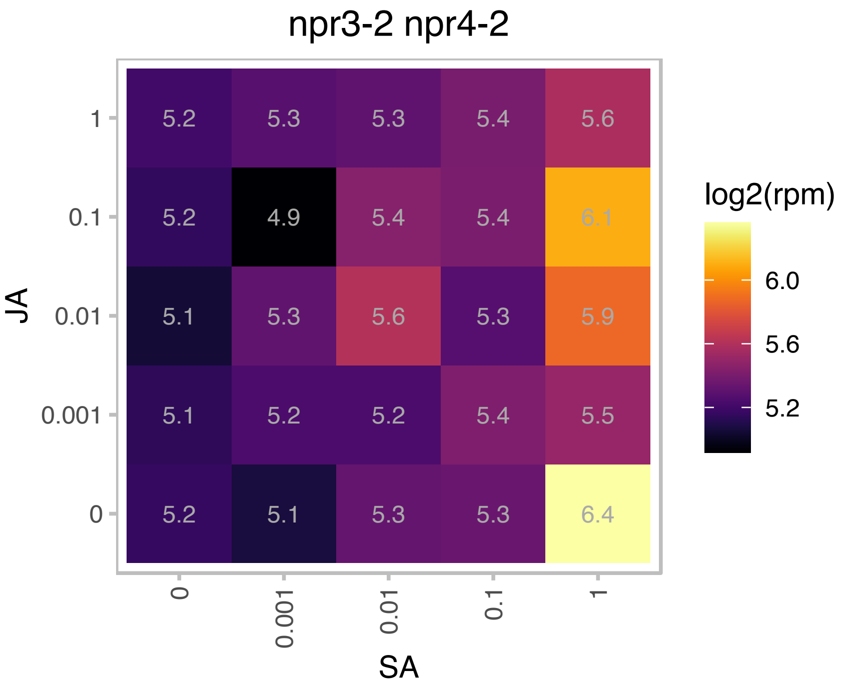


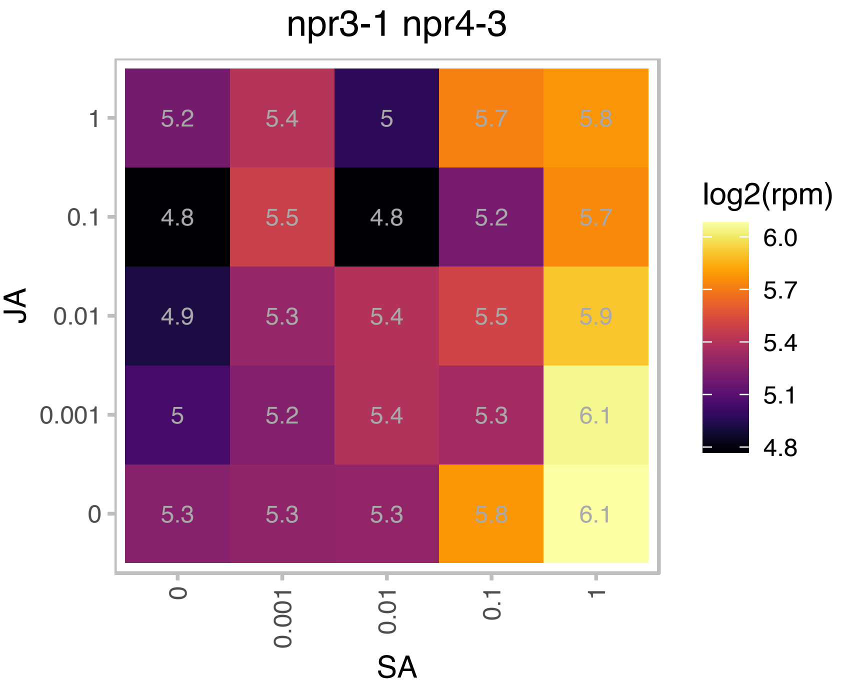

# AT3G22845.1 : NA

emp24/gp25L/p24 family/GOLD family protein

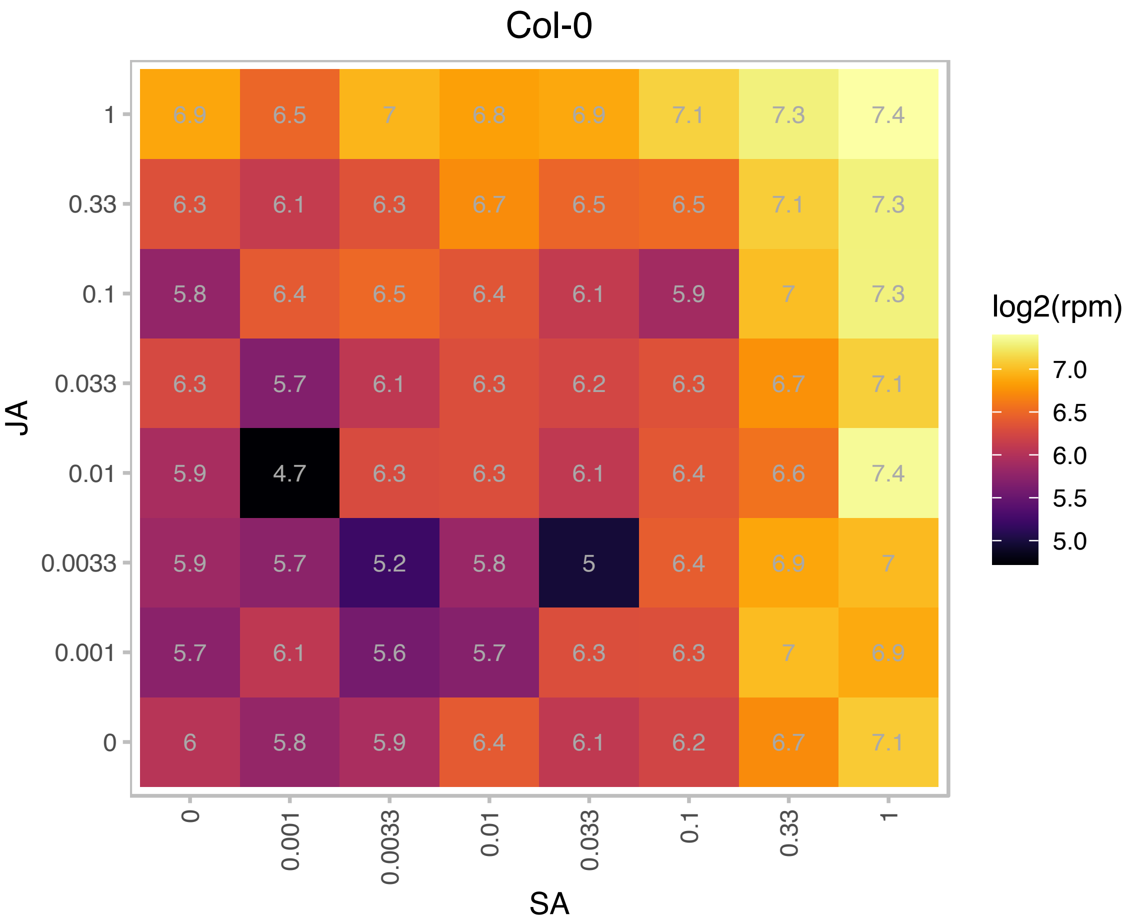


| | |
|---|---|
| Gene | AT3G22845 |
| is.expressed_genes | TRUE |
| is.responsive_genes | TRUE |
| is.apcluster | B-04 |

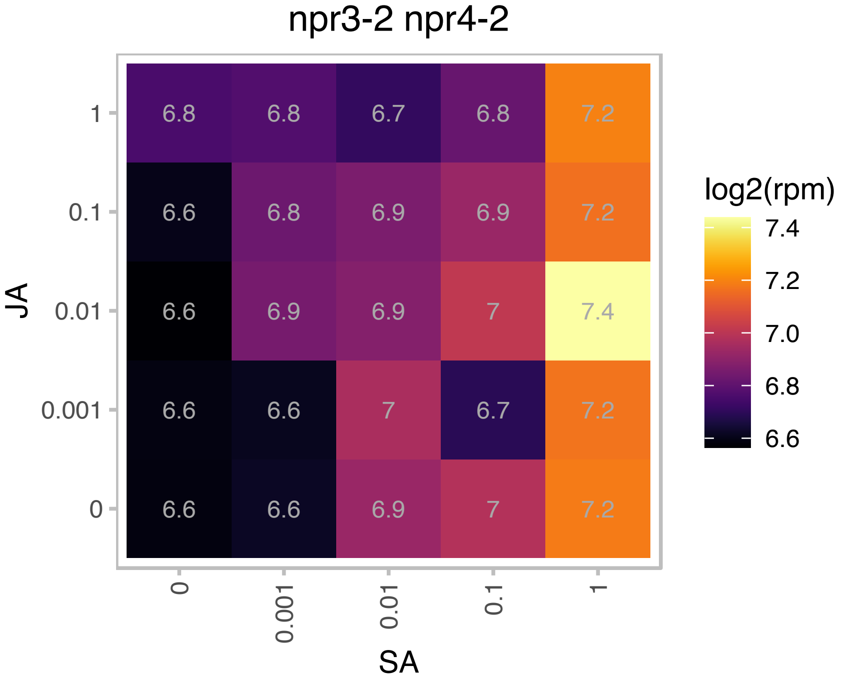


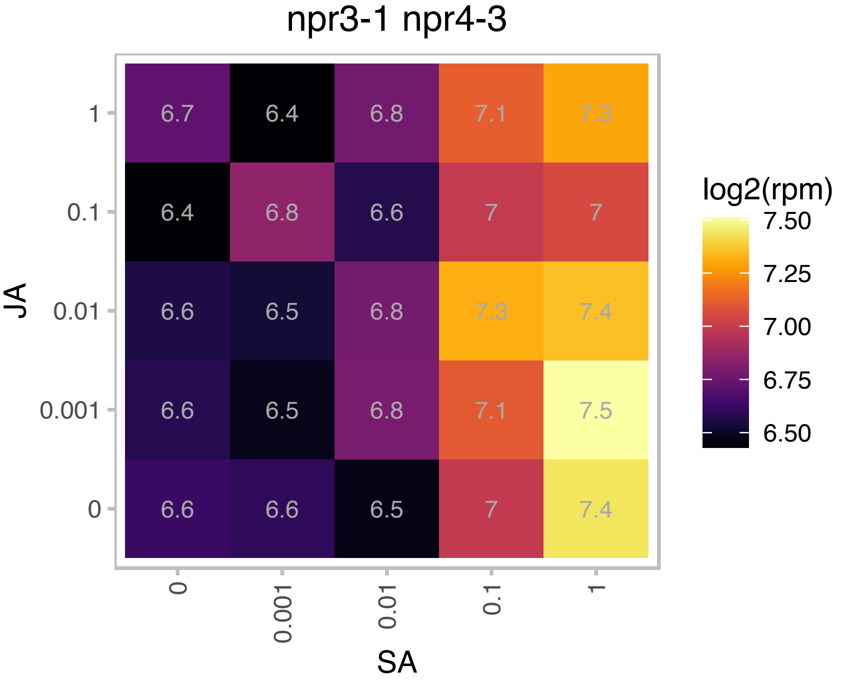

# AT3G27880.1 : NA

Protein of unknown function (DUF1645)

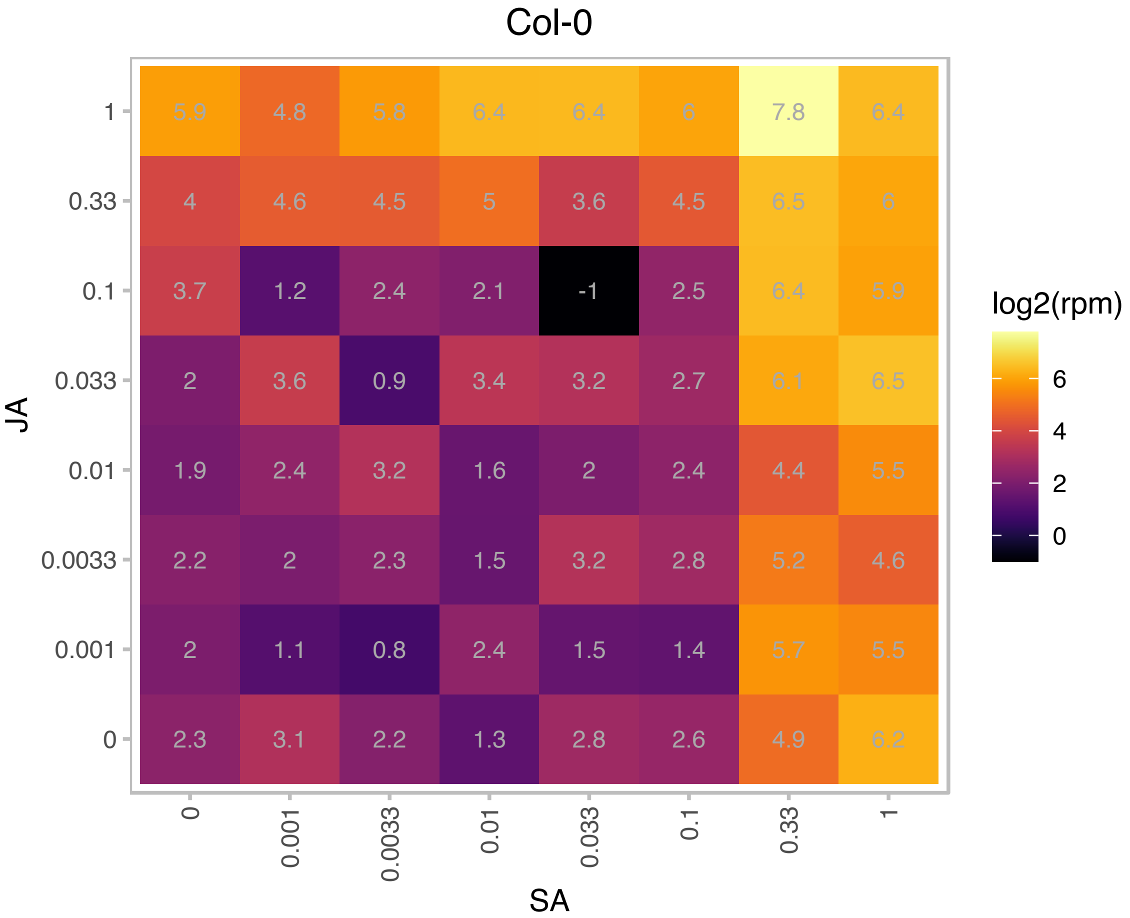


| | |
|---|---|
| Gene | AT3G27880 |
| is.expressed_genes | TRUE |
| is.responsive_genes | TRUE |
| is.apcluster | B-04 |

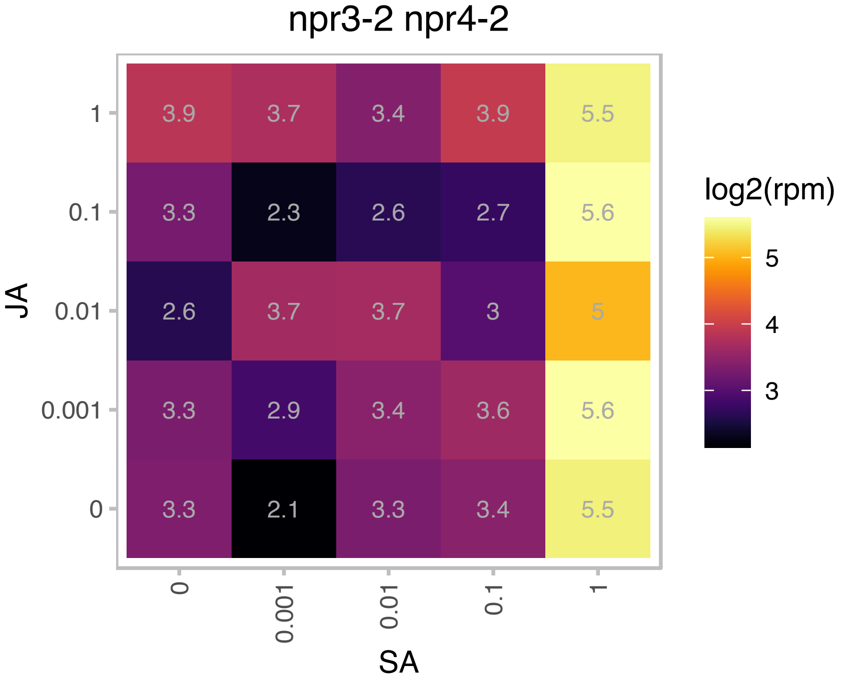


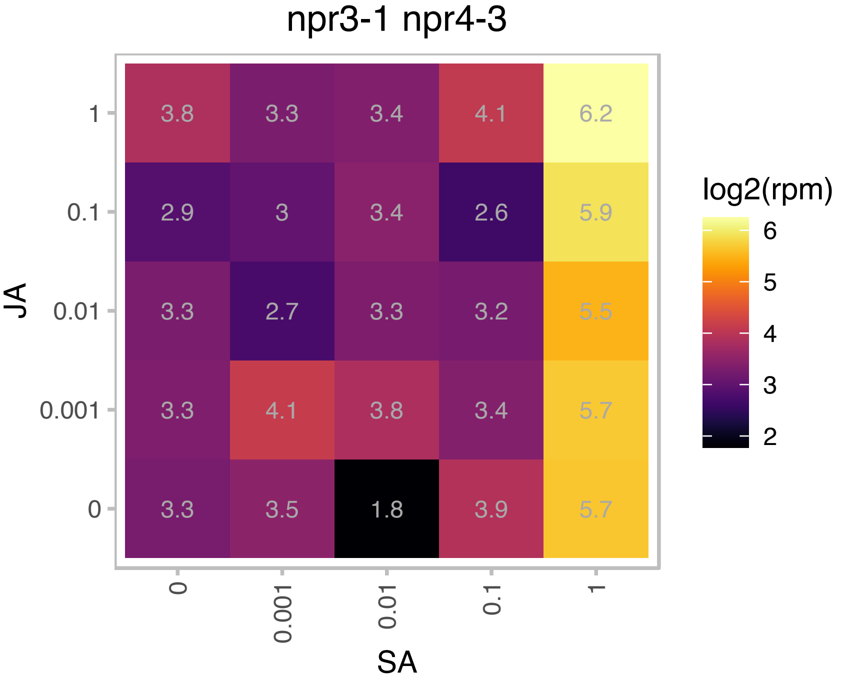

### AT3G28210.1 : PMZ, SAP12

zinc finger (AN1-like) family protein

Encodes a putative zinc finger protein (PMZ).

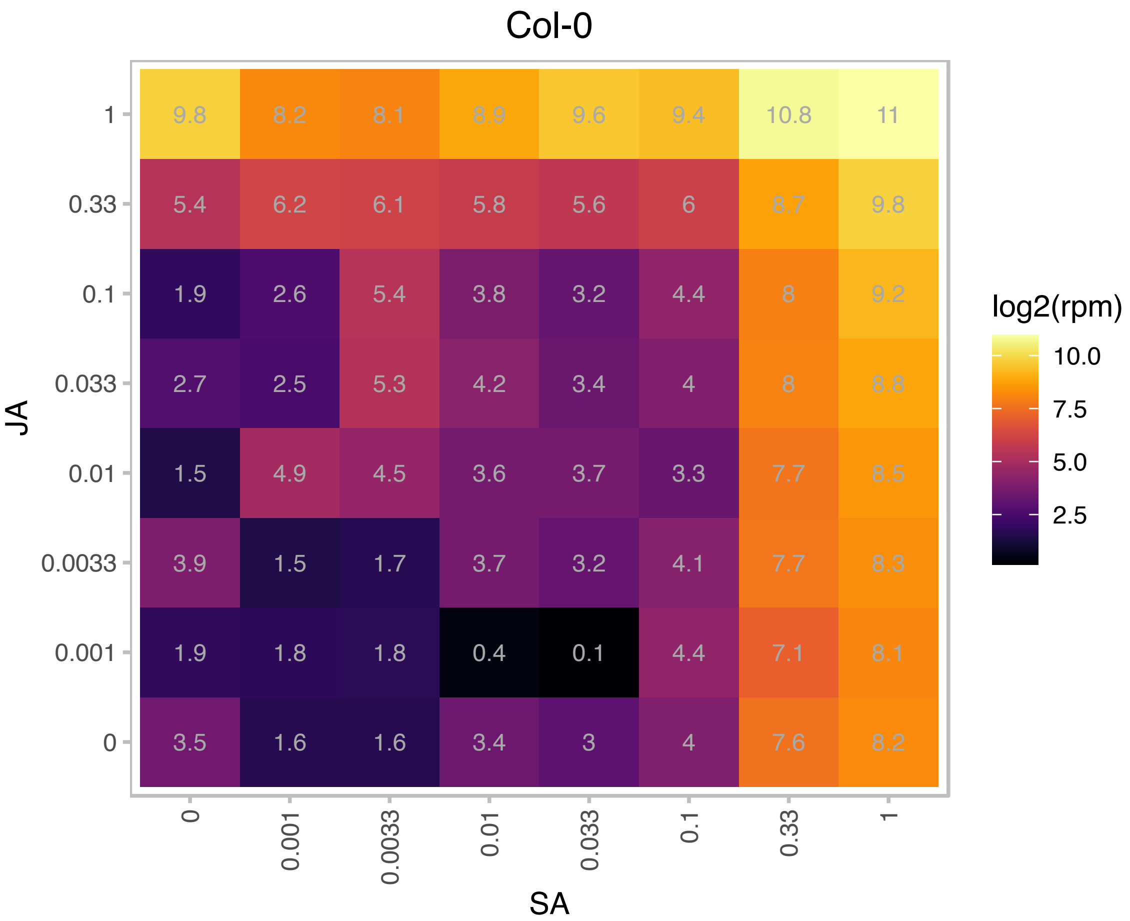


| | |
|---|---|
| Gene | AT3G28210 |
| is.expressed_genes | TRUE |
| is.responsive_genes | TRUE |
| is.apcluster | B-04 |

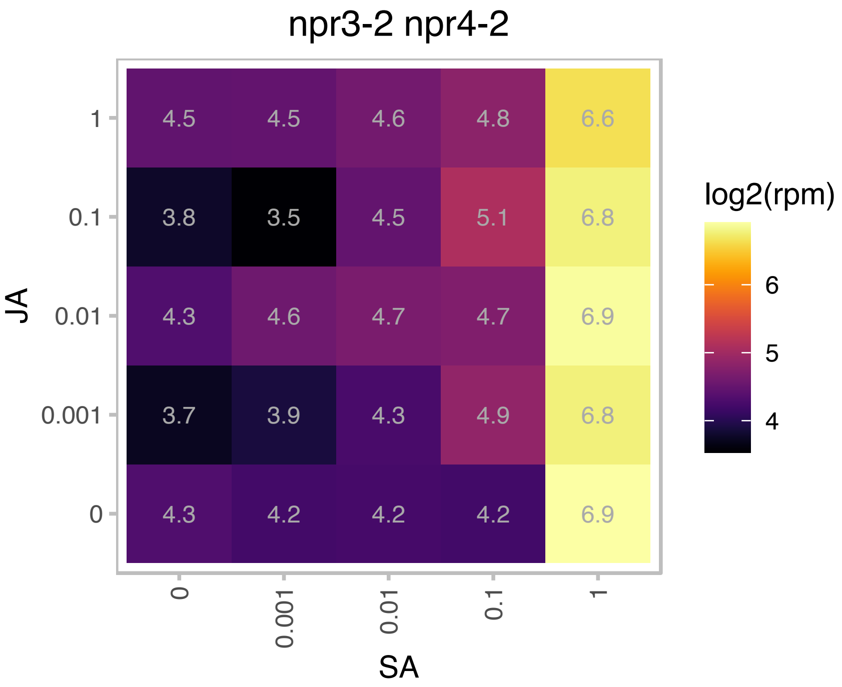


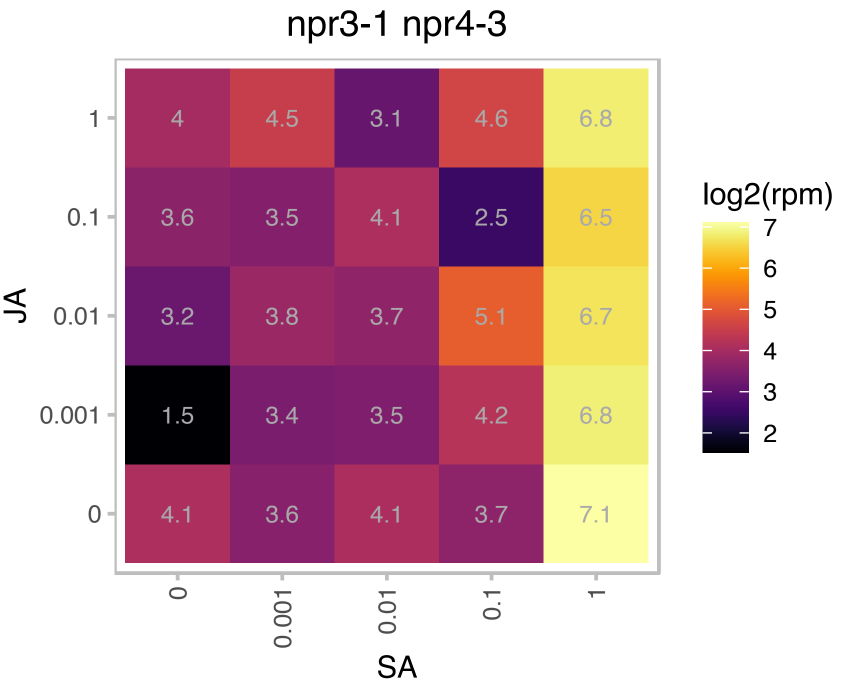

### AT3G44300.1 : AtNIT2, NIT2

nitrilase 2

Encodes an enzyme that catalyzes the hydrolysis of indole-3-acetonitrile (IAN) to indole-3-acetic acid (IAA) (nitrile aminohydrolase, EC 3.5.5.1) and IAN to indole-3-acetamide (IAM) at lower levels. Mutants have reduced sensitivity to IAN and are sensitive to IAA. This enzyme likely participates in other non-auxin-related metabolic pathways. The mRNA is cell-to-cell mobile.

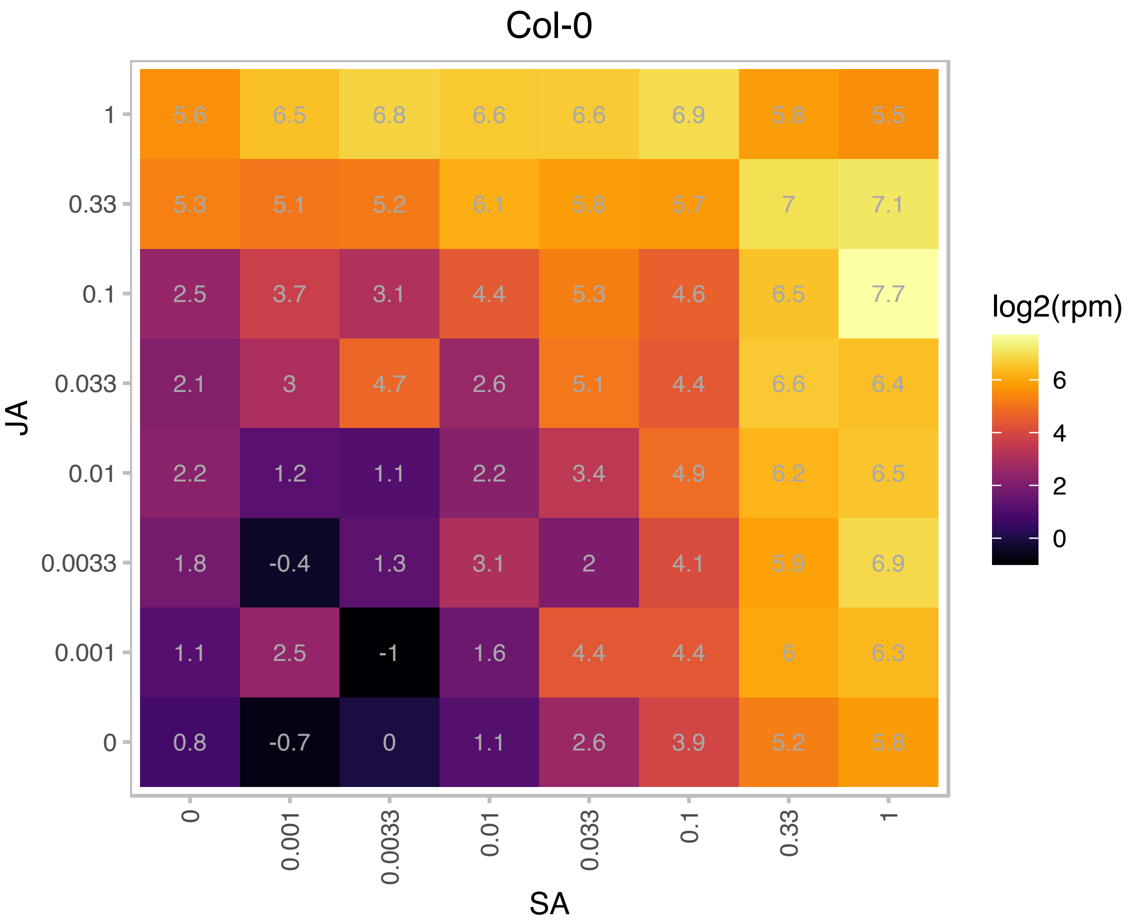


| | |
|---|---|
| Gene | AT3G44300 |
| is.expressed_genes | TRUE |
| is.responsive_genes | TRUE |
| is.apcluster | B-04 |

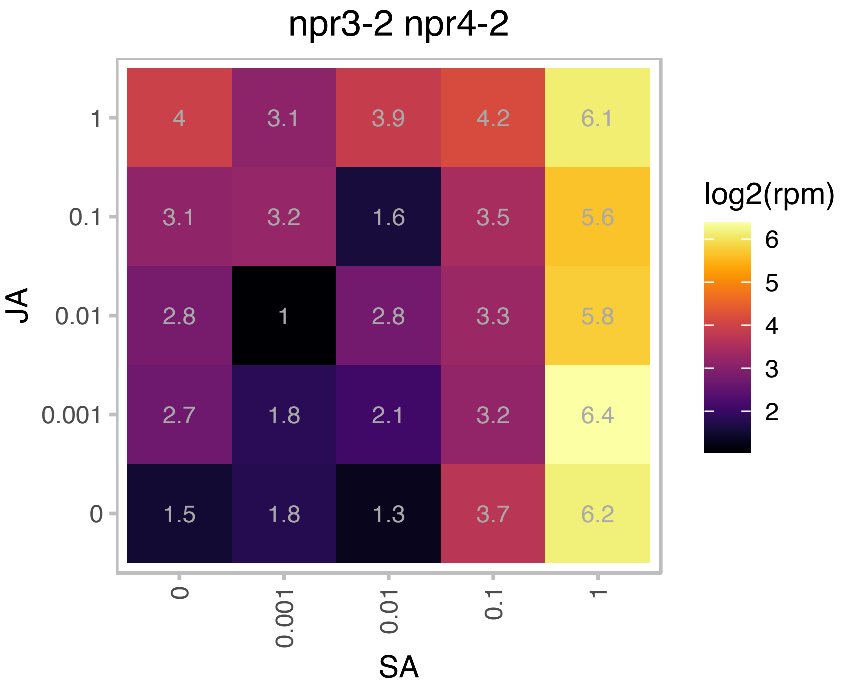


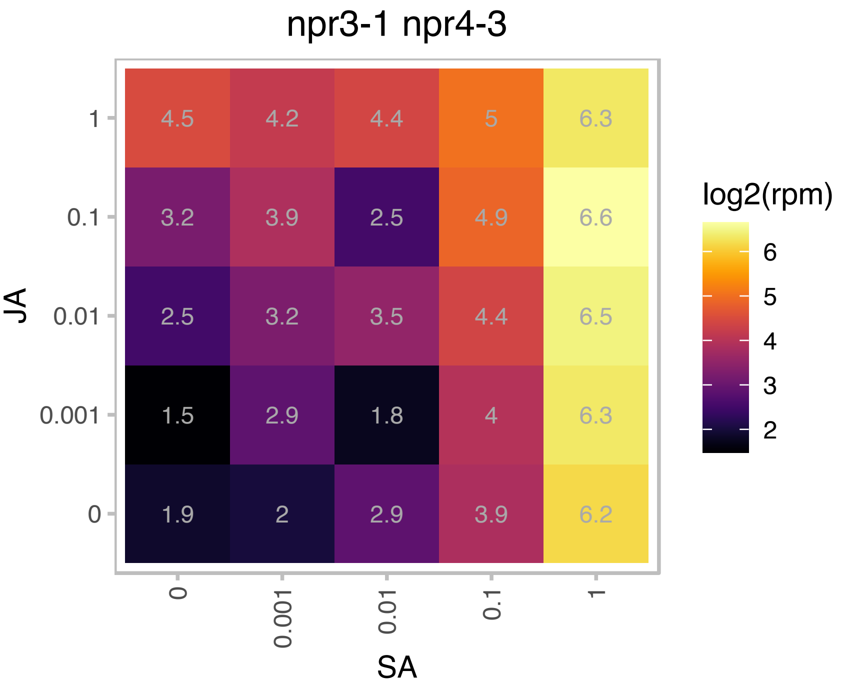

# AT3G46080.1 : NA

C2H2-type zinc finger family protein

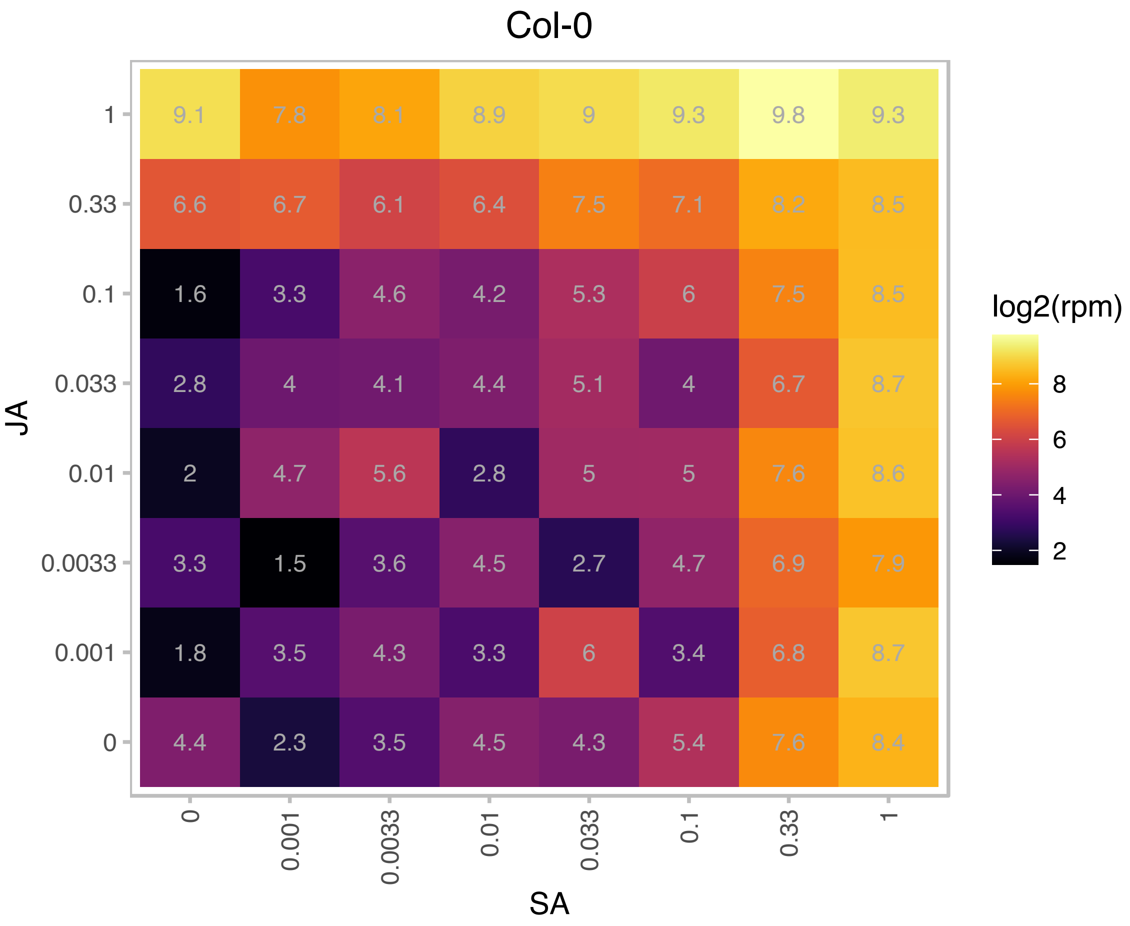


| | |
|---|---|
| Gene | AT3G46080 |
| is.expressed_genes | TRUE |
| is.responsive_genes | TRUE |
| is.apcluster | B-04 |

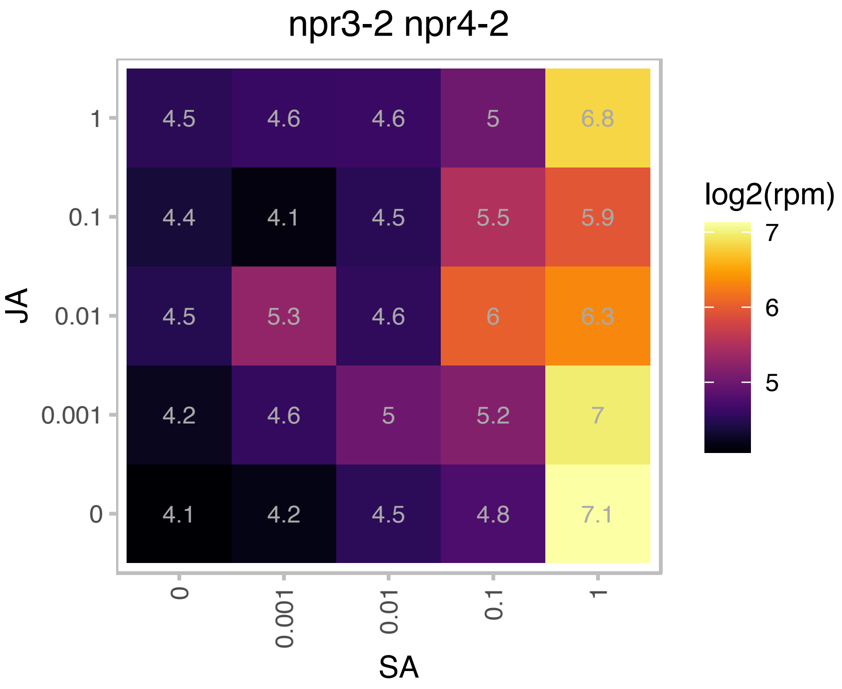


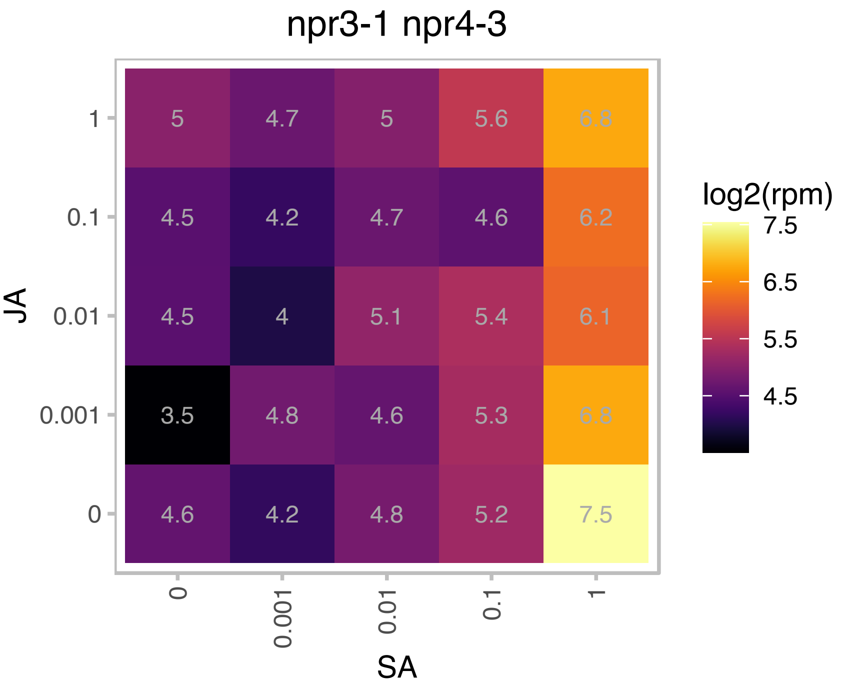

# AT3G46668.1 : NA

other RNA

Unknown gene

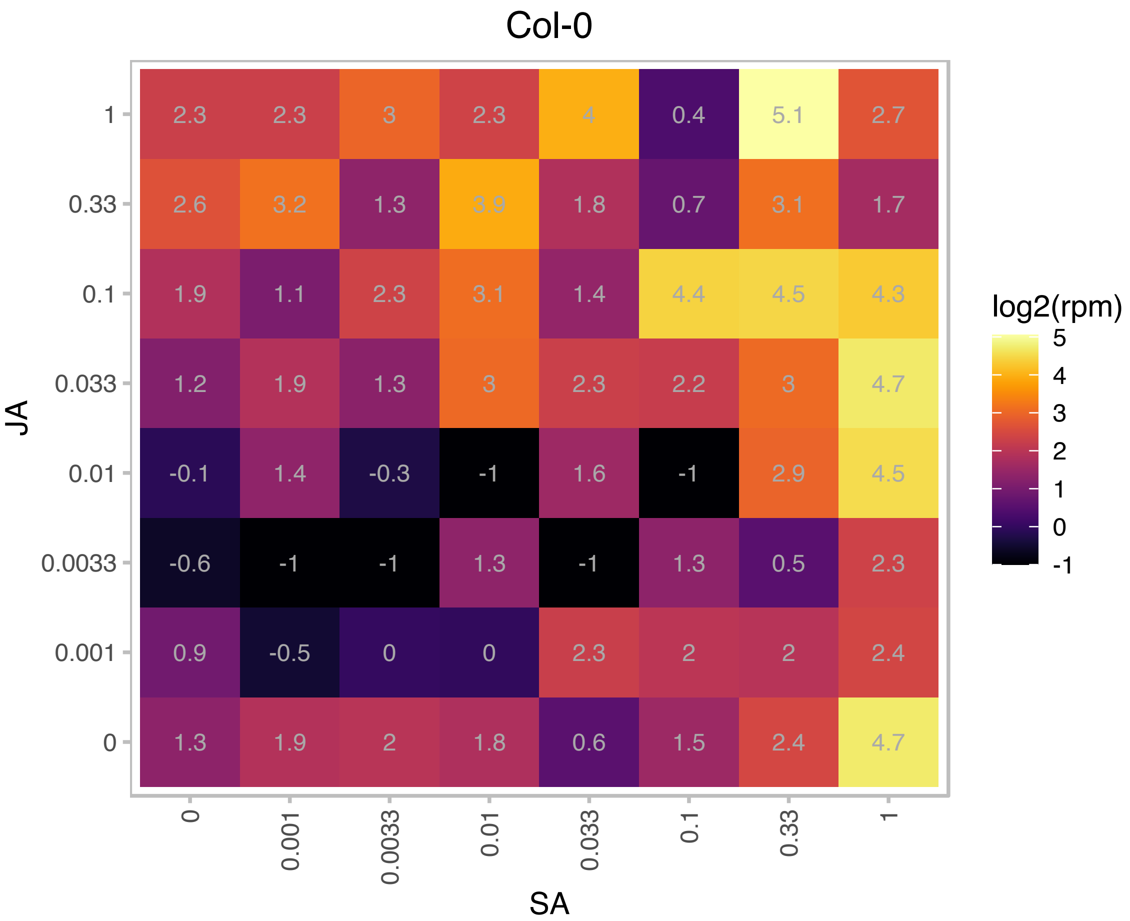


| Gene | AT3G46668 |
|---|---|
| is.expressed_genes | TRUE |
| is.responsive_genes | TRUE |
| is.apcluster | B-04 |

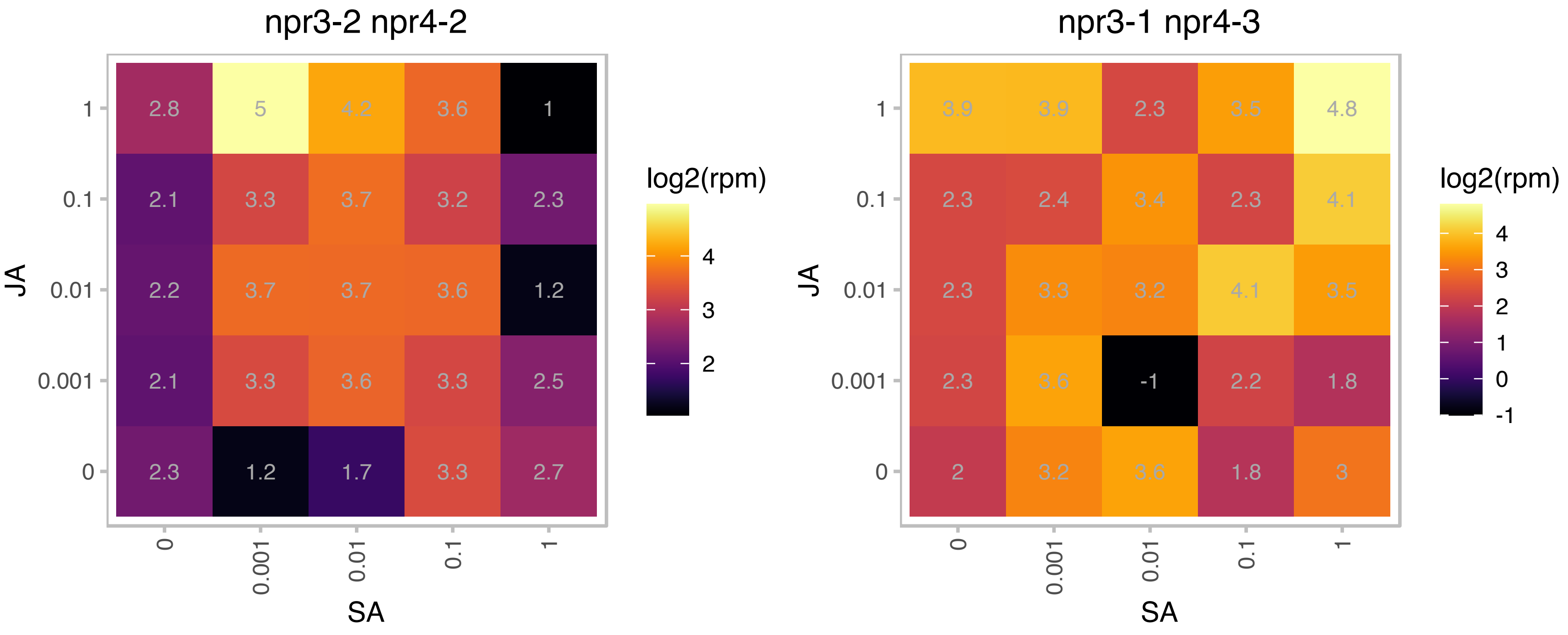

### AT3G47810.1 : ATVPS29, MAG1, VPS29

Calcineurin-like metallo-phosphoesterase superfamily protein

Homolog of yeast retromer subunit VPS29.  Part of a retromer-like protein complex involved in endosome to lysosome protein transport.

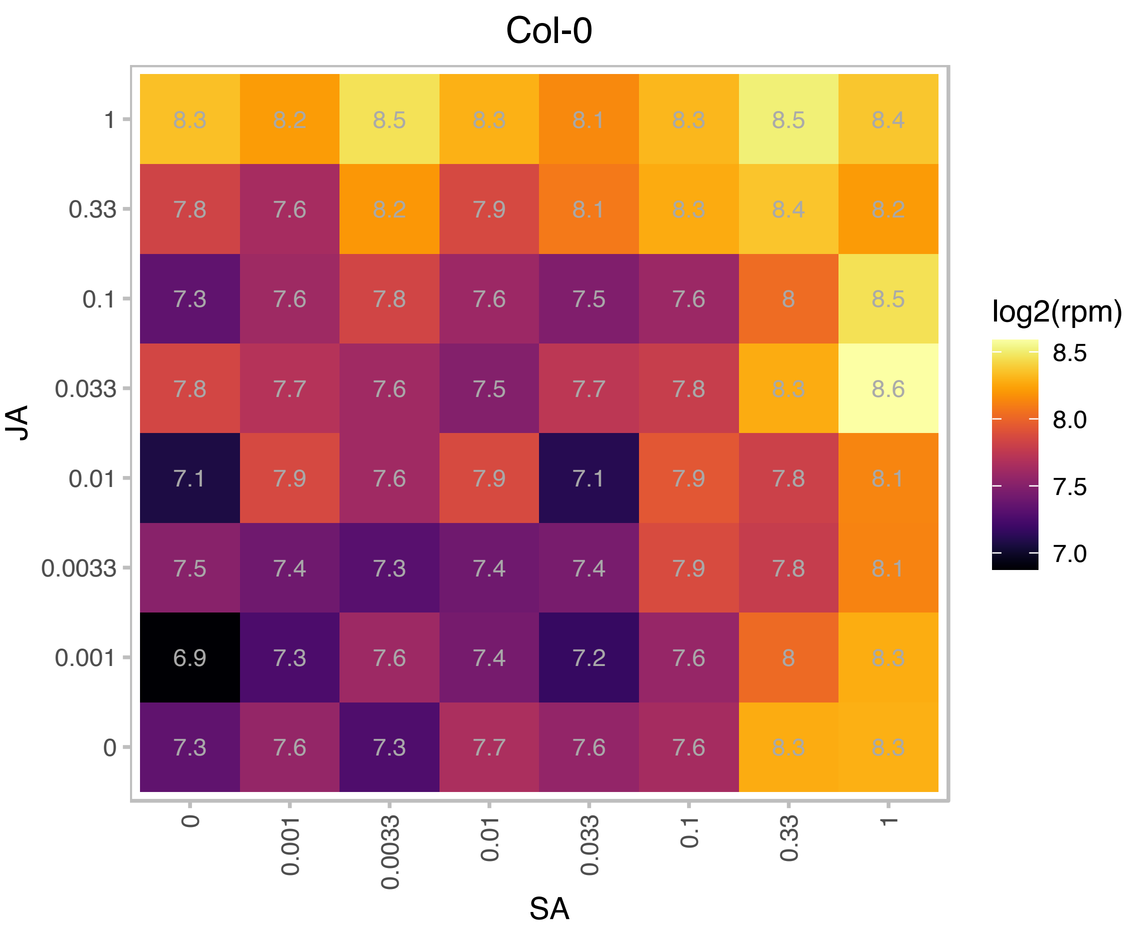


| | |
|---|---|
| Gene | AT3G47810 |
| is.expressed_genes | TRUE |
| is.responsive_genes | TRUE |
| is.apcluster | B-04 |

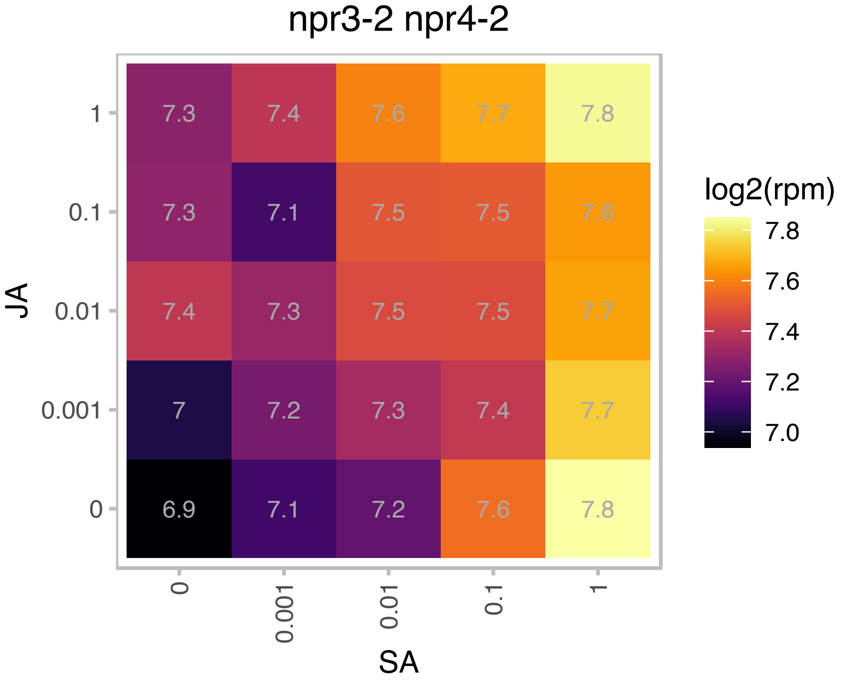


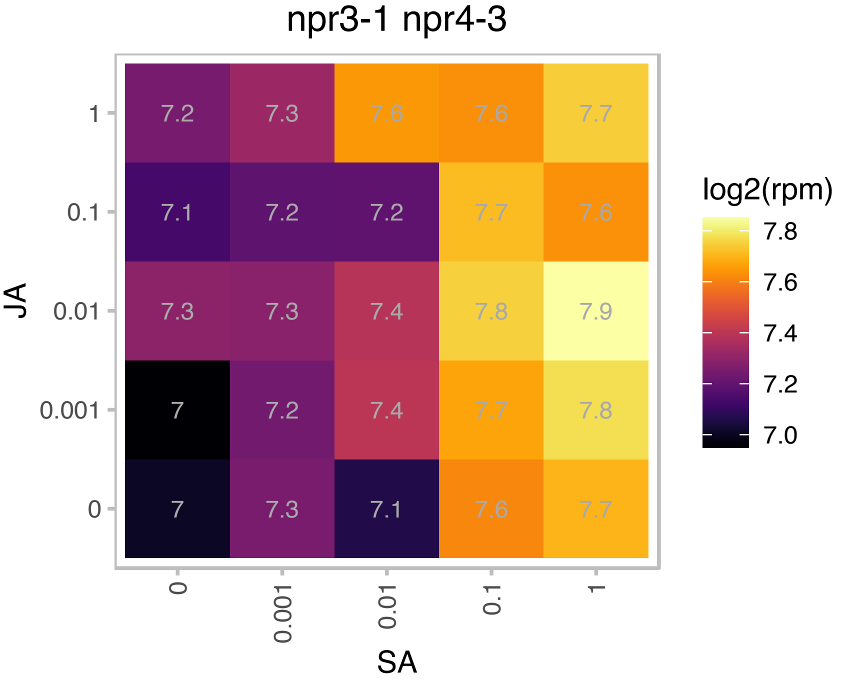

### AT3G50000.1 : ATCKA2, CKA2

casein kinase II, alpha chain 2

Encodes the casein kinase II (CK2) catalytic subunit (alpha).

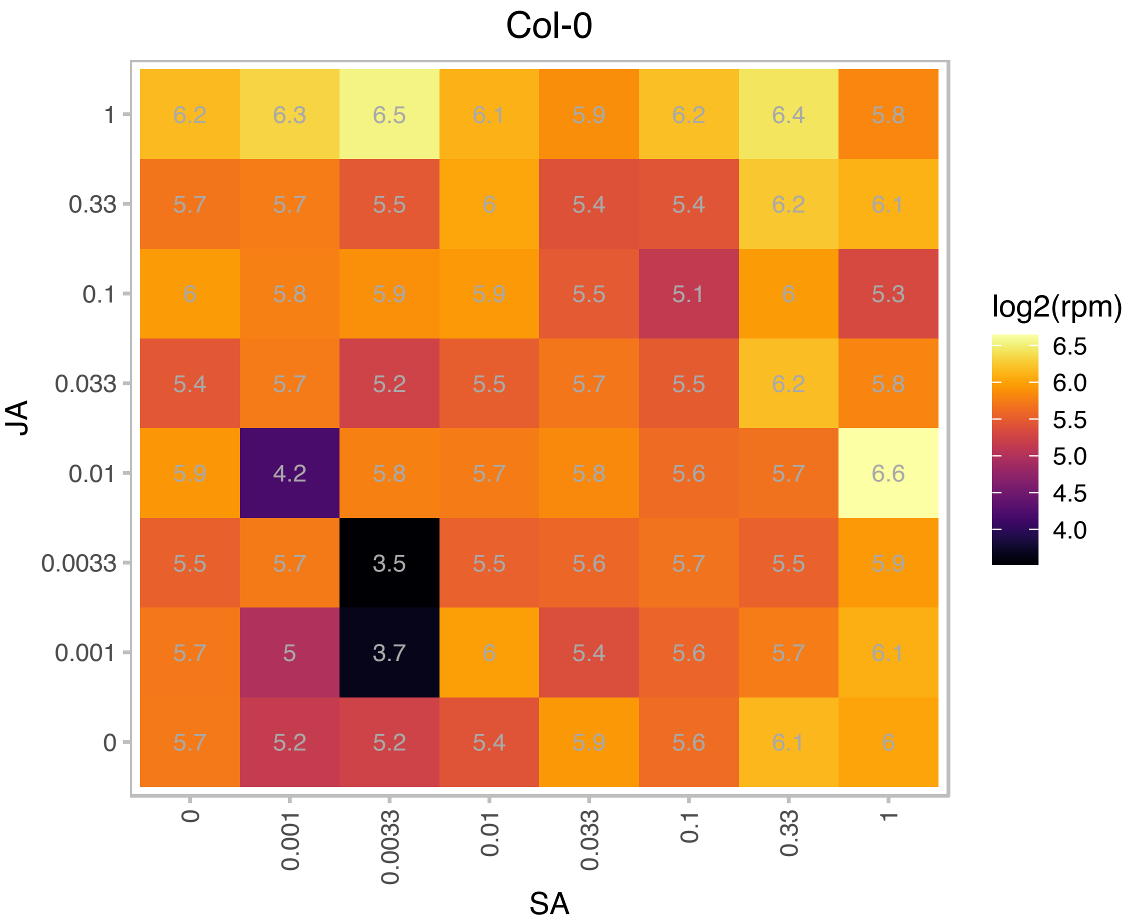


| | |
|---|---|
| Gene | AT3G50000 |
| is.expressed_genes | TRUE |
| is.responsive_genes | TRUE |
| is.apcluster | B-04 |

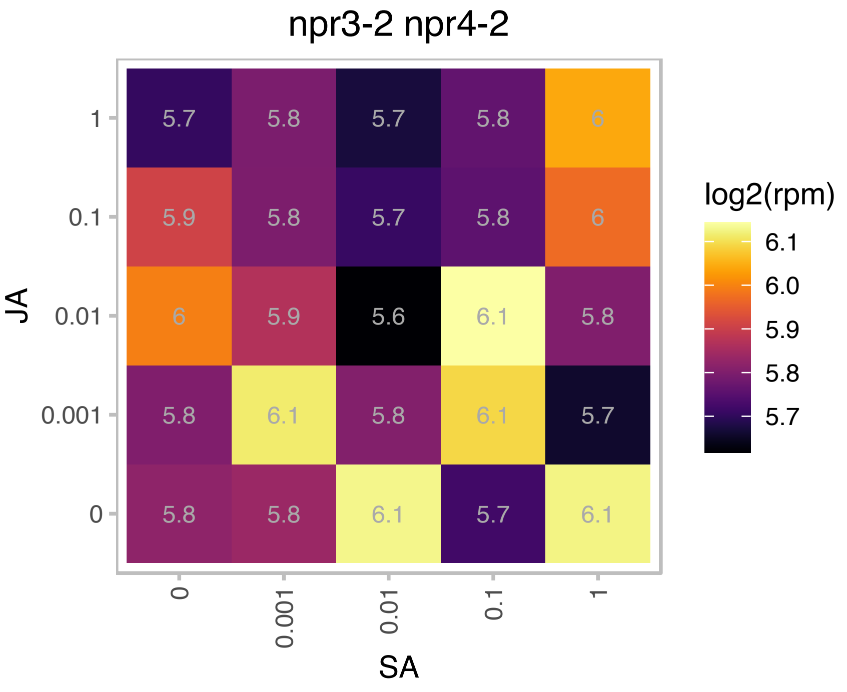


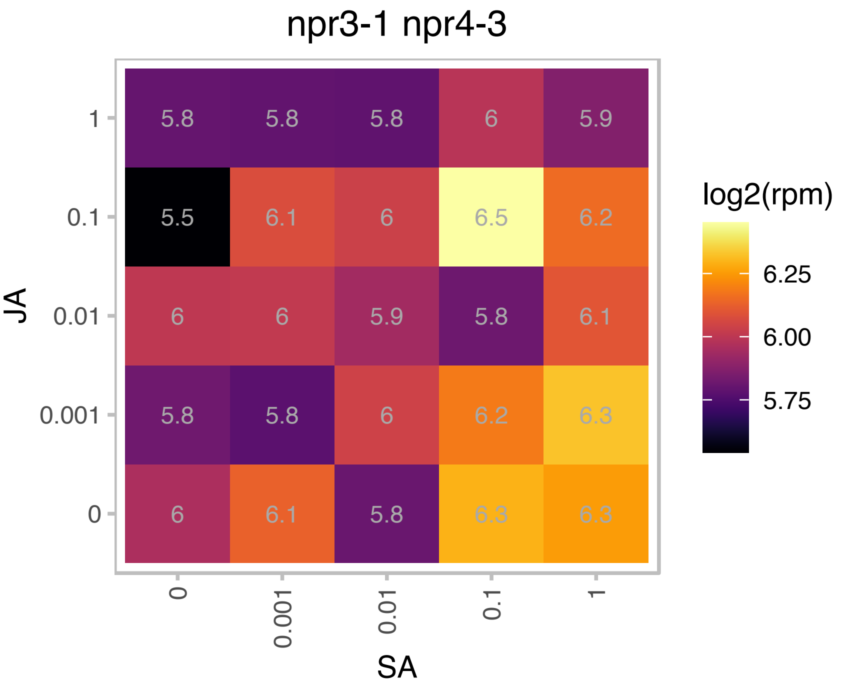

### AT3G50970.1 : LTI30, XERO2

dehydrin family protein

Belongs to the dehydrin protein family, which contains highly conserved stretches of 7-17 residues that are repetitively scattered in their sequences, the K-, S-, Y- and lysine rich segments.  LTI29 and LTI30 double overexpressors confer freeze tolerance. Located in membranes. mRNA upregulated by water deprivation and abscisic acid. The mRNA is cell-to-cell mobile.

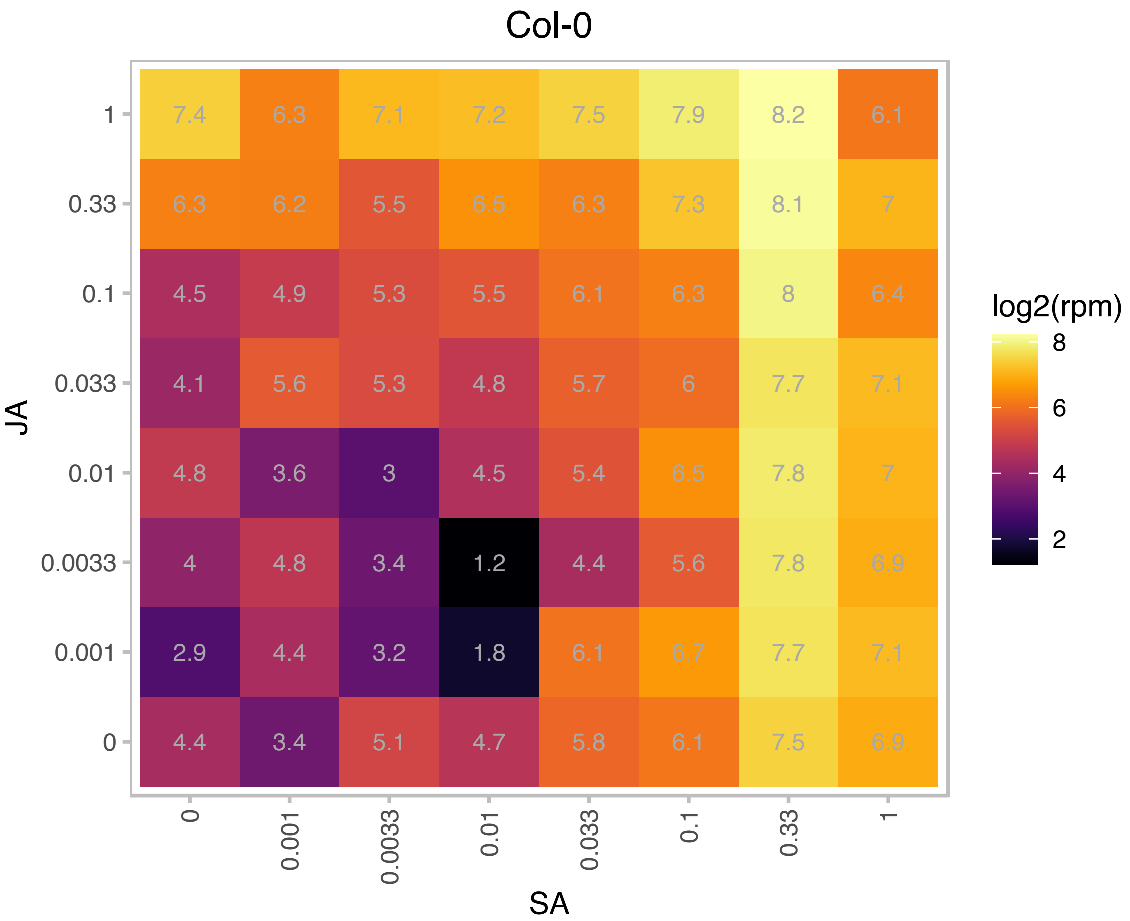


| | |
|---|---|
| Gene | AT3G50970 |
| is.expressed_genes | TRUE |
| is.responsive_genes | TRUE |
| is.apcluster | B-04 |

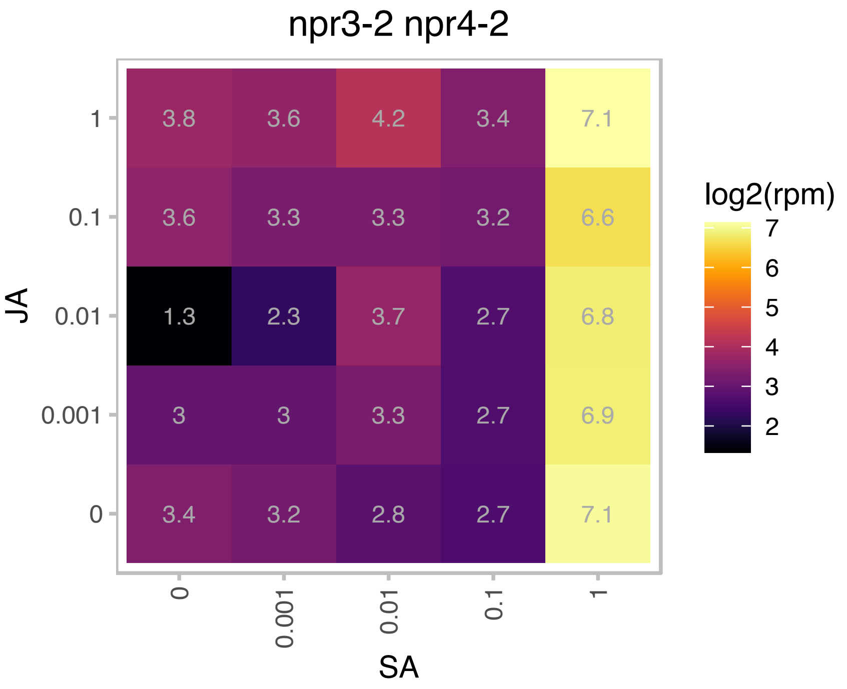


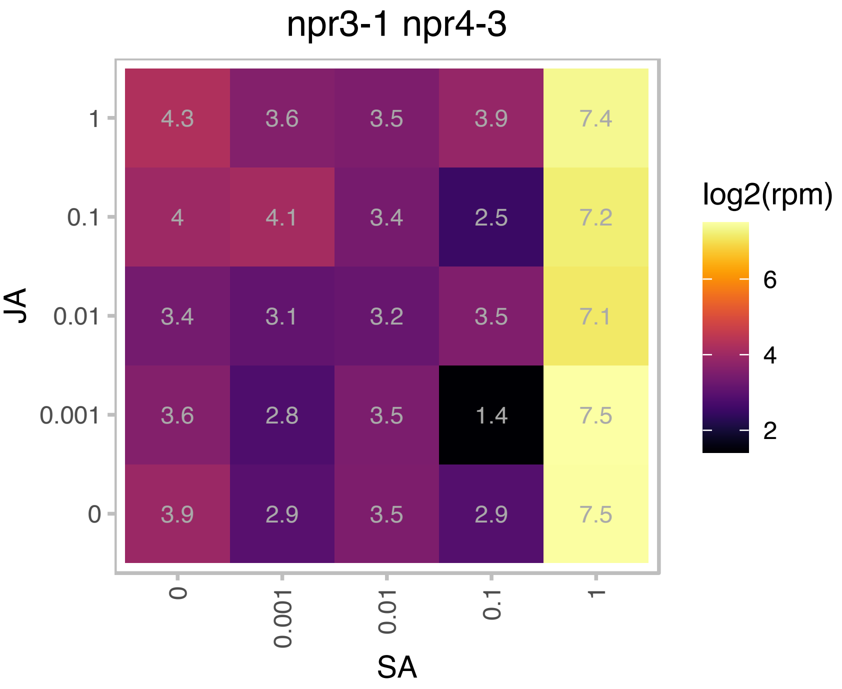

# AT3G57800.1 : NA

basic helix-loop-helix (bHLH) DNA-binding superfamily protein

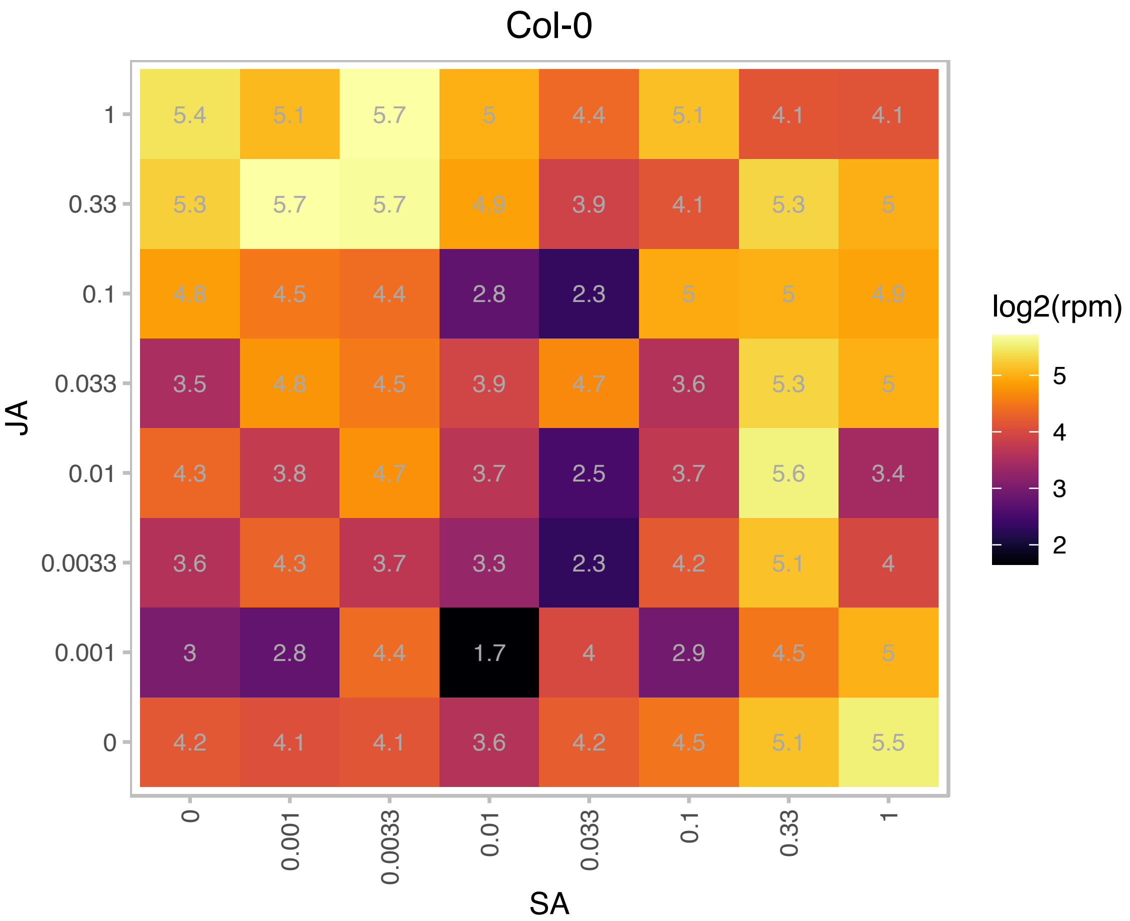


| | |
|---|---|
| Gene | AT3G57800 |
| is.expressed_genes | TRUE |
| is.responsive_genes | TRUE |
| is.apcluster | B-04 |

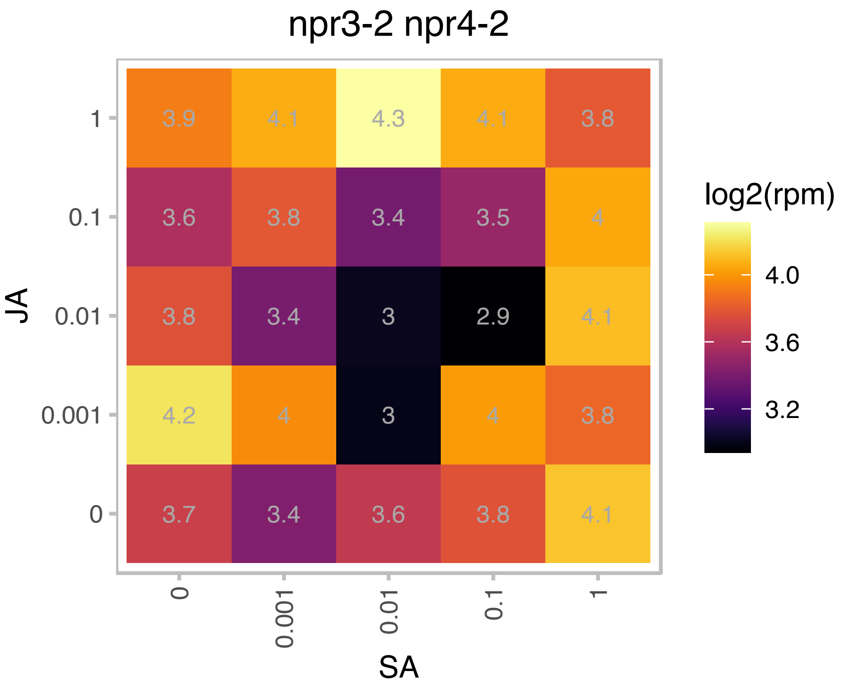


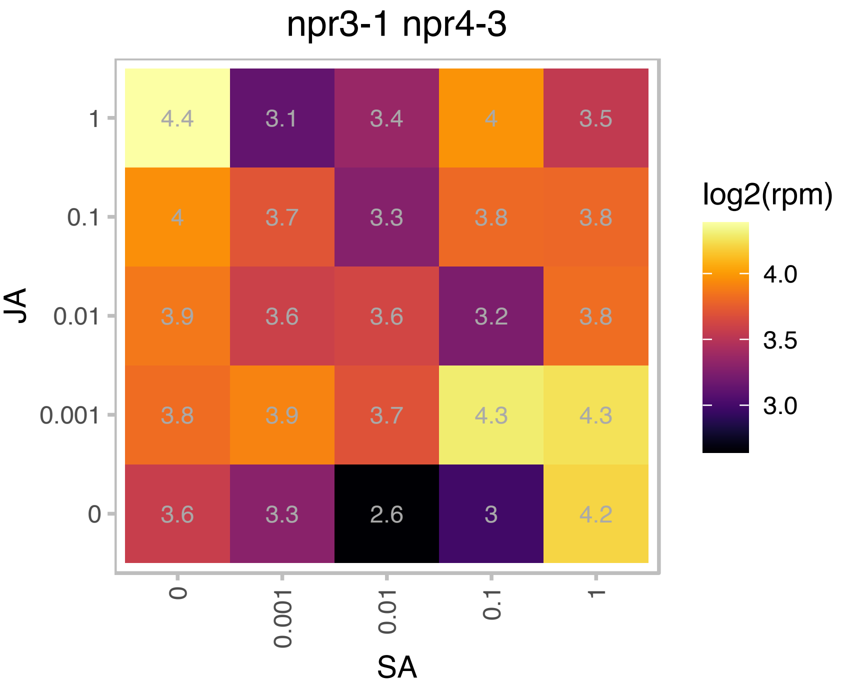

# AT3G60810.1 : NA

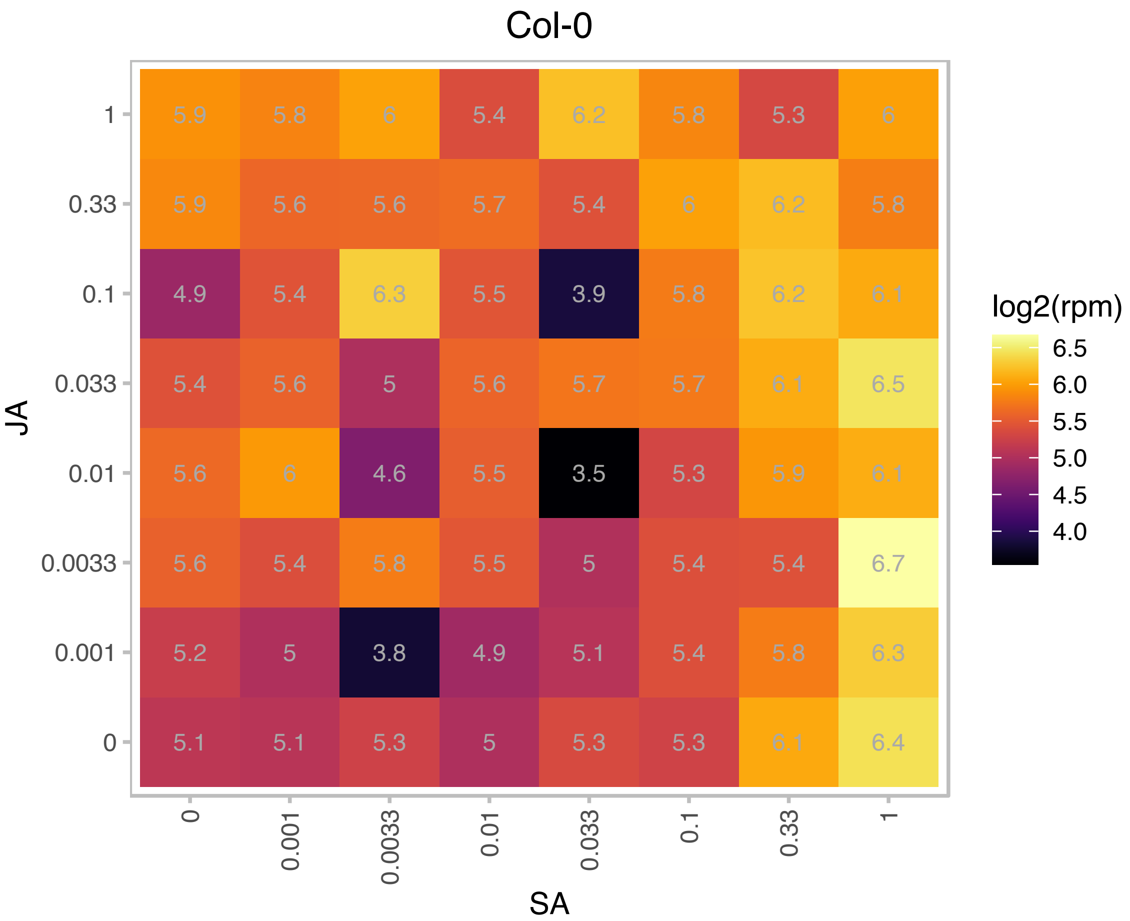


| | |
|---|---|
| Gene | AT3G60810 |
| is.expressed_genes | TRUE |
| is.responsive_genes | TRUE |
| is.apcluster | B-04 |

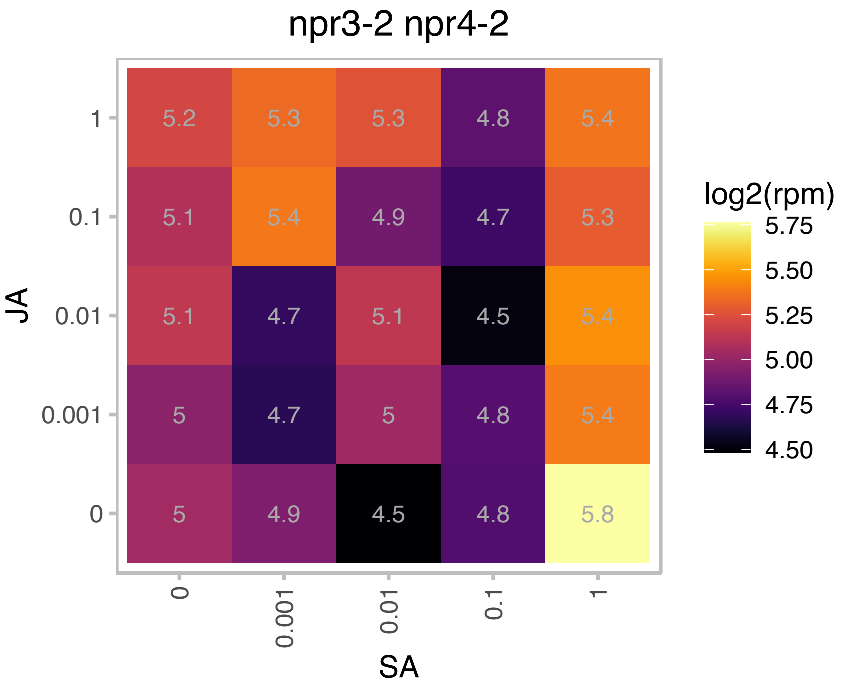


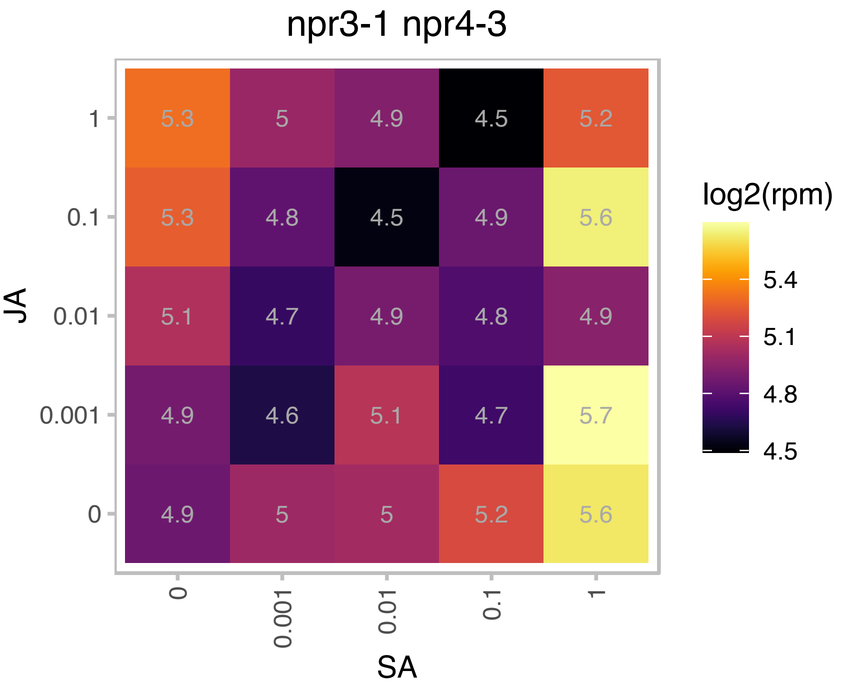

### AT4G02940.1 : ALKBH10B

oxidoreductase, 2OG-Fe(II) oxygenase family protein
ALKBH10B is a functional RNA N6-methyladenosine demethylase. Reduction in ALKBH10B decreases m6A levels, affects the stability of flowering time genes including FT, SPL3 and SPL9.  Mutant plants are early flowering.

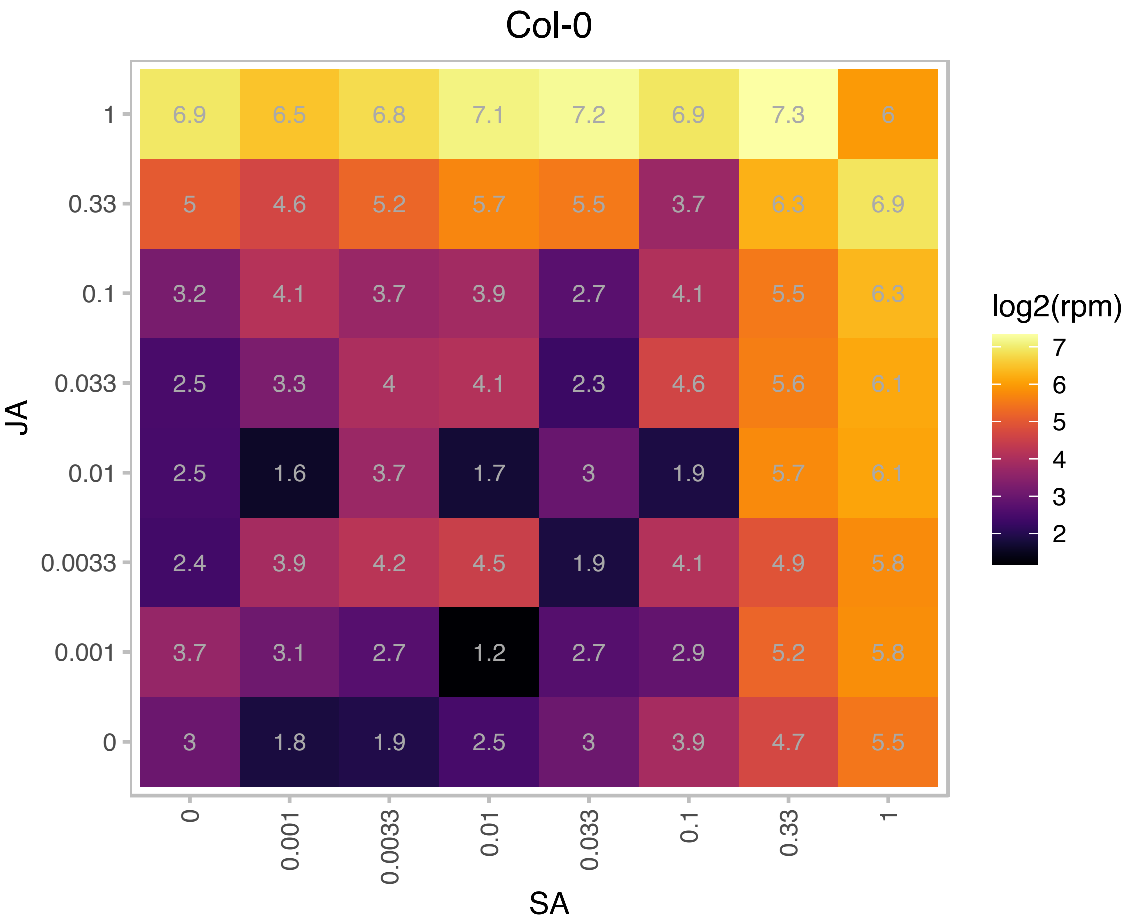


| | |
|---|---|
| Gene | AT4G02940 |
| is.expressed_genes | TRUE |
| is.responsive_genes | TRUE |
| is.apcluster | B-04 |

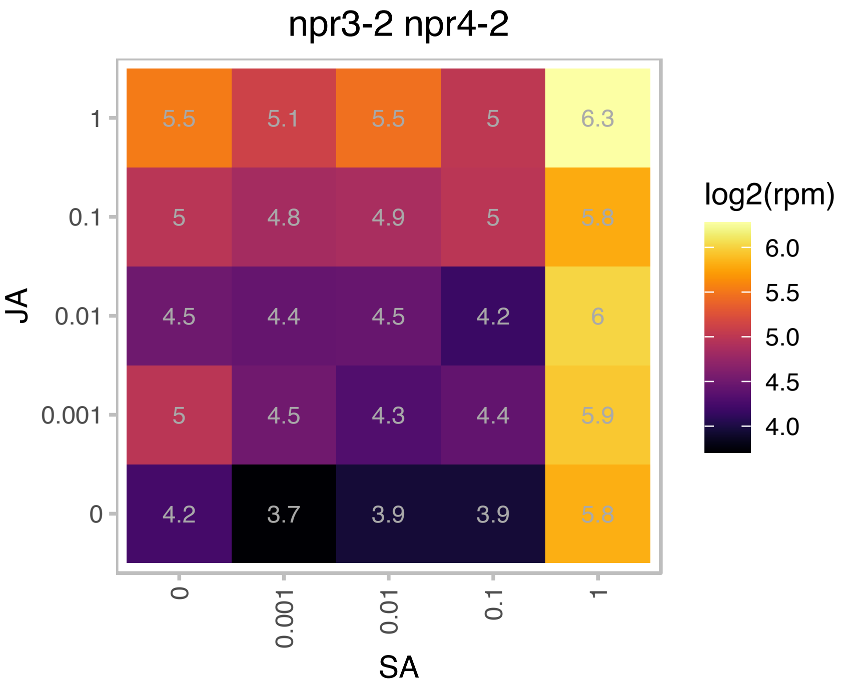


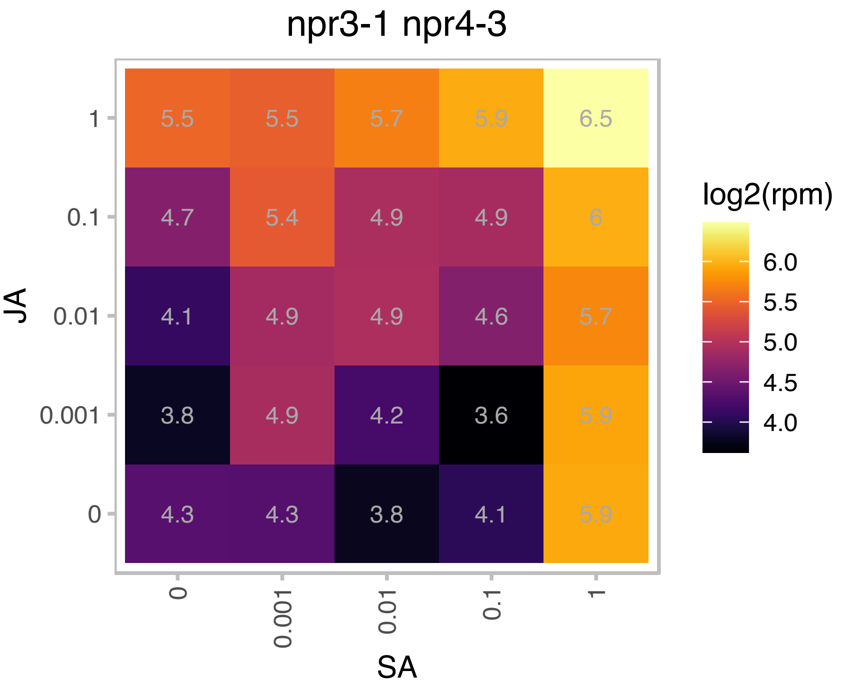

### AT4G05020.1 : NDB2

NAD(P)H dehydrogenase B2

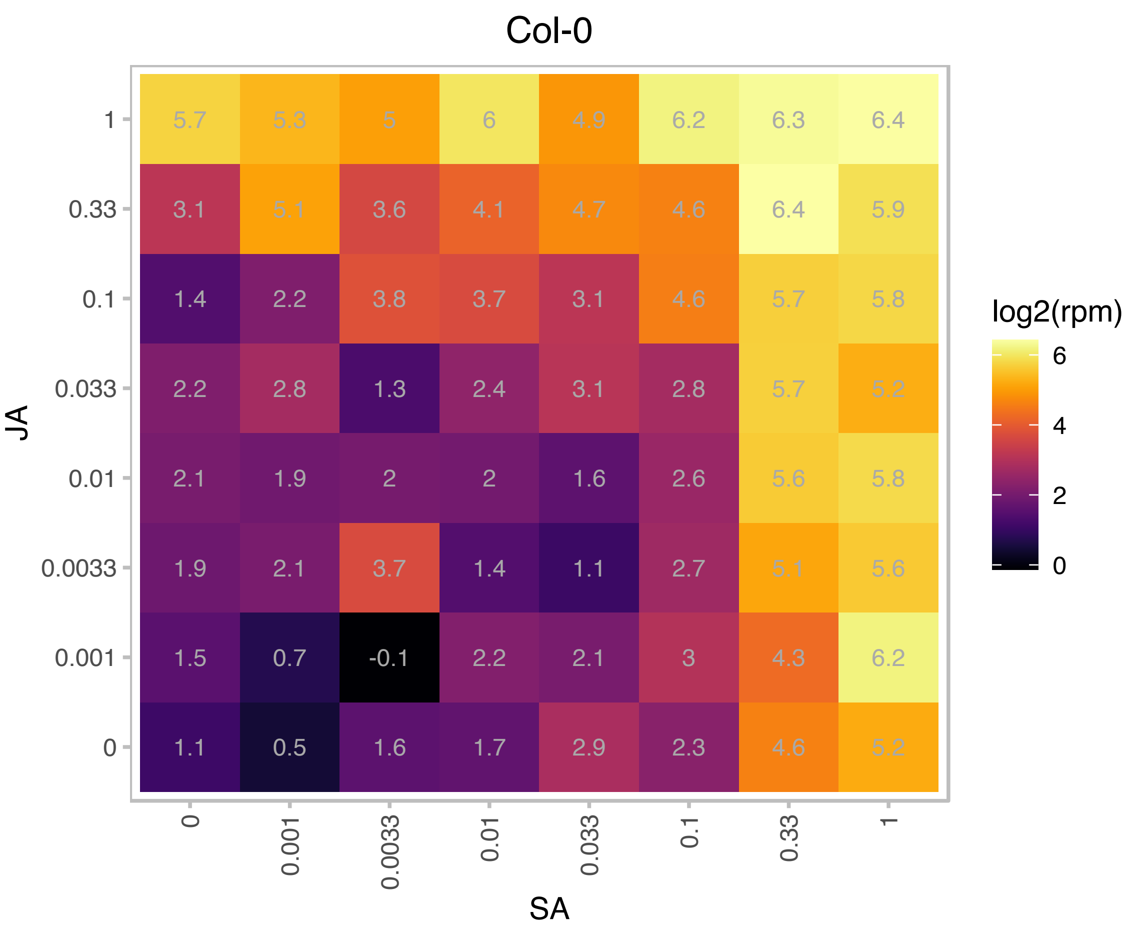


| | |
|---|---|
| Gene | AT4G05020 |
| is.expressed_genes | TRUE |
| is.responsive_genes | TRUE |
| is.apcluster | B-04 |

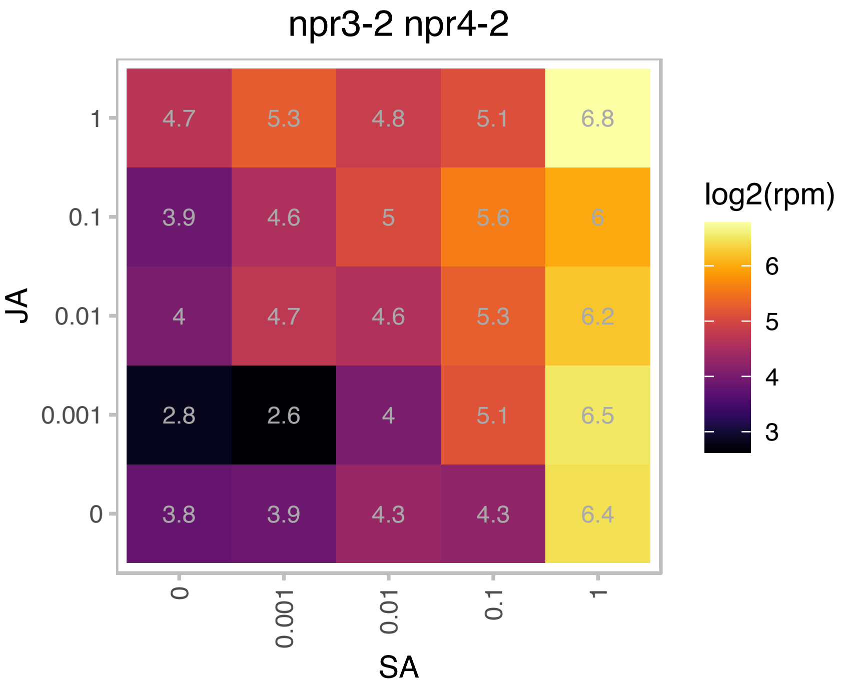


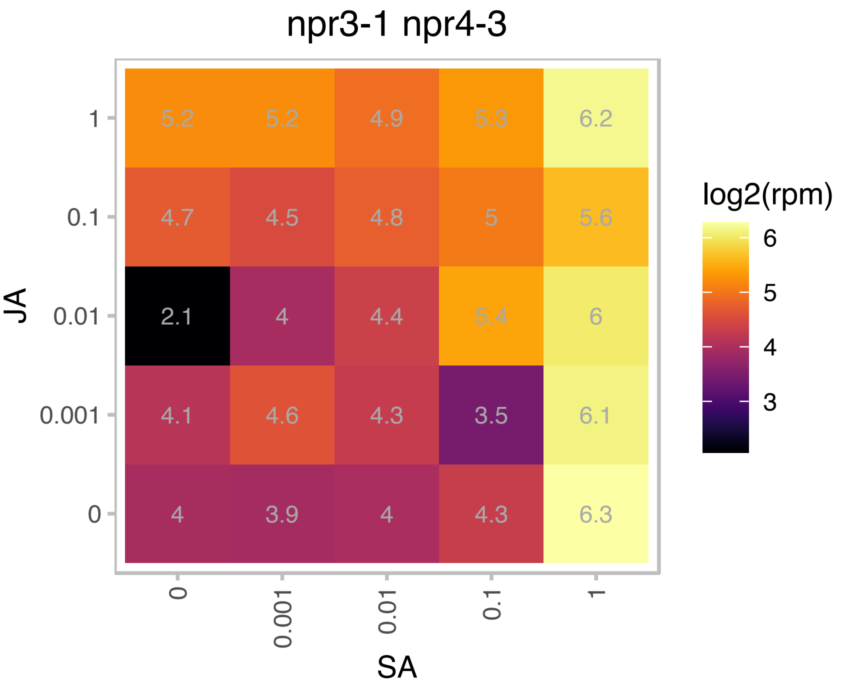

### AT4G05050.1 : UBQ11

ubiquitin 11

polyubiquitin gene, belongs to a subtype group with UBQ10 and UBQ14.  Various ecotypes of Arabidopsis have different numbers of ubiquitin repeats within this gene.

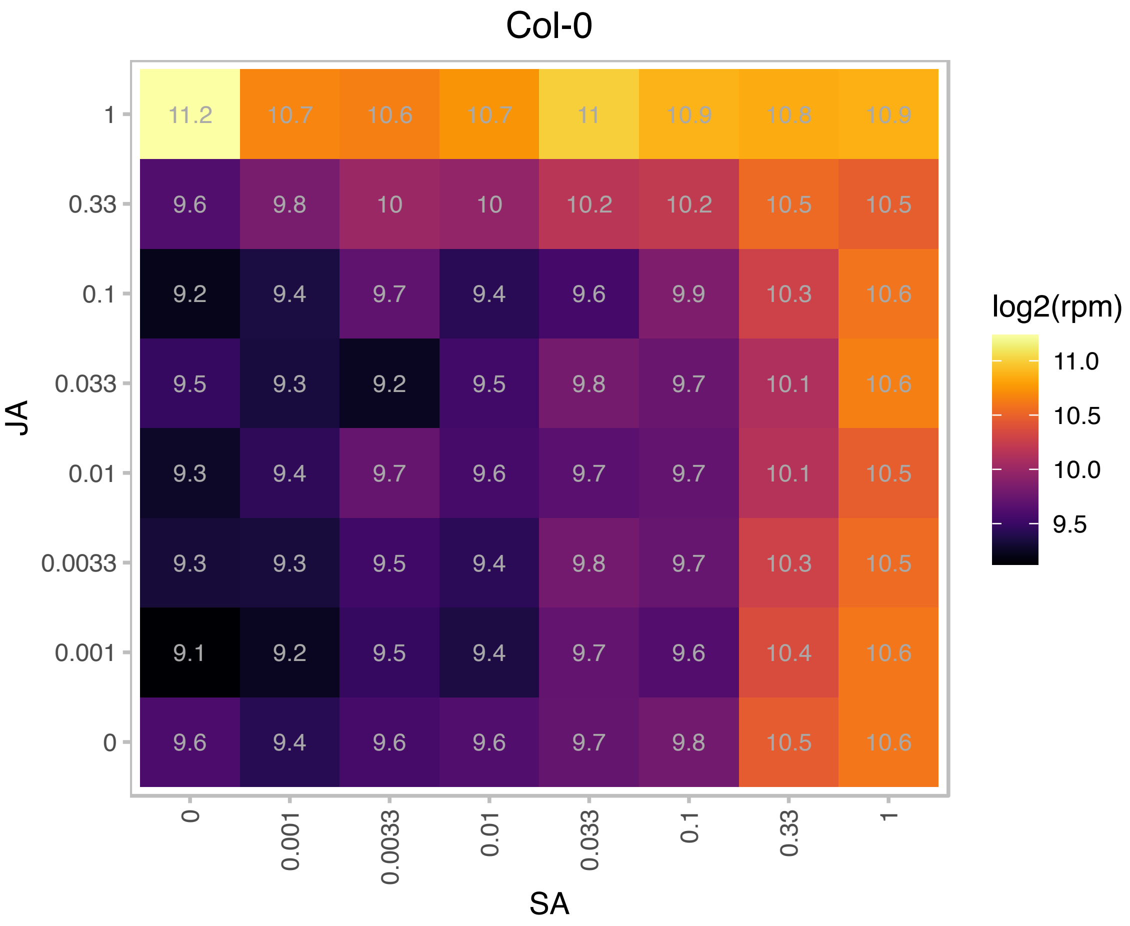


| | |
|---|---|
| Gene | AT4G05050 |
| is.expressed_genes | TRUE |
| is.responsive_genes | TRUE |
| is.apcluster | B-04 |

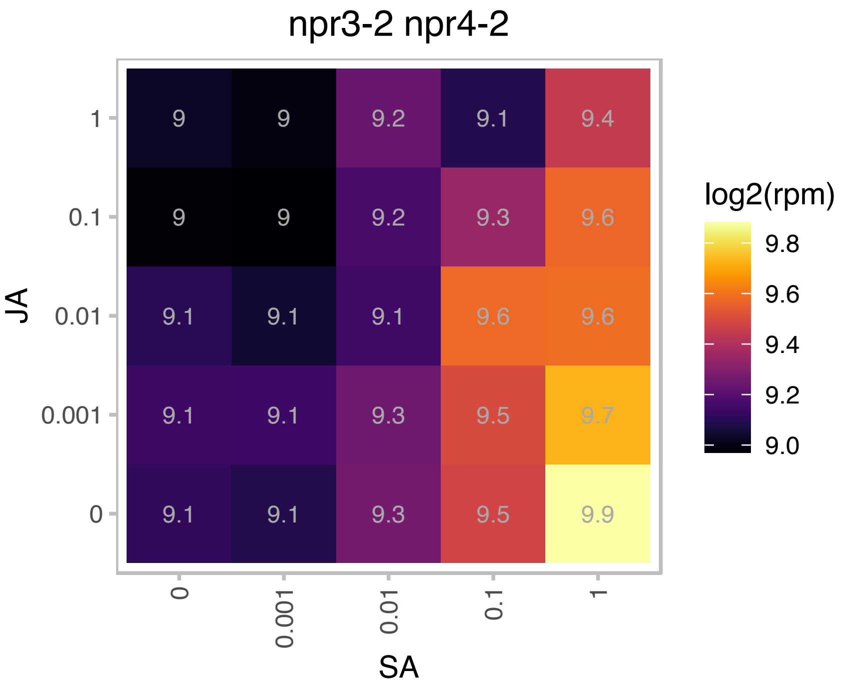


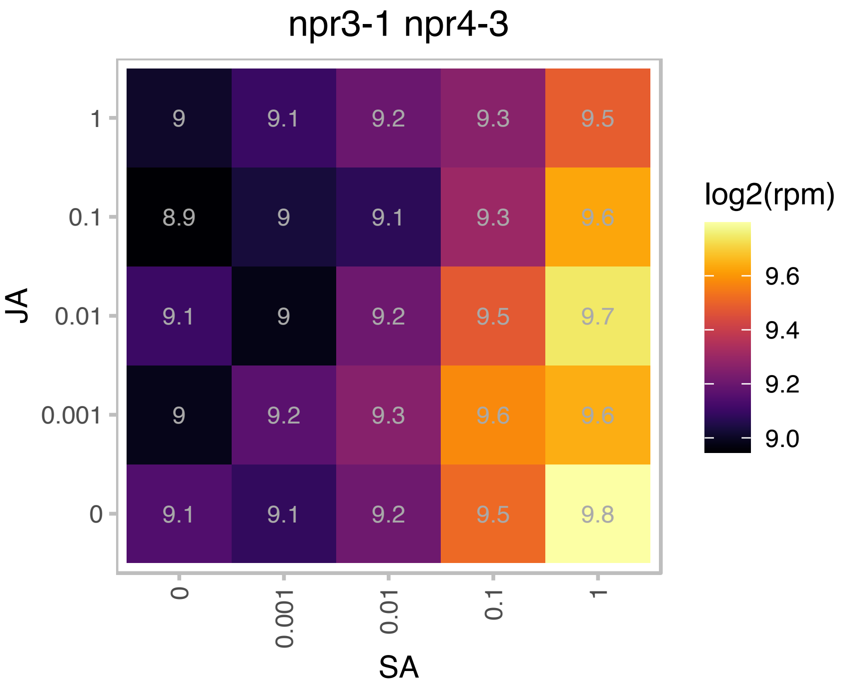

### AT4G05320.1 : UBI10, UBQ10

polyubiquitin 10

One of five polyubiquitin genes in A. thaliana.  These genes encode the highly conserved 76-amino acid protein ubiquitin that is covalently attached to substrate proteins targeting most for degradation. Polyubiquitin genes are characterized by the presence of tandem repeats of the 228 bp that encode a ubiquitin monomer. Induced by salicylic acid. Independent of NPR1 for their induction by salicylic acid. The mRNA is cell-to-cell mobile.

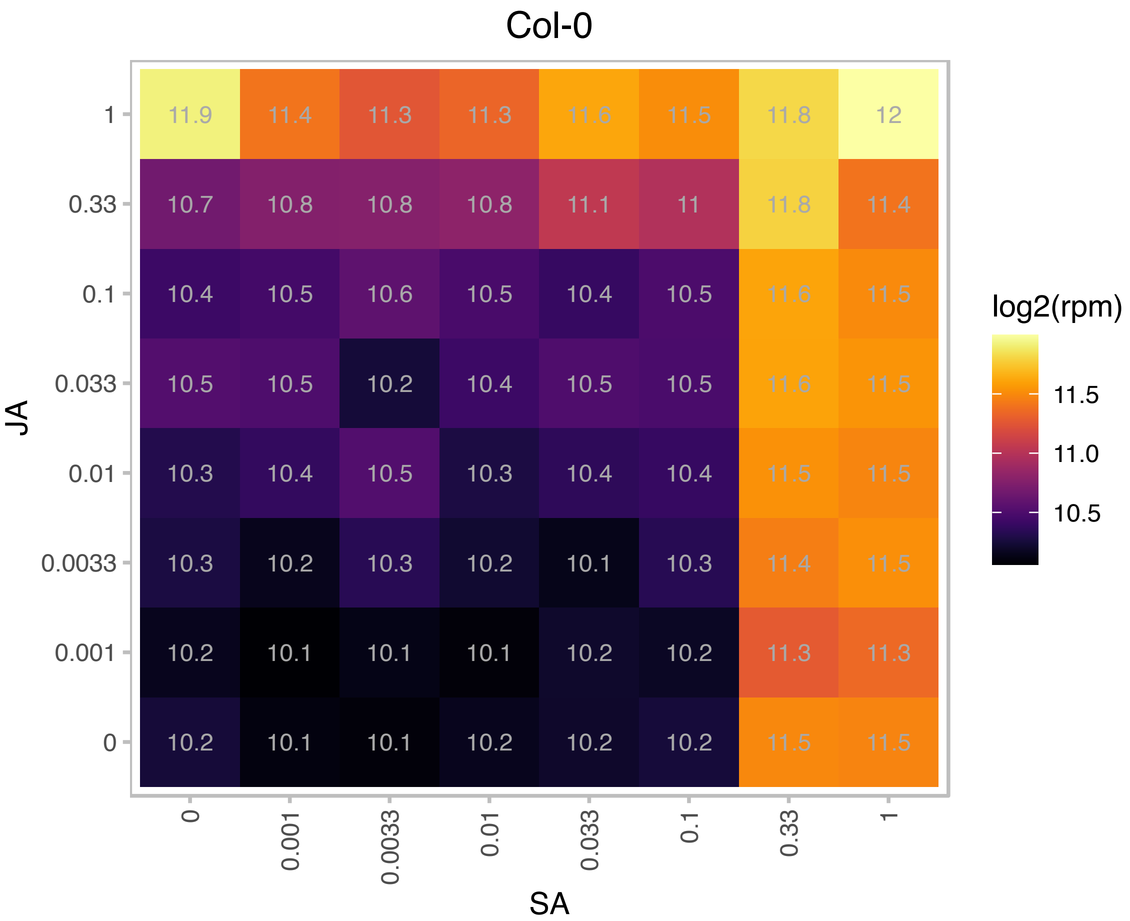


| | |
|---|---|
| Gene | AT4G05320 |
| is.expressed_genes | TRUE |
| is.responsive_genes | TRUE |
| is.apcluster | B-04 |

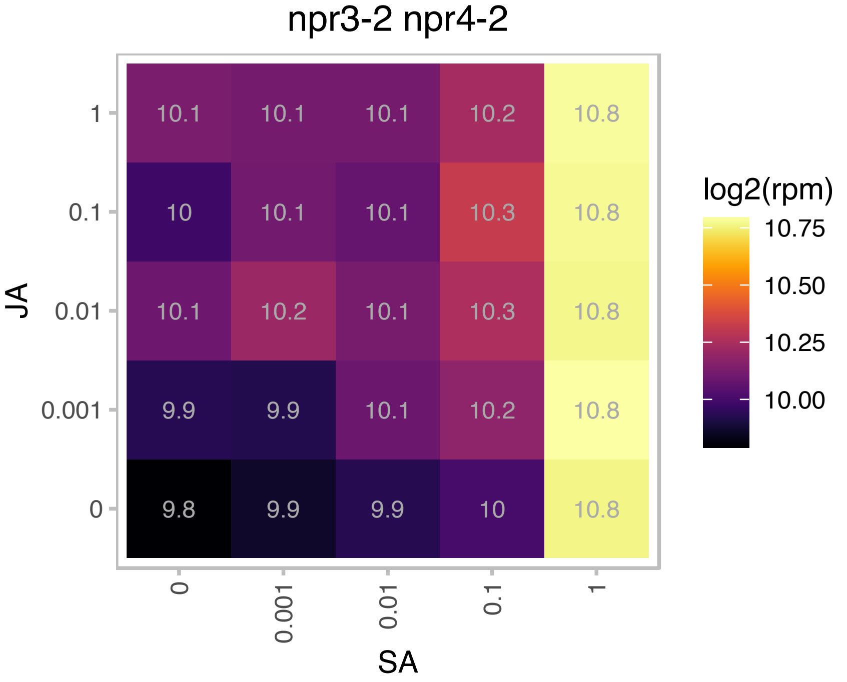


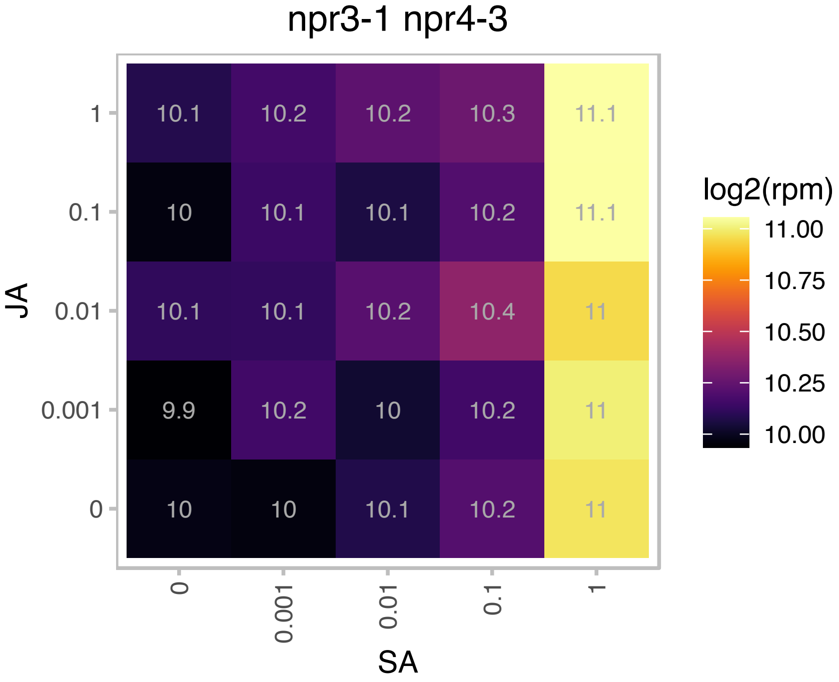

### AT4G11960.1 : PGRL1B

PGR5-like B
Encodes PGRL1B, a transmembrane protein present in thylakoids. PGRL1B has a highly homologous isoform PGF
encoded by At4g22890. Plants lacking PGRL1 show perturbation of cyclic electron flow, similar to PGR5-deficient plants. PGRL1 and PGR5 interact physically and associate with PSI (photosystem I).

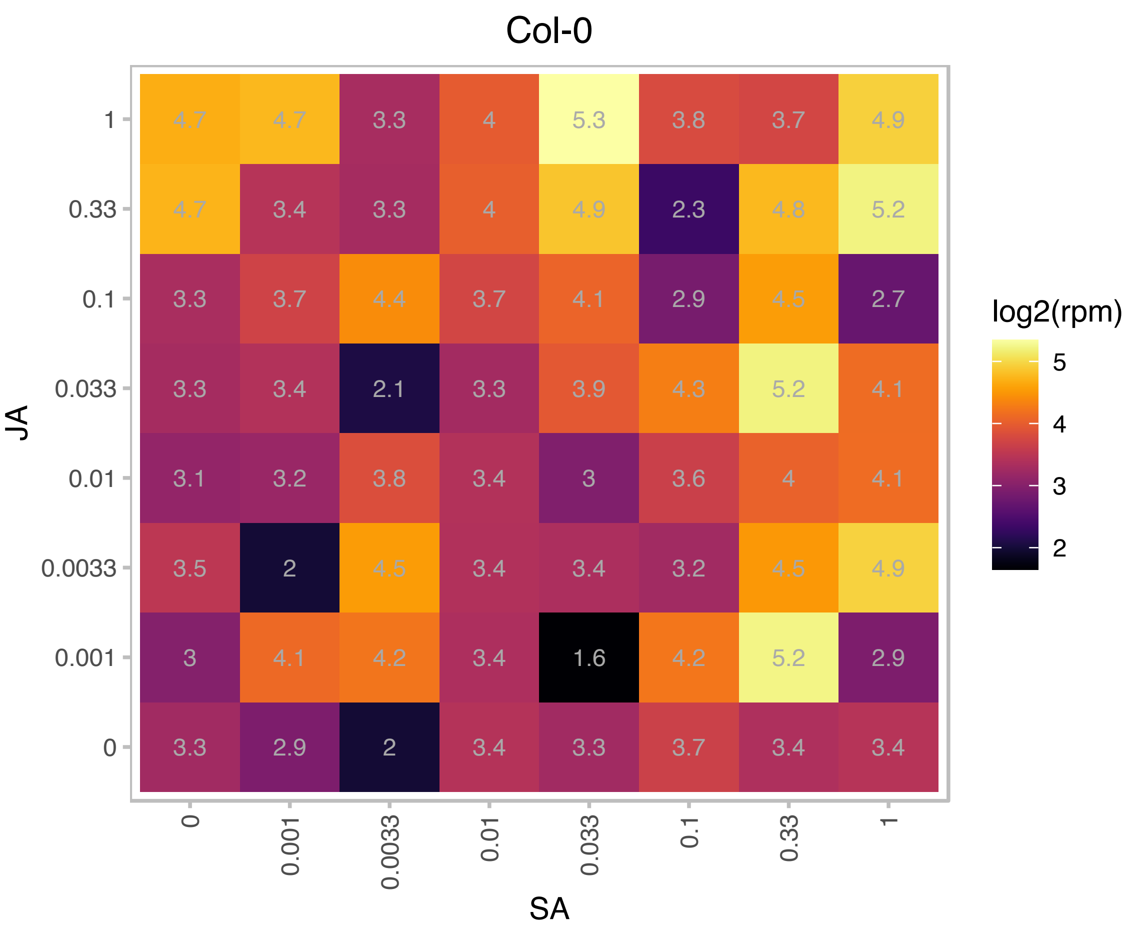


| | |
|---|---|
| Gene | AT4G11960 |
| is.expressed_genes | TRUE |
| is.responsive_genes | TRUE |
| is.apcluster | B-04 |

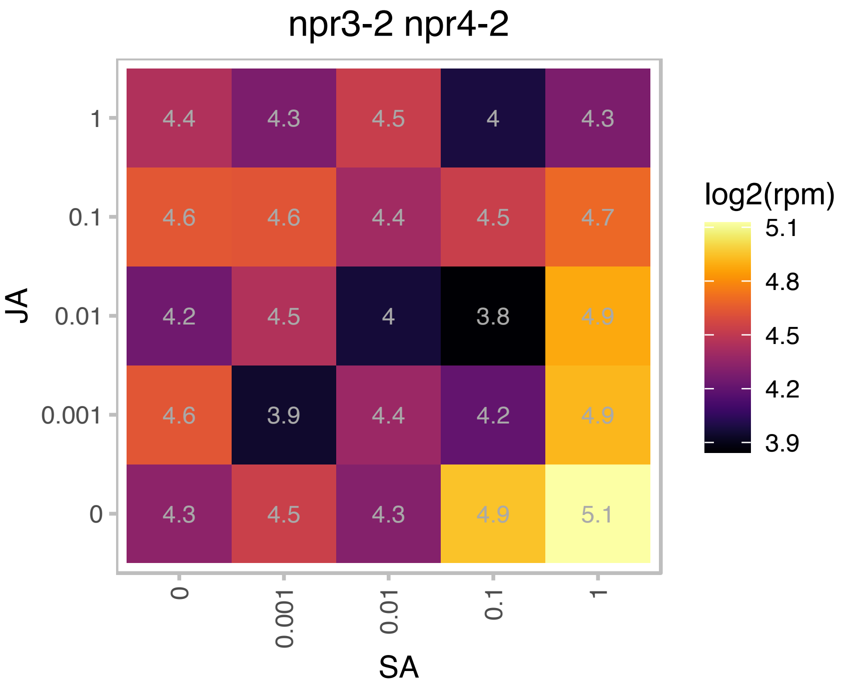


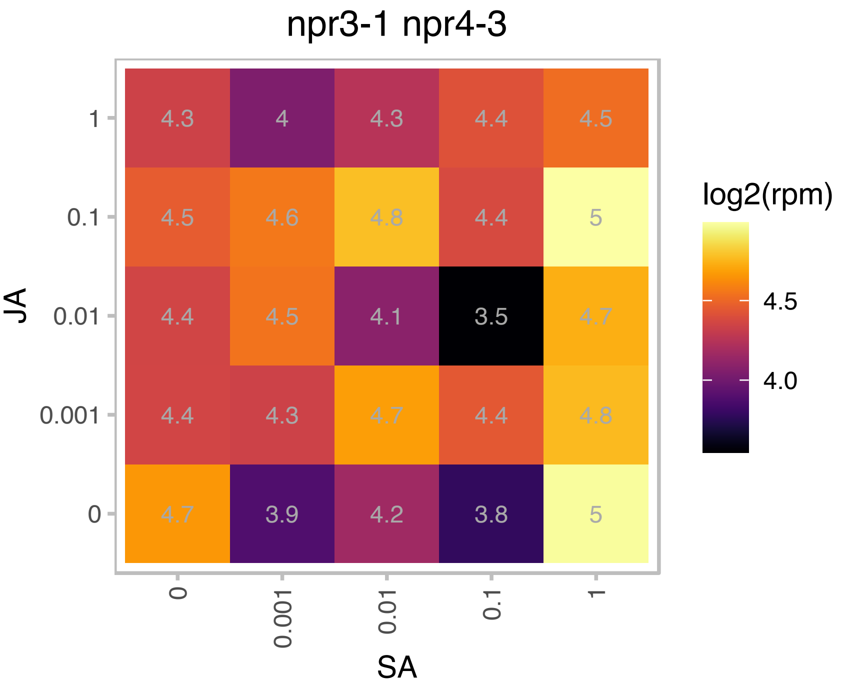

# AT4G12735.1 : NA

Encodes a peroxisomal protein.

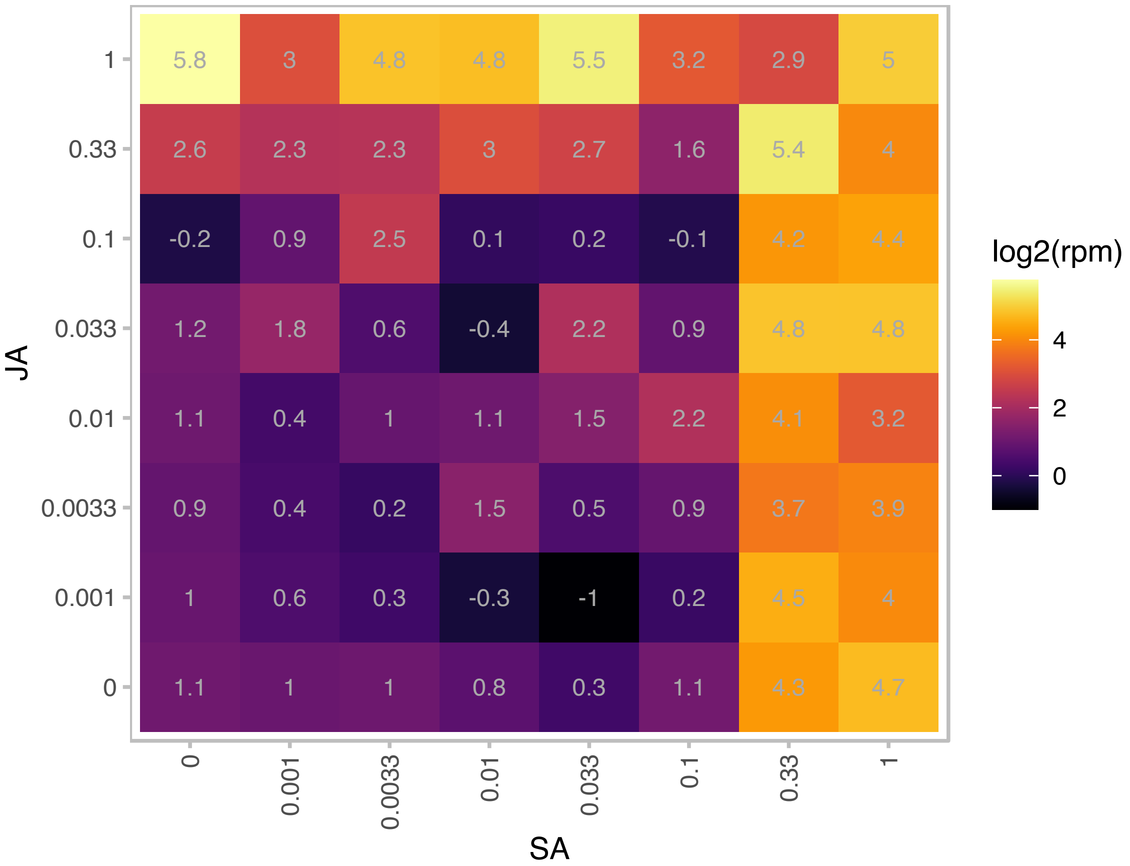


| Gene | AT4G12735 |
|---|---|
| is.expressed_genes | TRUE |
| is.responsive_genes | TRUE |
| is.apcluster | B-04 |

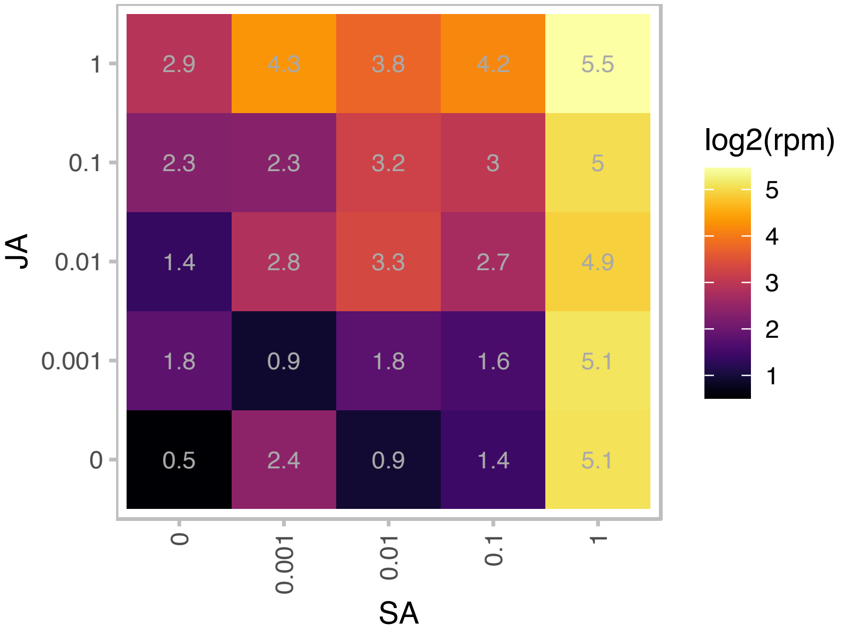


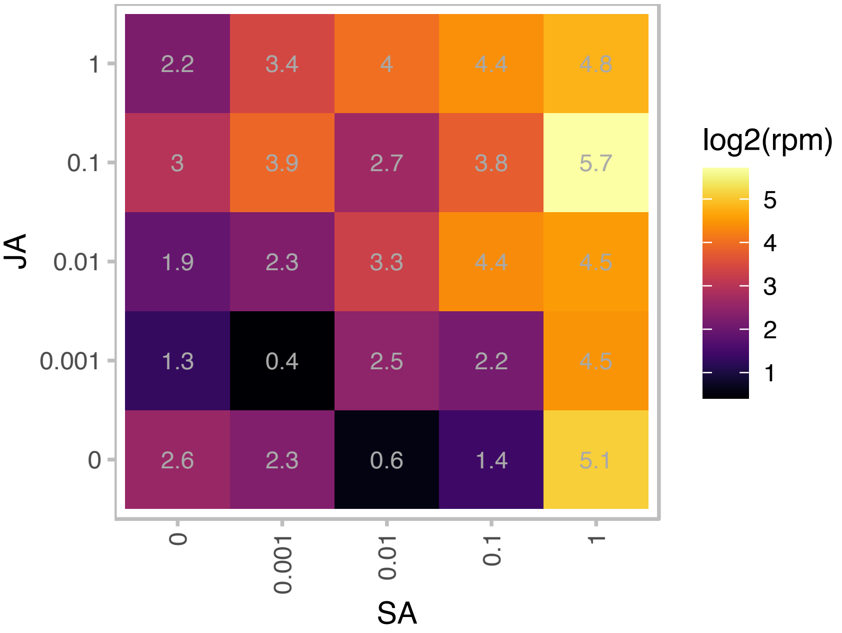

# AT4G15420.1 : NA

Ubiquitin fusion degradation UFD1 family protein

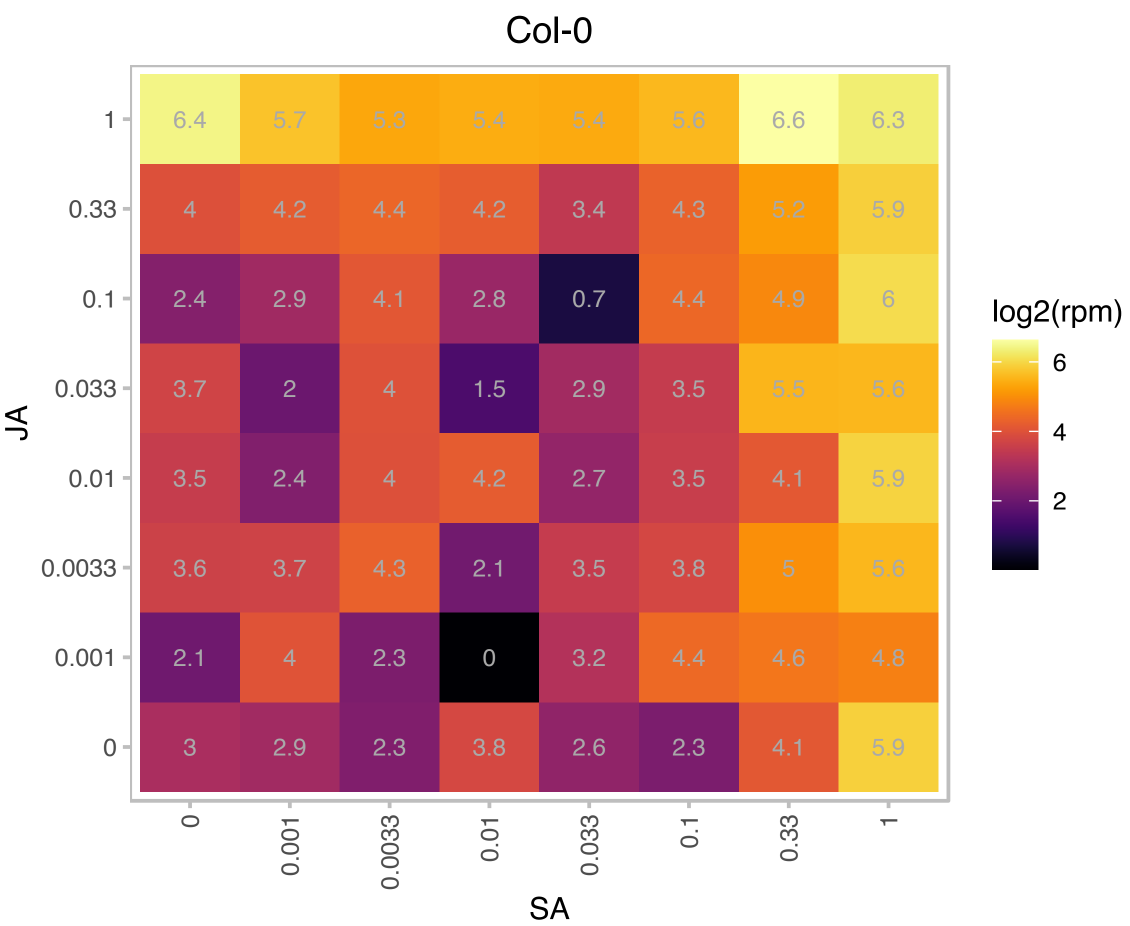


| | |
|---|---|
| Gene | AT4G15420 |
| is.expressed_genes | TRUE |
| is.responsive_genes | TRUE |
| is.apcluster | B-04 |

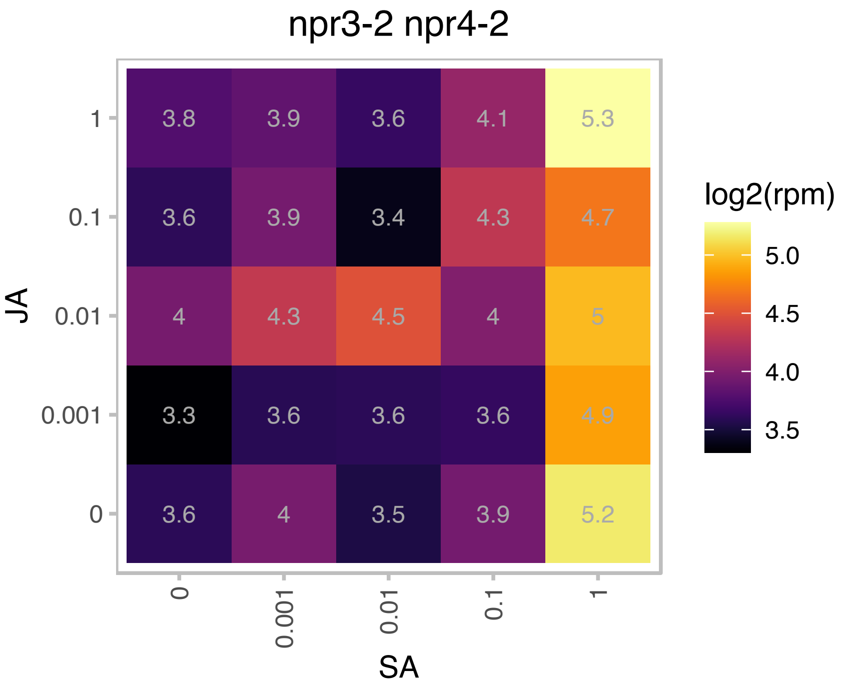


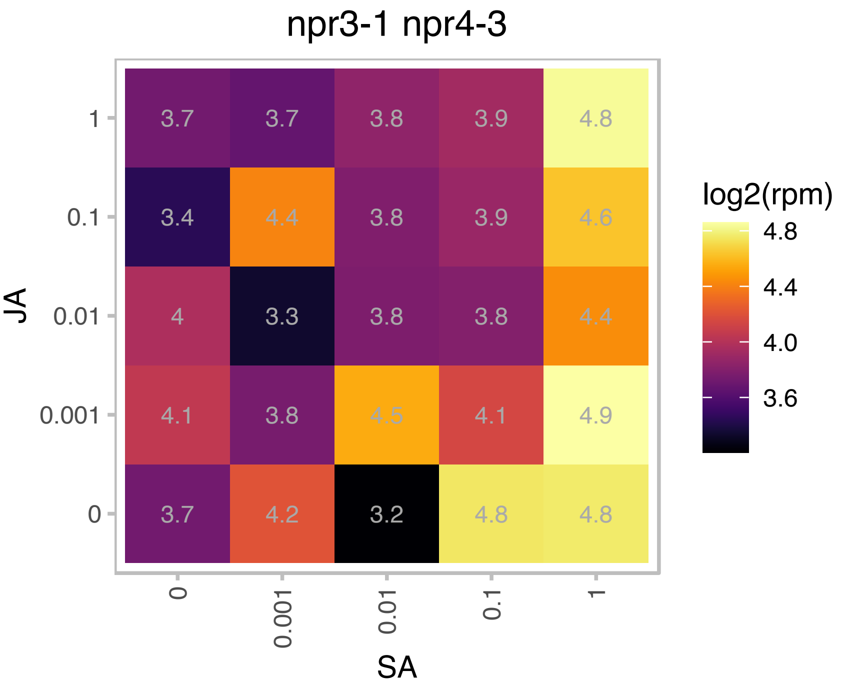

### AT4G21990.1 : APR3, ATAPR3, PRH-26, PRH26

APS reductase 3
Encodes a protein disulfide isomerase-like (PDIL) protein, a member of a multigene family within the thioredoxin (TRX) superfamily. This protein also belongs to the adenosine 5'-phosphosulfate reductase-like (APRL) group.

Col-0

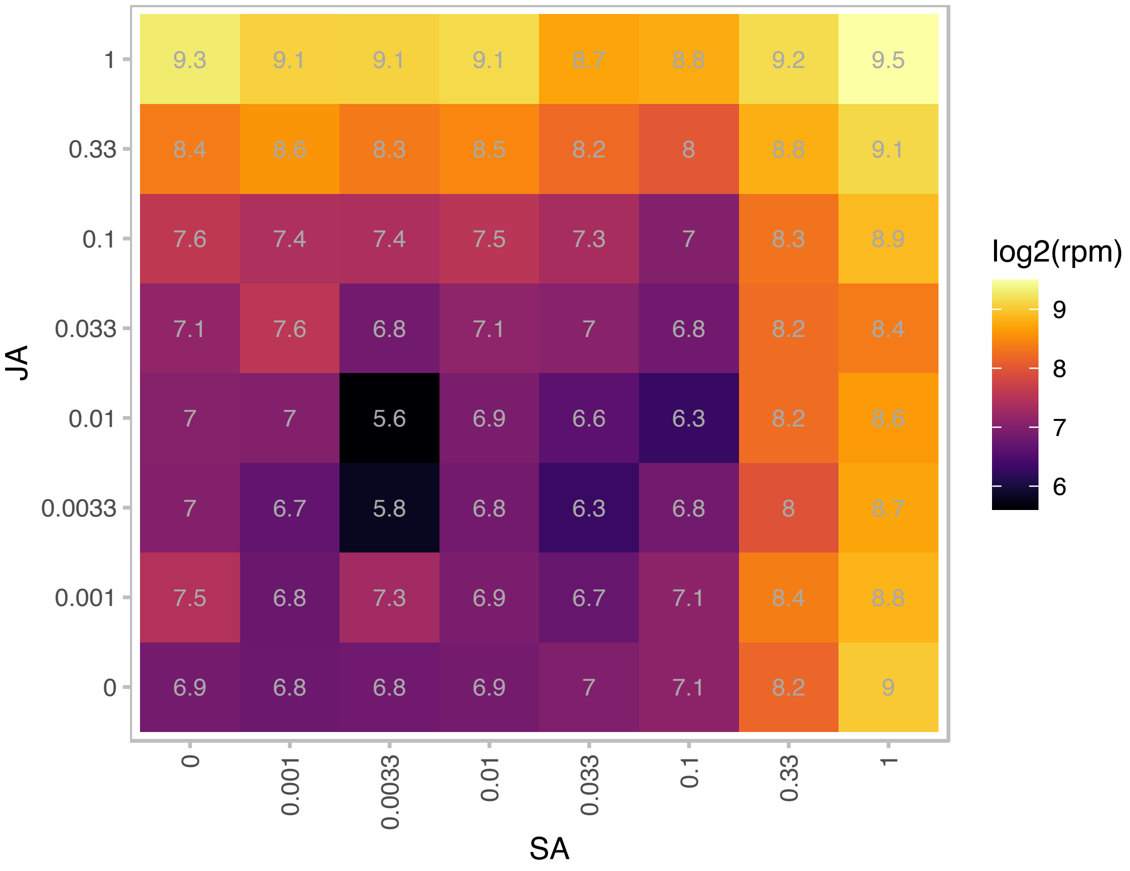


| | |
|---|---|
| Gene | AT4G21990 |
| is.expressed_genes | TRUE |
| is.responsive_genes | TRUE |
| is.apcluster | B-04 |

npr3-2 npr4-2

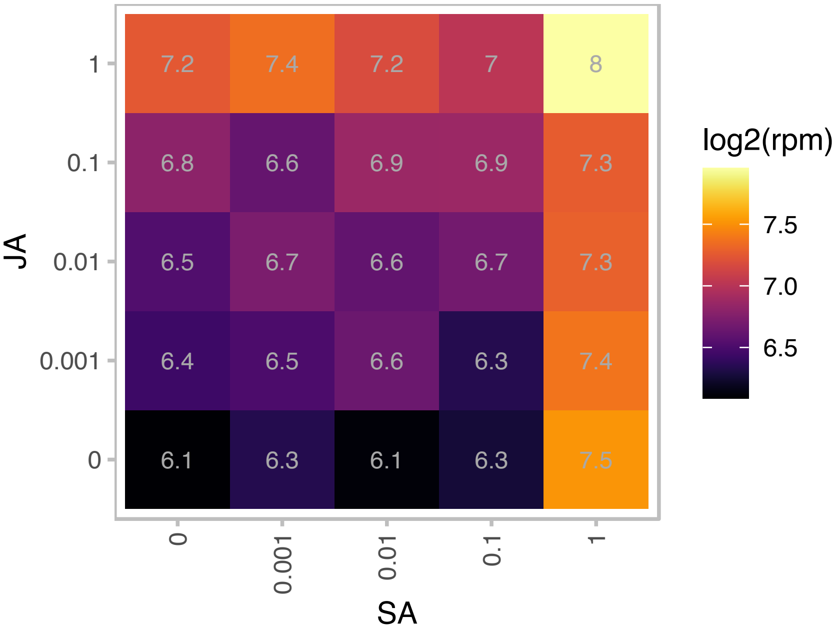


npr3-1 npr4-3

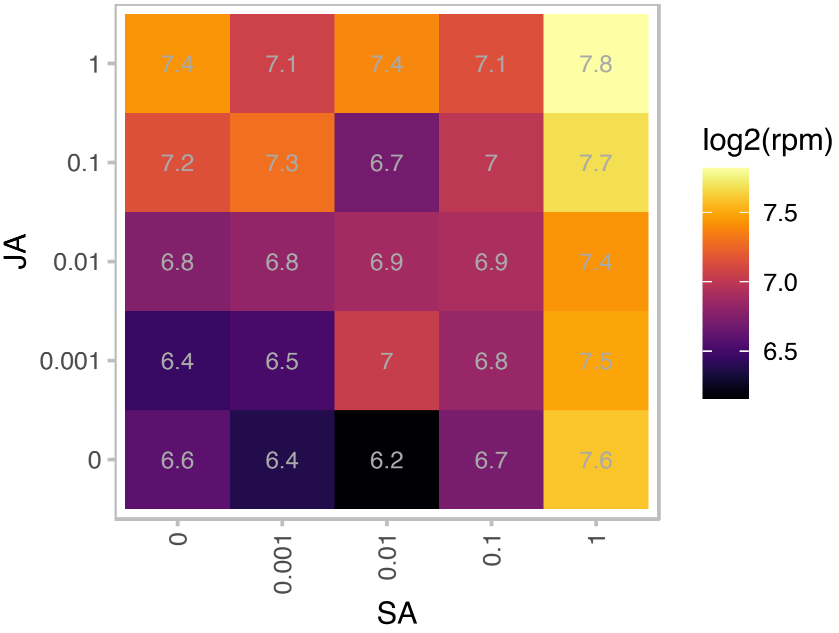

### AT4G29100.1 : bhlh68

basic helix-loop-helix (bHLH) DNA-binding superfamily protein

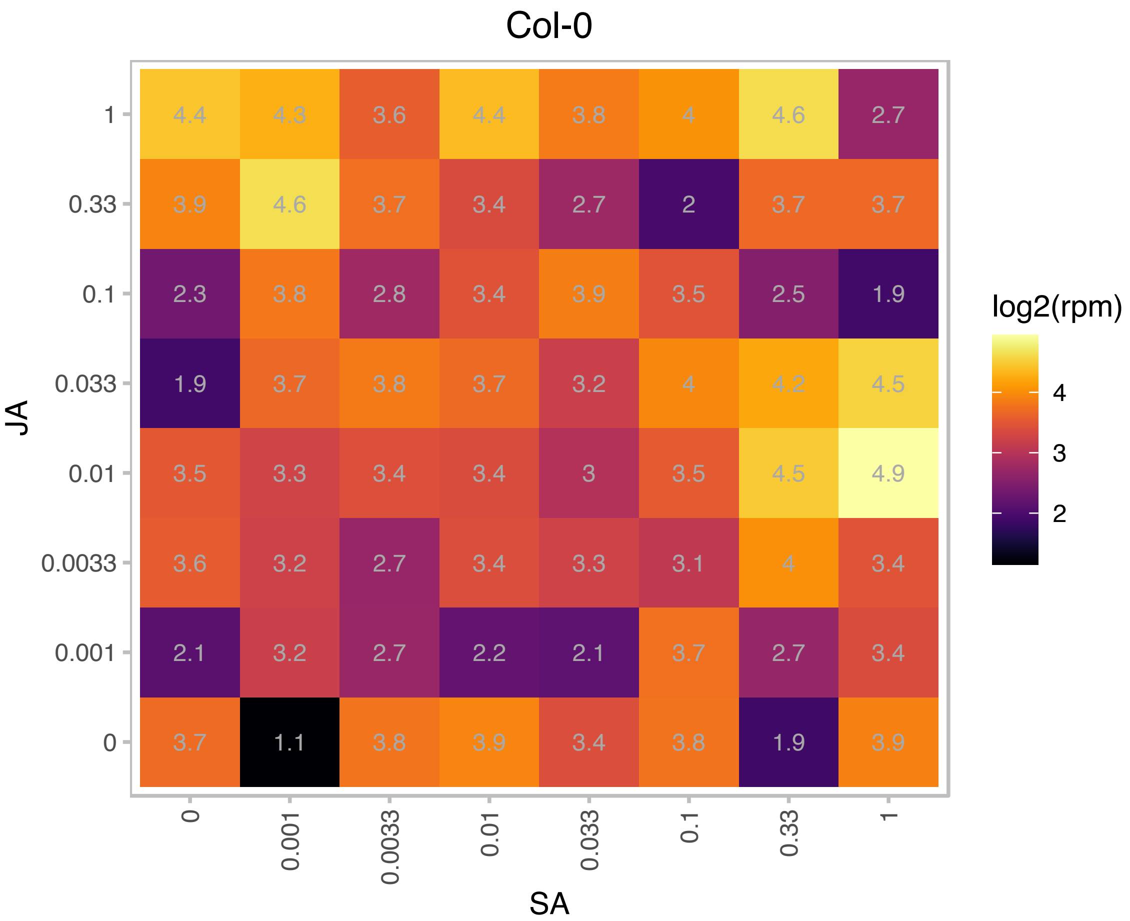


| Gene | AT4G29100 |
|---|---|
| is.expressed_genes | TRUE |
| is.responsive_genes | TRUE |
| is.apcluster | B-04 |

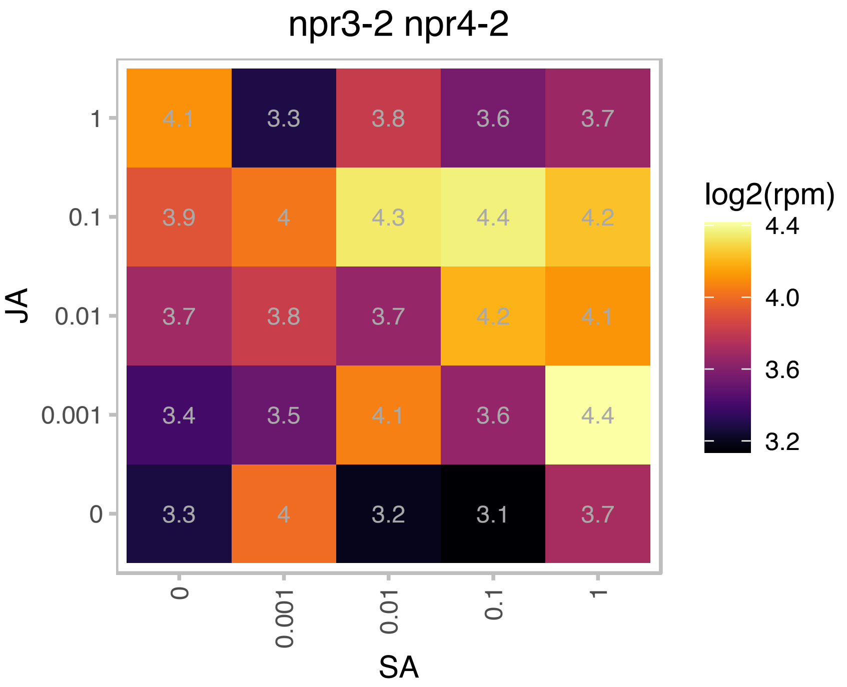


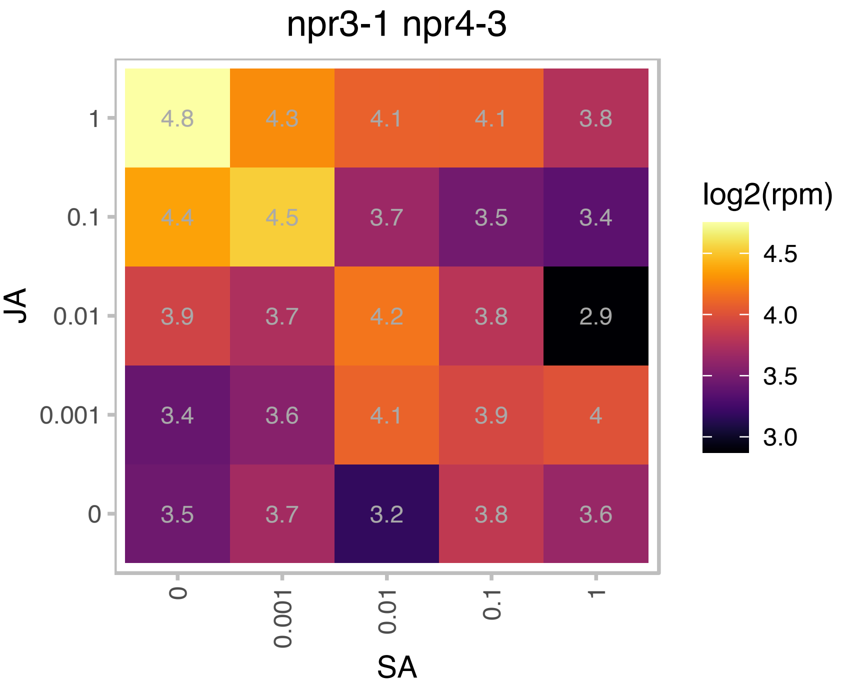

### AT4G30935.1 : ATWRKY32, WRKY32

WRKY DNA-binding protein 32

member of WRKY Transcription Factor; Group I

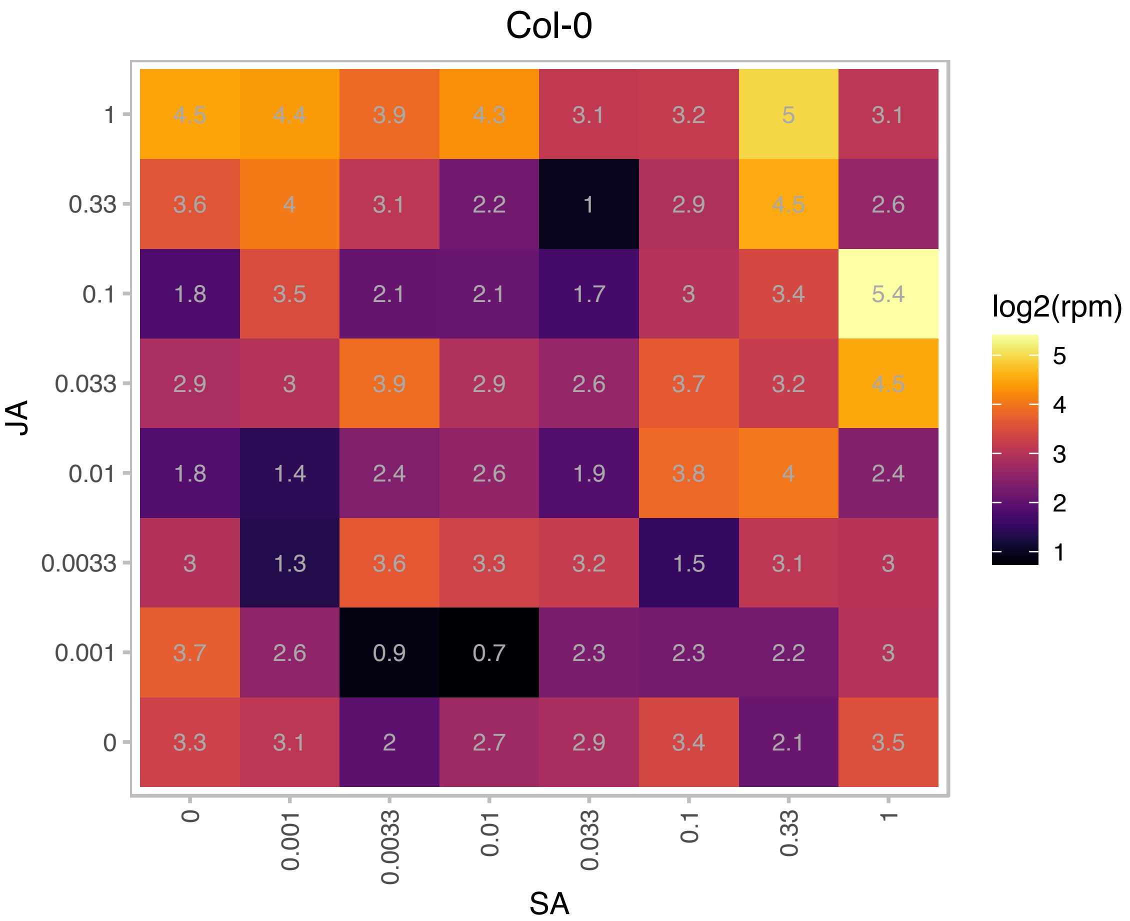


| | |
|---|---|
| Gene | AT4G30935 |
| is.expressed_genes | TRUE |
| is.responsive_genes | TRUE |
| is.apcluster | B-04 |

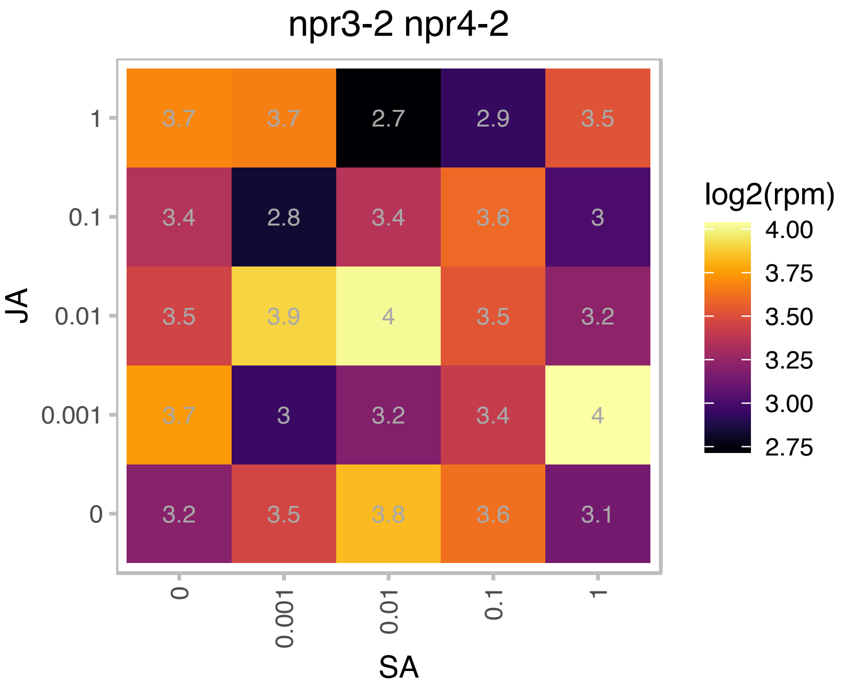


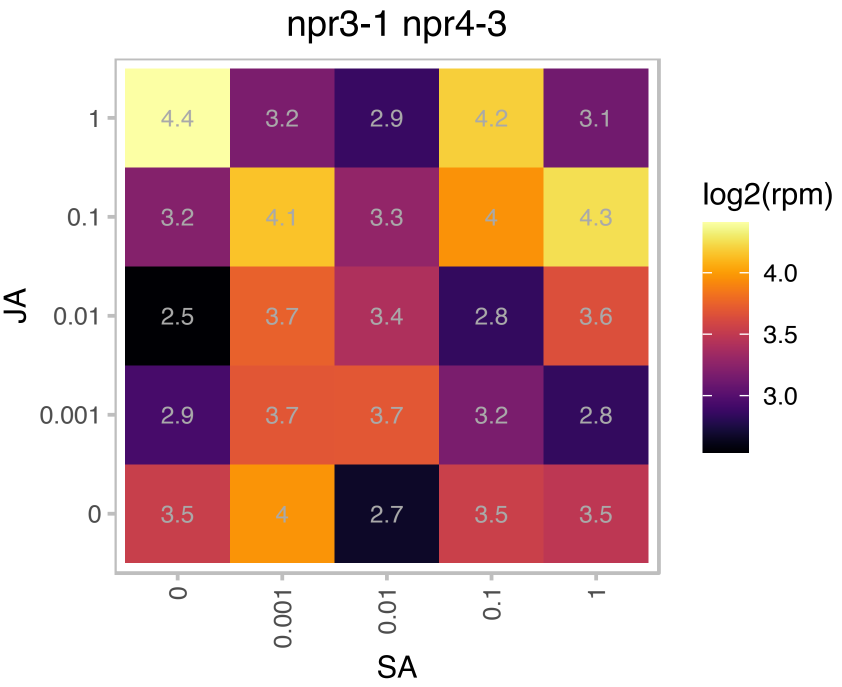

### AT4G34860.1 : A/N-InvB

Plant neutral invertase family protein

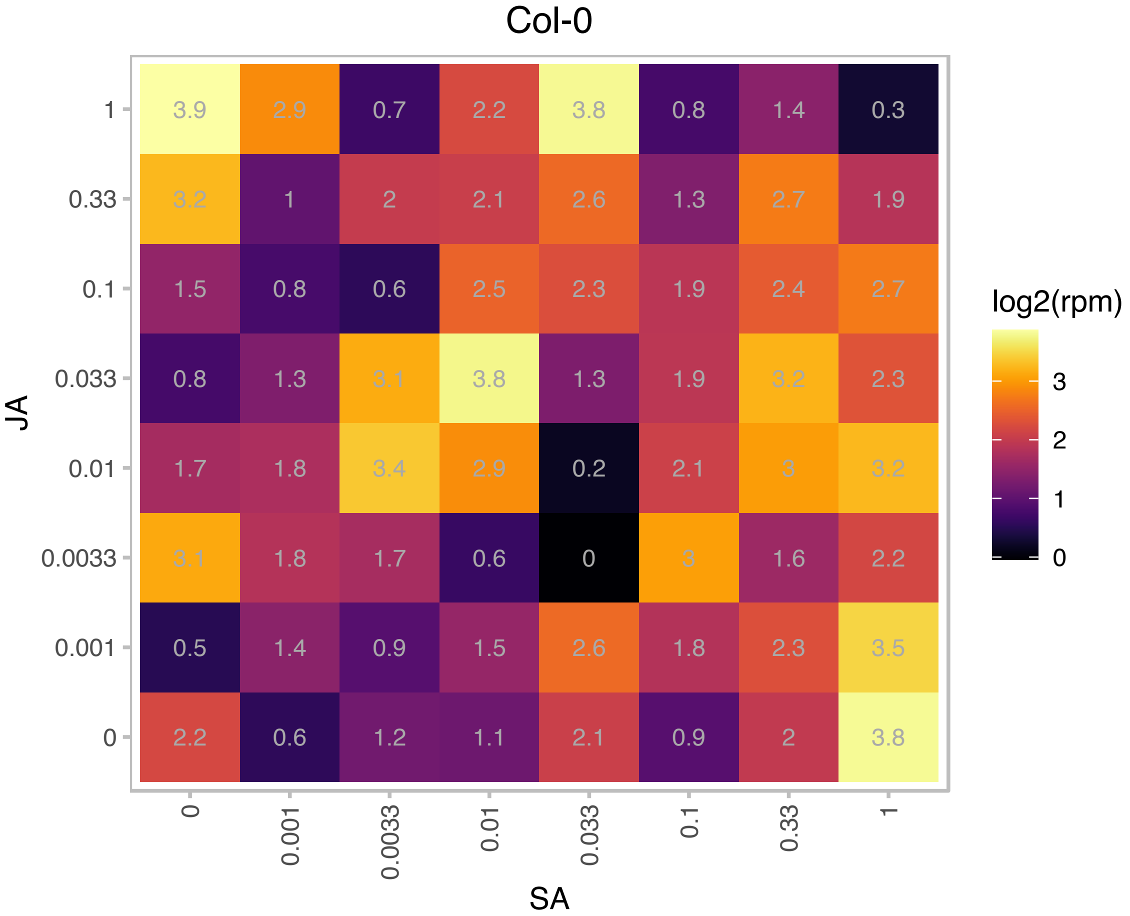


| | |
|---|---|
| Gene | AT4G34860 |
| is.expressed_genes | TRUE |
| is.responsive_genes | TRUE |
| is.apcluster | B-04 |

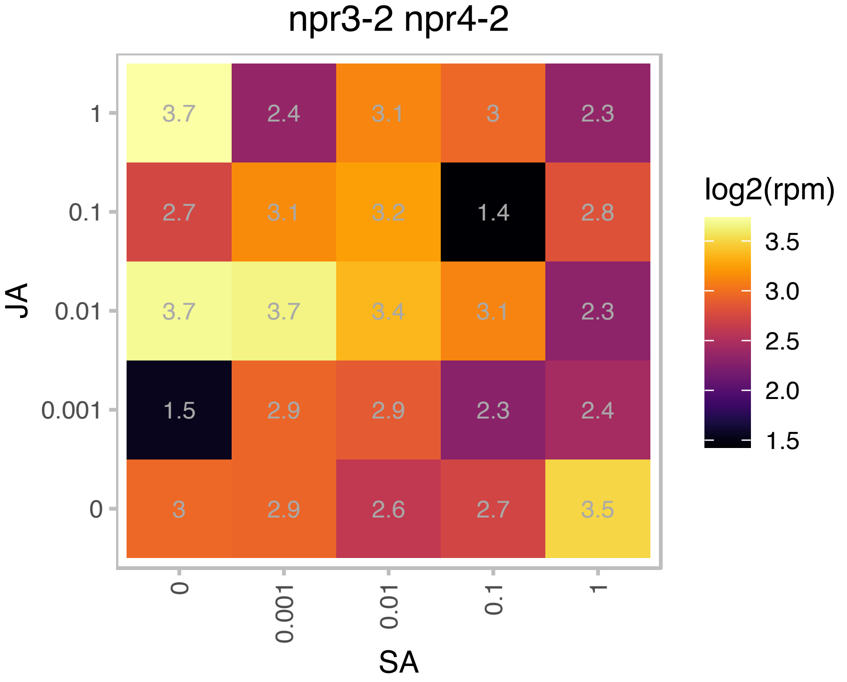


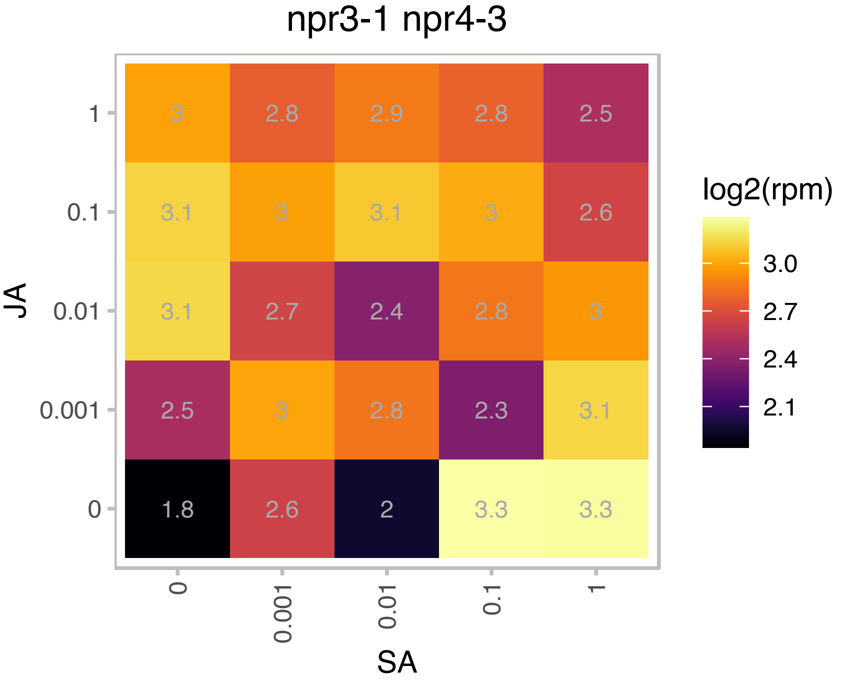

# AT4G36195.1 : NA

Serine carboxypeptidase S28 family protein

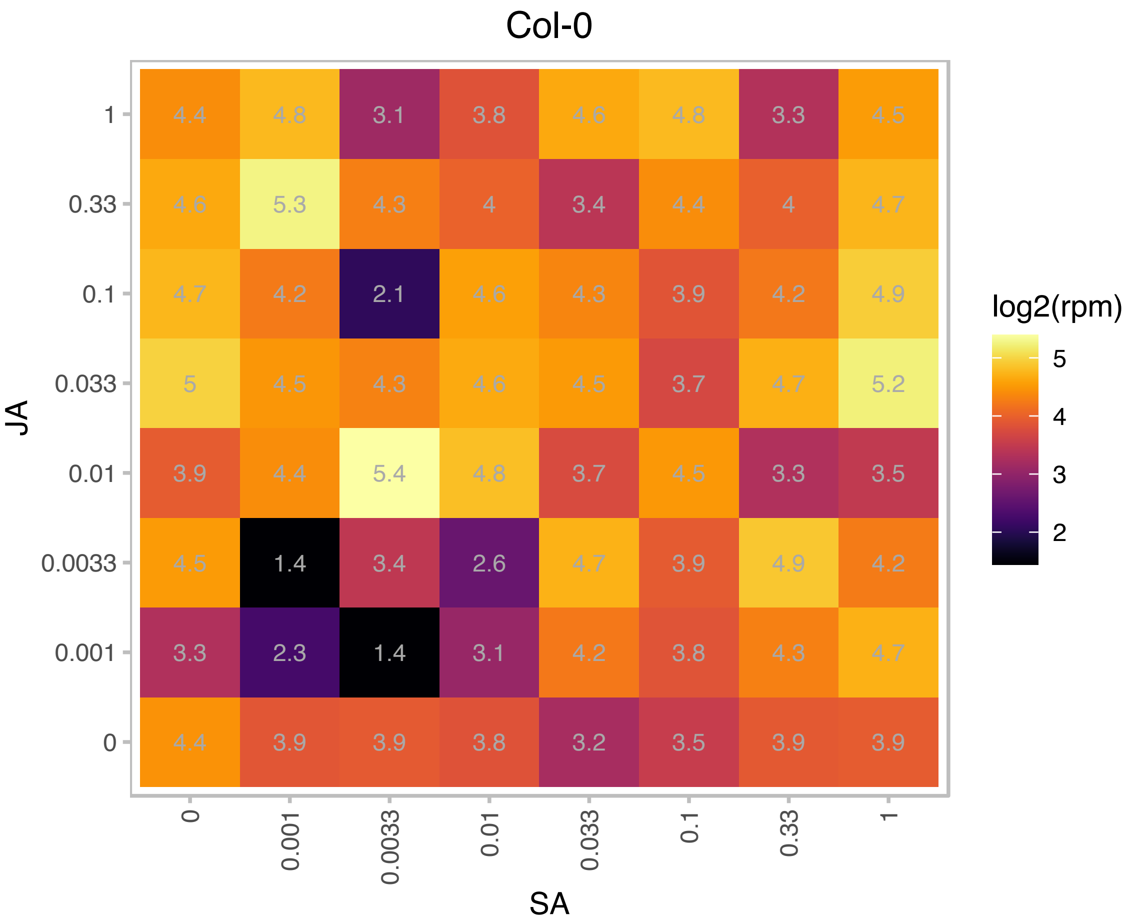


| | |
|---|---|
| Gene | AT4G36195 |
| is.expressed_genes | TRUE |
| is.responsive_genes | TRUE |
| is.apcluster | B-04 |

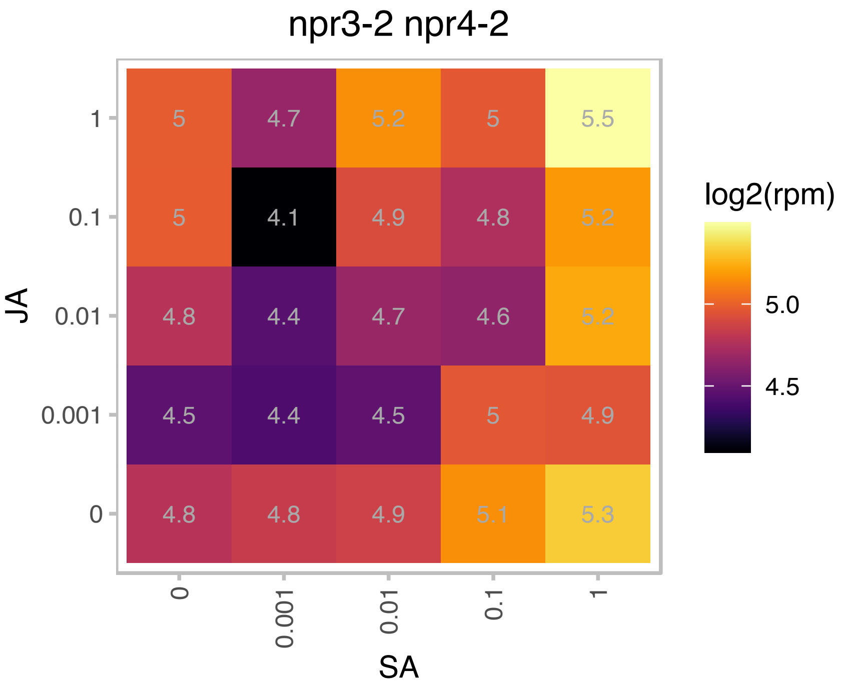


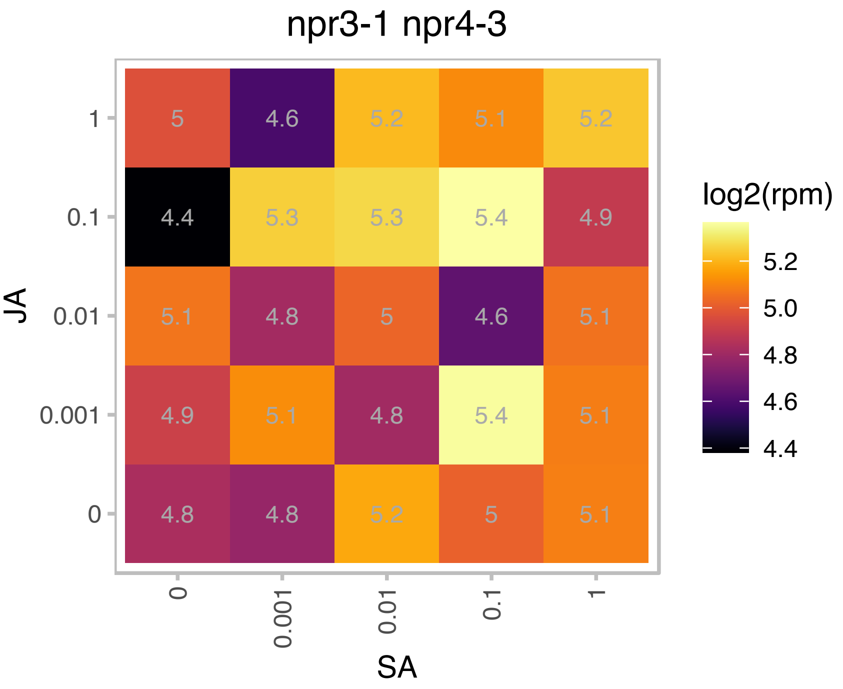

### AT5G08530.1 : CI51, NDUFV1

51 kDa subunit of complex I

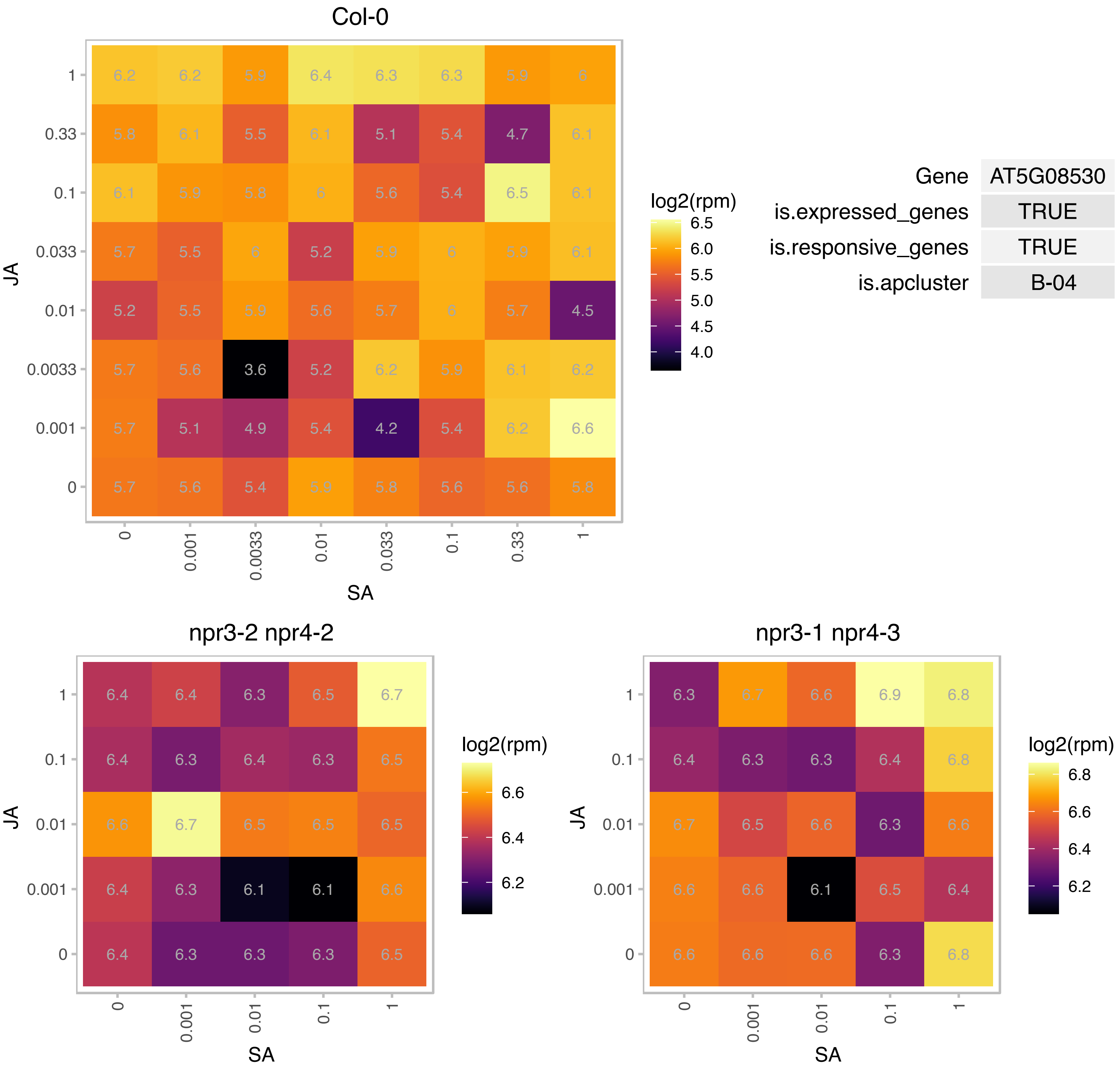

| | |
|---|---|
| Gene | AT5G08530 |
| is.expressed_genes | TRUE |
| is.responsive_genes | TRUE |
| is.apcluster | B-04 |

# AT5G10830.1 : NA

S-adenosyl-L-methionine-dependent methyltransferases superfamily protein

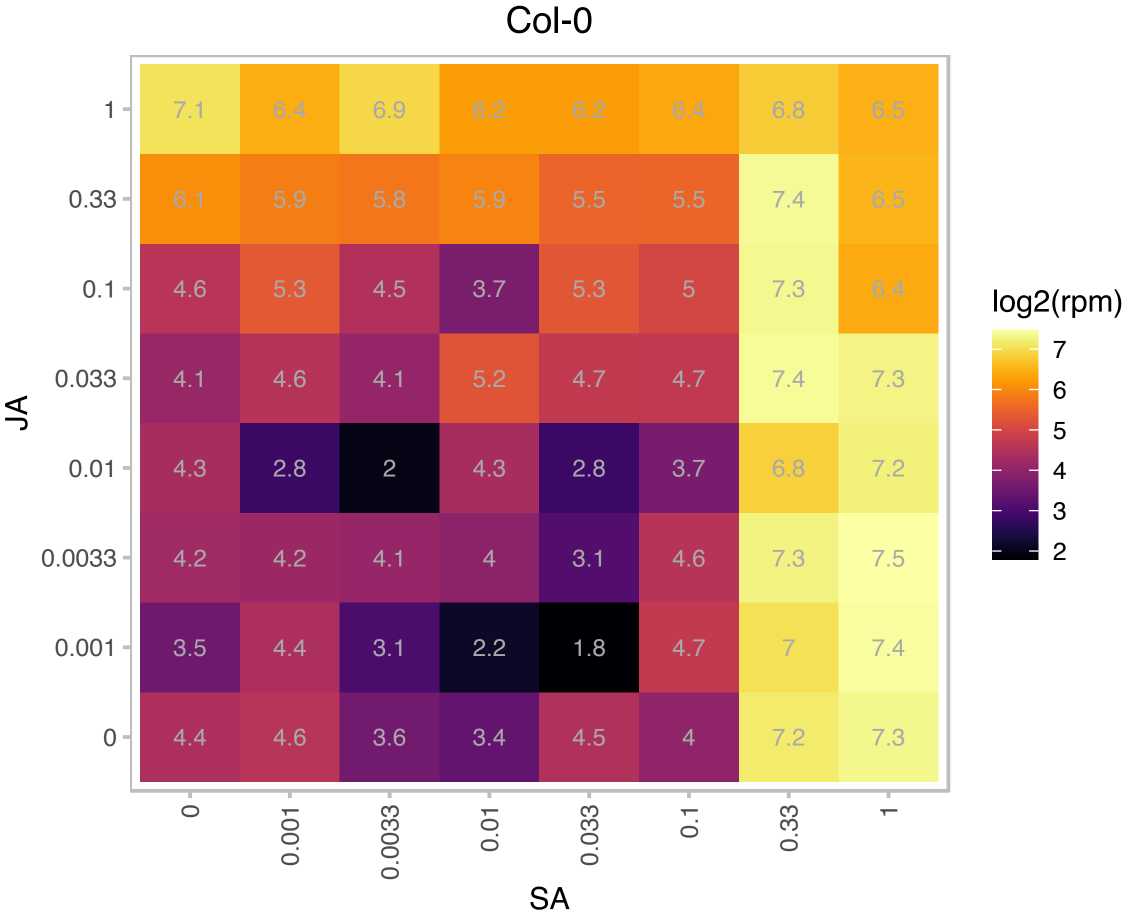


| | |
|---|---|
| Gene | AT5G10830 |
| is.expressed_genes | TRUE |
| is.responsive_genes | TRUE |
| is.apcluster | B-04 |

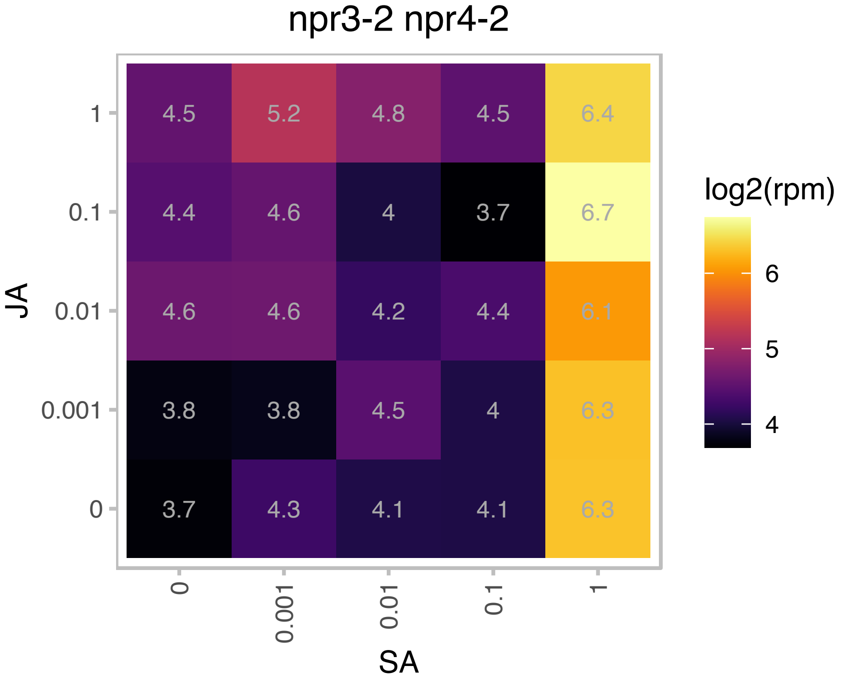


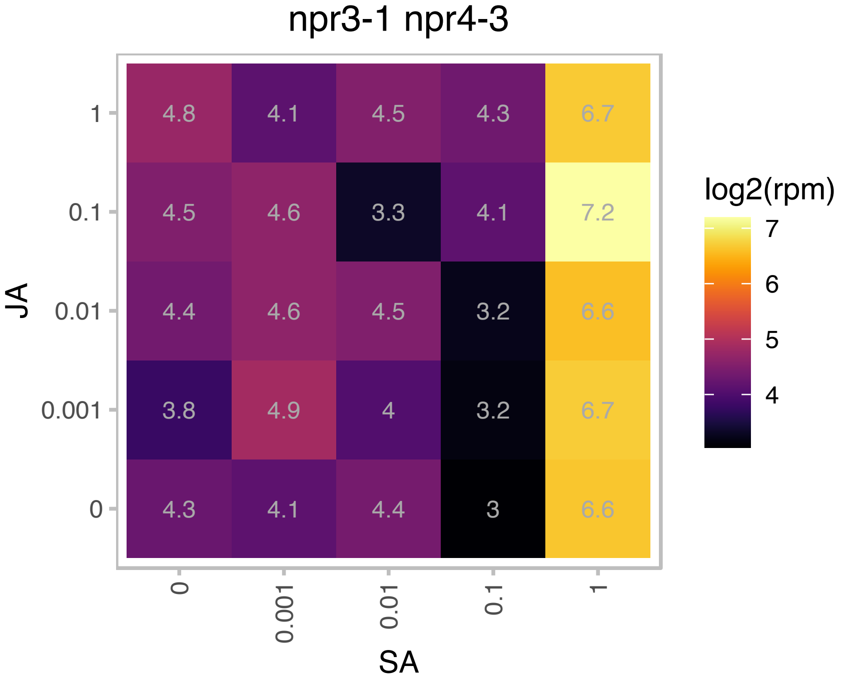

### AT5G13750.1 : ZIFL1

zinc induced facilitator-like 1

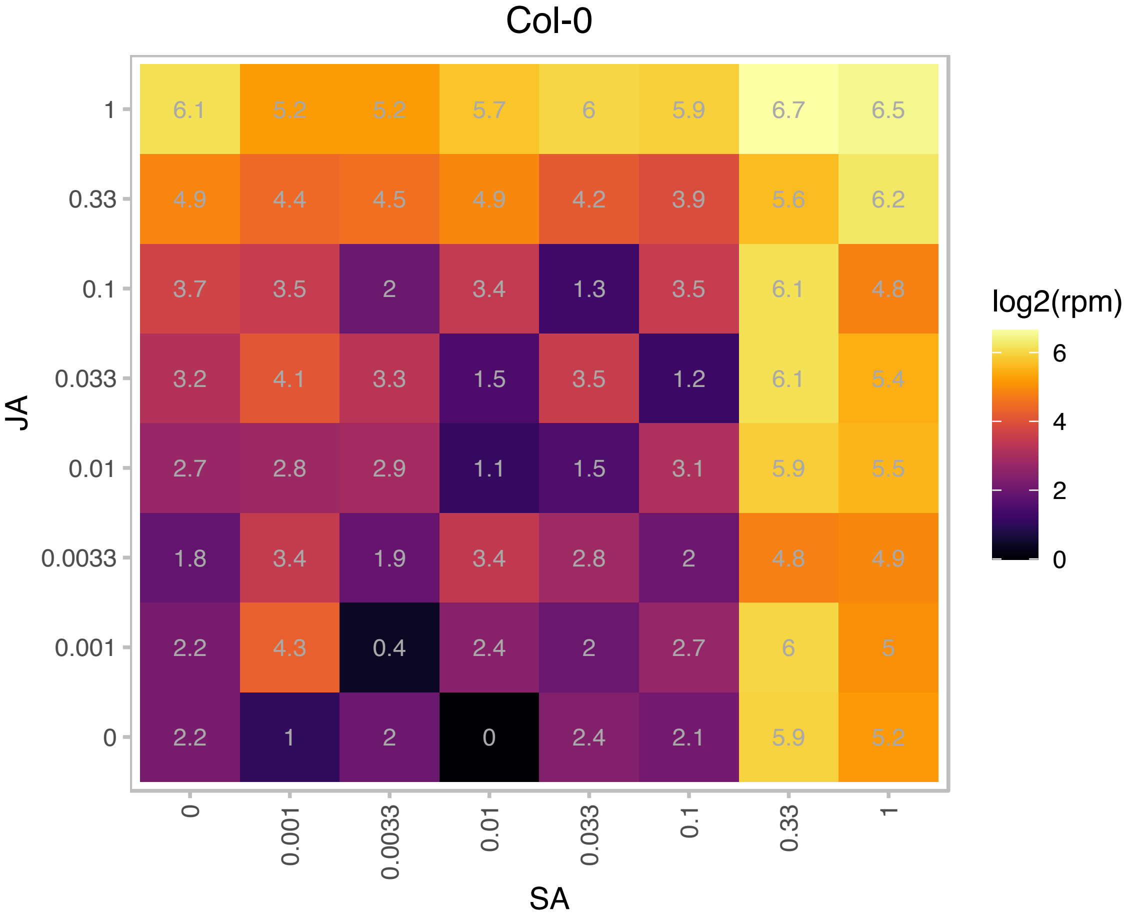


| | |
|---|---|
| Gene | AT5G13750 |
| is.expressed_genes | TRUE |
| is.responsive_genes | TRUE |
| is.apcluster | B-04 |

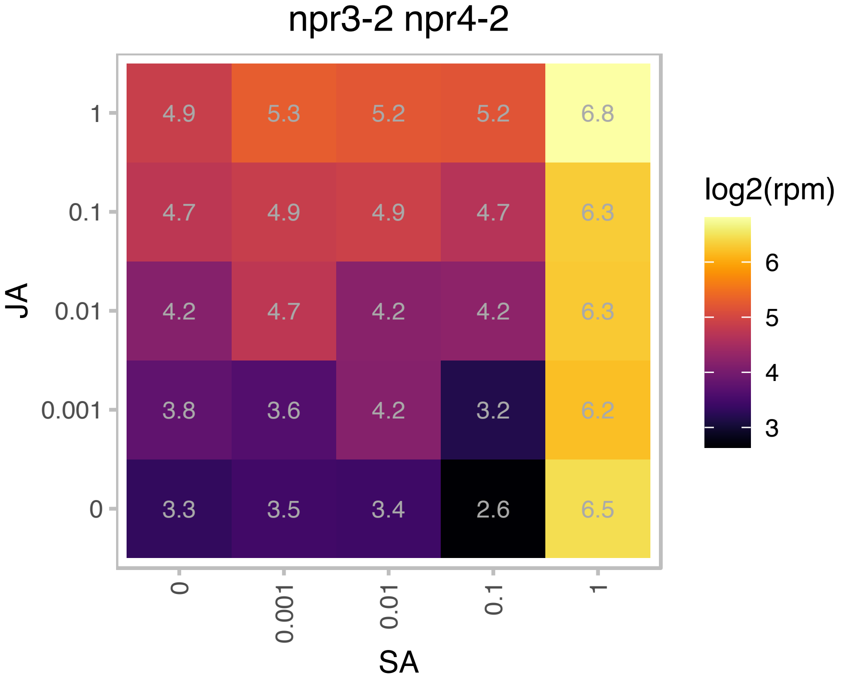


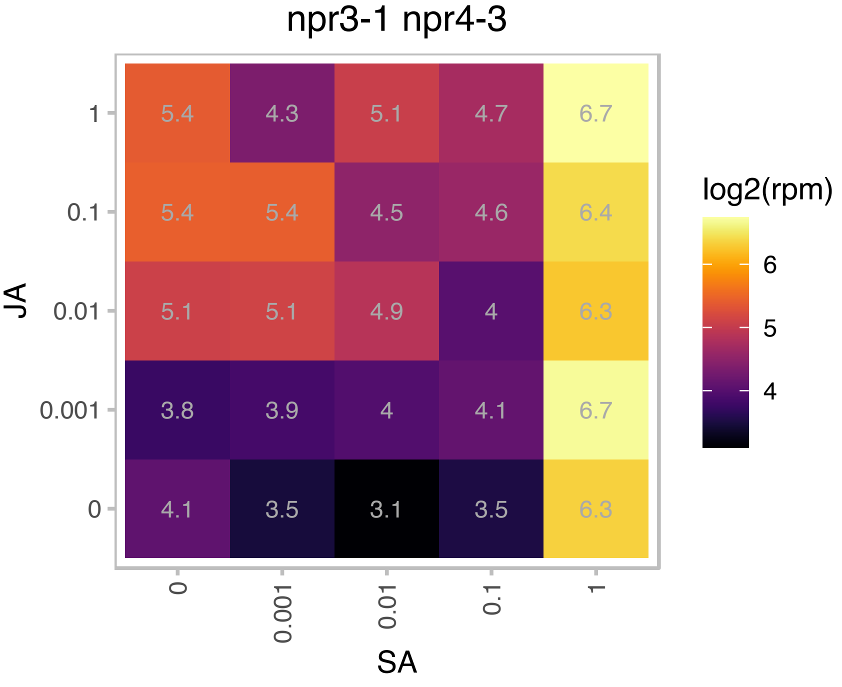

# AT5G16980.1 : NA

Zinc-binding dehydrogenase family protein

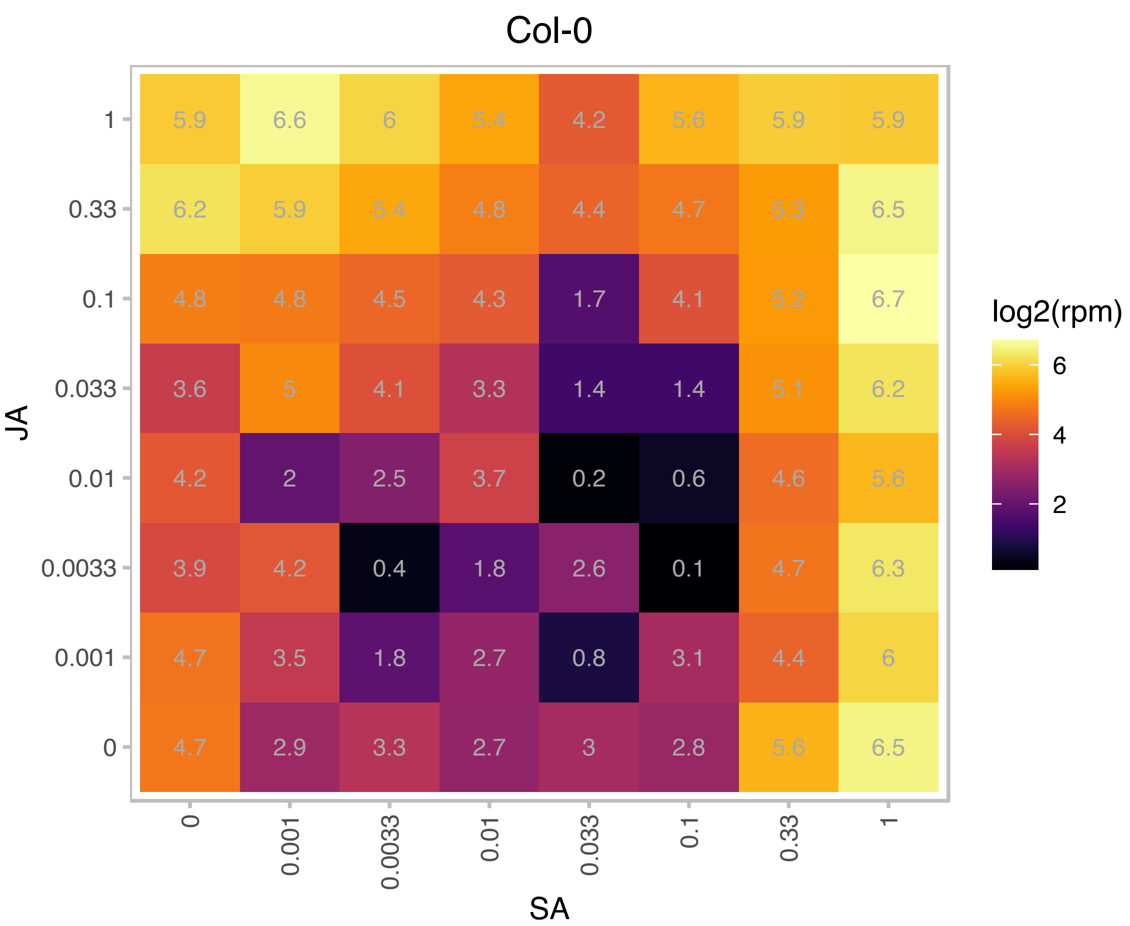


| Gene | AT5G16980 |
|---|---|
| is.expressed_genes | TRUE |
| is.responsive_genes | TRUE |
| is.apcluster | B-04 |

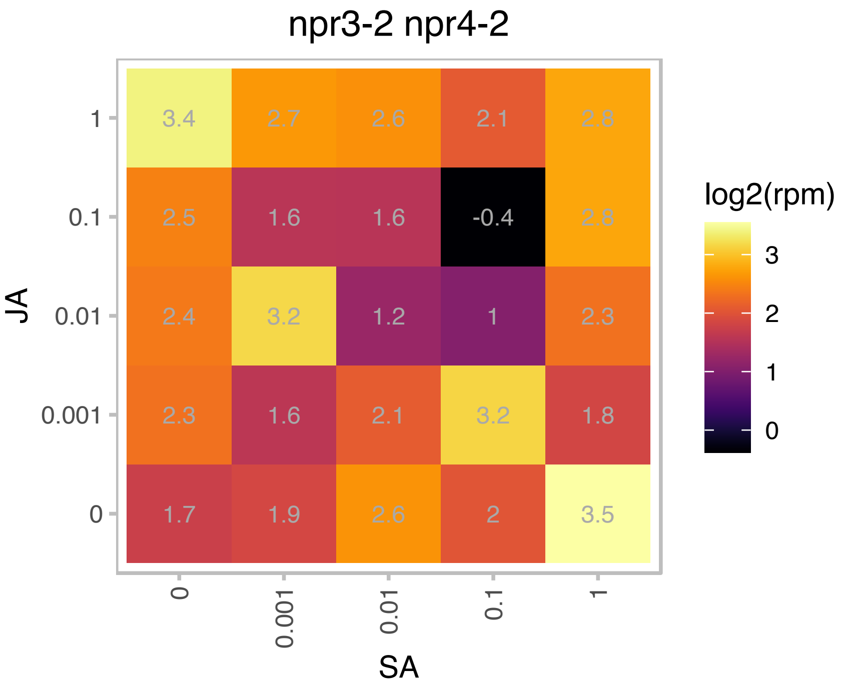


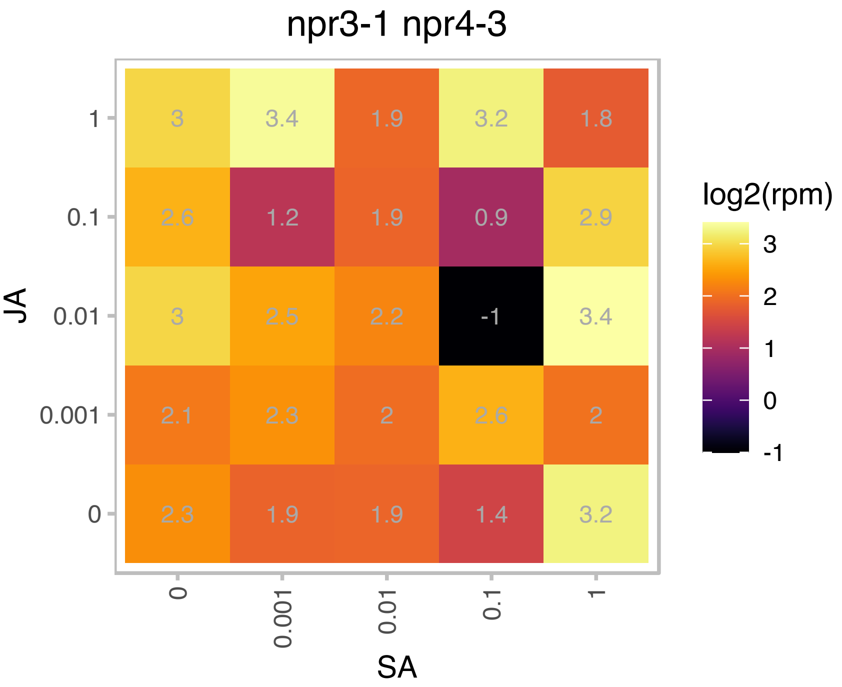

# AT5G17190.1 : NA

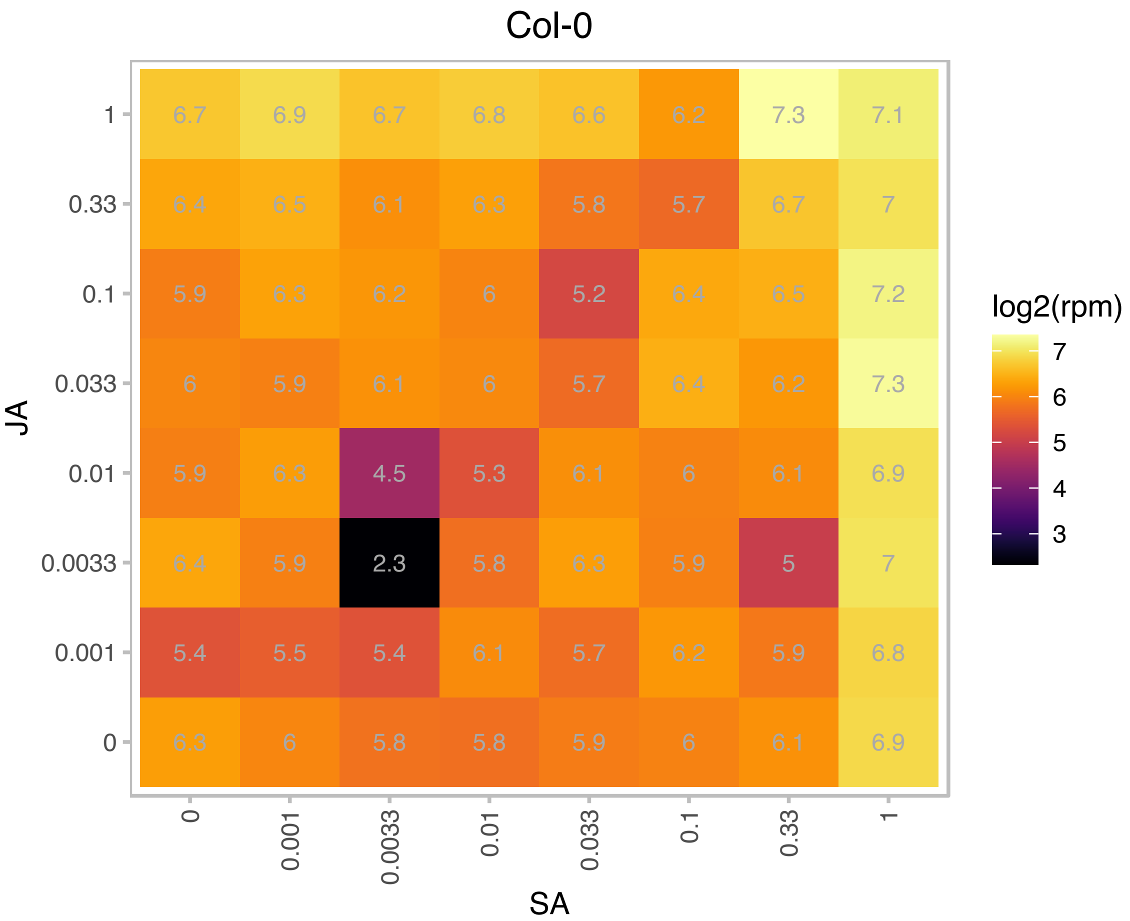


| | |
|---|---|
| Gene | AT5G17190 |
| is.expressed_genes | TRUE |
| is.responsive_genes | TRUE |
| is.apcluster | B-04 |

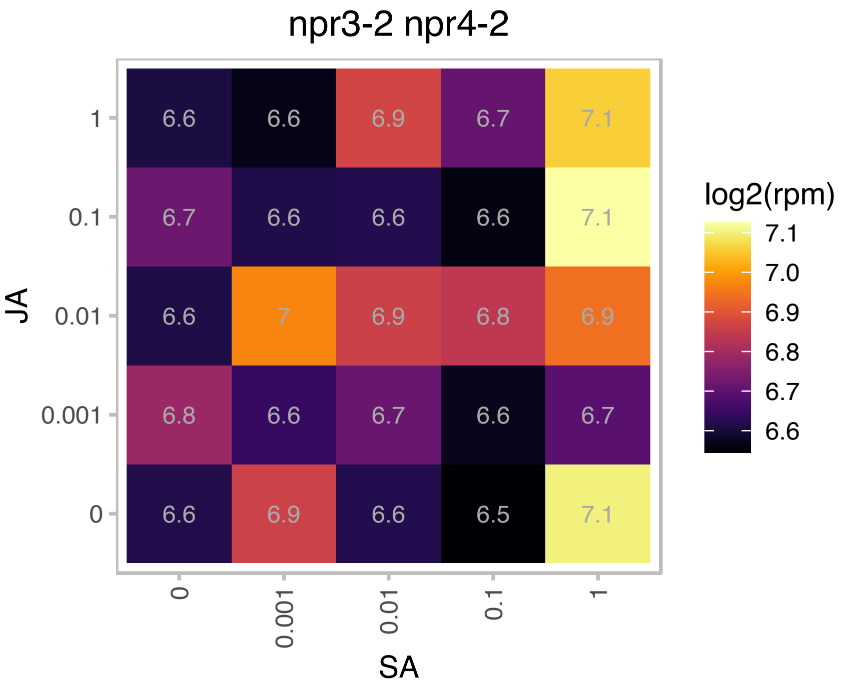


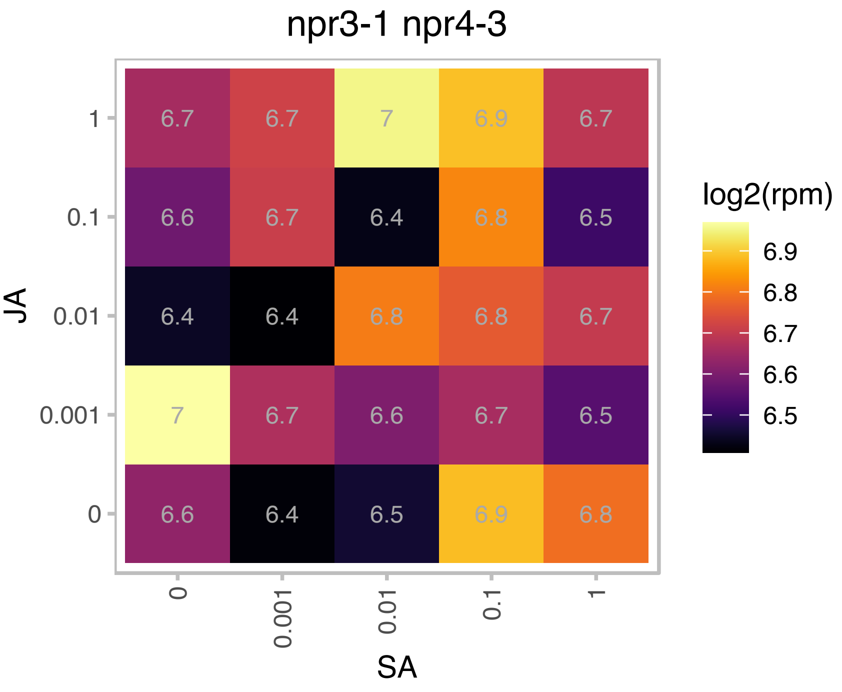

# AT5G22140.1 : NA

FAD/NAD(P)-binding oxidoreductase family protein

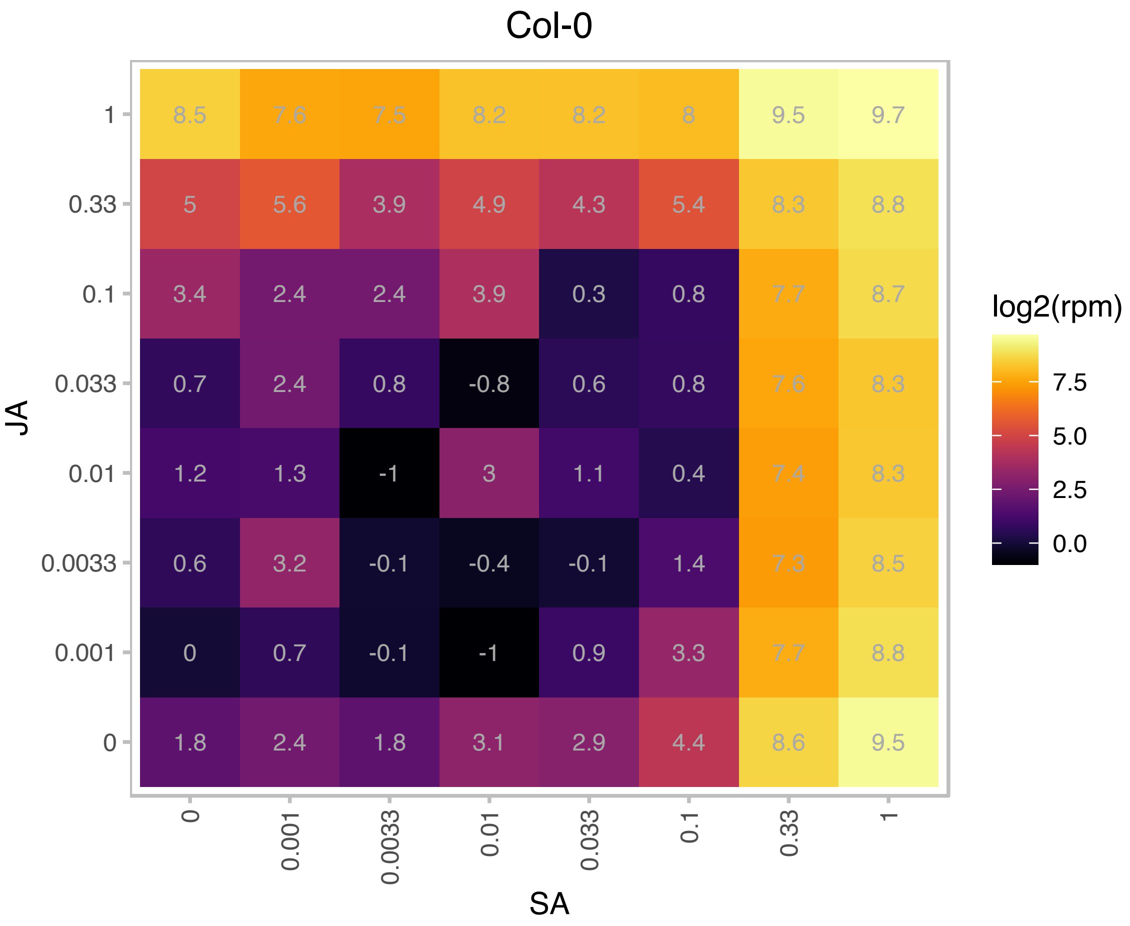


| | |
|---|---|
| Gene | AT5G22140 |
| is.expressed_genes | TRUE |
| is.responsive_genes | TRUE |
| is.apcluster | B-04 |

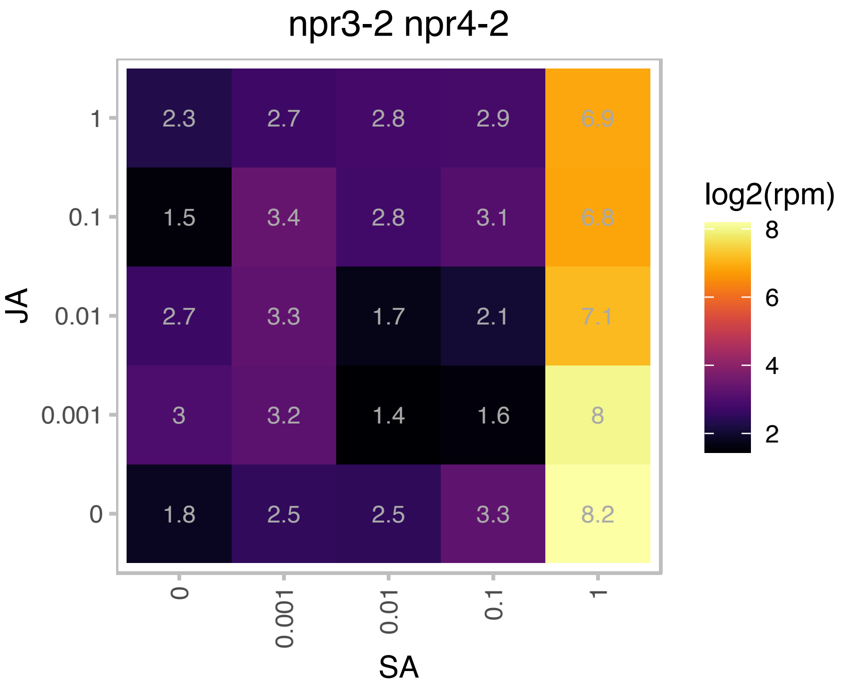


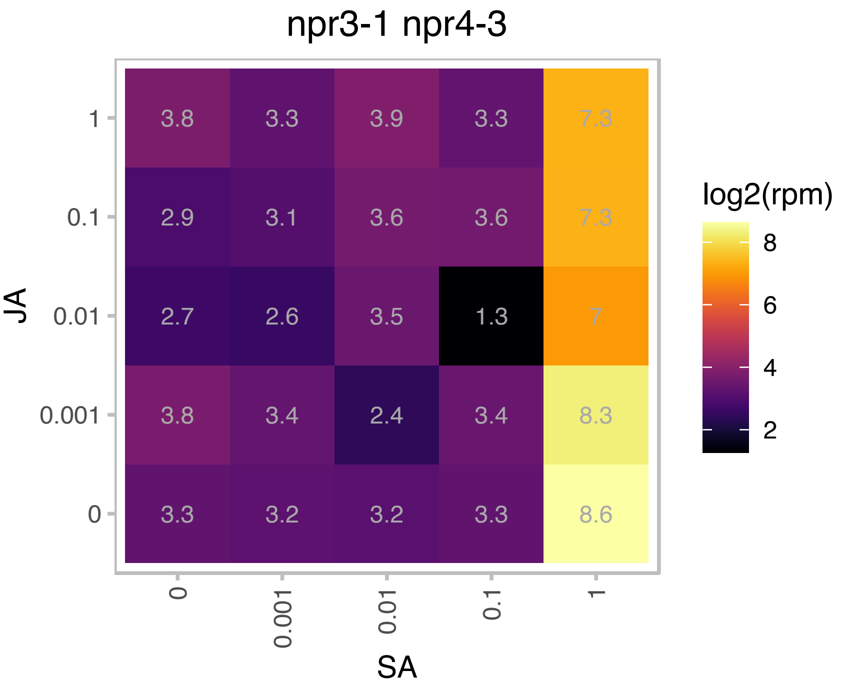

### AT5G33290.1 : XGD1

xylogalacturonan deficient 1

Acts as a xylogalacturonan xylosyltransferase within the XGA biosynthesis pathway. Involved in pectin biosynthesis.

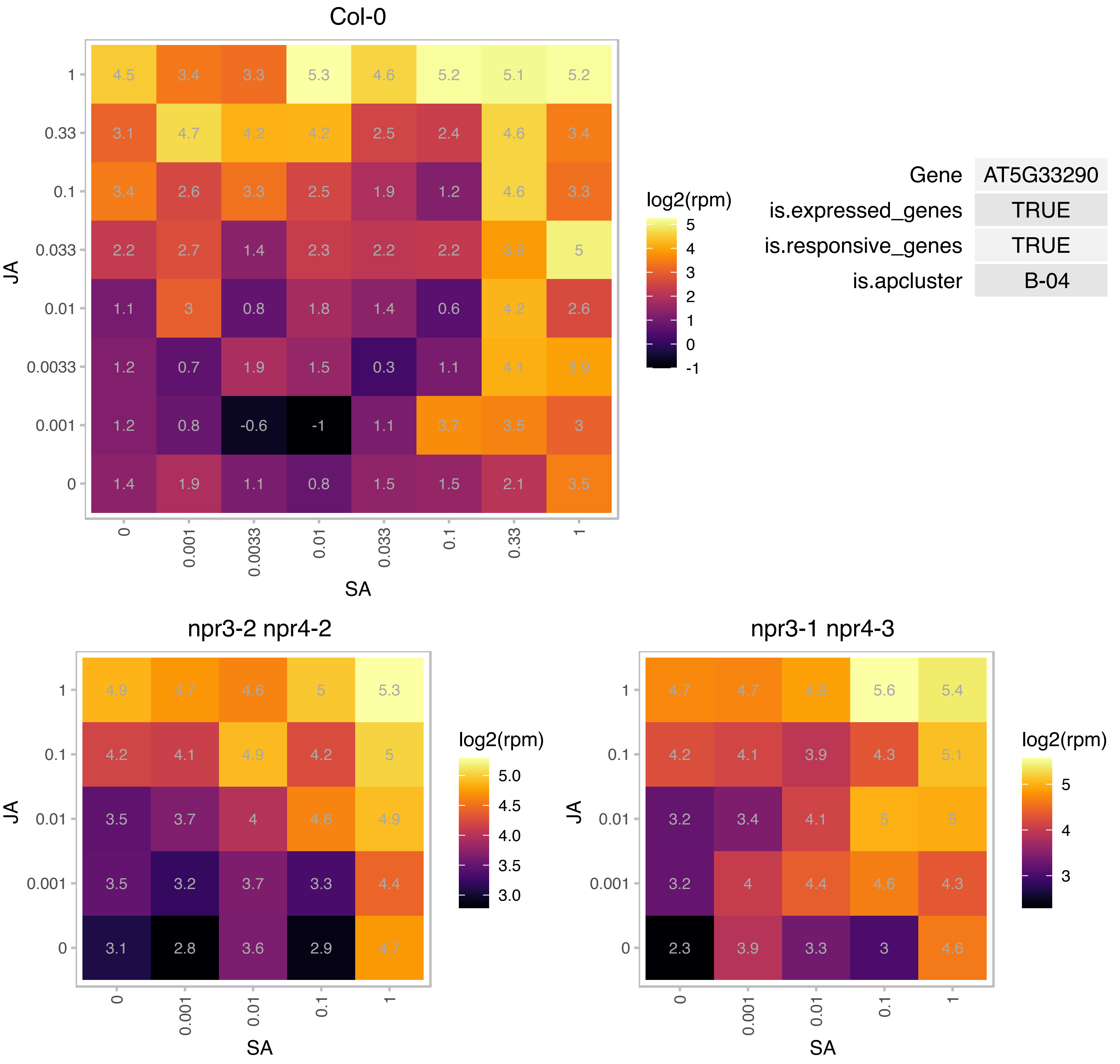


| | |
|---|---|
| Gene | AT5G33290 |
| is.expressed_genes | TRUE |
| is.responsive_genes | TRUE |
| is.apcluster | B-04 |

### AT5G40370.1 : AtGRXC2, GRX370, GRXC2

Glutaredoxin family protein

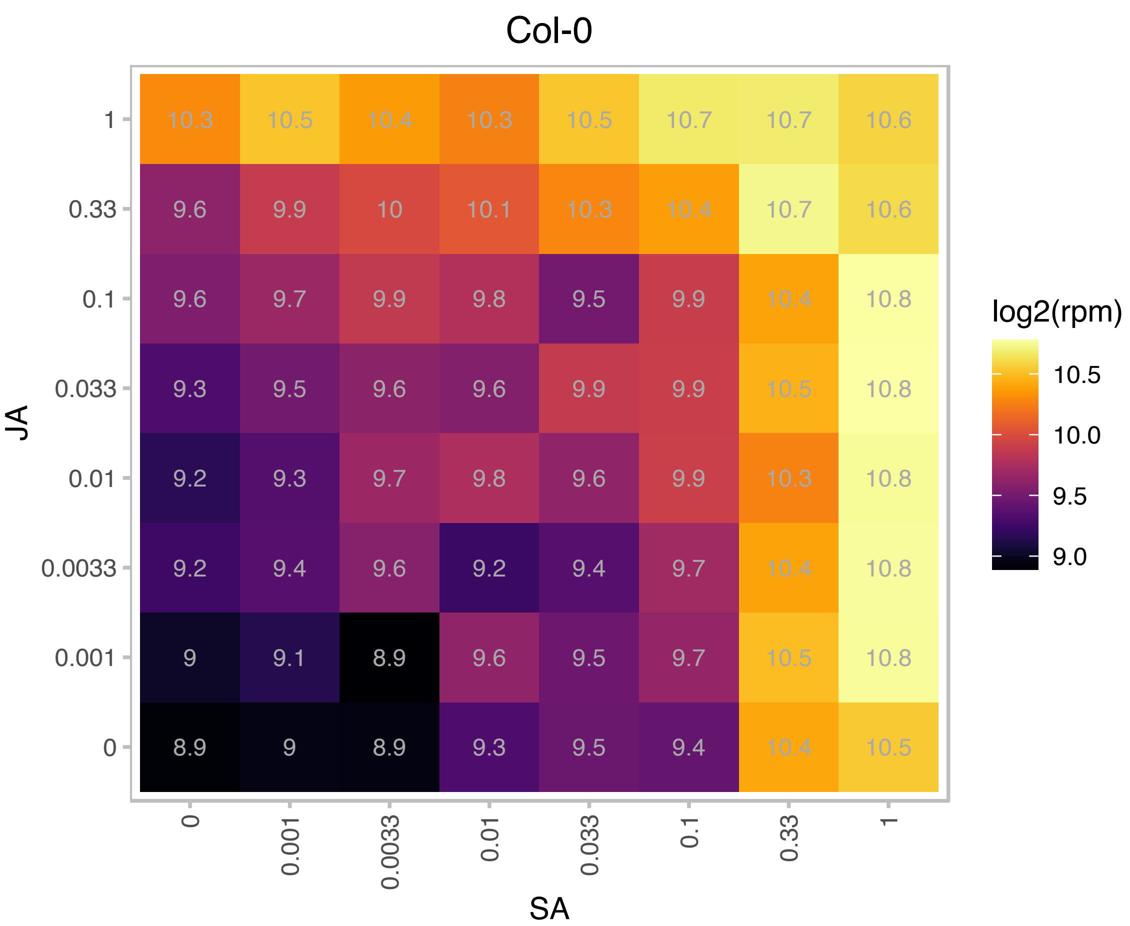


| | |
|---|---|
| Gene | AT5G40370 |
| is.expressed_genes | TRUE |
| is.responsive_genes | TRUE |
| is.apcluster | B-04 |

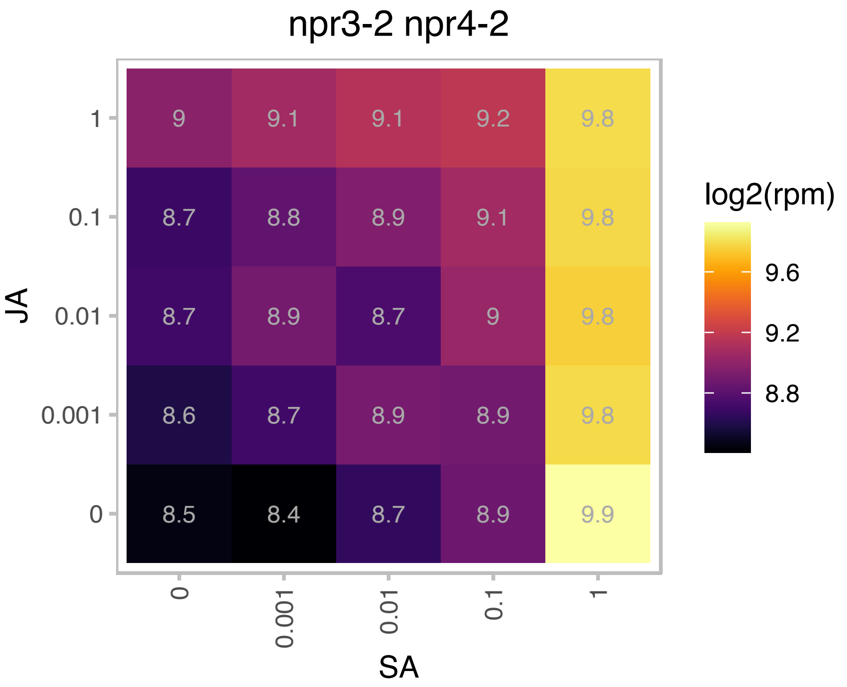


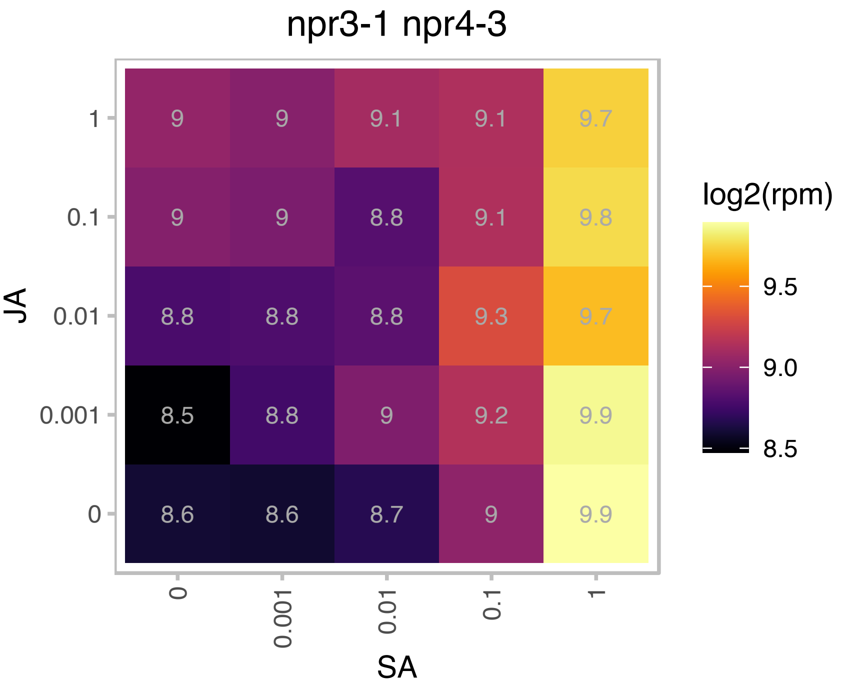

### AT5G49460.1 : ACLB-2

ATP citrate lyase subunit B 2

One of the two genes encoding subunit B of the cytosolic enzyme ATP Citrate Lyase (ACL)

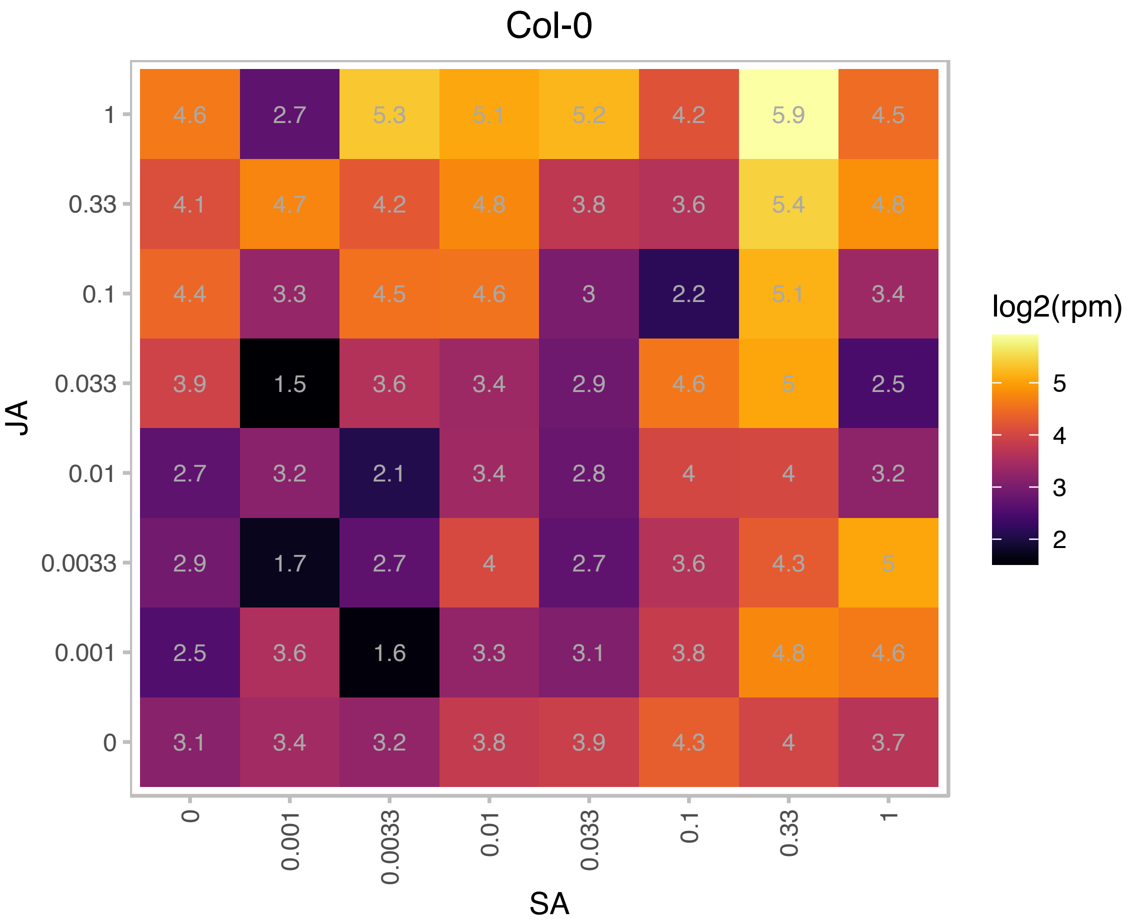


| | |
|---|---|
| Gene | AT5G49460 |
| is.expressed_genes | TRUE |
| is.responsive_genes | TRUE |
| is.apcluster | B-04 |

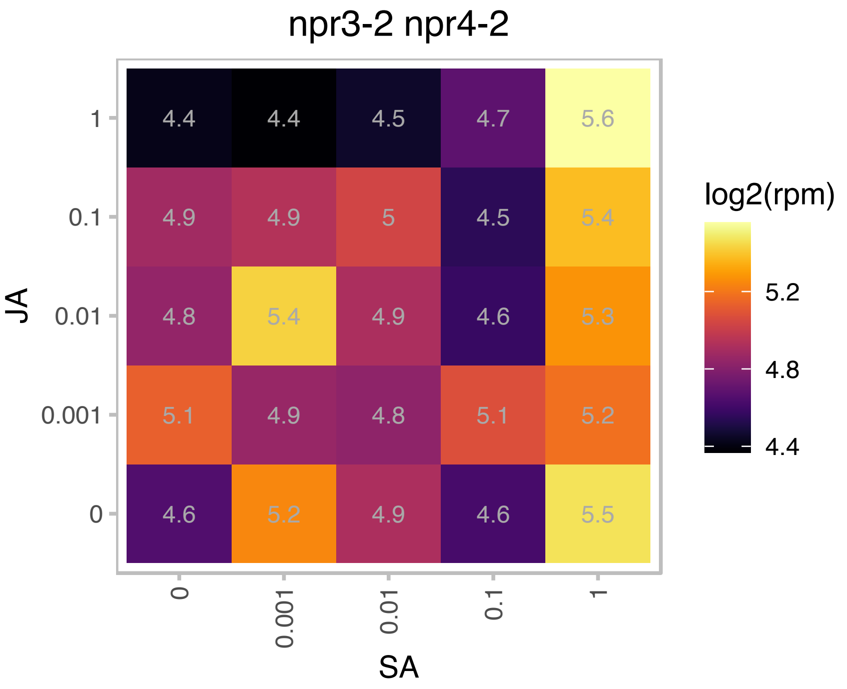


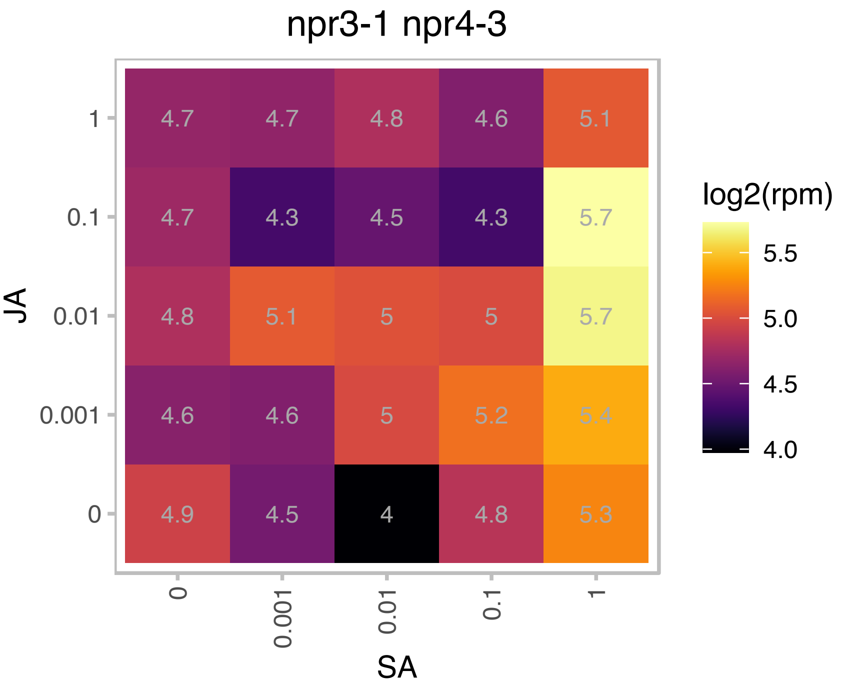

# AT5G54165.1 : NA

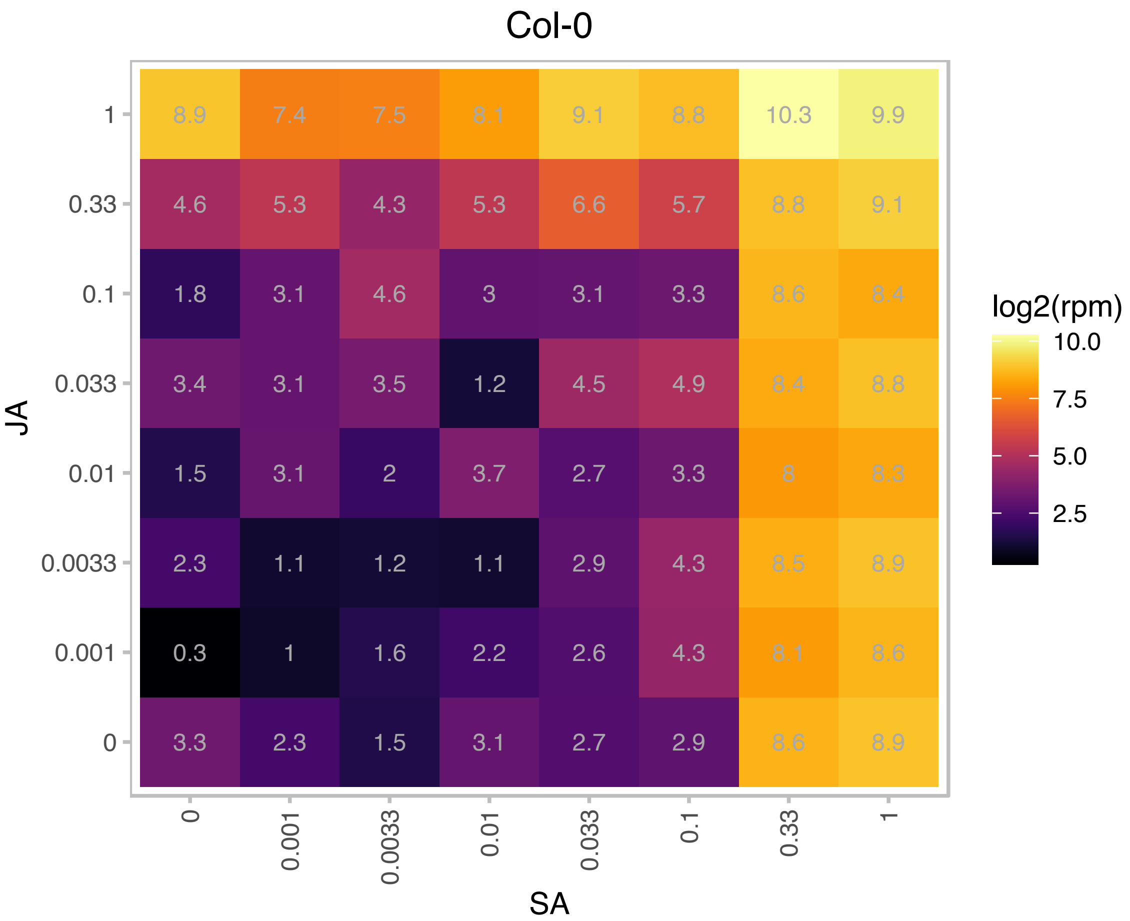


| | |
|---|---|
| Gene | AT5G54165 |
| is.expressed_genes | TRUE |
| is.responsive_genes | TRUE |
| is.apcluster | B-04 |

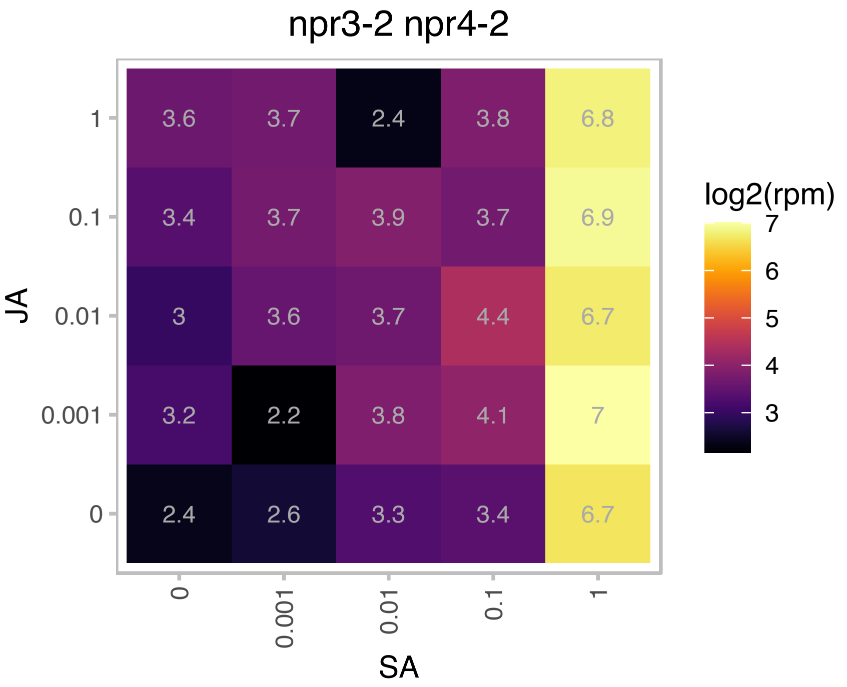


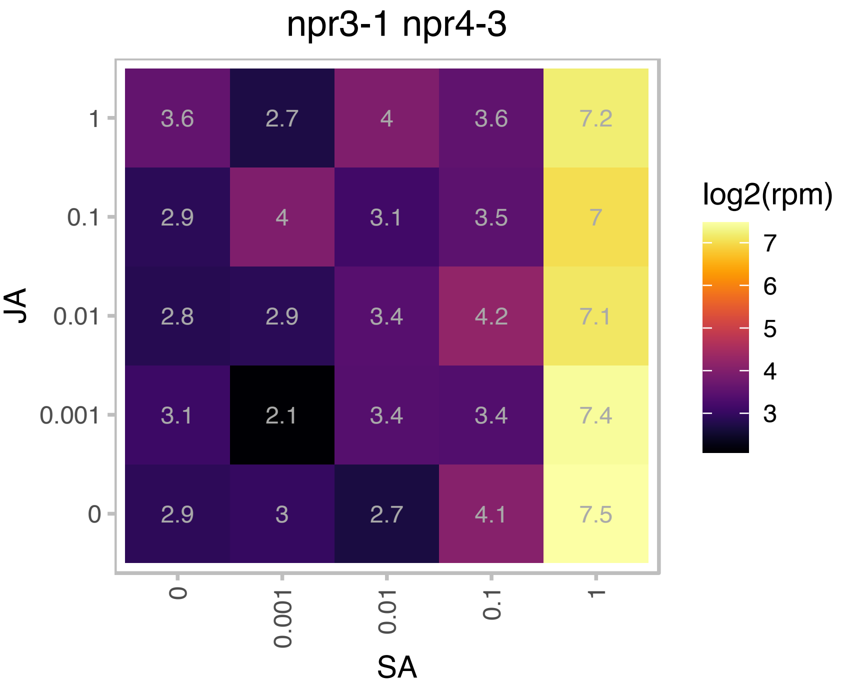

### AT5G54840.1 : ATSGP1, SGP1

Ras-related small GTP-binding family protein

Monomeric G protein. Expressed in the root quiescent center, flowers, and leaf guard cells and hydathodes.

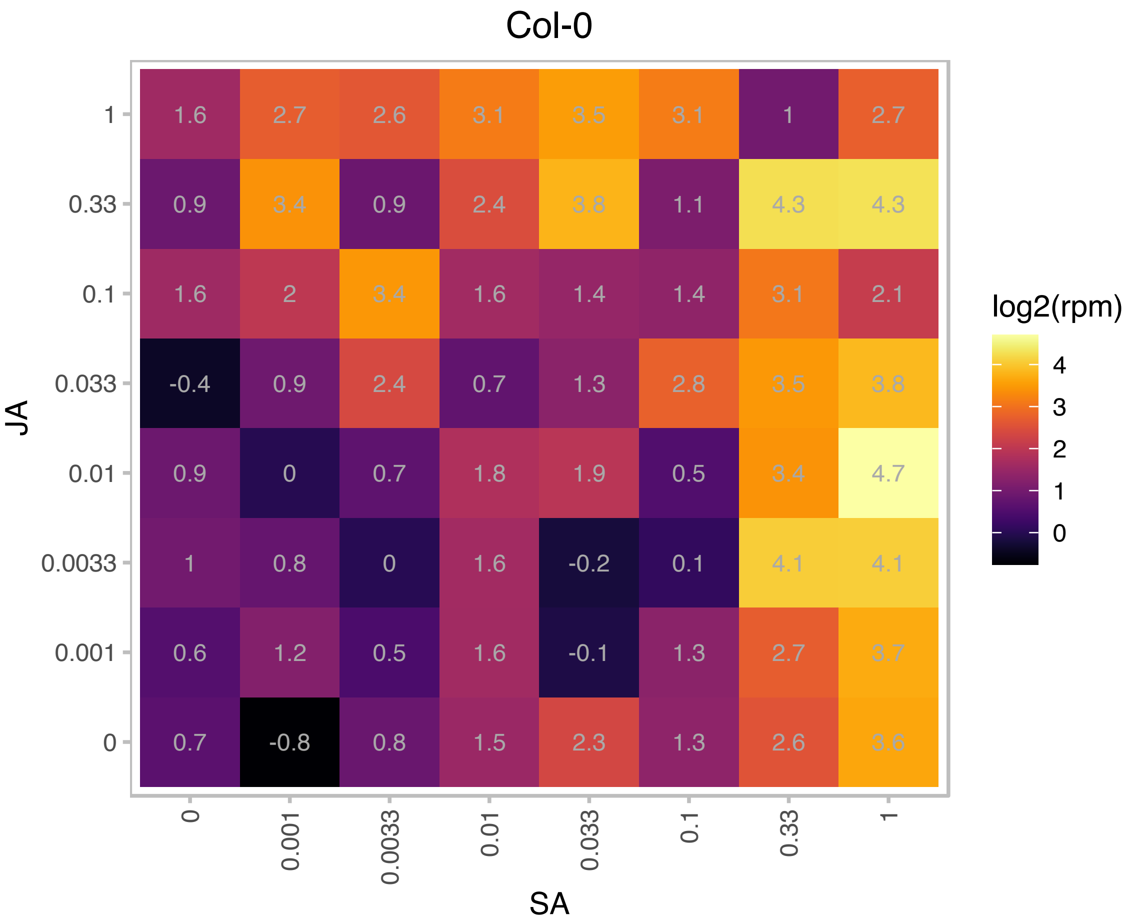


| | |
|---|---|
| Gene | AT5G54840 |
| is.expressed_genes | TRUE |
| is.responsive_genes | TRUE |
| is.apcluster | B-04 |

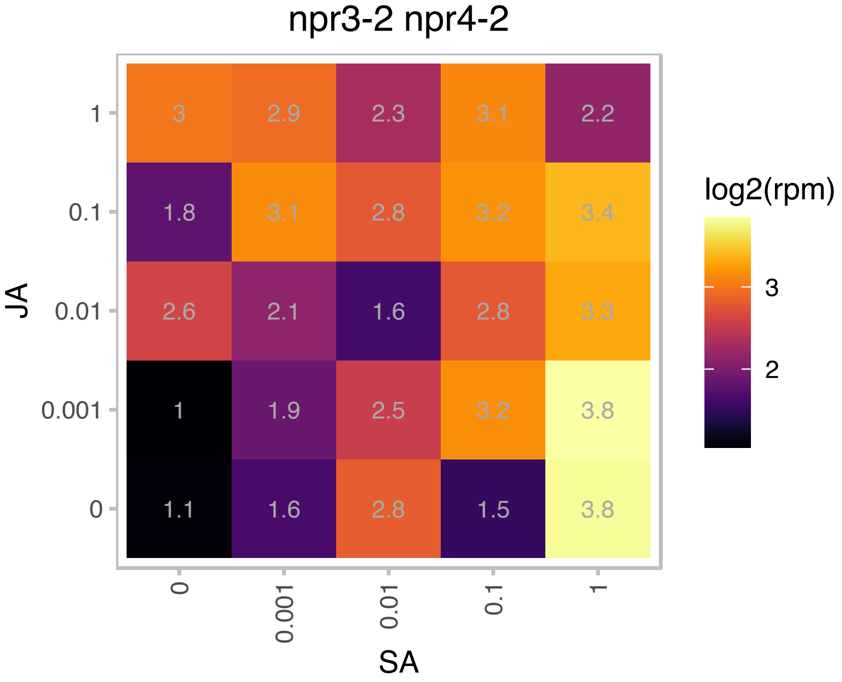


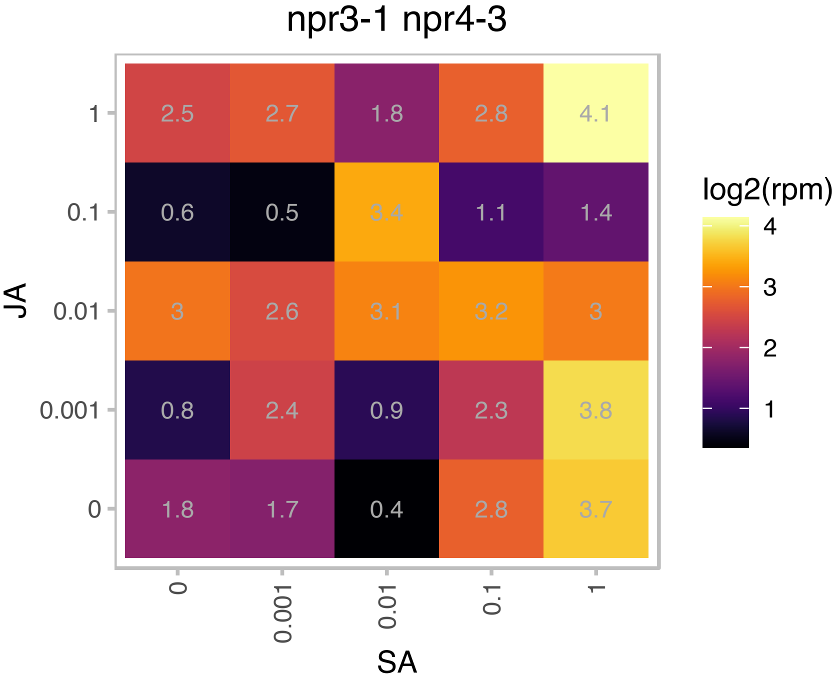

### AT5G59510.1 : DVL18, RTFL5

ROTUNDIFOLIA like 5

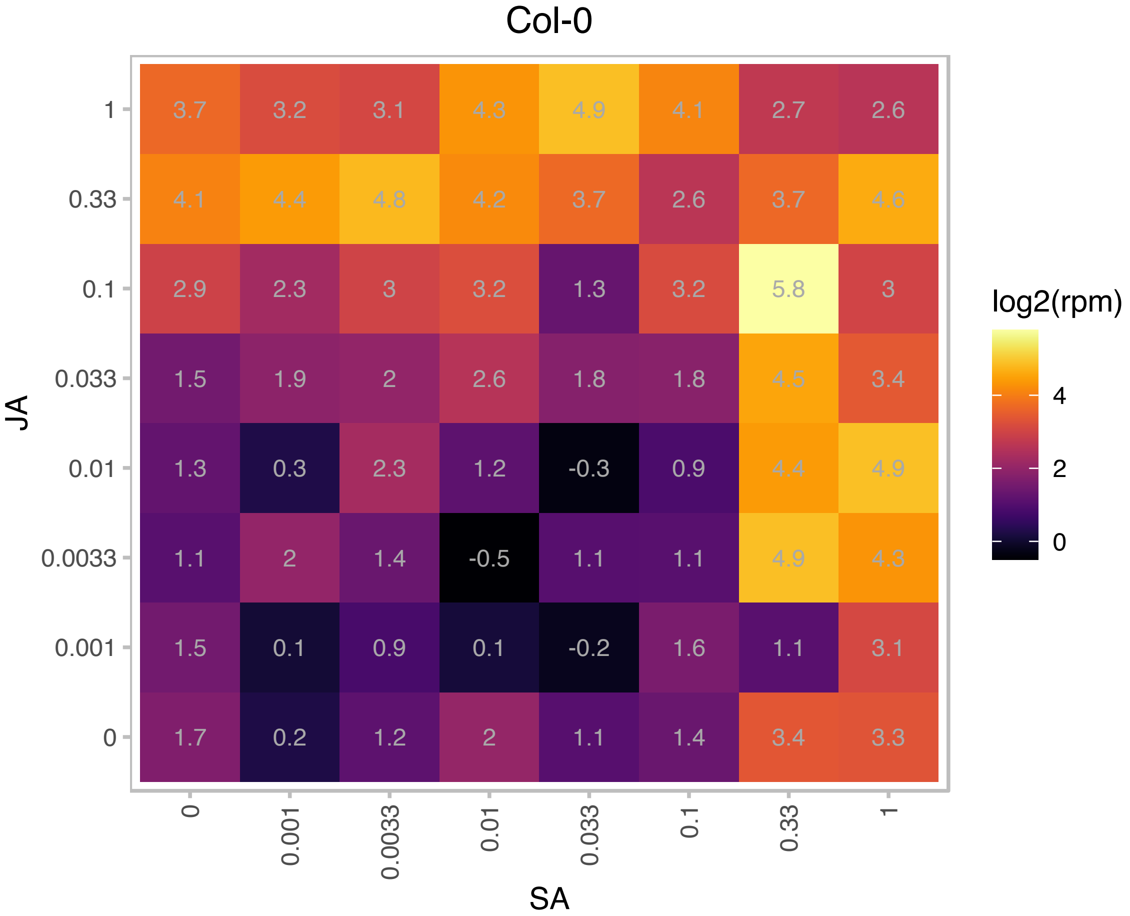


| | |
|---|---|
| Gene | AT5G59510 |
| is.expressed_genes | TRUE |
| is.responsive_genes | TRUE |
| is.apcluster | B-04 |

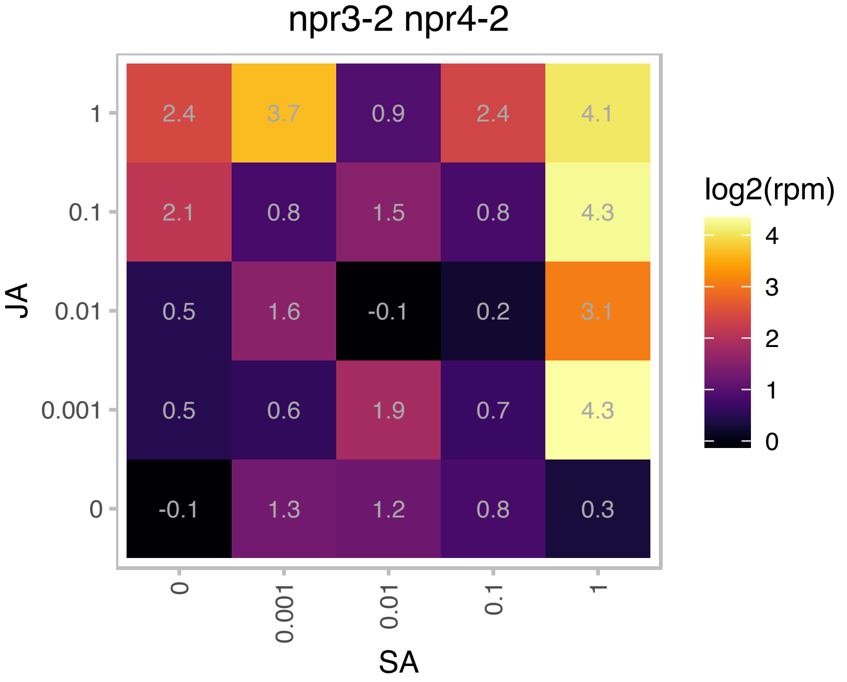


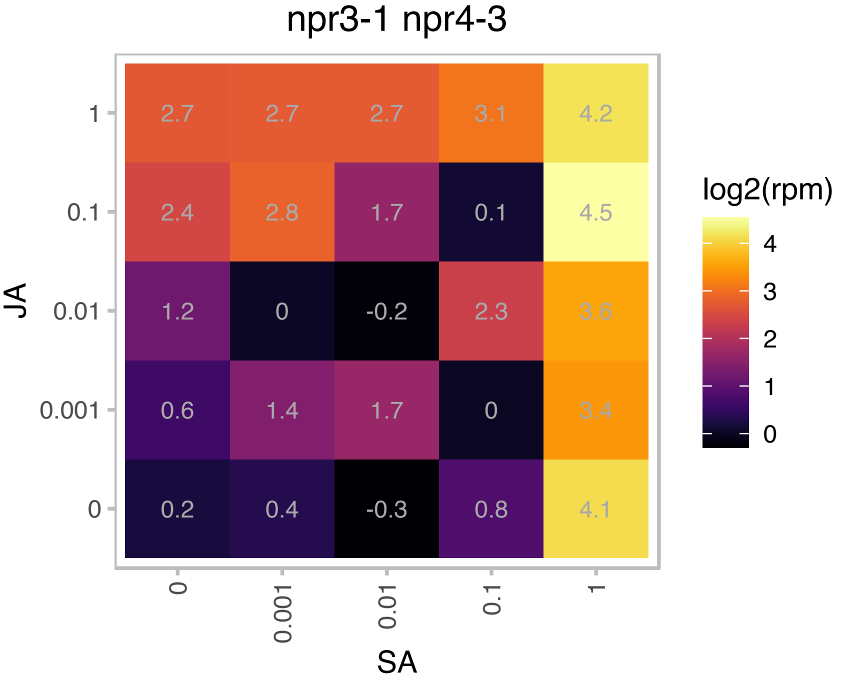

# AT5G67350.1 : NA

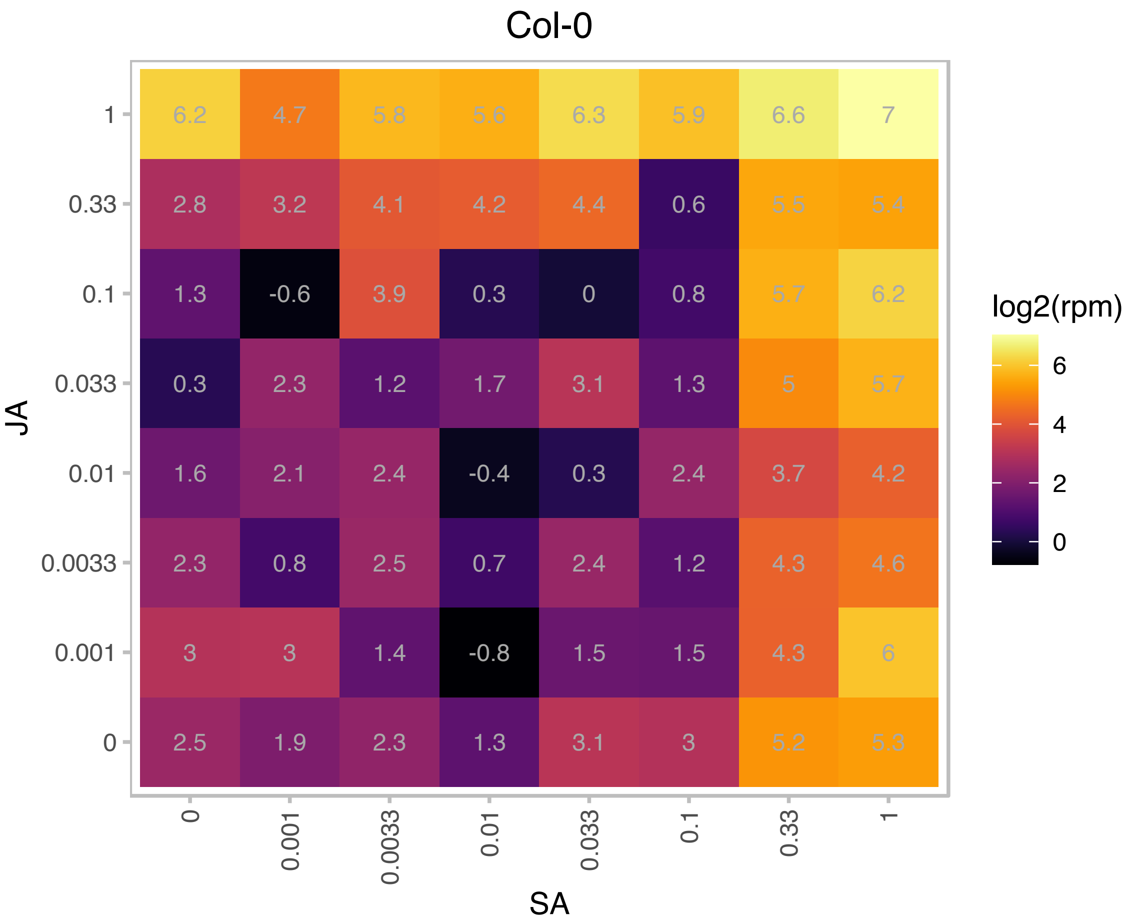


| | |
|---|---|
| Gene | AT5G67350 |
| is.expressed_genes | TRUE |
| is.responsive_genes | TRUE |
| is.apcluster | B-04 |

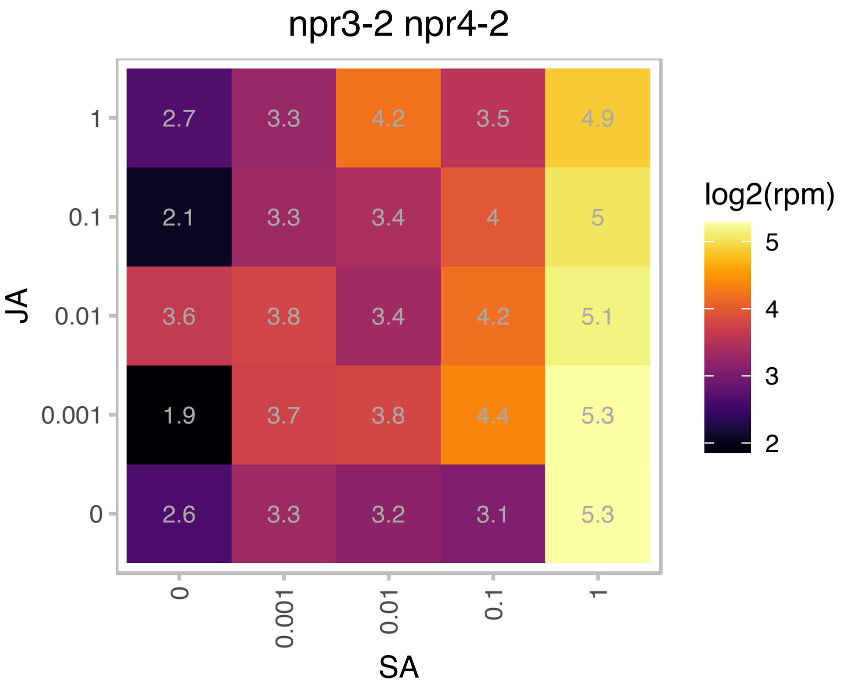


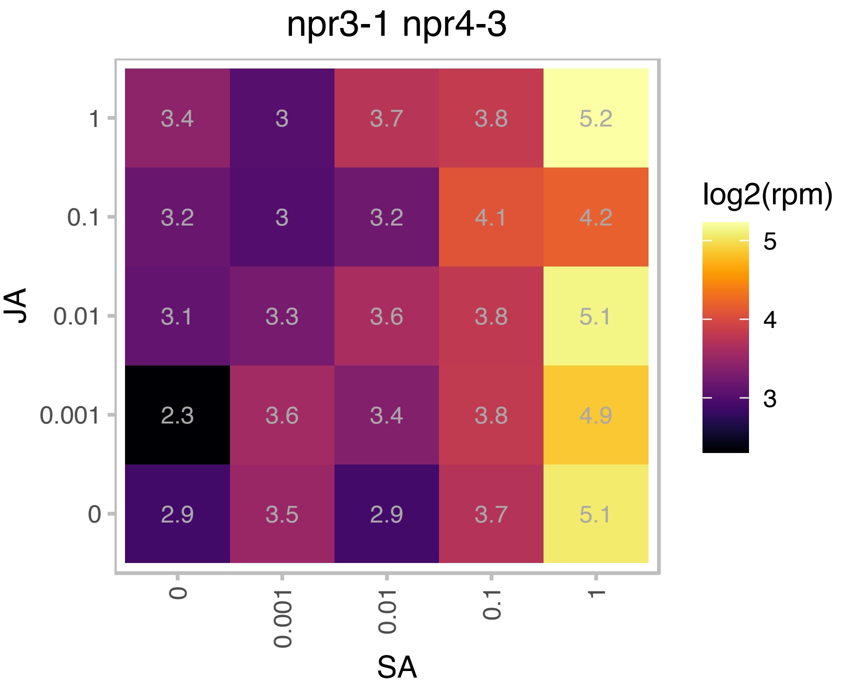

### AT1G02880.1 : TPK1

thiamin pyrophosphokinase1

Encodes a thiamine pyrophosphokinase capable of producing thiamine pyrophosphate from free thiamine.

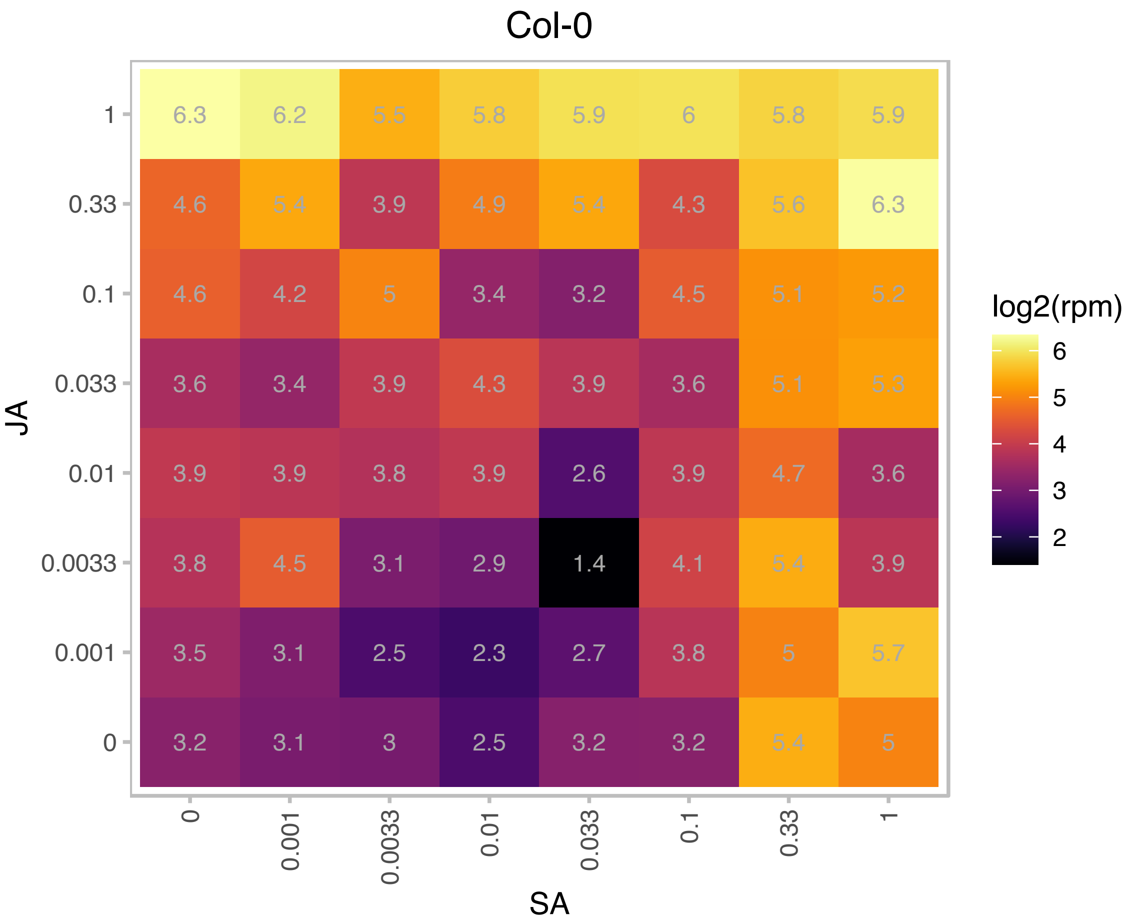


| | |
|---|---|
| Gene | AT1G02880 |
| is.expressed_genes | TRUE |
| is.responsive_genes | TRUE |
| is.apcluster | B-05 |

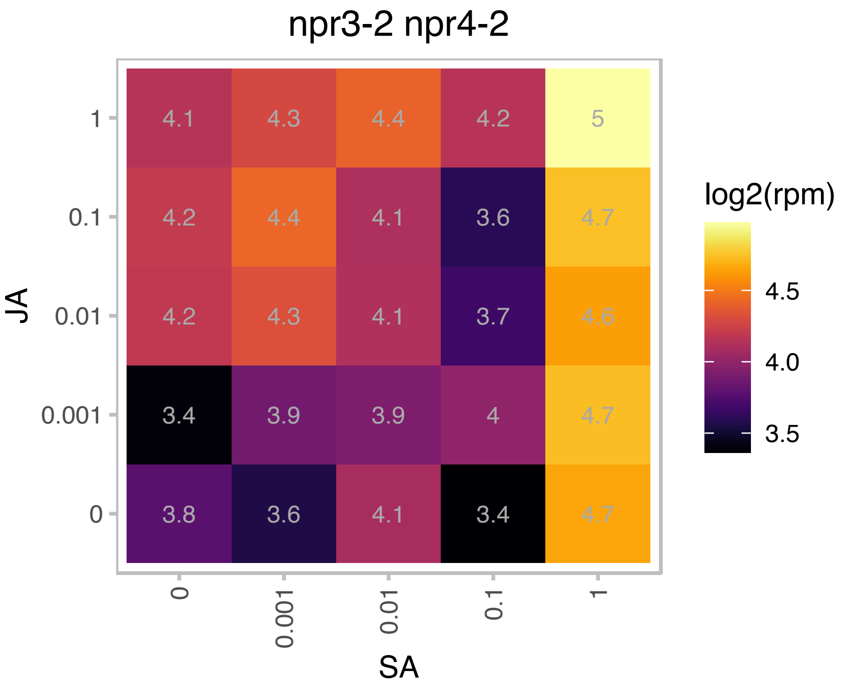


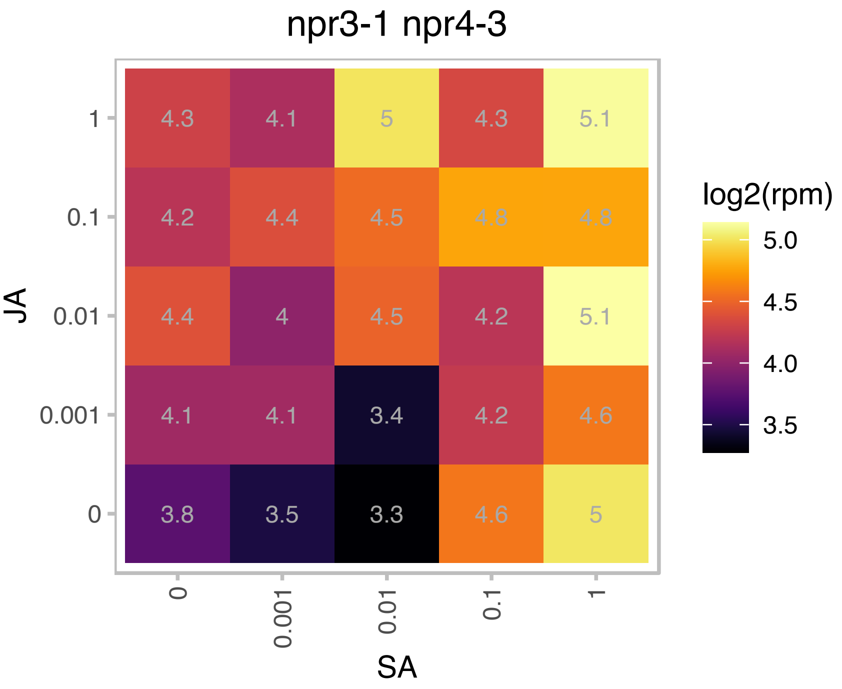

### AT1G05100.1 : MAPKKK18

mitogen-activated protein kinase kinase kinase 18

member of MEKK subfamily

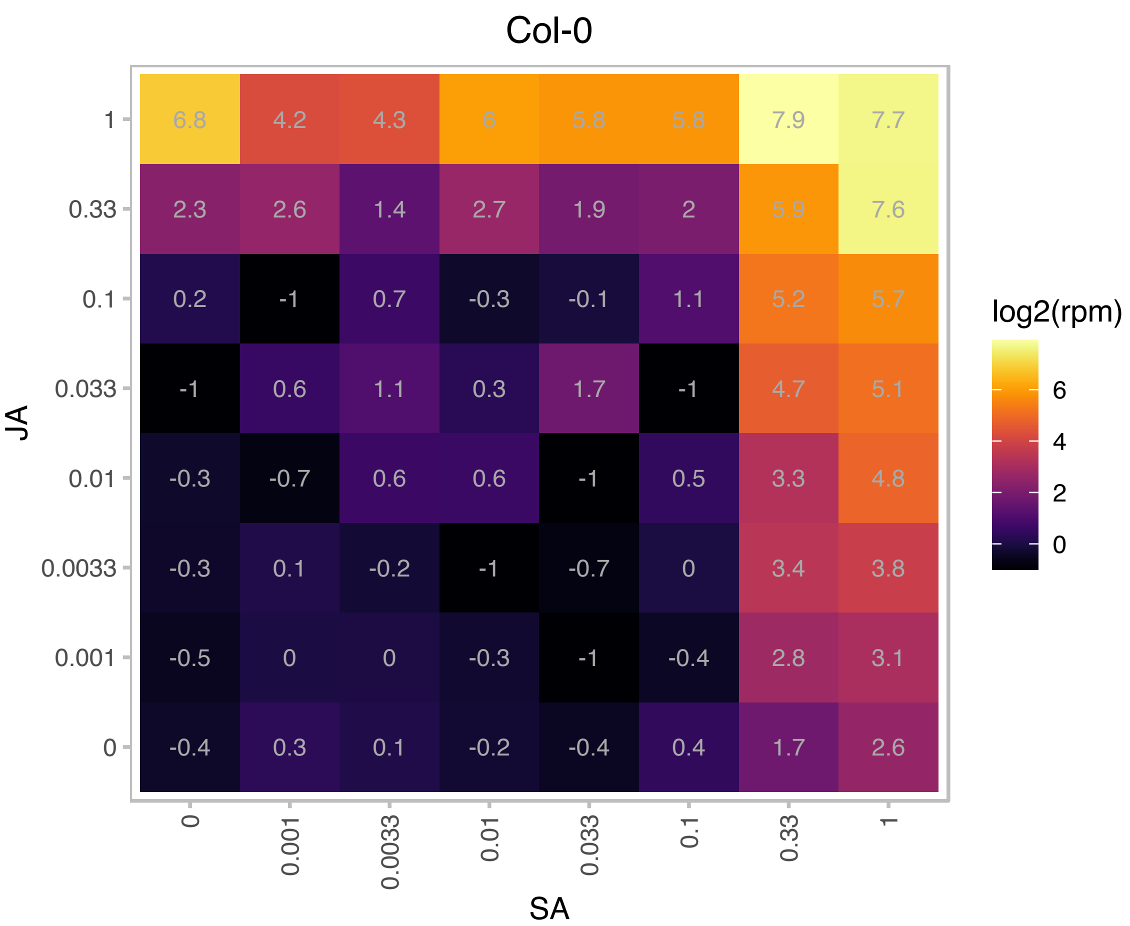


| | |
|---|---|
| Gene | AT1G05100 |
| is.expressed_genes | TRUE |
| is.responsive_genes | TRUE |
| is.apcluster | B-05 |

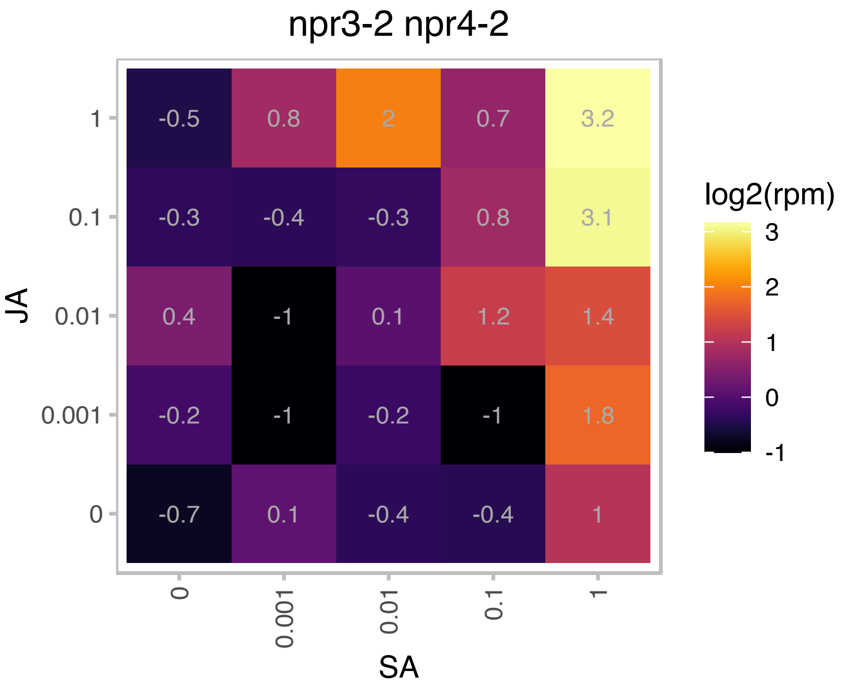


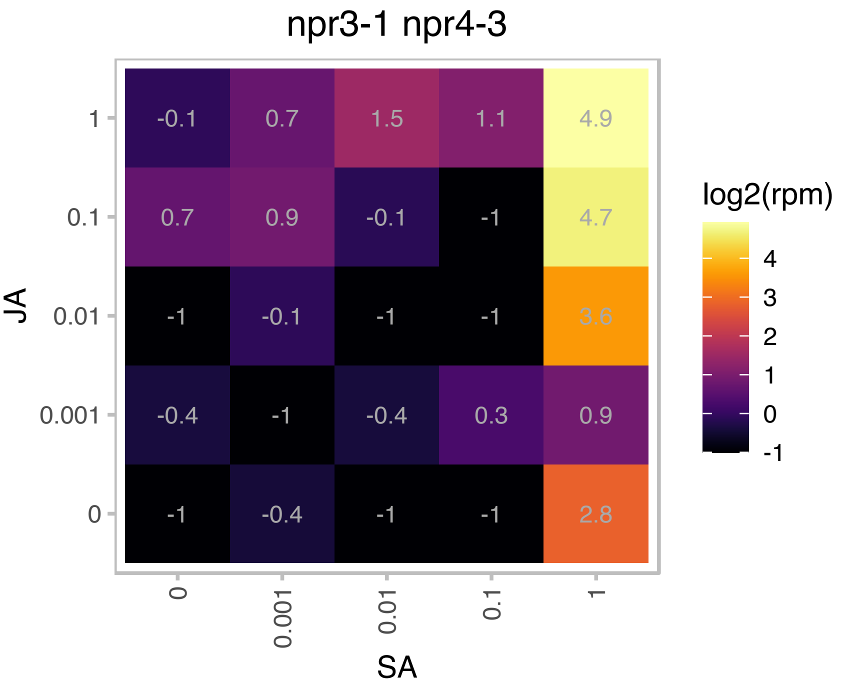

# AT1G05960.1 : NA

ARM repeat superfamily protein

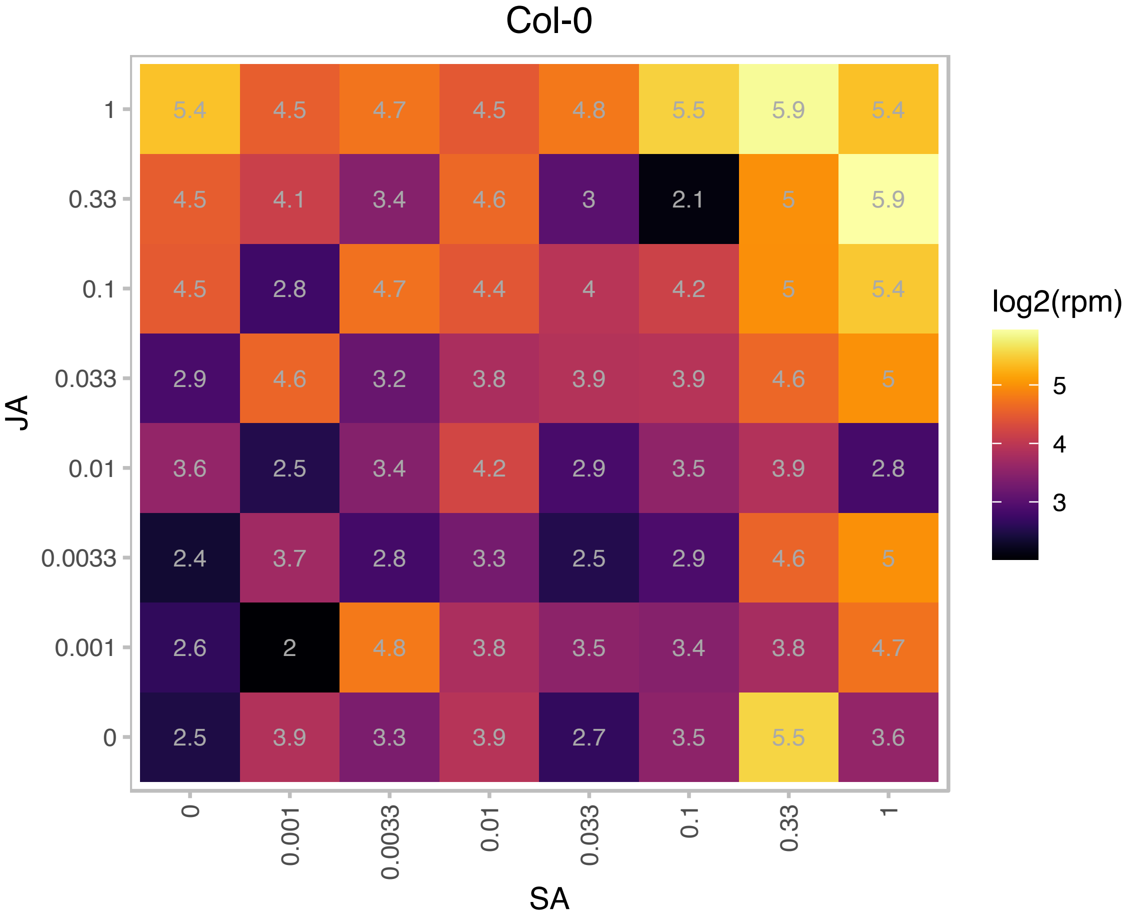


| | |
|---|---|
| Gene | AT1G05960 |
| is.expressed_genes | TRUE |
| is.responsive_genes | TRUE |
| is.apcluster | B-05 |

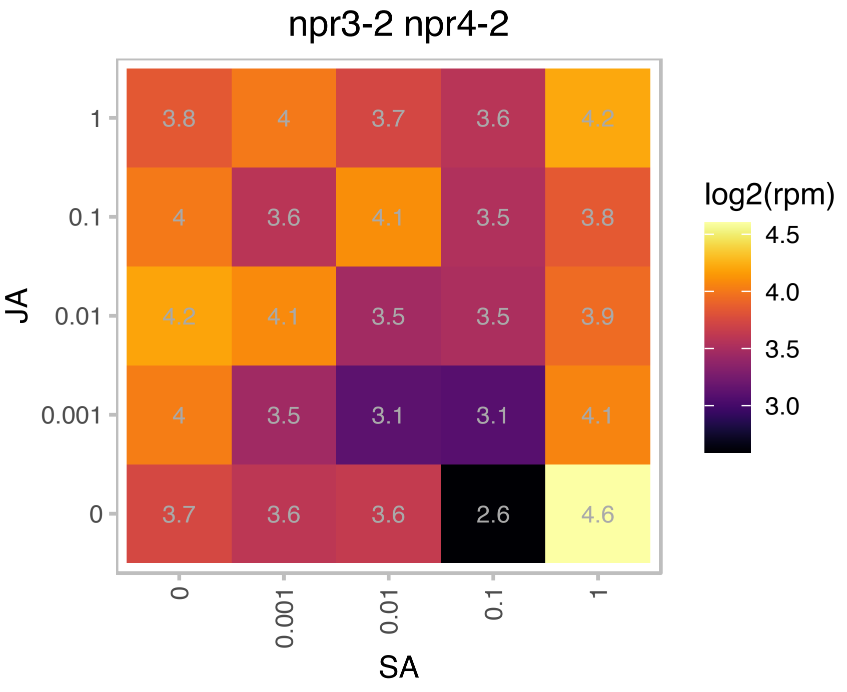


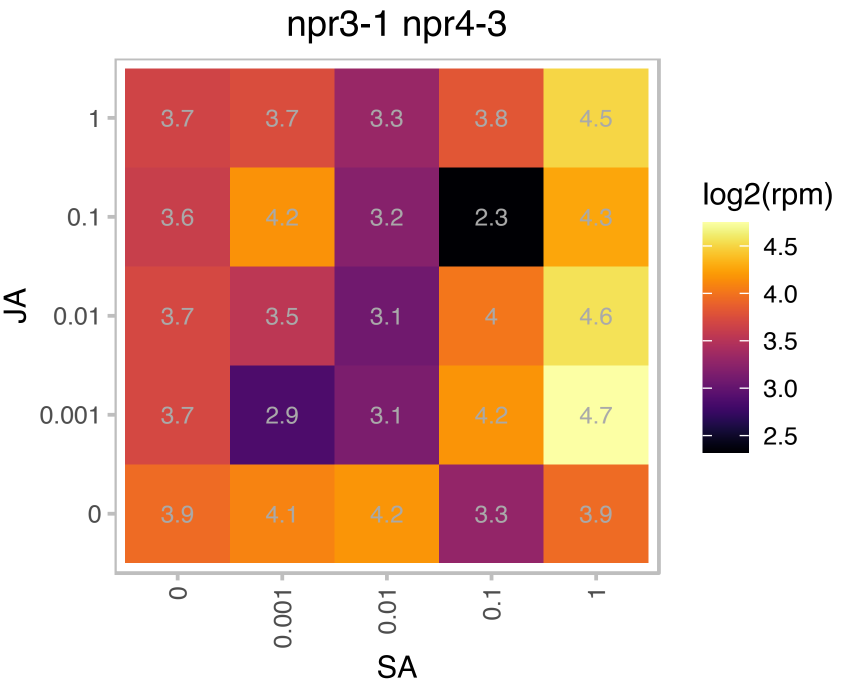

# AT1G07400.1 : NA

HSP20-like chaperones superfamily protein

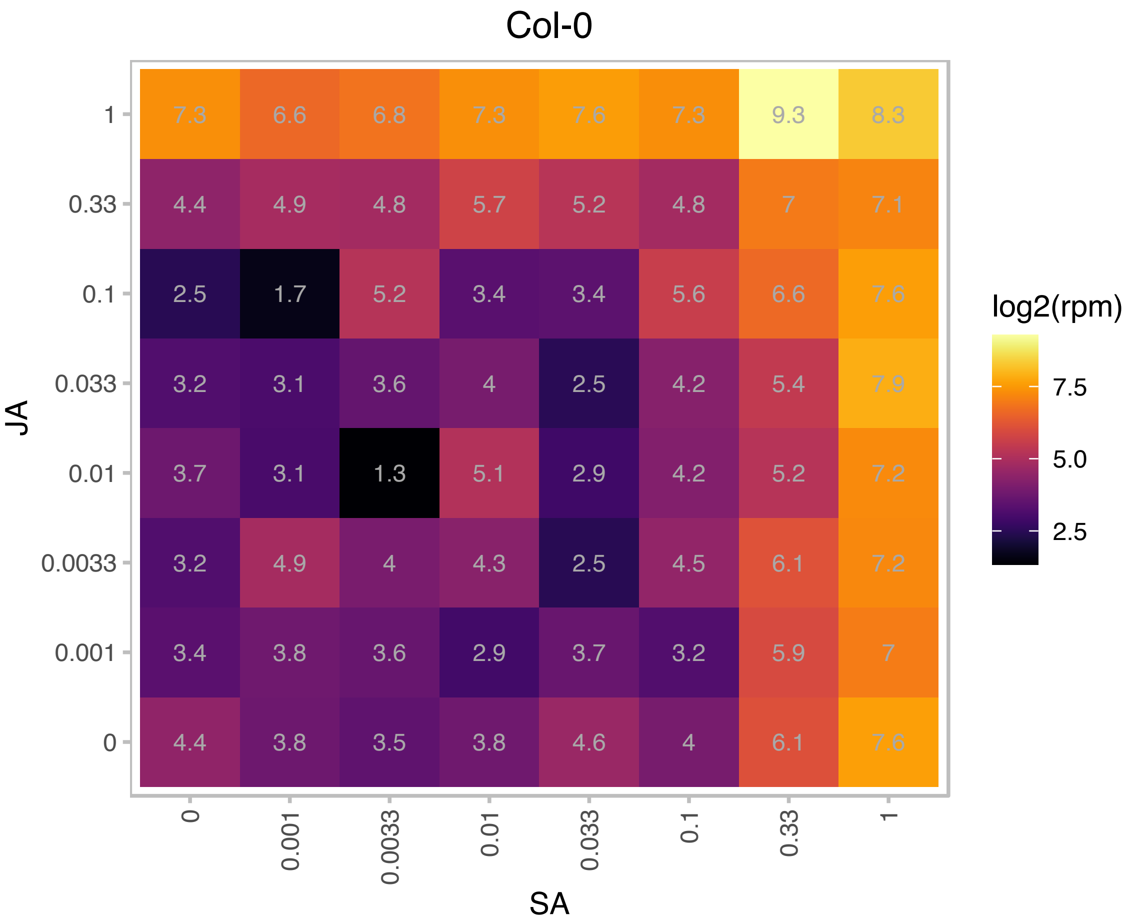


| | |
|---|---|
| Gene | AT1G07400 |
| is.expressed_genes | TRUE |
| is.responsive_genes | TRUE |
| is.apcluster | B-05 |

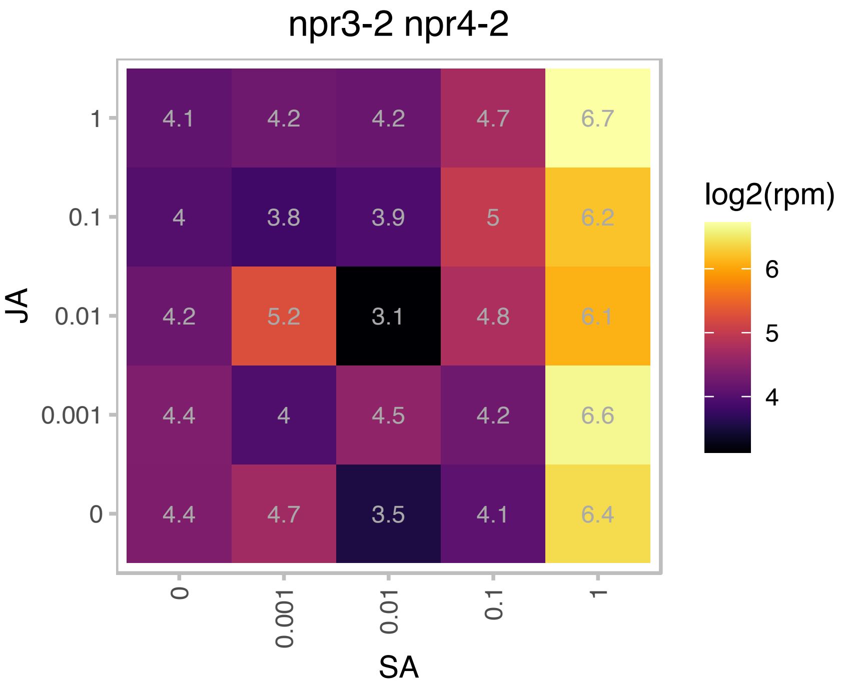


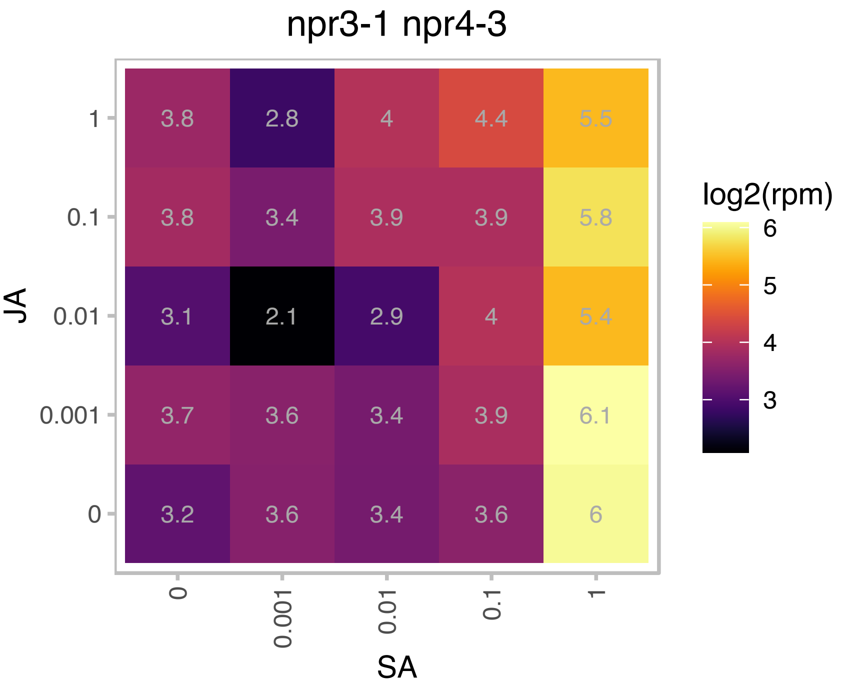

# AT1G07590.1 : NA

Tetratricopeptide repeat (TPR)-like superfamily protein

| | |
|---|---|
| Gene | AT1G07590 |
| is.expressed_genes | TRUE |
| is.responsive_genes | TRUE |
| is.apcluster | B-05 |

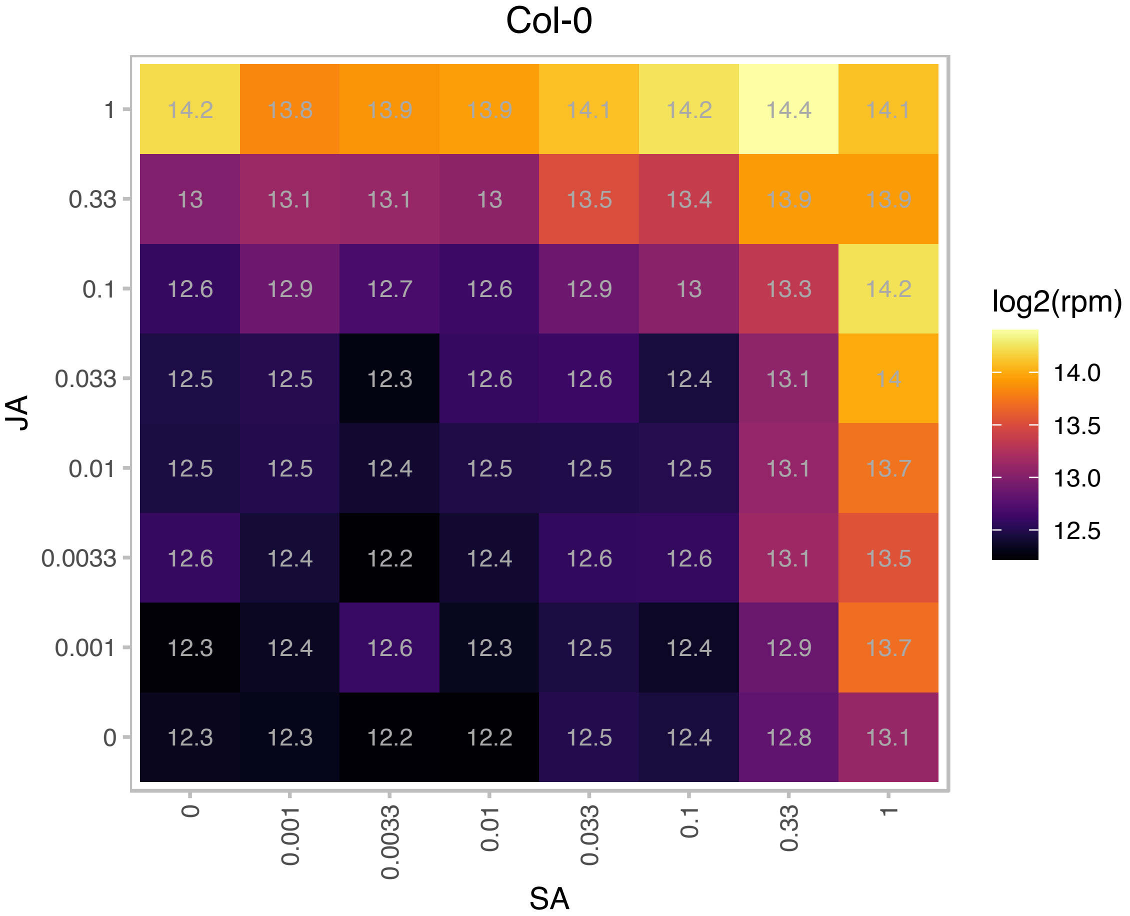


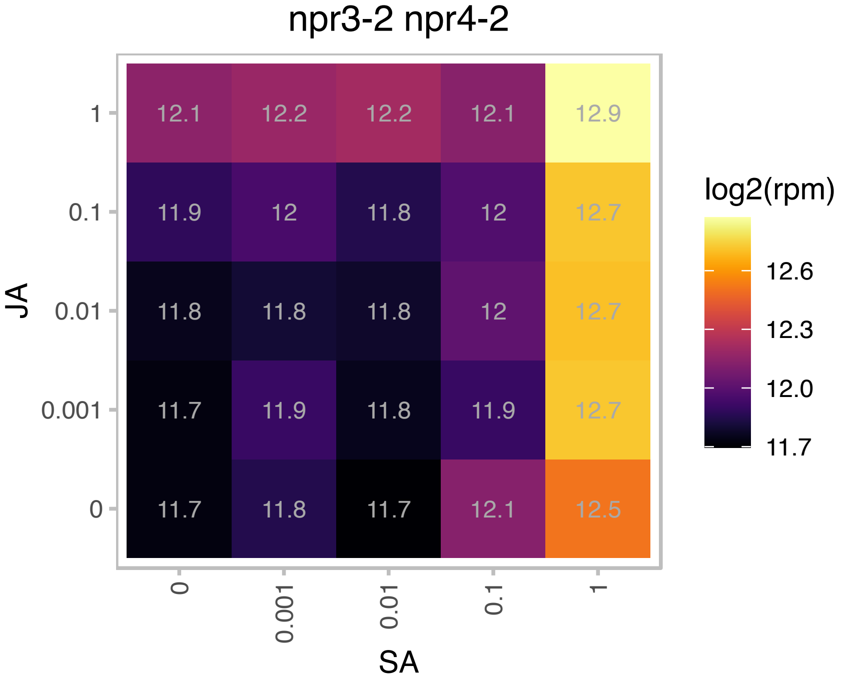


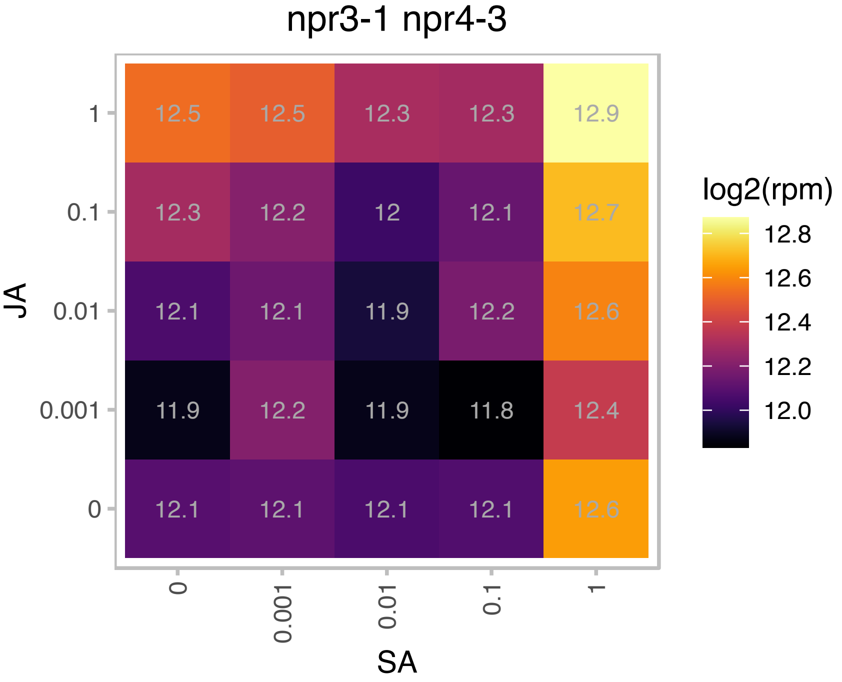

### AT1G09070.1 : (AT)SRC2, AtSRC2, SRC2

soybean gene regulated by cold-2

SRC2 specifically binds the peptide PIEPPPHH, and moves from ER to a vacuole fraction where it gets internalized. Involved in Protein Storage Vacuole targeting. The mRNA is cell-to-cell mobile.

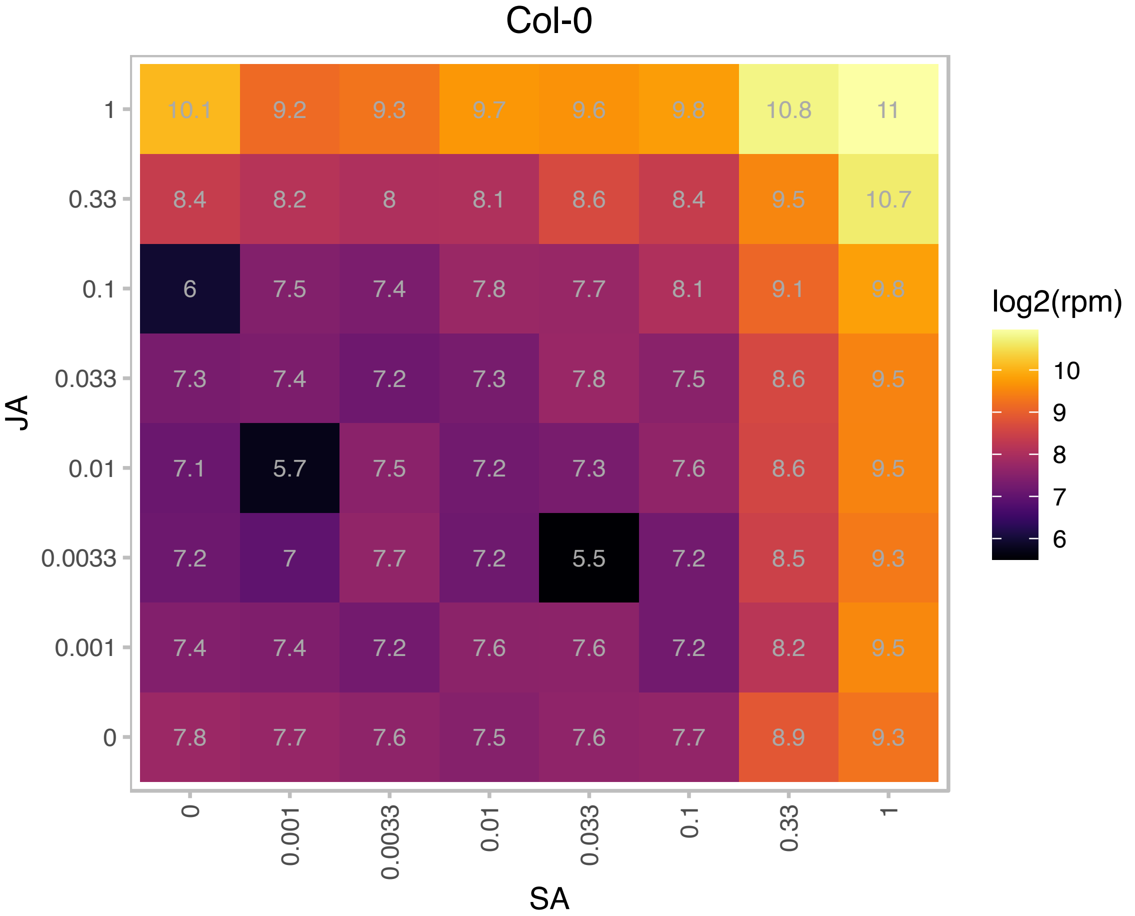


| | |
|---|---|
| Gene | AT1G09070 |
| is.expressed_genes | TRUE |
| is.responsive_genes | TRUE |
| is.apcluster | B-05 |

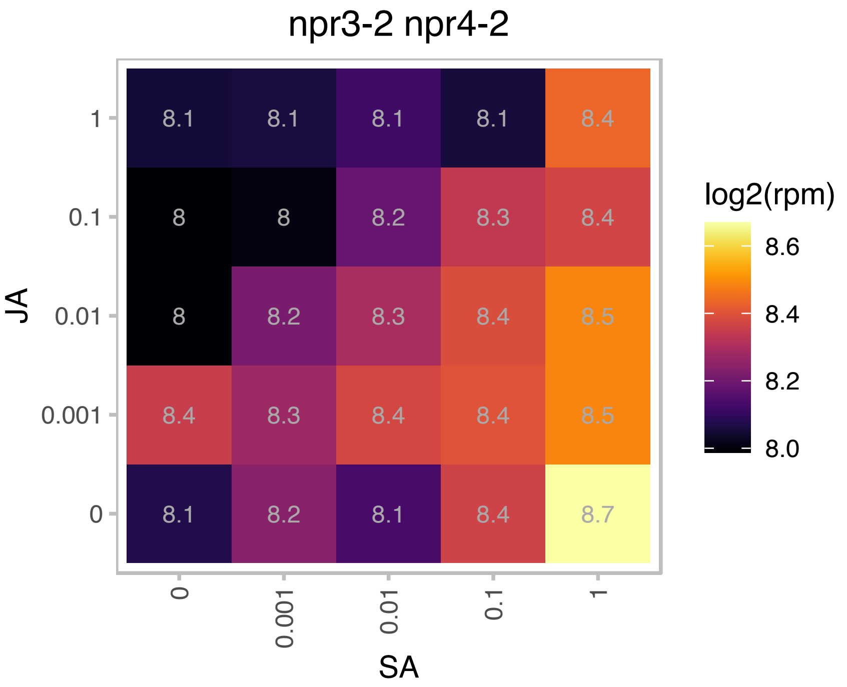


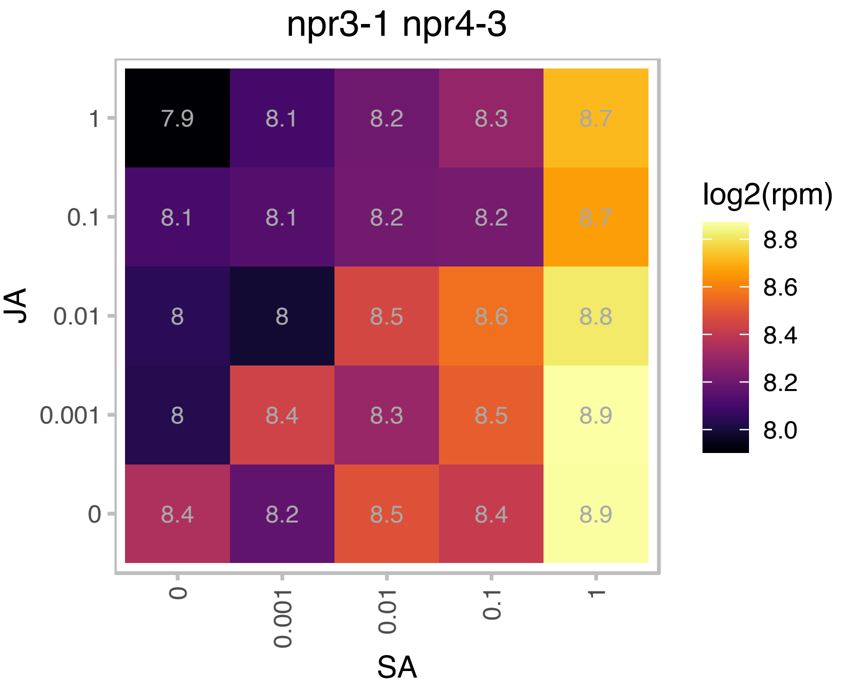

### AT1G09250.1 : AIF4

basic helix-loop-helix (bHLH) DNA-binding superfamily protein

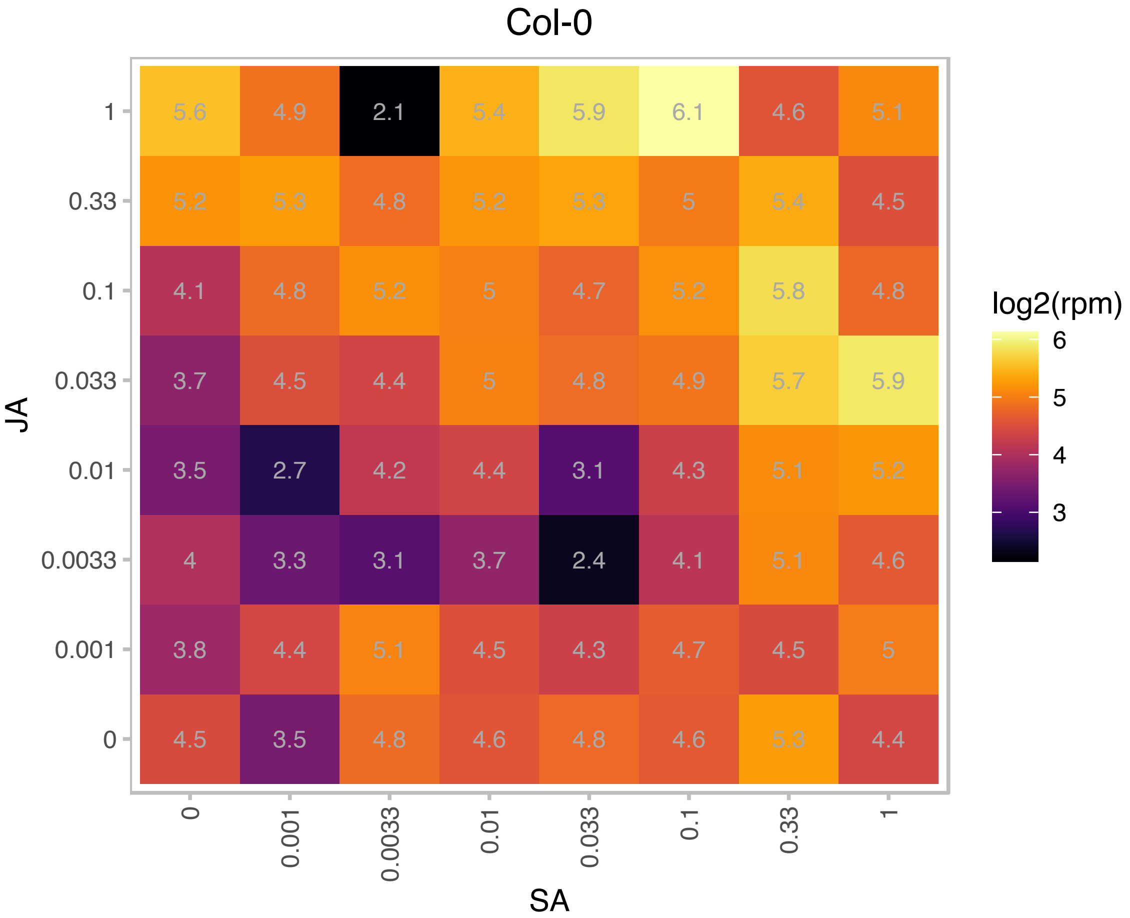


| | |
|---|---|
| Gene | AT1G09250 |
| is.expressed_genes | TRUE |
| is.responsive_genes | TRUE |
| is.apcluster | B-05 |

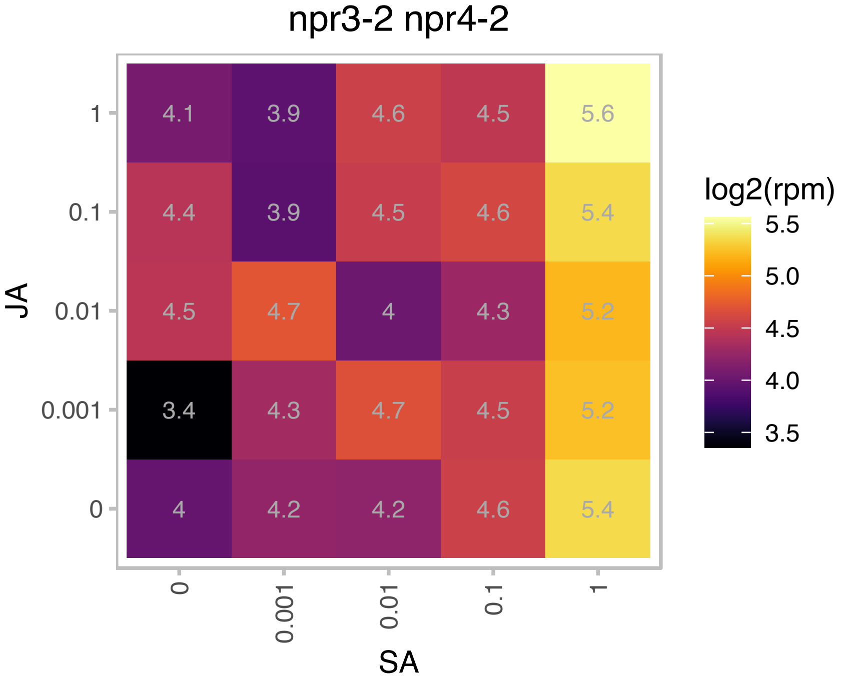


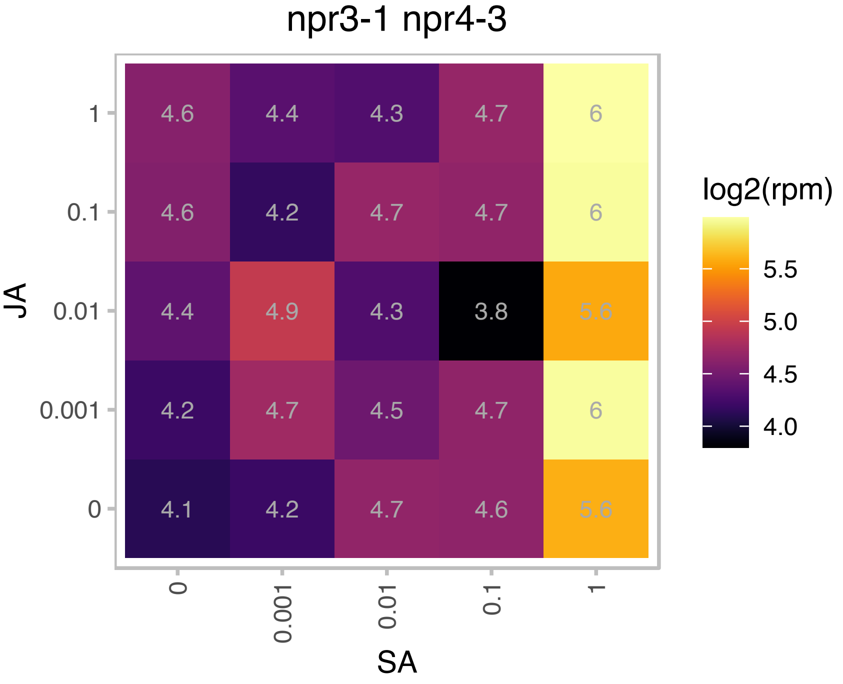

# AT1G12320.1 : NA

Protein of unknown function (DUF1442)

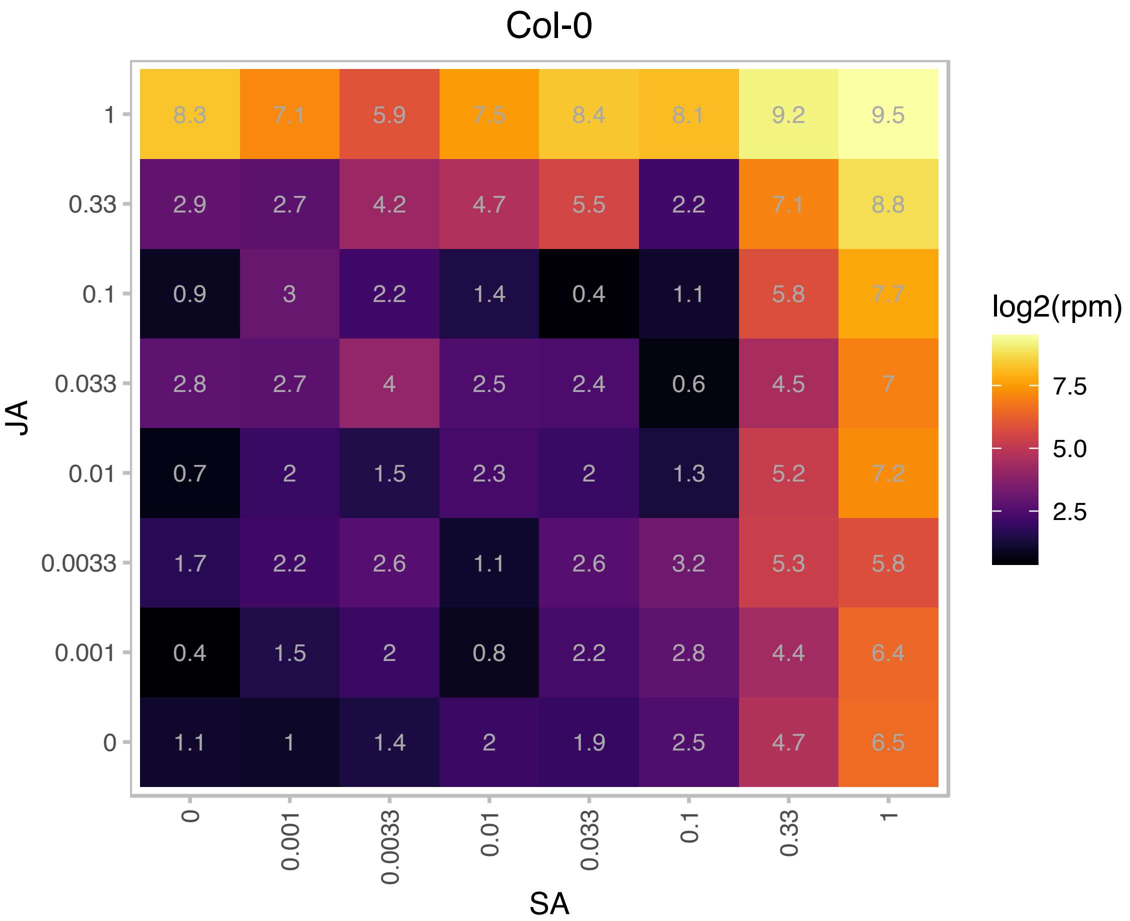


| | |
|---|---|
| Gene | AT1G12320 |
| is.expressed_genes | TRUE |
| is.responsive_genes | TRUE |
| is.apcluster | B-05 |

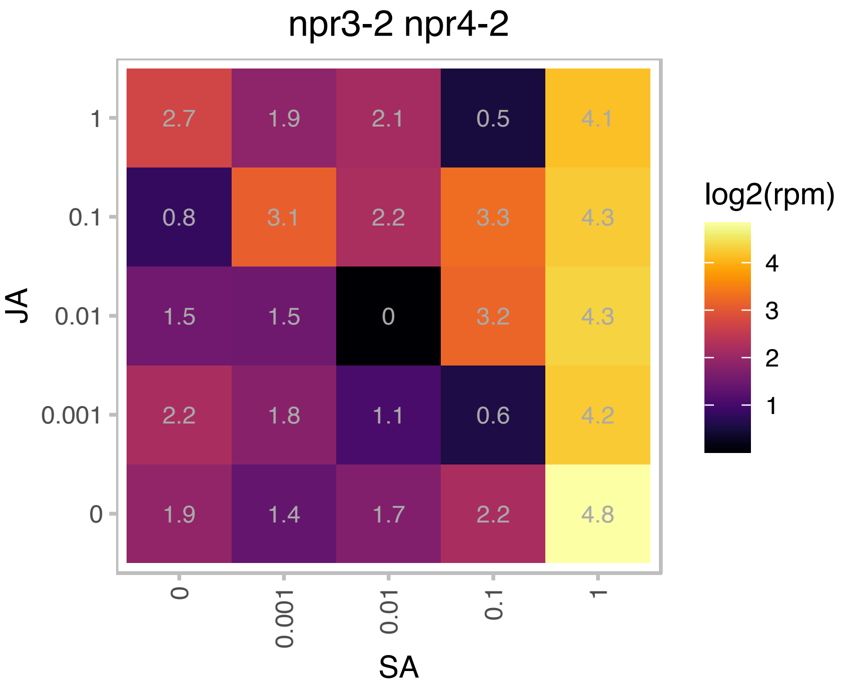


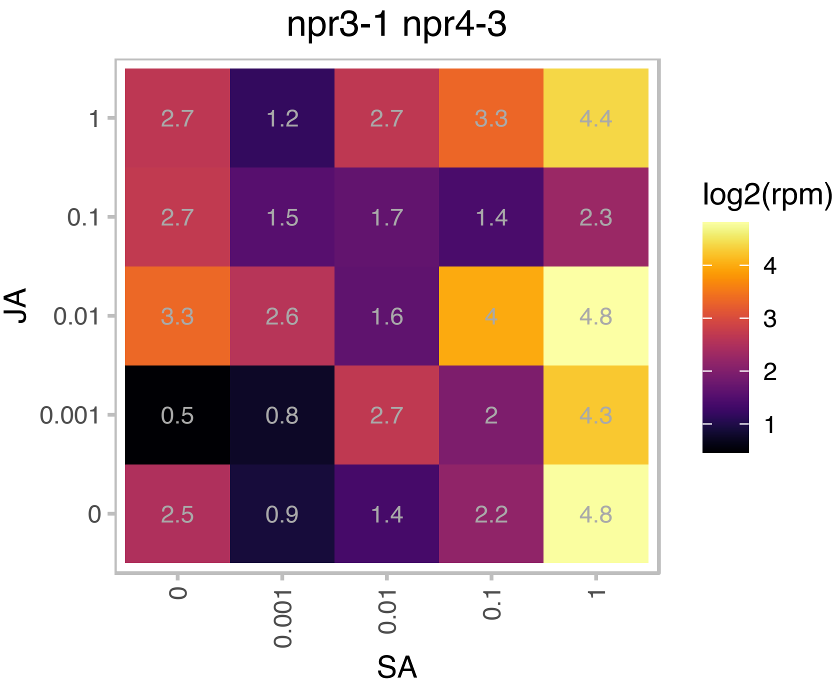

### AT1G13245.1 : DVL4, RTFL17

ROTUNDIFOLIA like 17

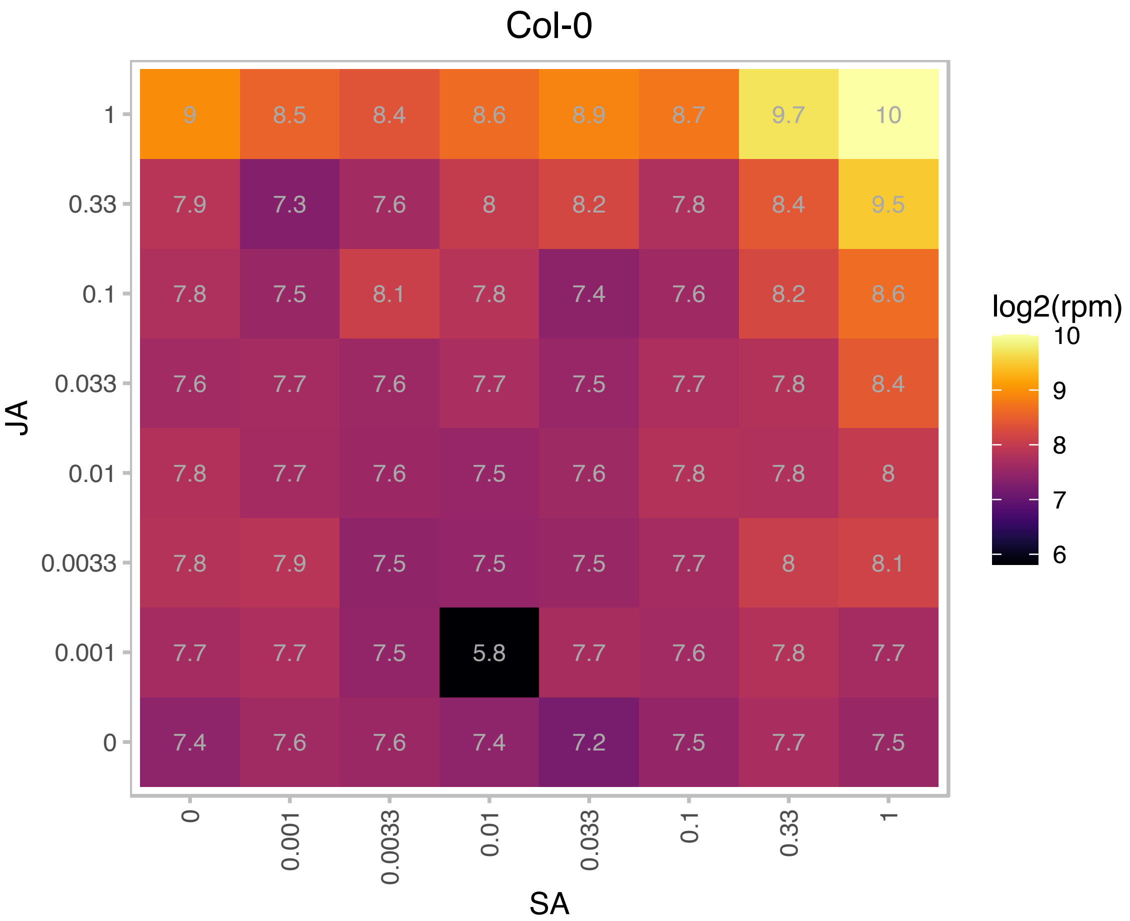


| | |
|---|---|
| Gene | AT1G13245 |
| is.expressed_genes | TRUE |
| is.responsive_genes | TRUE |
| is.apcluster | B-05 |

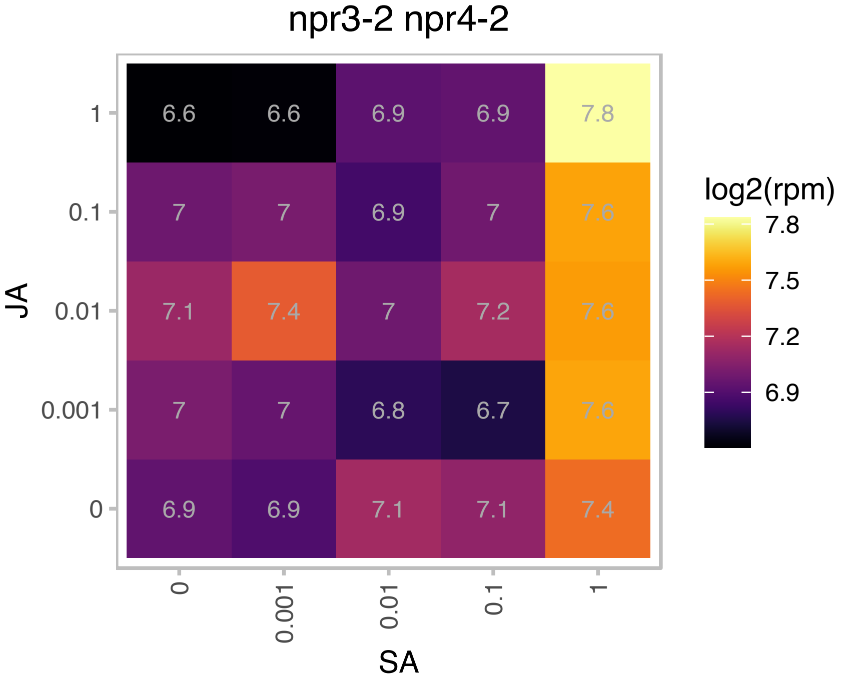


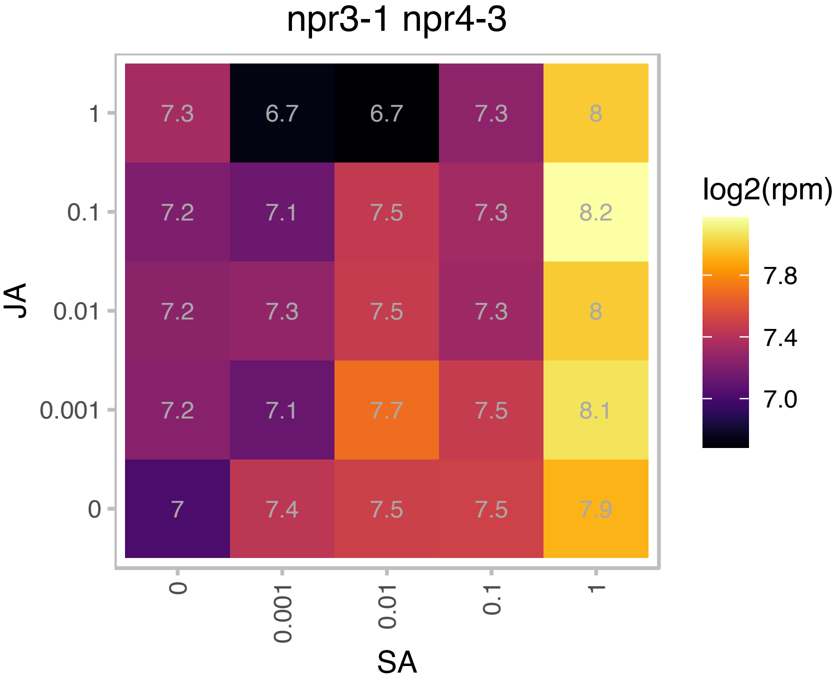

### AT1G13340.1 : ISTL6

Regulator of Vps4 activity in the MVB pathway protein

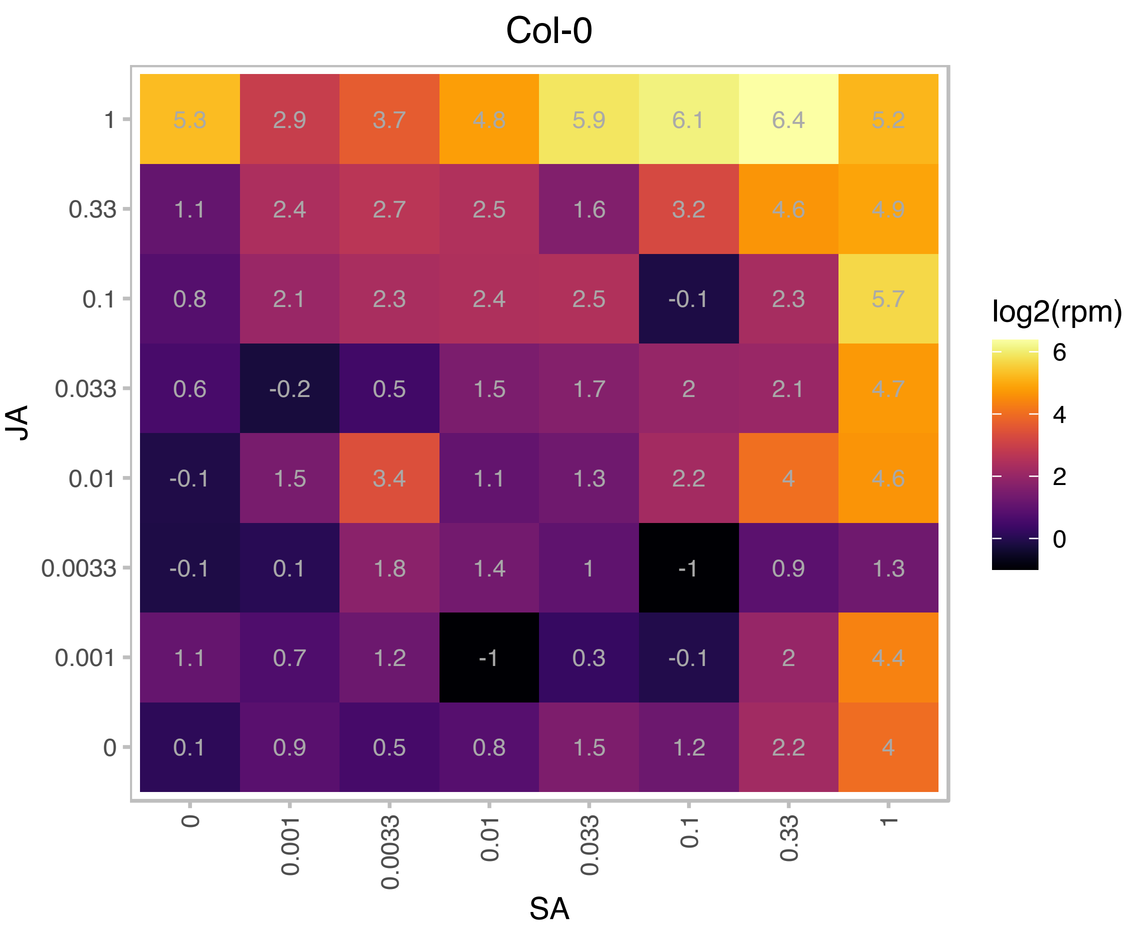


| | |
|---|---|
| Gene | AT1G13340 |
| is.expressed_genes | TRUE |
| is.responsive_genes | TRUE |
| is.apcluster | B-05 |

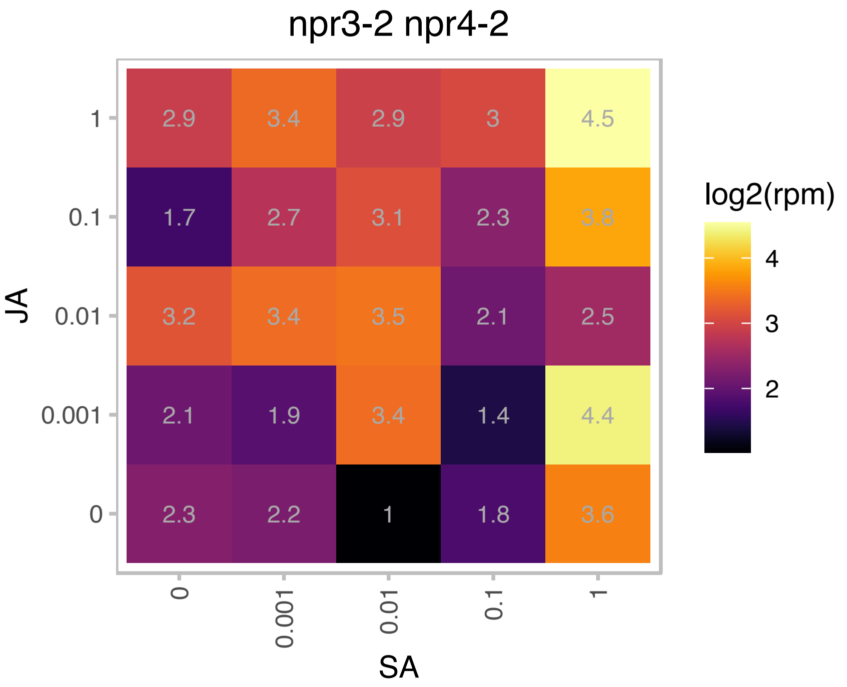


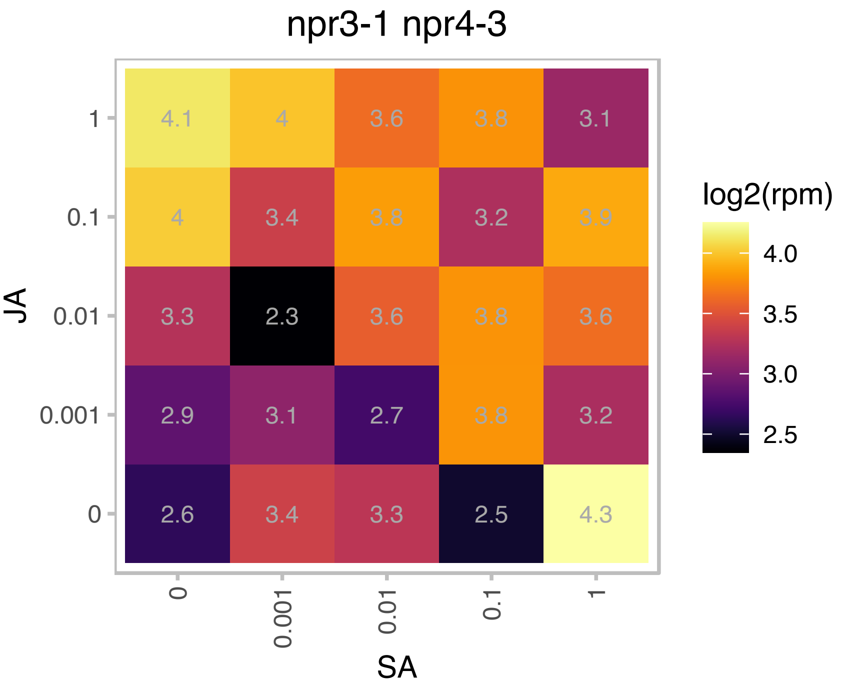

### AT1G15100.1 : RHA2A

RING-H2 finger A2A

Encodes a putative RING-H2 finger protein RHA2a.

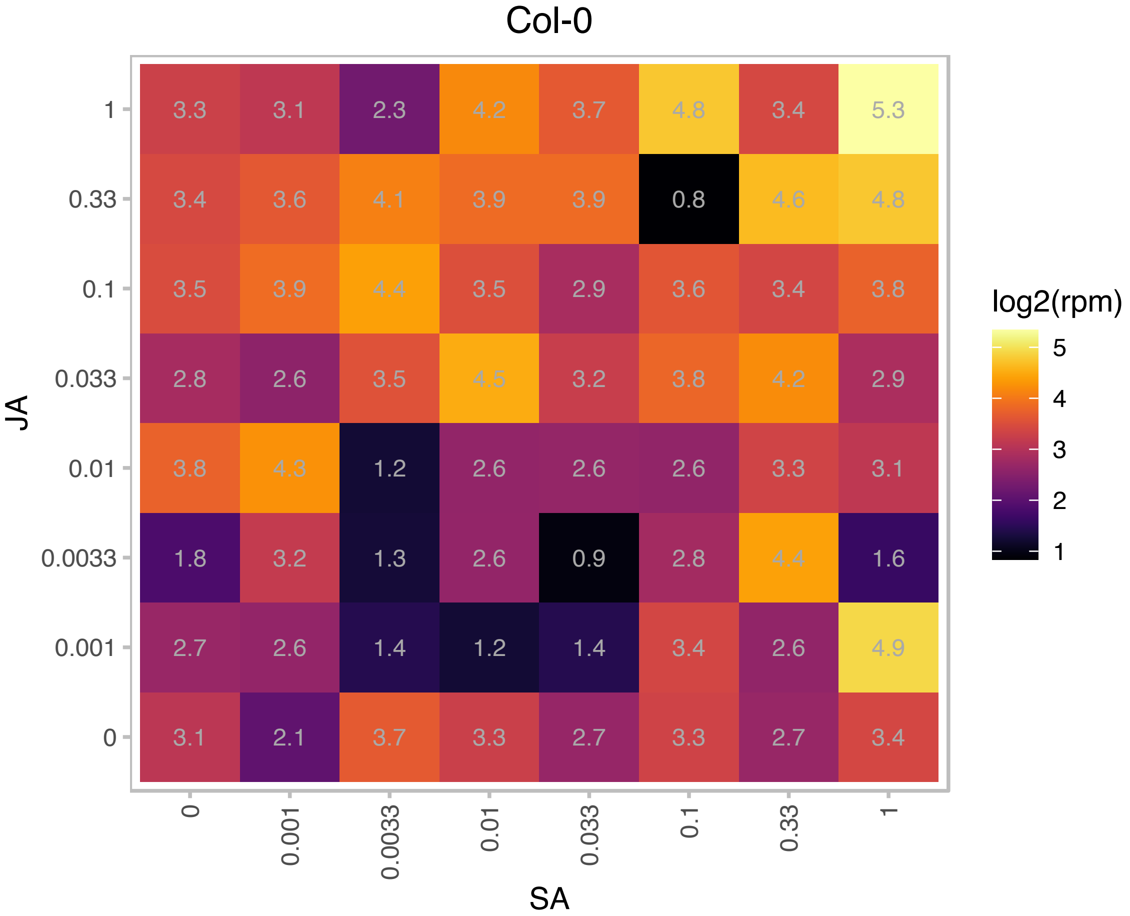


| | |
|---|---|
| Gene | AT1G15100 |
| is.expressed_genes | TRUE |
| is.responsive_genes | TRUE |
| is.apcluster | B-05 |

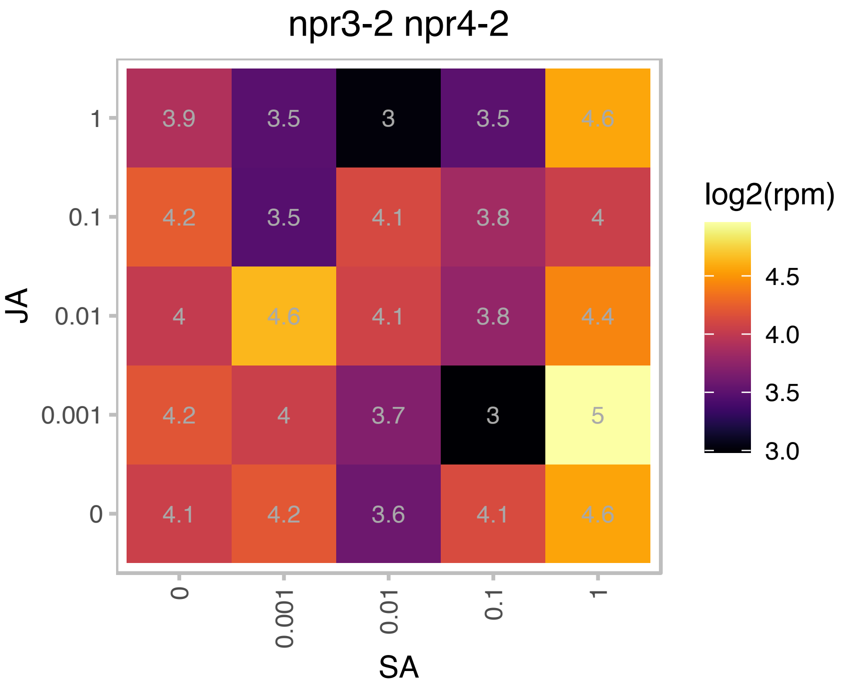


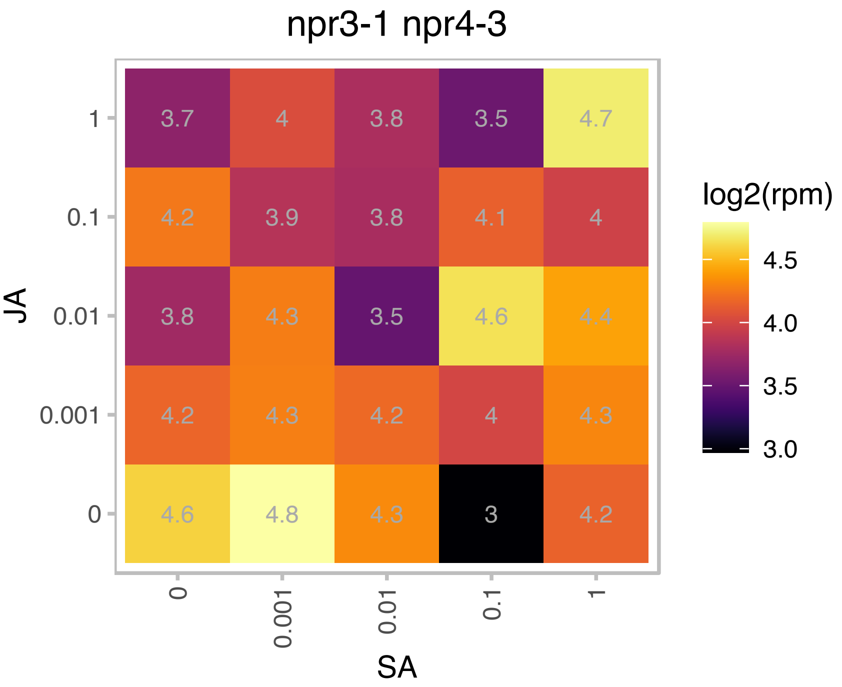

### AT1G15520.1 : ABCG40, ATABCG40, ATPDR12, PDR12

pleiotropic drug resistance 12

ABC transporter family involved in ABA transport and resistance to lead. Localizes to plasma membrane. Upregulated by lead. Expressed in leaves, flowers, stomata and roots.

Col-0

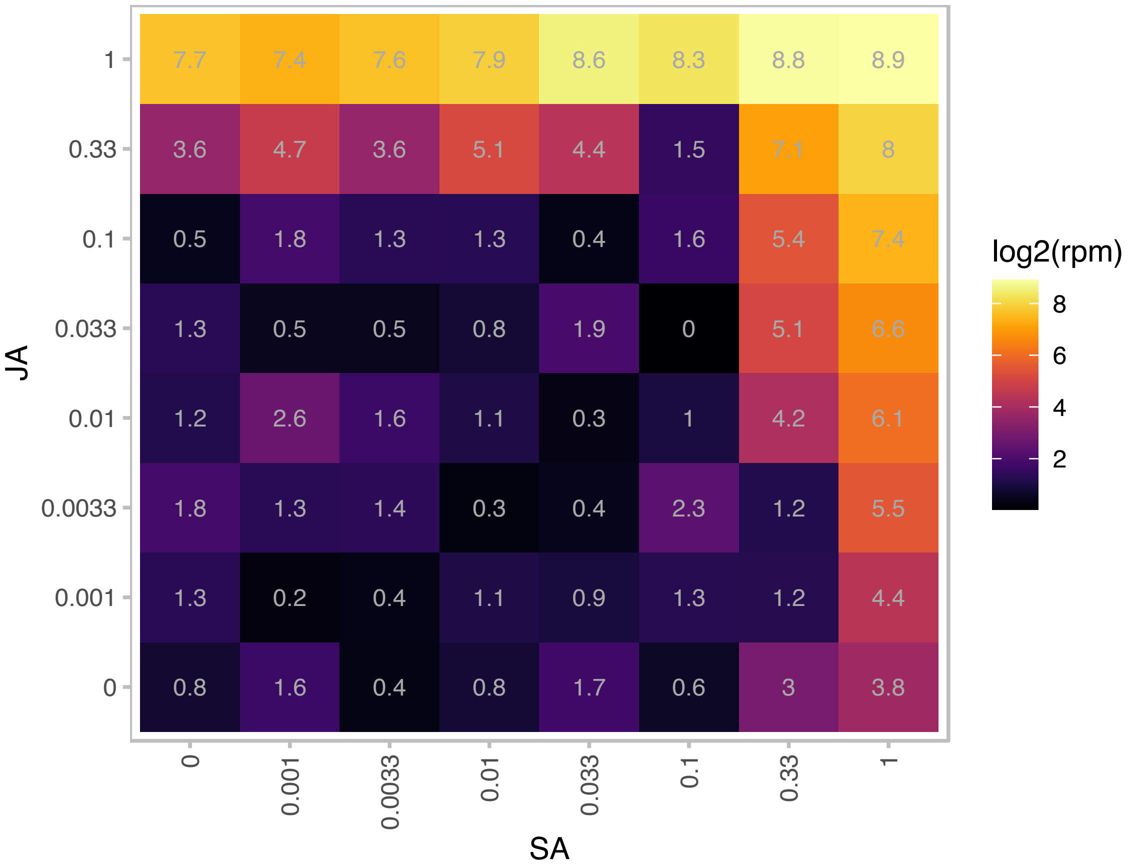


| | |
|---|---|
| Gene | AT1G15520 |
| is.expressed_genes | TRUE |
| is.responsive_genes | TRUE |
| is.apcluster | B-05 |

npr3-2 npr4-2

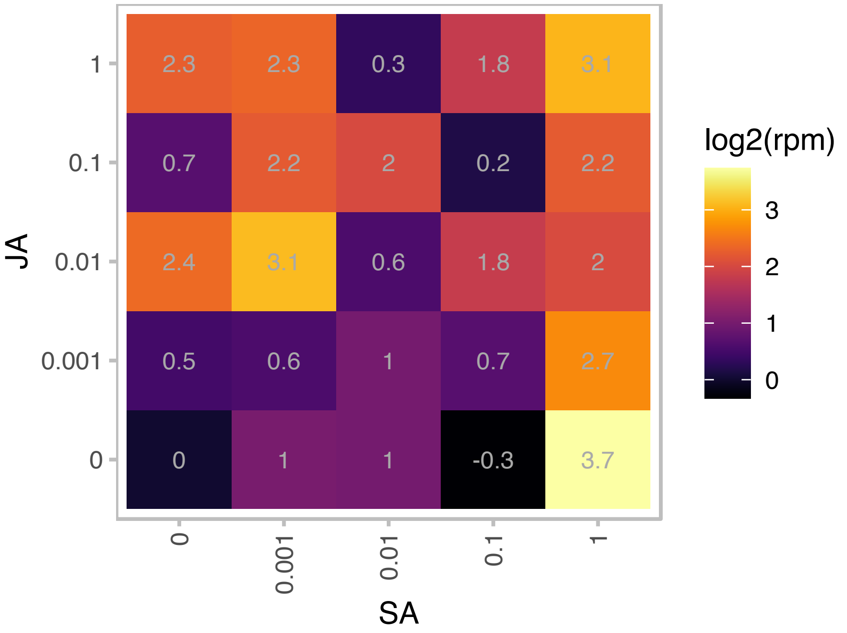


npr3-1 npr4-3

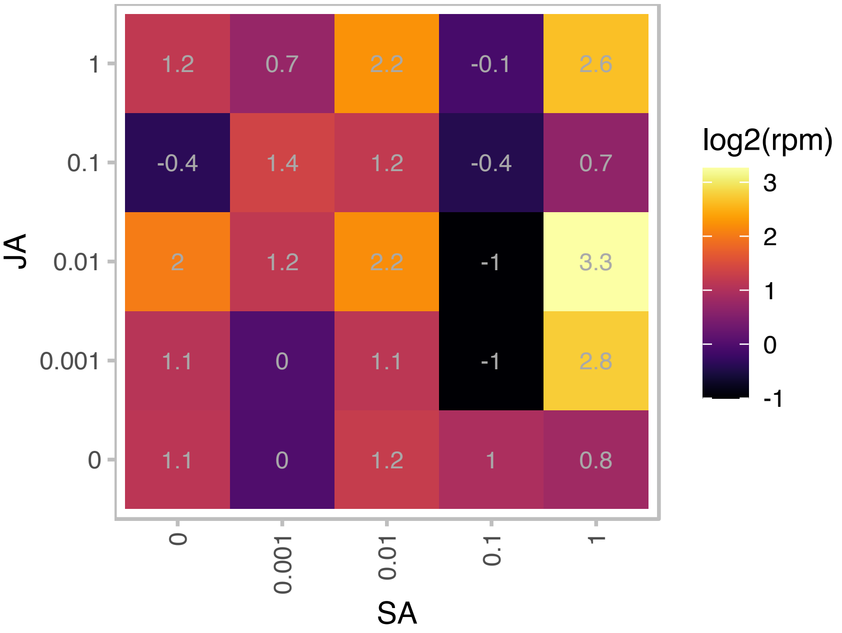

# AT1G25370.1 : NA

Protein of unknown function (DUF1639)

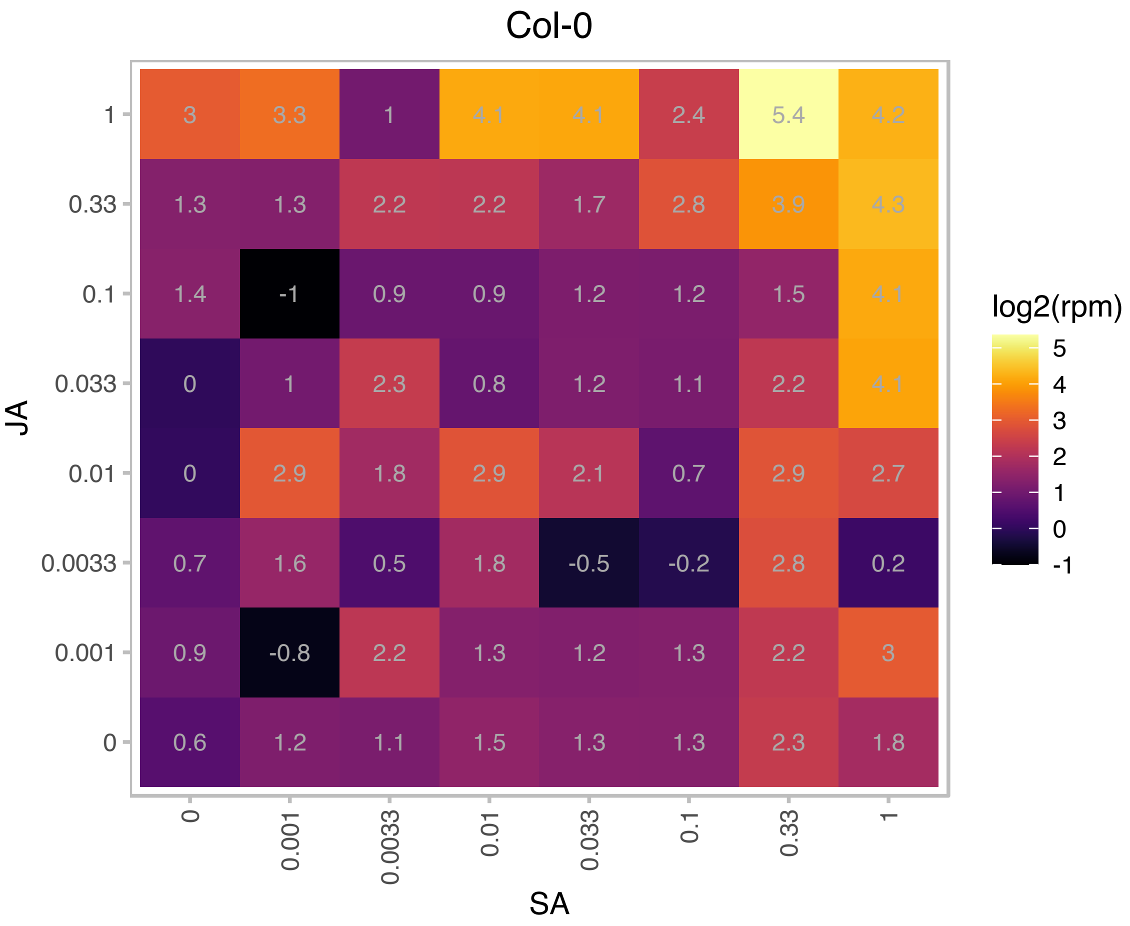


| | |
|---|---|
| Gene | AT1G25370 |
| is.expressed_genes | TRUE |
| is.responsive_genes | TRUE |
| is.apcluster | B-05 |

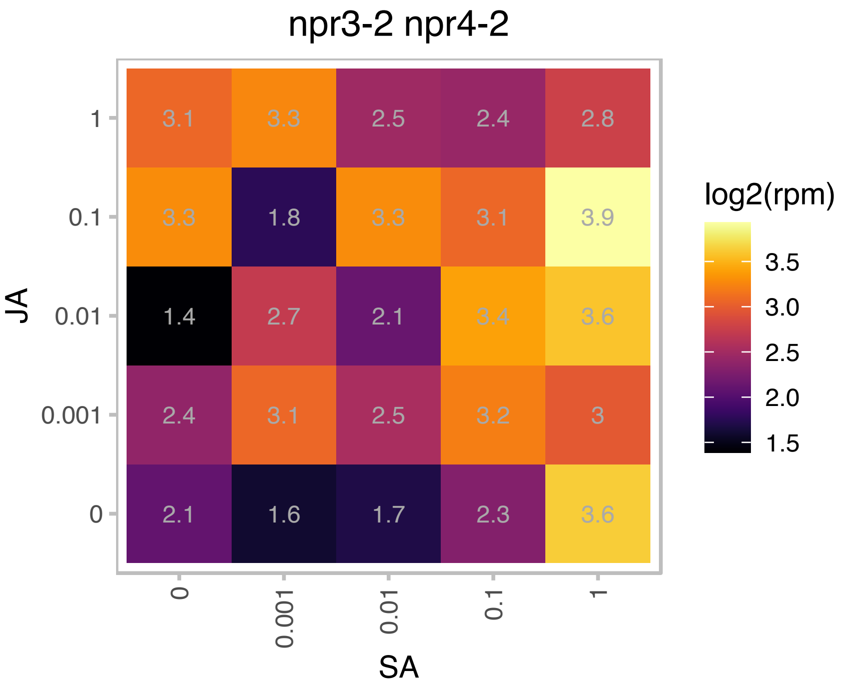


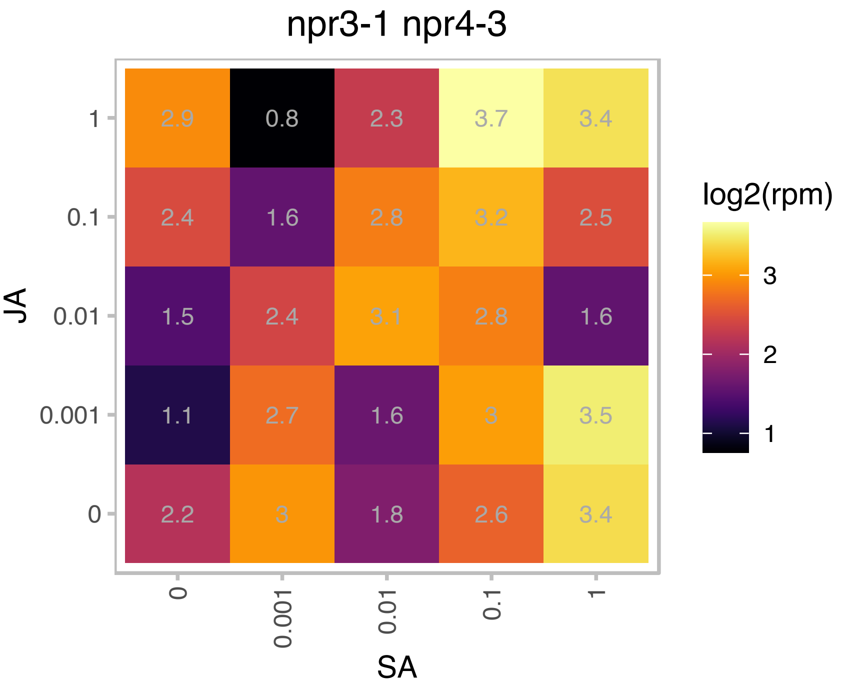

### AT1G29300.1 : UNE1

Plant protein of unknown function (DUF641)

| | |
|---|---|
| Gene | AT1G29300 |
| is.expressed_genes | TRUE |
| is.responsive_genes | TRUE |
| is.apcluster | B-05 |

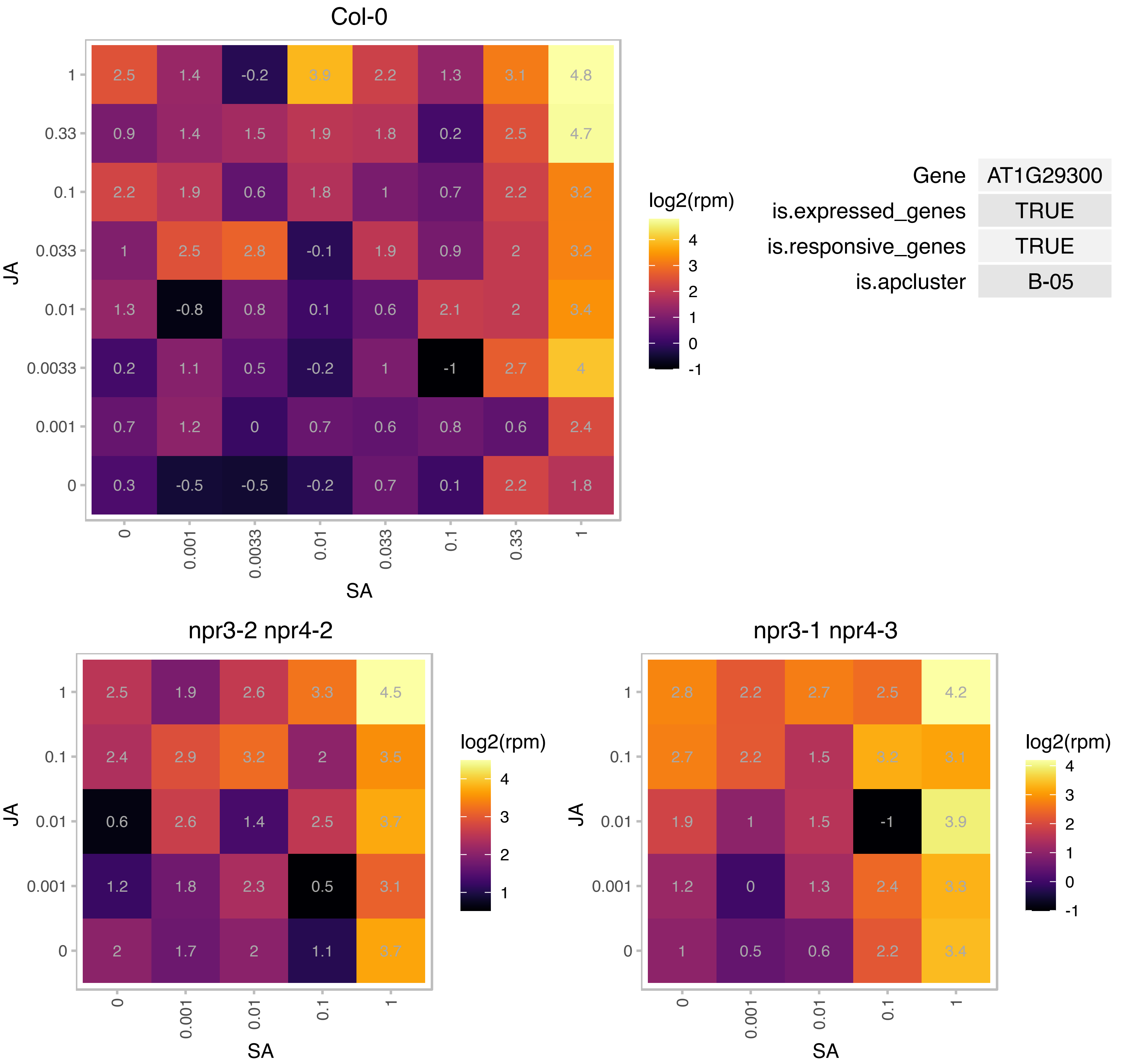

### AT1G29330.1 : AERD2, ATERD2, ERD2

ER lumen protein retaining receptor family protein
Encodes a protein similar in sequence to animal and yeast endoplasmic reticulum retention signal receptor. This protein can functionally complement the yeast homologue. Transcript is detected in flower buds, stems, root, and leaves.

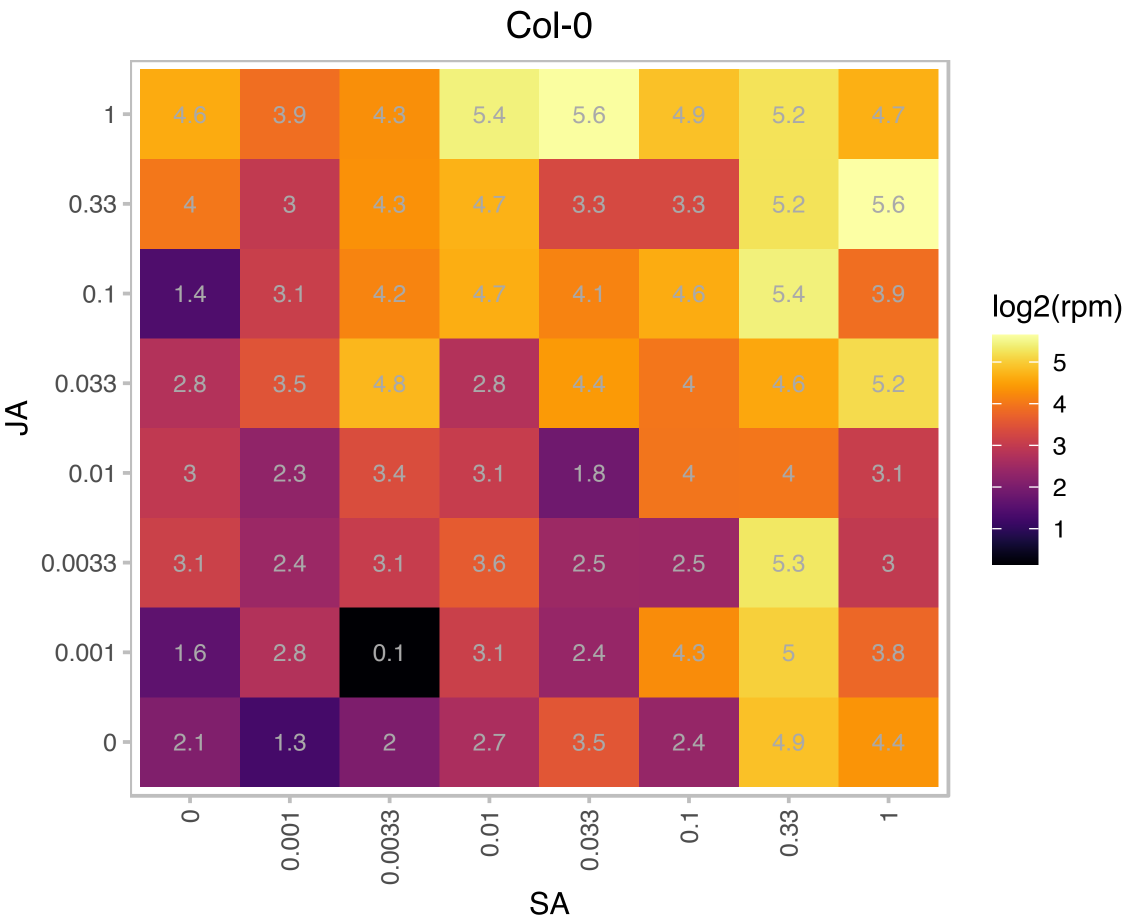


| | |
|---|---|
| Gene | AT1G29330 |
| is.expressed_genes | TRUE |
| is.responsive_genes | TRUE |
| is.apcluster | B-05 |

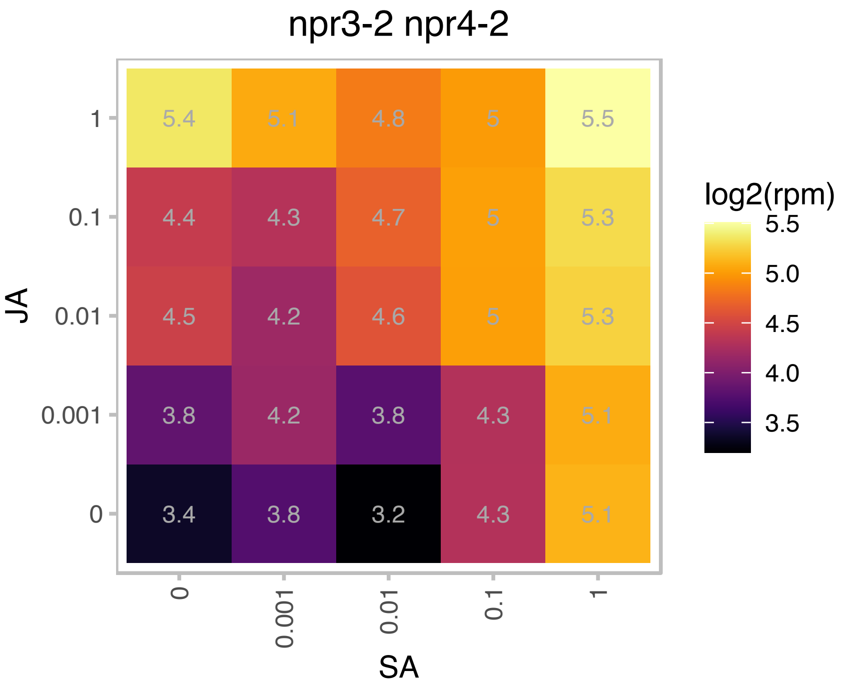


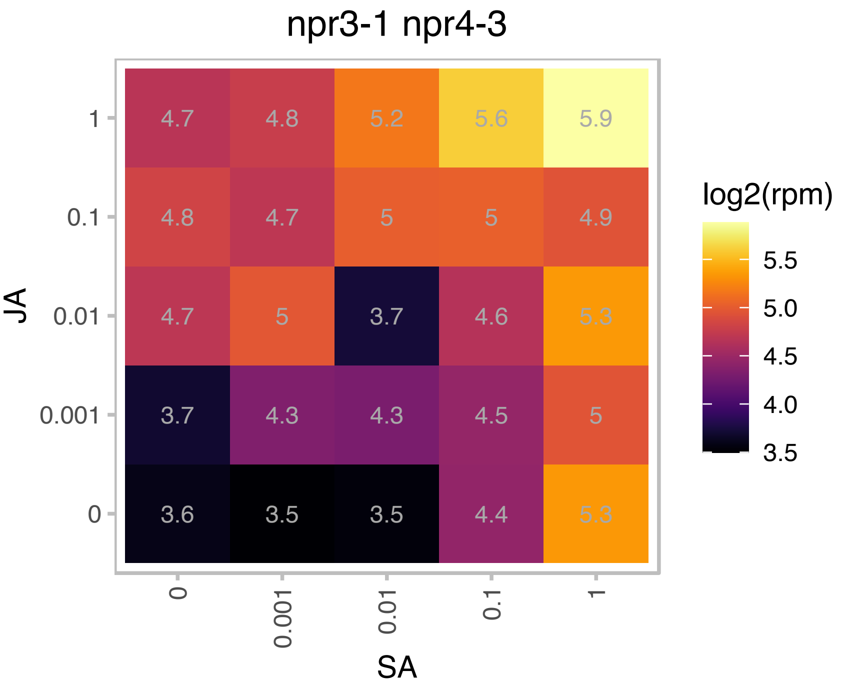

### AT1G29850.1 : AtPDCD5, PDCD5

double-stranded DNA-binding family protein

Encodes a protein that by its interaction with HAM acetyltransferases plays an important role during DNA damage responses induced by UV-B radiation and participates in programmed cell death programs.

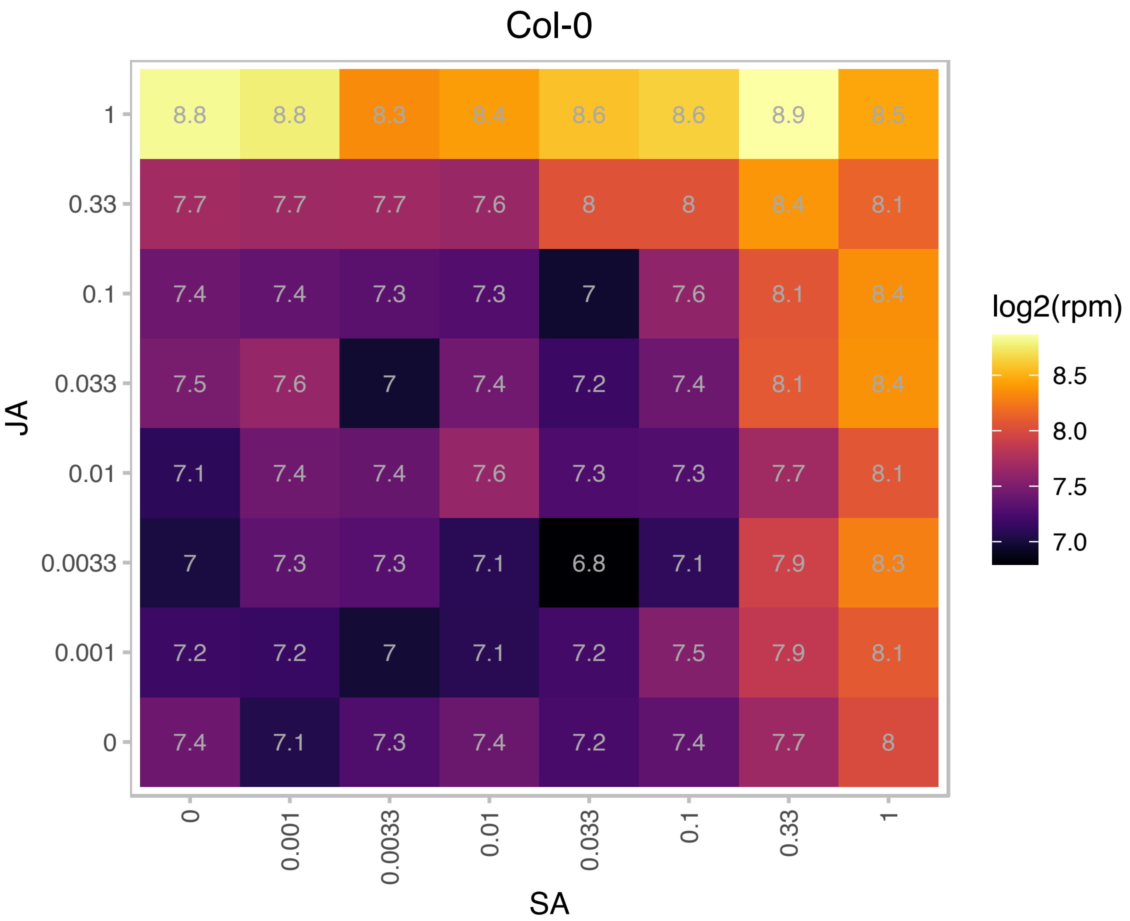


| | |
|---|---|
| Gene | AT1G29850 |
| is.expressed_genes | TRUE |
| is.responsive_genes | TRUE |
| is.apcluster | B-05 |

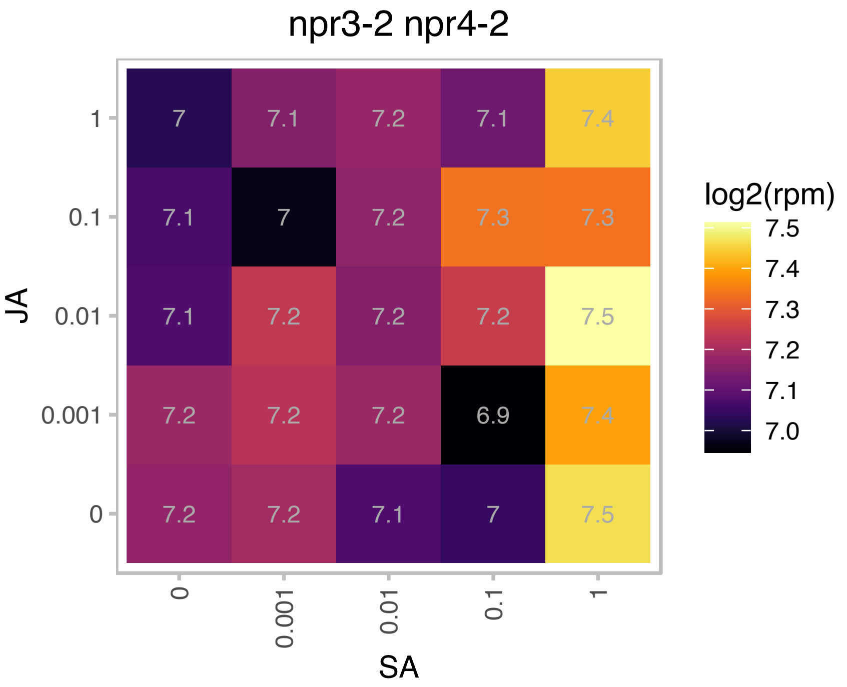


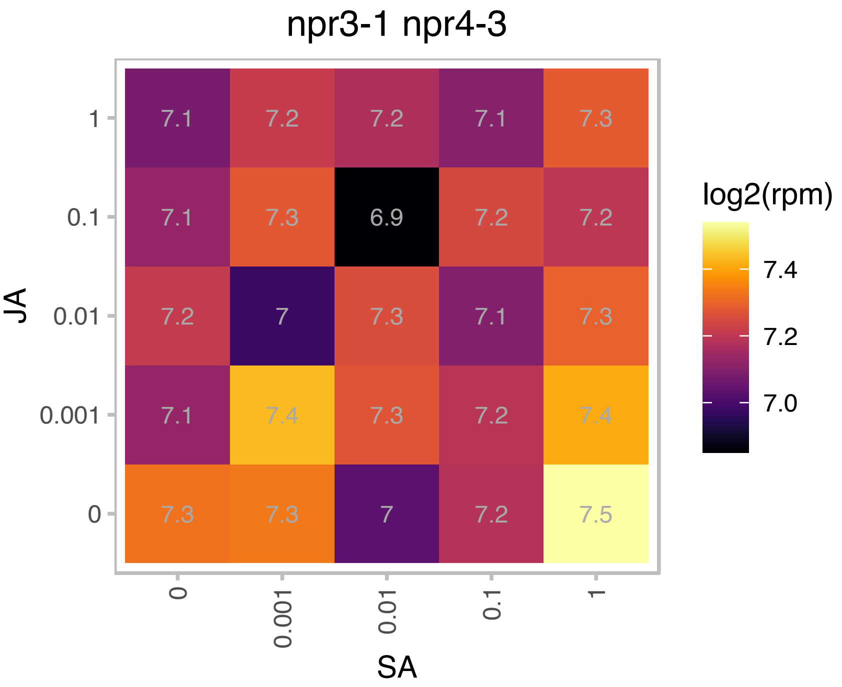

### AT1G31880.1 : BRX, NLM9

Belongs to five-member BRX gene family. Arabidopsis BRX genes share high levels of similarity among each others, with several conserved domains. The most distinct is BRX domain - highly conserved in all BRX genes among distantly related species. This protein-protein interaction domain is required and sufficient for BRX activity. BRX encodes a key regulator of cell proliferation and elongation in the root, which has been implicated in the brassinosteroid (BR) pathway as well as regulation of auxin-responsive gene expression. Also involved in cytokinin-mediated inhibition of lateral root initiation. A loss-of-function allele, named brx-2 in Rodrigues et al. (2009) Plant Physiol. but changed to brx-3 to resolve nomenclature conflict (Li et al. Planta 2009:229(3):593-603), shows enhanced response to ABA-mediated inhibition of root growth. [illegible]

DZC (Disease resistance/zinc finger/chromosome condensation-like region) domain containing protein

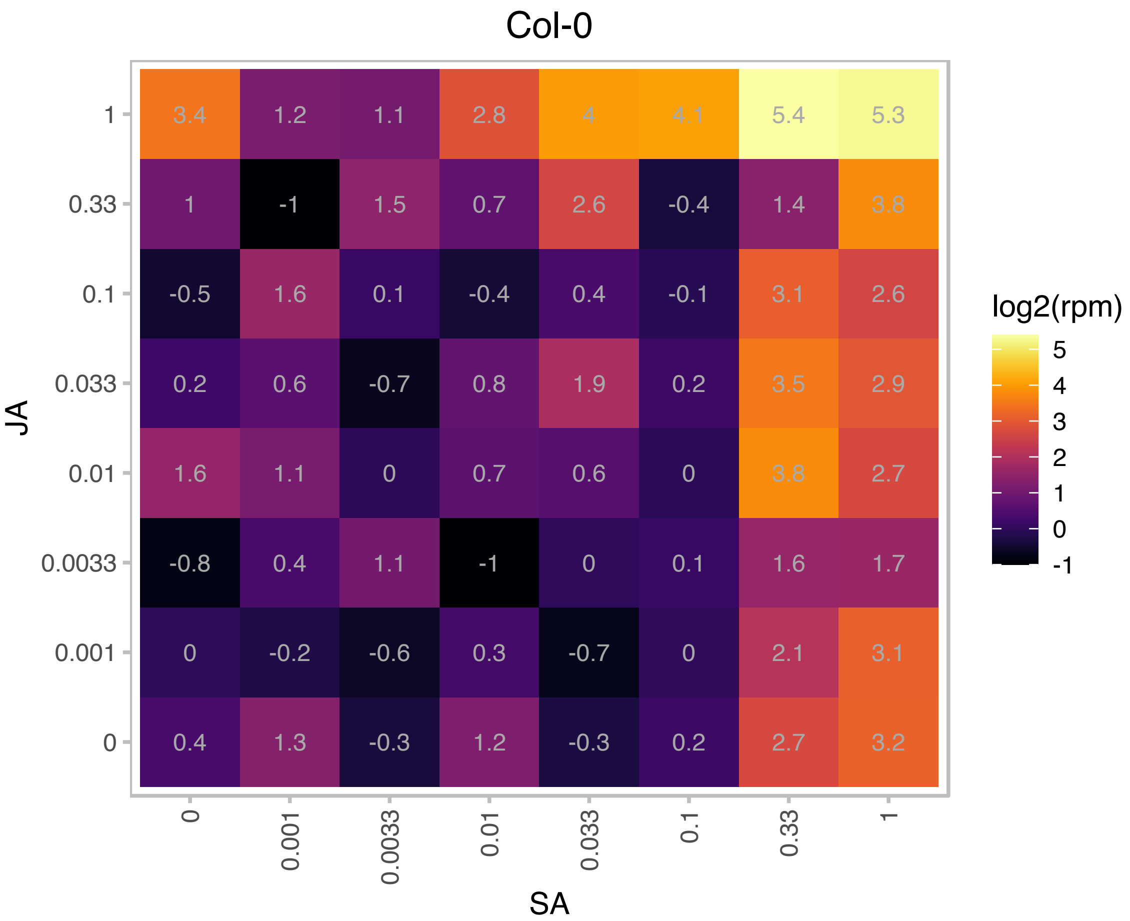


| | |
|---|---|
| Gene | AT1G31880 |
| is.expressed_genes | TRUE |
| is.responsive_genes | TRUE |
| is.apcluster | B-05 |

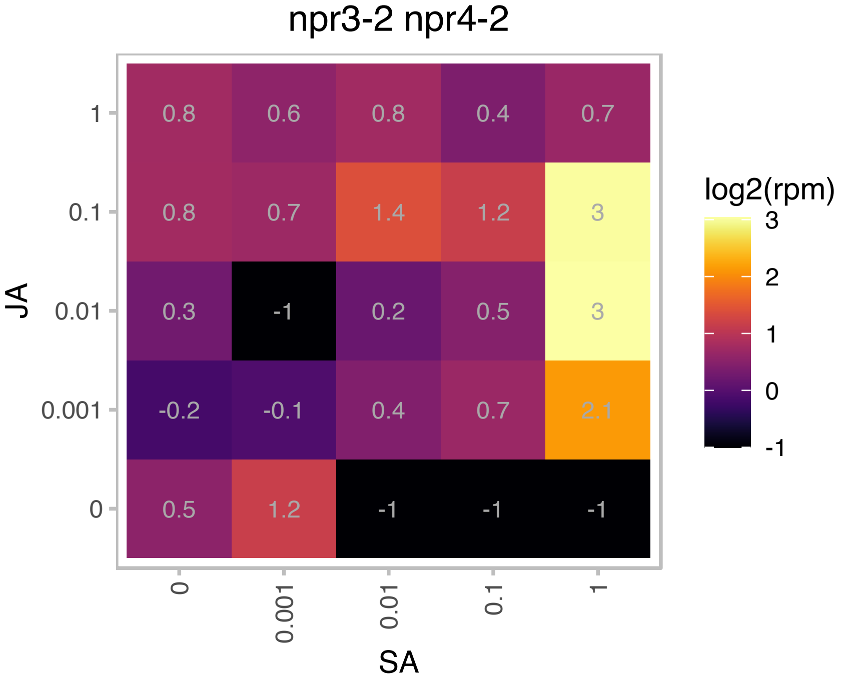


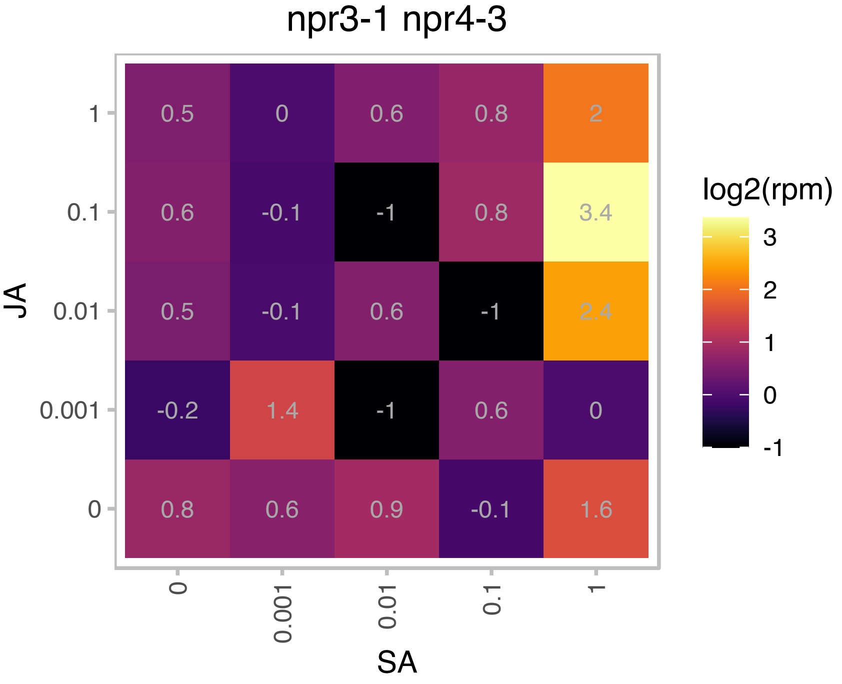

### AT1G32170.1 : XTH30, XTR4

xyloglucan endotransglucosylase/hydrolase 30

xyloglucan endotransglycosylase-related protein (XTR4) The mRNA is cell-to-cell mobile.

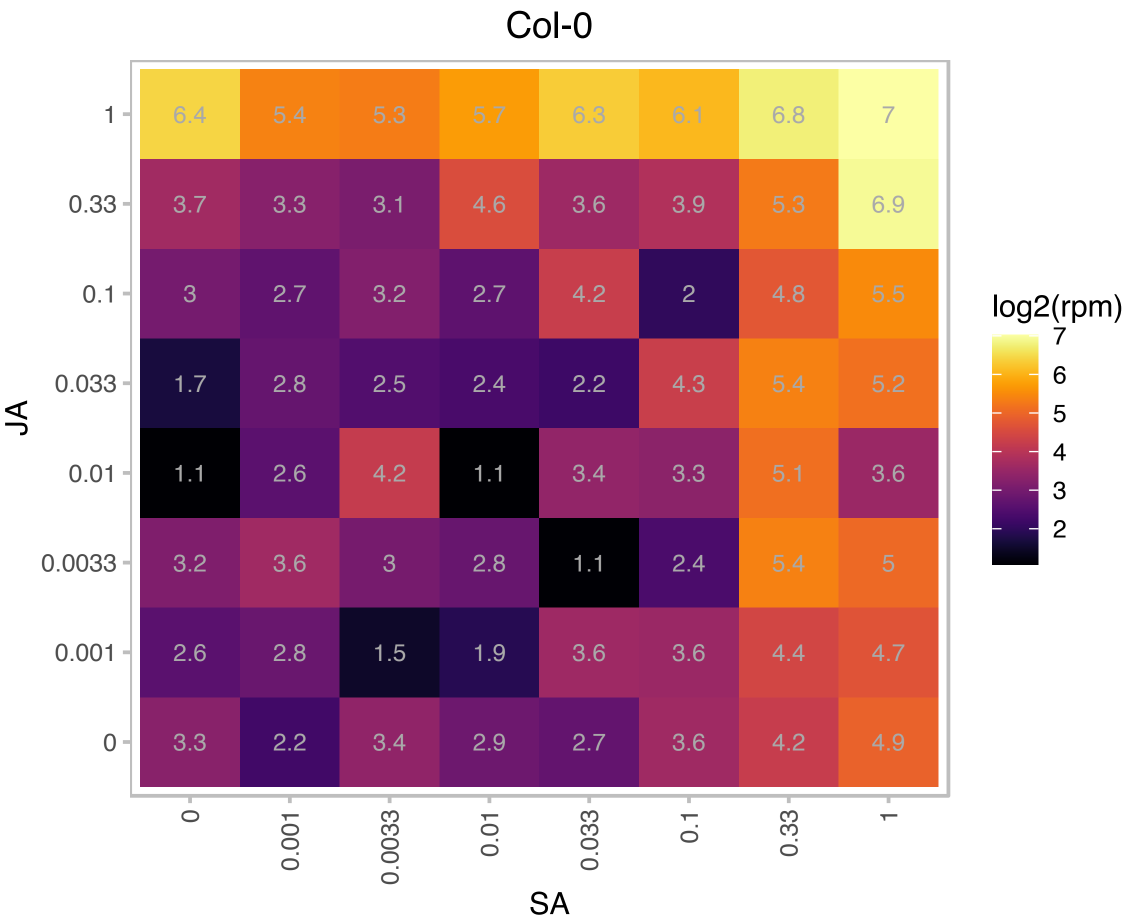


| | |
|---|---|
| Gene | AT1G32170 |
| is.expressed_genes | TRUE |
| is.responsive_genes | TRUE |
| is.apcluster | B-05 |

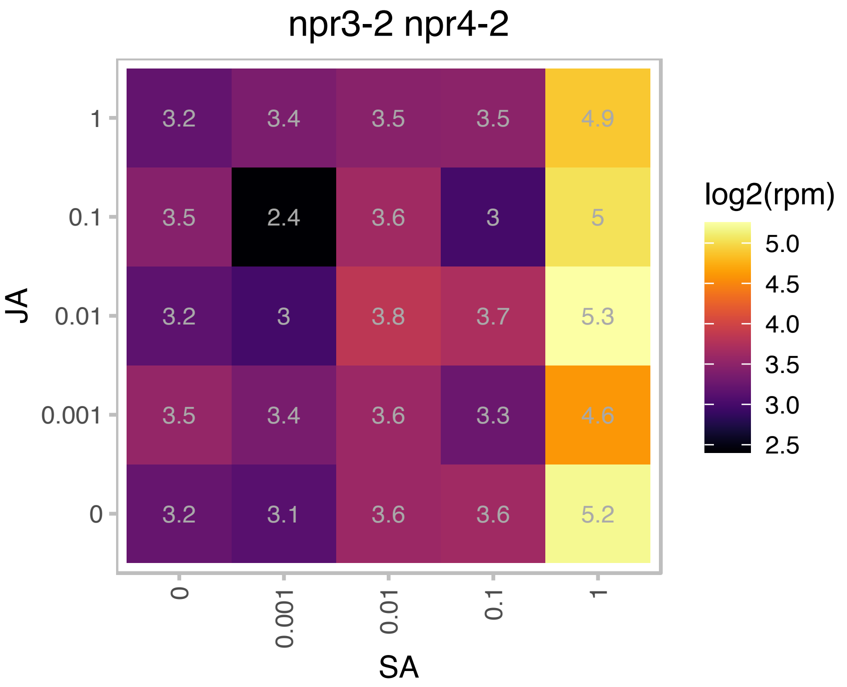


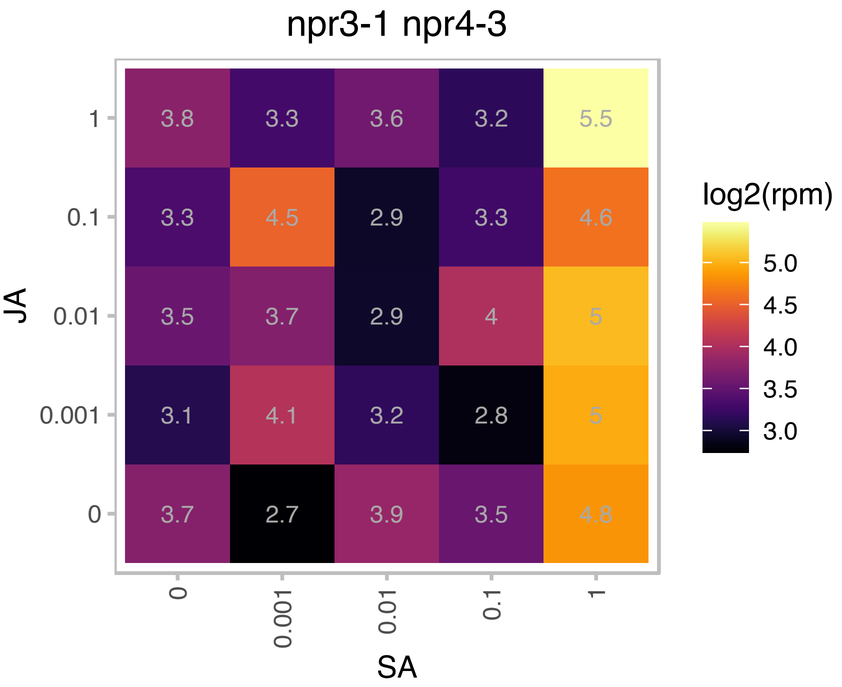

# AT1G32920.1 : NA

Col-0

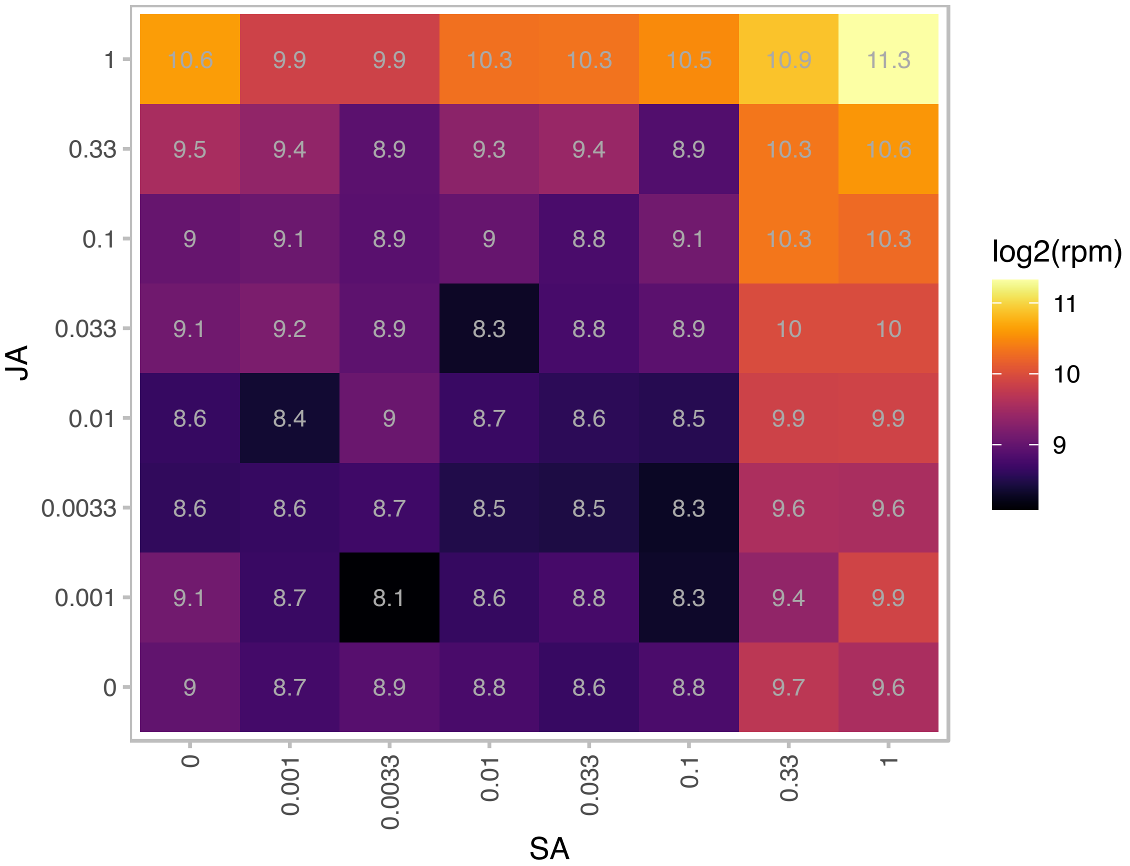


| | |
|---|---|
| Gene | AT1G32920 |
| is.expressed_genes | TRUE |
| is.responsive_genes | TRUE |
| is.apcluster | B-05 |

npr3-2 npr4-2

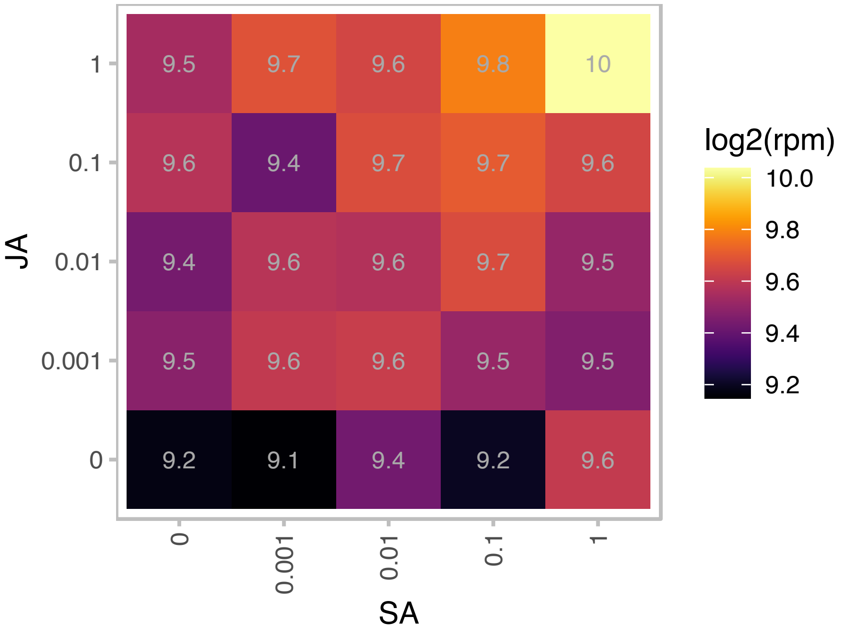


npr3-1 npr4-3

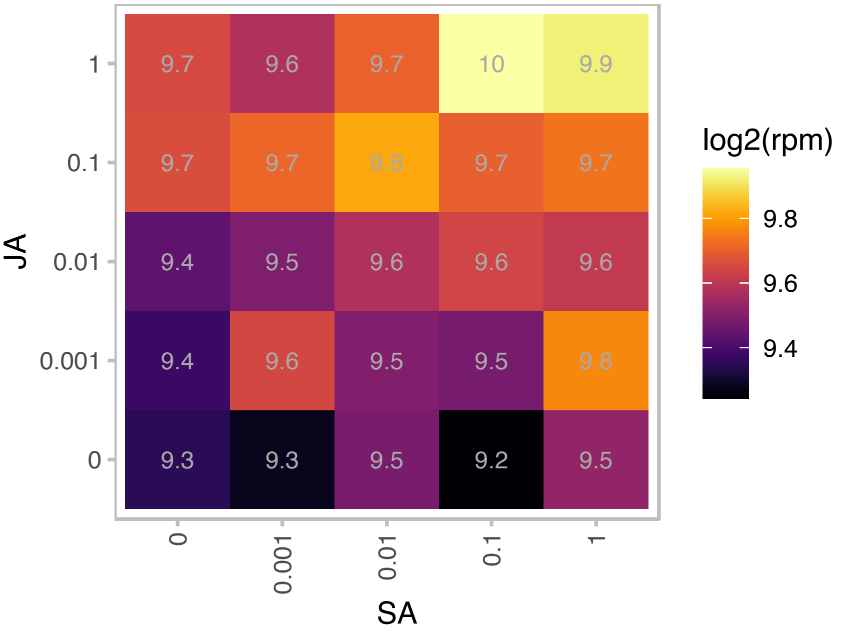

### AT1G53750.1 : RPT1A

regulatory particle triple-A 1A

26S proteasome AAA-ATPase subunit RPT1a (RPT1a) mRNA,

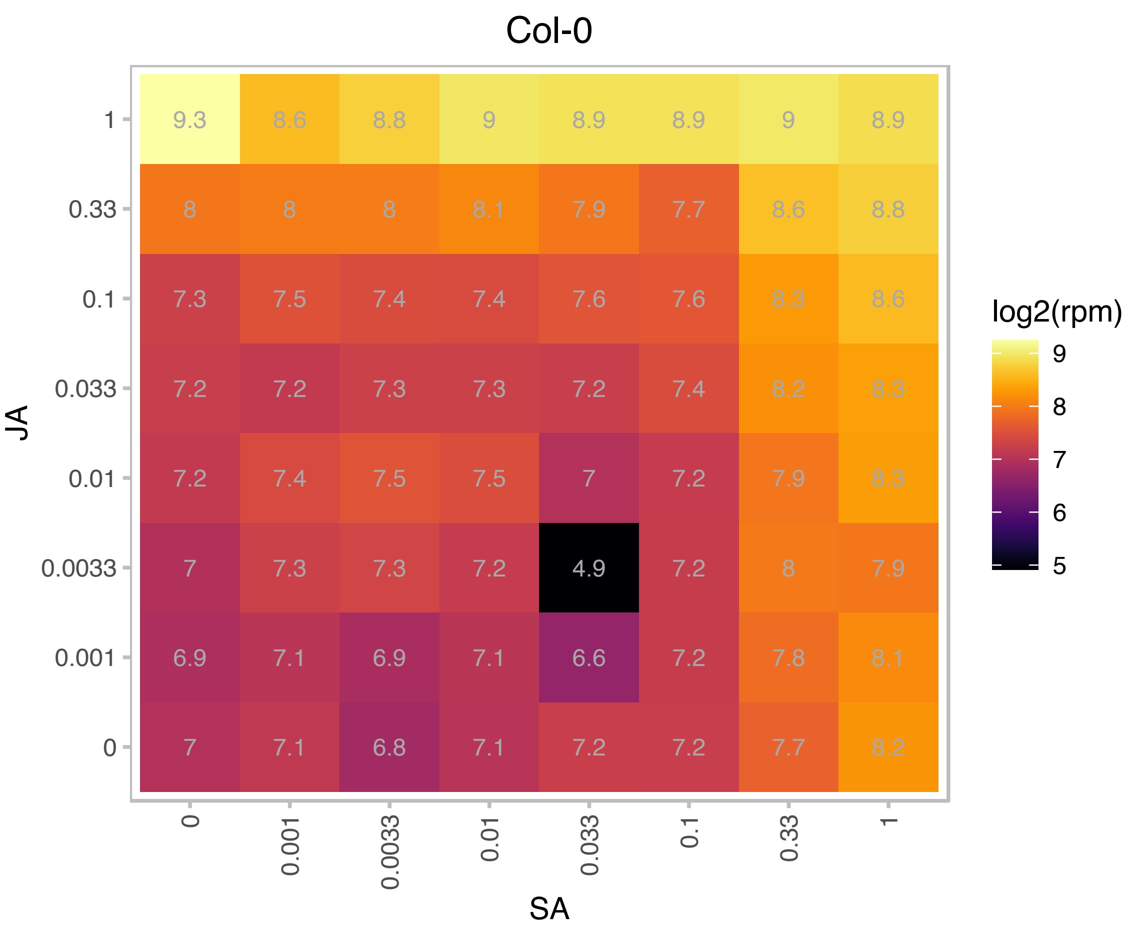


| | |
|---|---|
| Gene | AT1G53750 |
| is.expressed_genes | TRUE |
| is.responsive_genes | TRUE |
| is.apcluster | B-05 |

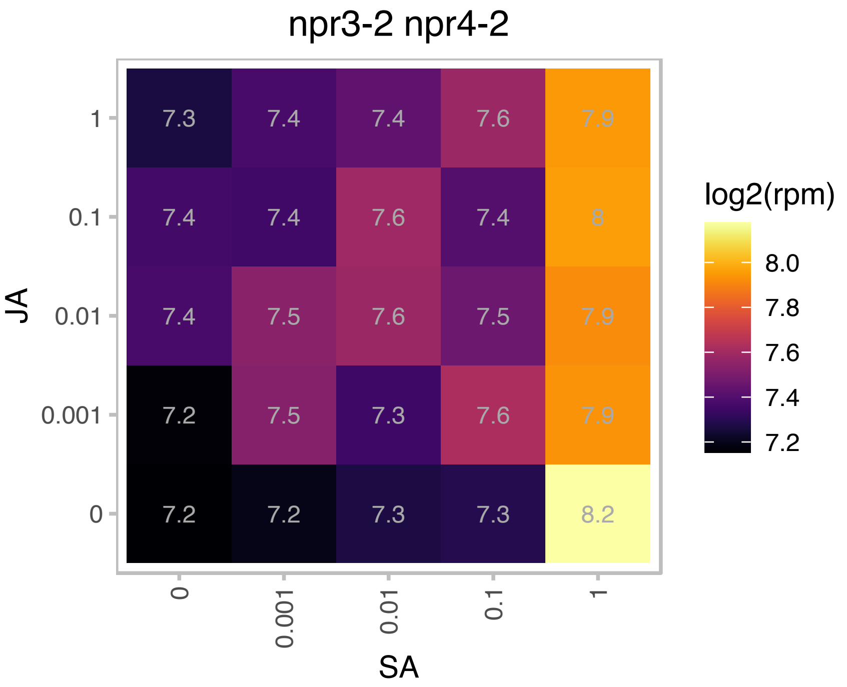


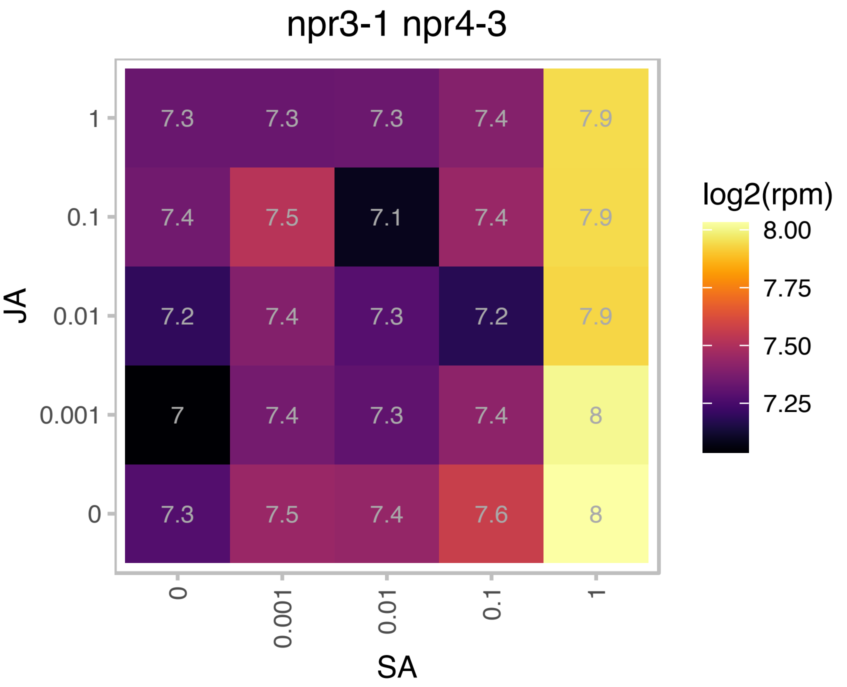

### AT1G55330.1 : AGP21, ATAGP21

arabinogalactan protein 21

Encodes a putative arabinogalactan-protein (AGP21).

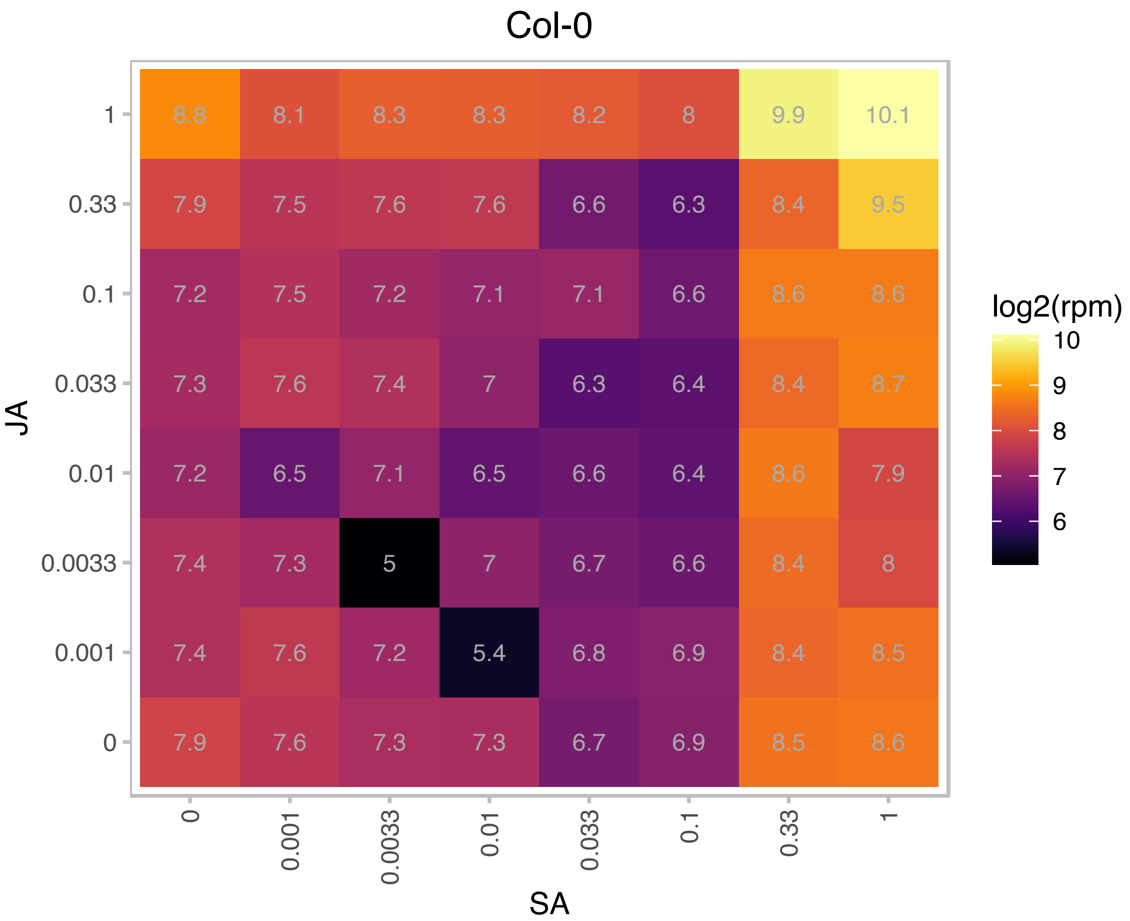


| | |
|---|---|
| Gene | AT1G55330 |
| is.expressed_genes | TRUE |
| is.responsive_genes | TRUE |
| is.apcluster | B-05 |

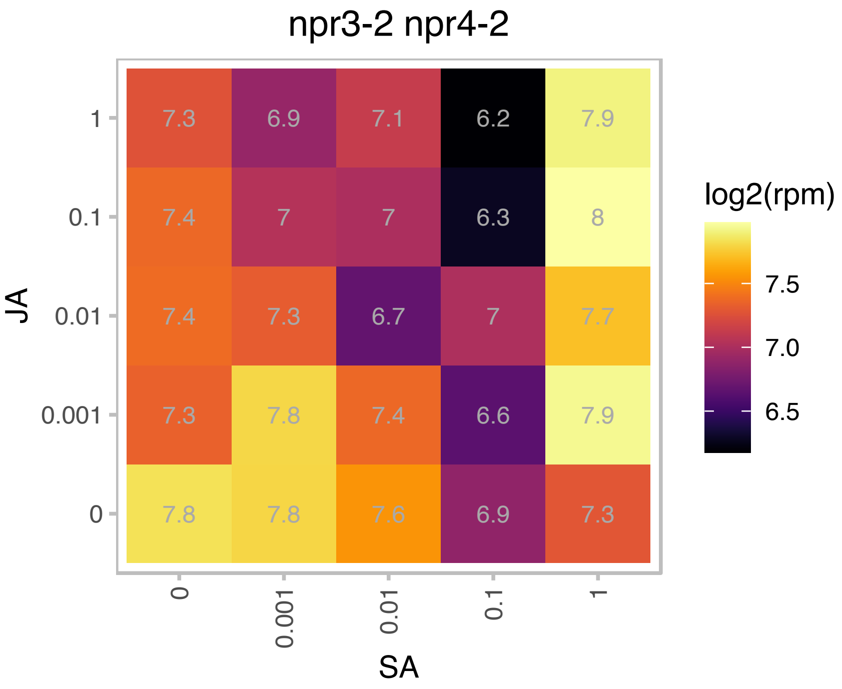


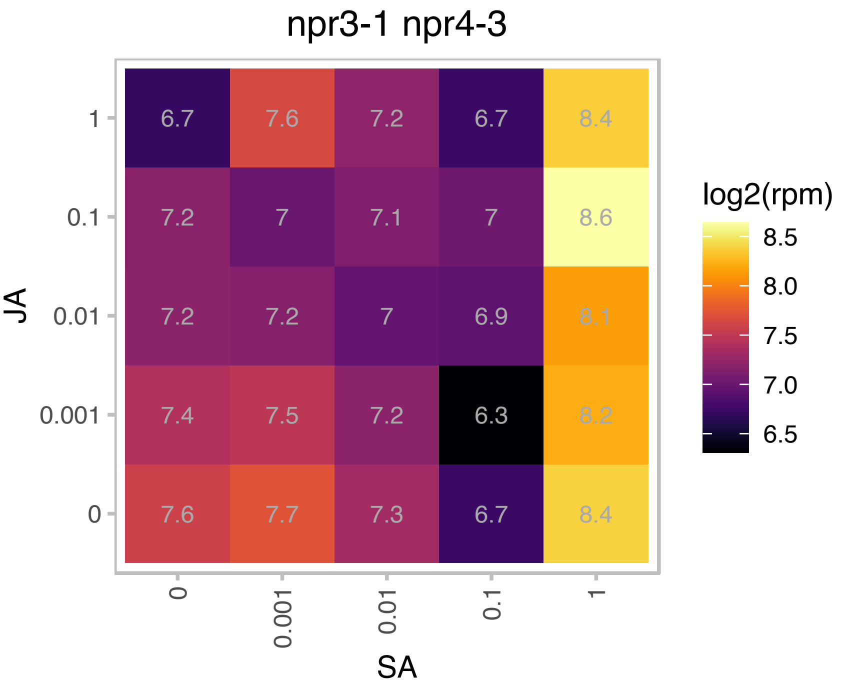

### AT1G59660.1 : DRAL

Nucleoporin autopeptidase

Encodes a protein with similarity to mammalian nucleoporin Nup98.Its expression is upregulated in mutants that are NUP deficient.

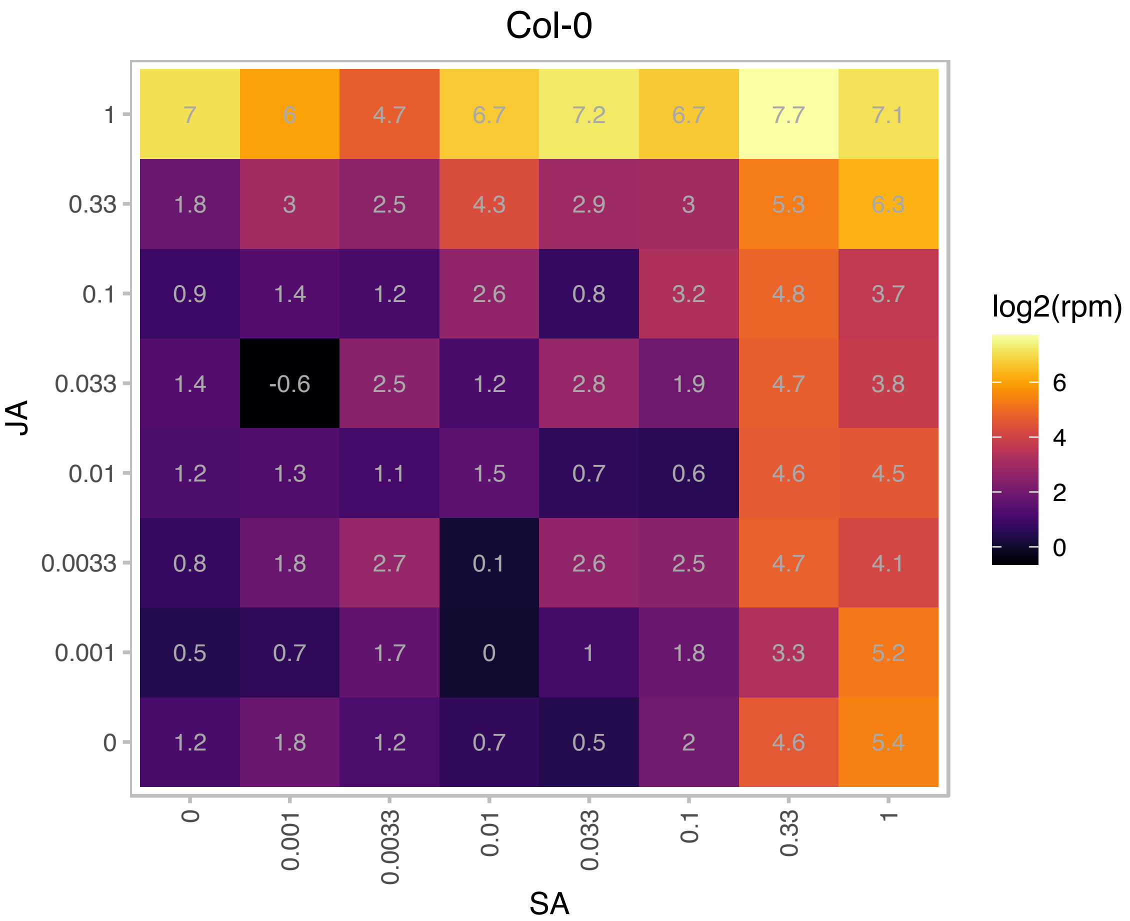


| | |
|---|---|
| Gene | AT1G59660 |
| is.expressed_genes | TRUE |
| is.responsive_genes | TRUE |
| is.apcluster | B-05 |

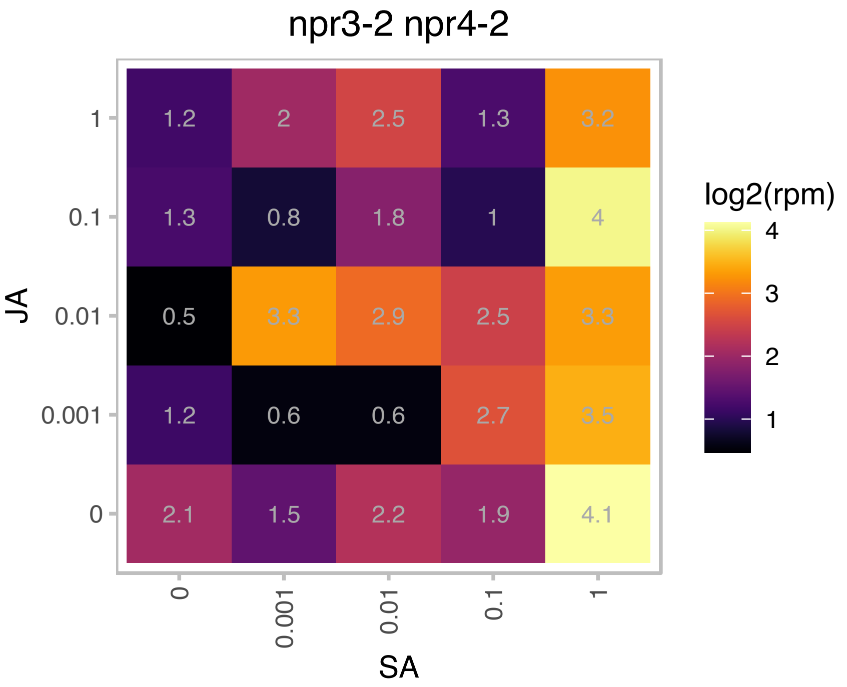


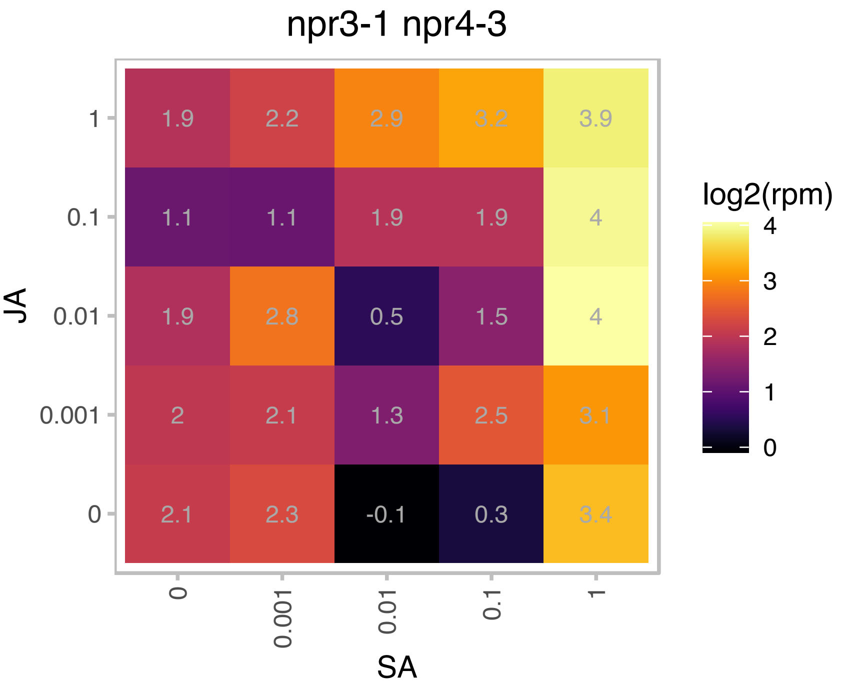

# AT1G59860.1 : NA

HSP20-like chaperones superfamily protein

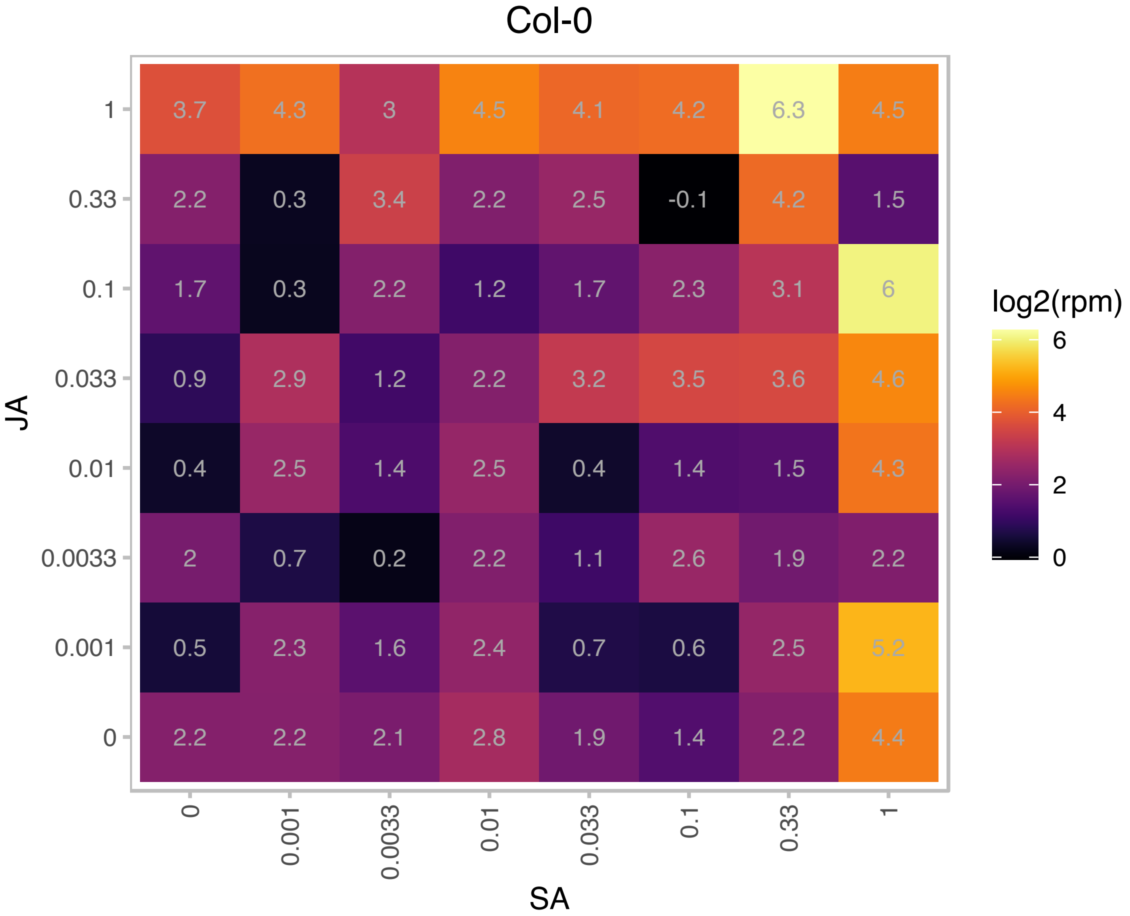


| | |
|---|---|
| Gene | AT1G59860 |
| is.expressed_genes | TRUE |
| is.responsive_genes | TRUE |
| is.apcluster | B-05 |

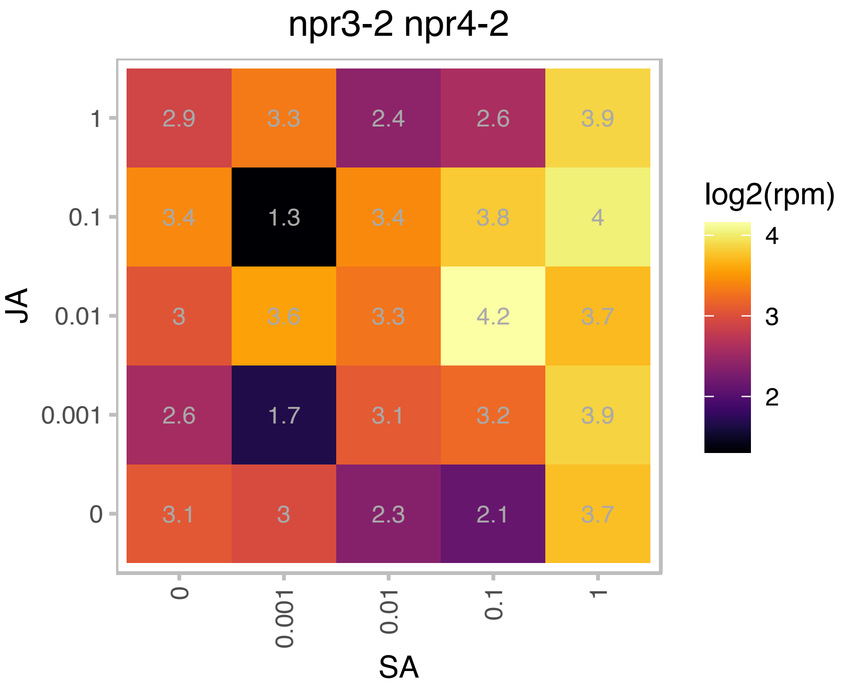


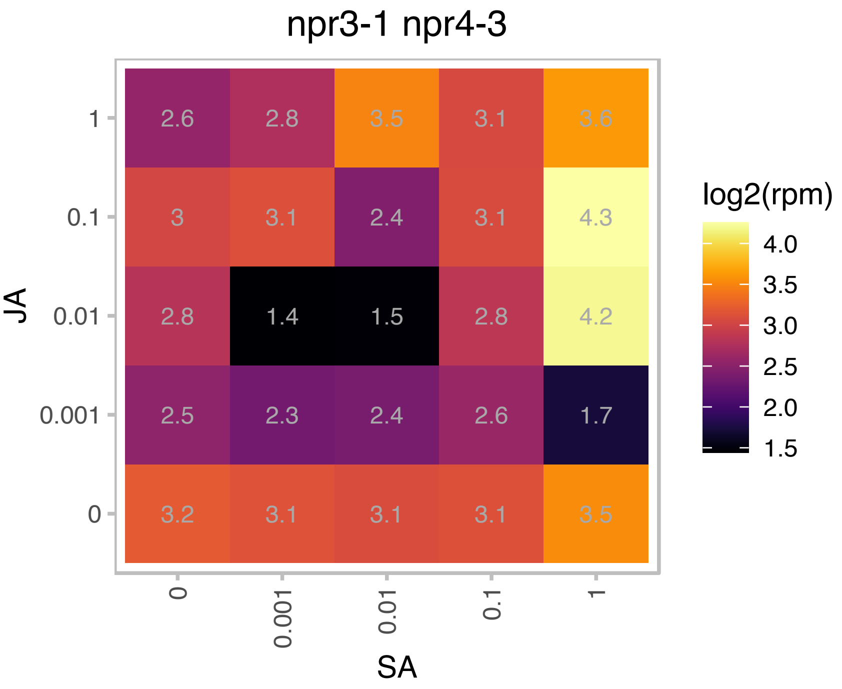

### AT1G63180.1 : UGE3

UDP-D-glucose/UDP-D-galactose 4-epimerase 3

Encodes a protein with UDP-D-glucose 4-epimerase activity. Involved in pollen development.

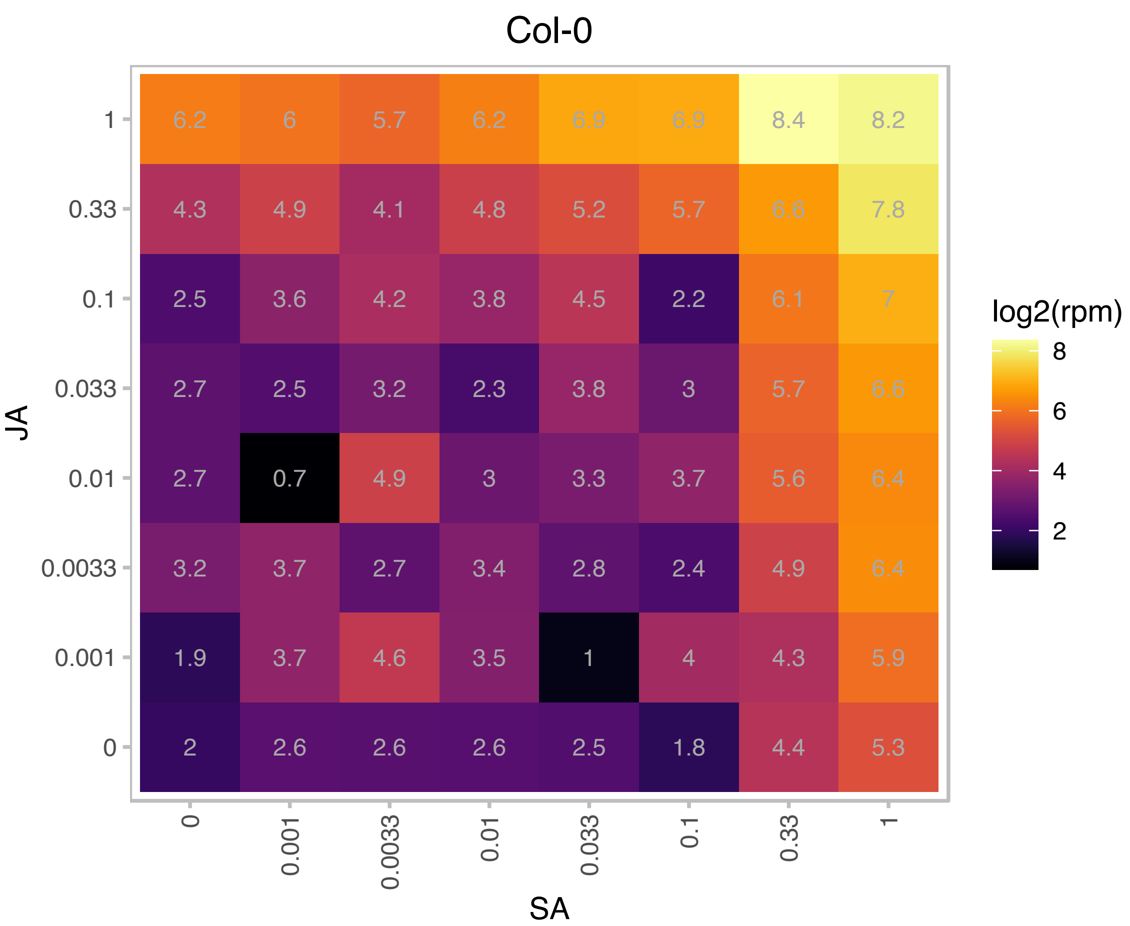


| | |
|---|---|
| Gene | AT1G63180 |
| is.expressed_genes | TRUE |
| is.responsive_genes | TRUE |
| is.apcluster | B-05 |

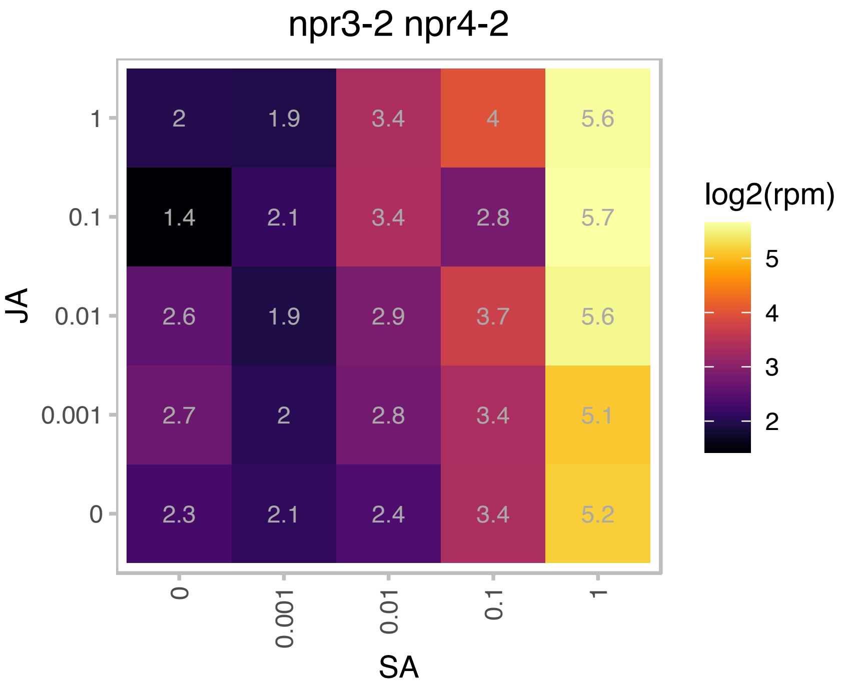


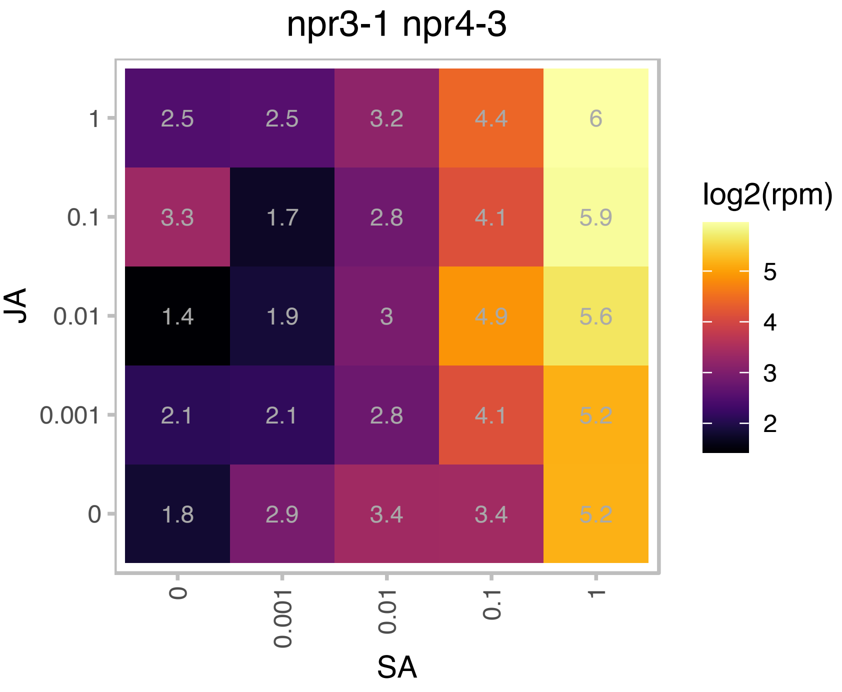

# AT1G65220.1 : NA

ARM repeat superfamily protein

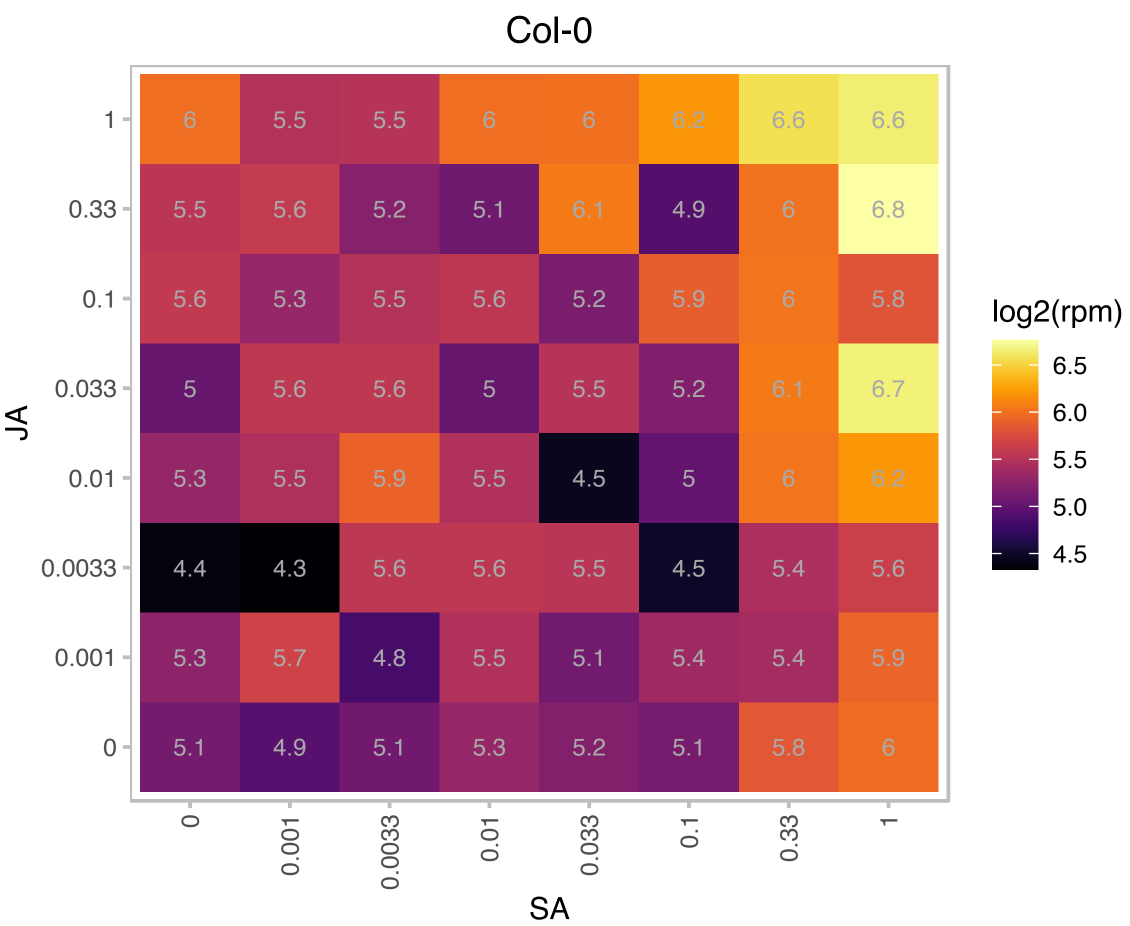


| | |
|---|---|
| Gene | AT1G65220 |
| is.expressed_genes | TRUE |
| is.responsive_genes | TRUE |
| is.apcluster | B-05 |

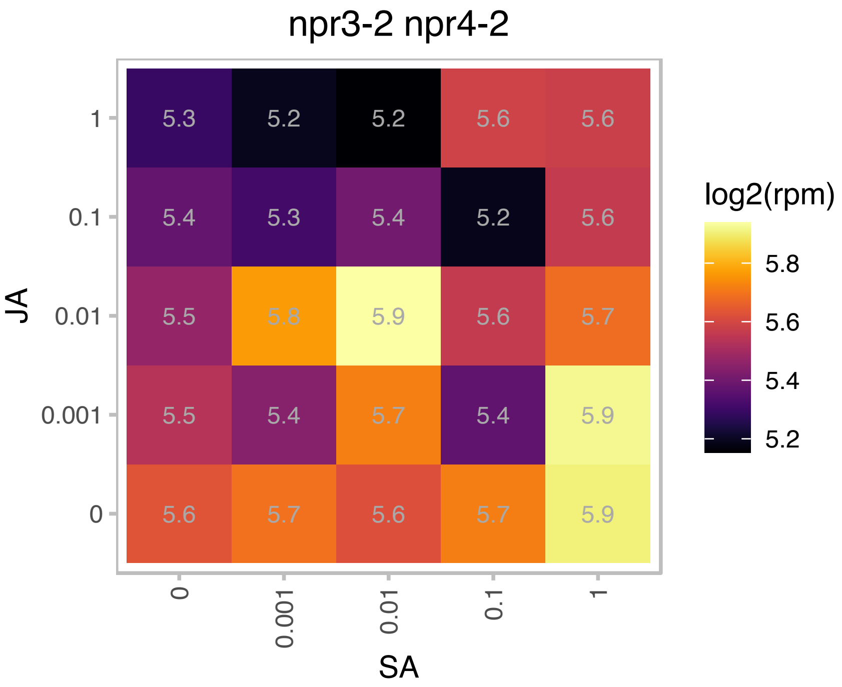


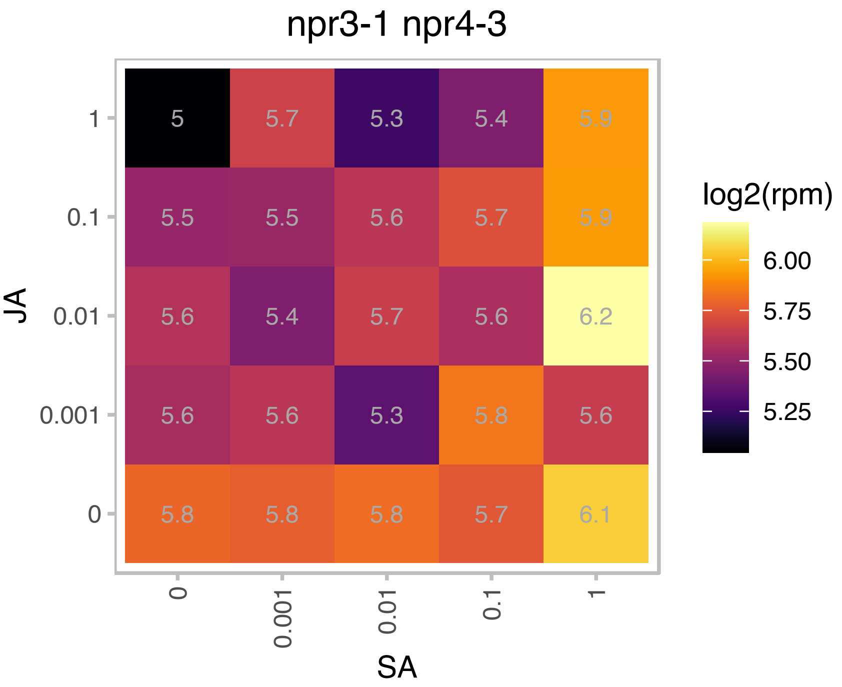

# AT1G66680.1 : AR401

S-adenosyl-L-methionine-dependent methyltransferases superfamily protein

unknown function

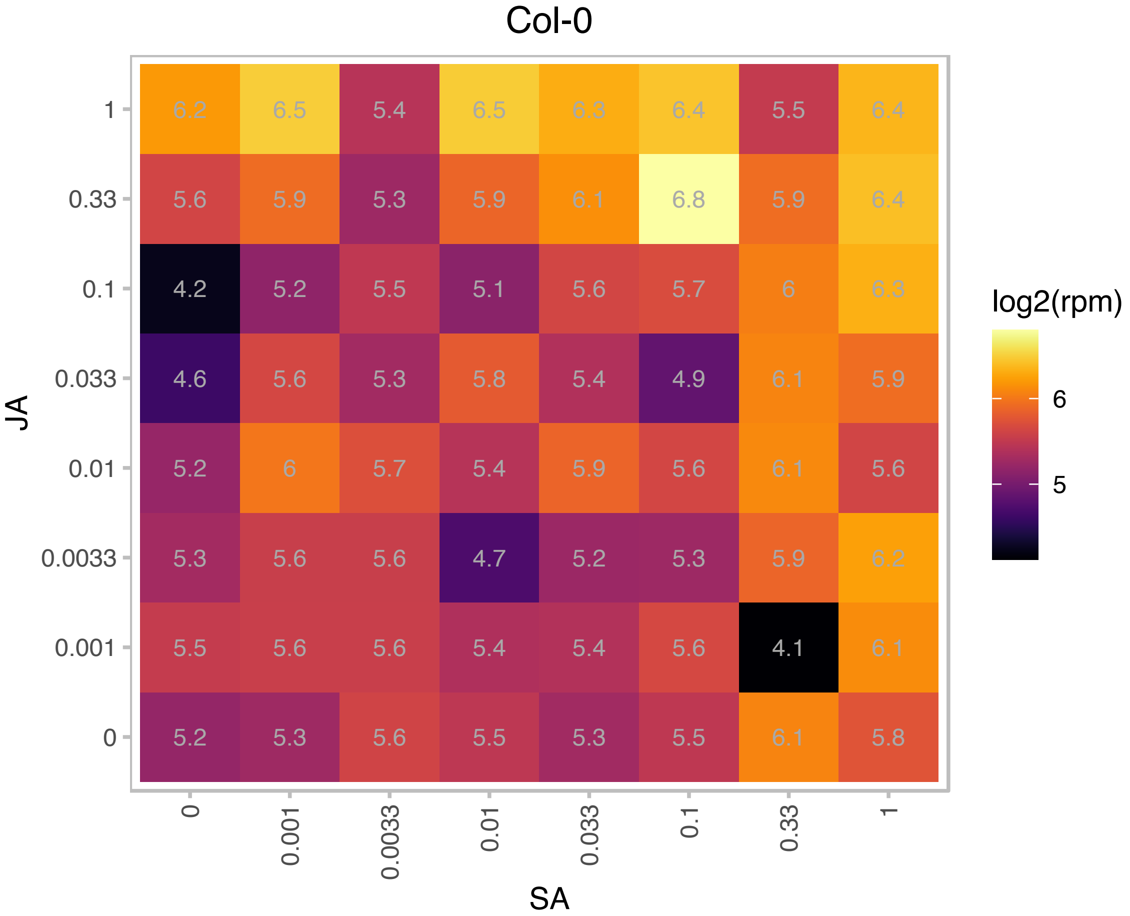


| | |
|---|---|
| Gene | AT1G66680 |
| is.expressed_genes | TRUE |
| is.responsive_genes | TRUE |
| is.apcluster | B-05 |

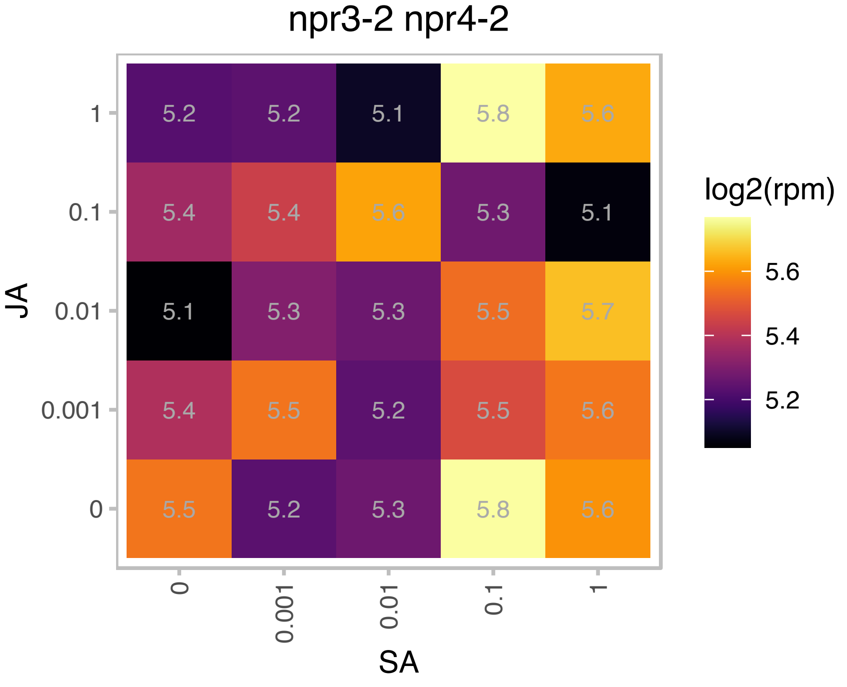


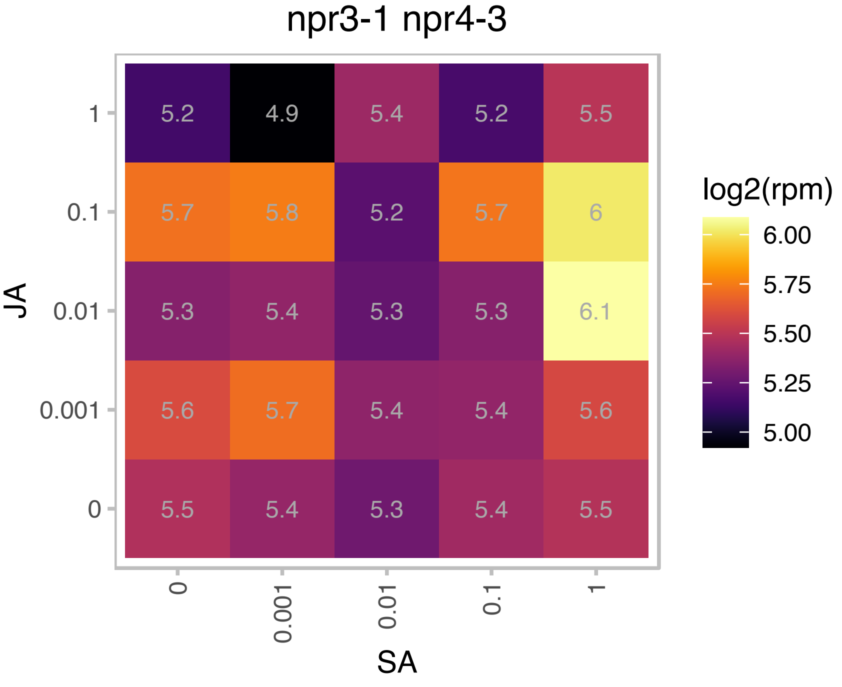

### AT1G67280.1 : AtGLYI6

Glyoxalase/Bleomycin resistance protein/Dioxygenase superfamily protein

Encodes a Ni+ dependent glyoxylase.

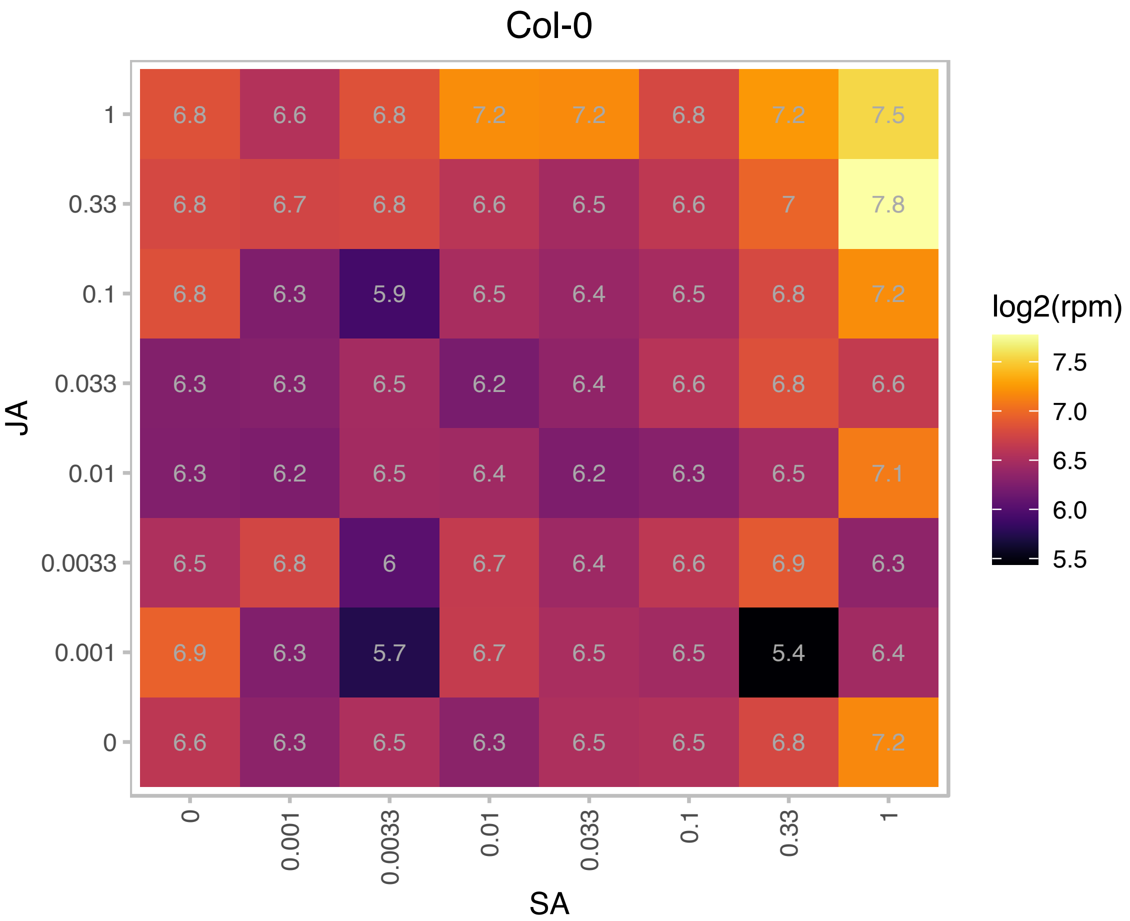


| | |
|---|---|
| Gene | AT1G67280 |
| is.expressed_genes | TRUE |
| is.responsive_genes | TRUE |
| is.apcluster | B-05 |

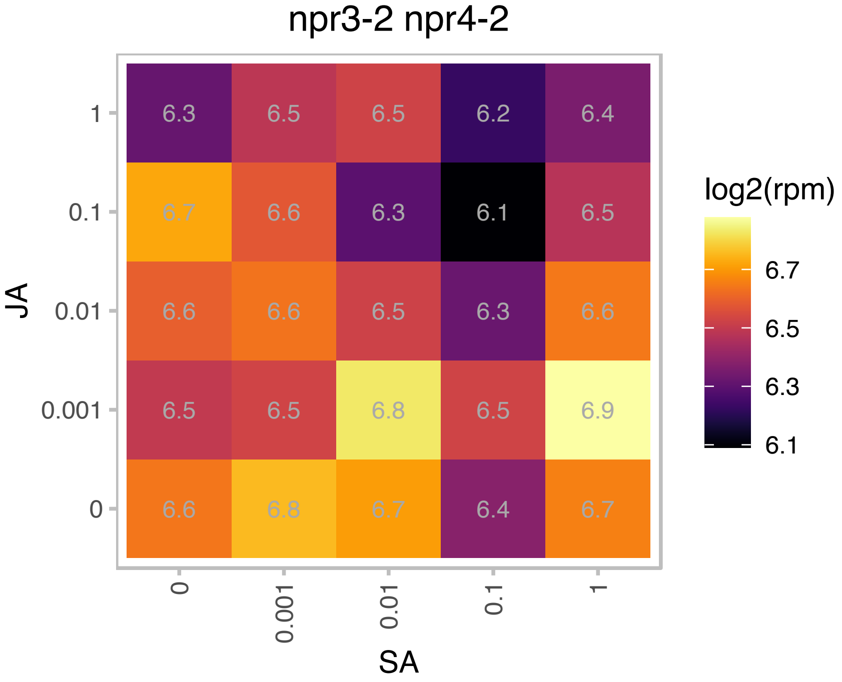


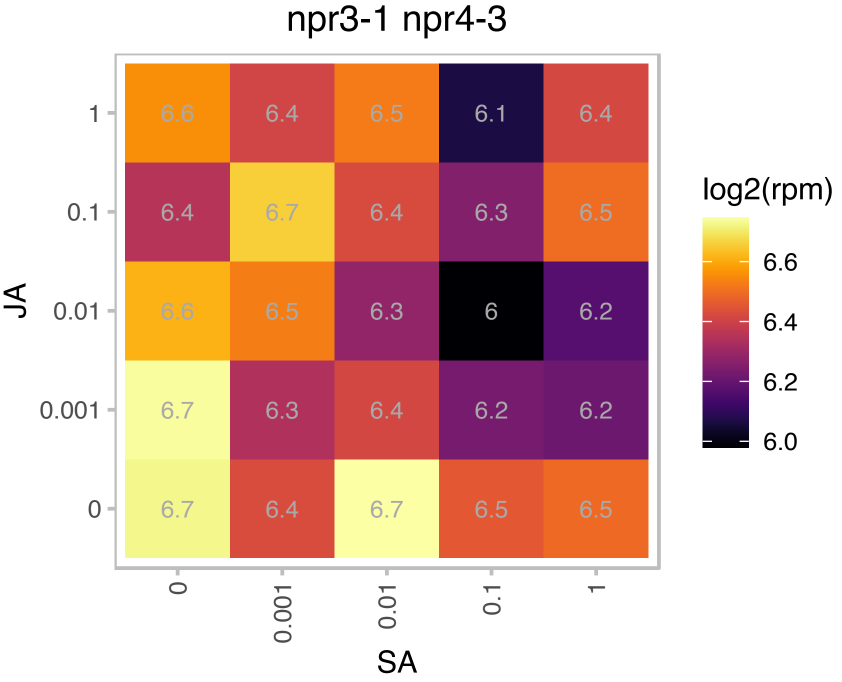

### AT1G67360.1 : LDAP1, SRP1

Rubber elongation factor protein (REF)

Encodes a small rubber particle protein homolog.  Plays dual roles as positive factors for tissue growth and development and in drought stress responses.

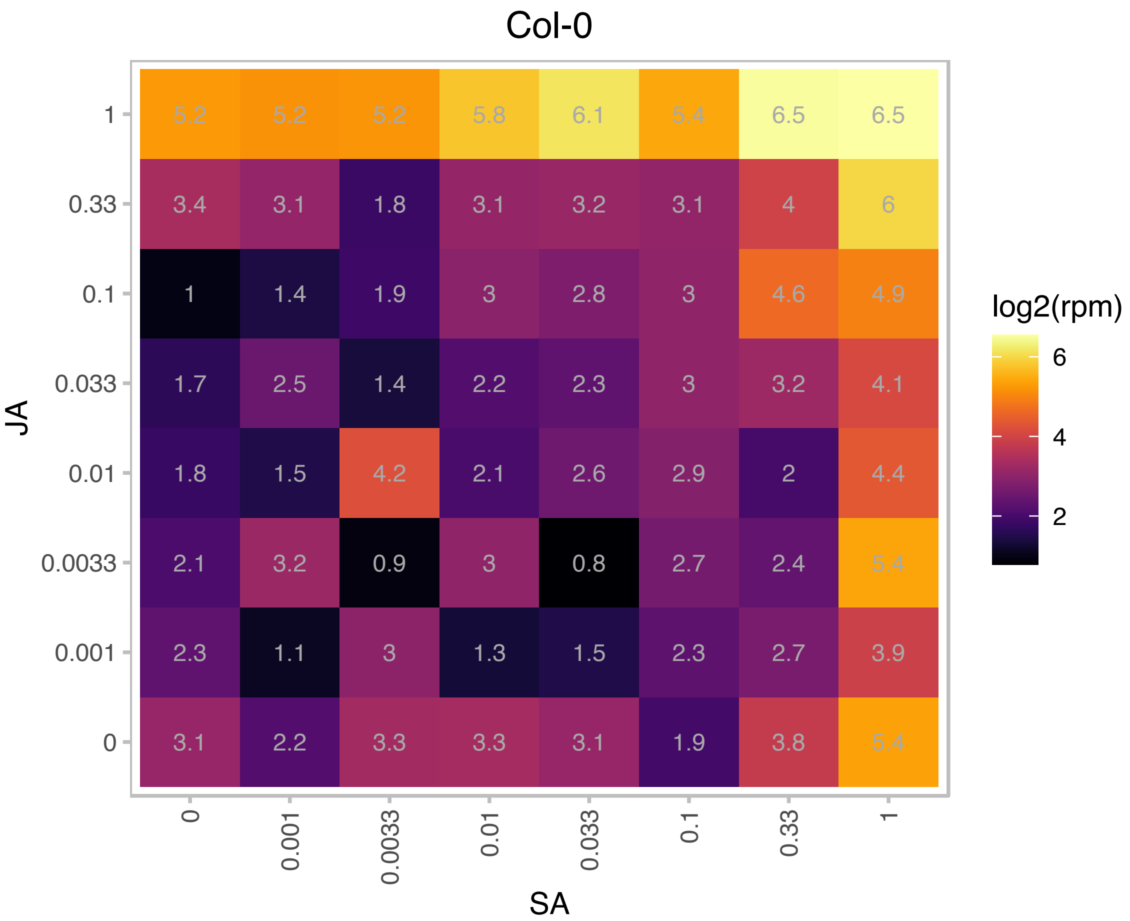


| | |
|---|---|
| Gene | AT1G67360 |
| is.expressed_genes | TRUE |
| is.responsive_genes | TRUE |
| is.apcluster | B-05 |

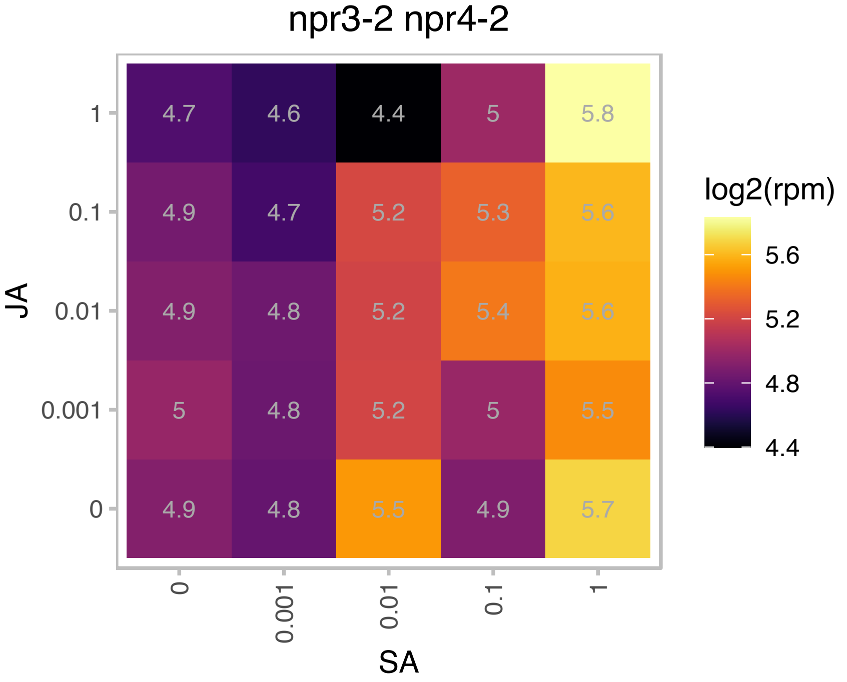


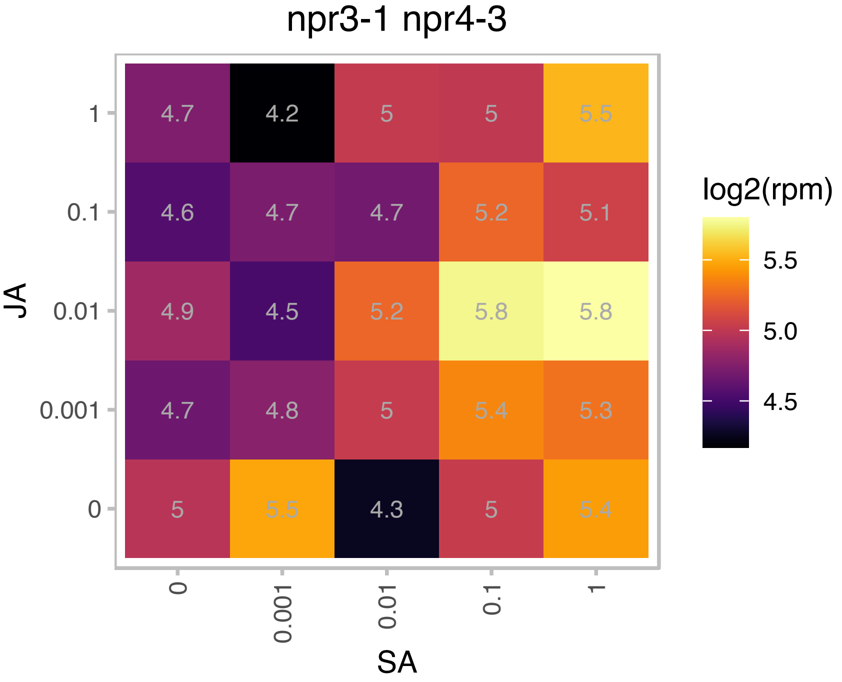

### AT1G67960.1 : POD1

POD1 is involved in pollen tube guidence and early embryo patterning.

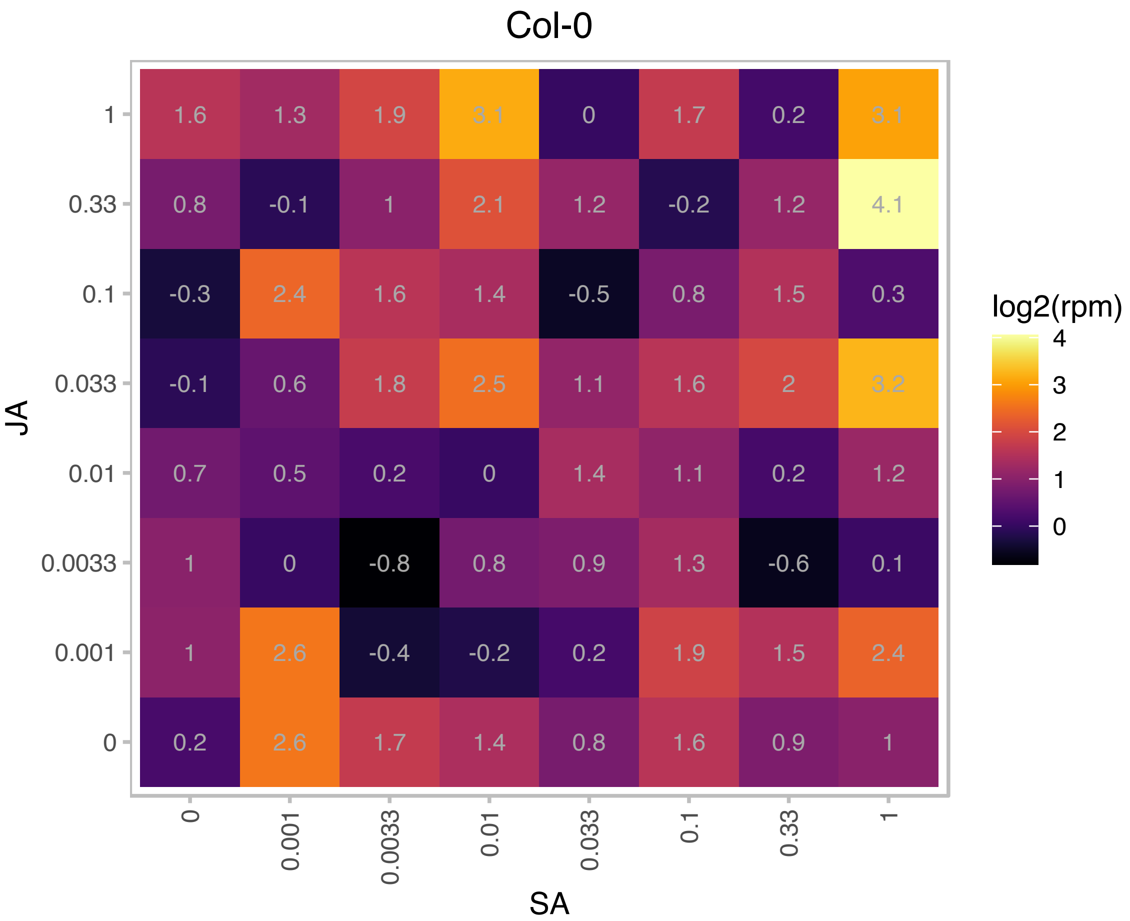


| | |
|---|---|
| Gene | AT1G67960 |
| is.expressed_genes | TRUE |
| is.responsive_genes | TRUE |
| is.apcluster | B-05 |

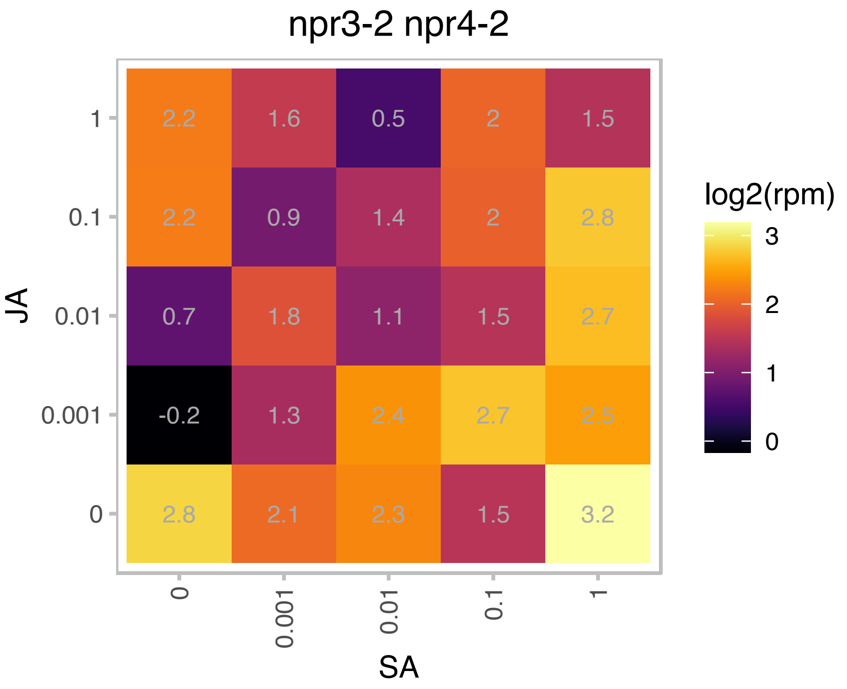


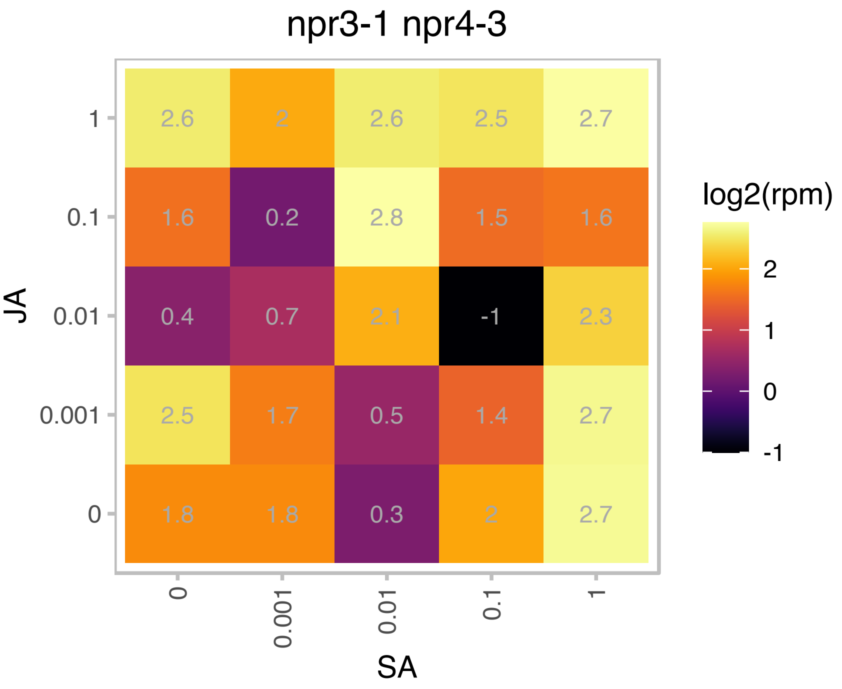

# AT1G68440.1 : NA

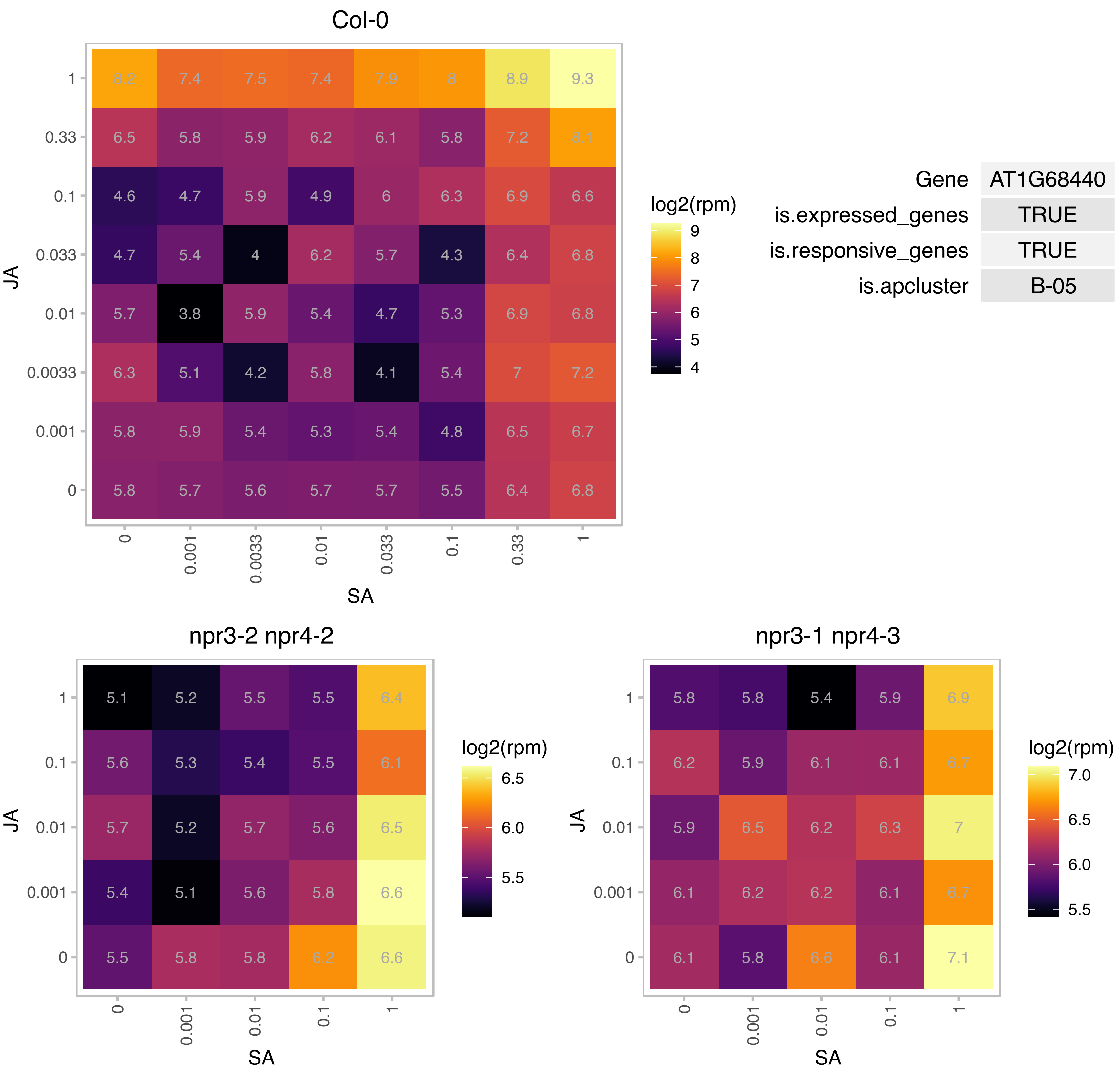

### AT1G71140.1 : DTX14

MATE efflux family protein

MATE transporter that can export the antibiotic norfloxacin.

Col-0

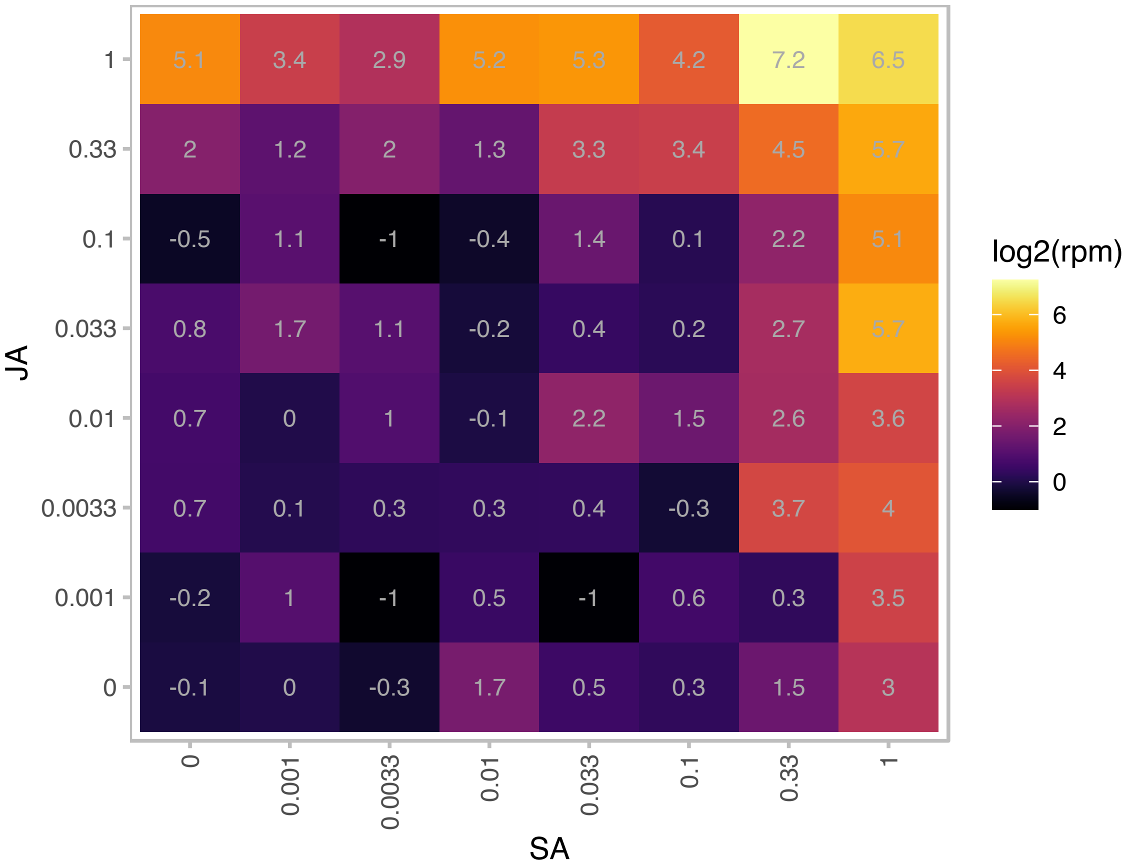


| | |
|---|---|
| Gene | AT1G71140 |
| is.expressed_genes | TRUE |
| is.responsive_genes | TRUE |
| is.apcluster | B-05 |

npr3-2 npr4-2

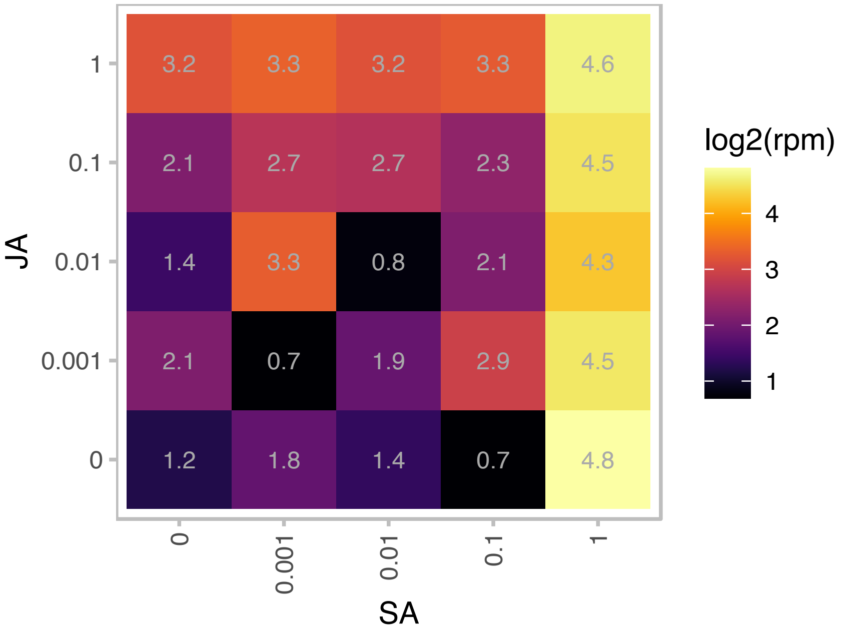


npr3-1 npr4-3

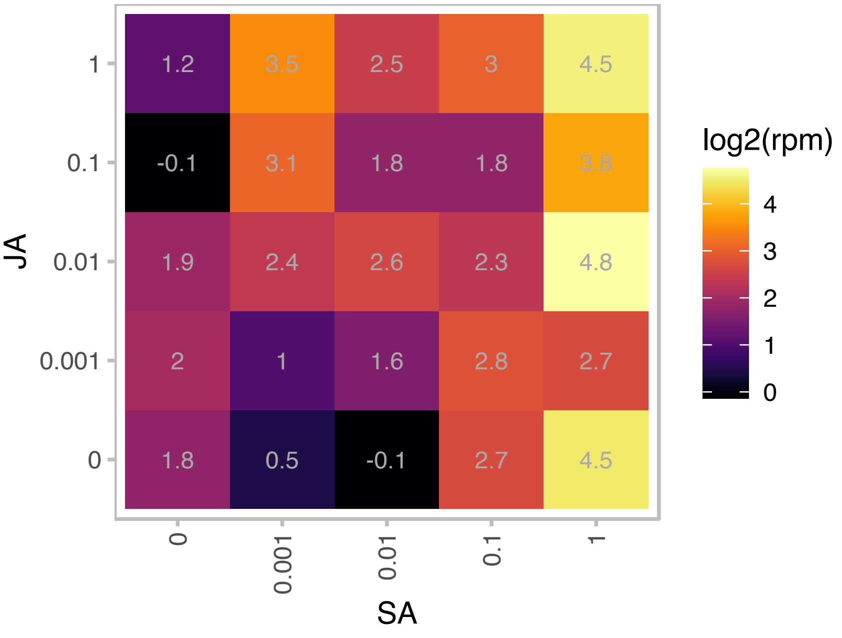

# AT1G72800.1 : NA

RNA-binding (RRM/RBD/RNP motifs) family protein

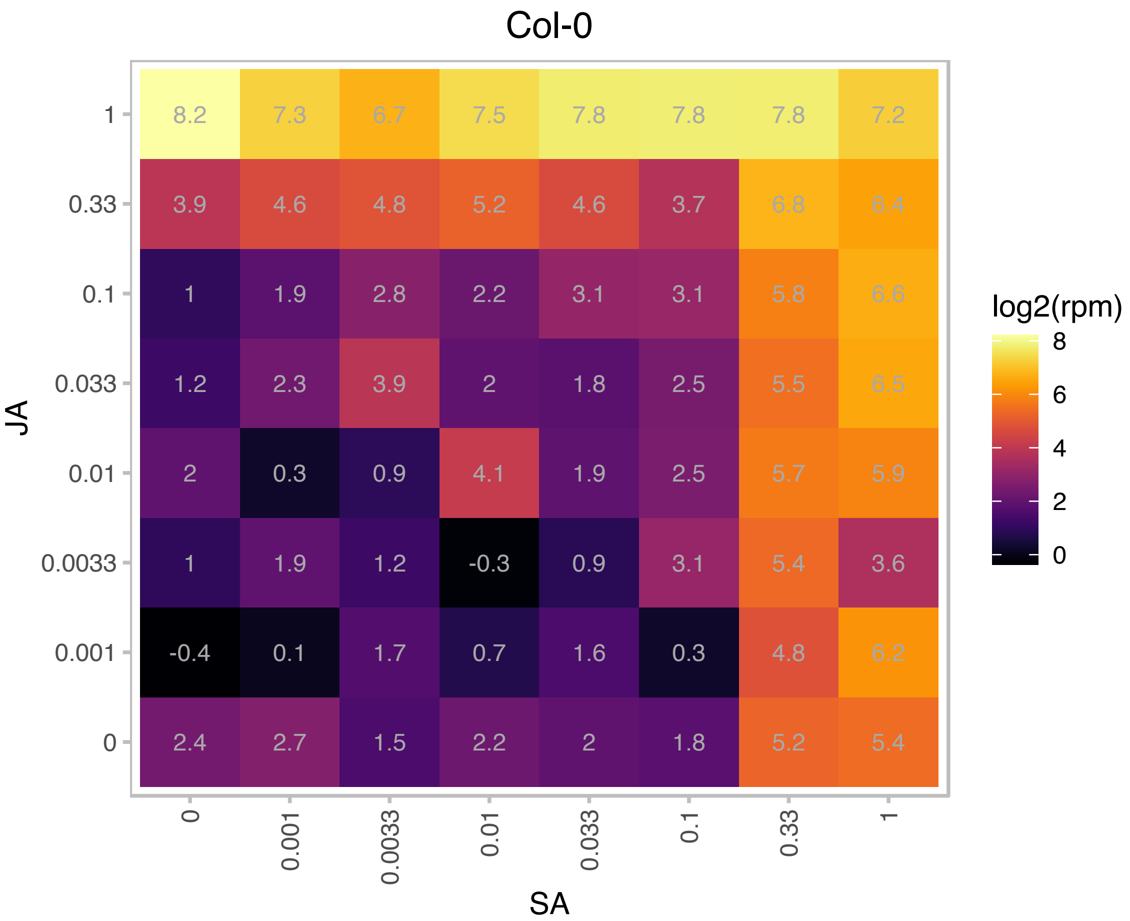


| | |
|---|---|
| Gene | AT1G72800 |
| is.expressed_genes | TRUE |
| is.responsive_genes | TRUE |
| is.apcluster | B-05 |

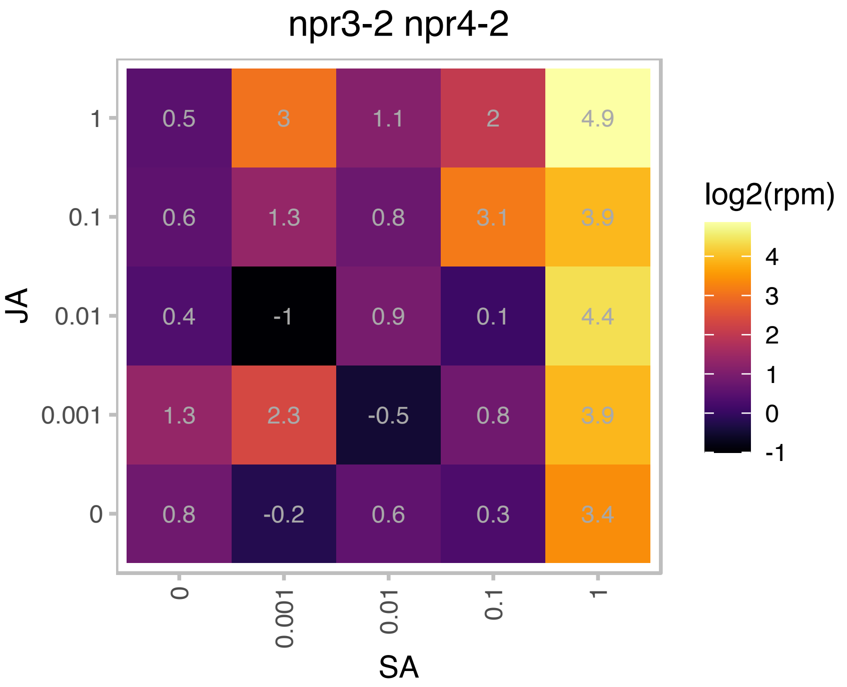


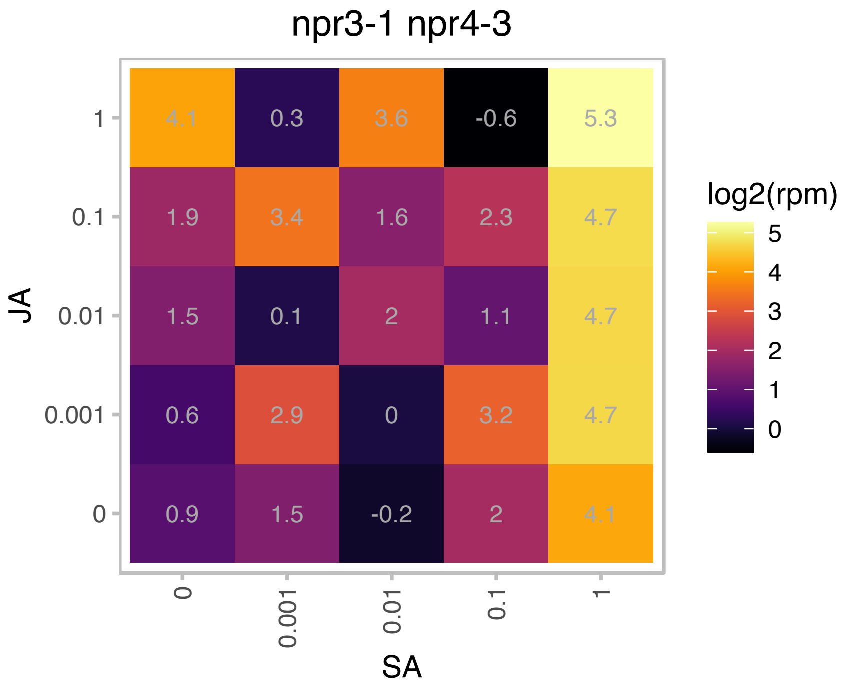

### AT1G73880.1 : UGT89B1

UDP-glucosyl transferase 89B1

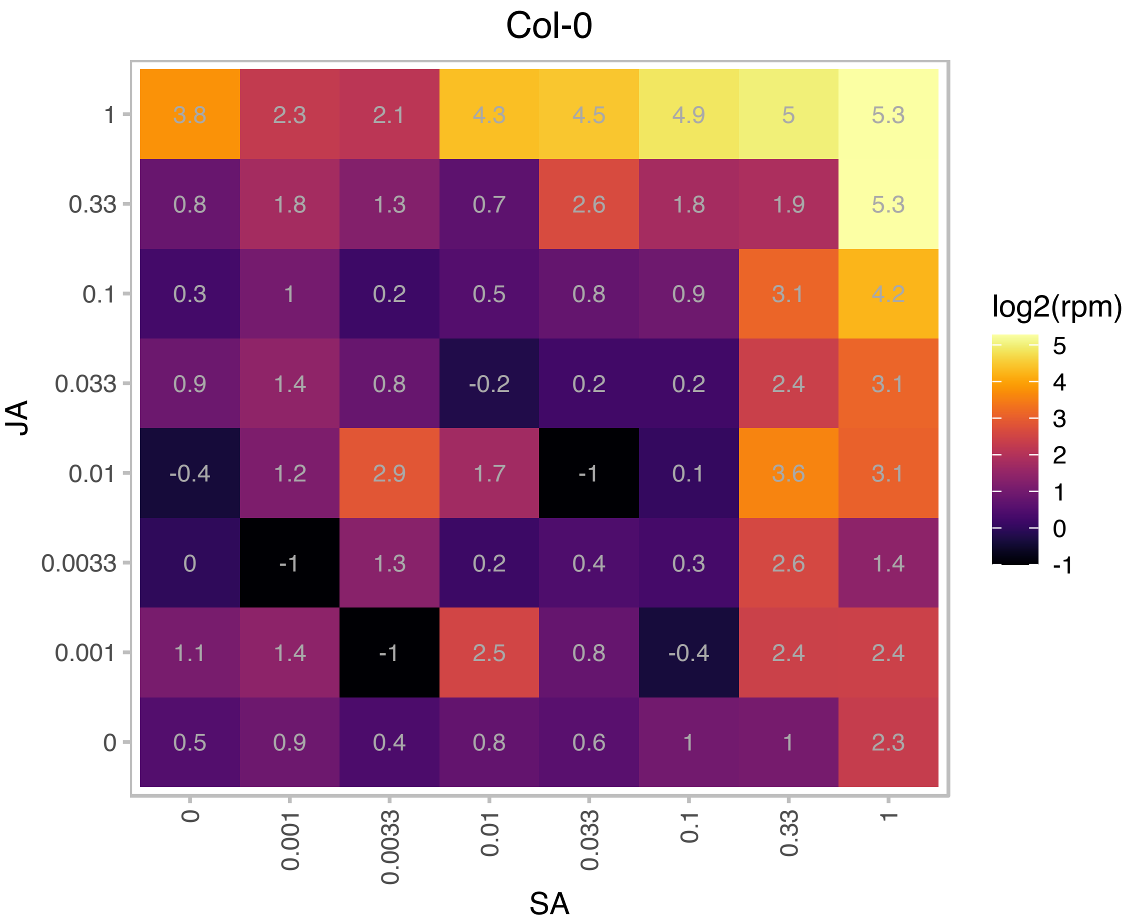


| | |
|---|---|
| Gene | AT1G73880 |
| is.expressed_genes | TRUE |
| is.responsive_genes | TRUE |
| is.apcluster | B-05 |

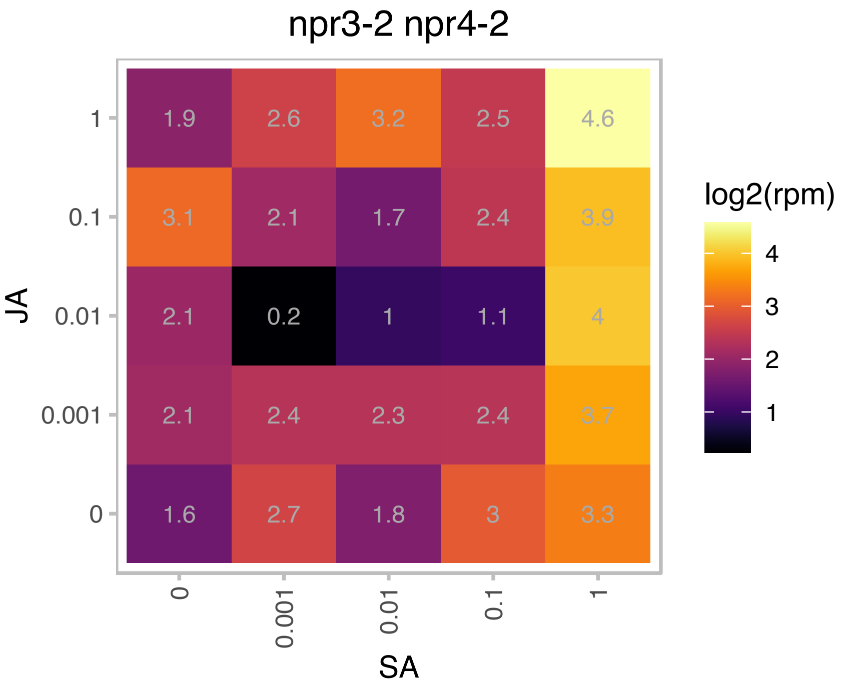


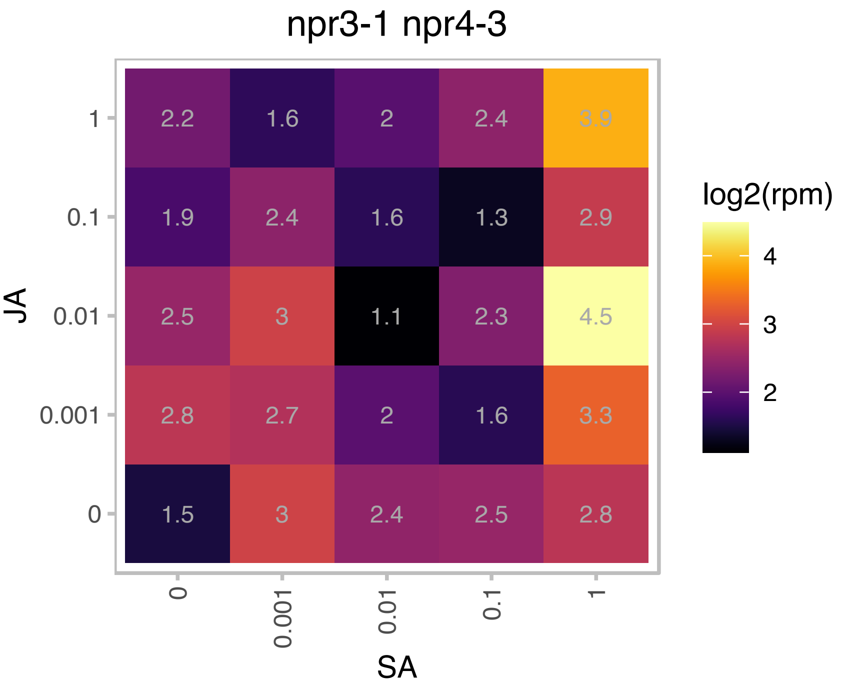

### AT1G80110.1 : ATPP2-B11, PP2-B11

phloem protein 2-B11

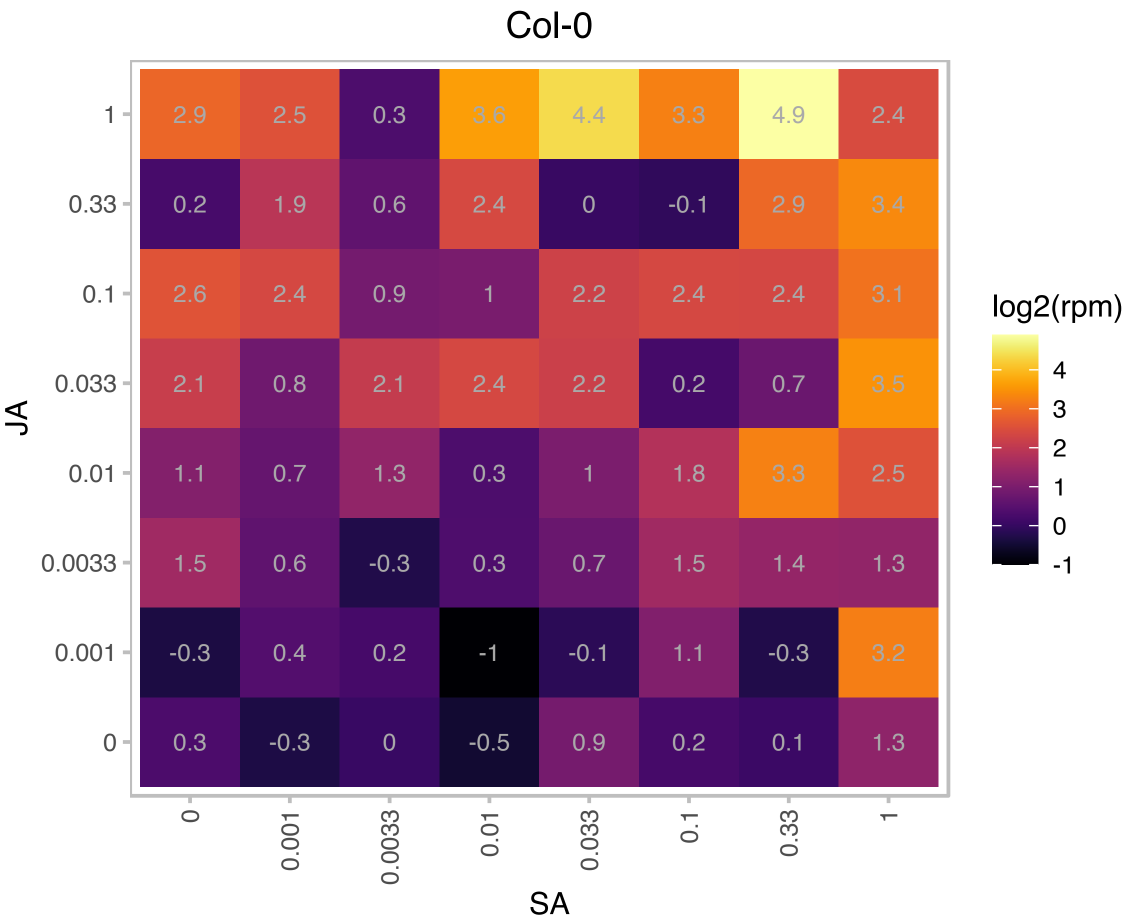


| | |
|---|---|
| Gene | AT1G80110 |
| is.expressed_genes | TRUE |
| is.responsive_genes | TRUE |
| is.apcluster | B-05 |

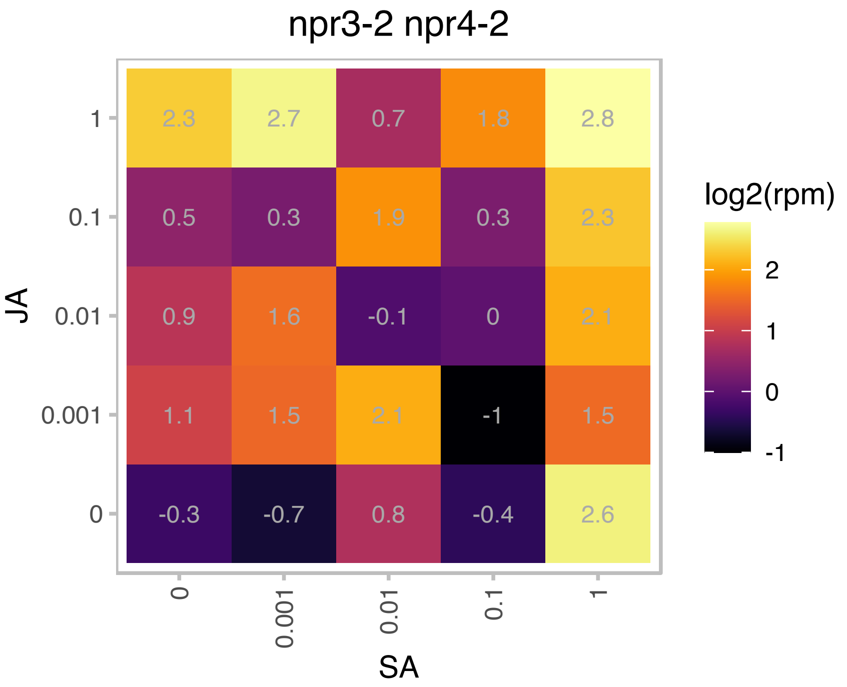


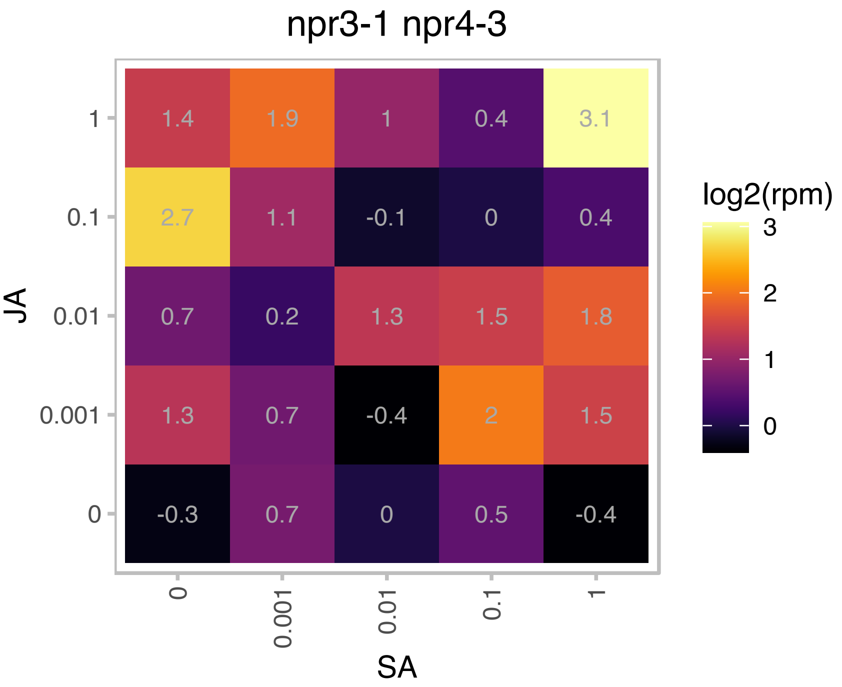

# AT2G16900.1 : NA

Arabidopsis phospholipase-like protein (PEARLI 4) family

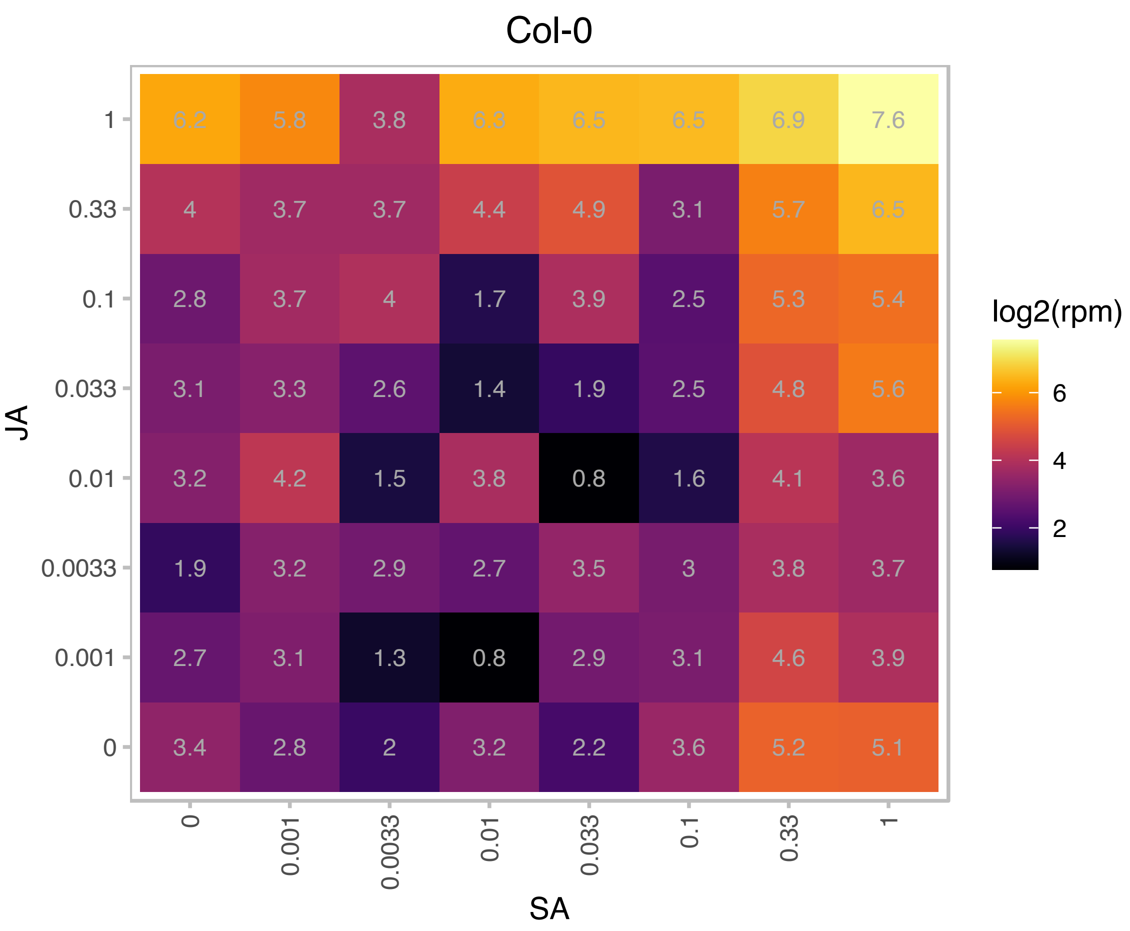


| | |
|---|---|
| Gene | AT2G16900 |
| is.expressed_genes | TRUE |
| is.responsive_genes | TRUE |
| is.apcluster | B-05 |

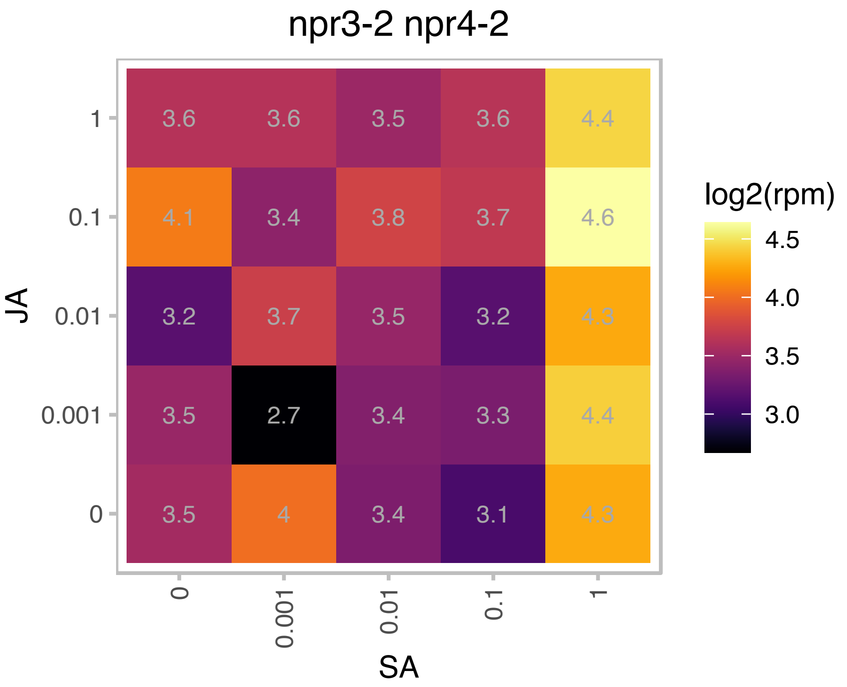


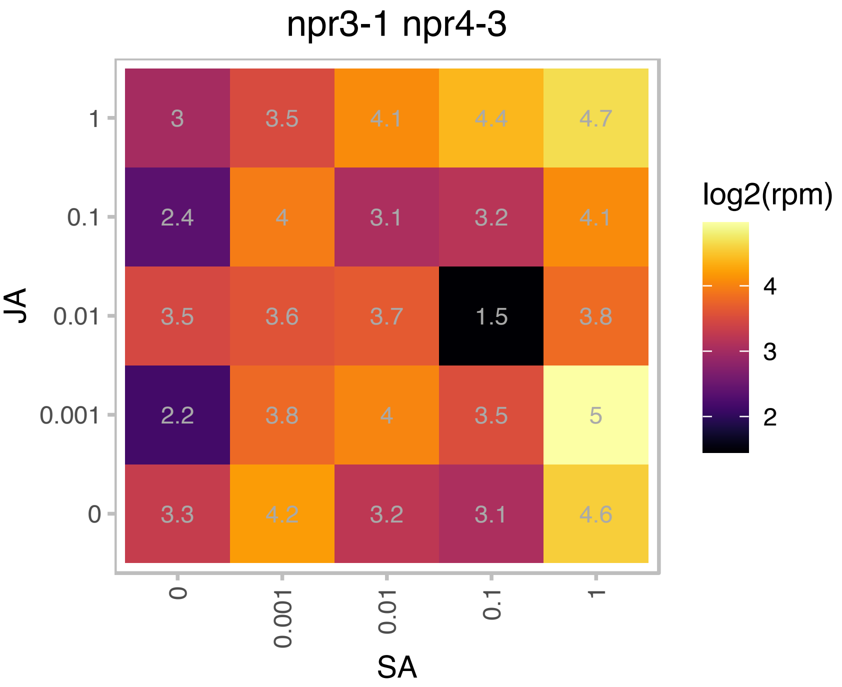

### AT2G17550.1 : TRM26

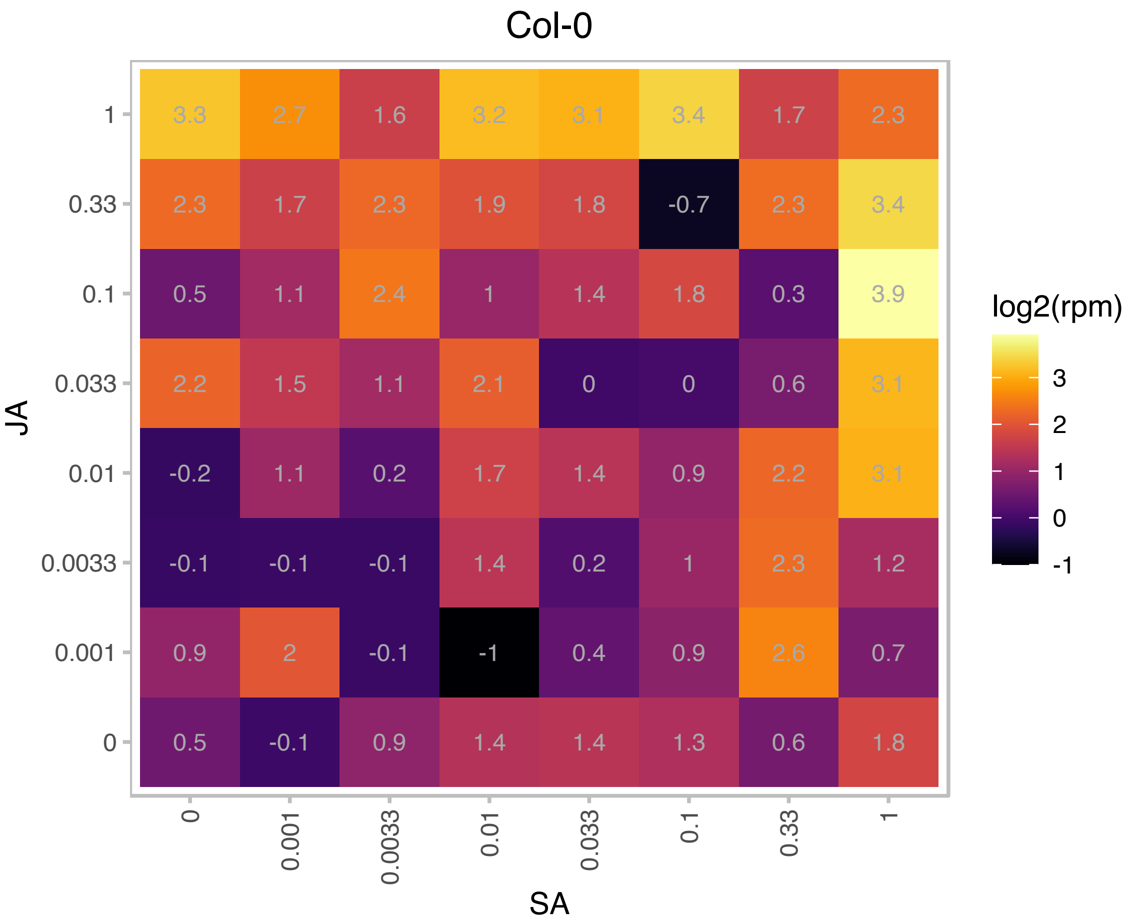


| | |
|---|---|
| Gene | AT2G17550 |
| is.expressed_genes | TRUE |
| is.responsive_genes | TRUE |
| is.apcluster | B-05 |

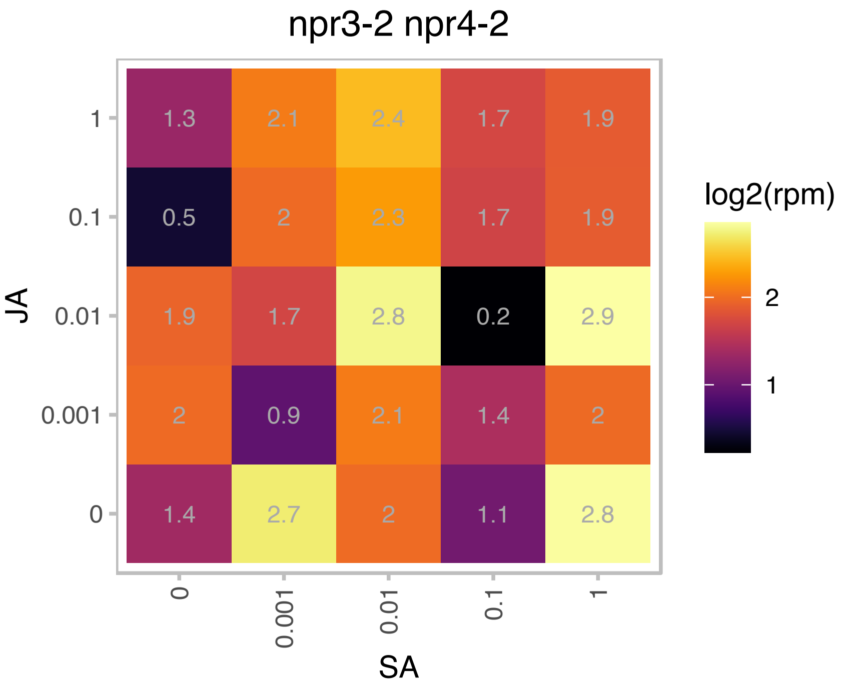


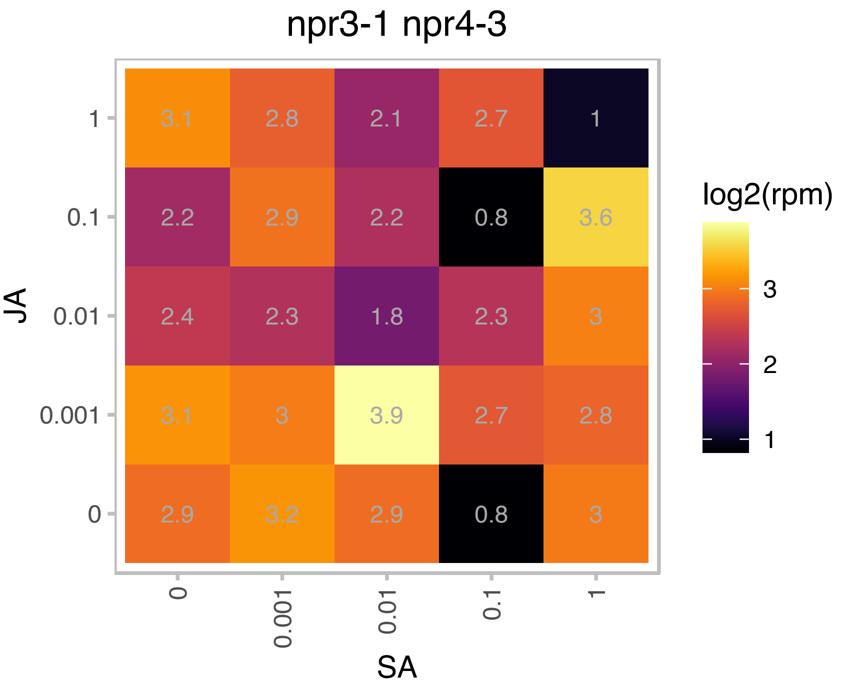

### AT2G17730.1 : NIP2

NEP-interacting protein 2

Intrinsic thylakoid membrane protein that fixes RPOTmp on the stromal side of the thylakoid membrane.

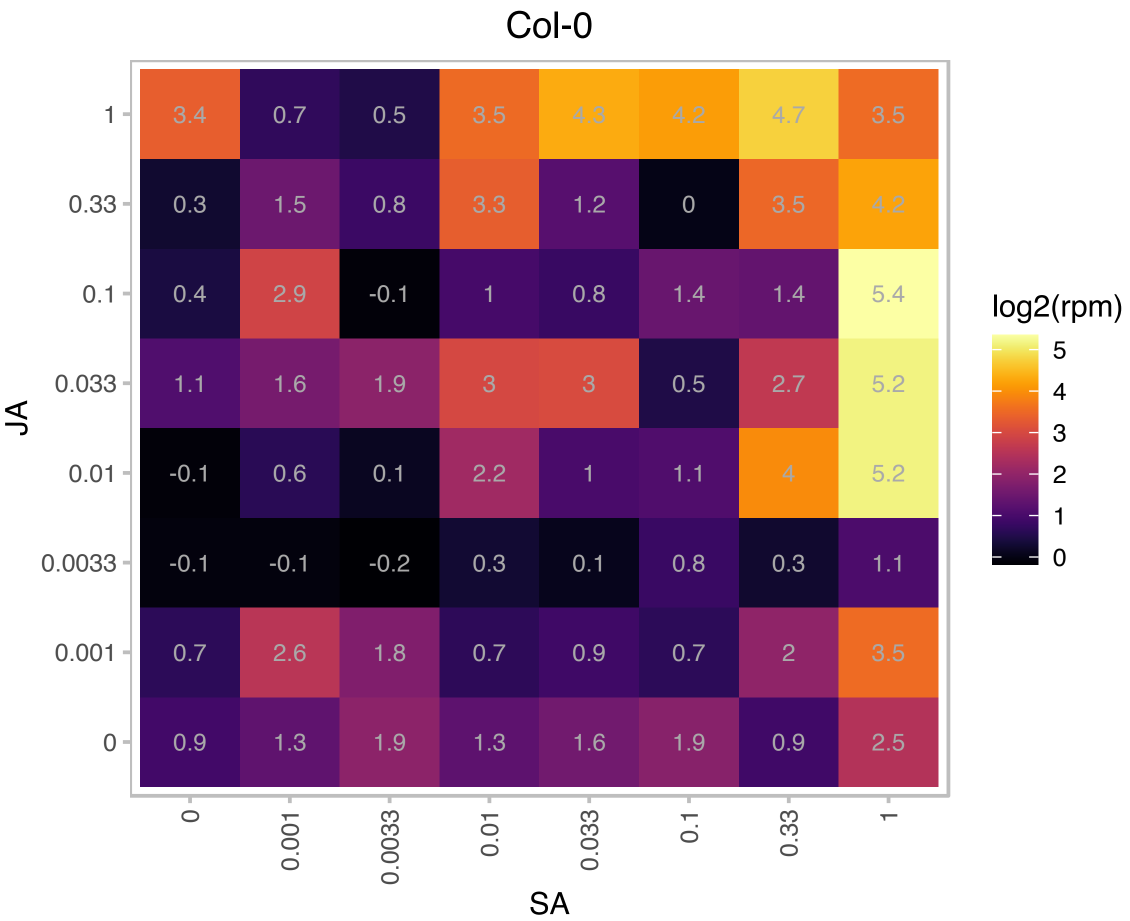


| | |
|---|---|
| Gene | AT2G17730 |
| is.expressed_genes | TRUE |
| is.responsive_genes | TRUE |
| is.apcluster | B-05 |

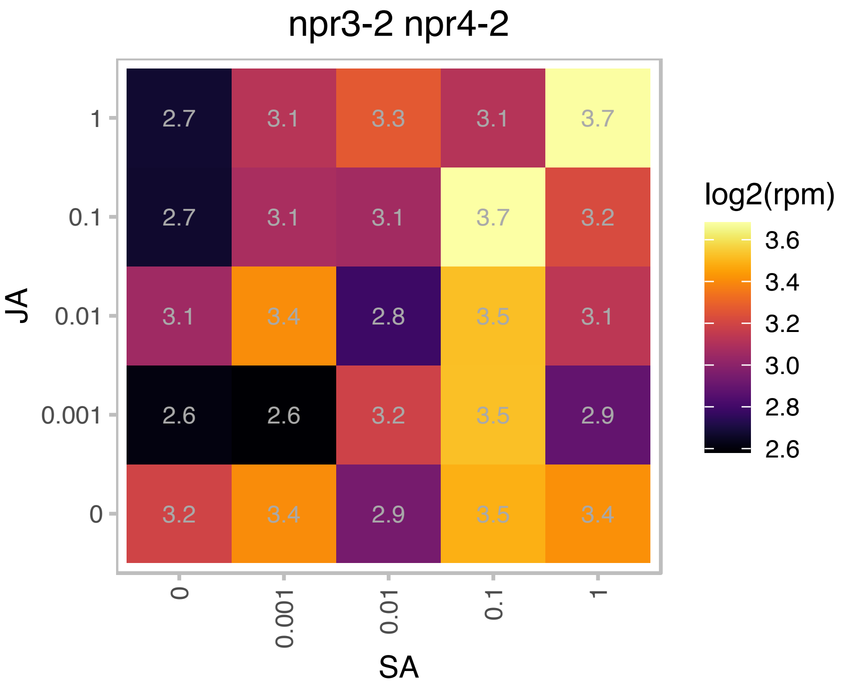


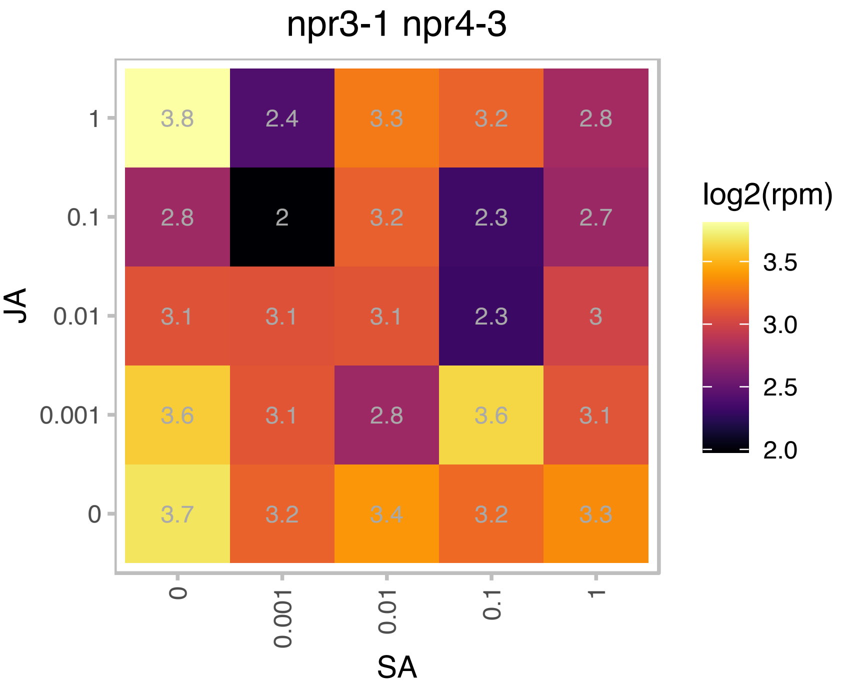

# AT2G18600.1 : NA

Ubiquitin-conjugating enzyme family protein

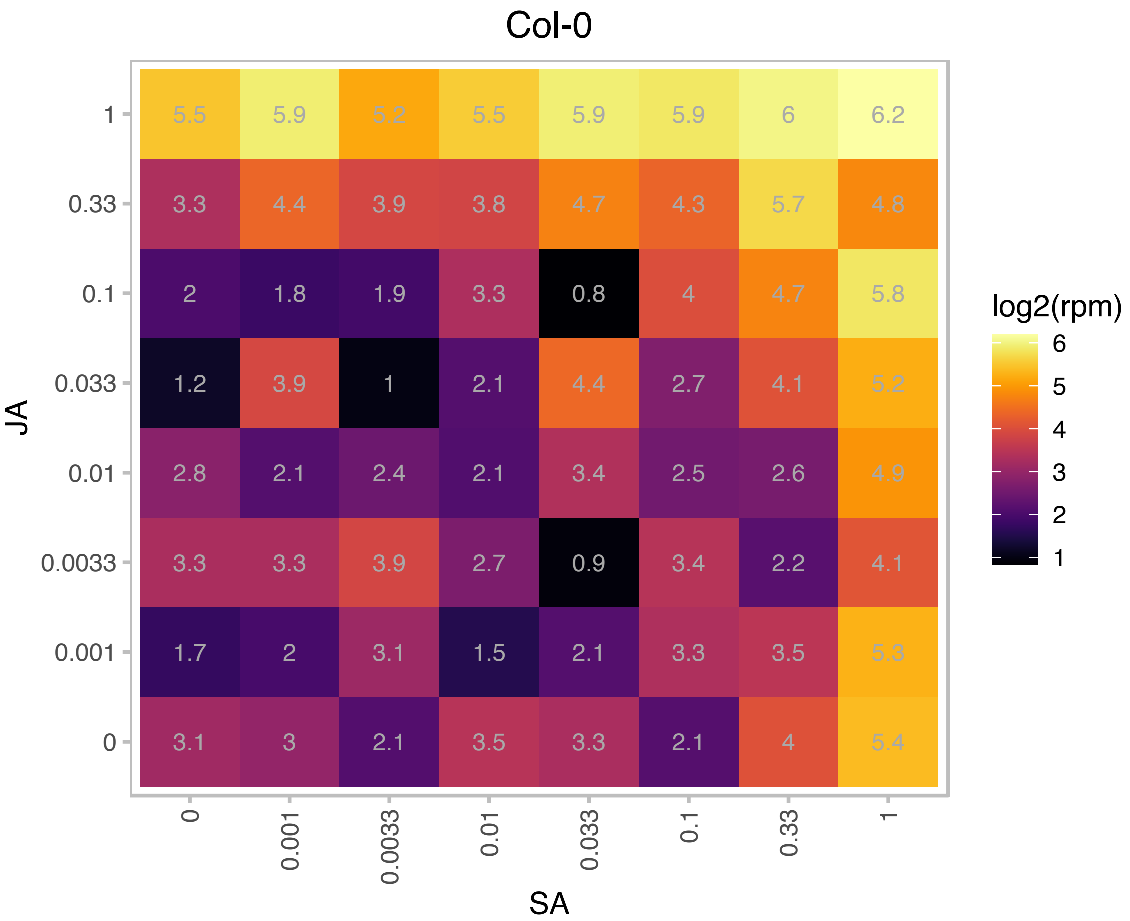


| | |
|---|---|
| Gene | AT2G18600 |
| is.expressed_genes | TRUE |
| is.responsive_genes | TRUE |
| is.apcluster | B-05 |

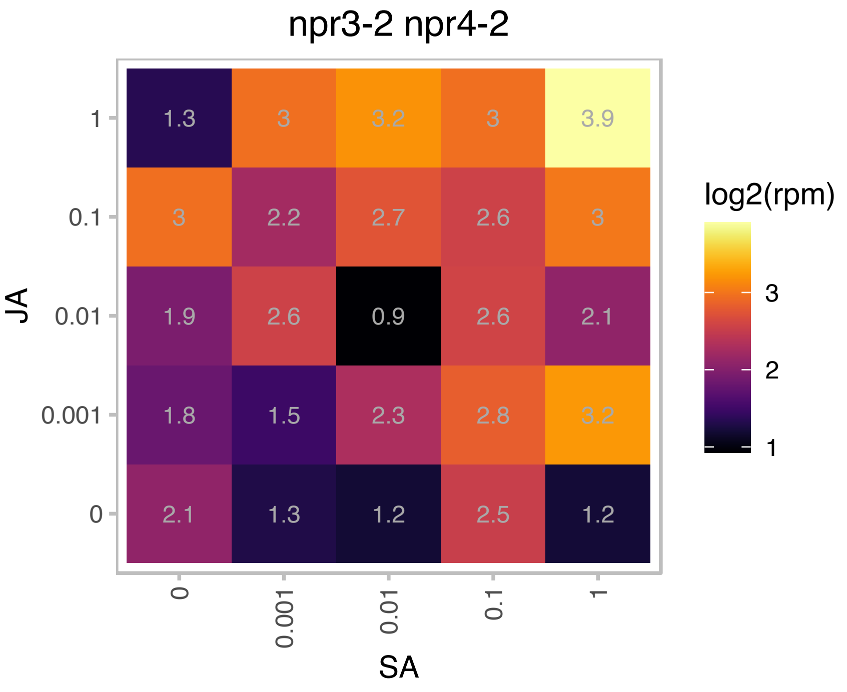


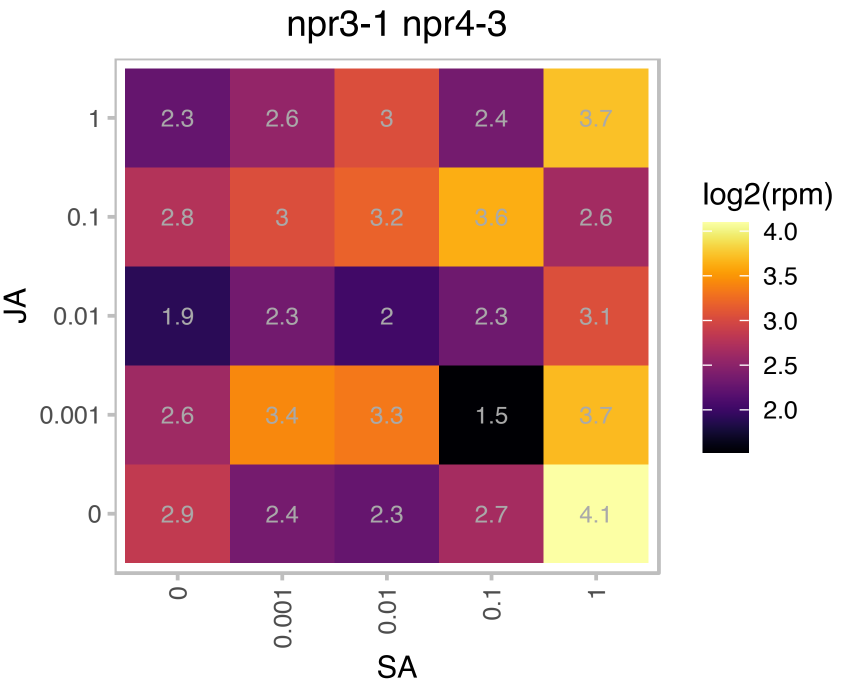

### AT2G26990.1 : ATCSN2, COP12, CSN2, FUS12

proteasome family protein

Represses photomorphogenesis and induces skotomorphogenesis in the dark.

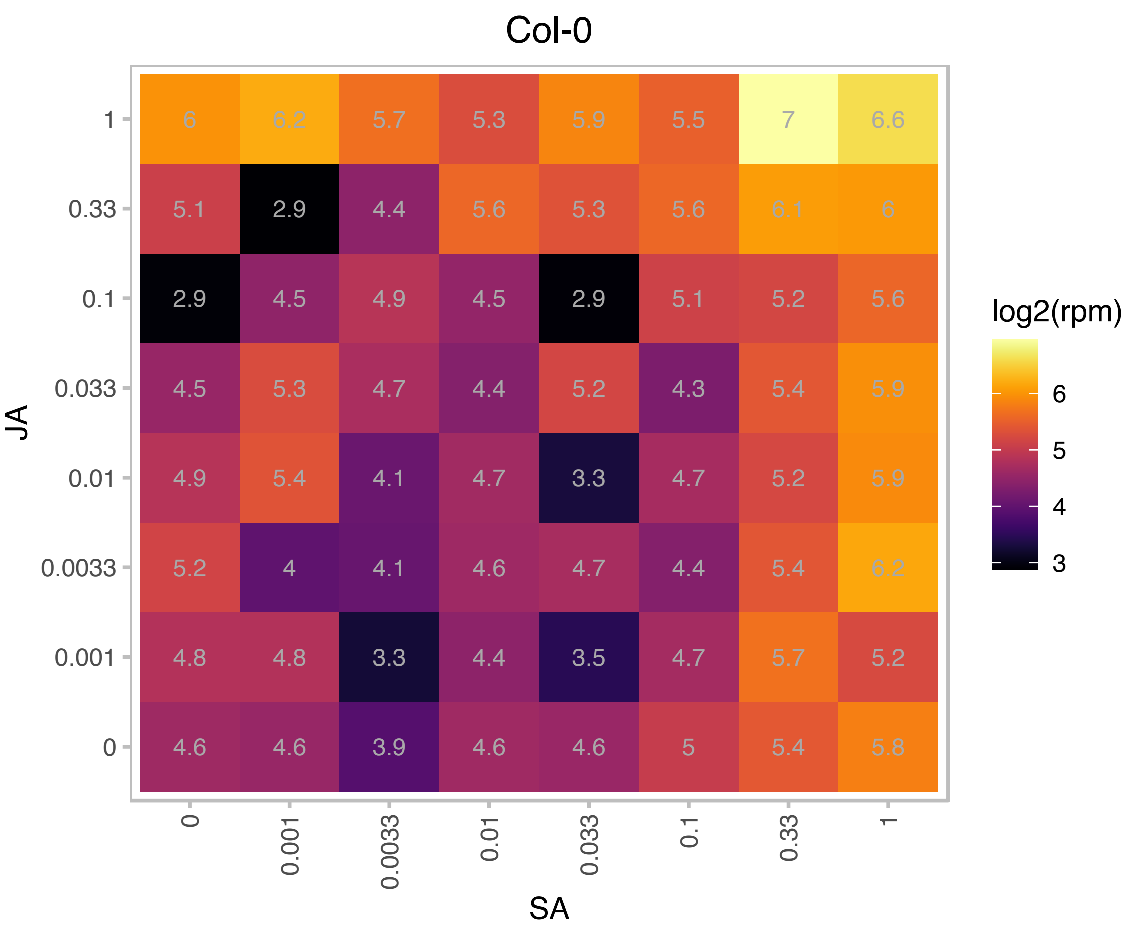


| | |
|---|---|
| Gene | AT2G26990 |
| is.expressed_genes | TRUE |
| is.responsive_genes | TRUE |
| is.apcluster | B-05 |

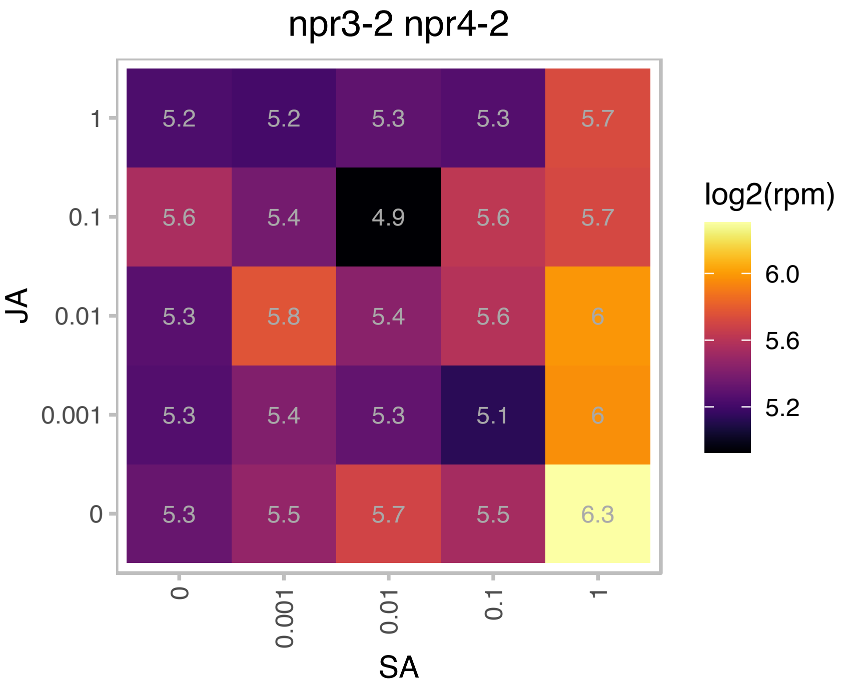


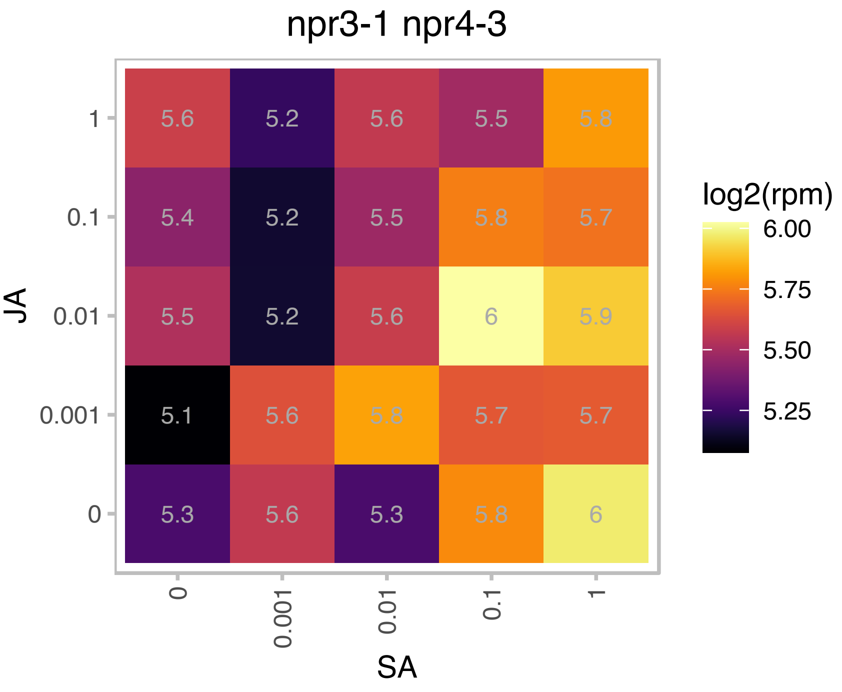

### AT2G31260.1 : APG9, ATAPG9, ATG9

autophagy 9 (APG9)

Involved in autophagy, the process of vacuolar bulk degradation of cytoplasmic components.  Mutant shows accelerated bolting and senescence.

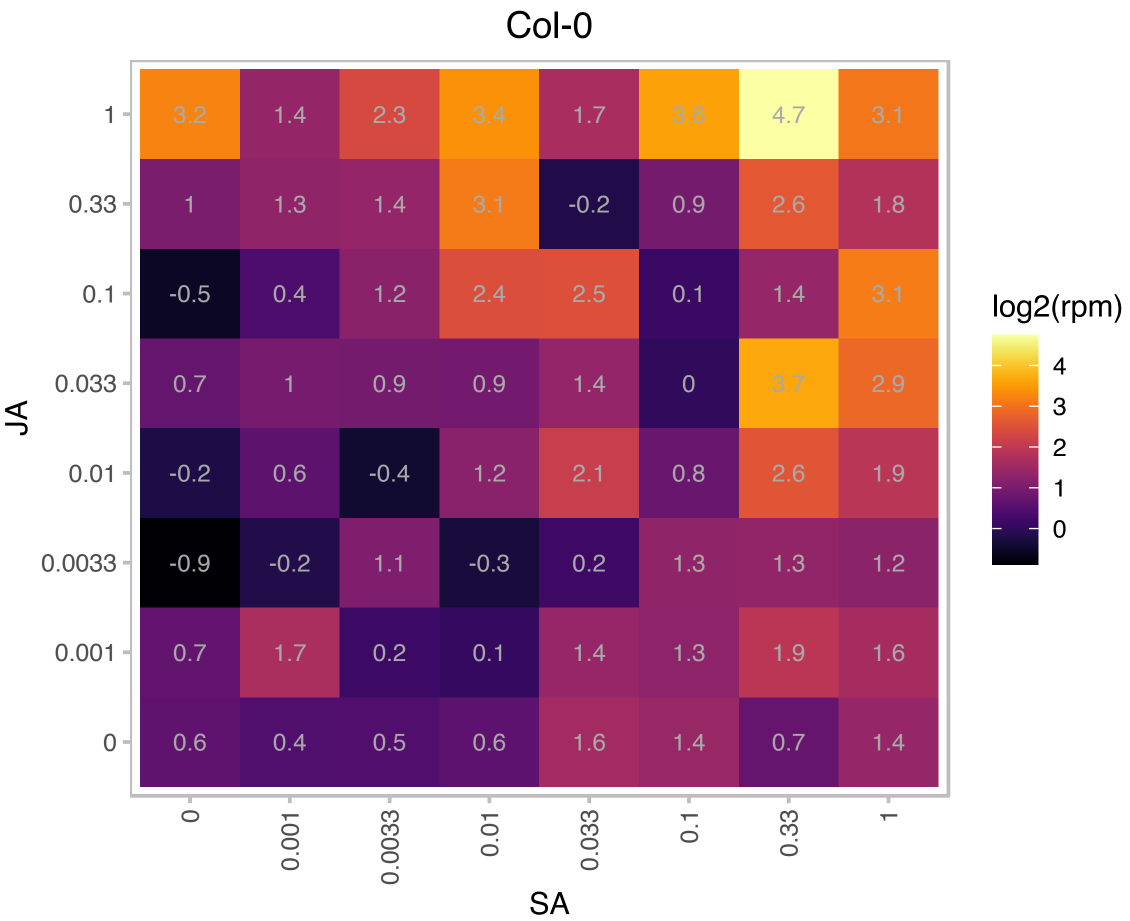


| | |
|---|---|
| Gene | AT2G31260 |
| is.expressed_genes | TRUE |
| is.responsive_genes | TRUE |
| is.apcluster | B-05 |

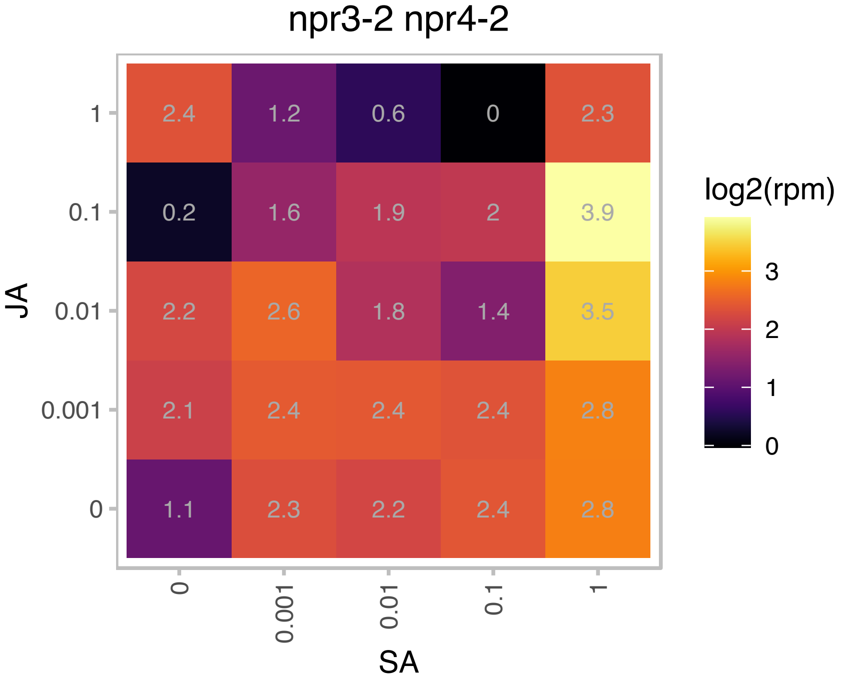


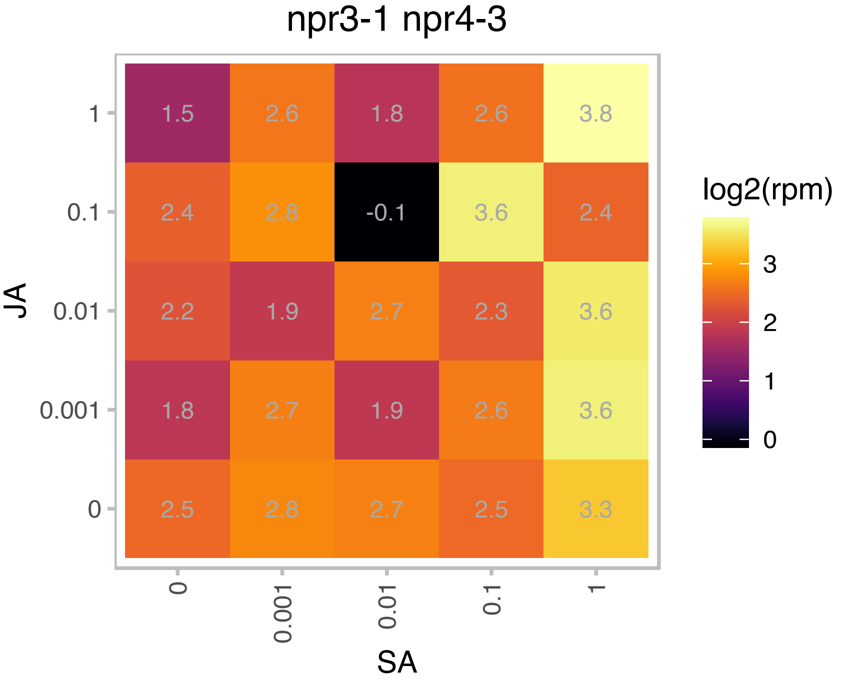

# AT2G31945.1 : NA

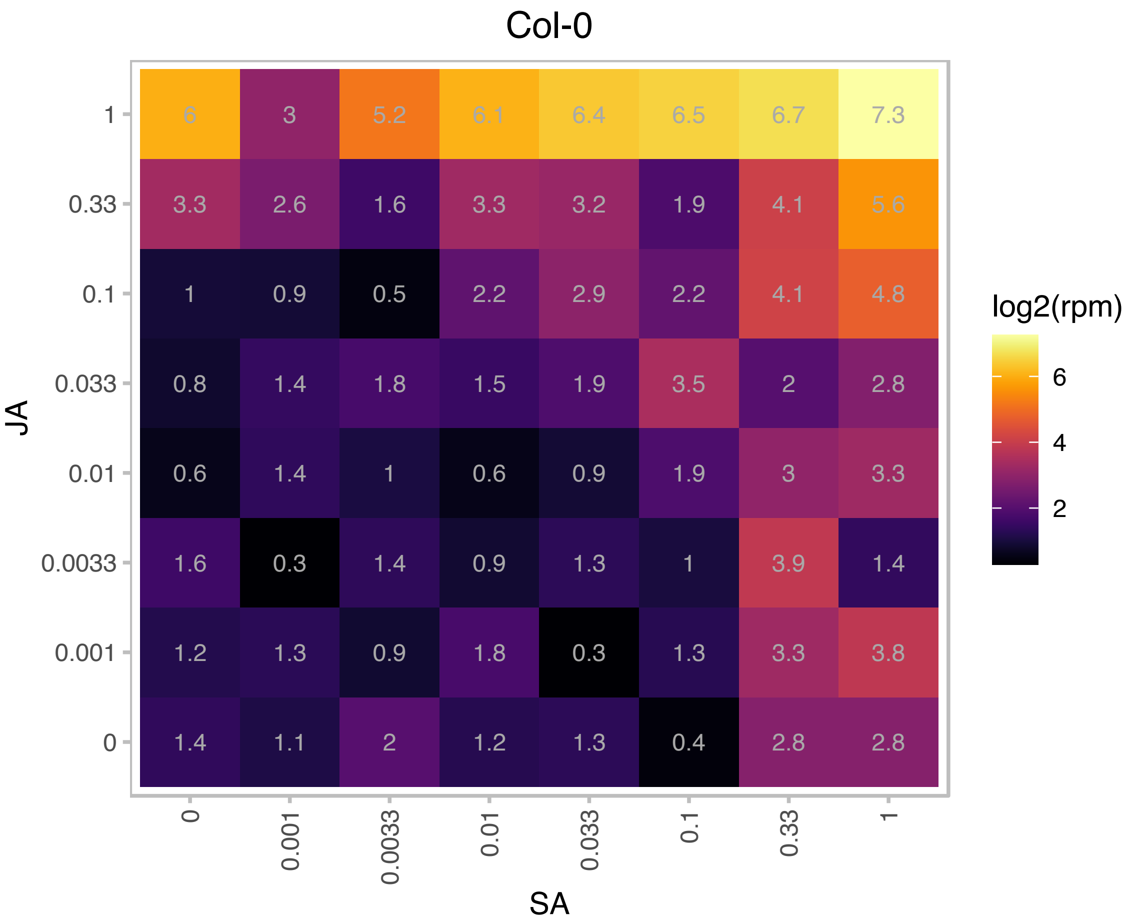


| | |
|---|---|
| Gene | AT2G31945 |
| is.expressed_genes | TRUE |
| is.responsive_genes | TRUE |
| is.apcluster | B-05 |

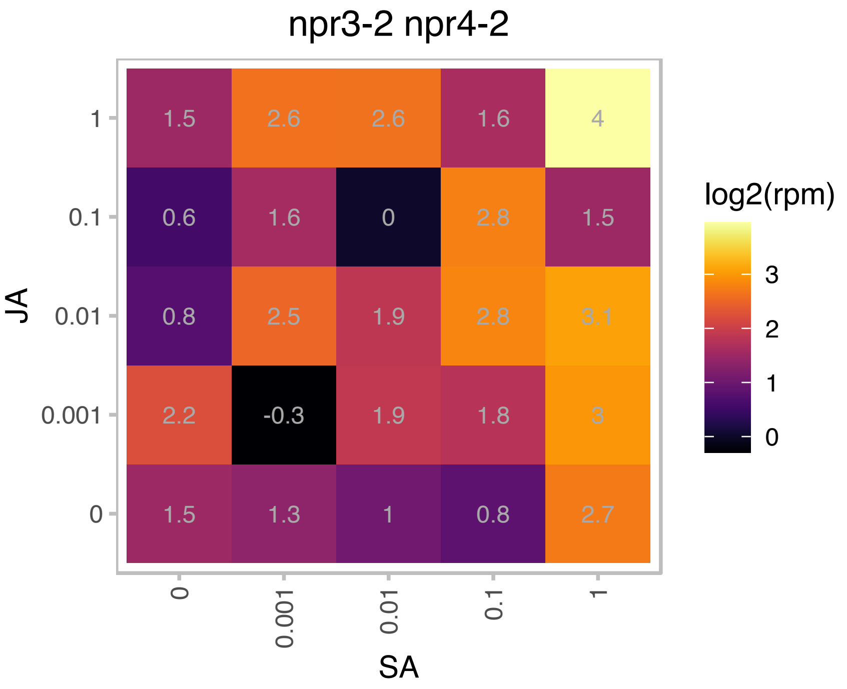


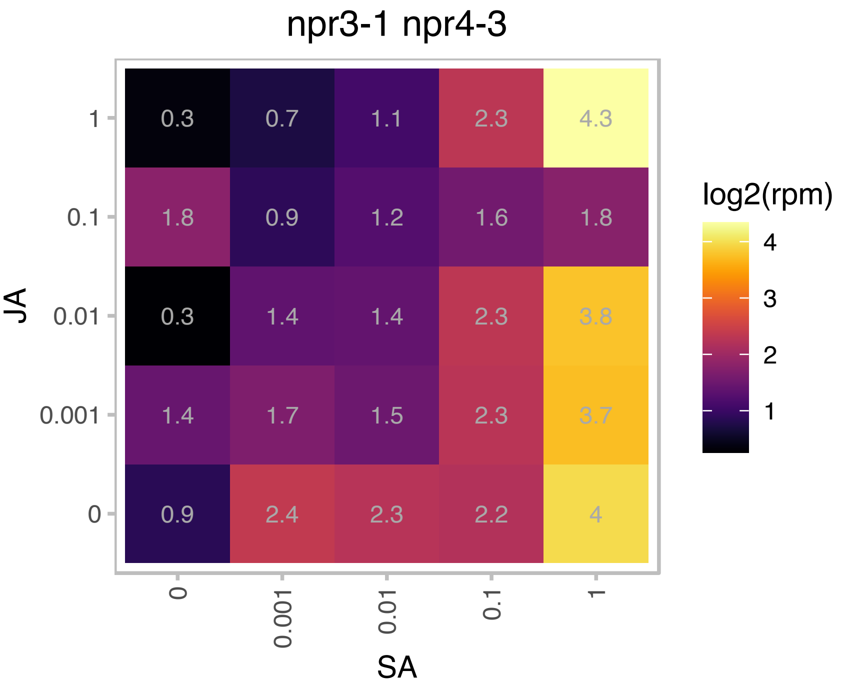

# AT2G36300.1 : NA

Integral membrane Yip1 family protein

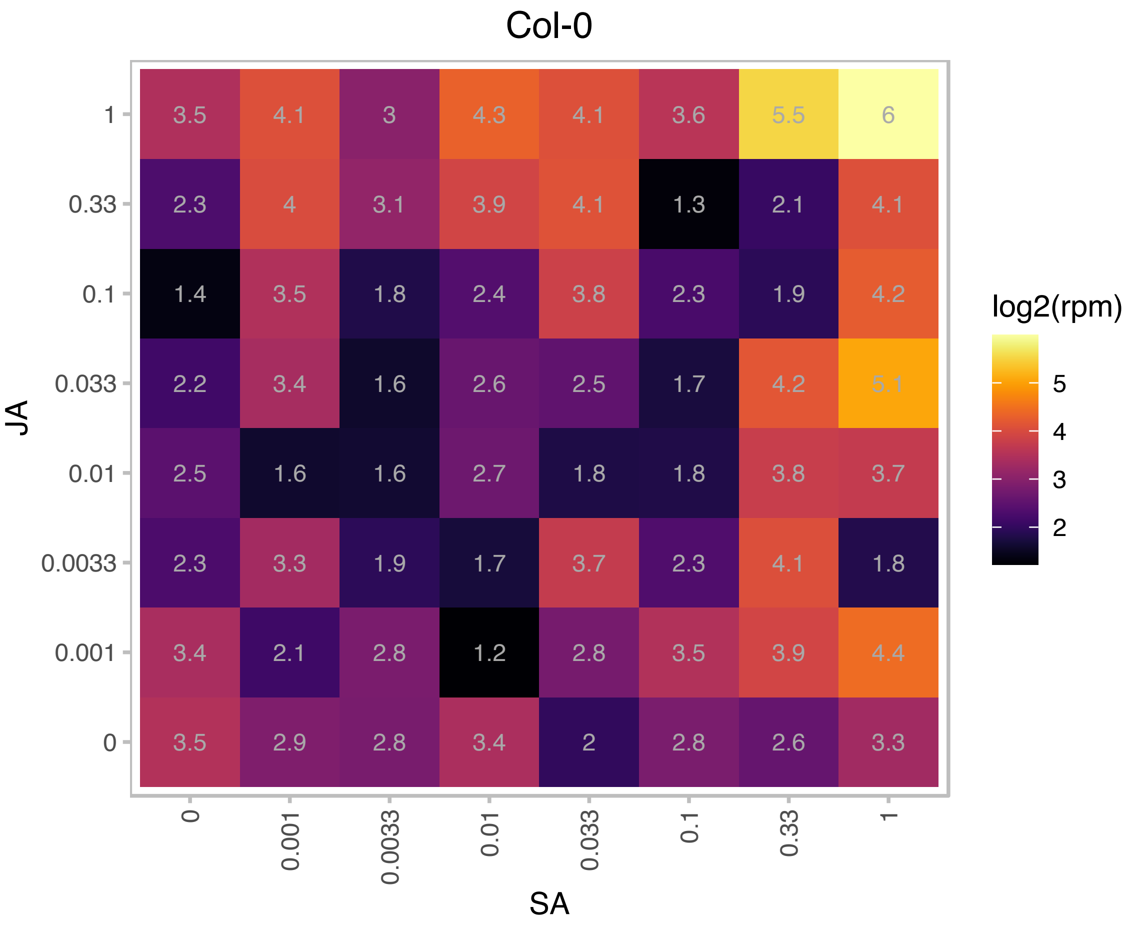


| | |
|---|---|
| Gene | AT2G36300 |
| is.expressed_genes | TRUE |
| is.responsive_genes | TRUE |
| is.apcluster | B-05 |

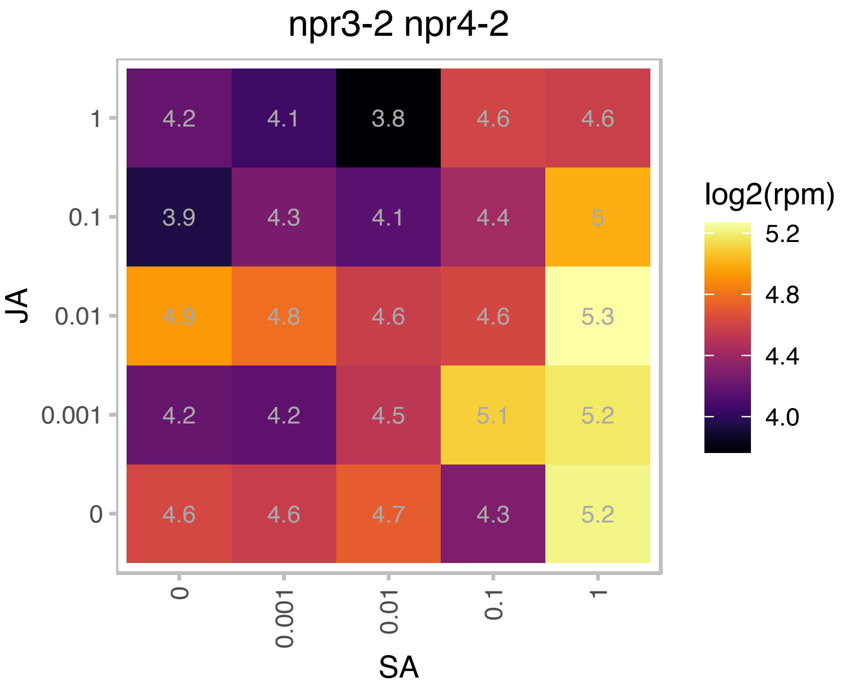


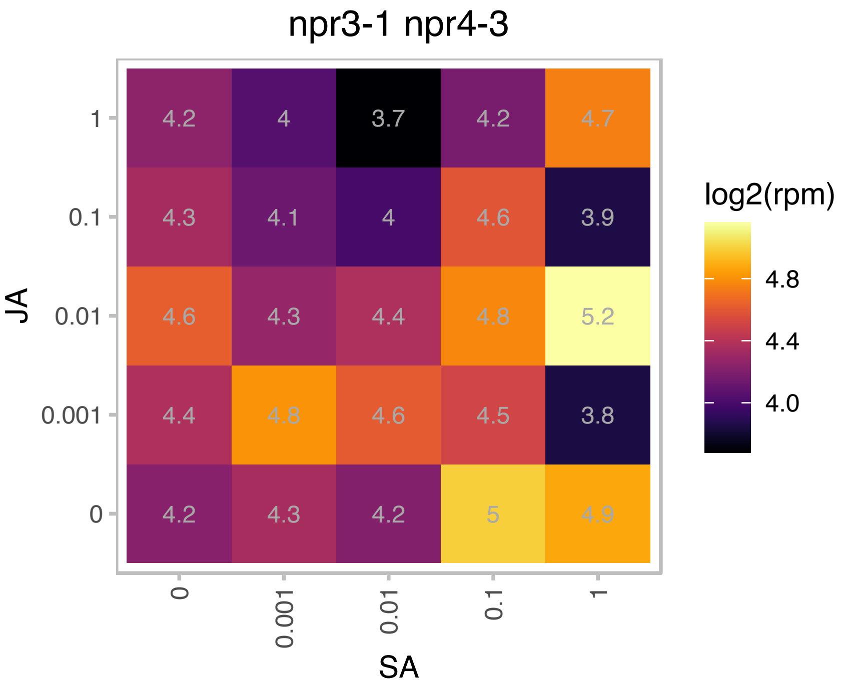

### AT2G36750.1 : UGT73C1

UDP-glucosyl transferase 73C1

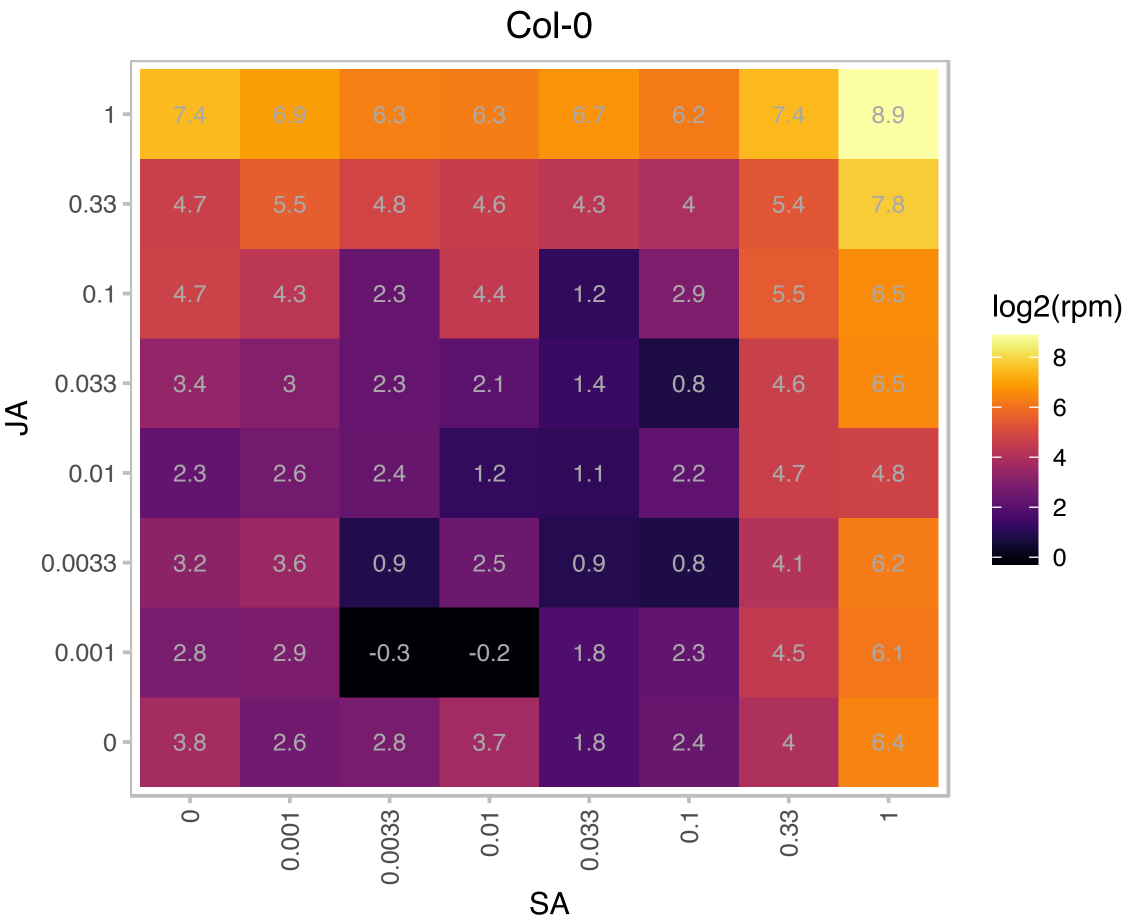


| | |
|---|---|
| Gene | AT2G36750 |
| is.expressed_genes | TRUE |
| is.responsive_genes | TRUE |
| is.apcluster | B-05 |

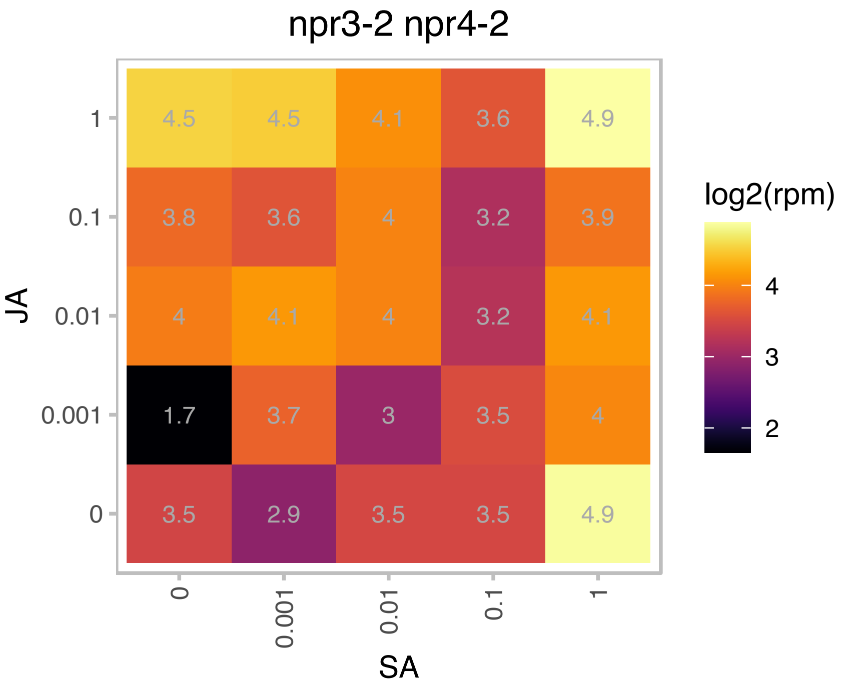


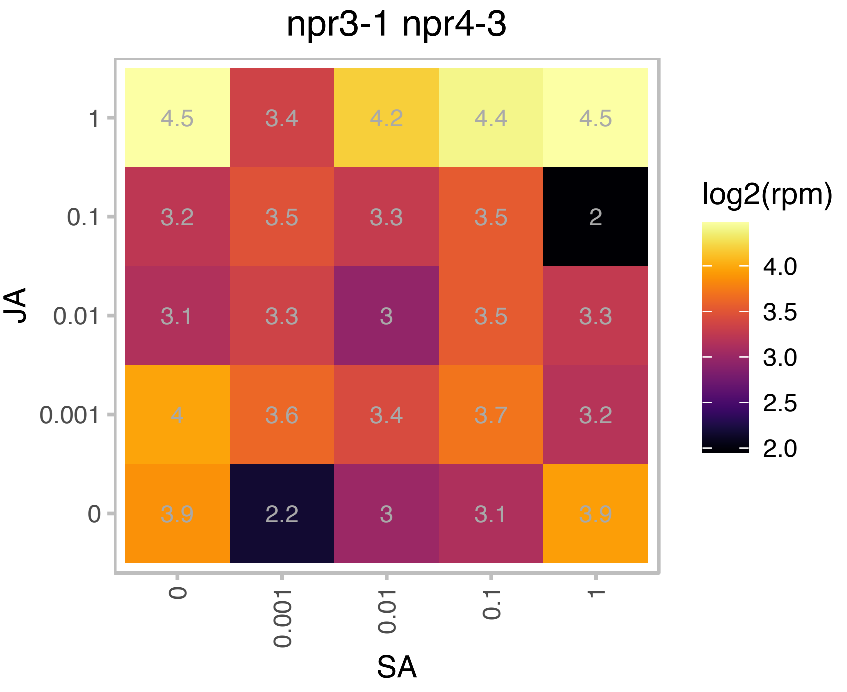

### AT2G36790.1 : UGT73C6

UDP-glucosyl transferase 73C6
The At2g36790 gene encodes a UDP-glucose:flavonol-3-O-glycoside-7-O-glucosyltransferase (UGT73C6)attaching glucosyl residue to the 7-O-position of the flavonols kaempferol, quercetin and their 3-O-glycoside derivatives.

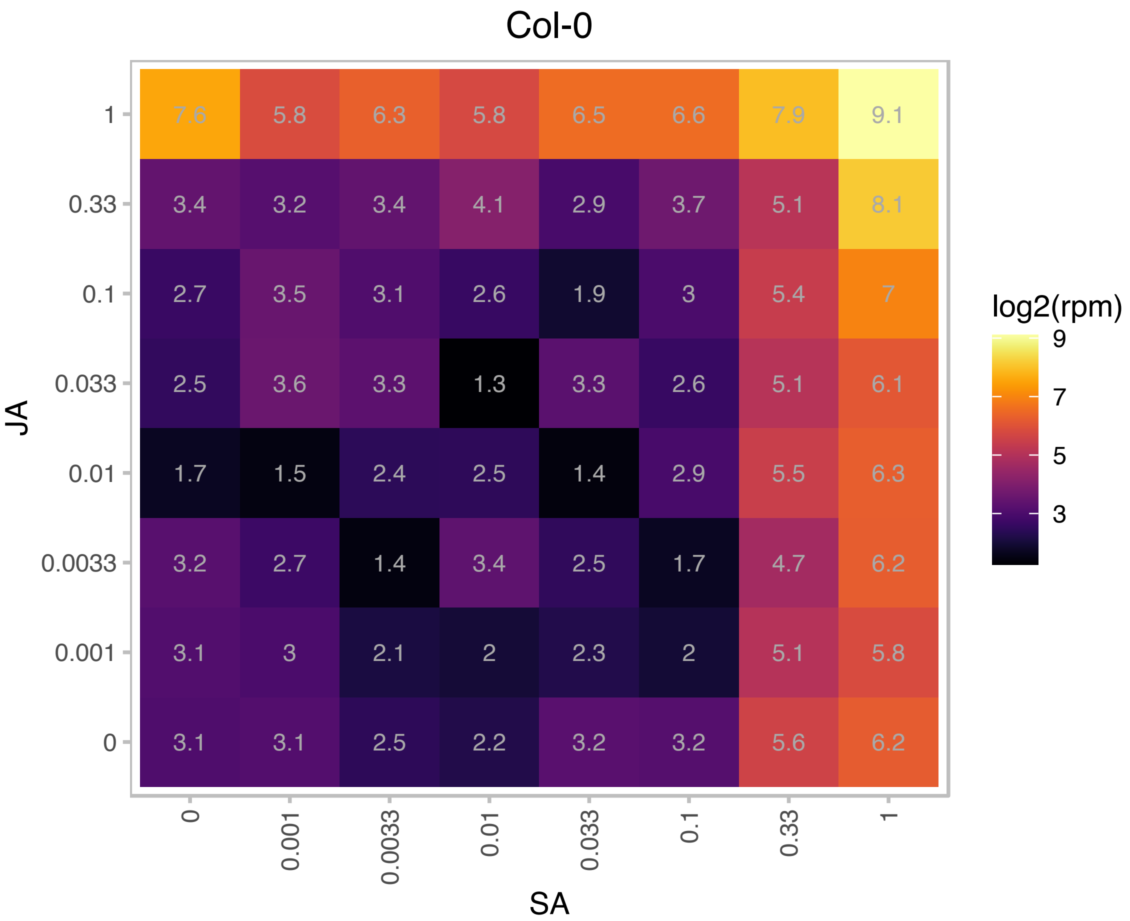


| | |
|---|---|
| Gene | AT2G36790 |
| is.expressed_genes | TRUE |
| is.responsive_genes | TRUE |
| is.apcluster | B-05 |

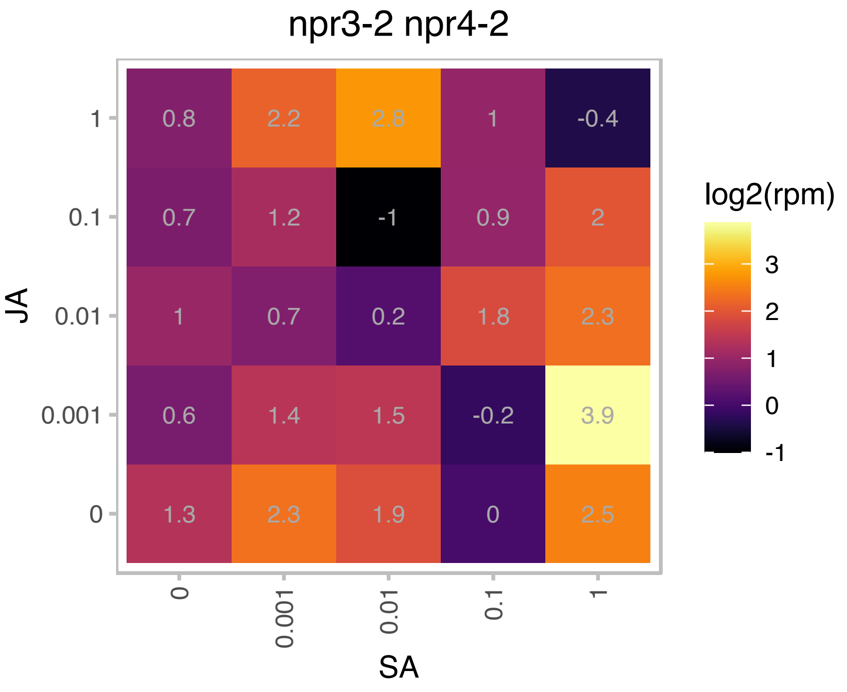


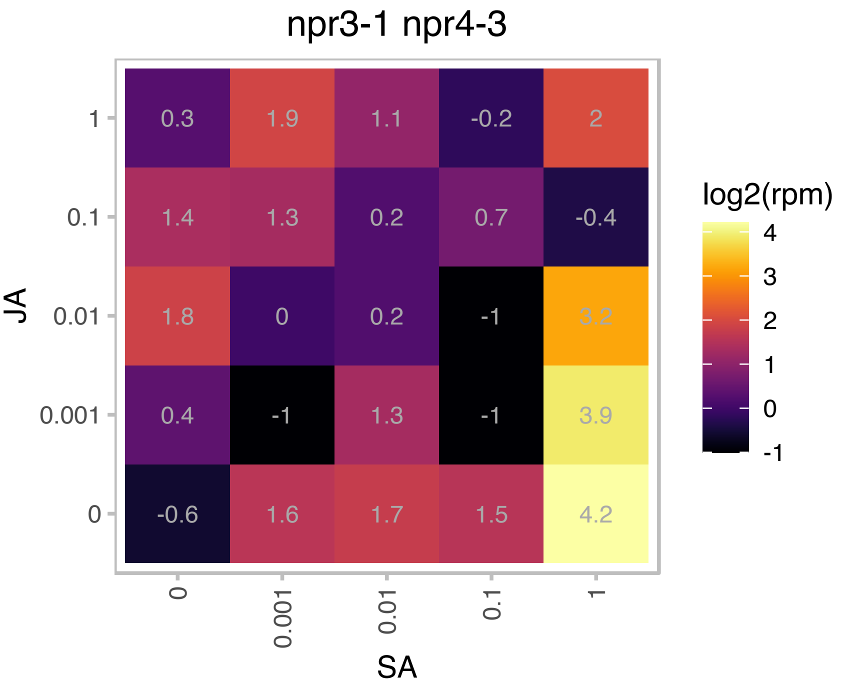

# AT2G38250.1 : NA

Homeodomain-like superfamily protein

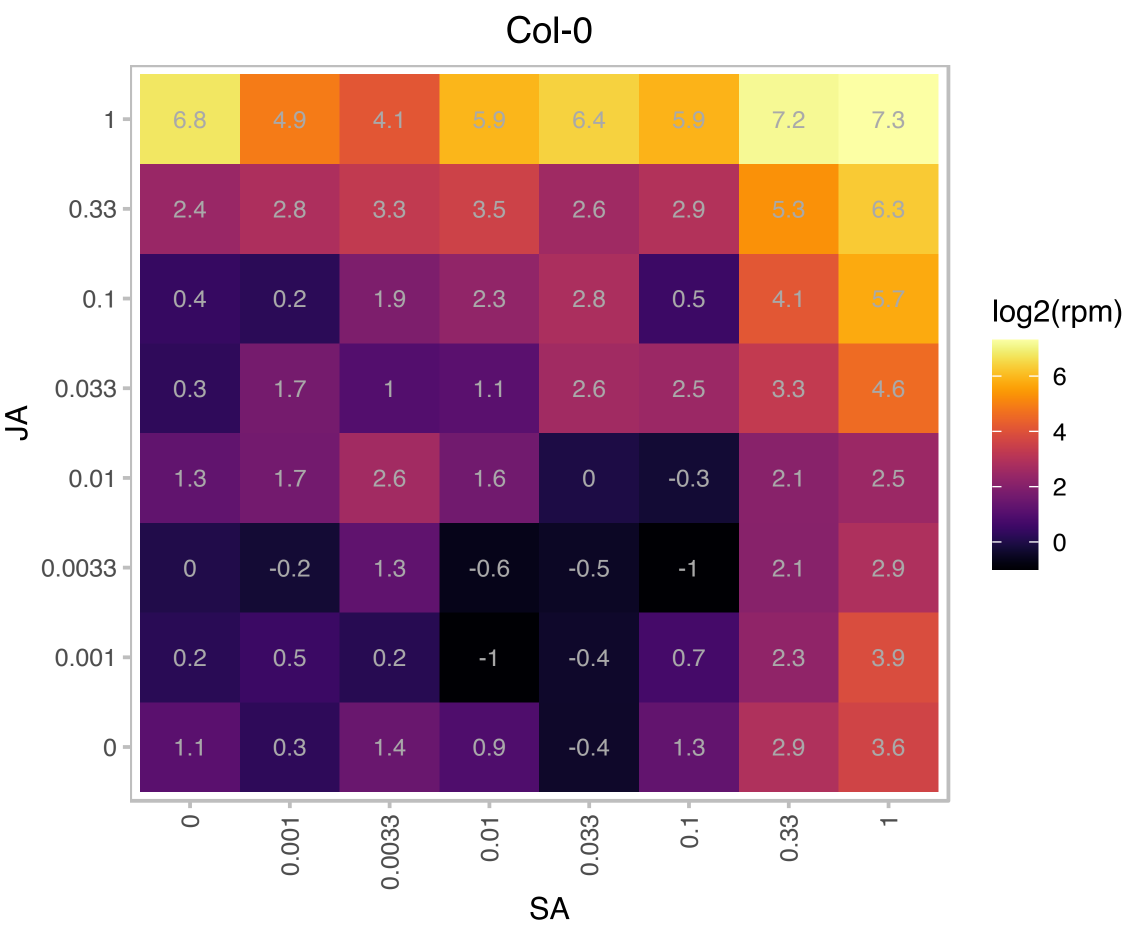


| | |
|---|---|
| Gene | AT2G38250 |
| is.expressed_genes | TRUE |
| is.responsive_genes | TRUE |
| is.apcluster | B-05 |

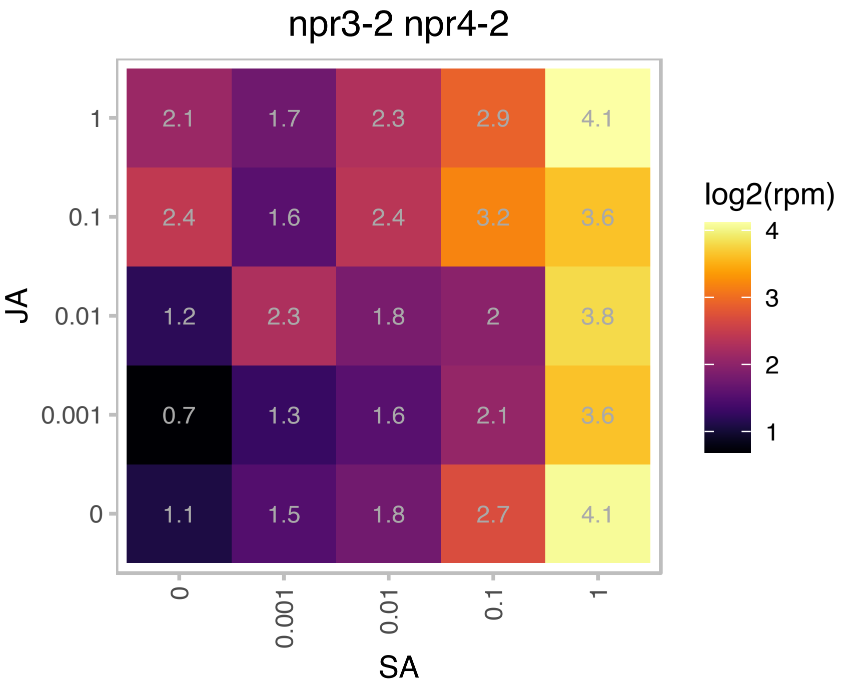


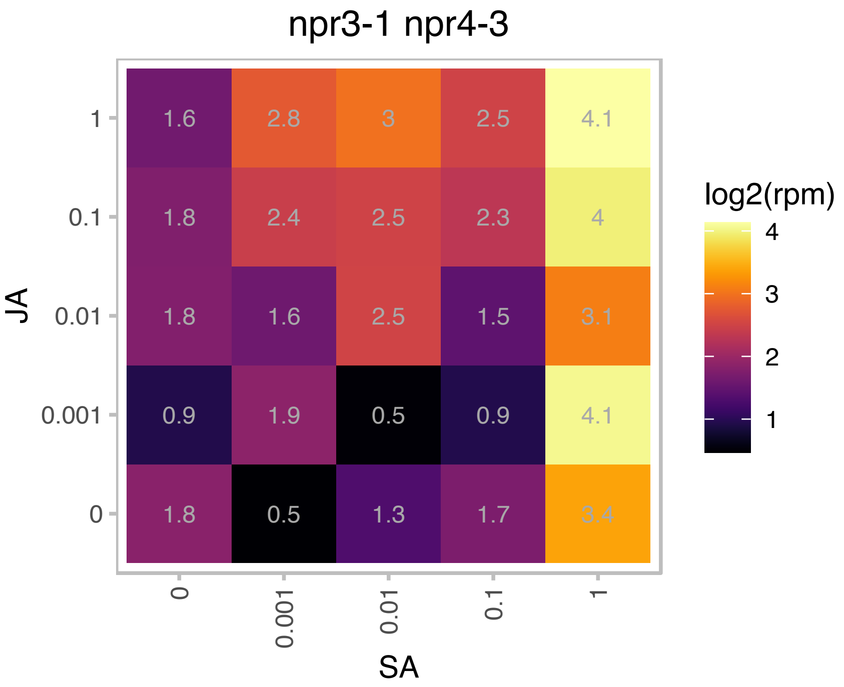

### AT2G40060.1 : CLC2

Clathrin light chain protein

Encodes a clathrin that is localized to the cortical division zone and the cell plate and colocalizes with TPLATE during cell plate anchoring. The mRNA is cell-to-cell mobile.

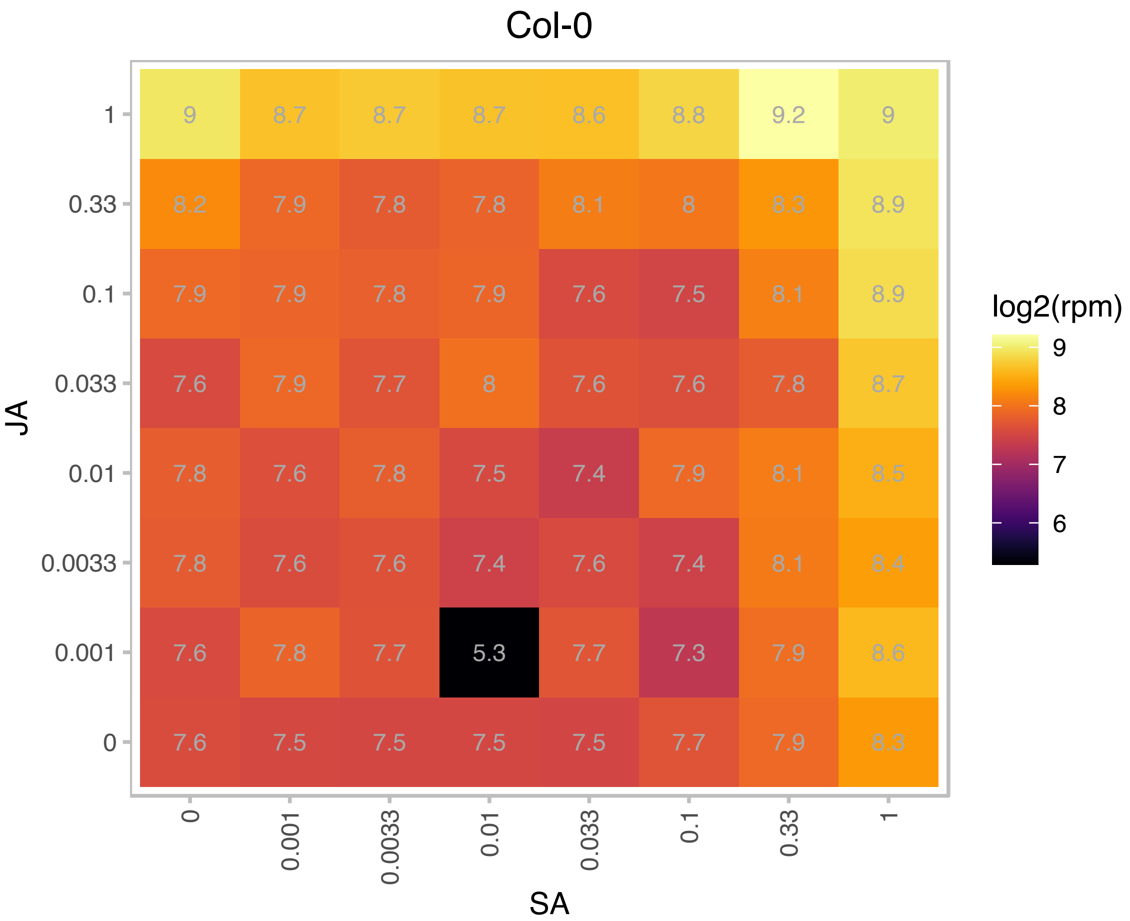


| | |
|---|---|
| Gene | AT2G40060 |
| is.expressed_genes | TRUE |
| is.responsive_genes | TRUE |
| is.apcluster | B-05 |

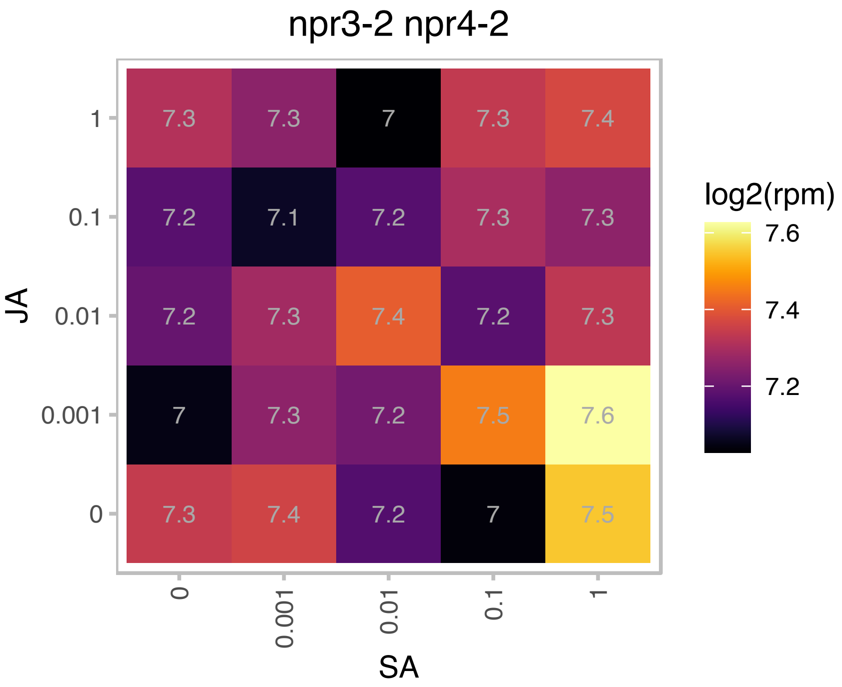


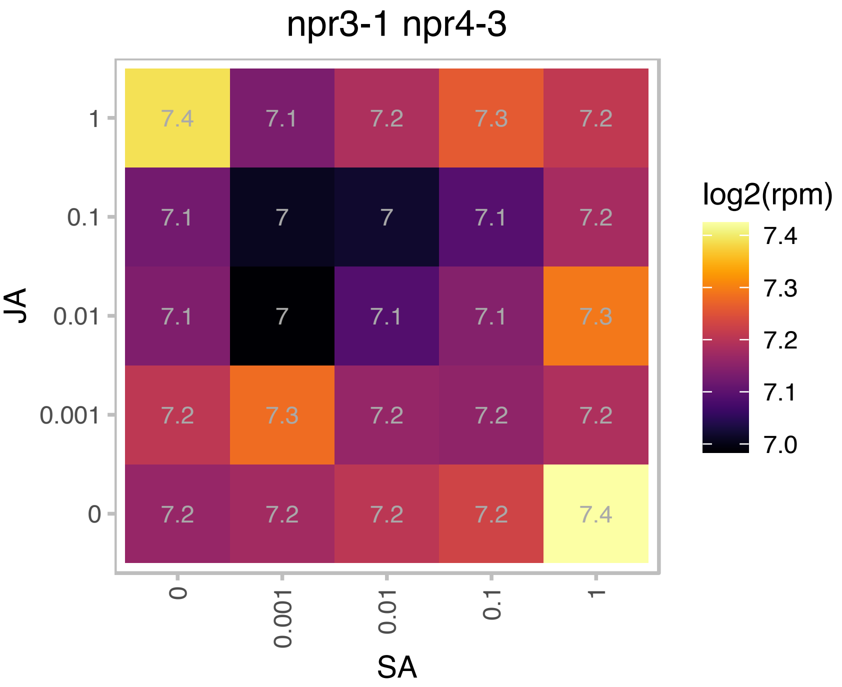

### AT2G42680.1 : ATMBF1A, MBF1A

multiprotein bridging factor 1A
One of three genes in A. thaliana encoding multiprotein bridging factor 1, a highly conserved transcriptional coactivator. May serve as a bridging factor between a bZIP factor and TBP. Its expression is developmentally regulated. The mRNA is cell-to-cell mobile.

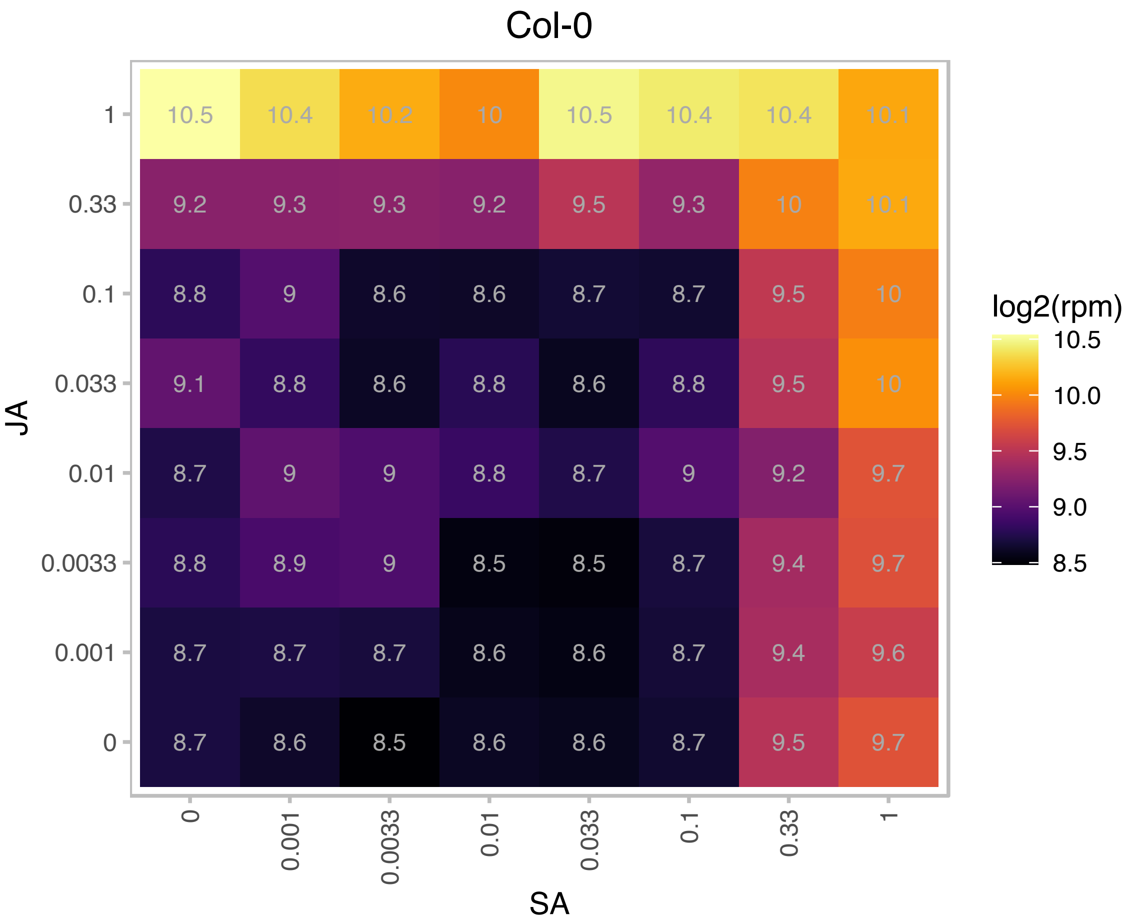


| | |
|---|---|
| Gene | AT2G42680 |
| is.expressed_genes | TRUE |
| is.responsive_genes | TRUE |
| is.apcluster | B-05 |

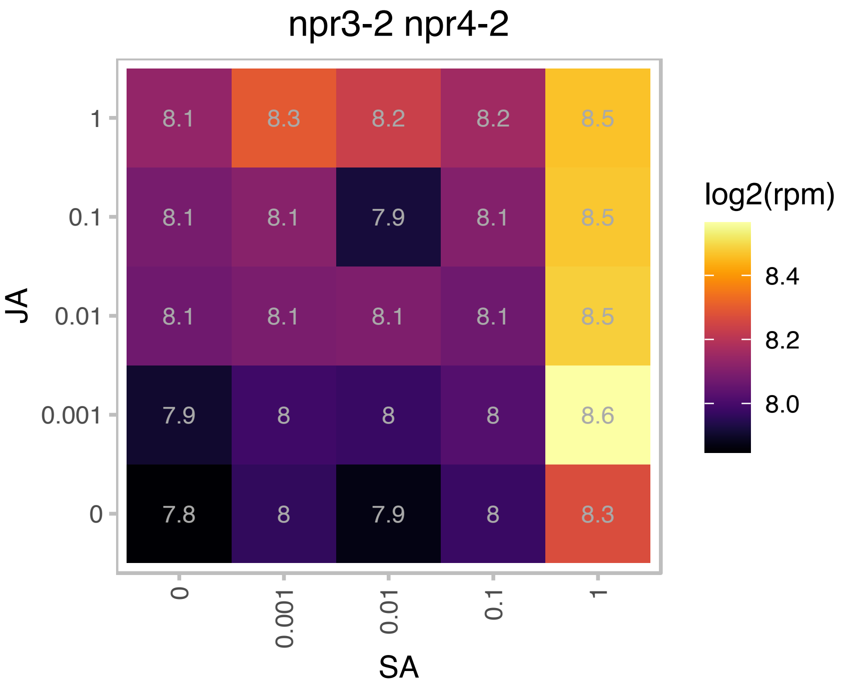


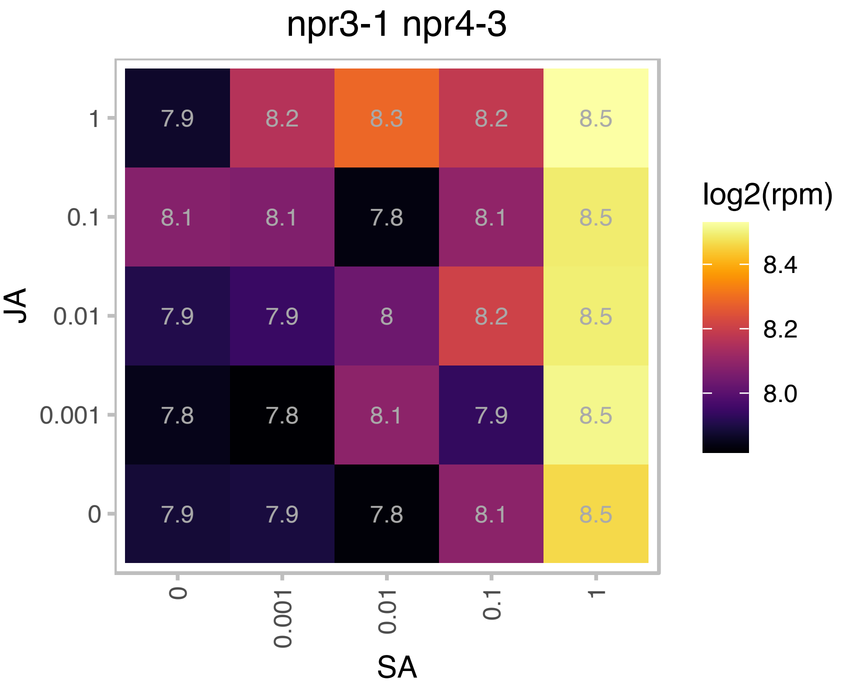

### AT2G42890.1 : AML2, ML2

MEI2-like 2

A member of mei2-like gene family, predominantly plant-based family of genes encoding RNA binding proteins with characteristic presence of a highly conserved RNA binding motif first described in the mei2 gene of the fission yeast S. pombe. In silico analyses reveal nine mei2 -like genes in A. thaliana. They were grouped into four distinct clades, based on overall sequence similarity and subfamily-specific sequence elements. AML2 is a member of two sister clades of mei2-like gene family, AML1 through AML5, and belongs to the clade named ALM235. AML2 is expressed during early embryo development (heart and torpedo stage) and predominantly [illegible]

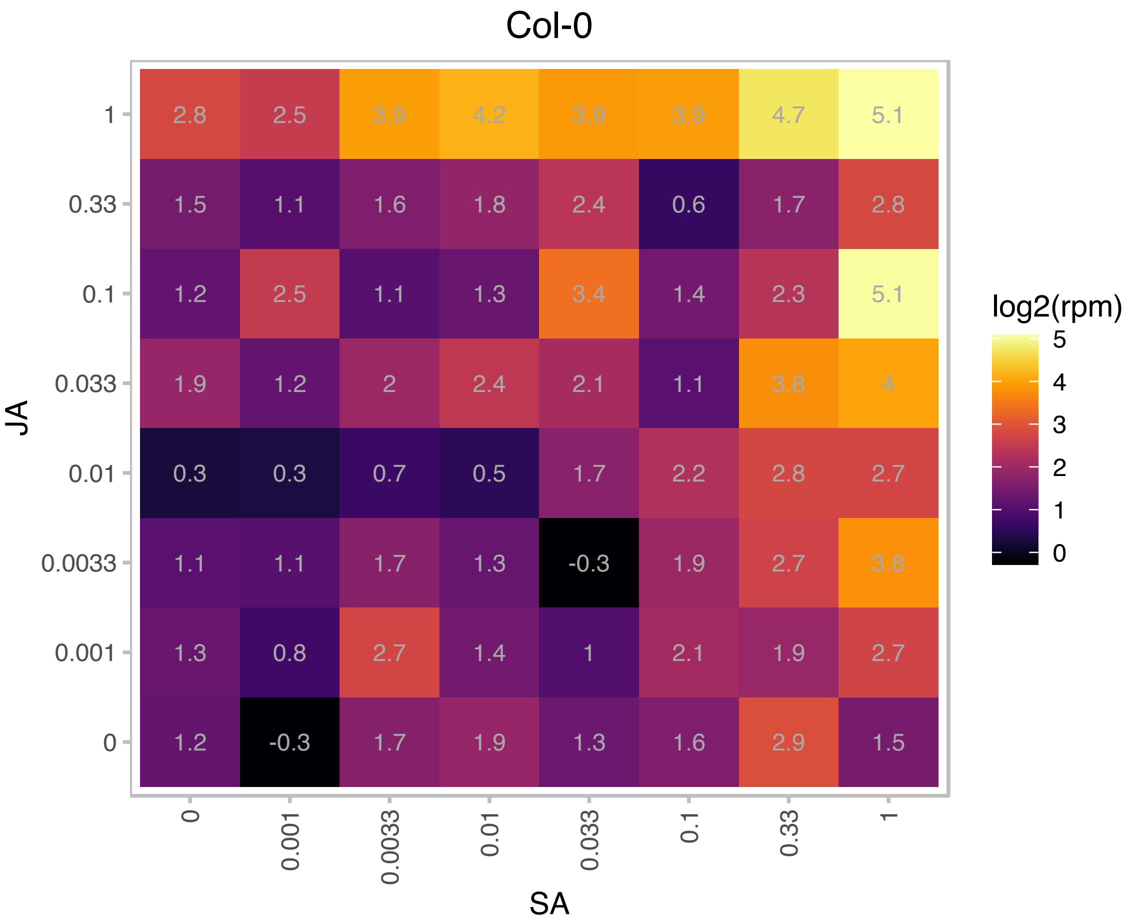


| | |
|---|---|
| Gene | AT2G42890 |
| is.expressed_genes | TRUE |
| is.responsive_genes | TRUE |
| is.apcluster | B-05 |

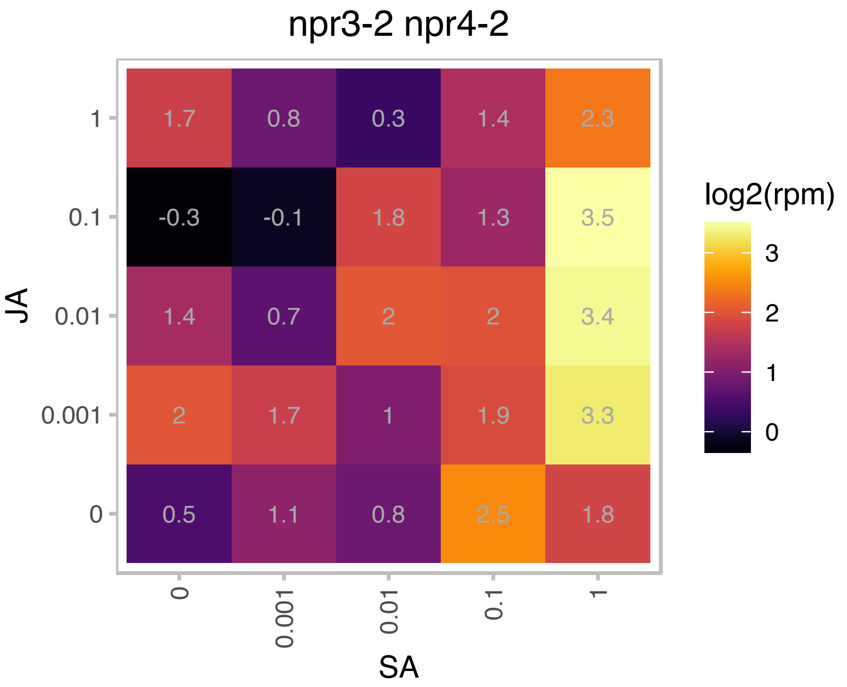


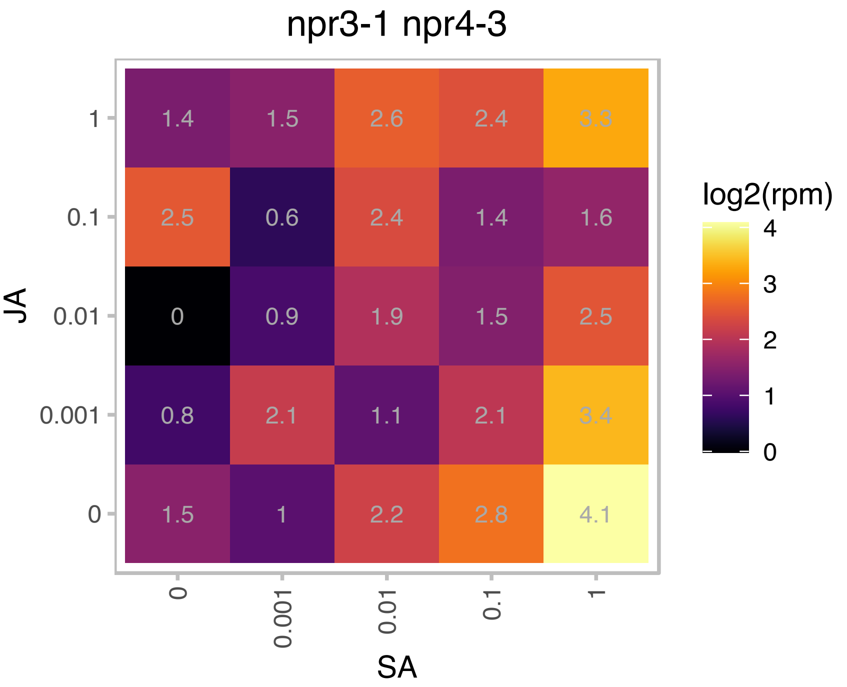

### AT3G01640.1 : ATGLCAK, GLCAK

glucuronokinase G

AtGlcAK is a sugar kinase able to phosphorylate D-GlcA to D-GlcA-1-phosphate in the presence of ATP.

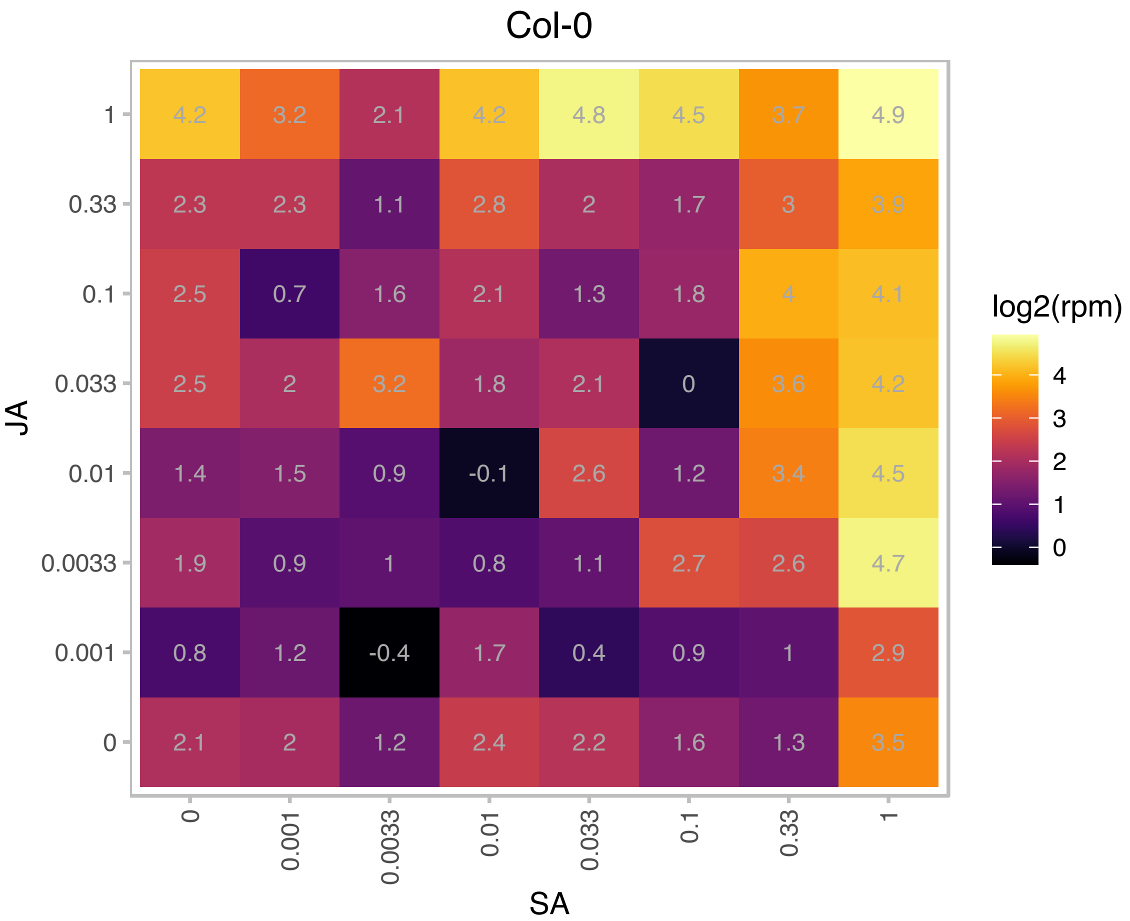


| Gene | AT3G01640 |
|---|---|
| is.expressed_genes | TRUE |
| is.responsive_genes | TRUE |
| is.apcluster | B-05 |

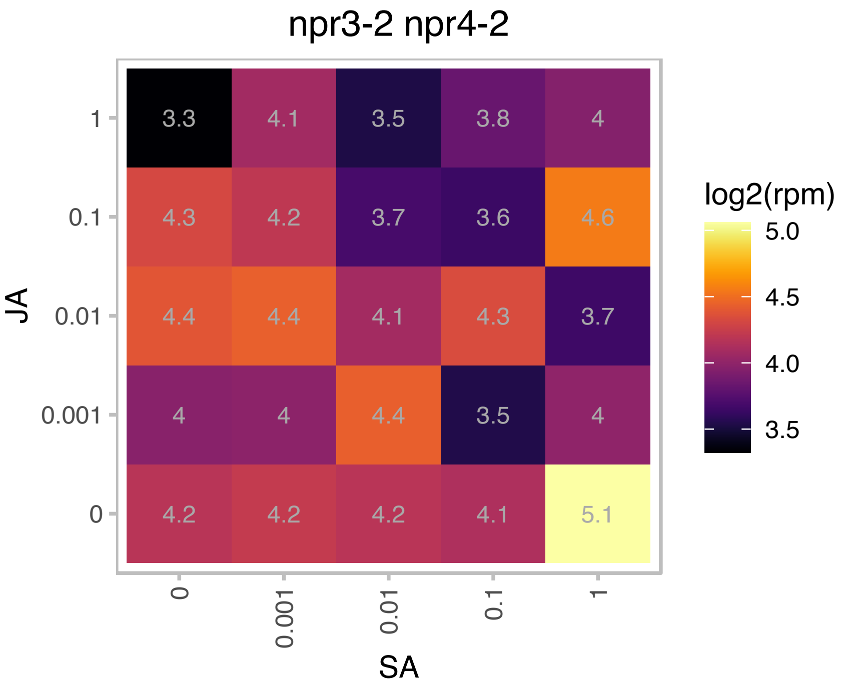


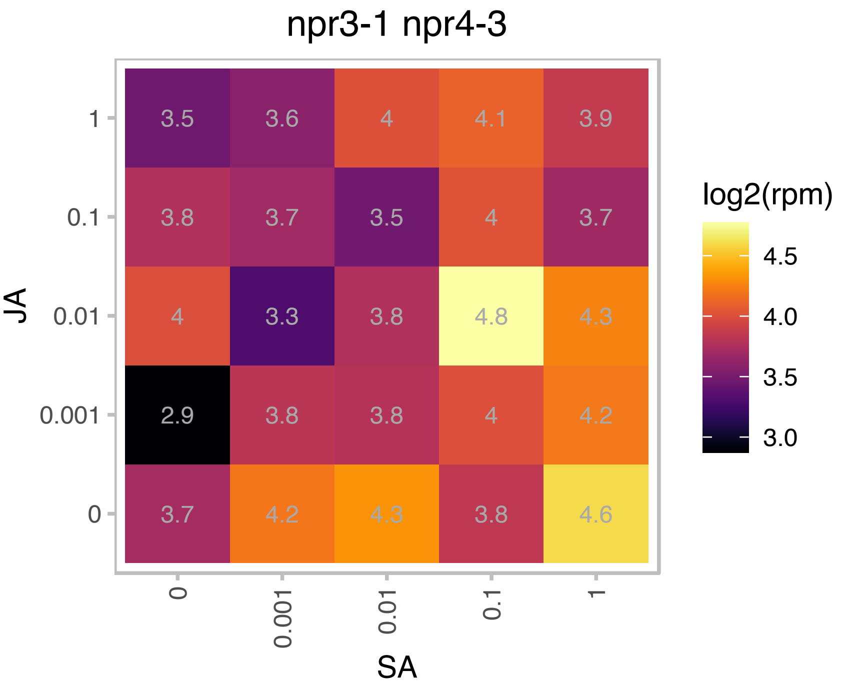

### AT3G01650.1 : RGLG1

RING domain ligase1

Encodes RGLG1 (RING domain ligase 1), a RING domain ubiquitin E3 ligase that negatively regulates the drought stress response by mediating ERF53 transcriptional activity.

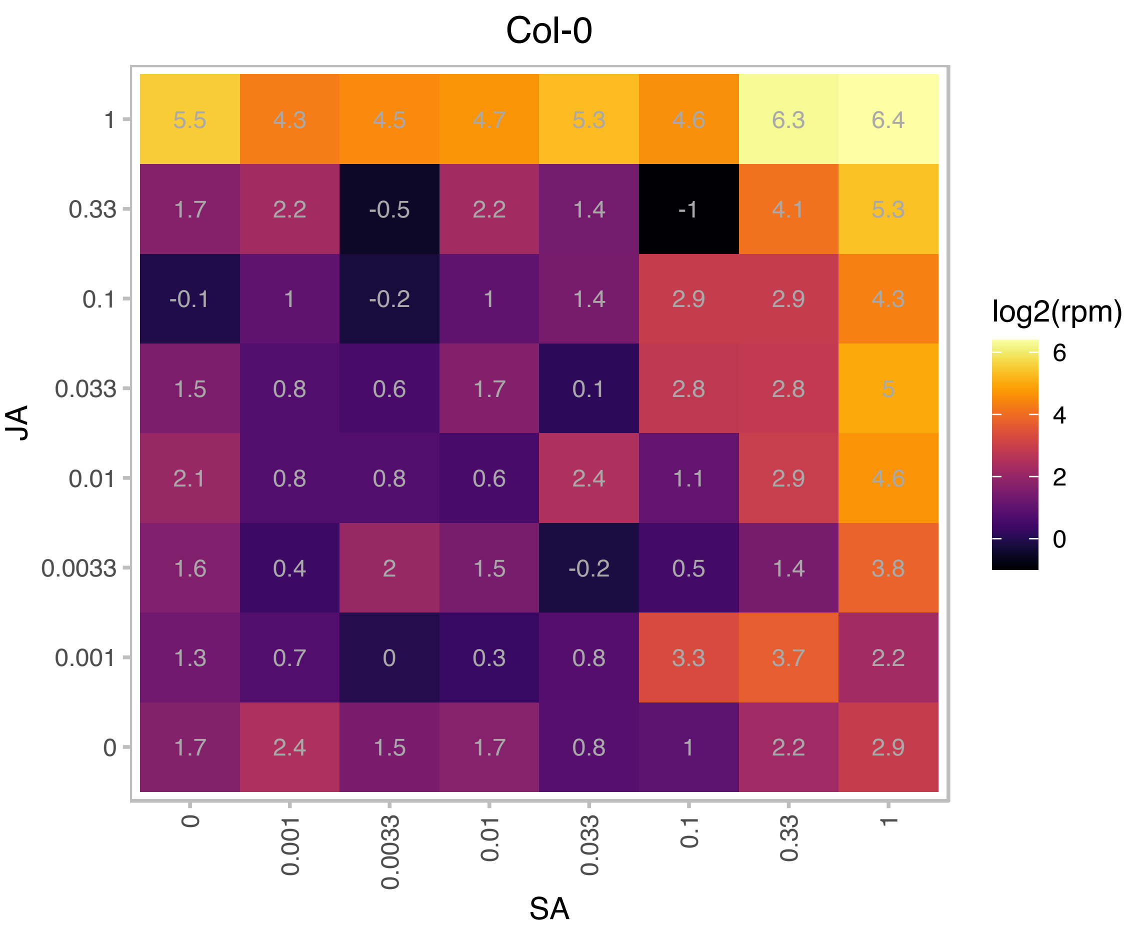


| | |
|---|---|
| Gene | AT3G01650 |
| is.expressed_genes | TRUE |
| is.responsive_genes | TRUE |
| is.apcluster | B-05 |

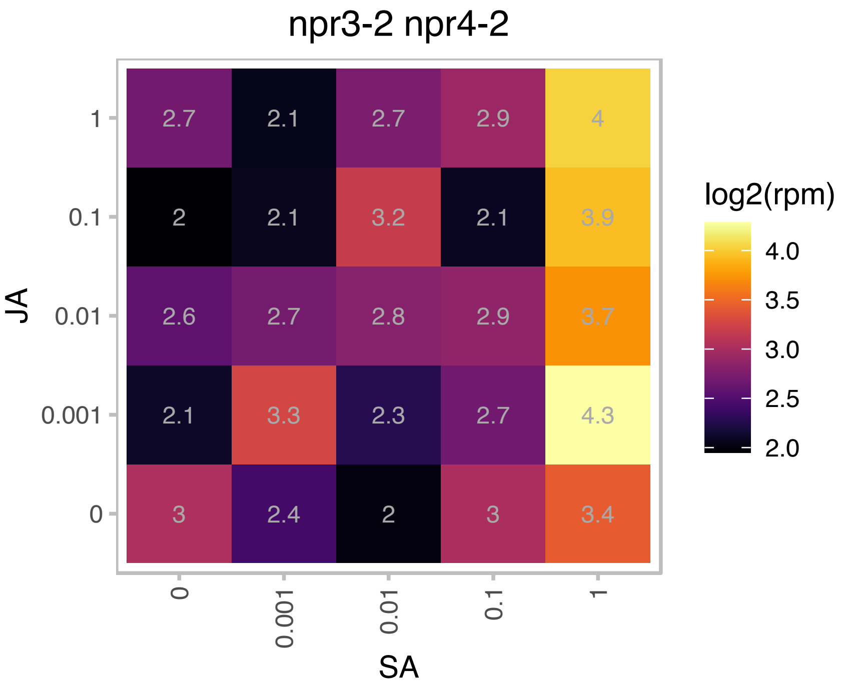


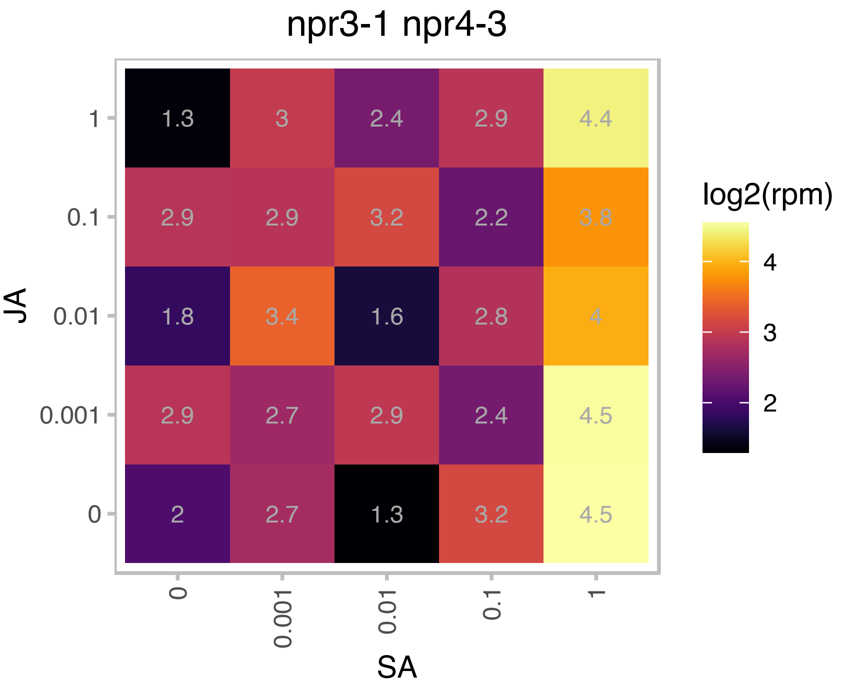

### AT3G03640.1 : BGLU25, GLUC

beta glucosidase 25

Encodes beta-glucosidase (GLUC).

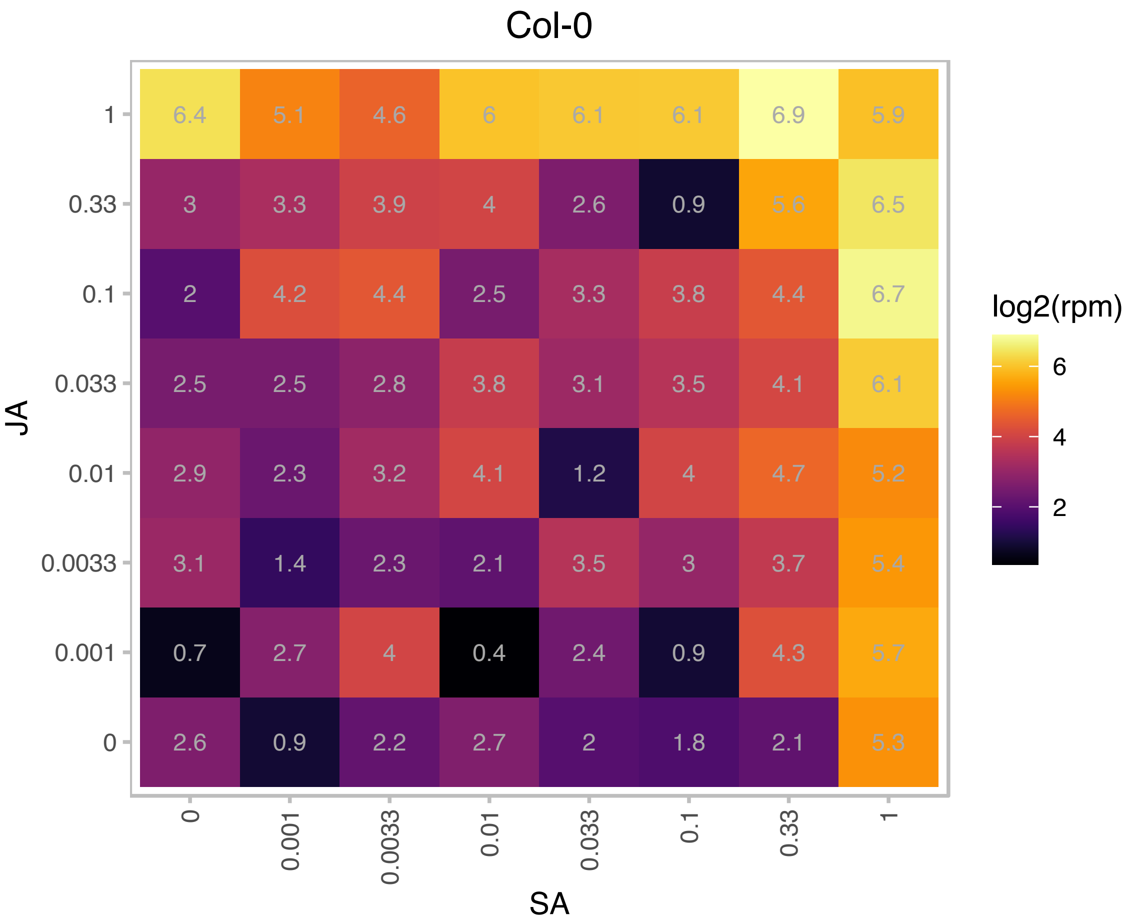


| Gene | AT3G03640 |
|---|---|
| is.expressed_genes | TRUE |
| is.responsive_genes | TRUE |
| is.apcluster | B-05 |

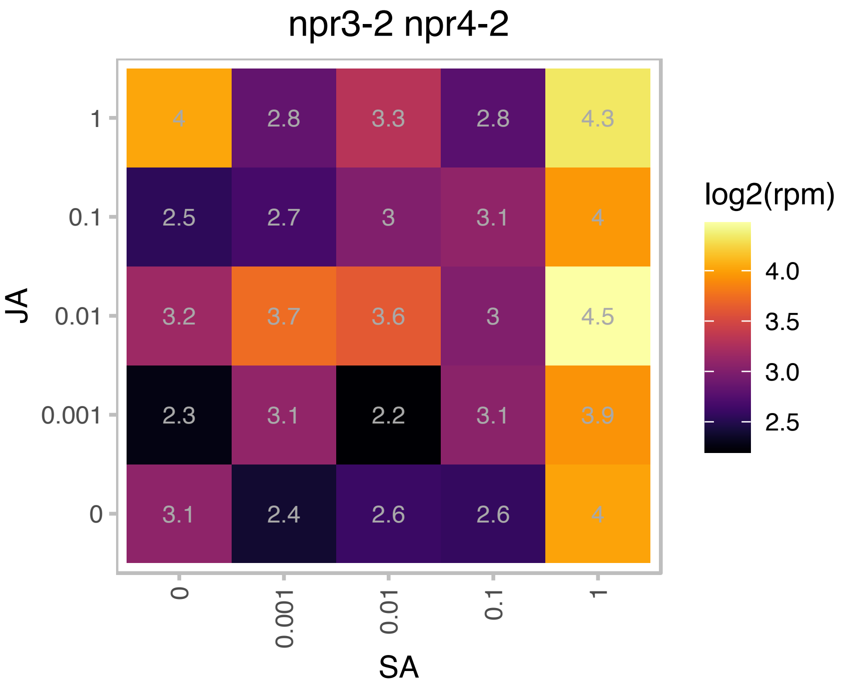


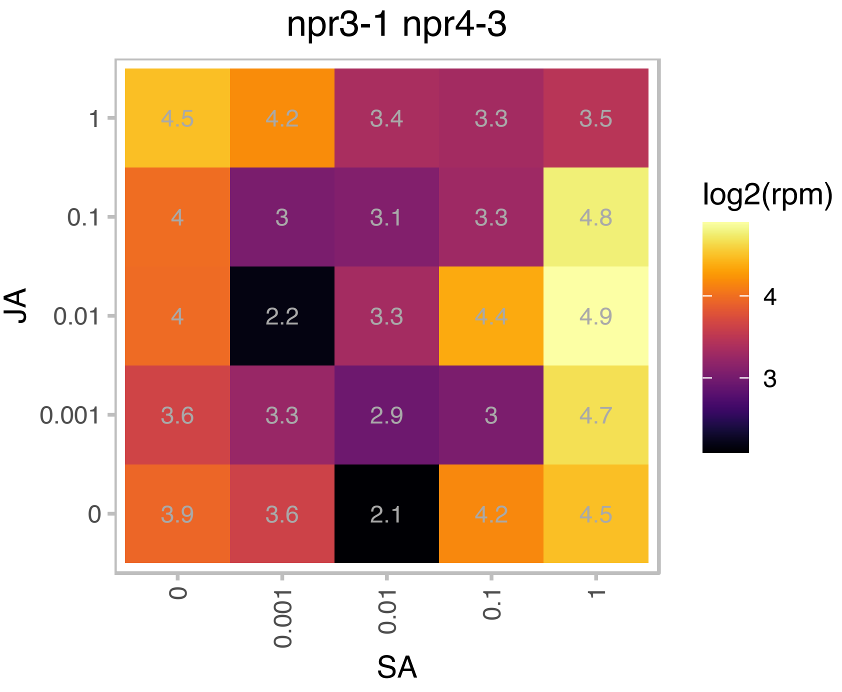

# AT3G04010.1 : NA

O-Glycosyl hydrolases family 17 protein

| | |
|---|---|
| Gene | AT3G04010 |
| is.expressed_genes | TRUE |
| is.responsive_genes | TRUE |
| is.apcluster | B-05 |

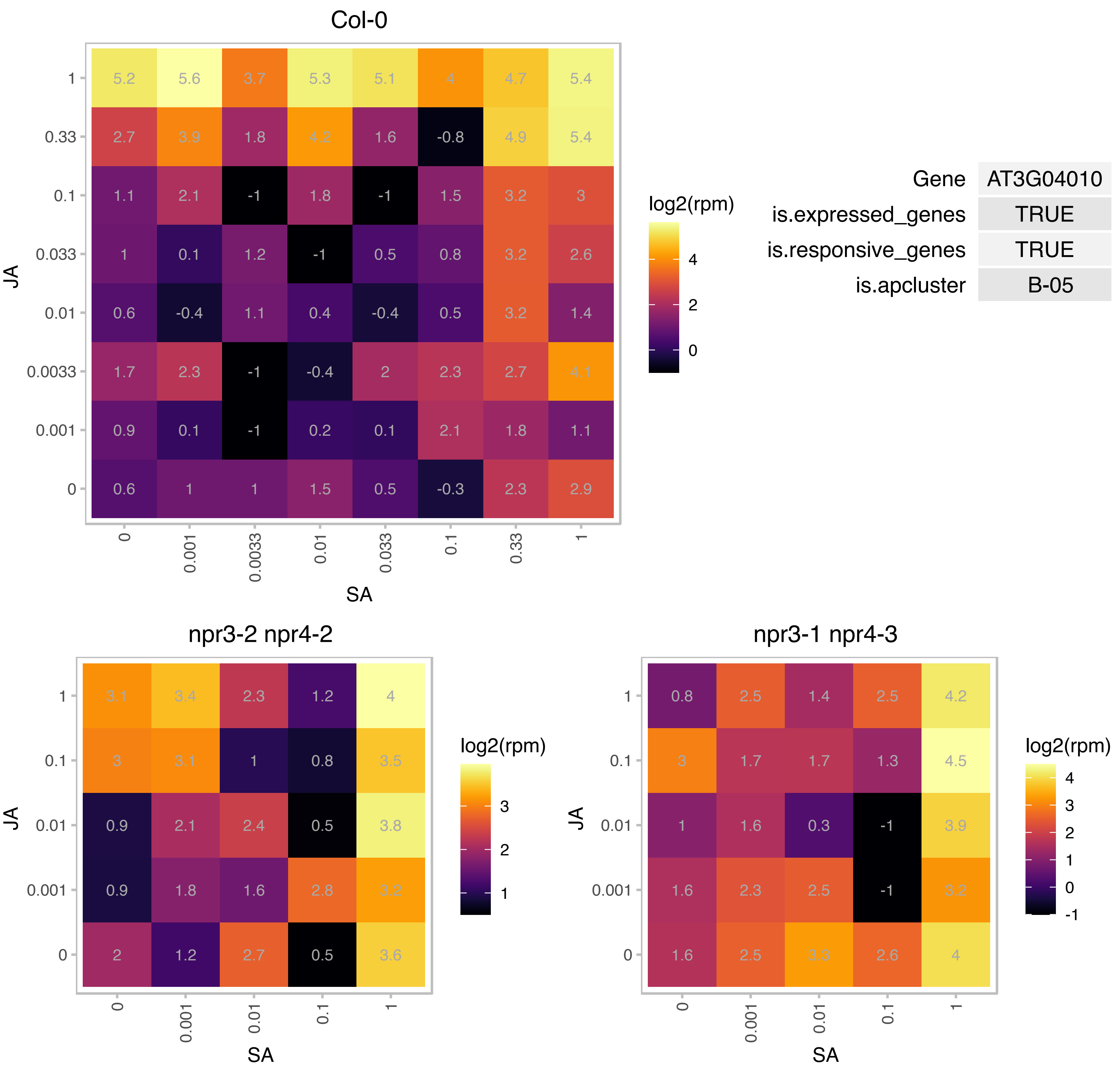

# AT3G05545.1 : NA

RING/U-box superfamily protein

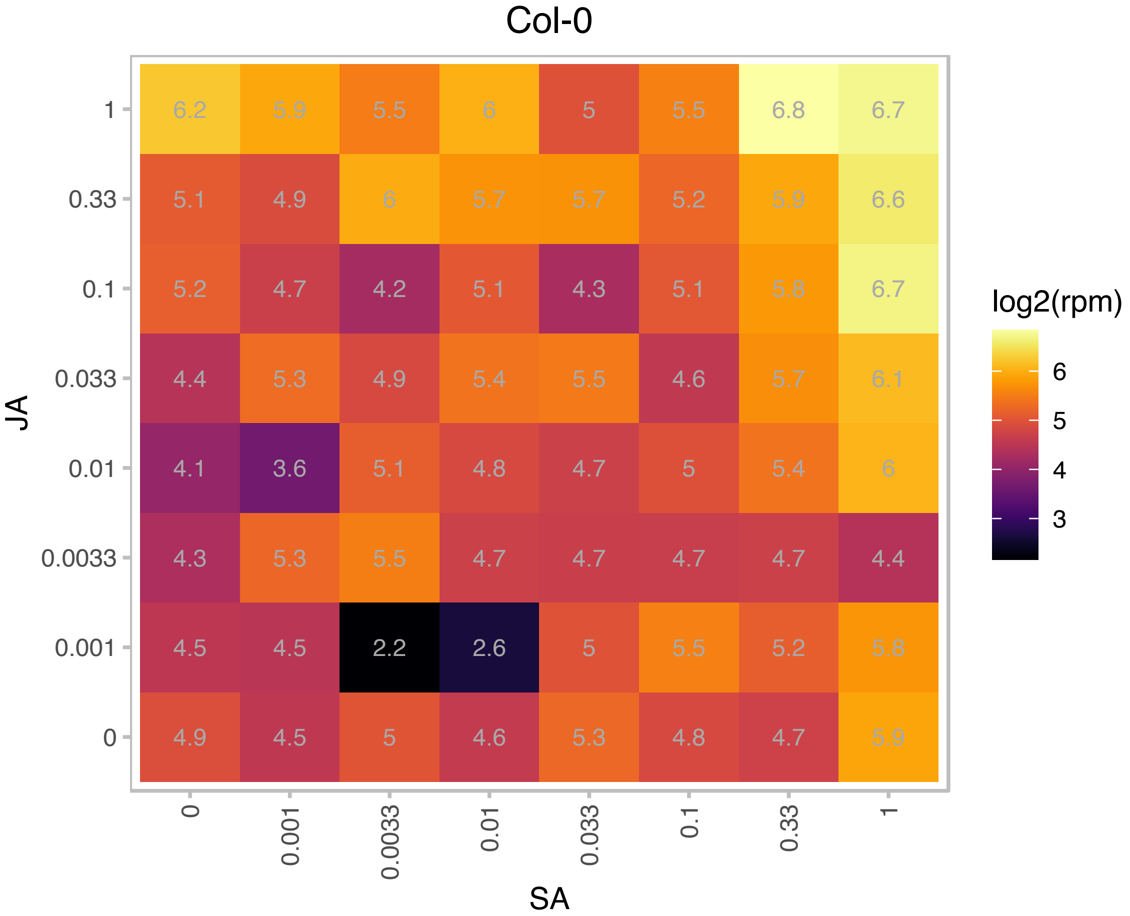


| | |
|---|---|
| Gene | AT3G05545 |
| is.expressed_genes | TRUE |
| is.responsive_genes | TRUE |
| is.apcluster | B-05 |

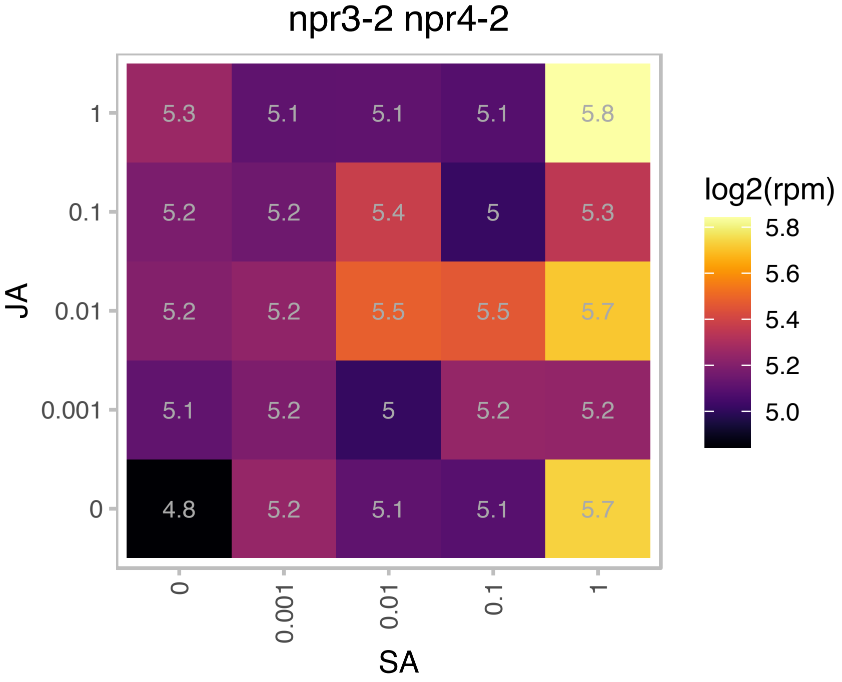


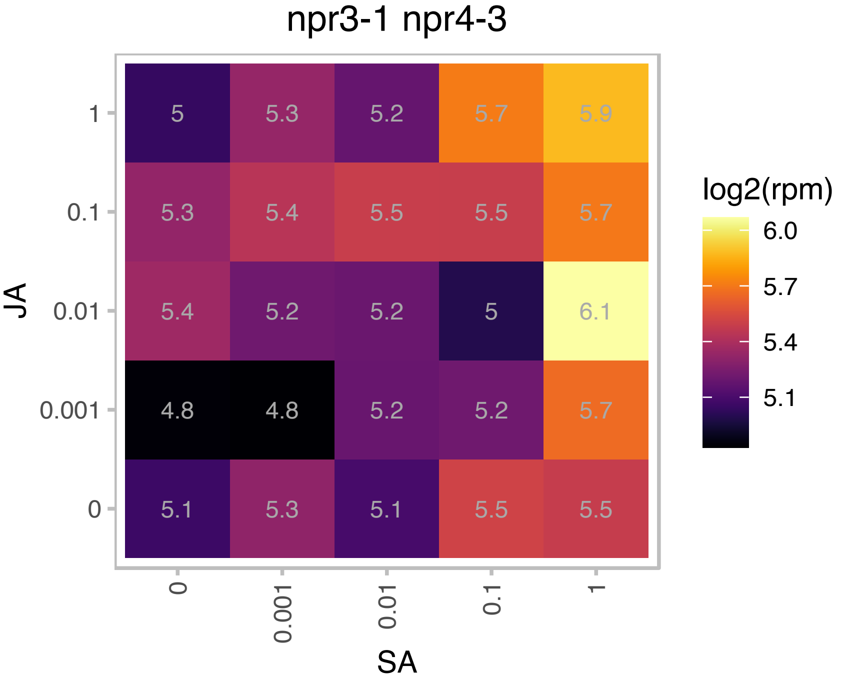

### AT3G08710.1 : ATH9, TH9, TRX H9

thioredoxin H-type 9

Associated to plasma membrane. Moves cell to cell, suggesting a role in intercellular communication.

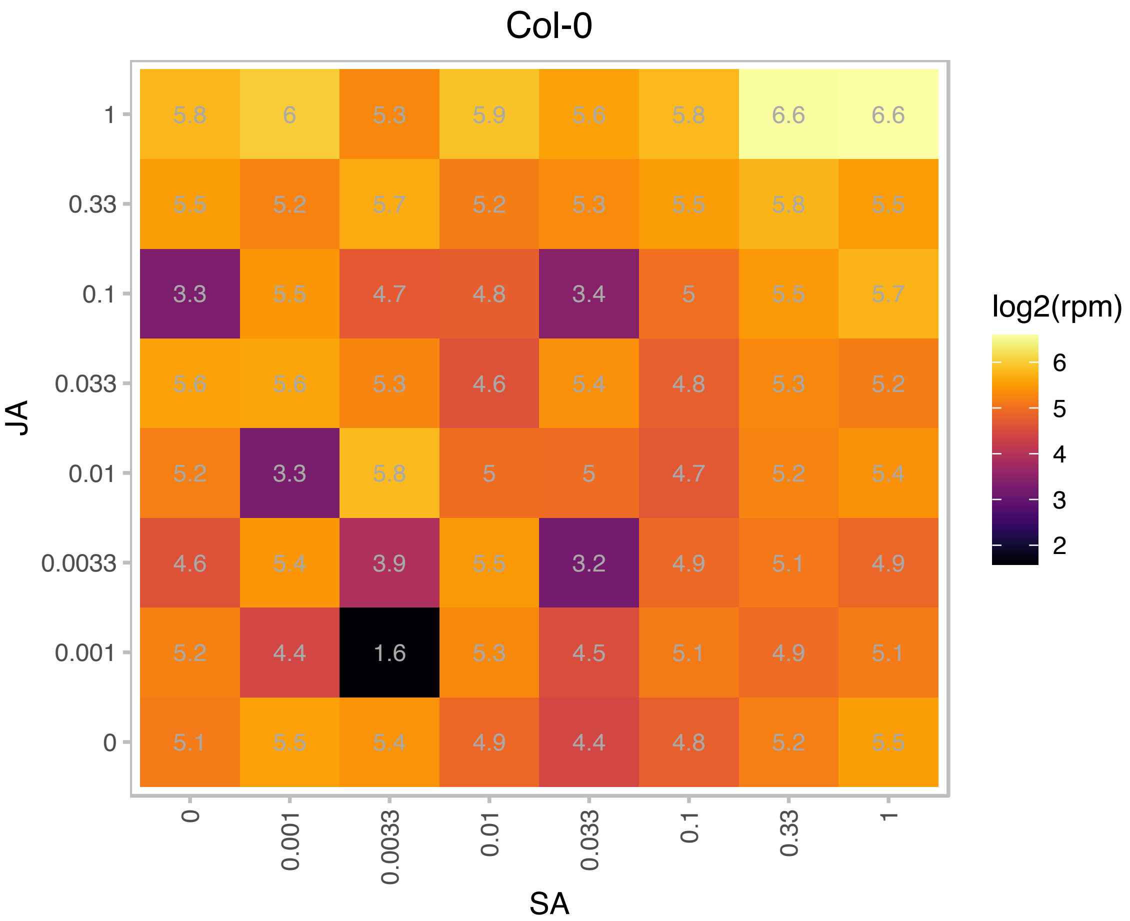


| | |
|---|---|
| Gene | AT3G08710 |
| is.expressed_genes | TRUE |
| is.responsive_genes | TRUE |
| is.apcluster | B-05 |

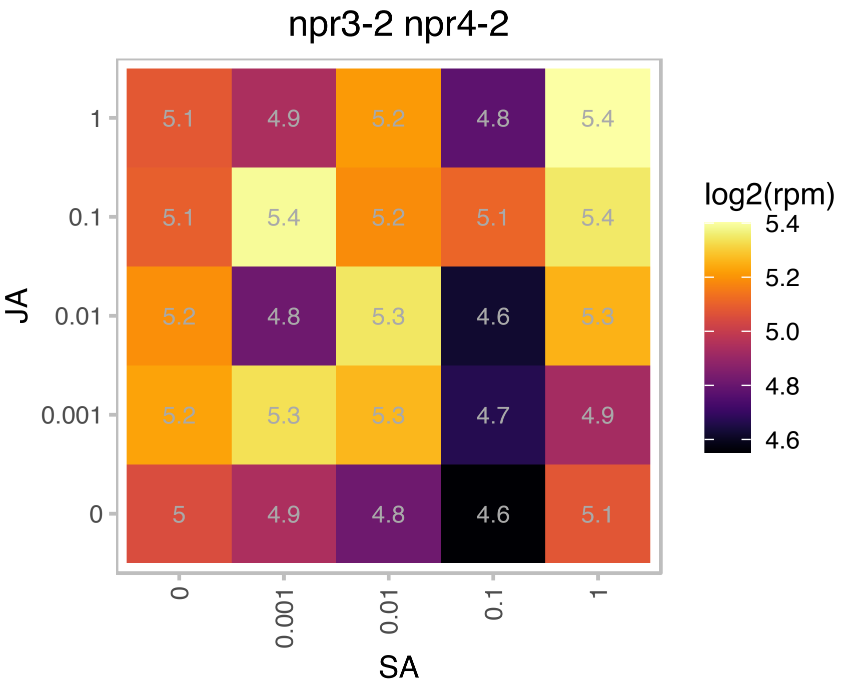


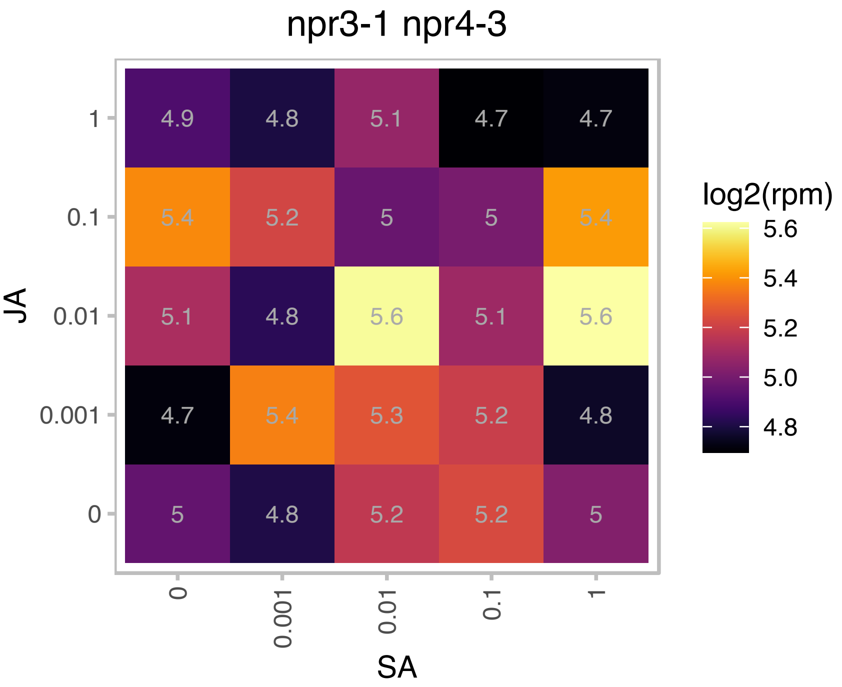

# AT3G09760.1 : NA

RING/U-box superfamily protein

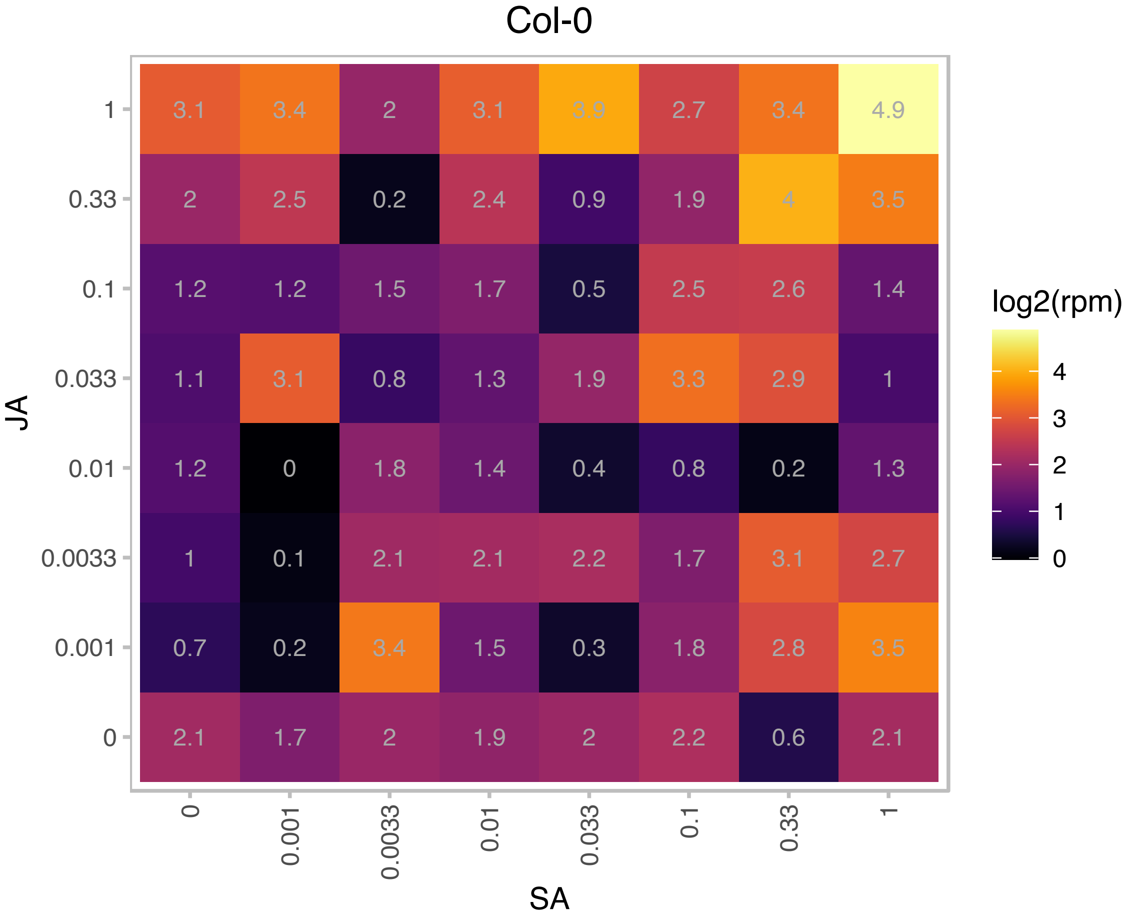


| | |
|---|---|
| Gene | AT3G09760 |
| is.expressed_genes | TRUE |
| is.responsive_genes | TRUE |
| is.apcluster | B-05 |

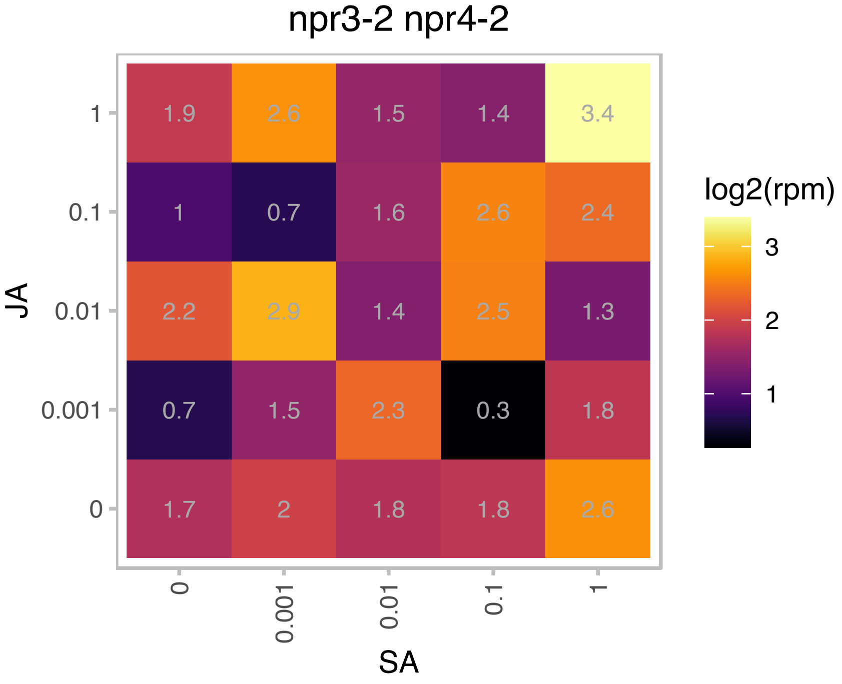


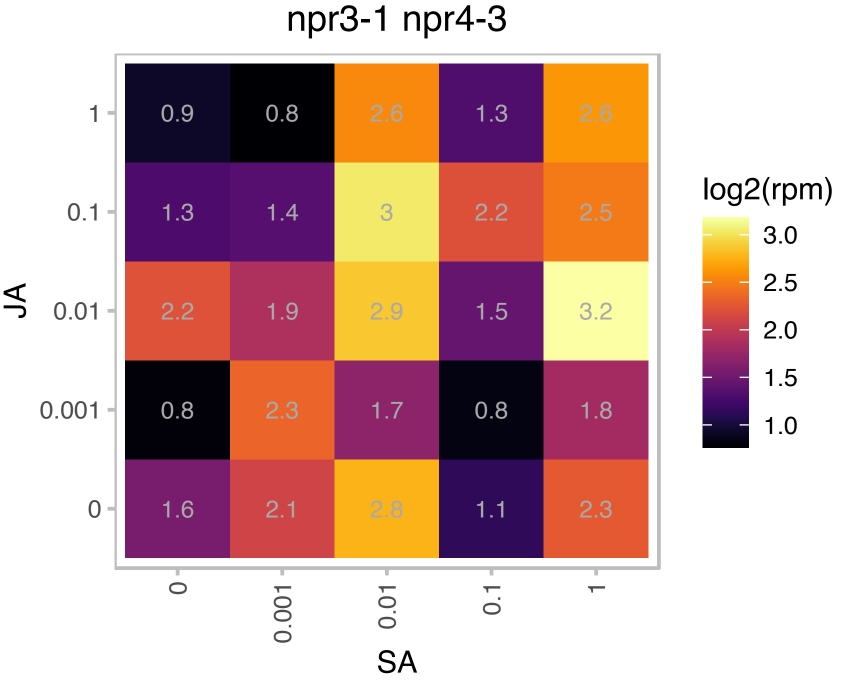

### AT3G12780.1 : PGK1

PGK1 was localized exclusively in the chloroplasts of photosynthetic tissues and is the photosynthetic phosphoglycerate kinase 1 isoform. The pgk1.1 knock-down mutant displayed reduced growth, lower photosynthetic capacity and starch content. Expression studies in PGK mutants showed that PGK1 and PGK3 were down-regulated in pgk3.2 and pgk1.1, respectively. These results indicate that the down-regulation of photosynthetic activity could be a plant strategy when glycolysis is impaired to achieve metabolic adjustment and optimize growth (DOI:10.1104/pp.17.01227

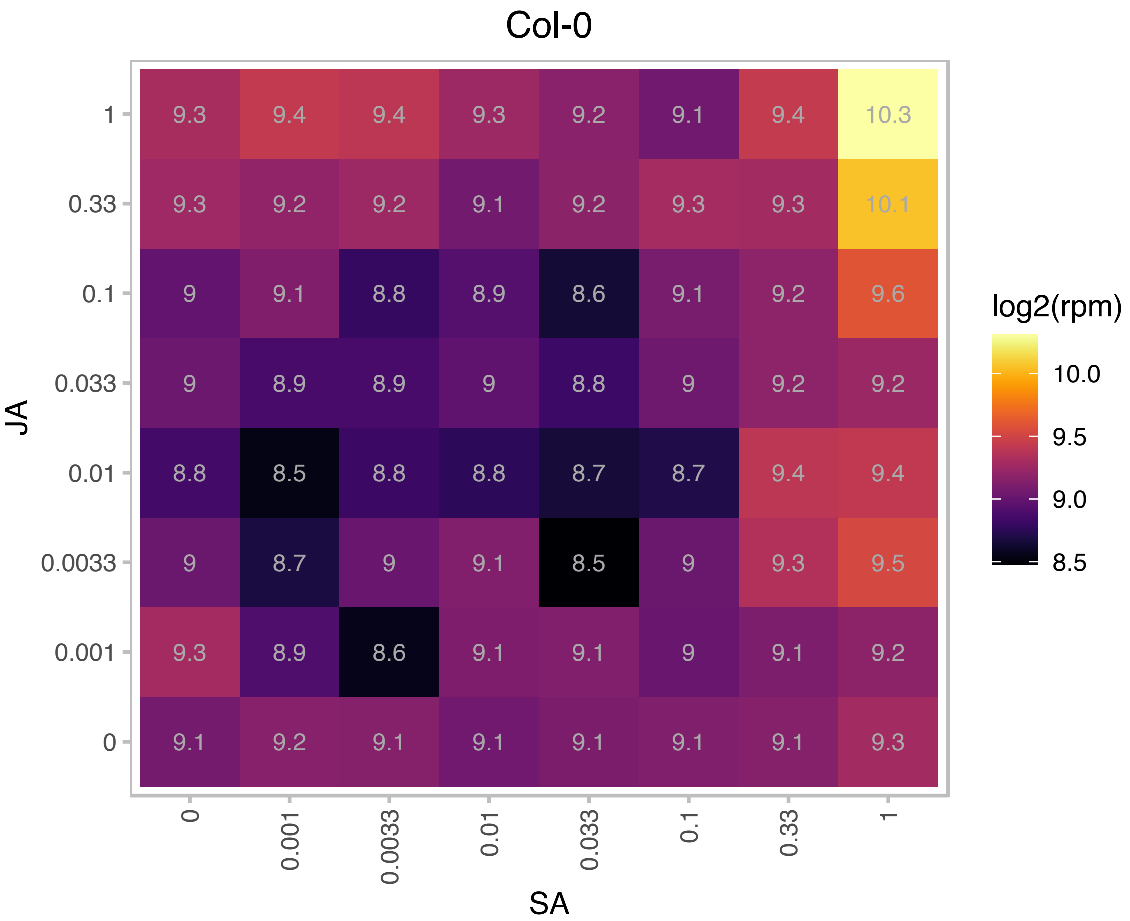


| | |
|---|---|
| Gene | AT3G12780 |
| is.expressed_genes | TRUE |
| is.responsive_genes | TRUE |
| is.apcluster | B-05 |

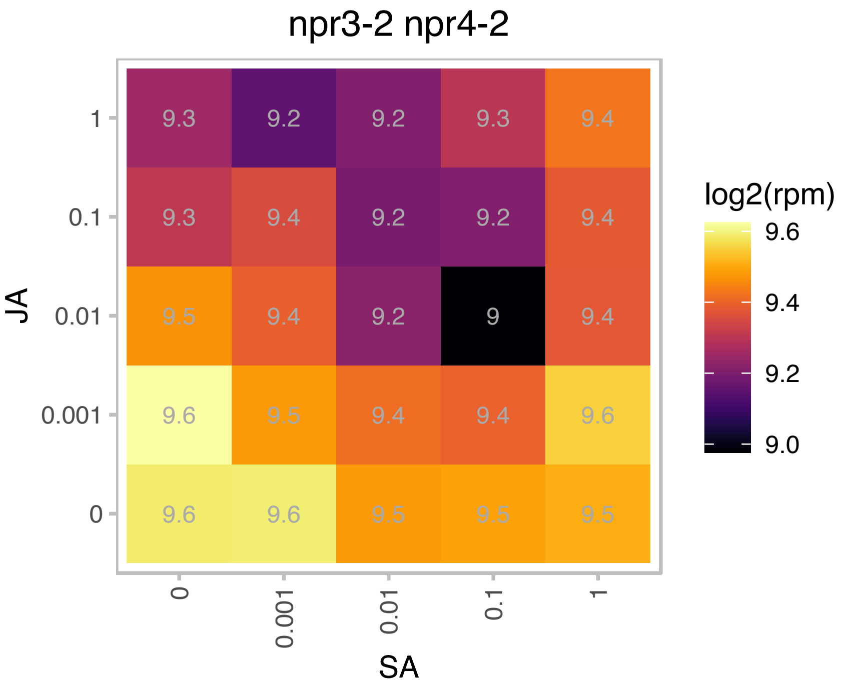


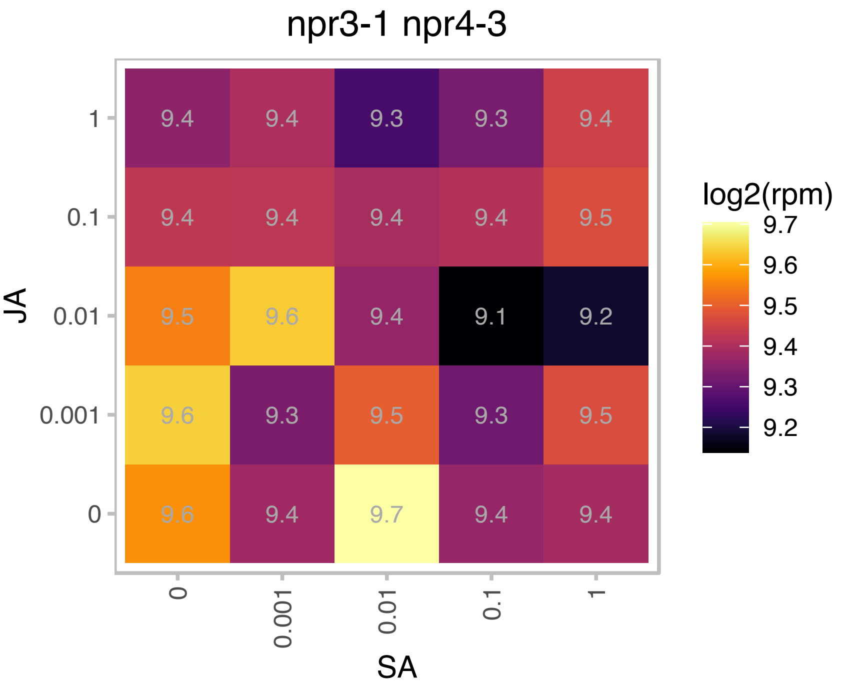

### AT3G13520.1 : AGP12, ATAGP12

arabinogalactan protein 12
Encodes a GPI-anchored arabinogalactan (AG) peptide with a short 'classical' backbone of 10 amino acids, seven of which are conserved among the 4 other Arabidopsis AG peptides.  These peptides may be involved in cell signaling.

Col-0

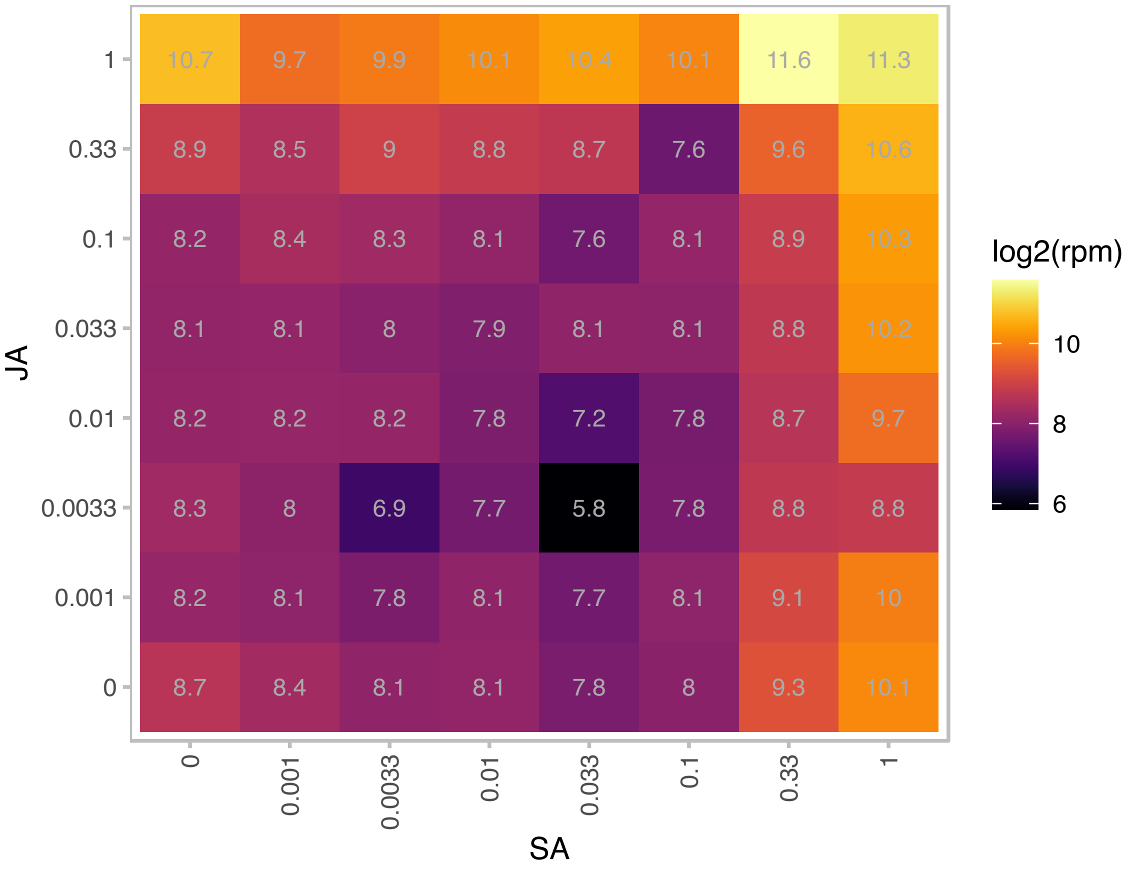


| | |
|---|---|
| Gene | AT3G13520 |
| is.expressed_genes | TRUE |
| is.responsive_genes | TRUE |
| is.apcluster | B-05 |

npr3-2 npr4-2

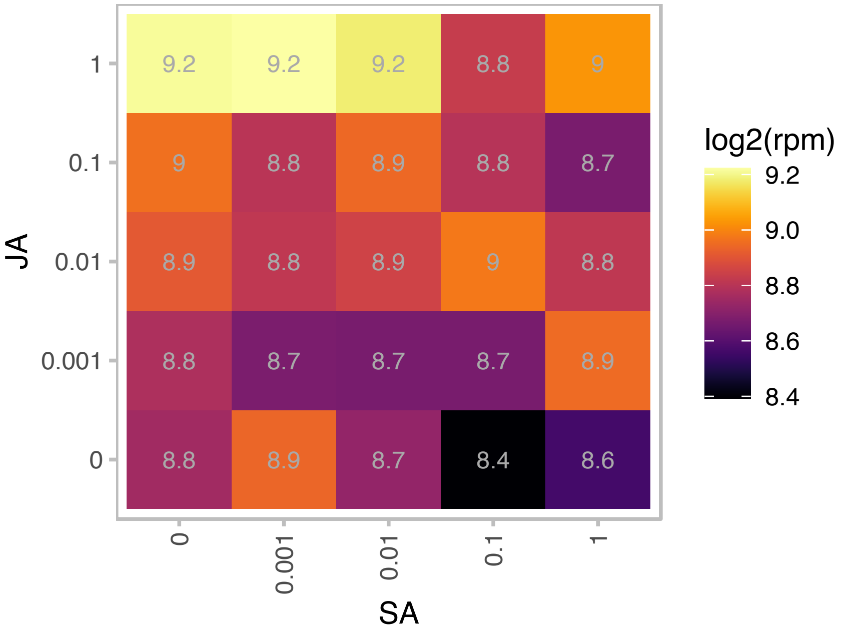


npr3-1 npr4-3

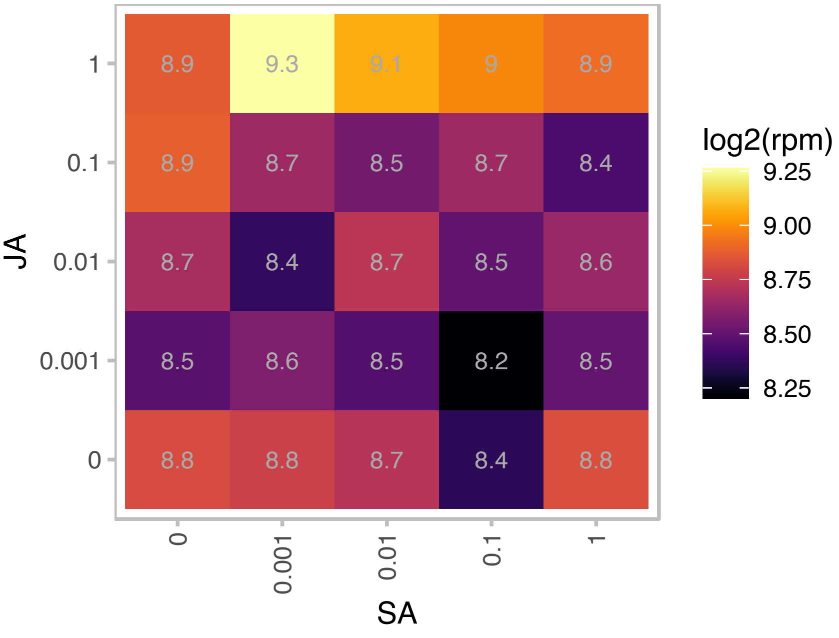

### AT3G20060.1 : UBC19

ubiquitin-conjugating enzyme19
Encodes one of two ubiquitin-conjugating enzymes belonging to the E2-C gene family (the other being UBC19). Transcript is always found in dividing cells, but also in other non-dividing cells. Protein is localized to the cytoplasm as well as to the nucleus.

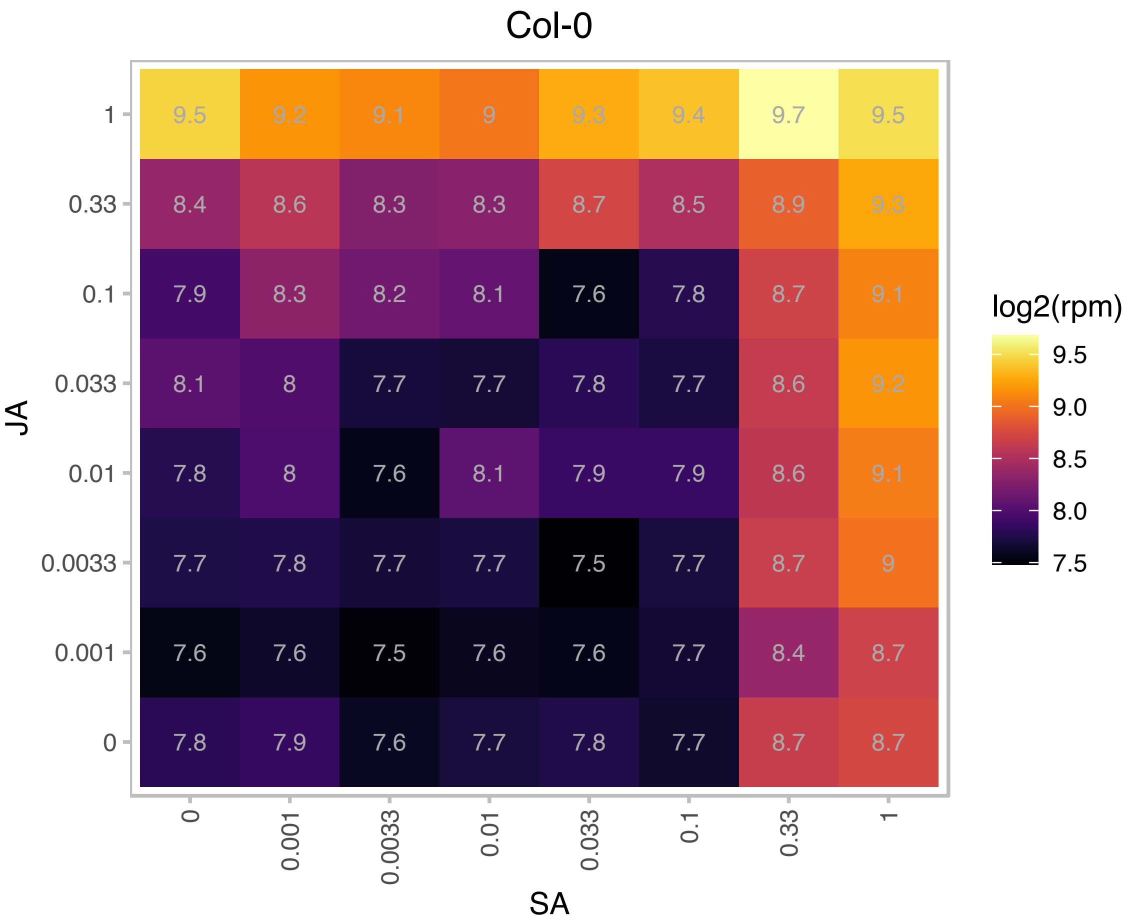


| | |
|---|---|
| Gene | AT3G20060 |
| is.expressed_genes | TRUE |
| is.responsive_genes | TRUE |
| is.apcluster | B-05 |

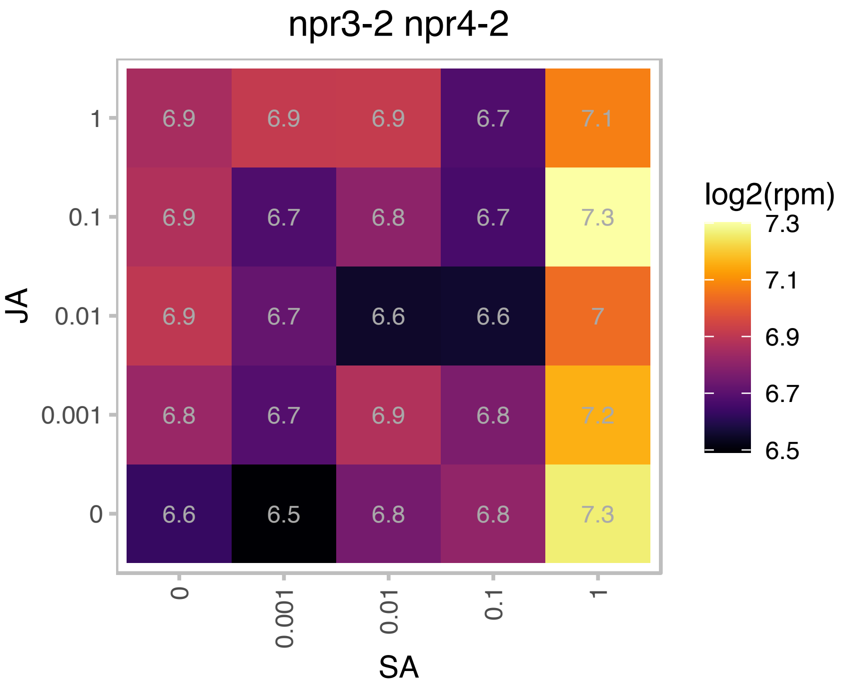


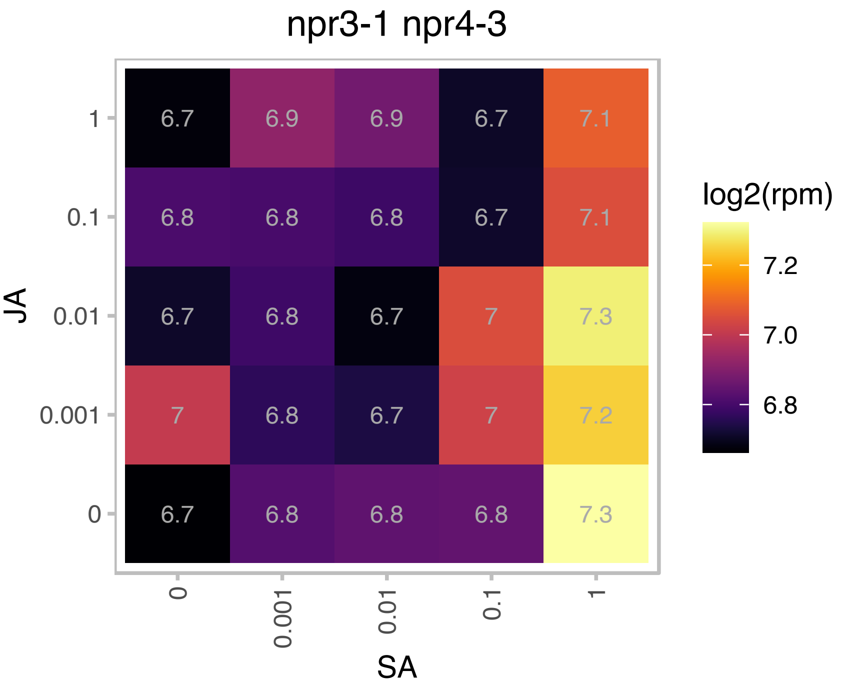

### AT3G21700.1 : ATSGP2, SGP2

Ras-related small GTP-binding family protein

Monomeric G protein. Expressed in root epidermal cells that are destined to become atrichoblasts. Also expressed during pollen development and in the pollen tube tip.

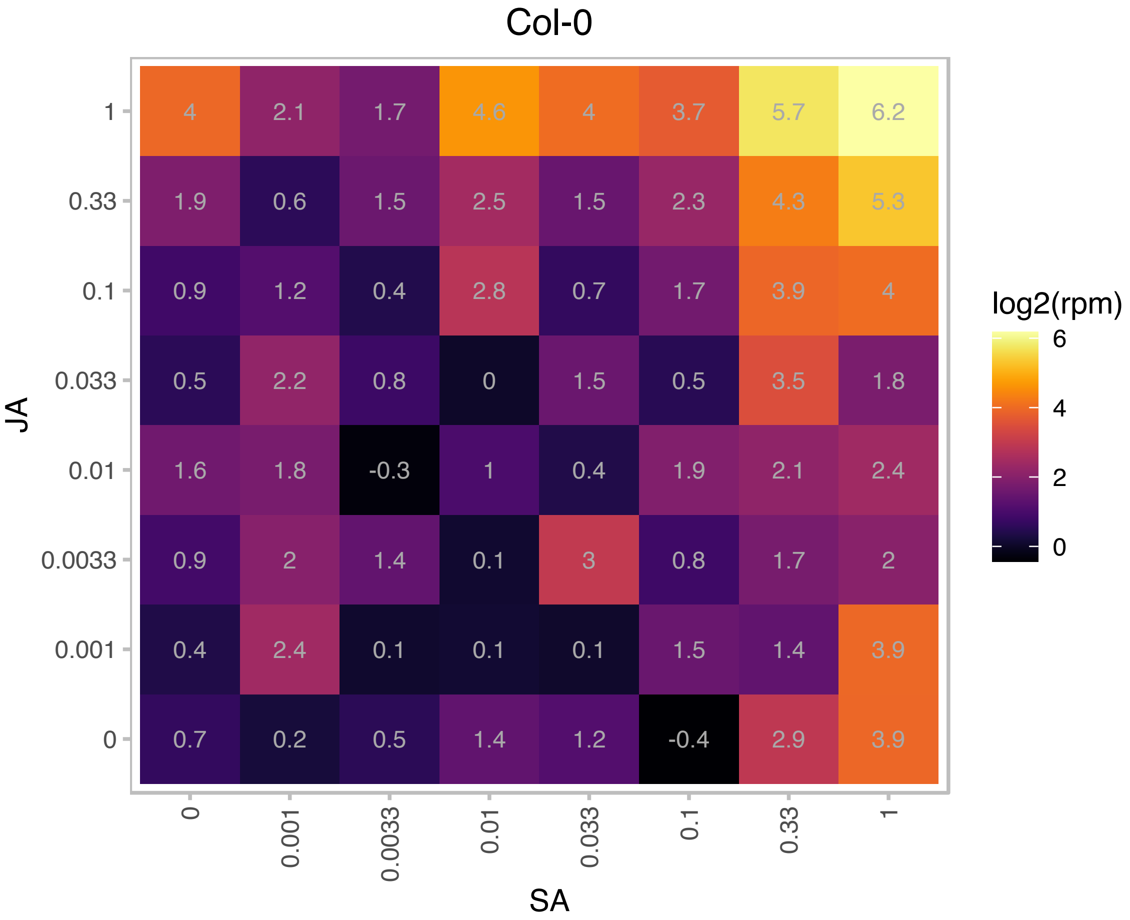


| | |
|---|---|
| Gene | AT3G21700 |
| is.expressed_genes | TRUE |
| is.responsive_genes | TRUE |
| is.apcluster | B-05 |

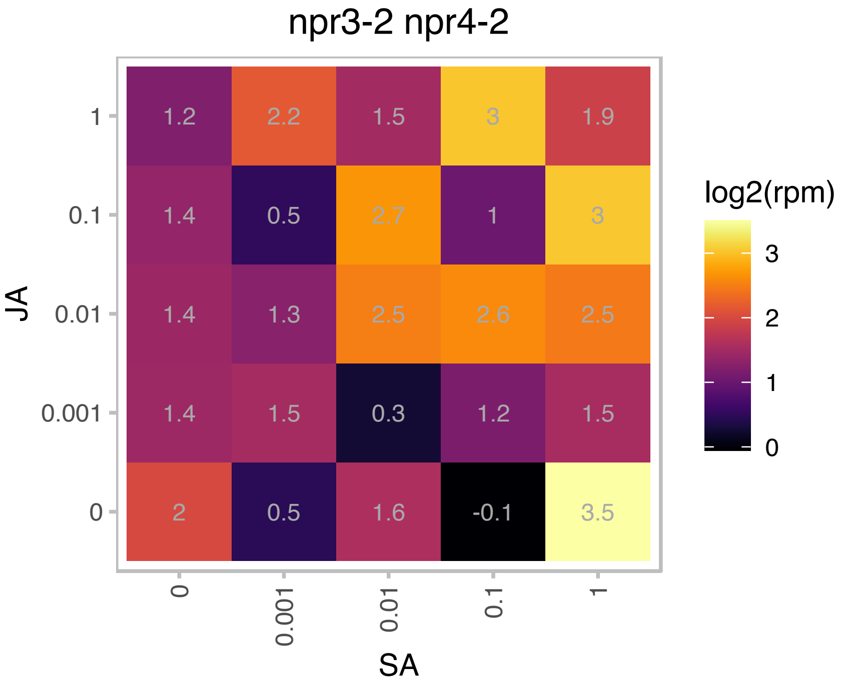


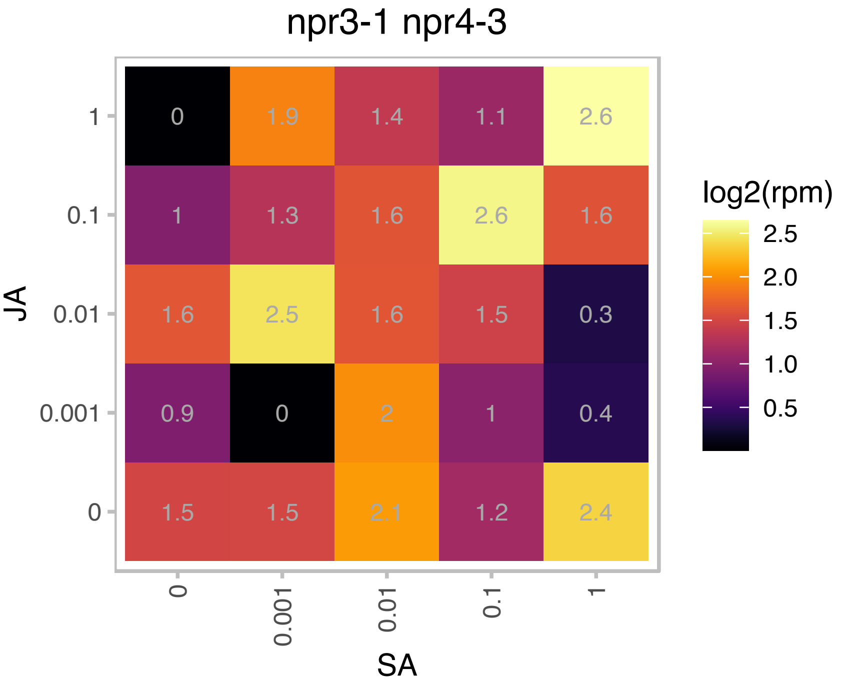

### AT3G25250.1 : AGC2, AGC2-1, AtOXI1, OXI1

AGC (cAMP-dependent, cGMP-dependent and protein kinase C) kinase family protein

Arabidopsis protein kinase The mRNA is cell-to-cell mobile.

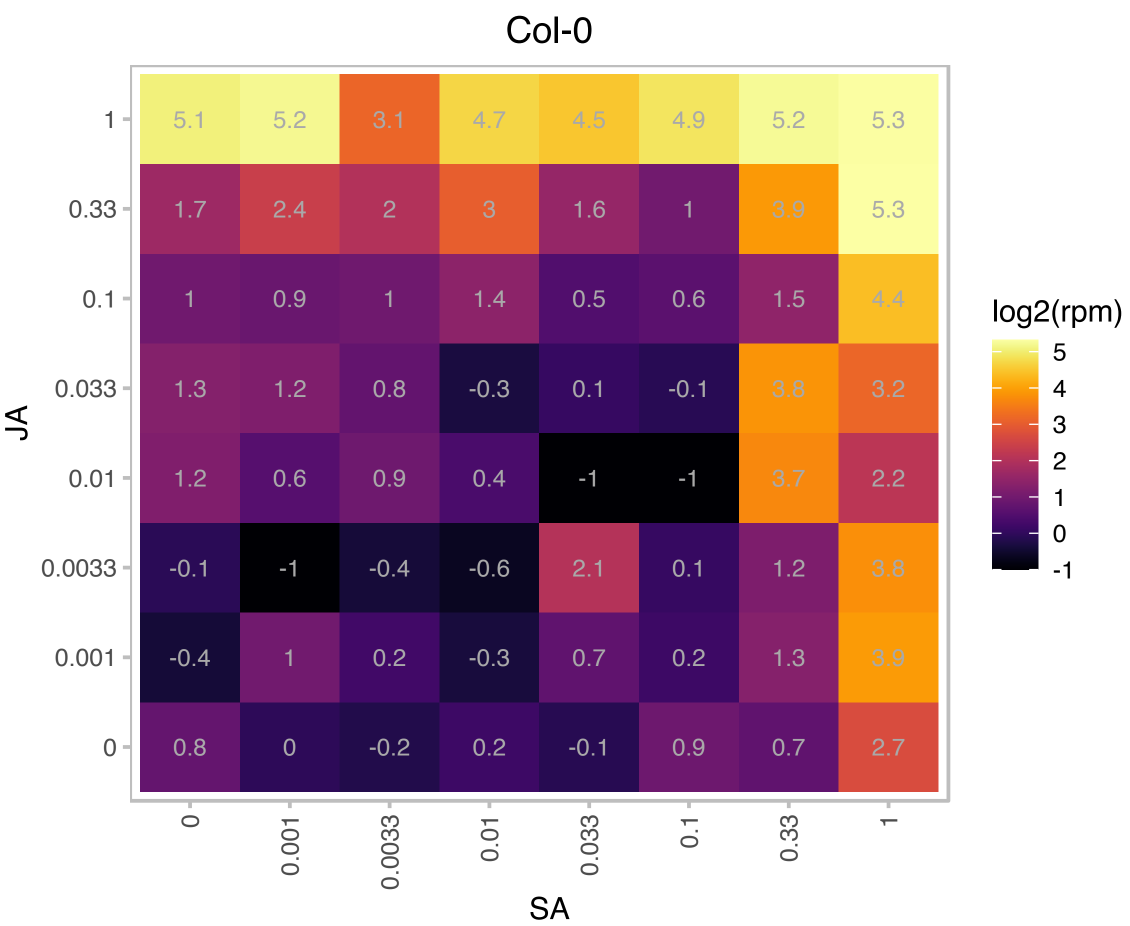


| | |
|---|---|
| Gene | AT3G25250 |
| is.expressed_genes | TRUE |
| is.responsive_genes | TRUE |
| is.apcluster | B-05 |

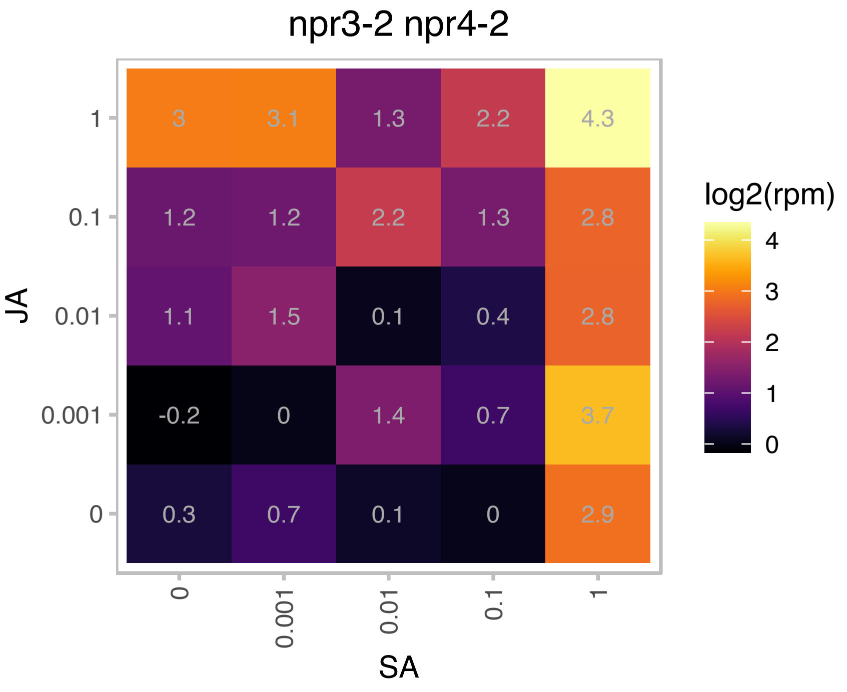


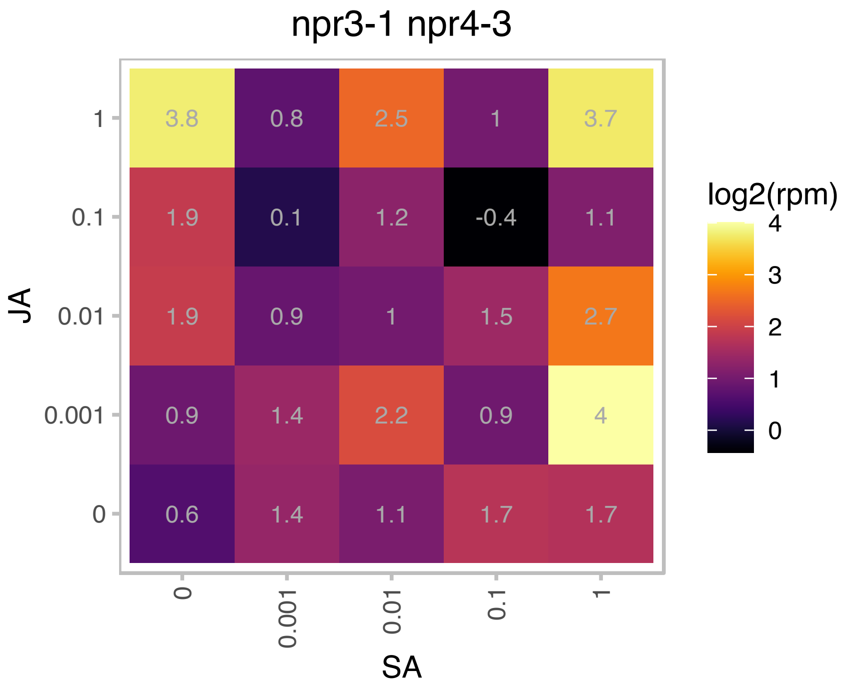

# AT3G25910.1 : NA

Protein of unknown function (DUF1644)

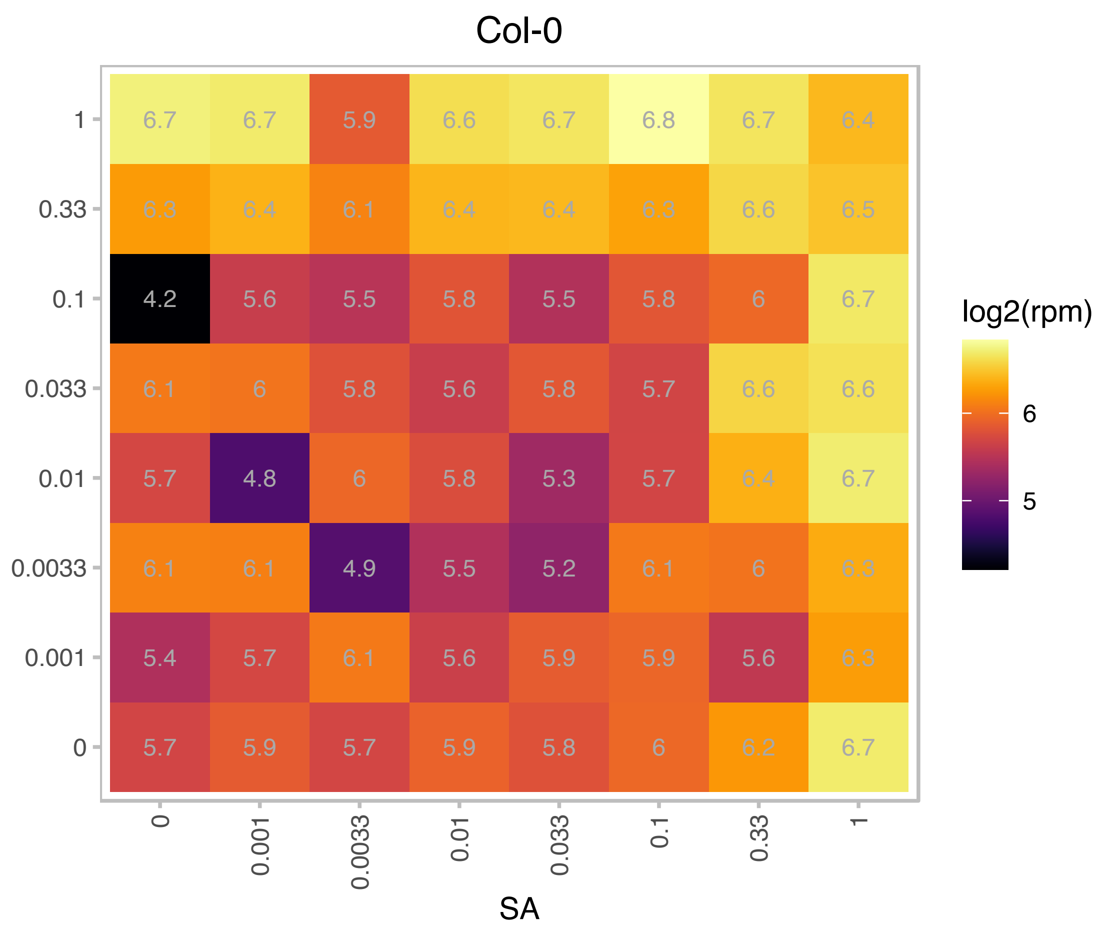


| | |
|---|---|
| Gene | AT3G25910 |
| is.expressed_genes | TRUE |
| is.responsive_genes | TRUE |
| is.apcluster | B-05 |

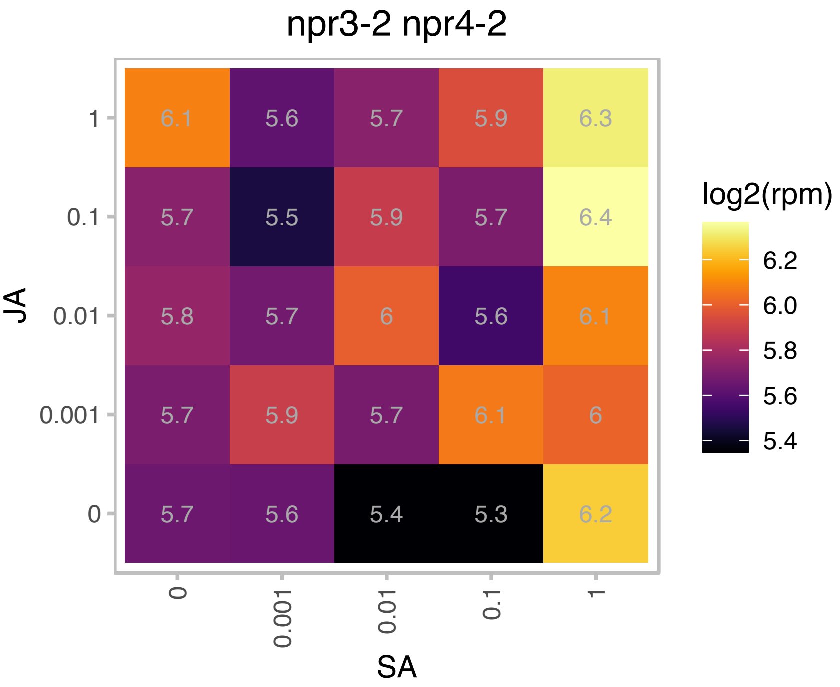


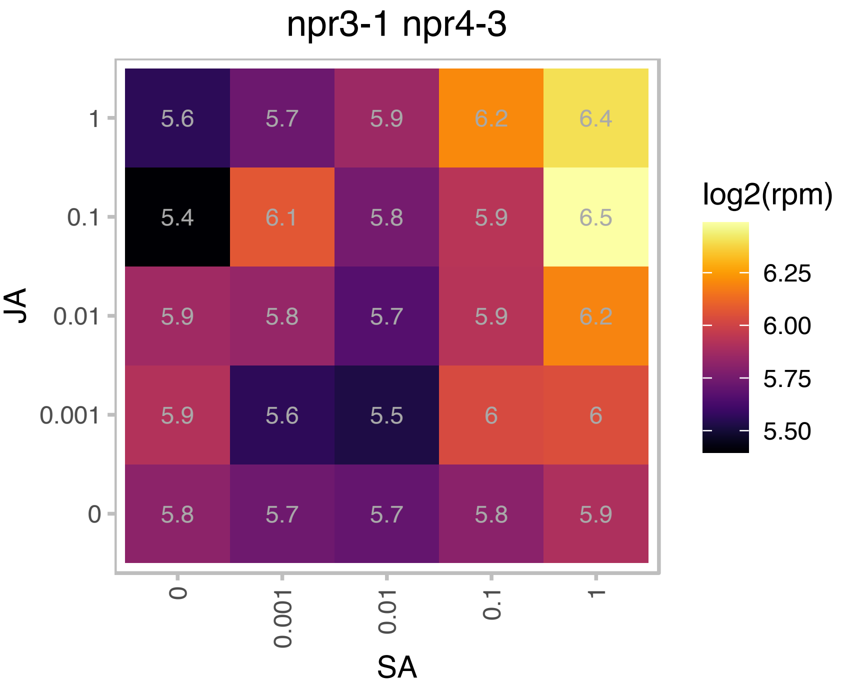

# AT3G51130.1 : NA

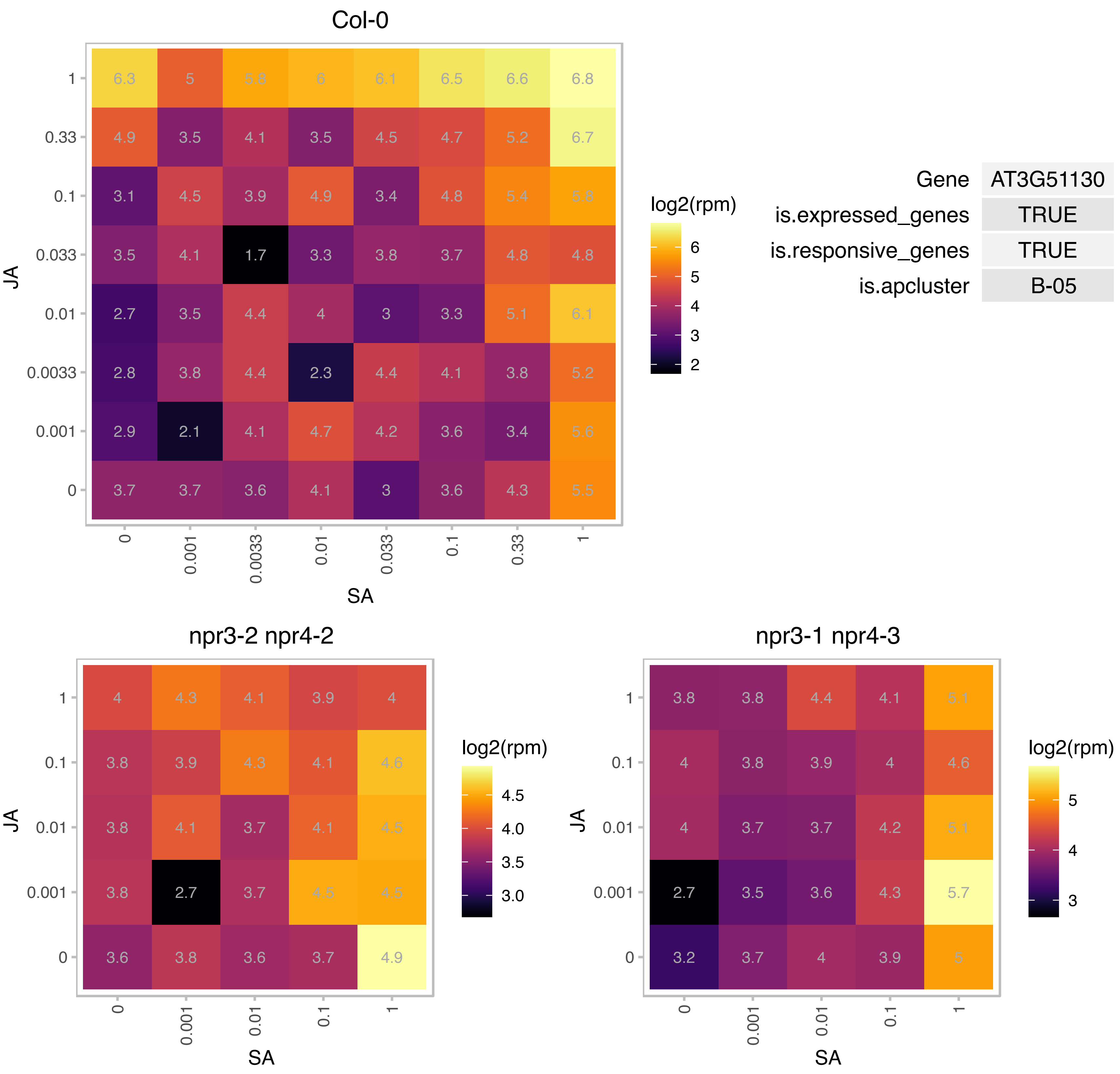

### AT3G52930.1 : AtFBA8, FBA8

Aldolase superfamily protein

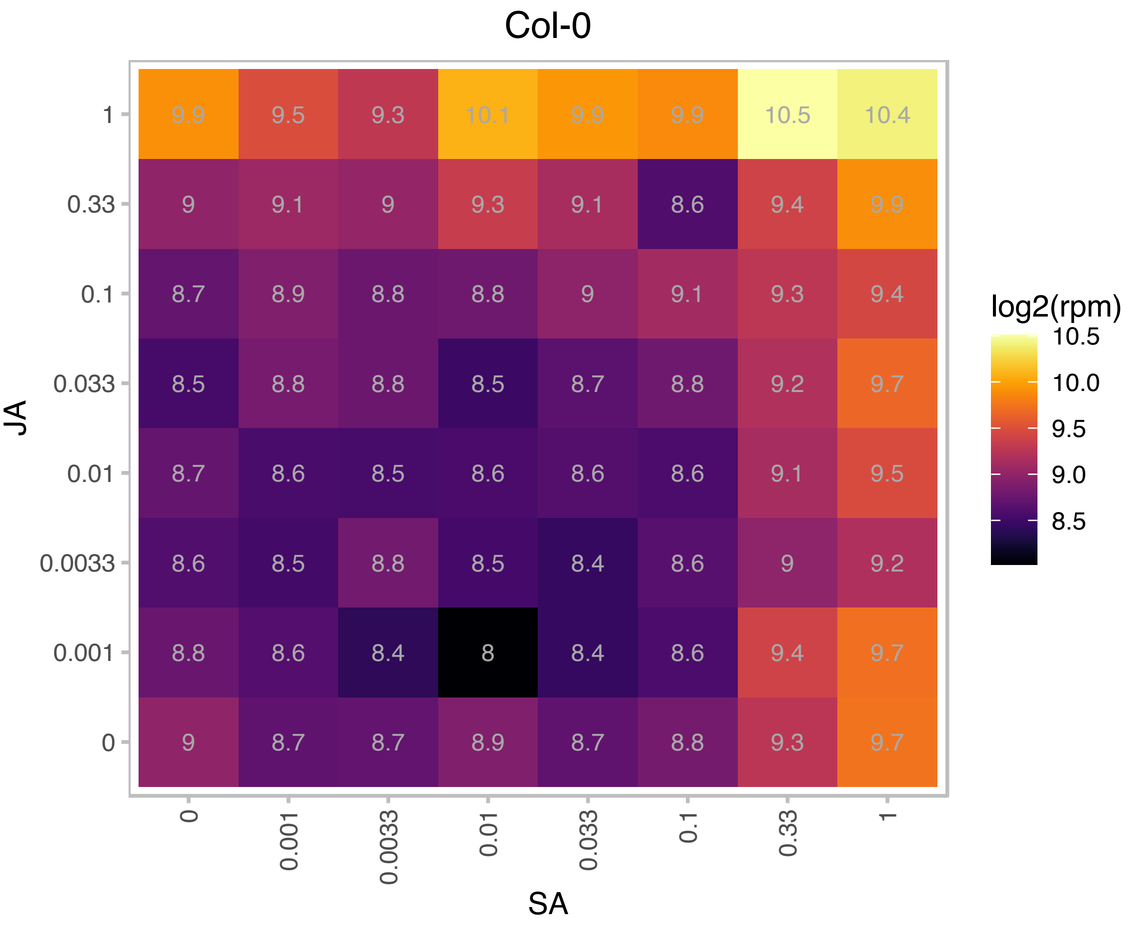


| | |
|---|---|
| Gene | AT3G52930 |
| is.expressed_genes | TRUE |
| is.responsive_genes | TRUE |
| is.apcluster | B-05 |

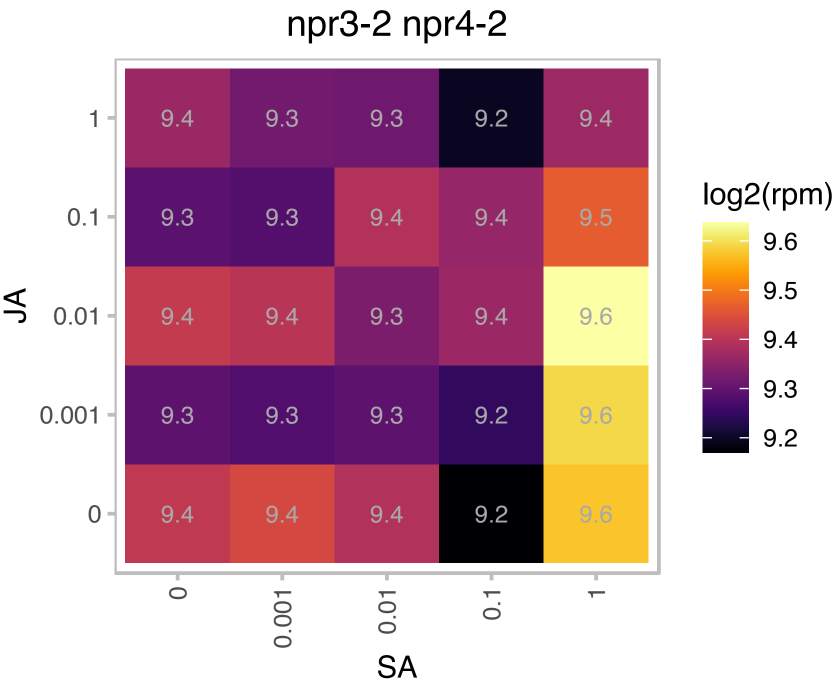


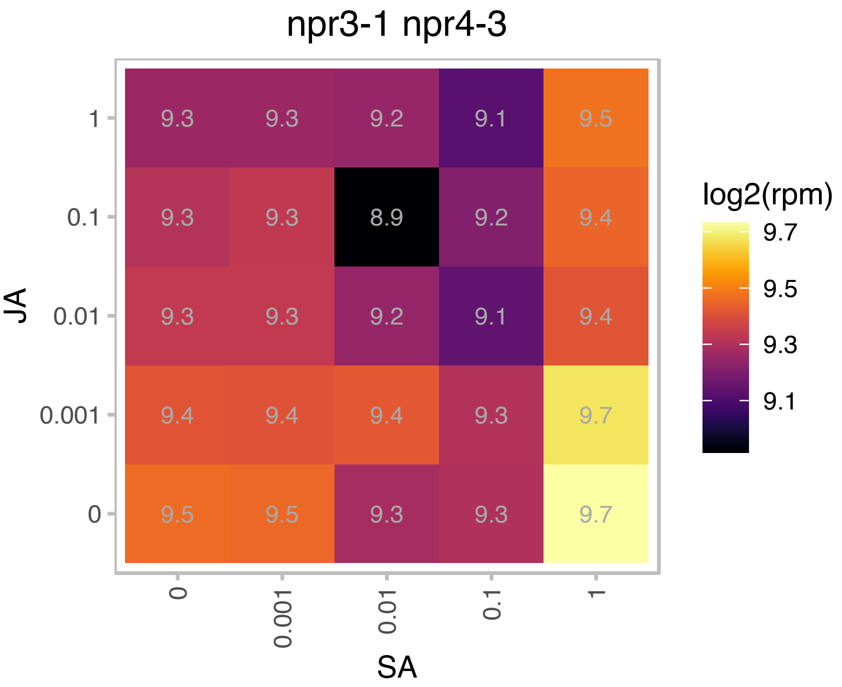

### AT3G53110.1 : LOS4

P-loop containing nucleoside triphosphate hydrolases superfamily protein

Encodes a putative DEAD-Box RNA Helicase and has RNA-dependent ATPase activity. Mutant is Sensitive to chilling stress and heat stress. Germination of the mutant is inhibited by ABA. LOS4 may be involved in temperature sensing. Is enriched in the nuclear envelope and also located in the cytoplasm. LOS4 is involved in export of poly A RNA. The mRNA is cell-to-cell mobile.

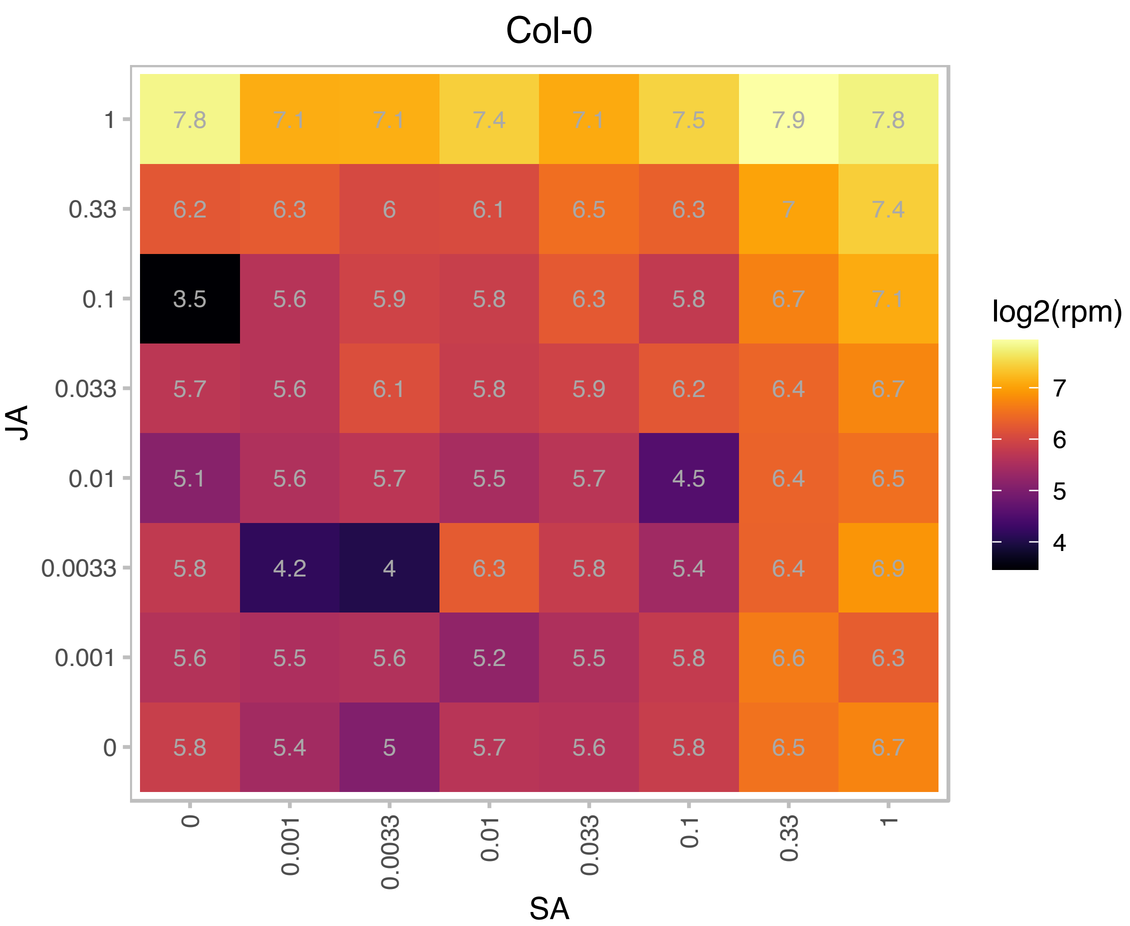


| | |
|---|---|
| Gene | AT3G53110 |
| is.expressed_genes | TRUE |
| is.responsive_genes | TRUE |
| is.apcluster | B-05 |

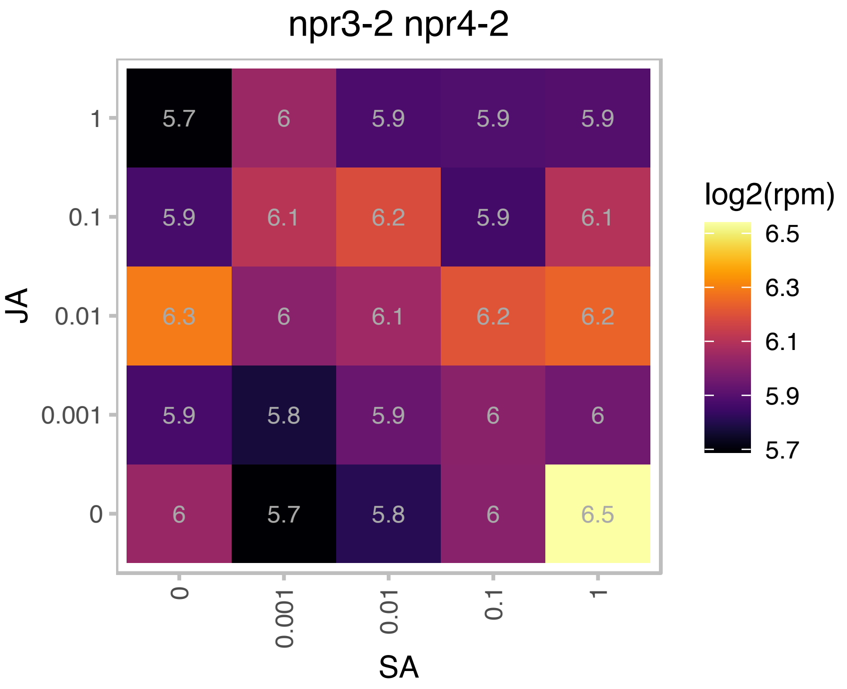


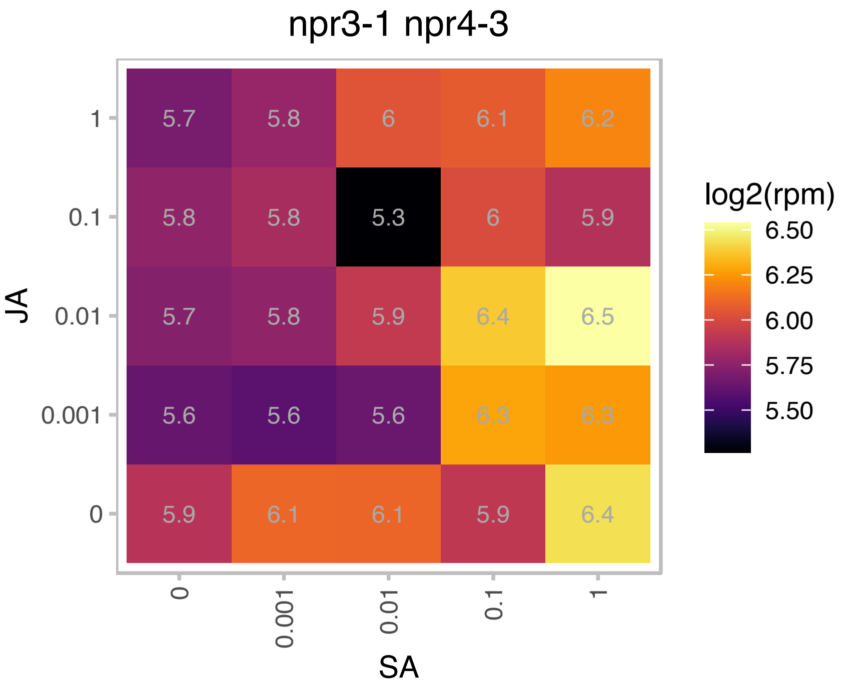

### AT3G53232.1 : DVL20, RTFL1

ROTUNDIFOLIA like 1

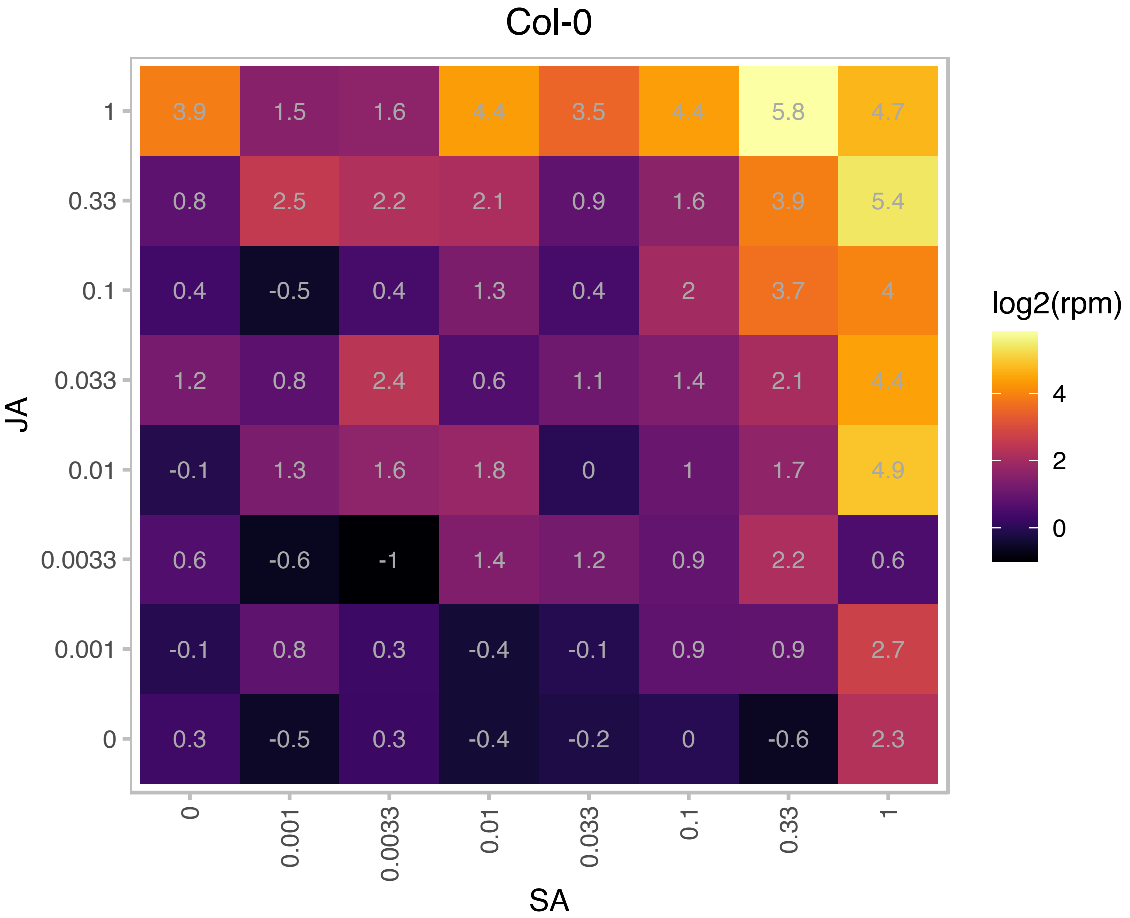


| | |
|---|---|
| Gene | AT3G53232 |
| is.expressed_genes | TRUE |
| is.responsive_genes | TRUE |
| is.apcluster | B-05 |

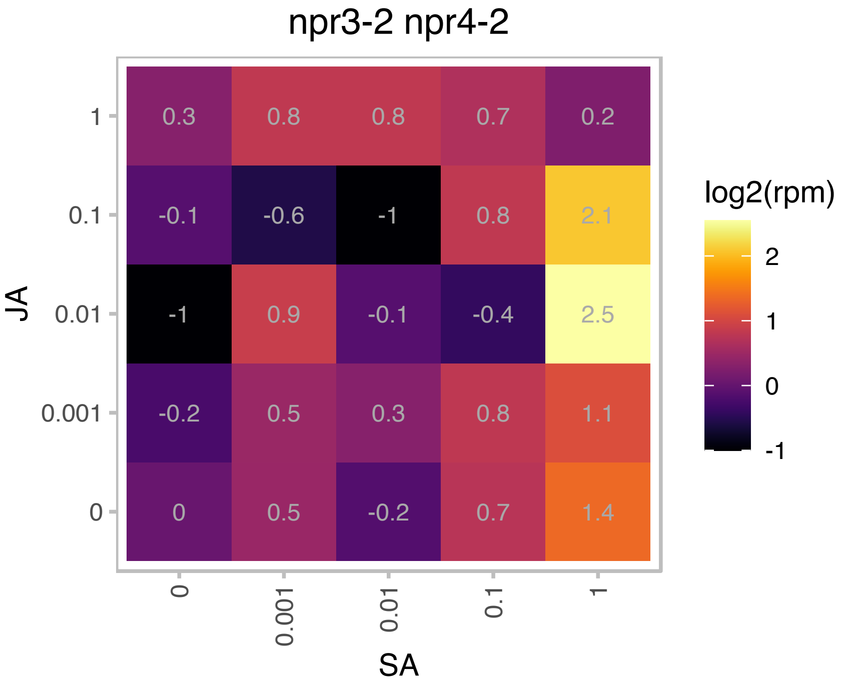


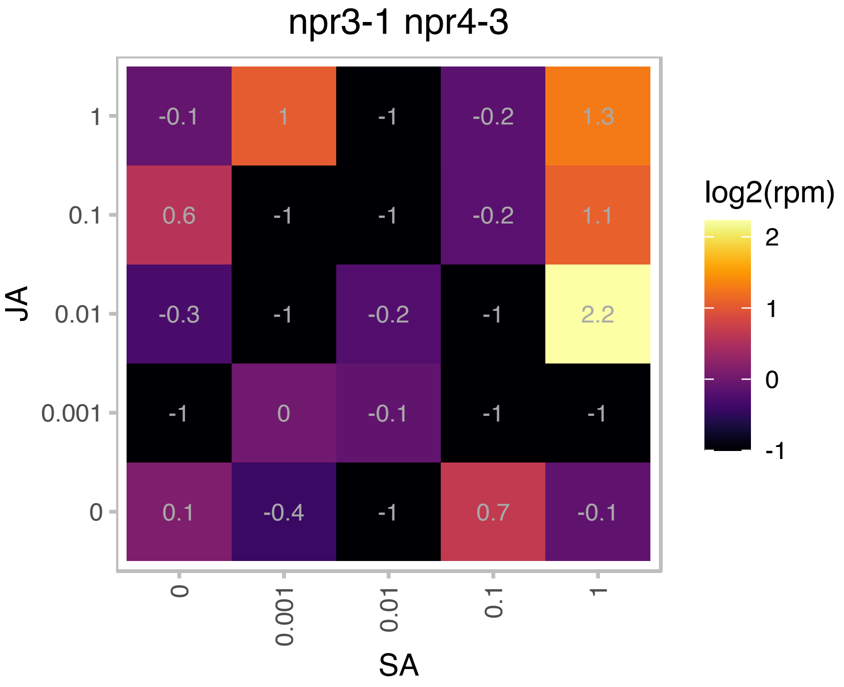

# AT3G55430.1 : NA

O-Glycosyl hydrolases family 17 protein

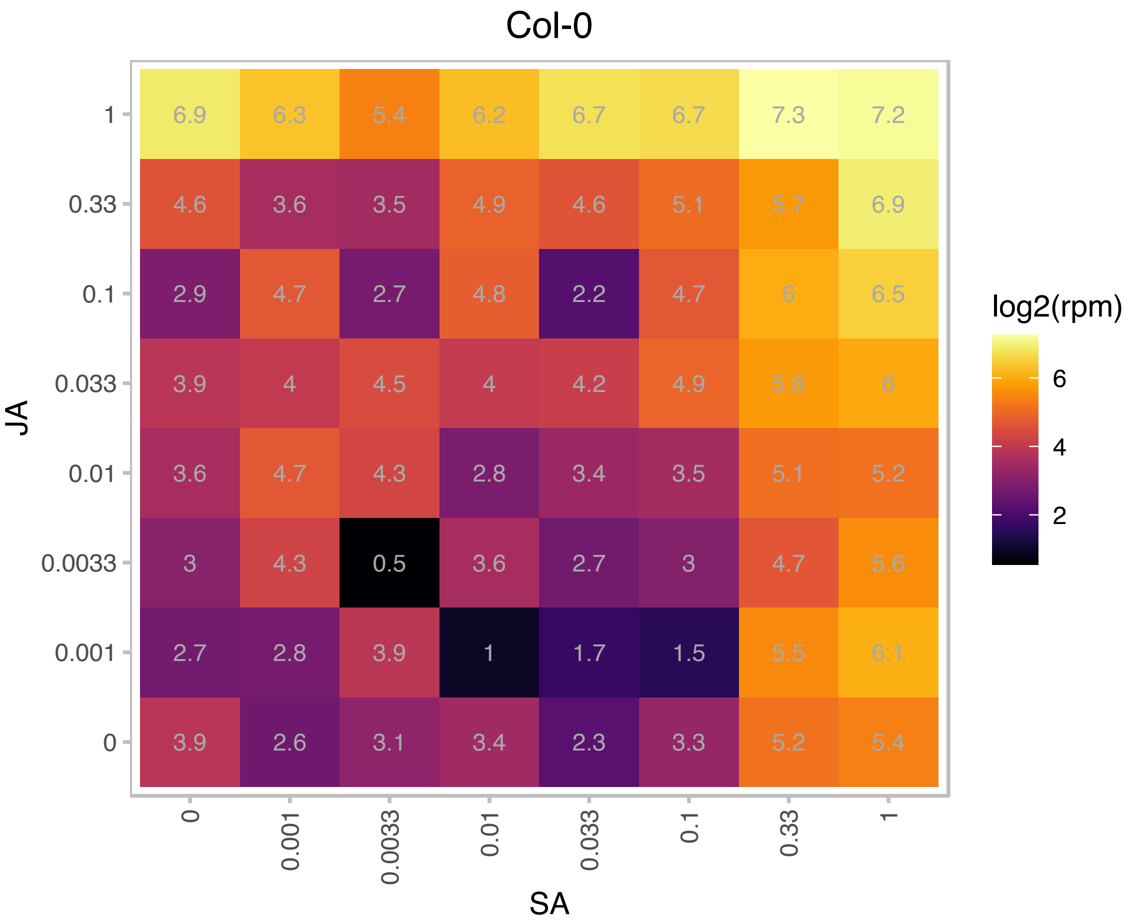


| | |
|---|---|
| Gene | AT3G55430 |
| is.expressed_genes | TRUE |
| is.responsive_genes | TRUE |
| is.apcluster | B-05 |

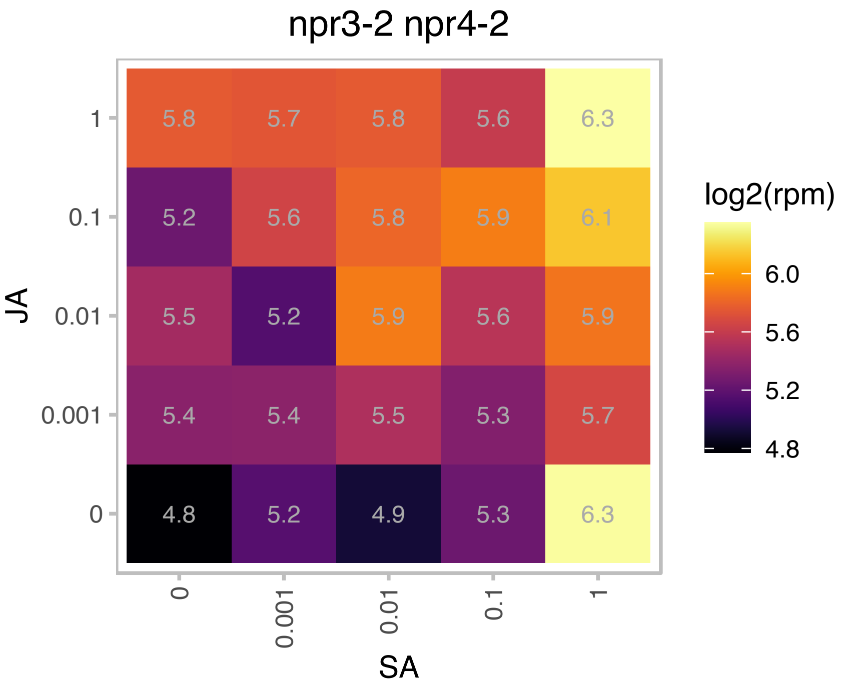


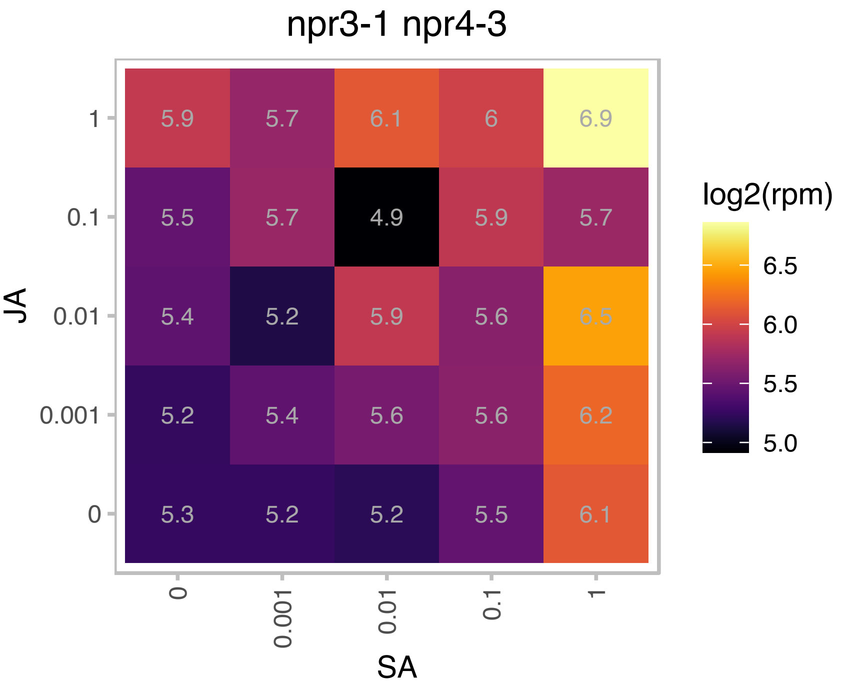

### AT4G02890.1 : UBQ14

Ubiquitin family protein

Polyubiquitin gene containing 4 ubiquitin repeats.

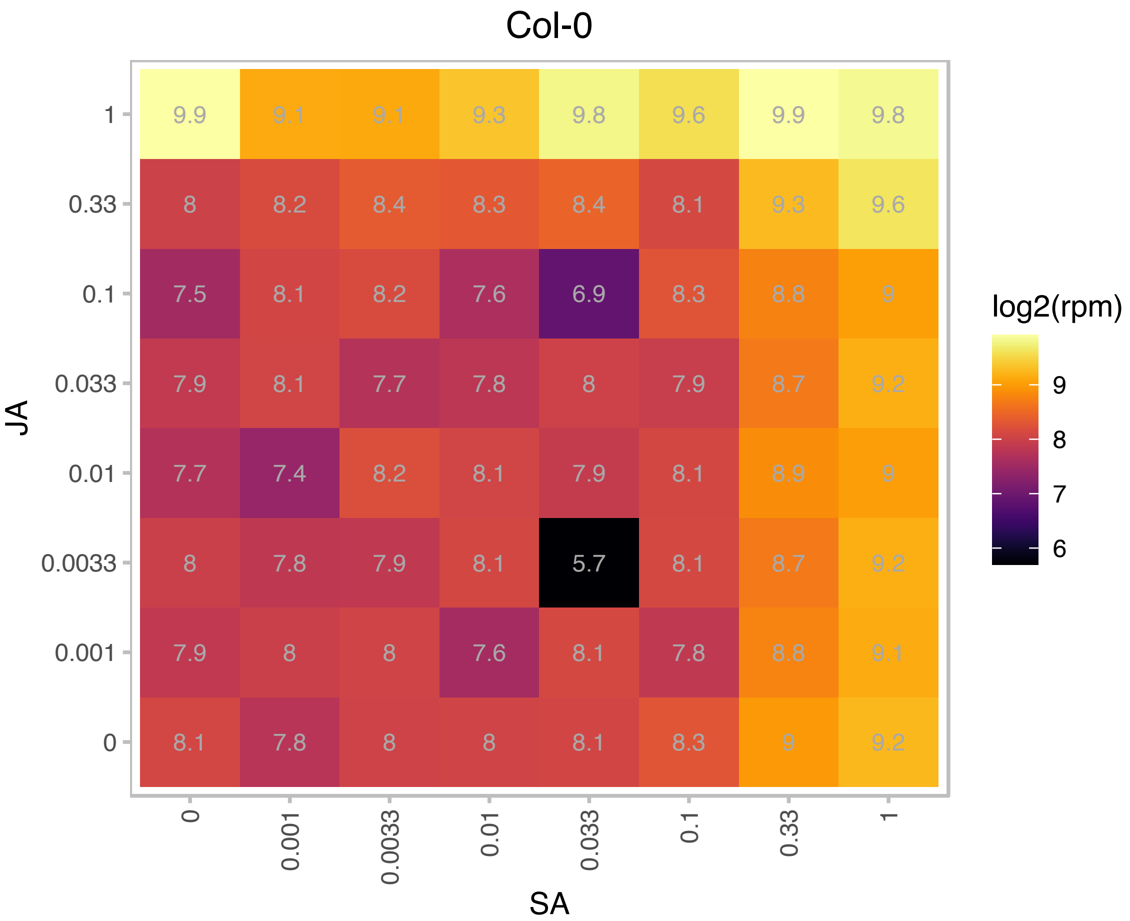


| | |
|---|---|
| Gene | AT4G02890 |
| is.expressed_genes | TRUE |
| is.responsive_genes | TRUE |
| is.apcluster | B-05 |

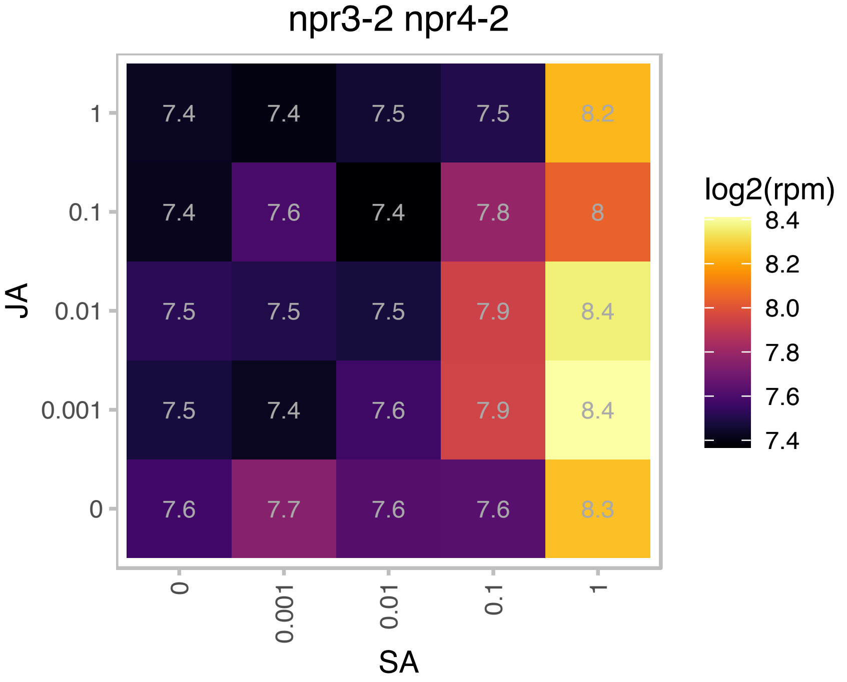


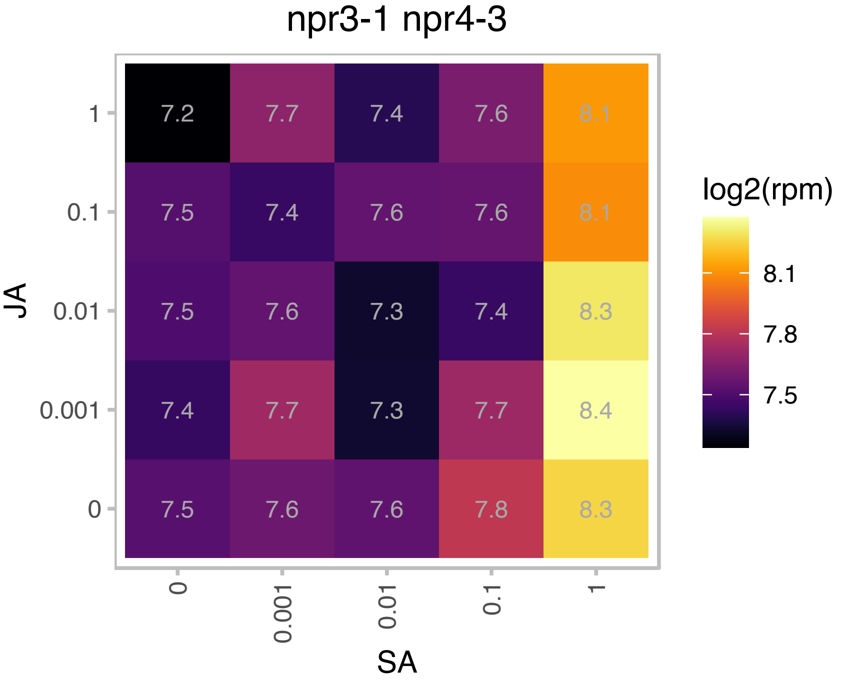

# AT4G09215.1 : NA

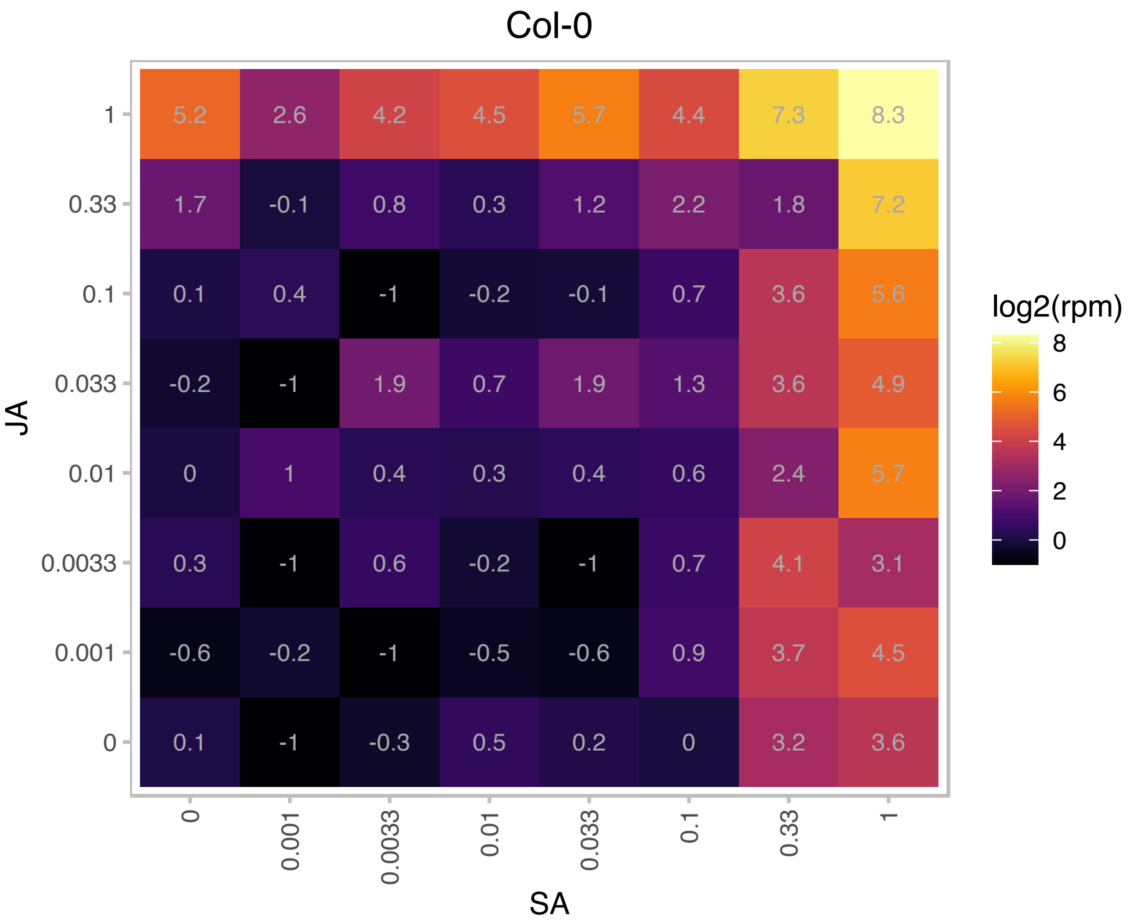


| | |
|---|---|
| Gene | AT4G09215 |
| is.expressed_genes | TRUE |
| is.responsive_genes | TRUE |
| is.apcluster | B-05 |

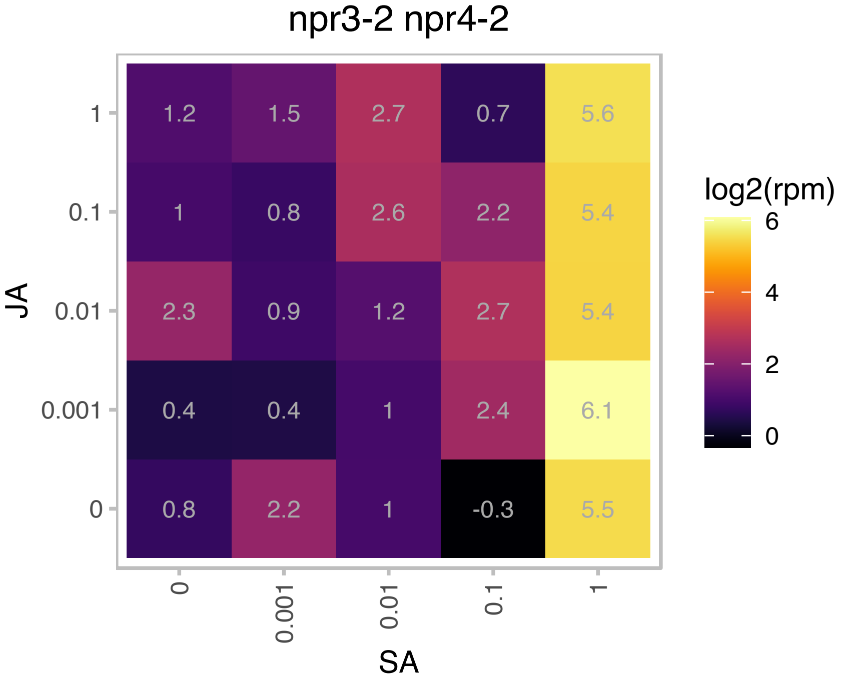


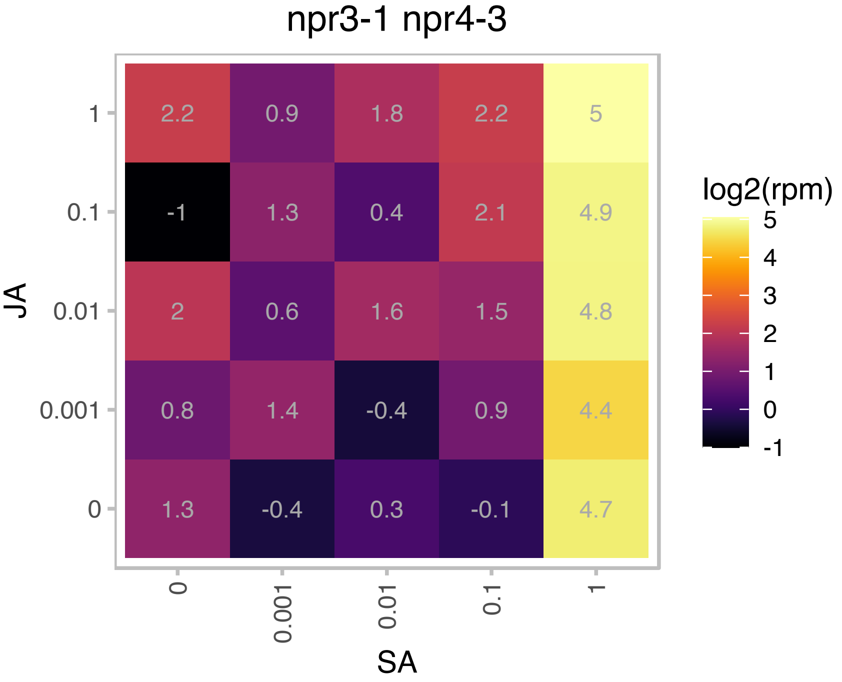

### AT4G14430.1 : ATECI2, ECHIB, ECI2, IBR10, PEC12

indole-3-butyric acid response 10
Encodes a peroxisomal delta3, delta2-enoyl CoA isomerase, involved in unsaturated fatty acid degradation. This enzyme might also be involved in the conversion of indole-3-butyric acid to indole-3-acetic acid via a beta-oxidation-like pathway.

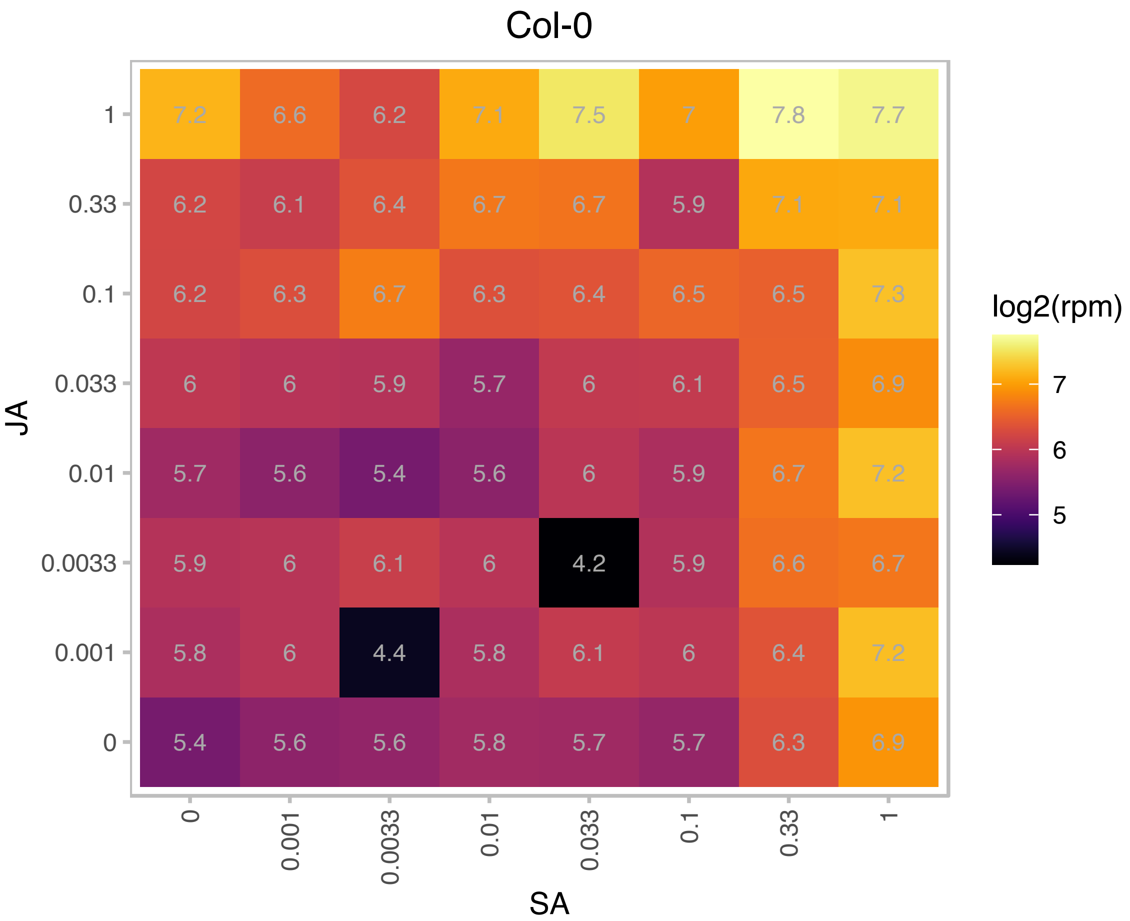


| | |
|---|---|
| Gene | AT4G14430 |
| is.expressed_genes | TRUE |
| is.responsive_genes | TRUE |
| is.apcluster | B-05 |

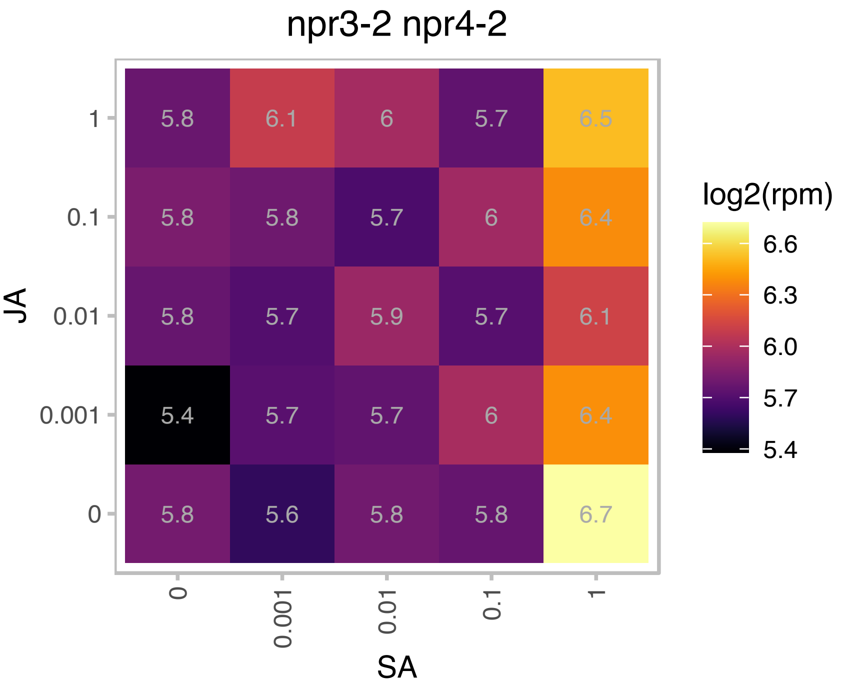


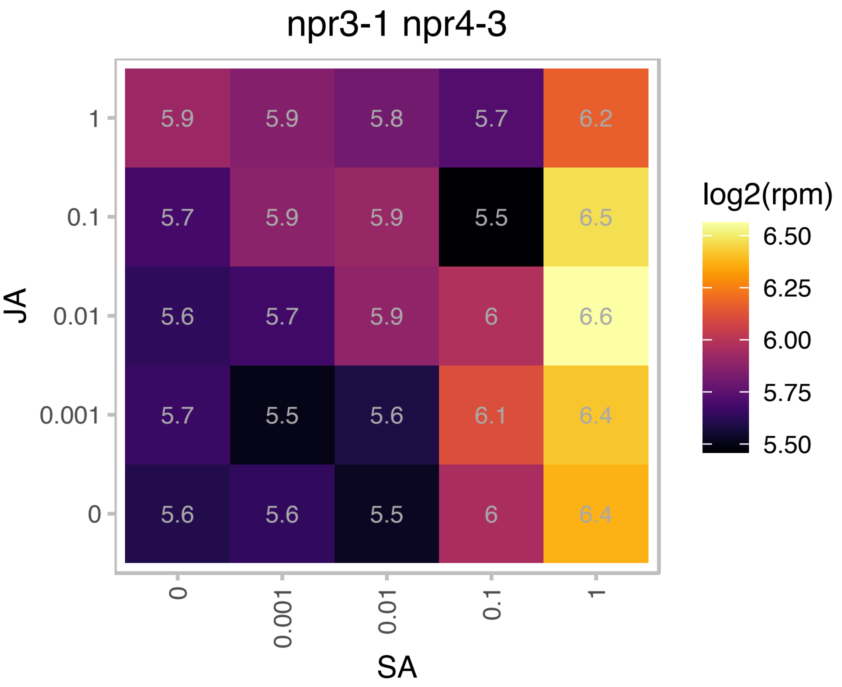

### AT4G15800.1 : RALFL33

ralf-like 33
Member of a diversely expressed predicted peptide family showing sequence similarity to tobacco Rapid Alkalinization Factor (RALF), and is believed to play an essential role in the physiology of Arabidopsis. Consists of a single exon and is characterized by a conserved C-terminal motif and N-terminal signal peptide. The mRNA is cell-to-cell mobile.

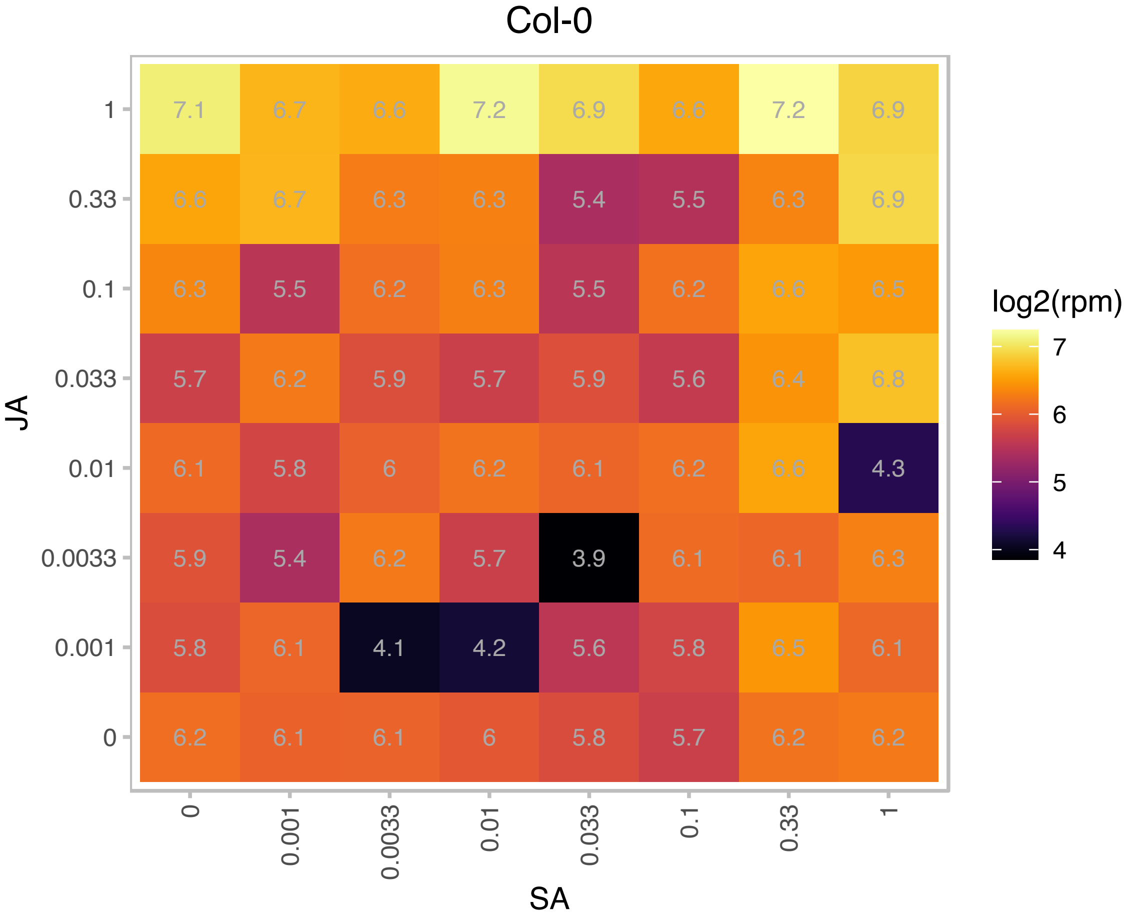


| | |
|---|---|
| Gene | AT4G15800 |
| is.expressed_genes | TRUE |
| is.responsive_genes | TRUE |
| is.apcluster | B-05 |

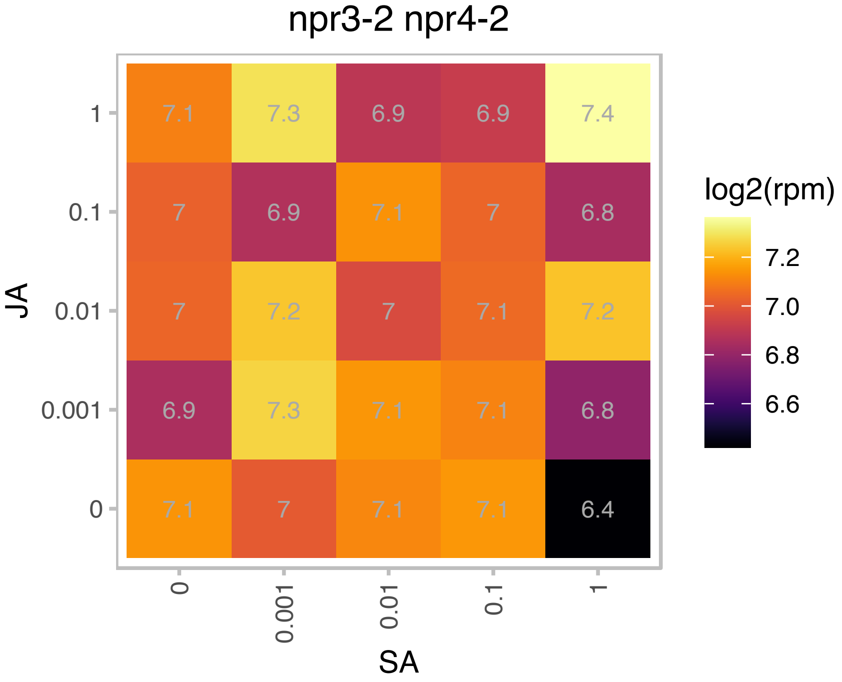


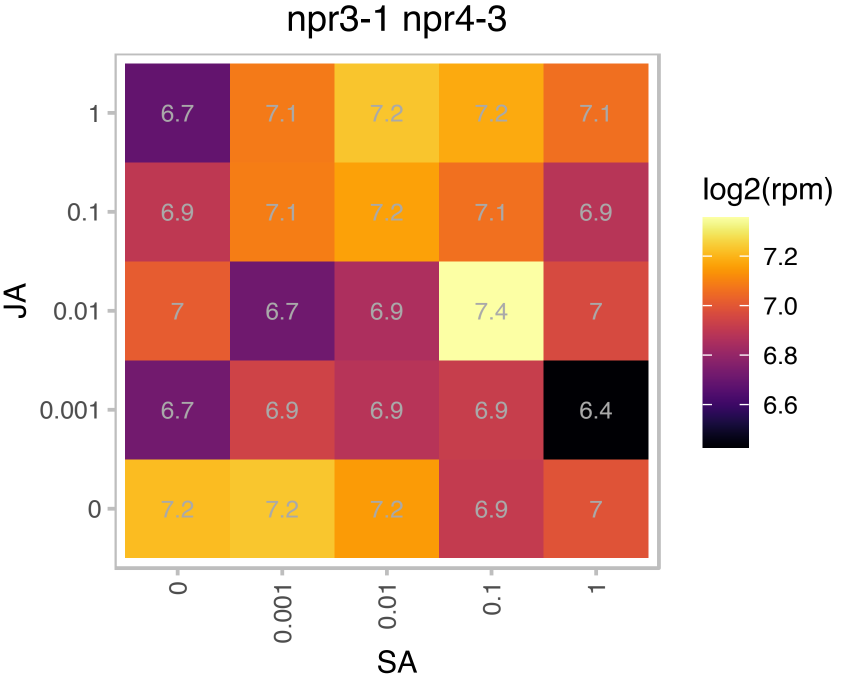

### AT4G17530.1 : ATRAB1C, ATRABD2C, RAB1C, RAB1C

RAB GTPase homolog 1C

AtRabD2c encodes a Rab GTPase, which plays important roles in pollen development, germination and tube elongation.

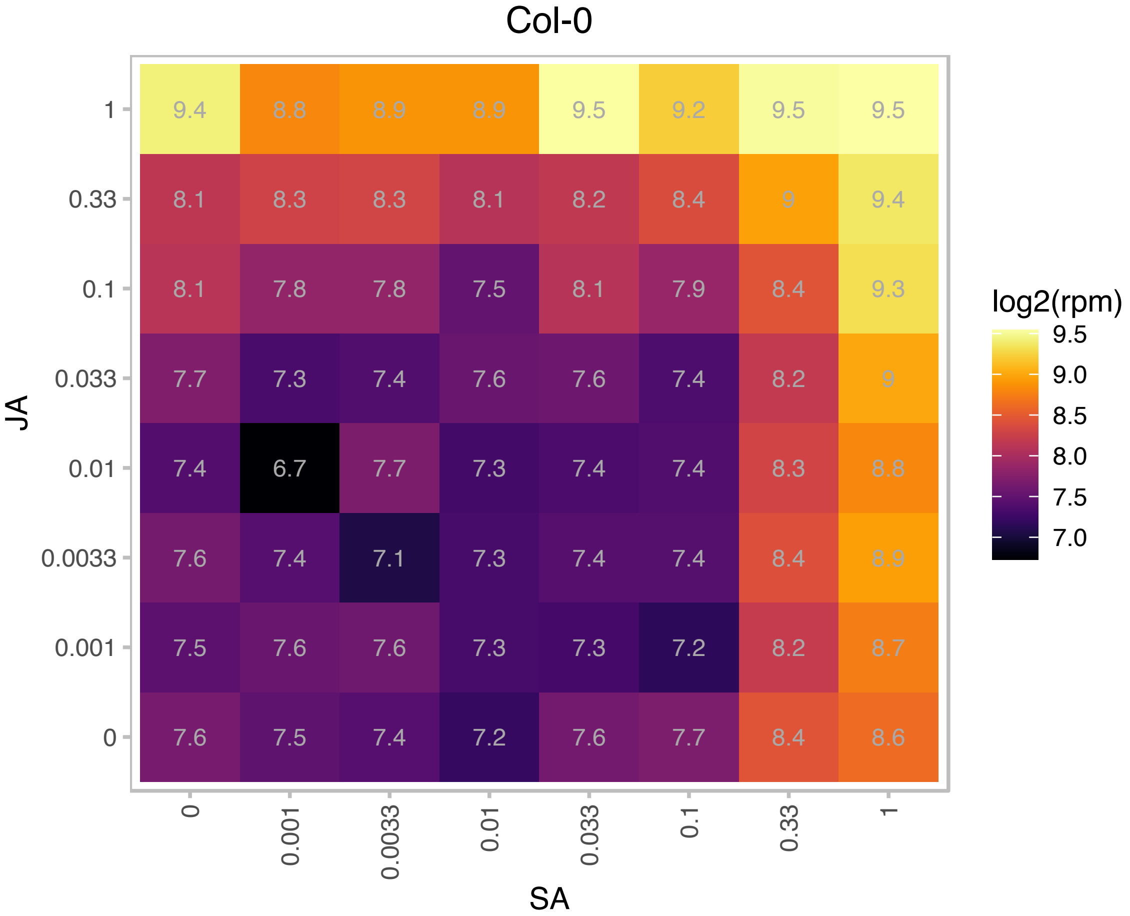


| | |
|---|---|
| Gene | AT4G17530 |
| is.expressed_genes | TRUE |
| is.responsive_genes | TRUE |
| is.apcluster | B-05 |

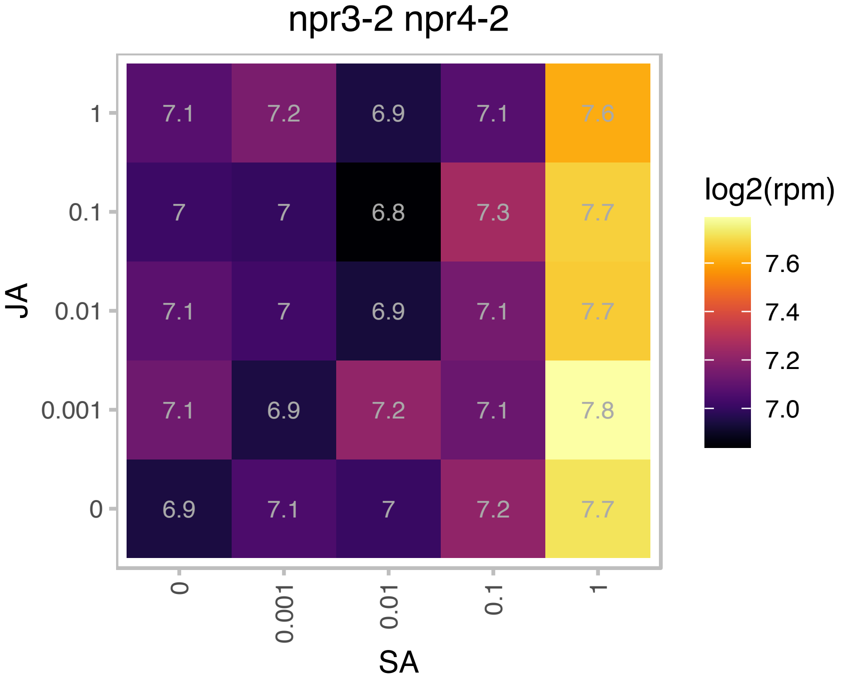


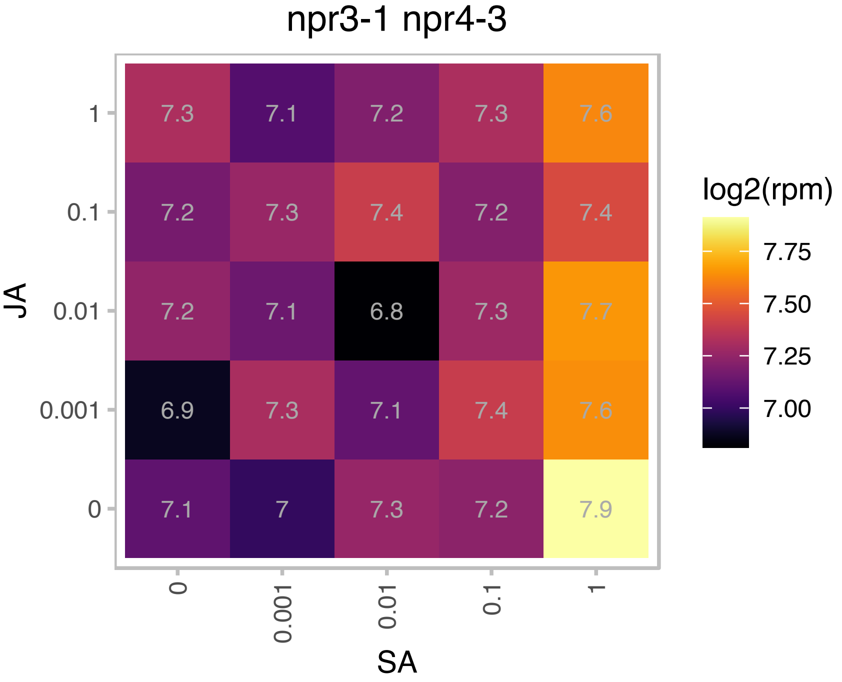

### AT4G18020.1 : APRR2, PRR2

CheY-like two-component responsive regulator family protein

Encodes pseudo-response regulator 2 (APRR2) that interacts with a calcium sensor (CML9).

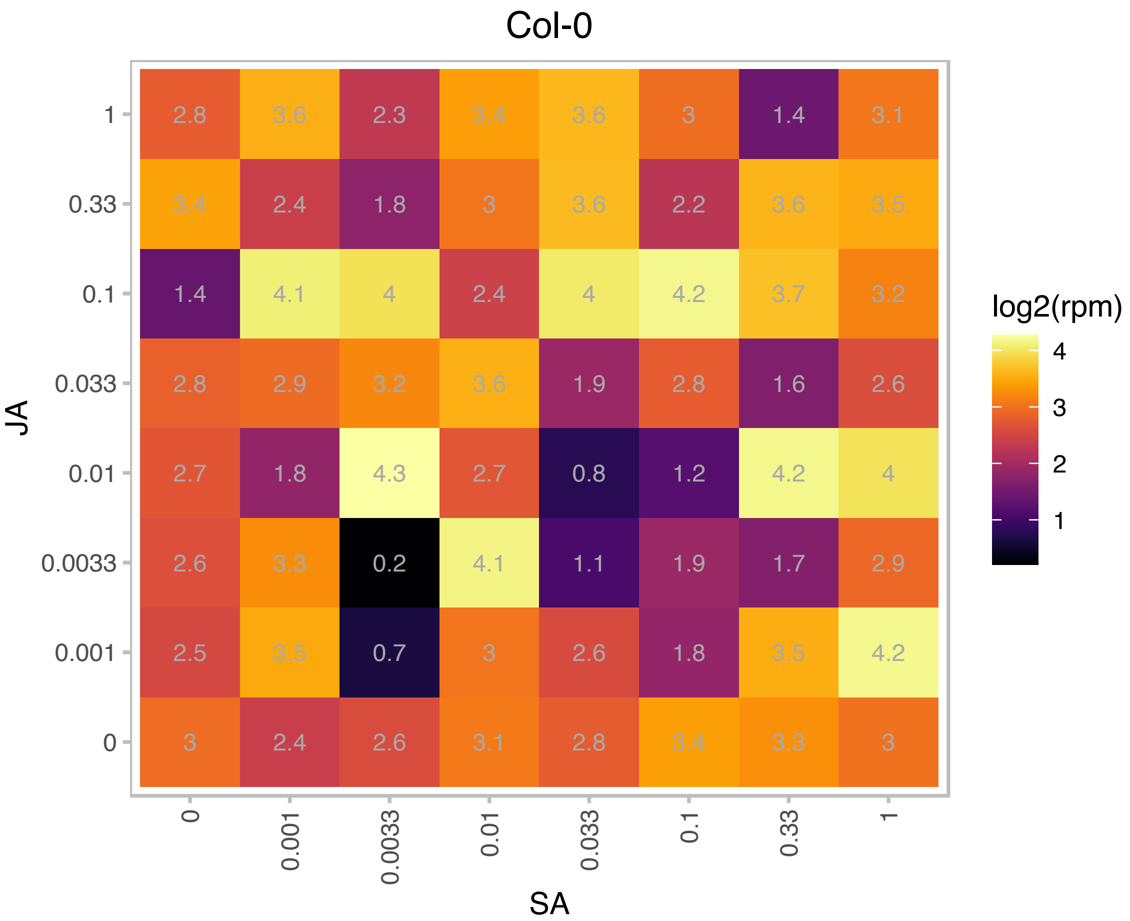


| | |
|---|---|
| Gene | AT4G18020 |
| is.expressed_genes | TRUE |
| is.responsive_genes | TRUE |
| is.apcluster | B-05 |

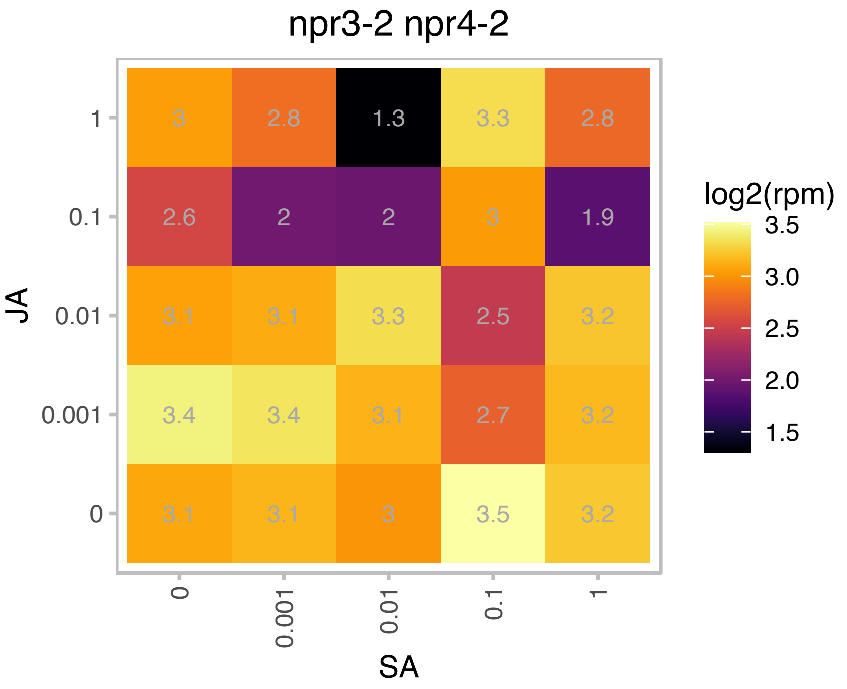


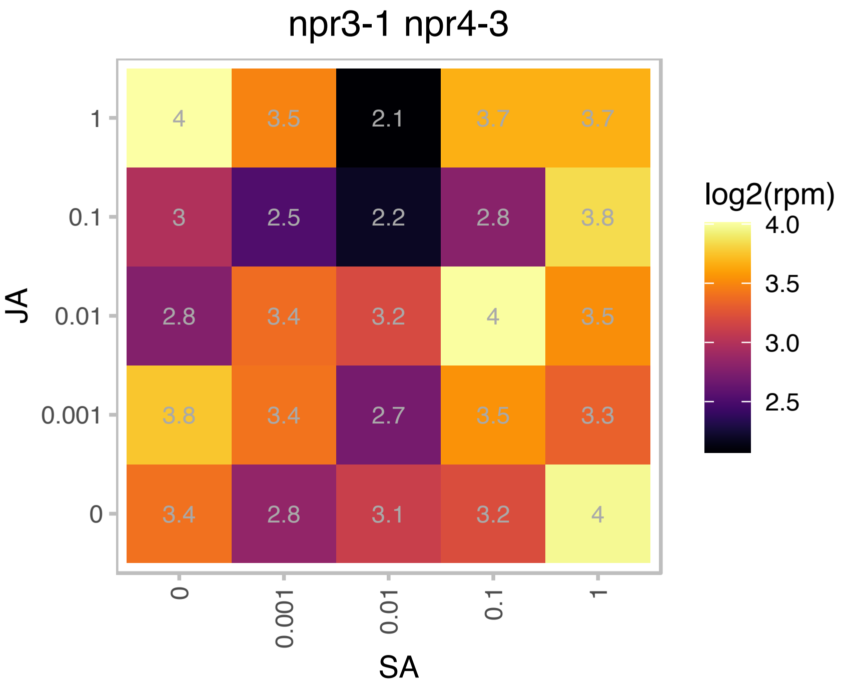

### AT4G18140.1 : SSP4b

SCP1-like small phosphatase 4b

Encodes a SCP1-like small phosphatase (SSP). Three SSPs form a unique group with long N-terminal extensions: AT5G46410 (SSP4), AT5G11860 (SSP5), AT4G18140 (SSP4b). SSP4 and SSP4b were localized exclusively in the nuclei, whereas SSP5 accumulated in both nuclei and cytoplasm. All three SSPs encodes active CTD phosphatases like animal SCP1 family proteins, with distinct substrate specificities: SSP4 and SSP4b could dephosphorylate both Ser2-PO(4) and Ser5-PO(4) of CTD, whereas SSP5 dephosphorylated only Ser5-PO(4).

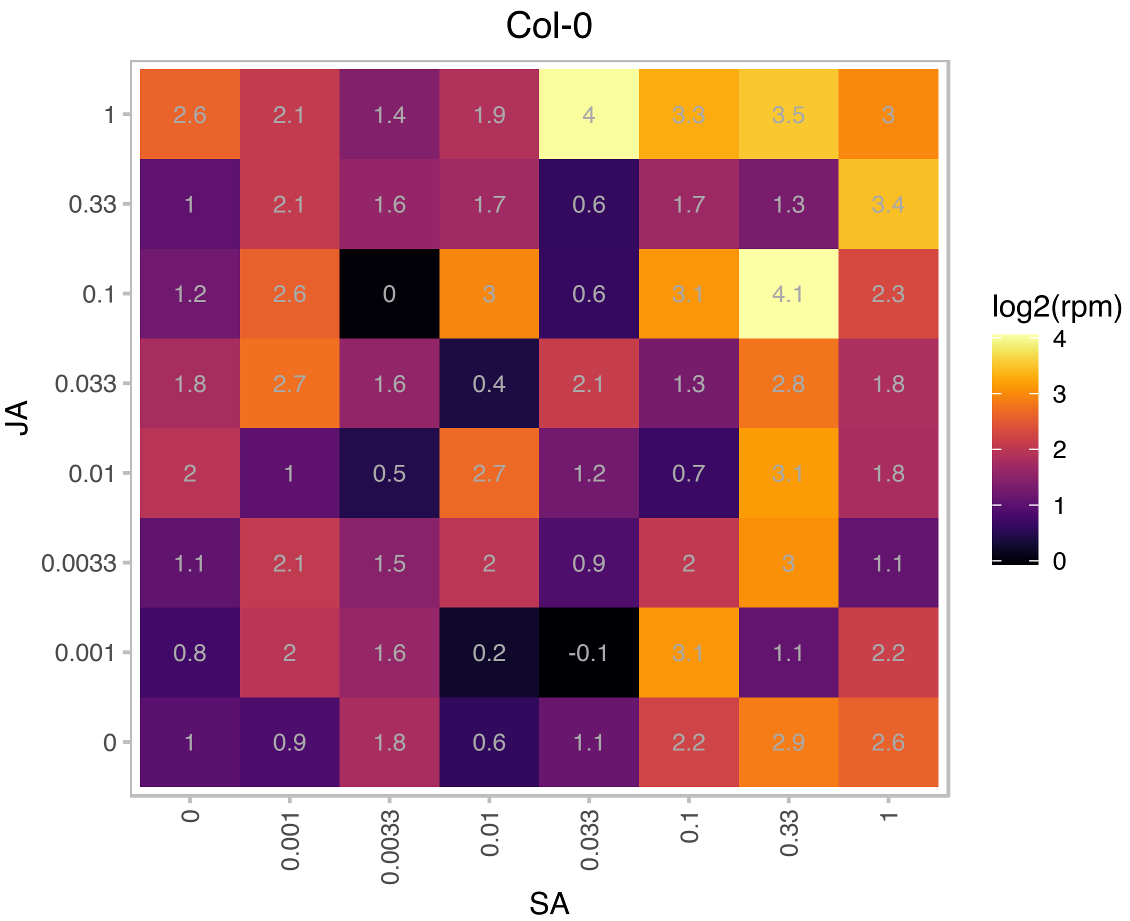


| | |
|---|---|
| Gene | AT4G18140 |
| is.expressed_genes | TRUE |
| is.responsive_genes | TRUE |
| is.apcluster | B-05 |

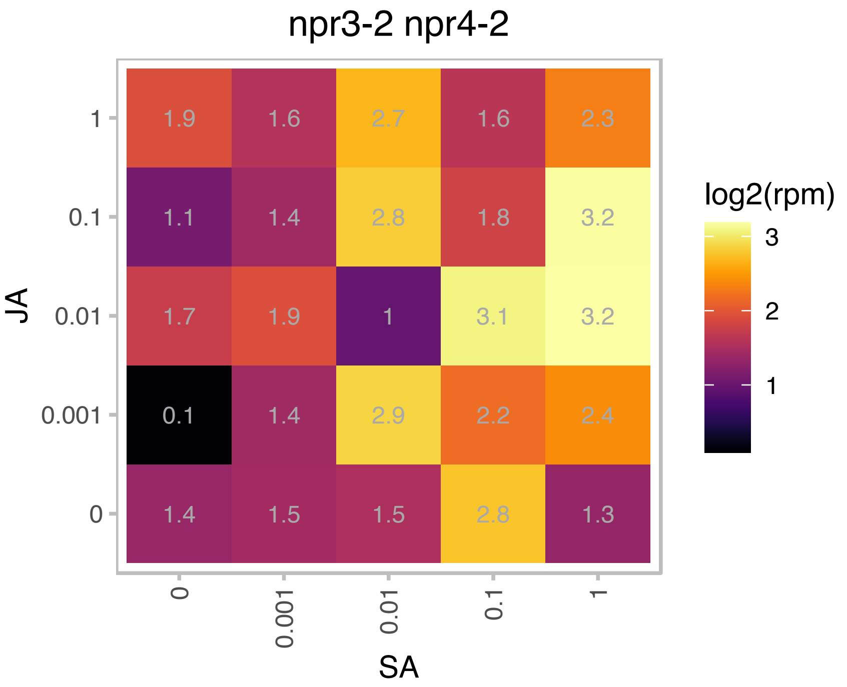


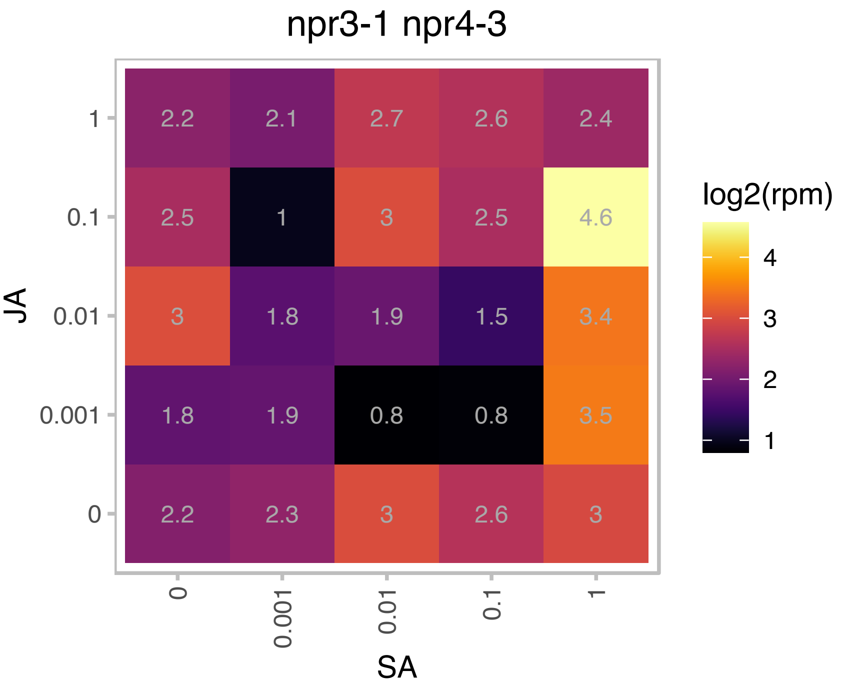

### AT4G19640.1 : ARA-7, ARA7, atARA7, ATRAB-F2B, ATRA

Ras-related small GTP-binding family protein

Encodes Ara7.

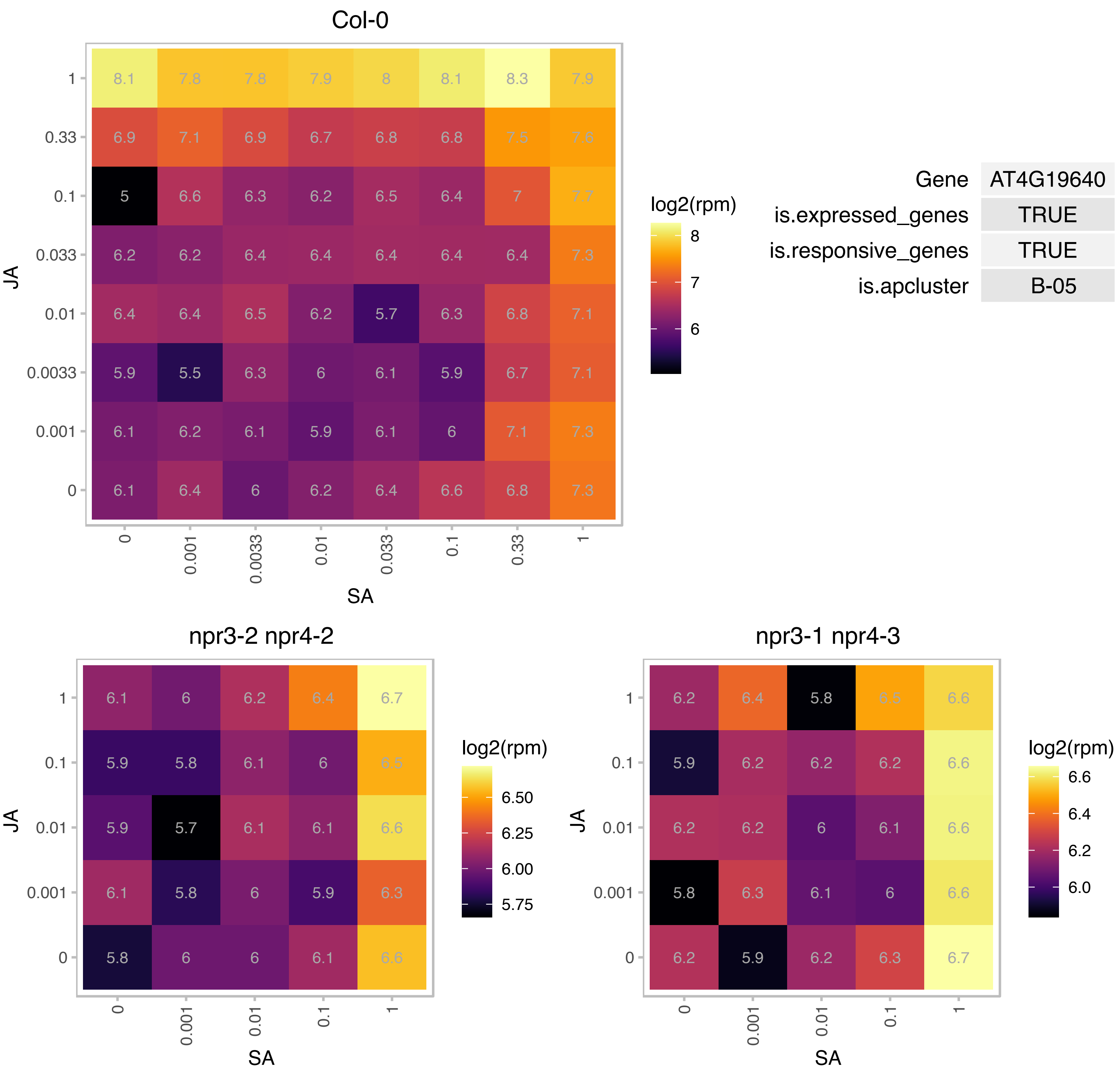


| | |
|---|---|
| Gene | AT4G19640 |
| is.expressed_genes | TRUE |
| is.responsive_genes | TRUE |
| is.apcluster | B-05 |

### AT4G30790.1 : ATG11

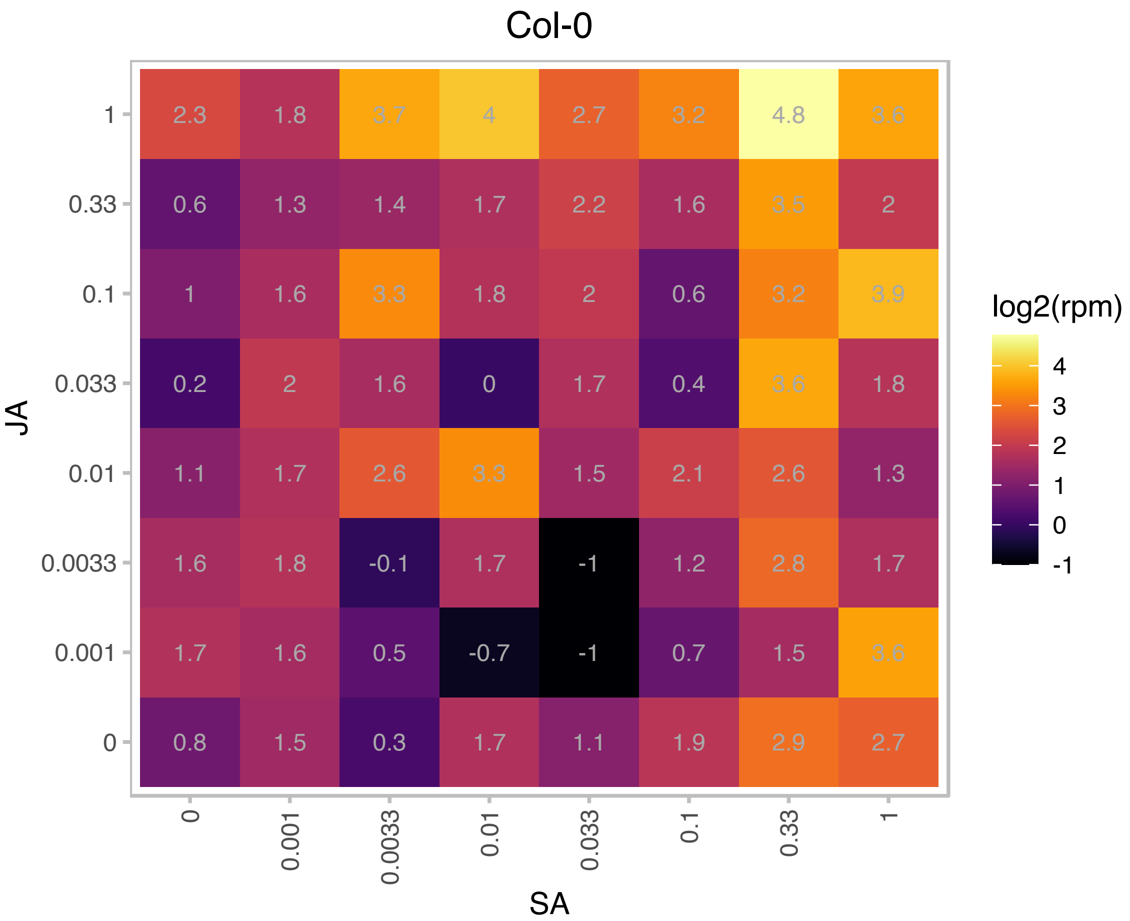


| | |
|---|---|
| Gene | AT4G30790 |
| is.expressed_genes | TRUE |
| is.responsive_genes | TRUE |
| is.apcluster | B-05 |

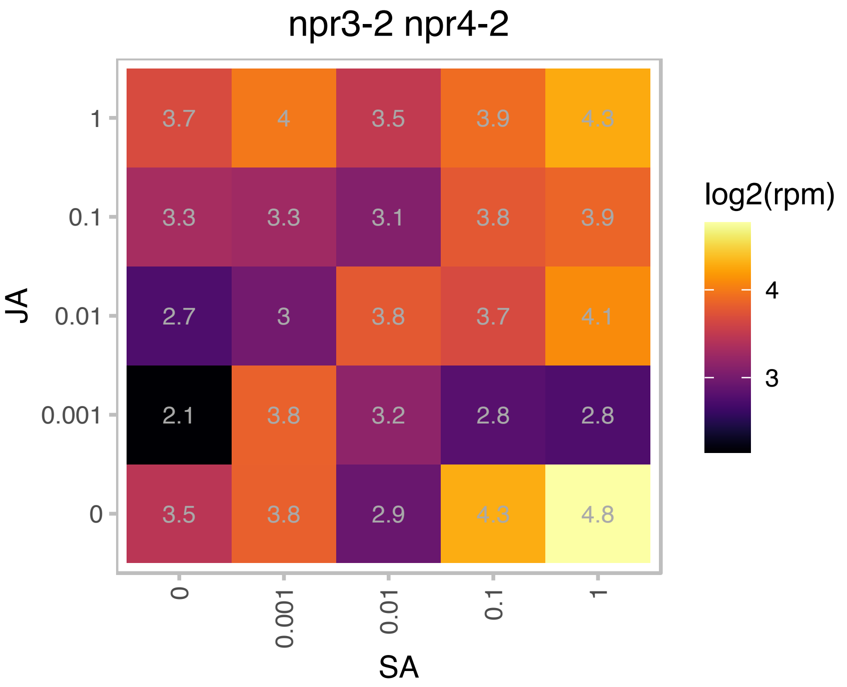


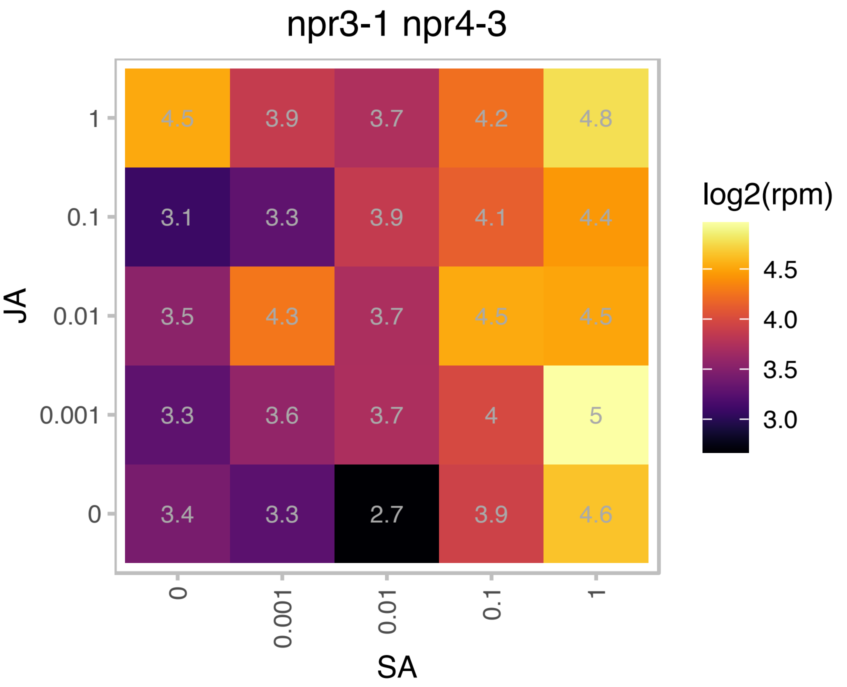

### AT4G33950.1 : ATOST1, OST1, P44, SNRK2-6, SNRK2.6,

Protein kinase superfamily protein

Encodes calcium-independent ABA-activated protein kinase, a member of SNF1-related protein kinases (SnRK2) whose activity is activated by ionic (salt) and non-ionic (mannitol) osmotic stress. Mutations disrupted ABA induction of stomatal closure as well as ABA inhibition of light-induced stomatal opening. However, regulation of stomatal opening/closing by light or CO(2) is not affected in these mutants. May act in the interval between ABA perception and reactive oxygen species production in the ABA signalling network.

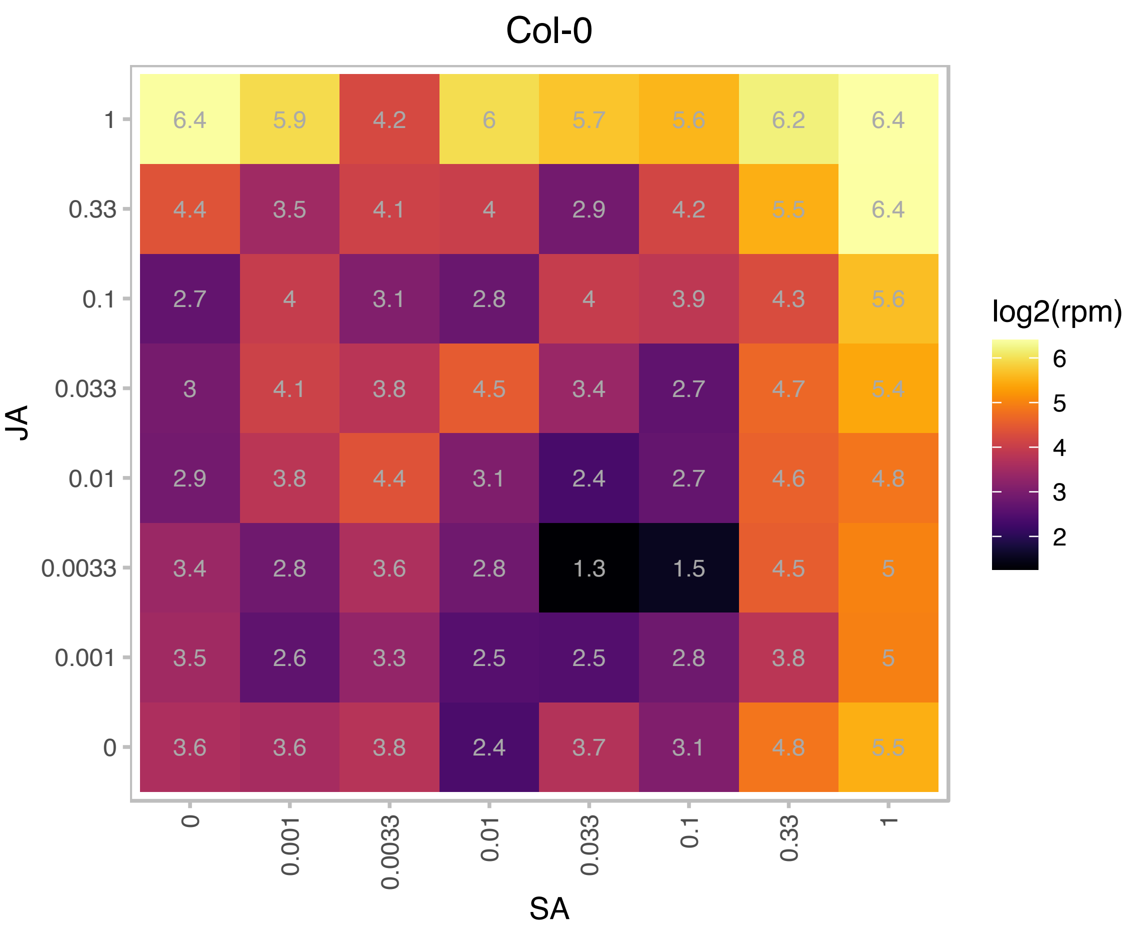


| | |
|---|---|
| Gene | AT4G33950 |
| is.expressed_genes | TRUE |
| is.responsive_genes | TRUE |
| is.apcluster | B-05 |

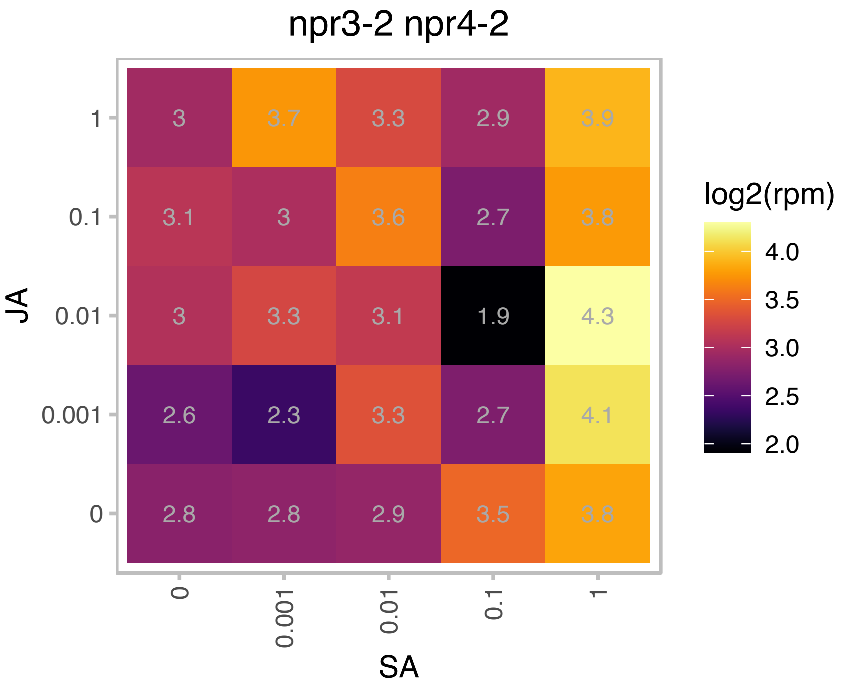


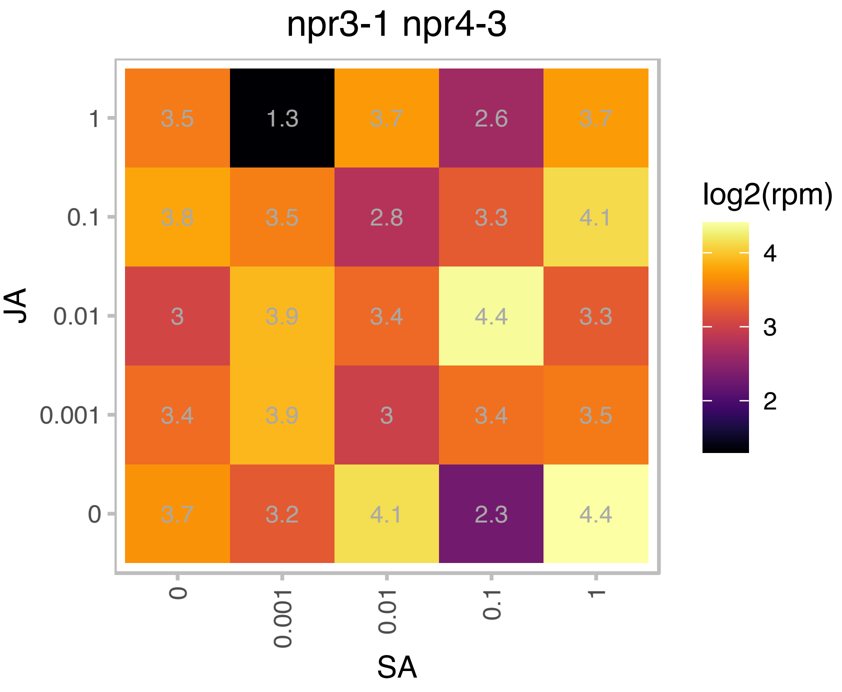

### AT4G34590.1 : ATB2, AtbZIP11, BZIP11, GBF6

G-box binding factor 6
Encodes a basic domain leucine zipper (bZip) transcription factor bZIP11. Translation is repressed by sucrose. Directly regulates gene expression of ASN1 and ProDH2, which are enzyme-coding genes involved in amino acid metabolism.

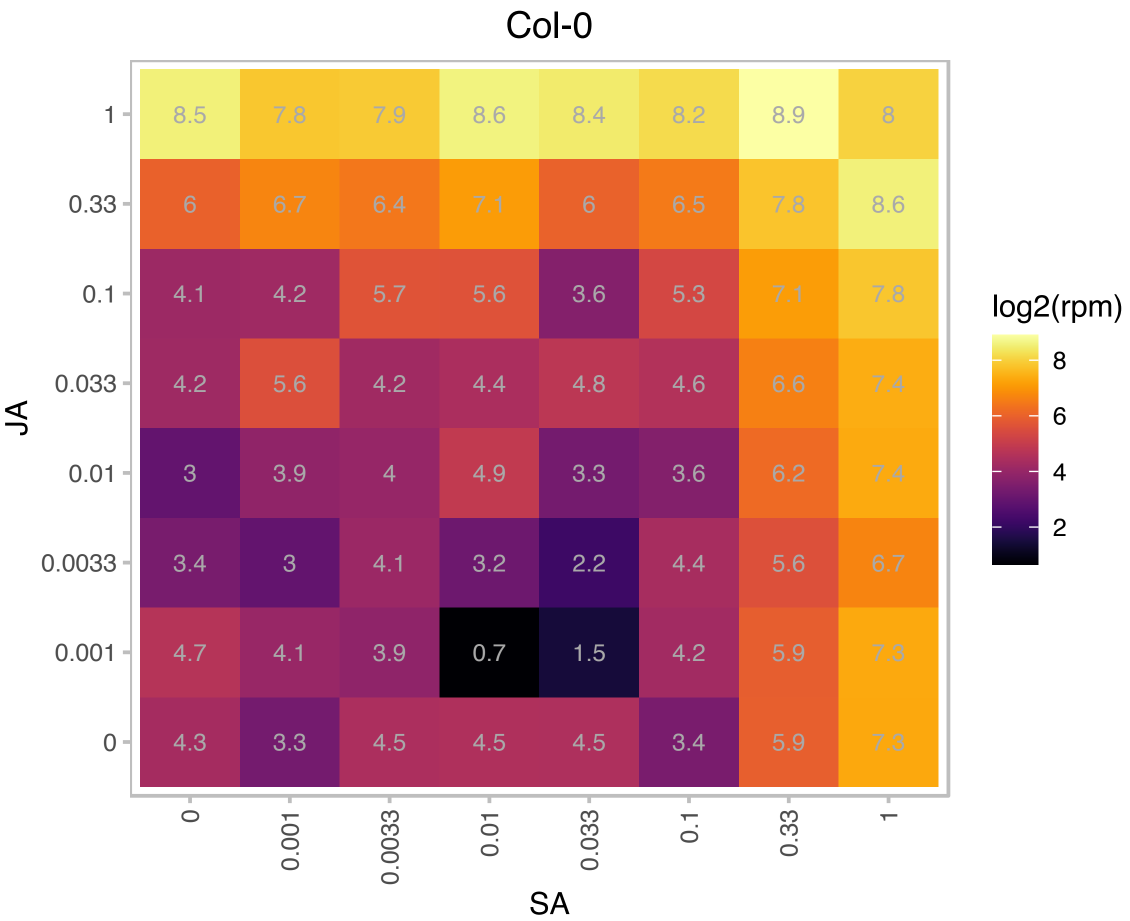


| | |
|---|---|
| Gene | AT4G34590 |
| is.expressed_genes | TRUE |
| is.responsive_genes | TRUE |
| is.apcluster | B-05 |

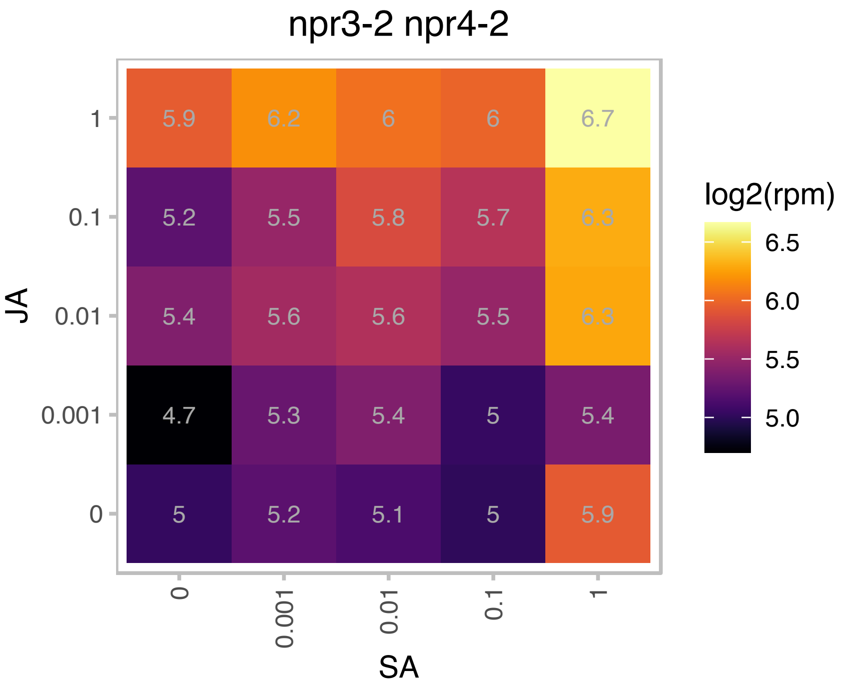


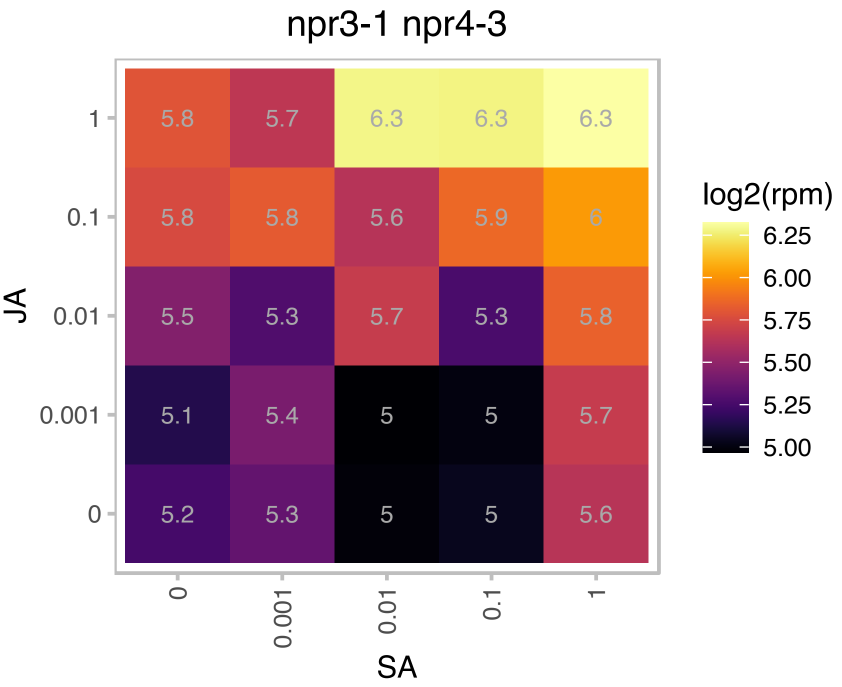

### AT4G36800.1 : RCE1

RUB1 conjugating enzyme 1

RUB1 conjugating enzyme that conjugates CUL1 and is involved in auxin response and embryogenesis. RCE1 protein physically interacts with RBX1, which may be the E3 for CUL1.

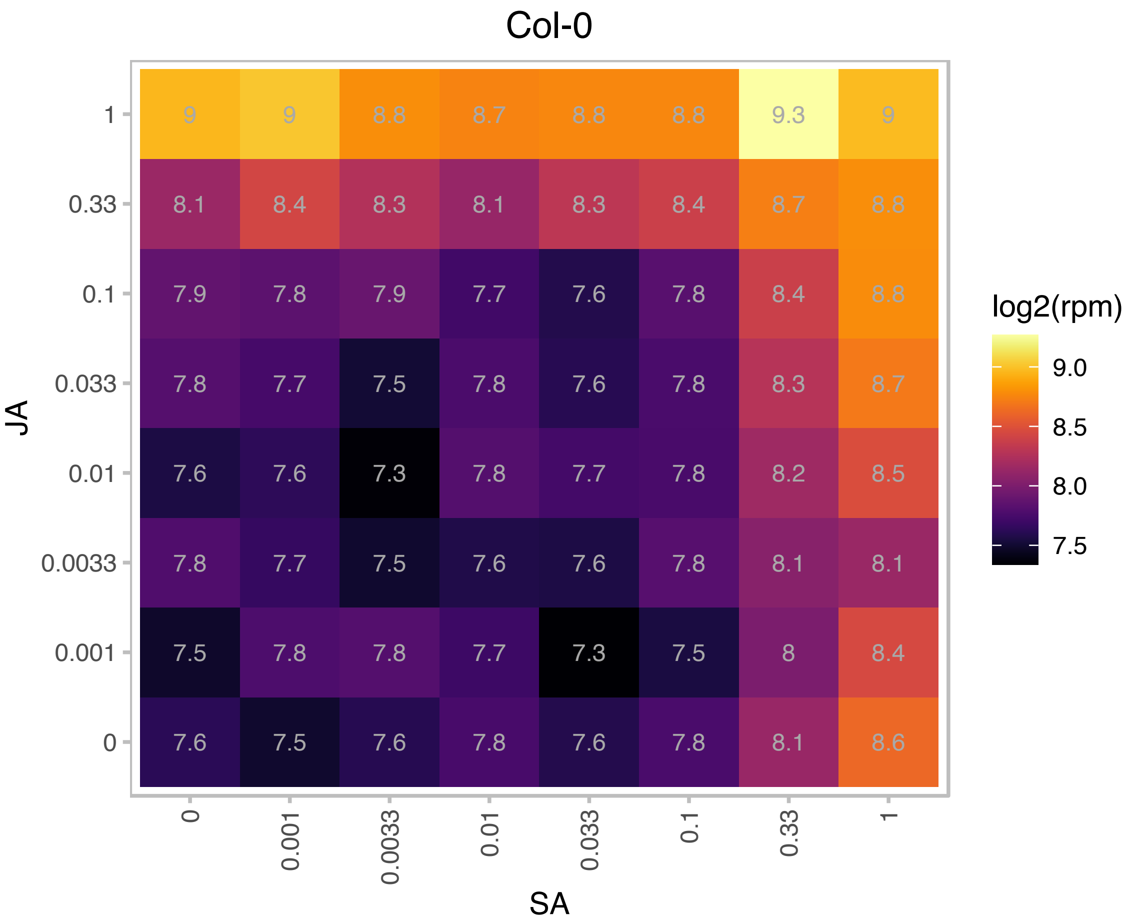


| | |
|---|---|
| Gene | AT4G36800 |
| is.expressed_genes | TRUE |
| is.responsive_genes | TRUE |
| is.apcluster | B-05 |

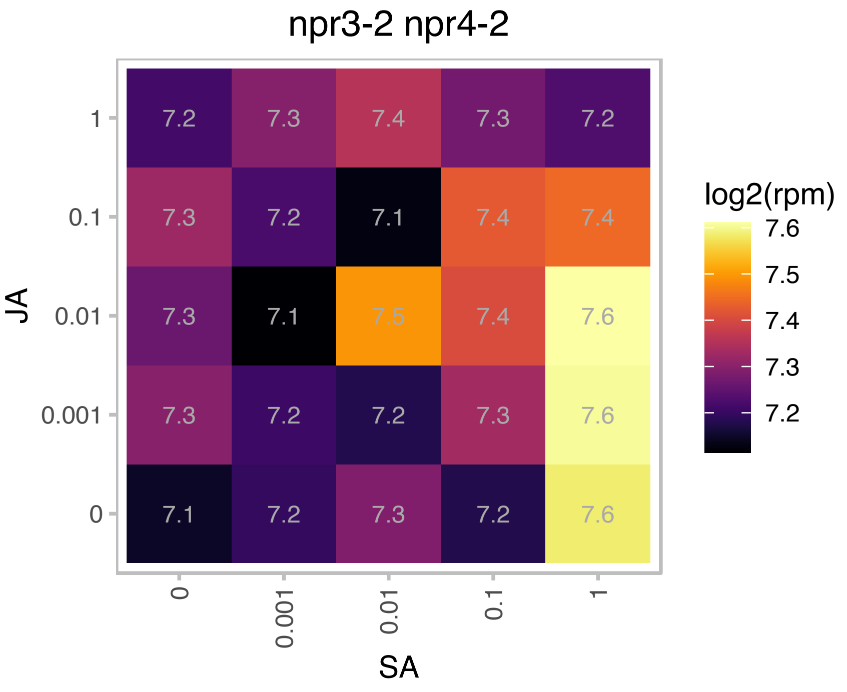


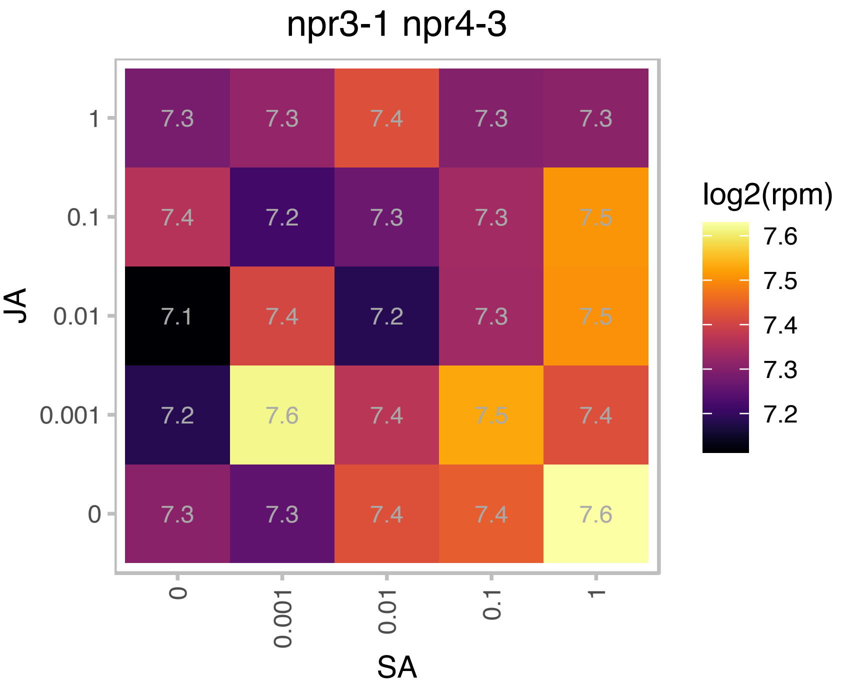

### AT4G37680.1 : HHP4

heptahelical protein 4

heptahelical transmembrane protein HHP4

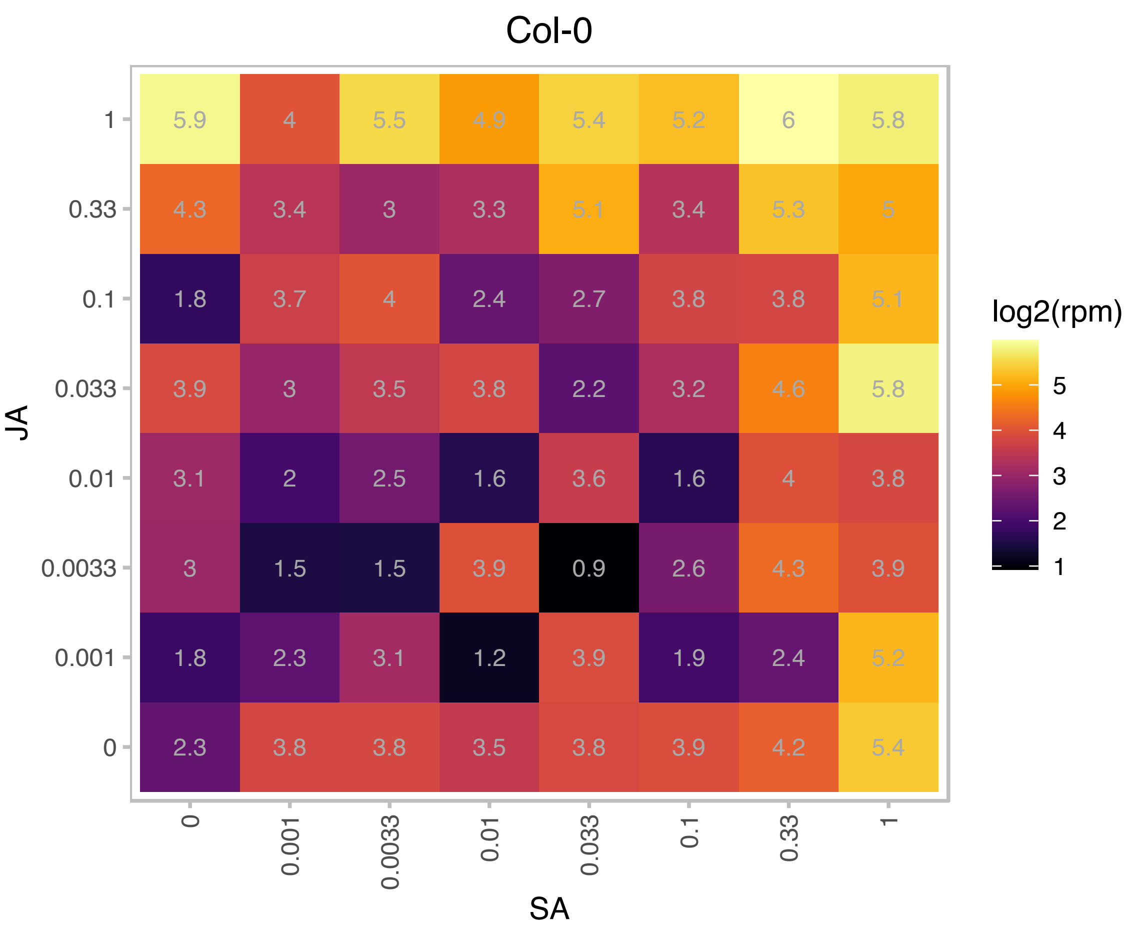


| | |
|---|---|
| Gene | AT4G37680 |
| is.expressed_genes | TRUE |
| is.responsive_genes | TRUE |
| is.apcluster | B-05 |

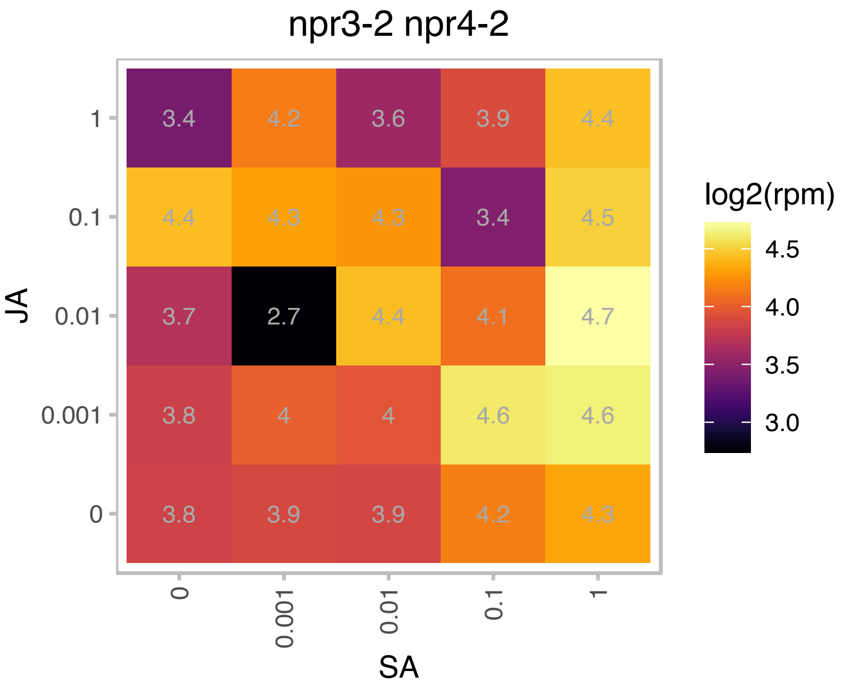


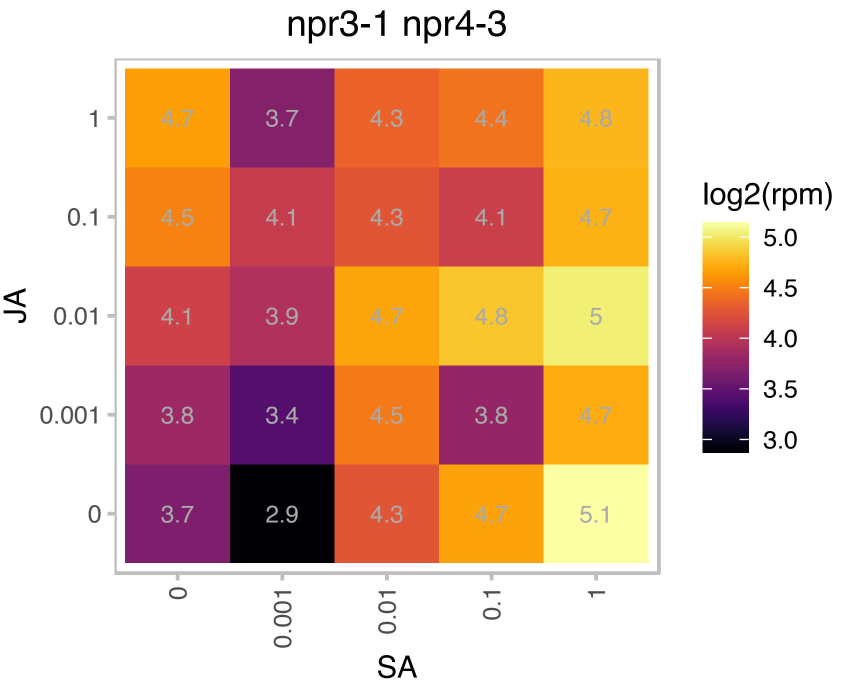

# AT4G39670.1 : NA

Glycolipid transfer protein (GLTP) family protein

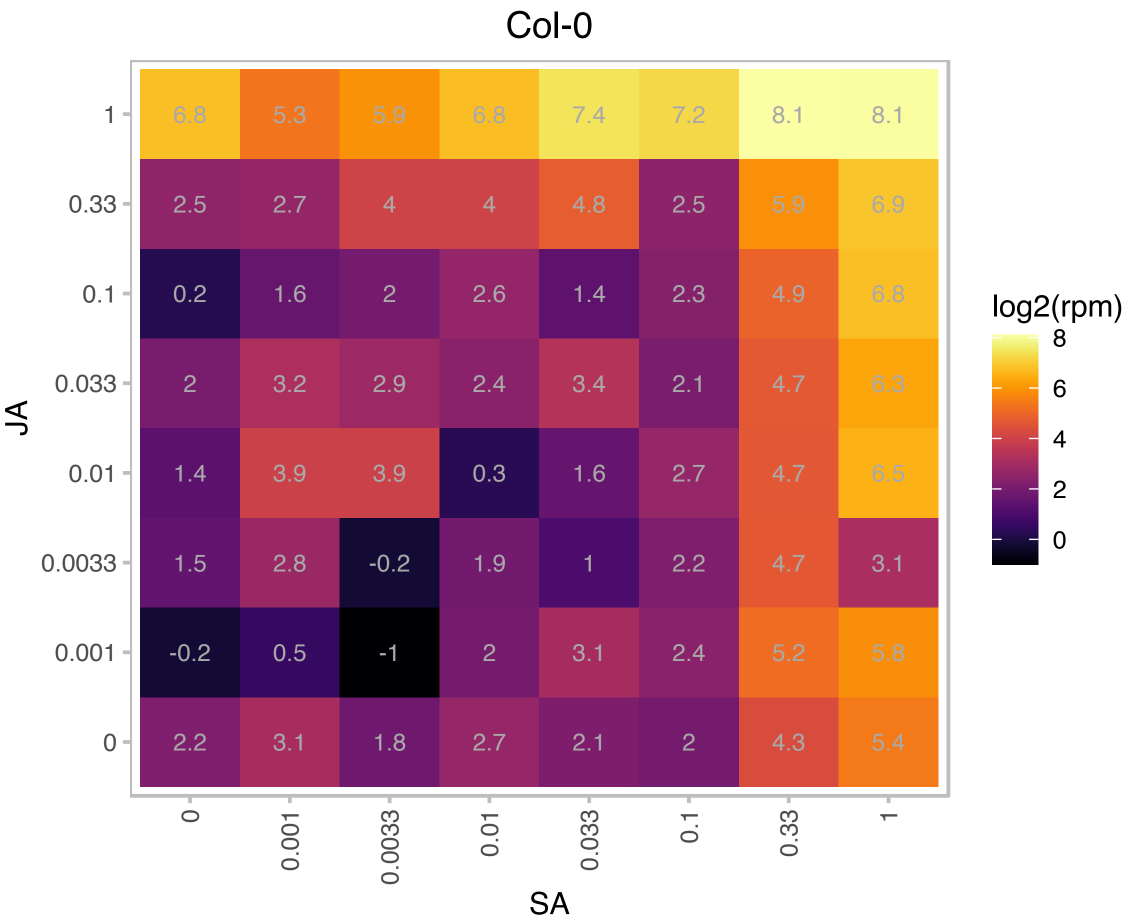


| | |
|---|---|
| Gene | AT4G39670 |
| is.expressed_genes | TRUE |
| is.responsive_genes | TRUE |
| is.apcluster | B-05 |

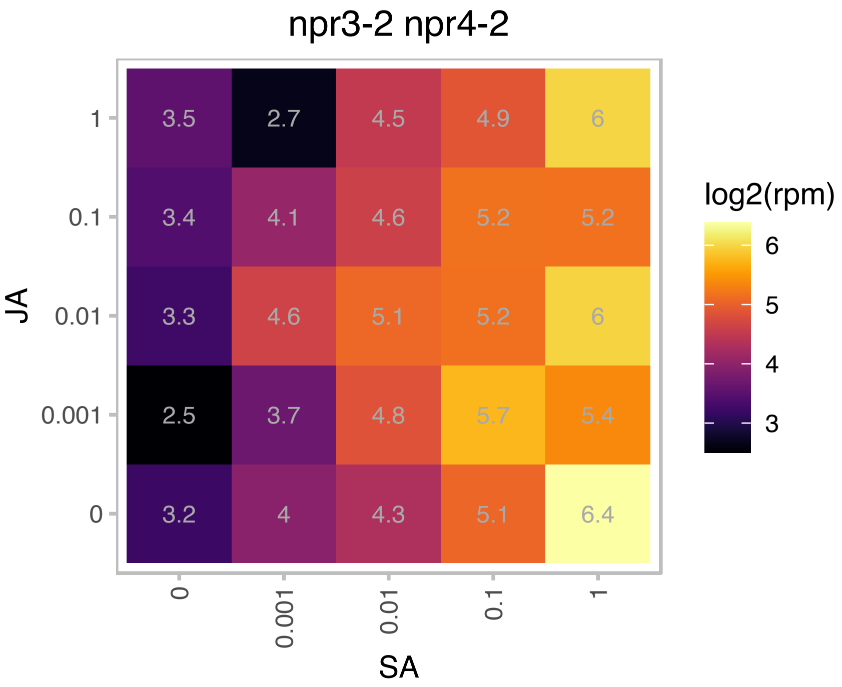


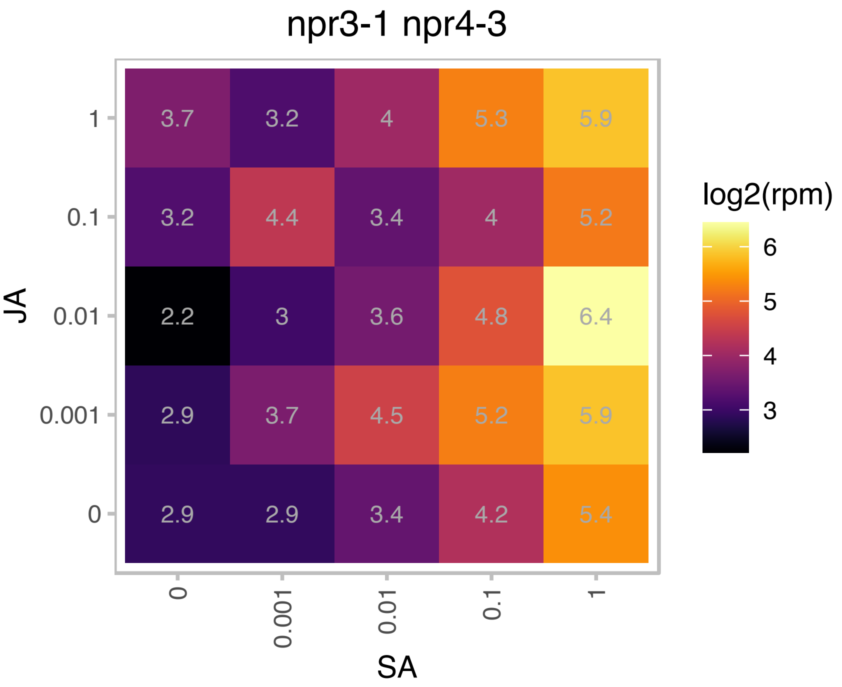

# AT5G03495.1 : NA

RNA-binding (RRM/RBD/RNP motifs) family protein

Col-0

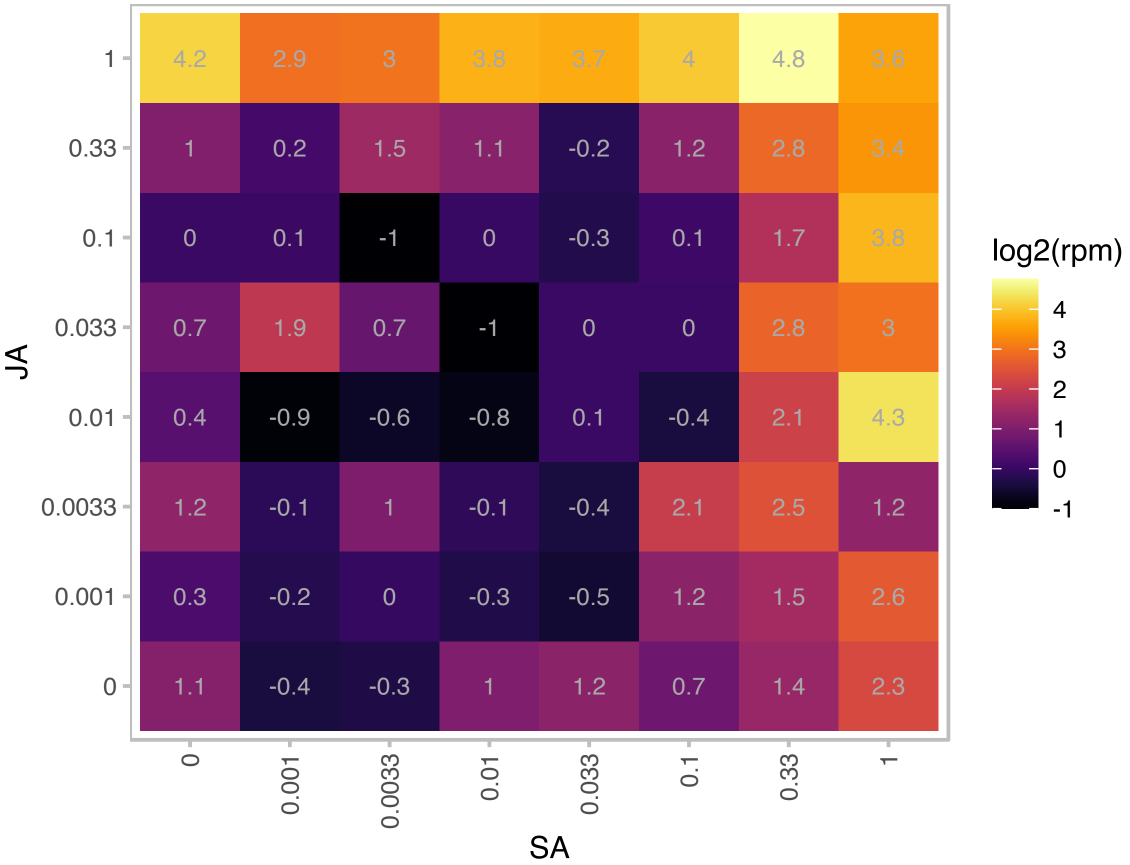


| | |
|---|---|
| Gene | AT5G03495 |
| is.expressed_genes | TRUE |
| is.responsive_genes | TRUE |
| is.apcluster | B-05 |

npr3-2 npr4-2

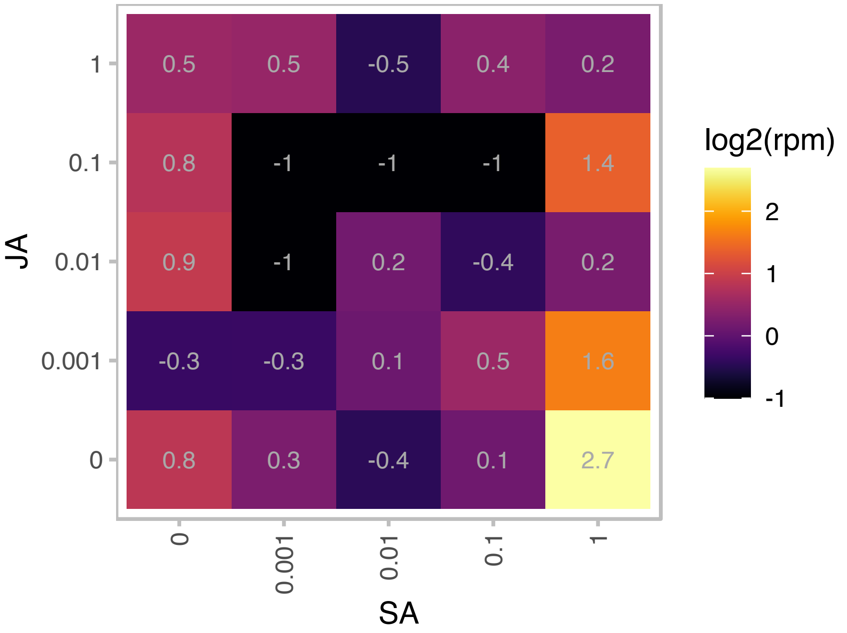


npr3-1 npr4-3

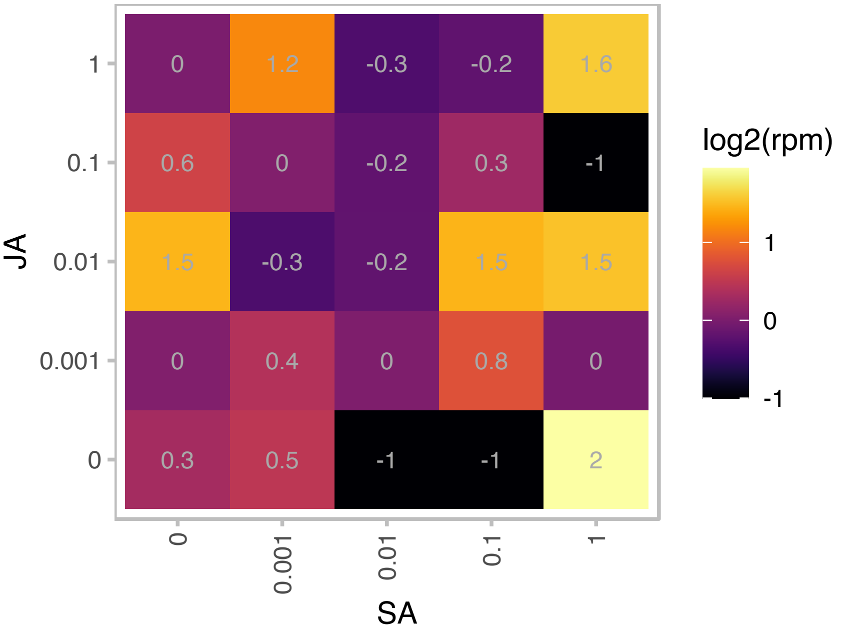

### AT5G04340.1 : AtZAT6, C2H2, CZF2, ZAT6

zinc finger of Arabidopsis thaliana 6
Encodes a C2H2 zinc finger transcription factor that coordinately activates phytochelatin-synthesis related gene expression and directly targets GSH1 by binding to its promoter to positively regulate Cd accumulation and tolerance.

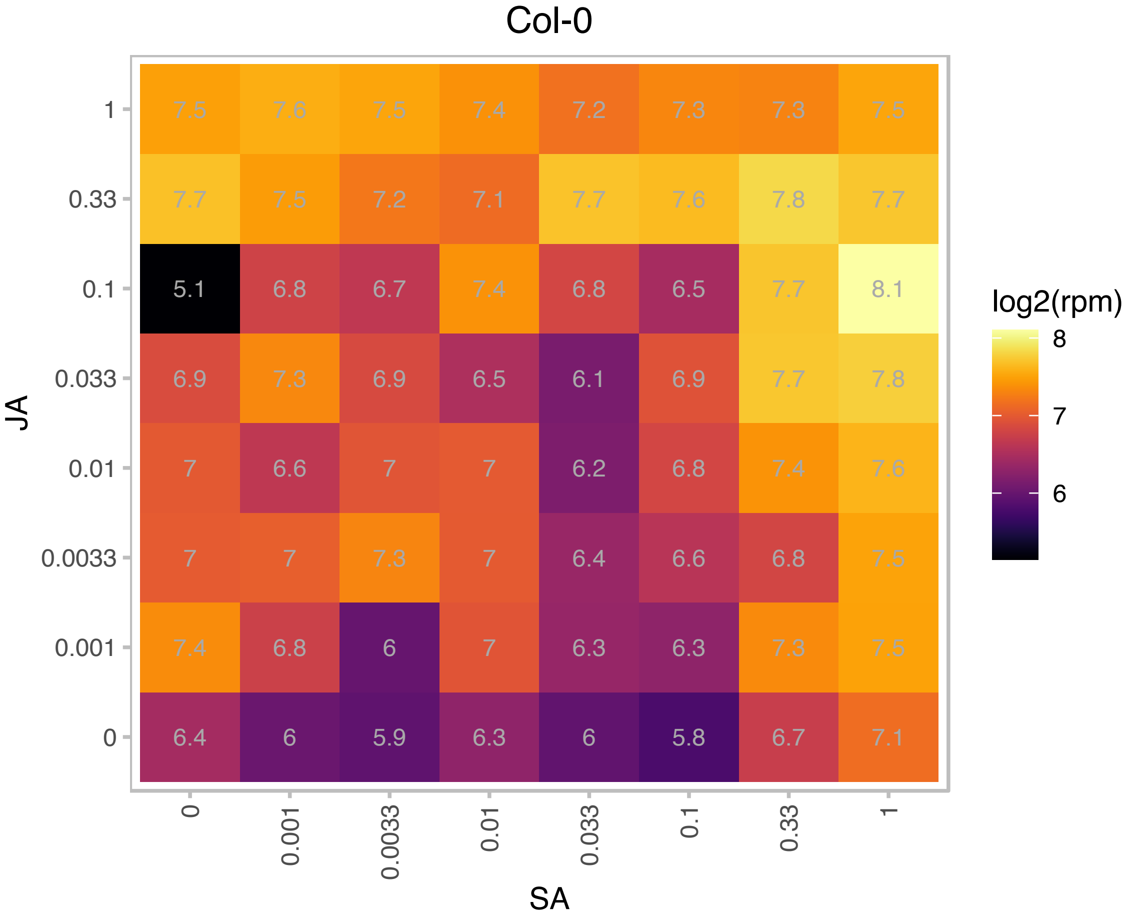


| | |
|---|---|
| Gene | AT5G04340 |
| is.expressed_genes | TRUE |
| is.responsive_genes | TRUE |
| is.apcluster | B-05 |

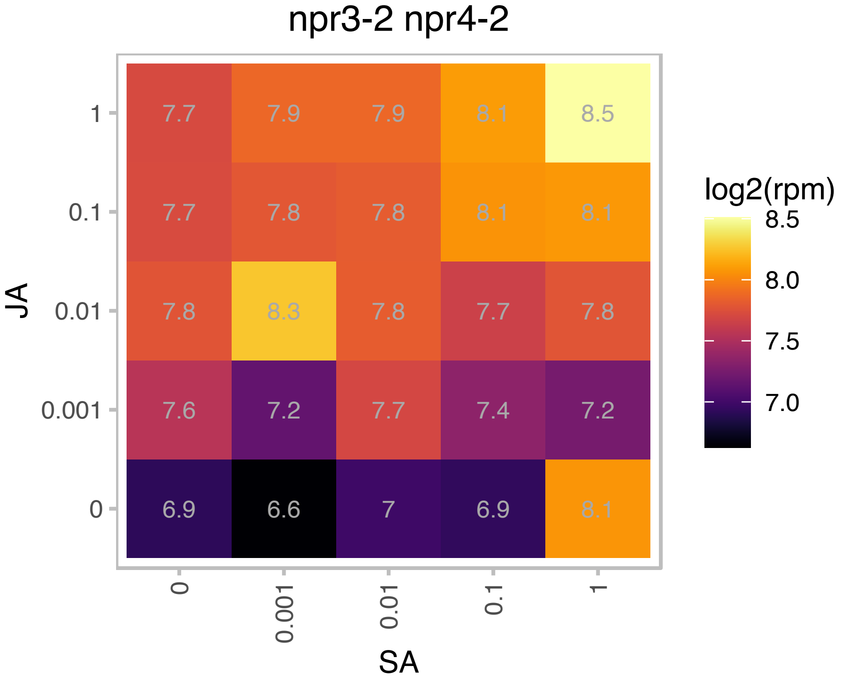


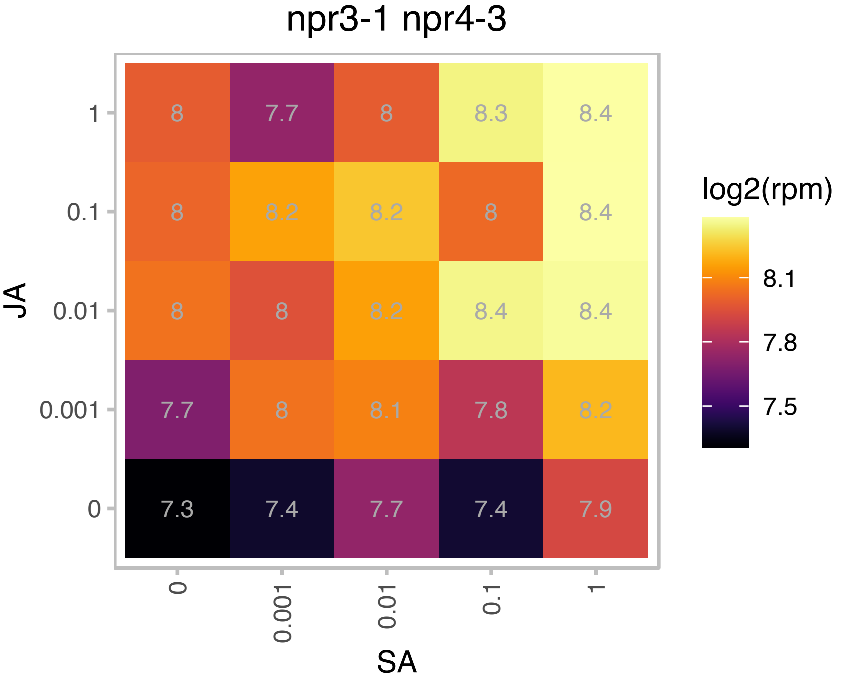

### AT5G05410.1 : DREB2, DREB2A

Encodes a transcription factor that specifically binds to DRE/CRT cis elements (responsive to drought and low-temperature stress). DRE-binding protein 2A. Belongs to the DREB subfamily A-2 of ERF/AP2 transcription factor family (DREB2A). There are eight members in this subfamily including DREB2B. The protein contains one AP2 domain. Overexpression of transcriptional activation domain of DREB2A resulted in significant drought stress tolerance but only slight freezing tolerance in transgenic Arabidopsis plants. Microarray and RNA gel blot analyses revealed that DREB2A regulates expression of many water stress□ □ nducible genes. The mRNA is

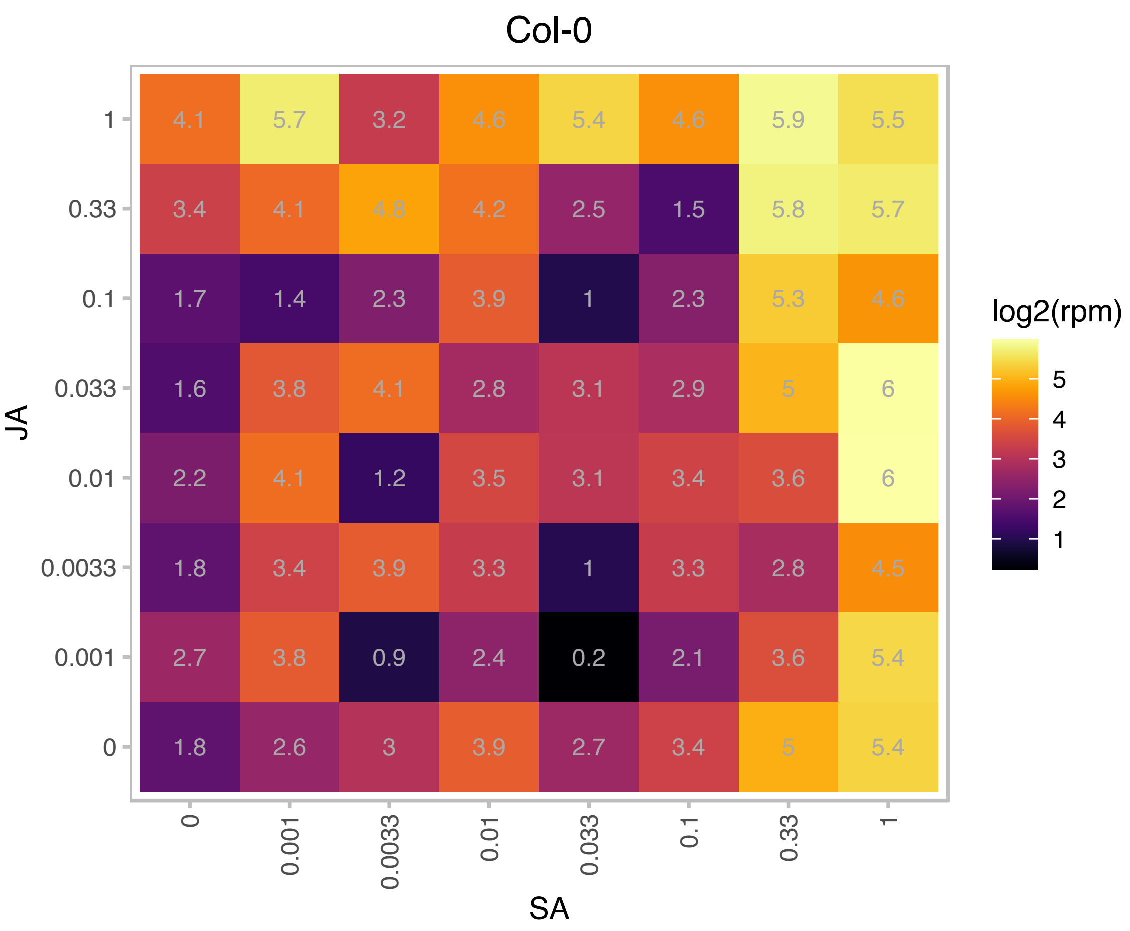


| | |
|---|---|
| Gene | AT5G05410 |
| is.expressed_genes | TRUE |
| is.responsive_genes | TRUE |
| is.apcluster | B-05 |

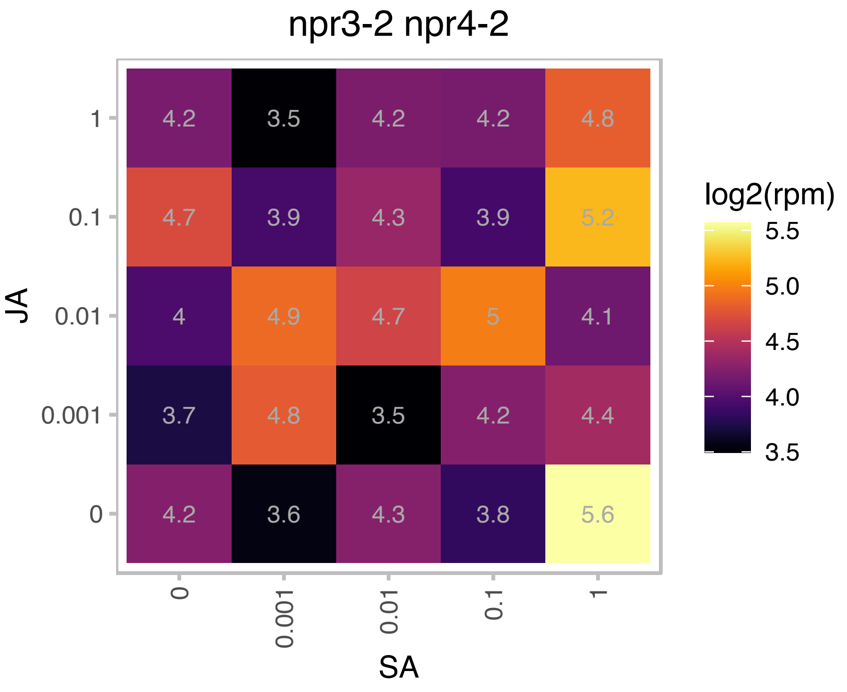


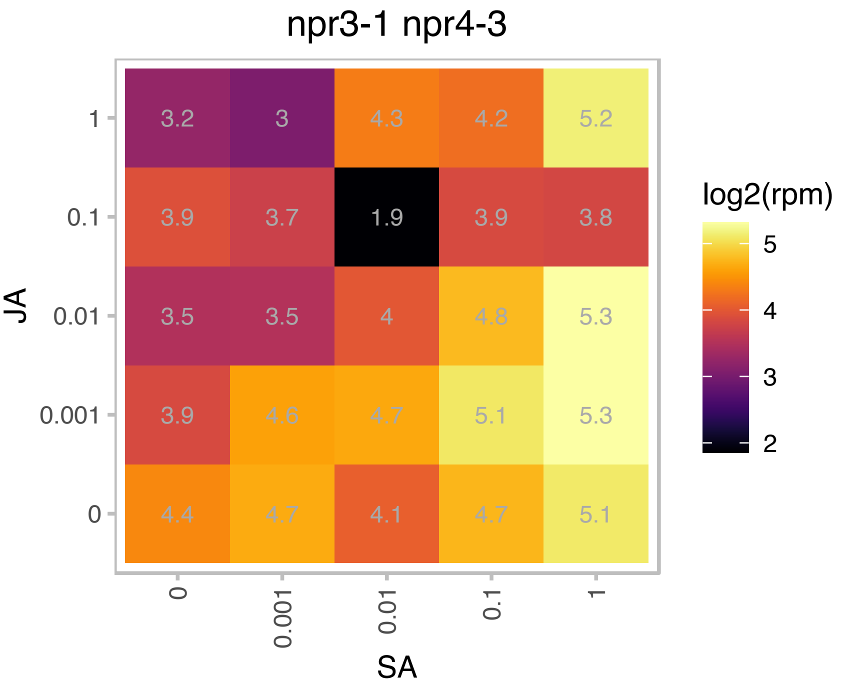

# AT5G08139.1 : NA

RING/U-box superfamily protein

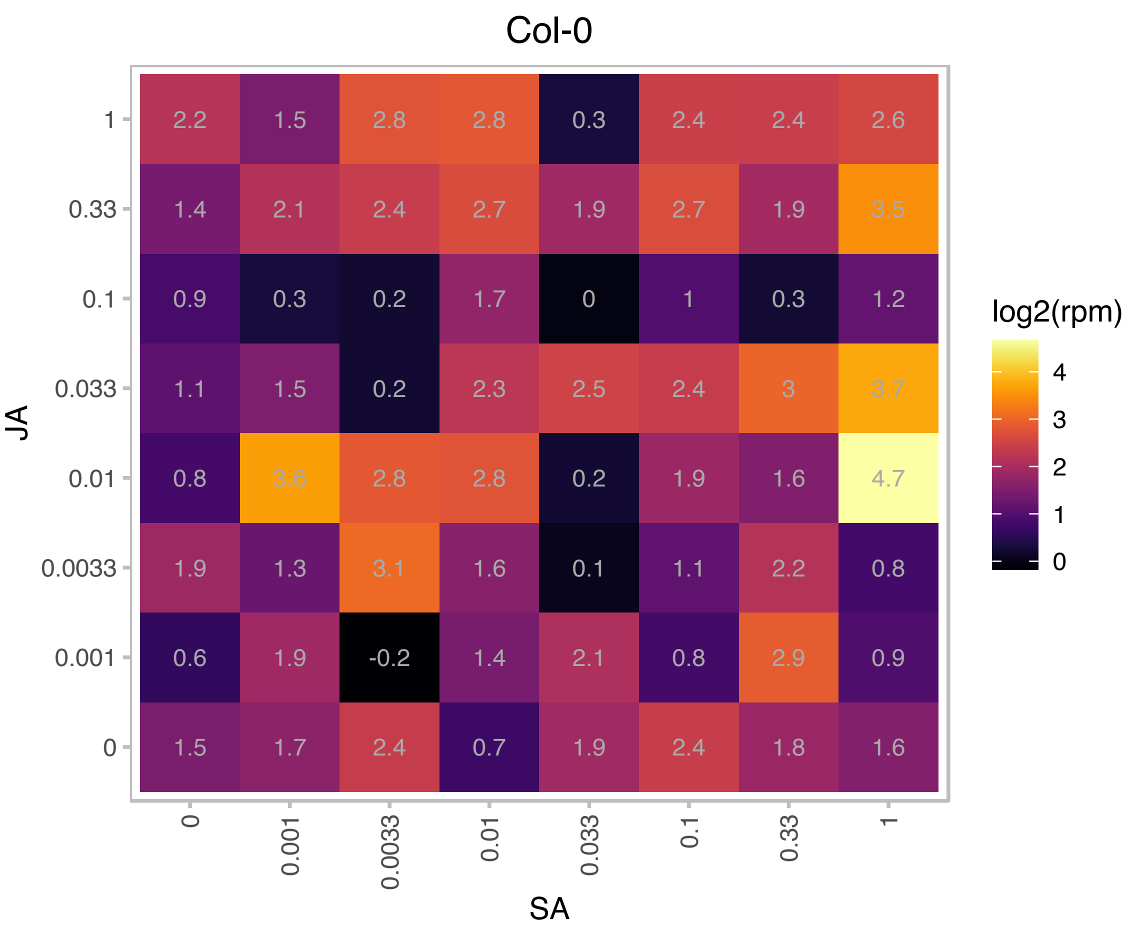


| | |
|---|---|
| Gene | AT5G08139 |
| is.expressed_genes | TRUE |
| is.responsive_genes | TRUE |
| is.apcluster | B-05 |

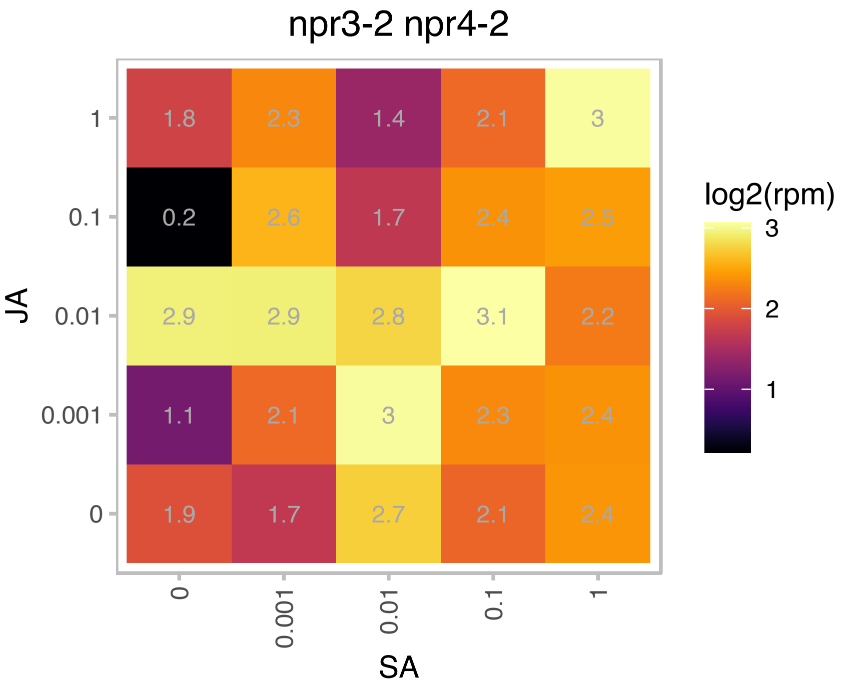


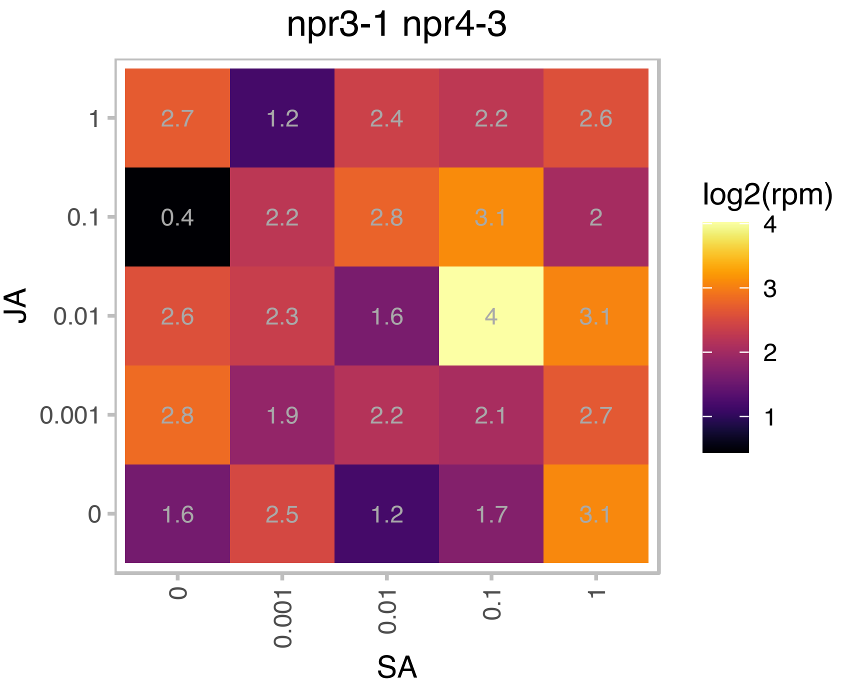

### AT5G13180.1 : ANAC083, NAC083, VNI2

NAC domain containing protein 83

Encodes a NAC domain transcription factor that interacts with VND7 and negatively regulates xylem vessel formation.

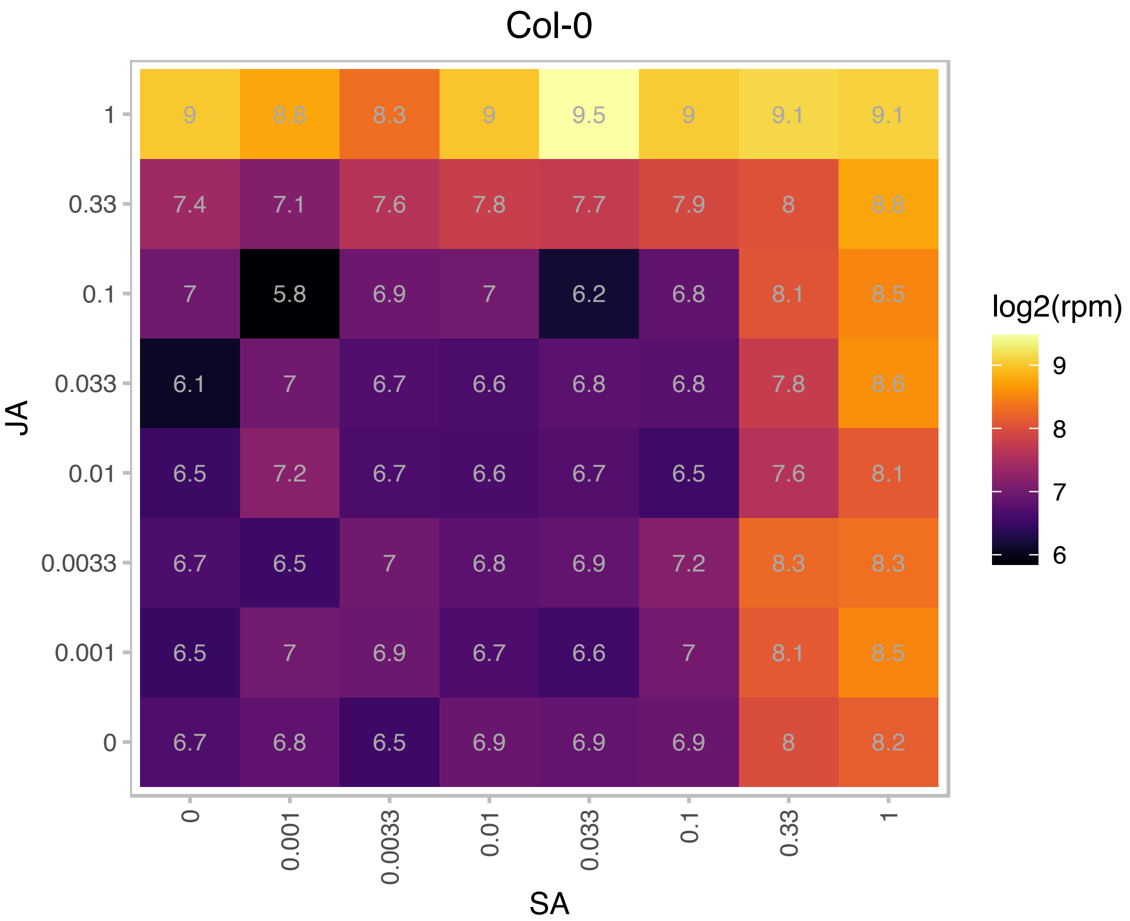


| | |
|---|---|
| Gene | AT5G13180 |
| is.expressed_genes | TRUE |
| is.responsive_genes | TRUE |
| is.apcluster | B-05 |

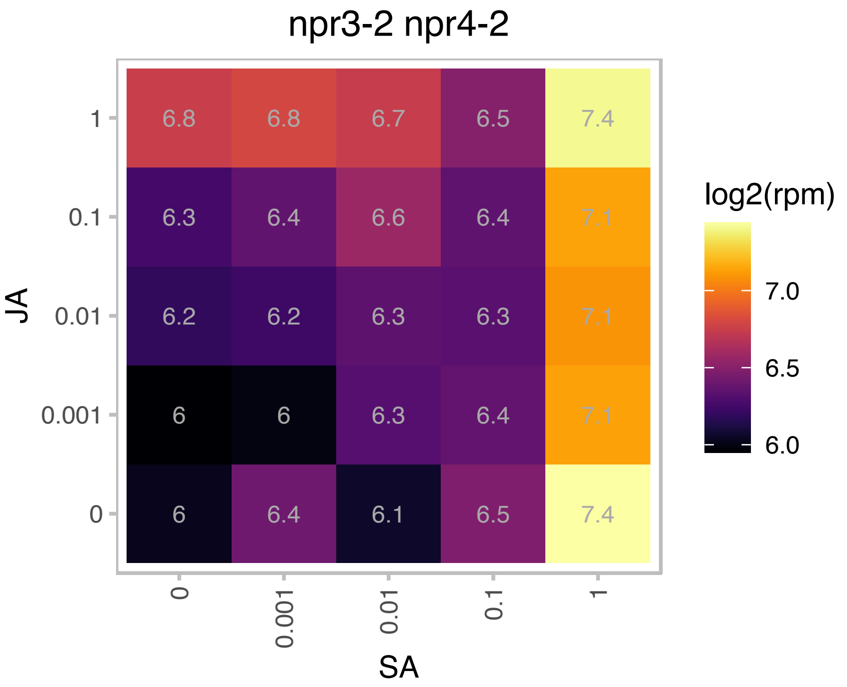


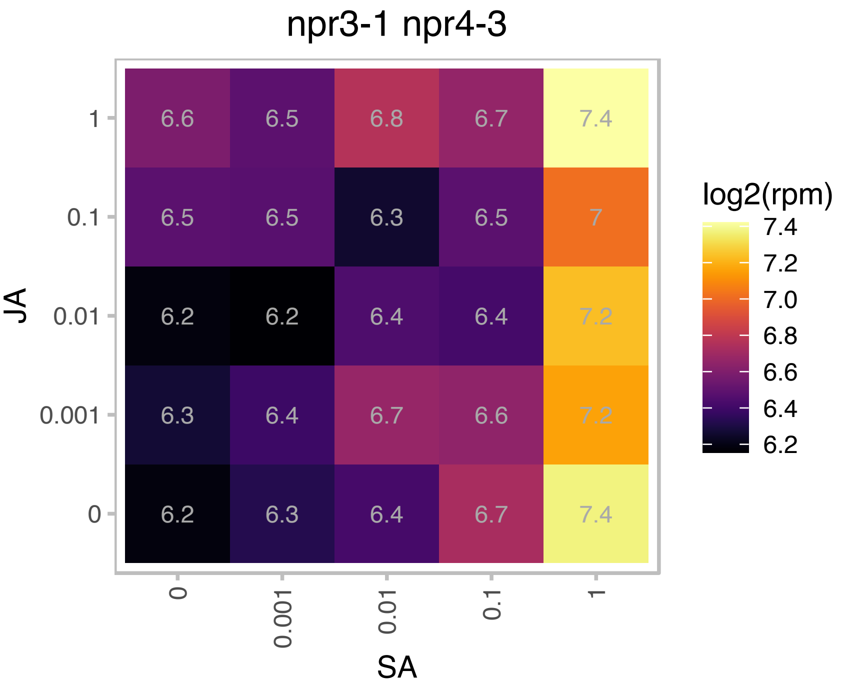

### AT5G13200.1 : GER5

GRAM domain family protein

Encodes a protein with unknown function that is involved in hormone mediated regulation of seed germination/dormancy.

Col-0

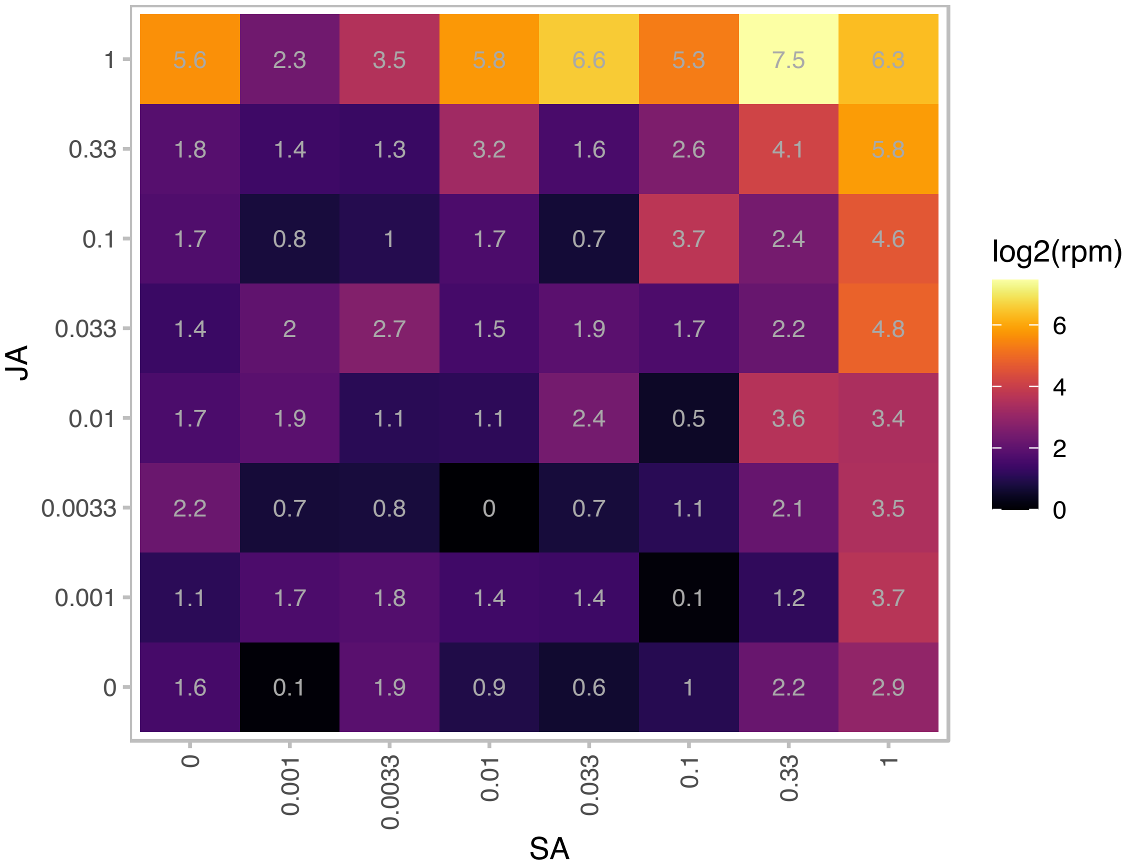


| | |
|---|---|
| Gene | AT5G13200 |
| is.expressed_genes | TRUE |
| is.responsive_genes | TRUE |
| is.apcluster | B-05 |

npr3-2 npr4-2

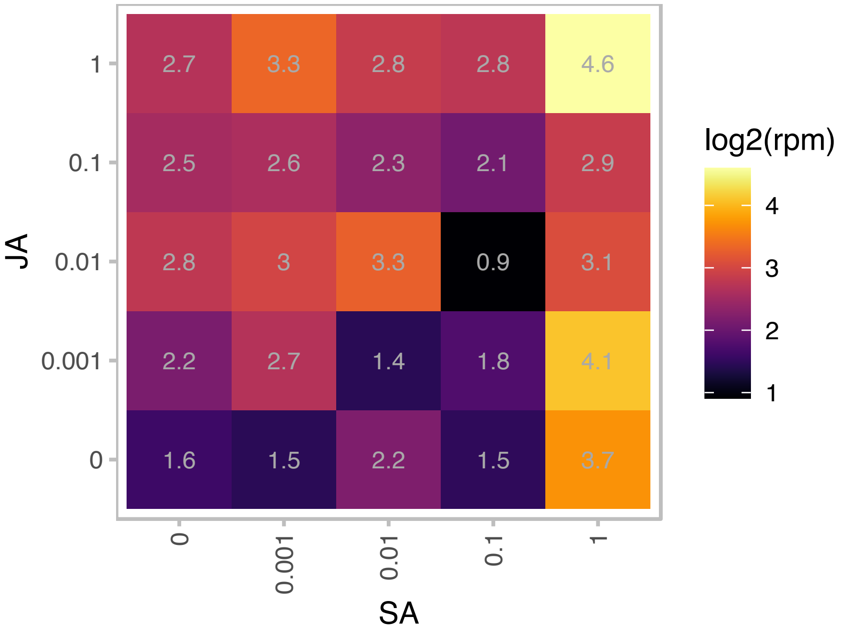


npr3-1 npr4-3

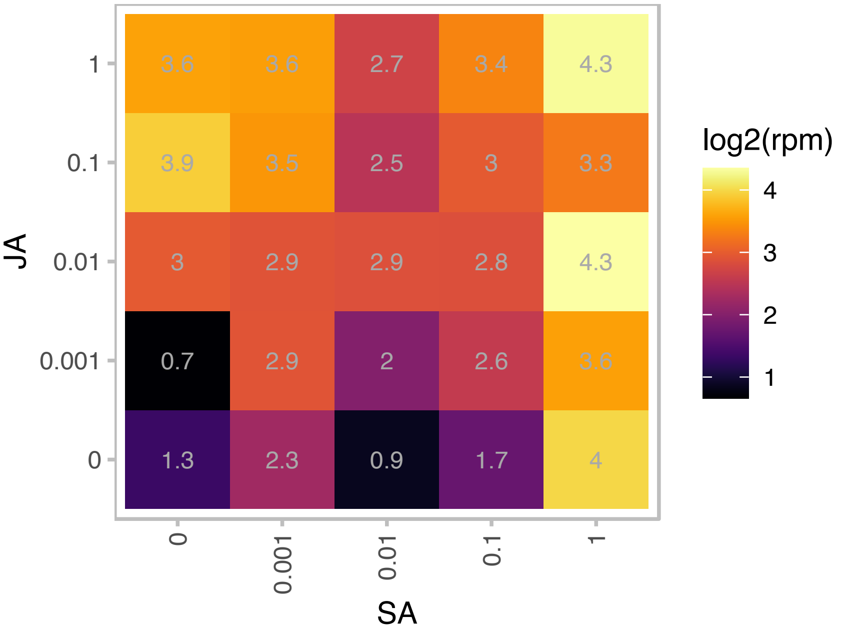

### AT5G16440.1 : IPP1

isopentenyl diphosphate isomerase 1

Encodes a protein with isopentenyl diphosphate:dimethylallyl diphosphate isomerase activity. There is genetic evidence that it functions in the mevalonate, but not the MEP biosynthetic pathway.

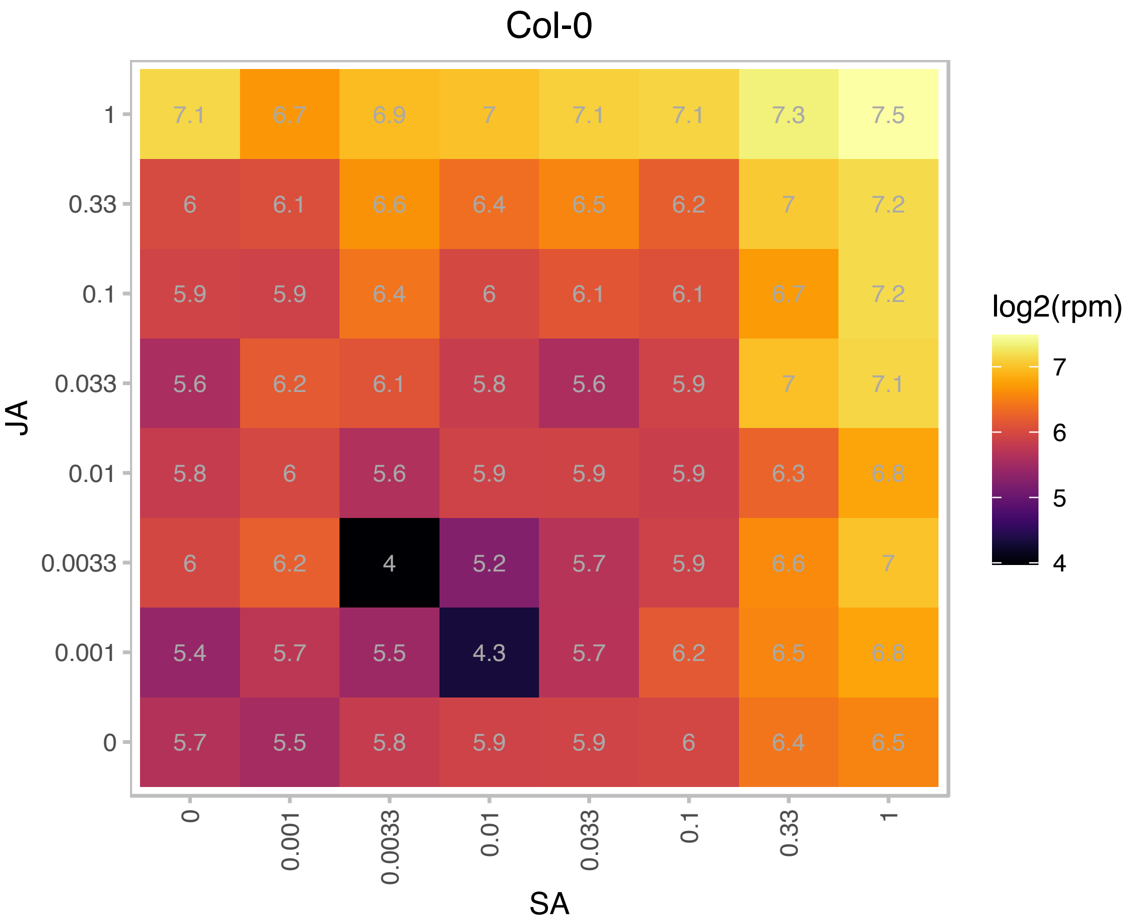


| | |
|---|---|
| Gene | AT5G16440 |
| is.expressed_genes | TRUE |
| is.responsive_genes | TRUE |
| is.apcluster | B-05 |

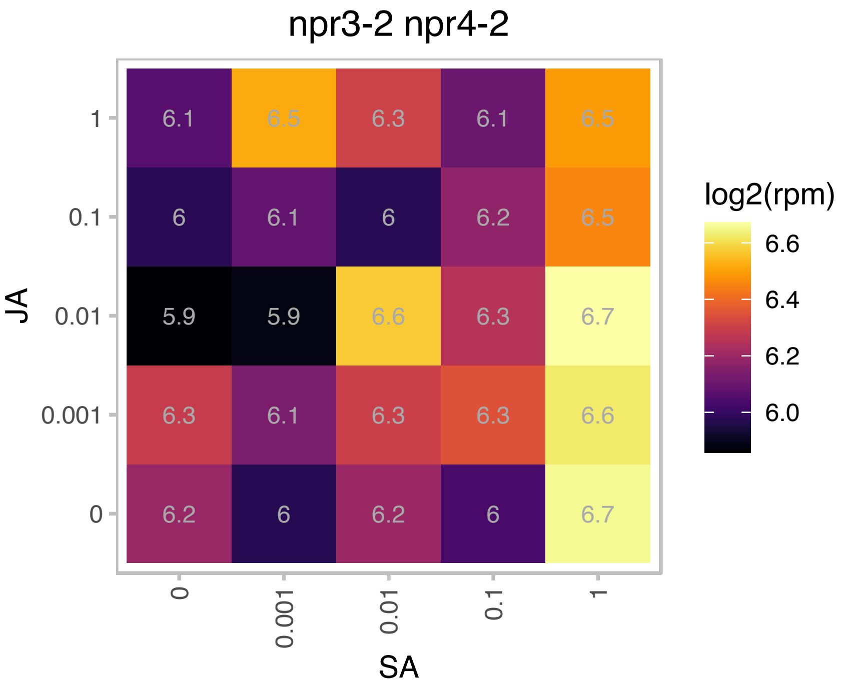


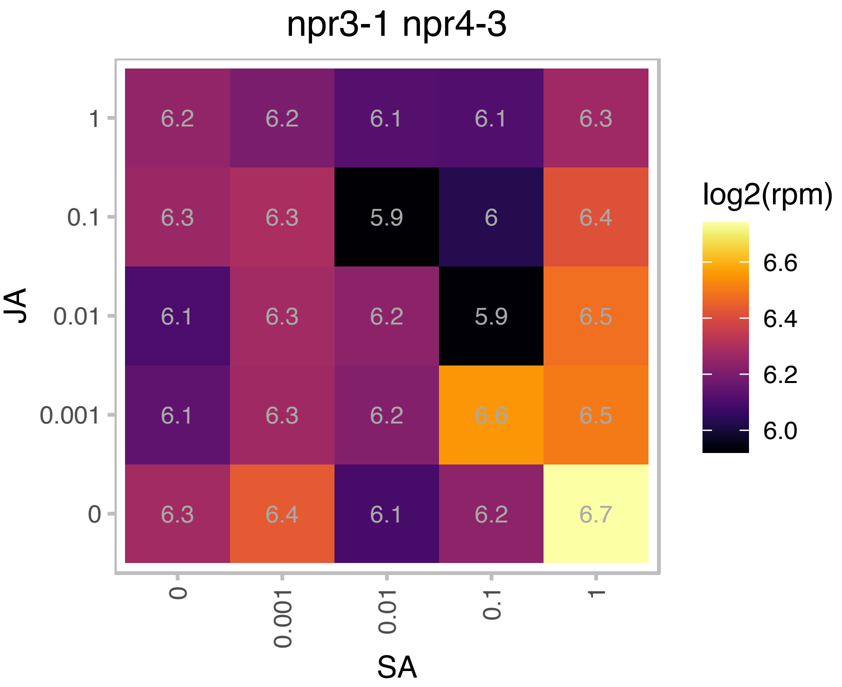

# AT5G17280.1 : NA

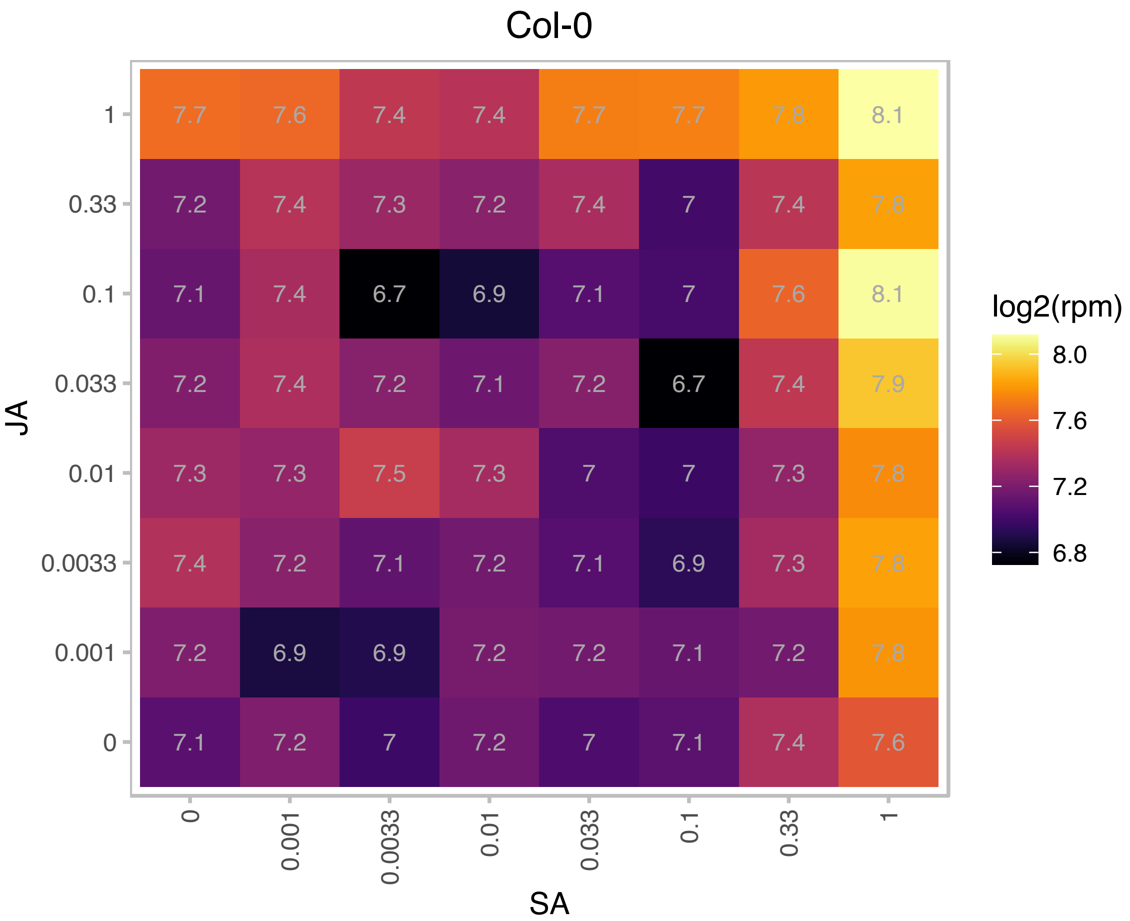


| | |
|---|---|
| Gene | AT5G17280 |
| is.expressed_genes | TRUE |
| is.responsive_genes | TRUE |
| is.apcluster | B-05 |

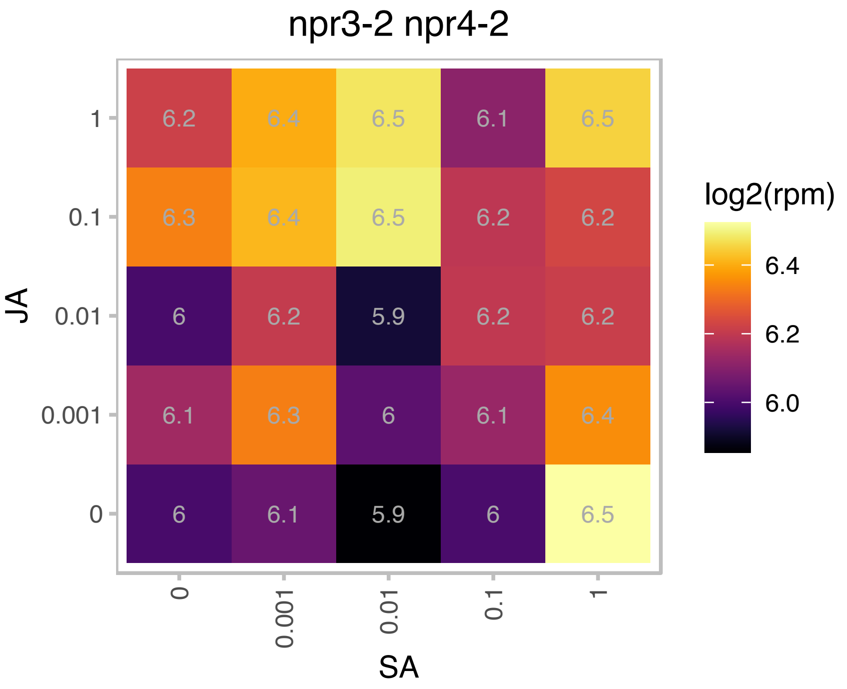


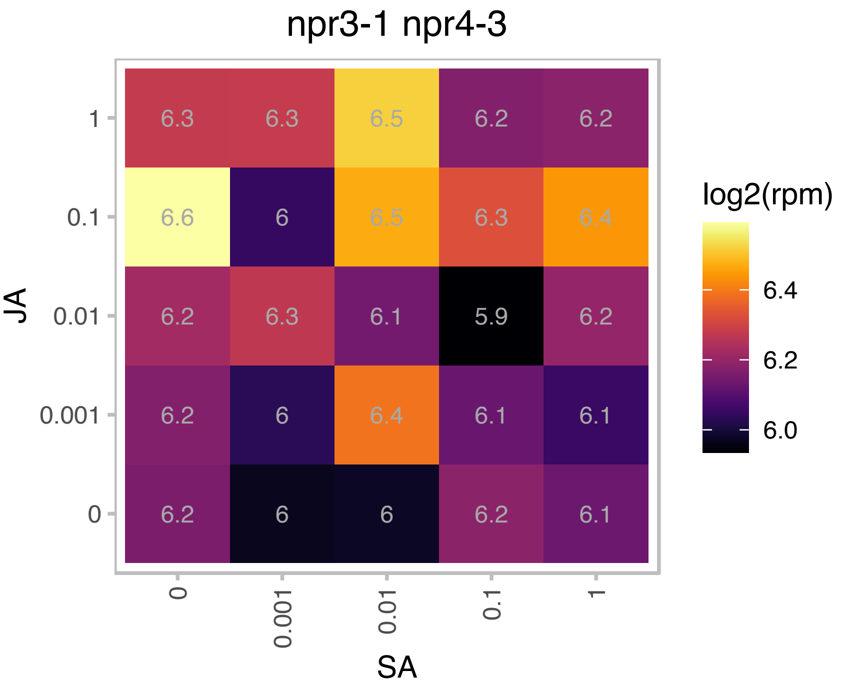

# AT5G17390.1 : NA

Adenine nucleotide alpha hydrolases-like superfamily protein

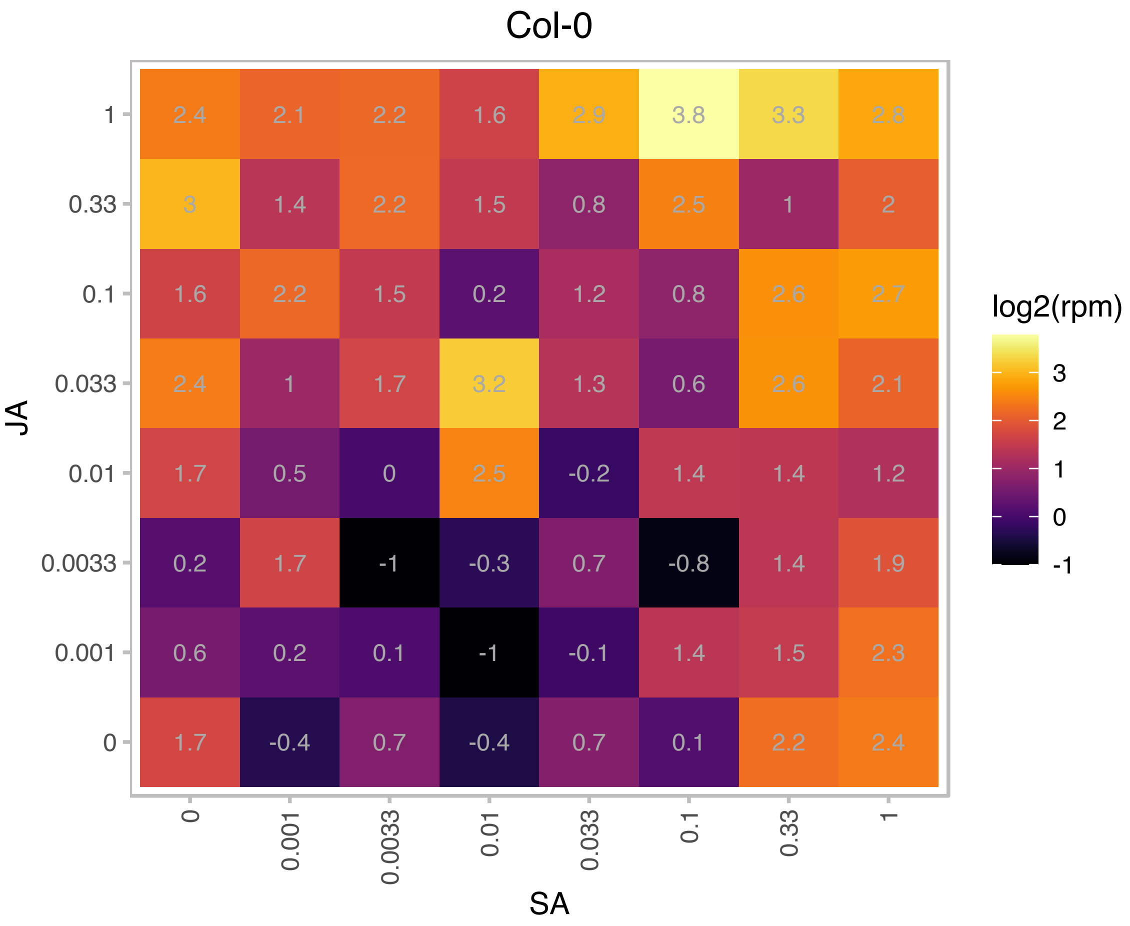


| | |
|---|---|
| Gene | AT5G17390 |
| is.expressed_genes | TRUE |
| is.responsive_genes | TRUE |
| is.apcluster | B-05 |

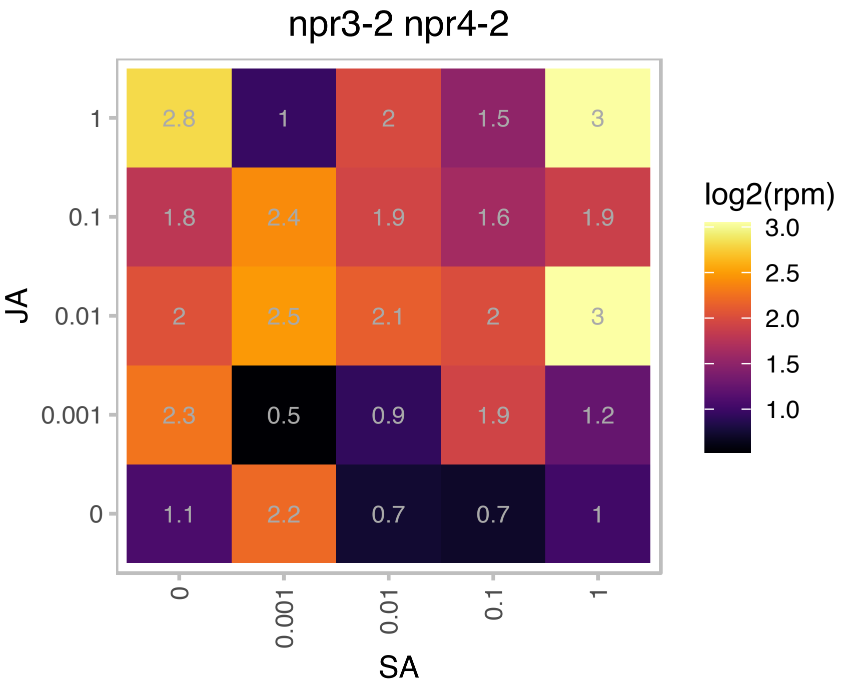


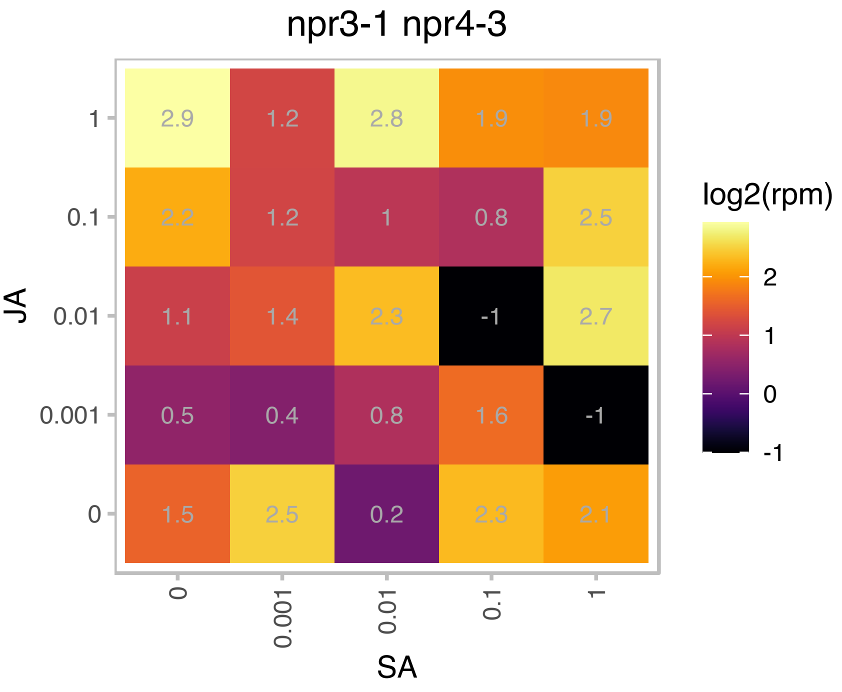

### AT5G20020.1 : RAN2

RAS-related GTP-binding nuclear protein 2

A member of RAN GTPase gene family. Encodes a small soluble GTP-binding protein. Likely to be involved in nuclear translocation of proteins.  May also be involved in cell cycle progression.

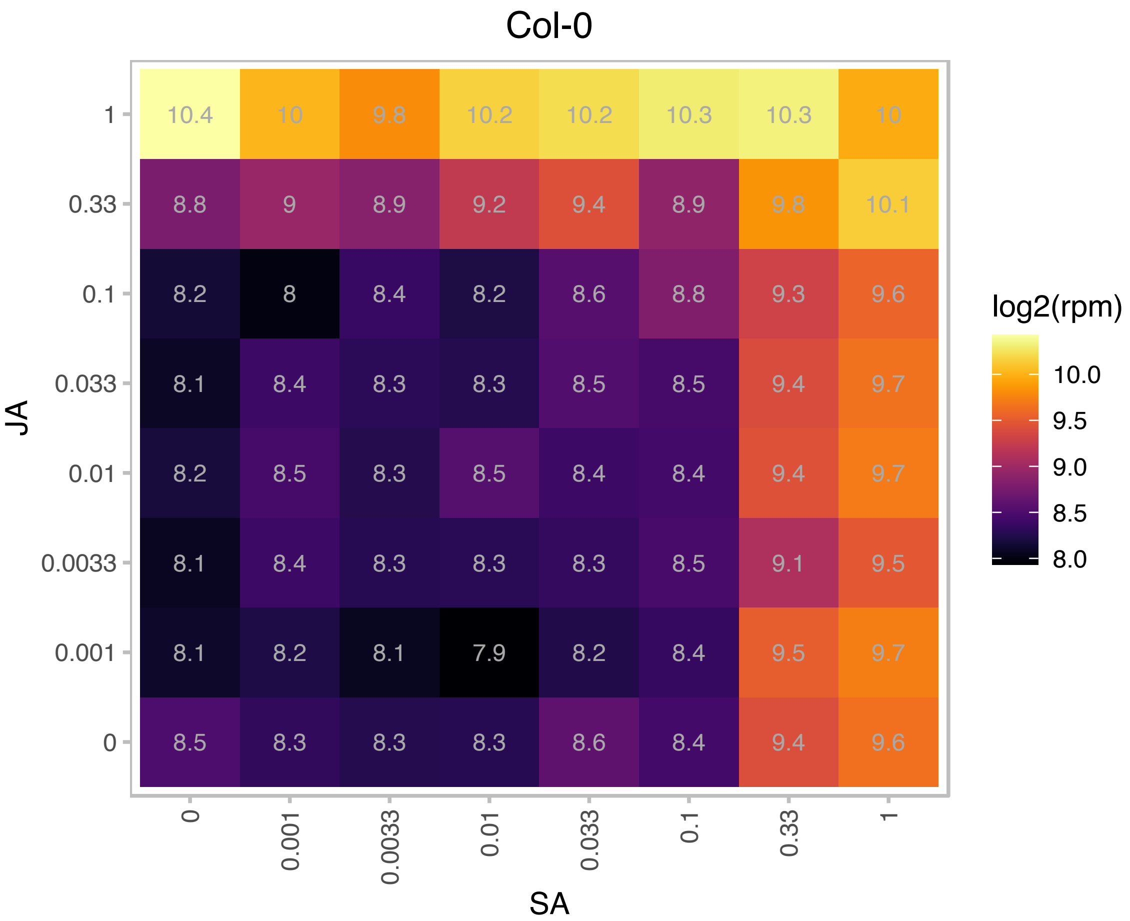


| | |
|---|---|
| Gene | AT5G20020 |
| is.expressed_genes | TRUE |
| is.responsive_genes | TRUE |
| is.apcluster | B-05 |

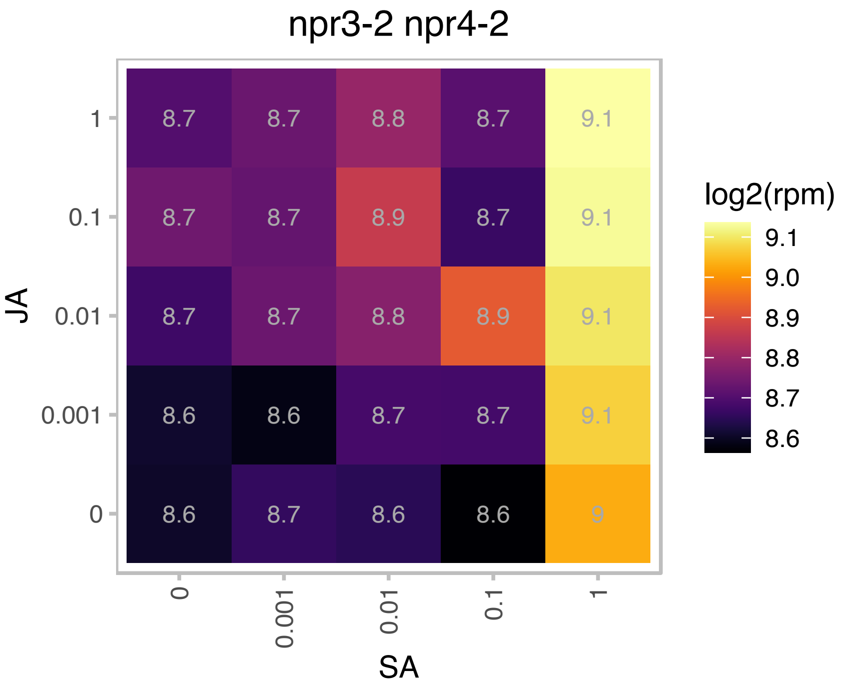


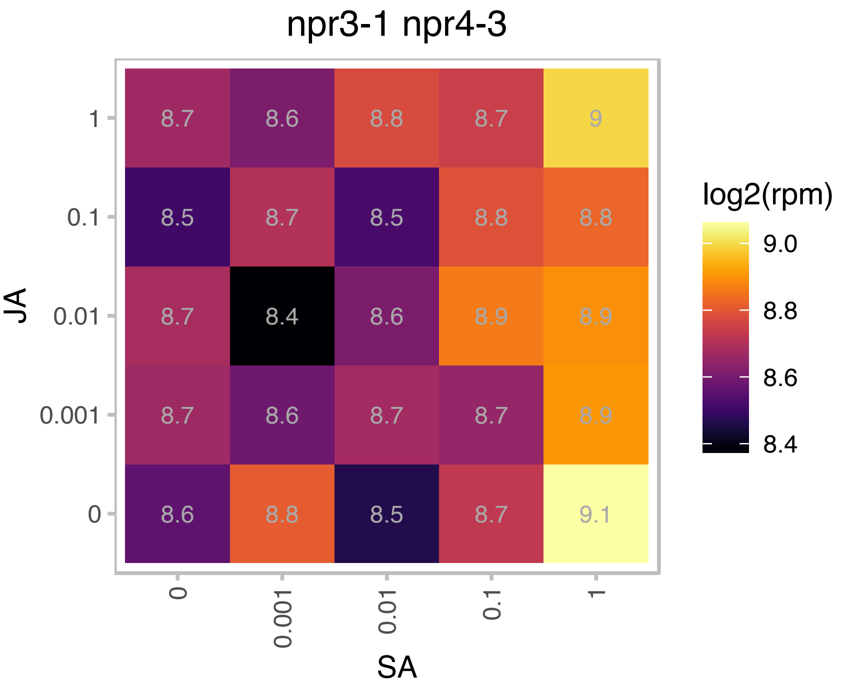

# AT5G22850.1 : NA

Eukaryotic aspartyl protease family protein

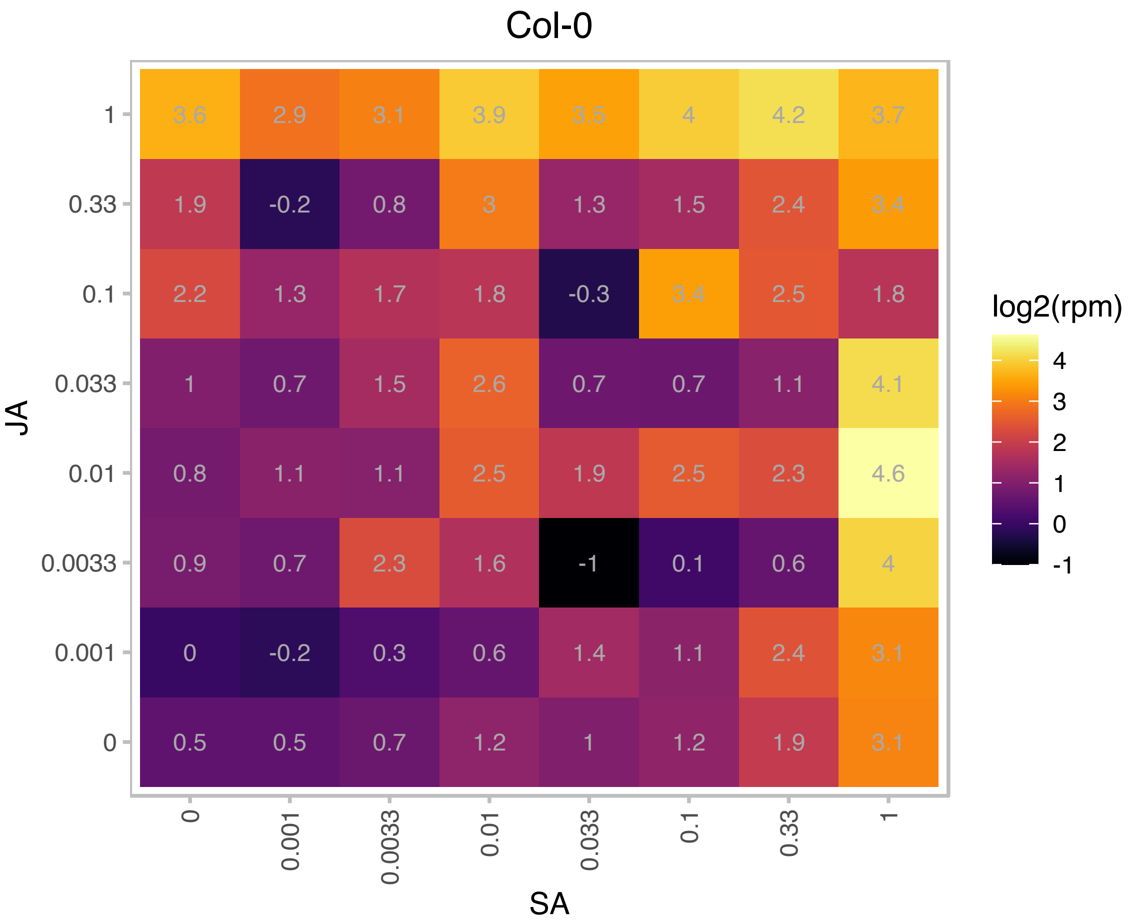


| | |
|---|---|
| Gene | AT5G22850 |
| is.expressed_genes | TRUE |
| is.responsive_genes | TRUE |
| is.apcluster | B-05 |

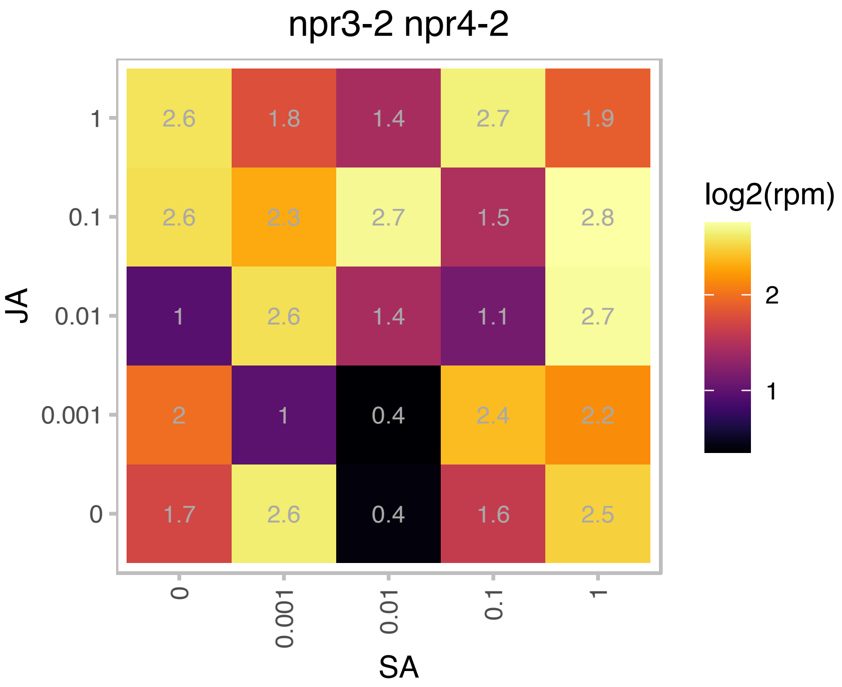


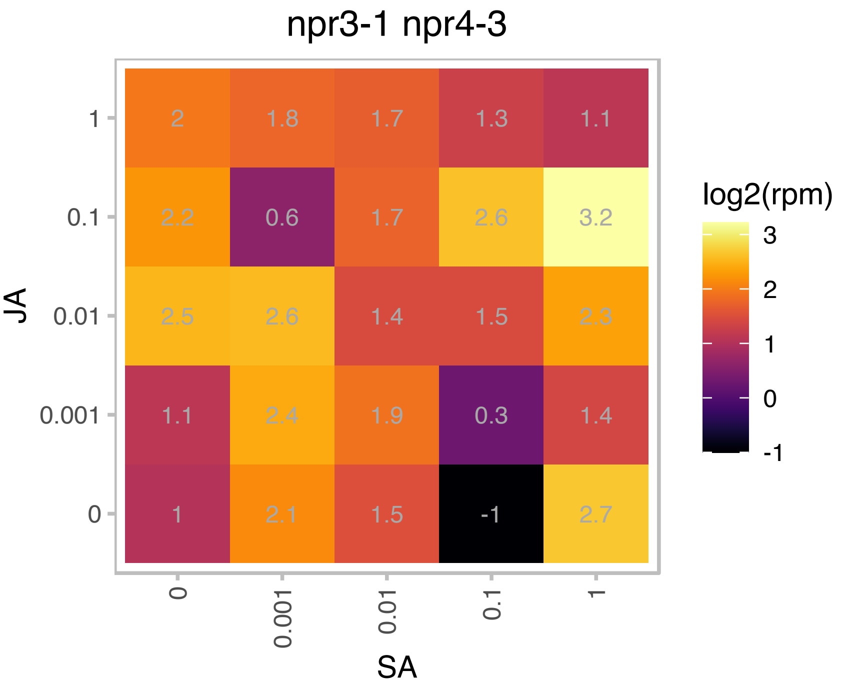

# AT5G25930.1 : NA

Protein kinase family protein with leucine-rich repeat domain

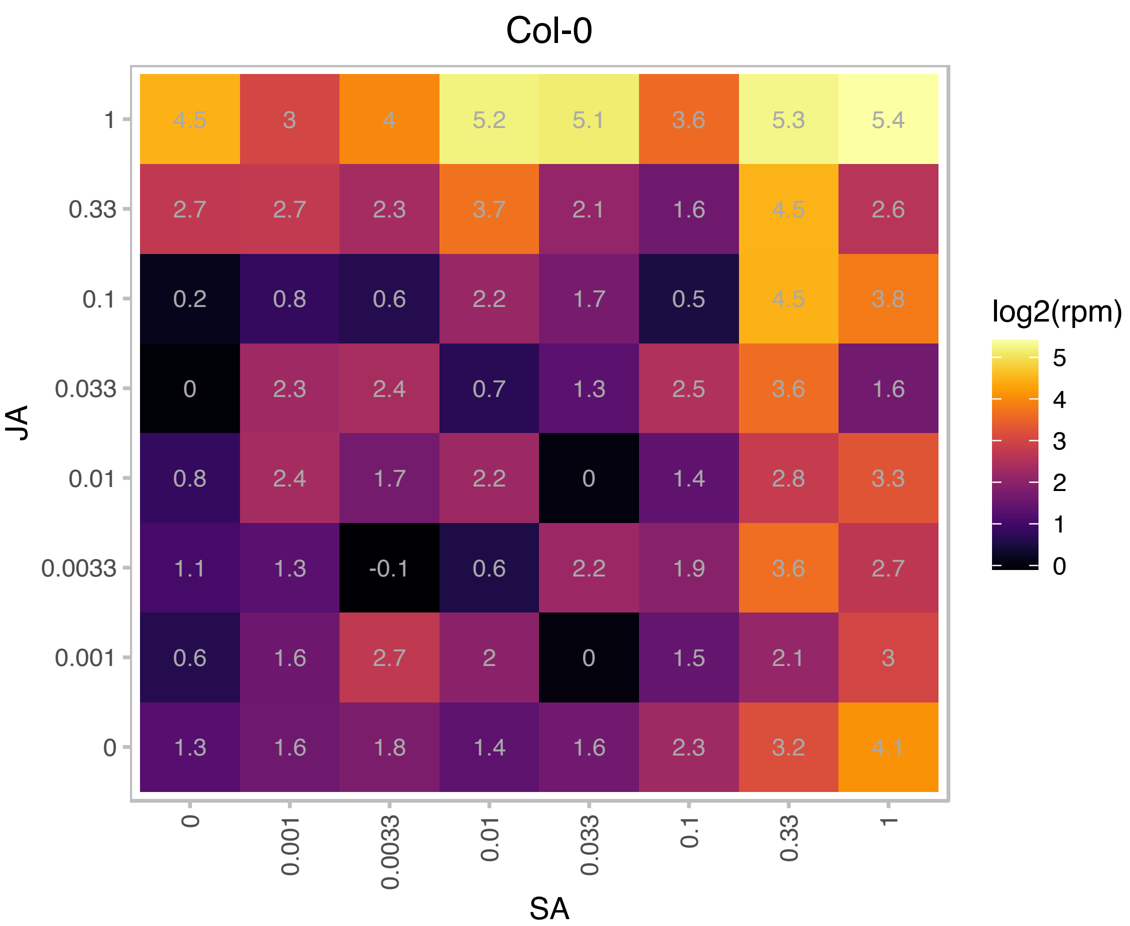


| Gene | AT5G25930 |
|---|---|
| is.expressed_genes | TRUE |
| is.responsive_genes | TRUE |
| is.apcluster | B-05 |

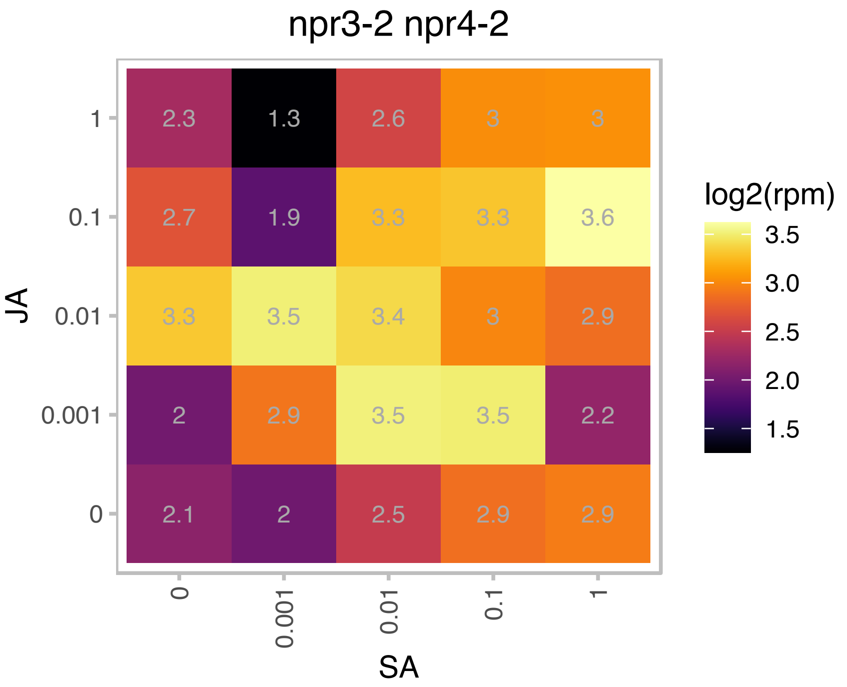


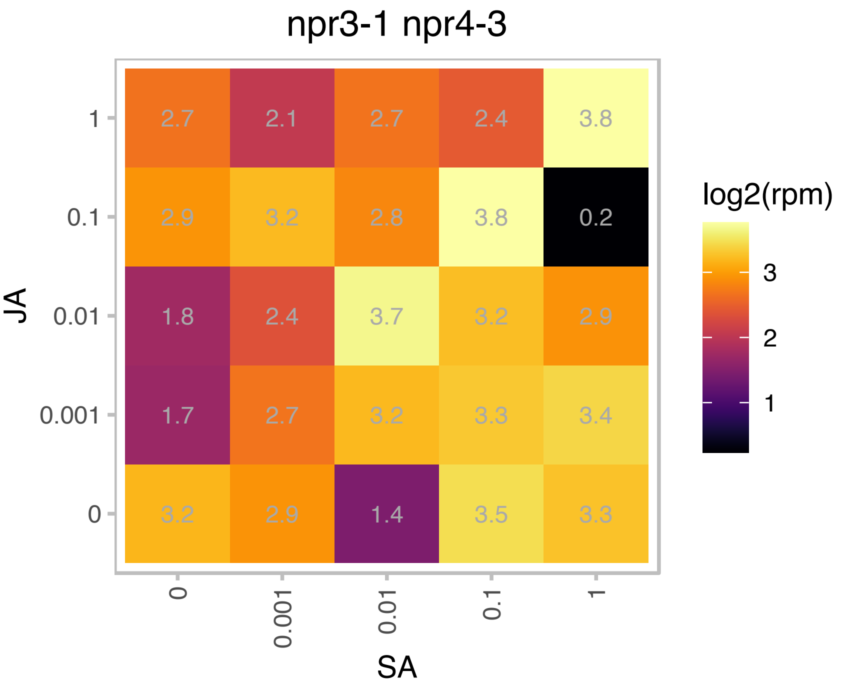

# AT5G35570.1 : NA

O-fucosyltransferase family protein

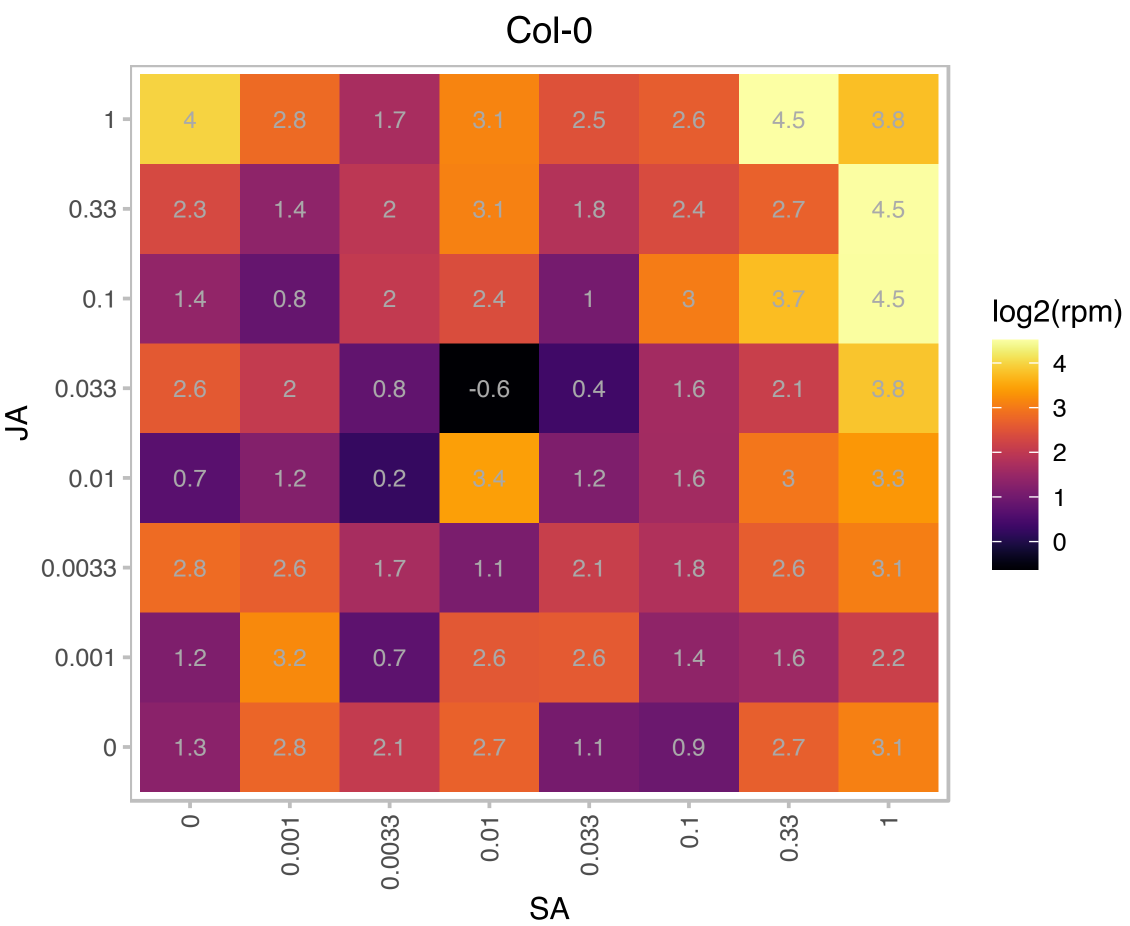


| | |
|---|---|
| Gene | AT5G35570 |
| is.expressed_genes | TRUE |
| is.responsive_genes | TRUE |
| is.apcluster | B-05 |

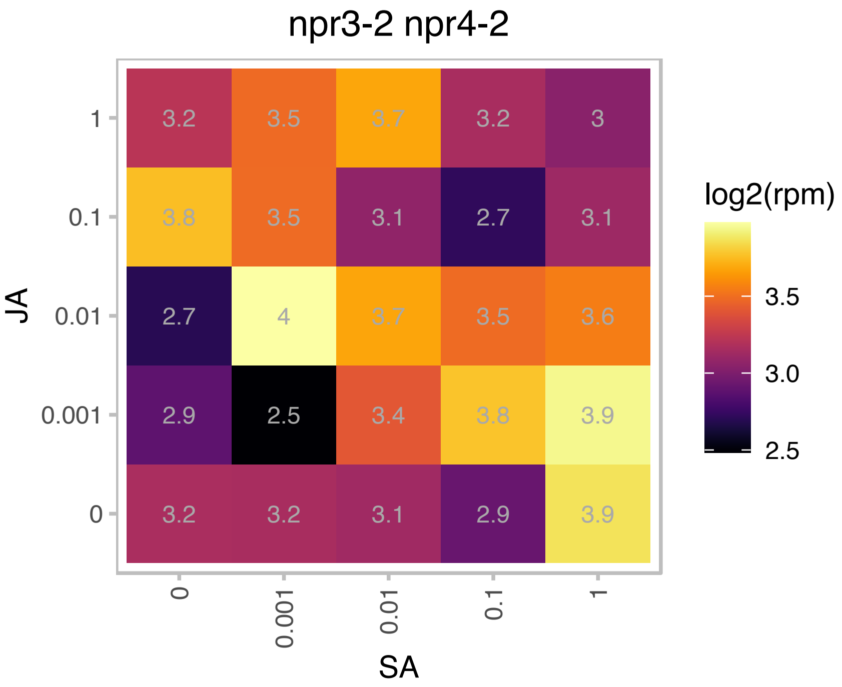


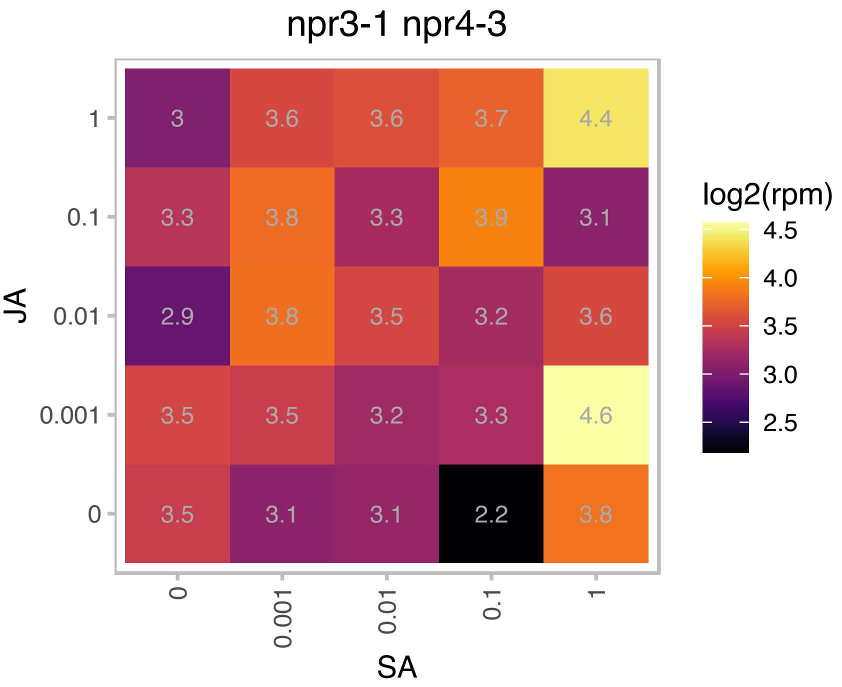

# AT5G38830.1 : NA

Cysteinyl-tRNA synthetase, class Ia family protein

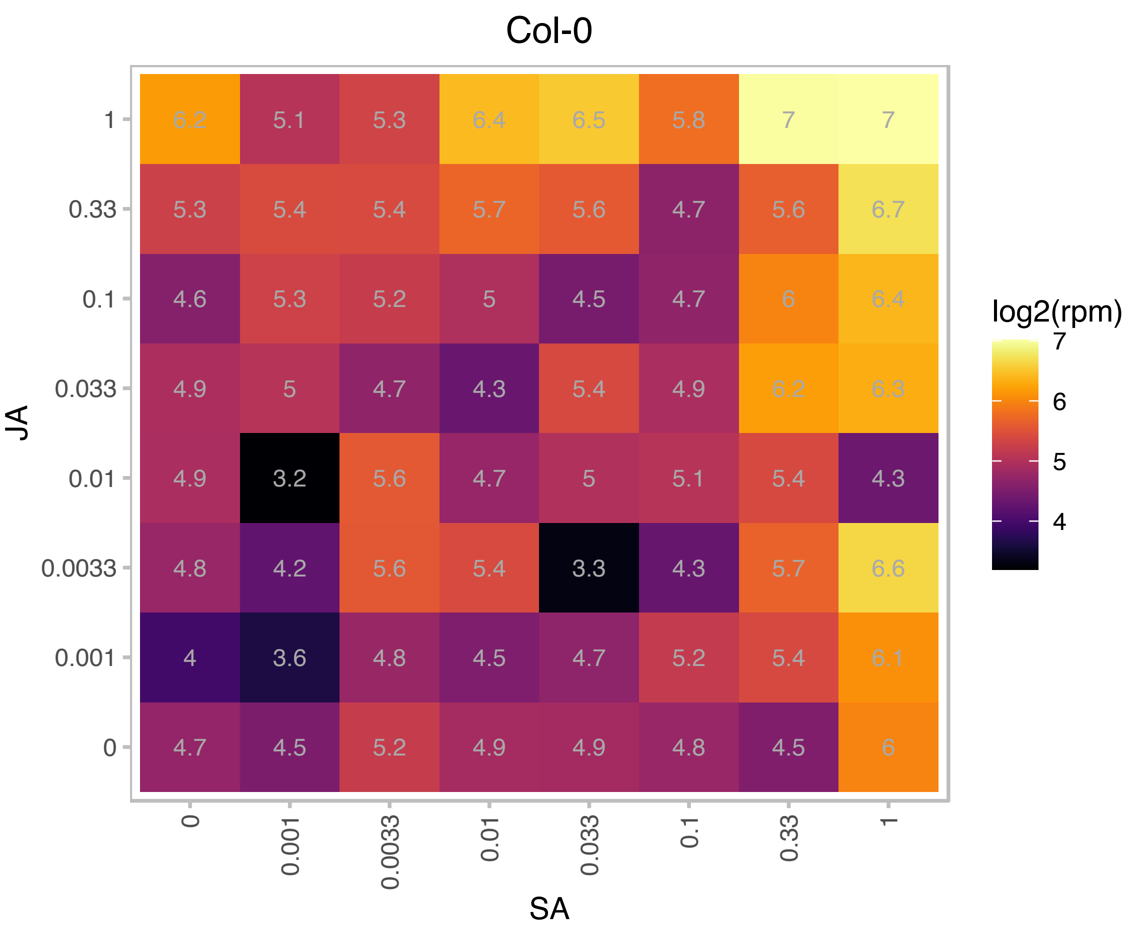


| | |
|---|---|
| Gene | AT5G38830 |
| is.expressed_genes | TRUE |
| is.responsive_genes | TRUE |
| is.apcluster | B-05 |

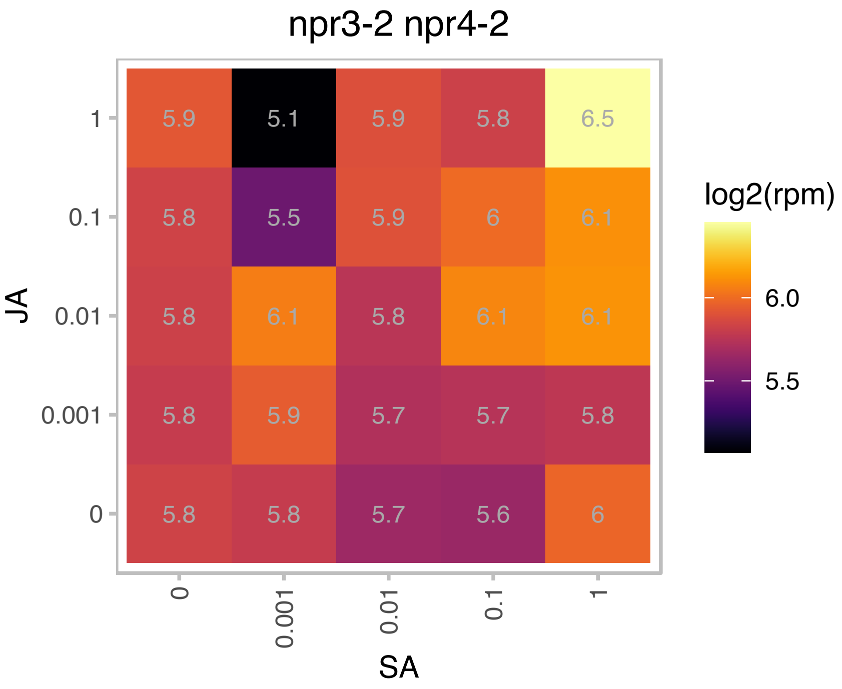


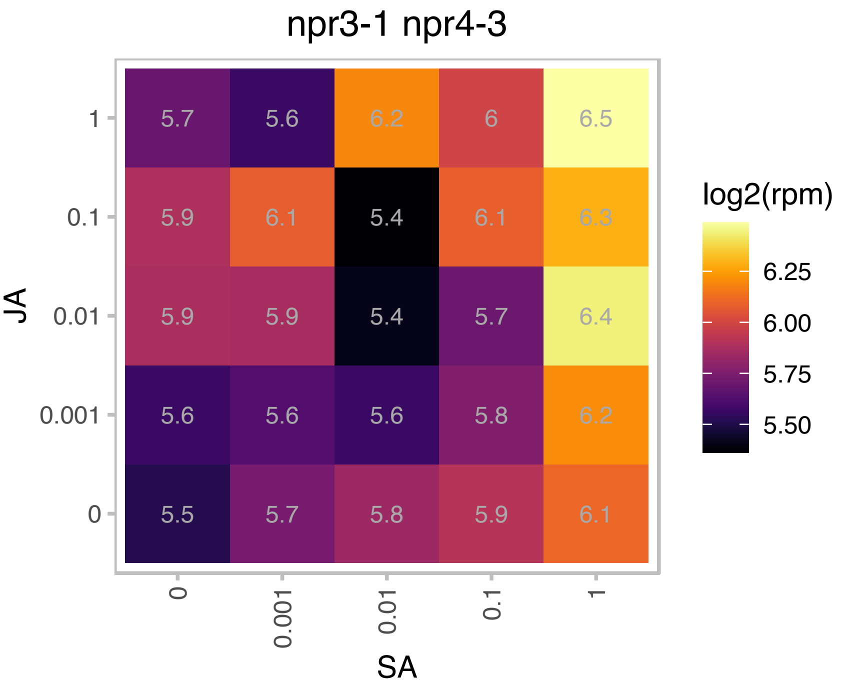

# AT5G43180.1 : NA

Protein of unknown function, DUF599

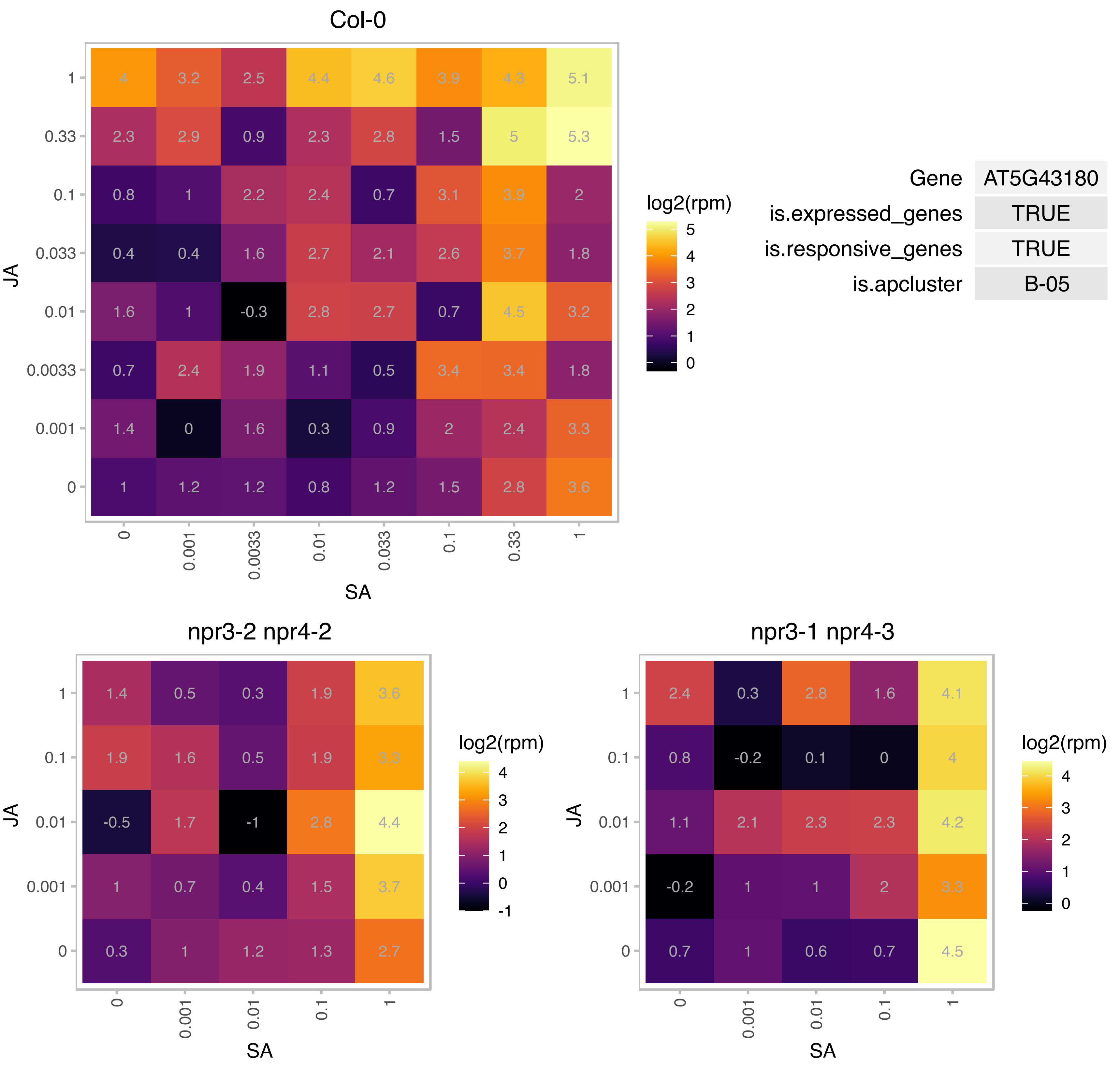


| | |
|---|---|
| Gene | AT5G43180 |
| is.expressed_genes | TRUE |
| is.responsive_genes | TRUE |
| is.apcluster | B-05 |

### AT5G43380.1 : TOPP6

type one serine/threonine protein phosphatase 6

encodes a type I serine/threonine protein phosphatase expressed in expressed in roots, rosettes and flowers.

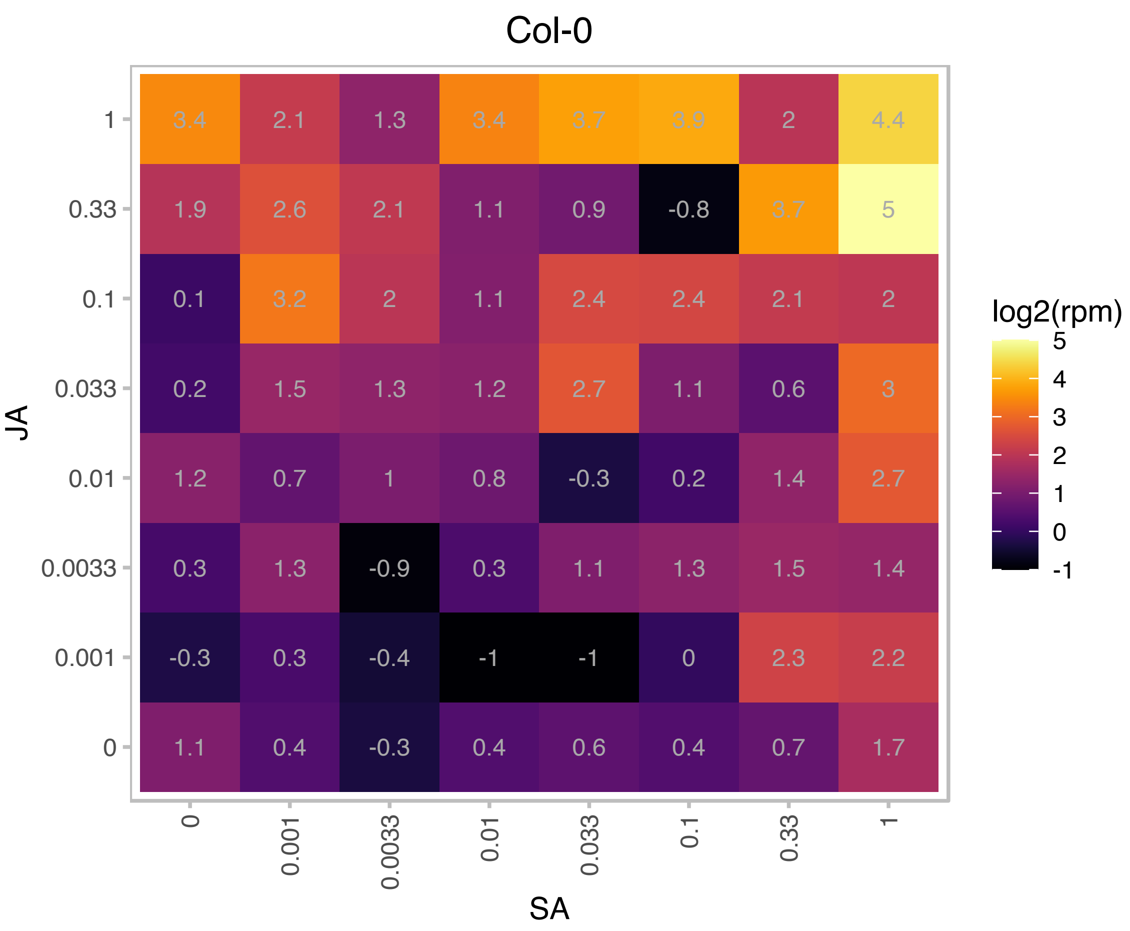


| | |
|---|---|
| Gene | AT5G43380 |
| is.expressed_genes | TRUE |
| is.responsive_genes | TRUE |
| is.apcluster | B-05 |

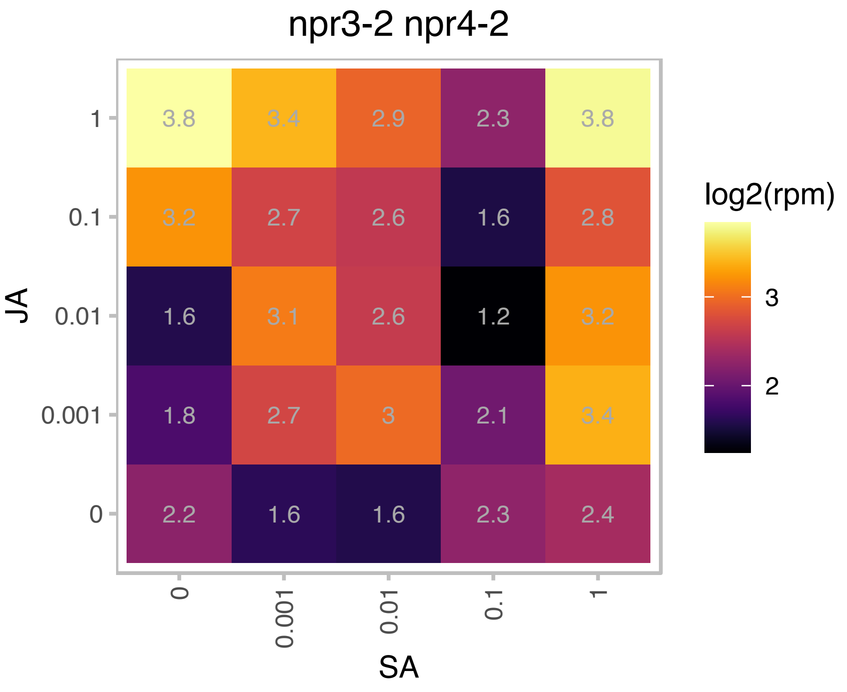


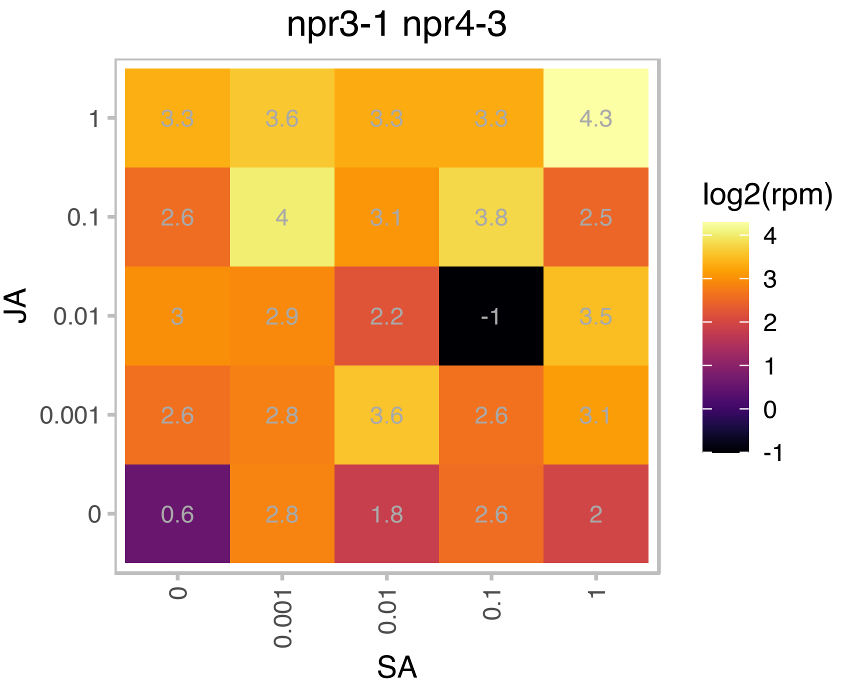

### AT5G47200.1 : ATRAB1A, ATRABD2B, RAB1A

RAB GTPase homolog 1A

AtRabD2b encodes a Rab GTPase, which plays important roles in pollen development, germination and tube elongation. The mRNA is cell-to-cell mobile.

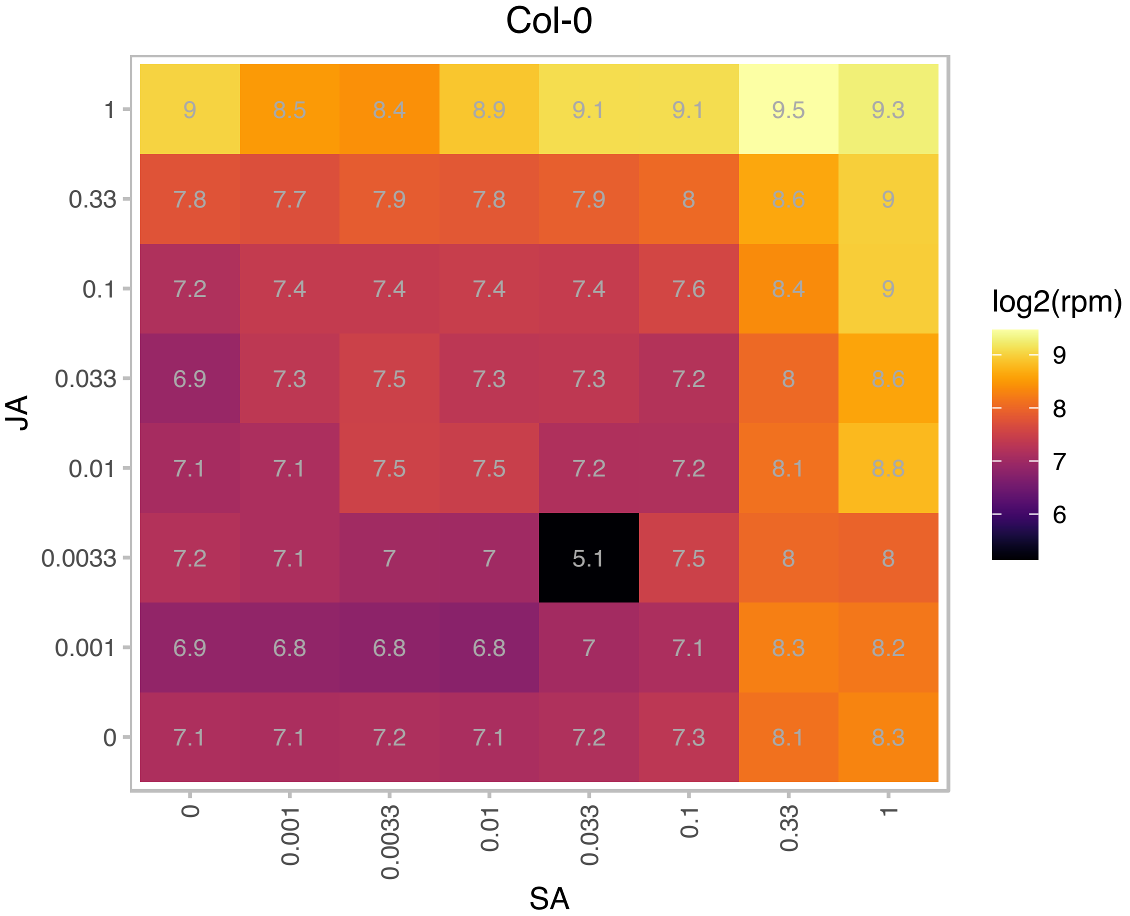


| | |
|---|---|
| Gene | AT5G47200 |
| is.expressed_genes | TRUE |
| is.responsive_genes | TRUE |
| is.apcluster | B-05 |

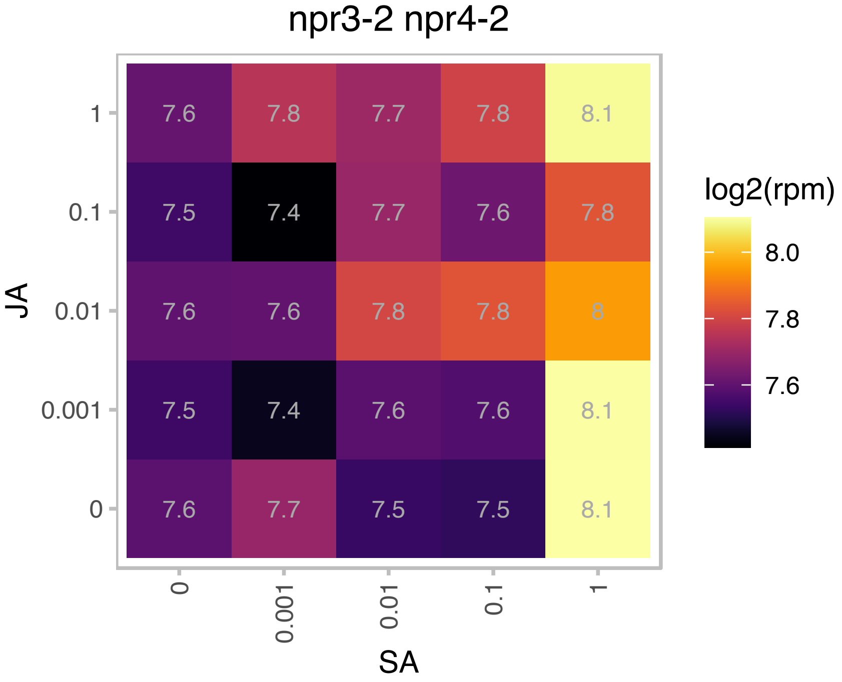


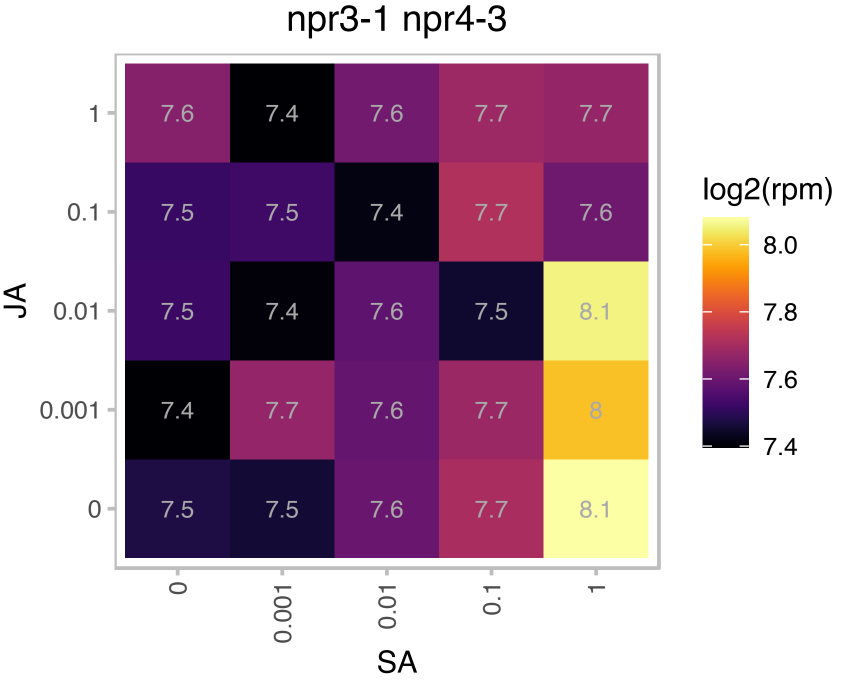

# AT5G49645.1 : NA

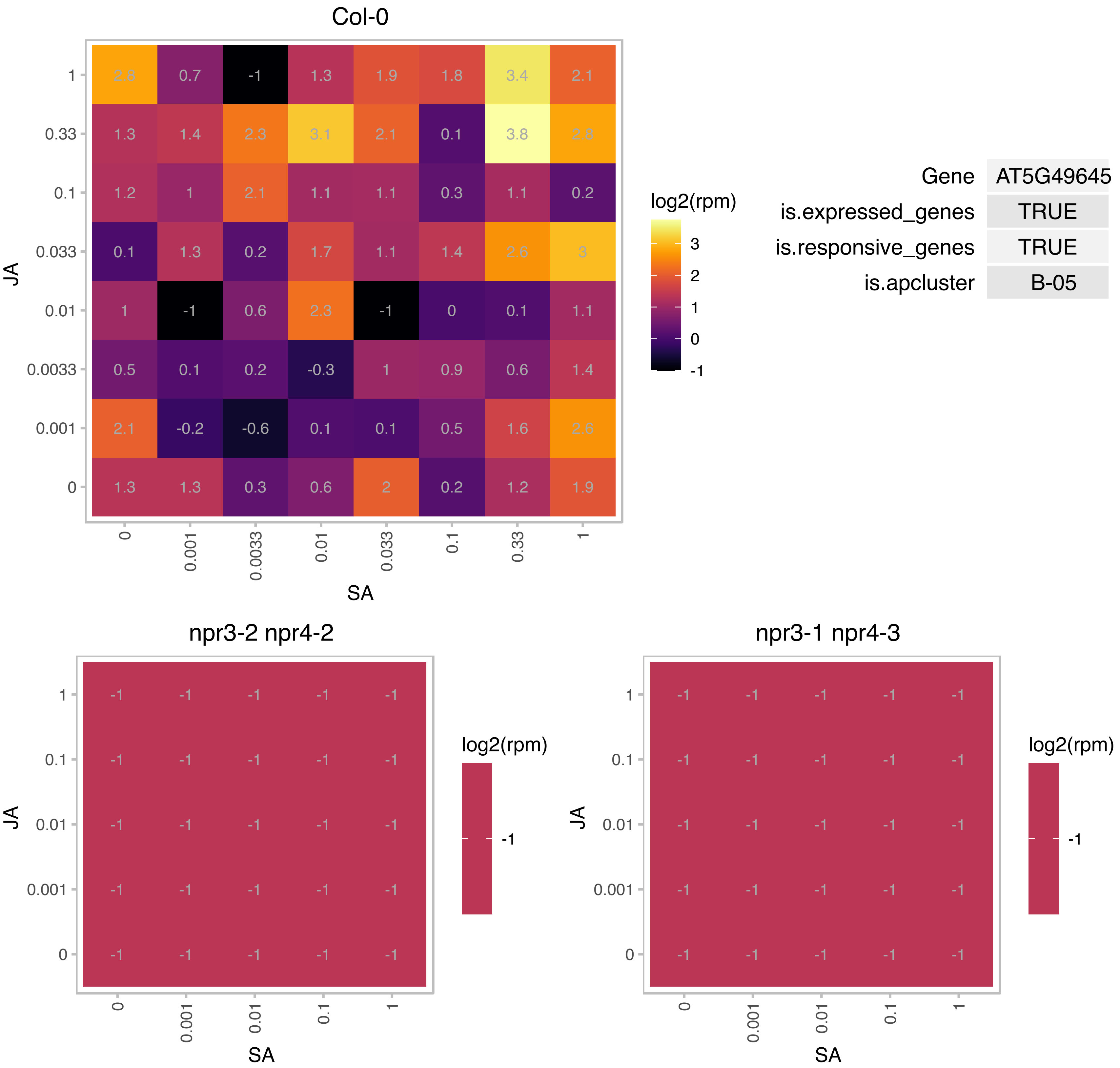


| | |
|---|---|
| Gene | AT5G49645 |
| is.expressed_genes | TRUE |
| is.responsive_genes | TRUE |
| is.apcluster | B-05 |

### AT5G50780.1 : AtMORC4

Histidine kinase-, DNA gyrase B-, and HSP90-like ATPase family protein

MORC4 is a member of a family of GHKL ATPases. It is localized in the nuceloplasm and adjacent to chromocenters. Along with MORC7, it appears to repress the expression

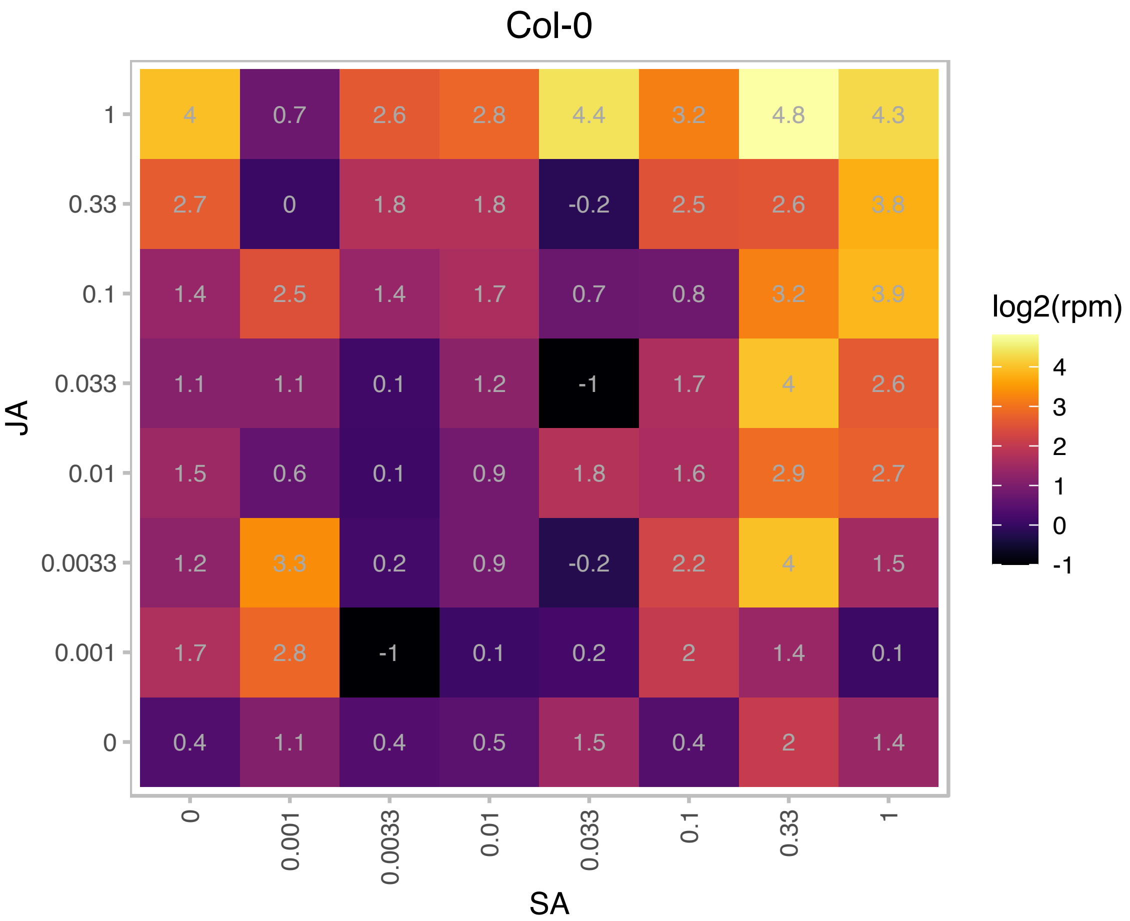


| | |
|---|---|
| Gene | AT5G50780 |
| is.expressed_genes | TRUE |
| is.responsive_genes | TRUE |
| is.apcluster | B-05 |

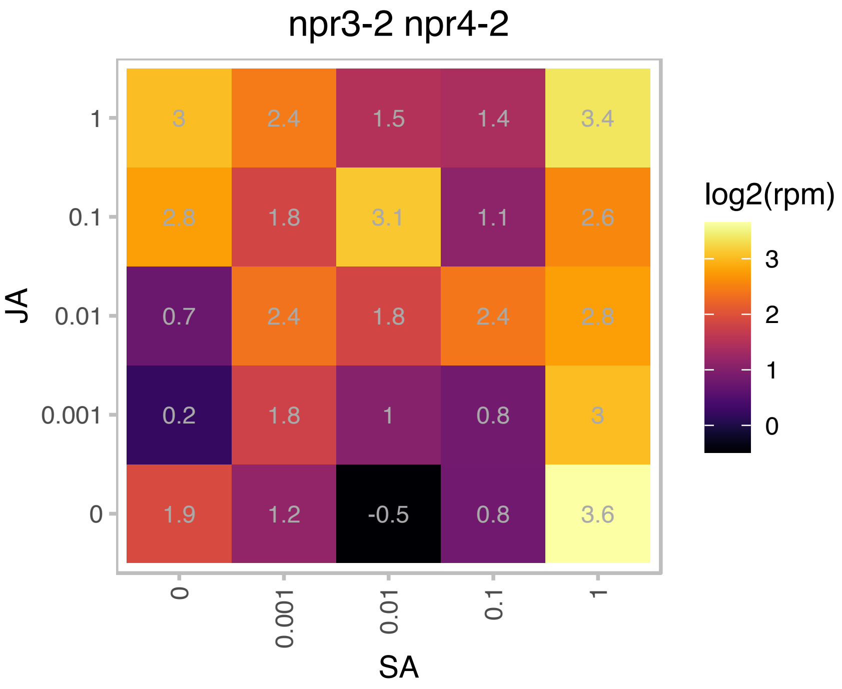


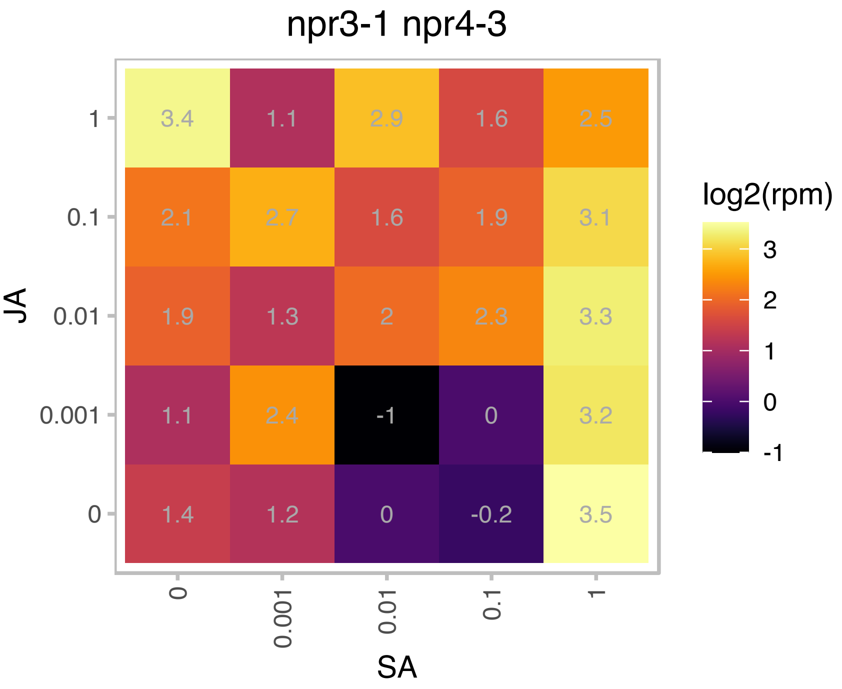

### AT5G50920.1 : ATHSP93-V, CLPC, CLPC1, DCA1, HSP93

CLPC homologue 1

Encodes a protein that is similar to ATP-dependent Clp protease ATP-binding subunit / ClpC. Involved in protein import into the chloroplast. May provide ATP source that drives the TIC (Translocon at the Inner envelope membrane of Chloroplasts) translocation machinery. Association of Hsp93 with the inner envelope membrane through its N domain is important for the functions of Hsp93 in vivo.

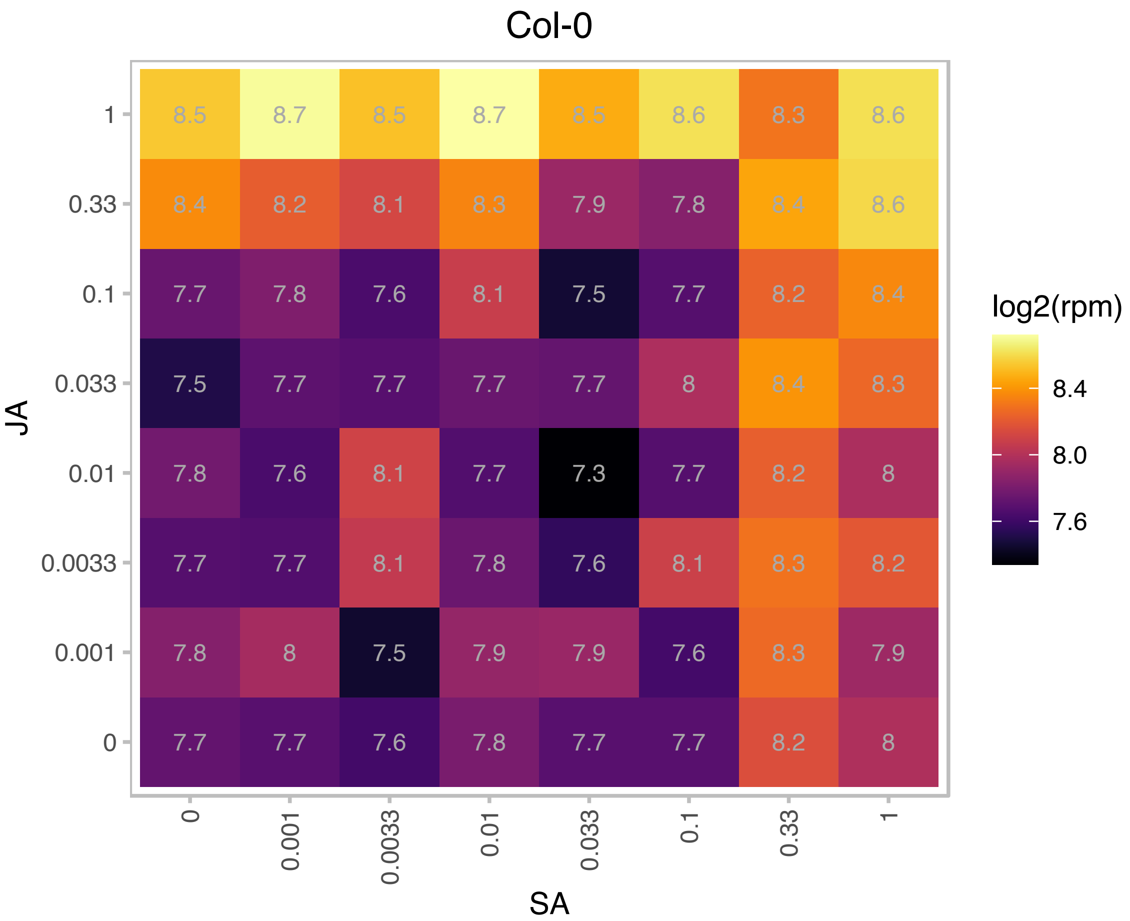


| | |
|---|---|
| Gene | AT5G50920 |
| is.expressed_genes | TRUE |
| is.responsive_genes | TRUE |
| is.apcluster | B-05 |

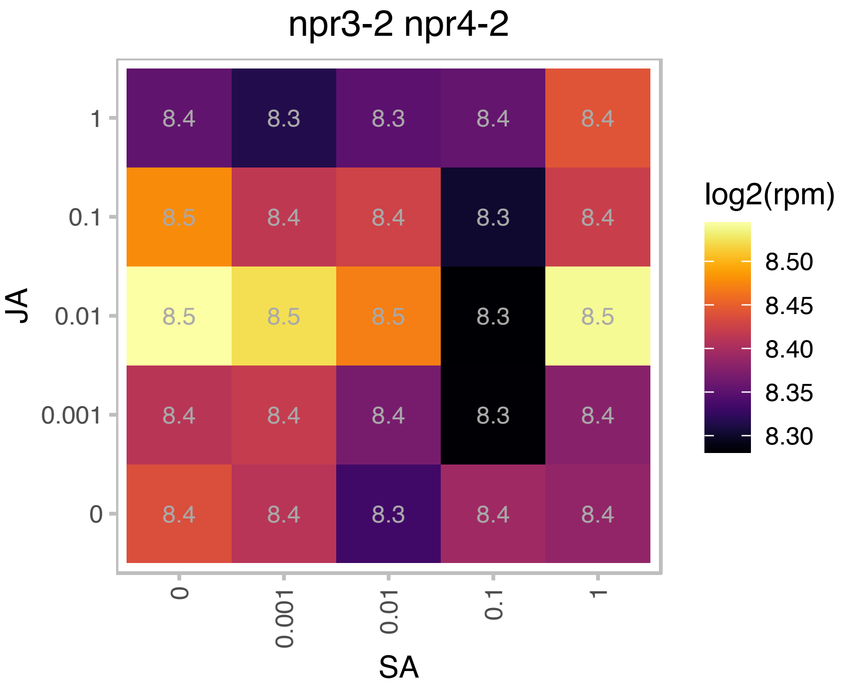


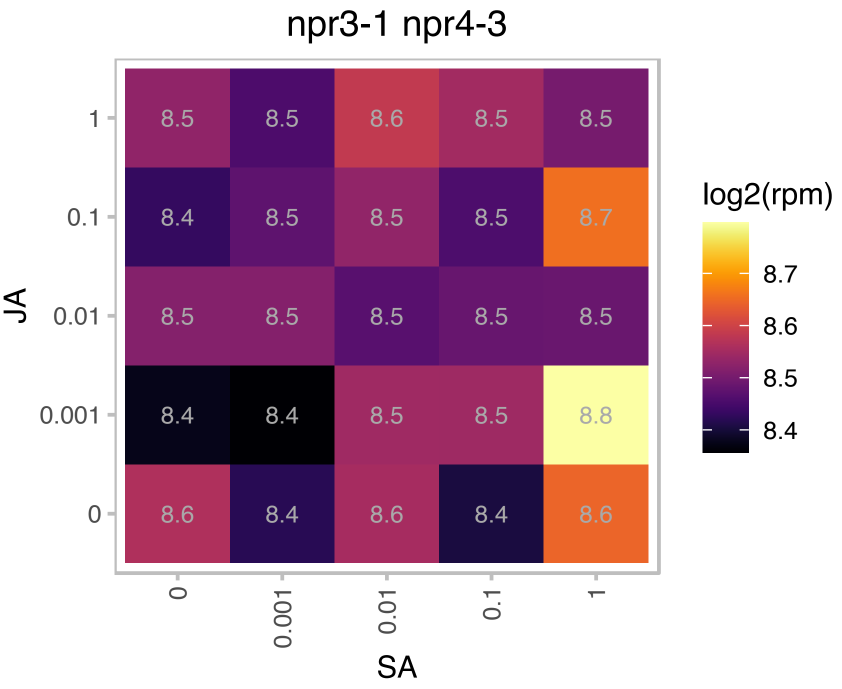

### AT5G52050.1 : AtDTX50, DTX50

MATE efflux family protein

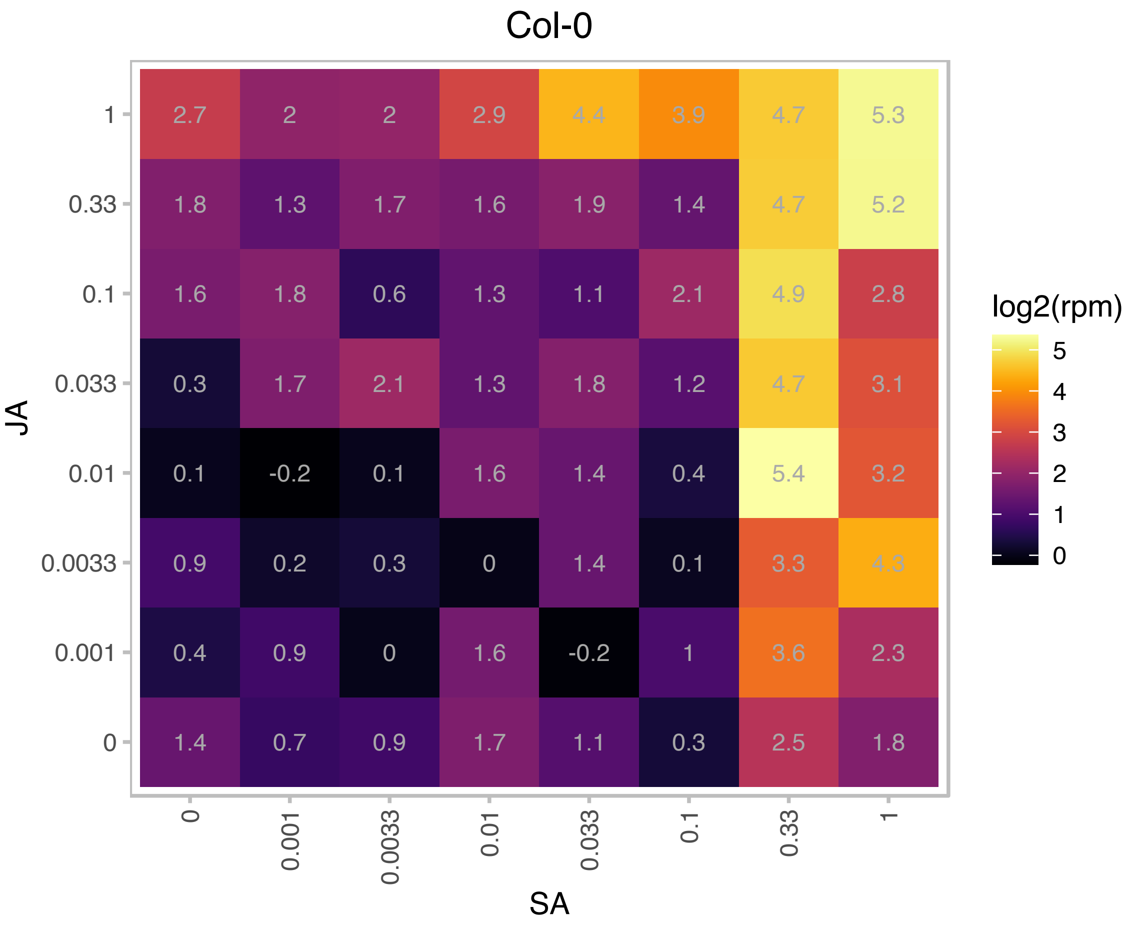


| | |
|---|---|
| Gene | AT5G52050 |
| is.expressed_genes | TRUE |
| is.responsive_genes | TRUE |
| is.apcluster | B-05 |

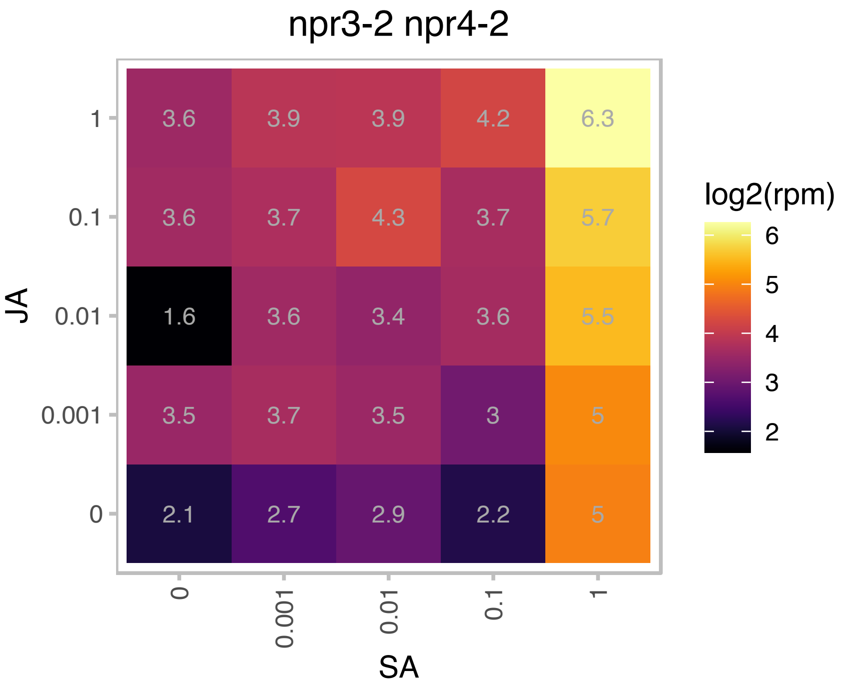


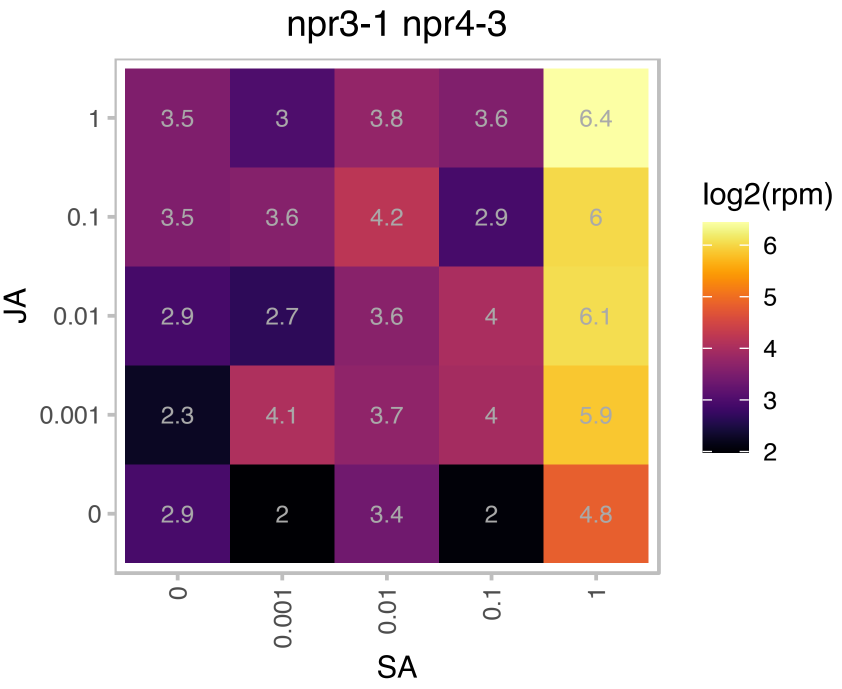

### AT5G52640.1 : ATHS83, AtHsp90-1, ATHSP90.1, HSP81-1

heat shock protein 90.1

Encodes a cytosolic heat shock protein AtHSP90.1.  AtHSP90.1 interacts with disease resistance signaling components SGT1b and RAR1 and is required for RPS2-mediated resistance. The mRNA is cell-to-cell mobile.

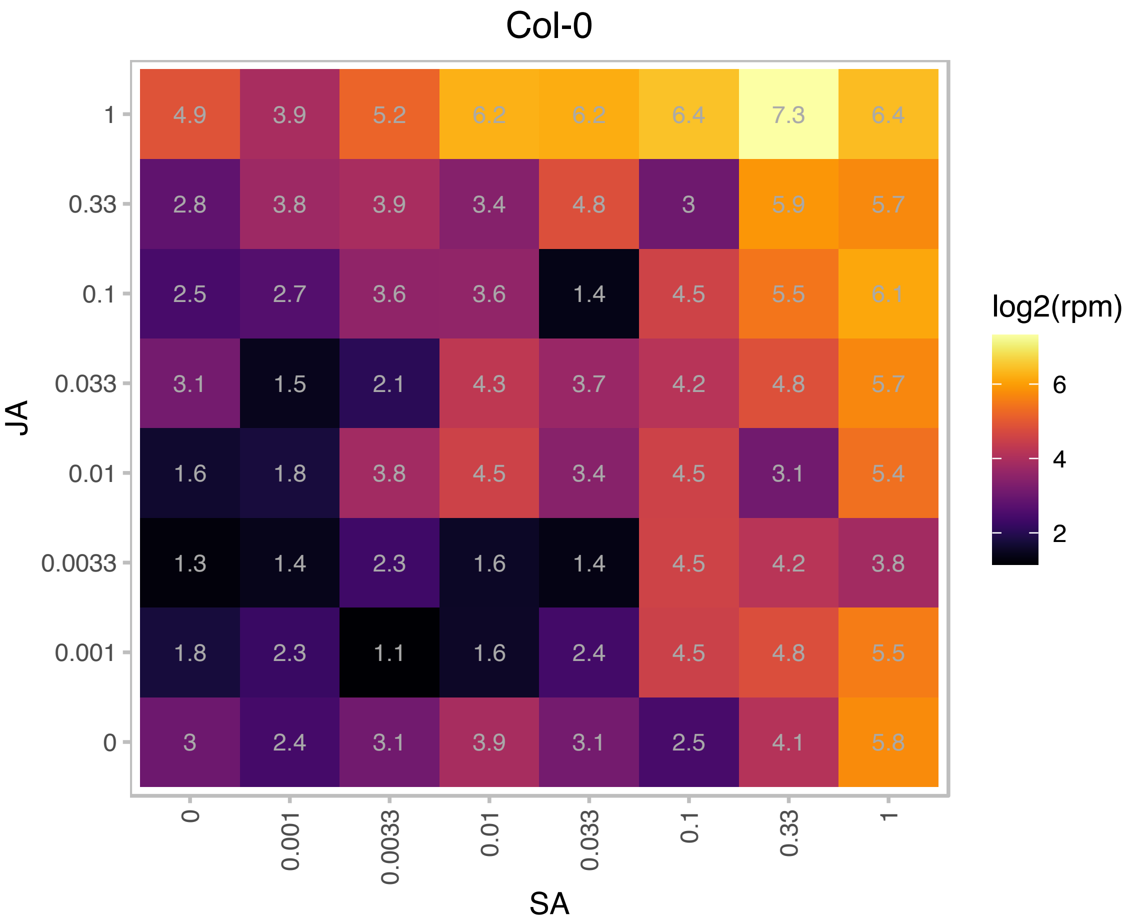


| | |
|---|---|
| Gene | AT5G52640 |
| is.expressed_genes | TRUE |
| is.responsive_genes | TRUE |
| is.apcluster | B-05 |

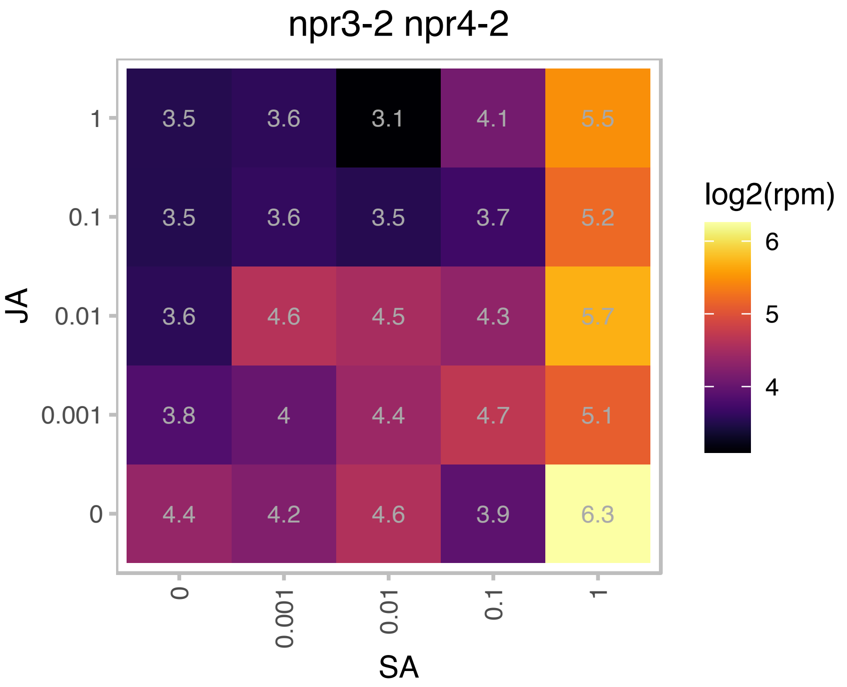


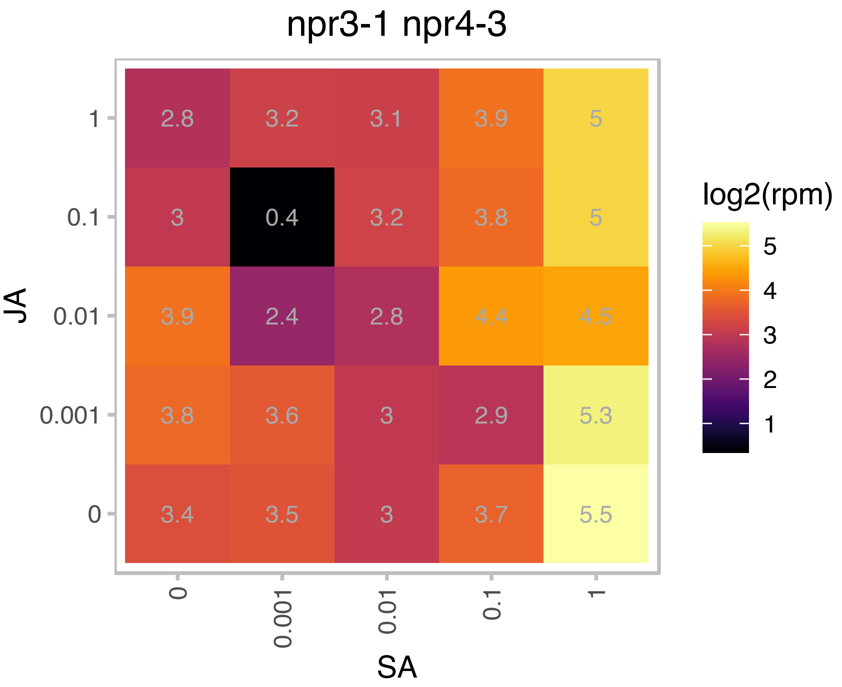

### AT5G53350.1 : CLPX

CLP protease regulatory subunit X

CLP protease regulatory subunit CLPX mRNA, nuclear gene

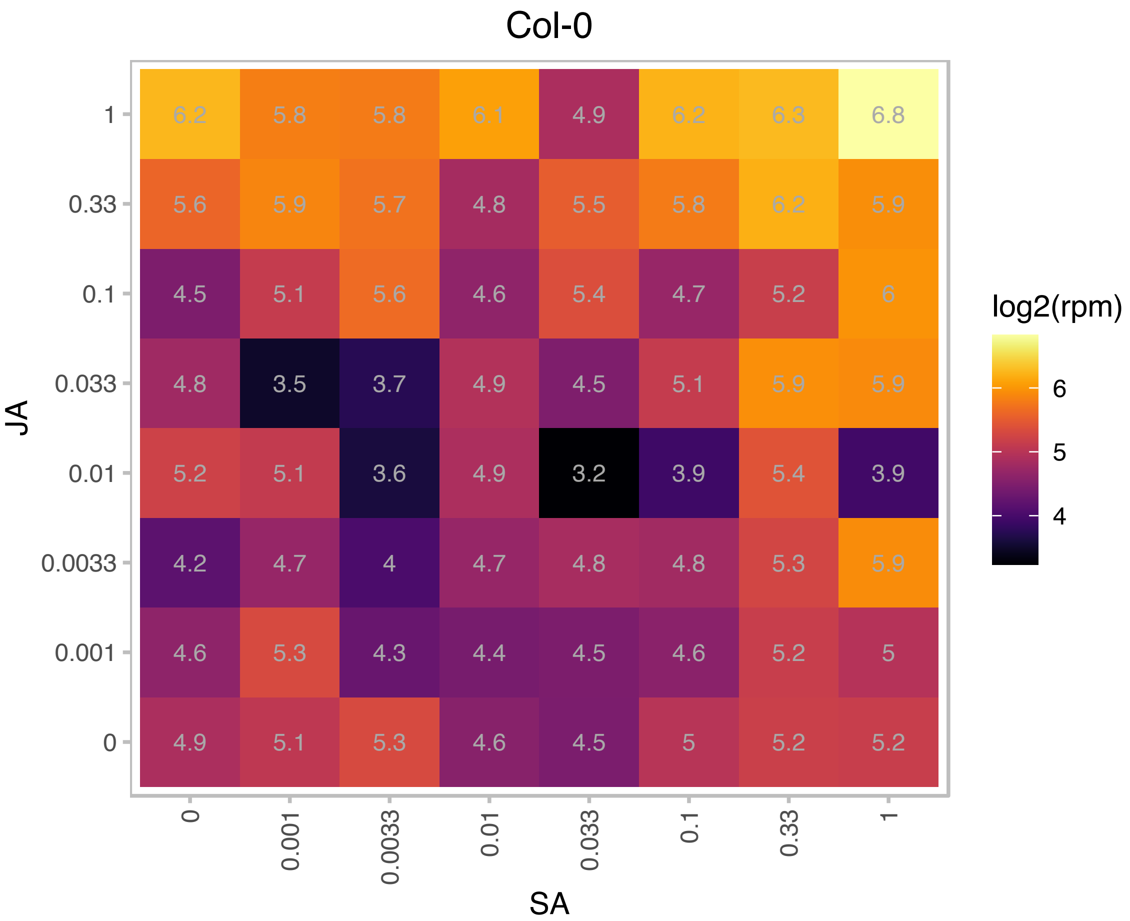


| | |
|---|---|
| Gene | AT5G53350 |
| is.expressed_genes | TRUE |
| is.responsive_genes | TRUE |
| is.apcluster | B-05 |

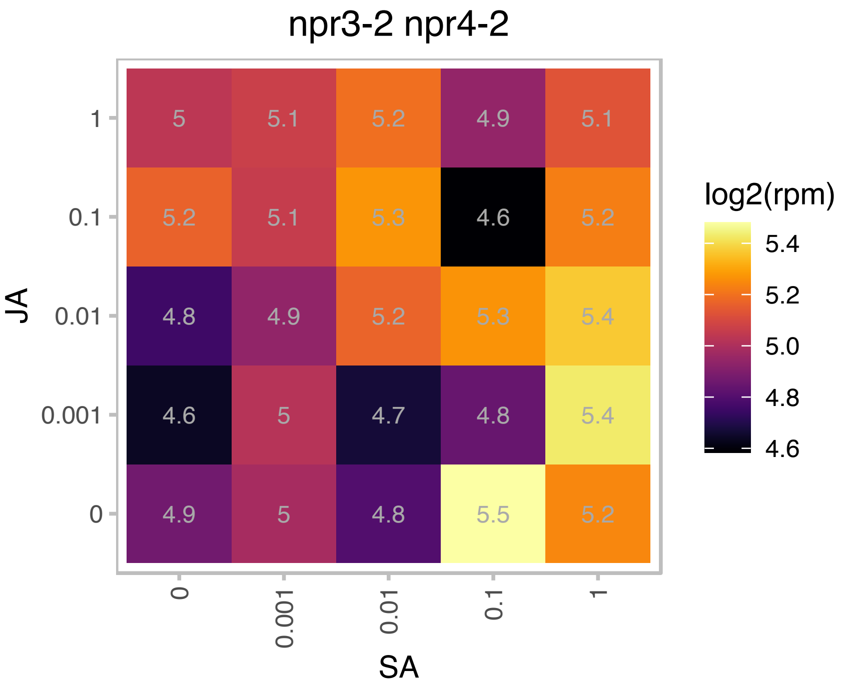


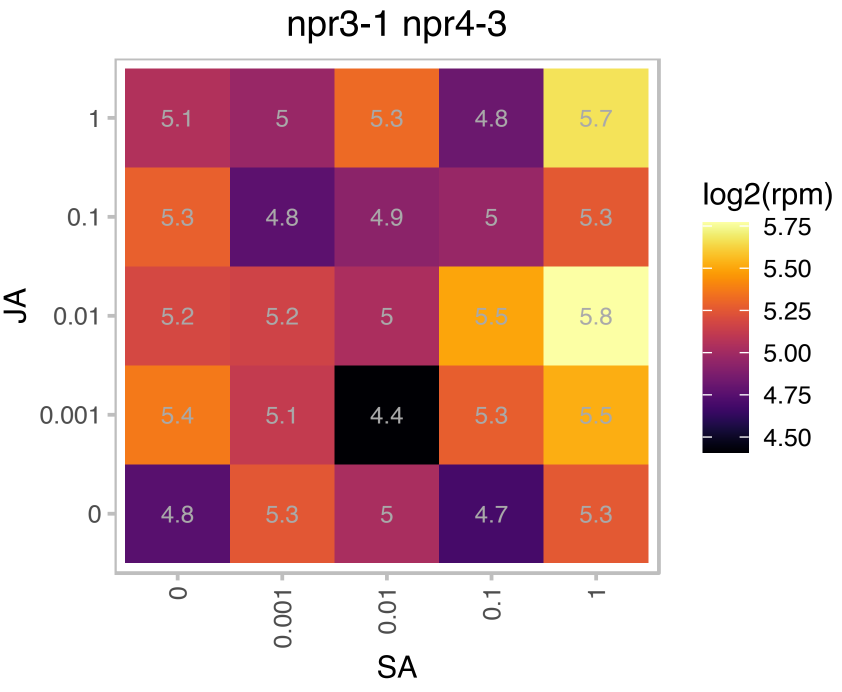

# AT5G54940.1 : NA

Translation initiation factor SUI1 family protein

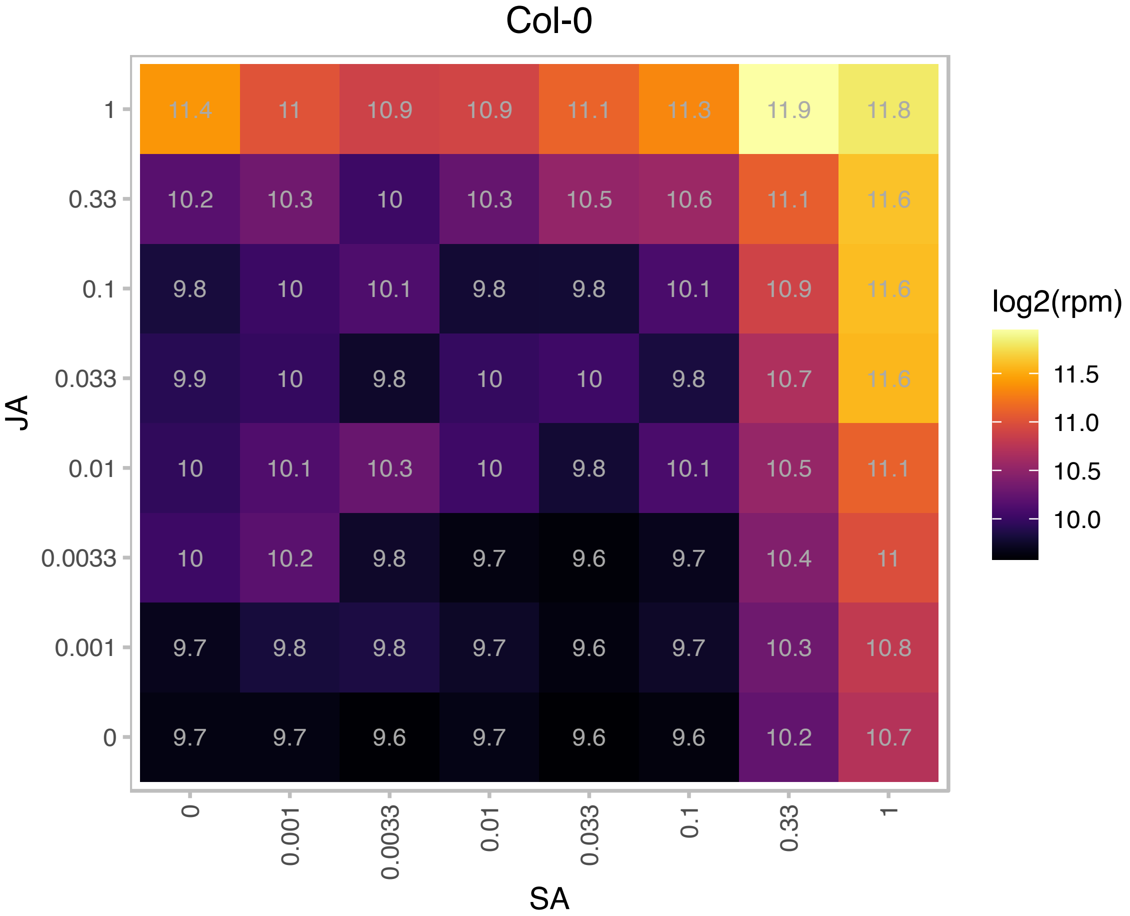


| | |
|---|---|
| Gene | AT5G54940 |
| is.expressed_genes | TRUE |
| is.responsive_genes | TRUE |
| is.apcluster | B-05 |

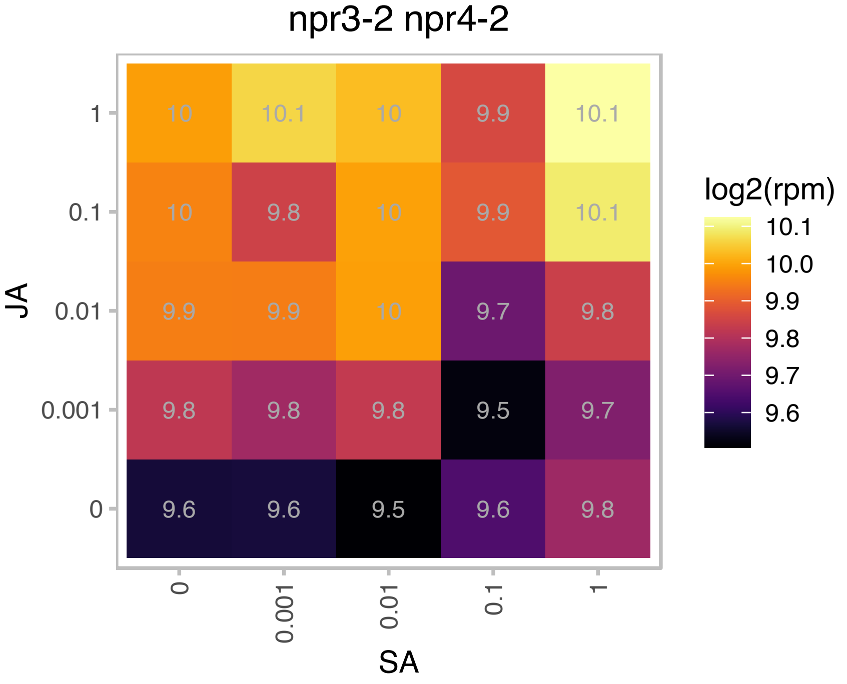


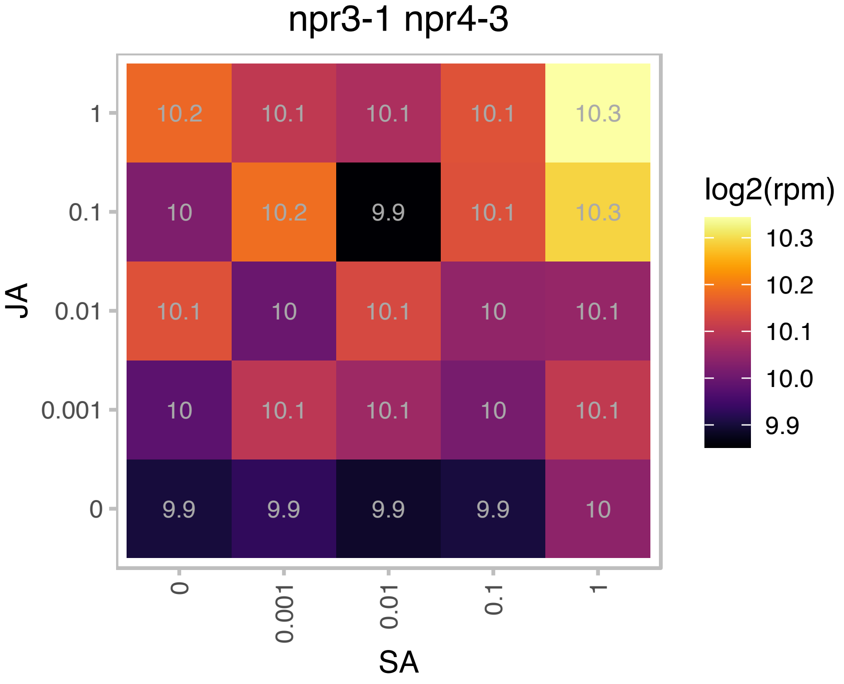

# AT5G56840.1 : NA

myb-like transcription factor family protein

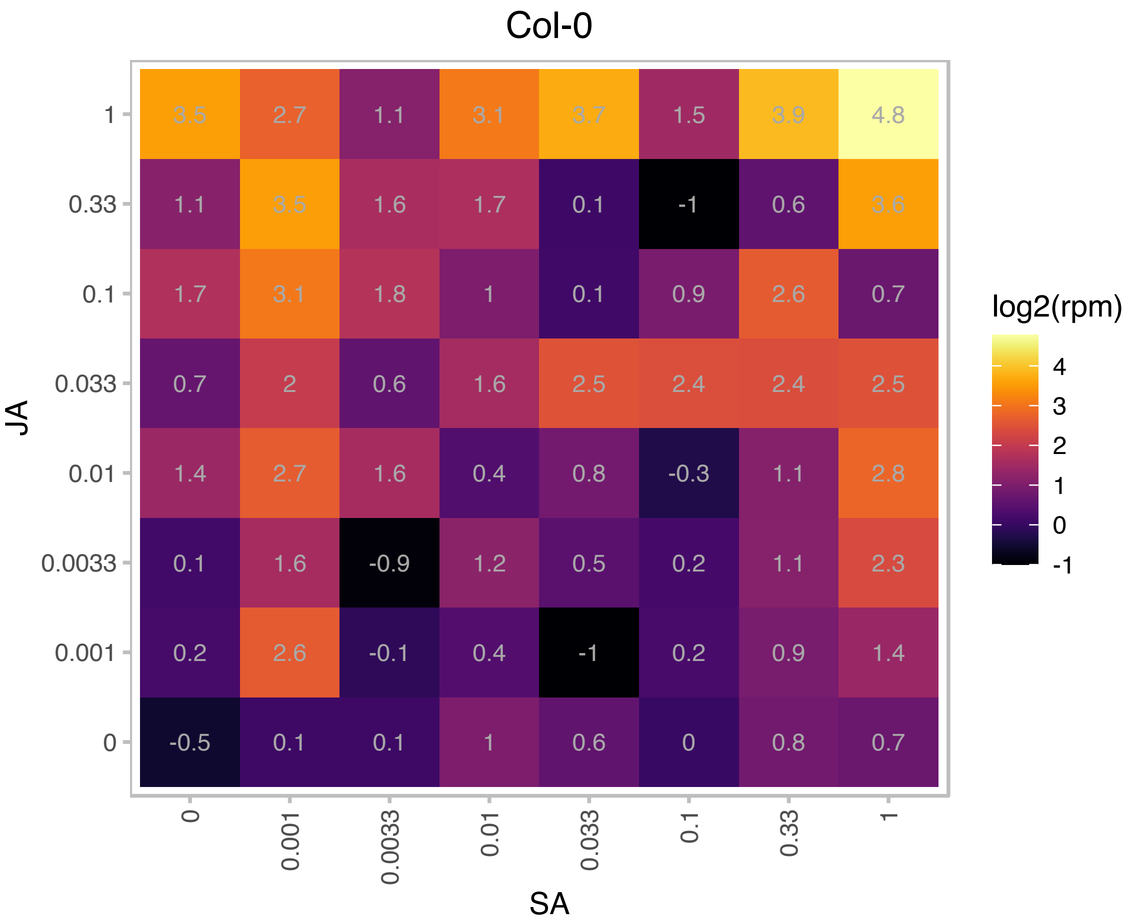


| | |
|---|---|
| Gene | AT5G56840 |
| is.expressed_genes | TRUE |
| is.responsive_genes | TRUE |
| is.apcluster | B-05 |

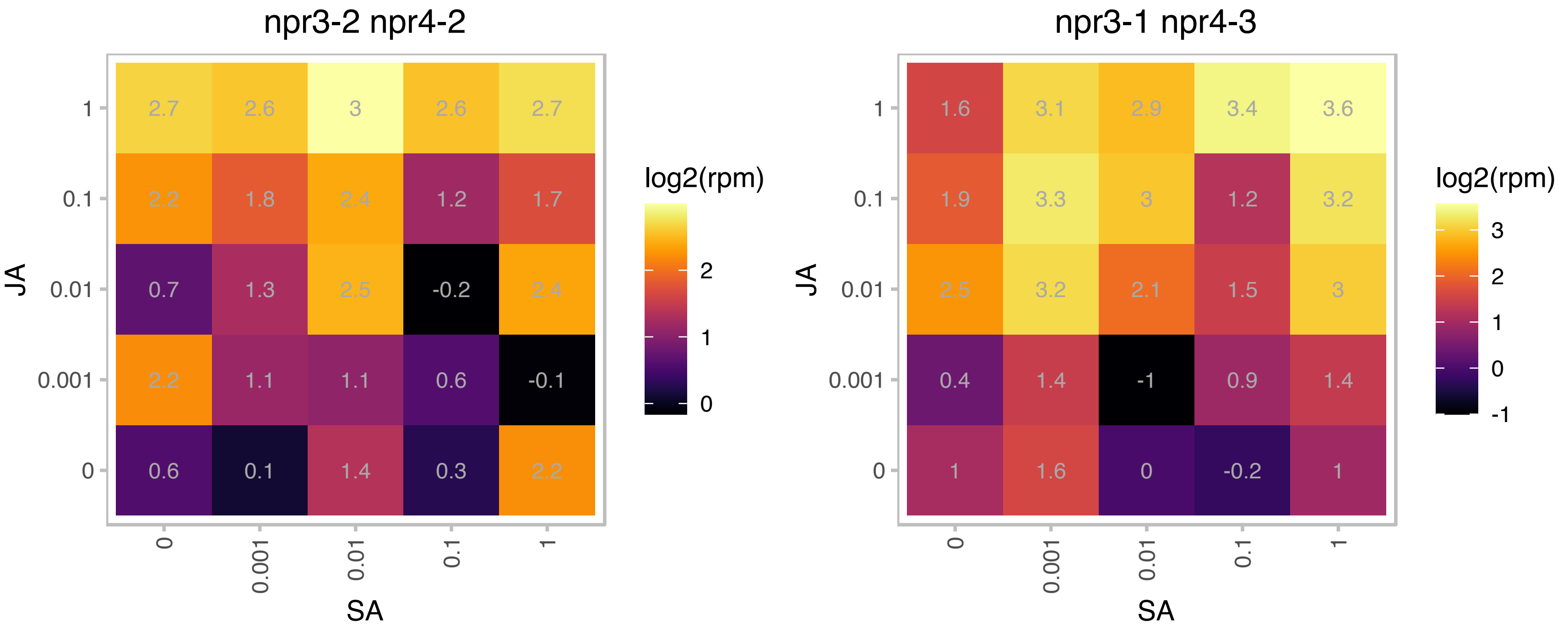

### AT5G58020.1 : AtRTF2, RTF2

Encodes a replication termination factor 2 domain containing protein involved in the regulation of pre-mRNA splicing.

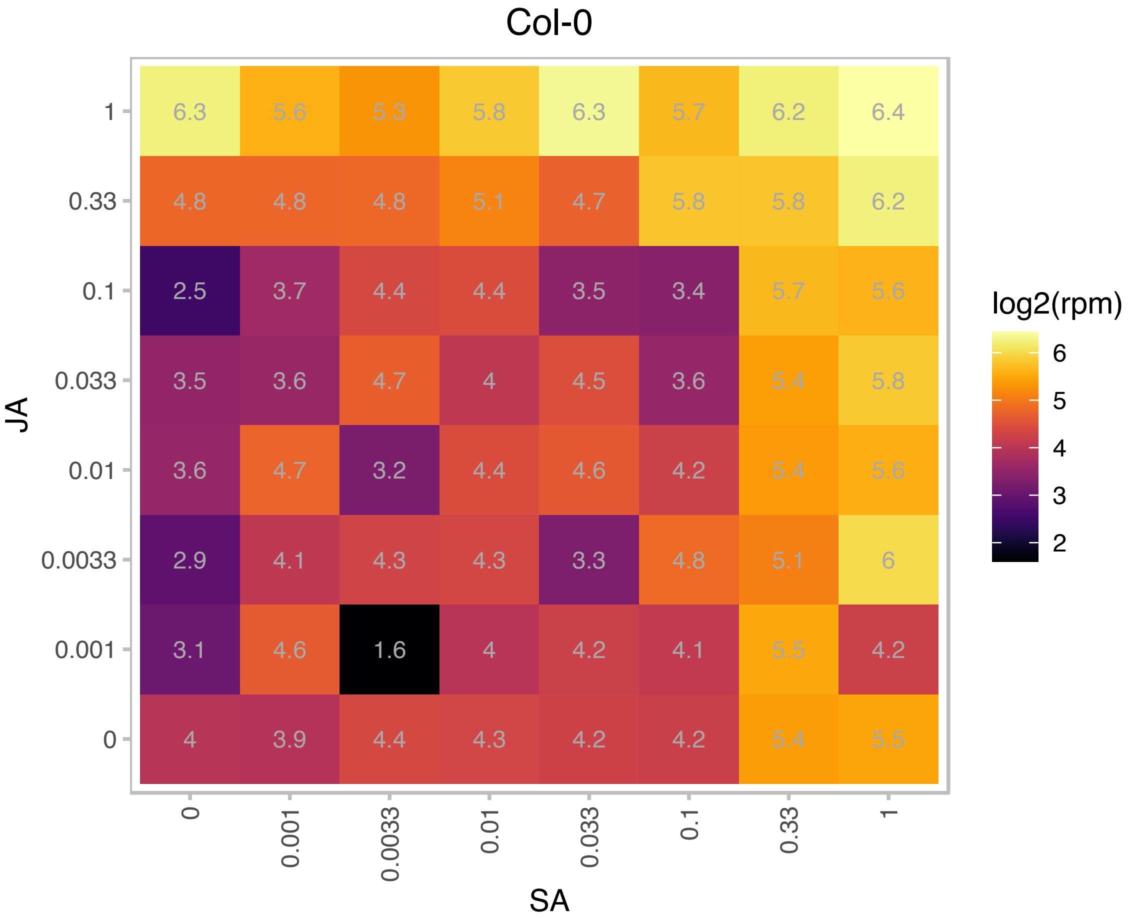


| | |
|---|---|
| Gene | AT5G58020 |
| is.expressed_genes | TRUE |
| is.responsive_genes | TRUE |
| is.apcluster | B-05 |

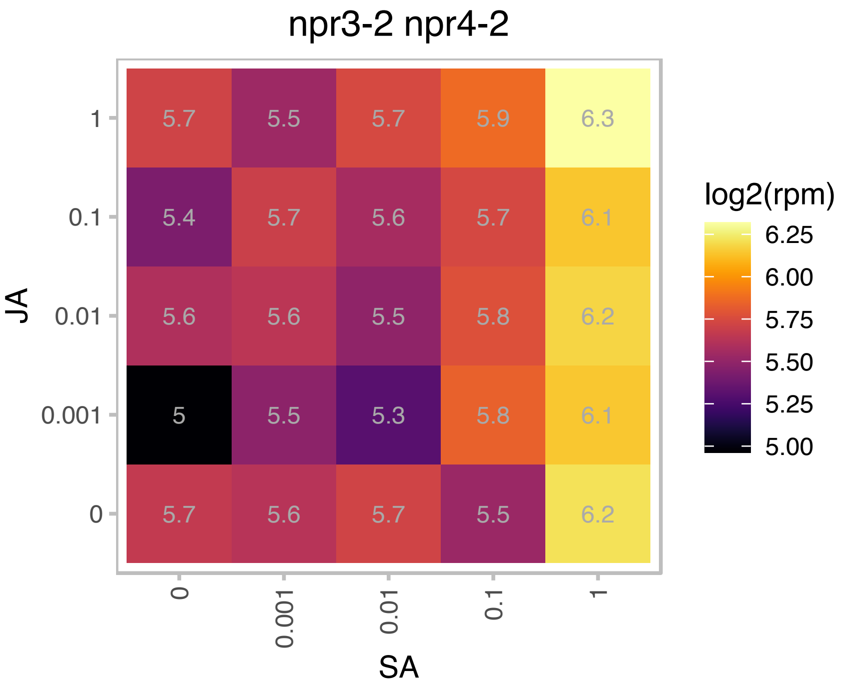


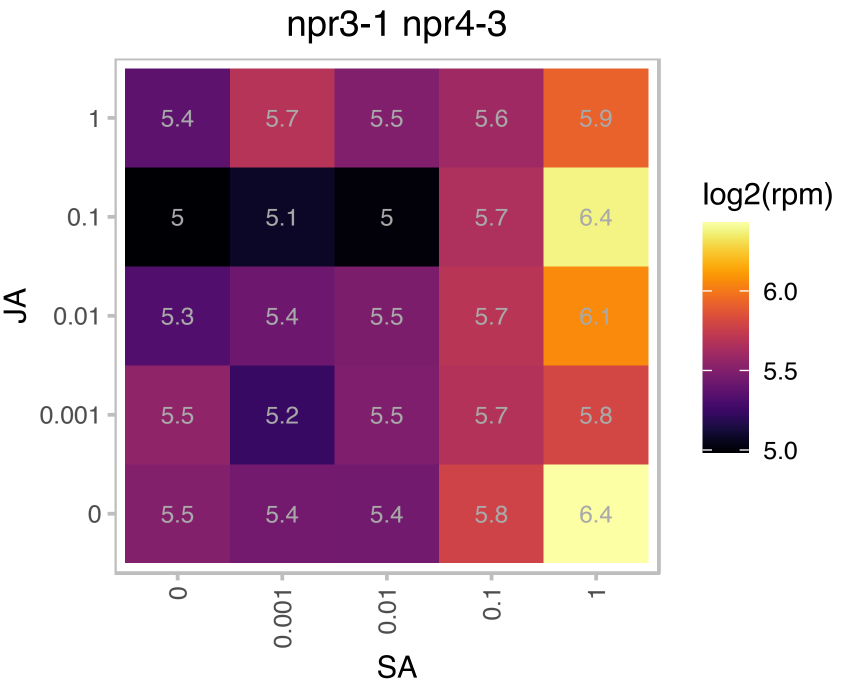

### AT5G58070.1 : ATTIL, TIL

temperature-induced lipocalin

Encodes a temperature-induced lipocalin TIL1.  Involved in thermotolerance.  Peripherally associated with plasma membrane.

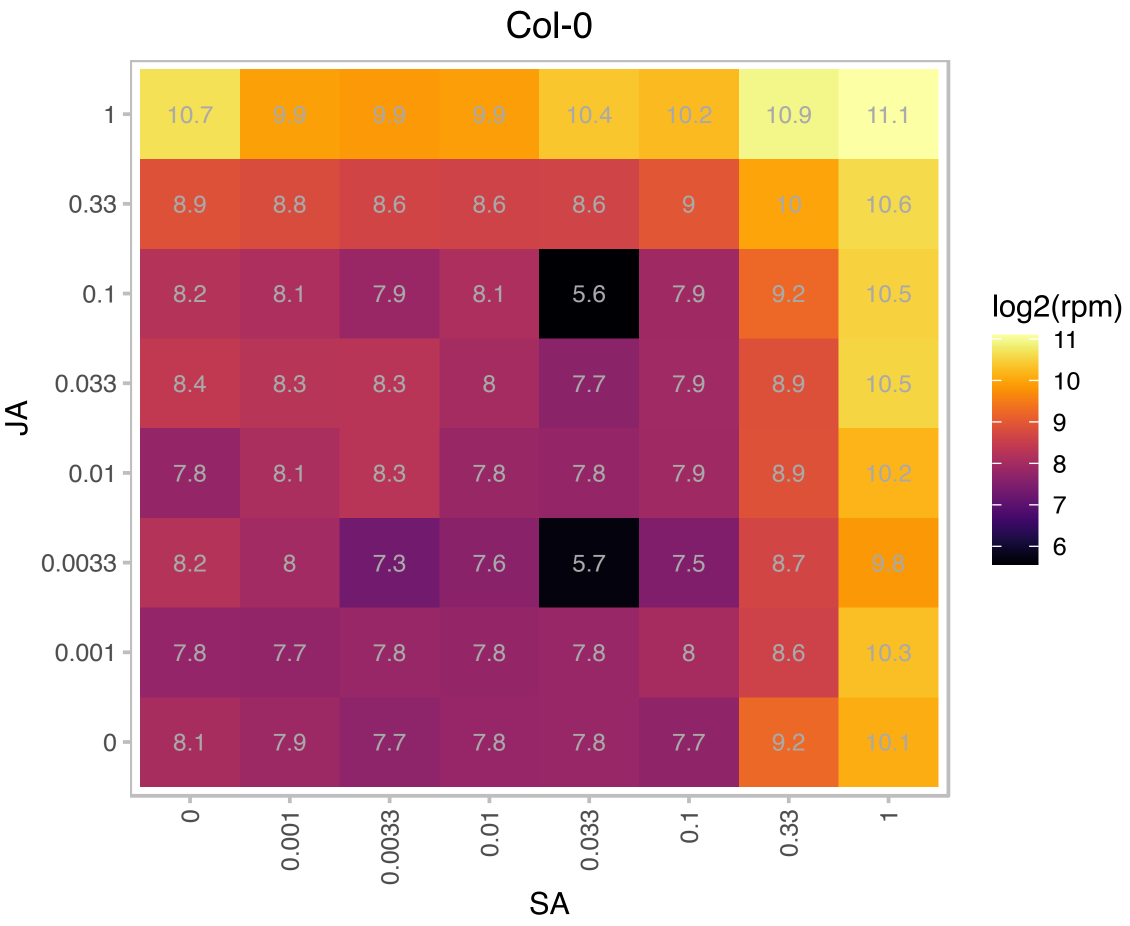


| | |
|---|---|
| Gene | AT5G58070 |
| is.expressed_genes | TRUE |
| is.responsive_genes | TRUE |
| is.apcluster | B-05 |

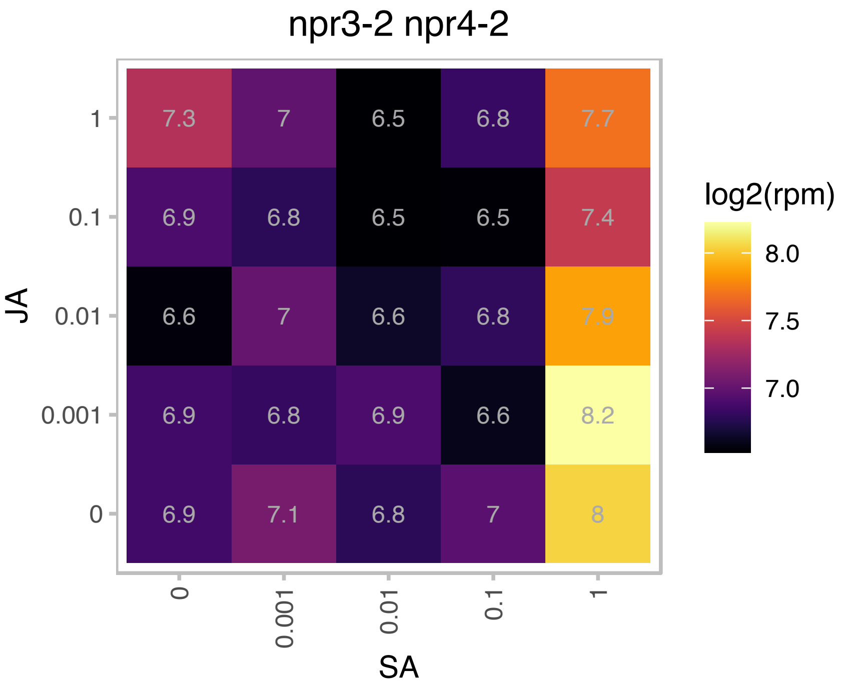


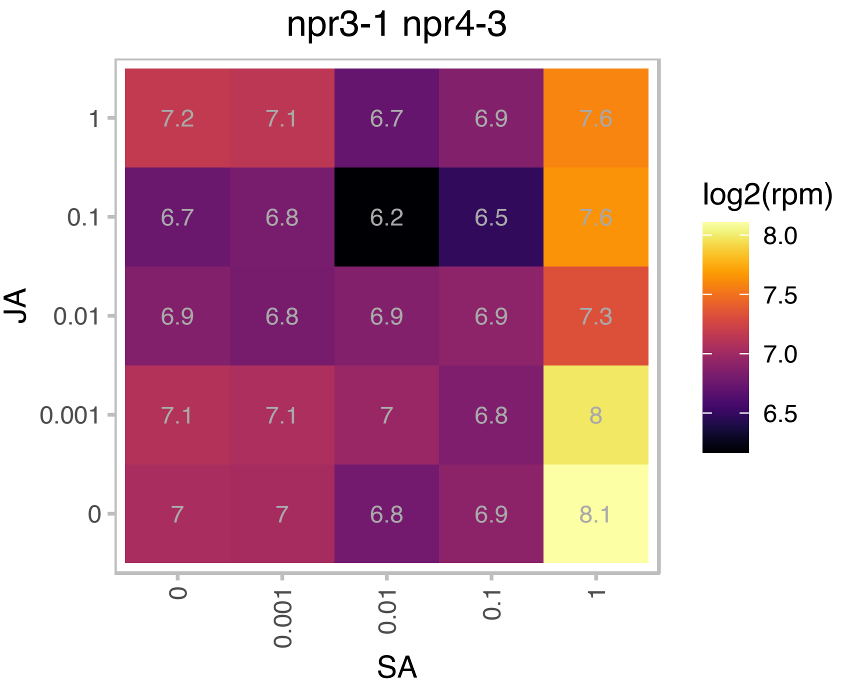

### AT5G60860.1 : AtRABA1f, RABA1f

RAB GTPase homolog A1F

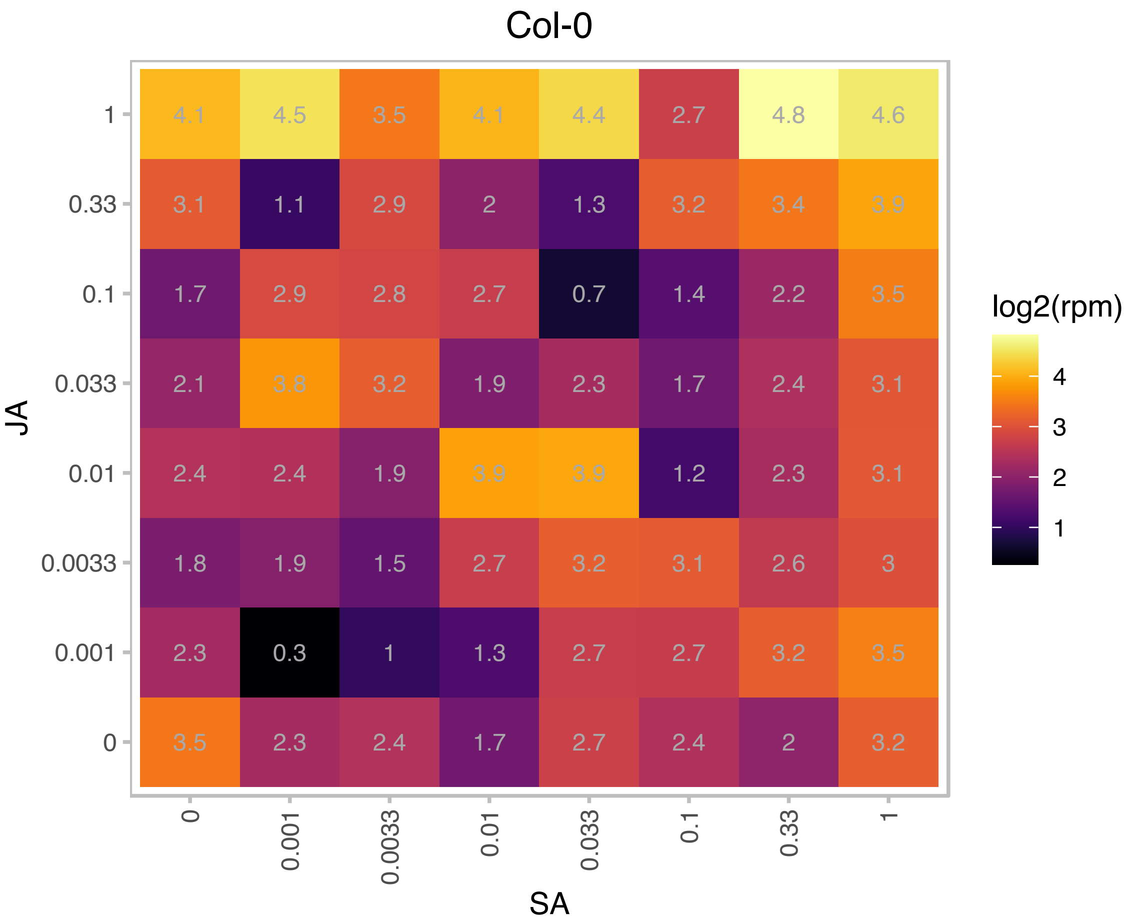


| | |
|---|---|
| Gene | AT5G60860 |
| is.expressed_genes | TRUE |
| is.responsive_genes | TRUE |
| is.apcluster | B-05 |

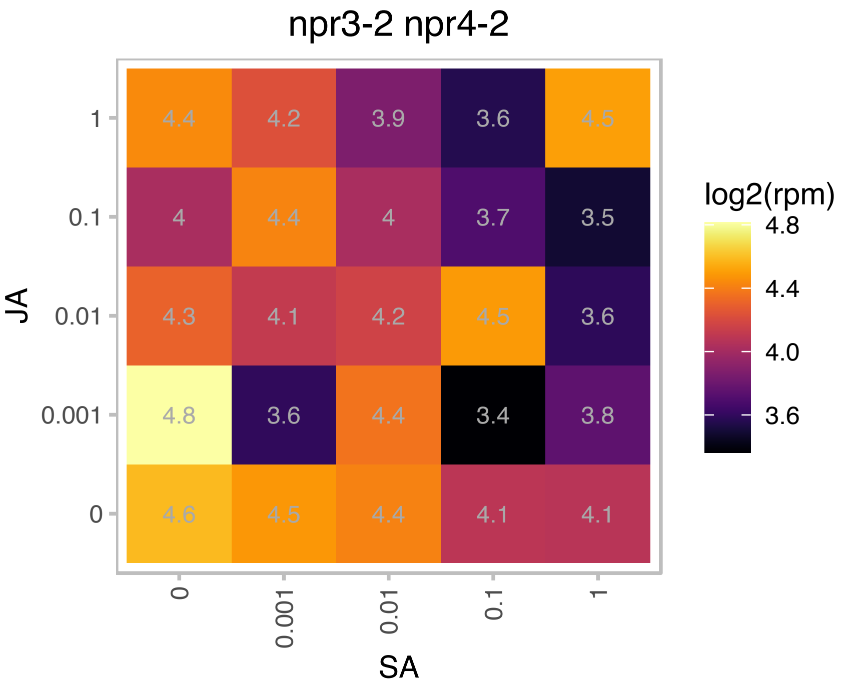


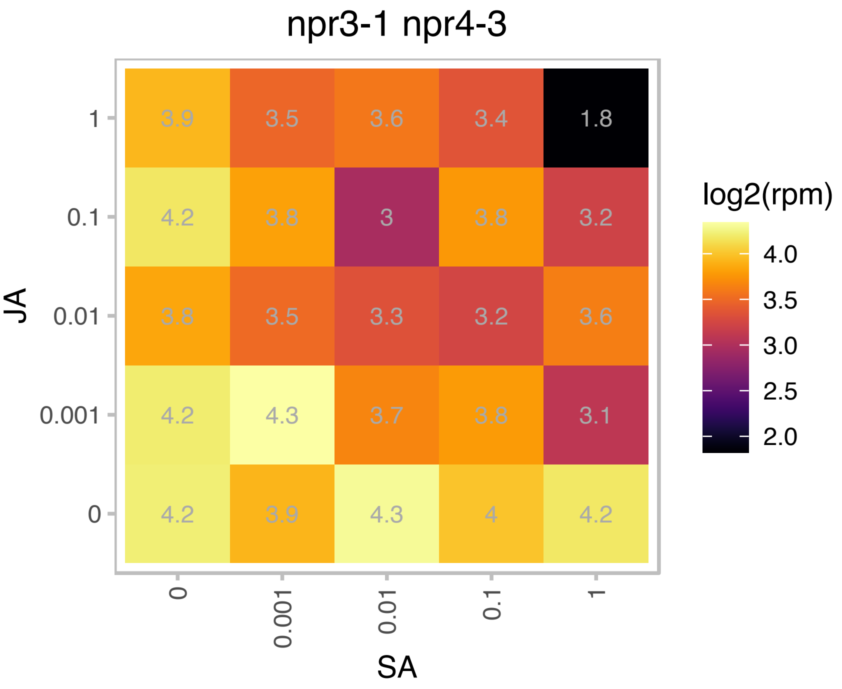

### AT5G62540.1 : UBC3

ubiquitin-conjugating enzyme 3
Encodes a protein predicted to be an E2 ubiquitin conjugating enzyme. It appears homologous to the RAD6 protein in yeast implicated in histone ubiquitination, but, UBC3 has not been experimentally associated with this process.

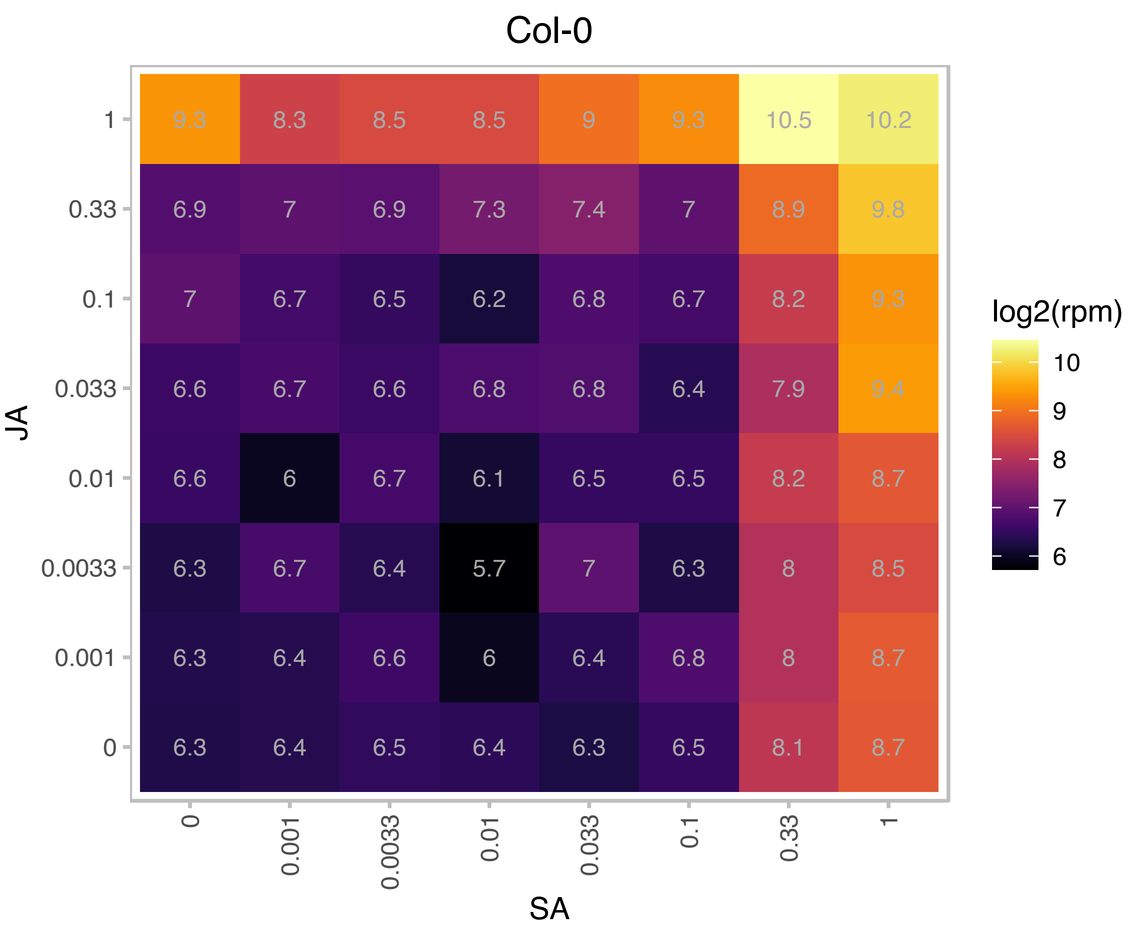


| | |
|---|---|
| Gene | AT5G62540 |
| is.expressed_genes | TRUE |
| is.responsive_genes | TRUE |
| is.apcluster | B-05 |

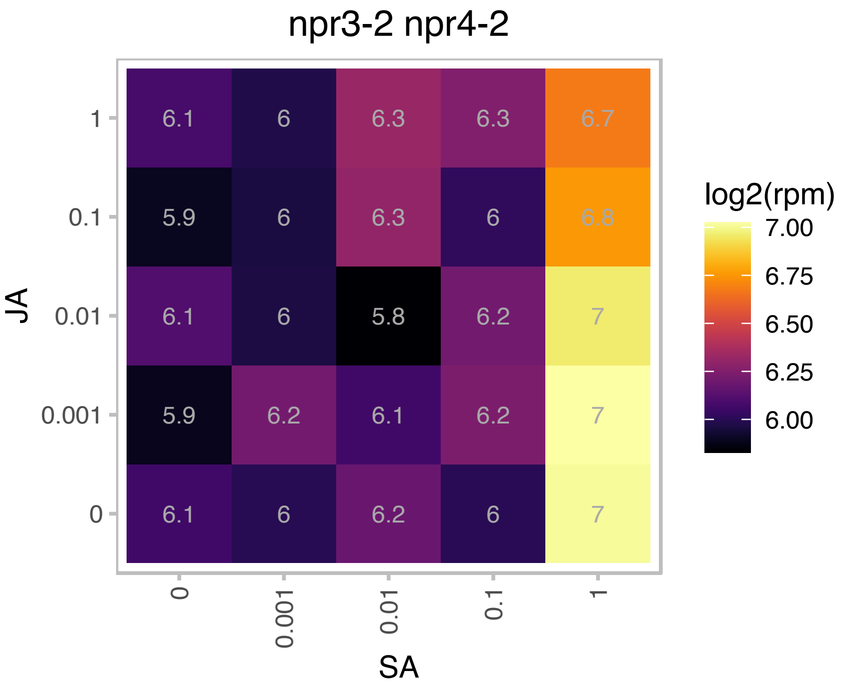


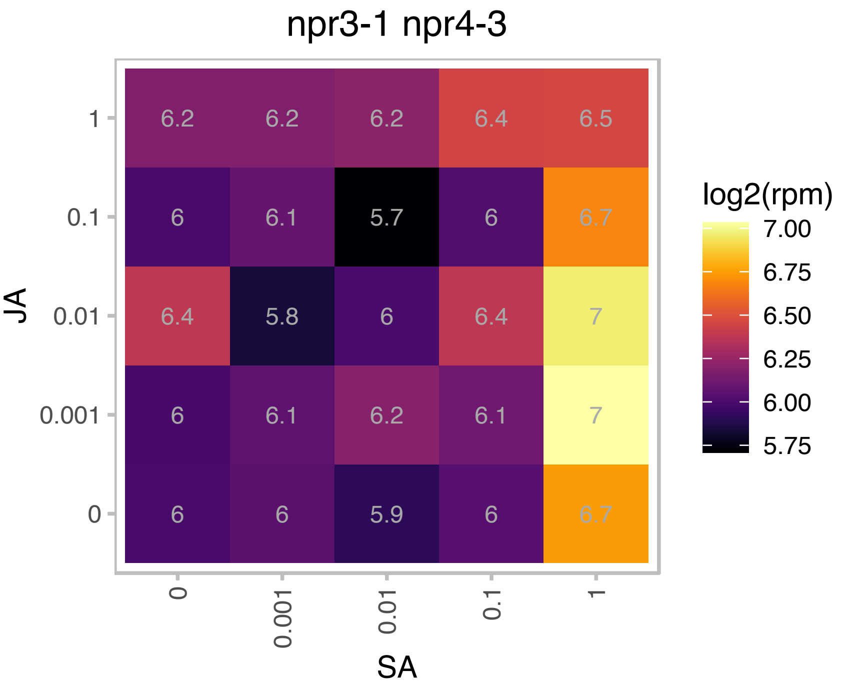

# AT5G64230.1 : NA

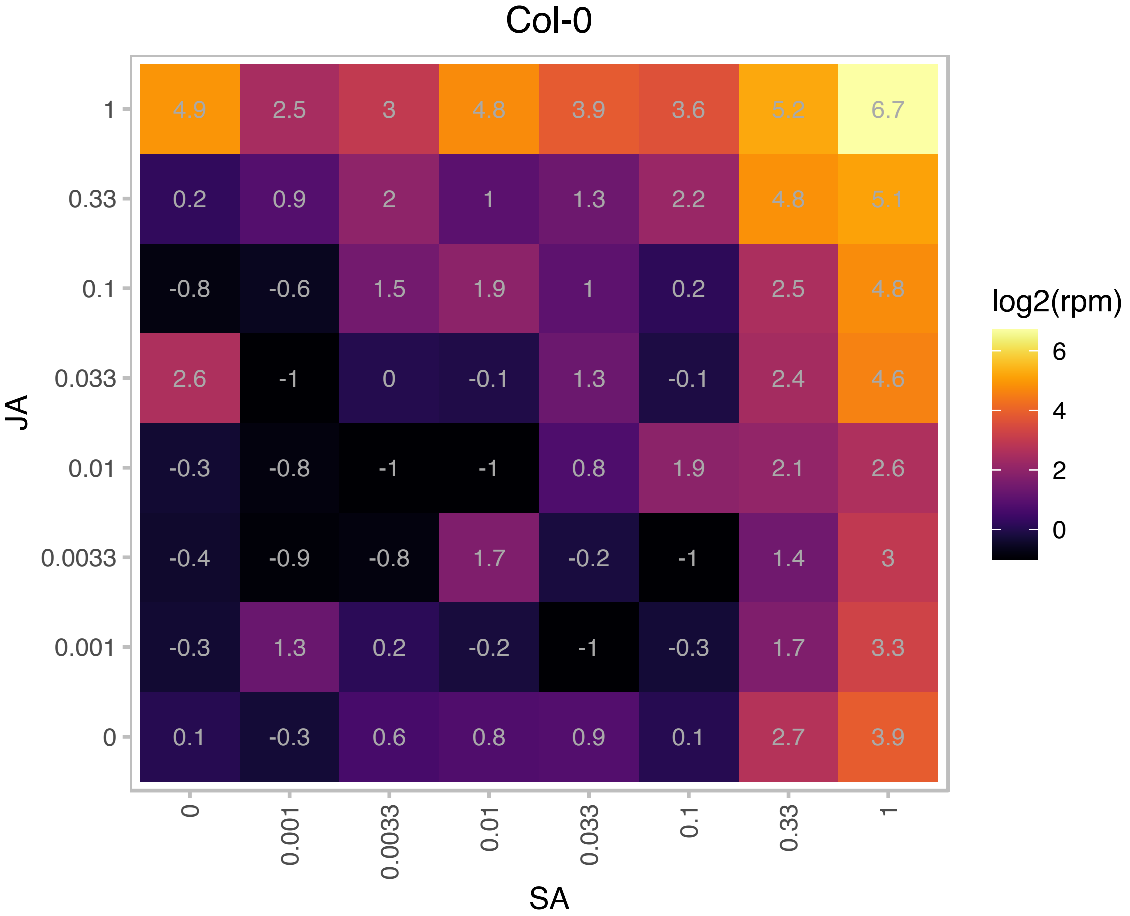


| | |
|---|---|
| Gene | AT5G64230 |
| is.expressed_genes | TRUE |
| is.responsive_genes | TRUE |
| is.apcluster | B-05 |

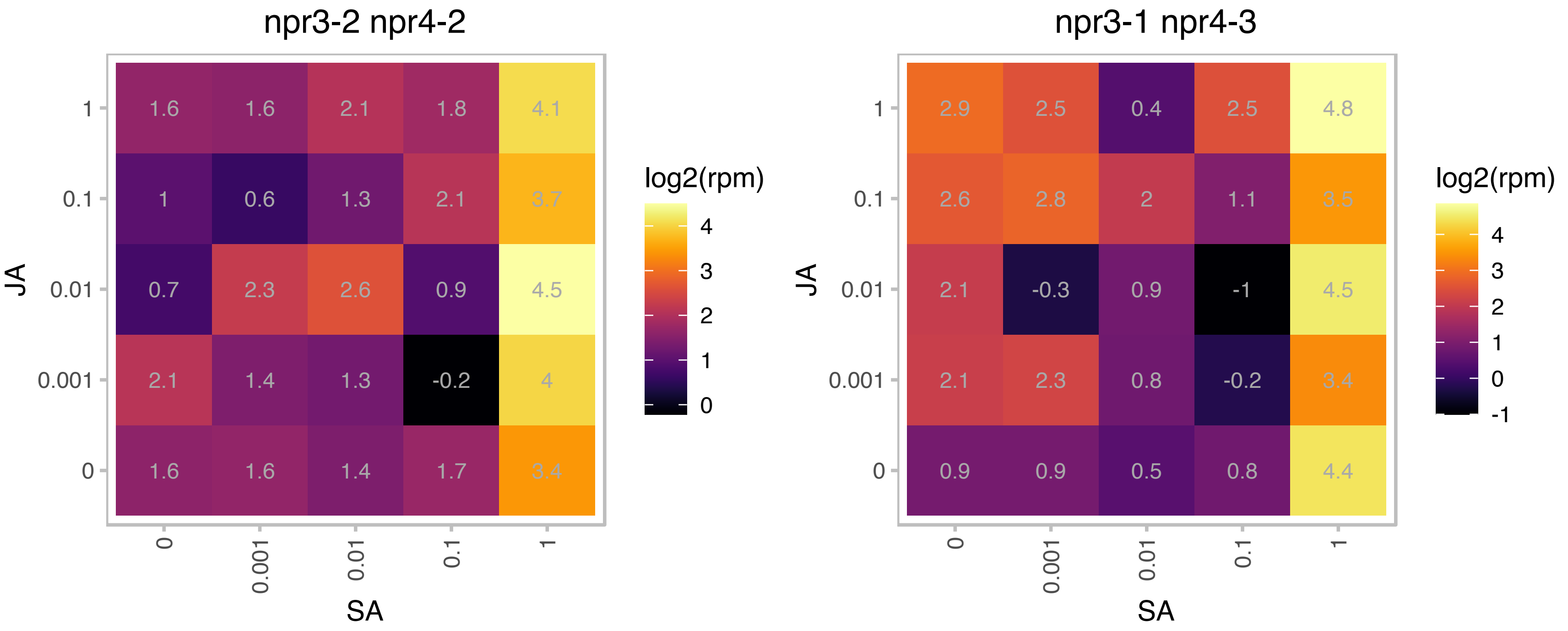

### AT5G66200.1 : ARO2

armadillo repeat only 2

Armadillo repeat protein. One of a family of four in Arabidopsis. Expressed in vegetative tissues, anthers and ovules.

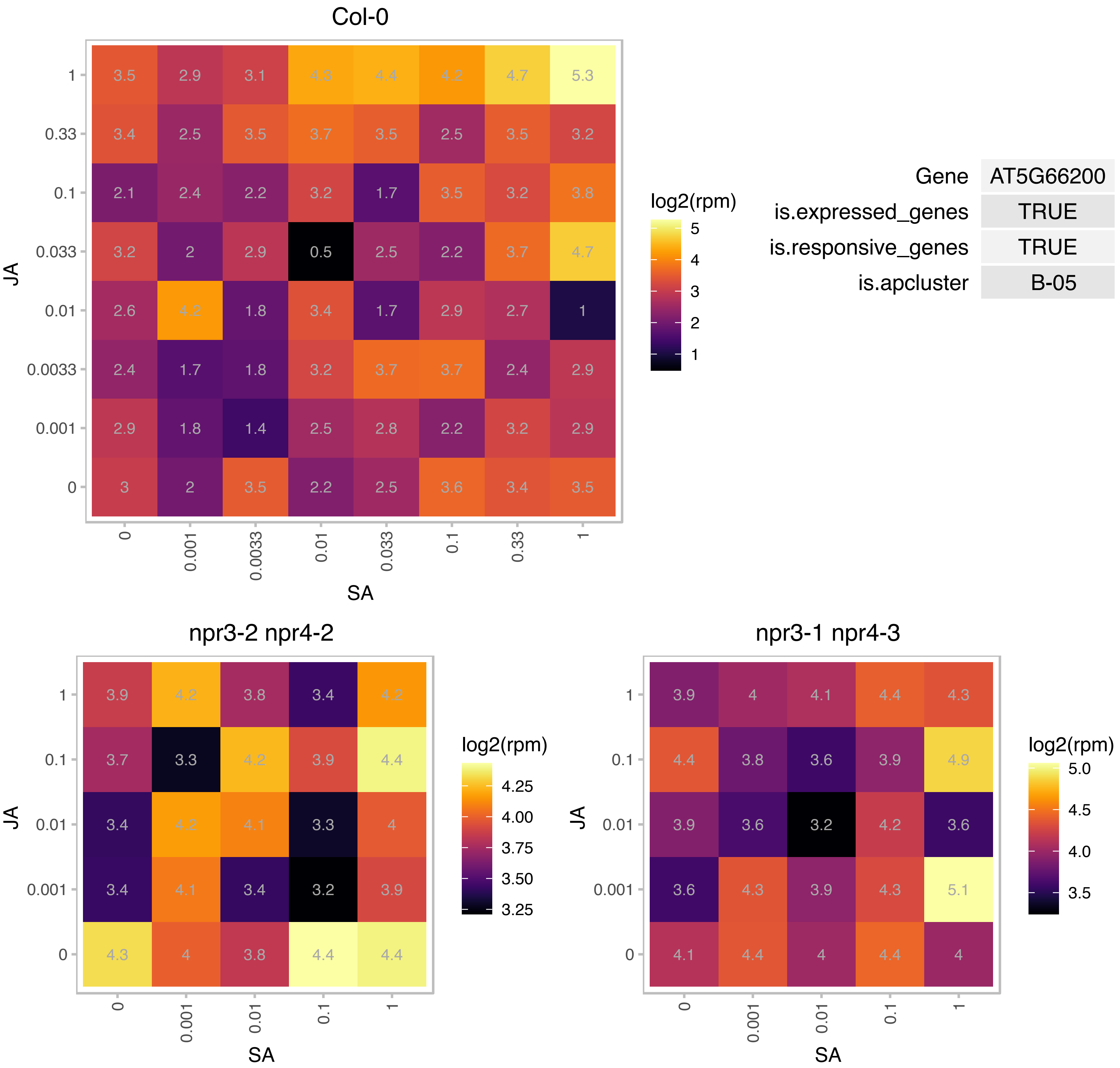


| | |
|---|---|
| Gene | AT5G66200 |
| is.expressed_genes | TRUE |
| is.responsive_genes | TRUE |
| is.apcluster | B-05 |

### AT1G02450.1 : NIMIN-1, NIMIN1

NIM1-interacting 1

NIMIN1 modulates PR gene expression according the following model: NPR1 forms a ternary complex with NIMIN1 and TGA factors upon SAR induction that binds to a positive regulatory cis-element of the PR-1 promoter, termed LS7. This leads to PR-1 gene induction. NIMIN1 decreases transcriptional activation, possibly through its EAR motif, which results in fine-tuning of PR-1 gene expression.

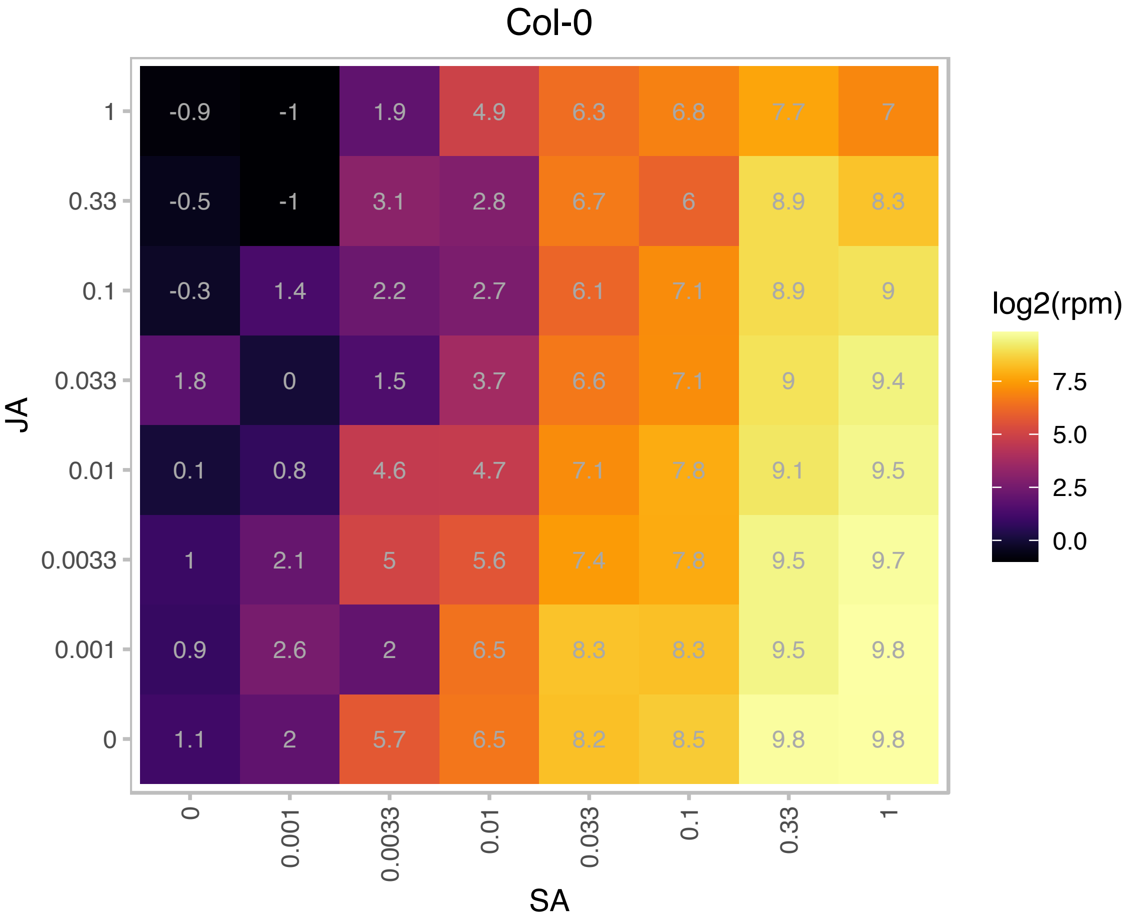


| | |
|---|---|
| Gene | AT1G02450 |
| is.expressed_genes | TRUE |
| is.responsive_genes | TRUE |
| is.apcluster | C-01 |

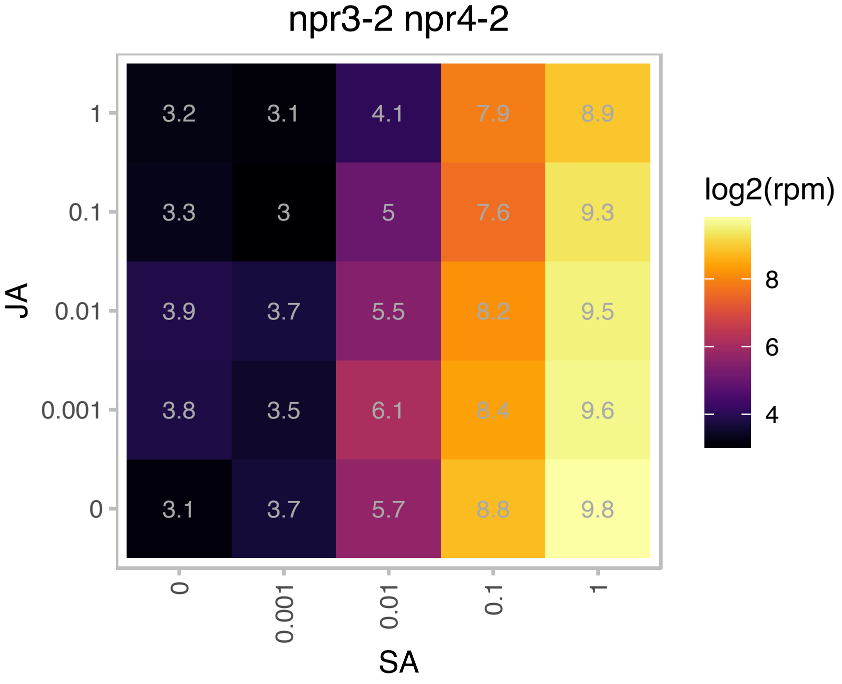


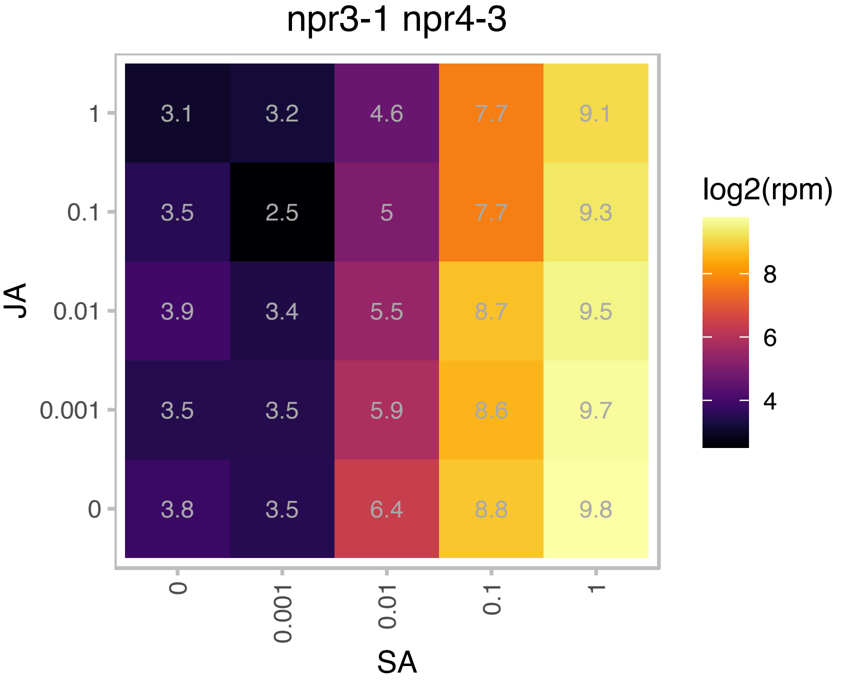

# AT1G02470.1 : NA

Polyketide cyclase/dehydrase and lipid transport superfamily protein

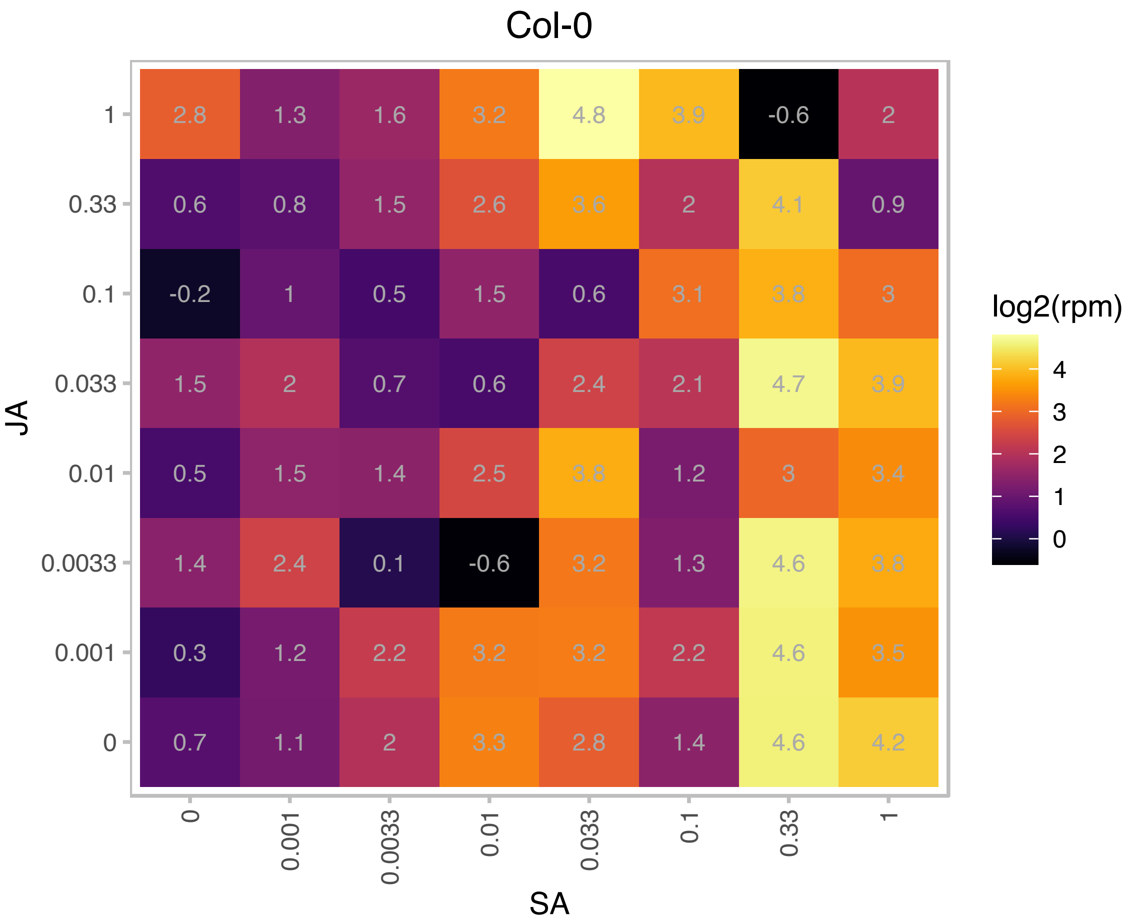


| | |
|---|---|
| Gene | AT1G02470 |
| is.expressed_genes | TRUE |
| is.responsive_genes | TRUE |
| is.apcluster | C-01 |

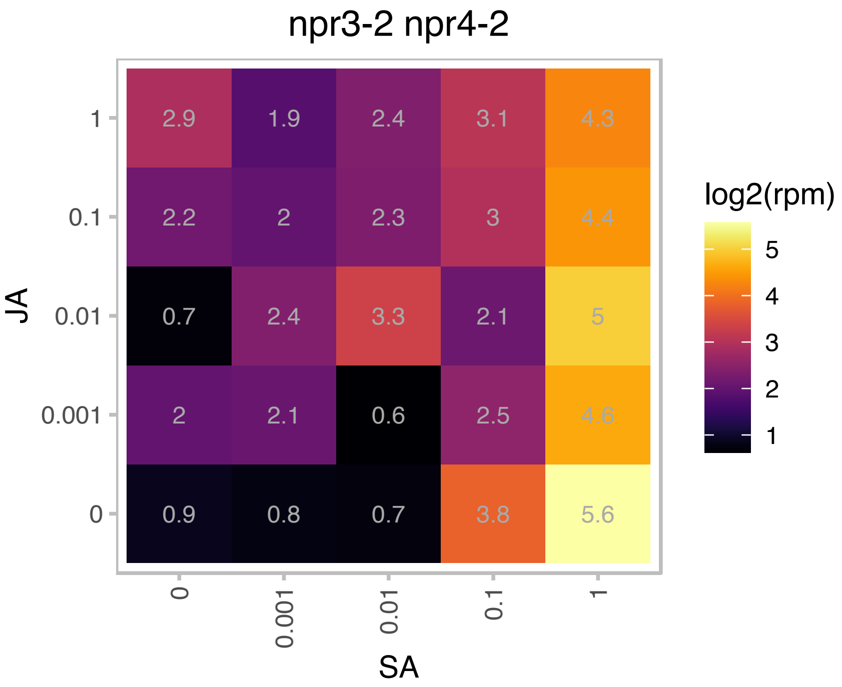


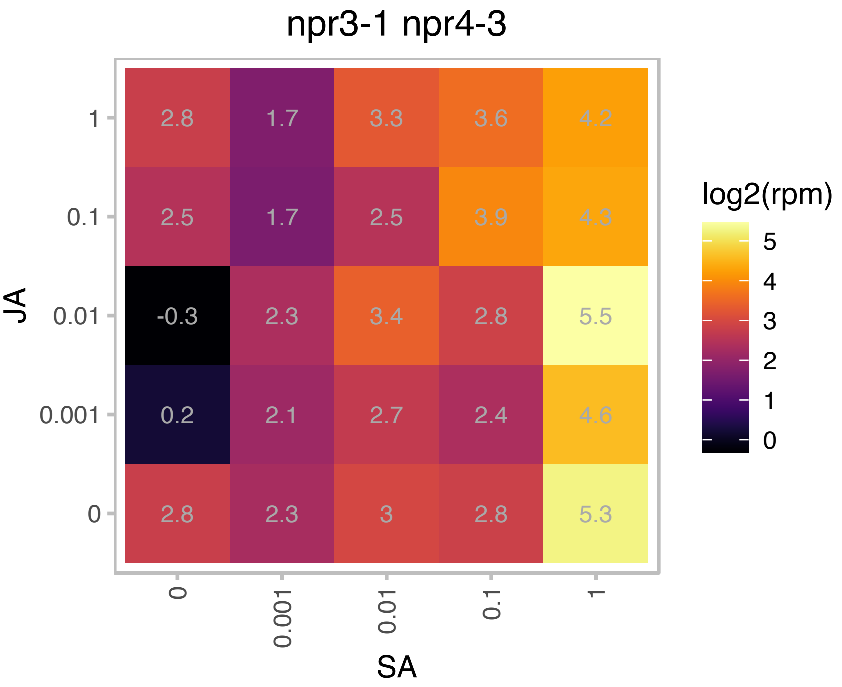

### AT1G04260.1 : MPI7, MPIP7, PRA1.D

CAMV movement protein interacting protein 7
Encodes protein that interacts with CaMV movement protein. Colocalizes in the cytoplasm with the movement protein. Has similarity to mammalian proteins (such as the rat PRA1) which have been described as rab acceptors.

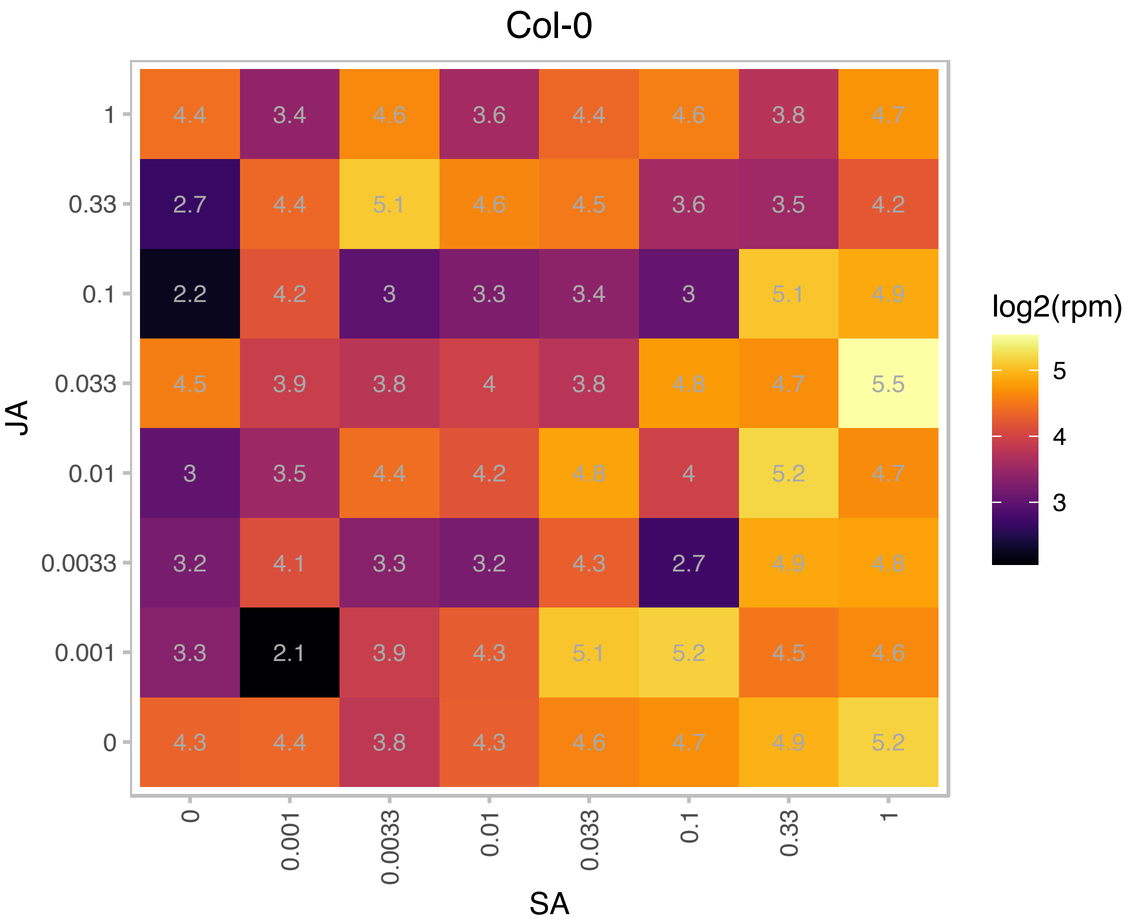


| | |
|---|---|
| Gene | AT1G04260 |
| is.expressed_genes | TRUE |
| is.responsive_genes | TRUE |
| is.apcluster | C-01 |

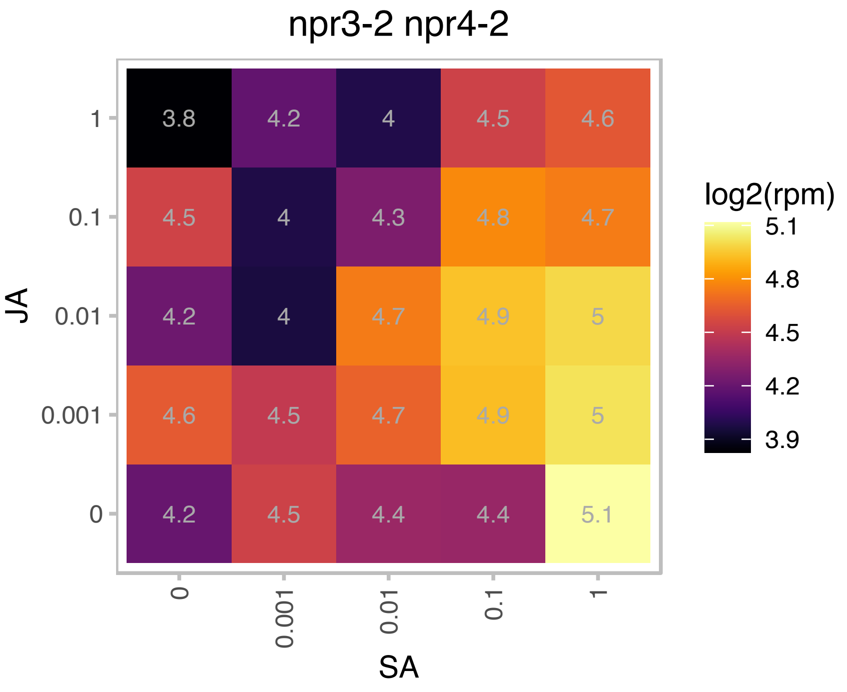


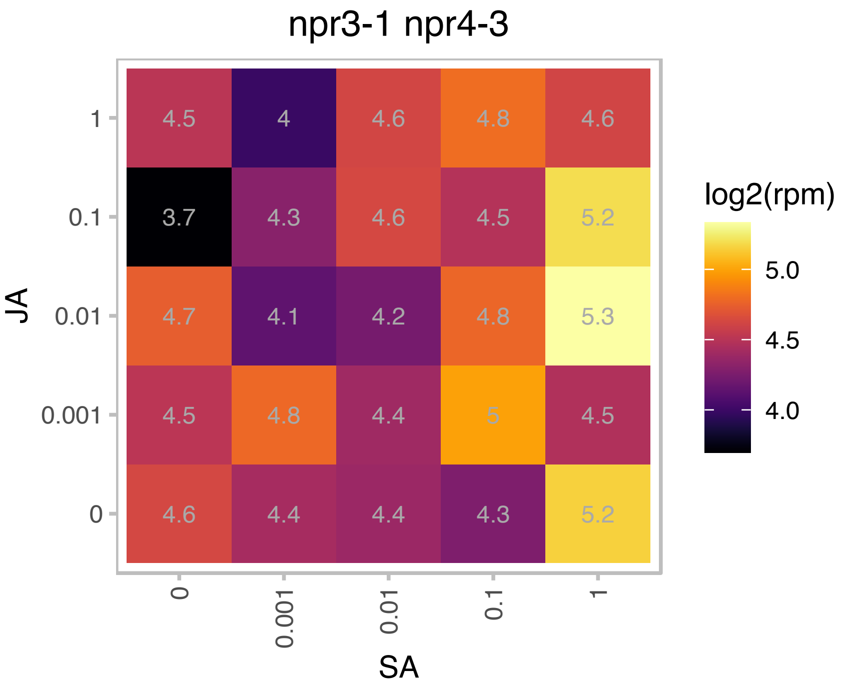

### AT1G07000.1 : ATEXO70B2, EXO70B2

exocyst subunit exo70 family protein B2

A member of EXO70 gene family, putative exocyst subunits, conserved in land plants. Arabidopsis thaliana contains 23 putative EXO70 genes, which can be classified into eight clusters on the phylogenetic tree.

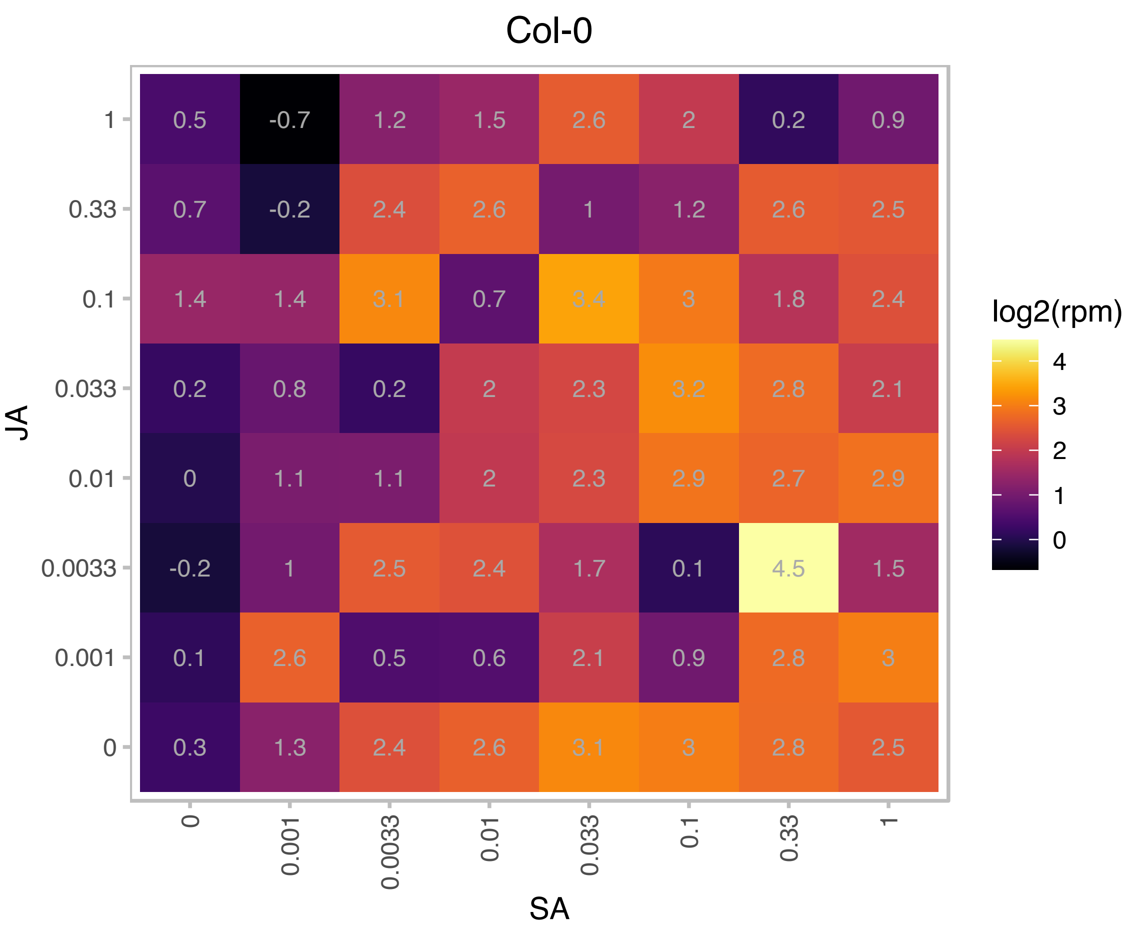


| | |
|---|---|
| Gene | AT1G07000 |
| is.expressed_genes | TRUE |
| is.responsive_genes | TRUE |
| is.apcluster | C-01 |

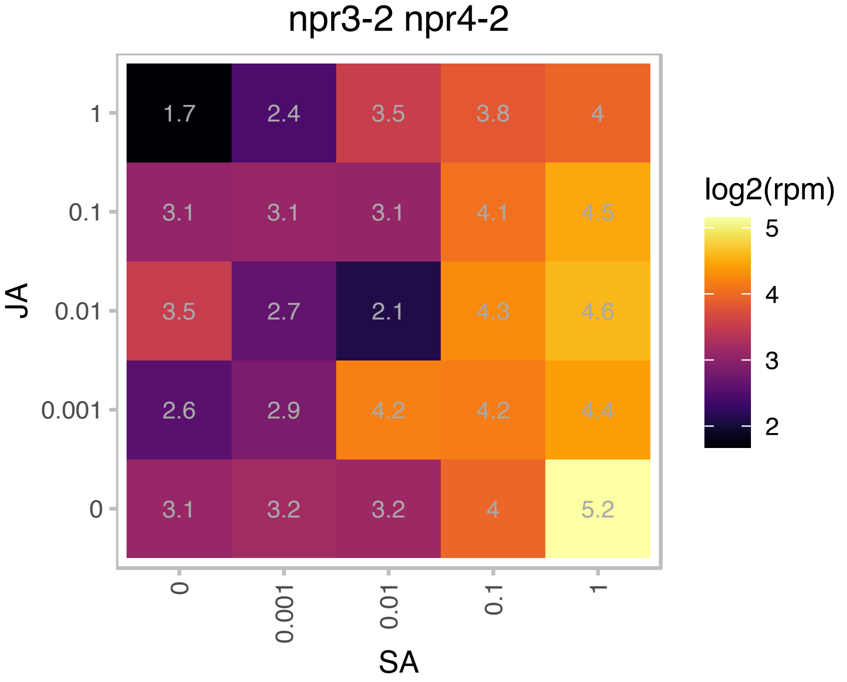


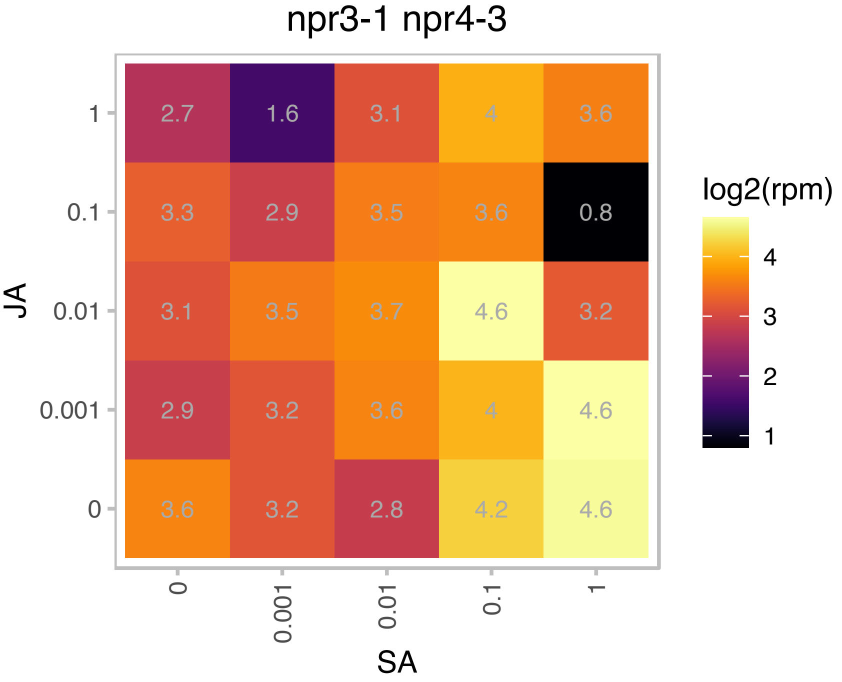

# AT1G09605.1 : NA

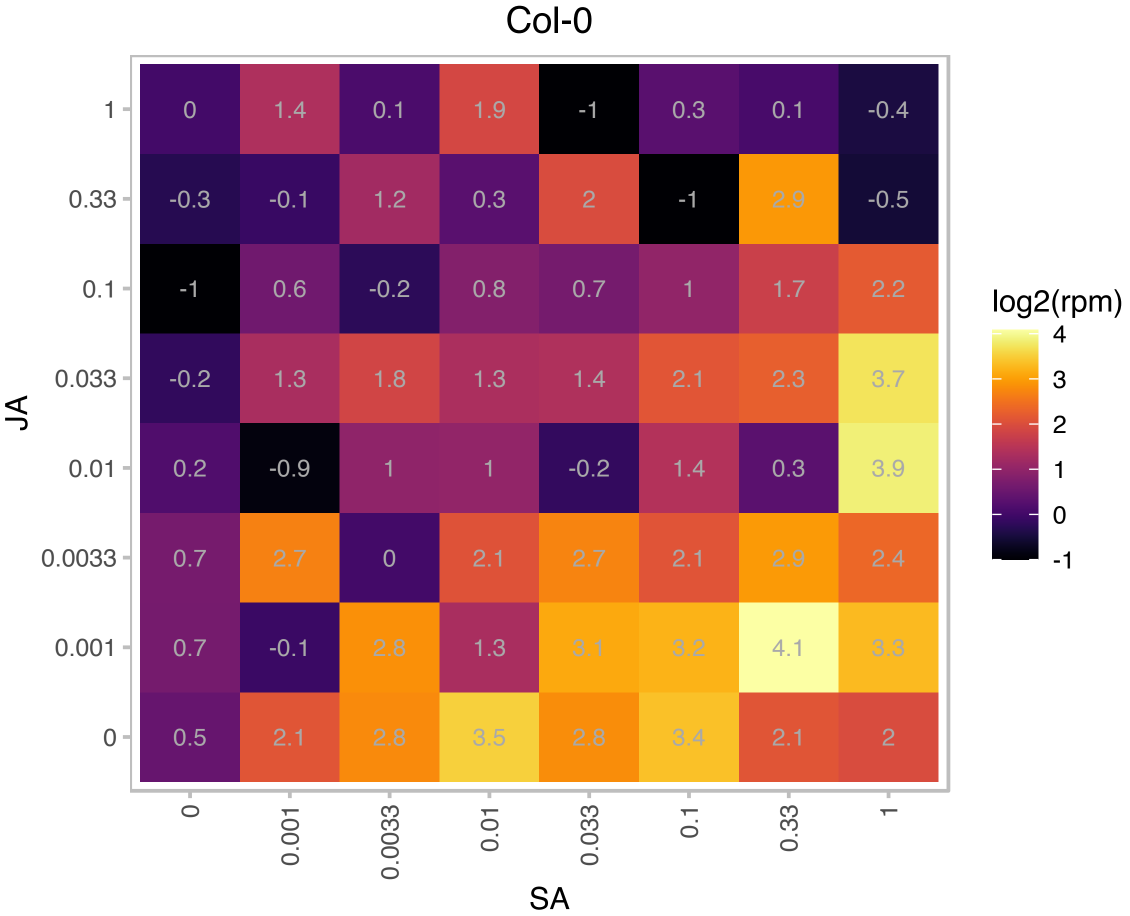


| | |
|---|---|
| Gene | AT1G09605 |
| is.expressed_genes | TRUE |
| is.responsive_genes | TRUE |
| is.apcluster | C-01 |

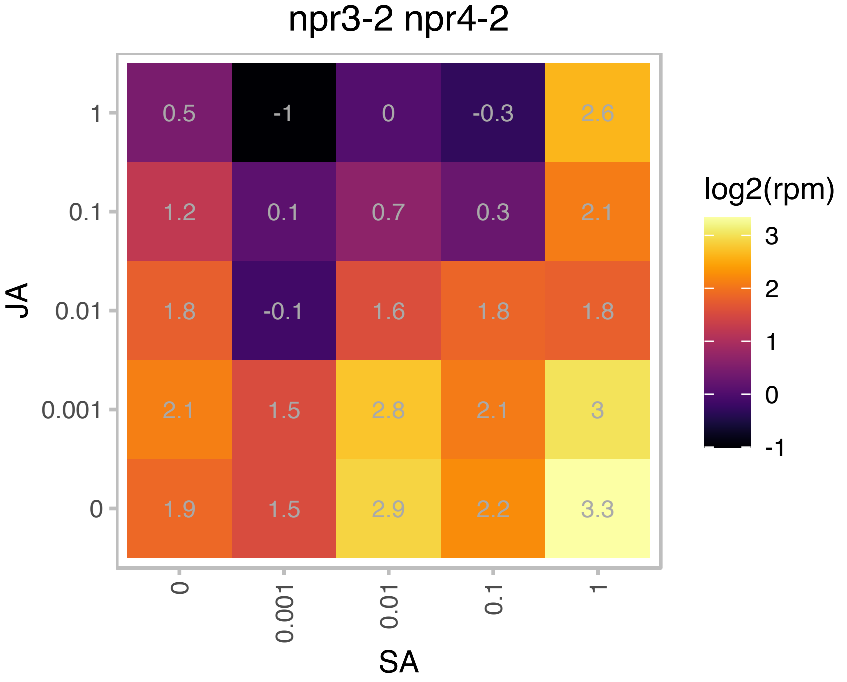


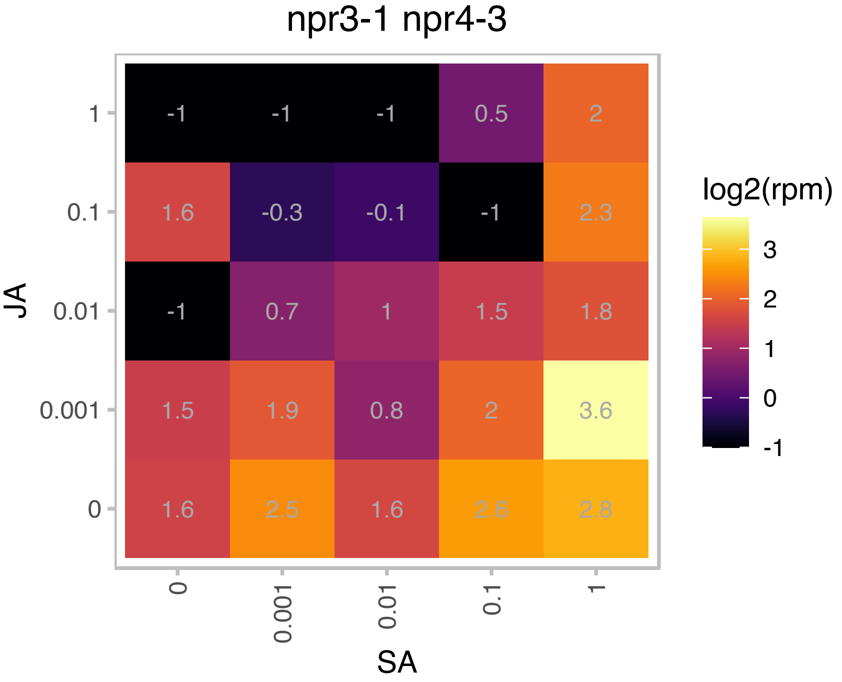

### AT1G14880.1 : AtPCR1, PCR1

PLANT CADMIUM RESISTANCE 1

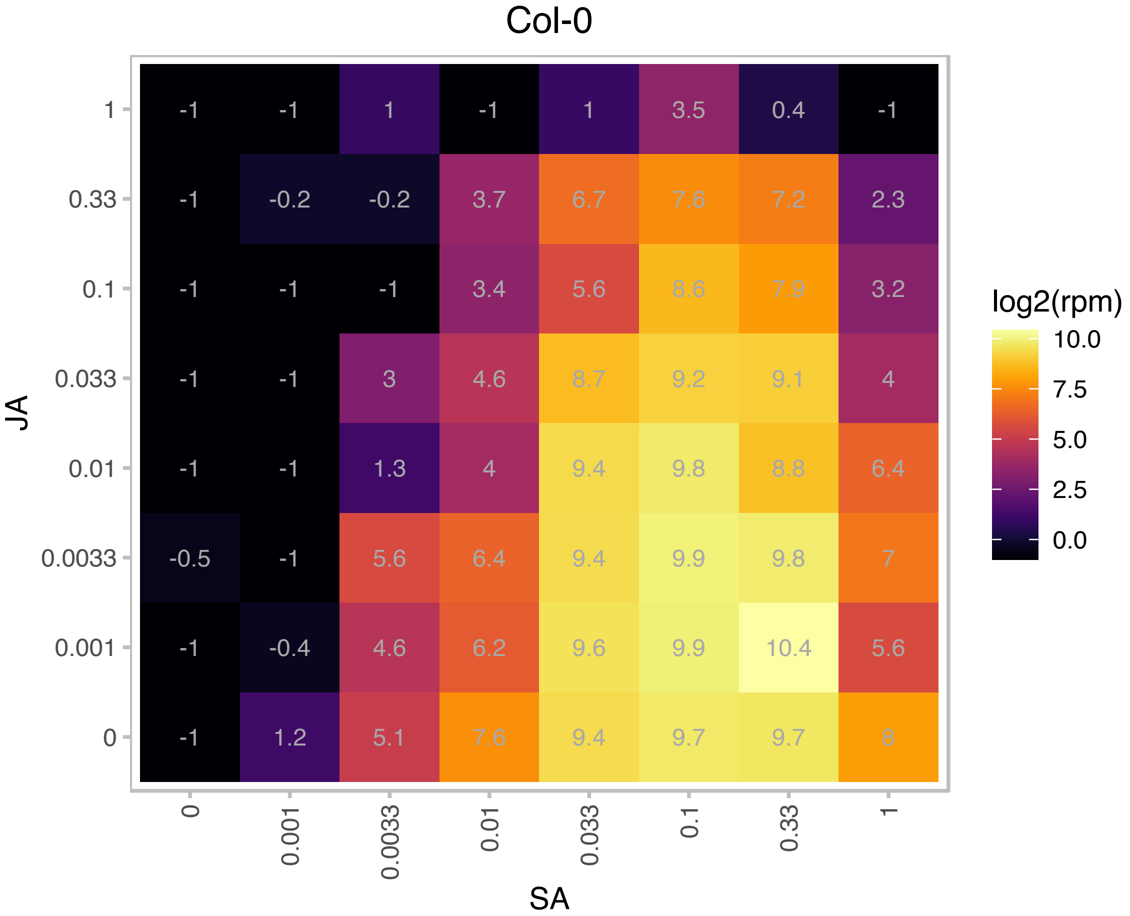


| | |
|---|---|
| Gene | AT1G14880 |
| is.expressed_genes | TRUE |
| is.responsive_genes | TRUE |
| is.apcluster | C-01 |

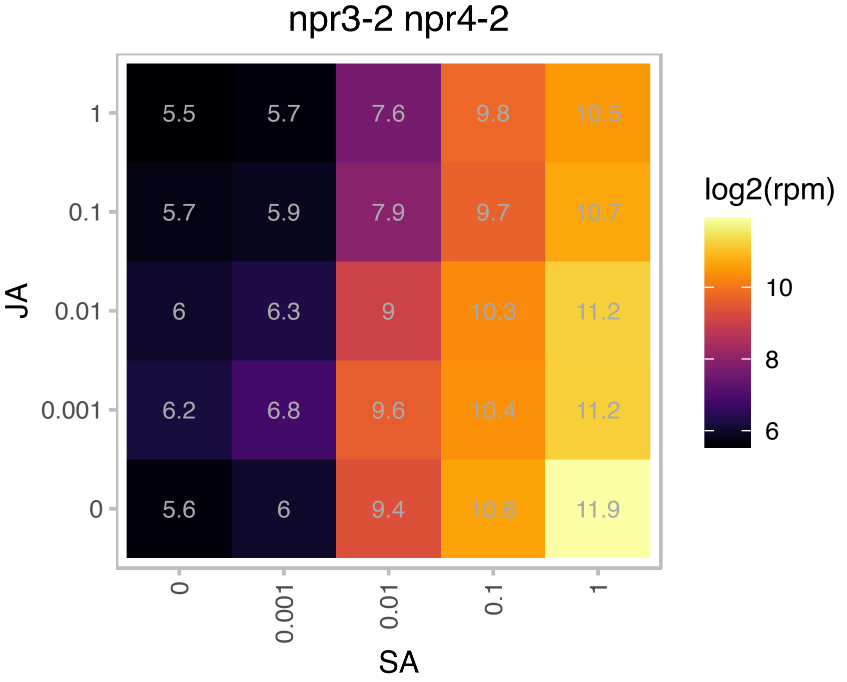


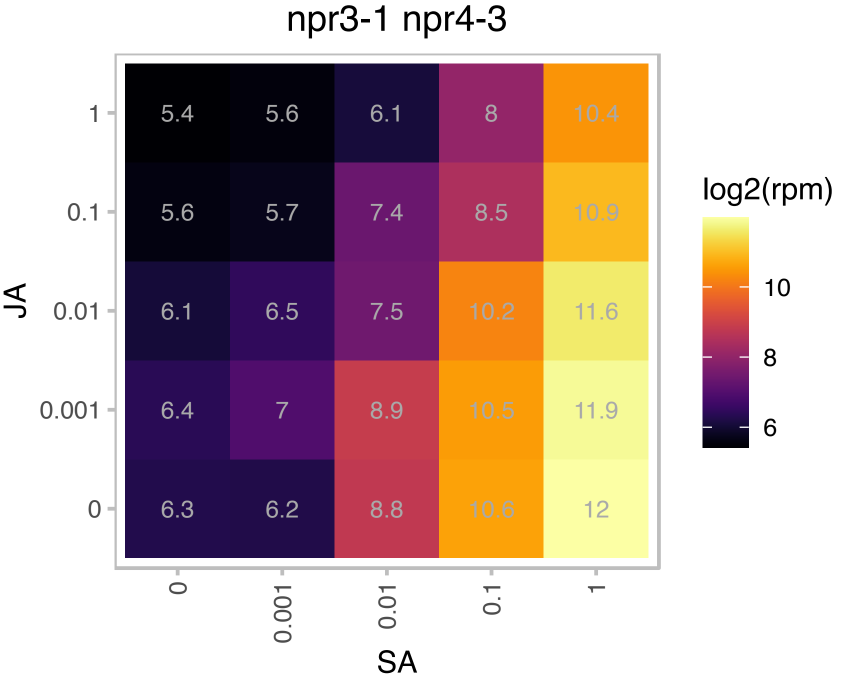

# AT1G16260.1 : NA

Wall-associated kinase family protein

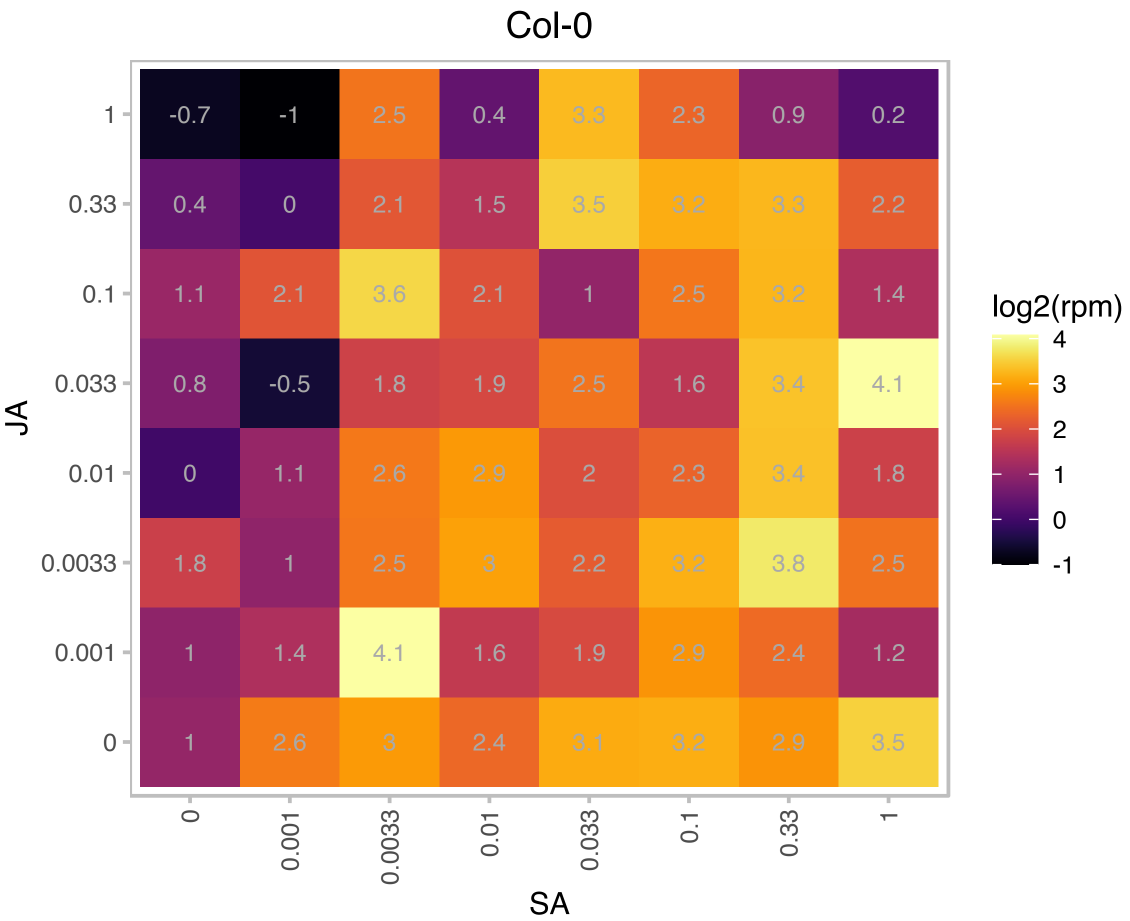


| | |
|---|---|
| Gene | AT1G16260 |
| is.expressed_genes | TRUE |
| is.responsive_genes | TRUE |
| is.apcluster | C-01 |

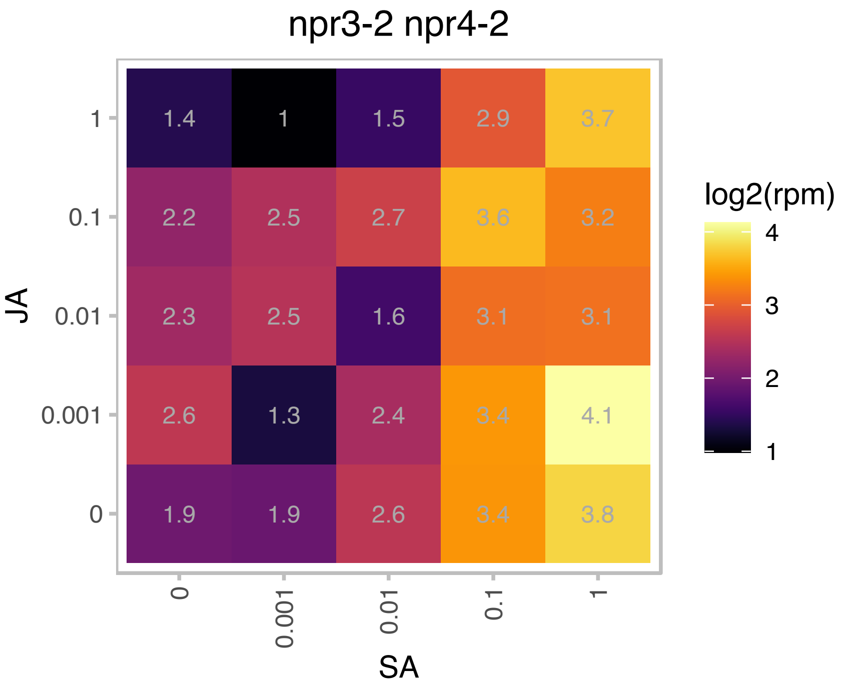


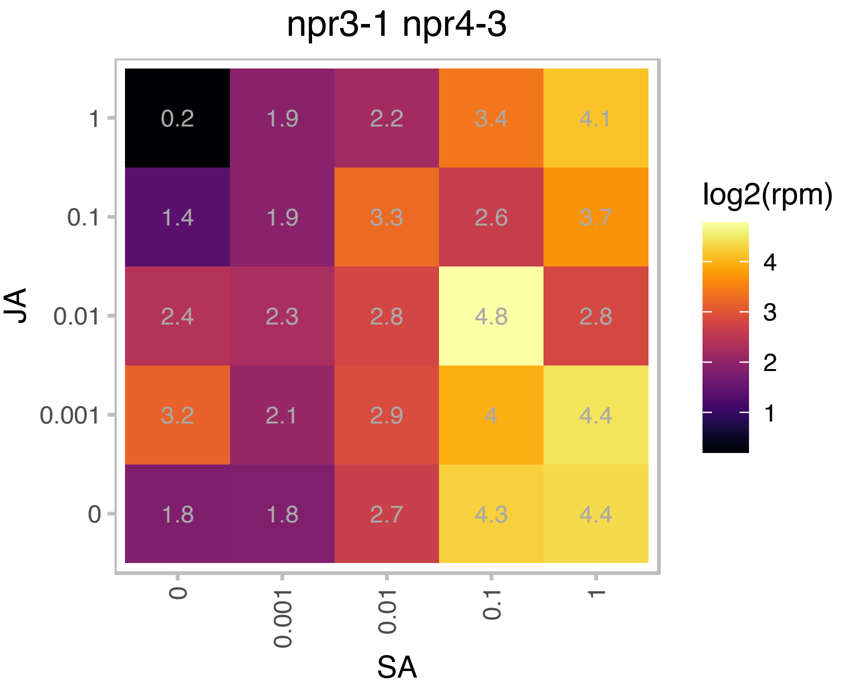

# AT1G24145.1 : NA

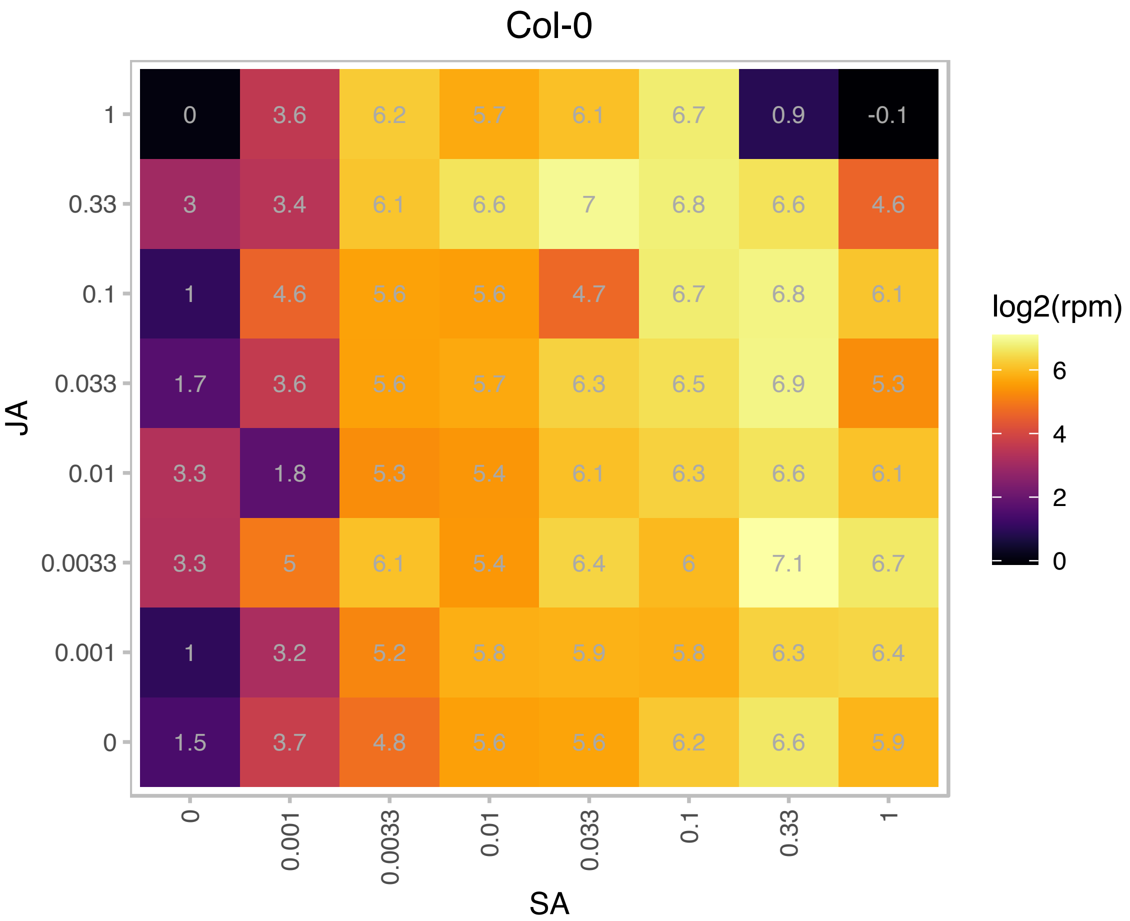


| | |
|---|---|
| Gene | AT1G24145 |
| is.expressed_genes | TRUE |
| is.responsive_genes | TRUE |
| is.apcluster | C-01 |

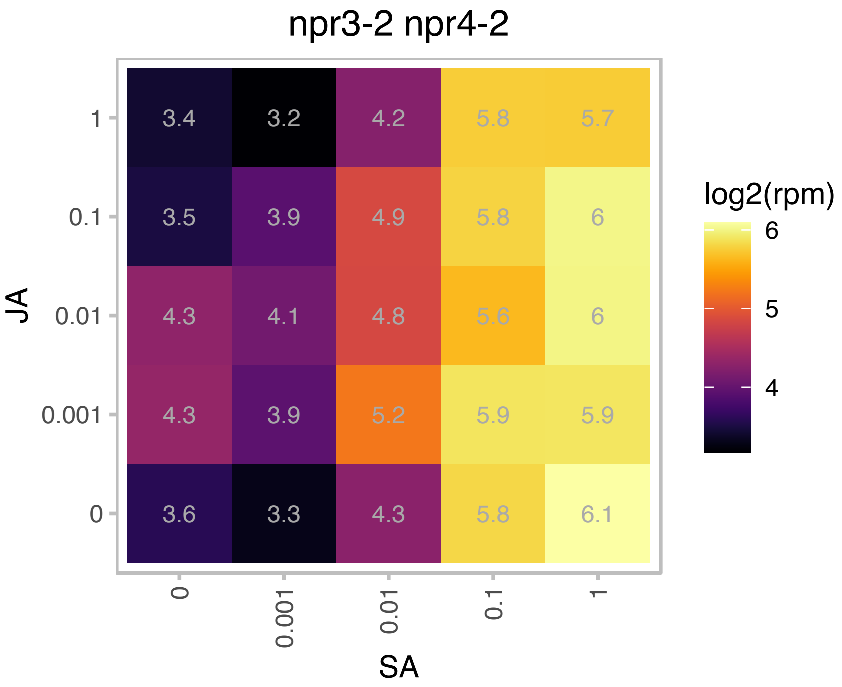


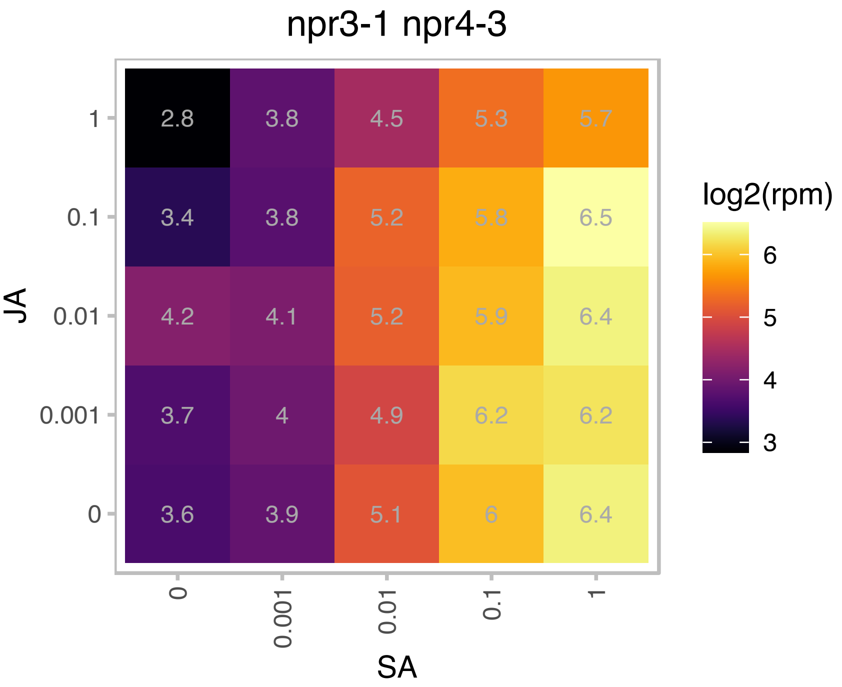

### AT1G33990.1 : ATMES14, MES14

methyl esterase 14

Encodes a protein predicted to act as a carboxylesterase. It has similarity to the SABP2 methyl salicylate esterase from tobacco. This protein does not act on methyl IAA, methyl JA, MeSA, MeGA4, or MEGA9 in vitro.

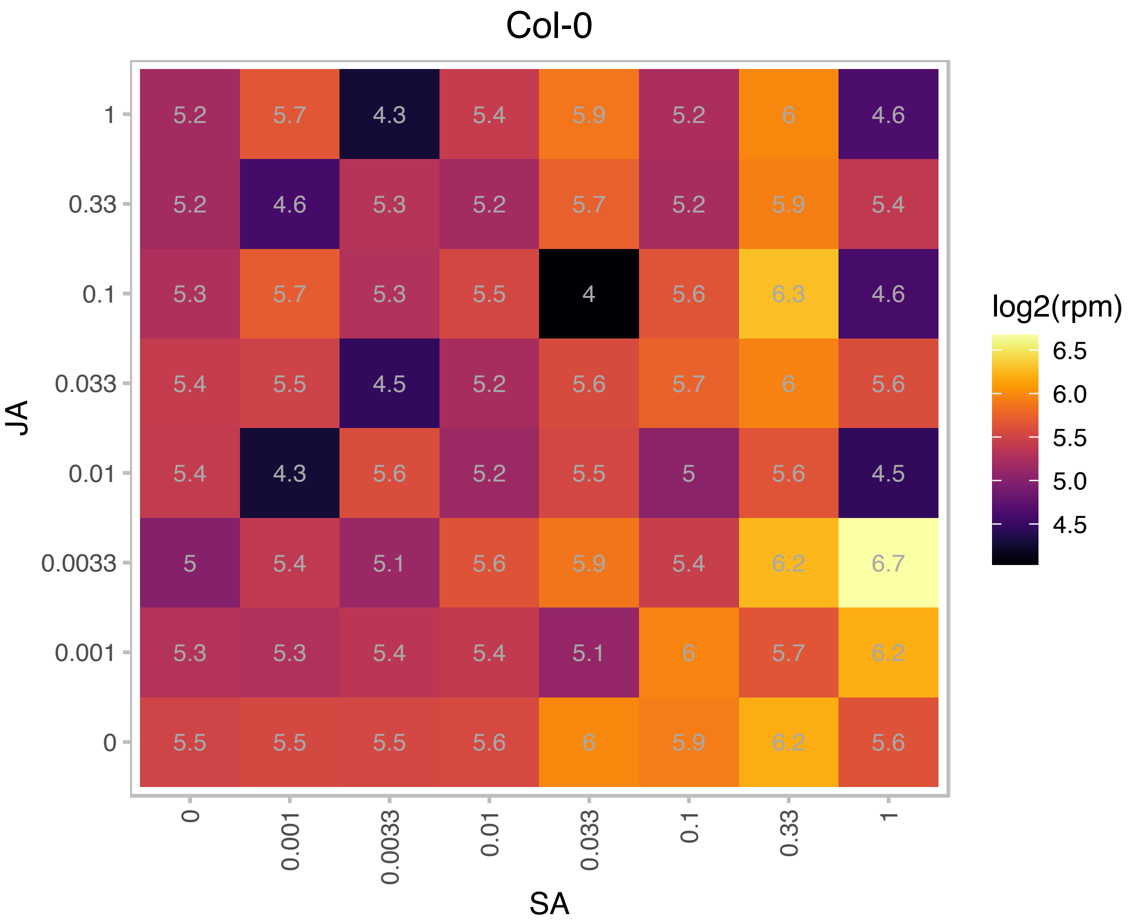


| | |
|---|---|
| Gene | AT1G33990 |
| is.expressed_genes | TRUE |
| is.responsive_genes | TRUE |
| is.apcluster | C-01 |

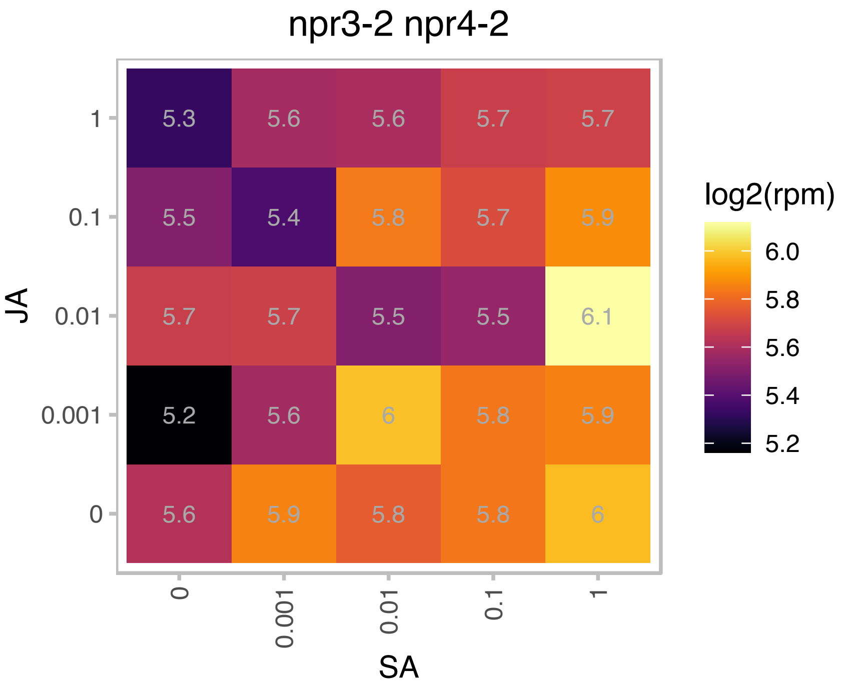


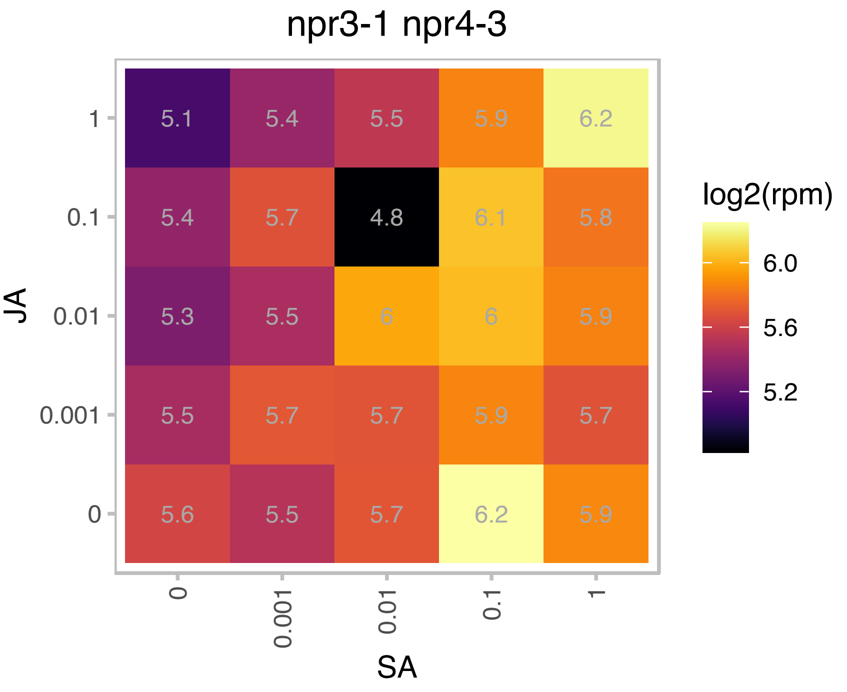

### AT1G35230.1 : AGP5

arabinogalactan protein 5

Encodes arabinogalactan-protein (AGP5). The mRNA is cell-to-cell mobile.

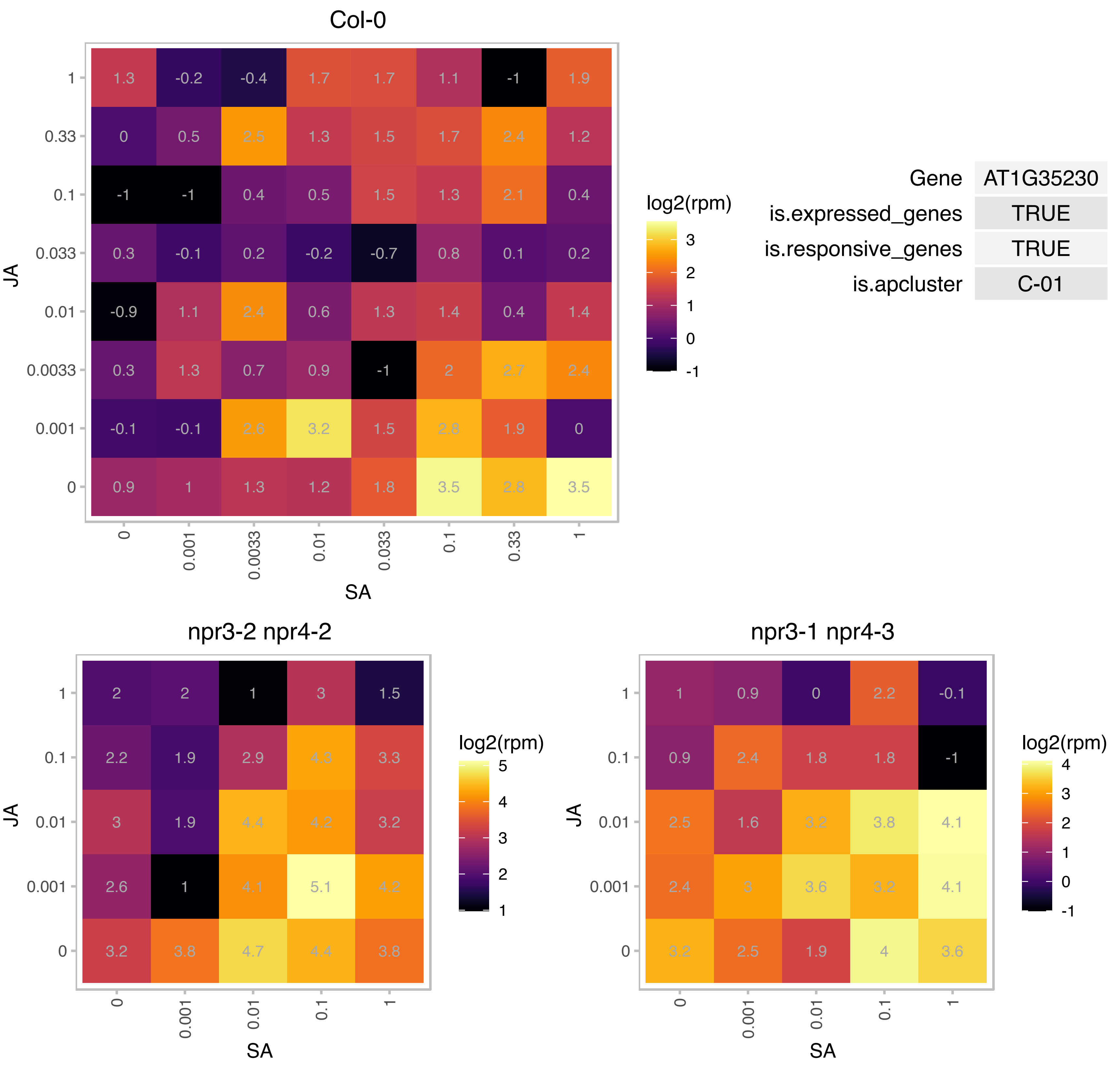


| | |
|---|---|
| Gene | AT1G35230 |
| is.expressed_genes | TRUE |
| is.responsive_genes | TRUE |
| is.apcluster | C-01 |

# AT1G53710.1 : NA

Calcineurin-like metallo-phosphoesterase superfamily protein

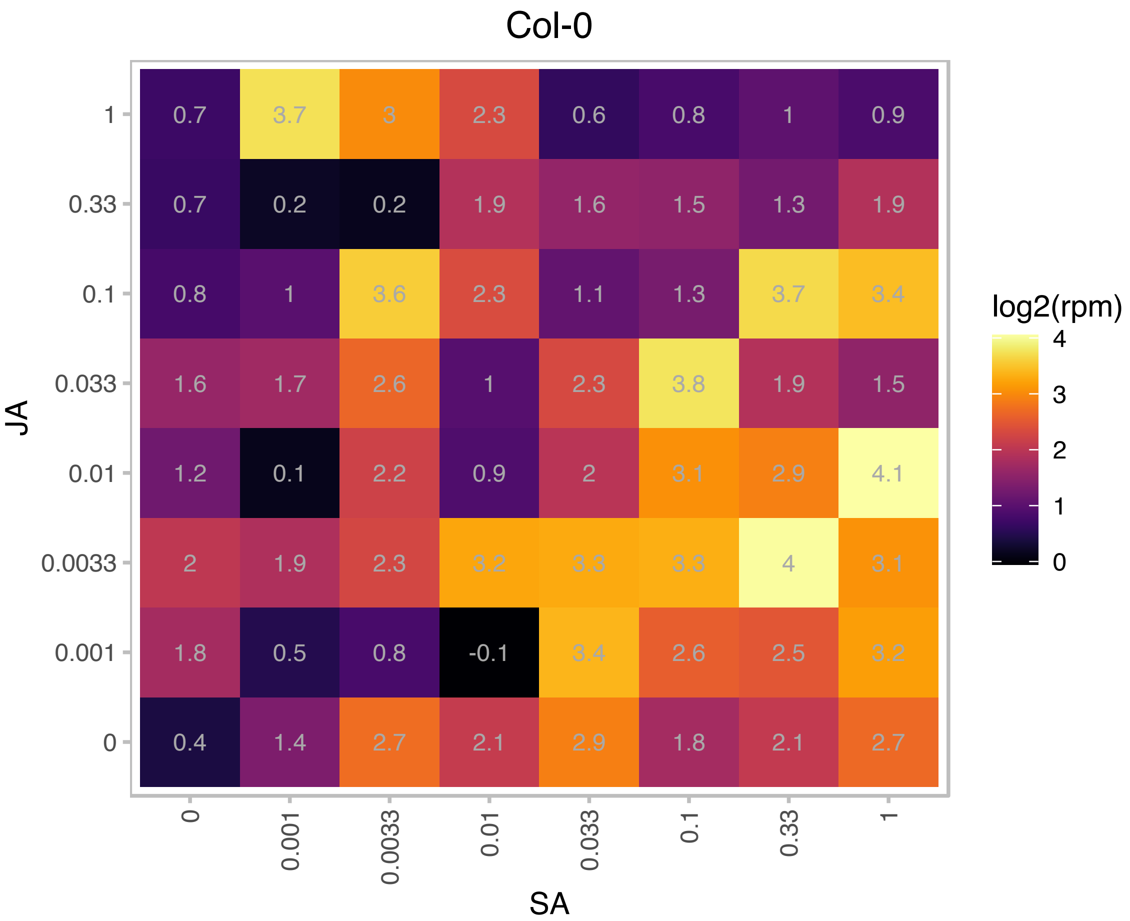


| | |
|---|---|
| Gene | AT1G53710 |
| is.expressed_genes | TRUE |
| is.responsive_genes | TRUE |
| is.apcluster | C-01 |

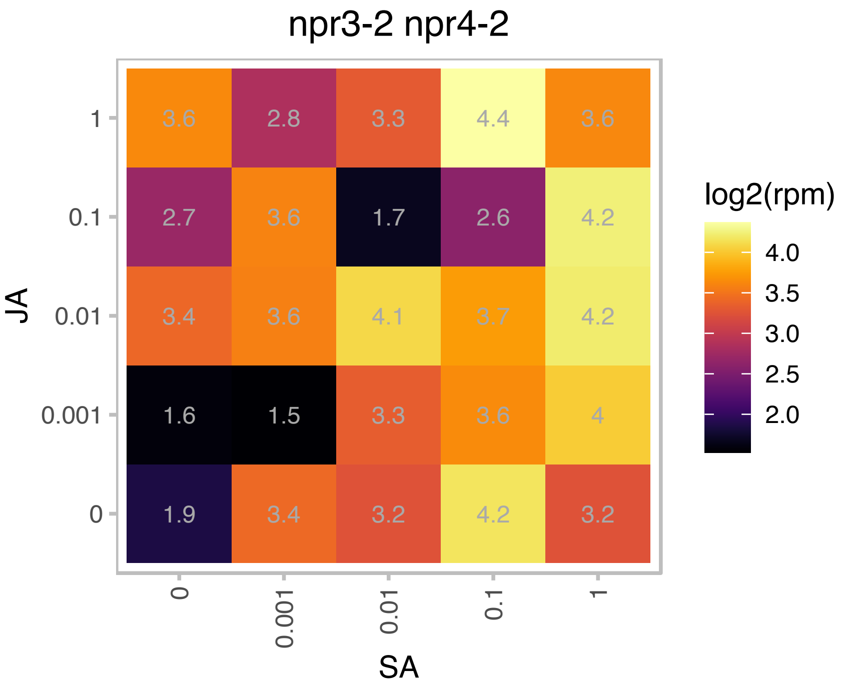


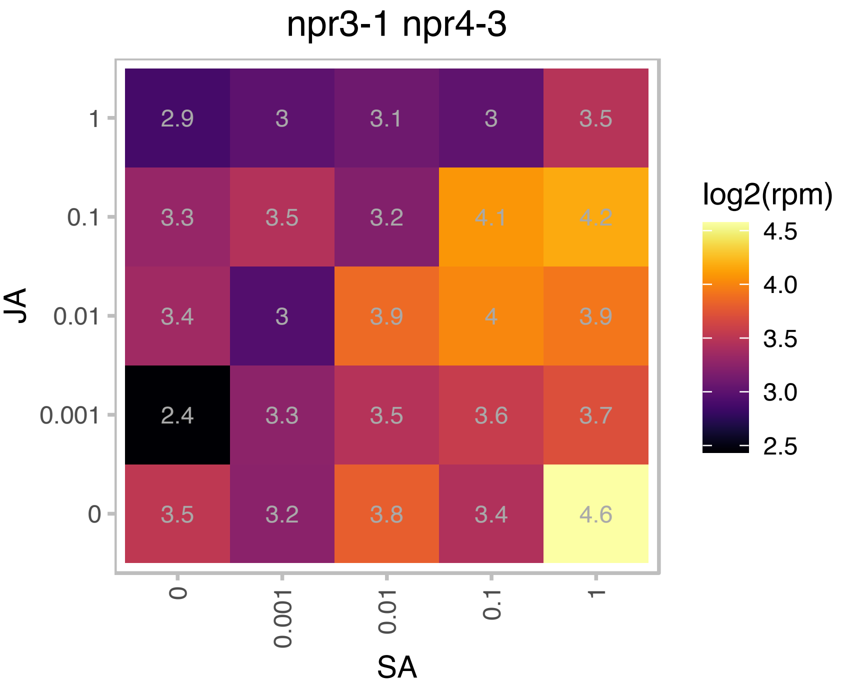

### AT1G55910.1 : ZIP11

zinc transporter 11 precursor

member of Putative zinc transporter ZIP2 - like family

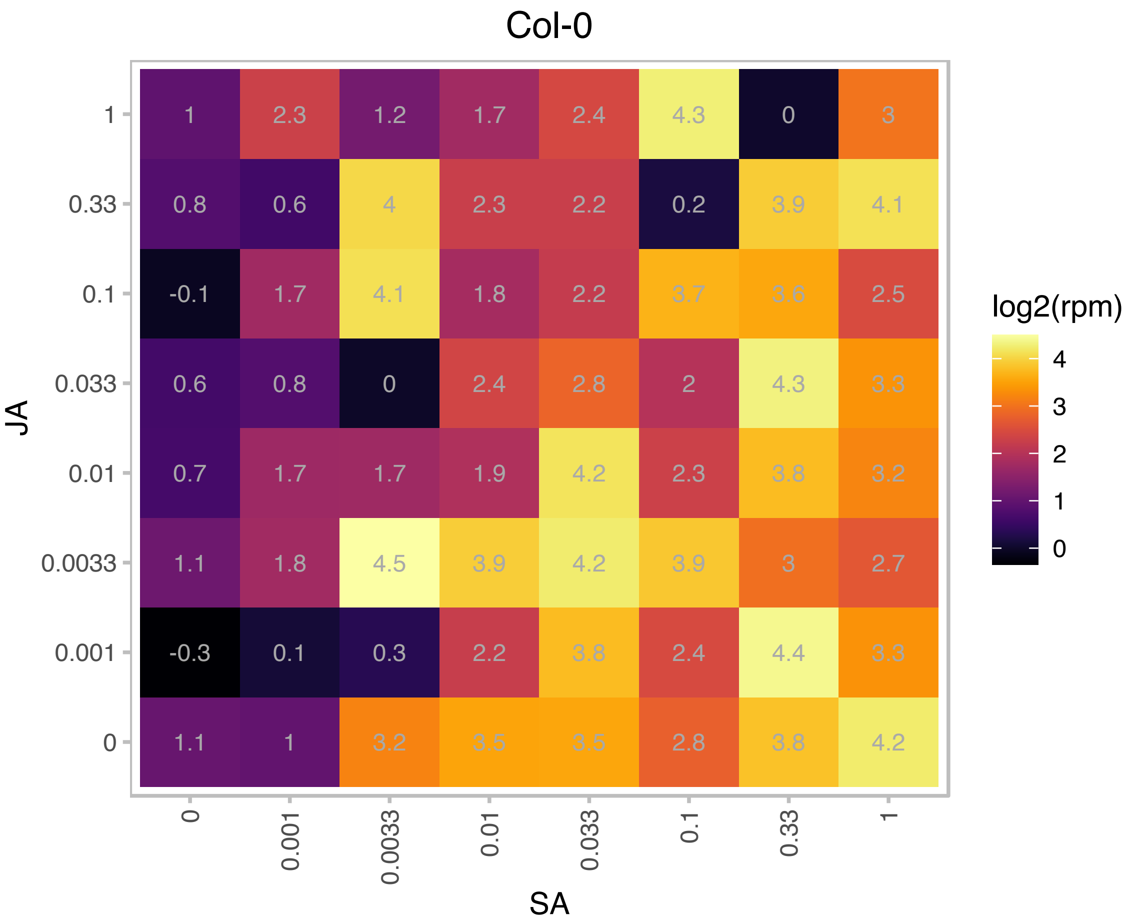


| | |
|---|---|
| Gene | AT1G55910 |
| is.expressed_genes | TRUE |
| is.responsive_genes | TRUE |
| is.apcluster | C-01 |

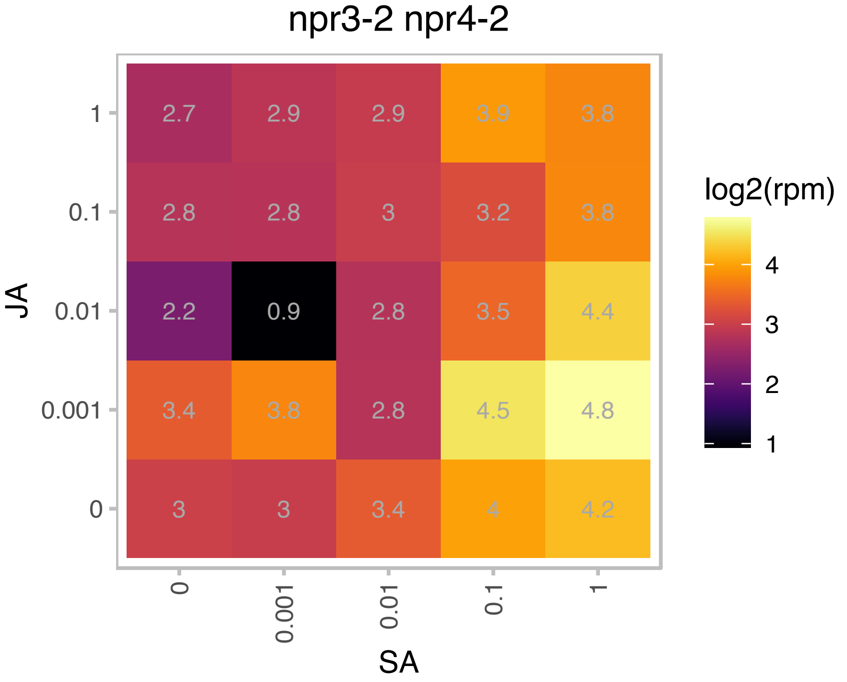


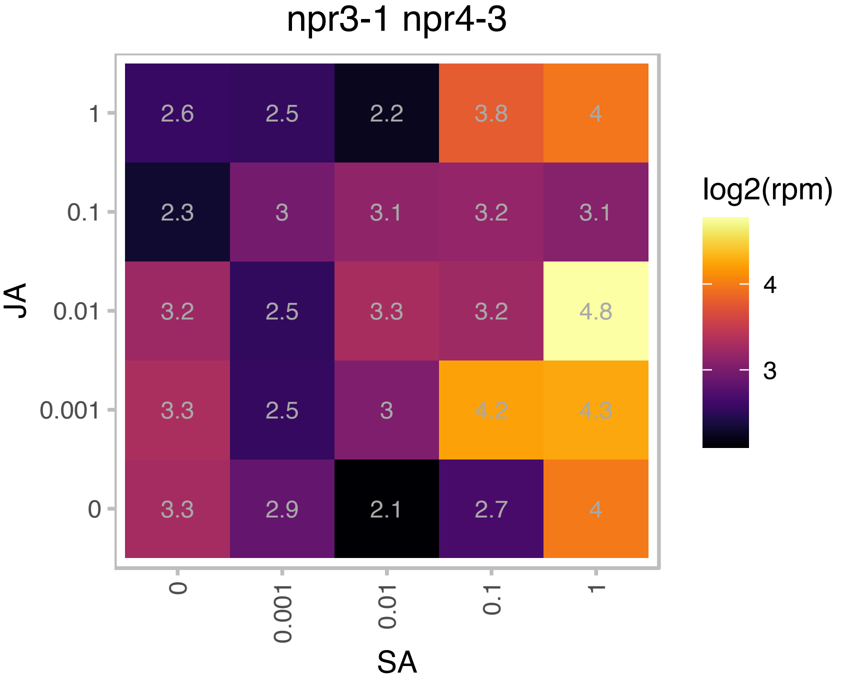

### AT1G64090.1 : RTN3, RTNLB3

Reticulan like protein B3

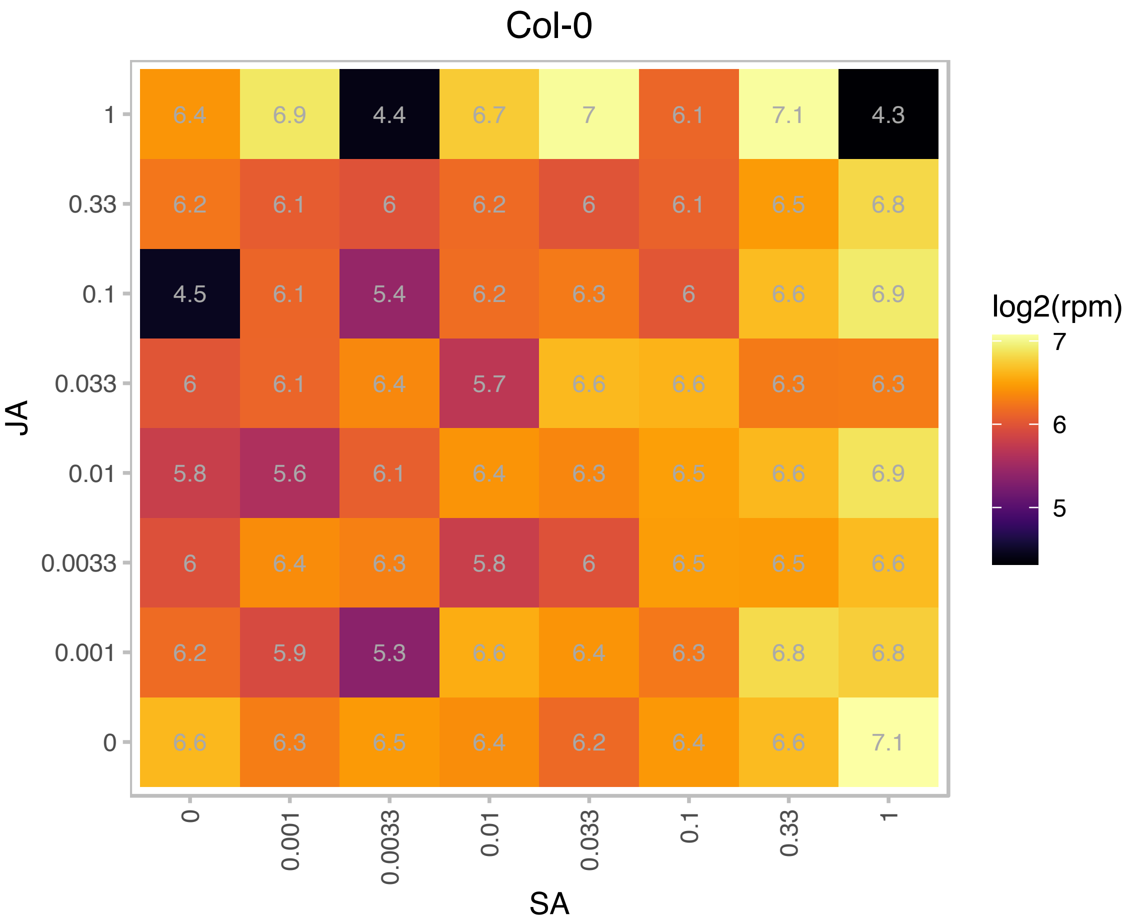


| | |
|---|---|
| Gene | AT1G64090 |
| is.expressed_genes | TRUE |
| is.responsive_genes | TRUE |
| is.apcluster | C-01 |

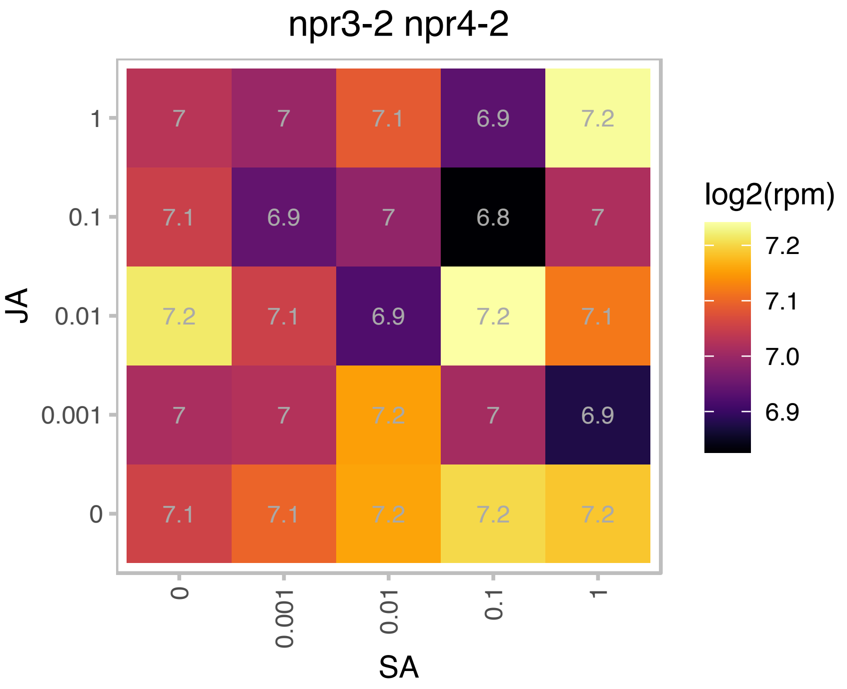


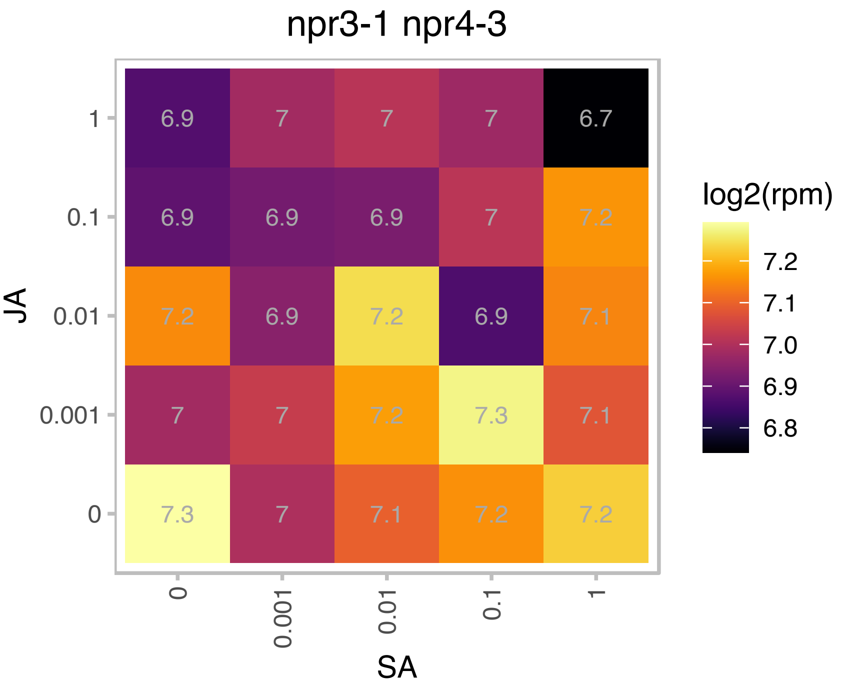

### AT1G73805.1 : SARD1

Calmodulin binding protein-like

Encodes SAR Deficient 1 (SARD1), a key regulator for ICS1 (Isochorismate Synthase 1) induction and salicylic acid (SA) synthesis.

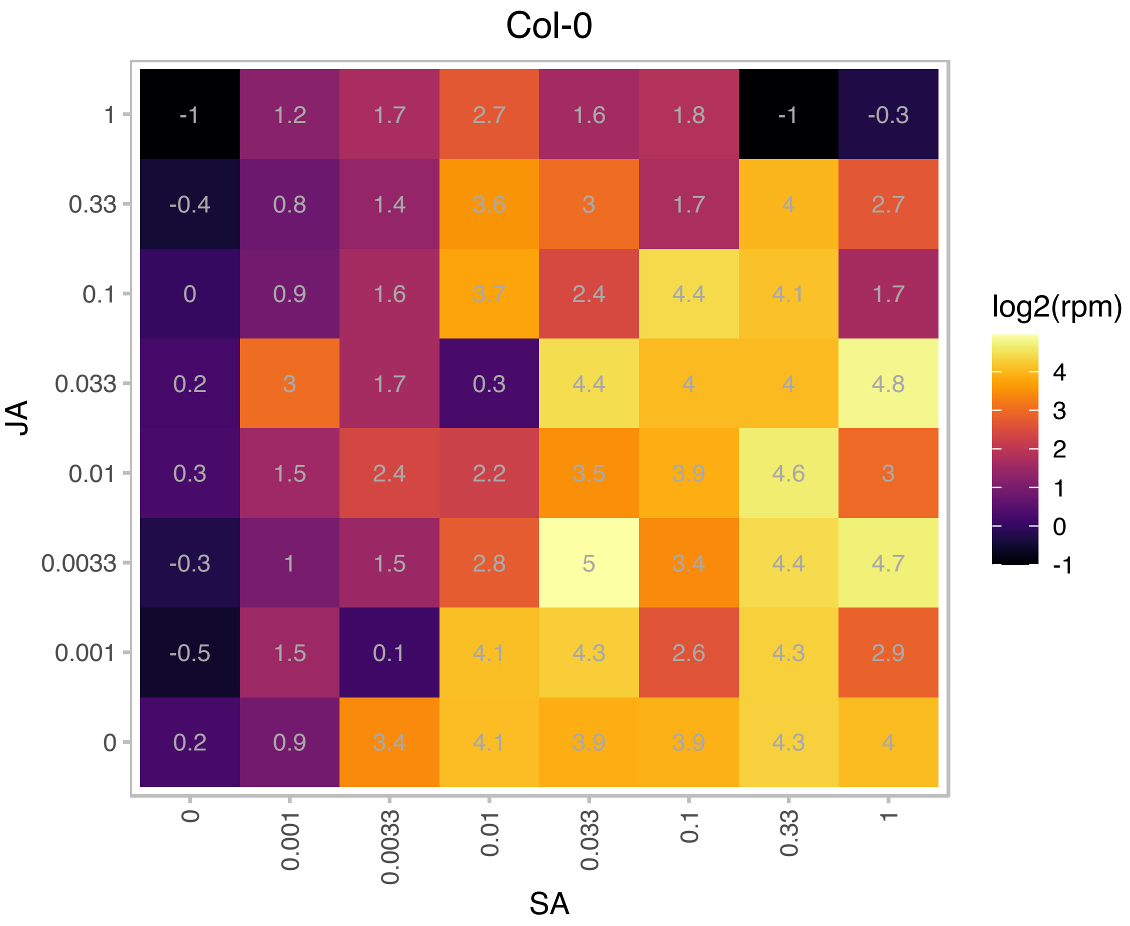


| | |
|---|---|
| Gene | AT1G73805 |
| is.expressed_genes | TRUE |
| is.responsive_genes | TRUE |
| is.apcluster | C-01 |

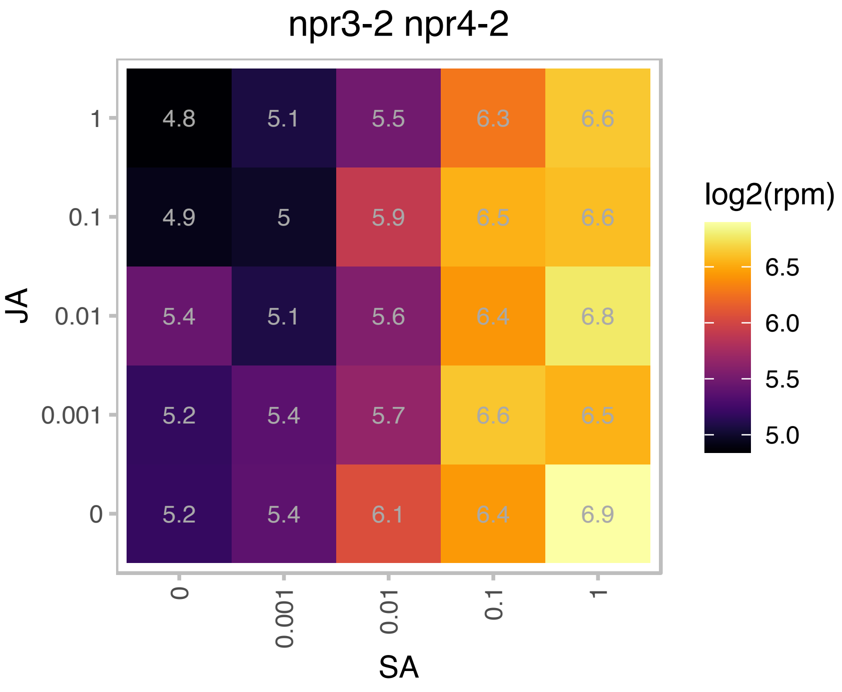


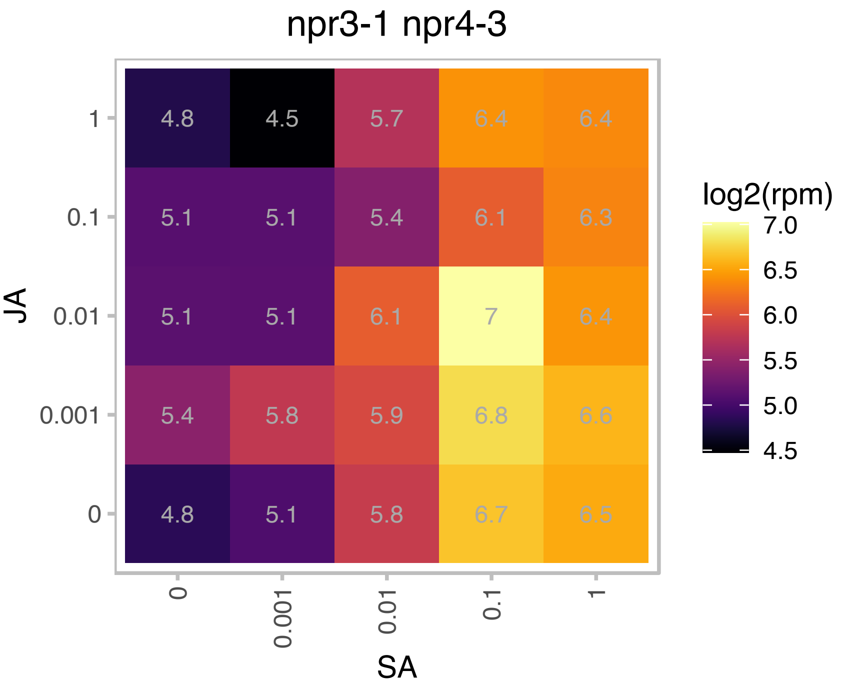

### AT2G14560.1 : LURP1

Protein of unknown function (DUF567)

Encodes LURP1, a member of the LURP cluster (late upregulated in response to Hyaloperonospora parasitica) which exhibits a pronounced upregulation after recognition of the pathogenic oomycte H. parasitica.  LURP1 is required for full basal defense to H. parasitica and resistance to this pathogen mediated by the R-proteins RPP4 and RPP5. The mRNA is cell-to-cell mobile.

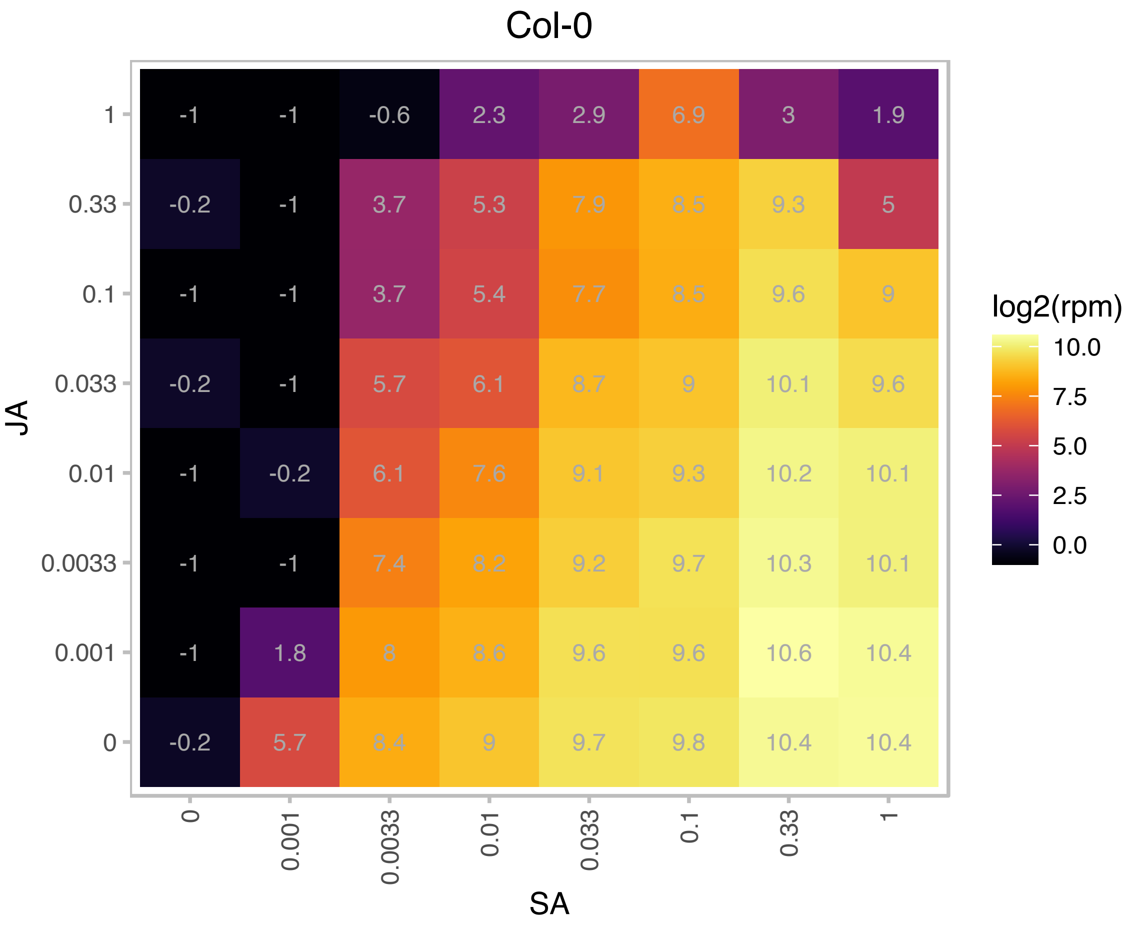


| | |
|---|---|
| Gene | AT2G14560 |
| is.expressed_genes | TRUE |
| is.responsive_genes | TRUE |
| is.apcluster | C-01 |

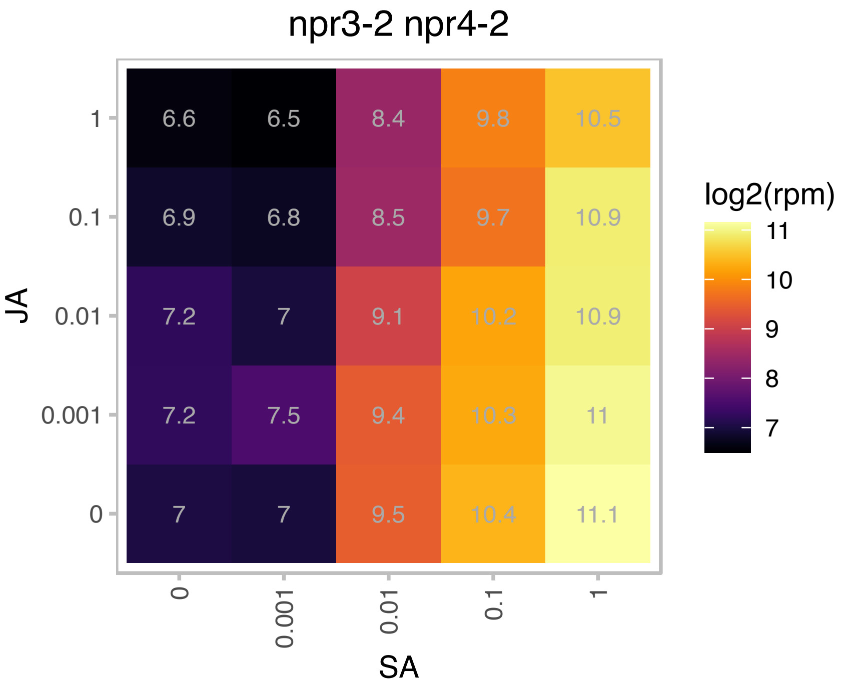


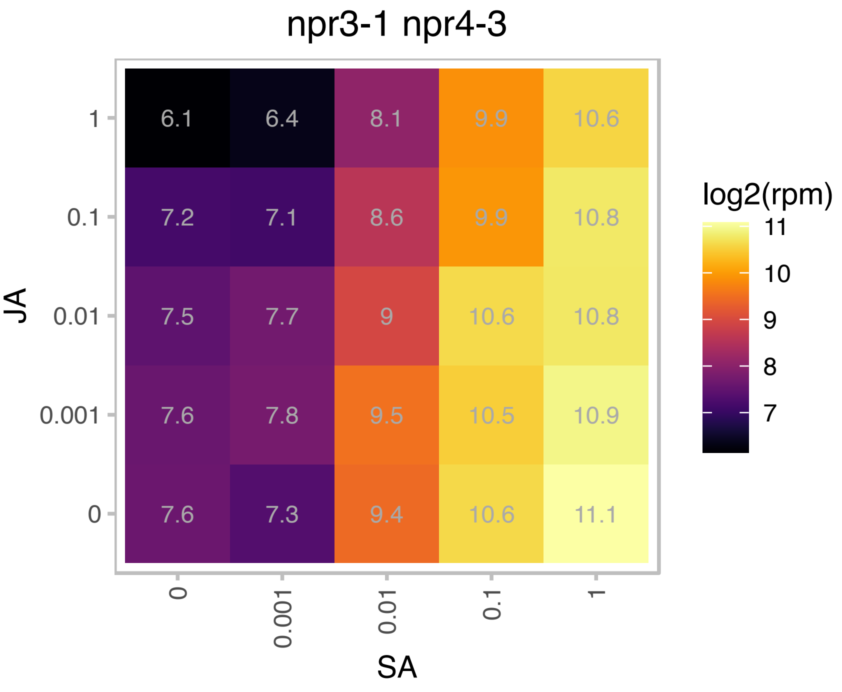

### AT2G14610.1 : AtCAPE9, ATPR1, PR 1, PR1

pathogenesis-related gene 1
PR1 gene expression is induced in response to a variety of pathogens. It is a useful molecular marker for the SAR response. Though the Genbank record for the cDNA associated to this gene is called 'PR-1-like', the sequence actually corresponds to PR1. Expression of this gene is salicylic-acid responsive.

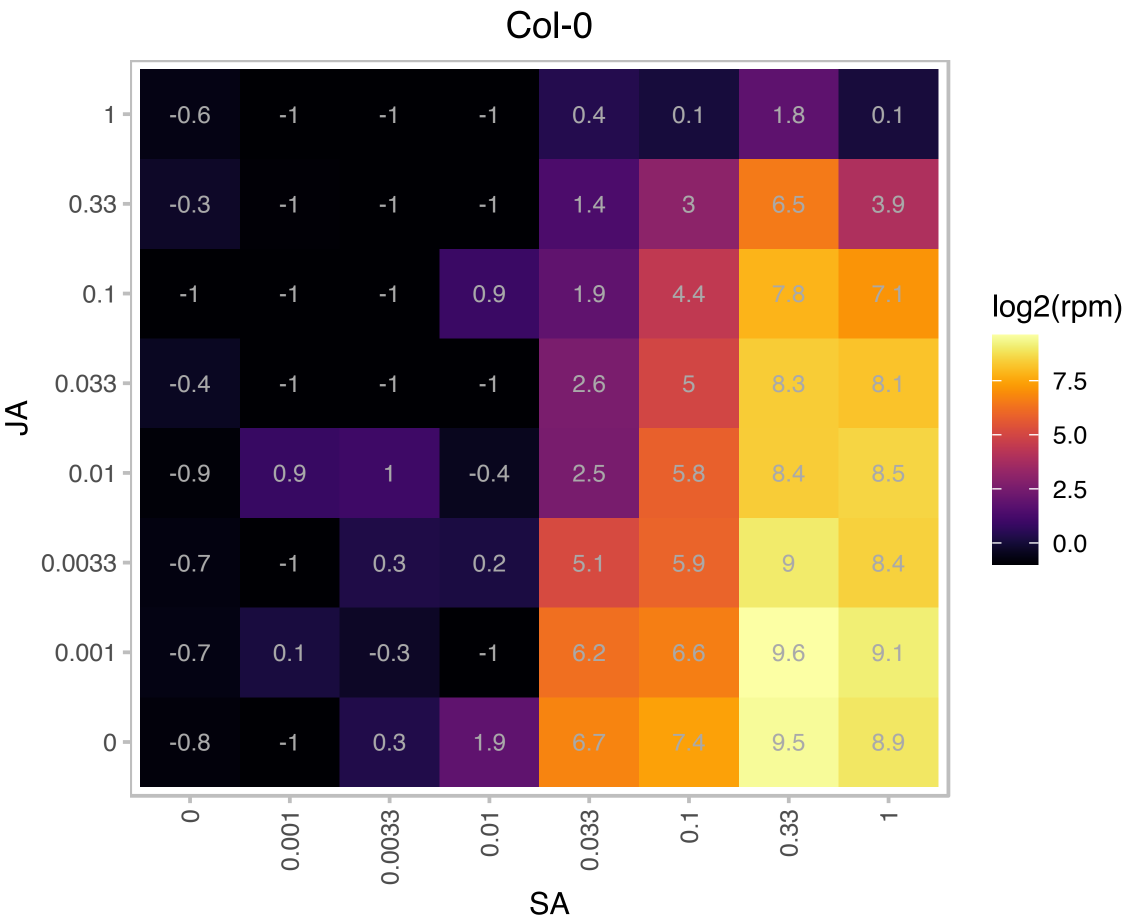


| | |
|---|---|
| Gene | AT2G14610 |
| is.expressed_genes | TRUE |
| is.responsive_genes | TRUE |
| is.apcluster | C-01 |

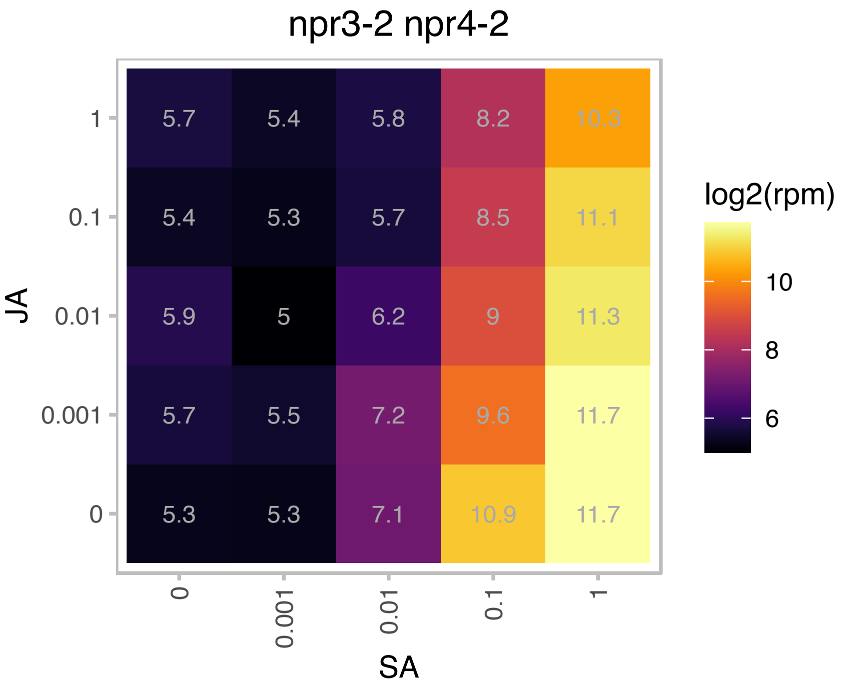


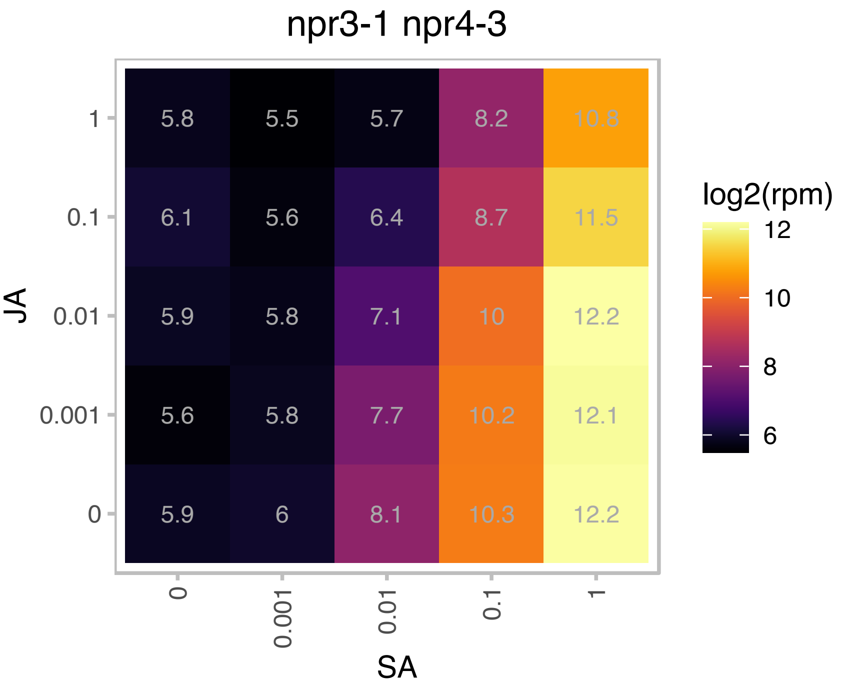

### AT2G19690.1 : PLA2-BETA

phospholipase A2-beta
Encodes one of the four Arabidopsis phospholipase PLA2 parologs: AT2G06925 (PLA2-ALPHA), AT2G19690 (PLA2-BETA), AT4G29460 (PLA2-GAMMA) and AT4G29470 (PLA2-DELTA). Involved in pollen development and germination and tube growth. Also involved in stomatal opening in response to light.

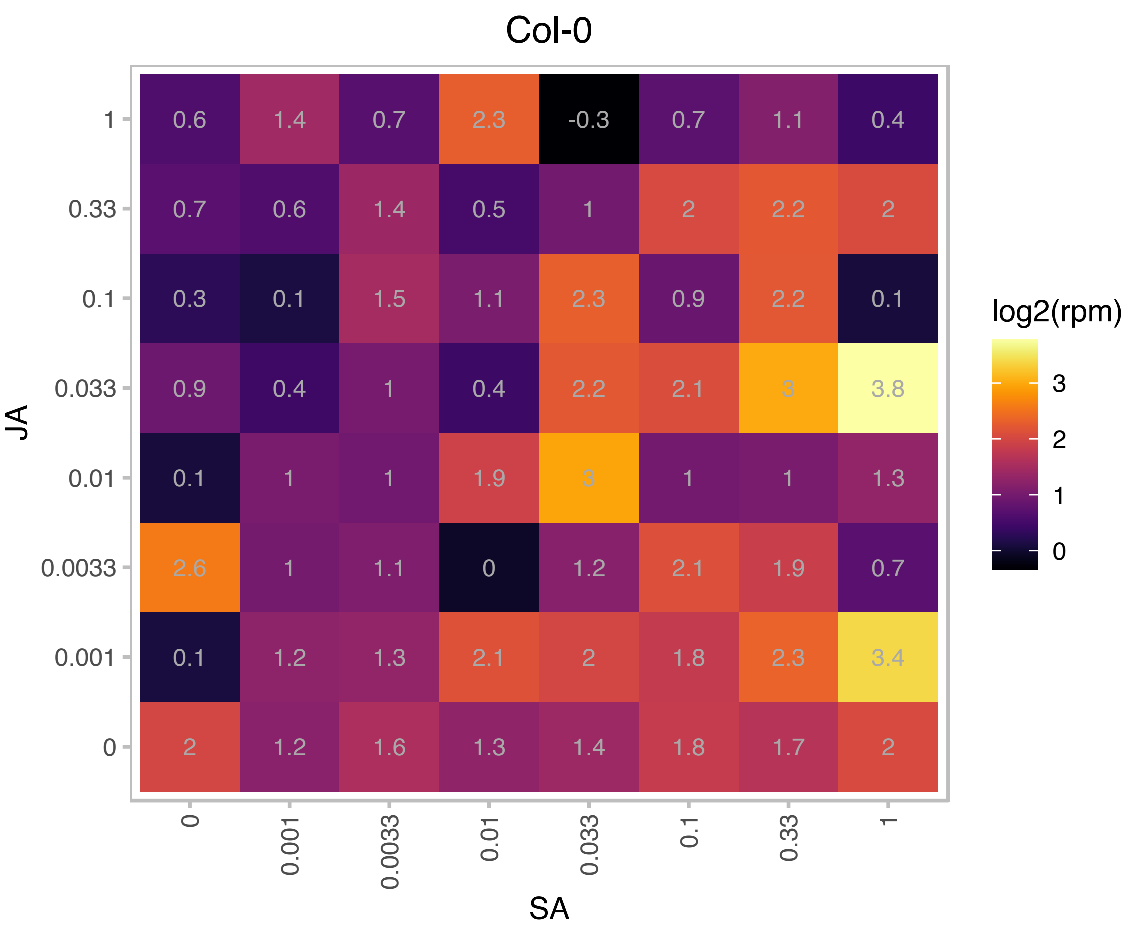


| | |
|---|---|
| Gene | AT2G19690 |
| is.expressed_genes | TRUE |
| is.responsive_genes | TRUE |
| is.apcluster | C-01 |

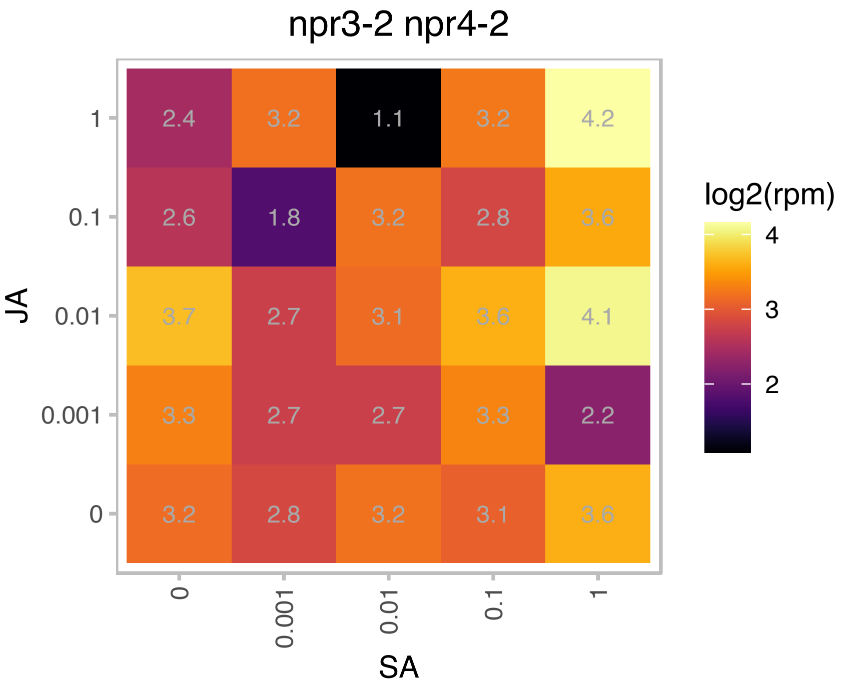


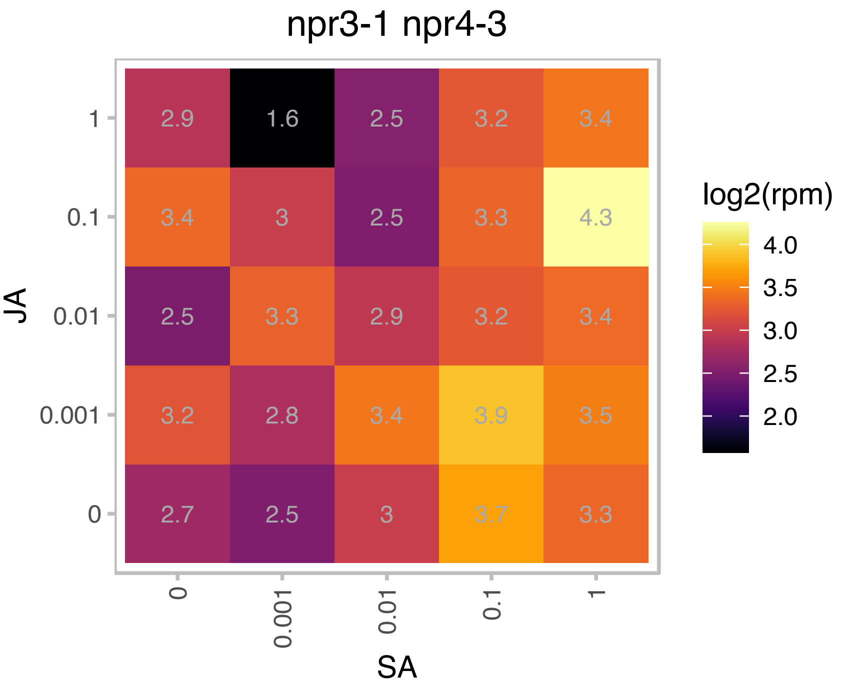

### AT2G21900.1 : ATWRKY59, WRKY59

WRKY DNA-binding protein 59

member of WRKY Transcription Factor; Group II-c

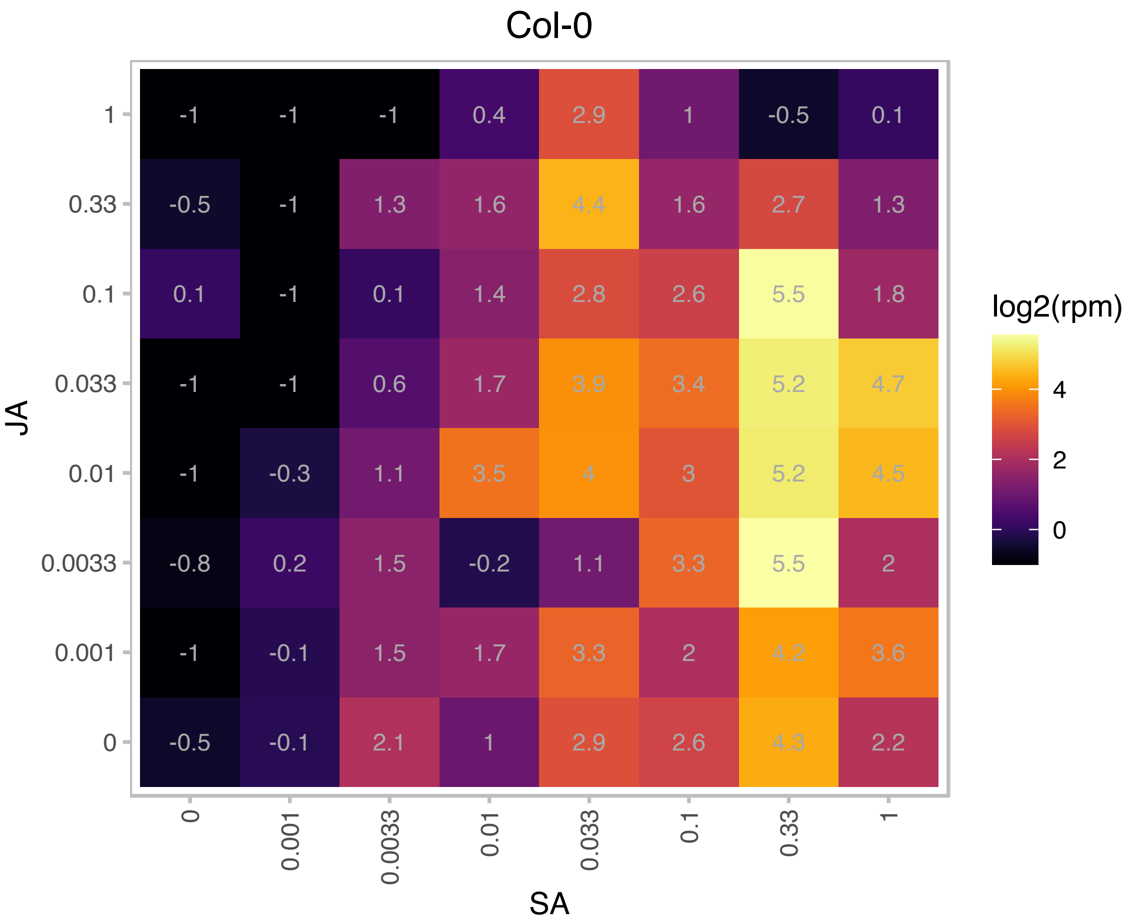


| | |
|---|---|
| Gene | AT2G21900 |
| is.expressed_genes | TRUE |
| is.responsive_genes | TRUE |
| is.apcluster | C-01 |

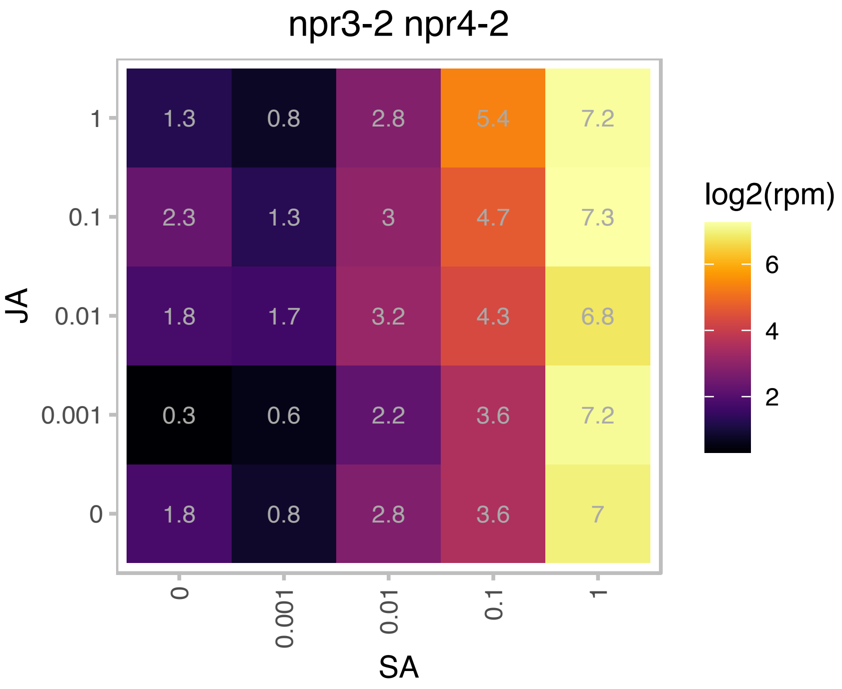


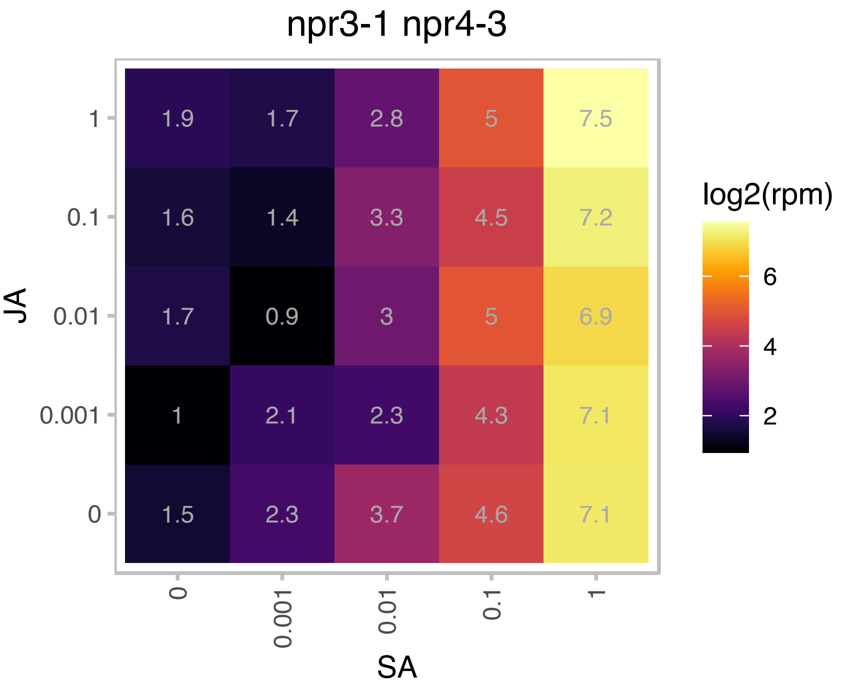

# AT2G23940.1 : NA

Protein of unknown function (DUF788)

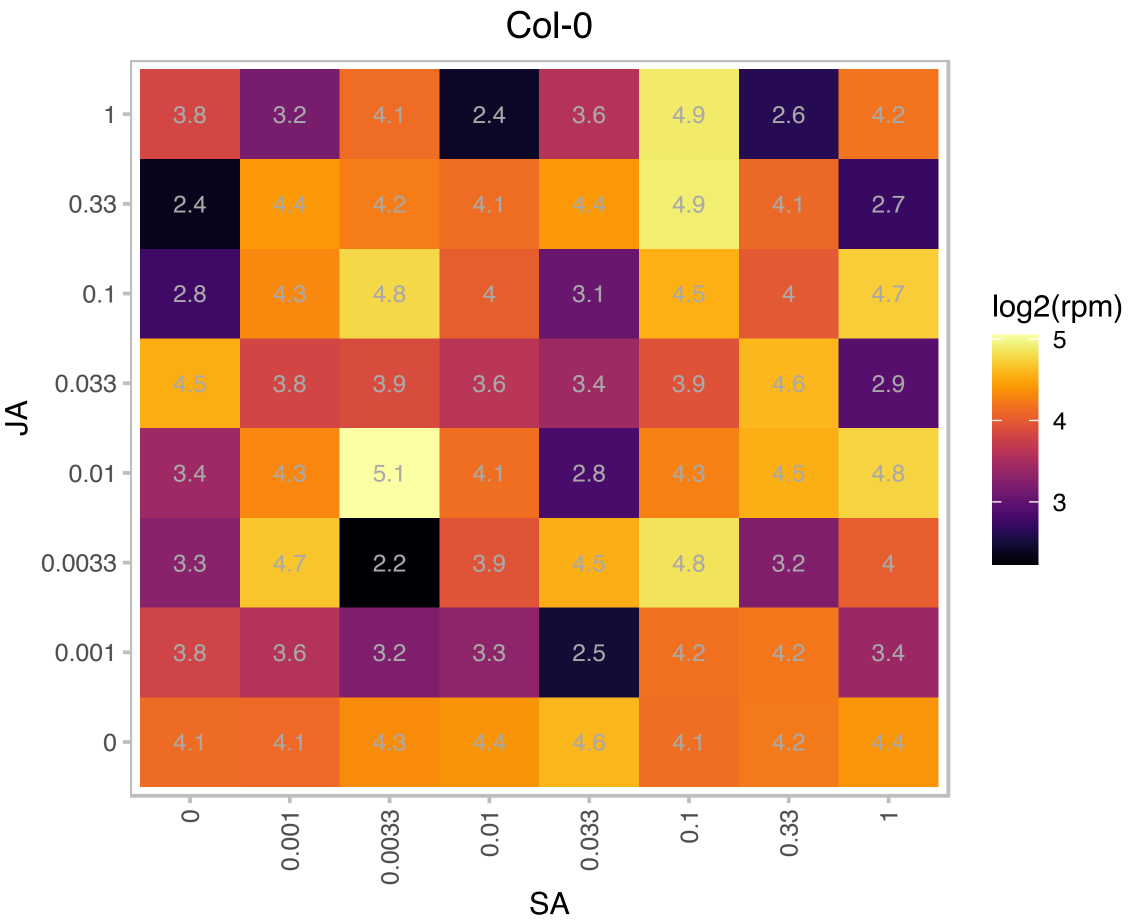


| | |
|---|---|
| Gene | AT2G23940 |
| is.expressed_genes | TRUE |
| is.responsive_genes | TRUE |
| is.apcluster | C-01 |

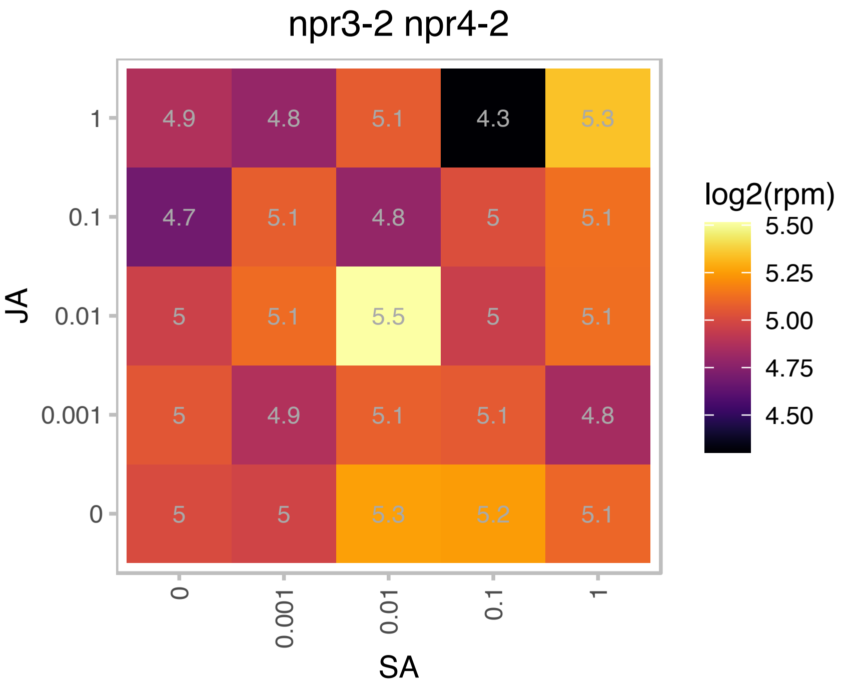


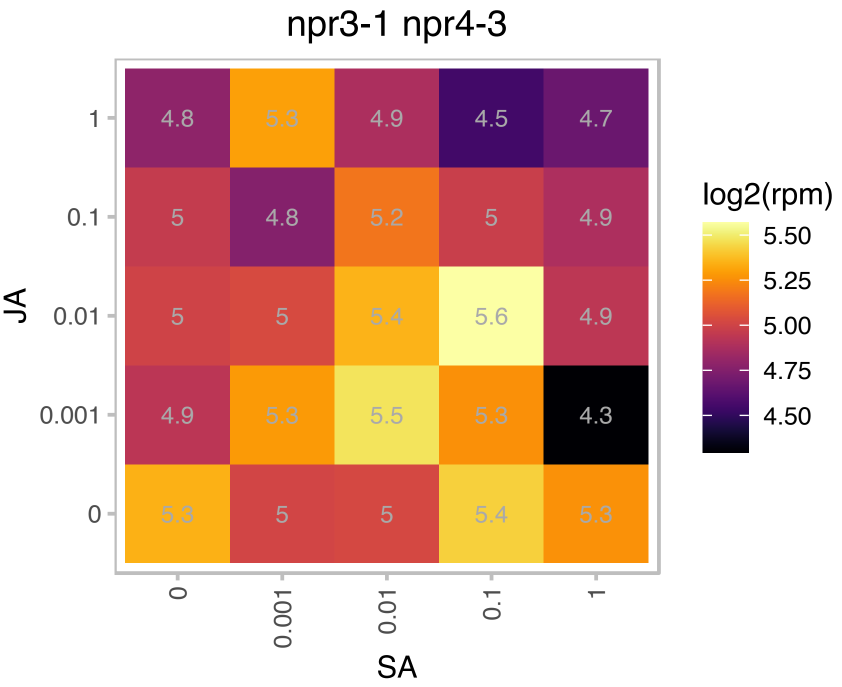

### AT2G26400.1 : ARD, ARD3, ATARD3

acireductone dioxygenase 3

Encodes a protein predicted to belong to the acireductone dioxygenase (ARD/ARD’)family.

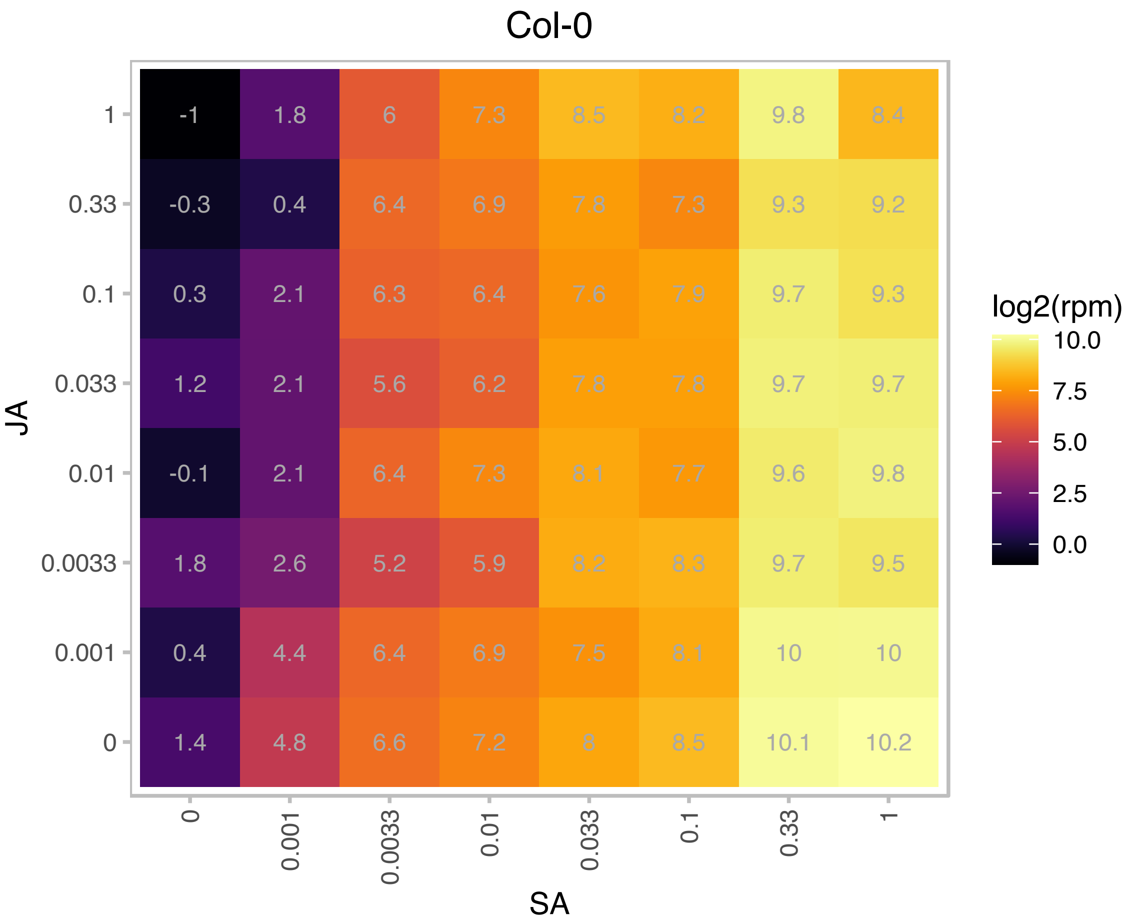


| | |
|---|---|
| Gene | AT2G26400 |
| is.expressed_genes | TRUE |
| is.responsive_genes | TRUE |
| is.apcluster | C-01 |

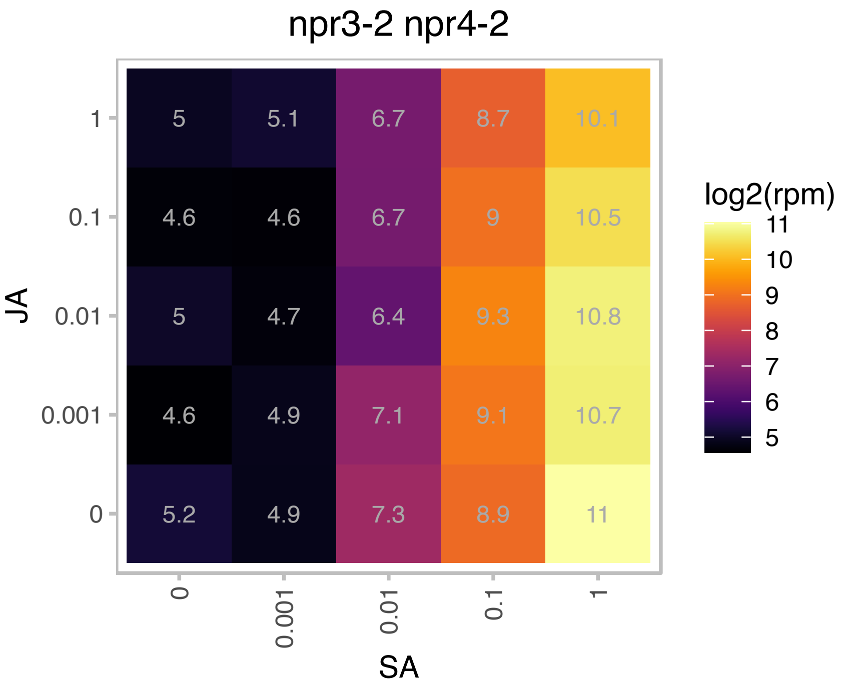


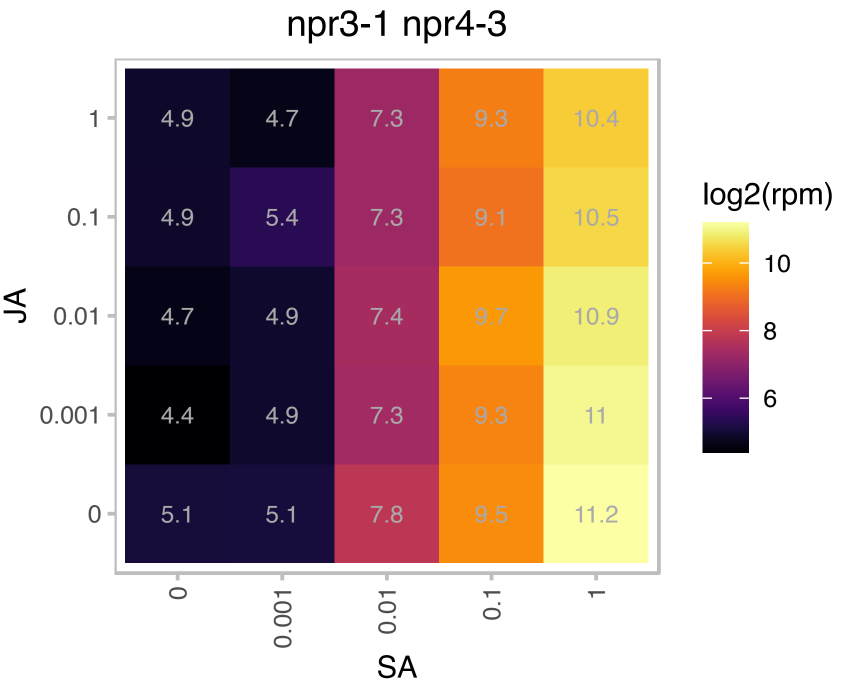

### AT2G28840.1 : XBAT31

XB3 ortholog 1 in Arabidopsis thaliana

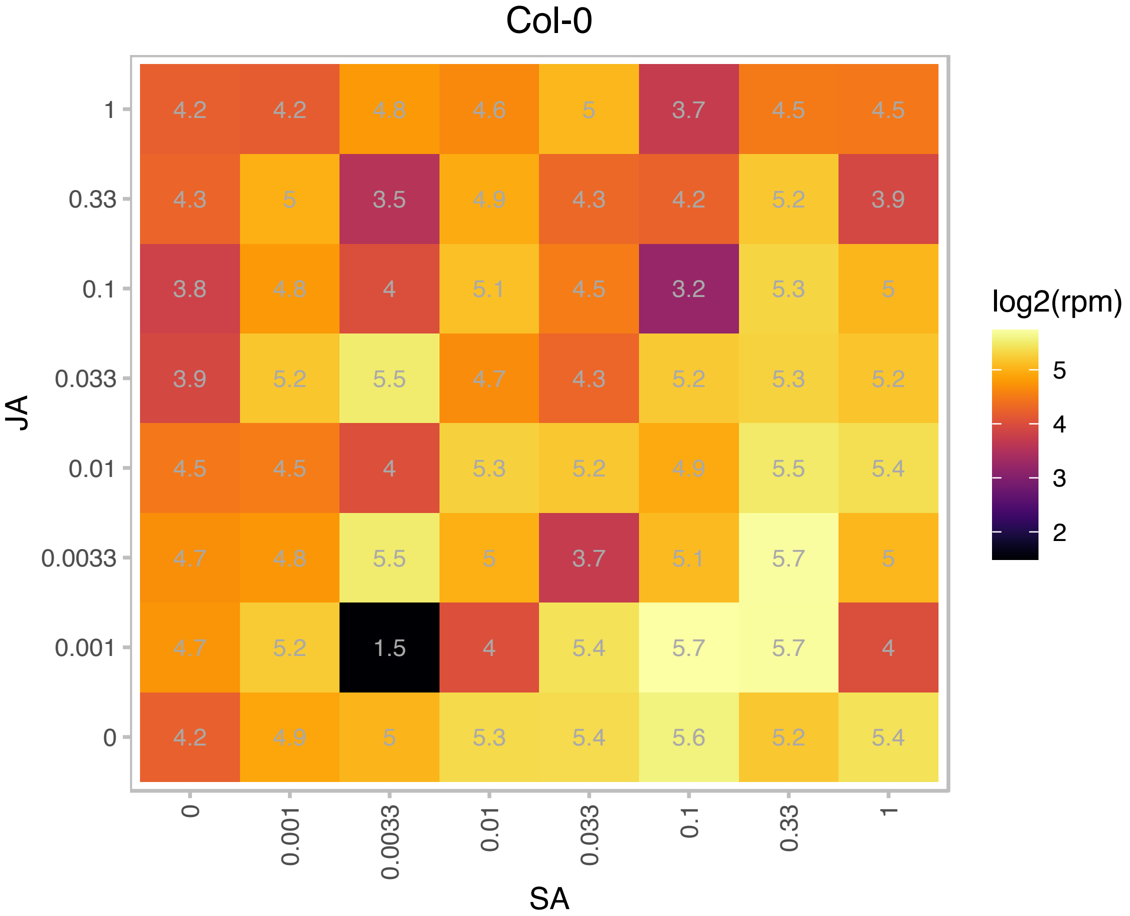


| | |
|---|---|
| Gene | AT2G28840 |
| is.expressed_genes | TRUE |
| is.responsive_genes | TRUE |
| is.apcluster | C-01 |

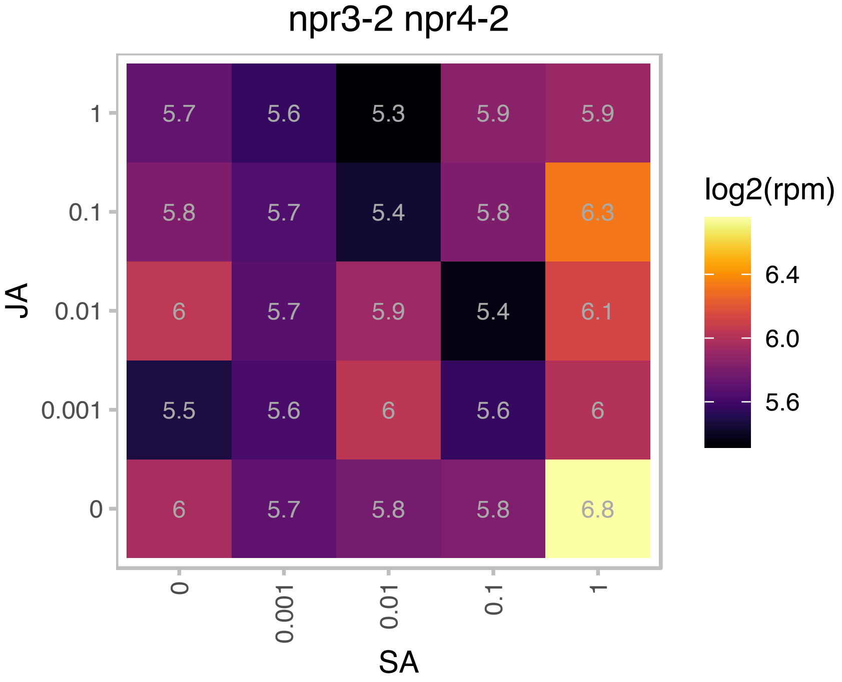


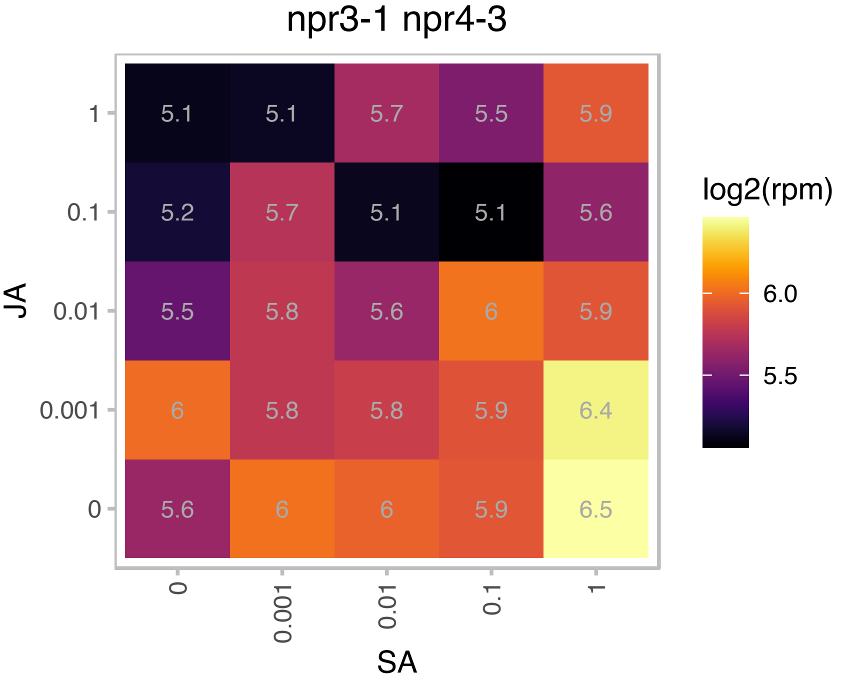

# AT2G32160.1 : NA

S-adenosyl-L-methionine-dependent methyltransferases superfamily protein

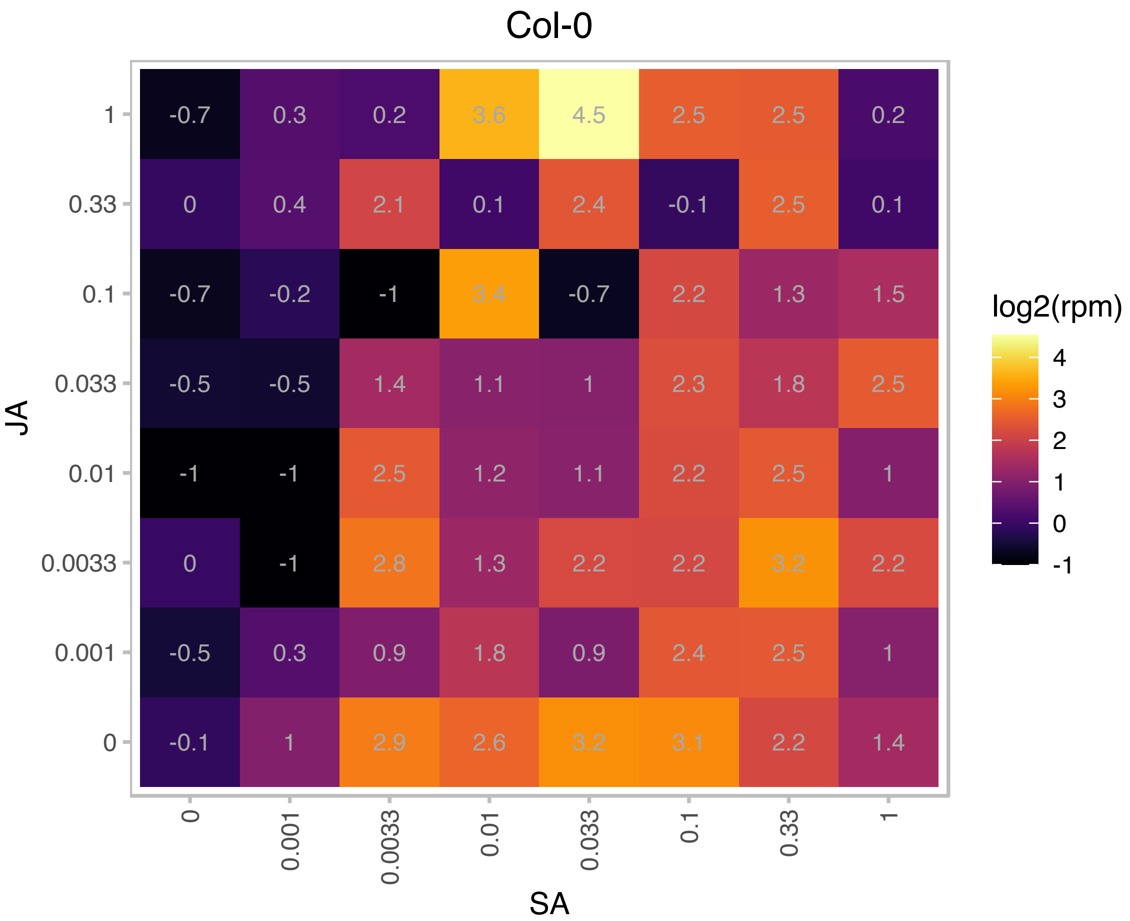


| | |
|---|---|
| Gene | AT2G32160 |
| is.expressed_genes | TRUE |
| is.responsive_genes | TRUE |
| is.apcluster | C-01 |

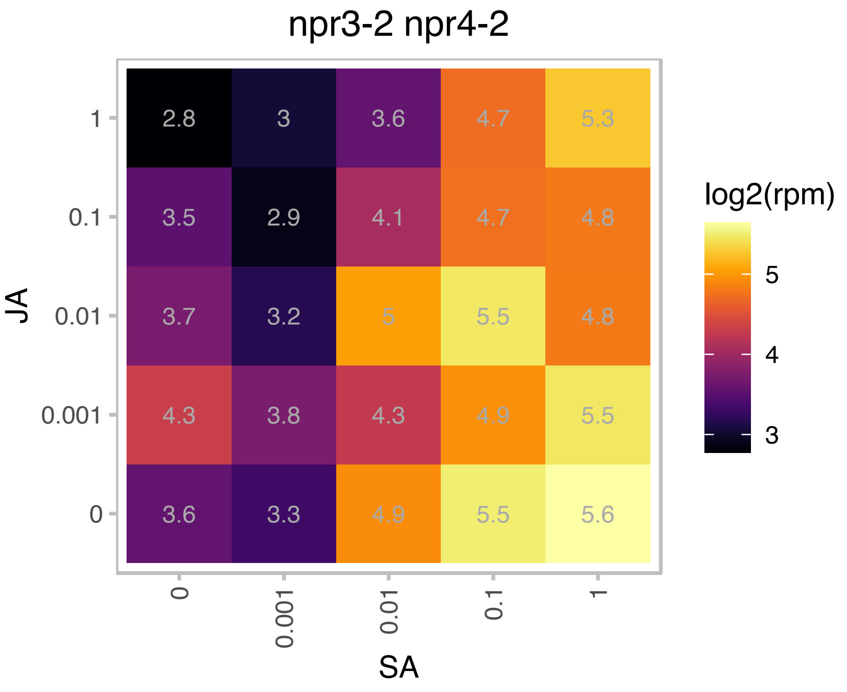


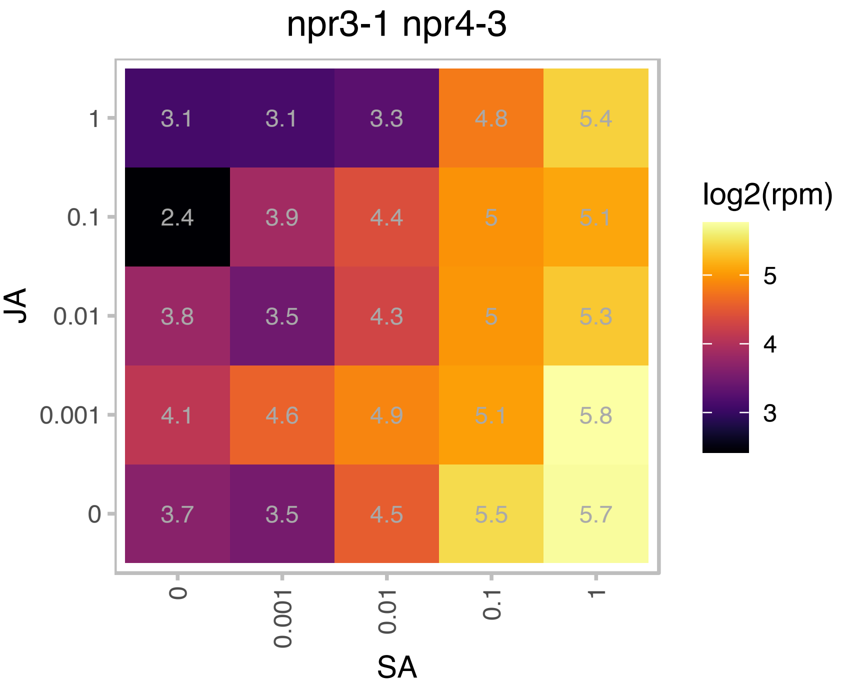

### AT2G37710.1 : LecRK-IV.1, RLK

receptor lectin kinase

Induced in response to Salicylic acid. The mRNA is cell-to-cell mobile.

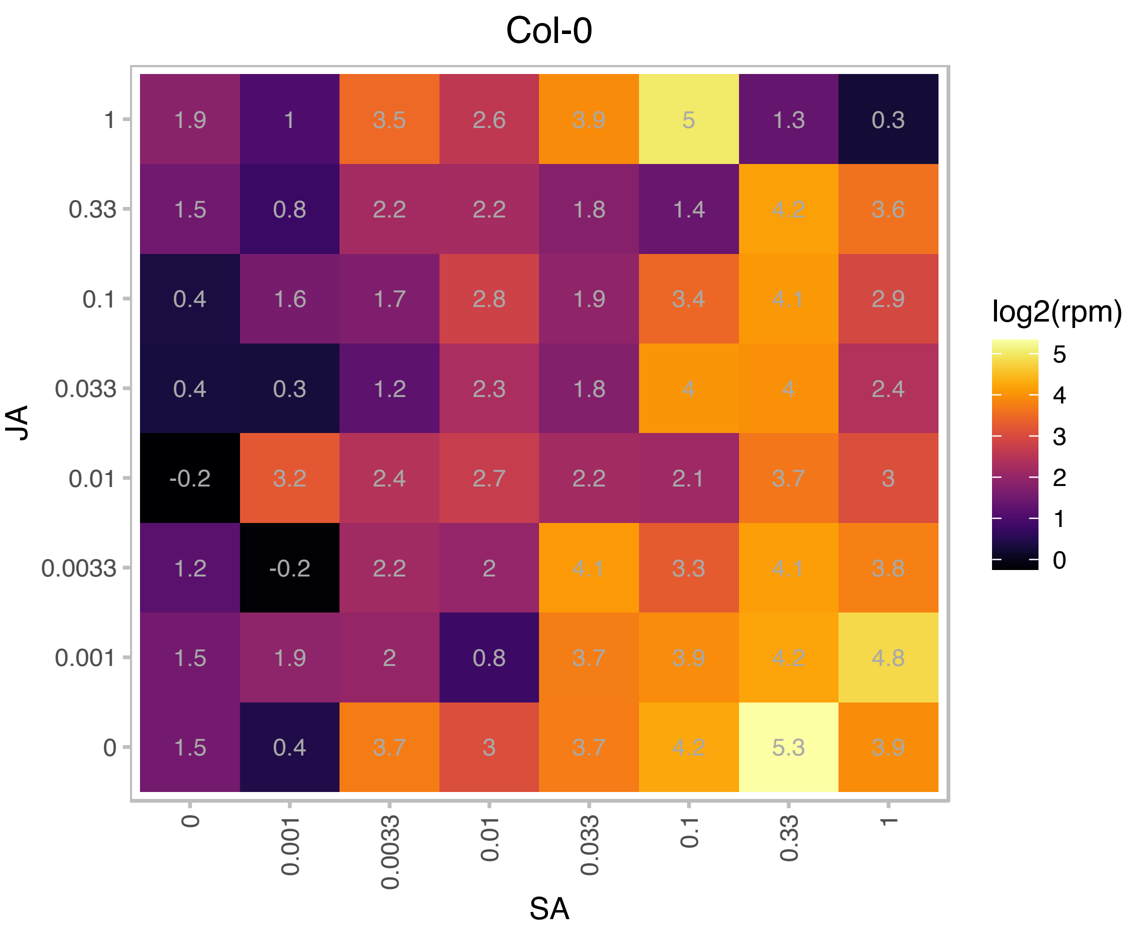


| | |
|---|---|
| Gene | AT2G37710 |
| is.expressed_genes | TRUE |
| is.responsive_genes | TRUE |
| is.apcluster | C-01 |

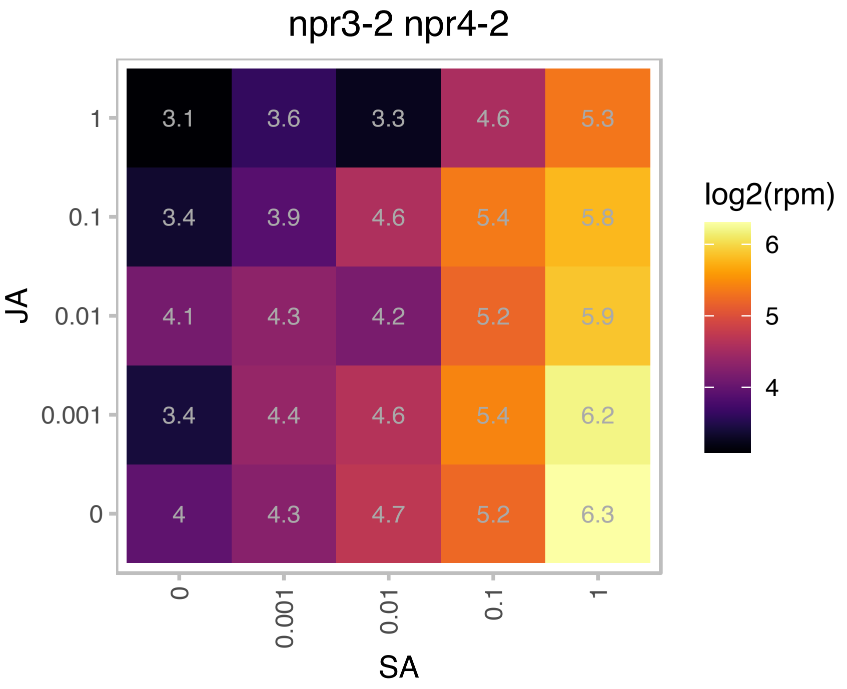


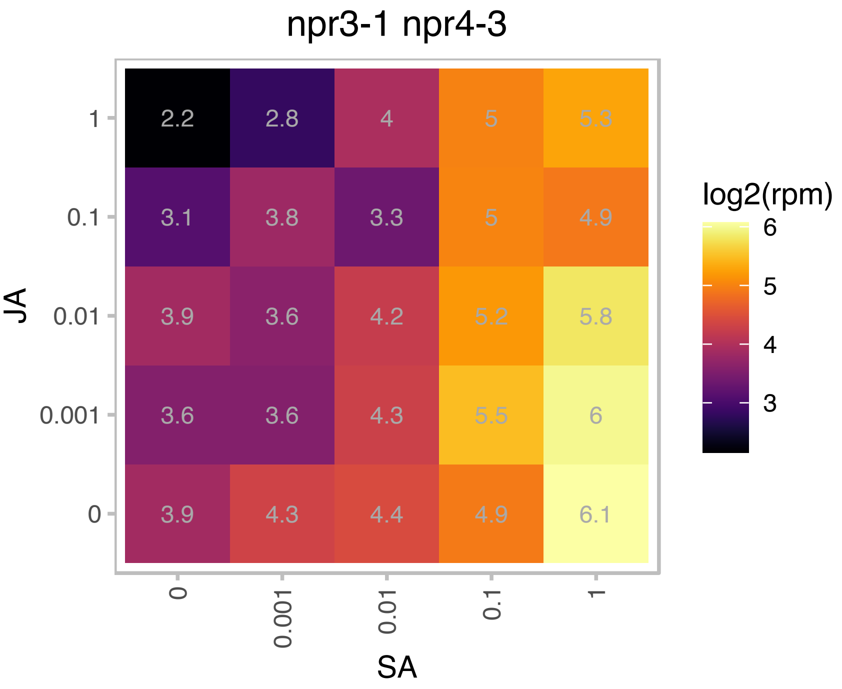

### AT2G40750.1 : ATWRKY54, WRKY54

WRKY DNA-binding protein 54

member of WRKY Transcription Factor; Group III

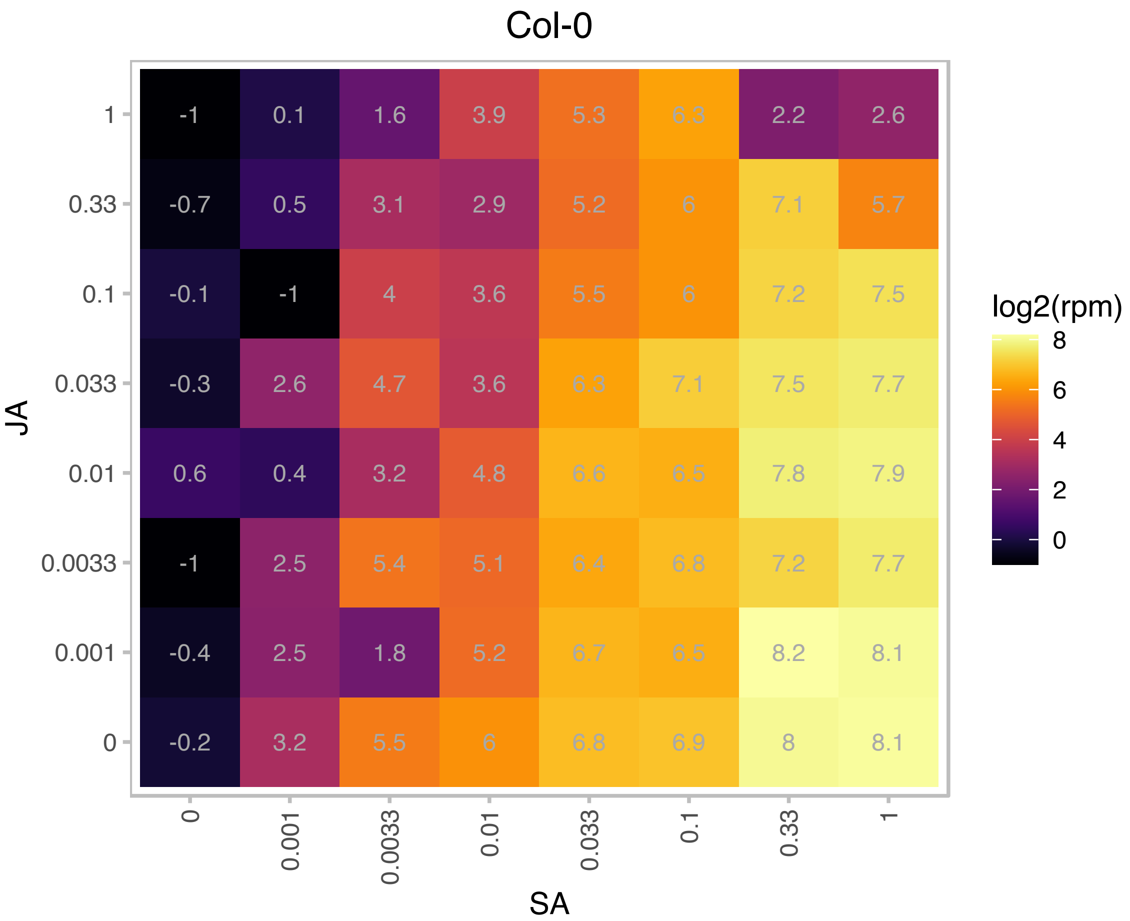


| | |
|---|---|
| Gene | AT2G40750 |
| is.expressed_genes | TRUE |
| is.responsive_genes | TRUE |
| is.apcluster | C-01 |

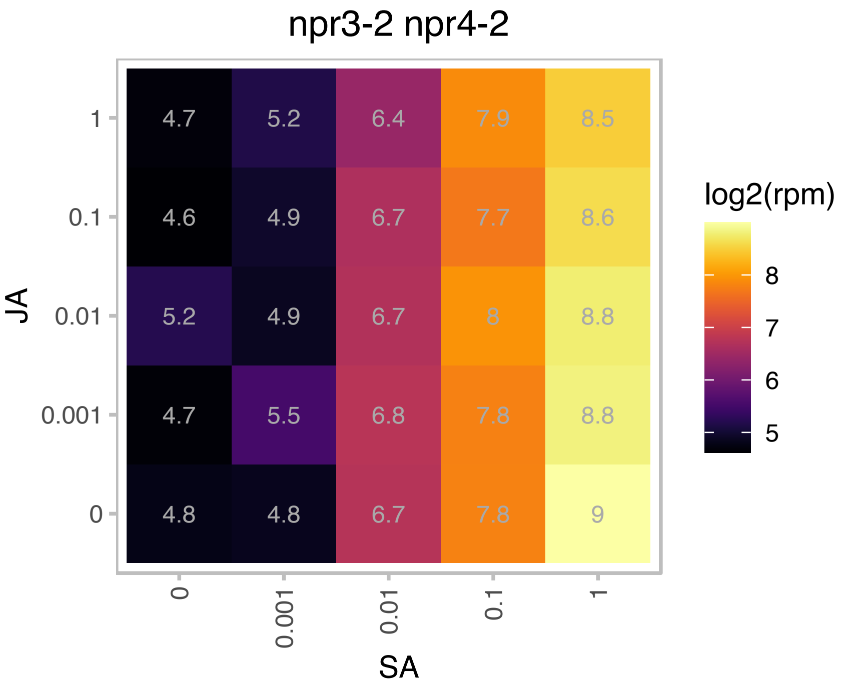


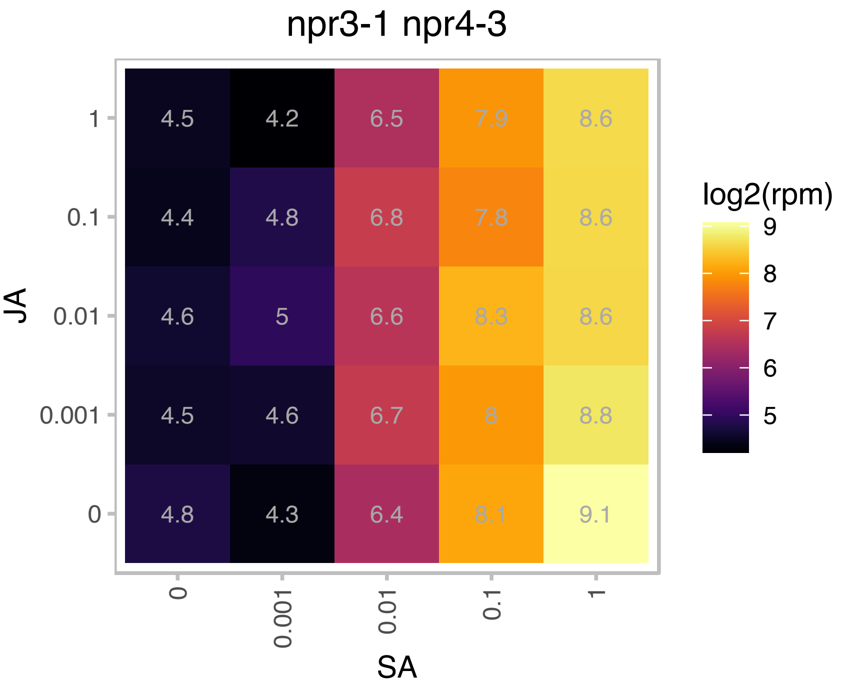

### AT2G46430.1 : ATCNGC3, CNGC3, CNGC3.C

cyclic nucleotide gated channel 3

Encodes a cyclic nucleotide gated channel, downstream component of the signaling pathways leading to hypersensitive response (HR) resistance.

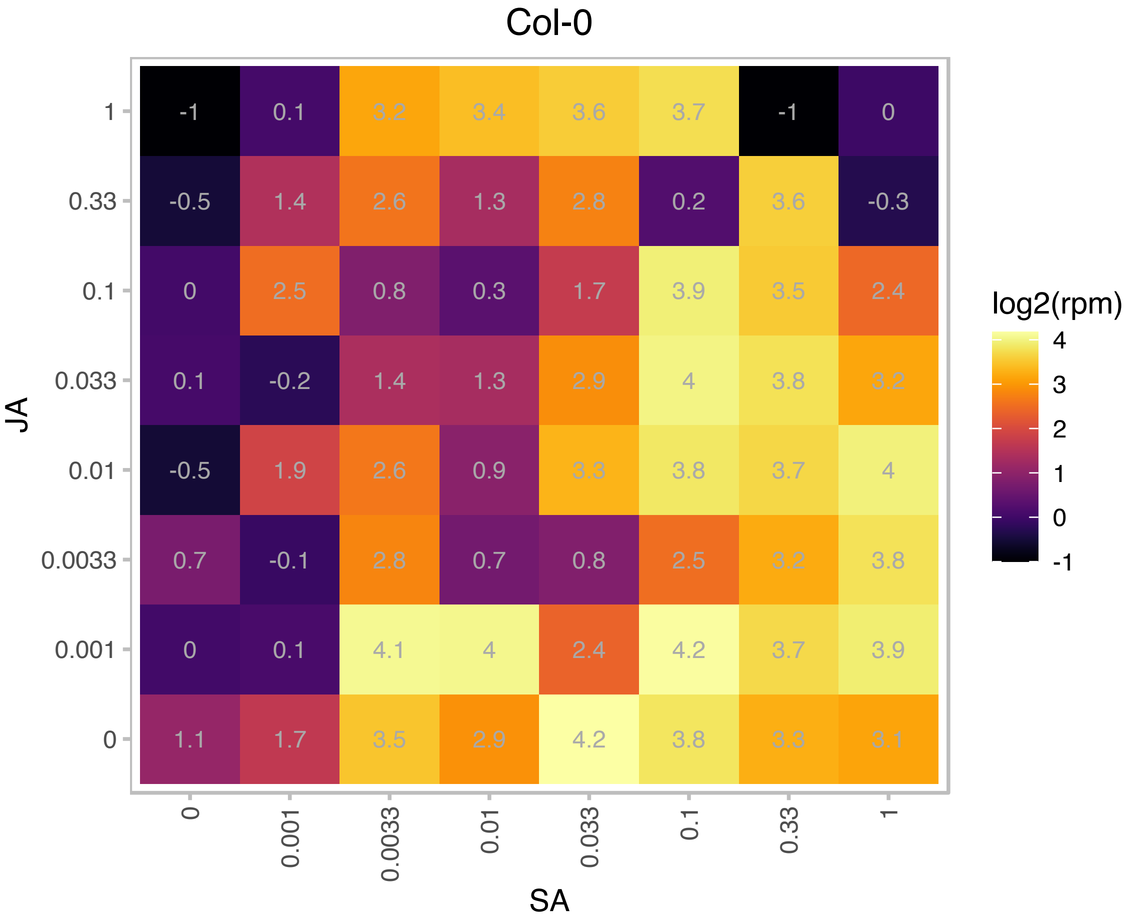


| | |
|---|---|
| Gene | AT2G46430 |
| is.expressed_genes | TRUE |
| is.responsive_genes | TRUE |
| is.apcluster | C-01 |

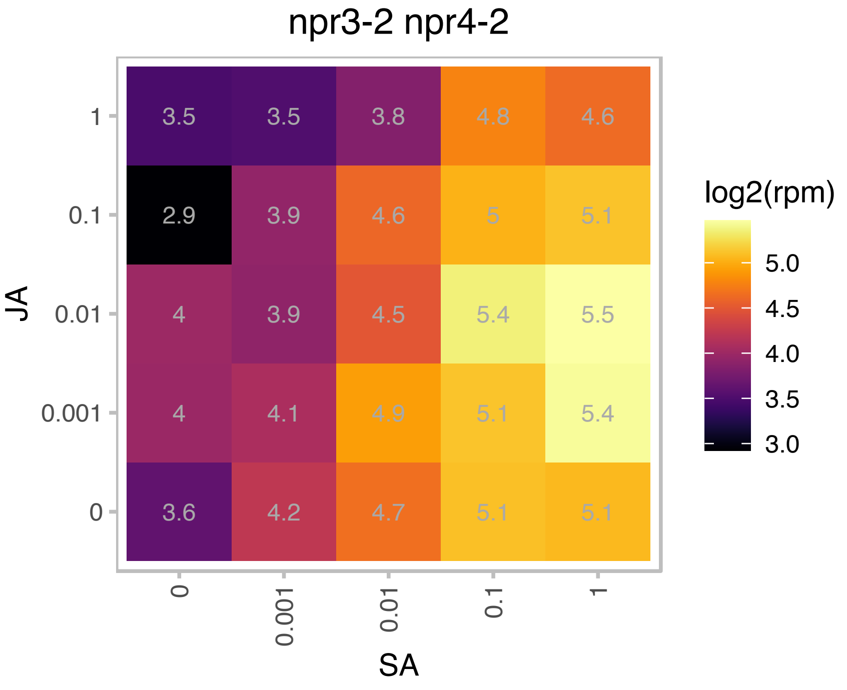


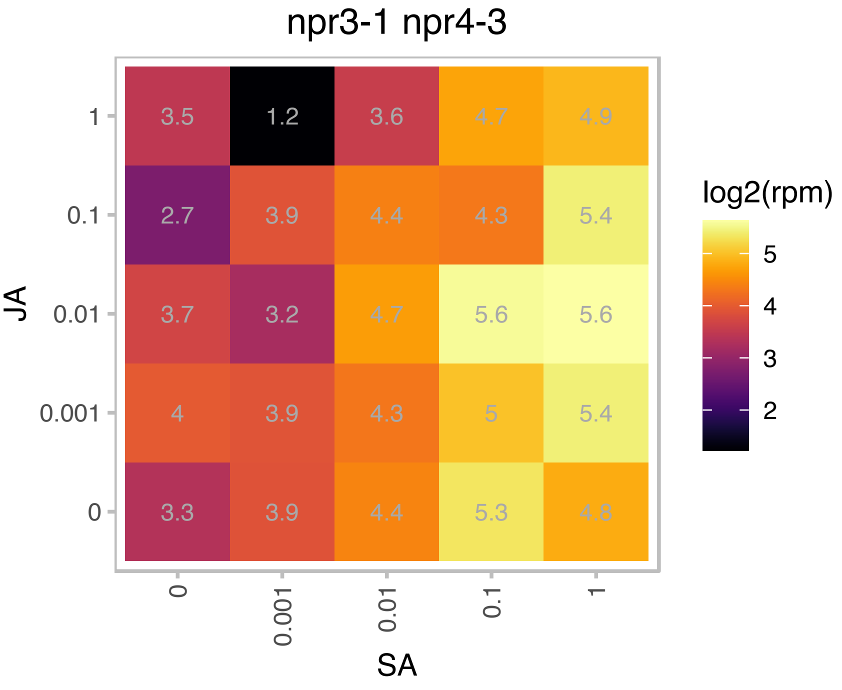

### AT2G46440.1 : ATCNGC11, CNGC11

cyclic nucleotide-gated channels

Member of Cyclic nucleotide gated channel family. Positive regulator of resistance against avirulent fungal pathogen. The mRNA is cell-to-cell mobile.

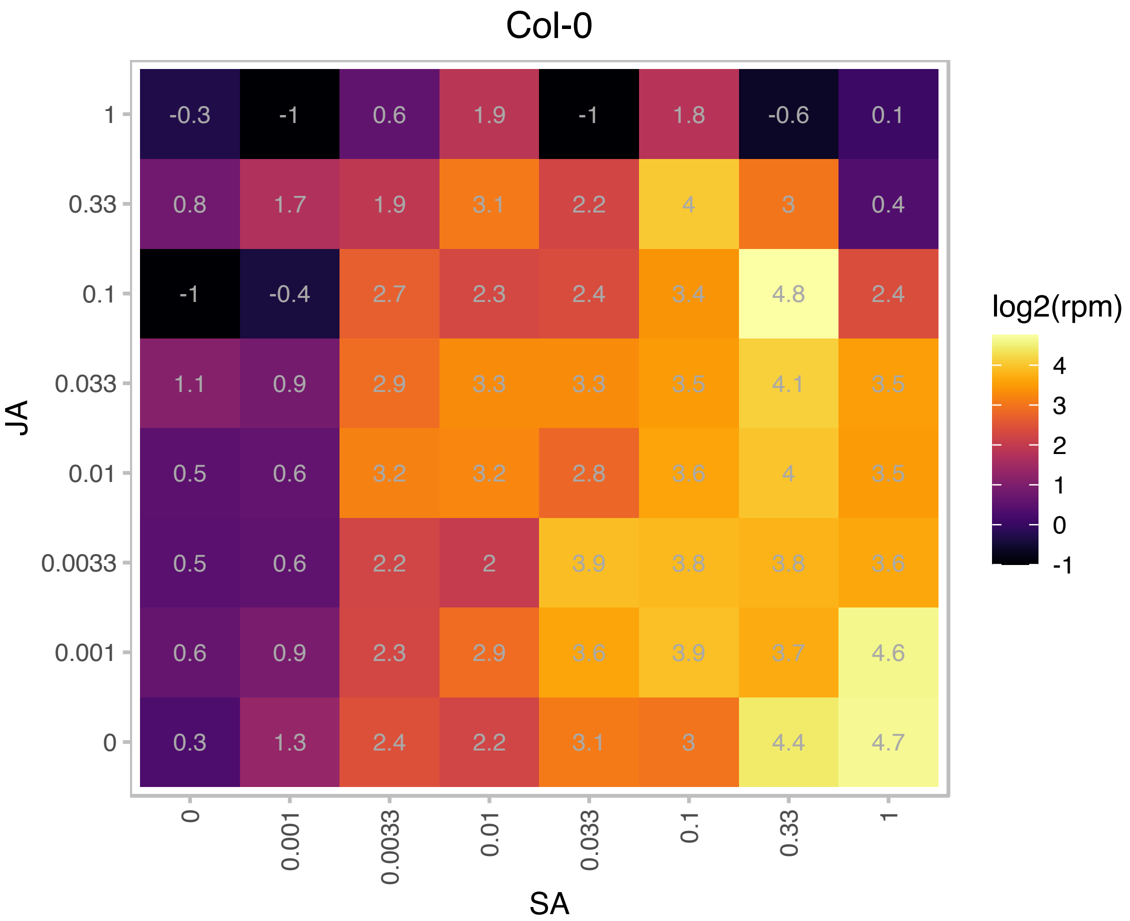


| | |
|---|---|
| Gene | AT2G46440 |
| is.expressed_genes | TRUE |
| is.responsive_genes | TRUE |
| is.apcluster | C-01 |

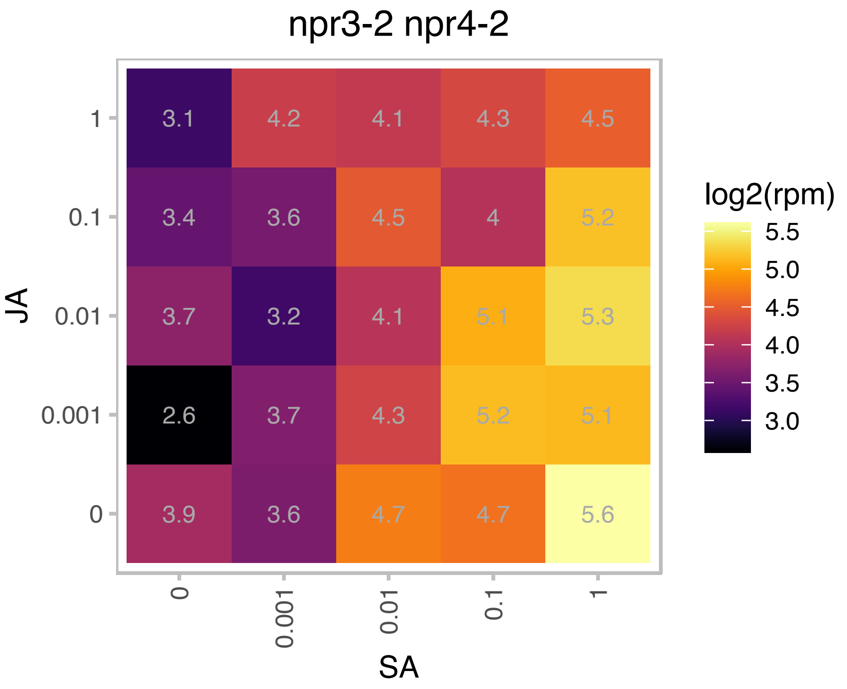


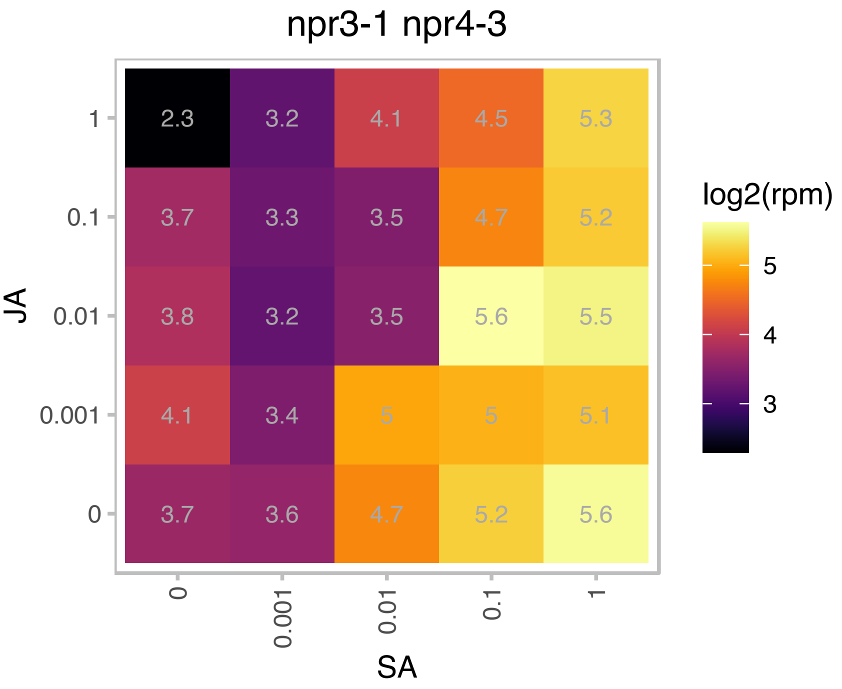

# AT2G47640.1 : NA

Small nuclear ribonucleoprotein family protein

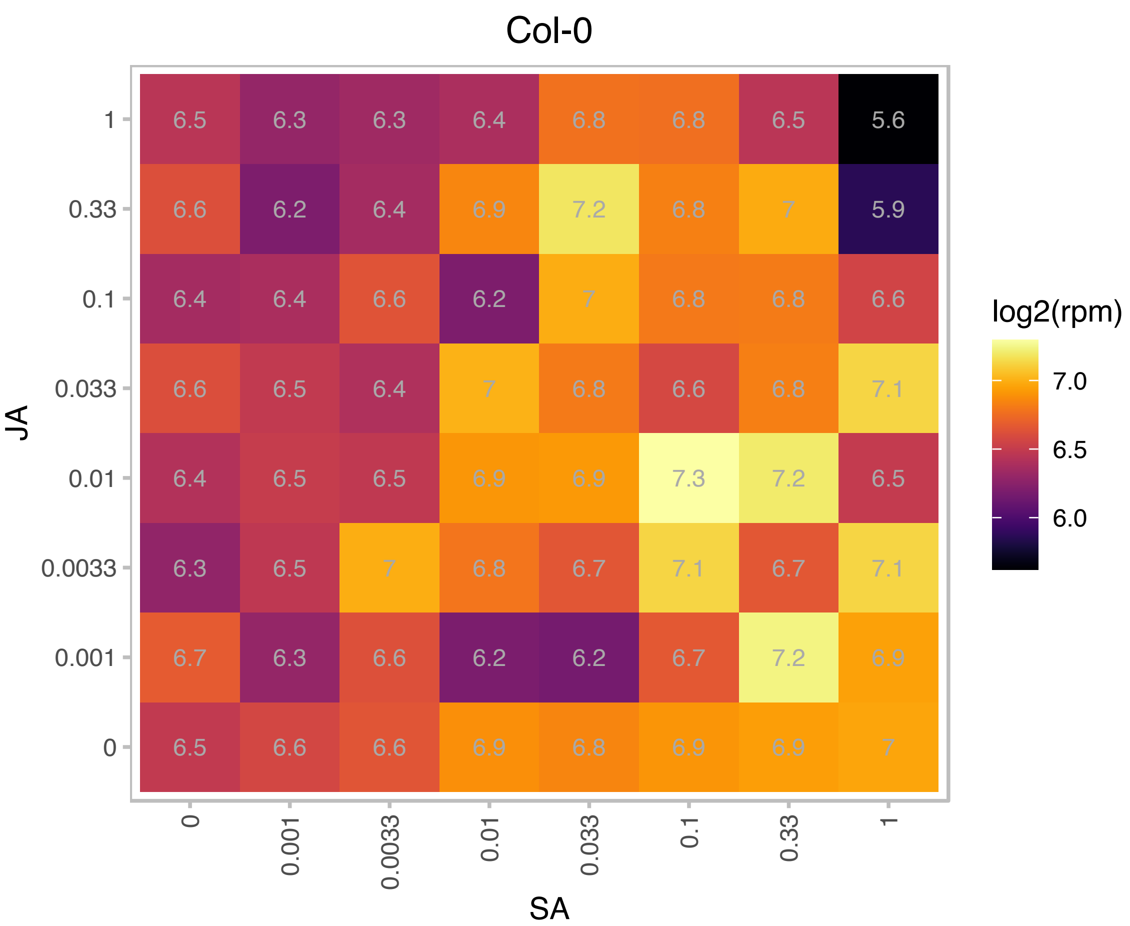


| | |
|---|---|
| Gene | AT2G47640 |
| is.expressed_genes | TRUE |
| is.responsive_genes | TRUE |
| is.apcluster | C-01 |

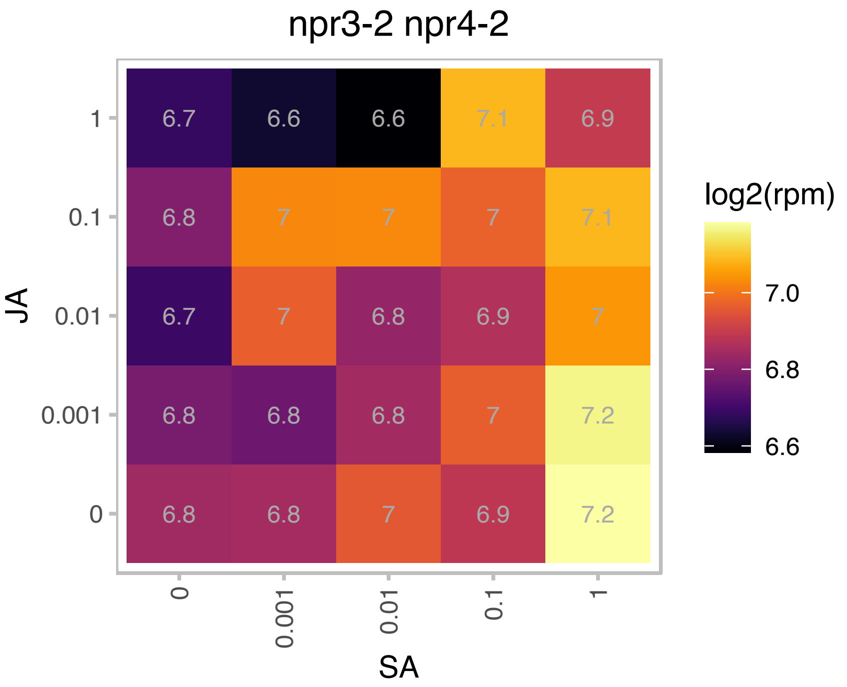


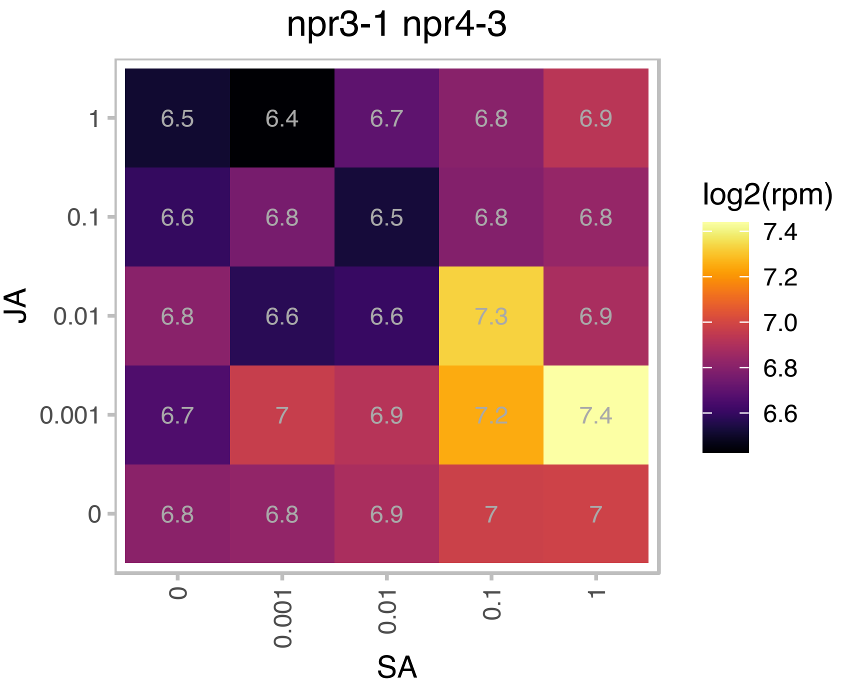

### AT3G22231.1 : PCC1

pathogen and circadian controlled 1
Encodes a member of a novel 6 member Arabidopsis gene family. Expression of PCC1 is regulated by the circadian clock and is upregulated in response to both virulent and avirulent strains of Pseudomonas syringae pv. tomato.

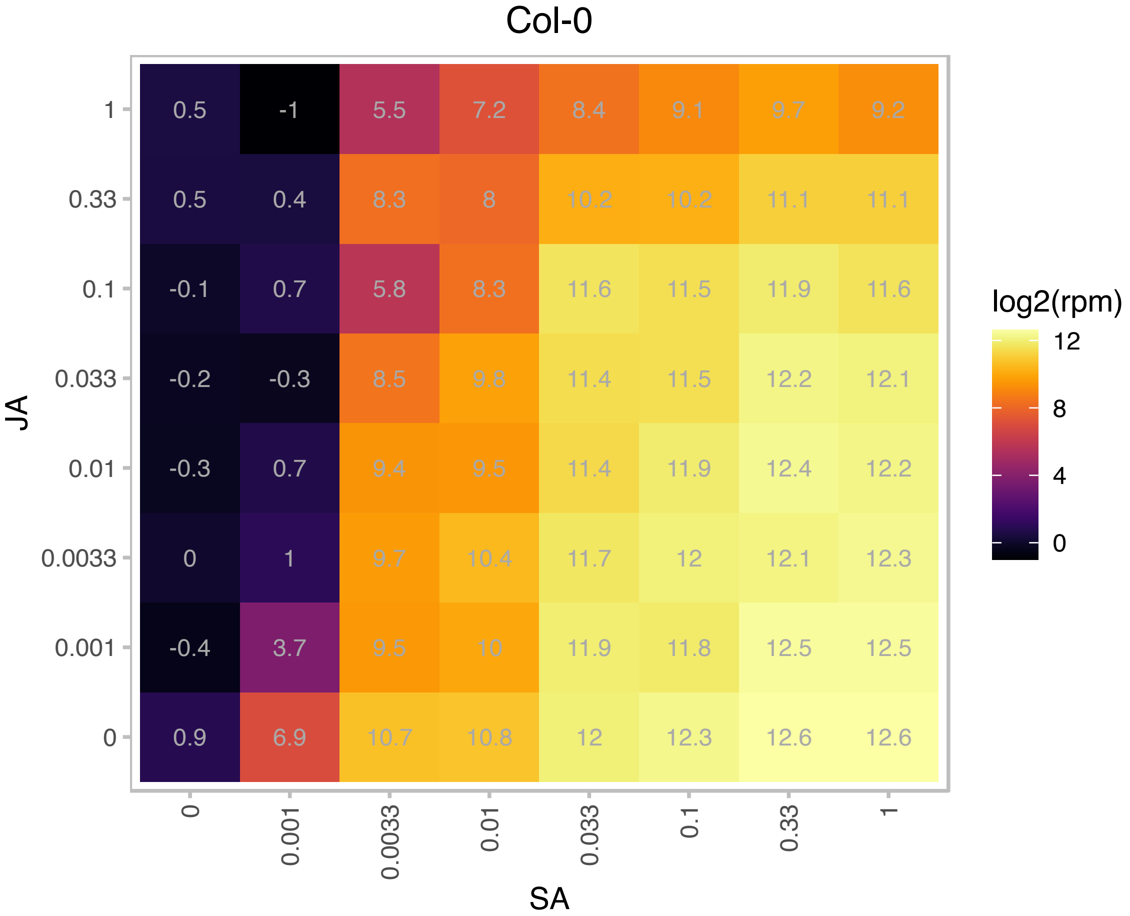


| Gene | AT3G22231 |
|---|---|
| is.expressed_genes | TRUE |
| is.responsive_genes | TRUE |
| is.apcluster | C-01 |

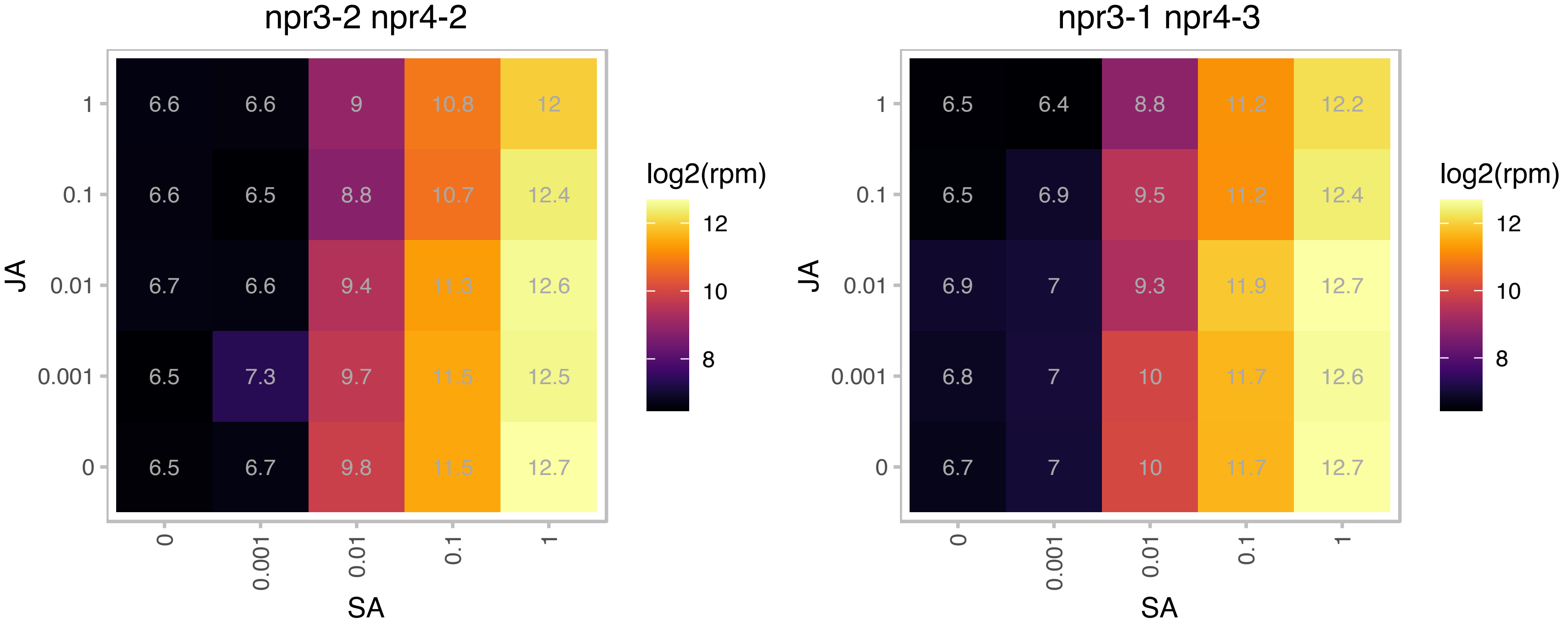

### AT3G22235.1 : AthCYSTM8

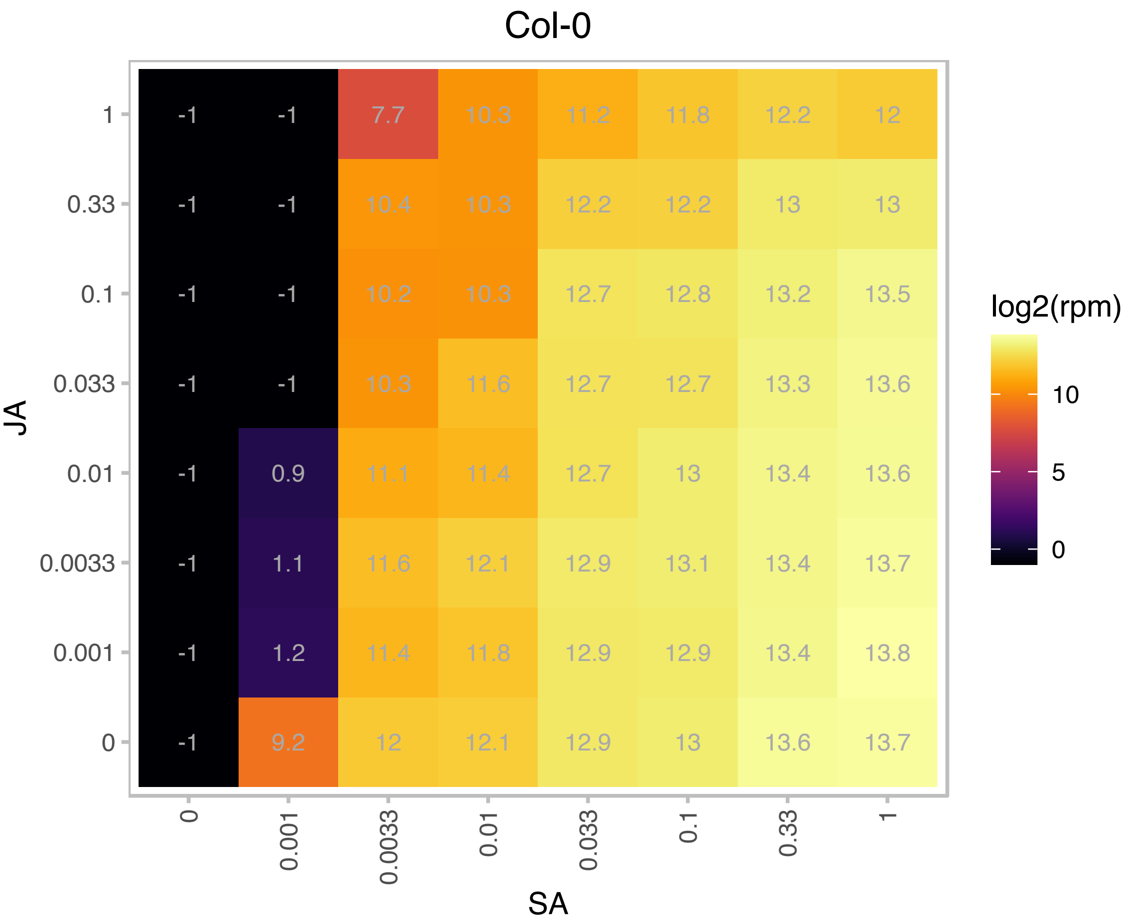


| | |
|---|---|
| Gene | AT3G22235 |
| is.expressed_genes | TRUE |
| is.responsive_genes | TRUE |
| is.apcluster | C-01 |

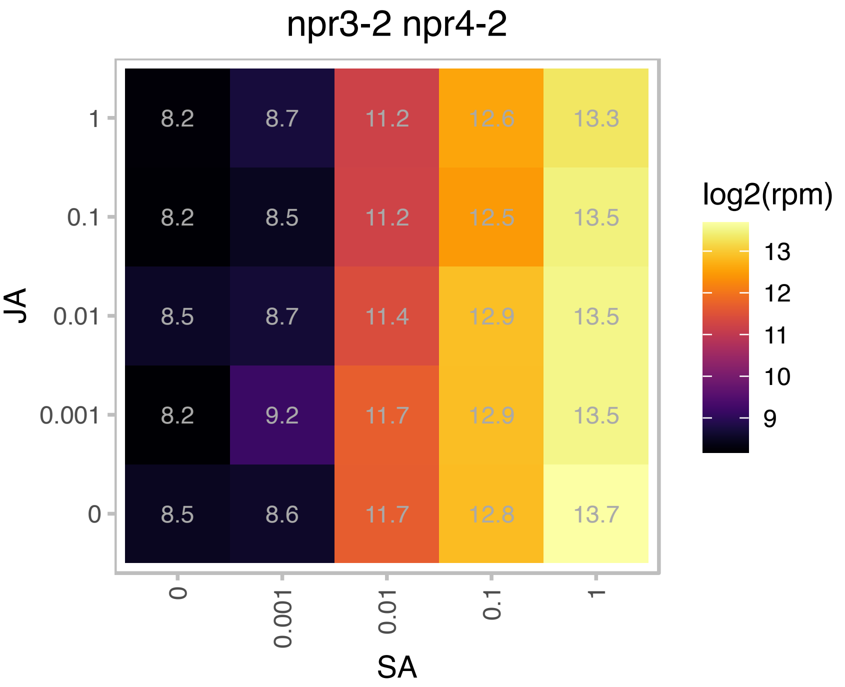


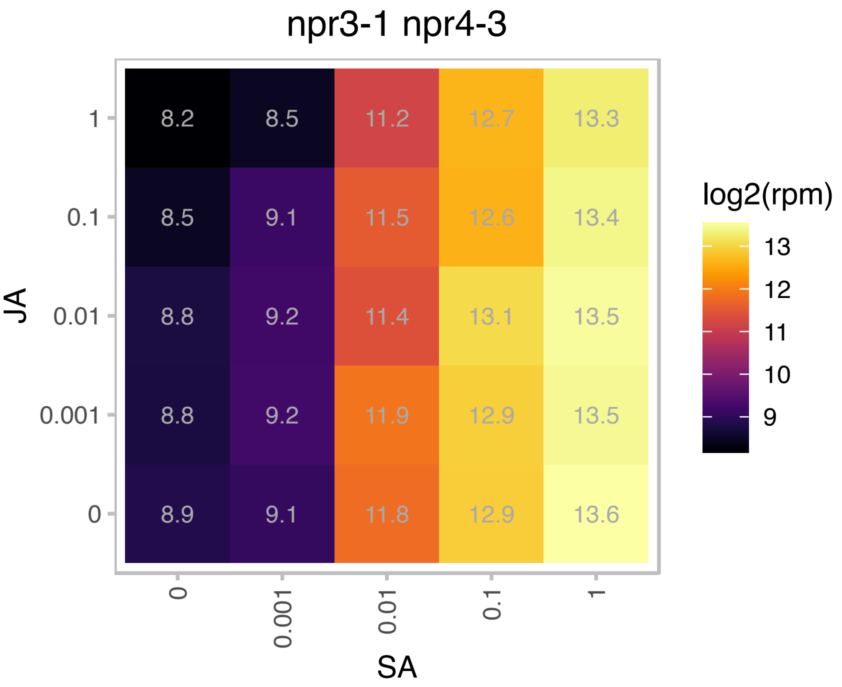

### AT3G22240.1 : AthCYSTM9

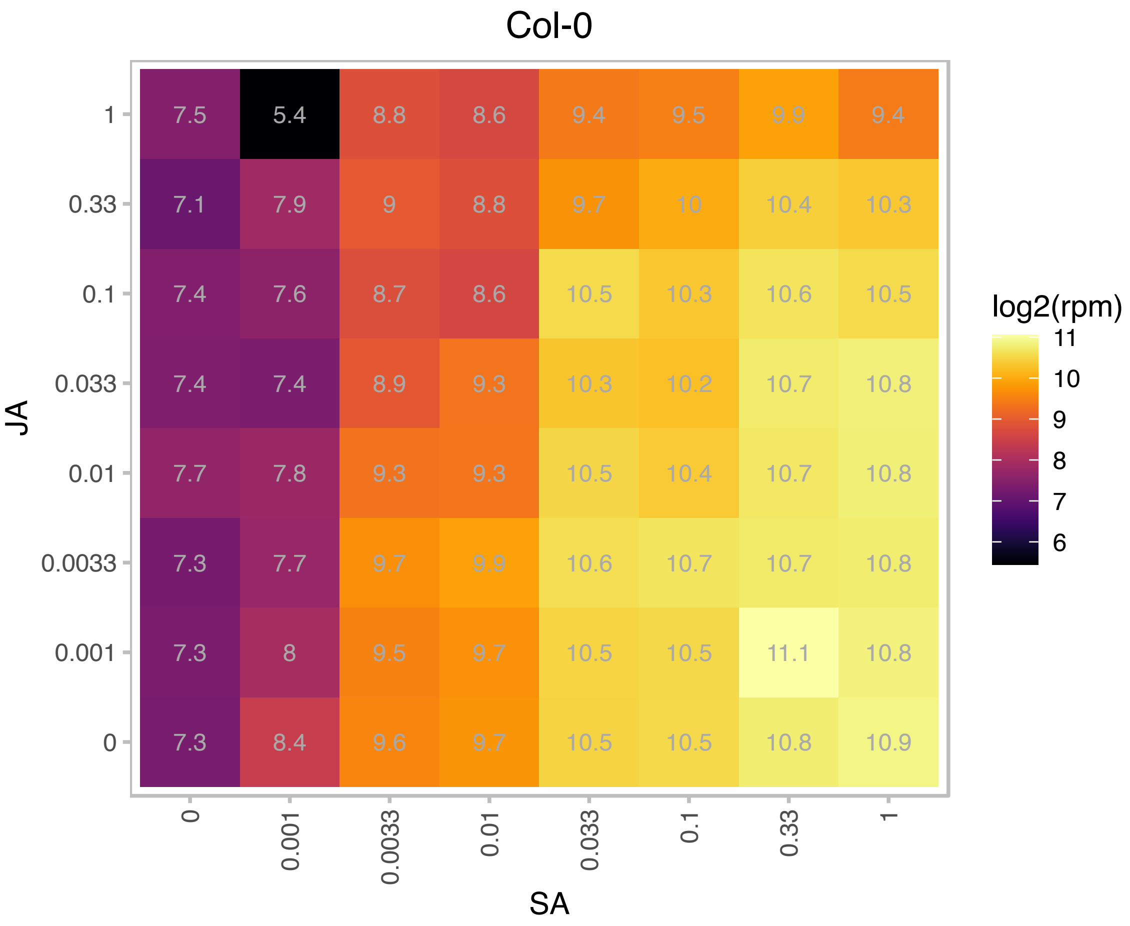


| | |
|---|---|
| Gene | AT3G22240 |
| is.expressed_genes | TRUE |
| is.responsive_genes | TRUE |
| is.apcluster | C-01 |

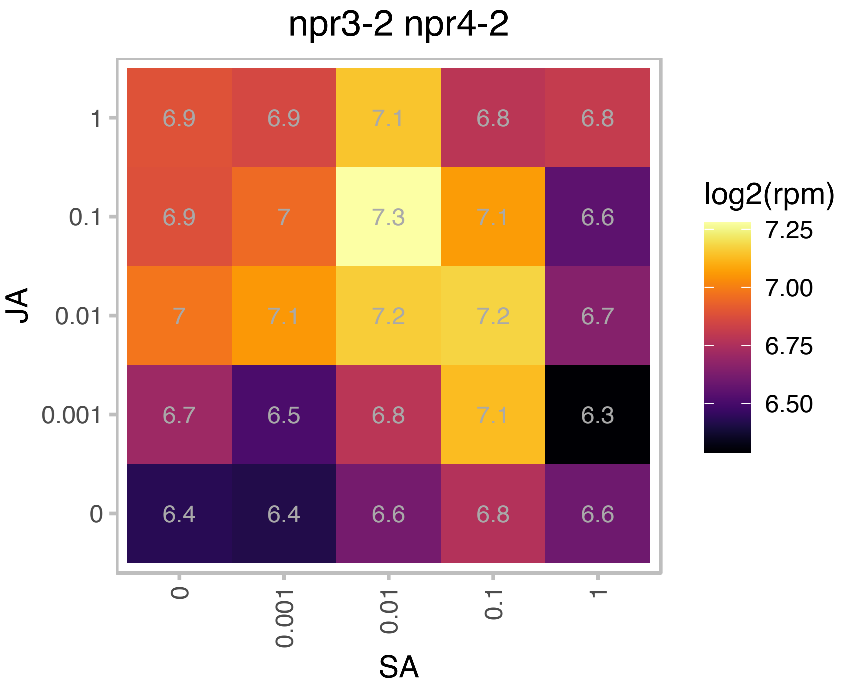


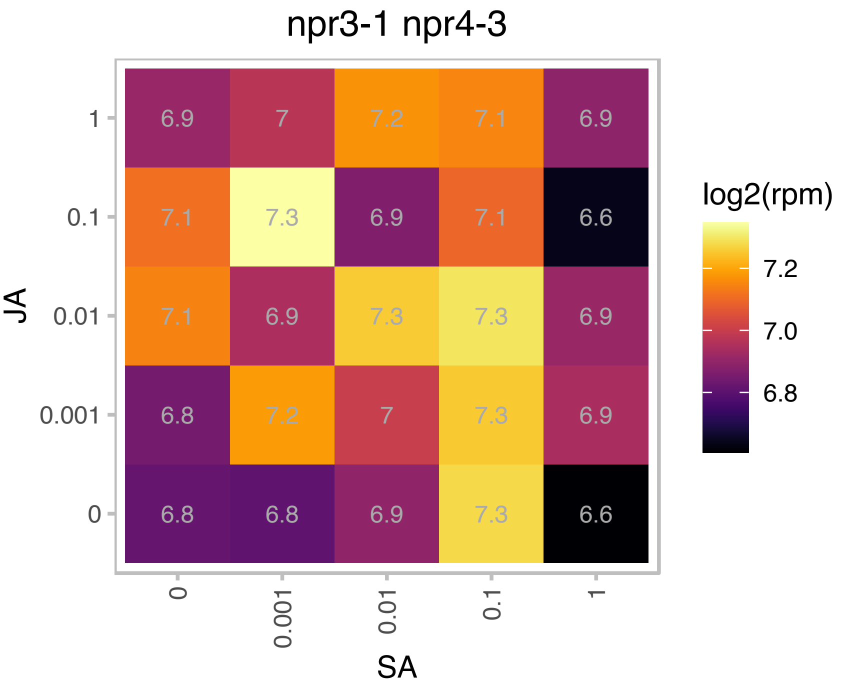

### AT3G25882.1 : NIMIN-2

NIM1-interacting 2

encodes a kinase that physically interacts with NPR1/NIM1

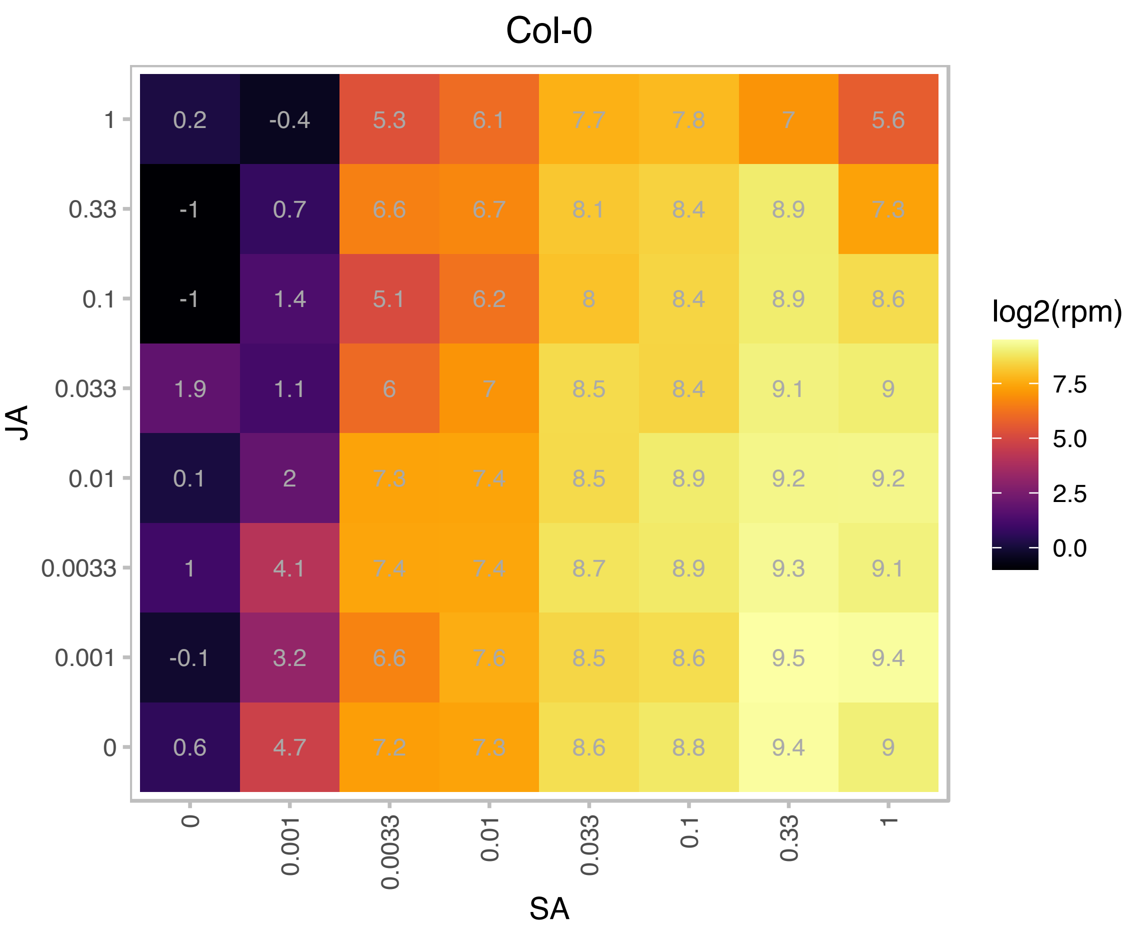


| | |
|---|---|
| Gene | AT3G25882 |
| is.expressed_genes | TRUE |
| is.responsive_genes | TRUE |
| is.apcluster | C-01 |

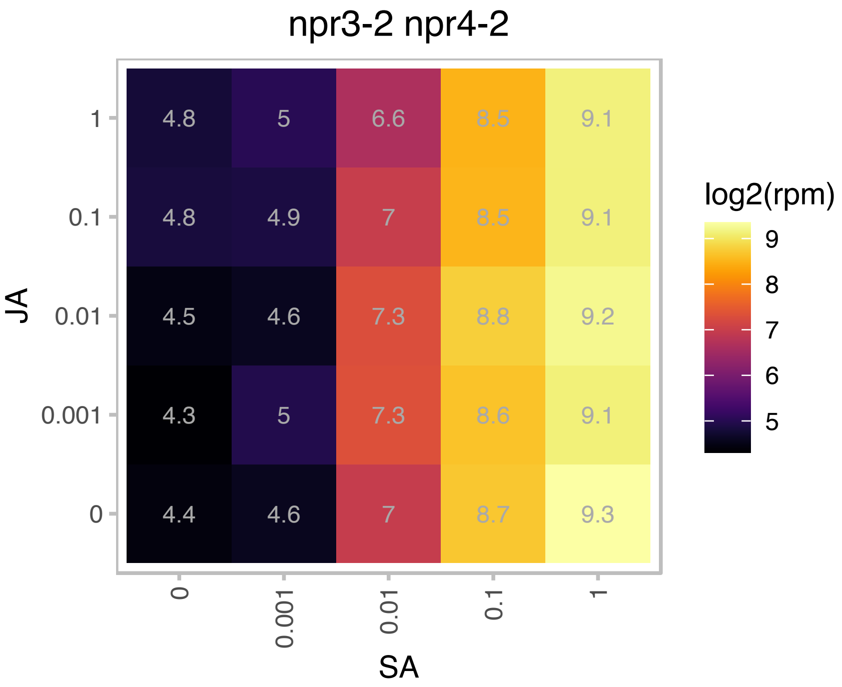


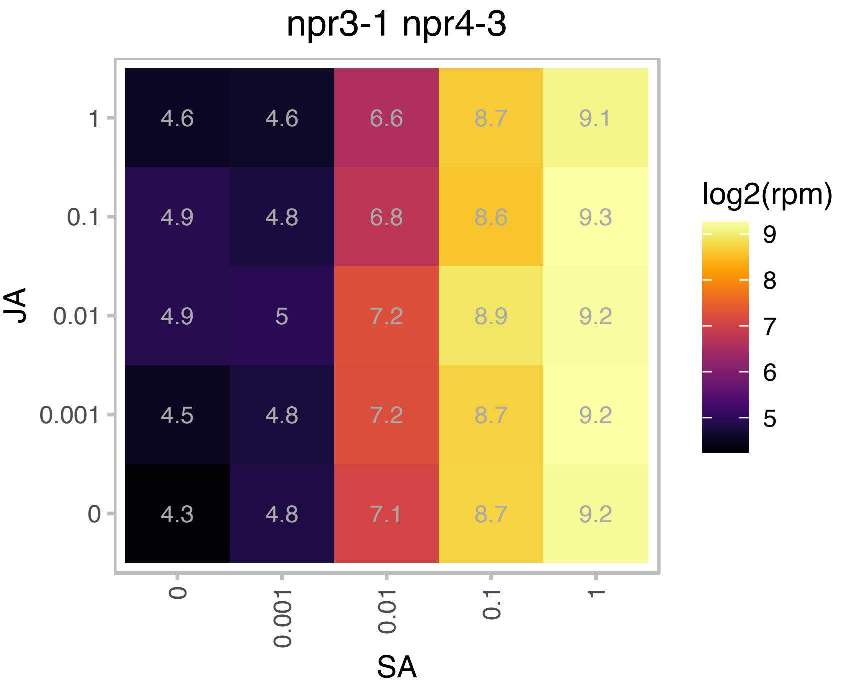

# AT3G27340.1 : NA

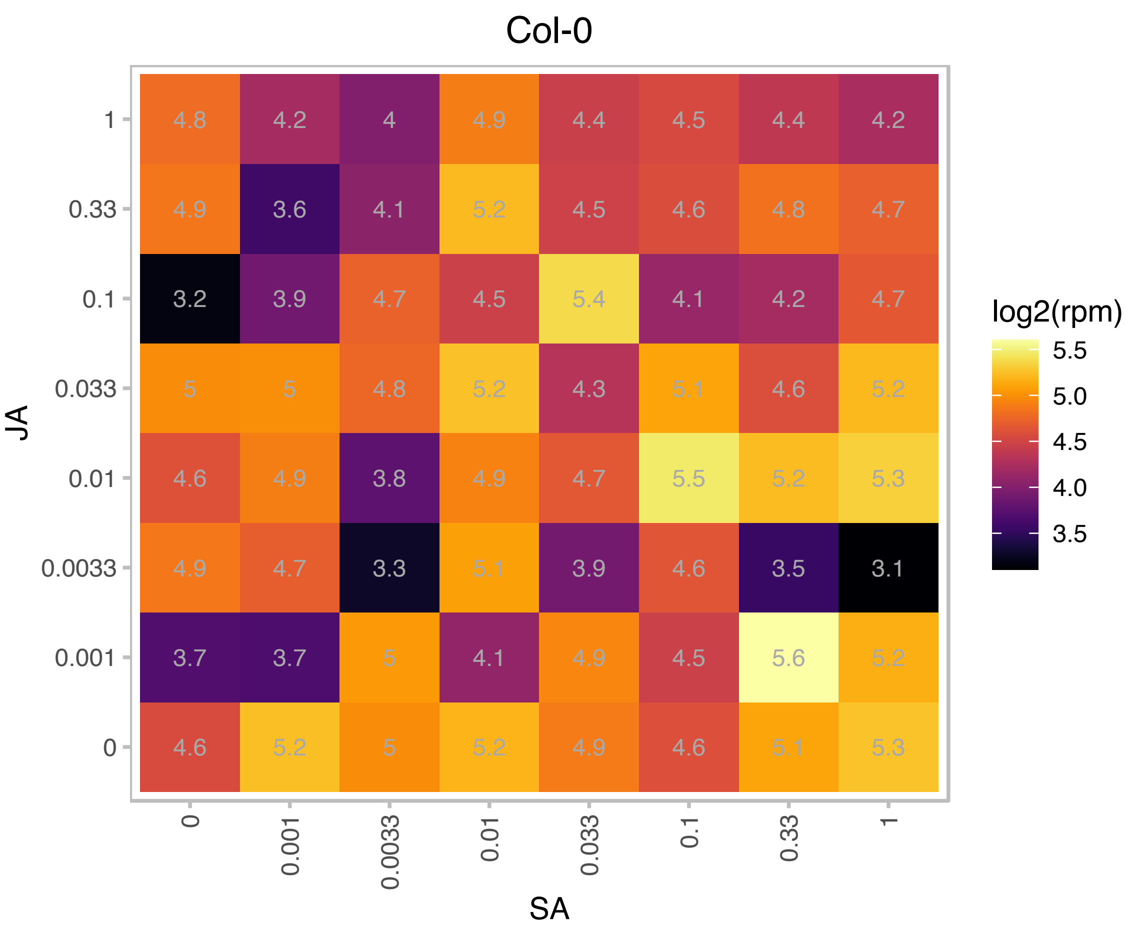


| | |
|---|---|
| Gene | AT3G27340 |
| is.expressed_genes | TRUE |
| is.responsive_genes | TRUE |
| is.apcluster | C-01 |

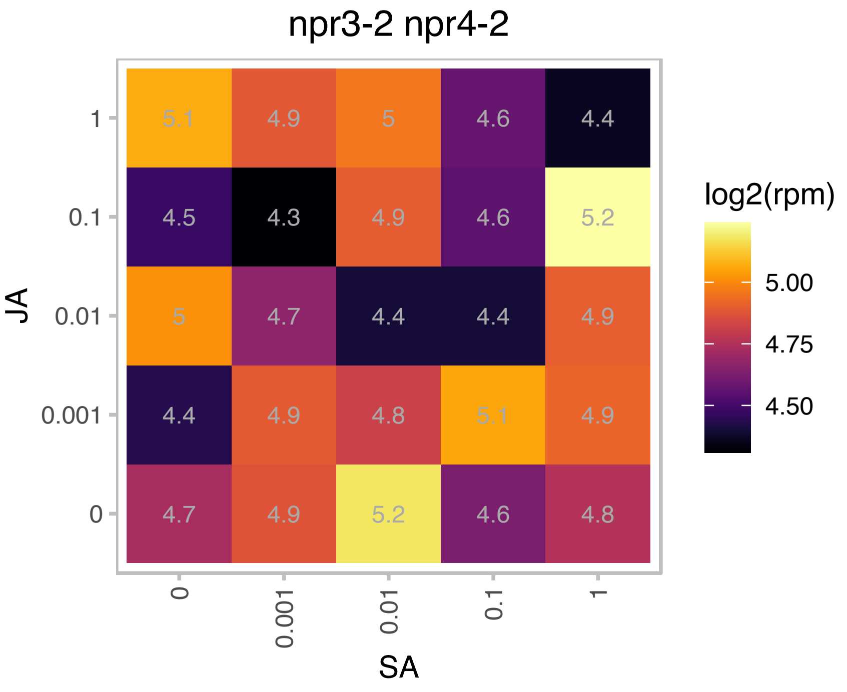


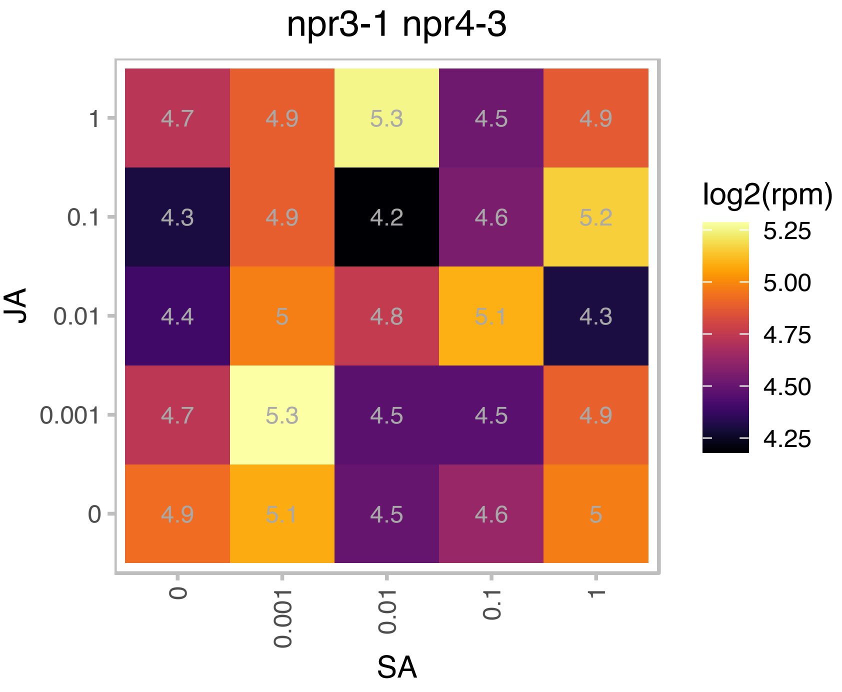

# AT3G28510.1 : NA

P-loop containing nucleoside triphosphate hydrolases superfamily protein

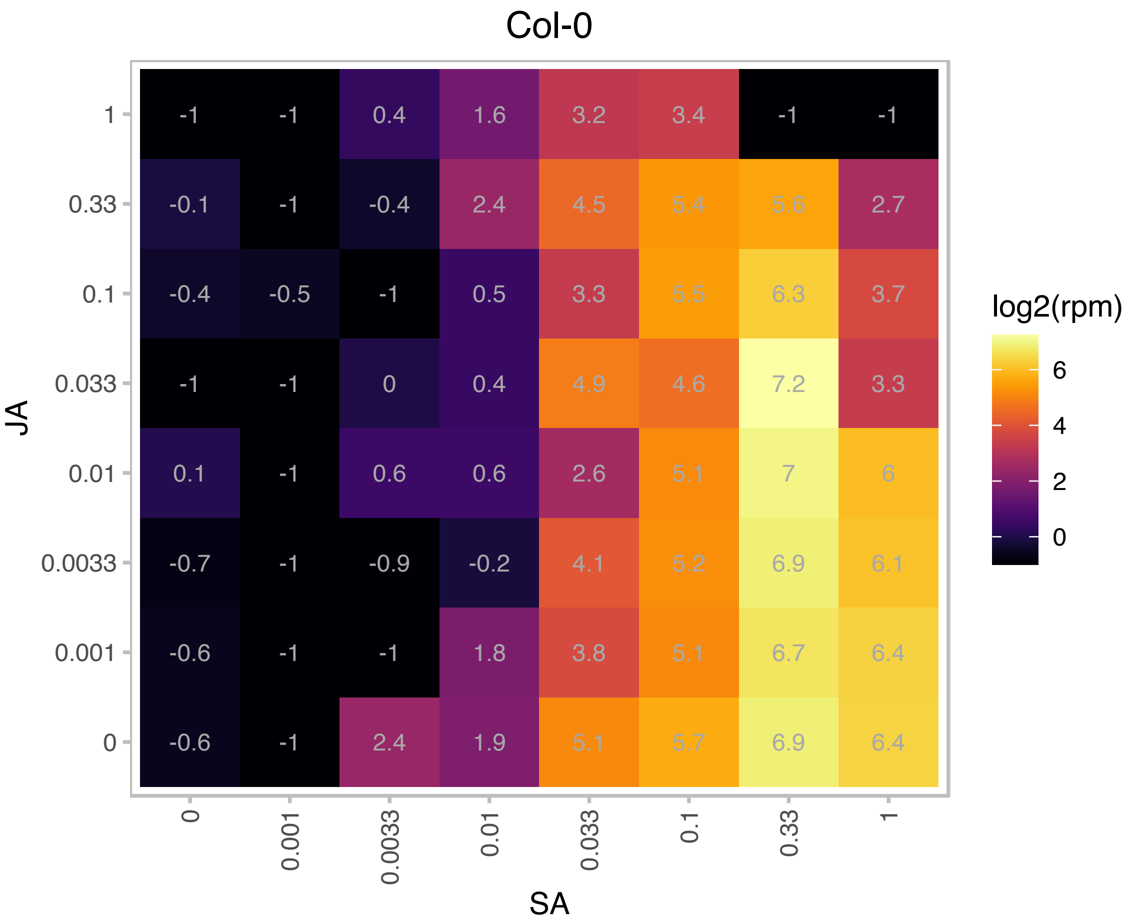


| | |
|---|---|
| Gene | AT3G28510 |
| is.expressed_genes | TRUE |
| is.responsive_genes | TRUE |
| is.apcluster | C-01 |

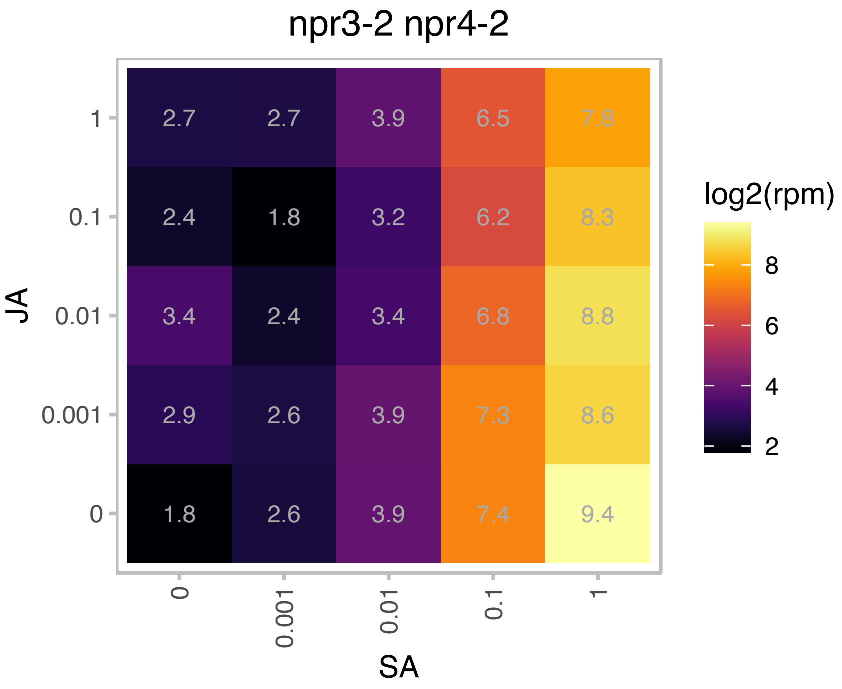


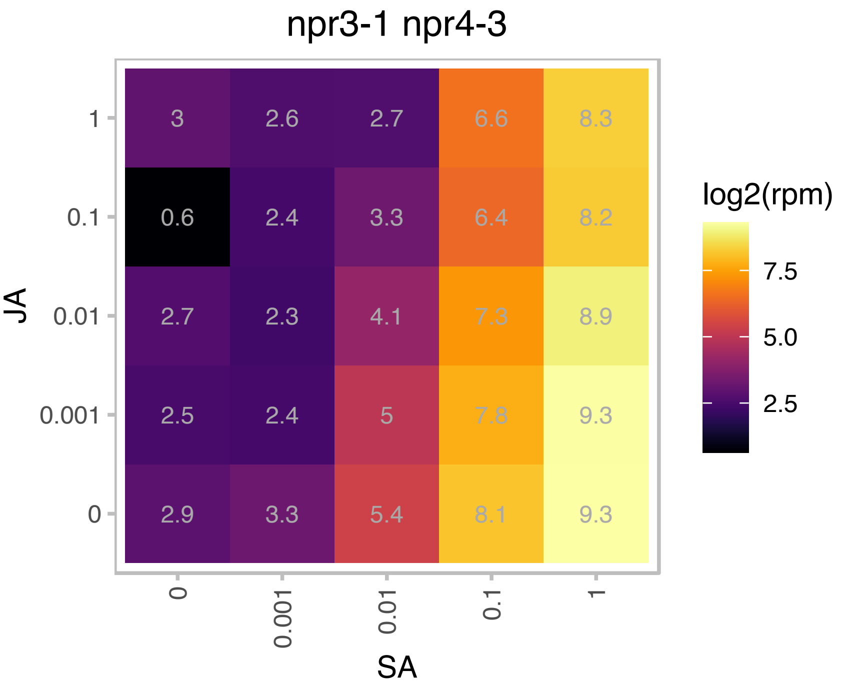

# AT3G28540.1 : NA

P-loop containing nucleoside triphosphate hydrolases superfamily protein

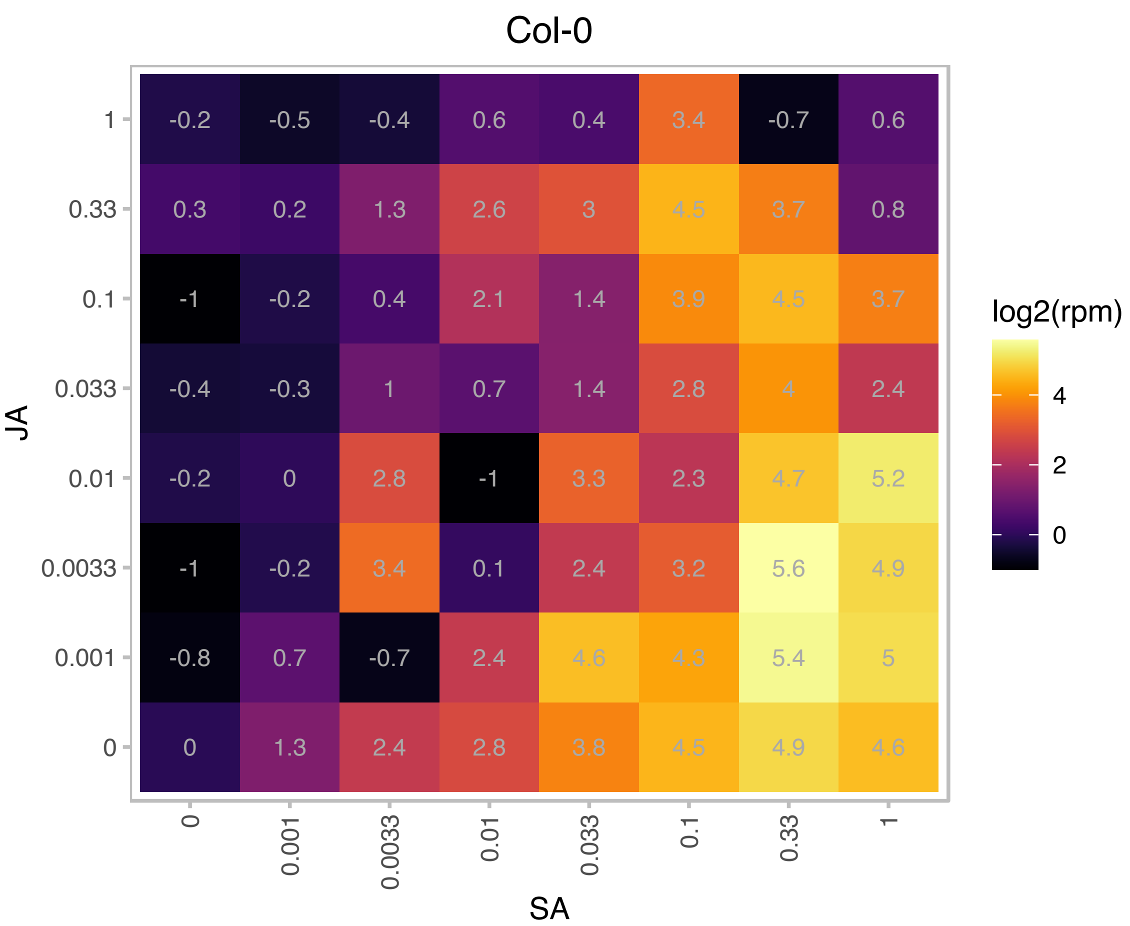


| | |
|---|---|
| Gene | AT3G28540 |
| is.expressed_genes | TRUE |
| is.responsive_genes | TRUE |
| is.apcluster | C-01 |

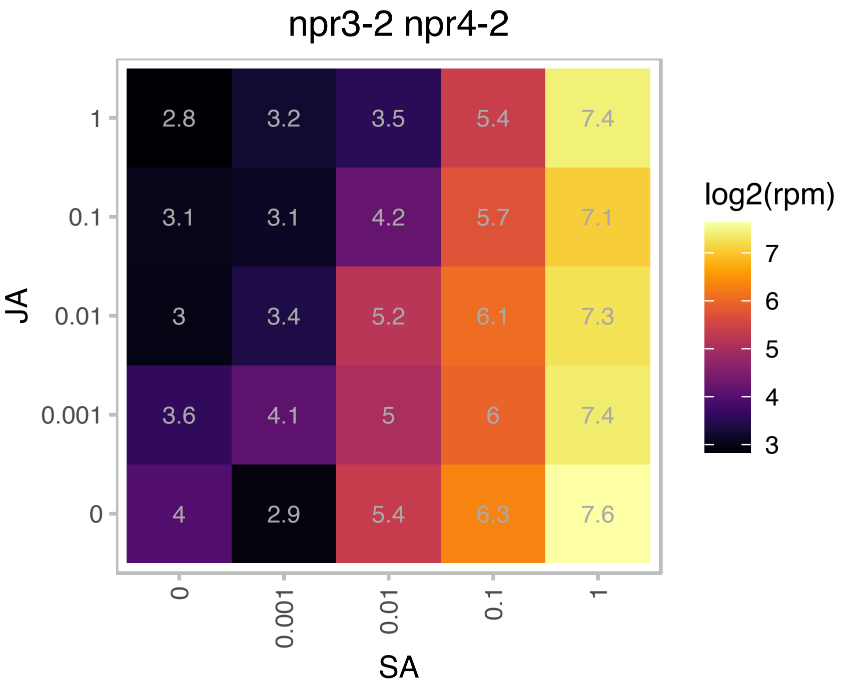


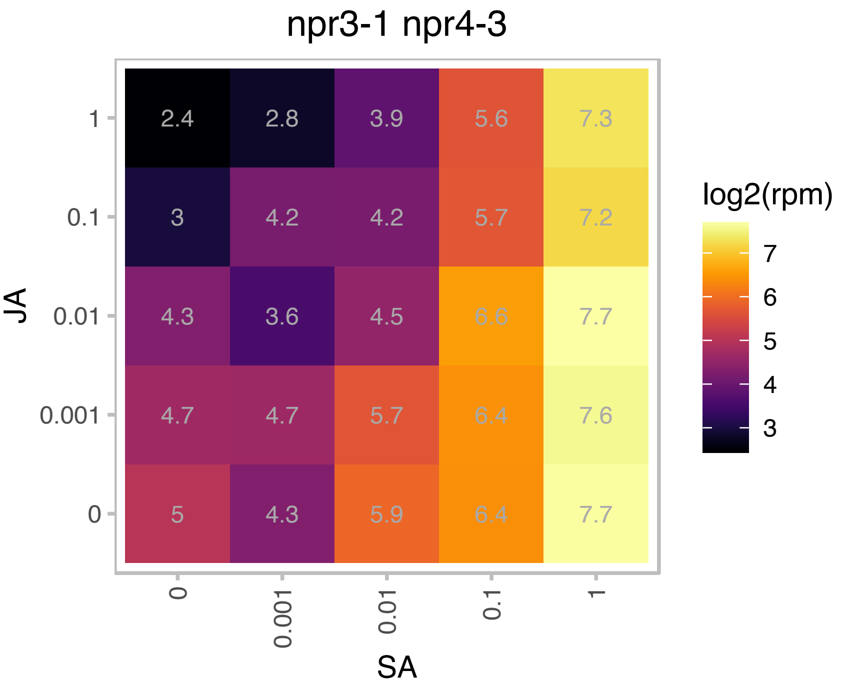

# AT3G29240.1 : NA

Protein of unknown function (DUF179)

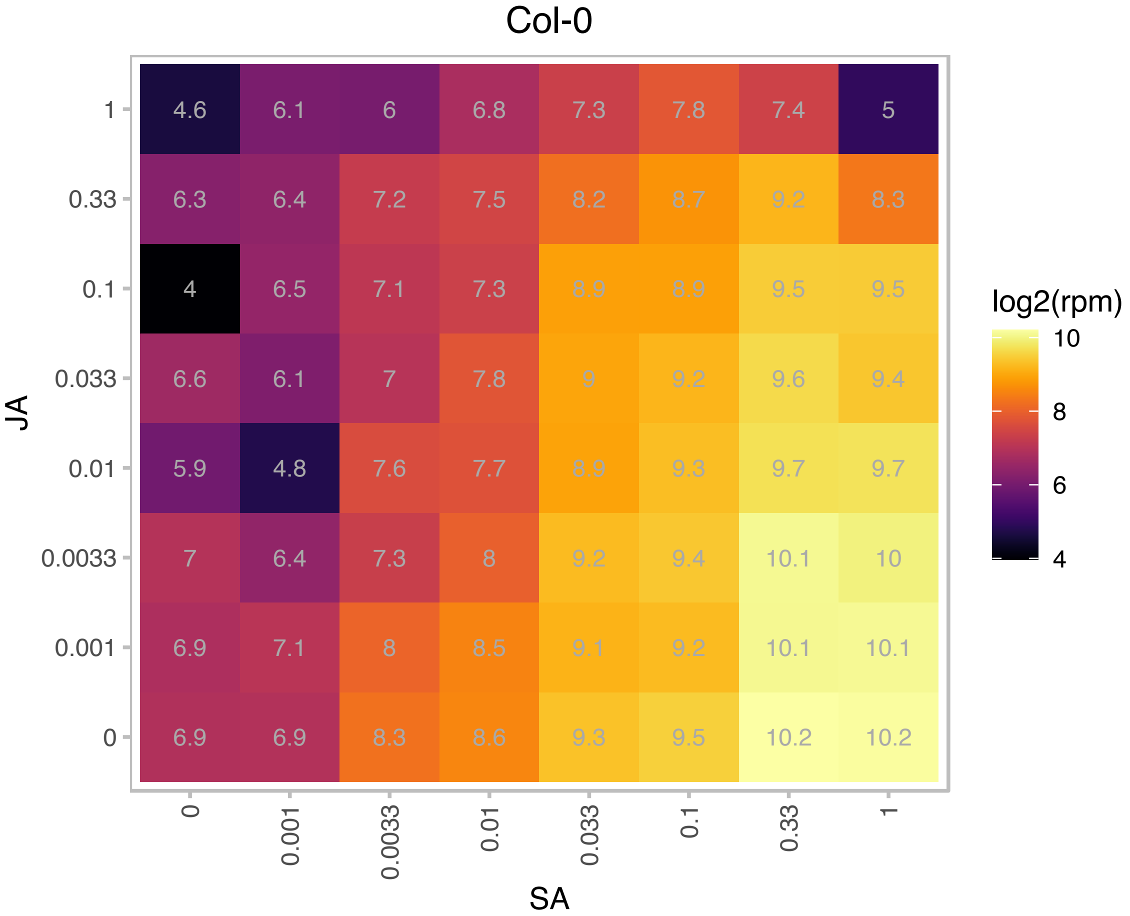


| | |
|---|---|
| Gene | AT3G29240 |
| is.expressed_genes | TRUE |
| is.responsive_genes | TRUE |
| is.apcluster | C-01 |

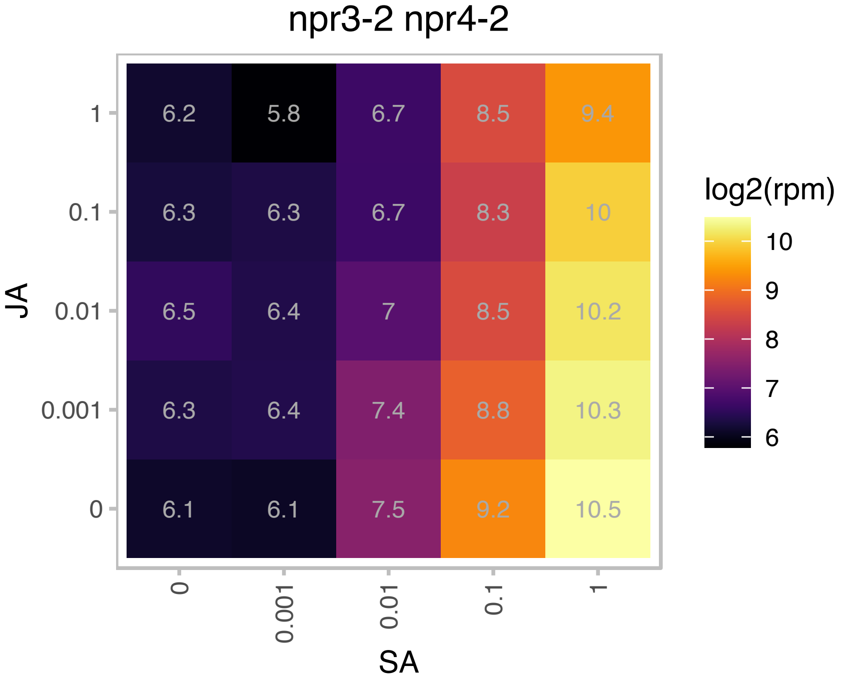


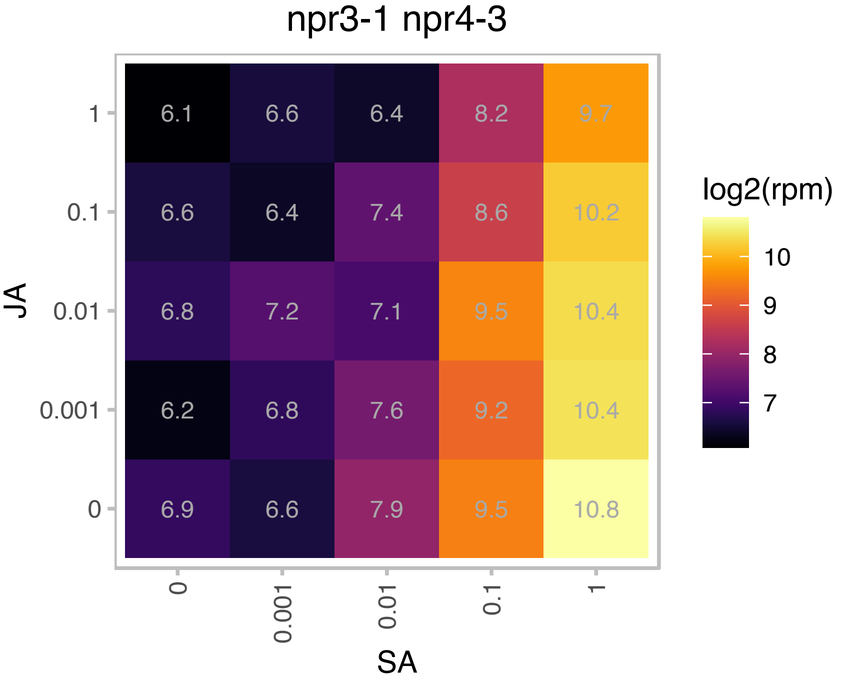

### AT3G50470.1 : HR3, MLA10

homolog of RPW8 3

Homolog of RPW8

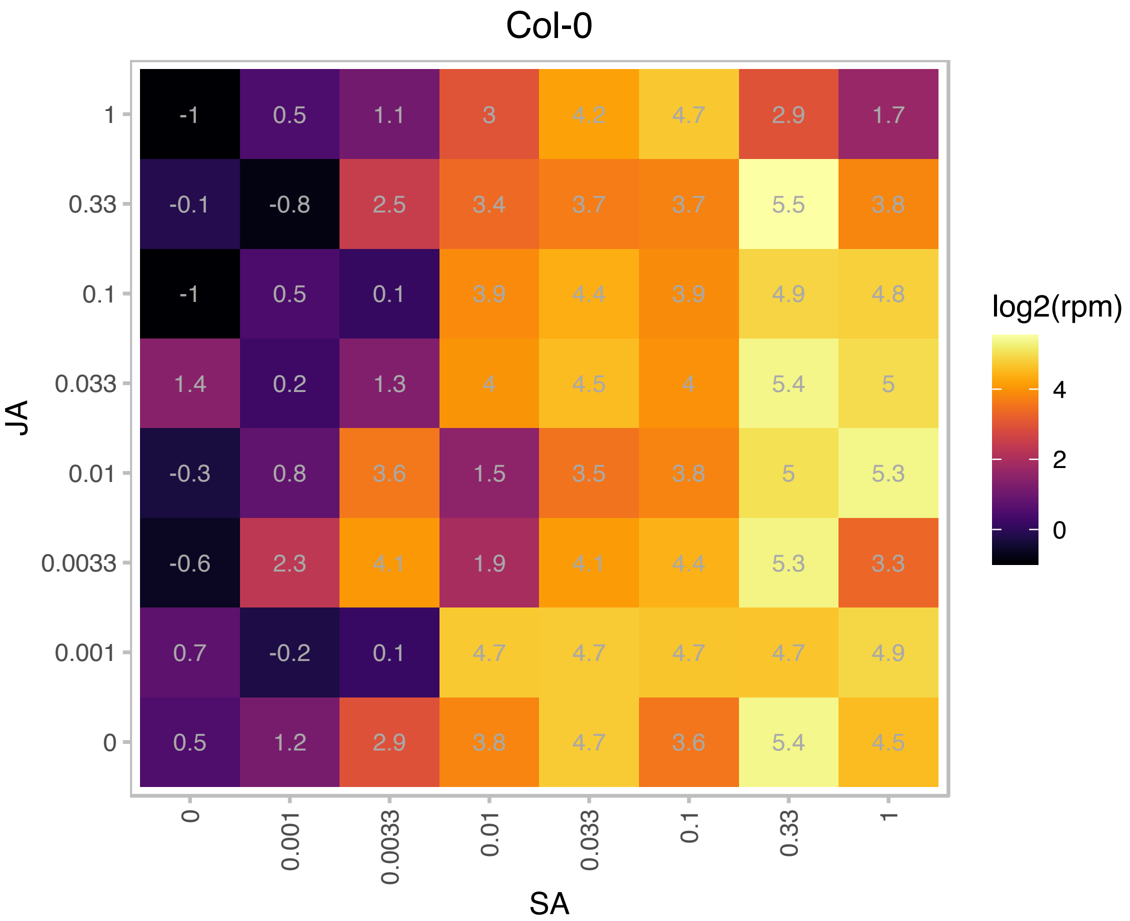


| Gene | AT3G50470 |
|---|---|
| is.expressed_genes | TRUE |
| is.responsive_genes | TRUE |
| is.apcluster | C-01 |

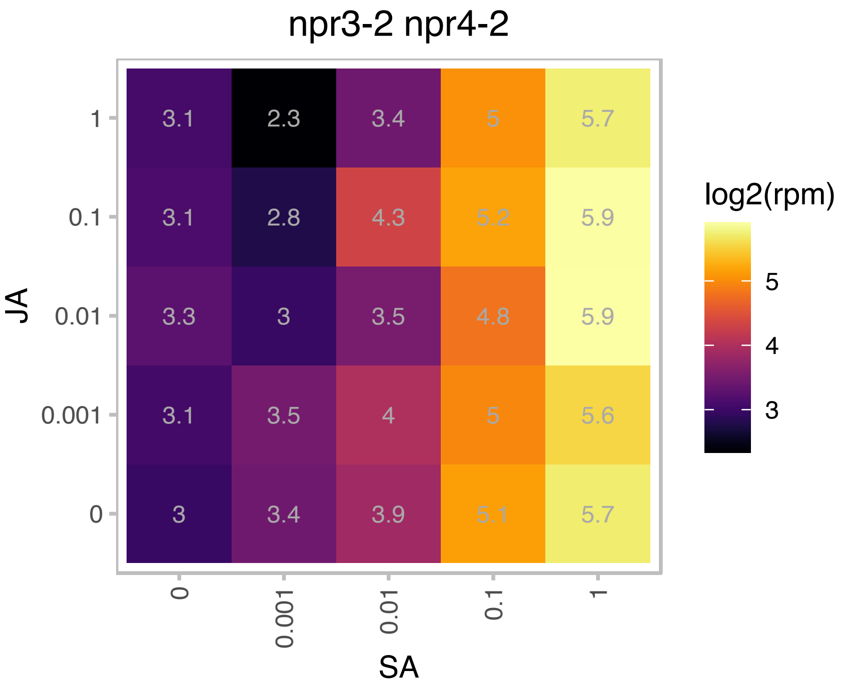


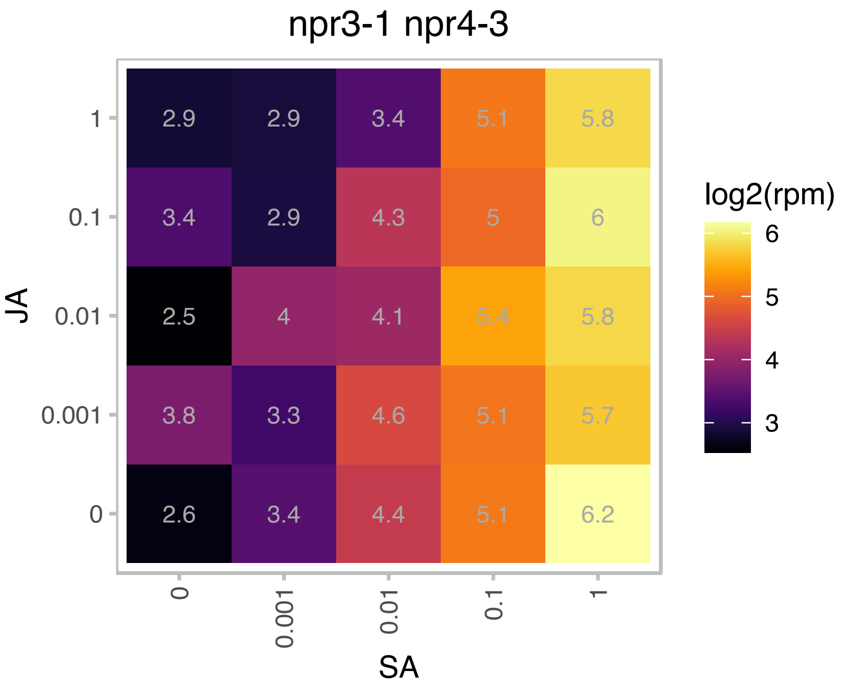

### AT3G56400.1 : ATWRKY70, WRKY70

WRKY DNA-binding protein 70

Member of WRKY Transcription Factor; Group III. Function as activator of SA-dependent defense genes and a repressor of JA-regulated genes. WRKY70-controlled suppression of JA-signaling is partly executed by NPR1.

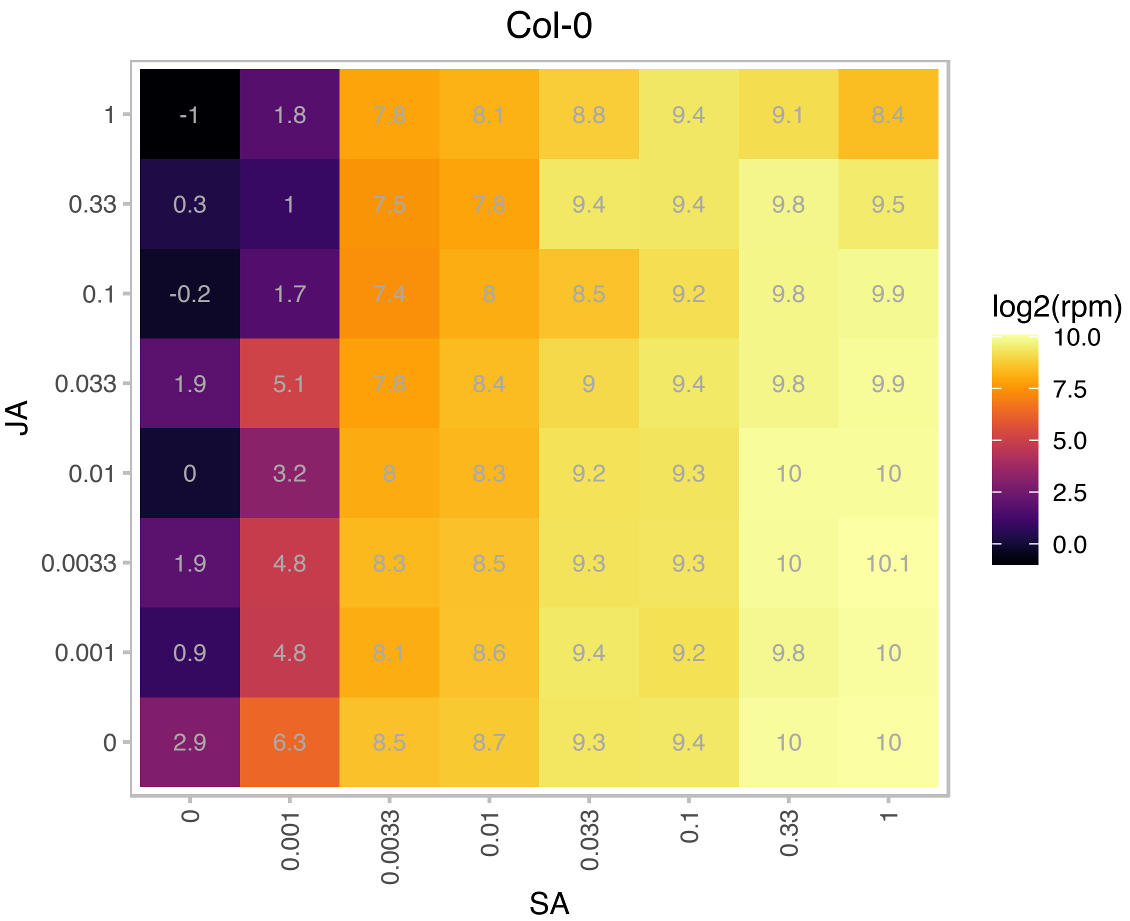


| | |
|---|---|
| Gene | AT3G56400 |
| is.expressed_genes | TRUE |
| is.responsive_genes | TRUE |
| is.apcluster | C-01 |

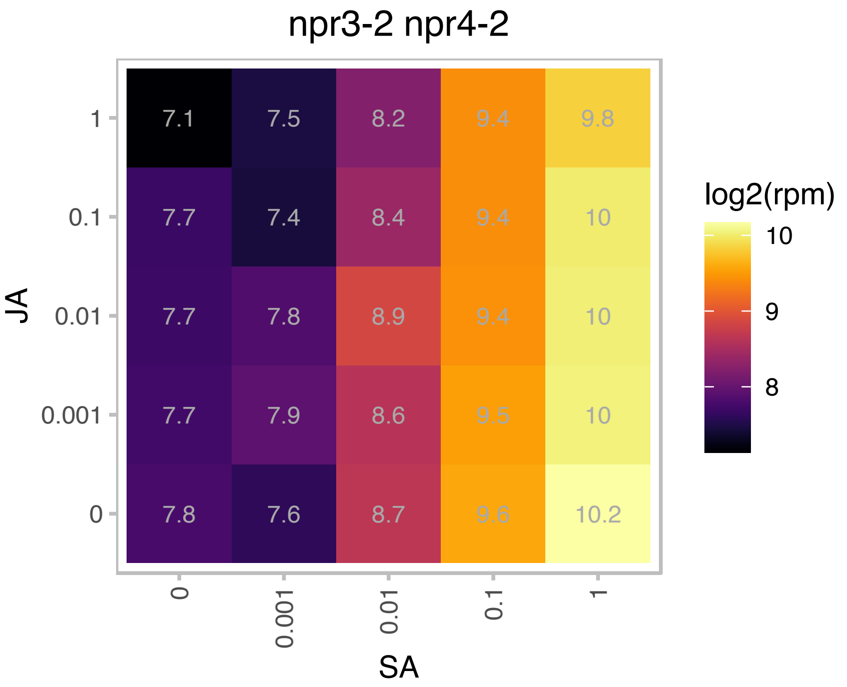


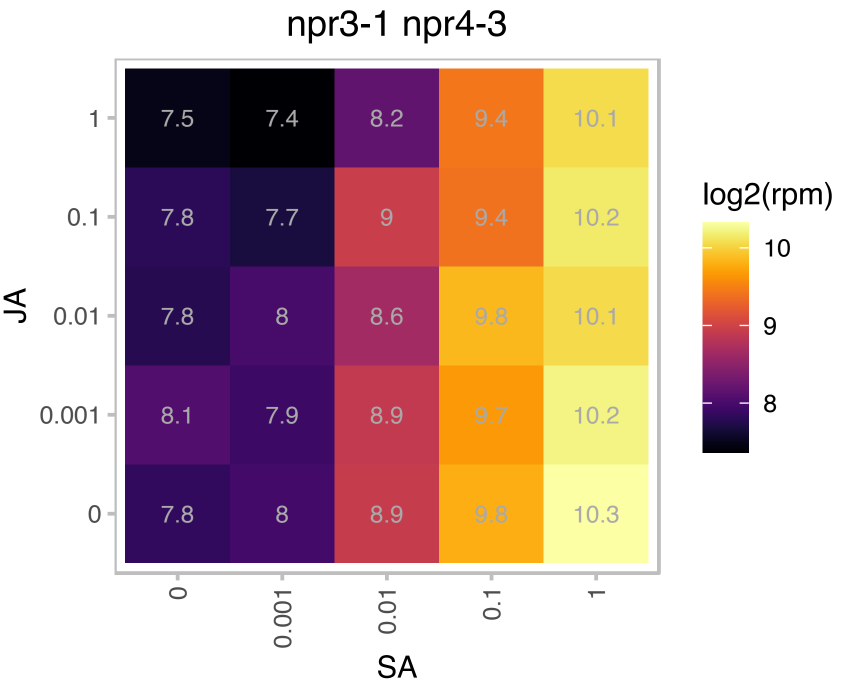

# AT3G60966.1 : NA

RING/U-box superfamily protein

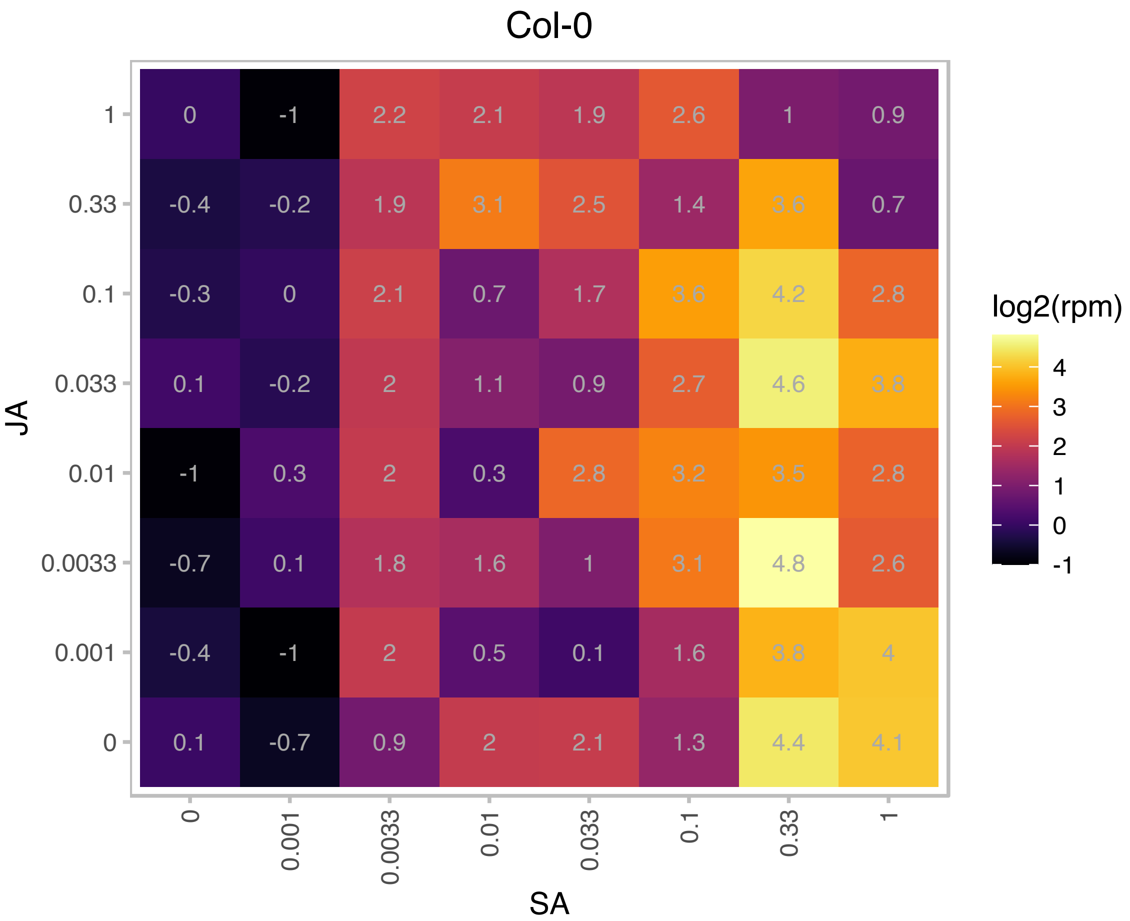


| | |
|---|---|
| Gene | AT3G60966 |
| is.expressed_genes | TRUE |
| is.responsive_genes | TRUE |
| is.apcluster | C-01 |

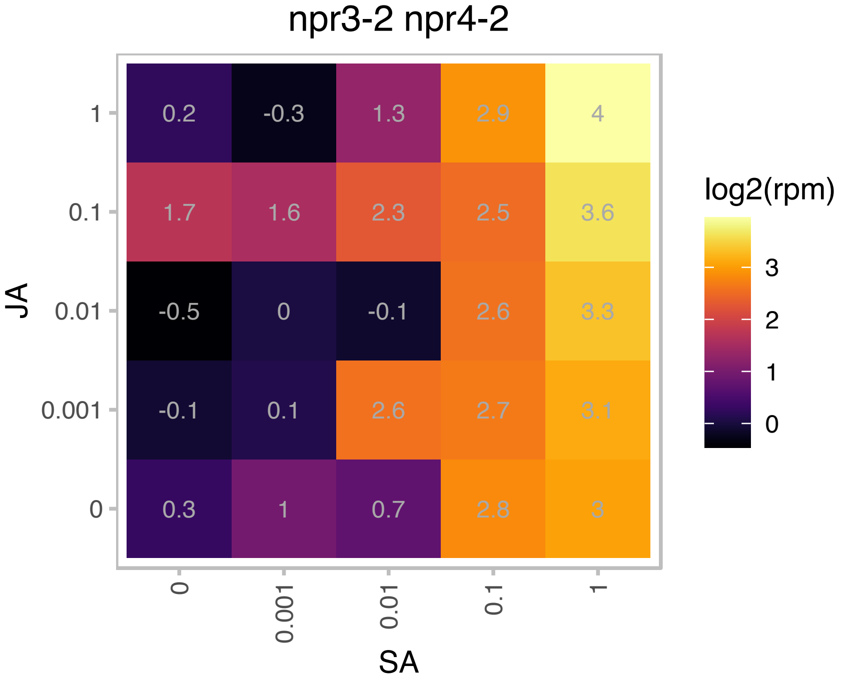


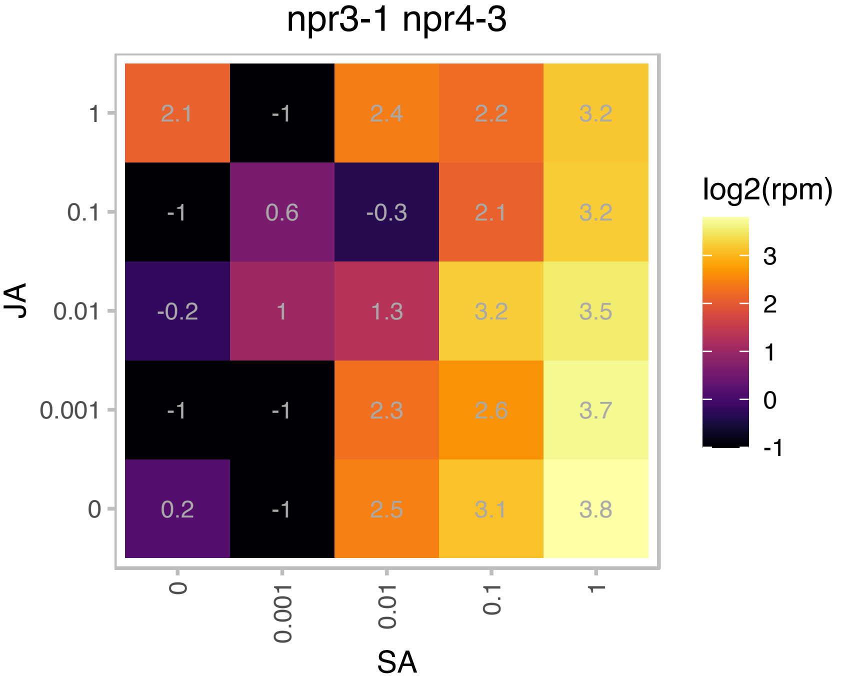

### AT3G63080.1 : ATGPX5, GPX5, GPXL5, MEE42

glutathione peroxidase 5

Encodes glutathione peroxidase.

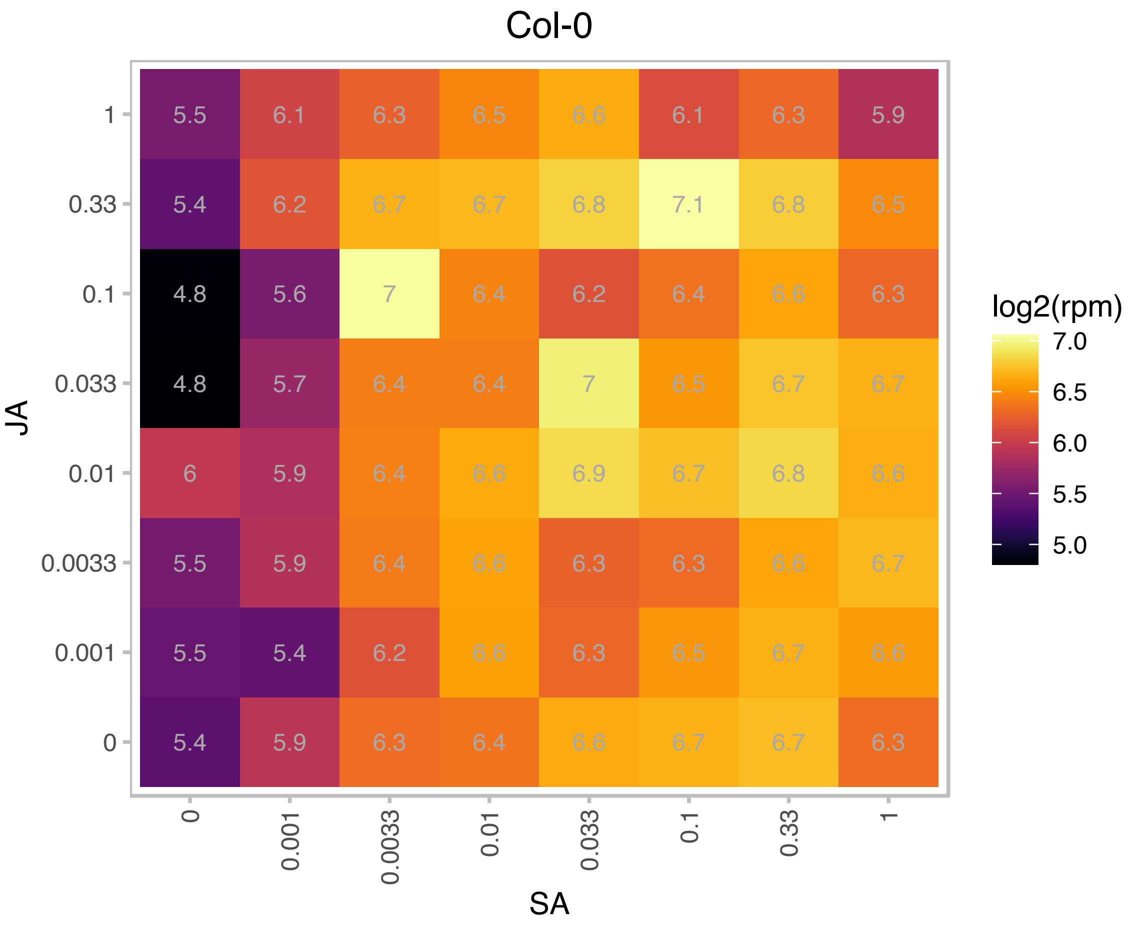


| | |
|---|---|
| Gene | AT3G63080 |
| is.expressed_genes | TRUE |
| is.responsive_genes | TRUE |
| is.apcluster | C-01 |

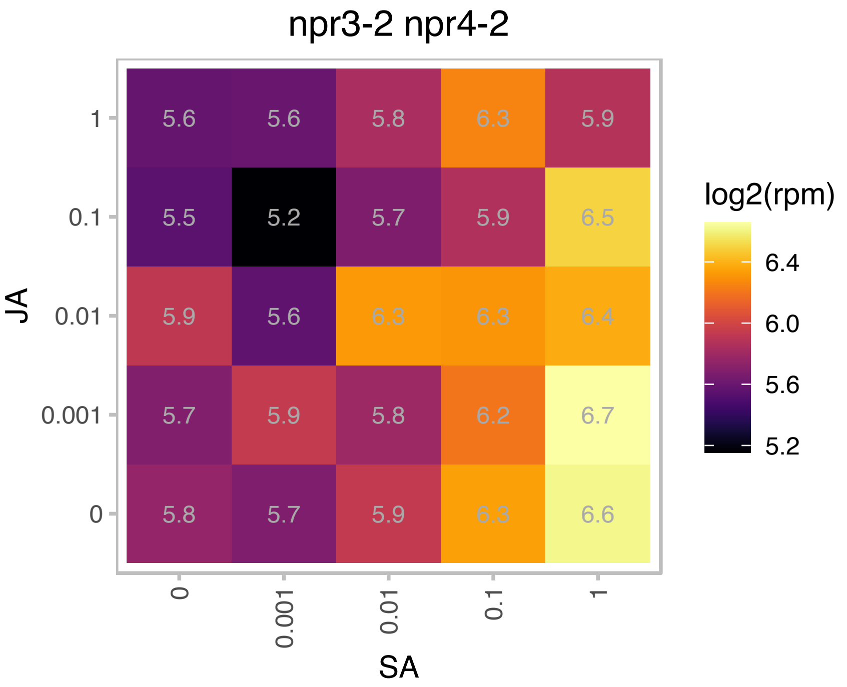


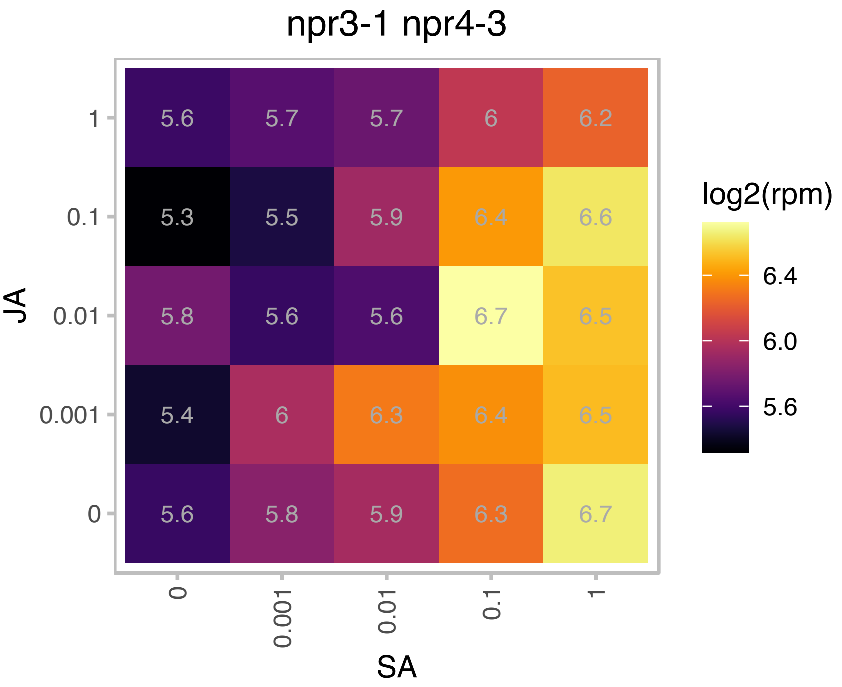

### AT4G23560.1 : AtGH9B15, GH9B15

glycosyl hydrolase 9B15

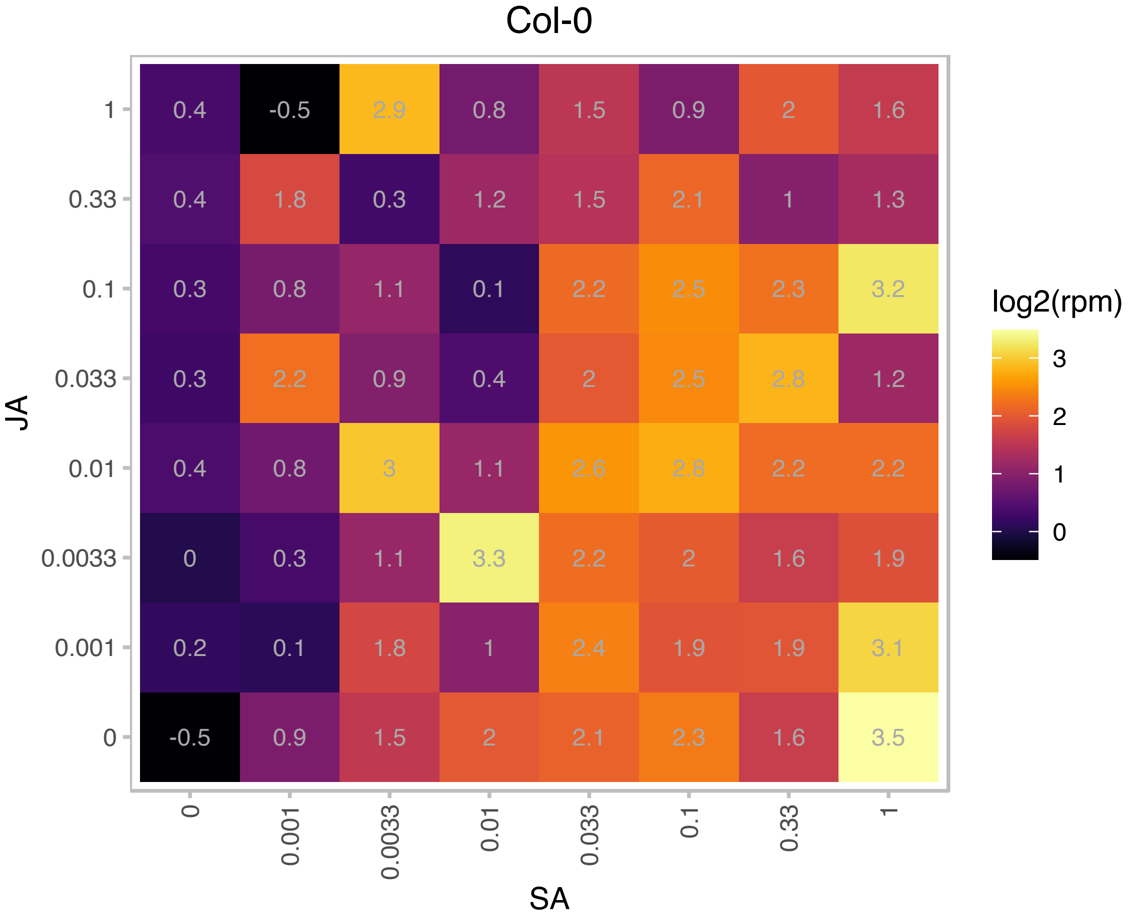


| | |
|---|---|
| Gene | AT4G23560 |
| is.expressed_genes | TRUE |
| is.responsive_genes | TRUE |
| is.apcluster | C-01 |

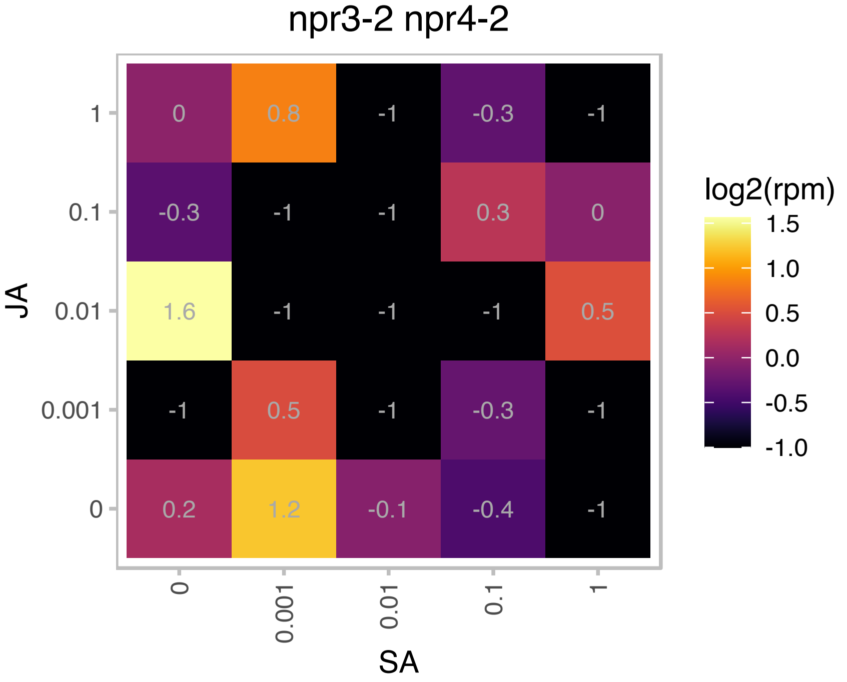


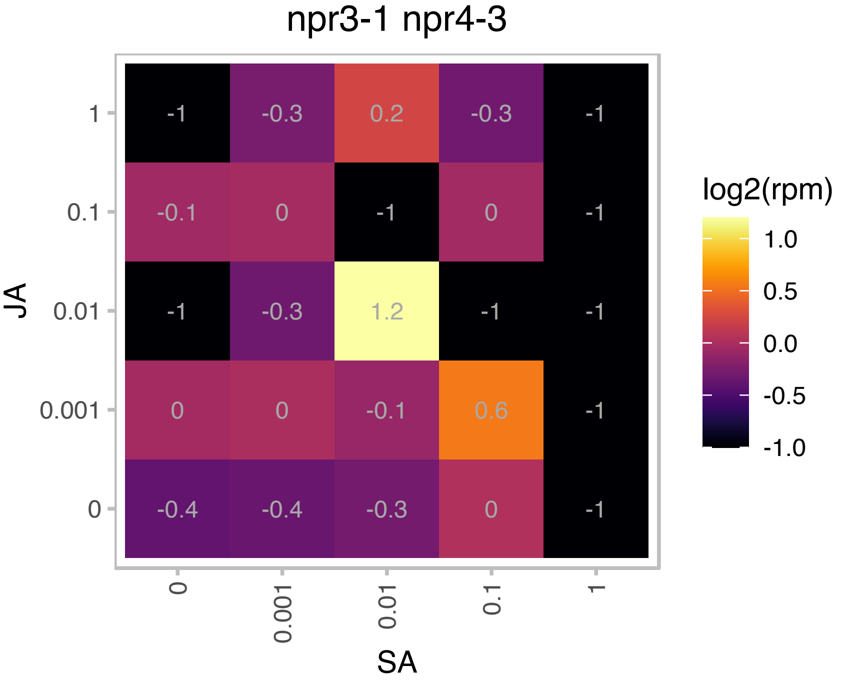

### AT4G24940.1 : AT-SAE1-1, ATSAE1A, SAE1A

SUMO-activating enzyme 1A
Encodes one of the two subunits of the SUMO activation enzyme required during sumolation. Sumolation is a post-translational protein modification process similar to ubiquitination during which a polypeptide (SUMO) is covalently attached to a target protein.

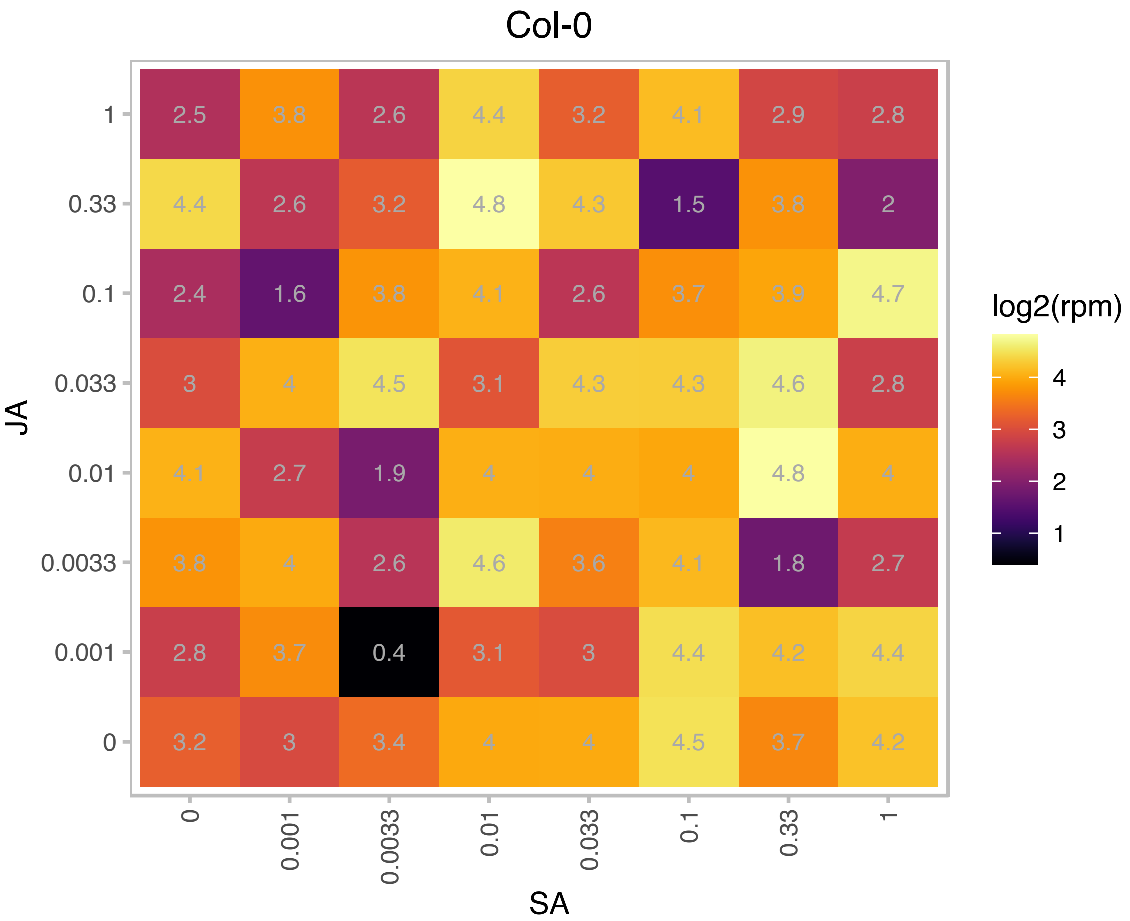


| | |
|---|---|
| Gene | AT4G24940 |
| is.expressed_genes | TRUE |
| is.responsive_genes | TRUE |
| is.apcluster | C-01 |

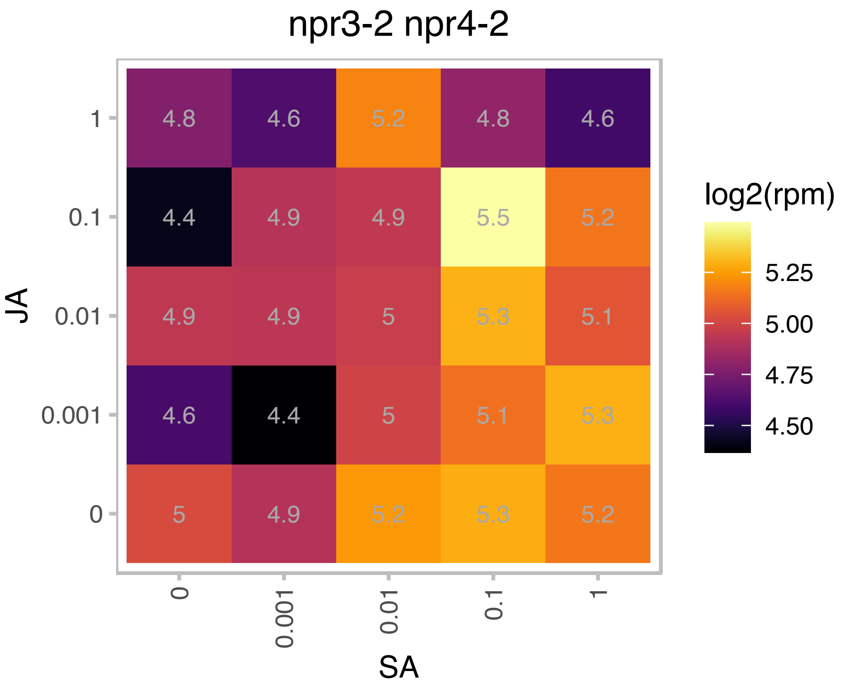


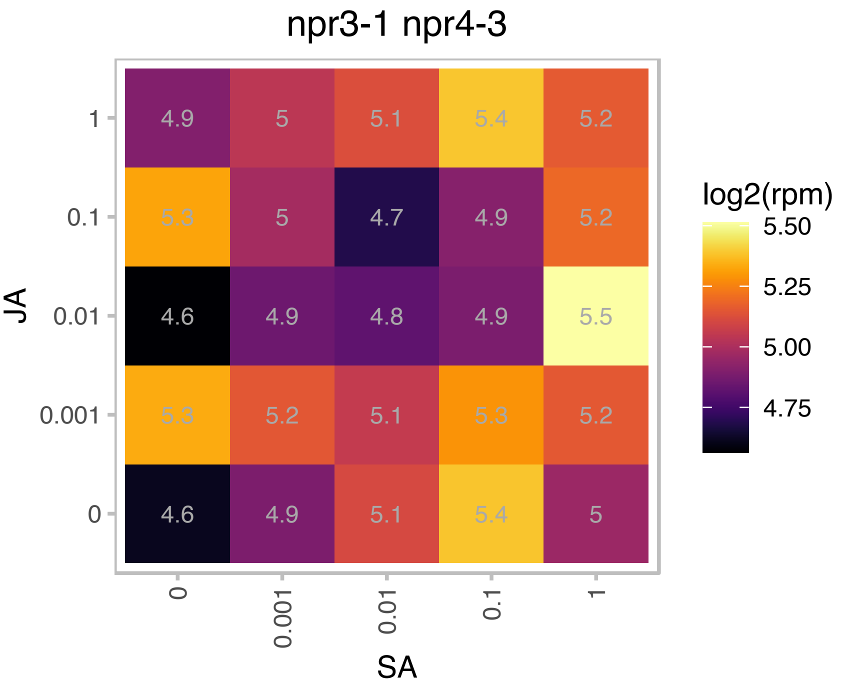

### AT4G26070.1 : ATMEK1, MEK1, MKK1, NMAPKK

MAP kinase/ ERK kinase 1

Member of MAP Kinase Kinase.  Likely functions in a stress-activated MAPK pathway. Can phosphorylate the MAPK
AtMPK4, in response to stress. Gets phosphorylated by MEKK1 in response to wounding.

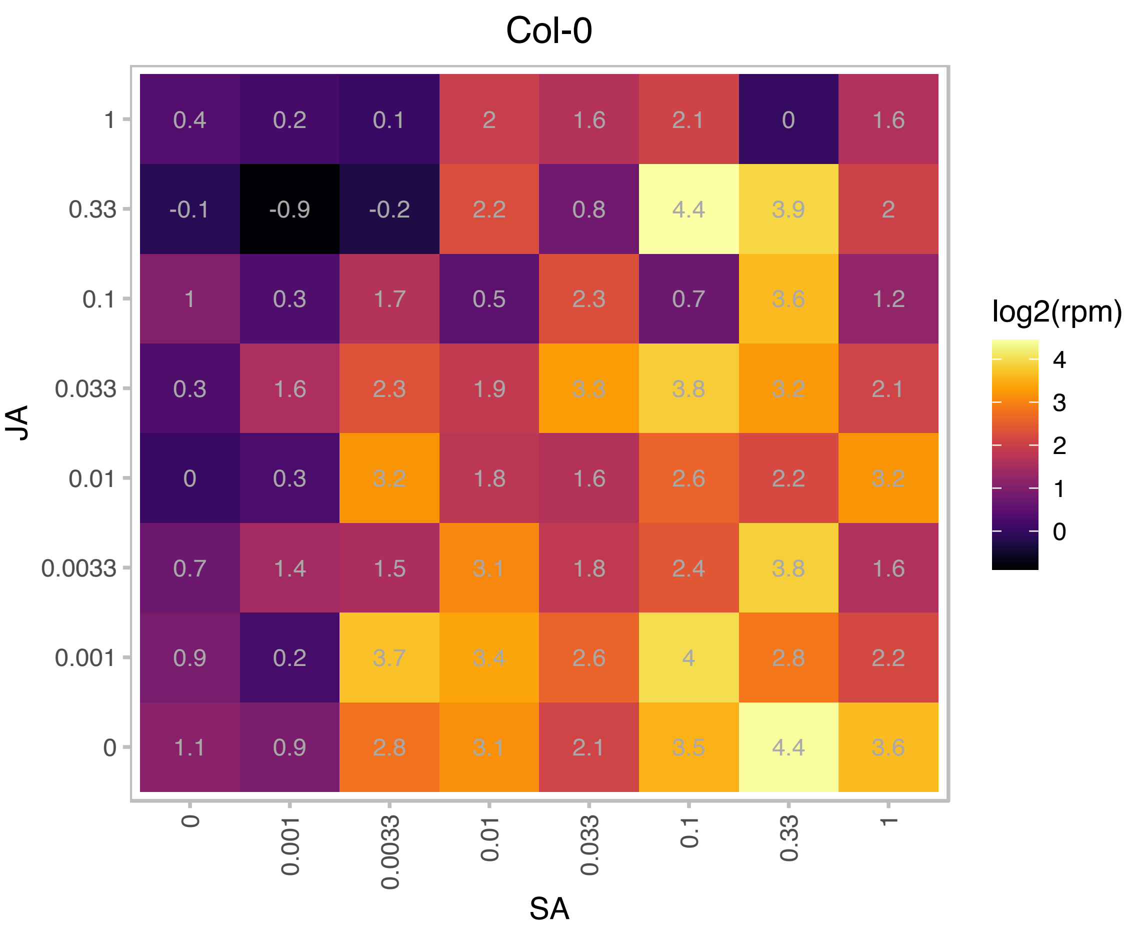


| | |
|---|---|
| Gene | AT4G26070 |
| is.expressed_genes | TRUE |
| is.responsive_genes | TRUE |
| is.apcluster | C-01 |

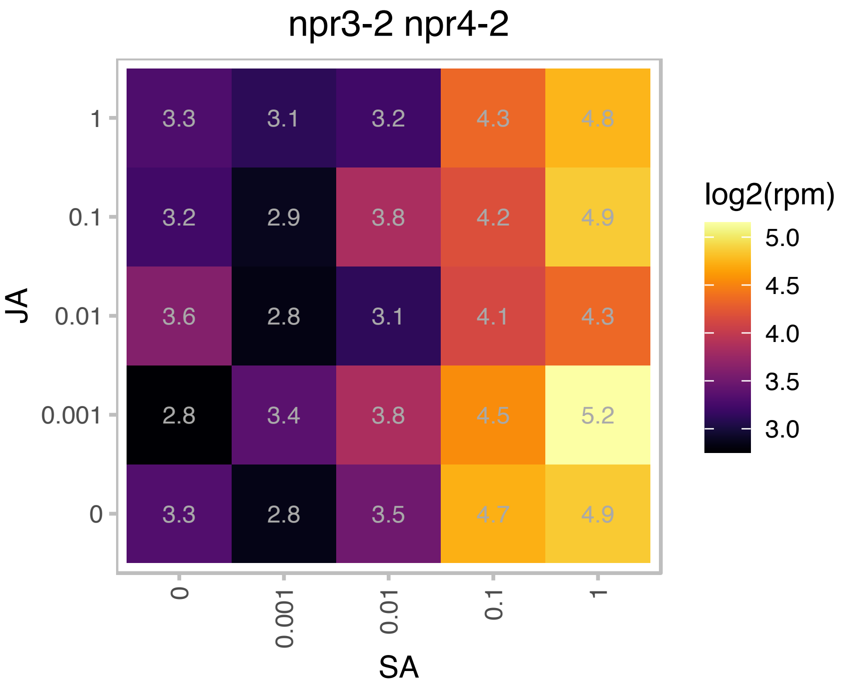


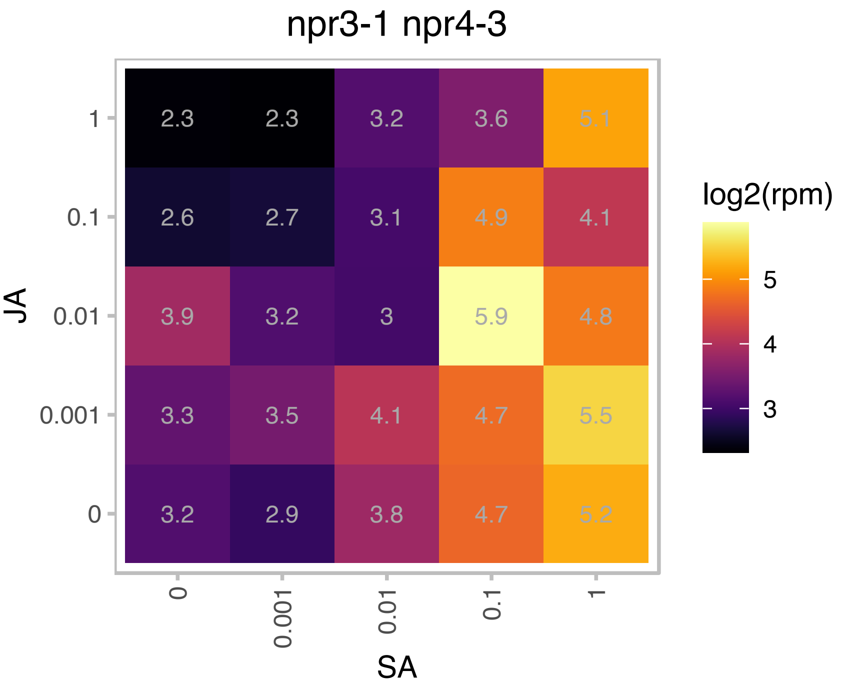

### AT4G28490.1 : HAE, RLK5

Leucine-rich receptor-like protein kinase family protein

Member of Receptor kinase-like protein family. Controls the separation step of floral organ abscission. The mRNA is cell-to-cell mobile.

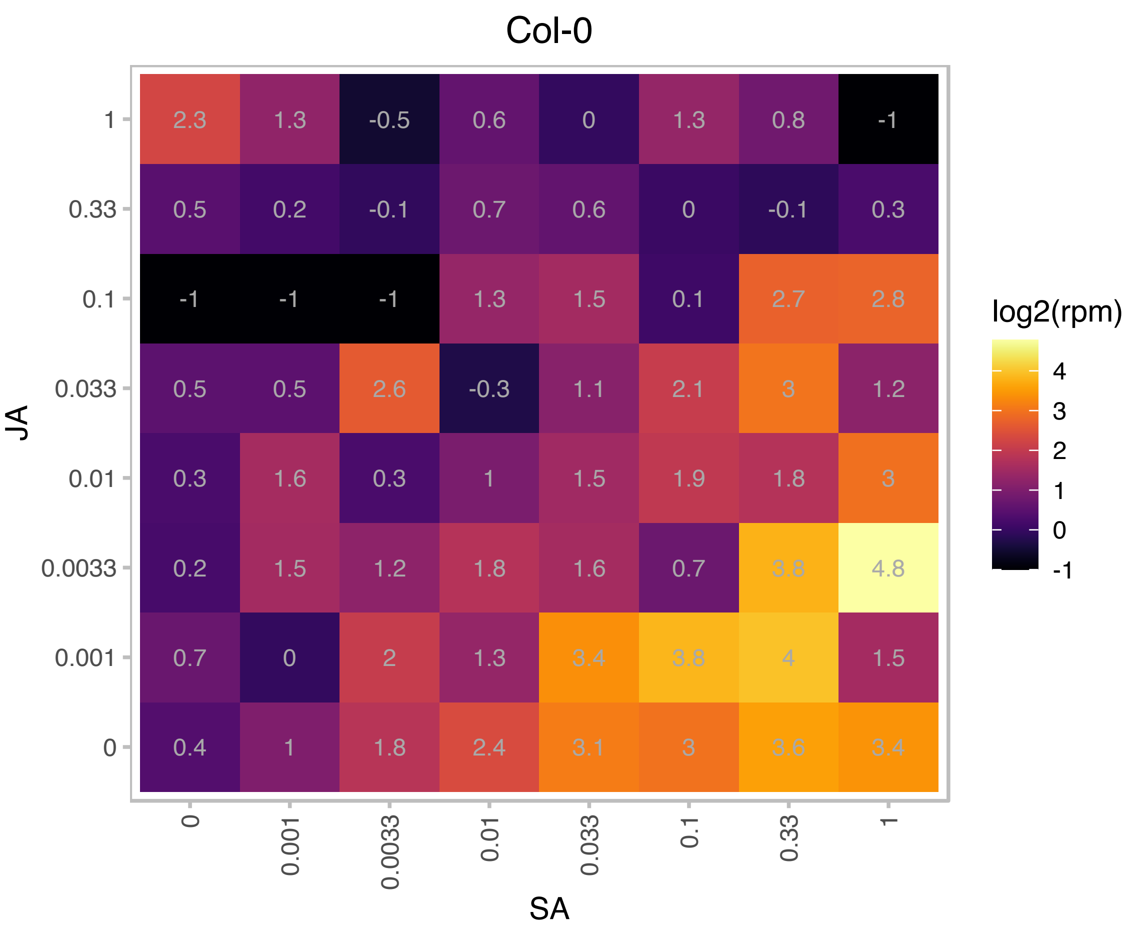


| | |
|---|---|
| Gene | AT4G28490 |
| is.expressed_genes | TRUE |
| is.responsive_genes | TRUE |
| is.apcluster | C-01 |

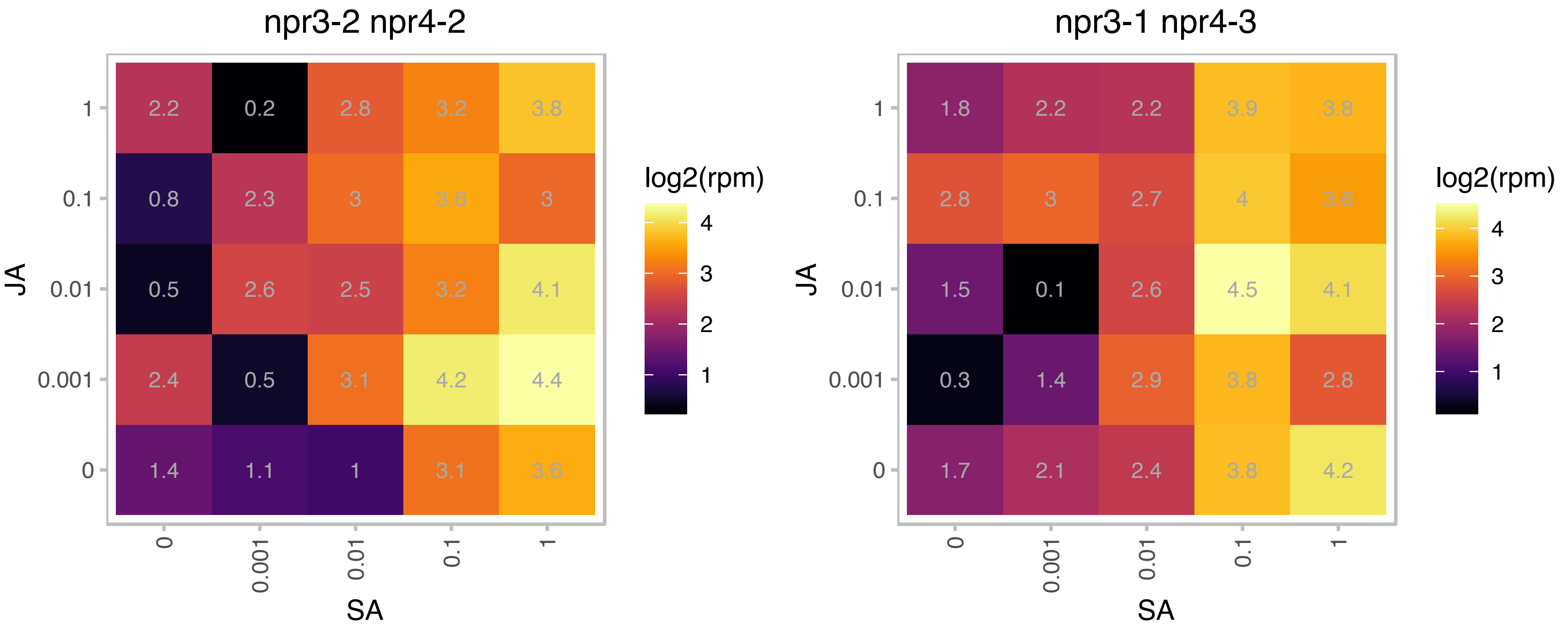

# AT4G32870.1 : NA

Polyketide cyclase/dehydrase and lipid transport superfamily protein

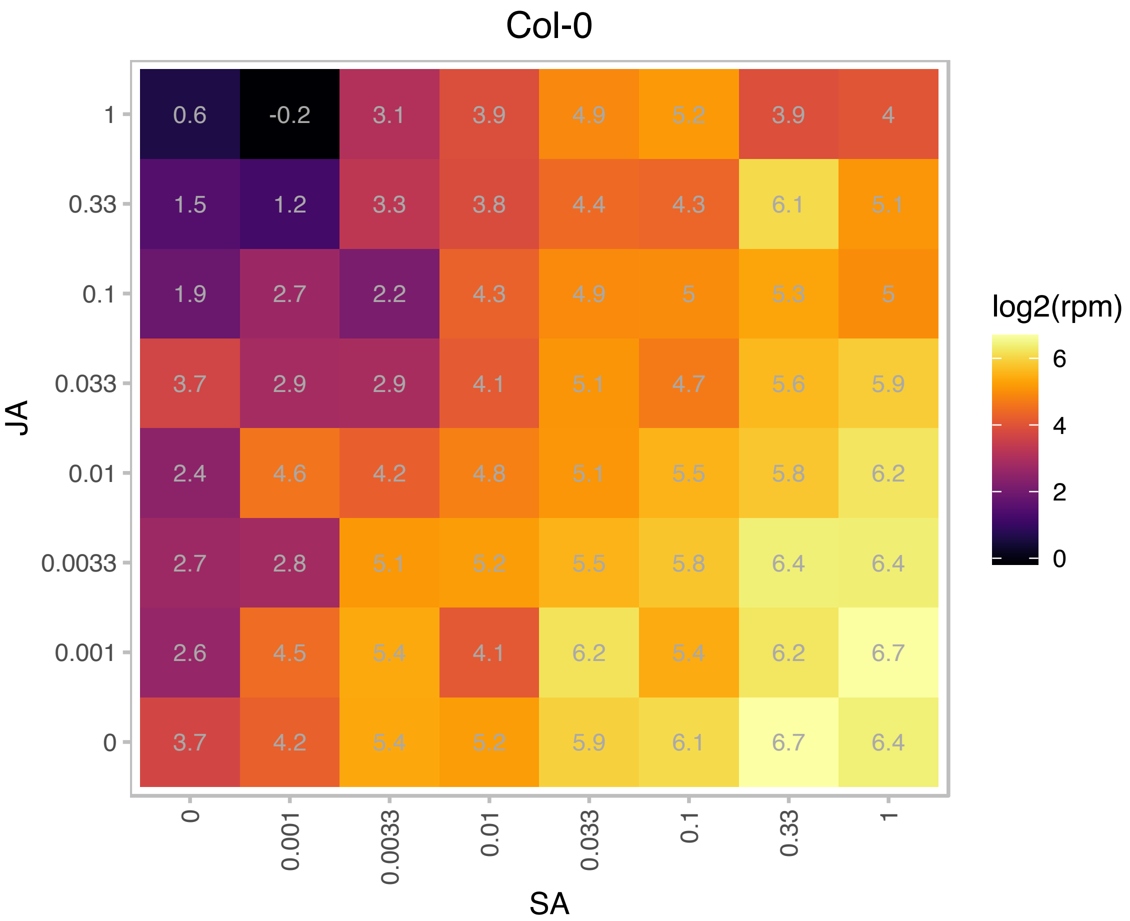


| | |
|---|---|
| Gene | AT4G32870 |
| is.expressed_genes | TRUE |
| is.responsive_genes | TRUE |
| is.apcluster | C-01 |

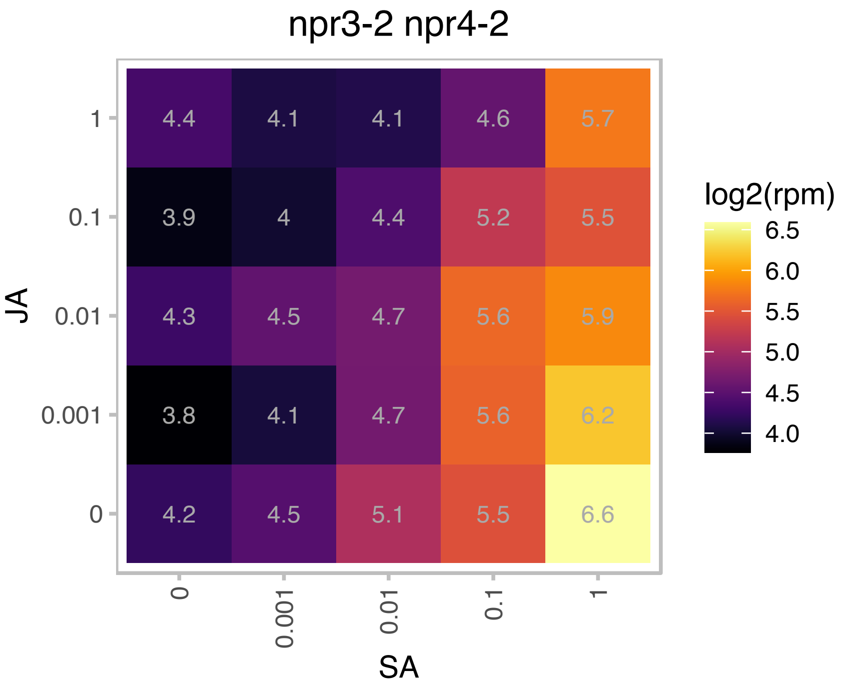


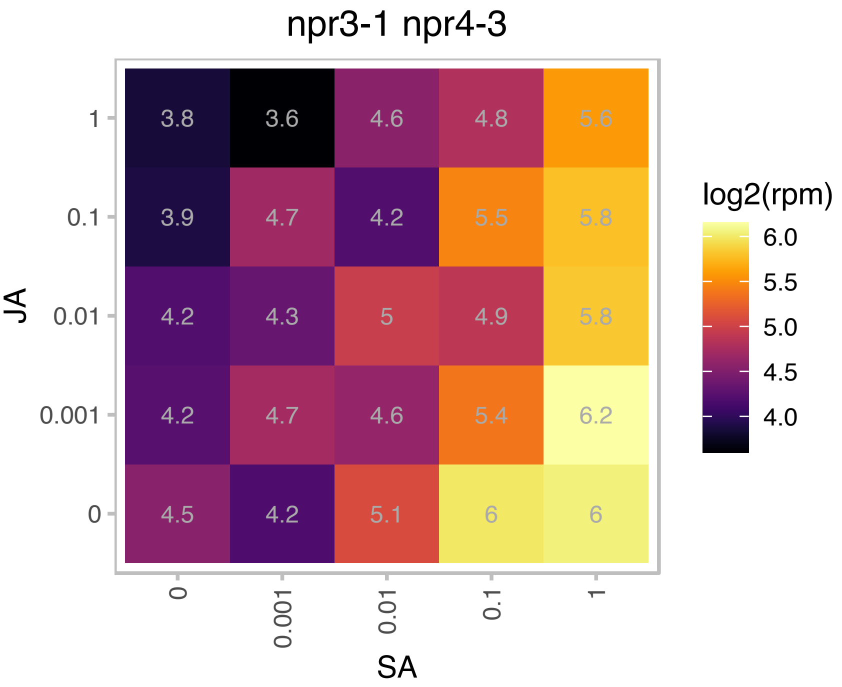

### AT4G34600.1 : CIF2

CAF2 is a peptide hormone expressed in the root stele that specifically binds the endodermis-expressed leucine-rich repeat receptor kinase GASSHO1 (GSO1)/SCHENGEN3 and its homolog, GSO2. Together with CAF1 i
required for formation of the casparian band.

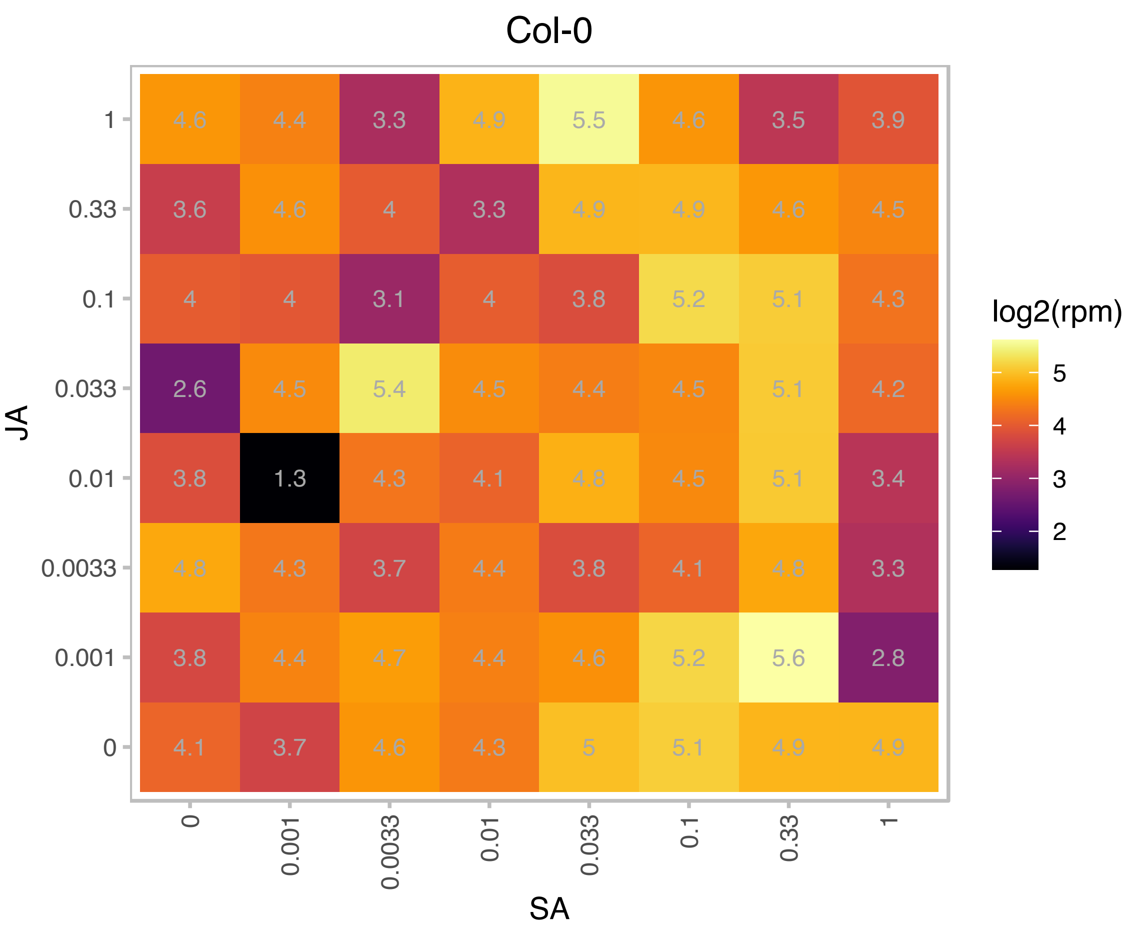


| | |
|---|---|
| Gene | AT4G34600 |
| is.expressed_genes | TRUE |
| is.responsive_genes | TRUE |
| is.apcluster | C-01 |

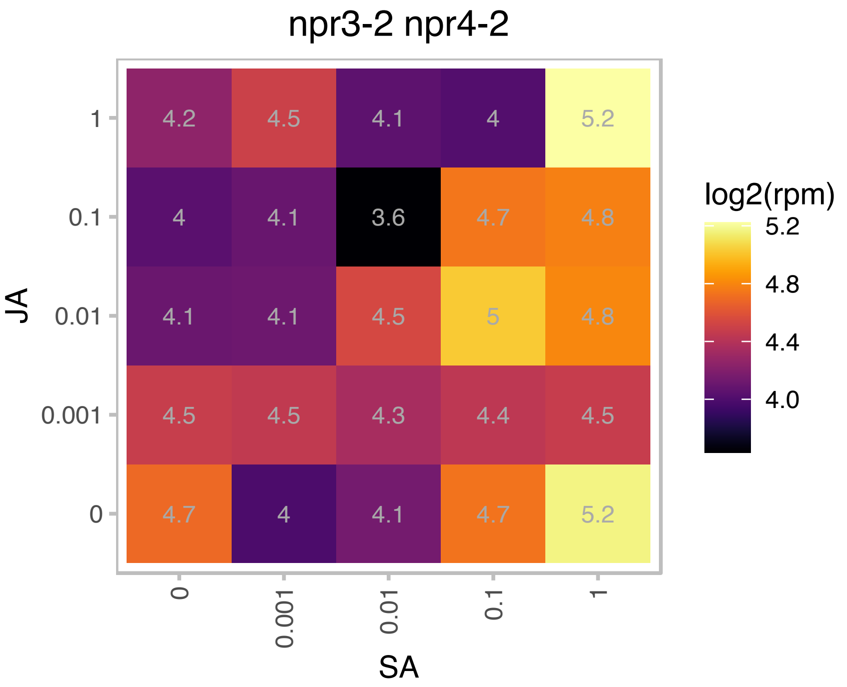


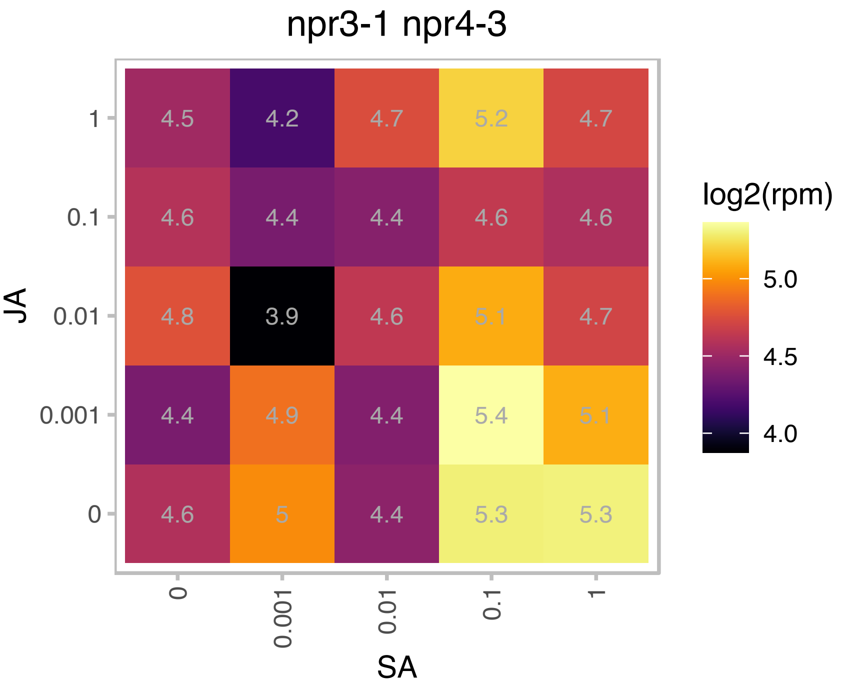

### AT4G35180.1 : LHT7

LYS/HIS transporter 7

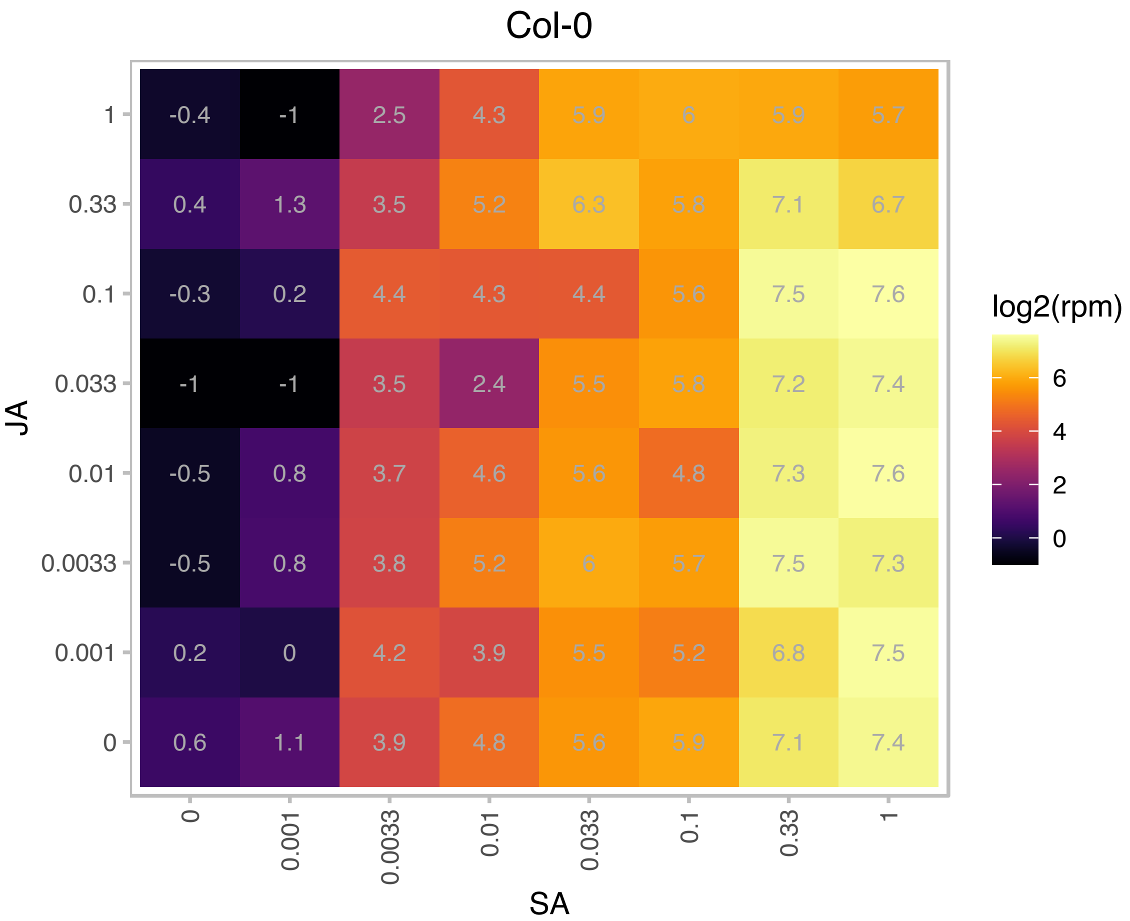


| | |
|---|---|
| Gene | AT4G35180 |
| is.expressed_genes | TRUE |
| is.responsive_genes | TRUE |
| is.apcluster | C-01 |

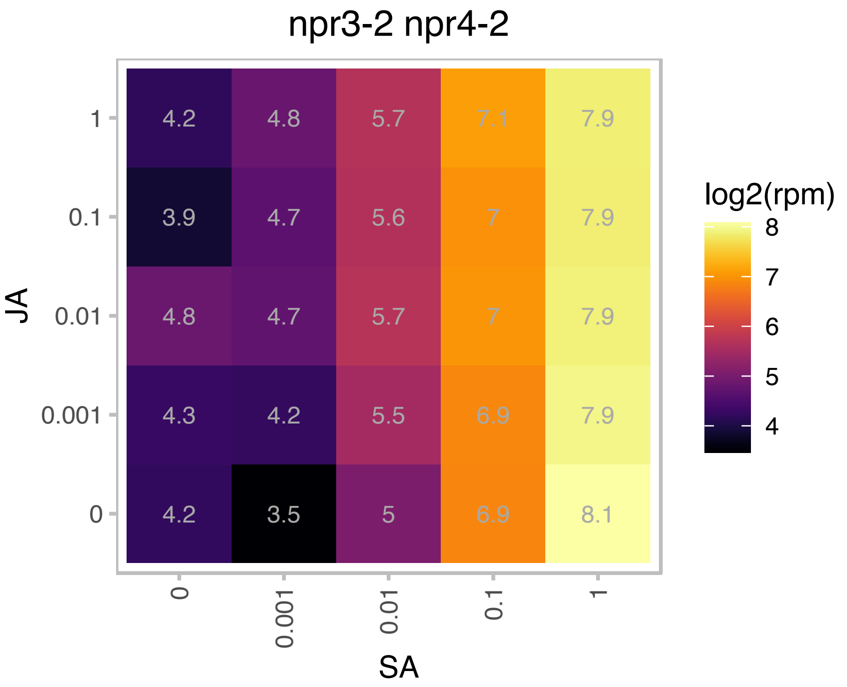


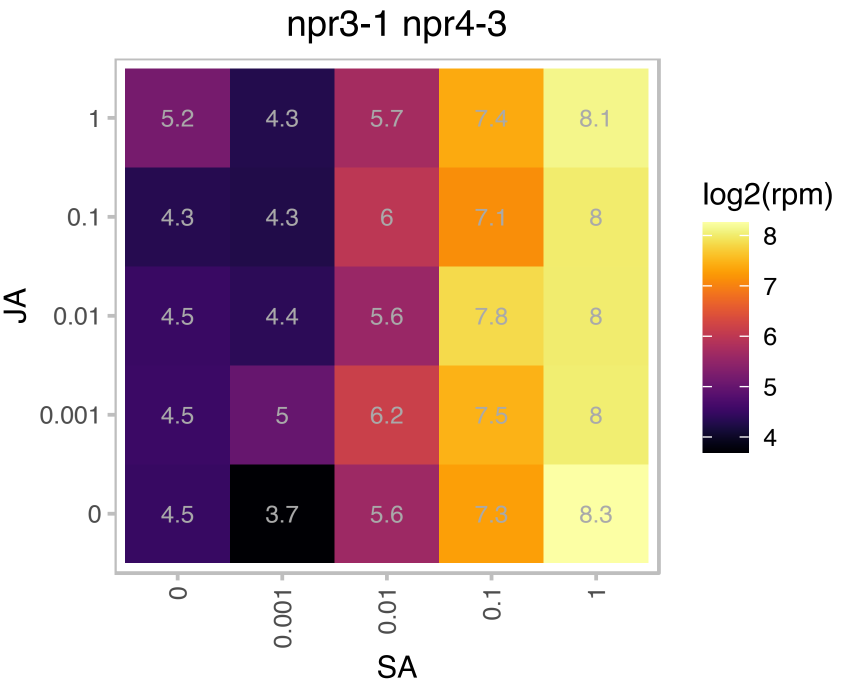

### AT5G01900.1 : ATWRKY62, WRKY62

WRKY DNA-binding protein 62

member of WRKY Transcription Factor; Group III

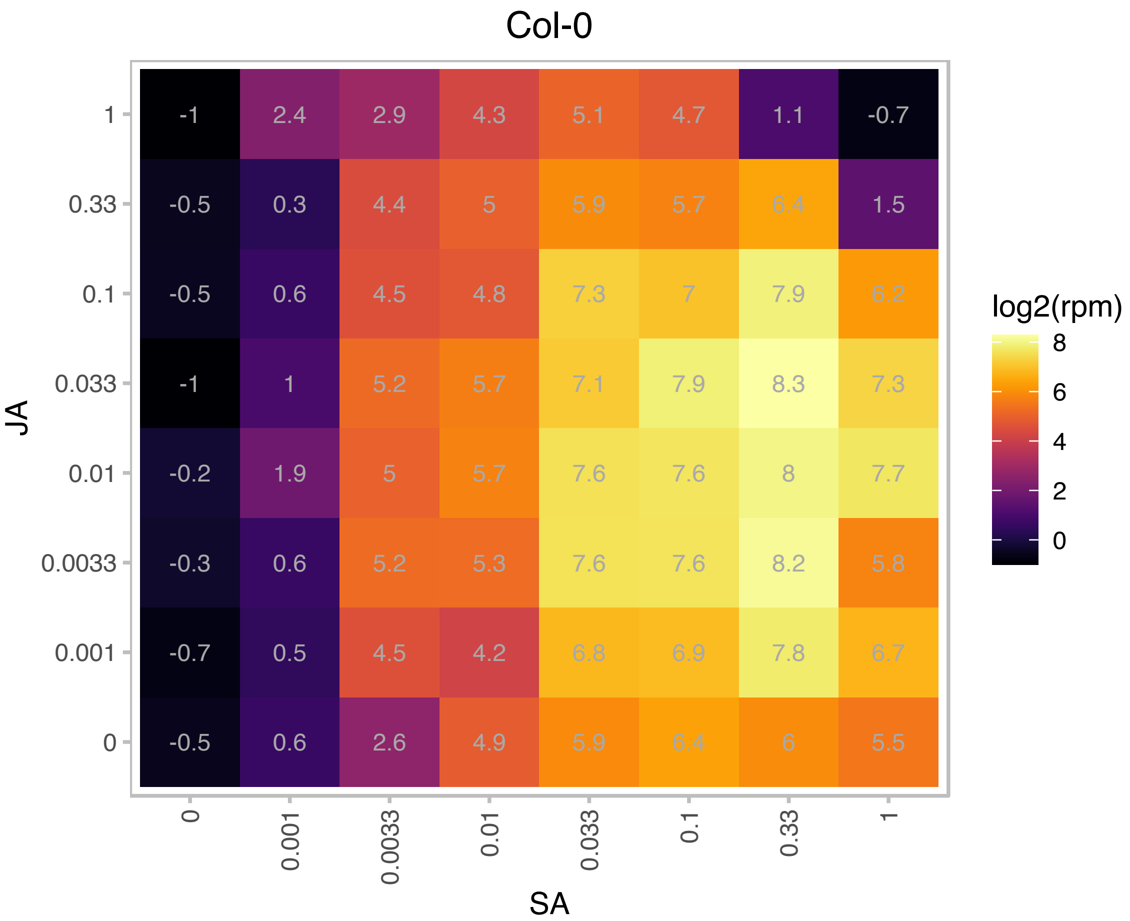


| | |
|---|---|
| Gene | AT5G01900 |
| is.expressed_genes | TRUE |
| is.responsive_genes | TRUE |
| is.apcluster | C-01 |

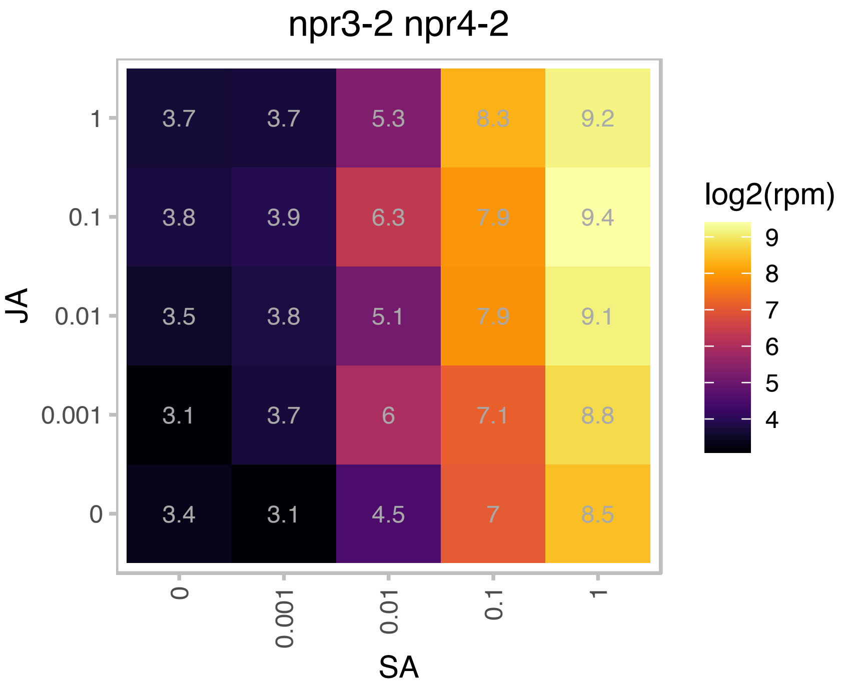


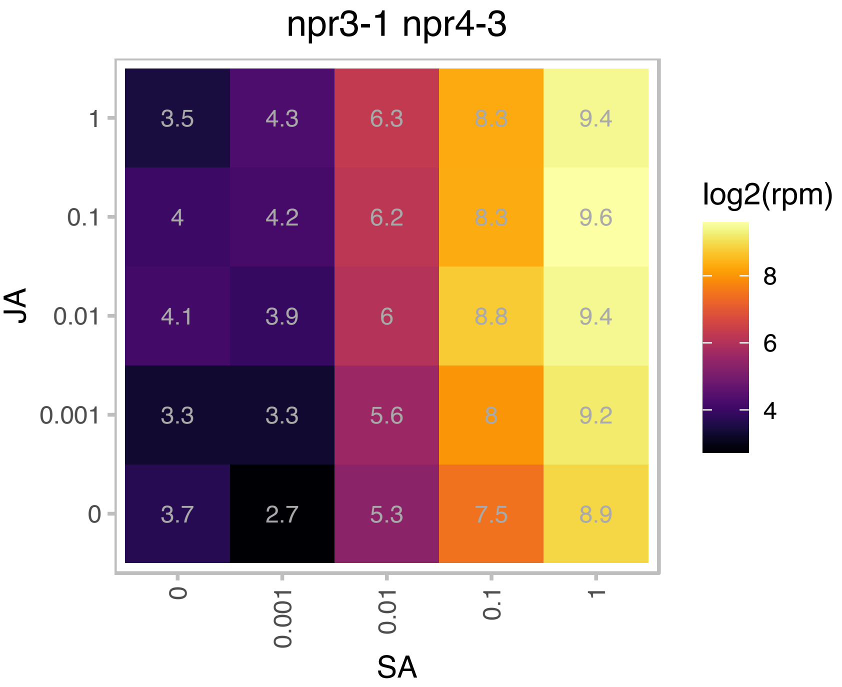

# AT5G03350.1 : NA

Legume lectin family protein

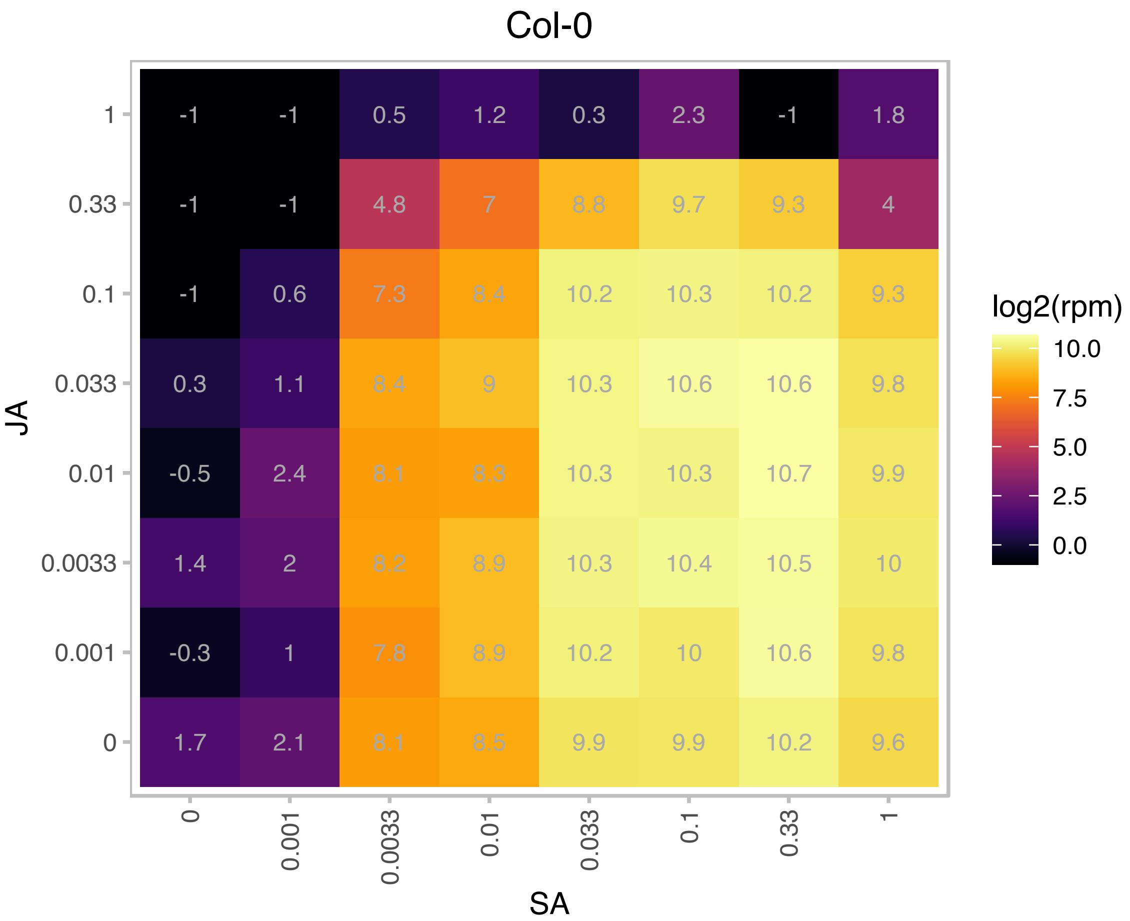


| | |
|---|---|
| Gene | AT5G03350 |
| is.expressed_genes | TRUE |
| is.responsive_genes | TRUE |
| is.apcluster | C-01 |

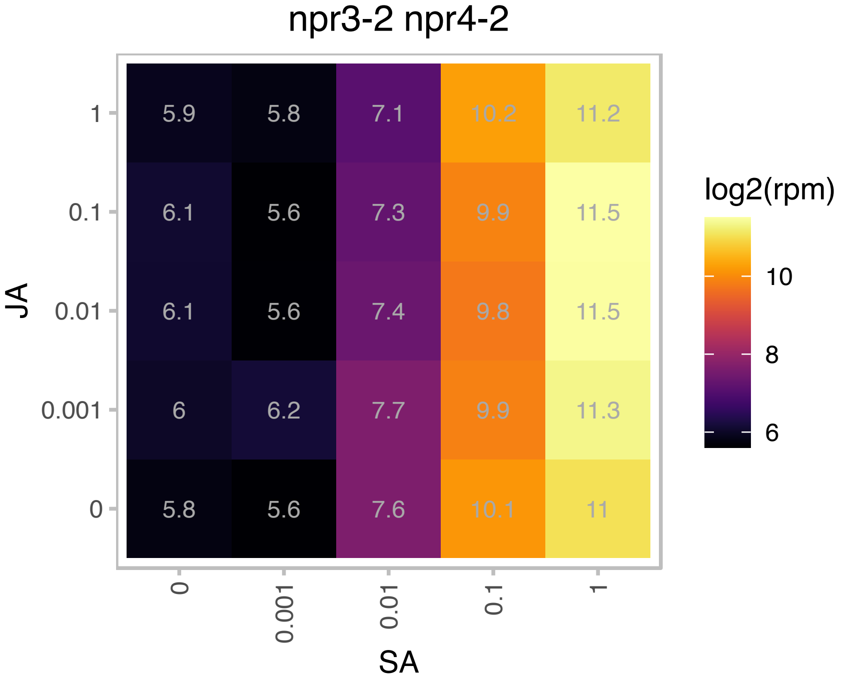


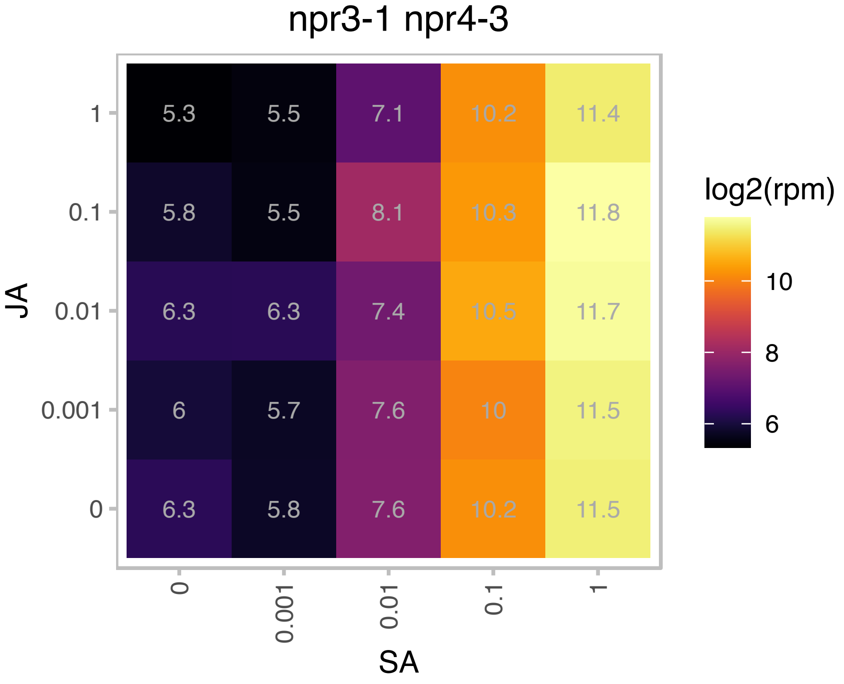

### AT5G05460.1 : AtENGase85A, ENGase85A

Glycosyl hydrolase family 85
Encodes a cytosolic beta-endo-N-acetylglucosaminidase (ENGase). ENGases N-glycans cleave the O-glycosidic linkage between the two GlcNAc residues of the N-glycan core structure and thus generate a protein with a single GlcNAc attached to asparagine.

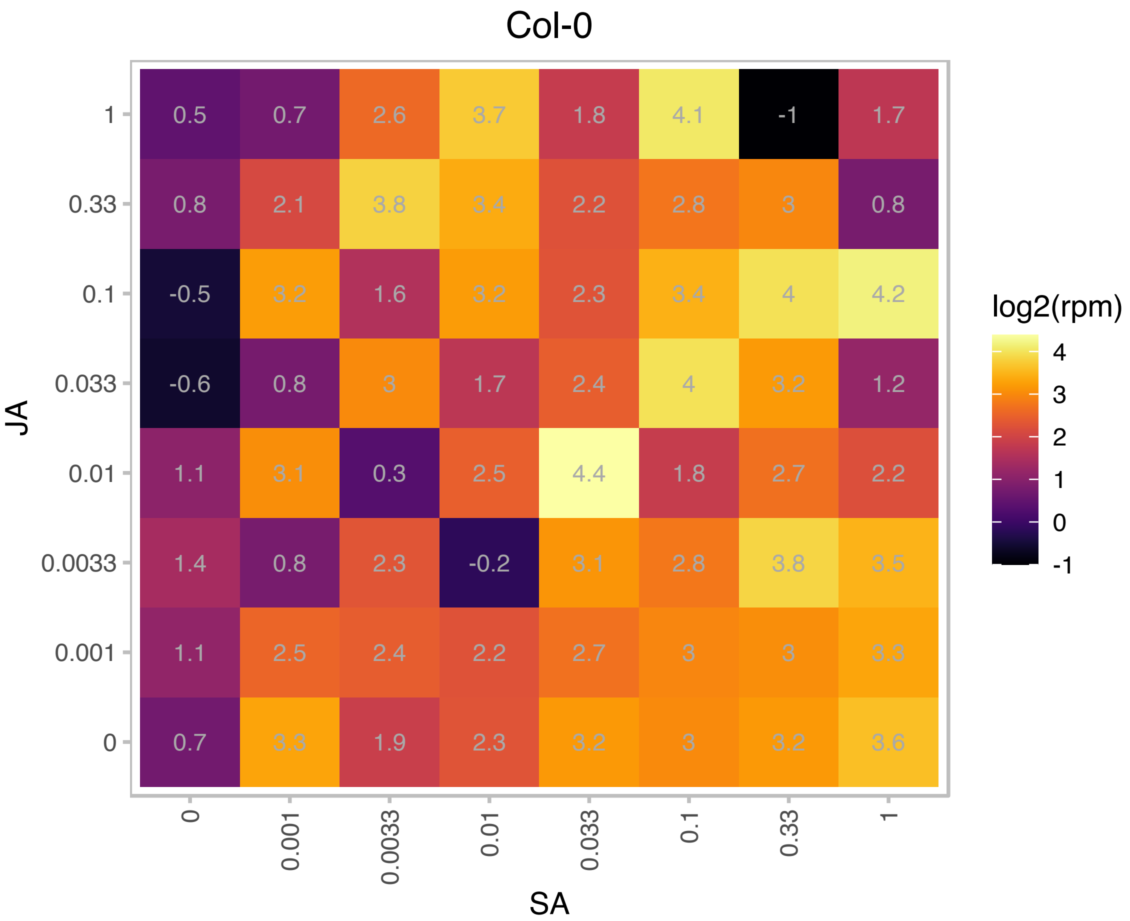


| | |
|---|---|
| Gene | AT5G05460 |
| is.expressed_genes | TRUE |
| is.responsive_genes | TRUE |
| is.apcluster | C-01 |

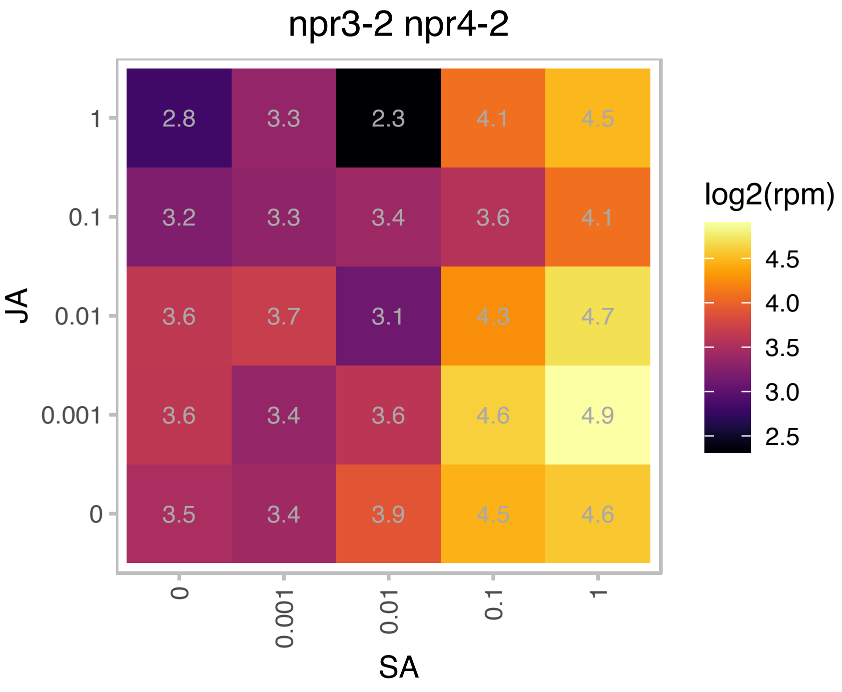


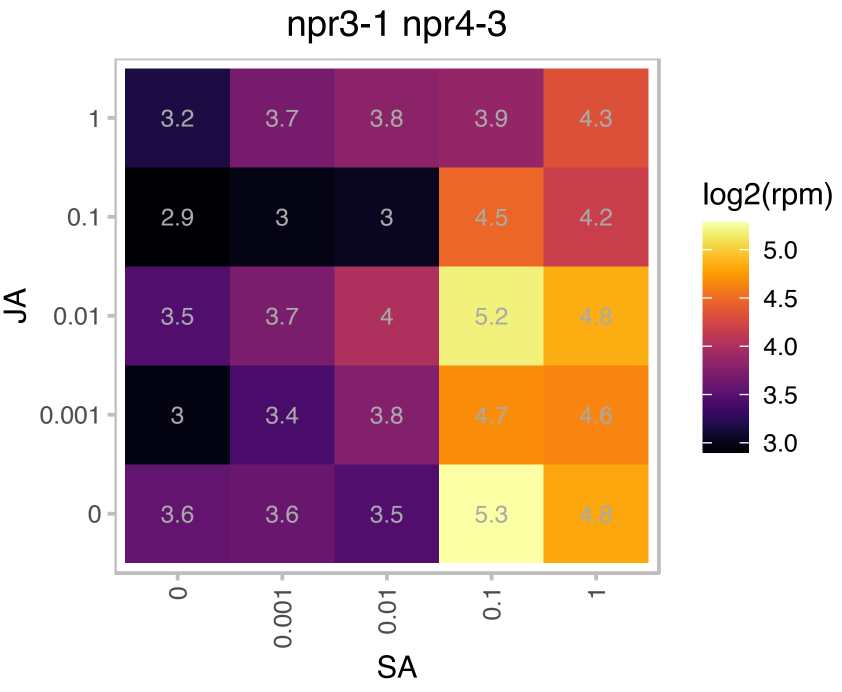

# AT5G12240.1 : NA

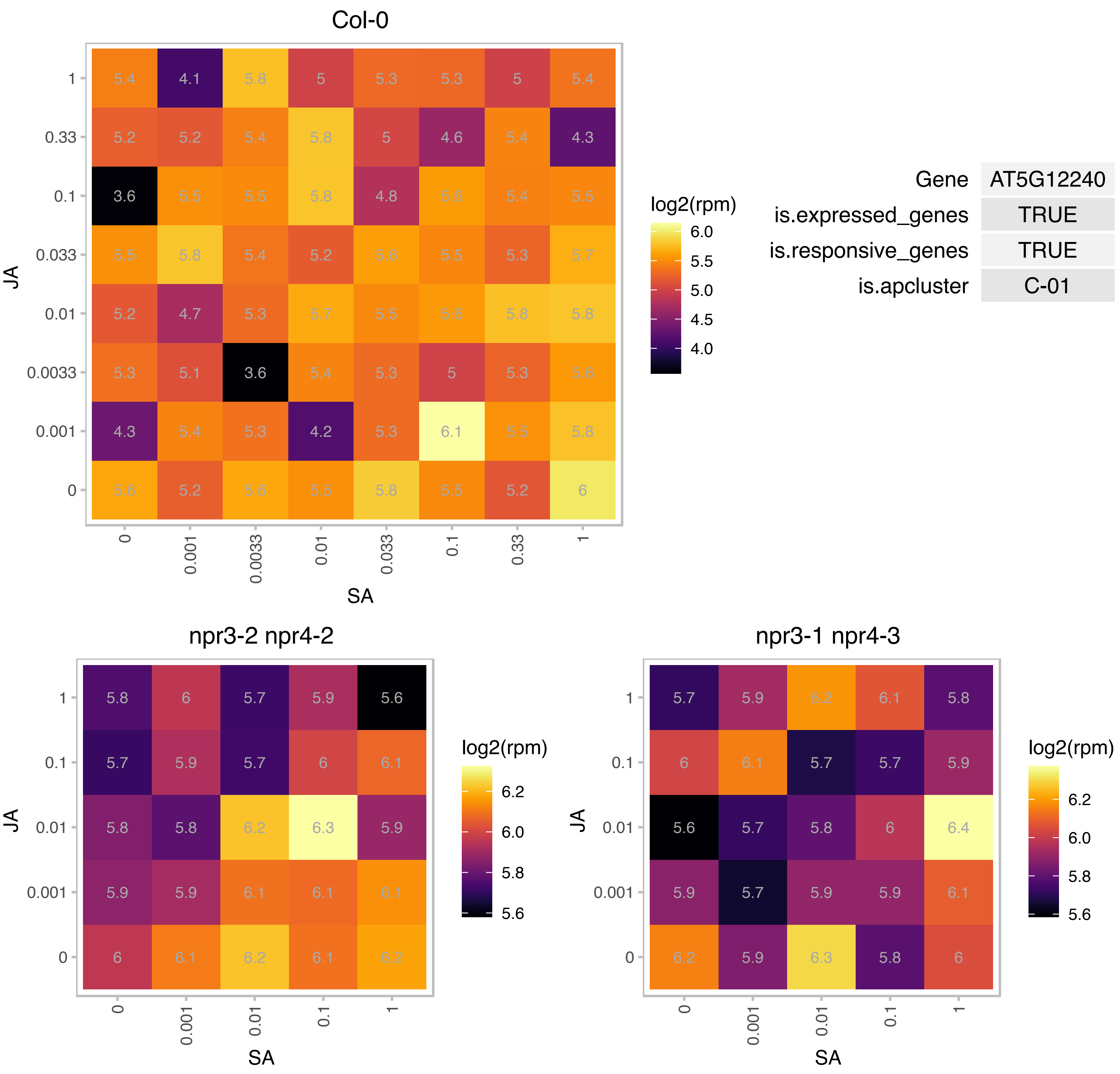

# AT5G19250.1 : NA

Glycoprotein membrane precursor GPI-anchored

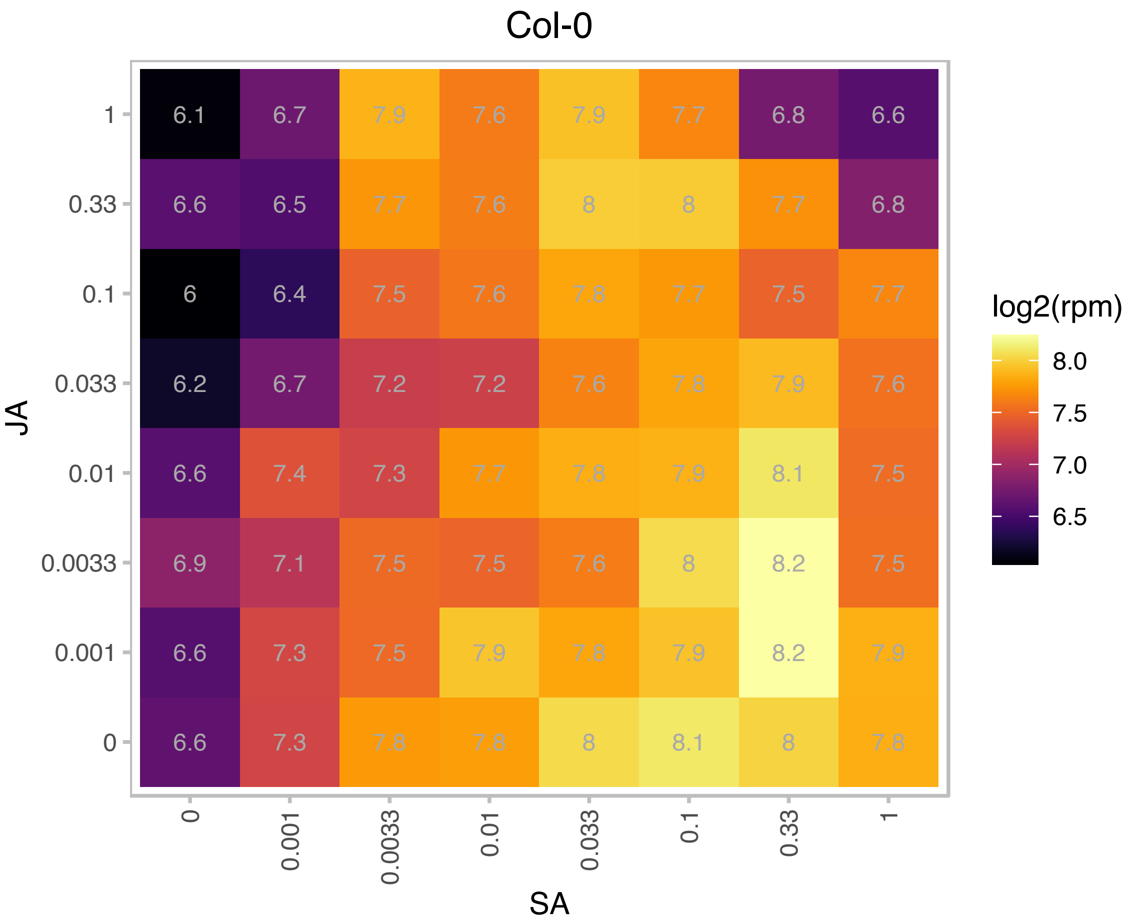


| Gene | AT5G19250 |
|---|---|
| is.expressed_genes | TRUE |
| is.responsive_genes | TRUE |
| is.apcluster | C-01 |

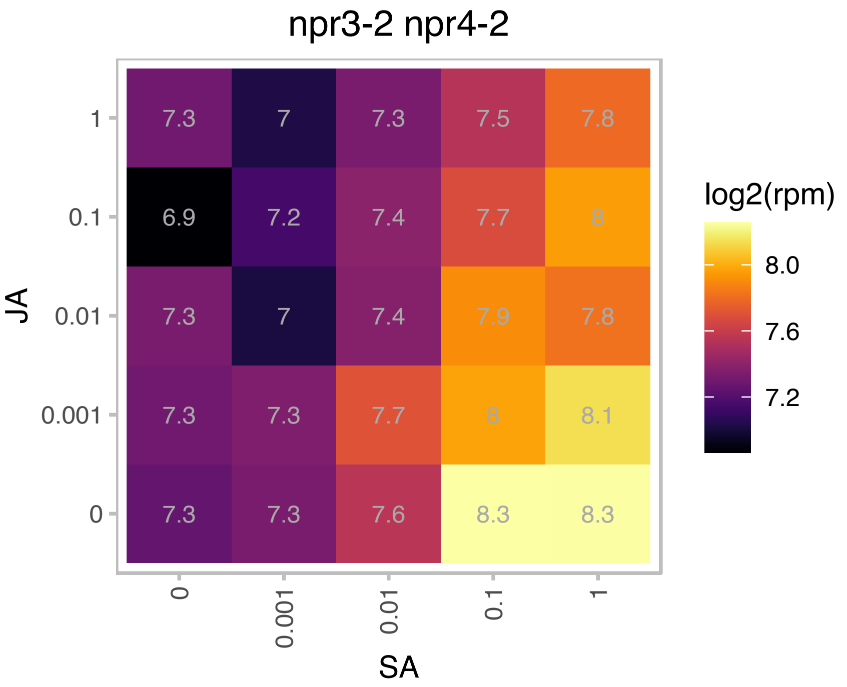


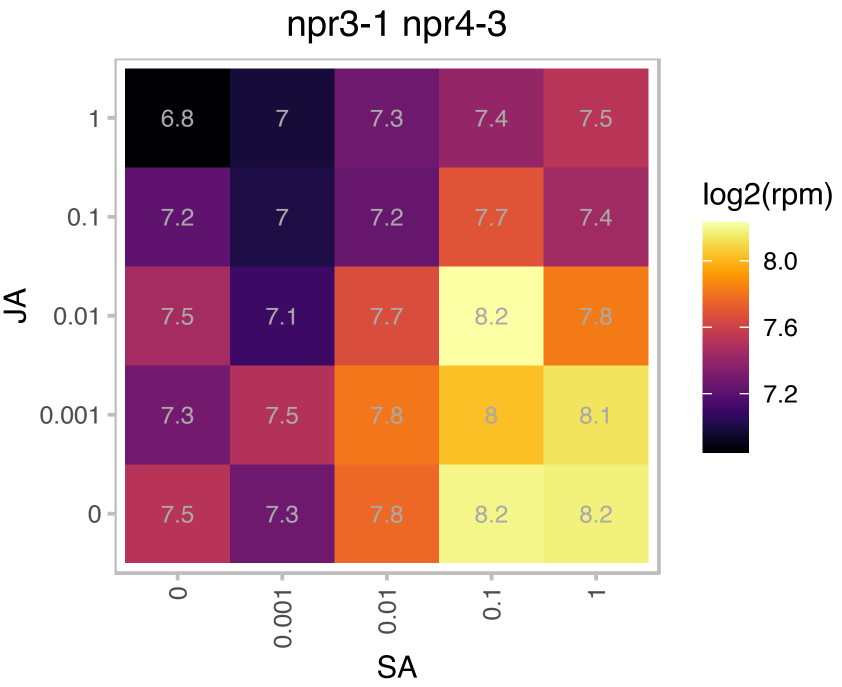

### AT5G19980.1 : GFT1, GONST4

golgi nucleotide sugar transporter 4

Encodes a Golgi-localized nucleotide-sugar transporter.

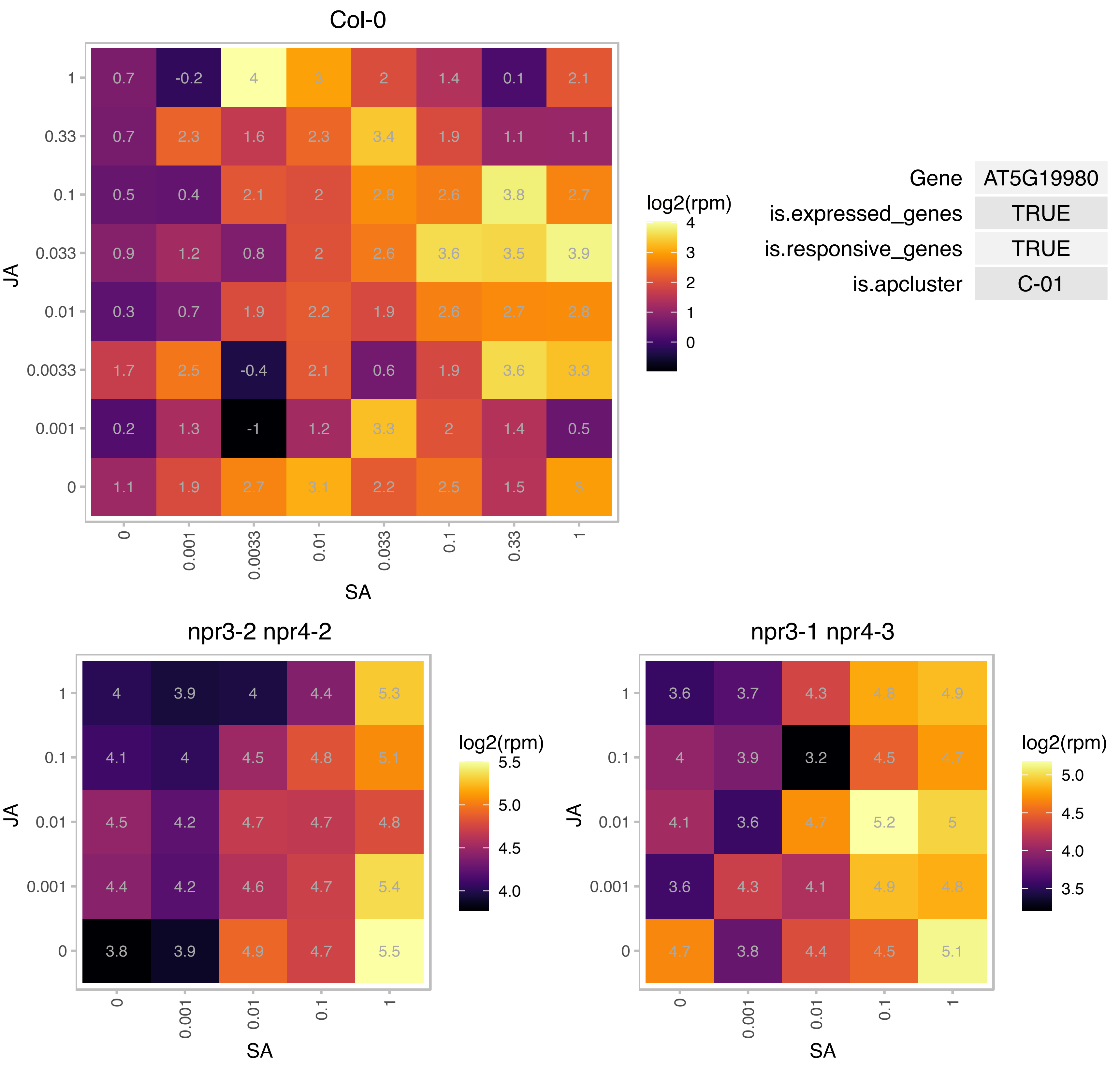


| | |
|---|---|
| Gene | AT5G19980 |
| is.expressed_genes | TRUE |
| is.responsive_genes | TRUE |
| is.apcluster | C-01 |

### AT5G22570.1 : ATWRKY38, WRKY38

WRKY DNA-binding protein 38

member of WRKY Transcription Factor; Group III

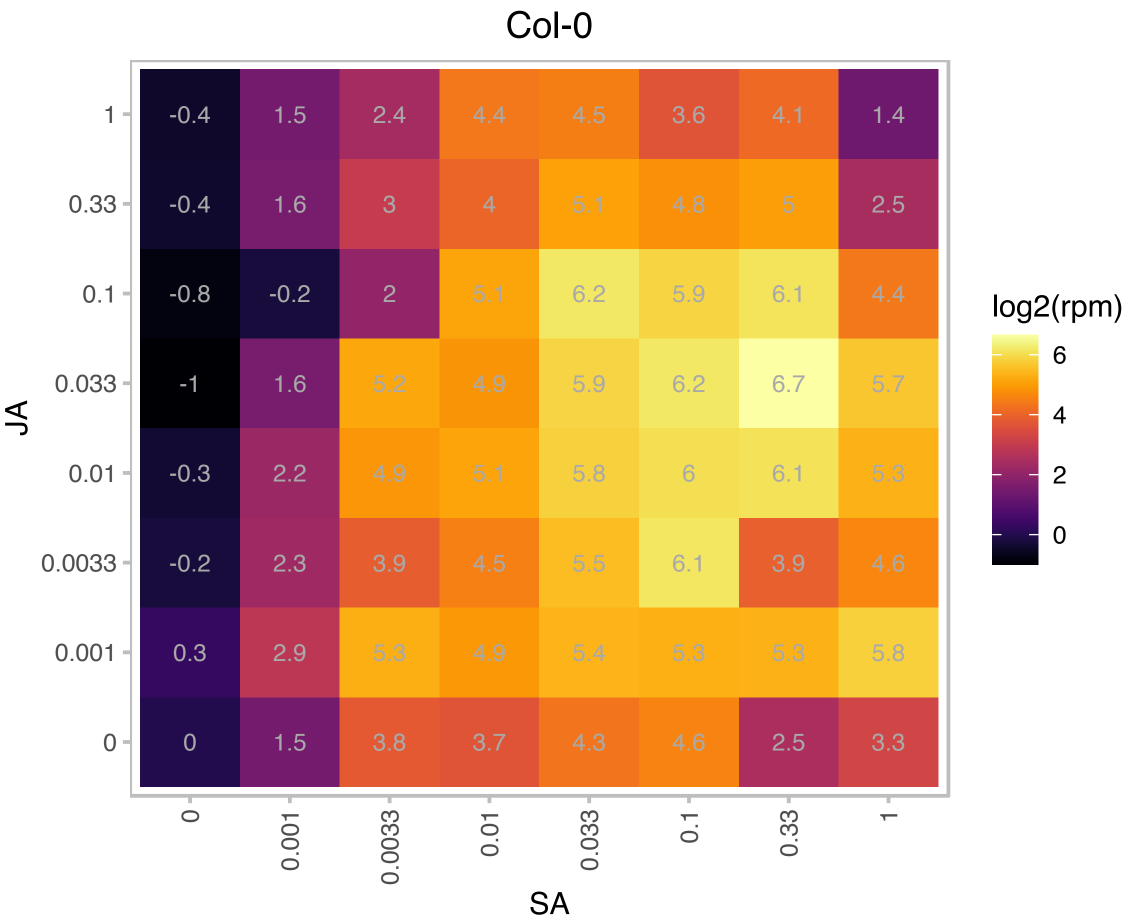


| | |
|---|---|
| Gene | AT5G22570 |
| is.expressed_genes | TRUE |
| is.responsive_genes | TRUE |
| is.apcluster | C-01 |

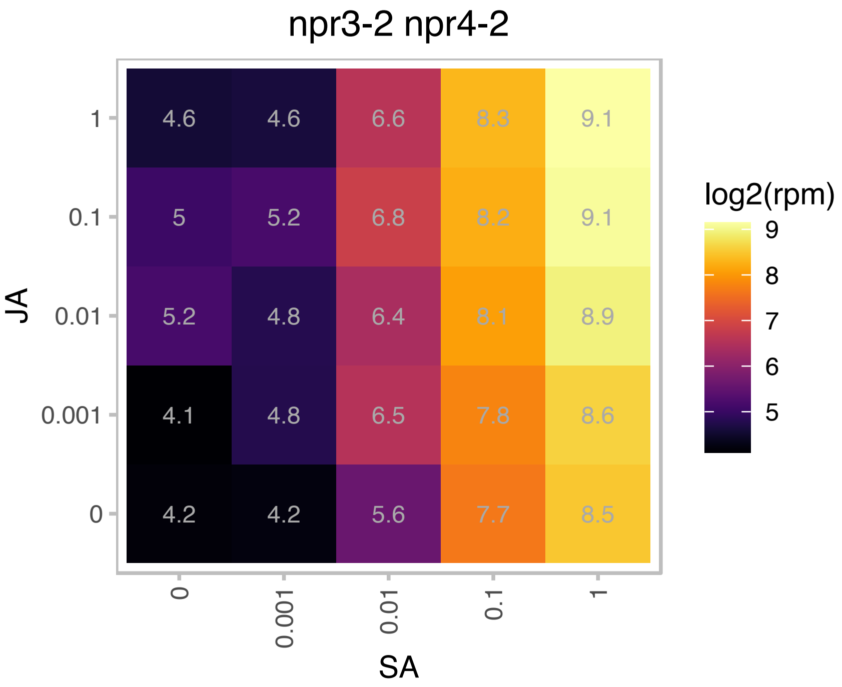


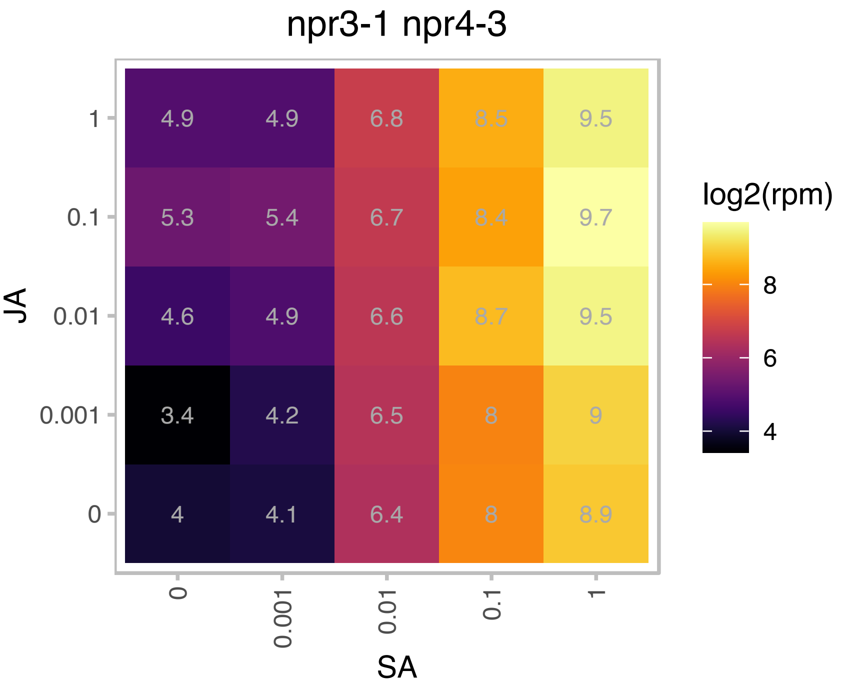

### AT5G24530.1 : AtDMR6, DMR6

2-oxoglutarate (2OG) and Fe(II)-dependent oxygenase superfamily protein

Encodes a putative 2OG-Fe(II) oxygenase that is defense-associated but required for susceptibility to downy mildew. The mRNA is cell-to-cell mobile.

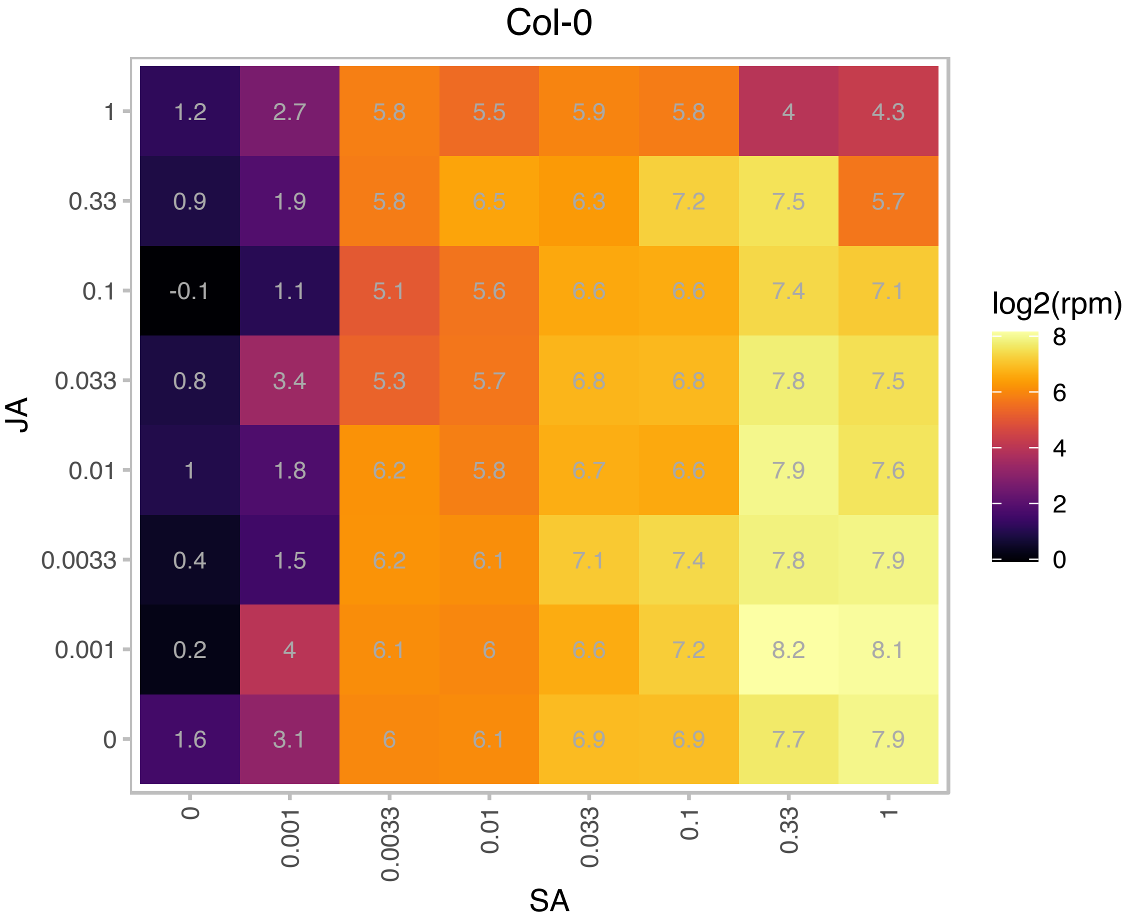


| | |
|---|---|
| Gene | AT5G24530 |
| is.expressed_genes | TRUE |
| is.responsive_genes | TRUE |
| is.apcluster | C-01 |

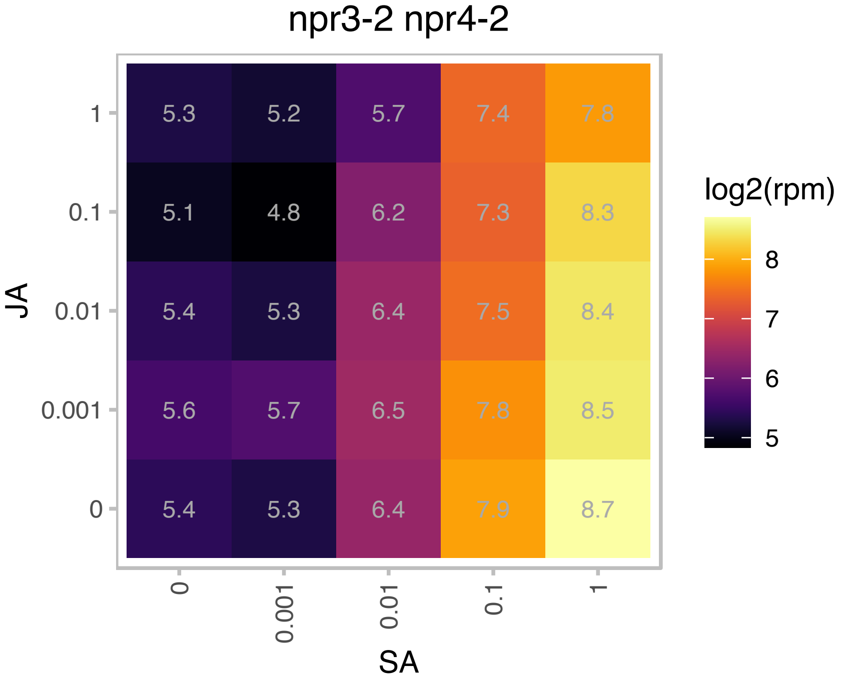


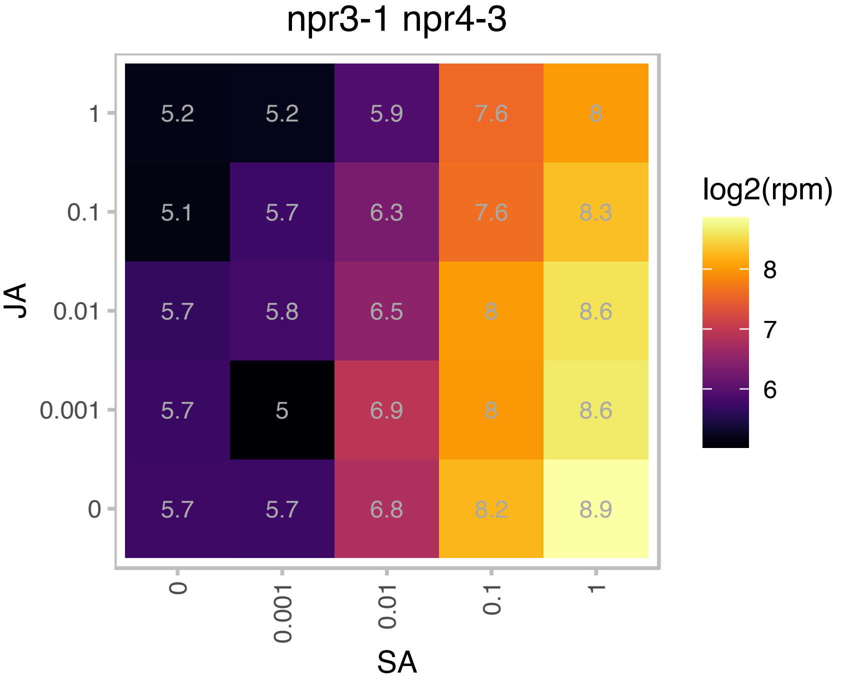

### AT5G26170.1 : ATWRKY50, WRKY50

WRKY DNA-binding protein 50

member of WRKY Transcription Factor; Group II-c.  Involved in jasmonic acid inducible defense responses.

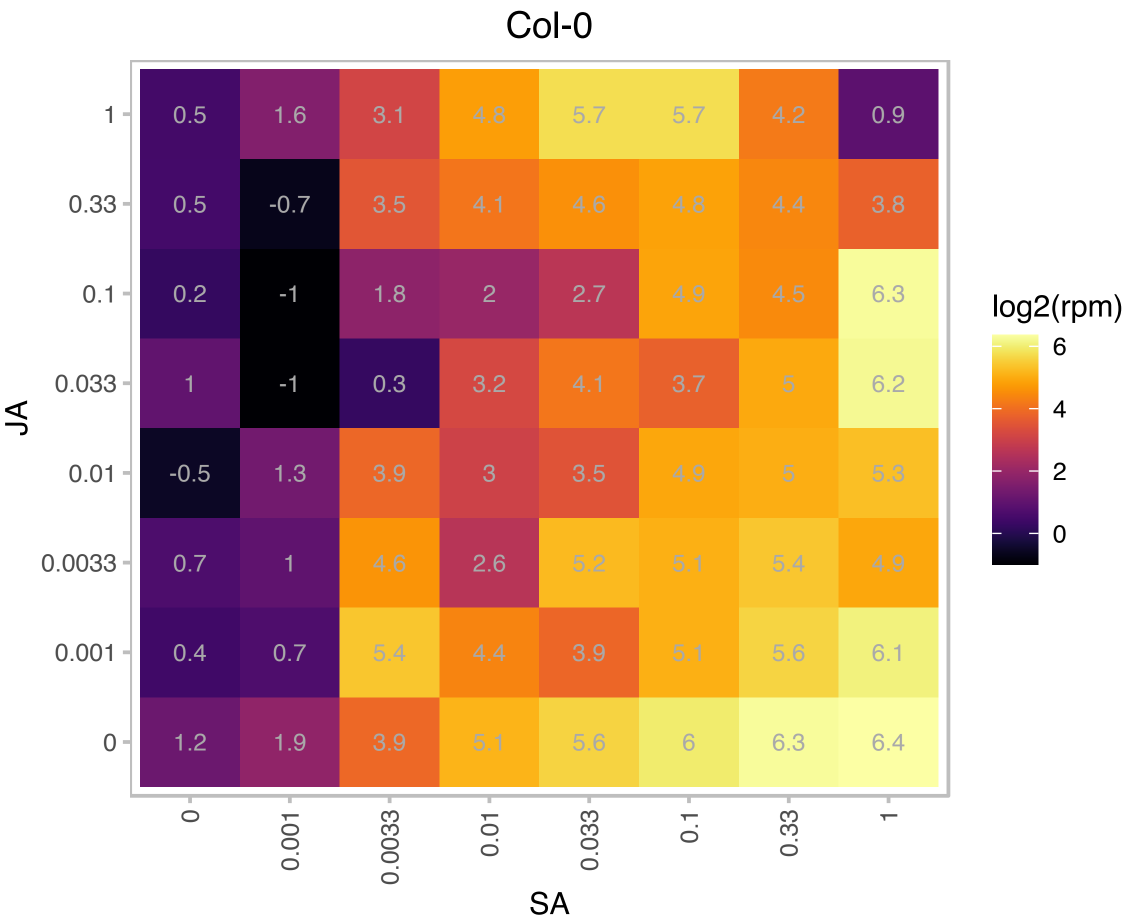


| | |
|---|---|
| Gene | AT5G26170 |
| is.expressed_genes | TRUE |
| is.responsive_genes | TRUE |
| is.apcluster | C-01 |

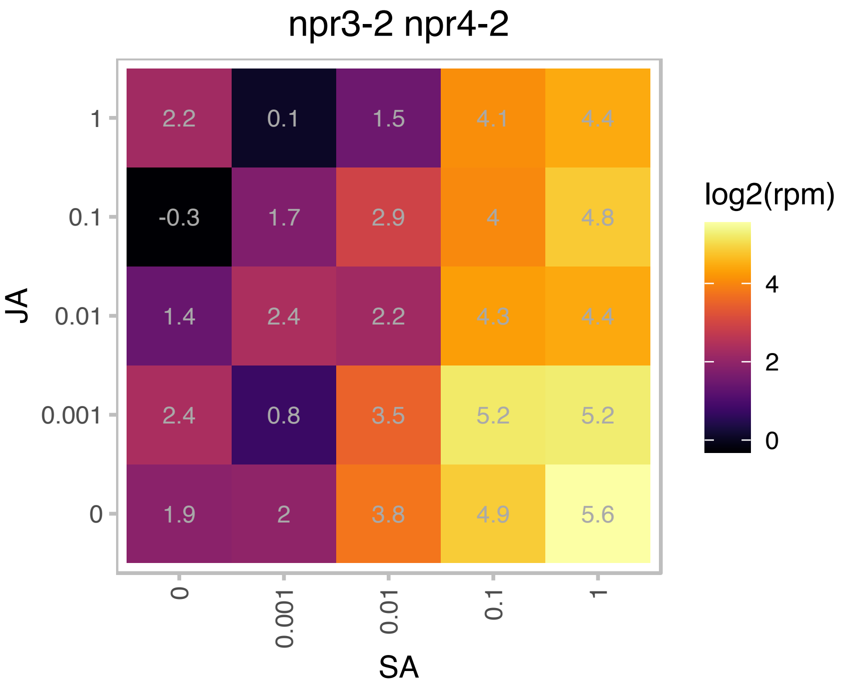


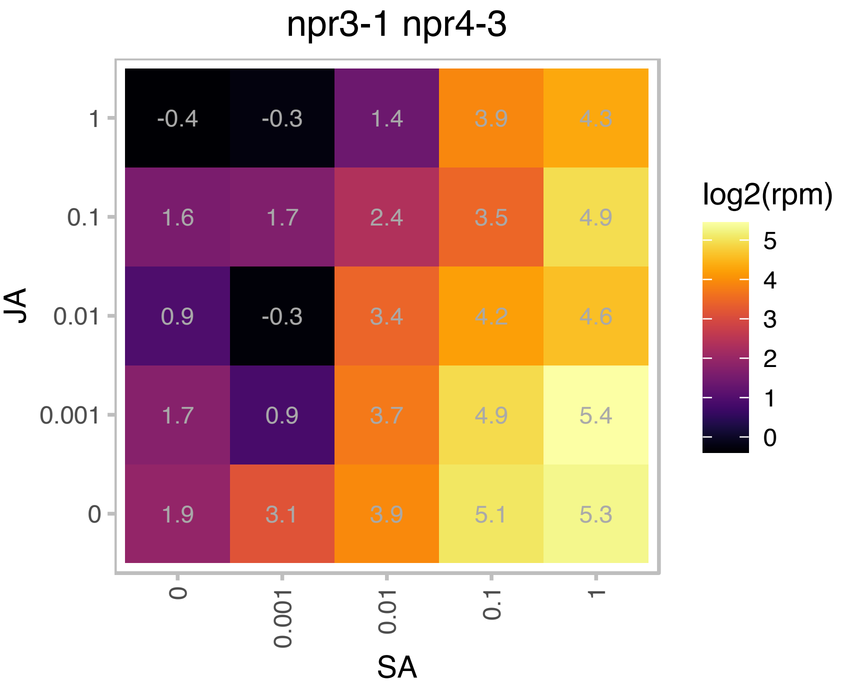

# AT5G26690.1 : NA

Heavy metal transport/detoxification superfamily protein

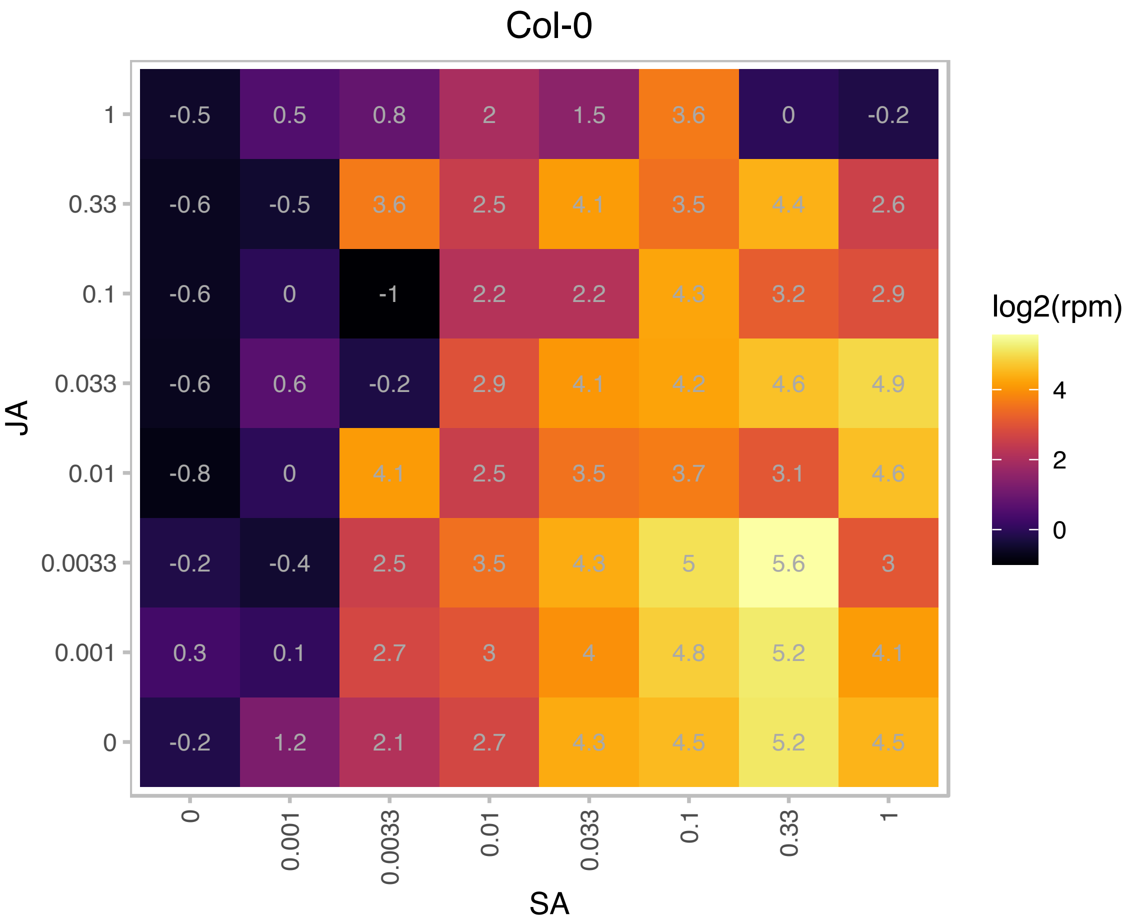


| | |
|---|---|
| Gene | AT5G26690 |
| is.expressed_genes | TRUE |
| is.responsive_genes | TRUE |
| is.apcluster | C-01 |

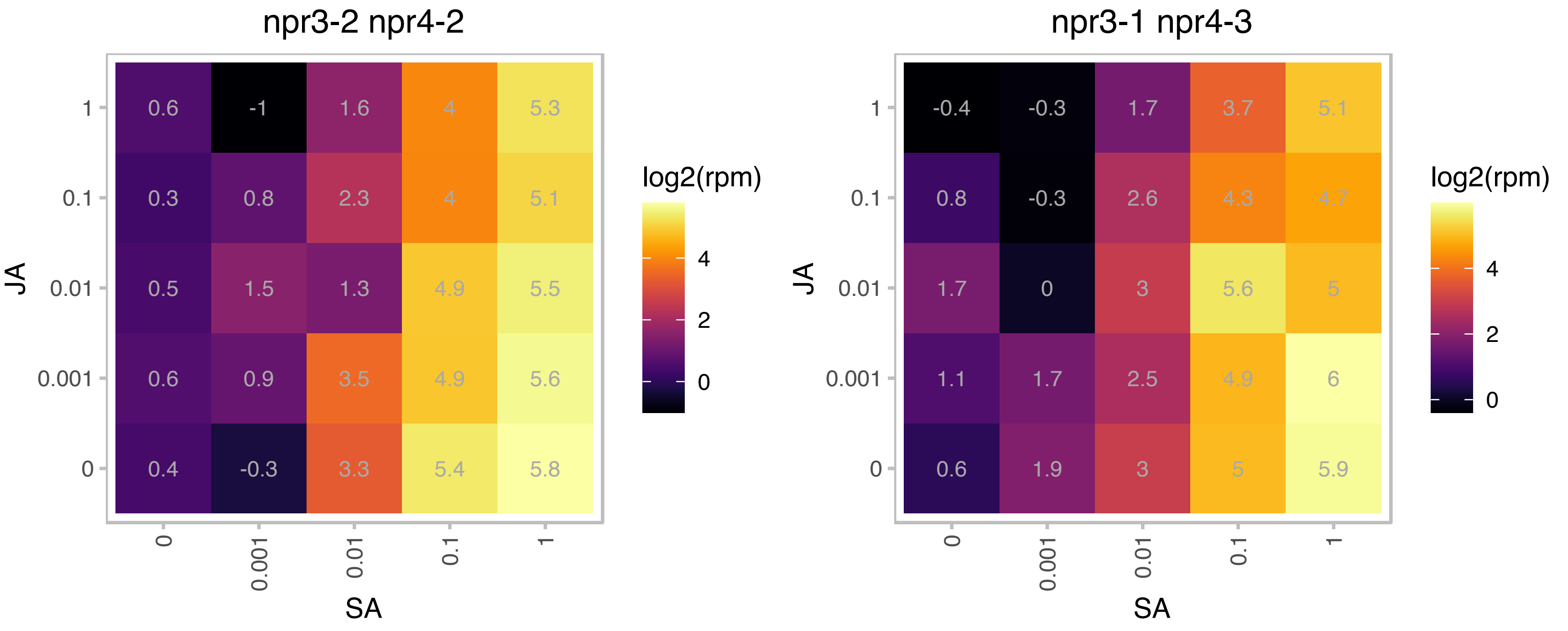

# AT5G37740.1 : NA

Calcium-dependent lipid-binding (CaLB domain) family protein

| | |
|---|---|
| Gene | AT5G37740 |
| is.expressed_genes | TRUE |
| is.responsive_genes | TRUE |
| is.apcluster | C-01 |

Col-0

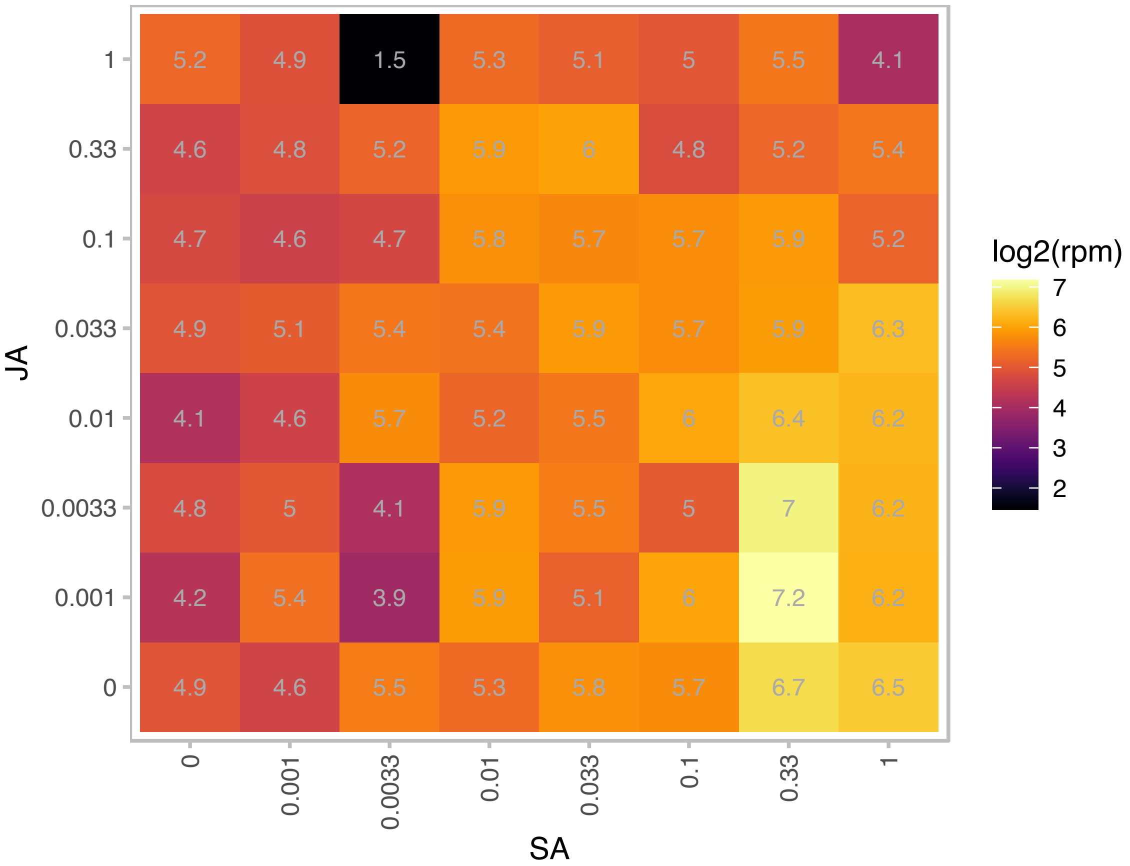


npr3-2 npr4-2

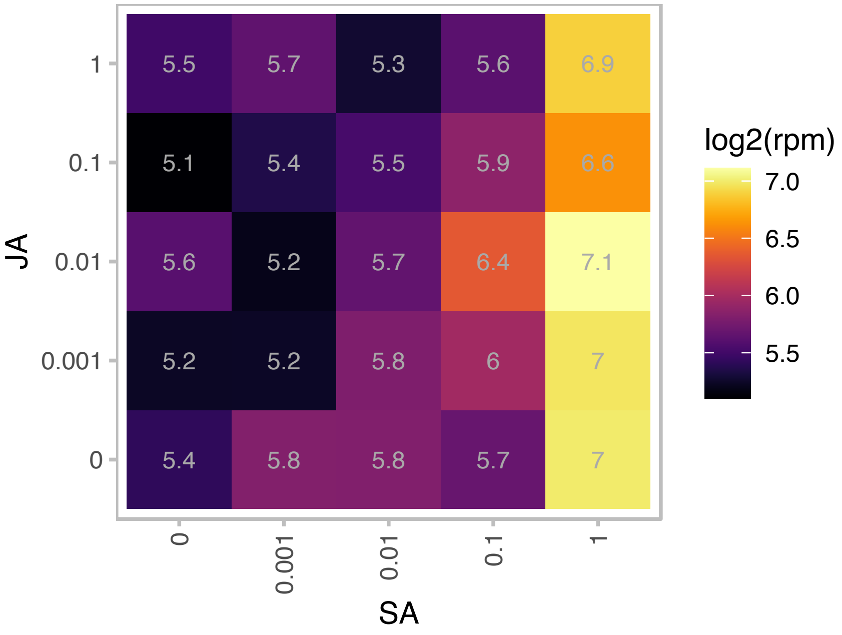


npr3-1 npr4-3

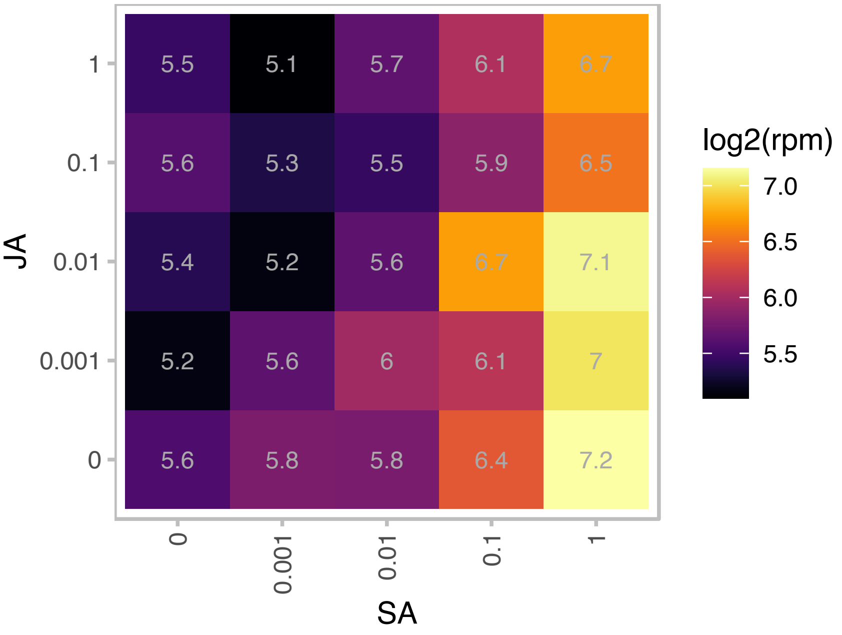

### AT5G44570.1 : CWM2

Encodes a PPR protein involved in mitochondrial functioning. Mutants suppress cell wall defects caused by C17 chemical inhibitor. Mutants are defective in cytochrome c maturation and activation of mitochondrial retrograde signalling.

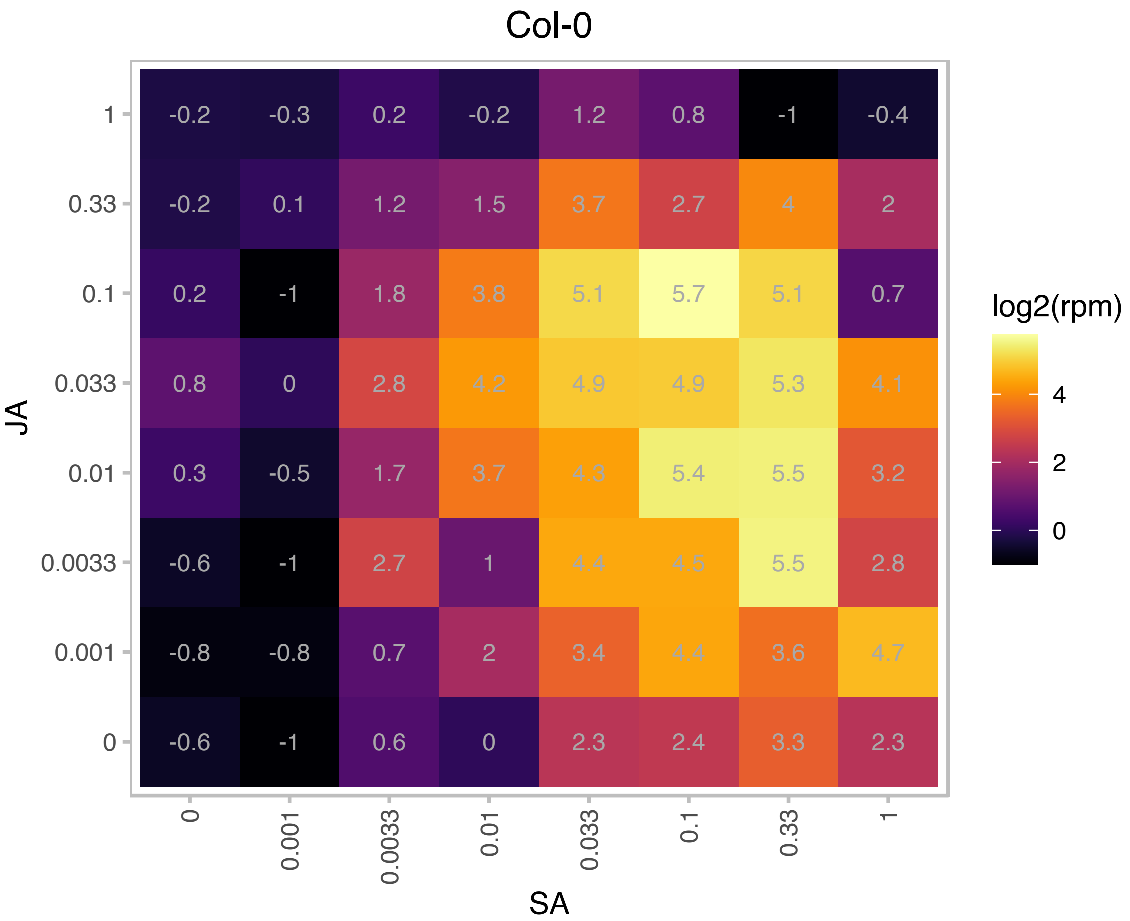


| | |
|---|---|
| Gene | AT5G44570 |
| is.expressed_genes | TRUE |
| is.responsive_genes | TRUE |
| is.apcluster | C-01 |

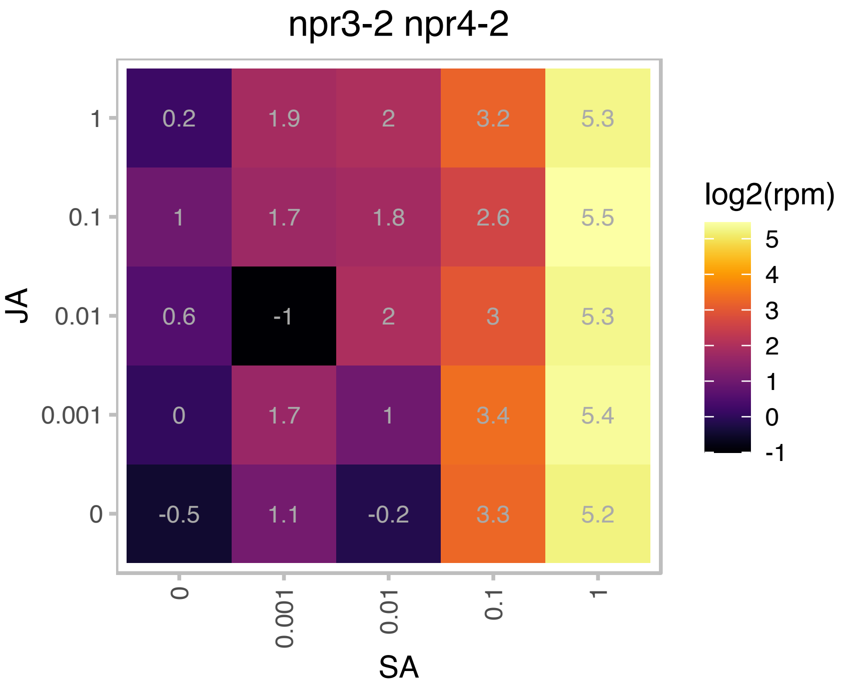


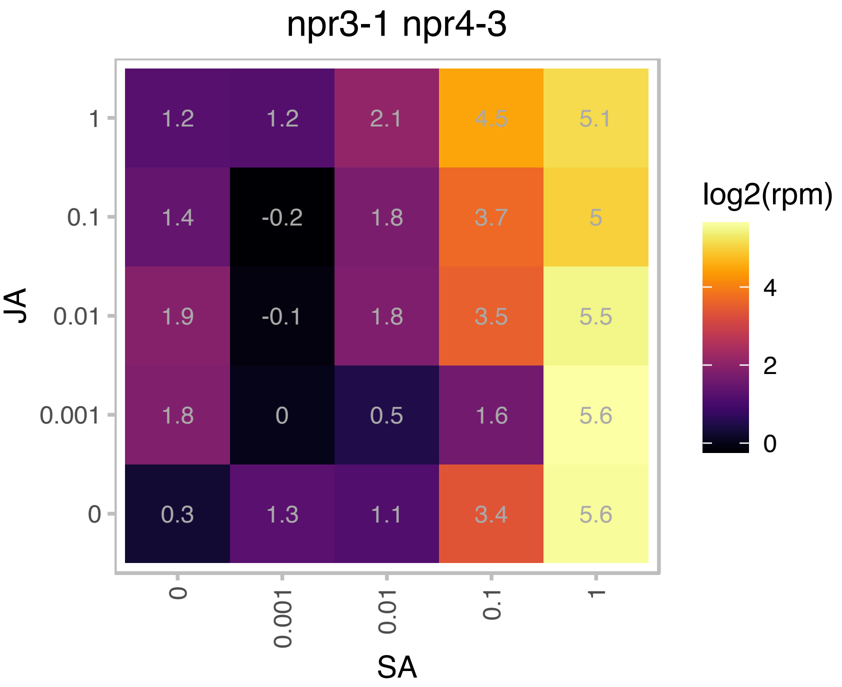

# AT5G44820.1 : NA

Nucleotide-diphospho-sugar transferase family protein

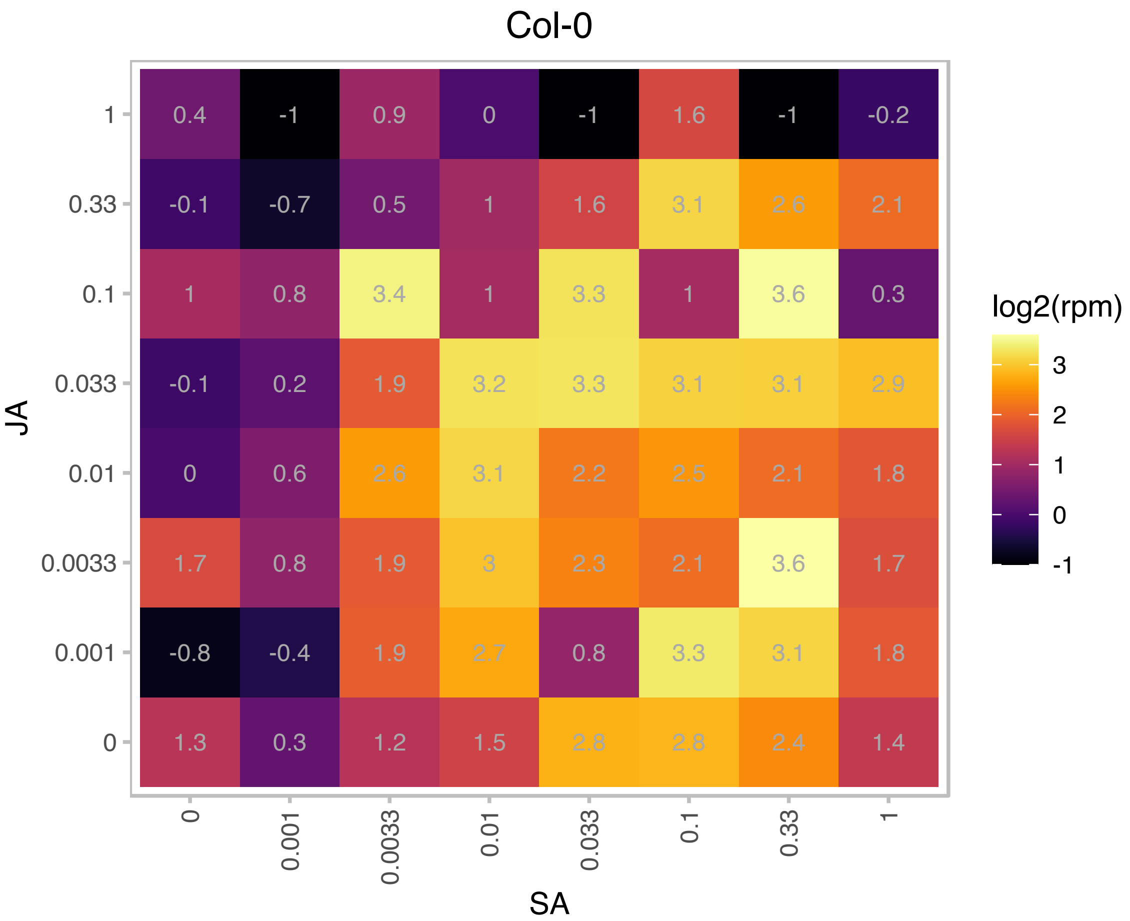


| | |
|---|---|
| Gene | AT5G44820 |
| is.expressed_genes | TRUE |
| is.responsive_genes | TRUE |
| is.apcluster | C-01 |

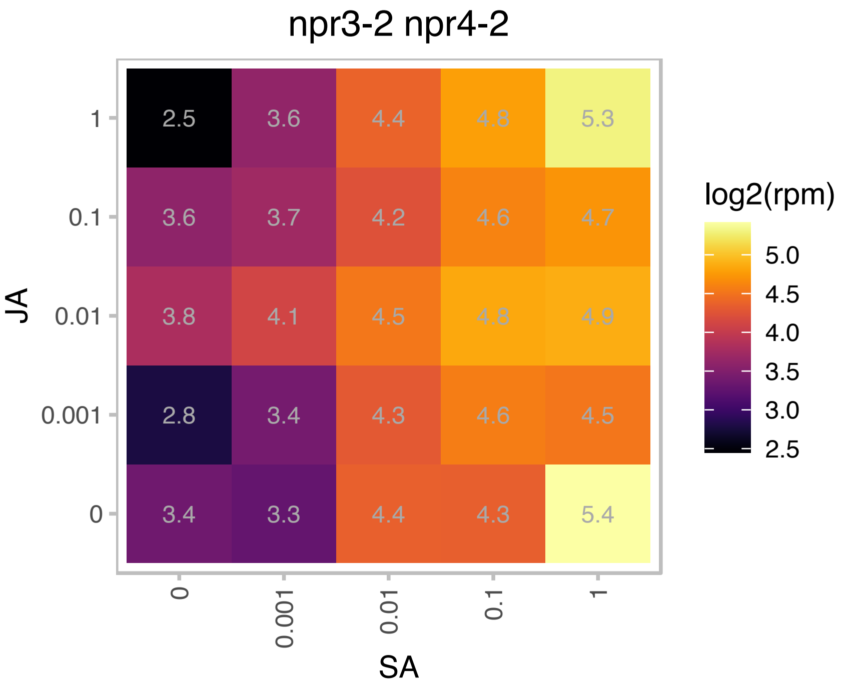


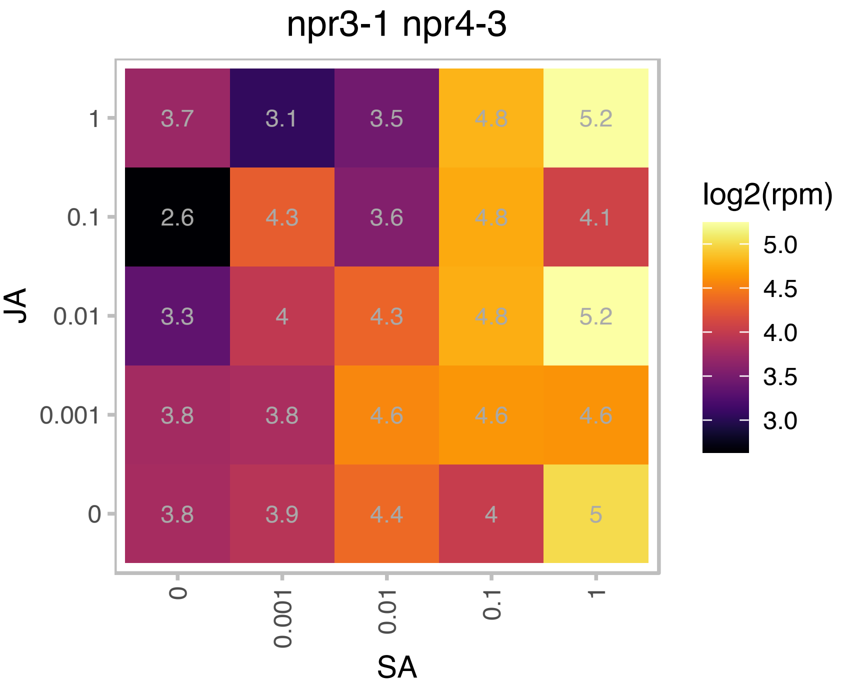

### AT5G54610.1 : ANK, BDA1

ankyrin

Induced in response to Salicylic acid. Belongs to the ankyrin repeat protein family.

Col-0

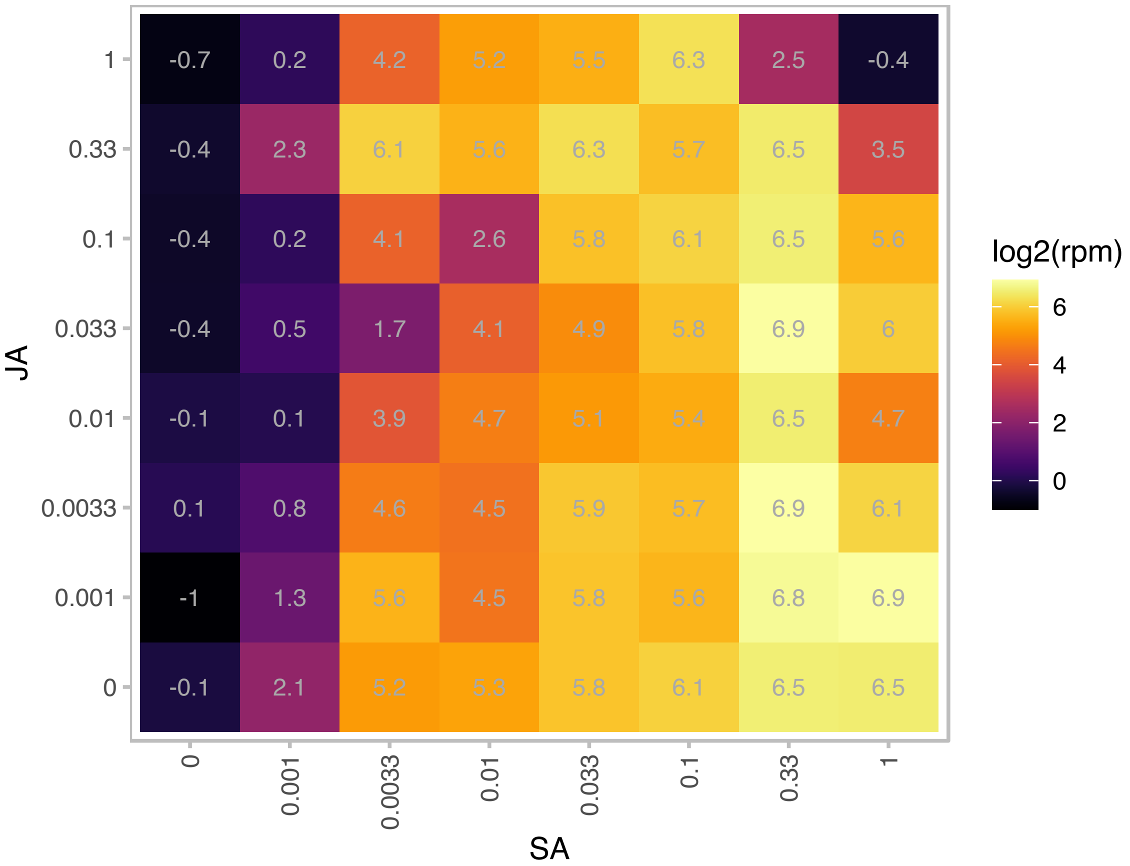


| | |
|---|---|
| Gene | AT5G54610 |
| is.expressed_genes | TRUE |
| is.responsive_genes | TRUE |
| is.apcluster | C-01 |

npr3-2 npr4-2

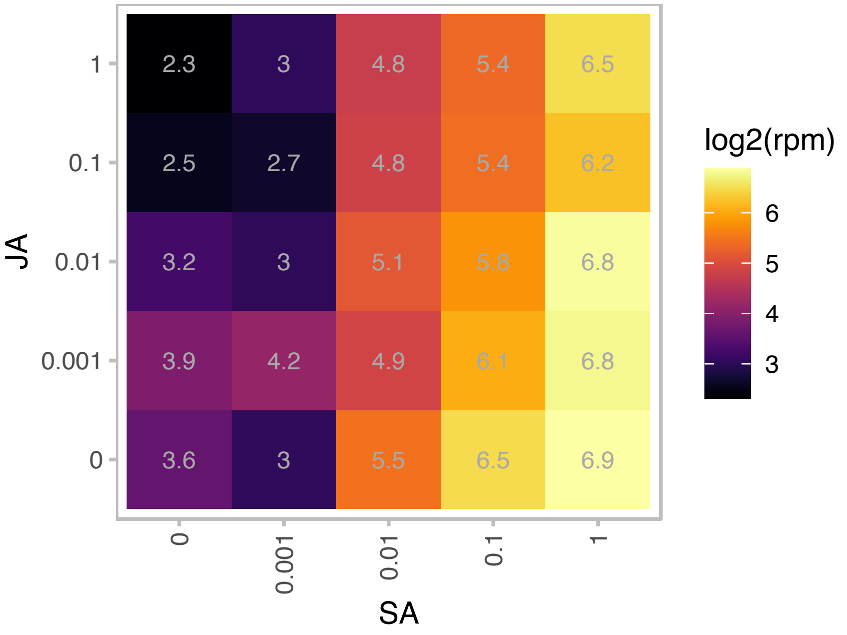


npr3-1 npr4-3

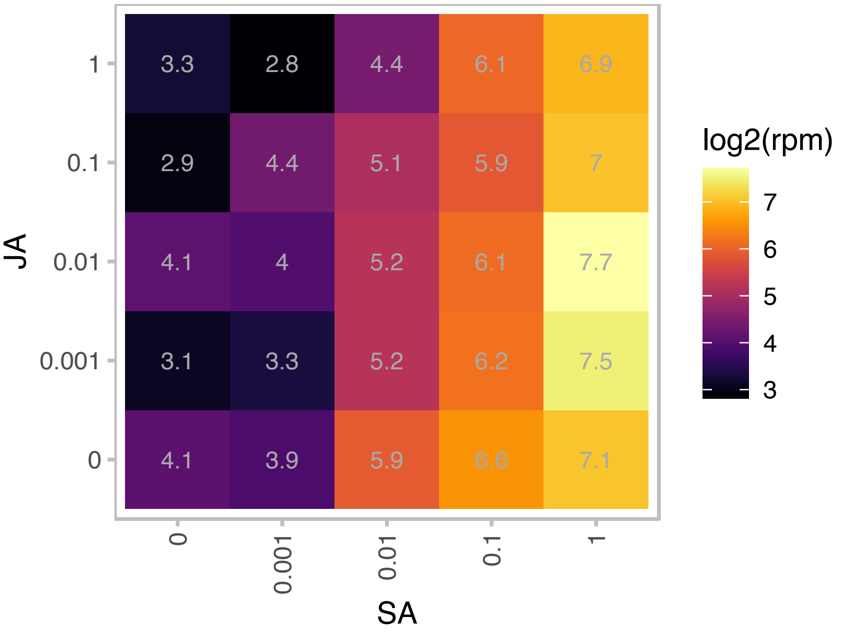

# AT5G62770.1 : NA

Protein of unknown function (DUF1645)

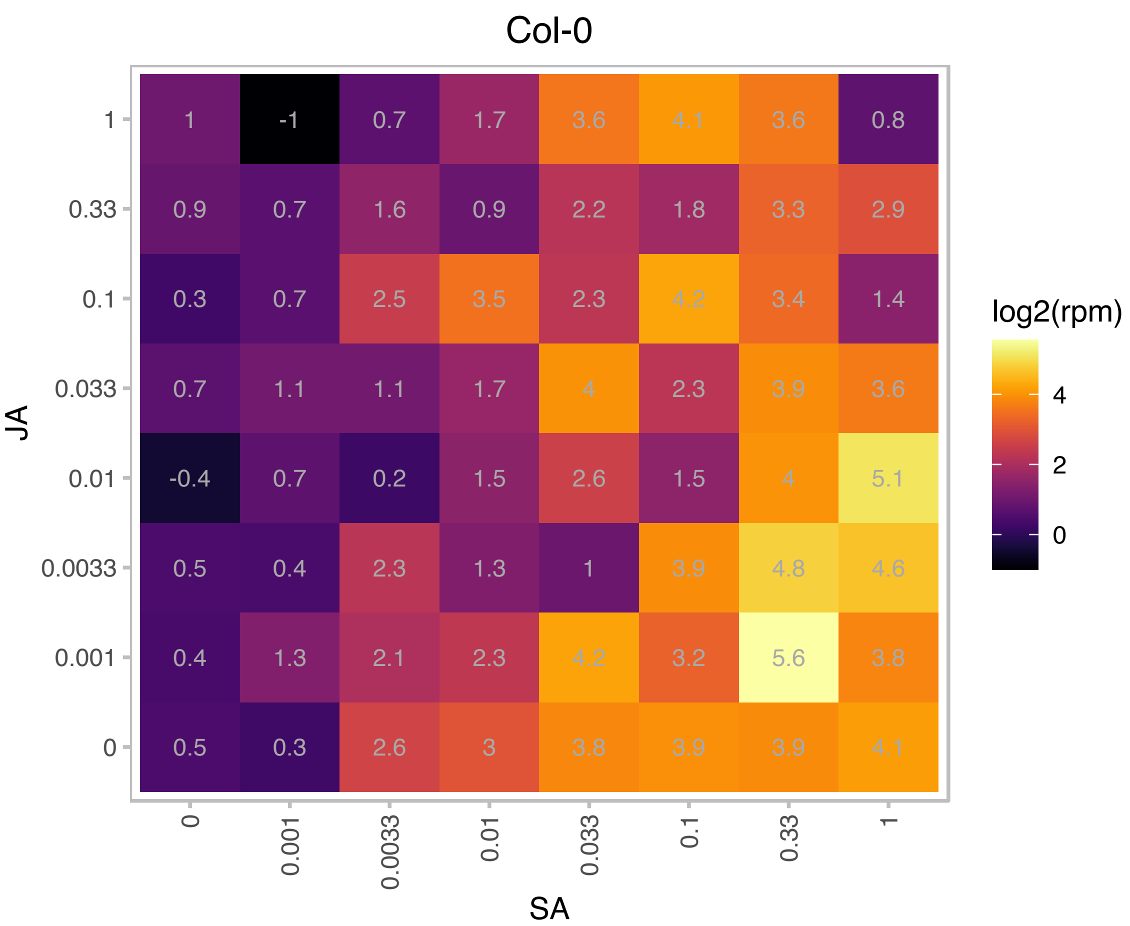


| Gene | AT5G62770 |
|---|---|
| is.expressed_genes | TRUE |
| is.responsive_genes | TRUE |
| is.apcluster | C-01 |

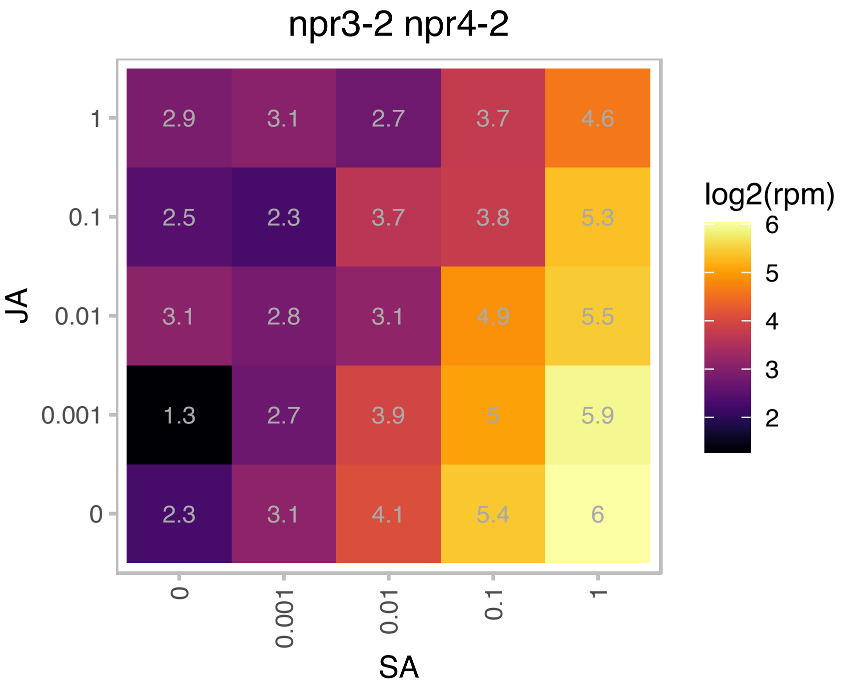


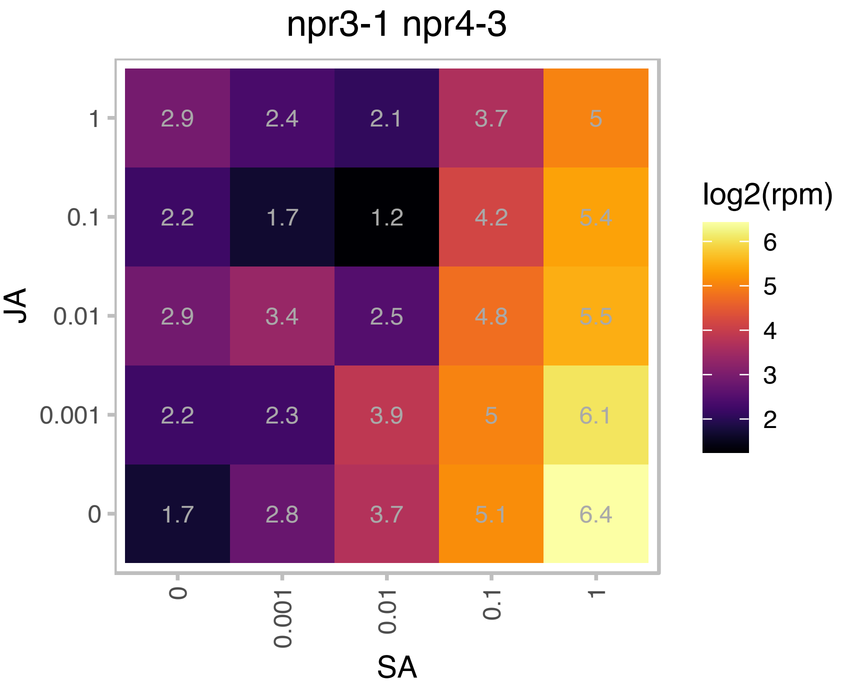

# AT5G63200.1 : NA

tetratricopeptide repeat (TPR)-containing protein

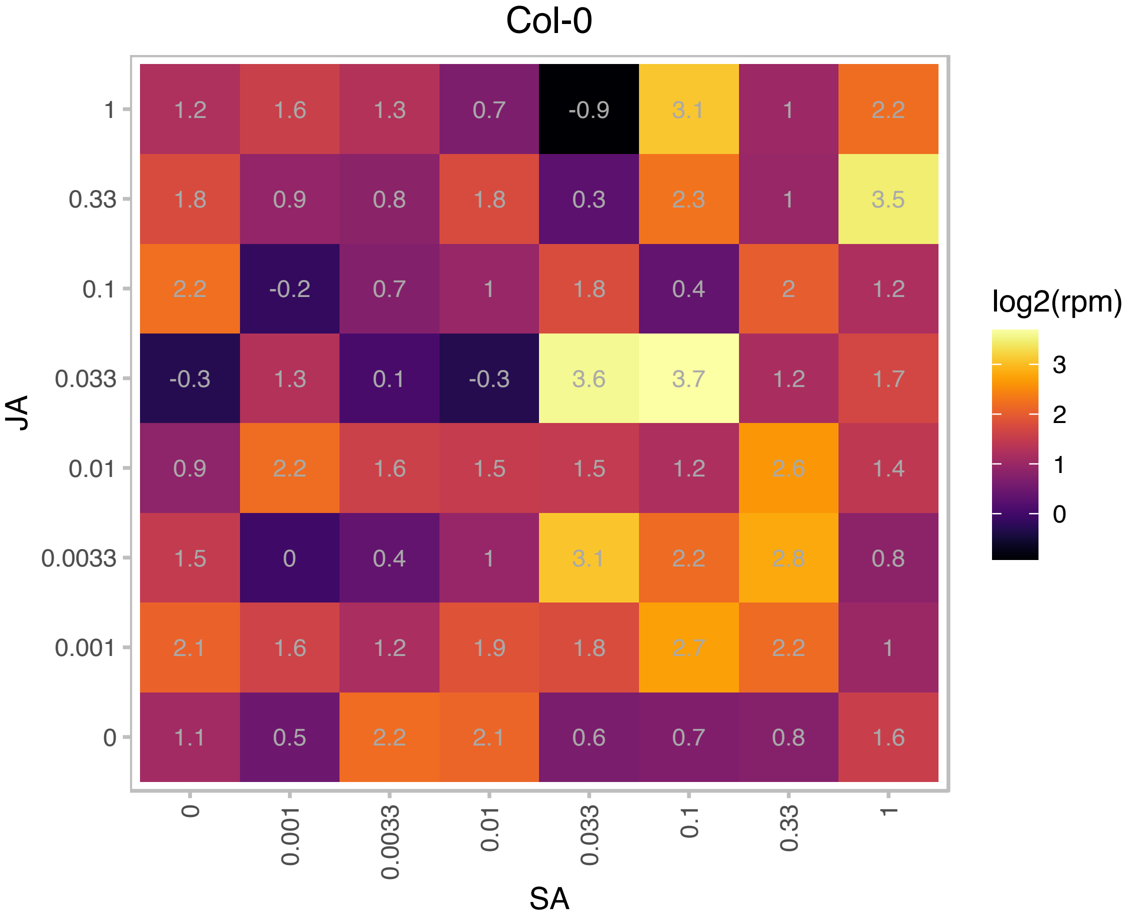


| | |
|---|---|
| Gene | AT5G63200 |
| is.expressed_genes | TRUE |
| is.responsive_genes | TRUE |
| is.apcluster | C-01 |

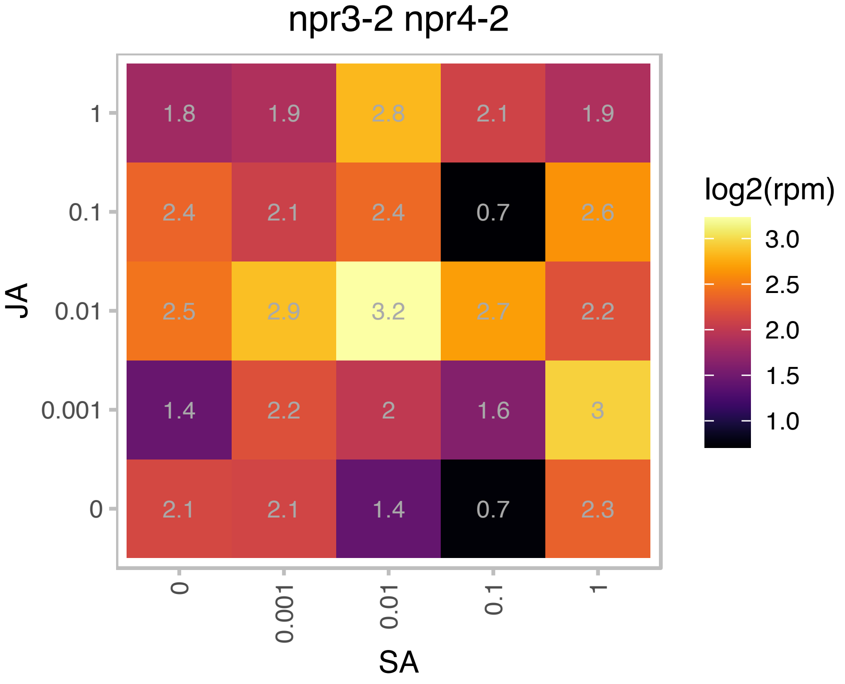


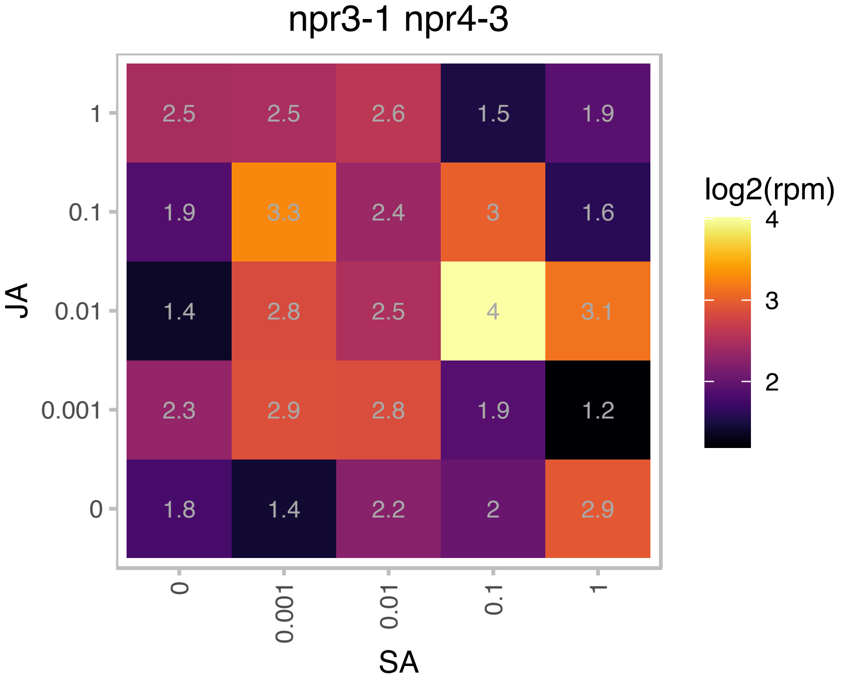

### AT5G64810.1 : ATWRKY51, WRKY51

WRKY DNA-binding protein 51

member of WRKY Transcription Factor; Group II-c.  Involved in jasmonic acid inducible defense responses.

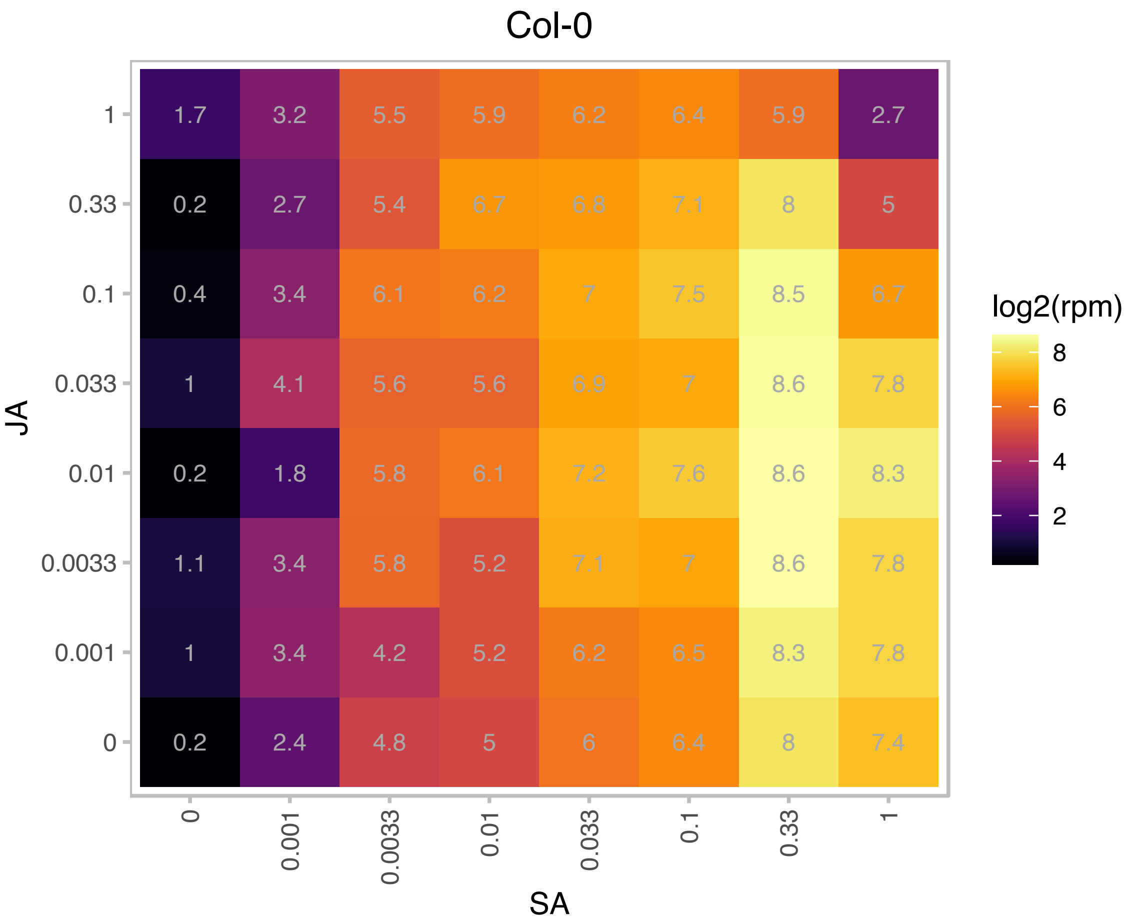


| | |
|---|---|
| Gene | AT5G64810 |
| is.expressed_genes | TRUE |
| is.responsive_genes | TRUE |
| is.apcluster | C-01 |

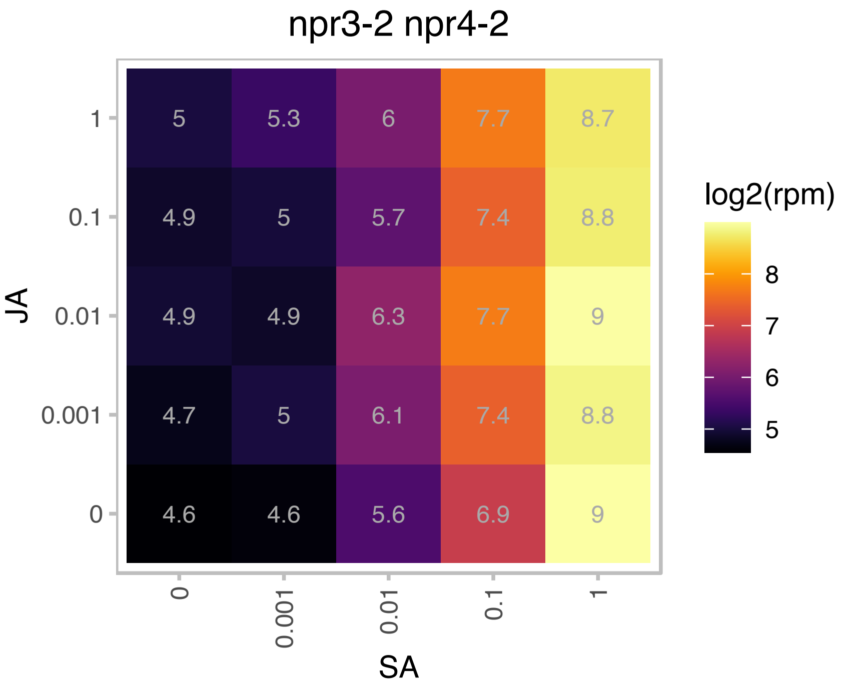


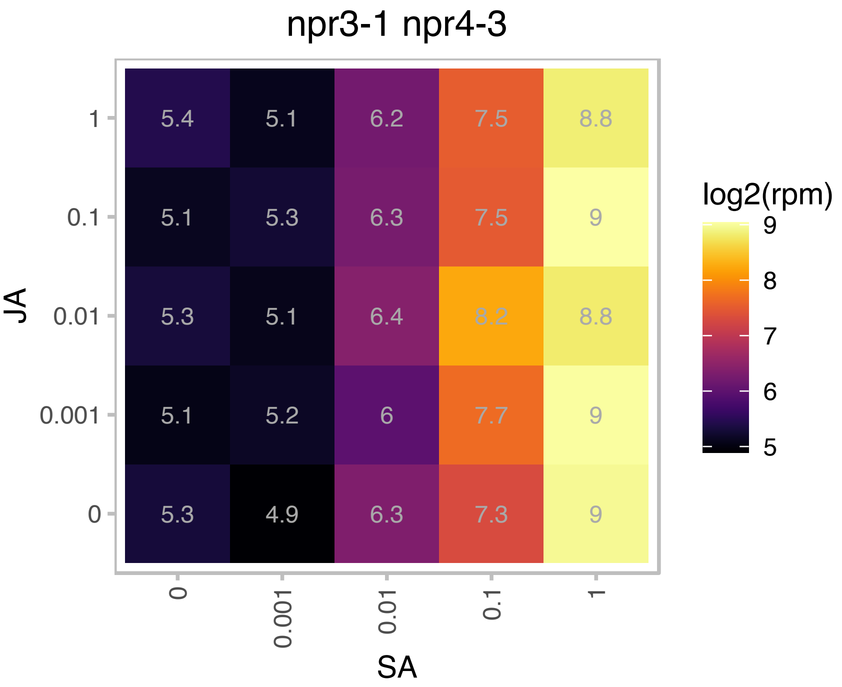

# AT5G66910.1 : NA

Disease resistance protein (CC-NBS-LRR class) family

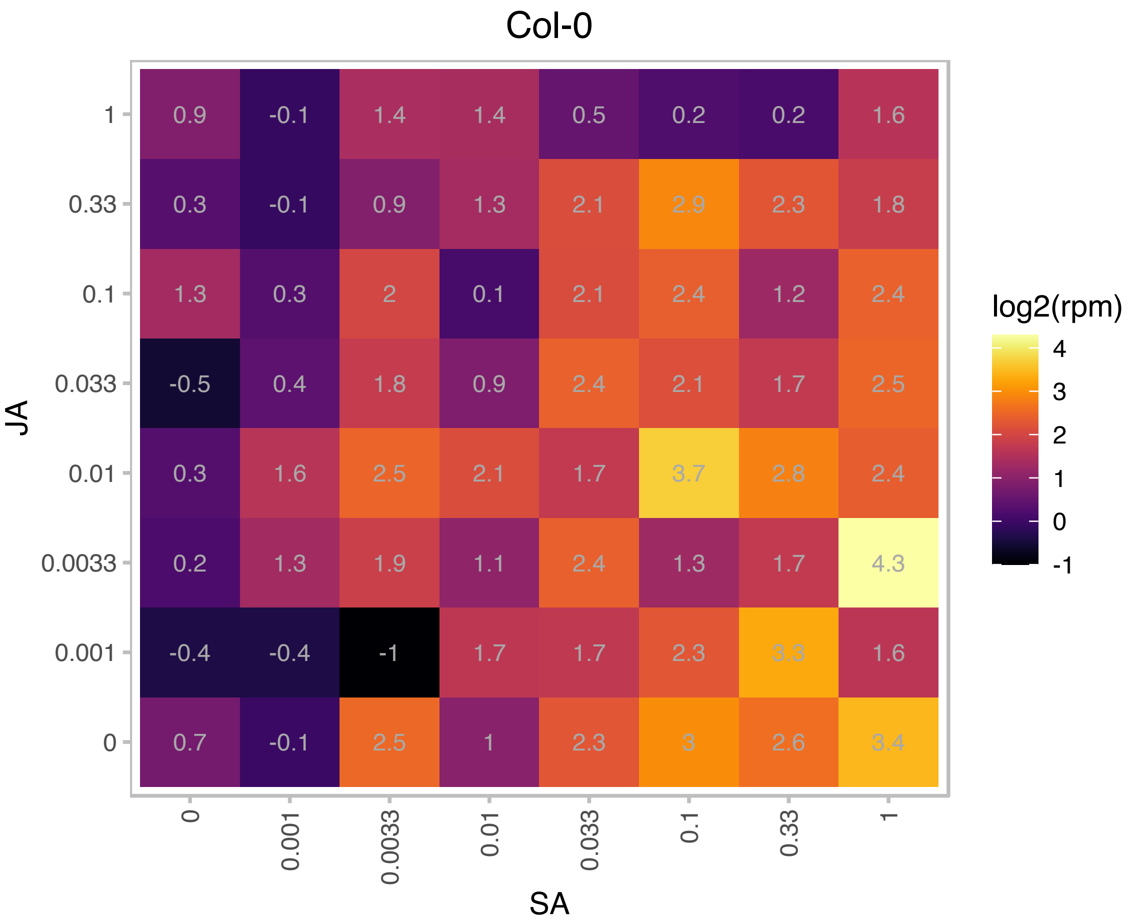


| | |
|---|---|
| Gene | AT5G66910 |
| is.expressed_genes | TRUE |
| is.responsive_genes | TRUE |
| is.apcluster | C-01 |

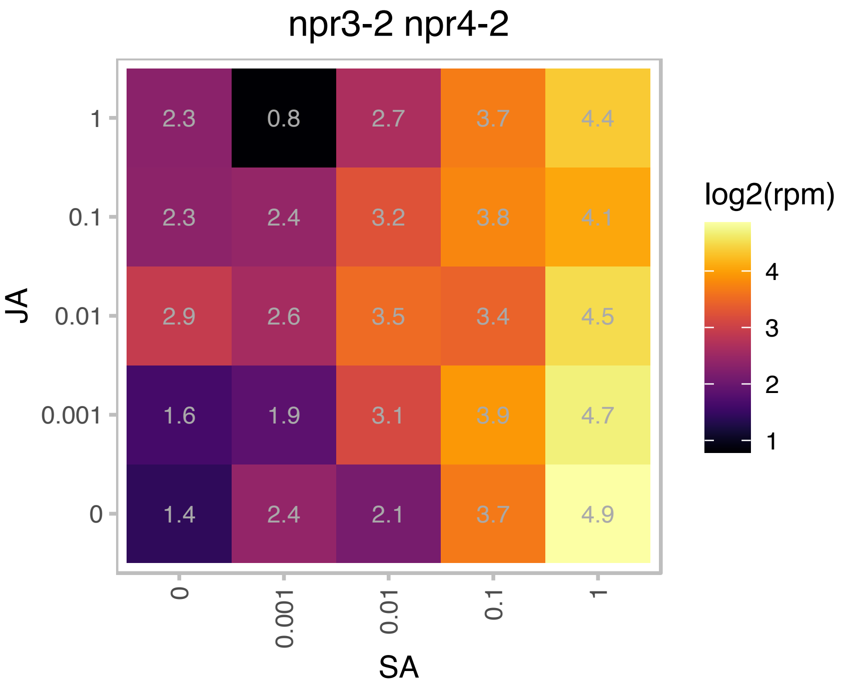


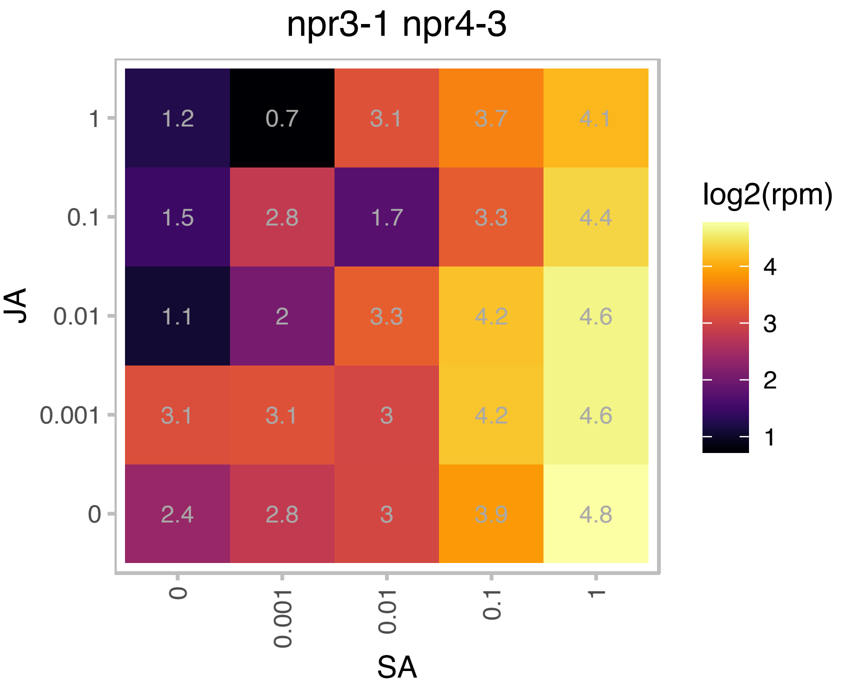

### AT1G03850.1 : ATGRXS13, GRXS13, ROXY18

Glutaredoxin family protein

Encodes glutaredoxin ATGRXS13, required to facilitate Botrytis cinerea infection of Arabidopsis thaliana plants. Sylvain La Camera et al (2011, PMID:21756272) reported a third splice variant in addition to the two annotated in TAIR10. It is a member of the CC-type glutaredoxin (ROXY) family that has been shown to interact with the transcription factor TGA2 and suppress ORA59 promoter activity.

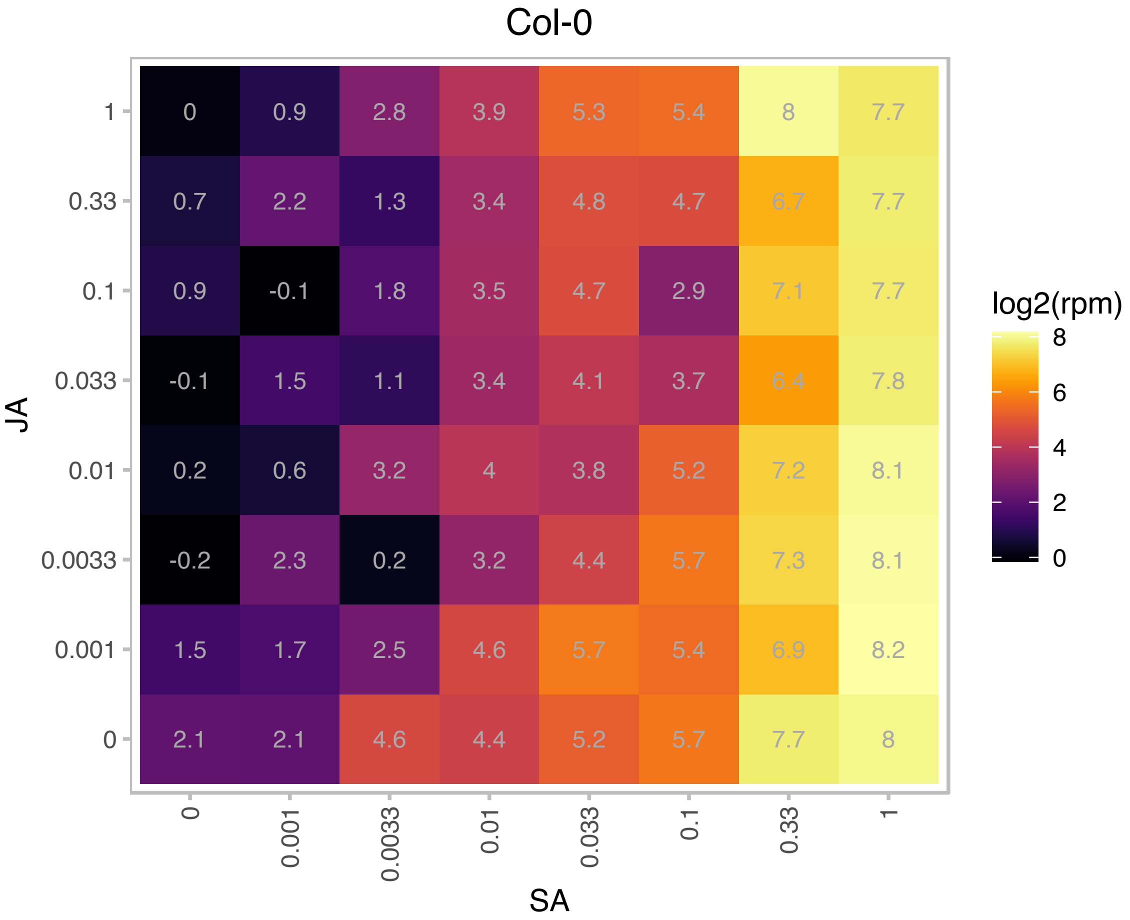


| | |
|---|---|
| Gene | AT1G03850 |
| is.expressed_genes | TRUE |
| is.responsive_genes | TRUE |
| is.apcluster | C-02 |

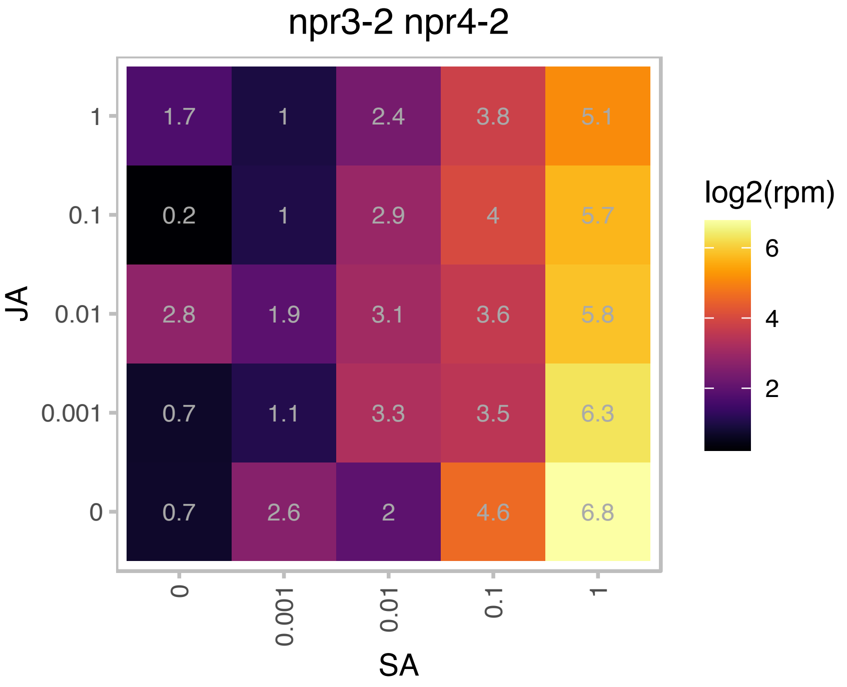


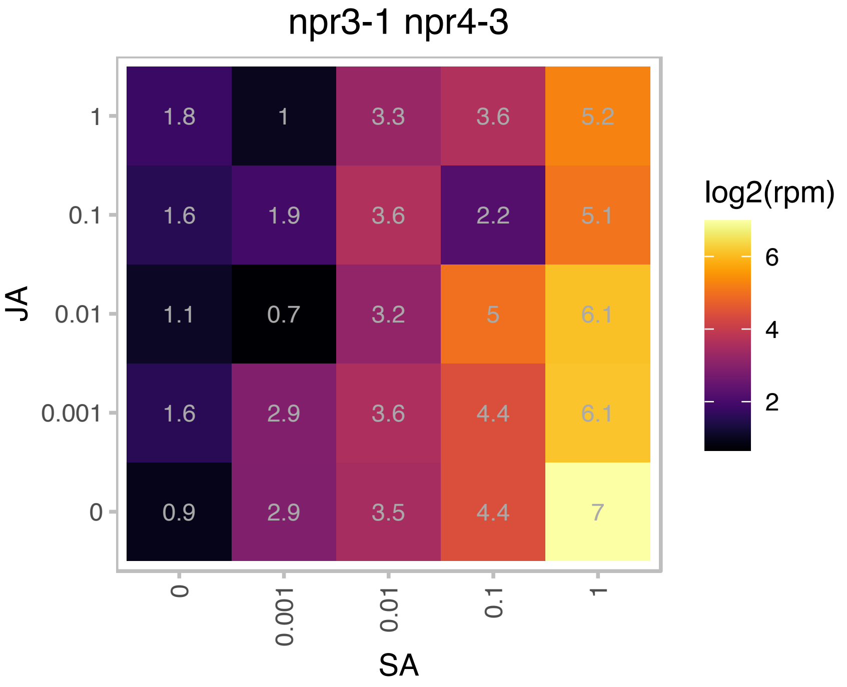

### AT1G17440.1 : CKH1, EER4, TAF12B

Encodes one of two Arabidopsis proteins with similarity to the TBP-associated factor TAF12. The gene product is an EIN3-interacting TFIID transcription factor required for proper ethylene response, including ERF1 induction. Loss of function mutants show enhanced response to ethylene. Located in nucleus and expressed throughout the plant. Required for ERF1 expression. Cytokinin-hypersensitive 1 (CKH1) mutants are characterized by rapidly growing calli with a green color at low levels of cytokinins, which are insufficient to induce such cytokinin responses in wild-type explants. It is hypothesized that CKH1 acts as a negative [illegible]

Transcription initiation factor TFIID subunit A

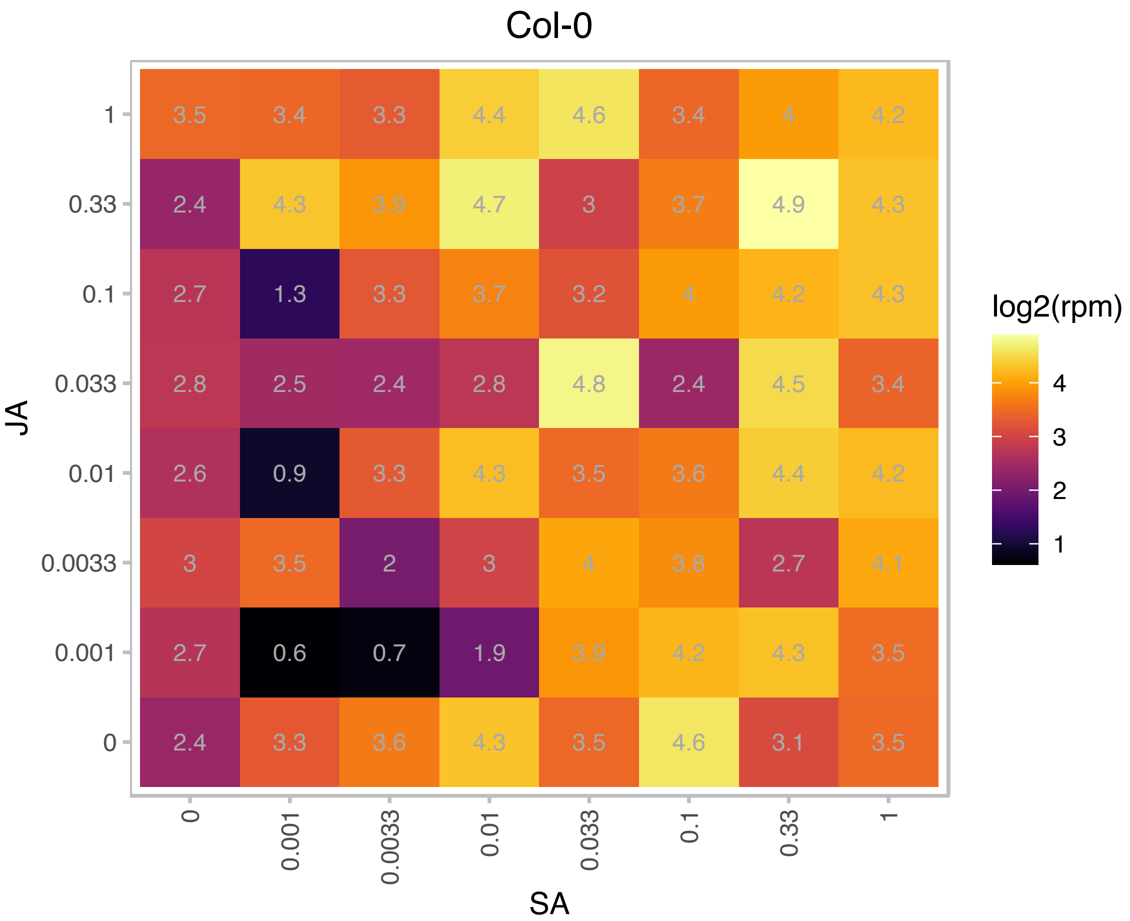


| | |
|---|---|
| Gene | AT1G17440 |
| is.expressed_genes | TRUE |
| is.responsive_genes | TRUE |
| is.apcluster | C-02 |

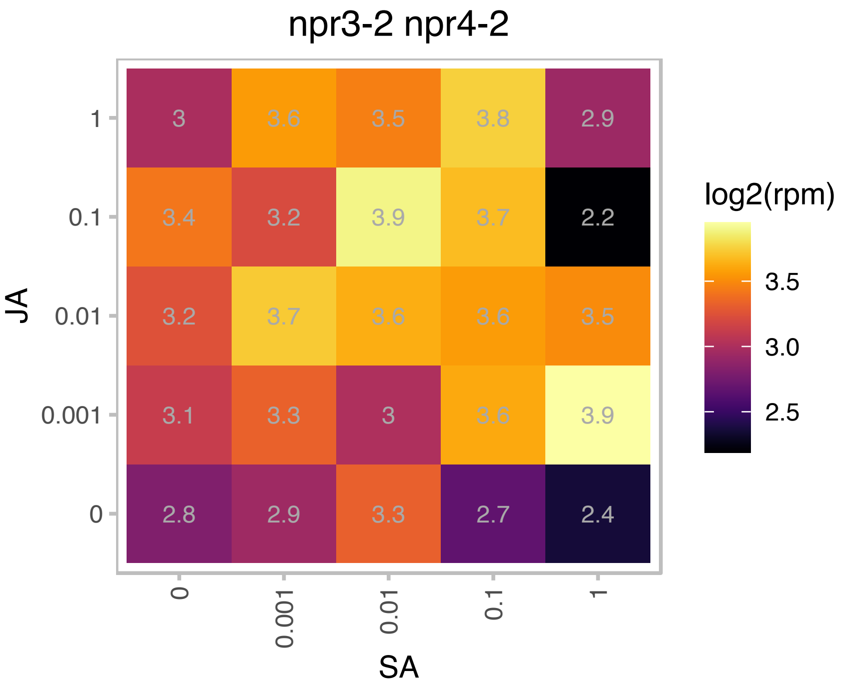


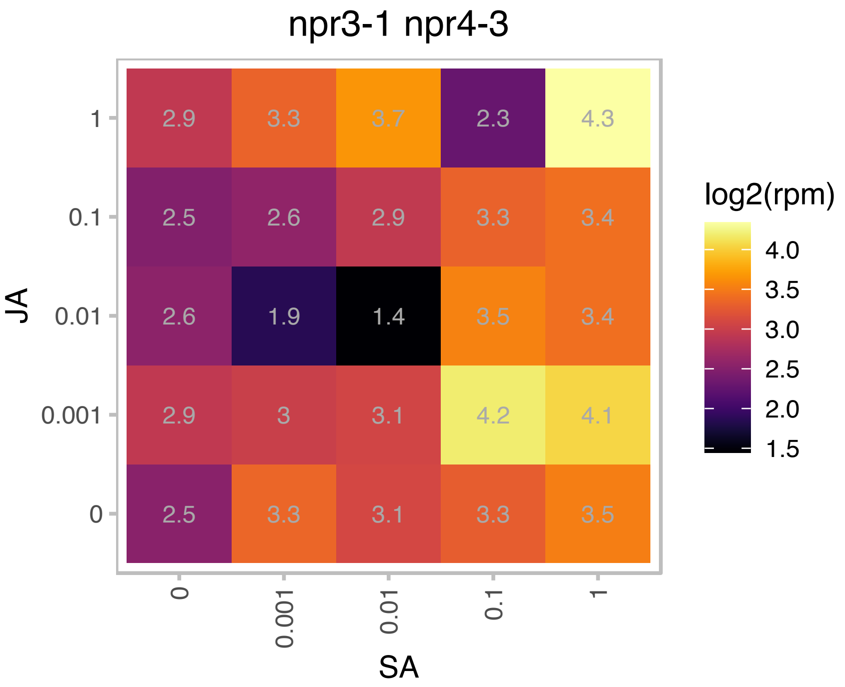

### AT1G28480.1 : GRX480, GRXC9, ROXY19

Thioredoxin superfamily protein

Encodes GRX480, a member of the glutaredoxin family that regulates protein redox state. GRX480 interacts with TGA factors and suppresses JA-responsive PDF1.2 transcription. GRX480 transcription is SA-inducible and requires NPR1. Maybe involved in SA/JA cross-talk. It has also been shown to interact with the transcription factor TGA2 and suppress ORA59 promoter activity.

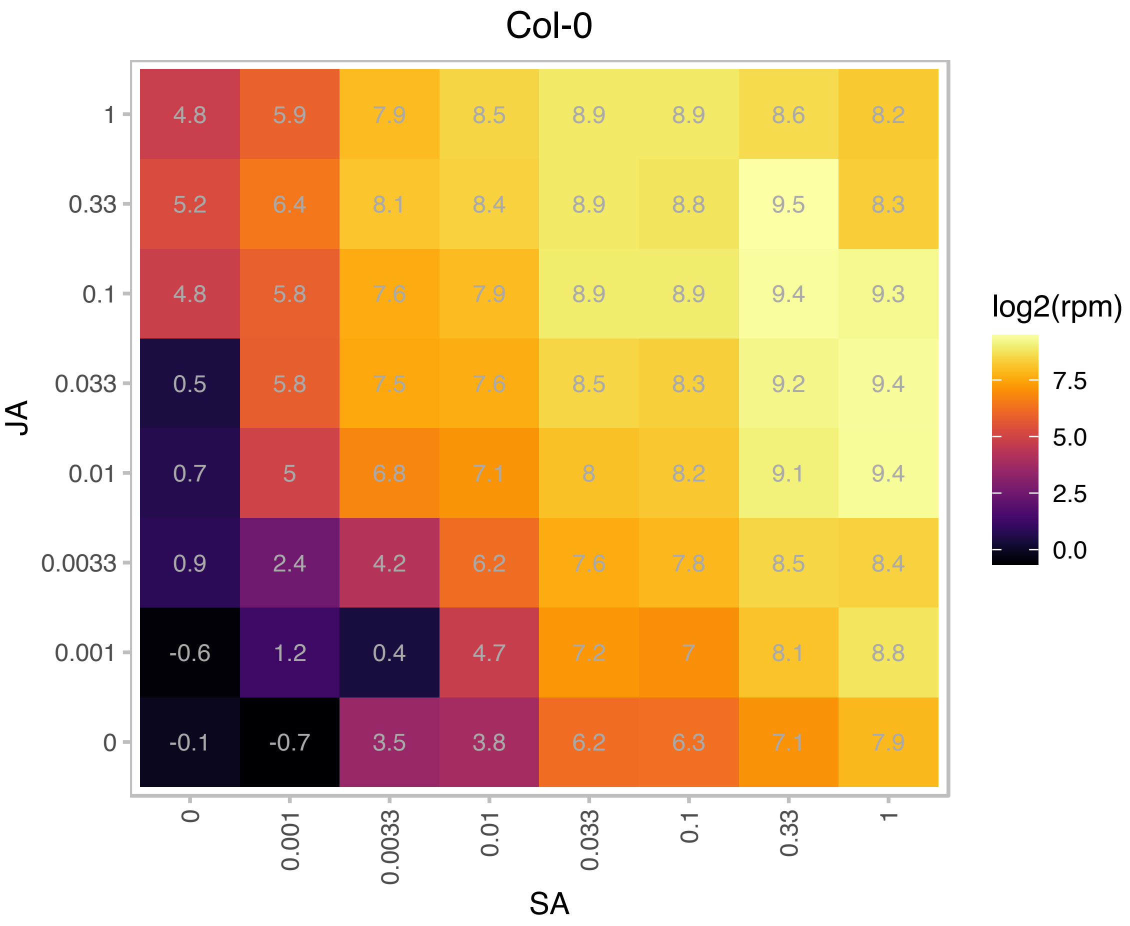


| | |
|---|---|
| Gene | AT1G28480 |
| is.expressed_genes | TRUE |
| is.responsive_genes | TRUE |
| is.apcluster | C-02 |

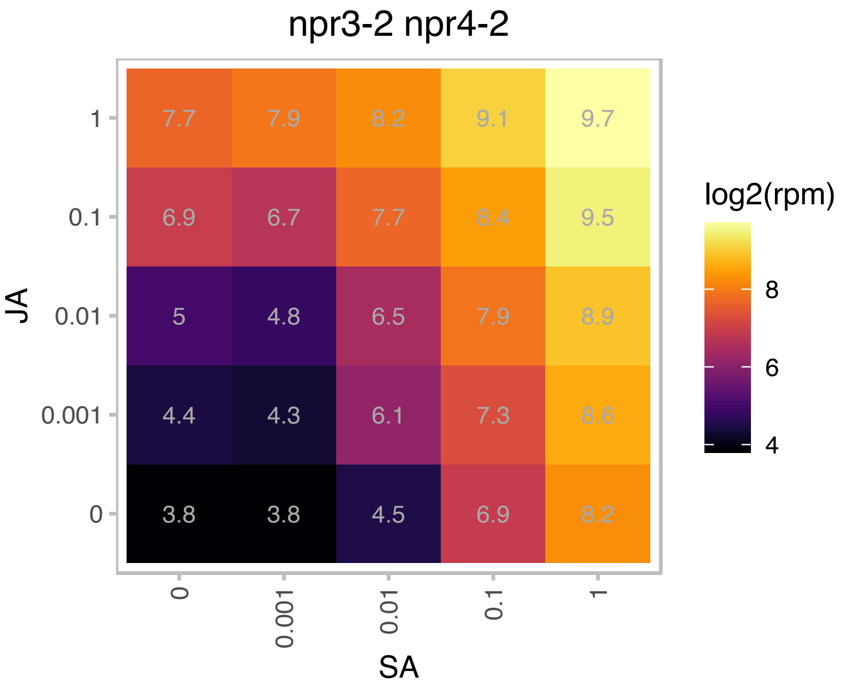


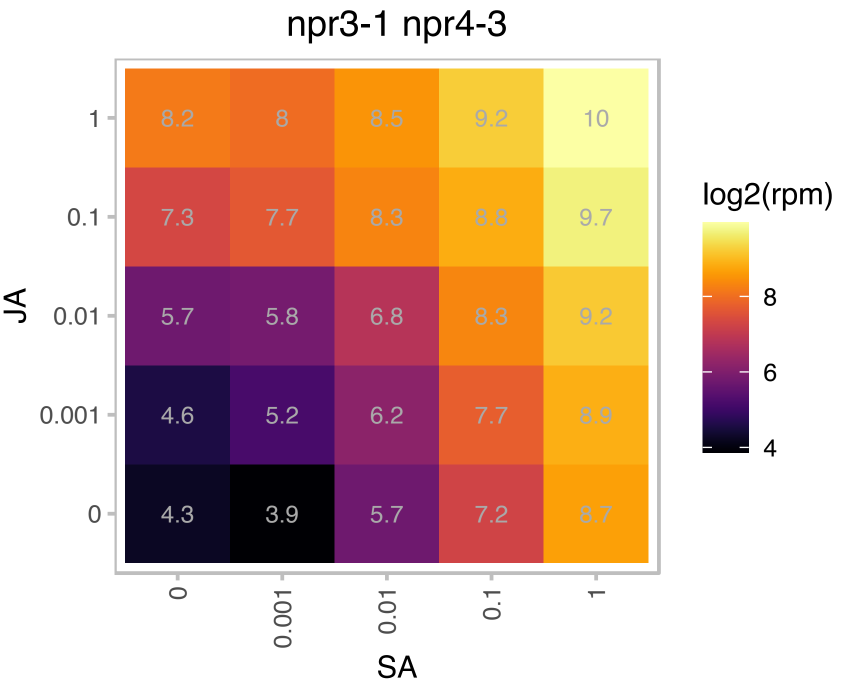

# AT1G29640.1 : NA

Protein of unknown function, DUF584

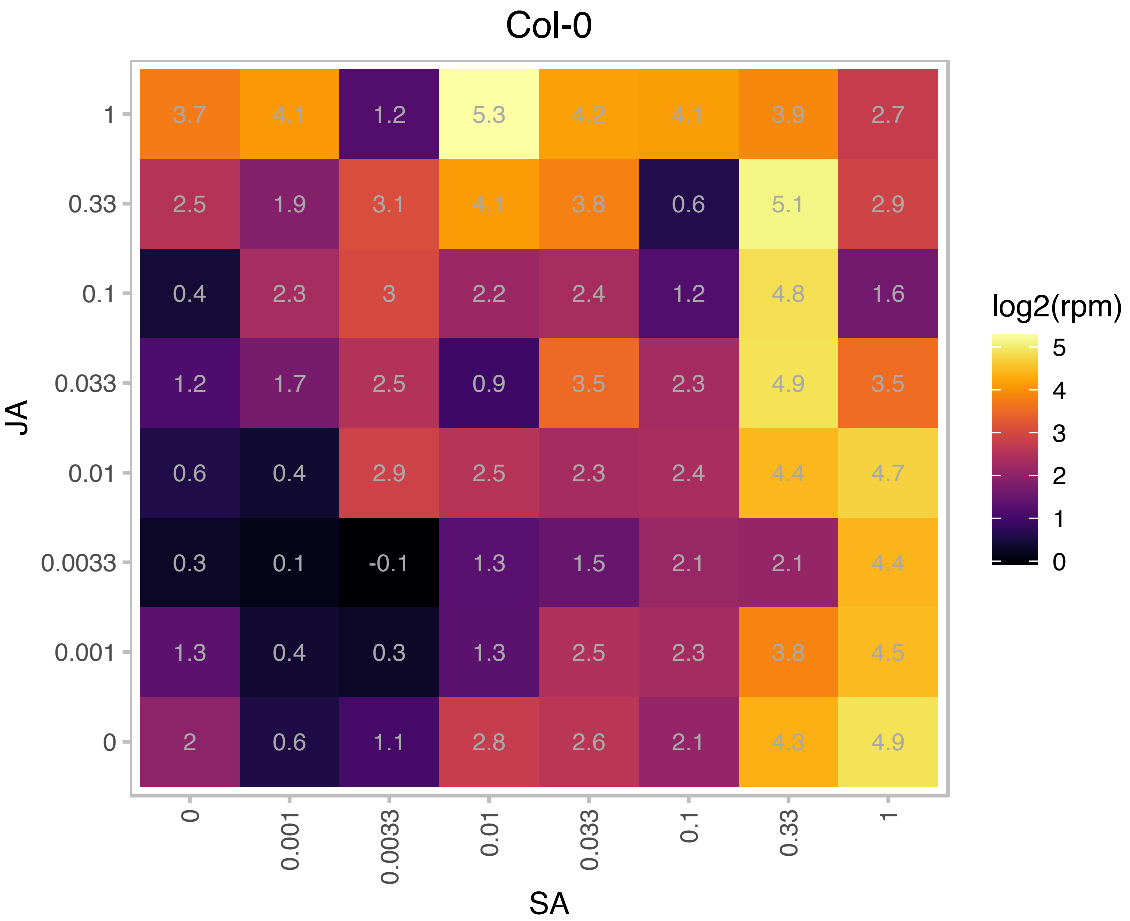


| | |
|---|---|
| Gene | AT1G29640 |
| is.expressed_genes | TRUE |
| is.responsive_genes | TRUE |
| is.apcluster | C-02 |

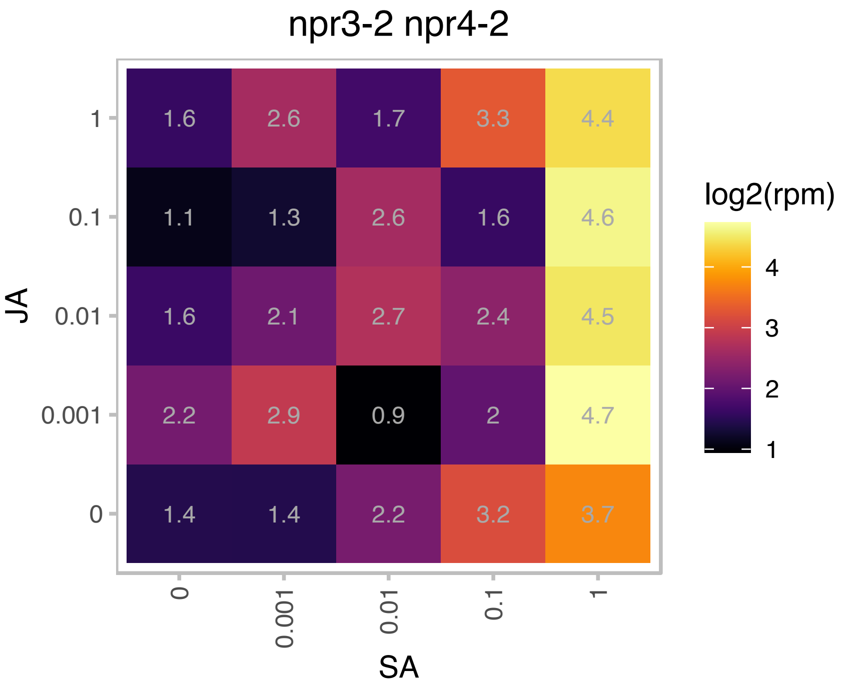


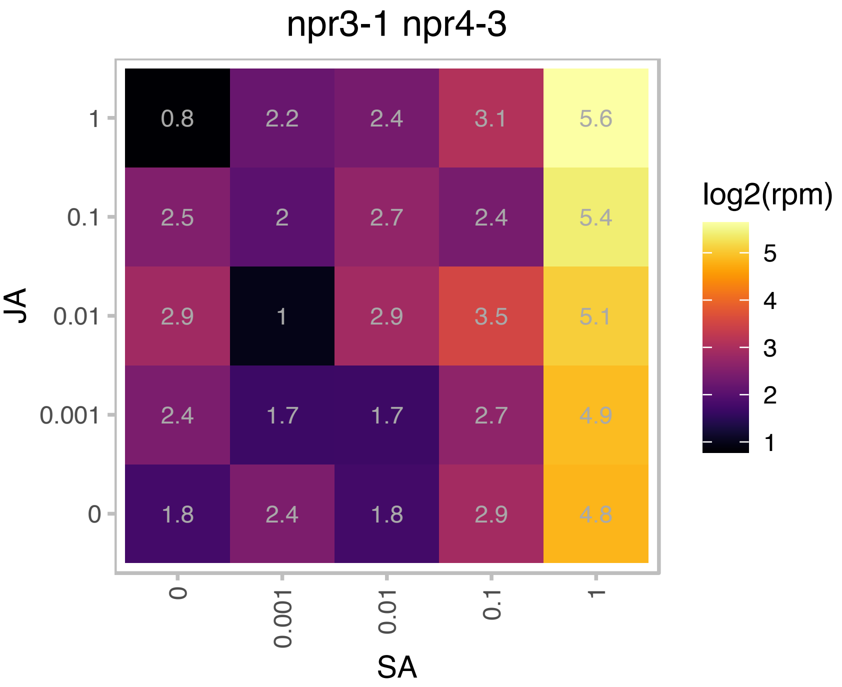

### AT1G66600.1 : ABO3, ATWRKY63, WRKY63

ABA overly sensitive mutant 3

A member of WRKY Transcription Factor; Group III. Involved in the regulation of plant responses to ABA and drought stress.

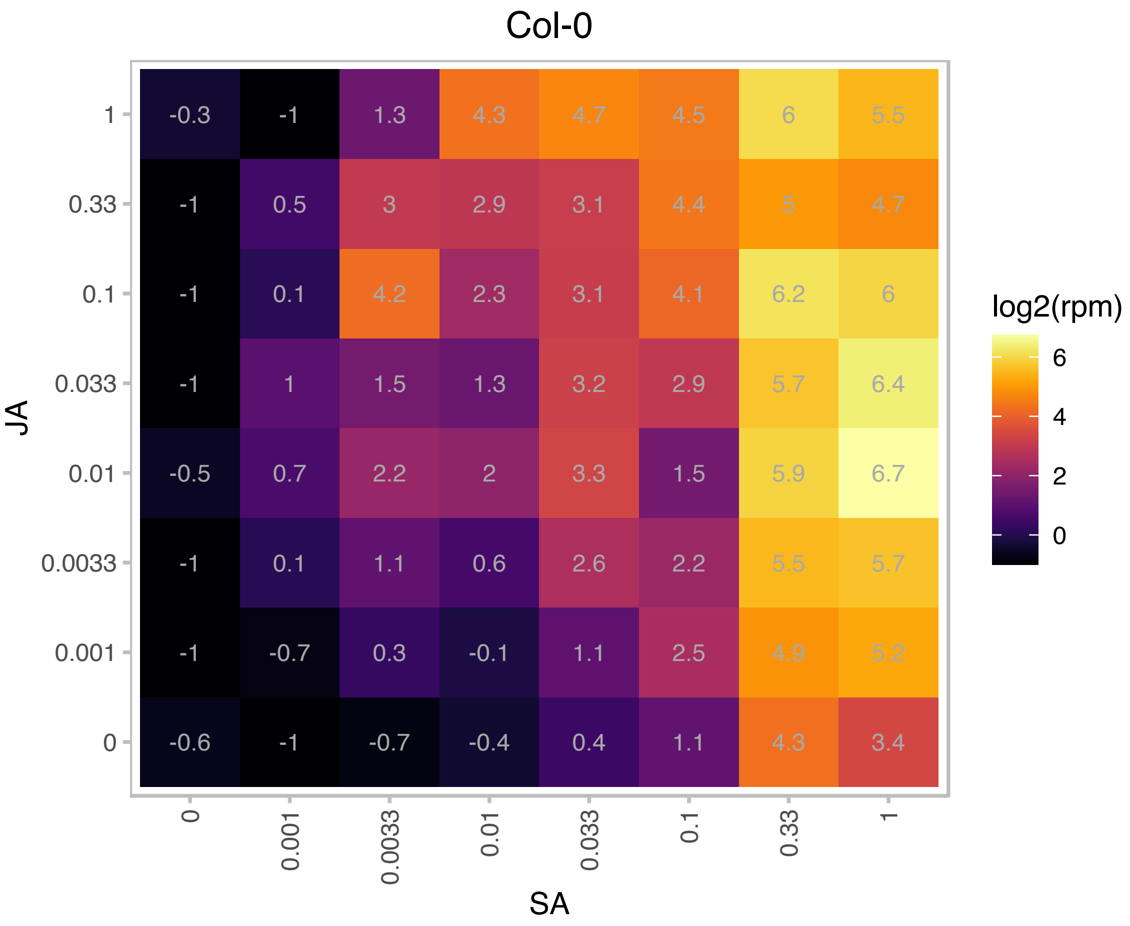


| | |
|---|---|
| Gene | AT1G66600 |
| is.expressed_genes | TRUE |
| is.responsive_genes | TRUE |
| is.apcluster | C-02 |

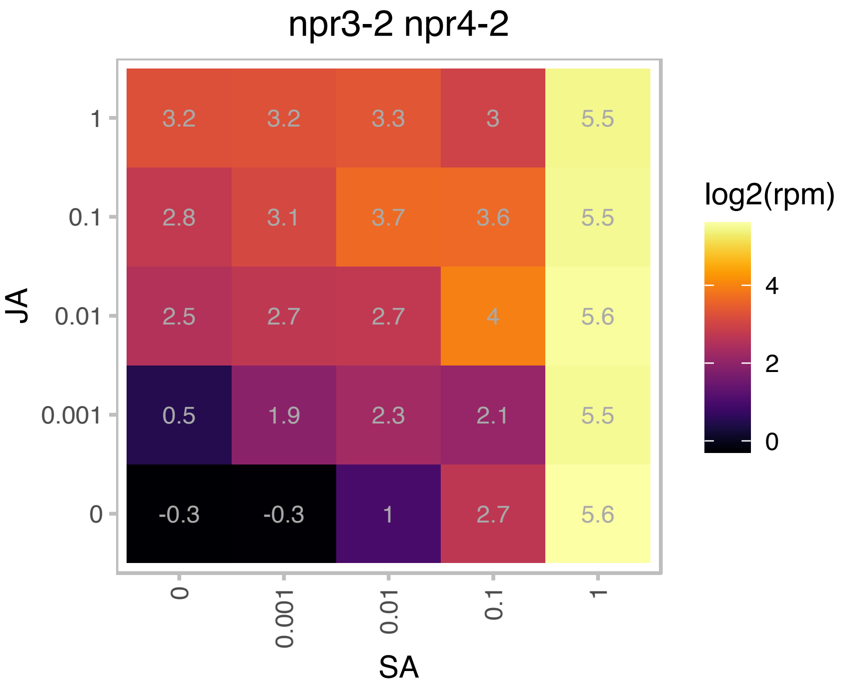


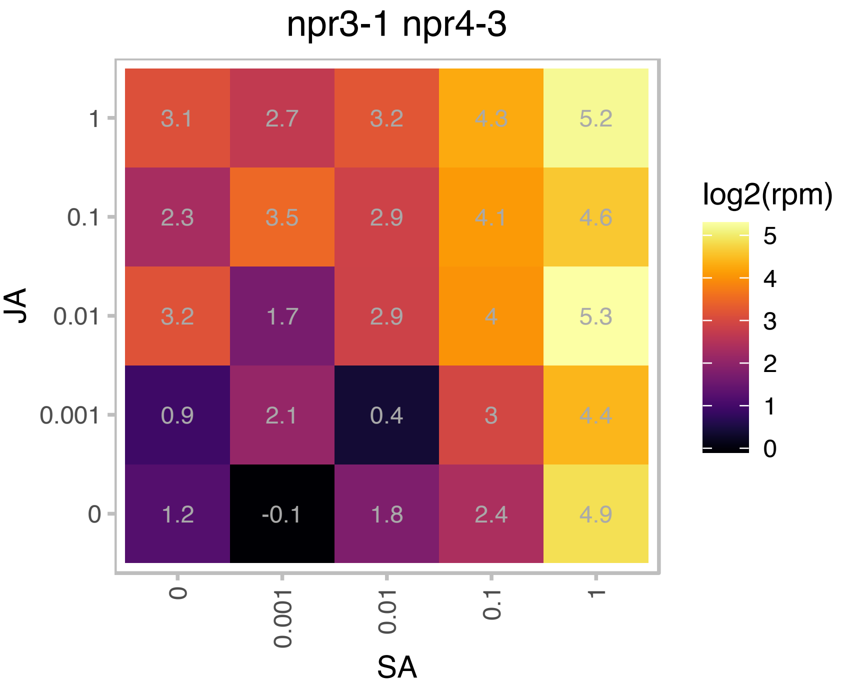

# AT1G66880.1 : NA

Protein kinase superfamily protein

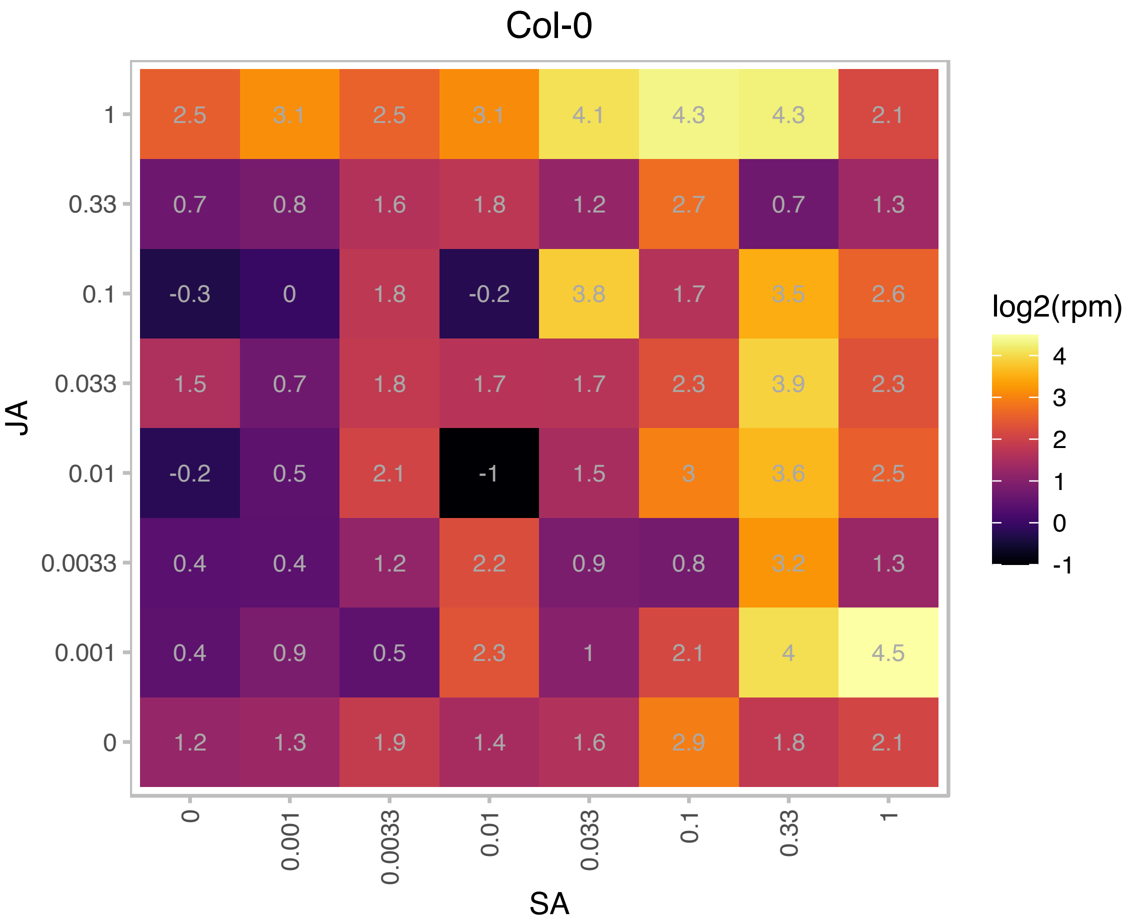


| | |
|---|---|
| Gene | AT1G66880 |
| is.expressed_genes | TRUE |
| is.responsive_genes | TRUE |
| is.apcluster | C-02 |

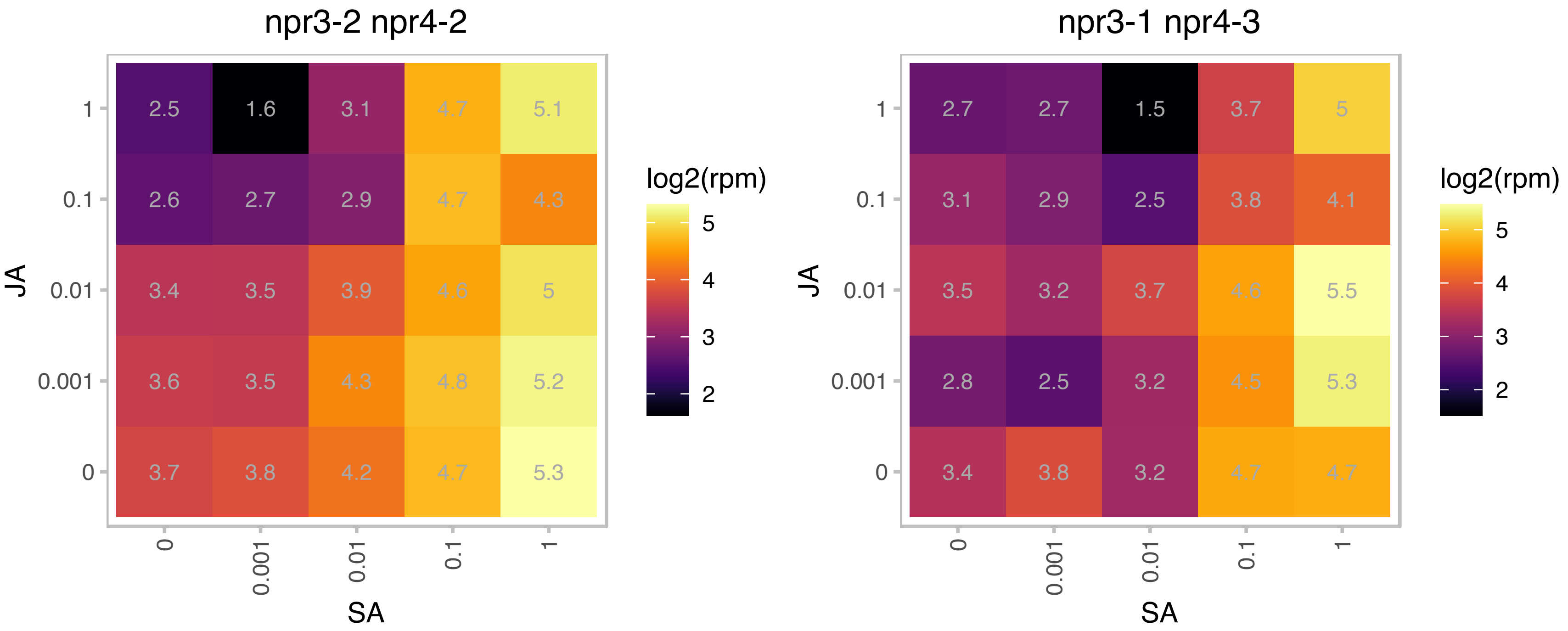

### AT1G69750.1 : ATCOX19-2, COX19-2

cytochrome c oxidase 19-2

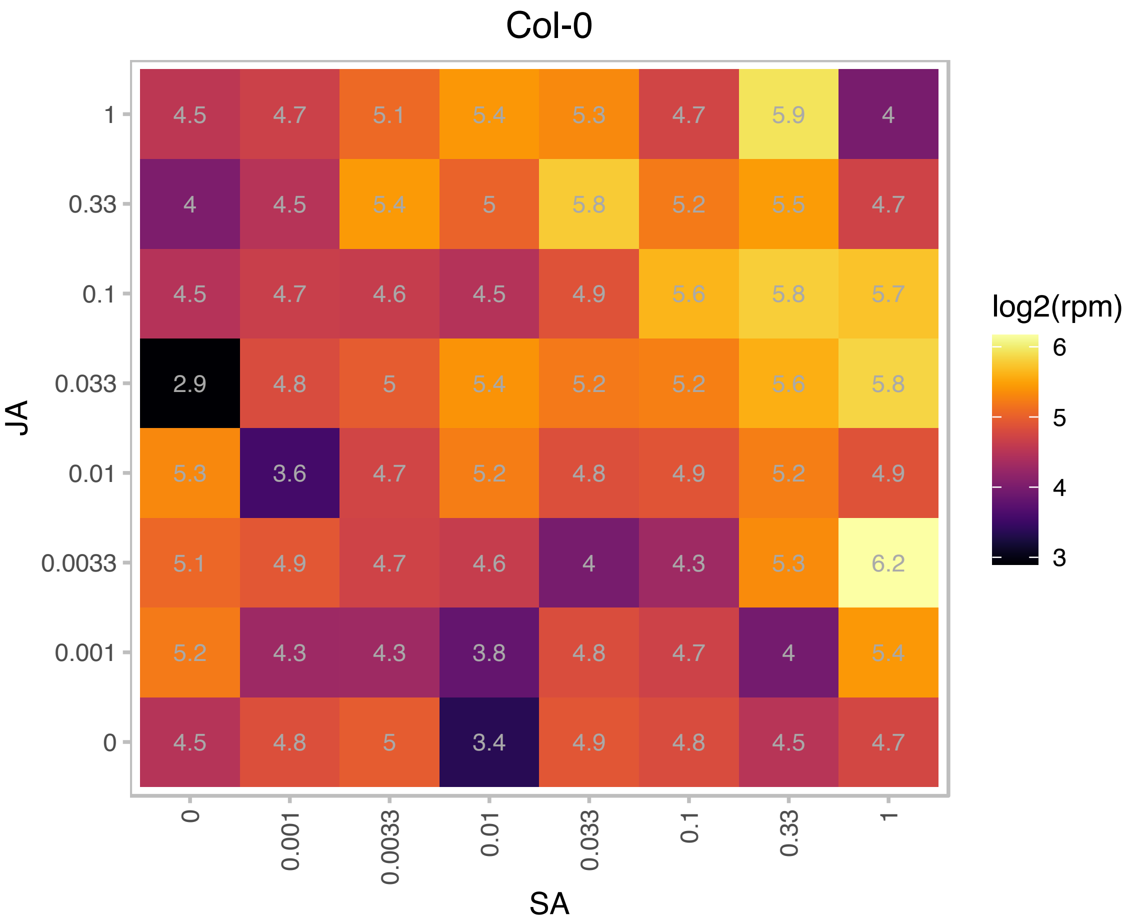


| | |
|---|---|
| Gene | AT1G69750 |
| is.expressed_genes | TRUE |
| is.responsive_genes | TRUE |
| is.apcluster | C-02 |

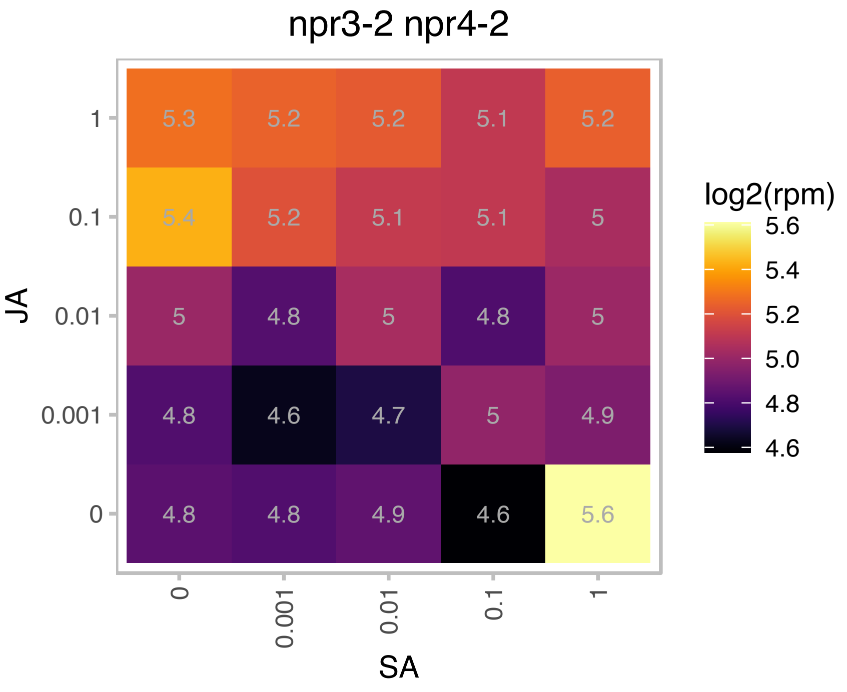


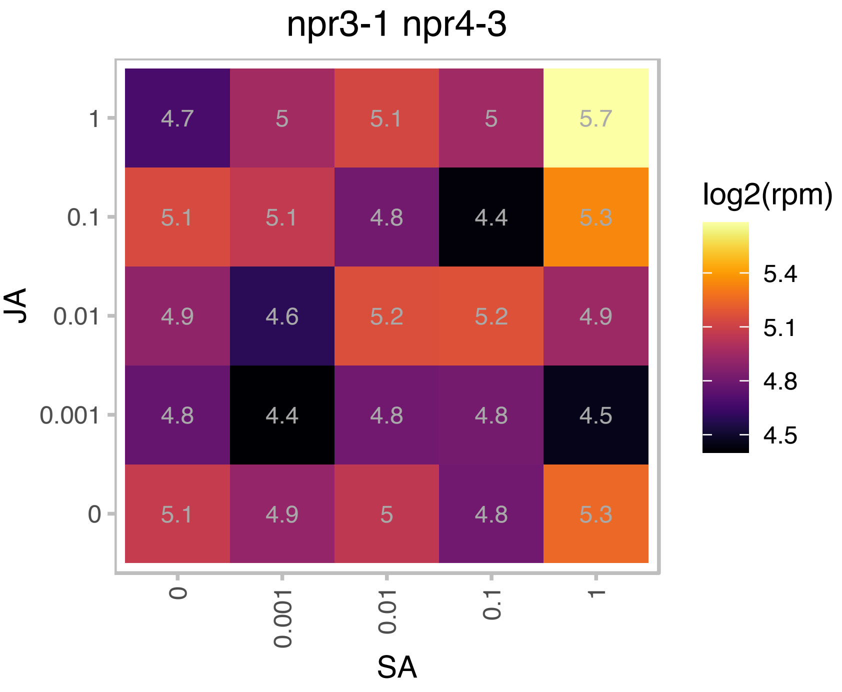

# AT1G71840.1 : NA

transducin family protein / WD-40 repeat family protein

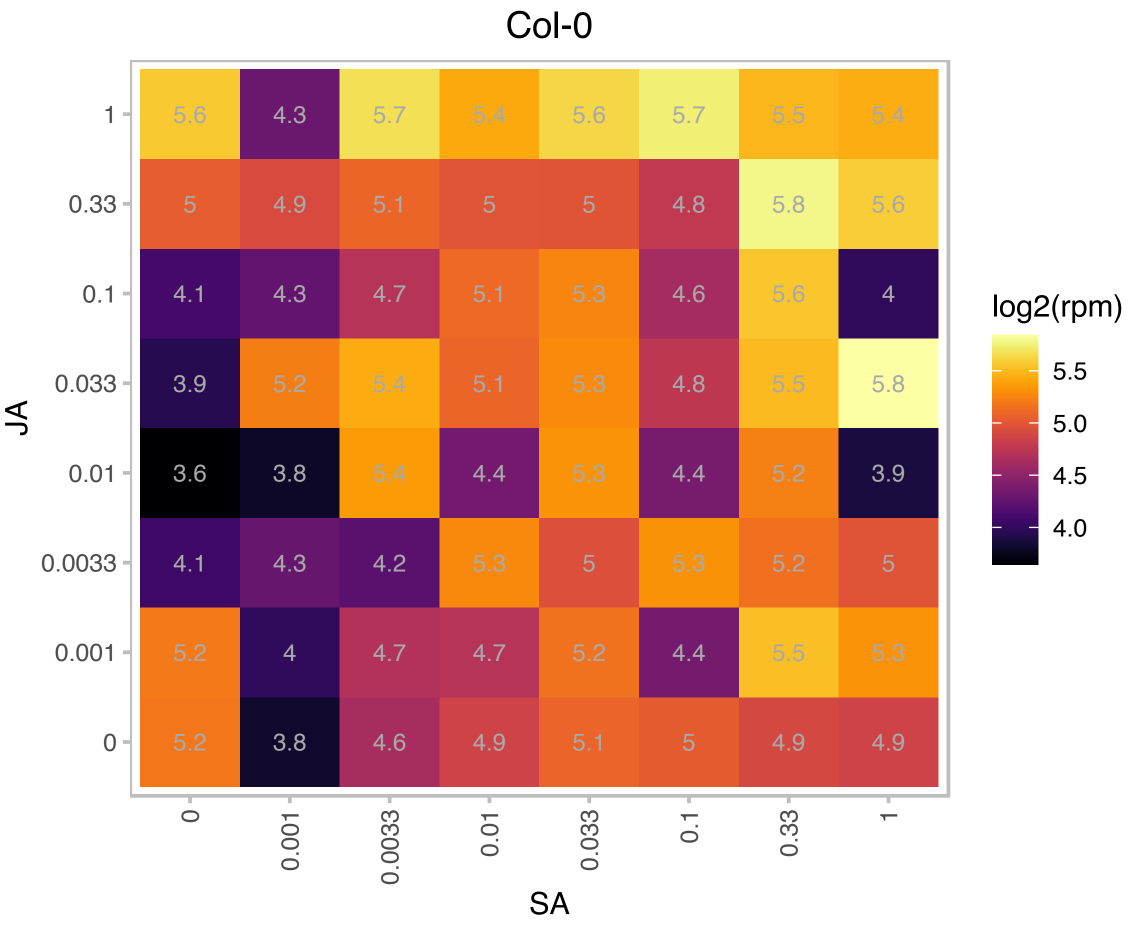


| | |
|---|---|
| Gene | AT1G71840 |
| is.expressed_genes | TRUE |
| is.responsive_genes | TRUE |
| is.apcluster | C-02 |

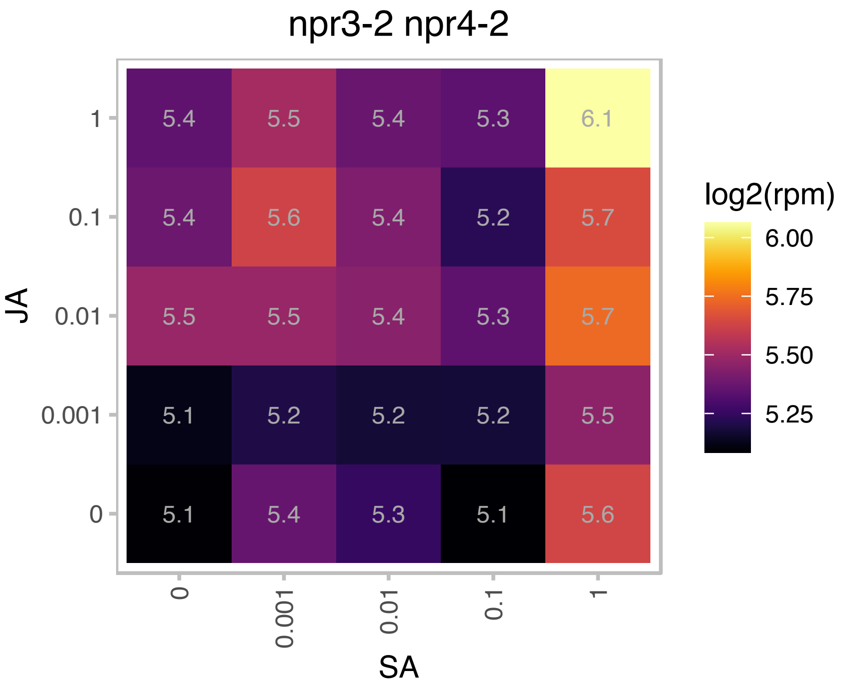


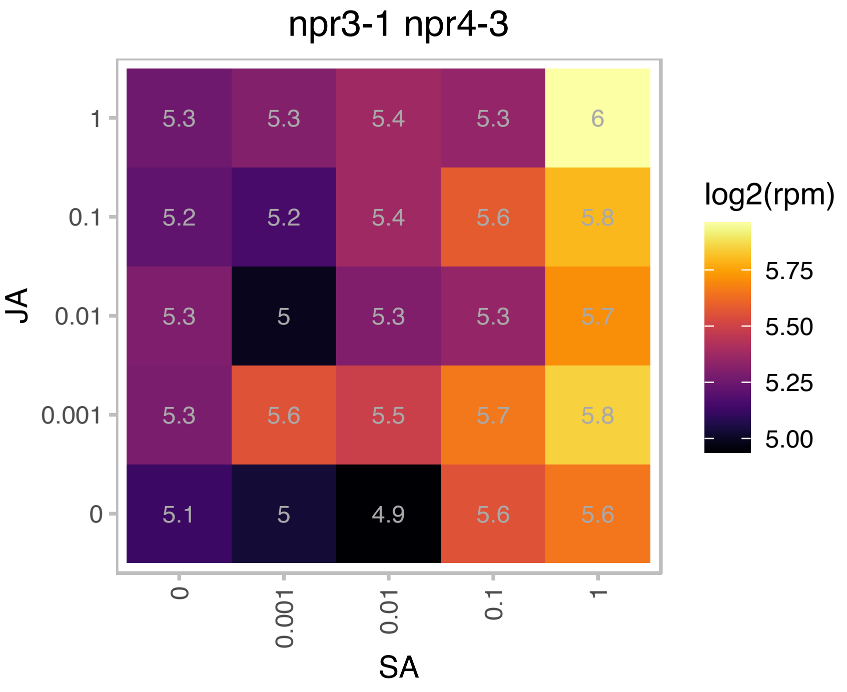

### AT2G30550.1 : DALL3

alpha/beta-Hydrolases superfamily protein

Encodes a lipase that hydrolyzes phosphatidylcholine, glycolipids as well as triacylglycerols.

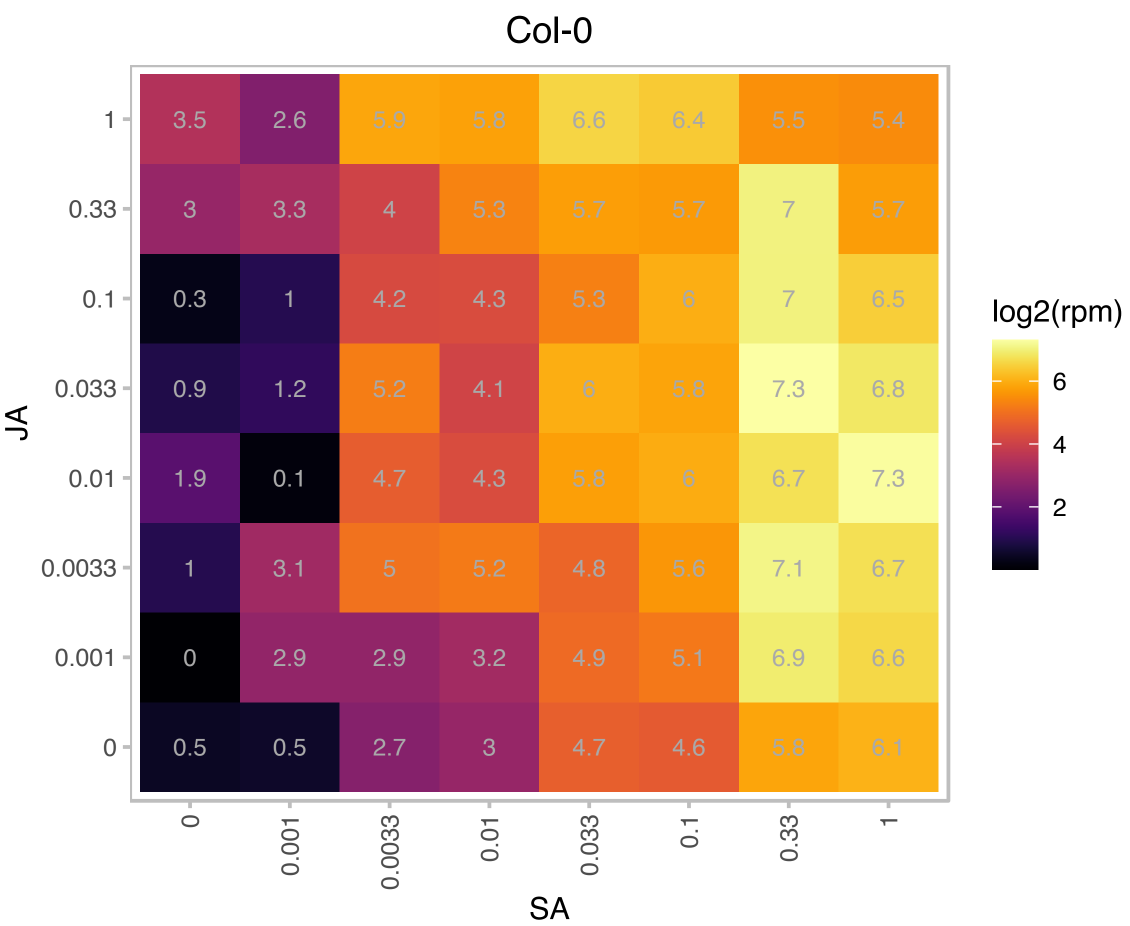


| | |
|---|---|
| Gene | AT2G30550 |
| is.expressed_genes | TRUE |
| is.responsive_genes | TRUE |
| is.apcluster | C-02 |

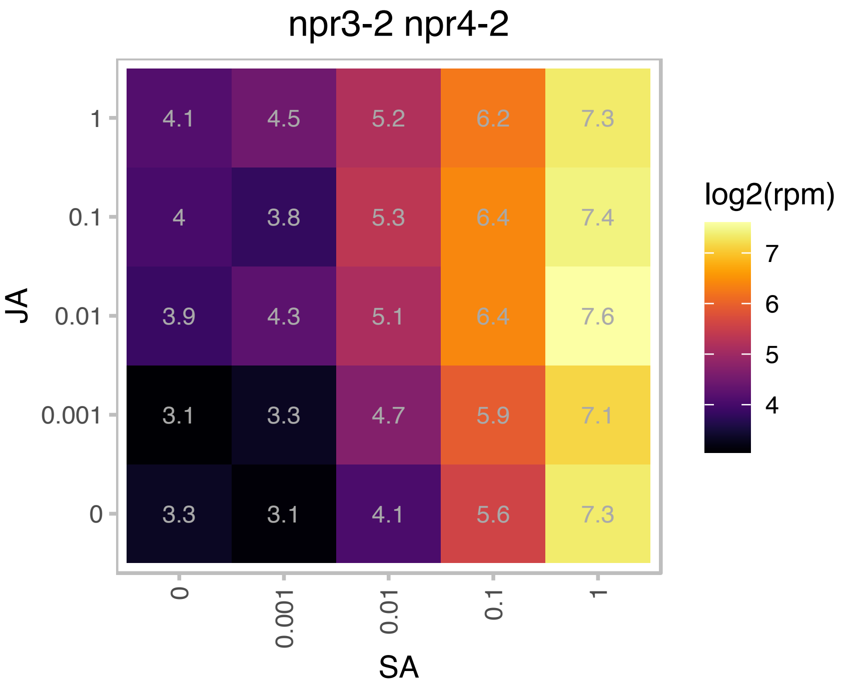


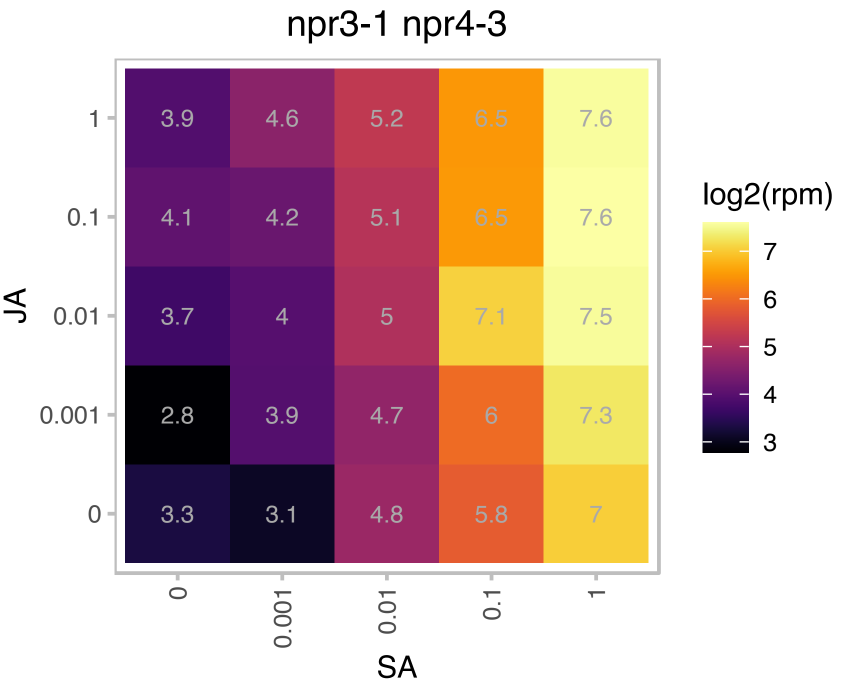

# AT3G04210.1 : TN13

Disease resistance protein (TIR-NBS class)

TN13 is a TIR-NBS protein involved in immune response. It co localizes with the ER and perinuclear membranes and interacts with MOS6.

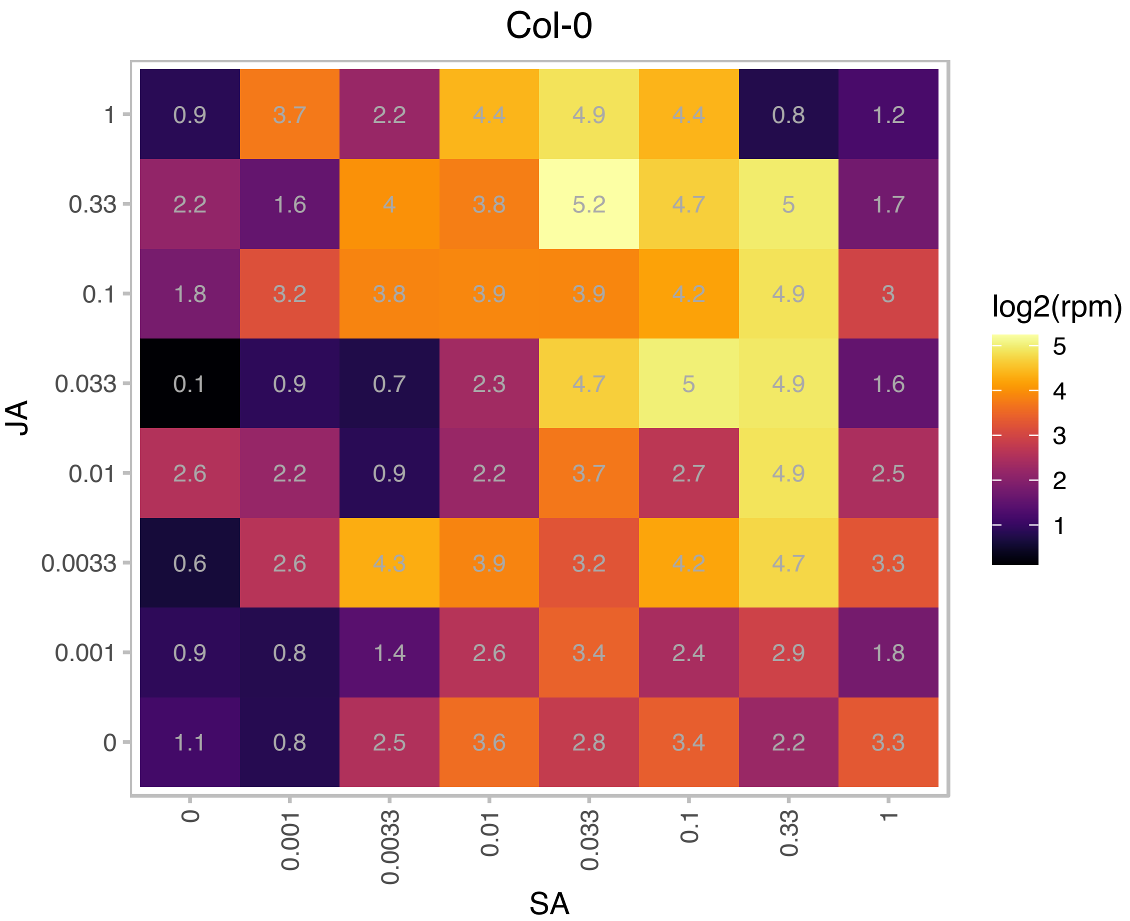


| | |
|---|---|
| Gene | AT3G04210 |
| is.expressed_genes | TRUE |
| is.responsive_genes | TRUE |
| is.apcluster | C-02 |

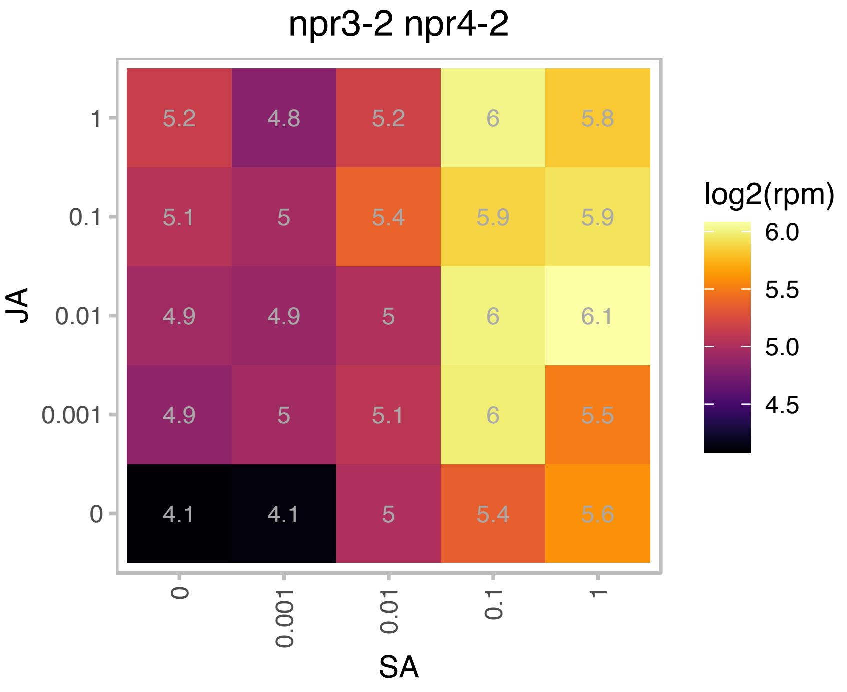


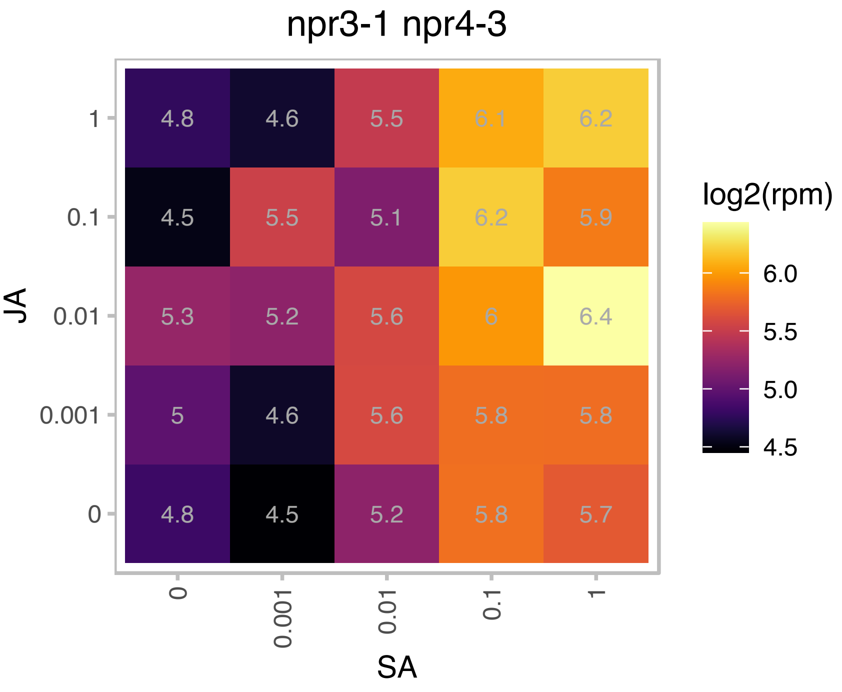

### AT3G11340.1 : UGT76B1

UDP-Glycosyltransferase superfamily protein

Encodes a glucosyltransferase that conjugates isoleucic acid and modulates plant defense and senescence.

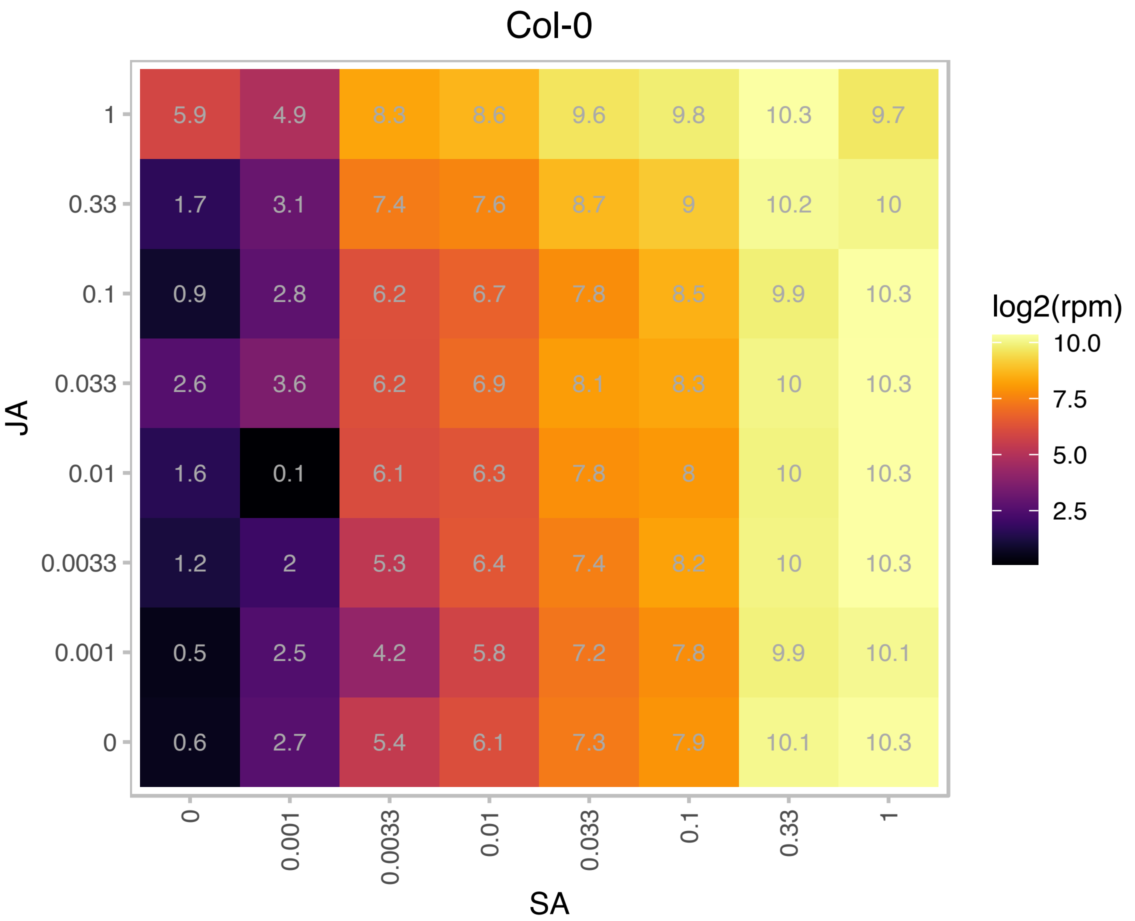


| | |
|---|---|
| Gene | AT3G11340 |
| is.expressed_genes | TRUE |
| is.responsive_genes | TRUE |
| is.apcluster | C-02 |

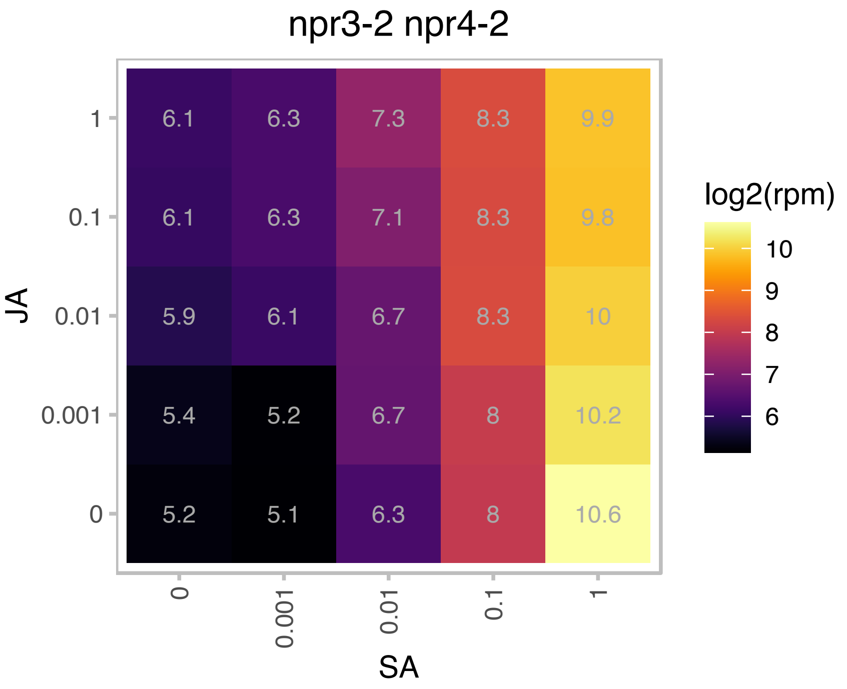


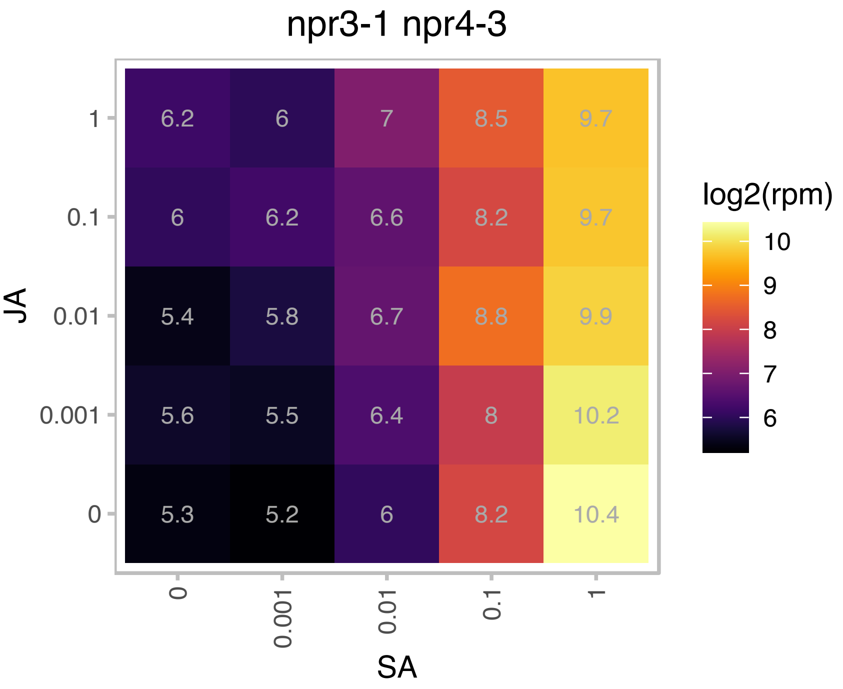

# AT3G13950.1 : NA

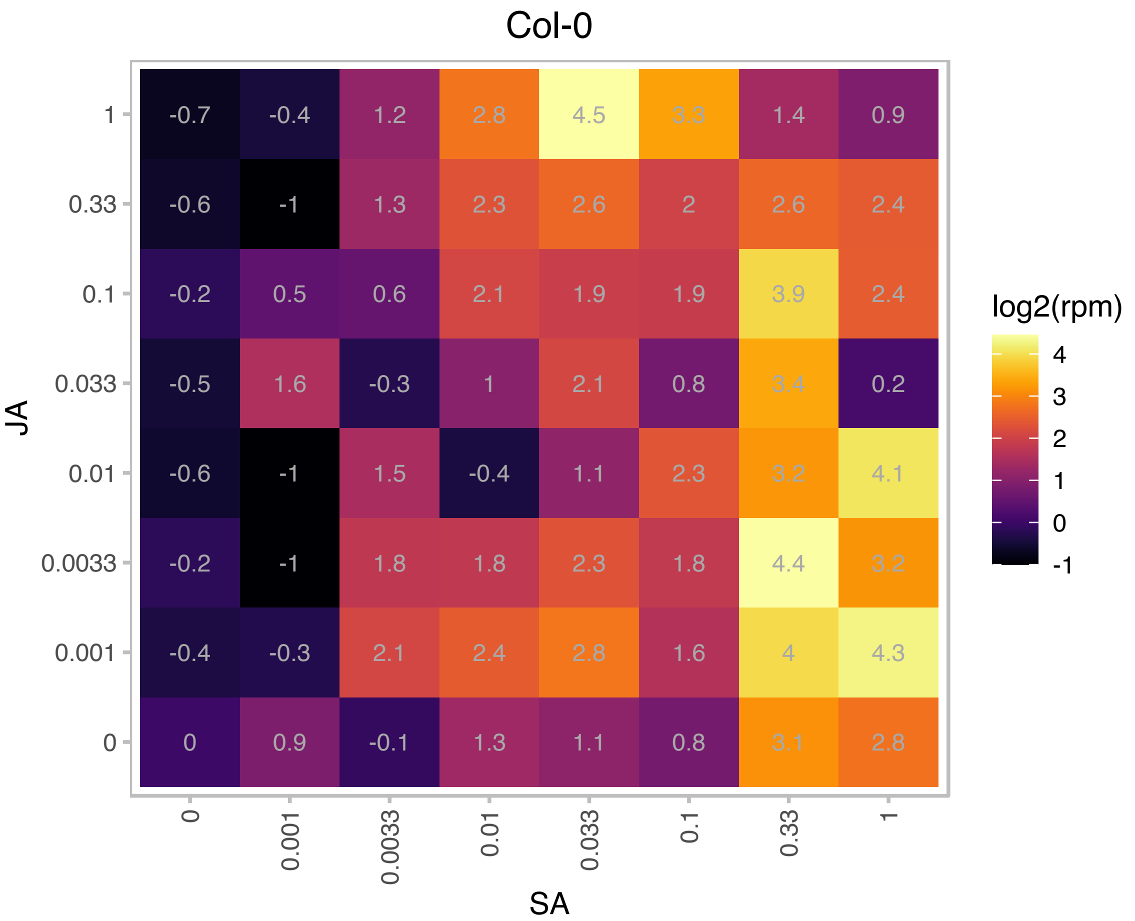


| | |
|---|---|
| Gene | AT3G13950 |
| is.expressed_genes | TRUE |
| is.responsive_genes | TRUE |
| is.apcluster | C-02 |

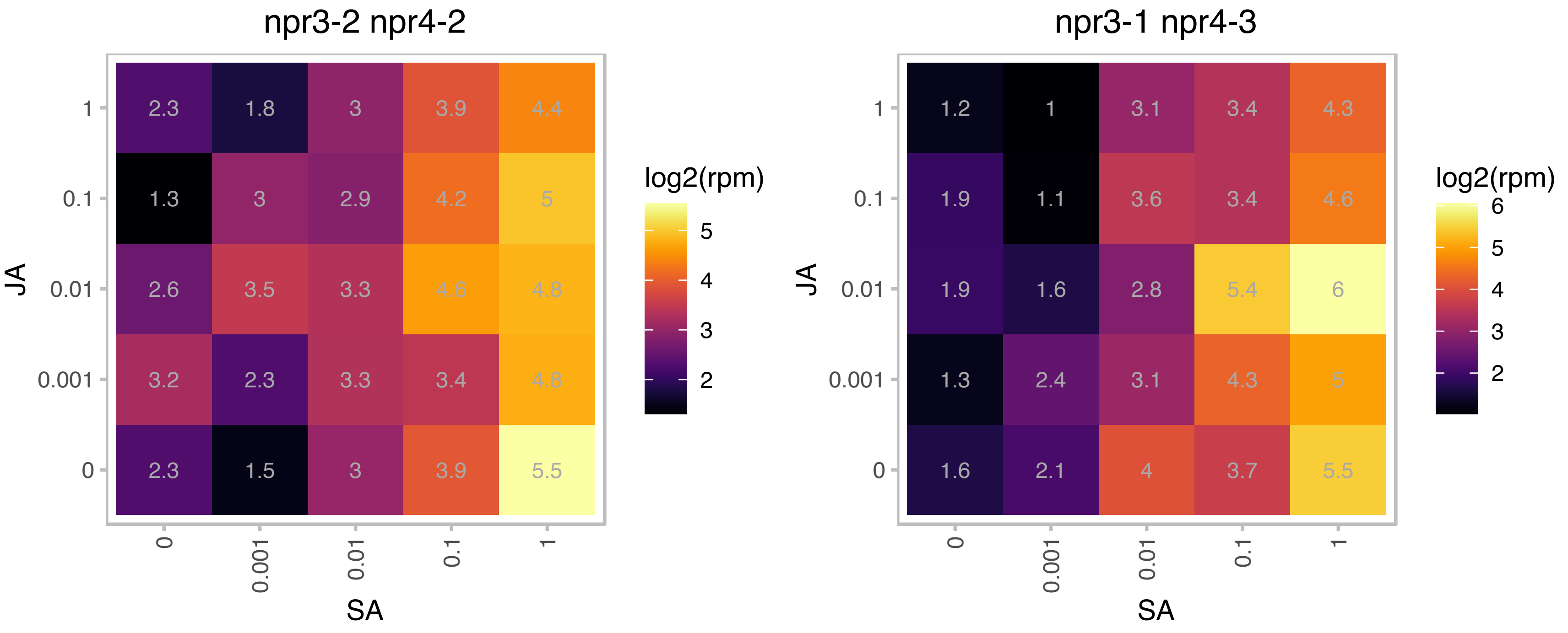

# AT3G14700.1 : NA

SART-1 family

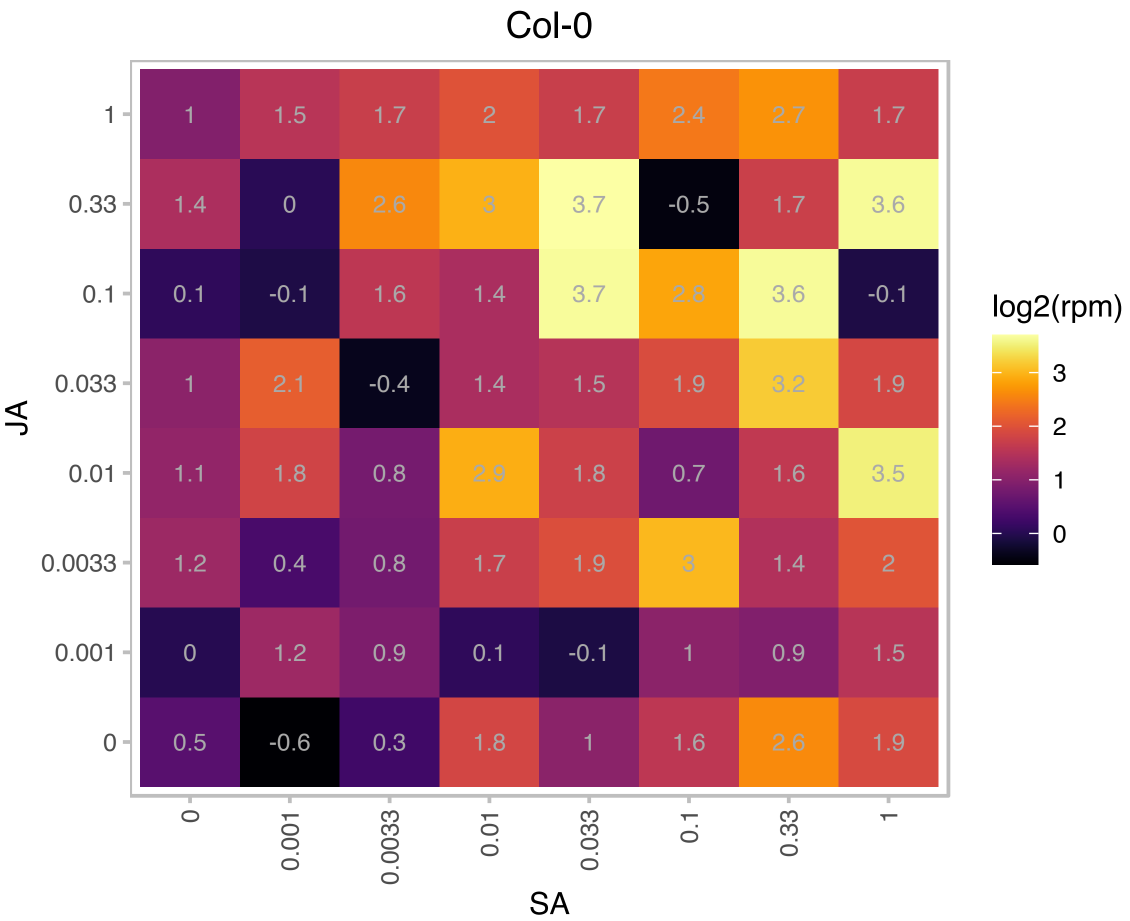


| | |
|---|---|
| Gene | AT3G14700 |
| is.expressed_genes | TRUE |
| is.responsive_genes | TRUE |
| is.apcluster | C-02 |

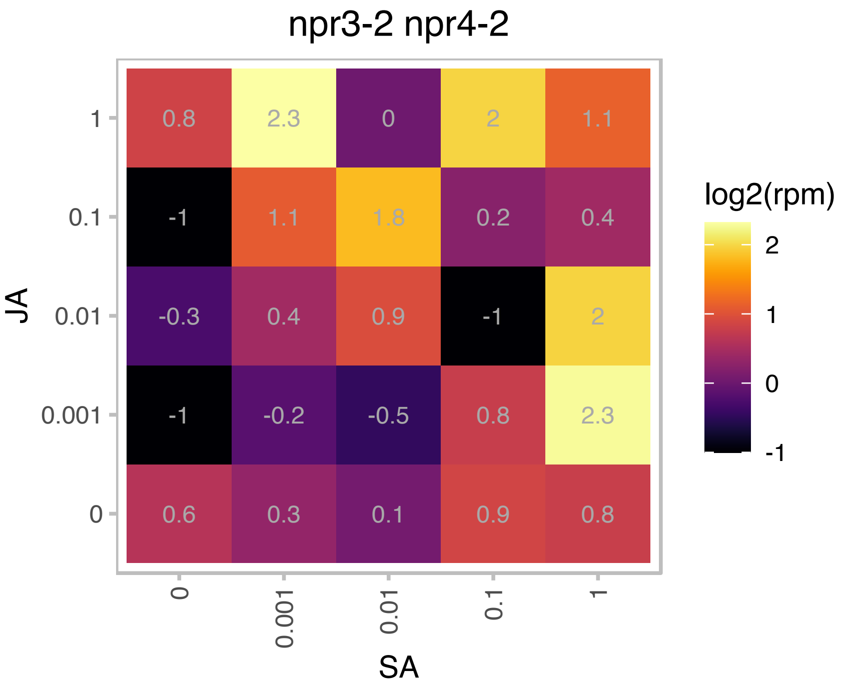


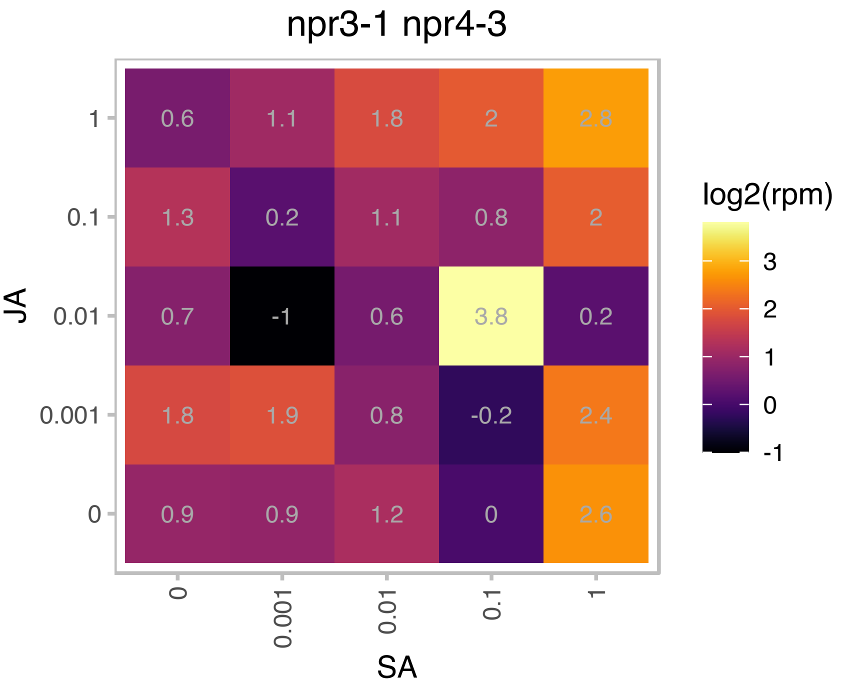

### AT3G18290.1 : BTS, EMB2454

zinc finger protein-related

Encodes BRUTUS (BTS), a putative E3 ligase protein with metal ion binding and DNA binding domains, which negatively regulates the response to iron deficiency. The mRNA is cell-to-cell mobile.

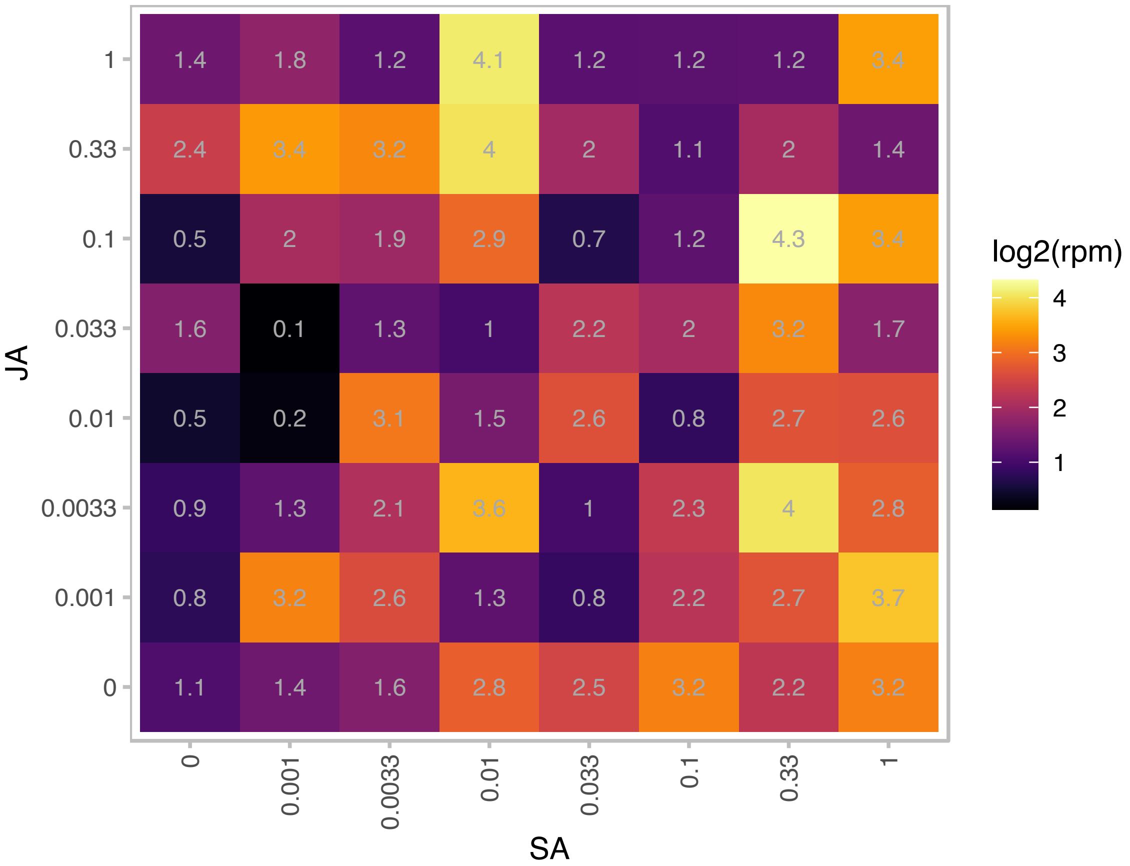


| | |
|---|---|
| Gene | AT3G18290 |
| is.expressed_genes | TRUE |
| is.responsive_genes | TRUE |
| is.apcluster | C-02 |

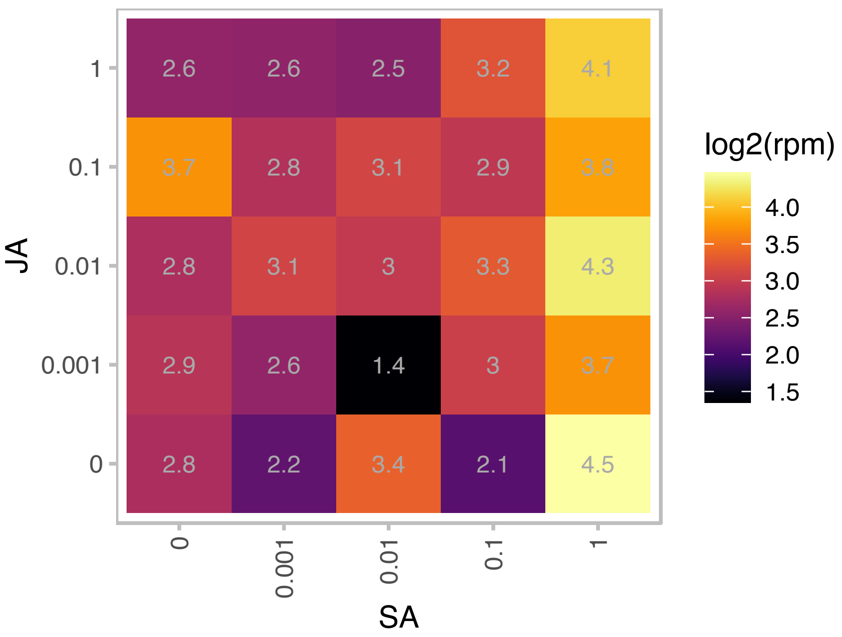


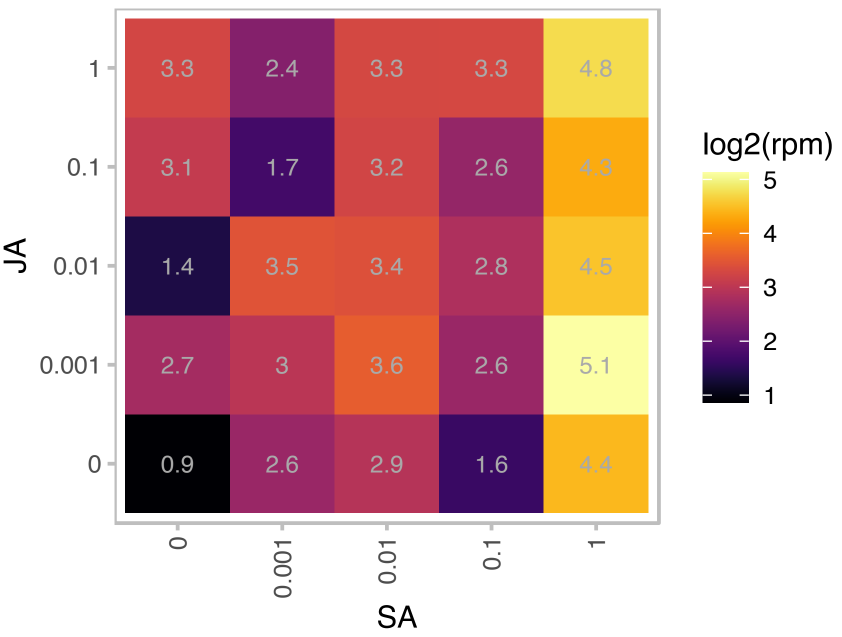

# AT3G43430.1 : NA

RING/U-box superfamily protein

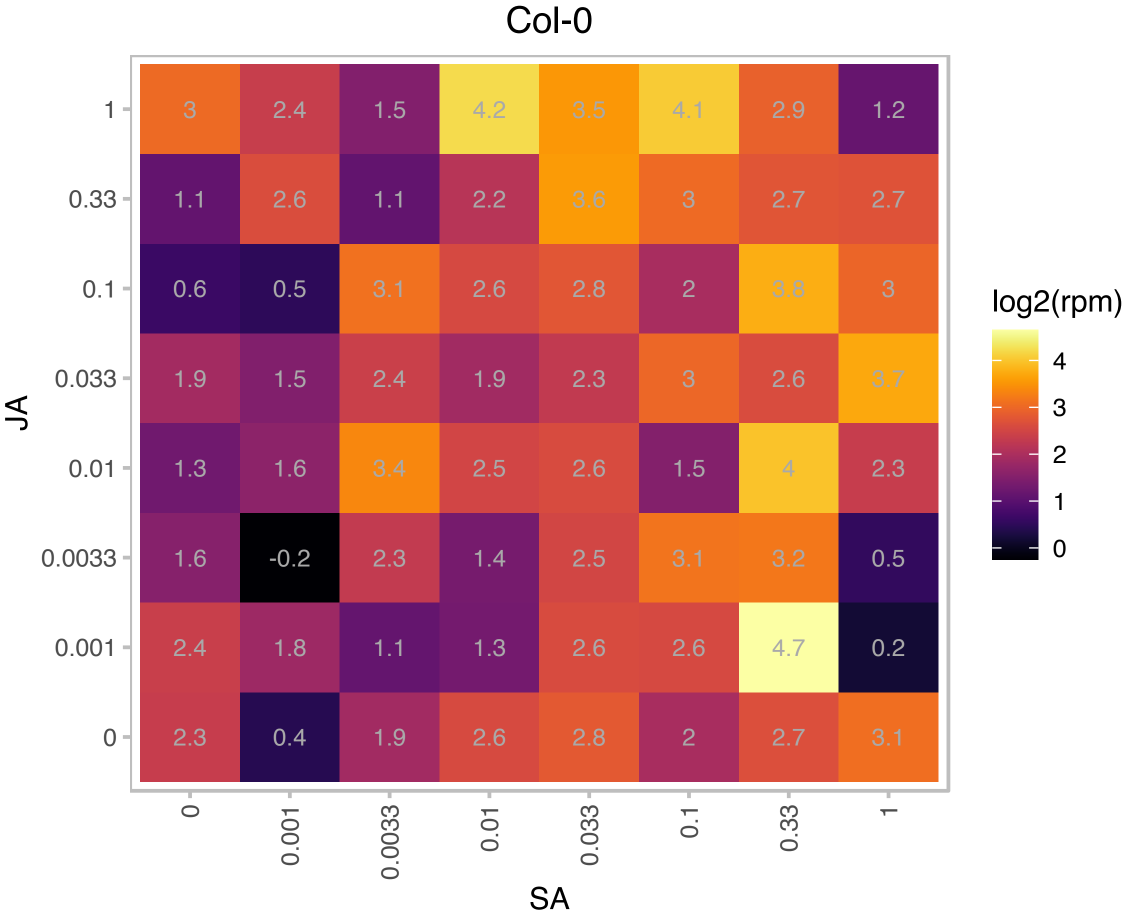


| | |
|---|---|
| Gene | AT3G43430 |
| is.expressed_genes | TRUE |
| is.responsive_genes | TRUE |
| is.apcluster | C-02 |

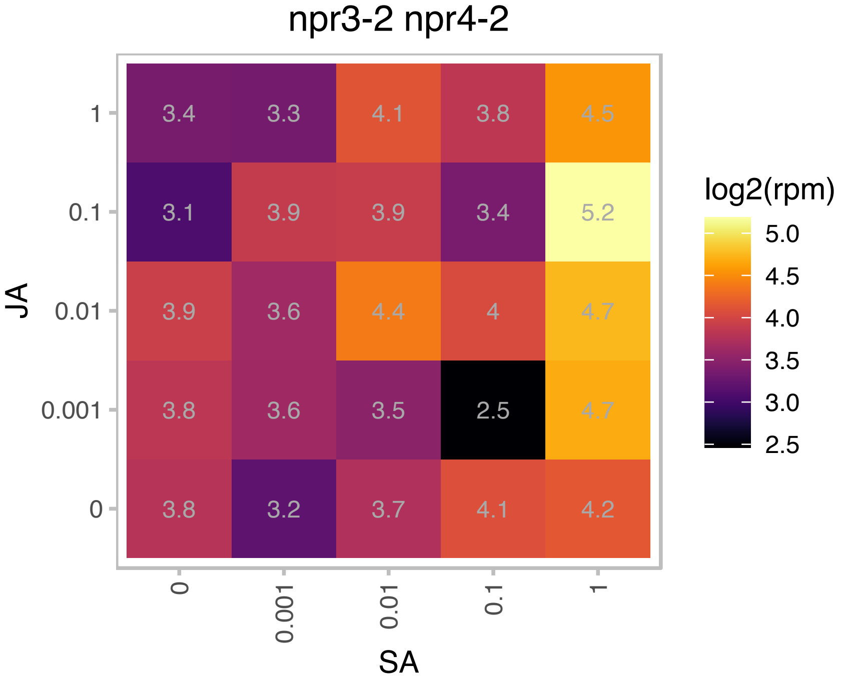


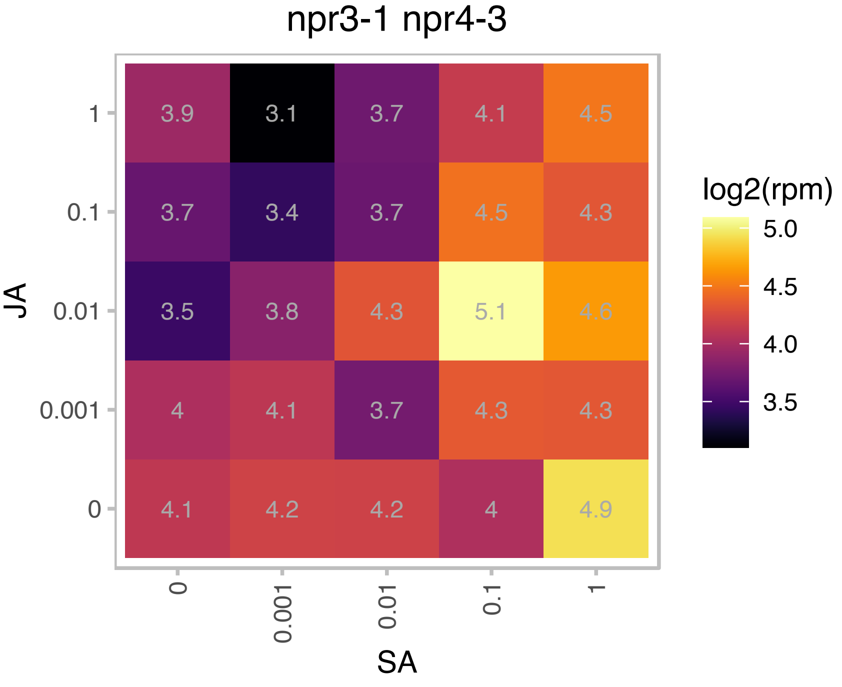

# AT3G50650.1 : NA

GRAS family transcription factor

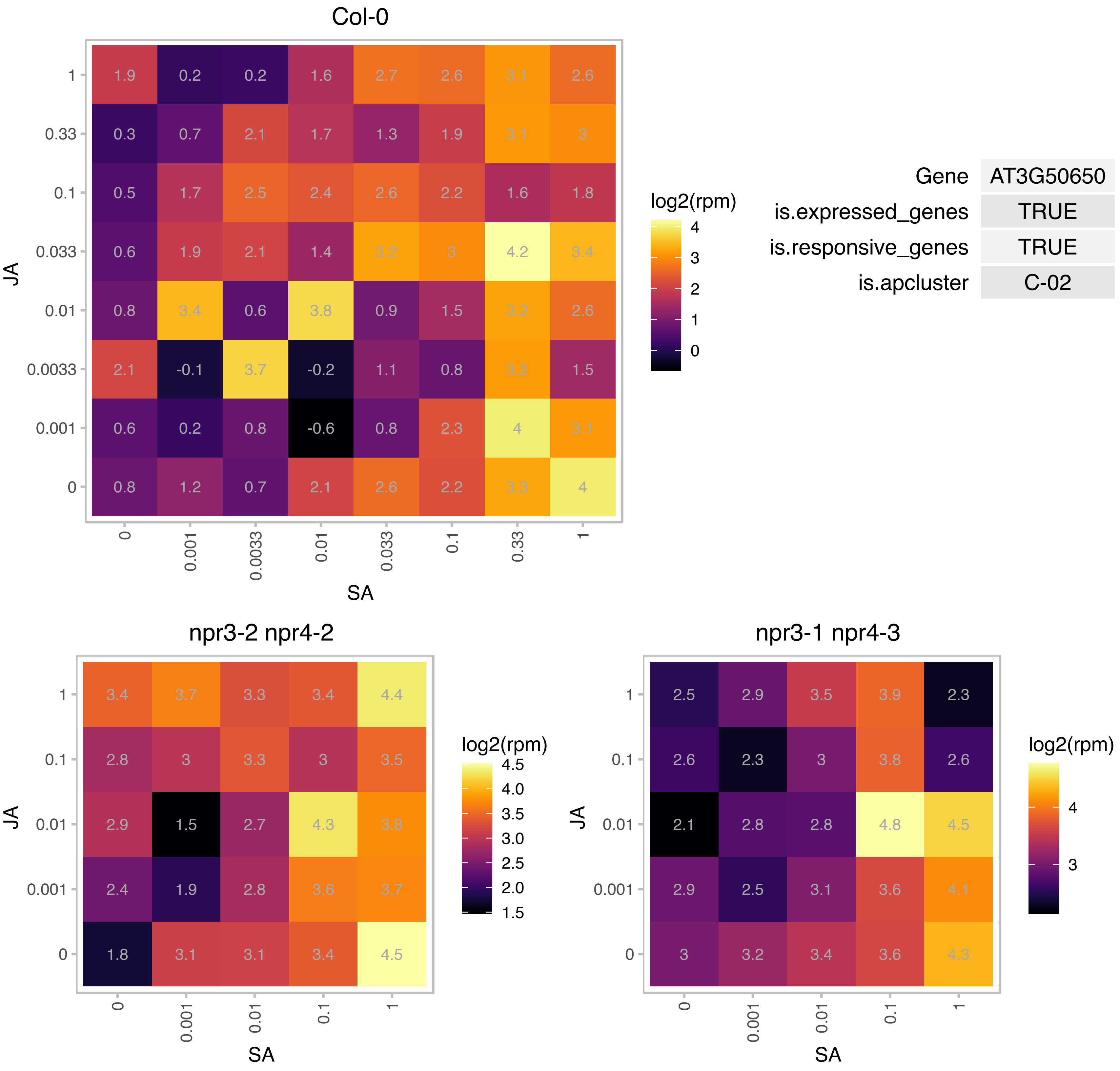

### AT3G56710.1 : SIB1

sigma factor binding protein 1

Sig1 binding protein; interacts with Sig1R4. As well as Sig1, SibI is imported into chloroplasts and its expression is light-dependent in mature chloroplasts.

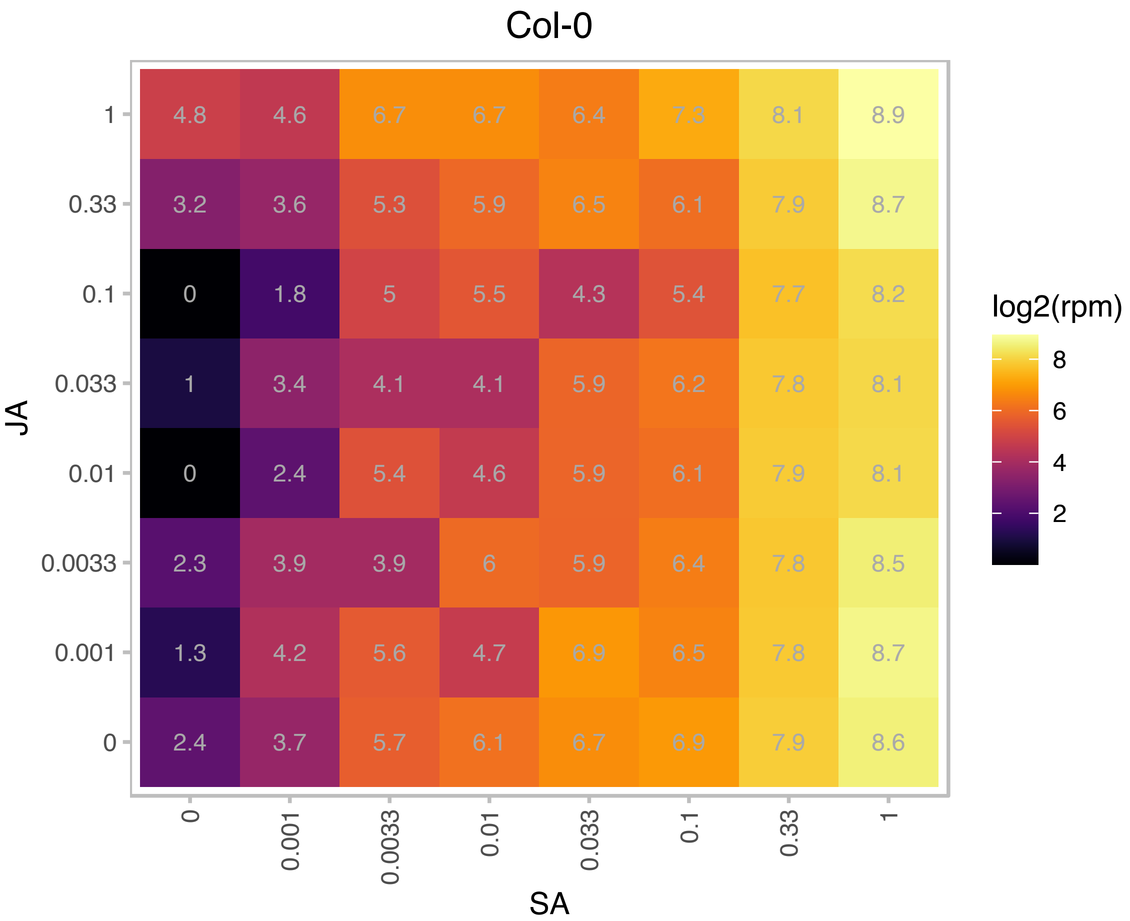


| | |
|---|---|
| Gene | AT3G56710 |
| is.expressed_genes | TRUE |
| is.responsive_genes | TRUE |
| is.apcluster | C-02 |

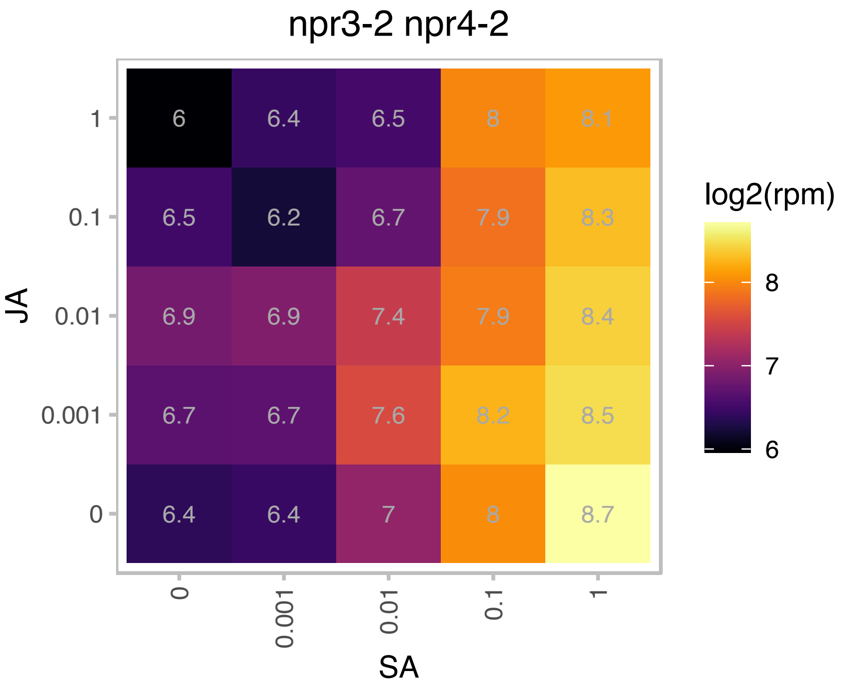


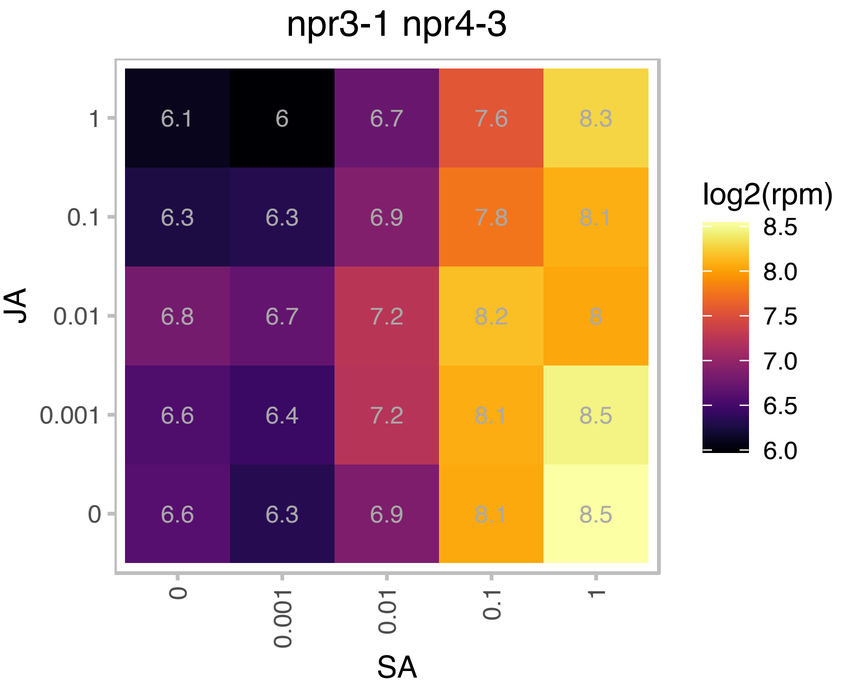

### AT3G59380.1 : ATFTA, FTA, PFT/PGGT-IALPHA, PLP

farnesyltransferase A

Encodes the alpha-subunit shared between protein farnesyltransferase and protein geranylgeranyltransferase-I. Involved in protein prenylation: covalent attachment of the C-15 isoprene farnesyl or the C-20 isoprene geranylgeranyl groups to the C-terminal end of some proteins. Involved in shoot and flower meristem homeostasis, and response to ABA and drought. Also regulates leaf cell shape. Mutant is epistatic to era1.

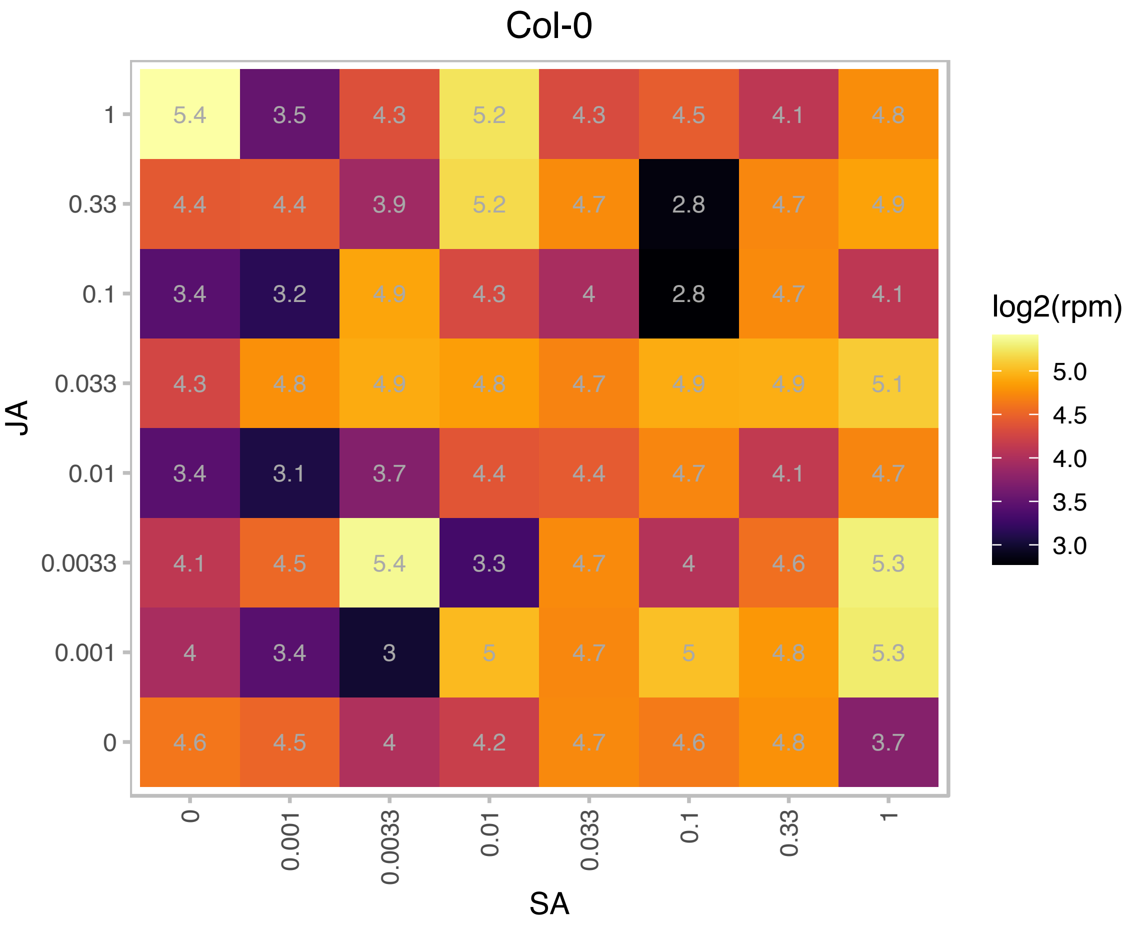


| | |
|---|---|
| Gene | AT3G59380 |
| is.expressed_genes | TRUE |
| is.responsive_genes | TRUE |
| is.apcluster | C-02 |

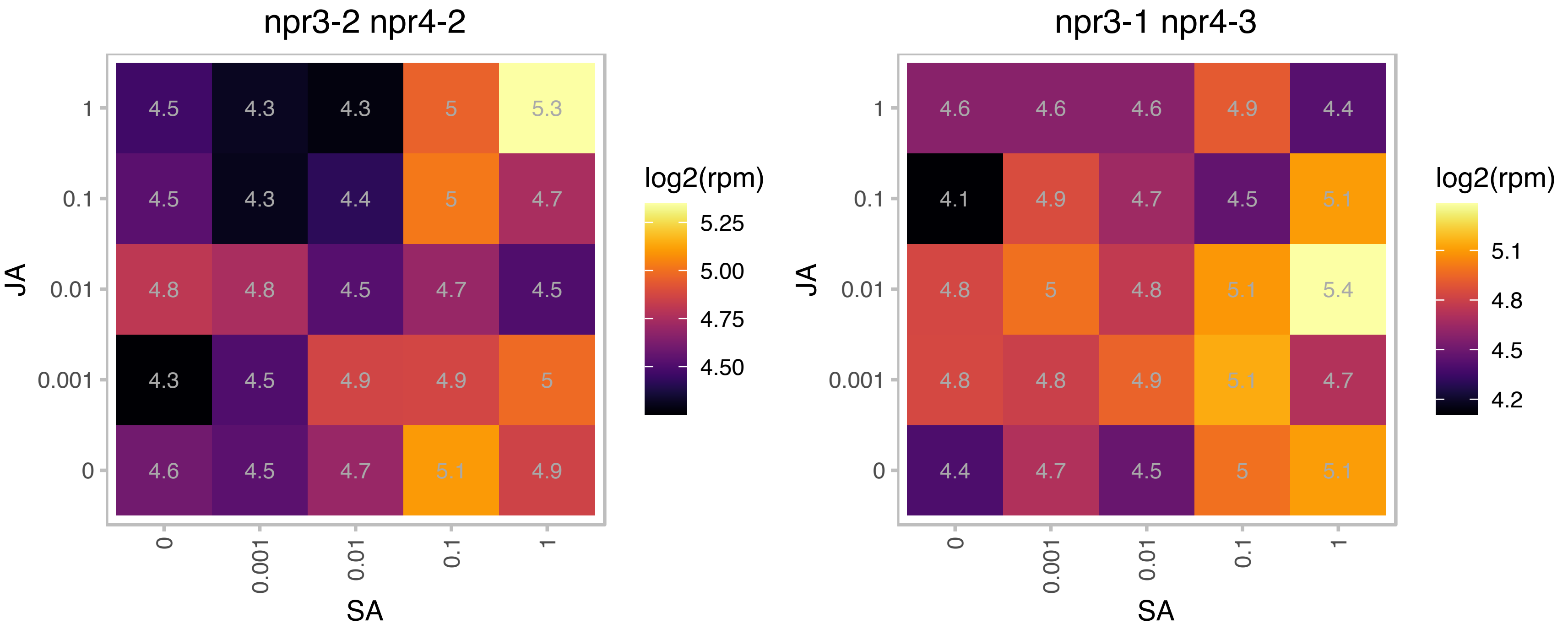

### AT3G63380.1 : ACA12

ATPase E1-E2 type family protein / haloacid dehalogenase-like hydrolase family protein

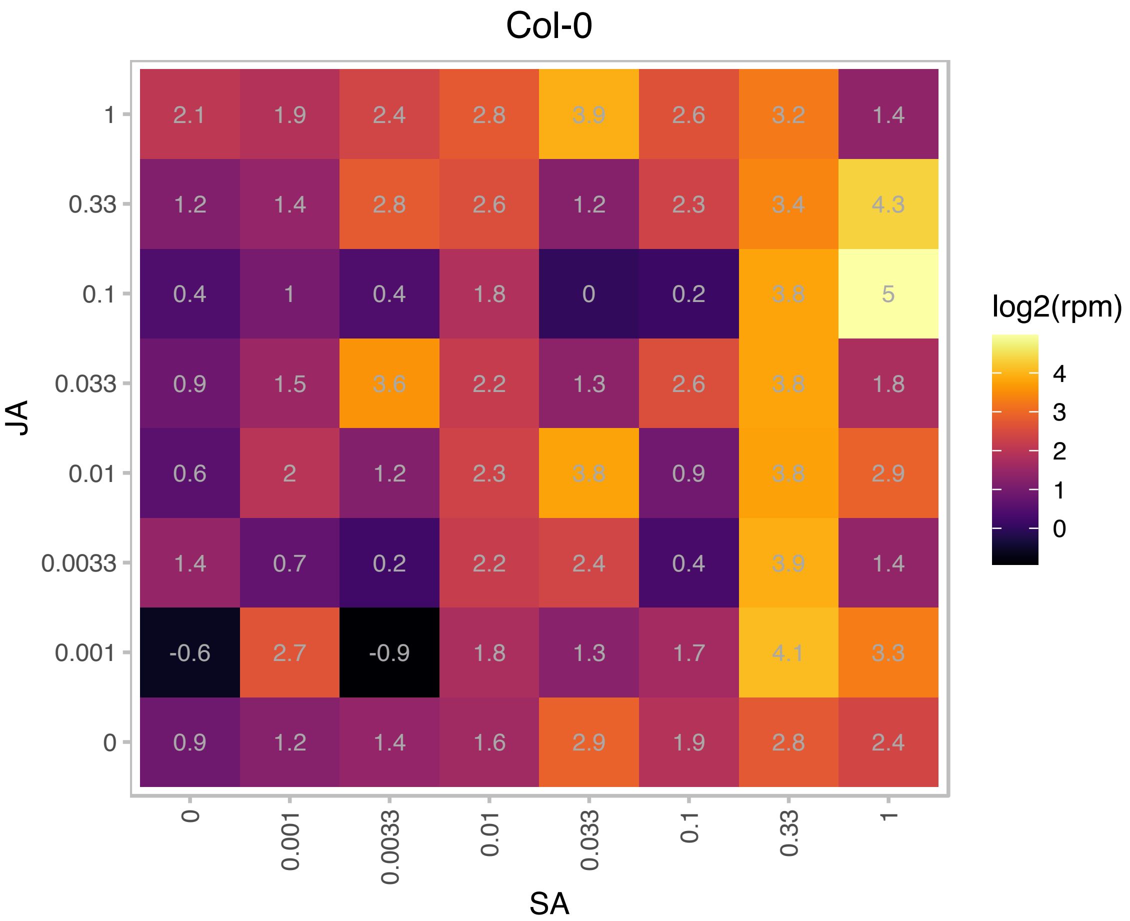


| | |
|---|---|
| Gene | AT3G63380 |
| is.expressed_genes | TRUE |
| is.responsive_genes | TRUE |
| is.apcluster | C-02 |

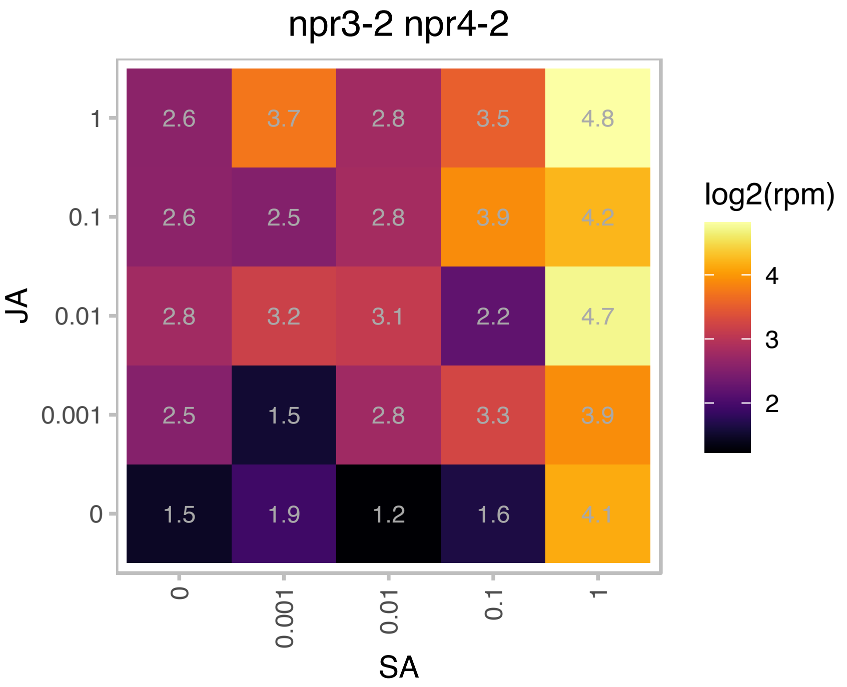


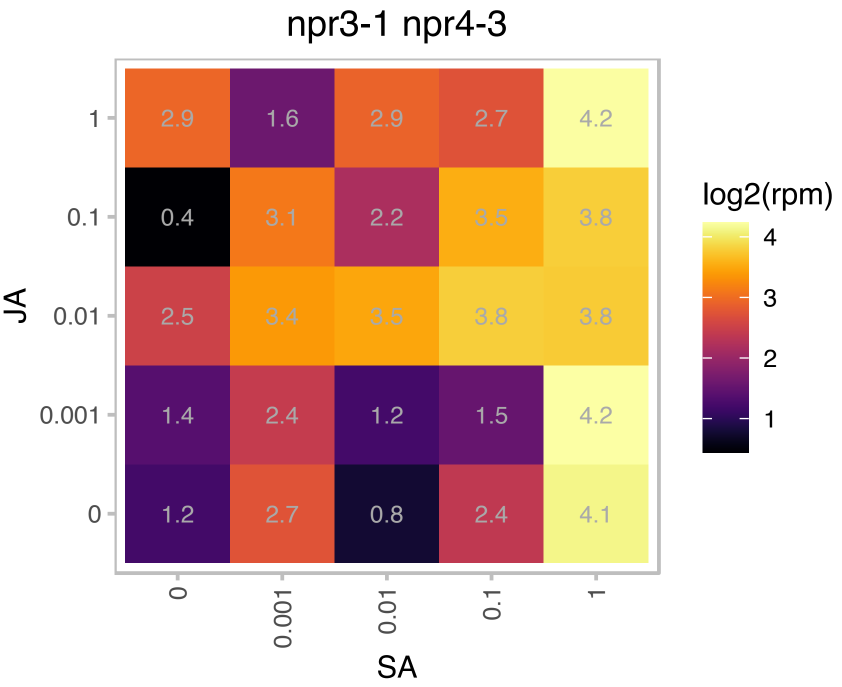

### AT4G13510.1 : AMT1;1, ATAMT1, ATAMT1;1

ammonium transporter 1;1

Encodes a plasma membrane localized ammonium transporter.  Contains a cytosolic trans-activation domain essential for ammonium uptake. The mRNA is cell-to-cell mobile.

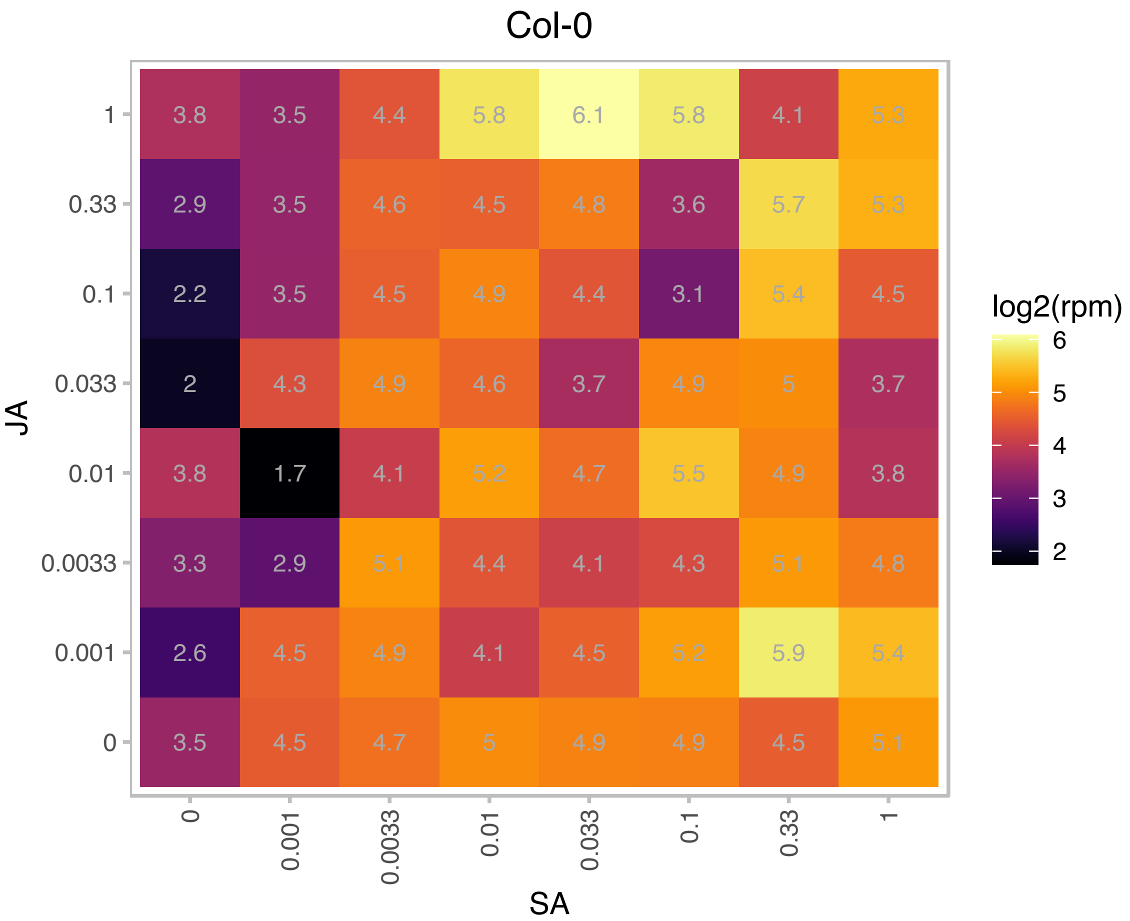


| | |
|---|---|
| Gene | AT4G13510 |
| is.expressed_genes | TRUE |
| is.responsive_genes | TRUE |
| is.apcluster | C-02 |

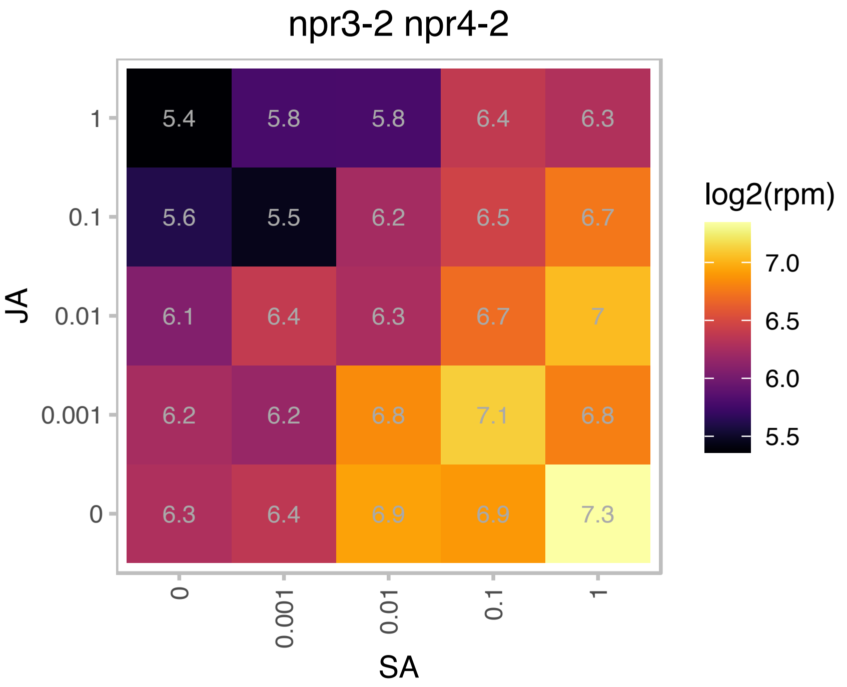


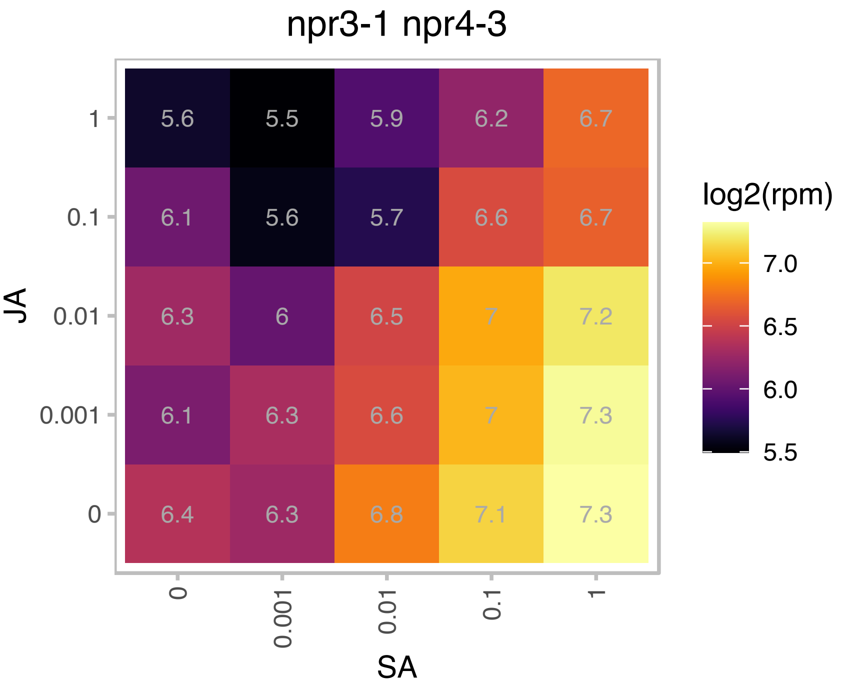

### AT4G23980.1 : ARF9

auxin response factor 9

Encodes auxin response factor 9 (ARF9). The mRNA is cell-to-cell mobile.

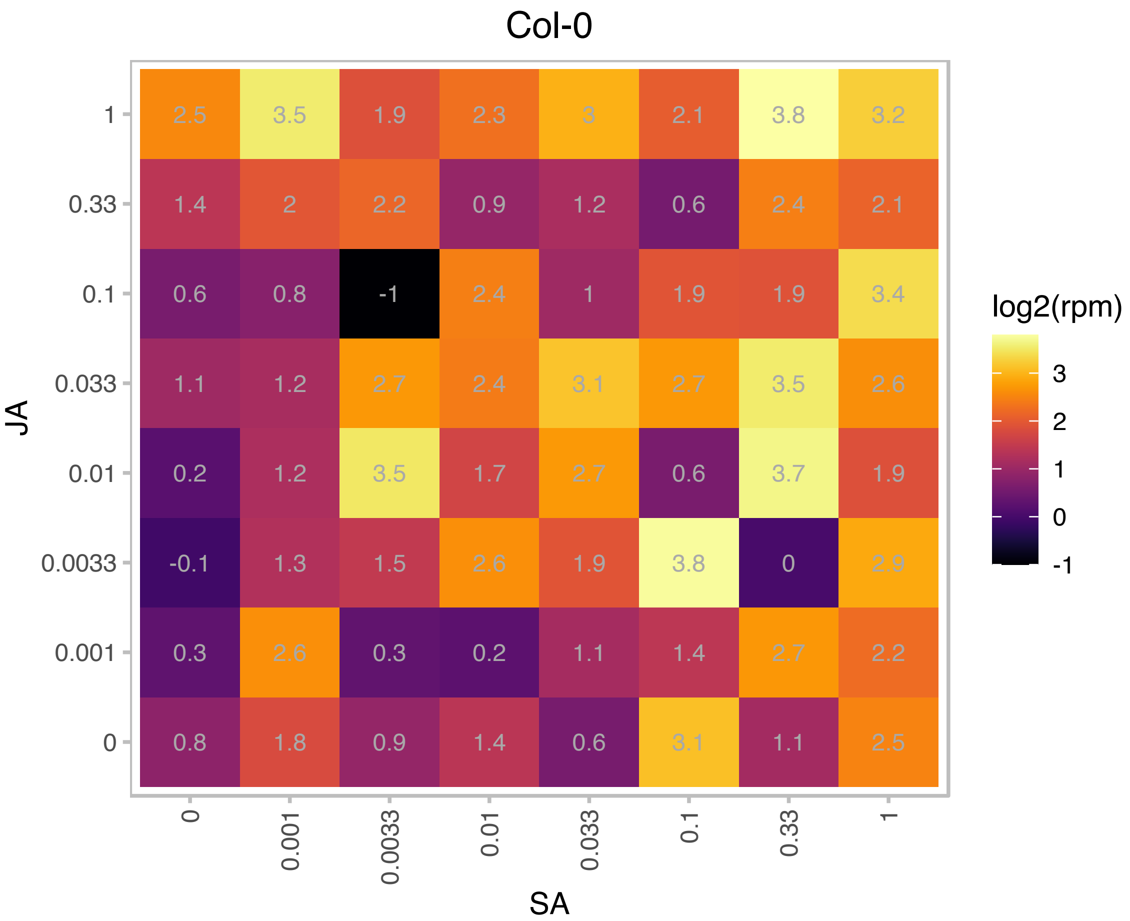


| | |
|---|---|
| Gene | AT4G23980 |
| is.expressed_genes | TRUE |
| is.responsive_genes | TRUE |
| is.apcluster | C-02 |

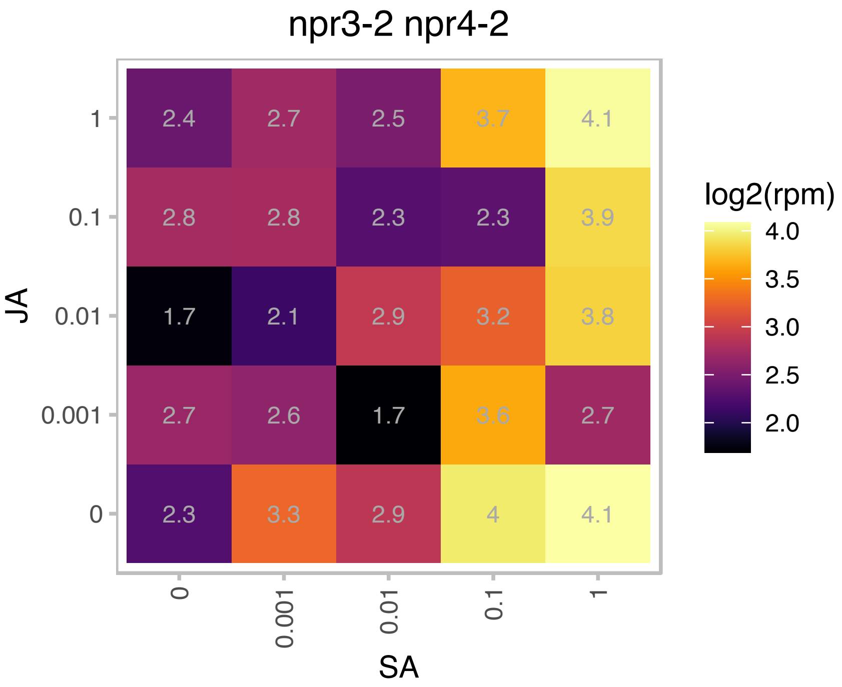


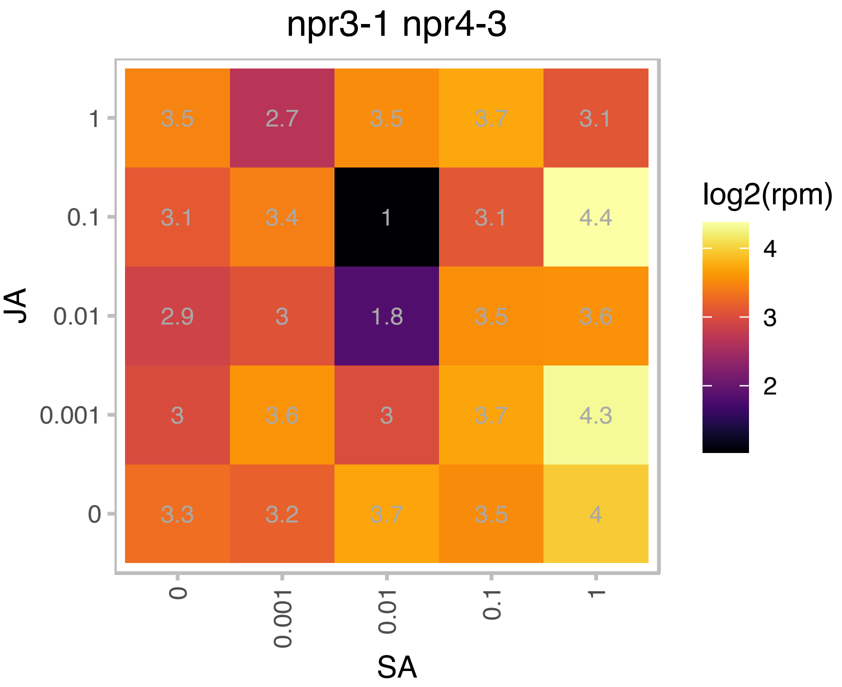

# AT5G01850.1 : NA

Protein kinase superfamily protein

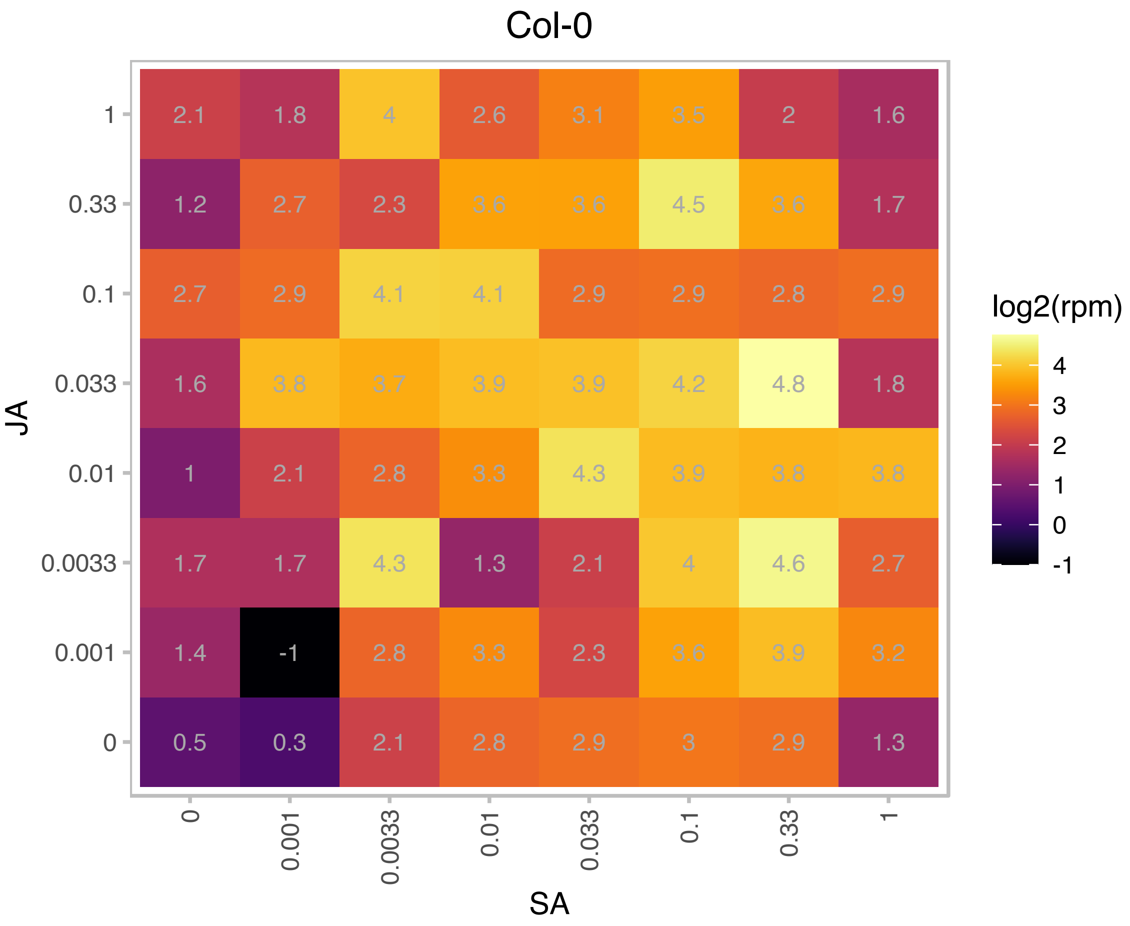


| | |
|---|---|
| Gene | AT5G01850 |
| is.expressed_genes | TRUE |
| is.responsive_genes | TRUE |
| is.apcluster | C-02 |

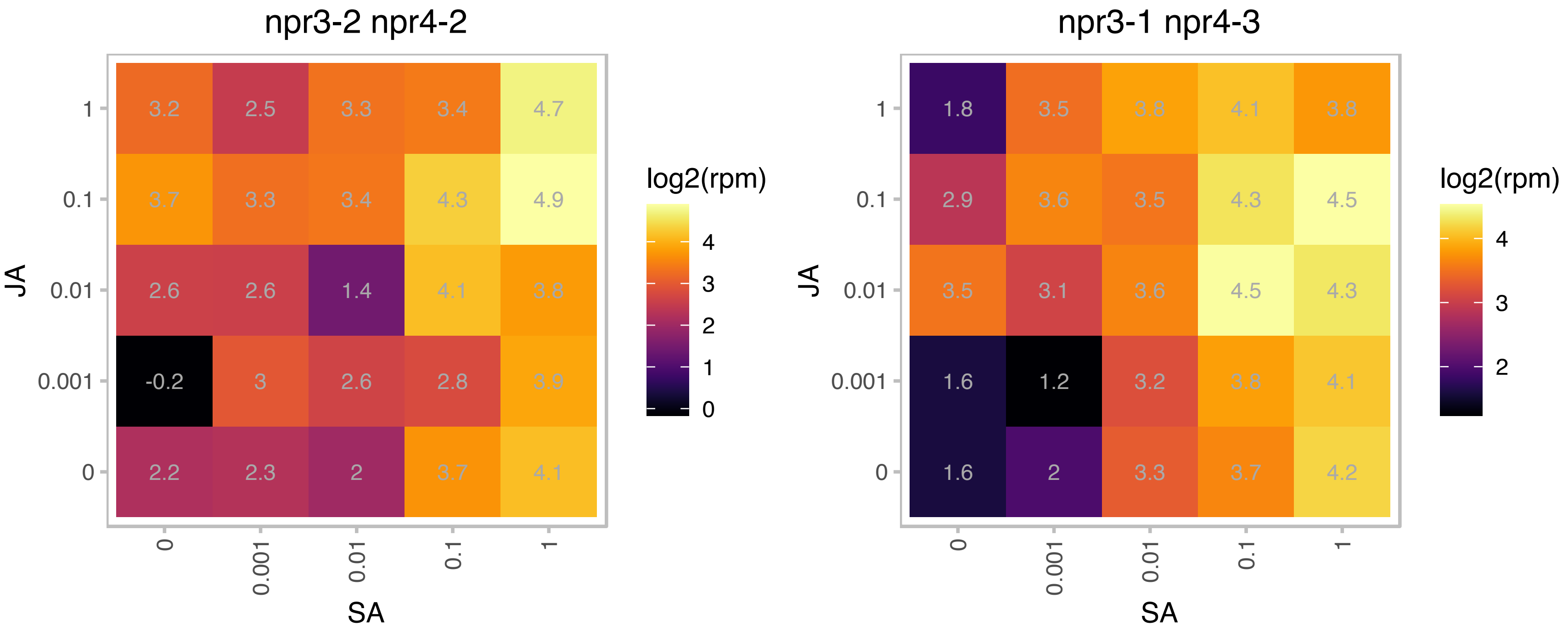

# AT5G01960.1 : NA

RING/U-box superfamily protein

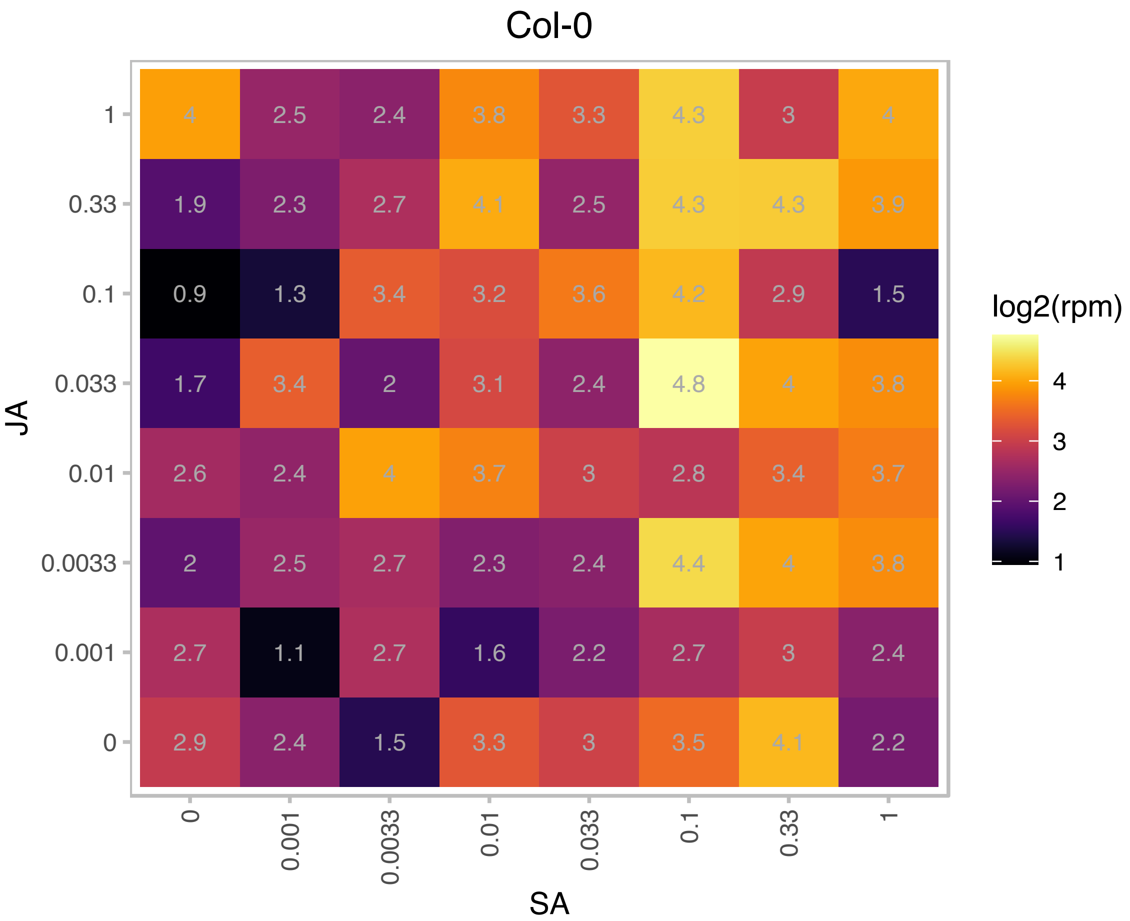


| | |
|---|---|
| Gene | AT5G01960 |
| is.expressed_genes | TRUE |
| is.responsive_genes | TRUE |
| is.apcluster | C-02 |

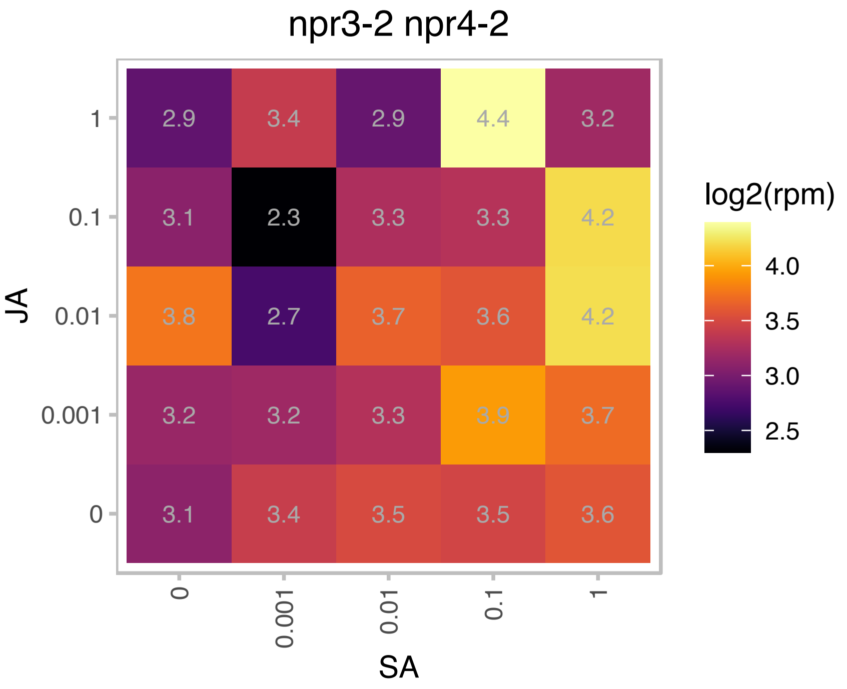


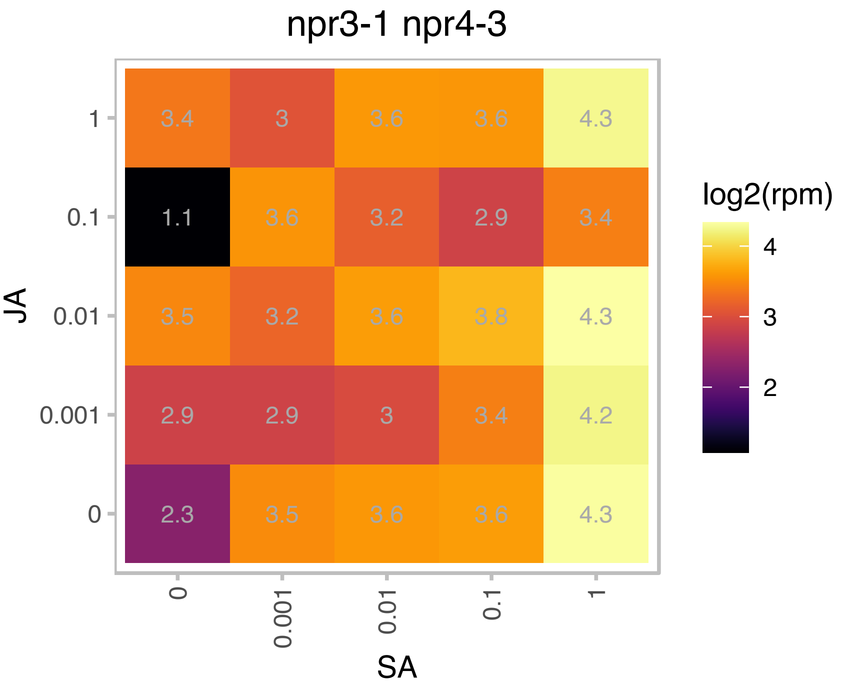

# AT5G07360.1 : NA

Amidase family protein

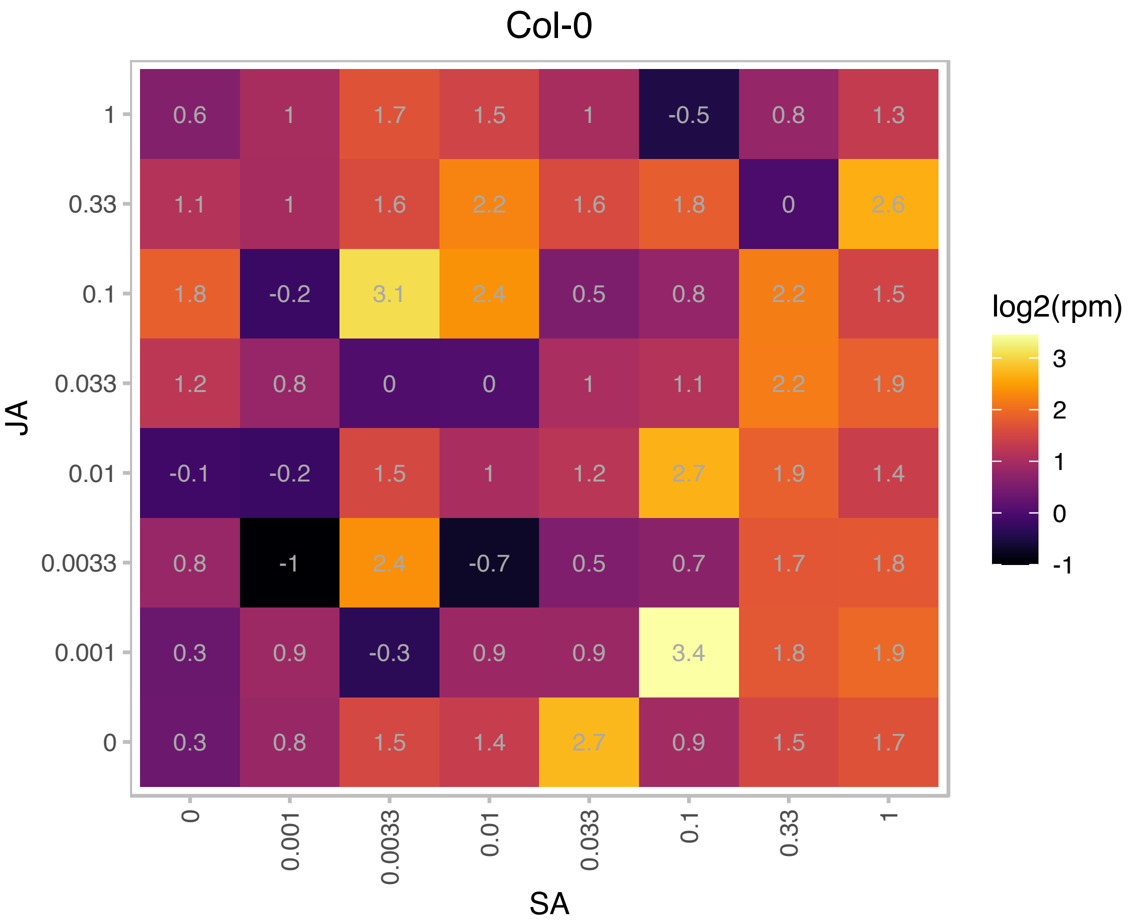


| | |
|---|---|
| Gene | AT5G07360 |
| is.expressed_genes | TRUE |
| is.responsive_genes | TRUE |
| is.apcluster | C-02 |

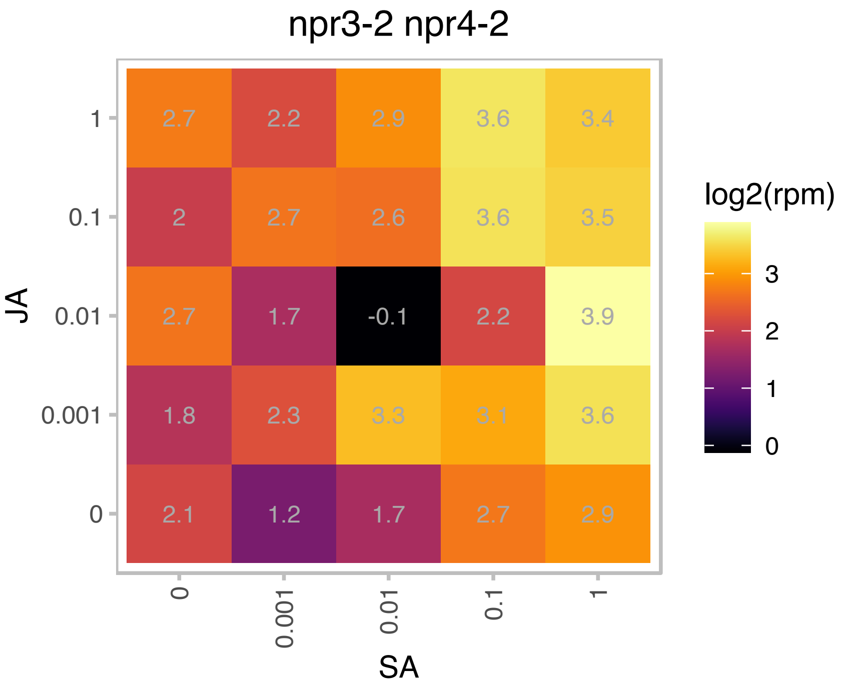


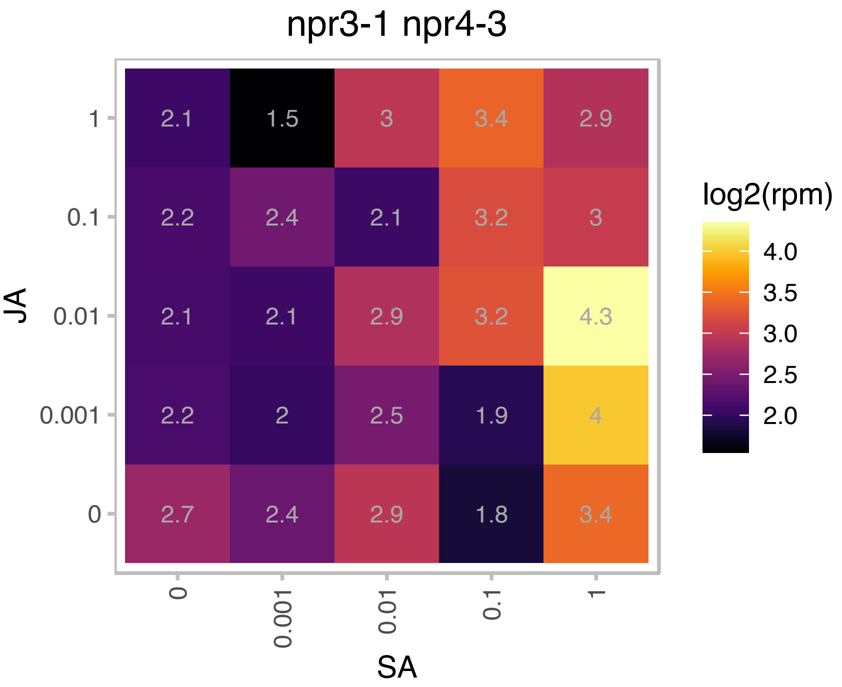

### AT5G10610.1 : CYP81K1

cytochrome P450, family 81, subfamily K, polypeptide 1

member of CYP81K The mRNA is cell-to-cell mobile.

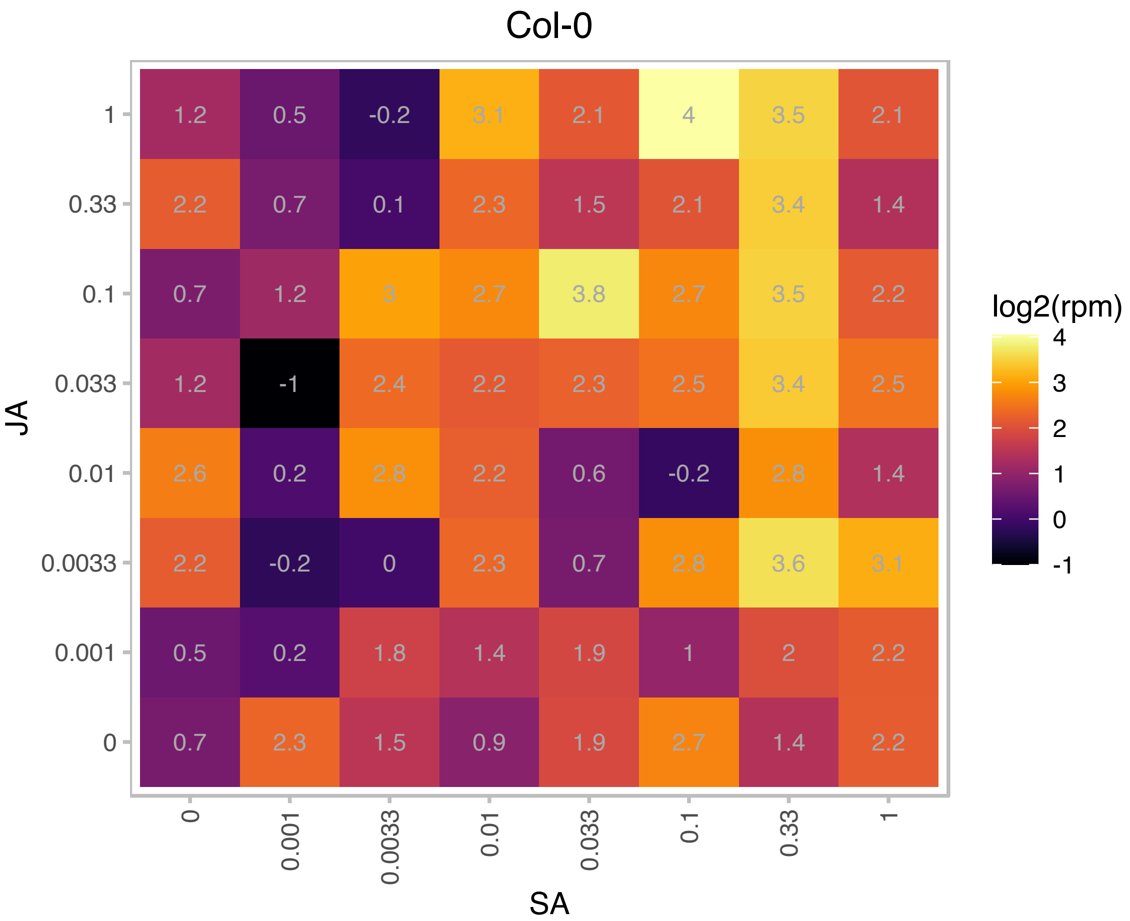


| | |
|---|---|
| Gene | AT5G10610 |
| is.expressed_genes | TRUE |
| is.responsive_genes | TRUE |
| is.apcluster | C-02 |

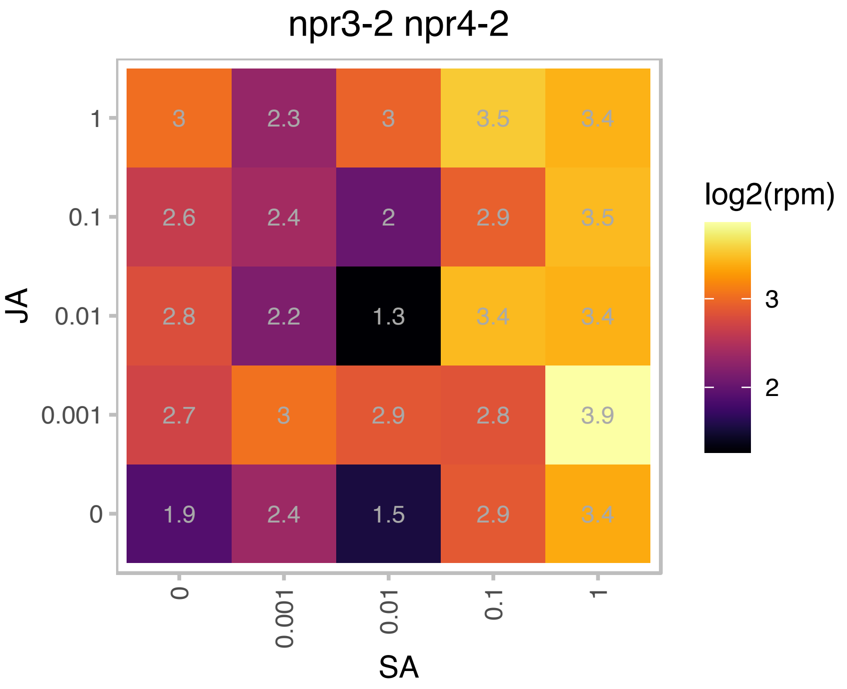


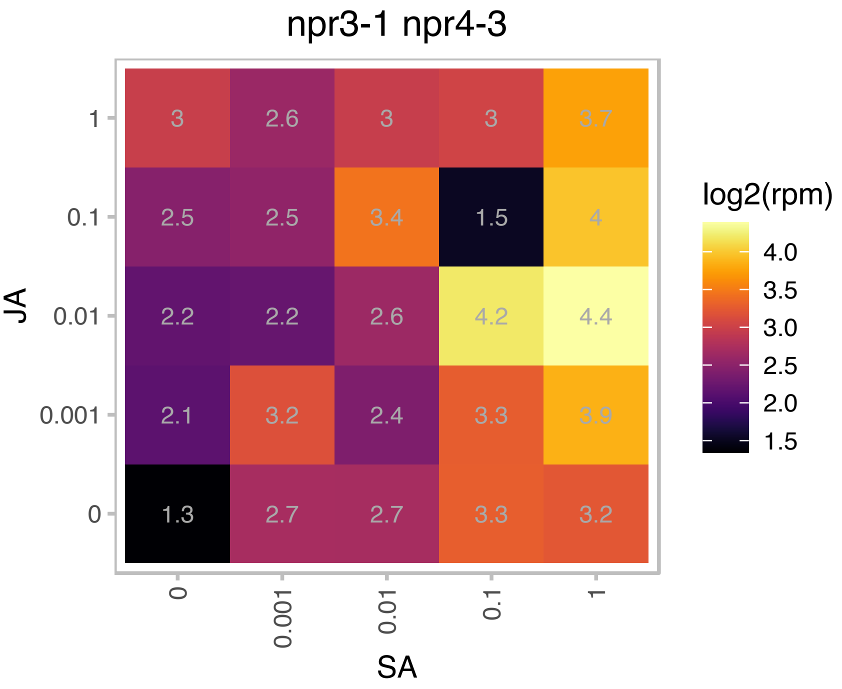

### AT5G11060.1 : KNAT4

KNOTTED1-like homeobox gene 4

A member of Class II KN1-like homeodomain transcription factors (together with KNAT3 and KNAT5), with greatest homology to the maize knox1 homeobox protein. Expression regulated by light. Detected in all tissues examined, but most prominent in leaves and young siliques. Transient expression of GFP translational fusion protein suggests bipartite localization in nucleus and cytoplasm. KNAT4 promoter activity showed cell-type specific pattern along longitudinal root axis; GUS expression pattern started at the elongation zone, predominantly in the phloem and pericycle cells, extending to endodermis toward the base of the root.

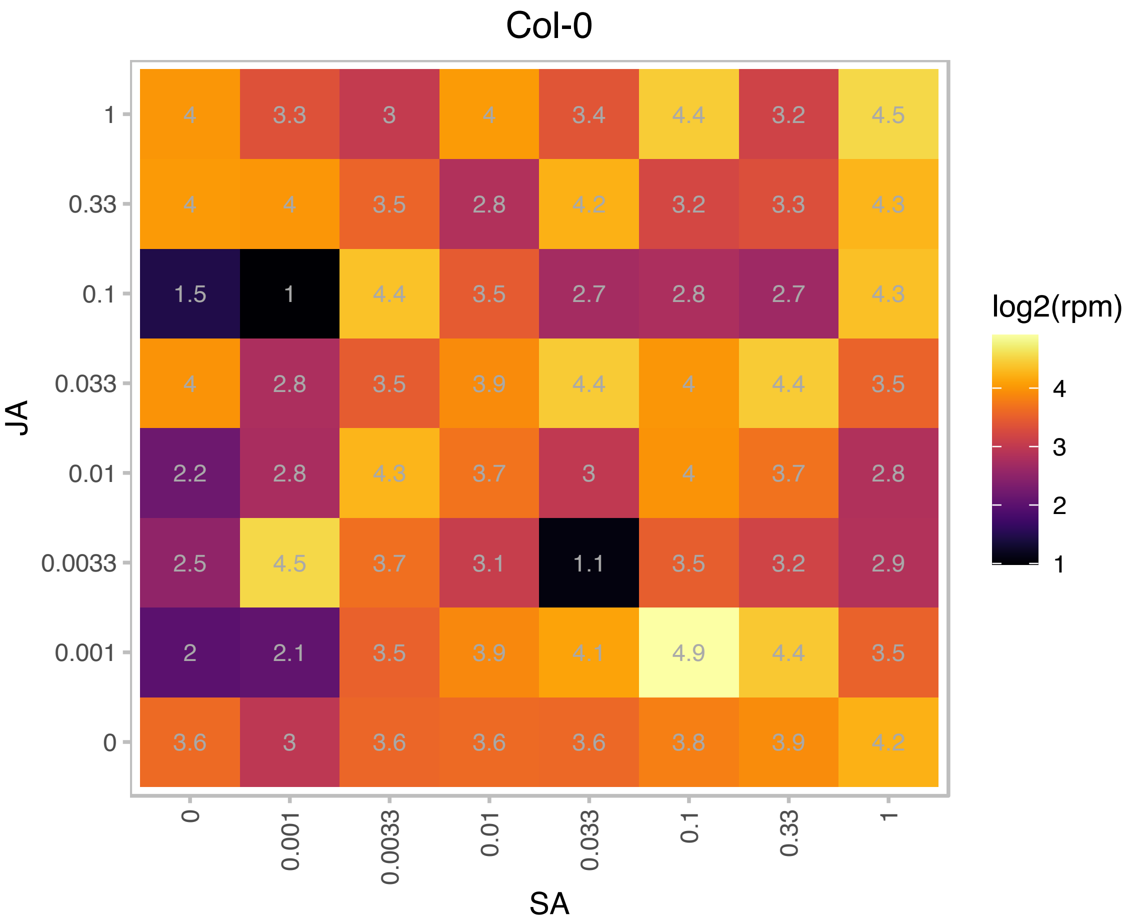


| | |
|---|---|
| Gene | AT5G11060 |
| is.expressed_genes | TRUE |
| is.responsive_genes | TRUE |
| is.apcluster | C-02 |

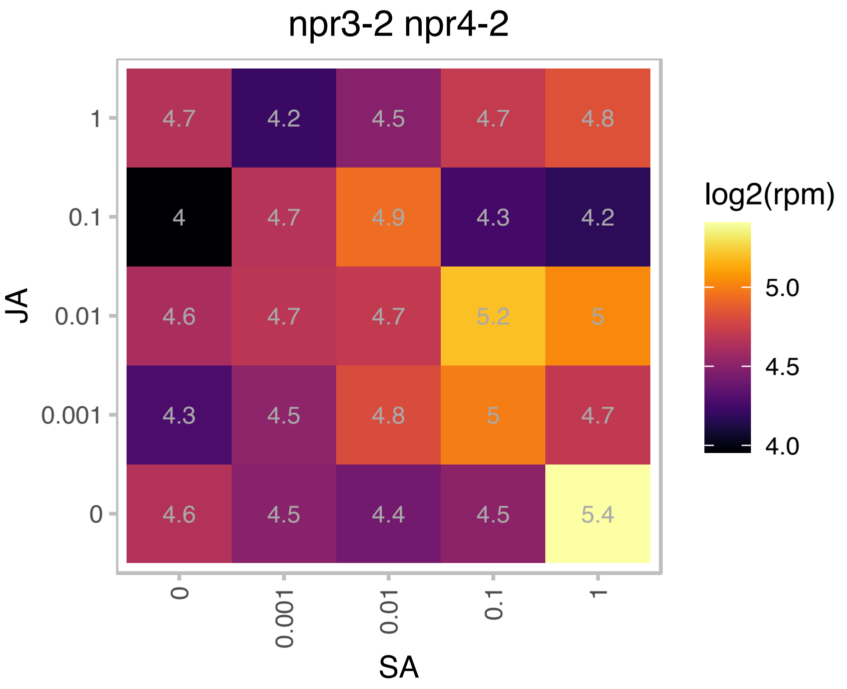


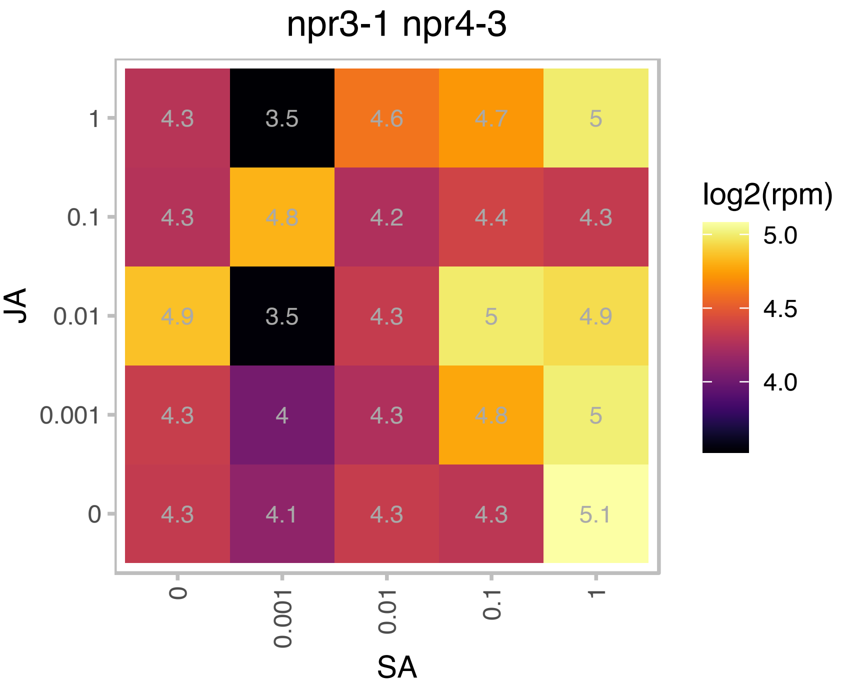

# AT5G15870.1 : NA

glycosyl hydrolase family 81 protein

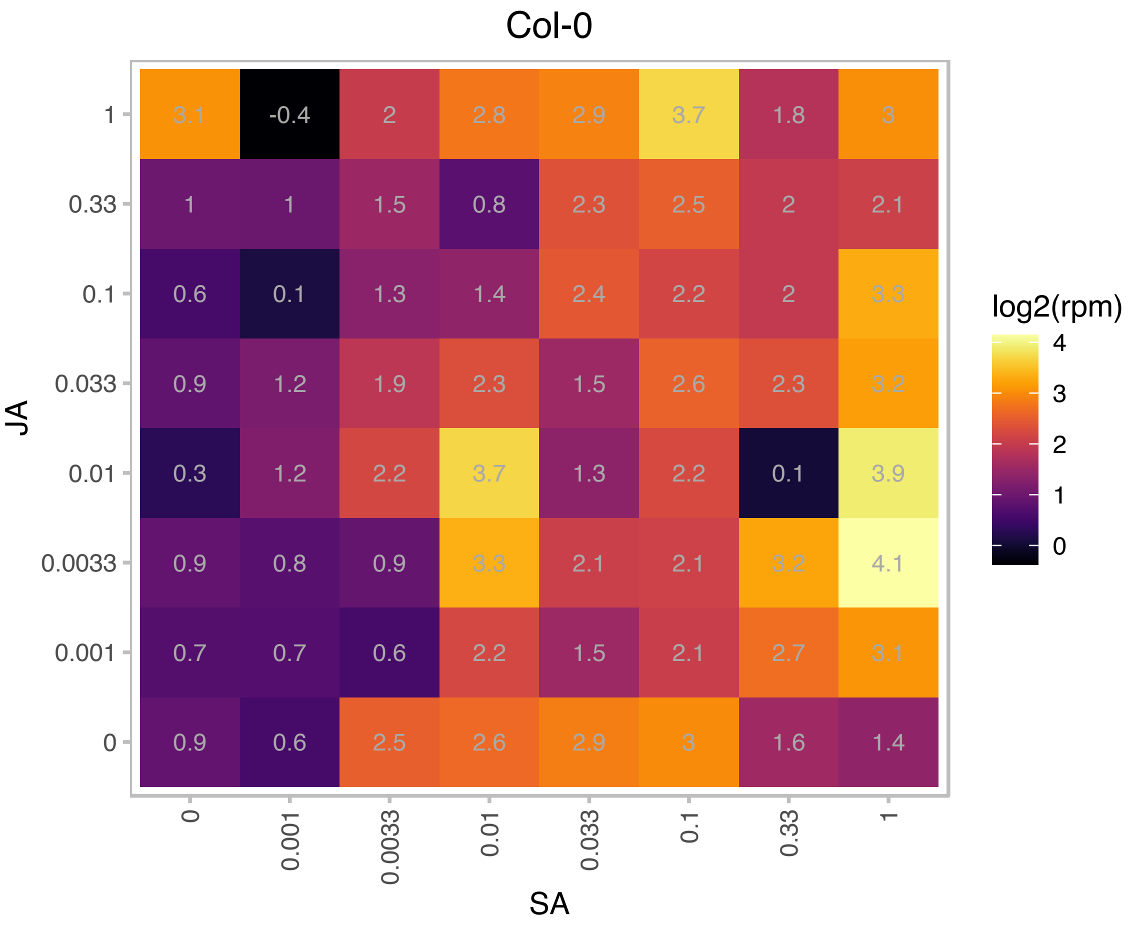


| | |
|---|---|
| Gene | AT5G15870 |
| is.expressed_genes | TRUE |
| is.responsive_genes | TRUE |
| is.apcluster | C-02 |

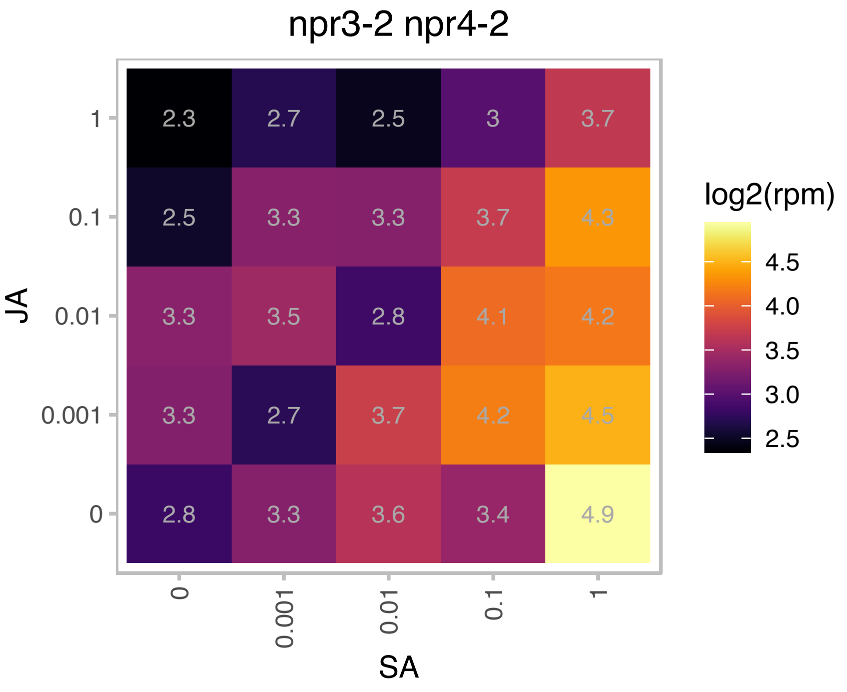


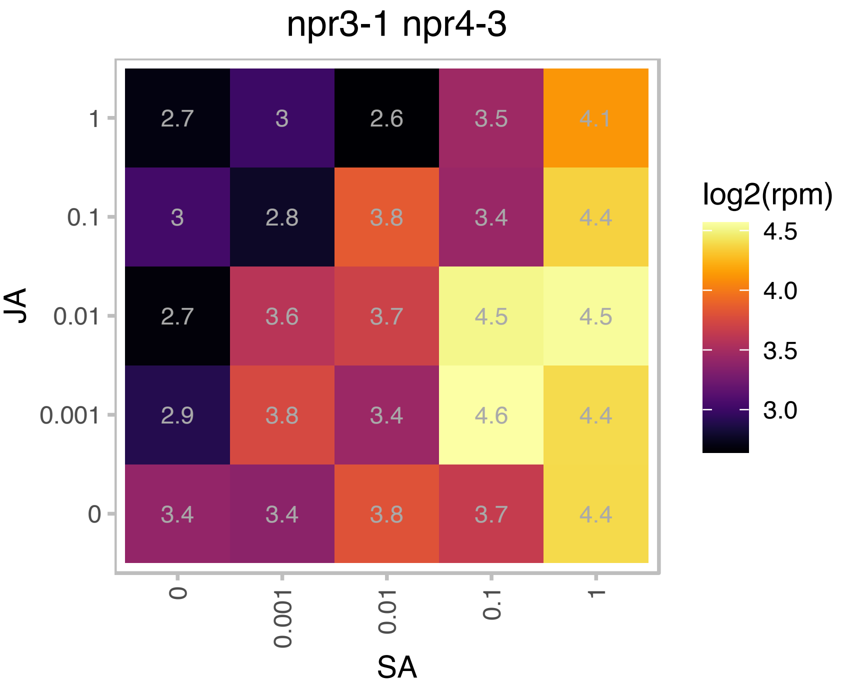

# AT5G18900.1 : NA

2-oxoglutarate (2OG) and Fe(II)-dependent oxygenase superfamily protein

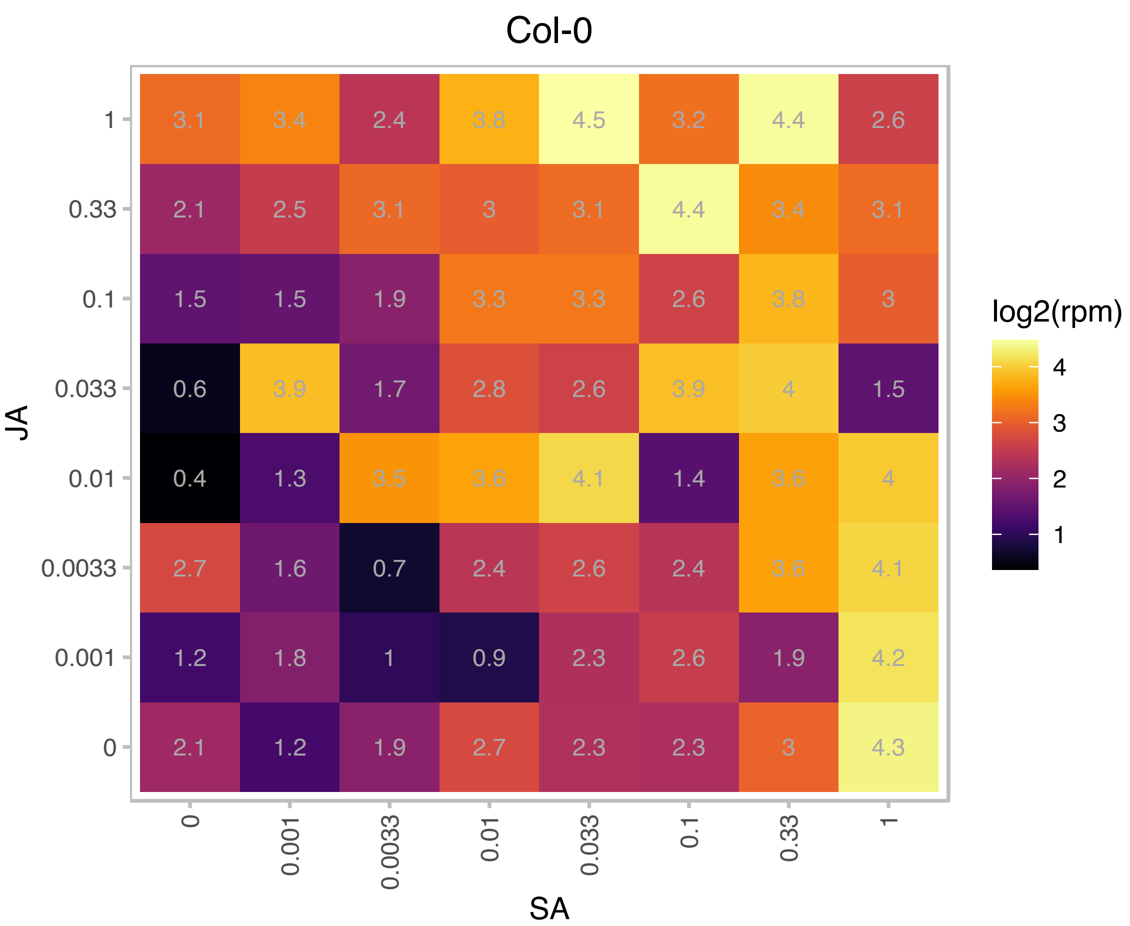


| | |
|---|---|
| Gene | AT5G18900 |
| is.expressed_genes | TRUE |
| is.responsive_genes | TRUE |
| is.apcluster | C-02 |

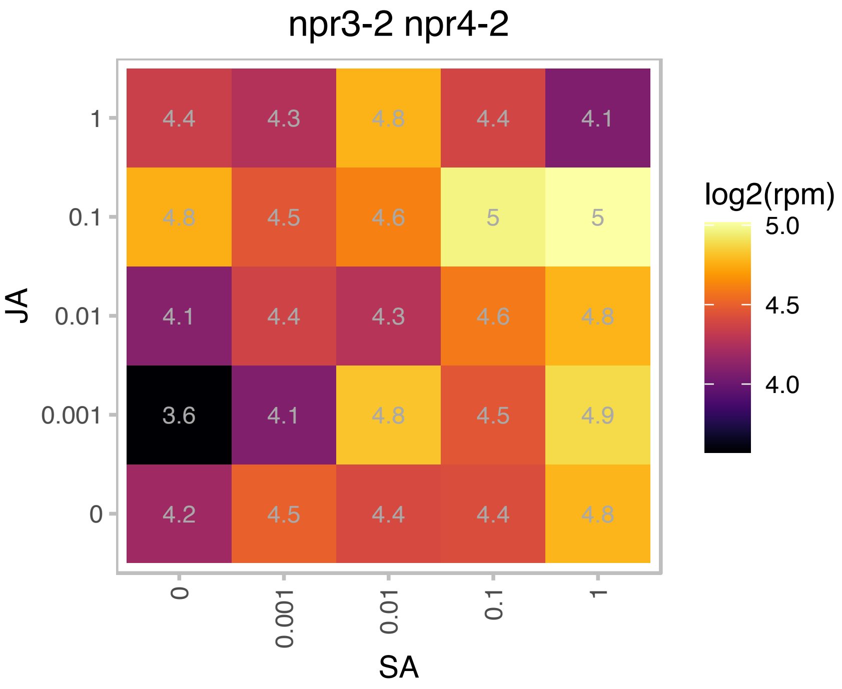


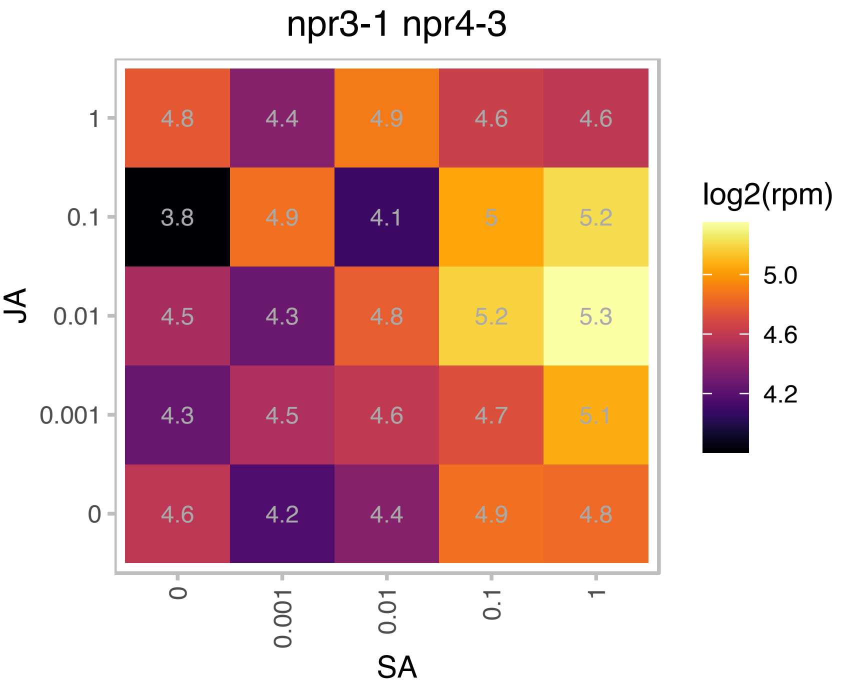

### AT5G22000.1 : RHF2A

RING-H2 group F2A

encodes a RING-type E3 ubiquitin ligase implicated in gametogenesis. Double mutant analyses with RHF1a suggests that RHF2a may be involved in targetting ICK4KRP6 for degradation following meiosis in order to allow the mitoses associated with megagametogenesis and microgametogenesis to occur. RHF2a is expressed in all four floral whorls and is present at ~8-fold higher levels than RHF1a in inflorescences by RT-PCR analyses.

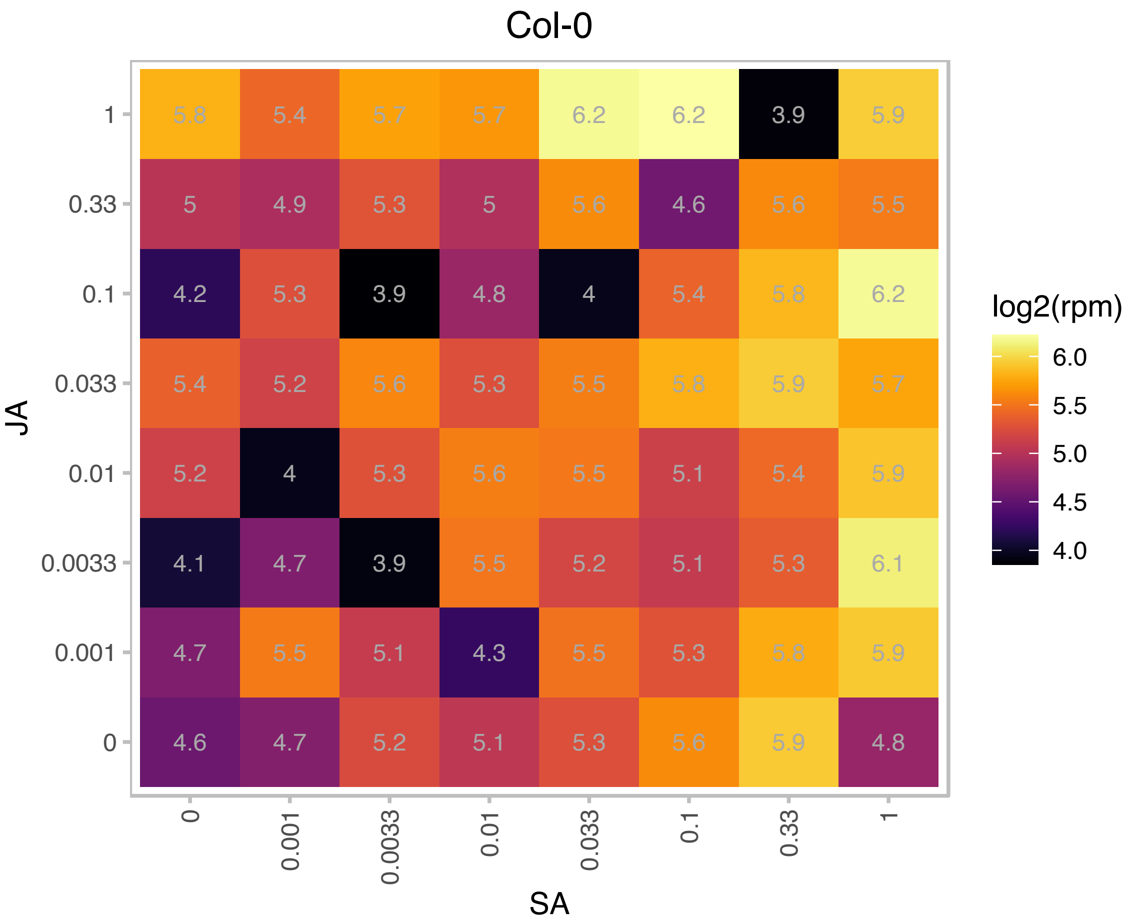


| | |
|---|---|
| Gene | AT5G22000 |
| is.expressed_genes | TRUE |
| is.responsive_genes | TRUE |
| is.apcluster | C-02 |

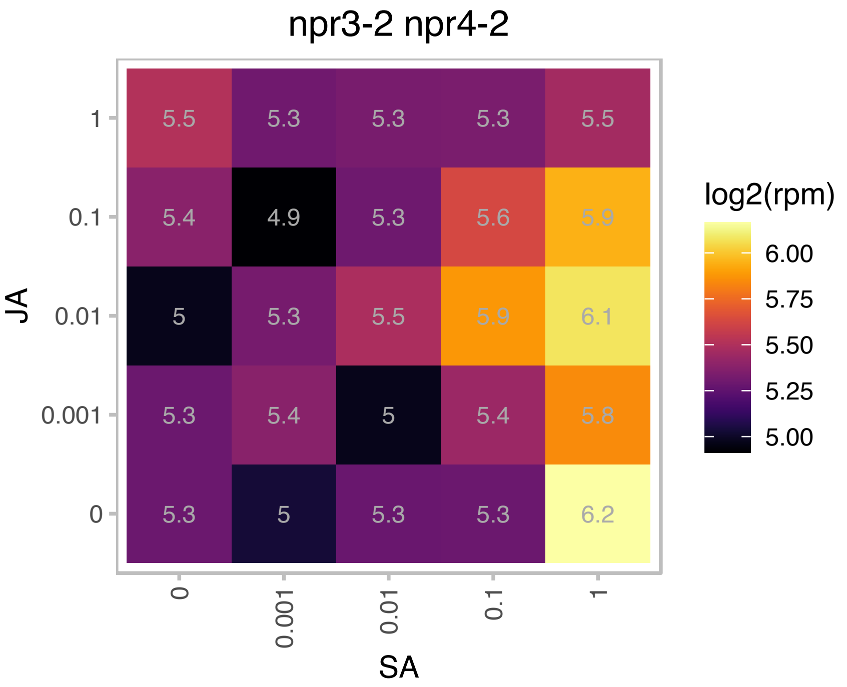


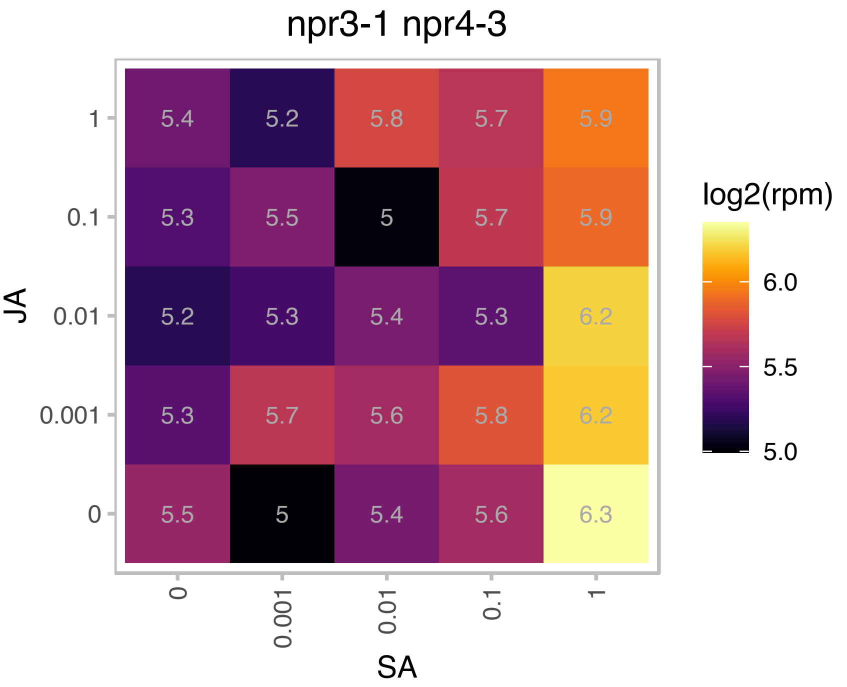

### AT5G45110.1 : ATNPR3, NPR3

NPR1-like protein 3

Encodes NPR3, a paralog of NPR1. Involved in negative regulation of defense responses against bacterial and oomycete pathogens. npr3 mutants has elevated level of PR1 expression. Interacts with TGA2, TGA3, TGA5 and TGA6 in yeast two hybrid assays. NPR3 and NPR4 are receptors for the immune signal salicylic acid. The mRNA is cell-to-cell mobile.

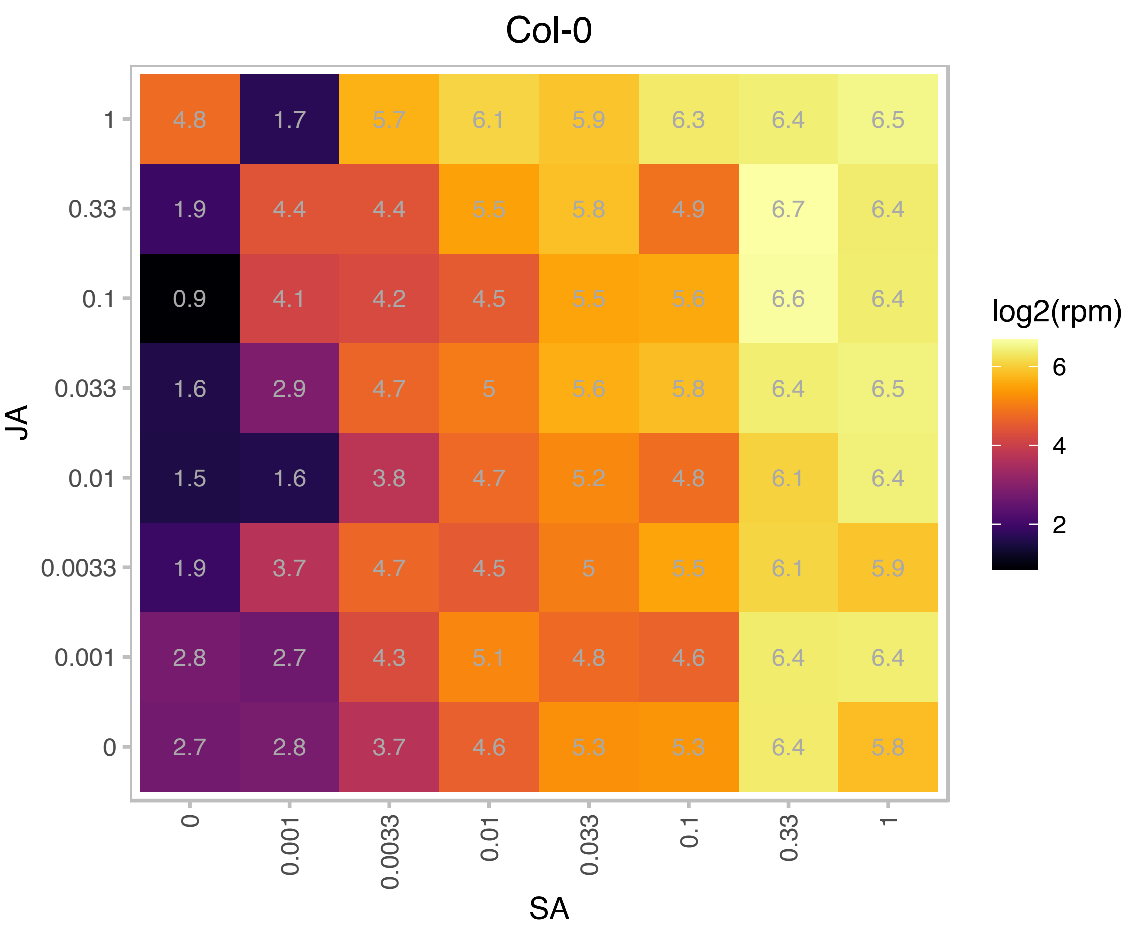


| | |
|---|---|
| Gene | AT5G45110 |
| is.expressed_genes | TRUE |
| is.responsive_genes | TRUE |
| is.apcluster | C-02 |

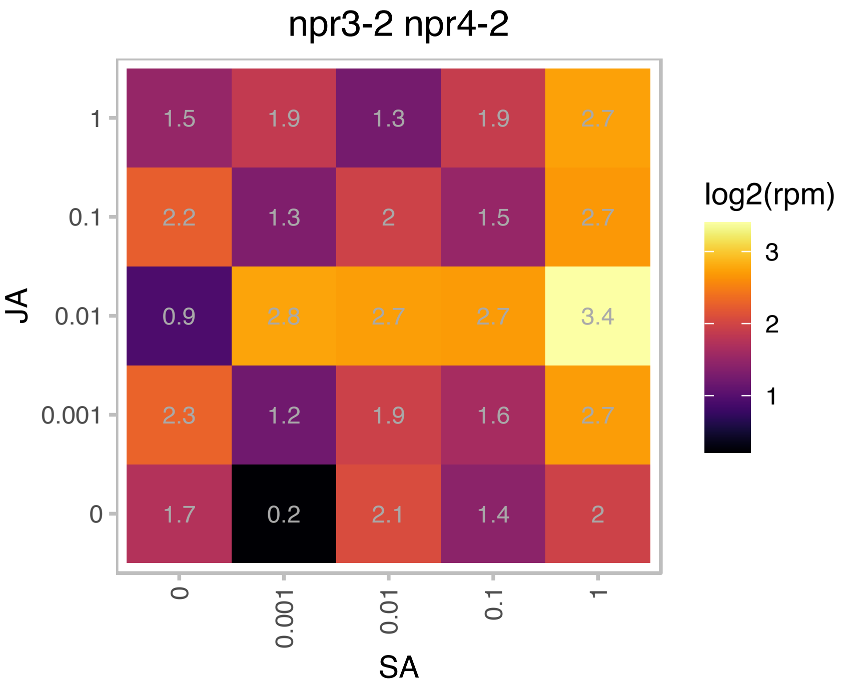


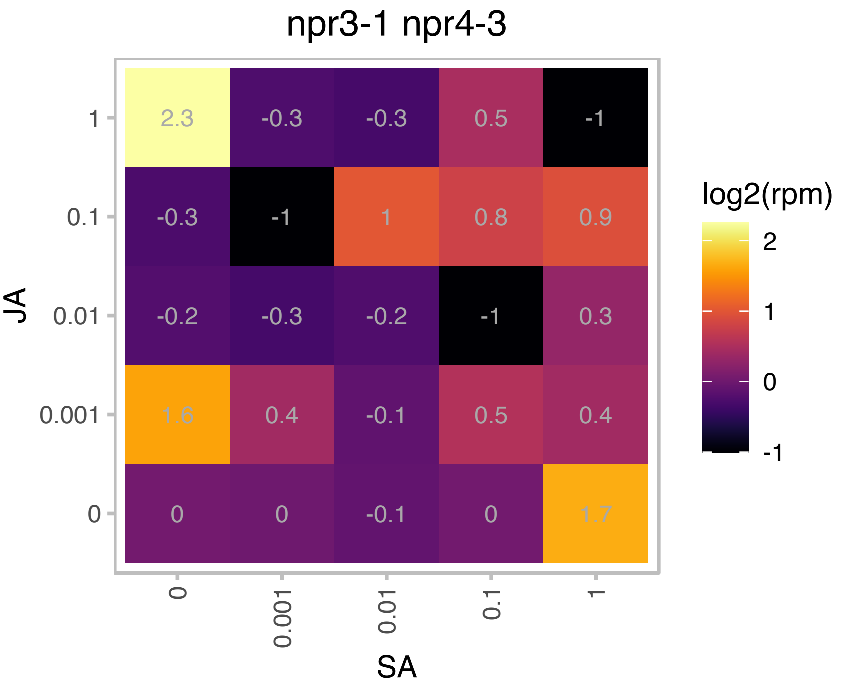

### AT5G55940.1 : emb2731

Uncharacterised protein family (UPF0172)

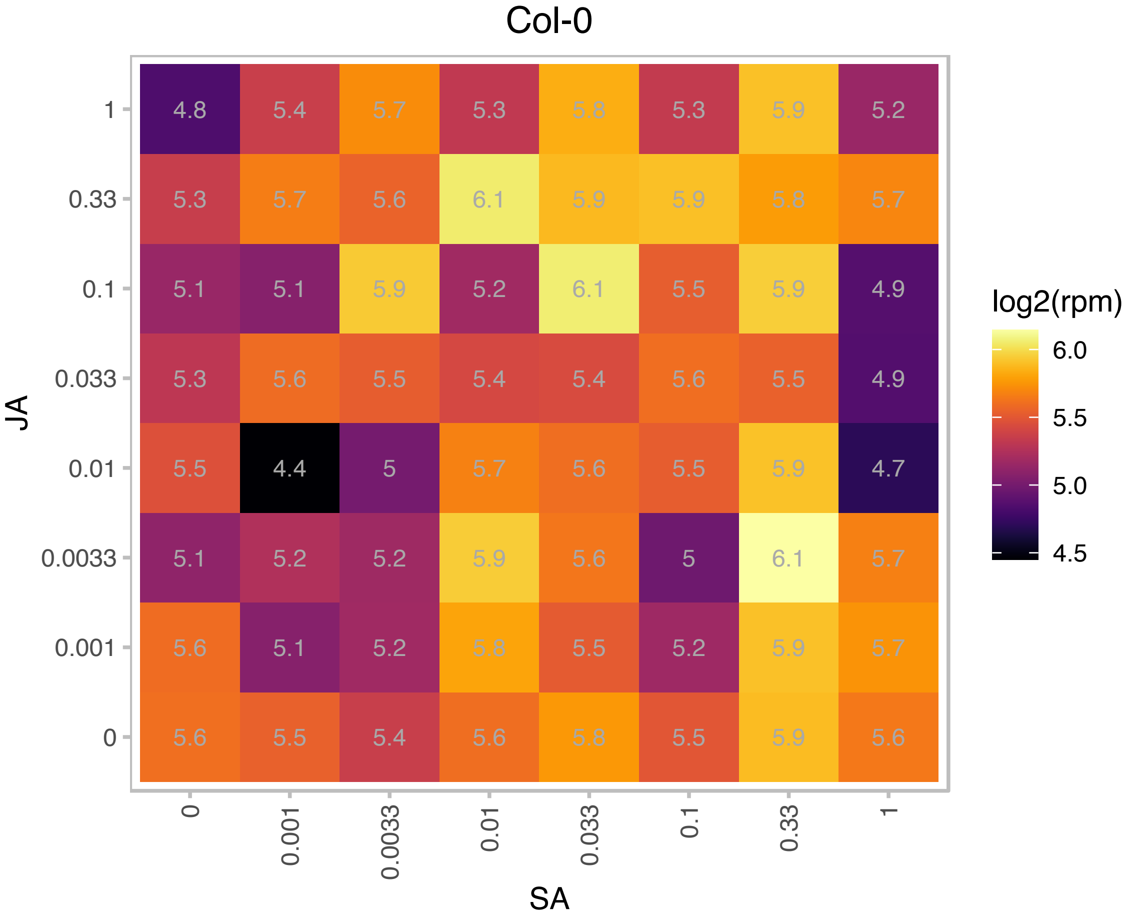


| | |
|---|---|
| Gene | AT5G55940 |
| is.expressed_genes | TRUE |
| is.responsive_genes | TRUE |
| is.apcluster | C-02 |

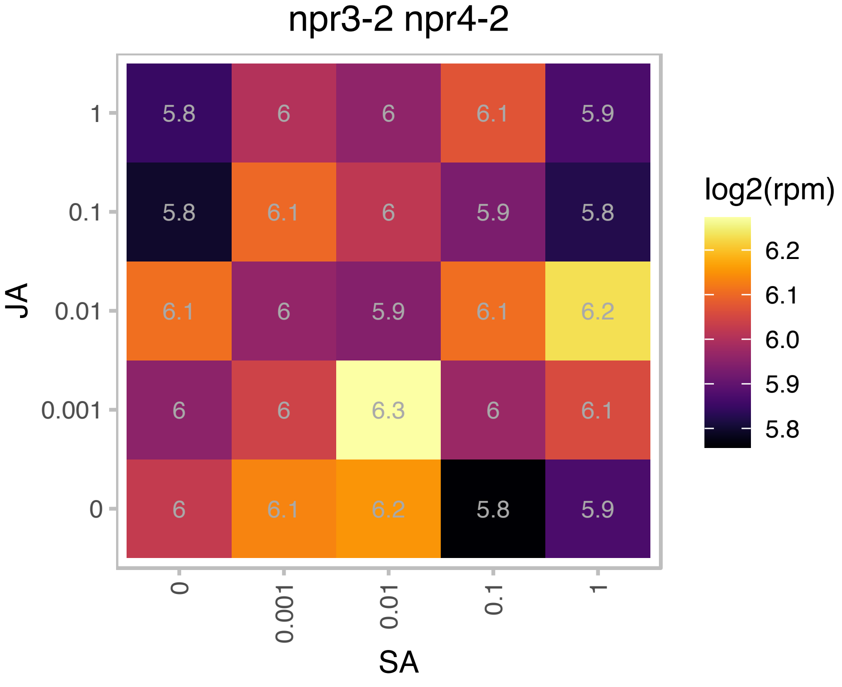


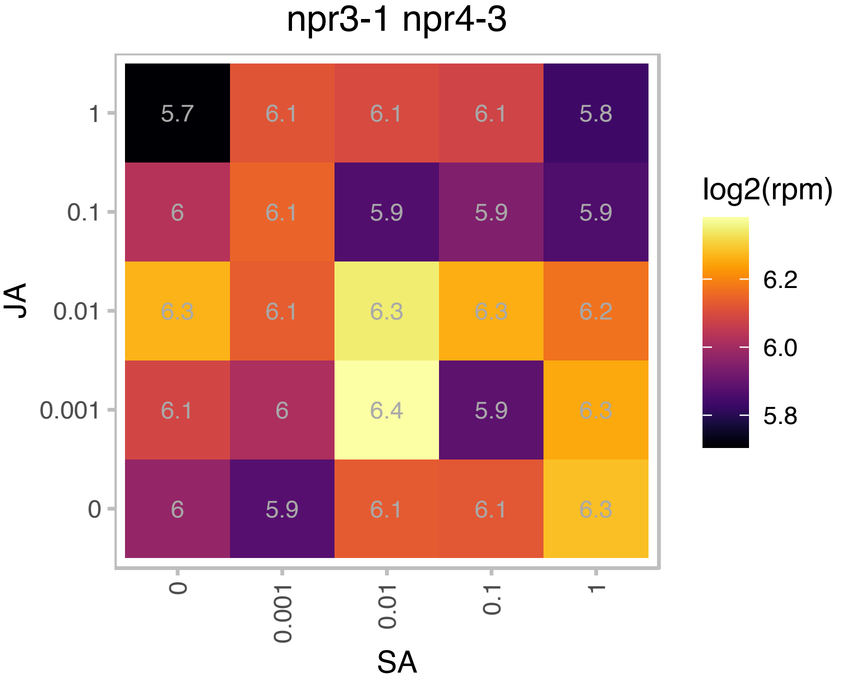

# AT5G57580.1 : NA

Calmodulin-binding protein

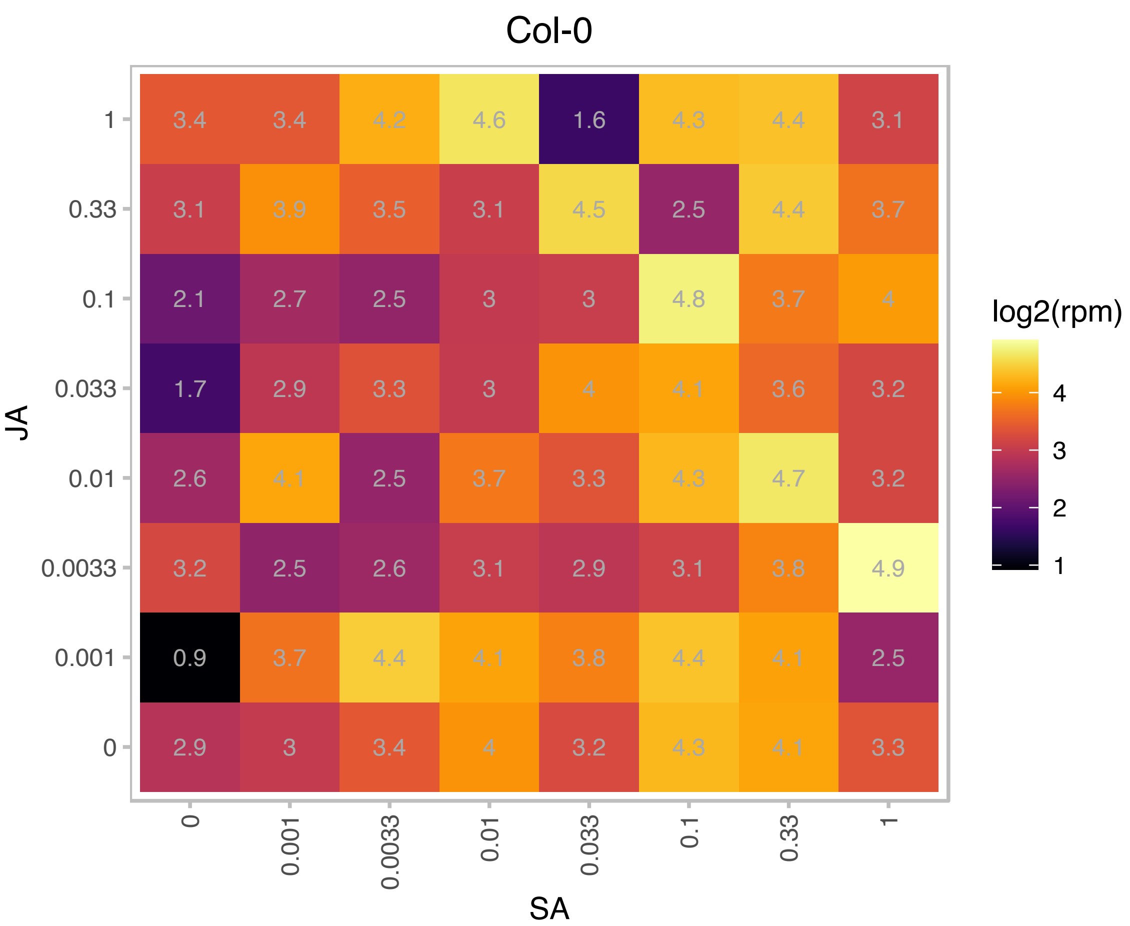


| | |
|---|---|
| Gene | AT5G57580 |
| is.expressed_genes | TRUE |
| is.responsive_genes | TRUE |
| is.apcluster | C-02 |

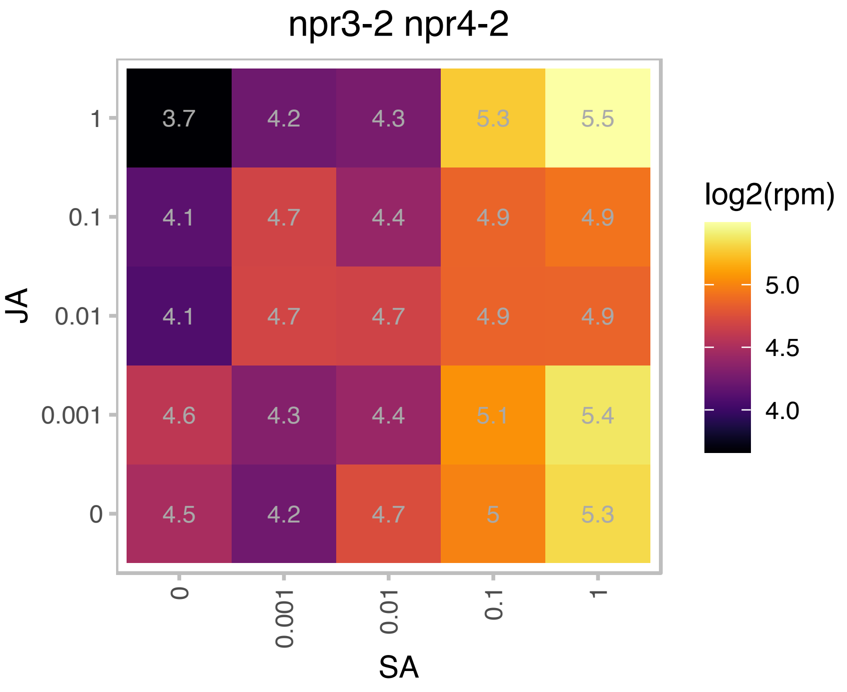


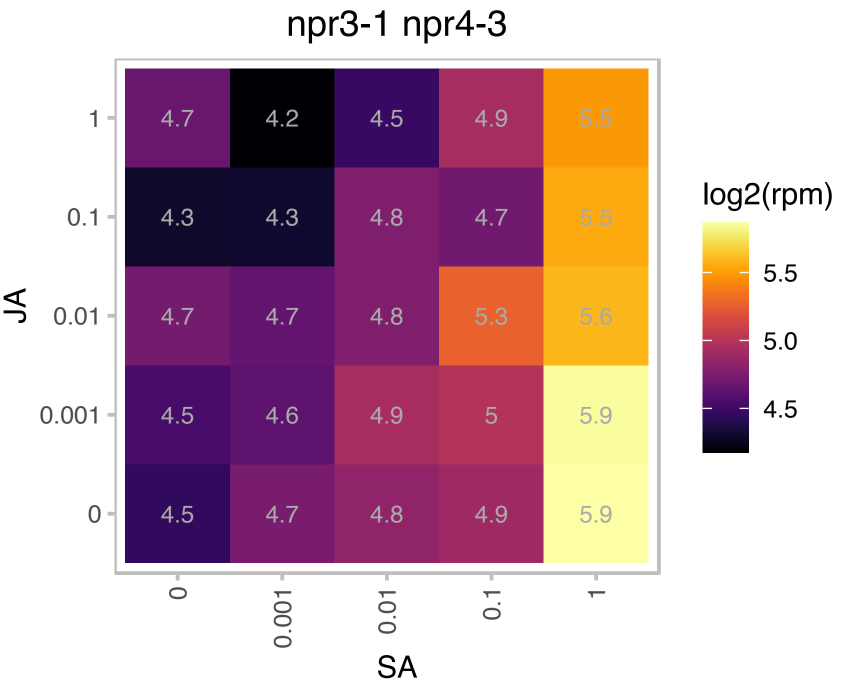

# AT5G65207.1 : NA

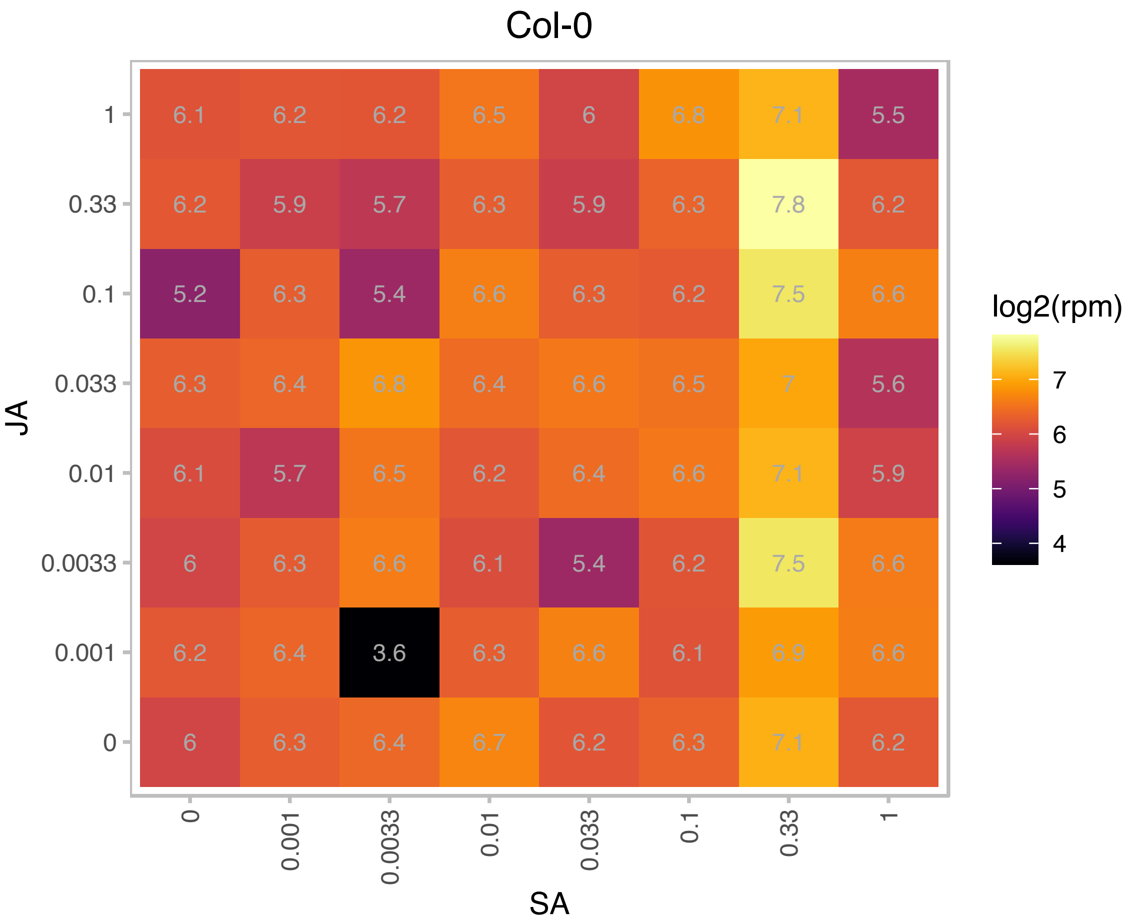


| | |
|---|---|
| Gene | AT5G65207 |
| is.expressed_genes | TRUE |
| is.responsive_genes | TRUE |
| is.apcluster | C-02 |

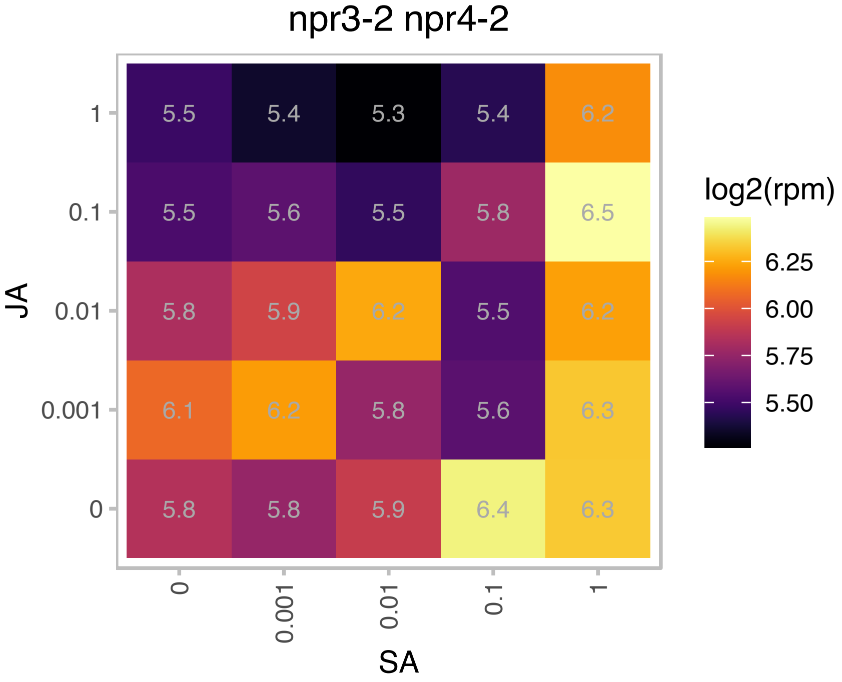


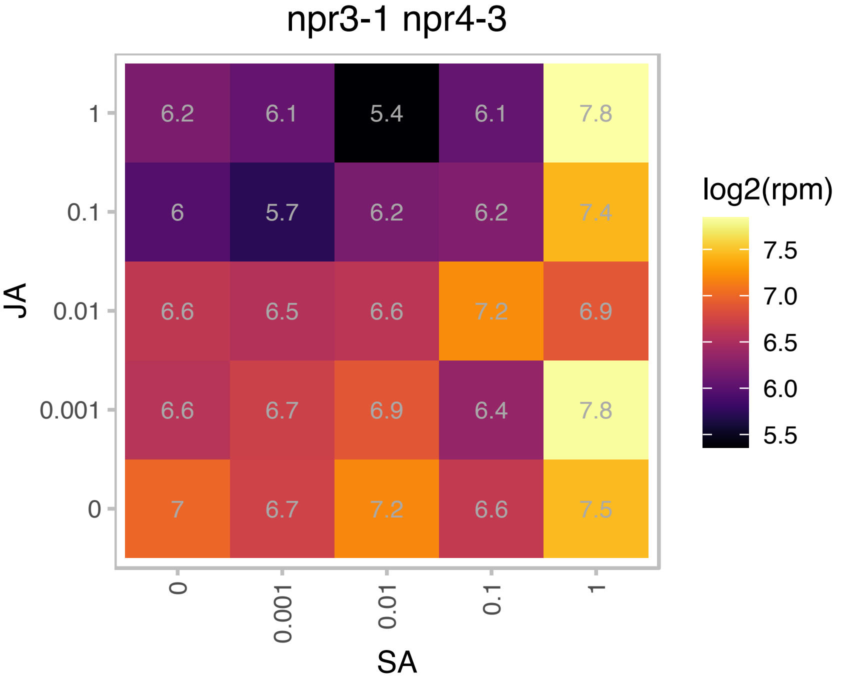

# AT1G05410.1 : NA

Protein of unknown function (DUF1423)

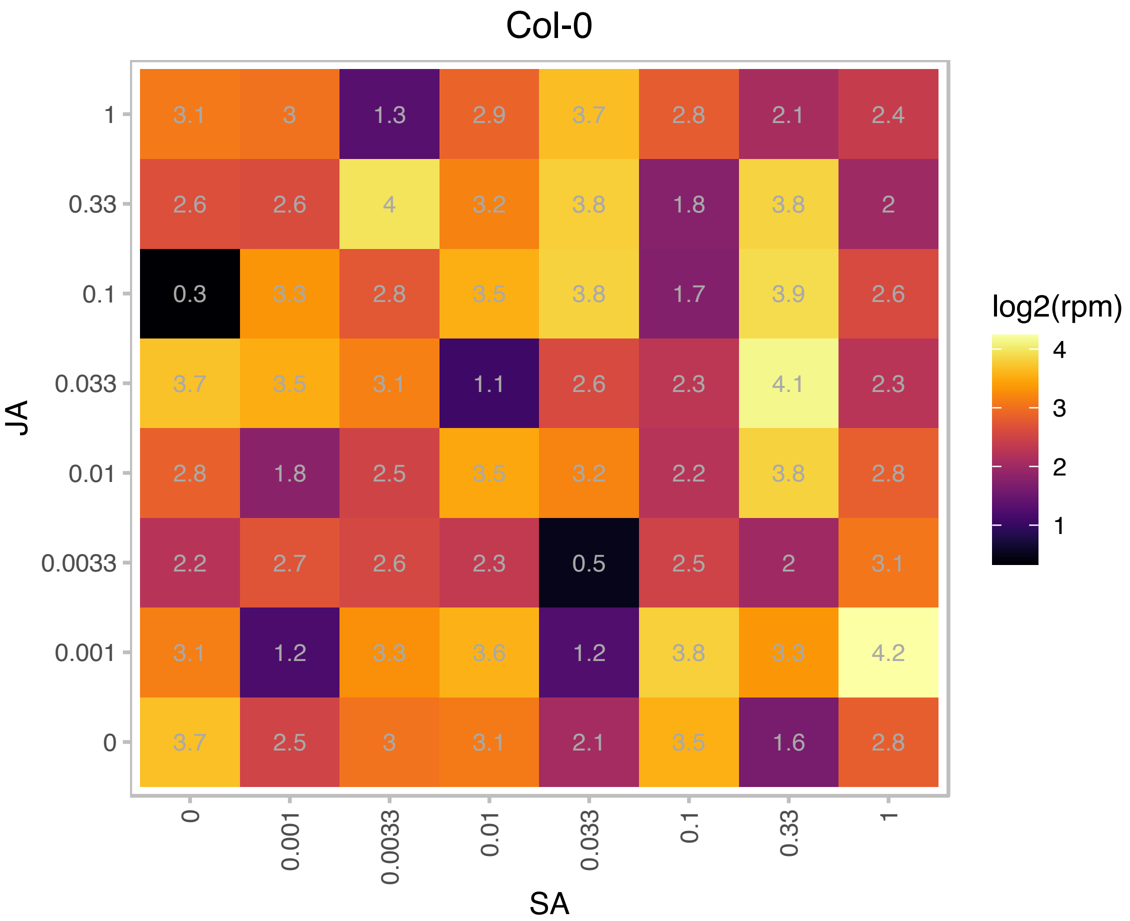


| | |
|---|---|
| Gene | AT1G05410 |
| is.expressed_genes | TRUE |
| is.responsive_genes | TRUE |
| is.apcluster | C-03 |

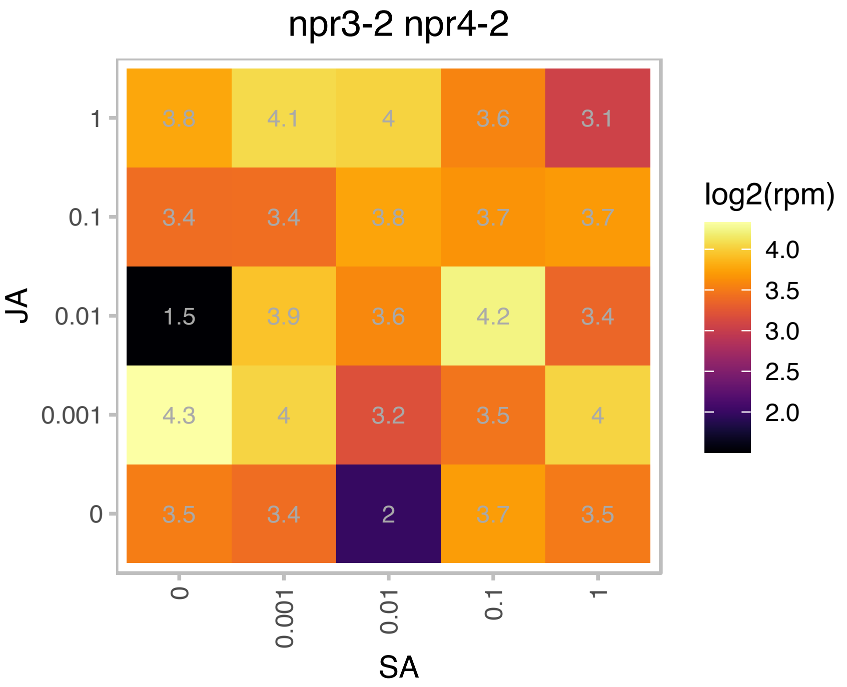


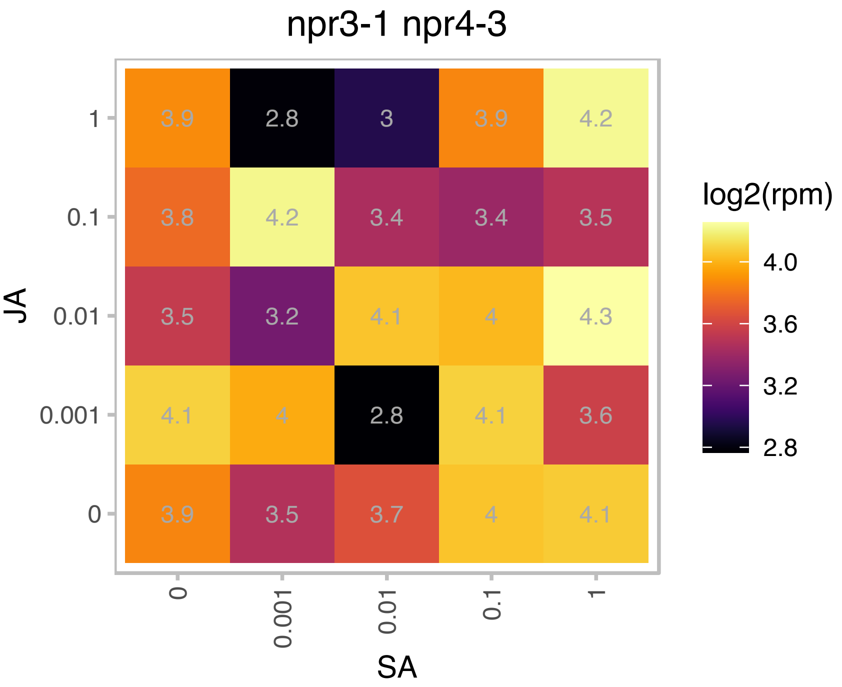

### AT1G06150.1 : EMB1444, LHL1

basic helix-loop-helix (bHLH) DNA-binding superfamily protein

Encodes a LHW-like protein with 79% amino acid identity to LHW.

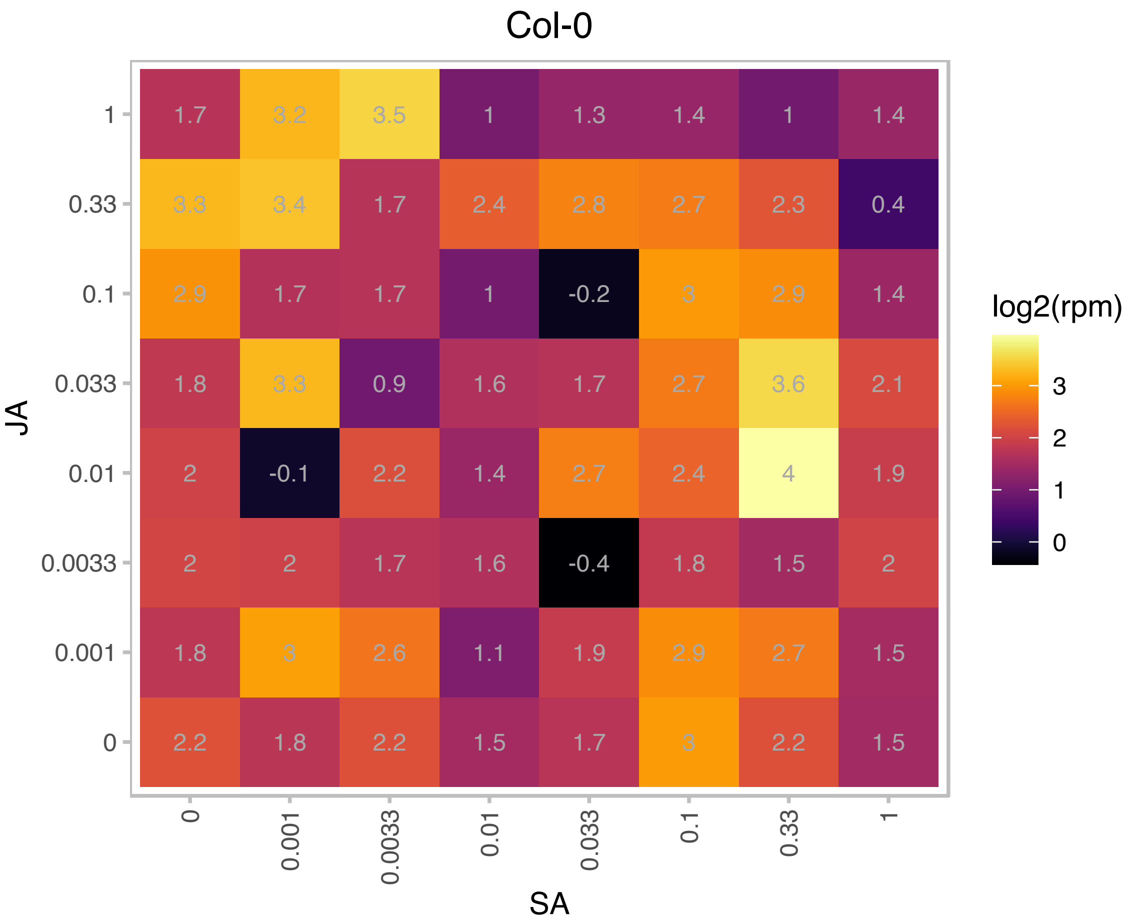


| | |
|---|---|
| Gene | AT1G06150 |
| is.expressed_genes | TRUE |
| is.responsive_genes | TRUE |
| is.apcluster | C-03 |

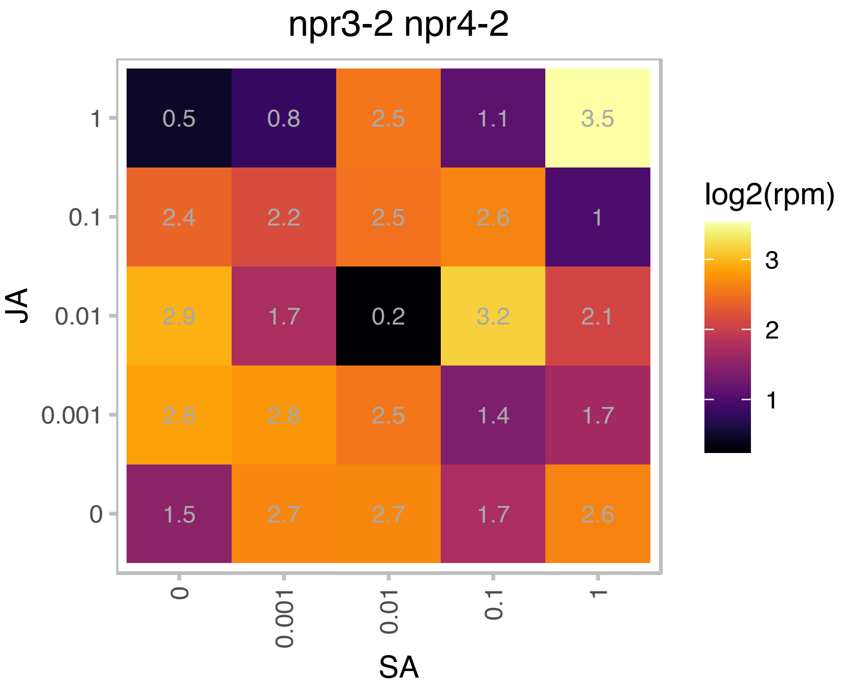


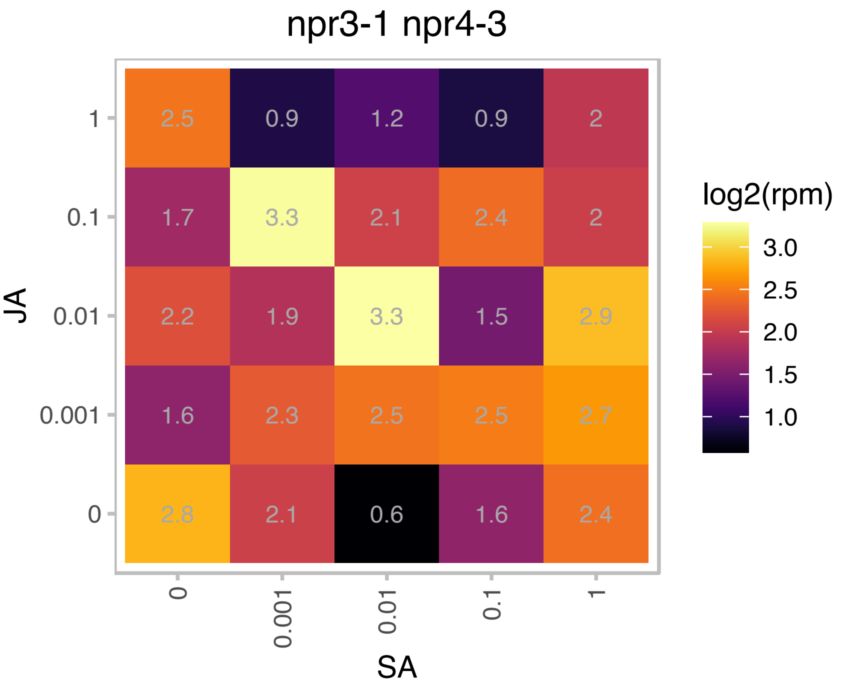

# AT1G06650.1 : NA

2-oxoglutarate (2OG) and Fe(II)-dependent oxygenase superfamily protein

encodes a protein whose sequence is similar to 2-oxoglutarate-dependent dioxygenase The mRNA is cell-to-cell mobile.

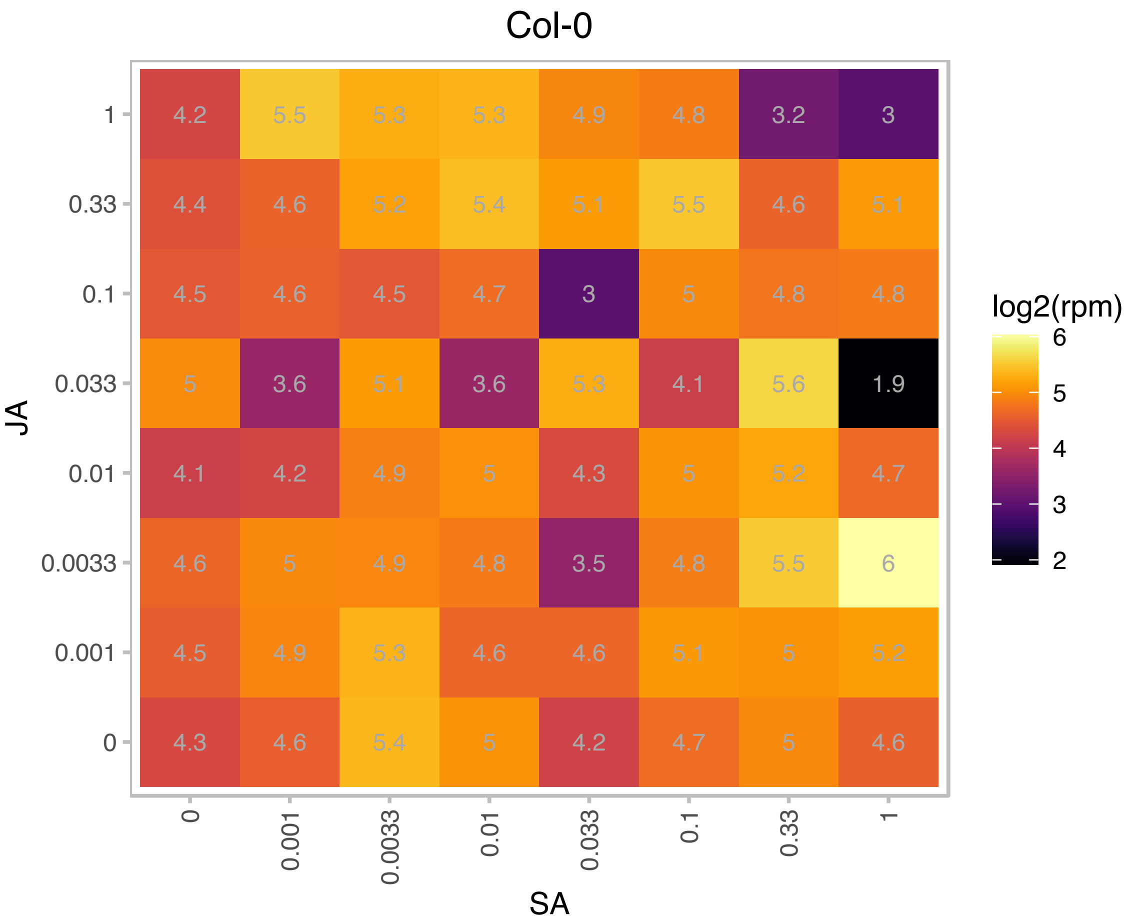


| | |
|---|---|
| Gene | AT1G06650 |
| is.expressed_genes | TRUE |
| is.responsive_genes | TRUE |
| is.apcluster | C-03 |

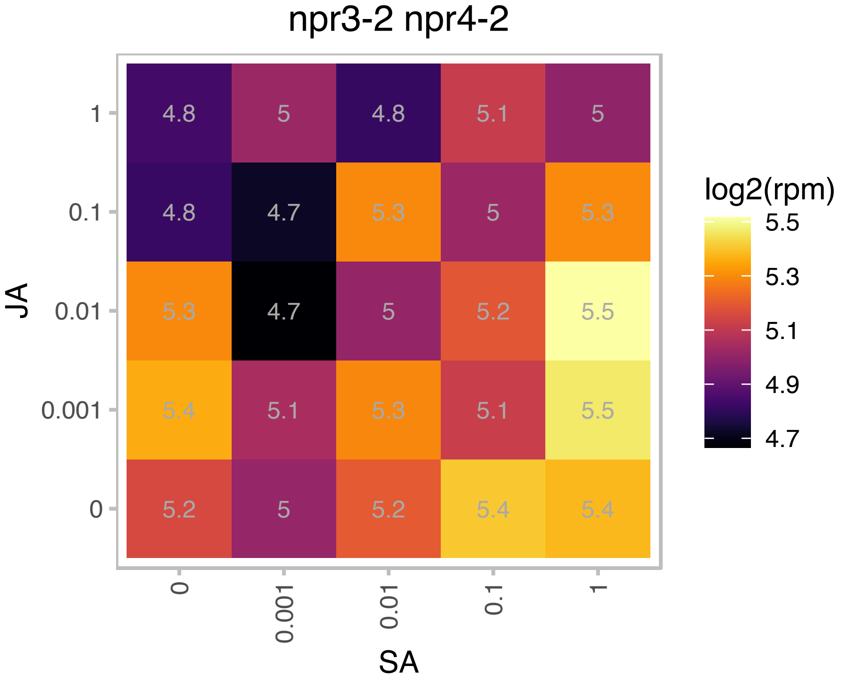


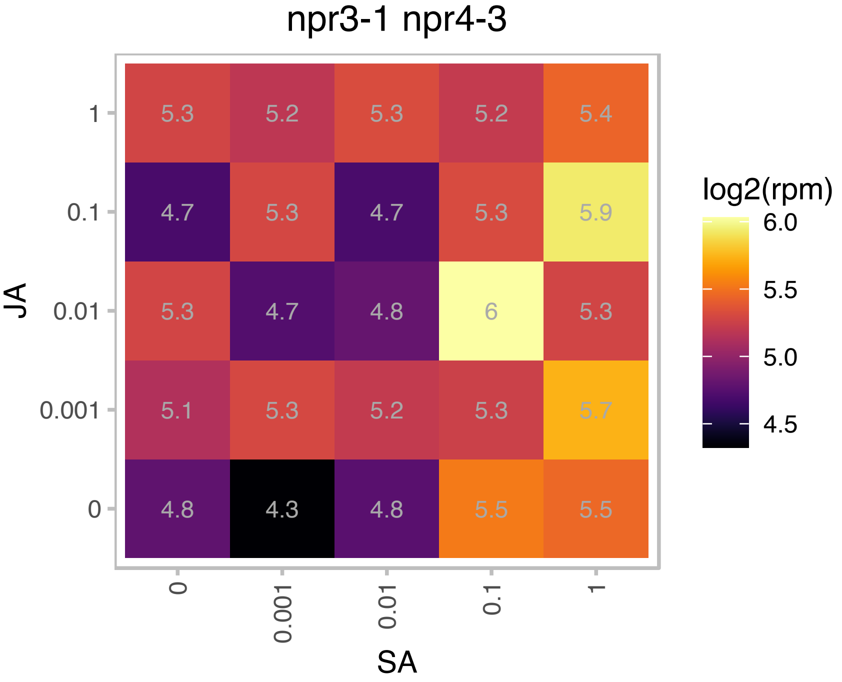

# AT1G18680.1 : NA

HNH endonuclease domain-containing protein

| | |
|---|---|
| Gene | AT1G18680 |
| is.expressed_genes | TRUE |
| is.responsive_genes | TRUE |
| is.apcluster | C-03 |

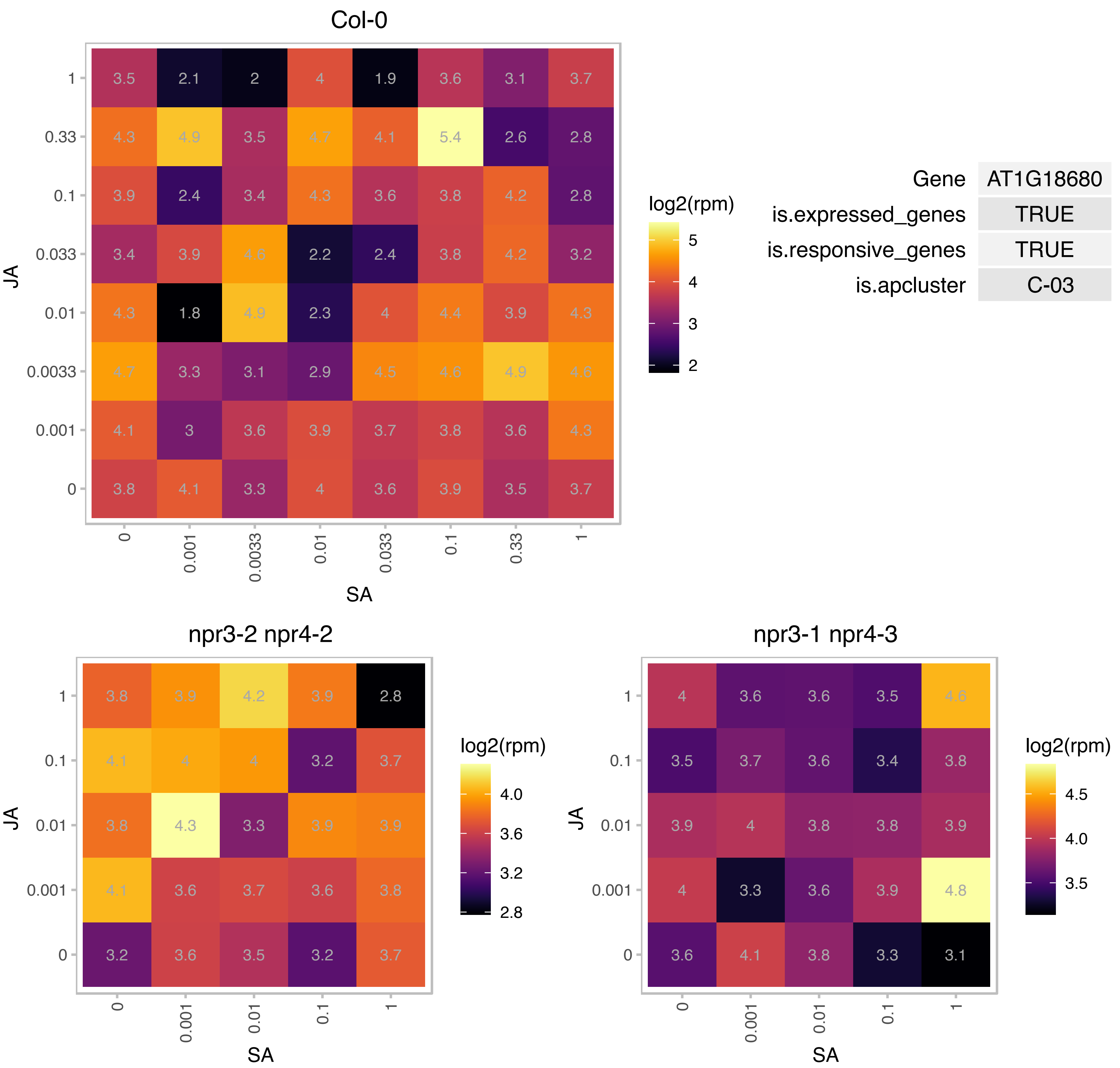

### AT1G20450.1 : ERD10, LTI29, LTI45

Dehydrin family protein

Encodes a gene induced by low temperature and dehydration. Inhibits e.coli growth while overexpressed. Belongs to the dehydrin protein family, which contains highly conserved stretches of 7-17 residues that are repetitively scattered in their sequences, the K-, S-, Y- and lysine rich segments. LTI29 and LTI30 double overexpressors confer cold tolerance. Localized to membranes and cytoplasm.

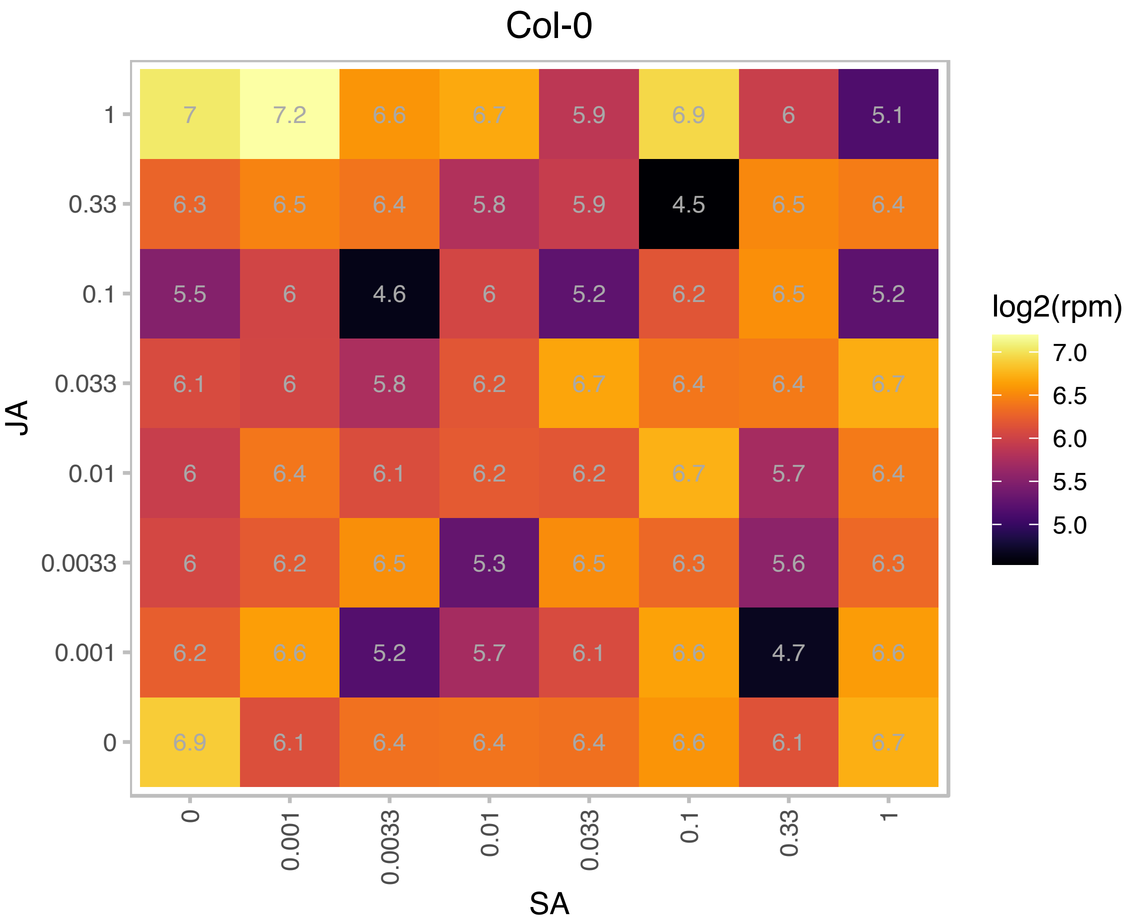


| | |
|---|---|
| Gene | AT1G20450 |
| is.expressed_genes | TRUE |
| is.responsive_genes | TRUE |
| is.apcluster | C-03 |

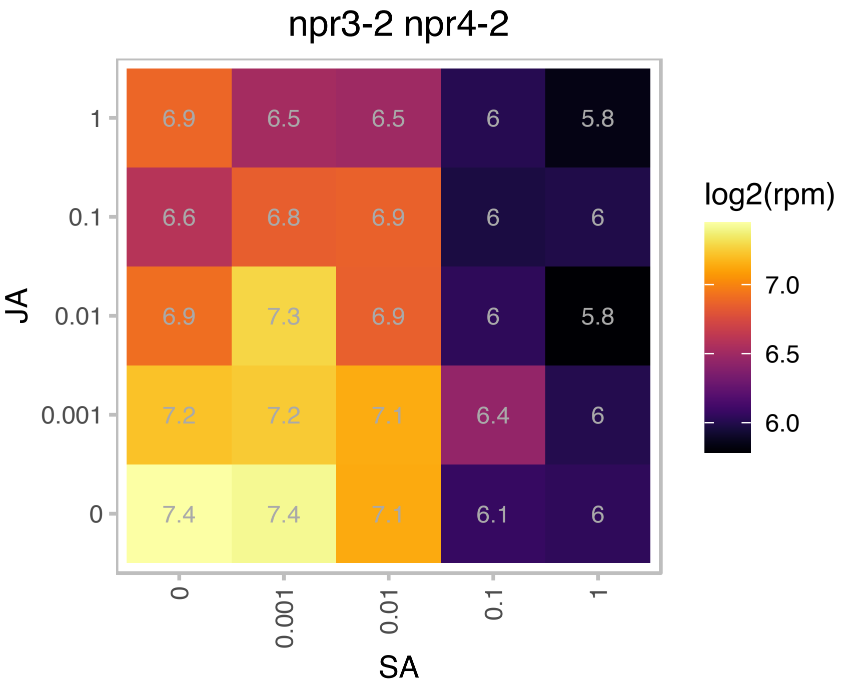


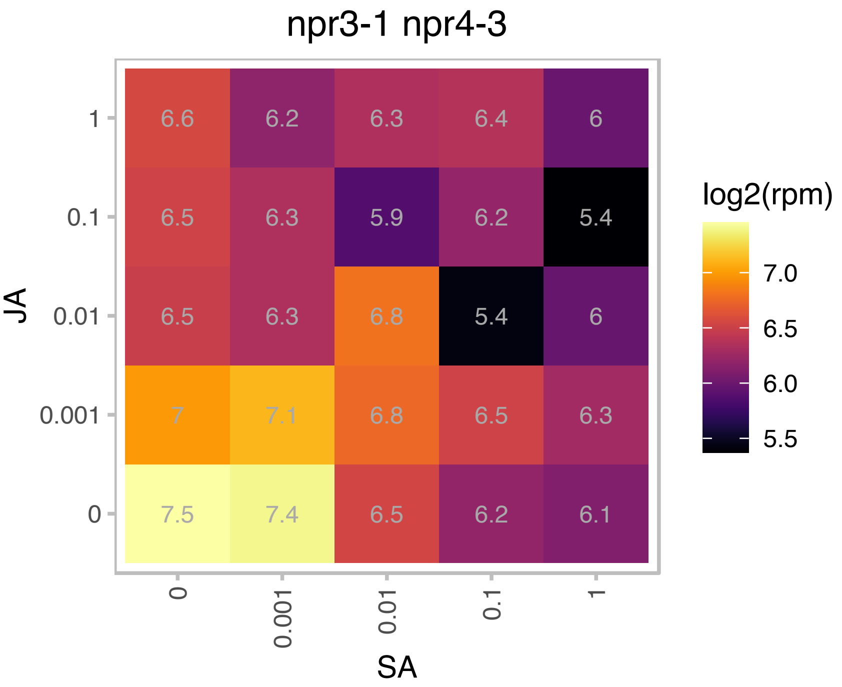

# AT1G52590.1 : NA

Putative thiol-disulphide oxidoreductase DCC

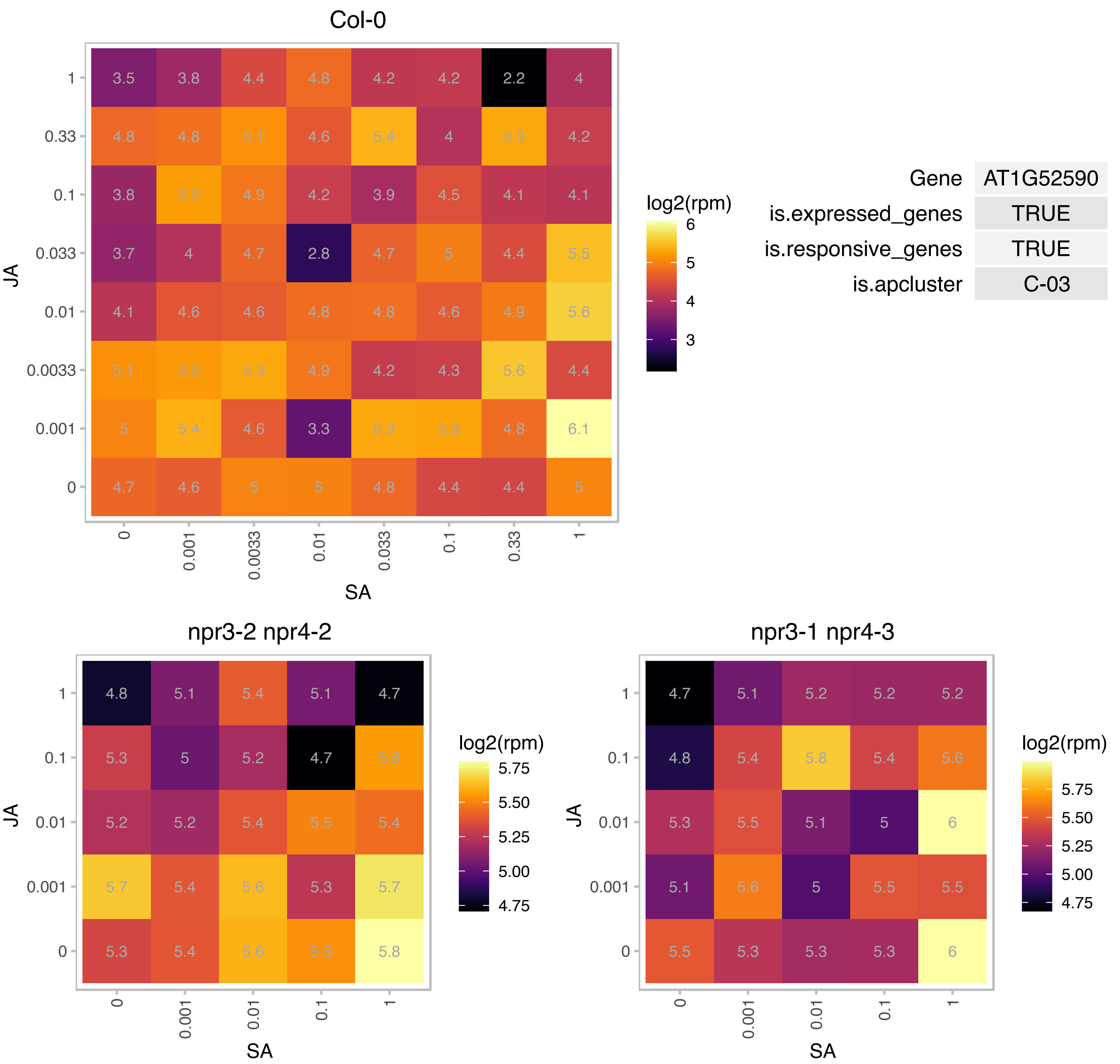


| | |
|---|---|
| Gene | AT1G52590 |
| is.expressed_genes | TRUE |
| is.responsive_genes | TRUE |
| is.apcluster | C-03 |

### AT1G63780.1 : IMP4

Ribosomal RNA processing Brix domain protein

Small nucleolar ribonucleoprotein protein involved in ribosomal RNA processing. Located in nucleolus and cajal bodies.

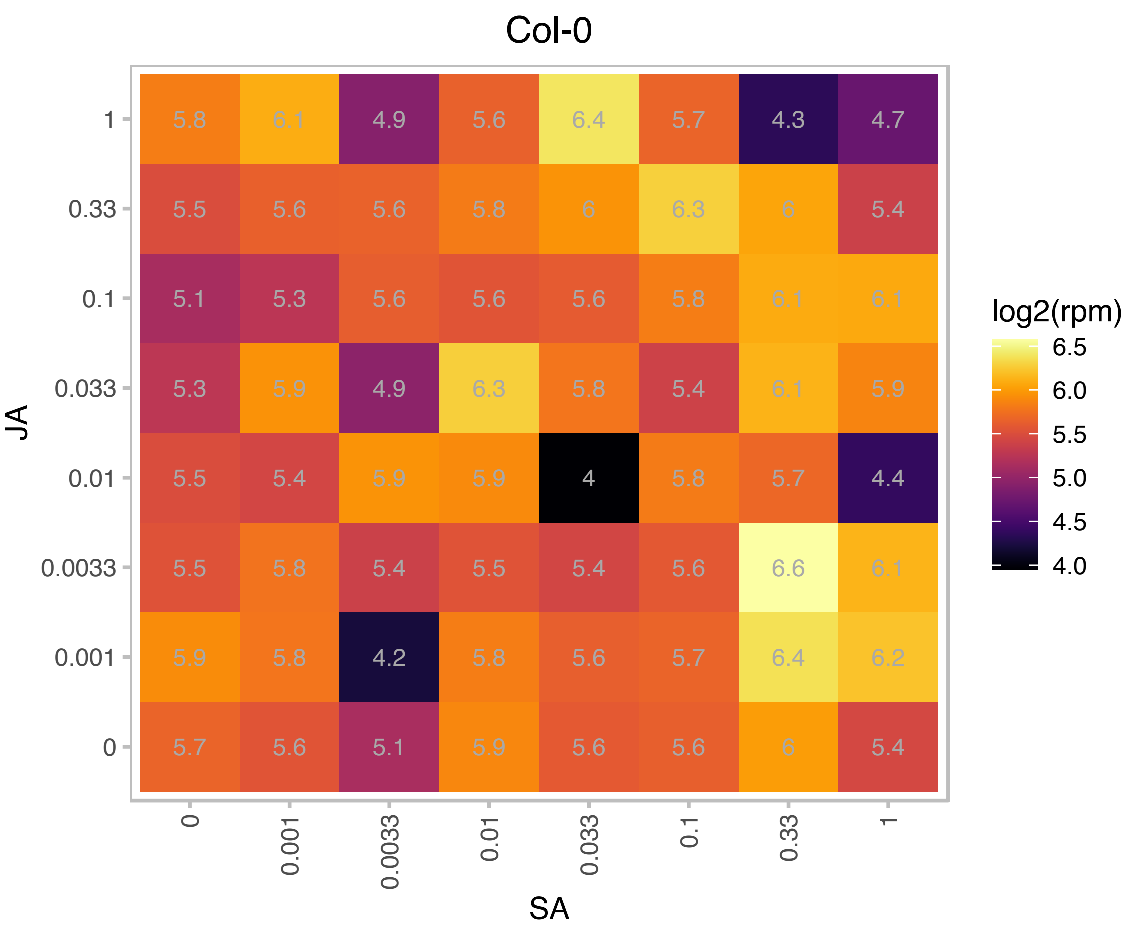


| | |
|---|---|
| Gene | AT1G63780 |
| is.expressed_genes | TRUE |
| is.responsive_genes | TRUE |
| is.apcluster | C-03 |

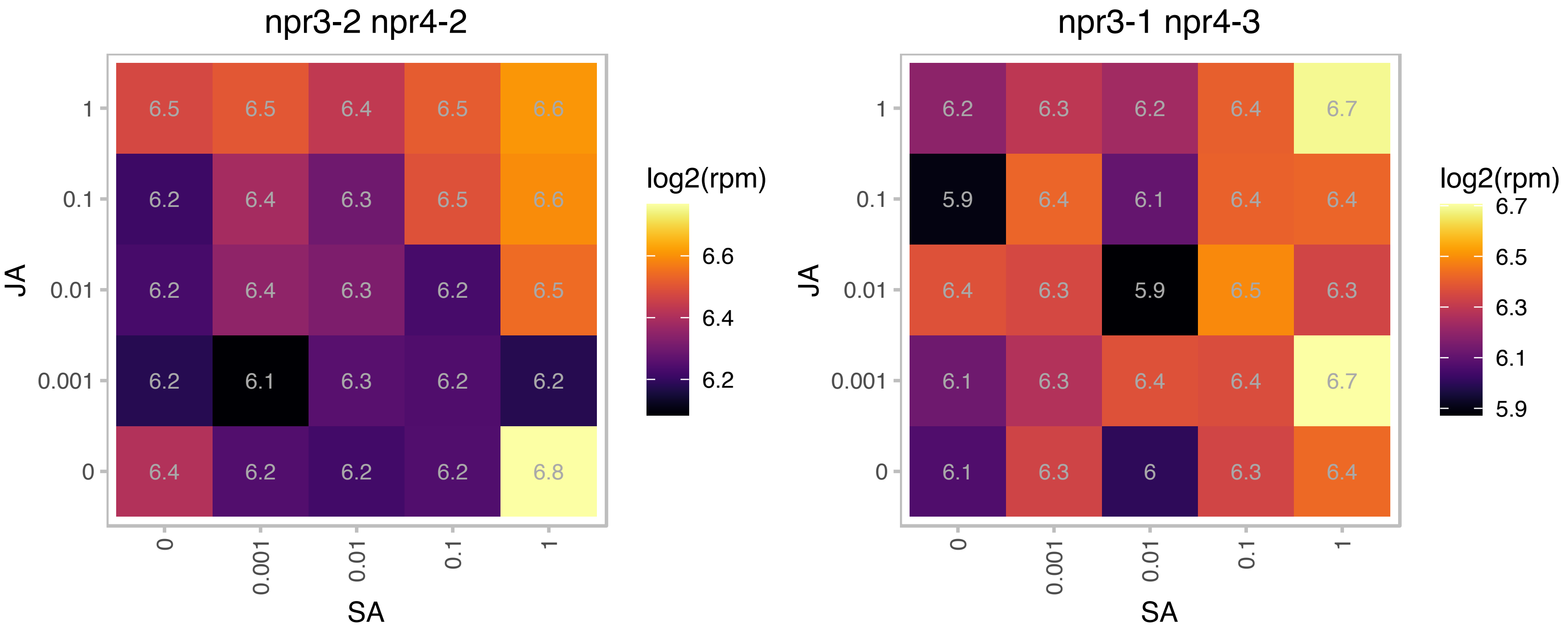

### AT1G65700.1 : AtLSM8, LSM8

Small nuclear ribonucleoprotein family protein

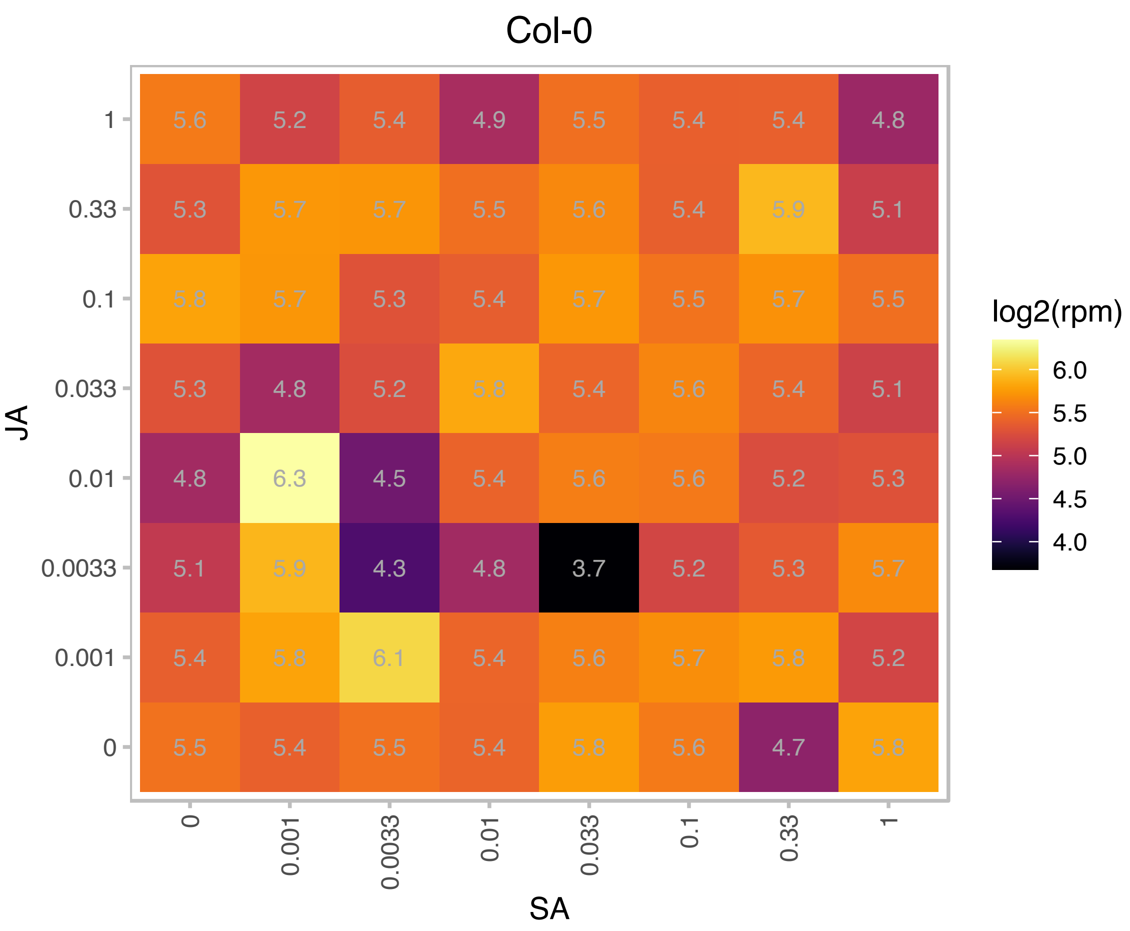


| | |
|---|---|
| Gene | AT1G65700 |
| is.expressed_genes | TRUE |
| is.responsive_genes | TRUE |
| is.apcluster | C-03 |

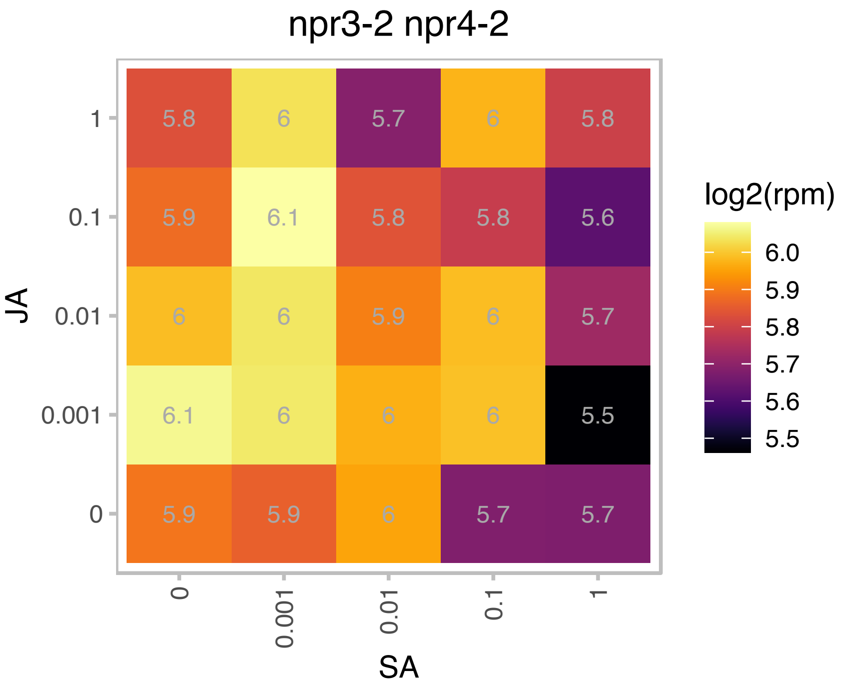


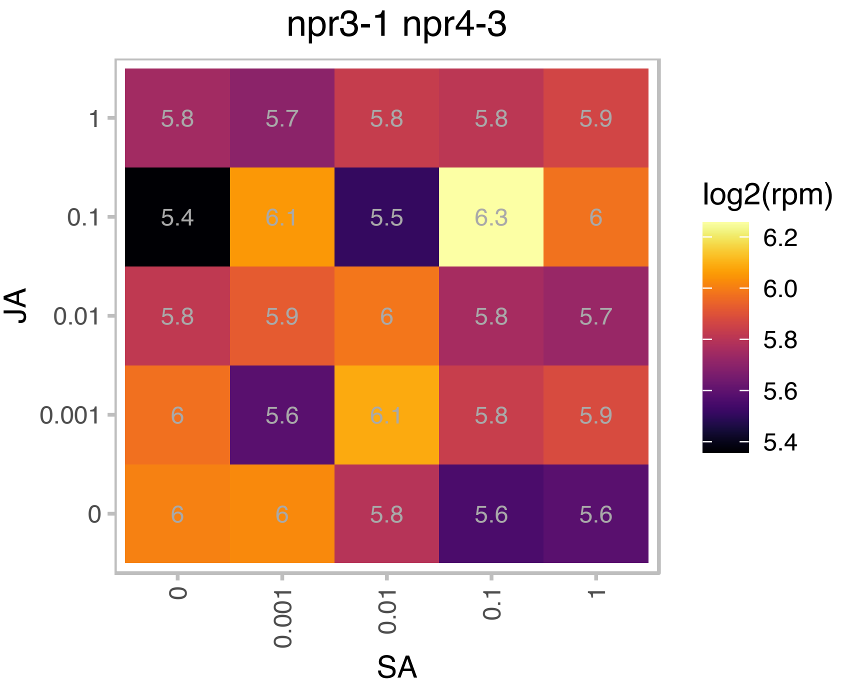

### AT2G02560.1 : ATCAND1, CAND1, ETA2, HVE, TIP120

Arabidopsis thaliana homolog of human CAND1 (cullin-associated and neddylation-dissociated). Putative cullin-associated and neddylation dissociated similarity to TBP-interacting protein TIP120. Ubiquitously expressed in plant tissues throughout development. T-DNA insertion mutant plants were completely sterile and resistant to sirtinol and auxin, but not to gibberellins or brassinolide. Displayed developmental phenotypes similar to those of axr1, namely, short petioles, downwardly curling leaves, shorter inflorescence. Required for SCF function and appears to modulate SCF complex cycling. Physically interacts with CUL1. The mRNA is cell-to-cell mobile.

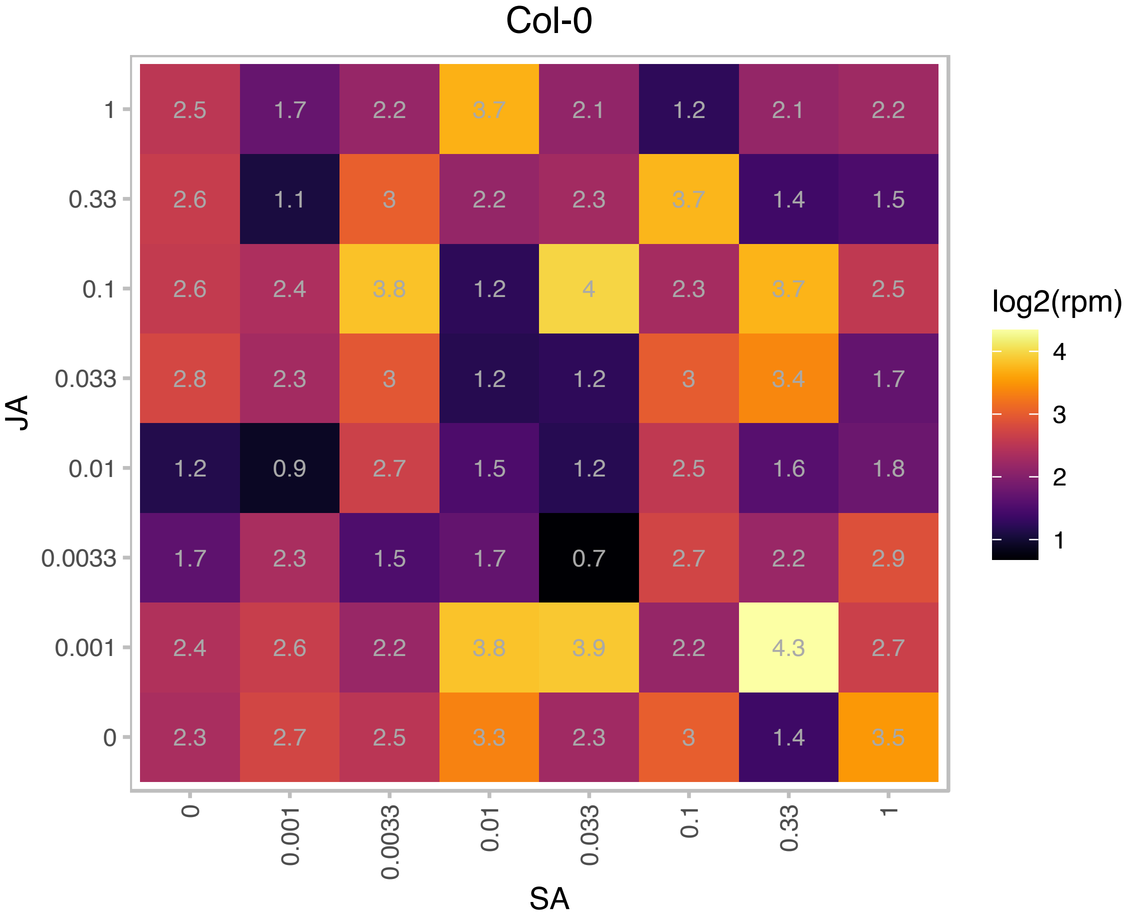


| | |
|---|---|
| Gene | AT2G02560 |
| is.expressed_genes | TRUE |
| is.responsive_genes | TRUE |
| is.apcluster | C-03 |

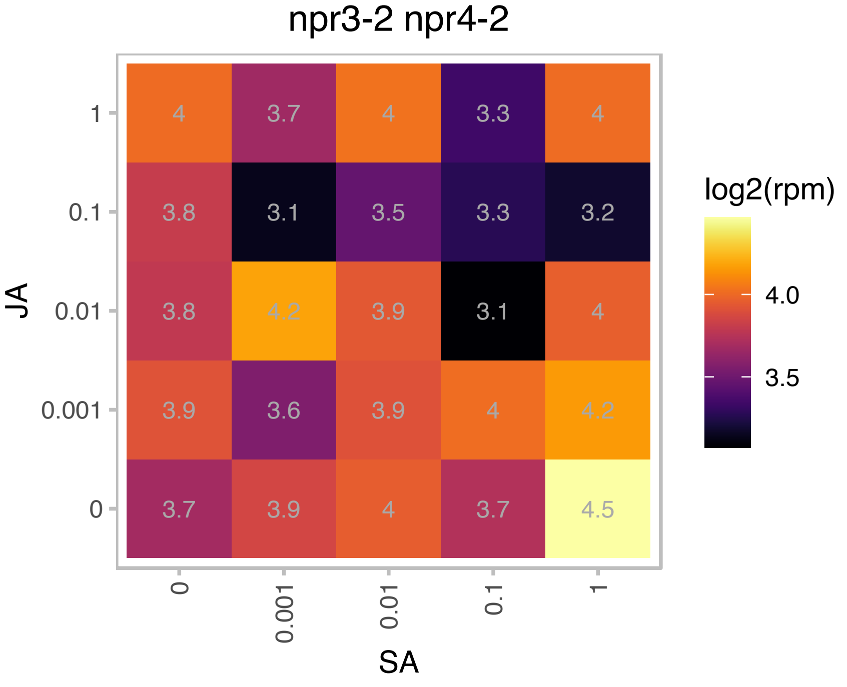


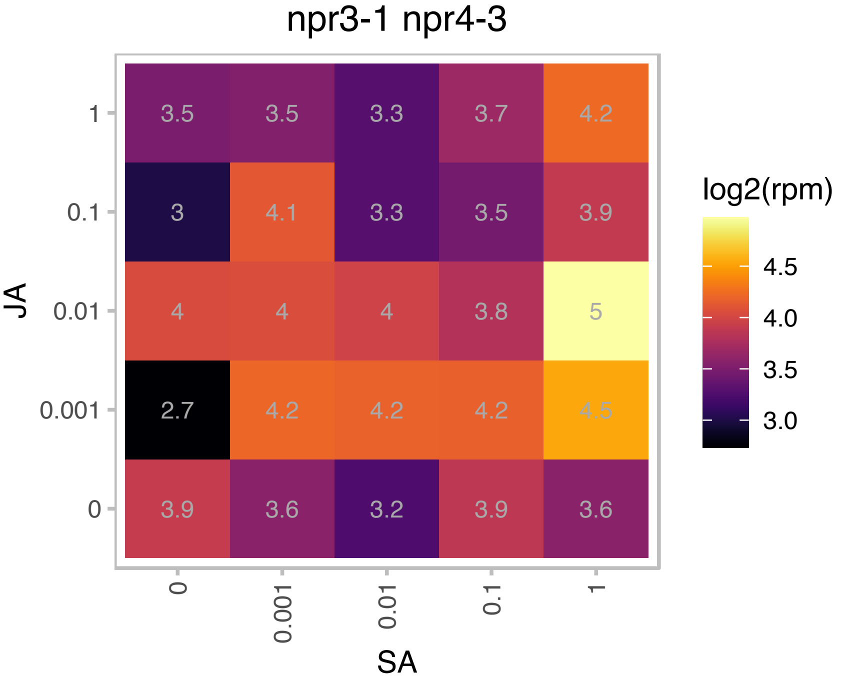

### AT2G21280.1 : ATSULA, GC1, SULA

NAD(P)-binding Rossmann-fold superfamily protein
A nuclear-encoded, plastid-targeted protein (AtSulA) whose overexpression causes severe yet stochastic plastid (shown in chloroplasts and leucoplasts) division defects. The protein does not appear to interact with either AtFtsZ proteins when studied in a yeast two-hybrid system.

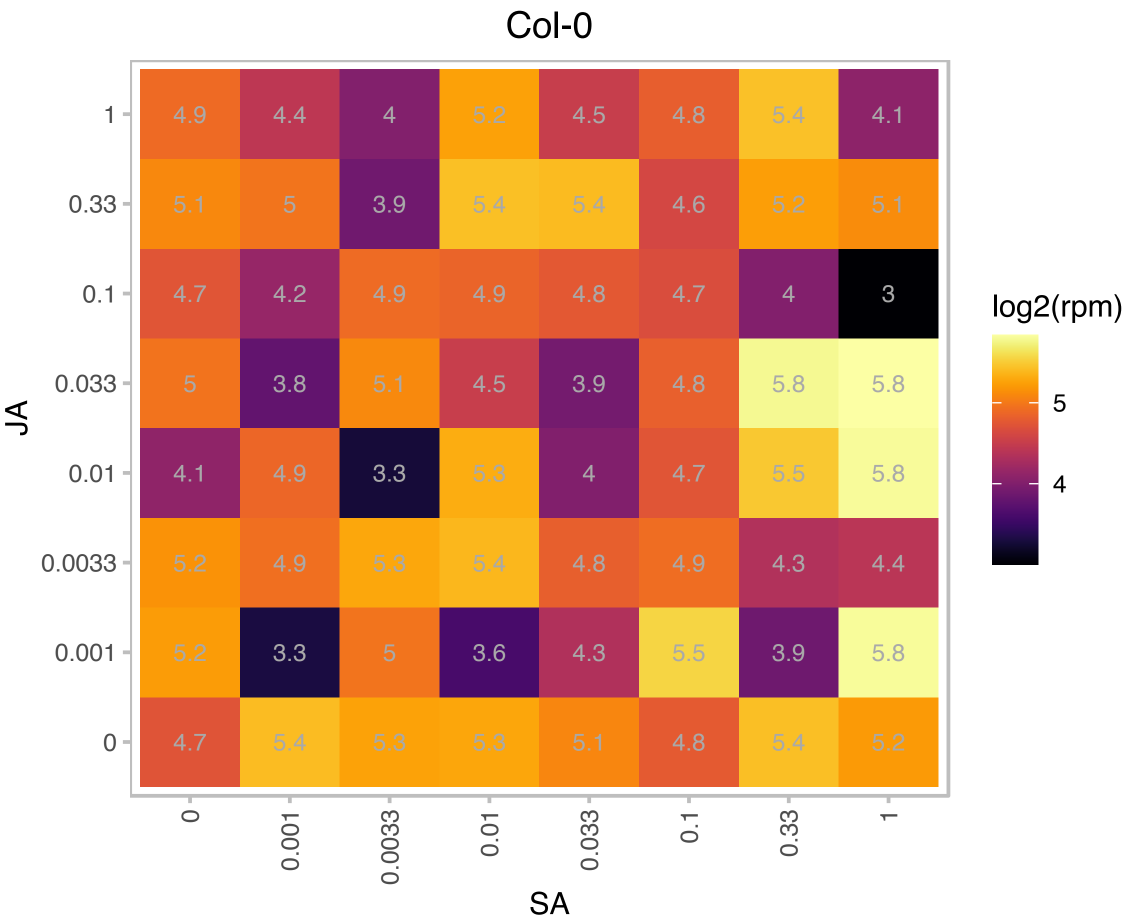


| | |
|---|---|
| Gene | AT2G21280 |
| is.expressed_genes | TRUE |
| is.responsive_genes | TRUE |
| is.apcluster | C-03 |

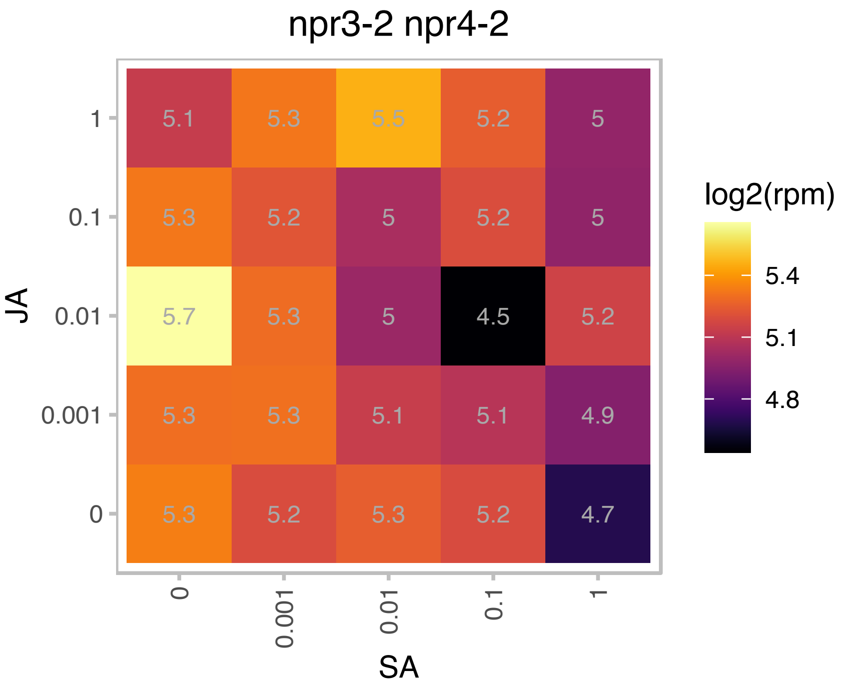


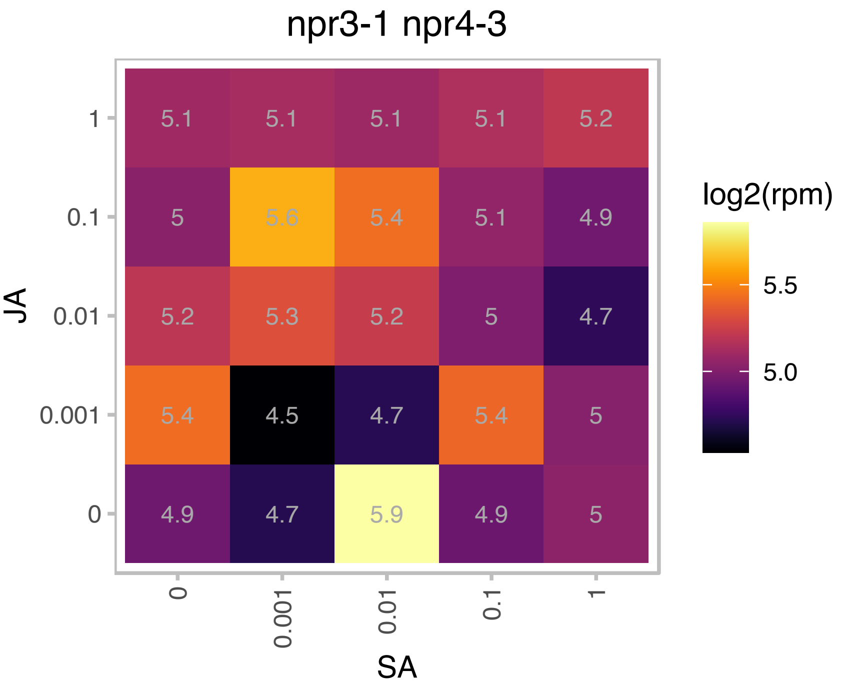

### AT2G22470.1 : AGP2, ATAGP2

arabinogalactan protein 2

Encodes arabinogalactan-protein (AGP2).

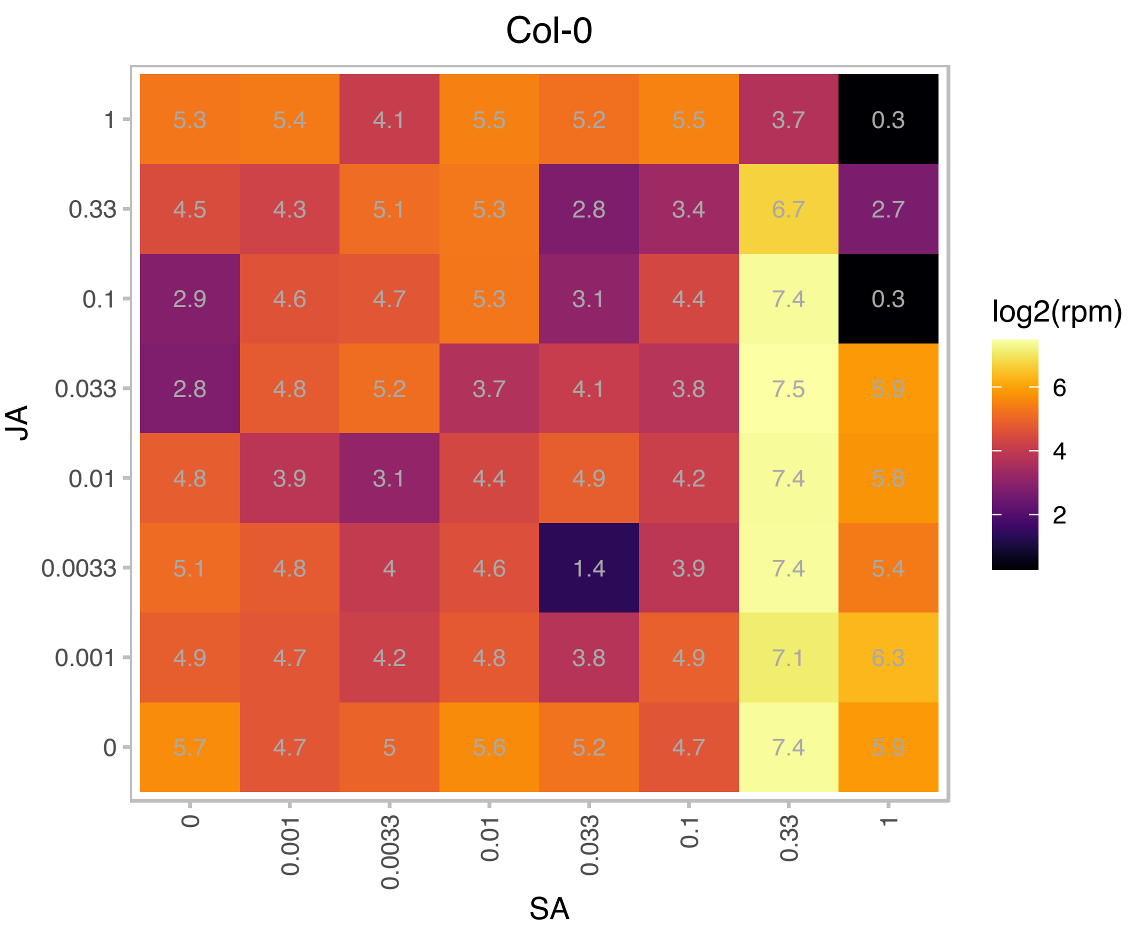


| | |
|---|---|
| Gene | AT2G22470 |
| is.expressed_genes | TRUE |
| is.responsive_genes | TRUE |
| is.apcluster | C-03 |

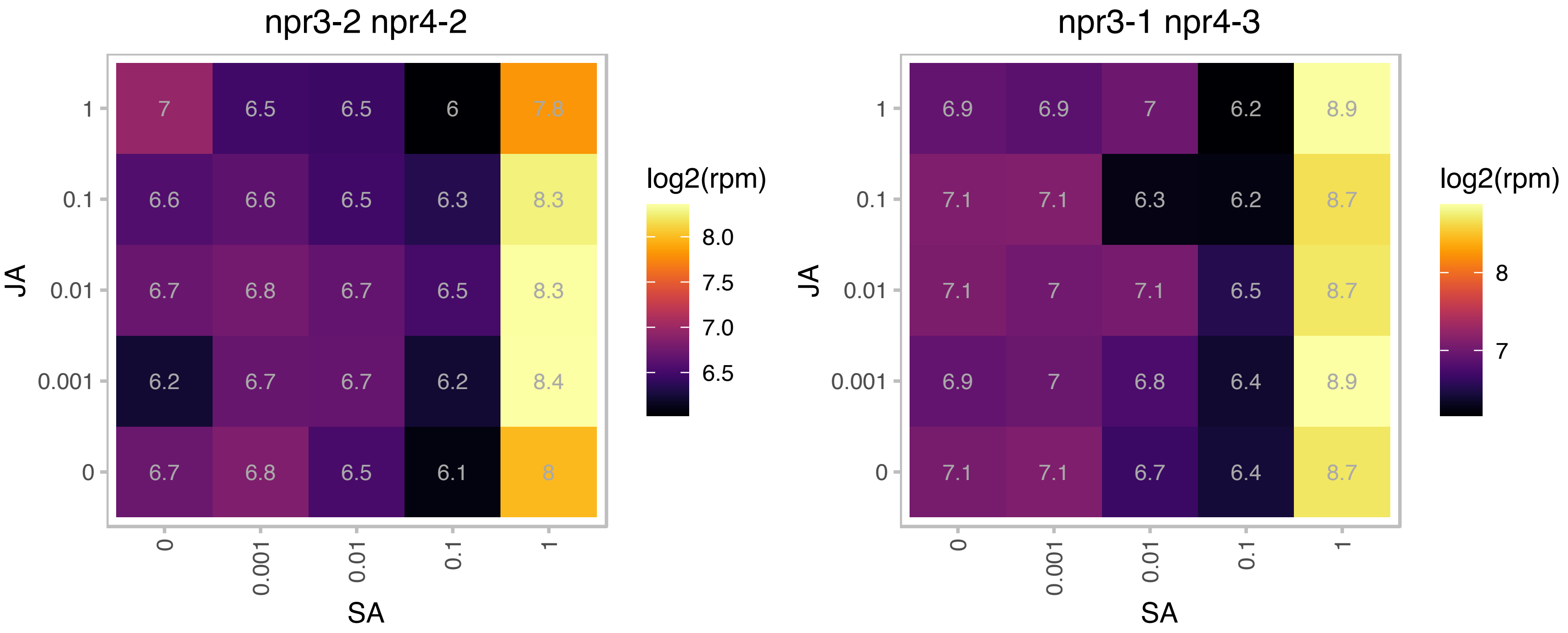

# AT2G25610.1 : NA

ATPase, F0/V0 complex, subunit C protein

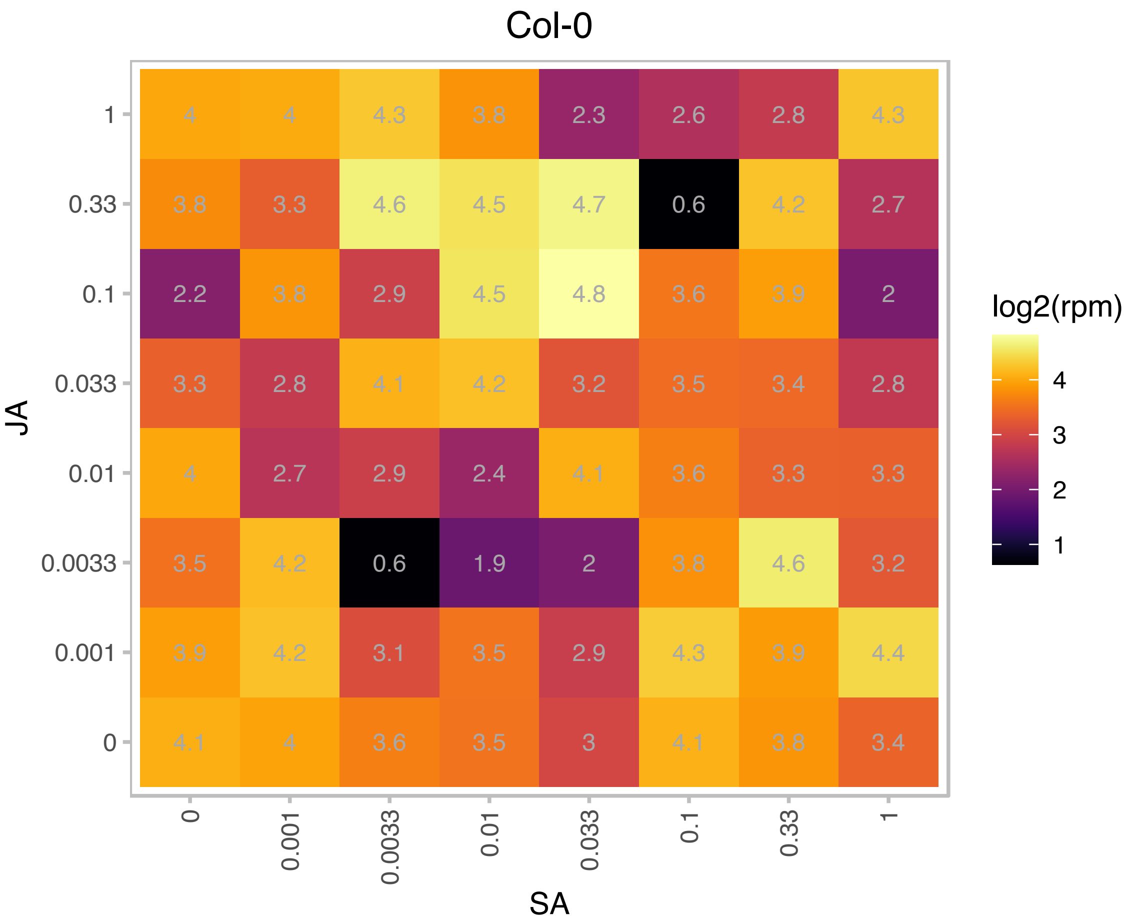


| | |
|---|---|
| Gene | AT2G25610 |
| is.expressed_genes | TRUE |
| is.responsive_genes | TRUE |
| is.apcluster | C-03 |

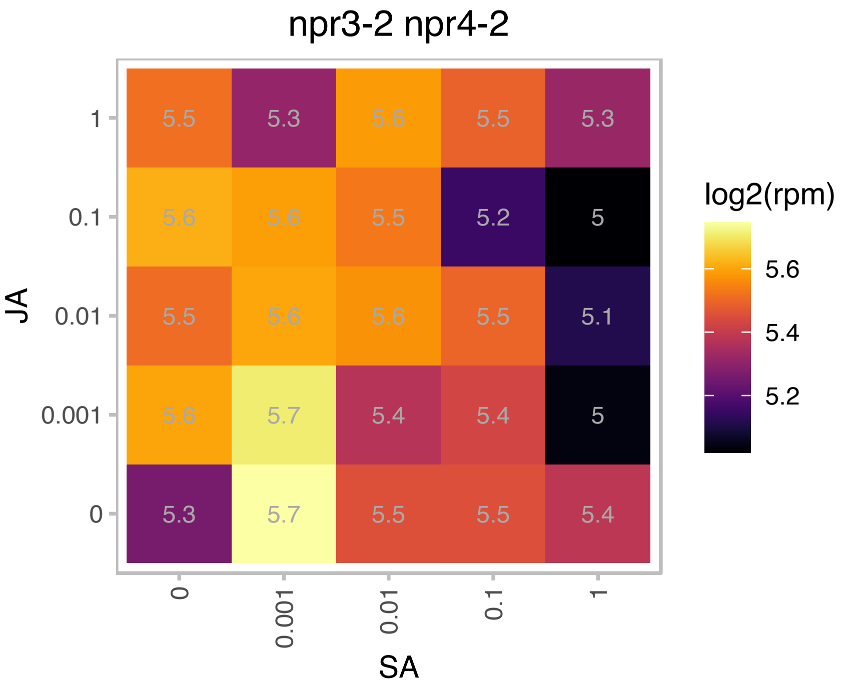


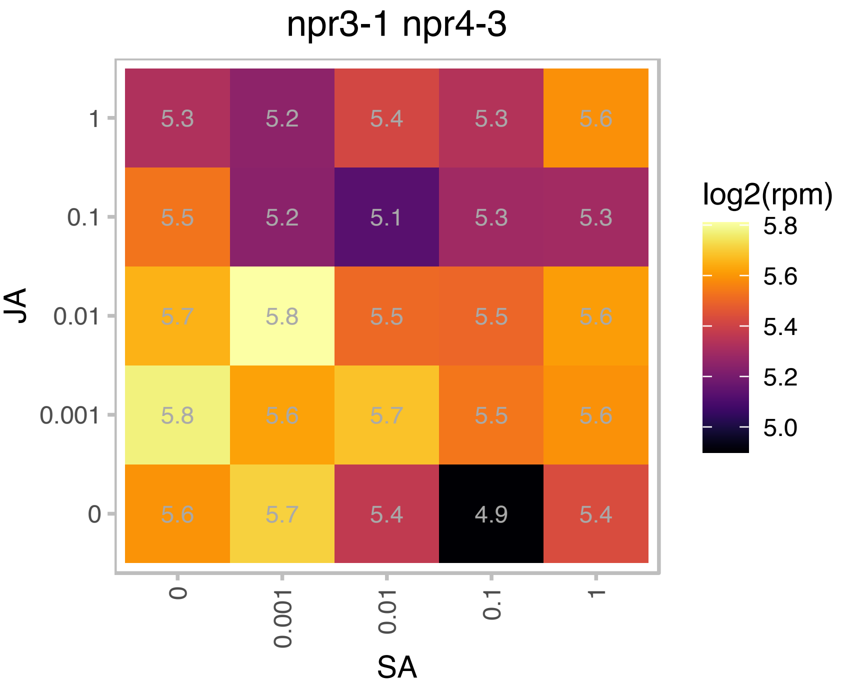

### AT2G33740.1 : CUTA

Nitrogen regulatory PII-like, alpha/beta

encodes a copper binding protein that forms tetramers in vitro. Gene is expressed in all tissues examined and protein is localized to the chloroplast. The mRNA is cell-to-cell mobile.

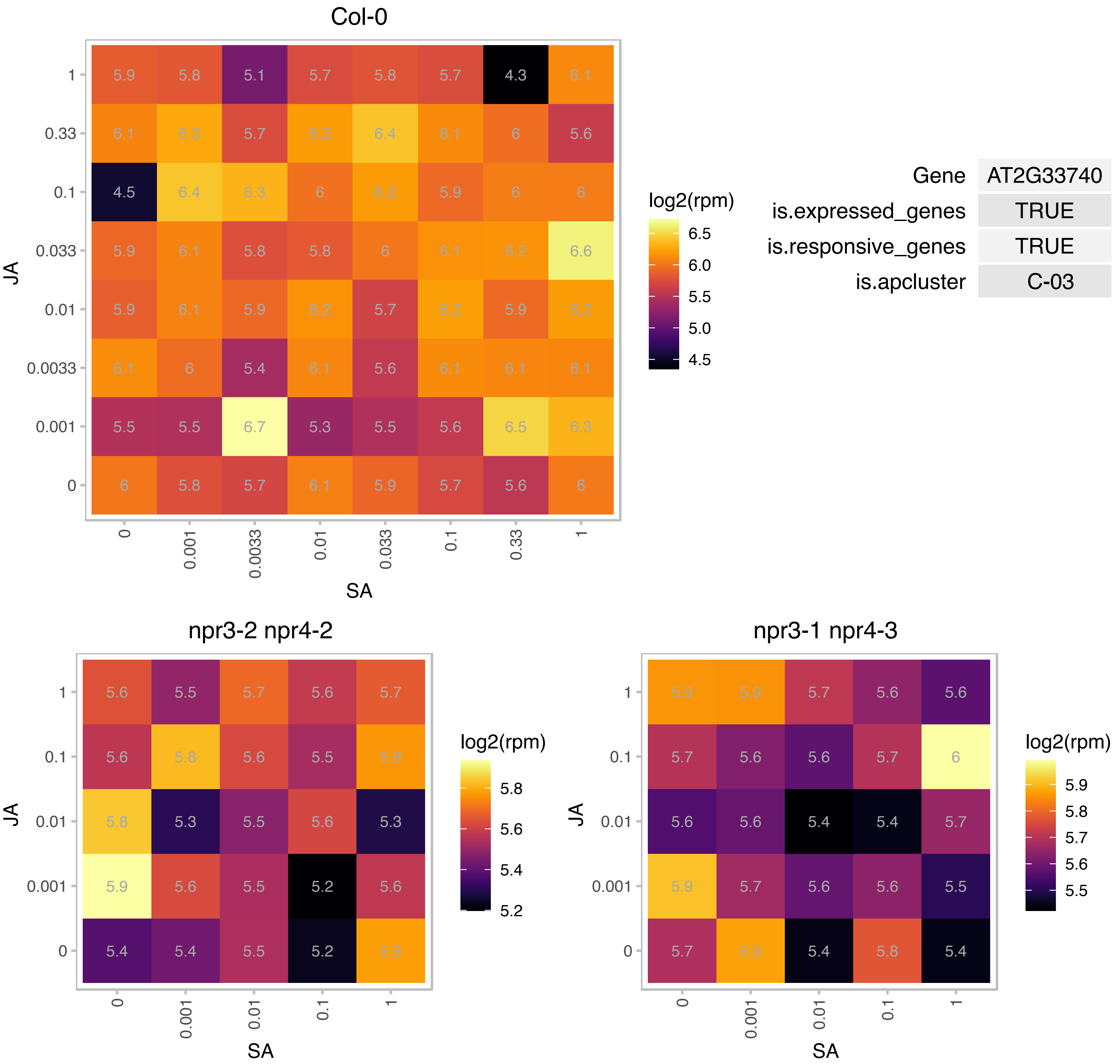


| | |
|---|---|
| Gene | AT2G33740 |
| is.expressed_genes | TRUE |
| is.responsive_genes | TRUE |
| is.apcluster | C-03 |

### AT2G42910.1 : AtPRS4, PRS4

Phosphoribosyltransferase family protein

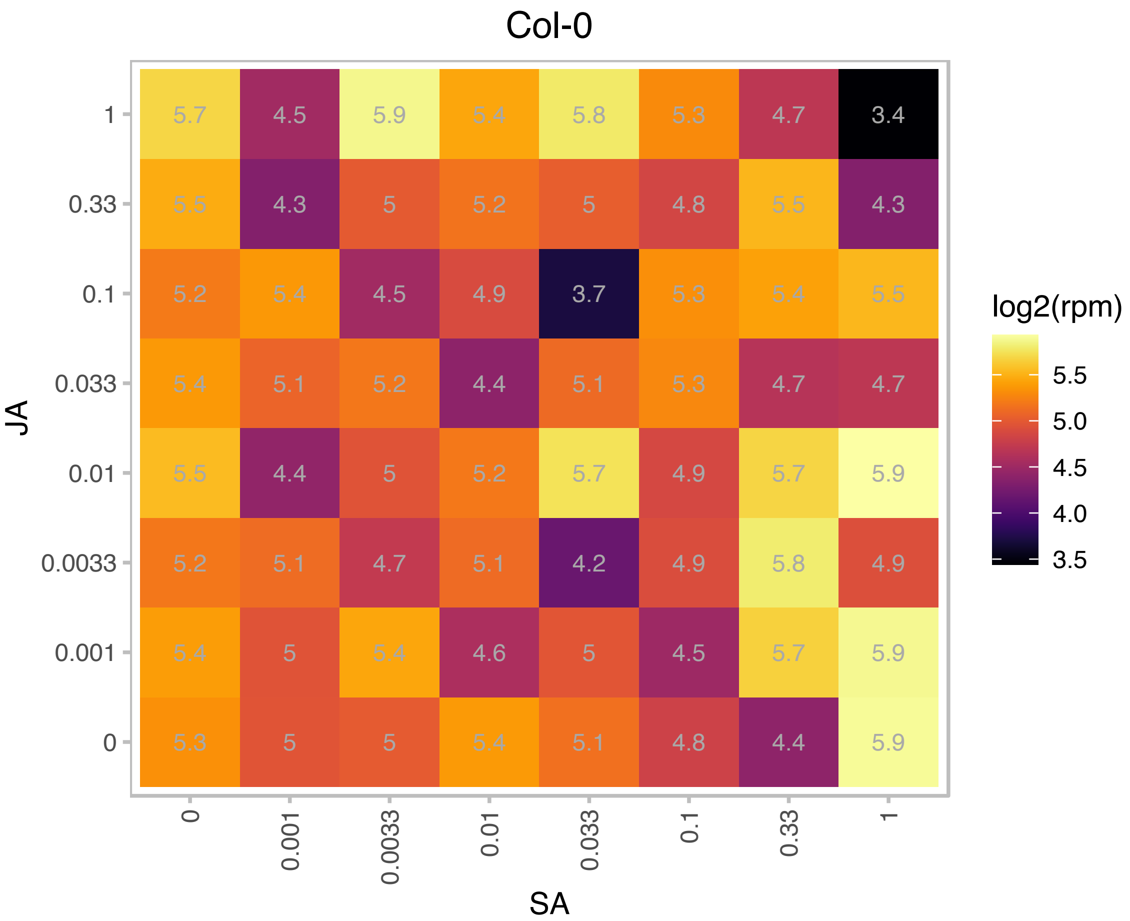


| | |
|---|---|
| Gene | AT2G42910 |
| is.expressed_genes | TRUE |
| is.responsive_genes | TRUE |
| is.apcluster | C-03 |

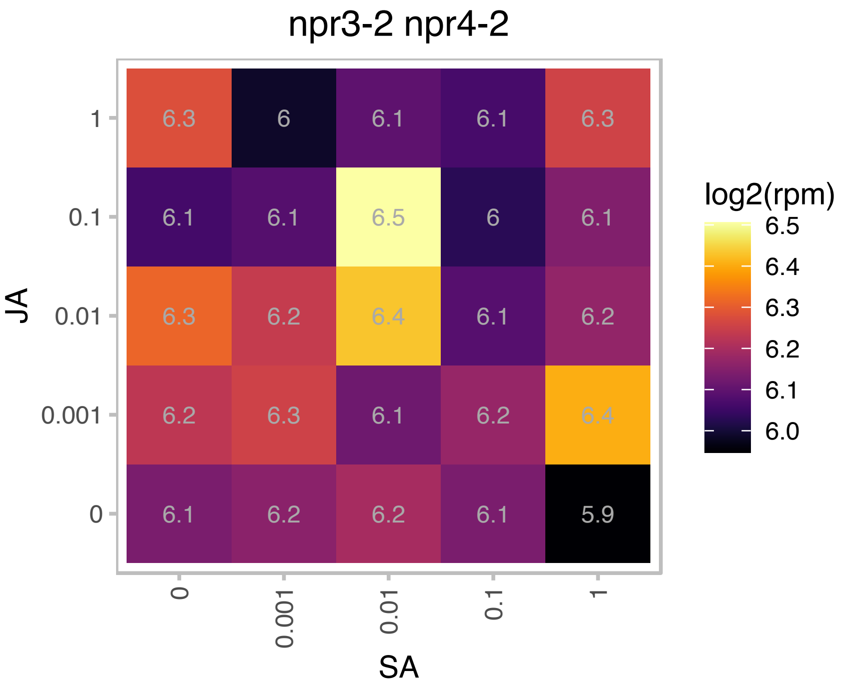


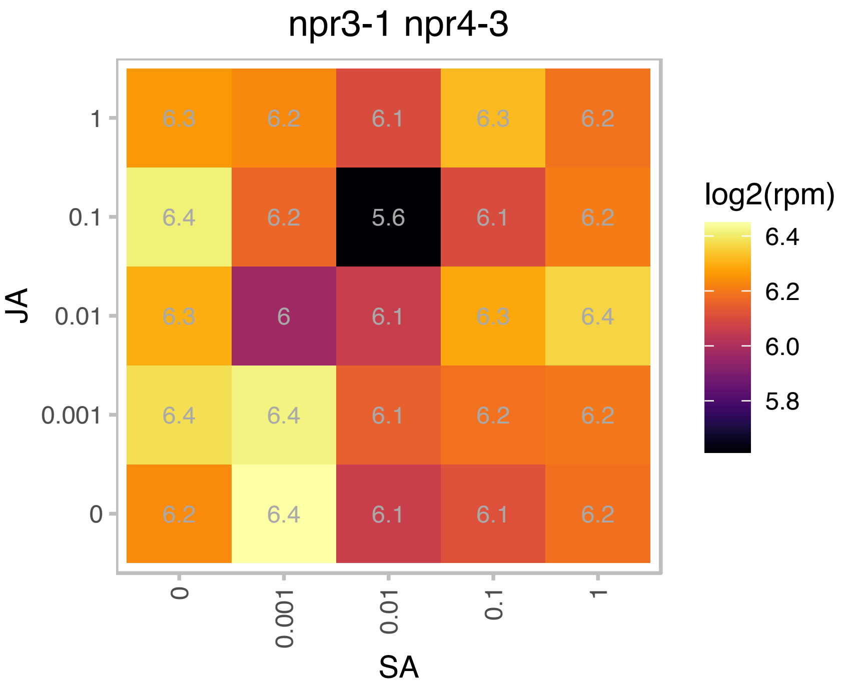

# AT3G02720.1 : DJ-1d, DJ1D

Class I glutamine amidotransferase-like superfamily protein
Encodes a glyoxalase with high activity towards glyoxal and methylglyoxal.  It differs from its animal and bacterial homologs with respect to the configuration of its catalytic residues and the oligomeric property of the enzyme. The mRNA is cell-to-cell mobile.

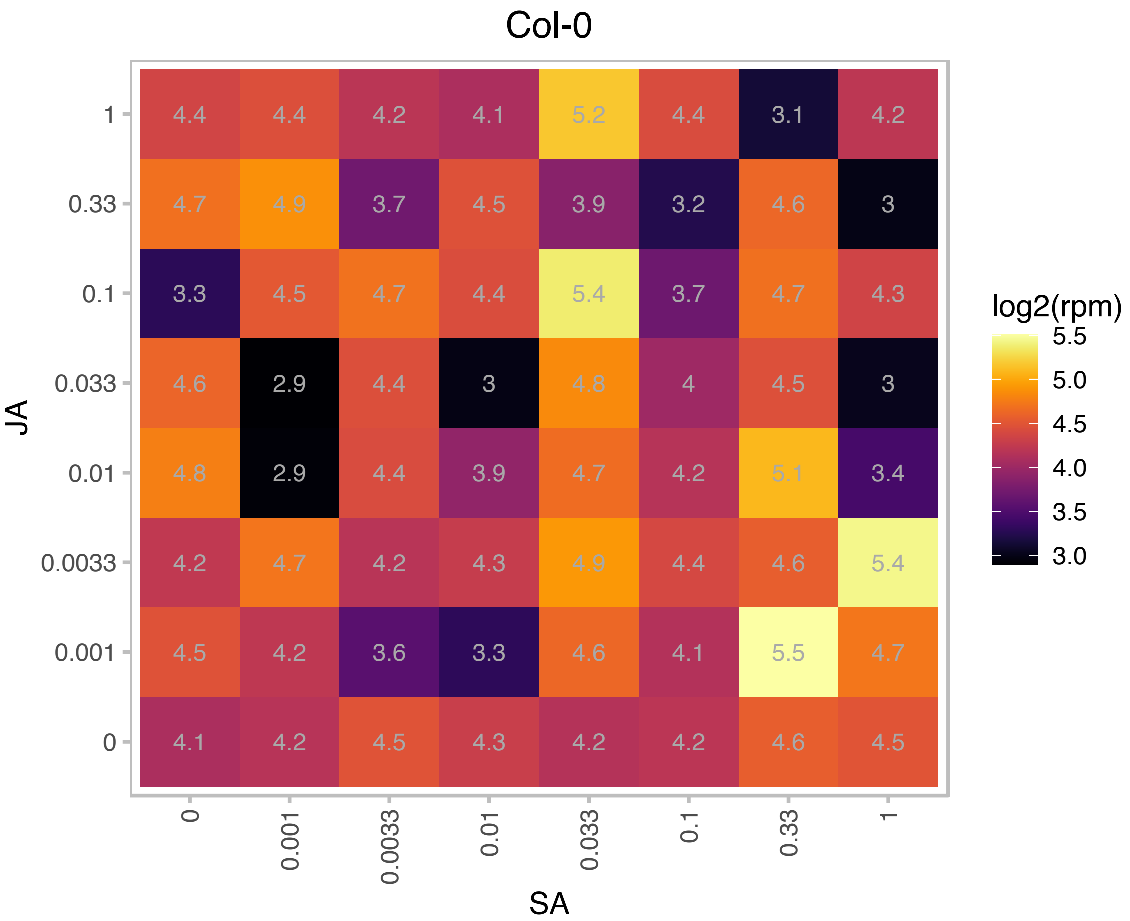


| | |
|---|---|
| Gene | AT3G02720 |
| is.expressed_genes | TRUE |
| is.responsive_genes | TRUE |
| is.apcluster | C-03 |

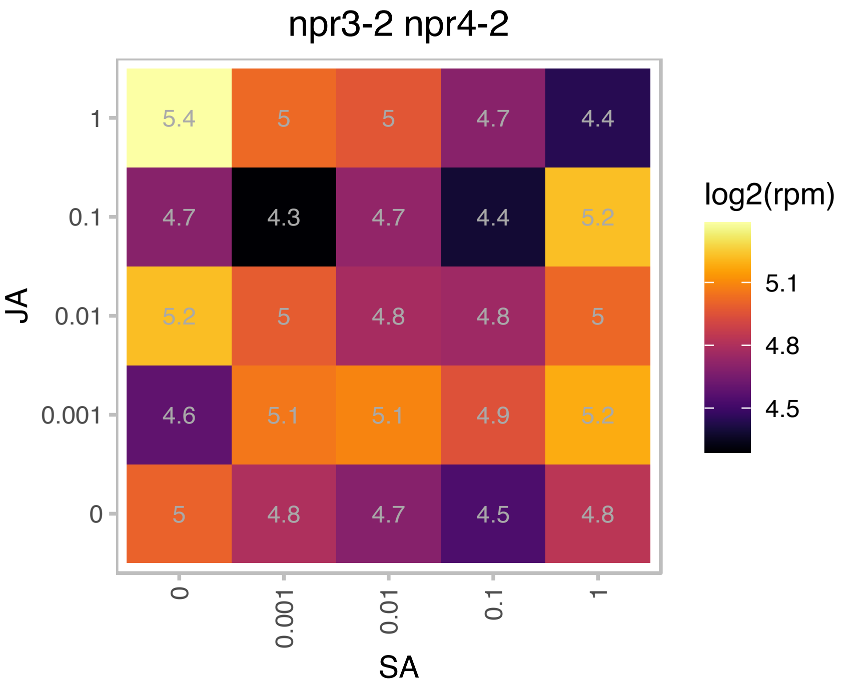


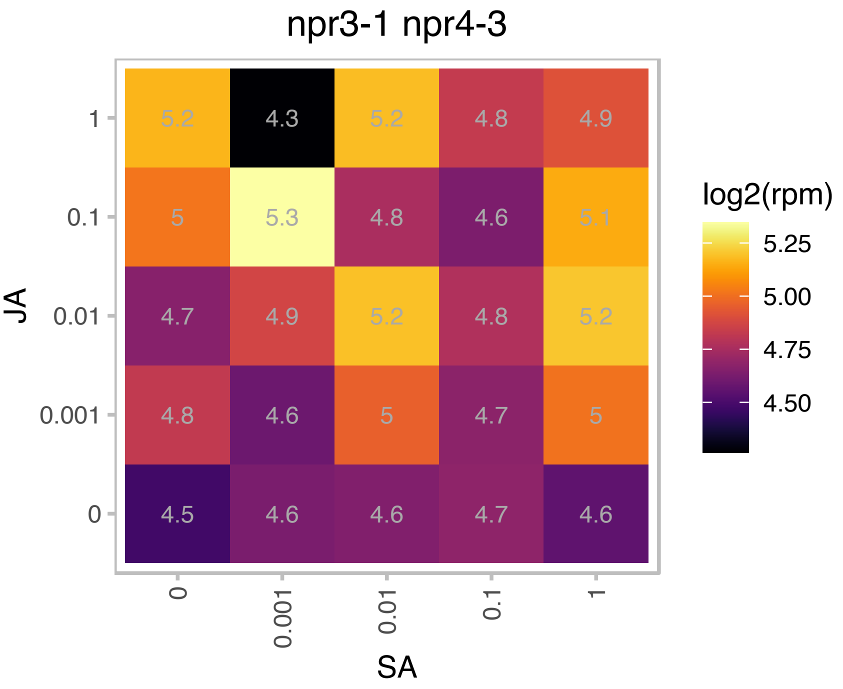

### AT3G09820.1 : ADK1, ATADK1

adenosine kinase 1

Involved in the salvage synthesis of adenylates and methyl recycling

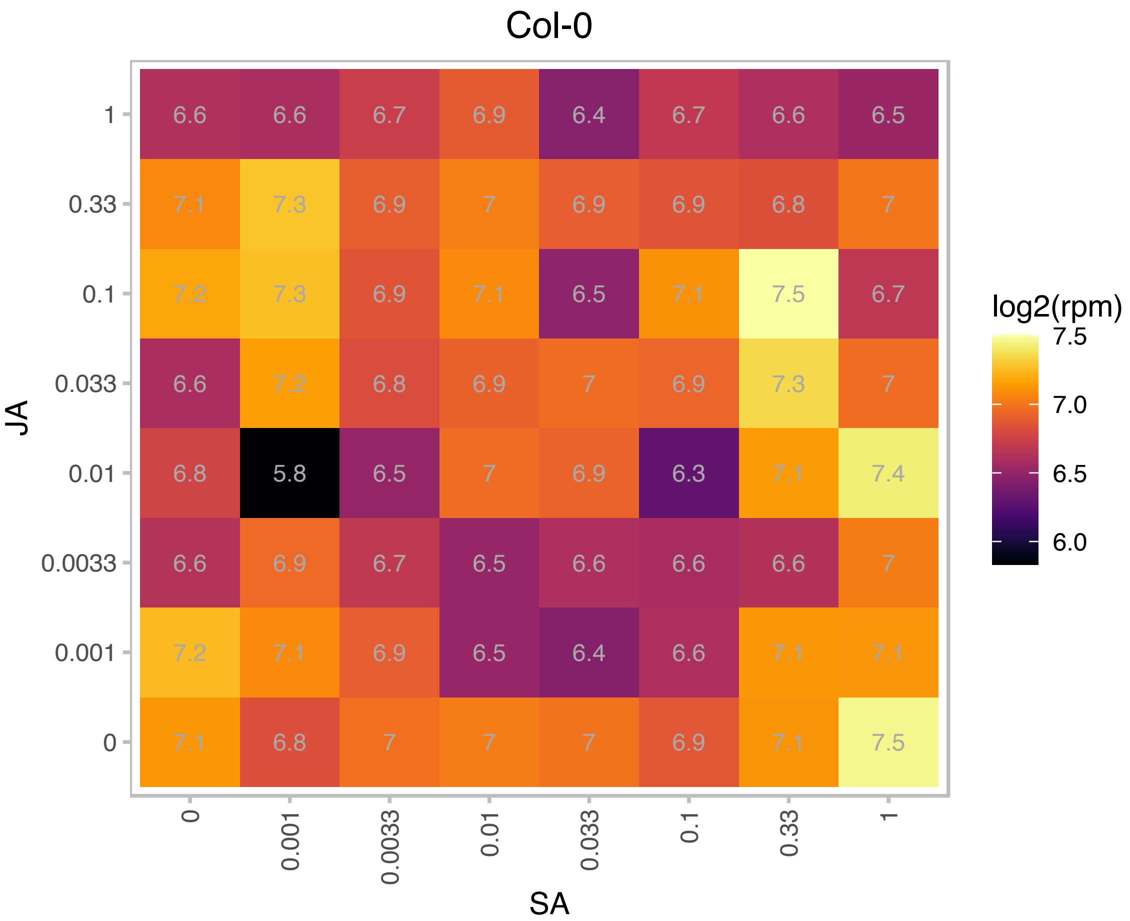


| | |
|---|---|
| Gene | AT3G09820 |
| is.expressed_genes | TRUE |
| is.responsive_genes | TRUE |
| is.apcluster | C-03 |

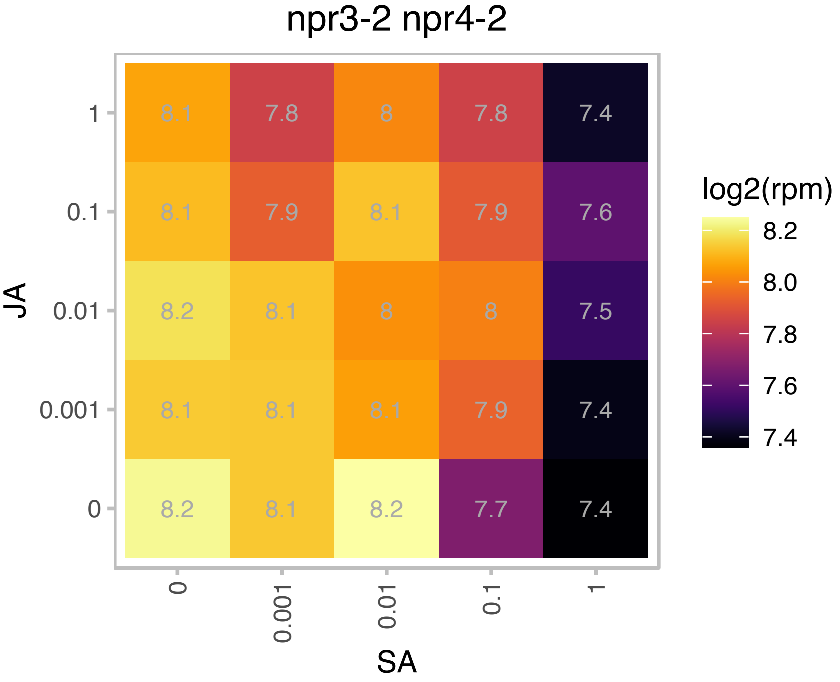


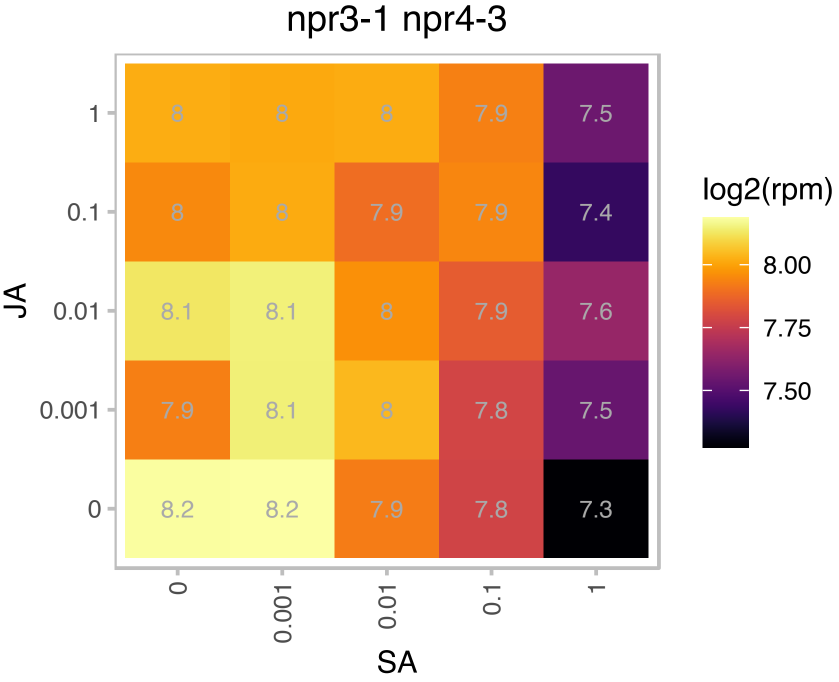

### AT3G22600.1 : LTPG5

Bifunctional inhibitor/lipid-transfer protein/seed storage 2S albumin superfamily protein

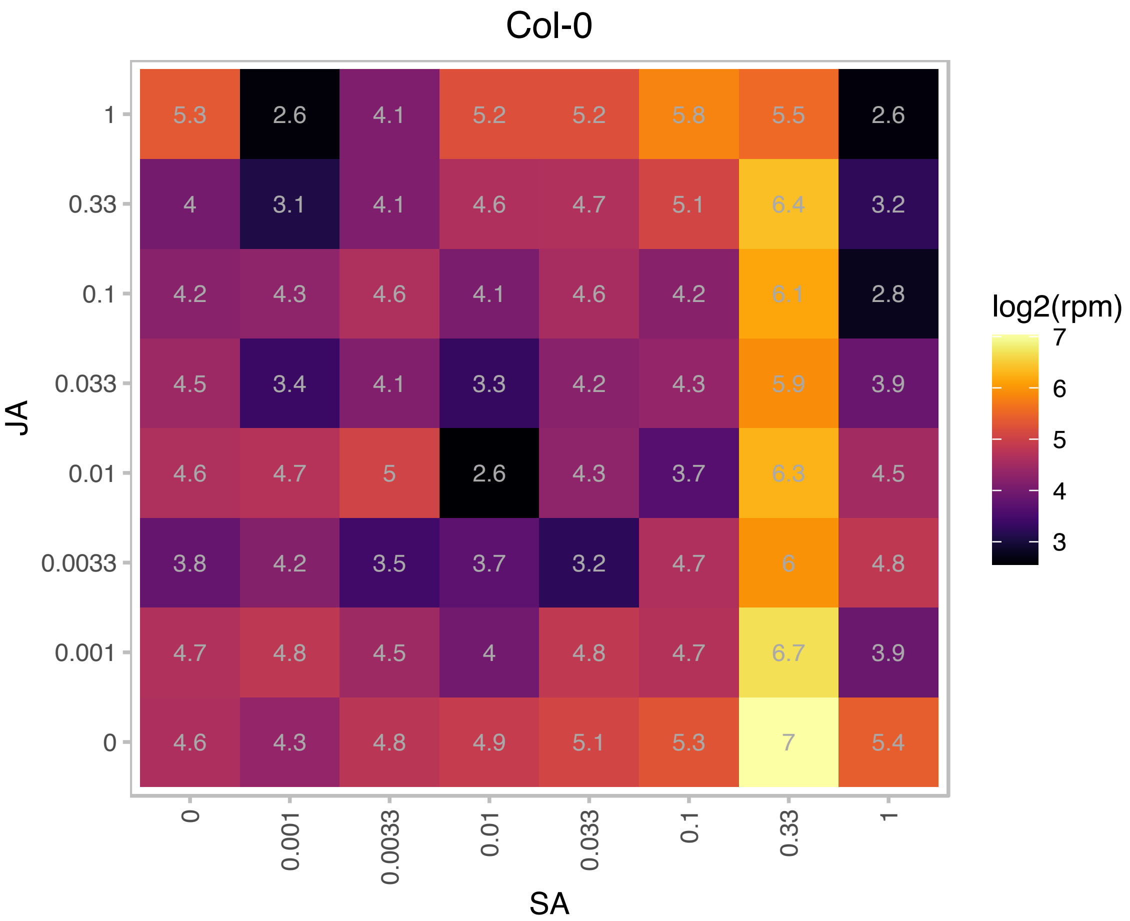


| | |
|---|---|
| Gene | AT3G22600 |
| is.expressed_genes | TRUE |
| is.responsive_genes | TRUE |
| is.apcluster | C-03 |

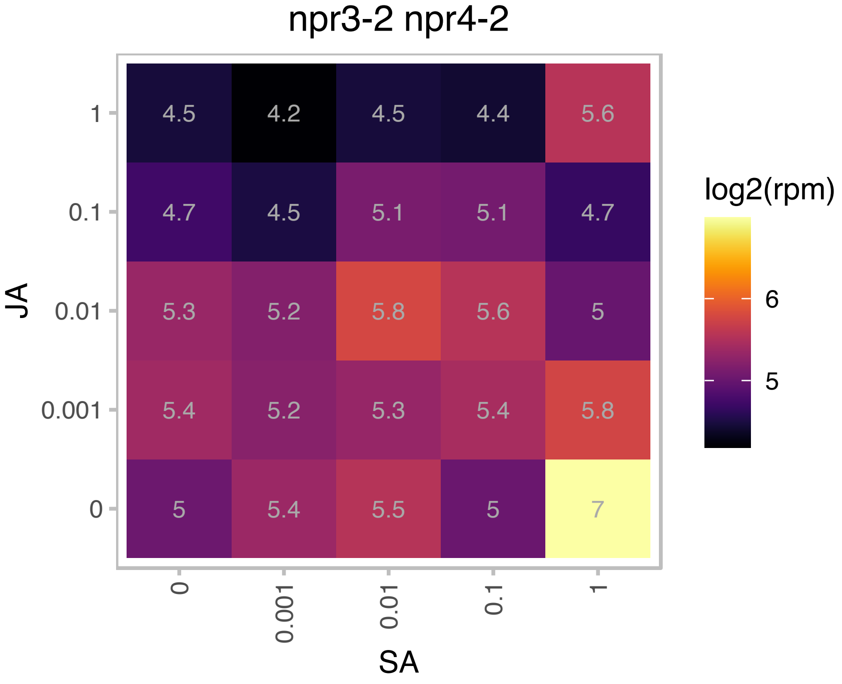


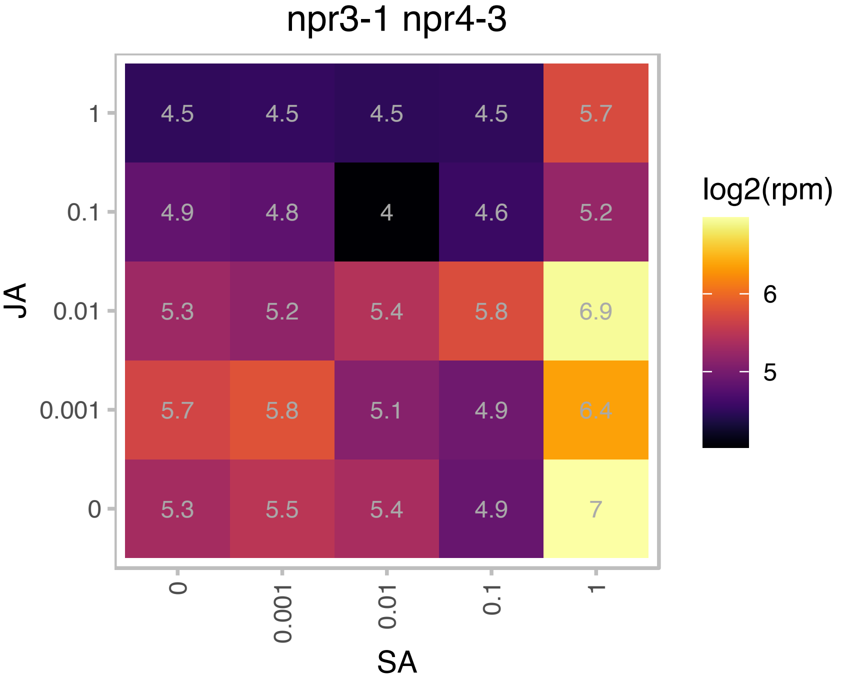

### AT3G56980.1 : BHLH039, bHLH39, ORG3

basic helix-loop-helix (bHLH) DNA-binding superfamily protein

Encodes a member of the basic helix-loop-helix transcription factor protein.

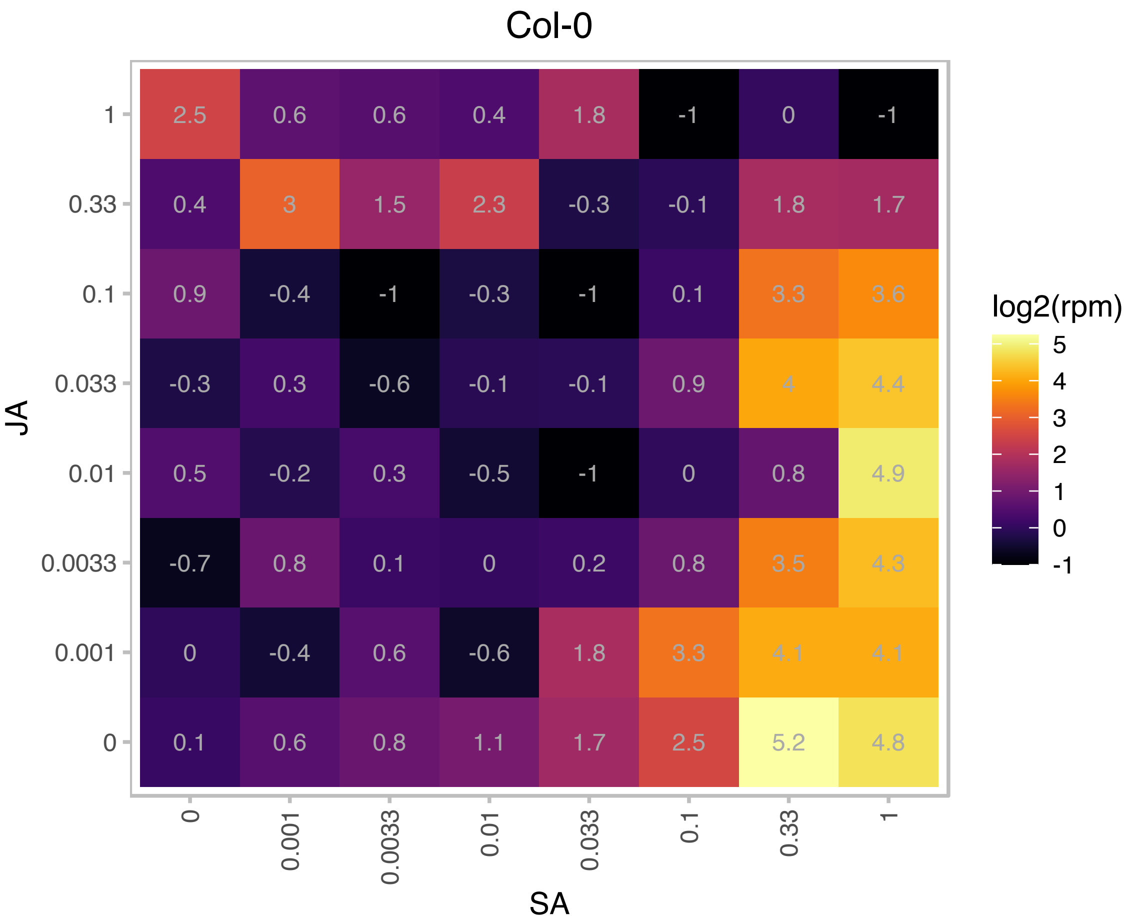


| | |
|---|---|
| Gene | AT3G56980 |
| is.expressed_genes | TRUE |
| is.responsive_genes | TRUE |
| is.apcluster | C-03 |

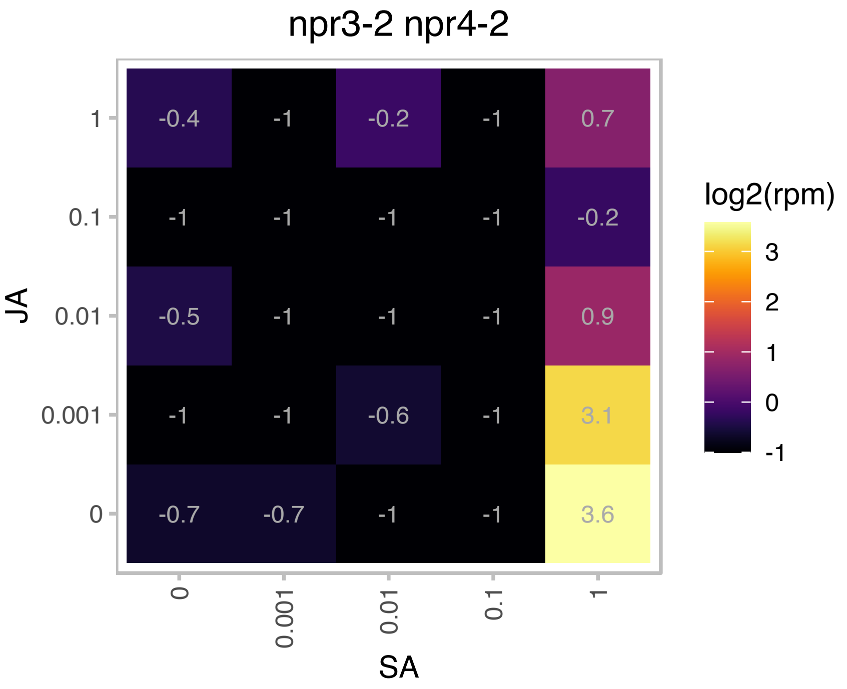


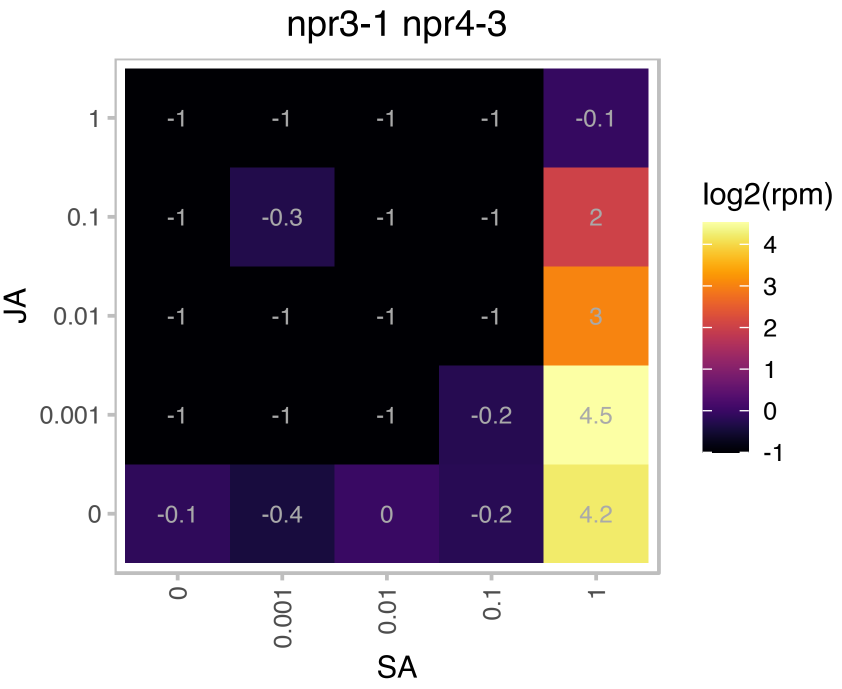

# AT4G26965.1 : NA

NADH:ubiquinone oxidoreductase, 17.2kDa subunit

| | |
|---|---|
| Gene | AT4G26965 |
| is.expressed_genes | TRUE |
| is.responsive_genes | TRUE |
| is.apcluster | C-03 |

Col-0

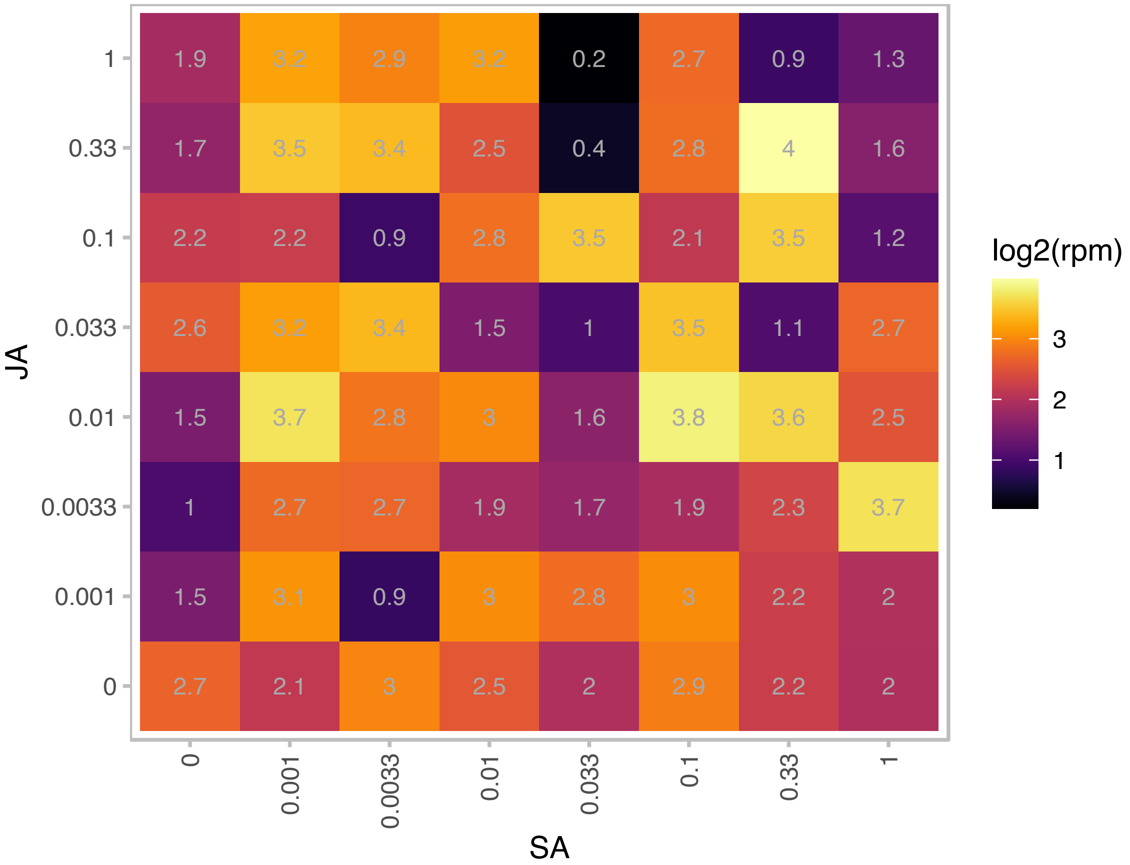


npr3-2 npr4-2

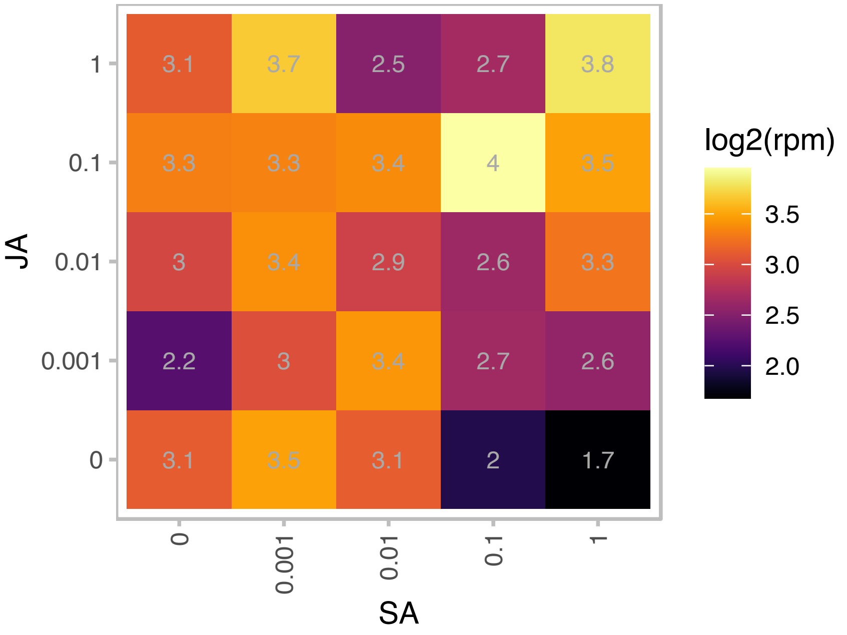


npr3-1 npr4-3

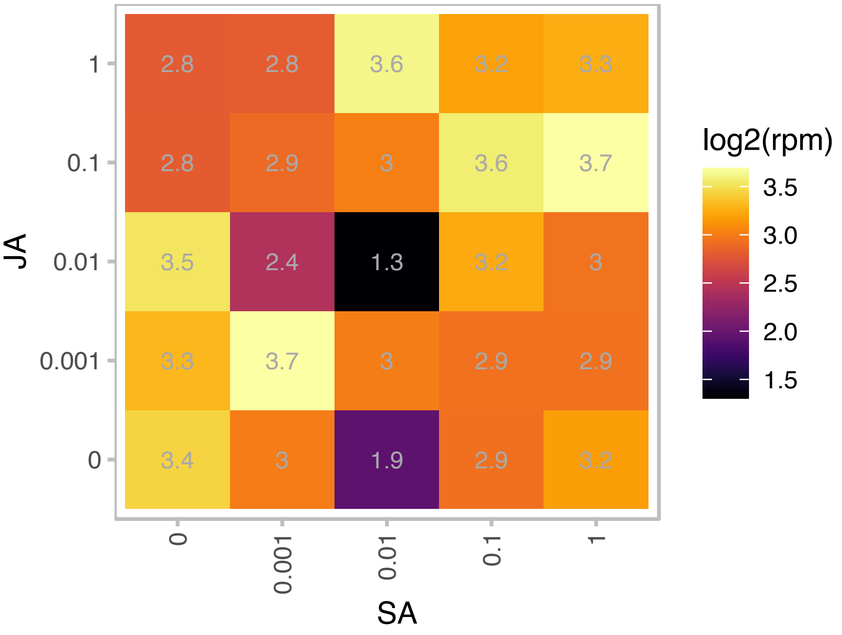

### AT4G38220.1 : AQI

Peptidase M20/M25/M40 family protein

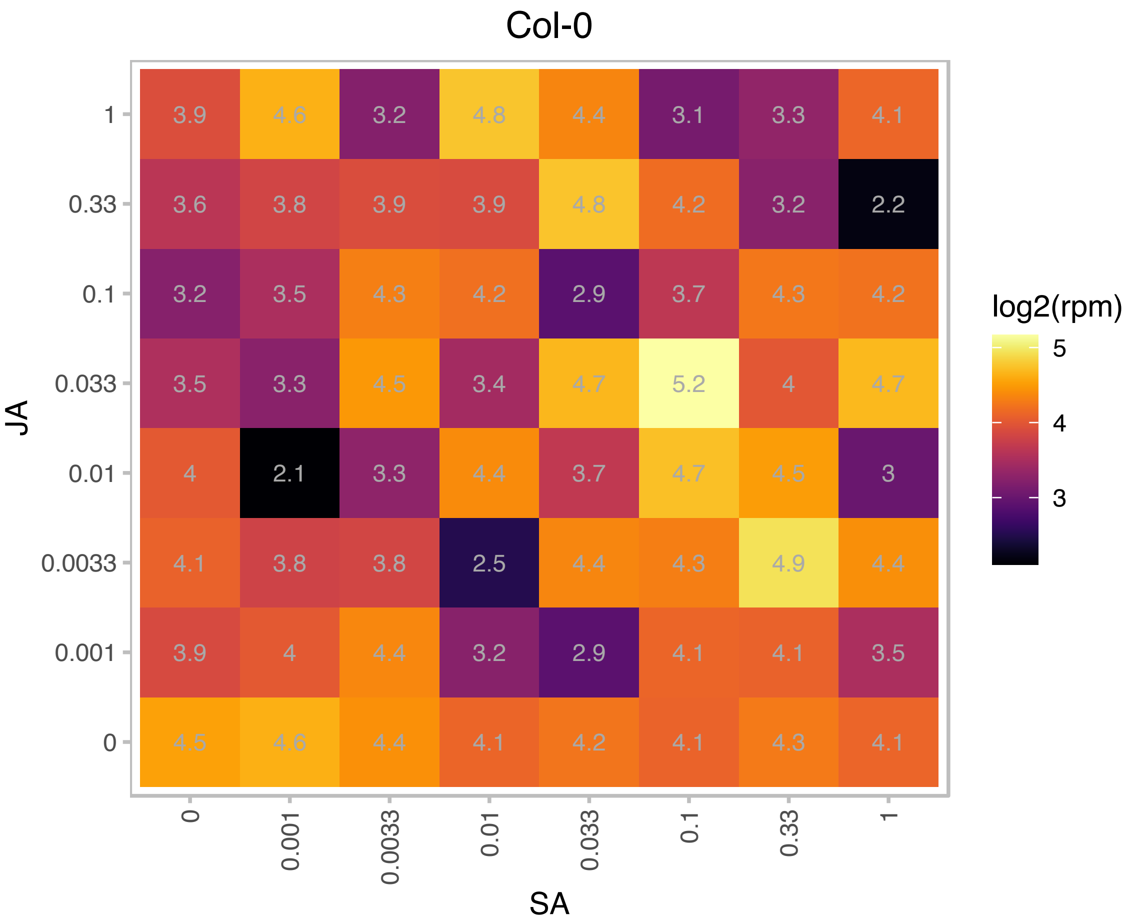


| | |
|---|---|
| Gene | AT4G38220 |
| is.expressed_genes | TRUE |
| is.responsive_genes | TRUE |
| is.apcluster | C-03 |

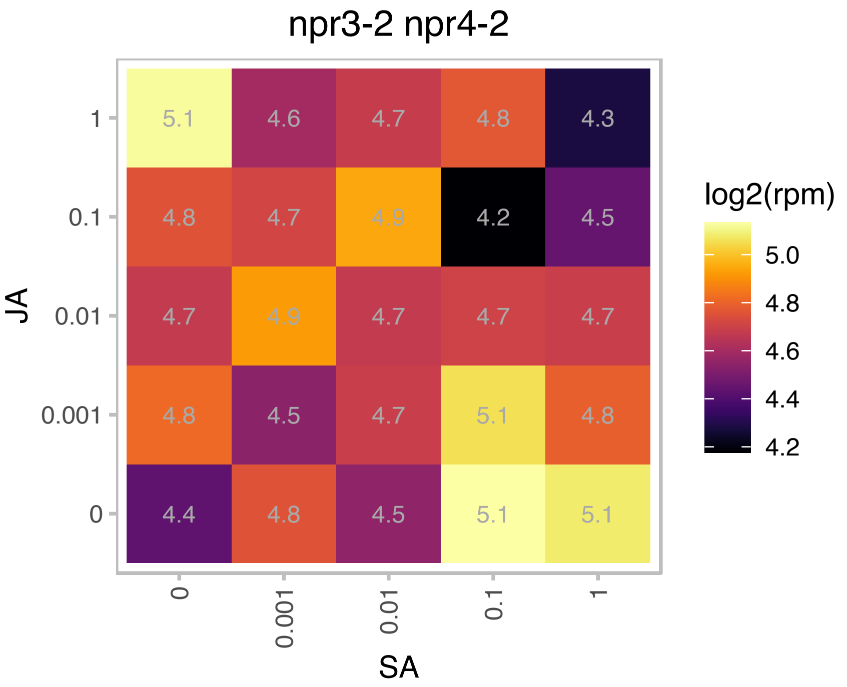


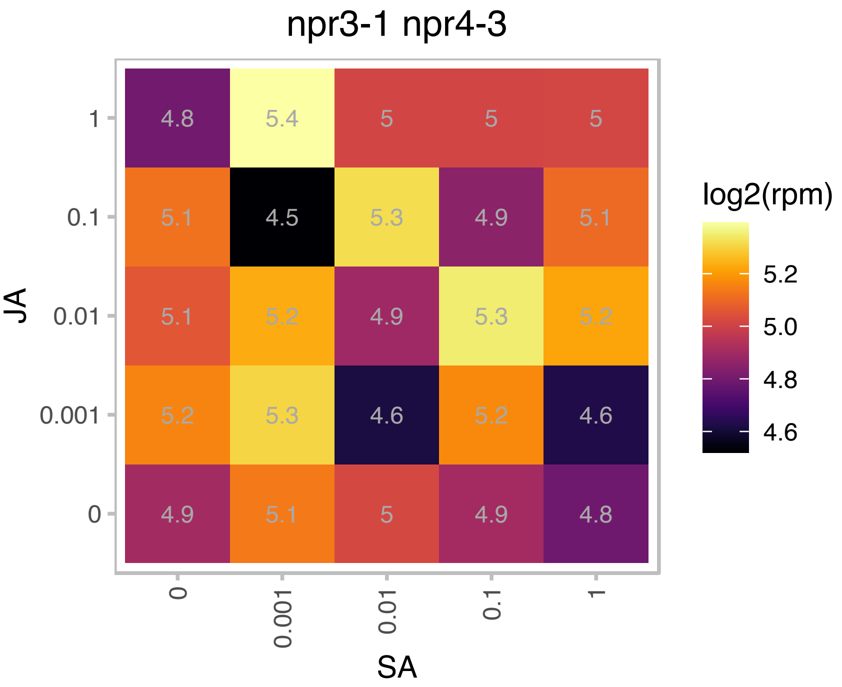

### AT5G09250.1 : KIWI

ssDNA-binding transcriptional regulator

putative transcriptional co-activator (KIWI) mRNA, complete

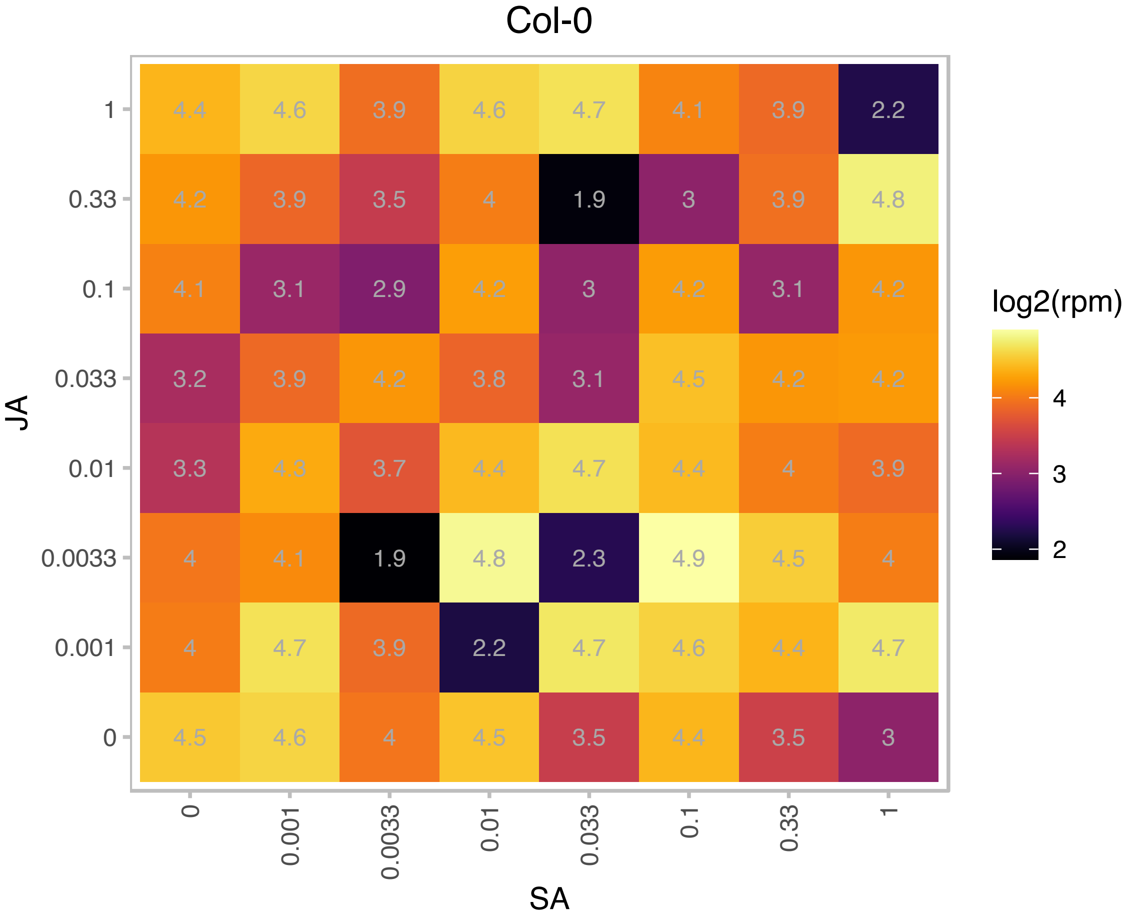


| | |
|---|---|
| Gene | AT5G09250 |
| is.expressed_genes | TRUE |
| is.responsive_genes | TRUE |
| is.apcluster | C-03 |

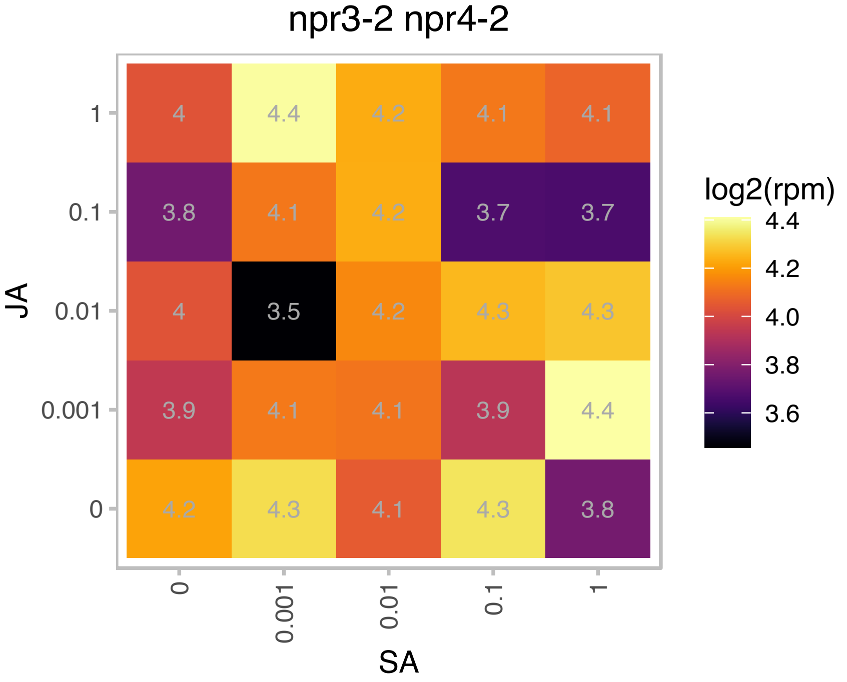


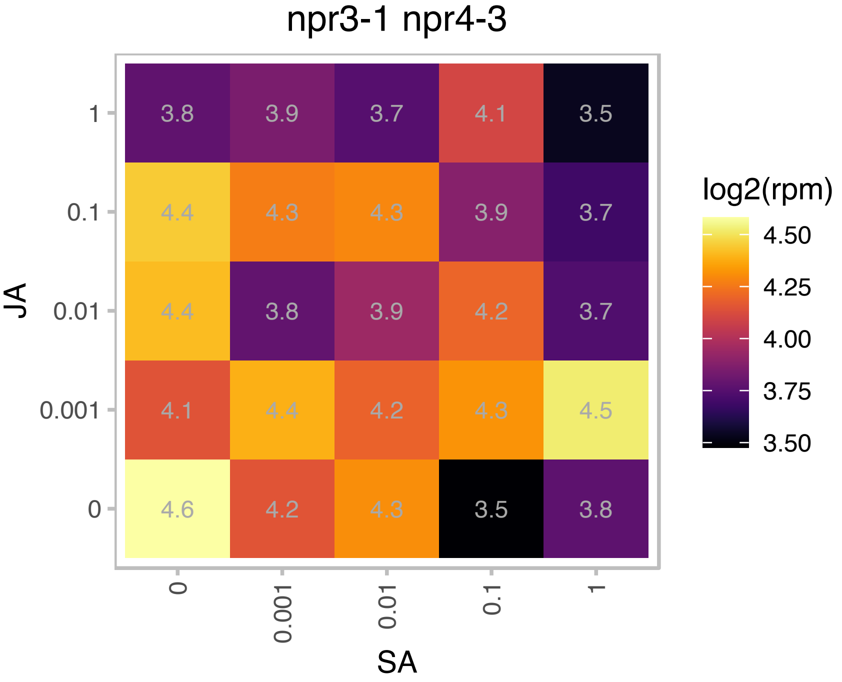

### AT5G10790.1 : UBP22

ubiquitin-specific protease 22

Encodes a ubiquitin-specific protease.

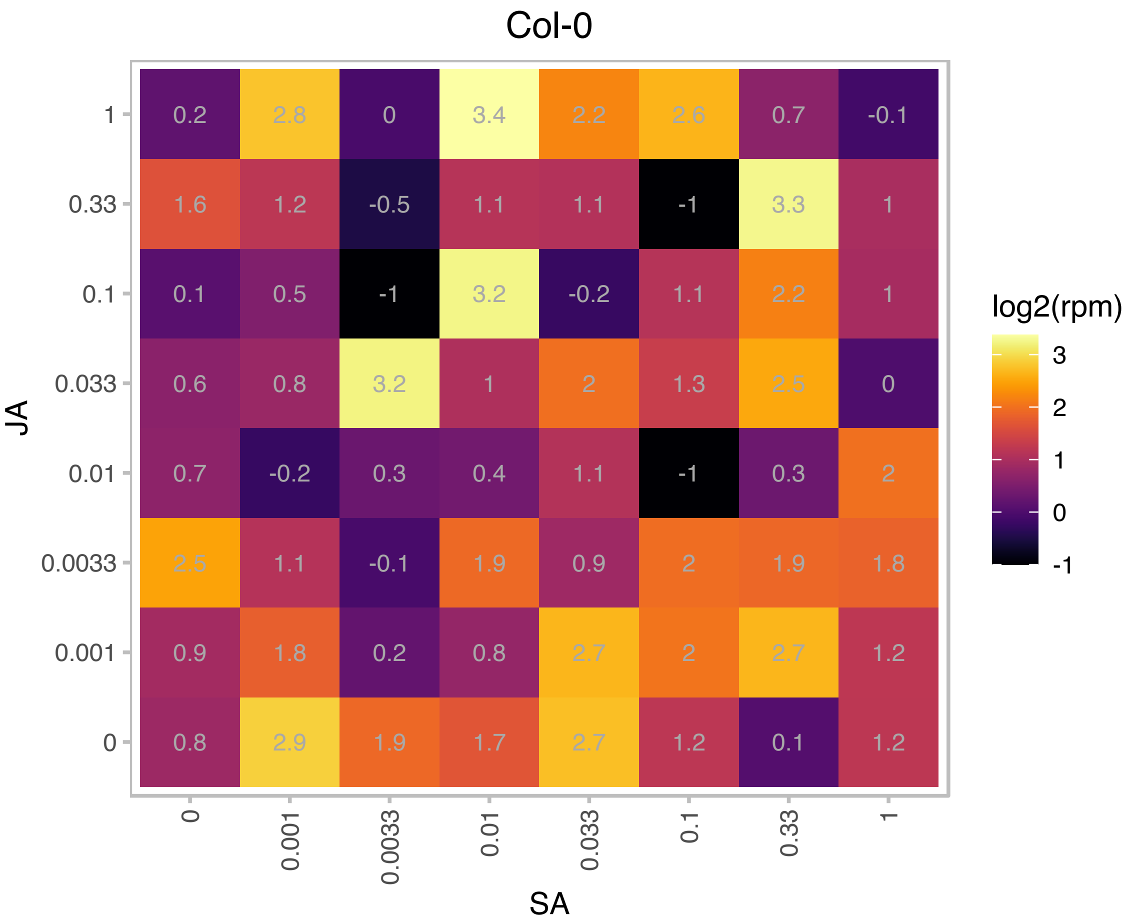


| | |
|---|---|
| Gene | AT5G10790 |
| is.expressed_genes | TRUE |
| is.responsive_genes | TRUE |
| is.apcluster | C-03 |

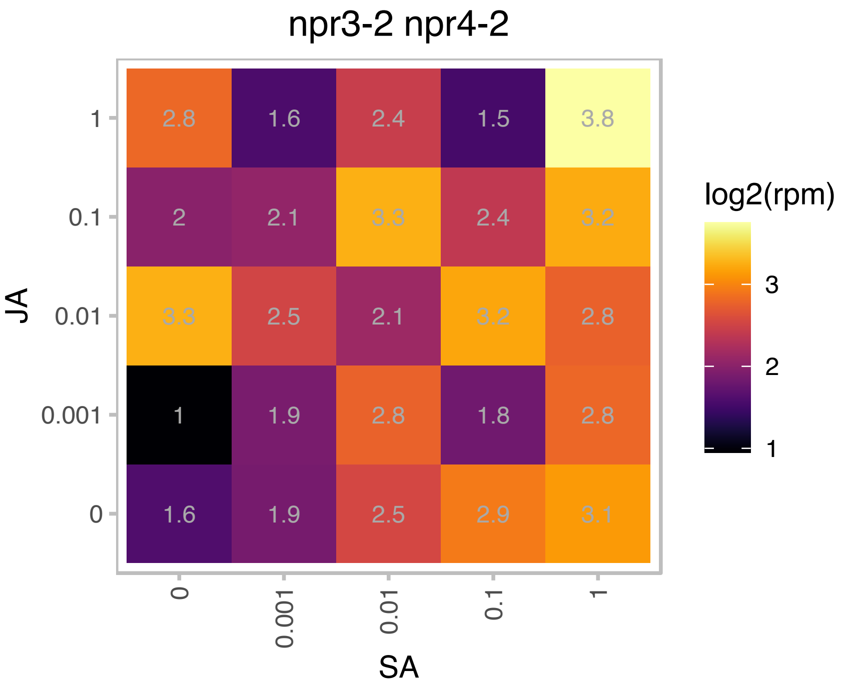


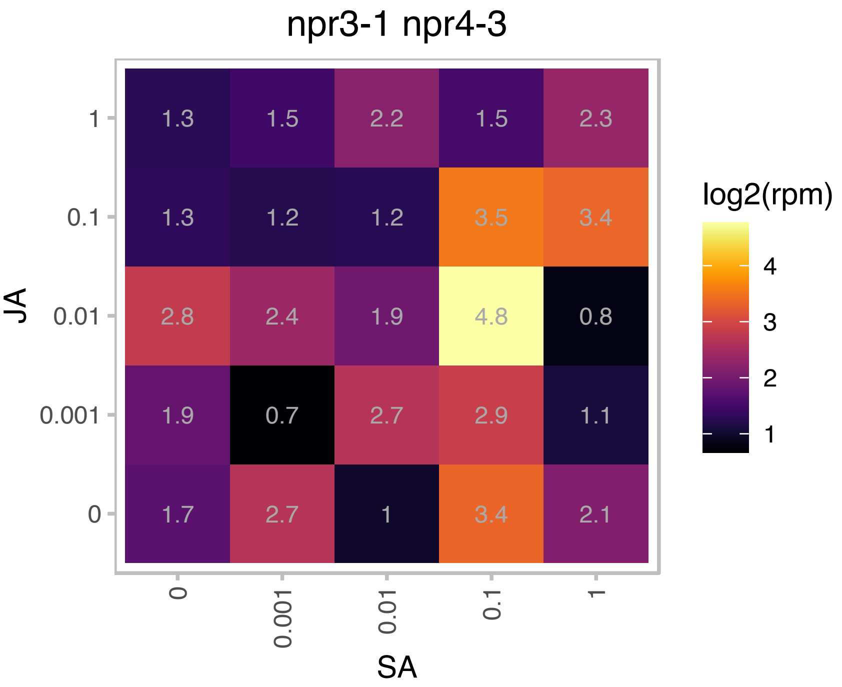

### AT5G12470.1 : RER4

Protein of unknown function (DUF3411)

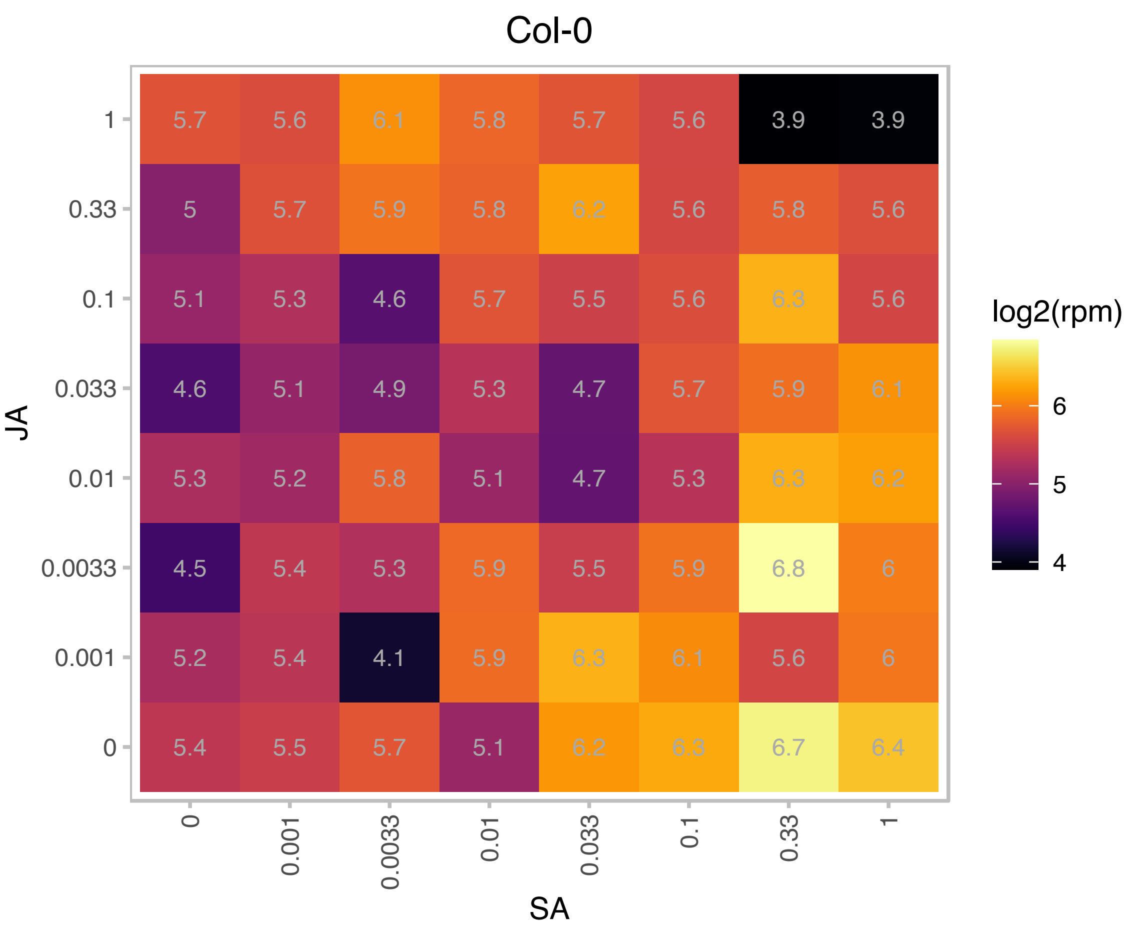


| | |
|---|---|
| Gene | AT5G12470 |
| is.expressed_genes | TRUE |
| is.responsive_genes | TRUE |
| is.apcluster | C-03 |

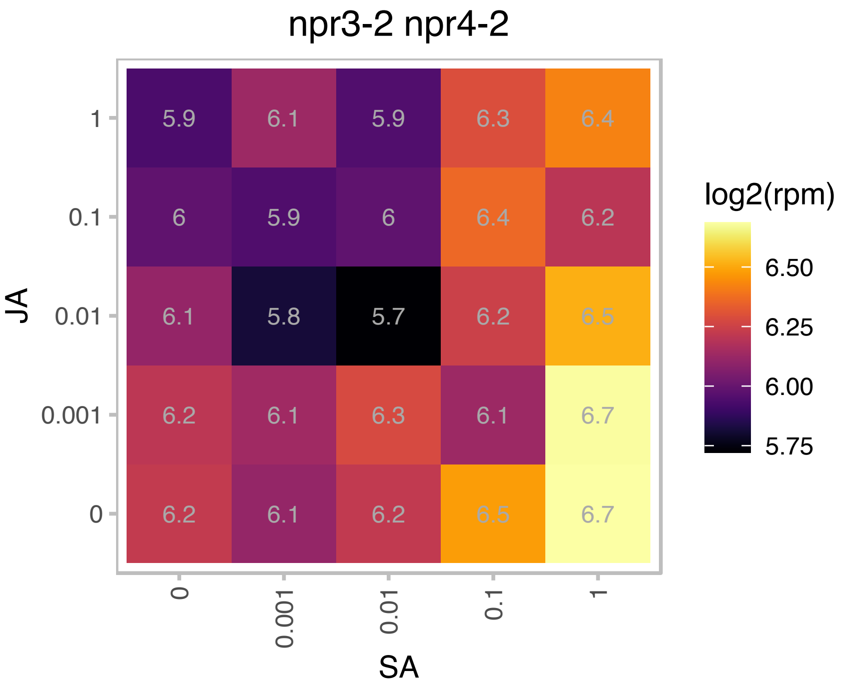


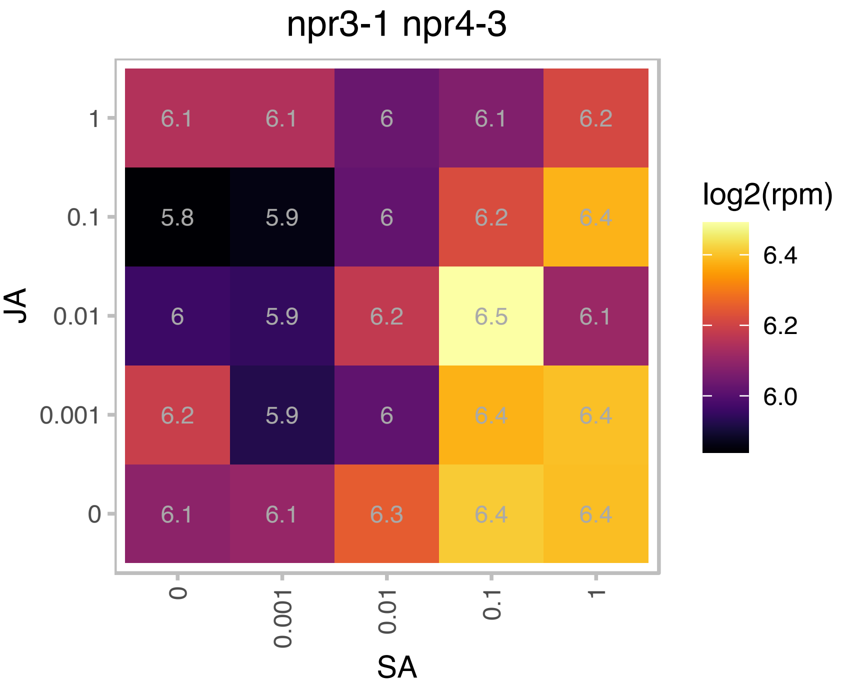

### AT5G17770.1 : ATCBR, CBR, CBR1

NADH:cytochrome B5 reductase 1
Encodes NADH:cytochrome (Cyt) b5 reductase that displayed strict specificity to NADH for the reduction of a recombinant Cyt b5 (AtB5-A), whereas no Cyt b5 reduction was observed when NADPH was used as the electron donor.

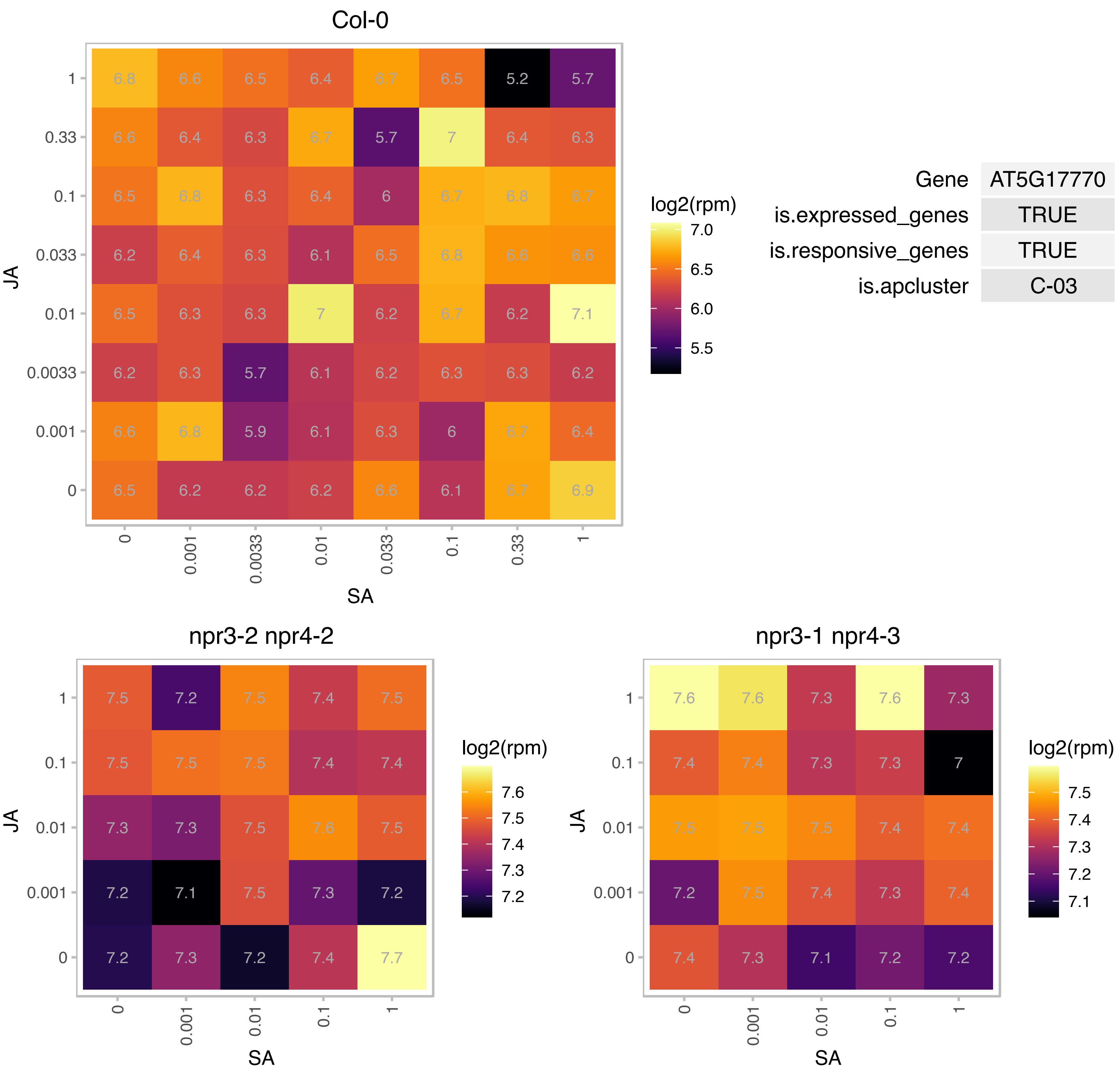


| | |
|---|---|
| Gene | AT5G17770 |
| is.expressed_genes | TRUE |
| is.responsive_genes | TRUE |
| is.apcluster | C-03 |

# AT5G47710.1 : NA

Calcium-dependent lipid-binding (CaLB domain) family protein

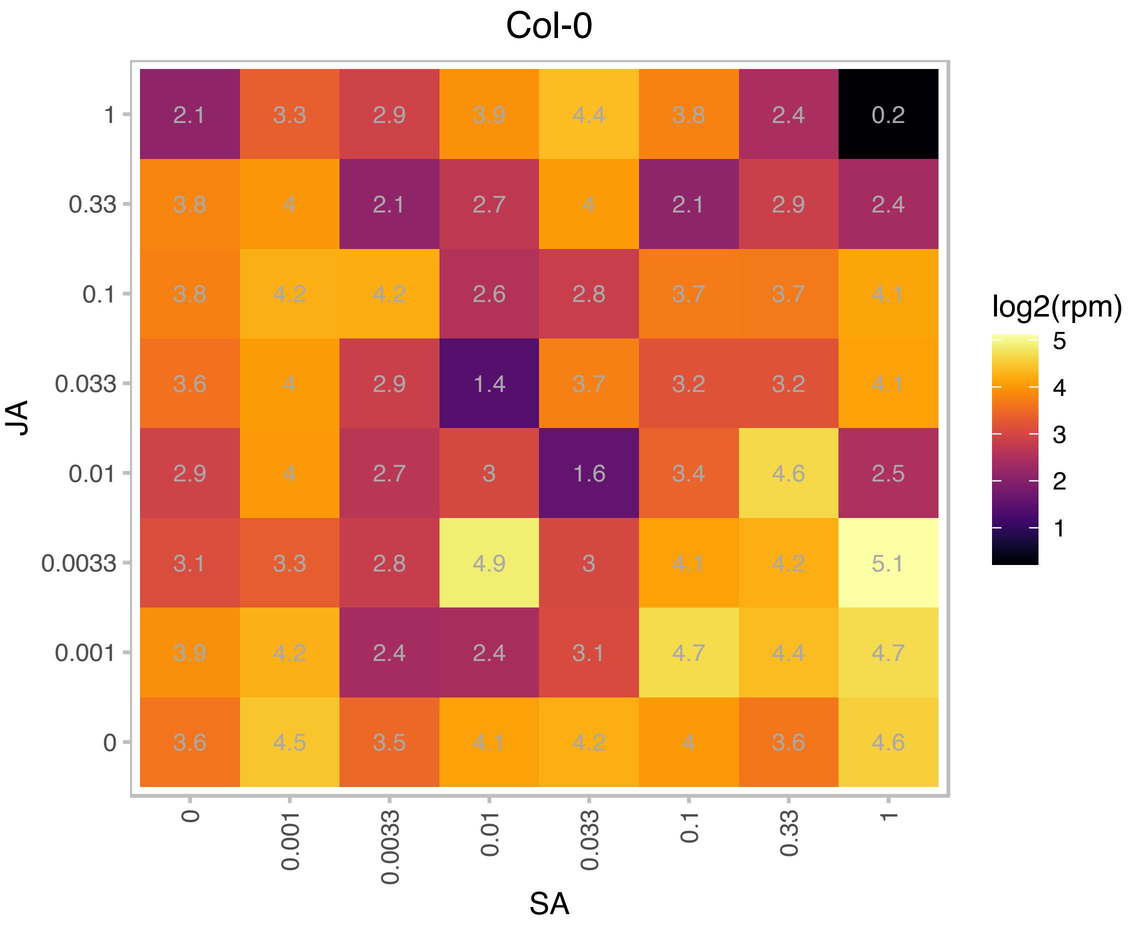


| | |
|---|---|
| Gene | AT5G47710 |
| is.expressed_genes | TRUE |
| is.responsive_genes | TRUE |
| is.apcluster | C-03 |

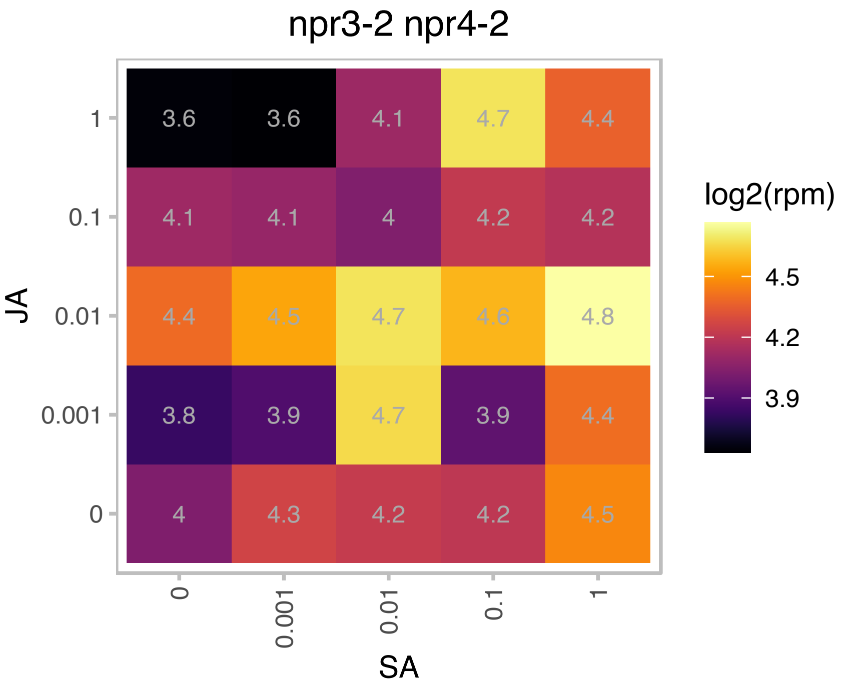


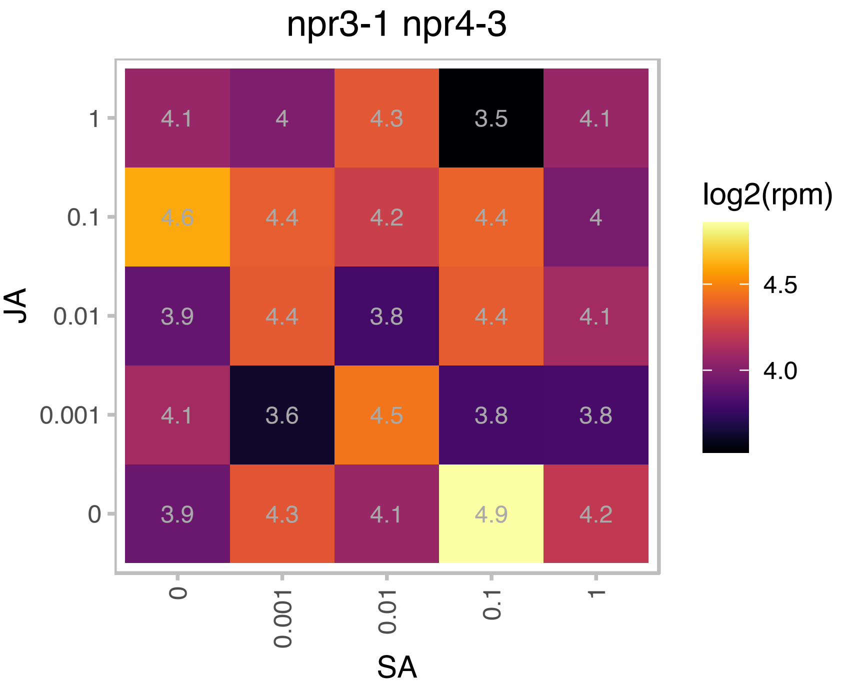

### AT5G55280.1 : ATFTSZ1-1, CPFTSZ, FTSZ1-1

homolog of bacterial cytokinesis Z-ring protein FTSZ 1-1

Encodes one of two FtsZ proteins, tubulin-like proteins, in Arabidopsis.  It is involved in chloroplast division.

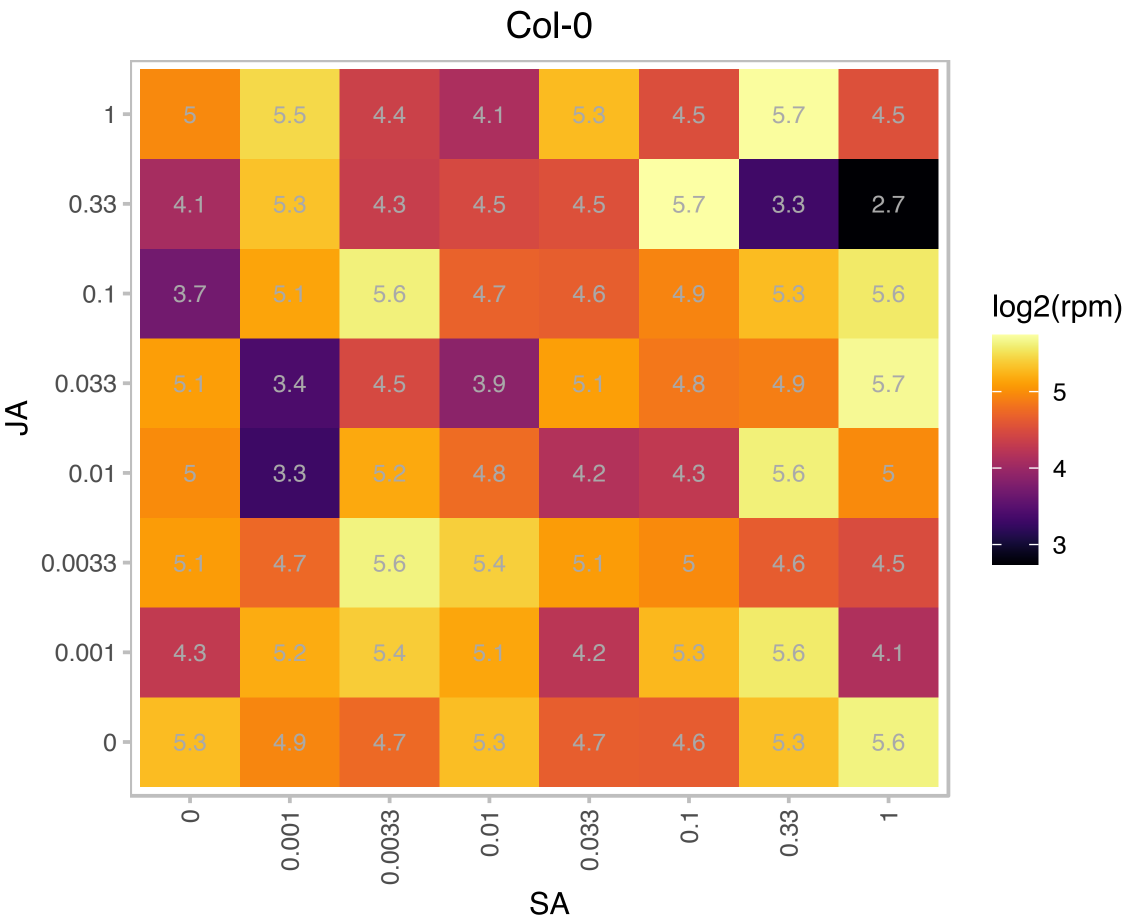


| | |
|---|---|
| Gene | AT5G55280 |
| is.expressed_genes | TRUE |
| is.responsive_genes | TRUE |
| is.apcluster | C-03 |

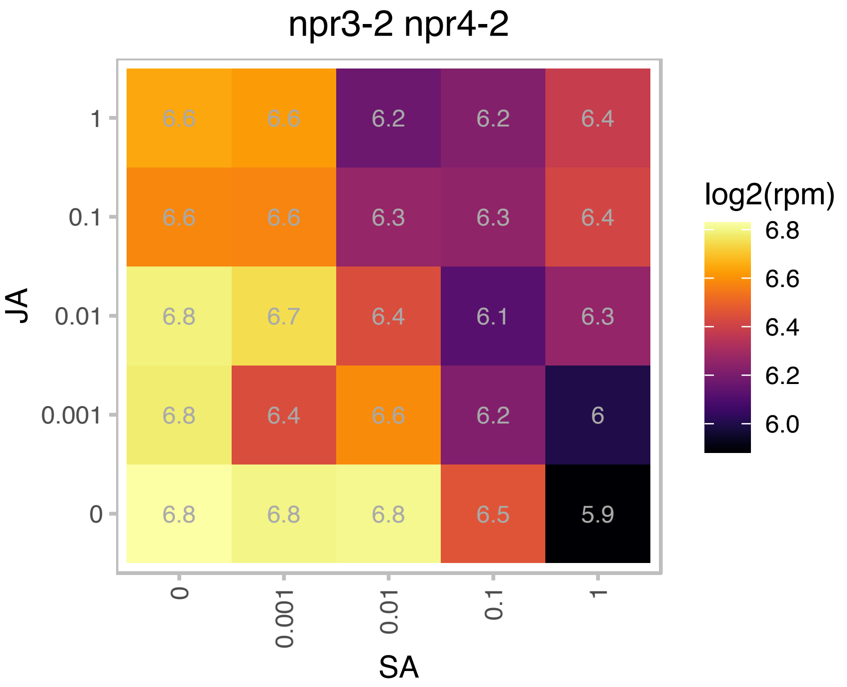


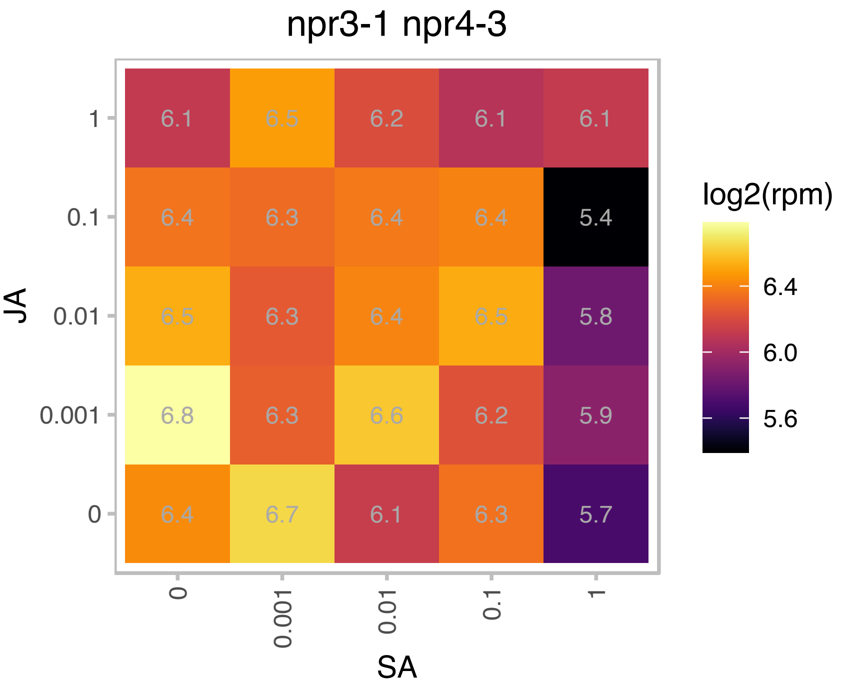

# AT1G15370.1 : NA

SNARE-like superfamily protein

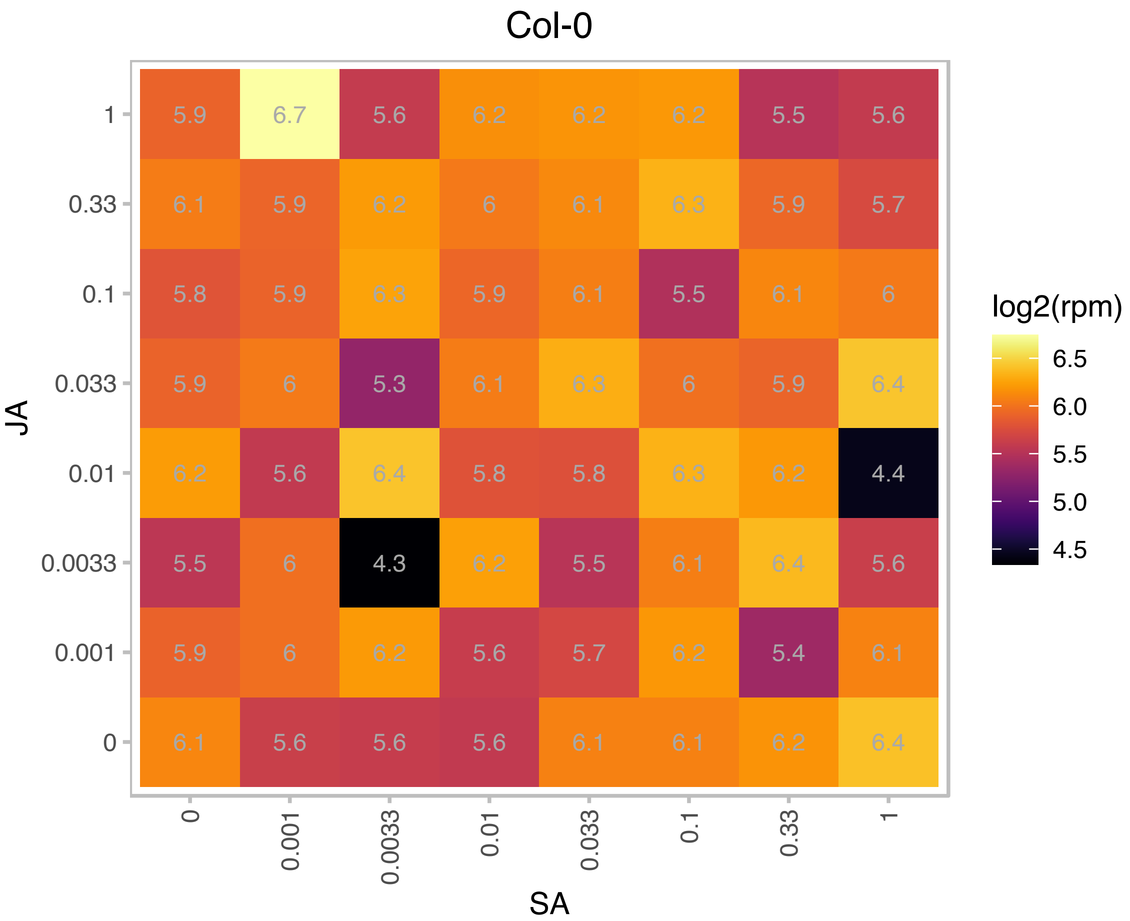


| | |
|---|---|
| Gene | AT1G15370 |
| is.expressed_genes | TRUE |
| is.responsive_genes | TRUE |
| is.apcluster | C-04 |

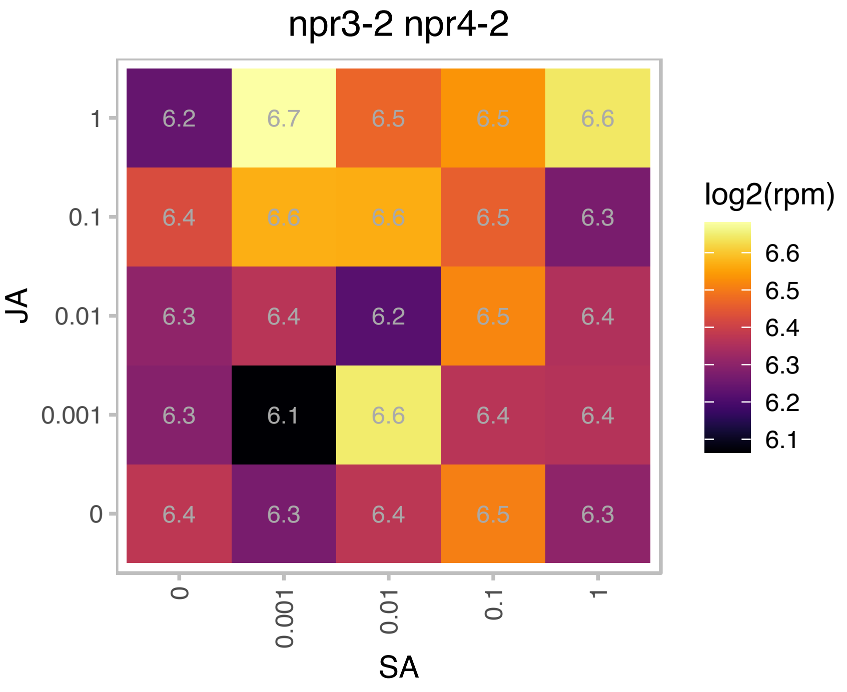


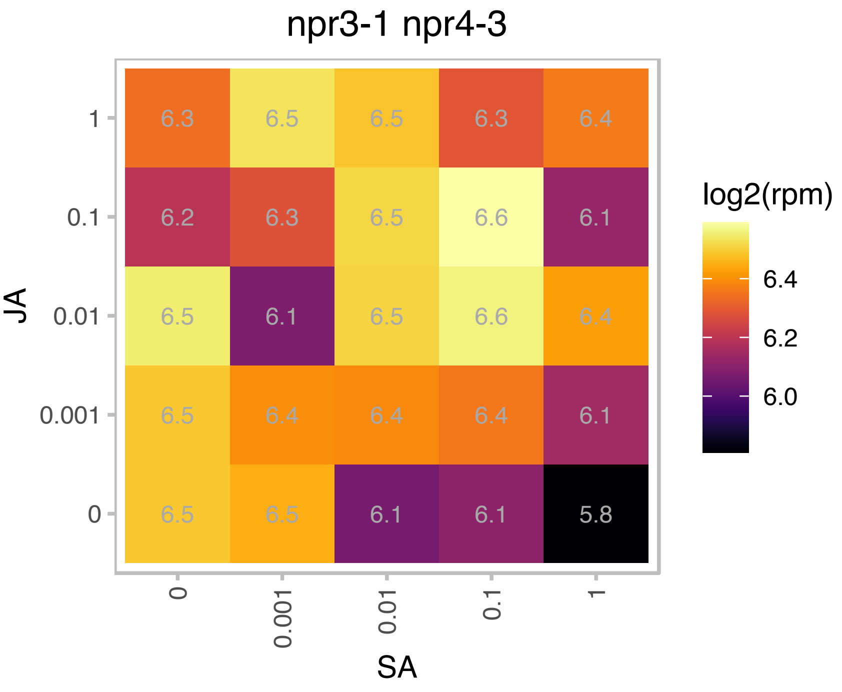

# AT1G17080.1 : NA

Ribosomal protein L18ae family

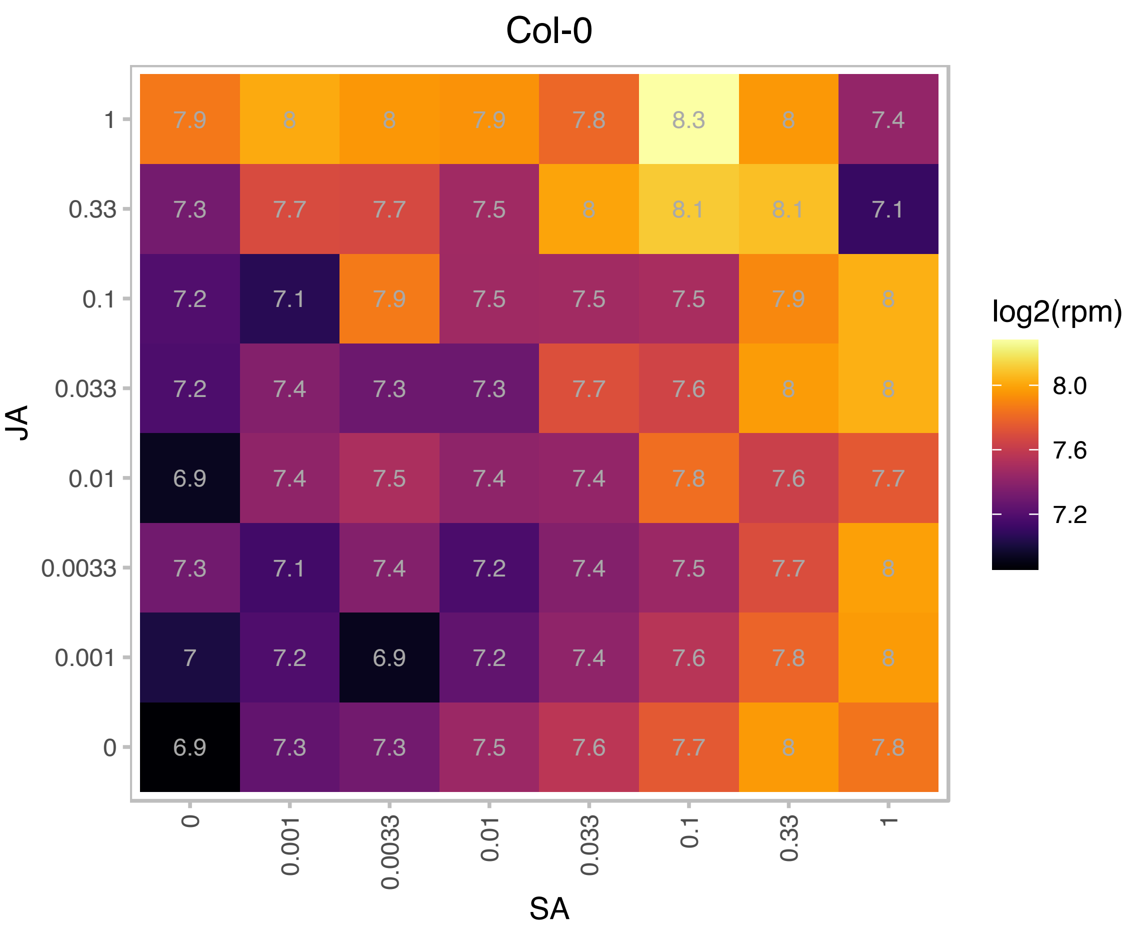


| | |
|---|---|
| Gene | AT1G17080 |
| is.expressed_genes | TRUE |
| is.responsive_genes | TRUE |
| is.apcluster | C-04 |

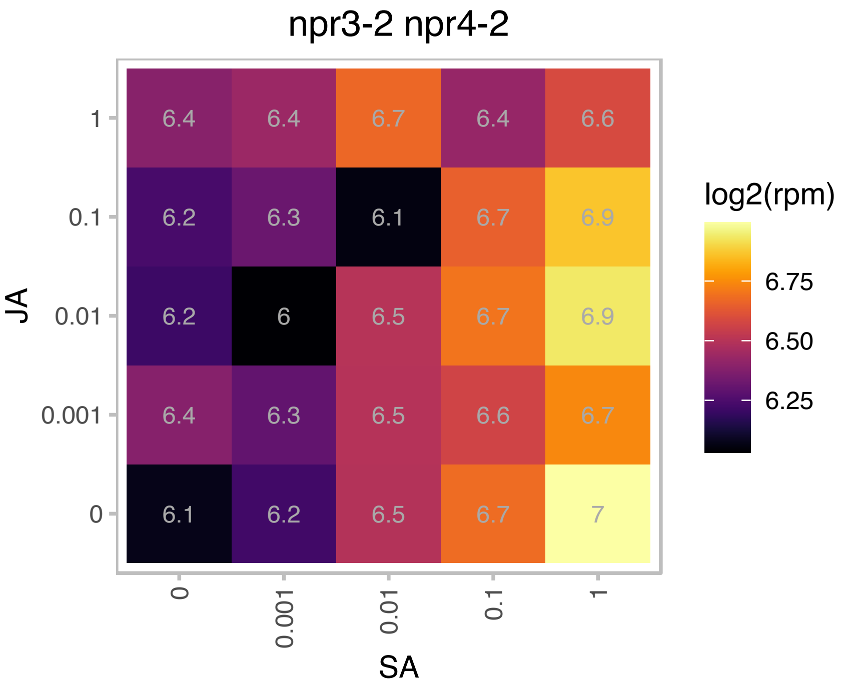


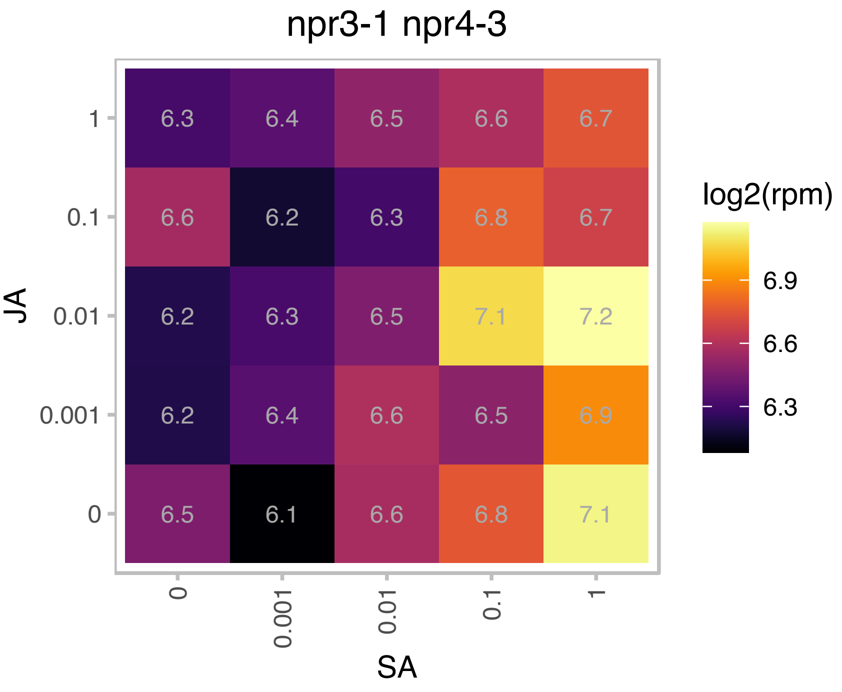

# AT1G21780.1 : NA

BTB/POZ domain-containing protein

BTB/POZ domain-containing protein. Contains similarity to gb:AJ000644 SPOP (speckle-type POZ protein) from Homo sapiens and contains a PF:00651 BTB/POZ domain. ESTs gb:T75841, gb:R89974, gb:R30221, gb:N96386, gb:T76457, gb:AI100013 and gb:T76456 come from this gene;supported by full-length. Interacts with CUL3A and CUL3B.

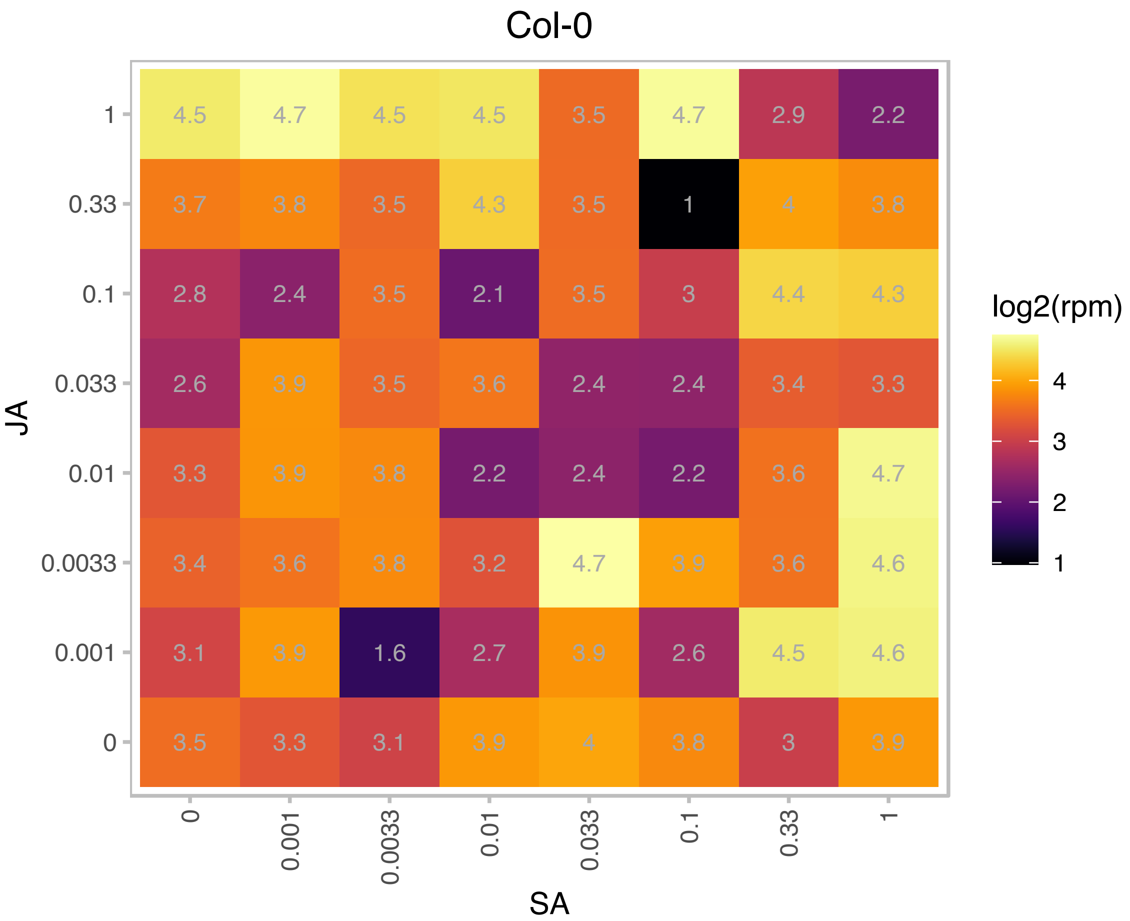


| | |
|---|---|
| Gene | AT1G21780 |
| is.expressed_genes | TRUE |
| is.responsive_genes | TRUE |
| is.apcluster | C-04 |

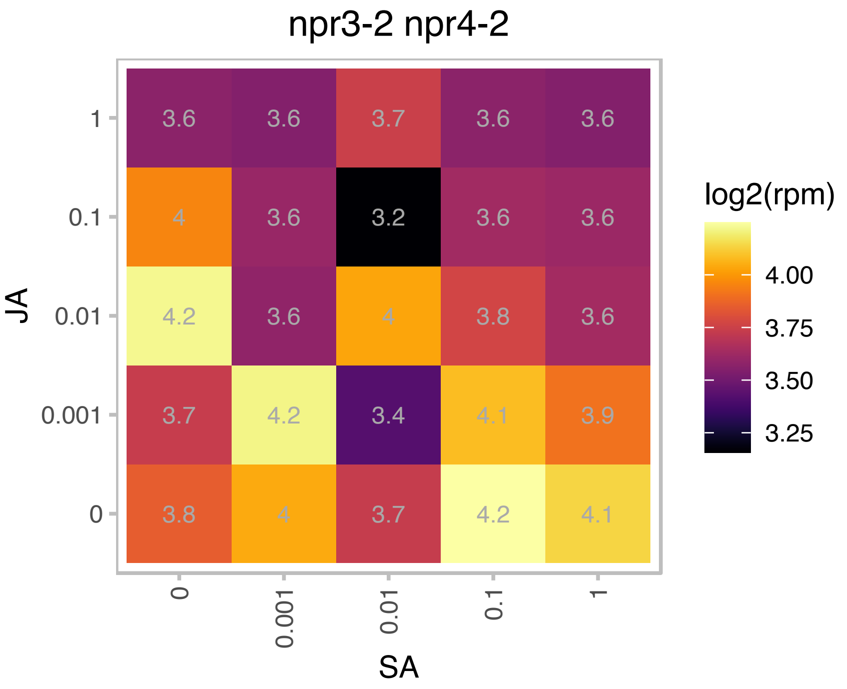


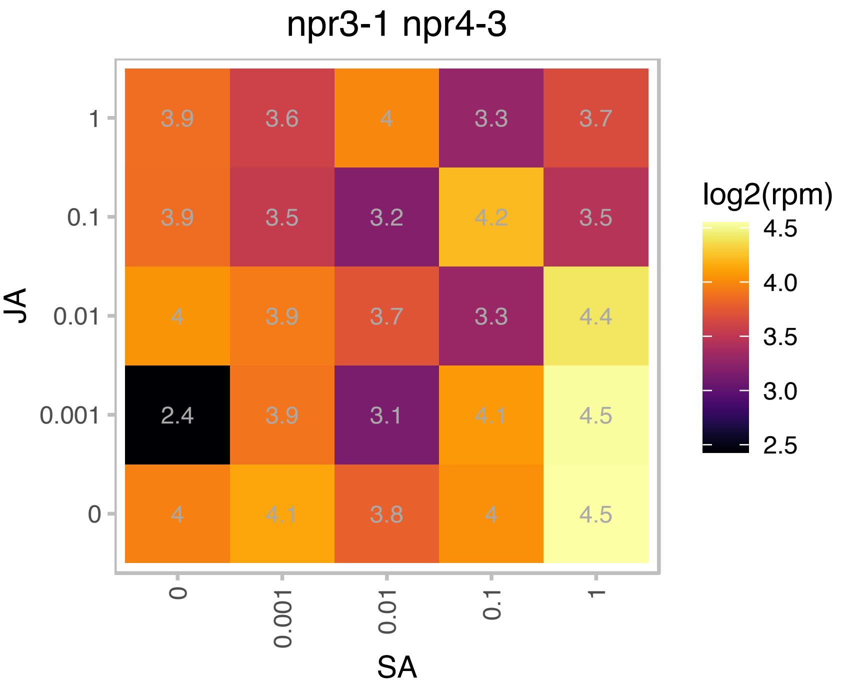

# AT1G30640.1 : NA

Protein kinase family protein

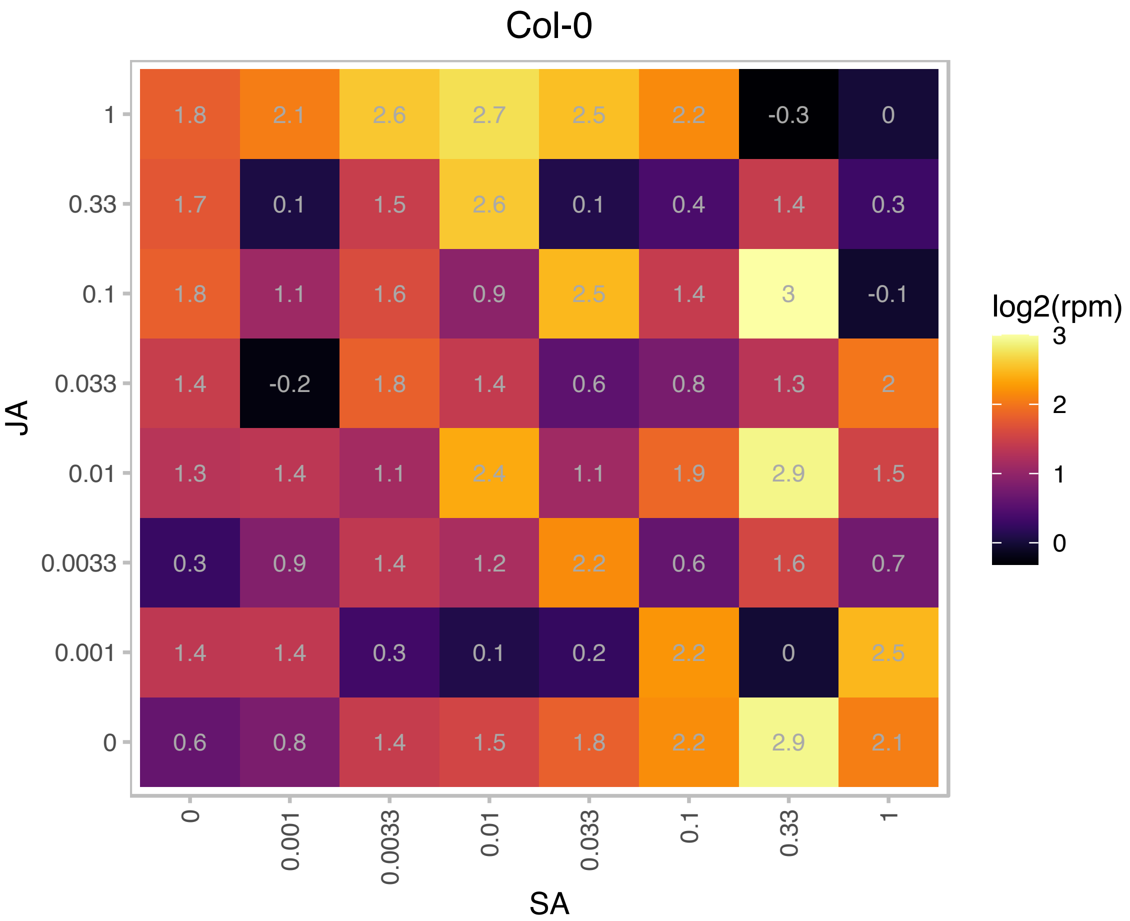


| | |
|---|---|
| Gene | AT1G30640 |
| is.expressed_genes | TRUE |
| is.responsive_genes | TRUE |
| is.apcluster | C-04 |

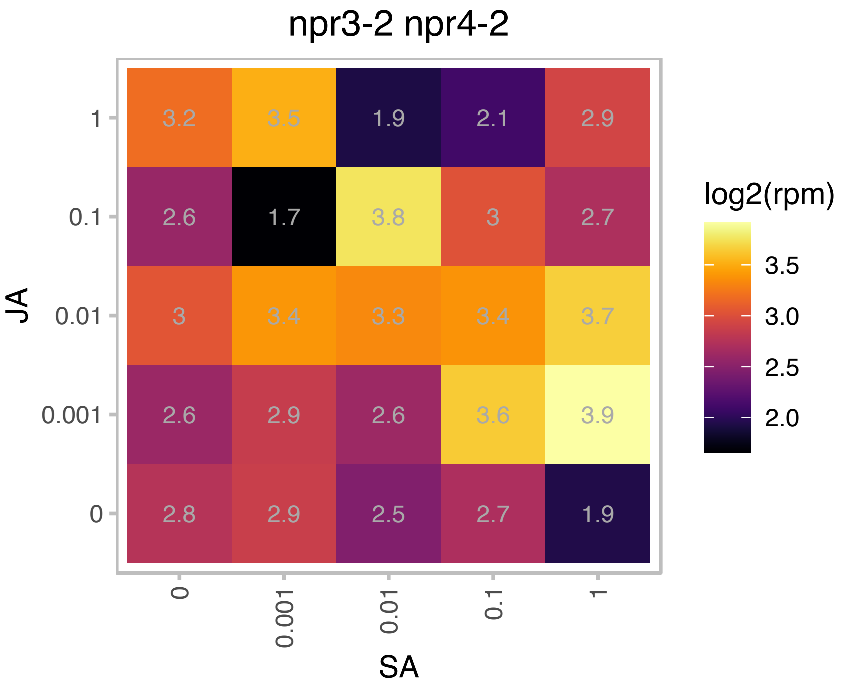


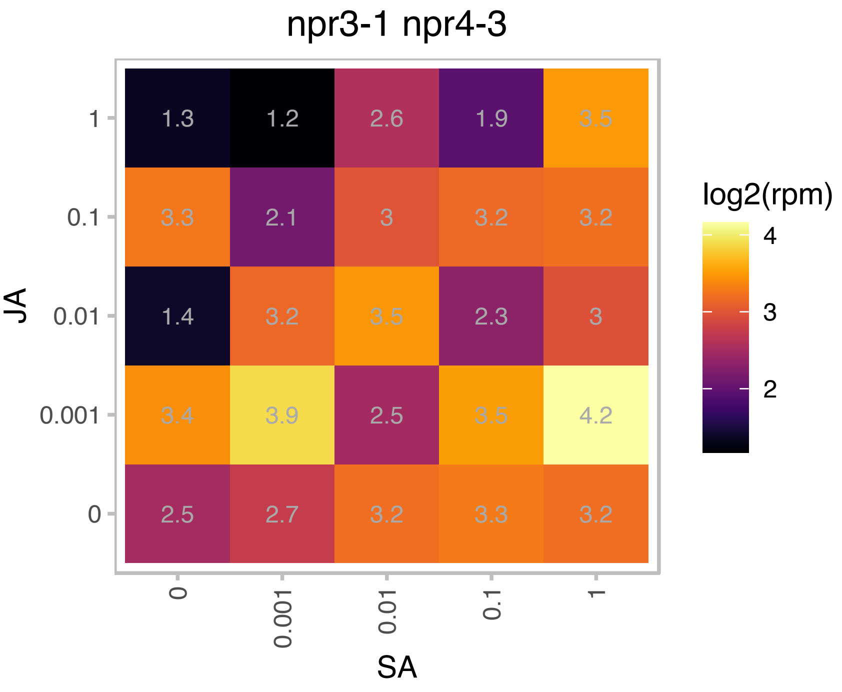

# AT1G49470.1 : NA

Family of unknown function (DUF716)

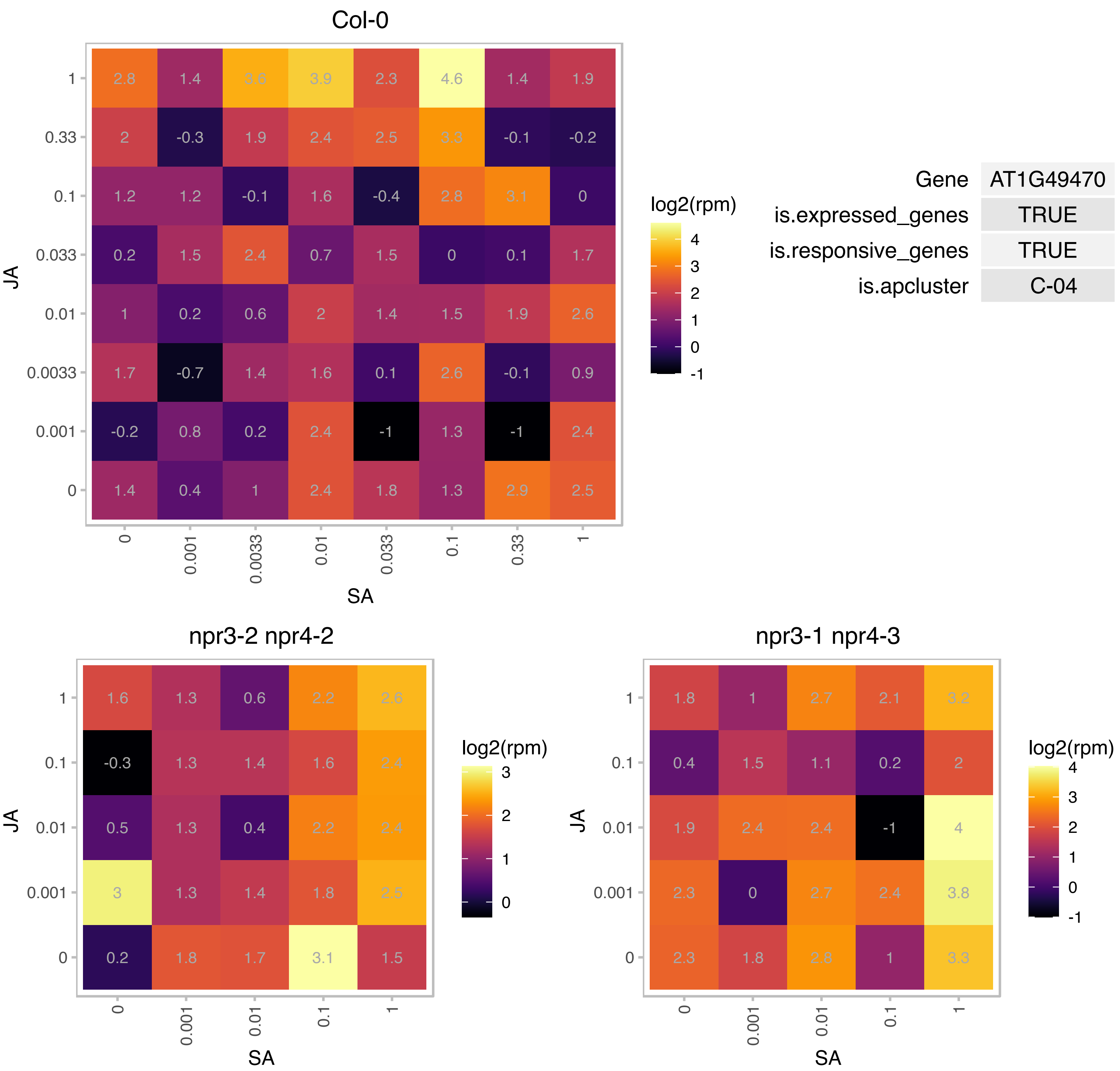


| | |
|---|---|
| Gene | AT1G49470 |
| is.expressed_genes | TRUE |
| is.responsive_genes | TRUE |
| is.apcluster | C-04 |

# AT1G63720.1 : NA

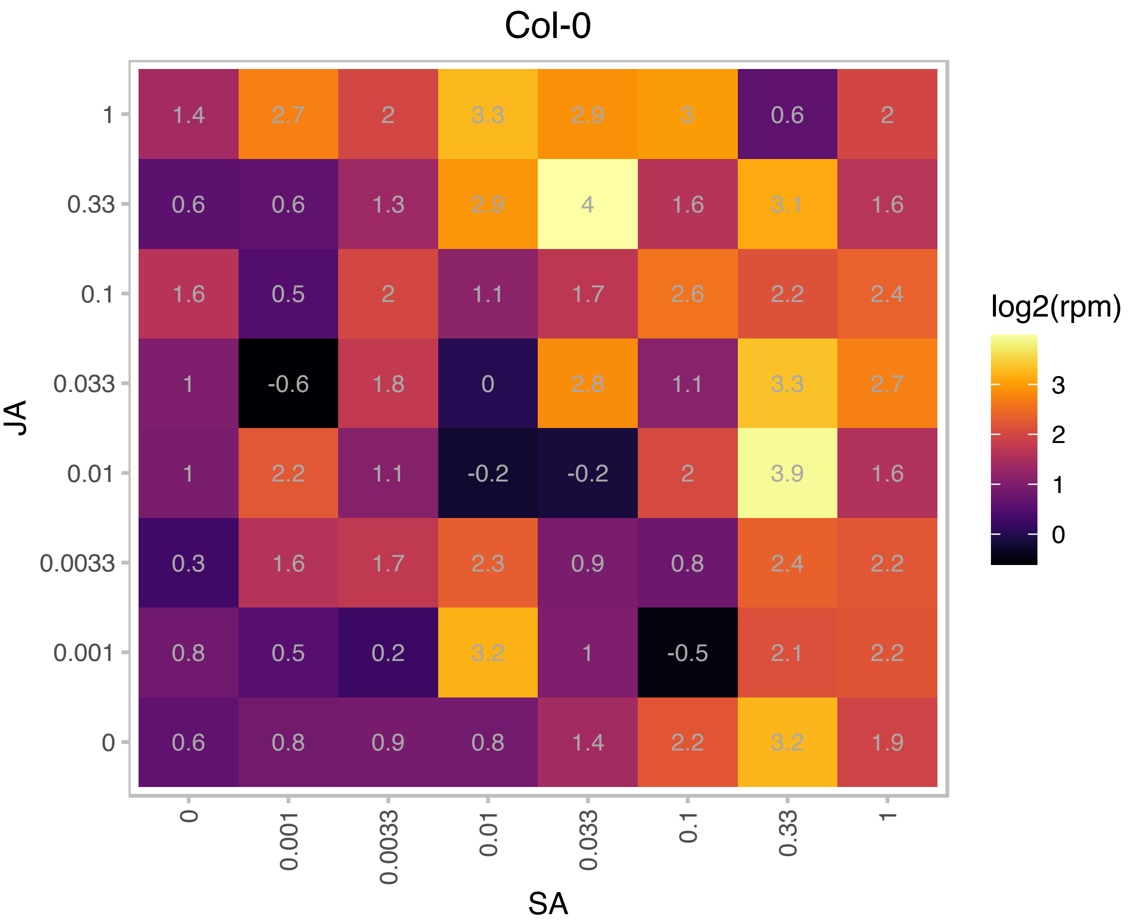


| | |
|---|---|
| Gene | AT1G63720 |
| is.expressed_genes | TRUE |
| is.responsive_genes | TRUE |
| is.apcluster | C-04 |

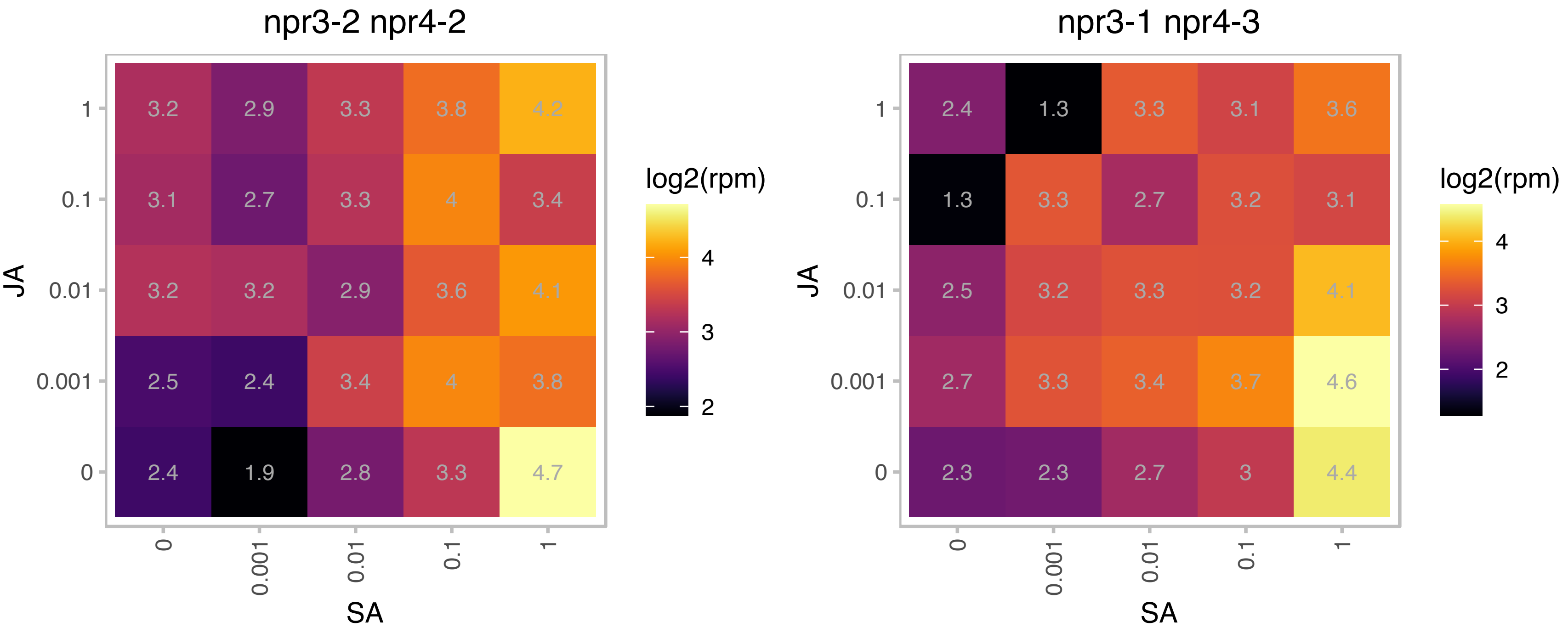

### AT1G64980.1 : CDI

Nucleotide-diphospho-sugar transferases superfamily protein

Encodes a putative nucleotide-diphospho-sugar transferase required for pollen germination and tube growth.

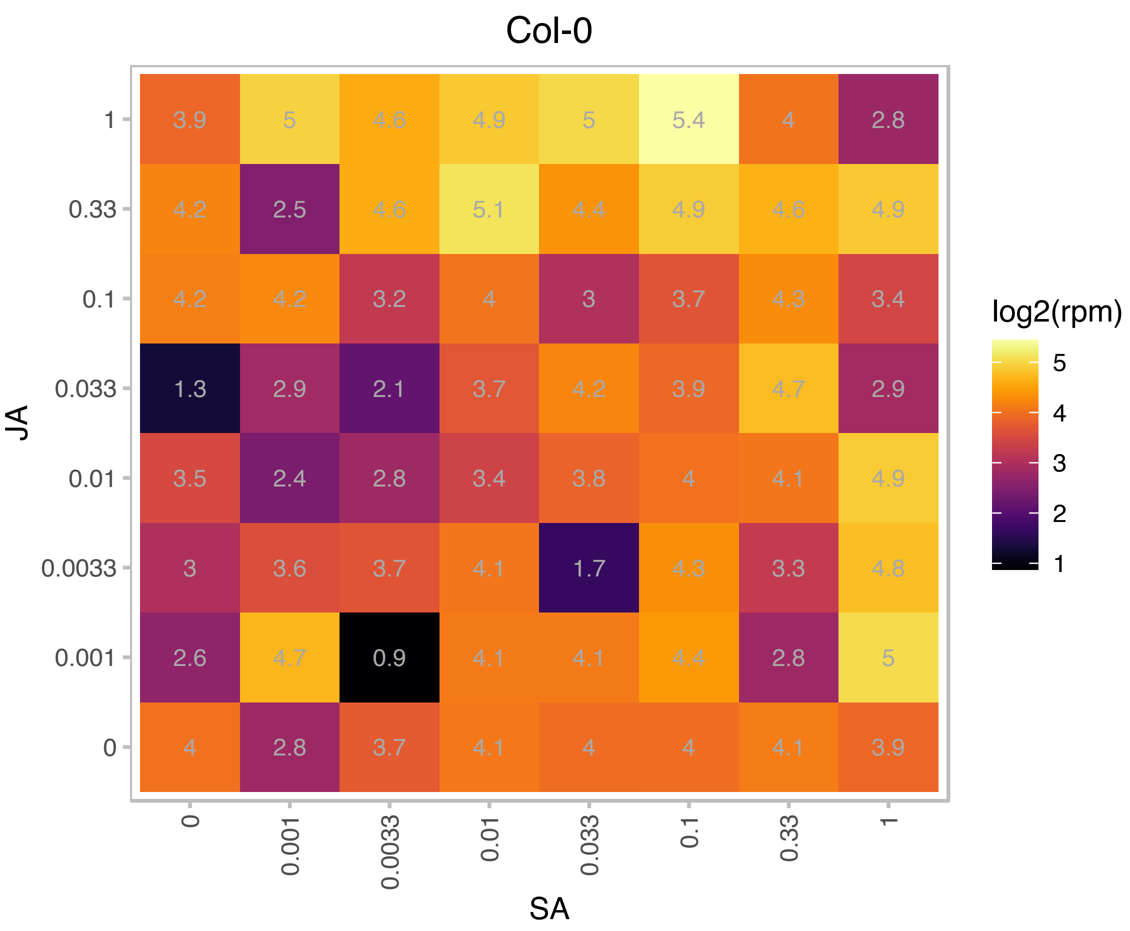


| | |
|---|---|
| Gene | AT1G64980 |
| is.expressed_genes | TRUE |
| is.responsive_genes | TRUE |
| is.apcluster | C-04 |

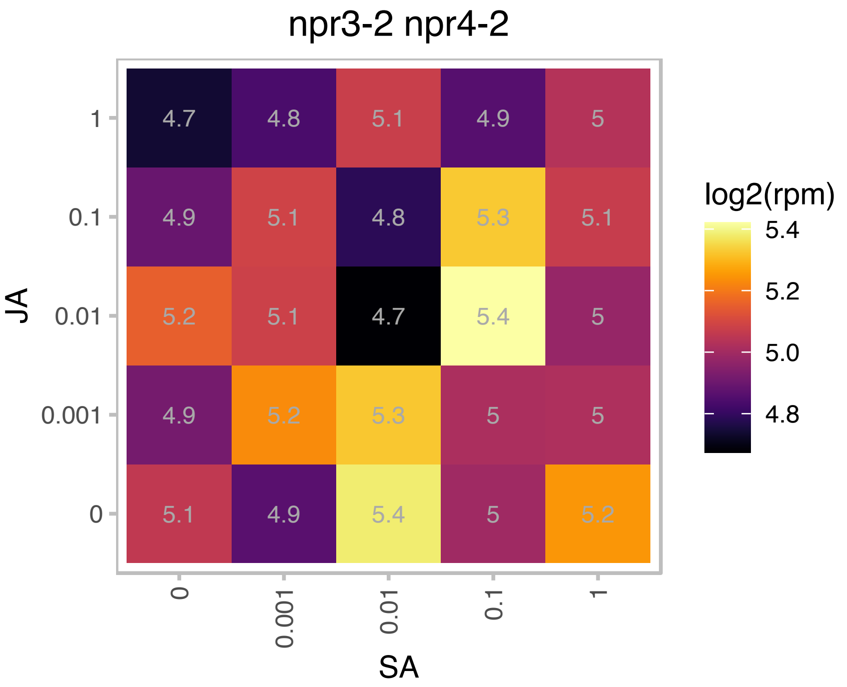


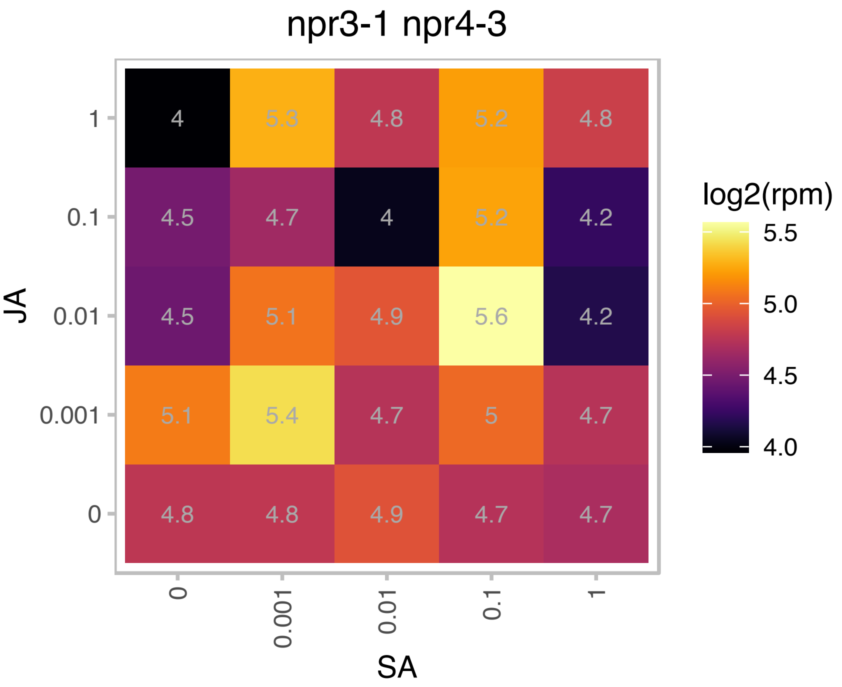

### AT1G67800.1 : RGLG5

Copine (Calcium-dependent phospholipid-binding protein) family

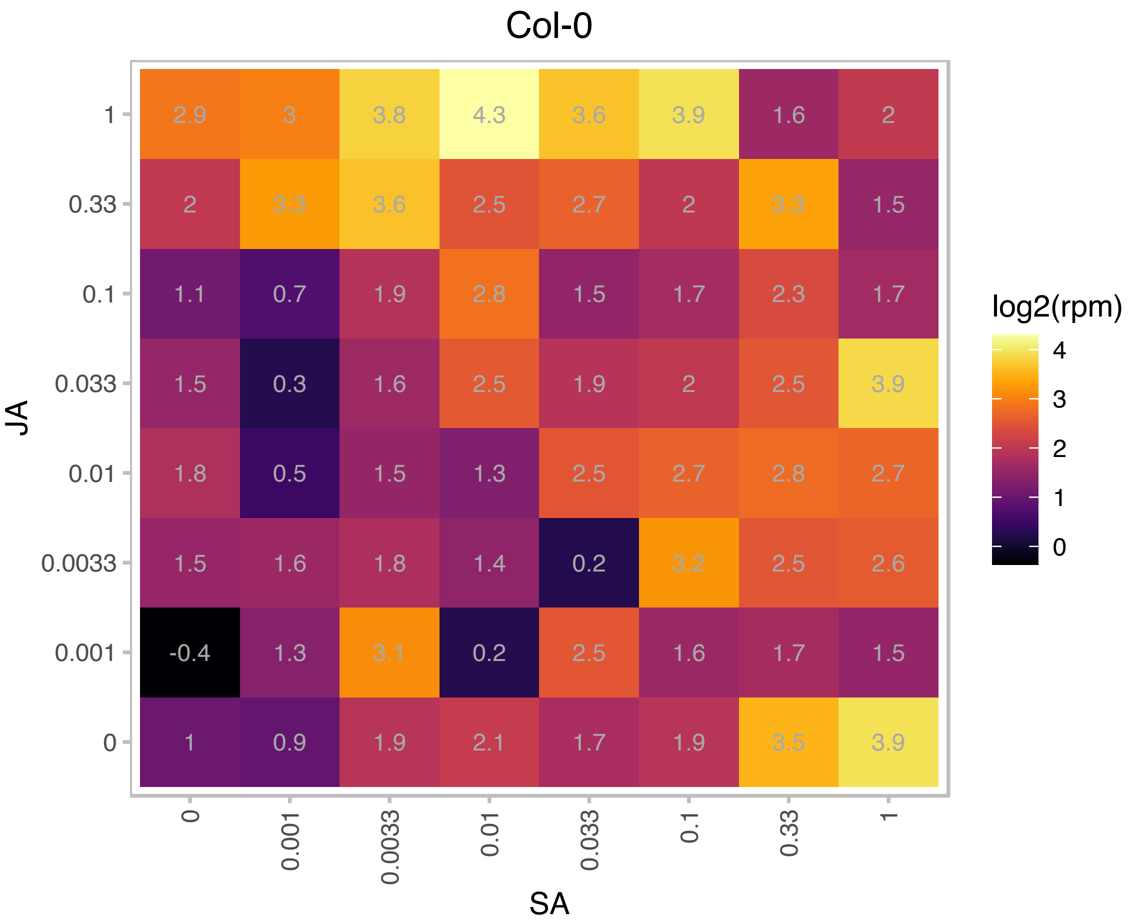


| | |
|---|---|
| Gene | AT1G67800 |
| is.expressed_genes | TRUE |
| is.responsive_genes | TRUE |
| is.apcluster | C-04 |

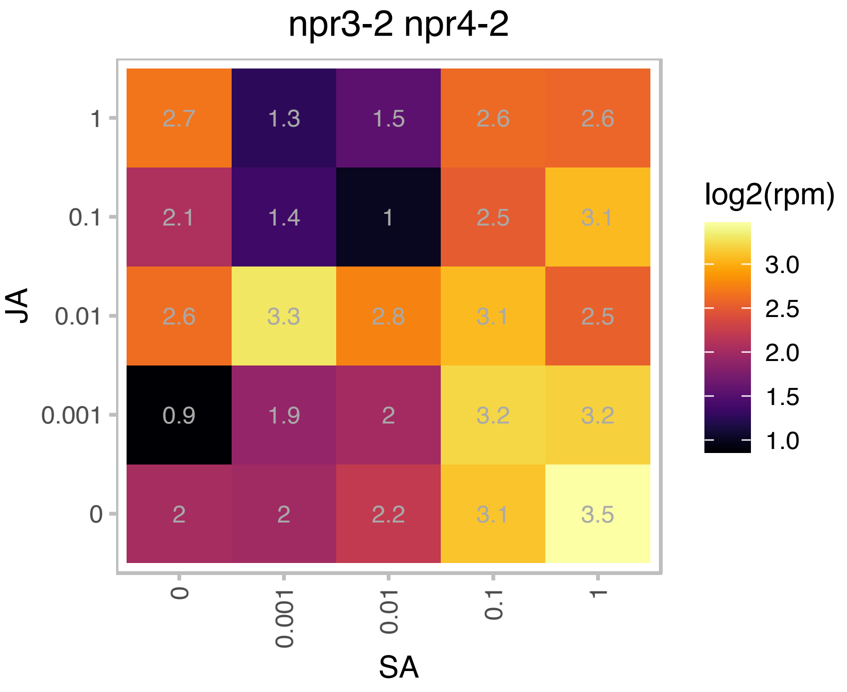


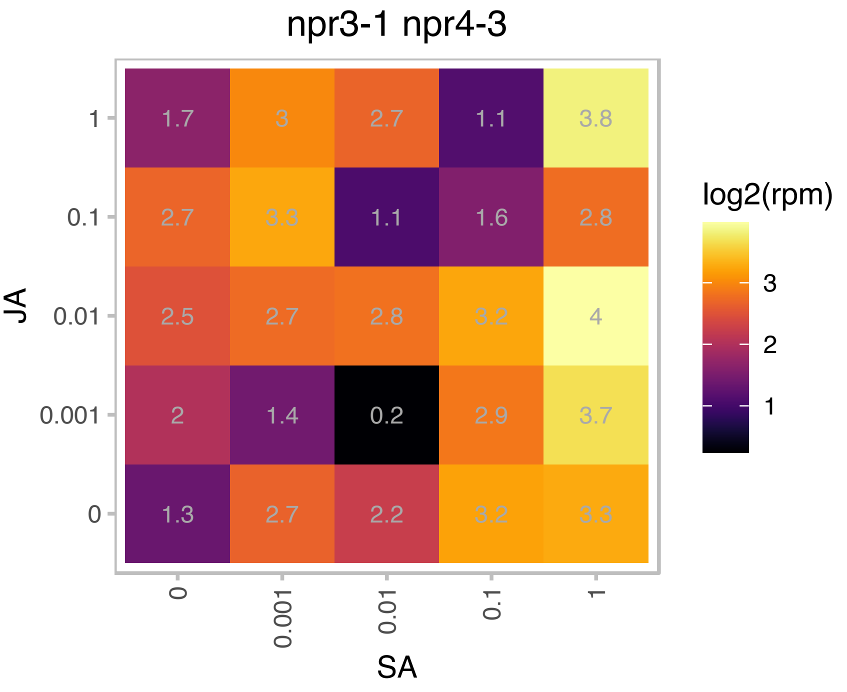

### AT1G72750.1 : ATTIM23-2, TIM23-2

translocase inner membrane subunit 23-2

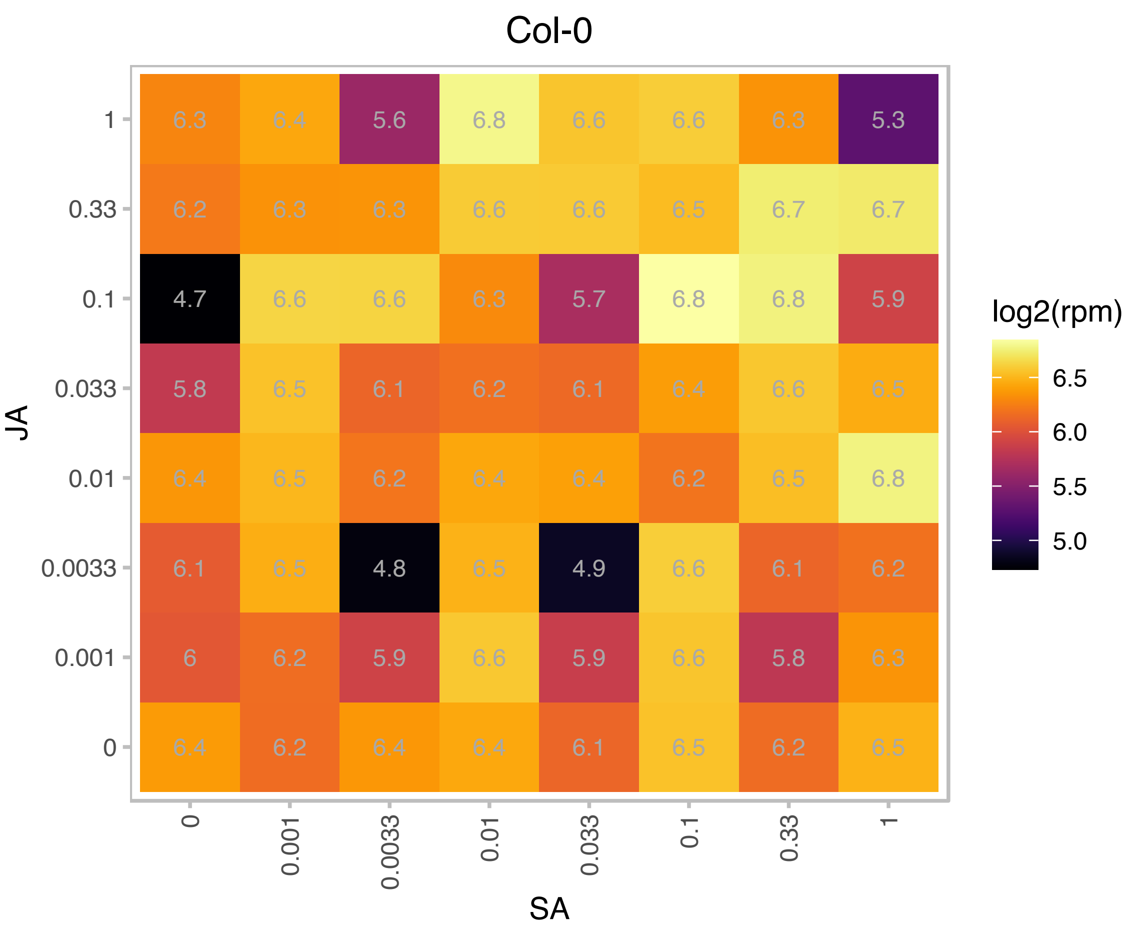


| | |
|---|---|
| Gene | AT1G72750 |
| is.expressed_genes | TRUE |
| is.responsive_genes | TRUE |
| is.apcluster | C-04 |

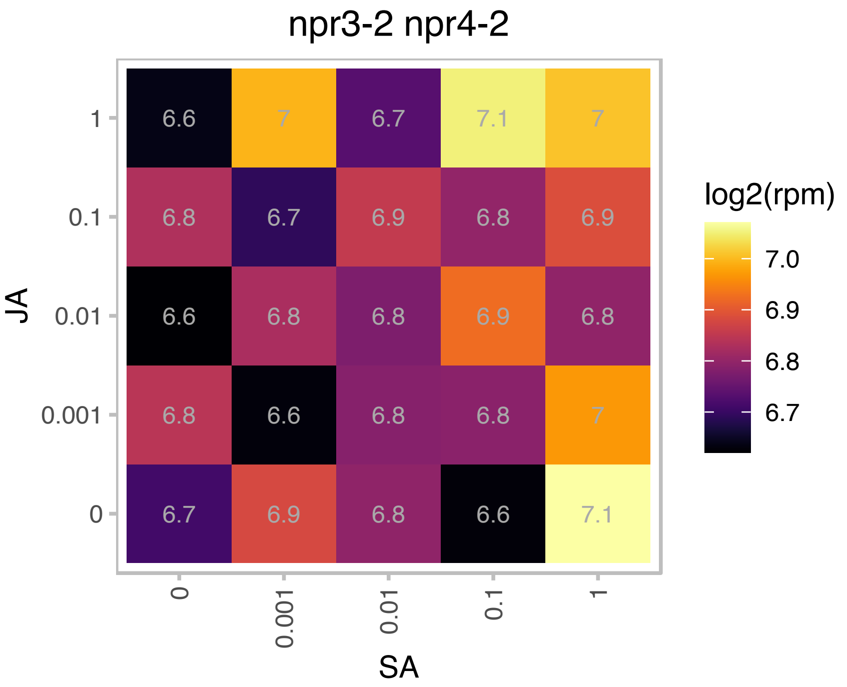


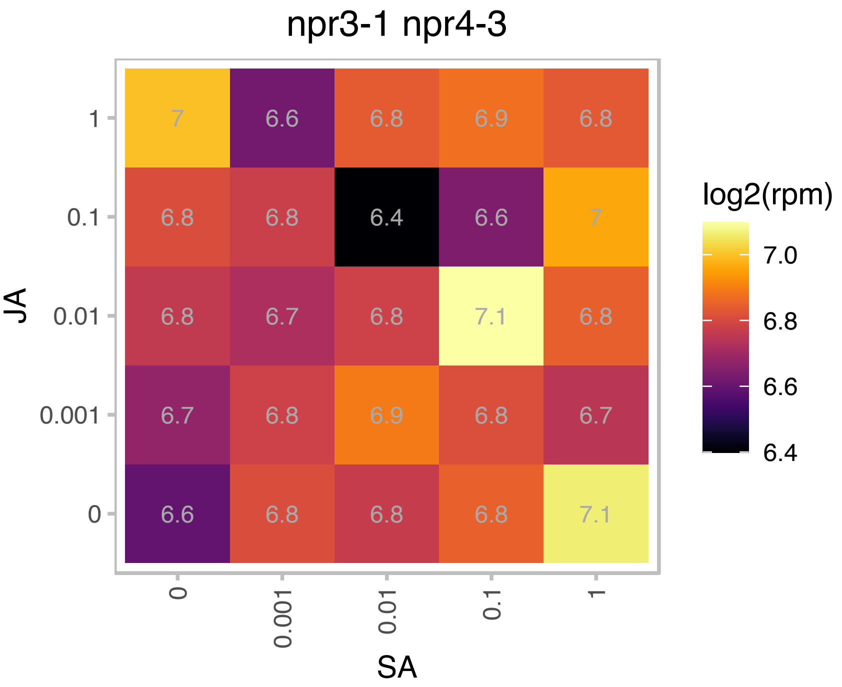

### AT1G73730.1 : AtEIL3, ATSLIM, EIL3, SLIM1

ETHYLENE-INSENSITIVE3-like 3

Encodes a putative transcription factor involved in ethylene signalling.  Isolated DNA binding domain has been shown to bind DNA in vitro.

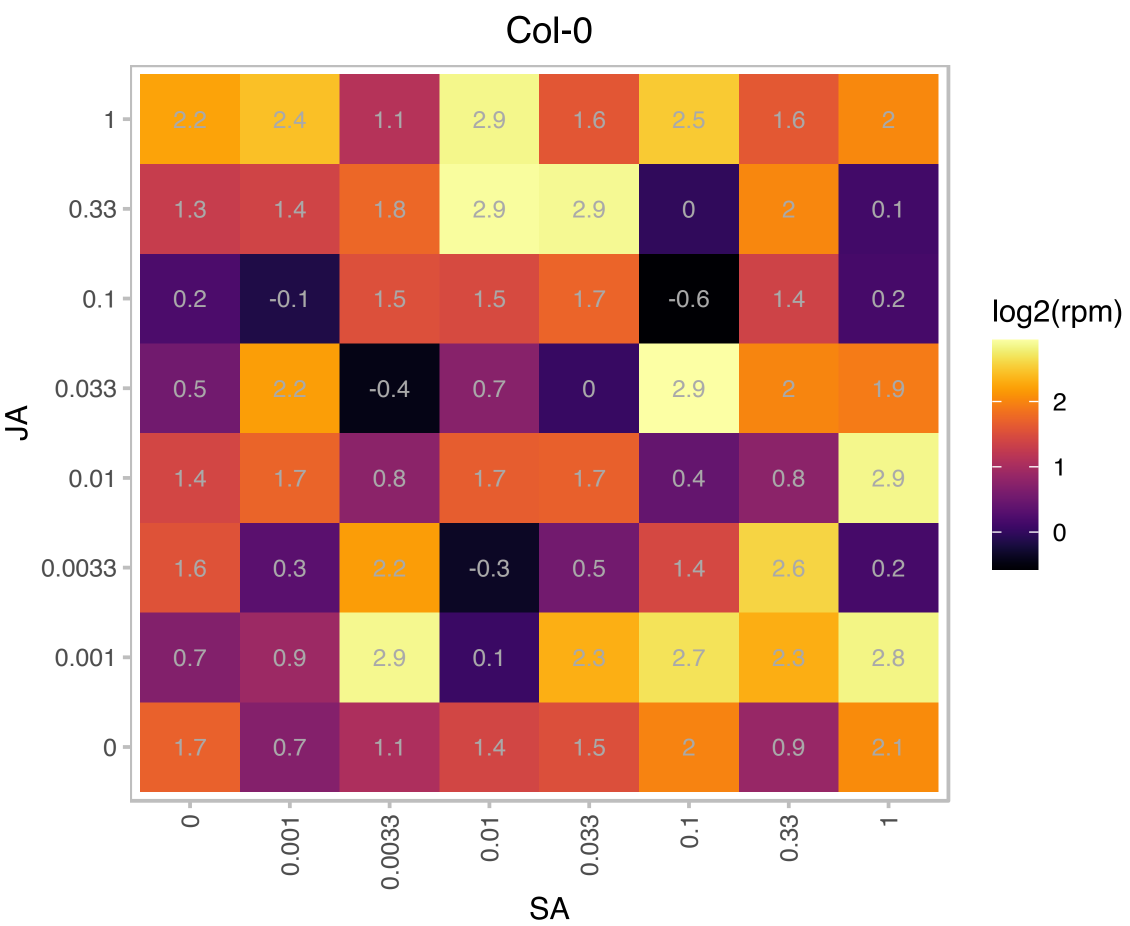


| | |
|---|---|
| Gene | AT1G73730 |
| is.expressed_genes | TRUE |
| is.responsive_genes | TRUE |
| is.apcluster | C-04 |

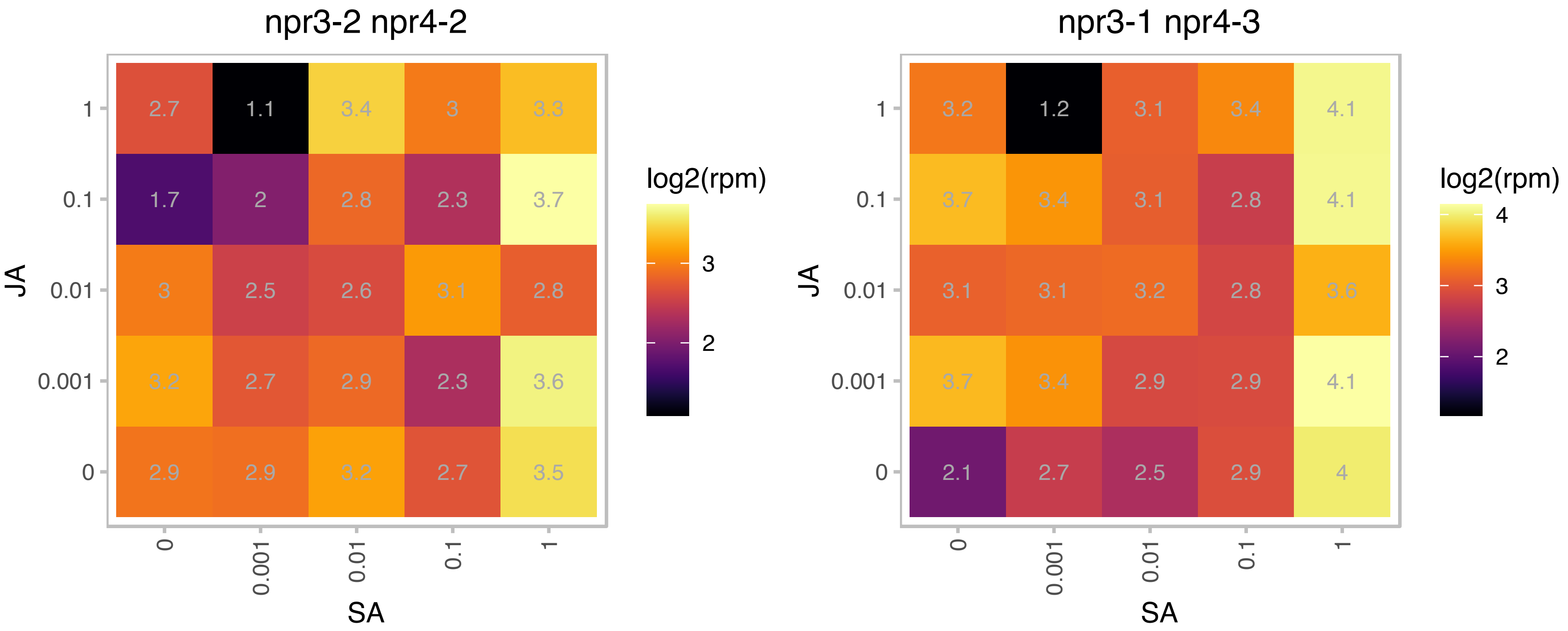

### AT1G78340.1 : ATGSTU22, GSTU22

glutathione S-transferase TAU 22

Encodes glutathione transferase belonging to the tau class of GSTs. Naming convention according to Wagner et al. (2002).

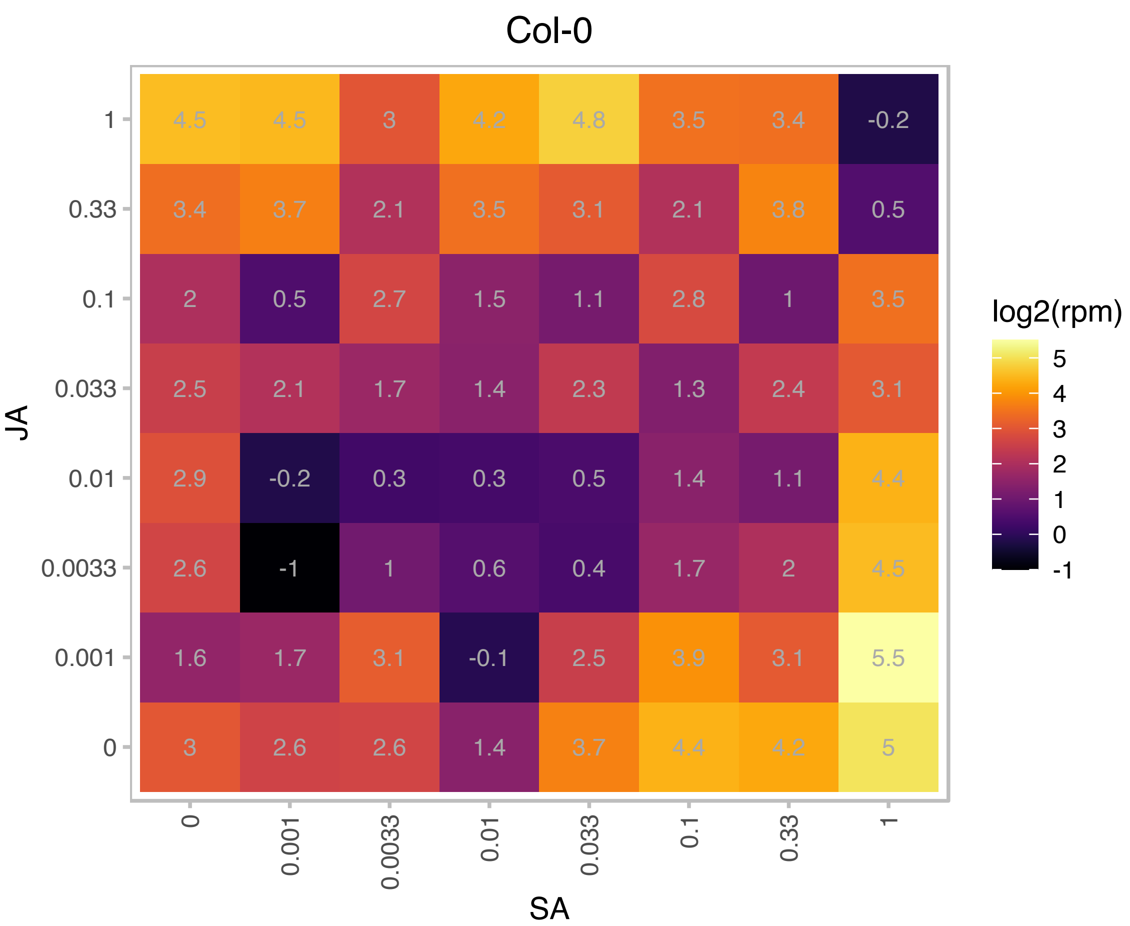


| | |
|---|---|
| Gene | AT1G78340 |
| is.expressed_genes | TRUE |
| is.responsive_genes | TRUE |
| is.apcluster | C-04 |

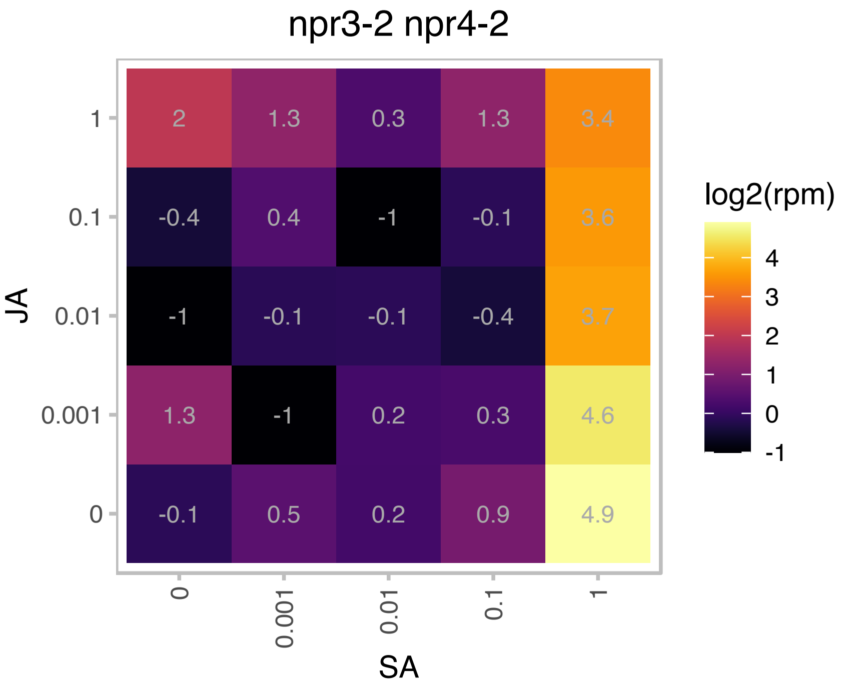


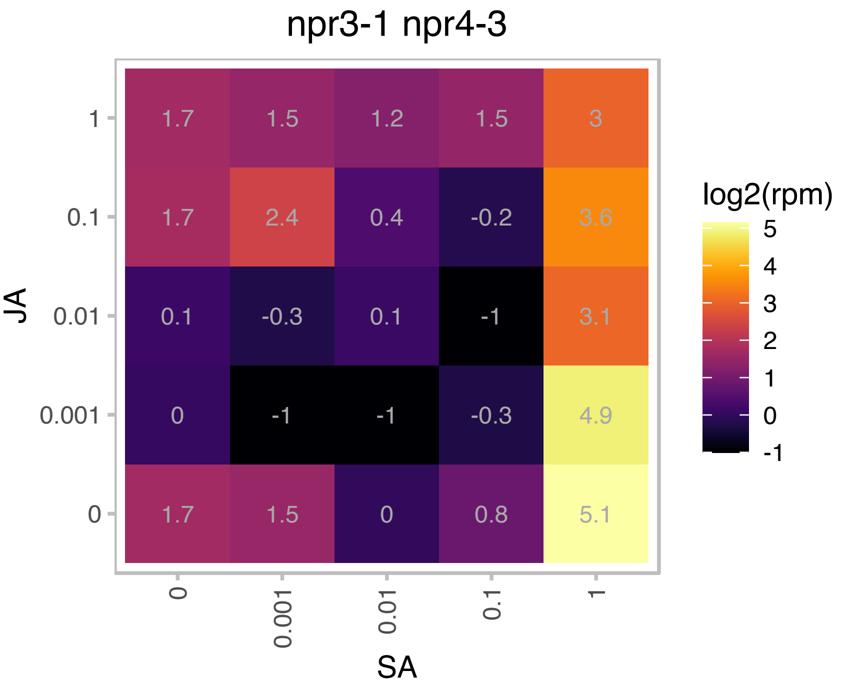

# AT2G14850.1 : NA

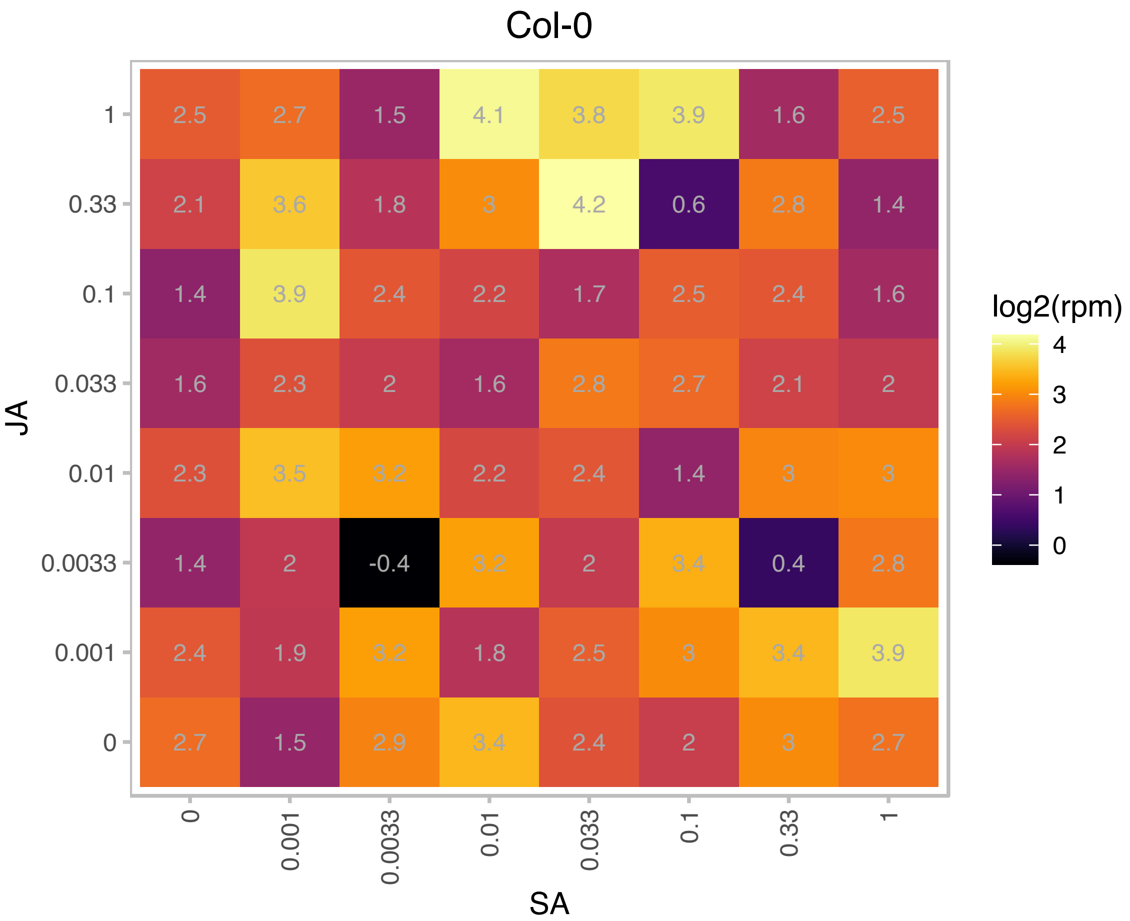


| | |
|---|---|
| Gene | AT2G14850 |
| is.expressed_genes | TRUE |
| is.responsive_genes | TRUE |
| is.apcluster | C-04 |

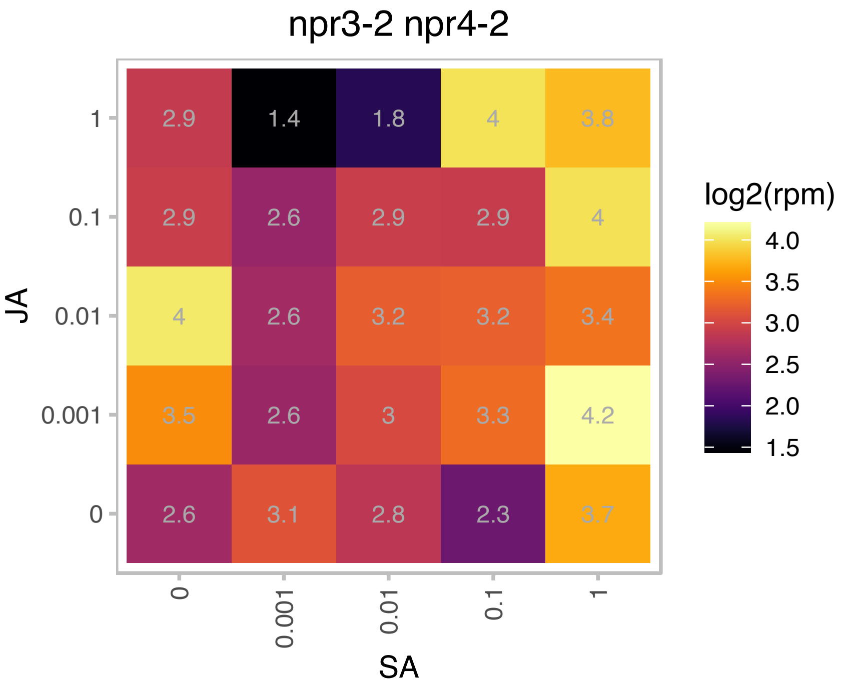


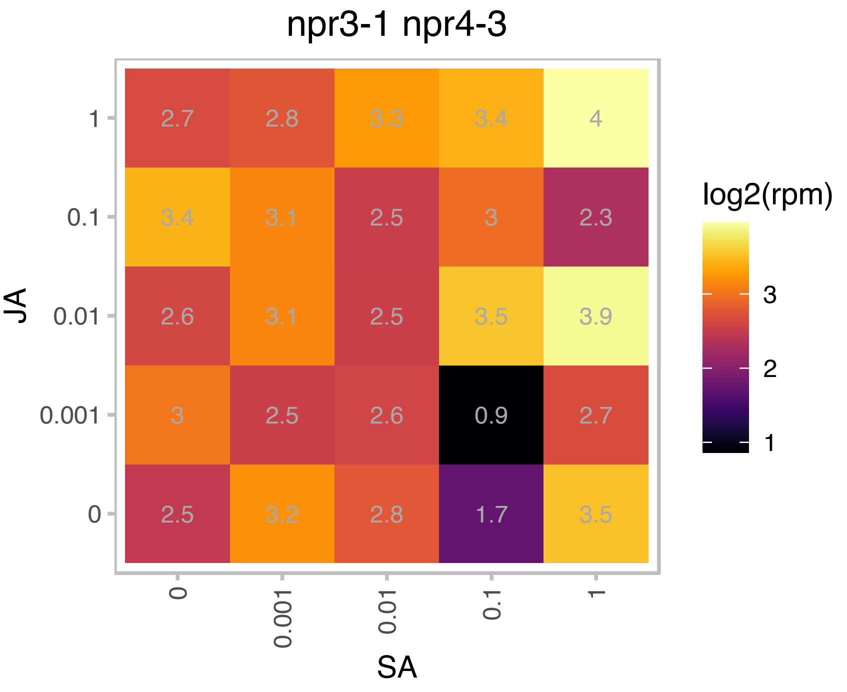

### AT2G31400.1 : GUN1

genomes uncoupled 1

Encodes a a chloroplast-localized pentatricopeptide-repeat protein involved in regulation of nuclear gene expression.

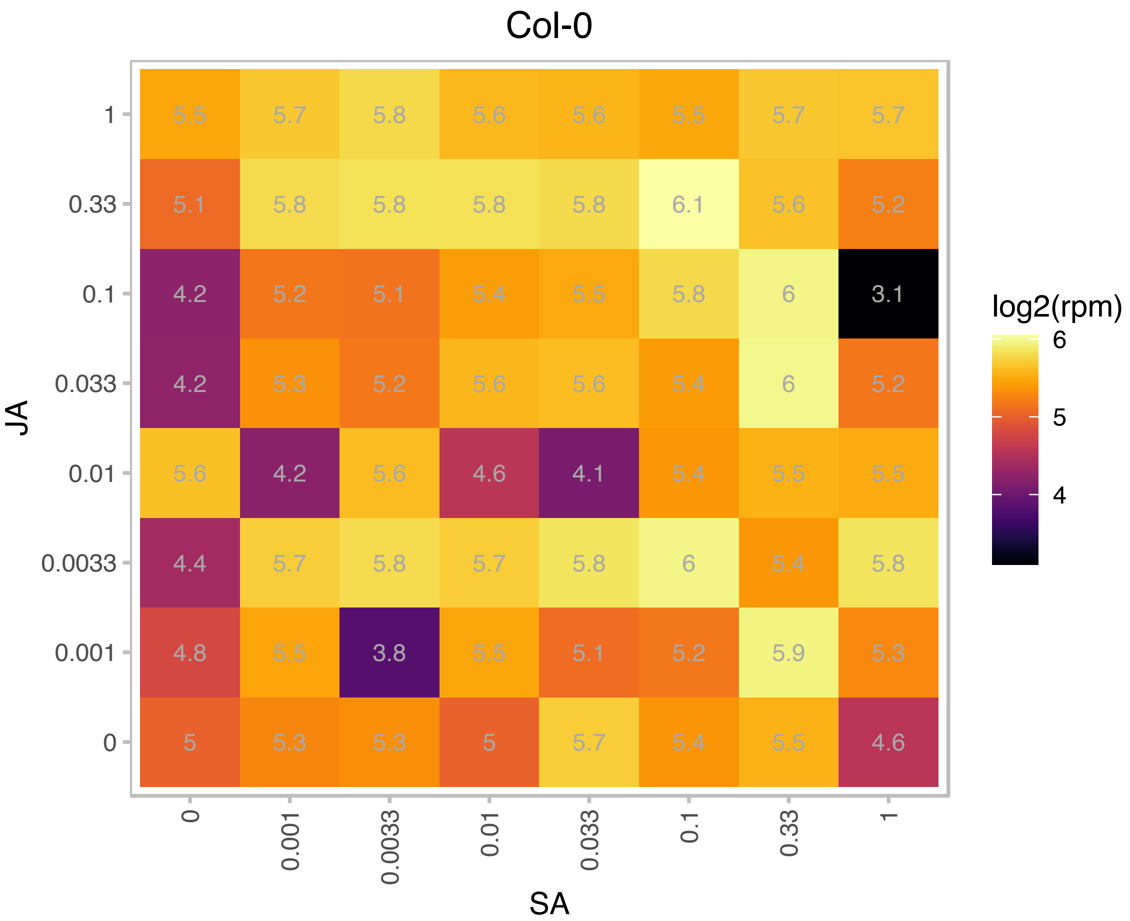


| | |
|---|---|
| Gene | AT2G31400 |
| is.expressed_genes | TRUE |
| is.responsive_genes | TRUE |
| is.apcluster | C-04 |

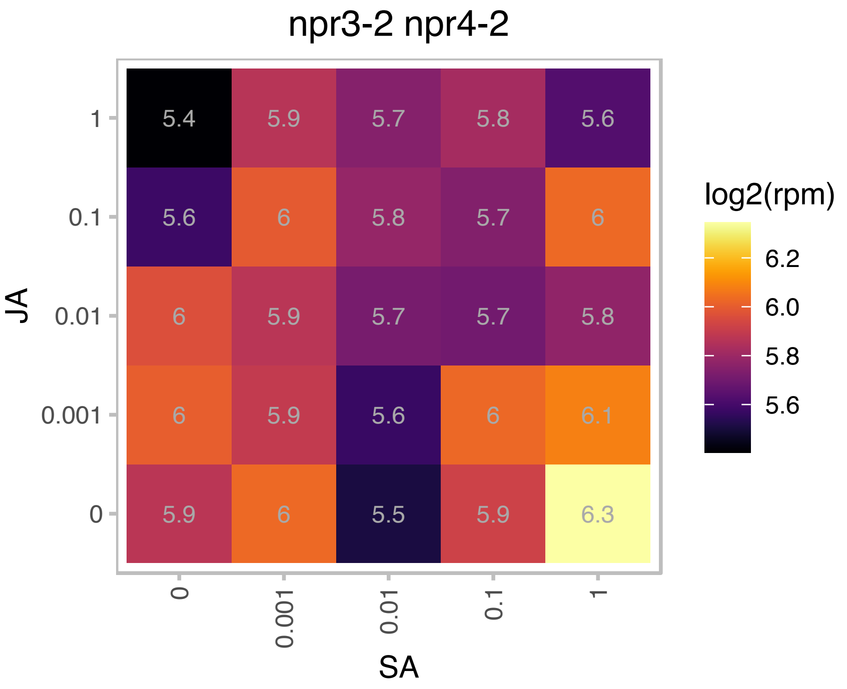


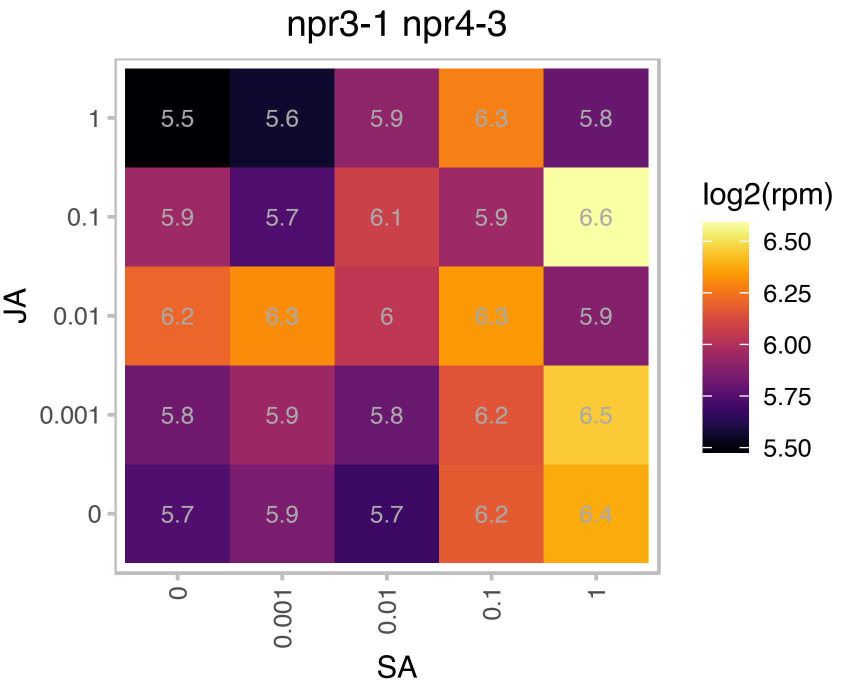

# AT2G35840.1 : NA

Sucrose-6F-phosphate phosphohydrolase family protein

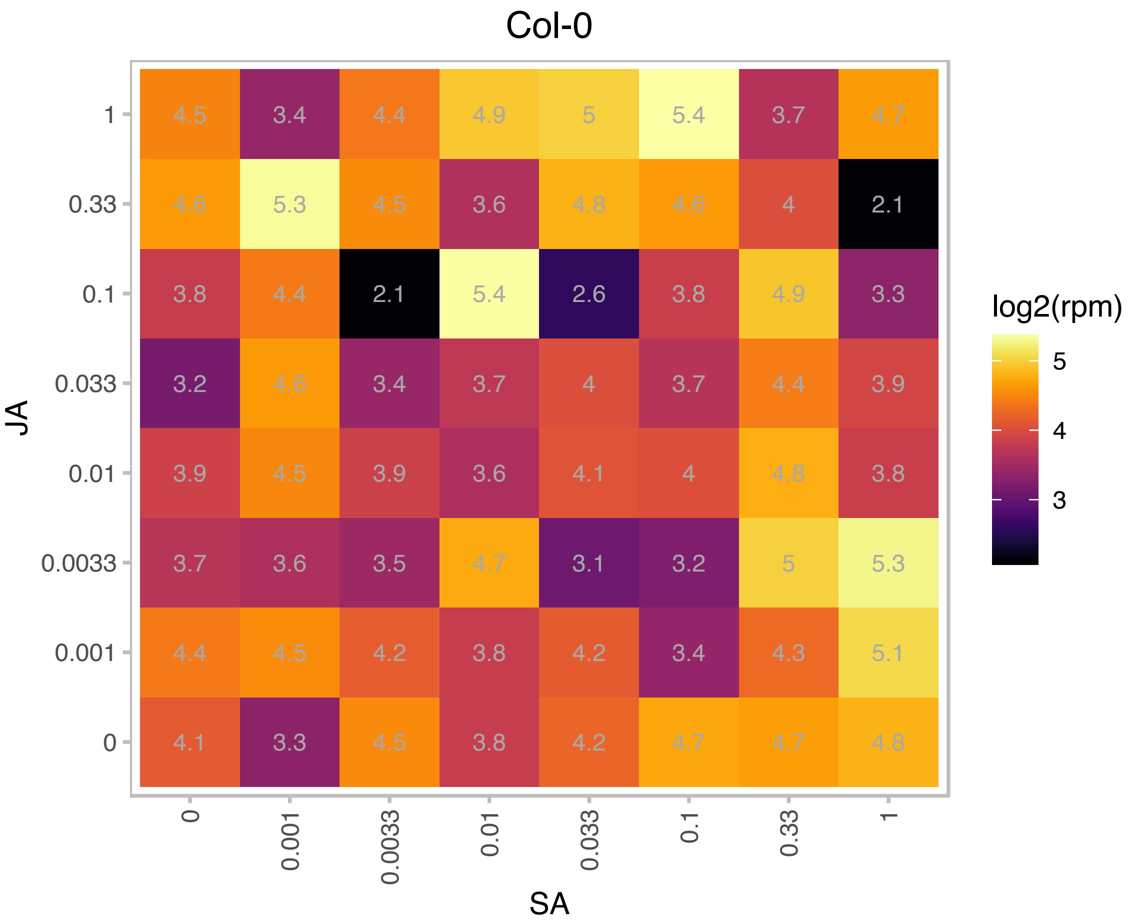


| | |
|---|---|
| Gene | AT2G35840 |
| is.expressed_genes | TRUE |
| is.responsive_genes | TRUE |
| is.apcluster | C-04 |

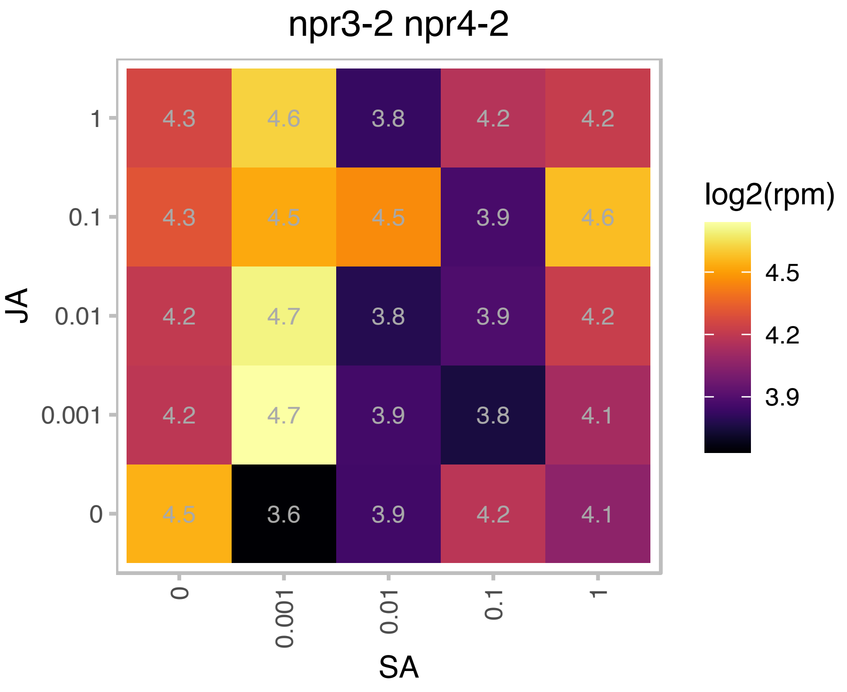


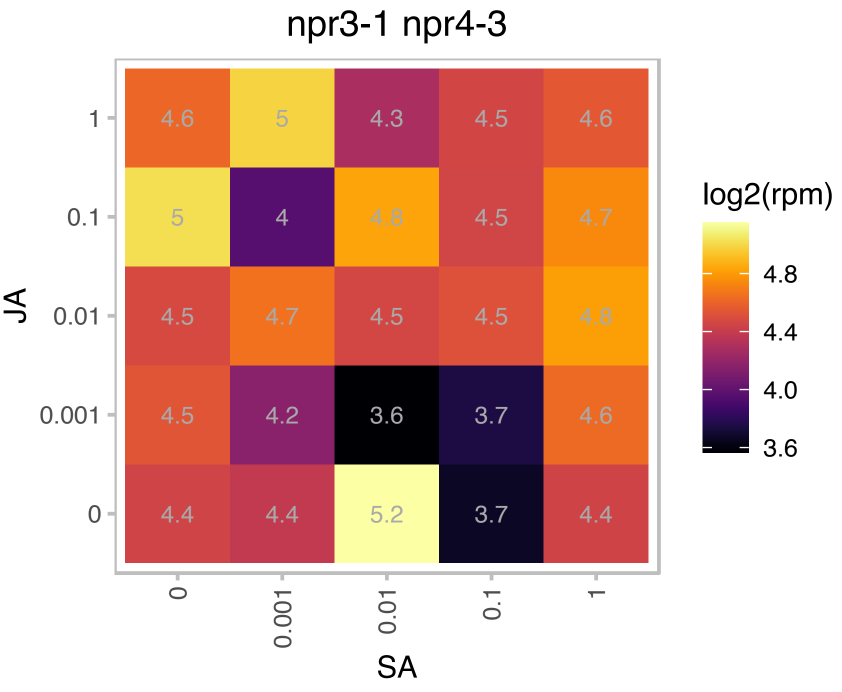

### AT2G38130.1 : ATMAK3

Acyl-CoA N-acyltransferases (NAT) superfamily protein
Encodes the Arabidopsis homolog of the yeast protein MAK3, a component of the N-terminal acetyltransferase complex C. In mutant plants, synthesis of plastome-encoded photosystem II core proteins D1 and CP47 is affected resulting in fewer thylakoid multiprotein complexes.

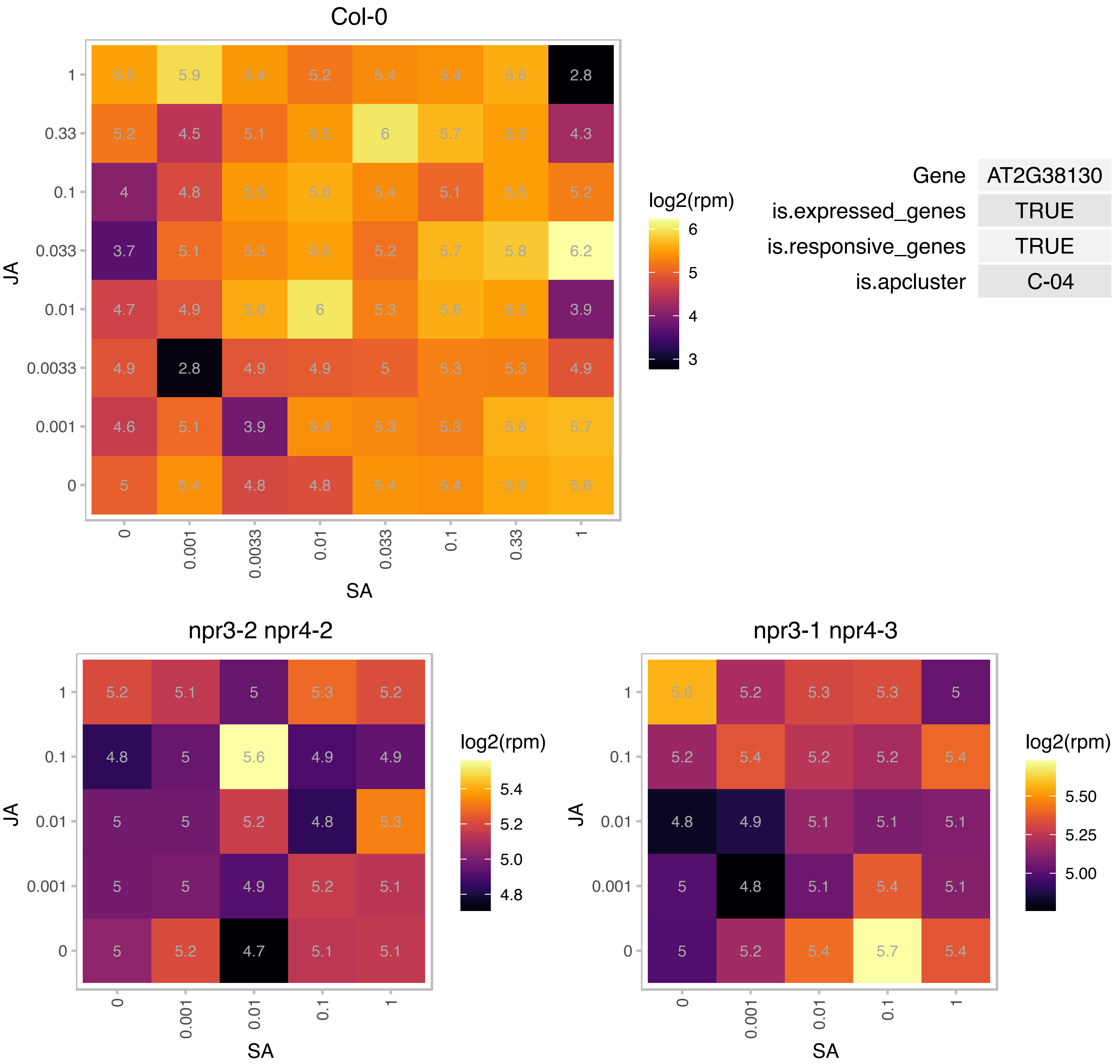


| | |
|---|---|
| Gene | AT2G38130 |
| is.expressed_genes | TRUE |
| is.responsive_genes | TRUE |
| is.apcluster | C-04 |

### AT2G41220.1 : GLU2

glutamate synthase 2

Encodes a gene whose sequence is similar to ferredoxin dependent glutamate synthase (Fd-GOGAT). Expression i
most abundant in root. The mRNA is cell-to-cell mobile.

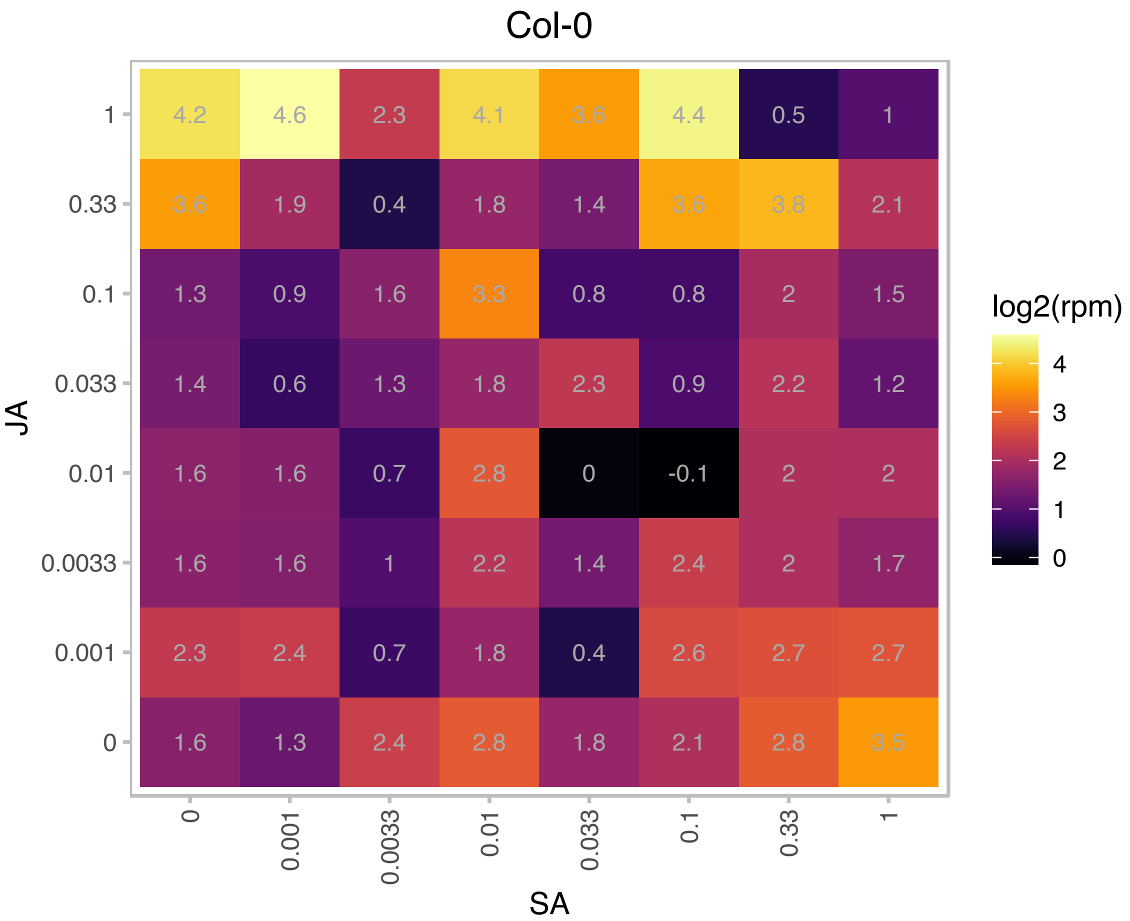


| | |
|---|---|
| Gene | AT2G41220 |
| is.expressed_genes | TRUE |
| is.responsive_genes | TRUE |
| is.apcluster | C-04 |

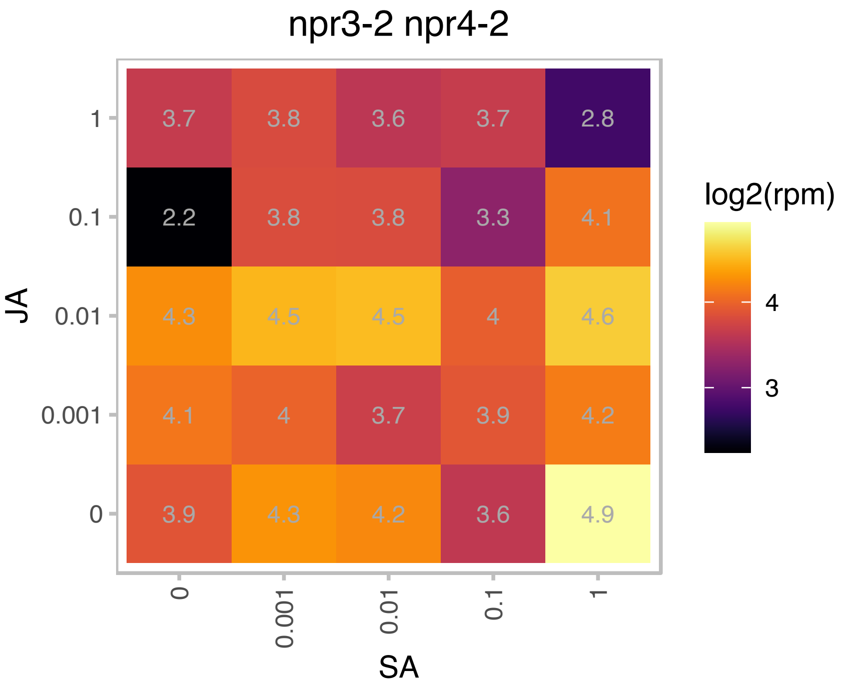


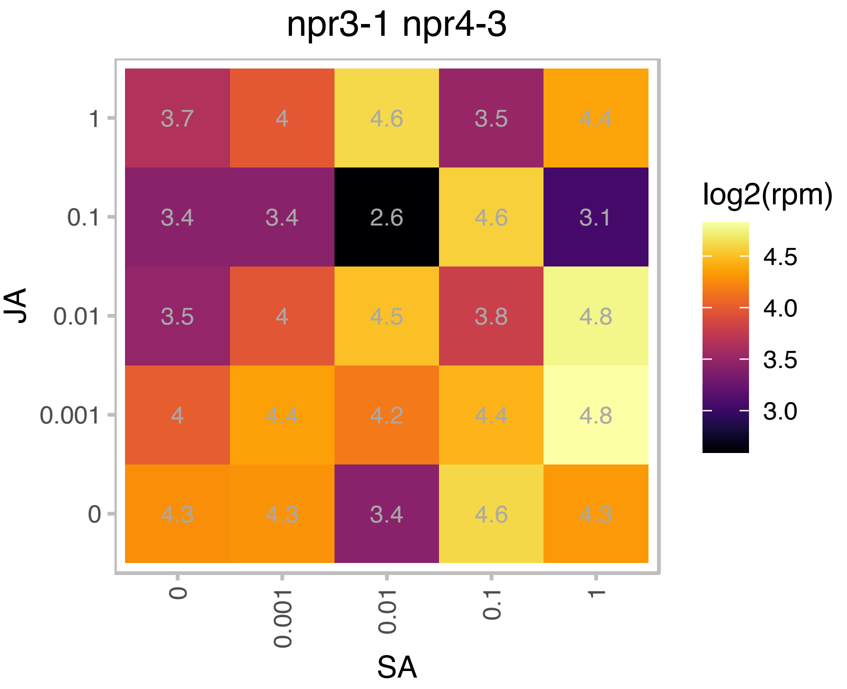

# AT2G45500.1 : NA

AAA-type ATPase family protein

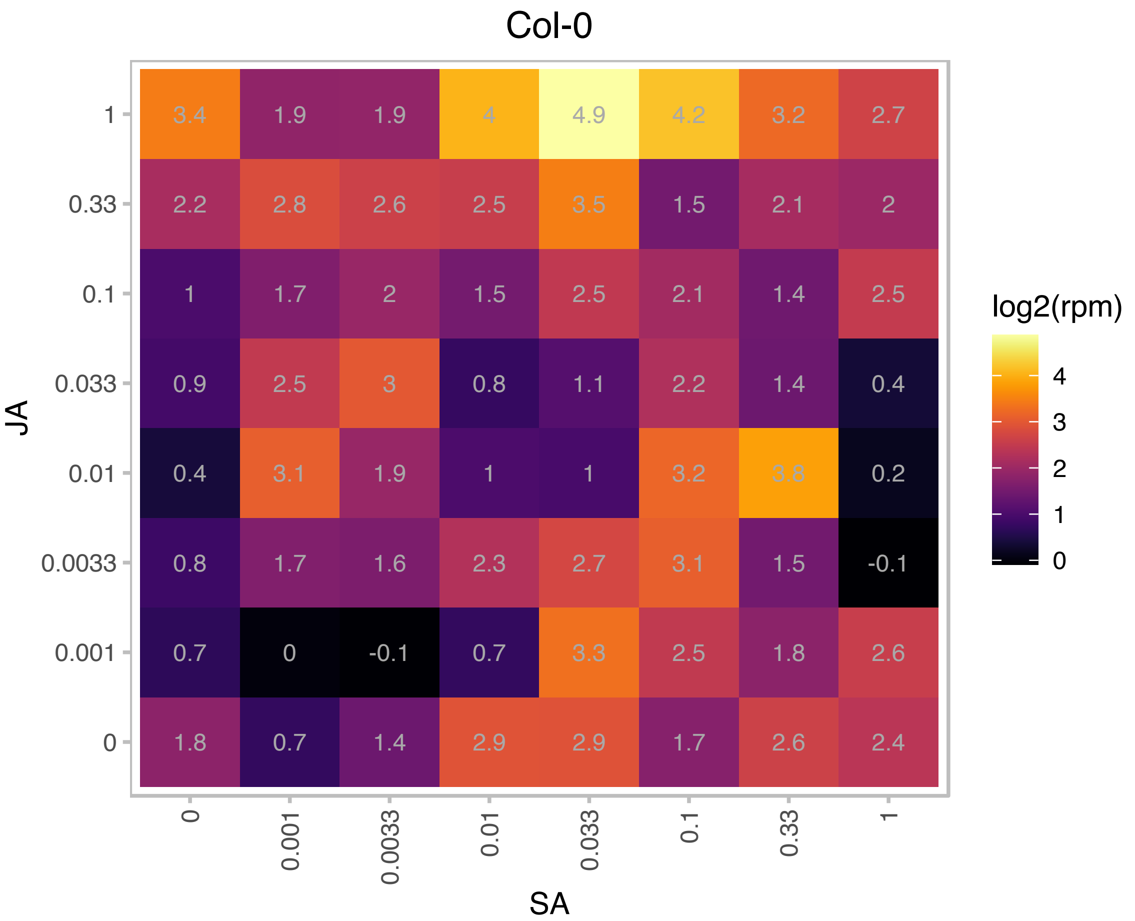


| | |
|---|---|
| Gene | AT2G45500 |
| is.expressed_genes | TRUE |
| is.responsive_genes | TRUE |
| is.apcluster | C-04 |

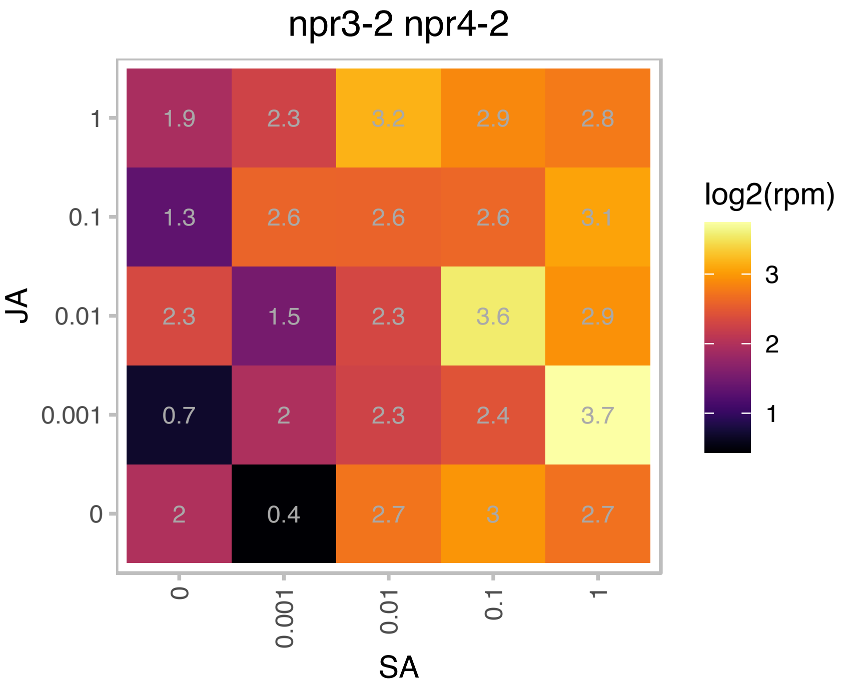


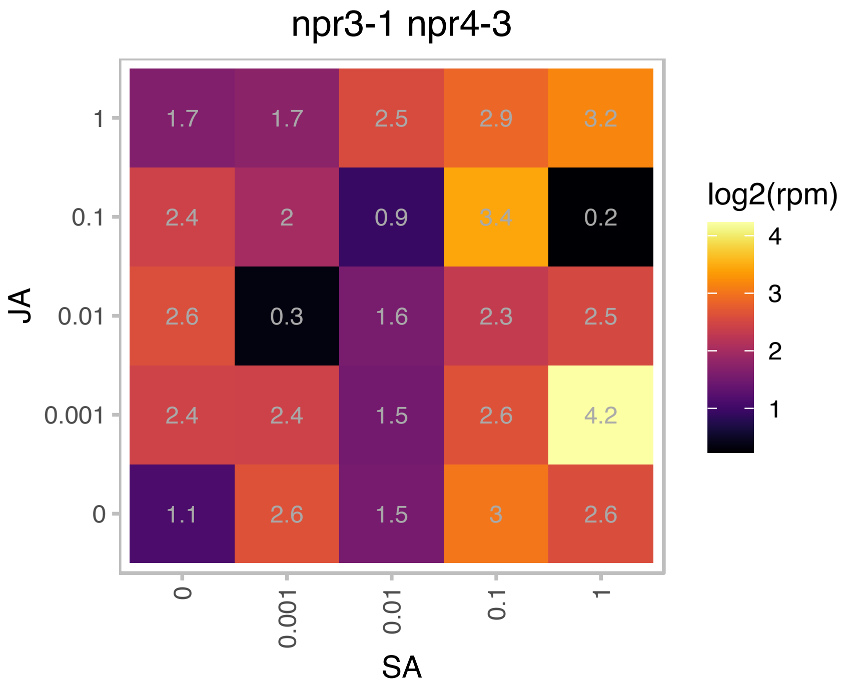

# AT2G46915.1 : NA

Protein of unknown function (DUF3754)

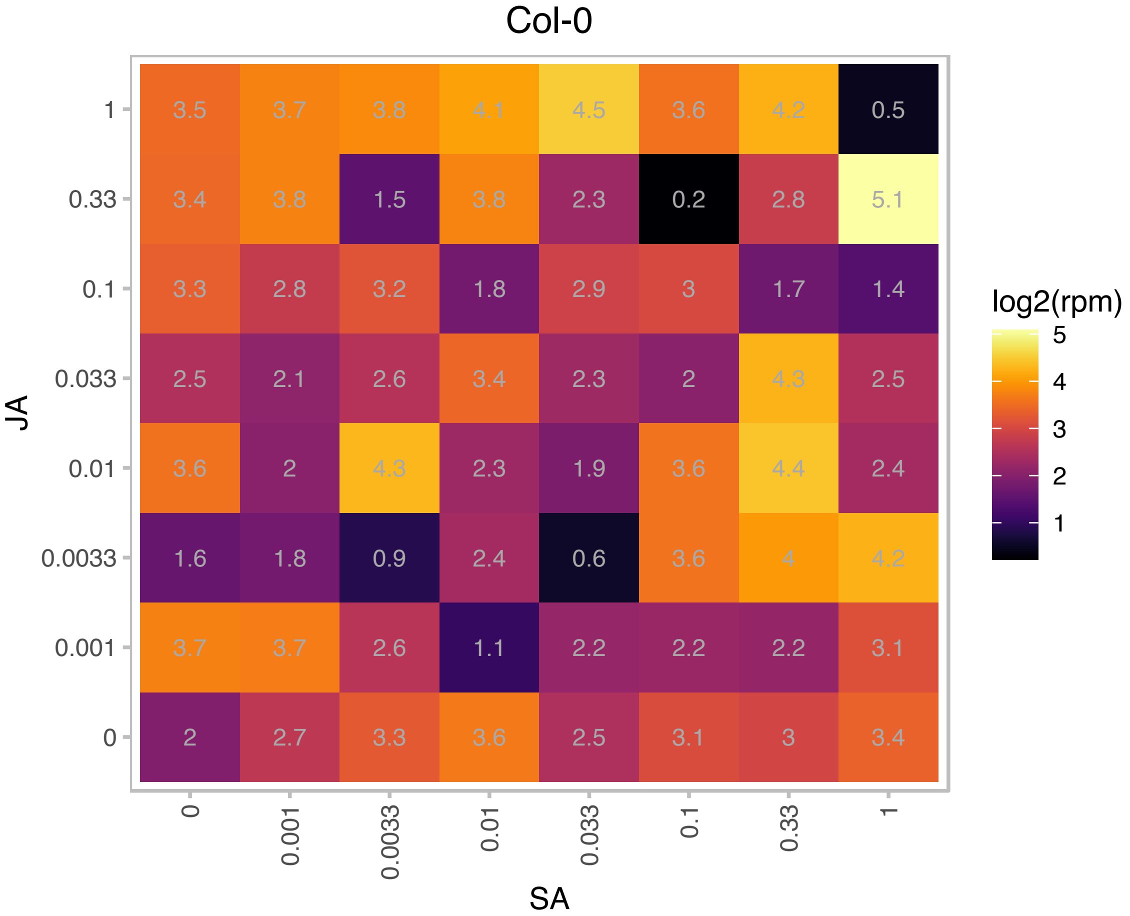


| | |
|---|---|
| Gene | AT2G46915 |
| is.expressed_genes | TRUE |
| is.responsive_genes | TRUE |
| is.apcluster | C-04 |

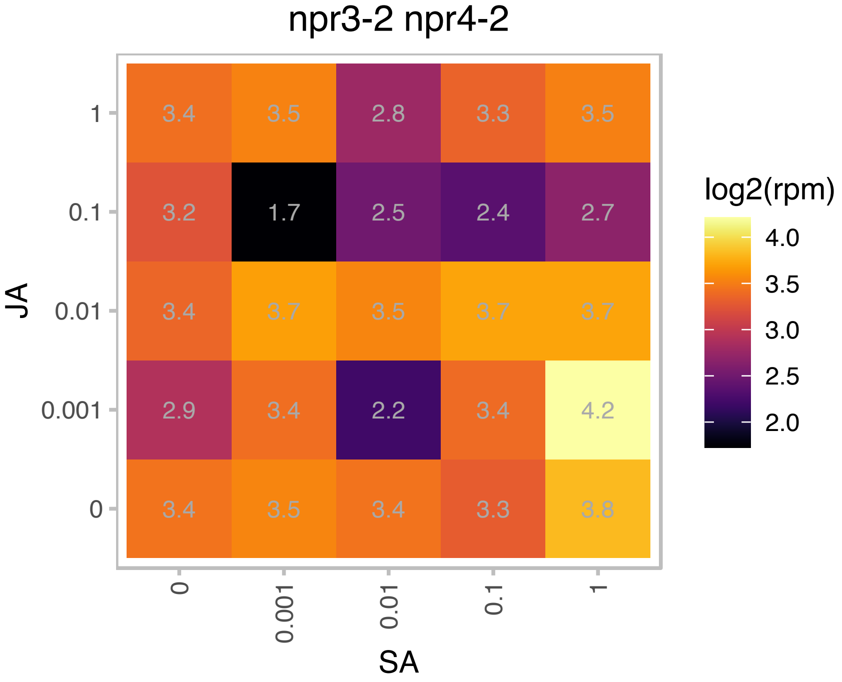


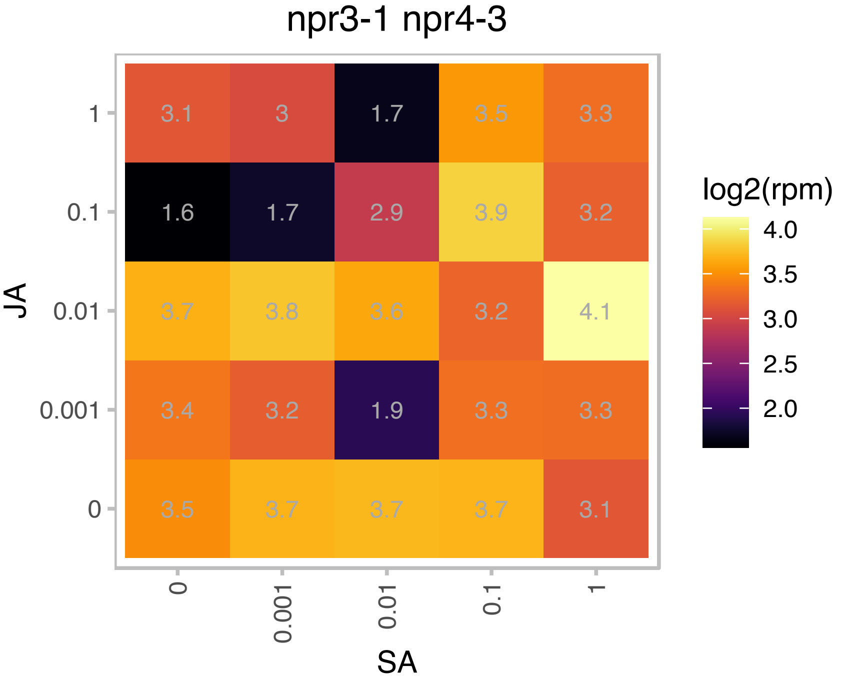

# AT3G05280.1 : NA

Integral membrane Yip1 family protein

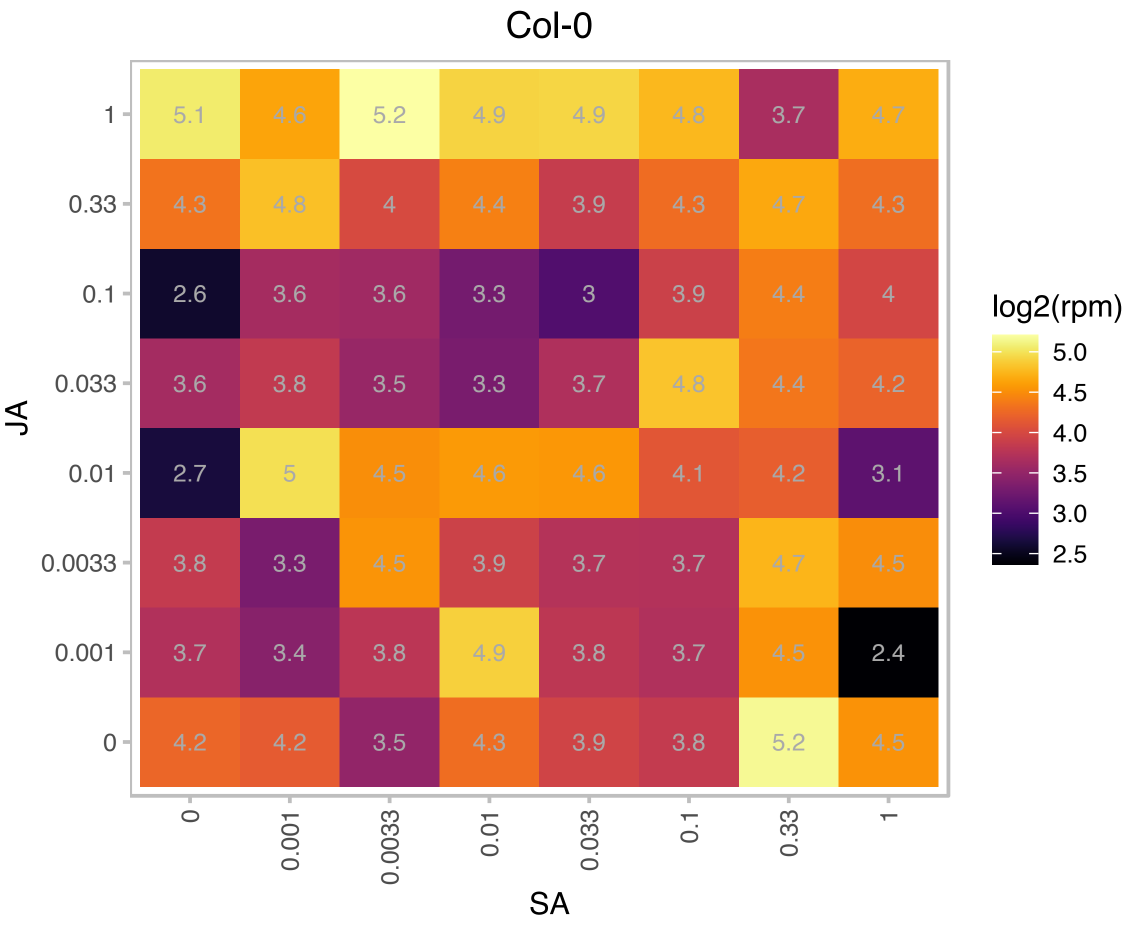


| | |
|---|---|
| Gene | AT3G05280 |
| is.expressed_genes | TRUE |
| is.responsive_genes | TRUE |
| is.apcluster | C-04 |

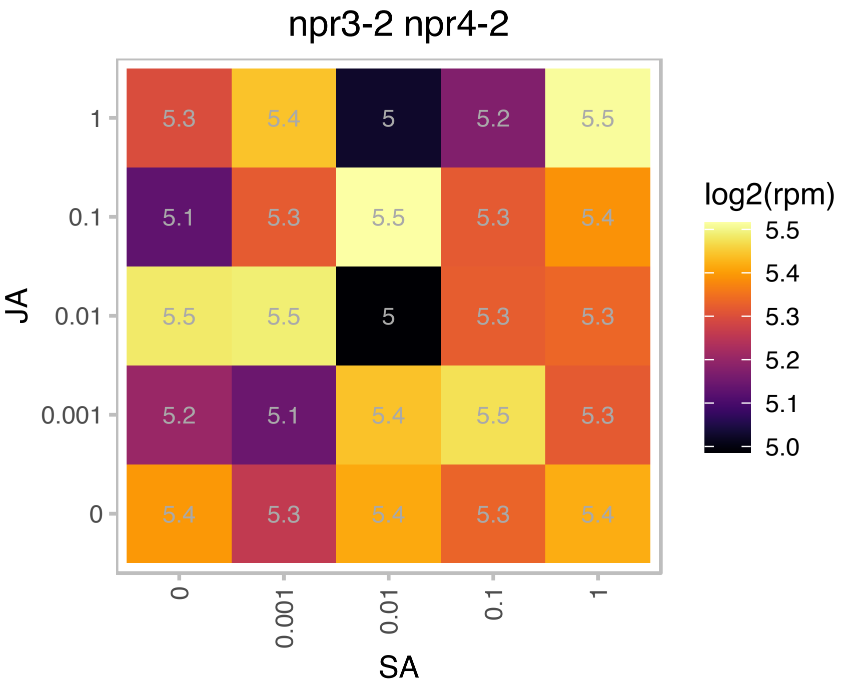


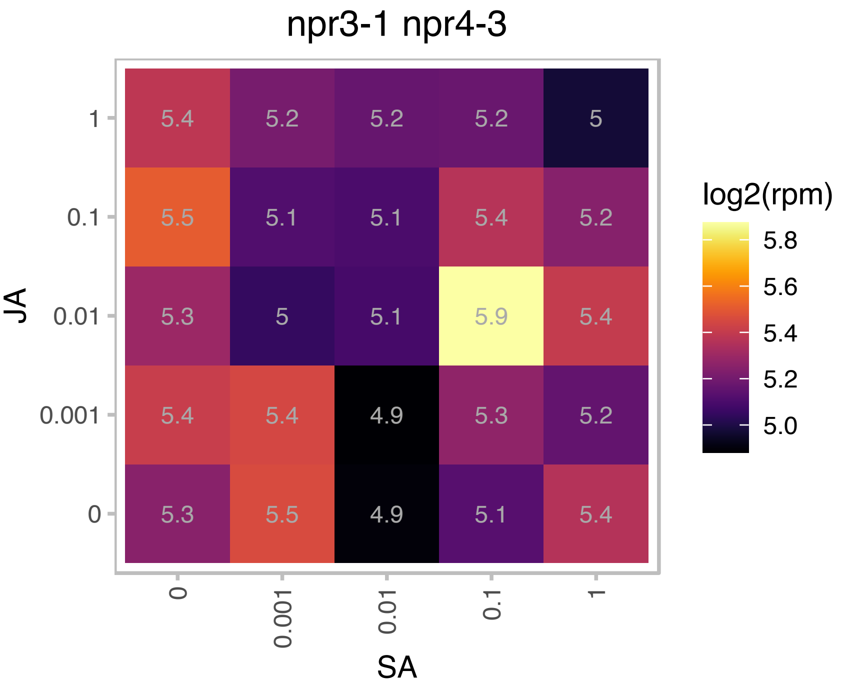

# AT3G22620.1 : NA

Bifunctional inhibitor/lipid-transfer protein/seed storage 2S albumin superfamily protein

| | |
|---|---|
| Gene | AT3G22620 |
| is.expressed_genes | TRUE |
| is.responsive_genes | TRUE |
| is.apcluster | C-04 |

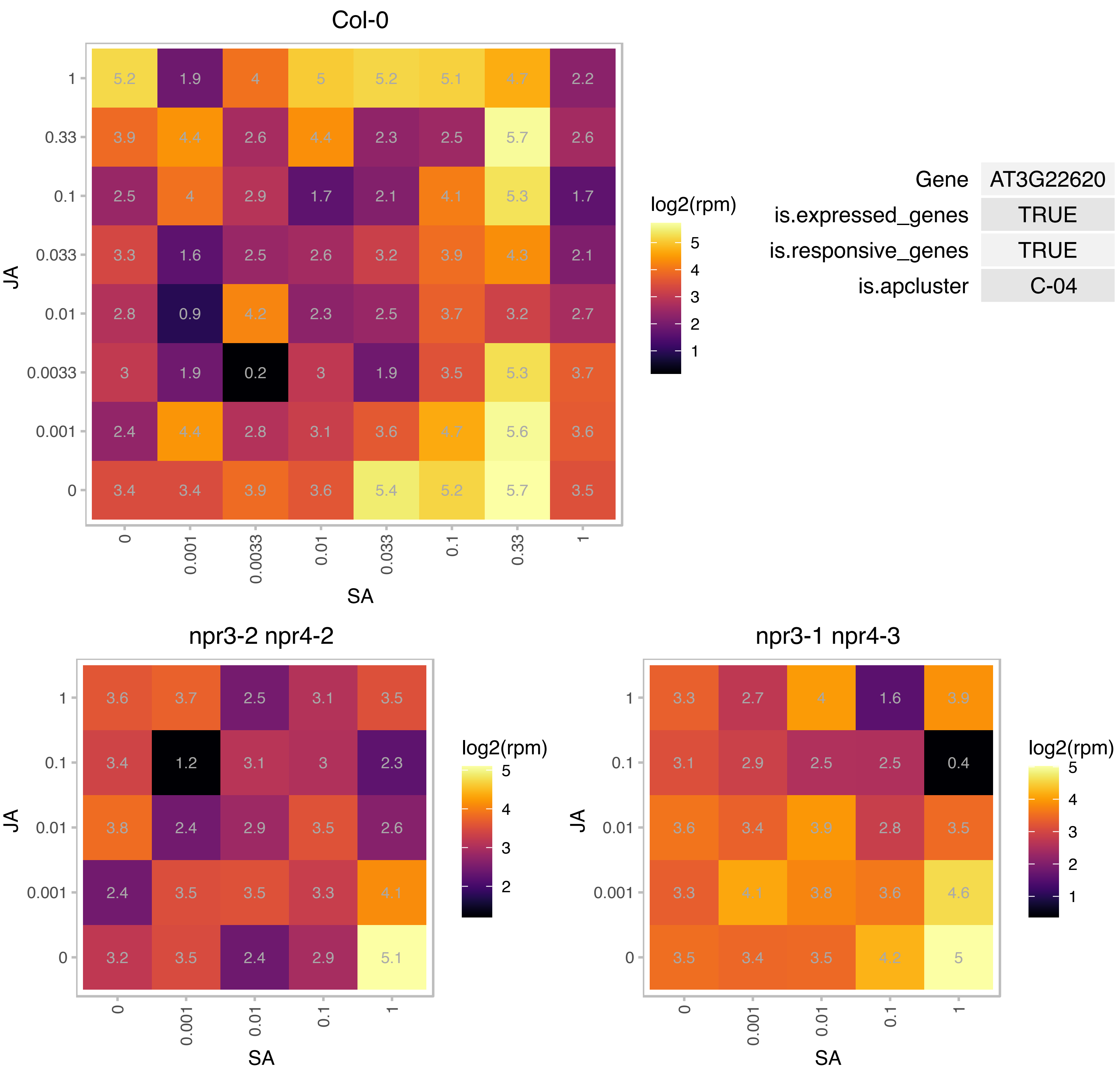

### AT3G23560.1 : ALF5

MATE efflux family protein

Member of the multidrug and toxic compound extrusion (MATE) family, protects roots from inhibitory compounds.

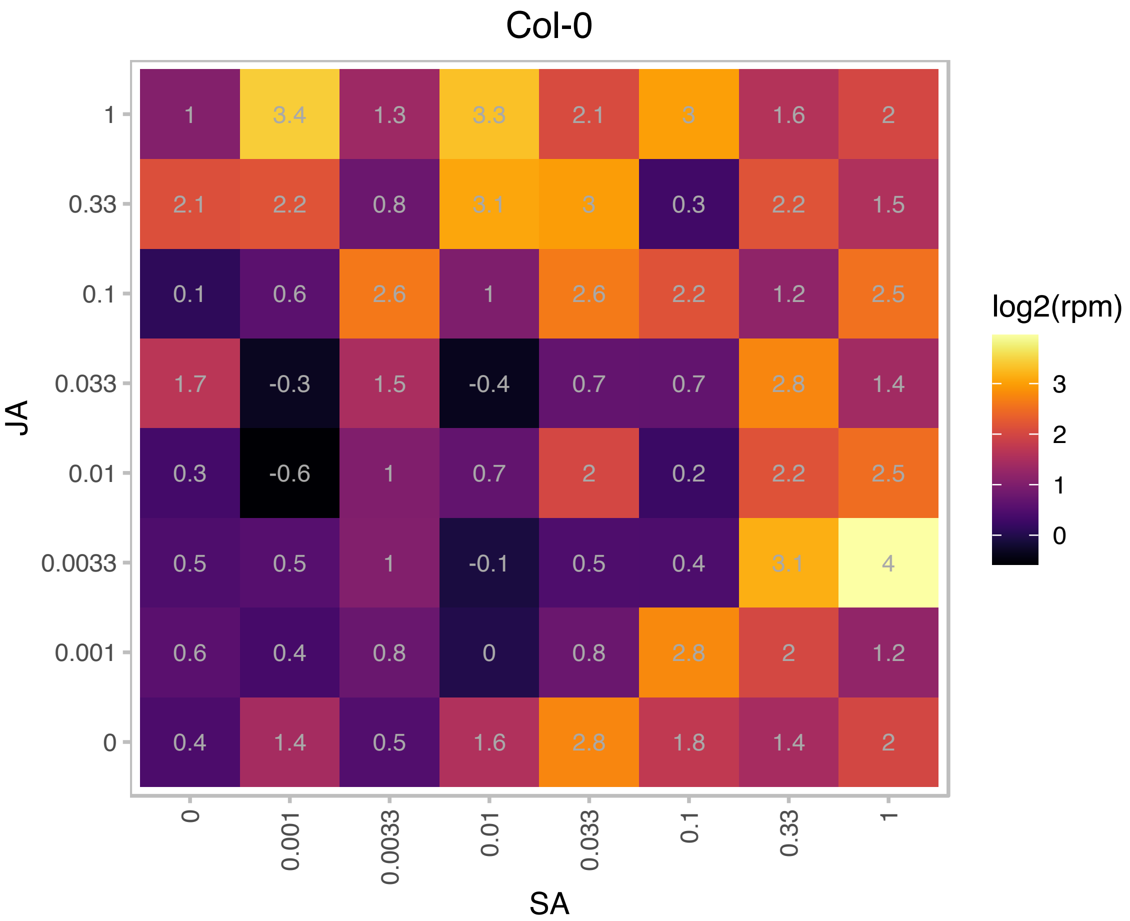


| | |
|---|---|
| Gene | AT3G23560 |
| is.expressed_genes | TRUE |
| is.responsive_genes | TRUE |
| is.apcluster | C-04 |

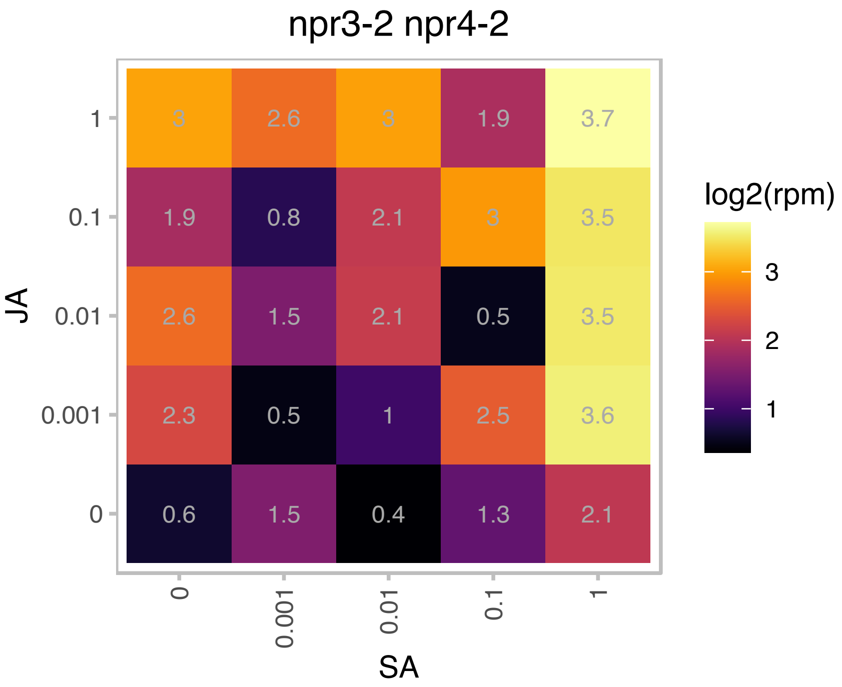


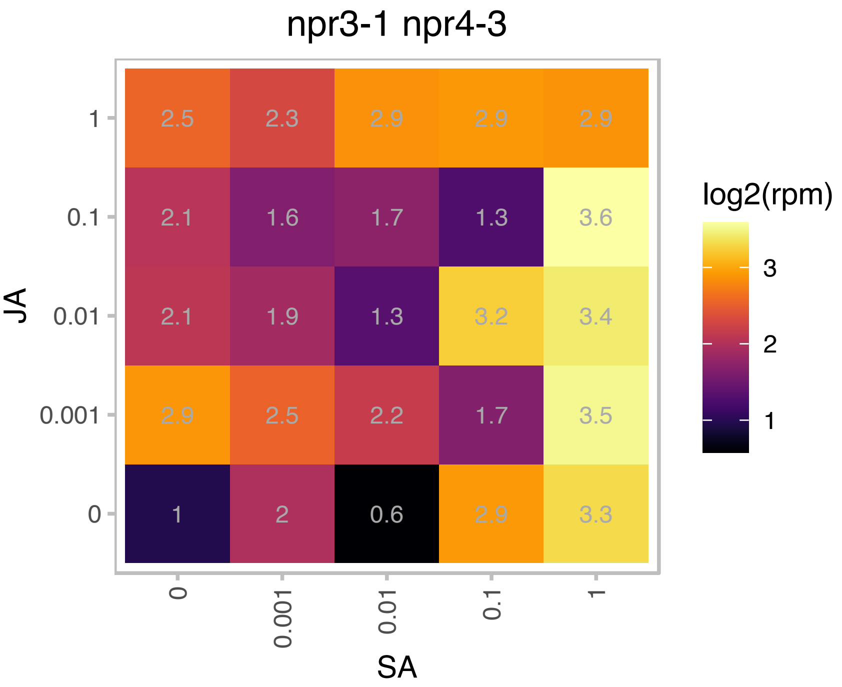

### AT3G47290.1 : ATPLC8, PLC8

phosphatidylinositol-speciwc phospholipase C8

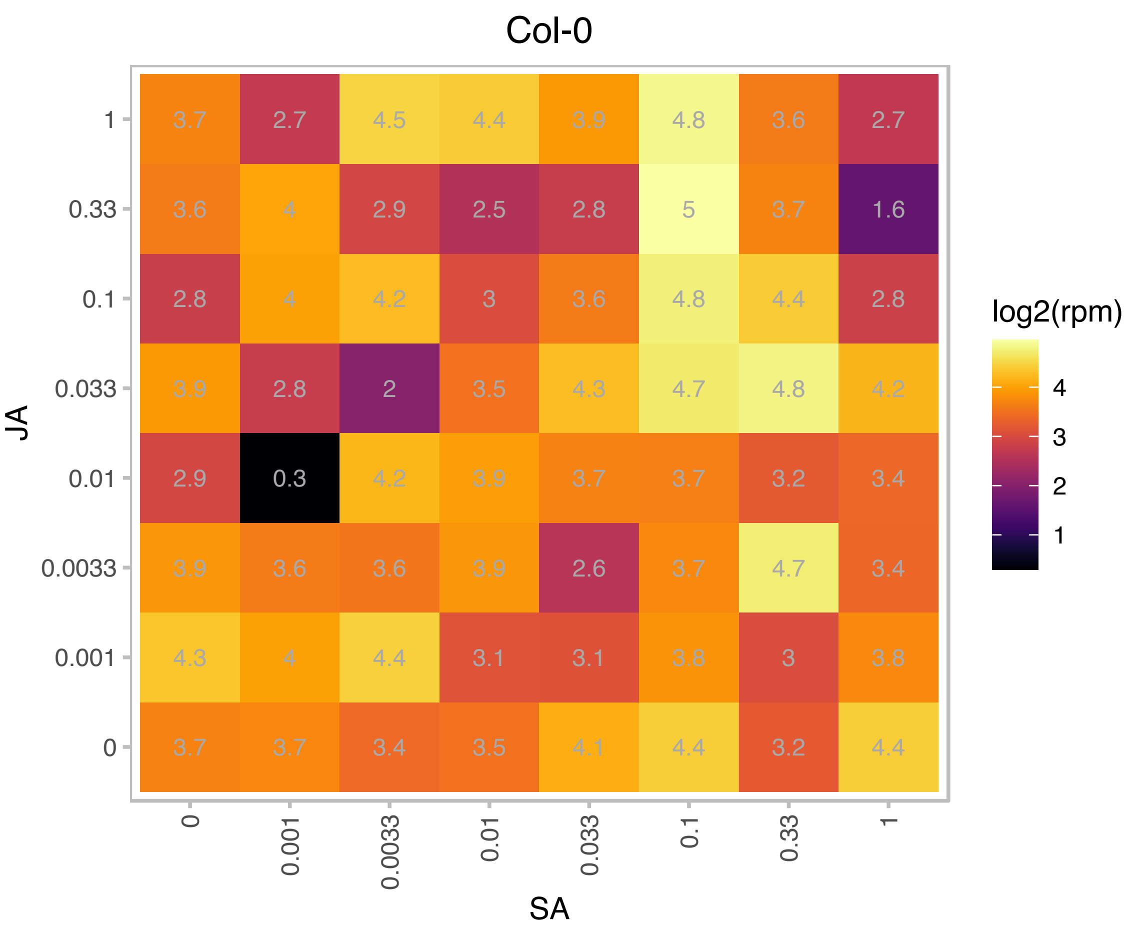


| | |
|---|---|
| Gene | AT3G47290 |
| is.expressed_genes | TRUE |
| is.responsive_genes | TRUE |
| is.apcluster | C-04 |

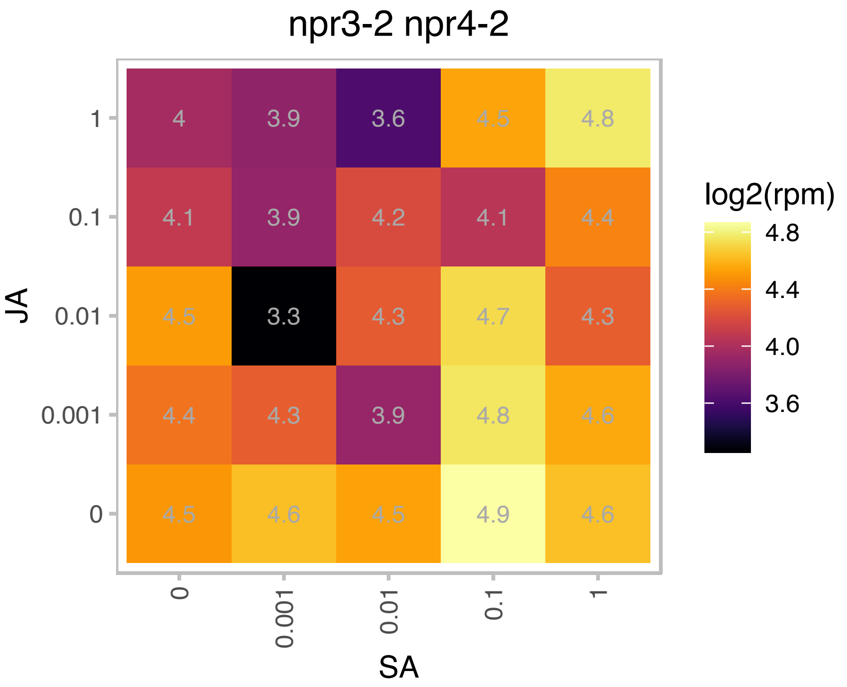


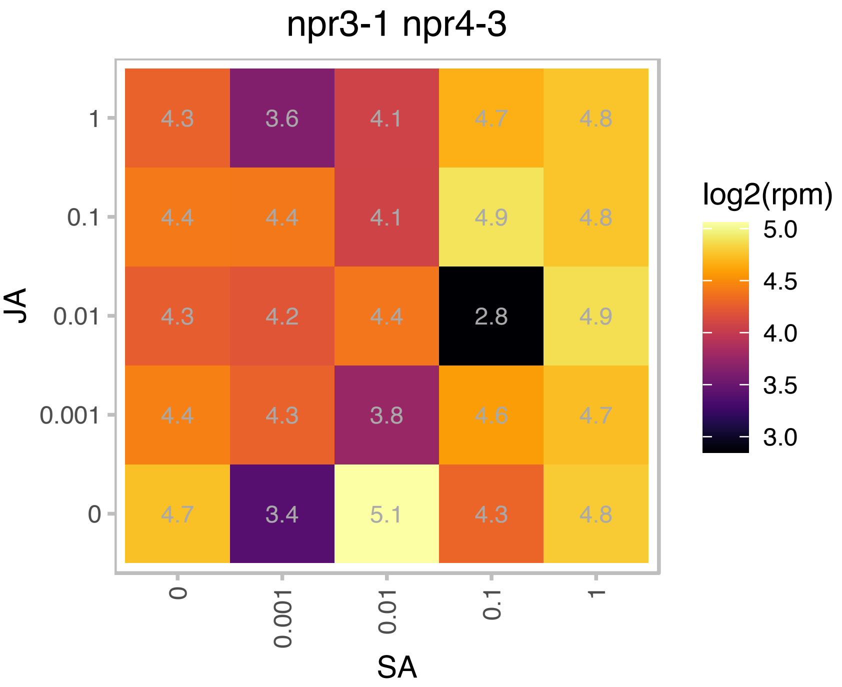

# AT3G50520.1 : NA

Phosphoglycerate mutase family protein

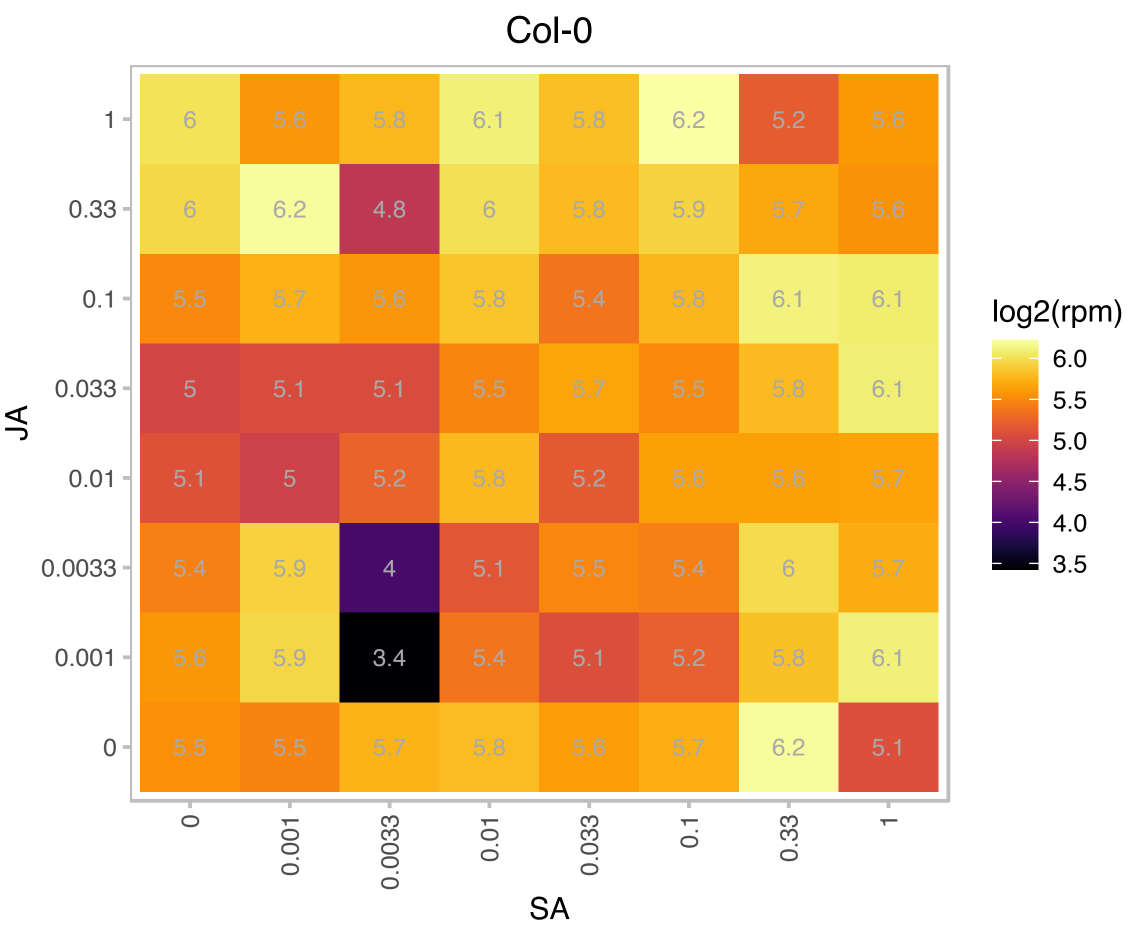


| | |
|---|---|
| Gene | AT3G50520 |
| is.expressed_genes | TRUE |
| is.responsive_genes | TRUE |
| is.apcluster | C-04 |

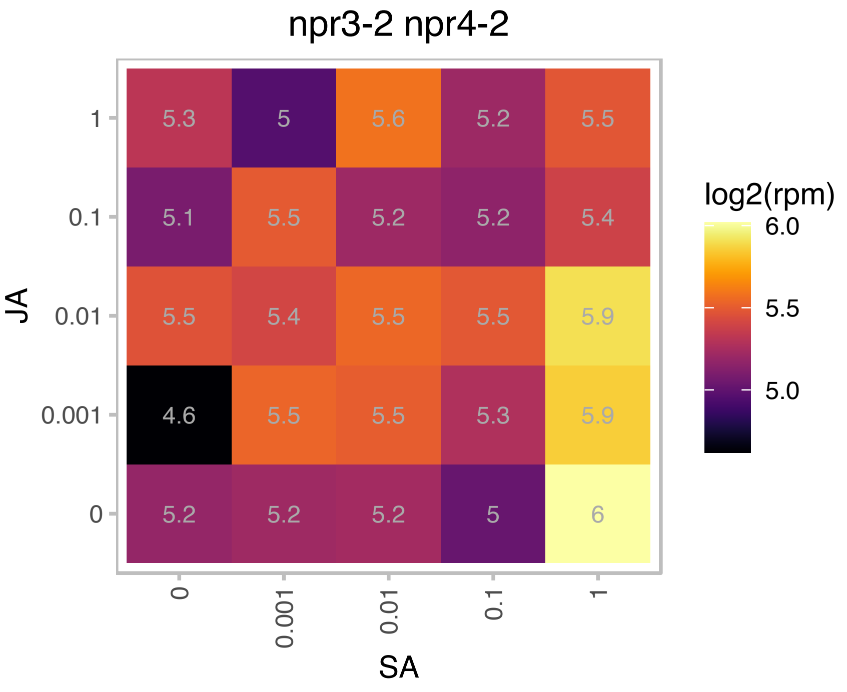


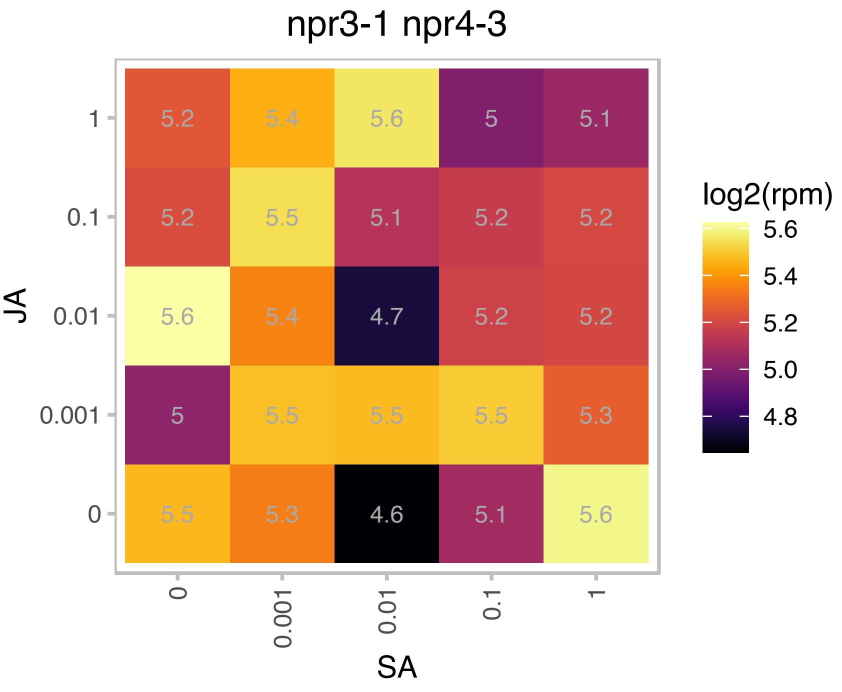

### AT3G56240.1 : CCH

copper chaperone

CCH protein belongs to a family of eukaryotic proteins that participate in intracellular copper homeostasis by delivering this metal to the secretory pathway; mainly located along the vascular bundles of senescing leaves and petioles as well as in stem sieve elements; hypothesized to have a role in copper mobilization from decaying organs towards reproductive structures, as a result of metalloprotein breakdown. The plant-specific C-terminal domain of the CCH protein forms amyloid-like fibrils in vitro.

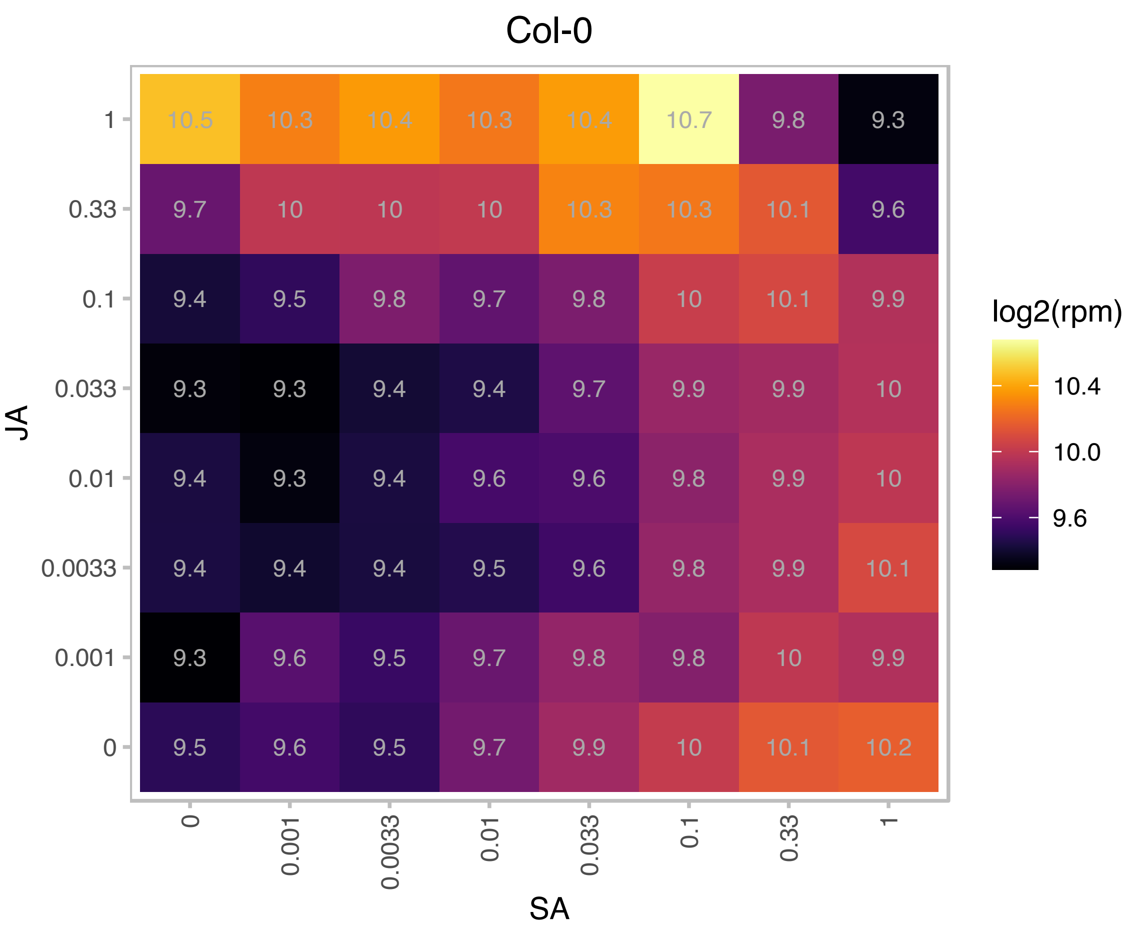


| | |
|---|---|
| Gene | AT3G56240 |
| is.expressed_genes | TRUE |
| is.responsive_genes | TRUE |
| is.apcluster | C-04 |

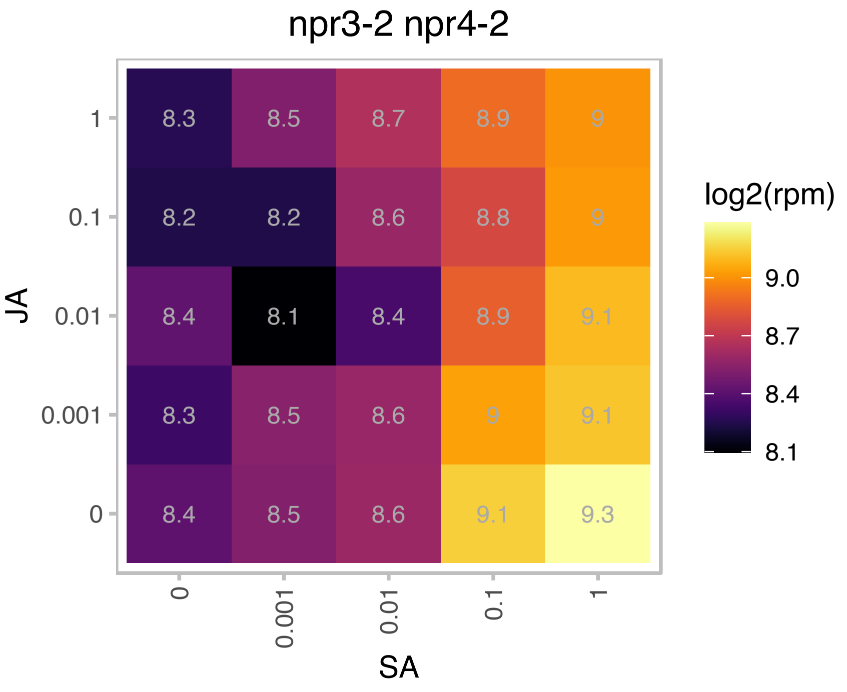


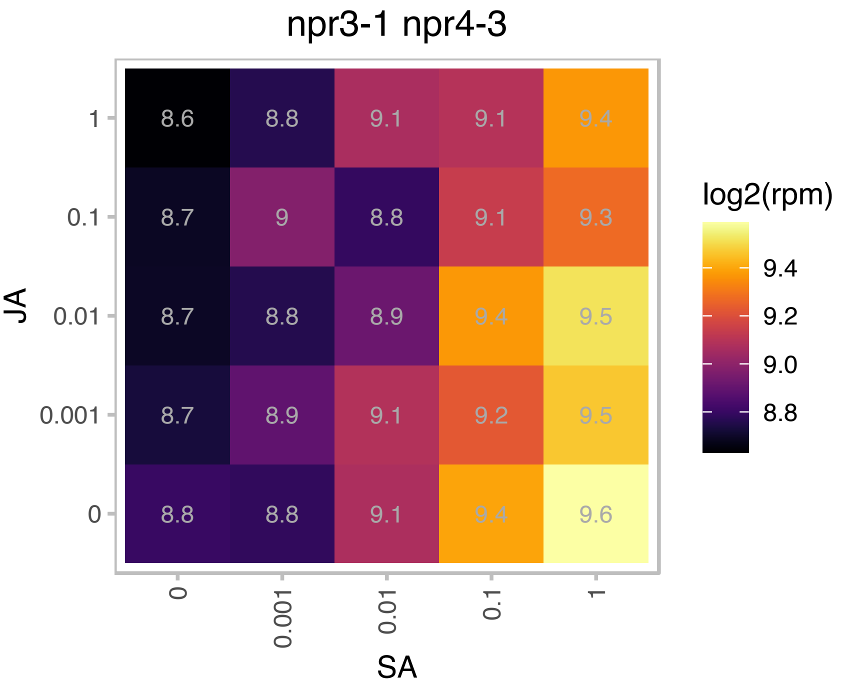

# AT3G57810.1 : NA

Cysteine proteinases superfamily protein

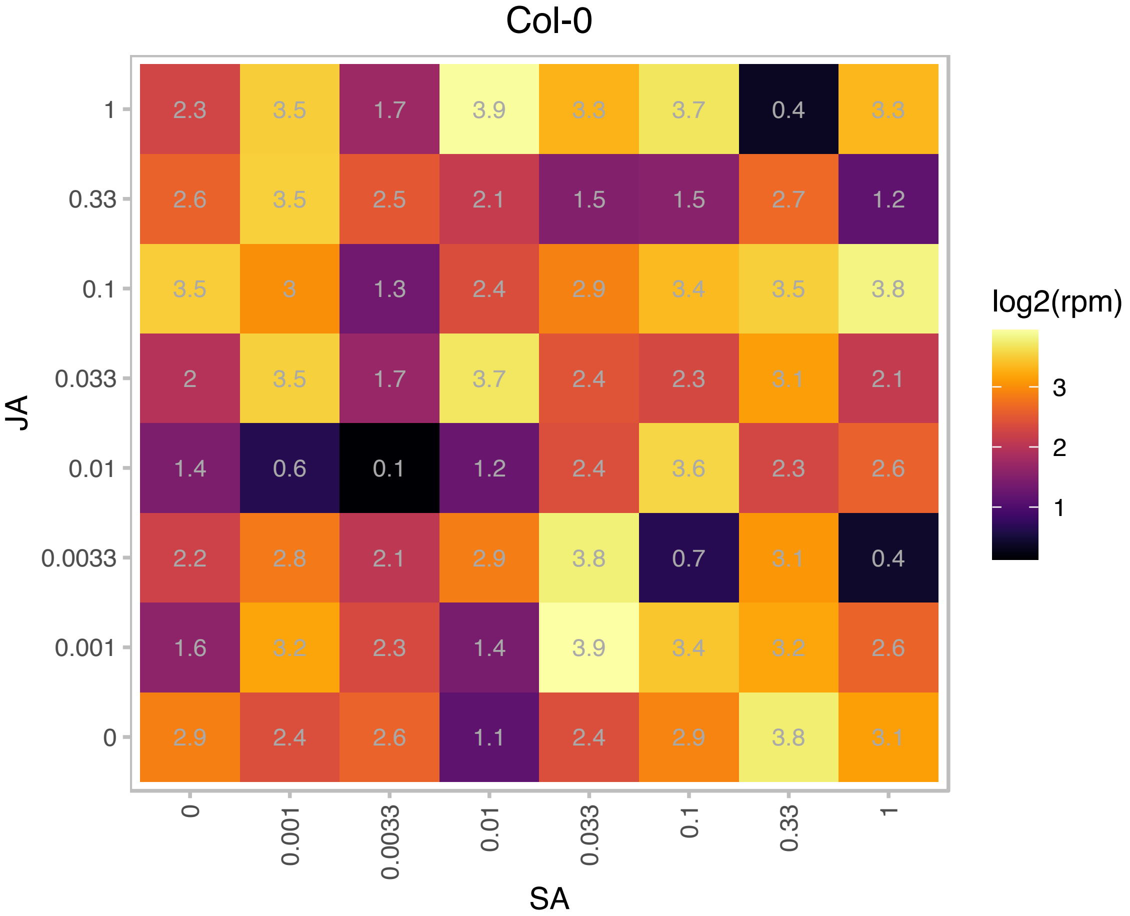


| | |
|---|---|
| Gene | AT3G57810 |
| is.expressed_genes | TRUE |
| is.responsive_genes | TRUE |
| is.apcluster | C-04 |

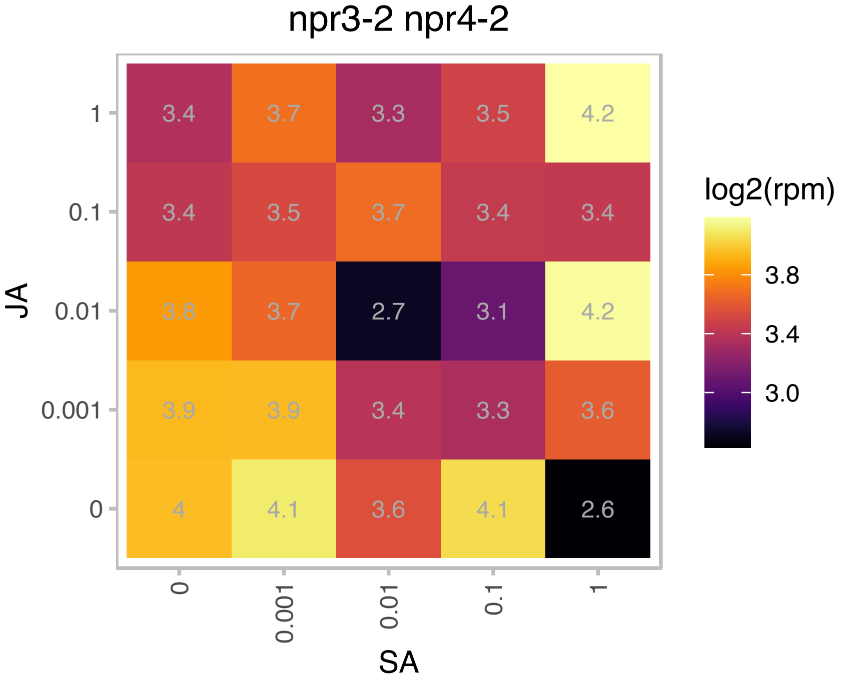


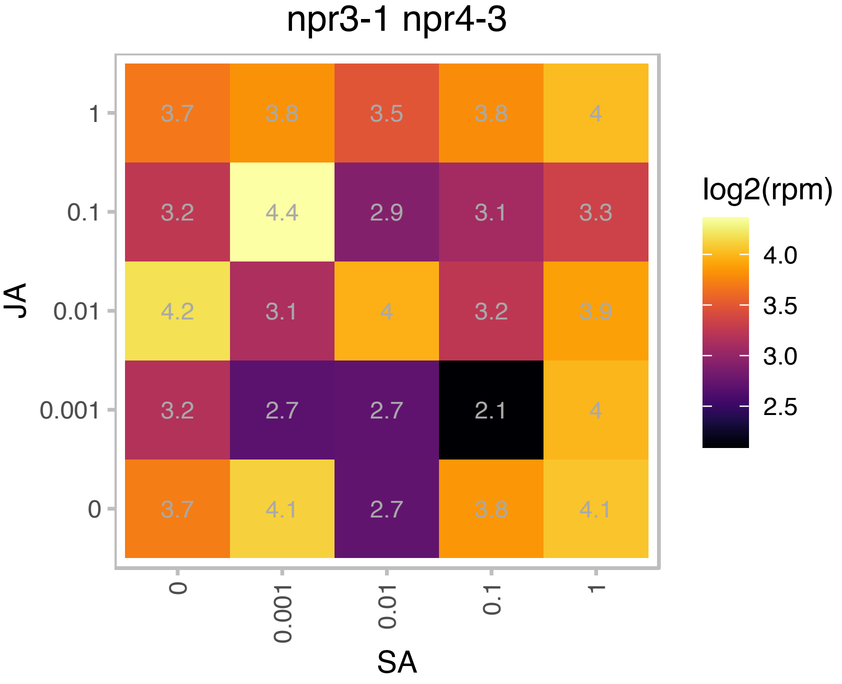

# AT3G58110.1 : NA

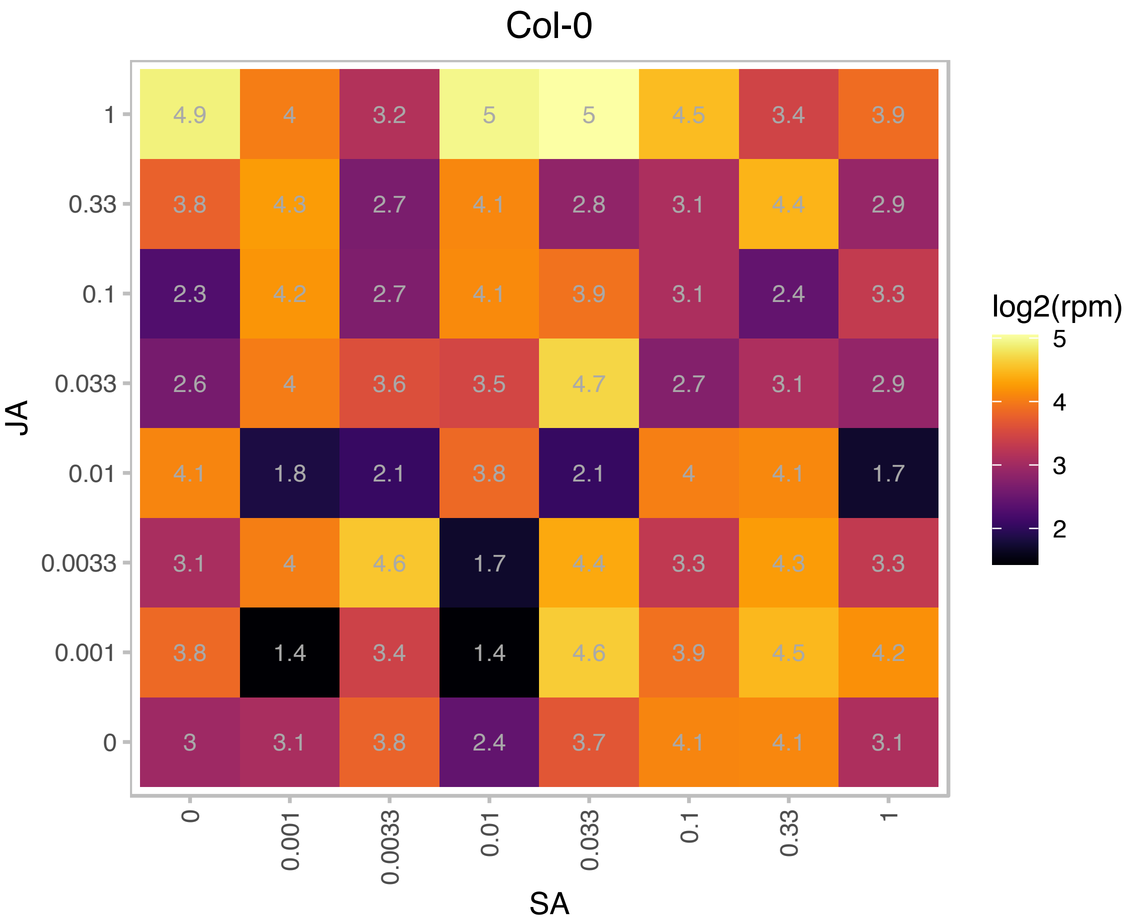


| | |
|---|---|
| Gene | AT3G58110 |
| is.expressed_genes | TRUE |
| is.responsive_genes | TRUE |
| is.apcluster | C-04 |

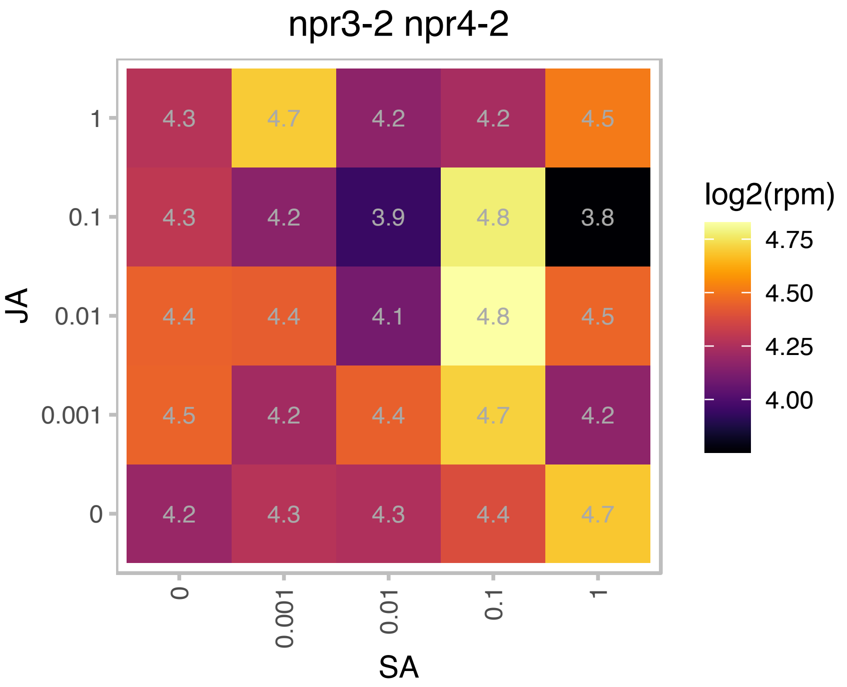


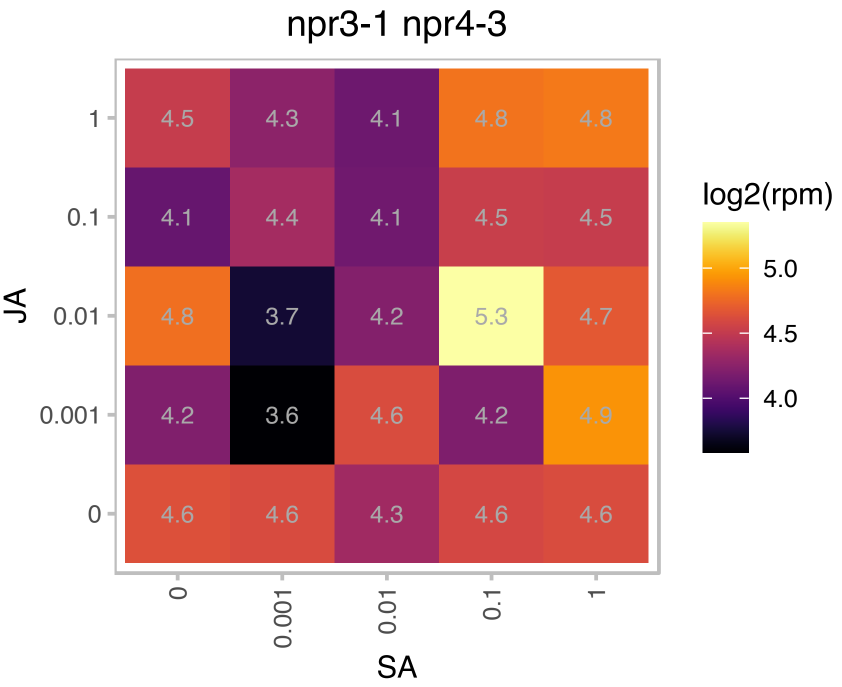

# AT4G02550.1 : NA

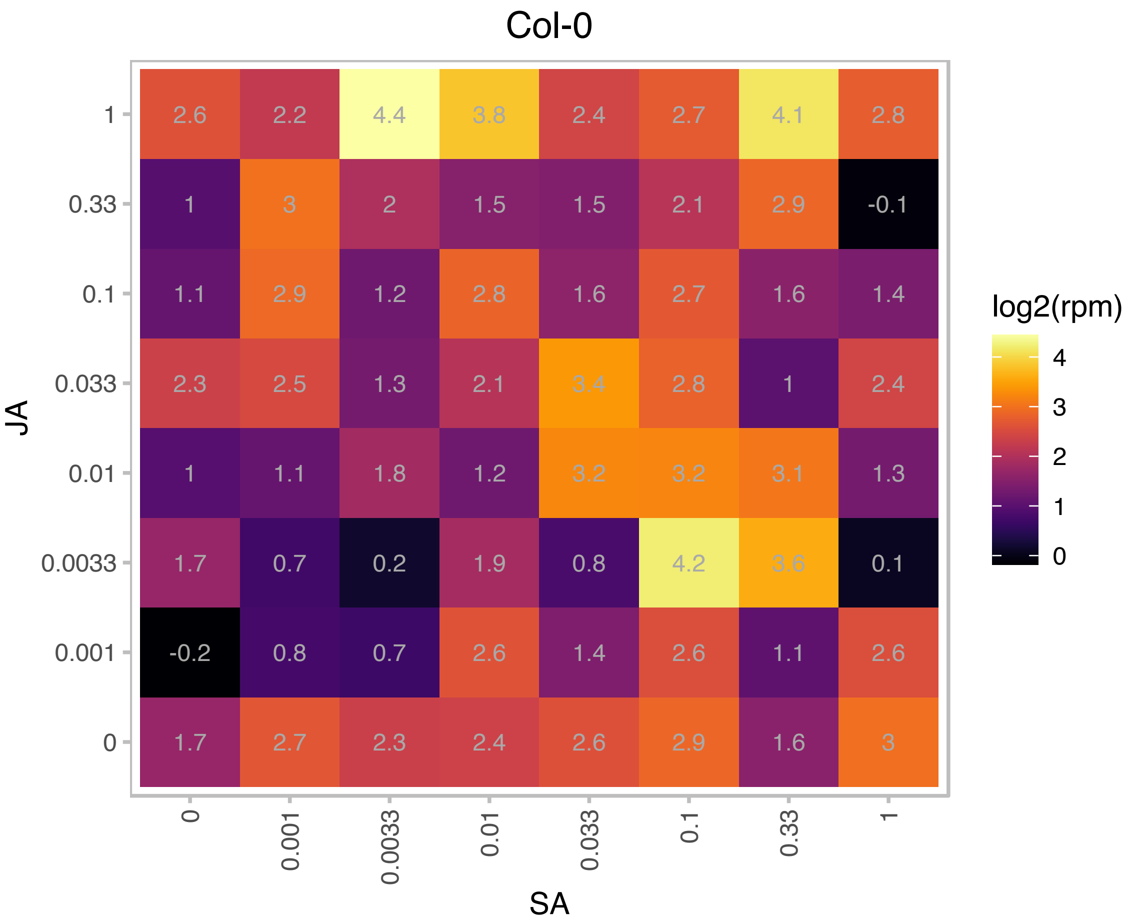


| | |
|---|---|
| Gene | AT4G02550 |
| is.expressed_genes | TRUE |
| is.responsive_genes | TRUE |
| is.apcluster | C-04 |

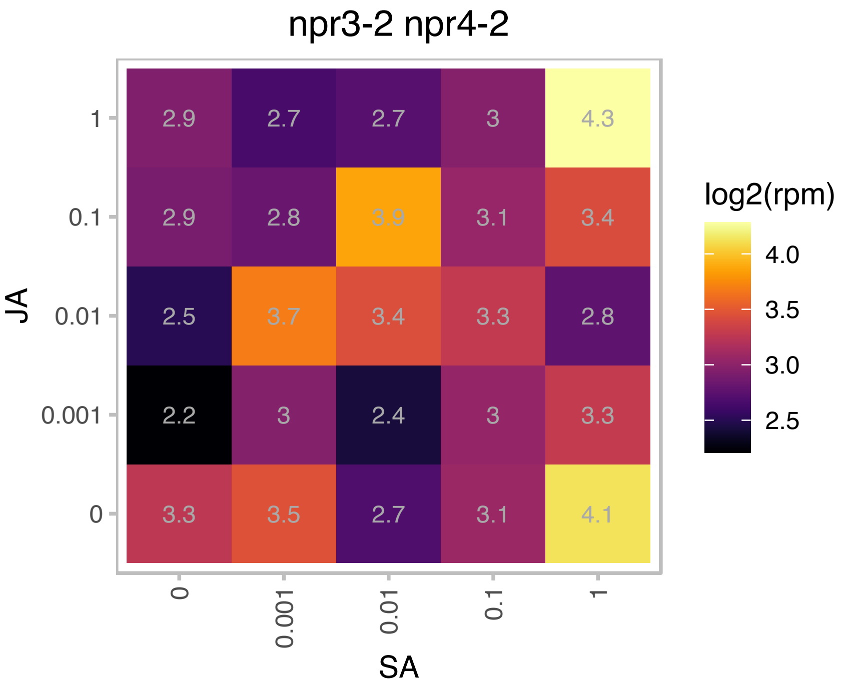


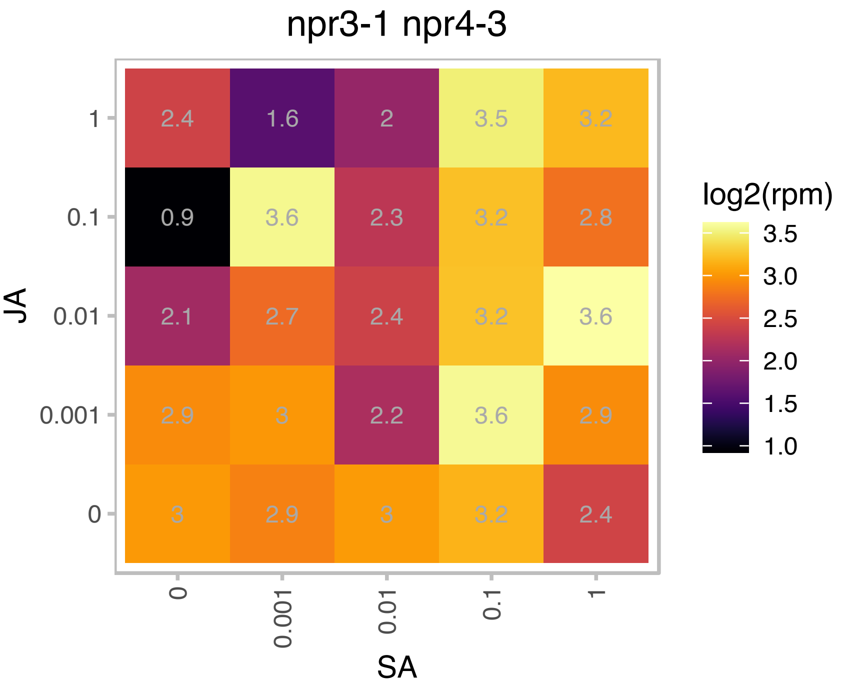

### AT4G03080.1 : BSL1

BRI1 suppressor 1 (BSU1)-like 1

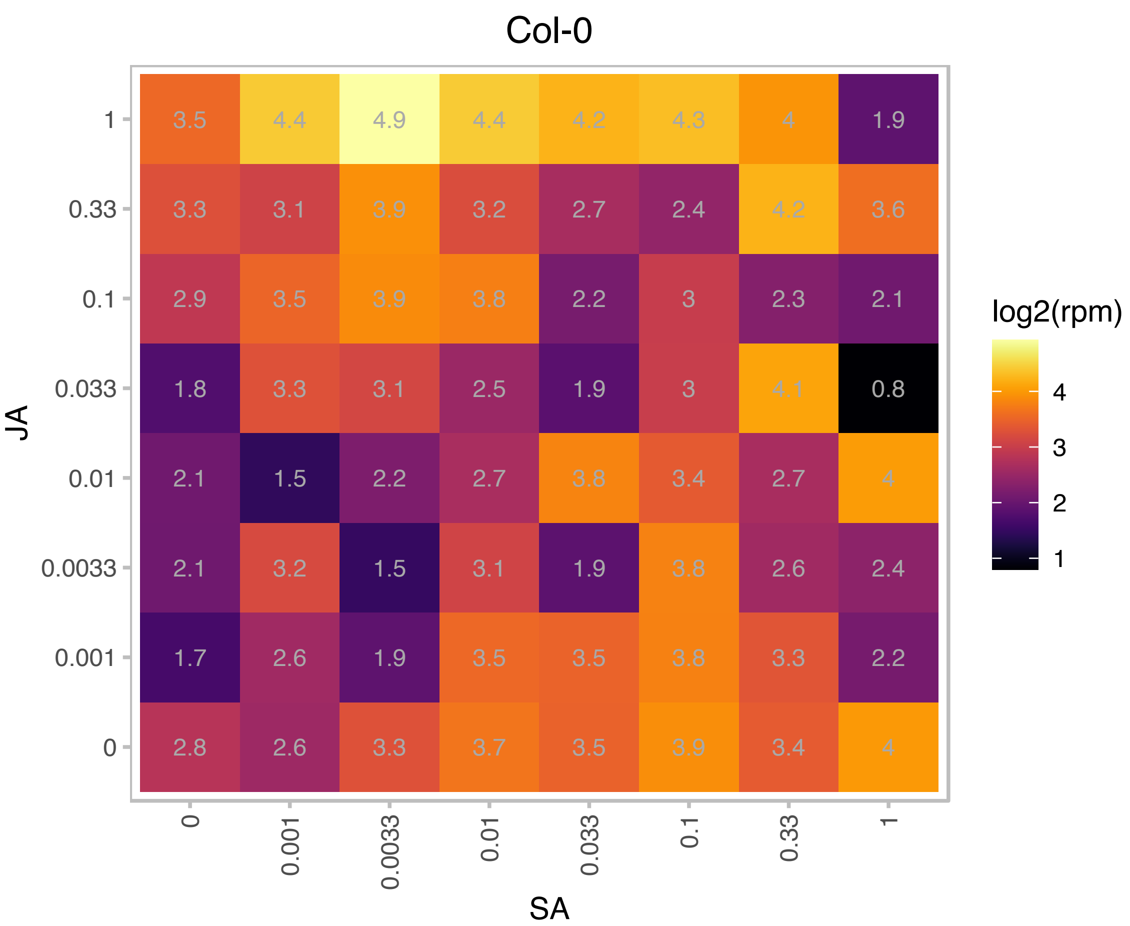


| | |
|---|---|
| Gene | AT4G03080 |
| is.expressed_genes | TRUE |
| is.responsive_genes | TRUE |
| is.apcluster | C-04 |

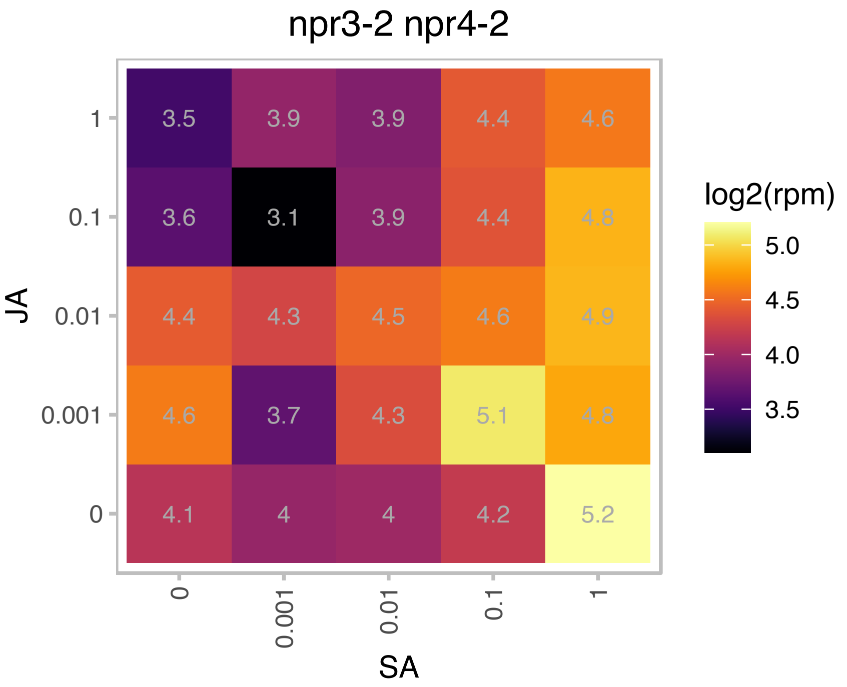


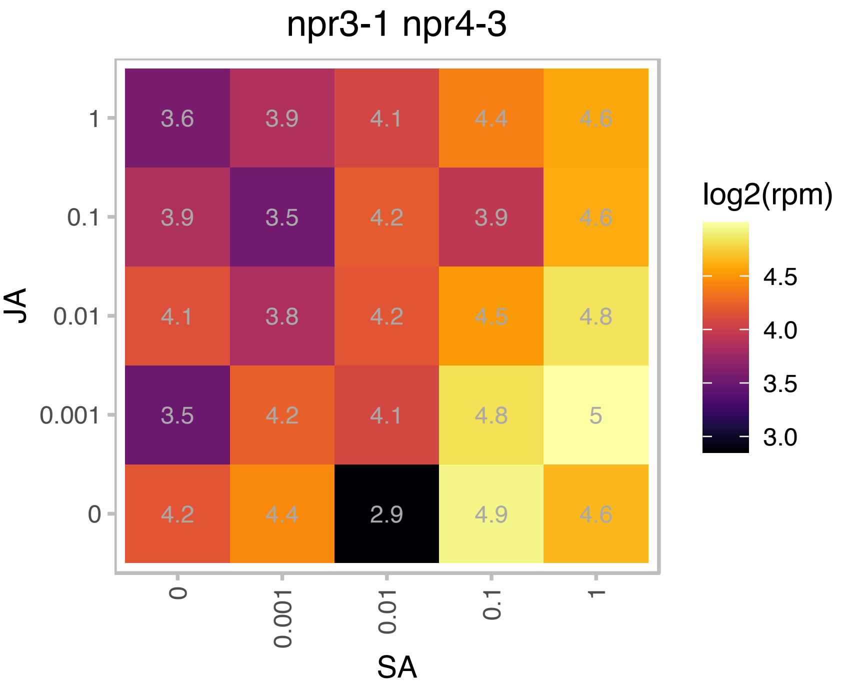

### AT4G18360.1 : GOX3

Aldolase-type TIM barrel family protein

Encodes a glycolate oxidase that modulates reactive oxygen species-mediated signal transduction during nonhost resistance.

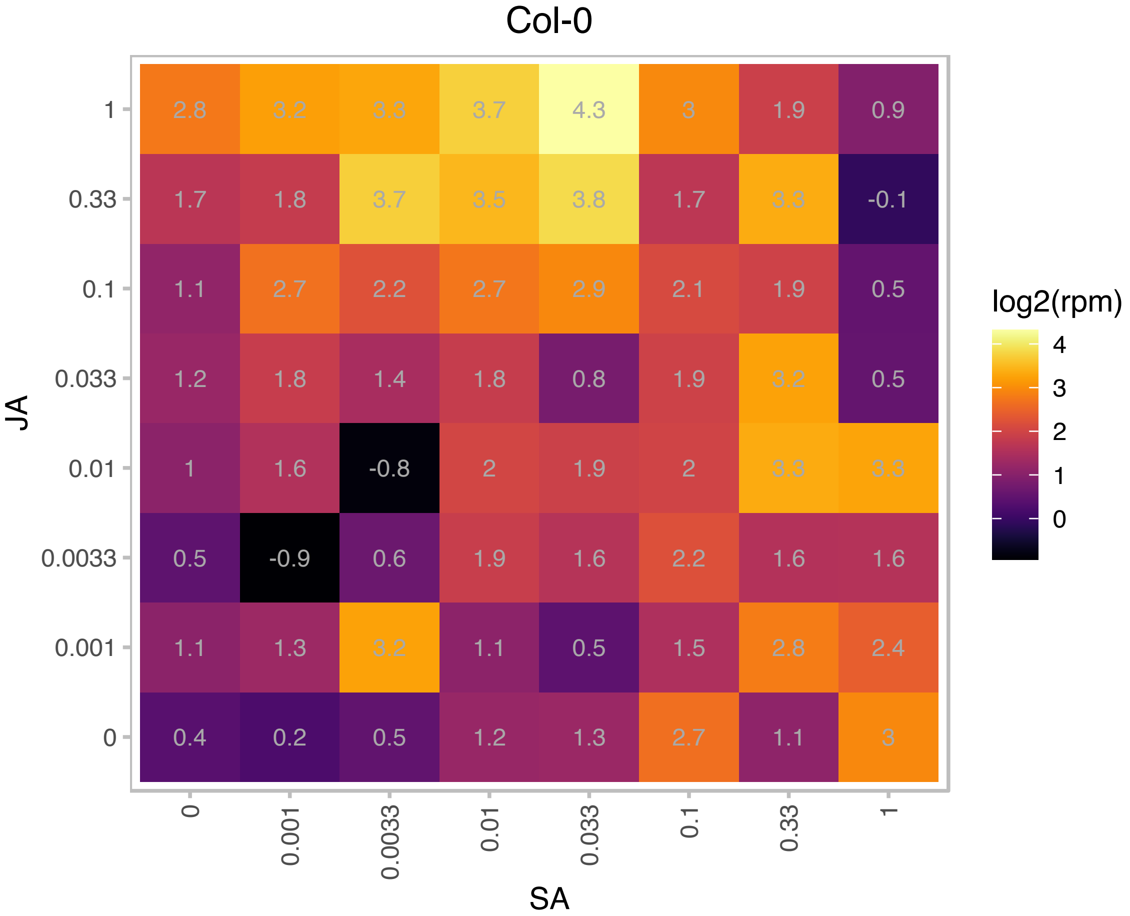


| JA \ SA | 0 | 0.001 | 0.0033 | 0.01 | 0.033 | 0.1 | 0.33 | 1 |
|---|---|---|---|---|---|---|---|---|
| 1 | 2.8 | 3.2 | 3.3 | 3.7 | 4.3 | 3 | 1.9 | 0.9 |
| 0.33 | 1.7 | 1.8 | 3.7 | 3.5 | 3.8 | 1.7 | 3.3 | -0.1 |
| 0.1 | 1.1 | 2.7 | 2.2 | 2.7 | 2.9 | 2.1 | 1.9 | 0.5 |
| 0.033 | 1.2 | 1.8 | 1.4 | 1.8 | 0.8 | 1.9 | 3.2 | 0.5 |
| 0.01 | 1 | 1.6 | -0.8 | 2 | 1.9 | 2 | 3.3 | 3.3 |
| 0.0033 | 0.5 | -0.9 | 0.6 | 1.9 | 1.6 | 2.2 | 1.6 | 1.6 |
| 0.001 | 1.1 | 1.3 | 3.2 | 1.1 | 0.5 | 1.5 | 2.8 | 2.4 |
| 0 | 0.4 | 0.2 | 0.5 | 1.2 | 1.3 | 2.7 | 1.1 | 3 |

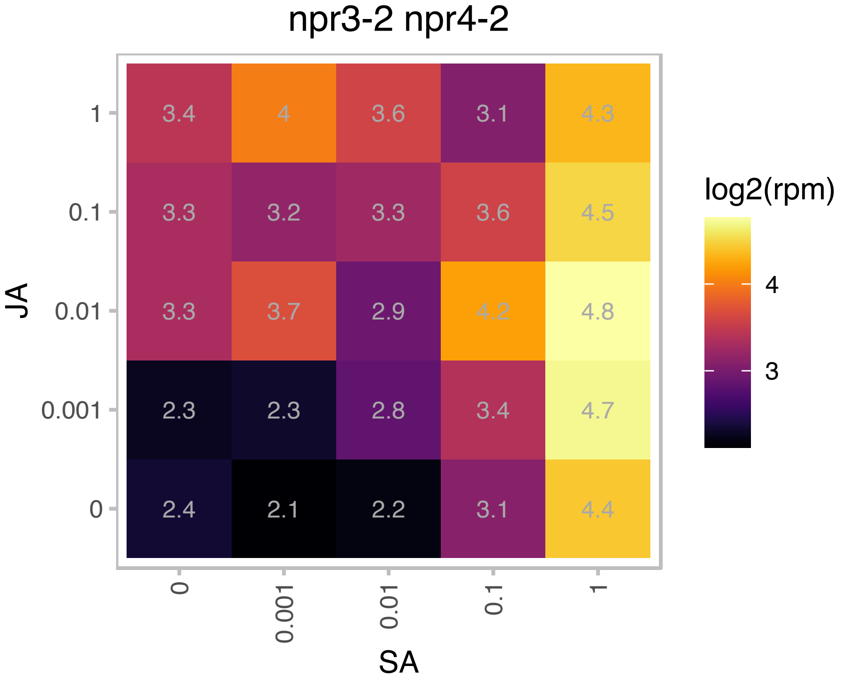


| JA \ SA | 0 | 0.001 | 0.01 | 0.1 | 1 |
|---|---|---|---|---|---|
| 1 | 3.4 | 4 | 3.6 | 3.1 | 4.3 |
| 0.1 | 3.3 | 3.2 | 3.3 | 3.6 | 4.5 |
| 0.01 | 3.3 | 3.7 | 2.9 | 4.2 | 4.8 |
| 0.001 | 2.3 | 2.3 | 2.8 | 3.4 | 4.7 |
| 0 | 2.4 | 2.1 | 2.2 | 3.1 | 4.4 |

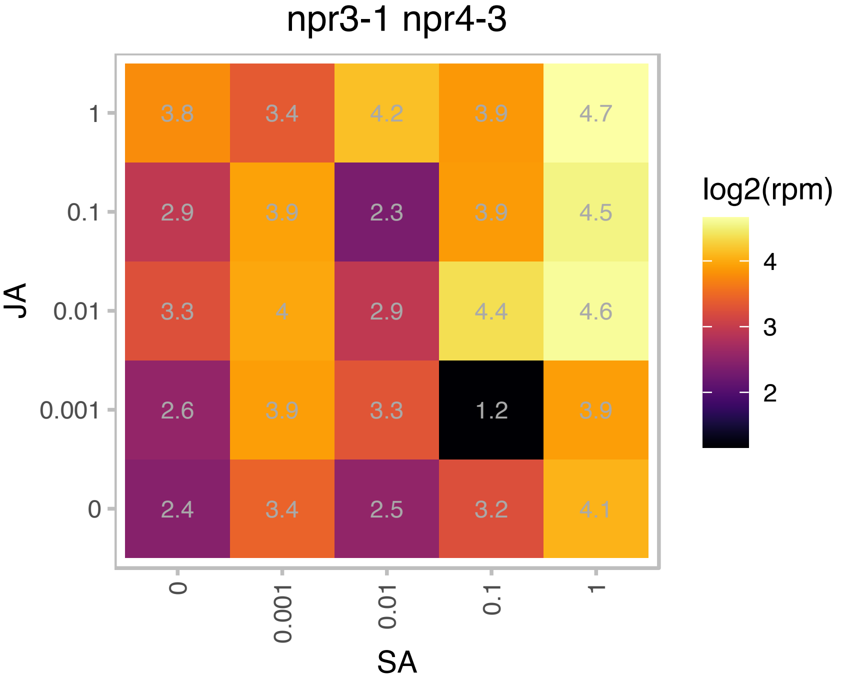


| JA \ SA | 0 | 0.001 | 0.01 | 0.1 | 1 |
|---|---|---|---|---|---|
| 1 | 3.8 | 3.4 | 4.2 | 3.9 | 4.7 |
| 0.1 | 2.9 | 3.9 | 2.3 | 3.9 | 4.5 |
| 0.01 | 3.3 | 4 | 2.9 | 4.4 | 4.6 |
| 0.001 | 2.6 | 3.9 | 3.3 | 1.2 | 3.9 |
| 0 | 2.4 | 3.4 | 2.5 | 3.2 | 4.1 |

# AT4G22740.1 : NA

glycine-rich protein

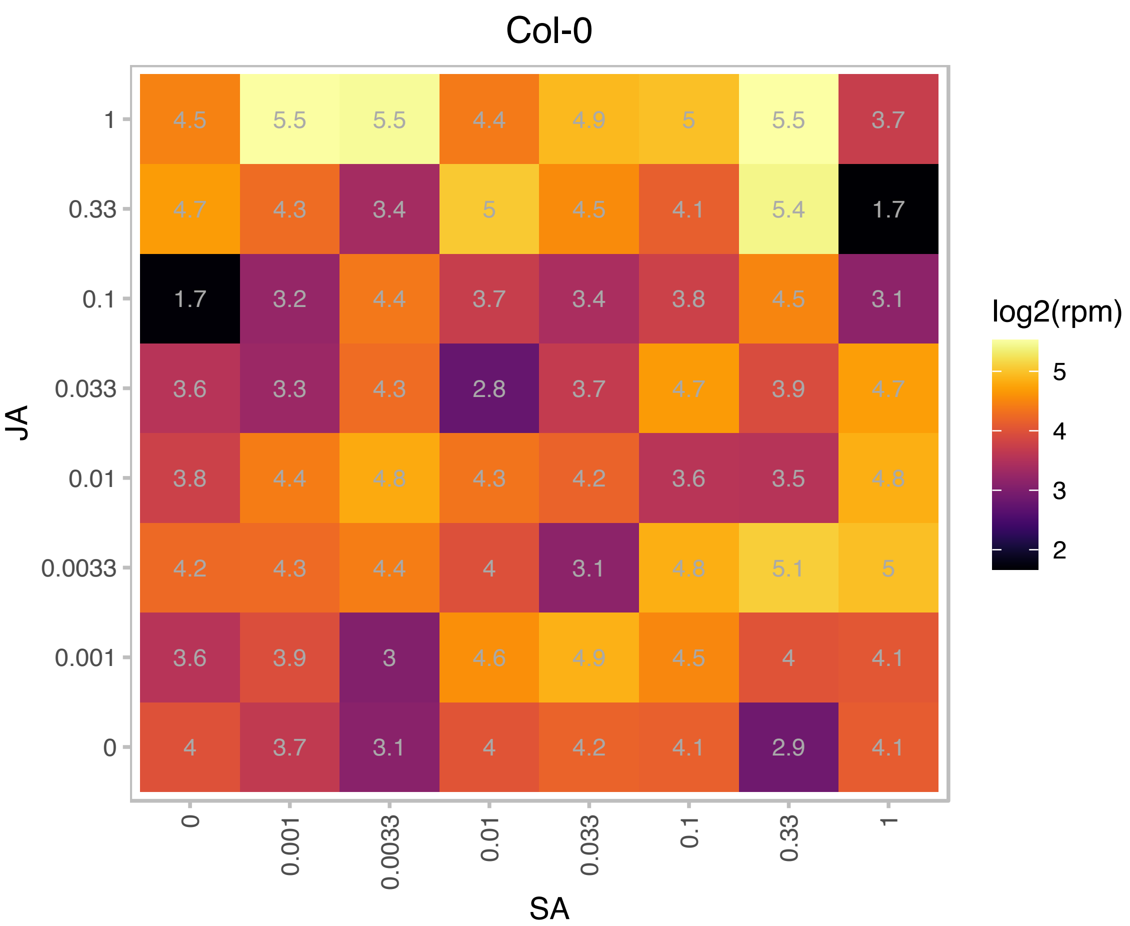


| | |
|---|---|
| Gene | AT4G22740 |
| is.expressed_genes | TRUE |
| is.responsive_genes | TRUE |
| is.apcluster | C-04 |

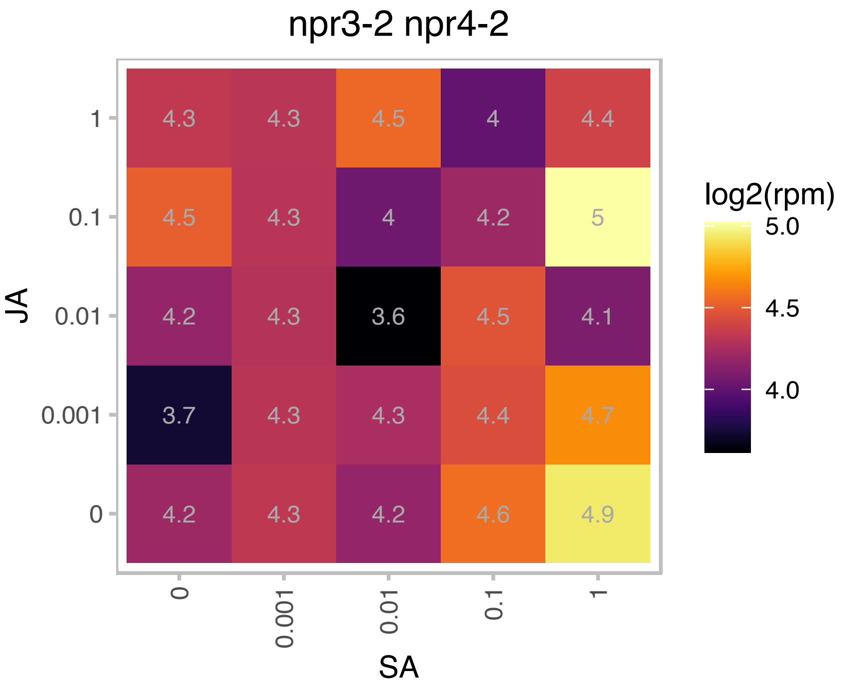


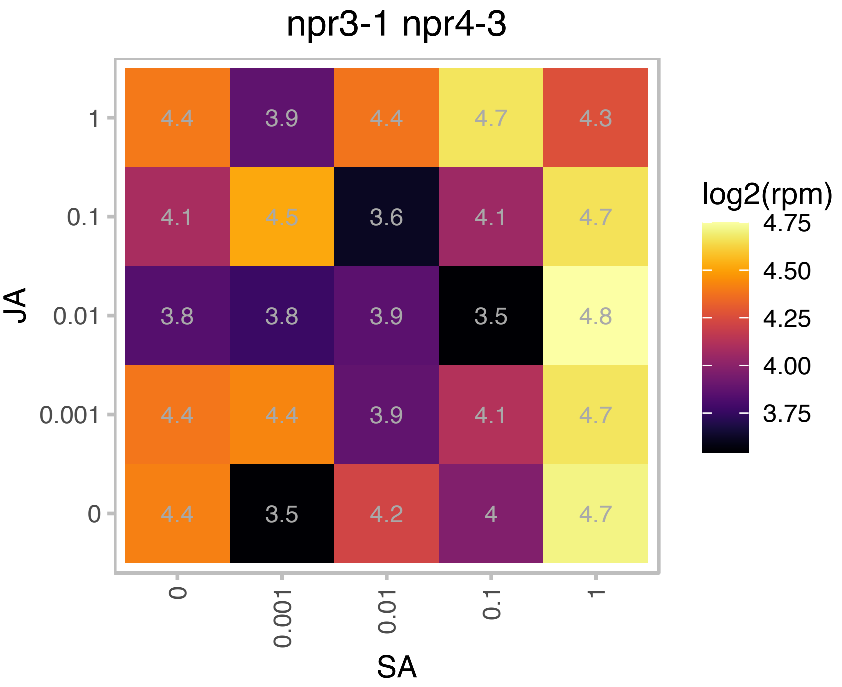

# AT4G24290.1 : NA

MAC/Perforin domain-containing protein

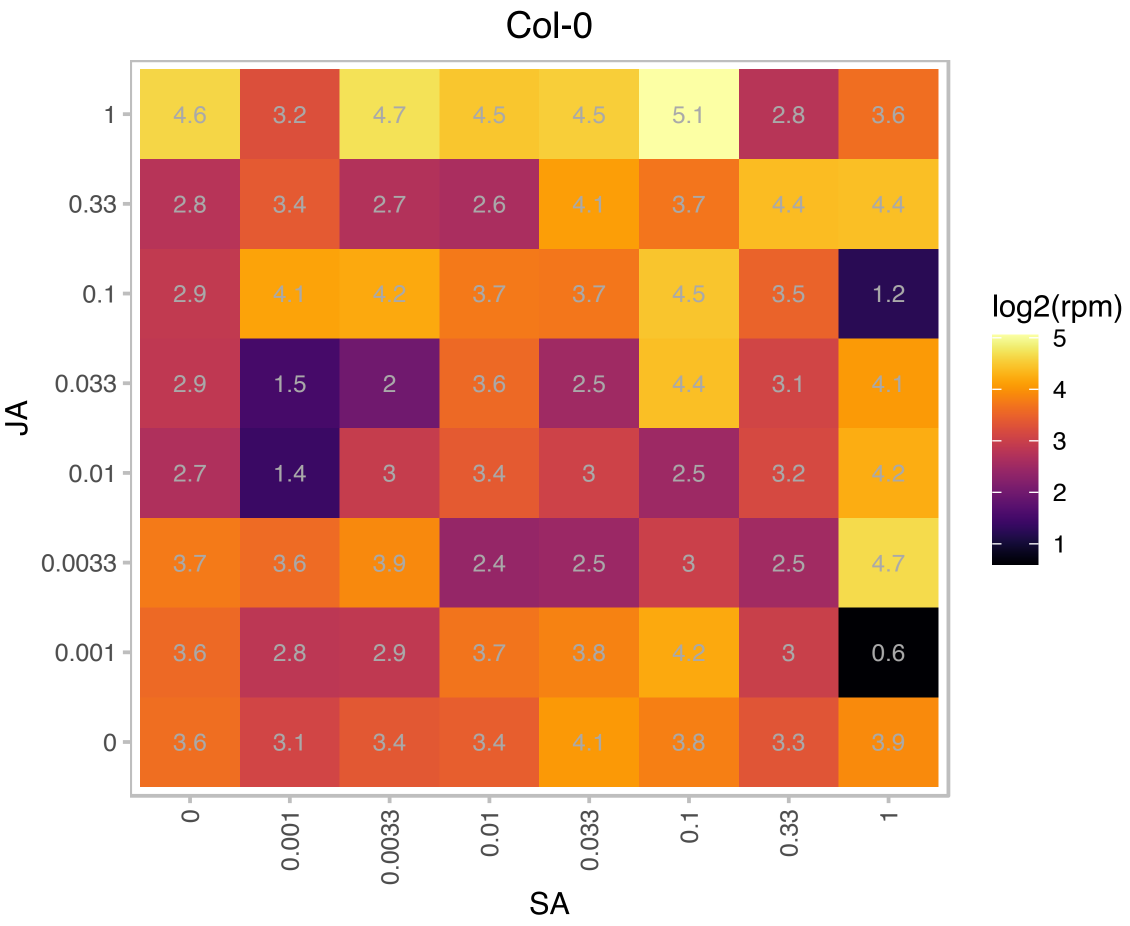


| | |
|---|---|
| Gene | AT4G24290 |
| is.expressed_genes | TRUE |
| is.responsive_genes | TRUE |
| is.apcluster | C-04 |

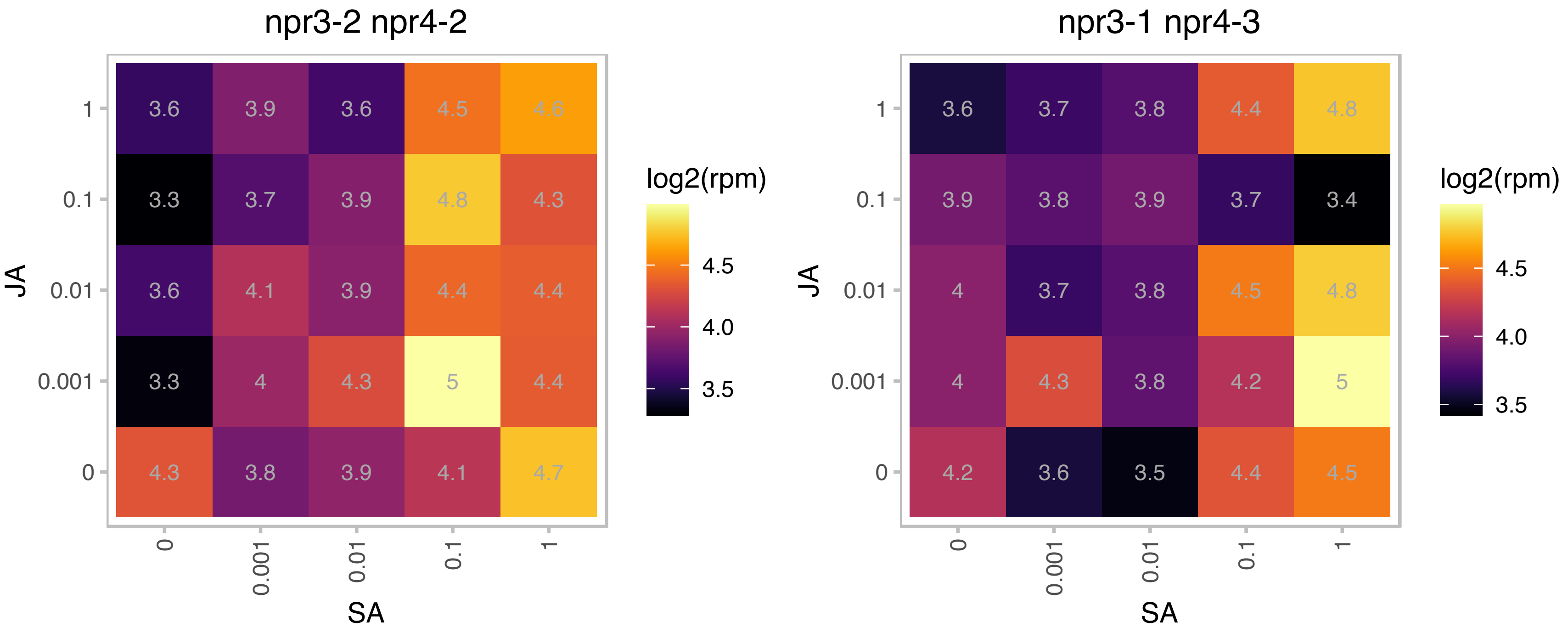

# AT4G25225.1 : NA

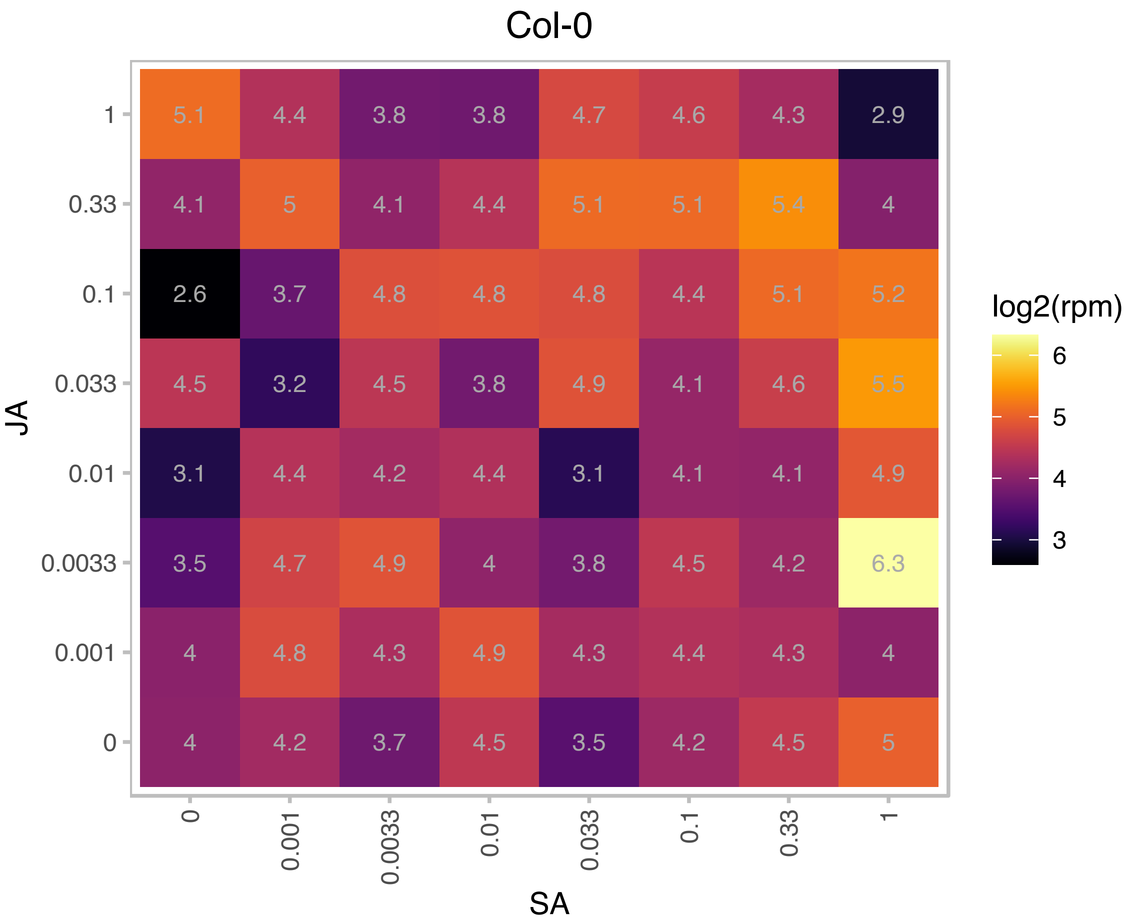


| | |
|---|---|
| Gene | AT4G25225 |
| is.expressed_genes | TRUE |
| is.responsive_genes | TRUE |
| is.apcluster | C-04 |

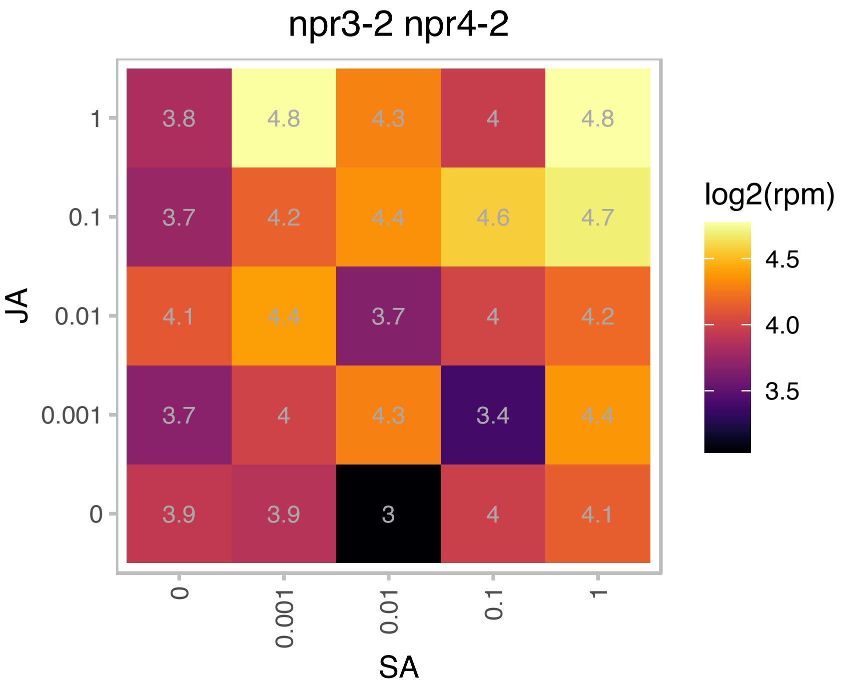


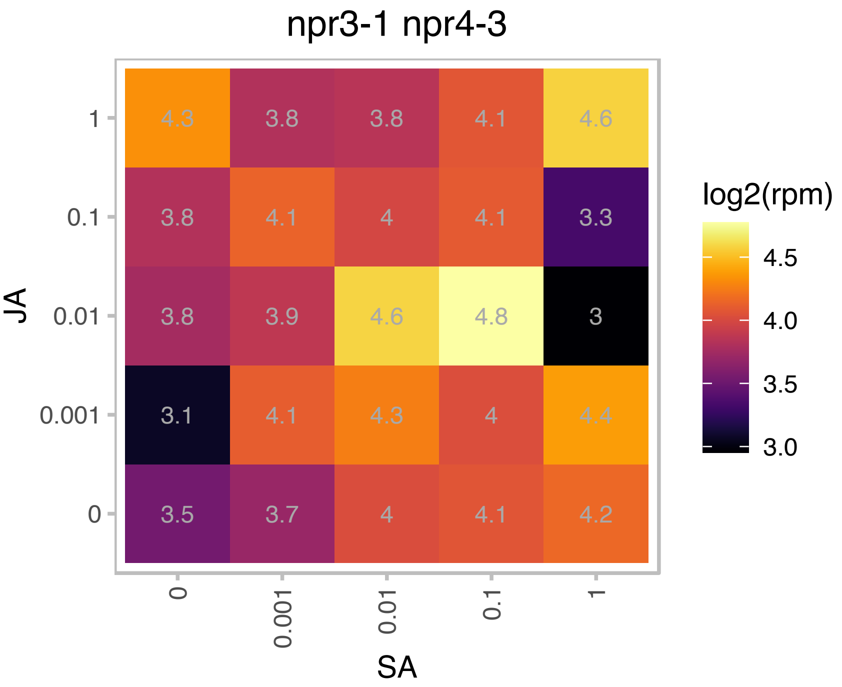

# AT4G27450.1 : NA

Aluminium induced protein with YGL and LRDR motifs

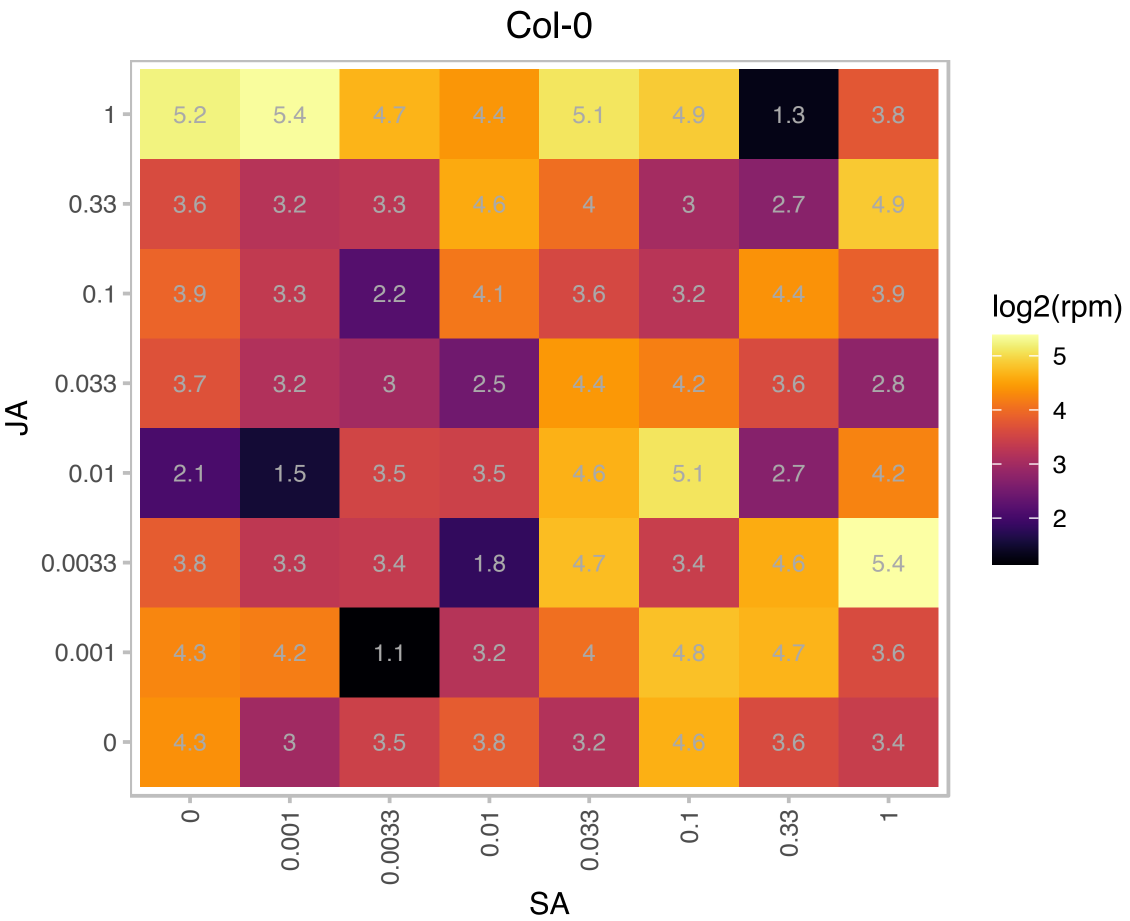


| | |
|---|---|
| Gene | AT4G27450 |
| is.expressed_genes | TRUE |
| is.responsive_genes | TRUE |
| is.apcluster | C-04 |

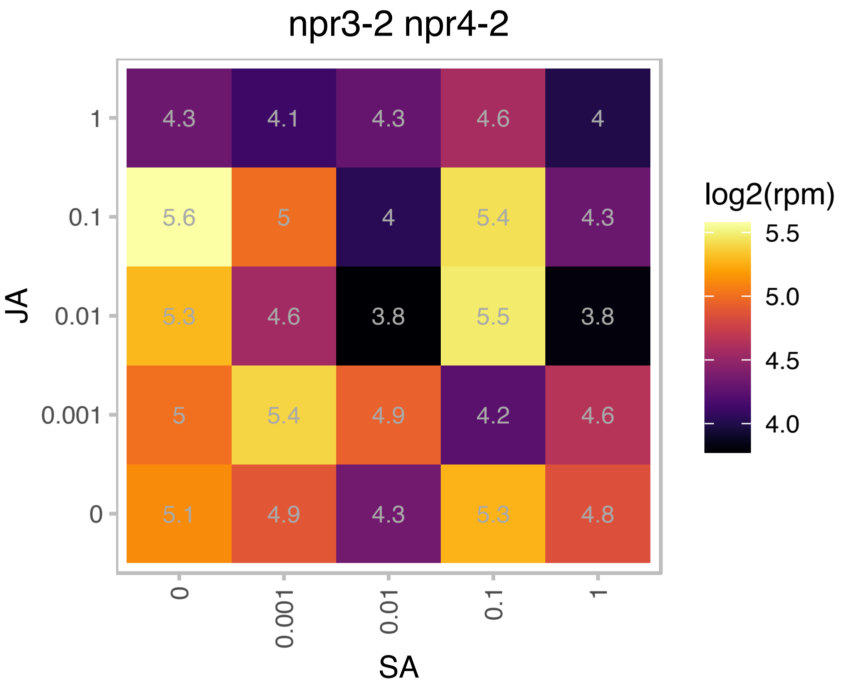


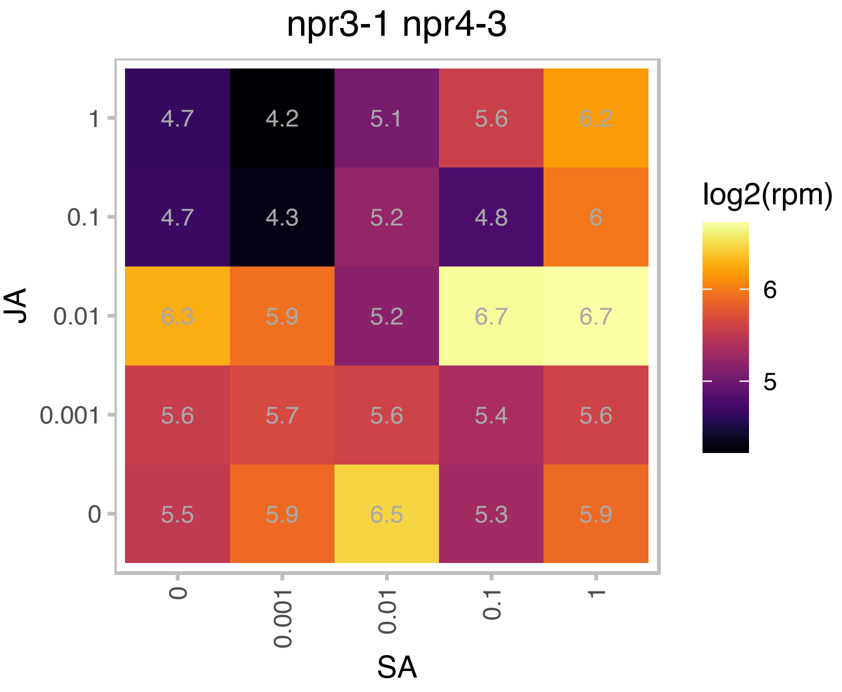

### AT4G31180.1 : IBI1

Class II aminoacyl-tRNA and biotin synthetases superfamily protein
The IBI1 gene encodes an aspartyl tRNA synthetase (AspRS). In addition, the IBI1 protein acts as a receptor protein of the chemical plant defence activator beta-aminobutyric acid (BABA). Binding of IBI1 to the active R-enantiomer of BABA primes non-canonical defence activity of the AspRS protein against pathogen attack.

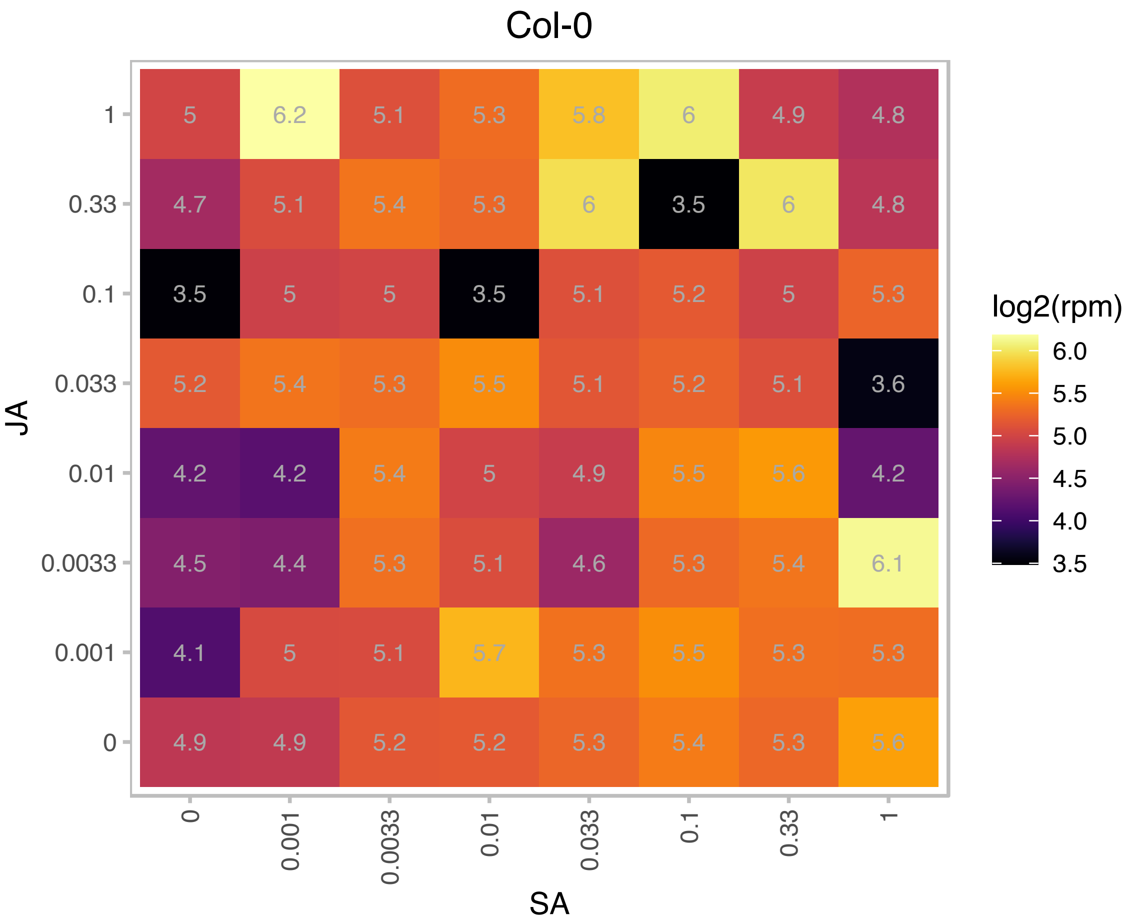


| | |
|---|---|
| Gene | AT4G31180 |
| is.expressed_genes | TRUE |
| is.responsive_genes | TRUE |
| is.apcluster | C-04 |

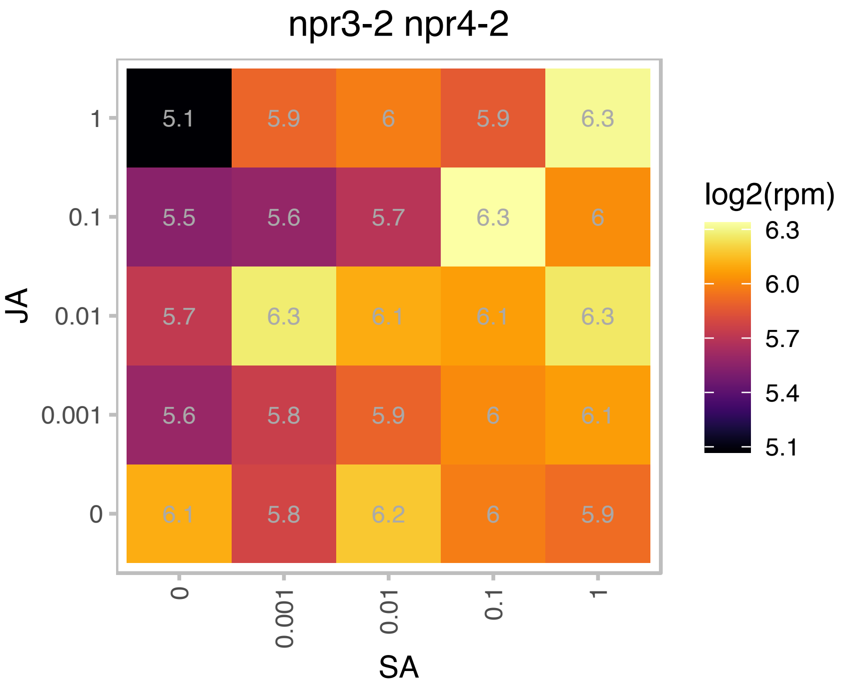


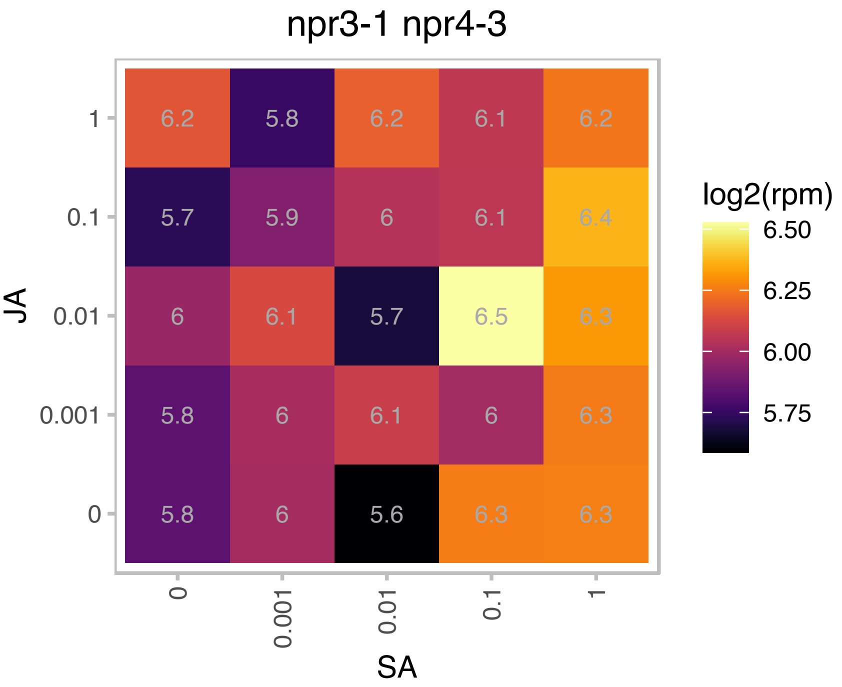

### AT4G39390.1 : ATNST-KT1, NST-K1, URGT4

nucleotide sugar transporter-KT 1

Encodes a golgi localized nucleotide sugar transporter.

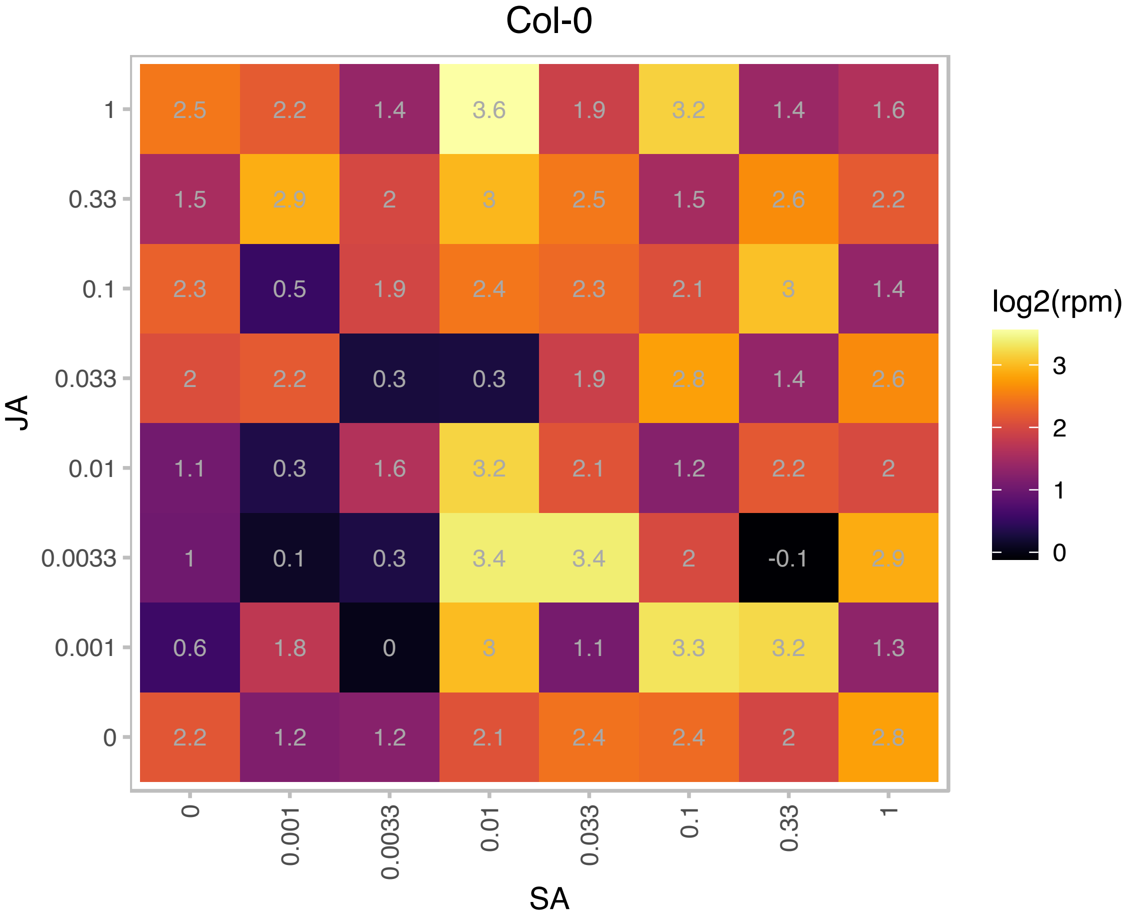


| | |
|---|---|
| Gene | AT4G39390 |
| is.expressed_genes | TRUE |
| is.responsive_genes | TRUE |
| is.apcluster | C-04 |

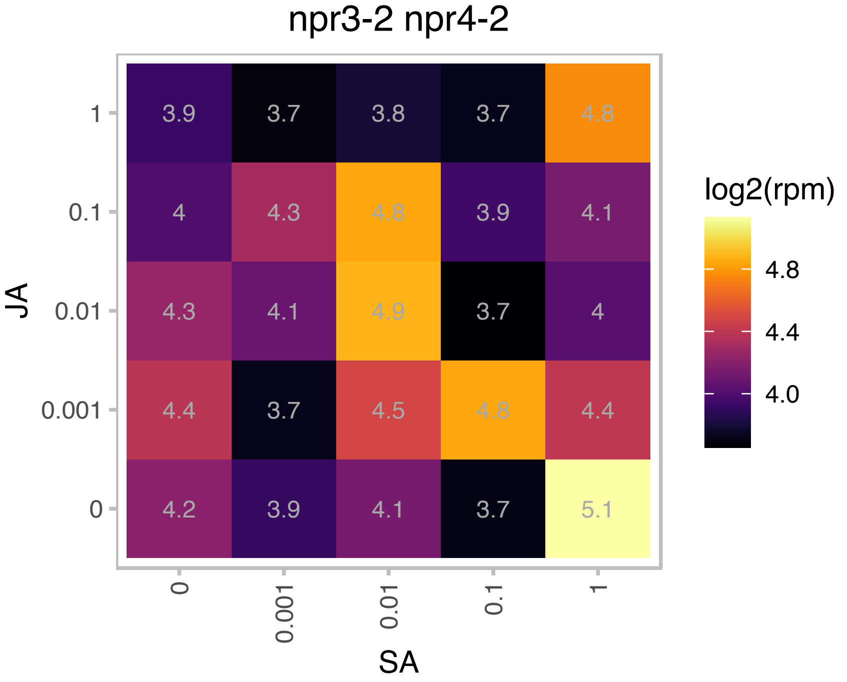


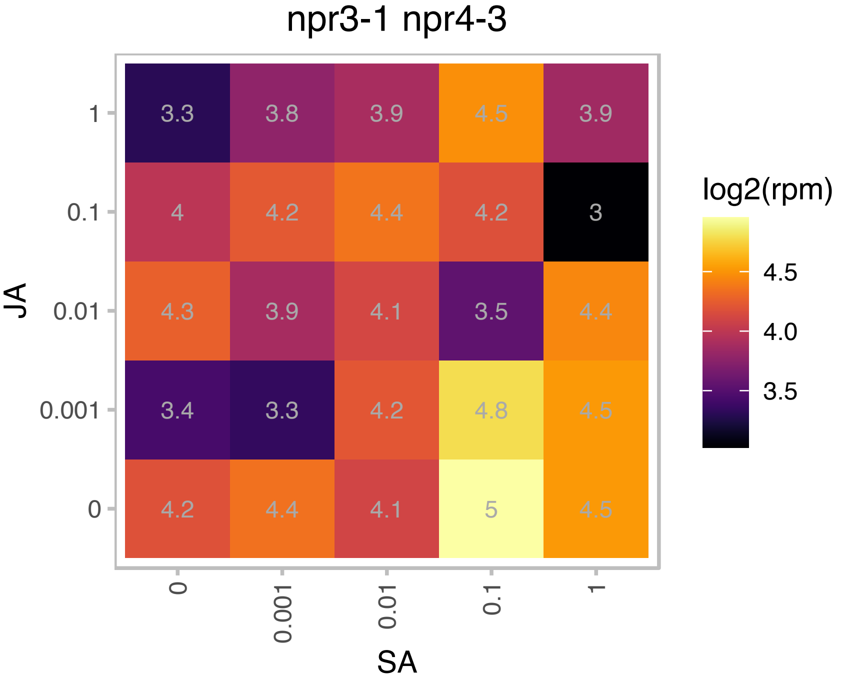

# AT5G05230.1 : NA

RING/U-box superfamily protein

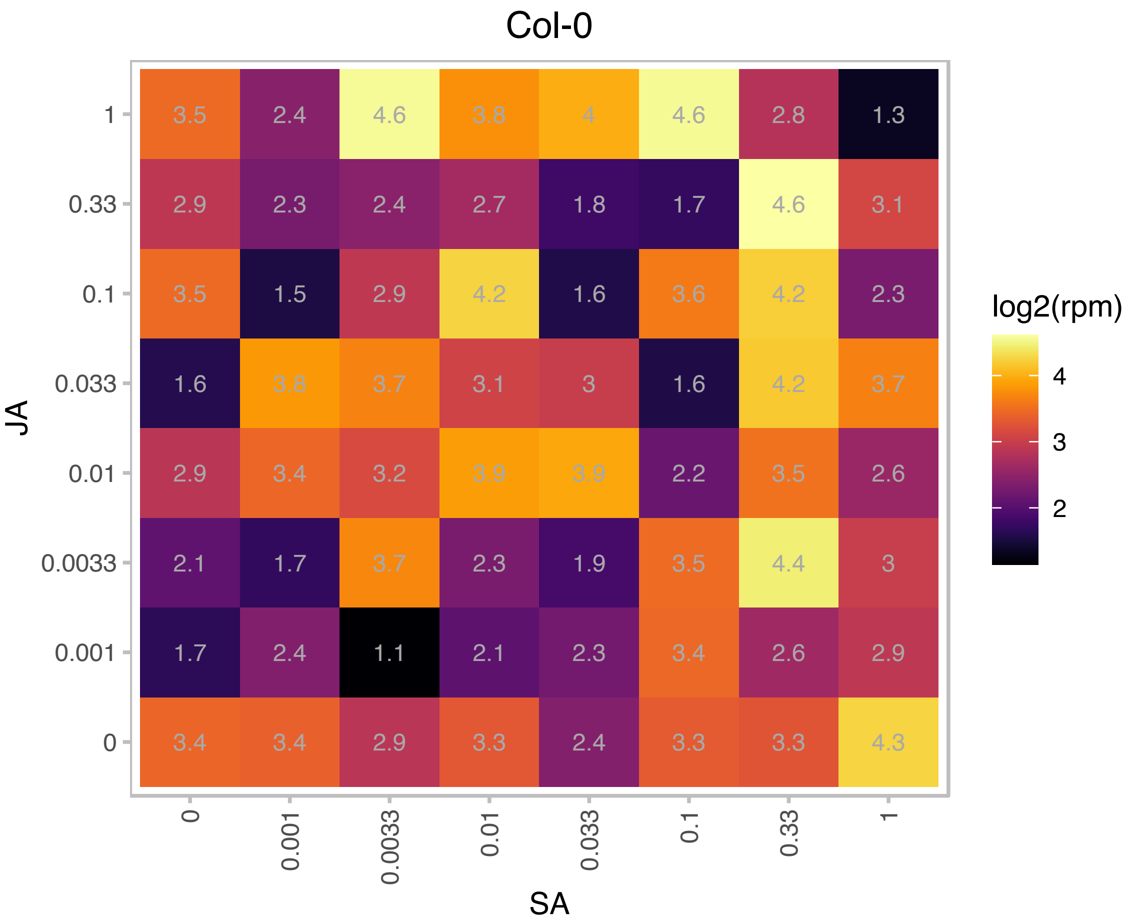


| | |
|---|---|
| Gene | AT5G05230 |
| is.expressed_genes | TRUE |
| is.responsive_genes | TRUE |
| is.apcluster | C-04 |

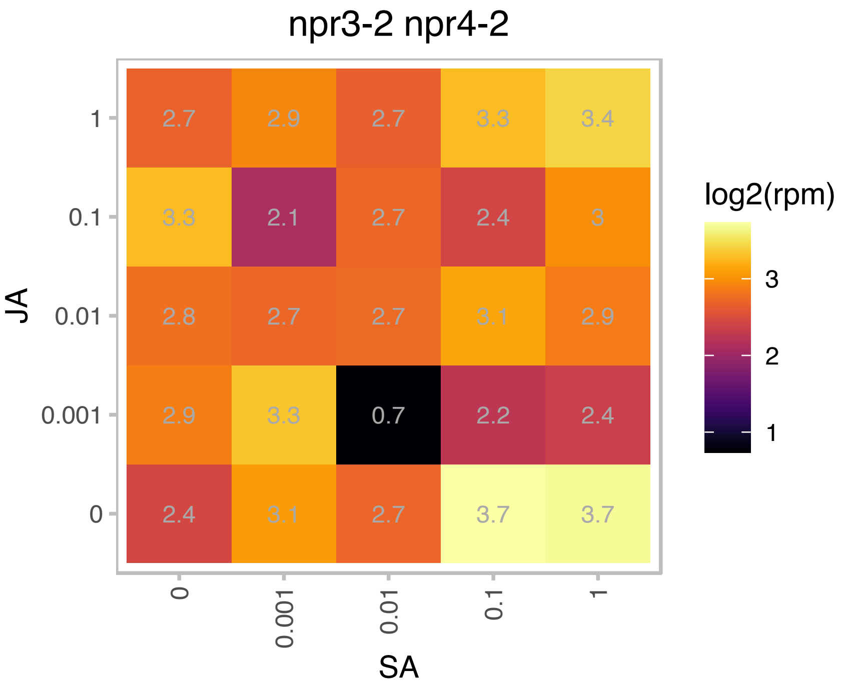


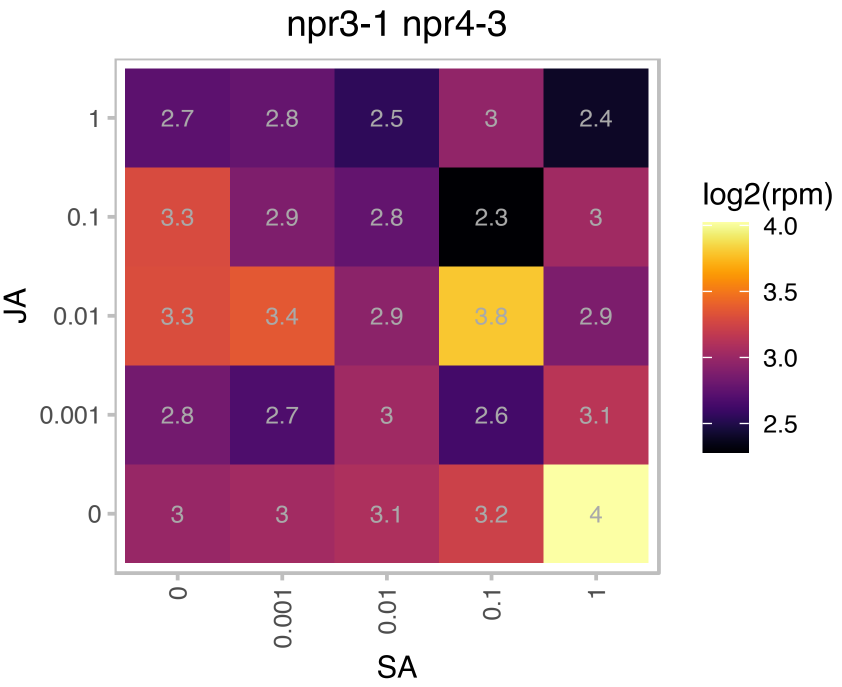

### AT5G08335.1 : ATICMTB, ATSTE14B, ICMTB

Isoprenylcysteine carboxyl methyltransferase (ICMT) family

Encodes an isoprenyl cysteine methylatransferase (ICMT) involved in the post-translational processing of proteins that have a C-terminal CaaX box. This protein appears to have higher catalytic activity and a higher transcript expression level than the other ICMT present in Arabidopsis (At5g23320). Analysis of ICMT RNAi lines suggests that this protein is involved in flower and stem development.

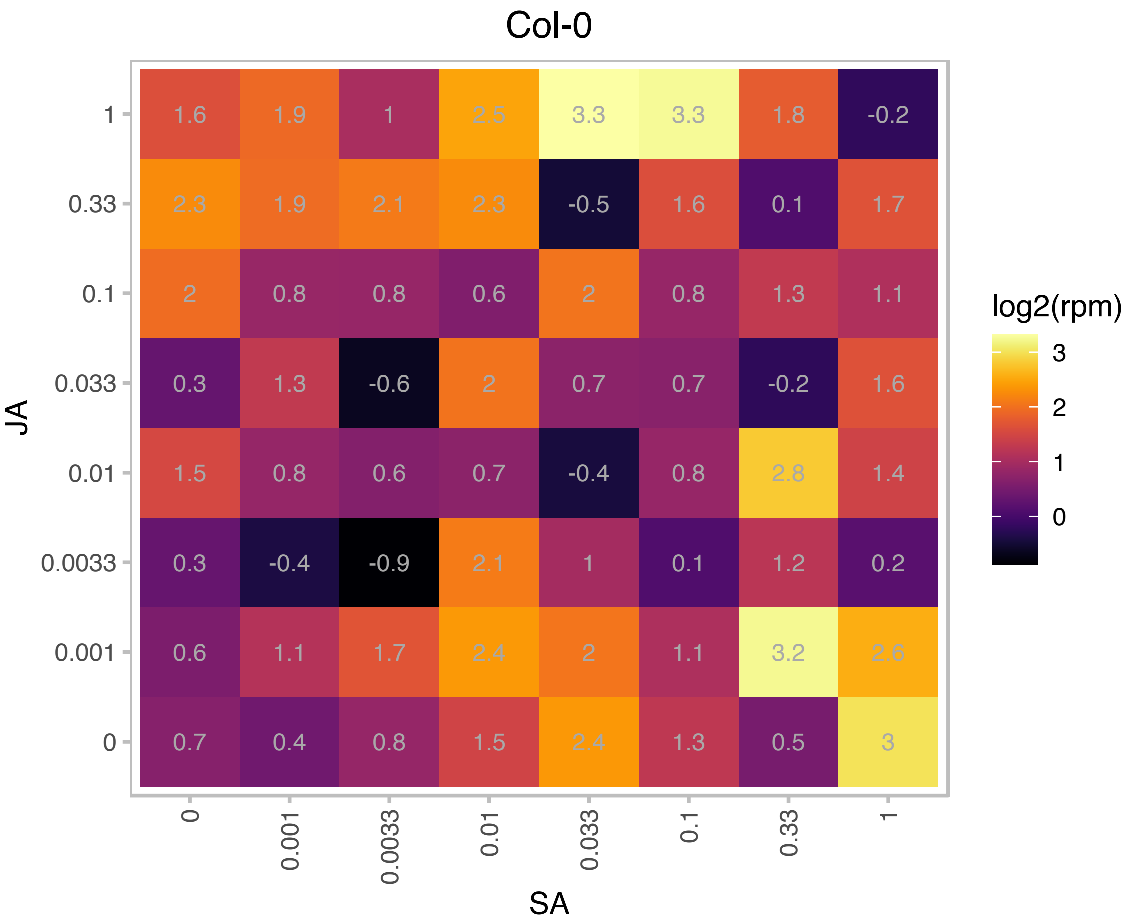


| | |
|---|---|
| Gene | AT5G08335 |
| is.expressed_genes | TRUE |
| is.responsive_genes | TRUE |
| is.apcluster | C-04 |

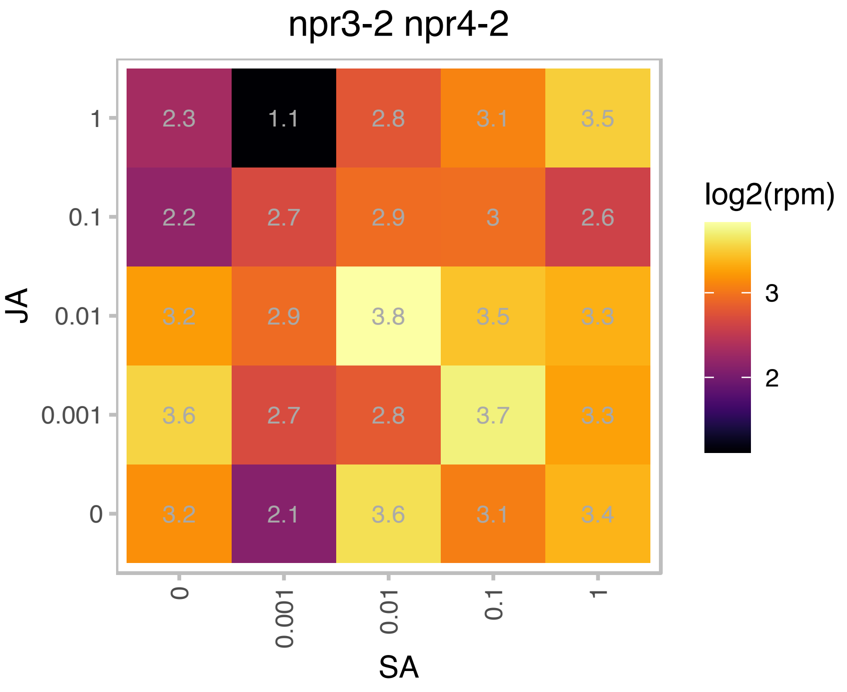


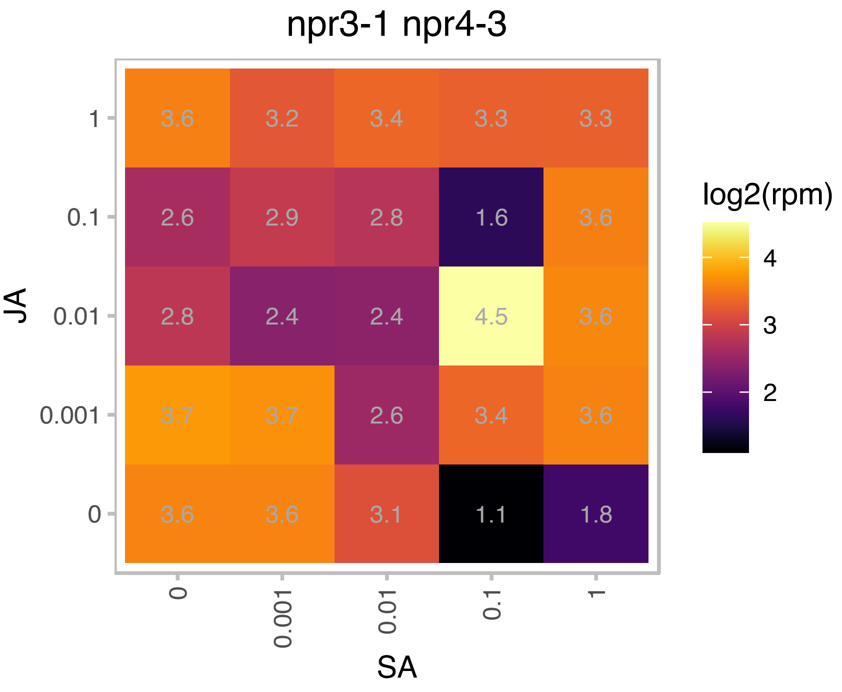

# AT5G11010.1 : NA

Pre-mRNA cleavage complex II protein family

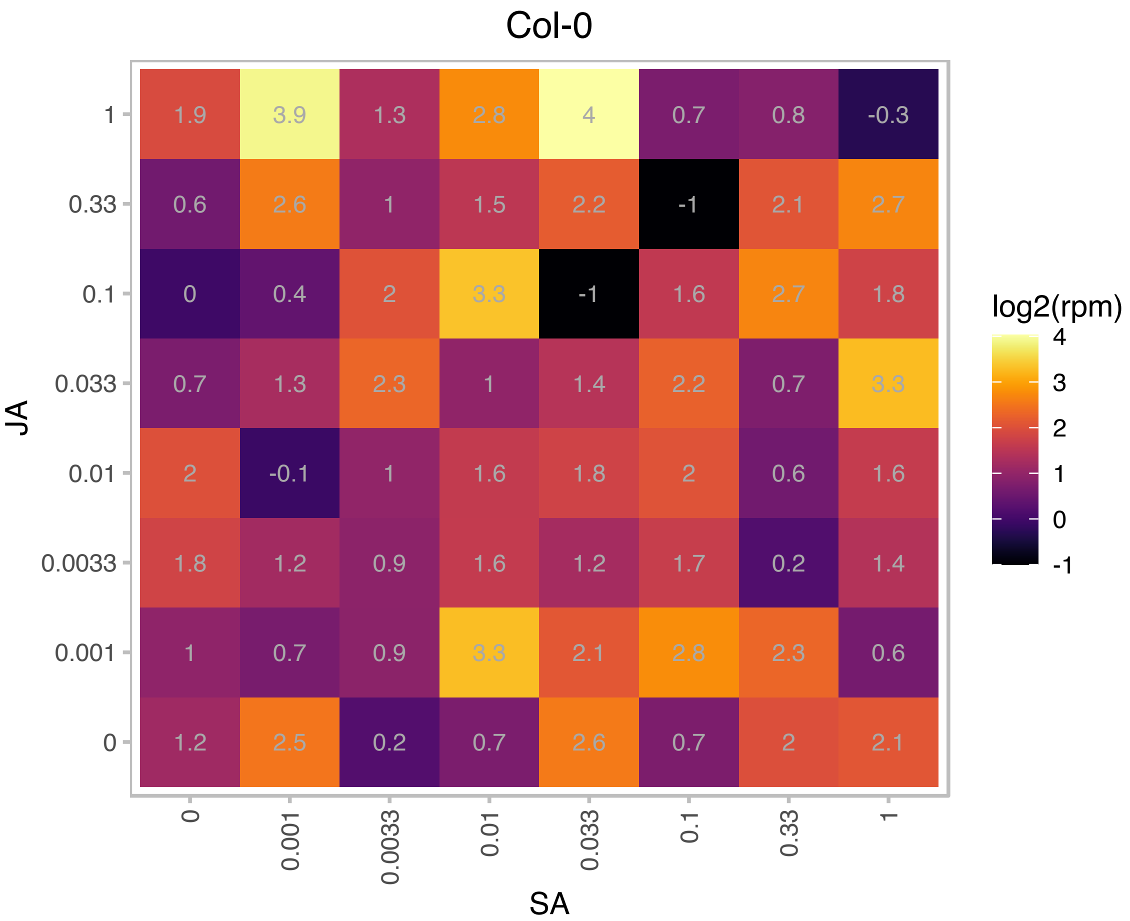


| | |
|---|---|
| Gene | AT5G11010 |
| is.expressed_genes | TRUE |
| is.responsive_genes | TRUE |
| is.apcluster | C-04 |

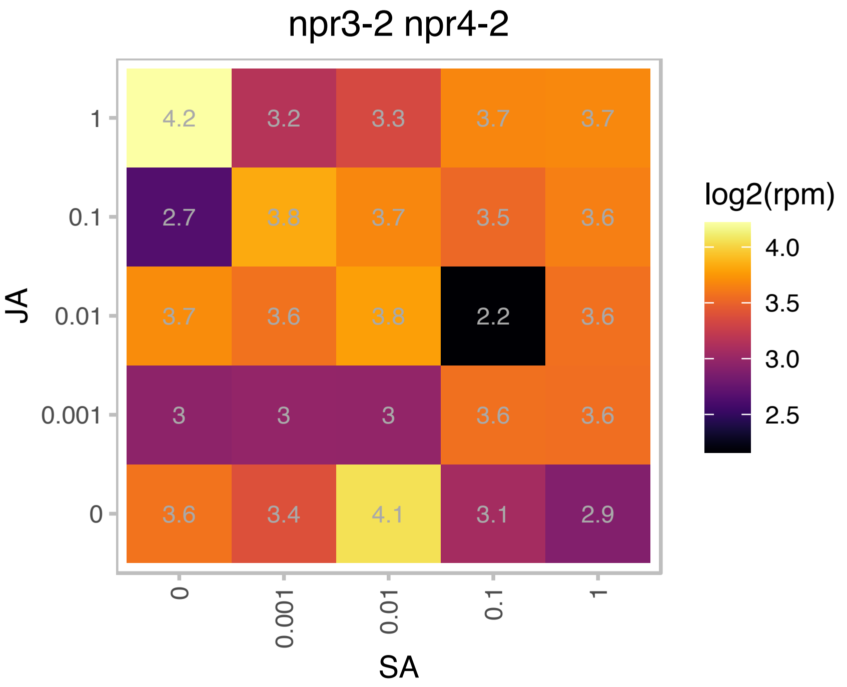


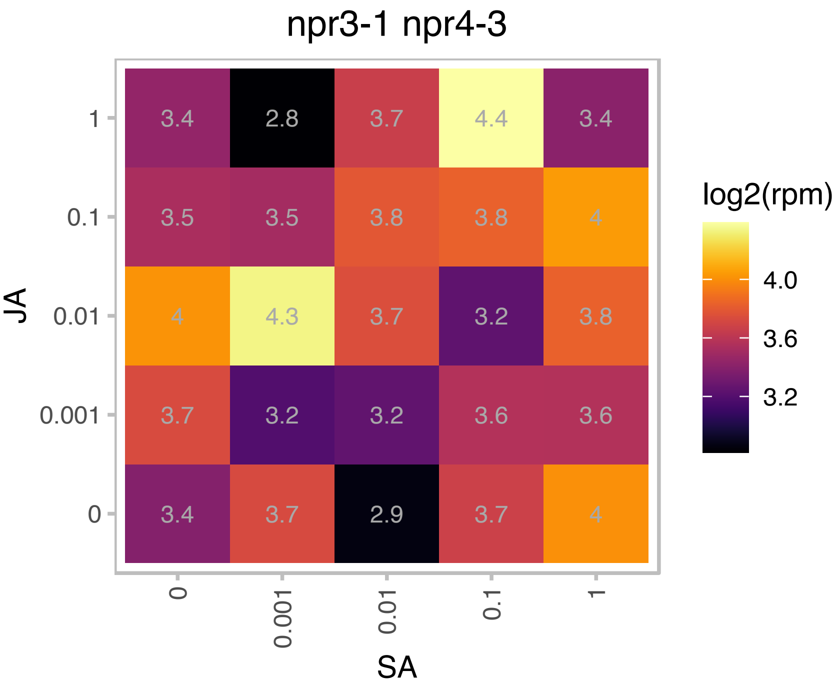

# AT5G19030.1 : NA

RNA-binding (RRM/RBD/RNP motifs) family protein

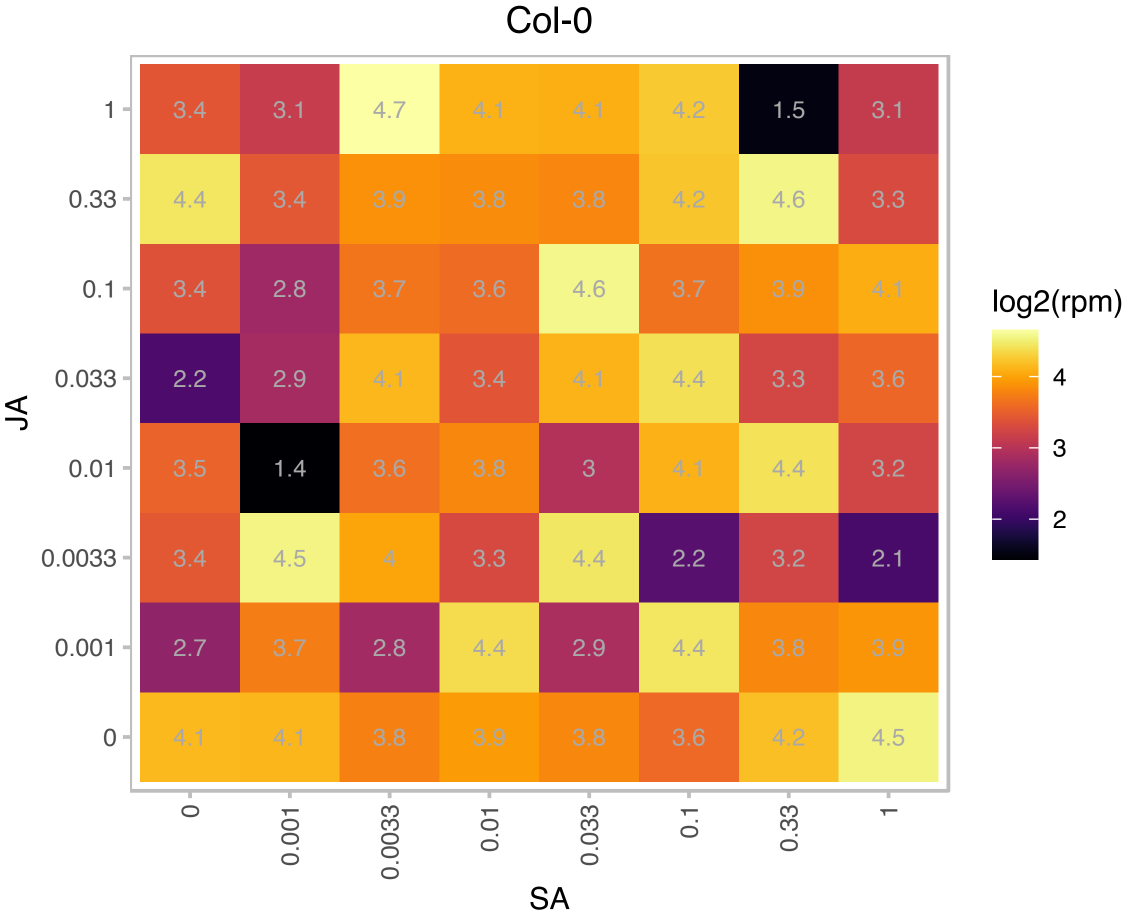


| | |
|---|---|
| Gene | AT5G19030 |
| is.expressed_genes | TRUE |
| is.responsive_genes | TRUE |
| is.apcluster | C-04 |

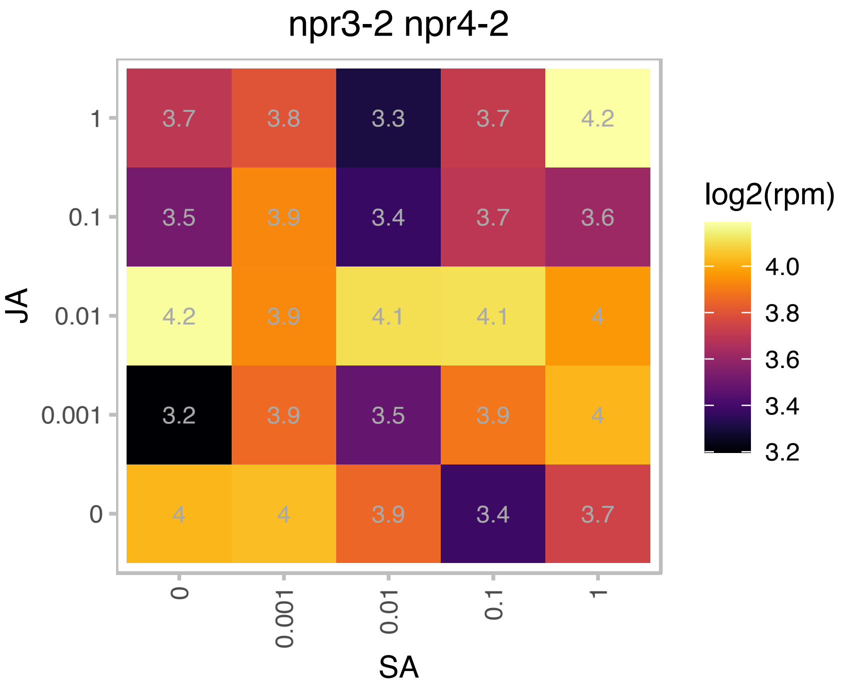


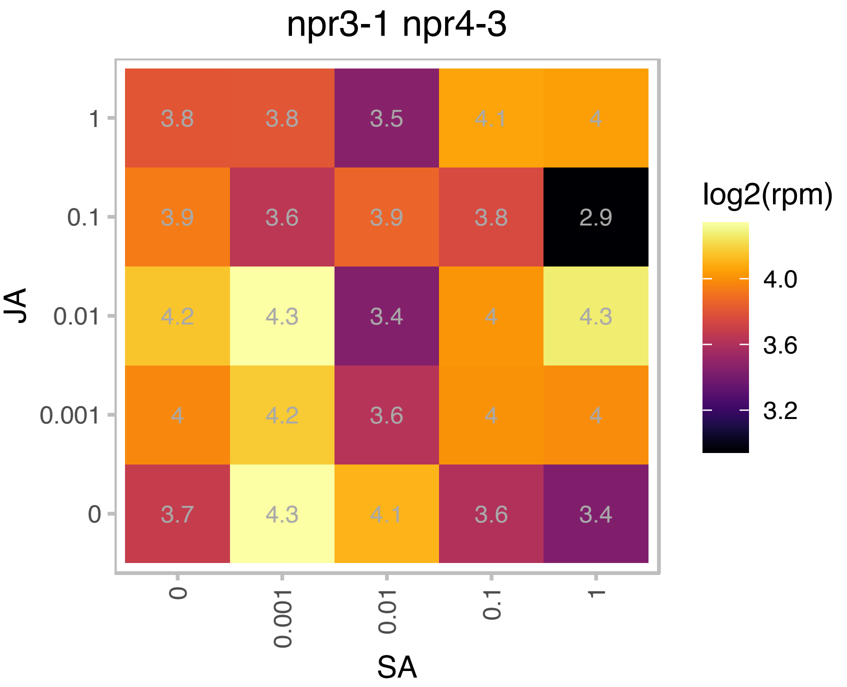

### AT5G34940.1 : AtGUS3, GUS3

glucuronidase 3

The protein is predicted (WoLF PSORT program) to be membrane-associated.

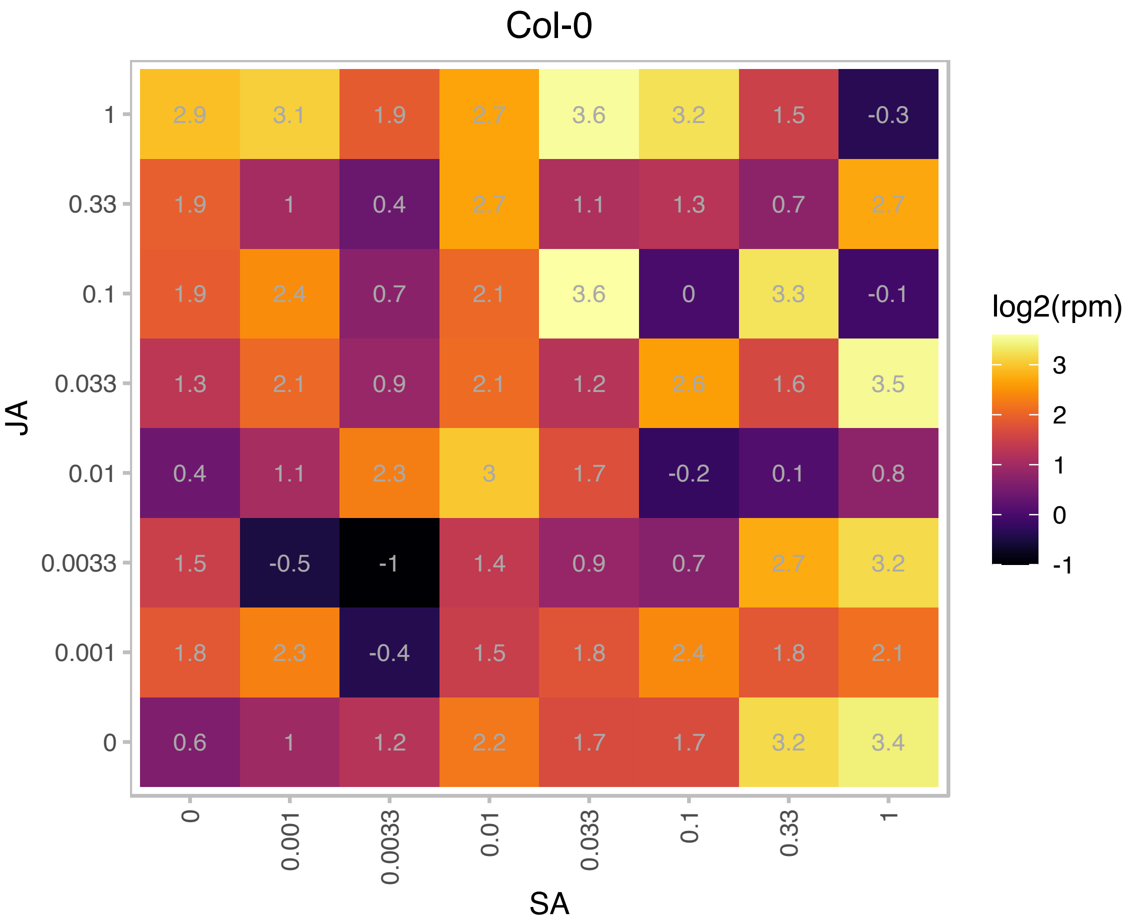


| | |
|---|---|
| Gene | AT5G34940 |
| is.expressed_genes | TRUE |
| is.responsive_genes | TRUE |
| is.apcluster | C-04 |

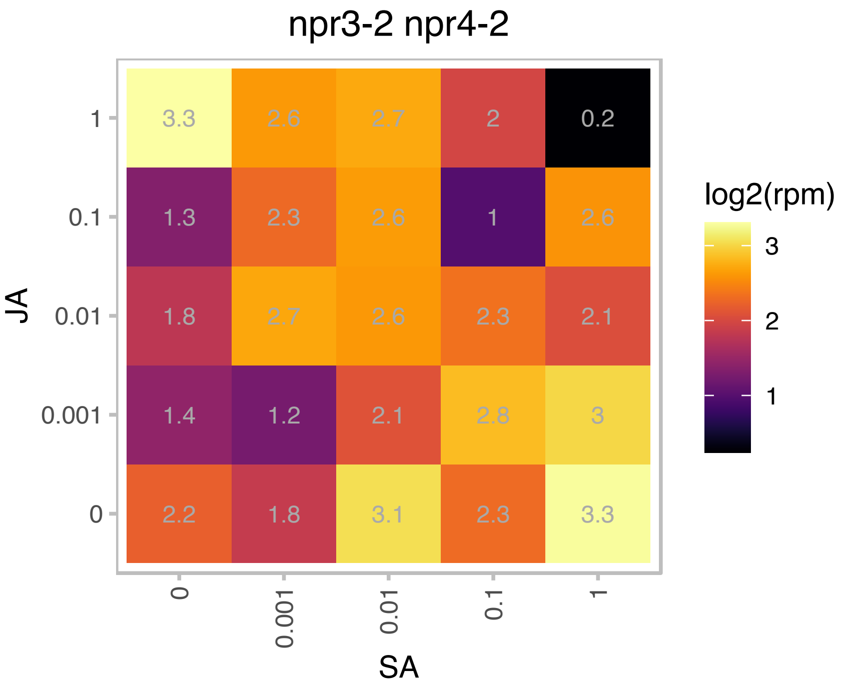


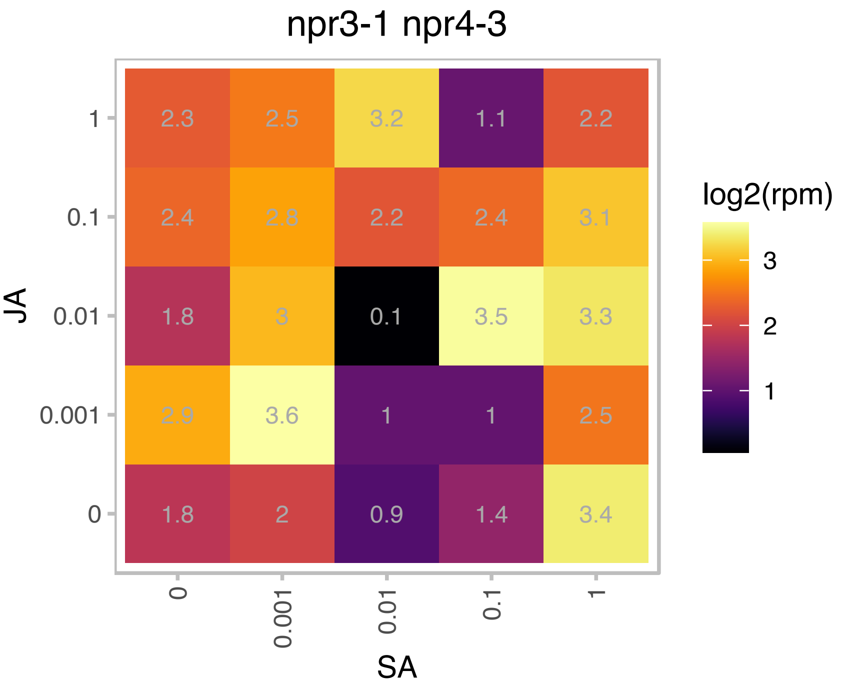

### AT5G37600.1 : ATGLN1;1, ATGSR1, GLN1;1, GSR 1

glutamine synthase clone R1

encodes a cytosolic glutamine synthetase, the enzyme has high affinity with substrate ammonium

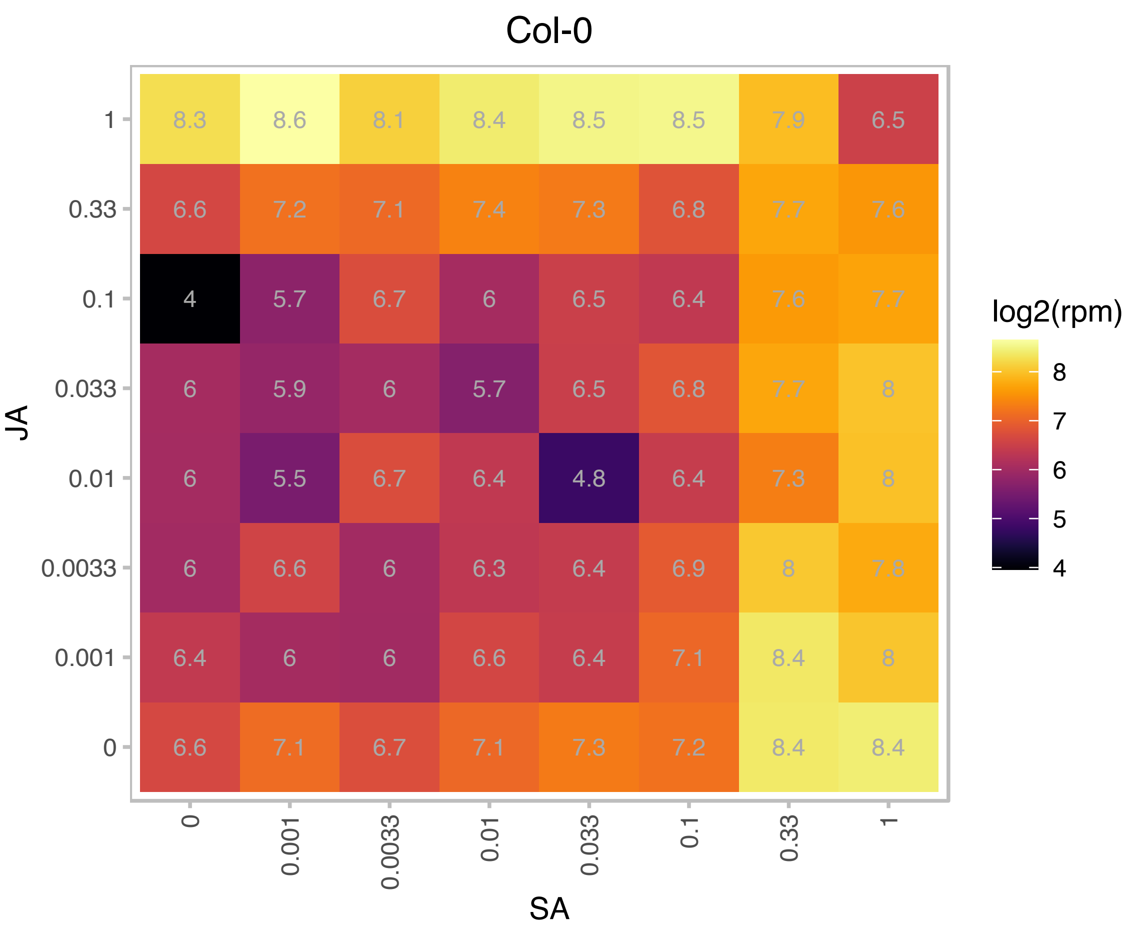


| | |
|---|---|
| Gene | AT5G37600 |
| is.expressed_genes | TRUE |
| is.responsive_genes | TRUE |
| is.apcluster | C-04 |

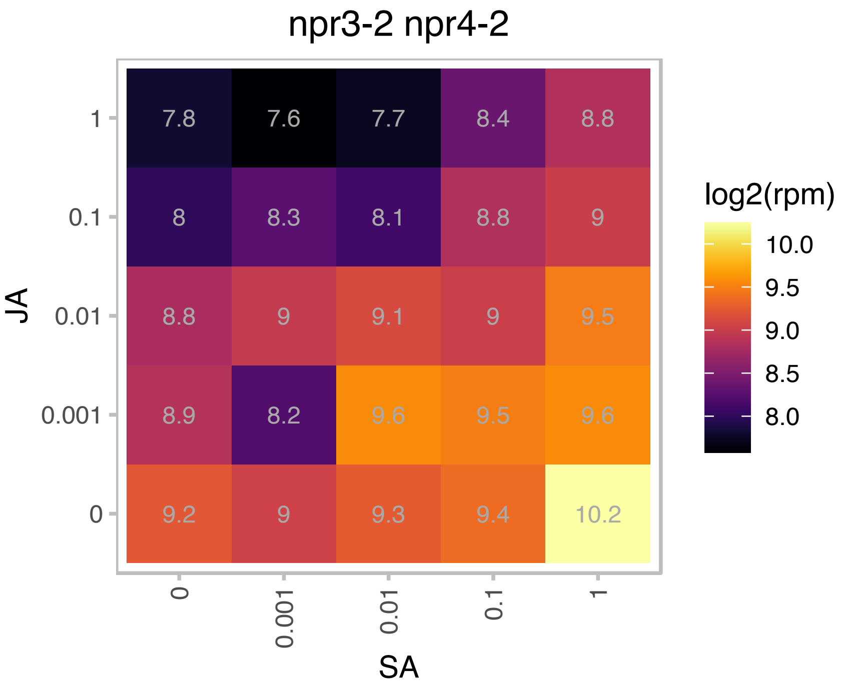


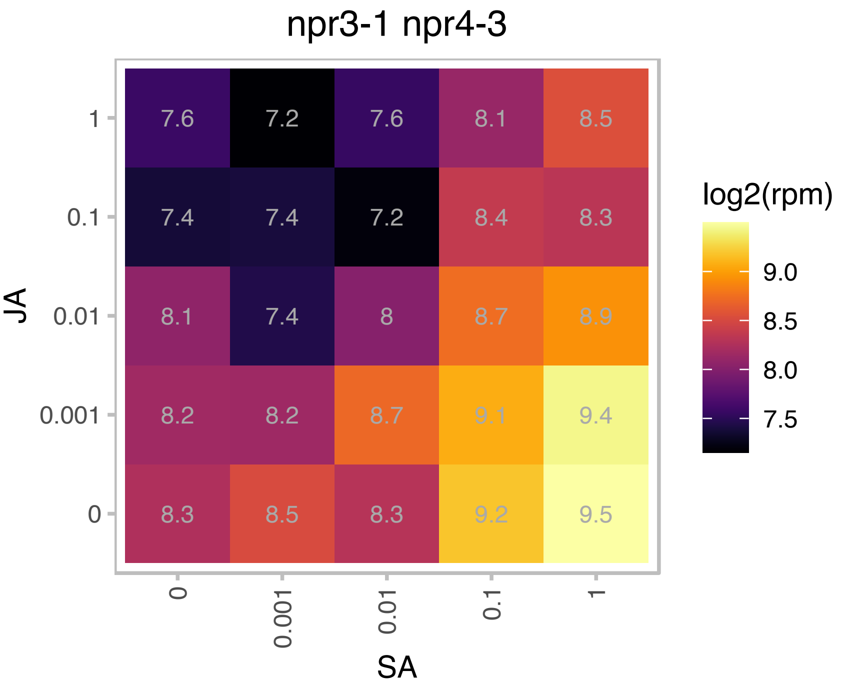

### AT5G41340.1 : ATUBC4, UBC4

ubiquitin conjugating enzyme 4

Belongs to Ubiquitin conjugating enzyme family. Gene expression is developmentally regulated.

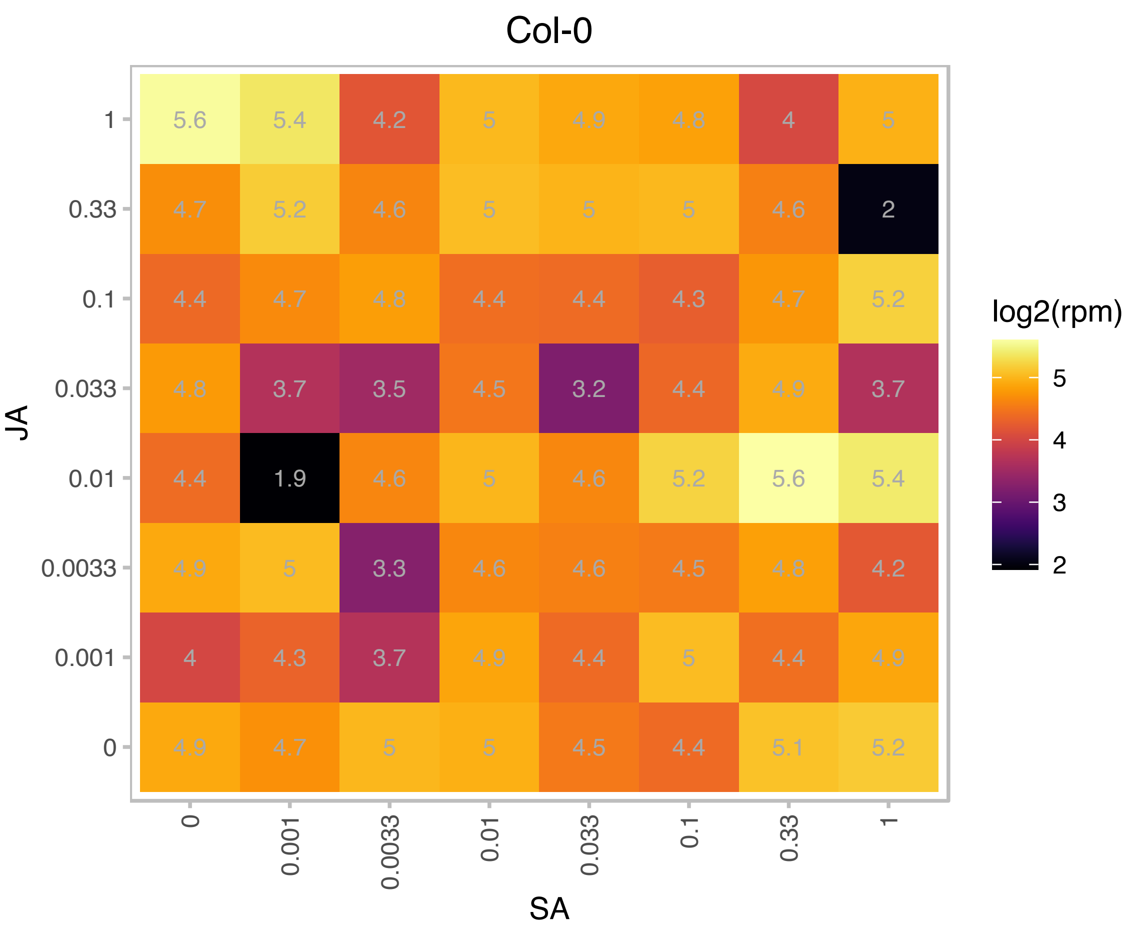


| | |
|---|---|
| Gene | AT5G41340 |
| is.expressed_genes | TRUE |
| is.responsive_genes | TRUE |
| is.apcluster | C-04 |

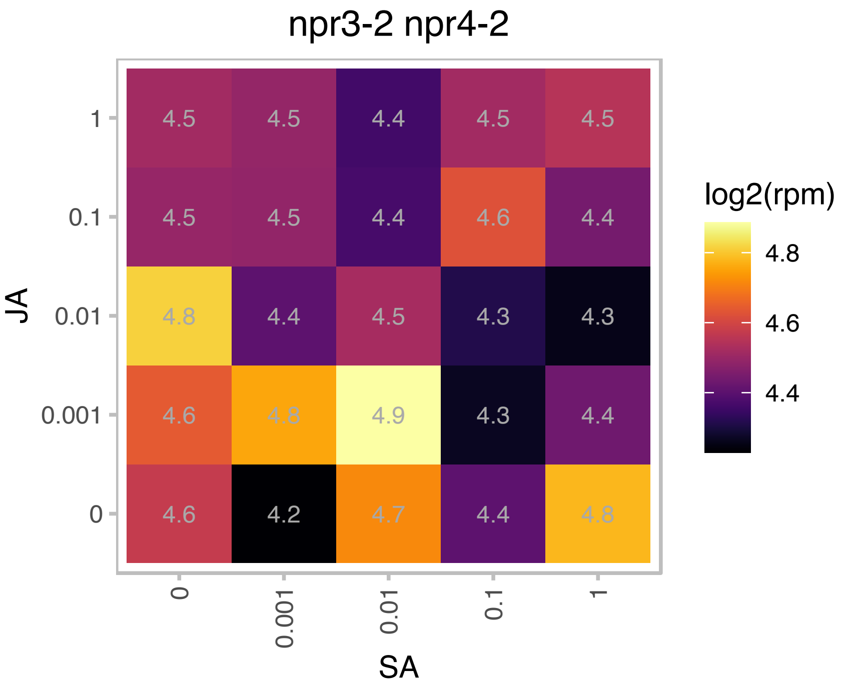


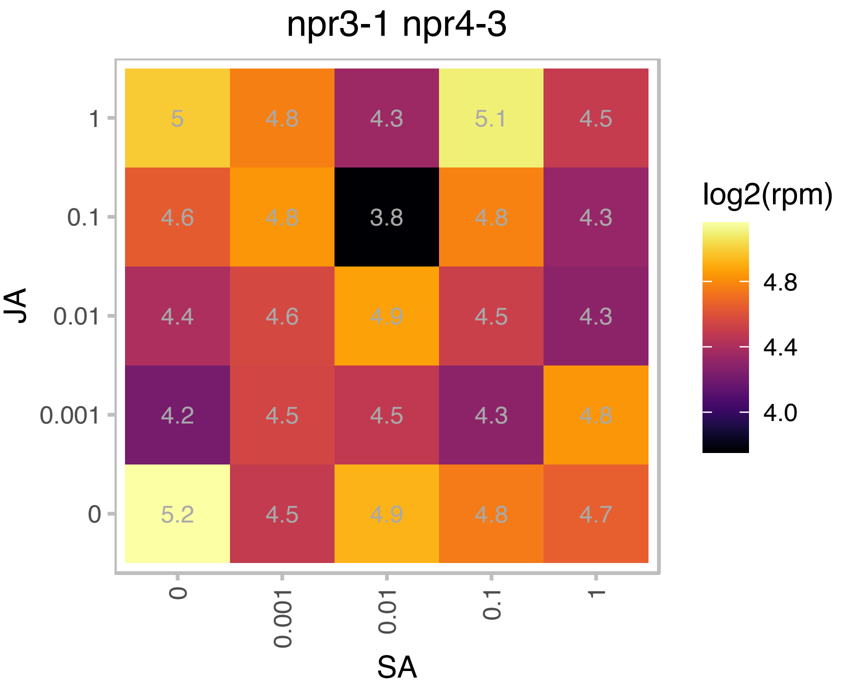

### AT5G58440.1 : SNX2a

sorting nexin 2A

Col-0

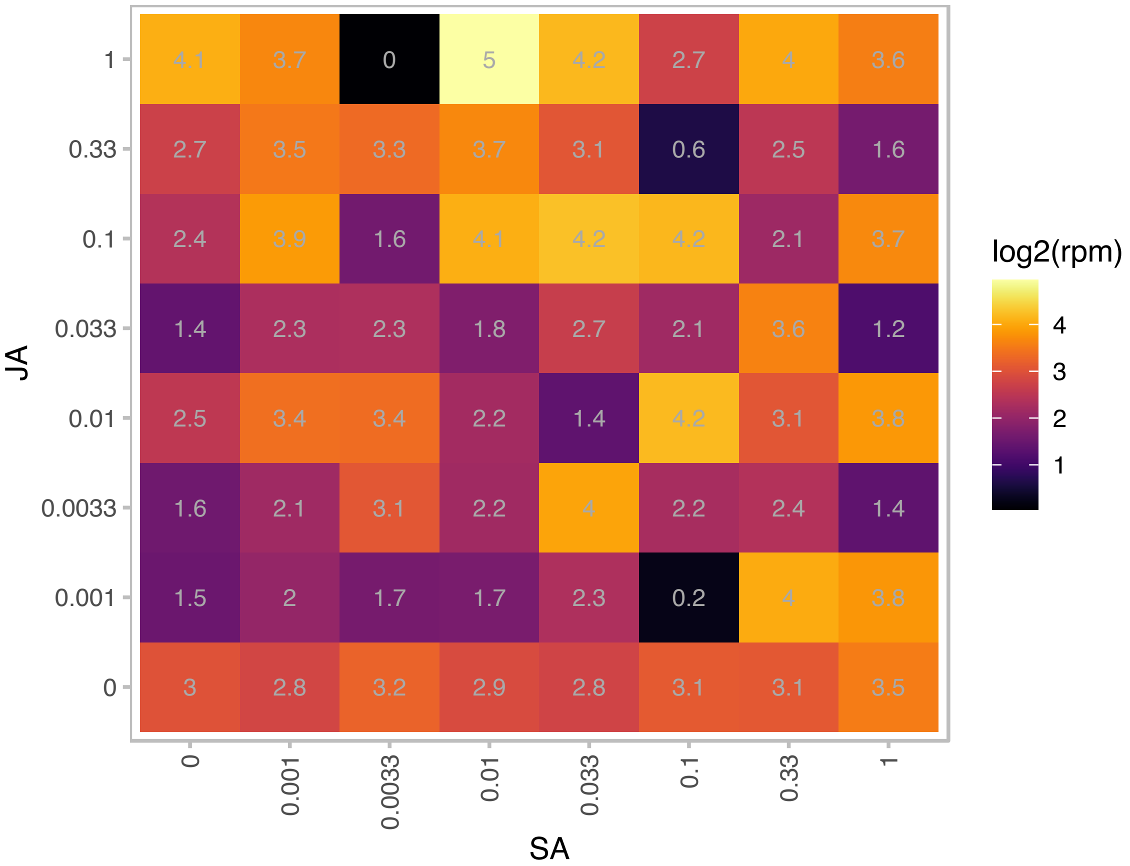


| | |
|---|---|
| Gene | AT5G58440 |
| is.expressed_genes | TRUE |
| is.responsive_genes | TRUE |
| is.apcluster | C-04 |

npr3-2 npr4-2

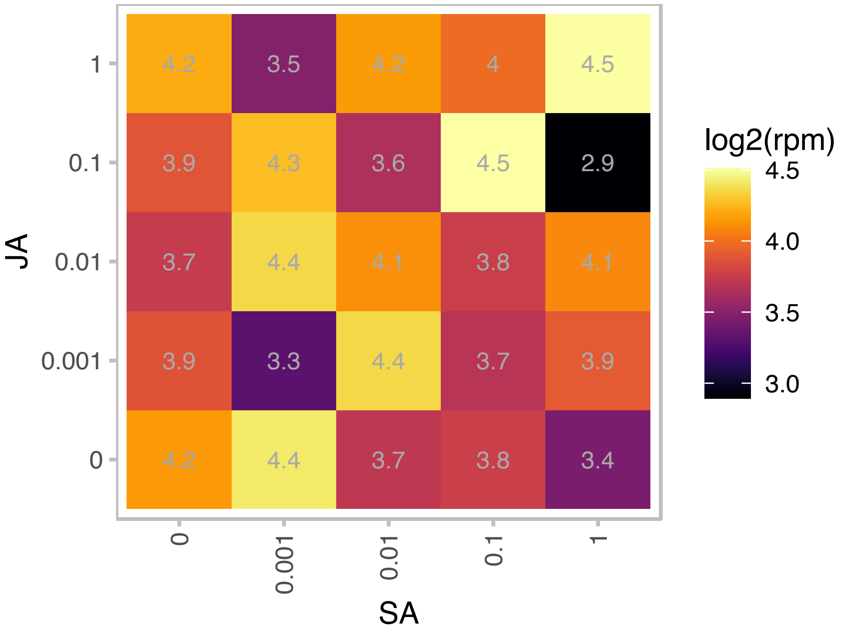


npr3-1 npr4-3

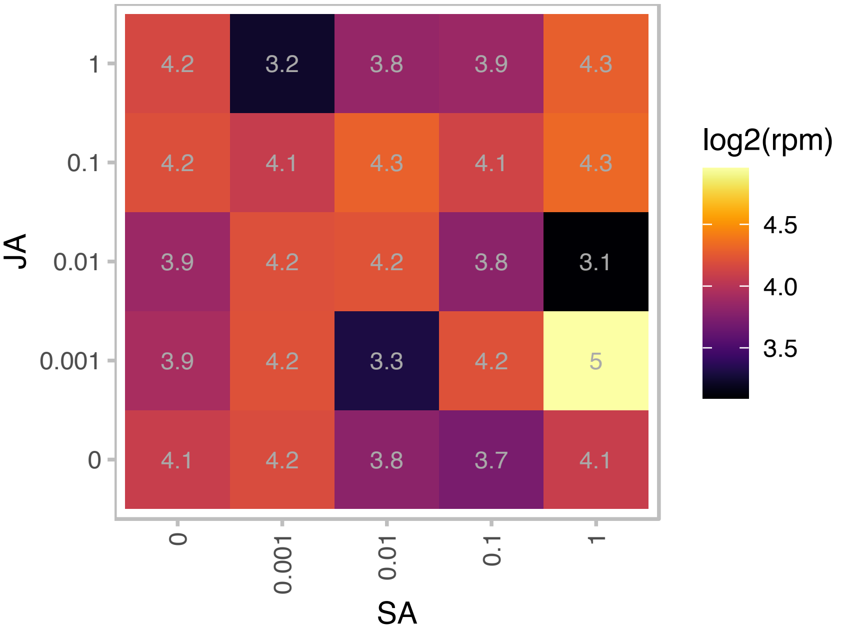

# AT5G67140.1 : NA

F-box/RNI-like superfamily protein

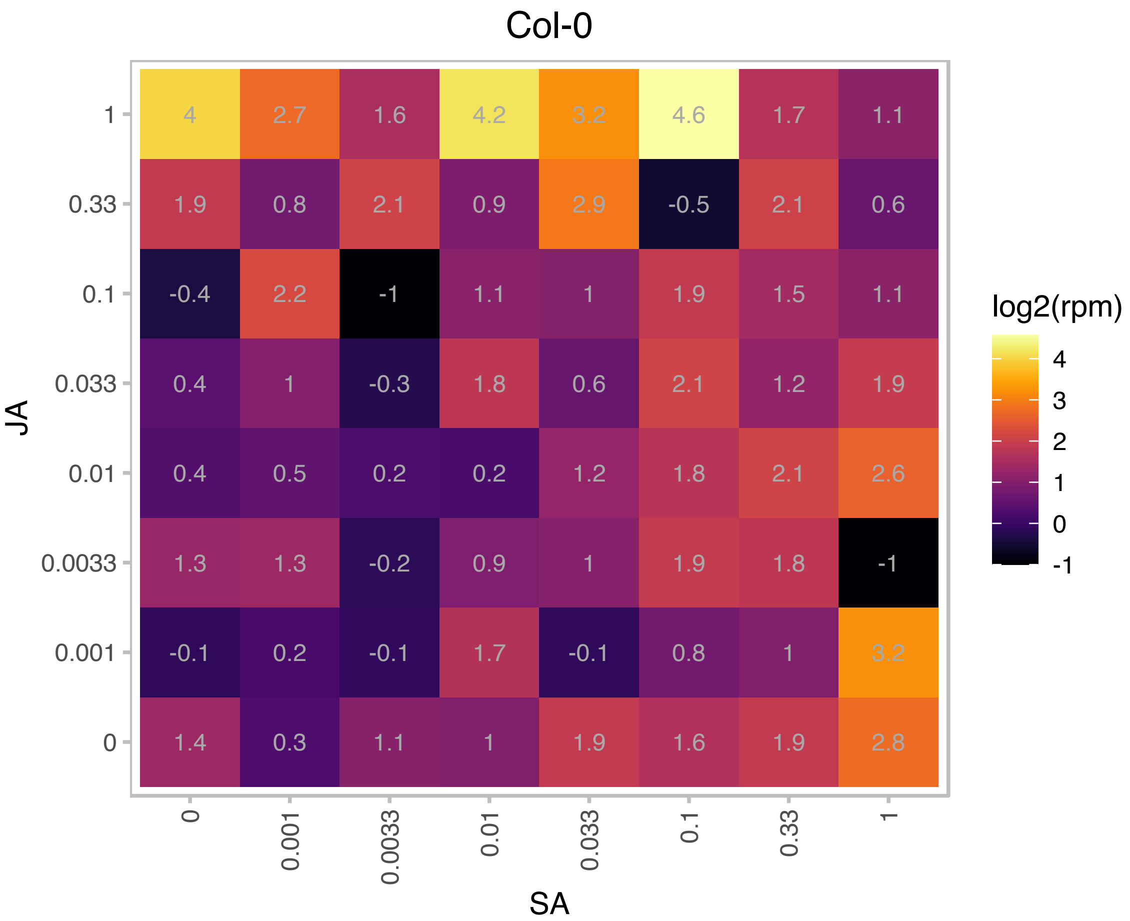


| | |
|---|---|
| Gene | AT5G67140 |
| is.expressed_genes | TRUE |
| is.responsive_genes | TRUE |
| is.apcluster | C-04 |

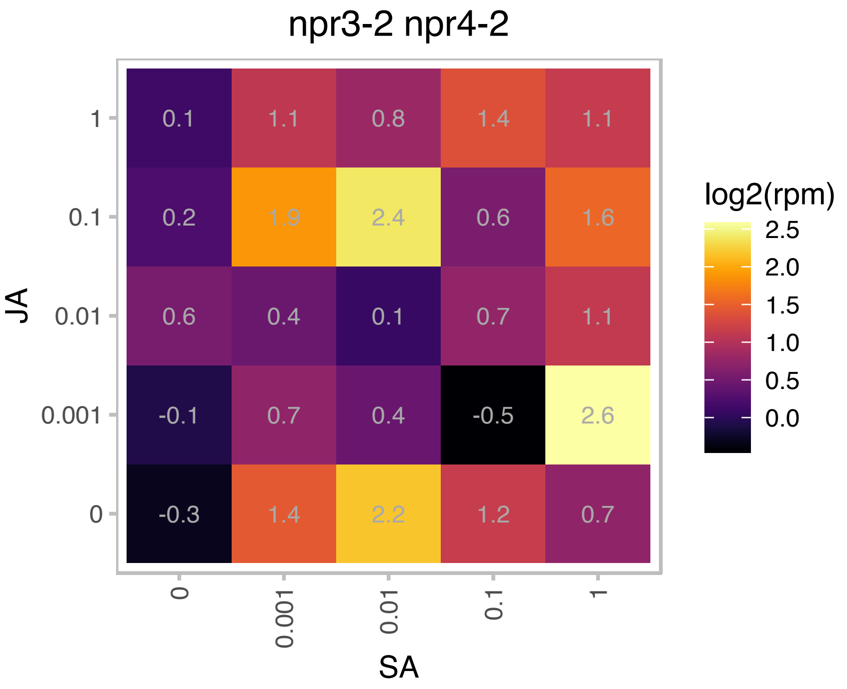


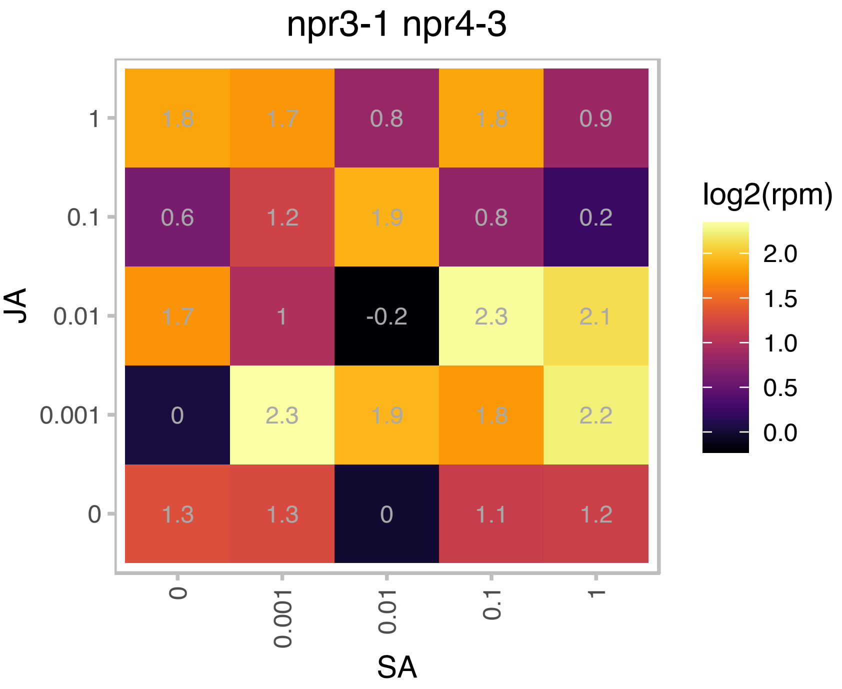

# AT1G30990.1 : NA

Polyketide cyclase/dehydrase and lipid transport superfamily protein

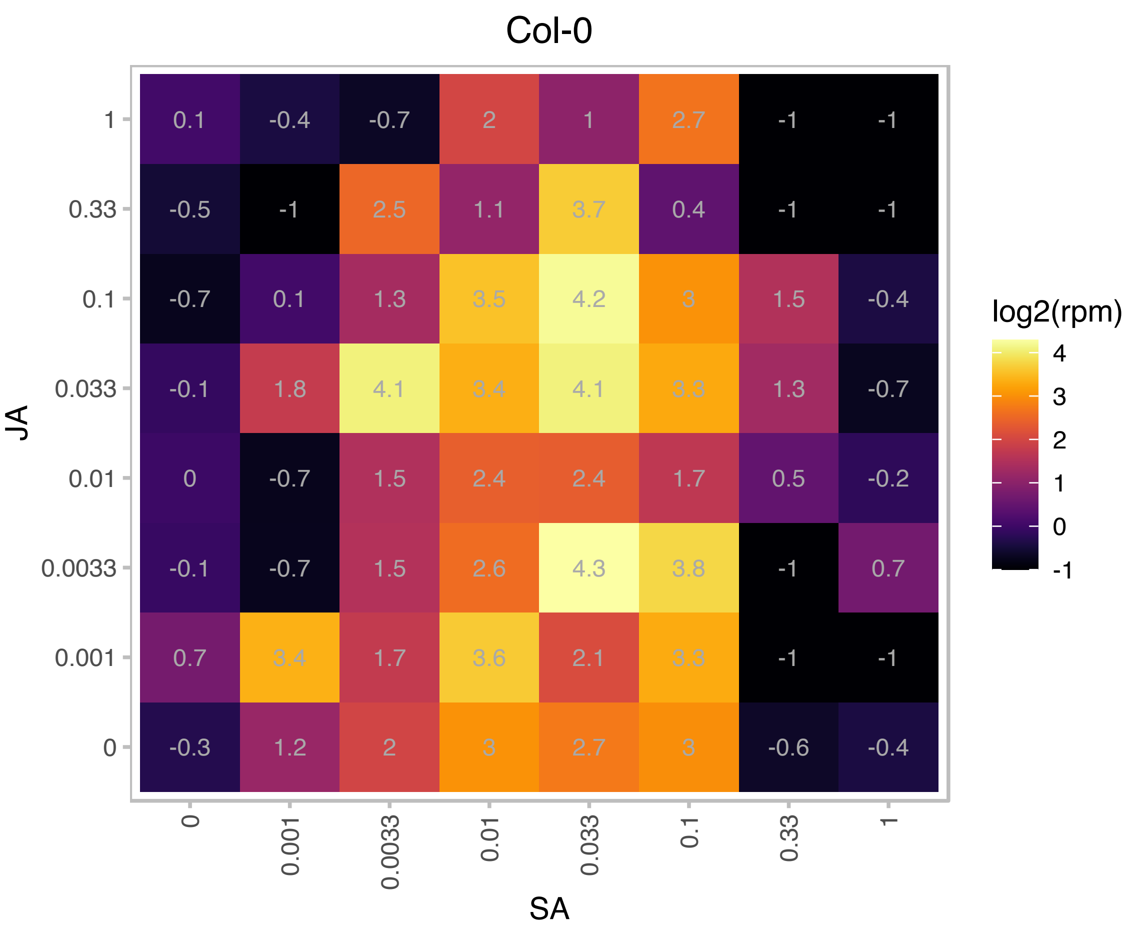


| | |
|---|---|
| Gene | AT1G30990 |
| is.expressed_genes | TRUE |
| is.responsive_genes | TRUE |
| is.apcluster | D-01 |

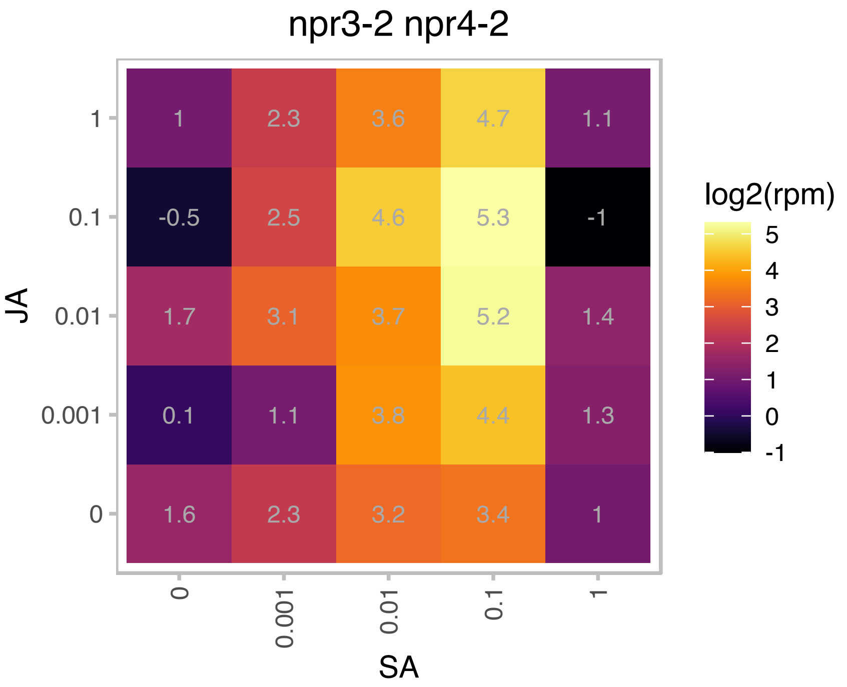


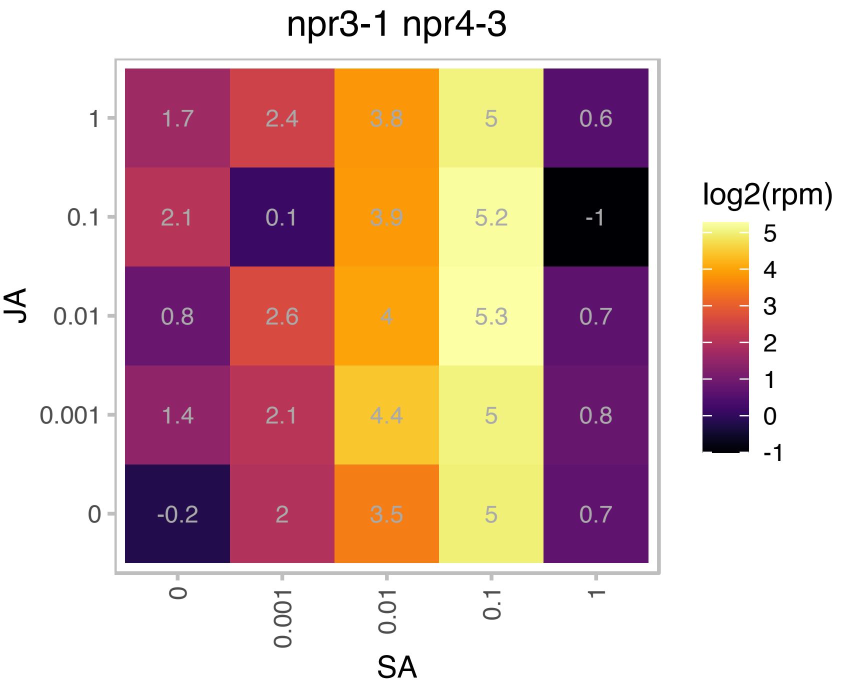

### AT1G51420.1 : ATSPP1, SPP1

sucrose-phosphatase 1

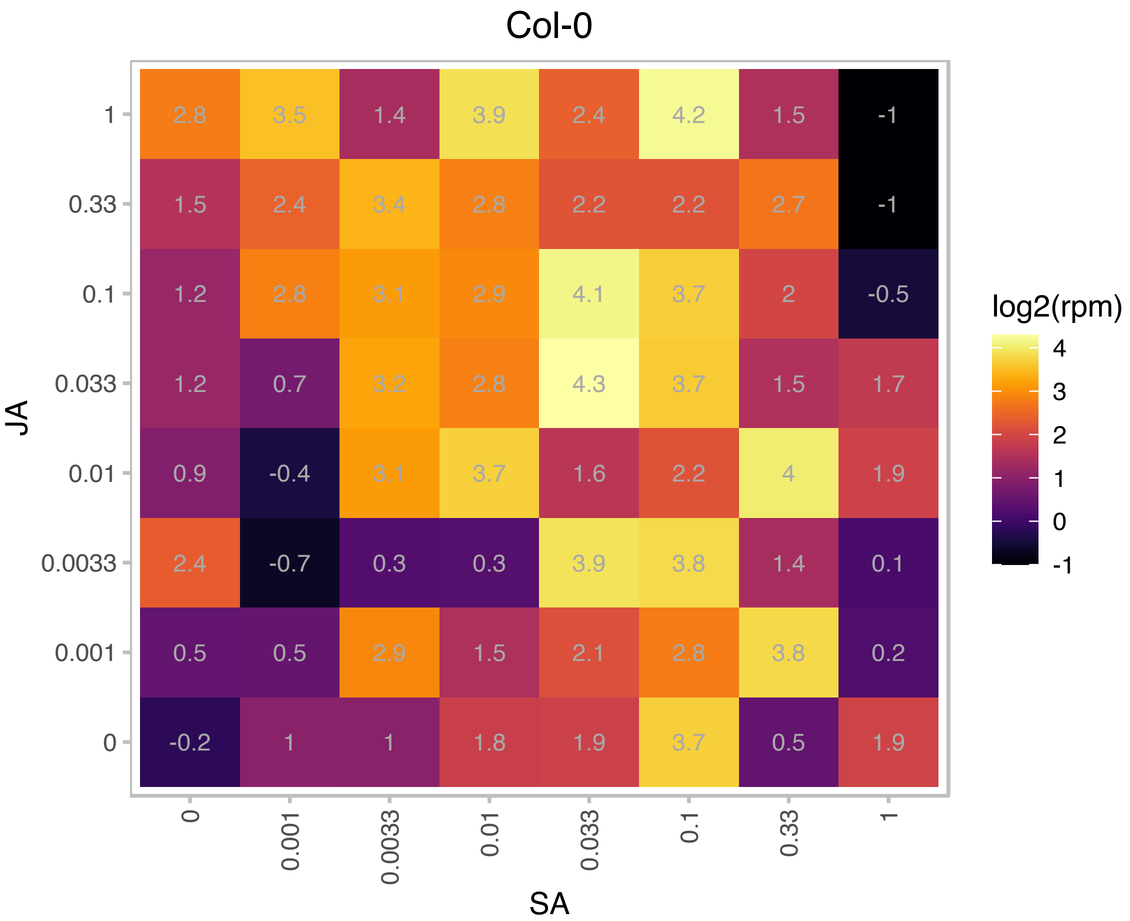


| | |
|---|---|
| Gene | AT1G51420 |
| is.expressed_genes | TRUE |
| is.responsive_genes | TRUE |
| is.apcluster | D-01 |

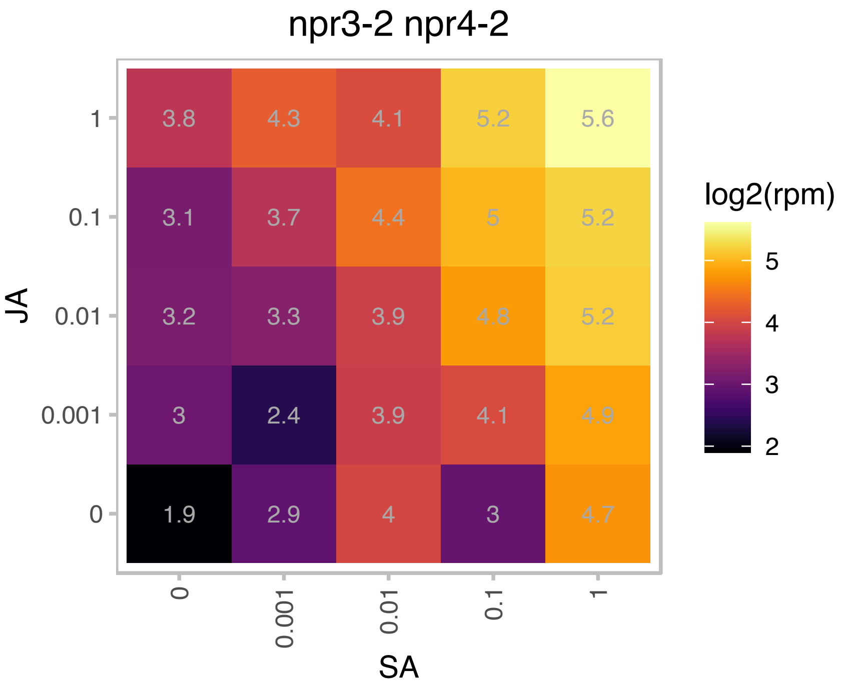


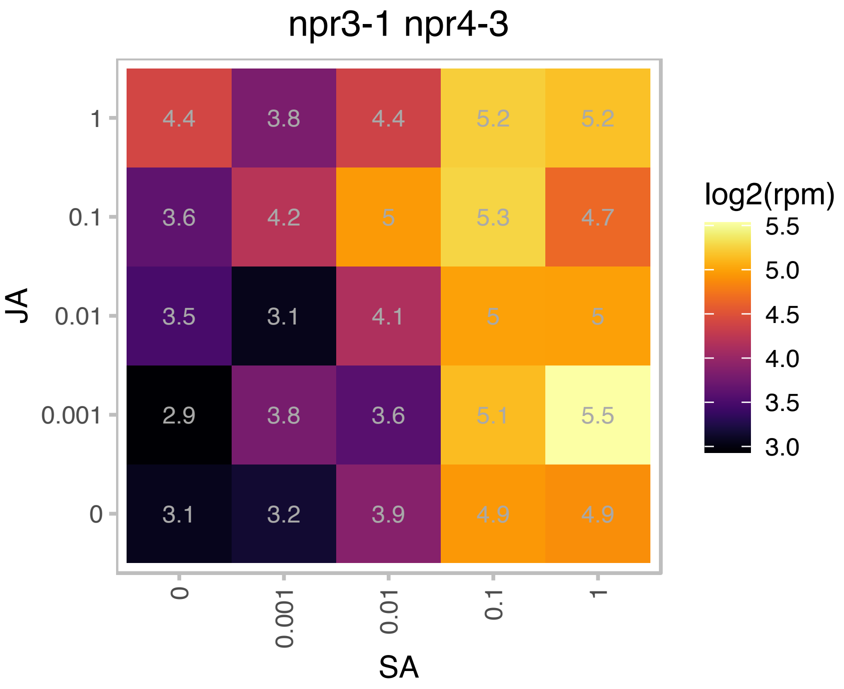

# AT2G04170.1 : NA

TRAF-like family protein

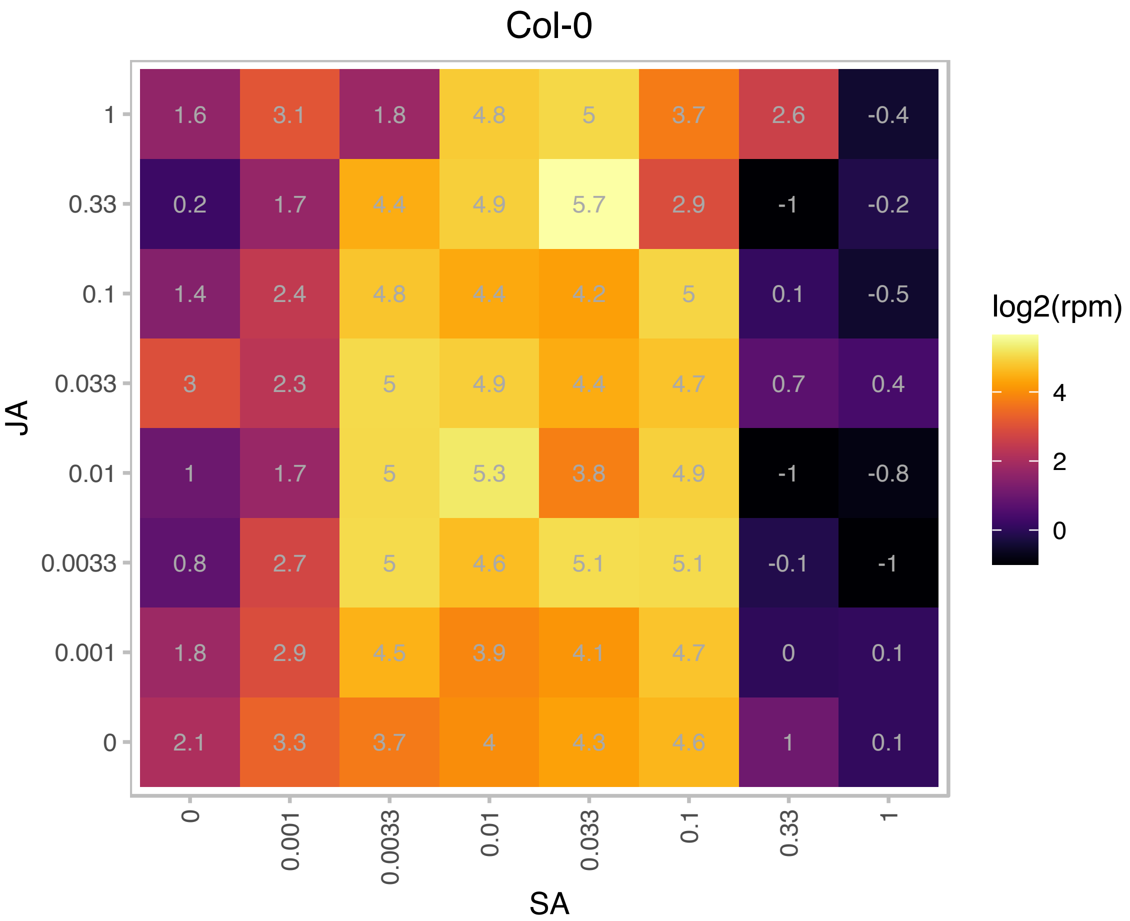


| Gene | AT2G04170 |
|---|---|
| is.expressed_genes | TRUE |
| is.responsive_genes | TRUE |
| is.apcluster | D-01 |

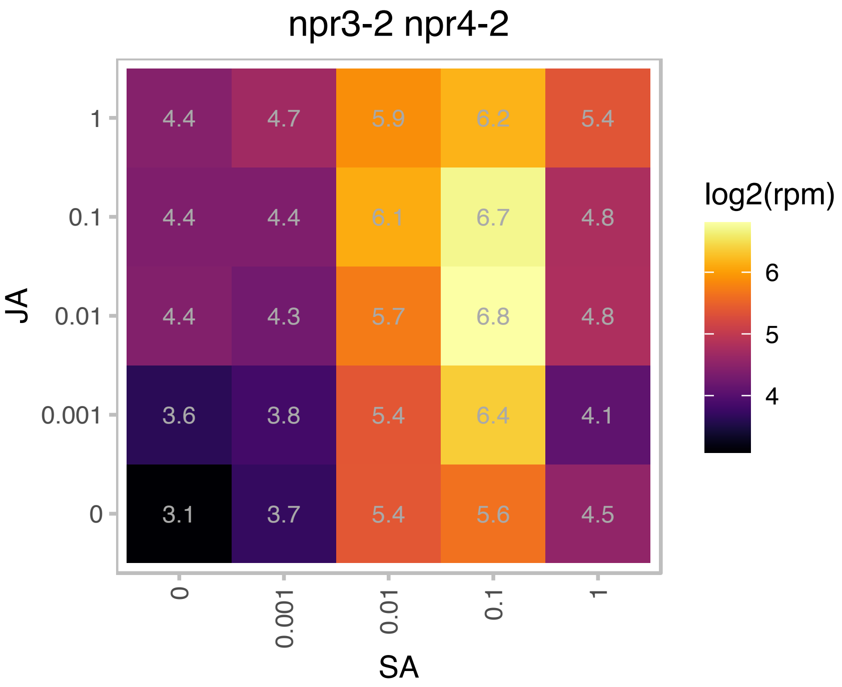


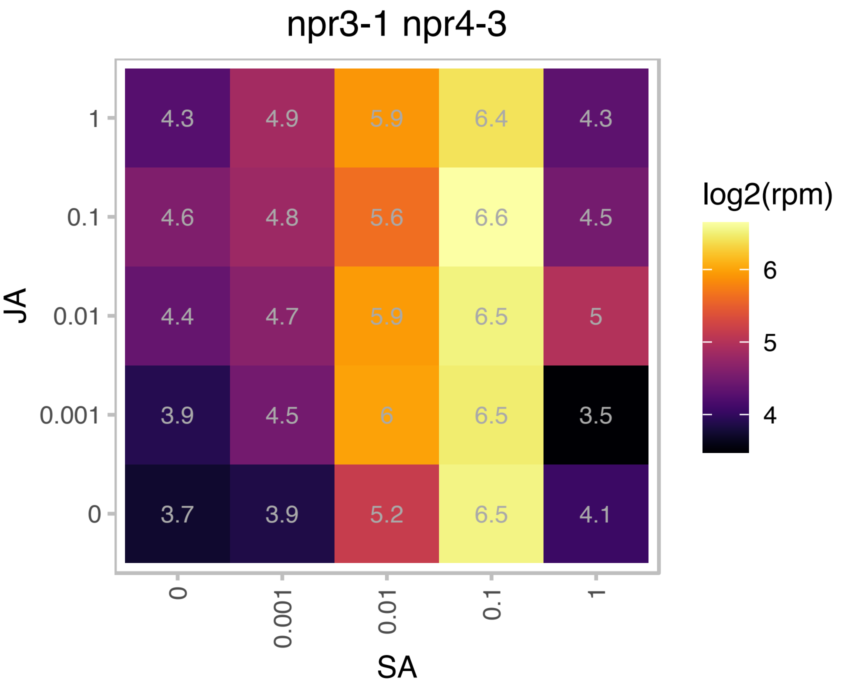

# AT2G04800.1 : NA

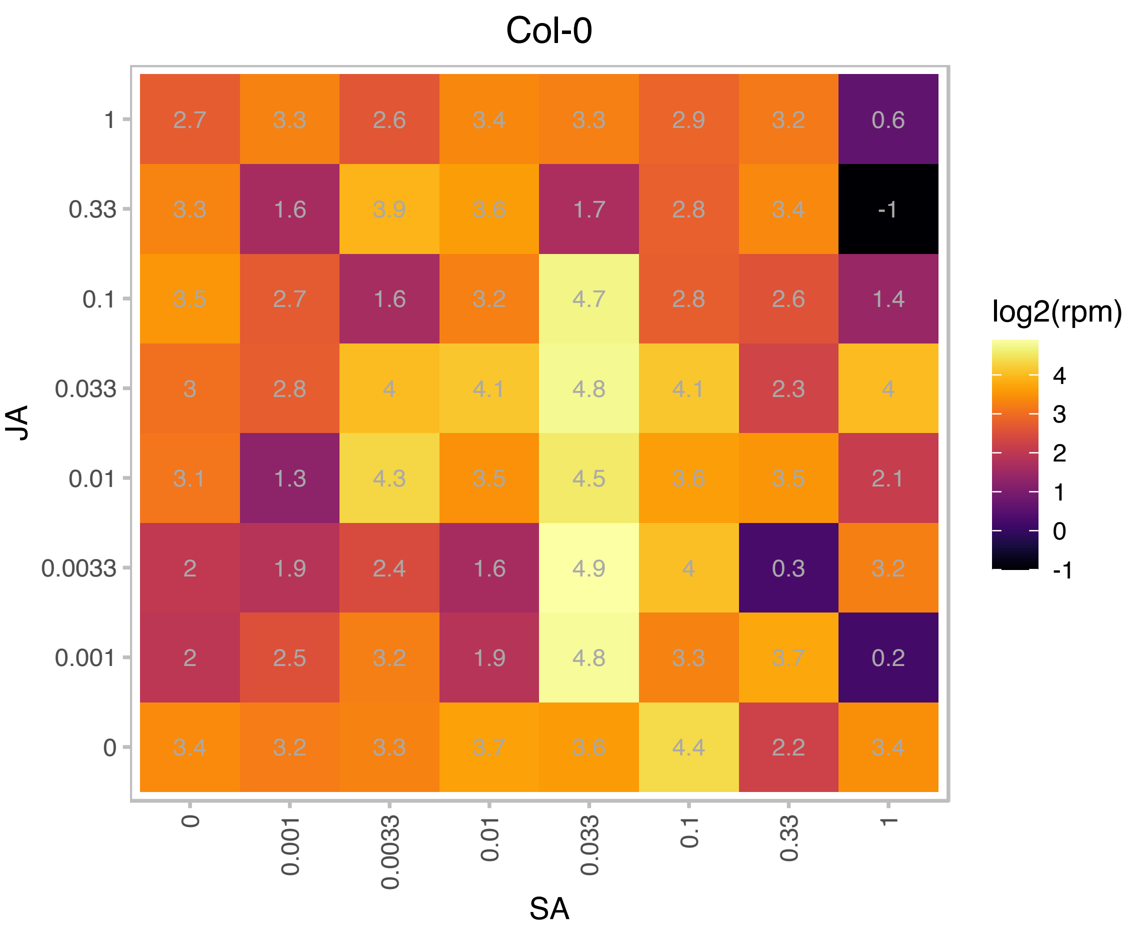


| Gene | AT2G04800 |
|---|---|
| is.expressed_genes | TRUE |
| is.responsive_genes | TRUE |
| is.apcluster | D-01 |

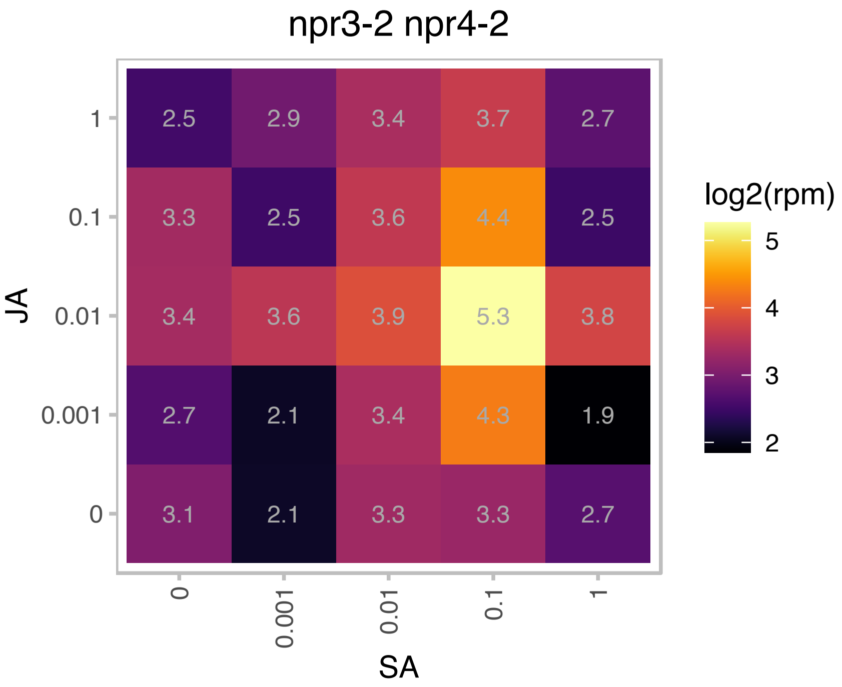


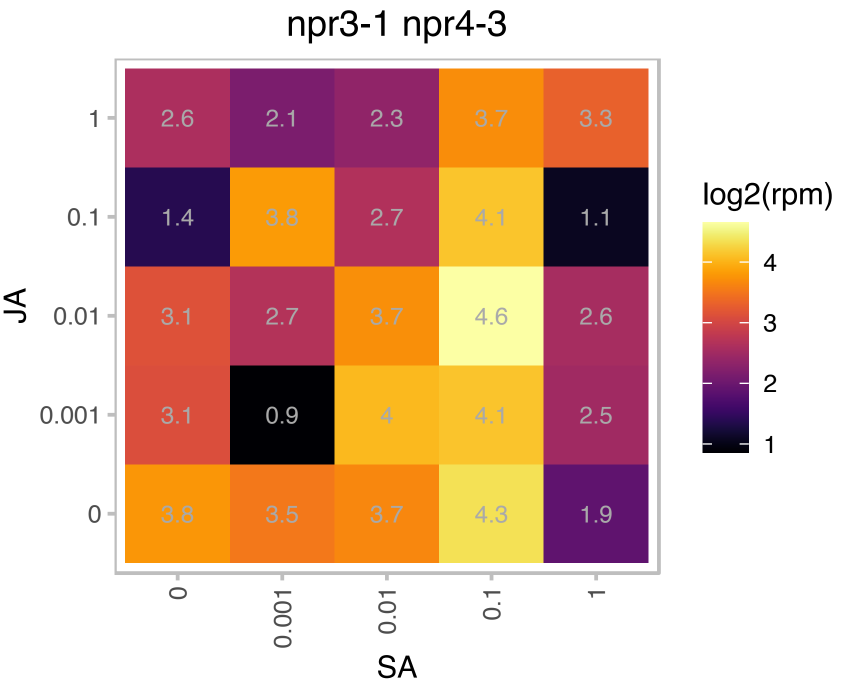

# AT2G30670.1 : NA

NAD(P)-binding Rossmann-fold superfamily protein

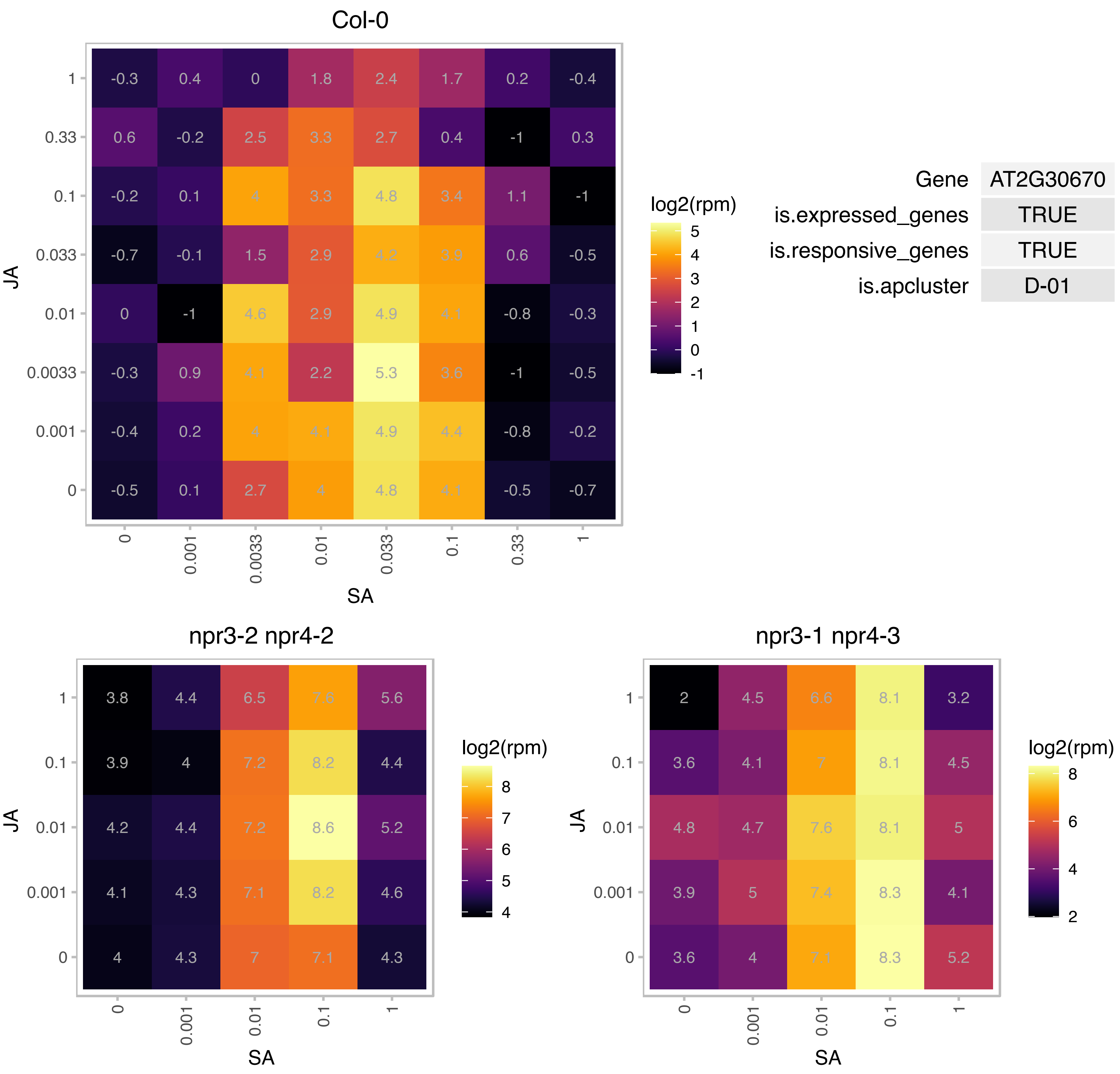

### AT2G38025.1 : OTU3

Cysteine proteinases superfamily protein

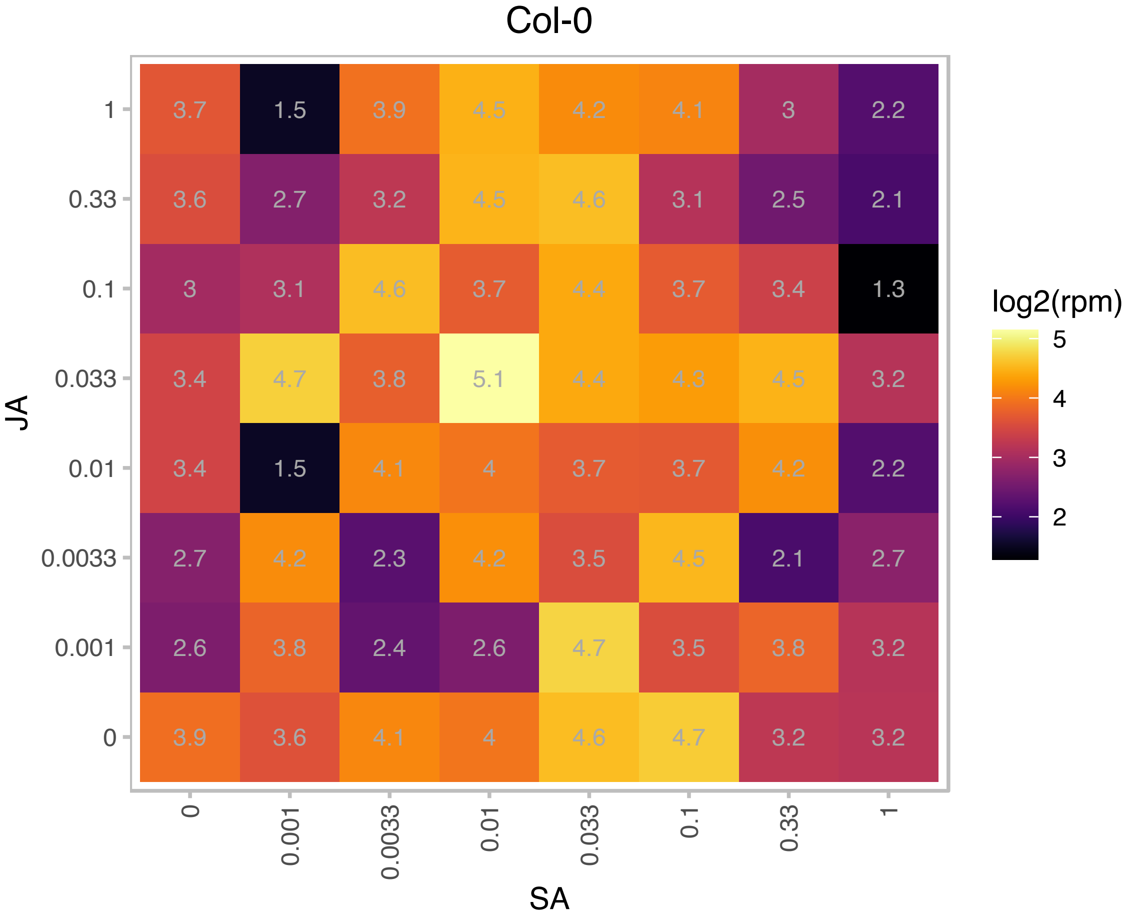


| | |
|---|---|
| Gene | AT2G38025 |
| is.expressed_genes | TRUE |
| is.responsive_genes | TRUE |
| is.apcluster | D-01 |

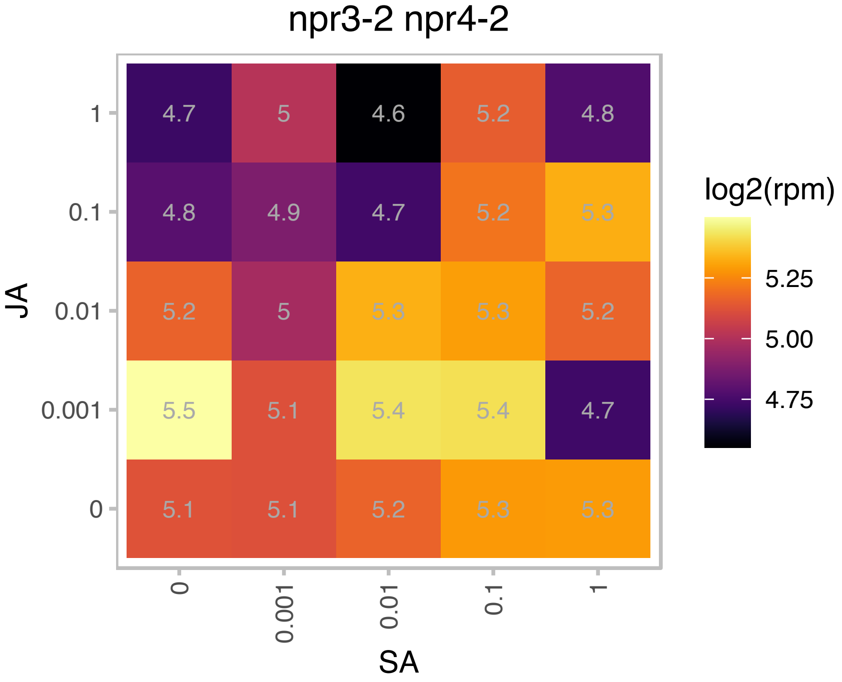


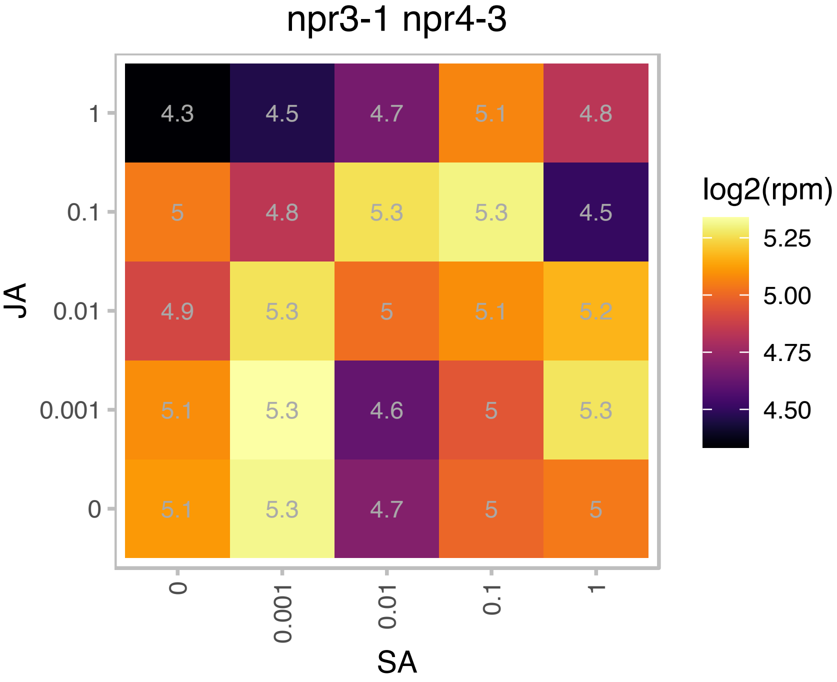

# AT2G41020.1 : NA

WW domain-containing protein

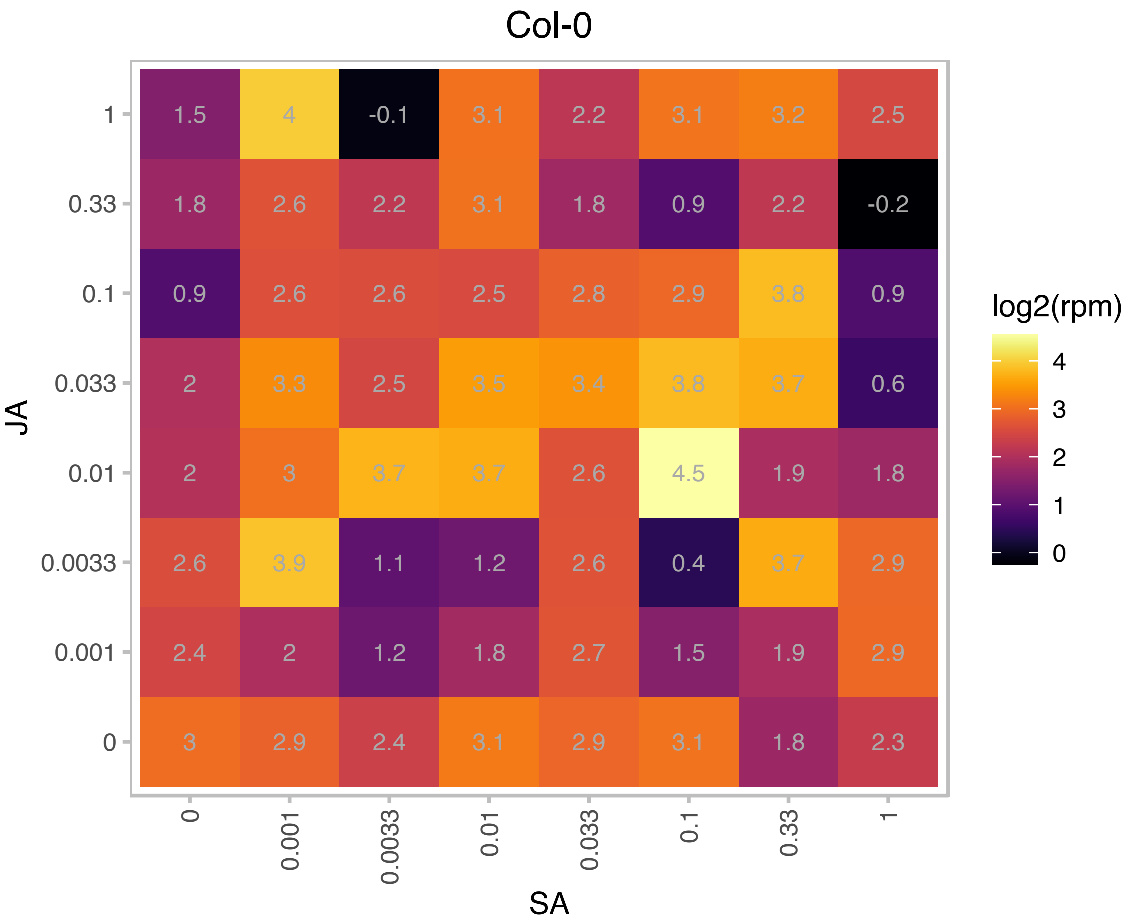


| | |
|---|---|
| Gene | AT2G41020 |
| is.expressed_genes | TRUE |
| is.responsive_genes | TRUE |
| is.apcluster | D-01 |

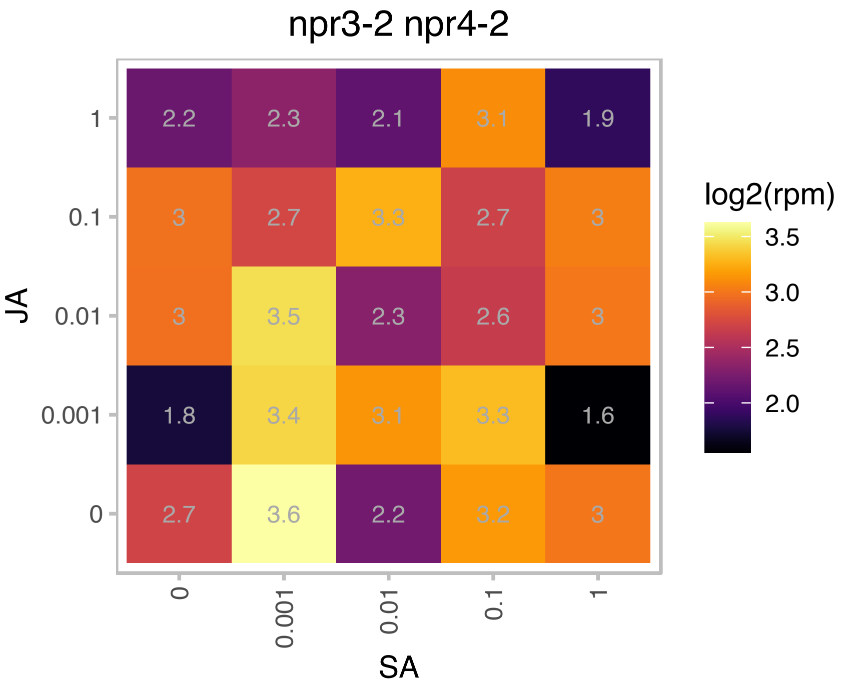


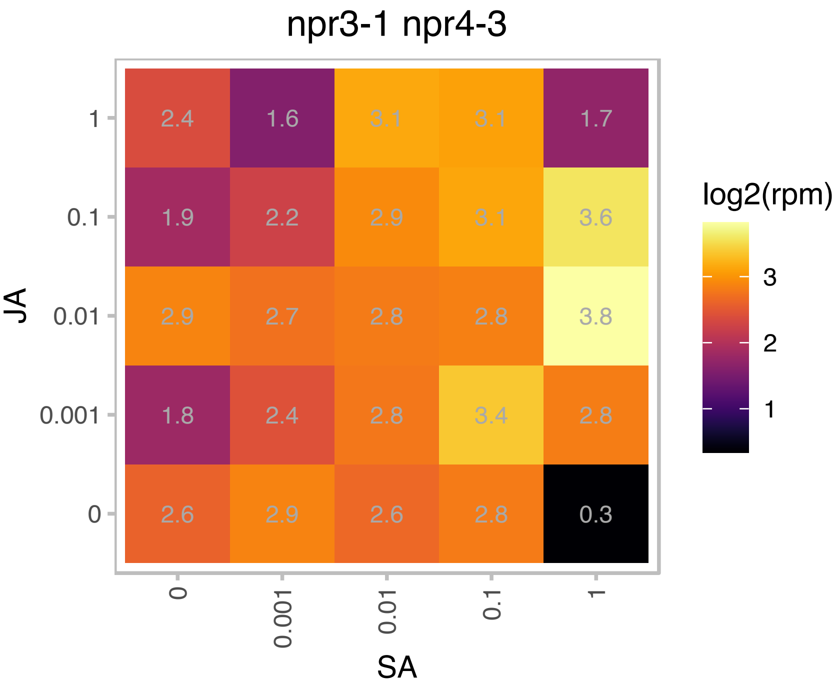

### AT3G20770.1 : AtEIN3, EIN3

Ethylene insensitive 3 family protein

Encodes EIN3 (ethylene-insensitive3), a nuclear transcription factor that initiates downstream transcriptional cascades for ethylene responses. EIN3 interacts with MYC2, MYC3 and MYC4 to inhibit jasmonate-induced expression of wound-responsive genes and herbivory-inducible genes, and plant defense against generalist herbivores.

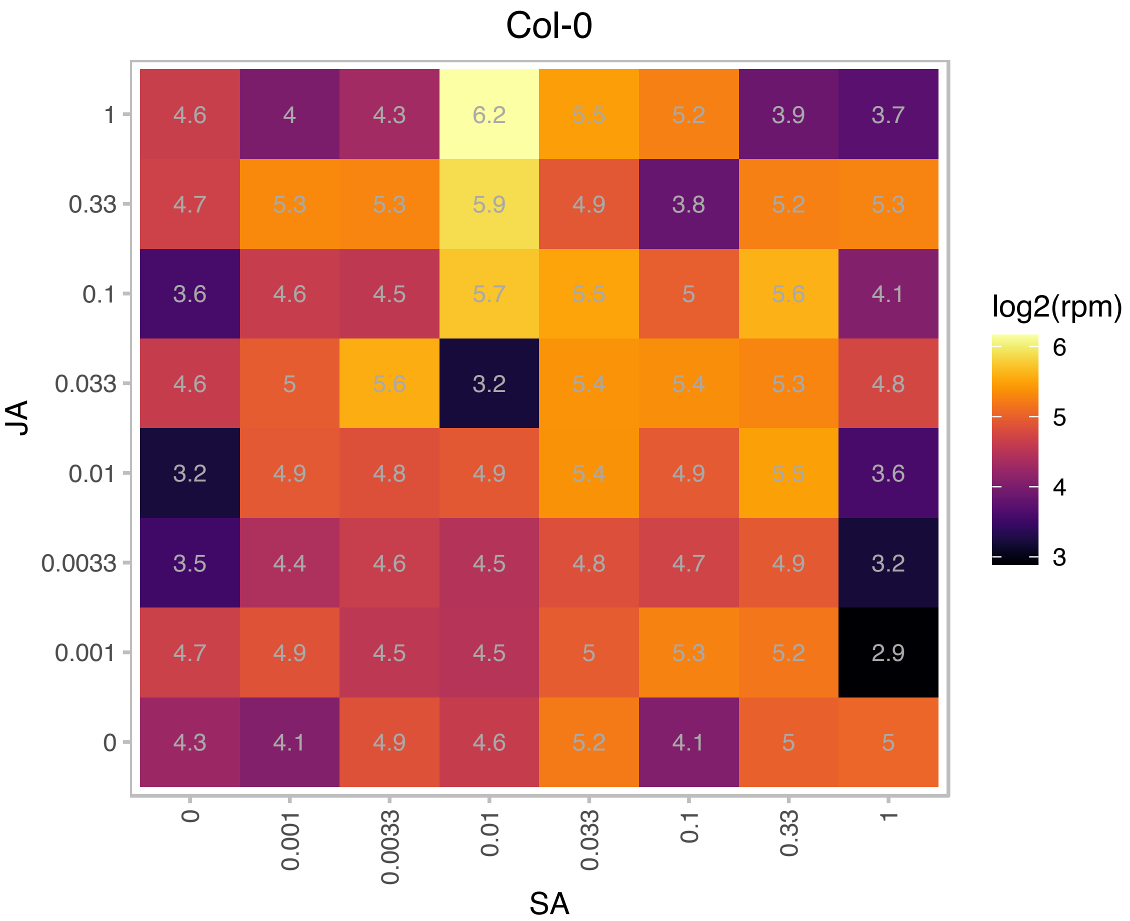


| | |
|---|---|
| Gene | AT3G20770 |
| is.expressed_genes | TRUE |
| is.responsive_genes | TRUE |
| is.apcluster | D-01 |

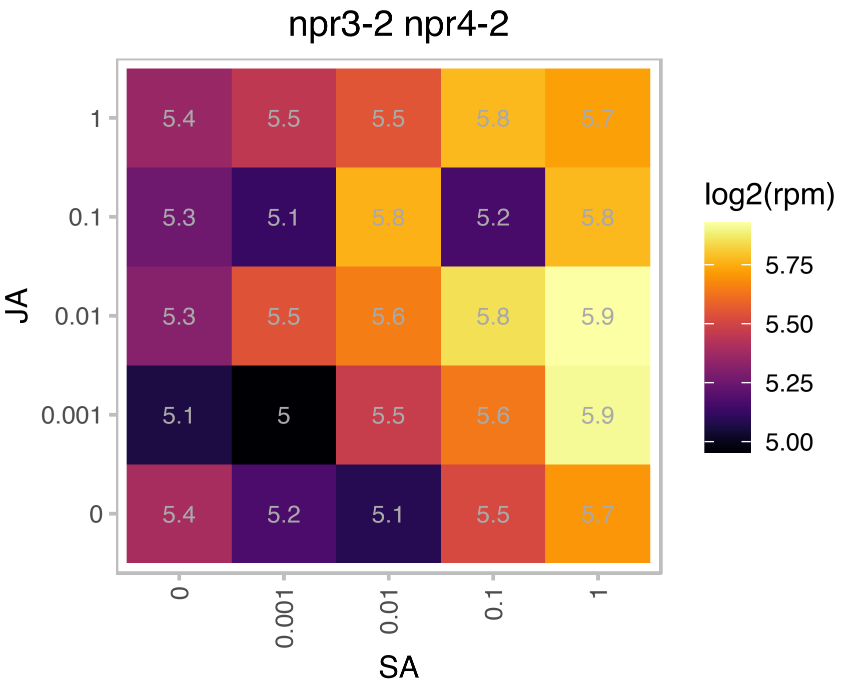


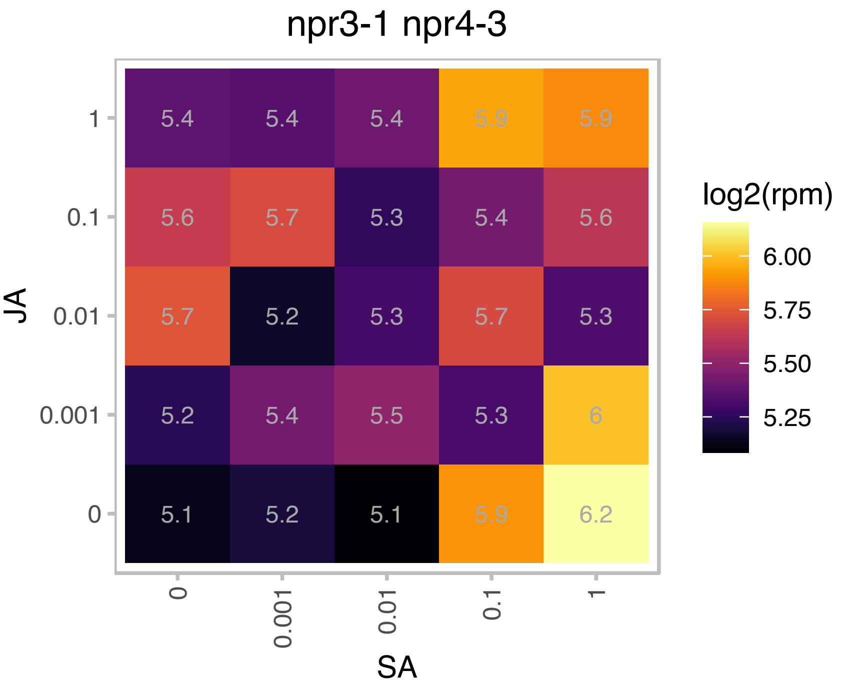

# AT3G23175.1 : NA

HR-like lesion-inducing protein-related

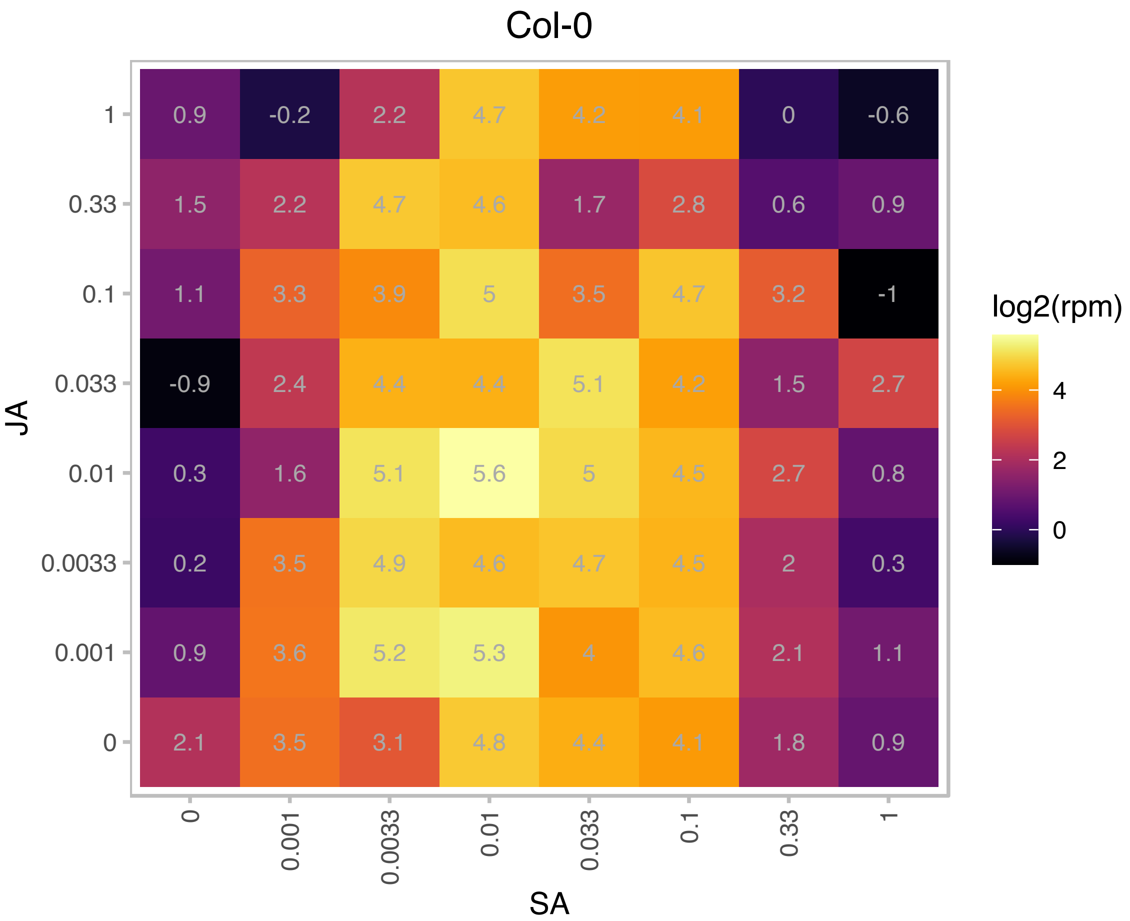


| | |
|---|---|
| Gene | AT3G23175 |
| is.expressed_genes | TRUE |
| is.responsive_genes | TRUE |
| is.apcluster | D-01 |

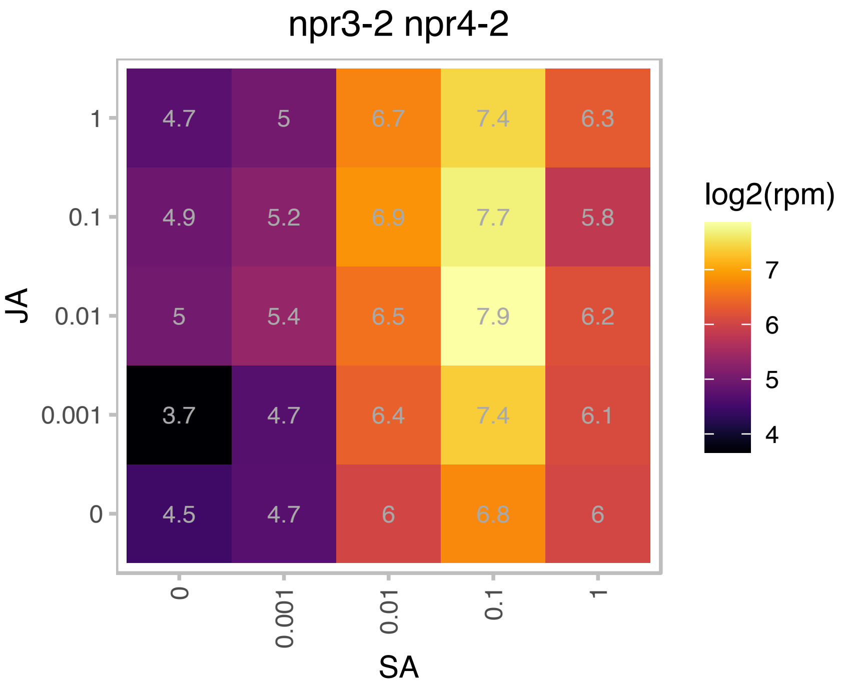


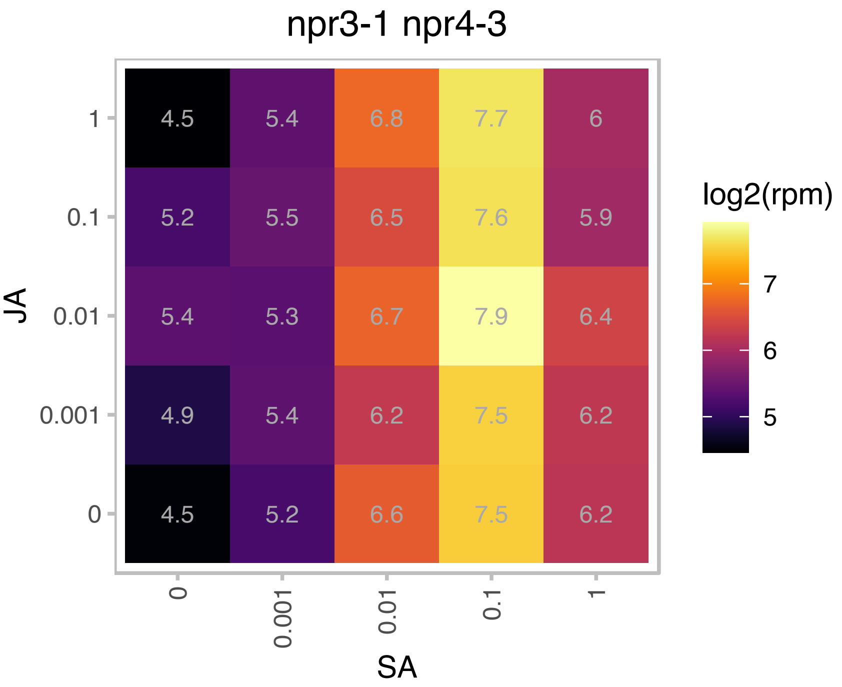

# AT3G23190.1 : NA

HR-like lesion-inducing protein-related

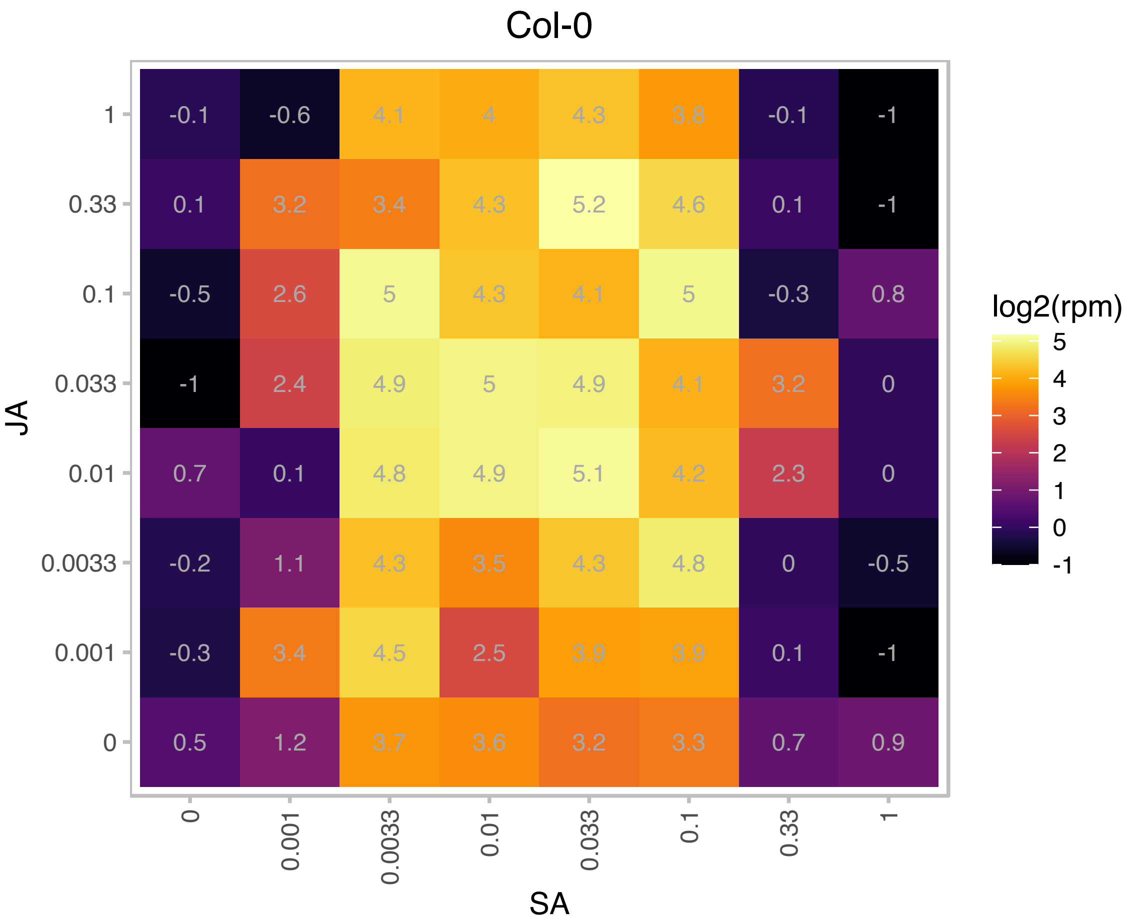


| | |
|---|---|
| Gene | AT3G23190 |
| is.expressed_genes | TRUE |
| is.responsive_genes | TRUE |
| is.apcluster | D-01 |

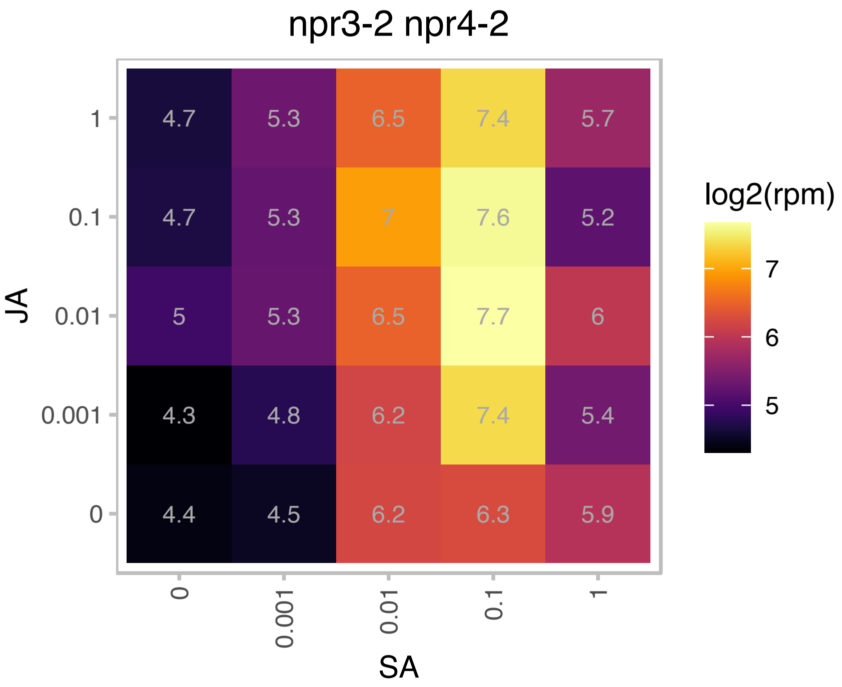


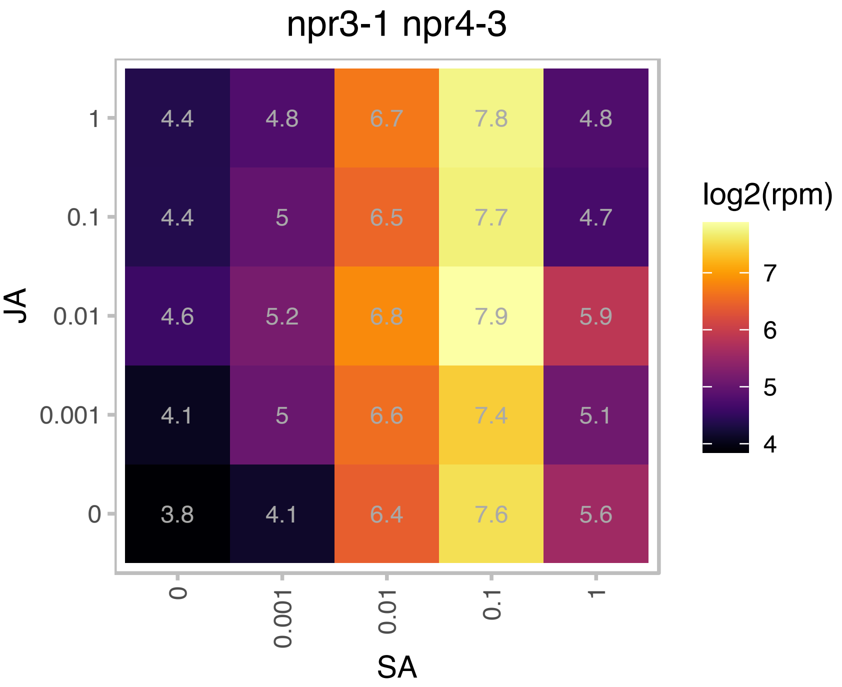

# AT3G23530.1 : NA

Cyclopropane-fatty-acyl-phospholipid synthase

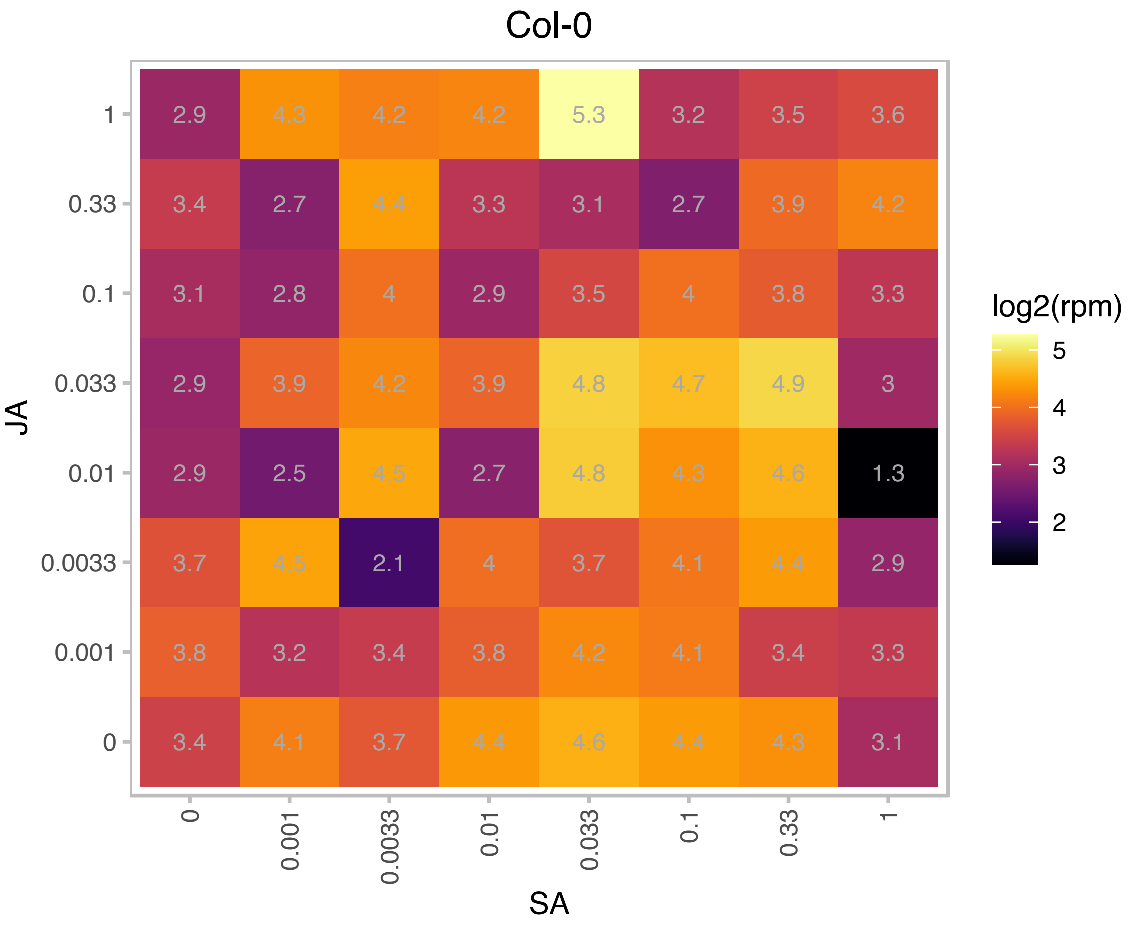


| | |
|---|---|
| Gene | AT3G23530 |
| is.expressed_genes | TRUE |
| is.responsive_genes | TRUE |
| is.apcluster | D-01 |

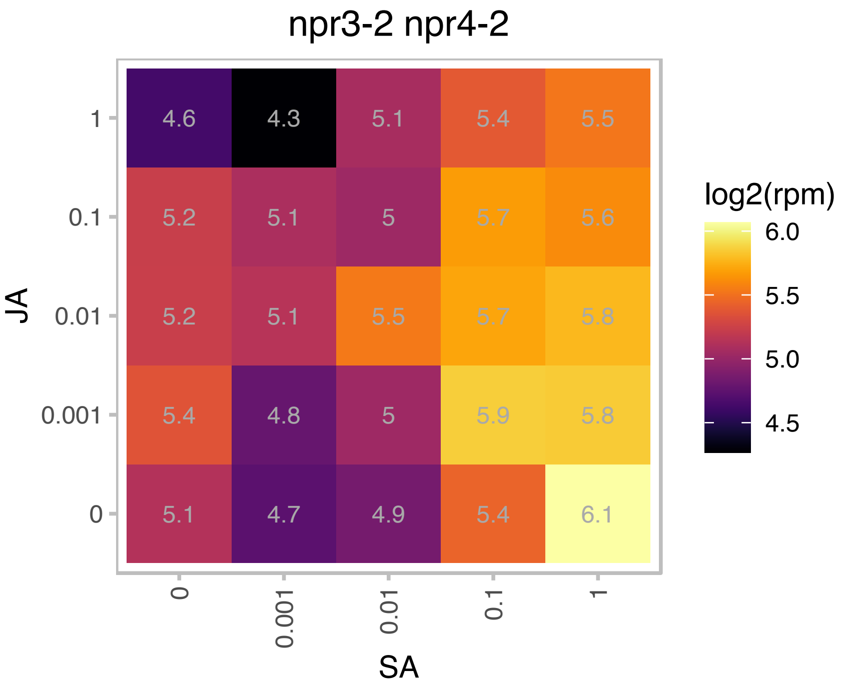


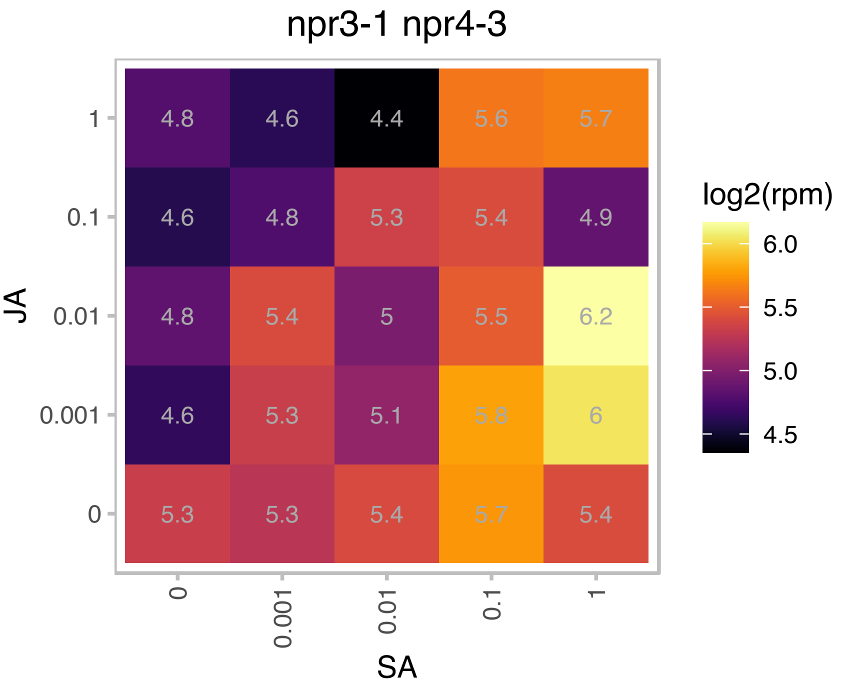

### AT3G27070.1 : TOM20-1

translocase outer membrane 20-1

Form of TOM20, which is a component of the TOM complex, involved in transport of nuclear-encoded mitochondrial proteins

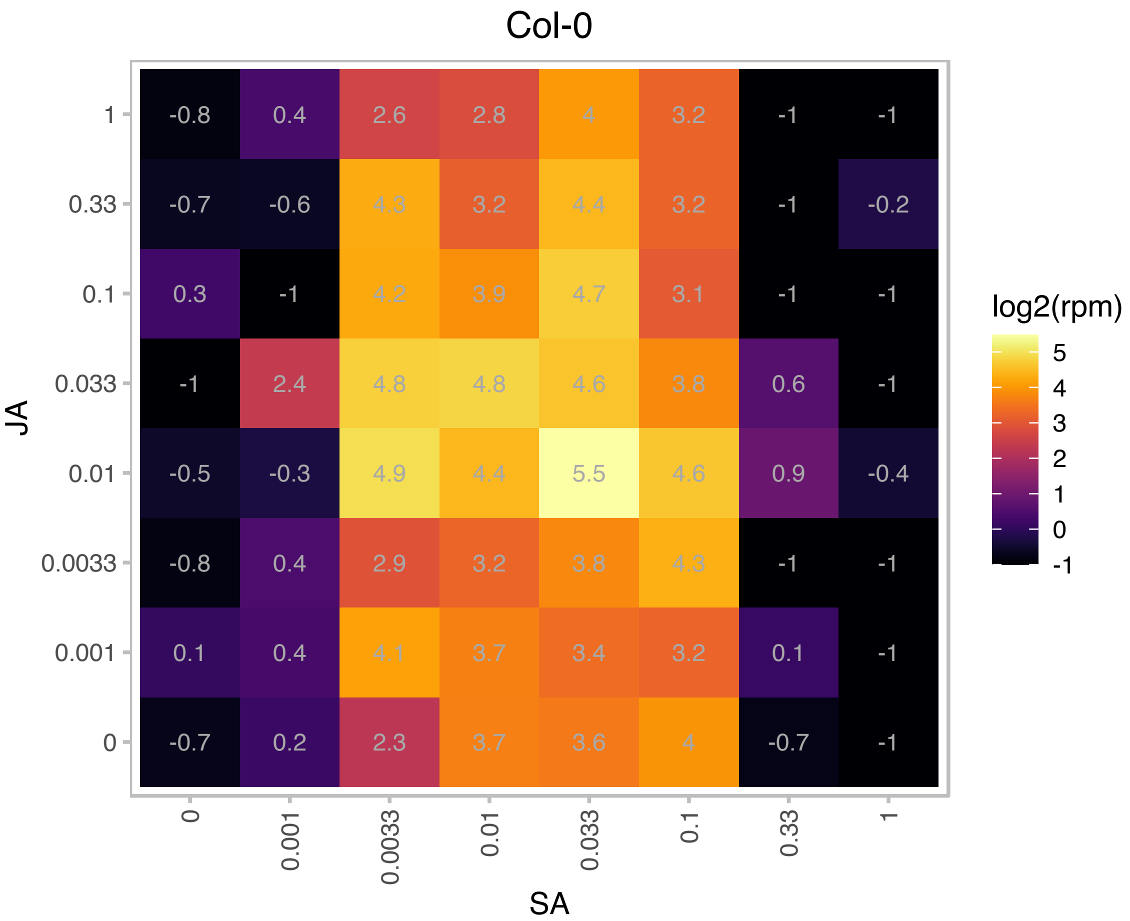


| | |
|---|---|
| Gene | AT3G27070 |
| is.expressed_genes | TRUE |
| is.responsive_genes | TRUE |
| is.apcluster | D-01 |

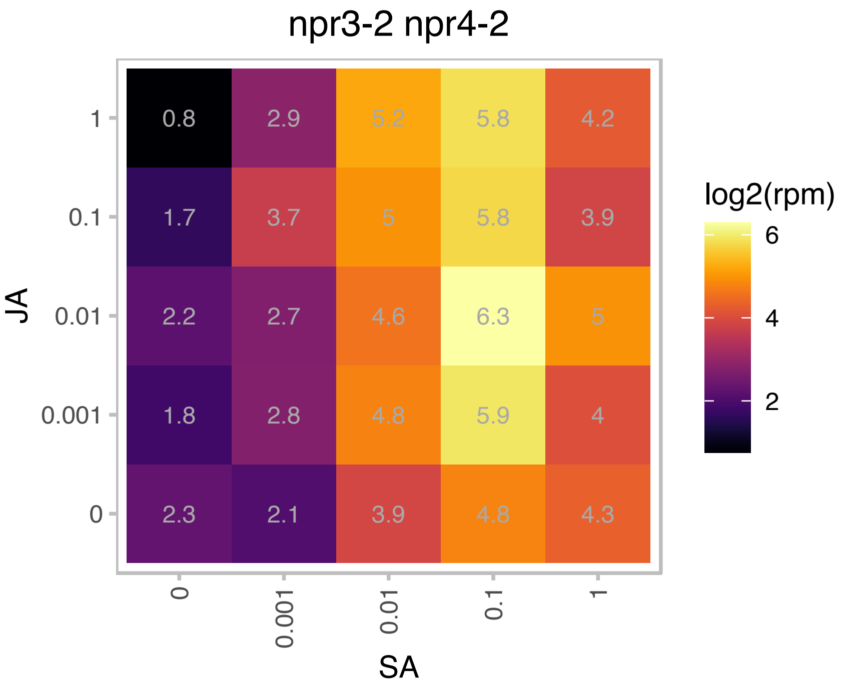


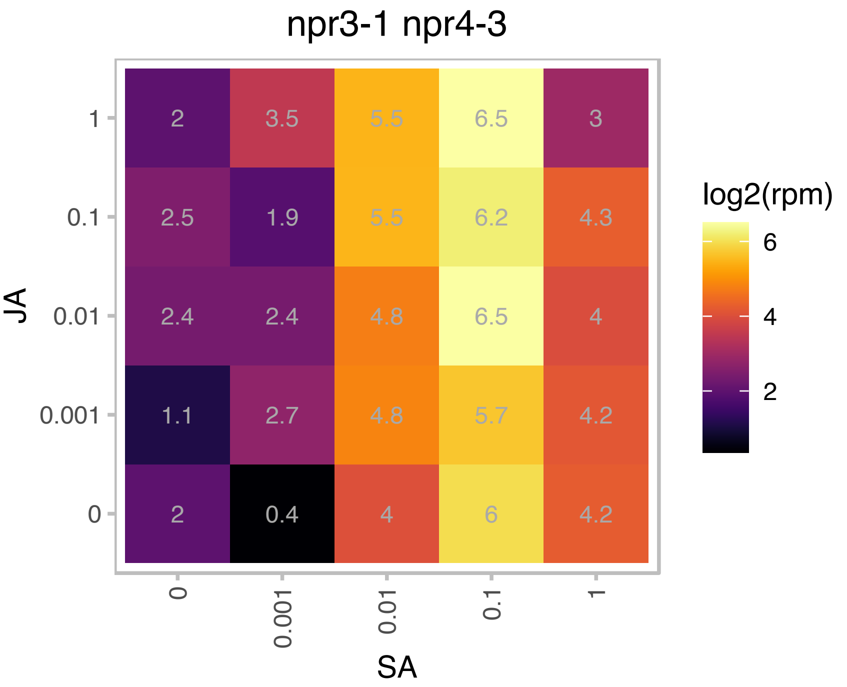

# AT3G46845.1 : NA

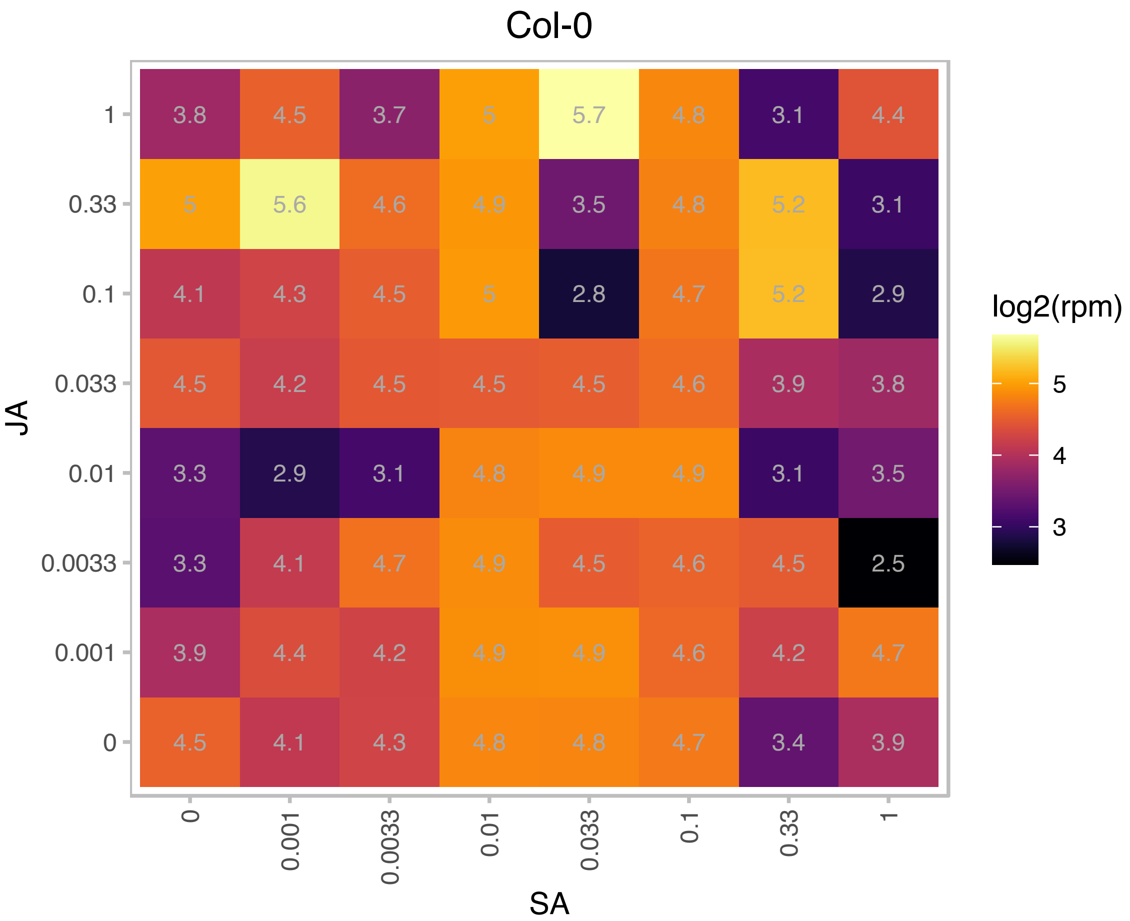


| | |
|---|---|
| Gene | AT3G46845 |
| is.expressed_genes | TRUE |
| is.responsive_genes | TRUE |
| is.apcluster | D-01 |

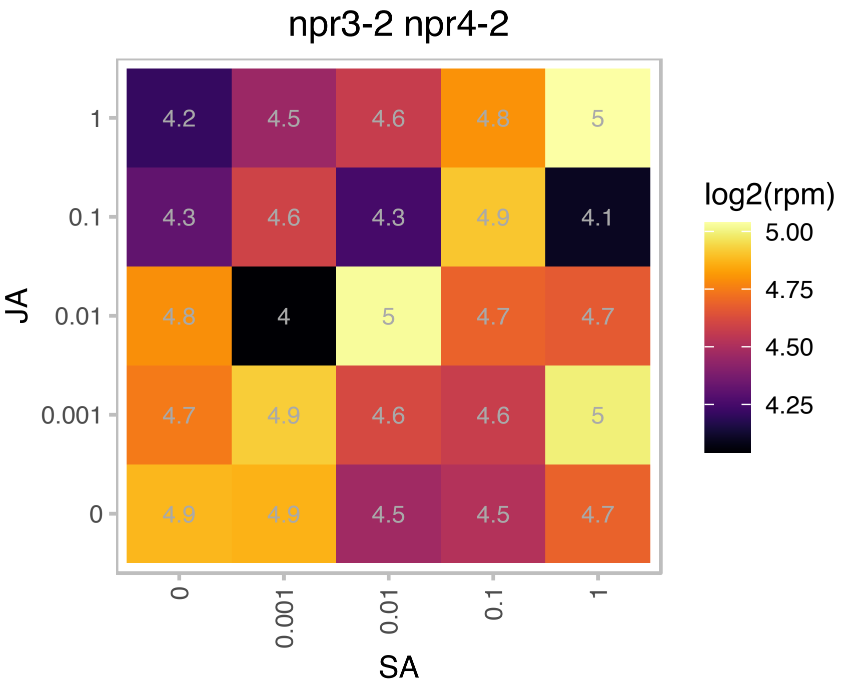


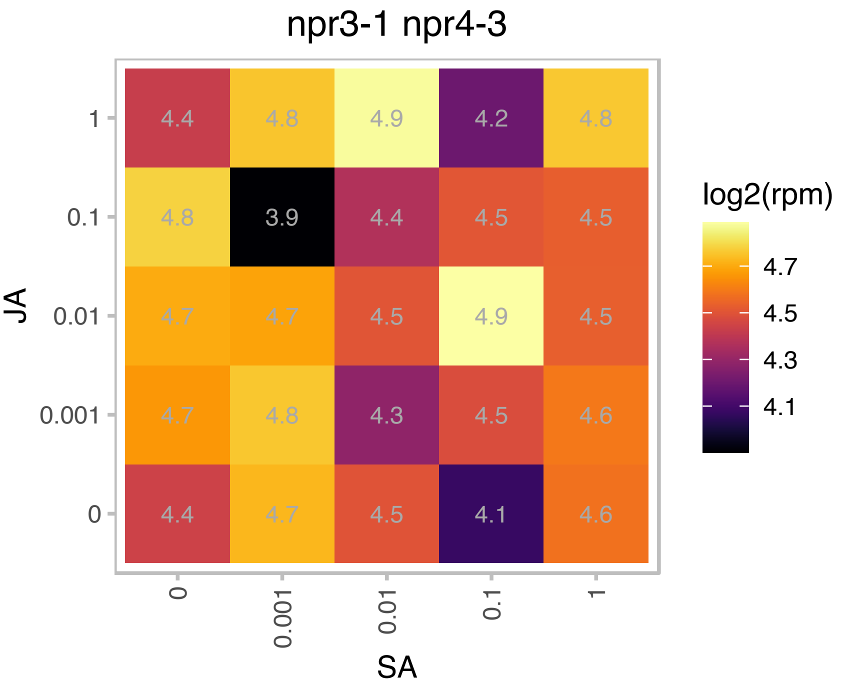

# AT3G62330.1 : NA

Zinc knuckle (CCHC-type) family protein

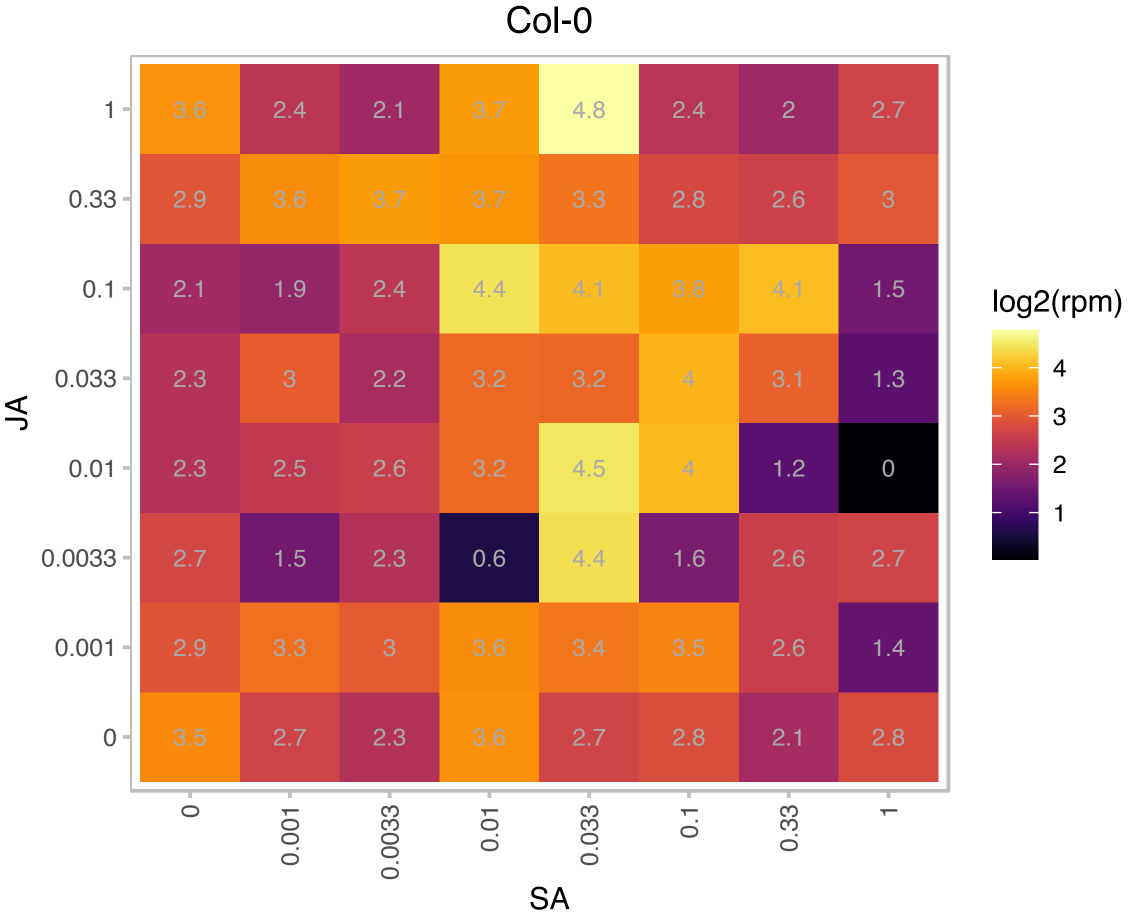


| | |
|---|---|
| Gene | AT3G62330 |
| is.expressed_genes | TRUE |
| is.responsive_genes | TRUE |
| is.apcluster | D-01 |

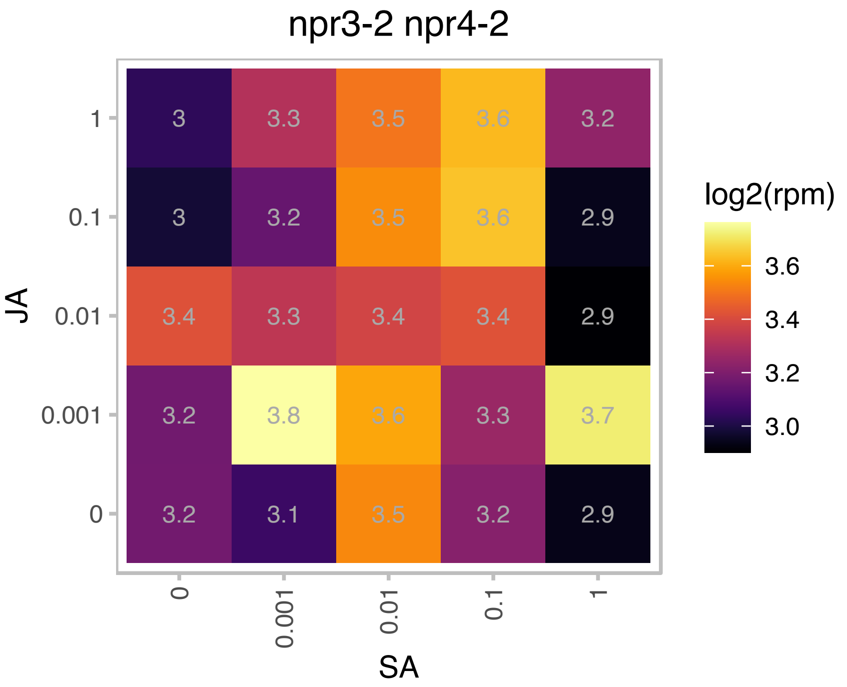


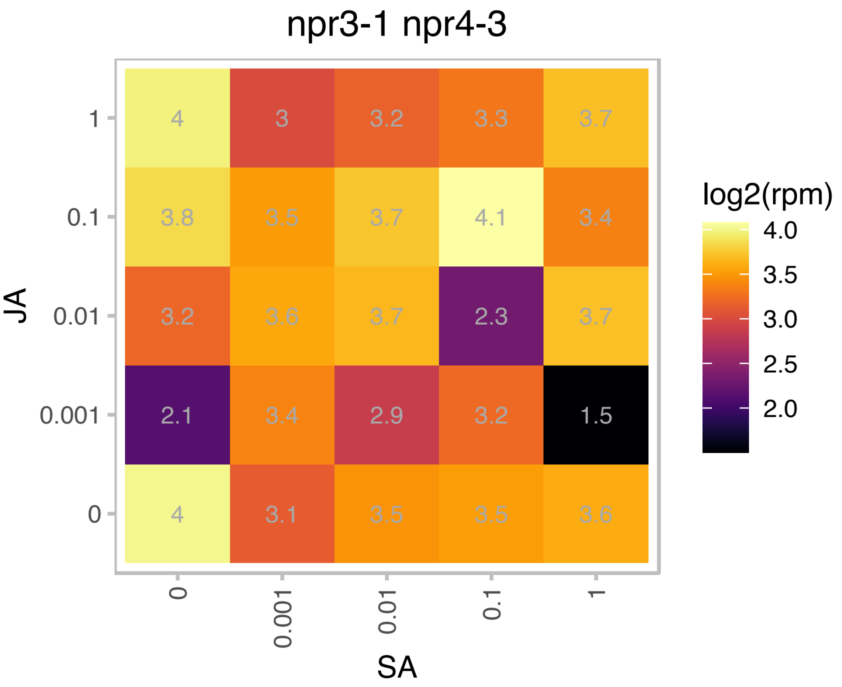

# AT3G62900.1 : NA

CW-type Zinc Finger

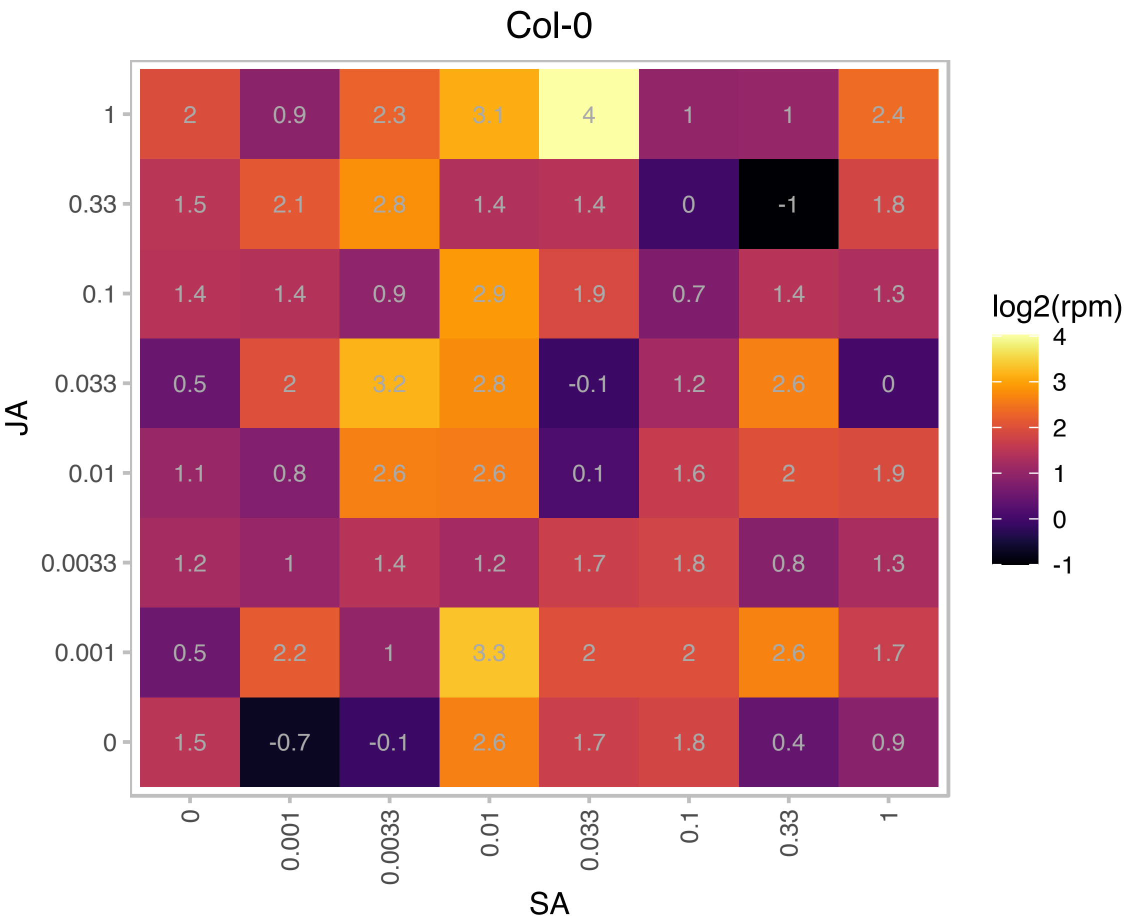


| | |
|---|---|
| Gene | AT3G62900 |
| is.expressed_genes | TRUE |
| is.responsive_genes | TRUE |
| is.apcluster | D-01 |

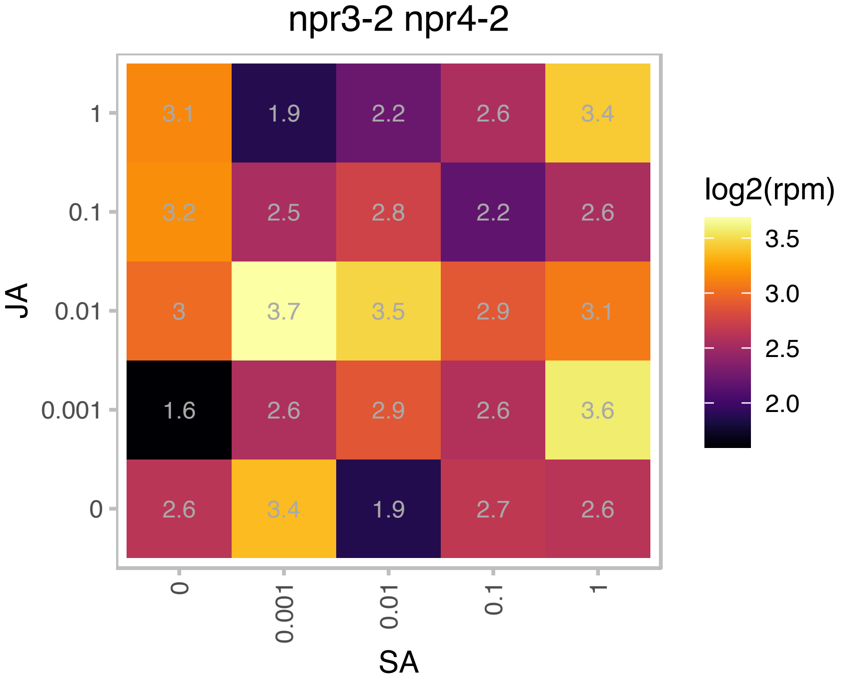


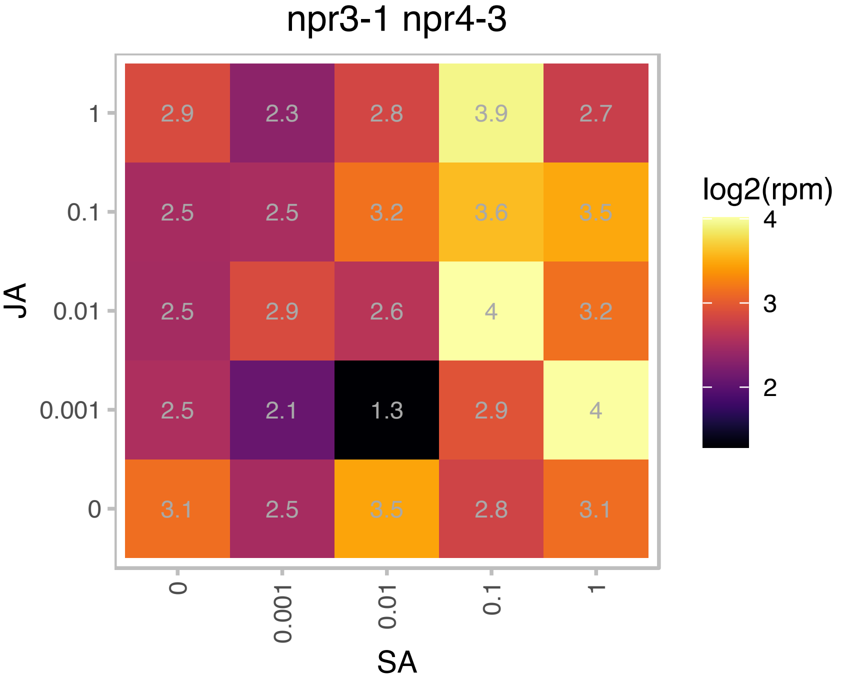

### AT4G04700.1 : CPK27

calcium-dependent protein kinase 27

member of Calcium Dependent Protein Kinase

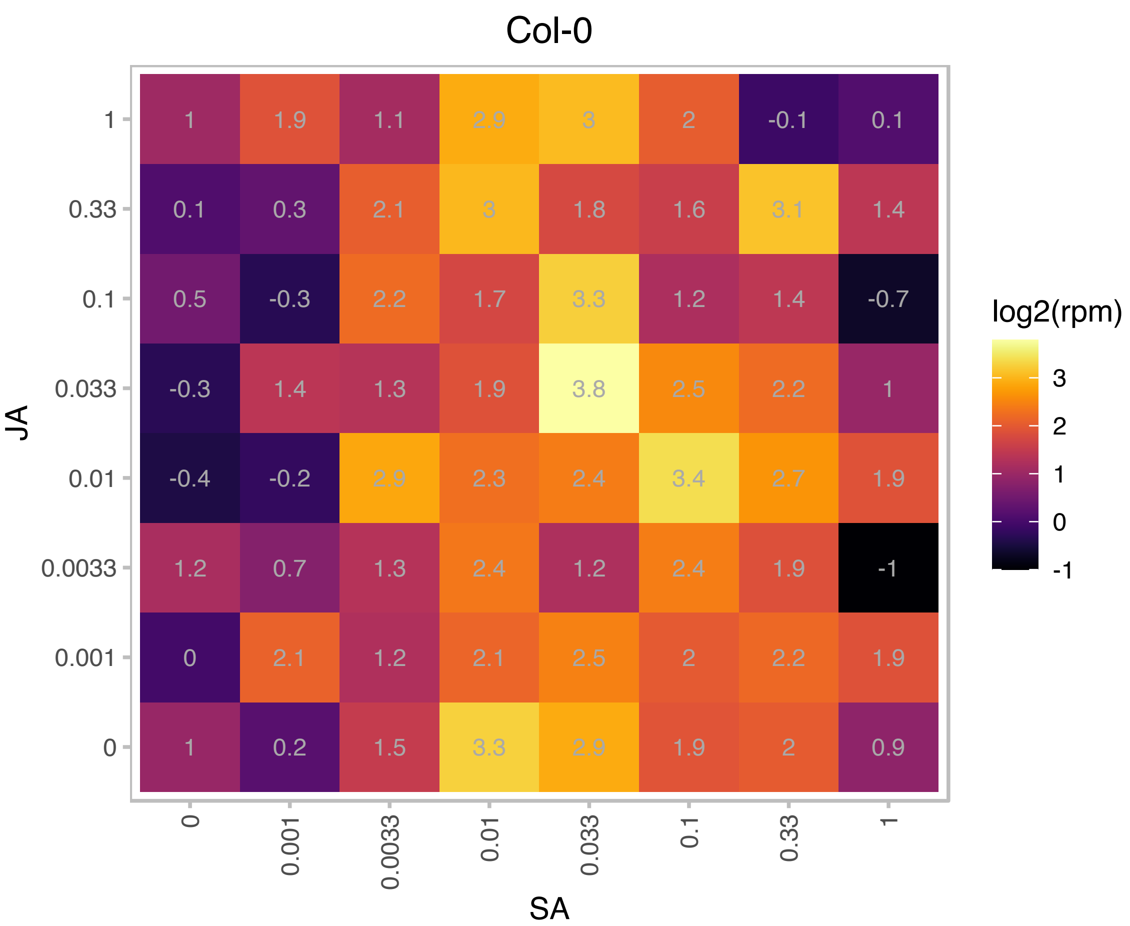


| | |
|---|---|
| Gene | AT4G04700 |
| is.expressed_genes | TRUE |
| is.responsive_genes | TRUE |
| is.apcluster | D-01 |

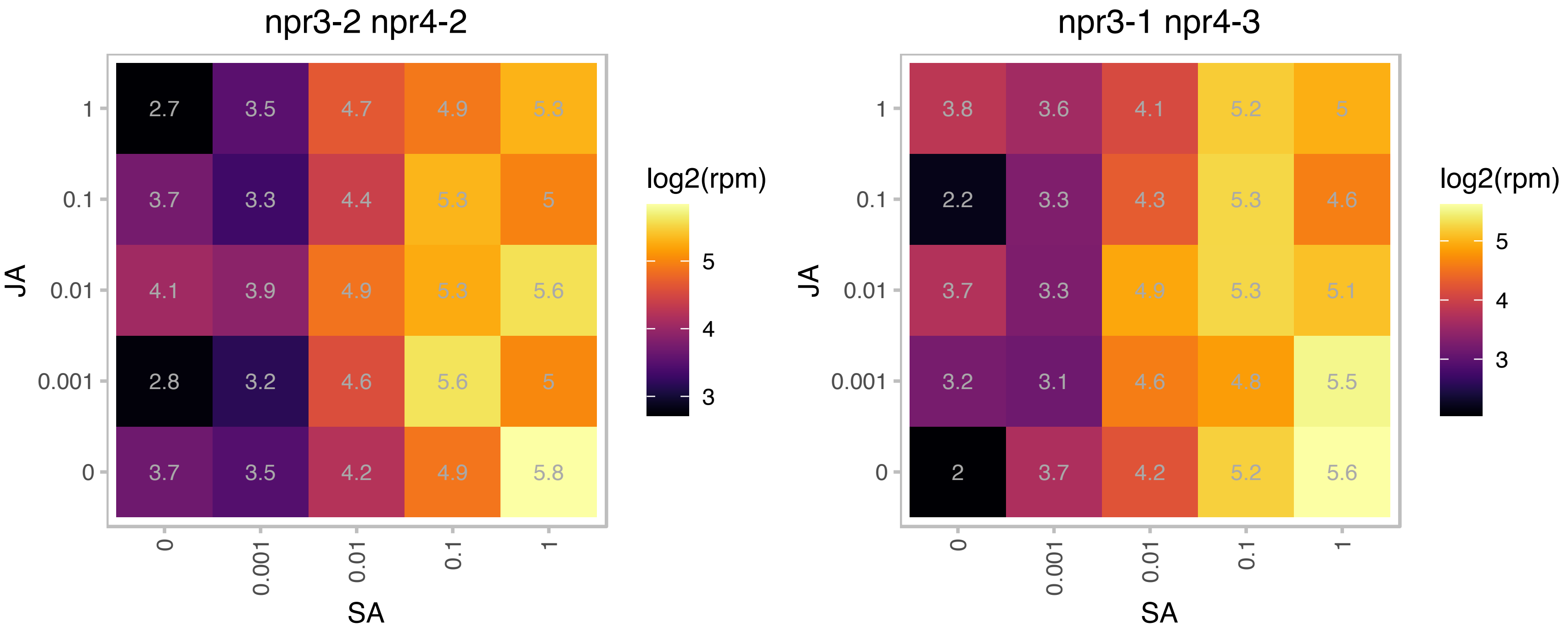

### AT4G05420.1 : DDB1A

damaged DNA binding protein 1A
Structurally similar to damaged DNA binding proteins. DDB1a is part of a 350 KDa nuclear localized DET1 protein complex. This complex may physically interact with histone tails and while bound to chromatin- repress transcription of genes involved in photomorphogenesis. DDB1a is shown to be RUB-modified.

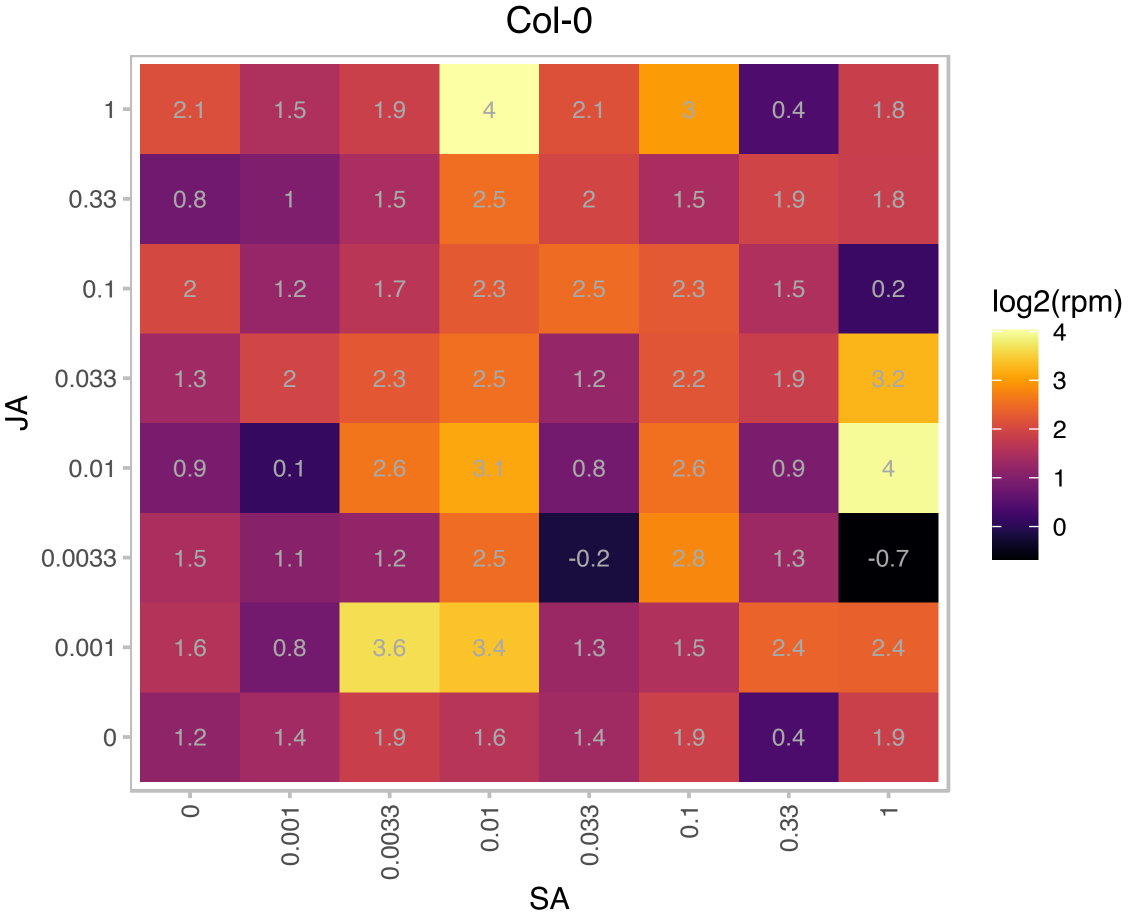


| | |
|---|---|
| Gene | AT4G05420 |
| is.expressed_genes | TRUE |
| is.responsive_genes | TRUE |
| is.apcluster | D-01 |

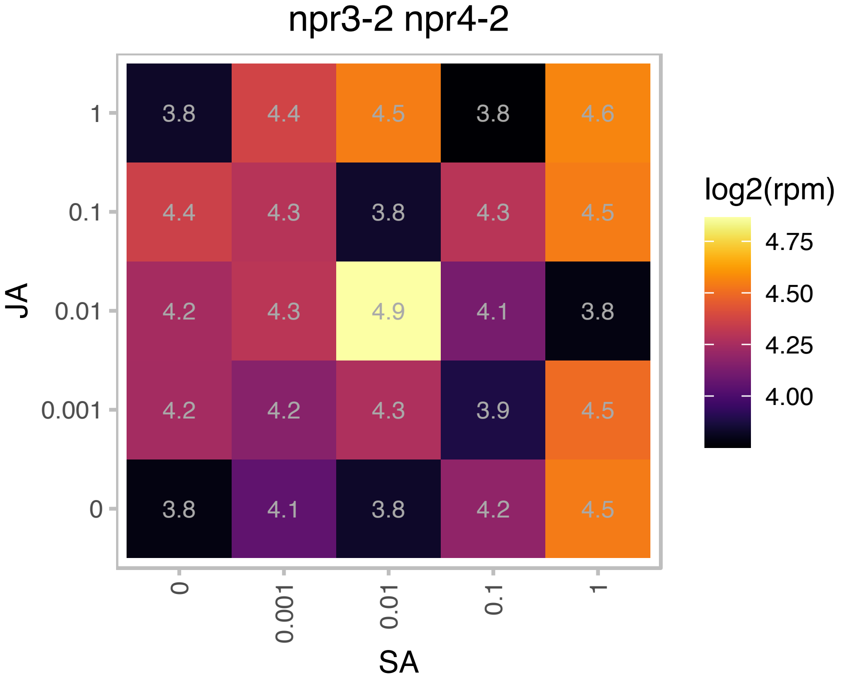


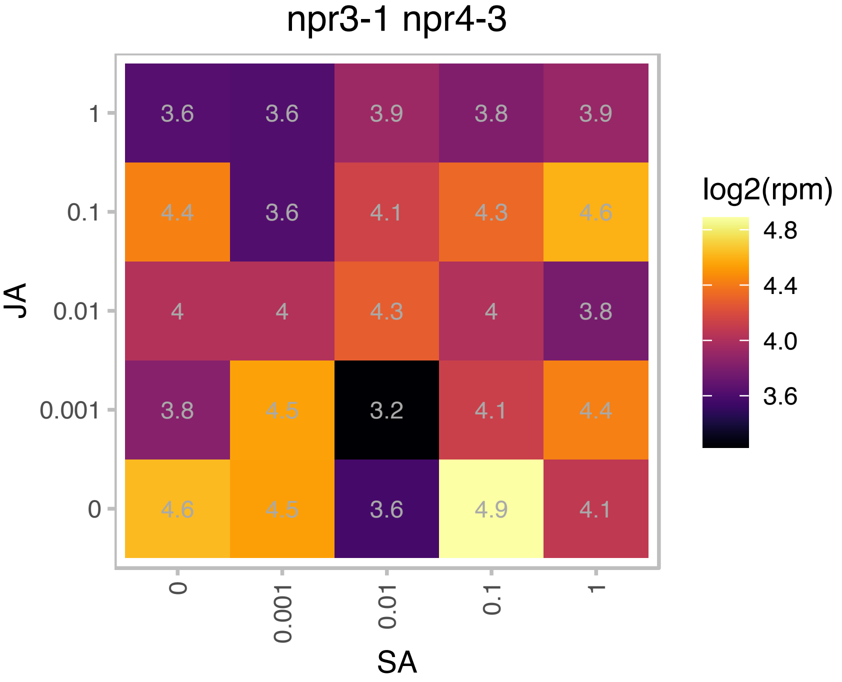

### AT4G10500.1 : DLO1

2-oxoglutarate (2OG) and Fe(II)-dependent oxygenase superfamily protein

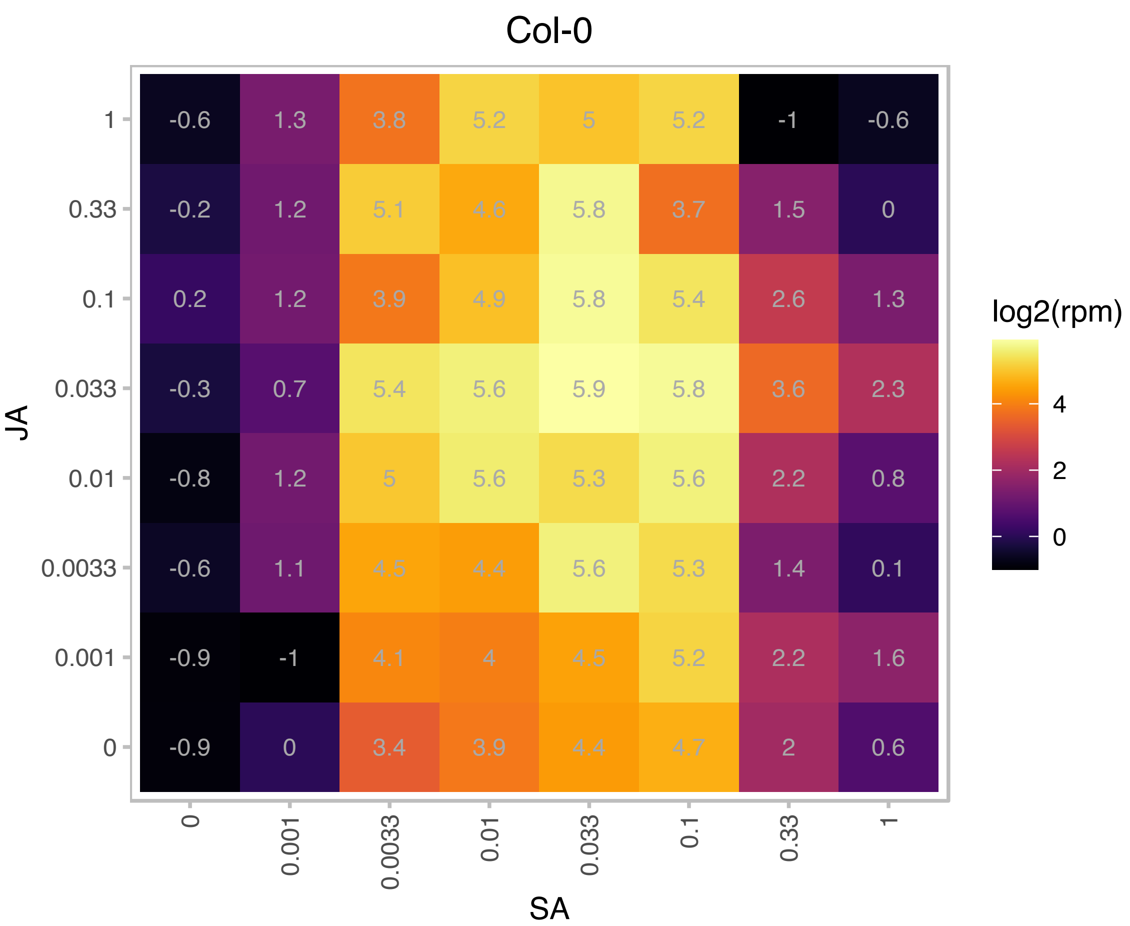


| | |
|---|---|
| Gene | AT4G10500 |
| is.expressed_genes | TRUE |
| is.responsive_genes | TRUE |
| is.apcluster | D-01 |

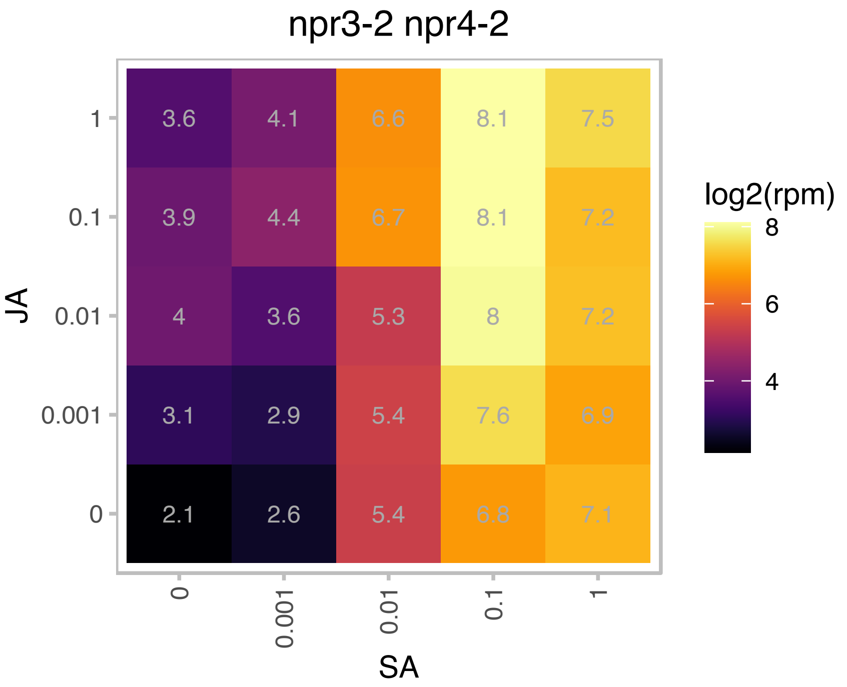


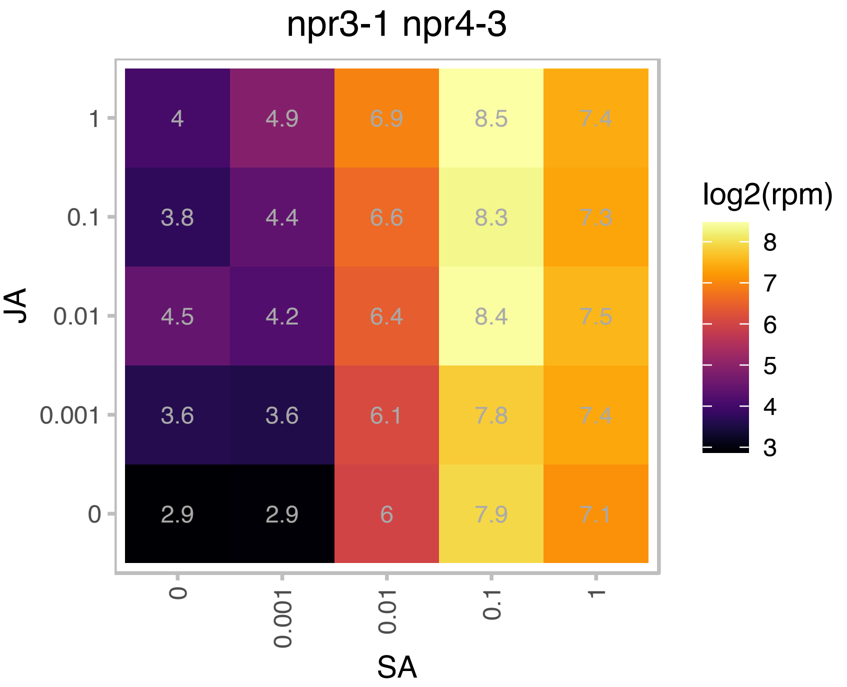

# AT4G18930.1 : NA

RNA ligase/cyclic nucleotide phosphodiesterase family protein

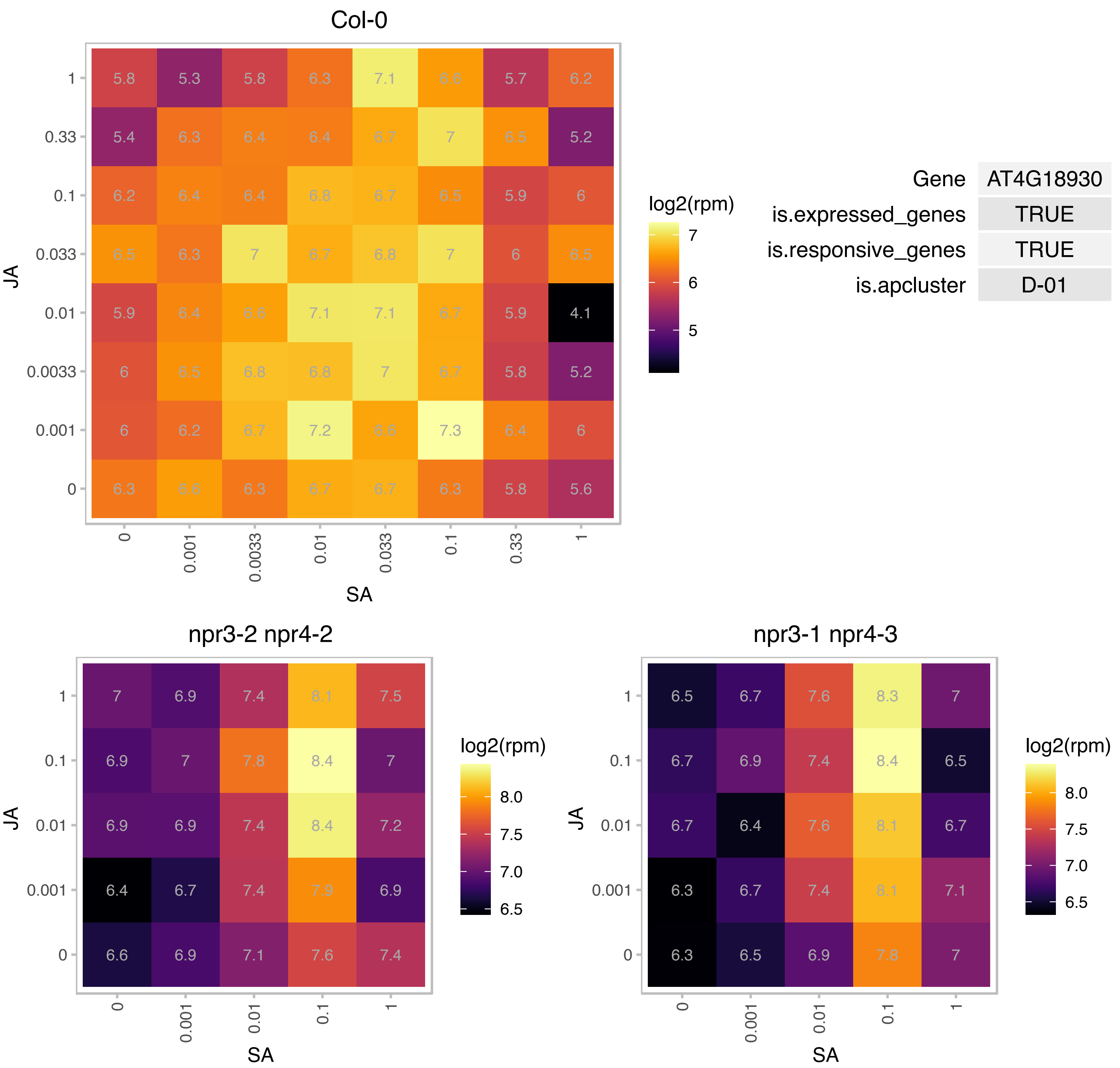


| Gene | AT4G18930 |
| --- | --- |
| is.expressed_genes | TRUE |
| is.responsive_genes | TRUE |
| is.apcluster | D-01 |

### AT4G32010.1 : HSI2-L1, HSL1, VAL2

HSI2-like 1

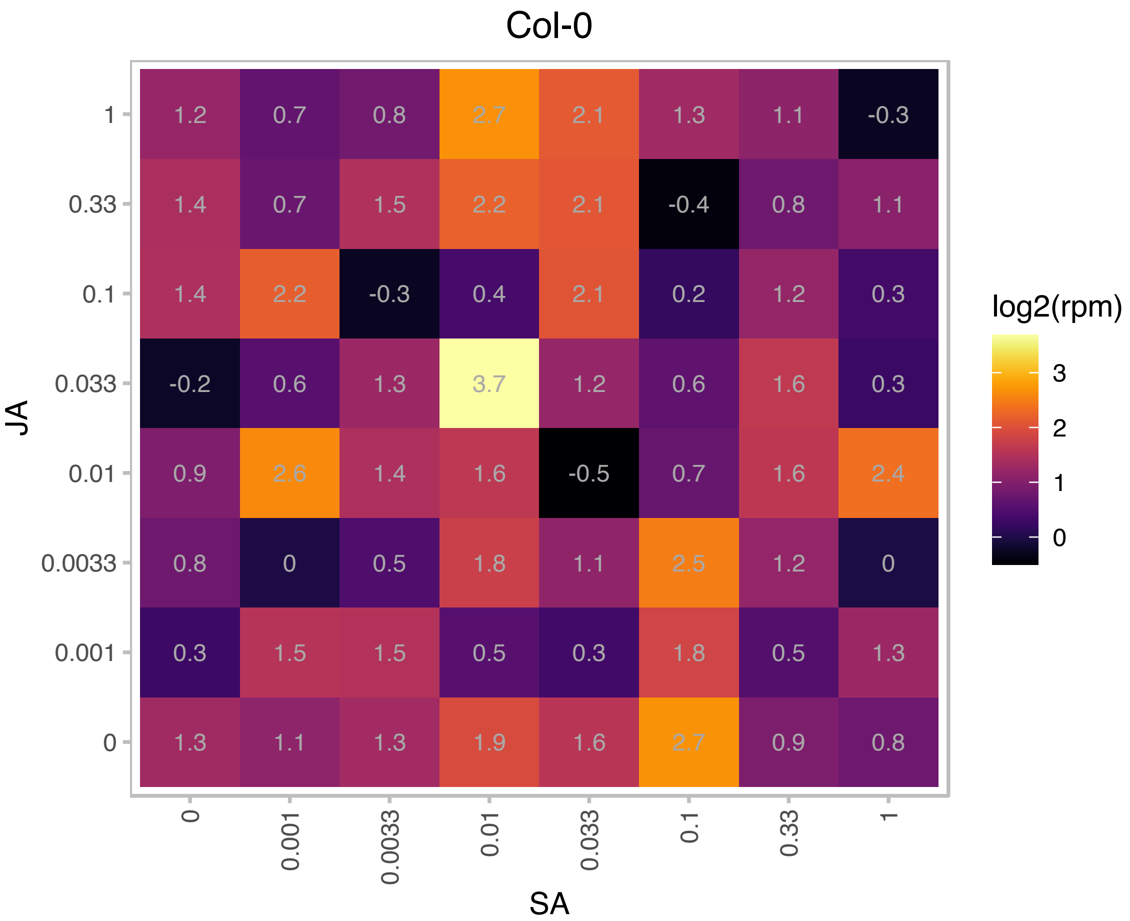


| | |
|---|---|
| Gene | AT4G32010 |
| is.expressed_genes | TRUE |
| is.responsive_genes | TRUE |
| is.apcluster | D-01 |

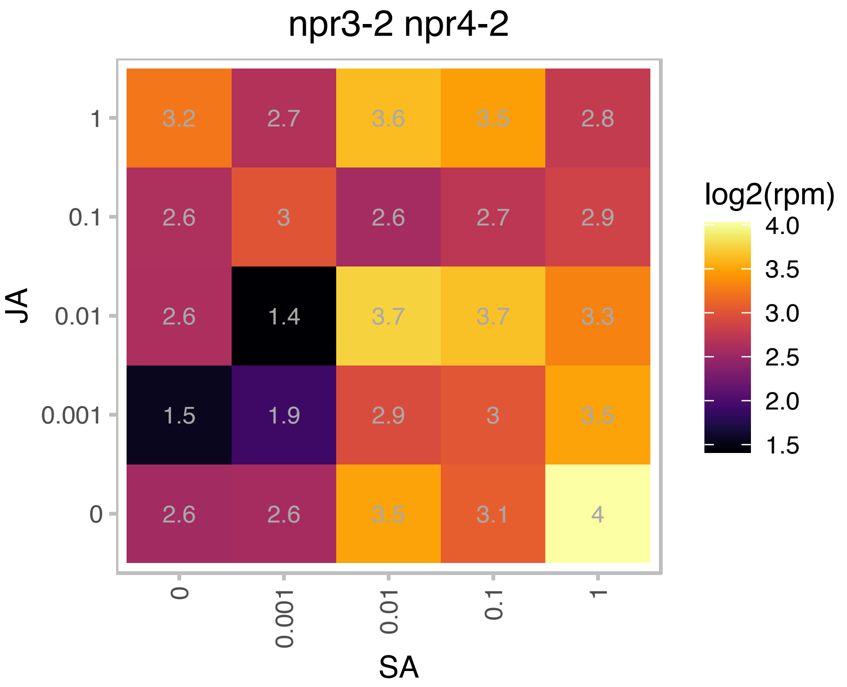


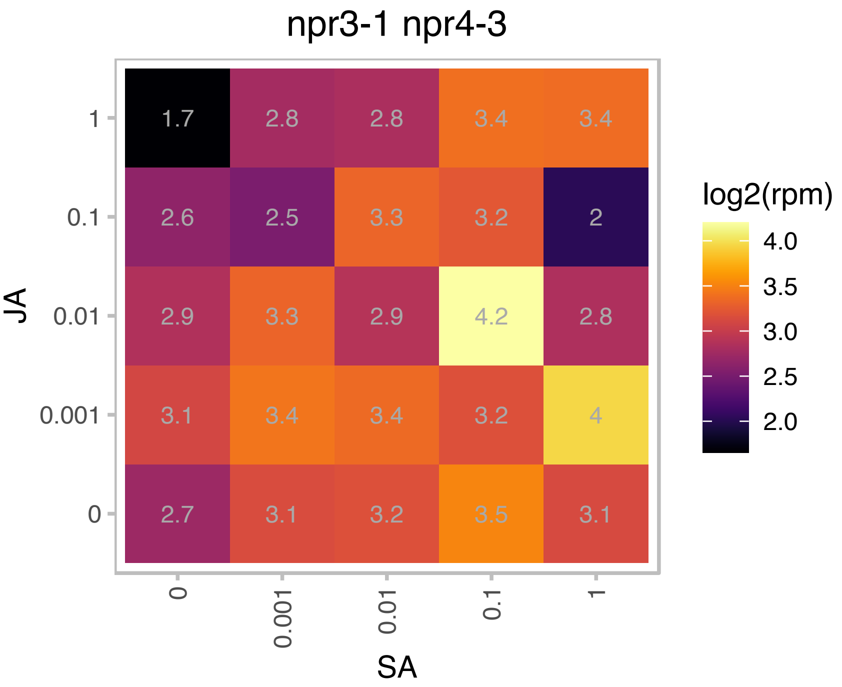

# AT4G32020.1 : NA

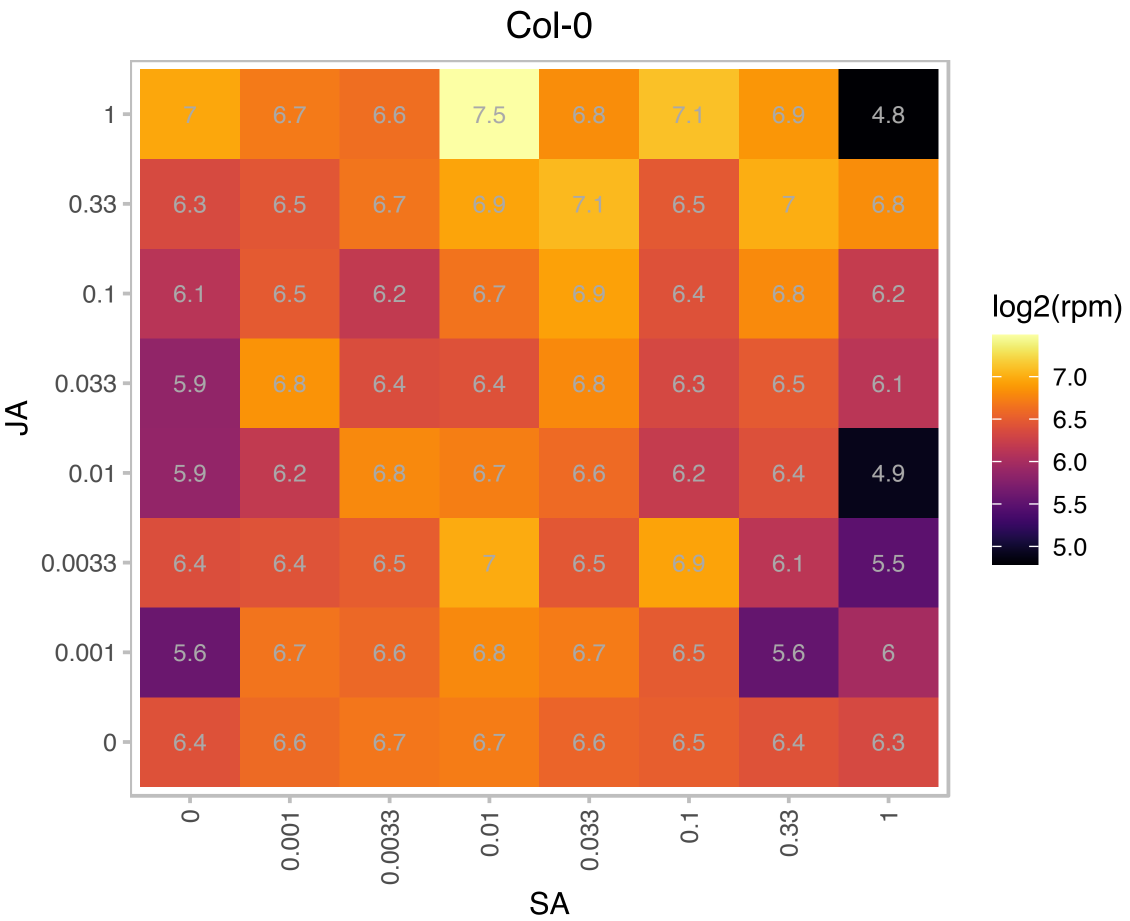


| | |
|---|---|
| Gene | AT4G32020 |
| is.expressed_genes | TRUE |
| is.responsive_genes | TRUE |
| is.apcluster | D-01 |

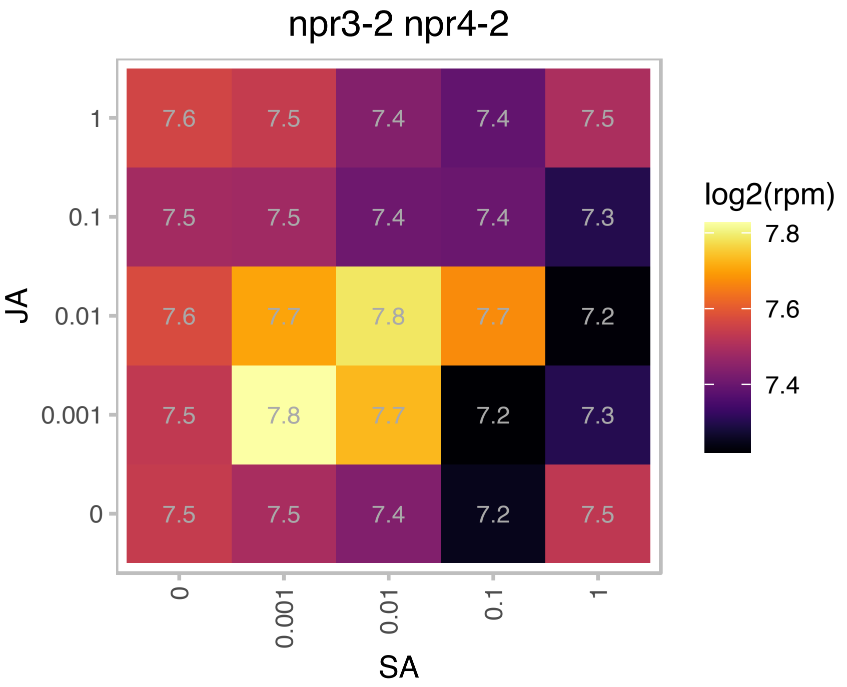


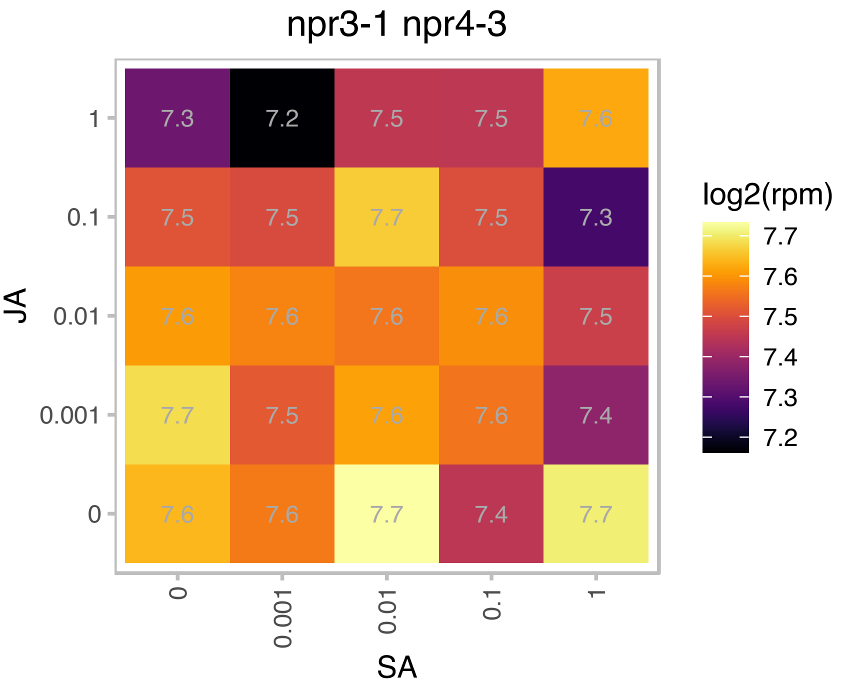

# AT5G12850.1 : NA

CCCH-type zinc finger protein with ARM repeat domain

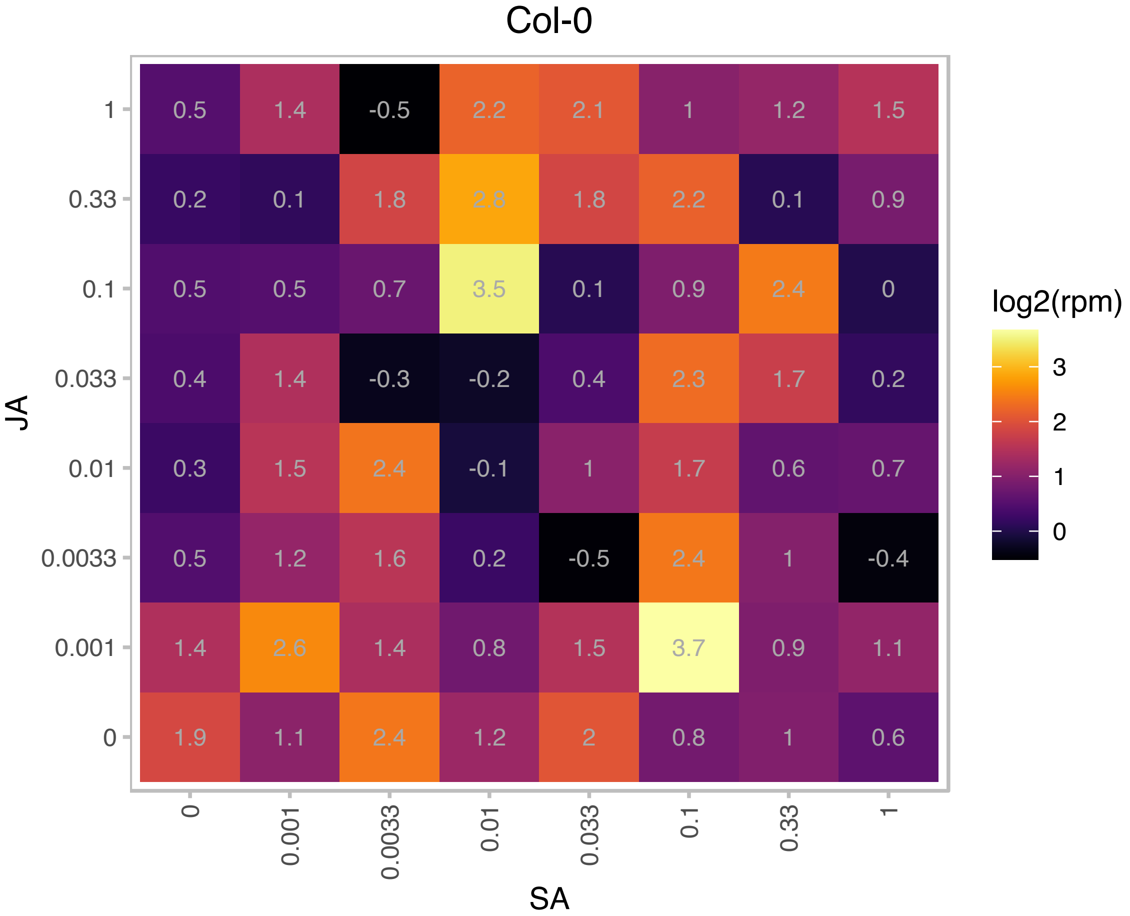


| | |
|---|---|
| Gene | AT5G12850 |
| is.expressed_genes | TRUE |
| is.responsive_genes | TRUE |
| is.apcluster | D-01 |

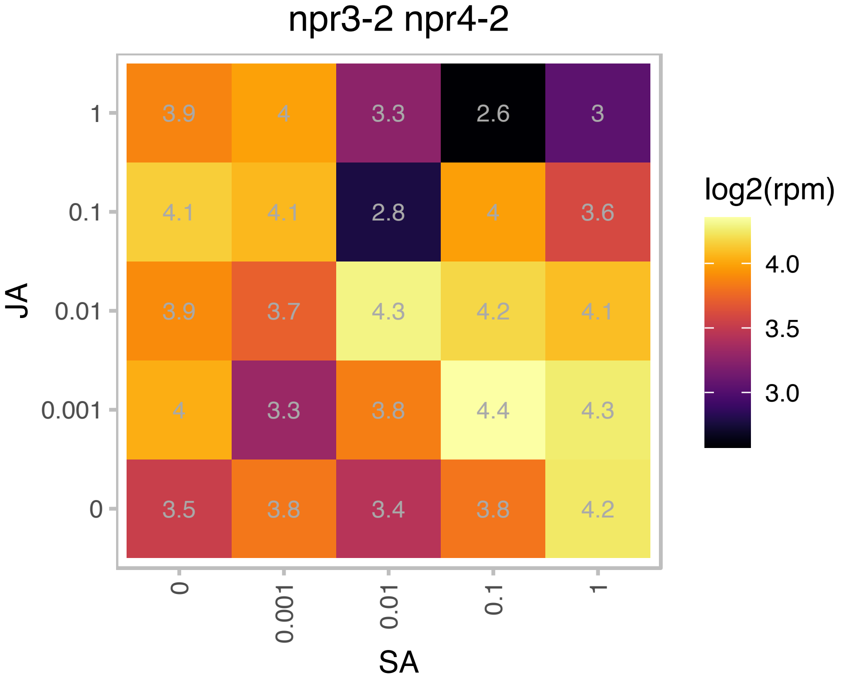


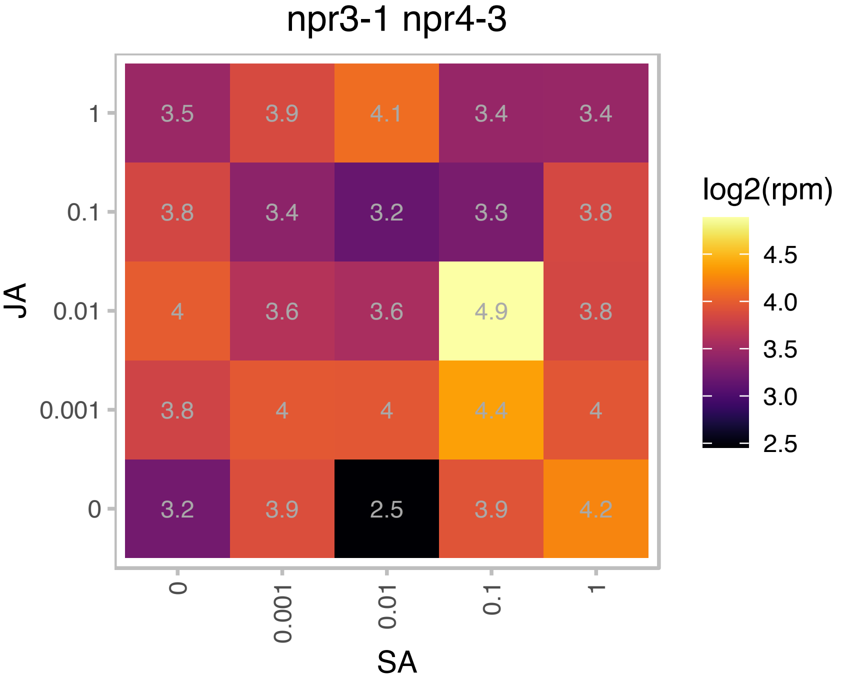

# AT5G22555.1 : NA

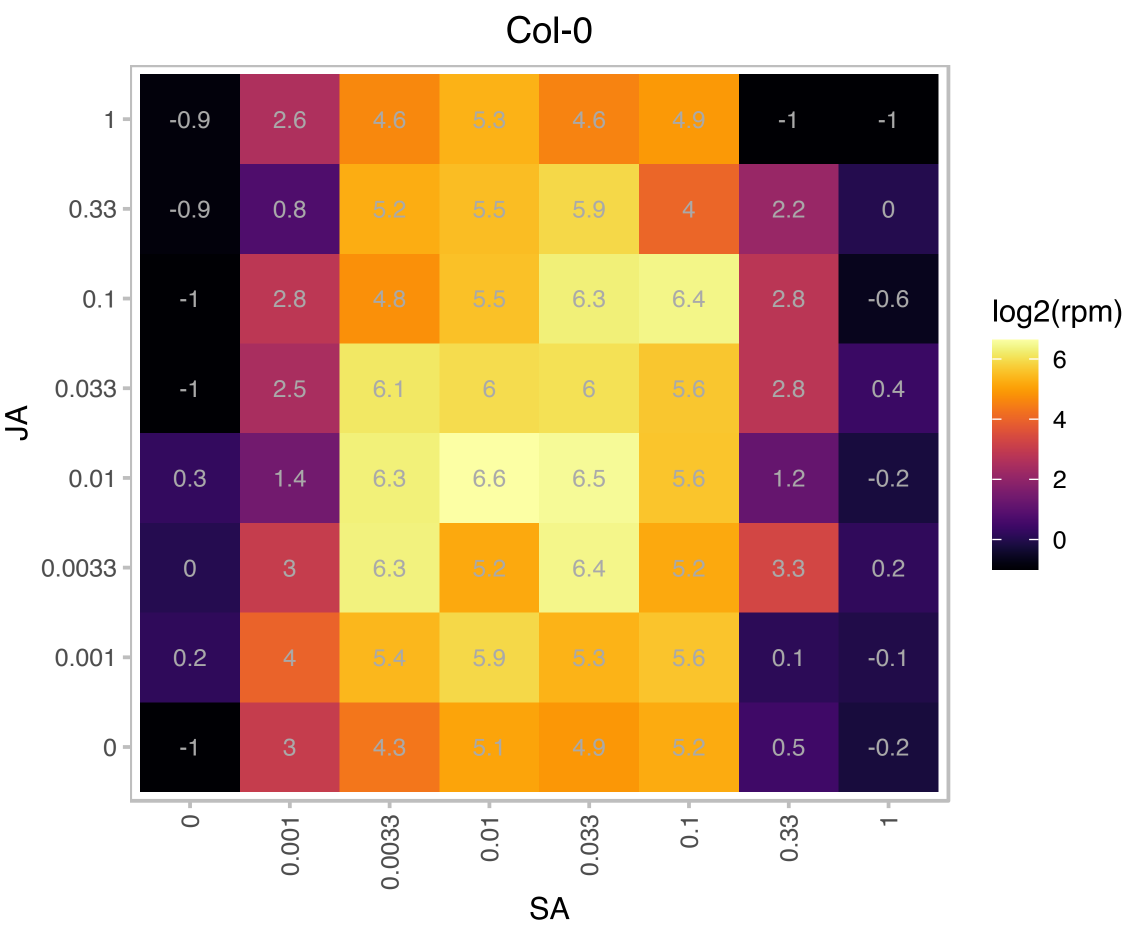


| | |
|---|---|
| Gene | AT5G22555 |
| is.expressed_genes | TRUE |
| is.responsive_genes | TRUE |
| is.apcluster | D-01 |

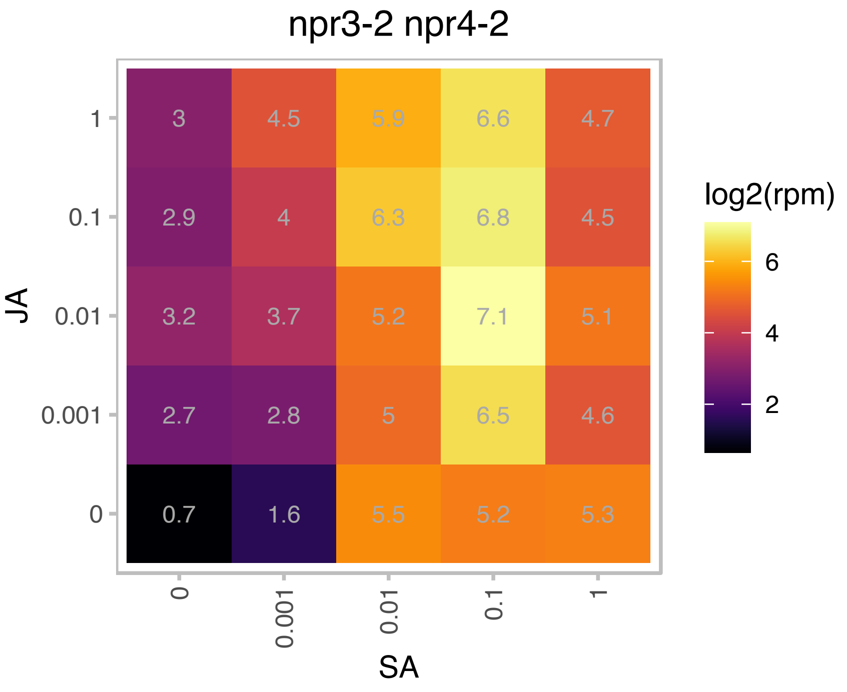


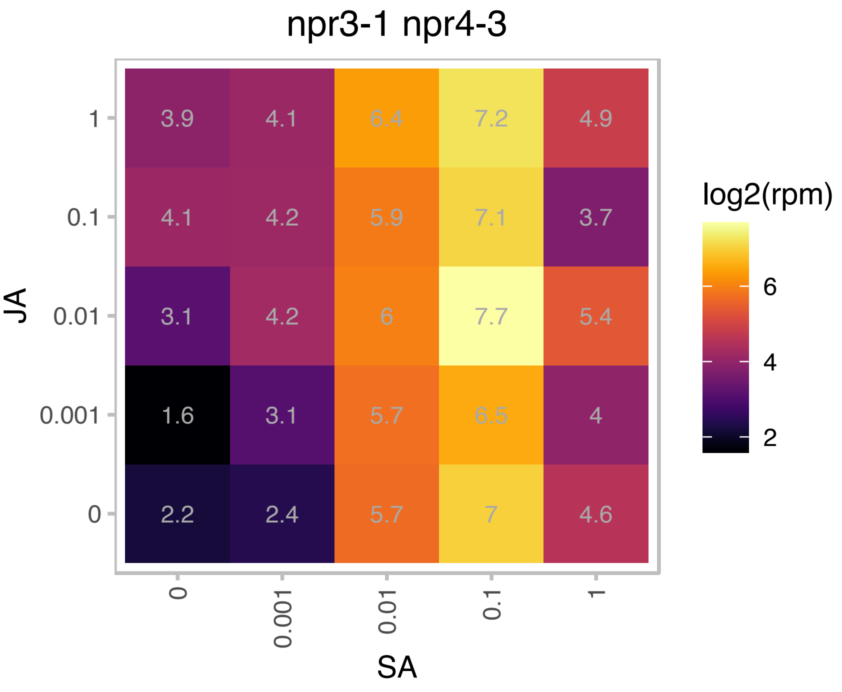

### AT5G23220.1 : NIC3

nicotinamidase 3

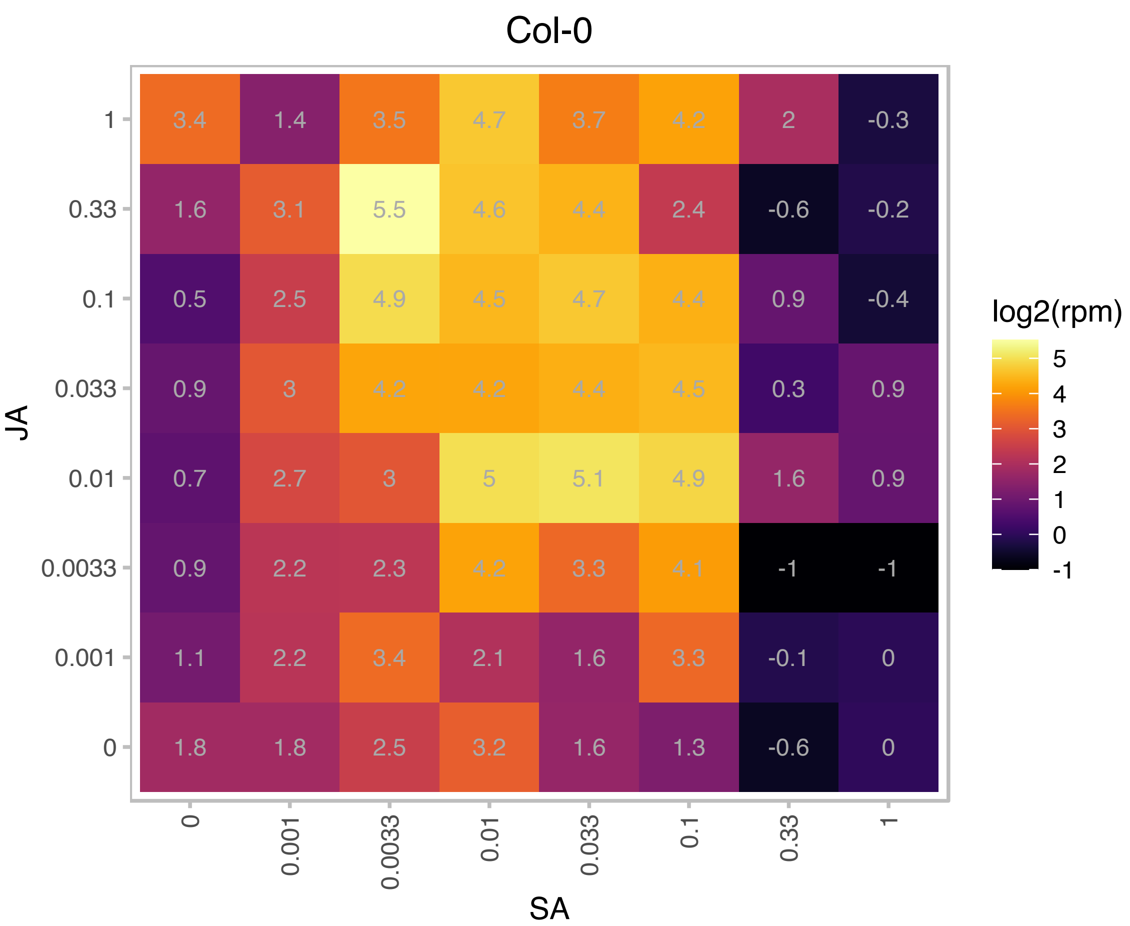


| | |
|---|---|
| Gene | AT5G23220 |
| is.expressed_genes | TRUE |
| is.responsive_genes | TRUE |
| is.apcluster | D-01 |

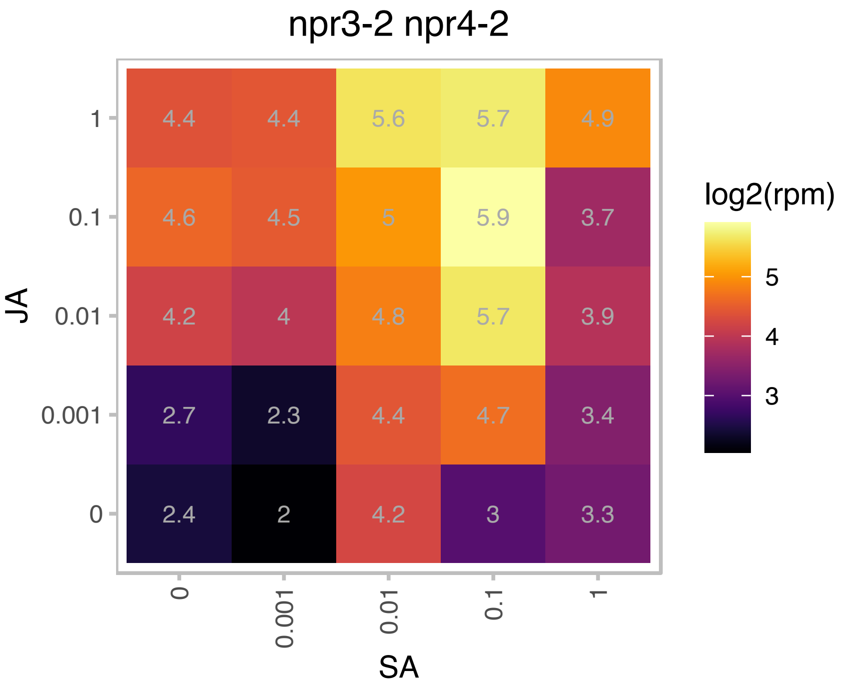


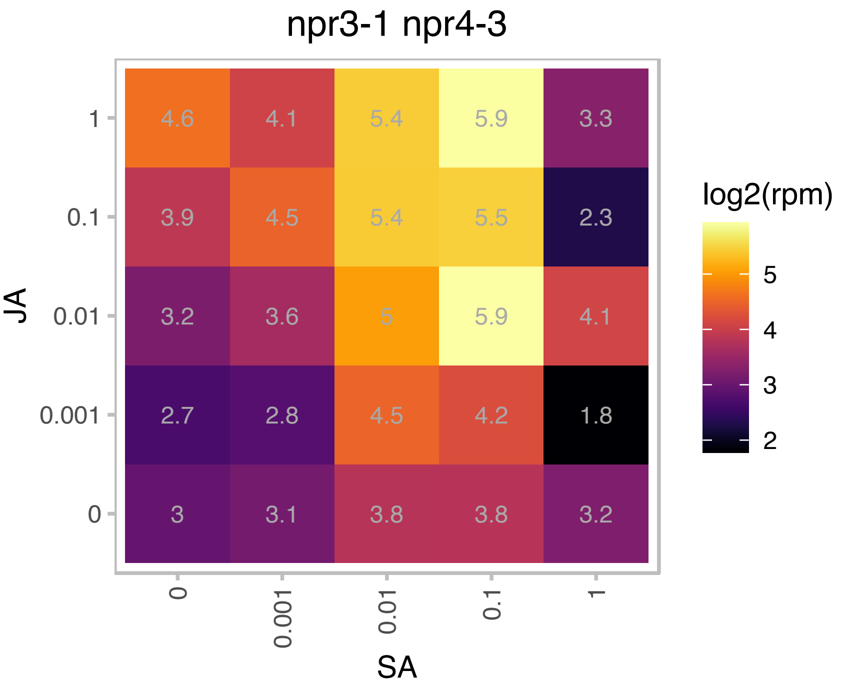

# AT5G45475.1 : NA

Potential natural antisense gene, locus overlaps with AT5G45480

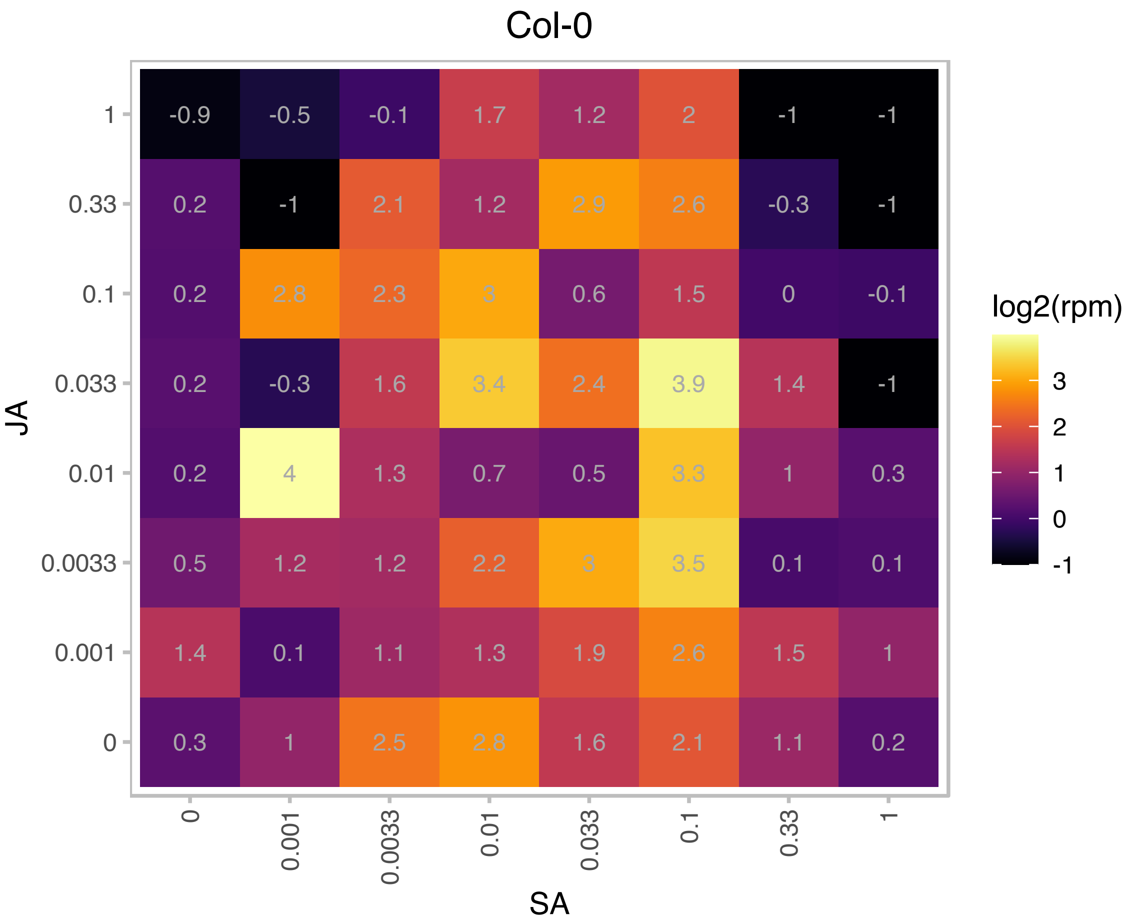


| | |
|---|---|
| Gene | AT5G45475 |
| is.expressed_genes | TRUE |
| is.responsive_genes | TRUE |
| is.apcluster | D-01 |

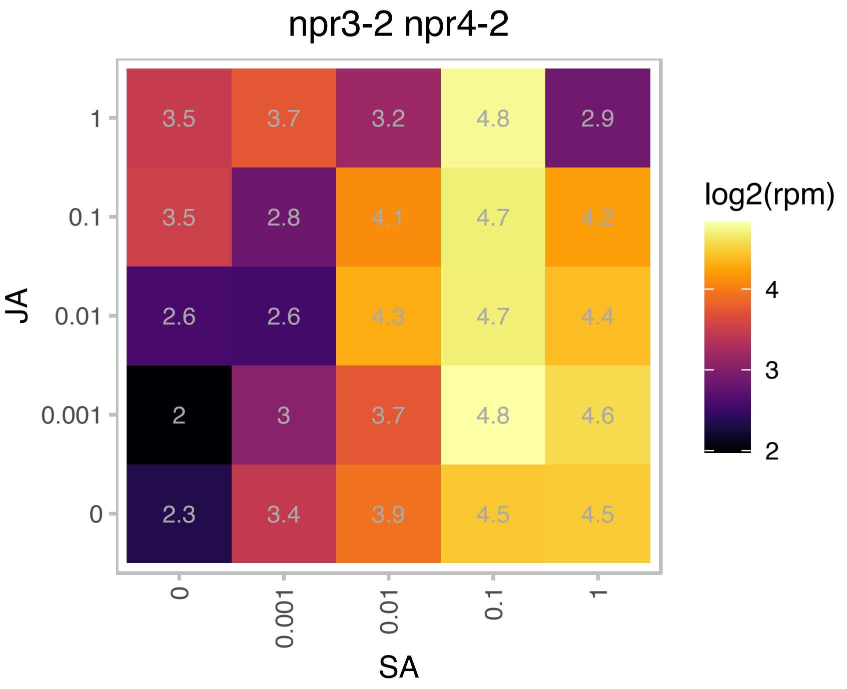


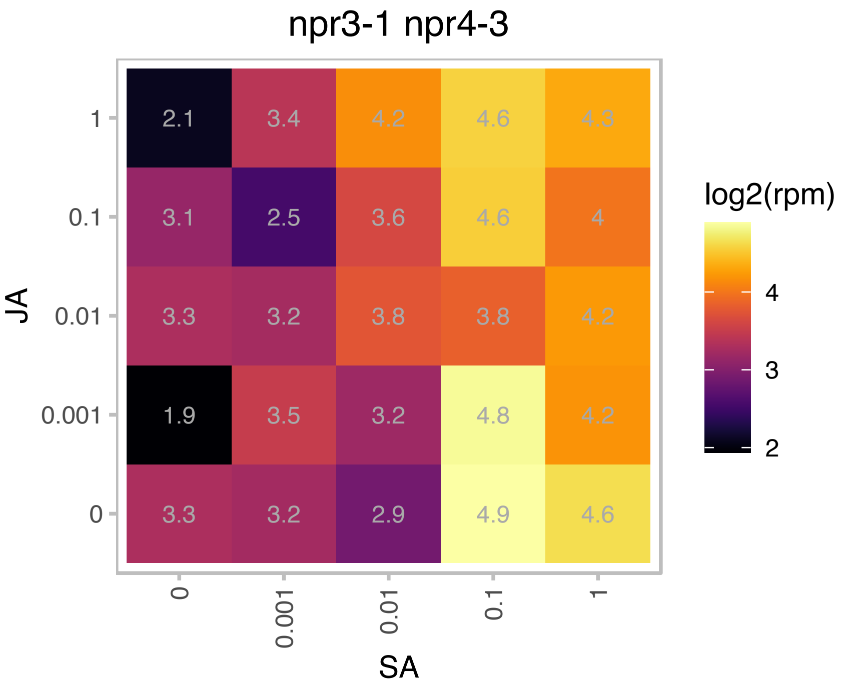

### AT5G54800.1 : ATGPT1, GPT1

glucose 6-phosphate/phosphate translocator 1

Encodes glucose6-Phosphate/phosphate transporter 1.  Essential for pollen maturation and embryo sac development. The mRNA is cell-to-cell mobile.

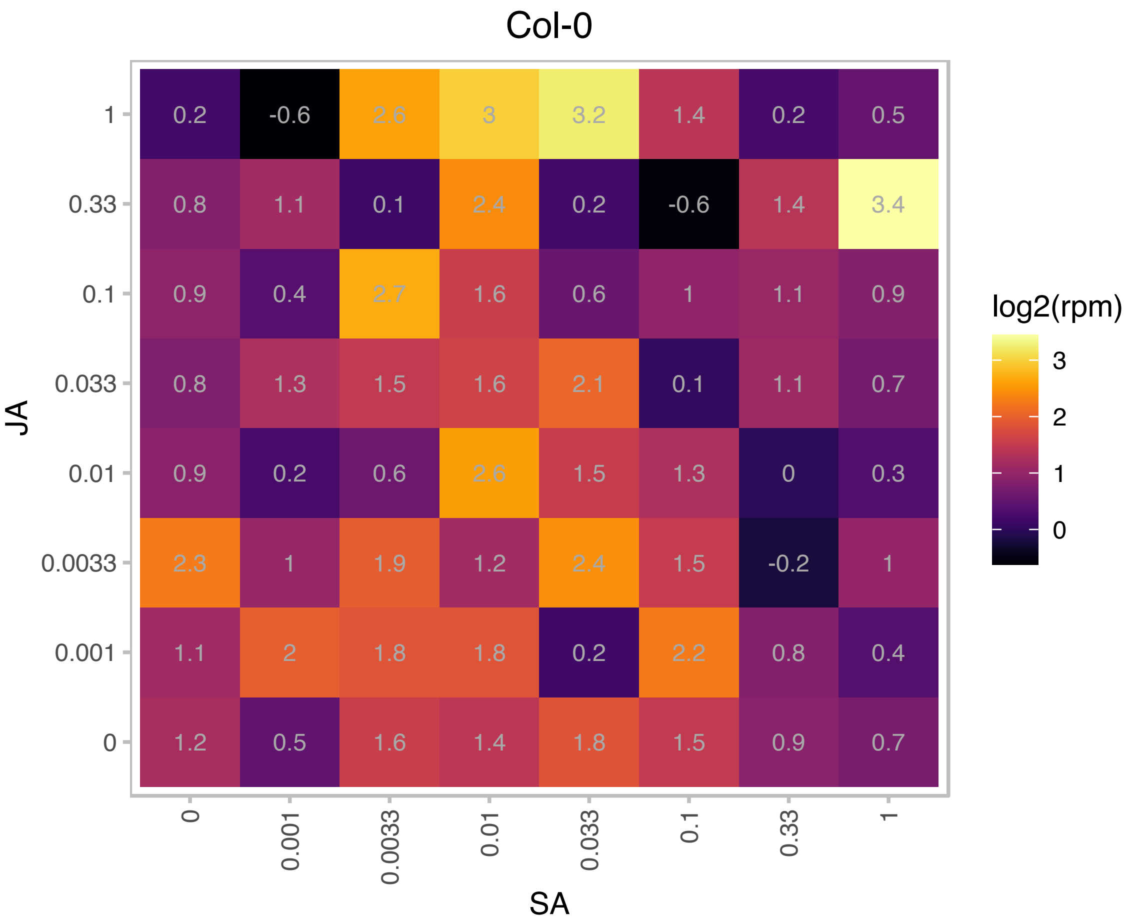


| | |
|---|---|
| Gene | AT5G54800 |
| is.expressed_genes | TRUE |
| is.responsive_genes | TRUE |
| is.apcluster | D-01 |

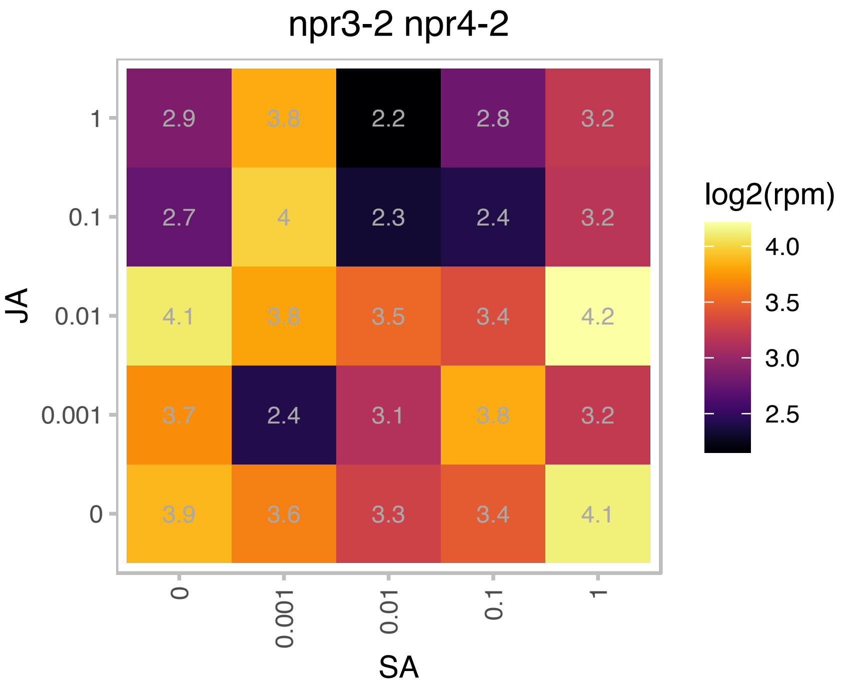


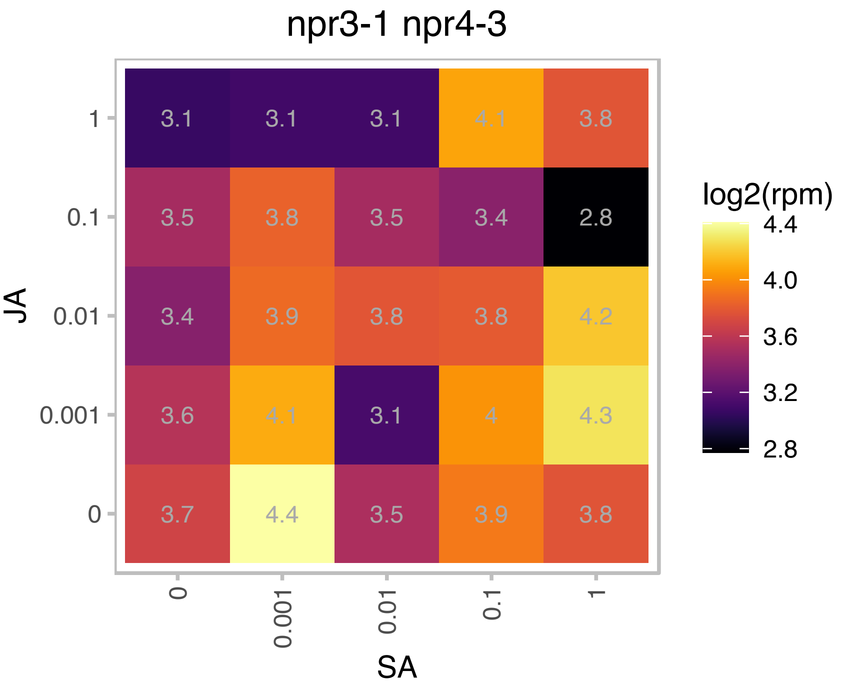

### AT5G65480.1 : CCI1

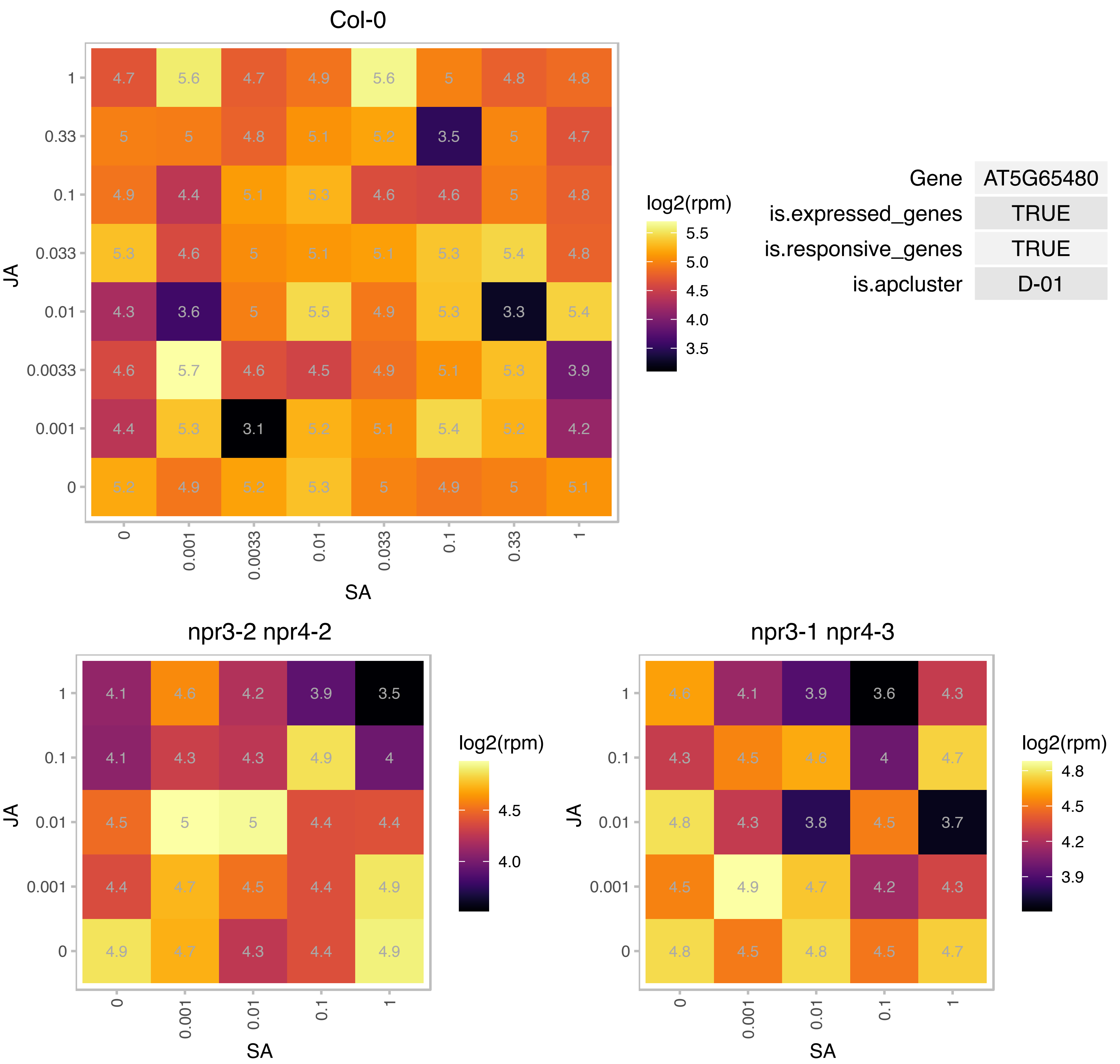


| | |
|---|---|
| Gene | AT5G65480 |
| is.expressed_genes | TRUE |
| is.responsive_genes | TRUE |
| is.apcluster | D-01 |

### AT5G66510.1 : GAMMA CA3

gamma carbonic anhydrase 3

Encodes mitochondrial gamma carbonic anhydrase. Component of the NADH dehydrogenase complex.

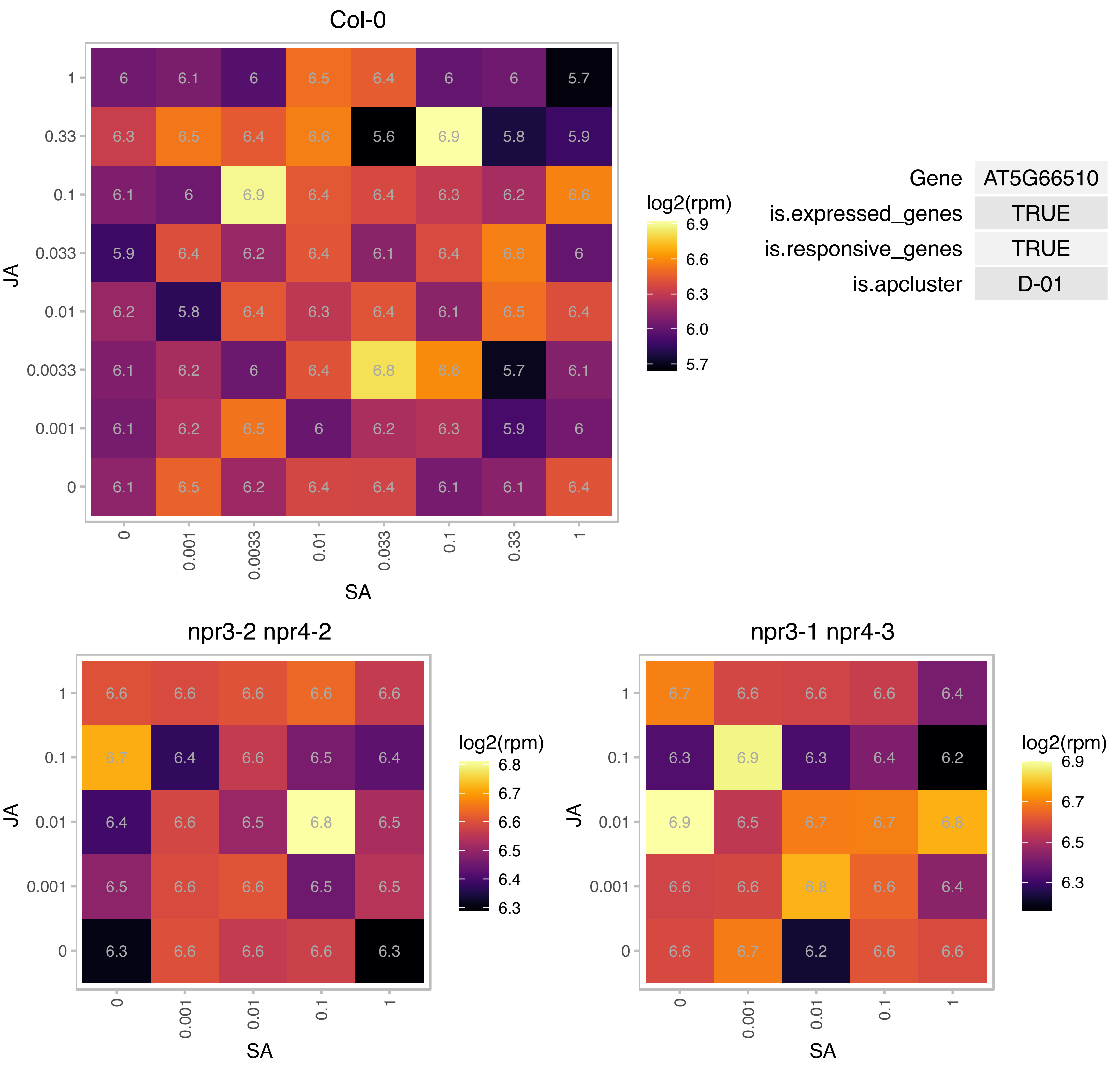


| | |
|---|---|
| Gene | AT5G66510 |
| is.expressed_genes | TRUE |
| is.responsive_genes | TRUE |
| is.apcluster | D-01 |

### ATCG00080.1 : PSBI

photosystem II reaction center protein I

PSII I protein

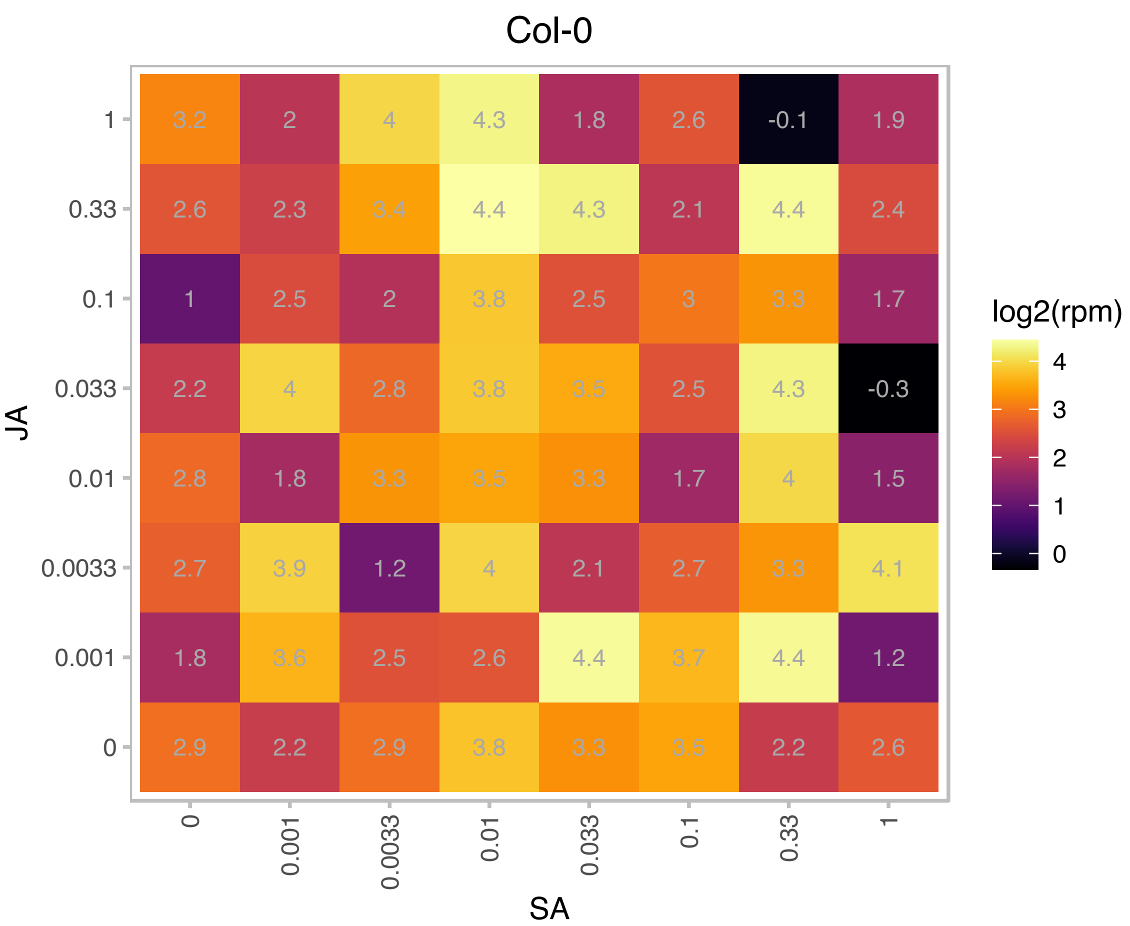


| | |
|---|---|
| Gene | ATCG00080 |
| is.expressed_genes | TRUE |
| is.responsive_genes | TRUE |
| is.apcluster | D-01 |

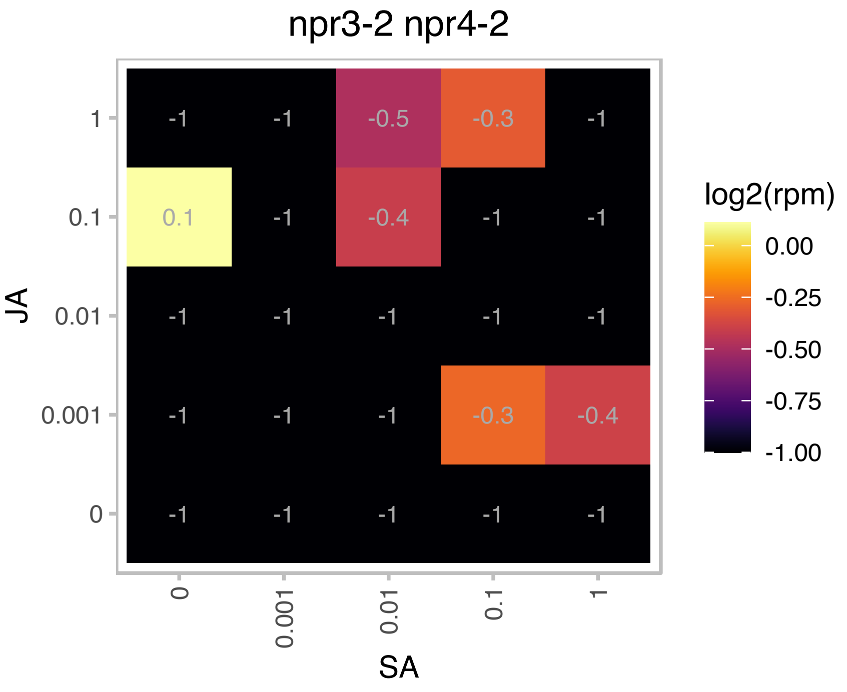


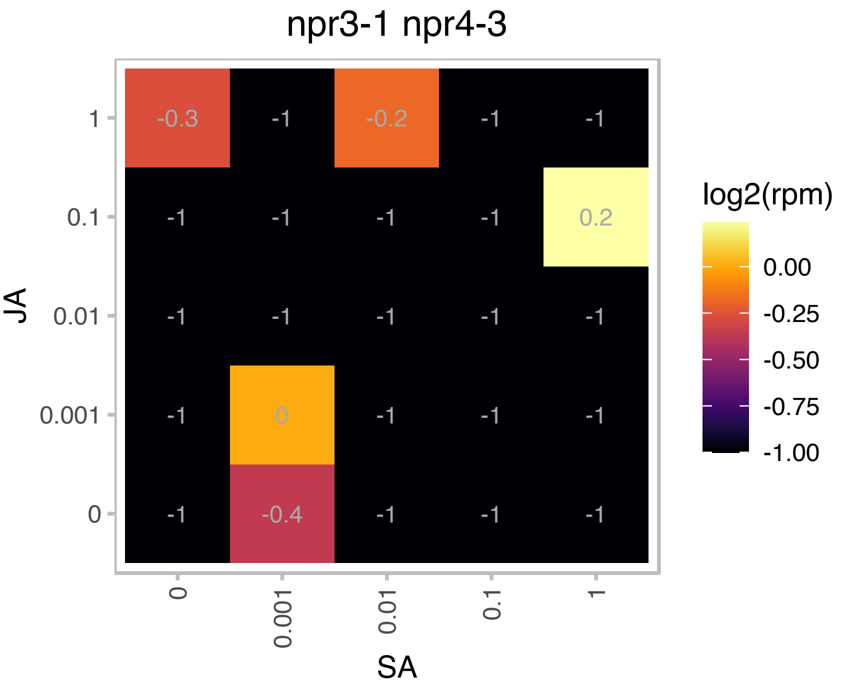

### ATCG01090.1 : NDHI

NADPH dehydrogenases

Encodes subunit of the chloroplast NAD(P)H dehydrogenase complex

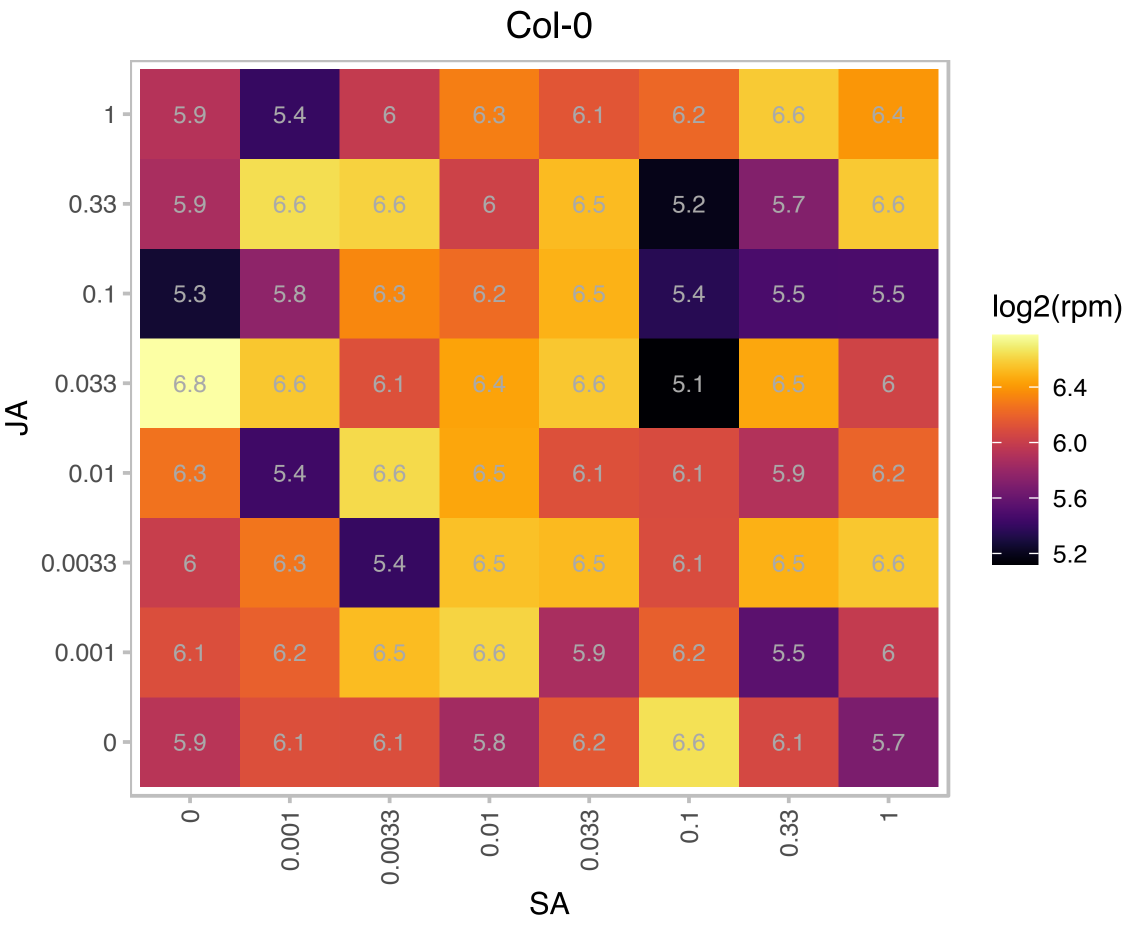


| | |
|---|---|
| Gene | ATCG01090 |
| is.expressed_genes | TRUE |
| is.responsive_genes | TRUE |
| is.apcluster | D-01 |

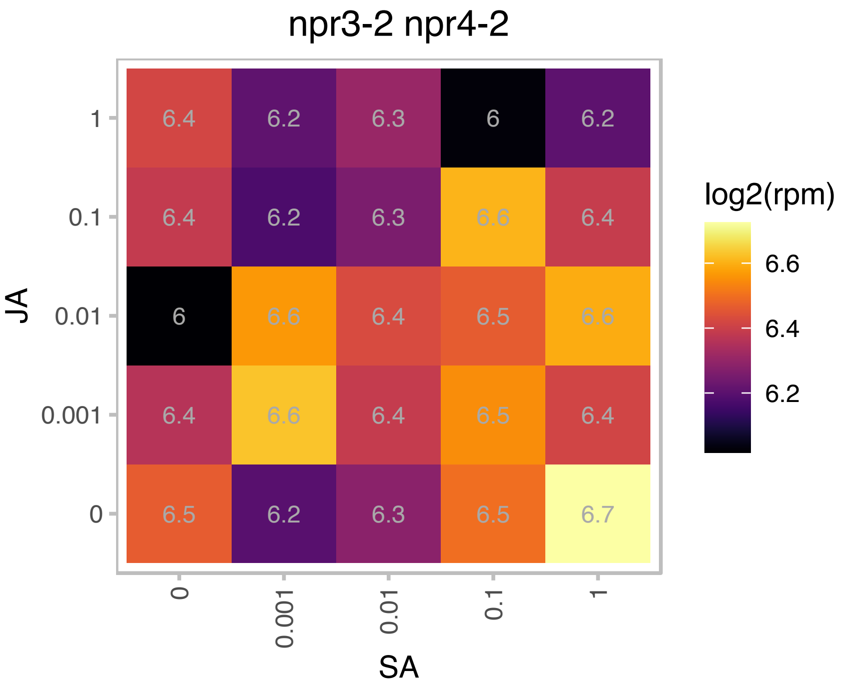


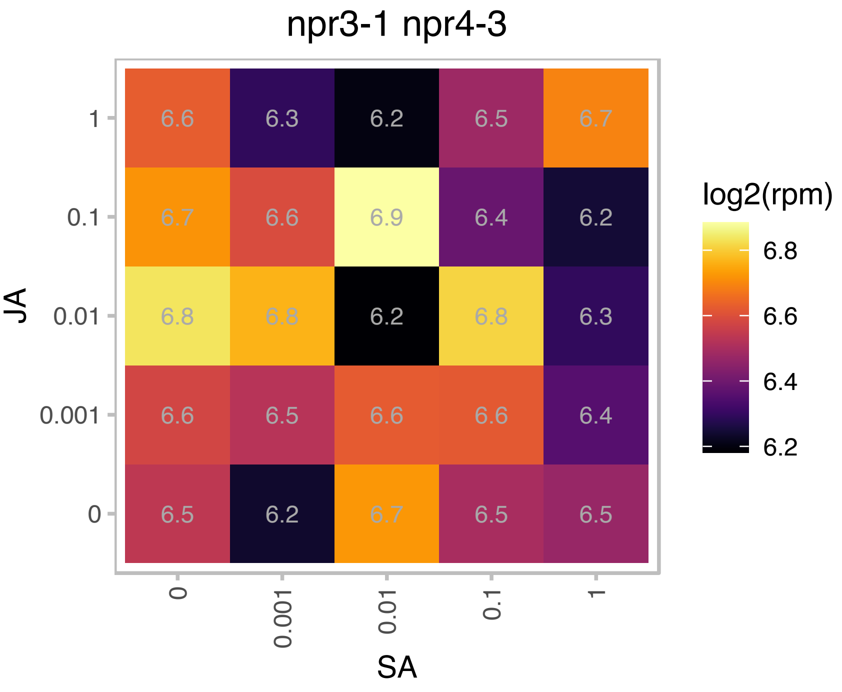

# AT1G29470.1 : NA

S-adenosyl-L-methionine-dependent methyltransferases superfamily protein

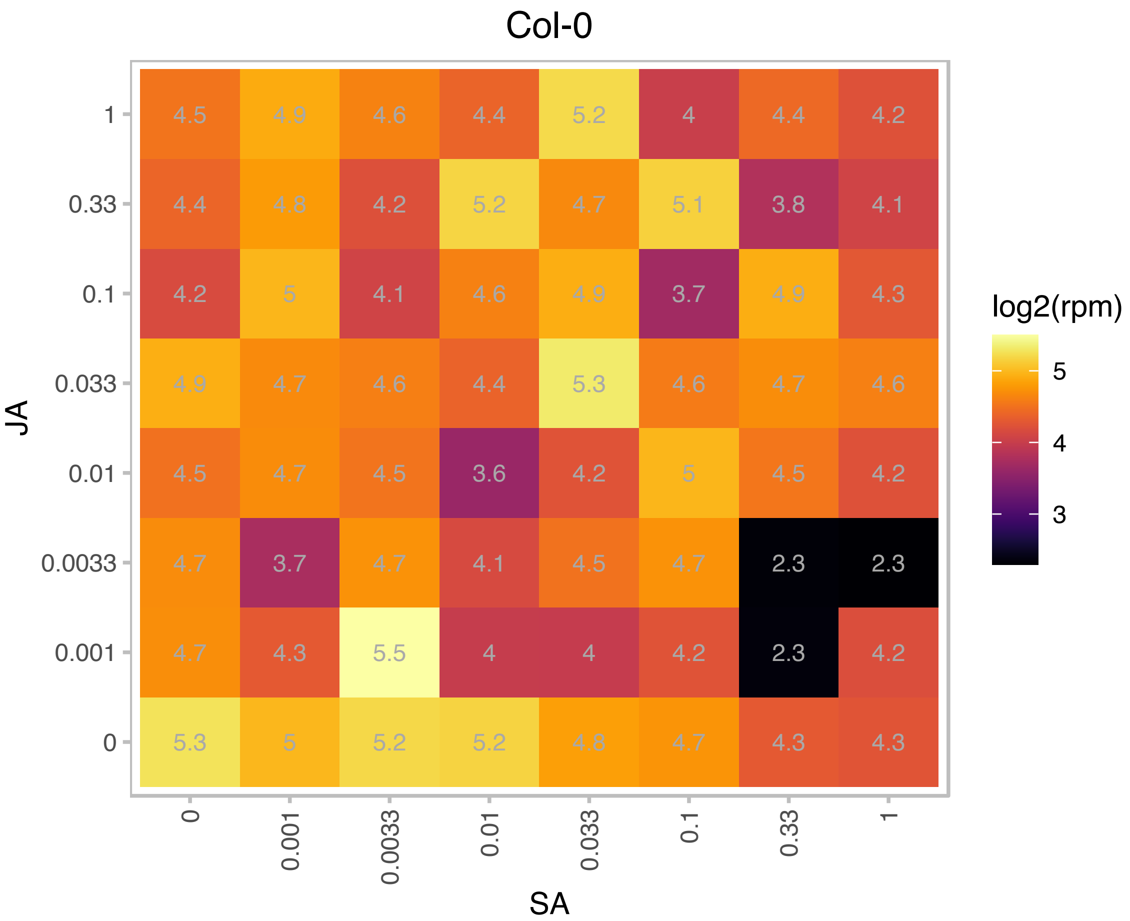


| | |
|---|---|
| Gene | AT1G29470 |
| is.expressed_genes | TRUE |
| is.responsive_genes | TRUE |
| is.apcluster | D-02 |

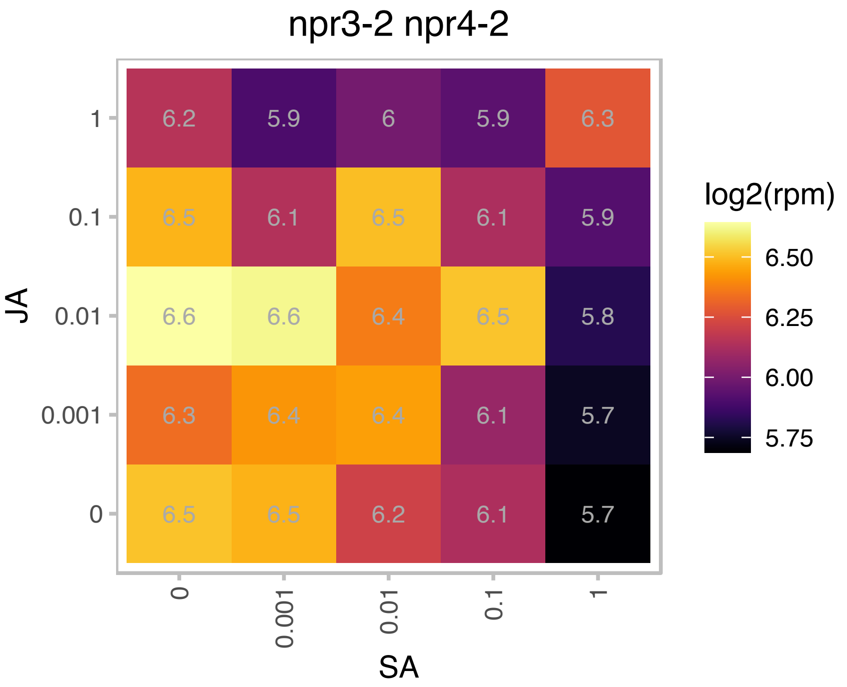


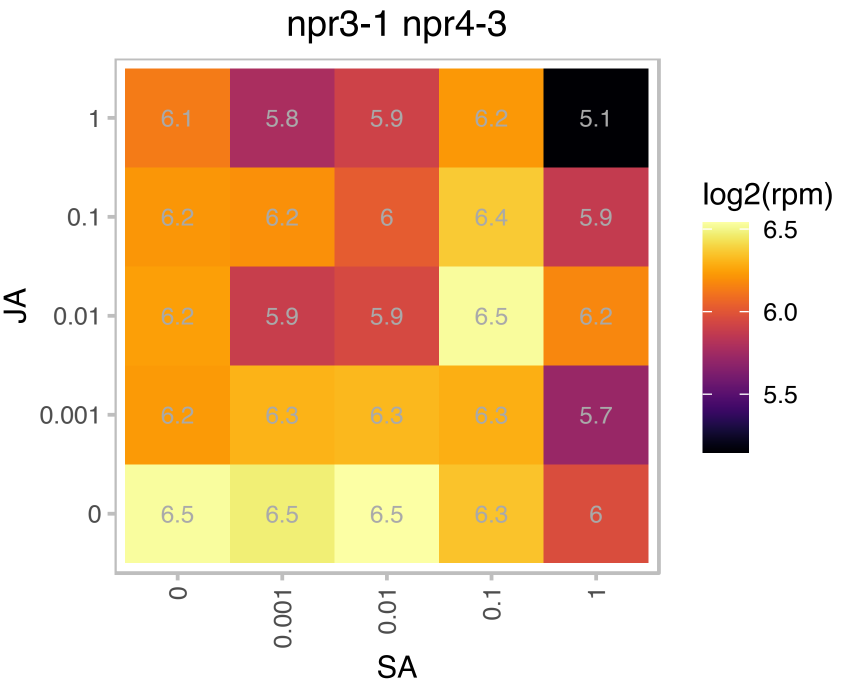

### AT1G47500.1 : ATRBP47C', RBP47C'

RNA-binding protein 47C'

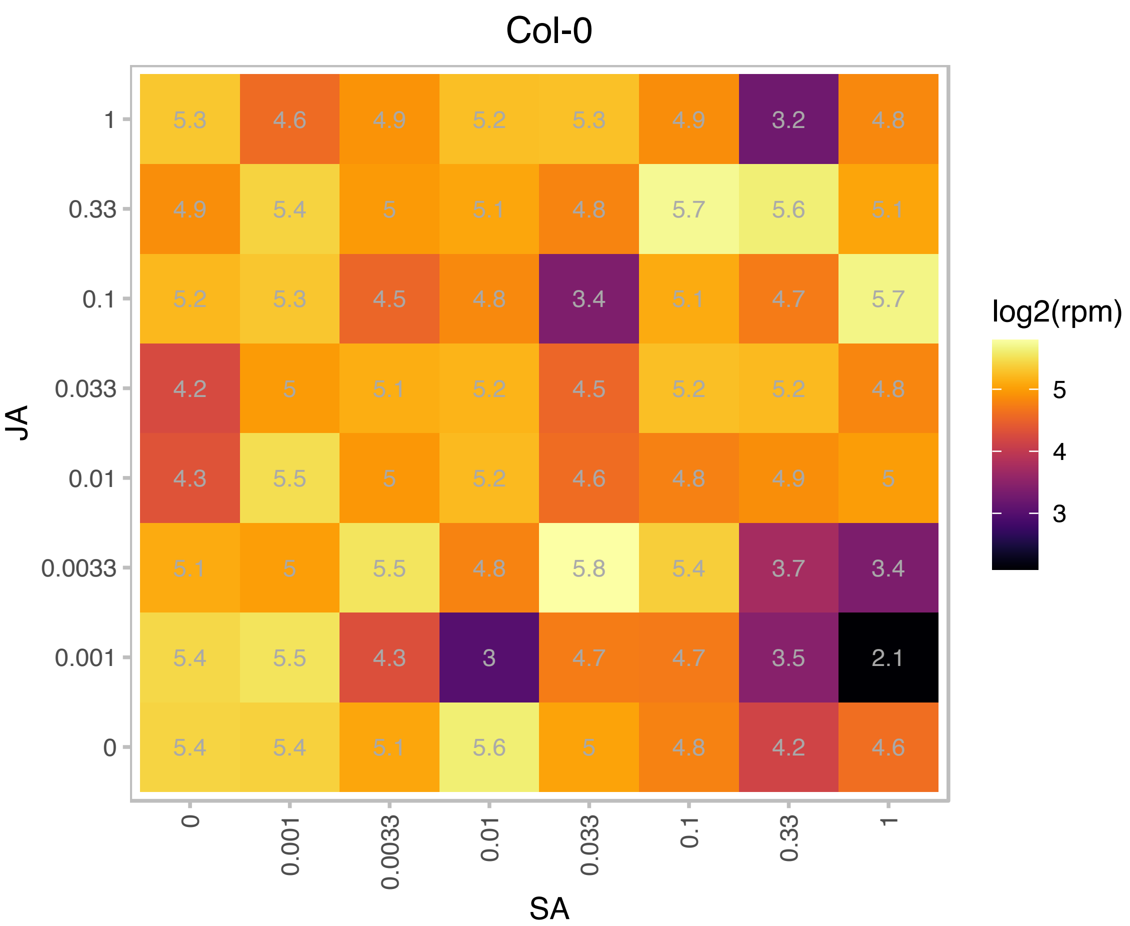


| | |
|---|---|
| Gene | AT1G47500 |
| is.expressed_genes | TRUE |
| is.responsive_genes | TRUE |
| is.apcluster | D-02 |

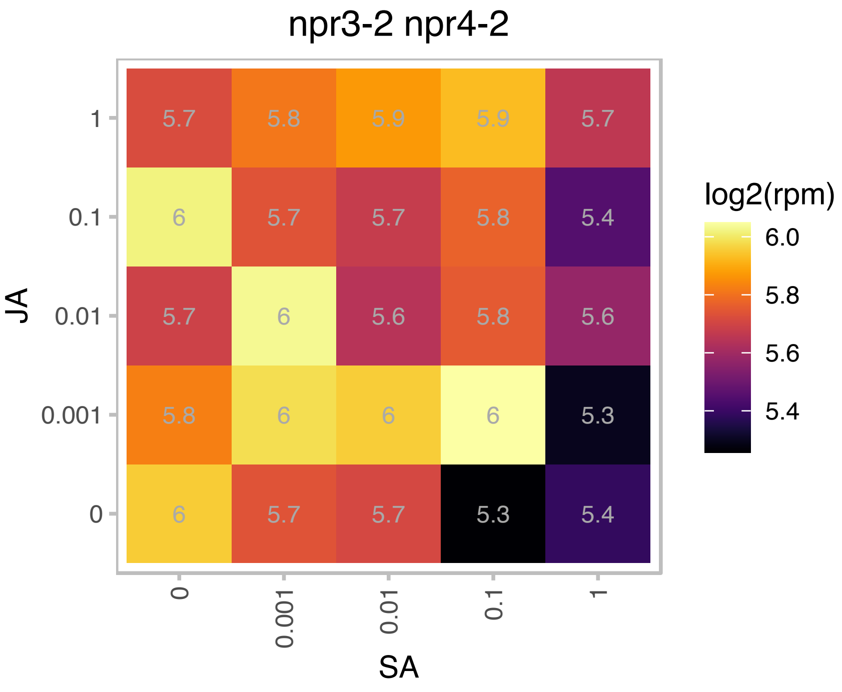


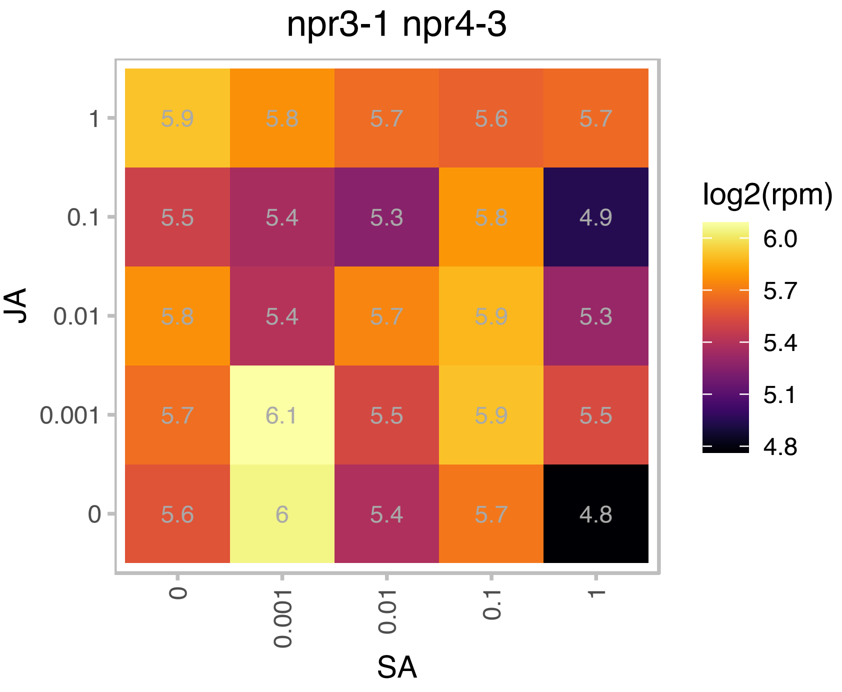

### AT1G56070.1 : LOS1

Ribosomal protein S5/Elongation factor G/III/V family protein
encodes a translation elongation factor 2-like protein that is involved in cold-induced translation. Mutations in this gene specifically blocks low temperature-induced transcription of cold-responsive genes but induces the expression of CBF genes and mutants carrying the recessive mutations fail to acclimate to cold and is freezing sensitive.

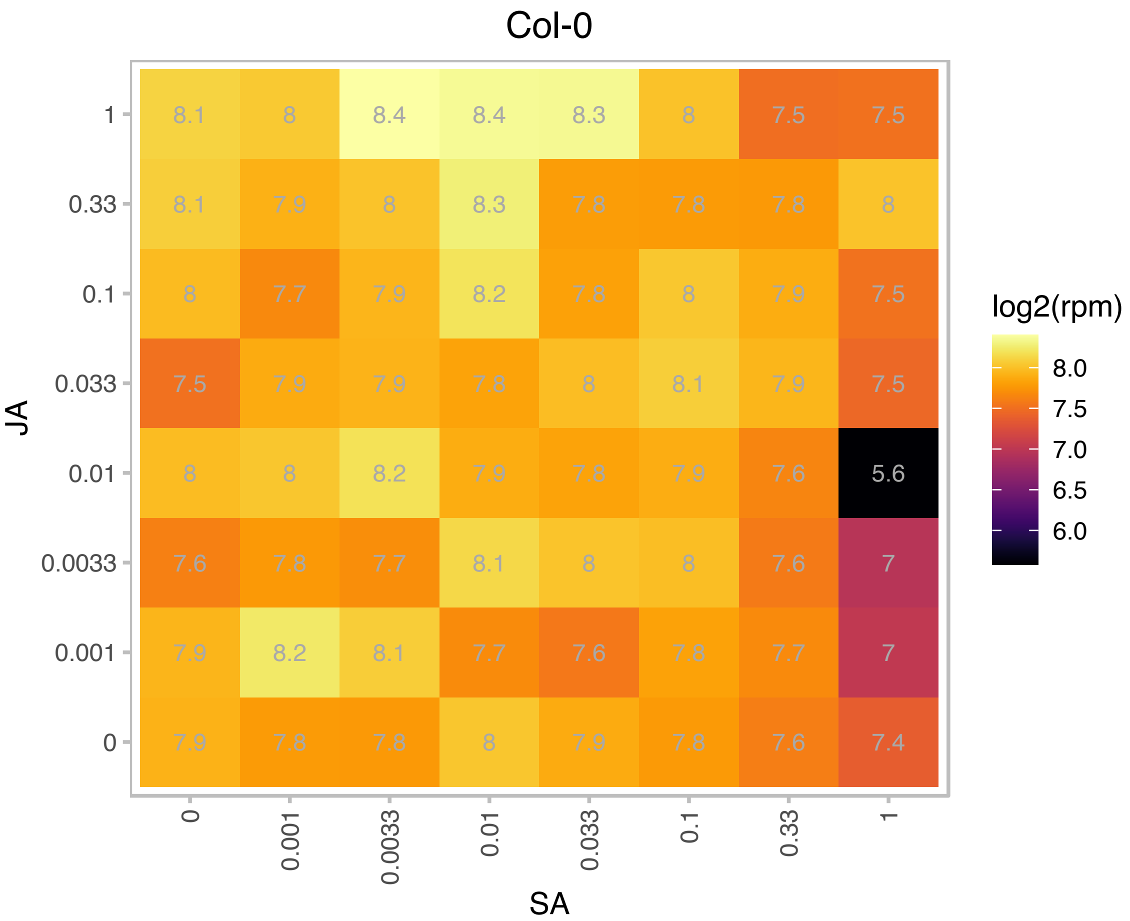


| | |
|---|---|
| Gene | AT1G56070 |
| is.expressed_genes | TRUE |
| is.responsive_genes | TRUE |
| is.apcluster | D-02 |

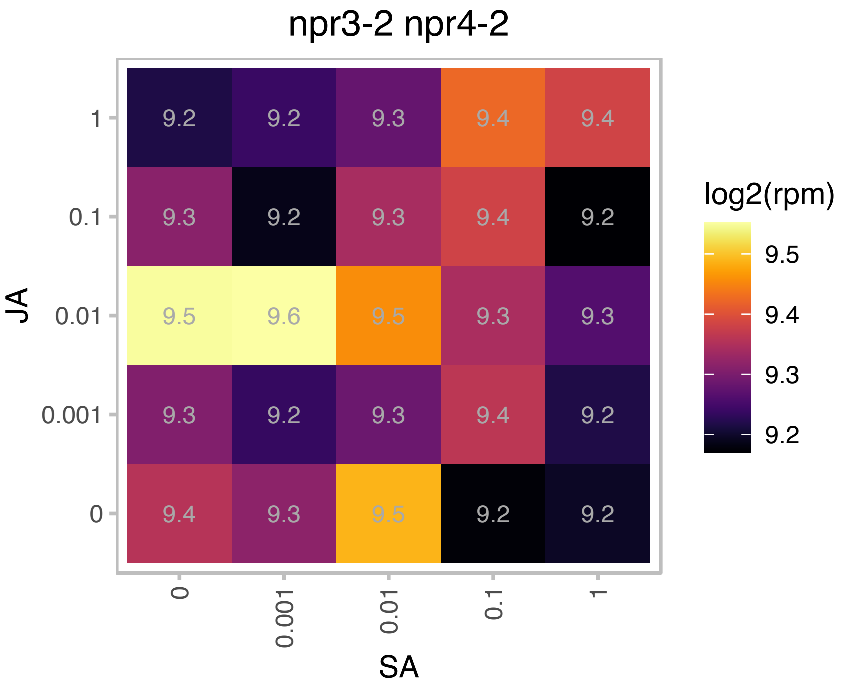


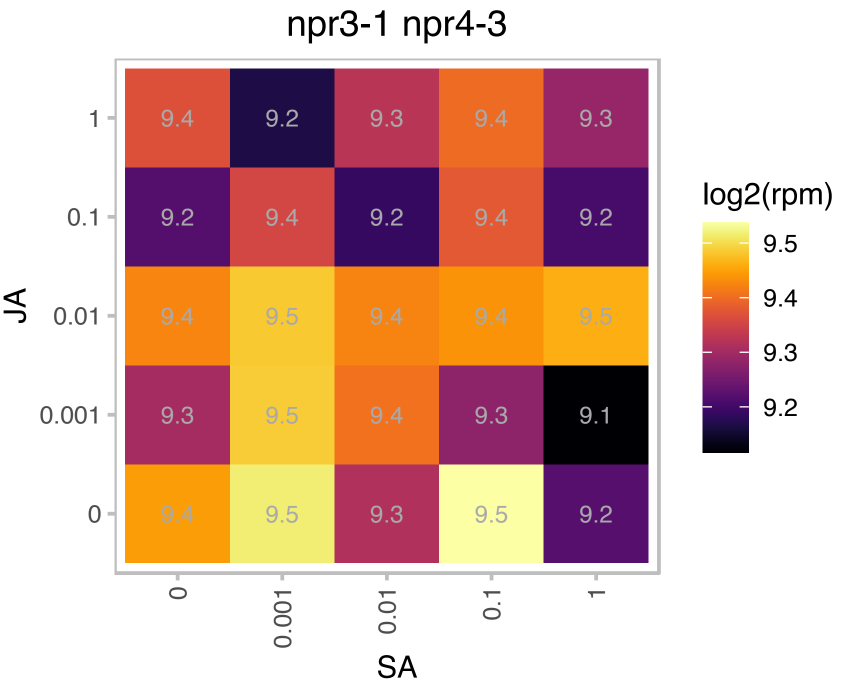

### AT2G05440.1 : ATGRP9, GRP9

GLYCINE RICH PROTEIN 9

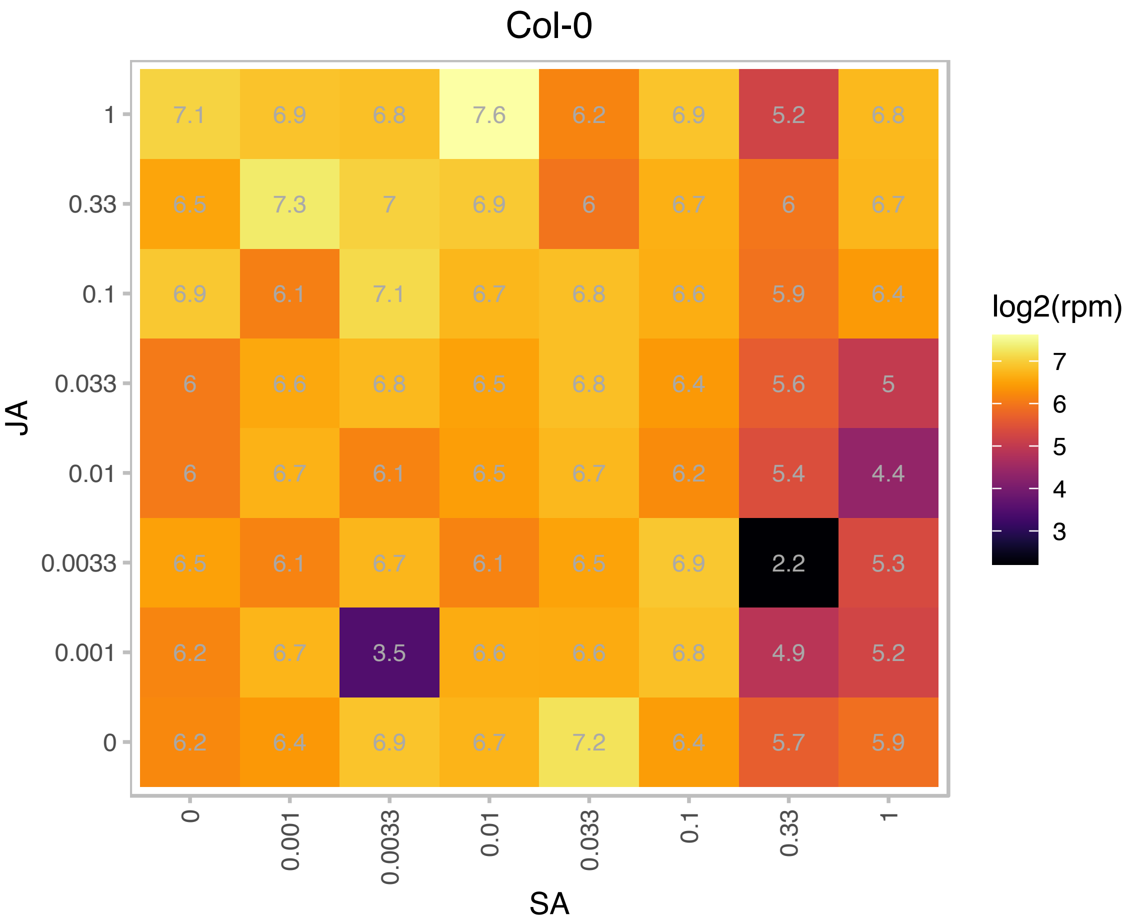


| | |
|---|---|
| Gene | AT2G05440 |
| is.expressed_genes | TRUE |
| is.responsive_genes | TRUE |
| is.apcluster | D-02 |

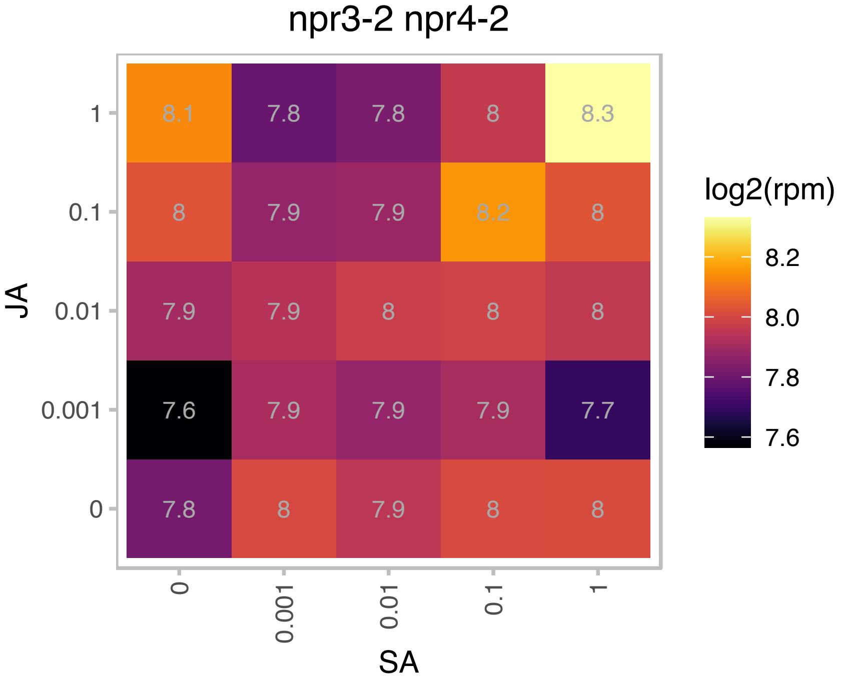


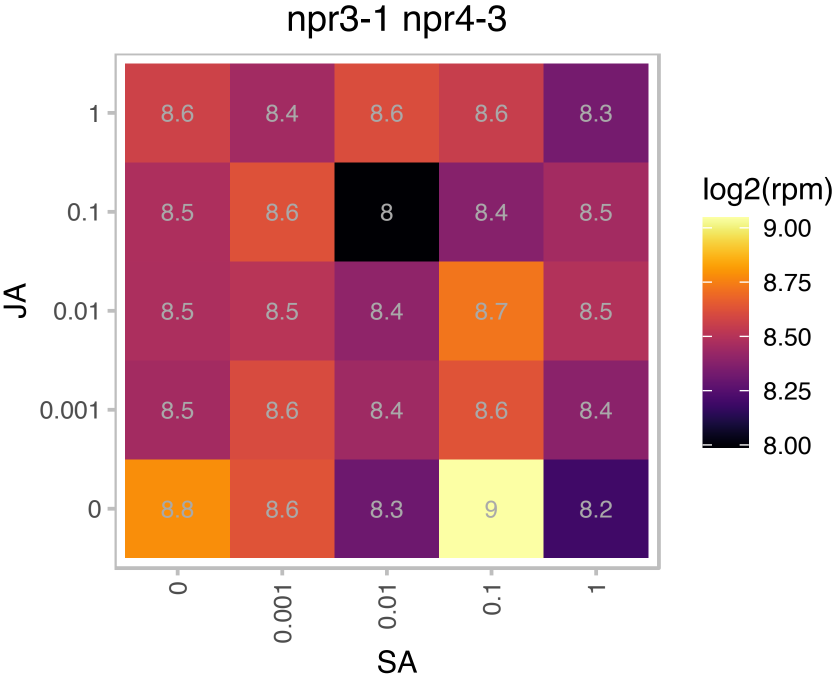

# AT2G23120.1 : NA

Late embryogenesis abundant protein, group 6

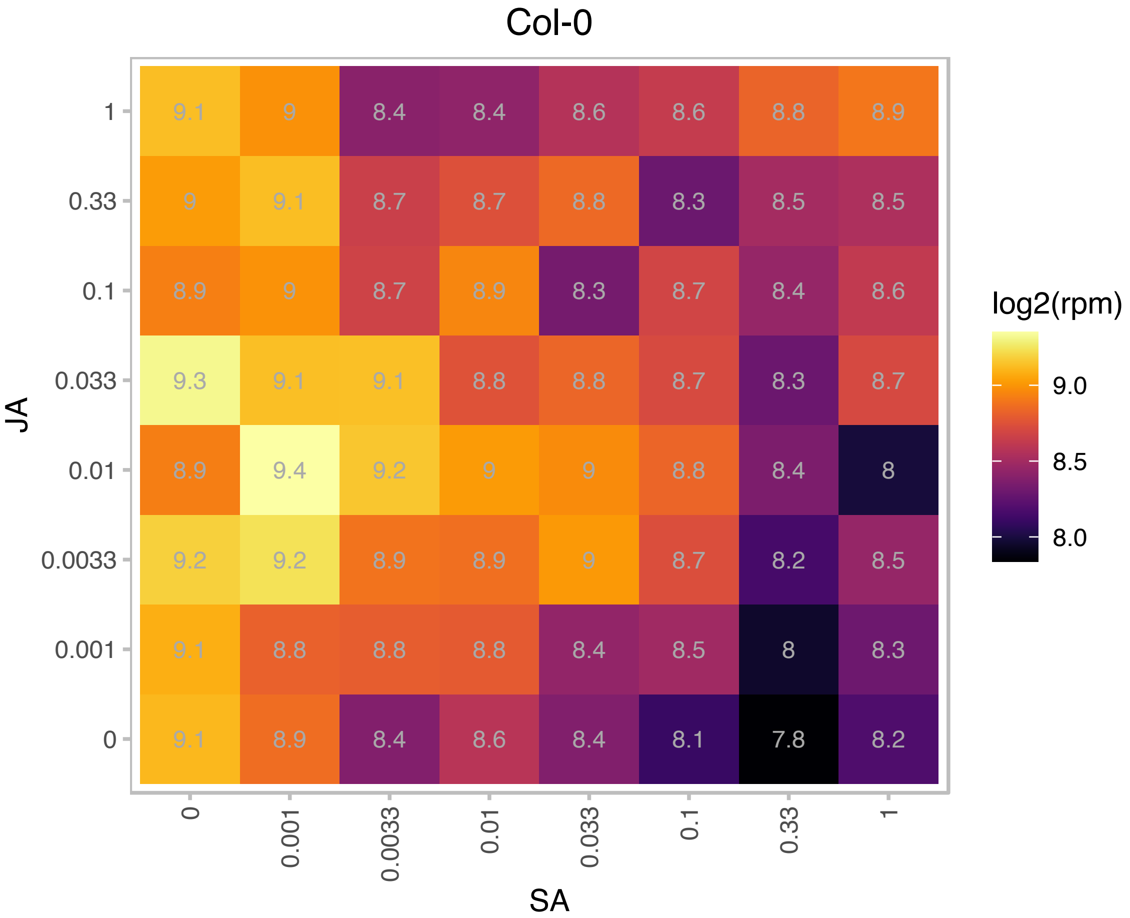


| | |
|---|---|
| Gene | AT2G23120 |
| is.expressed_genes | TRUE |
| is.responsive_genes | TRUE |
| is.apcluster | D-02 |

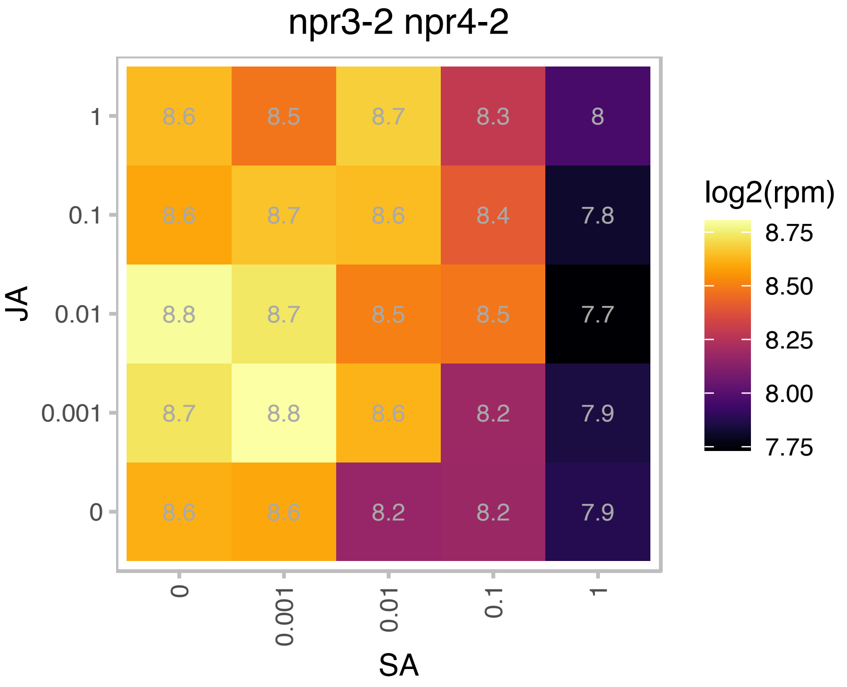


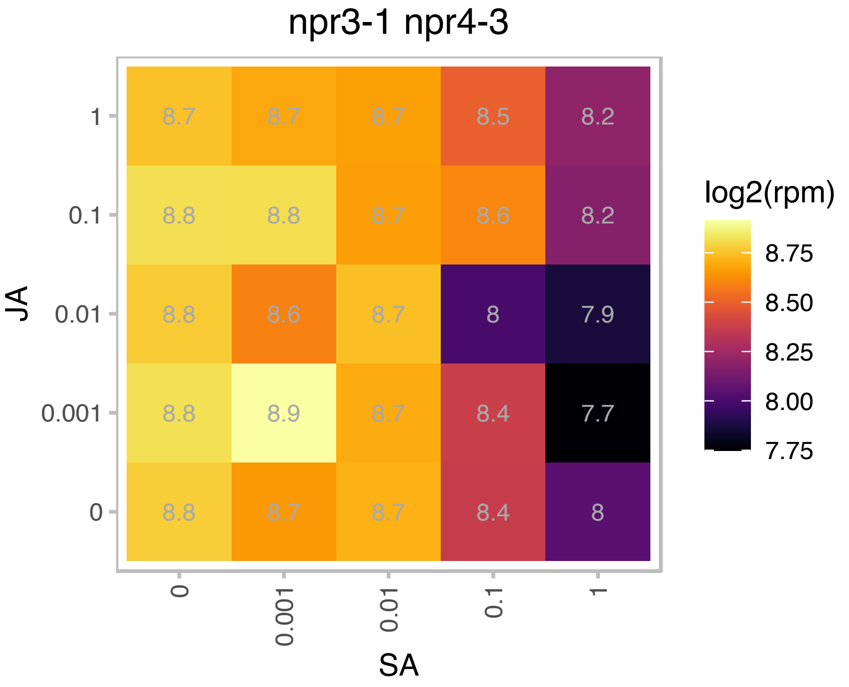

### AT2G29080.1 : AtFTSH3, FTSH3

FTSH protease 3

encodes an FtsH protease that is localized to the mitochondrion

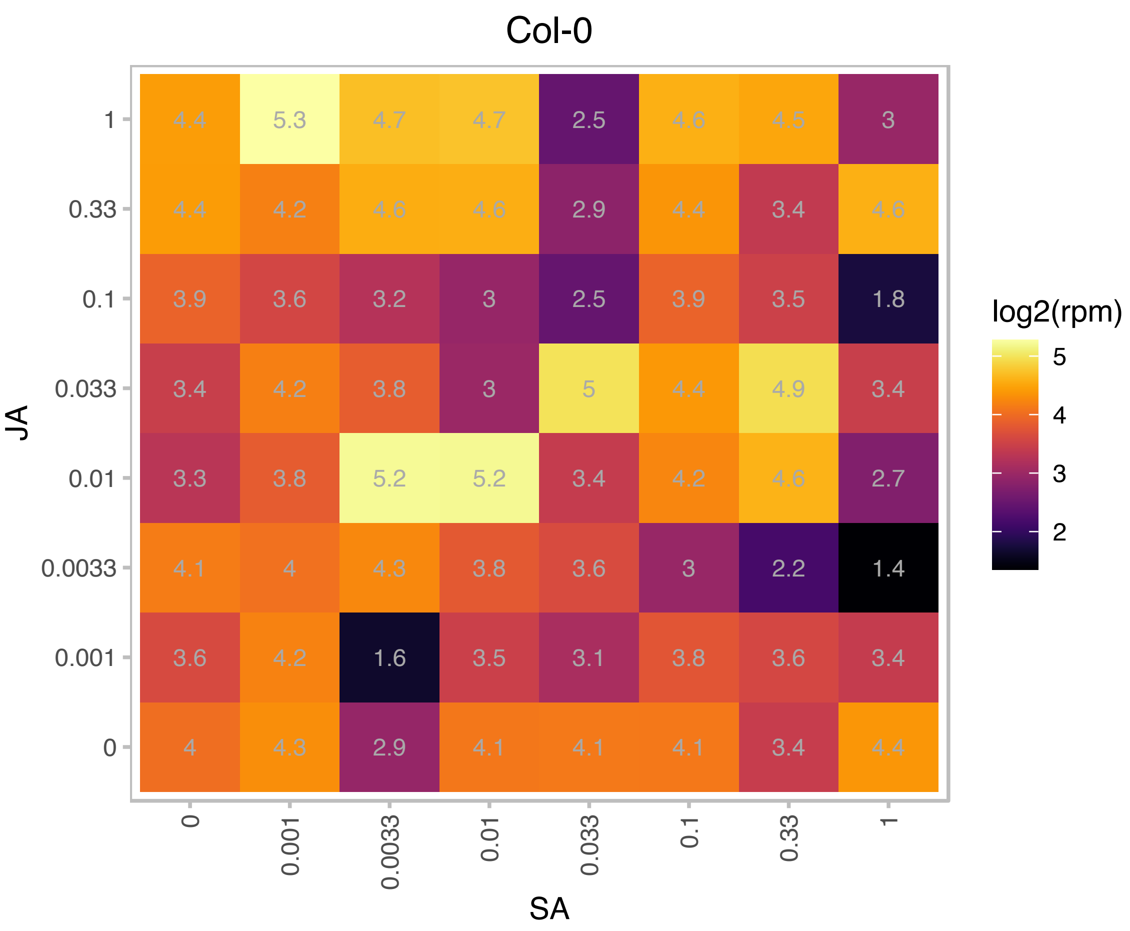


| | |
|---|---|
| Gene | AT2G29080 |
| is.expressed_genes | TRUE |
| is.responsive_genes | TRUE |
| is.apcluster | D-02 |

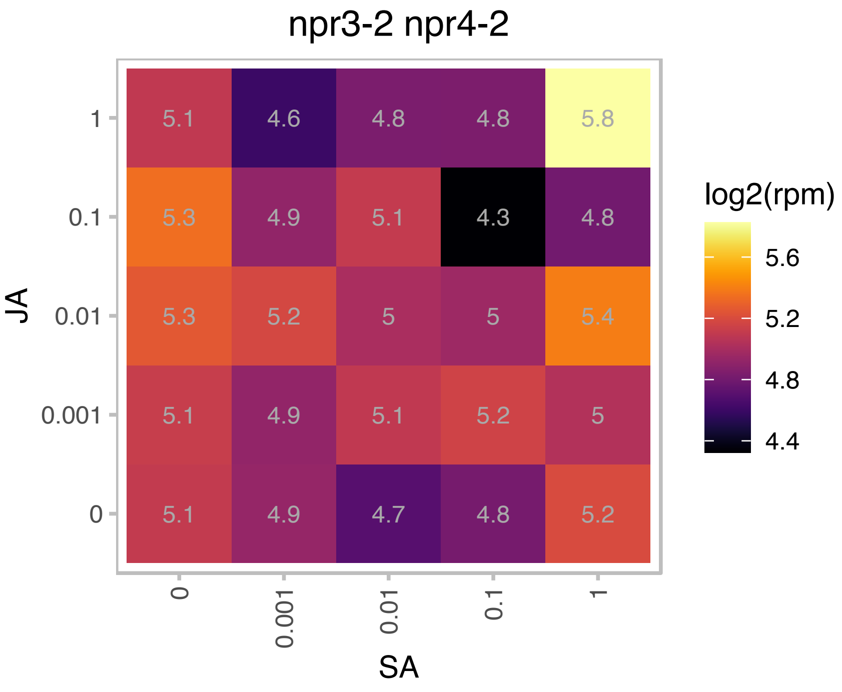


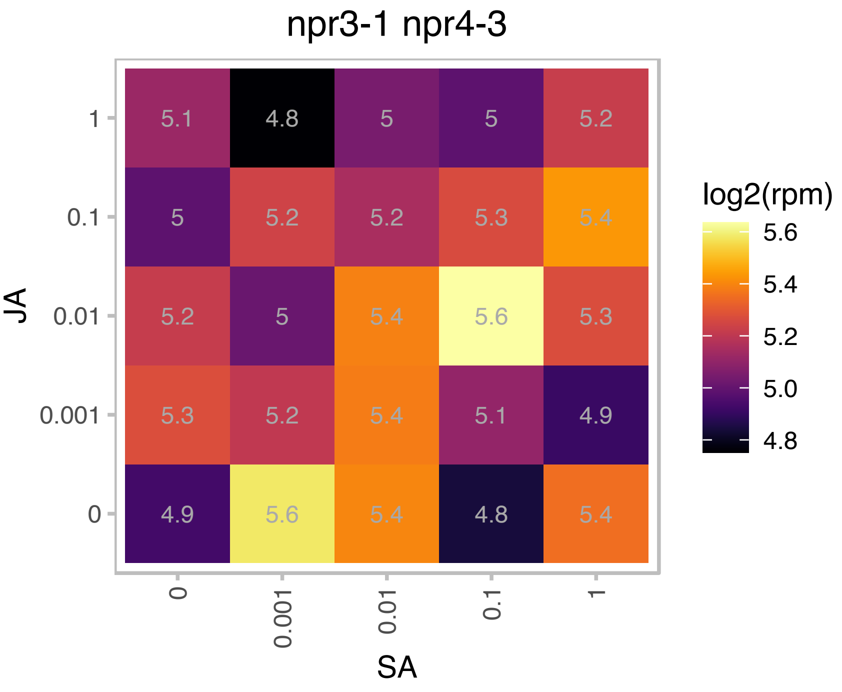

### AT2G31040.1 : AtCGL160, CGL160

ATP synthase protein I -related

Encodes an integral thylakoid protein that facilitates assembly of the membranous part of the chloroplast ATPase.

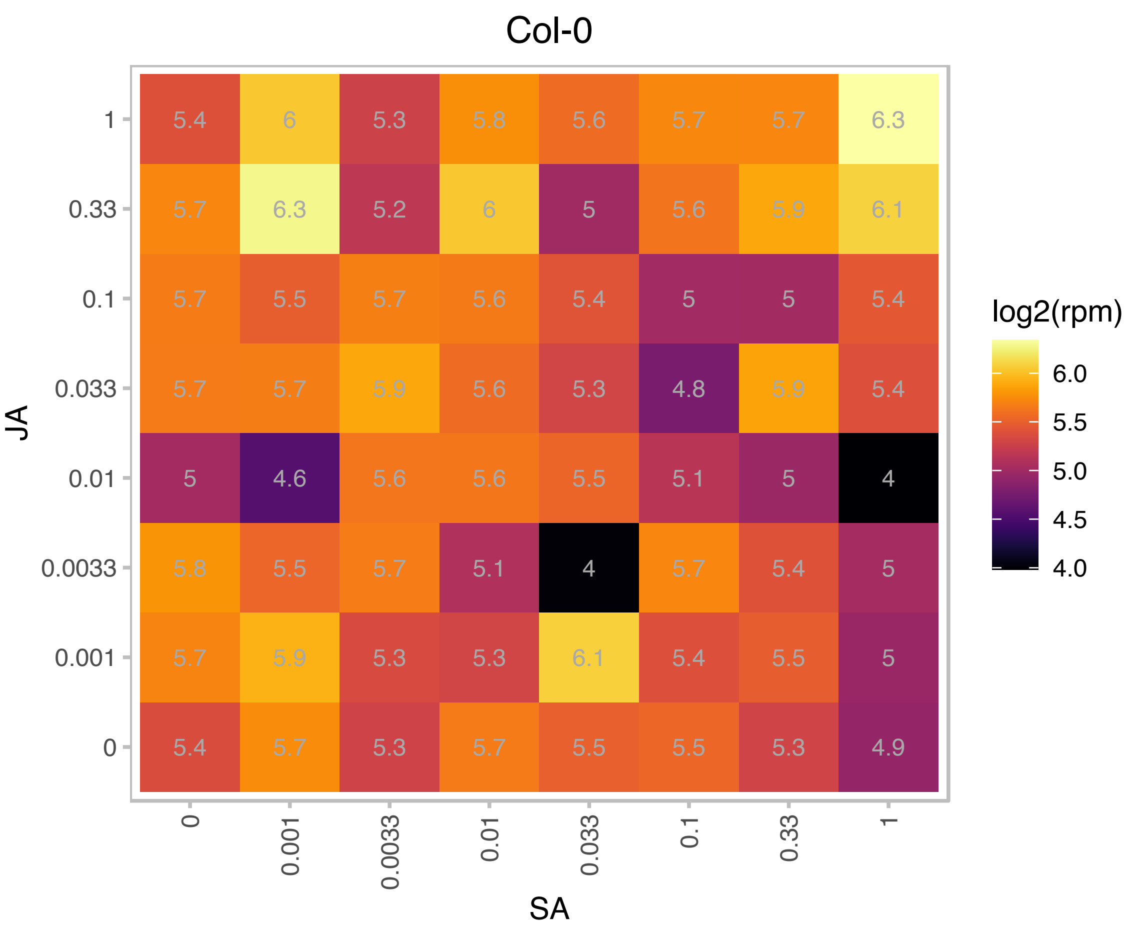


| | |
|---|---|
| Gene | AT2G31040 |
| is.expressed_genes | TRUE |
| is.responsive_genes | TRUE |
| is.apcluster | D-02 |

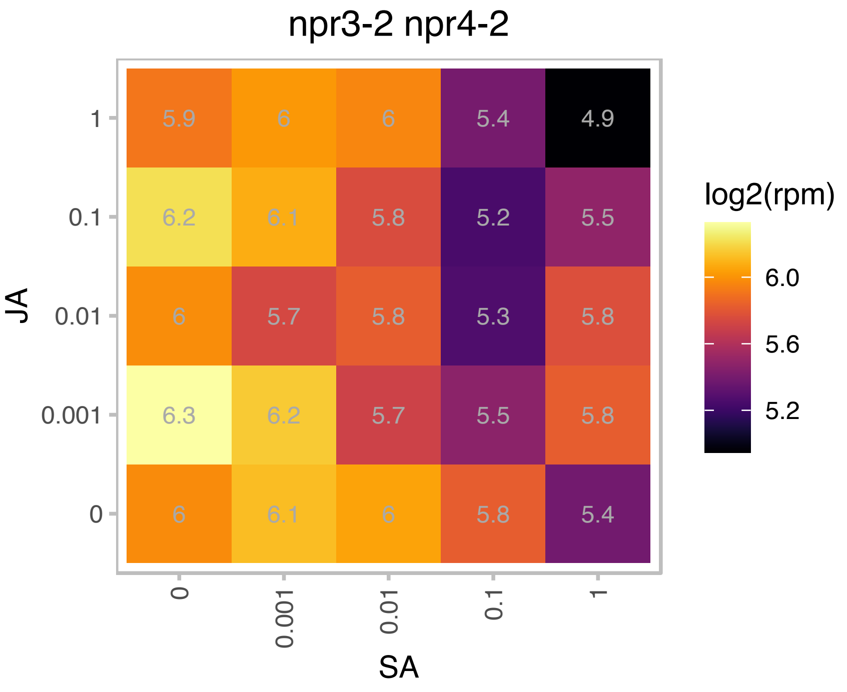


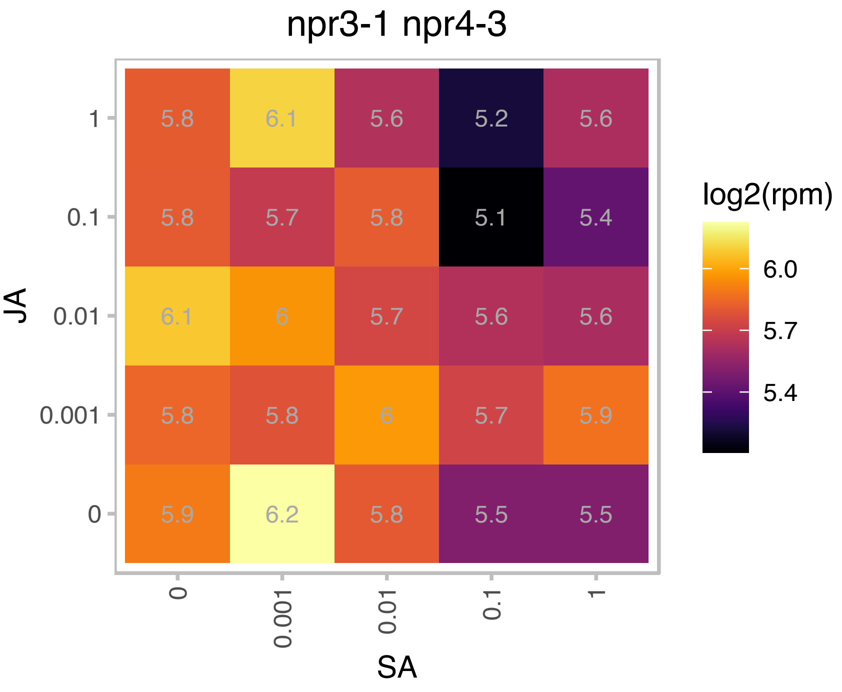

# AT2G34750.1 : NA

RNA polymerase I specific transcription initiation factor RRN3 protein

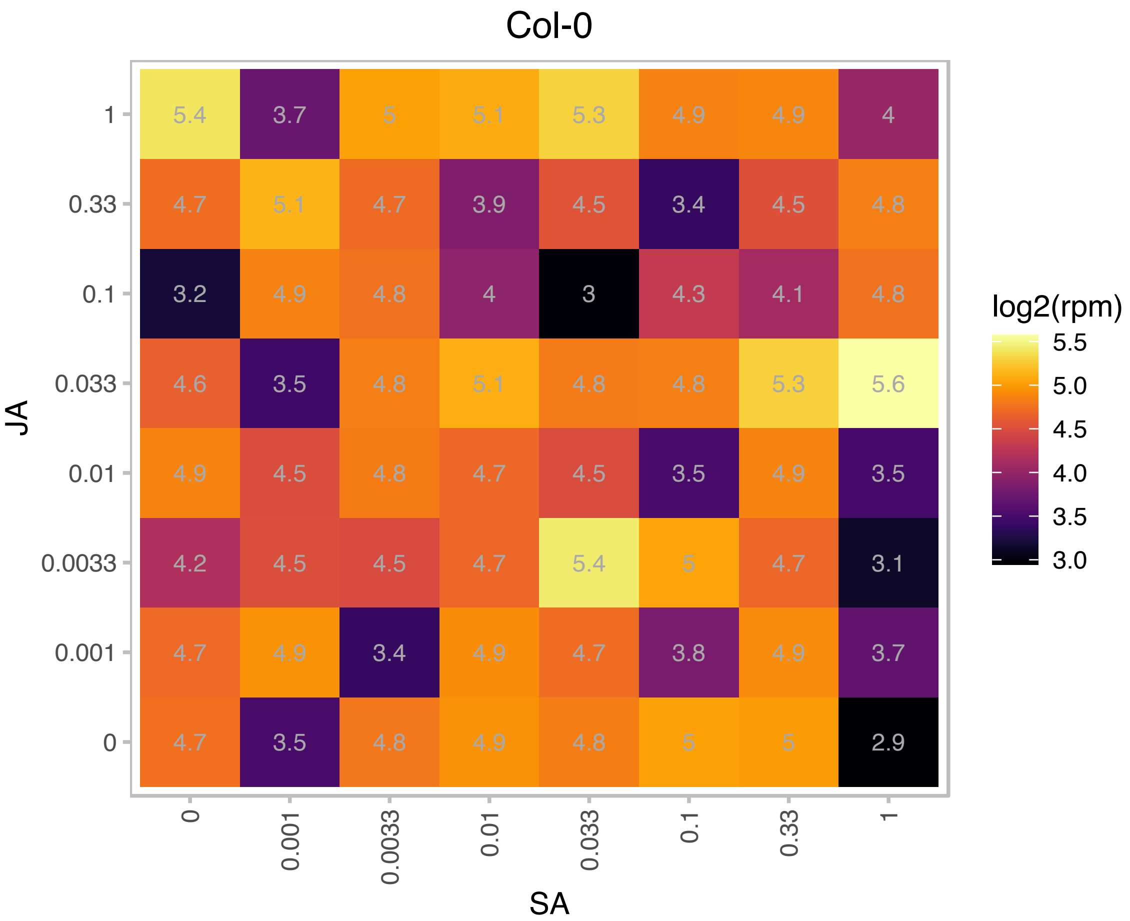


| | |
|---|---|
| Gene | AT2G34750 |
| is.expressed_genes | TRUE |
| is.responsive_genes | TRUE |
| is.apcluster | D-02 |

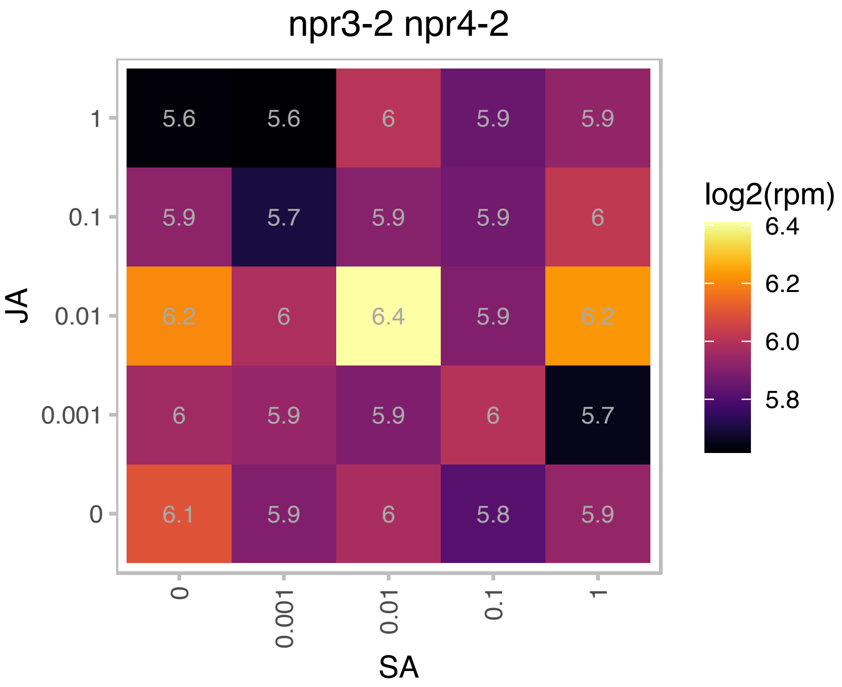


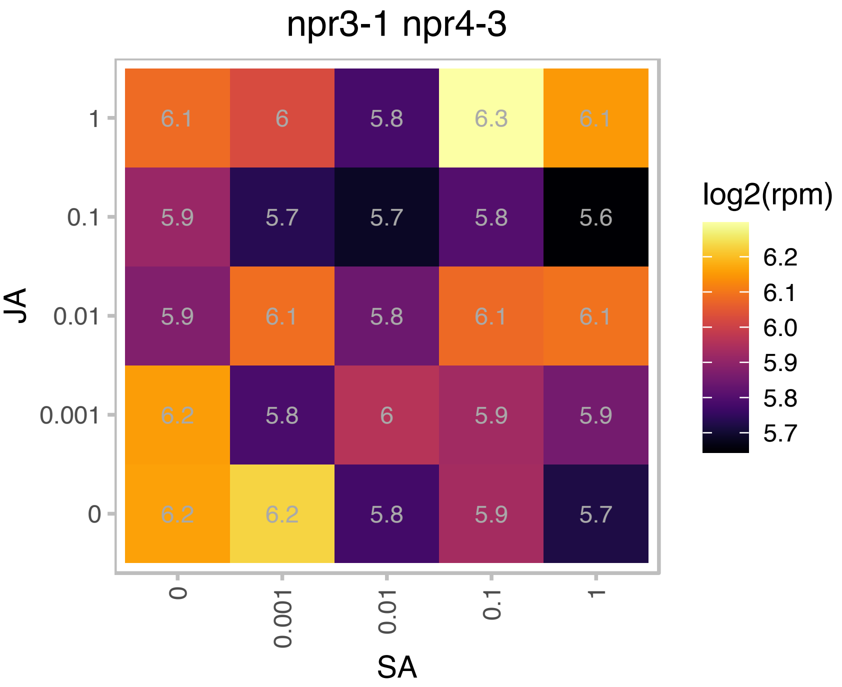

# AT2G37480.1 : NA

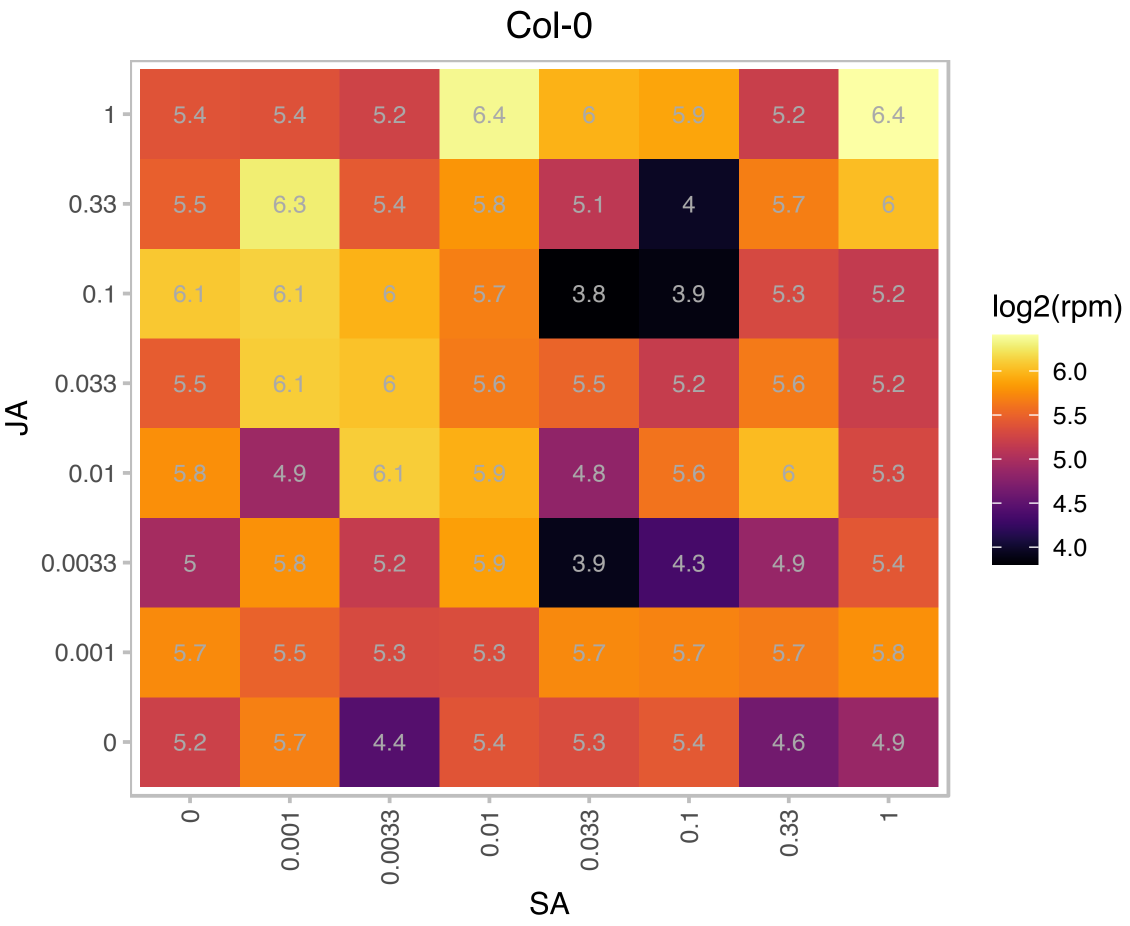


| | |
|---|---|
| Gene | AT2G37480 |
| is.expressed_genes | TRUE |
| is.responsive_genes | TRUE |
| is.apcluster | D-02 |

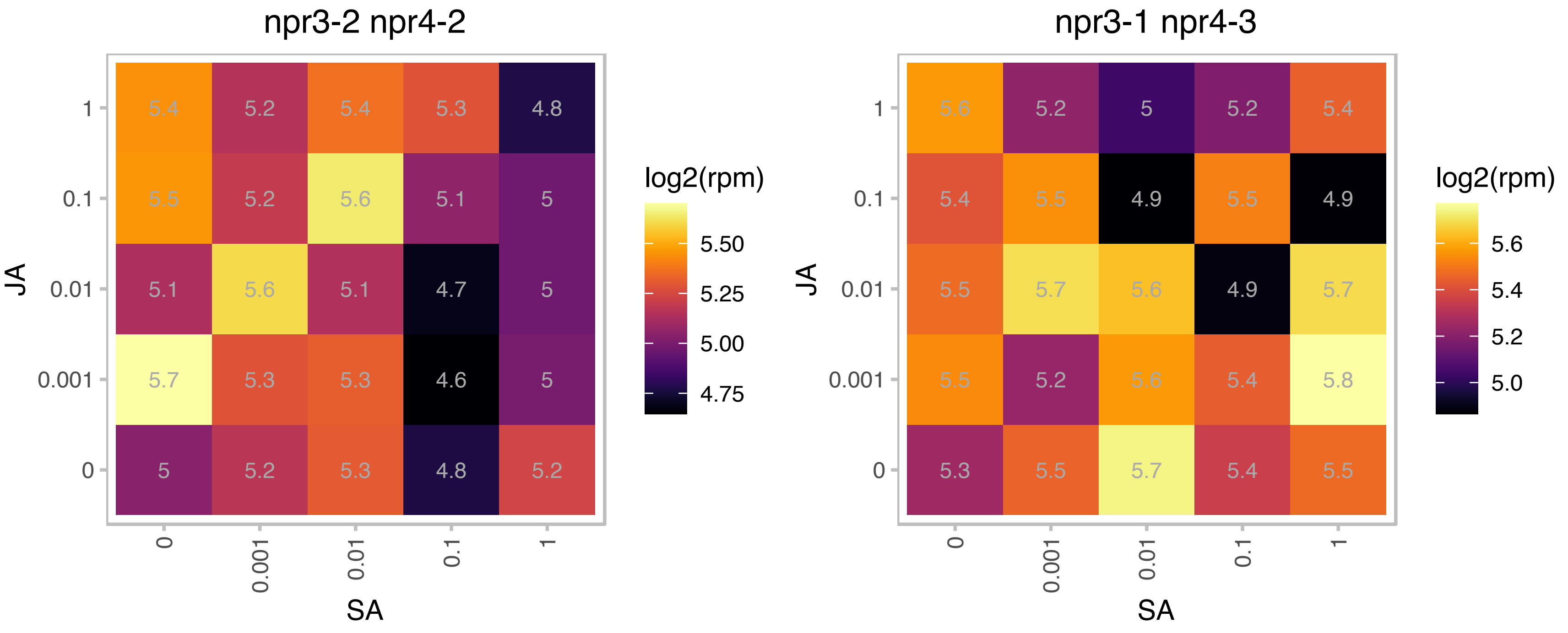

### AT3G04710.1 : TPR10

ankyrin repeat family protein

Encodes one of the 36 carboxylate clamp (CC)-tetratricopeptide repeat (TPR) proteins (Prasad 2010, Pubmed ID: 20856808) with potential to interact with Hsp90/Hsp70 as co-chaperones.

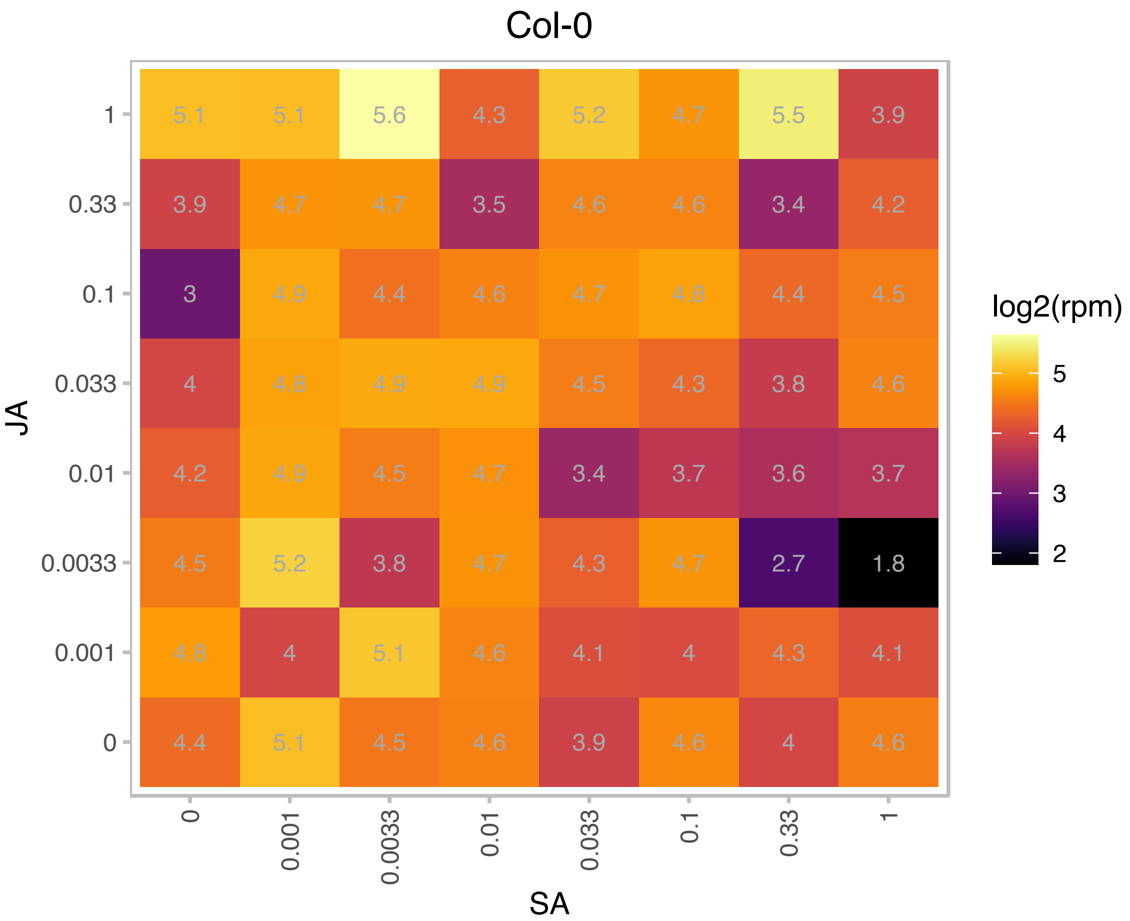


| | |
|---|---|
| Gene | AT3G04710 |
| is.expressed_genes | TRUE |
| is.responsive_genes | TRUE |
| is.apcluster | D-02 |

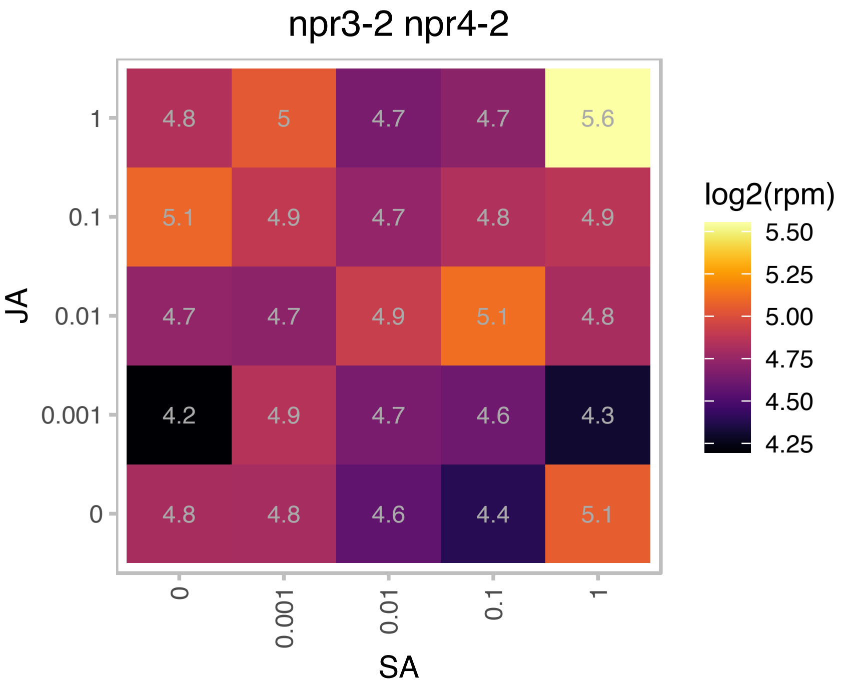


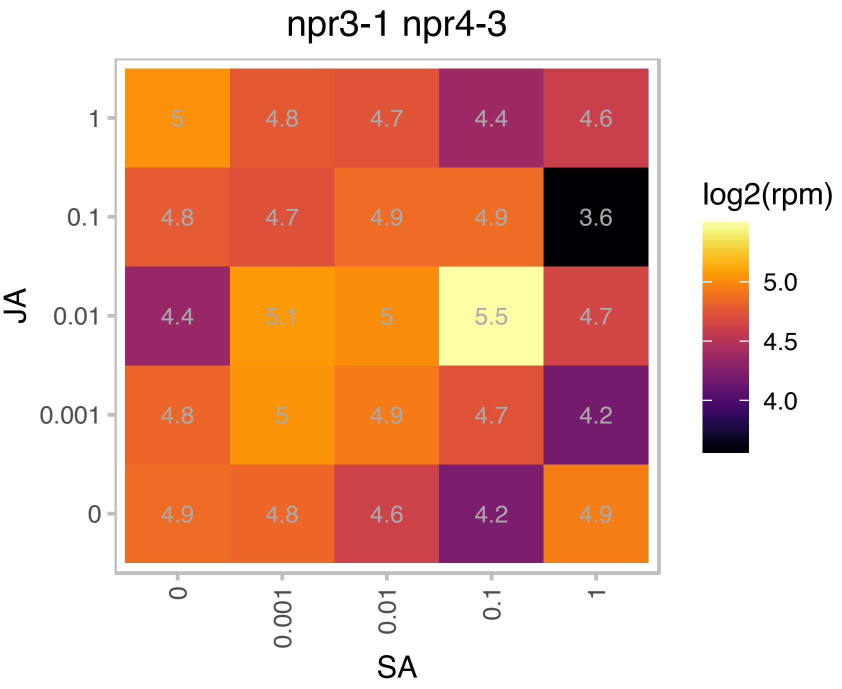

# AT3G32980.1 : NA

Peroxidase superfamily protein

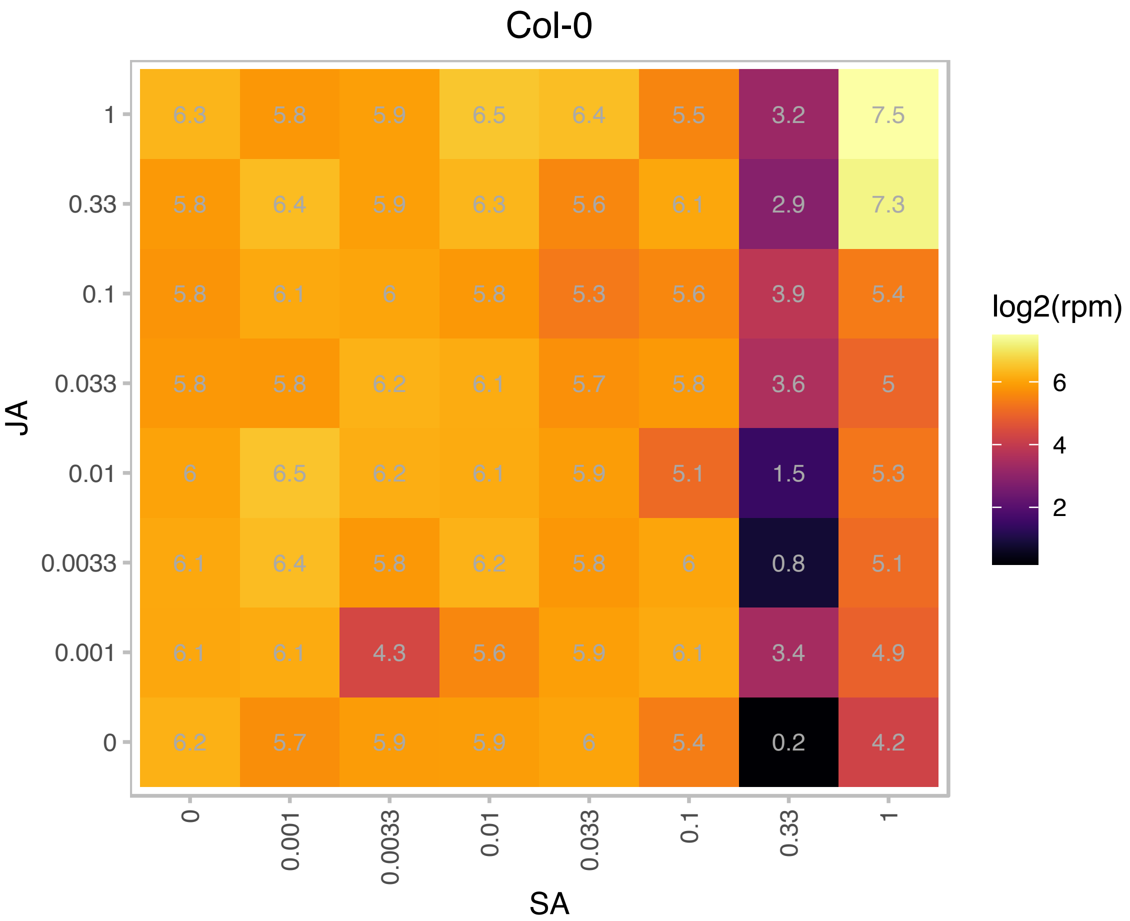


| | |
|---|---|
| Gene | AT3G32980 |
| is.expressed_genes | TRUE |
| is.responsive_genes | TRUE |
| is.apcluster | D-02 |

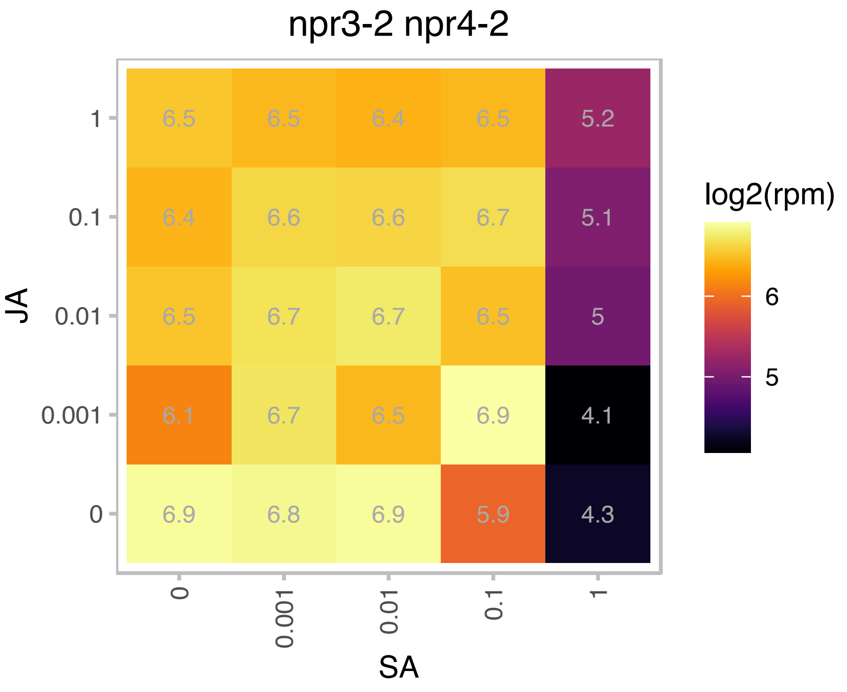


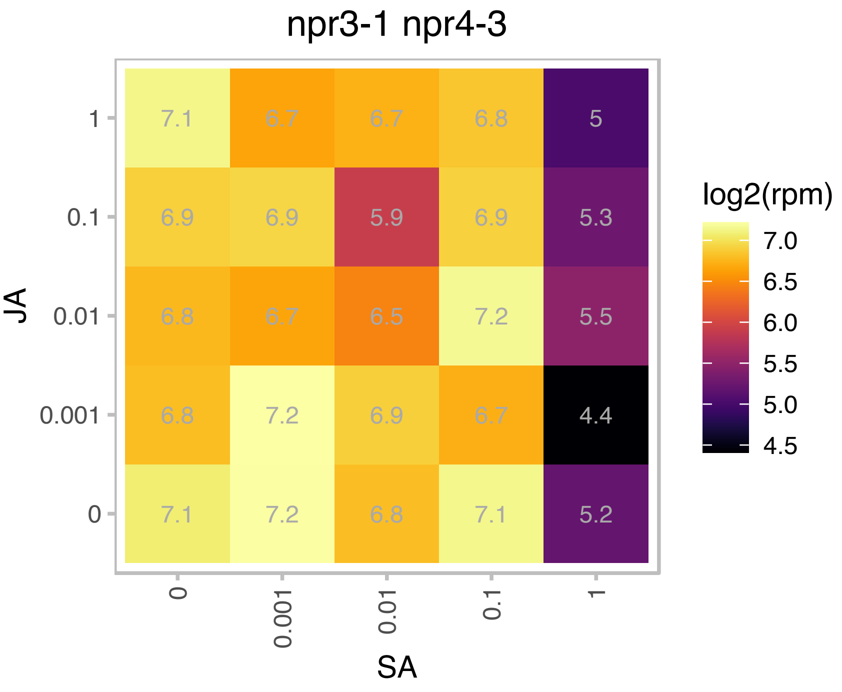

### AT3G48560.1 : AHAS, ALS, CSR1, IMR1, TZP5

chlorsulfuron/imidazolinone resistant 1

Catalyzes the formation of acetolactate from pyruvate, the first step in valine and isoleucine biosynthesis. Requires FAD, thiamine pyrophosphate and Mg. Inhibited by the sulphonylurea herbicide, chlorsulphuron, and the imidazolinone herbicide, imazapyr. The obtained crystal structure of acetohydroxyacid synthase AHAS, EC 2.2.1.6)in complex with herbicides of the sulphonylurea and imidazolinone family reveals the molecular basis for substrate/inhibitor binding.

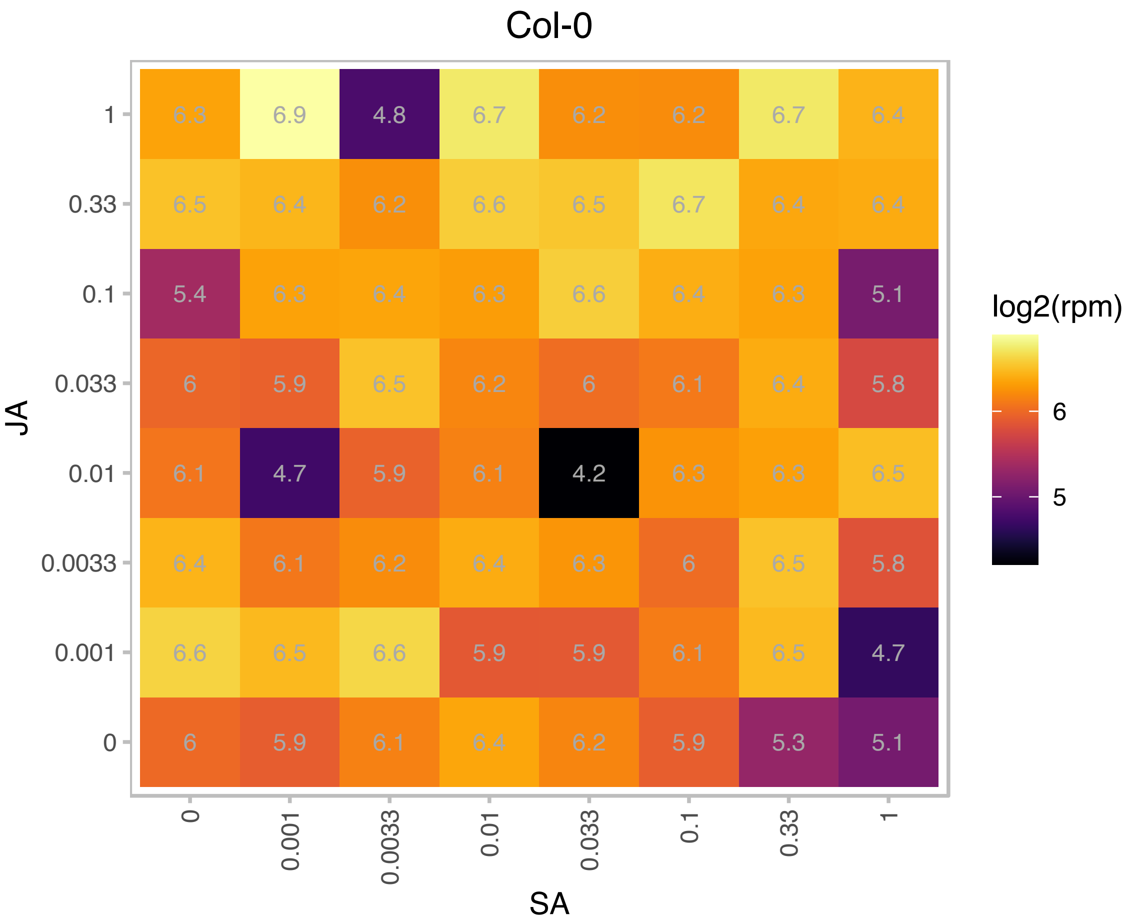


| | |
|---|---|
| Gene | AT3G48560 |
| is.expressed_genes | TRUE |
| is.responsive_genes | TRUE |
| is.apcluster | D-02 |

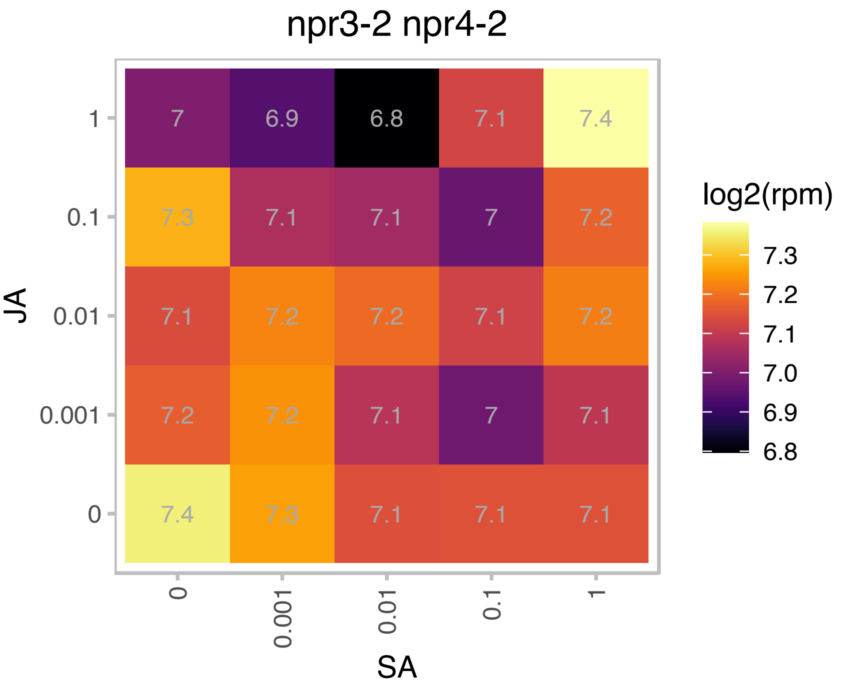


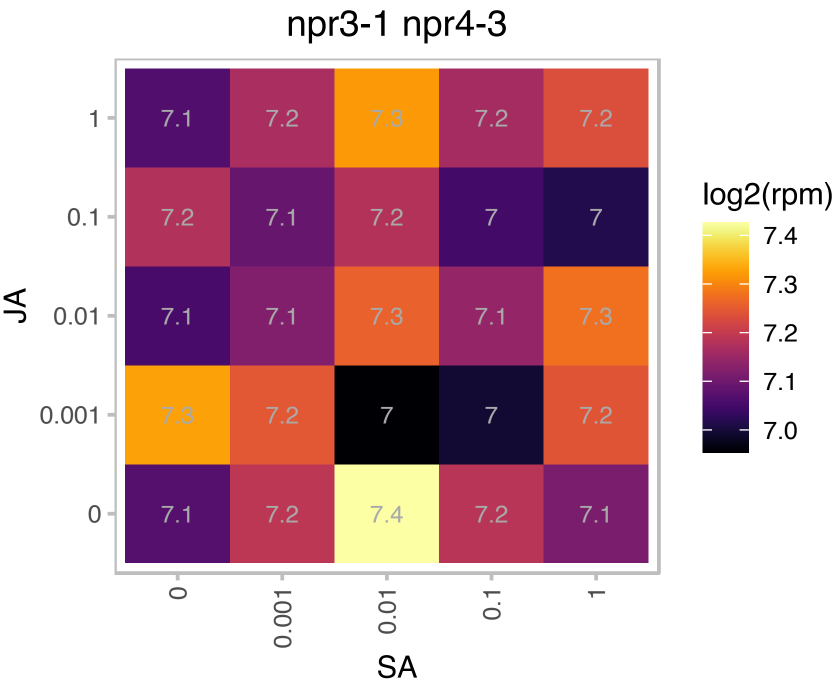

# AT3G56705.1 : U2.6

U2.6; snRNA

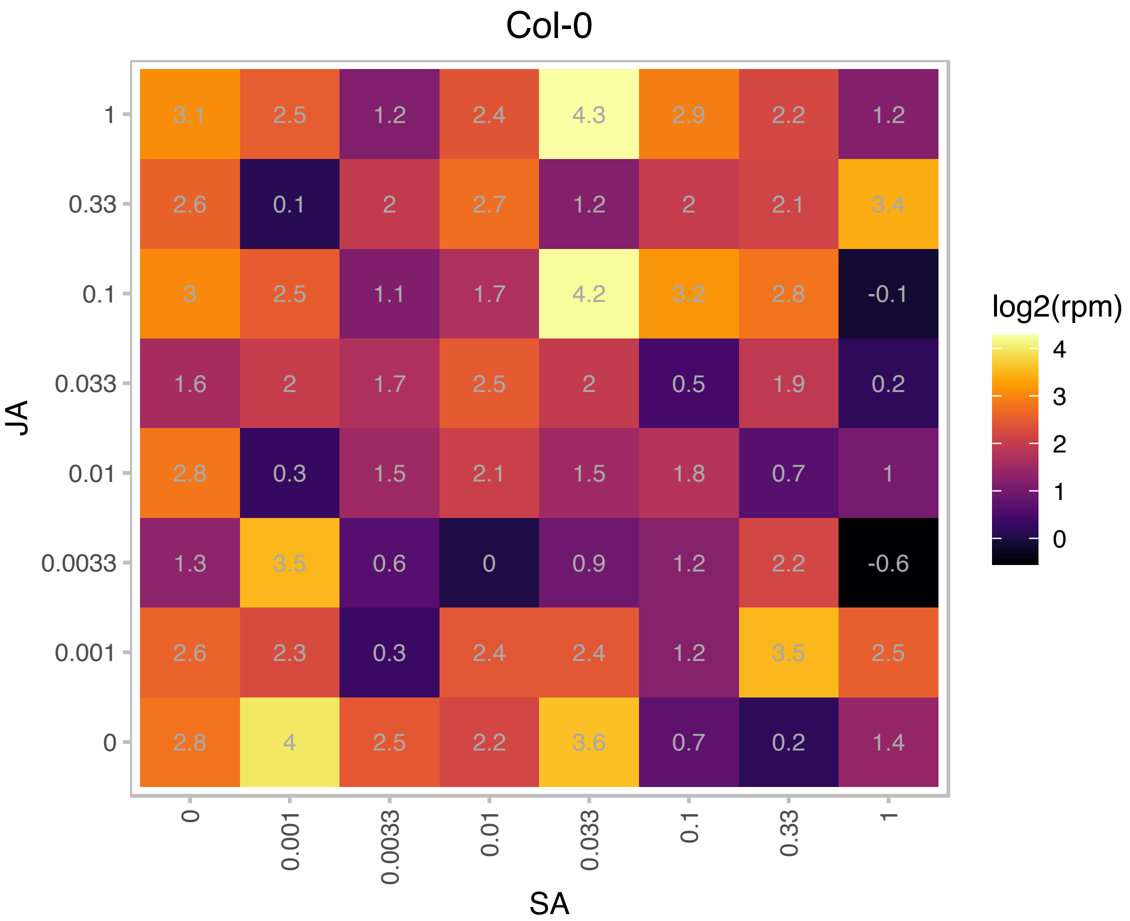


| | |
|---|---|
| Gene | AT3G56705 |
| is.expressed_genes | TRUE |
| is.responsive_genes | TRUE |
| is.apcluster | D-02 |

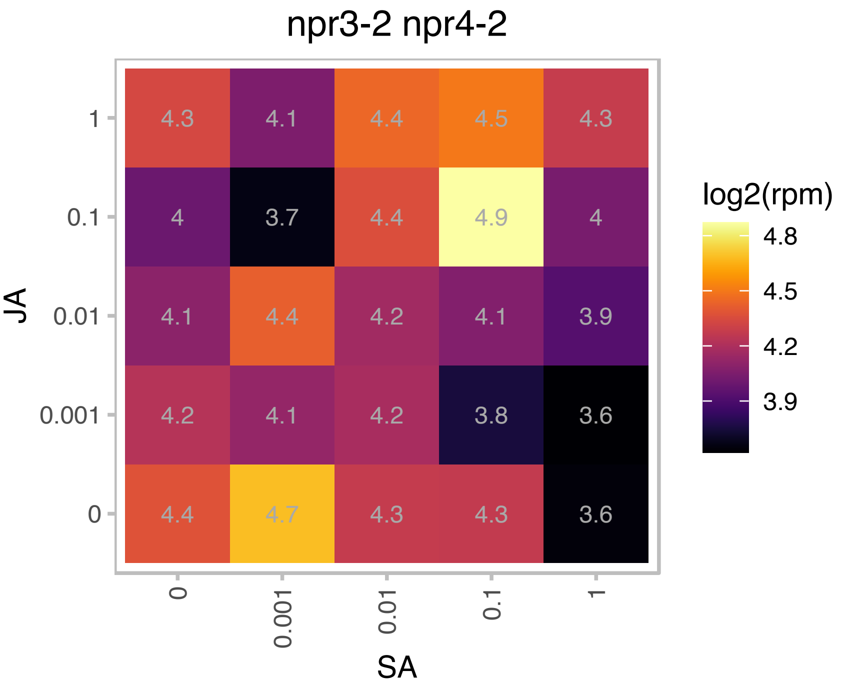


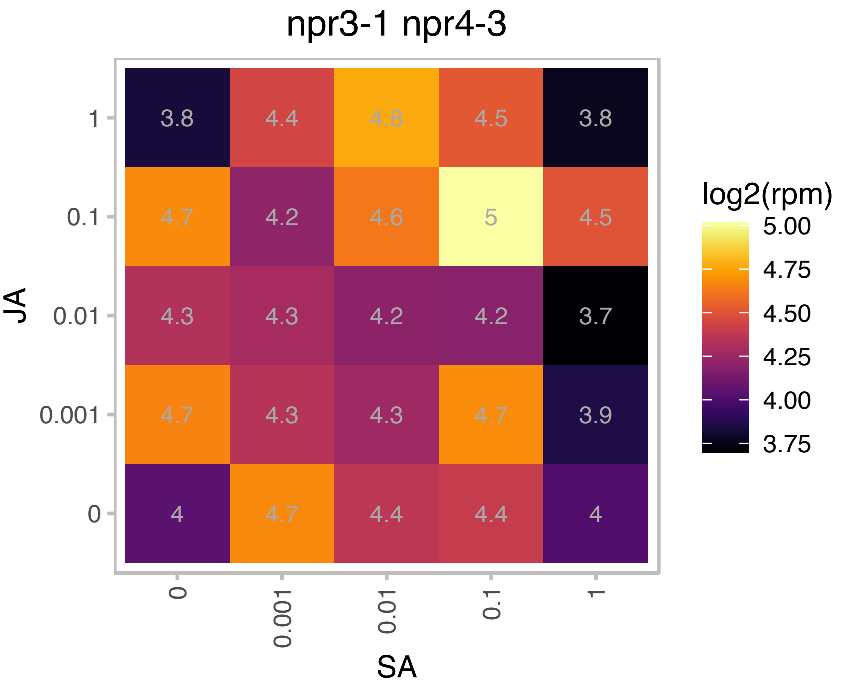

### AT3G59940.1 : AtKFB50, KFB50, KMD4

Encodes a member of a family of F-box proteins, called the KISS ME DEADLY (KMD) family, that targets type-B ARR proteins for degradation and is involved in the negative regulation of the cytokinin response. Also named as KFB50, a member of a group of Kelch repeat F-box proteins that negatively regulate phenylpropanoid biosynthesis by targeting the phenypropanoid biosynthesis enzyme phenylalanine ammonia-lyase. The mRNA is cell-to-cell mobile.

Galactose oxidase/kelch repeat superfamily protein

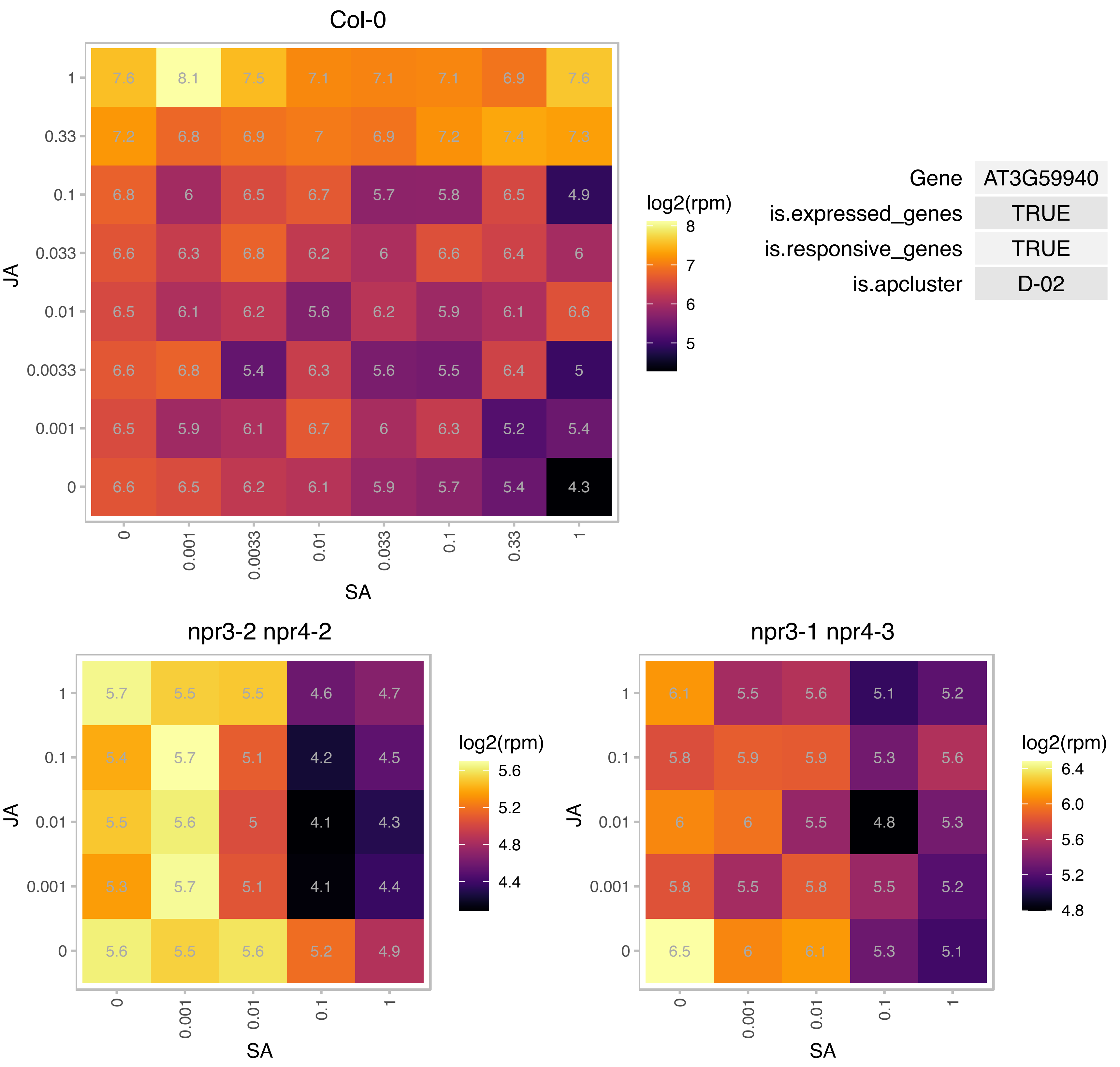


| | |
|---|---|
| Gene | AT3G59940 |
| is.expressed_genes | TRUE |
| is.responsive_genes | TRUE |
| is.apcluster | D-02 |

# AT4G02400.1 : NA

U3 ribonucleoprotein (Utp) family protein

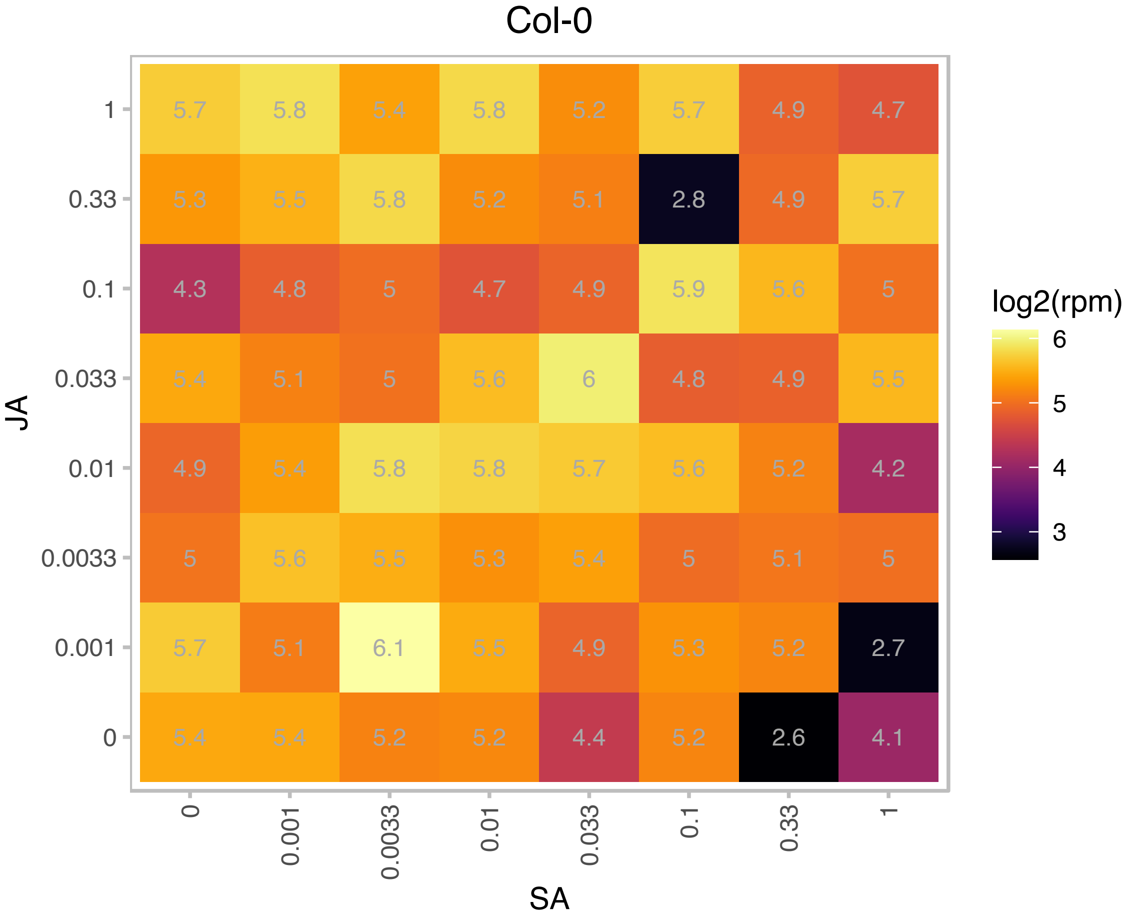


| | |
|---|---|
| Gene | AT4G02400 |
| is.expressed_genes | TRUE |
| is.responsive_genes | TRUE |
| is.apcluster | D-02 |

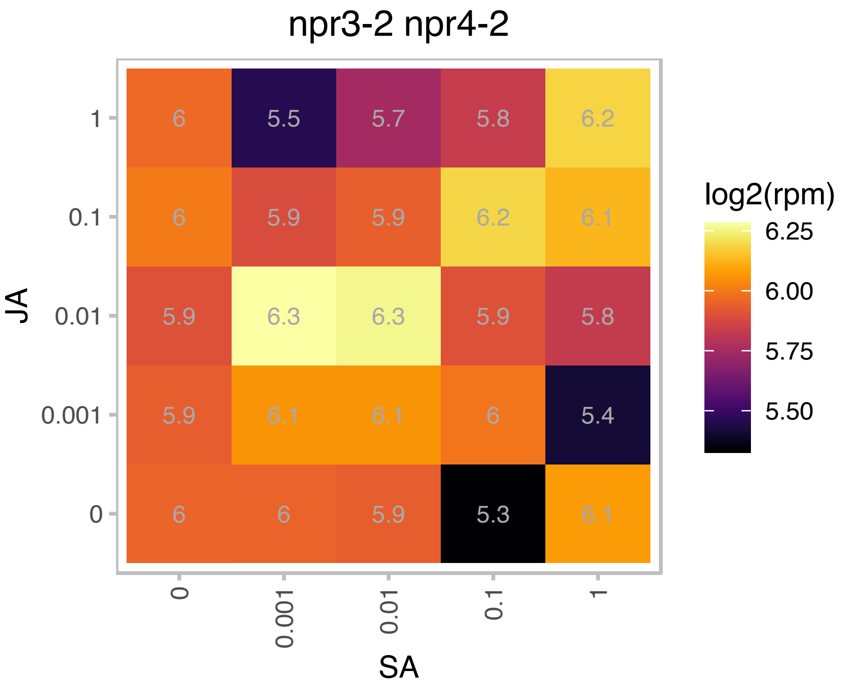


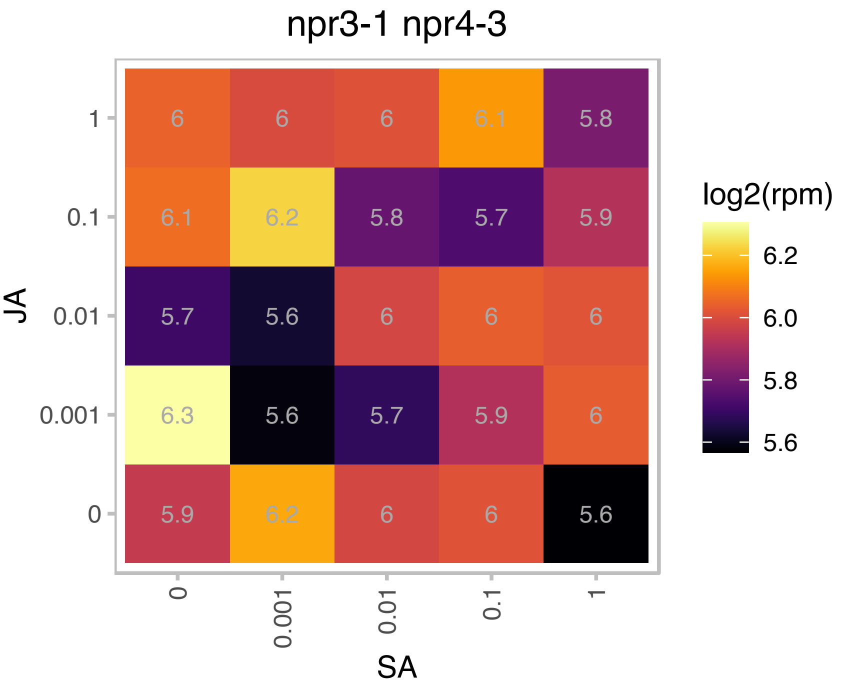

### AT4G13250.1 : NYC1

NAD(P)-binding Rossmann-fold superfamily protein

Encodes a chlorophyll b reductase involved in the degradation of chlorophyll b and LHCII (light harvesting complex II).

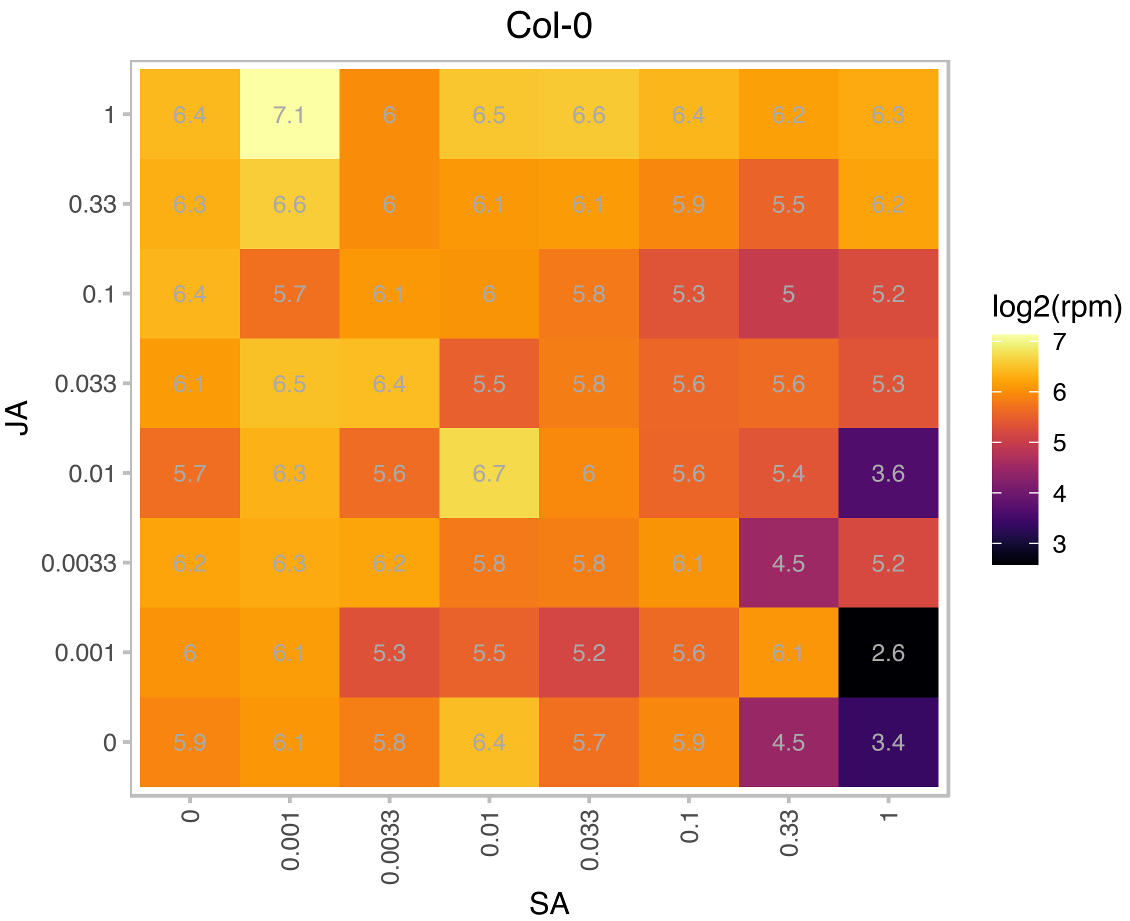


| | |
|---|---|
| Gene | AT4G13250 |
| is.expressed_genes | TRUE |
| is.responsive_genes | TRUE |
| is.apcluster | D-02 |

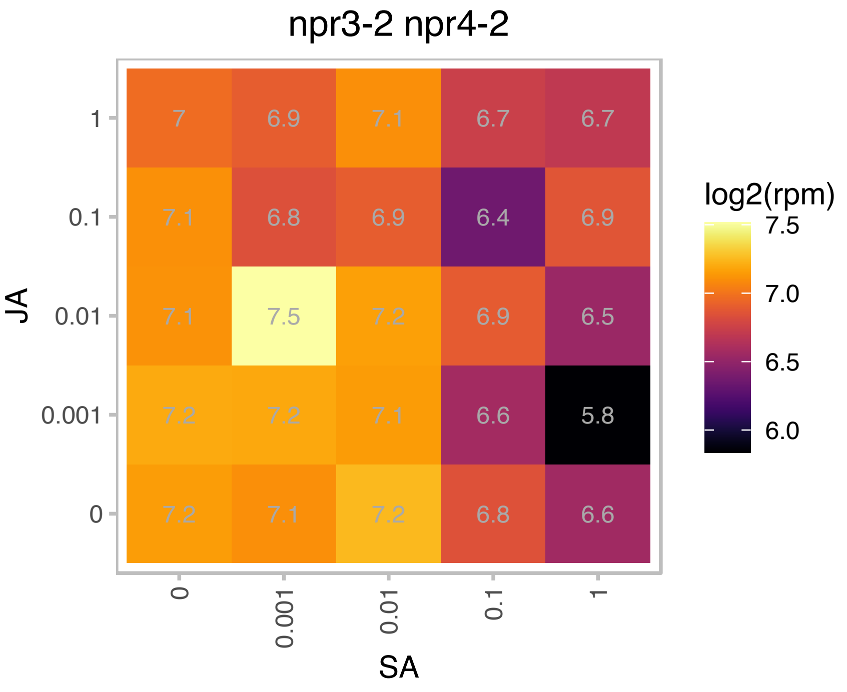


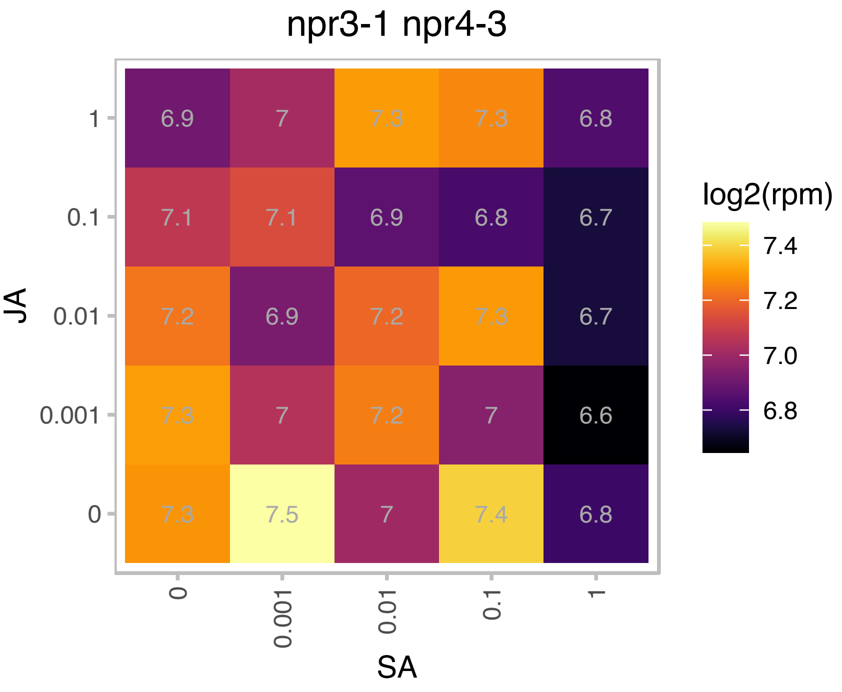

### AT4G30890.1 : UBP24

ubiquitin-specific protease 24

Encodes a ubiquitin-specific protease.

| | |
|---|---|
| Gene | AT4G30890 |
| is.expressed_genes | TRUE |
| is.responsive_genes | TRUE |
| is.apcluster | D-02 |

Col-0

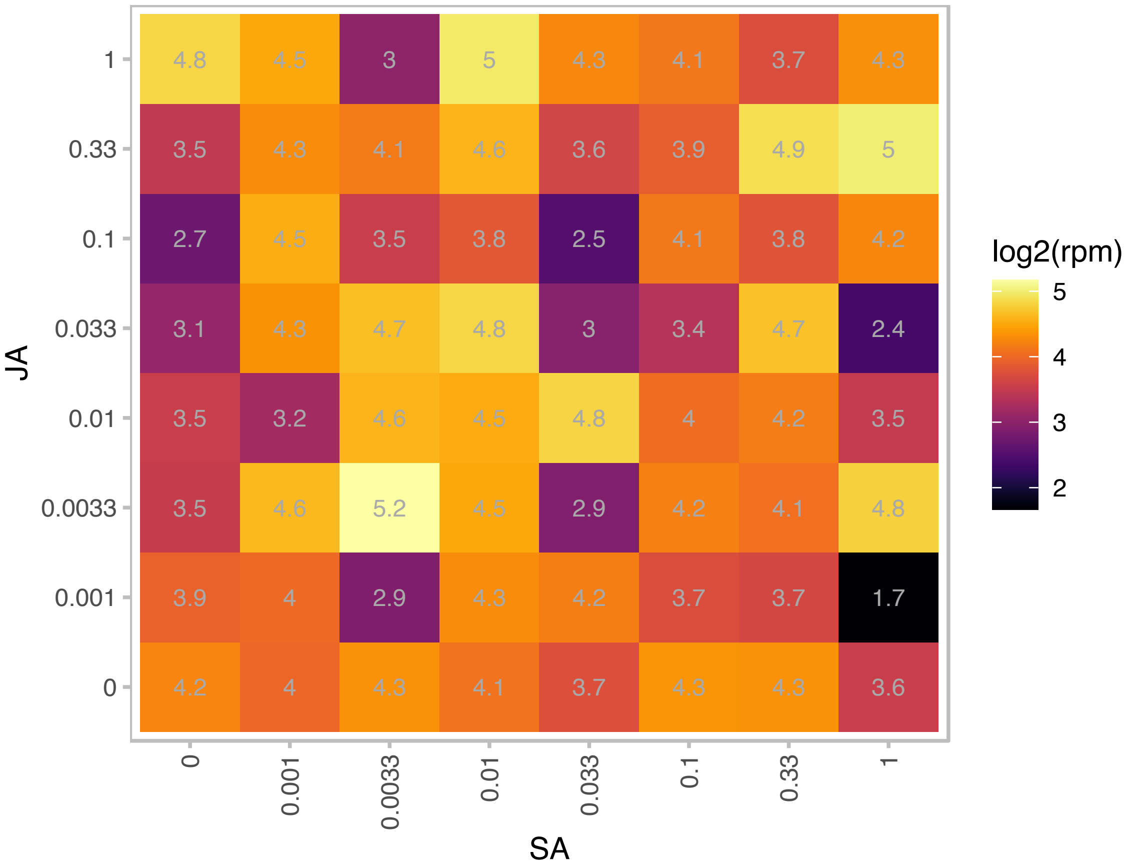


npr3-2 npr4-2

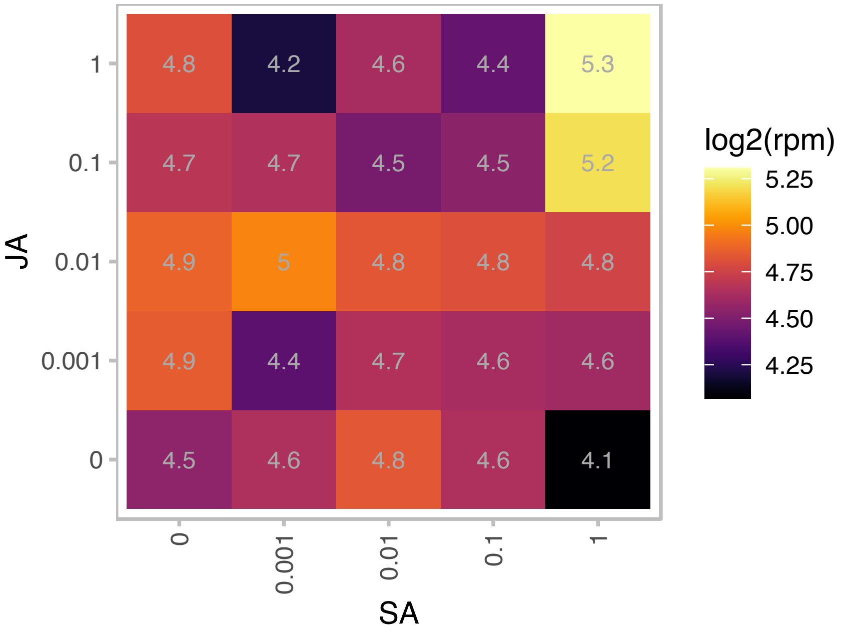


npr3-1 npr4-3

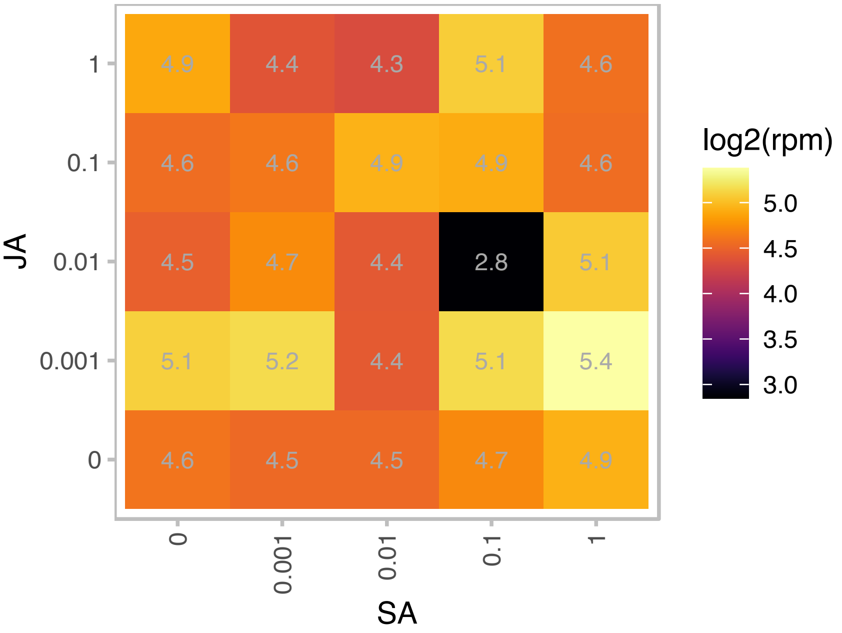

# AT4G37610.1 : BT5

BTB and TAZ domain protein 5

BTB and TAZ domain protein. Located in cytoplasm and expressed in fruit, flower and leaves.

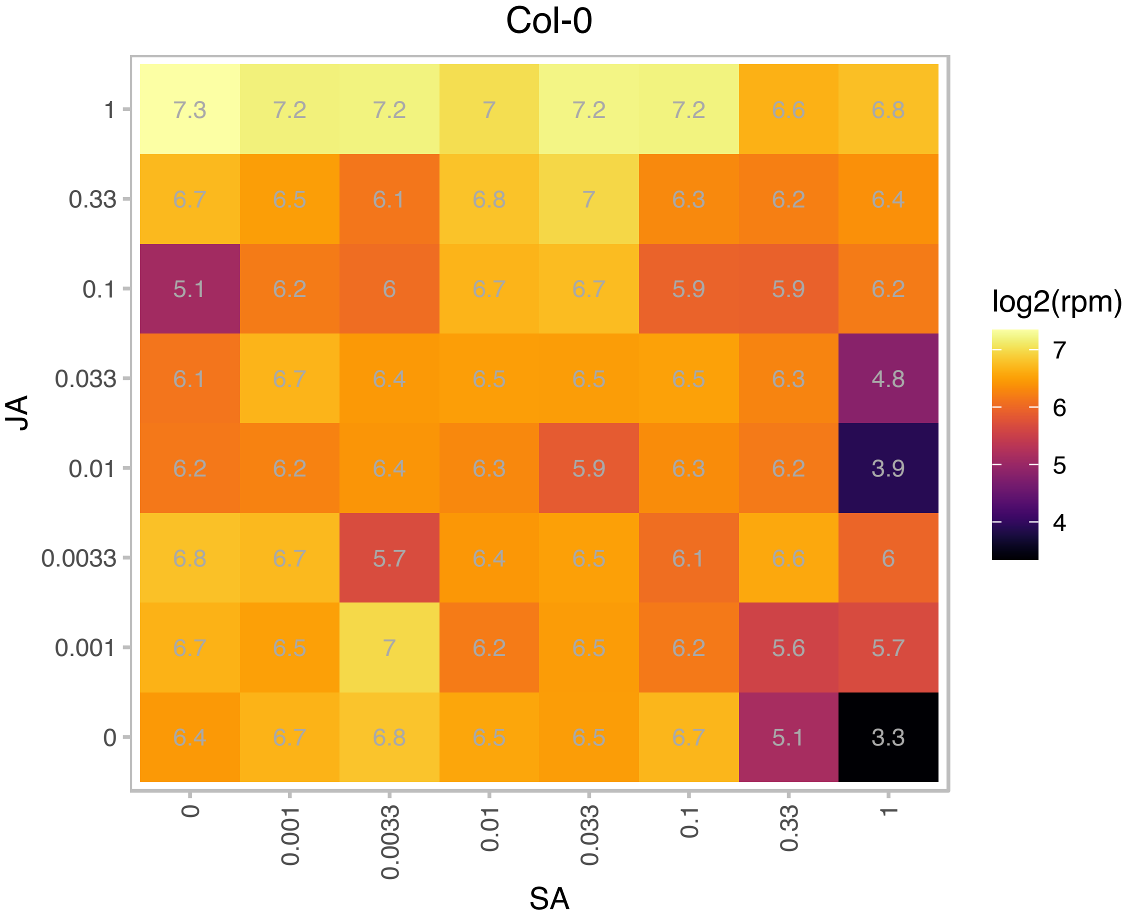


| | |
|---|---|
| Gene | AT4G37610 |
| is.expressed_genes | TRUE |
| is.responsive_genes | TRUE |
| is.apcluster | D-02 |

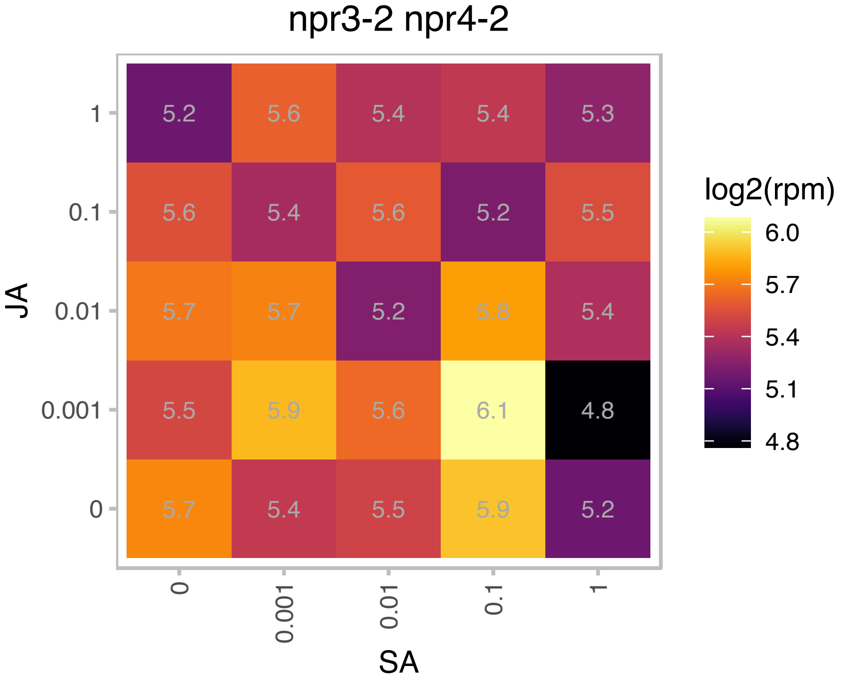


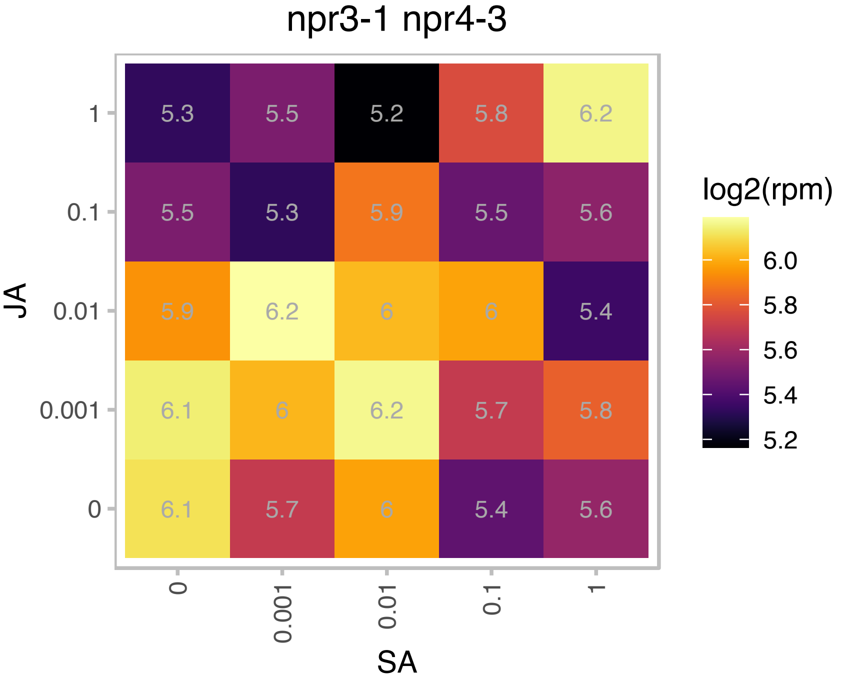

### AT5G05580.1 : AtFAD8, FAD8

fatty acid desaturase 8

Encodes a temperature sensitive plastidic fatty acid desaturase.

Col-0

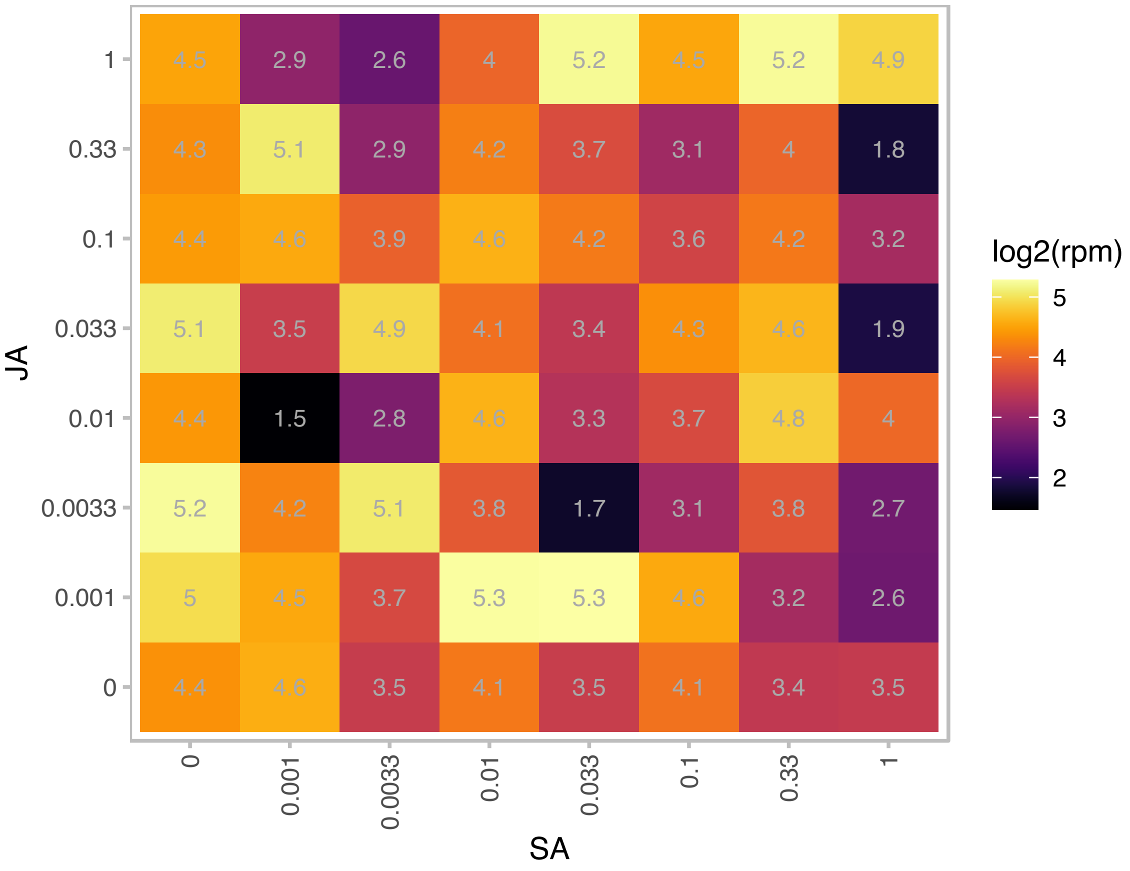


| | |
|---|---|
| Gene | AT5G05580 |
| is.expressed_genes | TRUE |
| is.responsive_genes | TRUE |
| is.apcluster | D-02 |

npr3-2 npr4-2

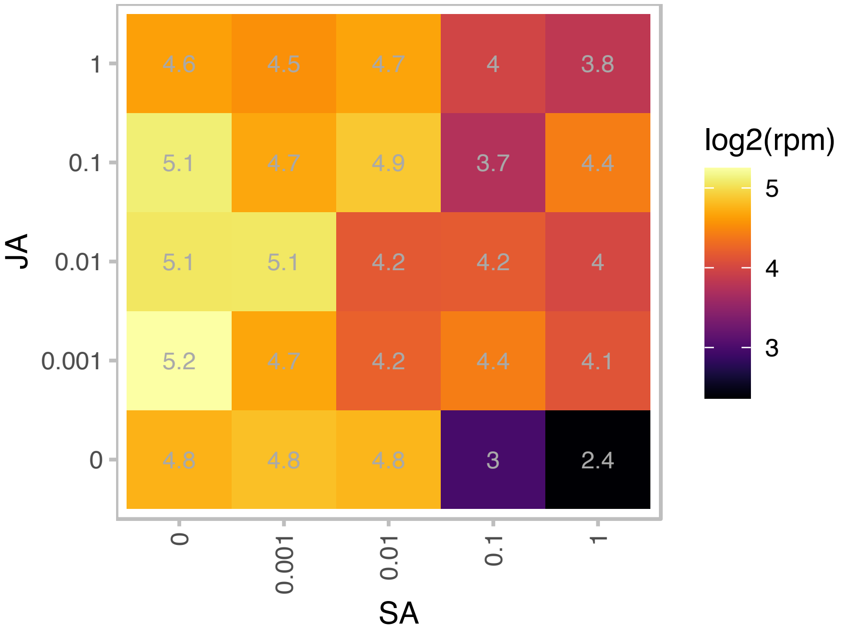


npr3-1 npr4-3

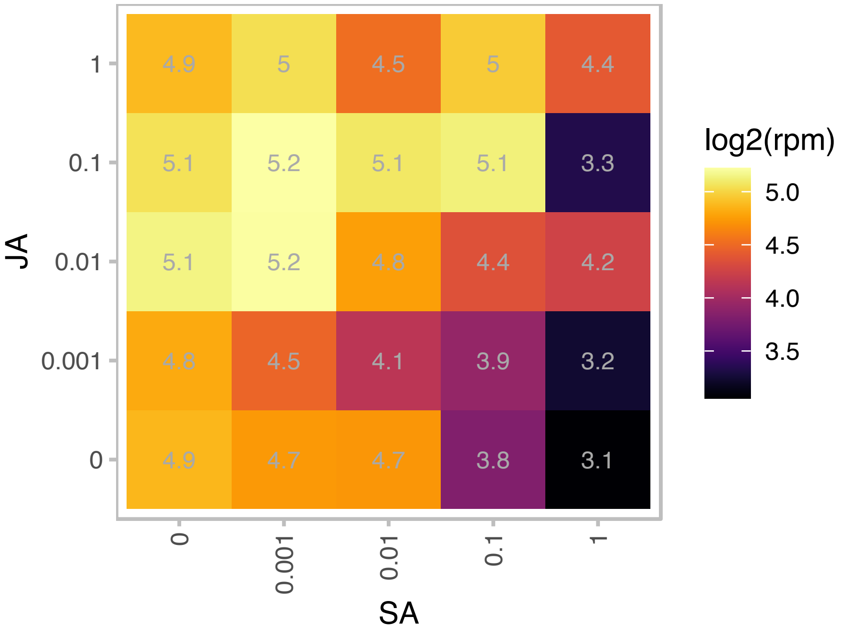

# AT5G09880.1 : NA

Splicing factor, CC1-like

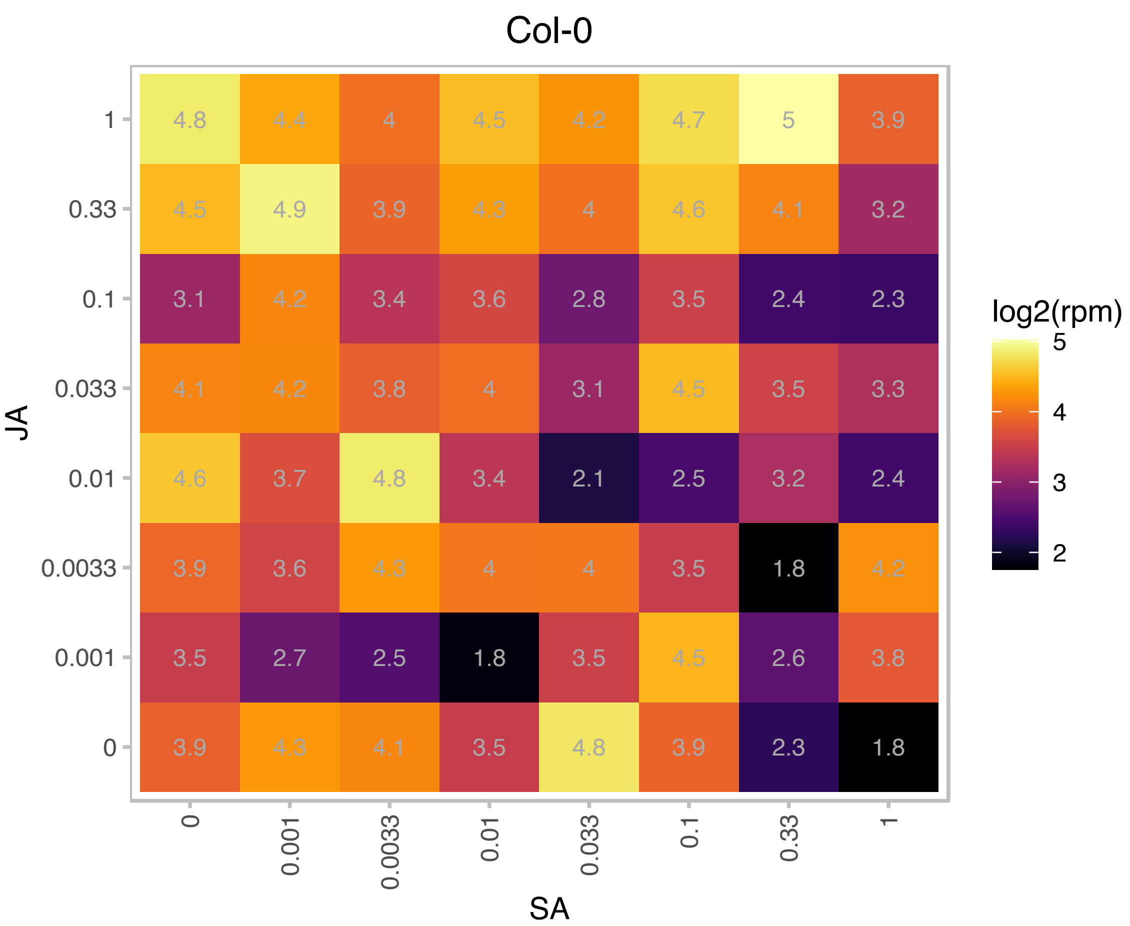


| | |
|---|---|
| Gene | AT5G09880 |
| is.expressed_genes | TRUE |
| is.responsive_genes | TRUE |
| is.apcluster | D-02 |

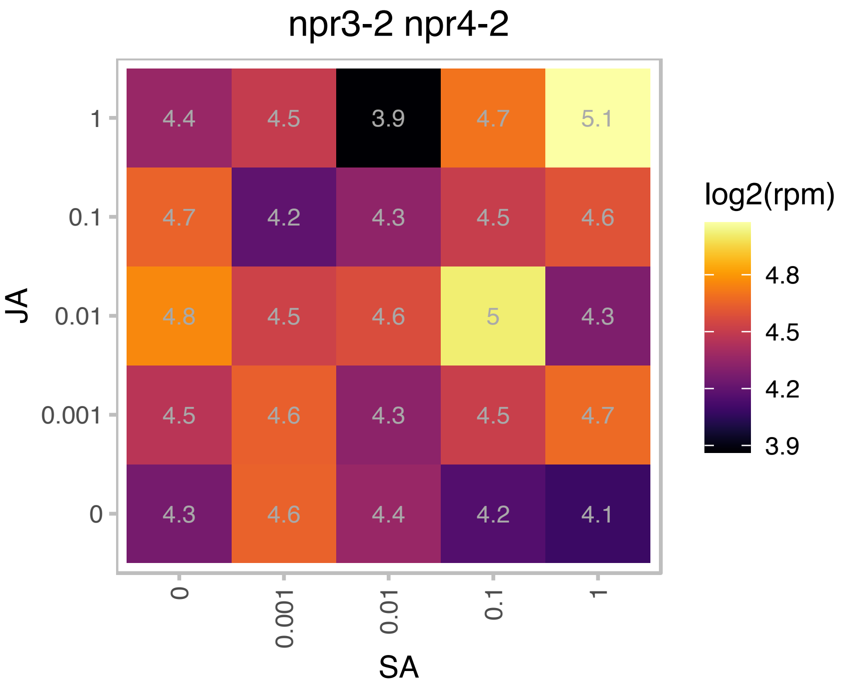


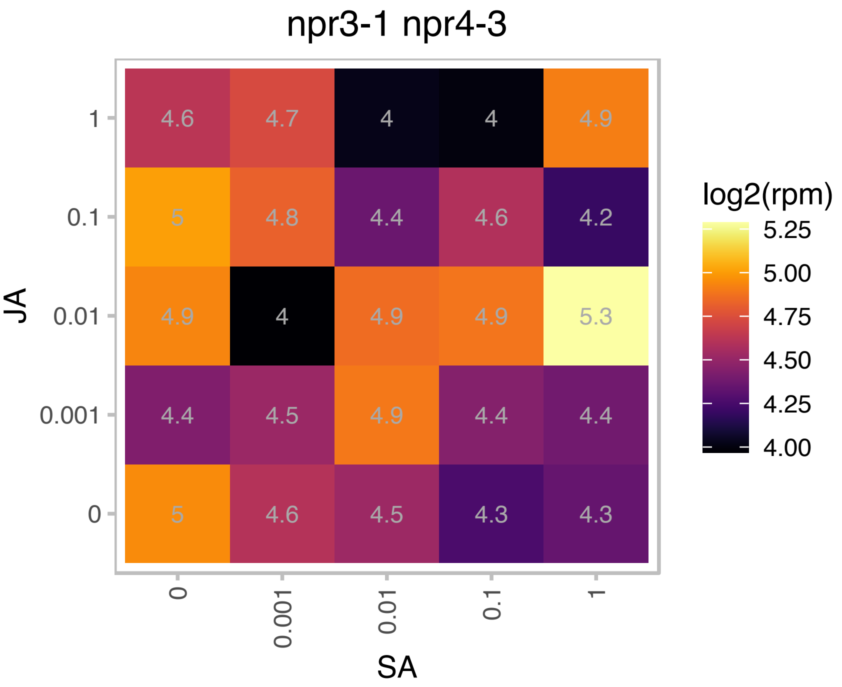

### AT5G11670.1 : ATNADP-ME2, NADP-ME2

NADP-malic enzyme 2
The malic enzyme (EC 1.1.1.40) encoded by AtNADP-ME2 is presumably a cytosolic enzyme involved in malate metabolism and possibly assisting the oxidative pentose phosphate pathway. AtNADP-ME2 counts for the major part of NADP-ME activity in mature tissues of Arabidopsis.

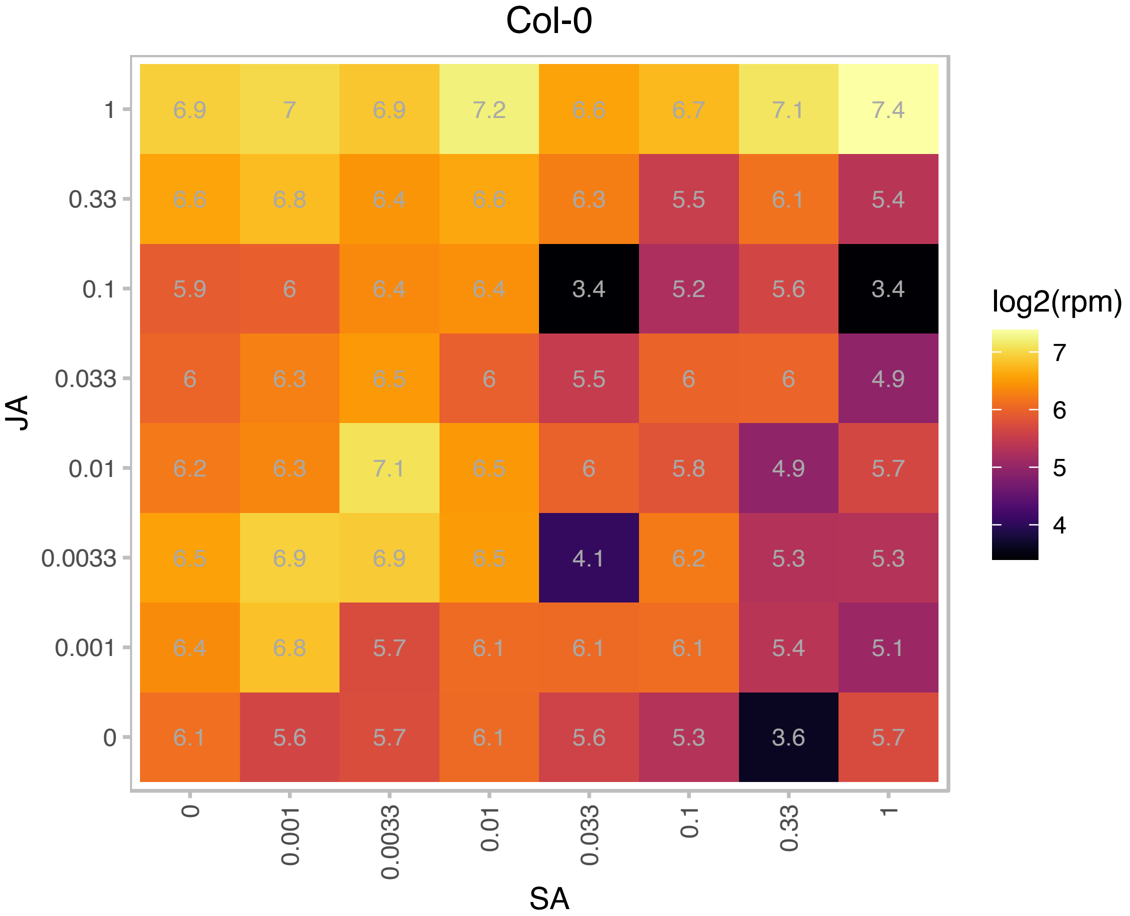


| JA \ SA | 0 | 0.001 | 0.0033 | 0.01 | 0.033 | 0.1 | 0.33 | 1 |
|---|---|---|---|---|---|---|---|---|
| 1 | 6.9 | 7 | 6.9 | 7.2 | 6.6 | 6.7 | 7.1 | 7.4 |
| 0.33 | 6.6 | 6.8 | 6.4 | 6.6 | 6.3 | 5.5 | 6.1 | 5.4 |
| 0.1 | 5.9 | 6 | 6.4 | 6.4 | 3.4 | 5.2 | 5.6 | 3.4 |
| 0.033 | 6 | 6.3 | 6.5 | 6 | 5.5 | 6 | 6 | 4.9 |
| 0.01 | 6.2 | 6.3 | 7.1 | 6.5 | 6 | 5.8 | 4.9 | 5.7 |
| 0.0033 | 6.5 | 6.9 | 6.9 | 6.5 | 4.1 | 6.2 | 5.3 | 5.3 |
| 0.001 | 6.4 | 6.8 | 5.7 | 6.1 | 6.1 | 6.1 | 5.4 | 5.1 |
| 0 | 6.1 | 5.6 | 5.7 | 6.1 | 5.6 | 5.3 | 3.6 | 5.7 |

| Gene | AT5G11670 |
|---|---|
| is.expressed_genes | TRUE |
| is.responsive_genes | TRUE |
| is.apcluster | D-02 |

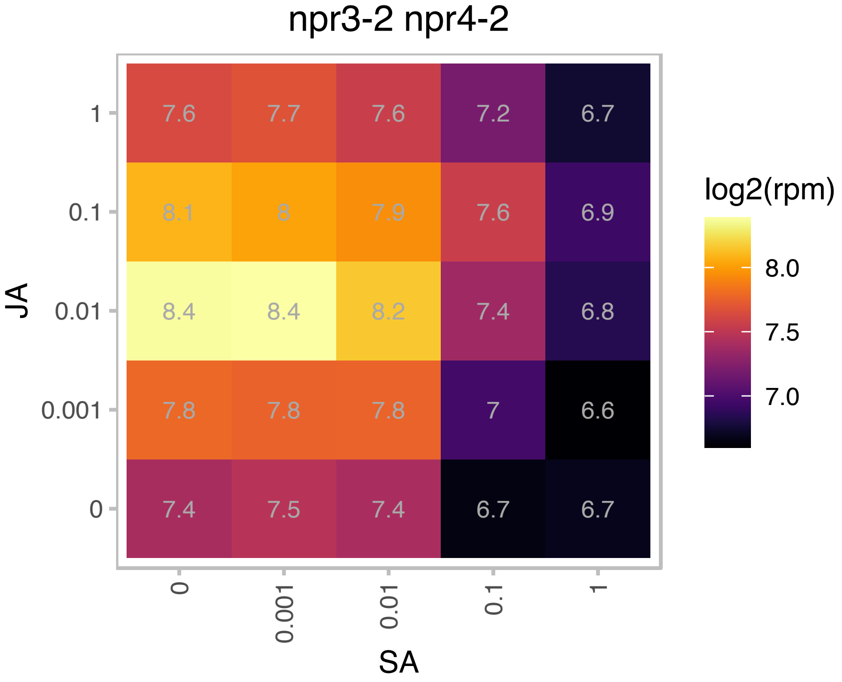


| JA \ SA | 0 | 0.001 | 0.01 | 0.1 | 1 |
|---|---|---|---|---|---|
| 1 | 7.6 | 7.7 | 7.6 | 7.2 | 6.7 |
| 0.1 | 8.1 | 8 | 7.9 | 7.6 | 6.9 |
| 0.01 | 8.4 | 8.4 | 8.2 | 7.4 | 6.8 |
| 0.001 | 7.8 | 7.8 | 7.8 | 7 | 6.6 |
| 0 | 7.4 | 7.5 | 7.4 | 6.7 | 6.7 |

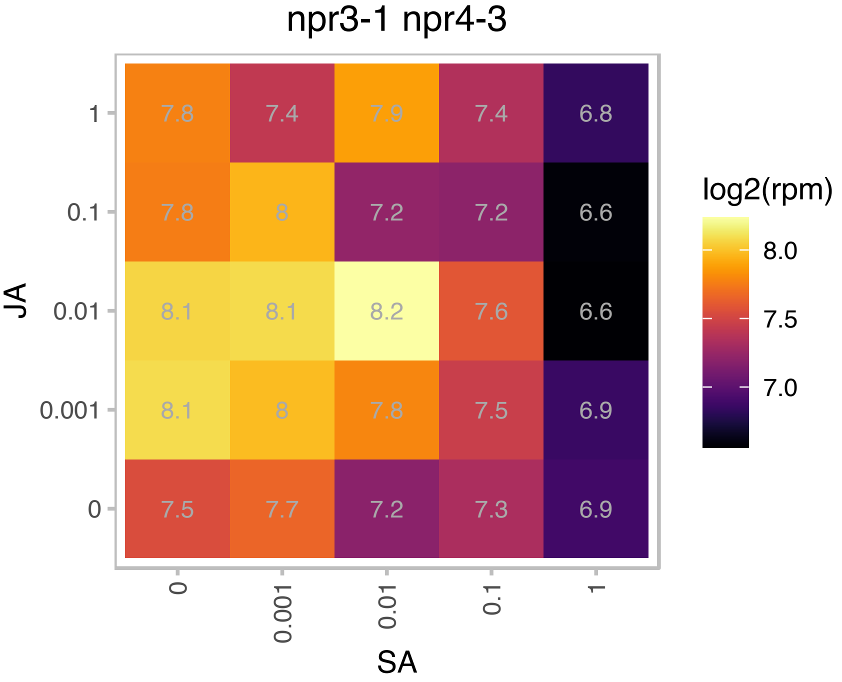


| JA \ SA | 0 | 0.001 | 0.01 | 0.1 | 1 |
|---|---|---|---|---|---|
| 1 | 7.8 | 7.4 | 7.9 | 7.4 | 6.8 |
| 0.1 | 7.8 | 8 | 7.2 | 7.2 | 6.6 |
| 0.01 | 8.1 | 8.1 | 8.2 | 7.6 | 6.6 |
| 0.001 | 8.1 | 8 | 7.8 | 7.5 | 6.9 |
| 0 | 7.5 | 7.7 | 7.2 | 7.3 | 6.9 |

### AT5G11740.1 : AGP15, ATAGP15

arabinogalactan protein 15

Encodes arabinogalactan protein (AGP15). The mRNA is cell-to-cell mobile.

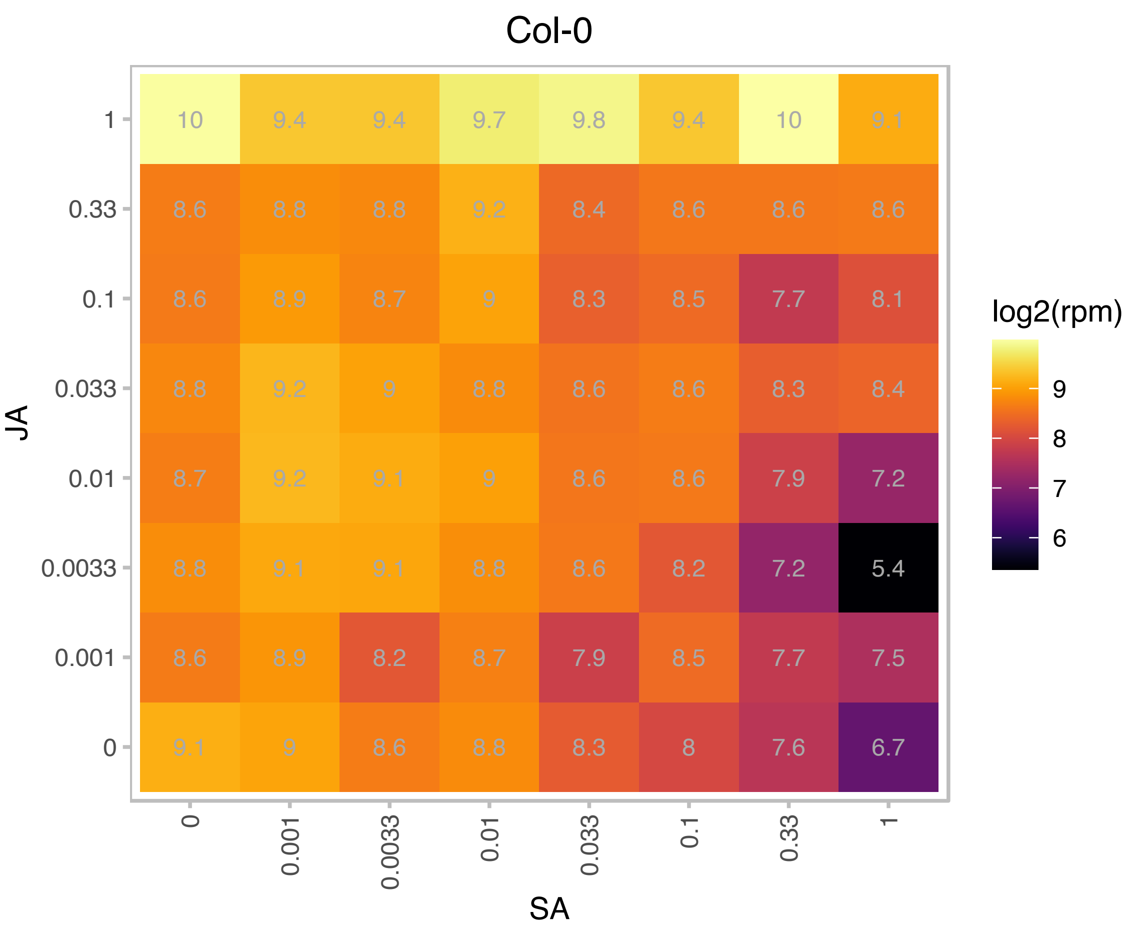


| | |
|---|---|
| Gene | AT5G11740 |
| is.expressed_genes | TRUE |
| is.responsive_genes | TRUE |
| is.apcluster | D-02 |

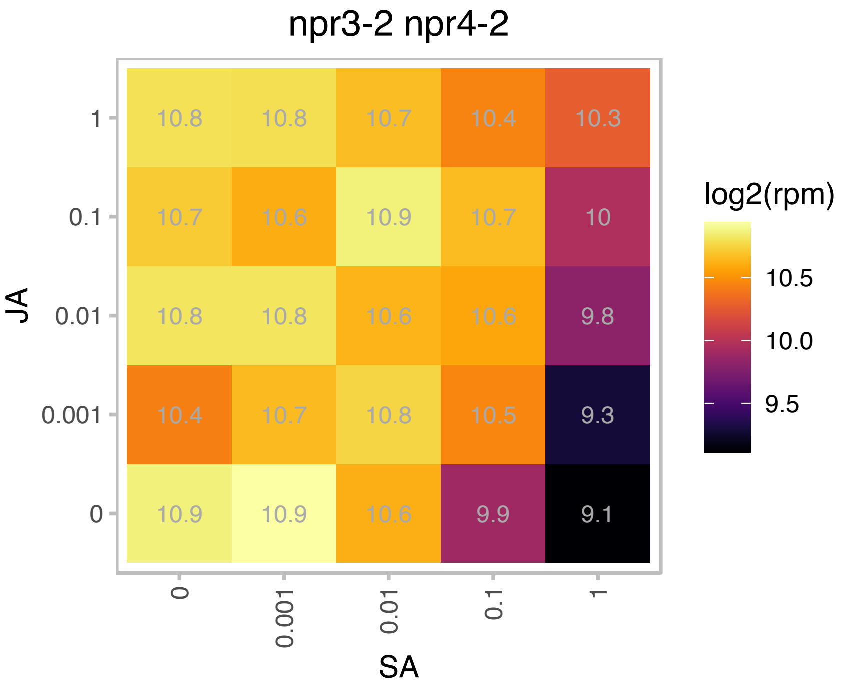


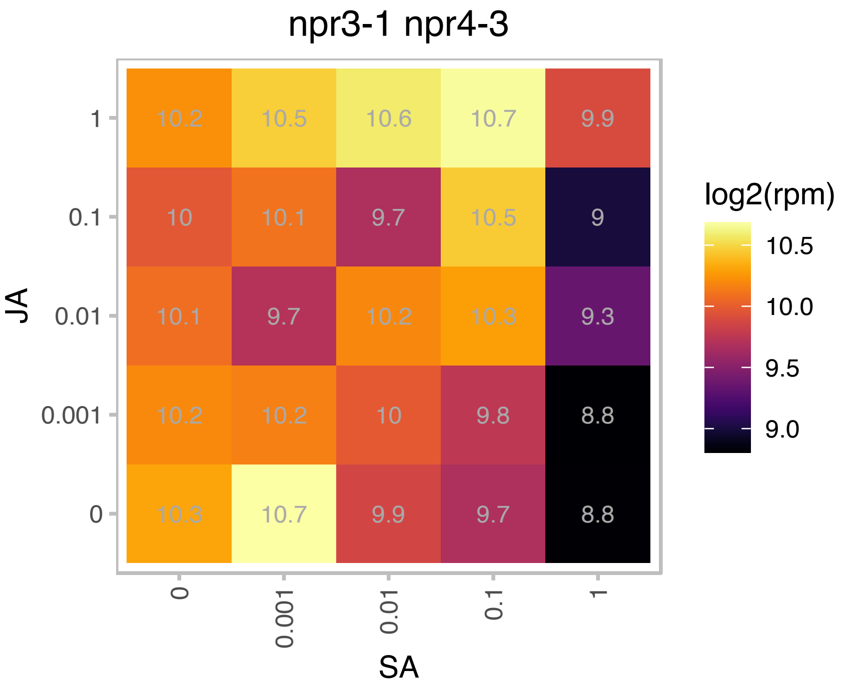

# AT5G32450.1 : NA

RNA binding (RRM/RBD/RNP motifs) family protein

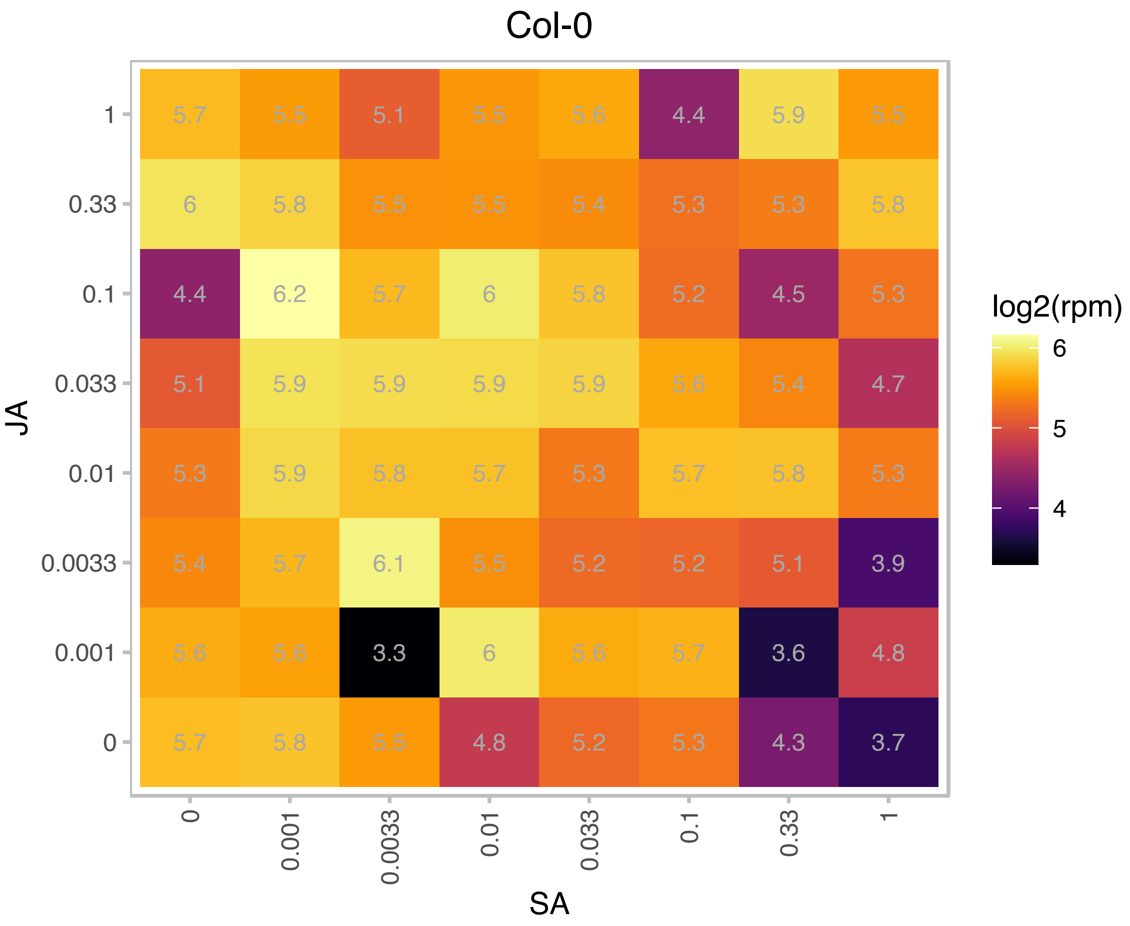


| | |
|---|---|
| Gene | AT5G32450 |
| is.expressed_genes | TRUE |
| is.responsive_genes | TRUE |
| is.apcluster | D-02 |

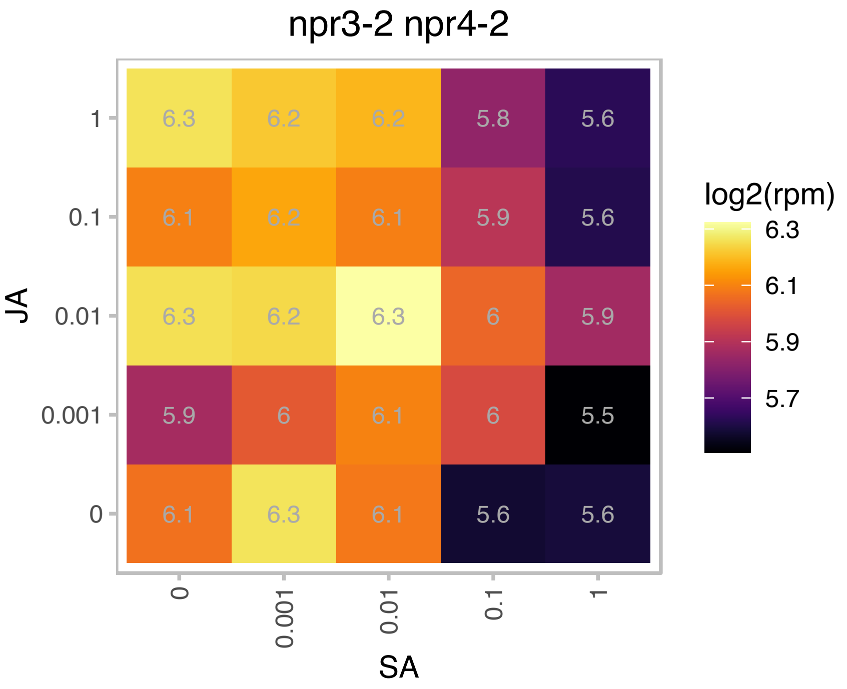


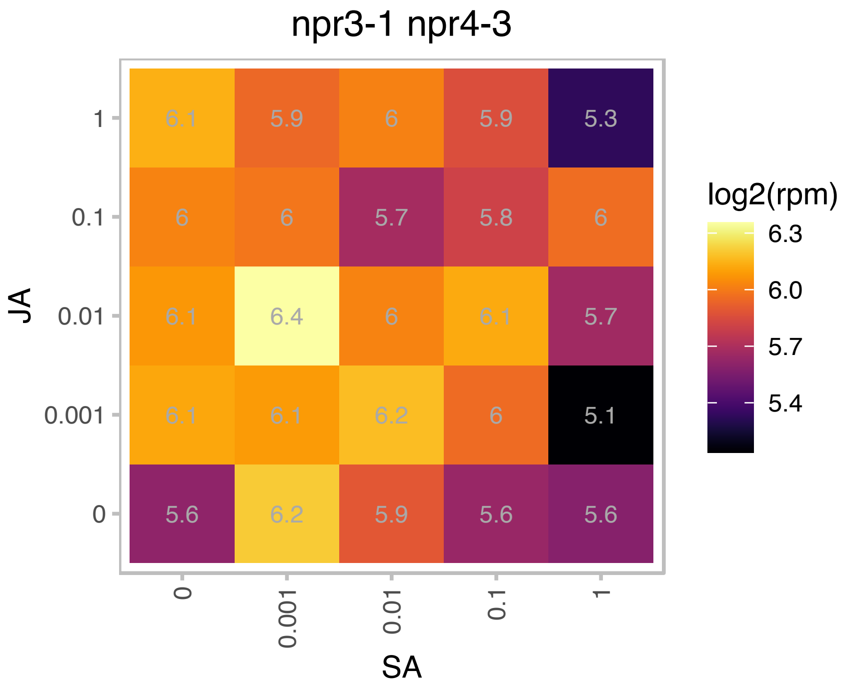

# AT5G43960.1 : NA

Nuclear transport factor 2 (NTF2) family protein with RNA binding (RRM-RBD-RNP motifs) domain

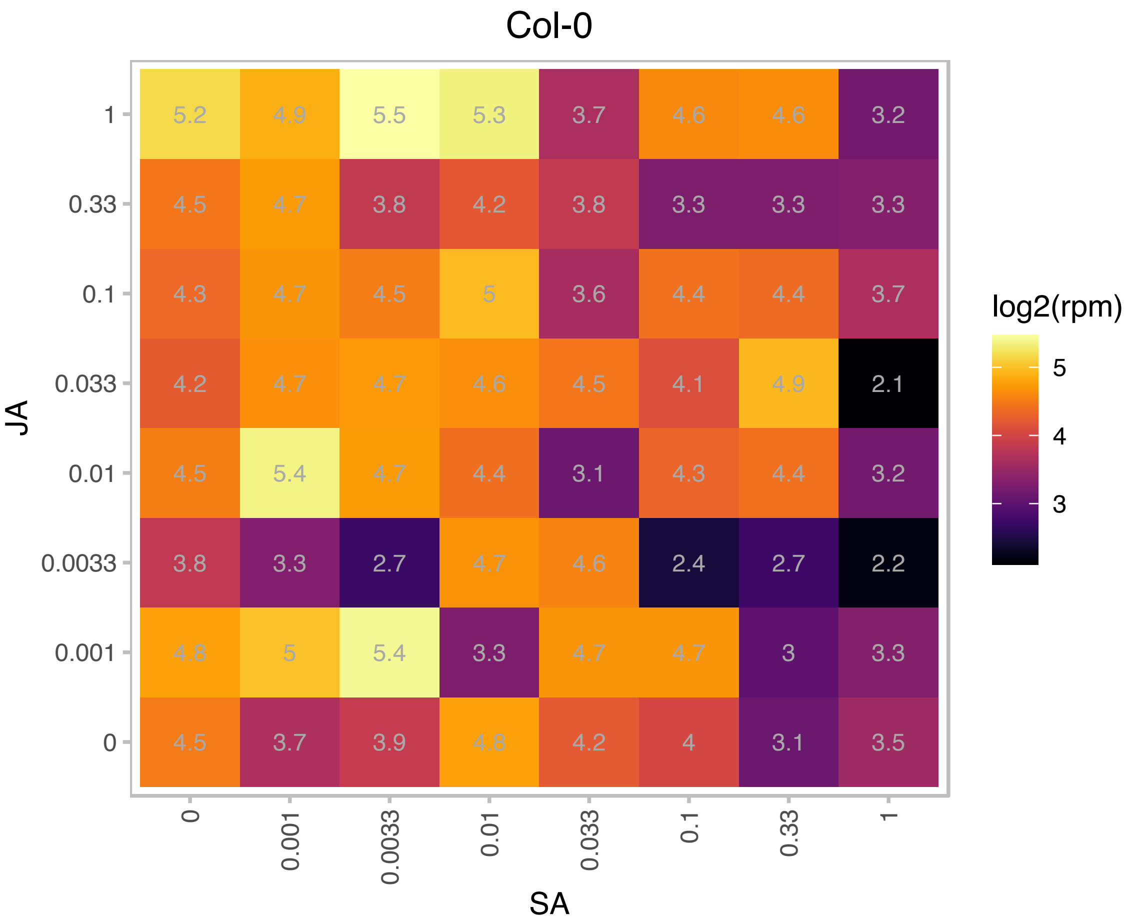


| | |
|---|---|
| Gene | AT5G43960 |
| is.expressed_genes | TRUE |
| is.responsive_genes | TRUE |
| is.apcluster | D-02 |

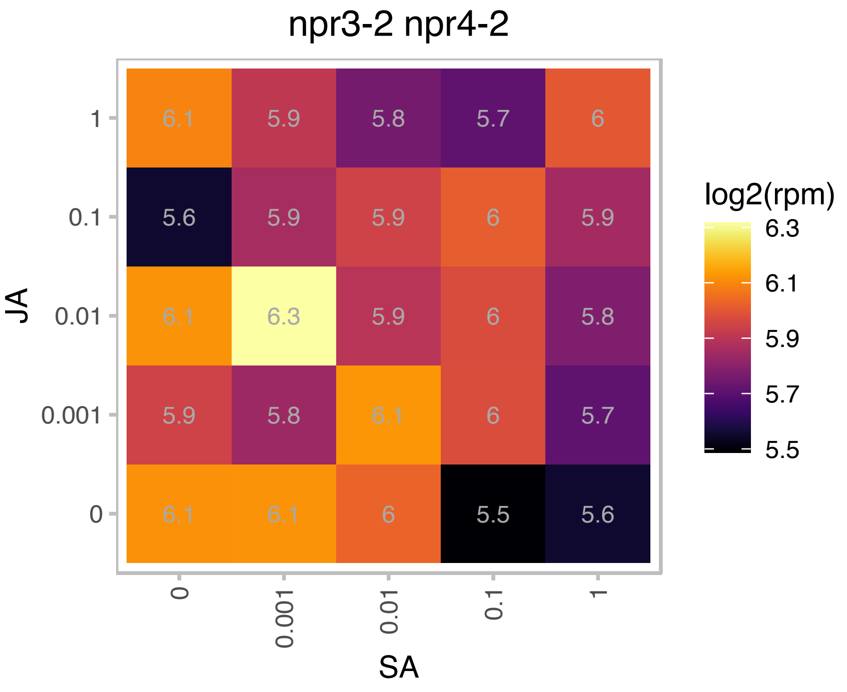


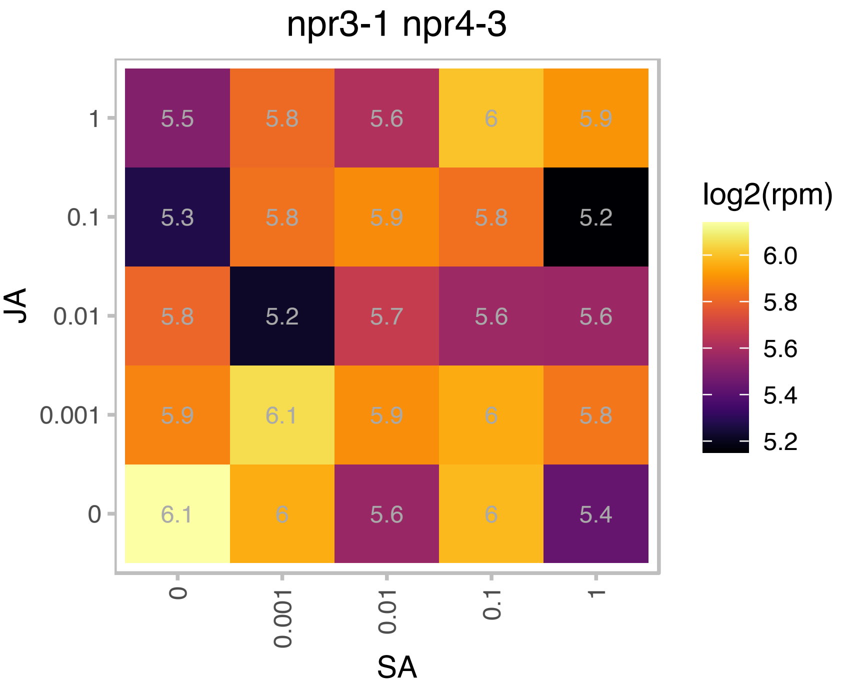

### AT5G47220.1 : ATERF-2, ATERF2, ERF2

ethylene responsive element binding factor 2

Encodes a member of the ERF (ethylene response factor) subfamily B-3 of ERF/AP2 transcription factor family (ATERF-2). The protein contains one AP2 domain. Functions as activator of GCC box□ □ ependent transcription. Positive regulator of JA-responsive defense genes and resistance to F. oxysporum and enhances JA inhibition of root elongation.

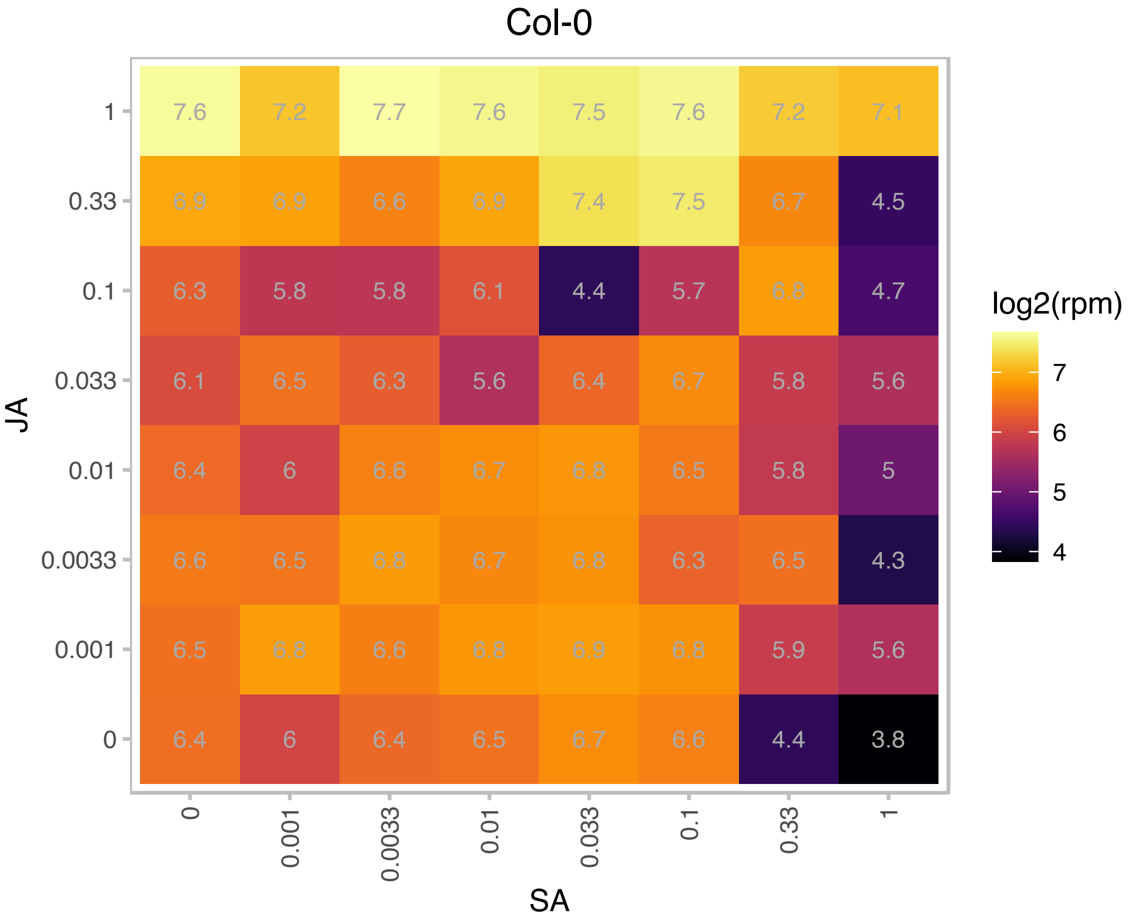


| | |
|---|---|
| Gene | AT5G47220 |
| is.expressed_genes | TRUE |
| is.responsive_genes | TRUE |
| is.apcluster | D-02 |

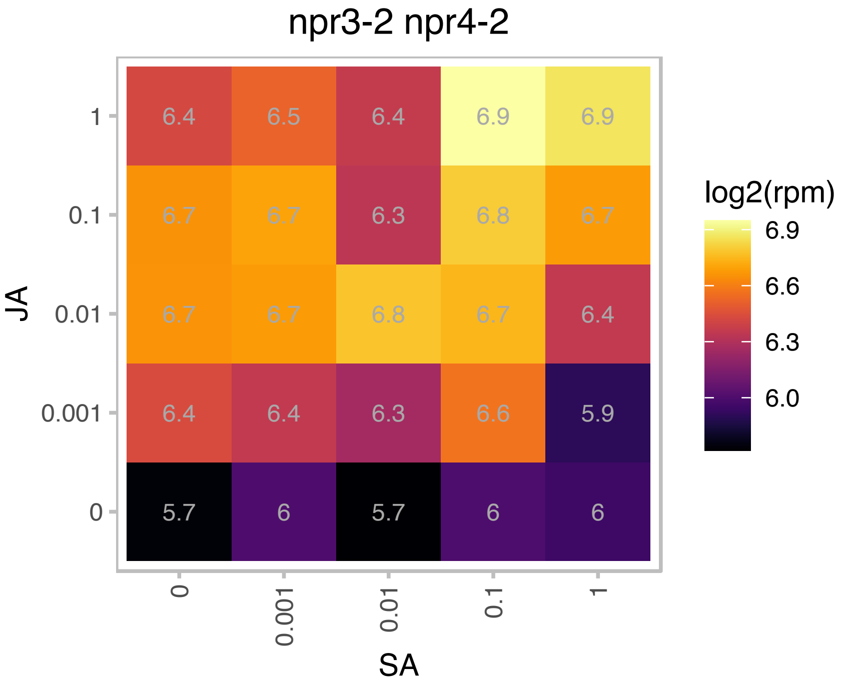


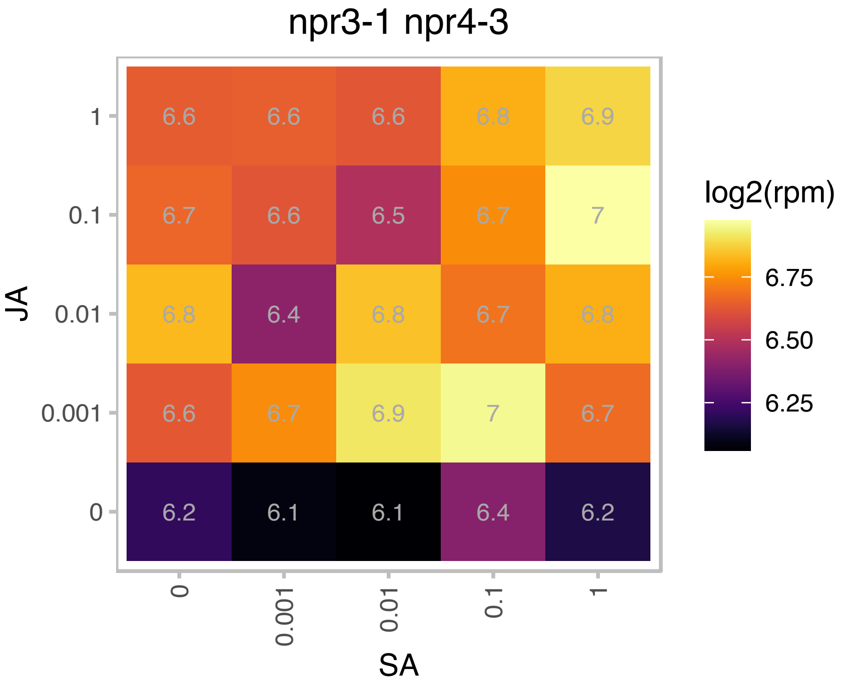

# AT5G55640.1 : NA

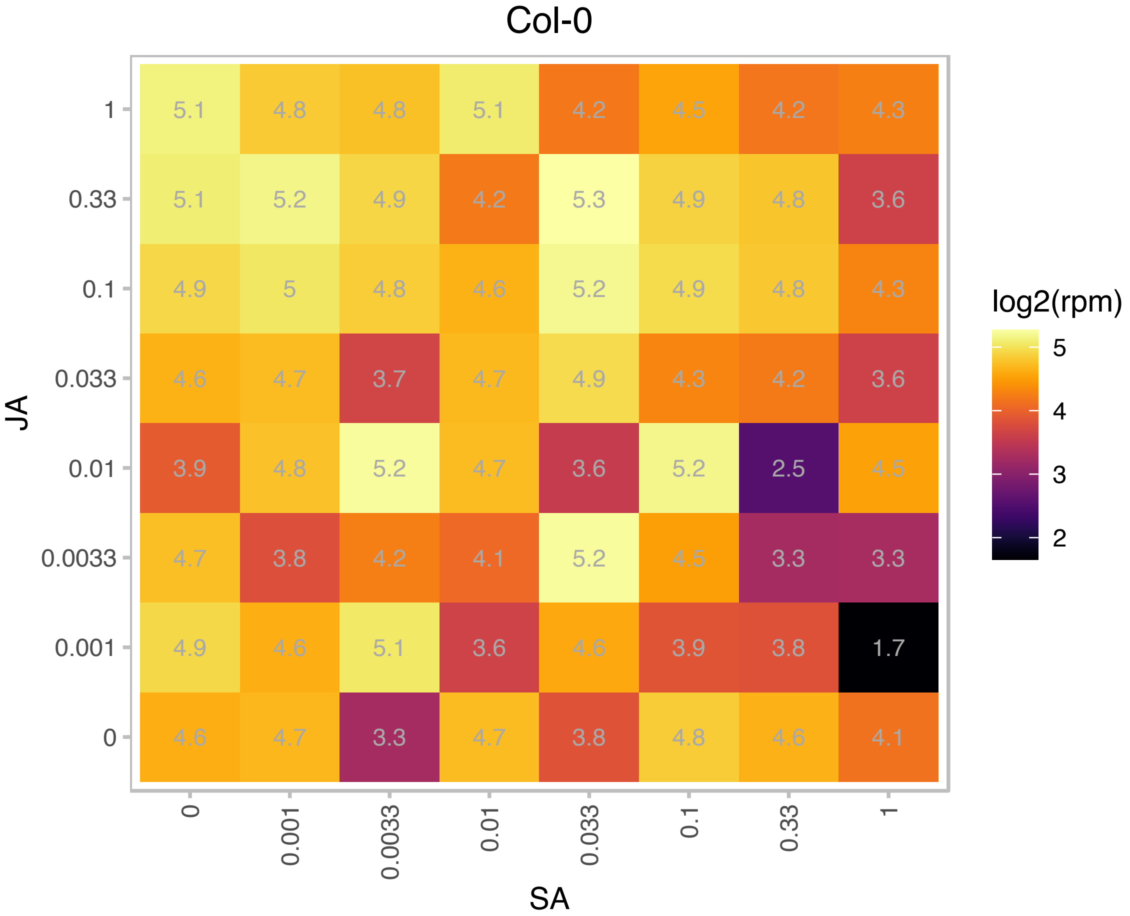


| | |
|---|---|
| Gene | AT5G55640 |
| is.expressed_genes | TRUE |
| is.responsive_genes | TRUE |
| is.apcluster | D-02 |

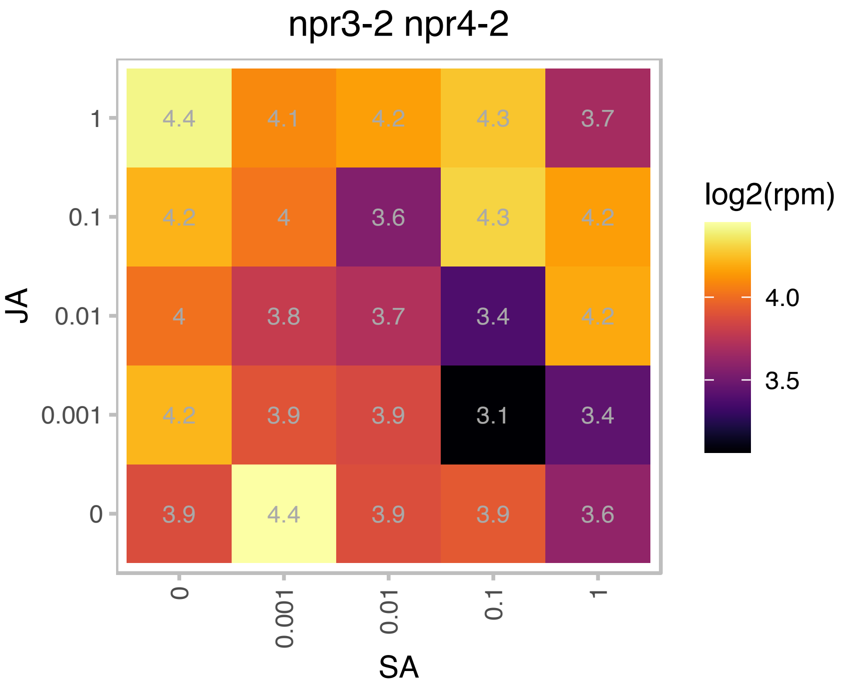


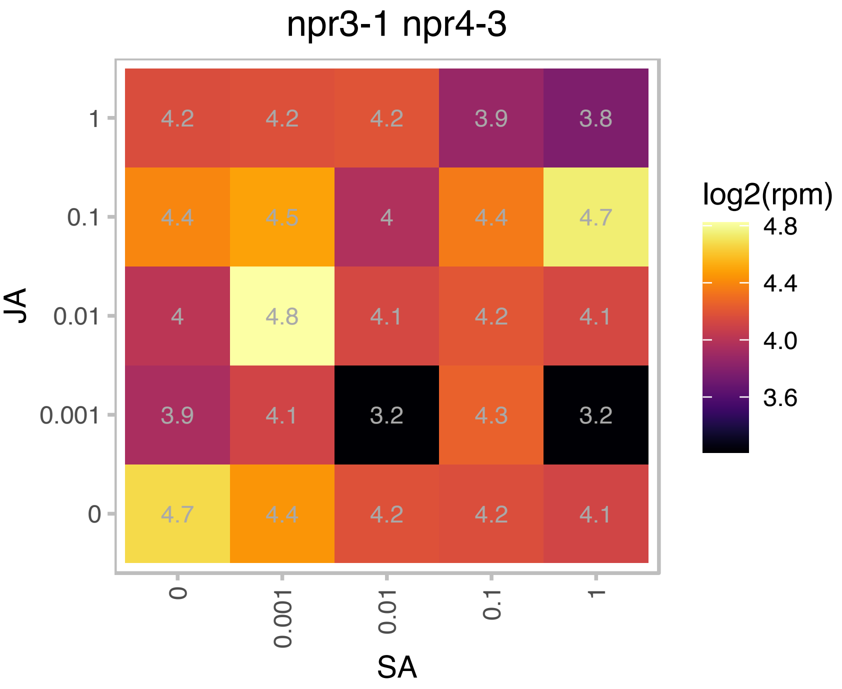

### AT5G56030.1 : AtHsp90.2, ERD8, HSP81-2, HSP81.2, HSI

heat shock protein 81-2

A member of heat shock protein 90 (HSP90) gene family. Expressed in all tissues and abundant in root apical meristem, pollen and tapetum. Expression is NOT heat-induced but induced by IAA and NaCl. Interacts with HsfA1d in the cytosol and the nucleus and negatively regulates HsfA1d. Did not bind to AtHsfA4c. The mRNA is cell-to-cell mobile.

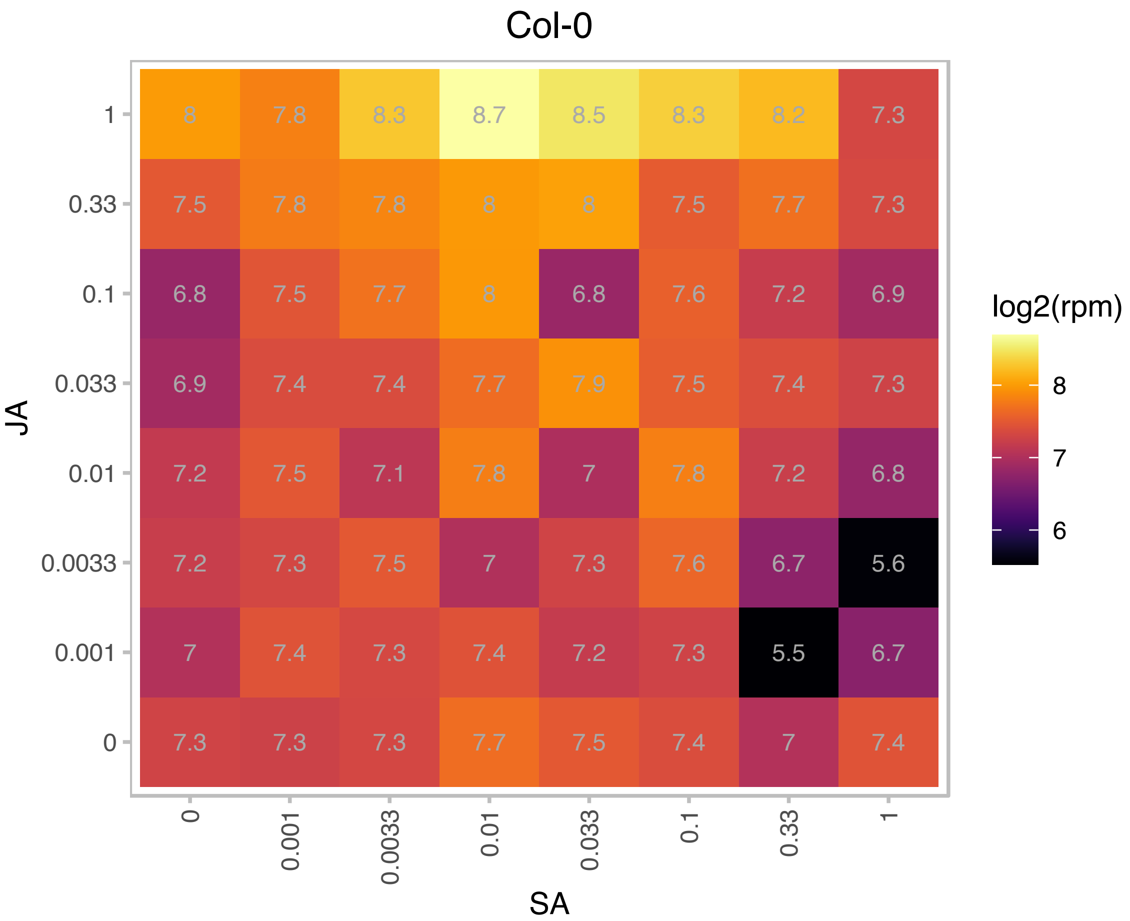


| | |
|---|---|
| Gene | AT5G56030 |
| is.expressed_genes | TRUE |
| is.responsive_genes | TRUE |
| is.apcluster | D-02 |

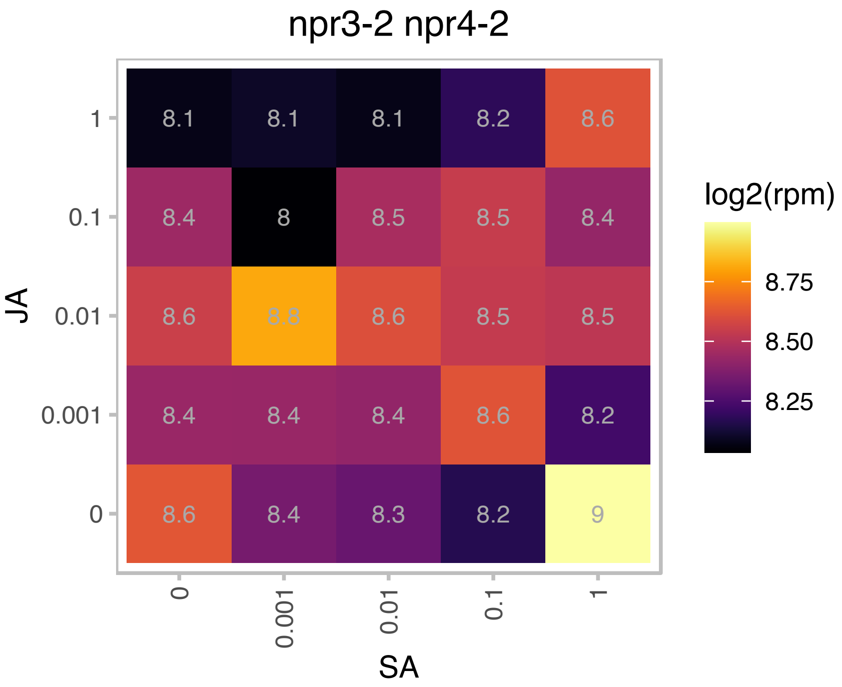


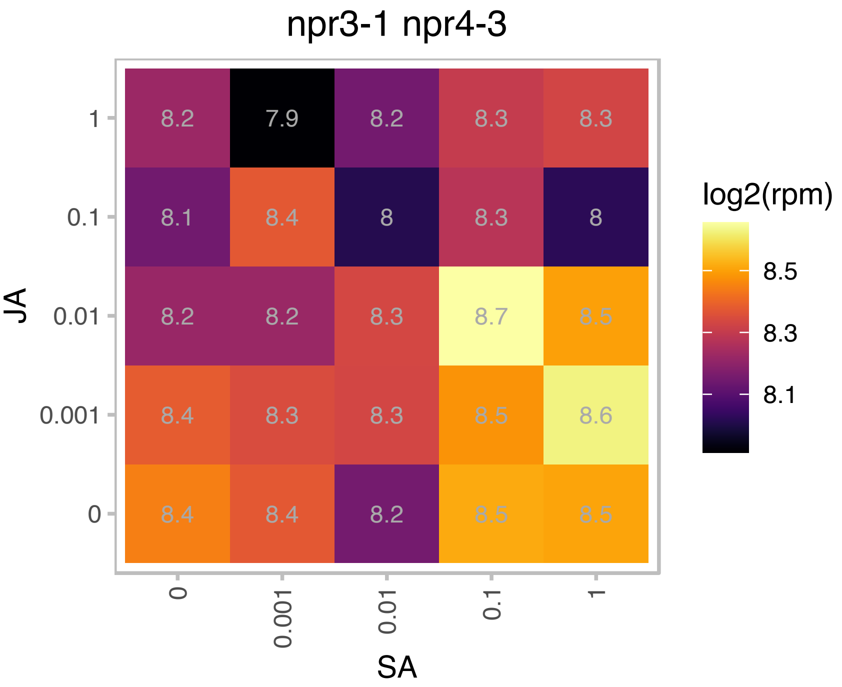

# AT5G58490.1 : NA

NAD(P)-binding Rossmann-fold superfamily protein

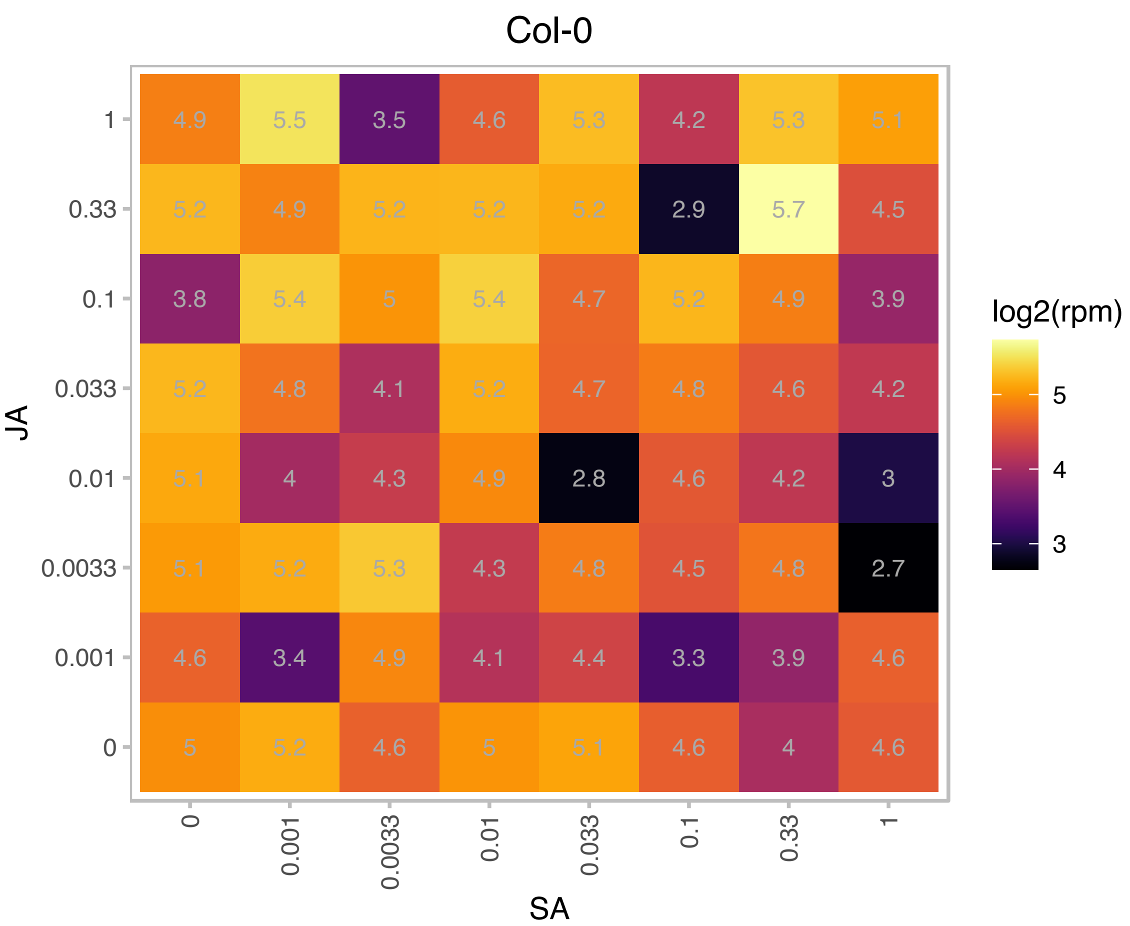


| | |
|---|---|
| Gene | AT5G58490 |
| is.expressed_genes | TRUE |
| is.responsive_genes | TRUE |
| is.apcluster | D-02 |

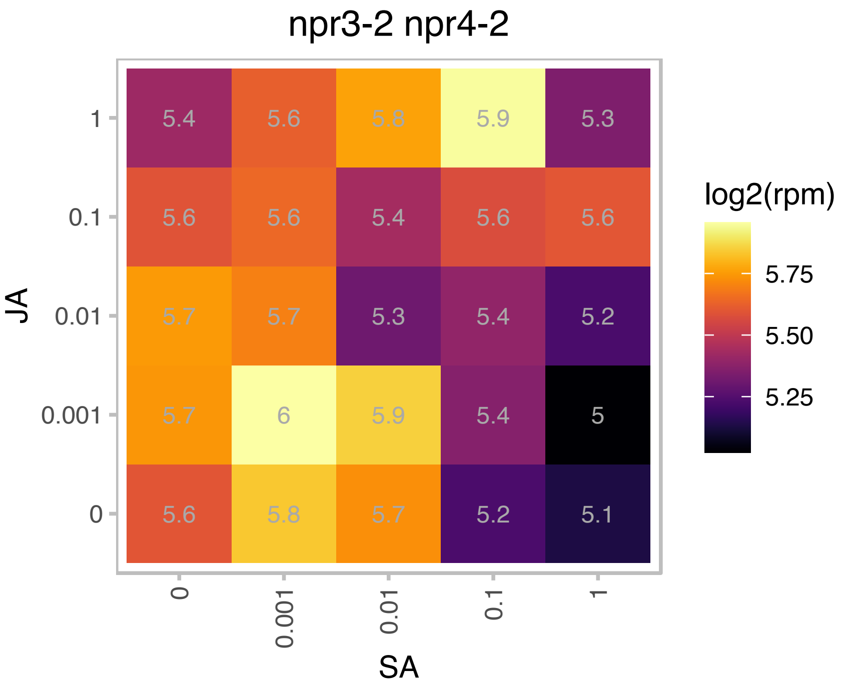


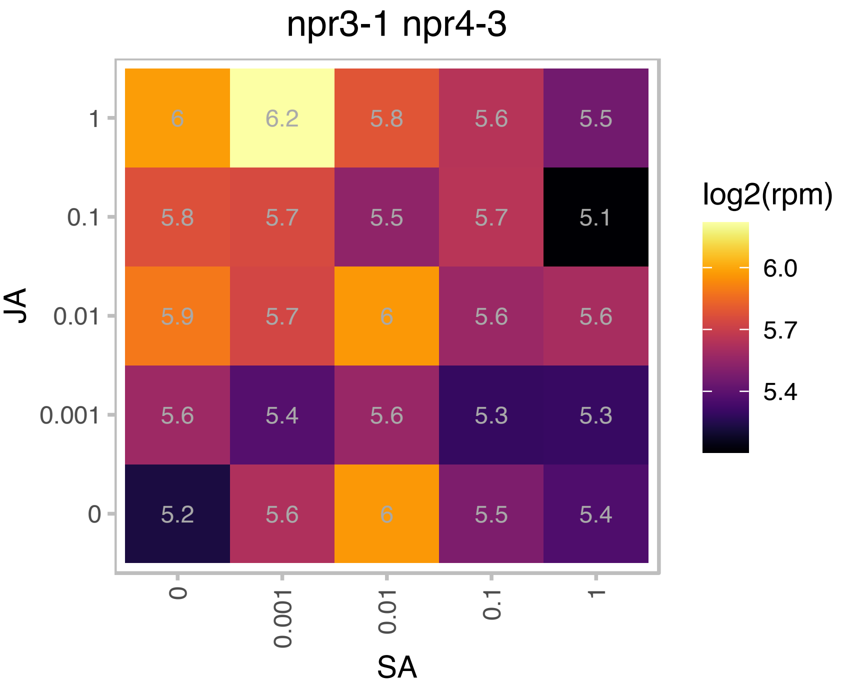

### AT5G62810.1 : AtPED2, ATPEX14, PED2, PEX14

peroxin 14

mutant has a defect in the intracellular transport of thiolase from the cytosol to glyoxysomes (formerly known as ped2) The mRNA is cell-to-cell mobile.

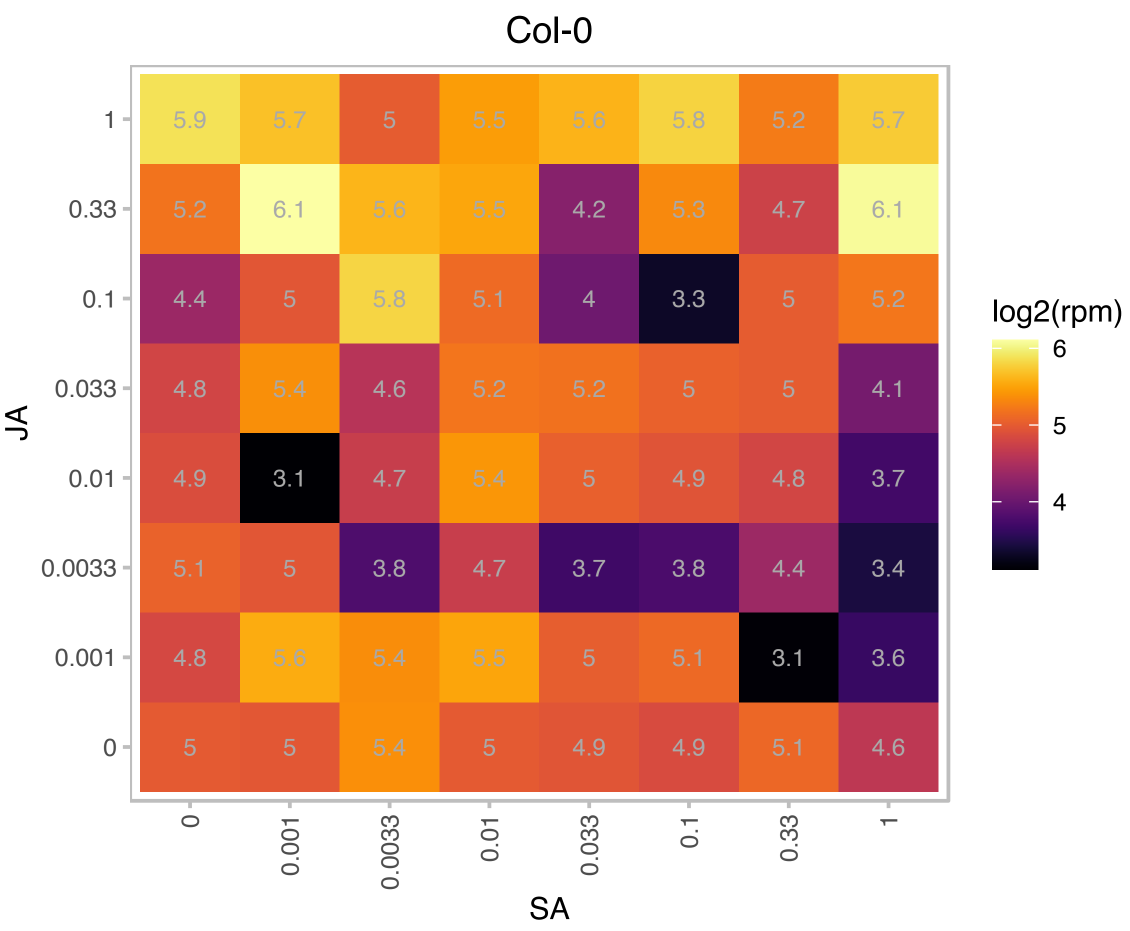


| | |
|---|---|
| Gene | AT5G62810 |
| is.expressed_genes | TRUE |
| is.responsive_genes | TRUE |
| is.apcluster | D-02 |

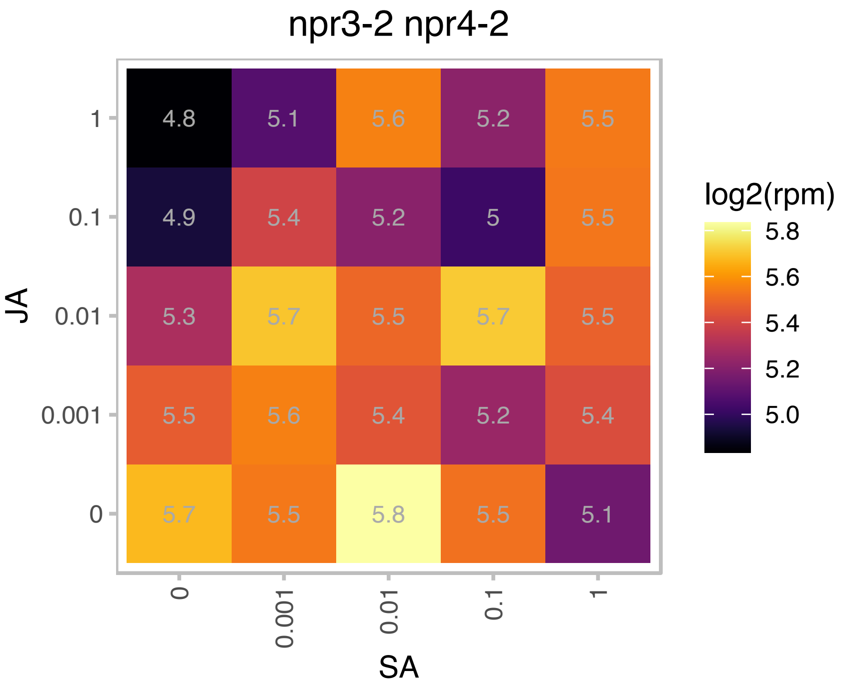


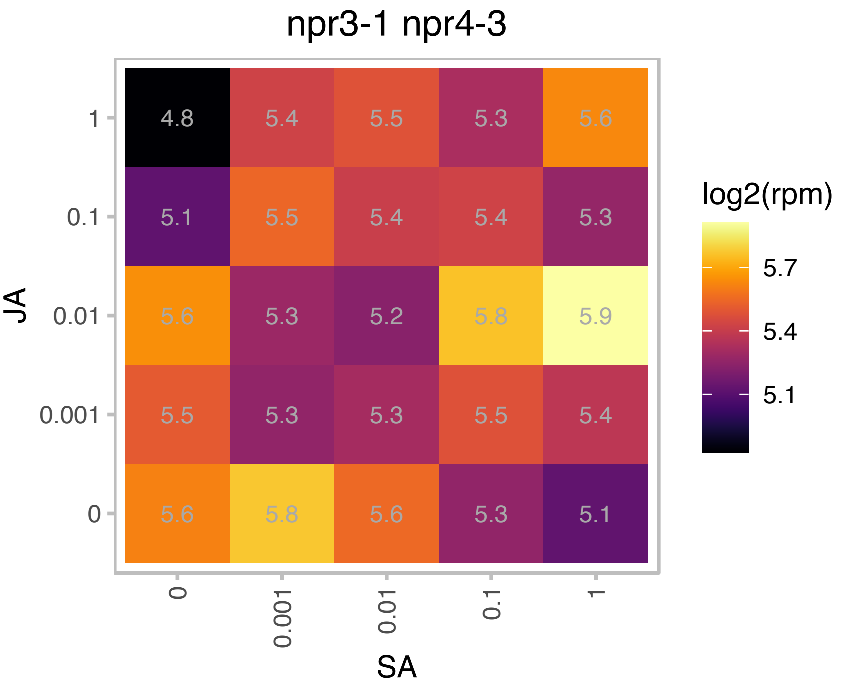

### AT5G66410.1 : PLP3b

phosducin-like protein 3 homolog
Encodes a protein that functions in microtubule assembly. Plants with reduced levels of both PLP3a (At3g50960) and PLP3b show defects in cytokinesis, cortical microtubule array formation, oriented cell growth, and maintenance of proper ploidy.

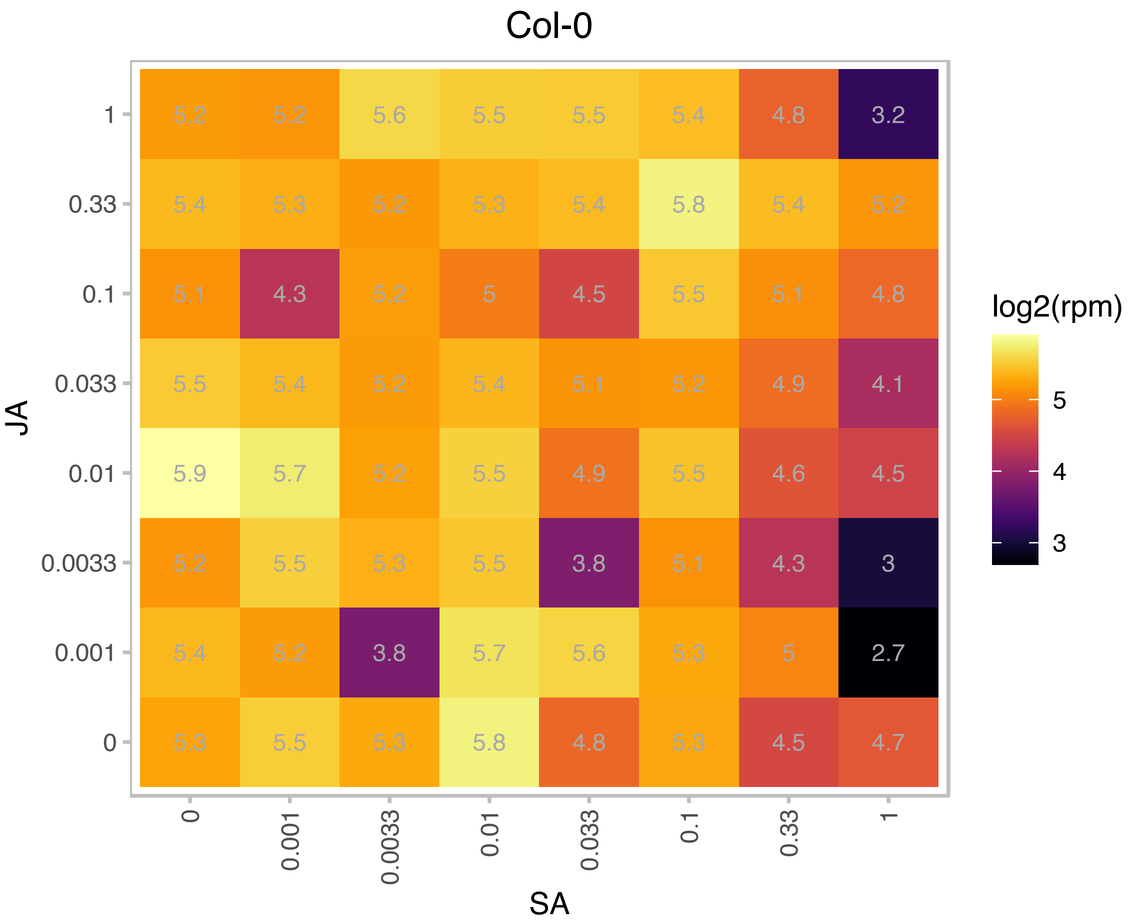


| | |
|---|---|
| Gene | AT5G66410 |
| is.expressed_genes | TRUE |
| is.responsive_genes | TRUE |
| is.apcluster | D-02 |

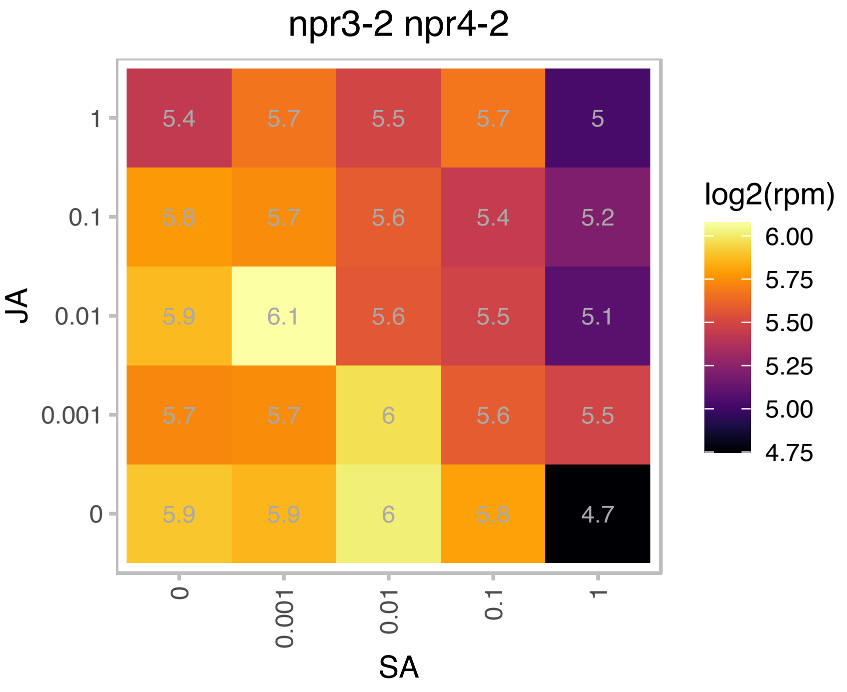


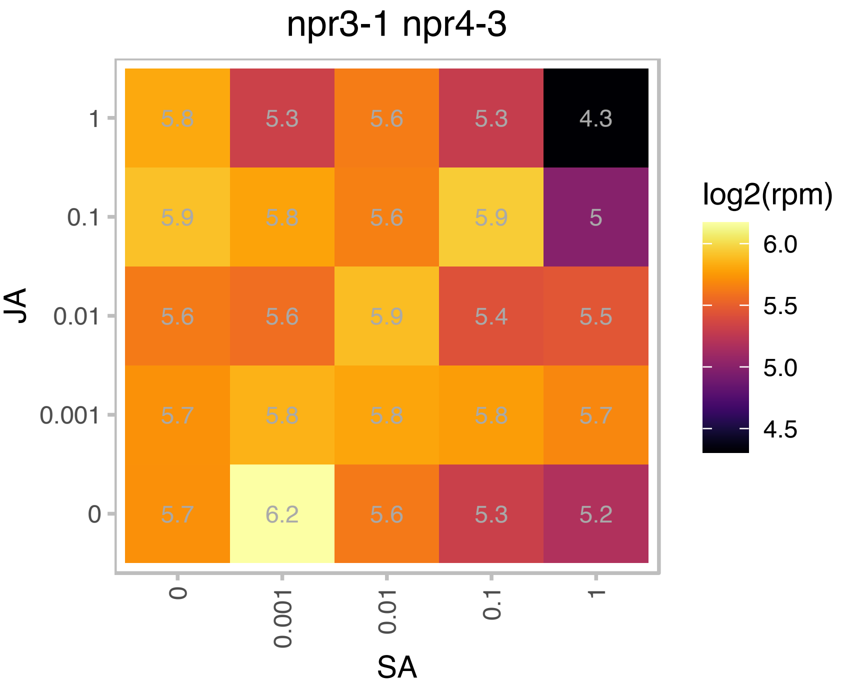

# AT1G01940.1 : NA

Cyclophilin-like peptidyl-prolyl cis-trans isomerase family protein

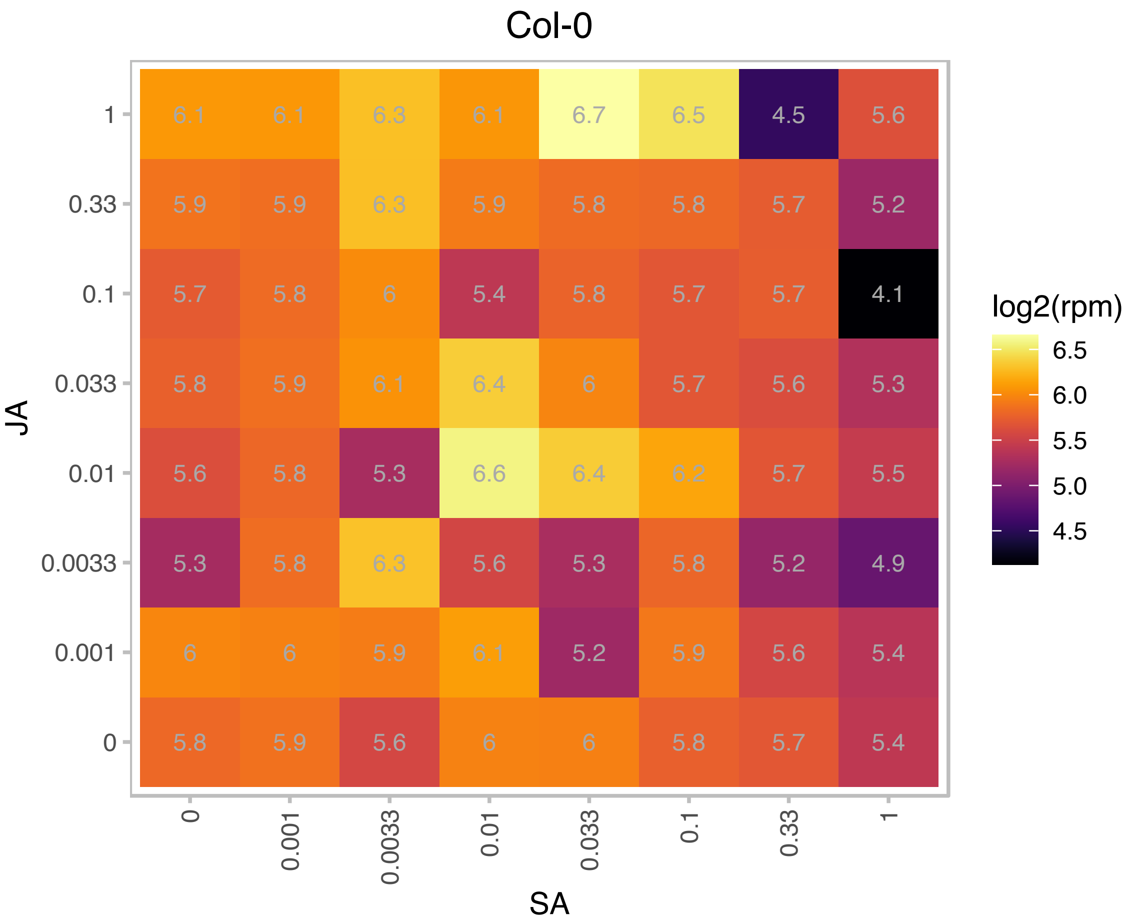


| | |
|---|---|
| Gene | AT1G01940 |
| is.expressed_genes | TRUE |
| is.responsive_genes | TRUE |
| is.apcluster | D-03 |

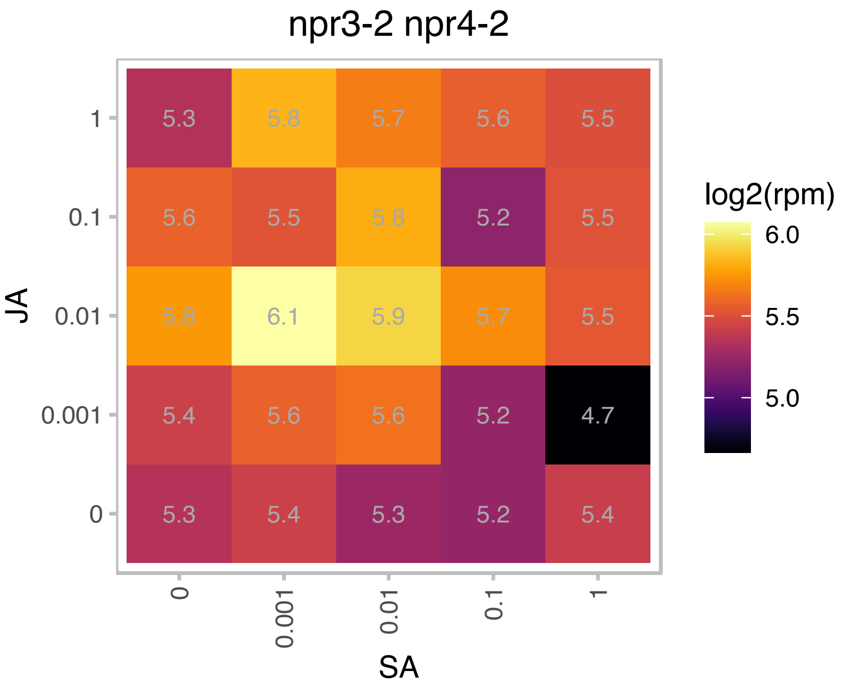


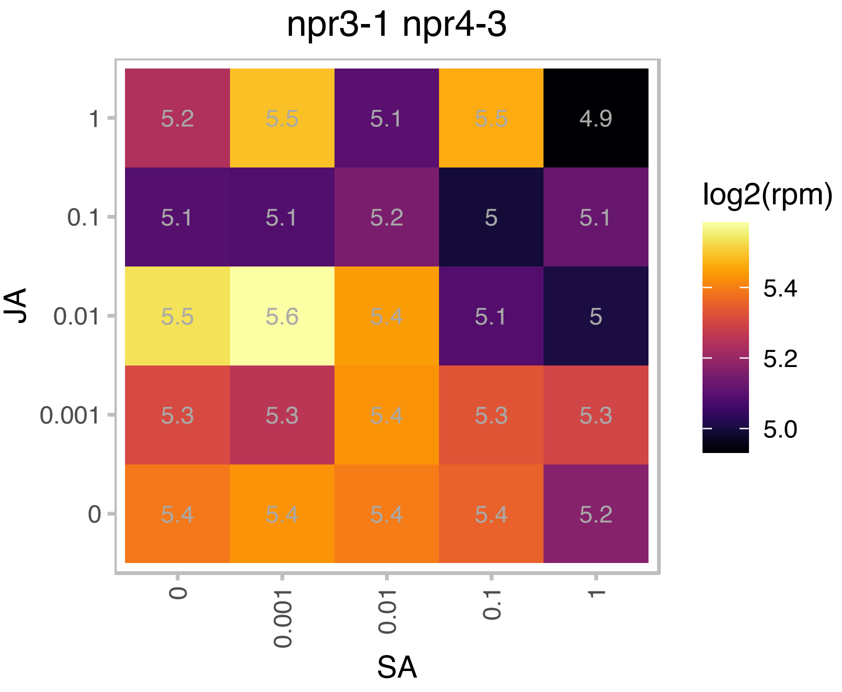

# AT1G05870.1 : NA

Protein of unknown function (DUF1685)

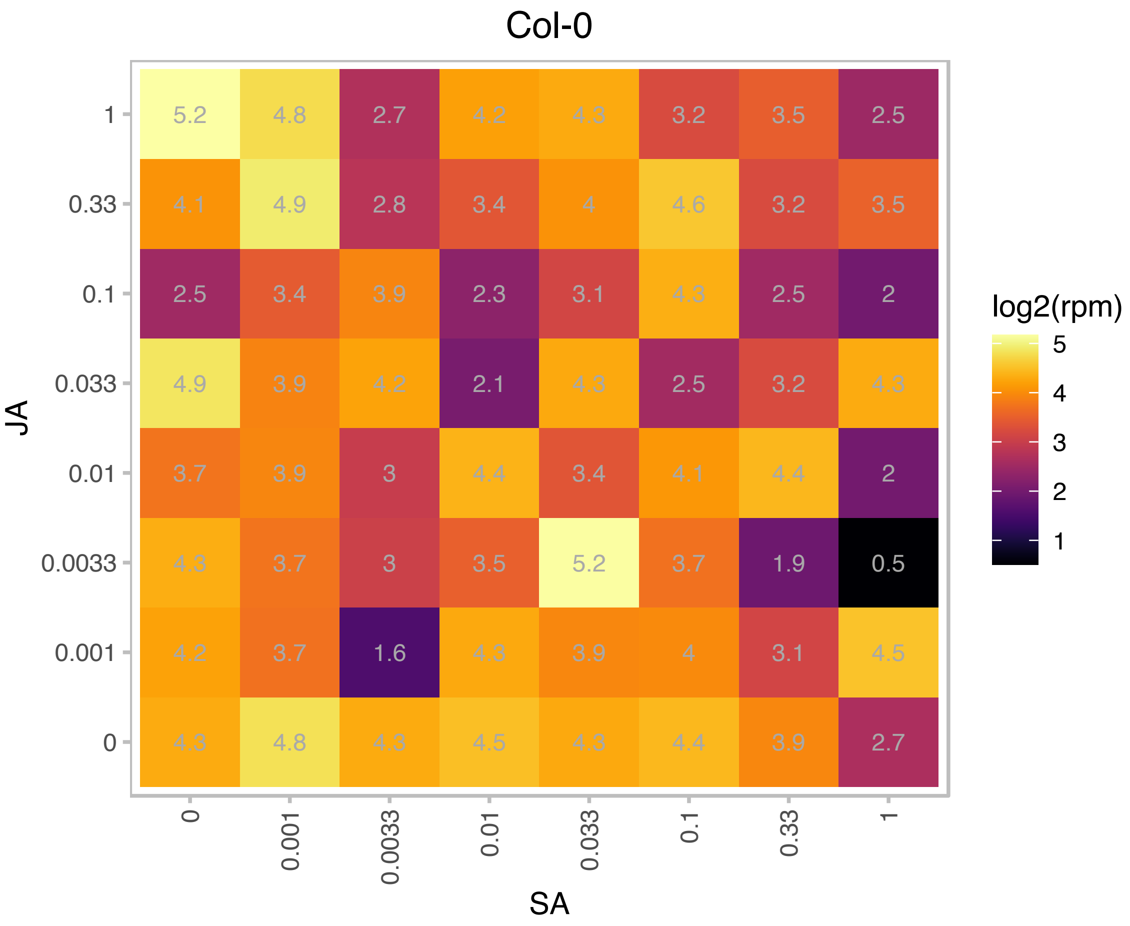


| | |
|---|---|
| Gene | AT1G05870 |
| is.expressed_genes | TRUE |
| is.responsive_genes | TRUE |
| is.apcluster | D-03 |

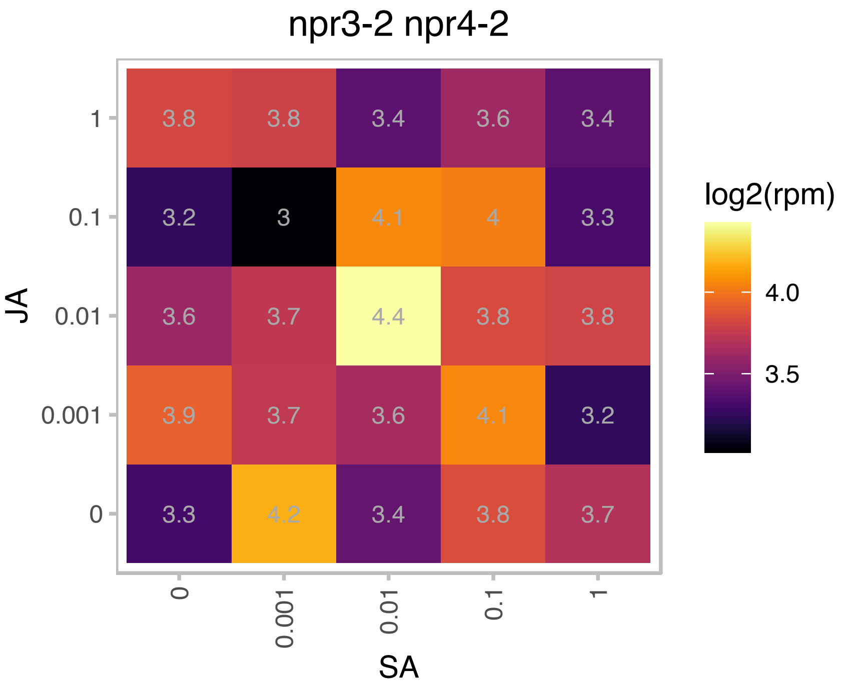


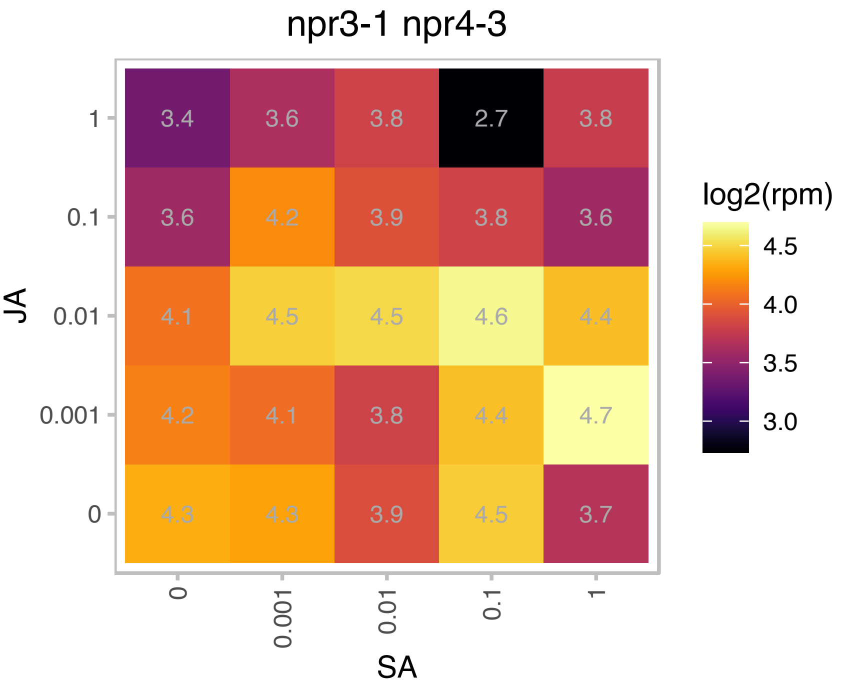

### AT1G18450.1 : ARP4, ATARP4

Encodes a gene similar to actin-related proteins in other organisms. Member of nuclear ARP family of genes. Component of chromatin remodeling complexes, involved in chromatin-mediated gene regulation. Phenotype of

actin-related protein 4

the arp4-1 mutant allele revealed partial sterility due to defects in anther development. Targeting the distinct, 3' UTR of AtARP4 transcripts with RNA interference caused a drastic reduction in the level of AtARP4 protein expression, and resulted in strong pleiotropic phenotypes such as altered organization of plant organs, early flowering, delayed flower senescence and high levels of sterility. Western blot analysis and immunolabelling demonstrated a clear correlation between reductions in the level of AtARP4 expression and

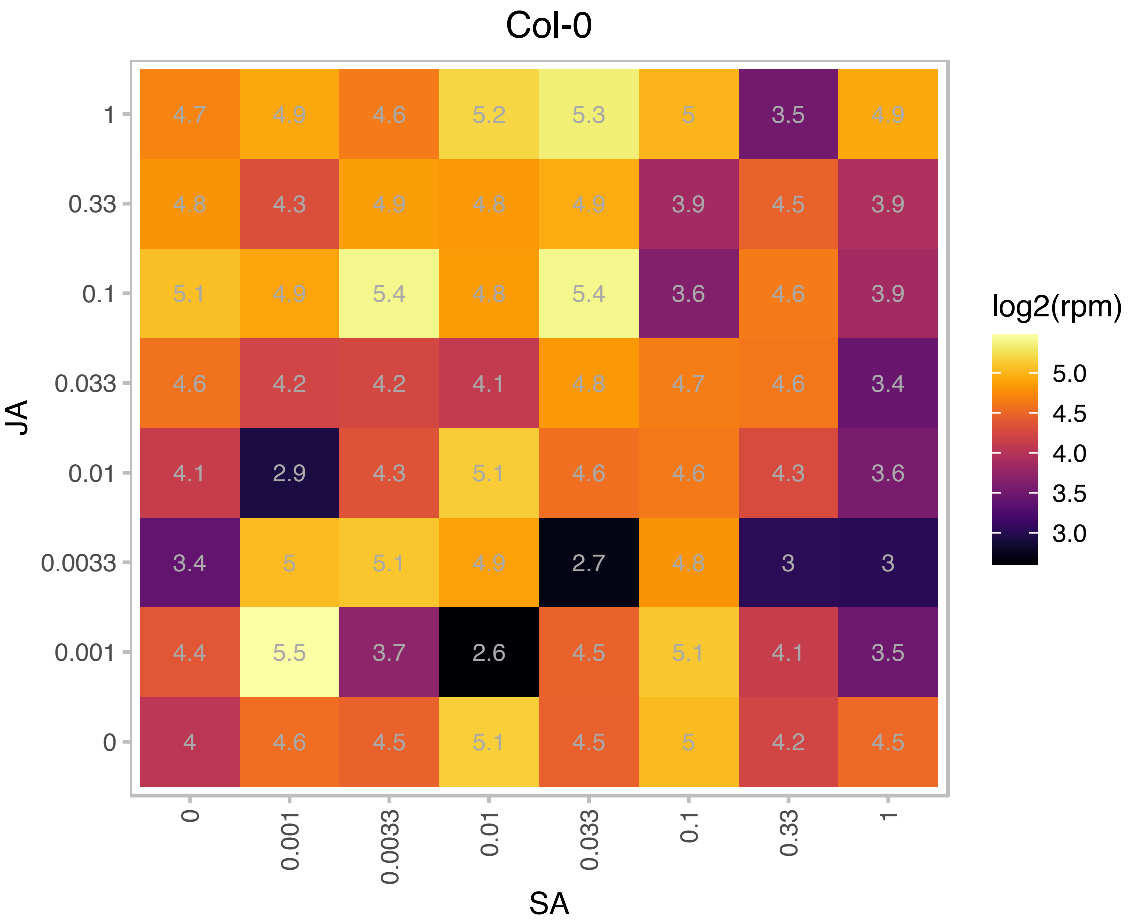


| | |
|---|---|
| Gene | AT1G18450 |
| is.expressed_genes | TRUE |
| is.responsive_genes | TRUE |
| is.apcluster | D-03 |

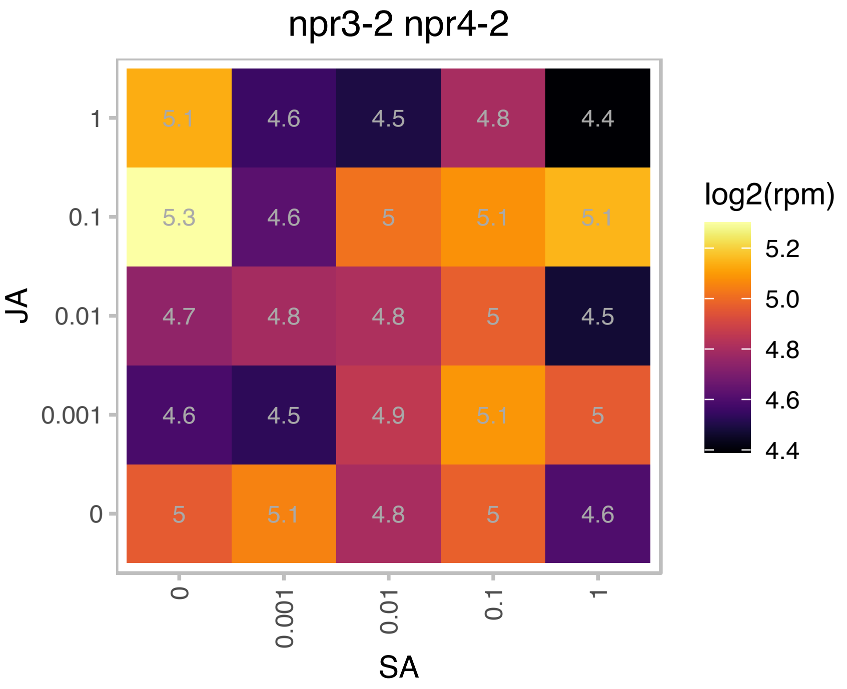


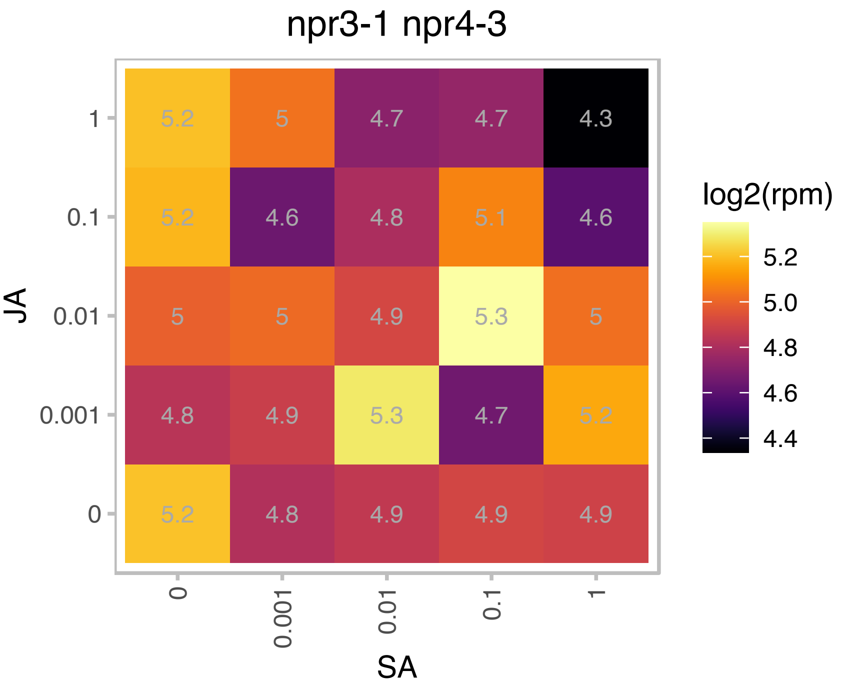

# AT1G20225.1 : NA

Thioredoxin superfamily protein

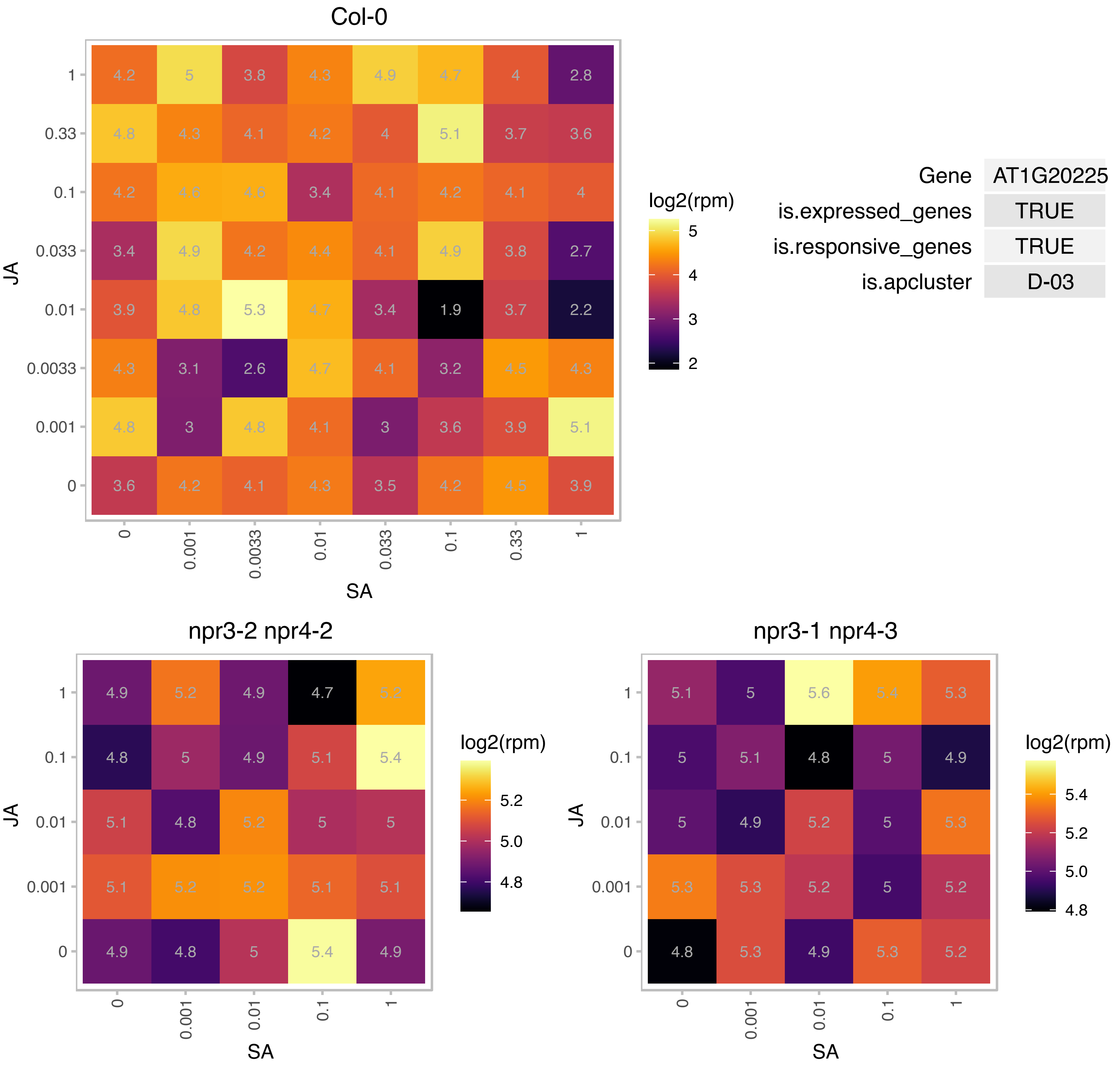


| | |
|---|---|
| Gene | AT1G20225 |
| is.expressed_genes | TRUE |
| is.responsive_genes | TRUE |
| is.apcluster | D-03 |

### AT1G22300.1 : 14-3-3EPSILON, GF14 EPSILON, GRF10

general regulatory factor 10

Encodes a 14-3-3 protein. This protein is reported to interact with the BZR1 transcription factor involved in brassinosteroid signaling and may affect the nucleocytoplasmic shuttling of BZR1

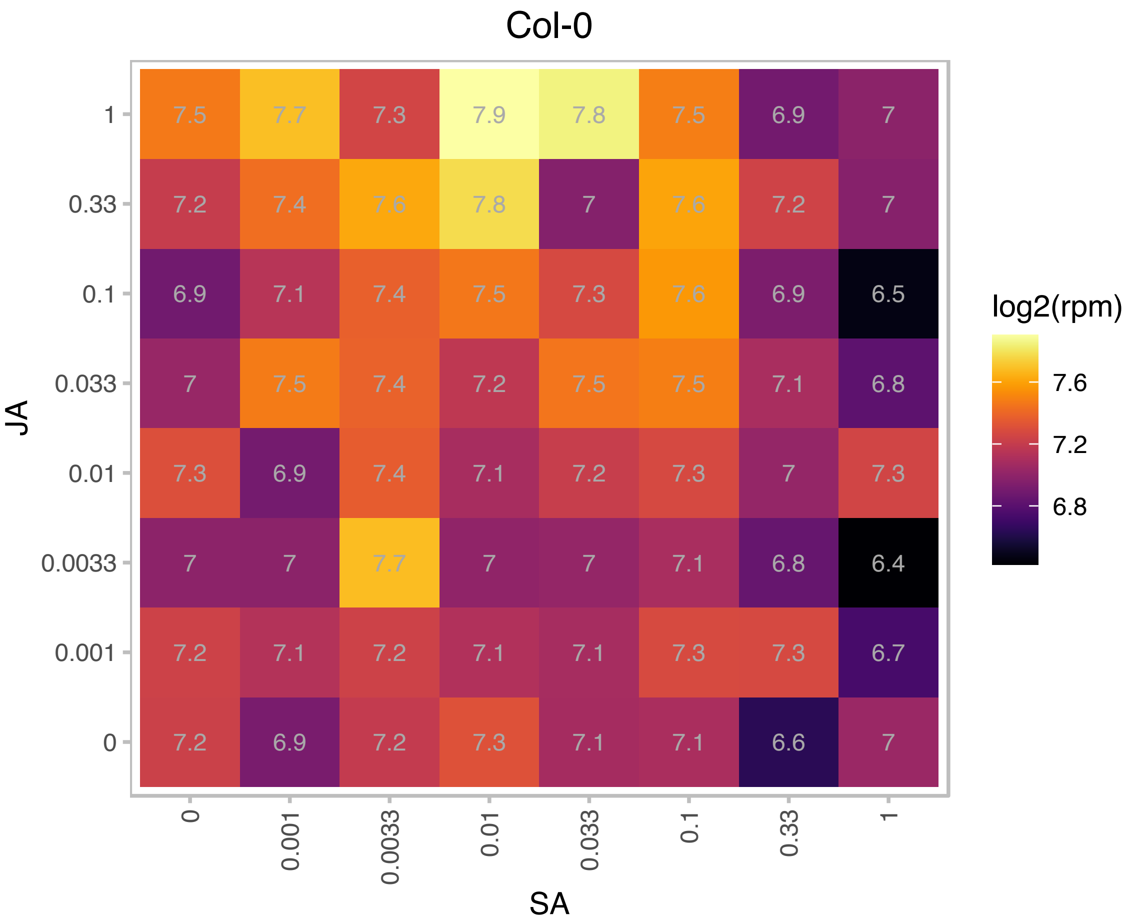


| | |
|---|---|
| Gene | AT1G22300 |
| is.expressed_genes | TRUE |
| is.responsive_genes | TRUE |
| is.apcluster | D-03 |

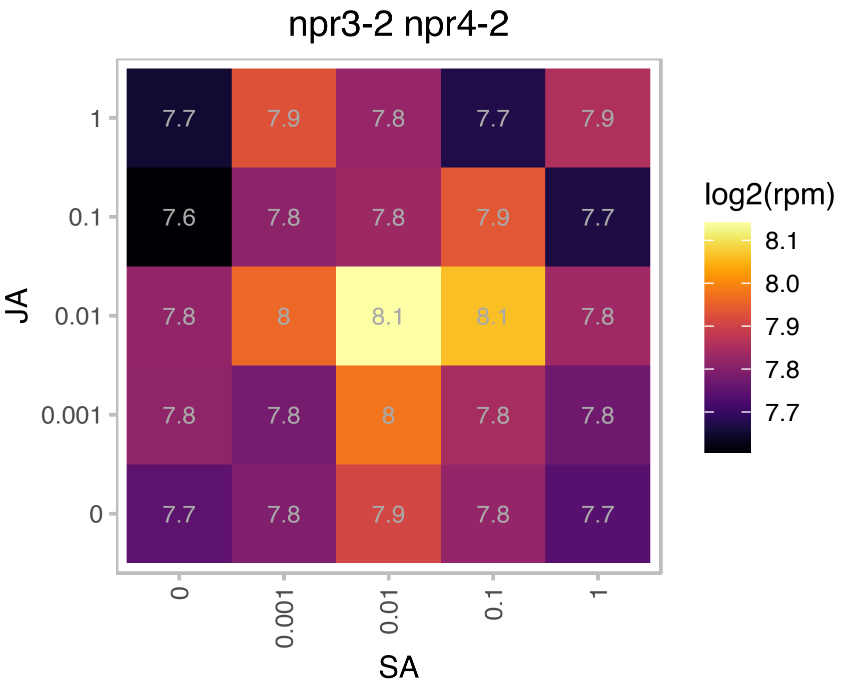


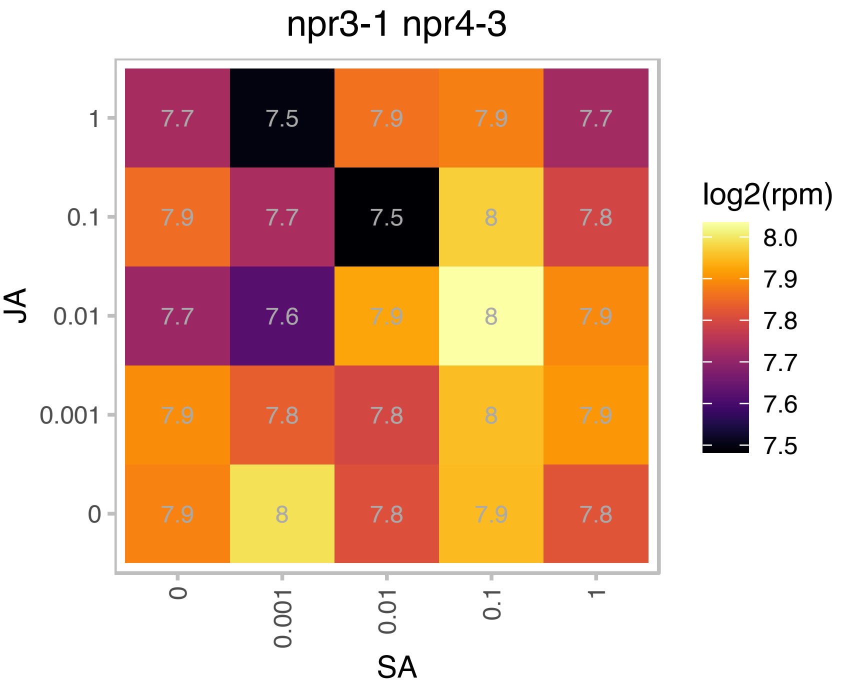

# AT1G50120.1 : NA

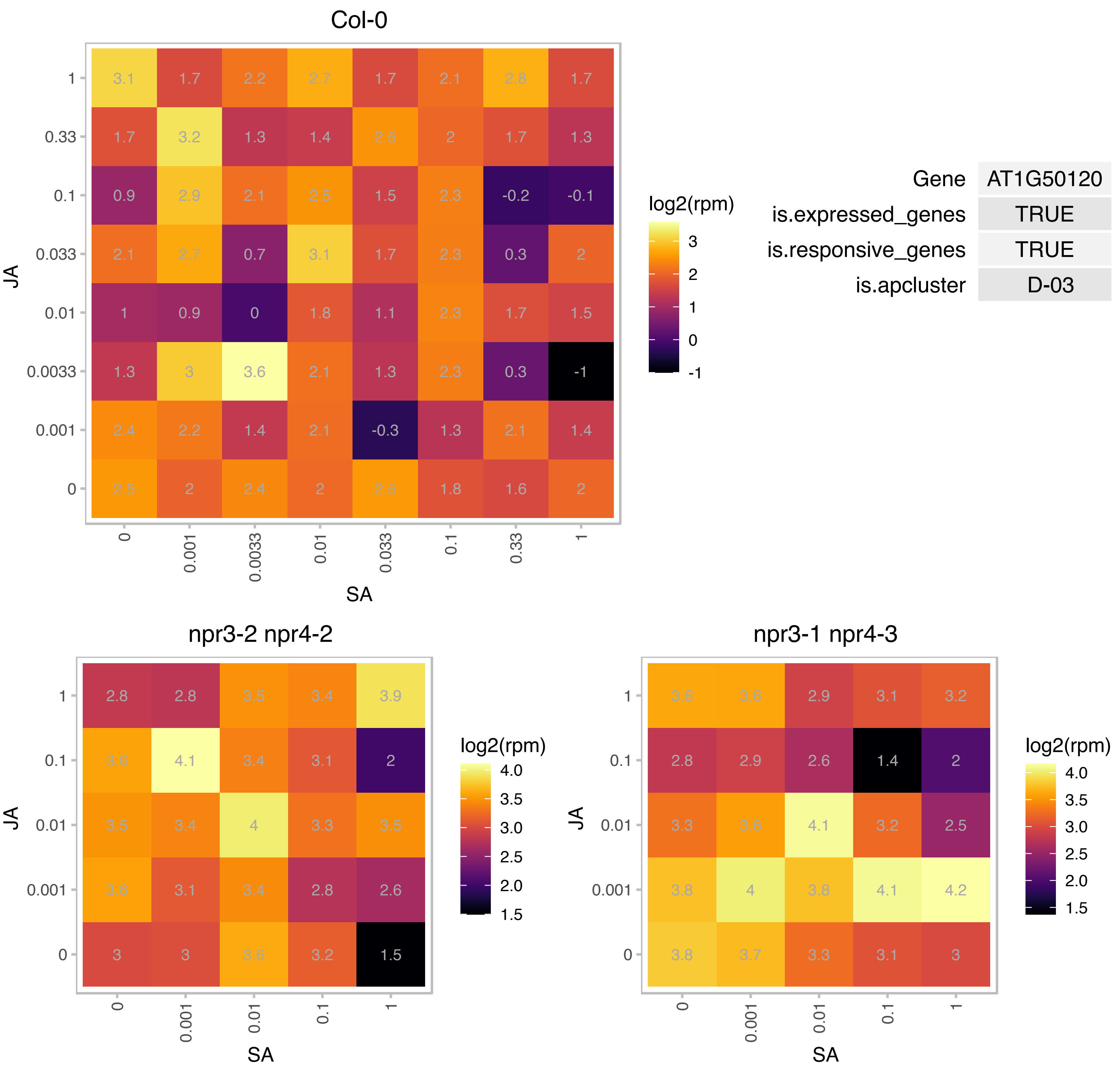


| | |
|---|---|
| Gene | AT1G50120 |
| is.expressed_genes | TRUE |
| is.responsive_genes | TRUE |
| is.apcluster | D-03 |

# AT1G64500.1 : NA

Glutaredoxin family protein
A member of a protein family found in plants and animals that contain conserved C-terminal glutaredoxin-like and putative zinc-binding cysteine-rich domains. It is involved in light stimulated actin bundling and chloroplast movement. The mRNA is cell-to-cell mobile.

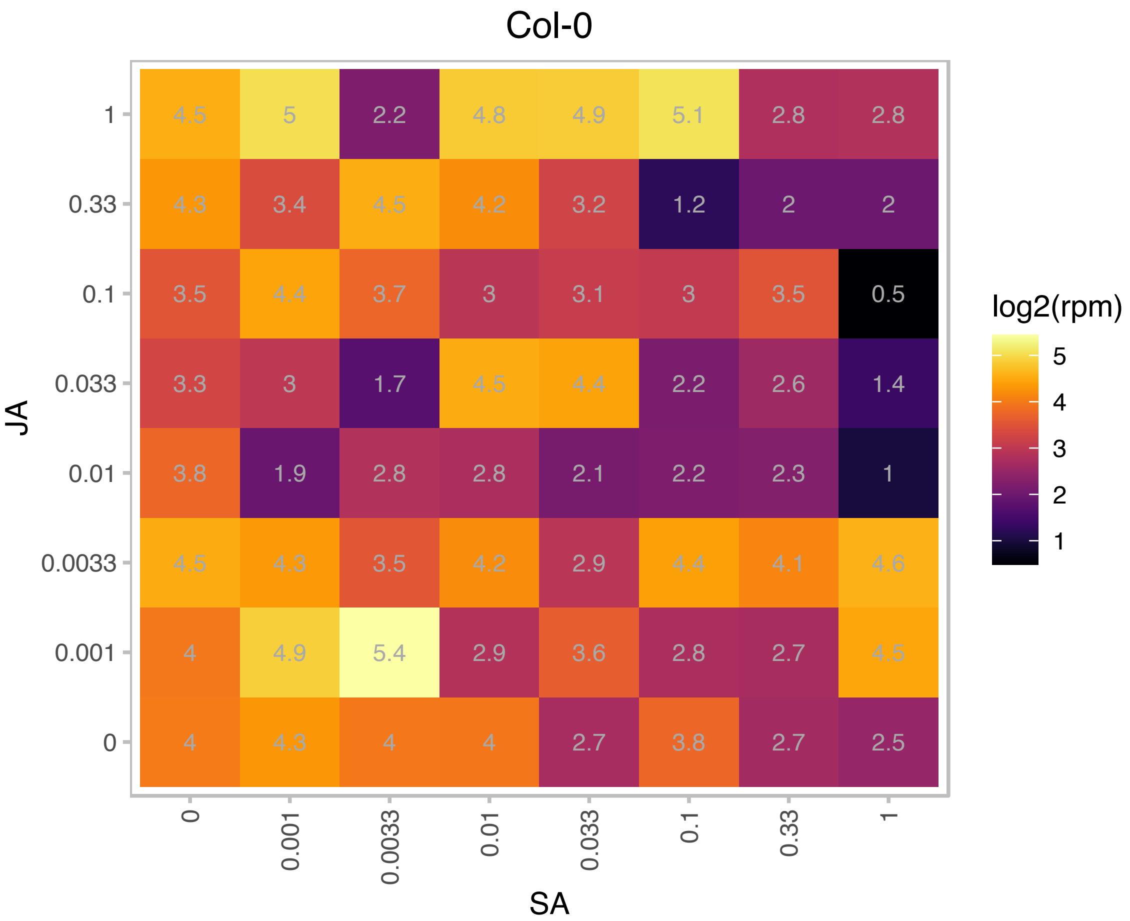


| | |
|---|---|
| Gene | AT1G64500 |
| is.expressed_genes | TRUE |
| is.responsive_genes | TRUE |
| is.apcluster | D-03 |

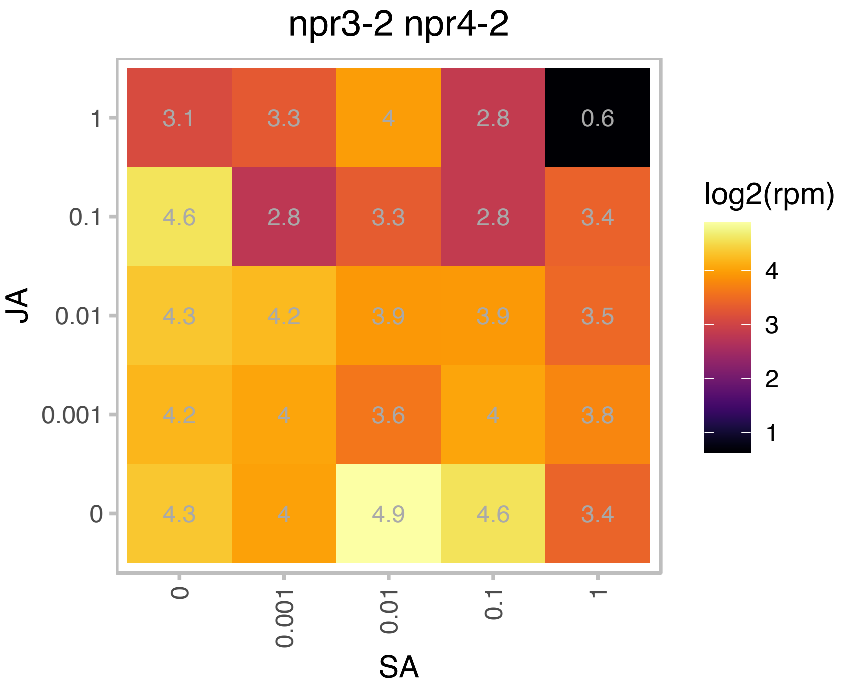


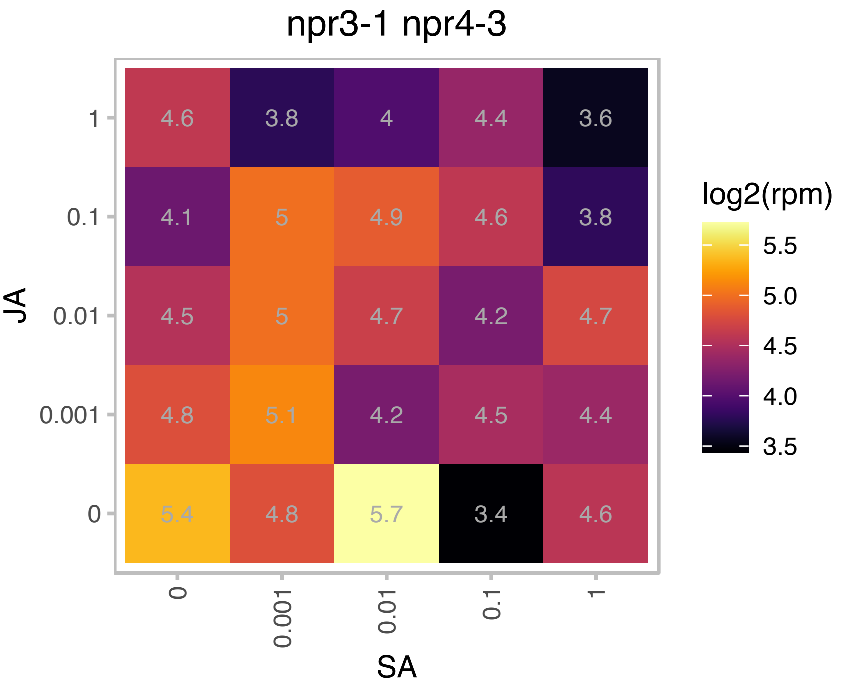

### AT1G65440.1 : GTB1, SPT6L

global transcription factor group B1
Related to yeast Spt6 protein, which functions as part of a protein complex in transcription initiation and also plays a role in chromatin structure / assembly. It encodes a putative WG/GW-repeat protein involved in the regulation of apical-basal polarity of embryo

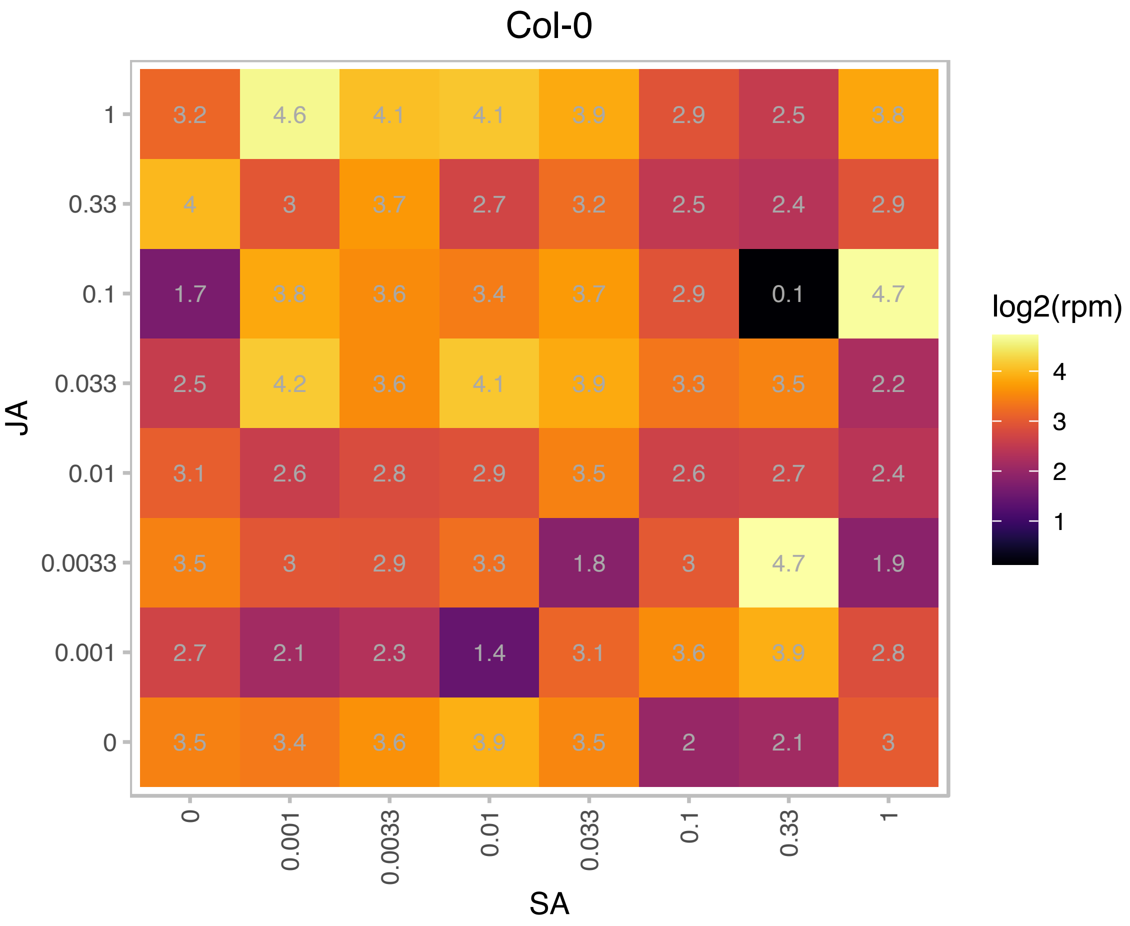


| | |
|---|---|
| Gene | AT1G65440 |
| is.expressed_genes | TRUE |
| is.responsive_genes | TRUE |
| is.apcluster | D-03 |

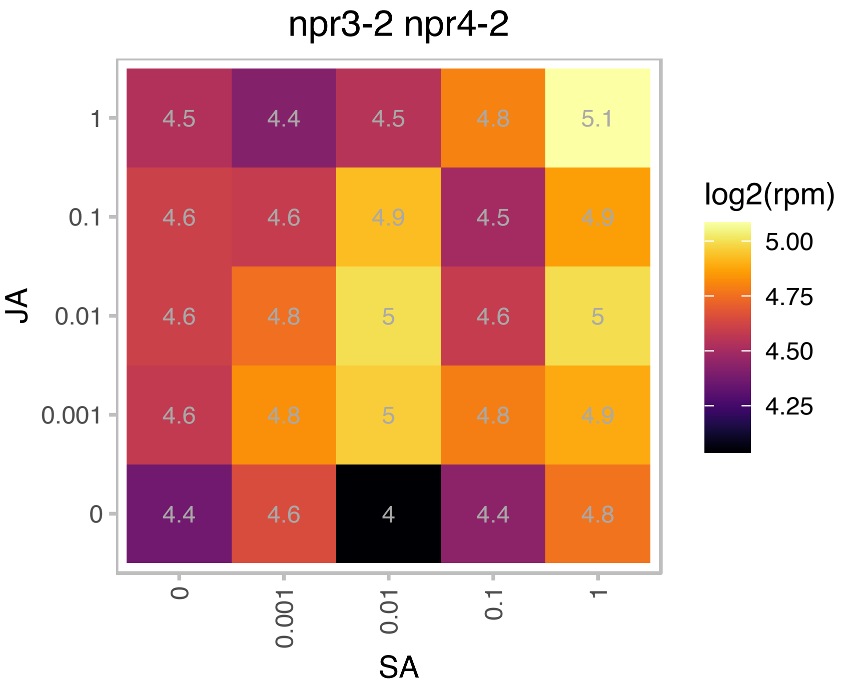


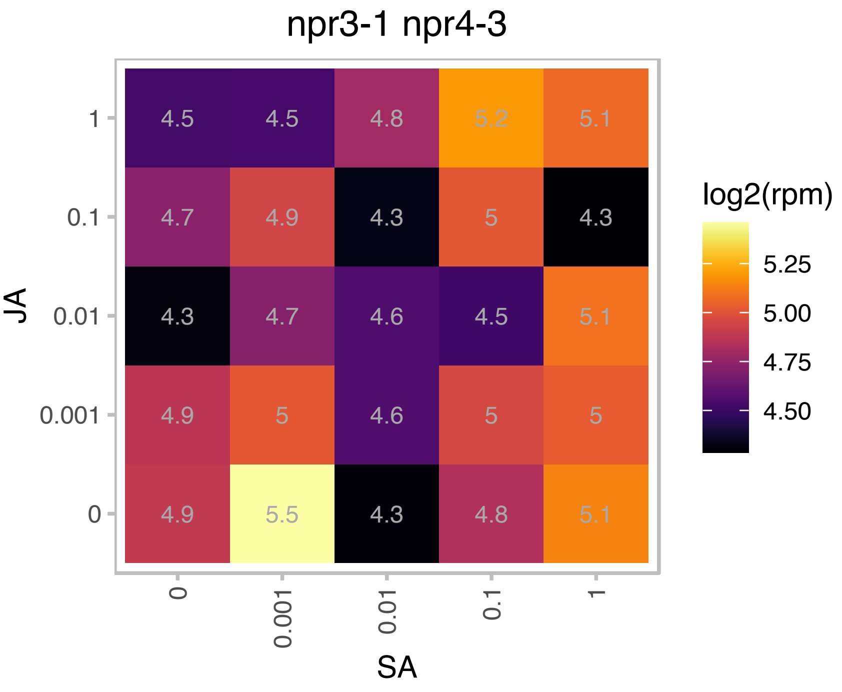

# AT1G72810.1 : NA

Pyridoxal-5'-phosphate-dependent enzyme family protein

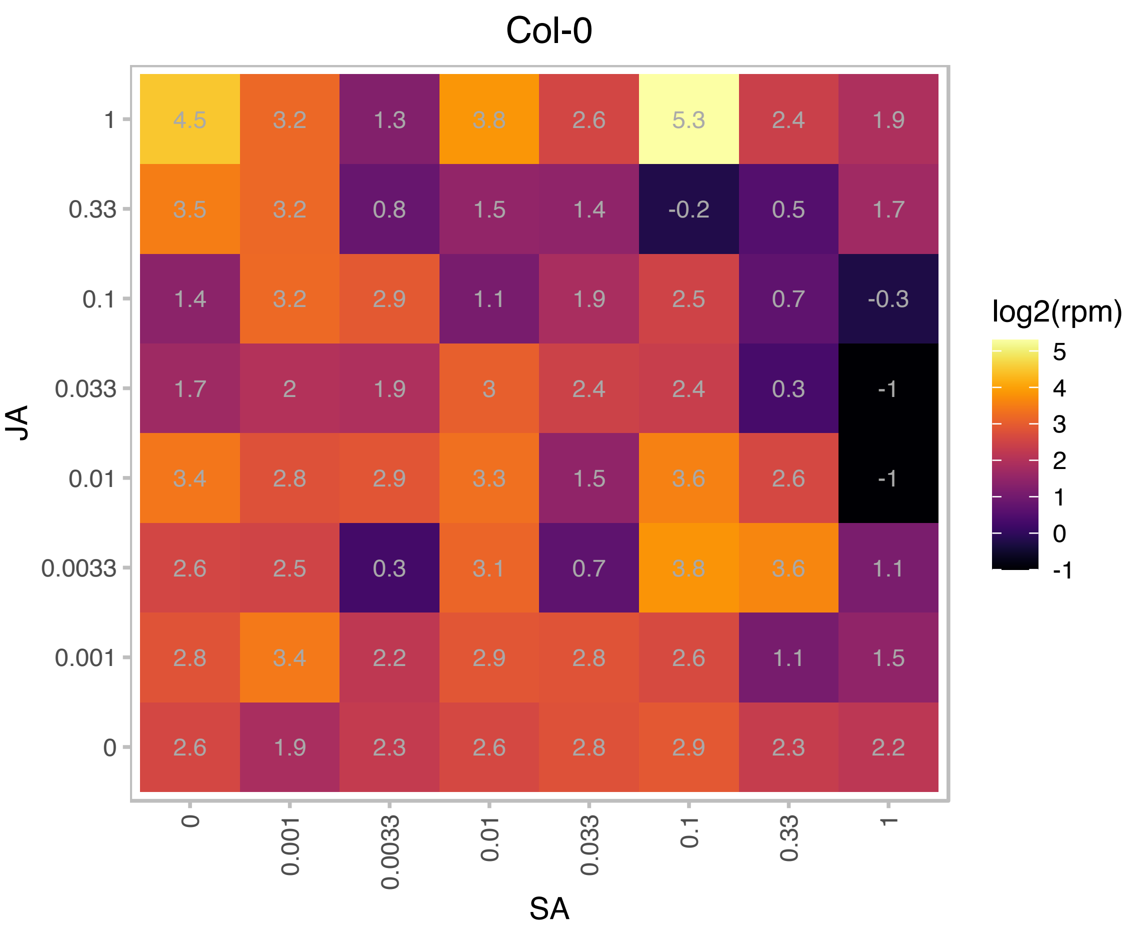


| | |
|---|---|
| Gene | AT1G72810 |
| is.expressed_genes | TRUE |
| is.responsive_genes | TRUE |
| is.apcluster | D-03 |

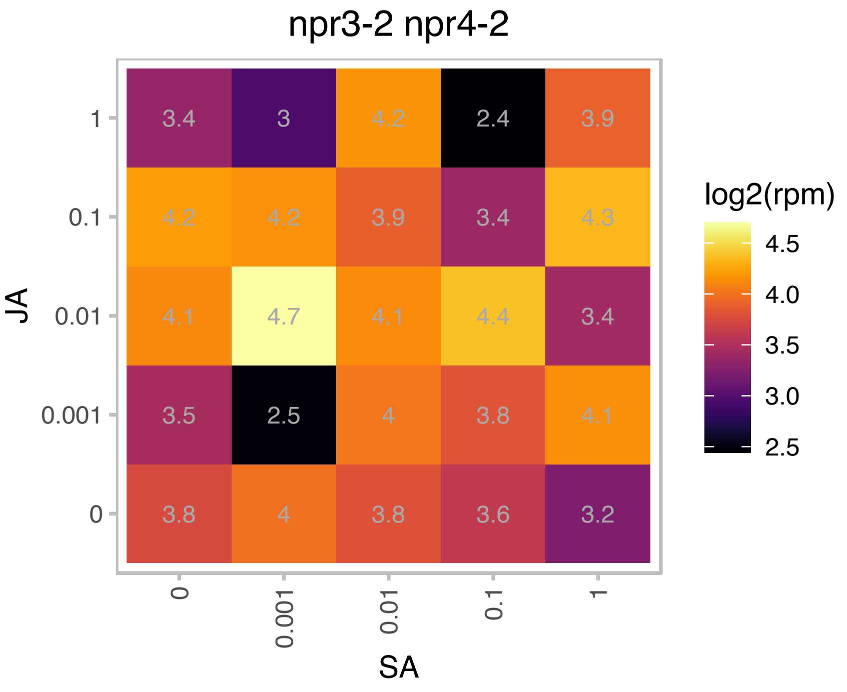


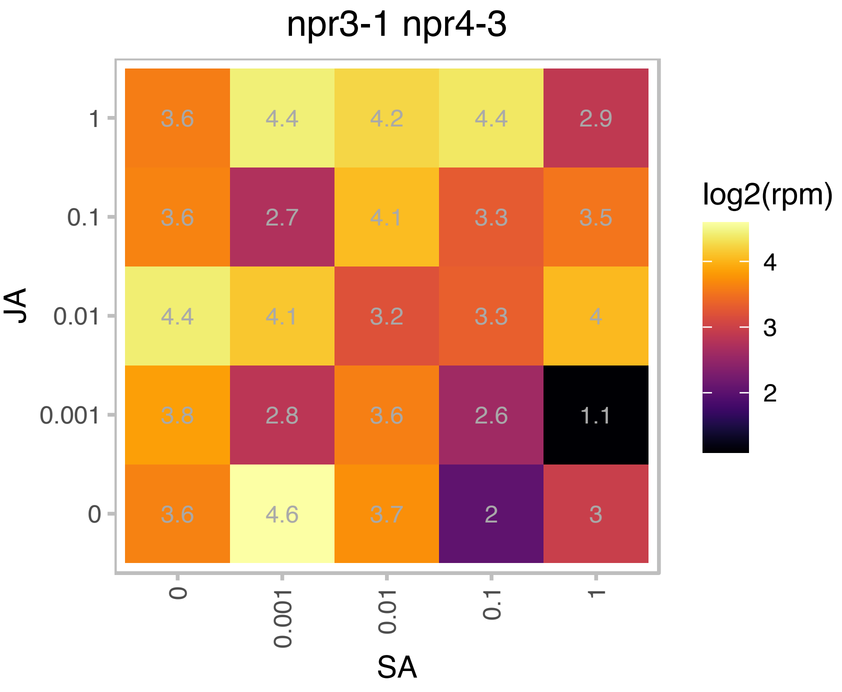

### AT2G01520.1 : MLP328, ZCE1

MLP-like protein 328

Encodes a cis-cinnamic acid responsive gene that is a member of the major latex protein-like gene family and plays a role in promoting vegetative growth and delaying ﬂowering. The mRNA is cell-to-cell mobile.

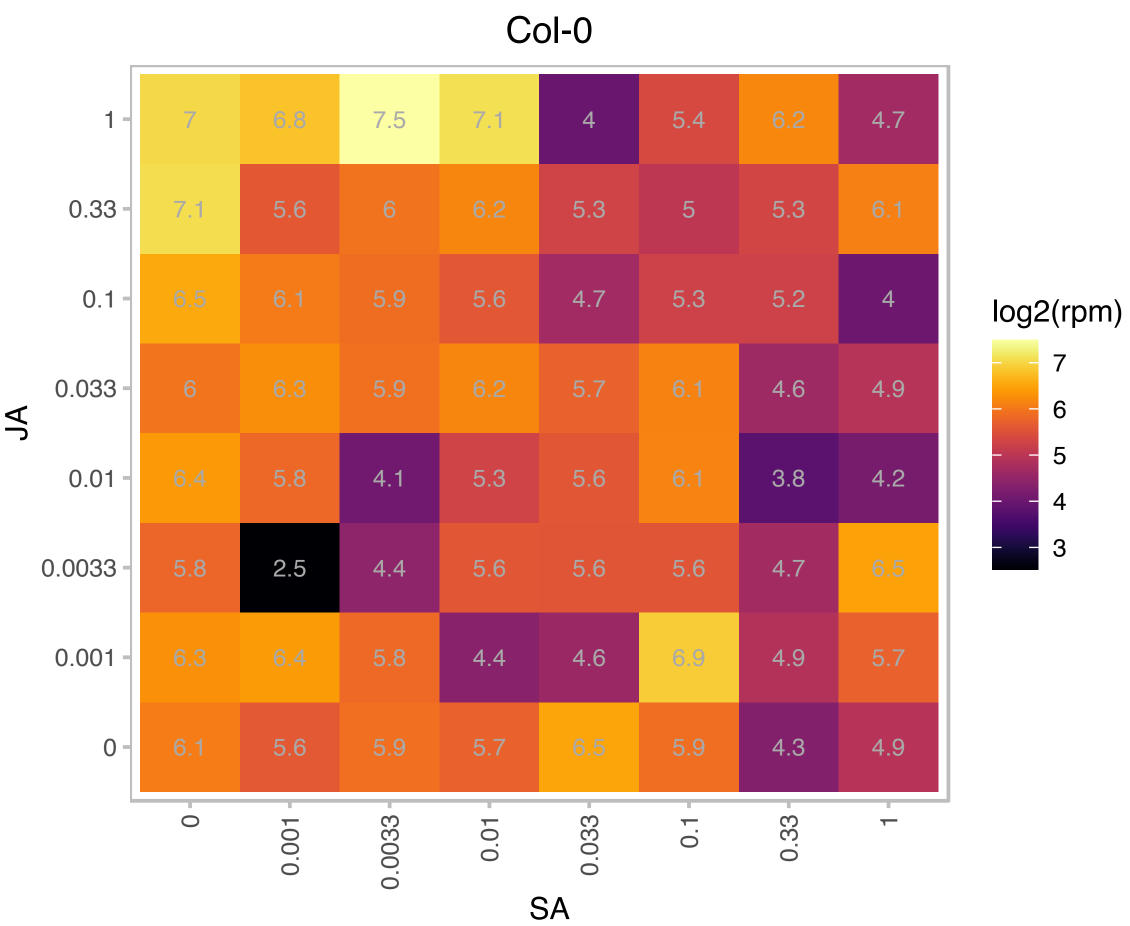


| | |
|---|---|
| Gene | AT2G01520 |
| is.expressed_genes | TRUE |
| is.responsive_genes | TRUE |
| is.apcluster | D-03 |

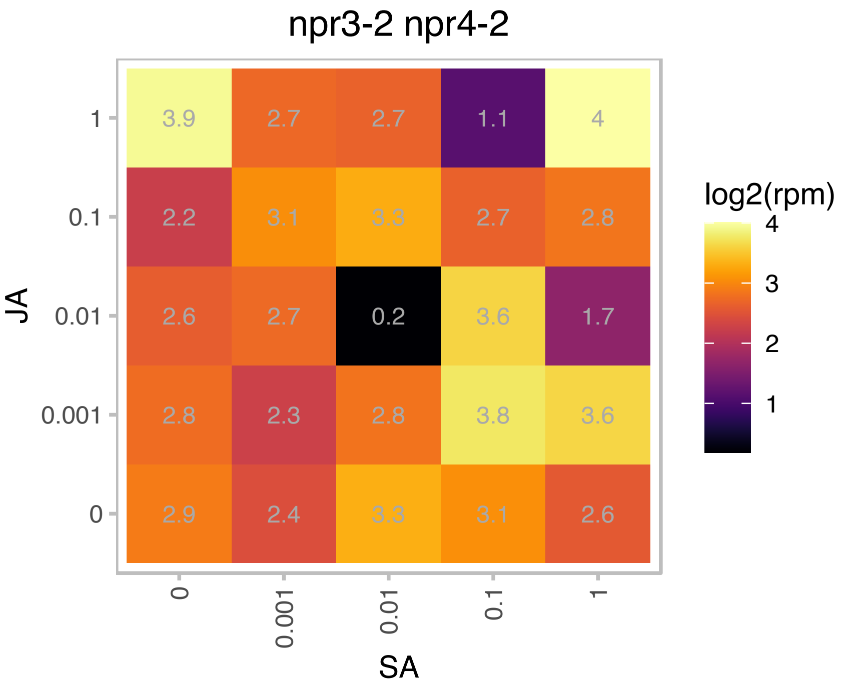


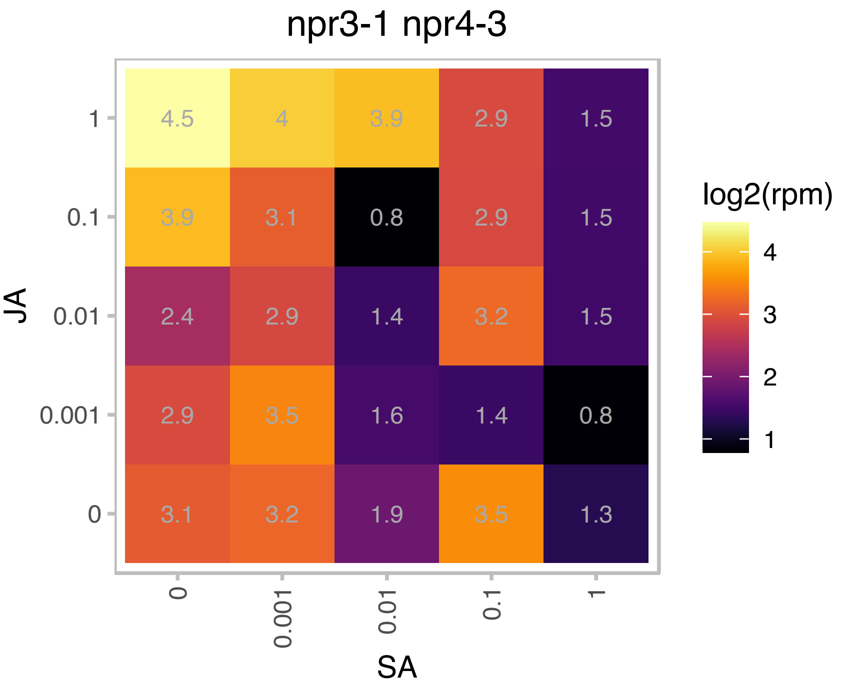

# AT2G08600.1 : NA

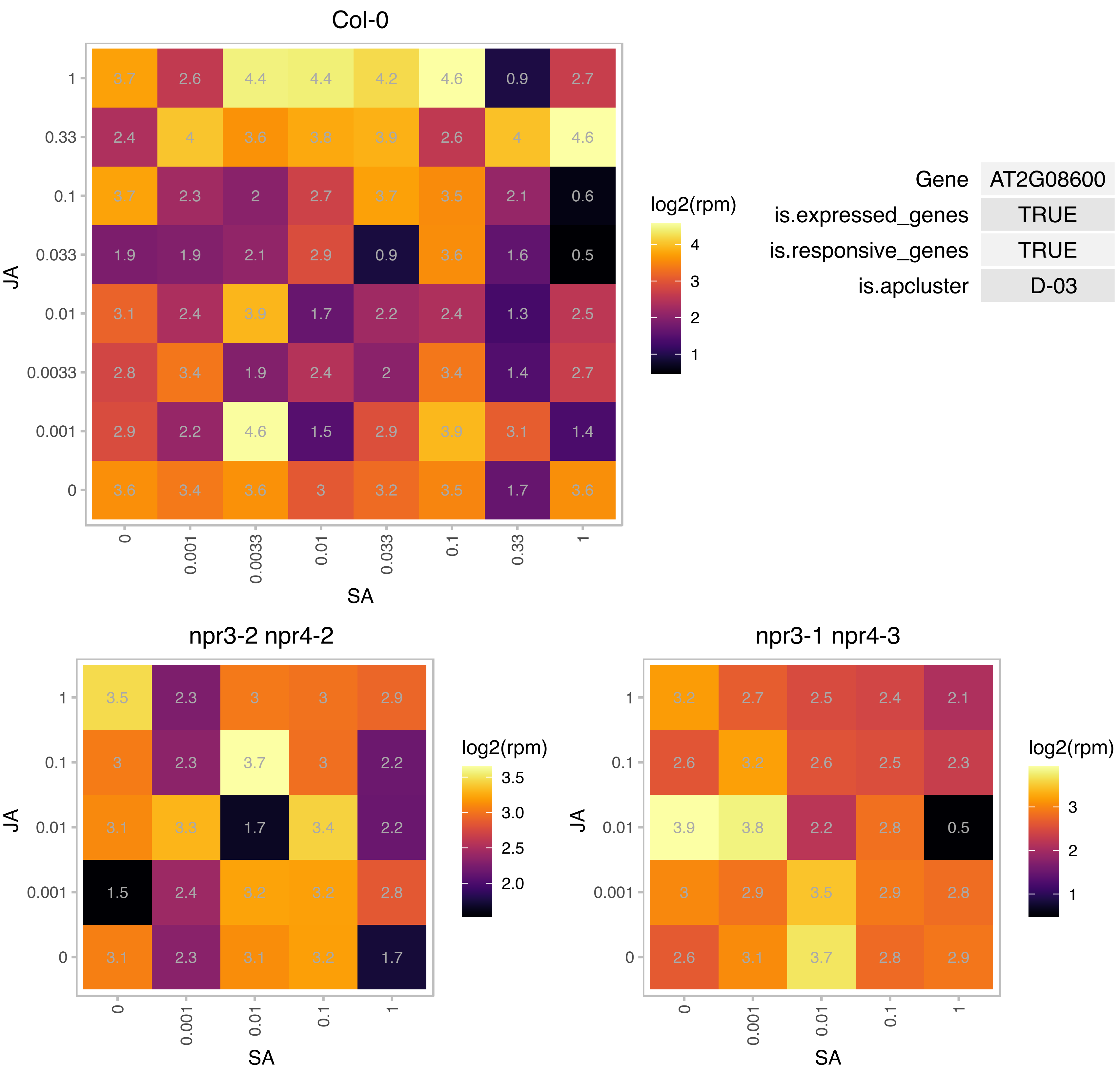

### AT2G22640.1 : ATBRK1, BRK1, HSPC300

BRICK1, putative
Component of the WAVE protein complex which act as activators of ARP2/3 complex involved in actin nucleation. Required for trichome morphogenesis. Required for accumulation of SCAR1 protein in vivo. Selectively stabilizes SCAR2.

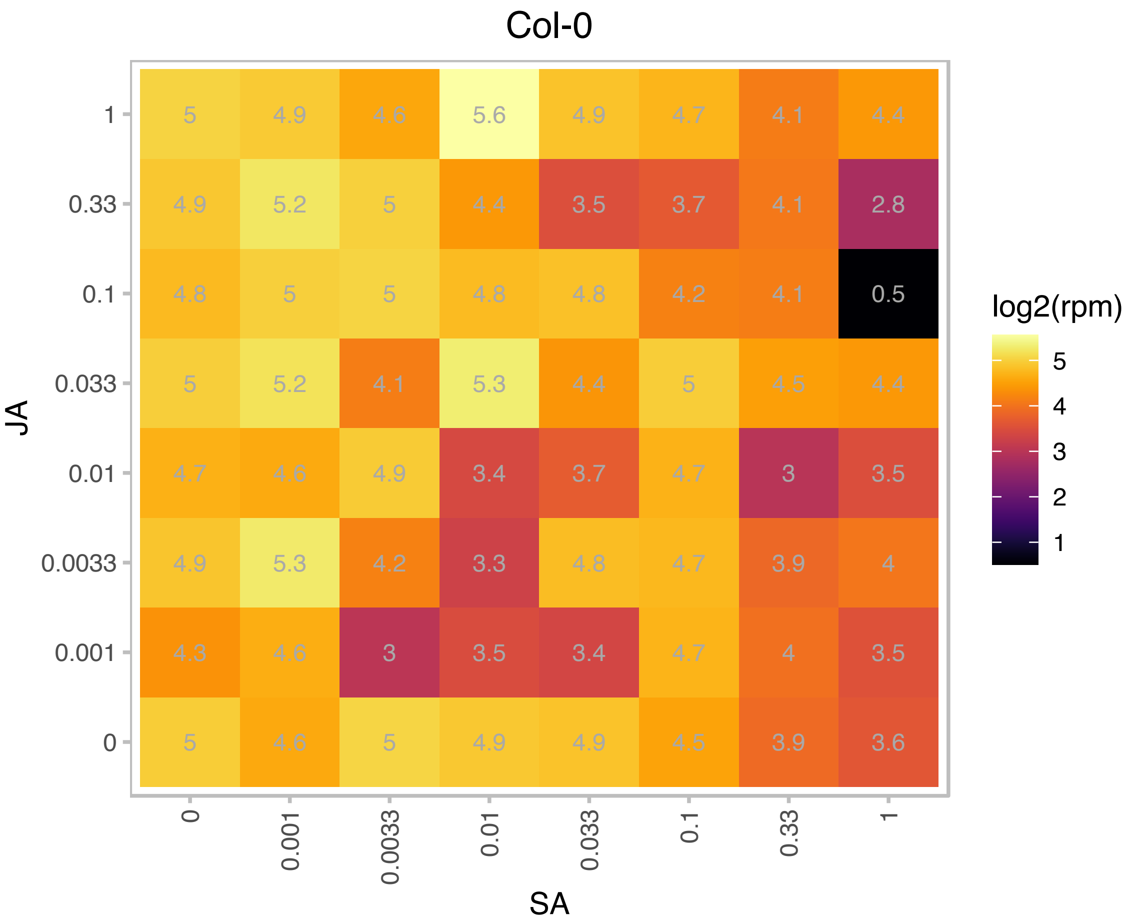


| | |
|---|---|
| Gene | AT2G22640 |
| is.expressed_genes | TRUE |
| is.responsive_genes | TRUE |
| is.apcluster | D-03 |

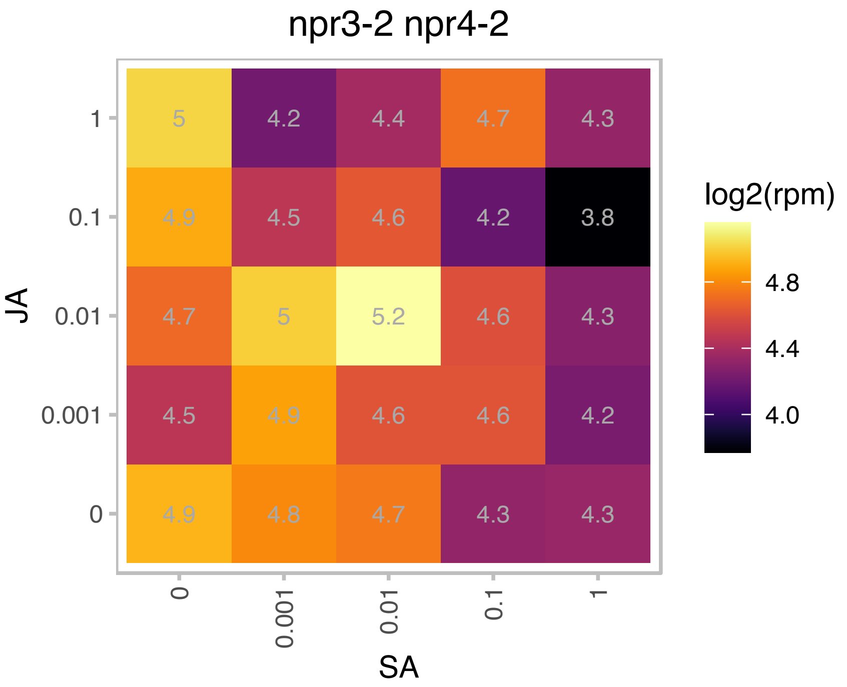


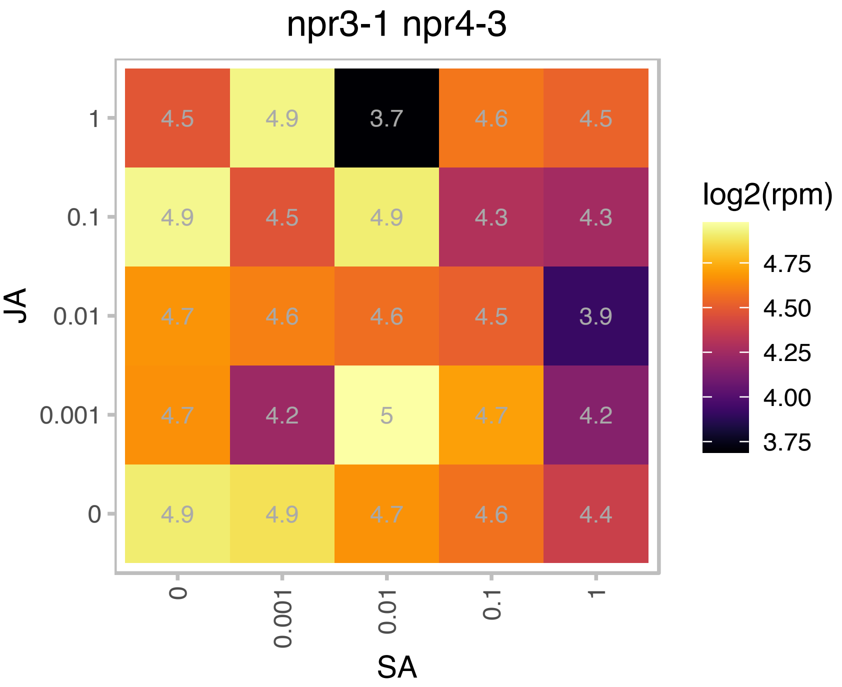

# AT2G23390.1 : NA

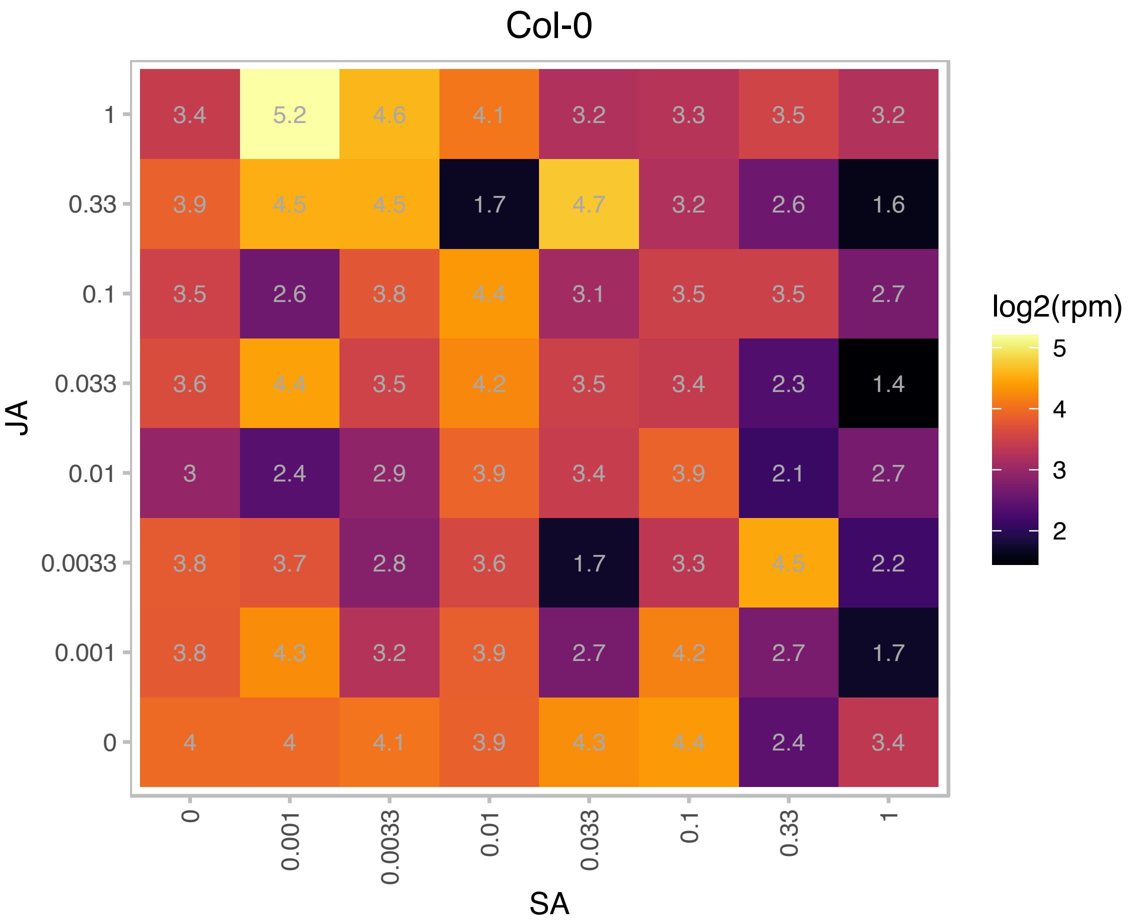


| | |
|---|---|
| Gene | AT2G23390 |
| is.expressed_genes | TRUE |
| is.responsive_genes | TRUE |
| is.apcluster | D-03 |

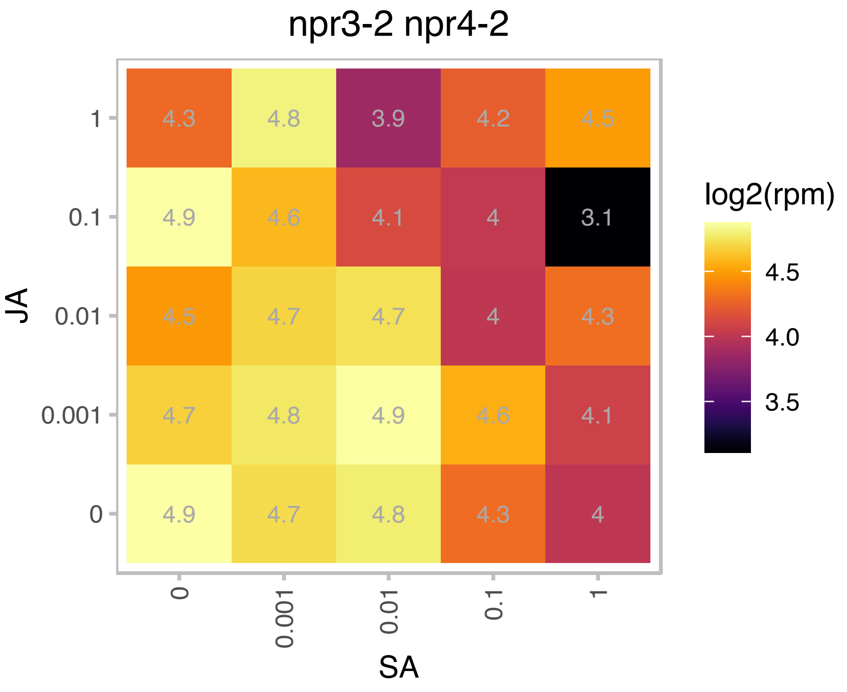


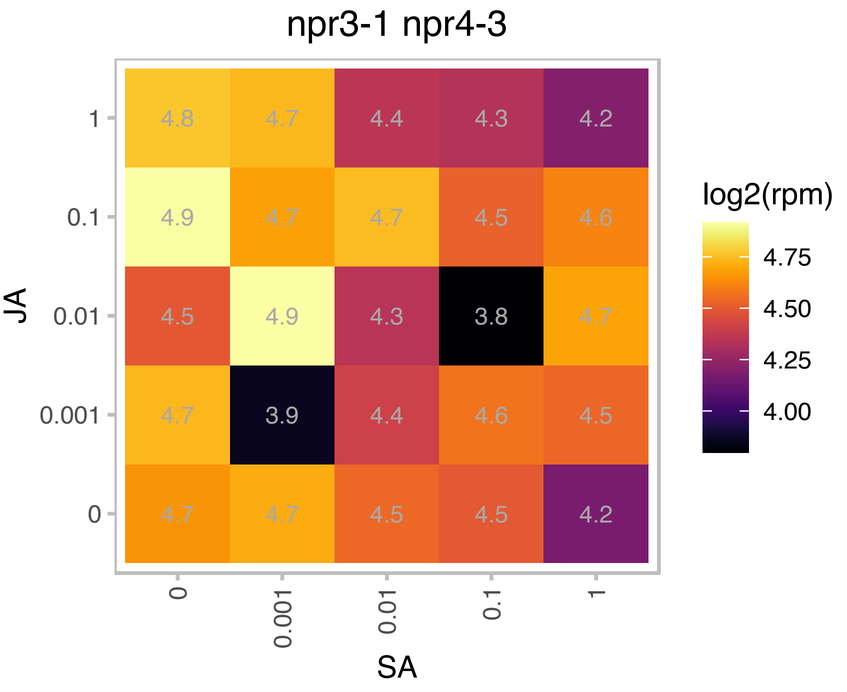

# AT2G28510.1 : NA

Dof-type zinc finger DNA-binding family protein

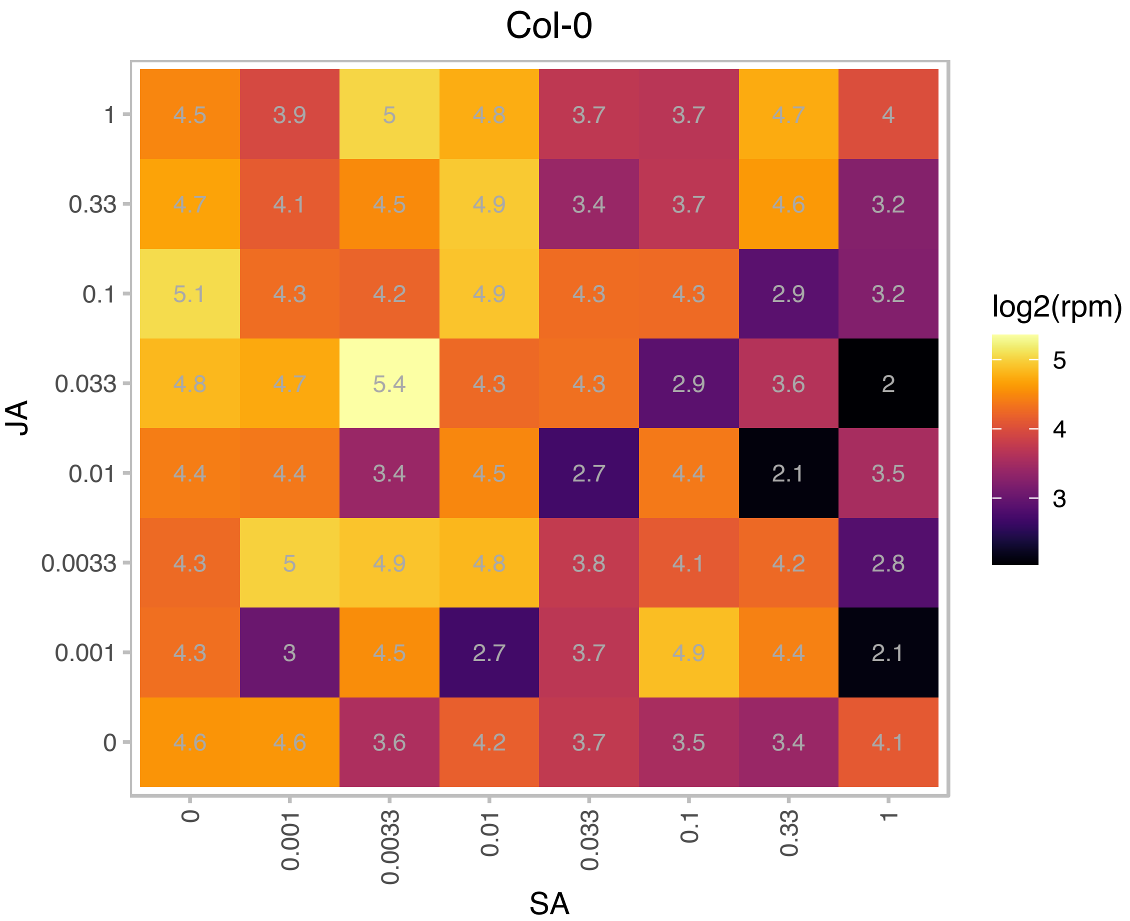


| | |
|---|---|
| Gene | AT2G28510 |
| is.expressed_genes | TRUE |
| is.responsive_genes | TRUE |
| is.apcluster | D-03 |

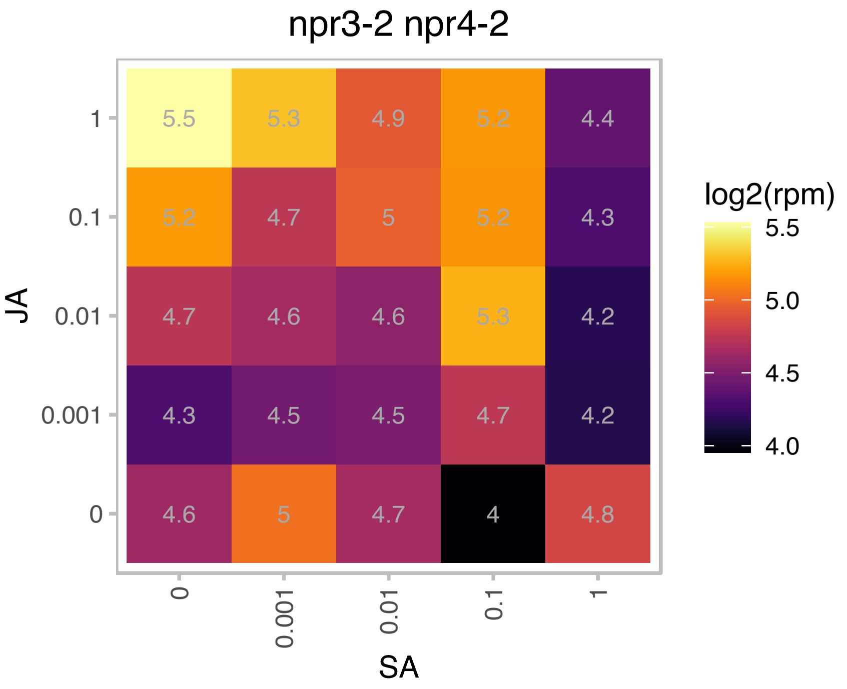


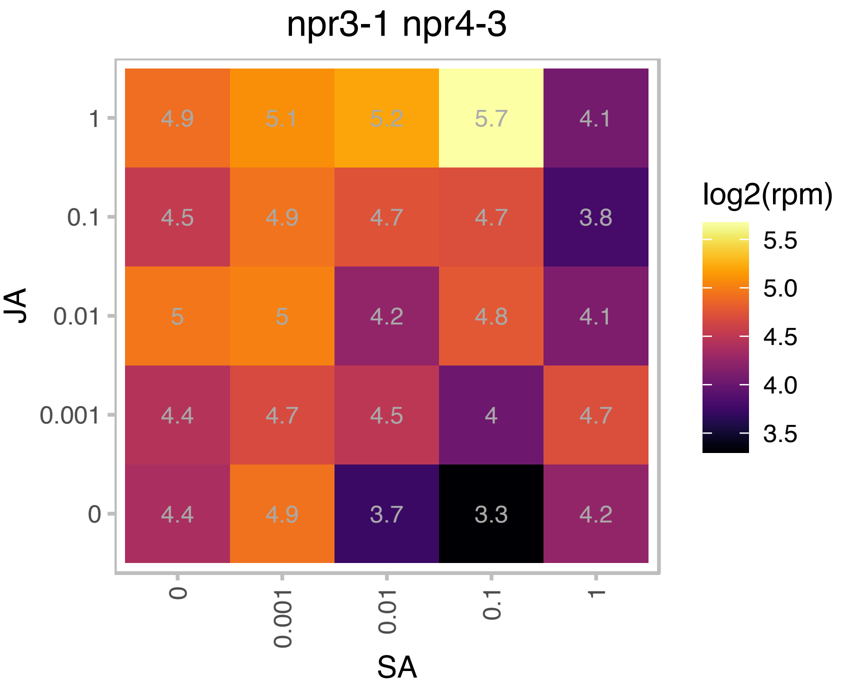

### AT2G29190.1 : APUM2, PUM2

pumilio 2
Encodes a member of the Arabidopsis Pumilio (APUM) proteins containing PUF domain (eight repeats of approximately 36 amino acids each). PUF proteins regulate both mRNA stability and translation through sequence-specific binding to the 3' UTR of target mRNA transcripts.

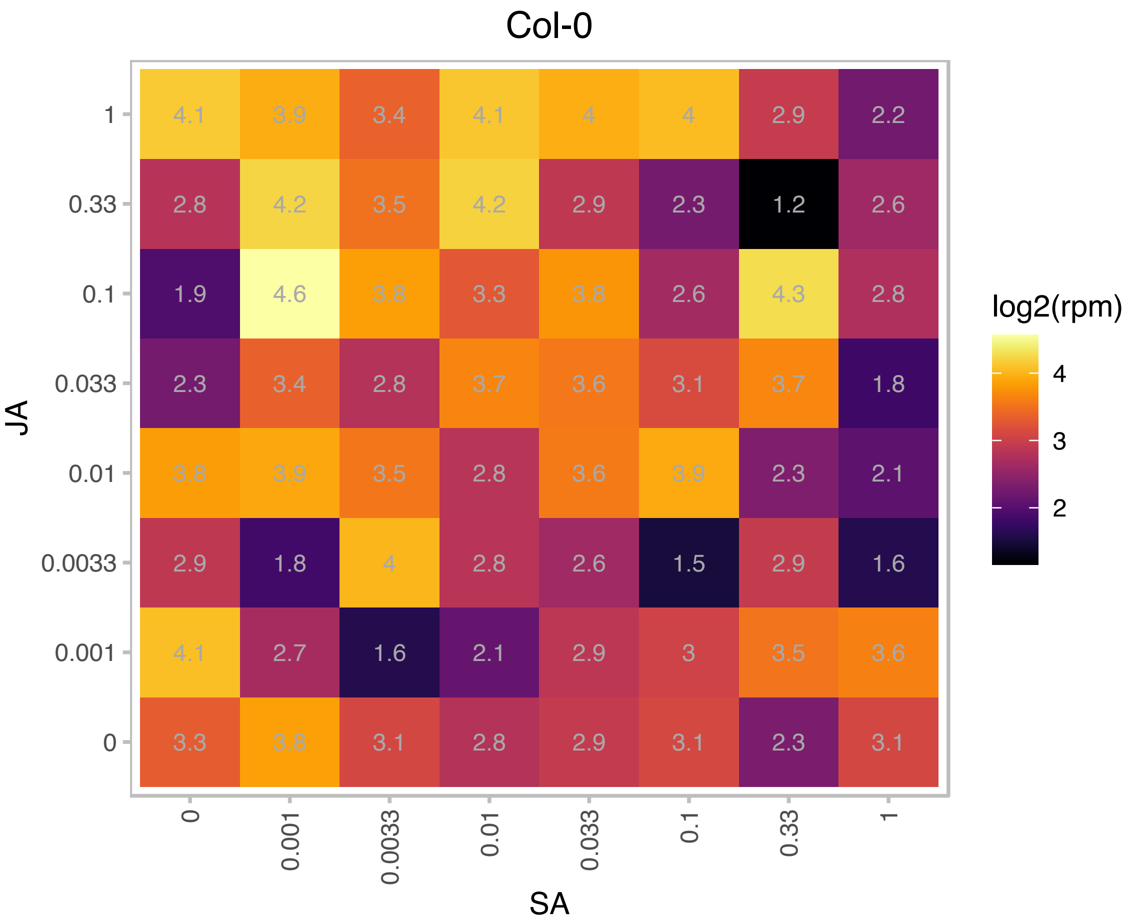


| | |
|---|---|
| Gene | AT2G29190 |
| is.expressed_genes | TRUE |
| is.responsive_genes | TRUE |
| is.apcluster | D-03 |

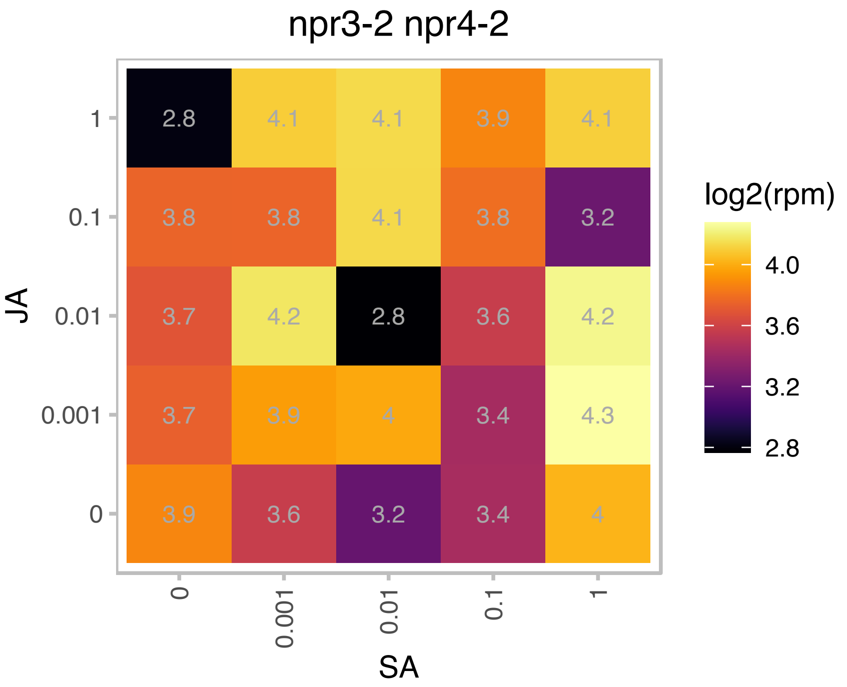


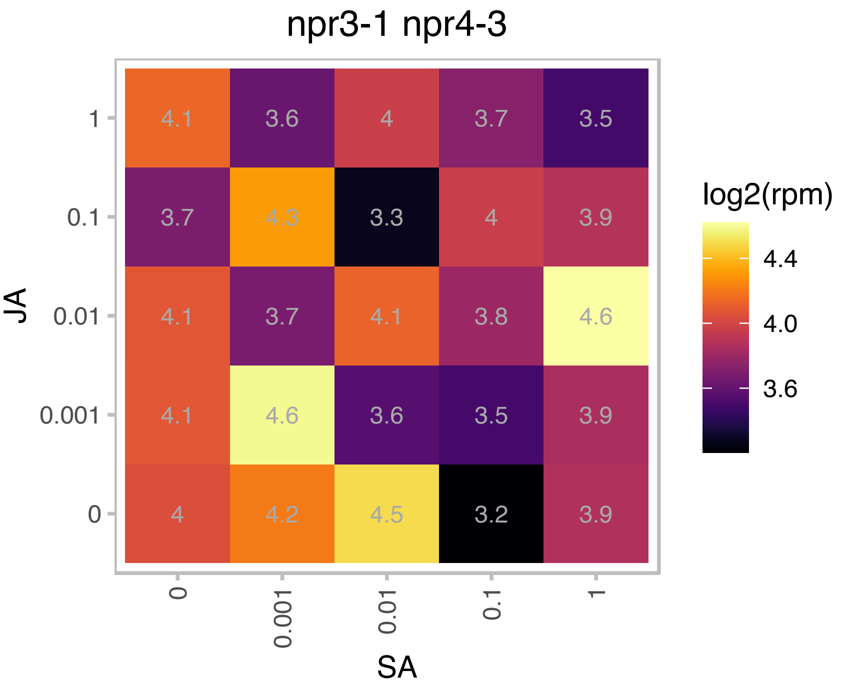

### AT2G30970.1 : ASP1

aspartate aminotransferase 1

ASPARTATE AMINOTRANSFERASE 1

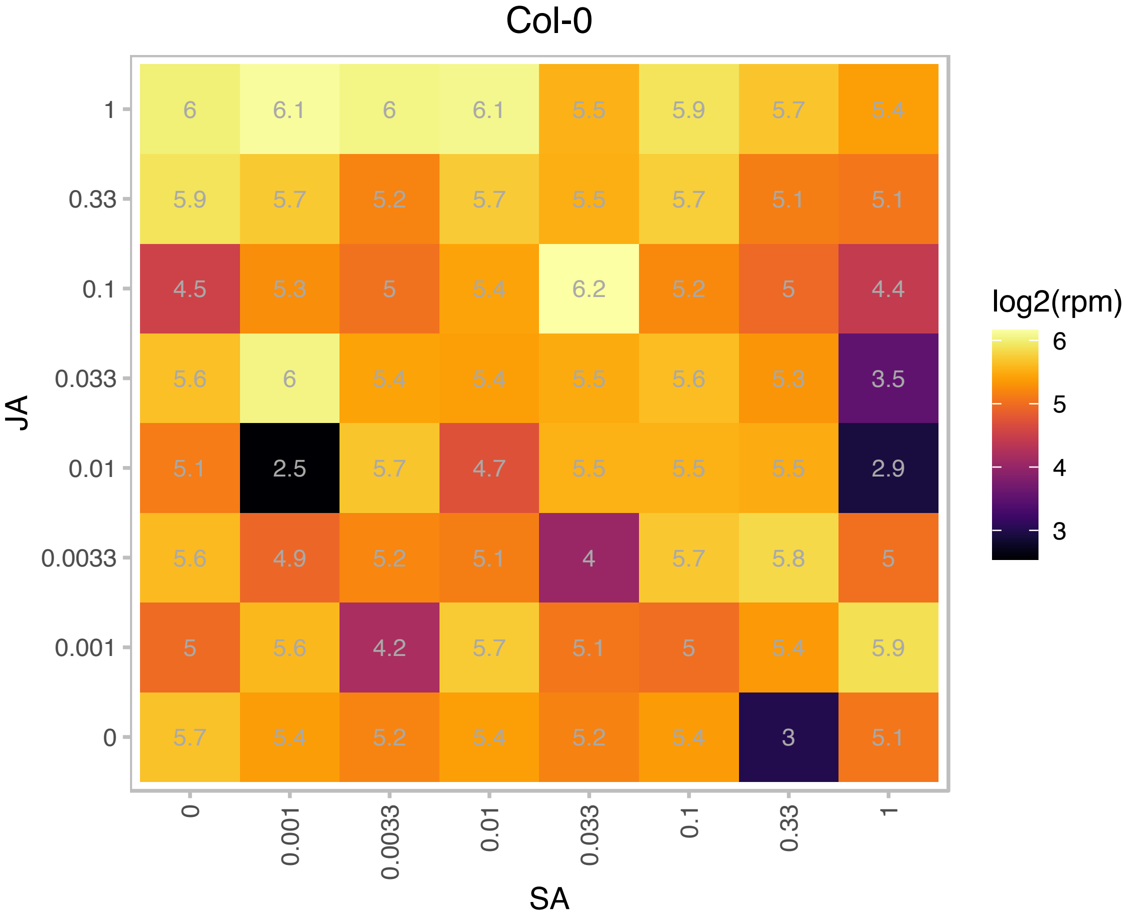


| Gene | AT2G30970 |
|---|---|
| is.expressed_genes | TRUE |
| is.responsive_genes | TRUE |
| is.apcluster | D-03 |

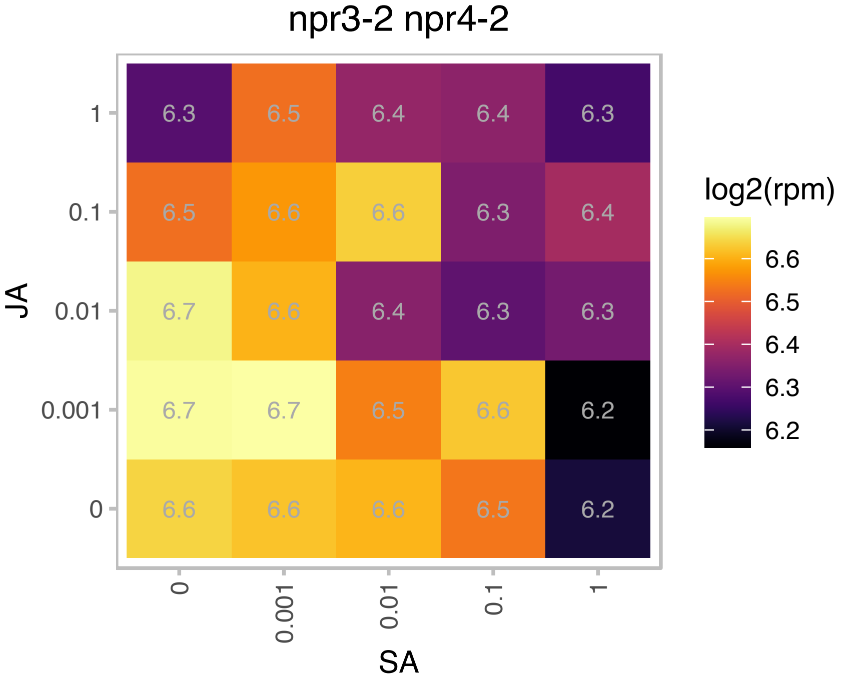


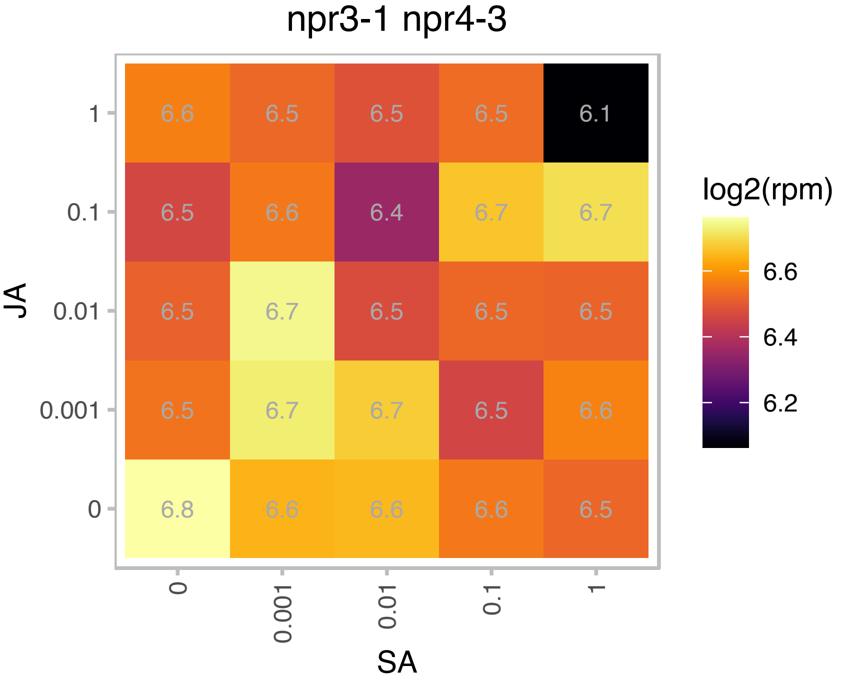

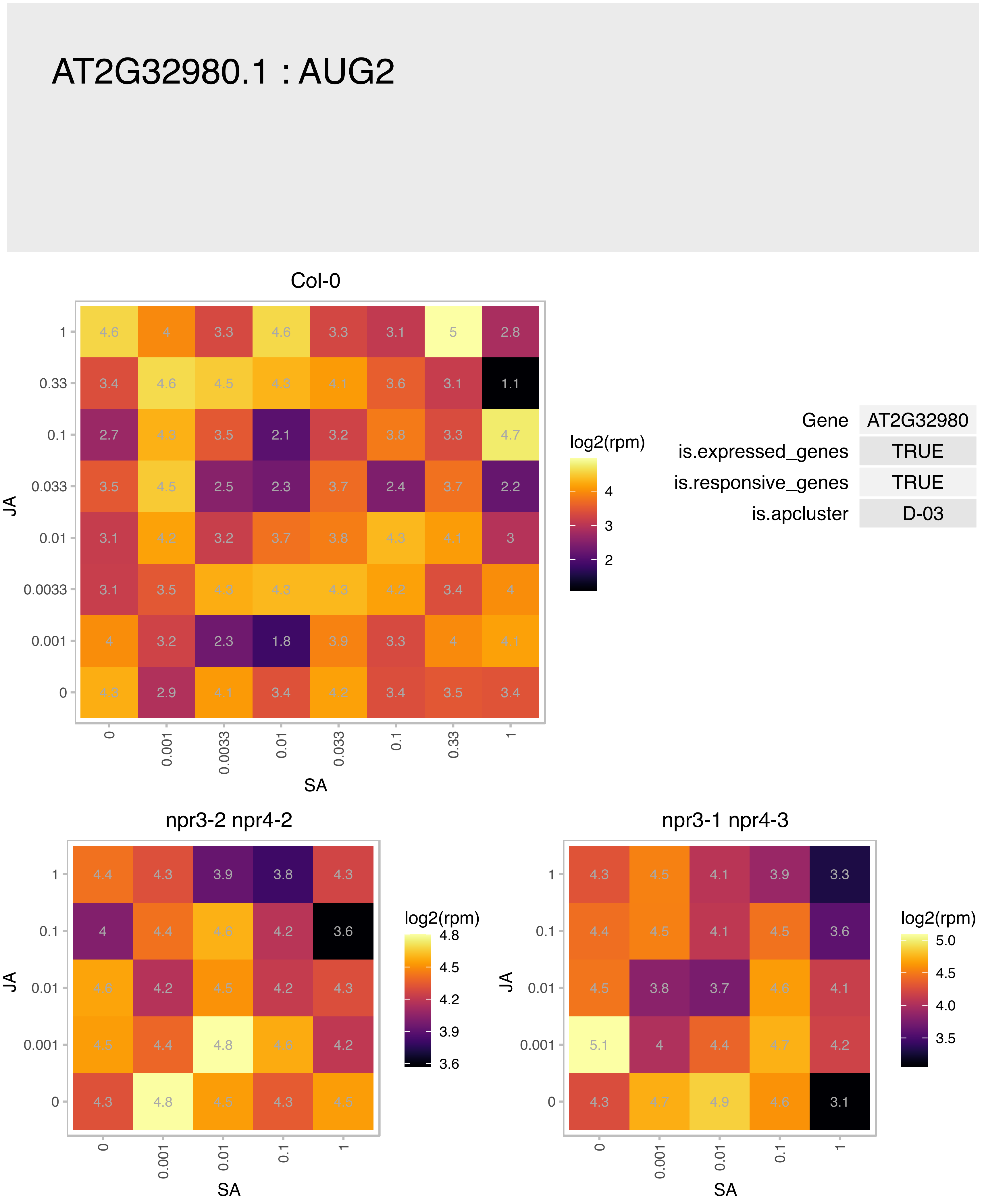
AT2G32980.1 : AUG2
Col-0
JA
SA
1 0.33 0.1 0.033 0.01 0.0033 0.001 0
0 0.001 0.0033 0.01 0.033 0.1 0.33 1
4.6 4 3.3 4.6 3.3 3.1 5 2.8
3.4 4.6 4.5 4.3 4.1 3.6 3.1 1.1
2.7 4.3 3.5 2.1 3.2 3.8 3.3 4.7
3.5 4.5 2.5 2.3 3.7 2.4 3.7 2.2
3.1 4.2 3.2 3.7 3.8 4.3 4.1 3
3.1 3.5 4.3 4.3 4.3 4.2 3.4 4
4 3.2 2.3 1.8 3.9 3.3 4 4.1
4.3 2.9 4.1 3.4 4.2 3.4 3.5 3.4
log2(rpm)
4
3
2
Gene AT2G32980
is.expressed_genes TRUE
is.responsive_genes TRUE
is.apcluster D-03
npr3-2 npr4-2
JA
SA
1 0.1 0.01 0.001 0
0 0.001 0.01 0.1 1
4.4 4.3 3.9 3.8 4.3
4 4.4 4.6 4.2 3.6
4.6 4.2 4.5 4.2 4.3
4.5 4.4 4.8 4.6 4.2
4.3 4.8 4.5 4.3 4.5
log2(rpm)
4.8
4.5
4.2
3.9
3.6
npr3-1 npr4-3
JA
SA
1 0.1 0.01 0.001 0
0 0.001 0.01 0.1 1
4.3 4.5 4.1 3.9 3.3
4.4 4.5 4.1 4.5 3.6
4.5 3.8 3.7 4.6 4.1
5.1 4 4.4 4.7 4.2
4.3 4.7 4.9 4.6 3.1
log2(rpm)
5.0
4.5
4.0
3.5

### AT2G34140.1 : CDF4

Dof-type zinc finger DNA-binding family protein

CDF4 is member of the group II DOF transcription factor family is involved in regulation of differentiation root columella cells. It is a direct target of the transcriptional repressor WOX5. CDF4 itself is a transcriptional repressor that appears to repress root columella stem cell identity. Ectopic expression of CDF leads to premature differentiation of root columella cells.

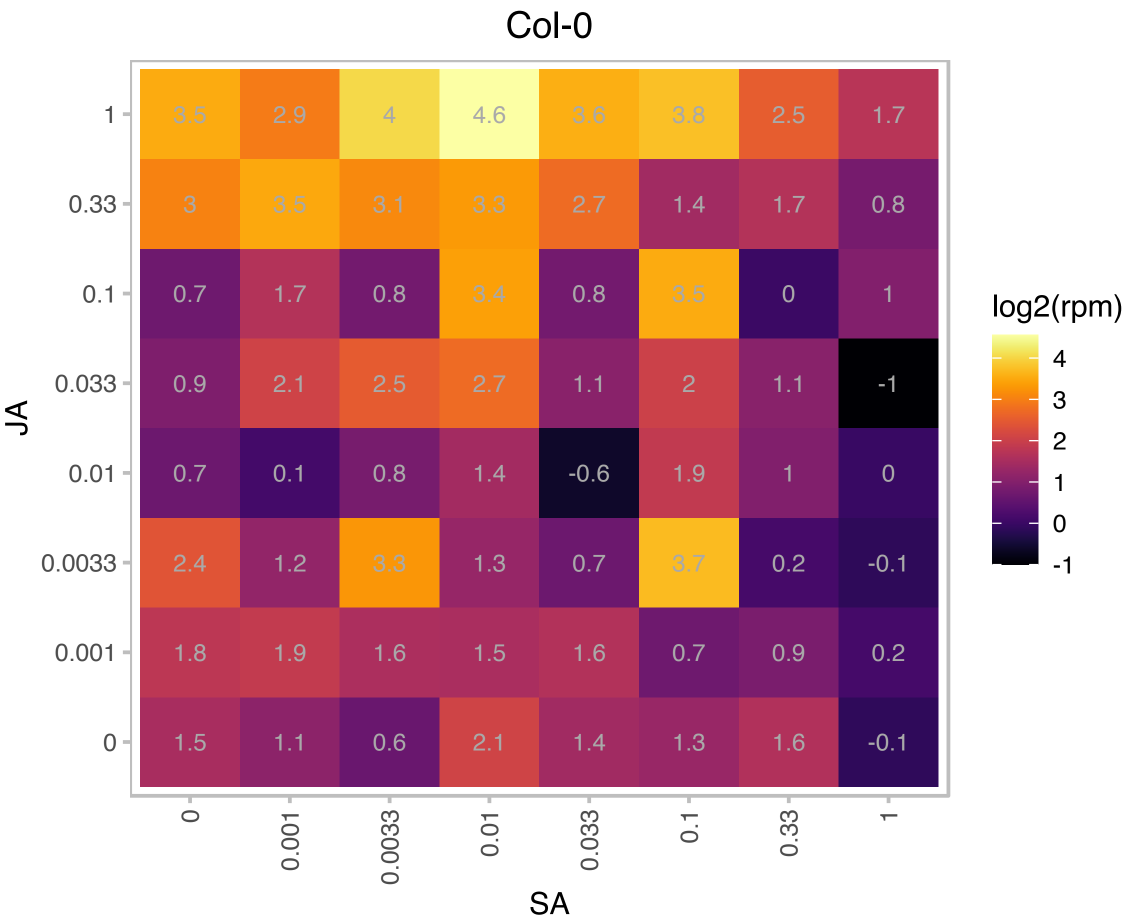


| | |
|---|---|
| Gene | AT2G34140 |
| is.expressed_genes | TRUE |
| is.responsive_genes | TRUE |
| is.apcluster | D-03 |

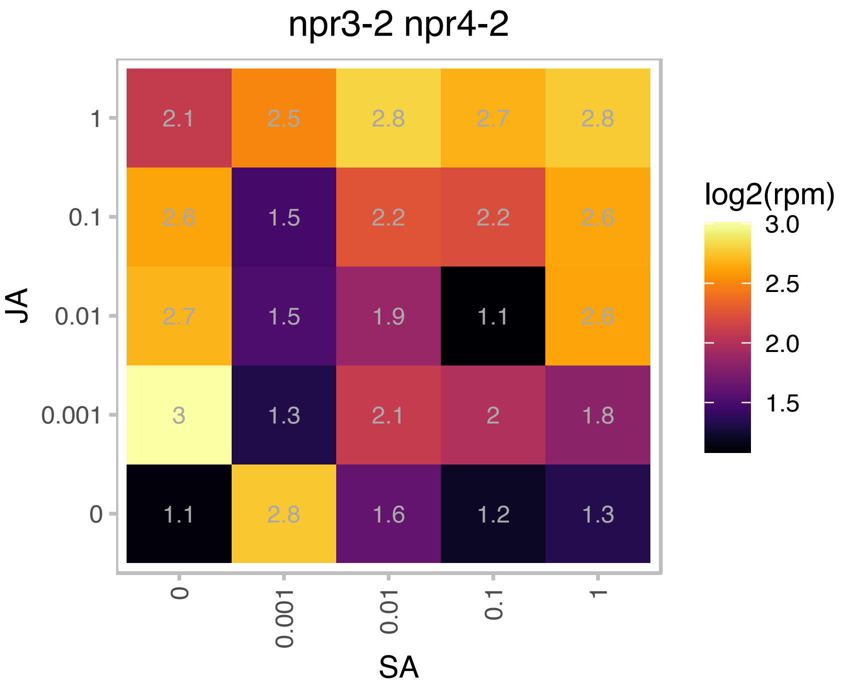


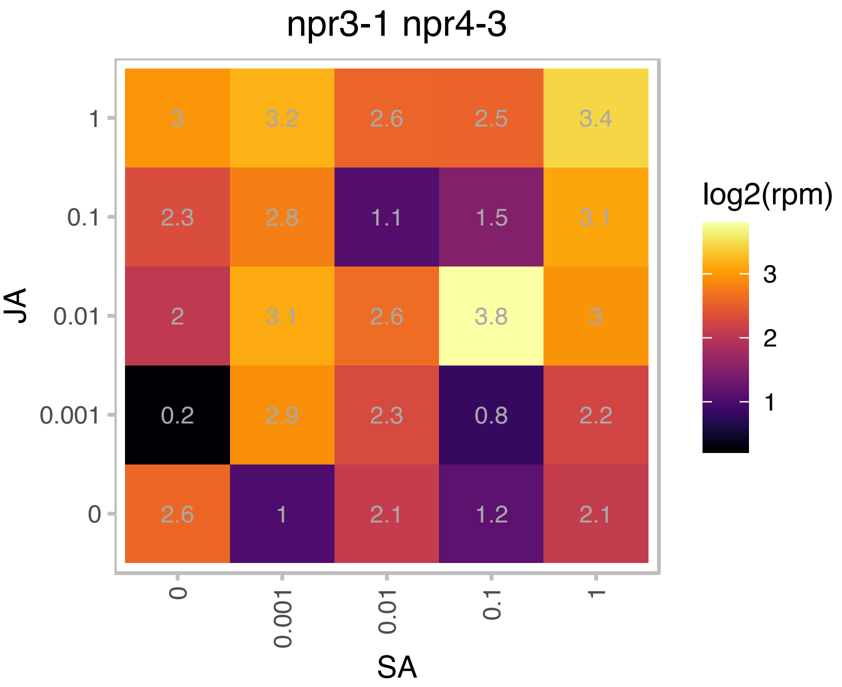

### AT2G46030.1 : UBC6

ubiquitin-conjugating enzyme 6

Ubiquitin conjugating enzyme E2

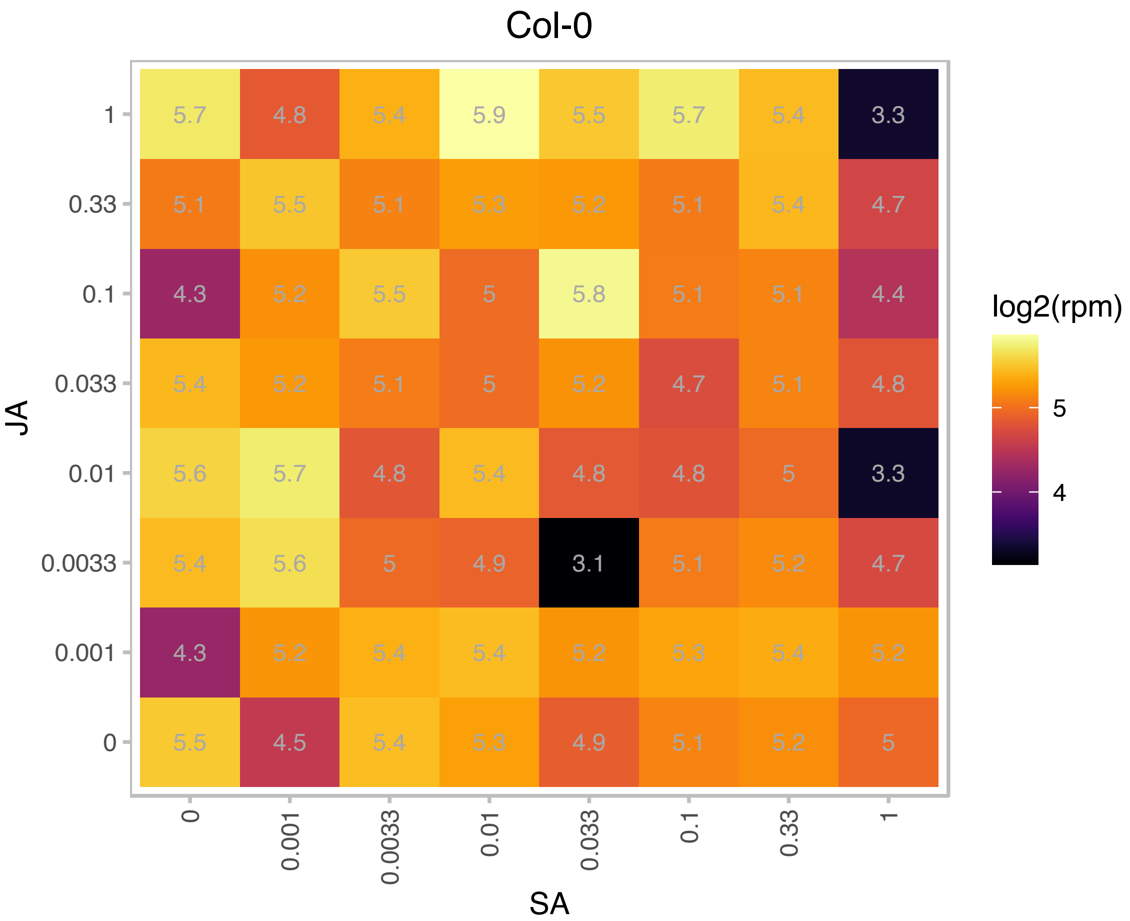


| | |
|---|---|
| Gene | AT2G46030 |
| is.expressed_genes | TRUE |
| is.responsive_genes | TRUE |
| is.apcluster | D-03 |

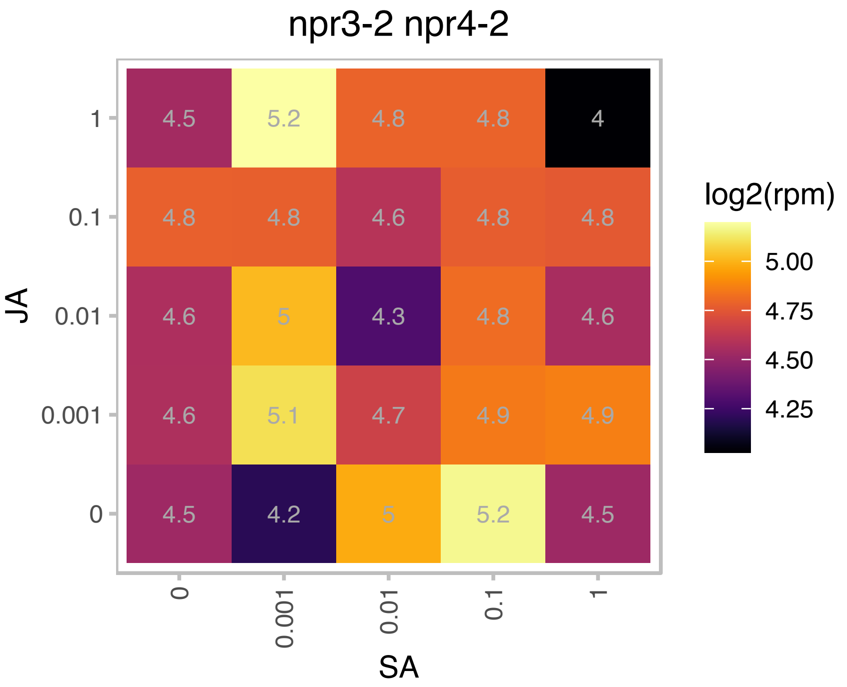


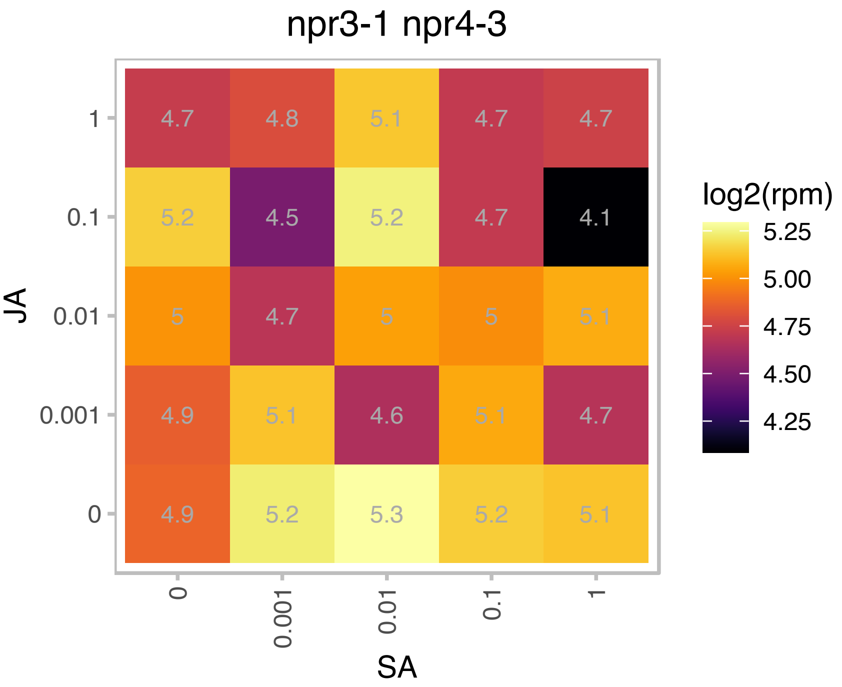

# AT2G48130.1 : NA

Bifunctional inhibitor/lipid-transfer protein/seed storage 2S albumin superfamily protein

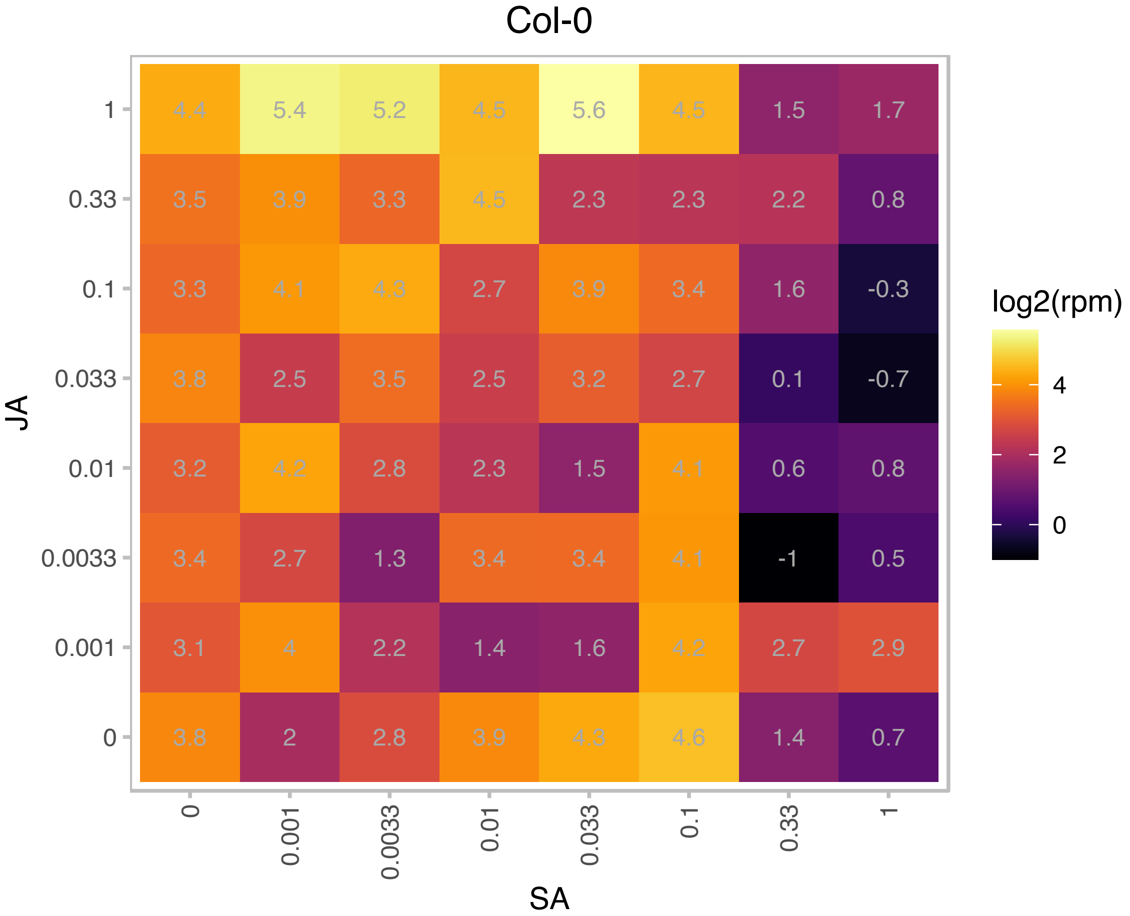


| | |
|---|---|
| Gene | AT2G48130 |
| is.expressed_genes | TRUE |
| is.responsive_genes | TRUE |
| is.apcluster | D-03 |

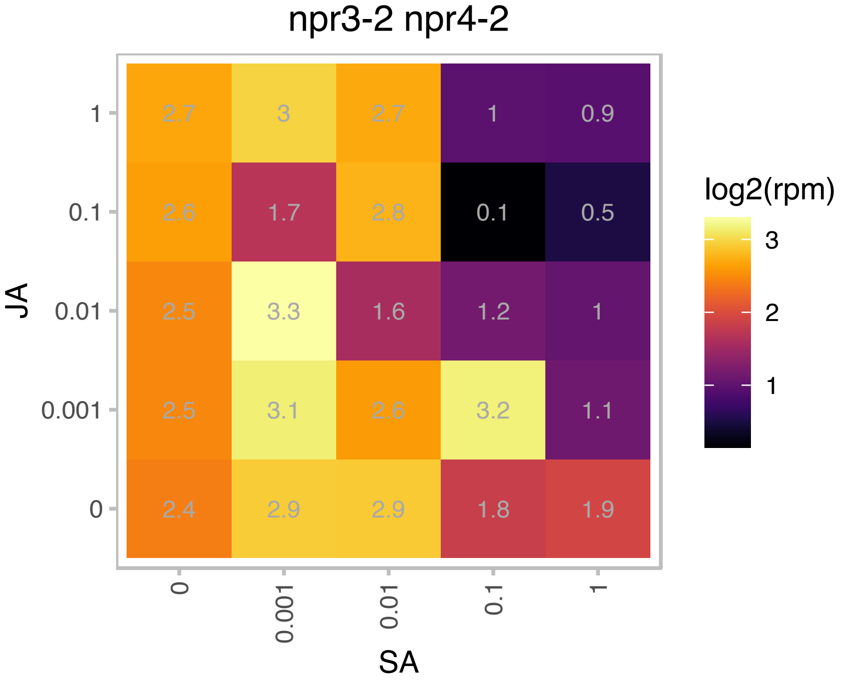


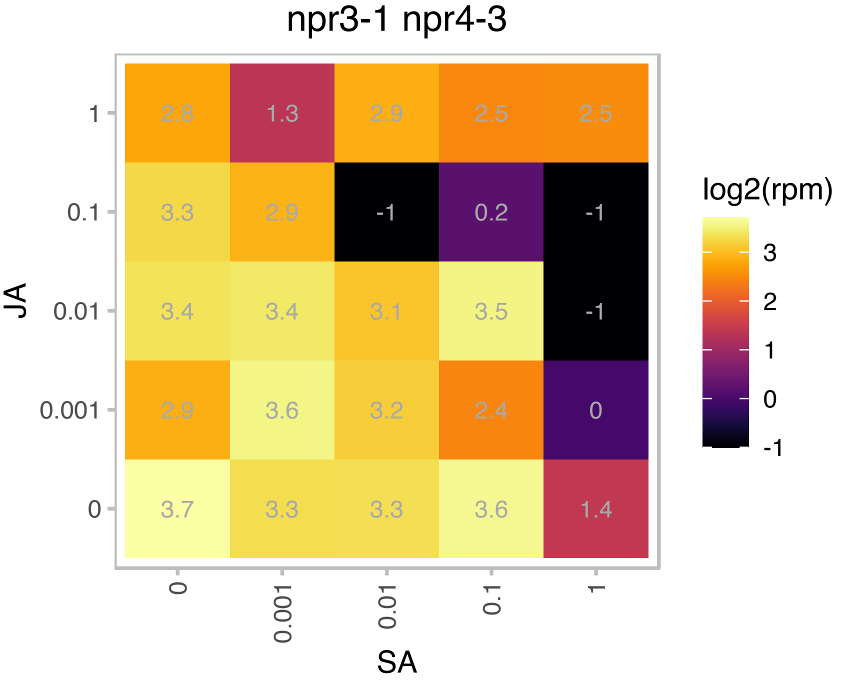

# AT3G01060.1 : NA

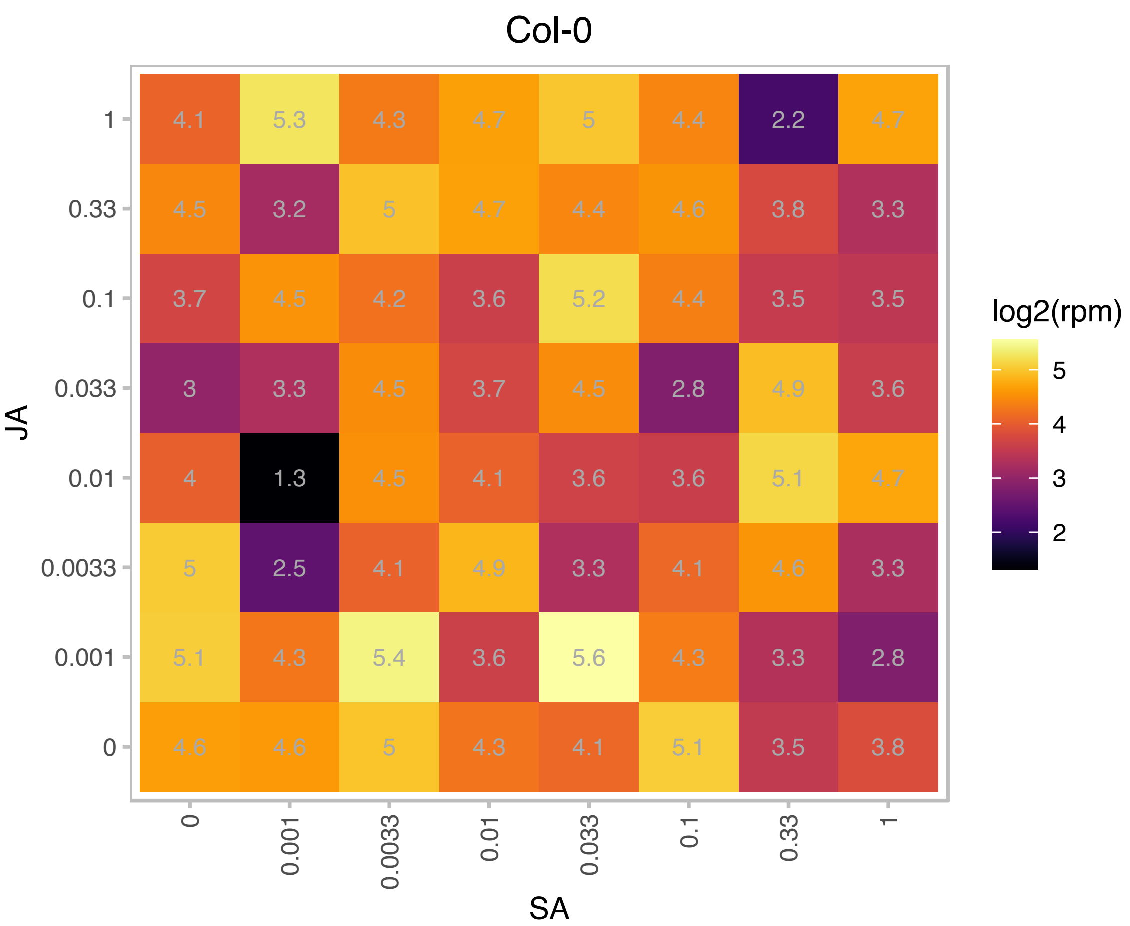


| | |
|---|---|
| Gene | AT3G01060 |
| is.expressed_genes | TRUE |
| is.responsive_genes | TRUE |
| is.apcluster | D-03 |

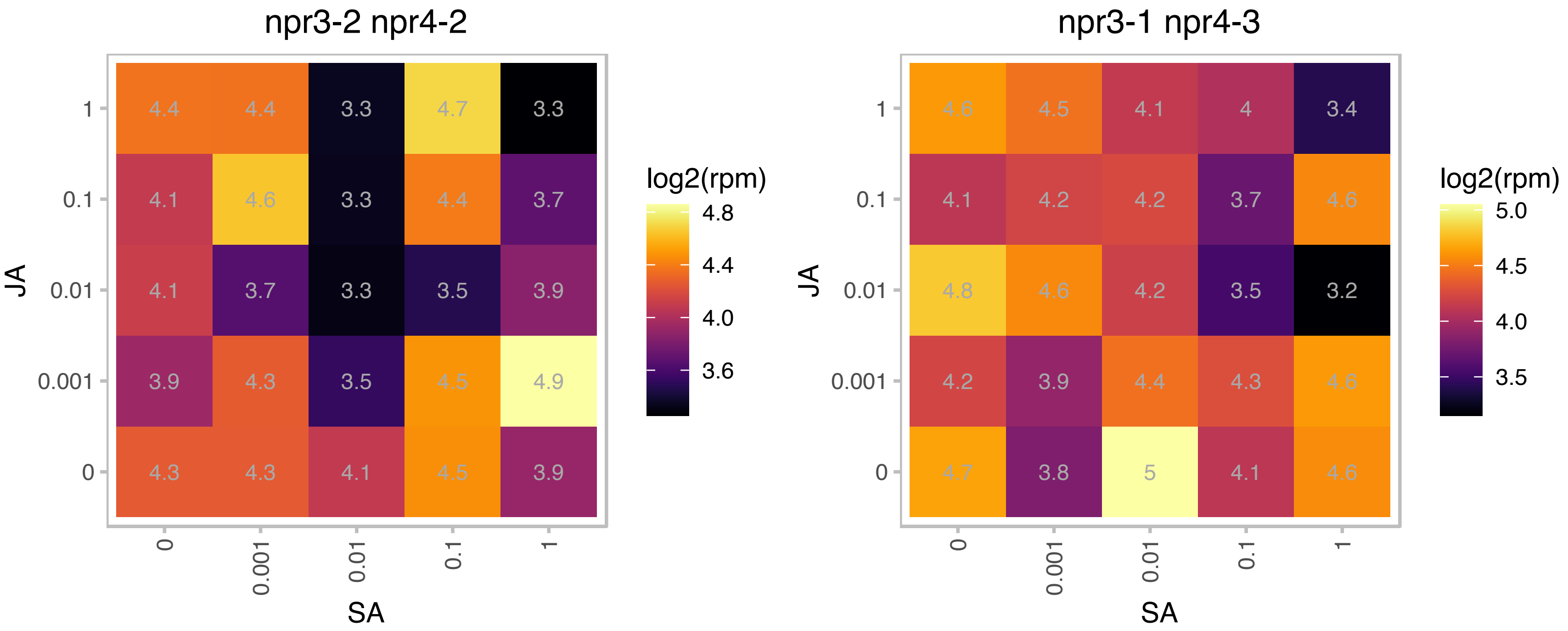

# AT3G04470.1 : NA

Ankyrin repeat family protein

Col-0

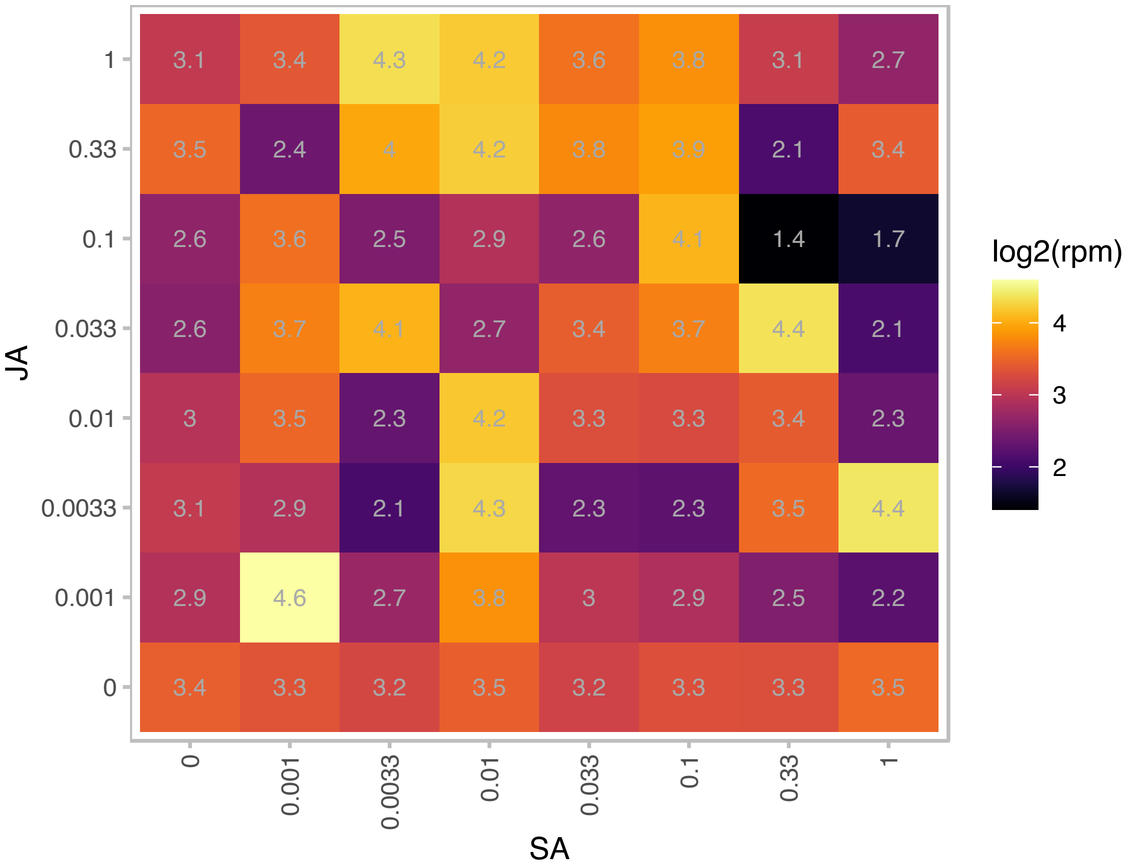


| | |
|---|---|
| Gene | AT3G04470 |
| is.expressed_genes | TRUE |
| is.responsive_genes | TRUE |
| is.apcluster | D-03 |

npr3-2 npr4-2

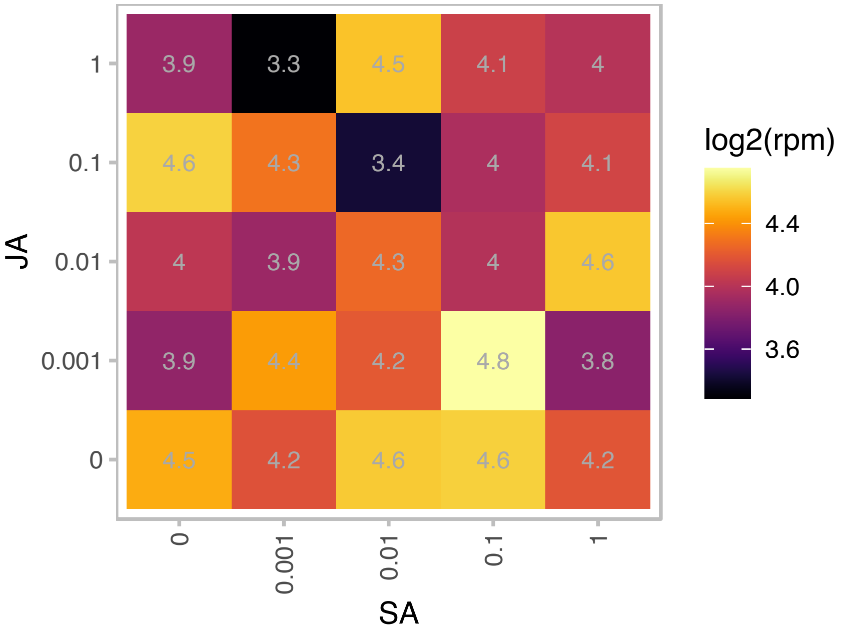


npr3-1 npr4-3

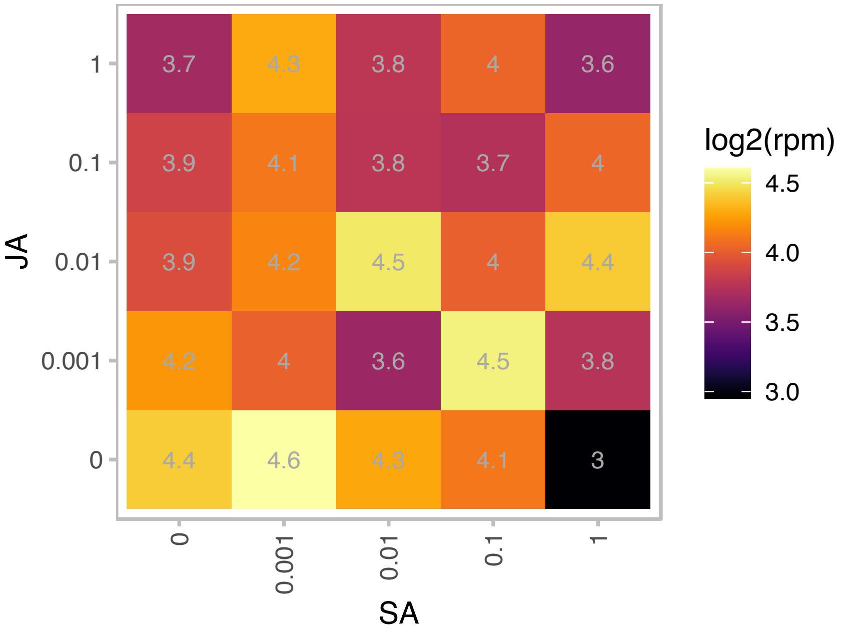

### AT3G11700.1 : FLA18

FASCICLIN-like arabinogalactan protein 18 precursor

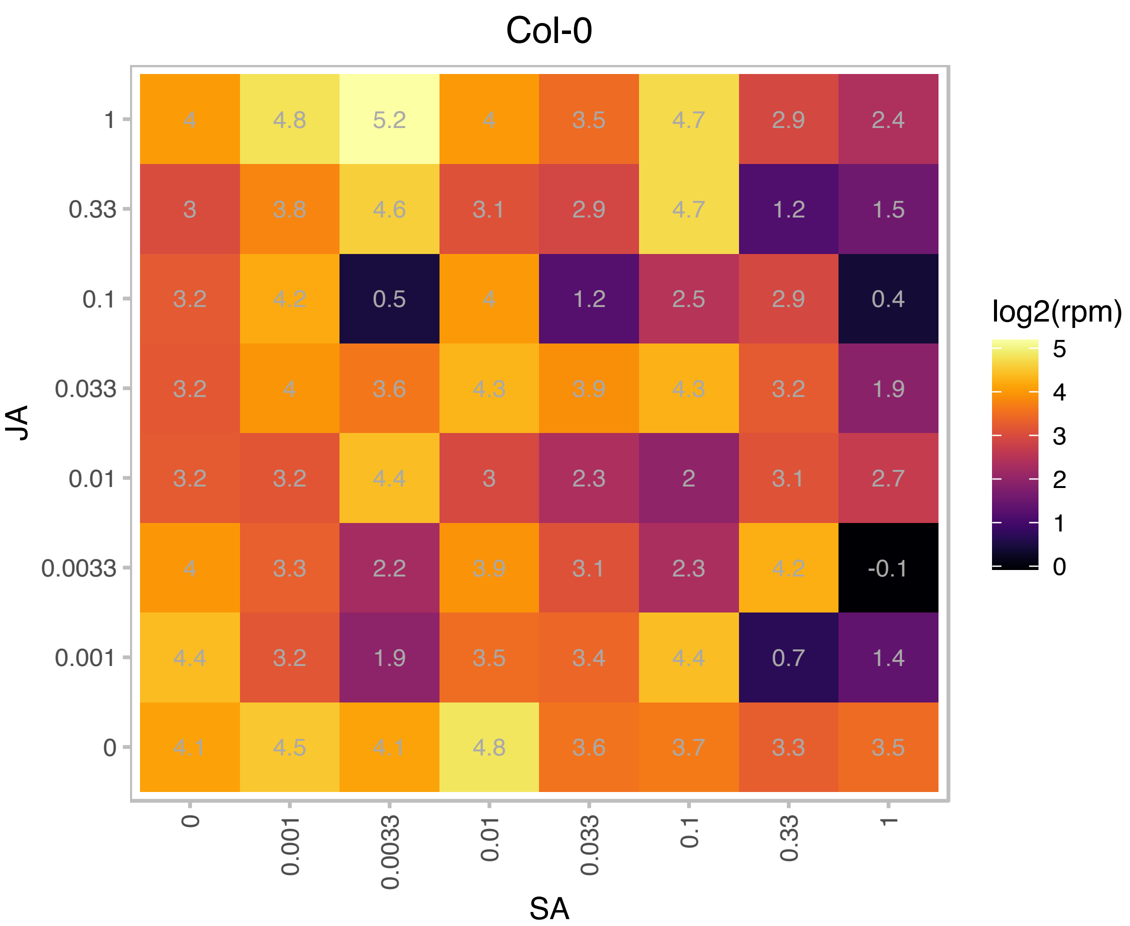


| | |
|---|---|
| Gene | AT3G11700 |
| is.expressed_genes | TRUE |
| is.responsive_genes | TRUE |
| is.apcluster | D-03 |

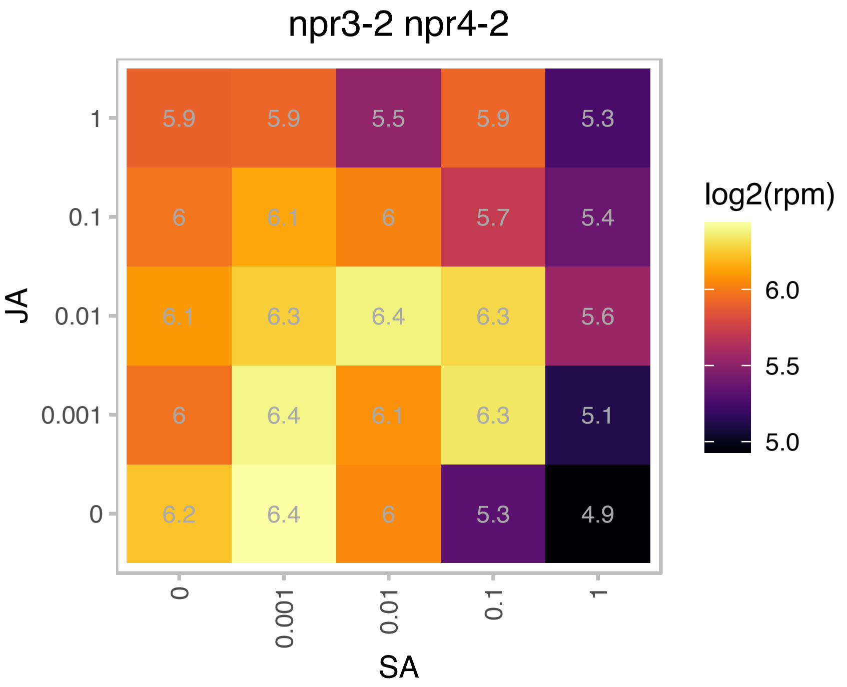


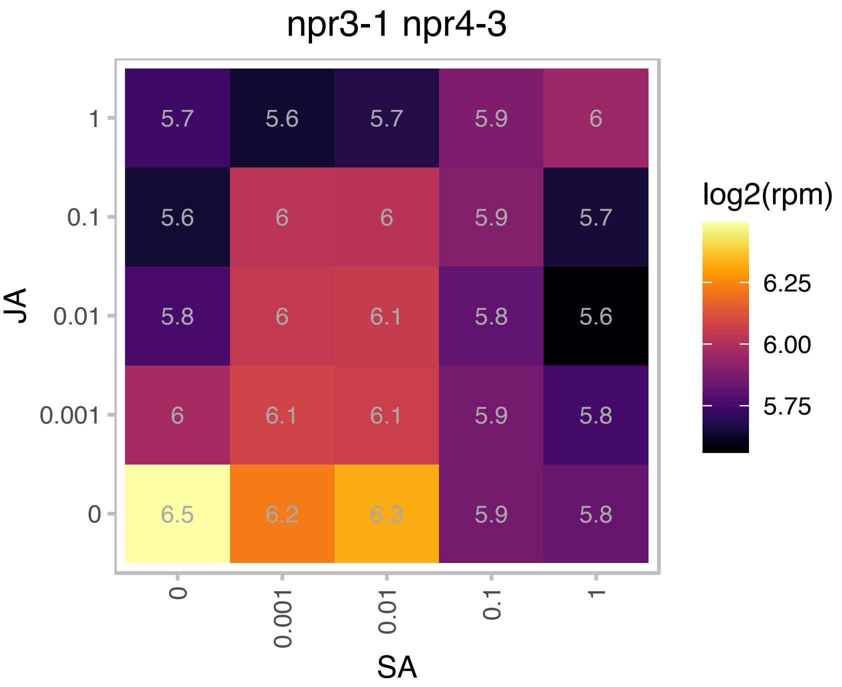

# AT3G11960.1 : NA

Cleavage and polyadenylation specificity factor (CPSF) A subunit protein

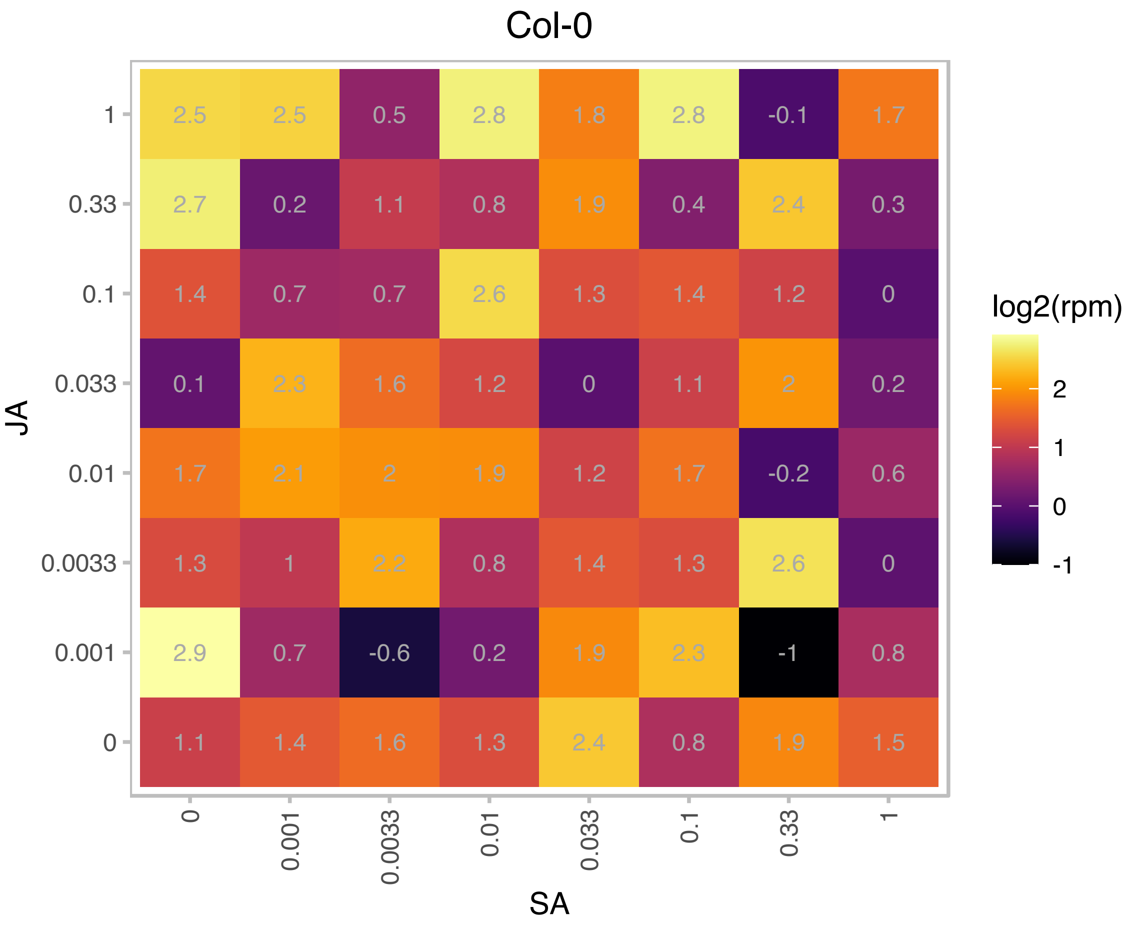


| | |
|---|---|
| Gene | AT3G11960 |
| is.expressed_genes | TRUE |
| is.responsive_genes | TRUE |
| is.apcluster | D-03 |

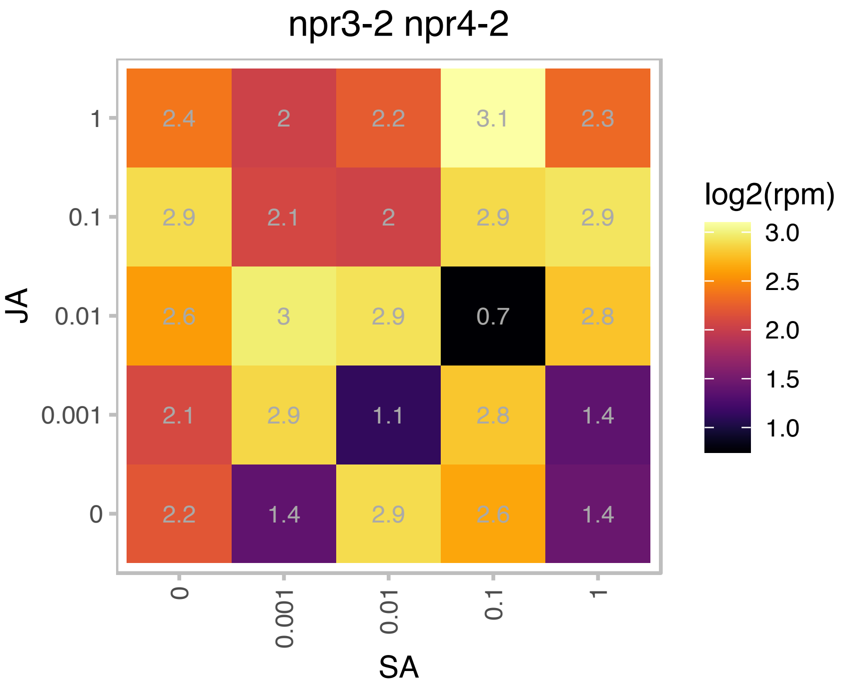


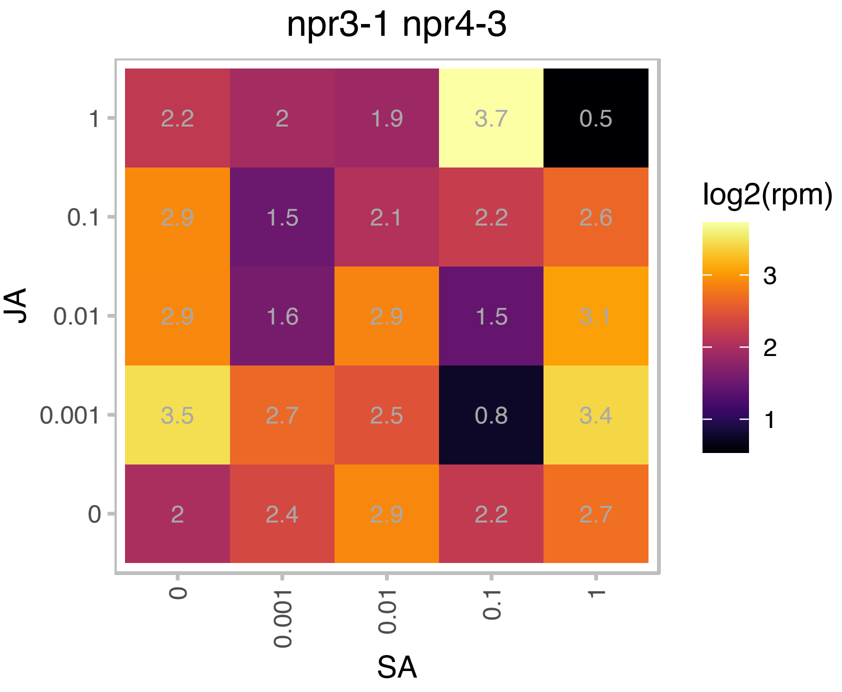

### AT3G13060.1 : ECT5

evolutionarily conserved C-terminal region 5

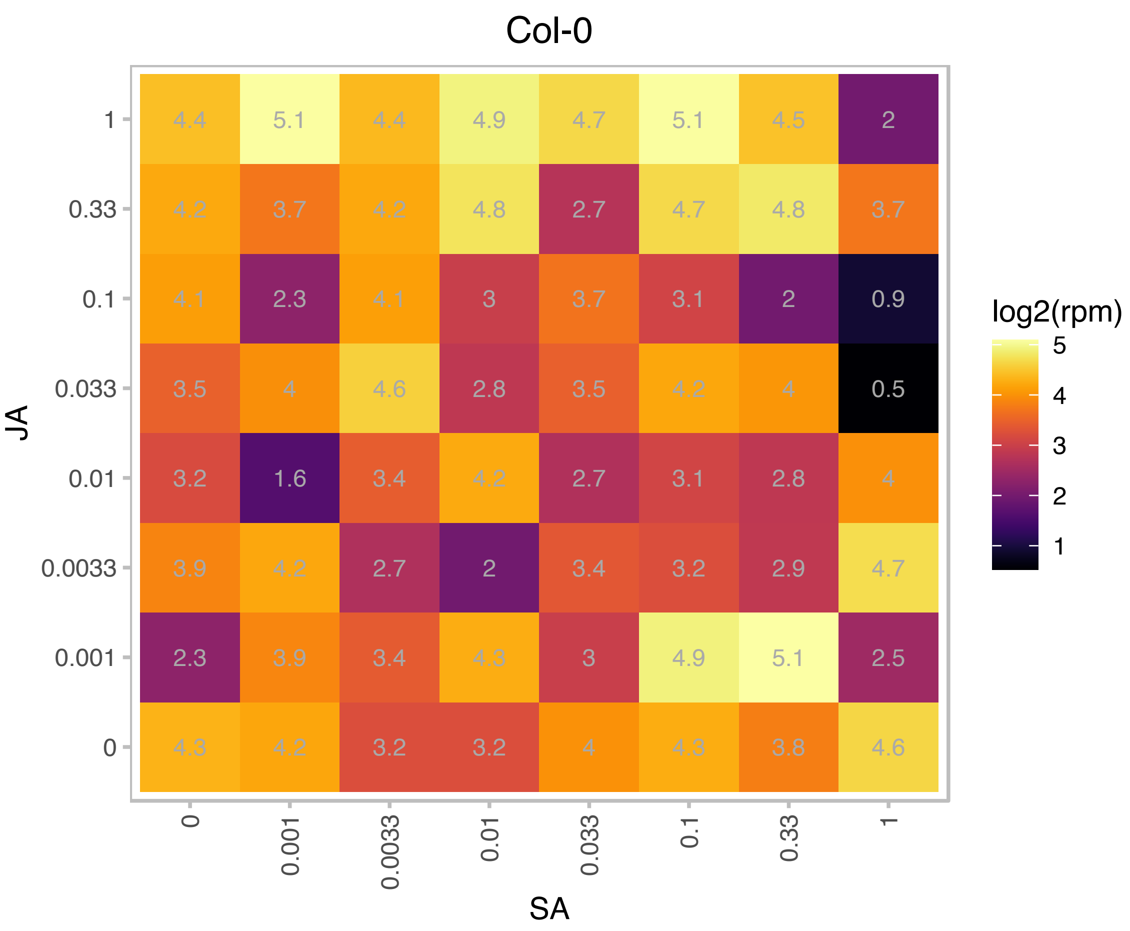


| | |
|---|---|
| Gene | AT3G13060 |
| is.expressed_genes | TRUE |
| is.responsive_genes | TRUE |
| is.apcluster | D-03 |

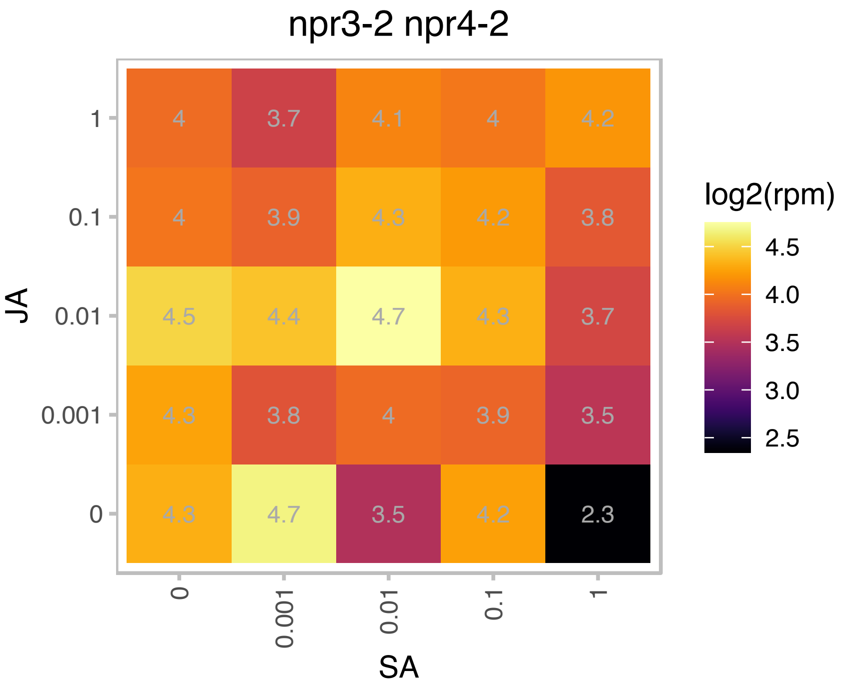


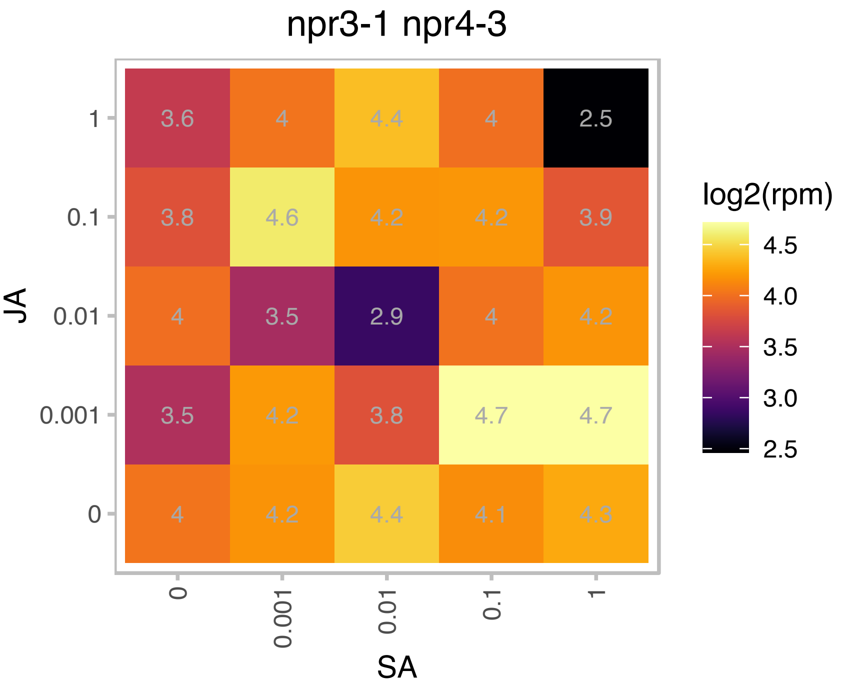

### AT3G48330.1 : ATPIMT1, PIMT1

protein-l-isoaspartate methyltransferase 1

Encodes protein-L-isoaspartate methyltransferase. Important for maintaining viability as the seed ages. Involved in germination.

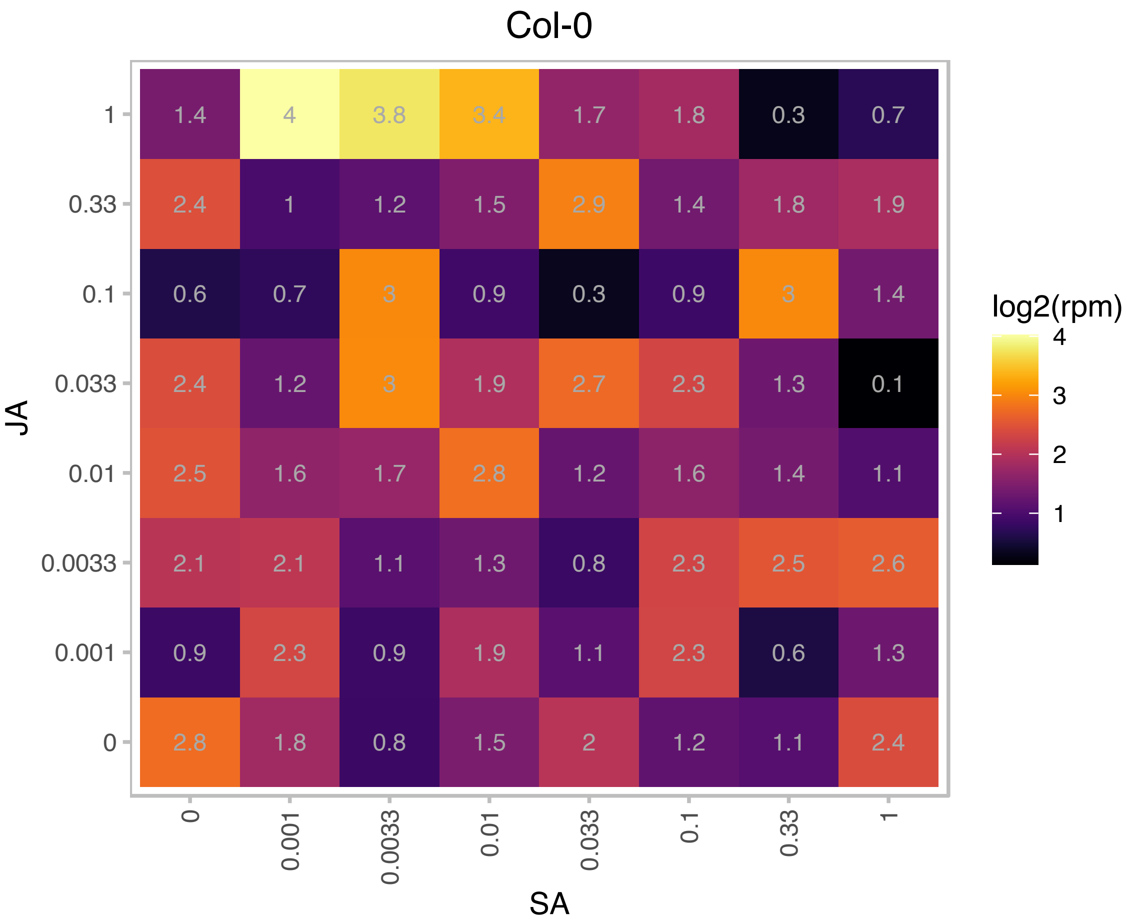


| | |
|---|---|
| Gene | AT3G48330 |
| is.expressed_genes | TRUE |
| is.responsive_genes | TRUE |
| is.apcluster | D-03 |

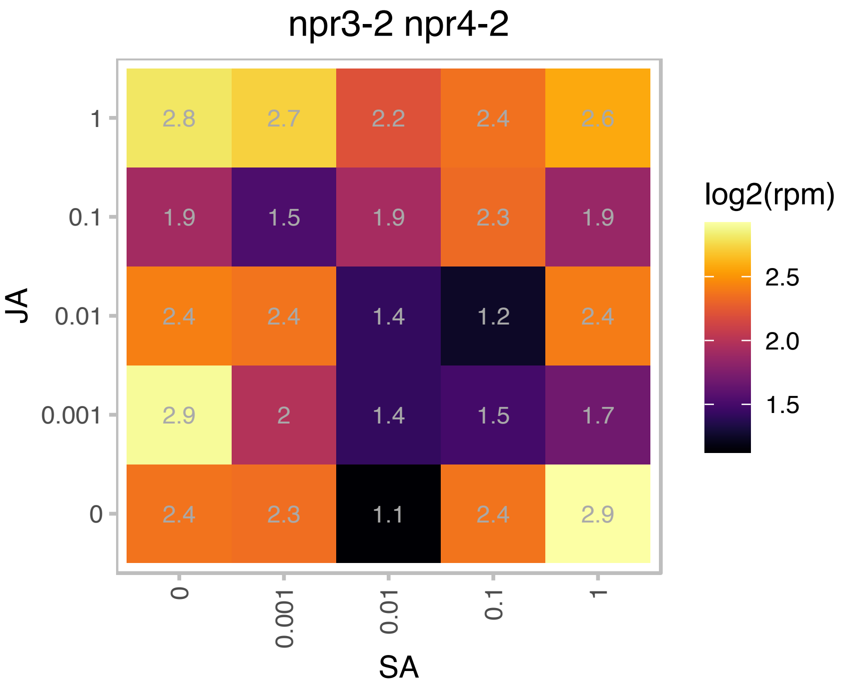


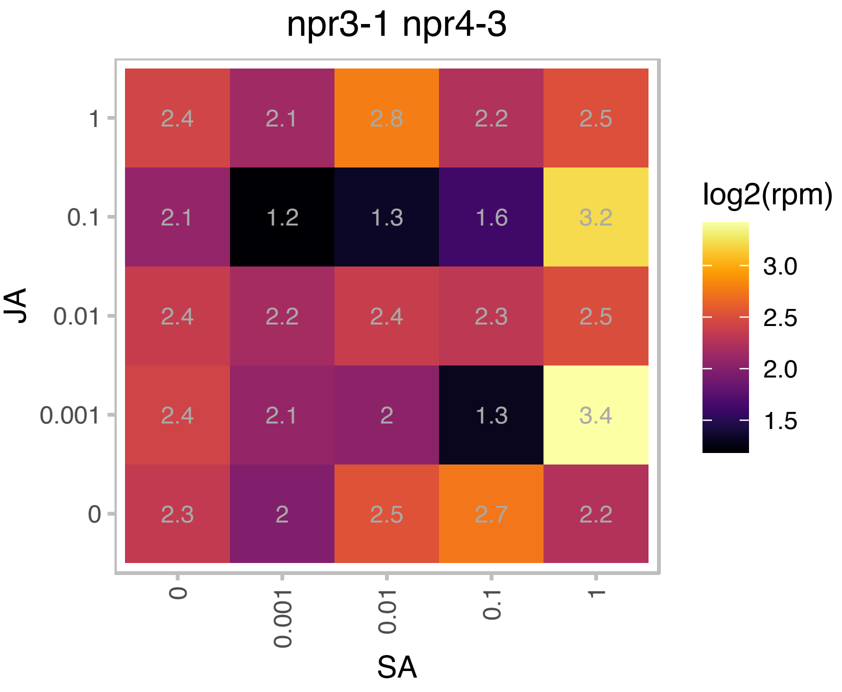

### AT4G12060.1 : ClpT2

Double Clp-N motif protein

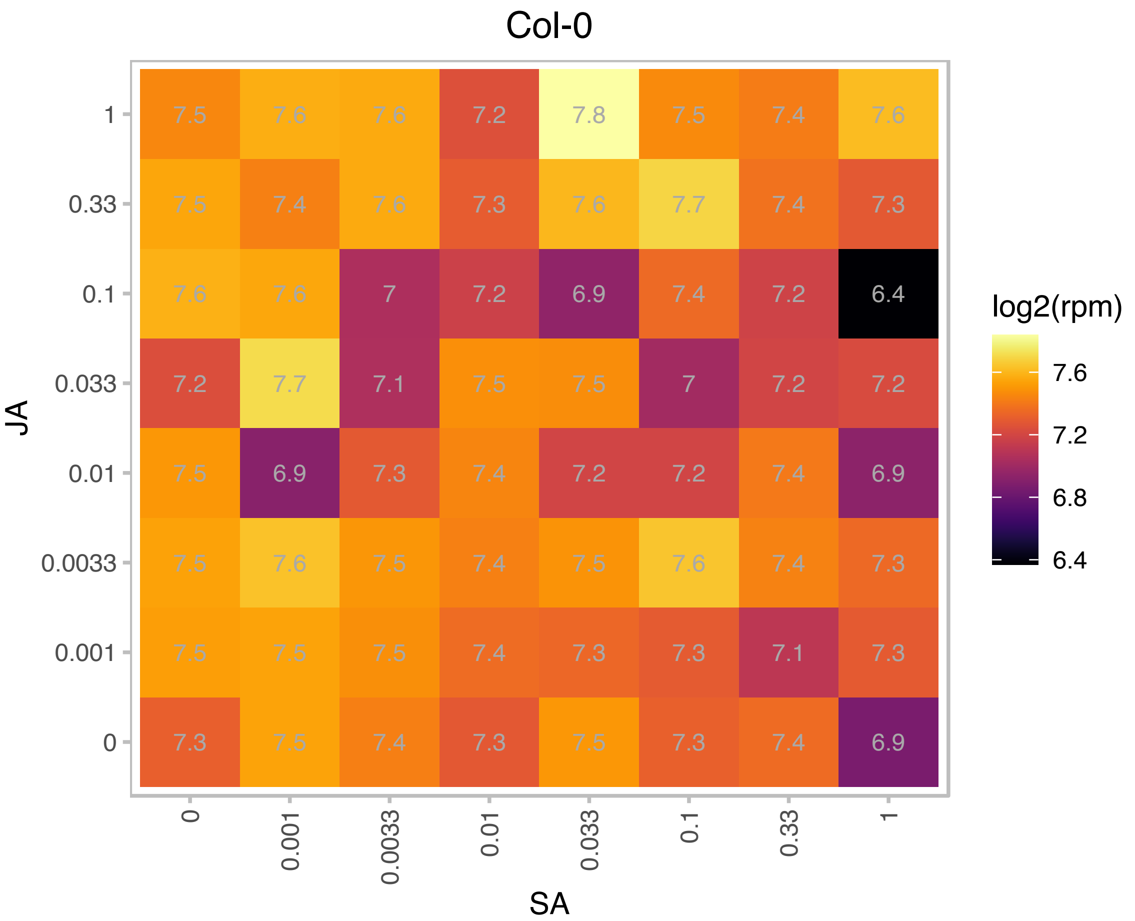


| | |
|---|---|
| Gene | AT4G12060 |
| is.expressed_genes | TRUE |
| is.responsive_genes | TRUE |
| is.apcluster | D-03 |

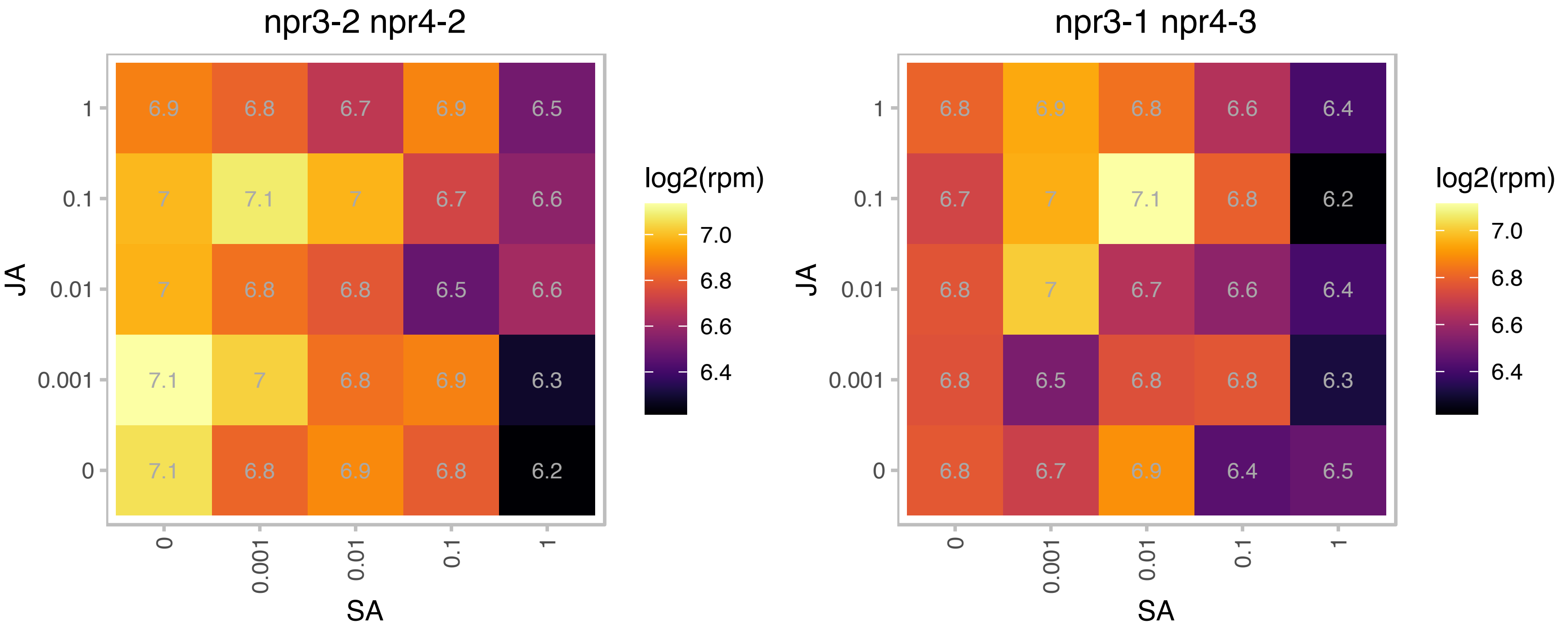

# AT4G17020.1 : NA

transcription factor-related

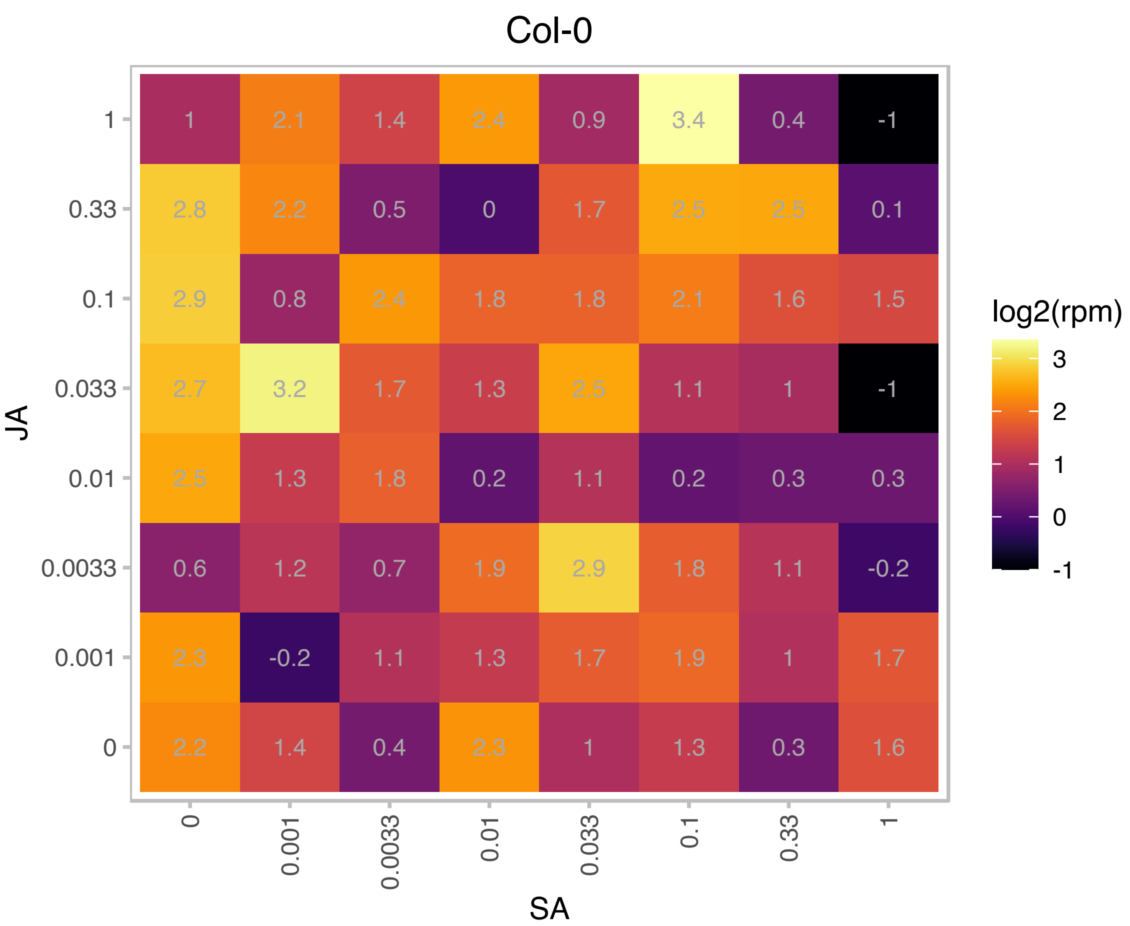


| | |
|---|---|
| Gene | AT4G17020 |
| is.expressed_genes | TRUE |
| is.responsive_genes | TRUE |
| is.apcluster | D-03 |

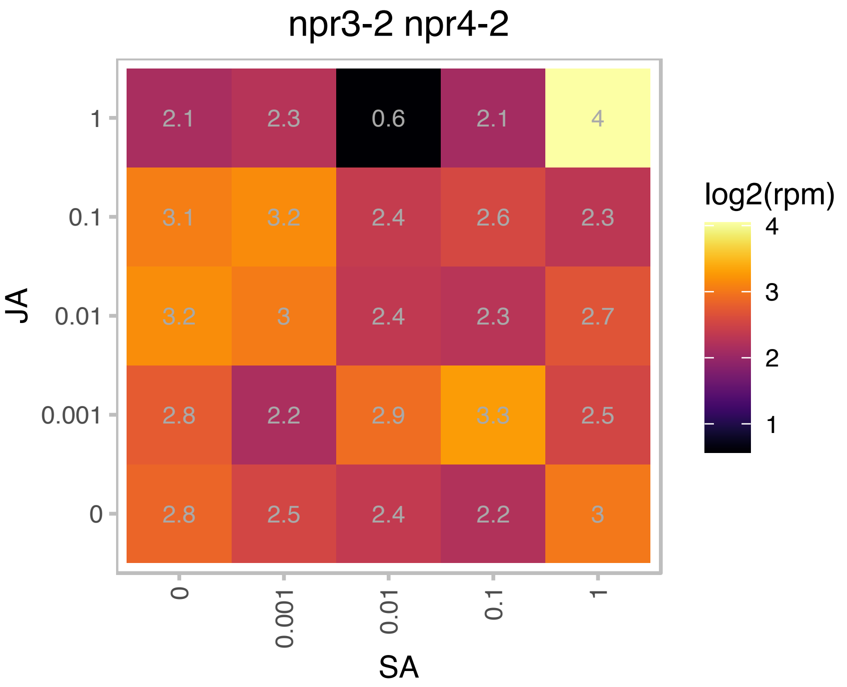


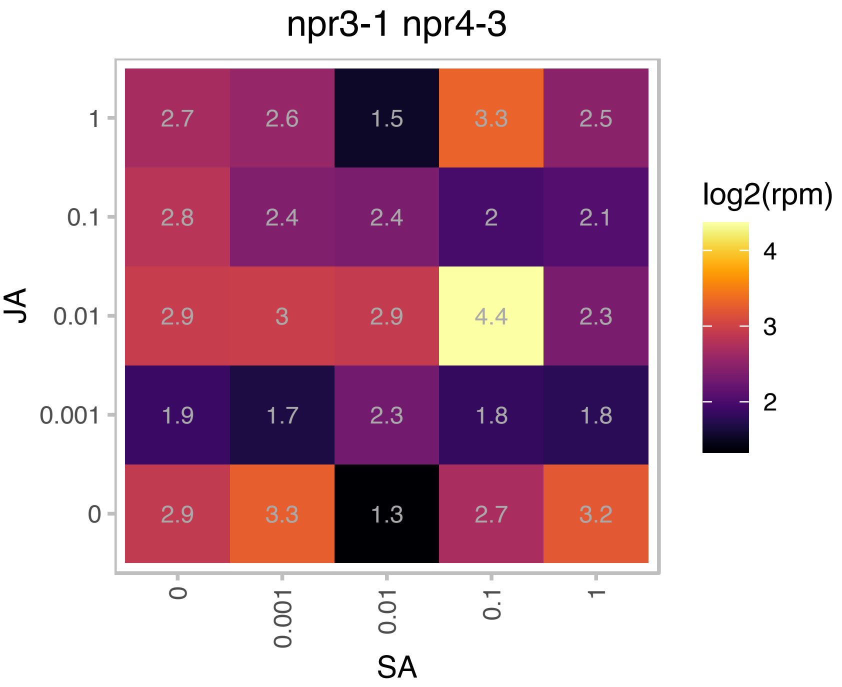

# AT4G18070.1 : NA

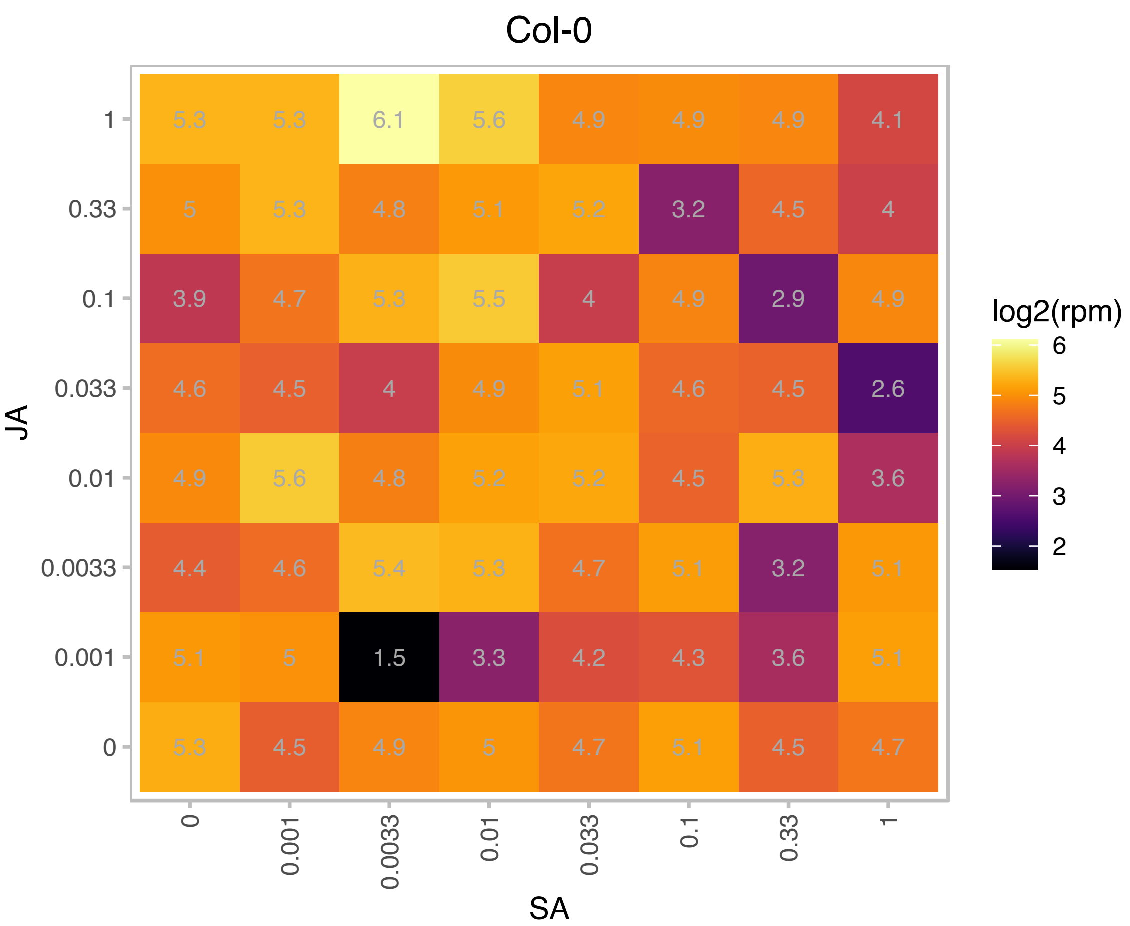


| | |
|---|---|
| Gene | AT4G18070 |
| is.expressed_genes | TRUE |
| is.responsive_genes | TRUE |
| is.apcluster | D-03 |

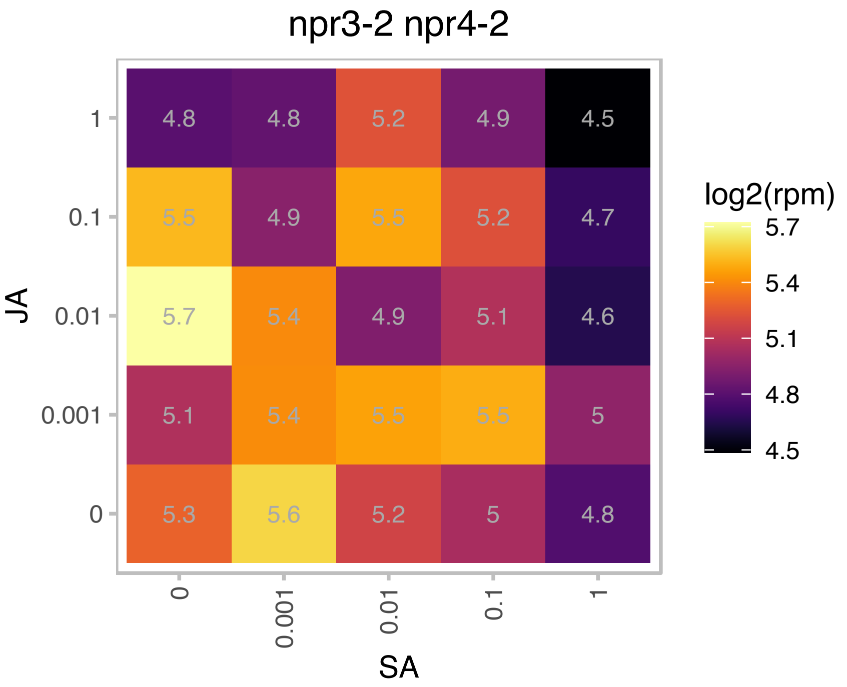


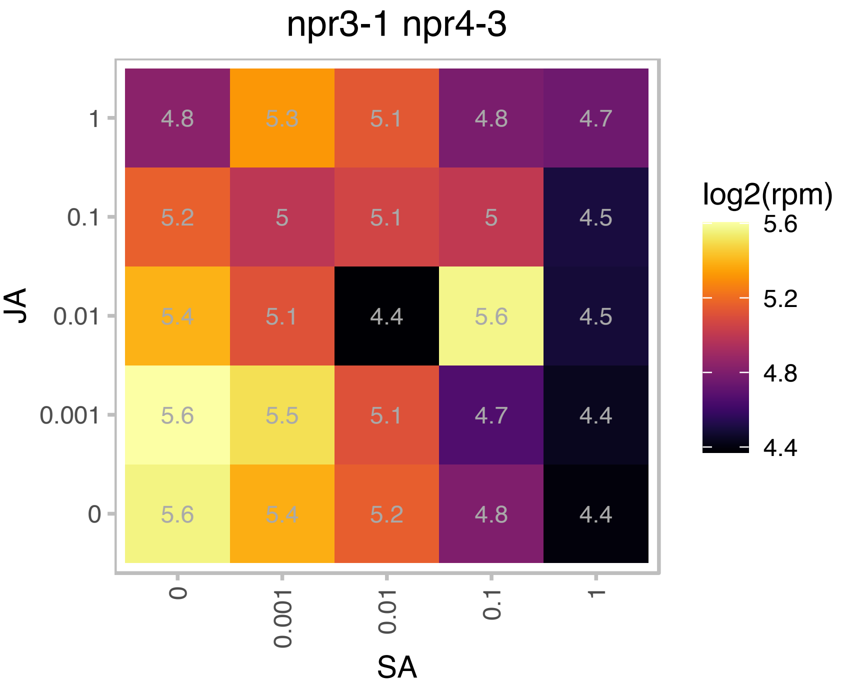

# AT4G36390.1 : NA

Methylthiotransferase

| | |
|---|---|
| Gene | AT4G36390 |
| is.expressed_genes | TRUE |
| is.responsive_genes | TRUE |
| is.apcluster | D-03 |

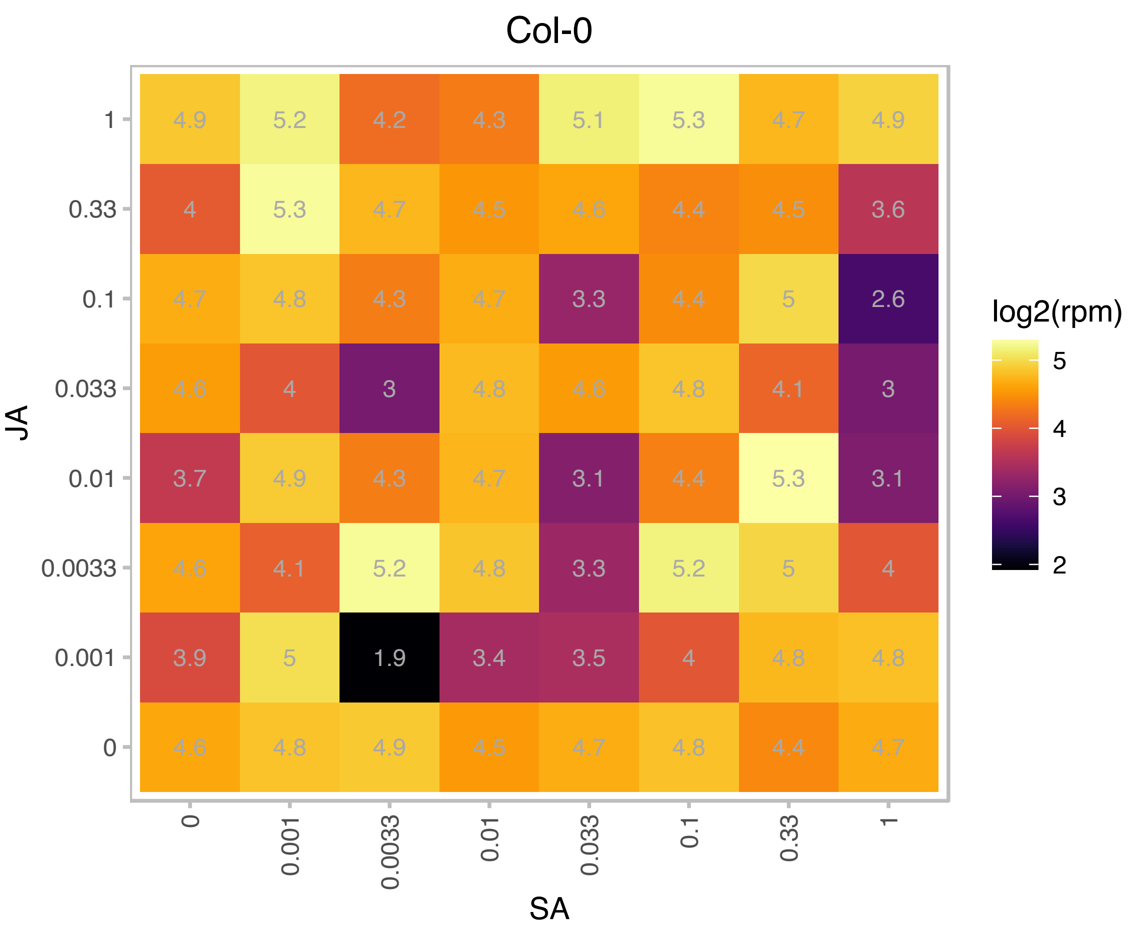


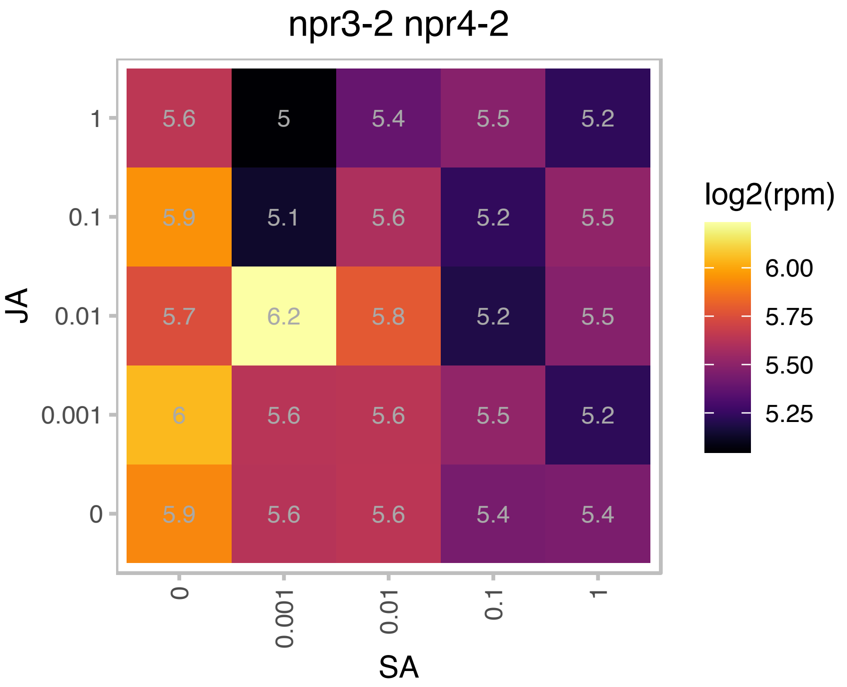


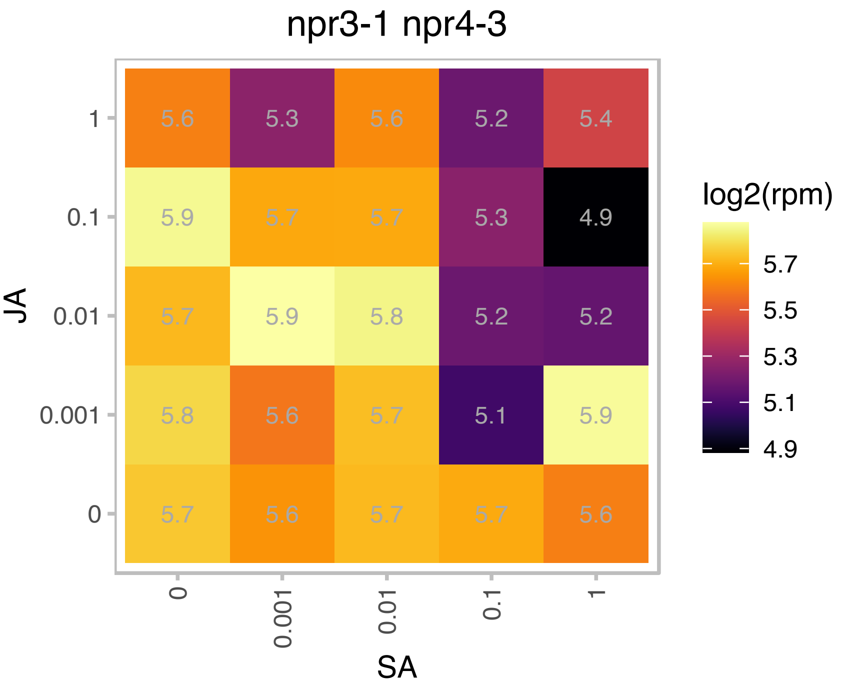

# AT5G42680.1 : NA

Protein of unknown function, DUF617

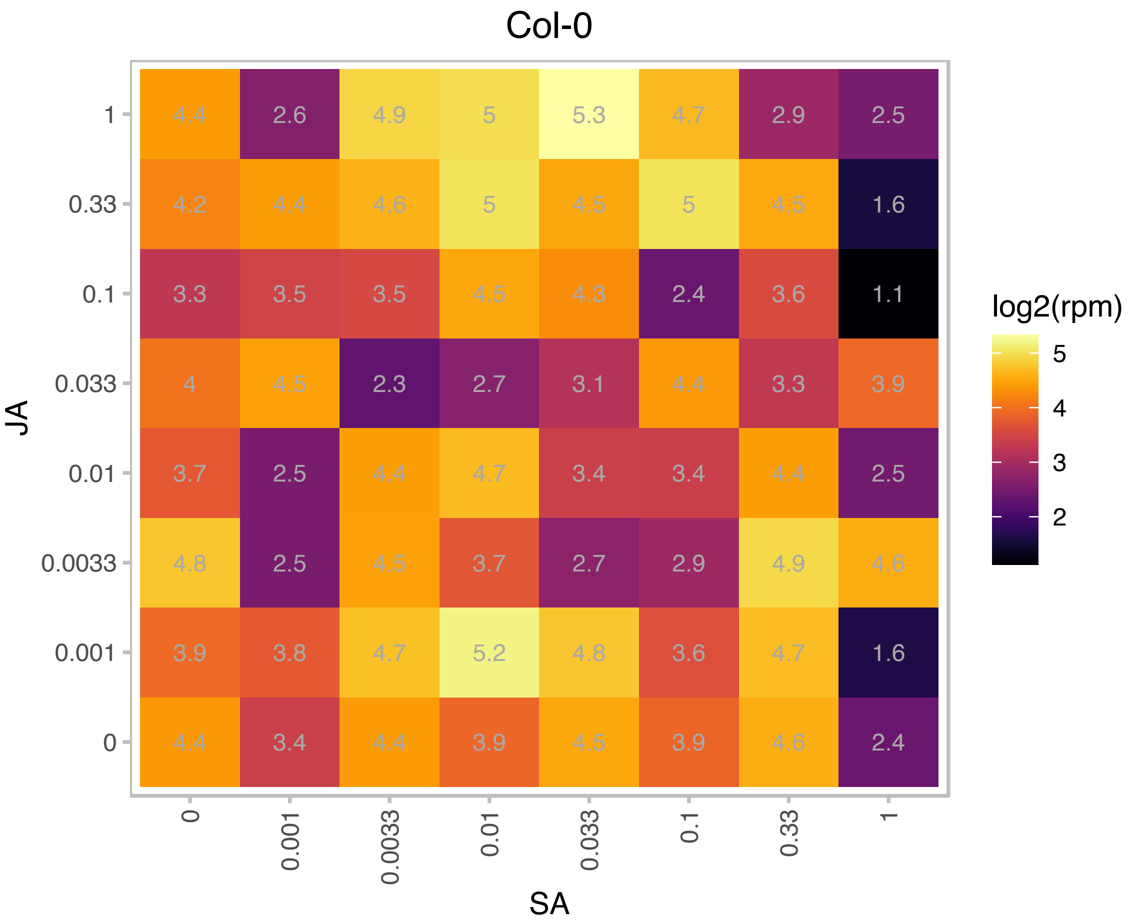


| | |
|---|---|
| Gene | AT5G42680 |
| is.expressed_genes | TRUE |
| is.responsive_genes | TRUE |
| is.apcluster | D-03 |

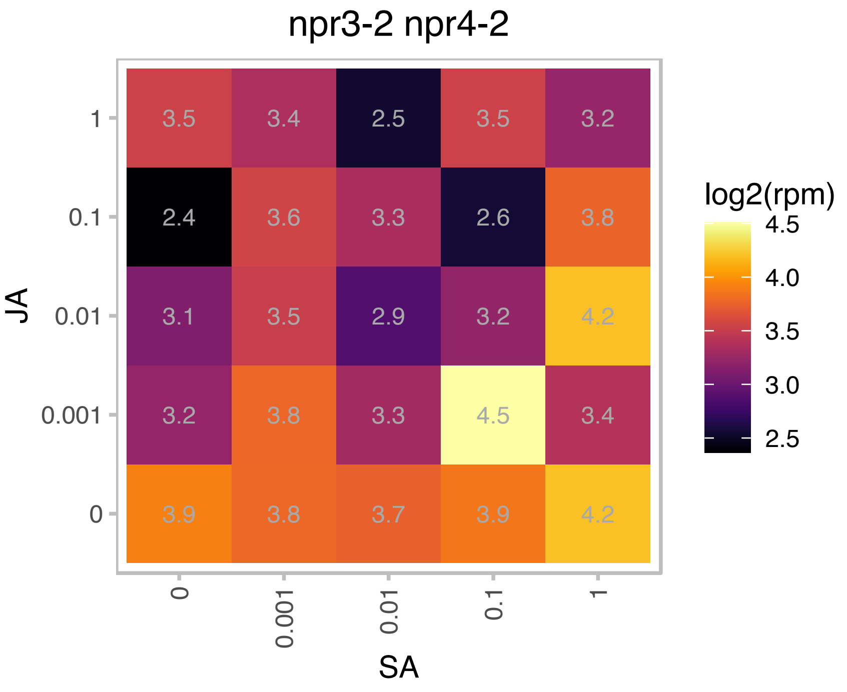


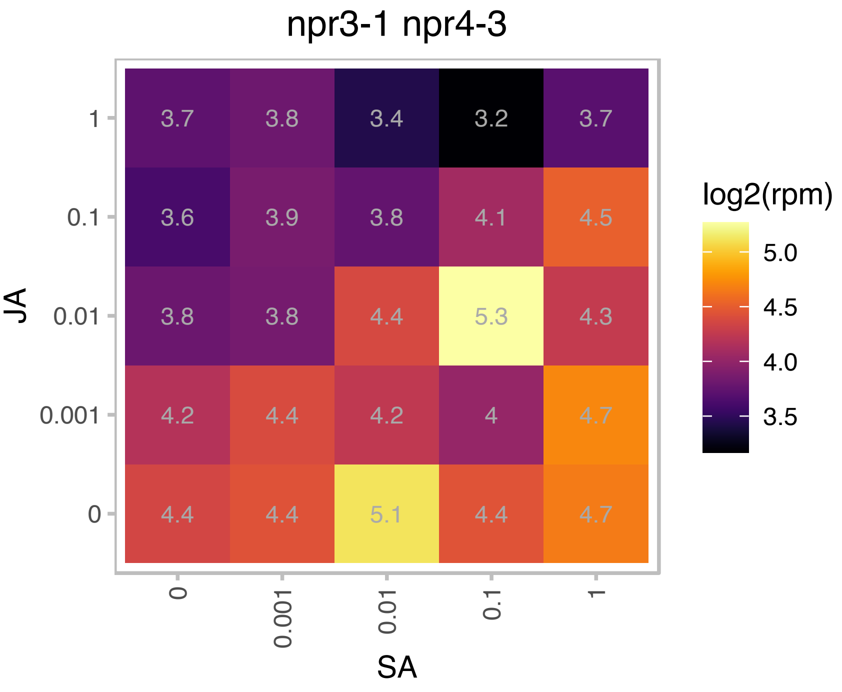

# AT5G43830.1 : NA

Aluminium induced protein with YGL and LRDR motifs

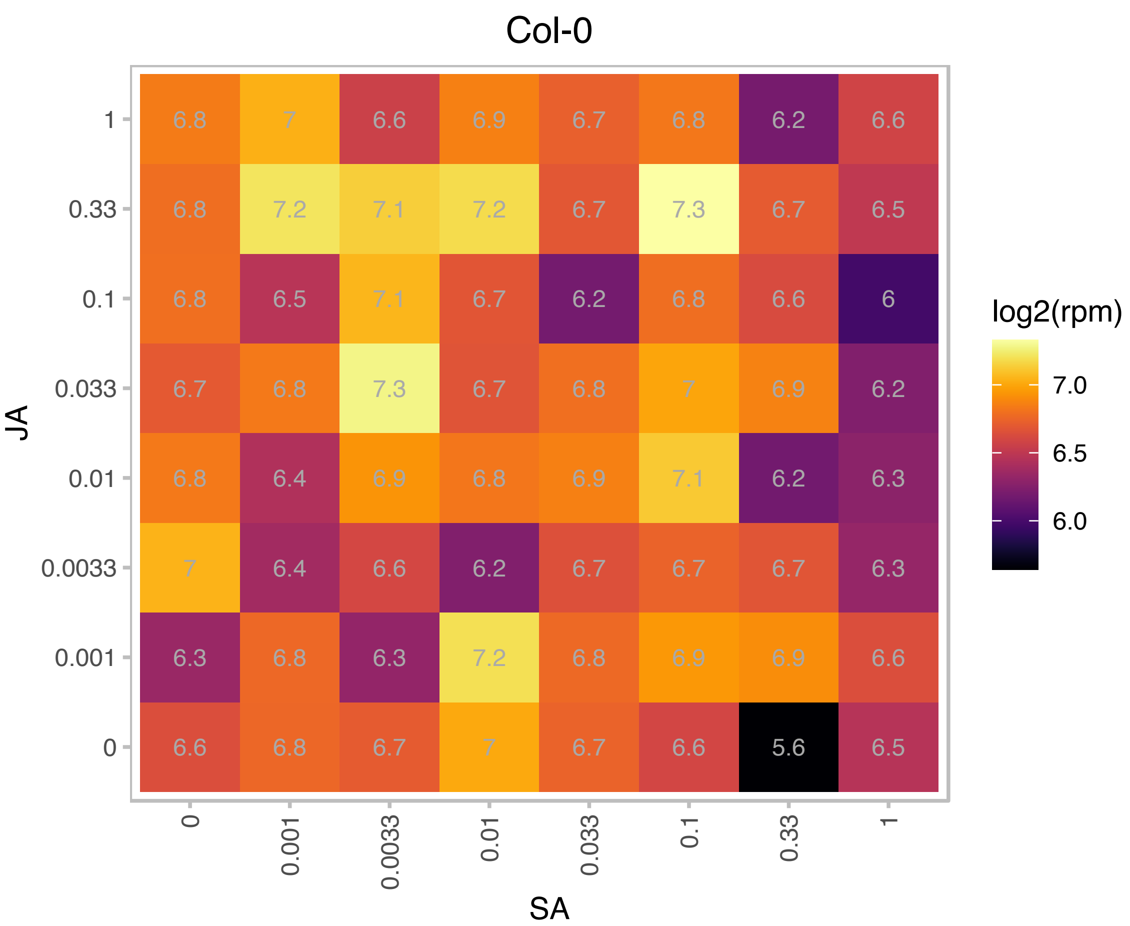


| | |
|---|---|
| Gene | AT5G43830 |
| is.expressed_genes | TRUE |
| is.responsive_genes | TRUE |
| is.apcluster | D-03 |

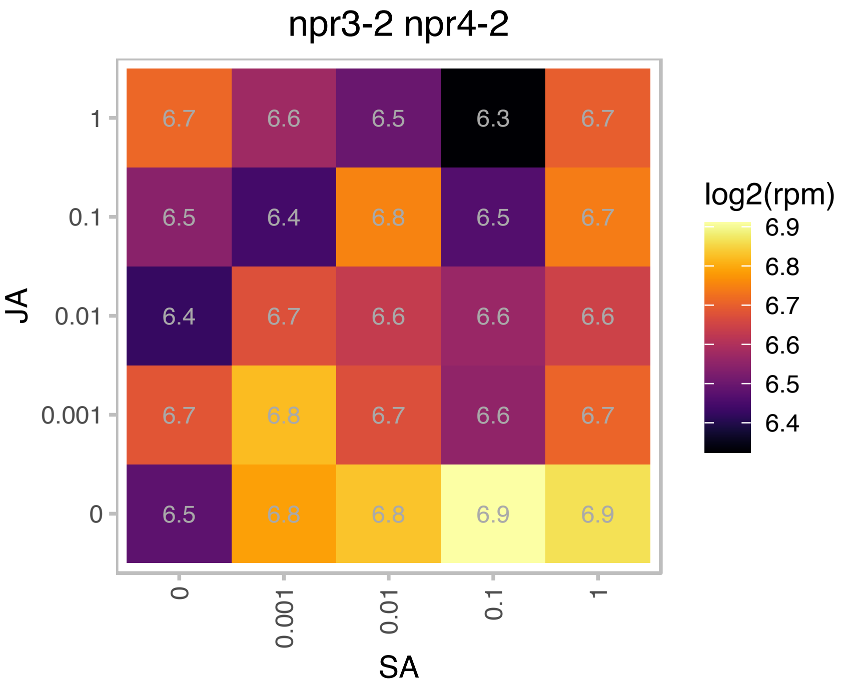


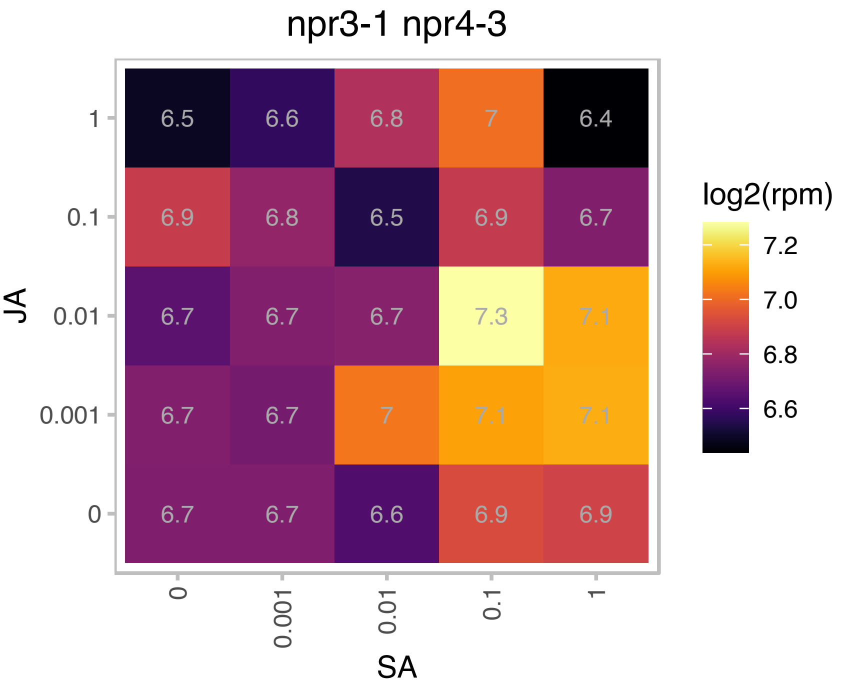

### AT5G49030.1 : OVA2

tRNA synthetase class I (I, L, M and V) family protein

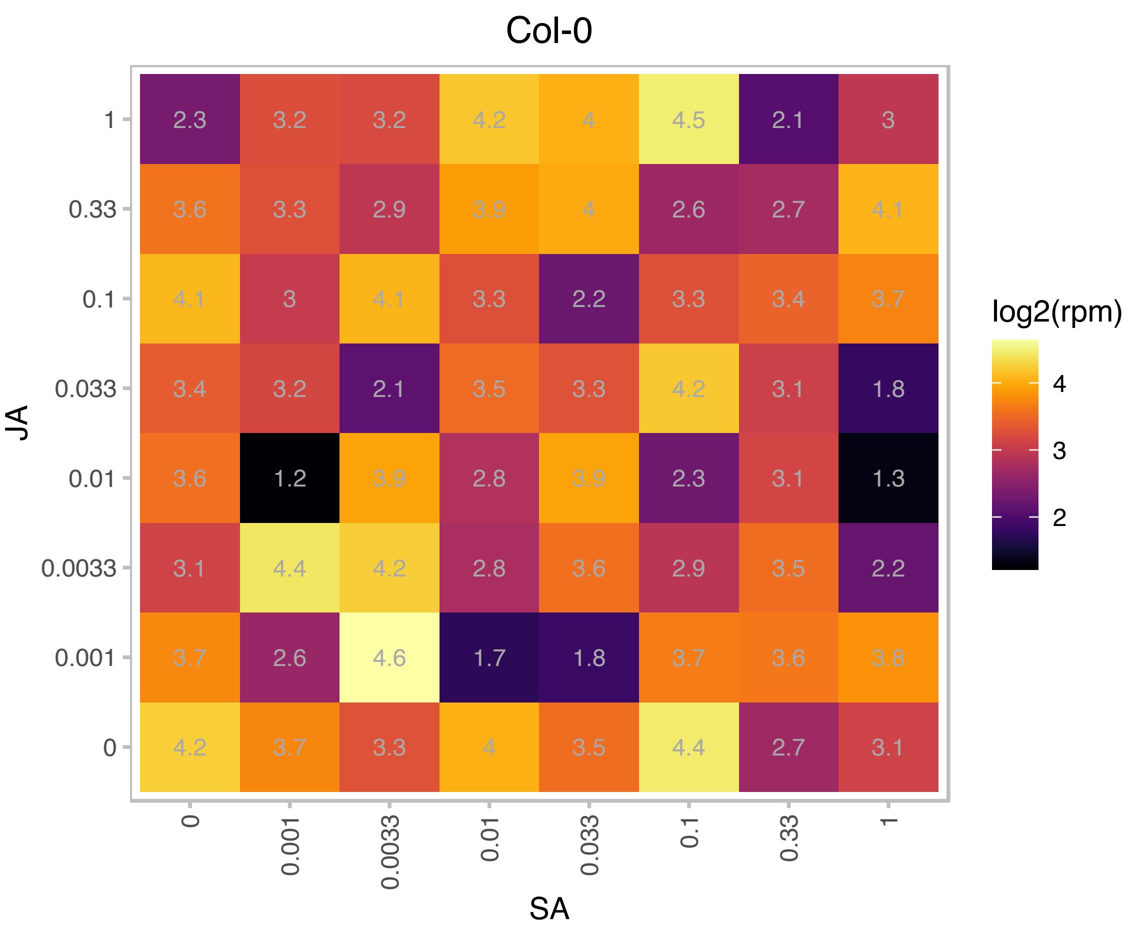


| | |
|---|---|
| Gene | AT5G49030 |
| is.expressed_genes | TRUE |
| is.responsive_genes | TRUE |
| is.apcluster | D-03 |

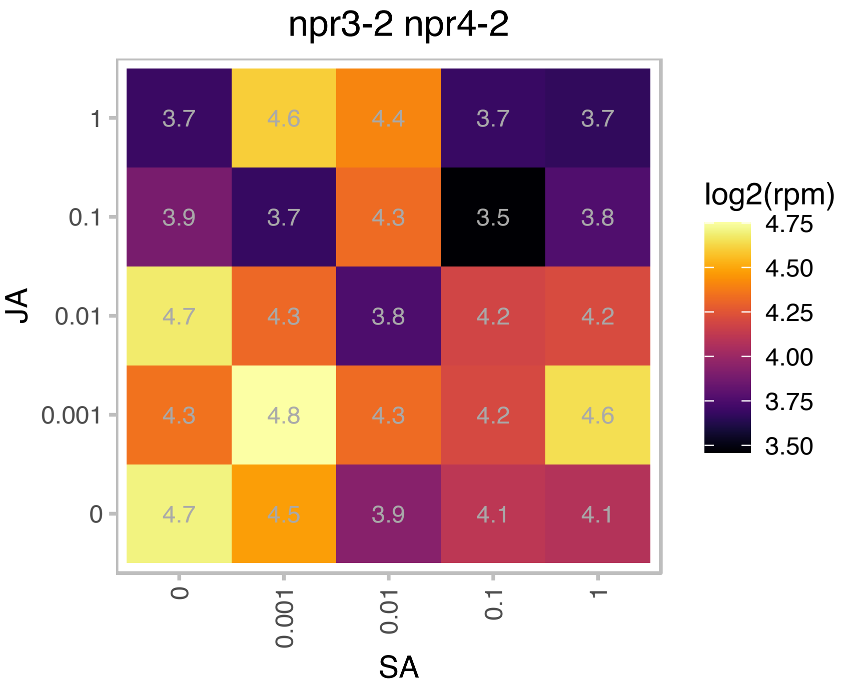


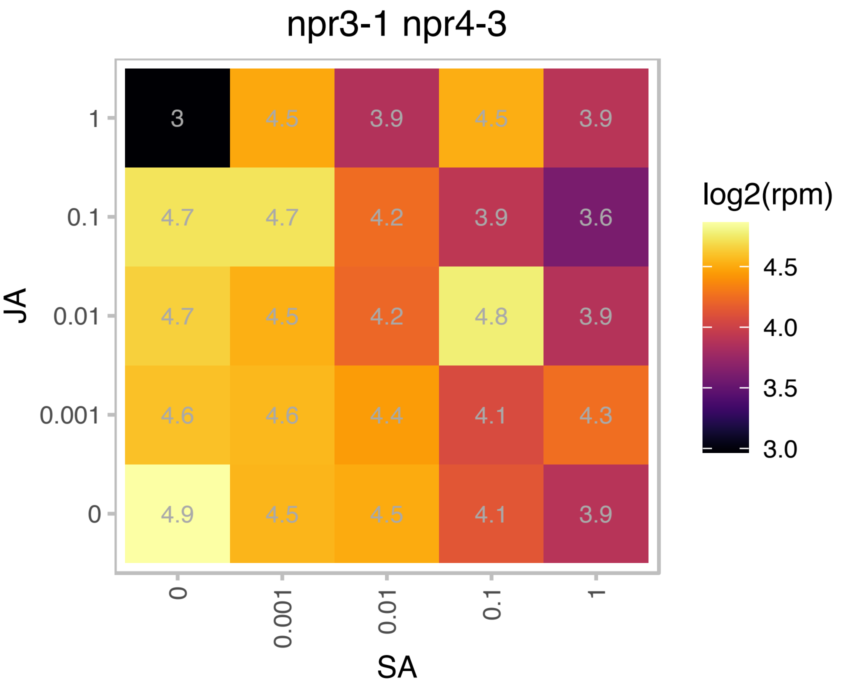

### AT5G54900.1 : ATRBP45A, RBP45A

RNA-binding protein 45A

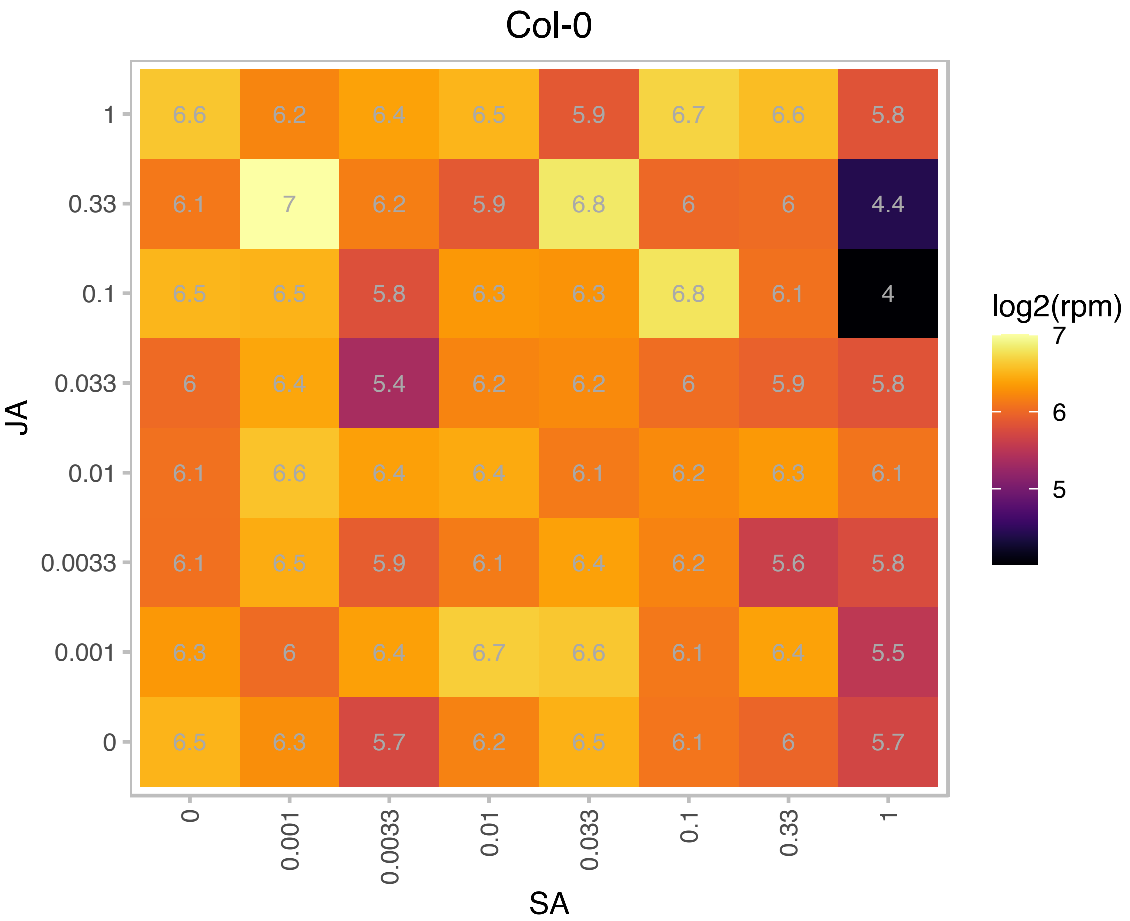


| | |
|---|---|
| Gene | AT5G54900 |
| is.expressed_genes | TRUE |
| is.responsive_genes | TRUE |
| is.apcluster | D-03 |

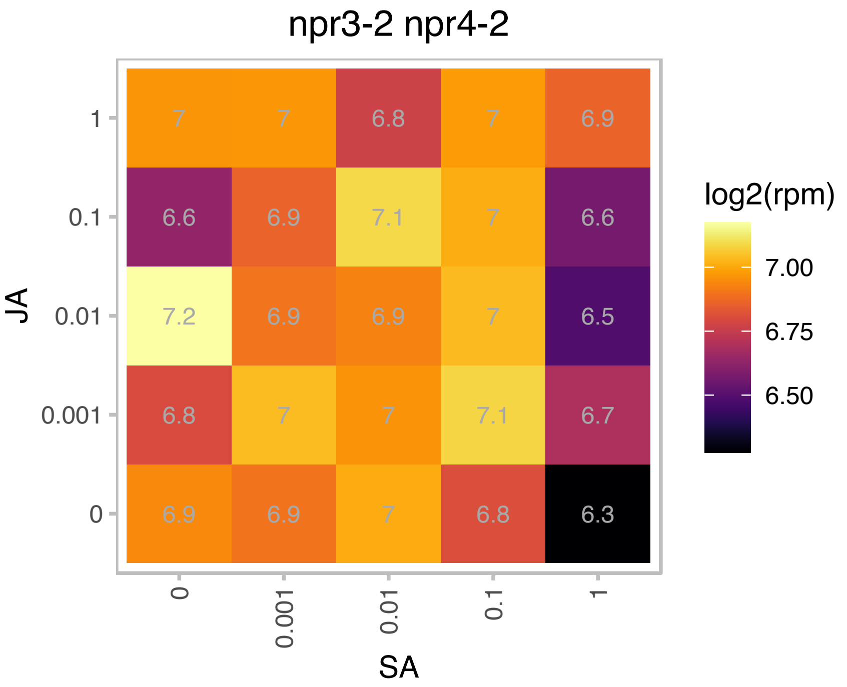


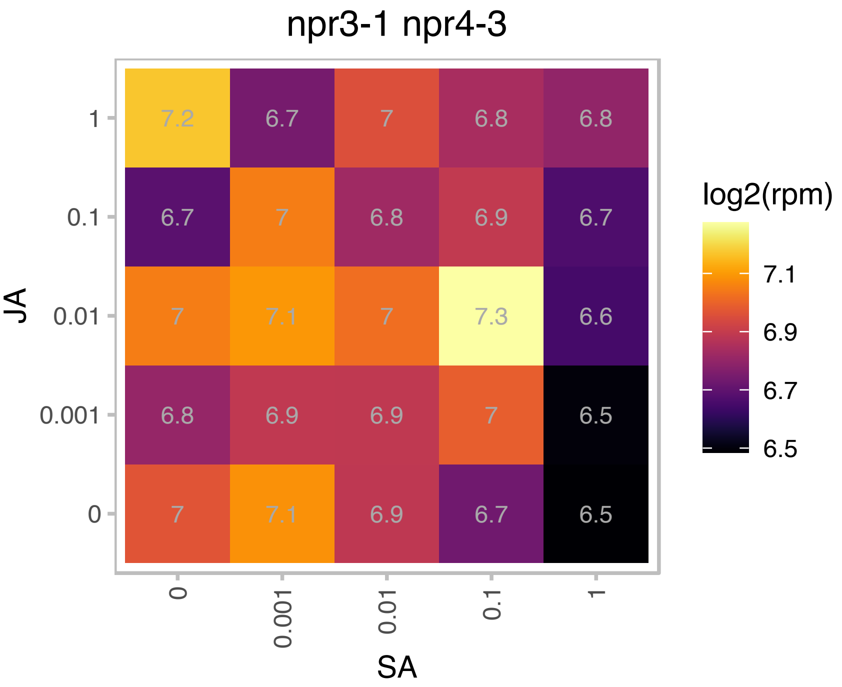

# AT5G56240.1 : NA

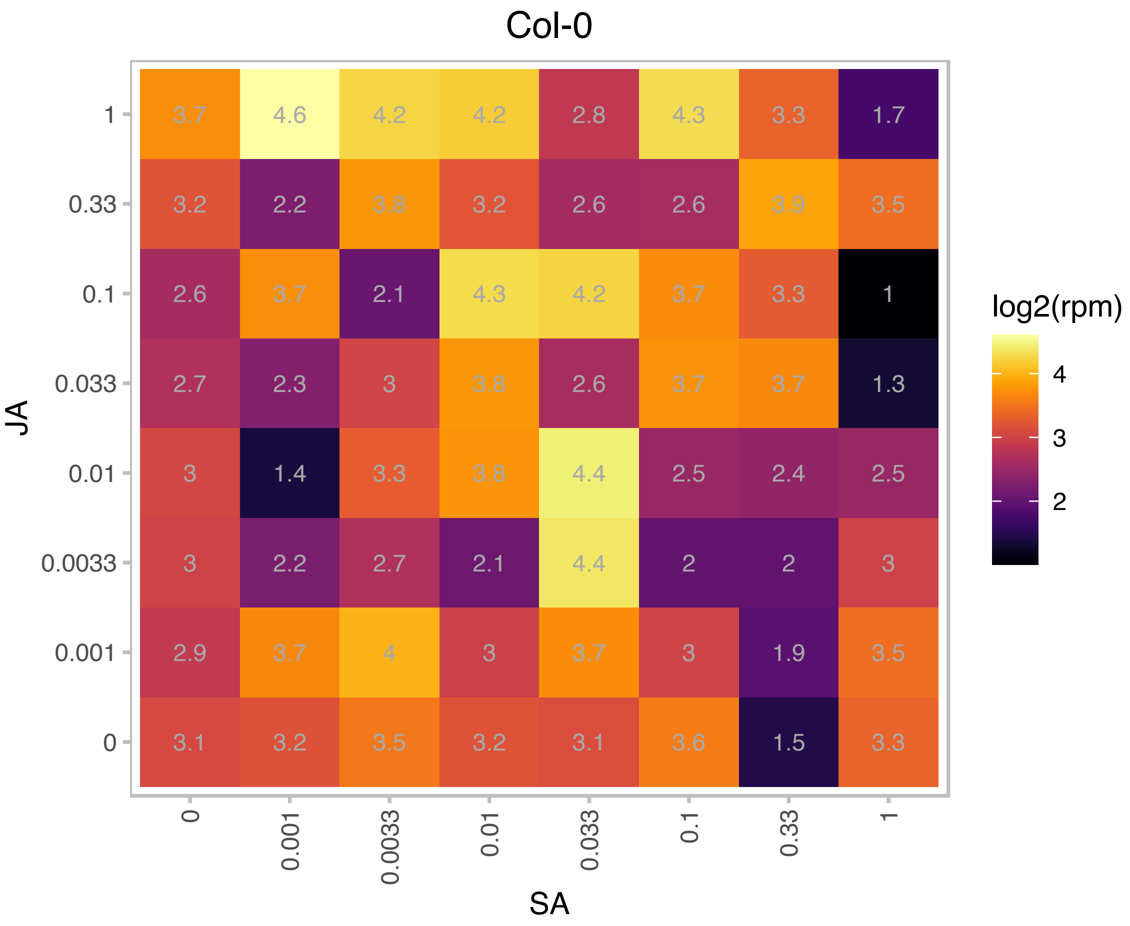


| | |
|---|---|
| Gene | AT5G56240 |
| is.expressed_genes | TRUE |
| is.responsive_genes | TRUE |
| is.apcluster | D-03 |

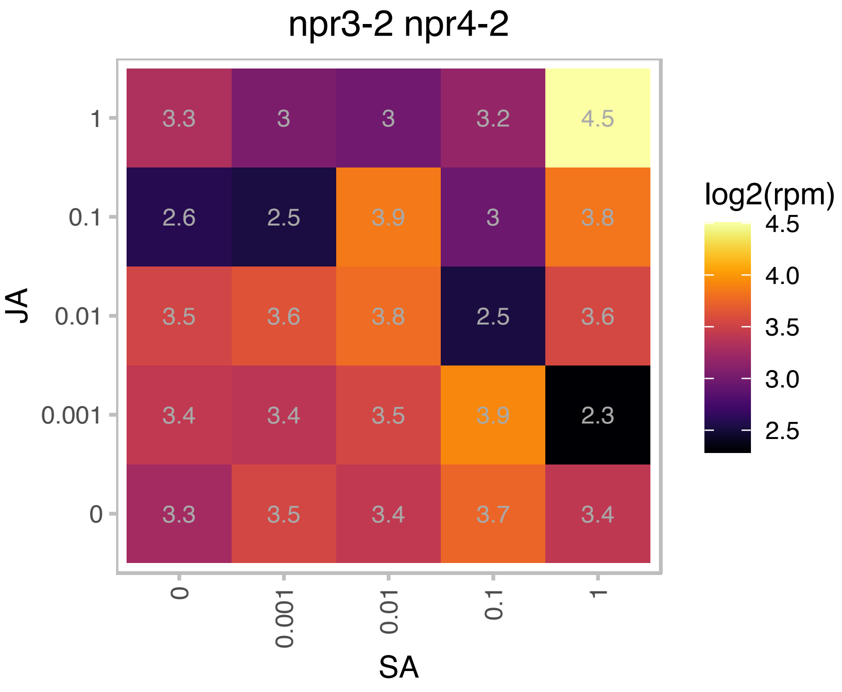


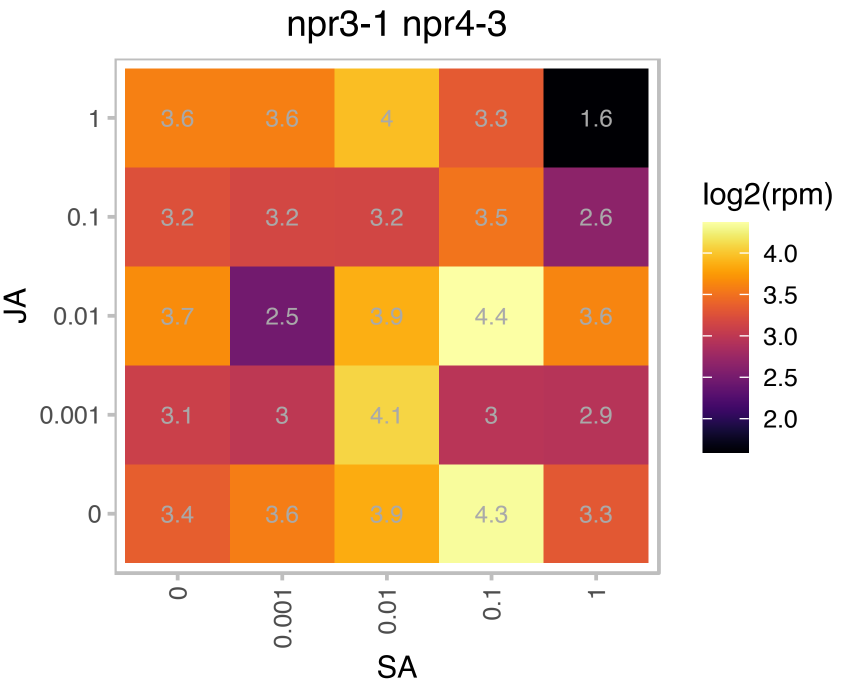

# AT5G58740.1 : NA

HSP20-like chaperones superfamily protein

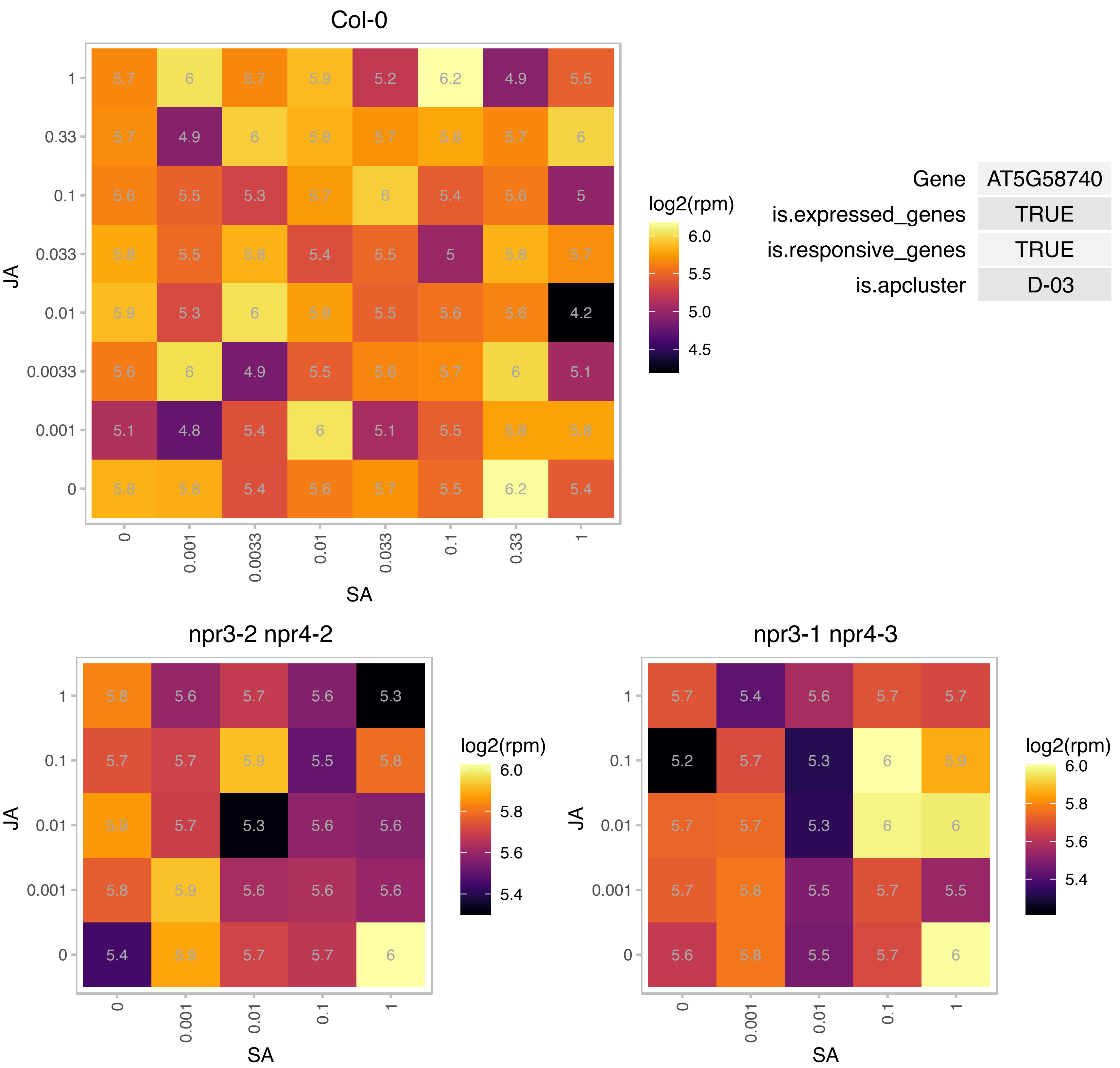


| | |
|---|---|
| Gene | AT5G58740 |
| is.expressed_genes | TRUE |
| is.responsive_genes | TRUE |
| is.apcluster | D-03 |

### AT5G63840.1 : PSL5, RSW3

Glycosyl hydrolases family 31 protein

radial swelling mutant shown to be specifically impaired in cellulose production. Encodes the alpha-subunit of a glucosidase II enzyme.

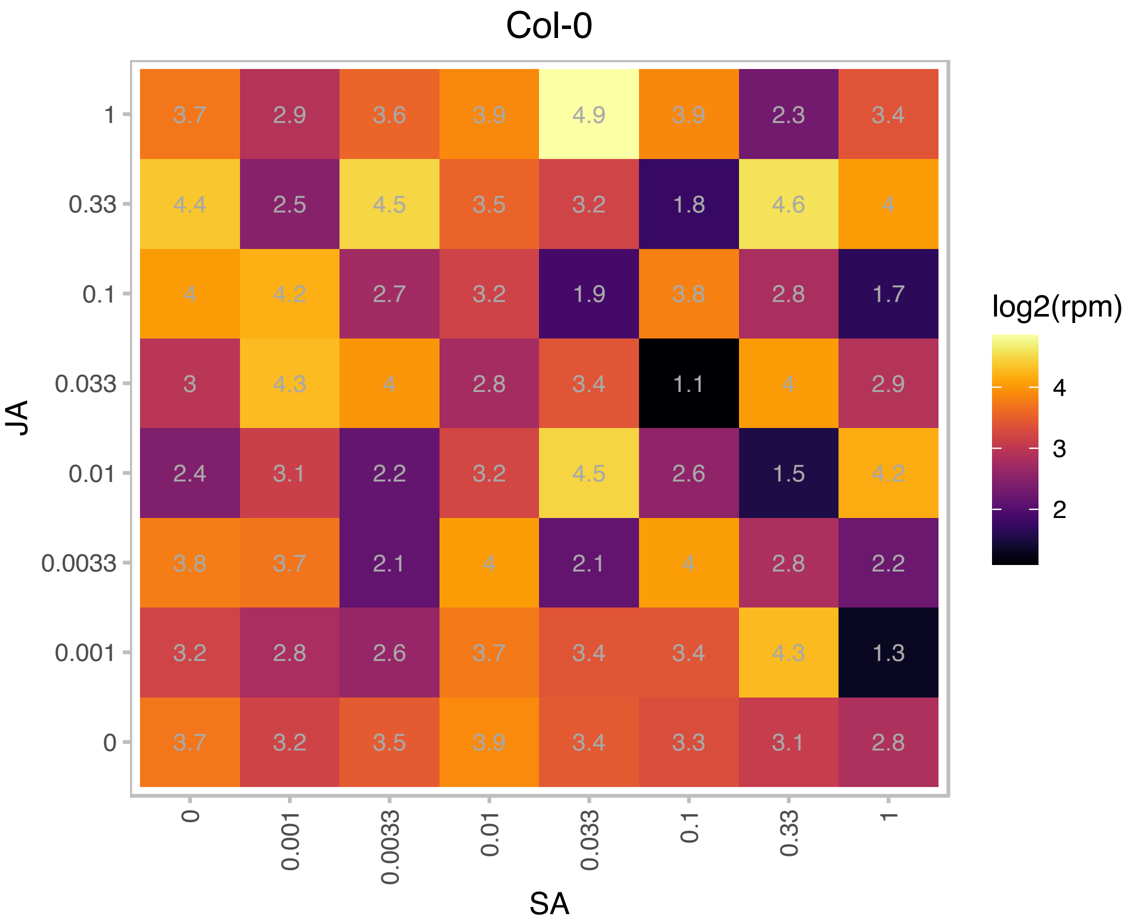


| | |
|---|---|
| Gene | AT5G63840 |
| is.expressed_genes | TRUE |
| is.responsive_genes | TRUE |
| is.apcluster | D-03 |

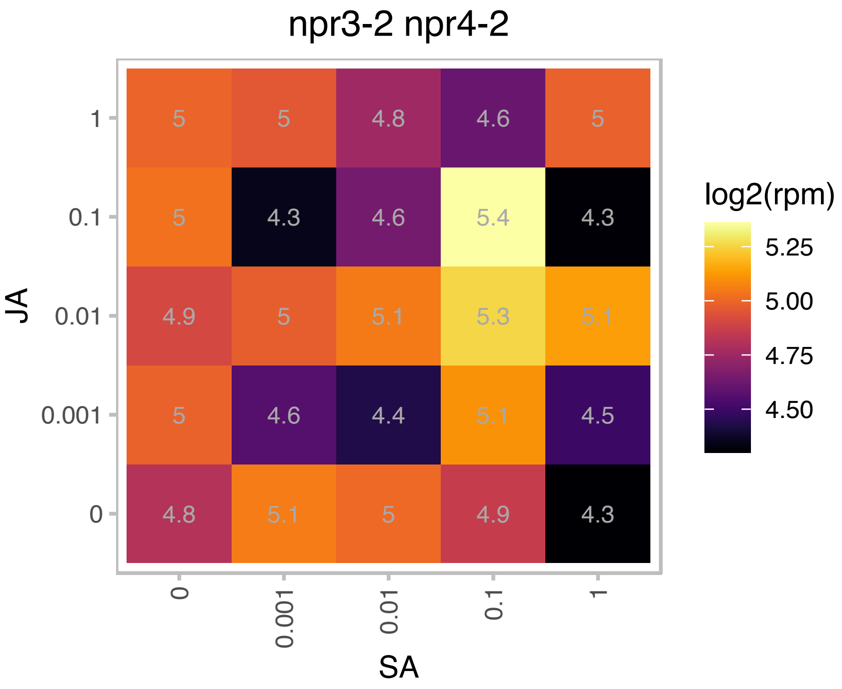


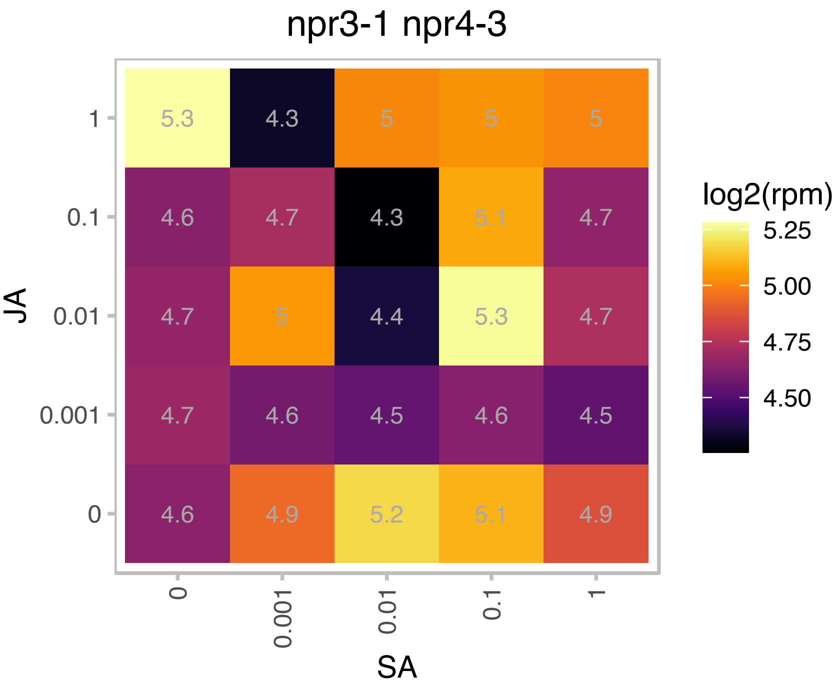

### AT1G09020.1 : ATSNF4, KINβγ, SNF4

homolog of yeast sucrose nonfermenting 4

putative activator subunit of SNF1-related protein kinase

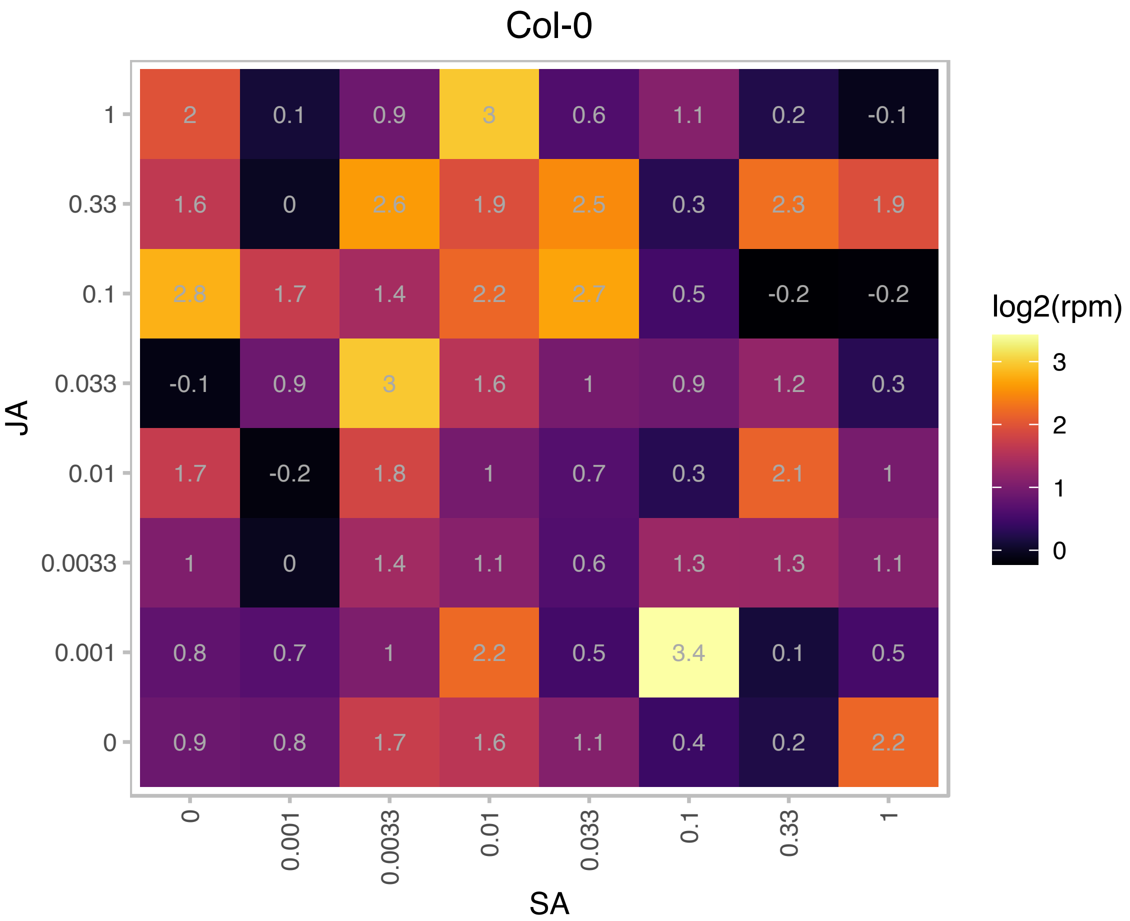


| | |
|---|---|
| Gene | AT1G09020 |
| is.expressed_genes | TRUE |
| is.responsive_genes | TRUE |
| is.apcluster | D-04 |

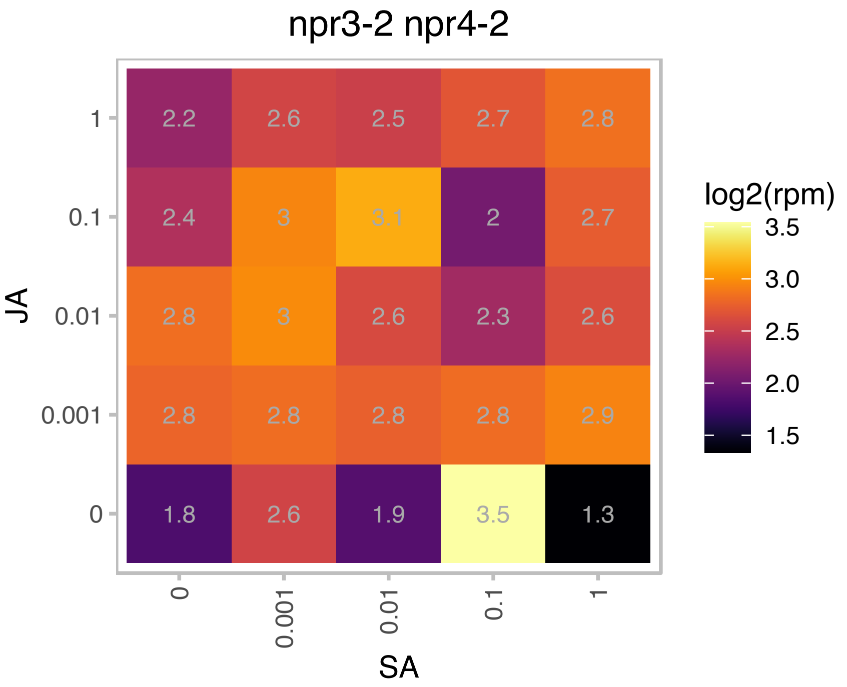


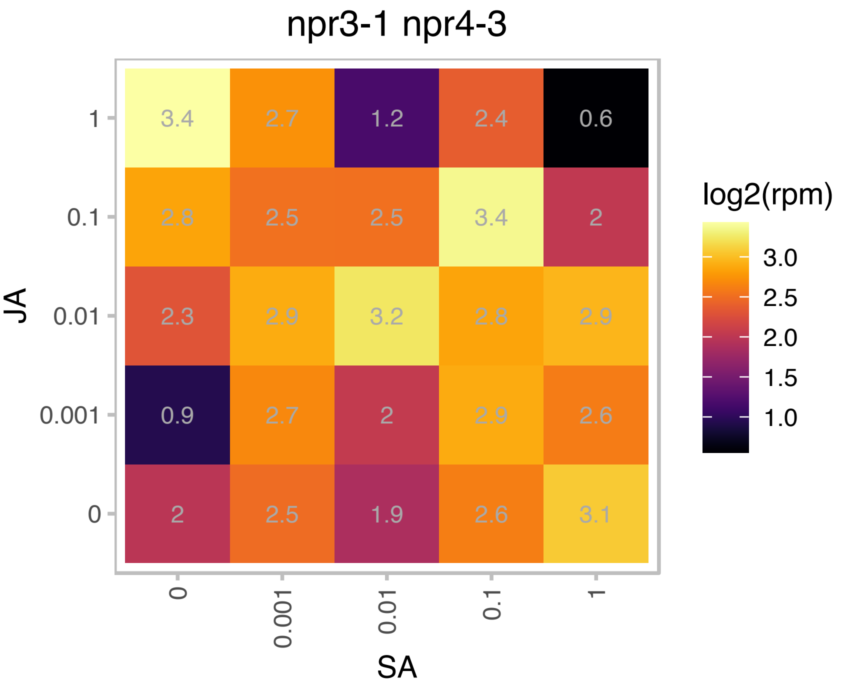

# AT1G11360.1 : NA

Adenine nucleotide alpha hydrolases-like superfamily protein

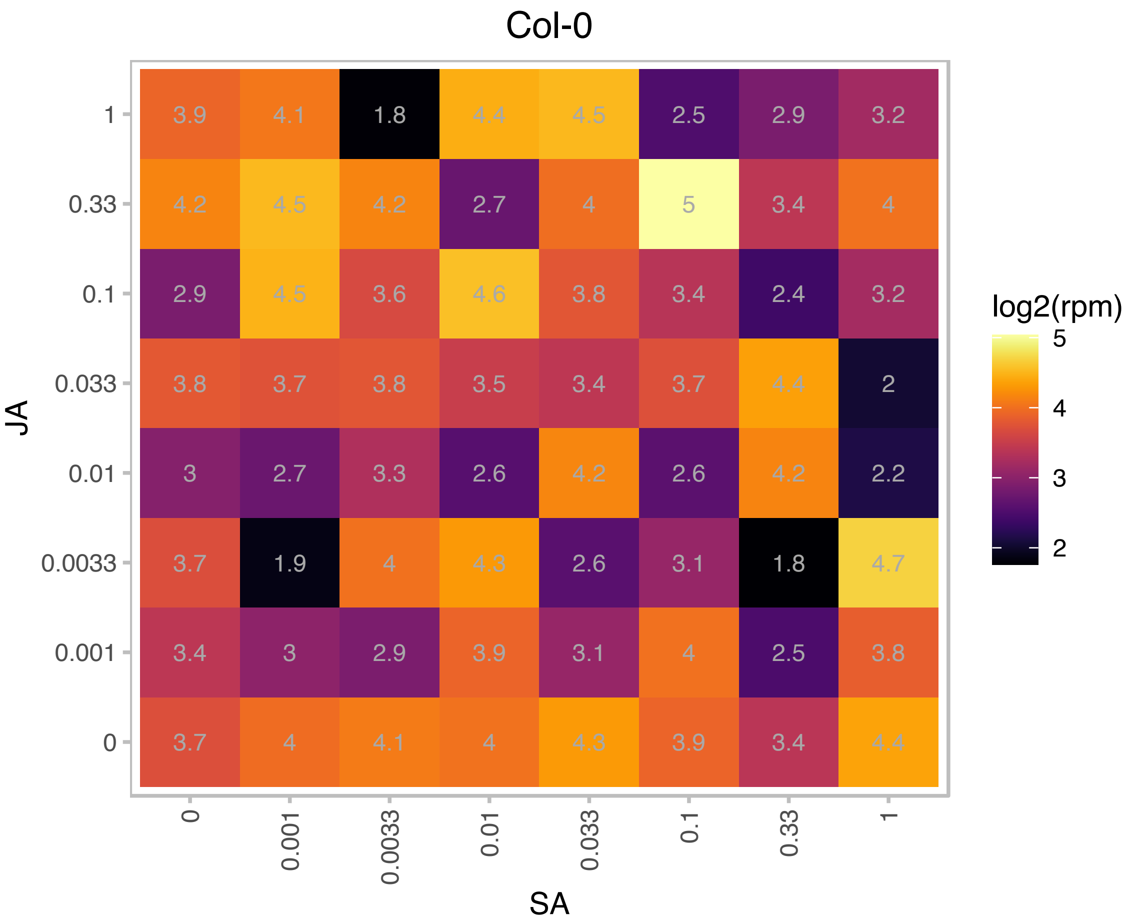


| | |
|---|---|
| Gene | AT1G11360 |
| is.expressed_genes | TRUE |
| is.responsive_genes | TRUE |
| is.apcluster | D-04 |

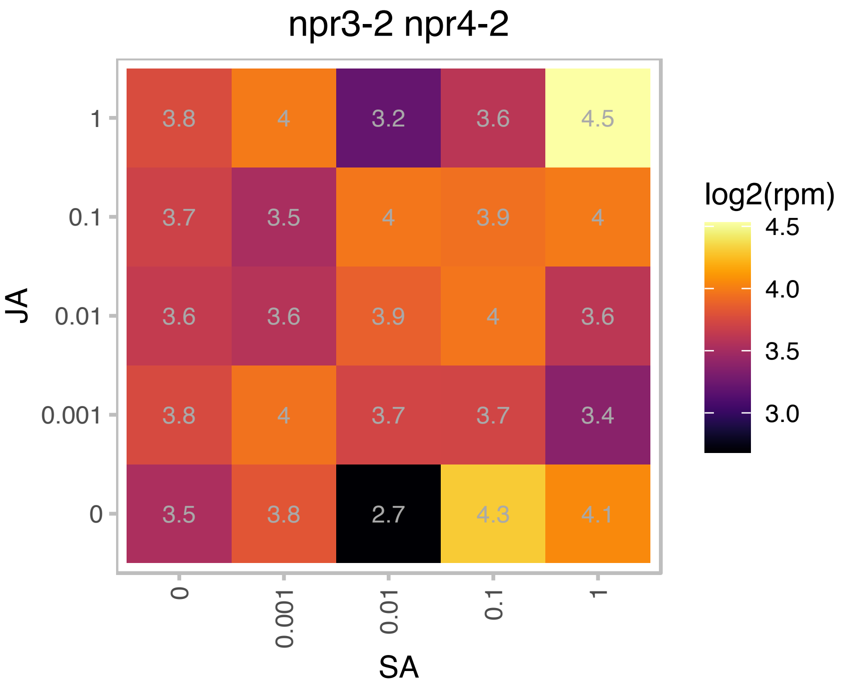


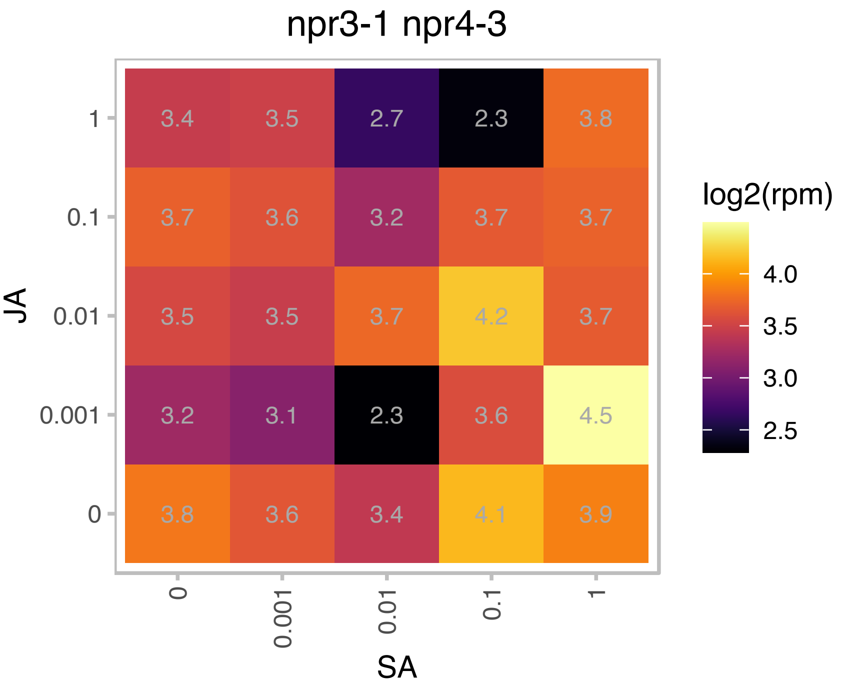

### AT1G12820.1 : AFB3

auxin signaling F-box 3

Auxin receptor involved in primary and lateral root growth inhibition in response to nitrate. Target of miR393. Induced by nitrate in primary roots.

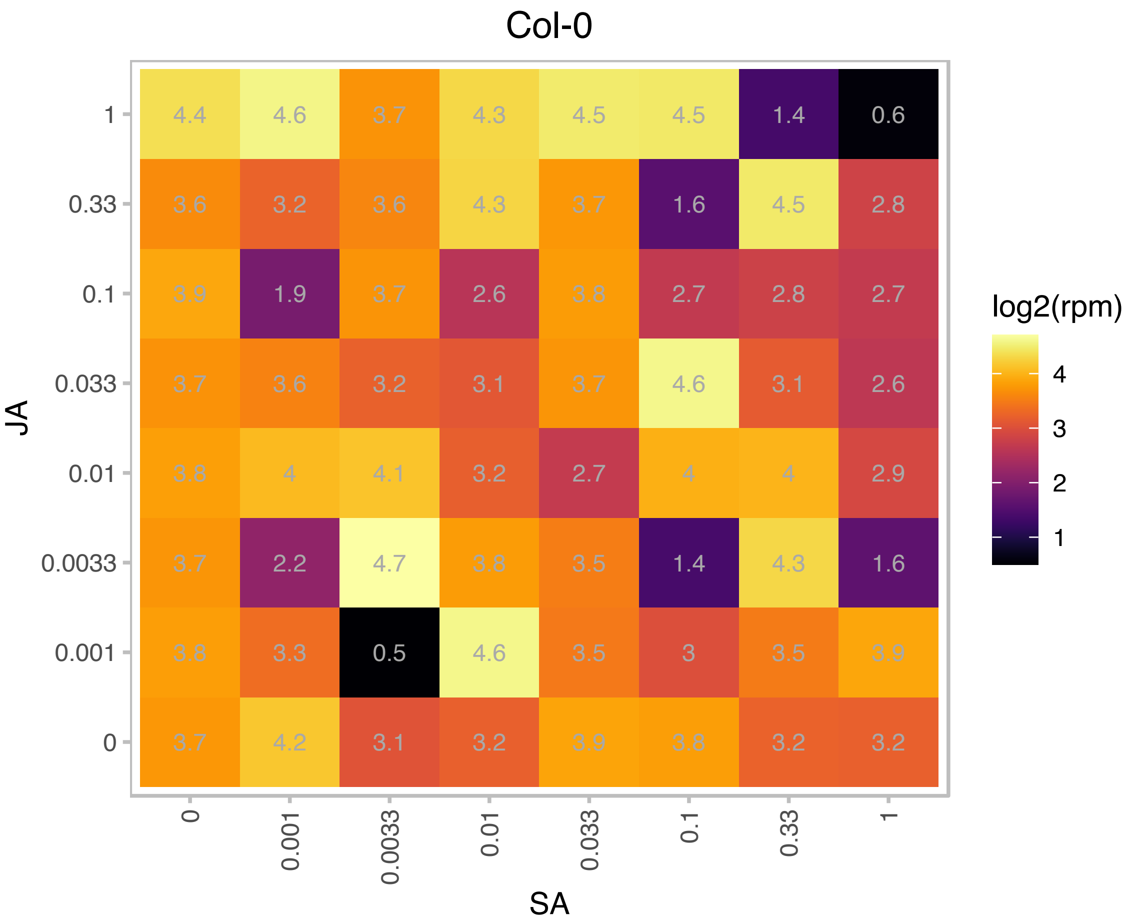


| | |
|---|---|
| Gene | AT1G12820 |
| is.expressed_genes | TRUE |
| is.responsive_genes | TRUE |
| is.apcluster | D-04 |

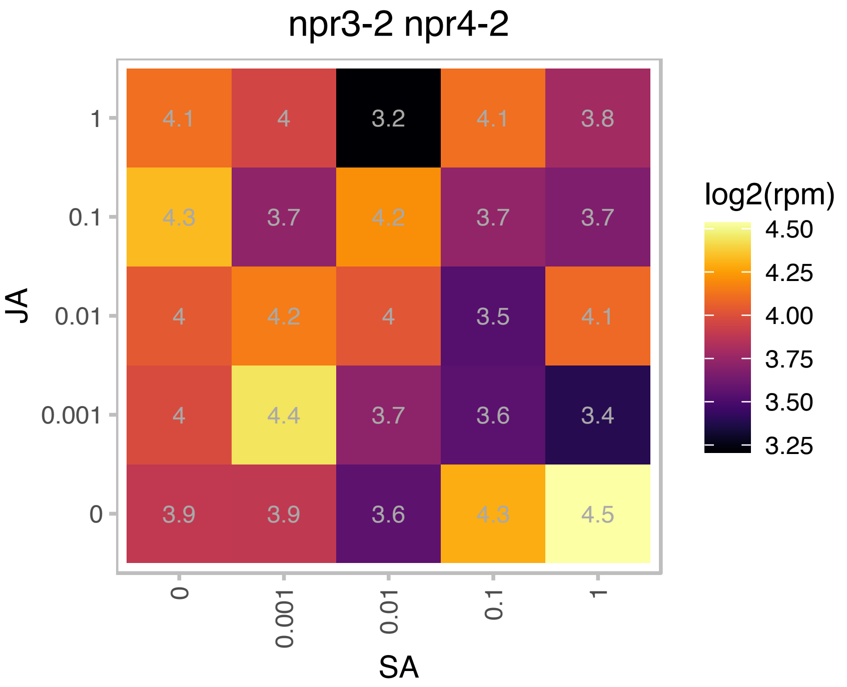


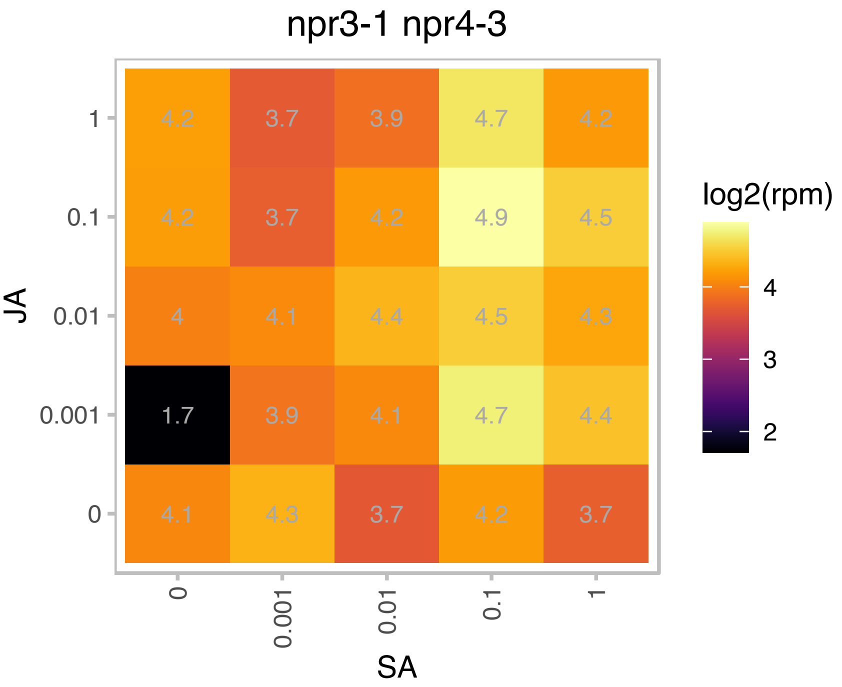

# AT1G26750.1 : NA

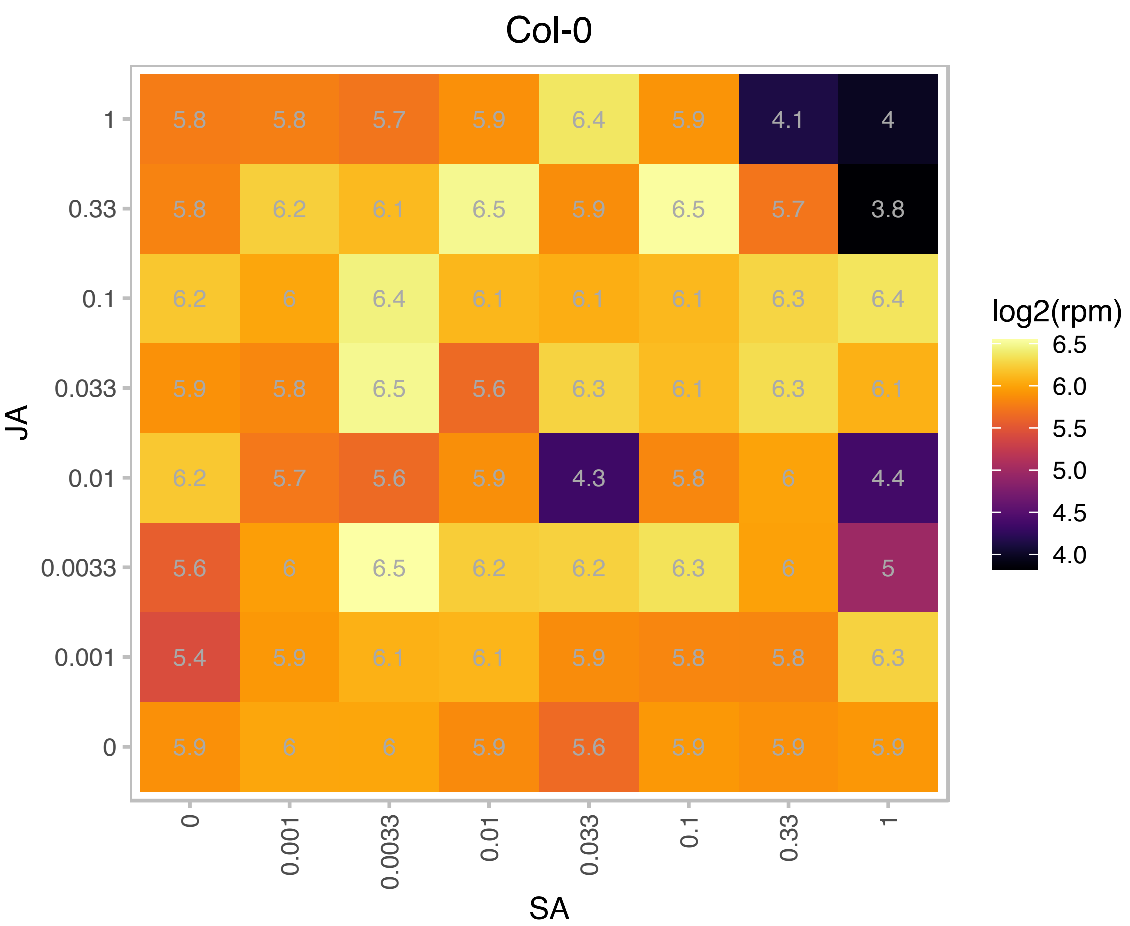


| | |
|---|---|
| Gene | AT1G26750 |
| is.expressed_genes | TRUE |
| is.responsive_genes | TRUE |
| is.apcluster | D-04 |

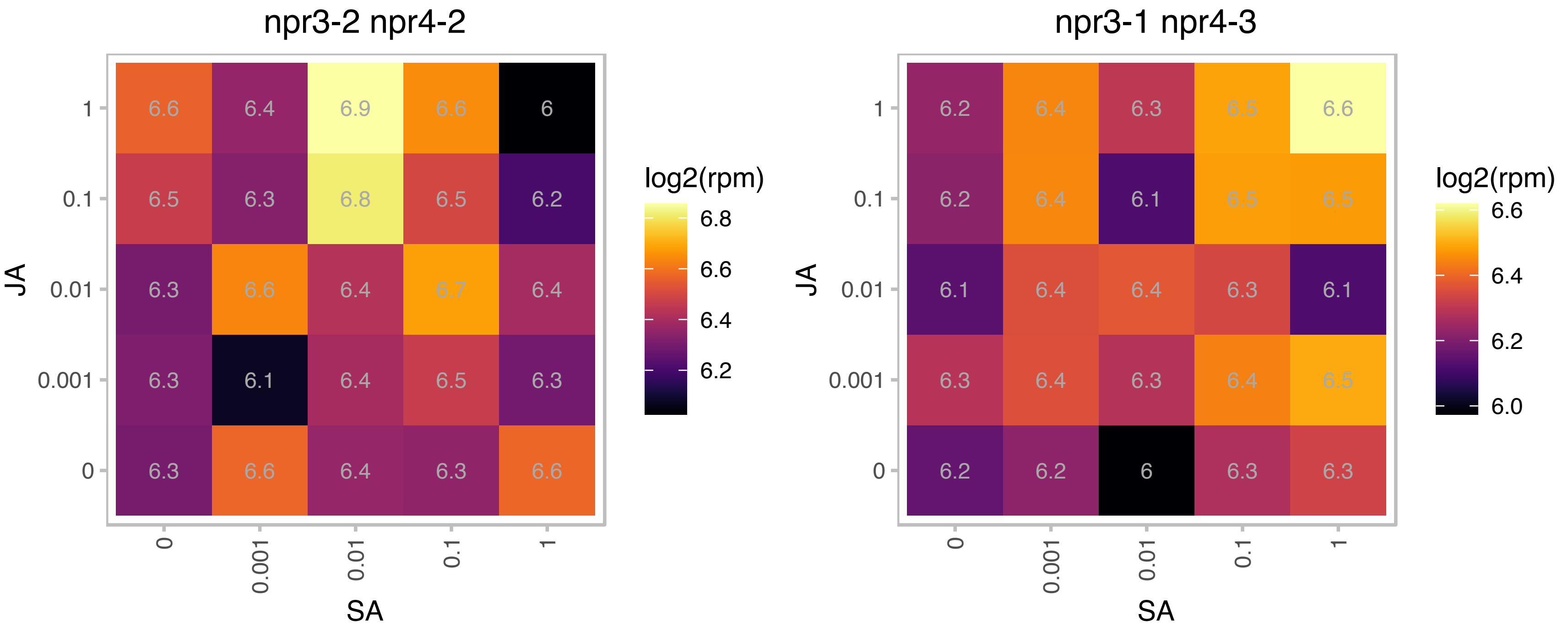

### AT1G53240.1 : mMDH1

Lactate/malate dehydrogenase family protein

mMDH1 encodes a mitochrondrial malate dehydrogenase. It is expressed at higher levels than the other mitochrondrial isoform mMDH2 (At3G15020) according to transcript and proteomic analyses.

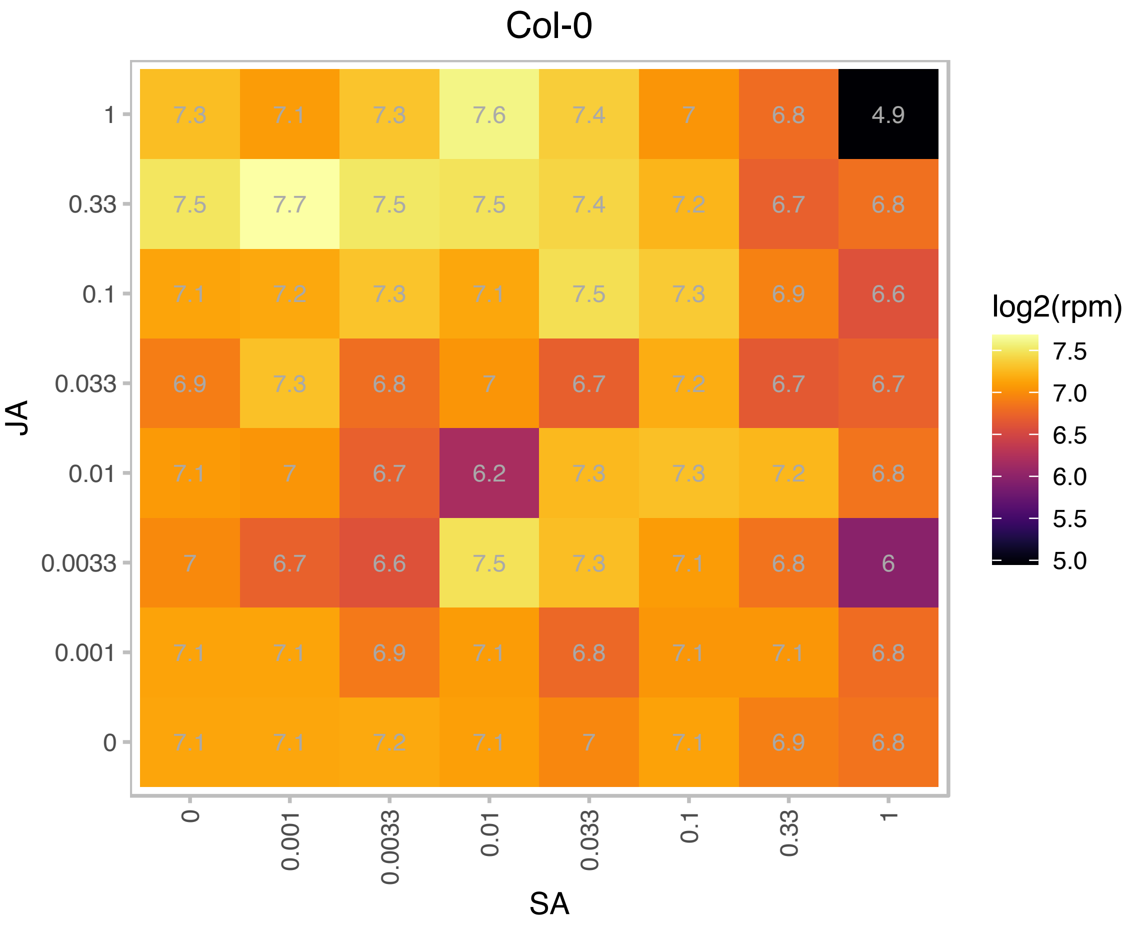


| | |
|---|---|
| Gene | AT1G53240 |
| is.expressed_genes | TRUE |
| is.responsive_genes | TRUE |
| is.apcluster | D-04 |

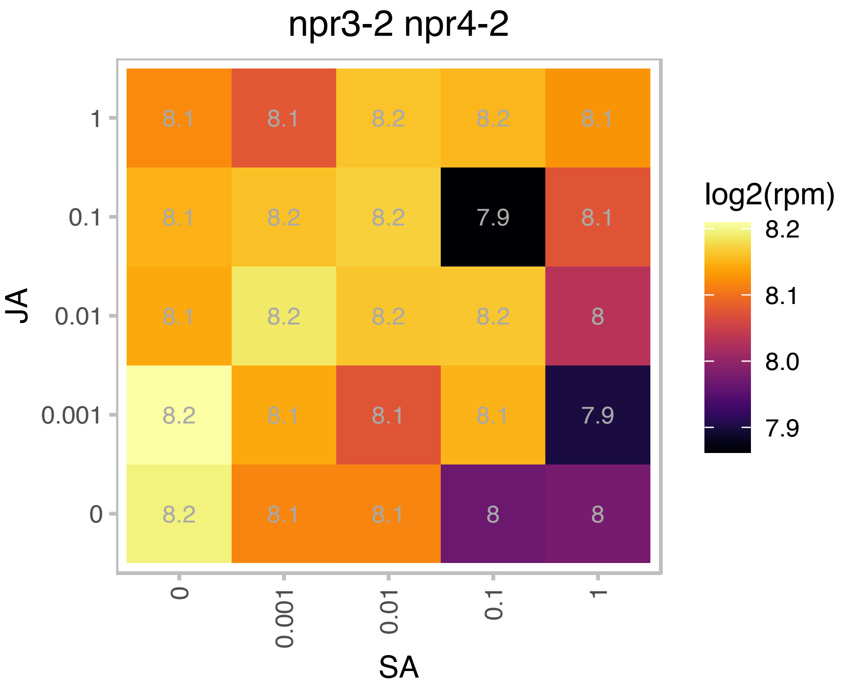


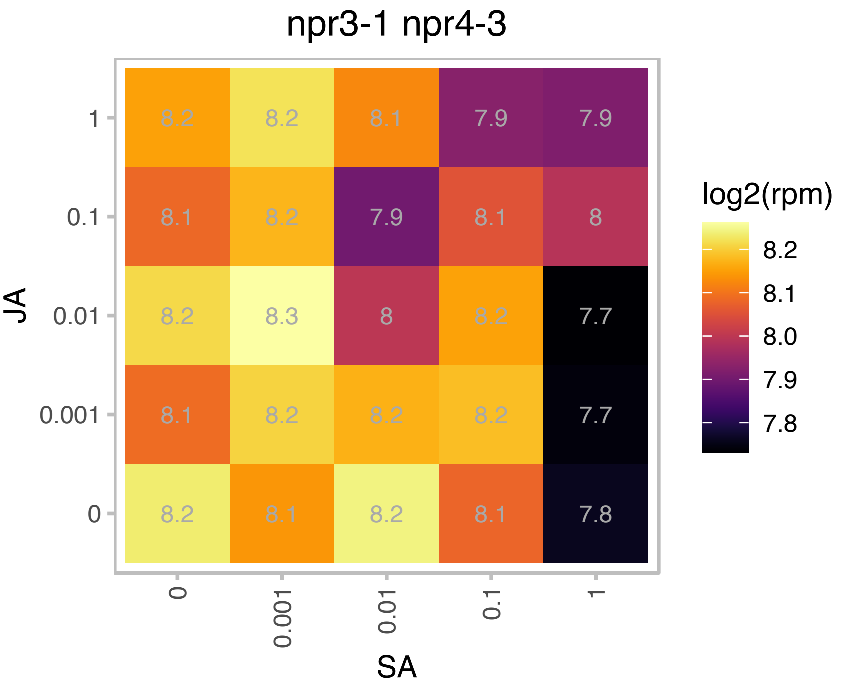

# AT1G63980.1 : NA

D111/G-patch domain-containing protein

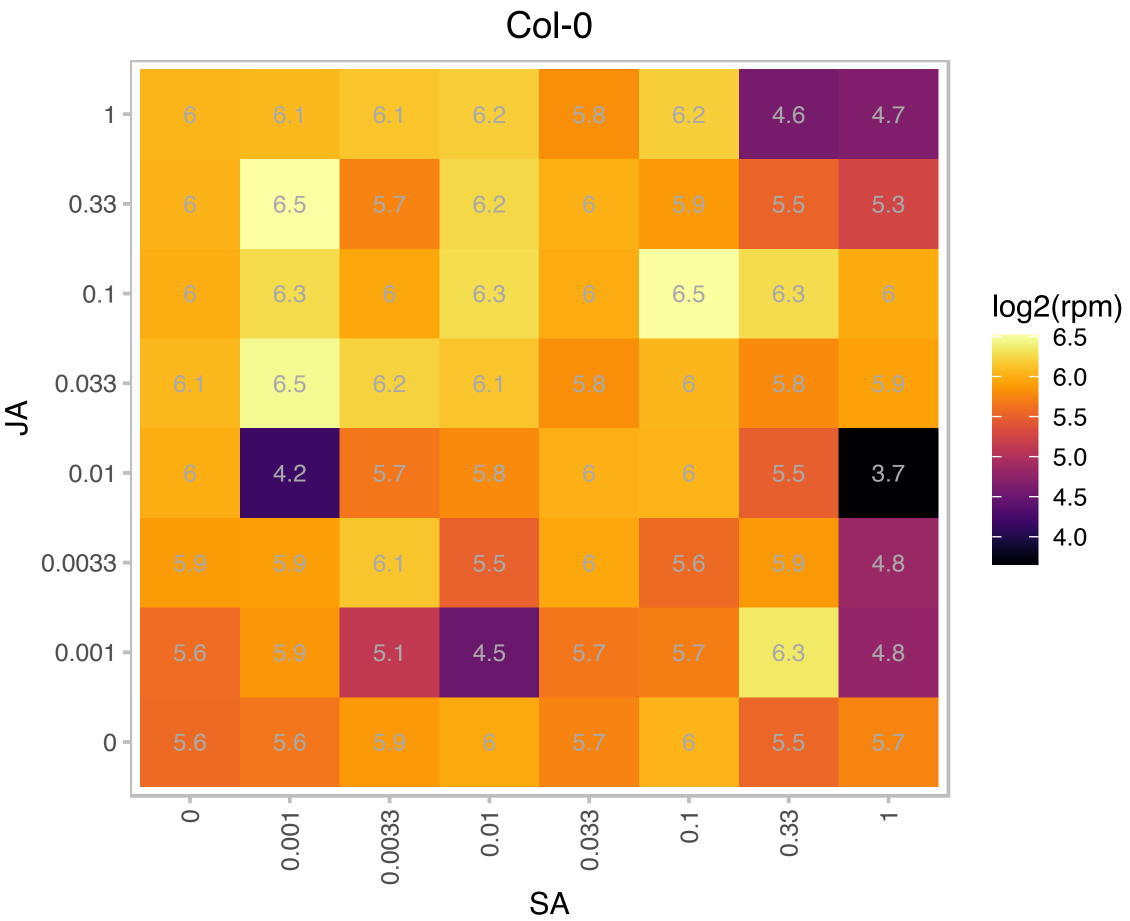


| | |
|---|---|
| Gene | AT1G63980 |
| is.expressed_genes | TRUE |
| is.responsive_genes | TRUE |
| is.apcluster | D-04 |

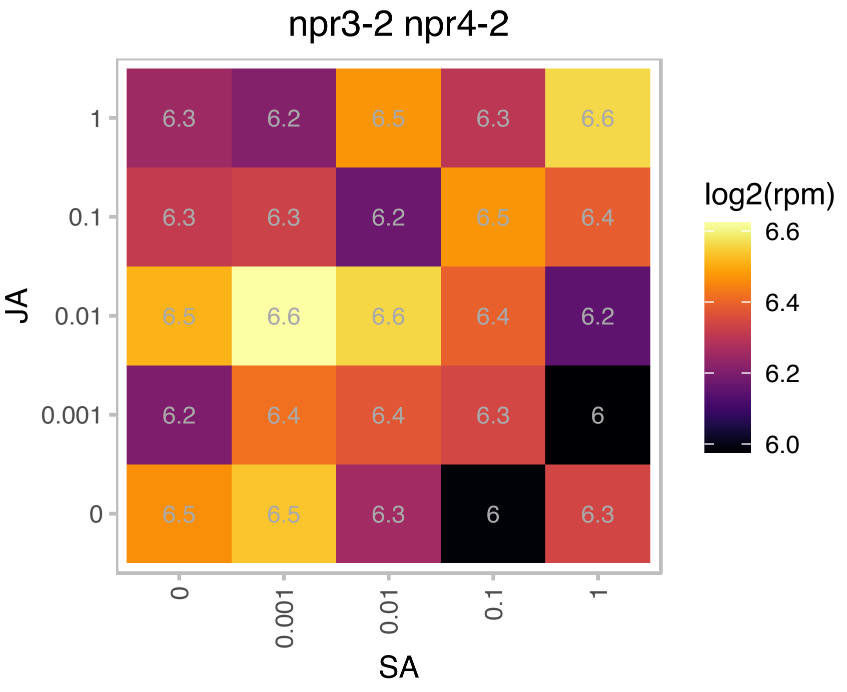


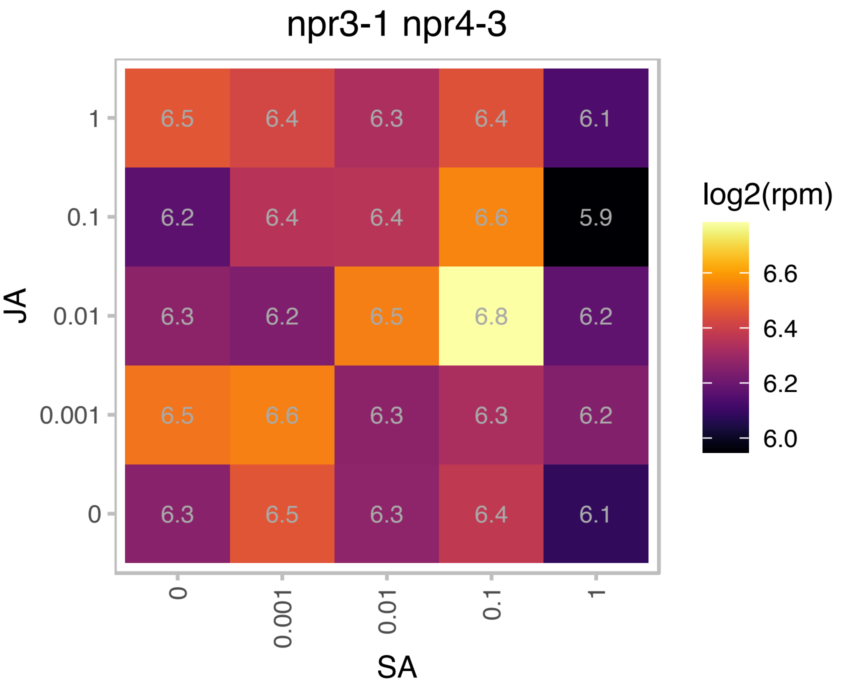

# AT1G72550.1 : NA

tRNA synthetase beta subunit family protein

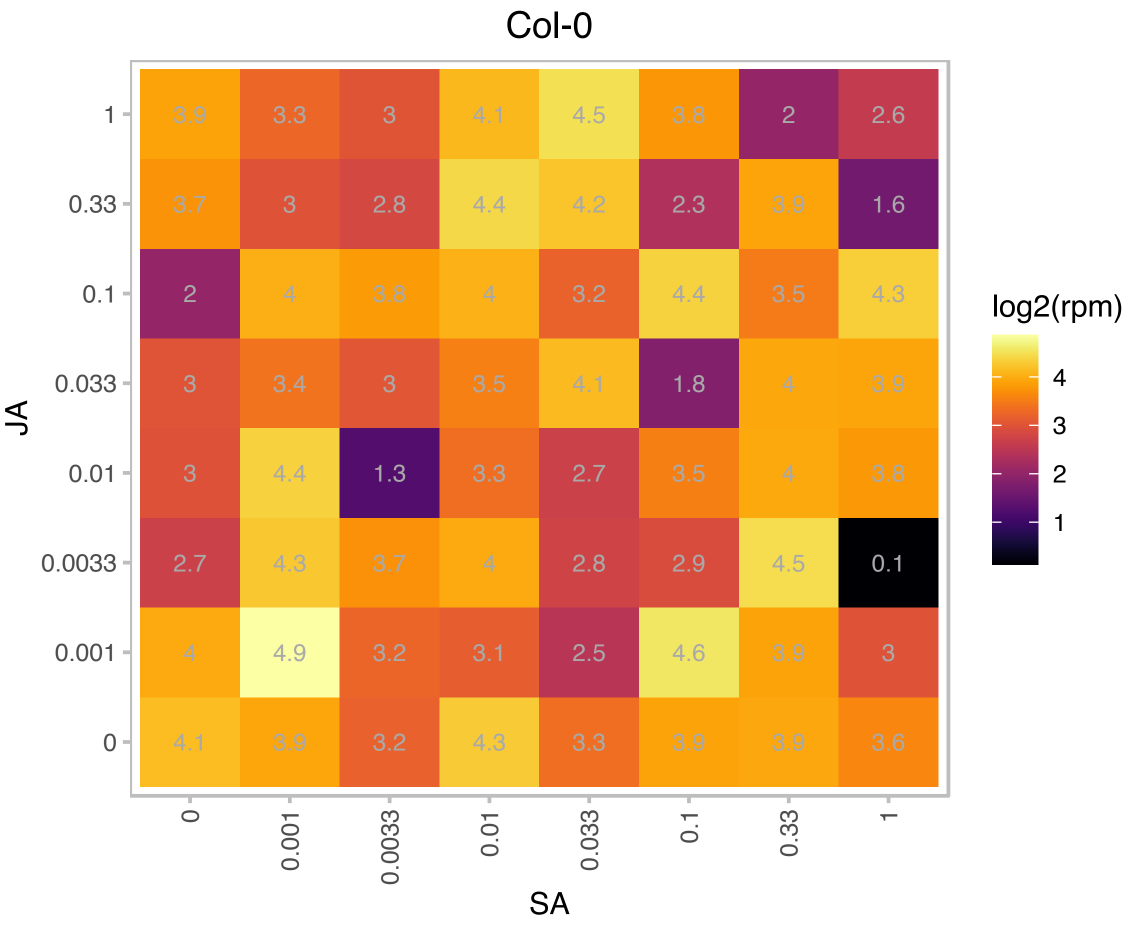


| | |
|---|---|
| Gene | AT1G72550 |
| is.expressed_genes | TRUE |
| is.responsive_genes | TRUE |
| is.apcluster | D-04 |

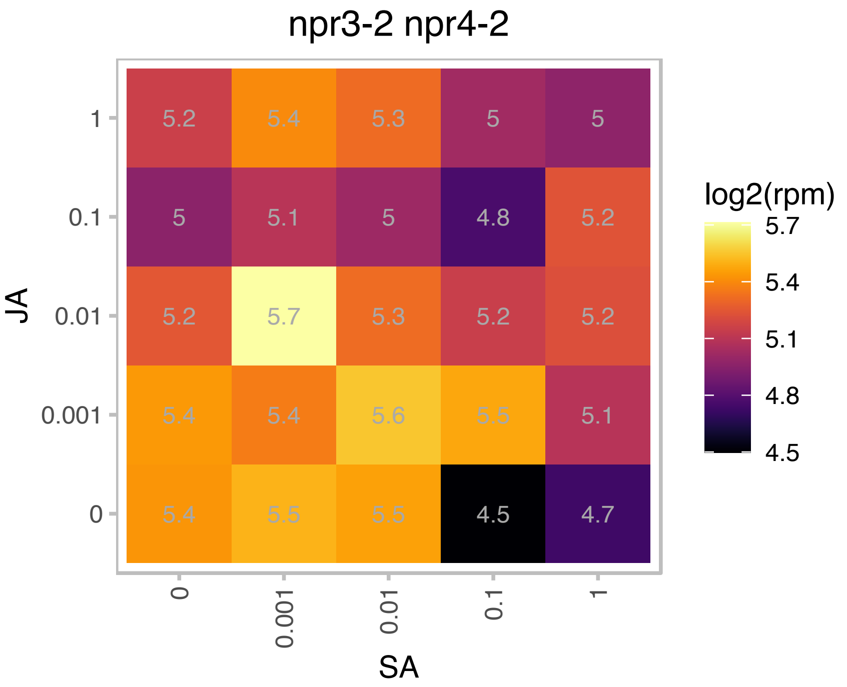


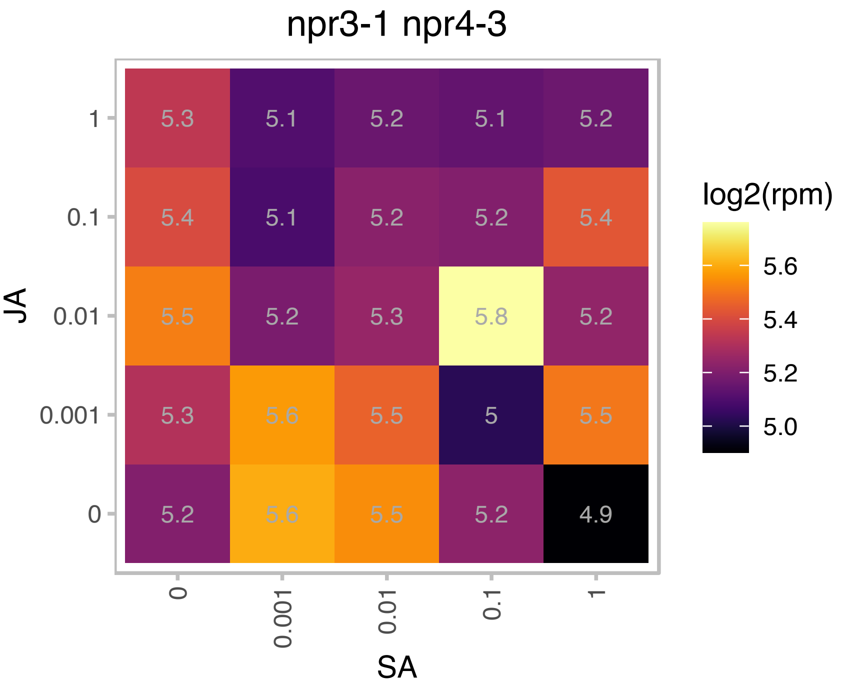

# AT1G78150.1 : NA

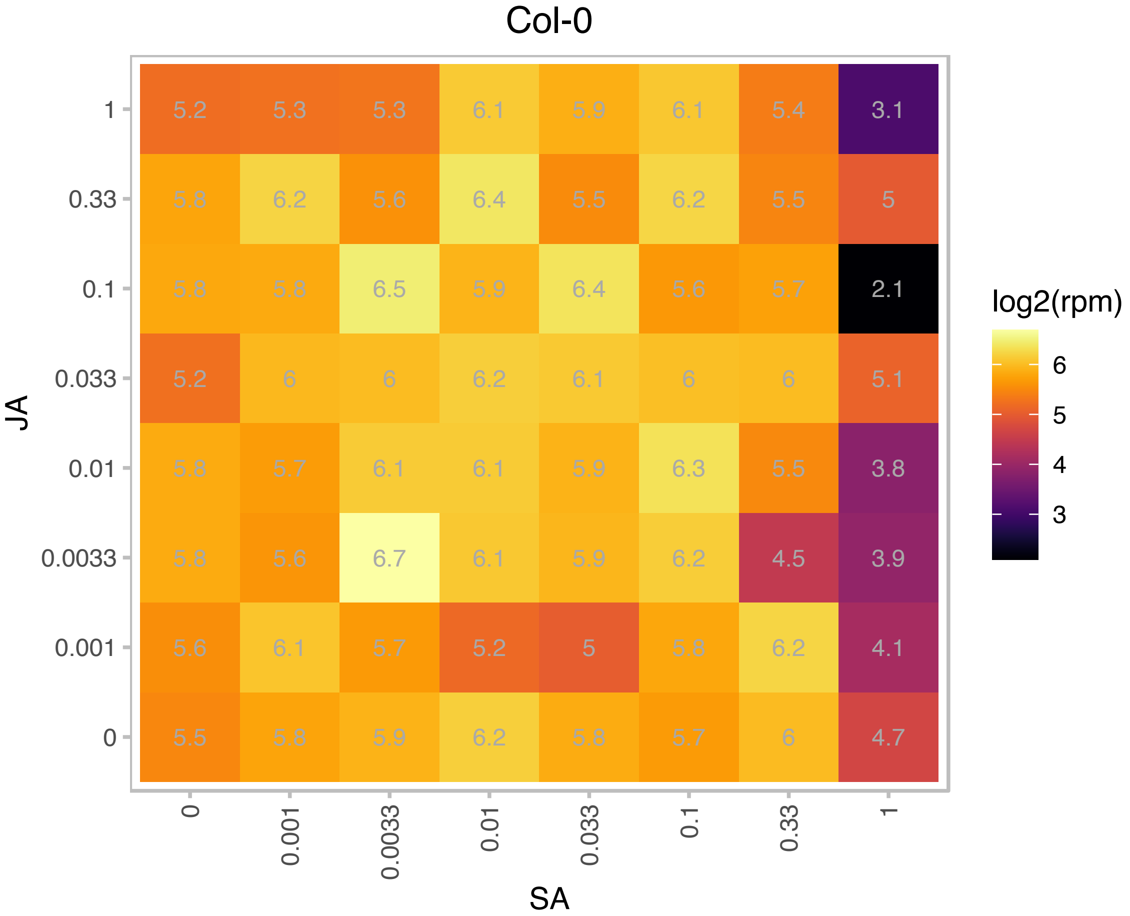


| | |
|---|---|
| Gene | AT1G78150 |
| is.expressed_genes | TRUE |
| is.responsive_genes | TRUE |
| is.apcluster | D-04 |

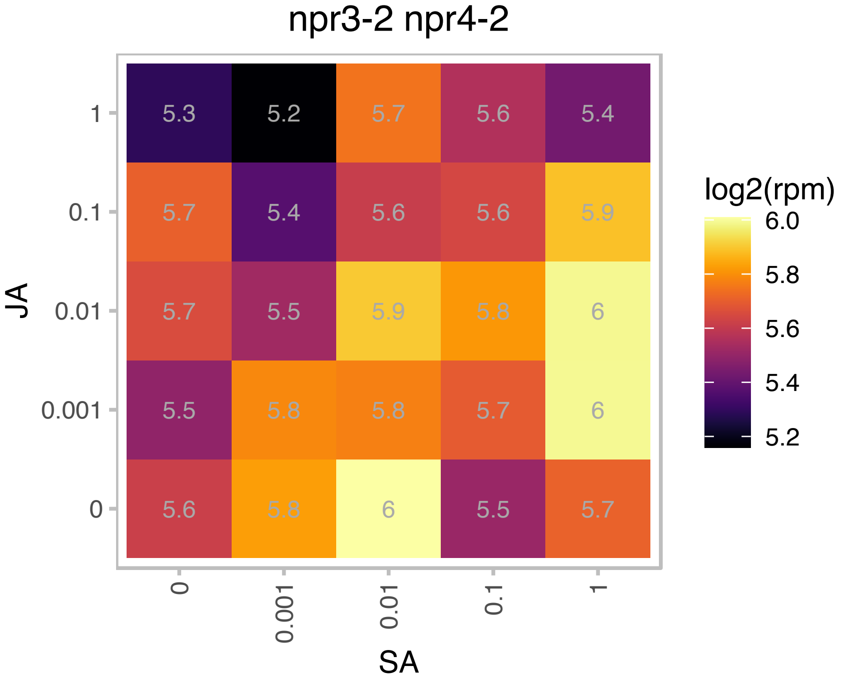


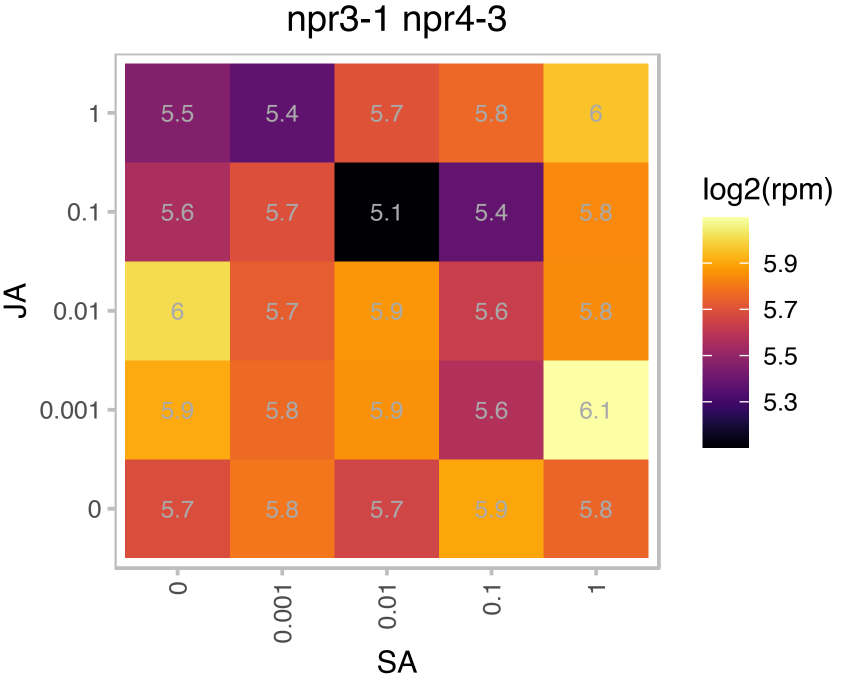

### AT2G01750.1 : ATMAP70-3, MAP70-3

microtubule-associated proteins 70-3

Encodes a microtubule associated protein (MAP70-3). Expressed in all tissues.

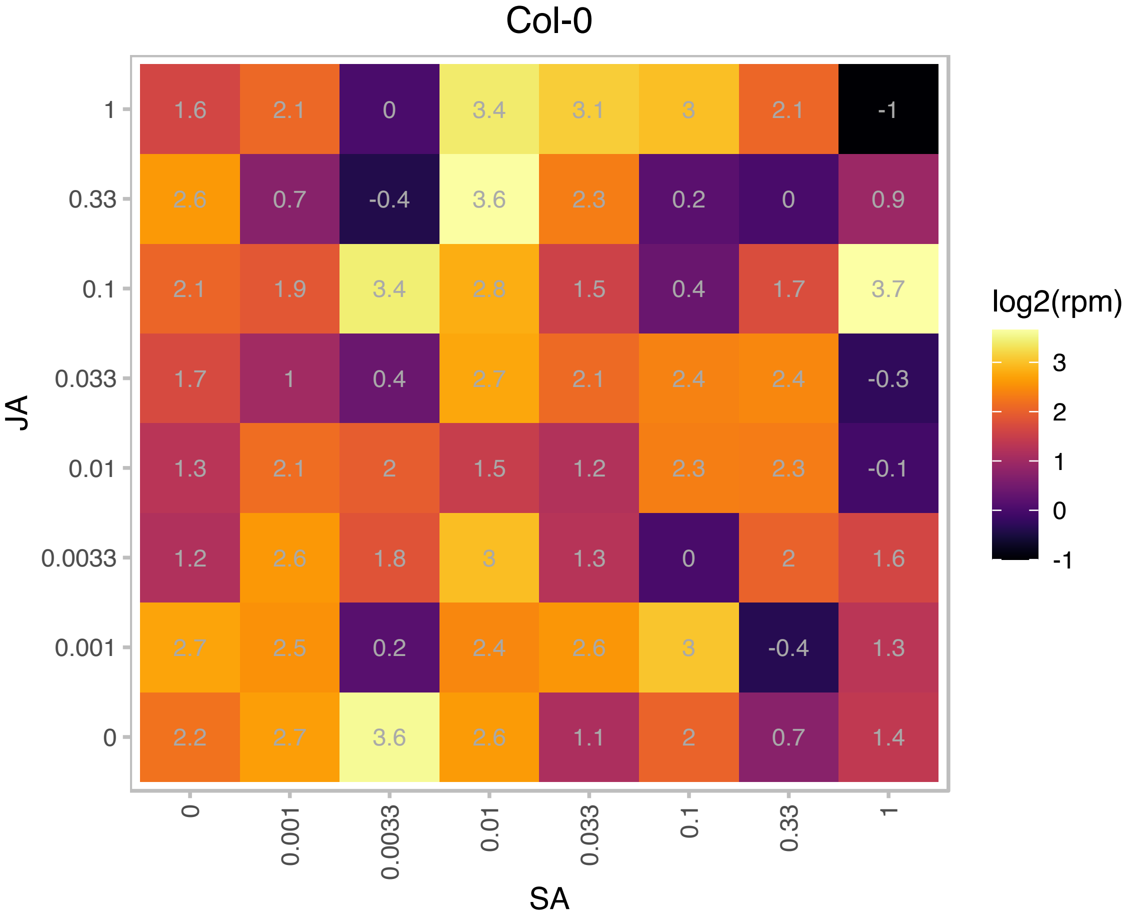


| | |
|---|---|
| Gene | AT2G01750 |
| is.expressed_genes | TRUE |
| is.responsive_genes | TRUE |
| is.apcluster | D-04 |

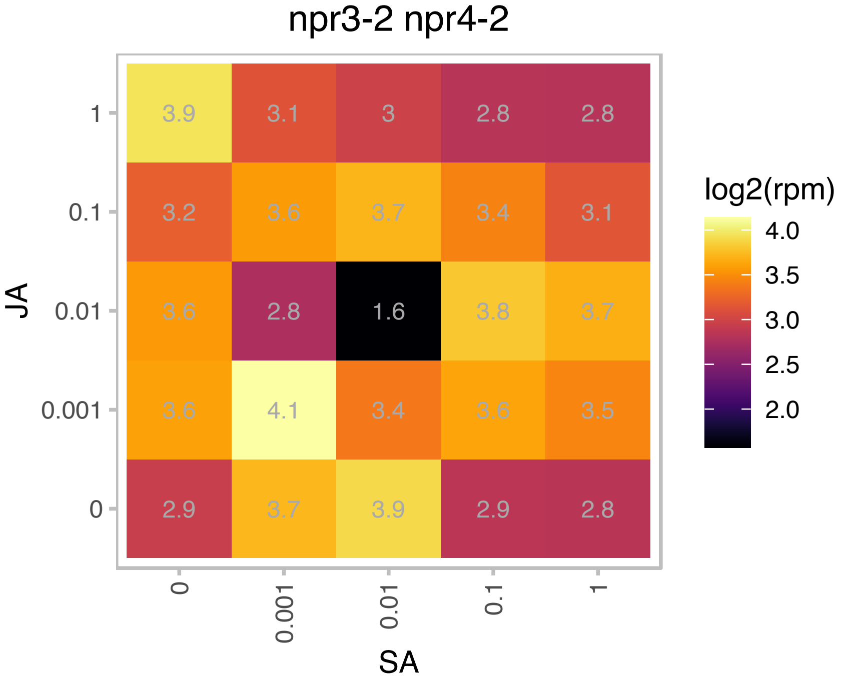


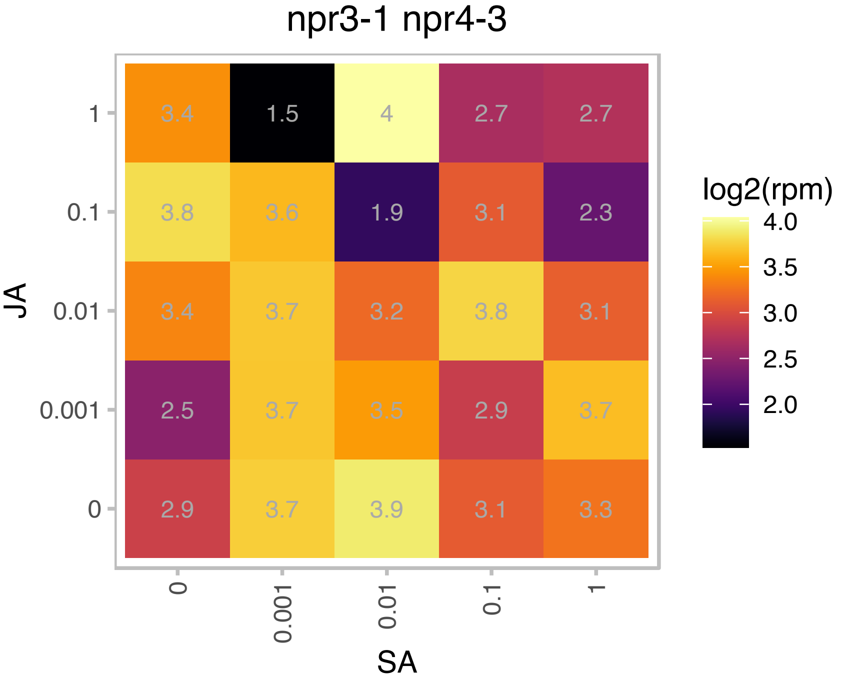

# AT2G02510.1 : NA

NADH dehydrogenase (ubiquinone)s

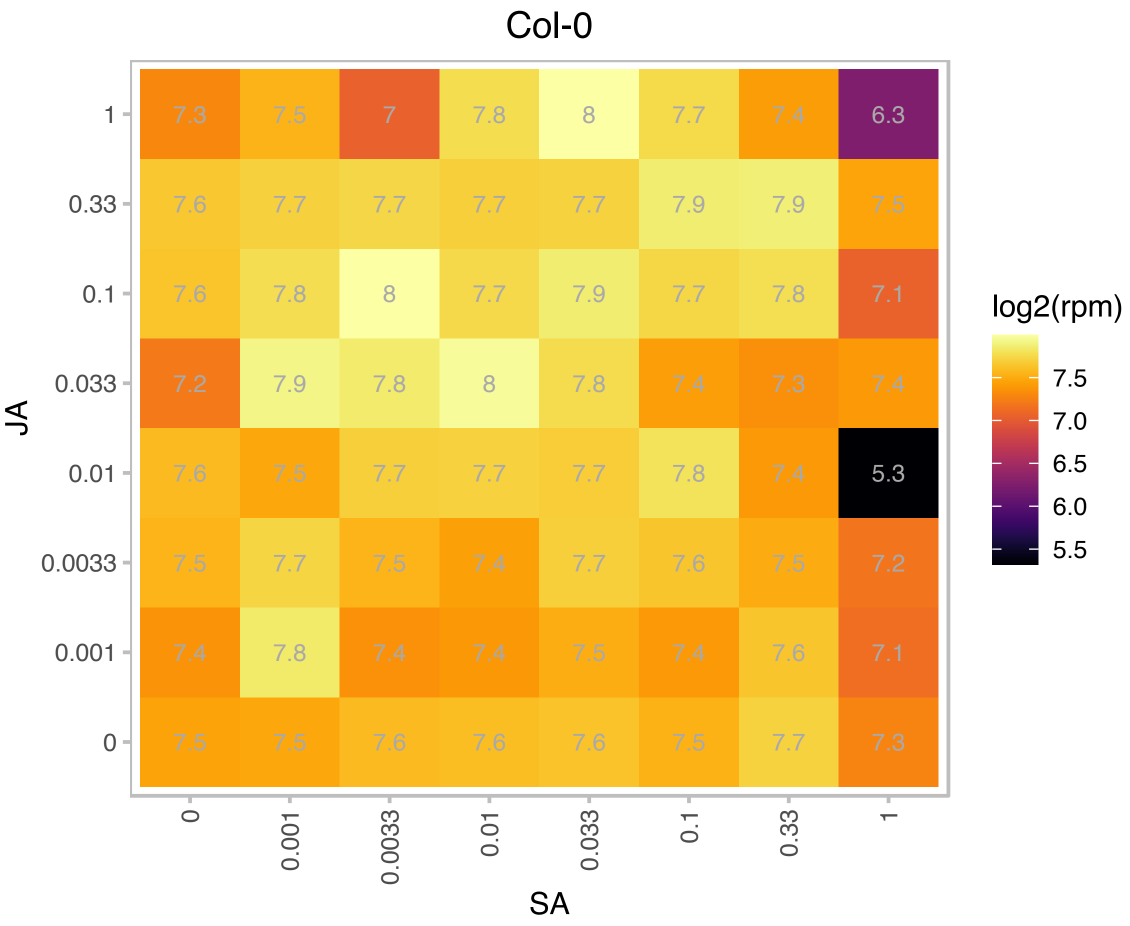


| | |
|---|---|
| Gene | AT2G02510 |
| is.expressed_genes | TRUE |
| is.responsive_genes | TRUE |
| is.apcluster | D-04 |

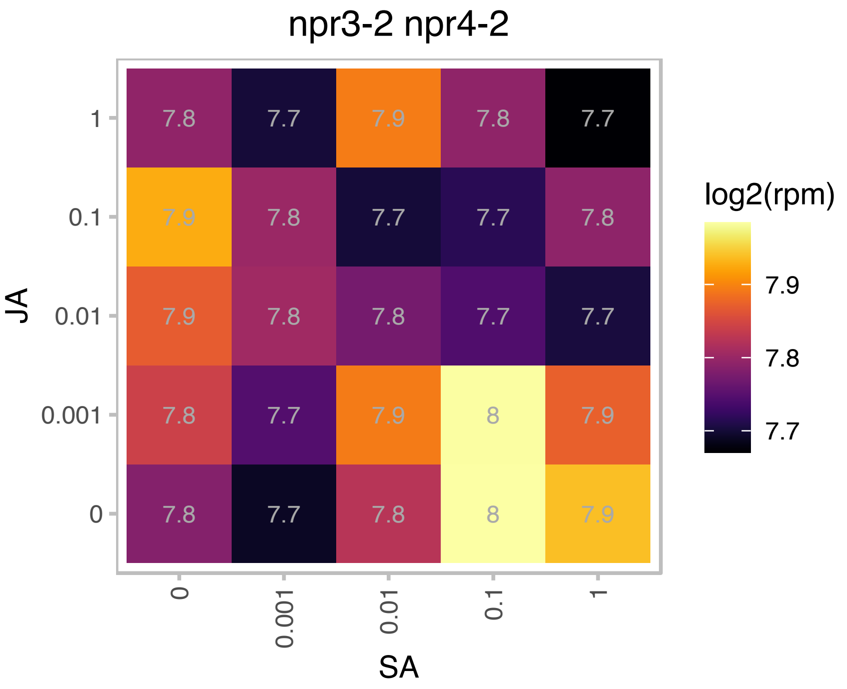


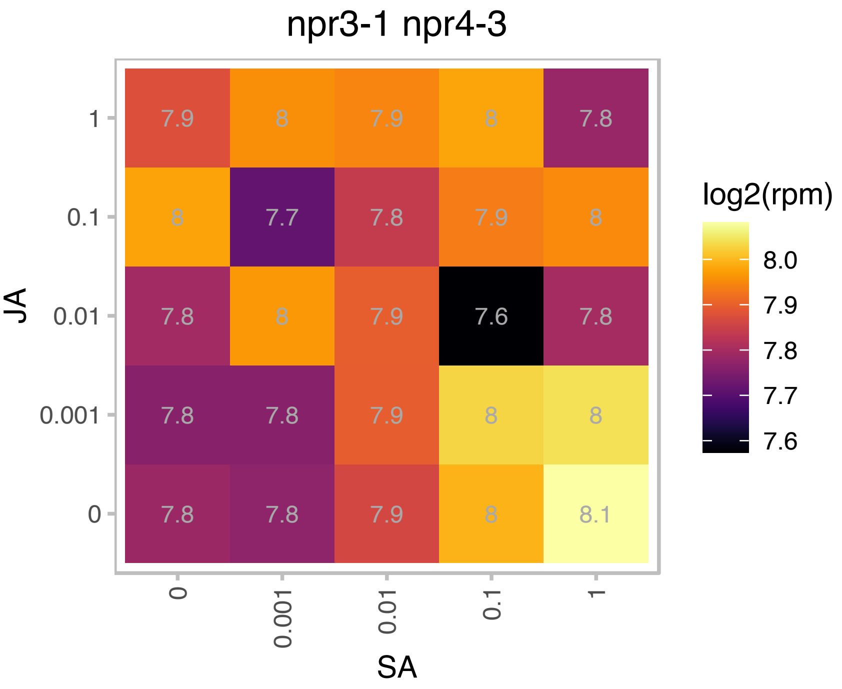

### AT2G24490.1 : ATRPA2, ATRPA32A, ROR1, RPA2, RPA32

replicon protein A2
Encodes a component of Replication Protein A. Component of transcriptional gene silencing which does not affect endogenous small RNA accumulation nor DNA methylation. Localized in the nucleus. Involved in DNA repair. Interacts physically with ROS1.

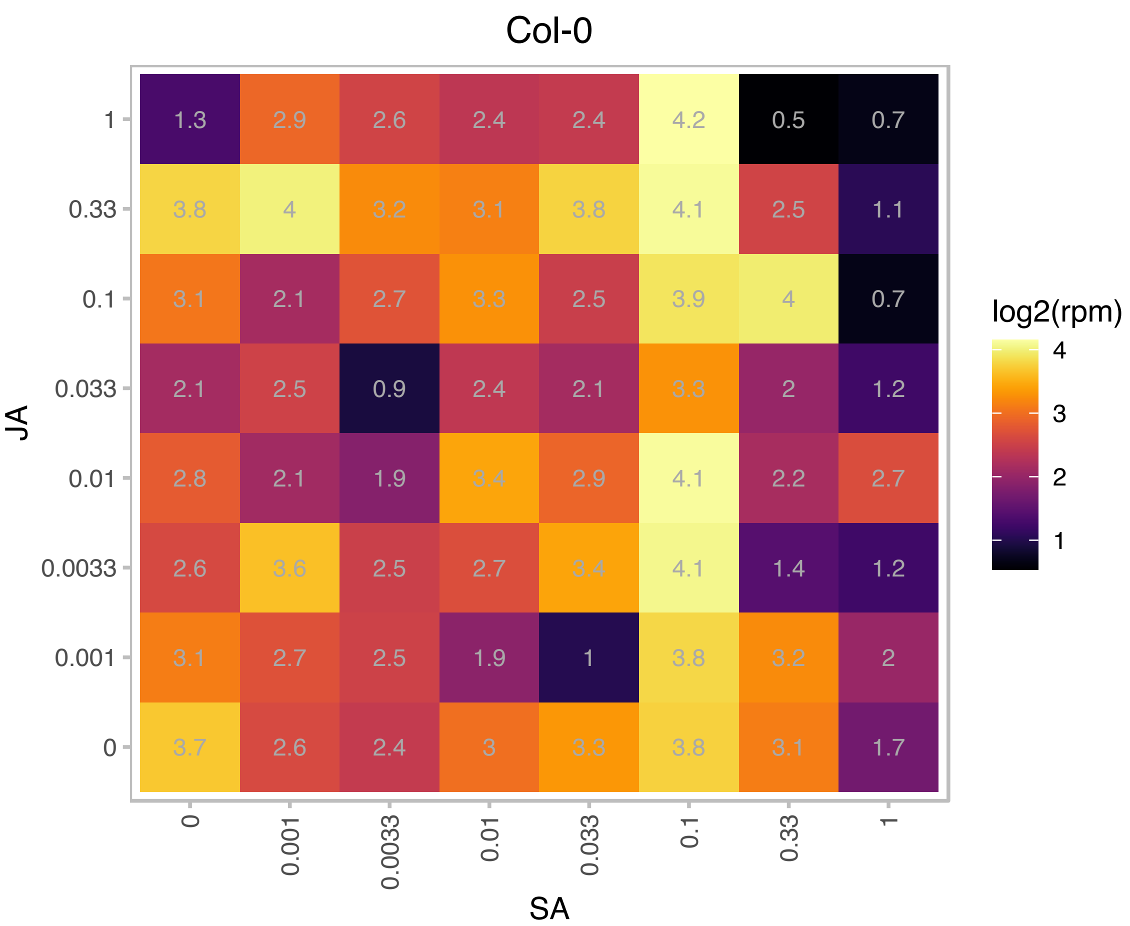


| | |
|---|---|
| Gene | AT2G24490 |
| is.expressed_genes | TRUE |
| is.responsive_genes | TRUE |
| is.apcluster | D-04 |

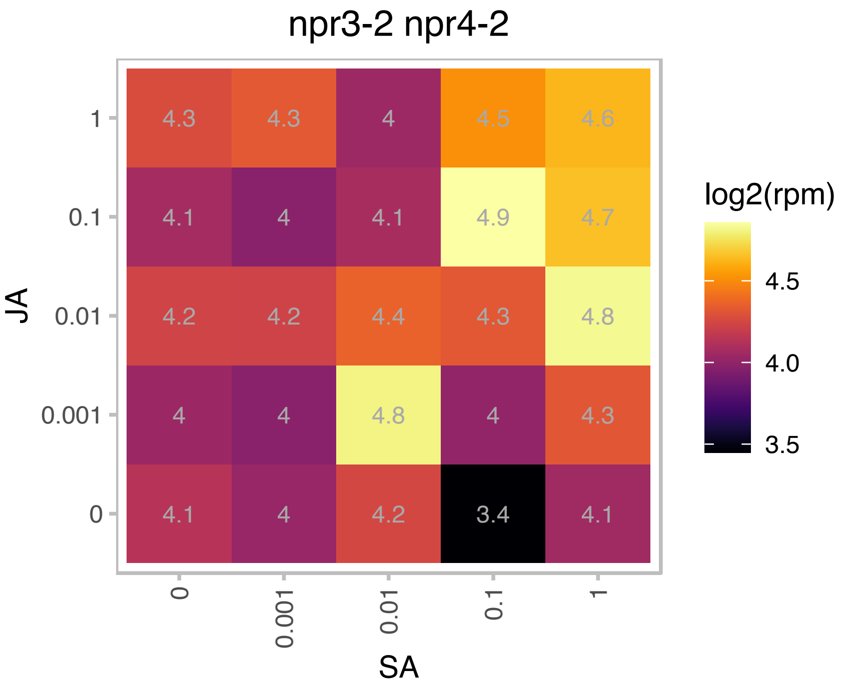


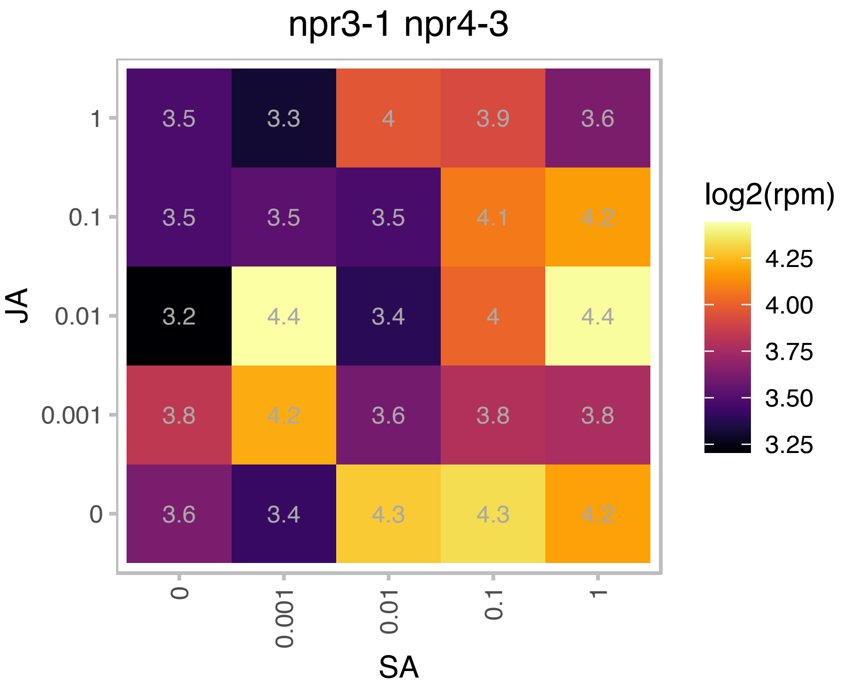

### AT2G38560.1 : RDO2, TFIIS

transcript elongation factor IIS
Encodes RNA polymerase II transcript elongation factor TFIIS.  Complements yeast TFIIS mutation.  Mutant plants display essentially normal development, but they flower slightly earlier than the wild type and show clearly reduced seed dormancy.

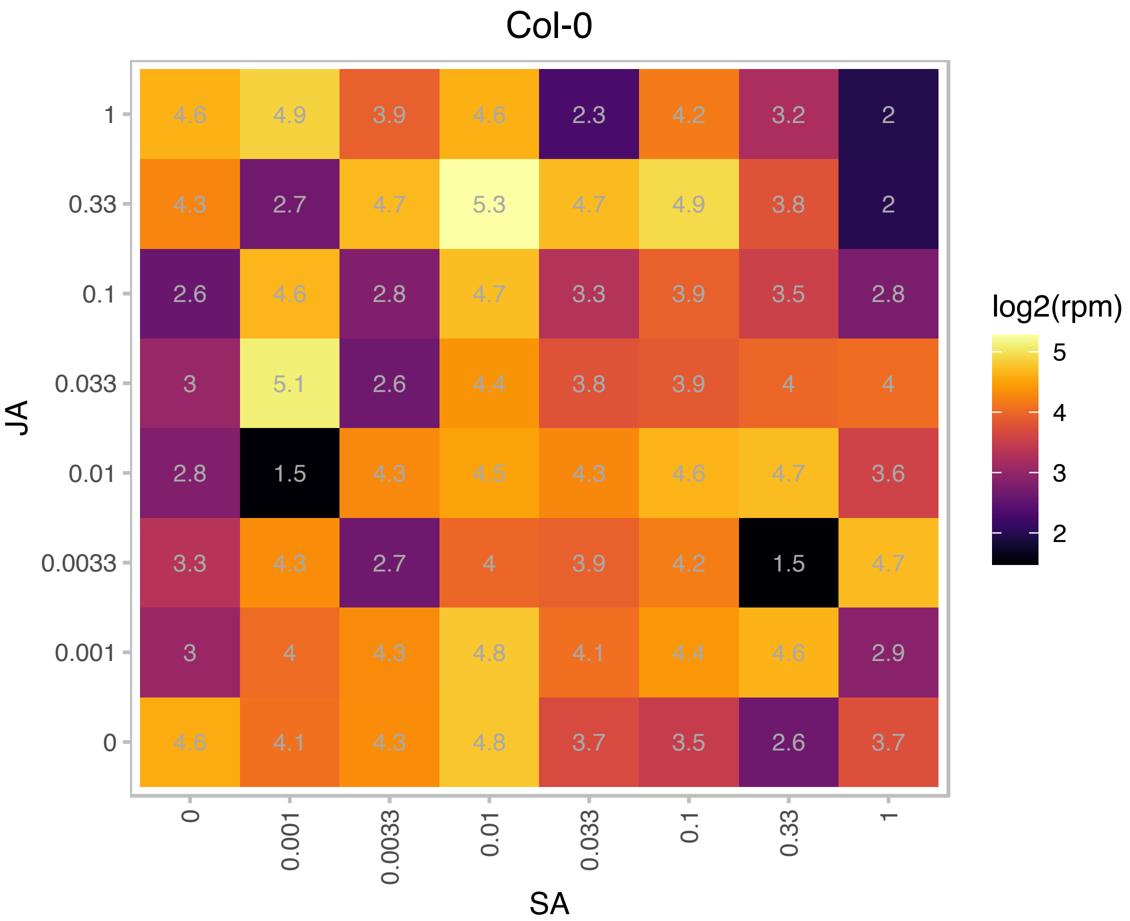


| | |
|---|---|
| Gene | AT2G38560 |
| is.expressed_genes | TRUE |
| is.responsive_genes | TRUE |
| is.apcluster | D-04 |

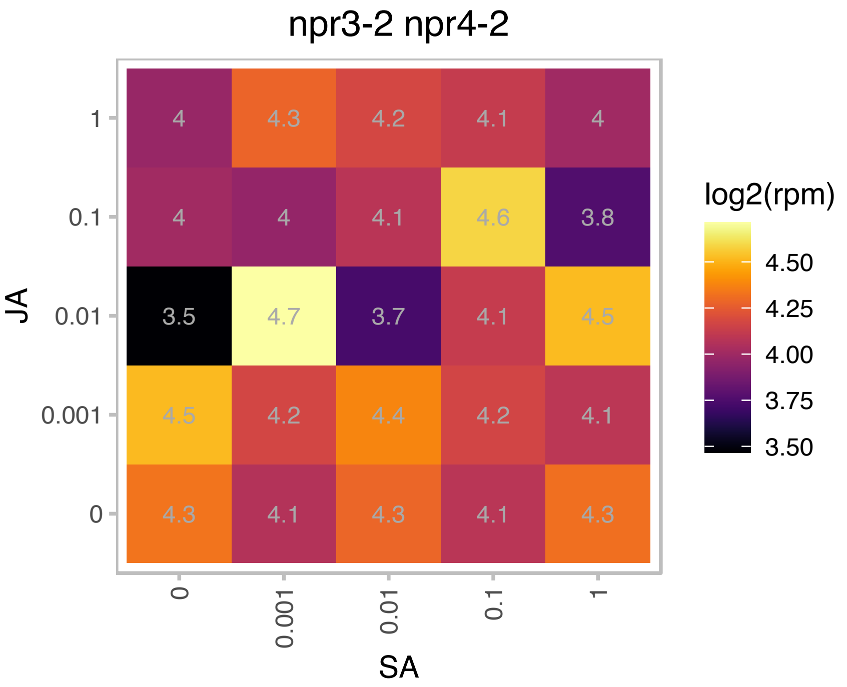


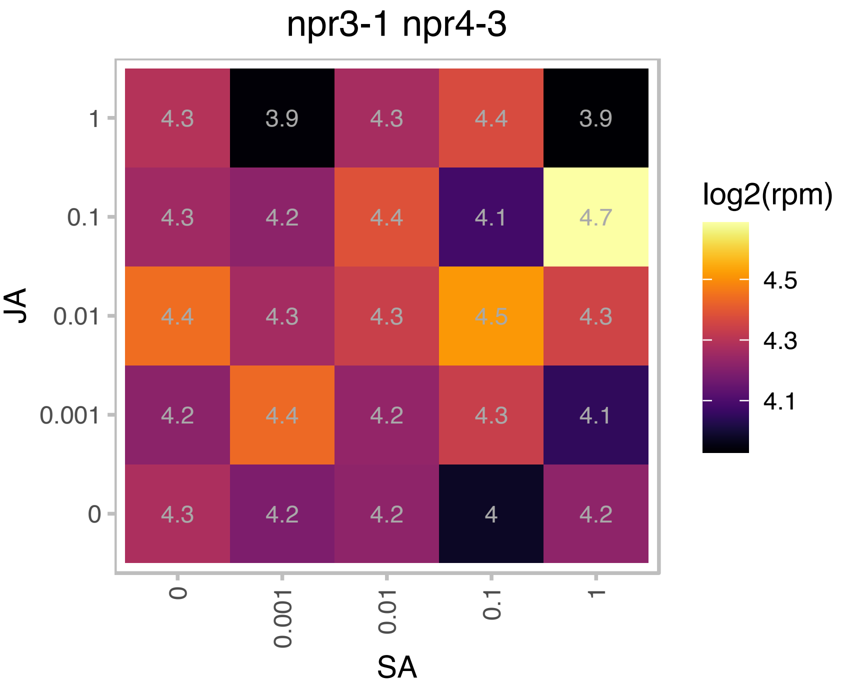

### AT3G05020.1 : ACP, ACP1, AtACP1

acyl carrier protein 1

encodes an acyl carrier protein expressed in leaves, roots, and dry seeds. Protein is not regulated by light.

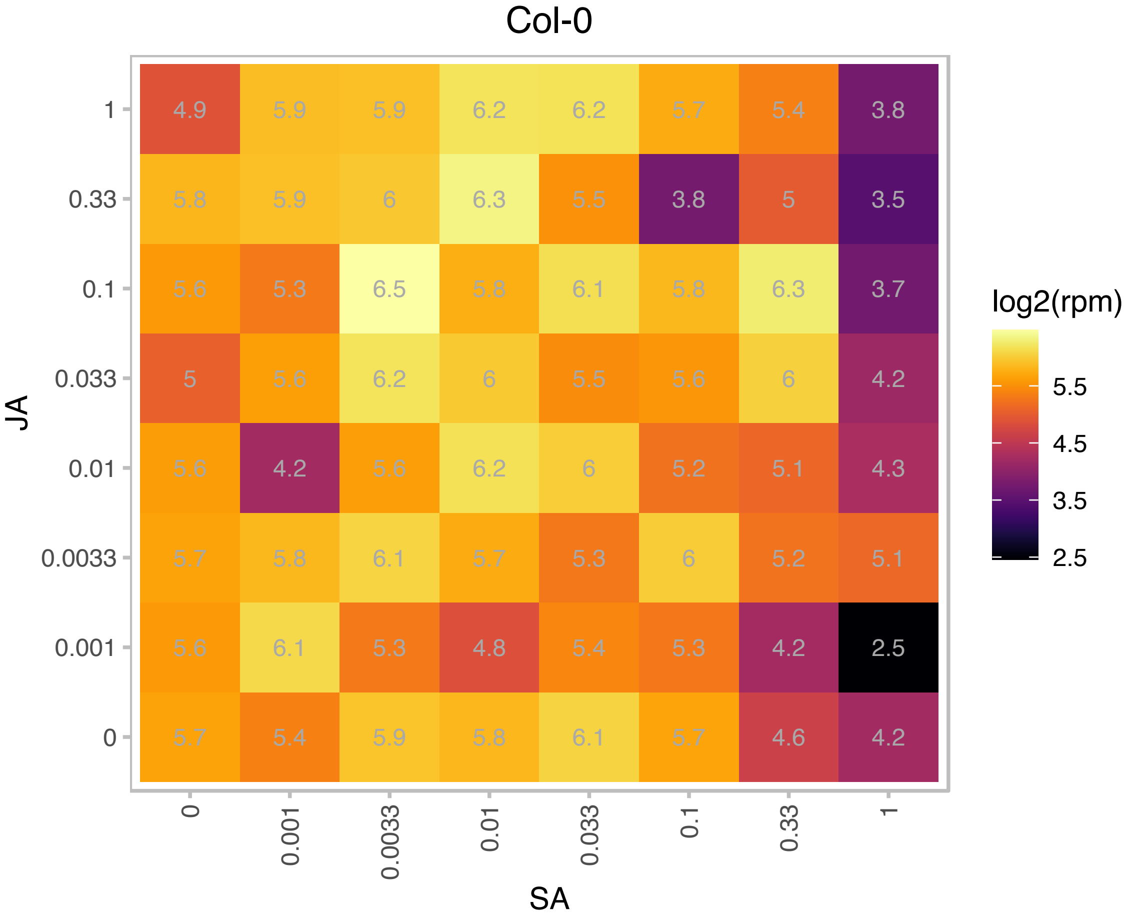


| | |
|---|---|
| Gene | AT3G05020 |
| is.expressed_genes | TRUE |
| is.responsive_genes | TRUE |
| is.apcluster | D-04 |

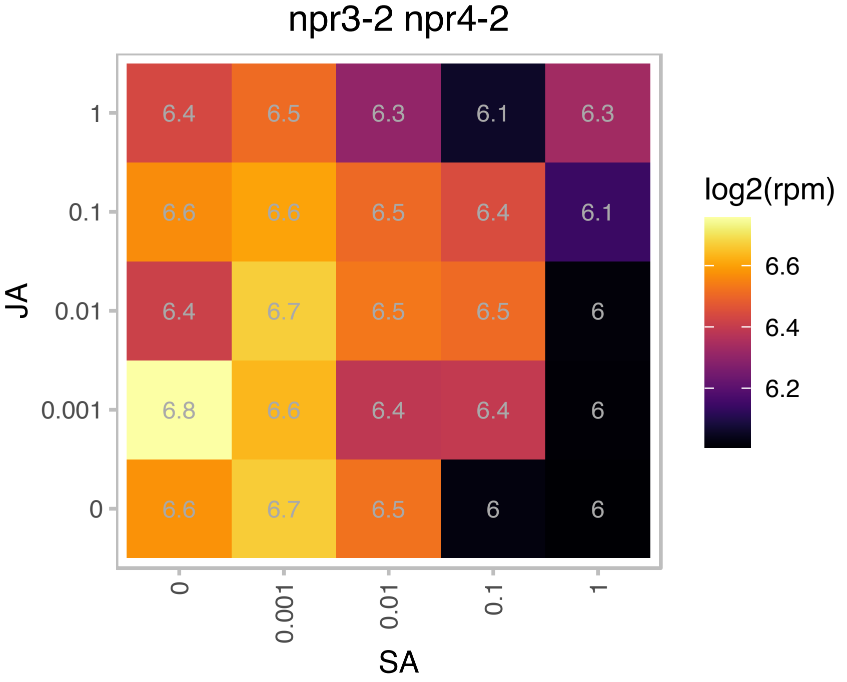


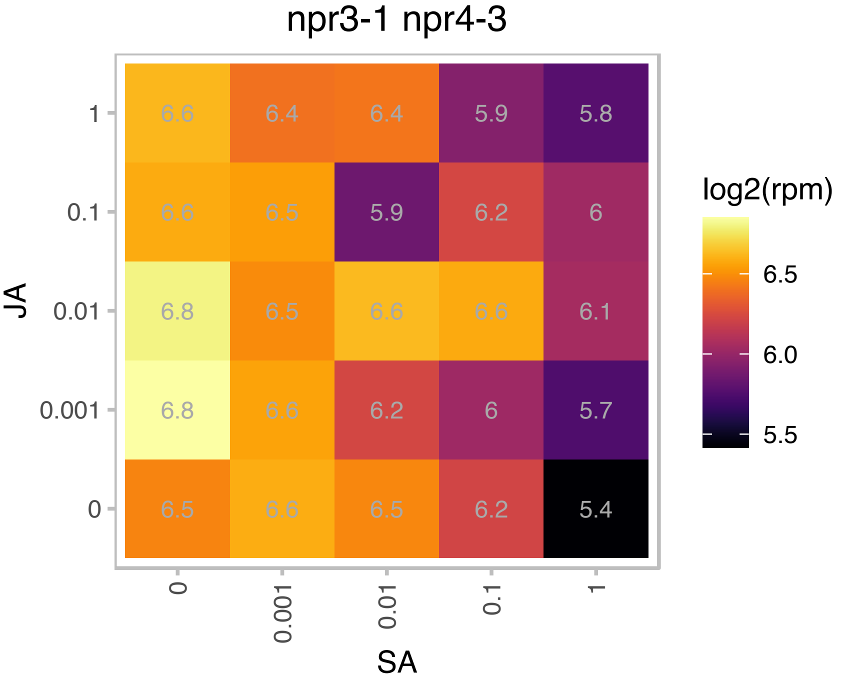

### AT3G15730.1 : PLD, PLDALPHA1

phospholipase D alpha 1
Encodes phospholipase D alpha 1 (PLD alpha 1). Positive regulator of abscisic acid (ABA) mediated stomatal movements.  PLD alpha 1 plays an important role in seed deterioration and aging in Arabidopsis. The mRNA is cell-to-cell mobile.

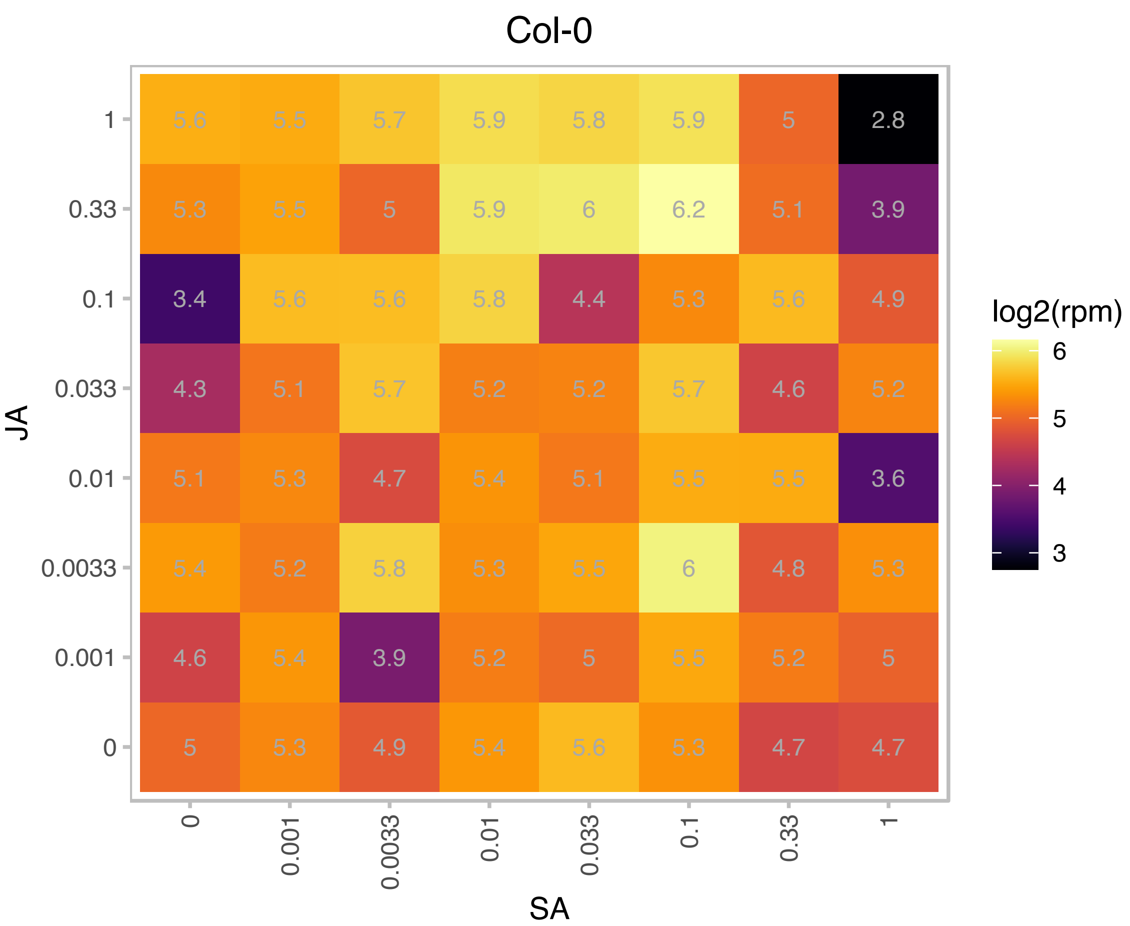


| | |
|---|---|
| Gene | AT3G15730 |
| is.expressed_genes | TRUE |
| is.responsive_genes | TRUE |
| is.apcluster | D-04 |

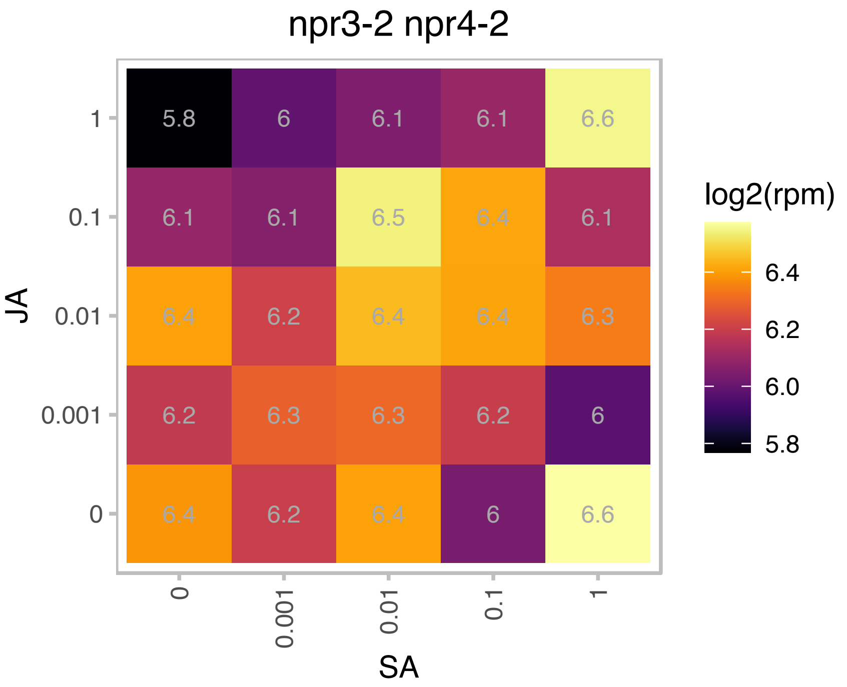


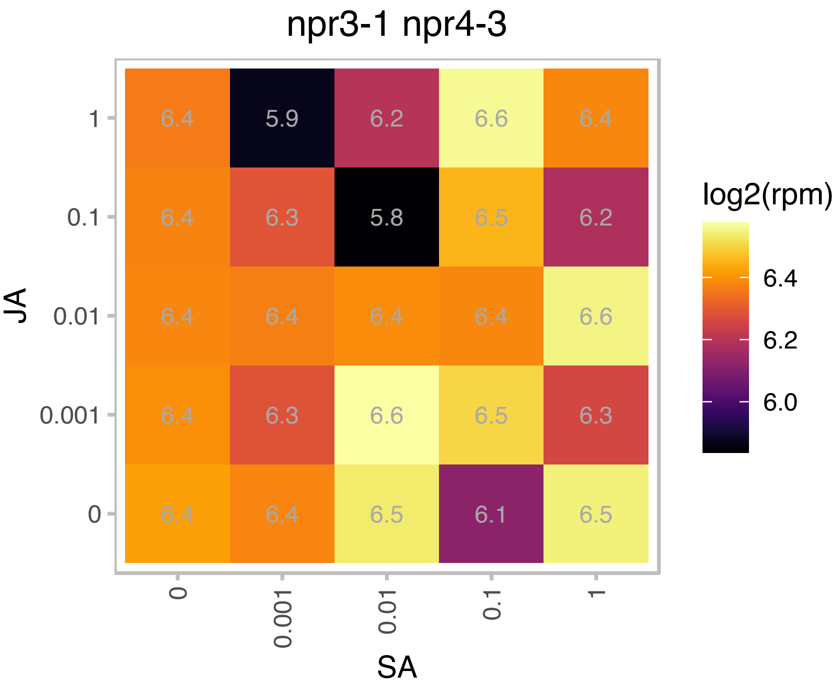

### AT3G20050.1 : ATTCP-1, TCP-1

T-complex protein 1 alpha subunit

Encodes a putative cytoplasmic chaperonin that is similar to mouse Tcp-1 (t complex polypeptide 1).

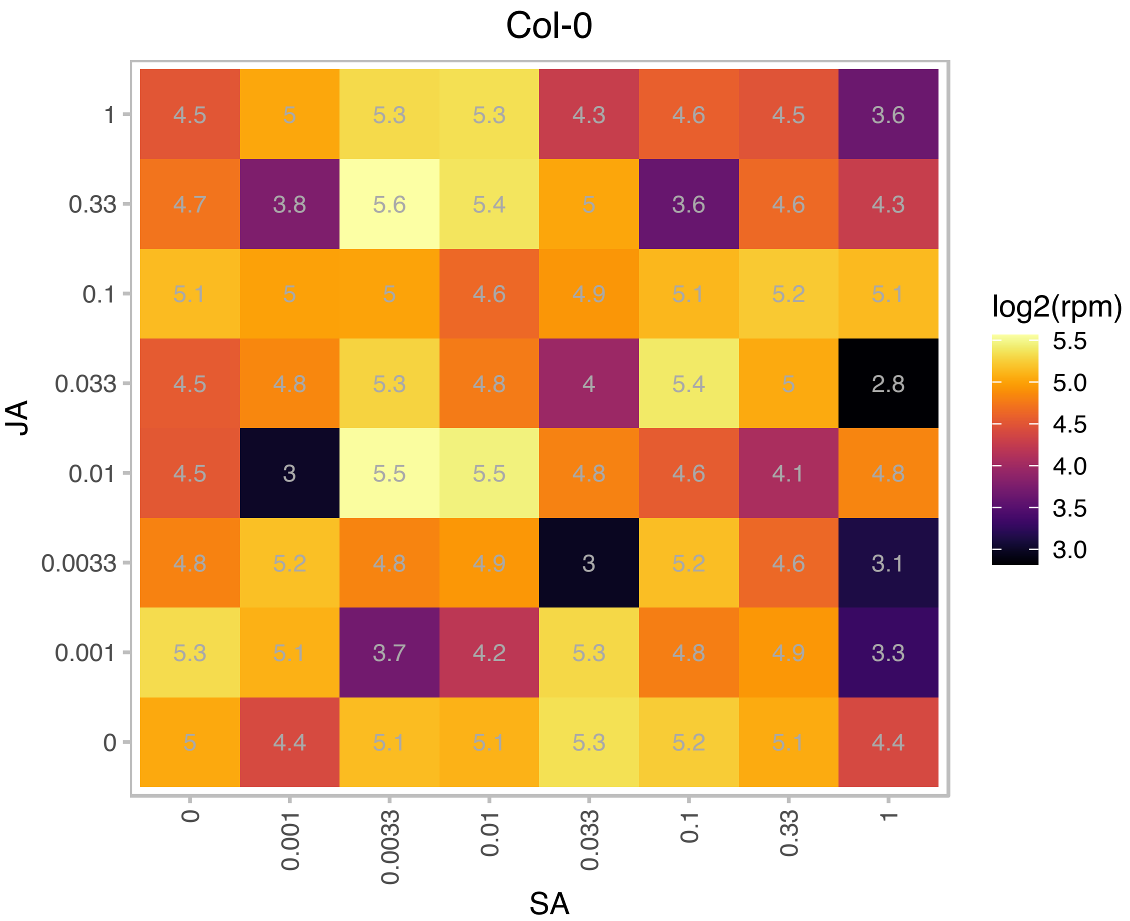


| | |
|---|---|
| Gene | AT3G20050 |
| is.expressed_genes | TRUE |
| is.responsive_genes | TRUE |
| is.apcluster | D-04 |

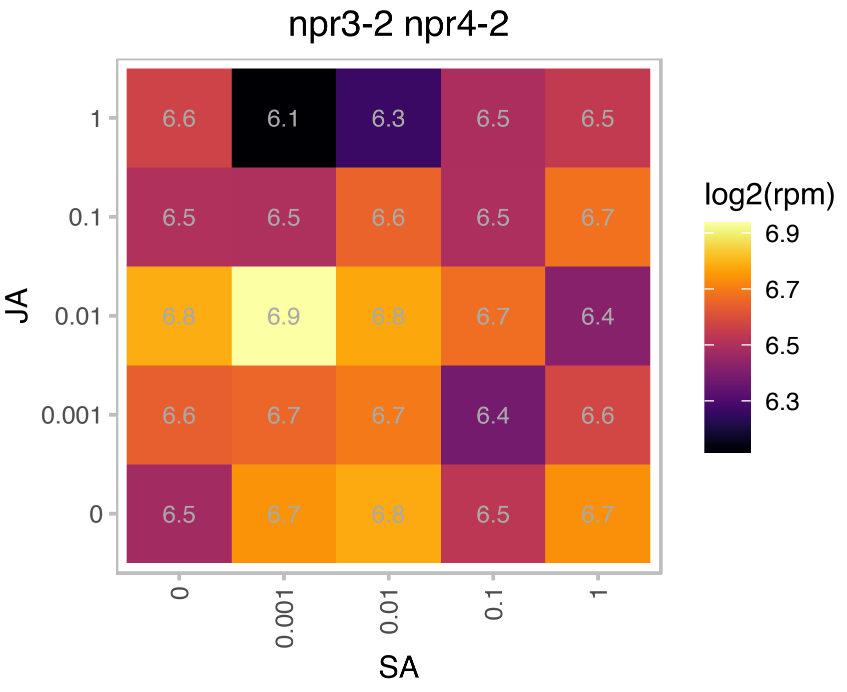


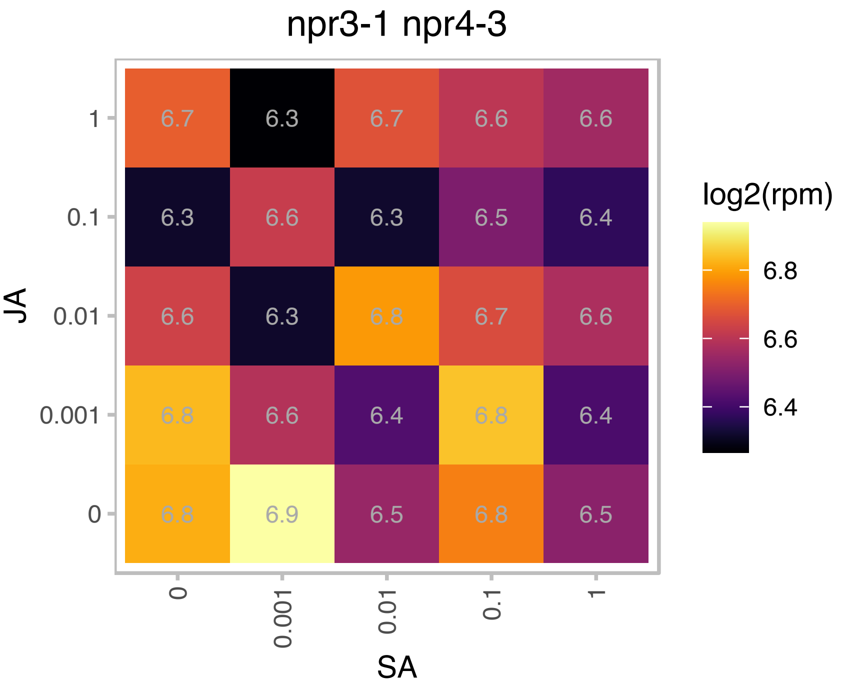

### AT3G57290.1 : ATEIF3E-1, ATINT6, EIF3E, INT-6, INT6, TI

eukaryotic translation initiation factor 3E
Encodes a protein that is found in not only the eif3 complex but also in association with subunits of the COP9 signalosome. eIF3e appears to be subjected to proteasome-dependent degradation that requires the PCI domain of eIF3e. The level of eIF3e present in cells appears to affect the rate of translation.

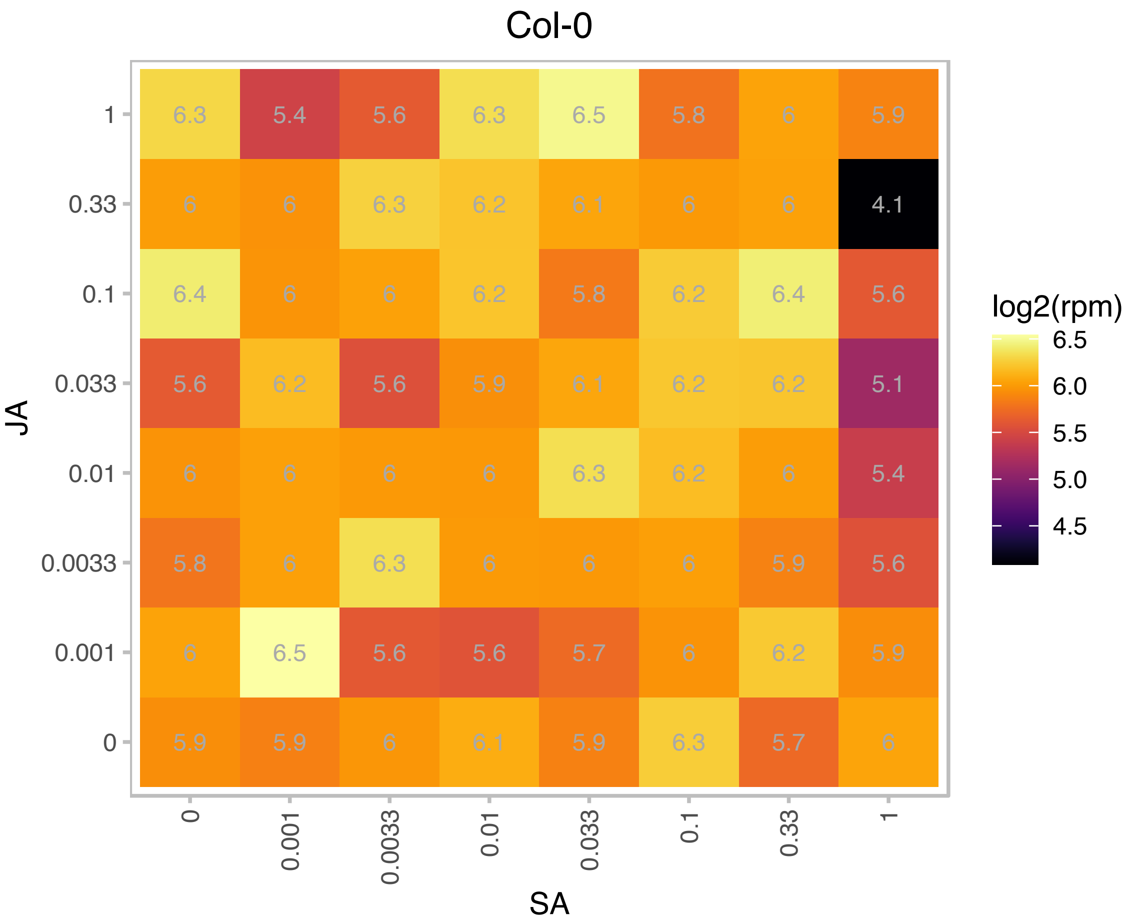


| | |
|---|---|
| Gene | AT3G57290 |
| is.expressed_genes | TRUE |
| is.responsive_genes | TRUE |
| is.apcluster | D-04 |

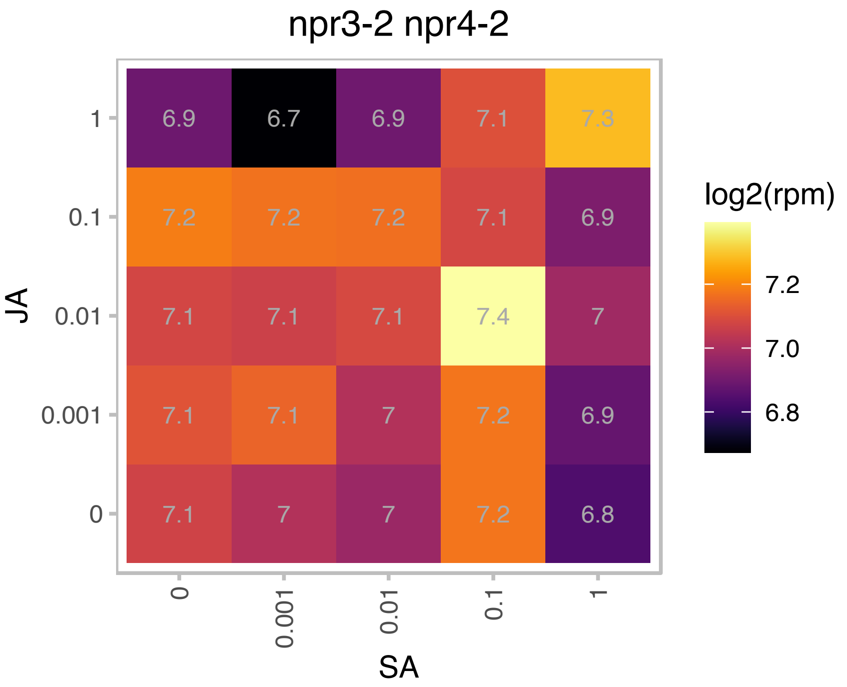


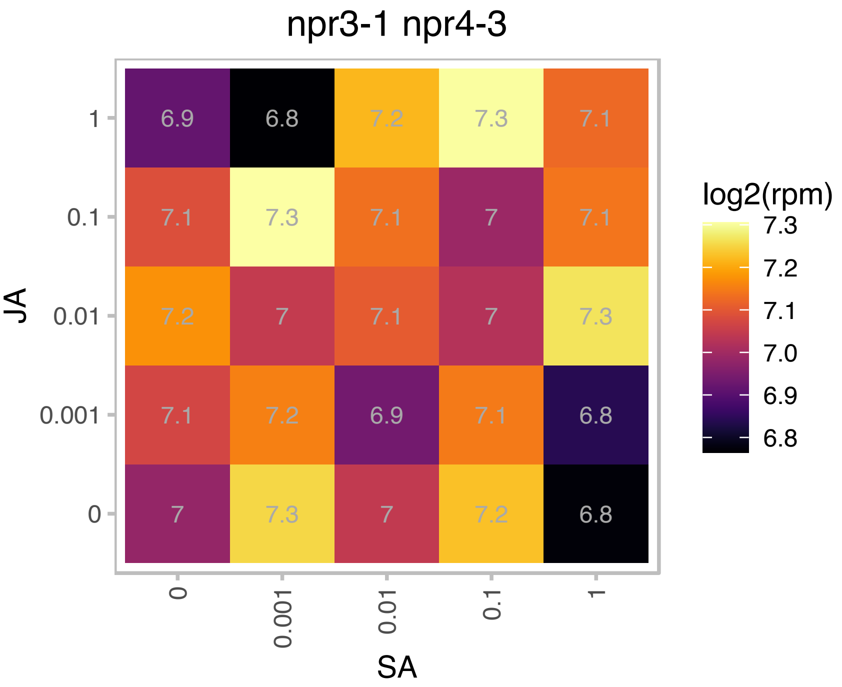

### AT3G62120.1 : AtProRS-Cyt, ProRS-Cyt

Class II aaRS and biotin synthetases superfamily protein

Encodes a cytosolic prolyl-tRNA synthetase.

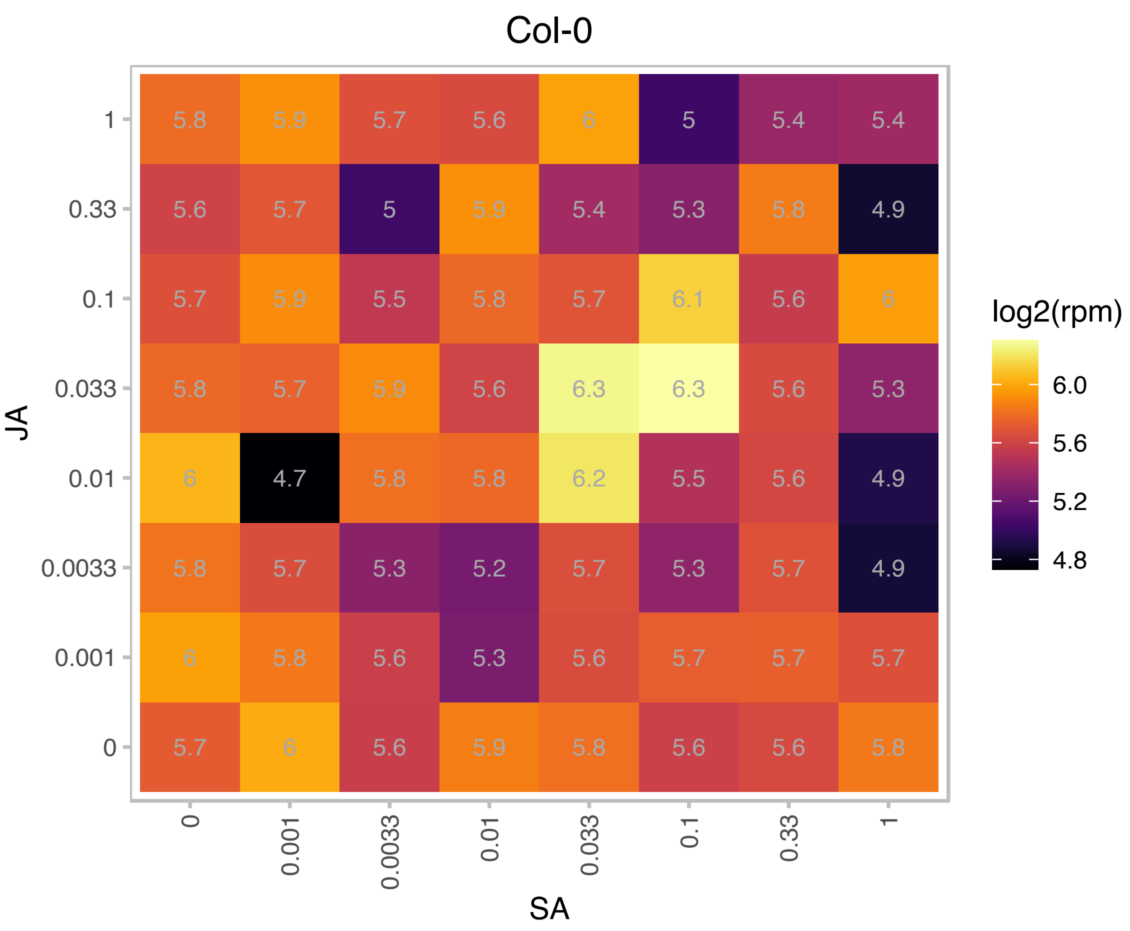


| | |
|---|---|
| Gene | AT3G62120 |
| is.expressed_genes | TRUE |
| is.responsive_genes | TRUE |
| is.apcluster | D-04 |

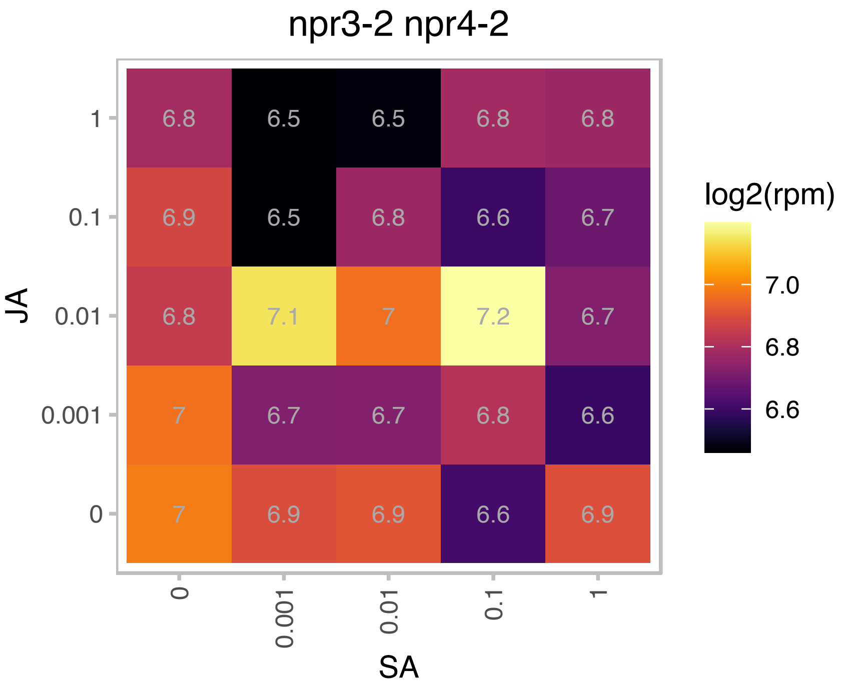


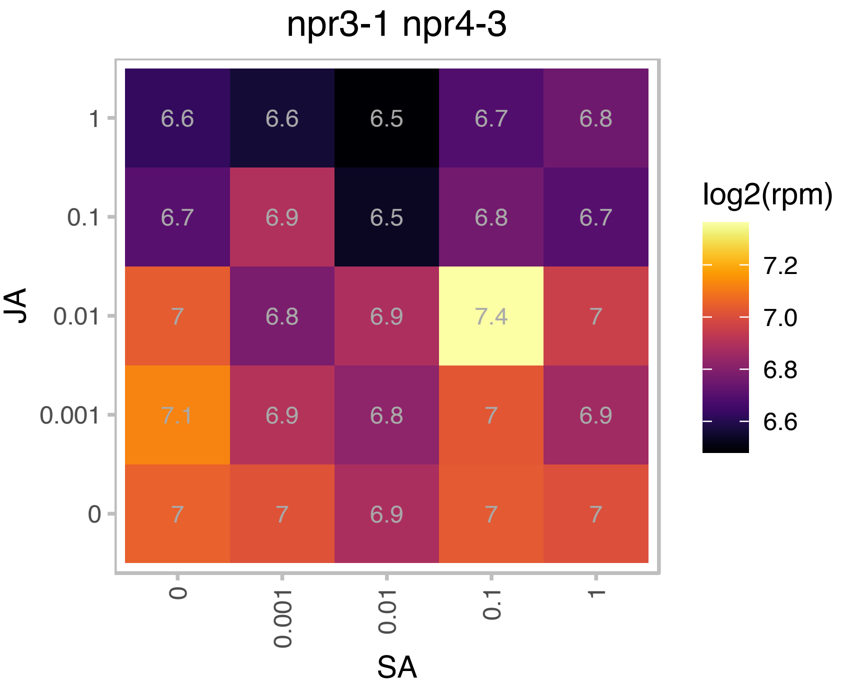

### AT3G62980.1 : AtTIR1, TIR1

F-box/RNI-like superfamily protein

Encodes an auxin receptor that mediates auxin-regulated transcription.  It contains leucine-rich repeats and an F-box and interacts with ASK1, ASK2 and AtCUL1 to form SCF-TIR1, an SCF ubiquitin ligase complex. Related to yeast Grr1p and human SKP2 proteins, involved in ubiquitin-mediated processes. Required for normal response to auxin and repressed in response to flagellin.  As part of the SCF complex and in the presence of auxin, TIR1 interacts with Aux/IAA transcriptional repressor proteins and mediates their degradation. Mutations in TIR1 block auxin stimulation of flavonoid synthesis.

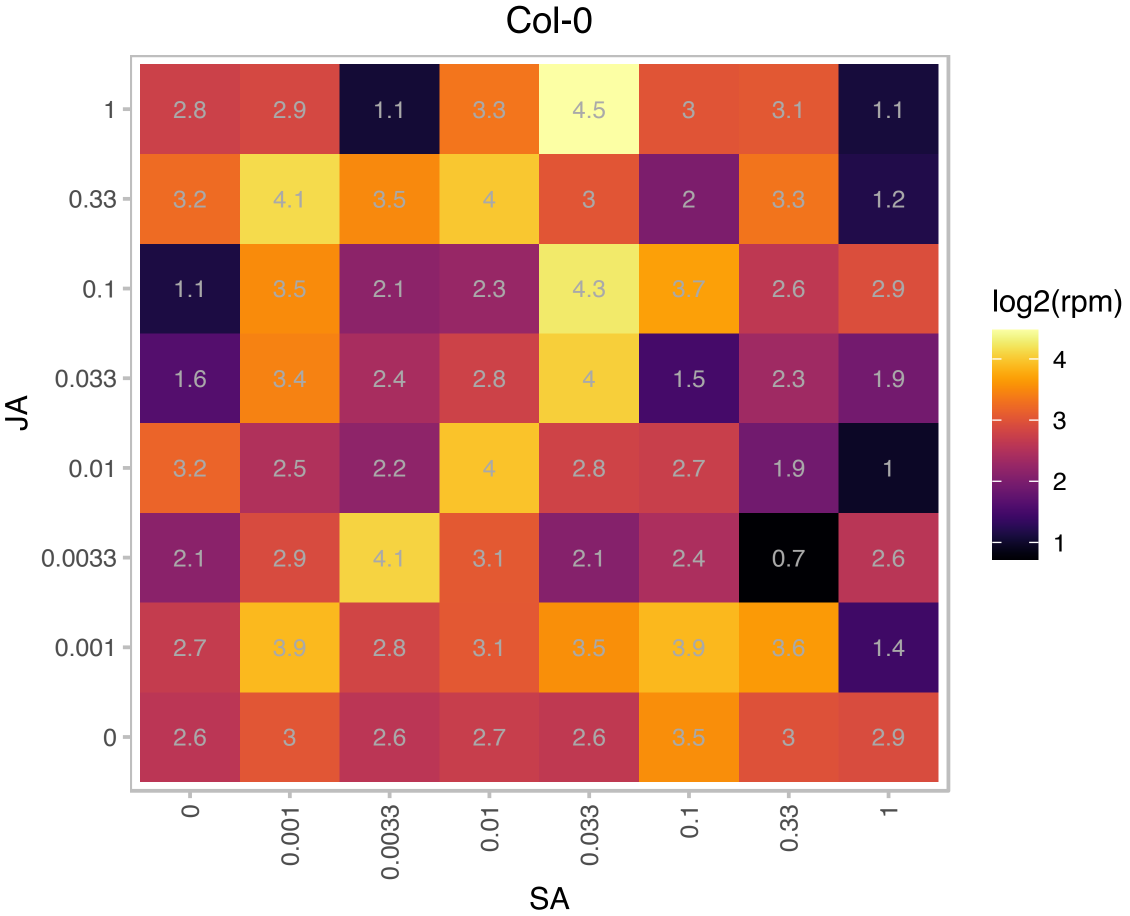


| | |
|---|---|
| Gene | AT3G62980 |
| is.expressed_genes | TRUE |
| is.responsive_genes | TRUE |
| is.apcluster | D-04 |

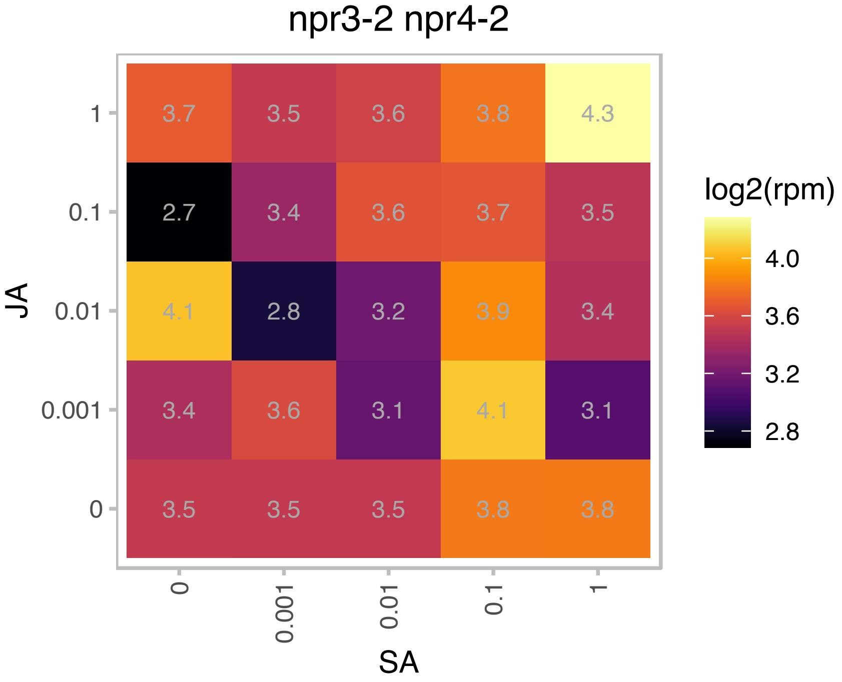


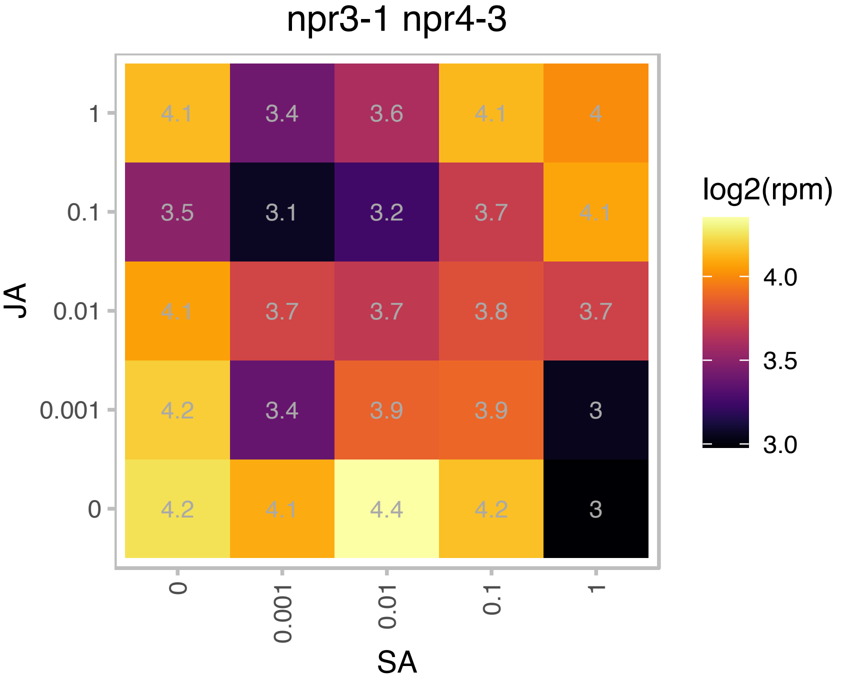

### AT4G03240.1 : ATFH, FH

frataxin homolog
Encodes AtFH, a frataxin homolog. Frataxin is required for the biogenesis of mitochondria in different organisms. AtFH knock-out mutants are lethal. Required for full activity of mitochondrial Fe-S proteins. Deficiency of AtFH induces oxidative stress.

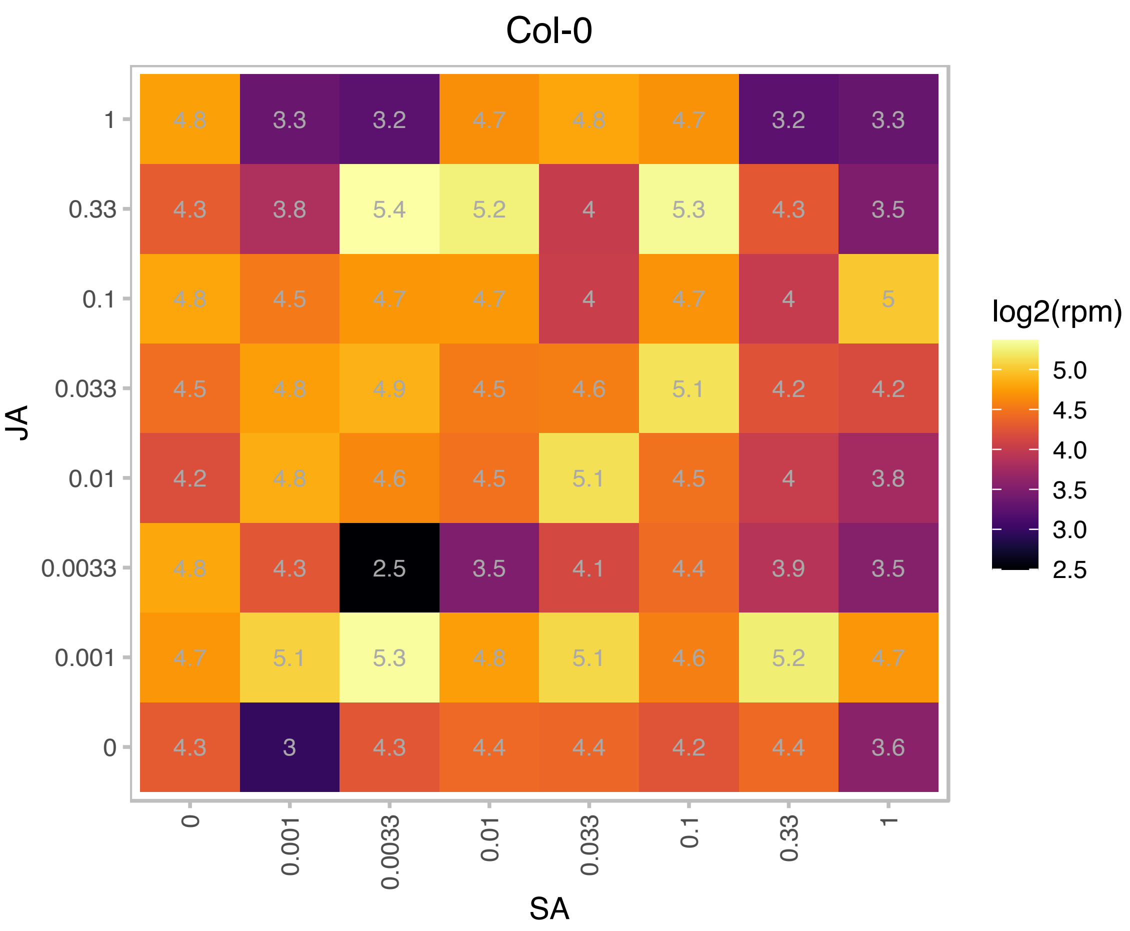


| | |
|---|---|
| Gene | AT4G03240 |
| is.expressed_genes | TRUE |
| is.responsive_genes | TRUE |
| is.apcluster | D-04 |

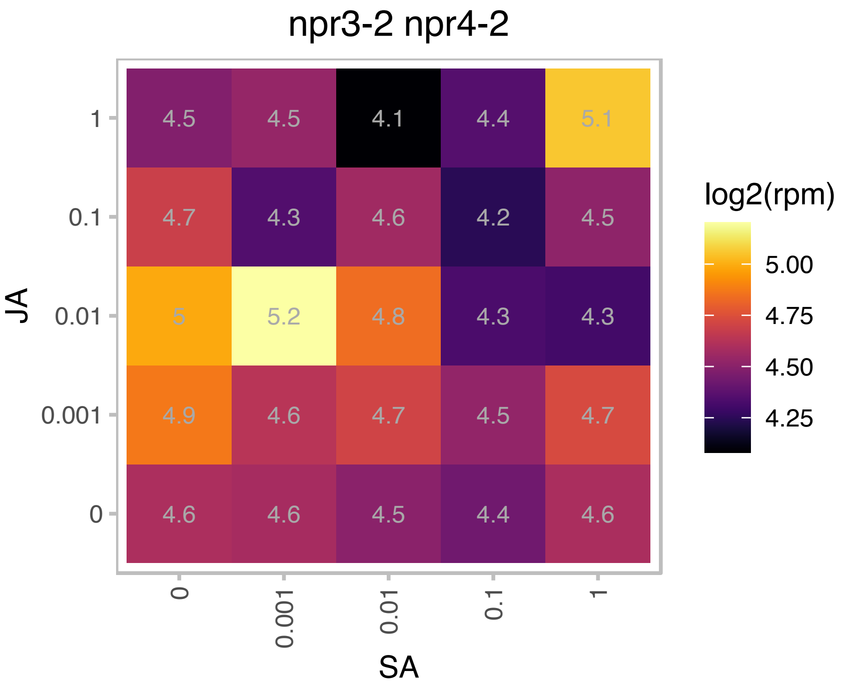


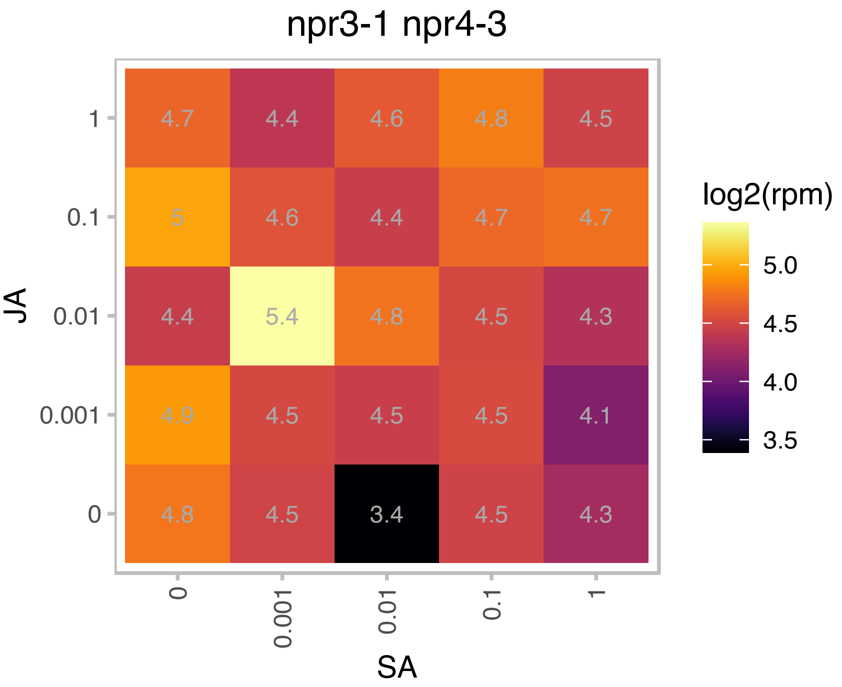

# AT4G11100.1 : NA

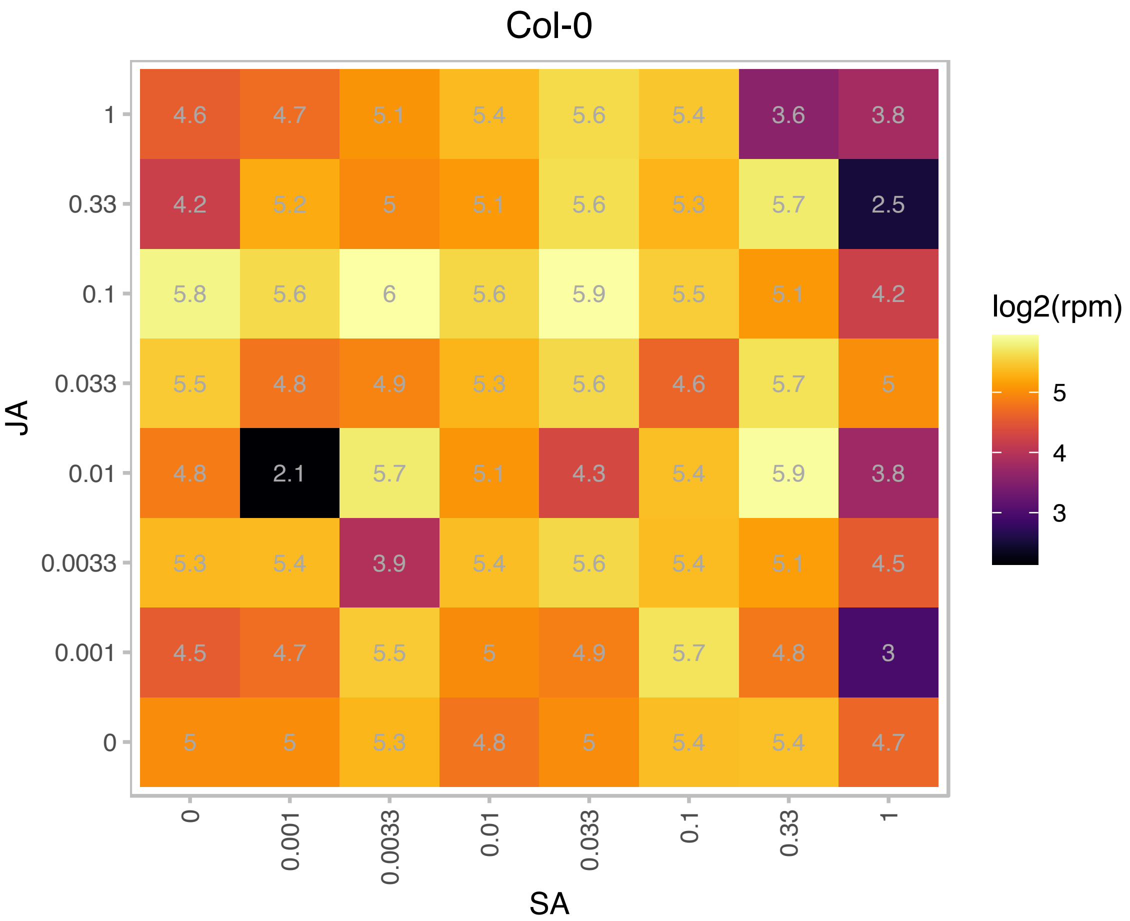


| | |
|---|---|
| Gene | AT4G11100 |
| is.expressed_genes | TRUE |
| is.responsive_genes | TRUE |
| is.apcluster | D-04 |

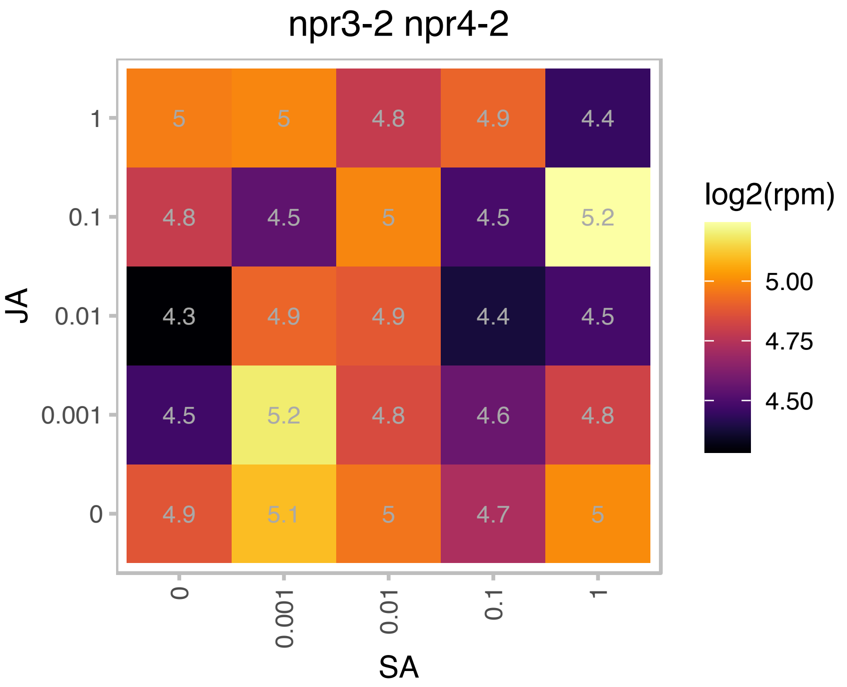


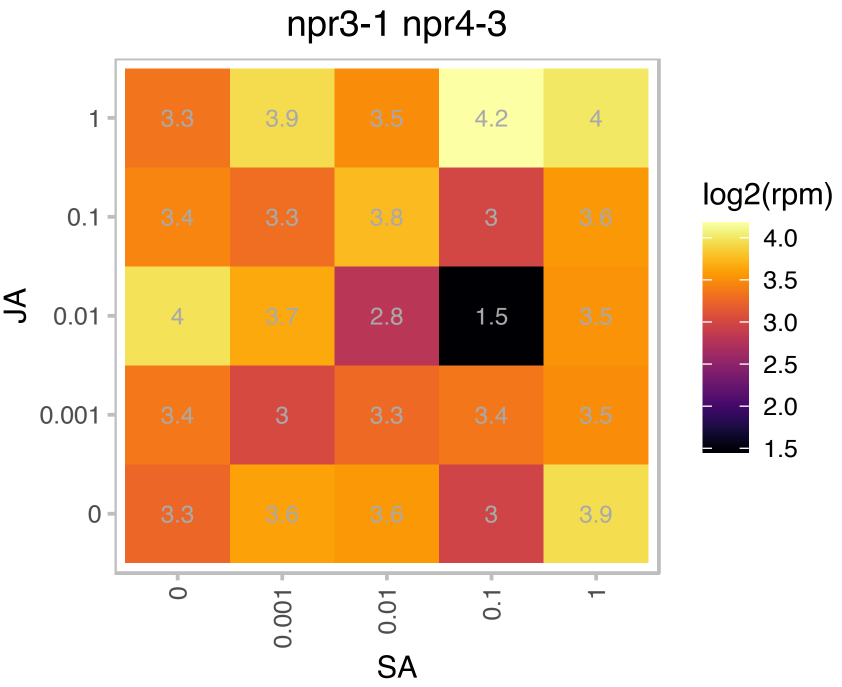

### AT4G11420.1 : ATEIF3A-1, ATTIF3A1, EIF3A, EIF3A-1, TIF

eukaryotic translation initiation factor 3A
Encodes a subunit of eukaryotic initiation factor 3 (eIF3), a multisubunit complex that is required for binding of mRNA to 40 S ribosomal subunits, stabilization of ternary complex binding to 40 S subunits, and dissociation of 40 and 60 S subunits.

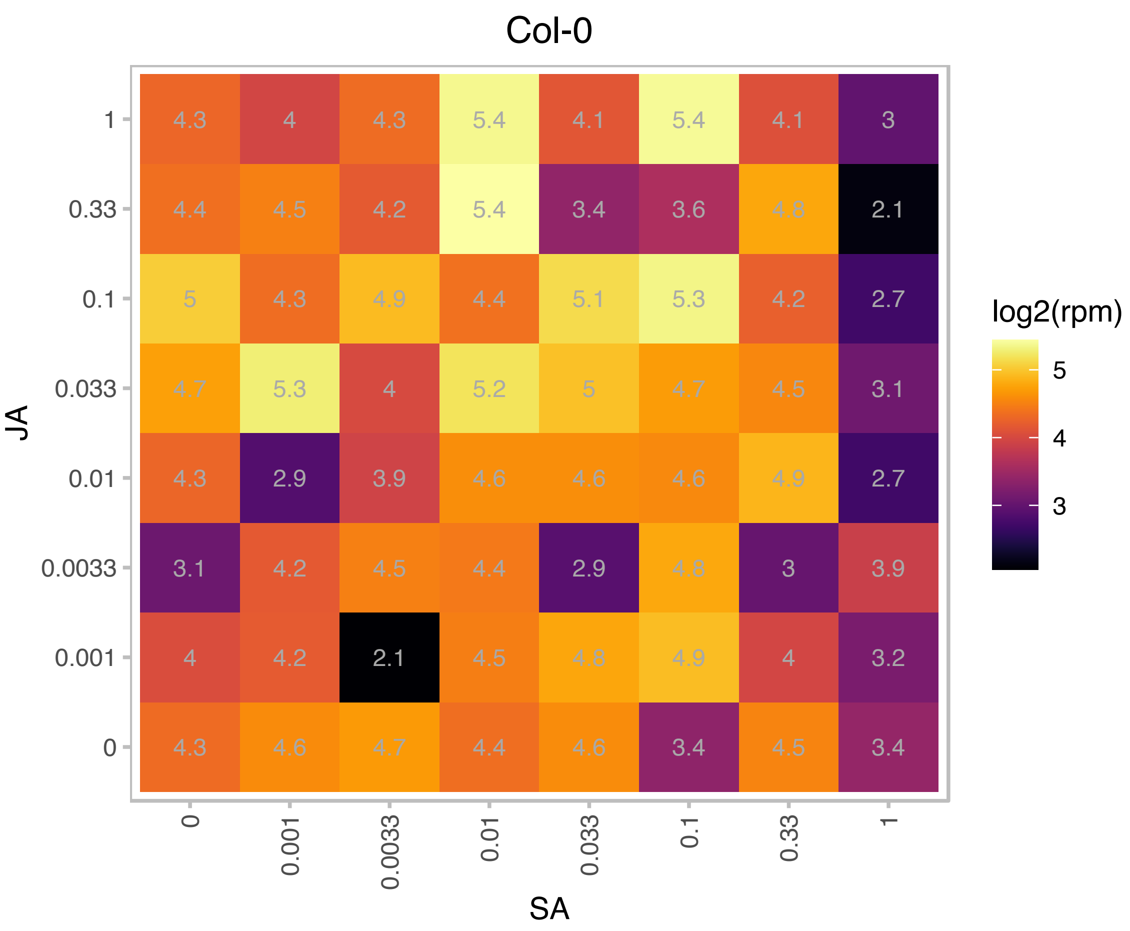


| | |
|---|---|
| Gene | AT4G11420 |
| is.expressed_genes | TRUE |
| is.responsive_genes | TRUE |
| is.apcluster | D-04 |

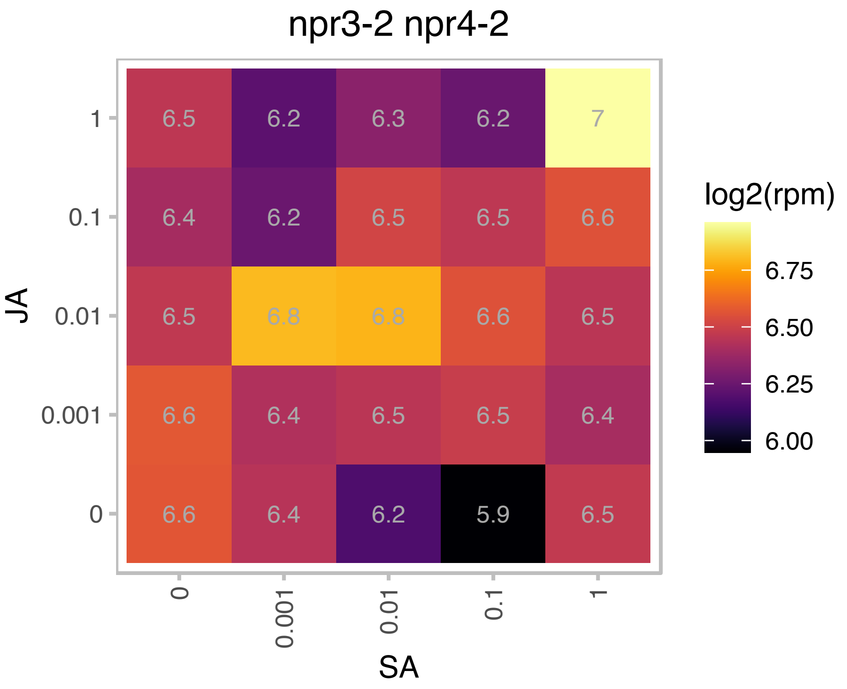


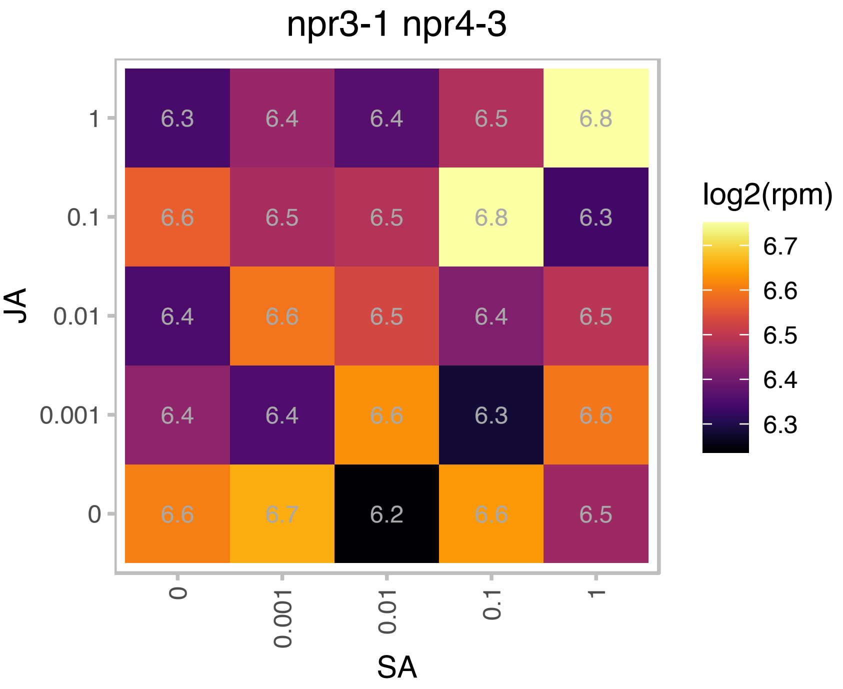

### AT4G22150.1 : PUX3

plant UBX domain-containing protein 3

Arabidopsis thaliana CDC48-interacting UBX-domain protein (PUX3)

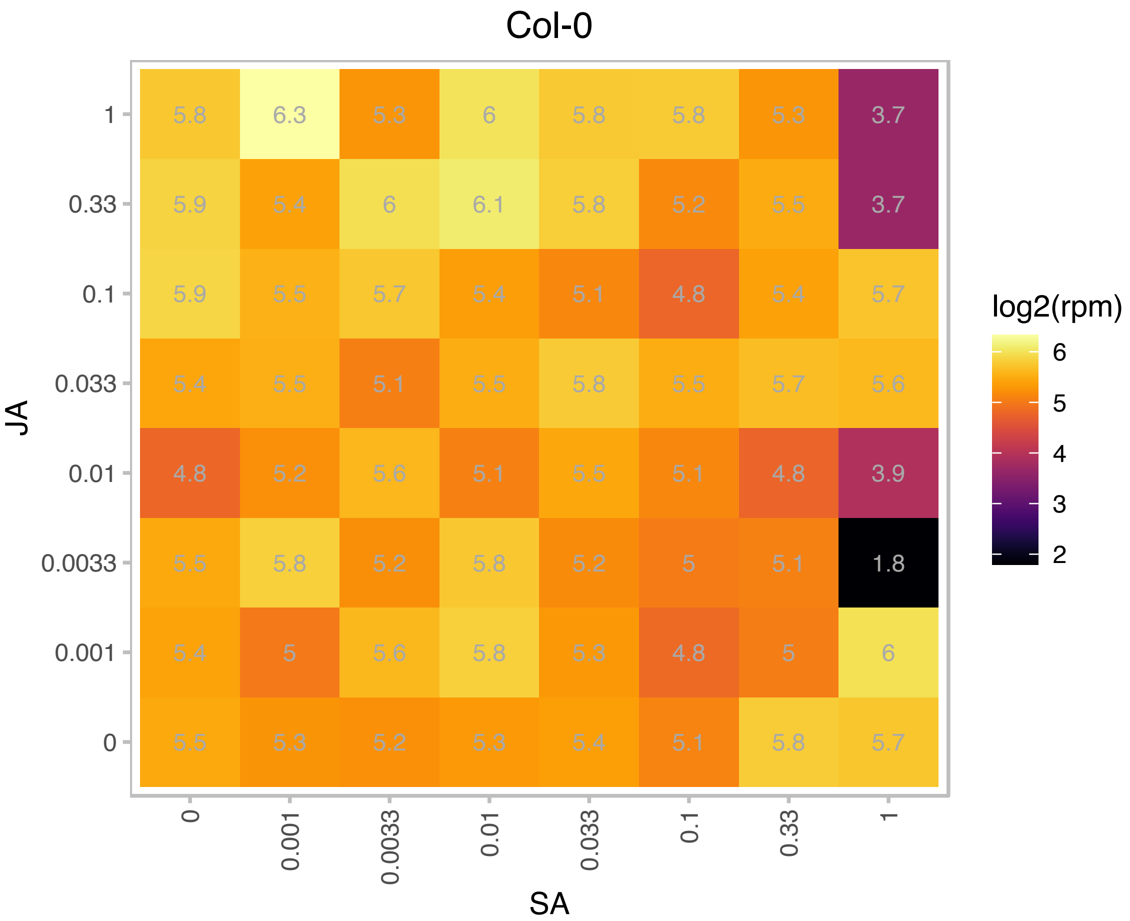


| | |
|---|---|
| Gene | AT4G22150 |
| is.expressed_genes | TRUE |
| is.responsive_genes | TRUE |
| is.apcluster | D-04 |

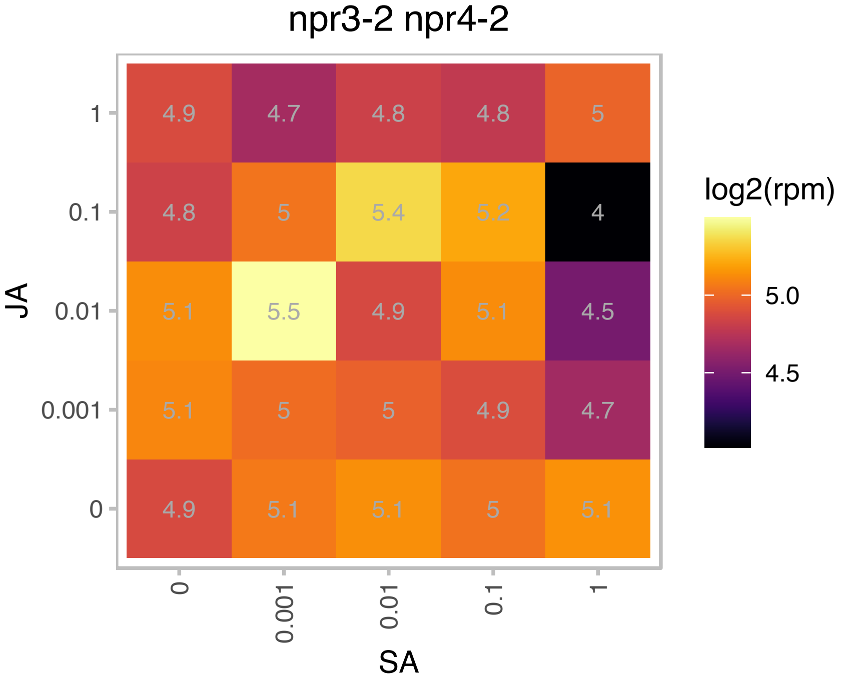


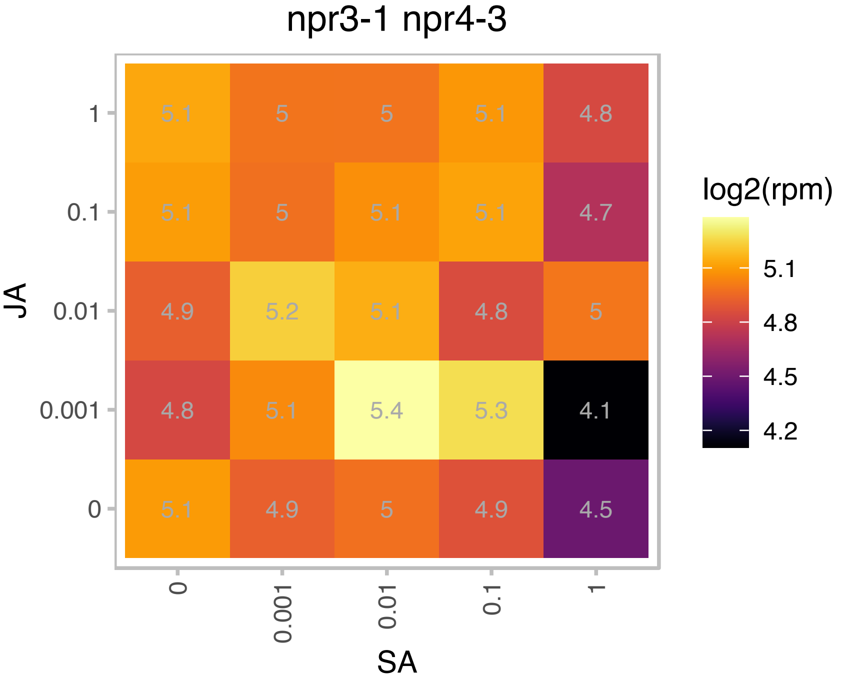

### AT4G25370.1 : ClpT1

Double Clp-N motif protein

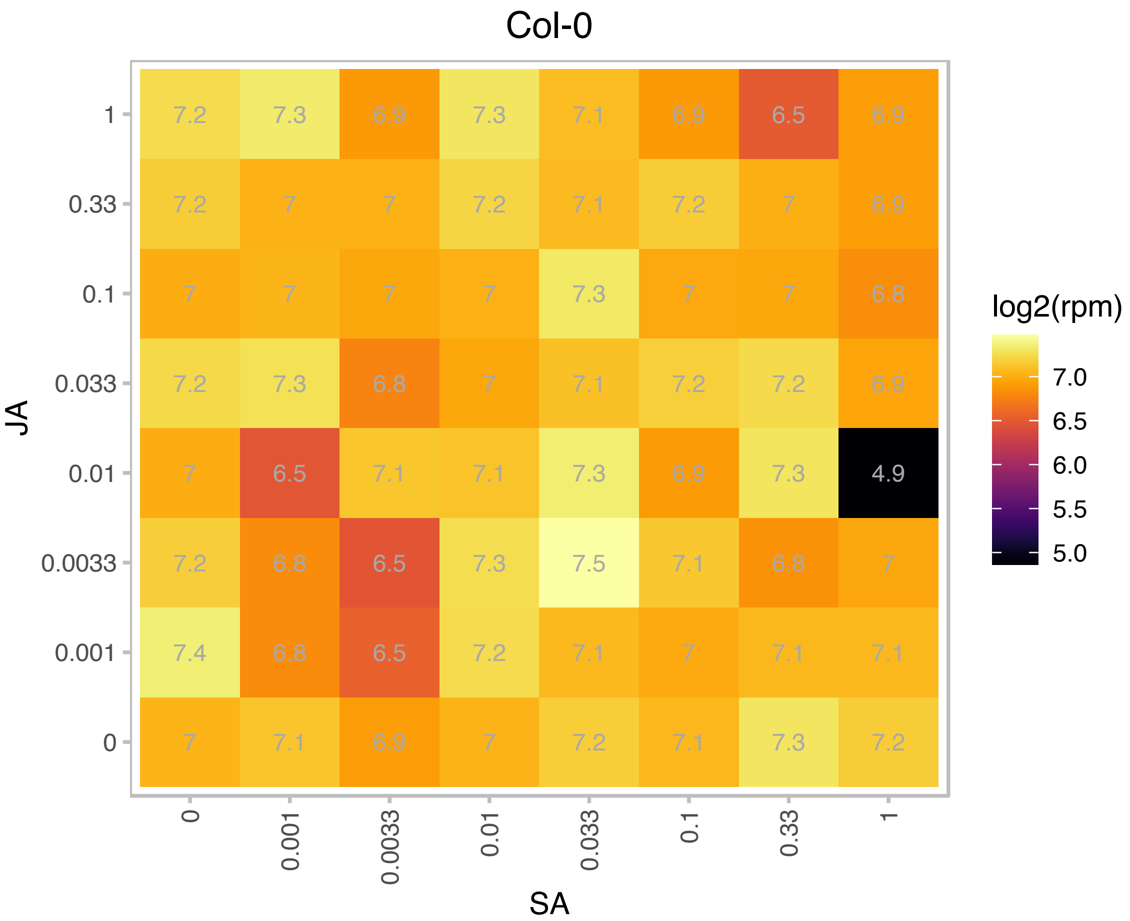


| | |
|---|---|
| Gene | AT4G25370 |
| is.expressed_genes | TRUE |
| is.responsive_genes | TRUE |
| is.apcluster | D-04 |

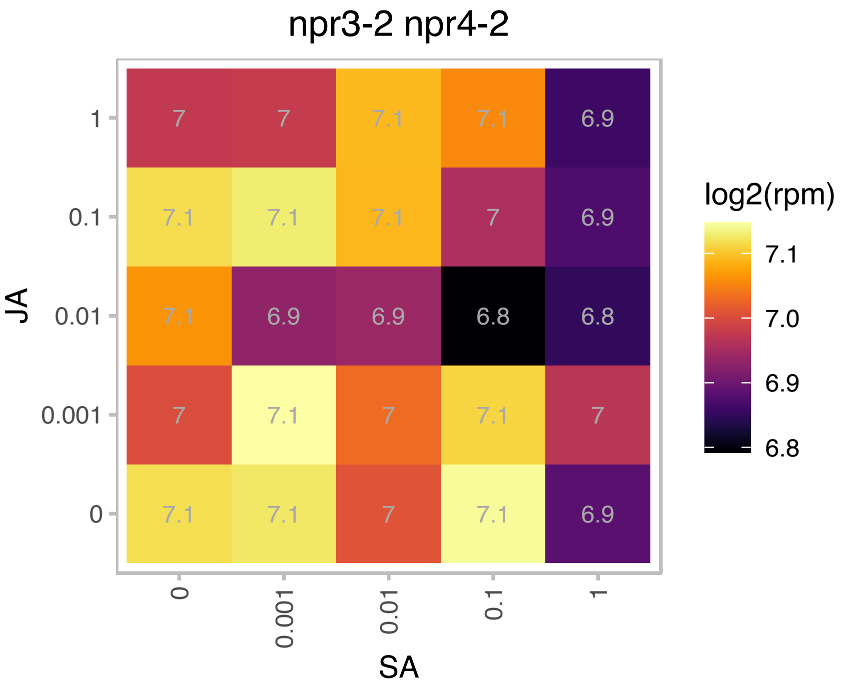


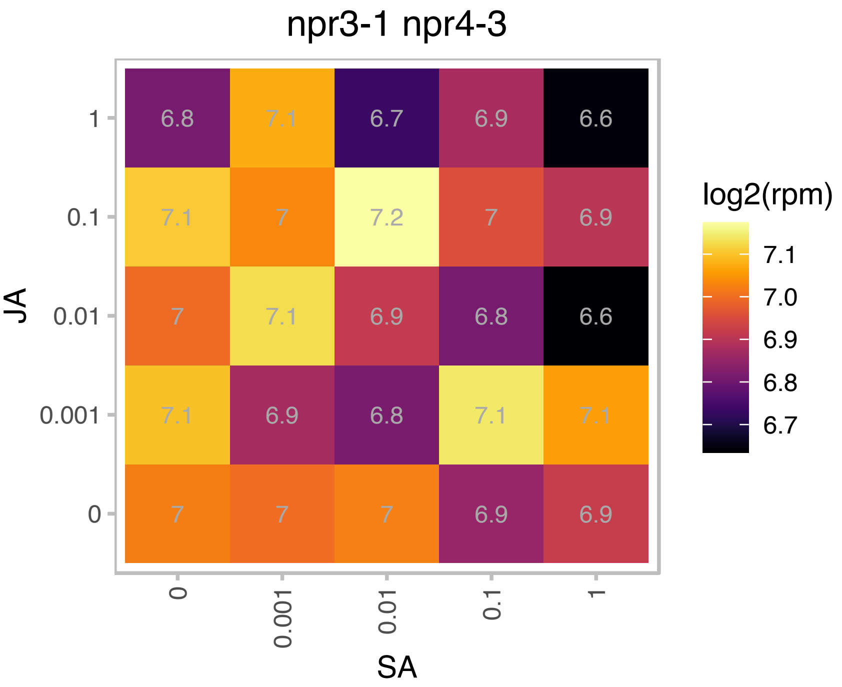

# AT5G01710.1 : NA

methyltransferases

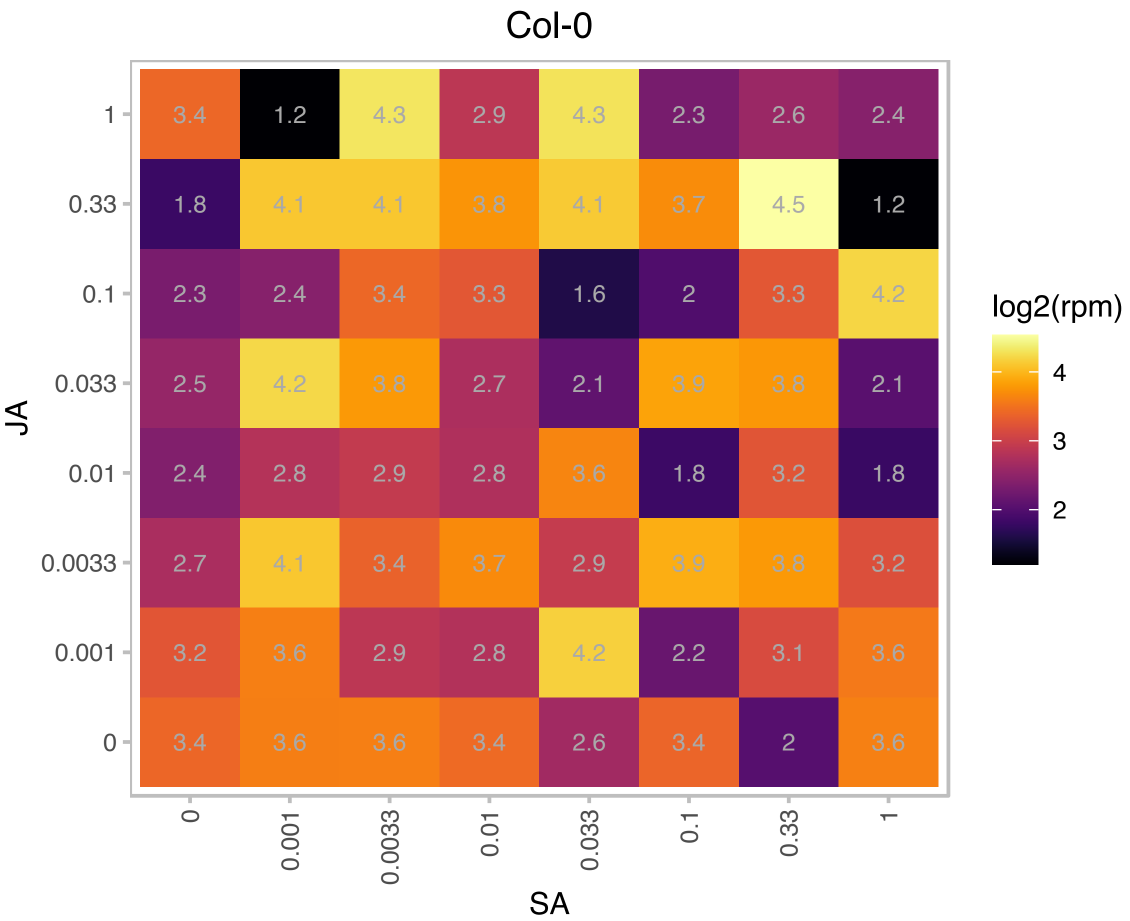


| | |
|---|---|
| Gene | AT5G01710 |
| is.expressed_genes | TRUE |
| is.responsive_genes | TRUE |
| is.apcluster | D-04 |

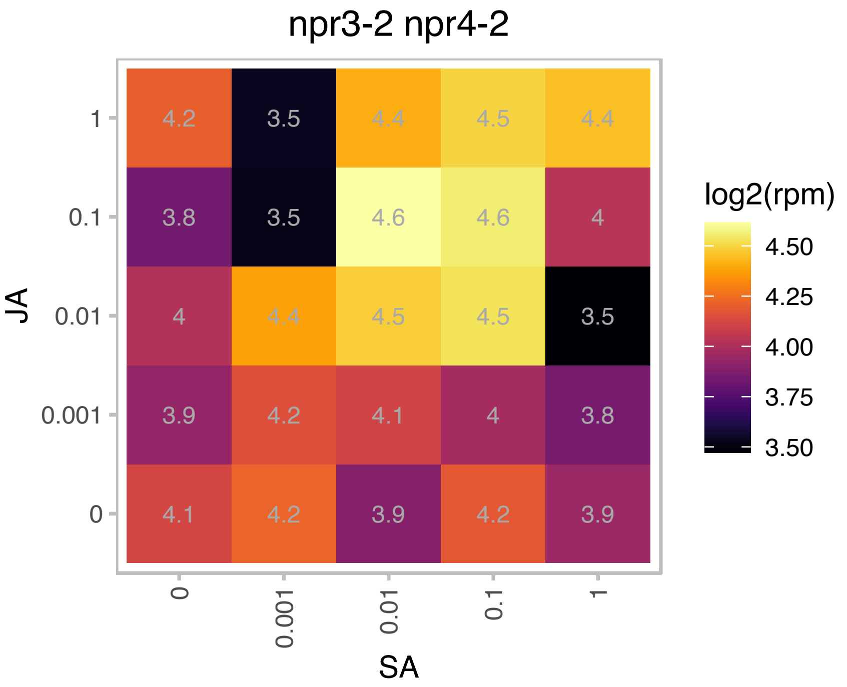


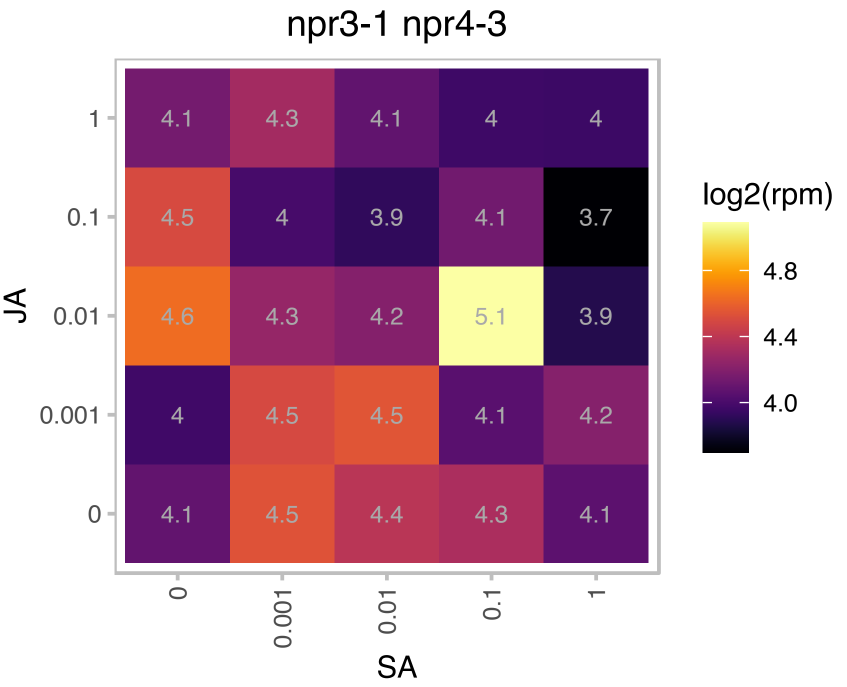

# AT5G08670.1 : NA

ATP synthase alpha/beta family protein

Encodes the mitochondrial ATP synthase beta-subunit.  This subunit is encoded by a multigene family of three members (At5g08670, At5g08680, At5g08690) that shared 98% sequence identity at the amino acid level.

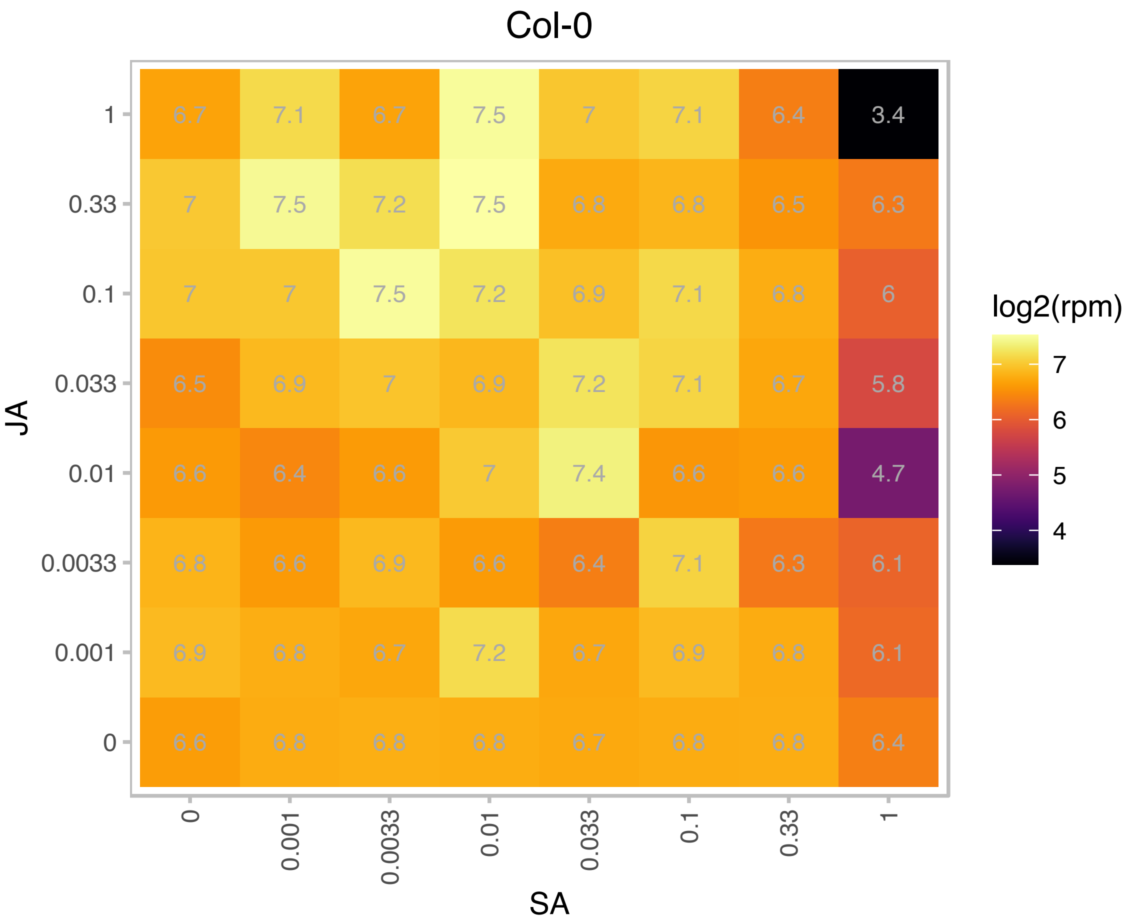


| | |
|---|---|
| Gene | AT5G08670 |
| is.expressed_genes | TRUE |
| is.responsive_genes | TRUE |
| is.apcluster | D-04 |

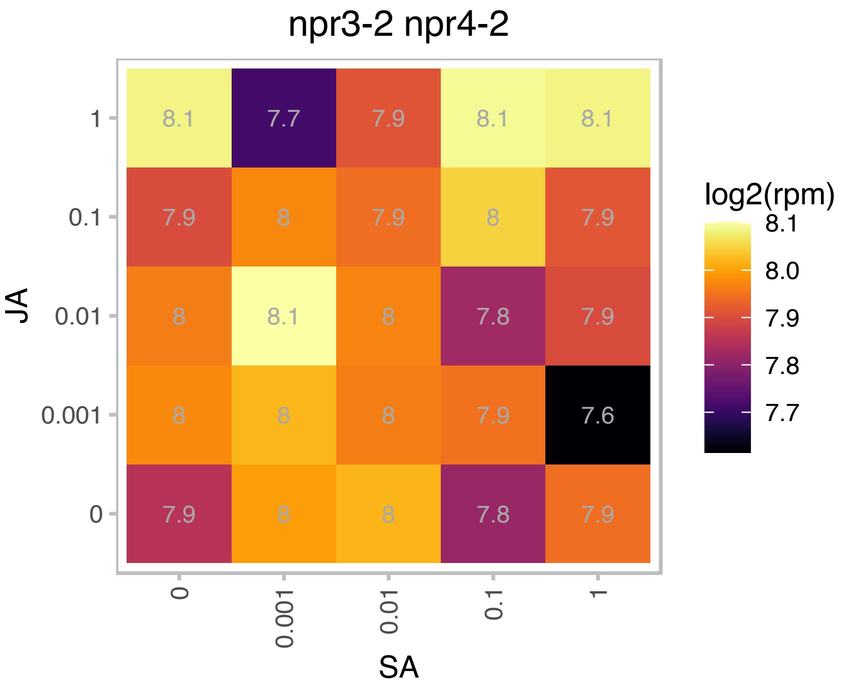


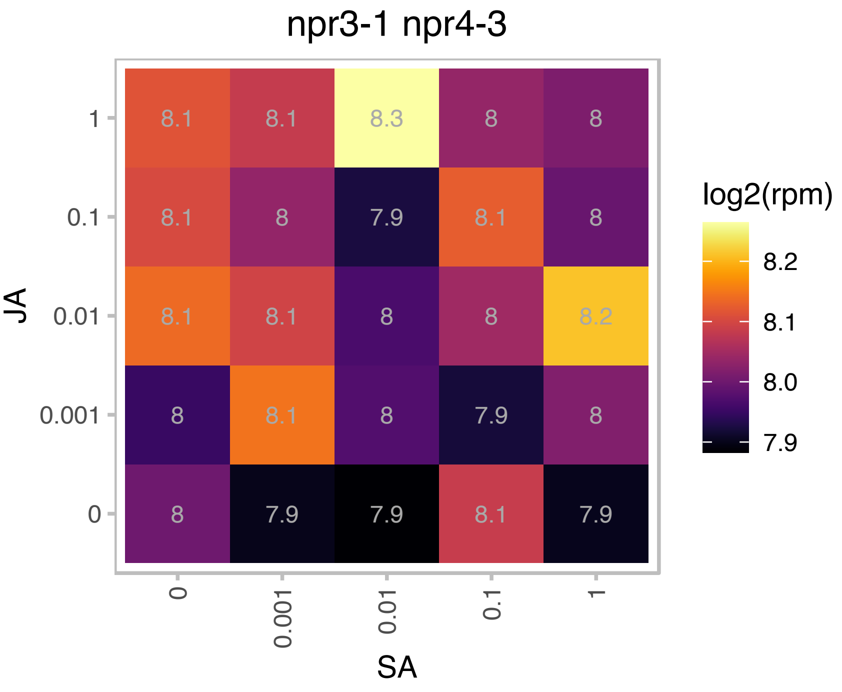

### AT5G18650.1 : MIEL1

CHY-type/CTCHY-type/RING-type Zinc finger protein

Encodes a RING-type E3 ubiquitin ligase that interacts with and ubiquitinates MYB30, leads to MYB30 proteasomal degradation and downregulation of its transcriptional activity. Since MYB30 is a positive regulator of Arabidopsis HR and defence responses, MIEL1 is involved in the negative regulation of these processes. The mRNA is cell-to-cell mobile.

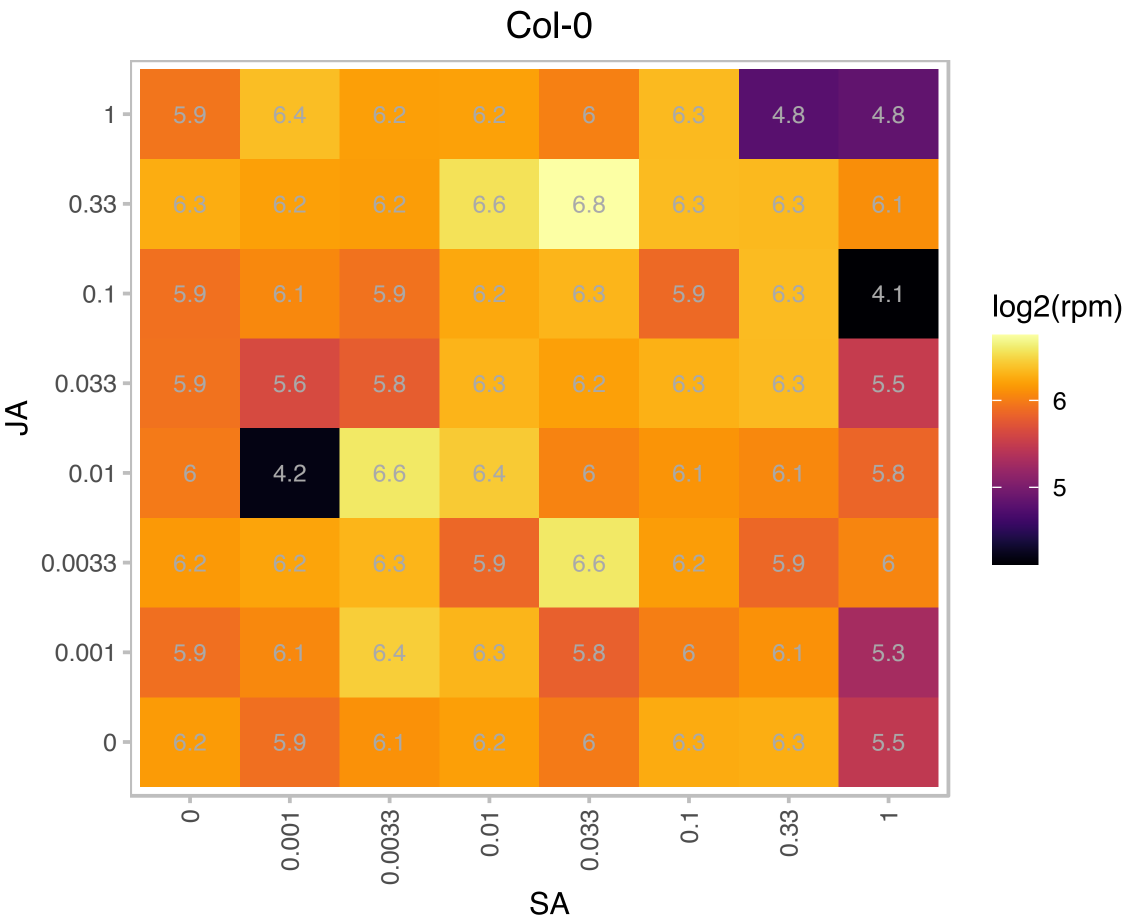


| | |
|---|---|
| Gene | AT5G18650 |
| is.expressed_genes | TRUE |
| is.responsive_genes | TRUE |
| is.apcluster | D-04 |

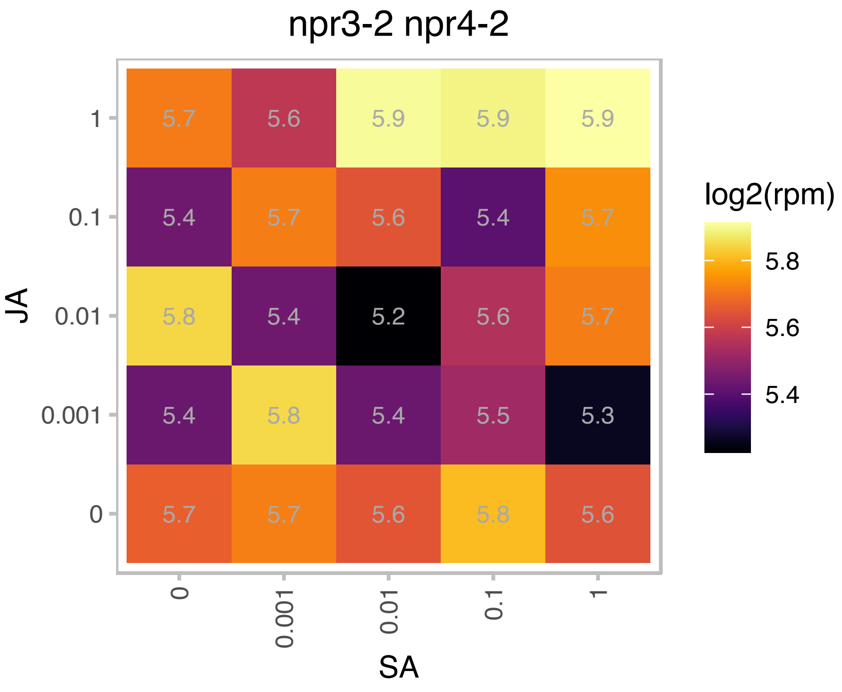


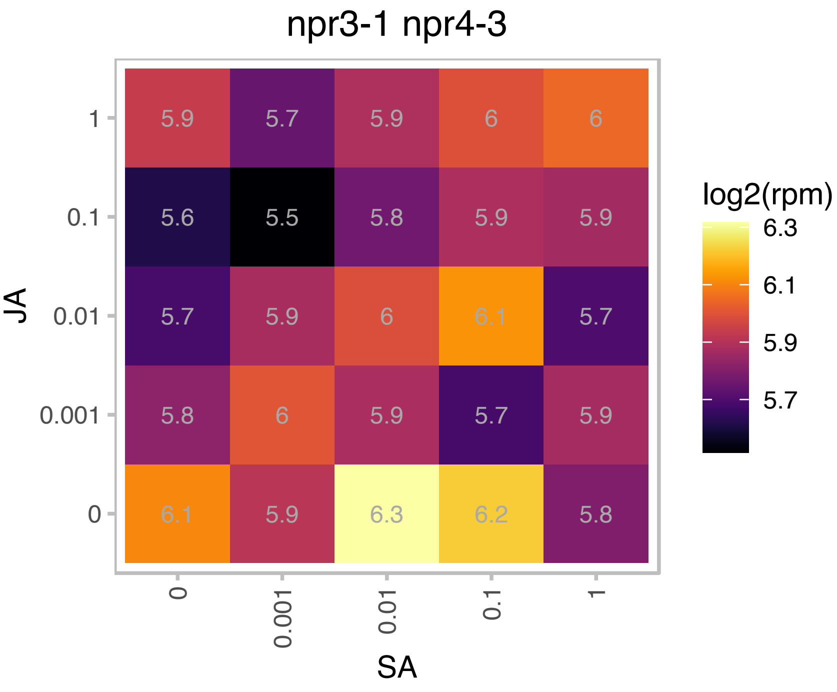

# AT5G19680.1 : NA

Leucine-rich repeat (LRR) family protein

Col-0

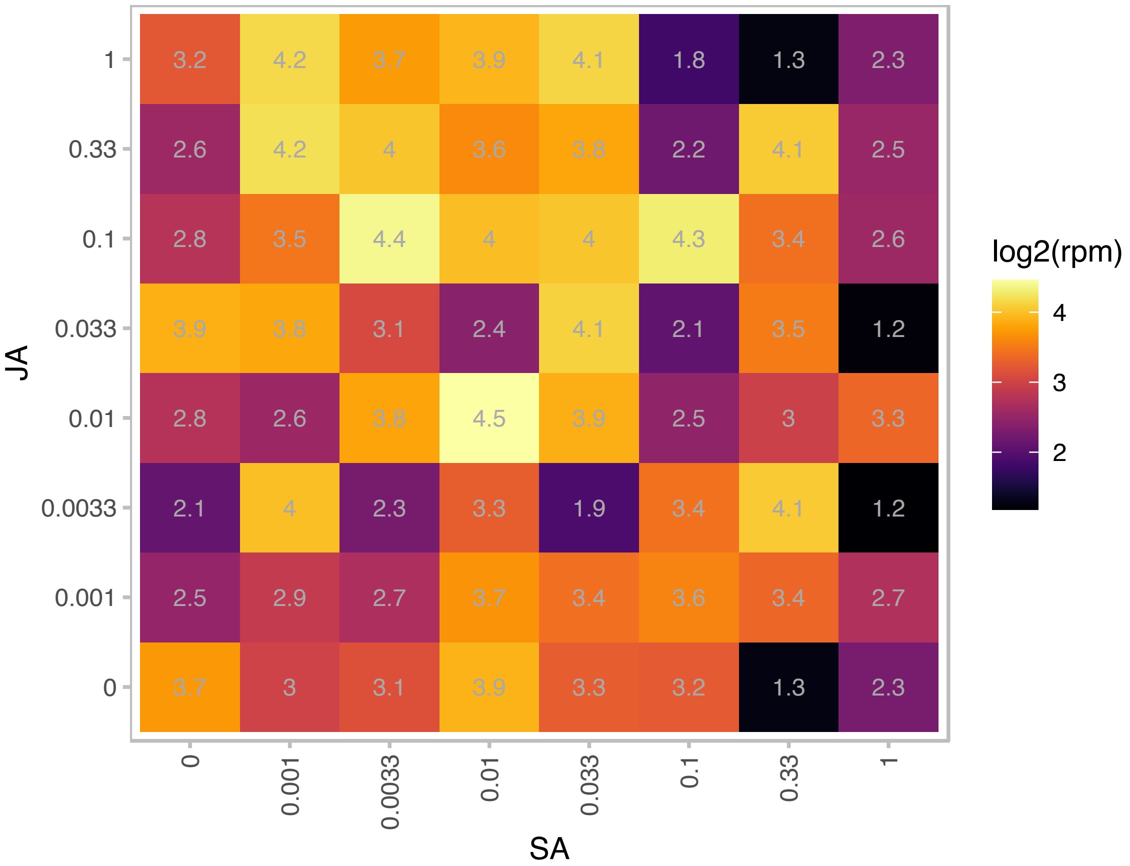


| | |
|---|---|
| Gene | AT5G19680 |
| is.expressed_genes | TRUE |
| is.responsive_genes | TRUE |
| is.apcluster | D-04 |

npr3-2 npr4-2

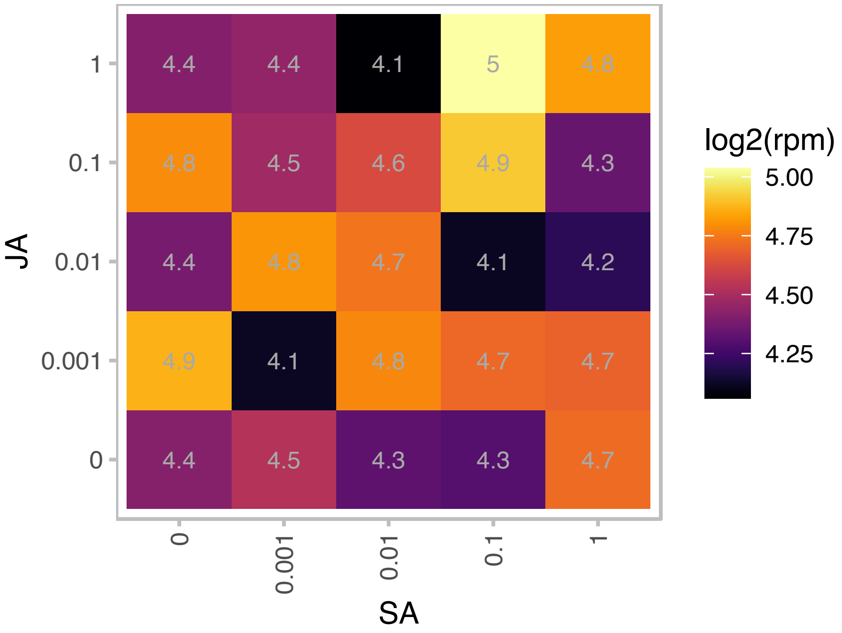


npr3-1 npr4-3

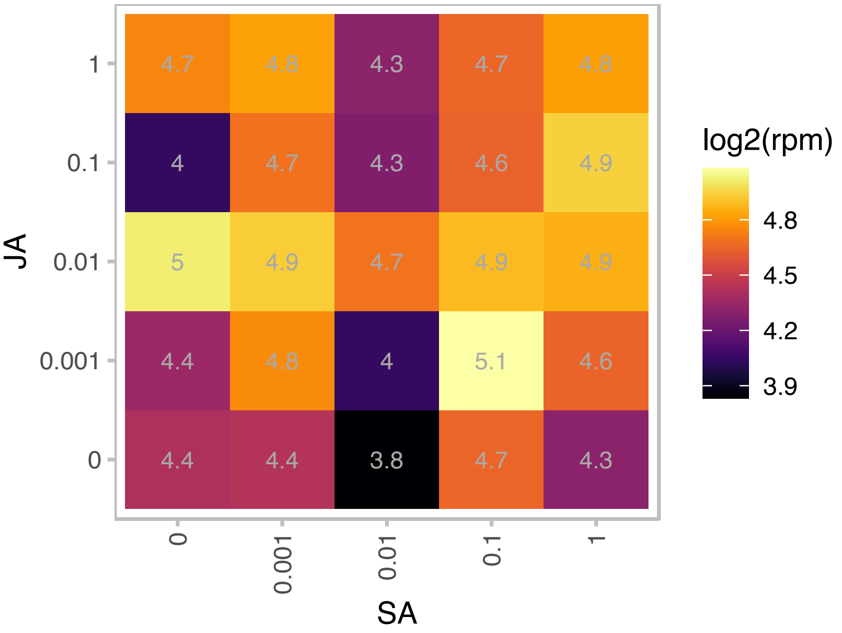

# AT5G20165.1 : NA

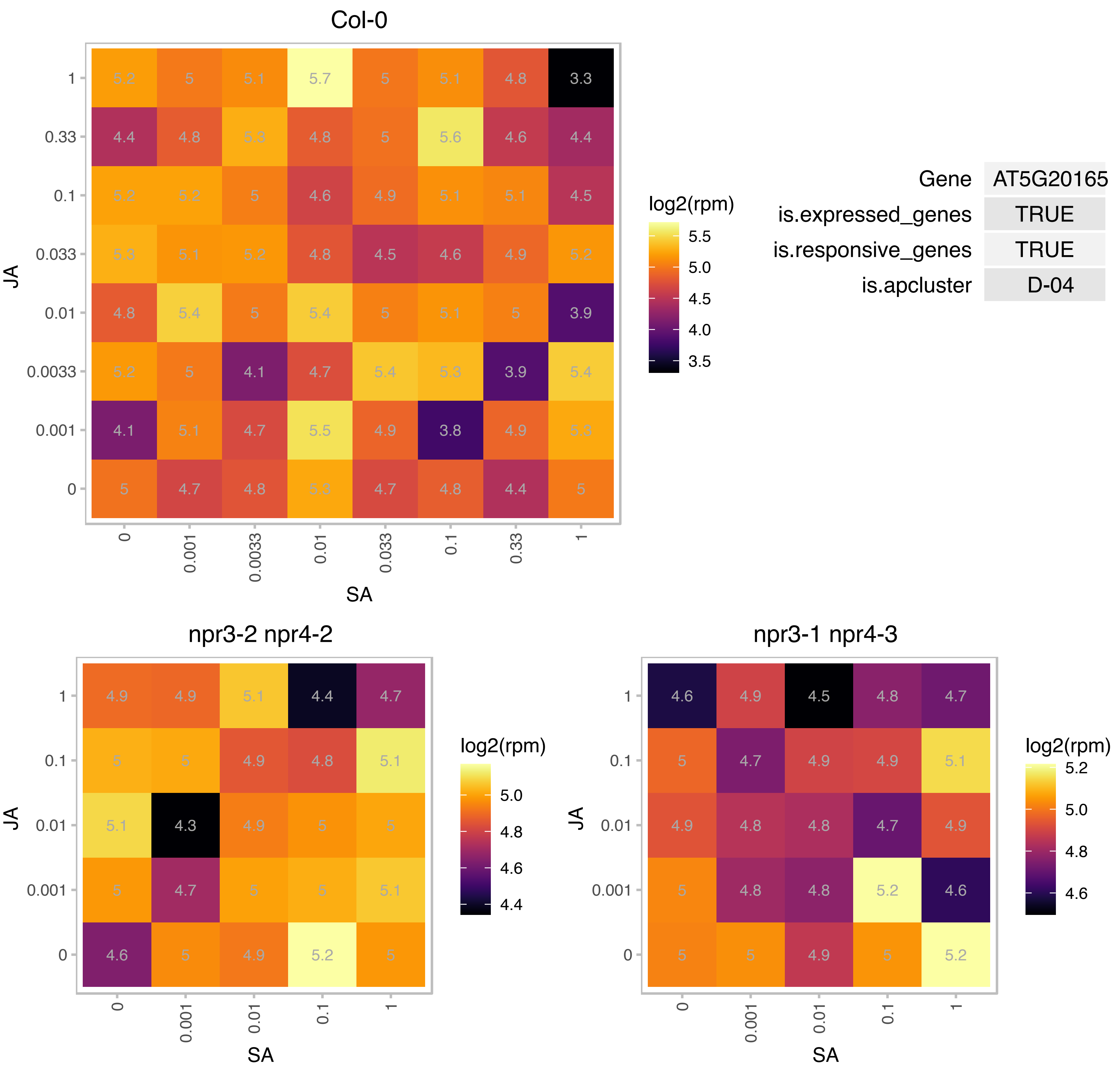

# AT5G20200.1 : NA

nucleoporin-related

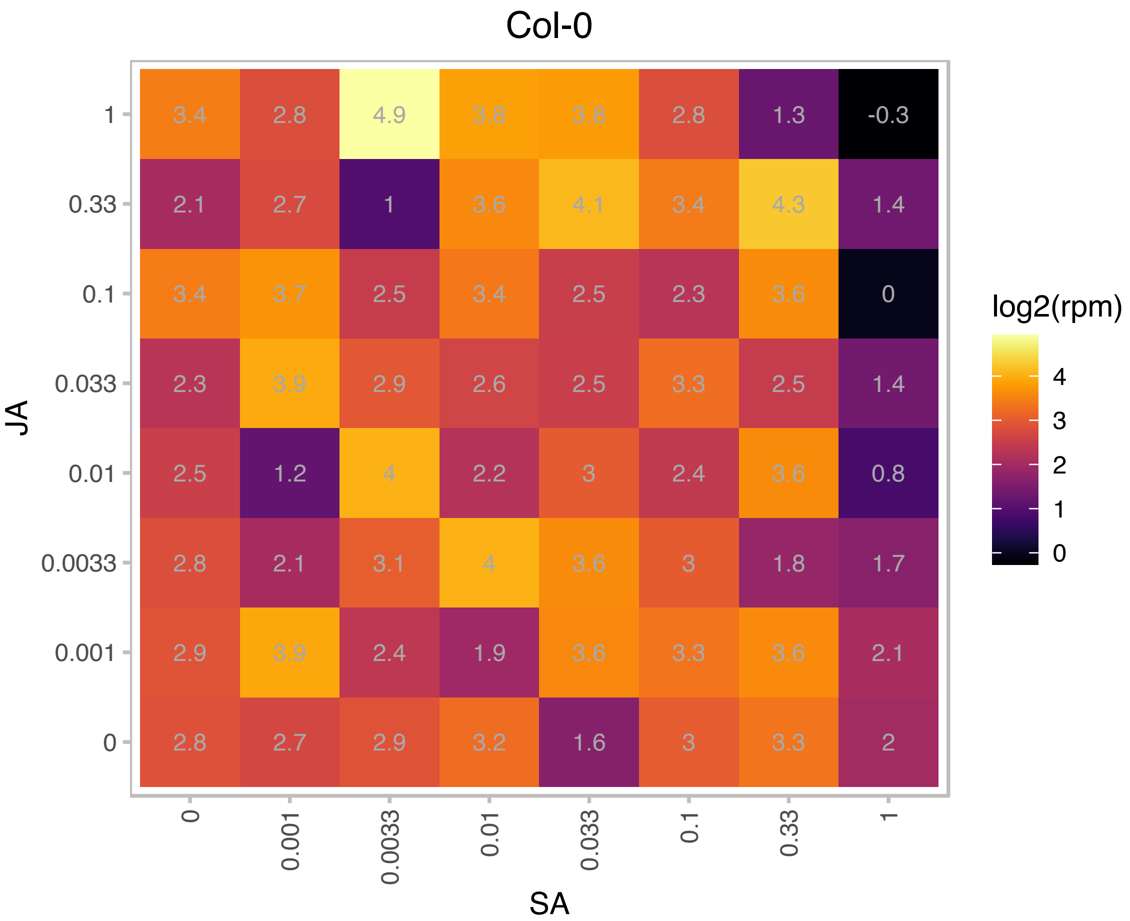


| | |
|---|---|
| Gene | AT5G20200 |
| is.expressed_genes | TRUE |
| is.responsive_genes | TRUE |
| is.apcluster | D-04 |

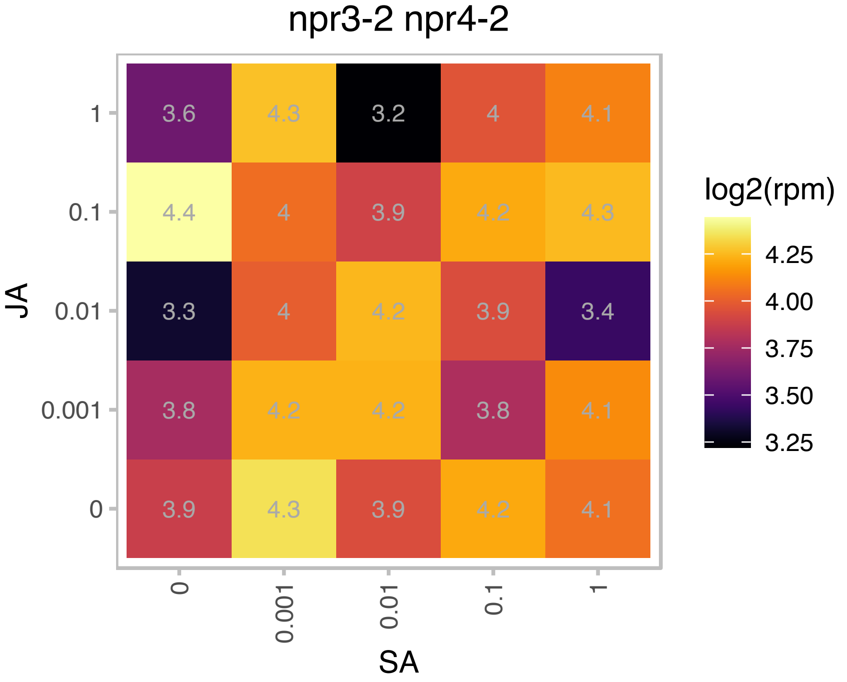


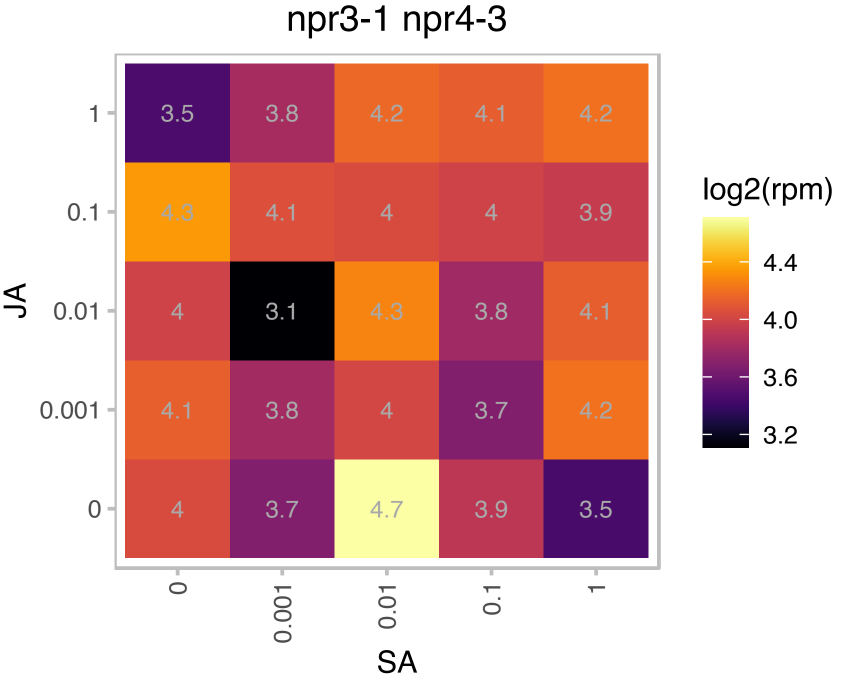

### AT5G22630.1 : ADT5

arogenate dehydratase 5
Encodes a plastid-localized arogenate dehydratase involved in phenylalanine biosynthesis.  Not less than six genes encoding ADT were identiﬁed in the Arabidopsis genome: ADT1 [At1g11790]; ADT2 [At3g07630]; AD [At2g27820]; ADT4 [At3g44720]; ADT5 [At5g22630]; and ADT6 [At1g08250]. The mRNA is cell-to-cell mobile.

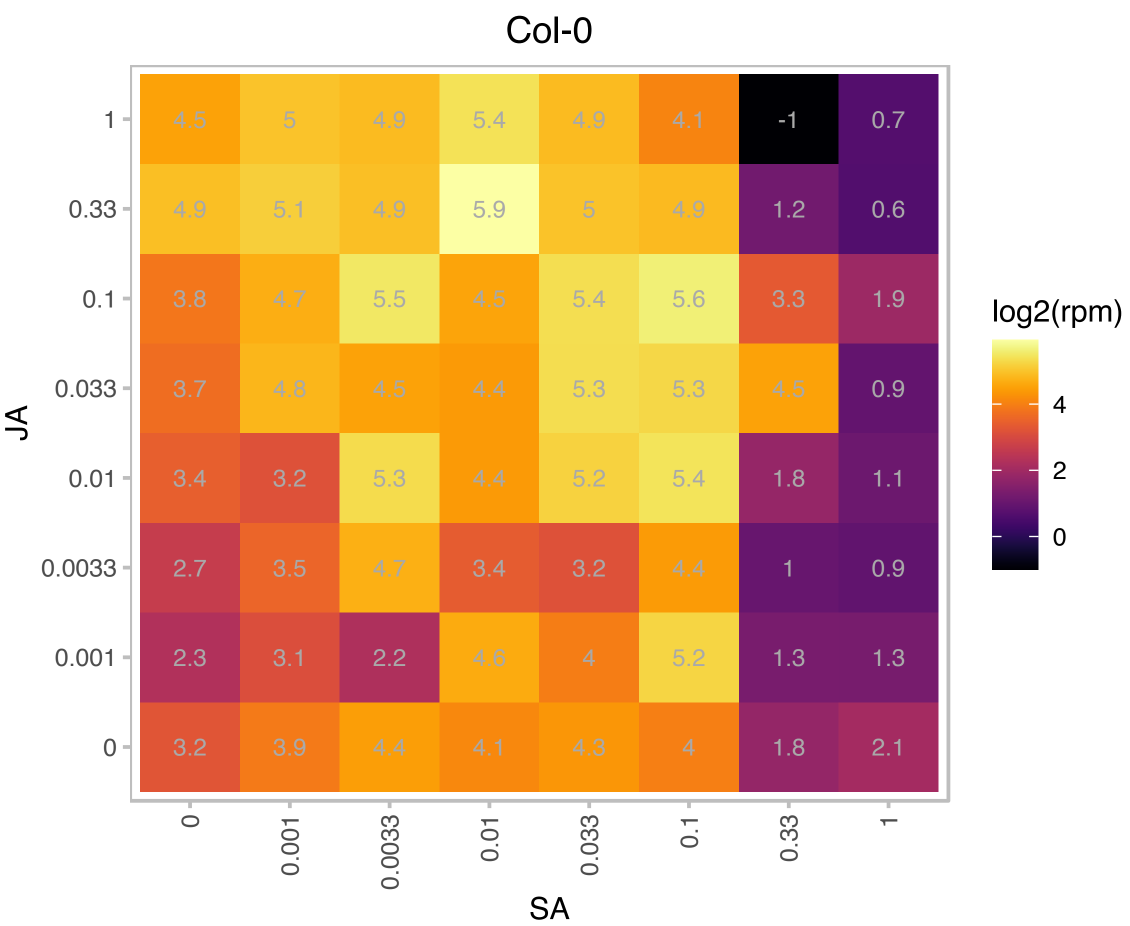


| | |
|---|---|
| Gene | AT5G22630 |
| is.expressed_genes | TRUE |
| is.responsive_genes | TRUE |
| is.apcluster | D-04 |

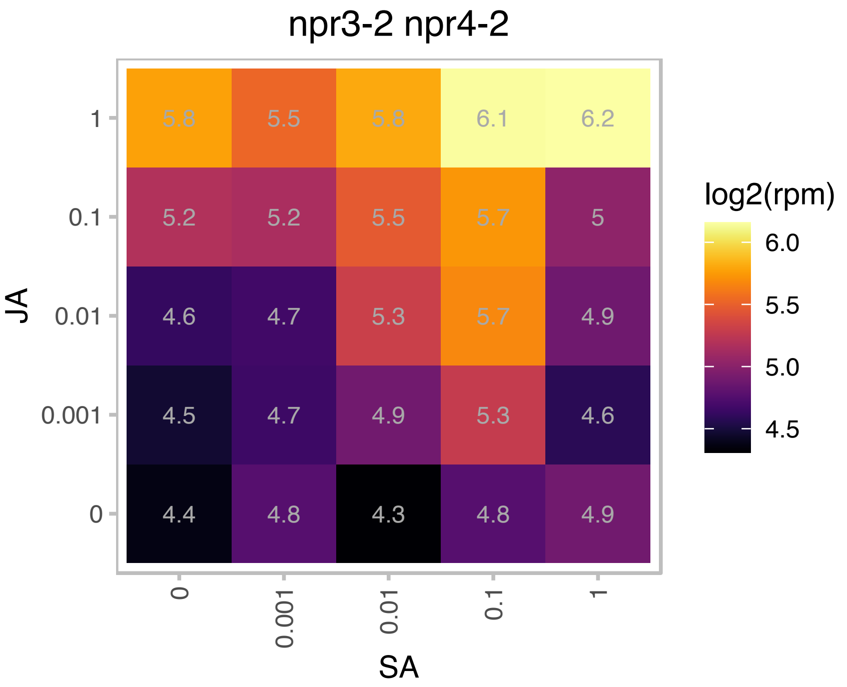


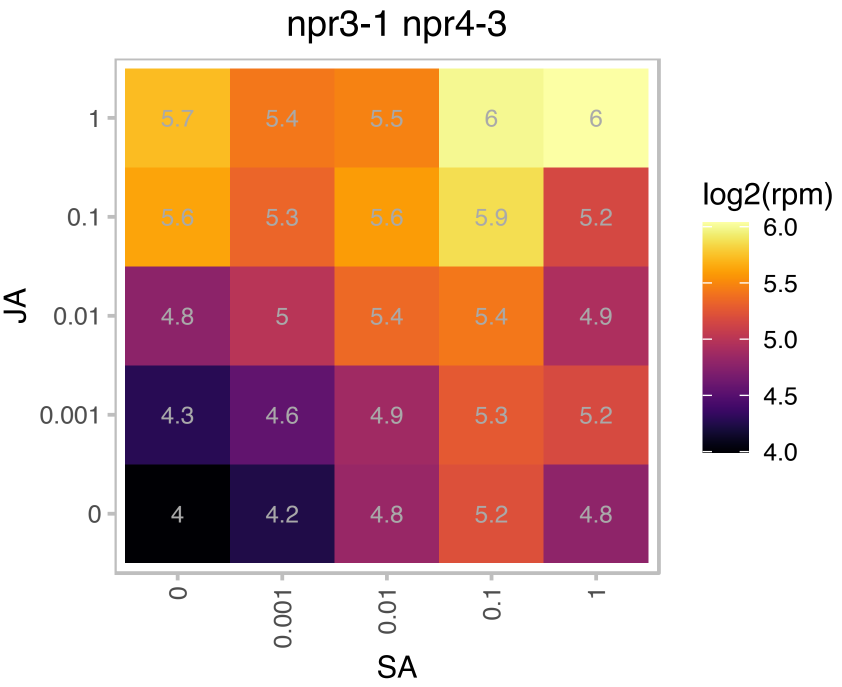

### AT5G51120.1 : ATPABN1, PABN1

polyadenylate-binding protein 1

Encodes a homolog of the protein PABN1, a polyadenylation factor subunit.

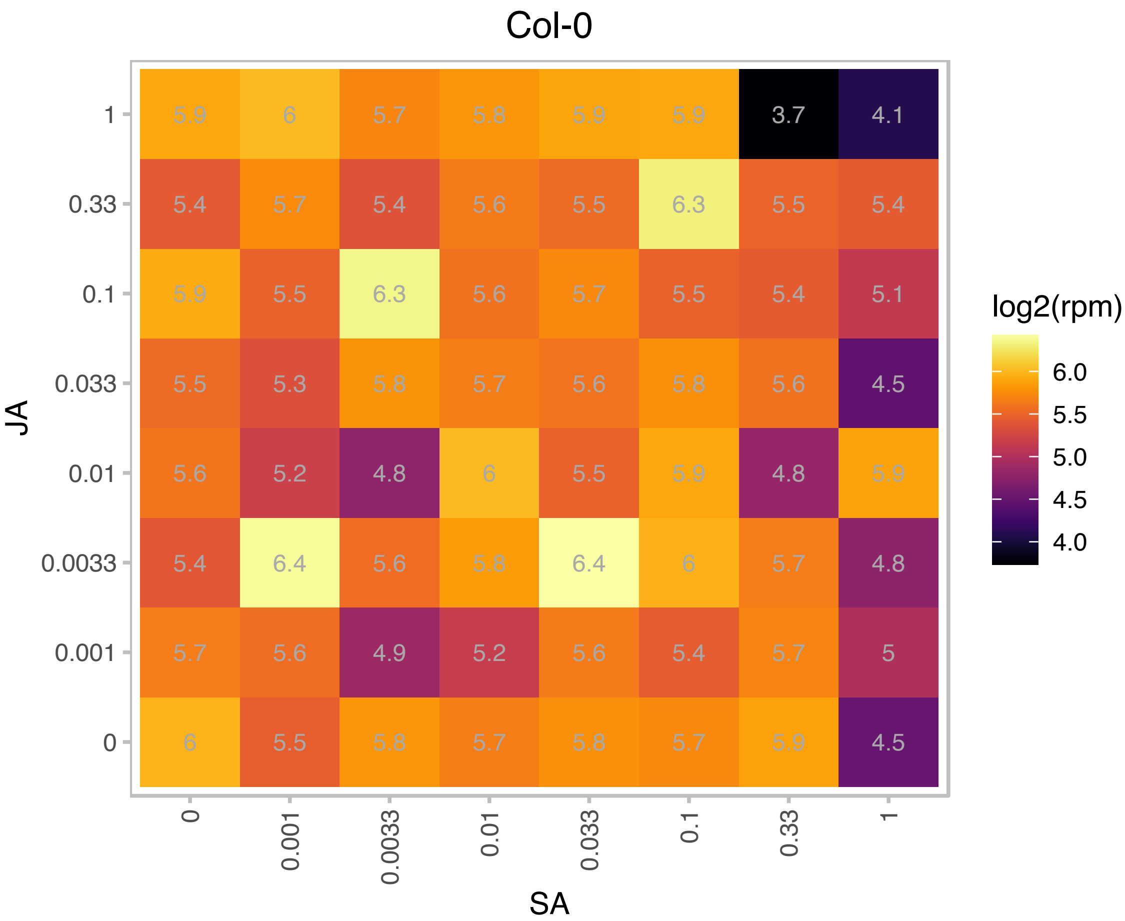


| | |
|---|---|
| Gene | AT5G51120 |
| is.expressed_genes | TRUE |
| is.responsive_genes | TRUE |
| is.apcluster | D-04 |

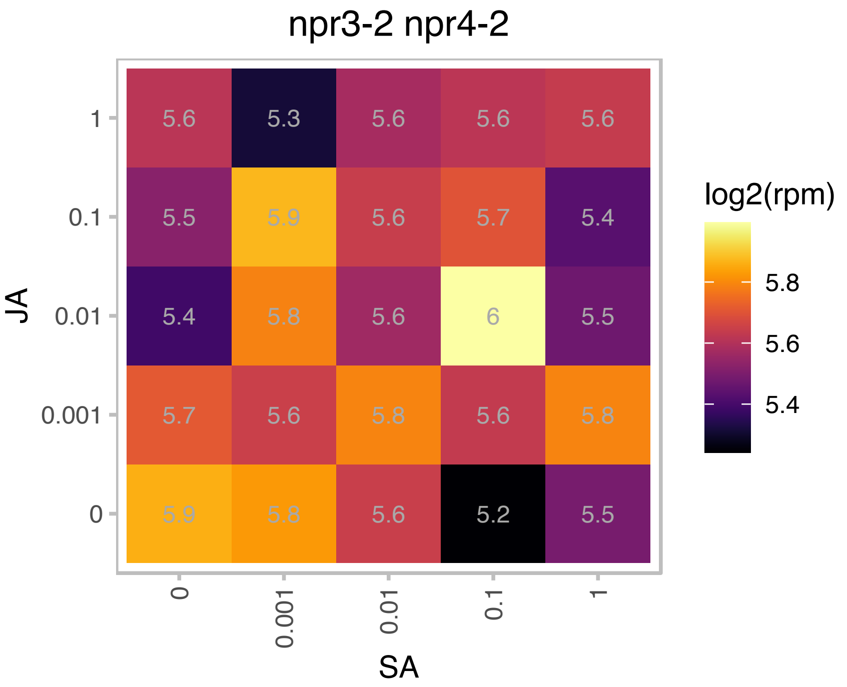


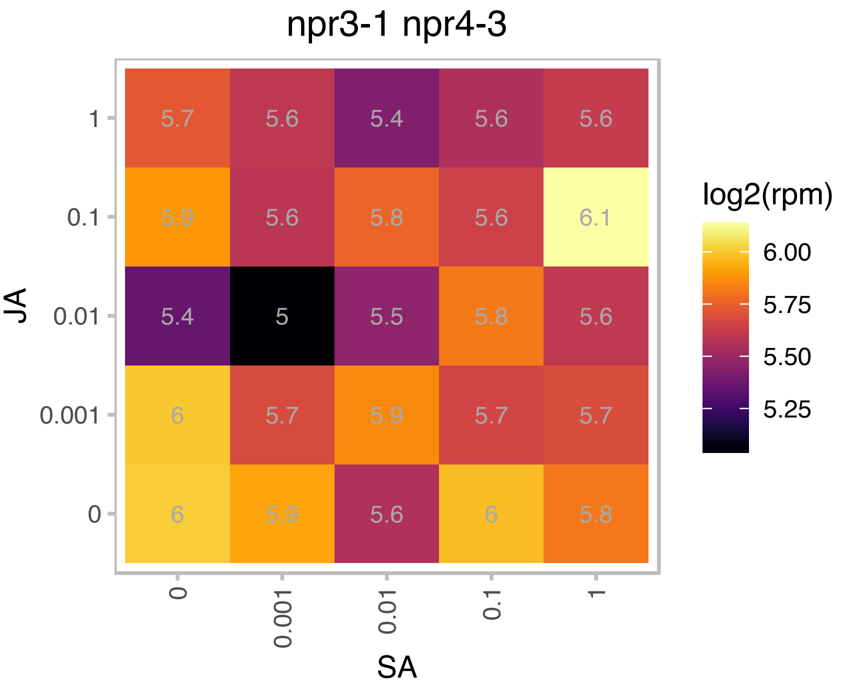

### AT5G53460.1 : GLT1

NADH-dependent glutamate synthase 1

NADH-dependent glutamate synthase The mRNA is cell-to-cell mobile.

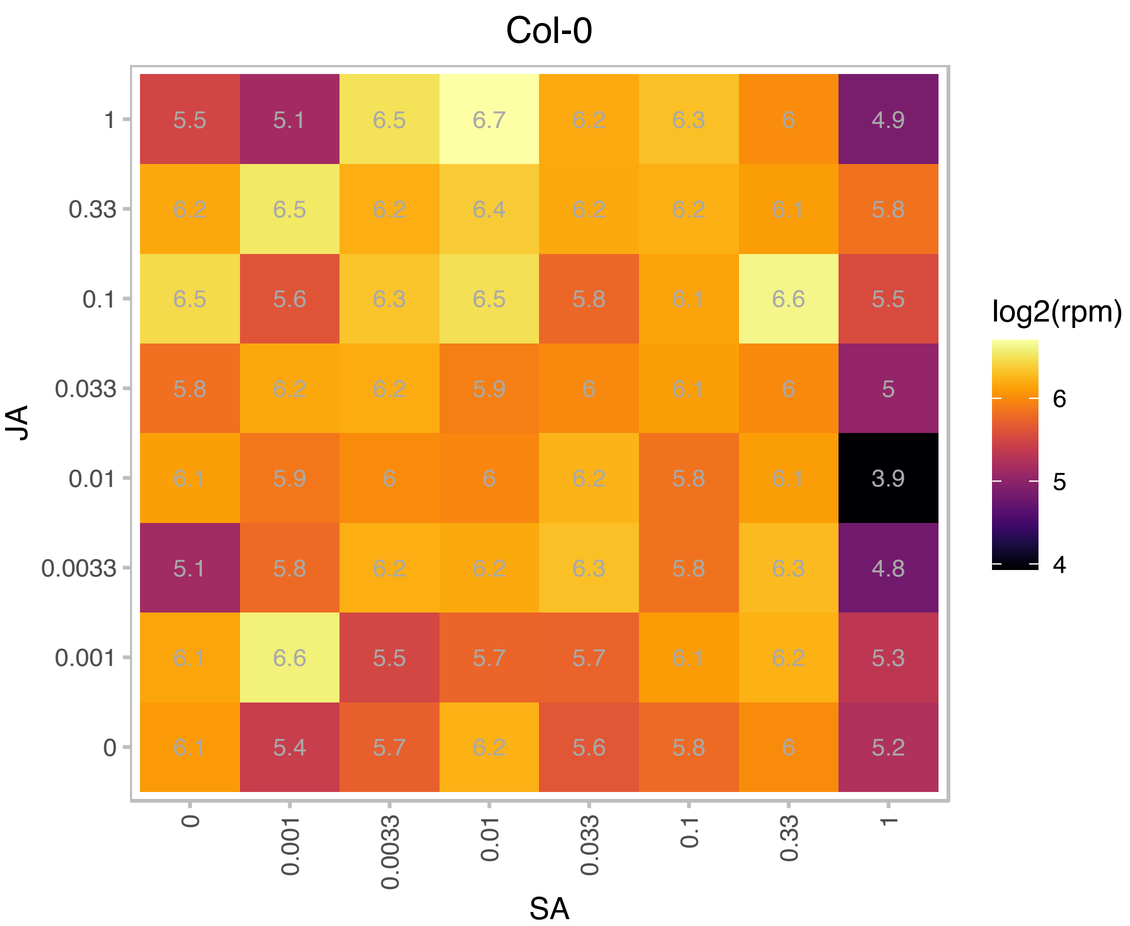


| | |
|---|---|
| Gene | AT5G53460 |
| is.expressed_genes | TRUE |
| is.responsive_genes | TRUE |
| is.apcluster | D-04 |

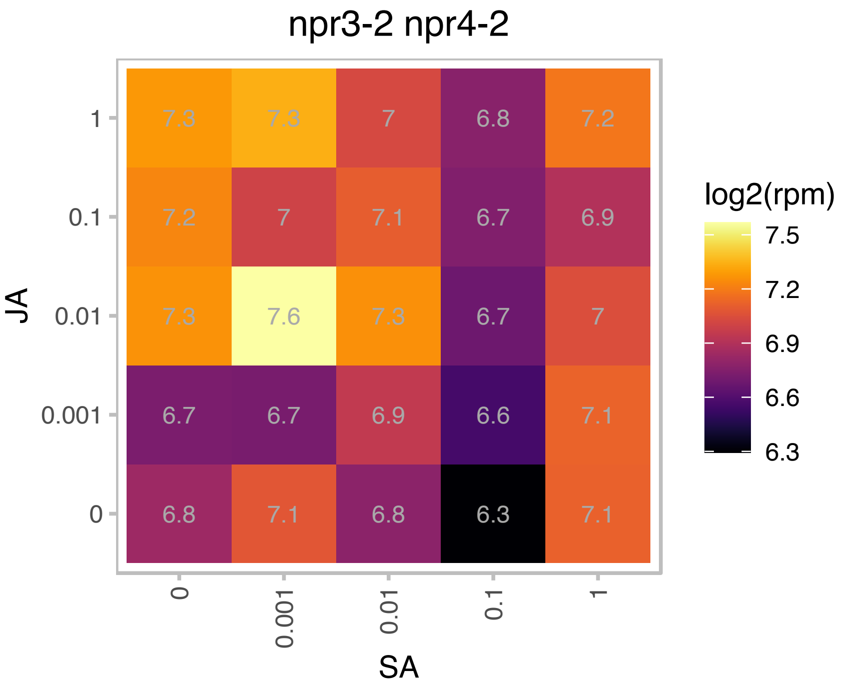


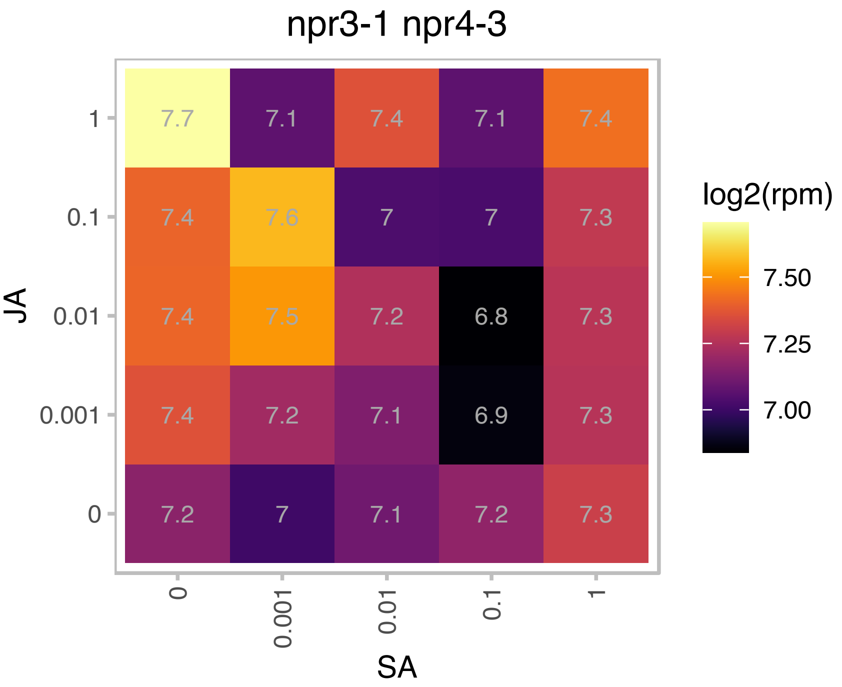

# AT5G54760.1 : NA

Translation initiation factor SUI1 family protein

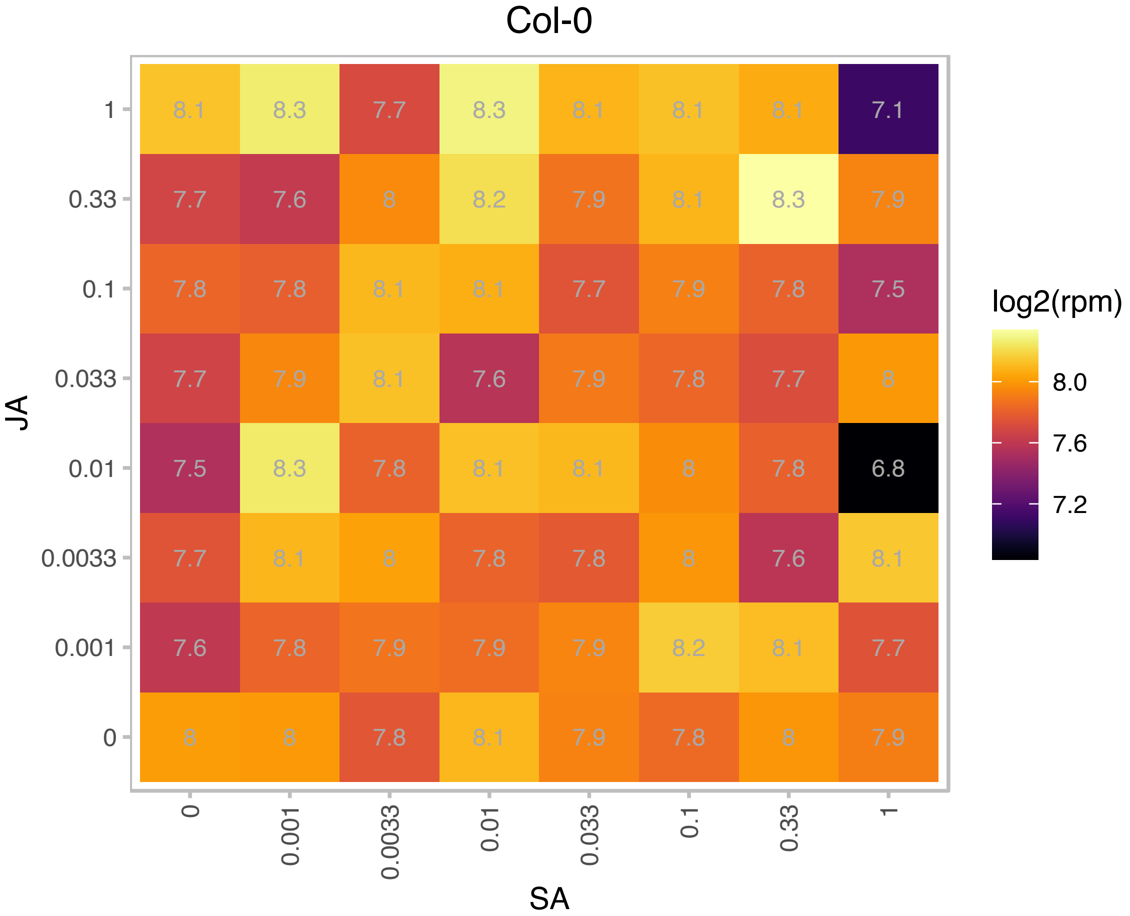


| | |
|---|---|
| Gene | AT5G54760 |
| is.expressed_genes | TRUE |
| is.responsive_genes | TRUE |
| is.apcluster | D-04 |

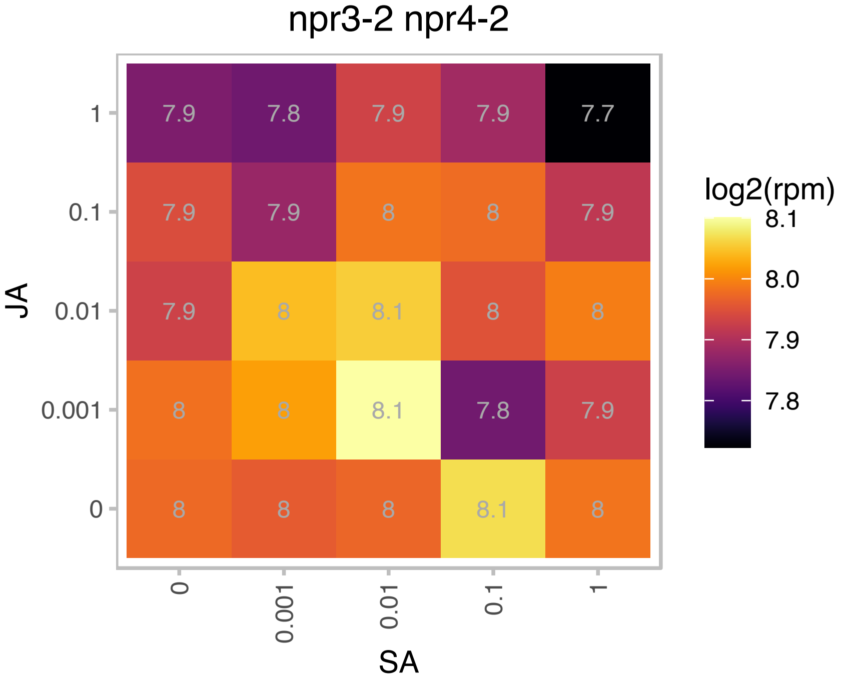


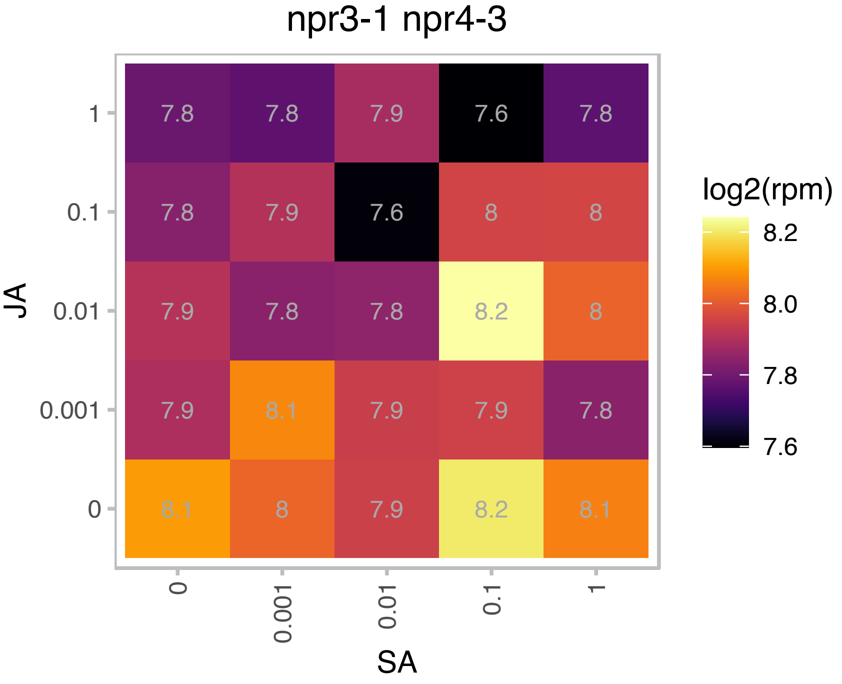

# AT5G57330.1 : NA

Galactose mutarotase-like superfamily protein

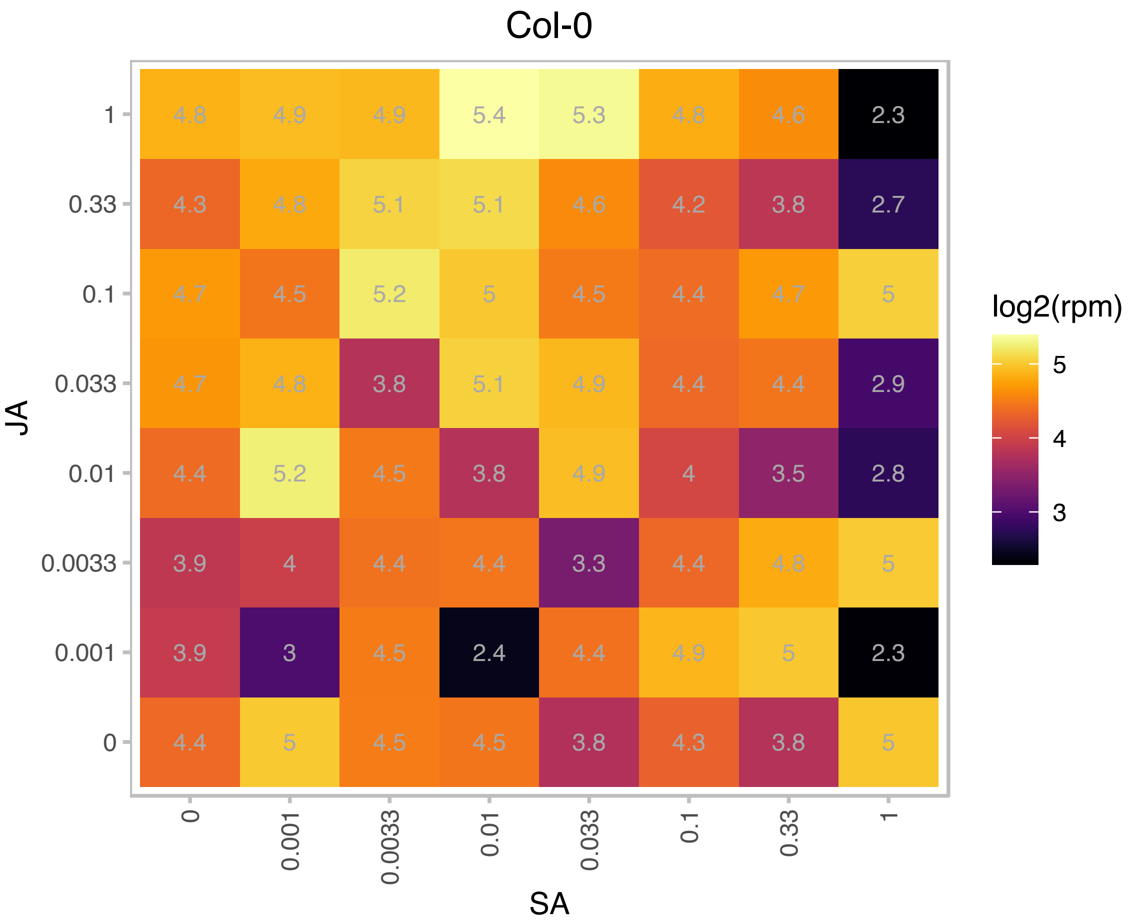


| | |
|---|---|
| Gene | AT5G57330 |
| is.expressed_genes | TRUE |
| is.responsive_genes | TRUE |
| is.apcluster | D-04 |

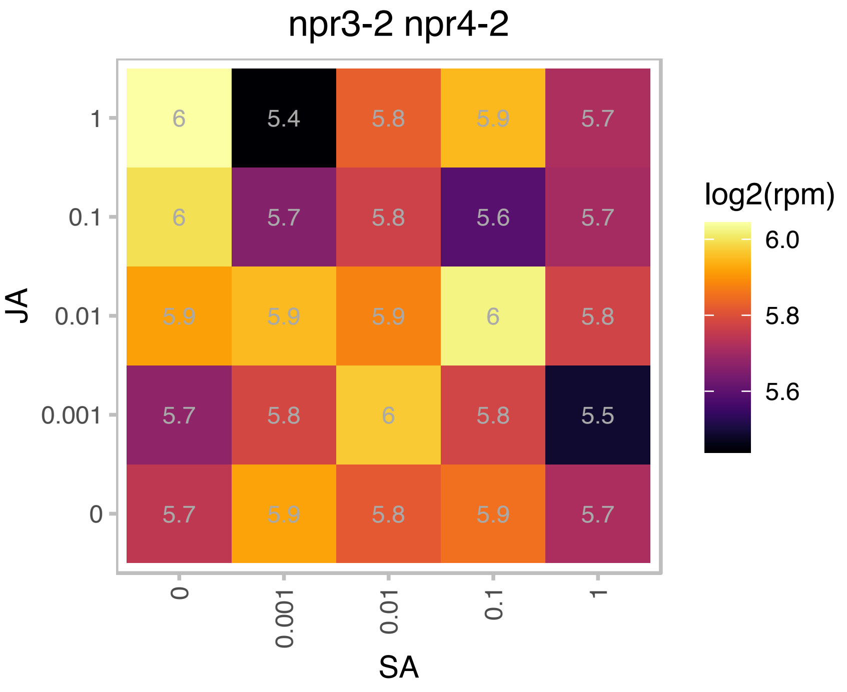


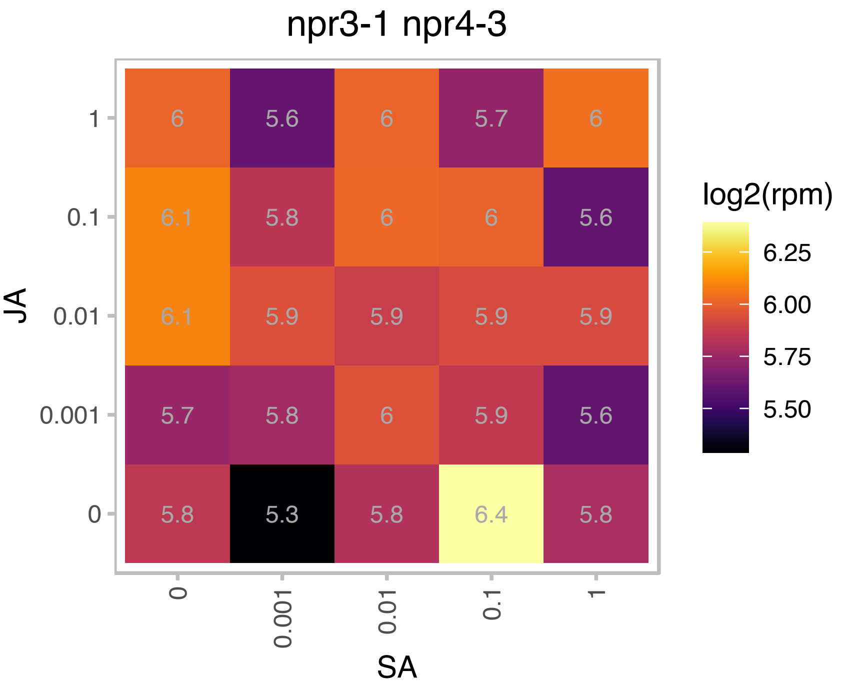

# AT5G59830.1 : NA

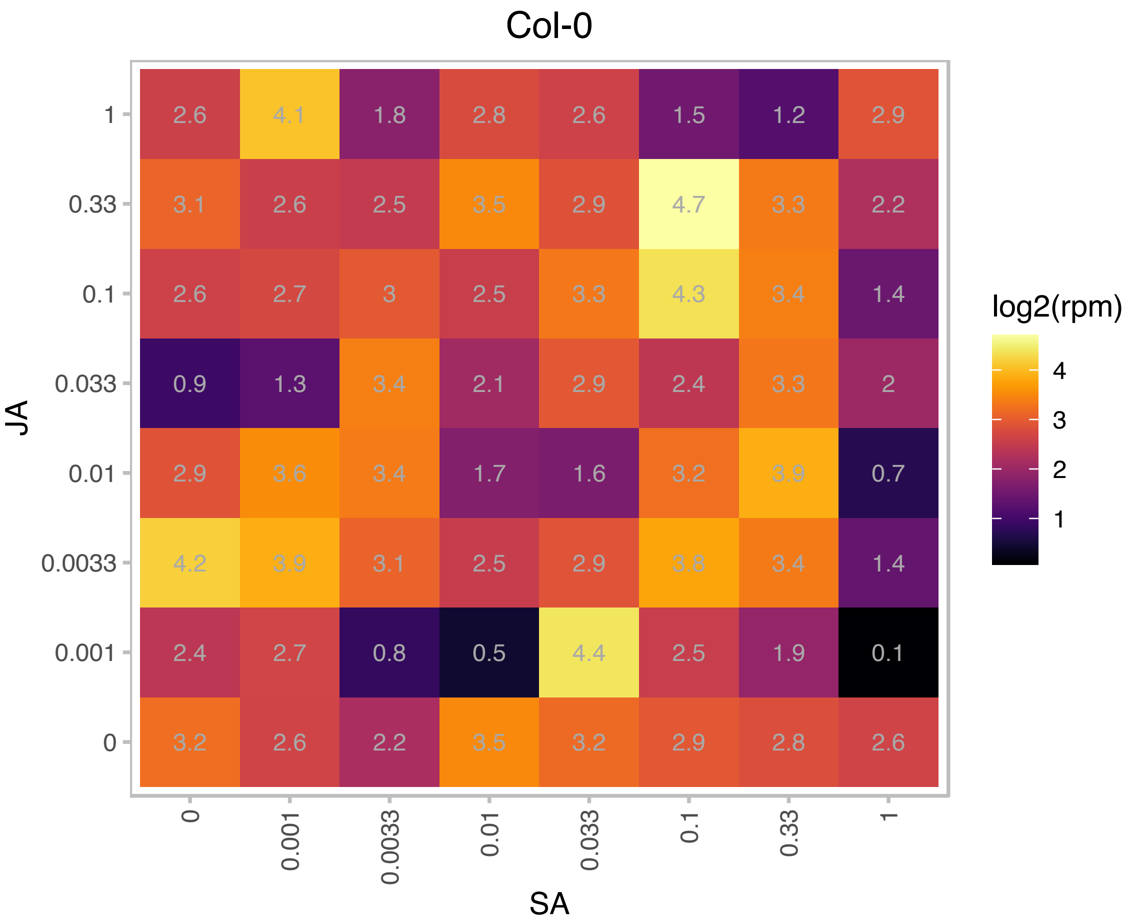


| | |
|---|---|
| Gene | AT5G59830 |
| is.expressed_genes | TRUE |
| is.responsive_genes | TRUE |
| is.apcluster | D-04 |

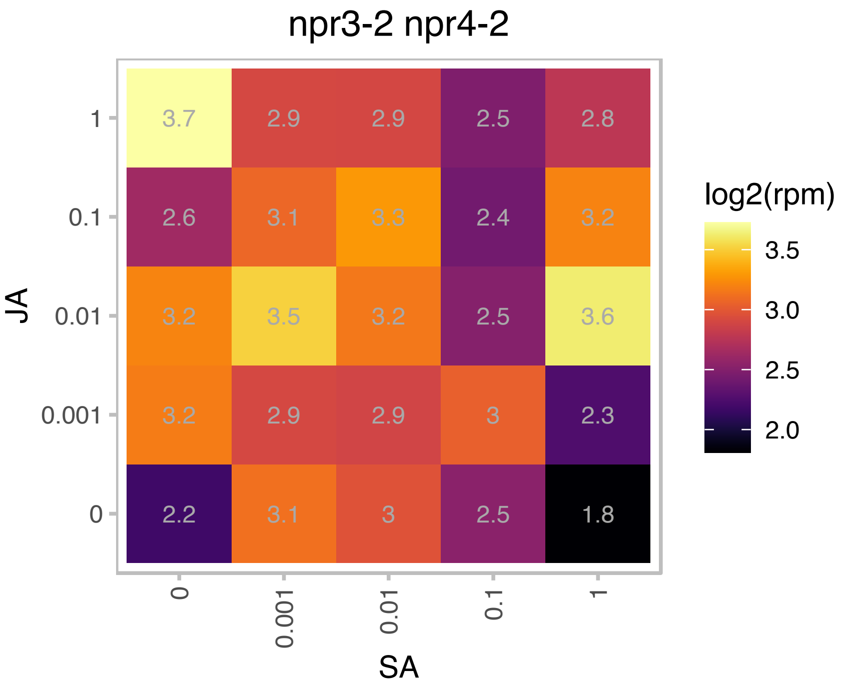


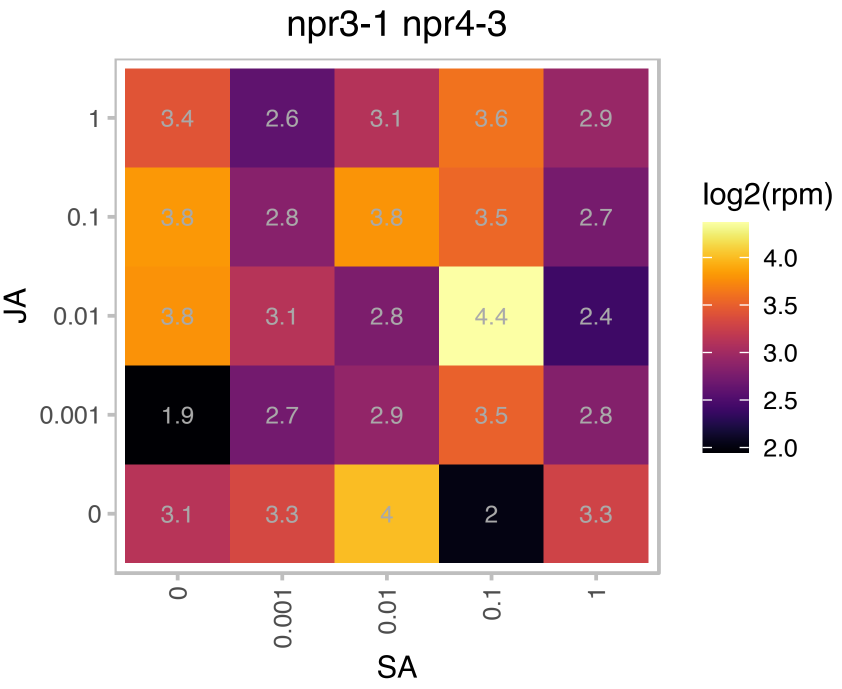

### AT5G66030.1 : ATGRIP, GRIP

Golgi-localized GRIP domain-containing protein

Involved in golgi protein trafficking. AtARL1 binds directly to the GRIP domain of AtGRIP in a GTP-dependent manner. Localized to the golgi apparatus, tyrosine 717 in AtGRIP is crucial for Golgi localization.

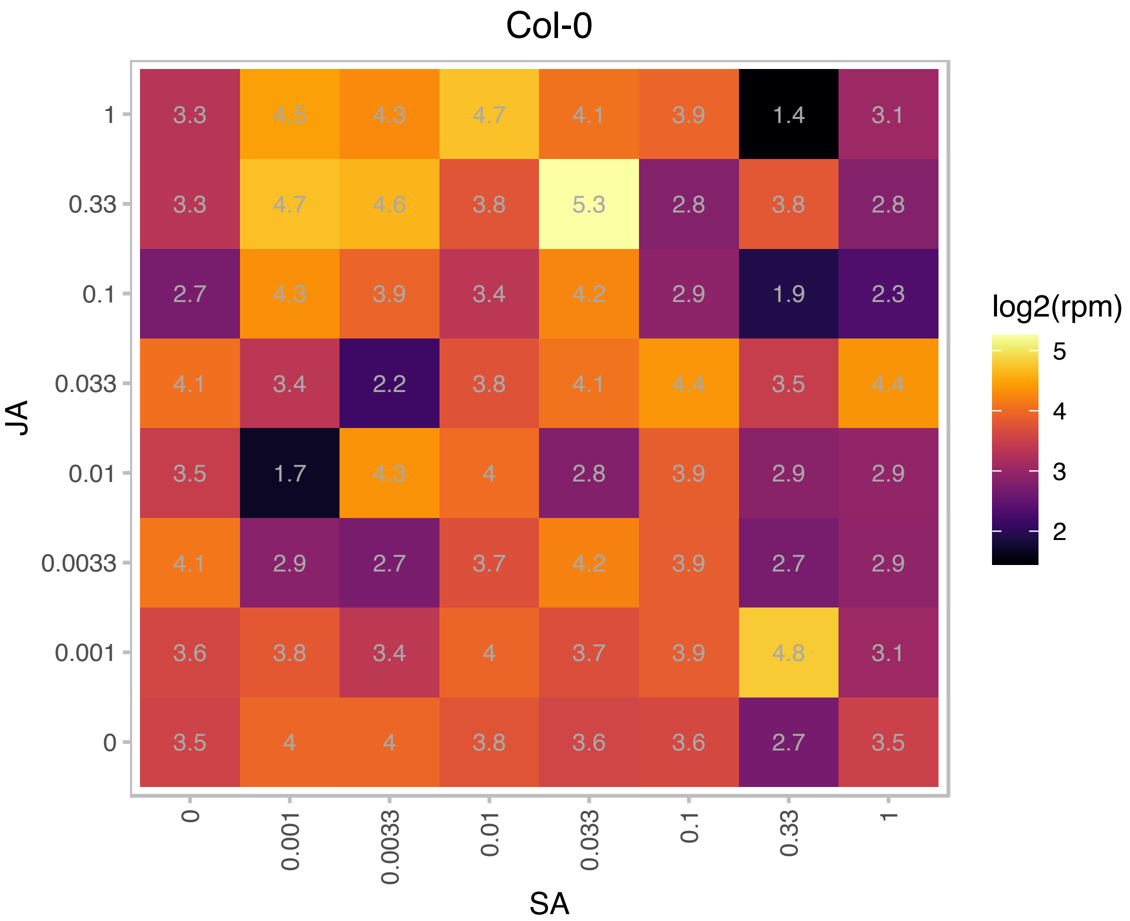


| | |
|---|---|
| Gene | AT5G66030 |
| is.expressed_genes | TRUE |
| is.responsive_genes | TRUE |
| is.apcluster | D-04 |

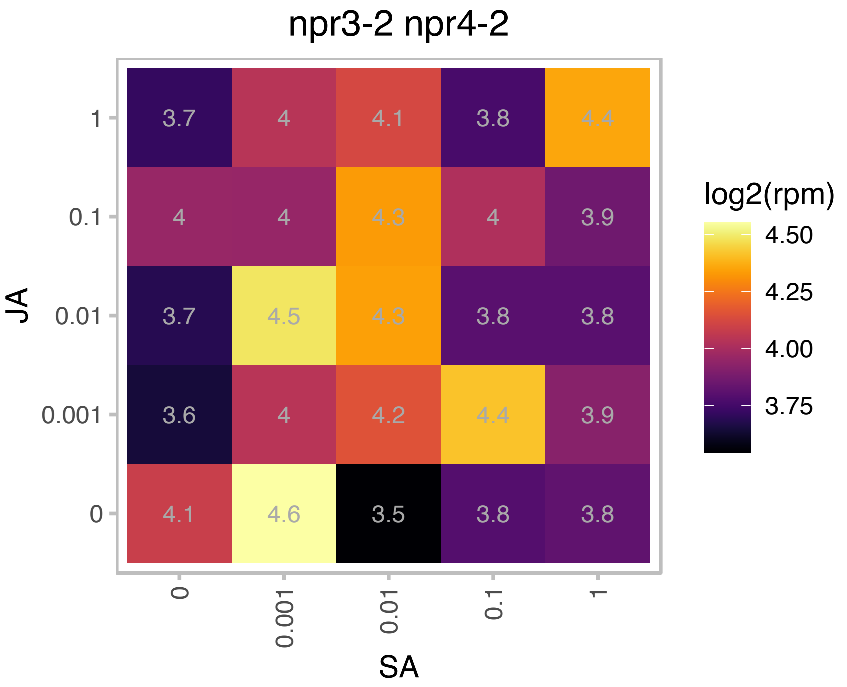


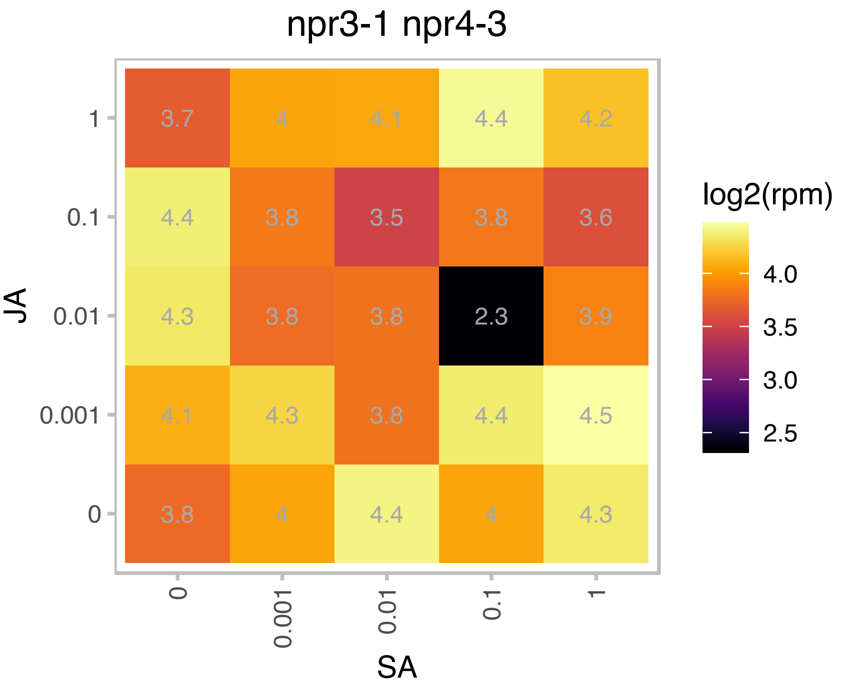

# AT1G03940.1 : NA

HXXXD-type acyl-transferase family protein

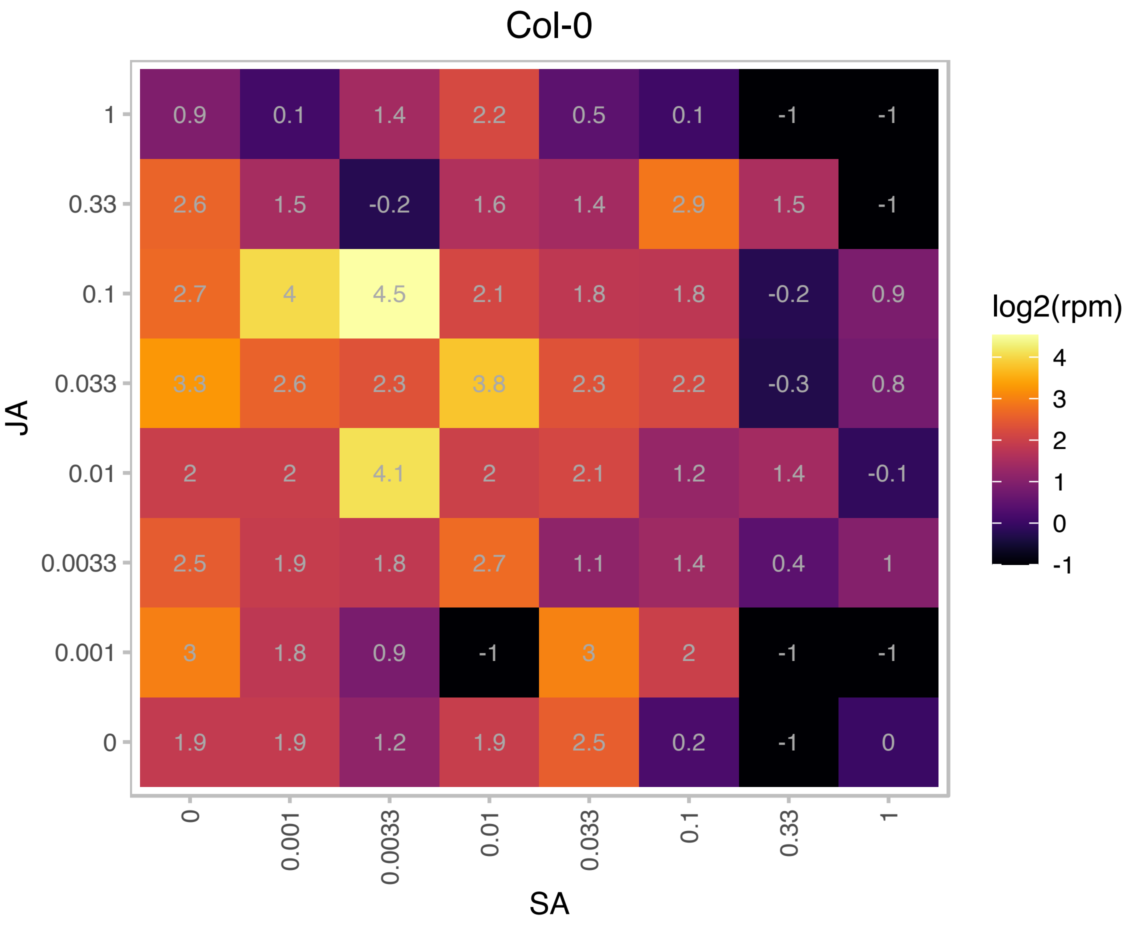


| | |
|---|---|
| Gene | AT1G03940 |
| is.expressed_genes | TRUE |
| is.responsive_genes | TRUE |
| is.apcluster | E-01 |

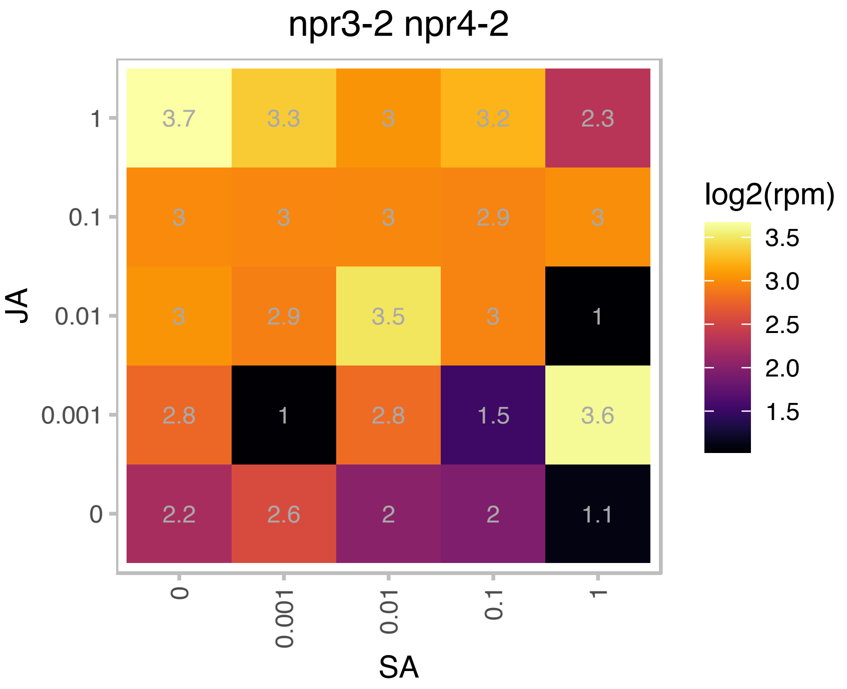


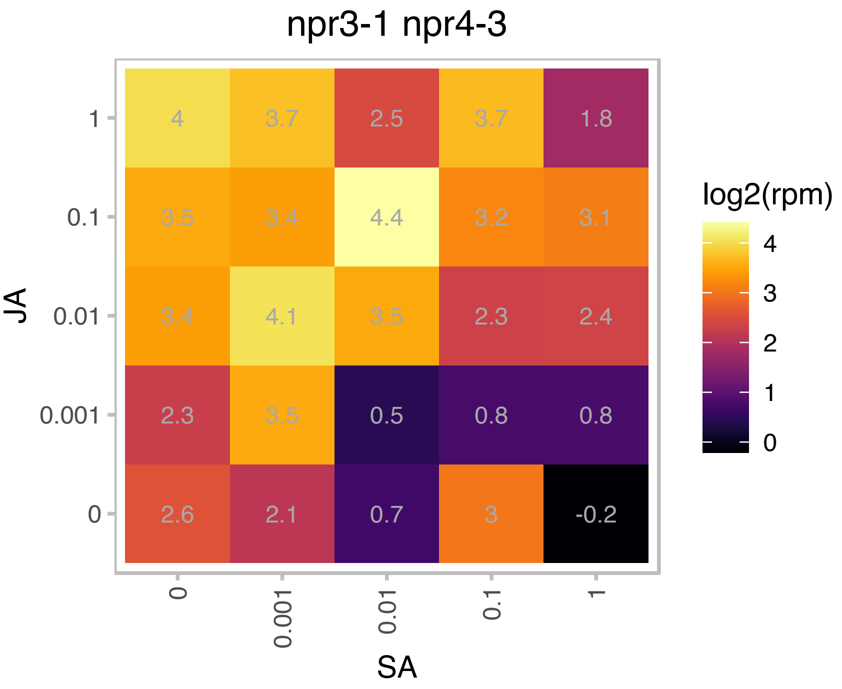

### AT1G04190.1 : TPR3

Tetratricopeptide repeat (TPR)-like superfamily protein

Encodes one of the 36 carboxylate clamp (CC)-tetratricopeptide repeat (TPR) proteins (Prasad 2010, Pubmed ID: 20856808) with potential to interact with Hsp90/Hsp70 as co-chaperones.

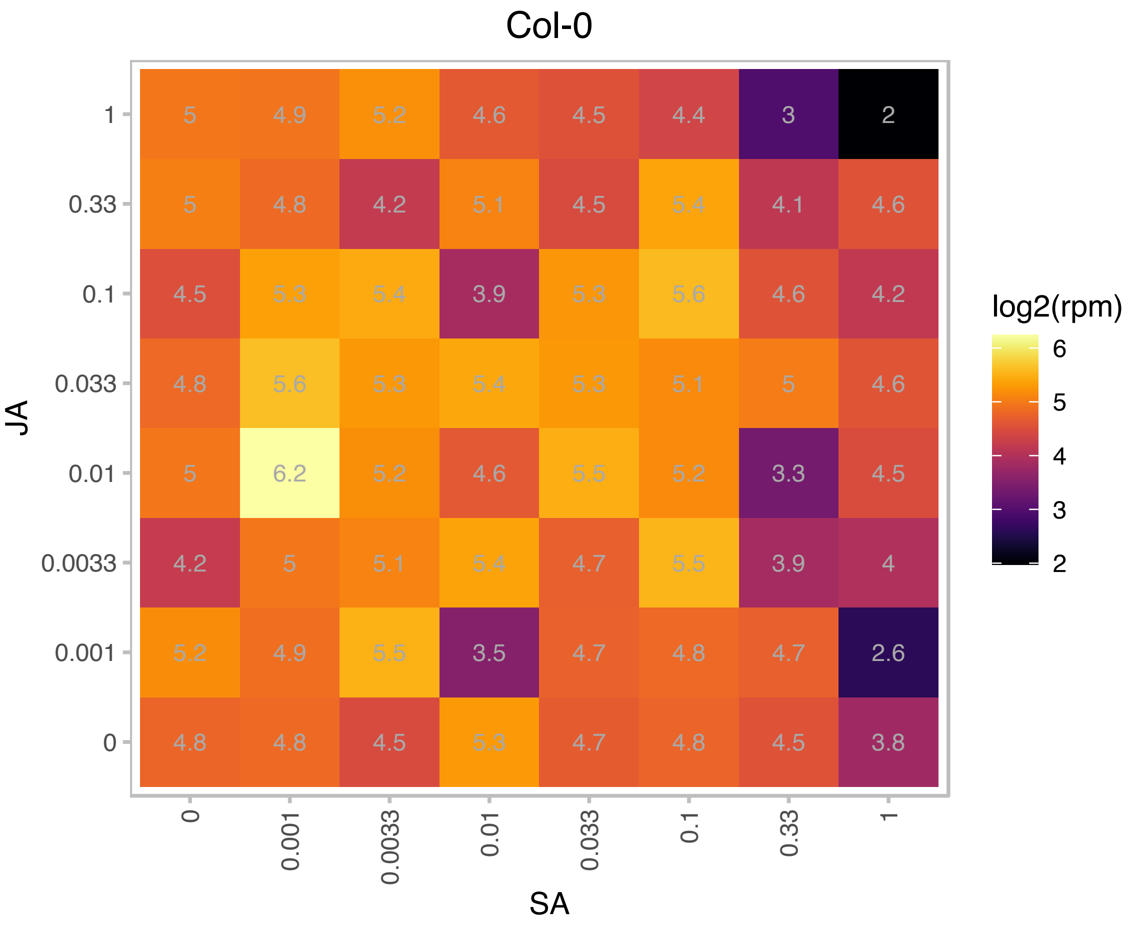


| | |
|---|---|
| Gene | AT1G04190 |
| is.expressed_genes | TRUE |
| is.responsive_genes | TRUE |
| is.apcluster | E-01 |

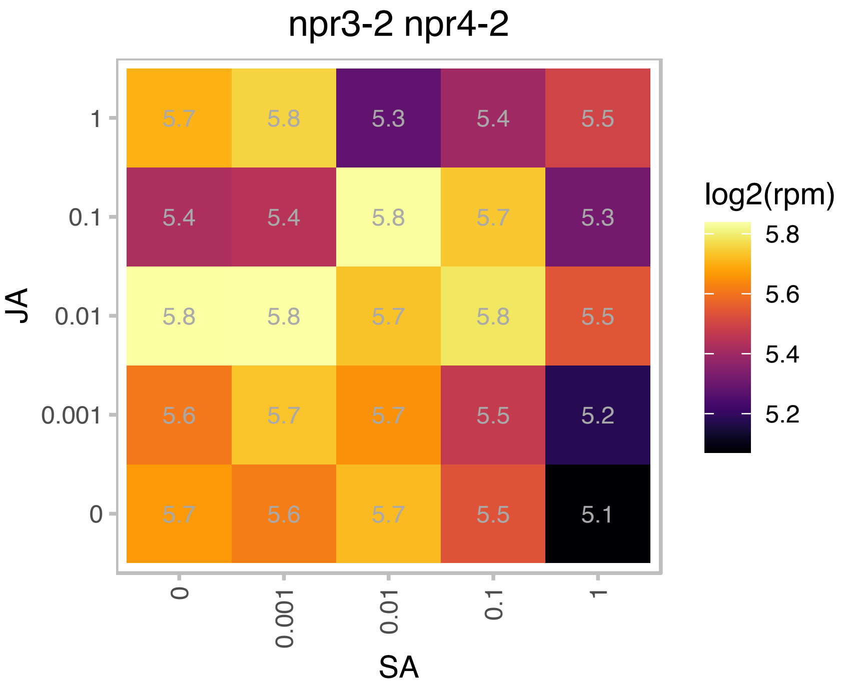


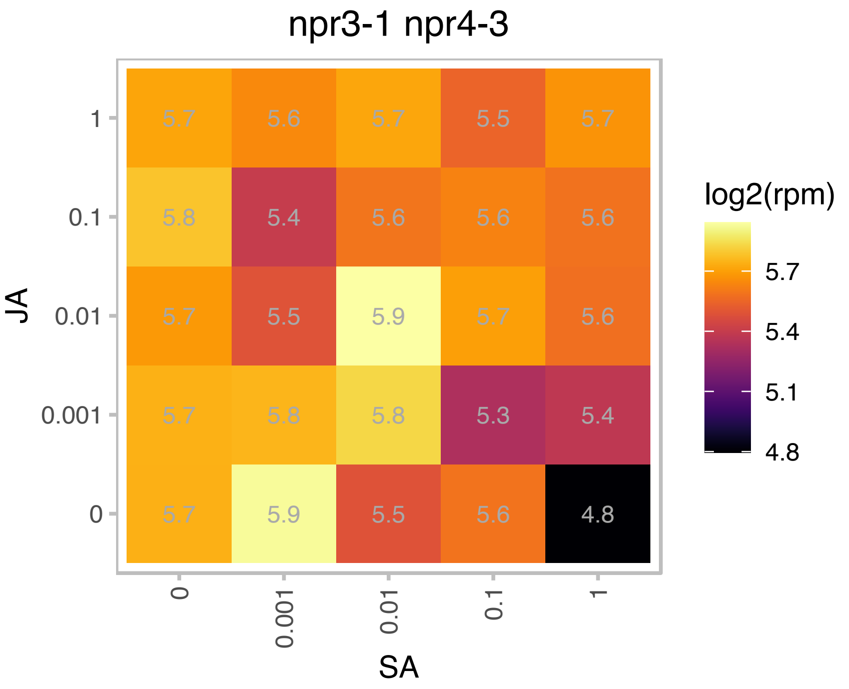

### AT1G07050.1 : FITNESS

CCT motif family protein

FITNESS encodes a protein with a single CCT domain and belongs to the CCT motif family genes (CMF). FITNESS acts upstream JUB1 thereby controlling H2O2 levels. FITNESS has a role in cellular redox homeostasis controlling H2O2 levels, due to changes in enzymes, metabolites and transcripts related to ROS detoxification.

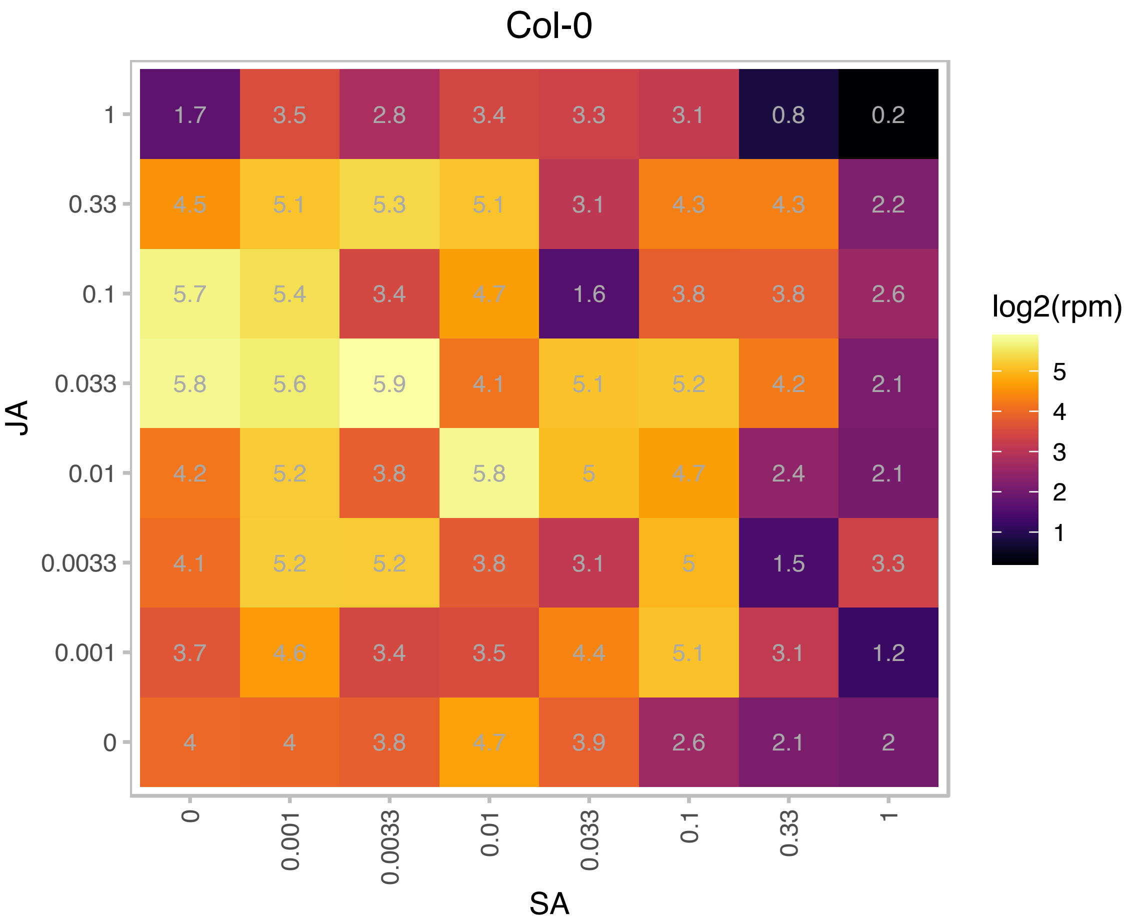


| | |
|---|---|
| Gene | AT1G07050 |
| is.expressed_genes | TRUE |
| is.responsive_genes | TRUE |
| is.apcluster | E-01 |

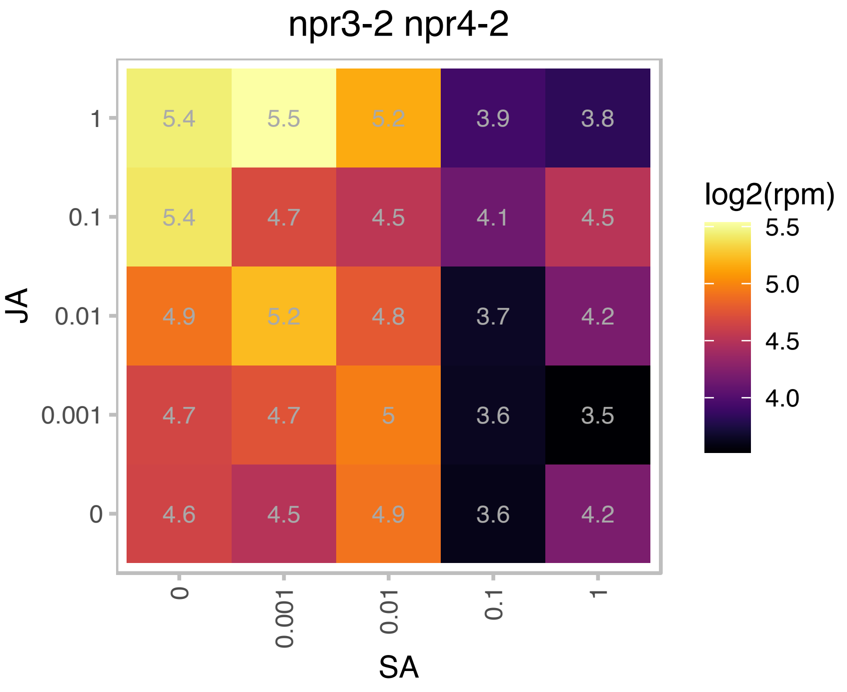


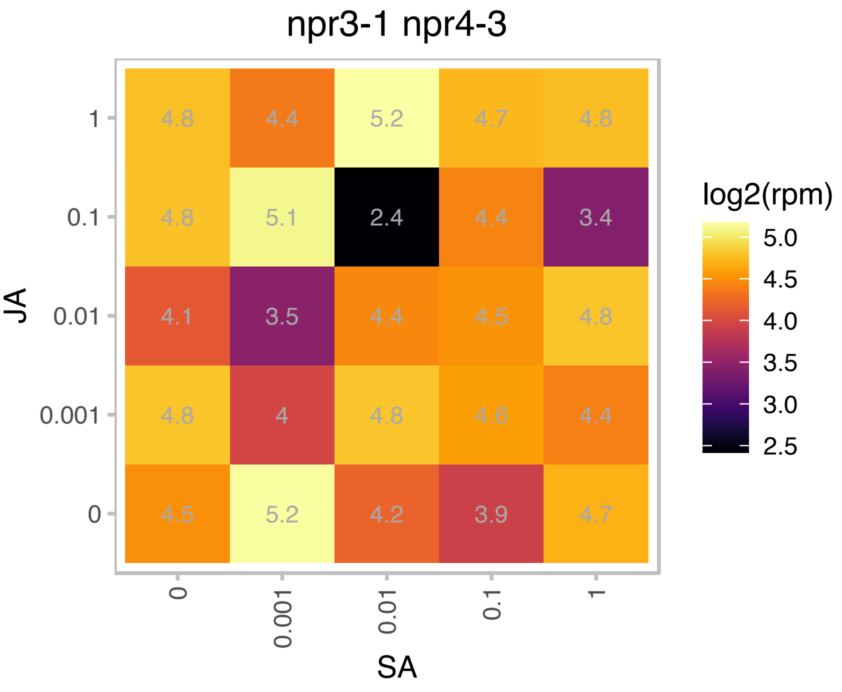

### AT1G13420.1 : ATST4B, ST4B

sulfotransferase 4B
Encodes a sulfotransferase. Unlike the related ST4A protein (At2g14920), in vitro experiments show that this enzyme does not act brassinosteroids. ST4B is expressed in the roots and transcript levels rise in response to cytokinin treatment.

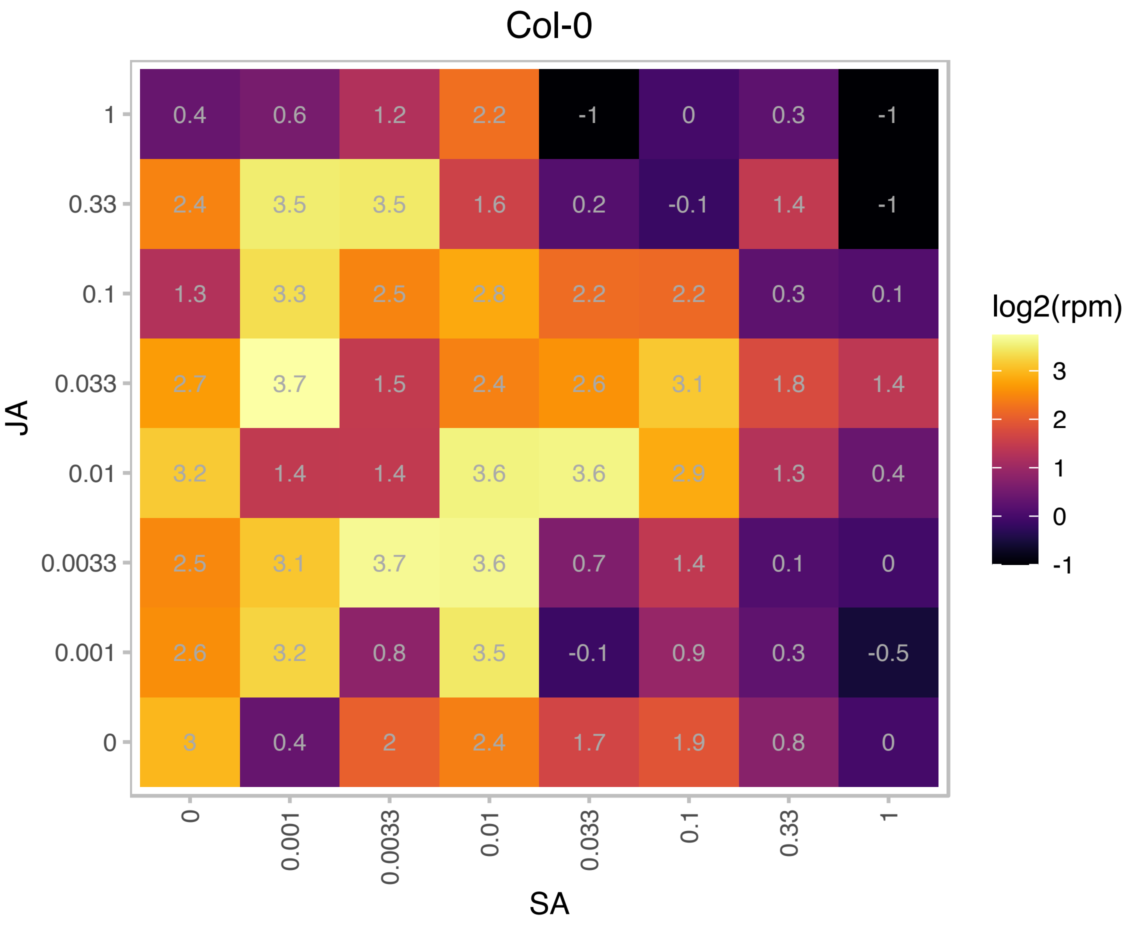


| | |
|---|---|
| Gene | AT1G13420 |
| is.expressed_genes | TRUE |
| is.responsive_genes | TRUE |
| is.apcluster | E-01 |

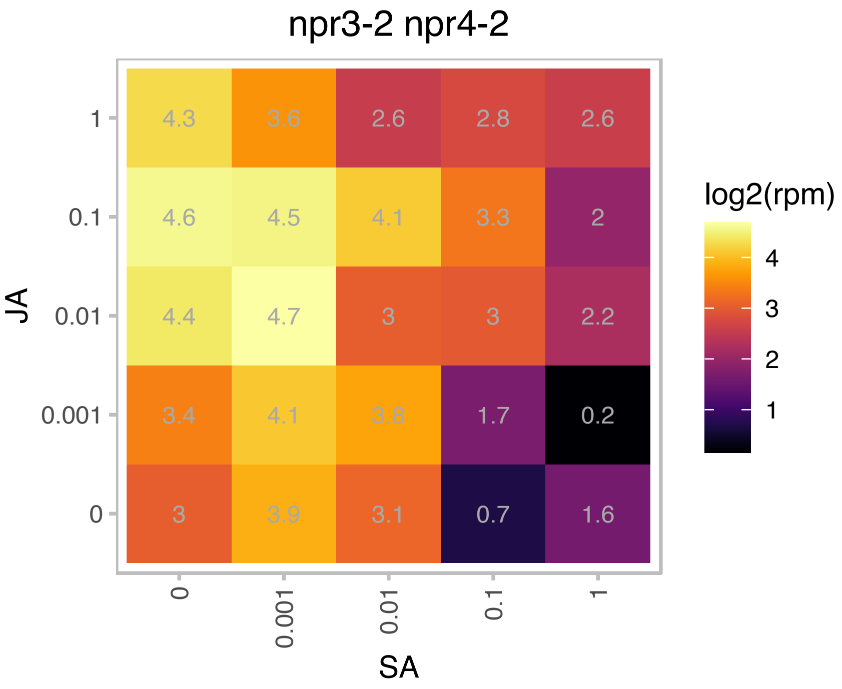


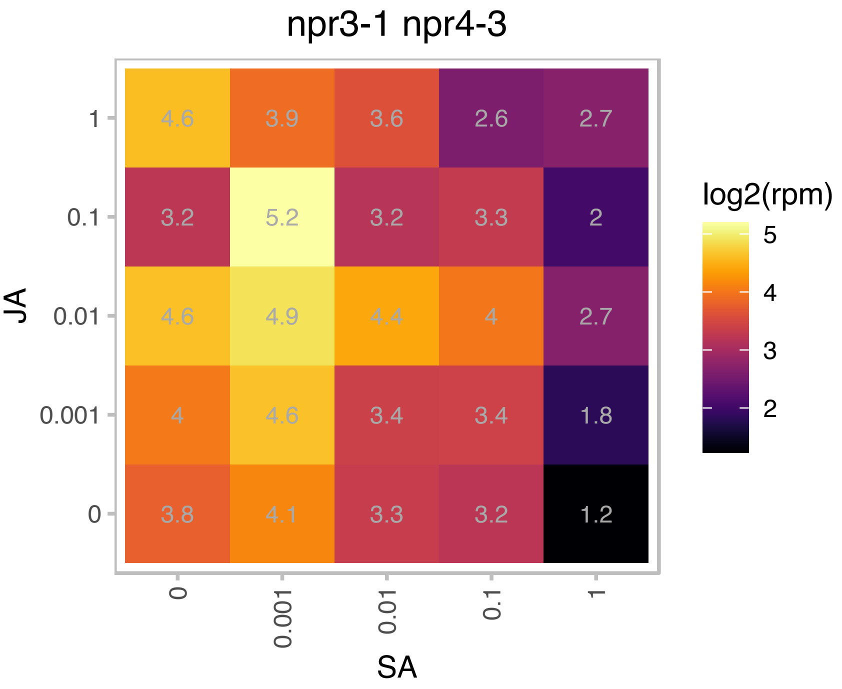

# AT1G23120.1 : NA

Polyketide cyclase/dehydrase and lipid transport superfamily protein

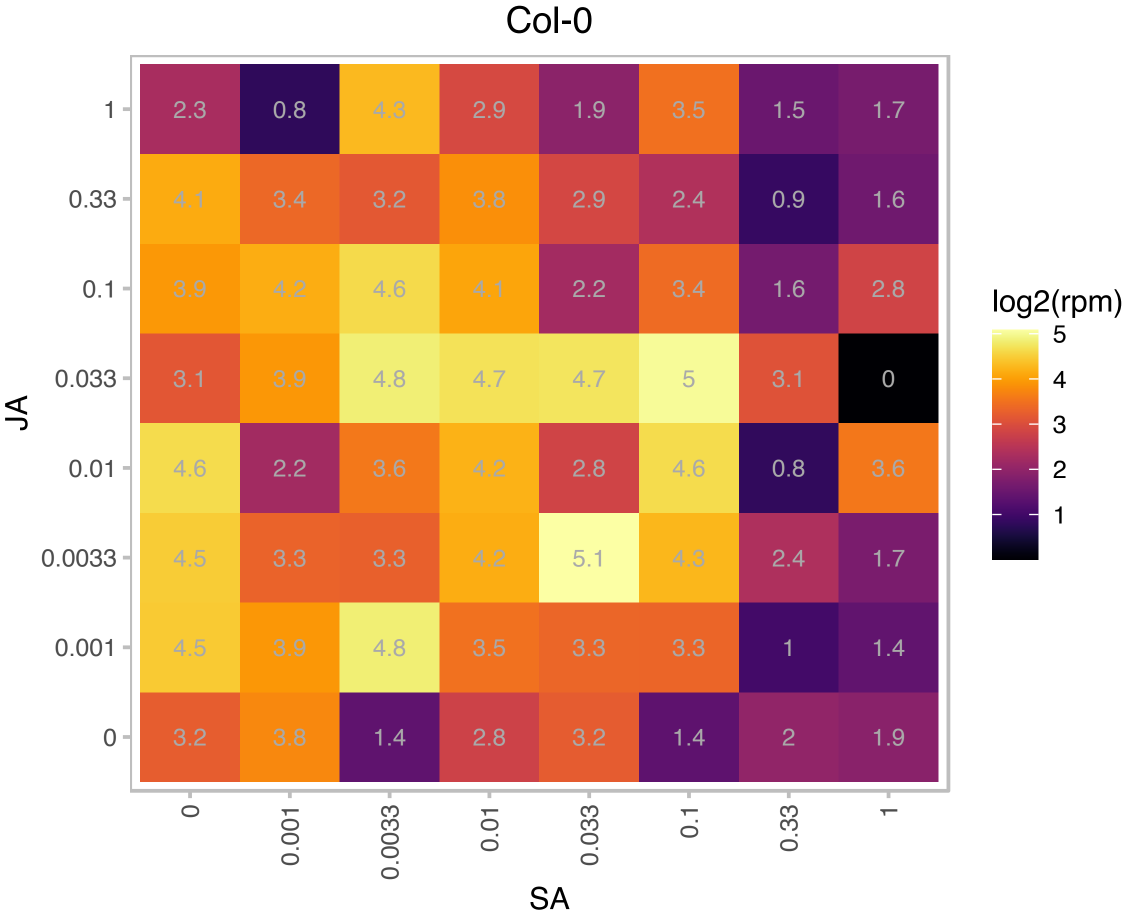


| | |
|---|---|
| Gene | AT1G23120 |
| is.expressed_genes | TRUE |
| is.responsive_genes | TRUE |
| is.apcluster | E-01 |

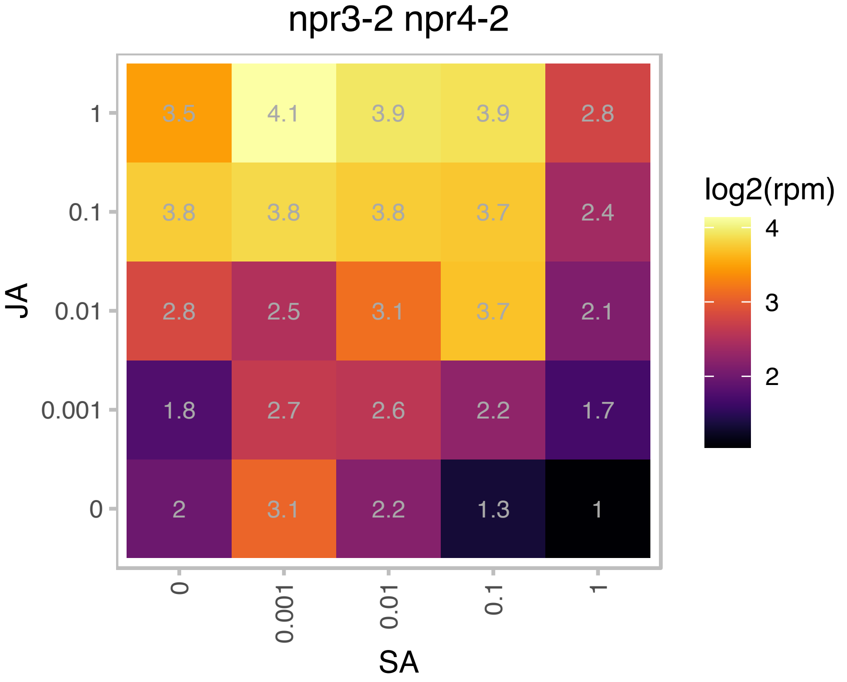


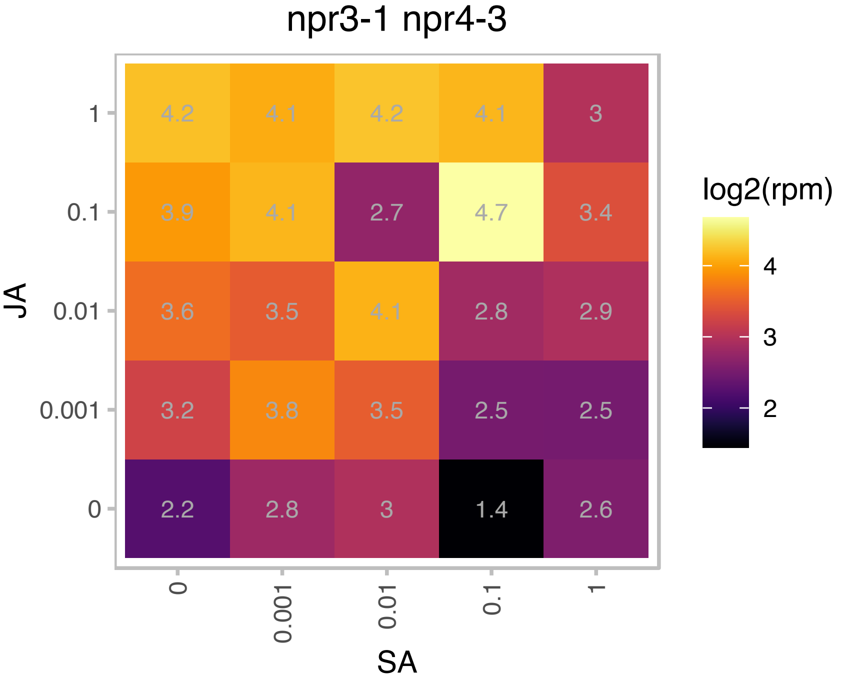

### AT1G25220.1 : ASB1, TRP4, WEI7

anthranilate synthase beta subunit 1

Catalyzes the first step of tryptophan biosynthesis: Chorismate L-Glutamine = Anthranilate Pyruvate L-Glutamate. Functions as a heterocomplex with anthranilate synthase alpha subunit (ASA1 or ASA2).

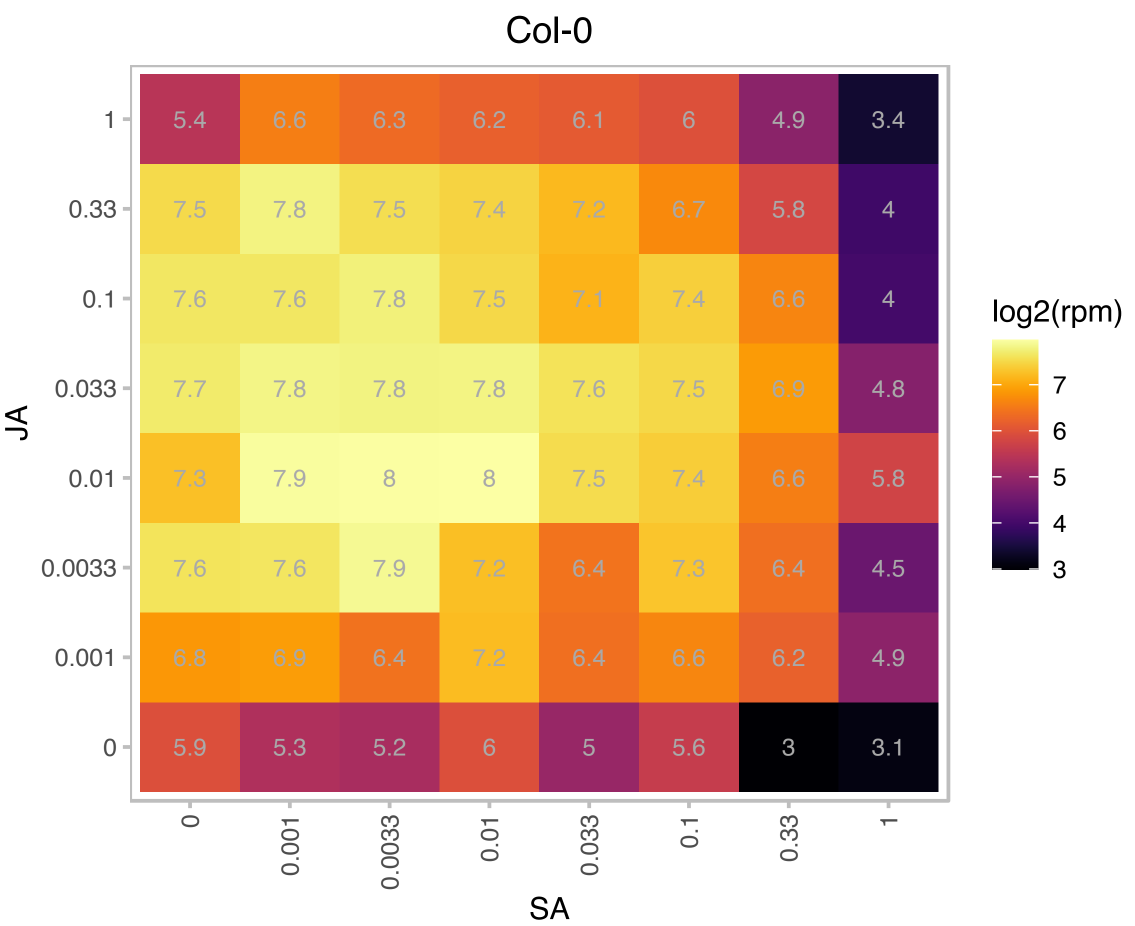


| | |
|---|---|
| Gene | AT1G25220 |
| is.expressed_genes | TRUE |
| is.responsive_genes | TRUE |
| is.apcluster | E-01 |

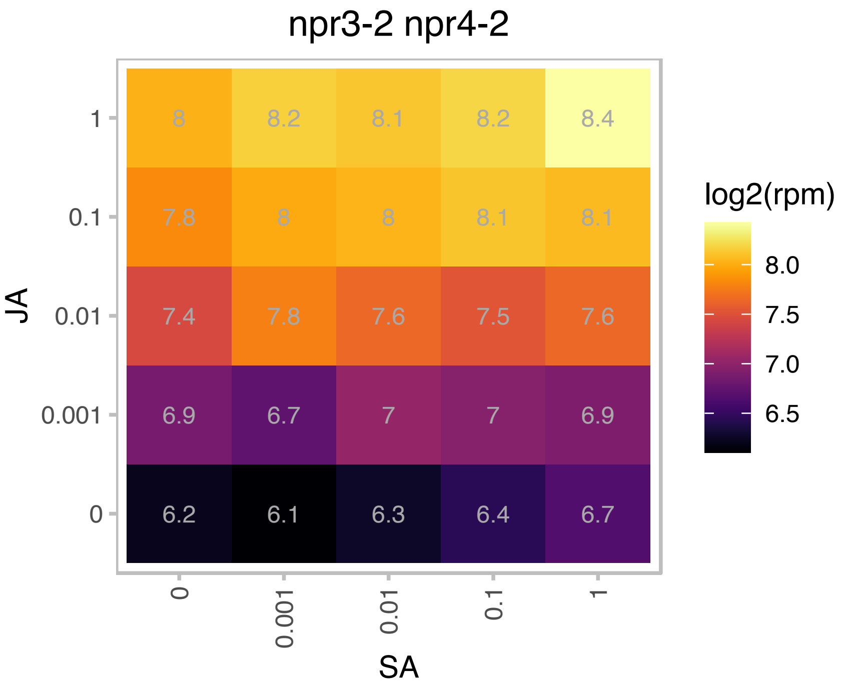


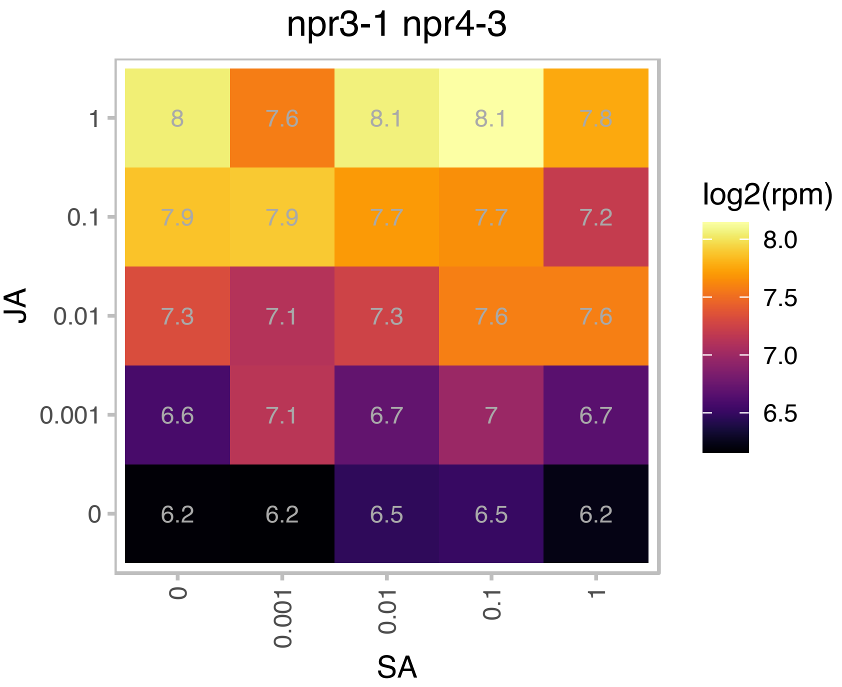

# AT1G26460.1 : NA

Tetratricopeptide repeat (TPR)-like superfamily protein

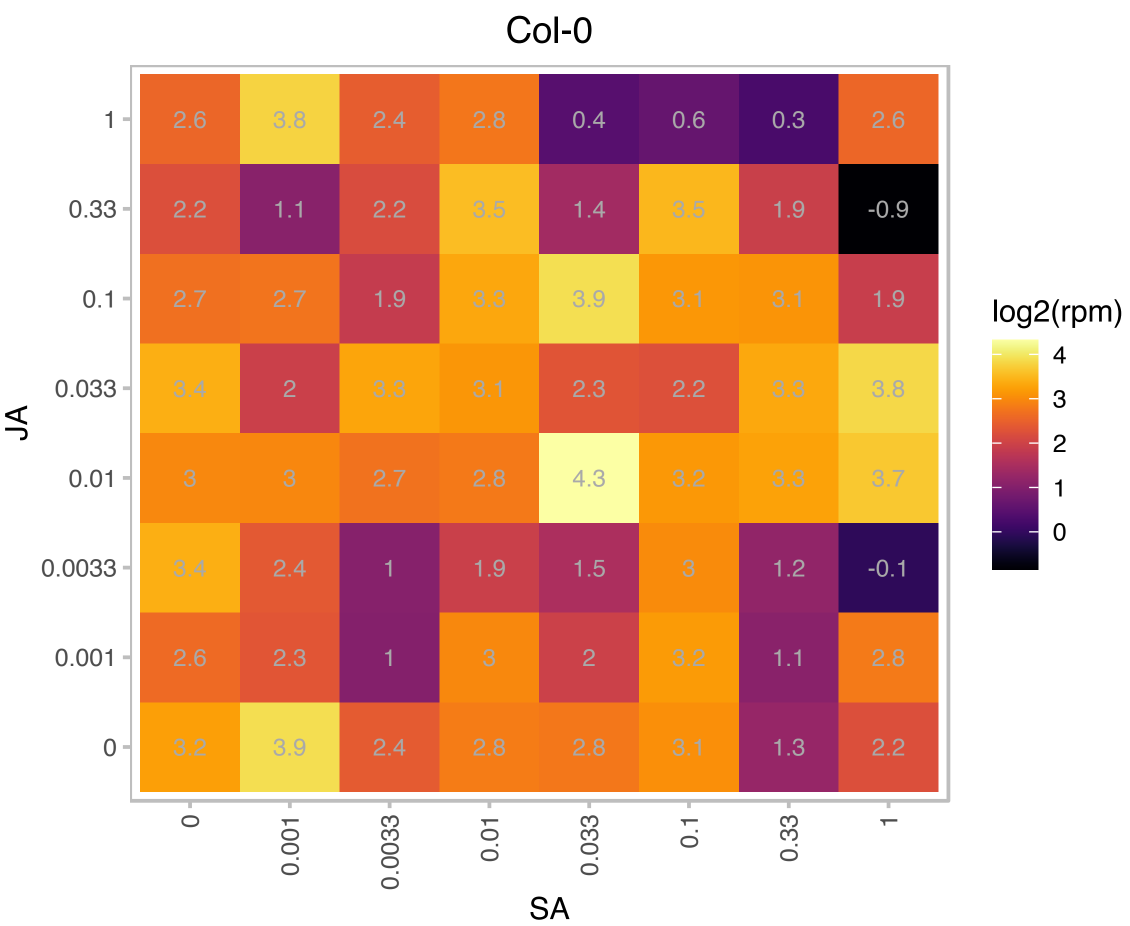


| | |
|---|---|
| Gene | AT1G26460 |
| is.expressed_genes | TRUE |
| is.responsive_genes | TRUE |
| is.apcluster | E-01 |

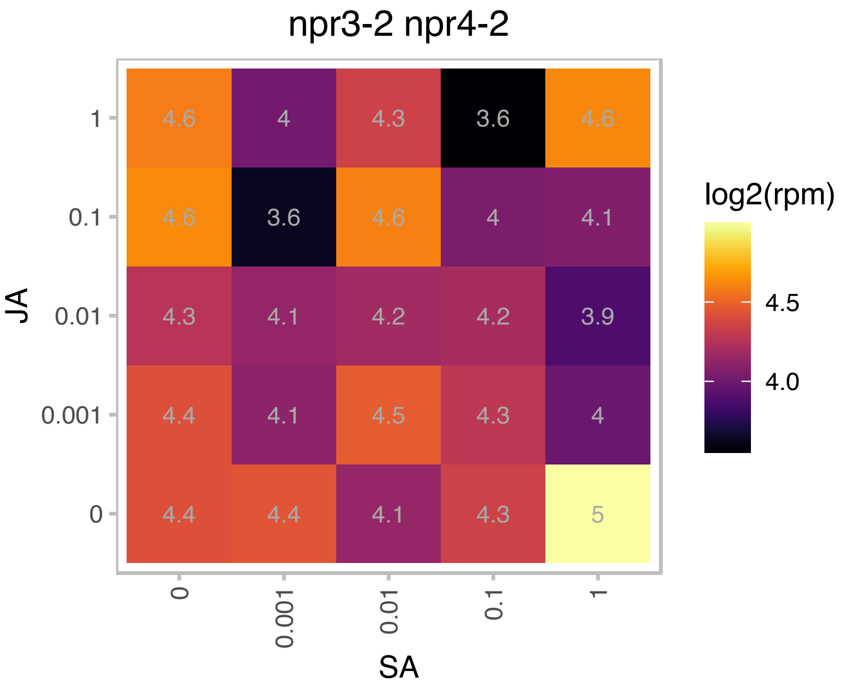


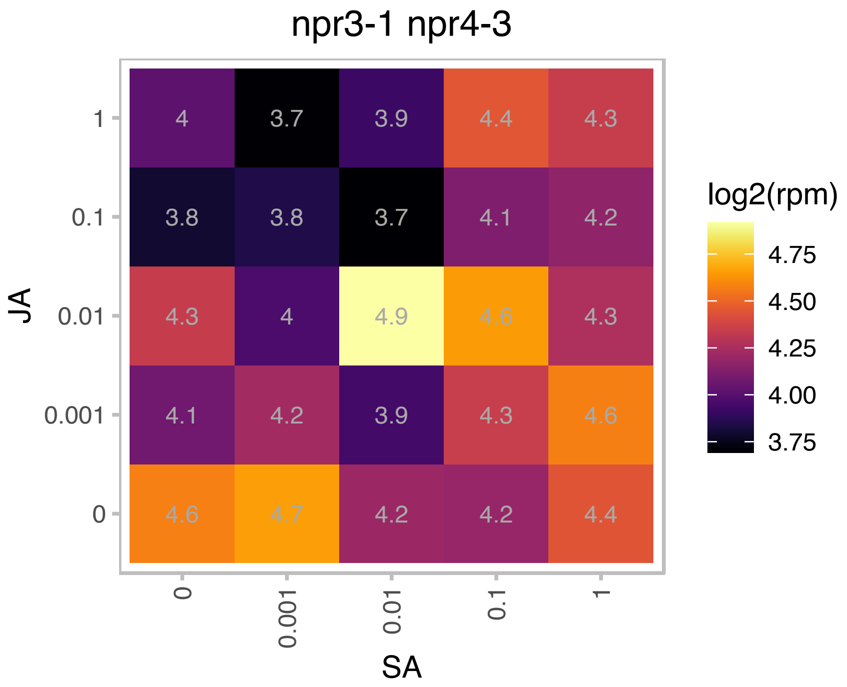

### AT1G31660.1 : atENP1, ENP1

Encodes a protein that is a ribosome biogenesis co-factor. Mutants display aberrant RNA processing and male and female gametophyte development.

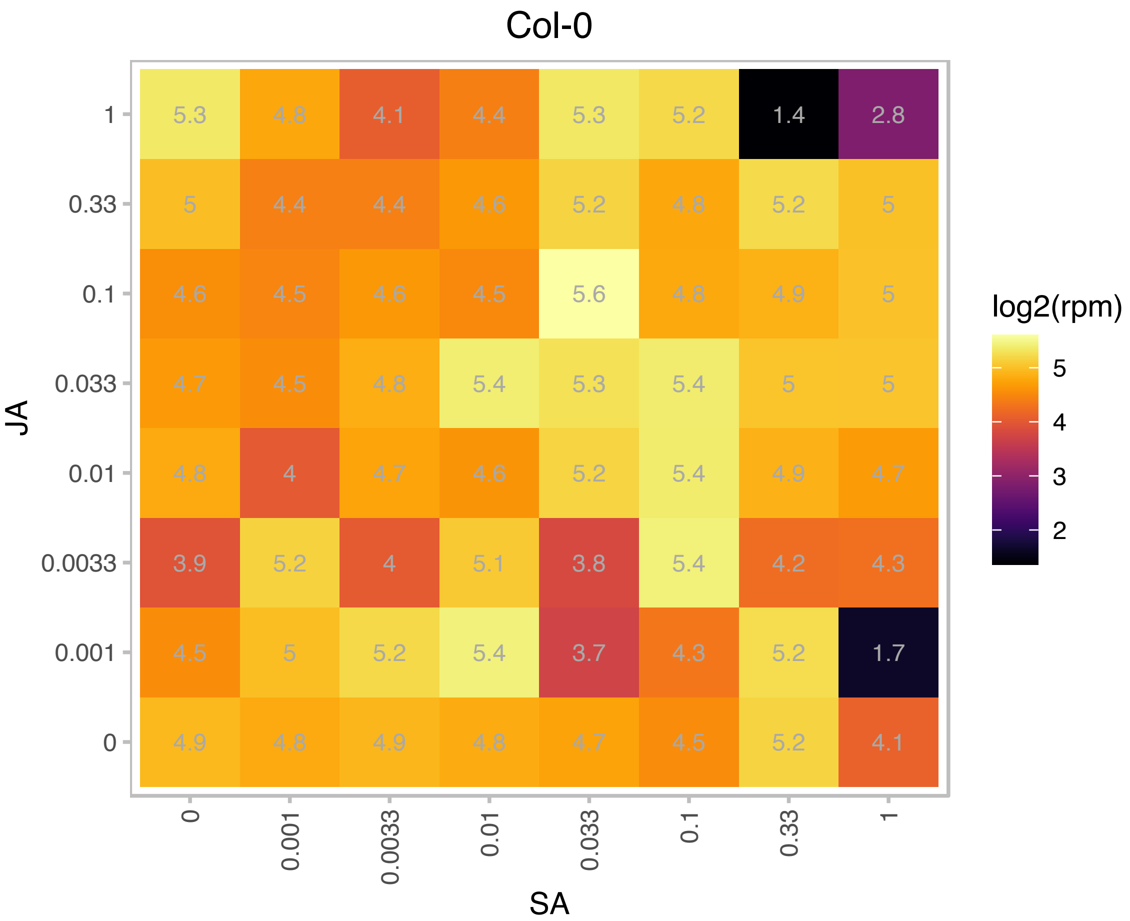


| | |
|---|---|
| Gene | AT1G31660 |
| is.expressed_genes | TRUE |
| is.responsive_genes | TRUE |
| is.apcluster | E-01 |

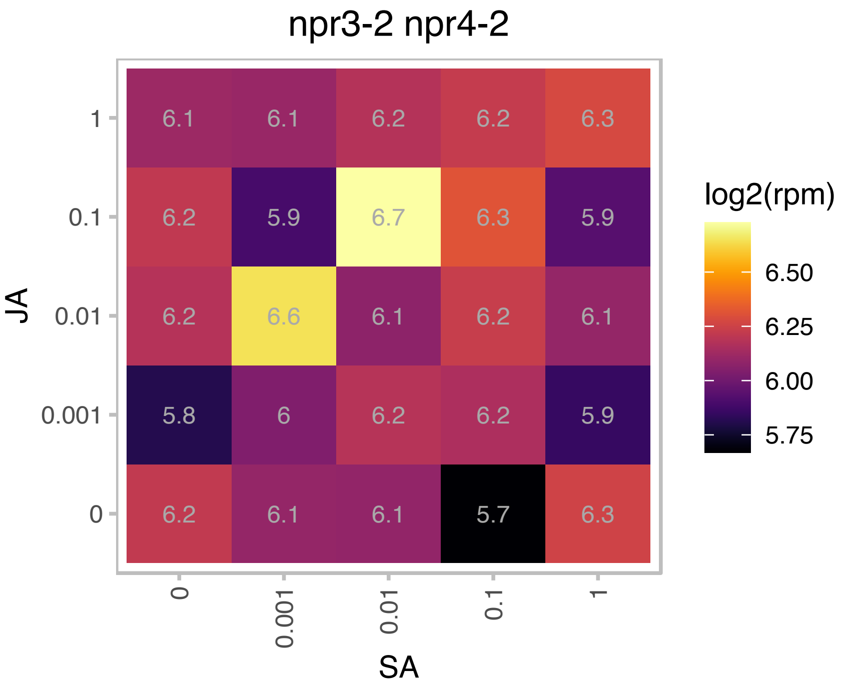


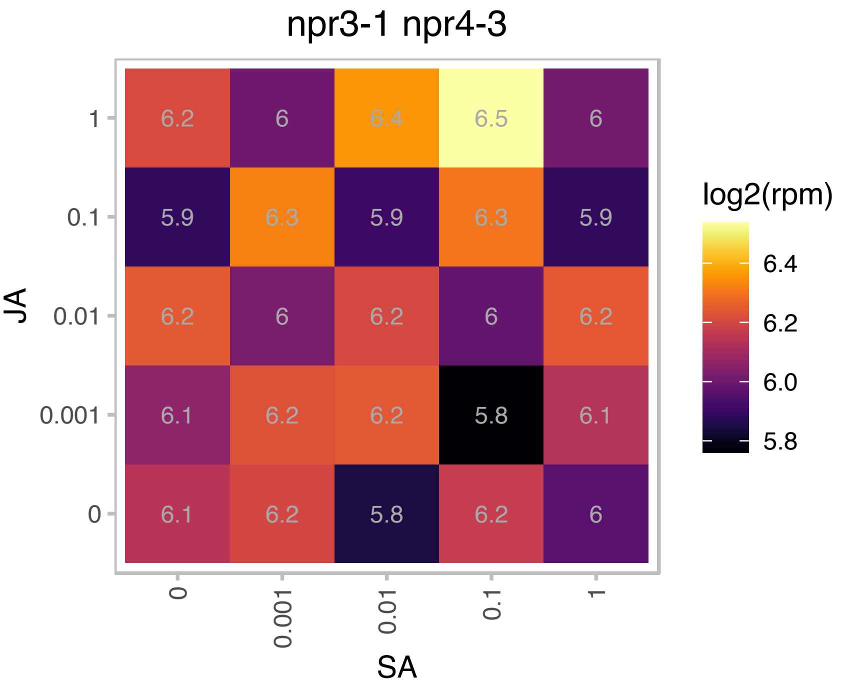

### AT1G32540.1 : LOL1

lsd one like 1

Encodes a protein with 3 plant-specific zinc finger domains that acts as a positive regulator of cell death.

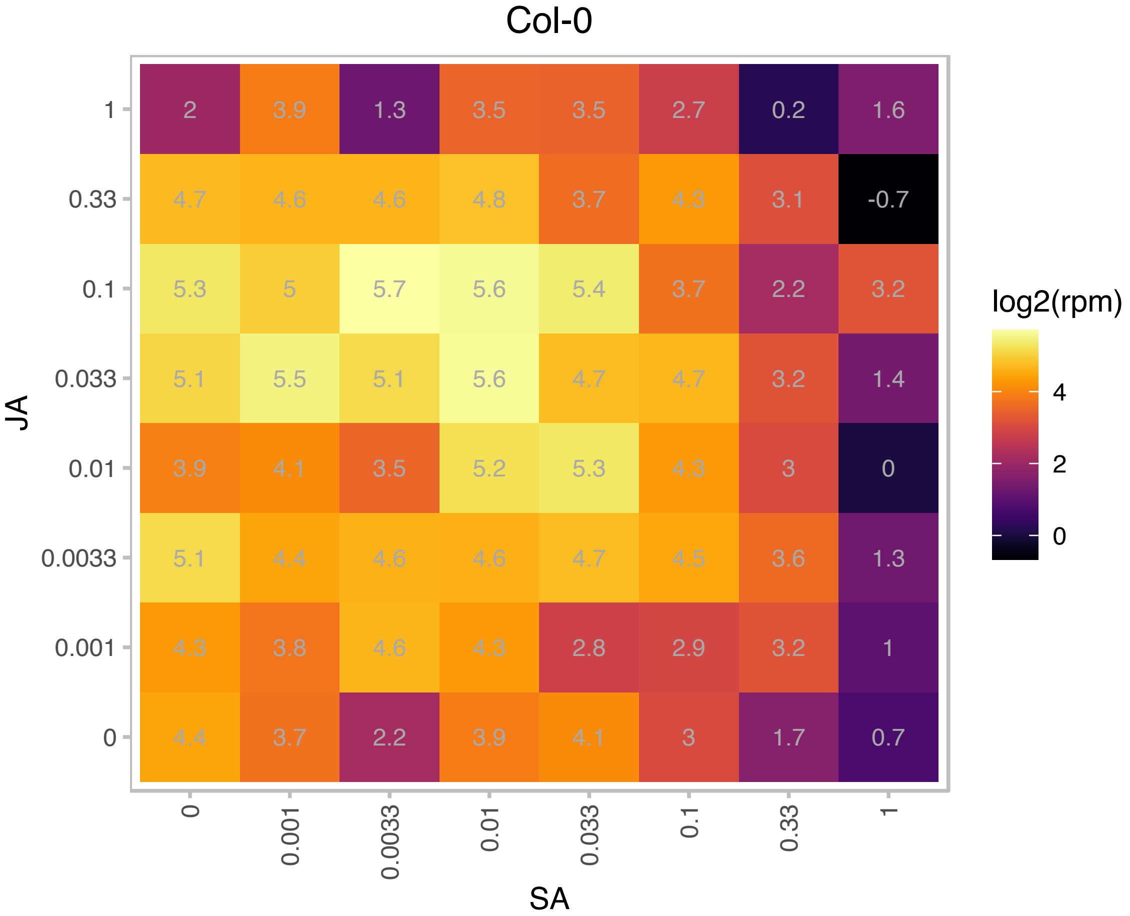


| | |
|---|---|
| Gene | AT1G32540 |
| is.expressed_genes | TRUE |
| is.responsive_genes | TRUE |
| is.apcluster | E-01 |

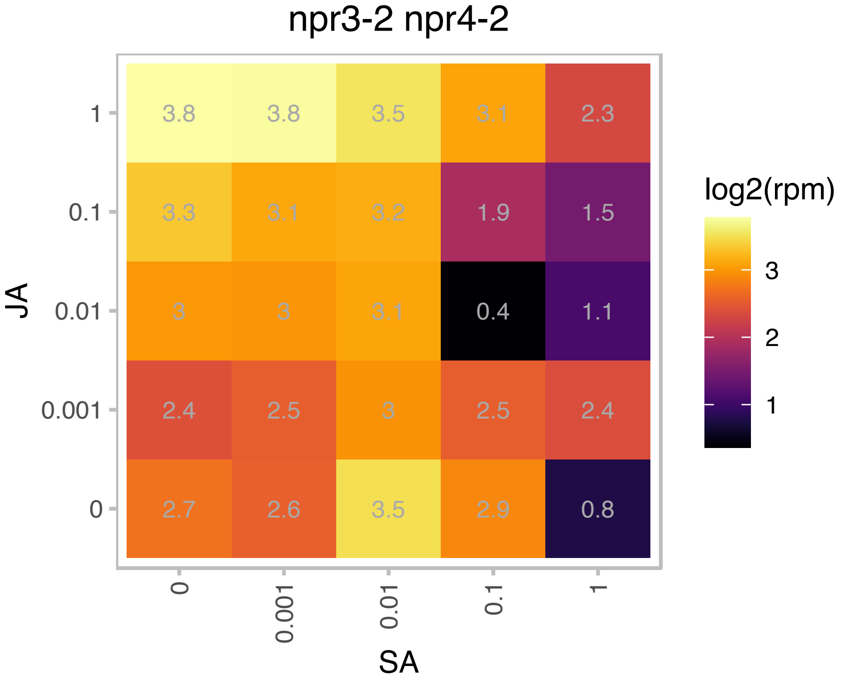


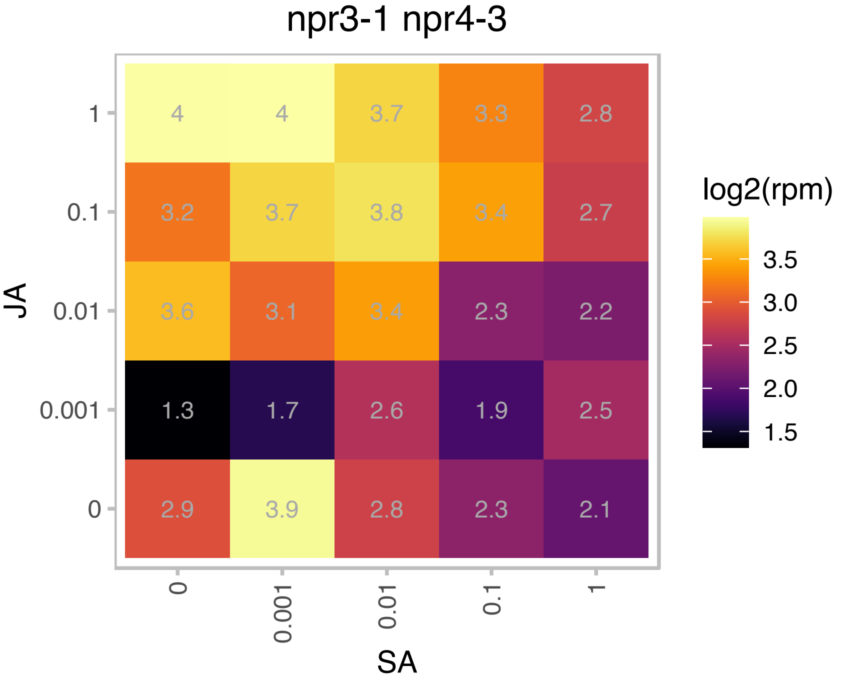

# AT1G52100.1 : NA

Mannose-binding lectin superfamily protein

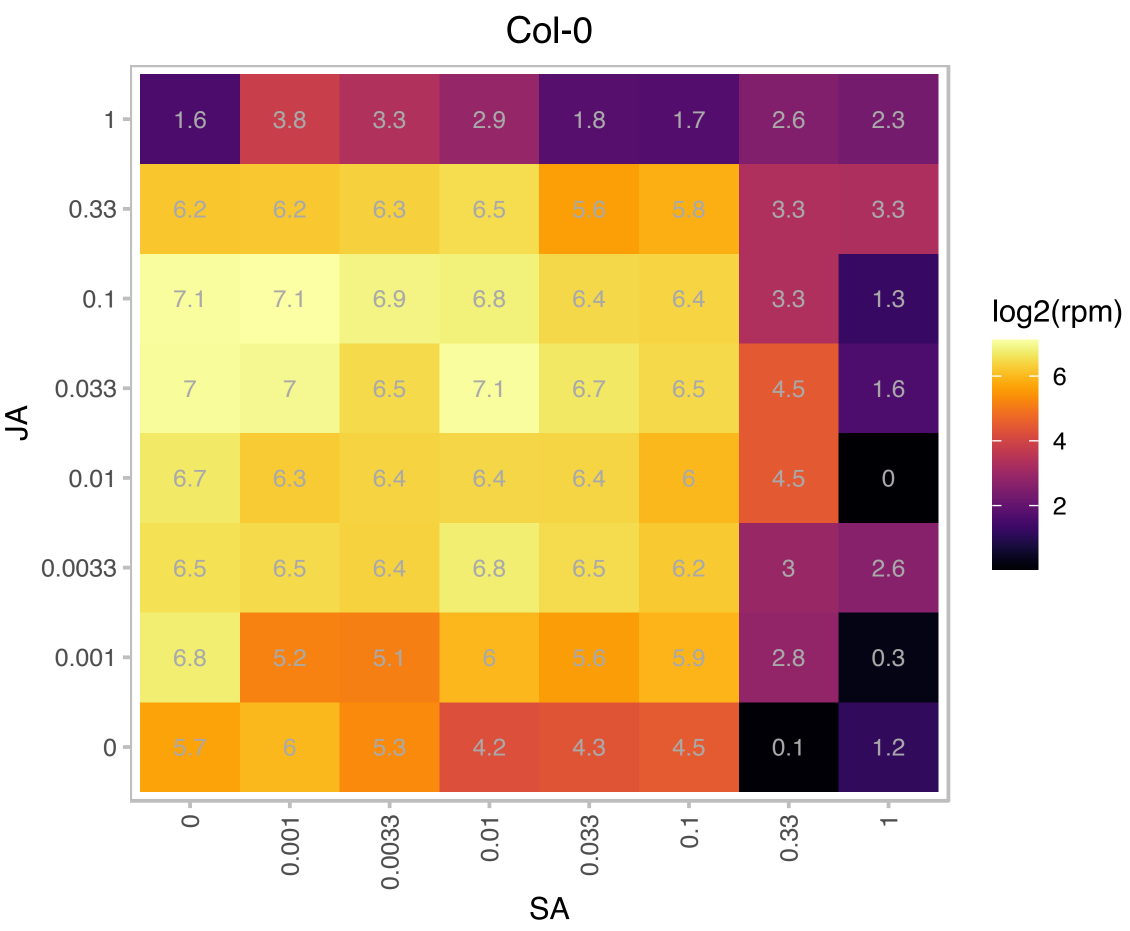


| | |
|---|---|
| Gene | AT1G52100 |
| is.expressed_genes | TRUE |
| is.responsive_genes | TRUE |
| is.apcluster | E-01 |

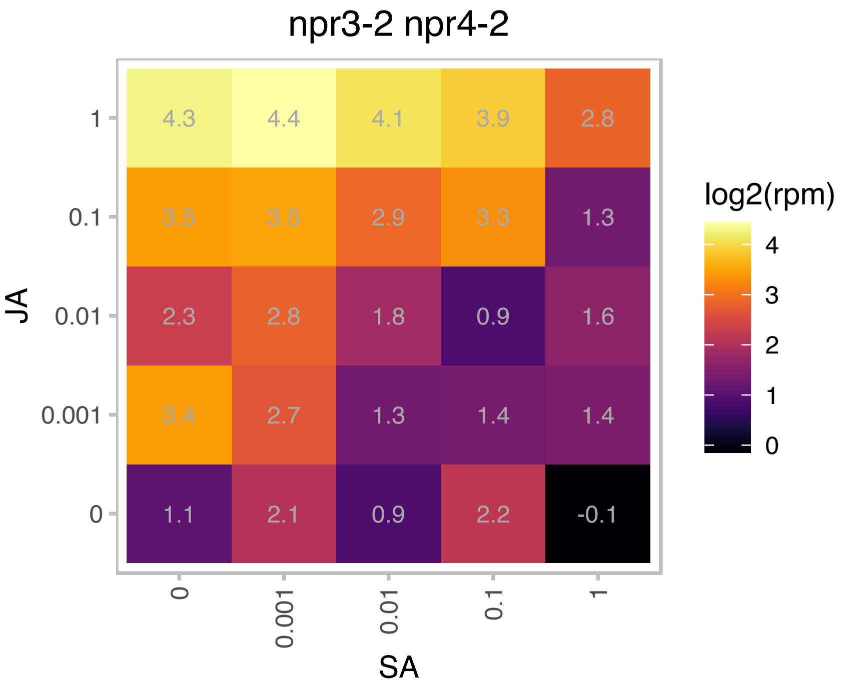


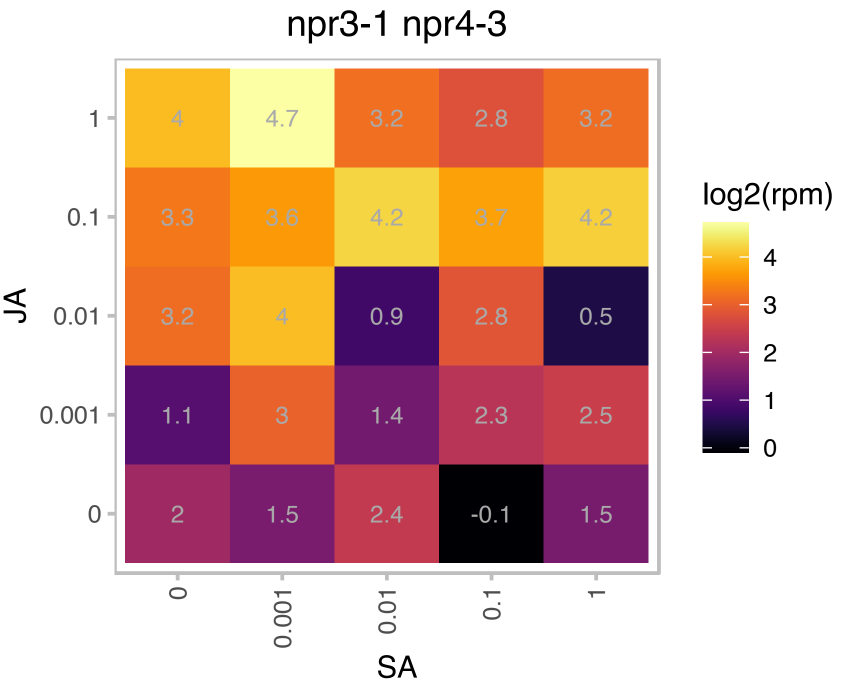

### AT1G54010.1 : GLL23

GDSL-like Lipase/Acylhydrolase superfamily protein

GDSL-motif esterase/acyltransferase/lipase. Enzyme group with broad substrate specificity that may catalyze acyltransfer or hydrolase reactions with lipid and non-lipid substrates.

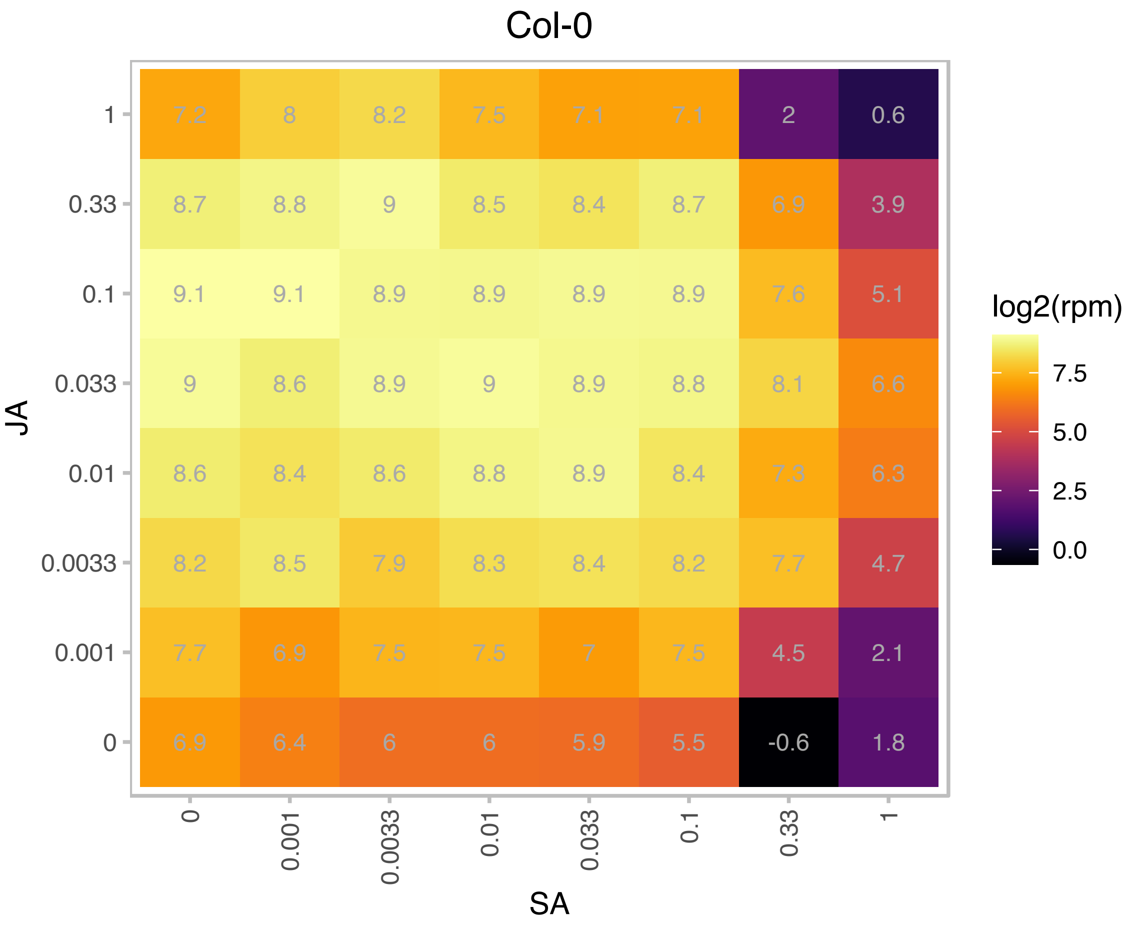


| | |
|---|---|
| Gene | AT1G54010 |
| is.expressed_genes | TRUE |
| is.responsive_genes | TRUE |
| is.apcluster | E-01 |

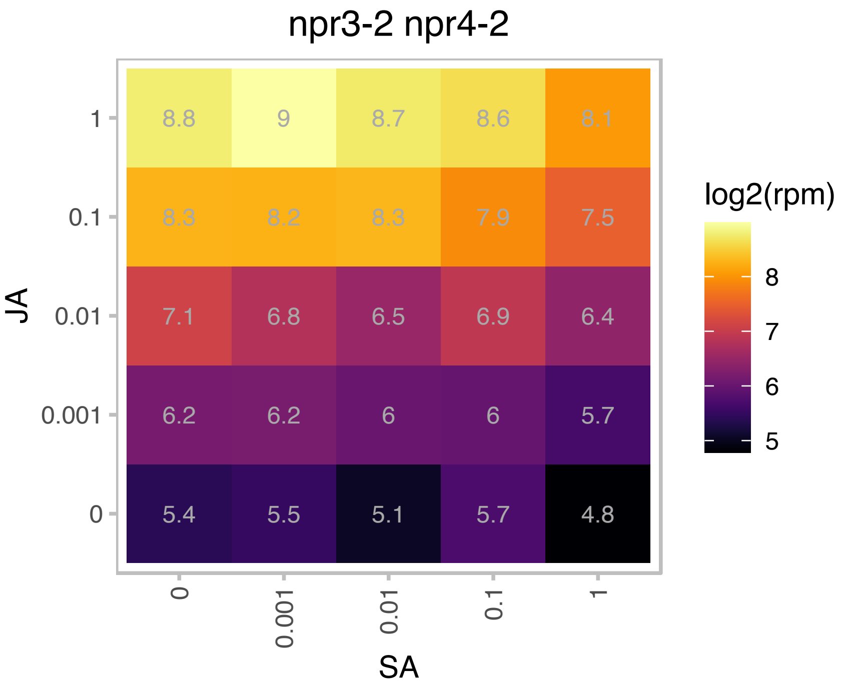


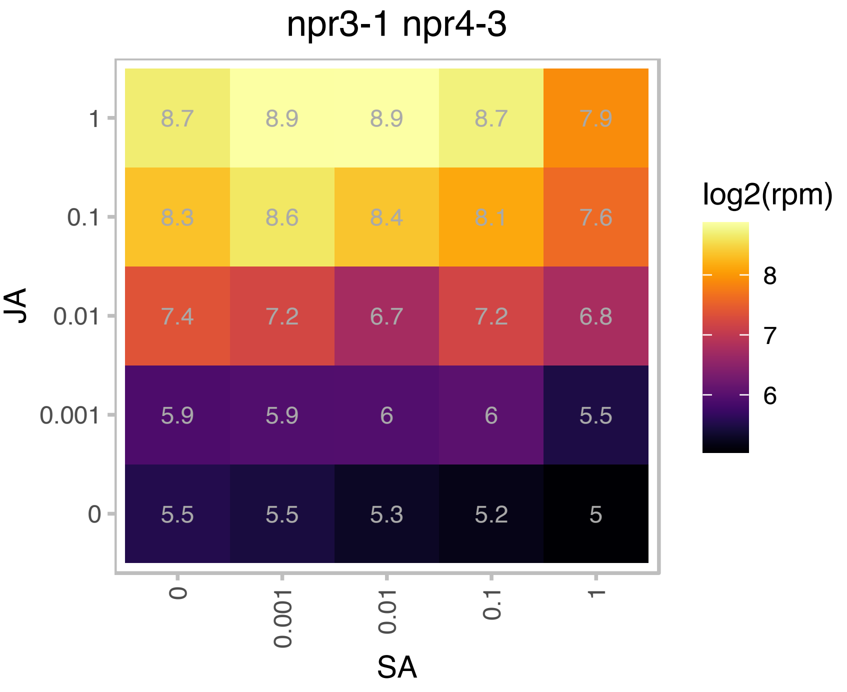

# AT1G54770.1 : NA

Fcf2 pre-rRNA processing protein

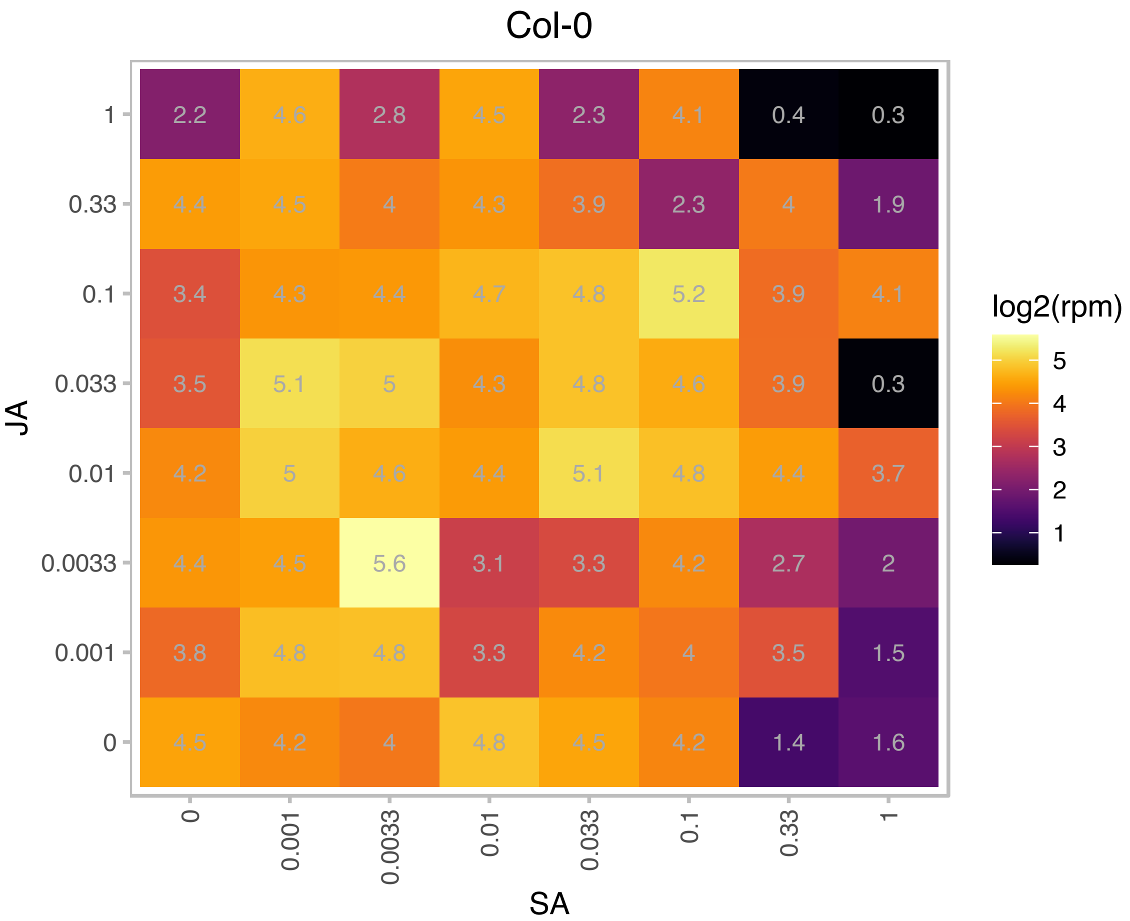


| | |
|---|---|
| Gene | AT1G54770 |
| is.expressed_genes | TRUE |
| is.responsive_genes | TRUE |
| is.apcluster | E-01 |

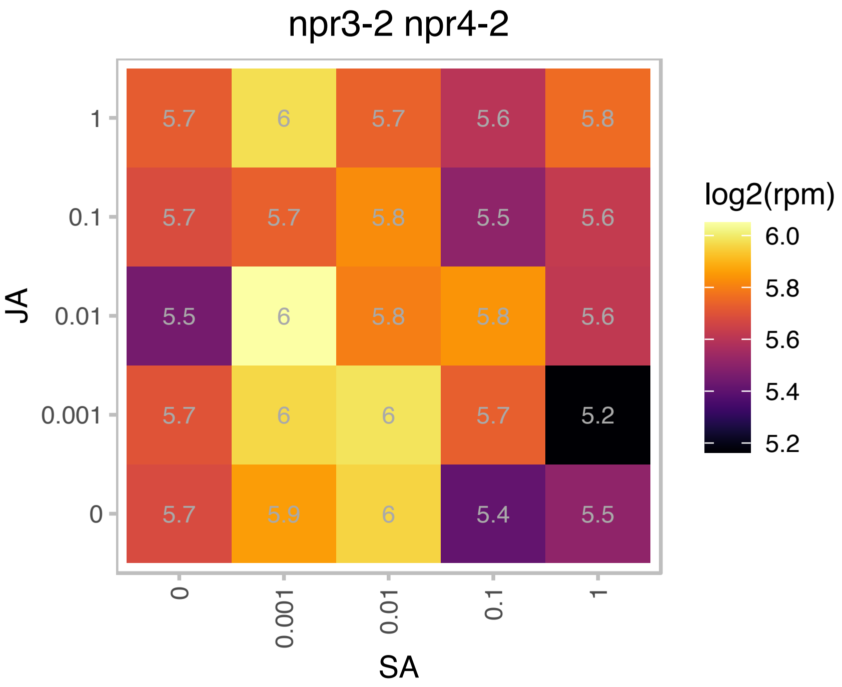


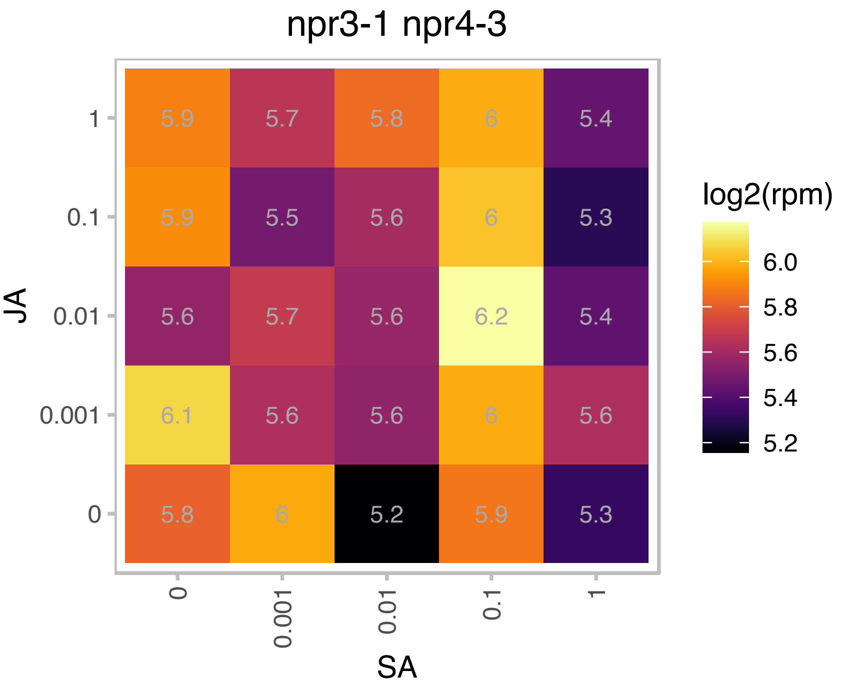

# AT1G69800.1 : NA

Cystathionine beta-synthase (CBS) protein

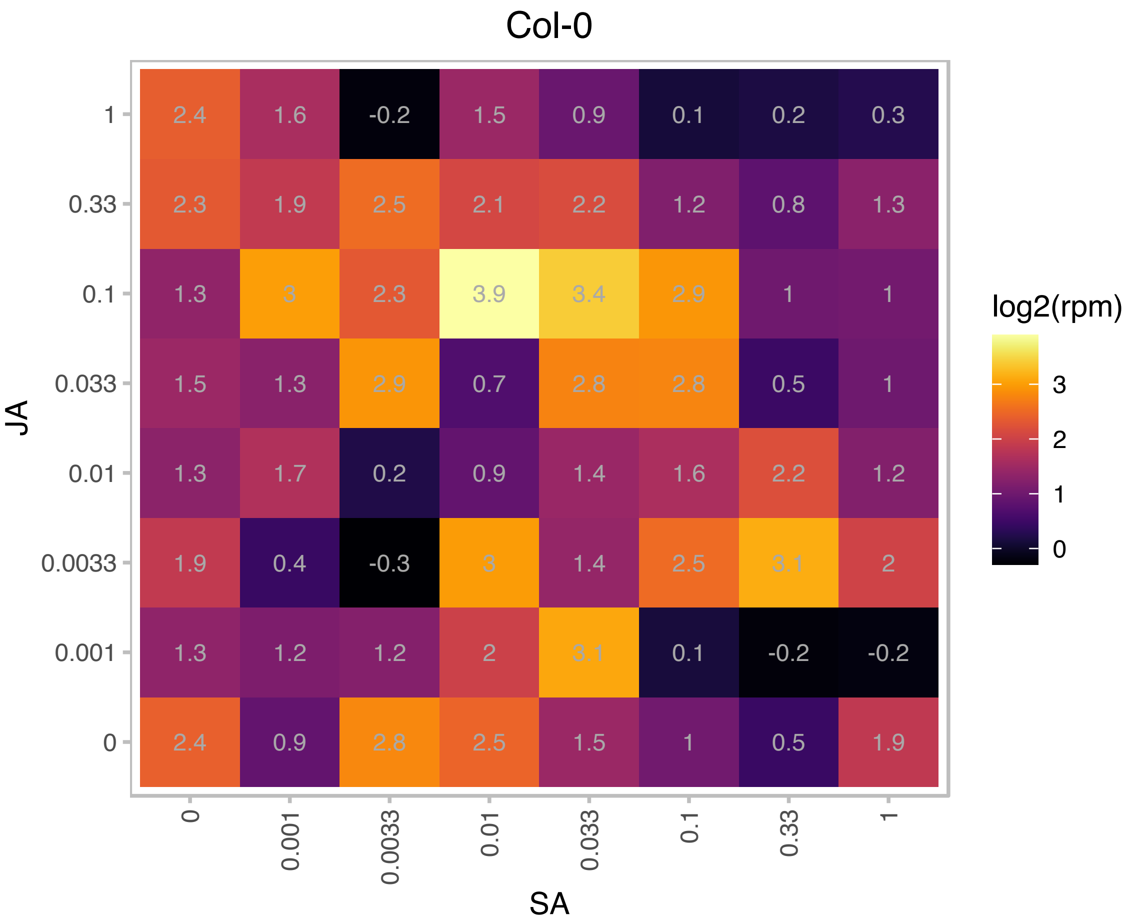


| | |
|---|---|
| Gene | AT1G69800 |
| is.expressed_genes | TRUE |
| is.responsive_genes | TRUE |
| is.apcluster | E-01 |

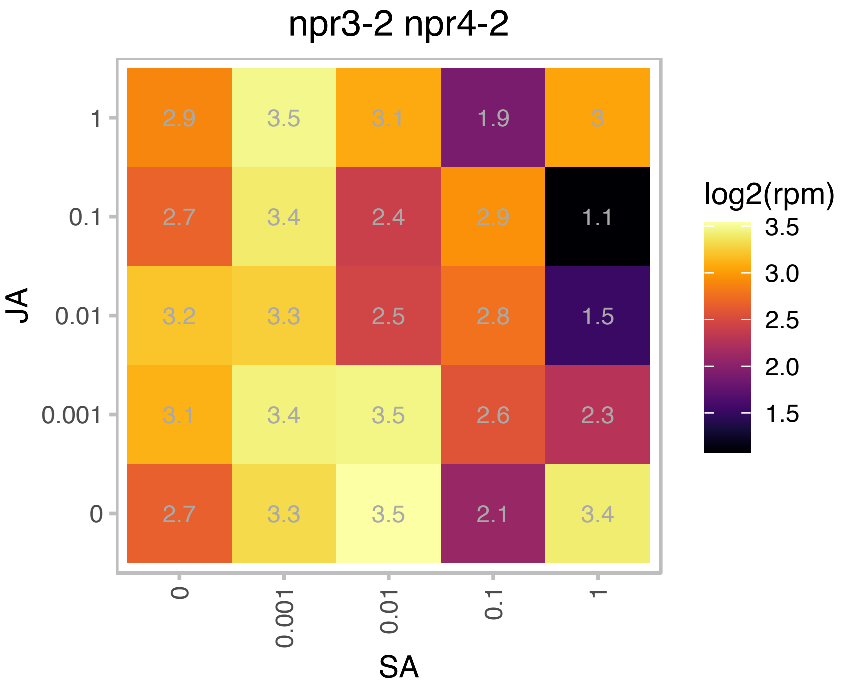


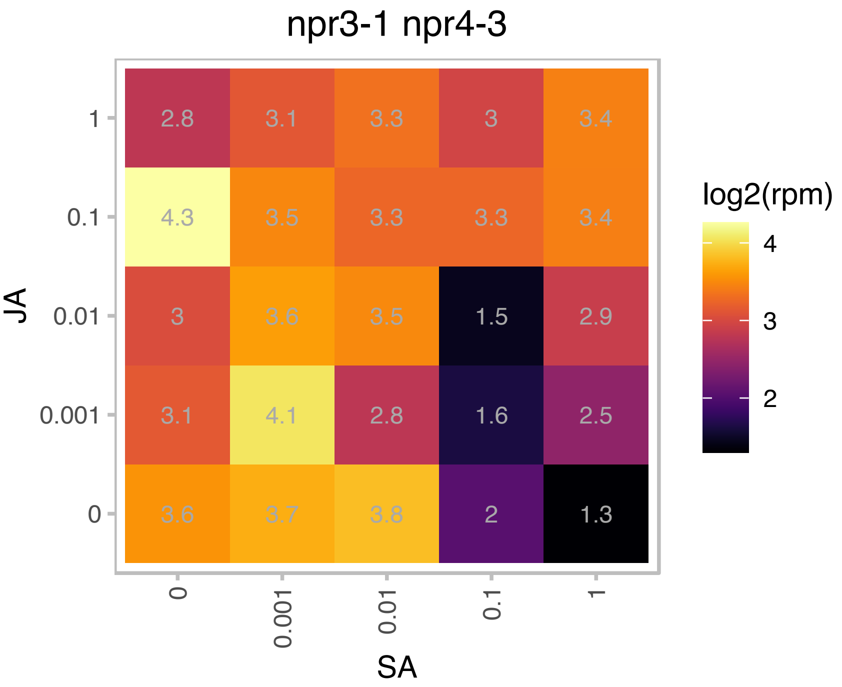

### AT1G72630.1 : ELF4-L2

ELF4-like 2

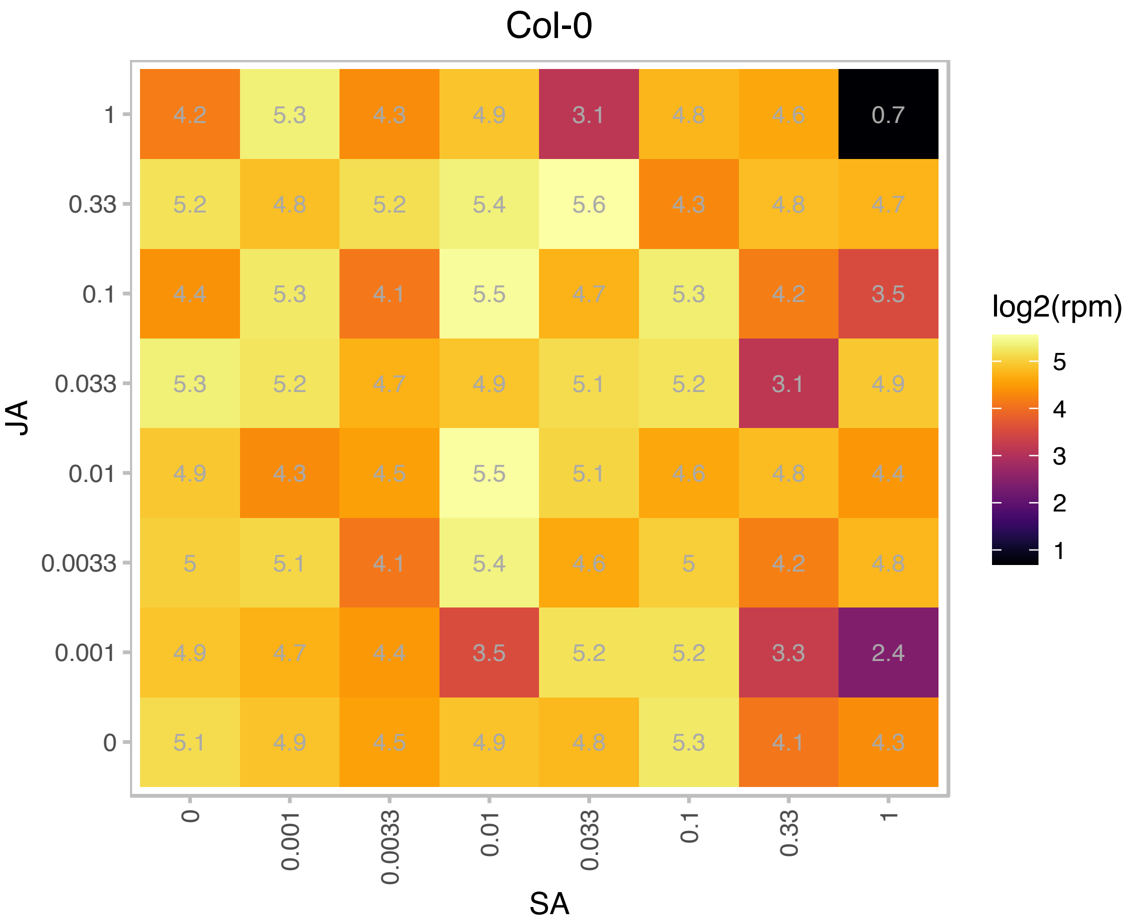


| | |
|---|---|
| Gene | AT1G72630 |
| is.expressed_genes | TRUE |
| is.responsive_genes | TRUE |
| is.apcluster | E-01 |

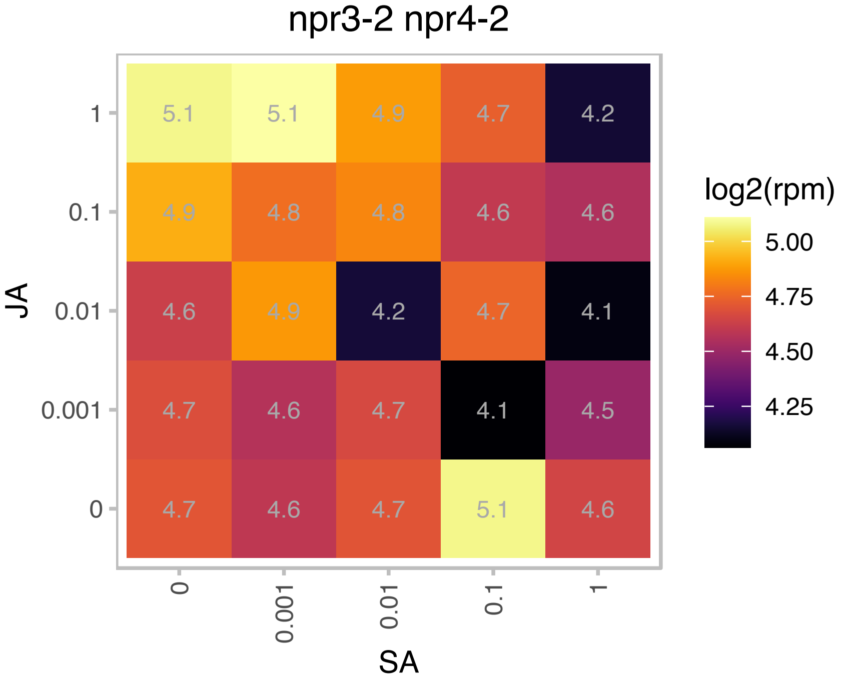


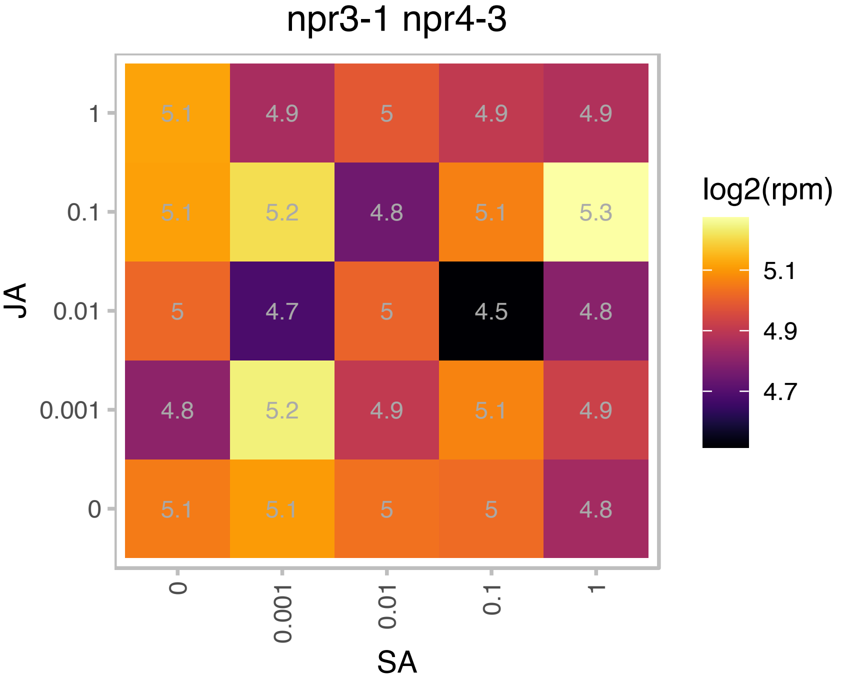

### AT1G74030.1 : ENO1

enolase 1

Encodes the plastid-localized phosphoenolpyruvate enolase. Mutant plants have abnormal trichomes.

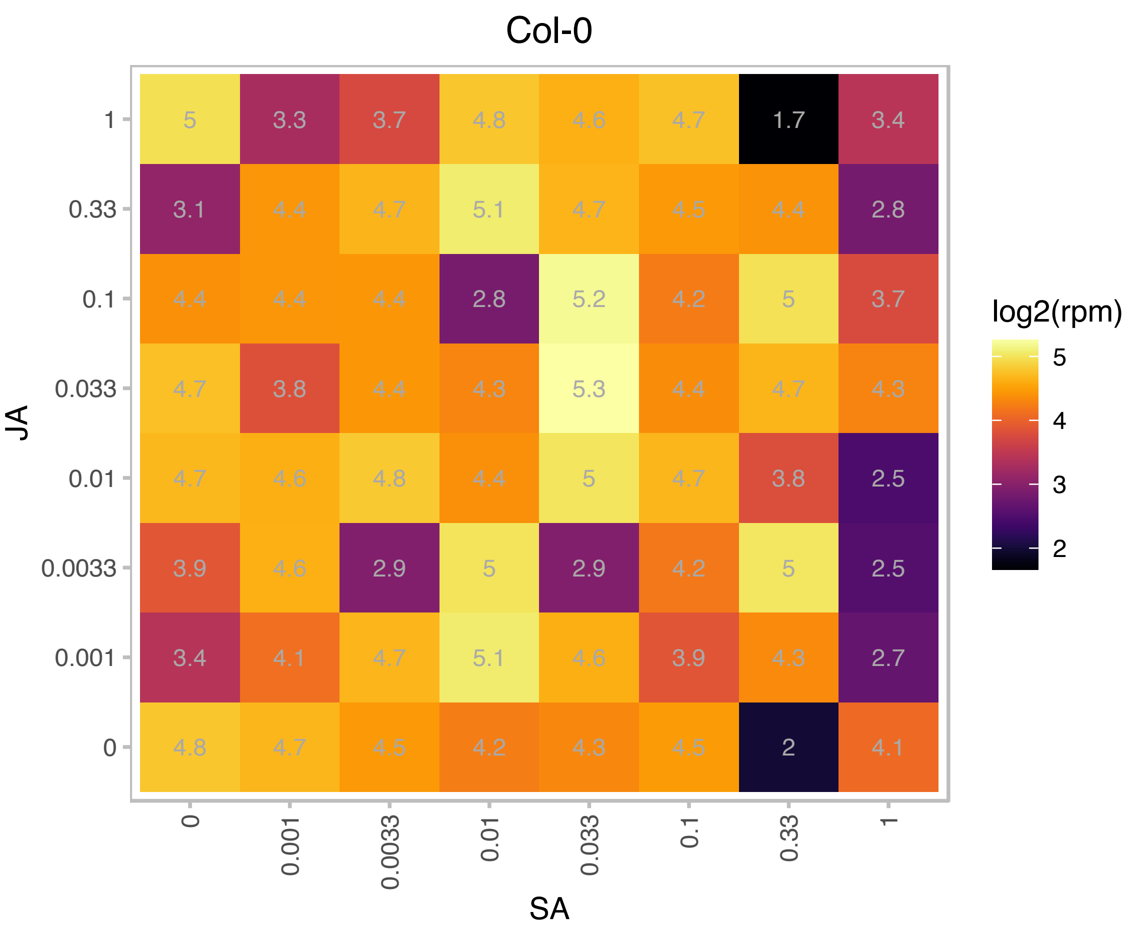


| | |
|---|---|
| Gene | AT1G74030 |
| is.expressed_genes | TRUE |
| is.responsive_genes | TRUE |
| is.apcluster | E-01 |

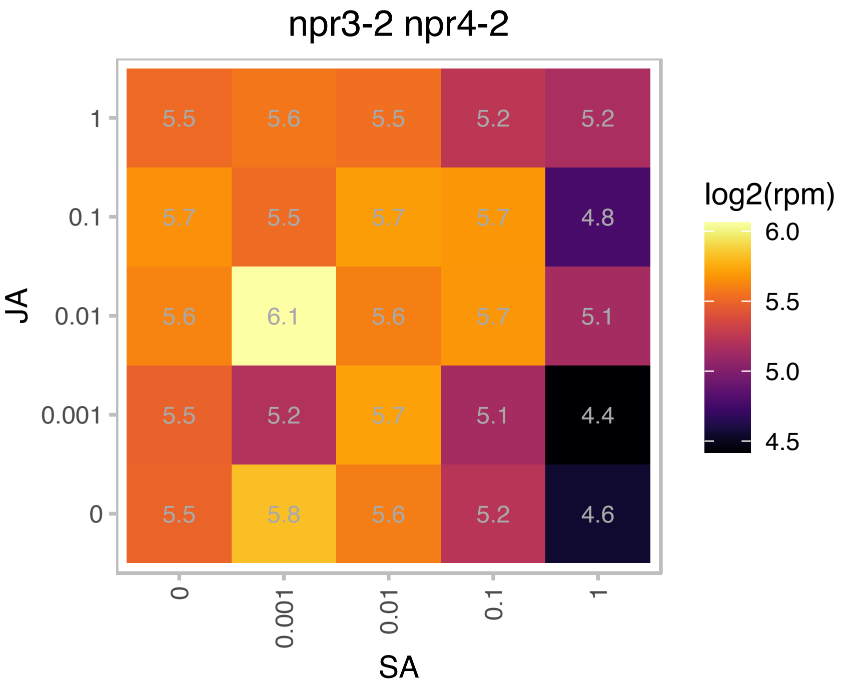


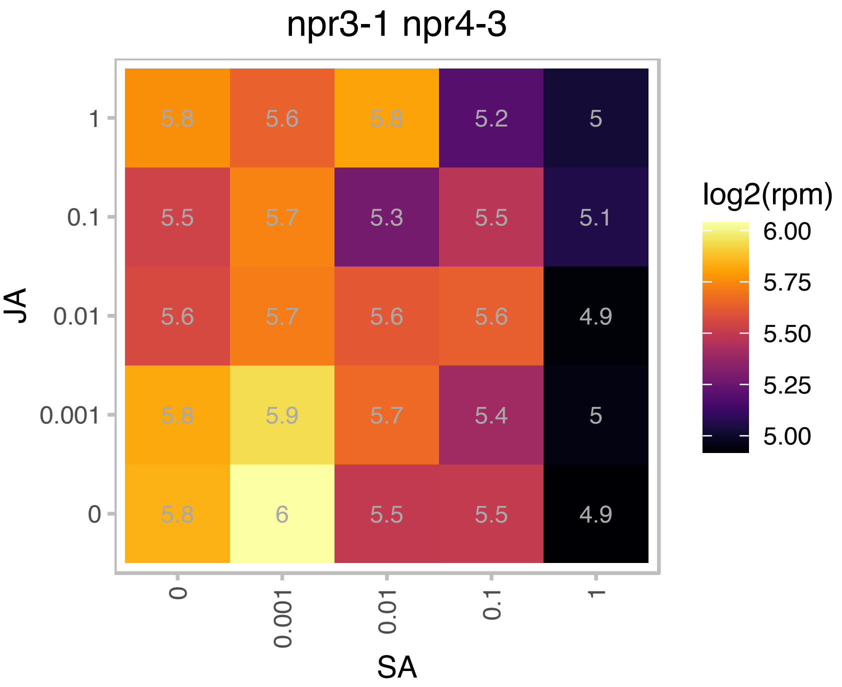

# AT1G77060.1 : NA

Phosphoenolpyruvate carboxylase family protein

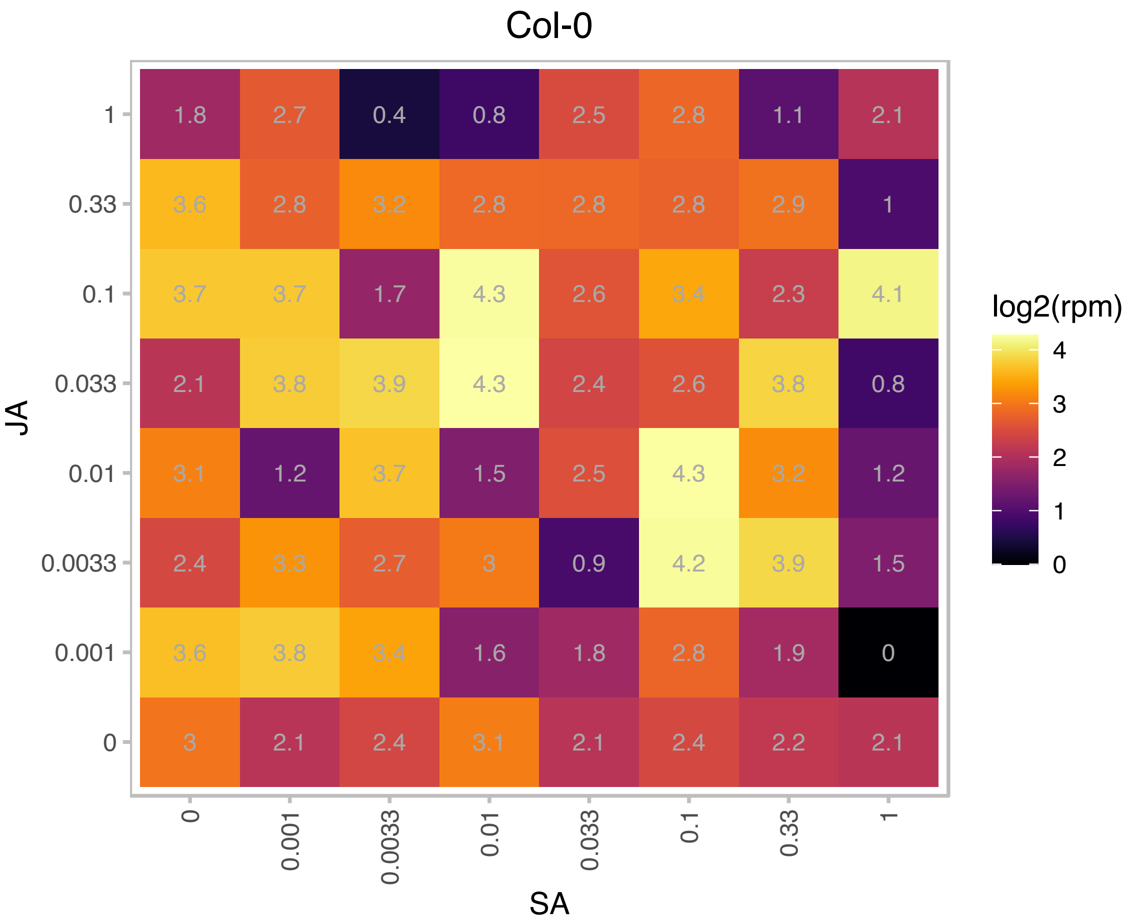


| | |
|---|---|
| Gene | AT1G77060 |
| is.expressed_genes | TRUE |
| is.responsive_genes | TRUE |
| is.apcluster | E-01 |

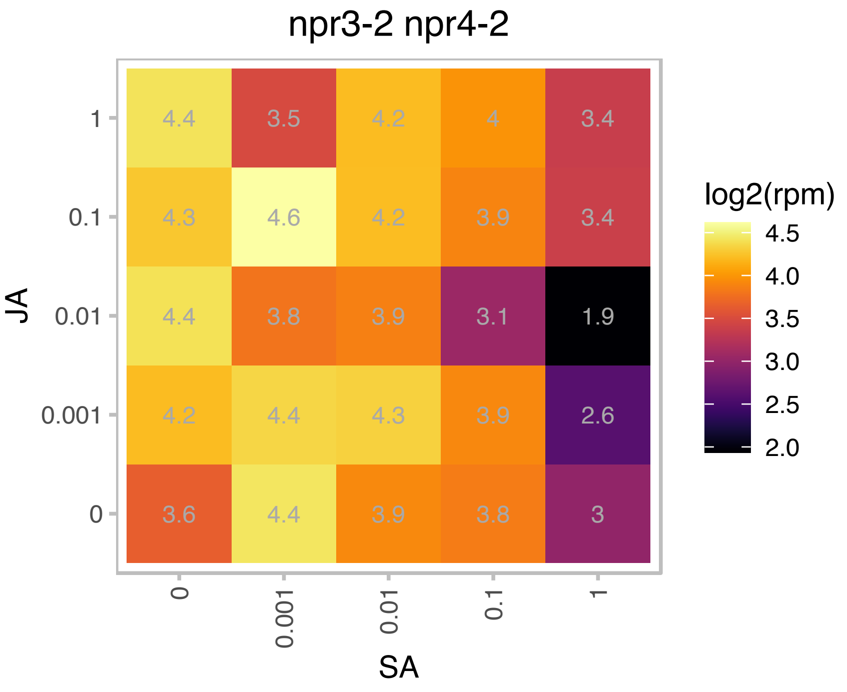


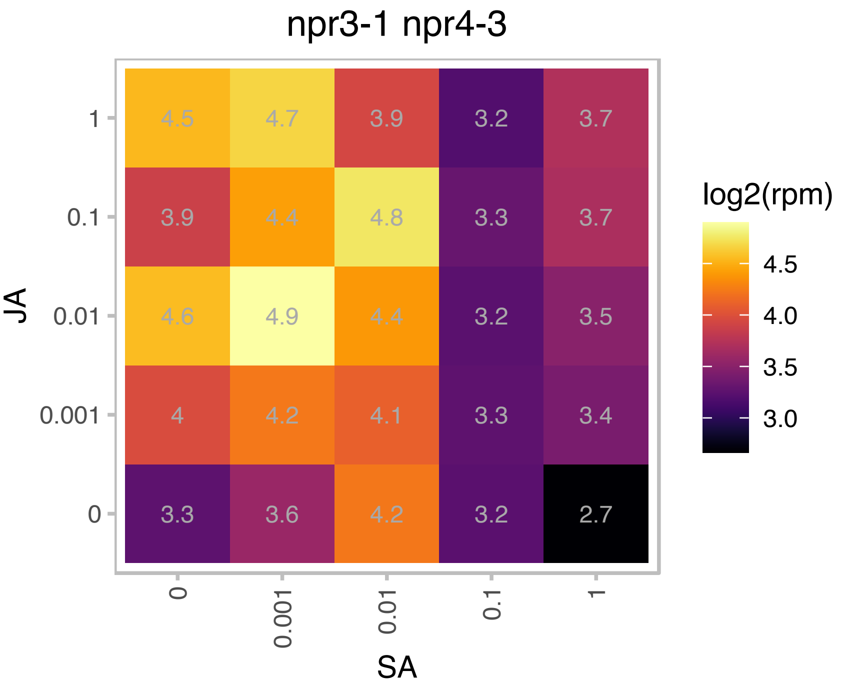

### AT2G20610.1 : ALF1, HLS3, RTY, RTY1, SUR1

Tyrosine transaminase family protein

Confers auxin overproduction. Mutants have an over-proliferation of lateral roots. Encodes a C-S lyase involved in converting S-alkylthiohydroximate to thiohydroximate in glucosinolate biosynthesis.

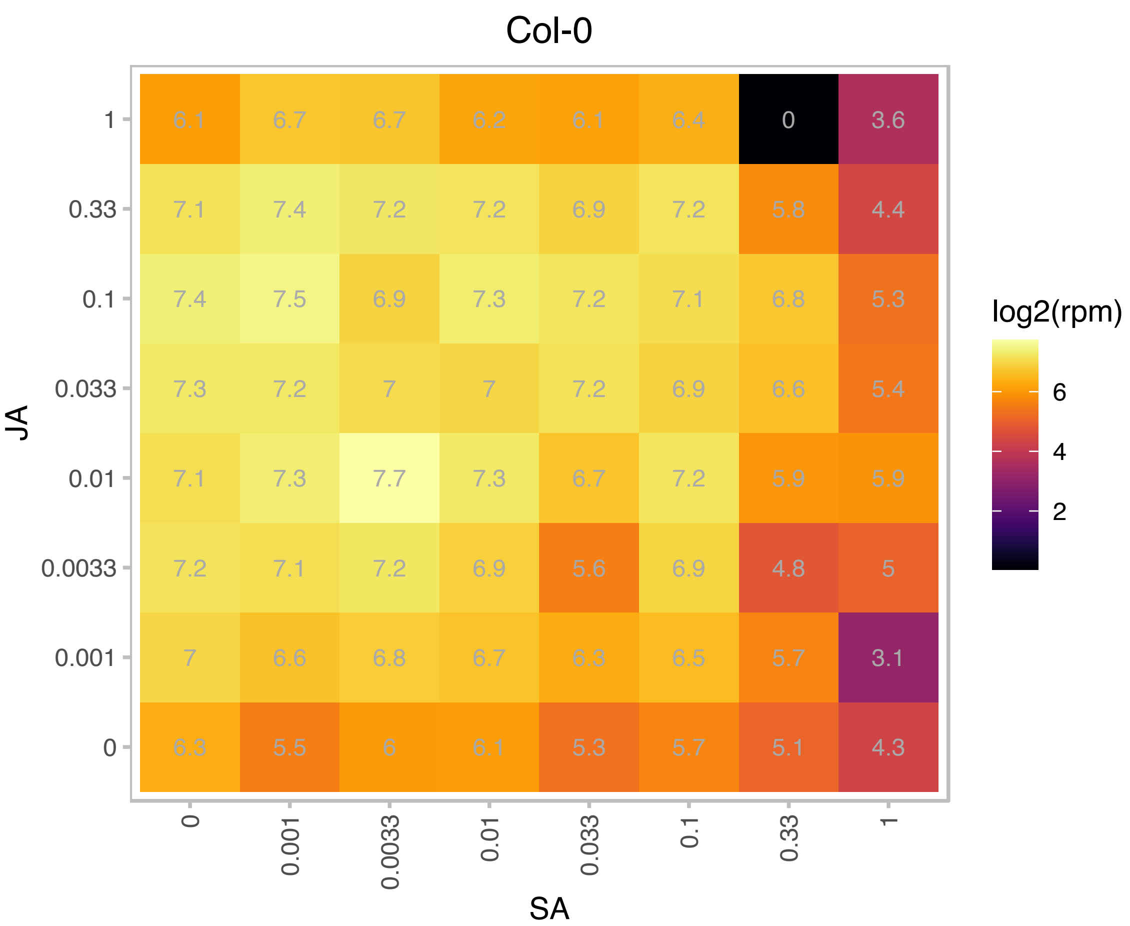


| | |
|---|---|
| Gene | AT2G20610 |
| is.expressed_genes | TRUE |
| is.responsive_genes | TRUE |
| is.apcluster | E-01 |

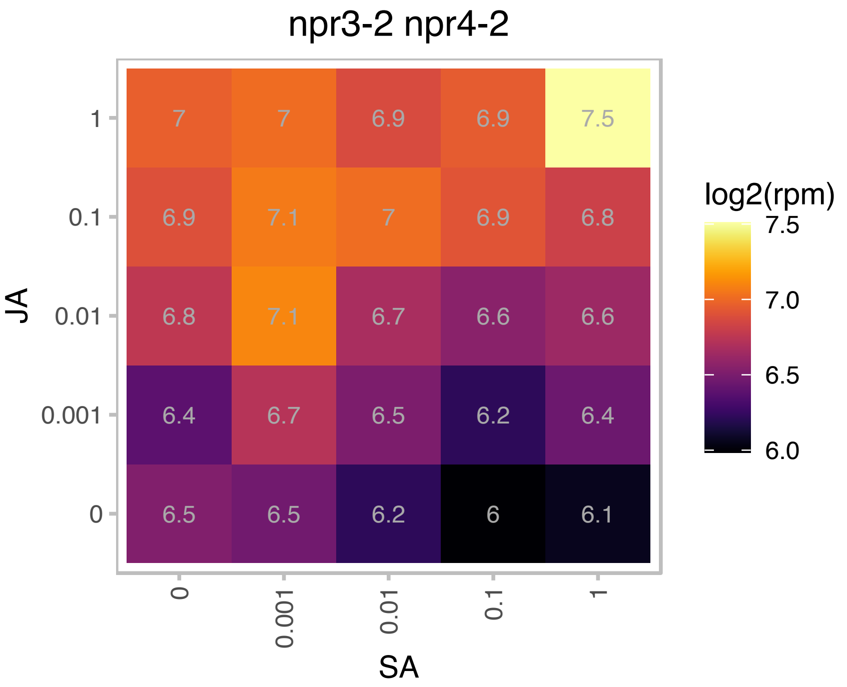


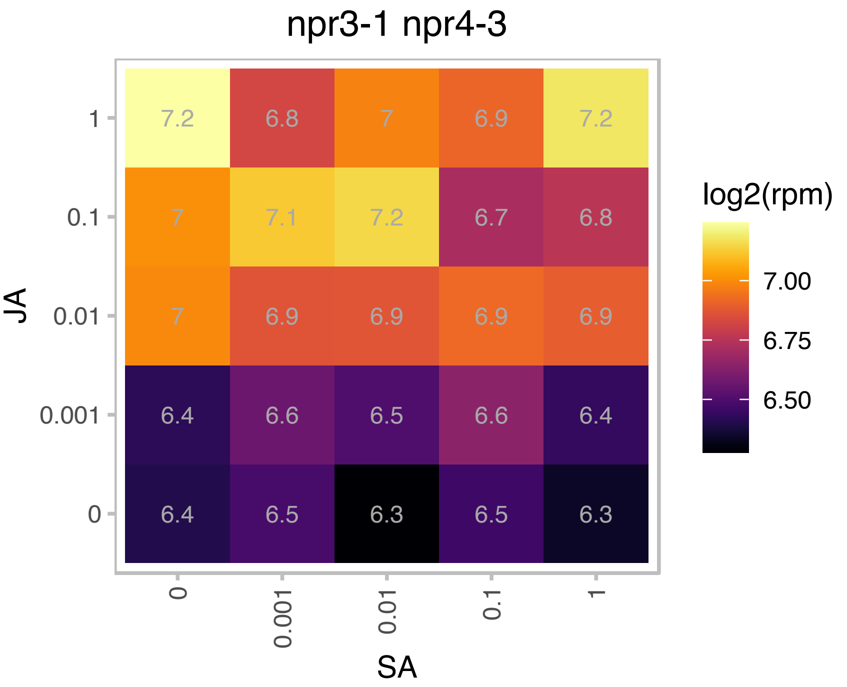

### AT2G22330.1 : CYP79B3

cytochrome P450, family 79, subfamily B, polypeptide 3

Encodes a cytochrome P450. Involved in tryptophan metabolism. Converts Trp to indole-3-acetaldoxime (IAOx), a precursor to IAA and indole glucosinolates. The mRNA is cell-to-cell mobile.

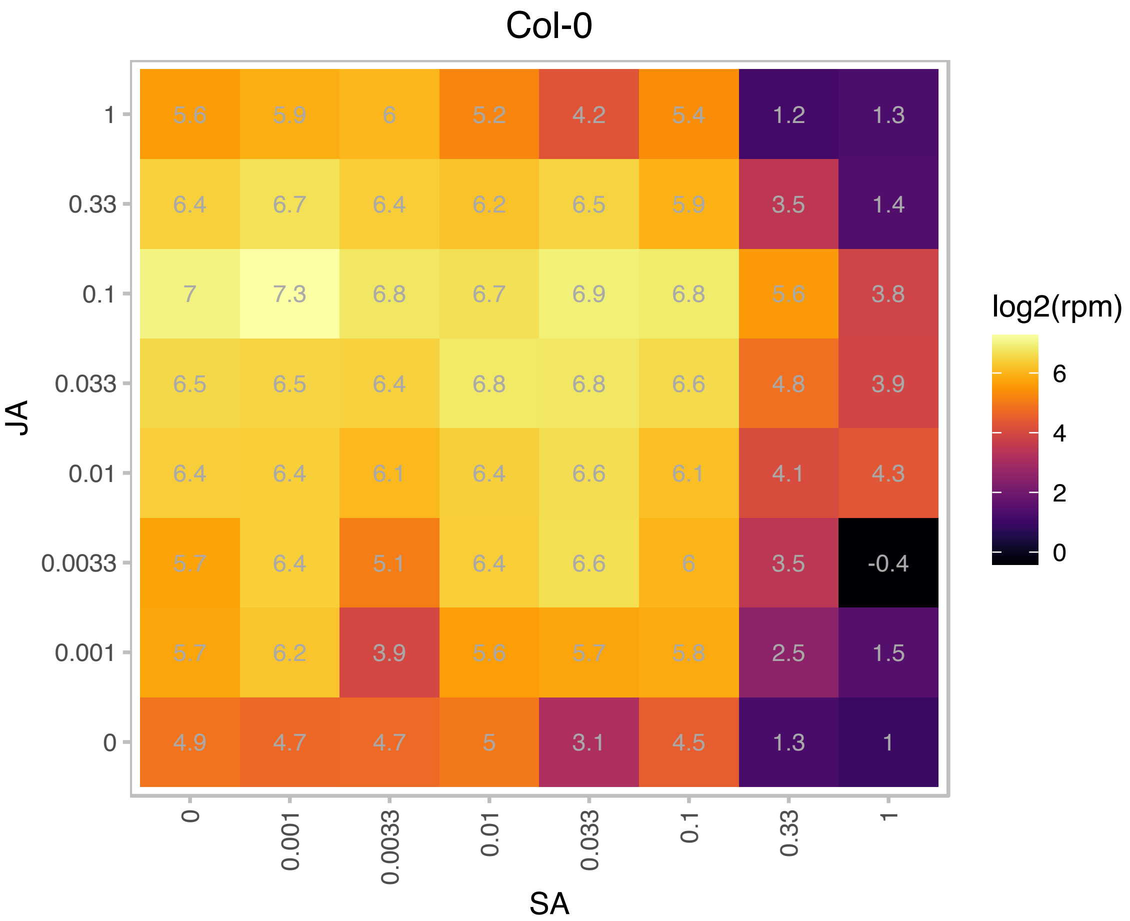


| | |
|---|---|
| Gene | AT2G22330 |
| is.expressed_genes | TRUE |
| is.responsive_genes | TRUE |
| is.apcluster | E-01 |

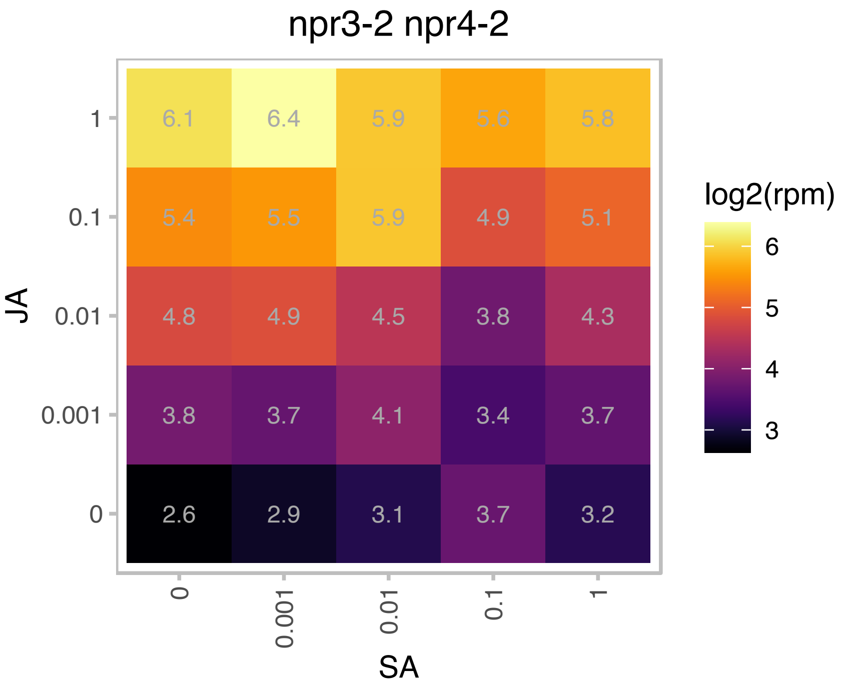


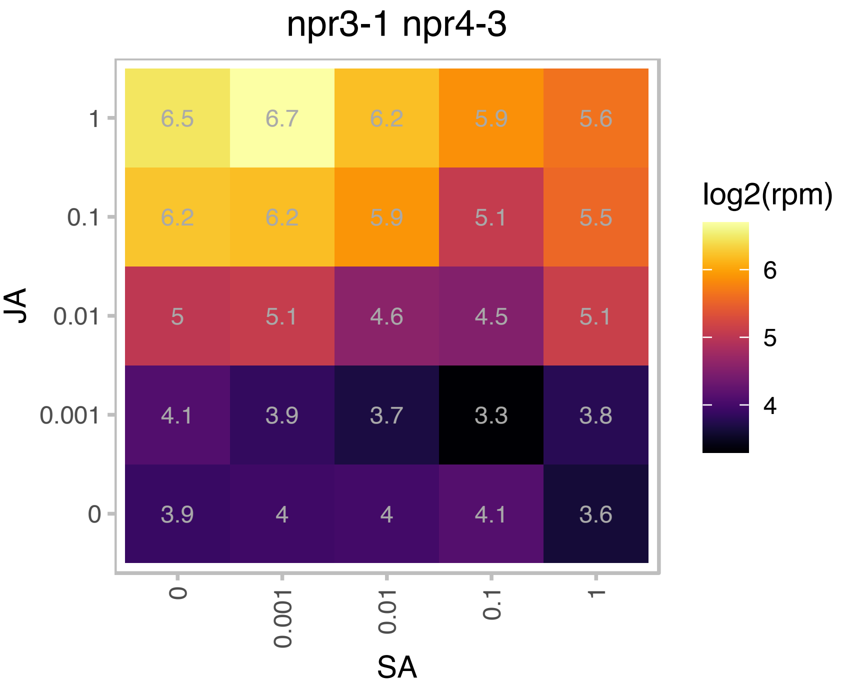

# AT2G22880.1 : VQ12

VQ motif-containing protein

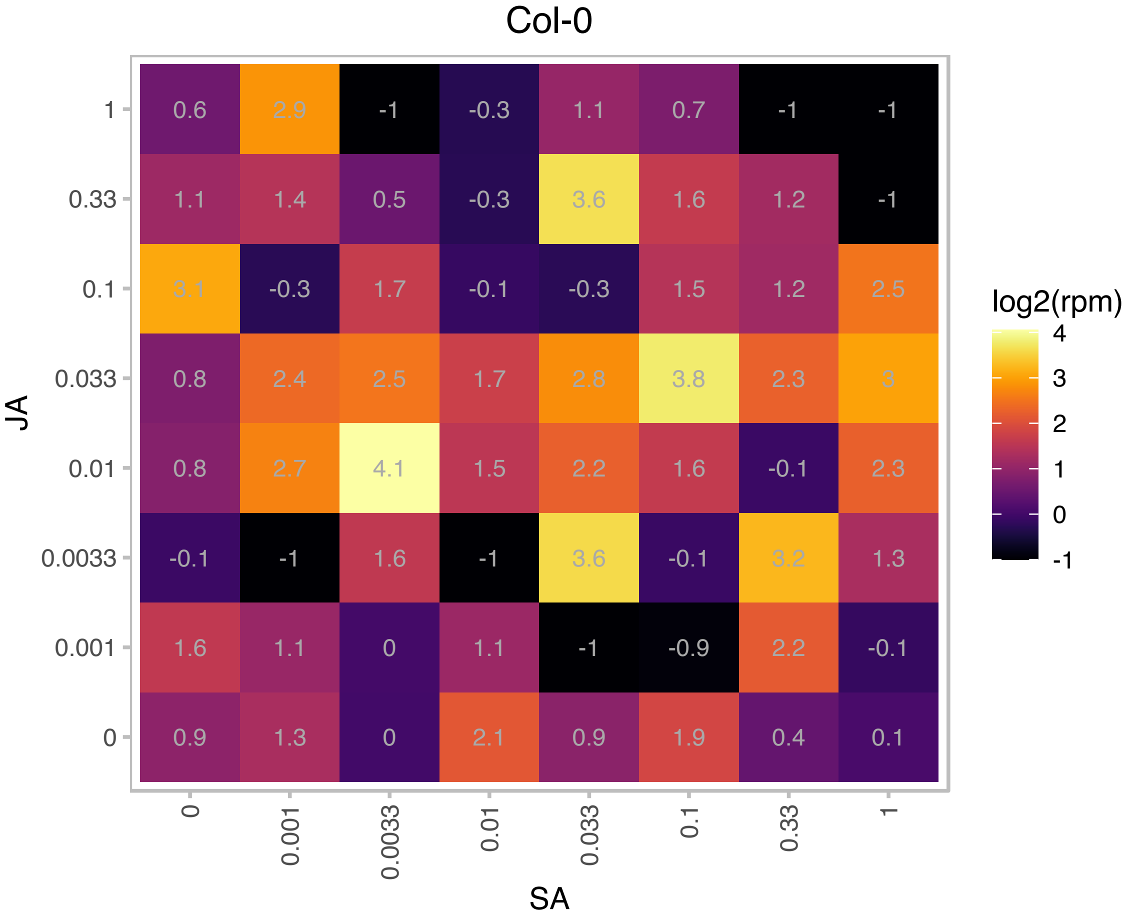


| | |
|---|---|
| Gene | AT2G22880 |
| is.expressed_genes | TRUE |
| is.responsive_genes | TRUE |
| is.apcluster | E-01 |

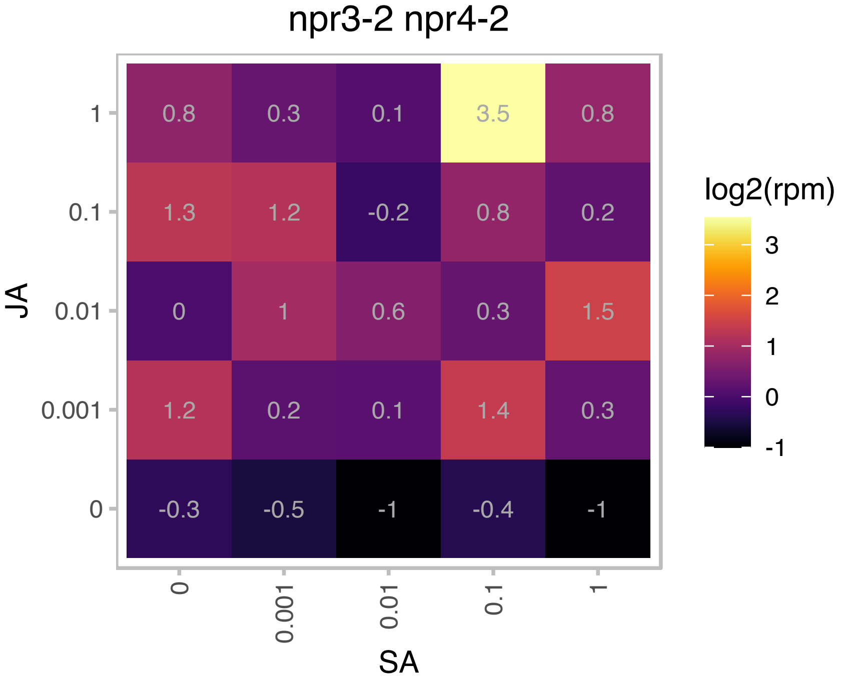


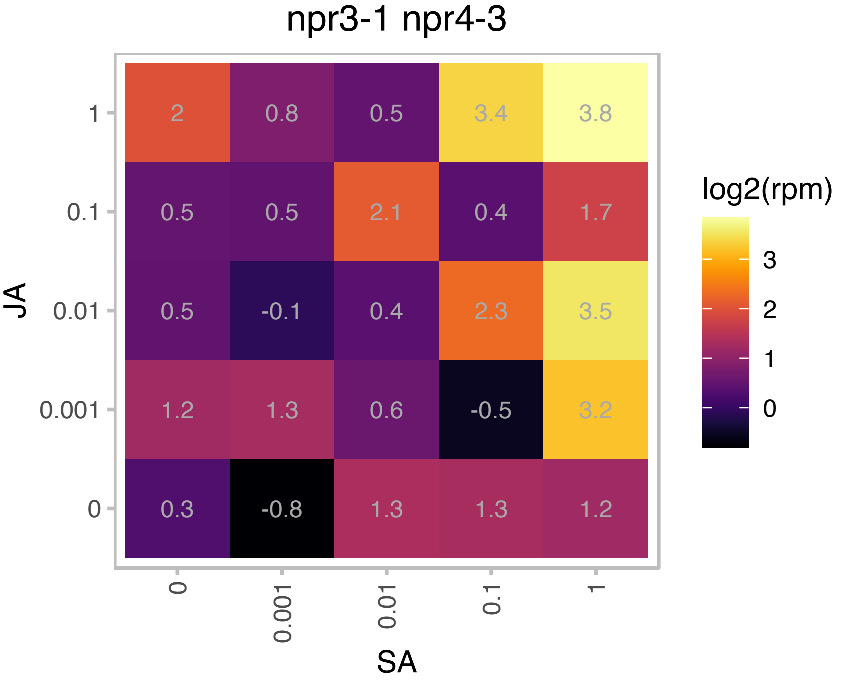

# AT2G25735.1 : NA

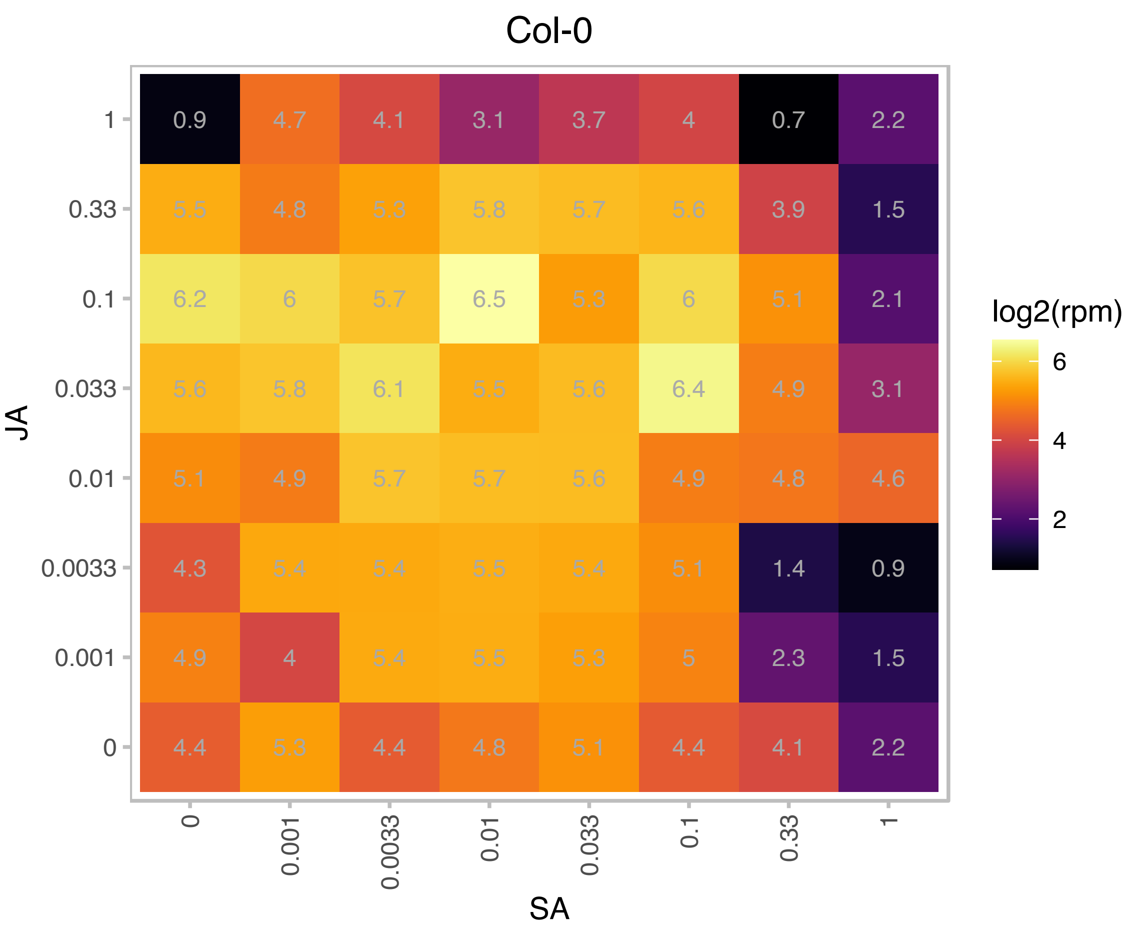


| | |
|---|---|
| Gene | AT2G25735 |
| is.expressed_genes | TRUE |
| is.responsive_genes | TRUE |
| is.apcluster | E-01 |

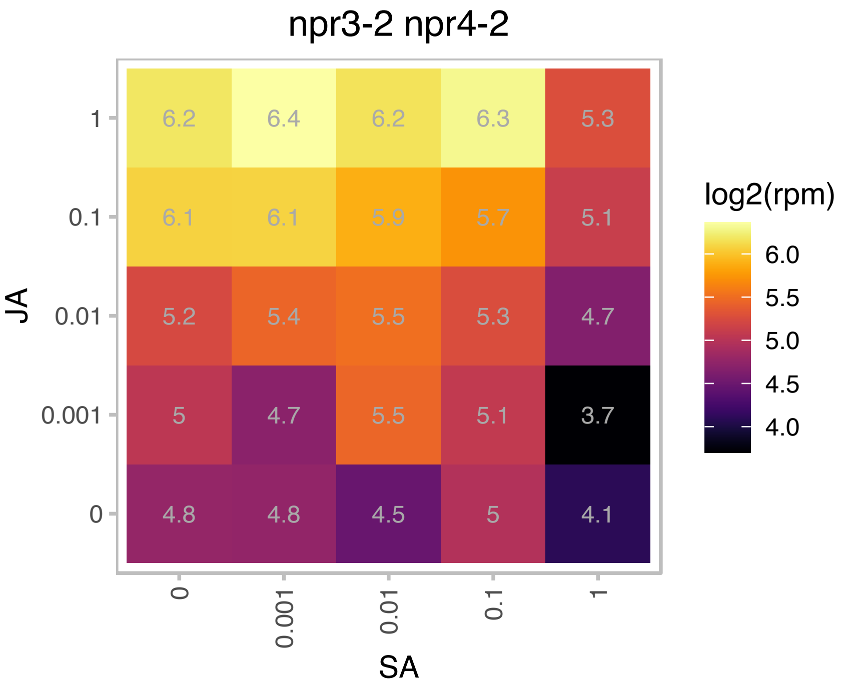


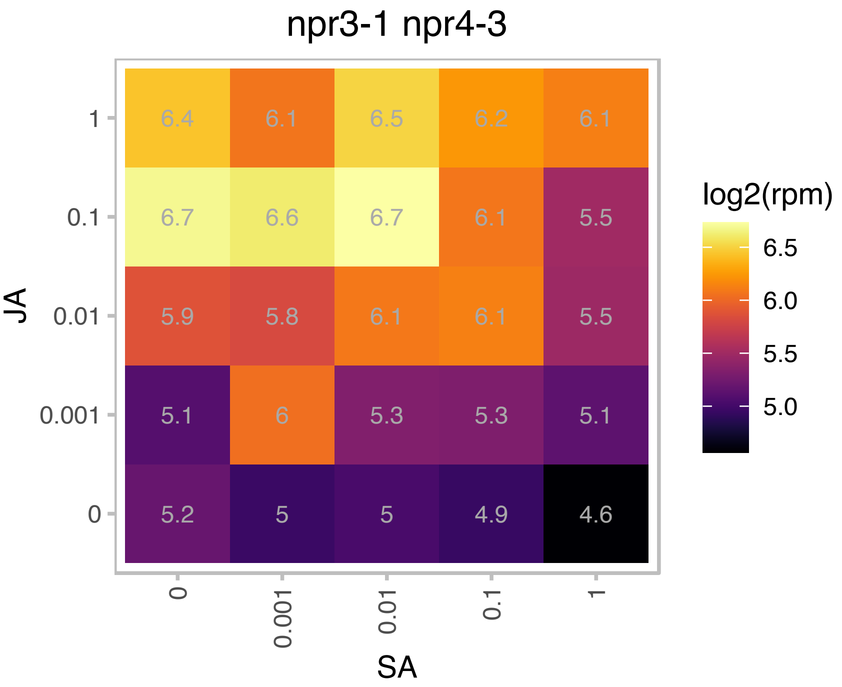

### AT2G29530.1 : AtTIM10, TIM10

Tim10/DDP family zinc finger protein
Encodes a small zinc finger-like protein that is a component of the mitochondrial protein import apparatus. Together with AtTIM9, AtTIM10 is non-redundantly essential for maintaining mitochondrial function of early embryo proper cells and endosperm free-nuclei.

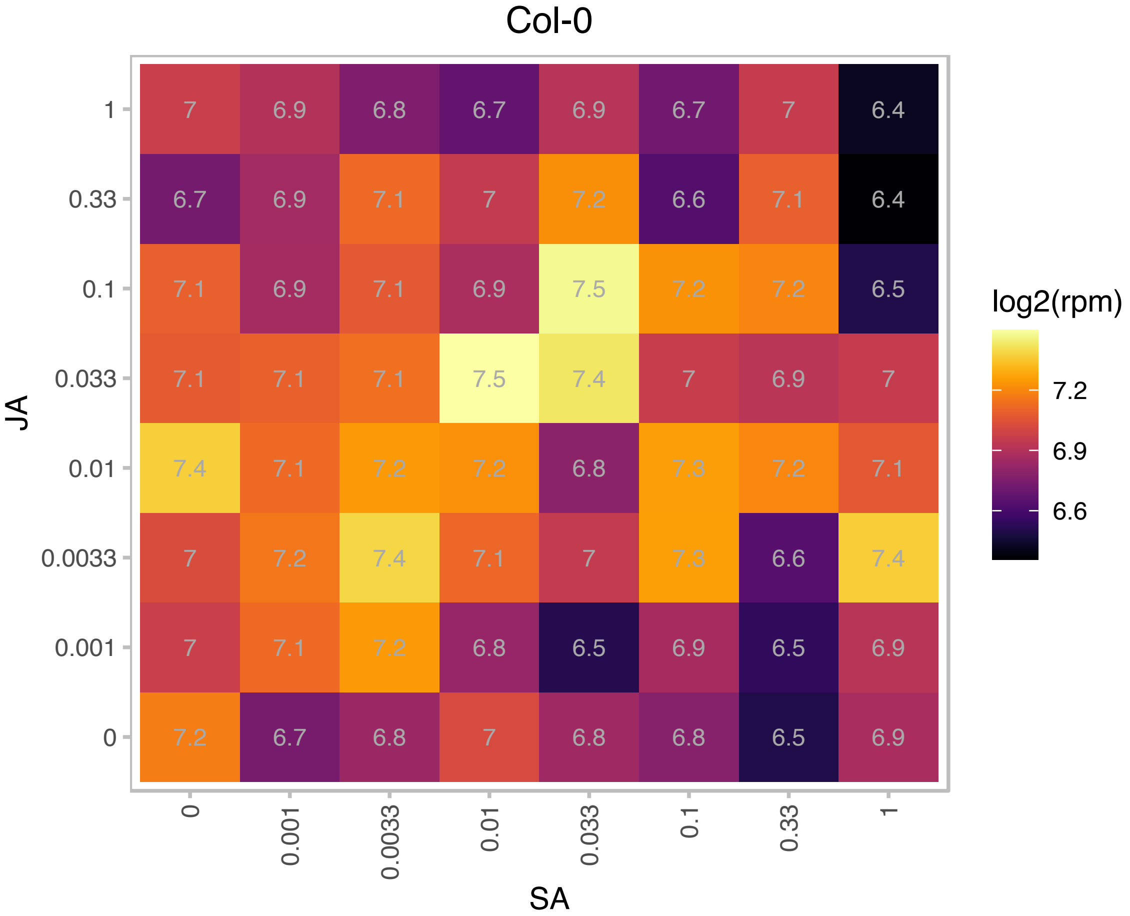


| | |
|---|---|
| Gene | AT2G29530 |
| is.expressed_genes | TRUE |
| is.responsive_genes | TRUE |
| is.apcluster | E-01 |

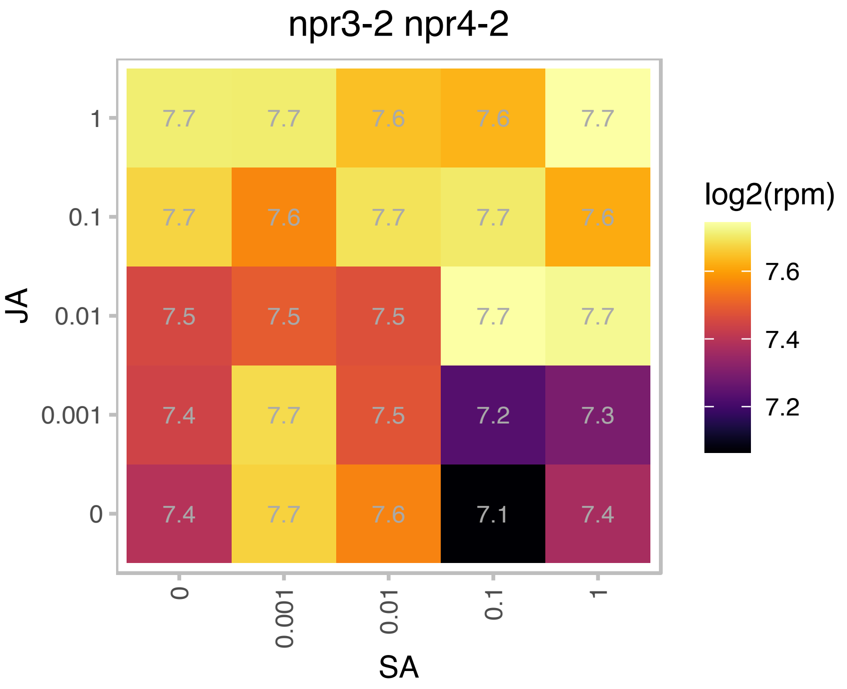


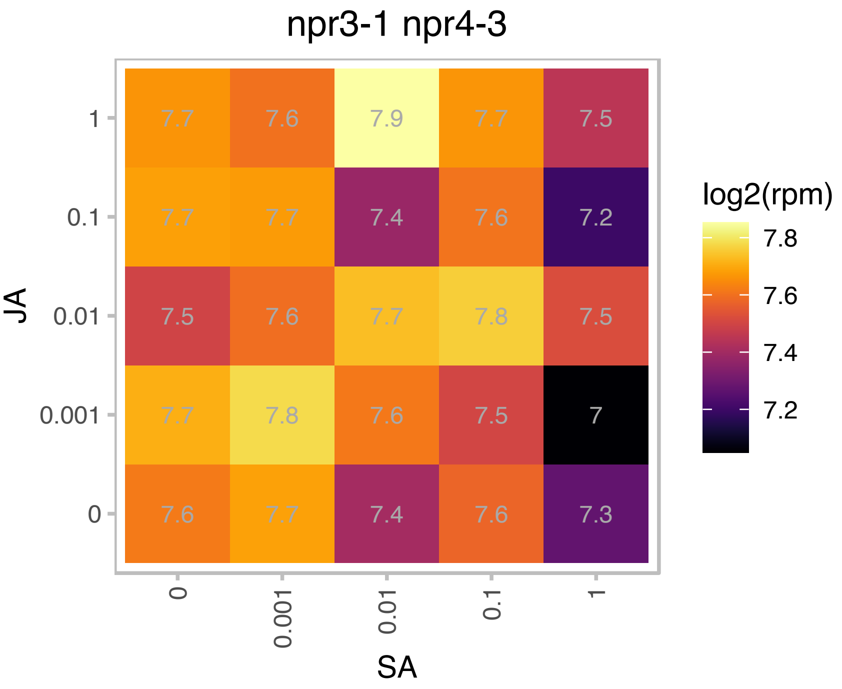

### AT2G30540.1 : ROXY7

Thioredoxin superfamily protein

Encodes a member of the CC-type glutaredoxin (ROXY) family that has been shown to interact with the transcription factor TGA2.

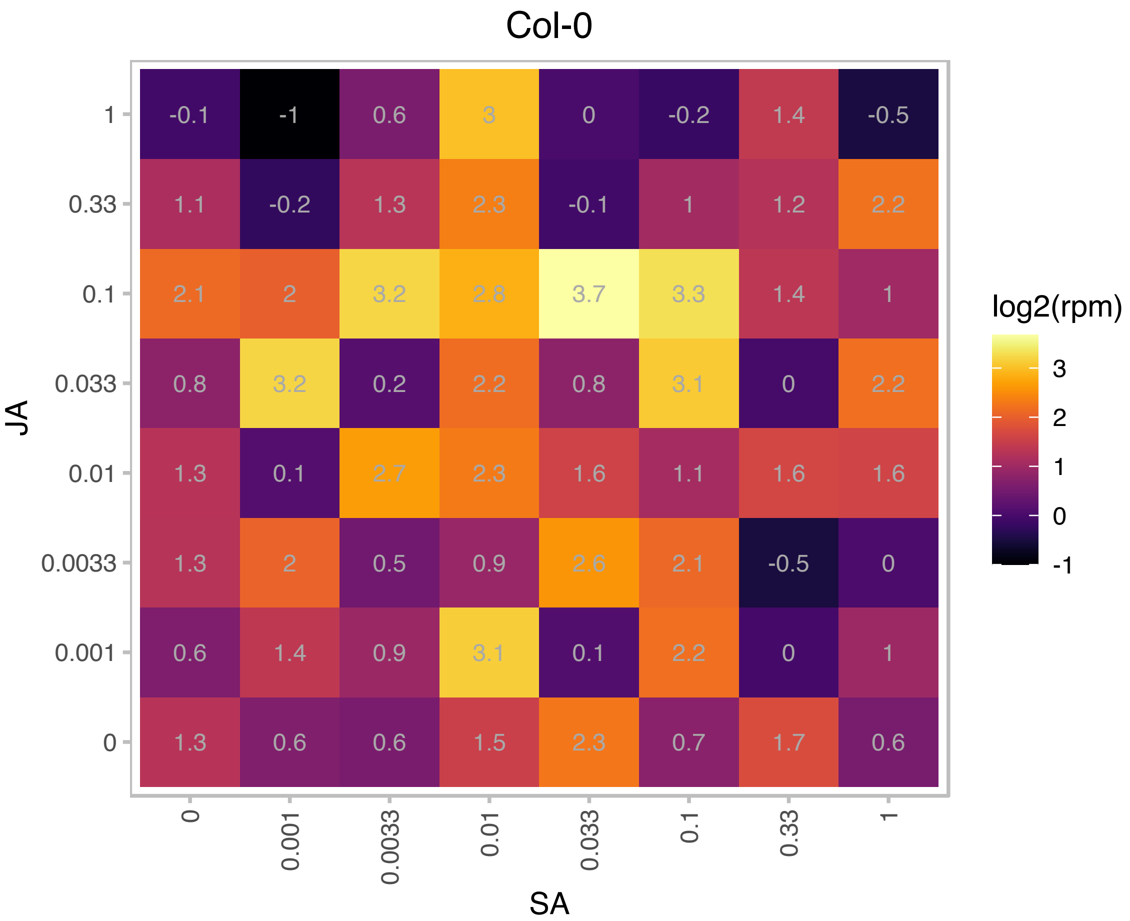


| | |
|---|---|
| Gene | AT2G30540 |
| is.expressed_genes | TRUE |
| is.responsive_genes | TRUE |
| is.apcluster | E-01 |

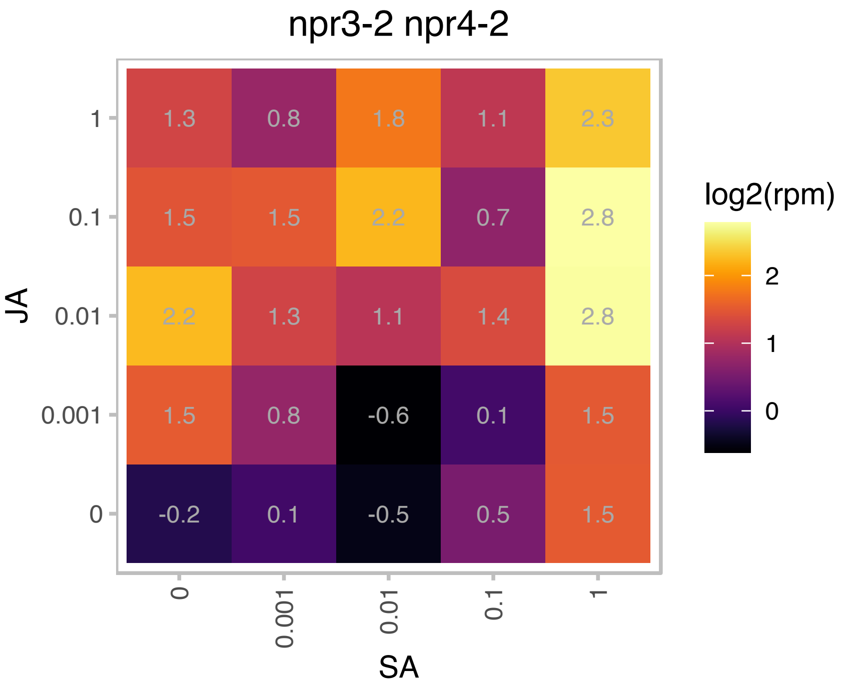


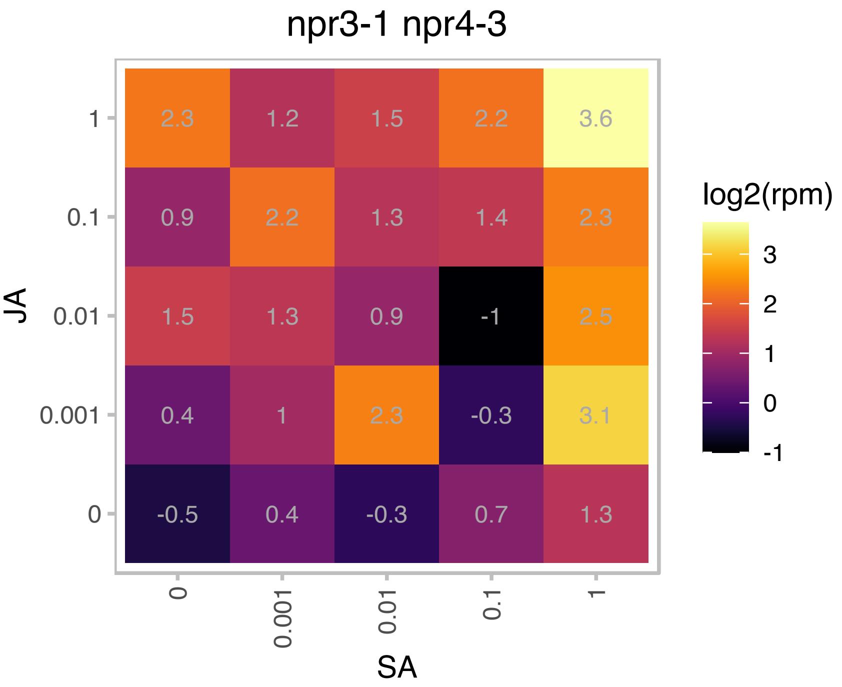

### AT2G30860.1 : ATGSTF7, ATGSTF9, GLUTTR, GSTF9

glutathione S-transferase PHI 9

Encodes glutathione transferase belonging to the phi class of GSTs. Naming convention according to Wagner et al. (2002).

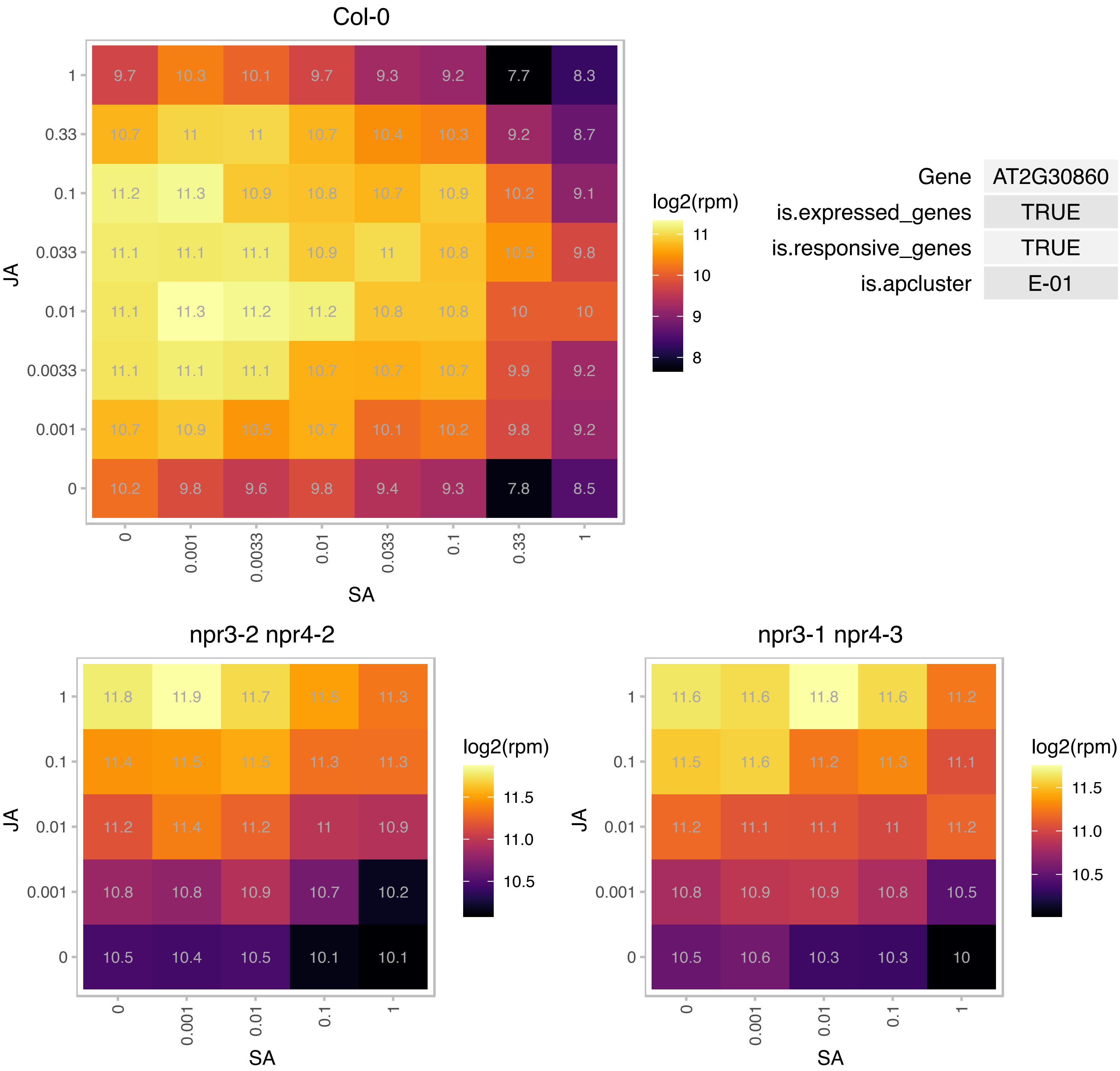


| | |
|---|---|
| Gene | AT2G30860 |
| is.expressed_genes | TRUE |
| is.responsive_genes | TRUE |
| is.apcluster | E-01 |

### AT2G30870.1 : ATGSTF10, ATGSTF4, ERD13, GSTF10

glutathione S-transferase PHI 10
early dehydration-induced gene ERD13 homologous to tobacco and maize glutathione S-transferases. Encodes glutathione transferase belonging to the phi class of GSTs. Naming convention according to Wagner et al. (2002)

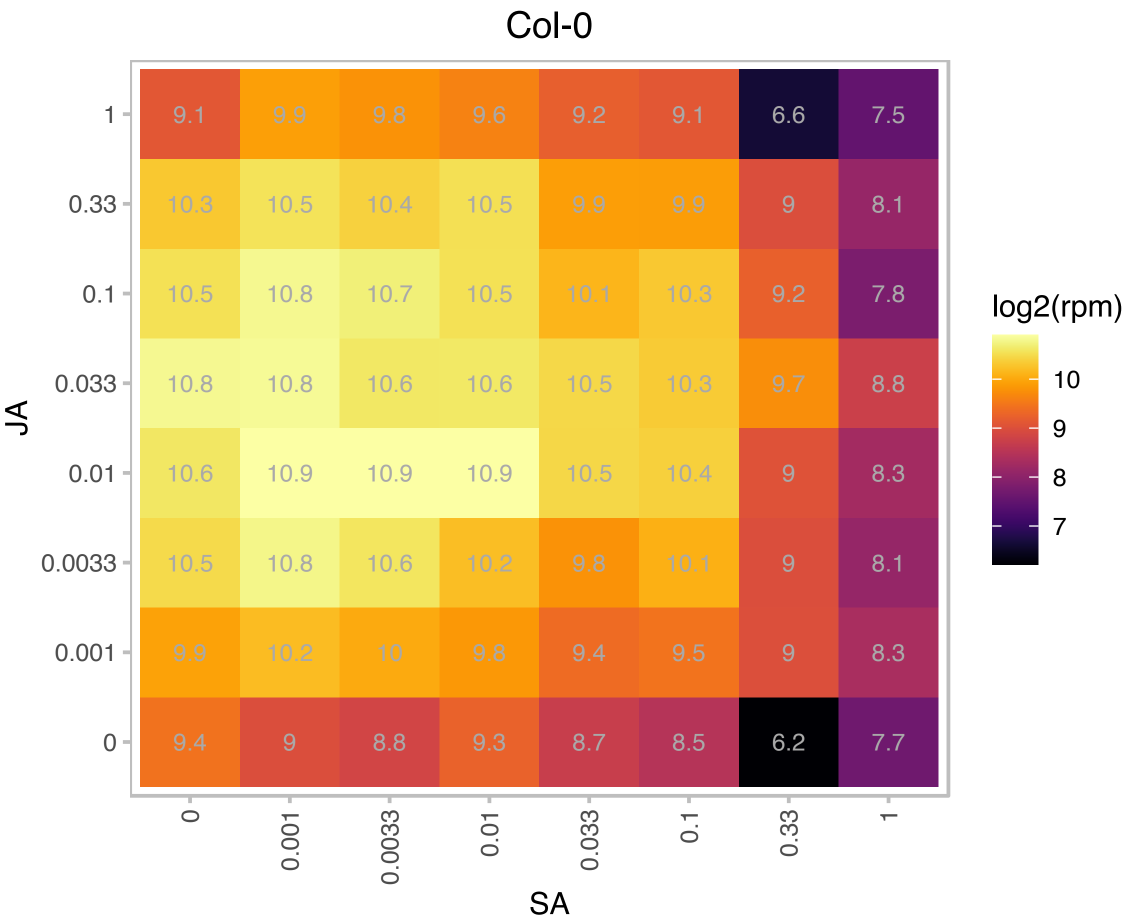


| | |
|---|---|
| Gene | AT2G30870 |
| is.expressed_genes | TRUE |
| is.responsive_genes | TRUE |
| is.apcluster | E-01 |

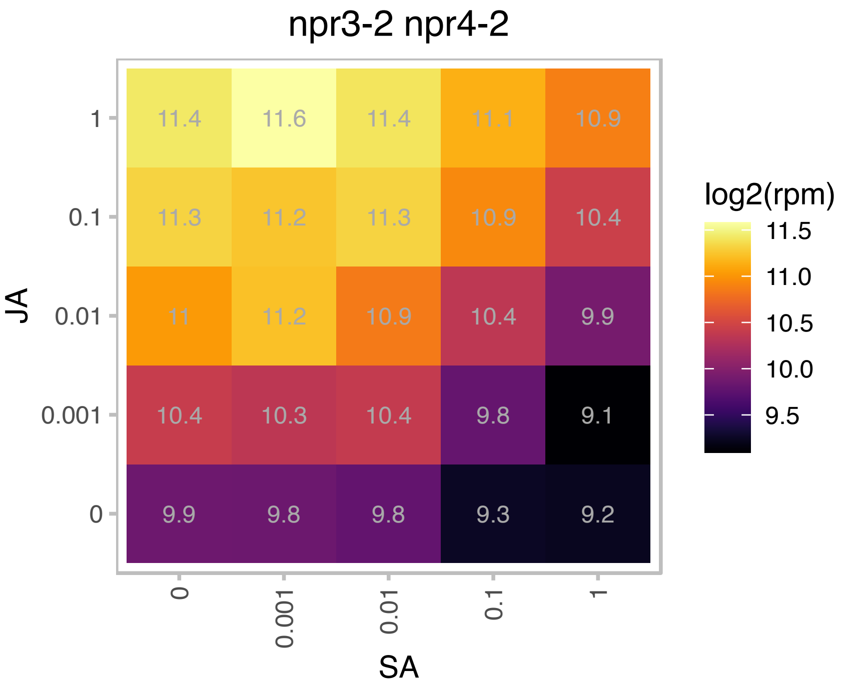


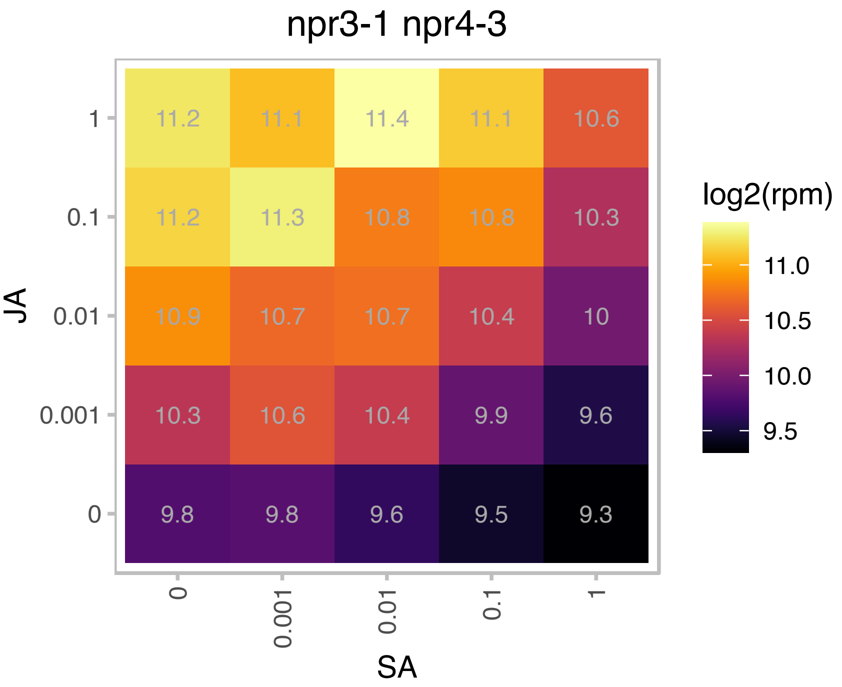

# AT2G30990.1 : NA

Protein of unknown function (DUF688)

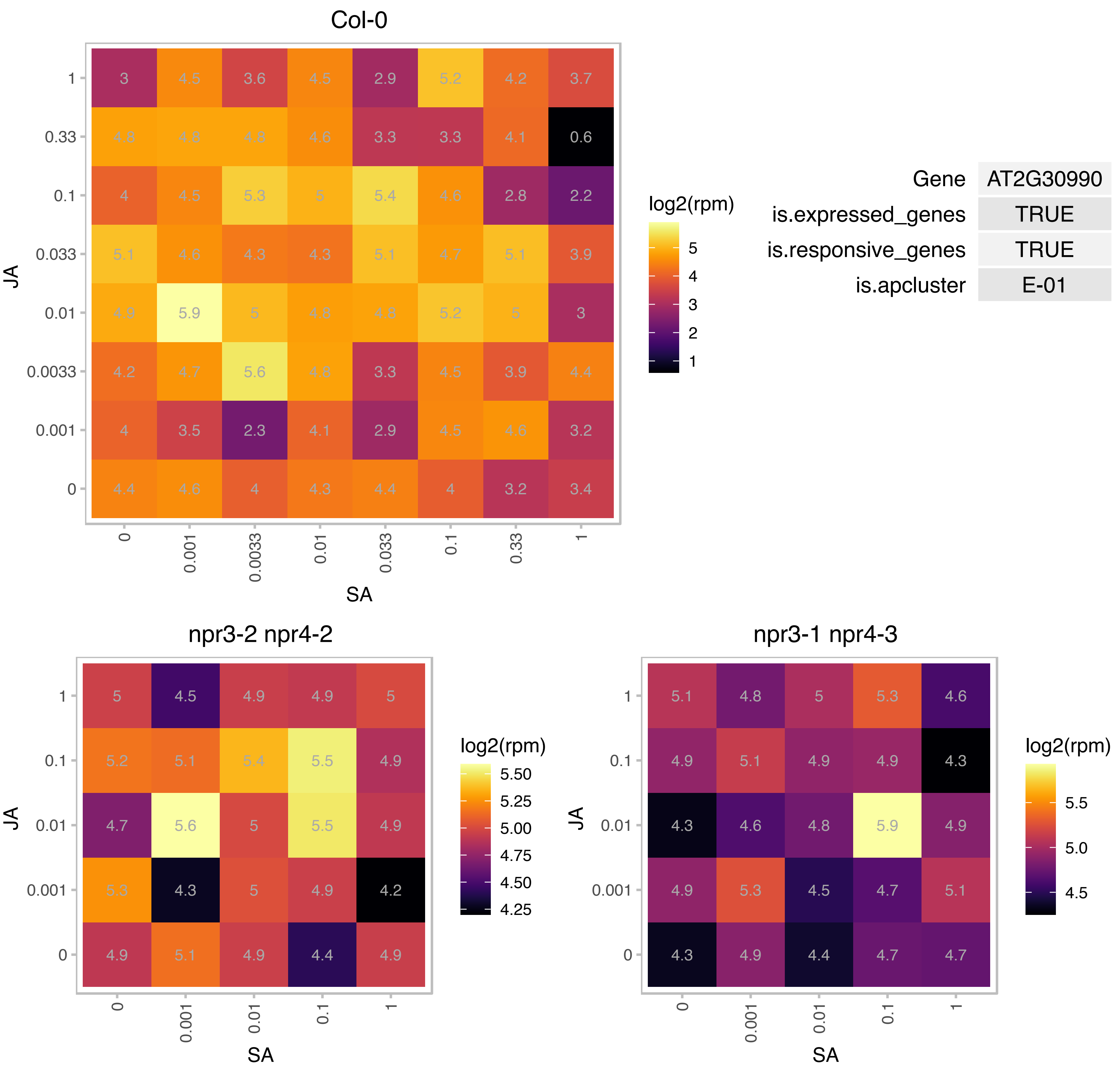


| Gene | AT2G30990 |
|---|---|
| is.expressed_genes | TRUE |
| is.responsive_genes | TRUE |
| is.apcluster | E-01 |

### AT2G31370.1 : AtbZIP59, PosF21

Basic-leucine zipper (bZIP) transcription factor family protein

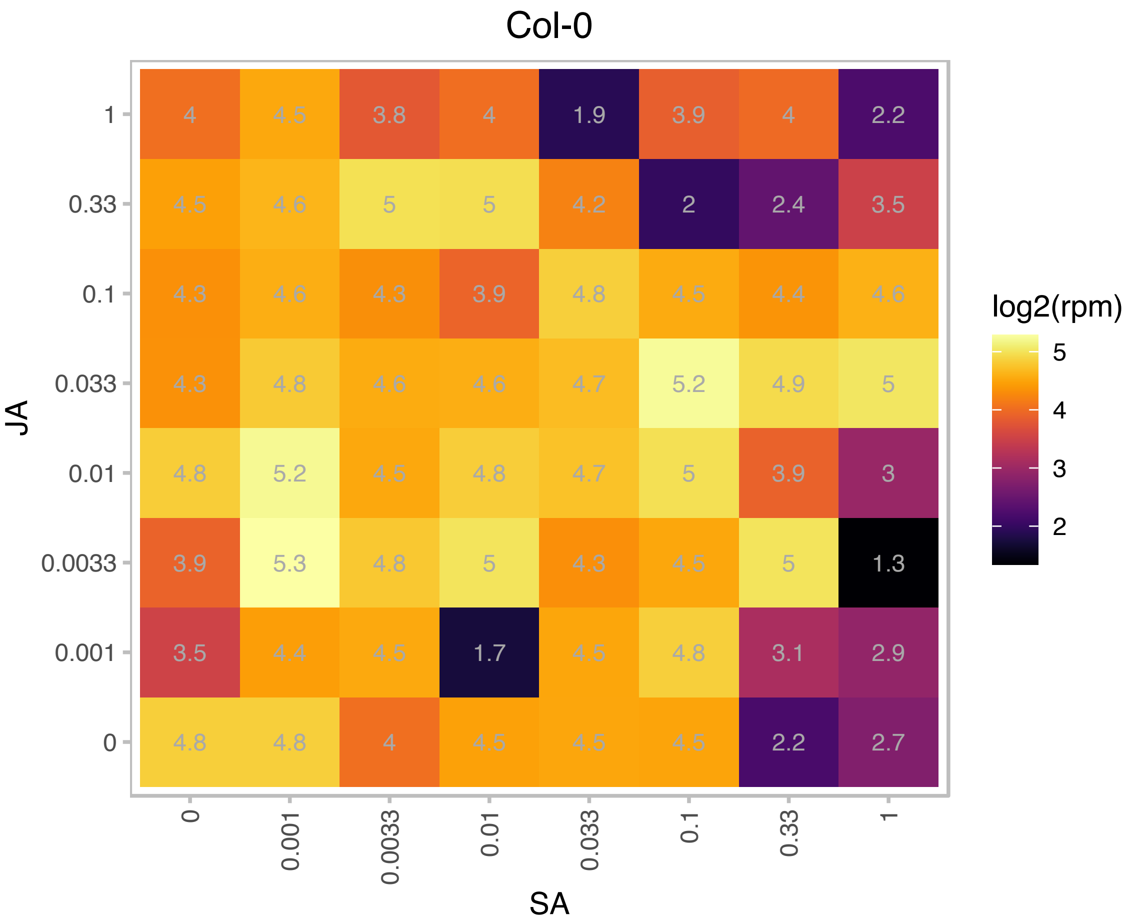


| Gene | AT2G31370 |
| --- | --- |
| is.expressed_genes | TRUE |
| is.responsive_genes | TRUE |
| is.apcluster | E-01 |

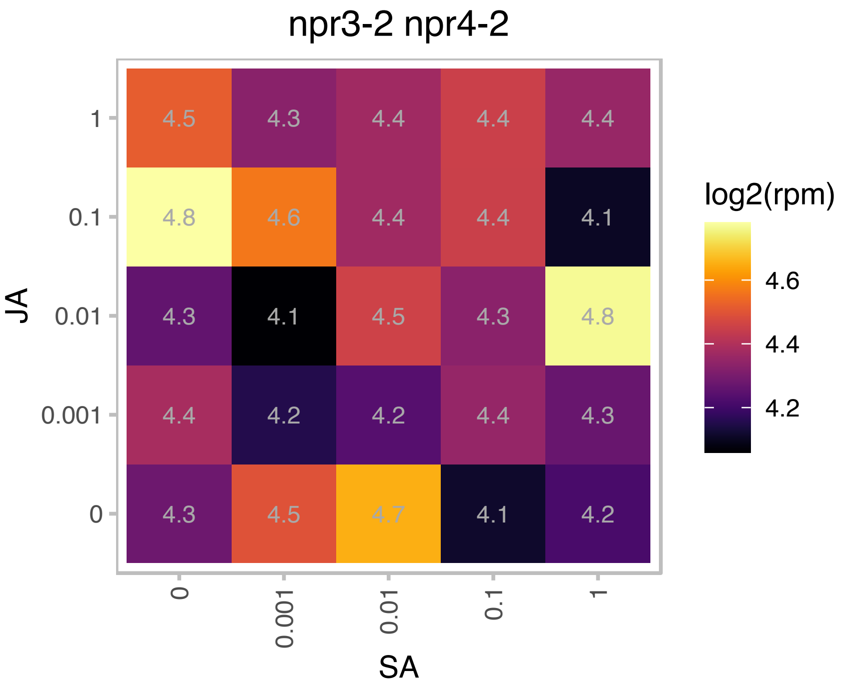


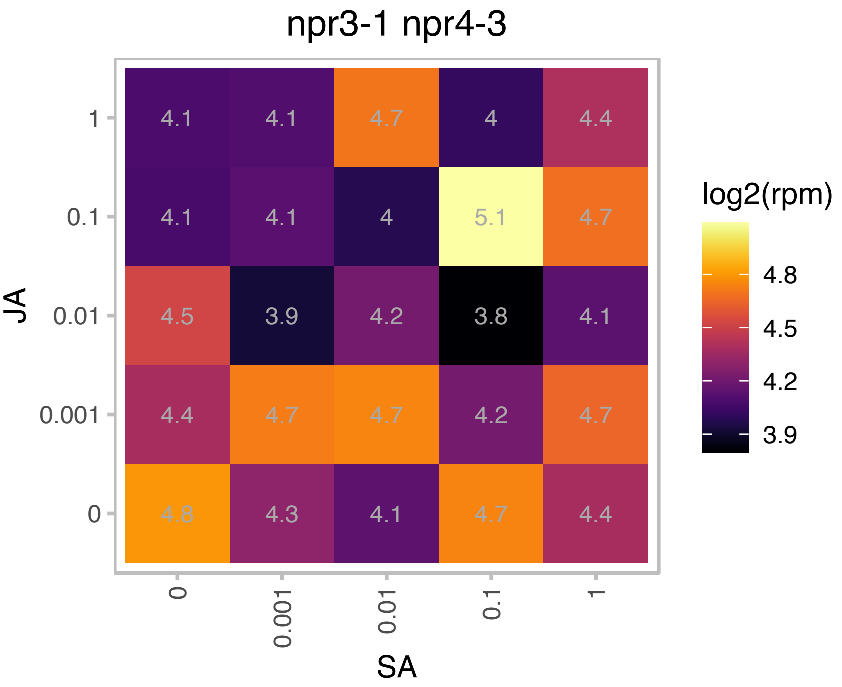

### AT2G36080.1 : ABS2, NGAL1

AP2/B3-like transcriptional factor family protein

Encodes a plant-specific B3 DNA-binding domain transcription factor.  Has transcription repressor activity.

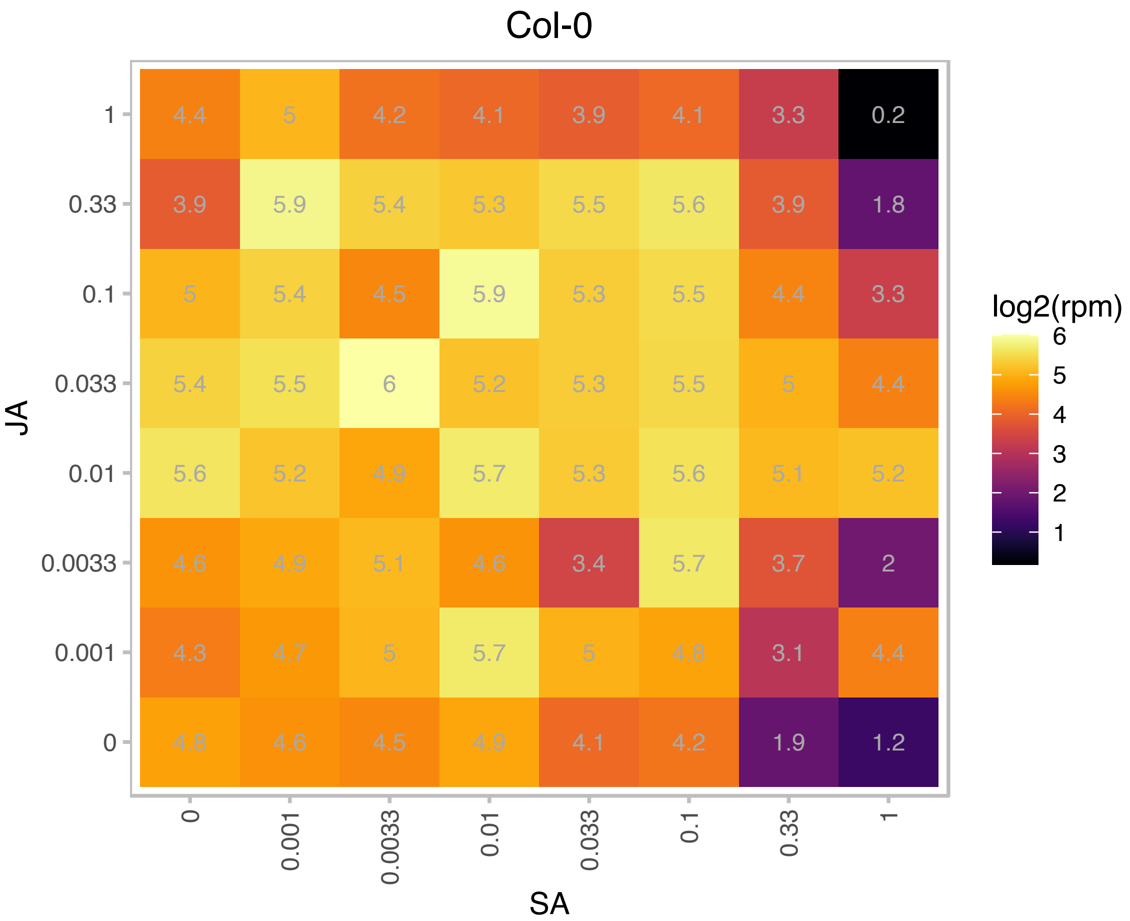


| | |
|---|---|
| Gene | AT2G36080 |
| is.expressed_genes | TRUE |
| is.responsive_genes | TRUE |
| is.apcluster | E-01 |

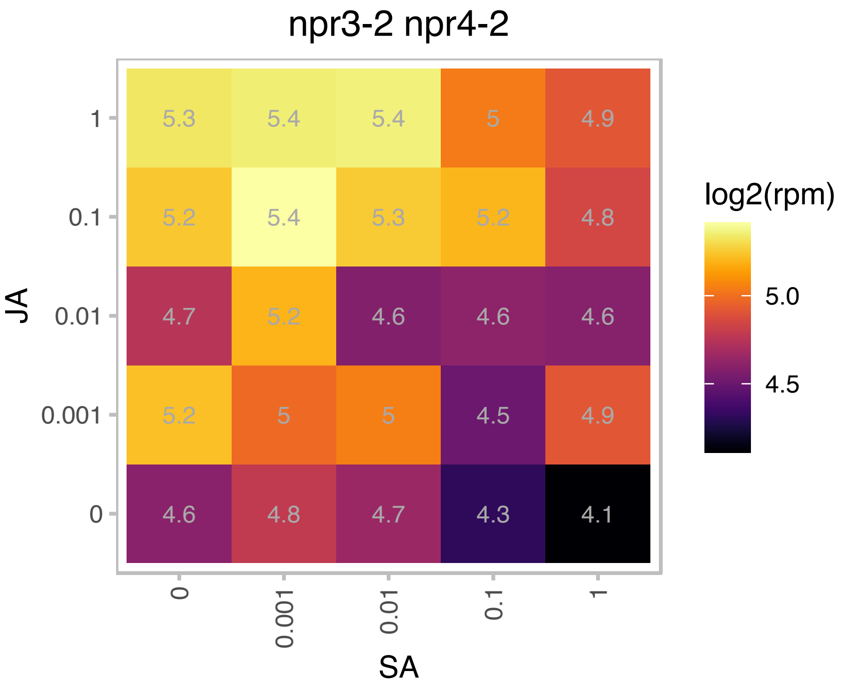


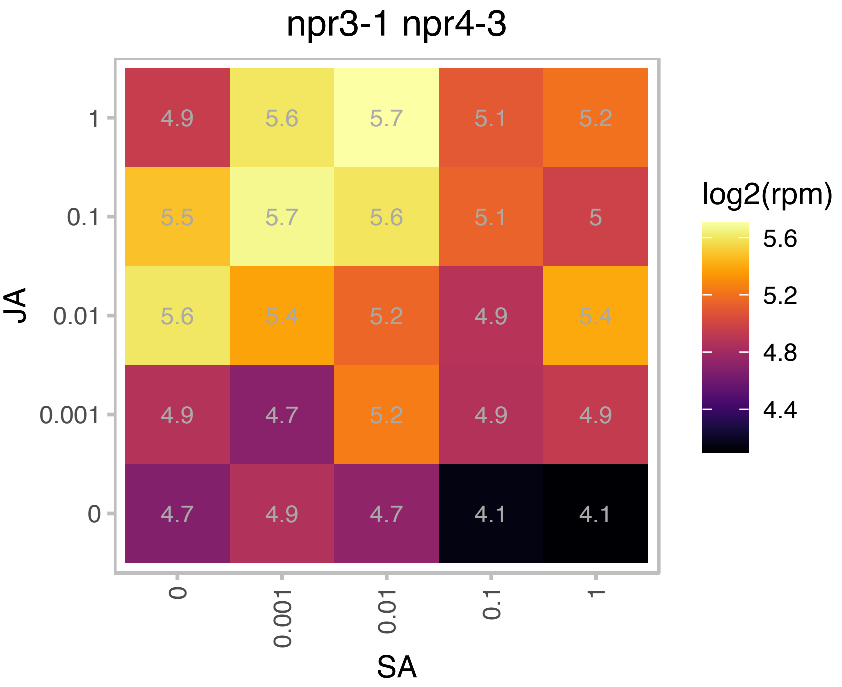

### AT2G42540.1 : AtCOR15A, COR15, COR15A

cold-regulated 15a

A cold-regulated gene whose product is targeted to the chloroplast. Cor15am protects stromal proteins from aggregation under various stress conditions. Constitutive expression increases freezing tolerance in protoplasts in vitro and chloroplasts in vivo. NMR and x-ray diffraction studies suggest that COR15a alters the intrinsic curvature of the inner membrane of chloroplast envelope. Late Embryogenesis abundant protein (LEA). Protects chloroplast membranes during freezing.

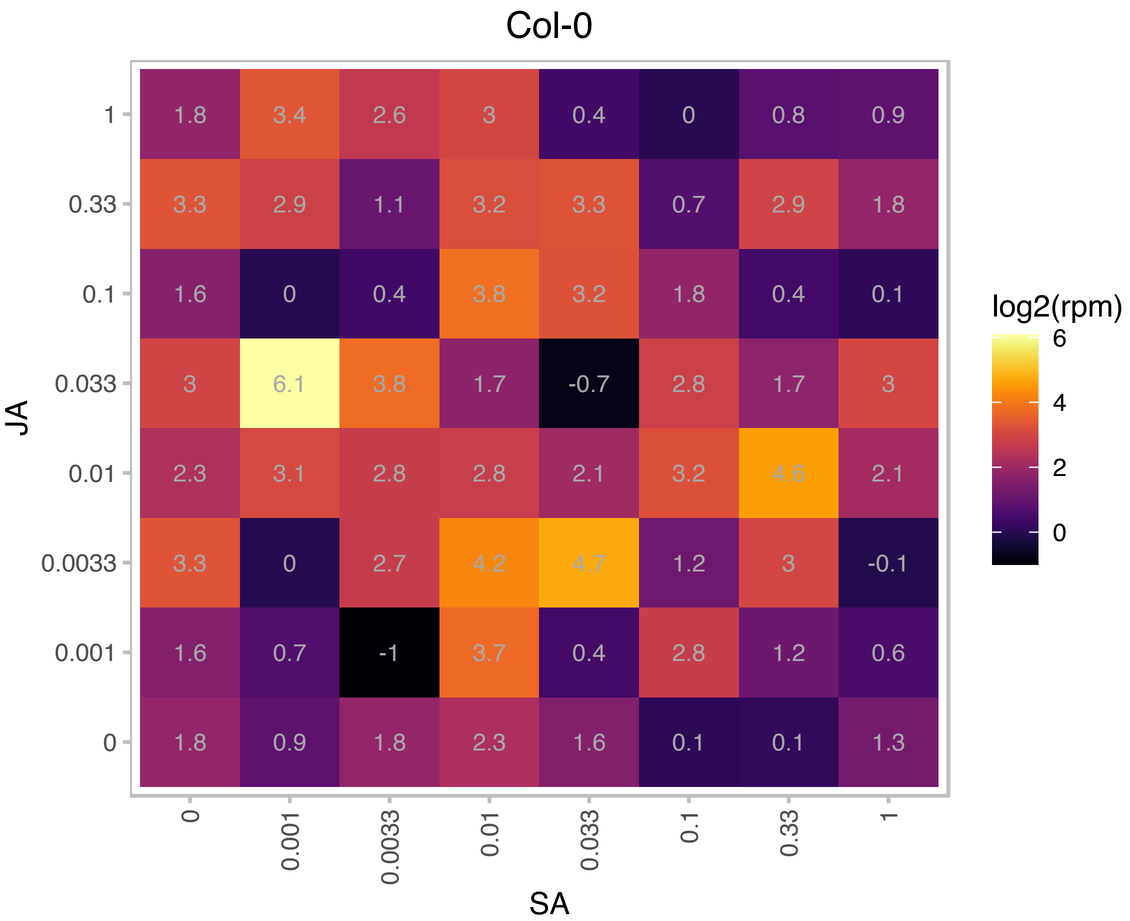


| | |
|---|---|
| Gene | AT2G42540 |
| is.expressed_genes | TRUE |
| is.responsive_genes | TRUE |
| is.apcluster | E-01 |

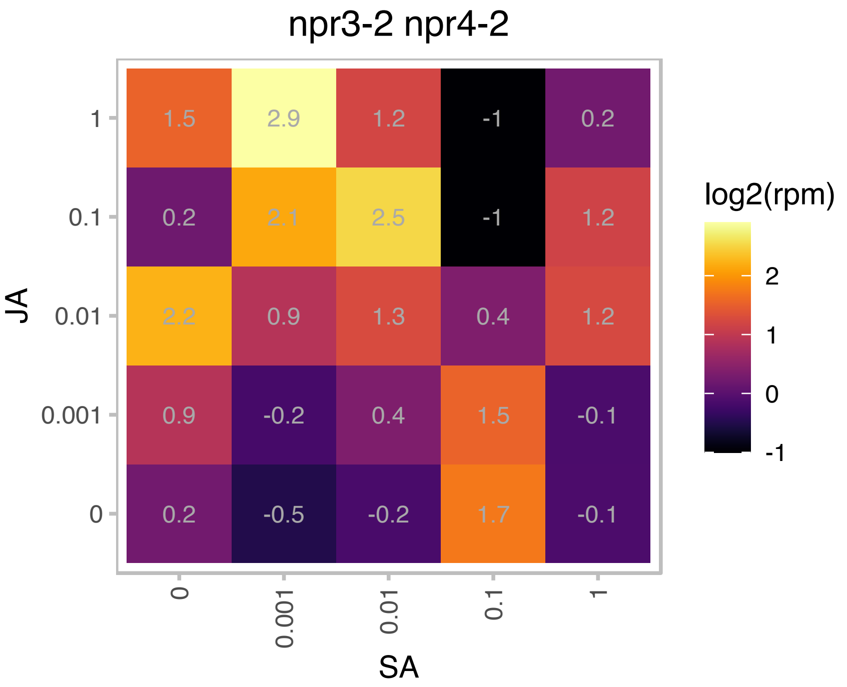


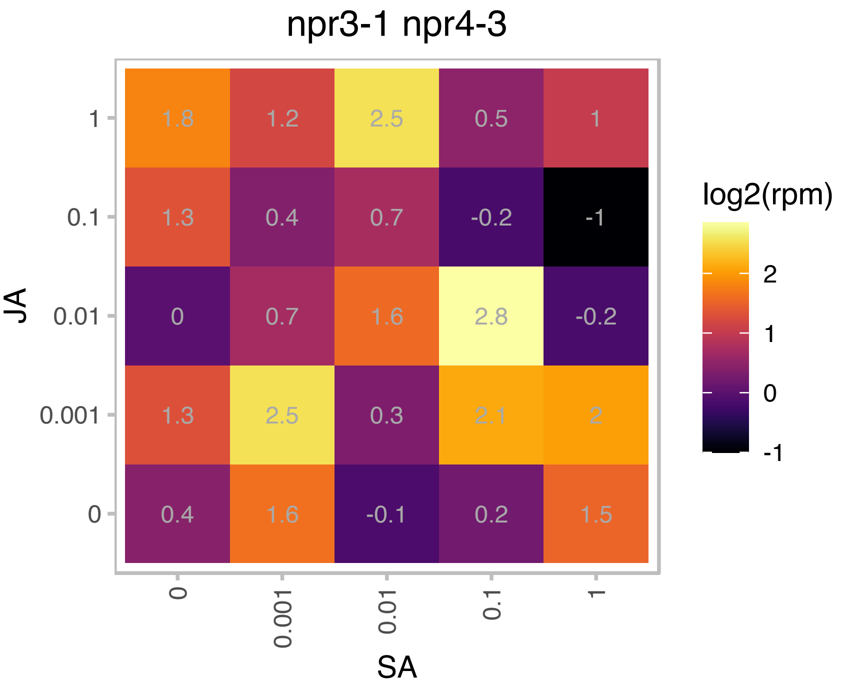

# AT2G43540.1 : NA

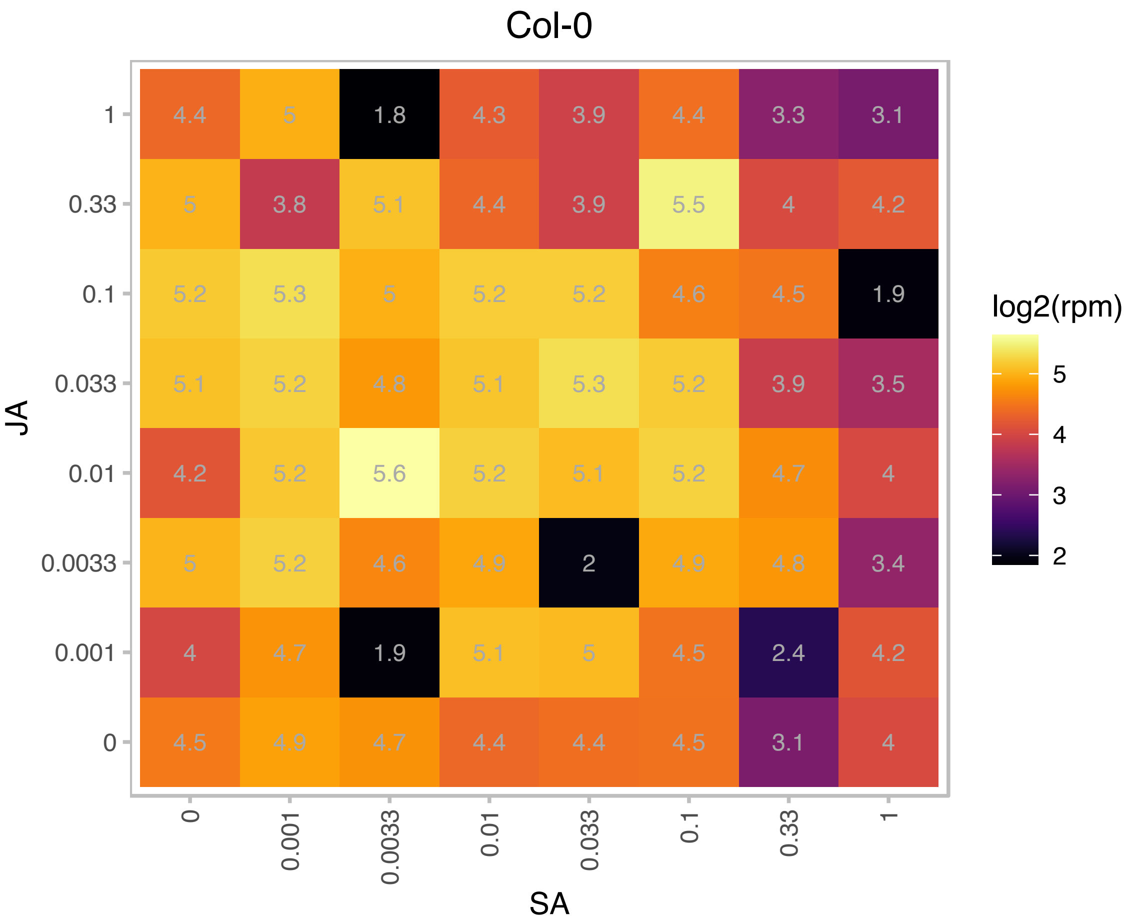


| | |
|---|---|
| Gene | AT2G43540 |
| is.expressed_genes | TRUE |
| is.responsive_genes | TRUE |
| is.apcluster | E-01 |

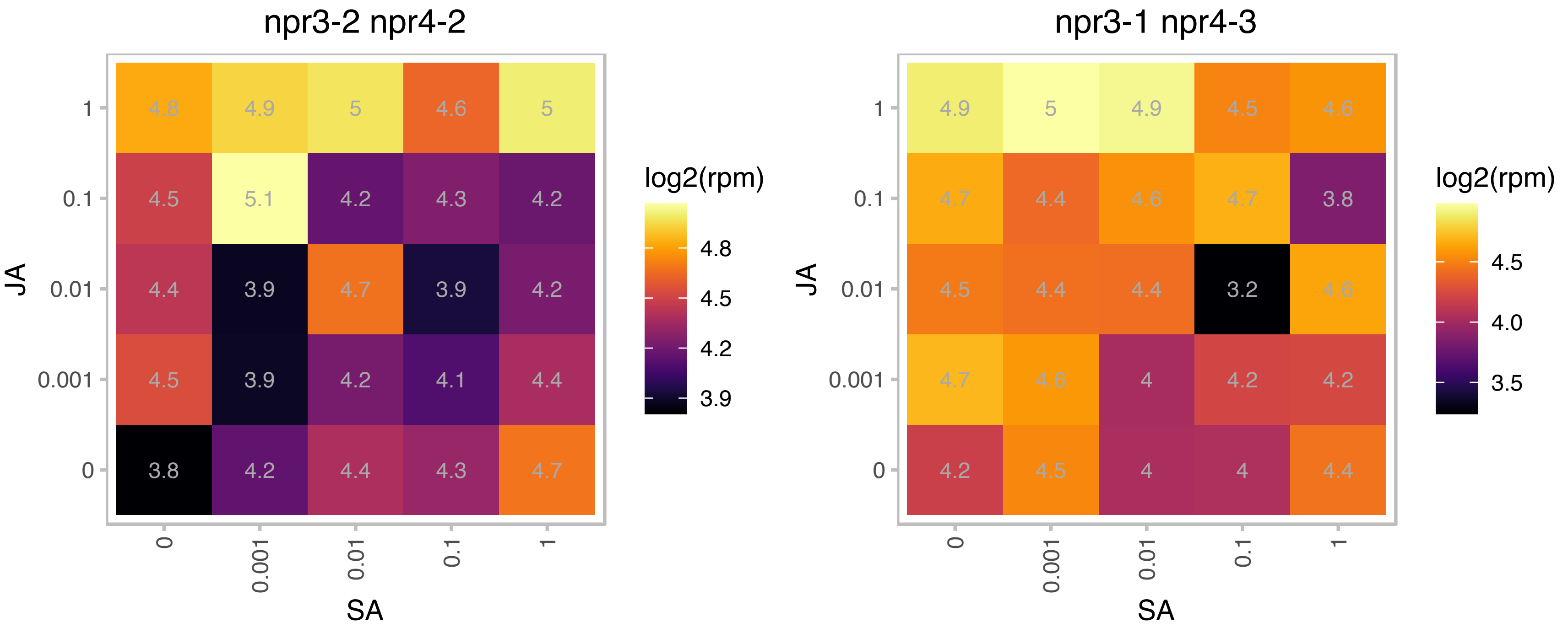

# AT2G44120.1 : NA

Ribosomal protein L30/L7 family protein

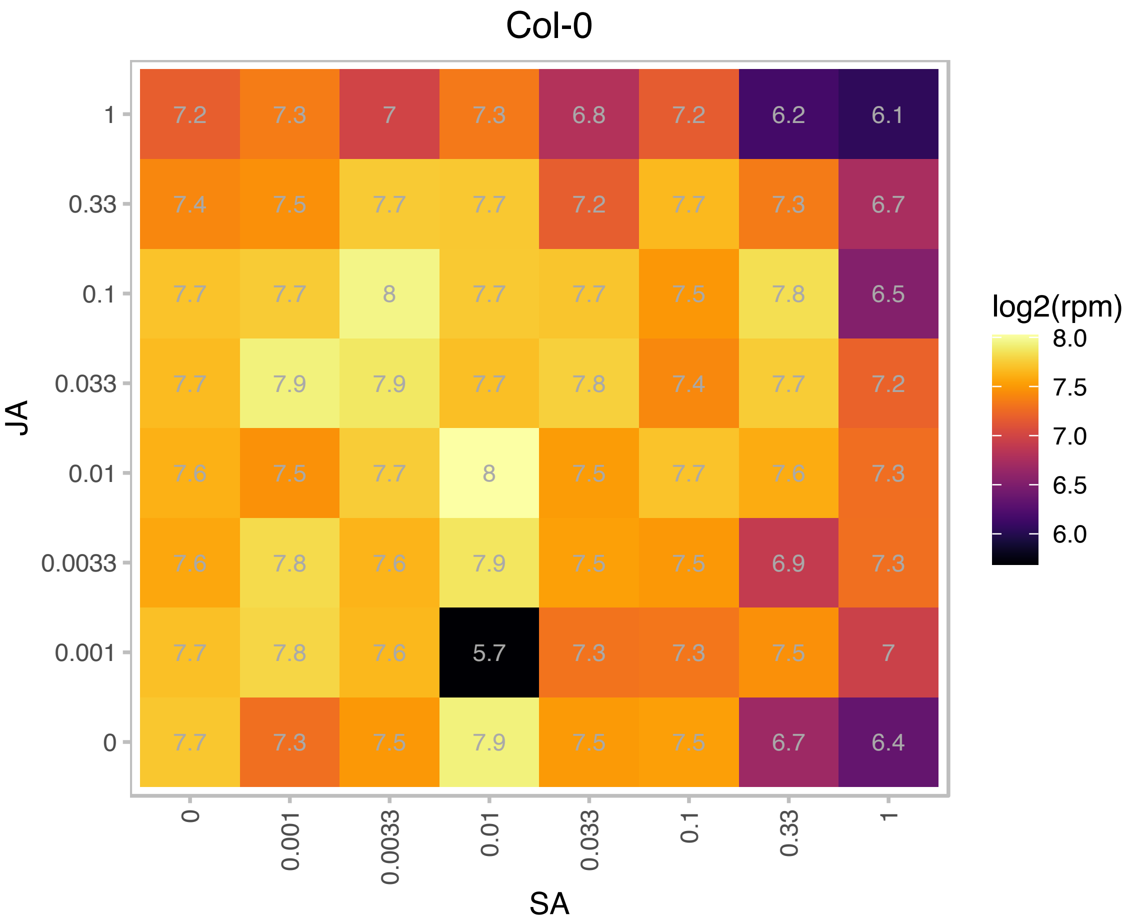


| | |
|---|---|
| Gene | AT2G44120 |
| is.expressed_genes | TRUE |
| is.responsive_genes | TRUE |
| is.apcluster | E-01 |

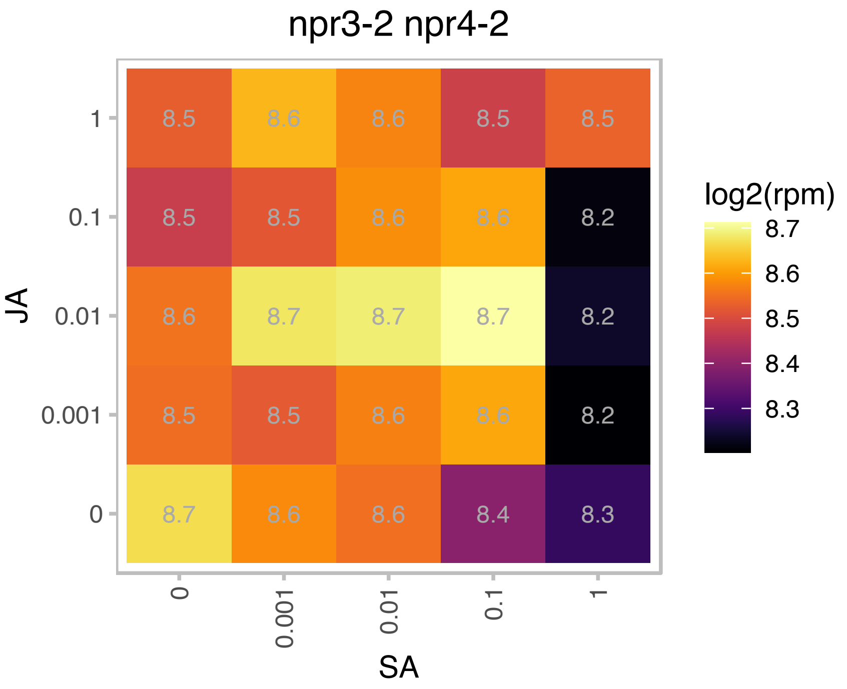


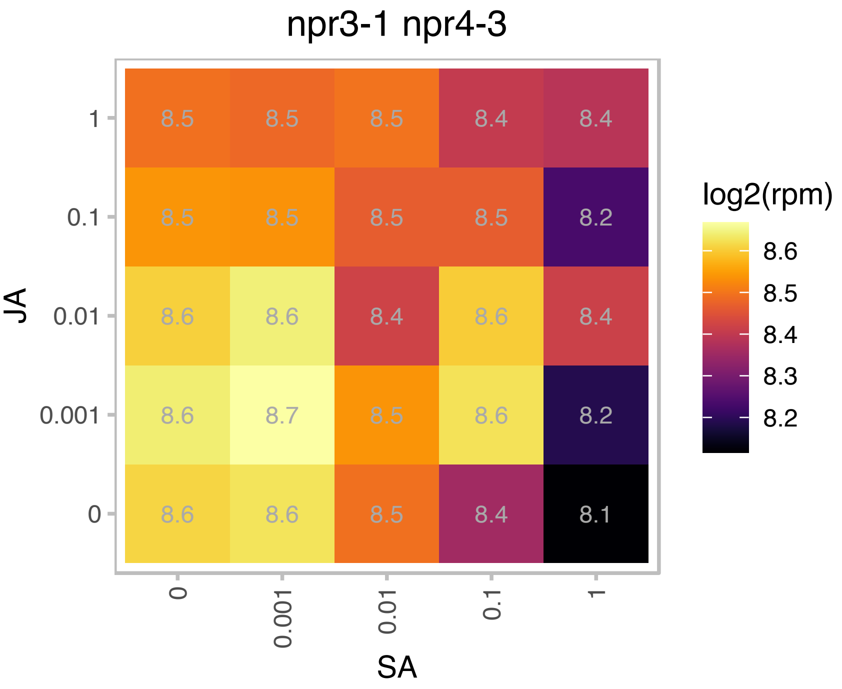

### AT2G46650.1 : ATCB5-C, B5 #1, CB5-C, CYTB5-C

cytochrome B5 isoform C

member of Cytochromes b5 The mRNA is cell-to-cell mobile.

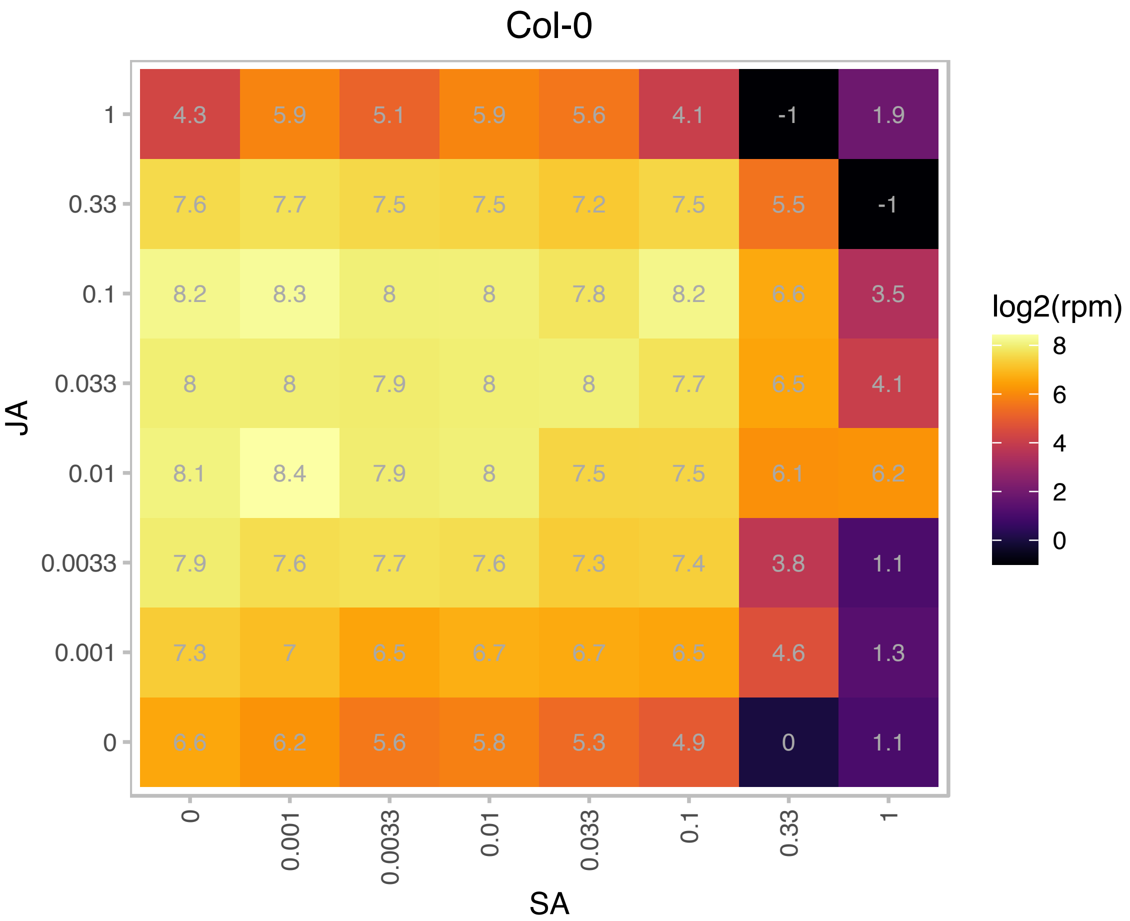


| | |
|---|---|
| Gene | AT2G46650 |
| is.expressed_genes | TRUE |
| is.responsive_genes | TRUE |
| is.apcluster | E-01 |

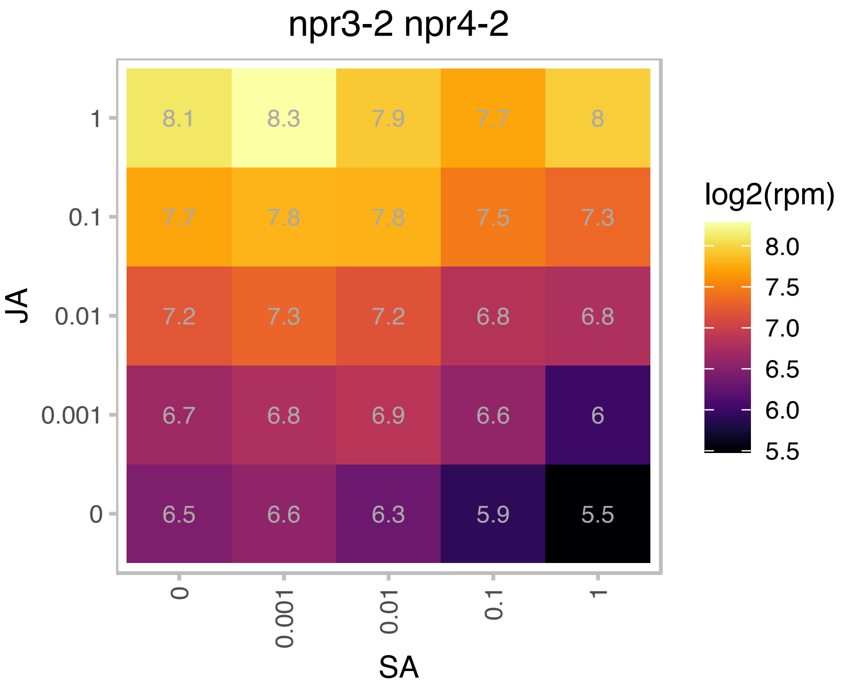


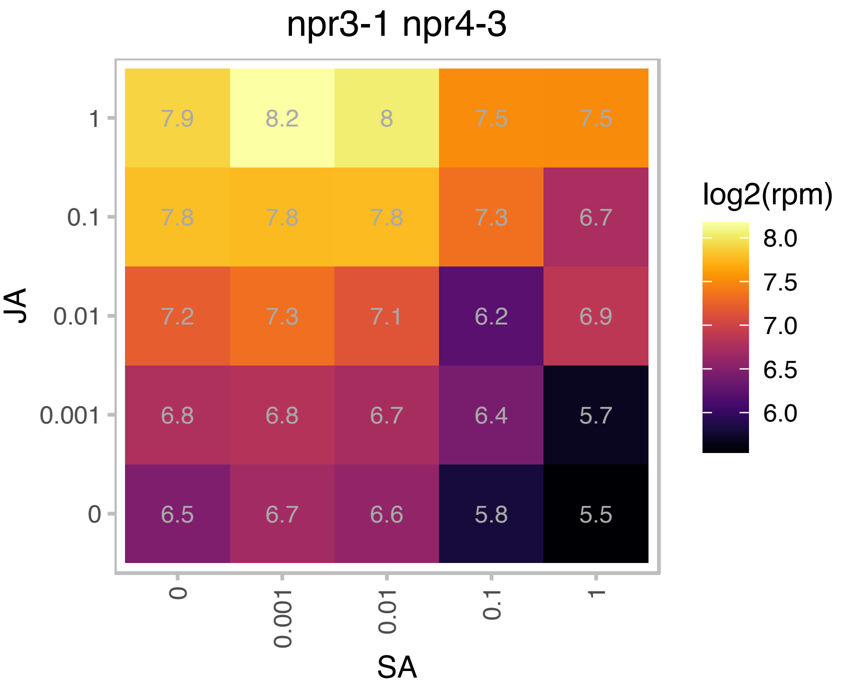

### AT3G06510.1 : ATSFR2, SFR2

Glycosyl hydrolase superfamily protein
Encodes a protein with beta-glucosidase and galactosyltransferase activity, mutants show increased sensitivity to freezing.  Though it is classified as a family I glycosyl hydrolase, it has no hydrolase activity in vitro.

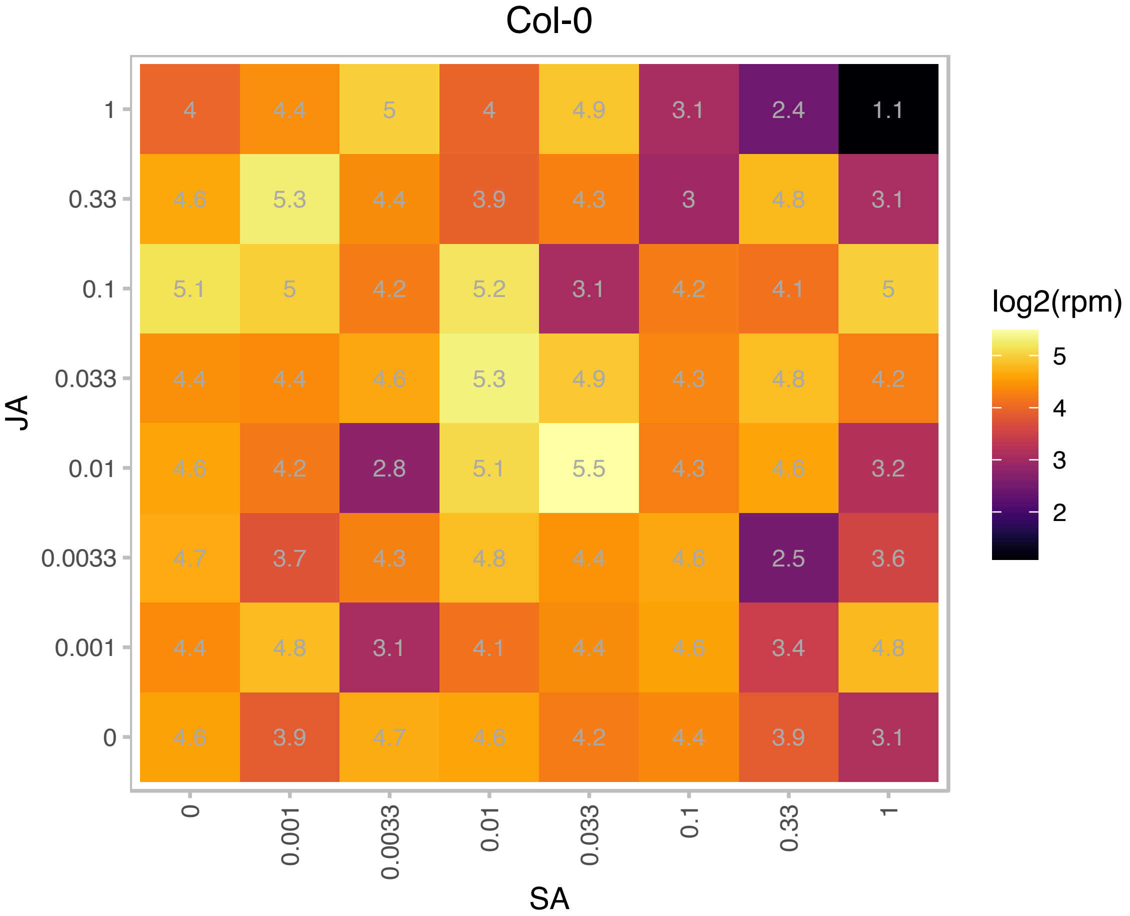


| | |
|---|---|
| Gene | AT3G06510 |
| is.expressed_genes | TRUE |
| is.responsive_genes | TRUE |
| is.apcluster | E-01 |

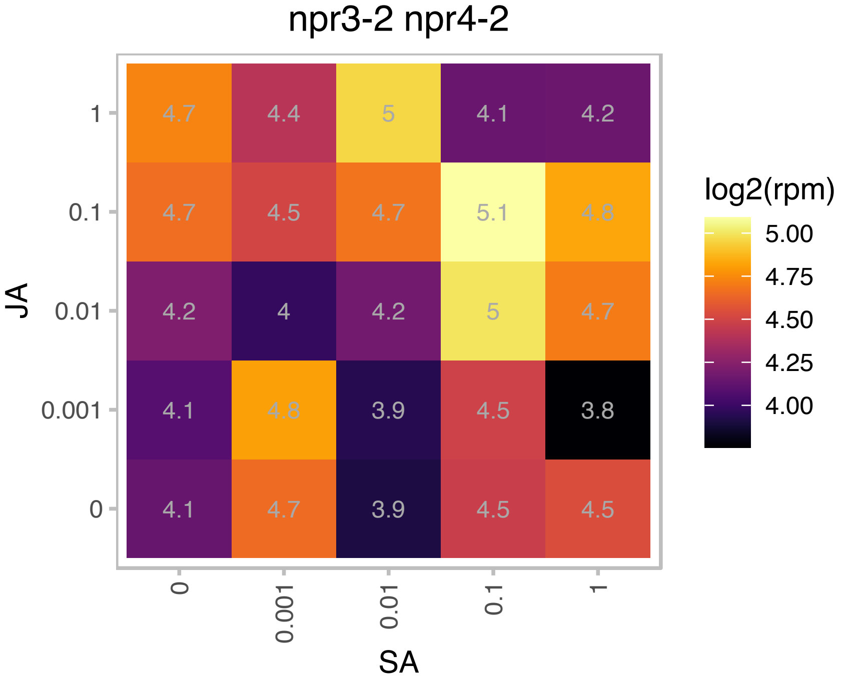


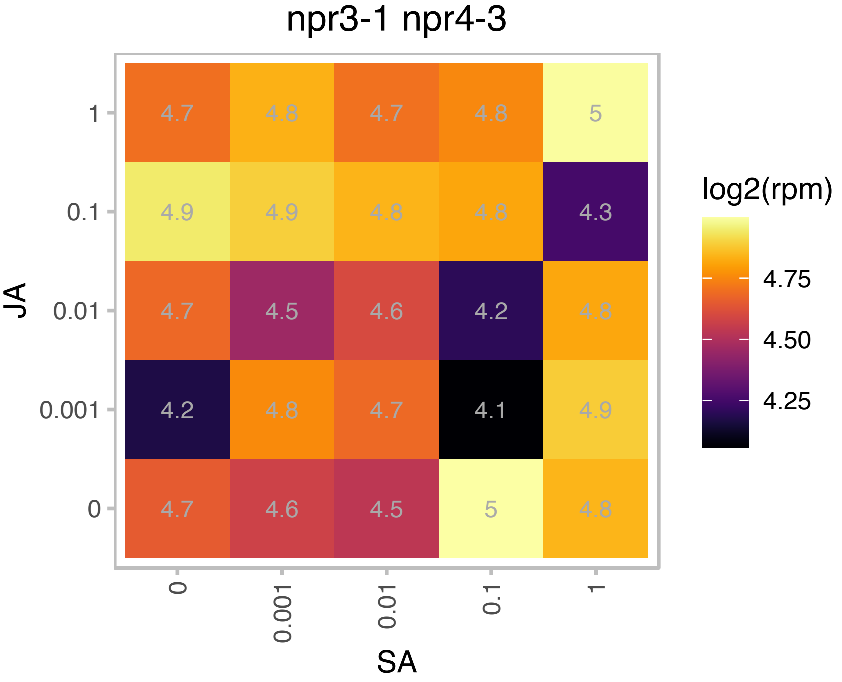

# AT3G07350.1 : NA

Protein of unknown function (DUF506)

| | |
|---|---|
| Gene | AT3G07350 |
| is.expressed_genes | TRUE |
| is.responsive_genes | TRUE |
| is.apcluster | E-01 |

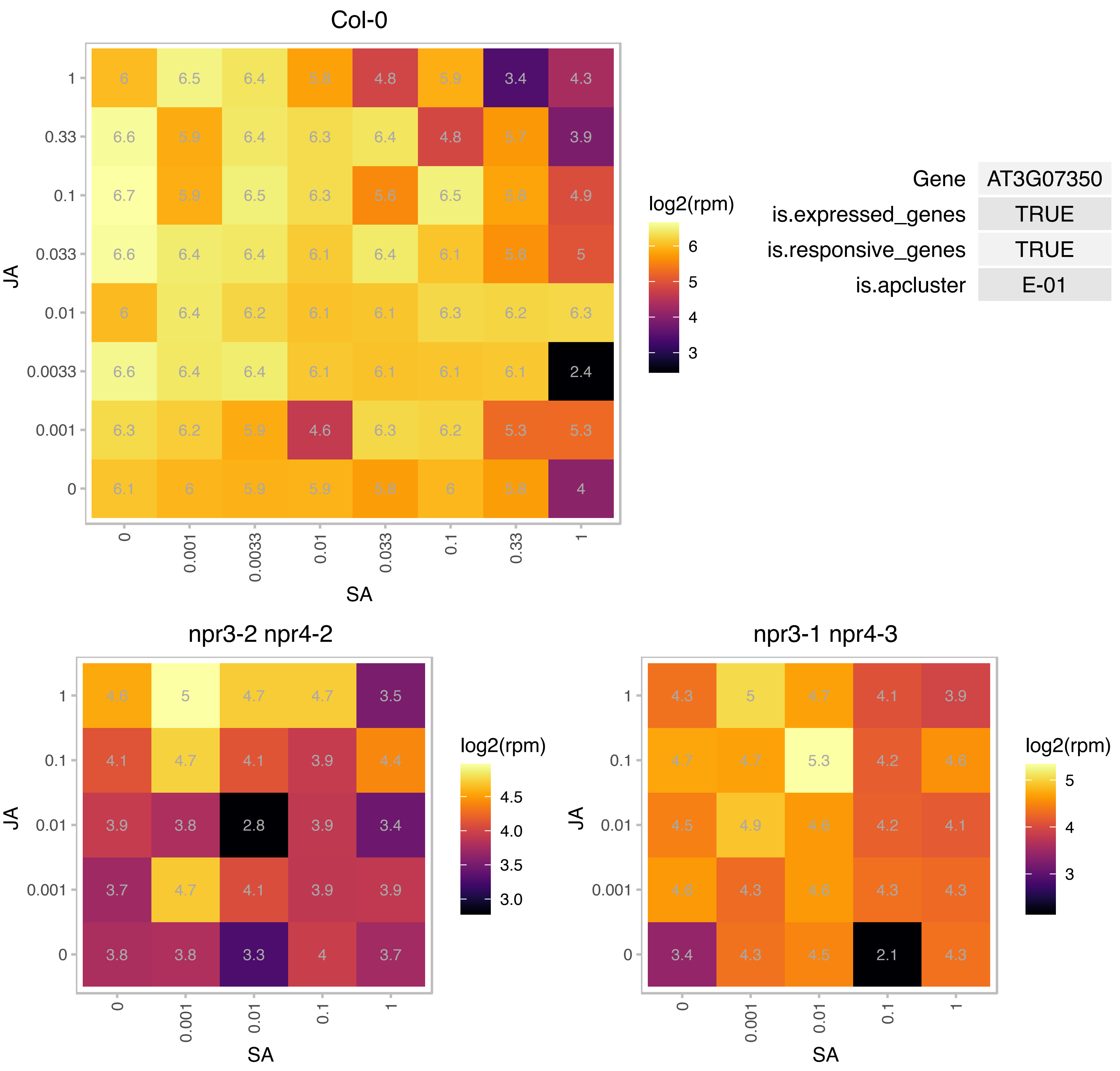

### AT3G12600.1 : atnudt16, NUDT16

nudix hydrolase homolog 16

Col-0

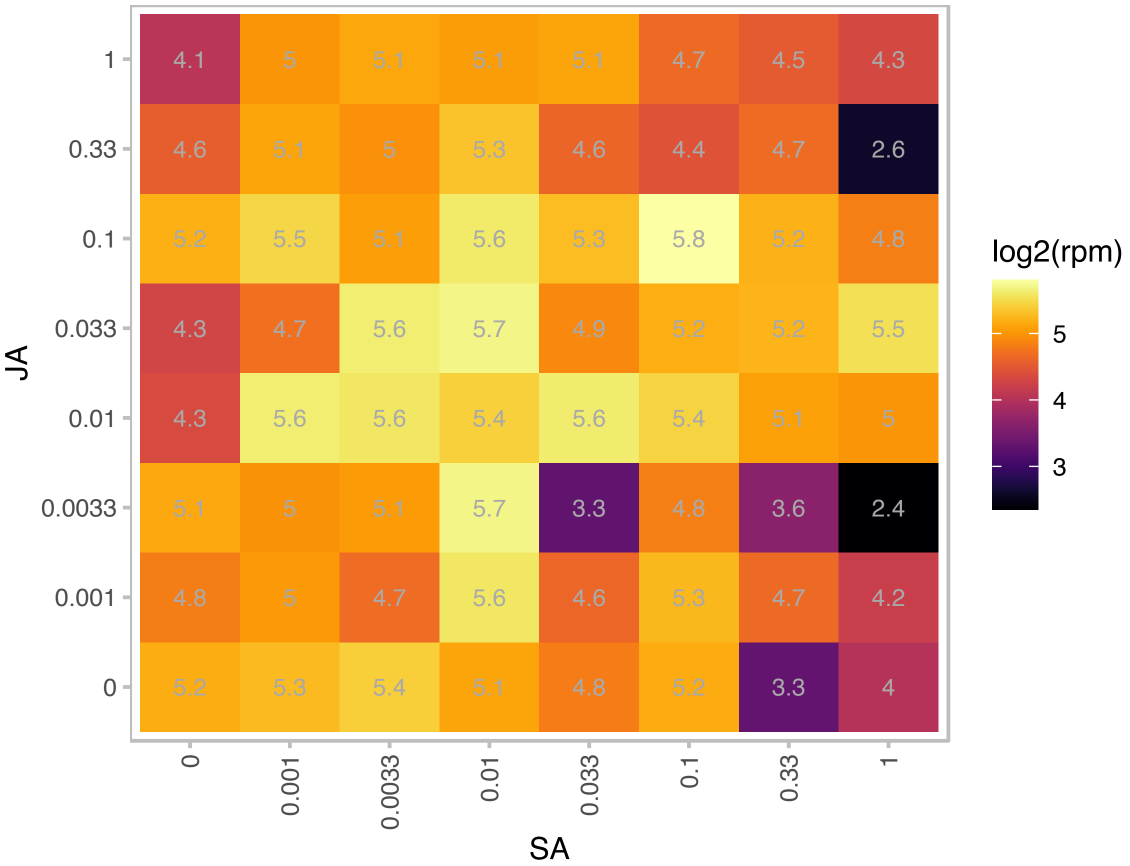


| | |
|---|---|
| Gene | AT3G12600 |
| is.expressed_genes | TRUE |
| is.responsive_genes | TRUE |
| is.apcluster | E-01 |

npr3-2 npr4-2

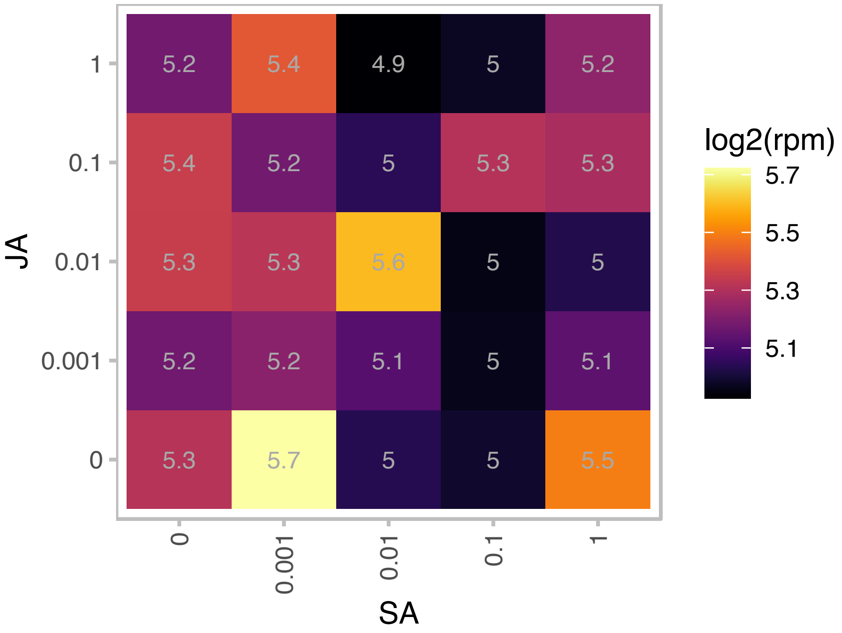


npr3-1 npr4-3

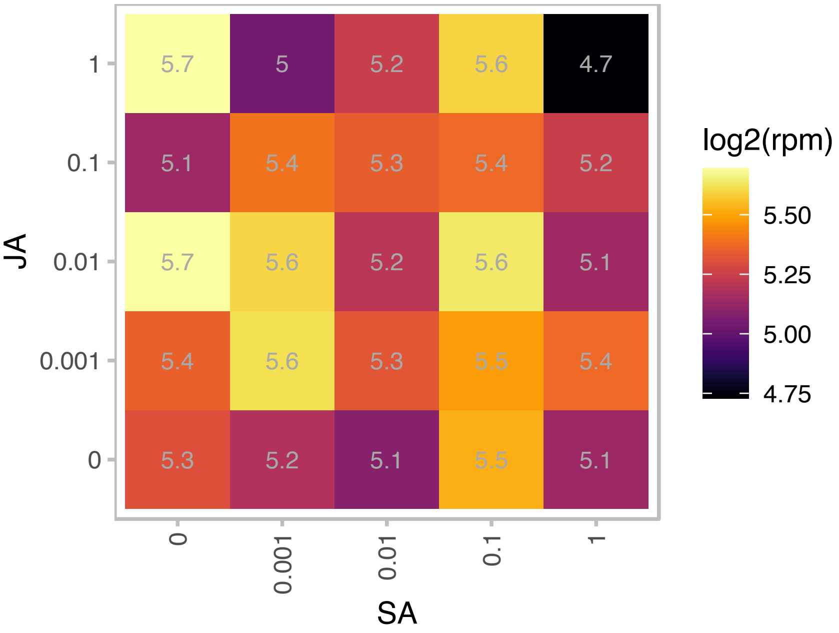

# AT3G15080.1 : NA

Polynucleotidyl transferase, ribonuclease H-like superfamily protein

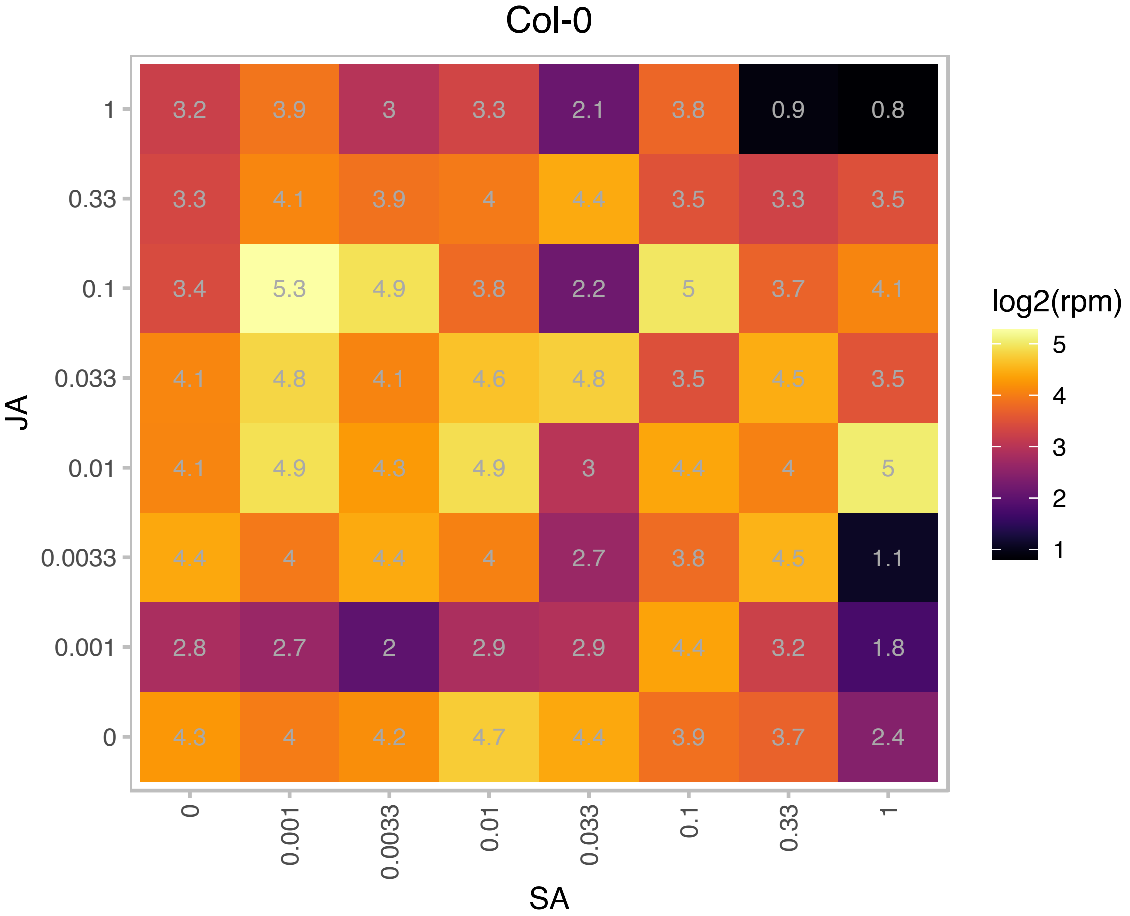


| Gene | AT3G15080 |
| --- | --- |
| is.expressed_genes | TRUE |
| is.responsive_genes | TRUE |
| is.apcluster | E-01 |

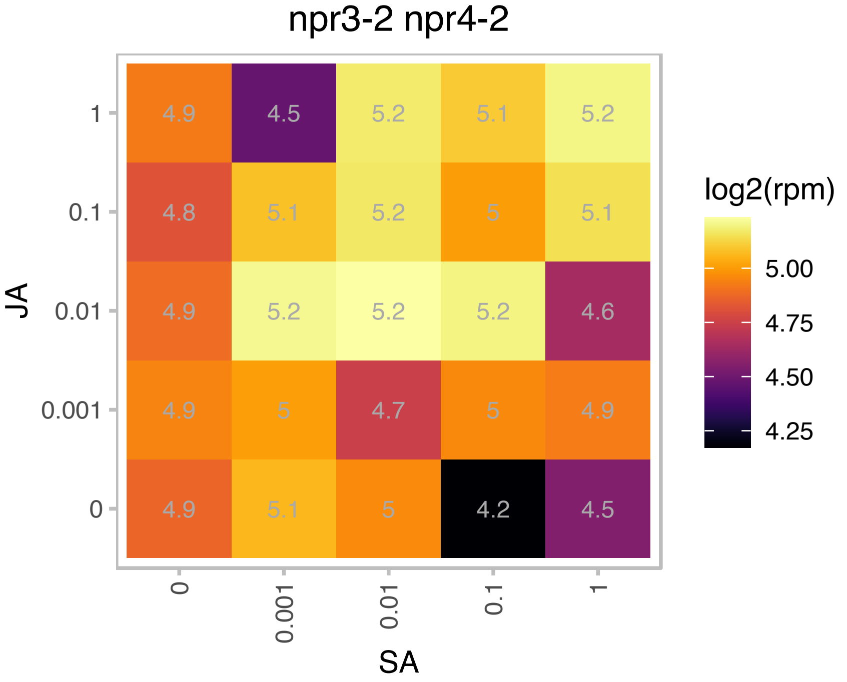


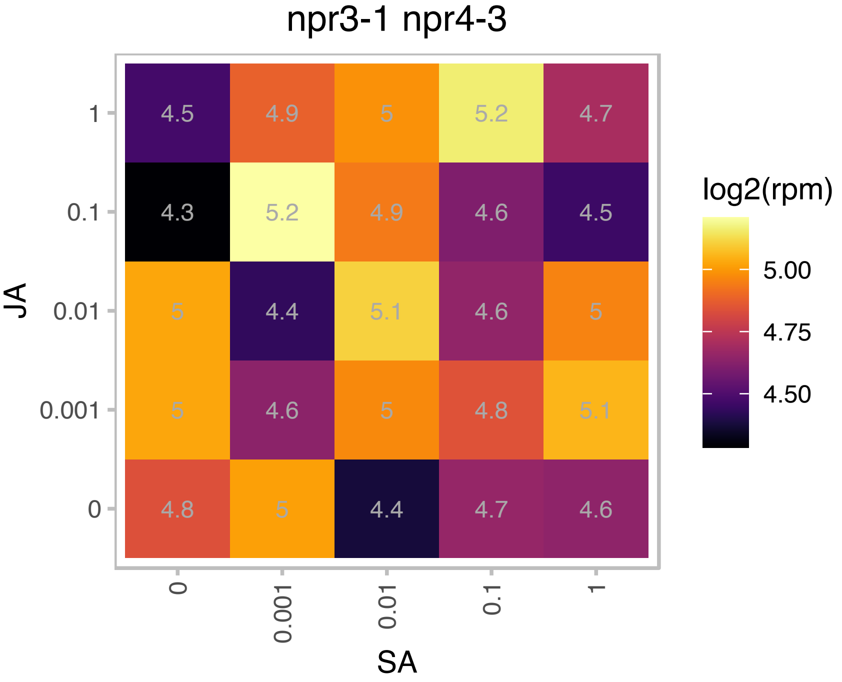

### AT3G15790.1 : ATMBD11, MBD11

methyl-CPG-binding domain 11

Protein containing methyl-CpG-binding domain.Has sequence similarity to human MBD proteins.

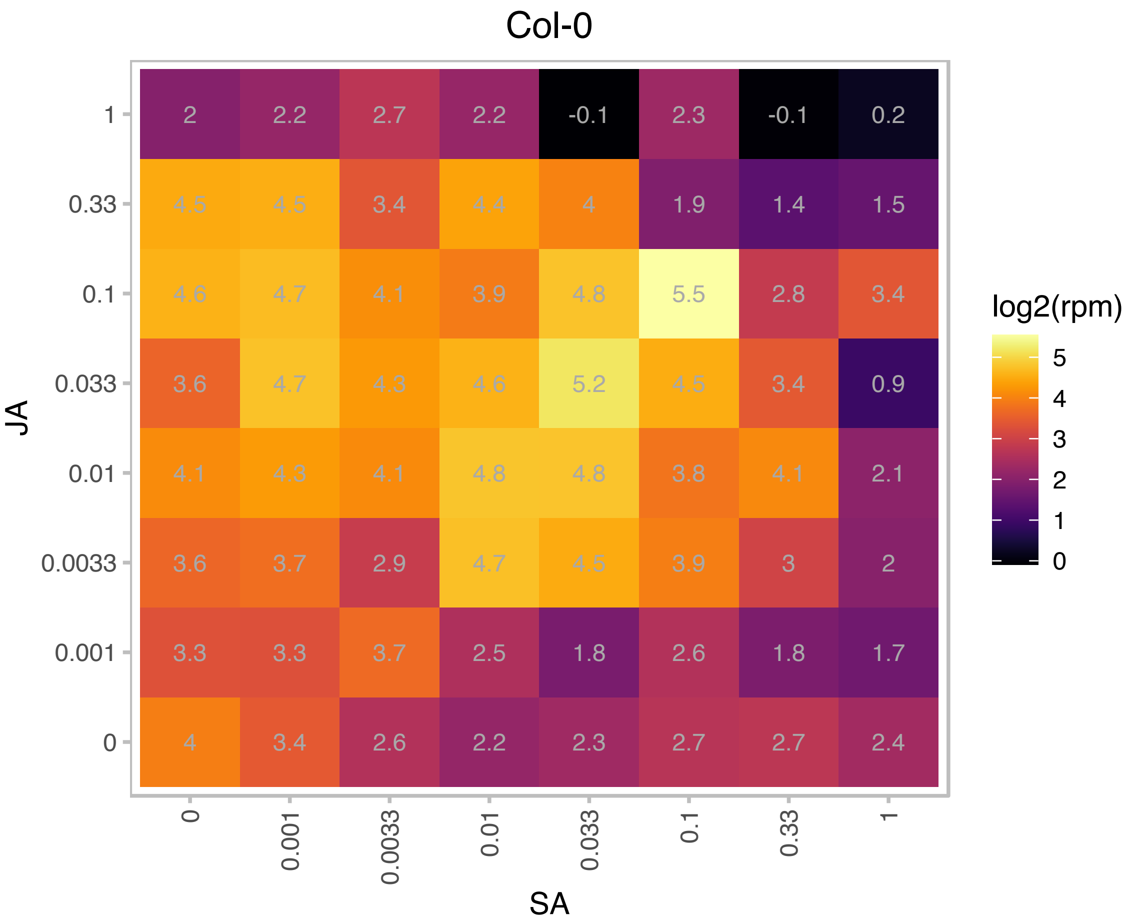


| | |
|---|---|
| Gene | AT3G15790 |
| is.expressed_genes | TRUE |
| is.responsive_genes | TRUE |
| is.apcluster | E-01 |

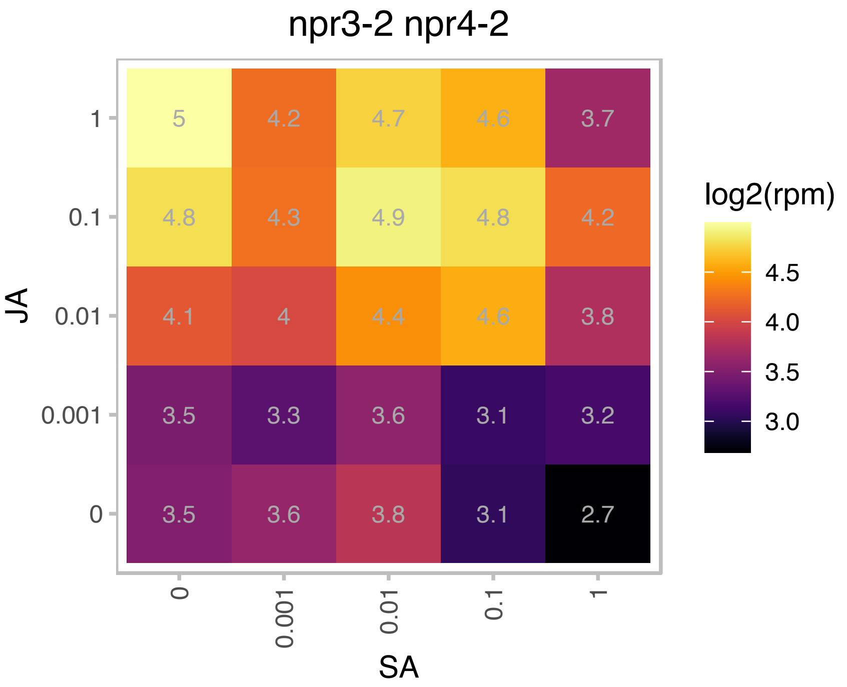


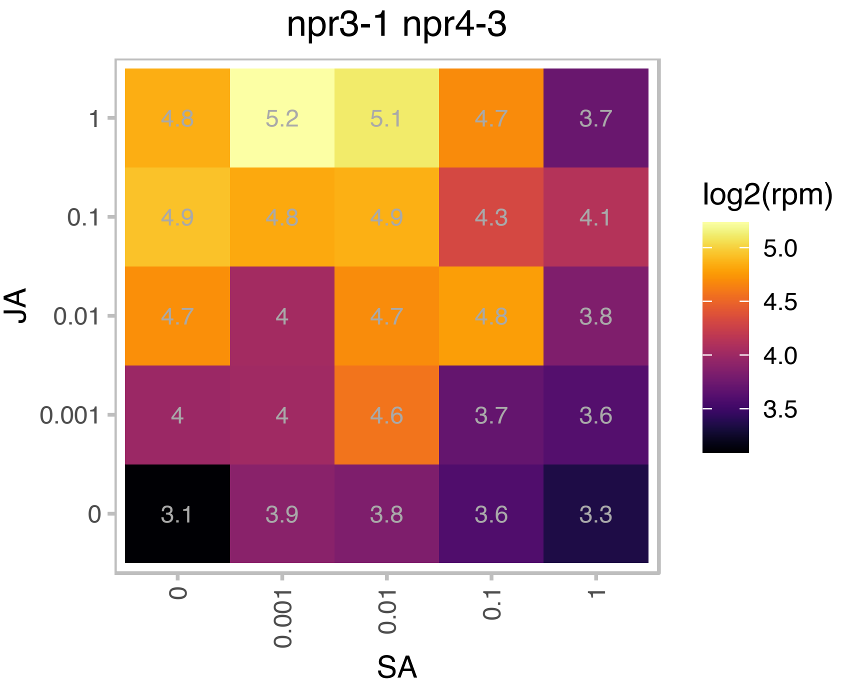

# AT3G17780.1 : NA

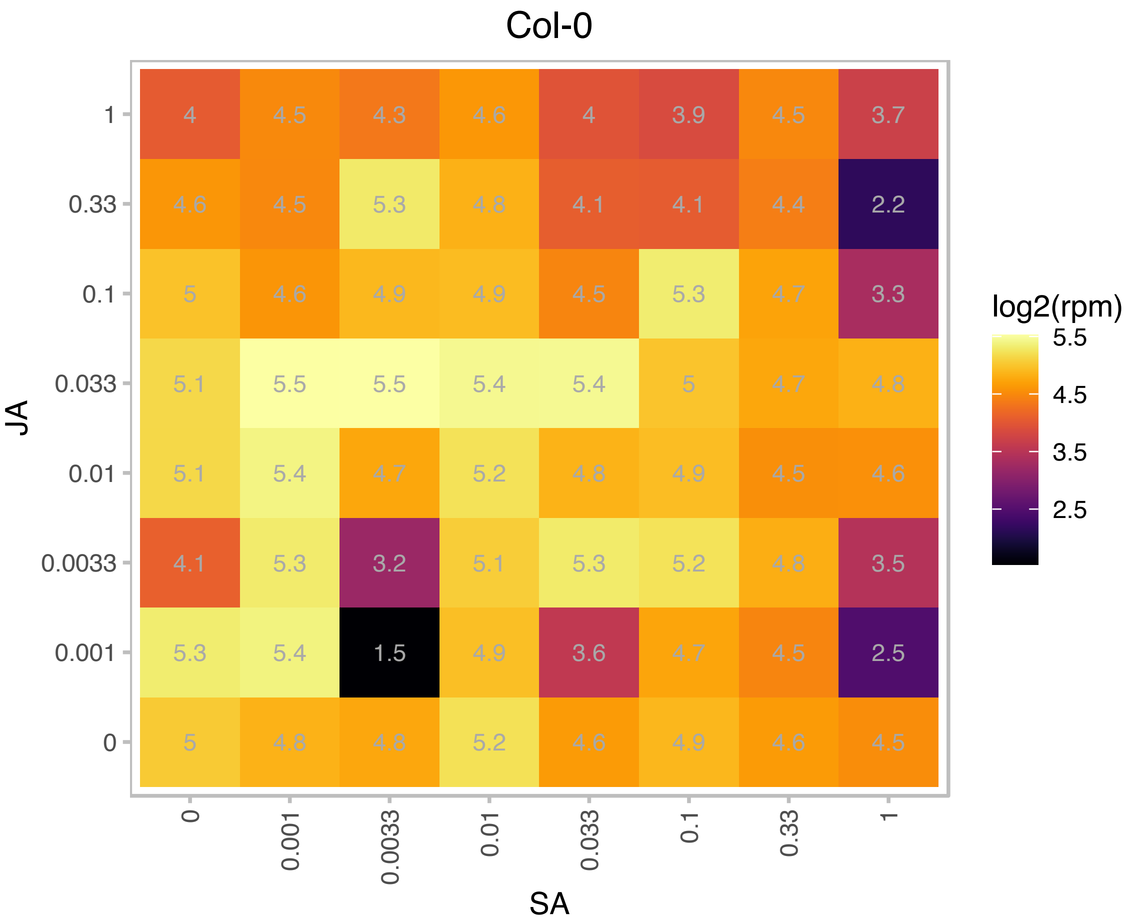


| | |
|---|---|
| Gene | AT3G17780 |
| is.expressed_genes | TRUE |
| is.responsive_genes | TRUE |
| is.apcluster | E-01 |

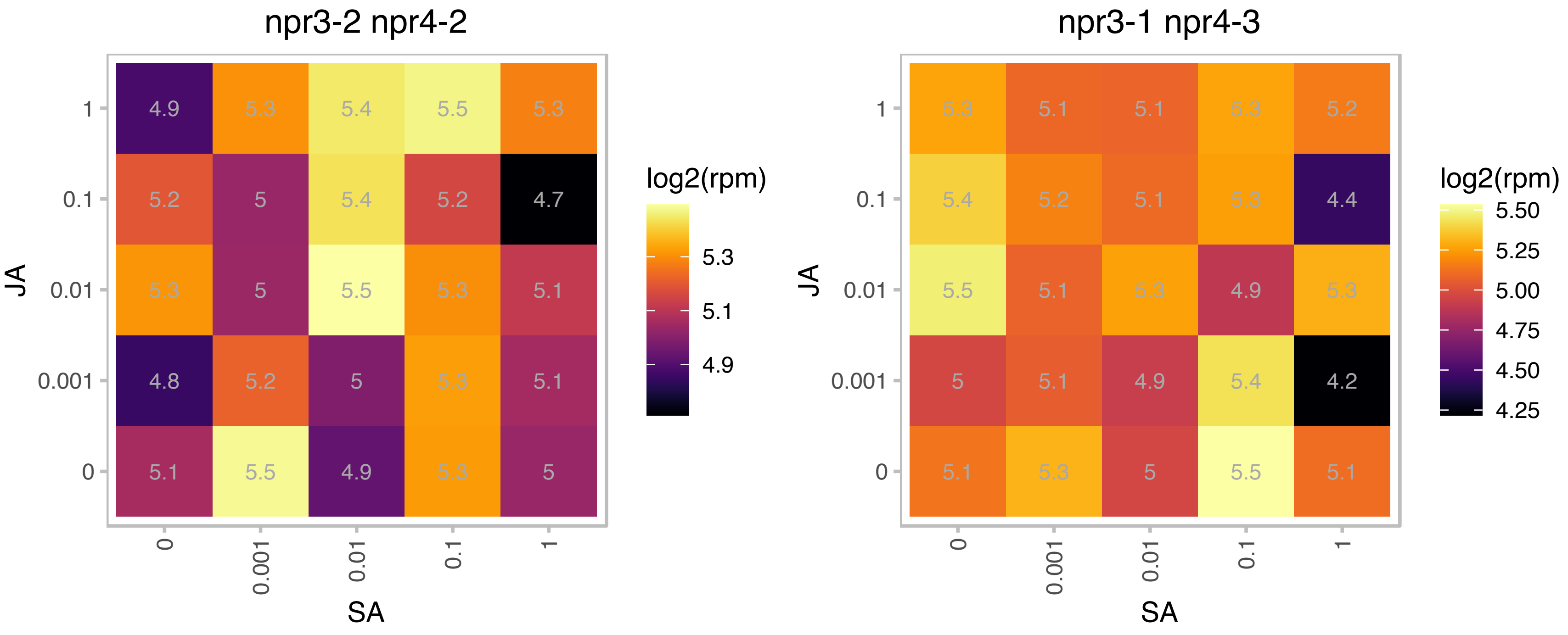

# AT3G20370.1 : NA

TRAF-like family protein

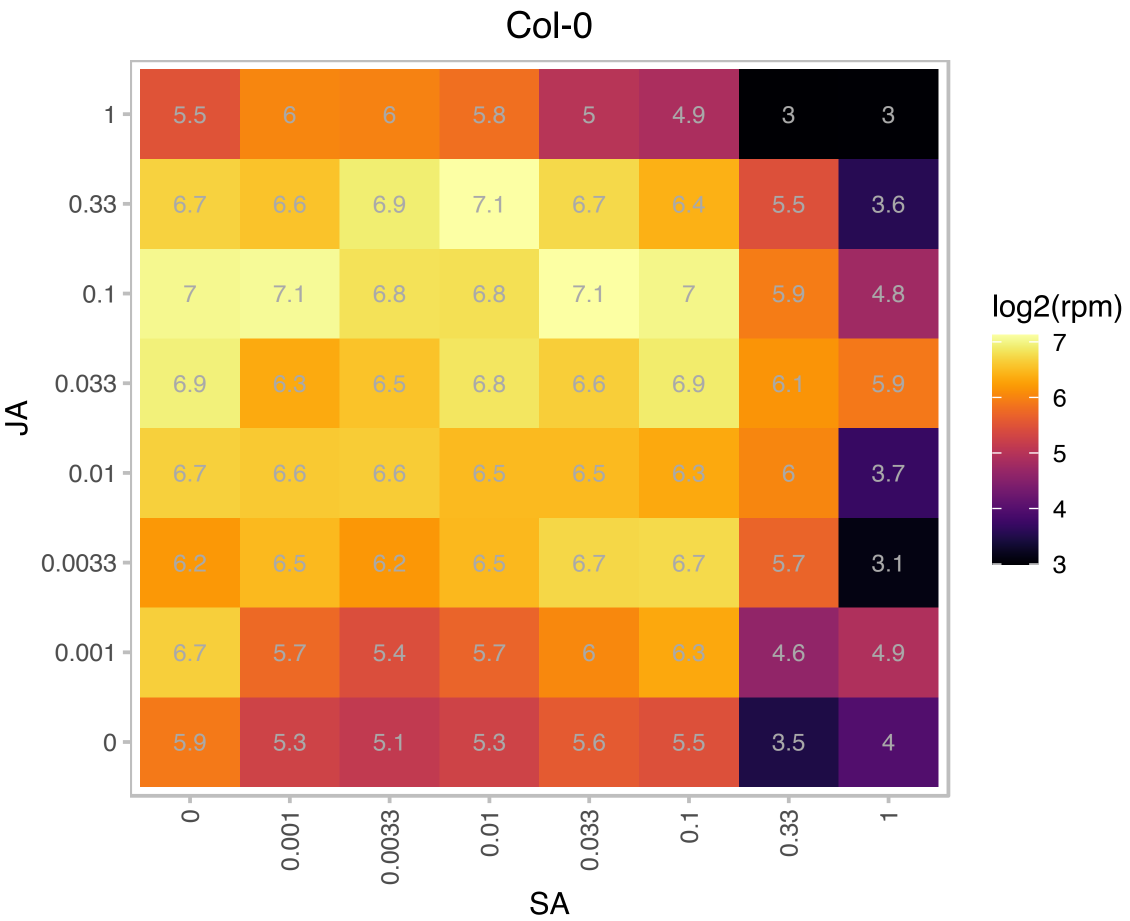


| | |
|---|---|
| Gene | AT3G20370 |
| is.expressed_genes | TRUE |
| is.responsive_genes | TRUE |
| is.apcluster | E-01 |

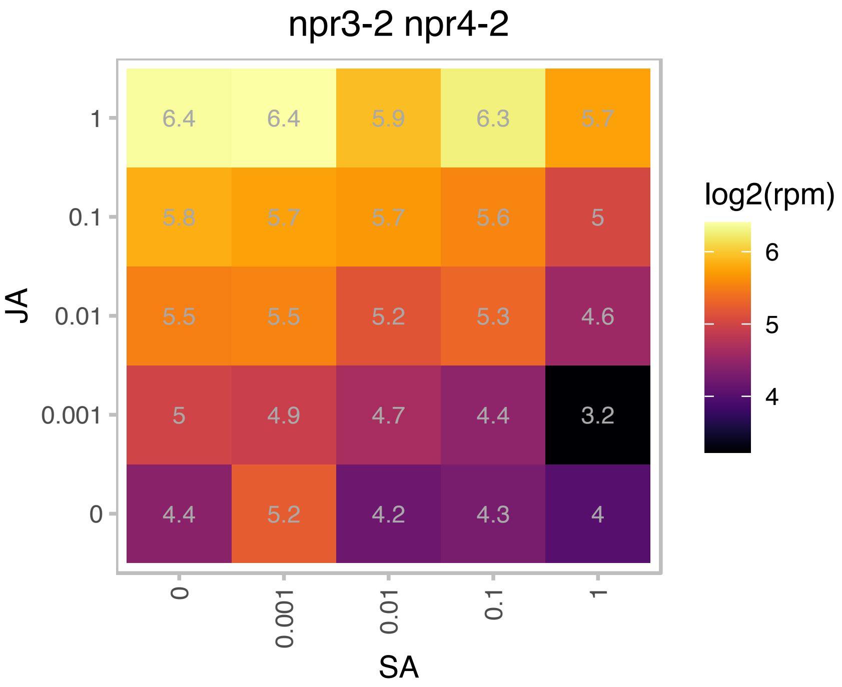


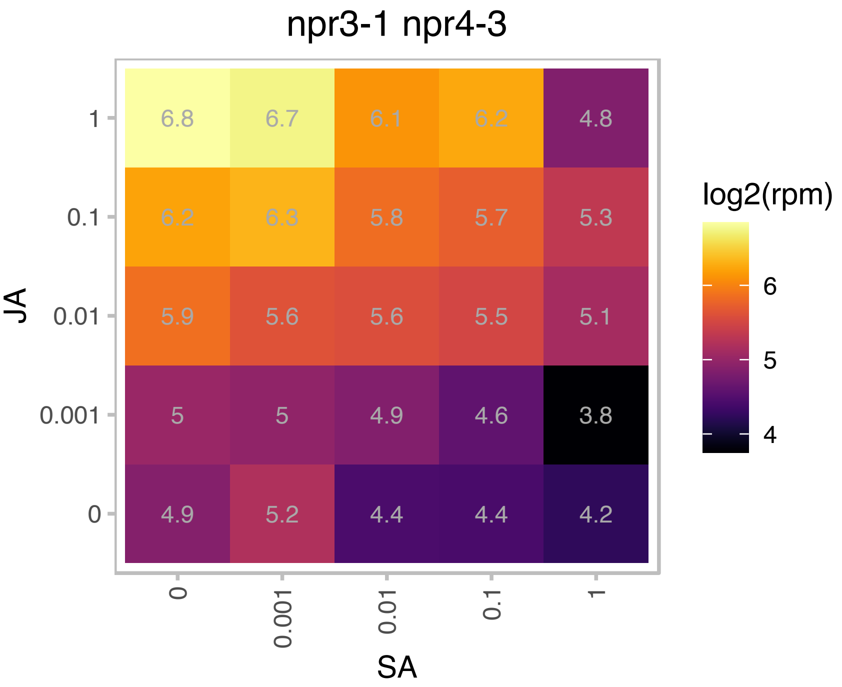

### AT3G26520.1 : GAMMA-TIP2, SITIP, TIP1;2, TIP2

tonoplast intrinsic protein 2
gamma tonoplast intrinsic protein 2 (TIP2). expressed throughout the plant and transcript level is increased upon NaCl or ABA treatments. NaCl stress-sensitive yeast mutant strains exhibit more resistance to salt when expressing this protein.

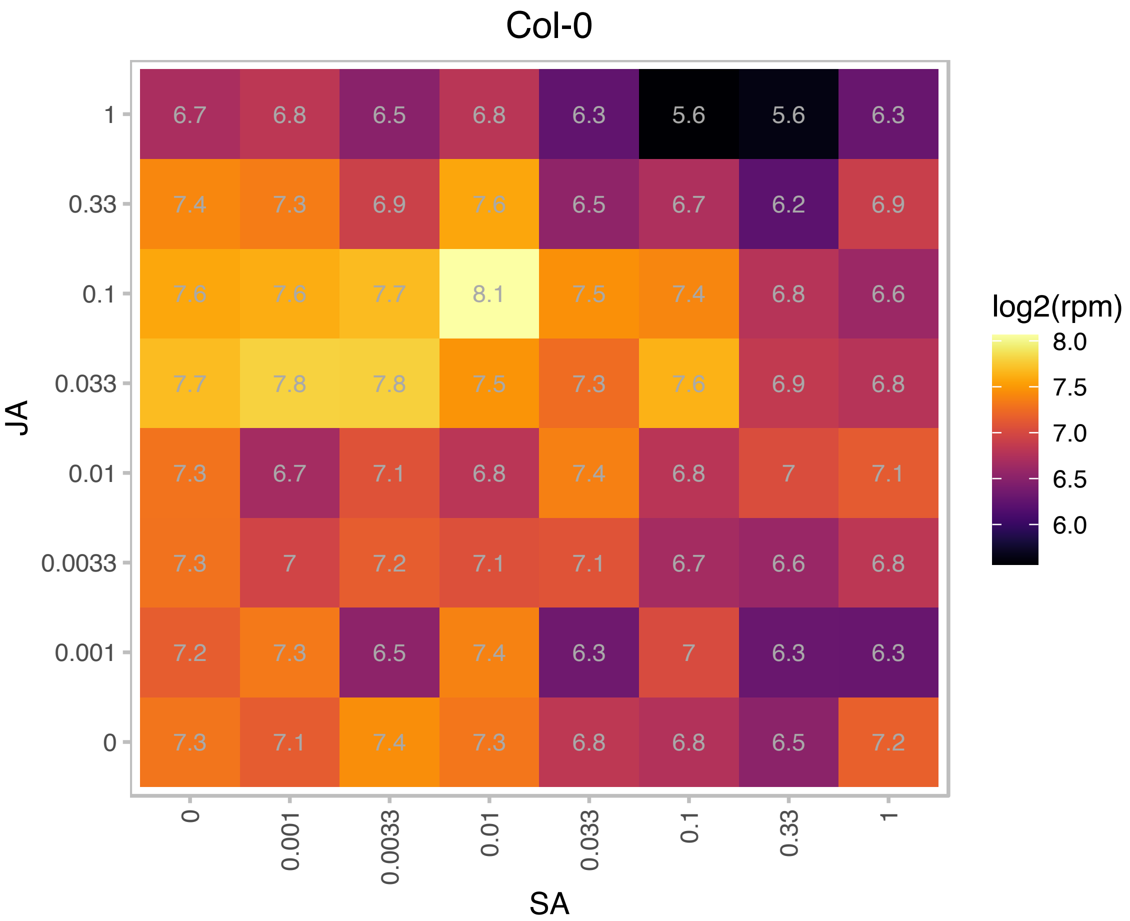


| | |
|---|---|
| Gene | AT3G26520 |
| is.expressed_genes | TRUE |
| is.responsive_genes | TRUE |
| is.apcluster | E-01 |

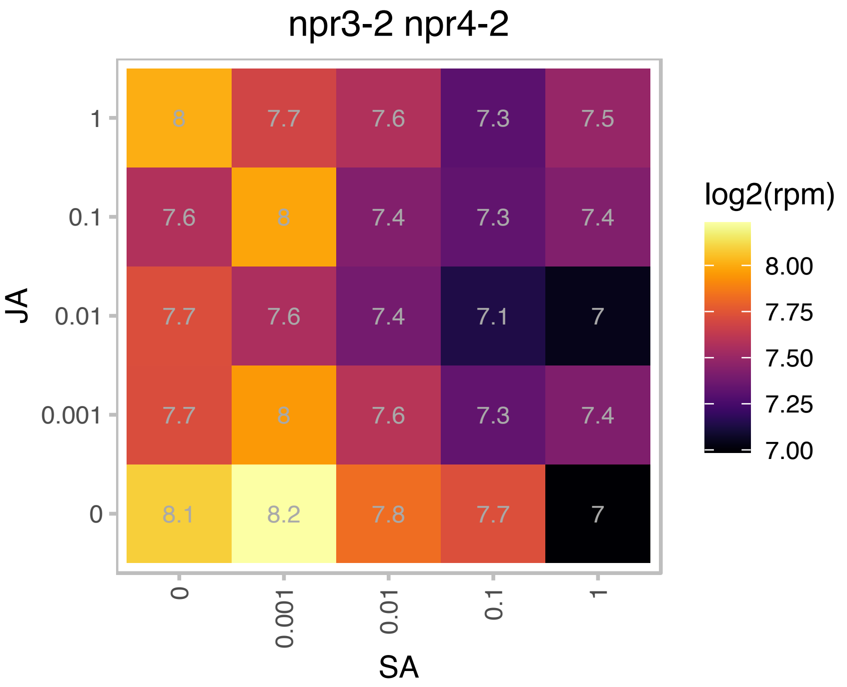


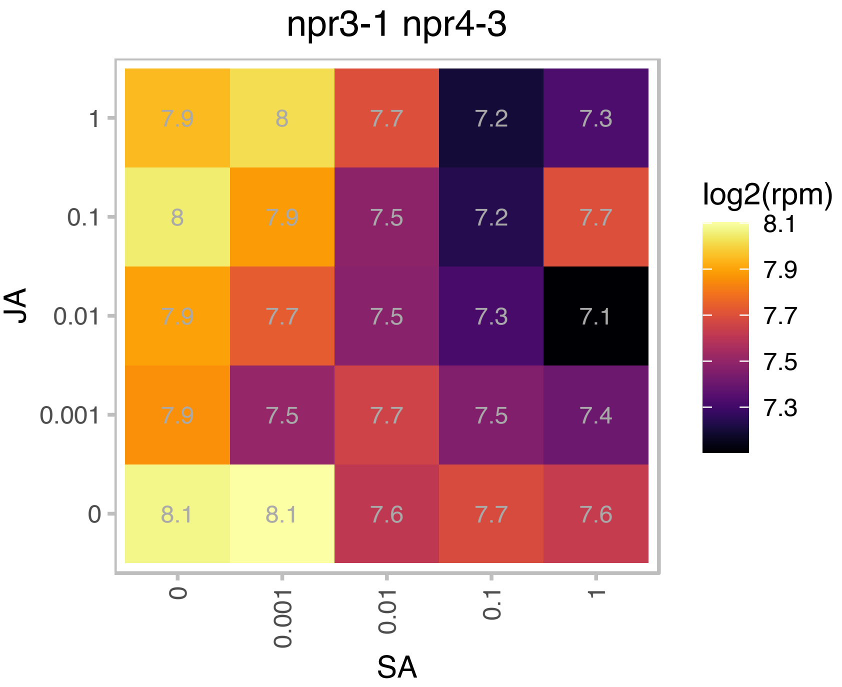

# AT3G45310.1 : NA

Cysteine proteinases superfamily protein

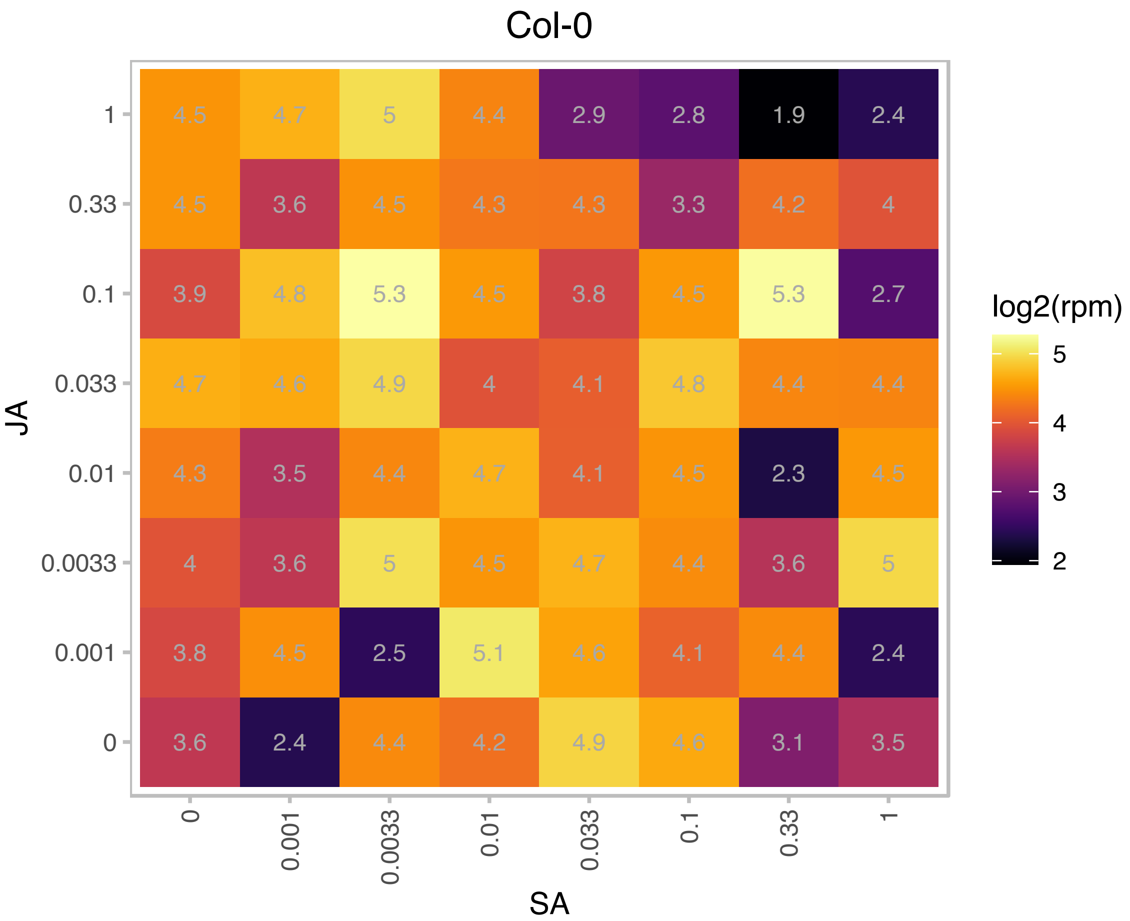


| | |
|---|---|
| Gene | AT3G45310 |
| is.expressed_genes | TRUE |
| is.responsive_genes | TRUE |
| is.apcluster | E-01 |

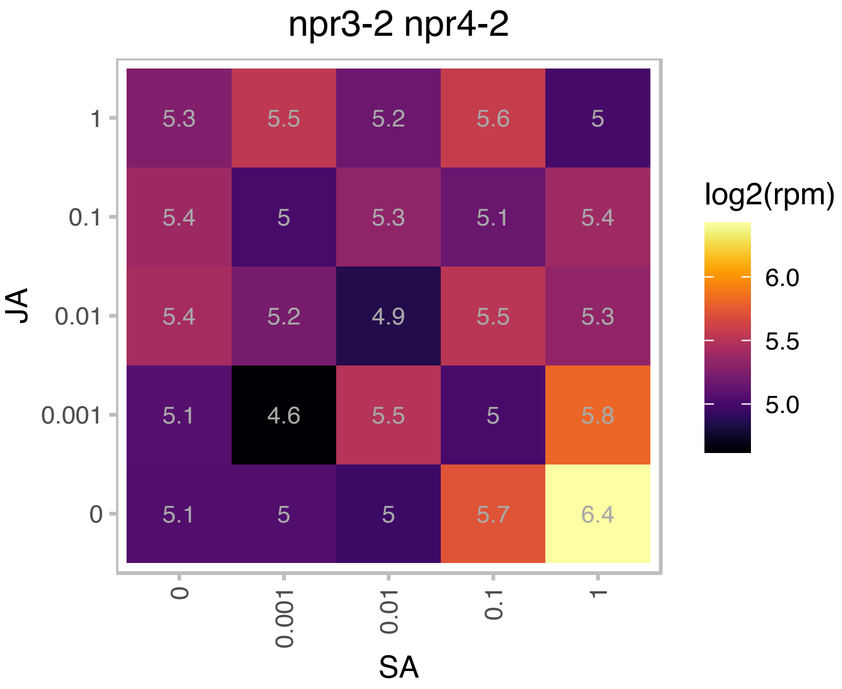


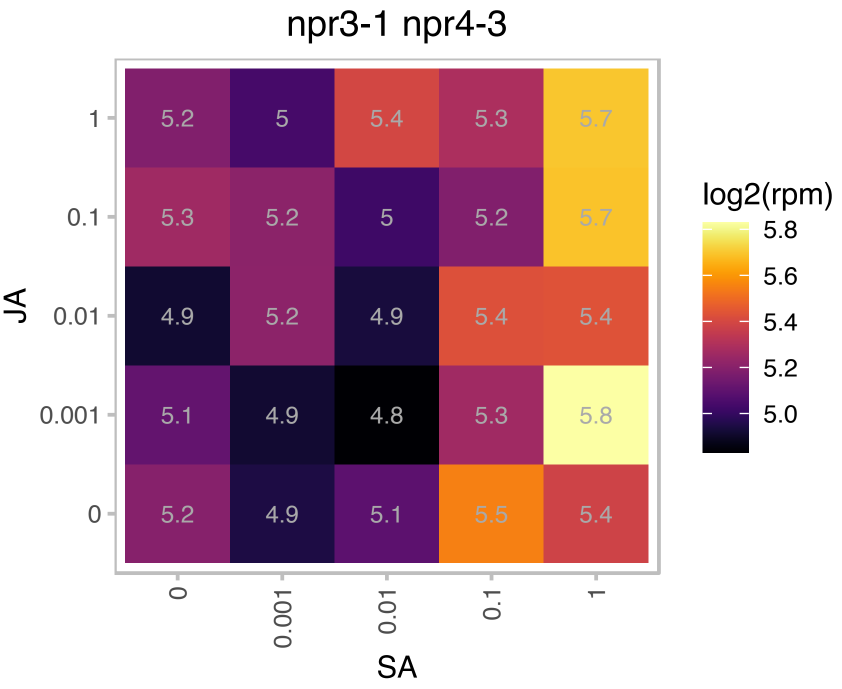

# AT3G52500.1 : NA

Eukaryotic aspartyl protease family protein

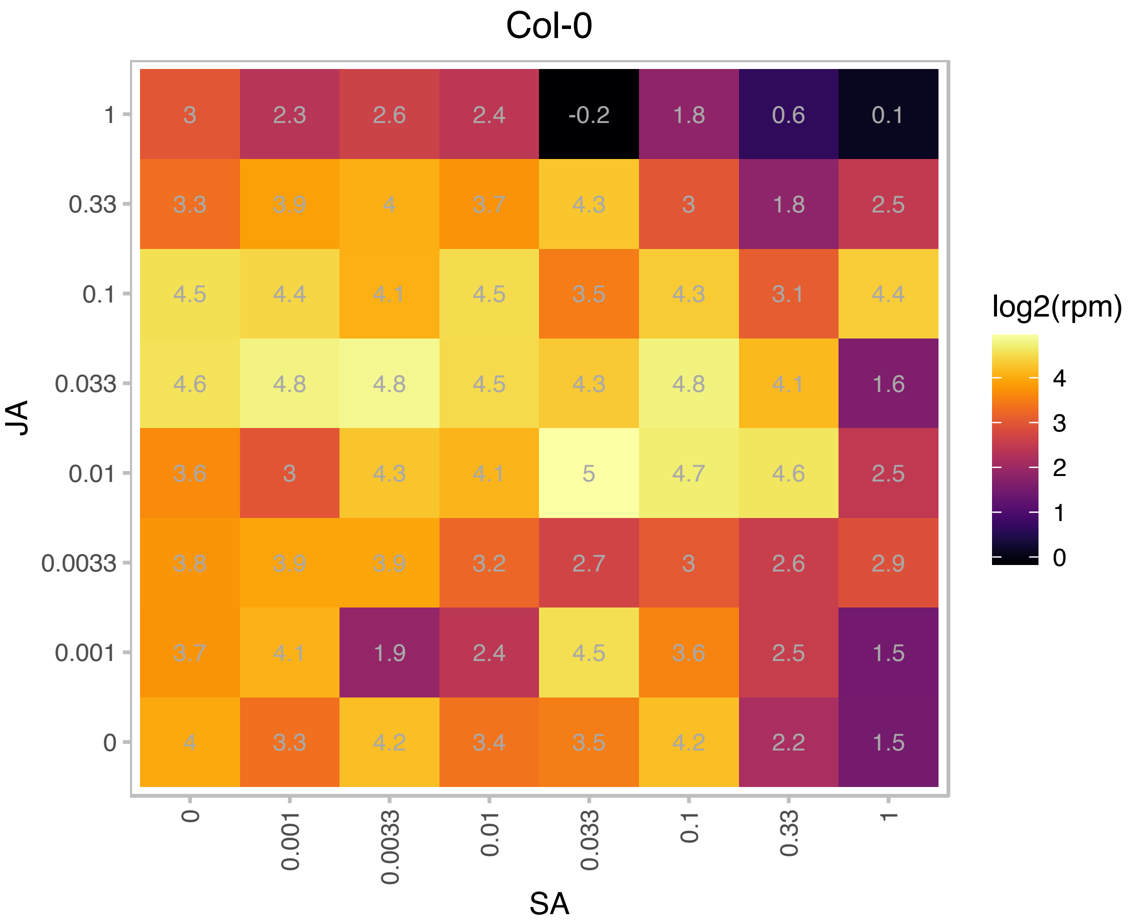


| | |
|---|---|
| Gene | AT3G52500 |
| is.expressed_genes | TRUE |
| is.responsive_genes | TRUE |
| is.apcluster | E-01 |

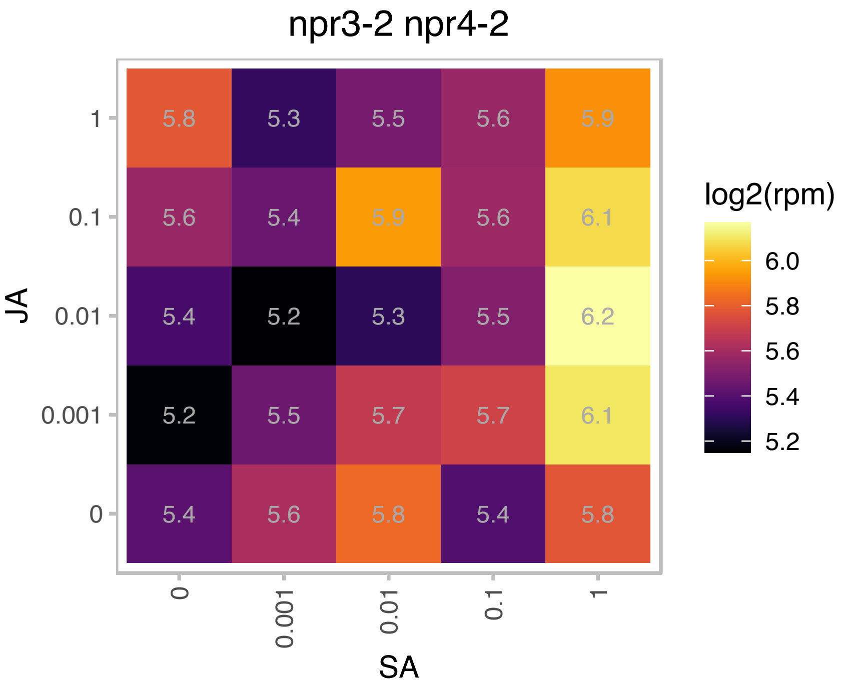


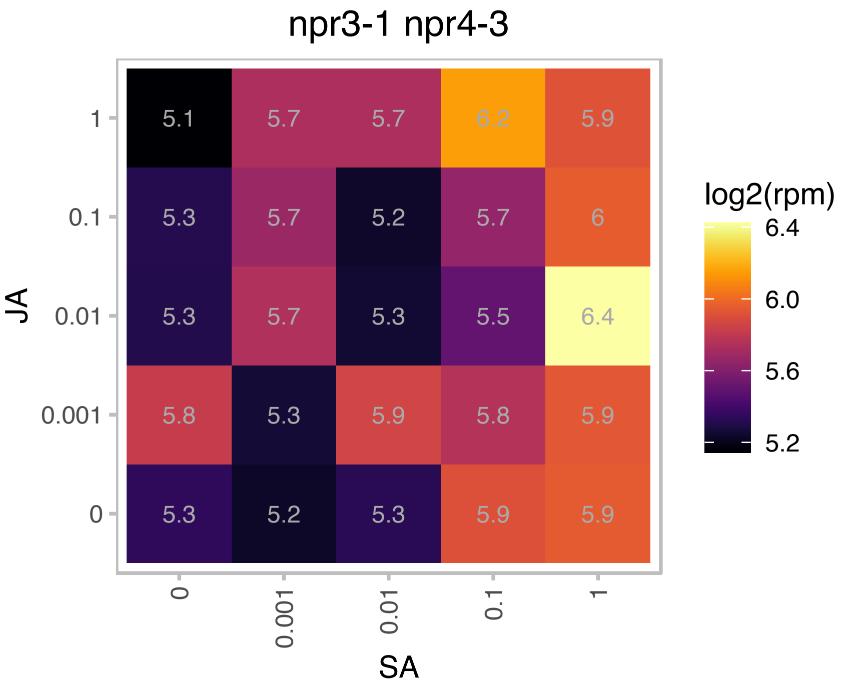

# AT3G52870.1 : NA

IQ calmodulin-binding motif family protein

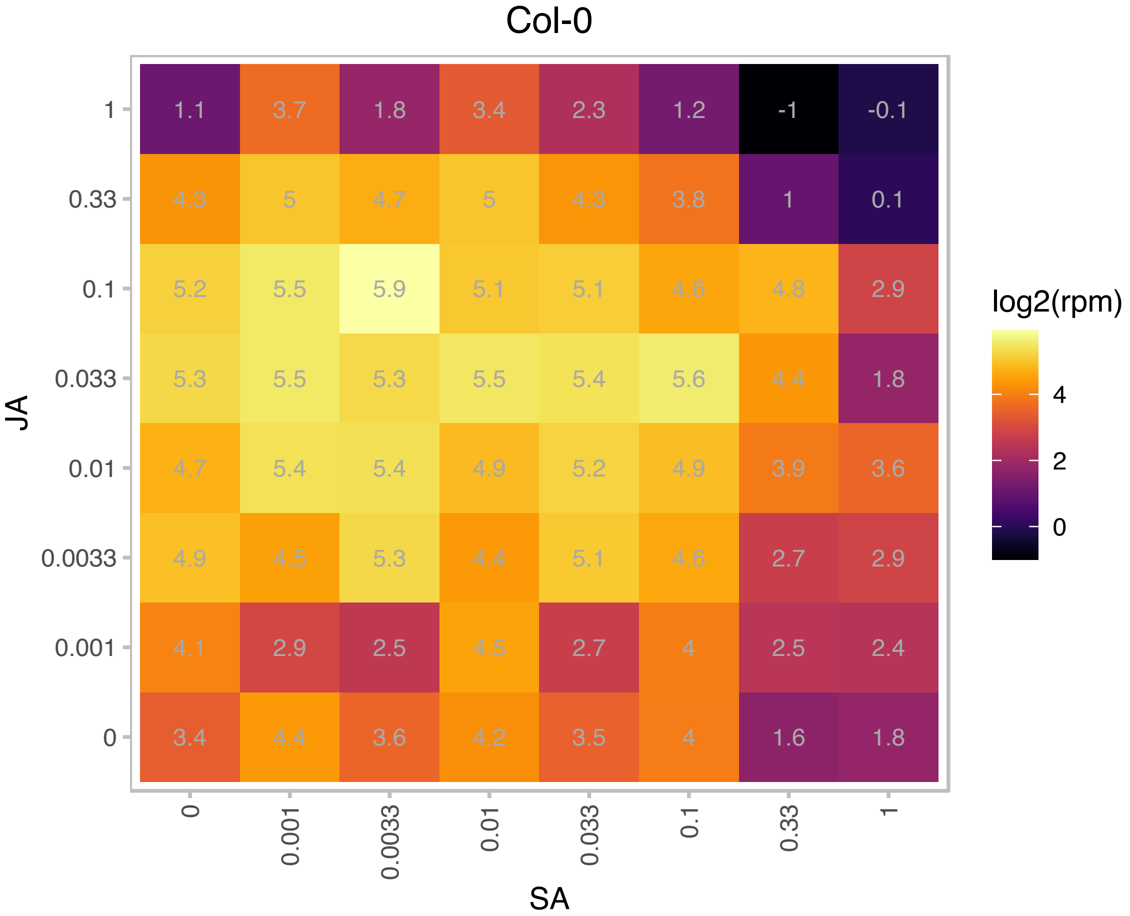


| | |
|---|---|
| Gene | AT3G52870 |
| is.expressed_genes | TRUE |
| is.responsive_genes | TRUE |
| is.apcluster | E-01 |

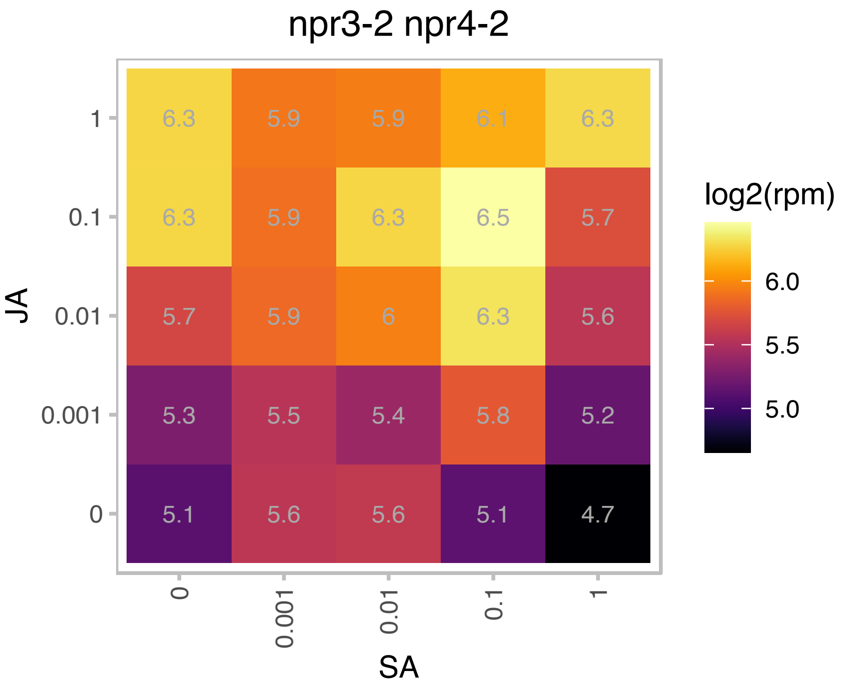


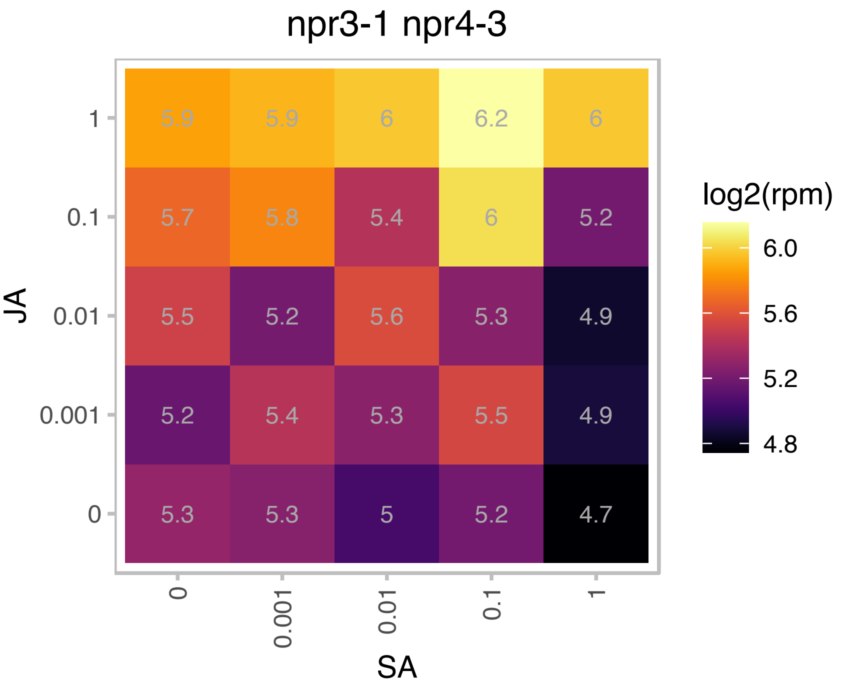

### AT3G54640.1 : TRP3, TSA1

tryptophan synthase alpha chain

Catalyzes the conversion of indole-3-glycerolphosphate to indole, the penultimate reaction in the biosynthesis of tryptophan. Functions as a heterocomplex with tryptophan synthase beta subunit (TSA2).

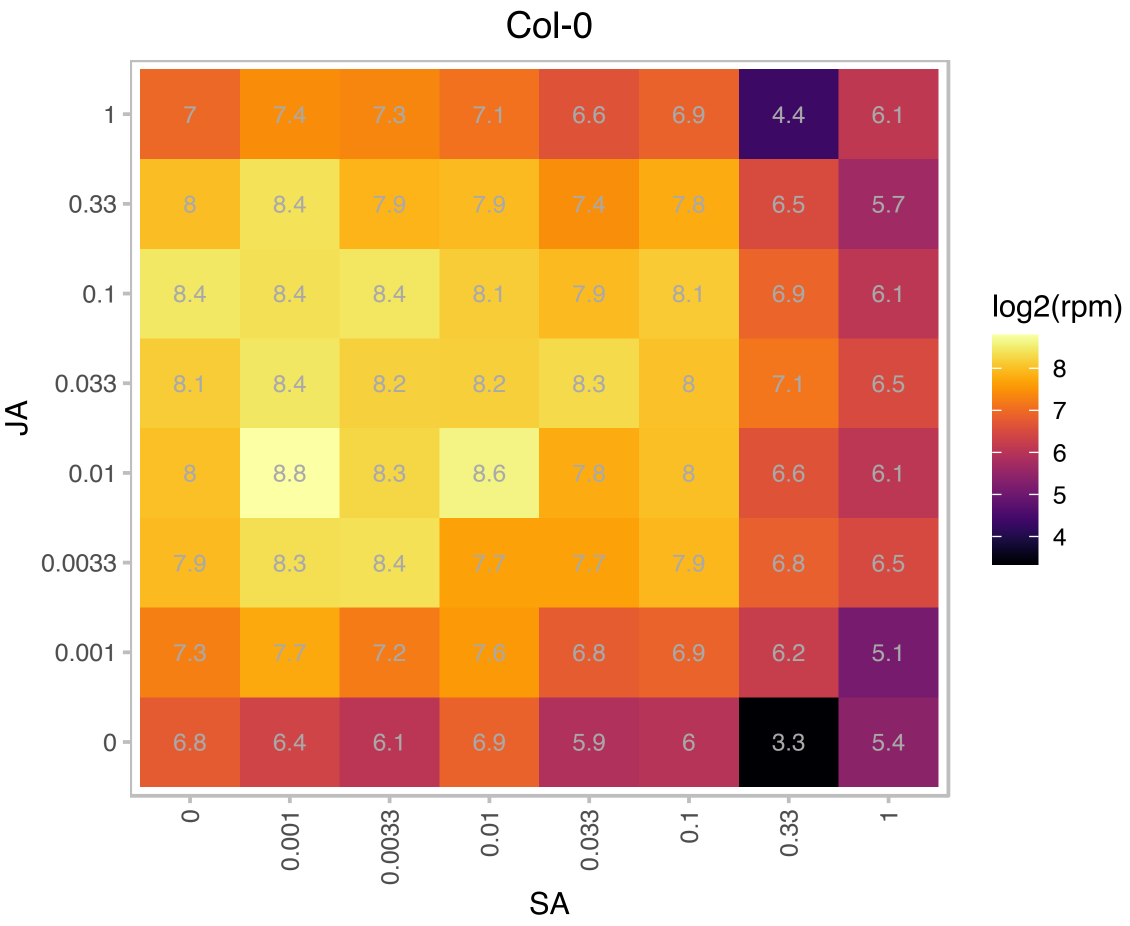


| | |
|---|---|
| Gene | AT3G54640 |
| is.expressed_genes | TRUE |
| is.responsive_genes | TRUE |
| is.apcluster | E-01 |

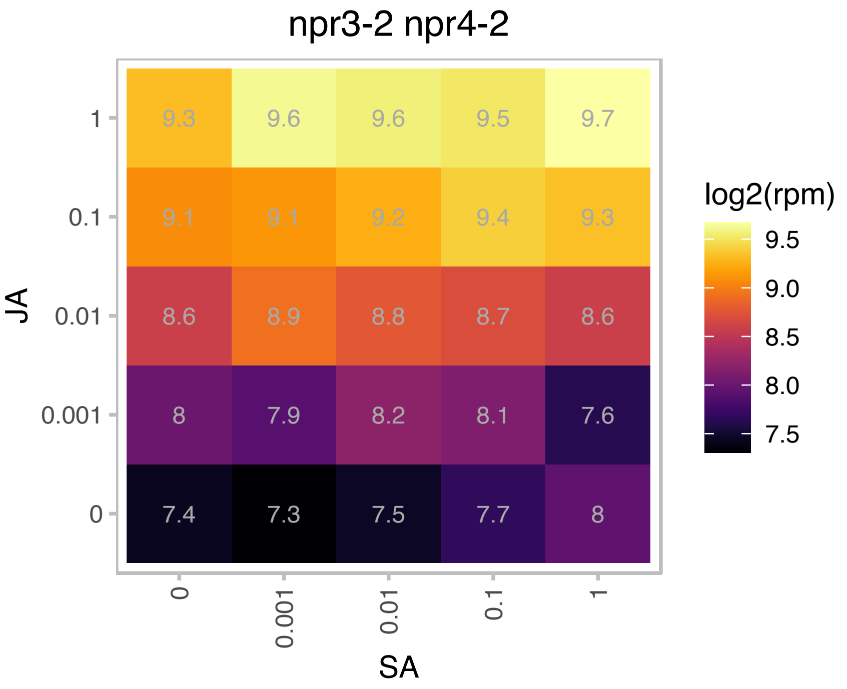


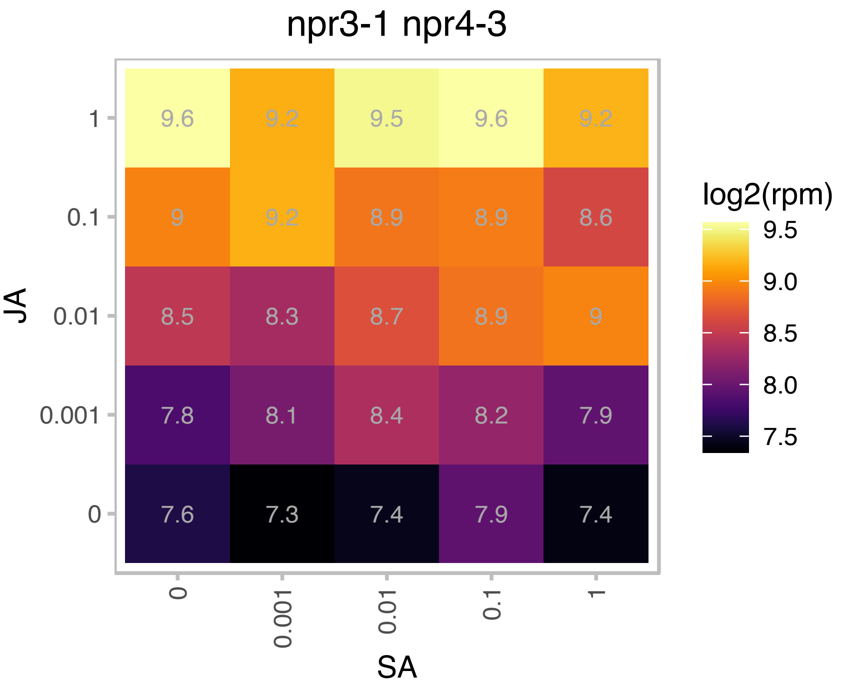

# AT3G57785.1 : NA

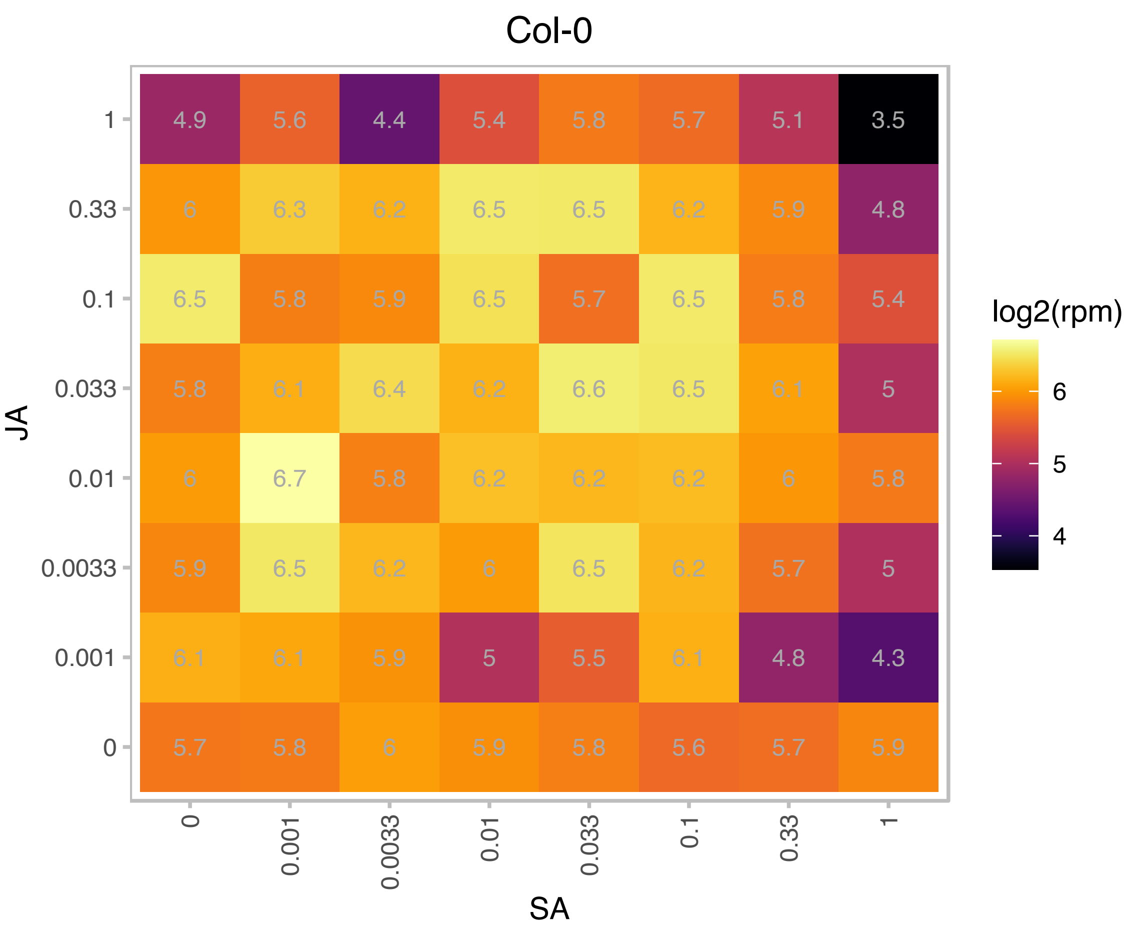


| | |
|---|---|
| Gene | AT3G57785 |
| is.expressed_genes | TRUE |
| is.responsive_genes | TRUE |
| is.apcluster | E-01 |

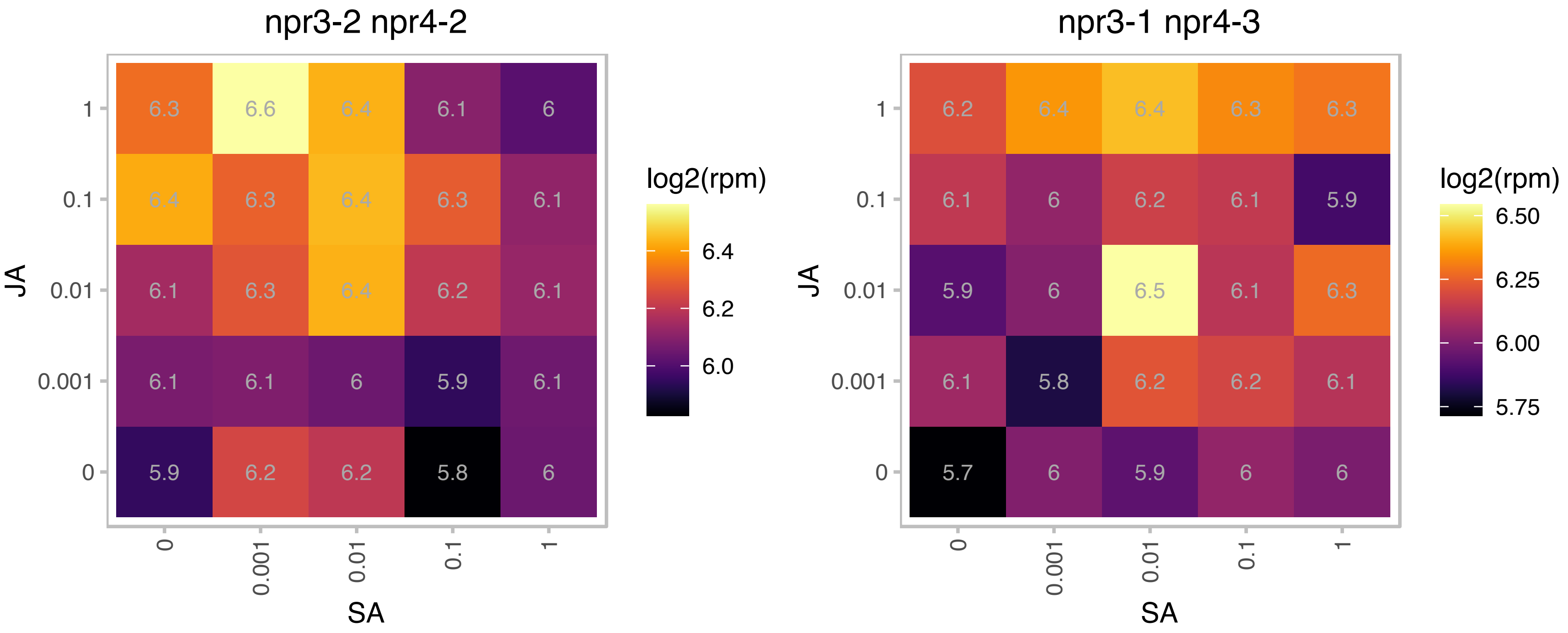

### AT3G58560.1 : AtCCR4a

DNAse I-like superfamily protein
Encodes a protein that is involved in mRNA processing and localized to cytoplasmic p-bodies. Double mutants with CCR4b show decreased sensitivity to high concentrations of sucrose. Involved in starch and sucrose metabolism.

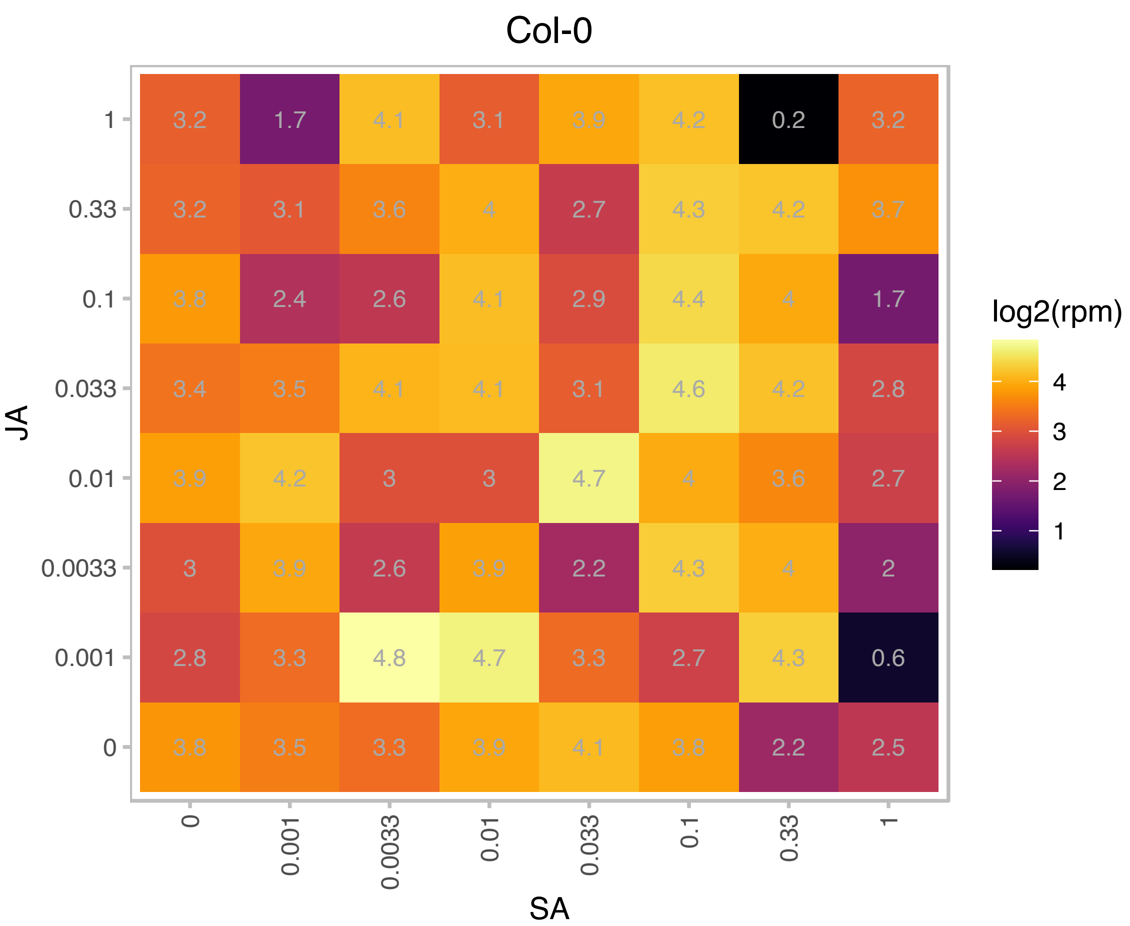


| | |
|---|---|
| Gene | AT3G58560 |
| is.expressed_genes | TRUE |
| is.responsive_genes | TRUE |
| is.apcluster | E-01 |

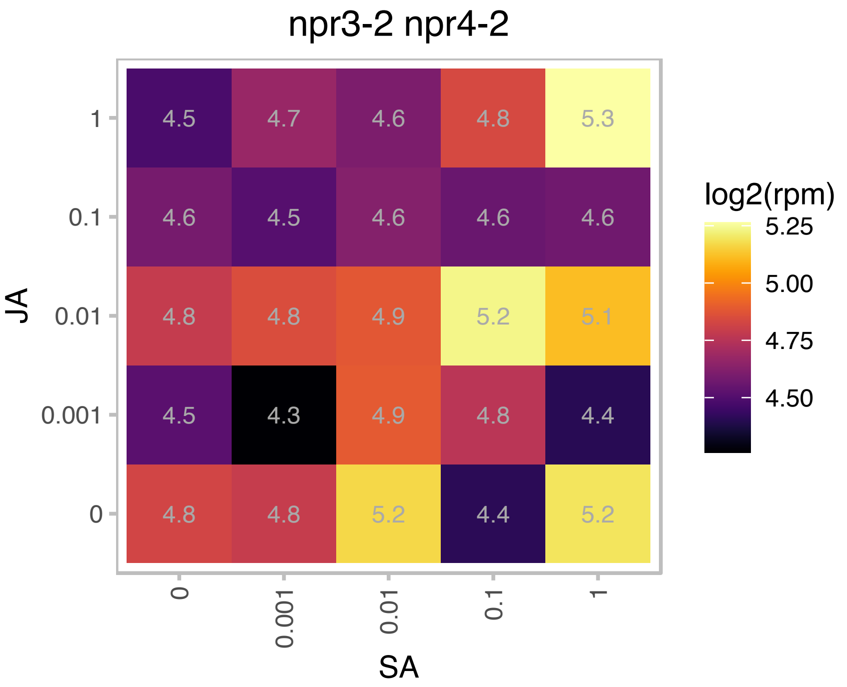


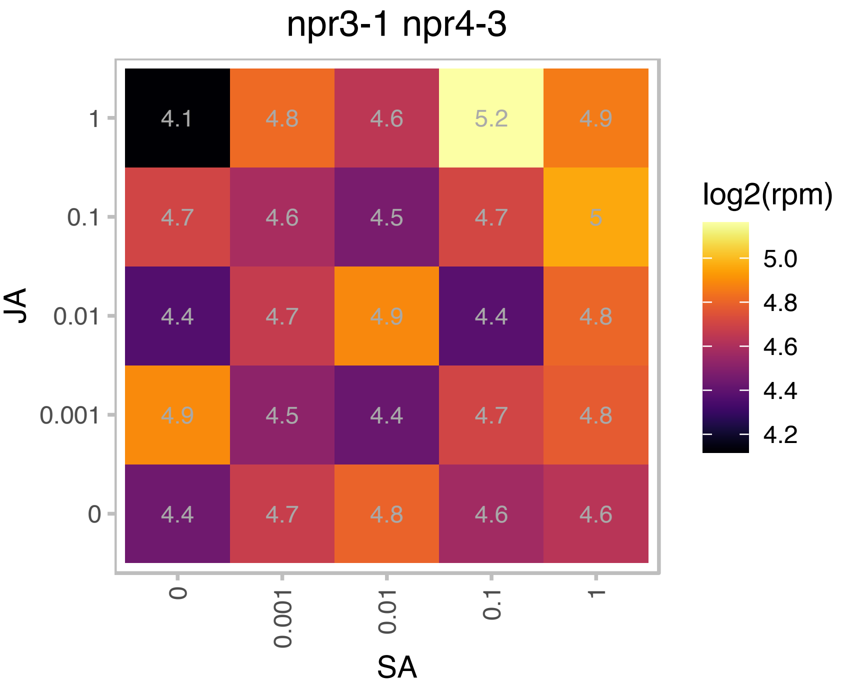

# AT4G00420.1 : NA

Double-stranded RNA-binding domain (DsRBD)-containing protein

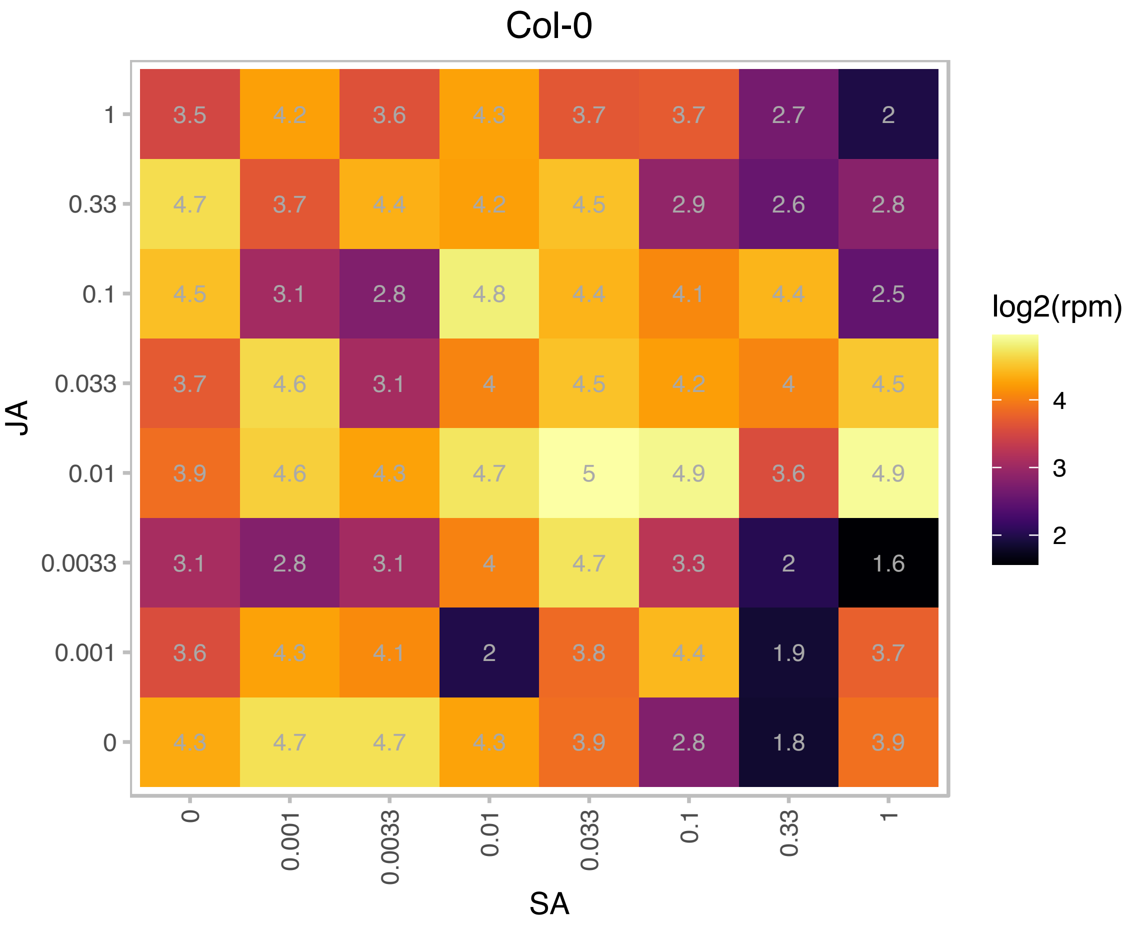


| | |
|---|---|
| Gene | AT4G00420 |
| is.expressed_genes | TRUE |
| is.responsive_genes | TRUE |
| is.apcluster | E-01 |

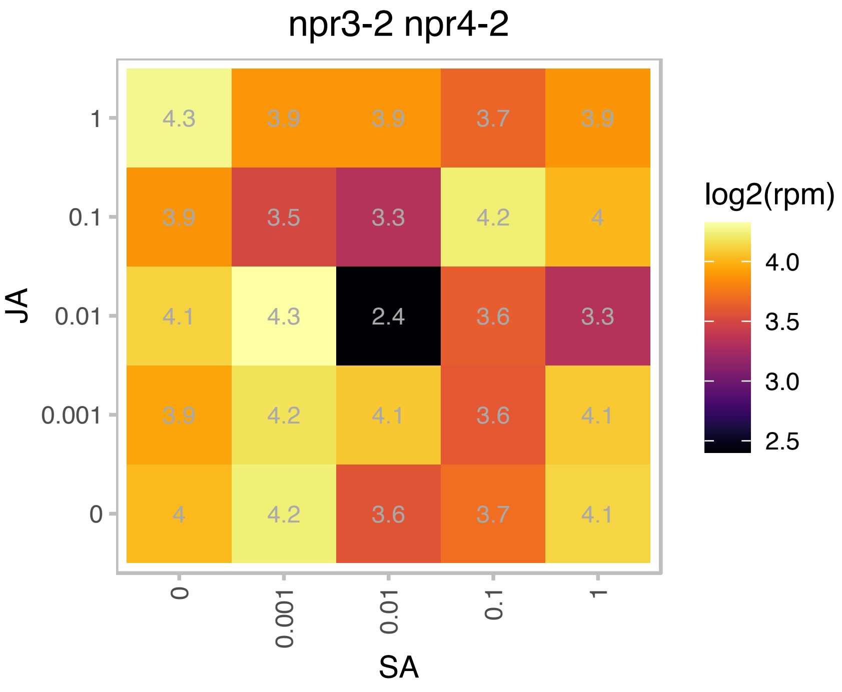


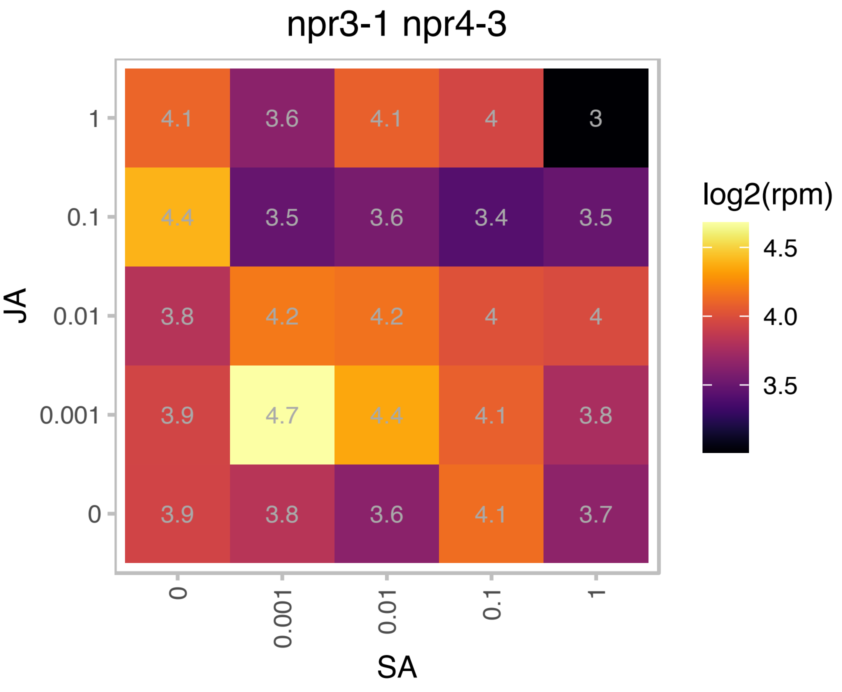

### AT4G02060.1 : MCM7, PRL

Minichromosome maintenance (MCM2/3/5) family protein
Member of the minichromosome maintenance complex, involved in DNA replication initiation. Abundant in proliferating and endocycling tissues. Localized in the nucleus during G1, S and G2 phases of the cell cycle, and are released into the cytoplasmic compartment during mitosis. Binds chromatin.

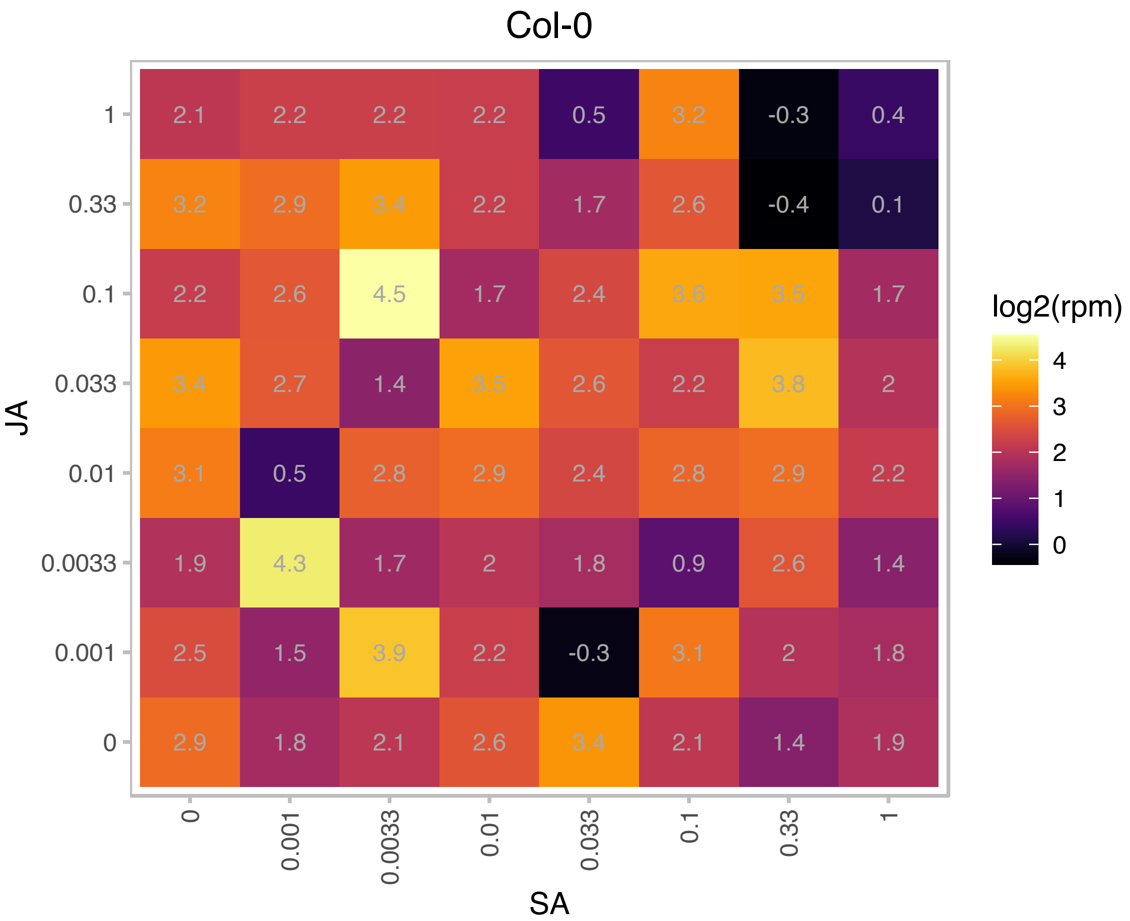


| | |
|---|---|
| Gene | AT4G02060 |
| is.expressed_genes | TRUE |
| is.responsive_genes | TRUE |
| is.apcluster | E-01 |

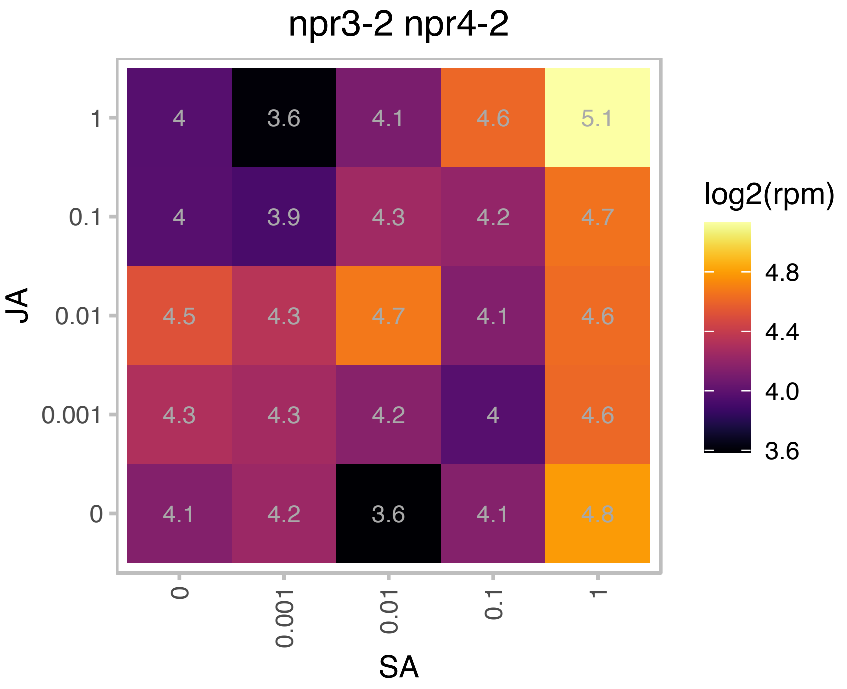


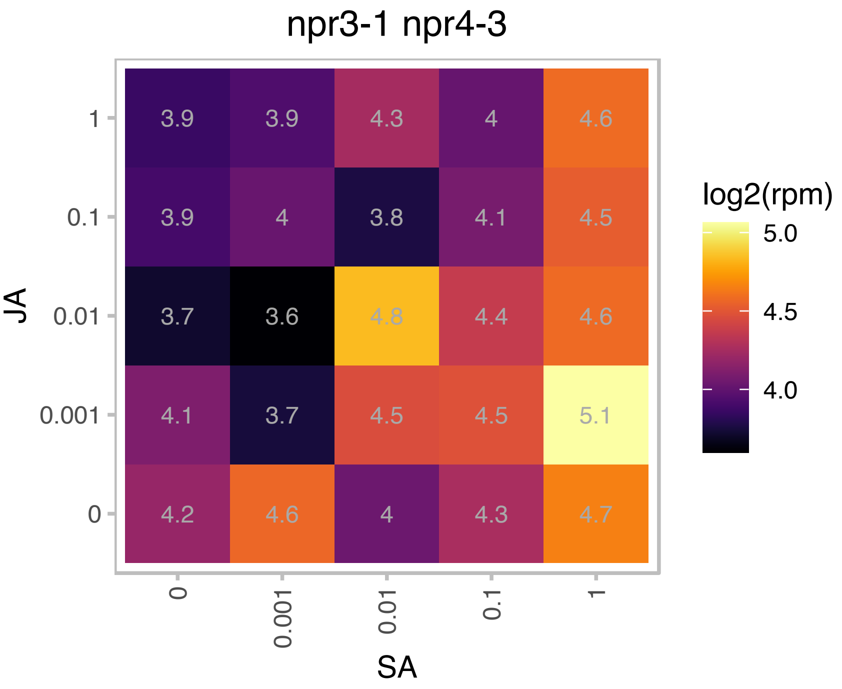

### AT4G11010.1 : NDPK3

nucleoside diphosphate kinase 3

nucleoside diphosphate kinase 3 (ndpk3), located to the inter-membrane space in mitochondria

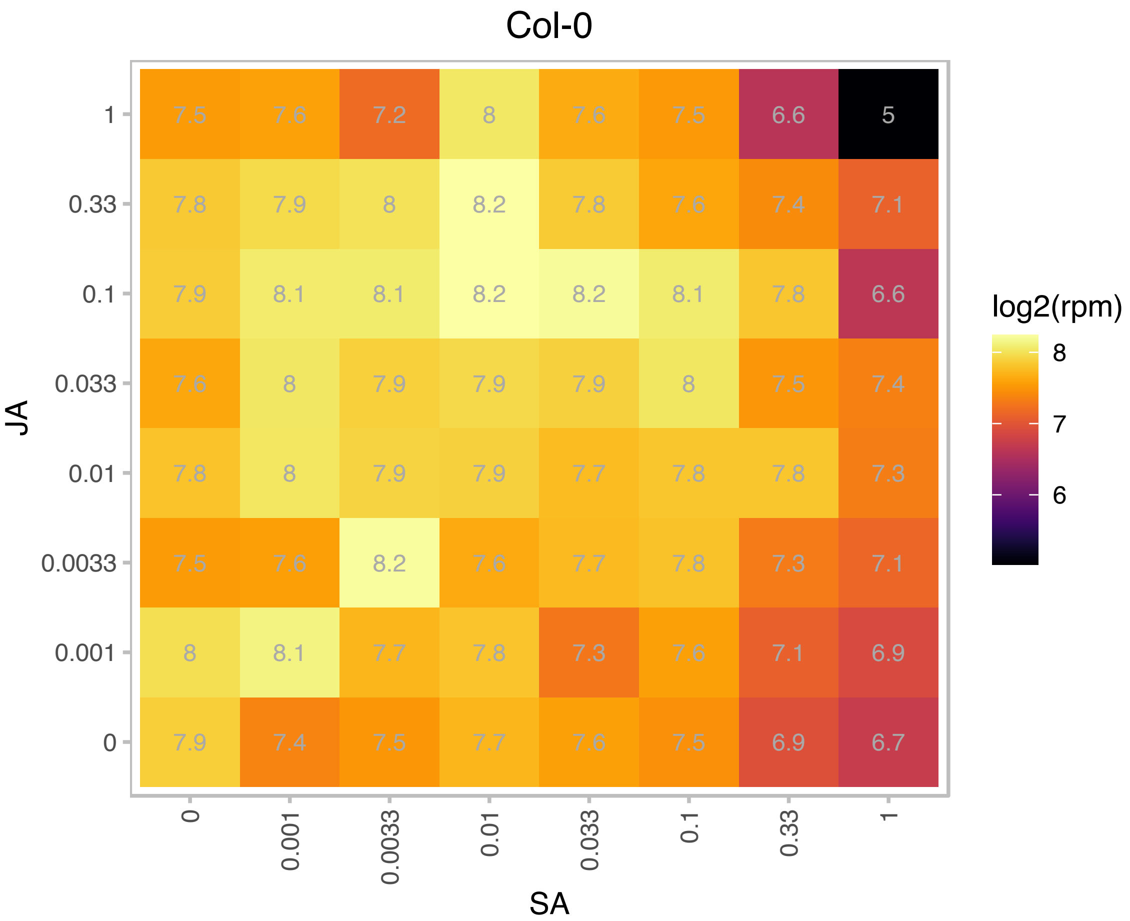


| | |
|---|---|
| Gene | AT4G11010 |
| is.expressed_genes | TRUE |
| is.responsive_genes | TRUE |
| is.apcluster | E-01 |

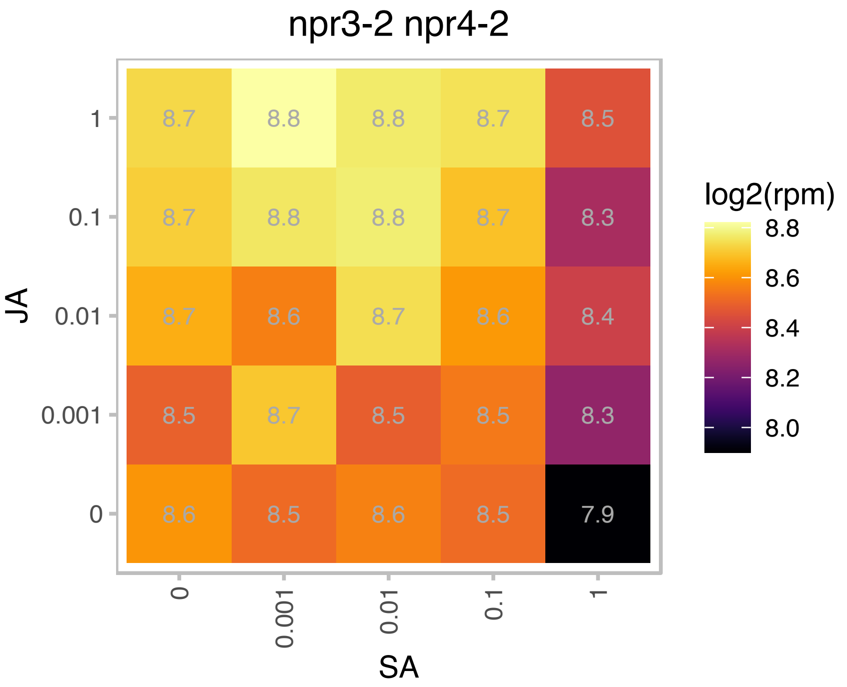


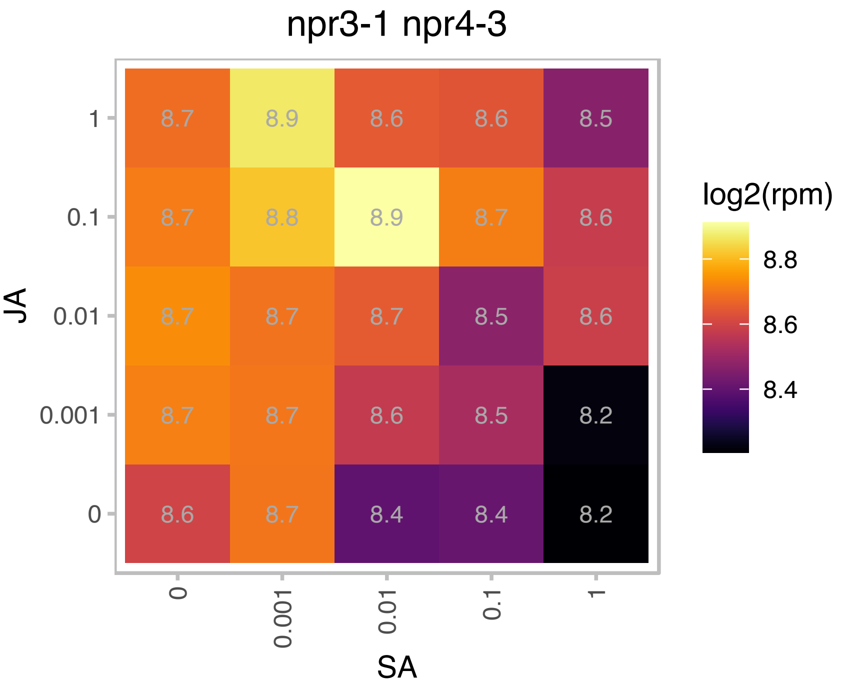

### AT4G13410.1 : ATCSLA15, CSLA15

Nucleotide-diphospho-sugar transferases superfamily protein

encodes a gene similar to cellulose synthase

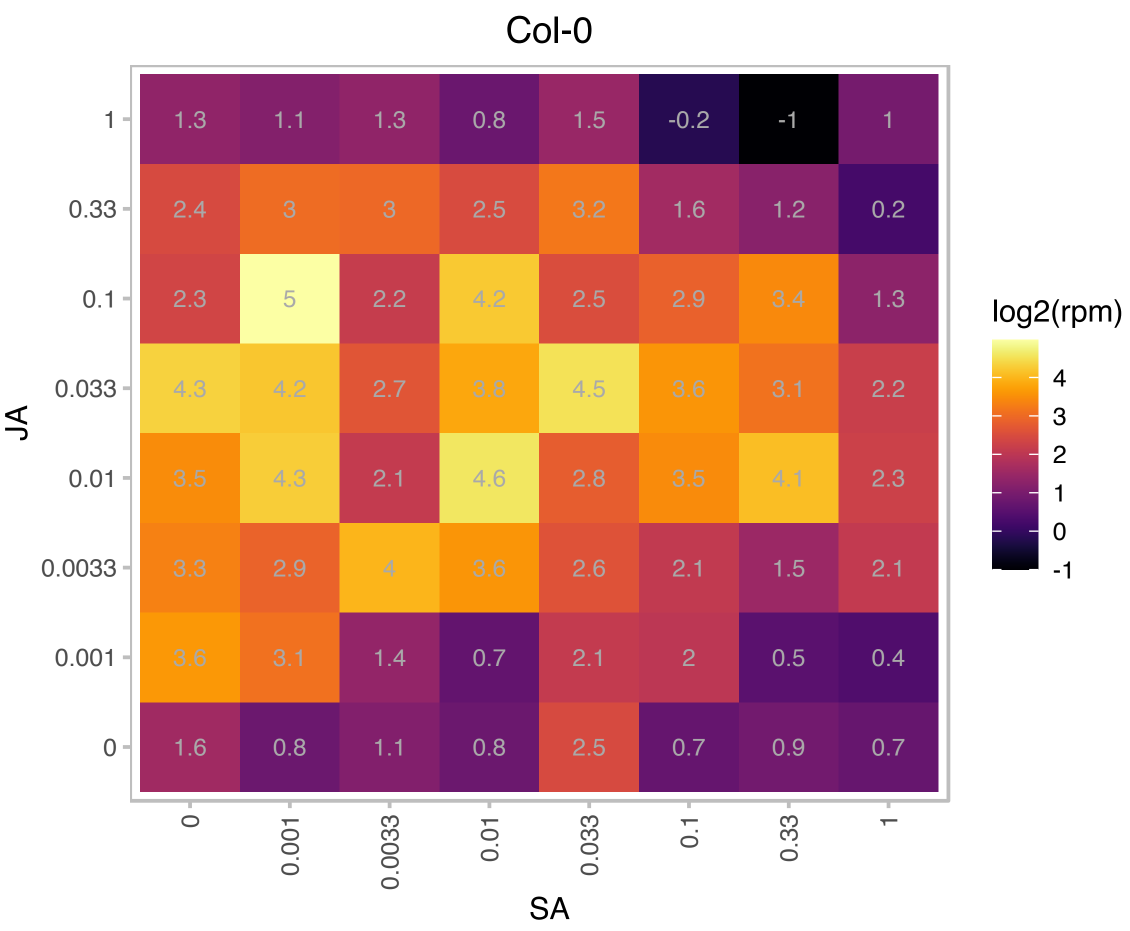


| | |
|---|---|
| Gene | AT4G13410 |
| is.expressed_genes | TRUE |
| is.responsive_genes | TRUE |
| is.apcluster | E-01 |

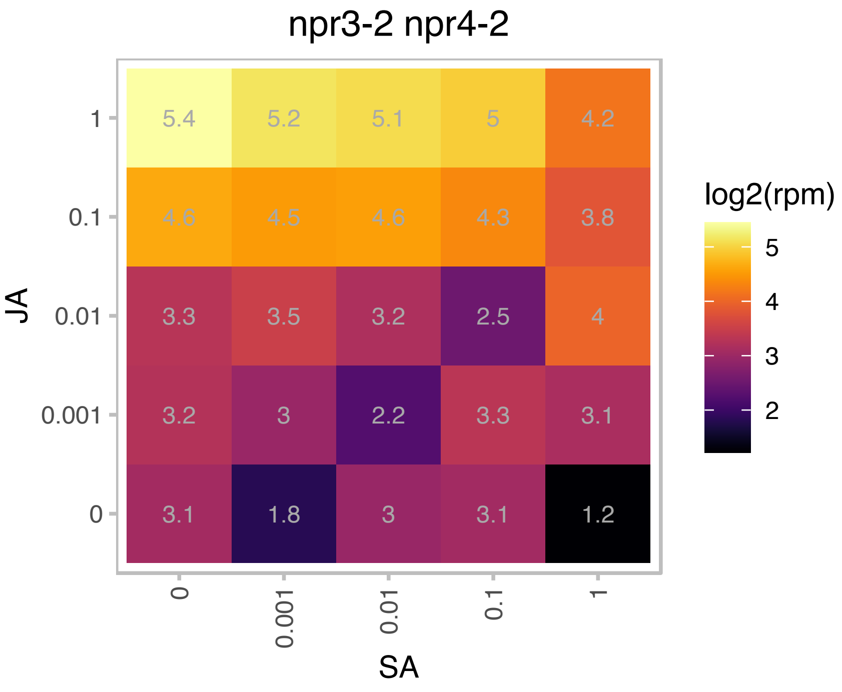


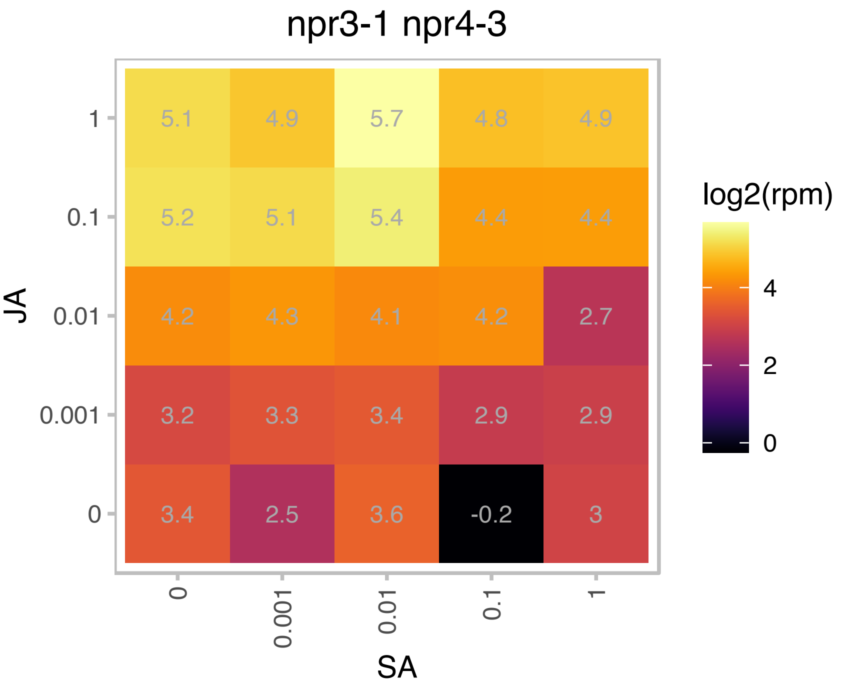

# AT4G19160.1 : NA

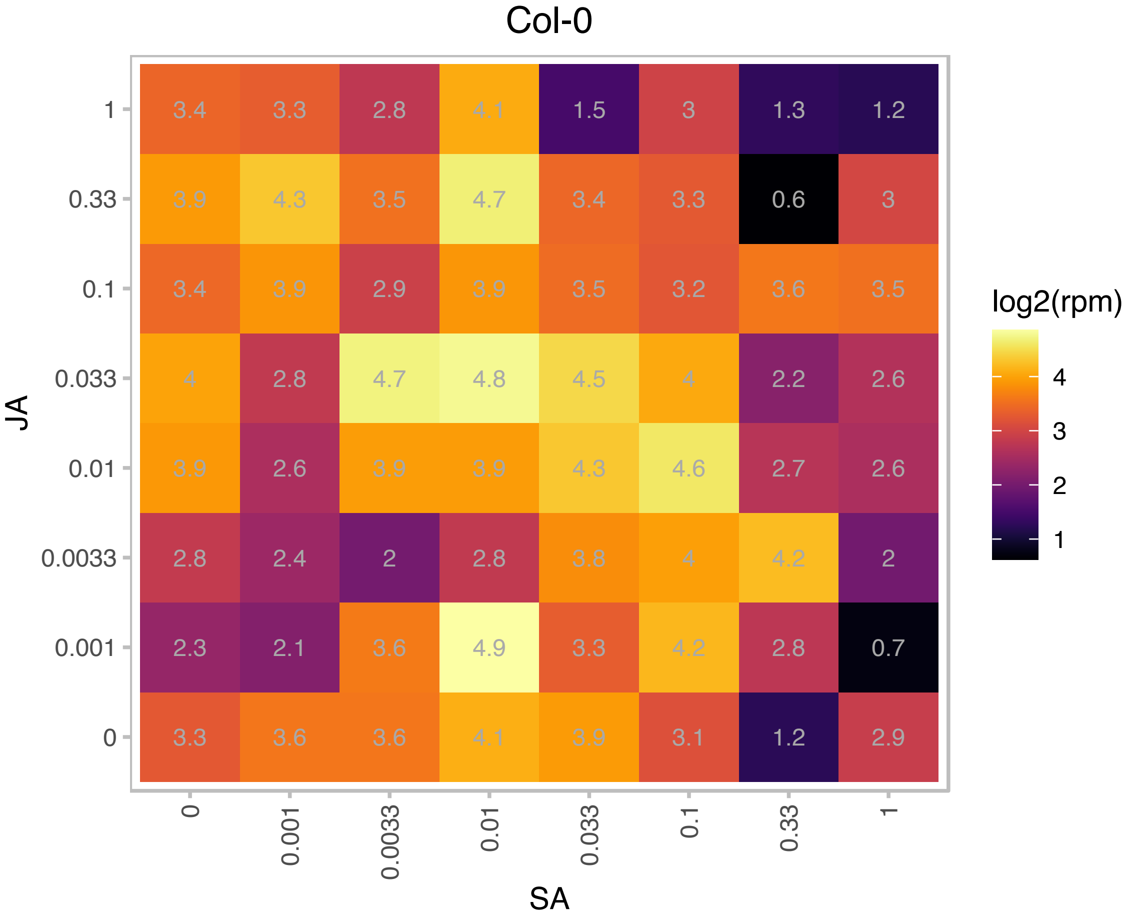


| | |
|---|---|
| Gene | AT4G19160 |
| is.expressed_genes | TRUE |
| is.responsive_genes | TRUE |
| is.apcluster | E-01 |

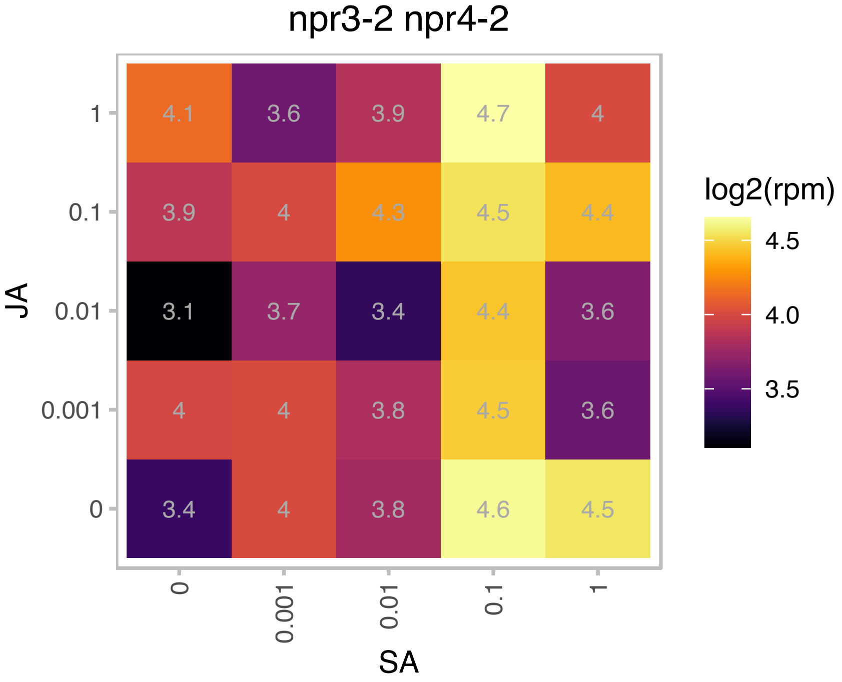


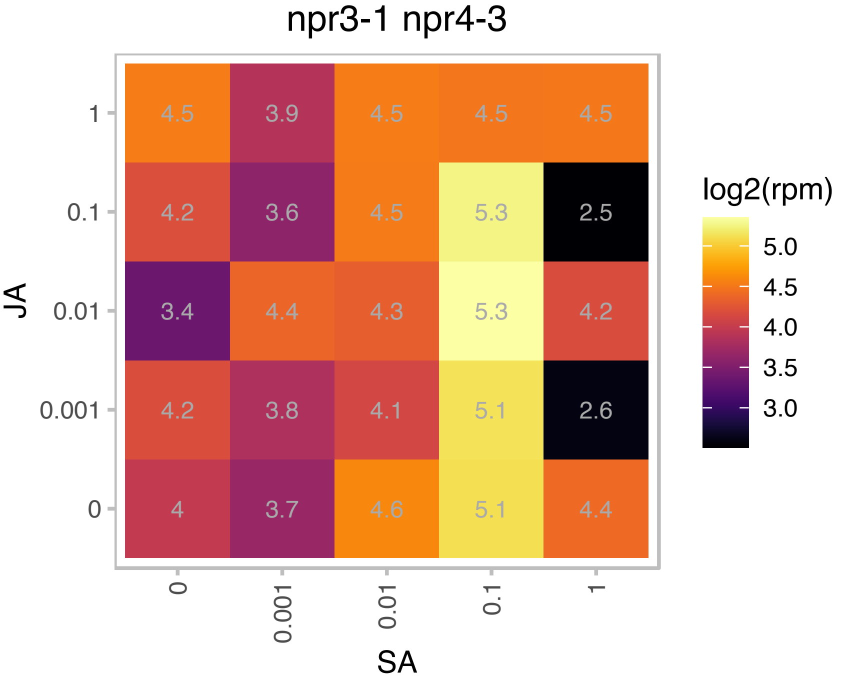

# AT4G21105.1 : NA

cytochrome-c oxidases;electron carriers

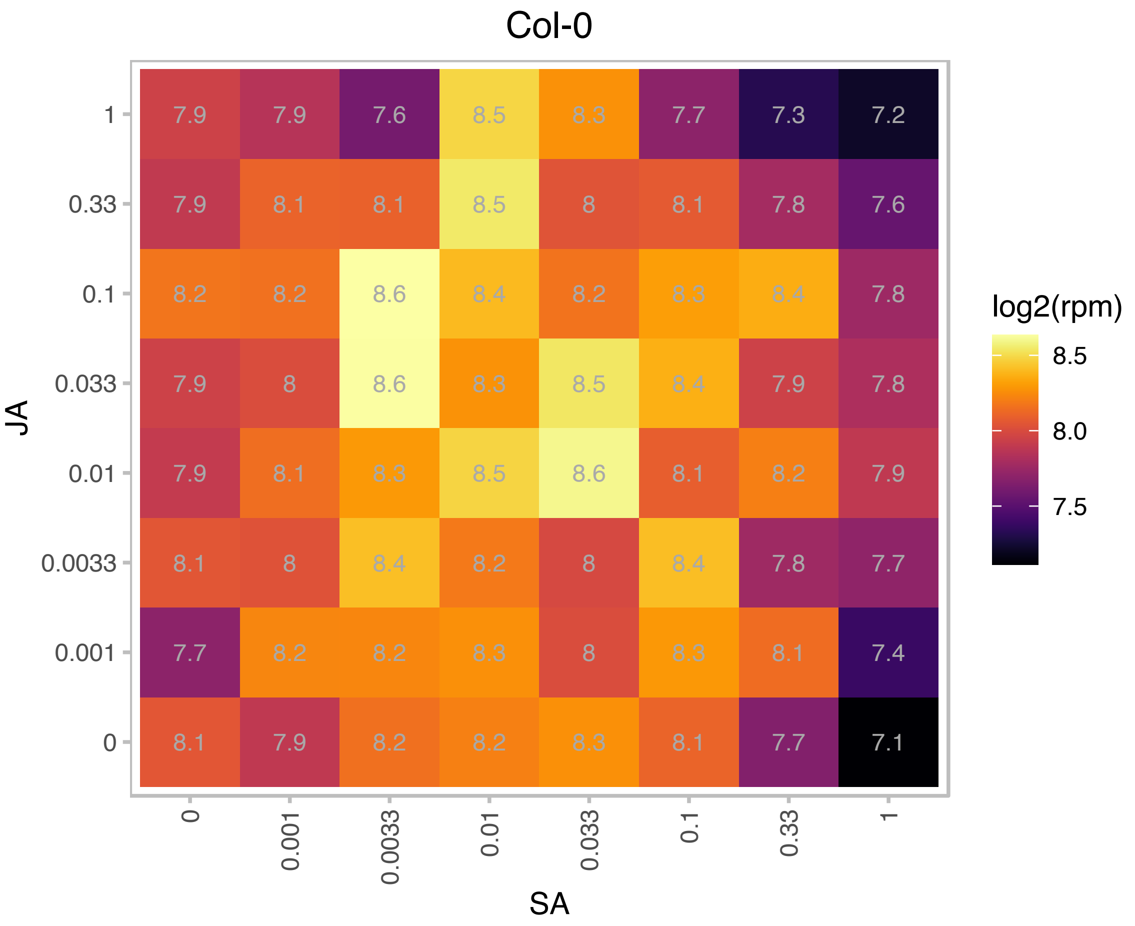


| | |
|---|---|
| Gene | AT4G21105 |
| is.expressed_genes | TRUE |
| is.responsive_genes | TRUE |
| is.apcluster | E-01 |

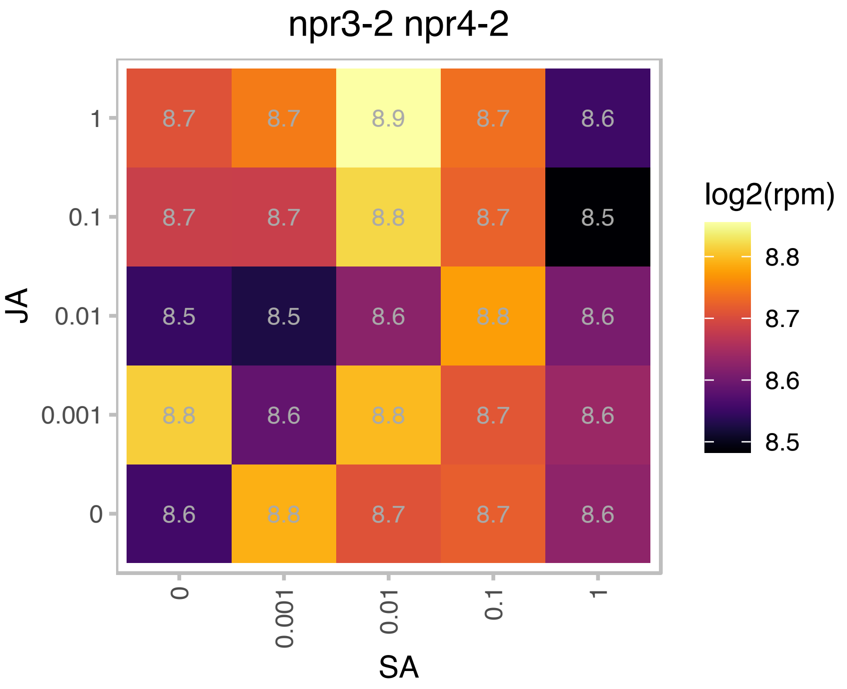


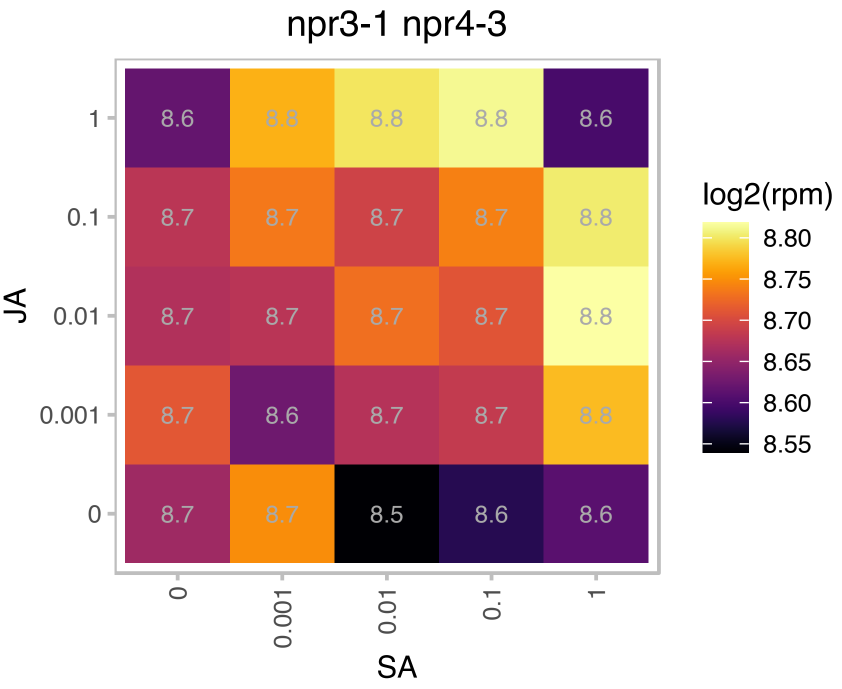

### AT4G23700.1 : ATCHX17, CHX17

cation/H+ exchanger 17

member of Putative Na+/H+ antiporter family

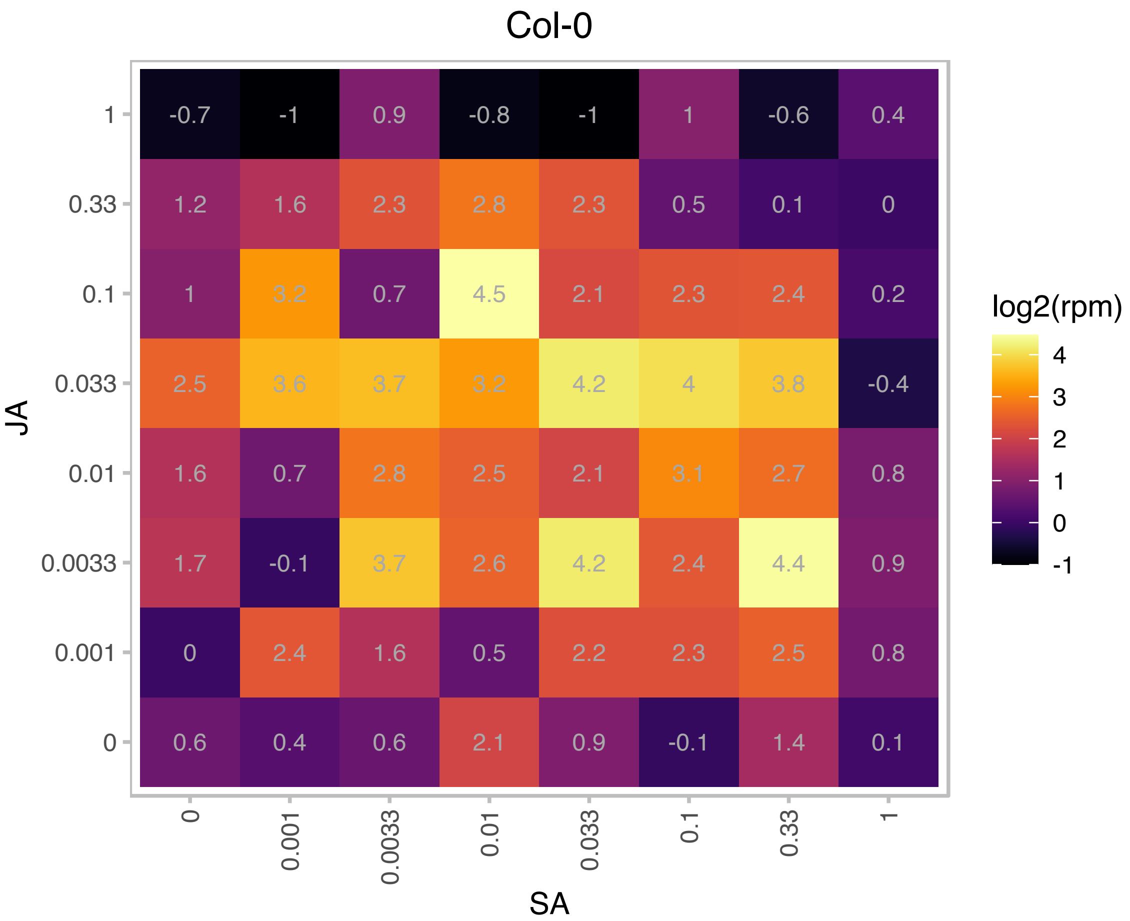


| Gene | AT4G23700 |
| --- | --- |
| is.expressed_genes | TRUE |
| is.responsive_genes | TRUE |
| is.apcluster | E-01 |

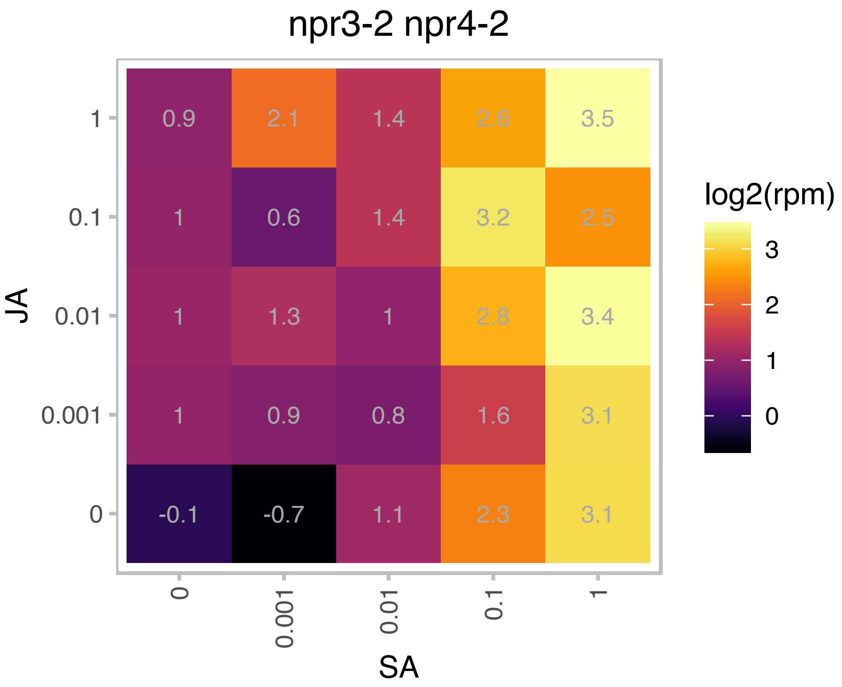


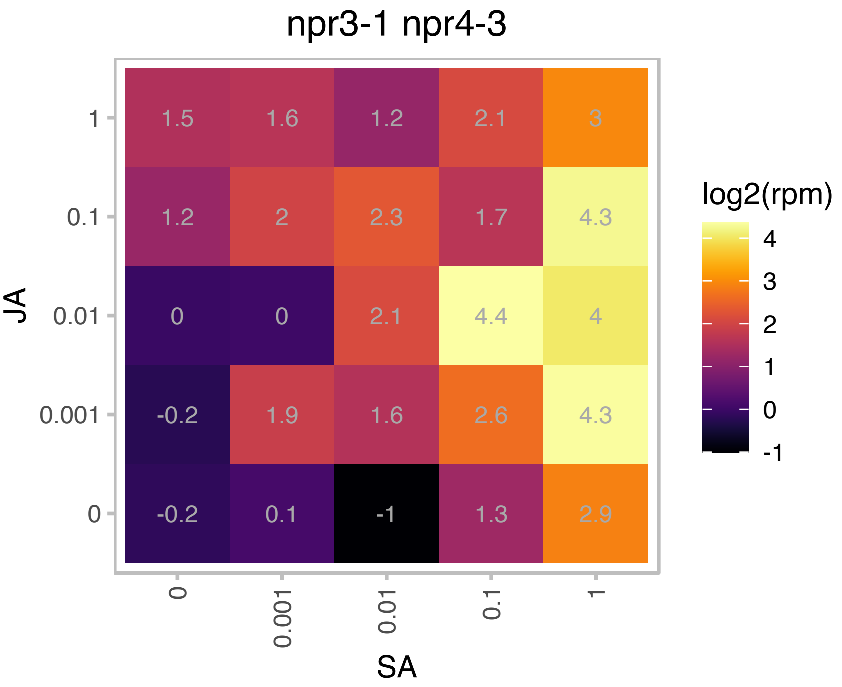

# AT4G24830.1 : NA

arginosuccinate synthase family

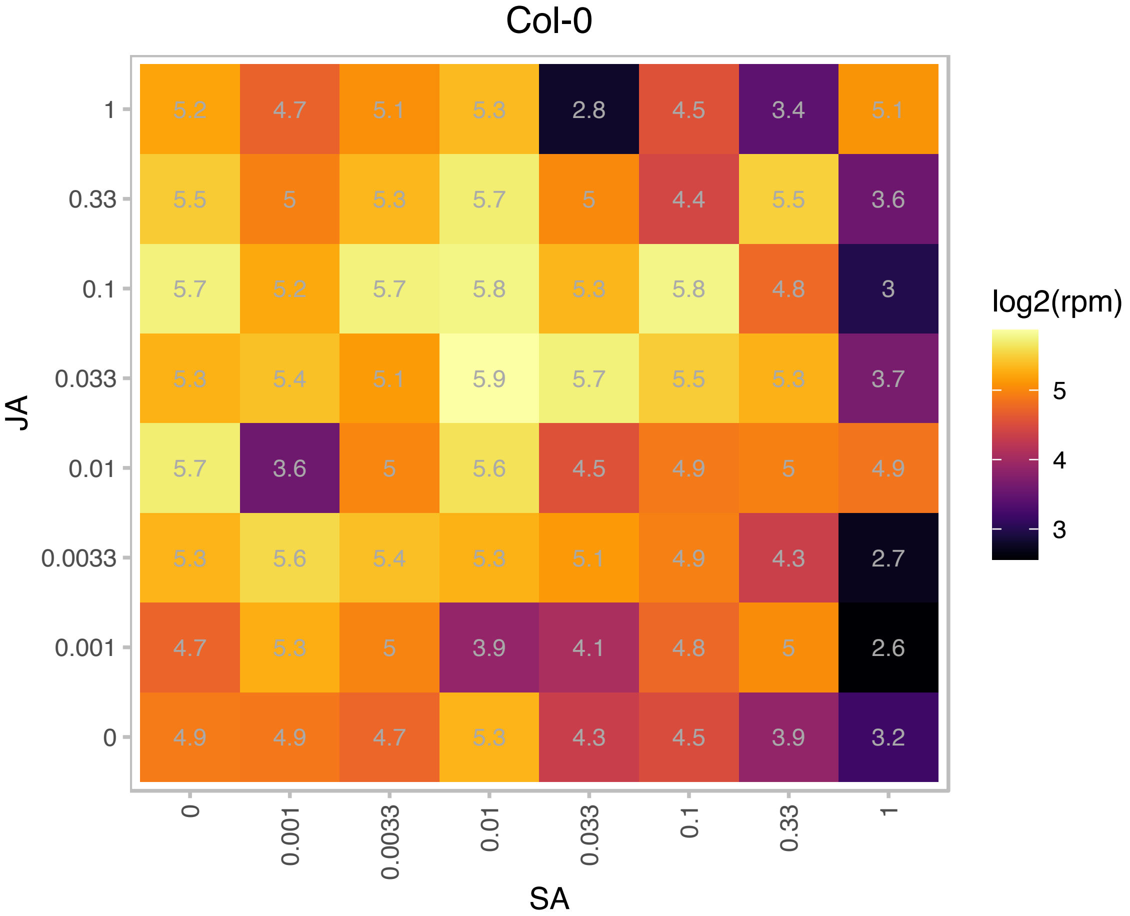


| Gene | AT4G24830 |
|---|---|
| is.expressed_genes | TRUE |
| is.responsive_genes | TRUE |
| is.apcluster | E-01 |

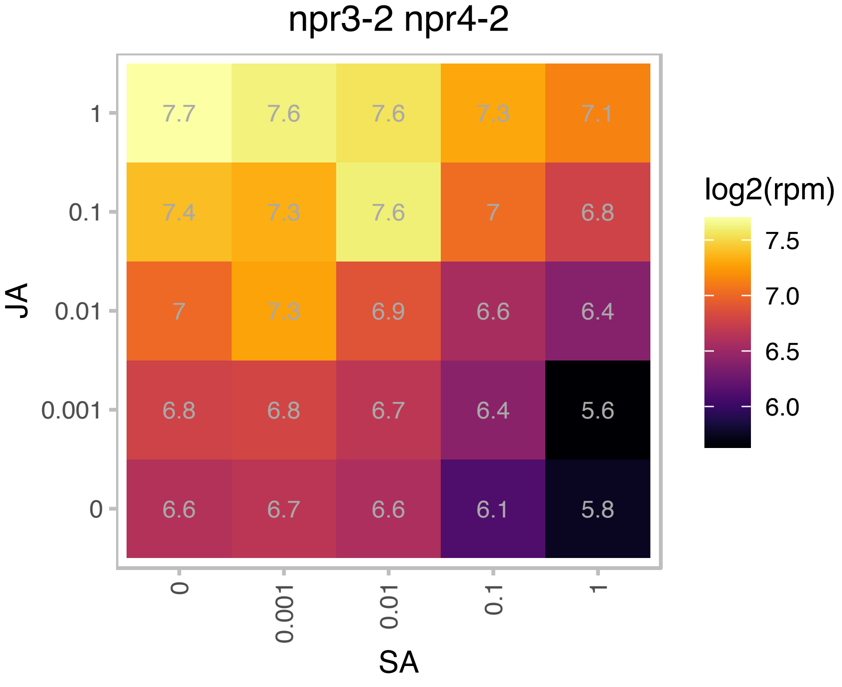


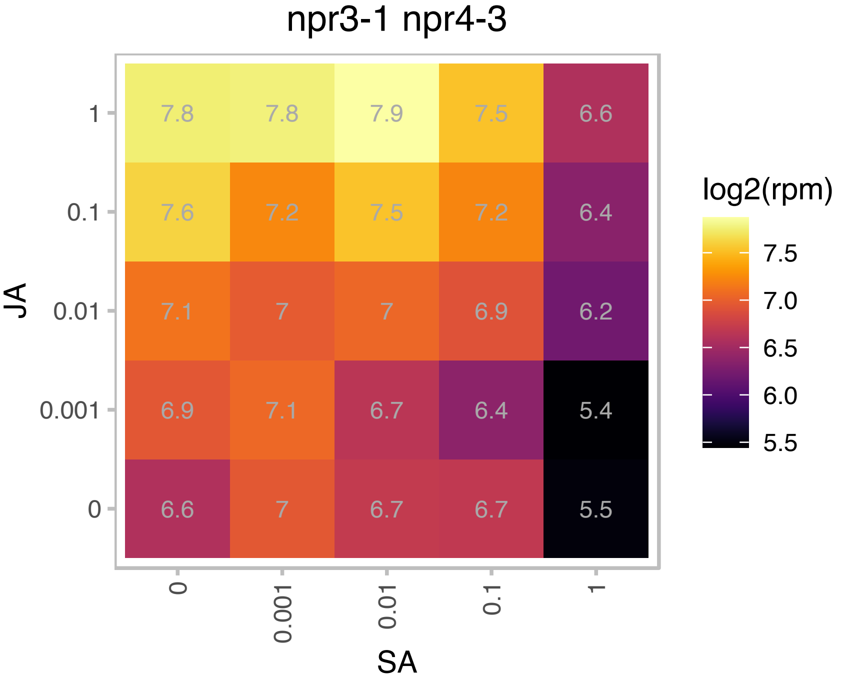

### AT4G26300.1 : emb1027

Arginyl-tRNA synthetase, class Ic

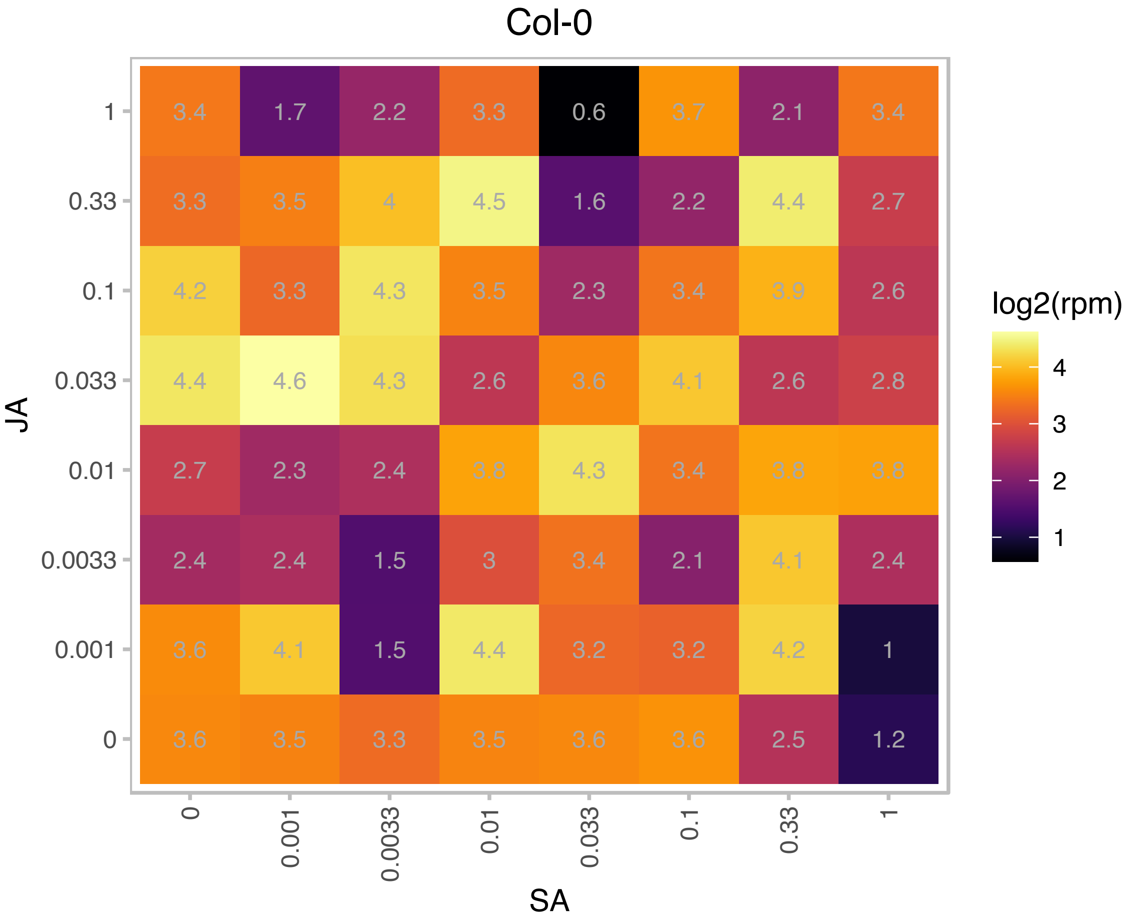


| | |
|---|---|
| Gene | AT4G26300 |
| is.expressed_genes | TRUE |
| is.responsive_genes | TRUE |
| is.apcluster | E-01 |

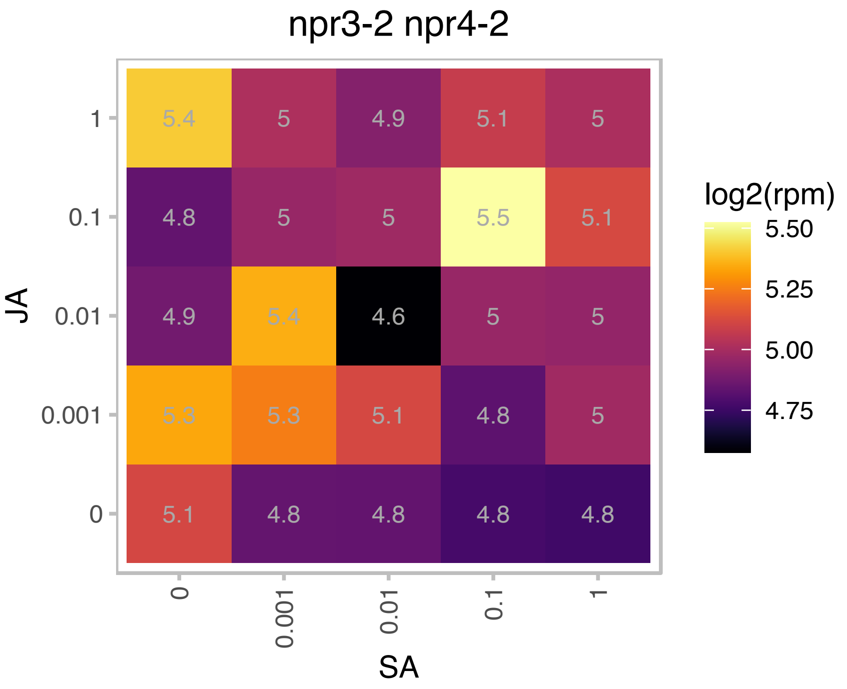


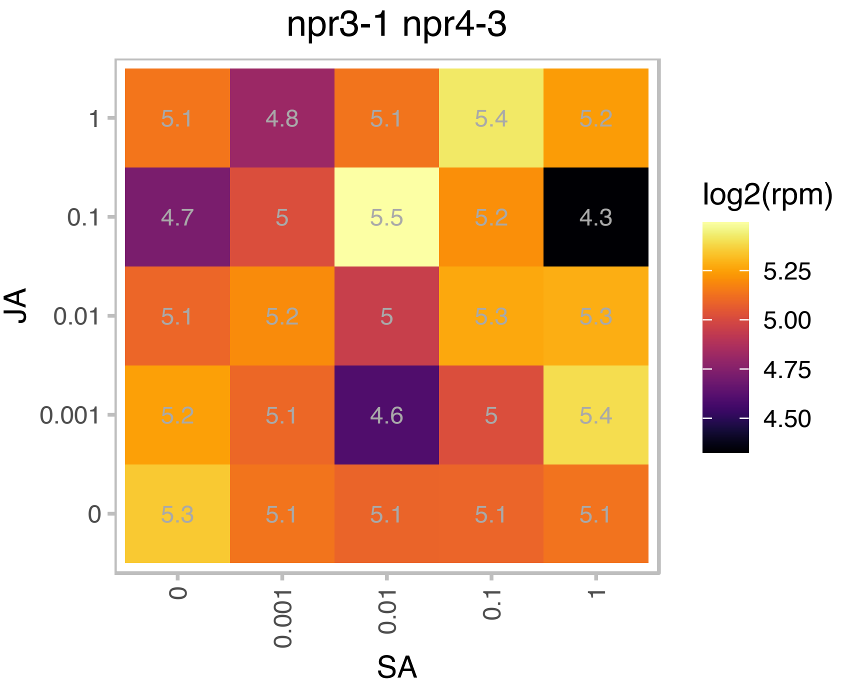

### AT4G31500.1 : ATR4, CYP83B1, RED1, RNT1, SUR2

cytochrome P450, family 83, subfamily B, polypeptide 1

Encodes an oxime-metabolizing enzyme in the biosynthetic pathway of glucosinolates.  Is required for phytochrome signal transduction in red light.  Mutation confers auxin overproduction.

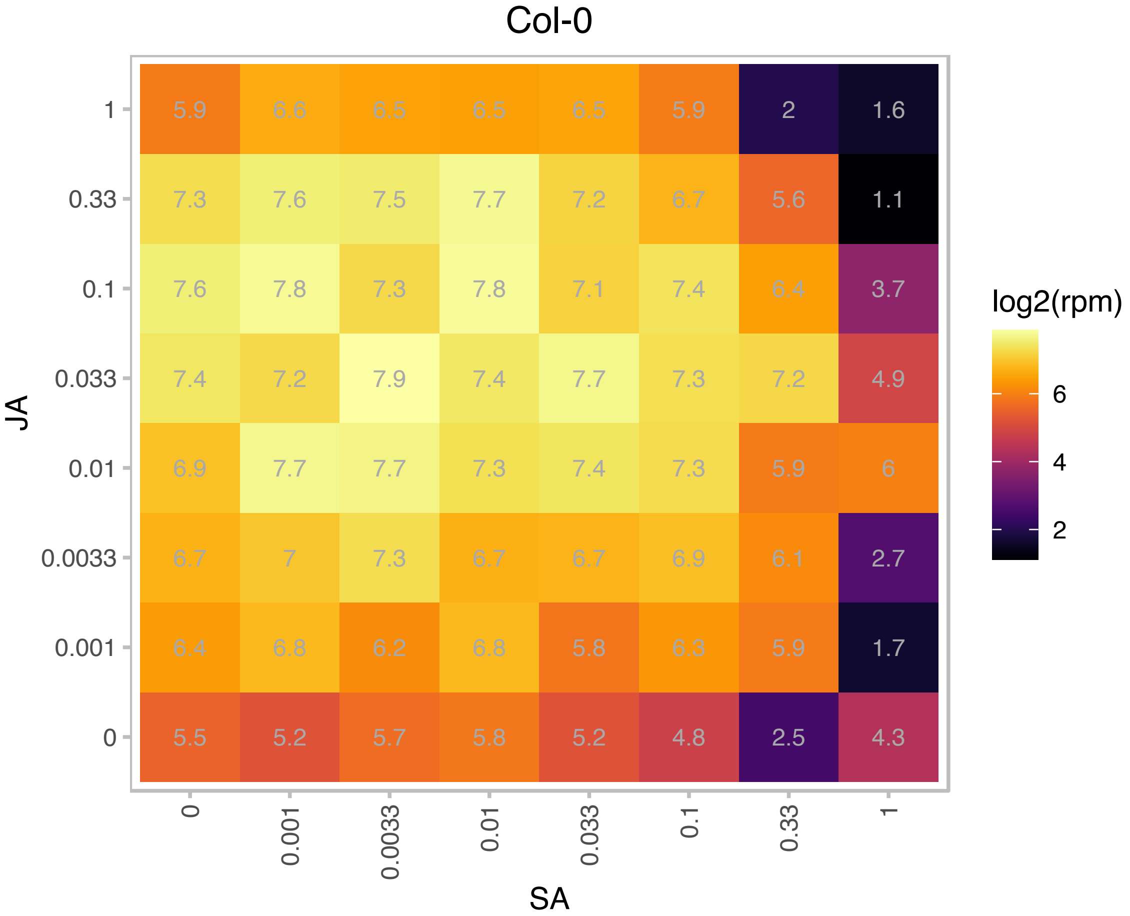


| | |
|---|---|
| Gene | AT4G31500 |
| is.expressed_genes | TRUE |
| is.responsive_genes | TRUE |
| is.apcluster | E-01 |

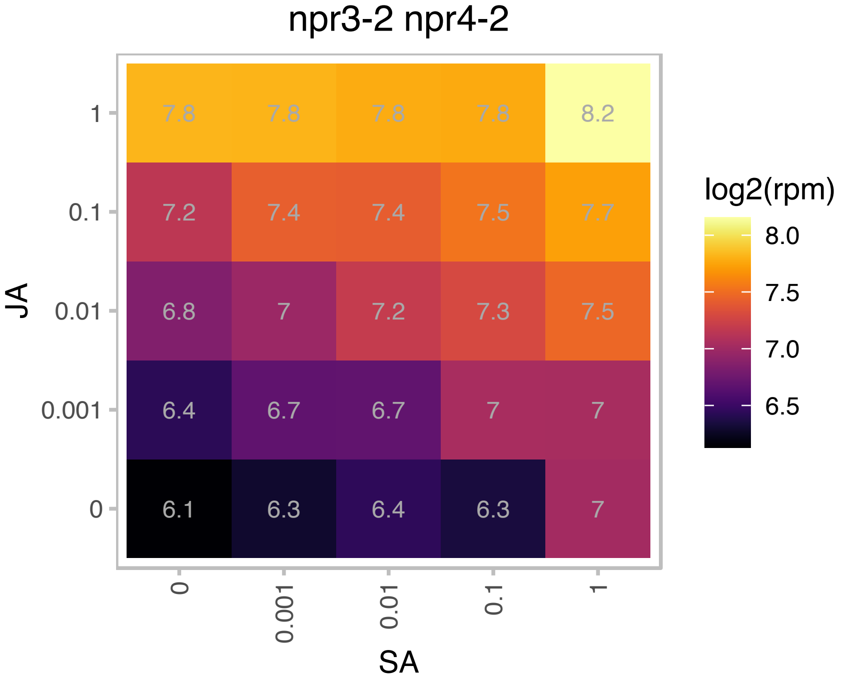


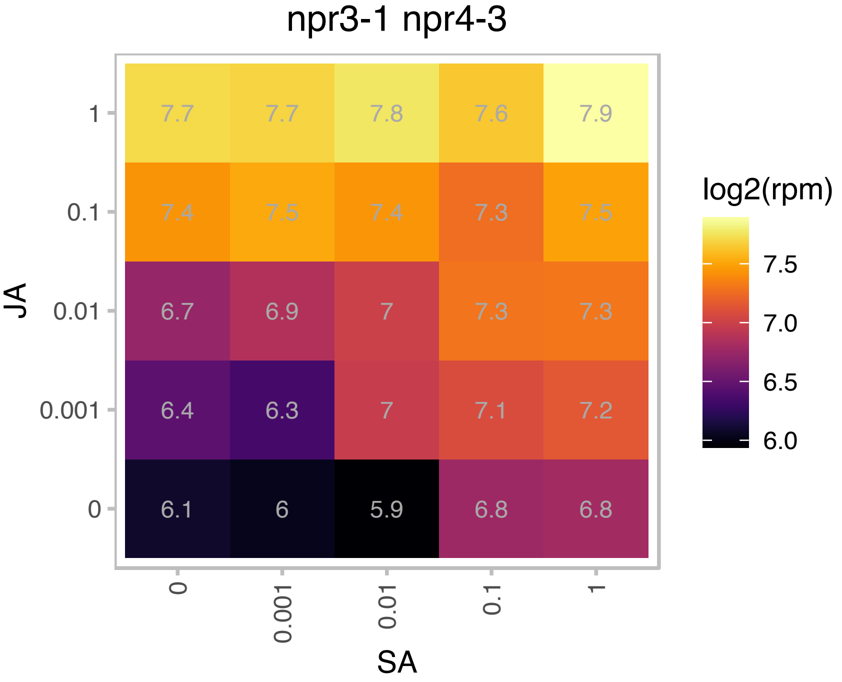

# AT4G37090.1 : NA

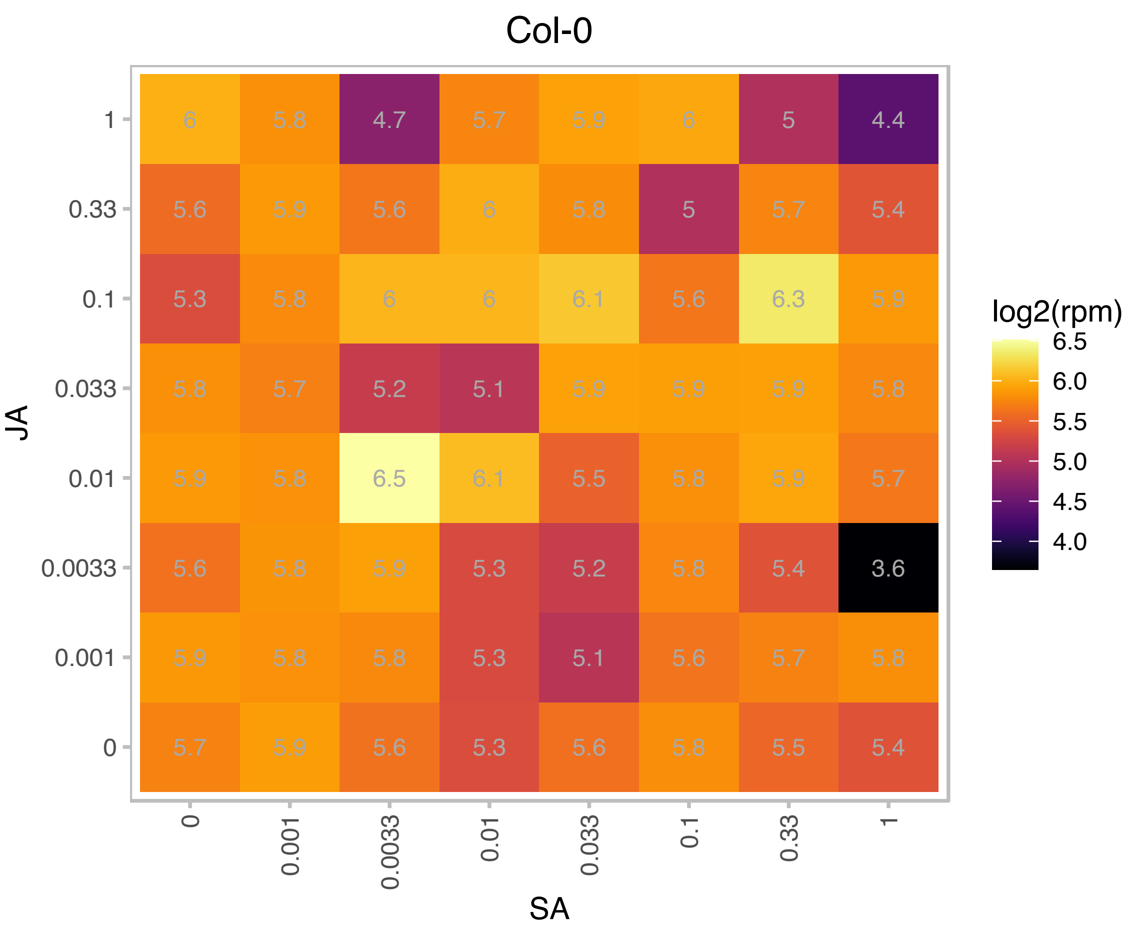


| | |
|---|---|
| Gene | AT4G37090 |
| is.expressed_genes | TRUE |
| is.responsive_genes | TRUE |
| is.apcluster | E-01 |

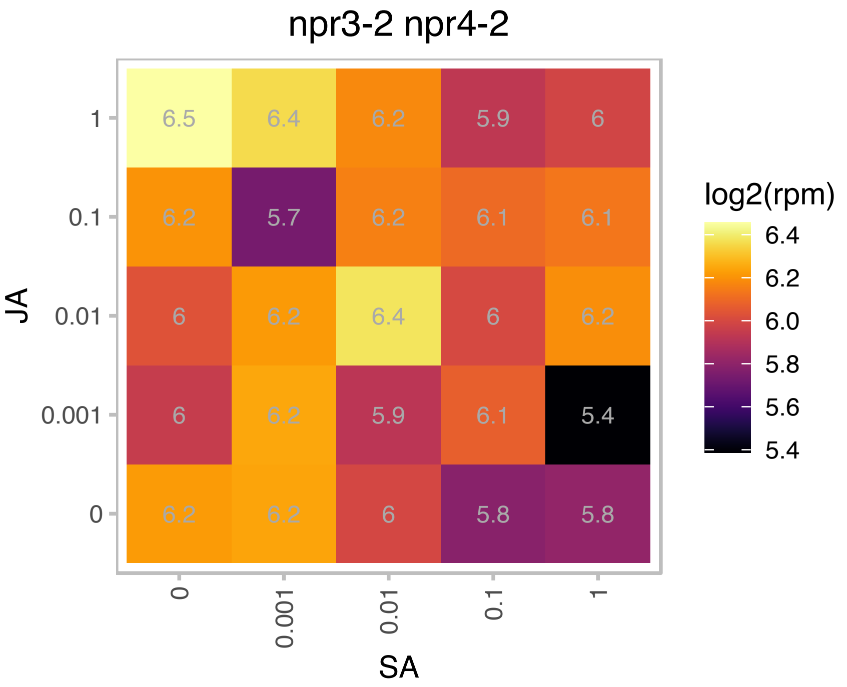


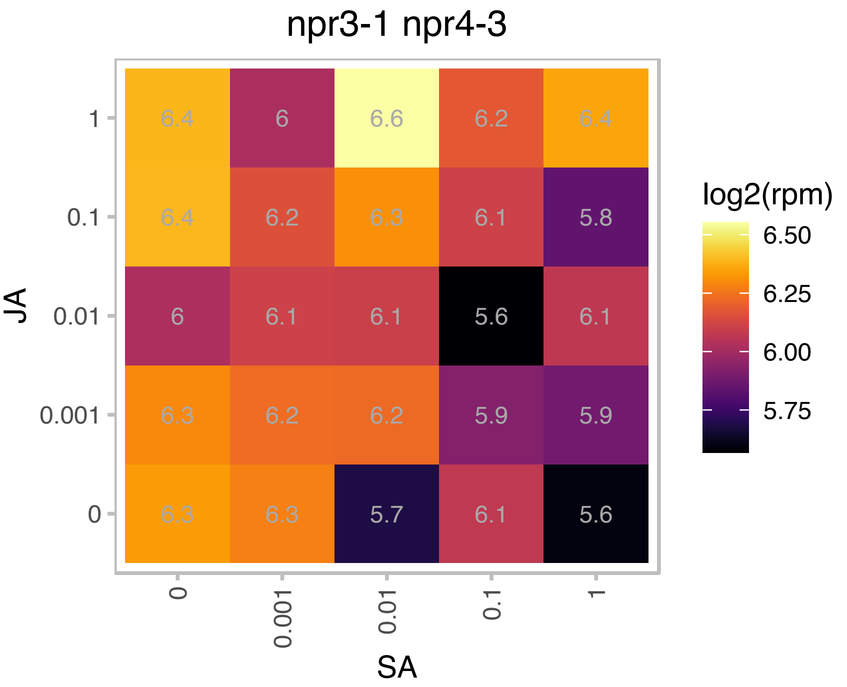

### AT4G38740.1 : ROC1

rotamase CYP 1

Encodes cytosolic cyclophilin ROC1.

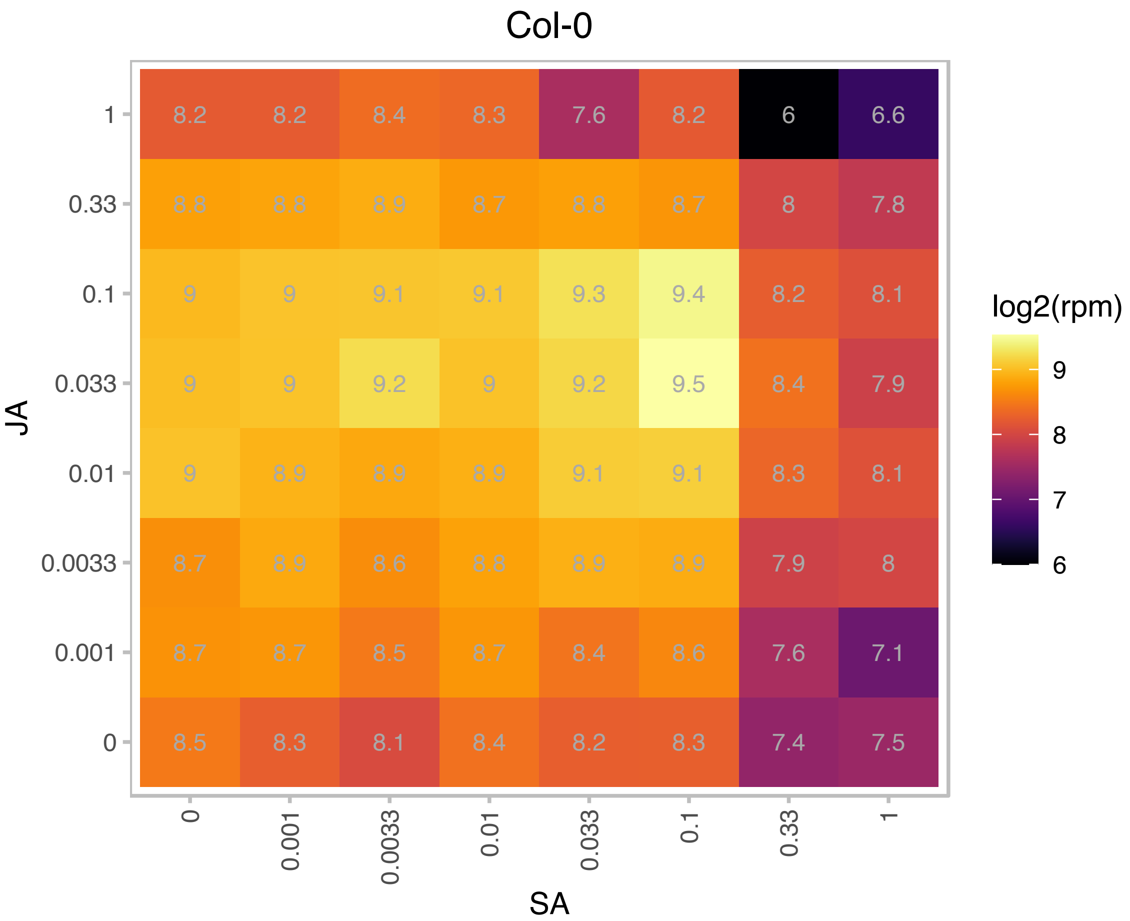


| | |
|---|---|
| Gene | AT4G38740 |
| is.expressed_genes | TRUE |
| is.responsive_genes | TRUE |
| is.apcluster | E-01 |

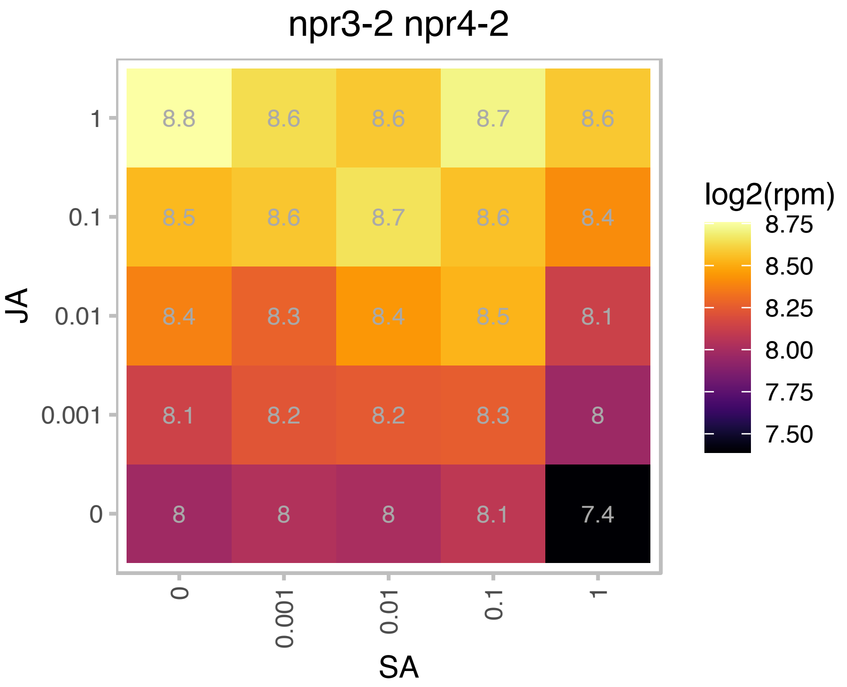


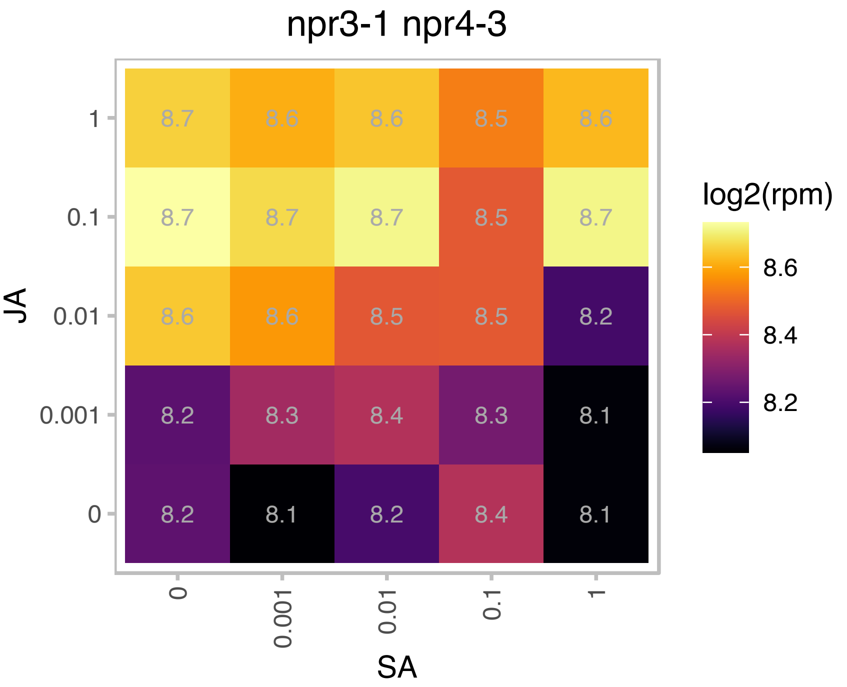

### AT4G39940.1 : AKN2, APK2

APS-kinase 2

adenosine-5'-phosphosulfate-kinase (akn2) mRNA, complete The mRNA is cell-to-cell mobile.

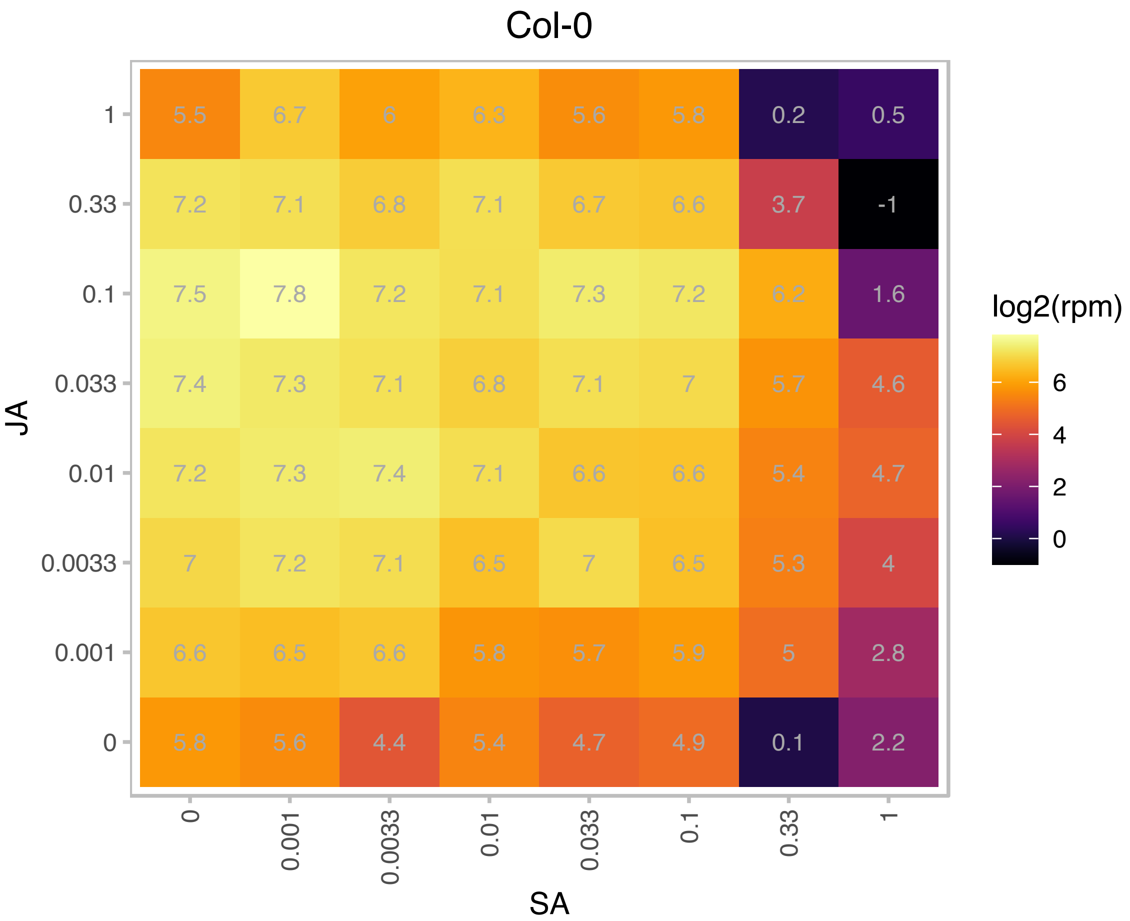


| | |
|---|---|
| Gene | AT4G39940 |
| is.expressed_genes | TRUE |
| is.responsive_genes | TRUE |
| is.apcluster | E-01 |

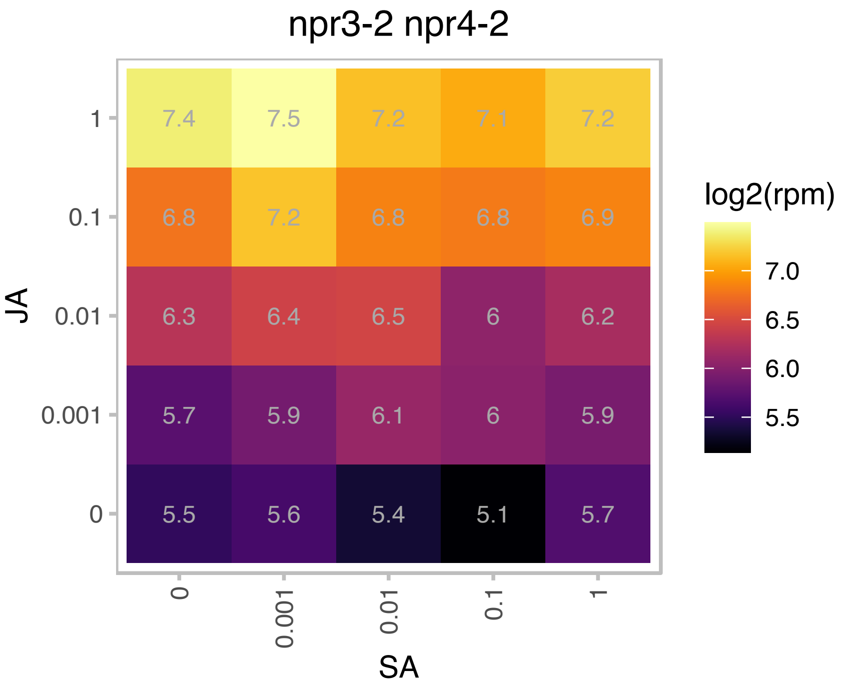


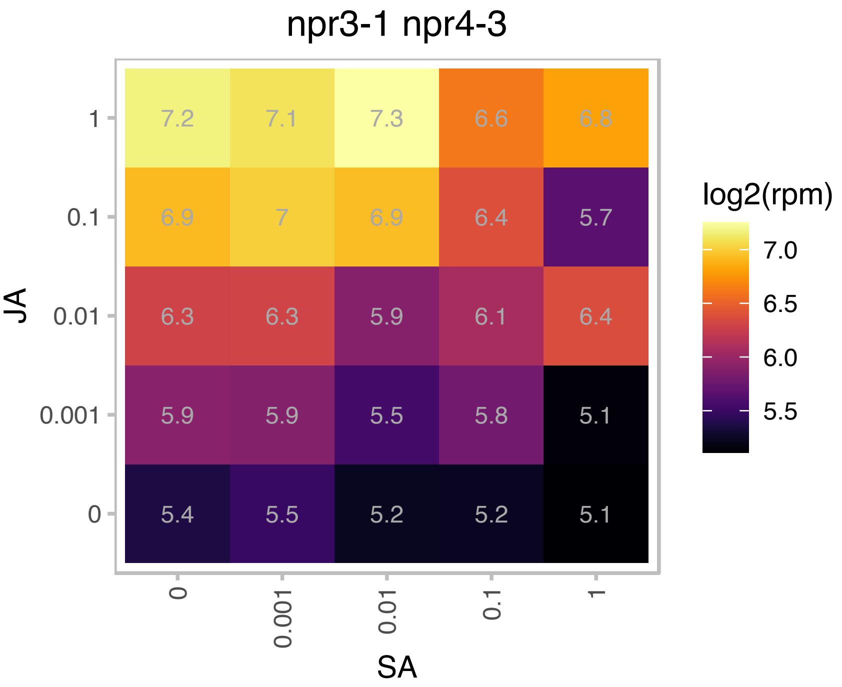

### AT5G02260.1 : ATEXP9, ATEXPA9, ATHEXP ALPHA 1.10,

expansin A9

member of Alpha-Expansin Gene Family. Naming convention from the Expansin Working Group (Kende et al, 2004. Plant Mol Bio)

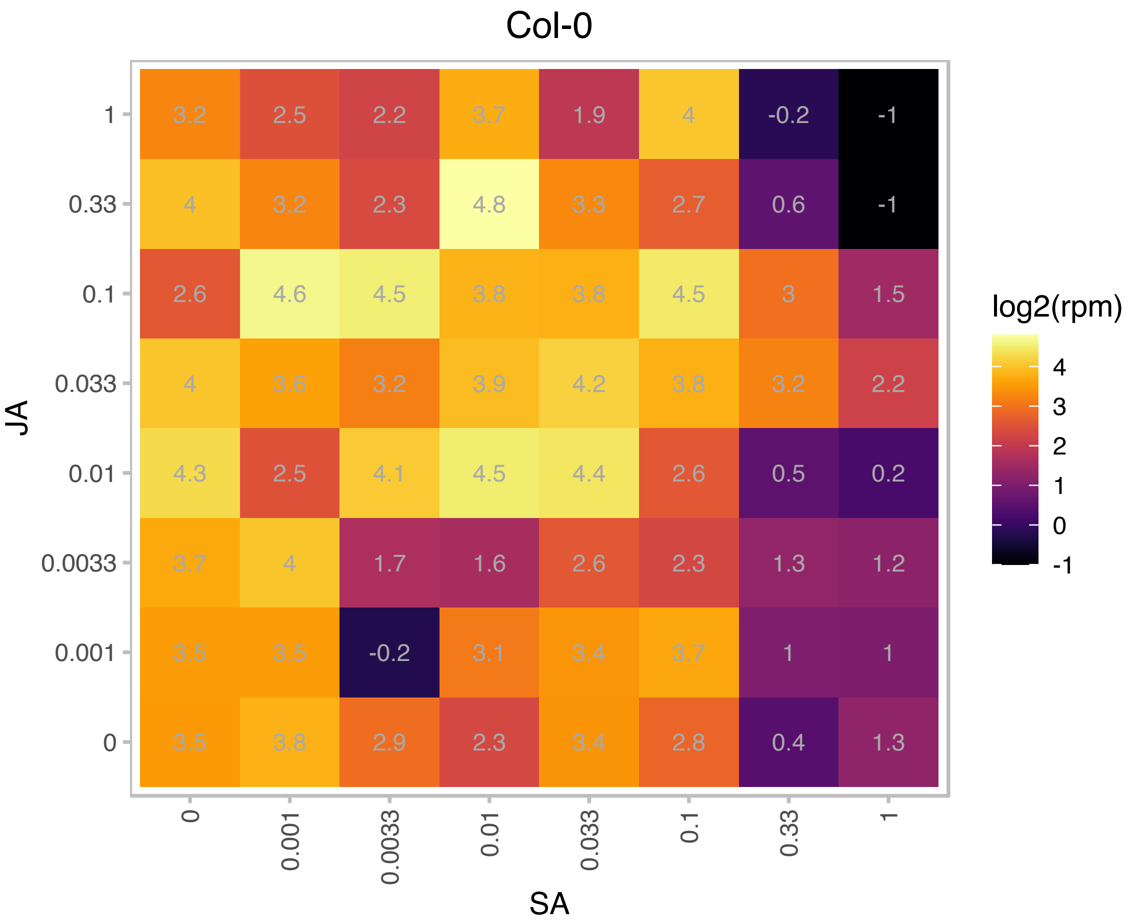


| | |
|---|---|
| Gene | AT5G02260 |
| is.expressed_genes | TRUE |
| is.responsive_genes | TRUE |
| is.apcluster | E-01 |

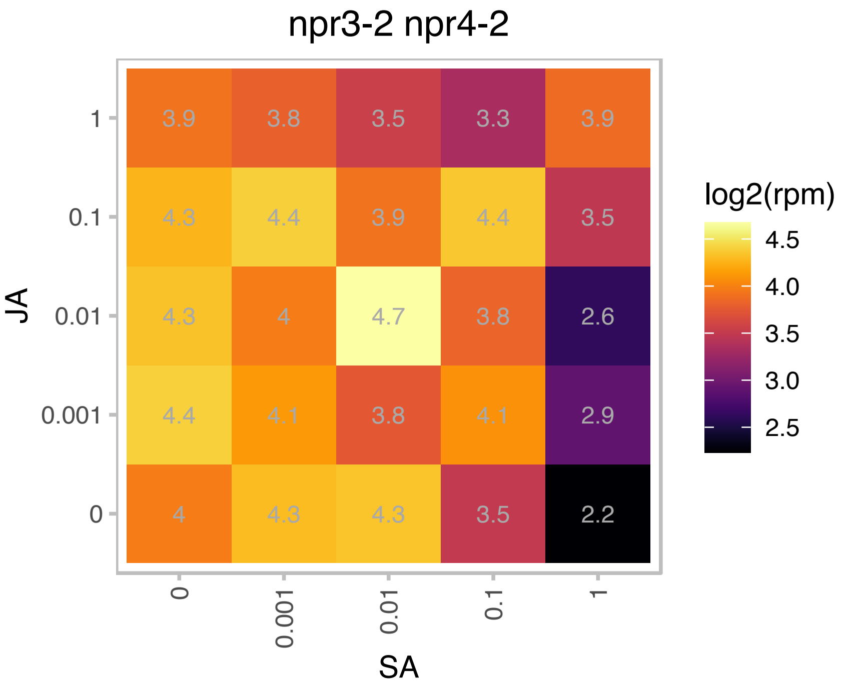


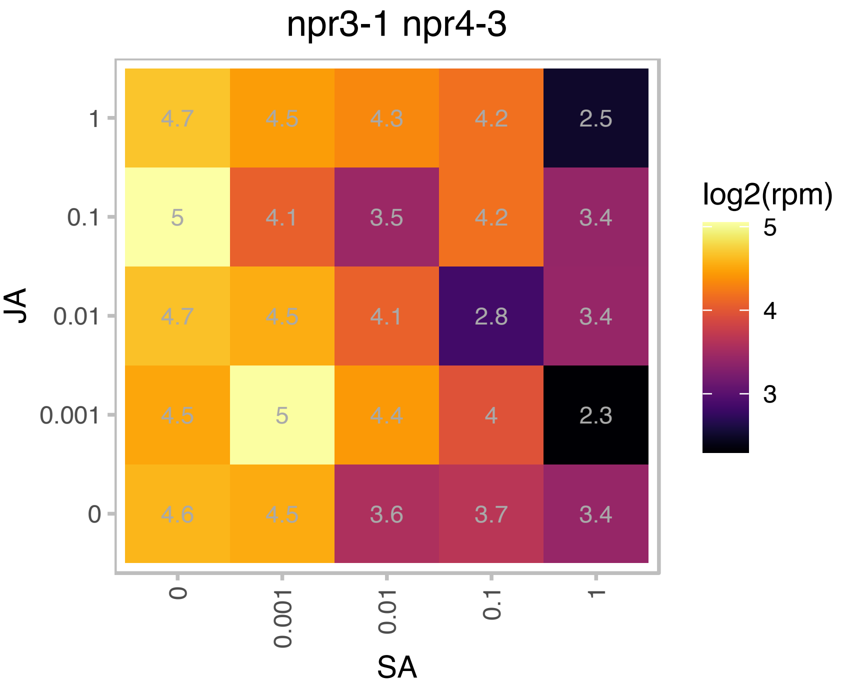

### AT5G07460.1 : ATMSRA2, PMSR2

peptidemethionine sulfoxide reductase 2
ubiquitous enzyme that repairs oxidatively damaged proteins. Methionine sulfoxide reductase activity. Mutant lacking reductase activity showed increased protein oxidation, nitration and glycation of specific amino acid residues during darkness.

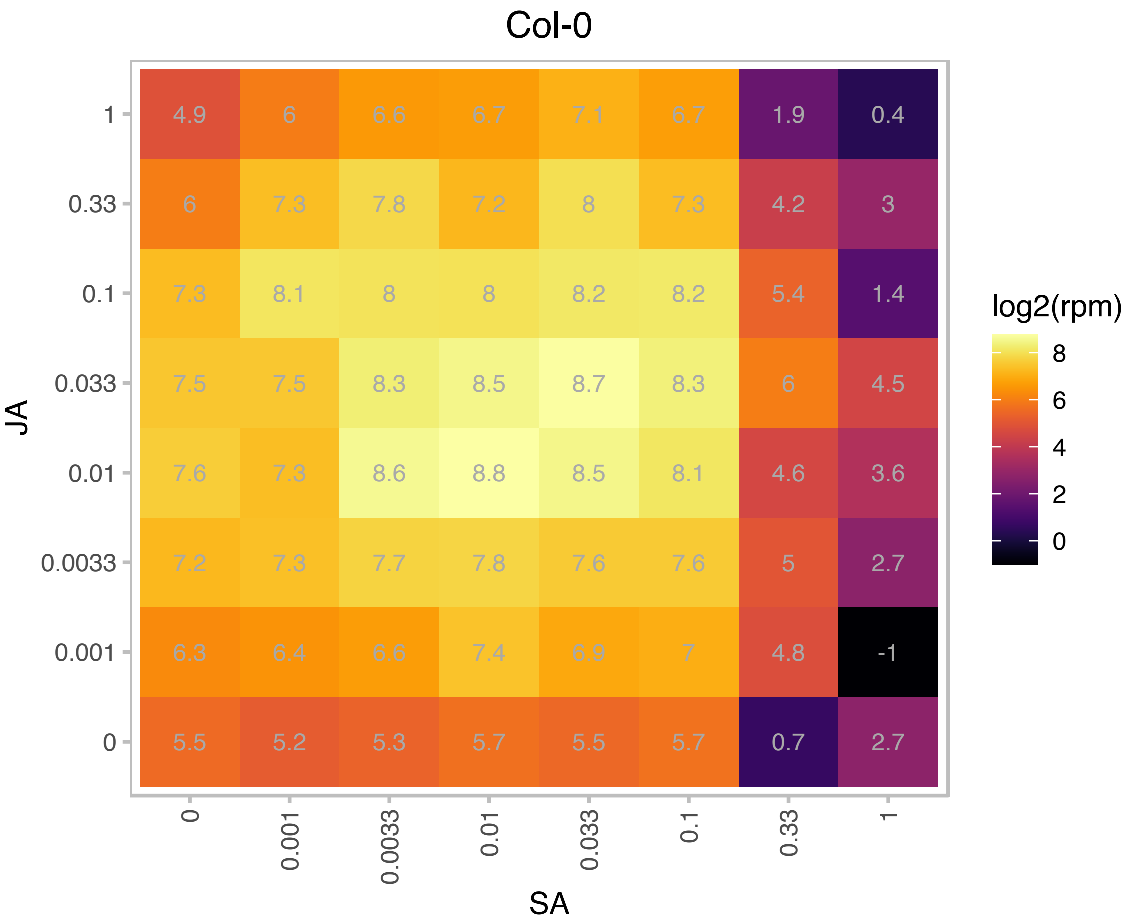


| | |
|---|---|
| Gene | AT5G07460 |
| is.expressed_genes | TRUE |
| is.responsive_genes | TRUE |
| is.apcluster | E-01 |

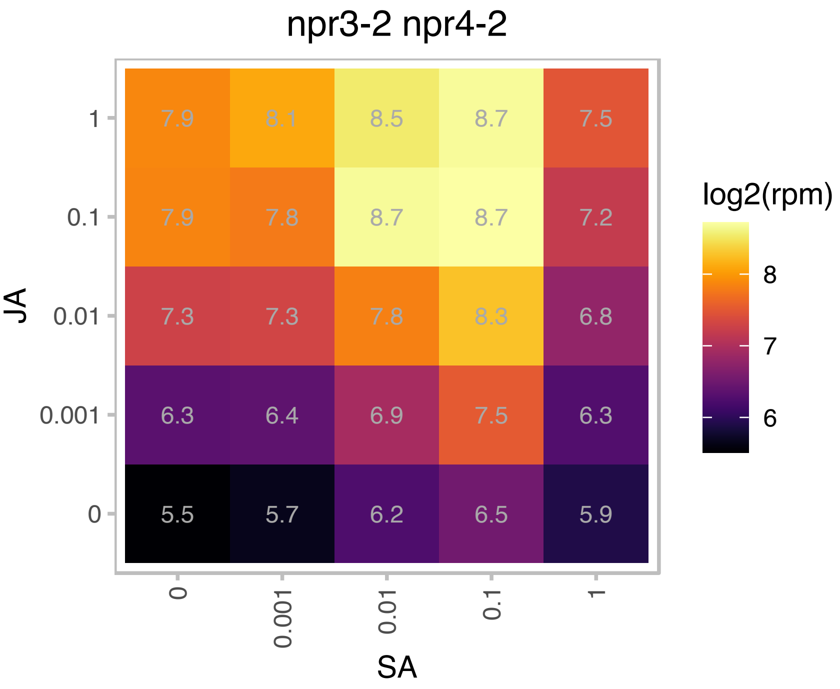


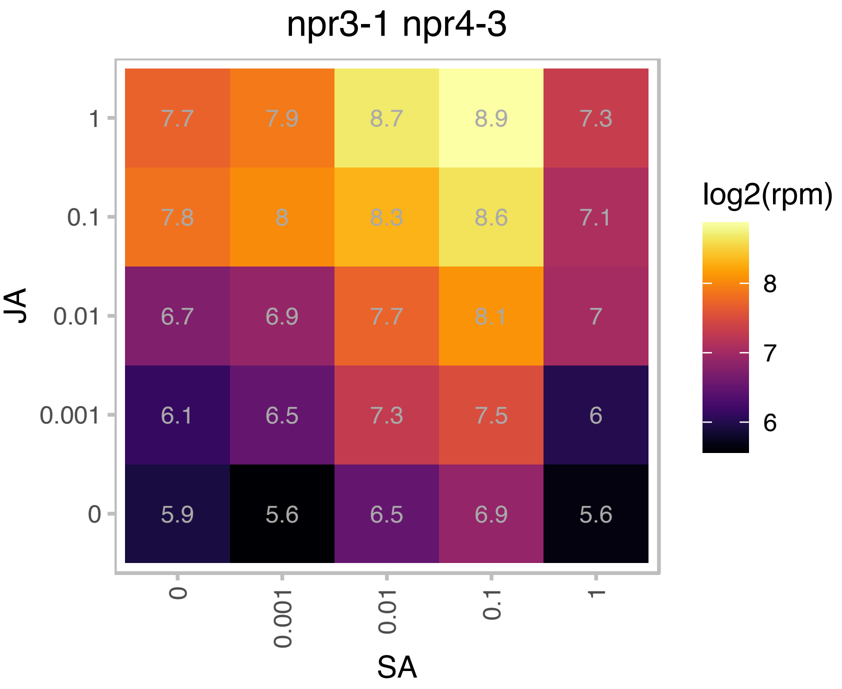

### AT5G09978.1 : PEP7

elicitor peptide 7 precursor

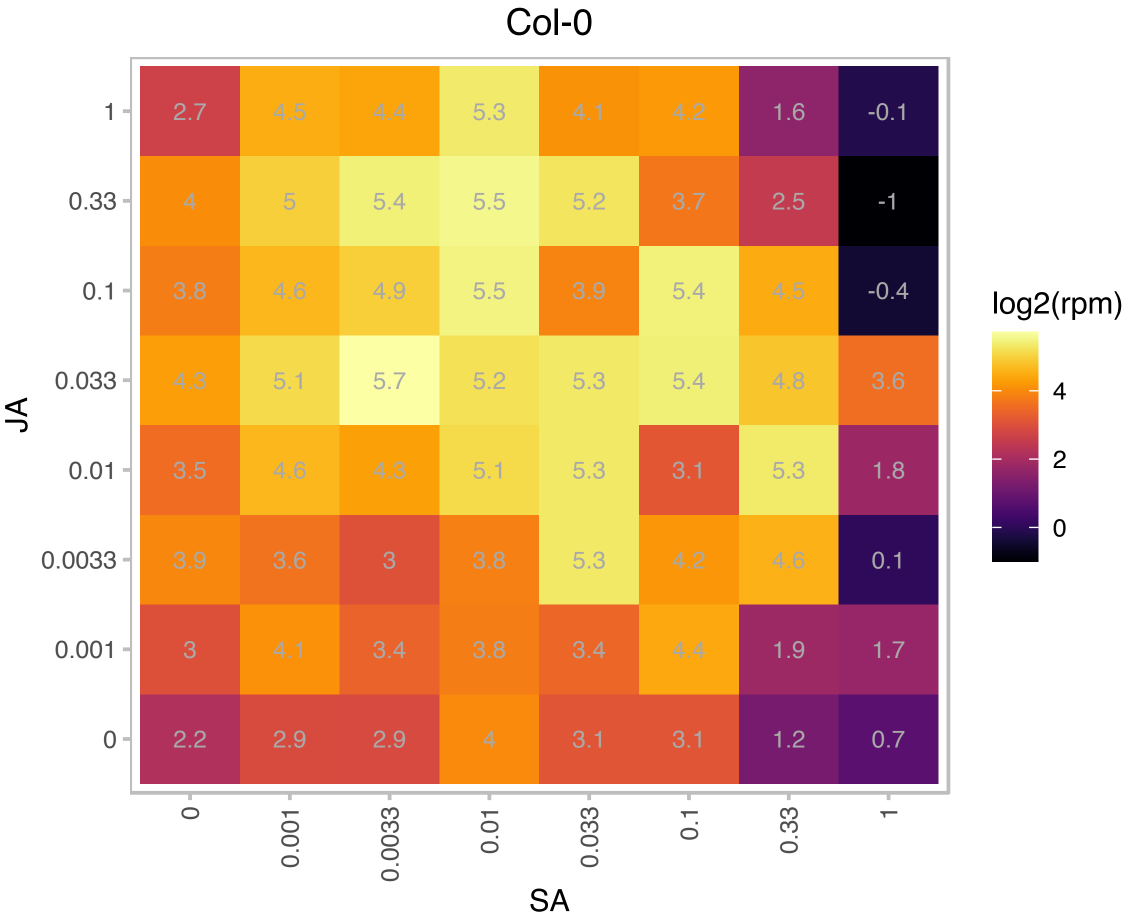


| Gene | AT5G09978 |
|---|---|
| is.expressed_genes | TRUE |
| is.responsive_genes | TRUE |
| is.apcluster | E-01 |

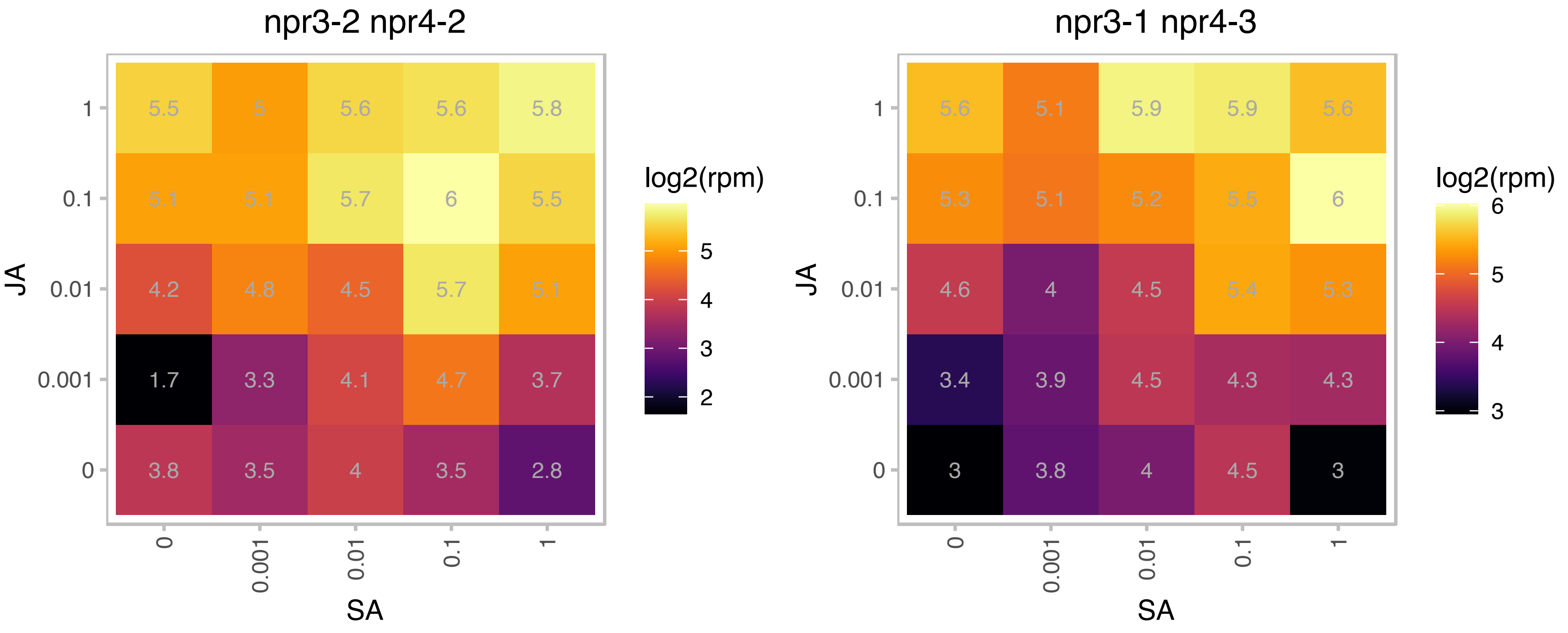

### AT5G23010.1 : GSM1, IMS3, MAM1

methylthioalkylmalate synthase 1
Encodes a methylthioalkylmalate synthase, catalyzes the condensation reactions of the first two rounds of methionine chain elongation in the biosynthesis of methionine-derived glucosinolates. The mRNA is cell-to-cell mobile.

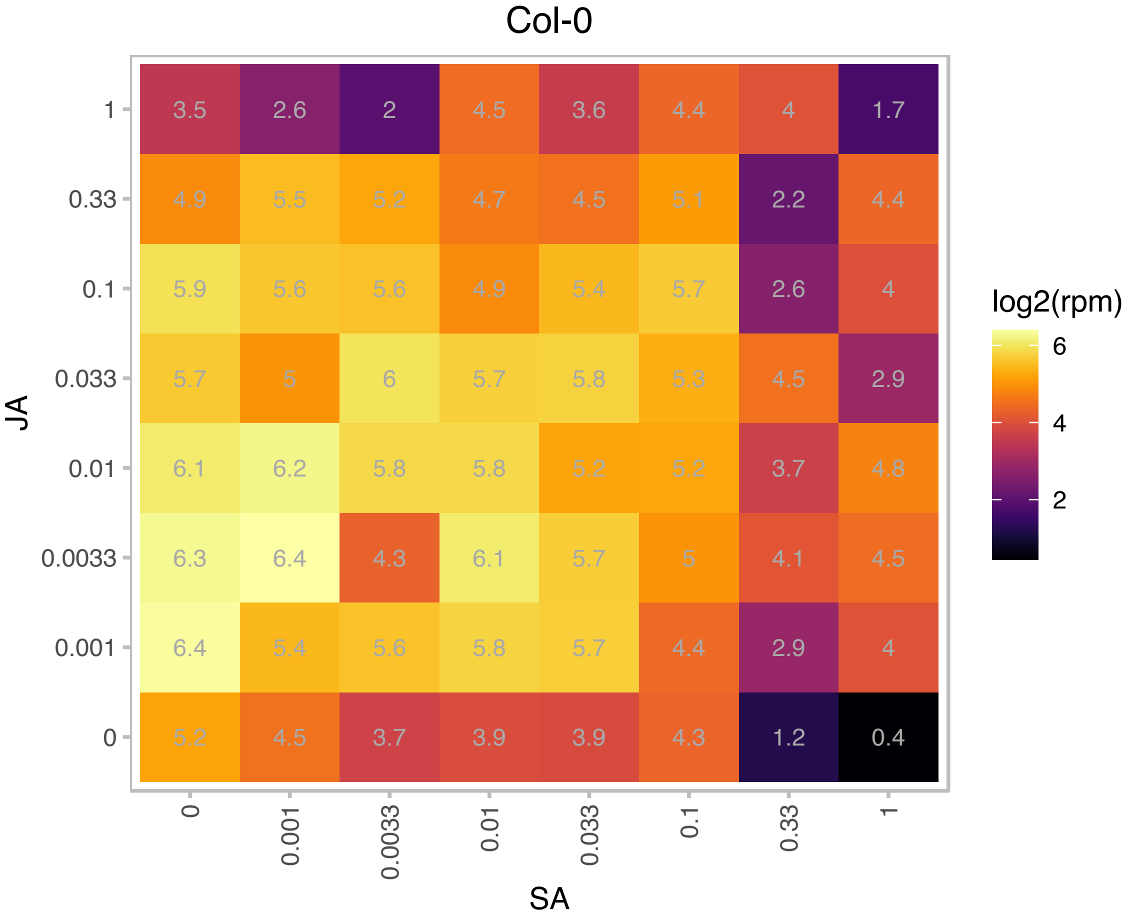


| | |
|---|---|
| Gene | AT5G23010 |
| is.expressed_genes | TRUE |
| is.responsive_genes | TRUE |
| is.apcluster | E-01 |

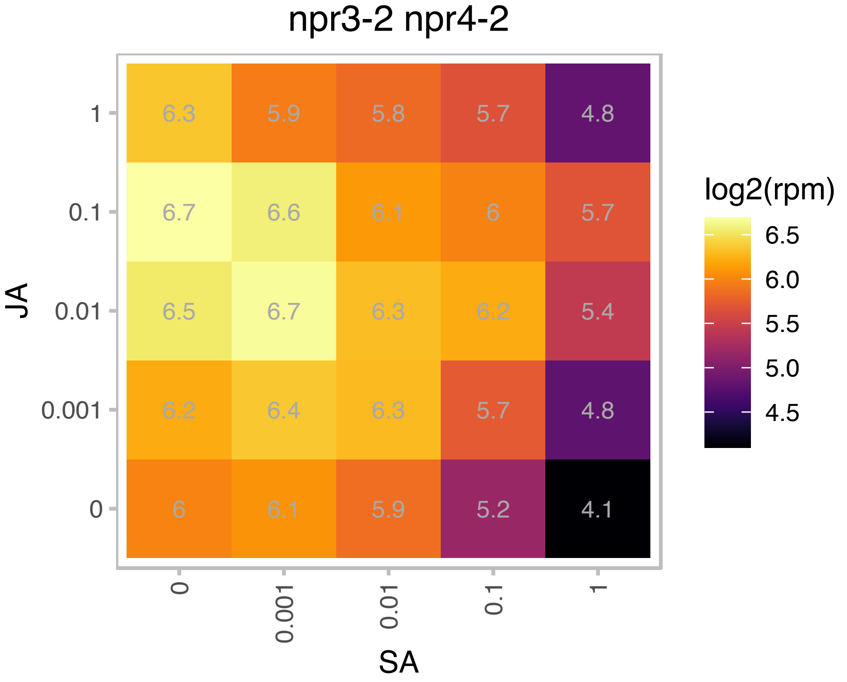


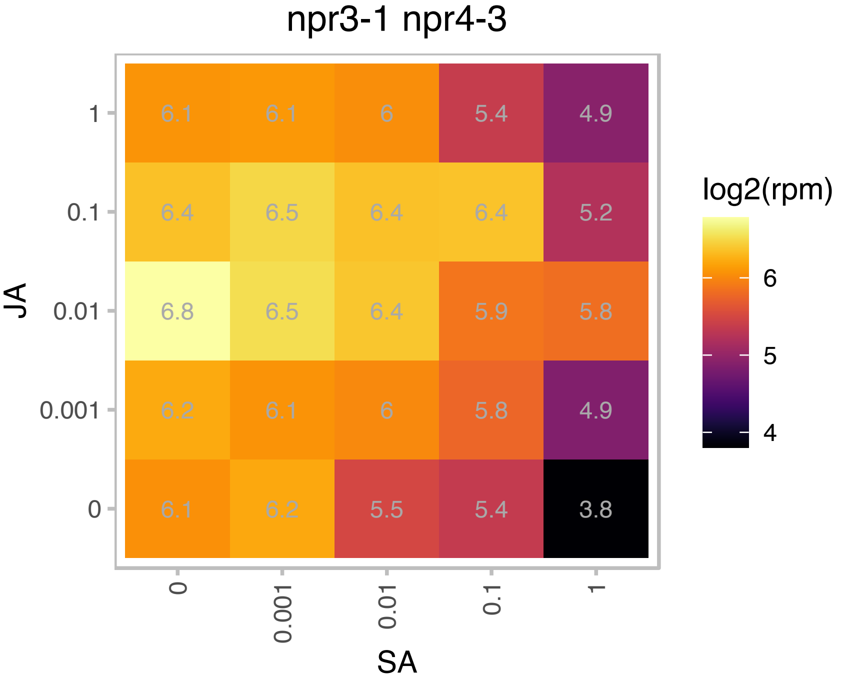

# AT5G24610.1 : NA

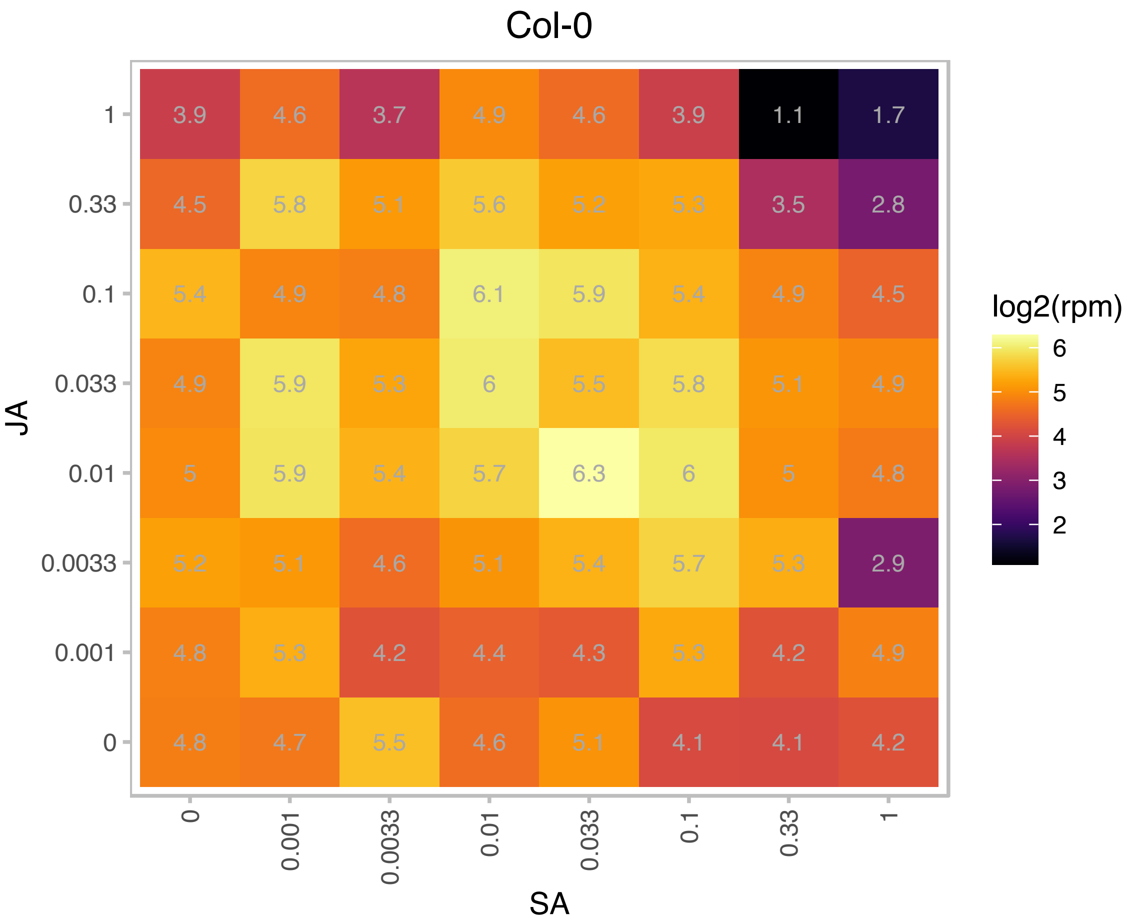


| | |
|---|---|
| Gene | AT5G24610 |
| is.expressed_genes | TRUE |
| is.responsive_genes | TRUE |
| is.apcluster | E-01 |

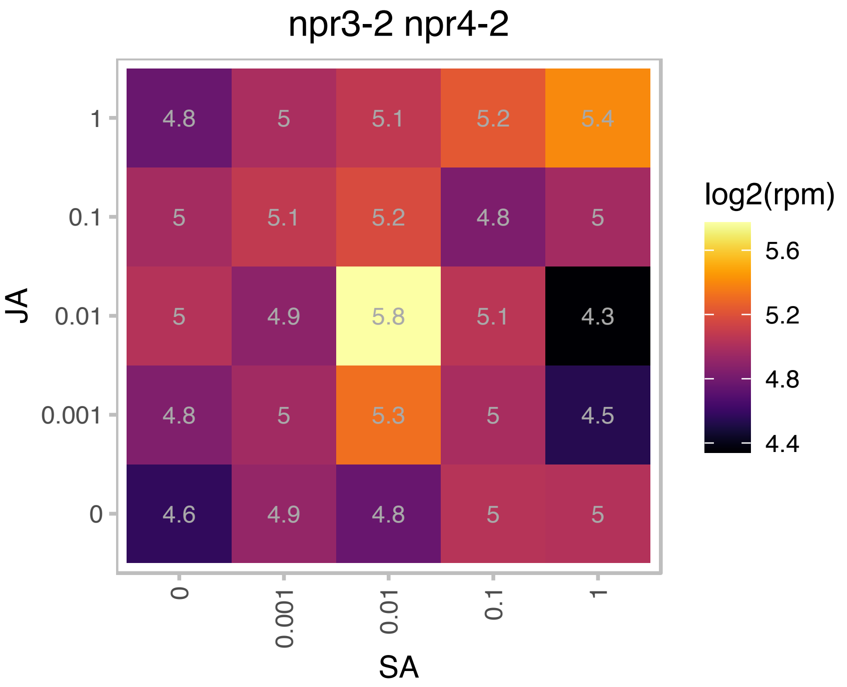


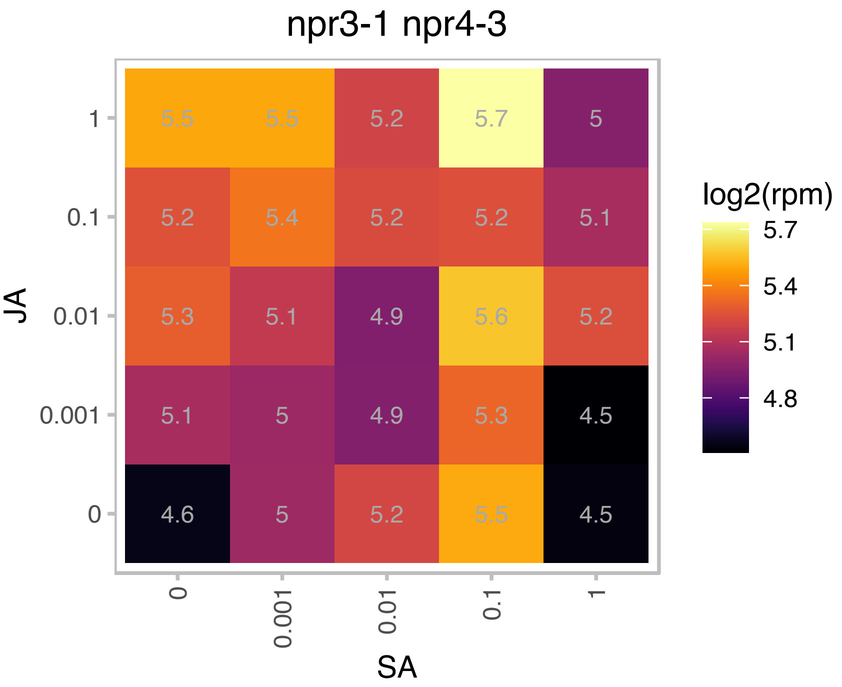

# AT5G36907.1 : NA

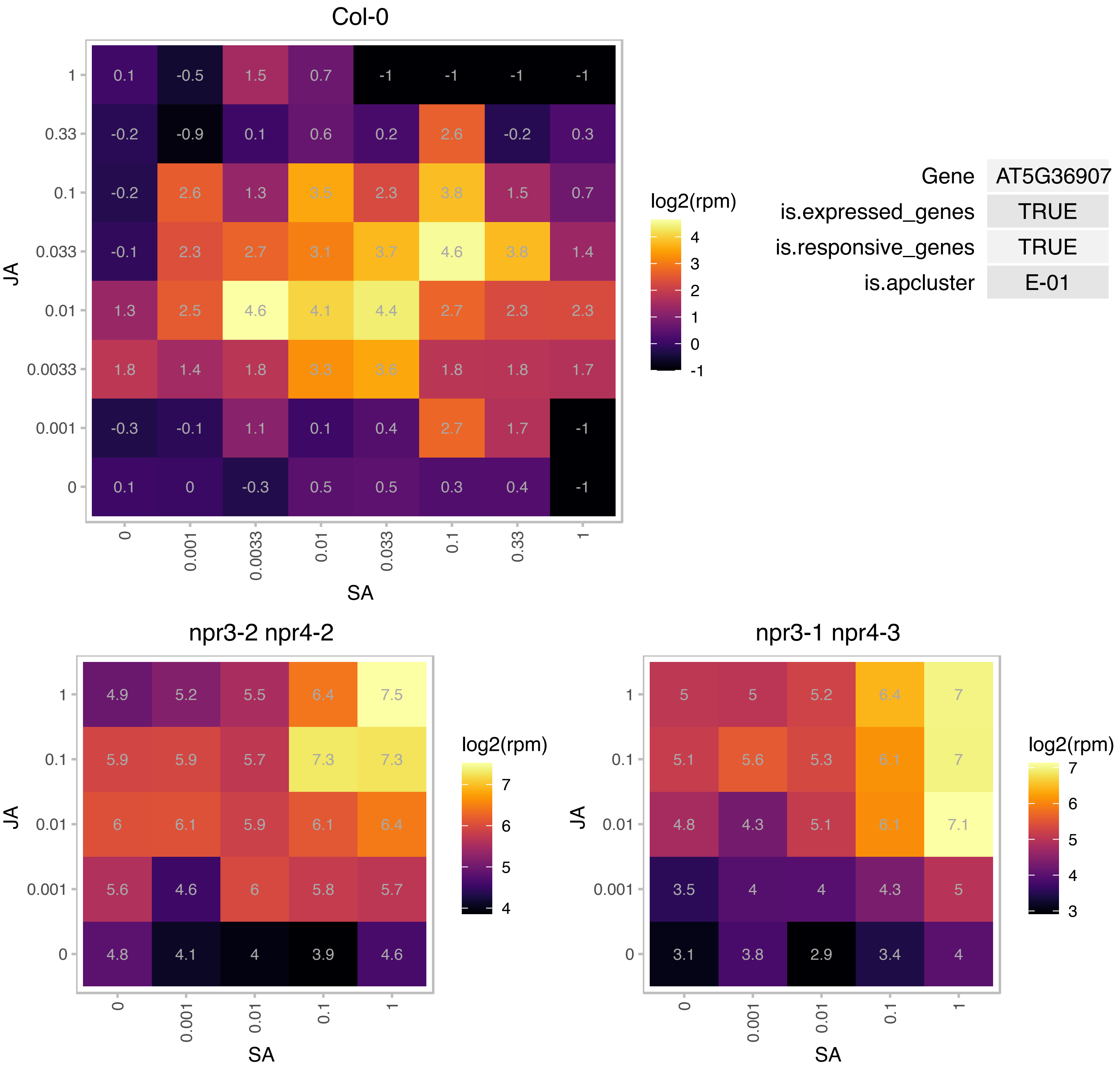

# AT5G40530.1 : NA

S-adenosyl-L-methionine-dependent methyltransferases superfamily protein

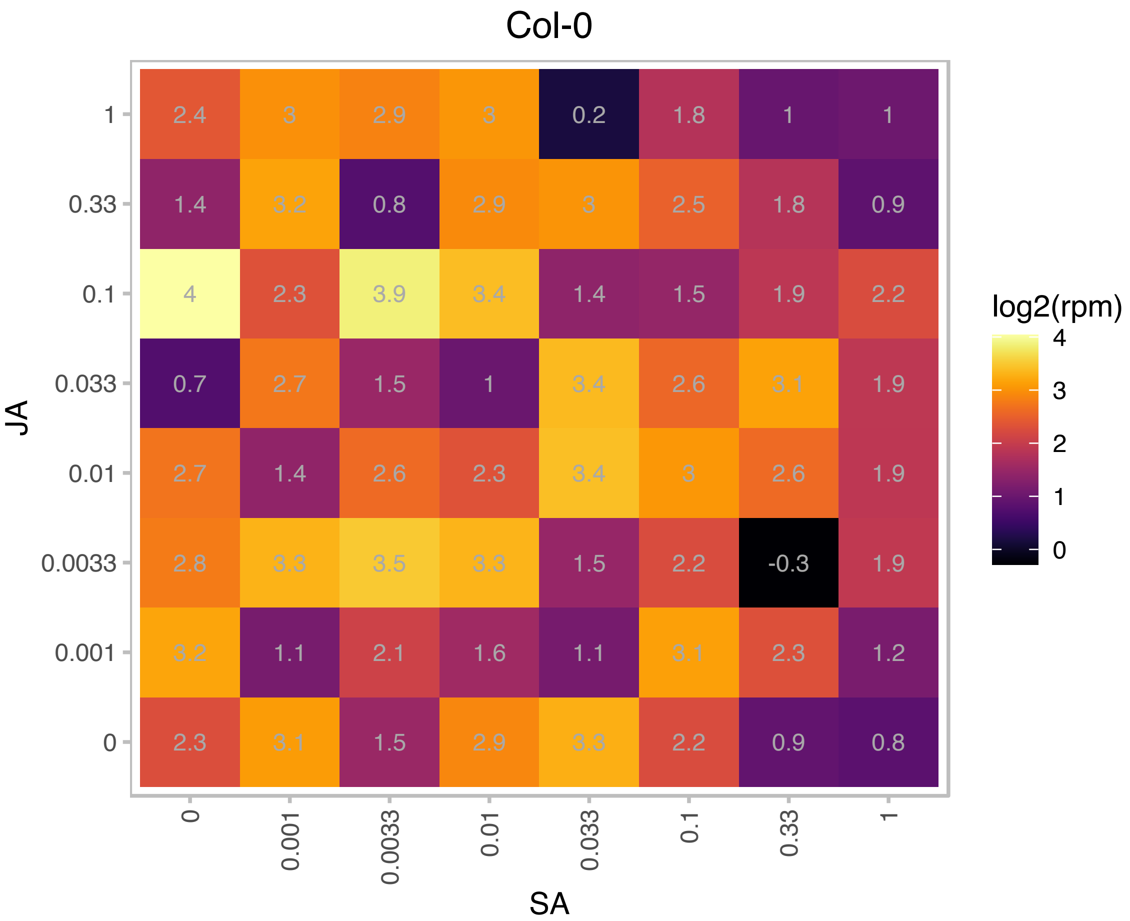


| | |
|---|---|
| Gene | AT5G40530 |
| is.expressed_genes | TRUE |
| is.responsive_genes | TRUE |
| is.apcluster | E-01 |

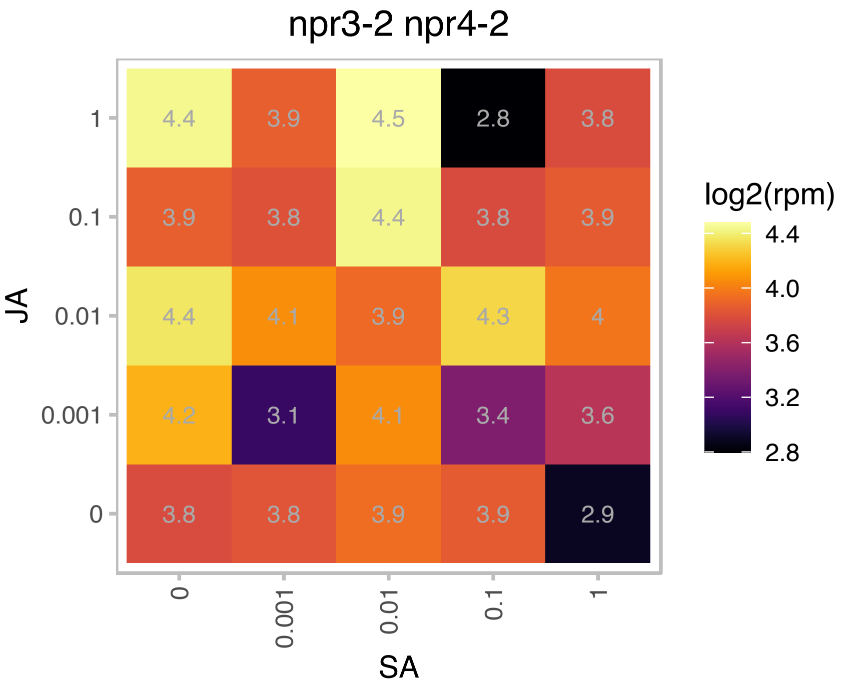


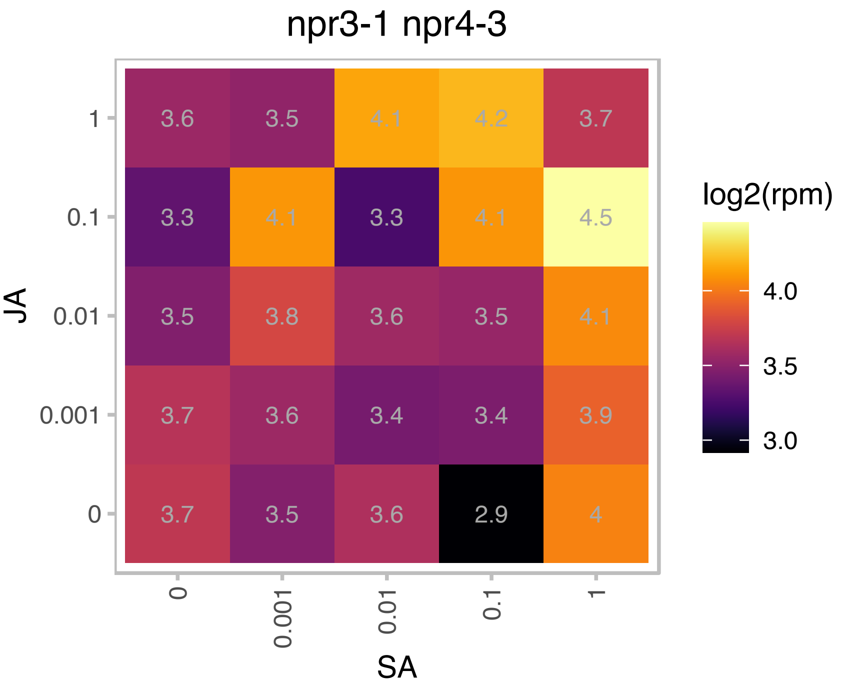

### AT5G48880.1 : KAT5, PKT1, PKT2

peroxisomal 3-keto-acyl-CoA thiolase 2

Encodes a peroxisomal 3-keto-acyl-CoA thiolase 2 precursor. EC2.3.1.16 thiolases. AT5G48880.1 is named PKT1 and AT5G48880.2 is named PKT2.

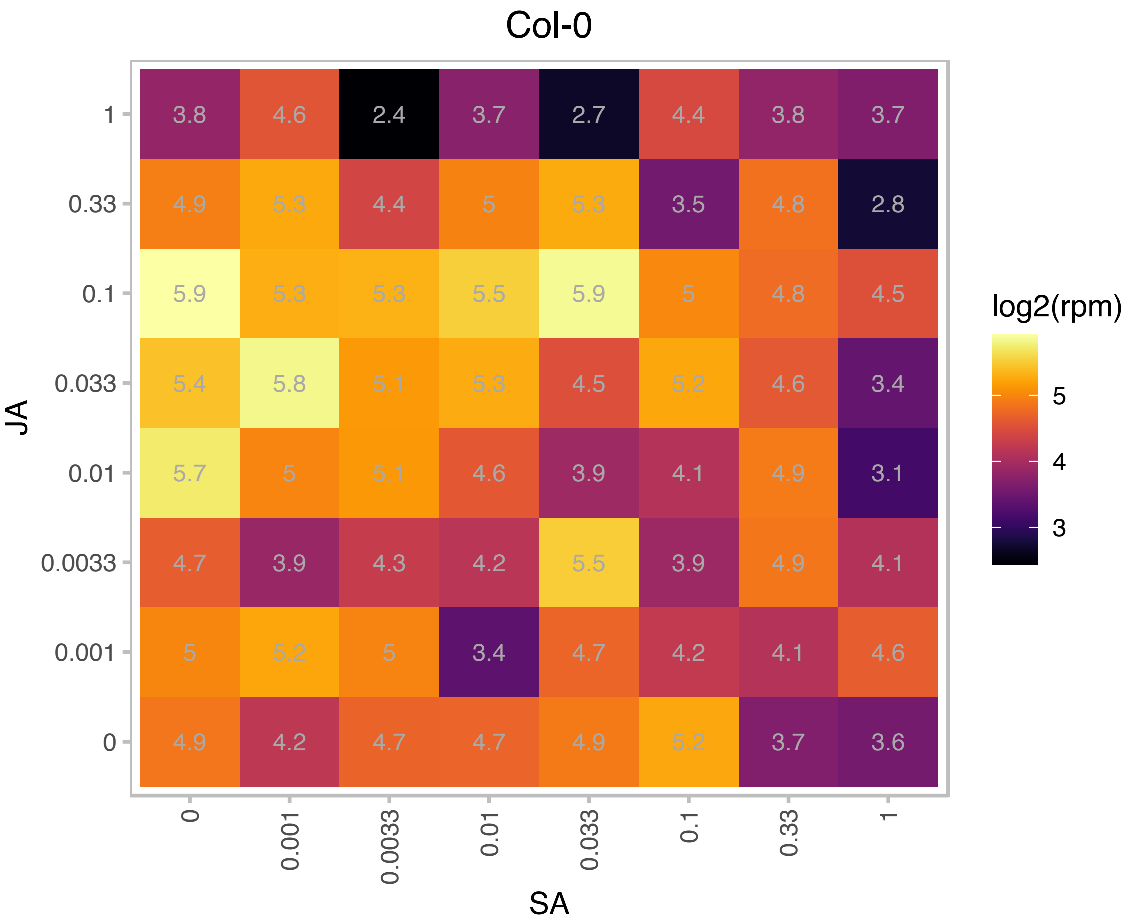


| | |
|---|---|
| Gene | AT5G48880 |
| is.expressed_genes | TRUE |
| is.responsive_genes | TRUE |
| is.apcluster | E-01 |

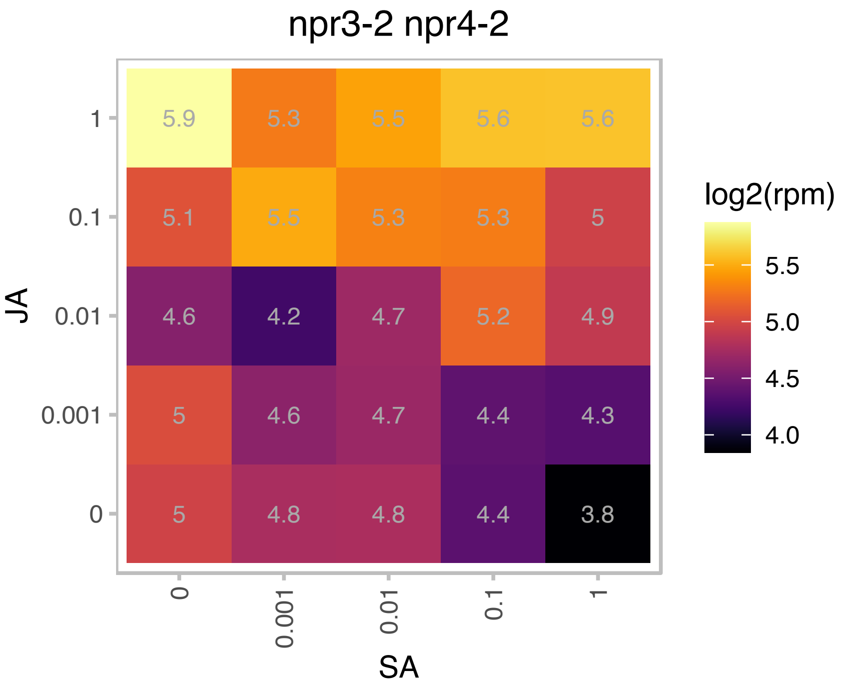


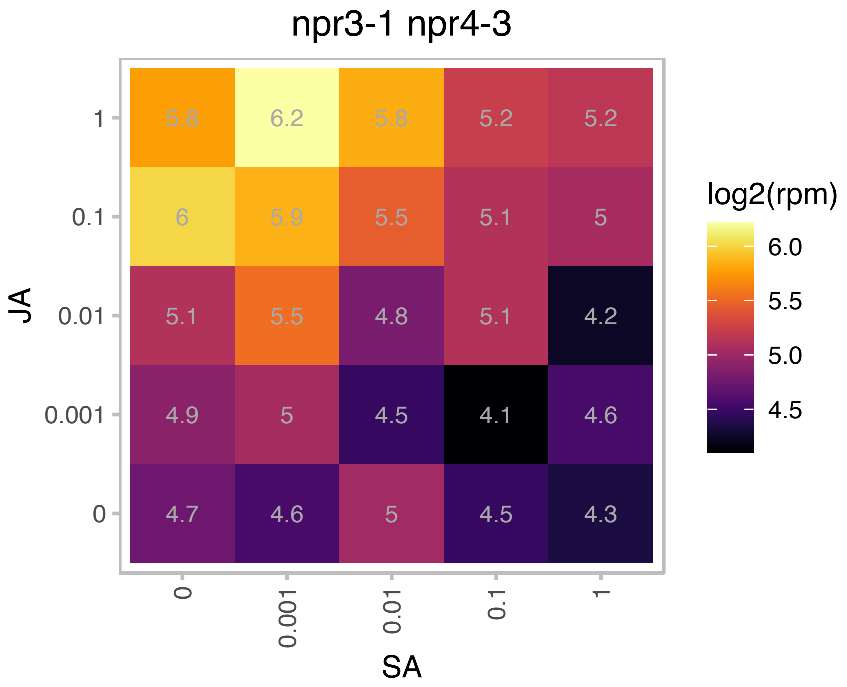

# AT5G49945.1 : NA

Protein of unknown function (DUF1682)

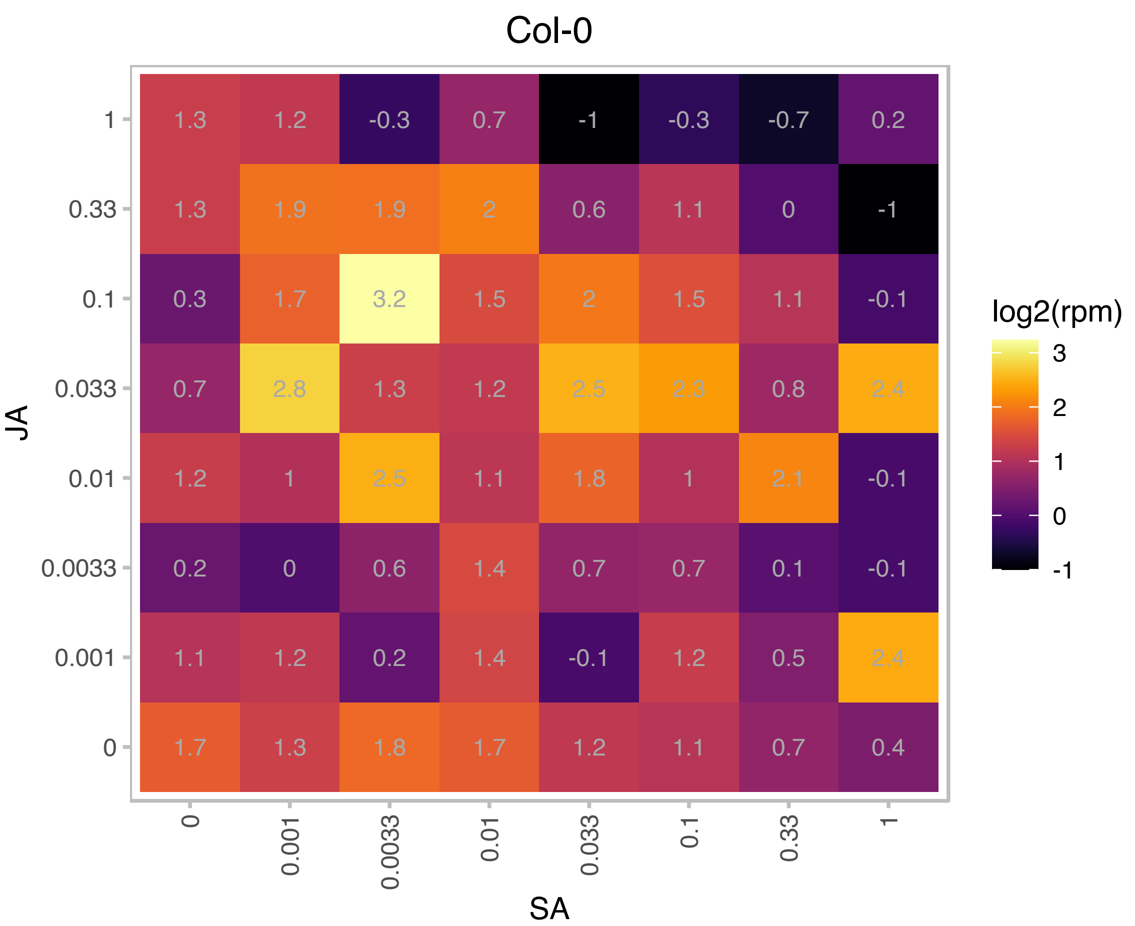


| | |
|---|---|
| Gene | AT5G49945 |
| is.expressed_genes | TRUE |
| is.responsive_genes | TRUE |
| is.apcluster | E-01 |

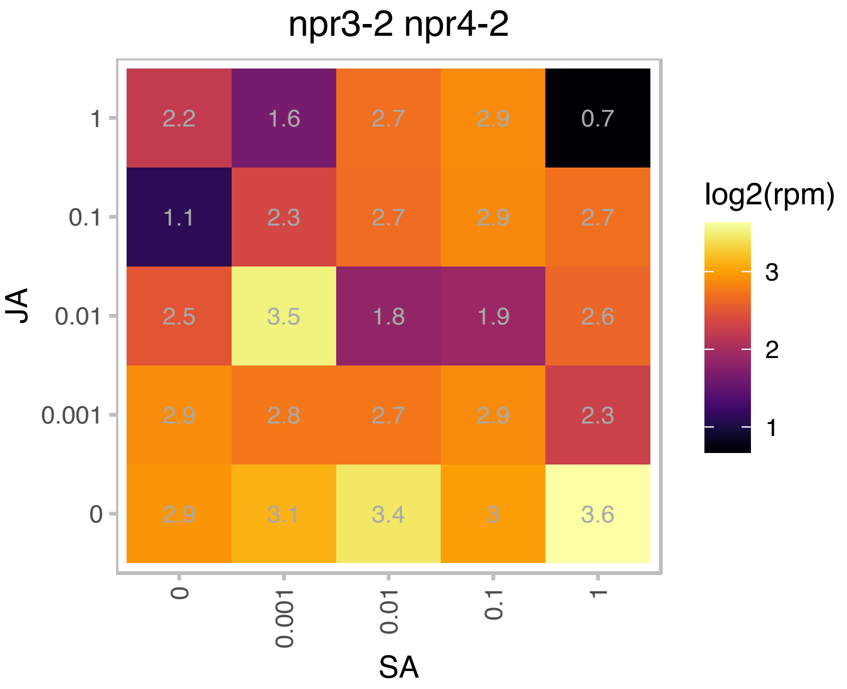


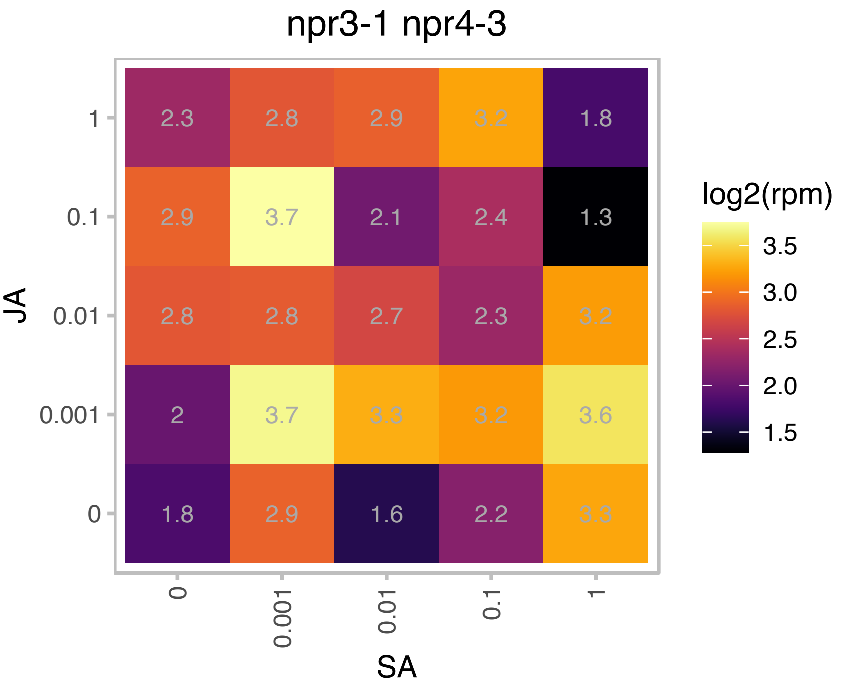

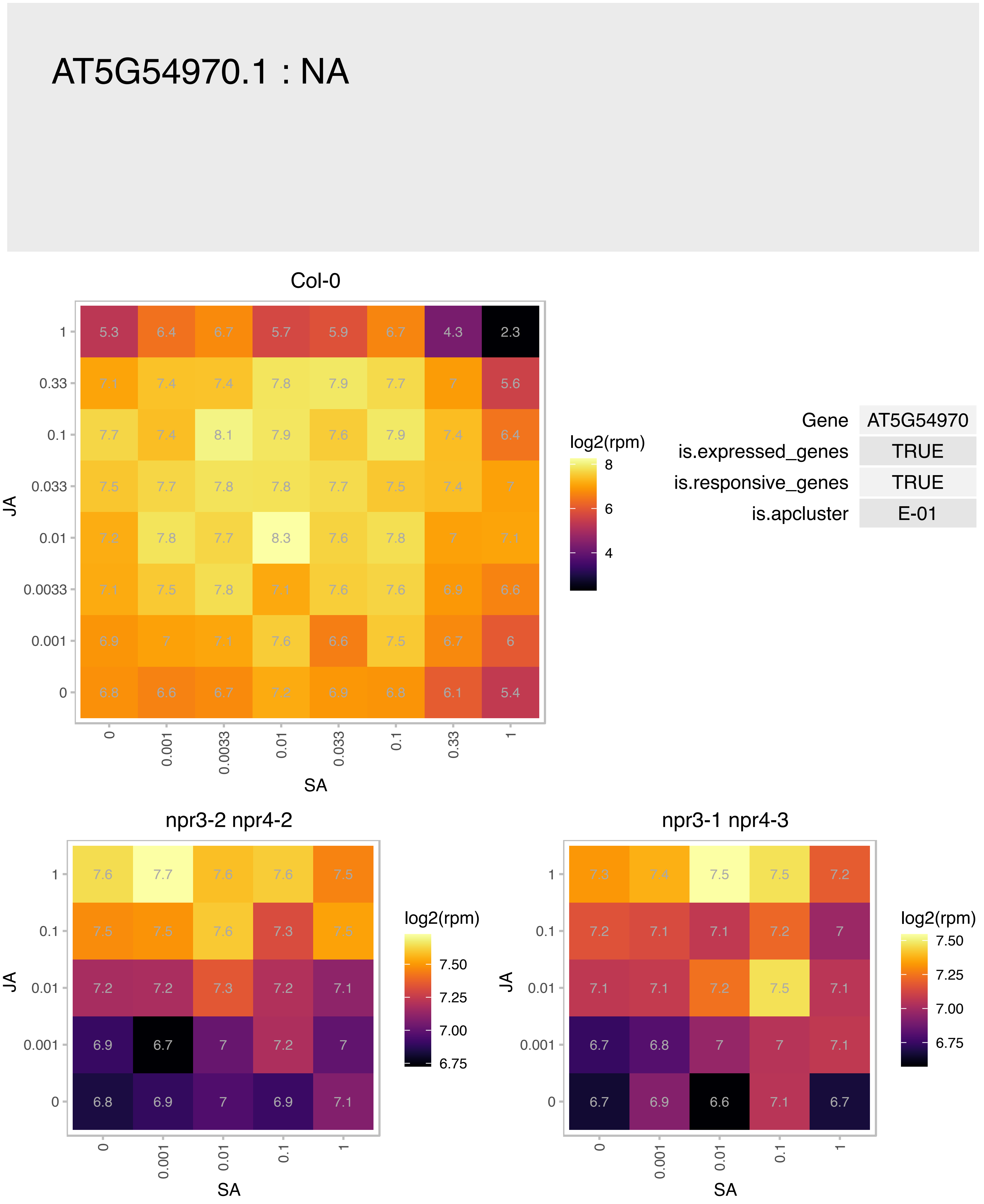
AT5G54970.1 : NA
Col-0
JA
SA
1 0.33 0.1 0.033 0.01 0.0033 0.001 0
0 0.001 0.0033 0.01 0.033 0.1 0.33 1
5.3 6.4 6.7 5.7 5.9 6.7 4.3 2.3
7.1 7.4 7.4 7.8 7.9 7.7 7 5.6
7.7 7.4 8.1 7.9 7.6 7.9 7.4 6.4
7.5 7.7 7.8 7.8 7.7 7.5 7.4 7
7.2 7.8 7.7 8.3 7.6 7.8 7 7.1
7.1 7.5 7.8 7.1 7.6 7.6 6.9 6.6
6.9 7 7.1 7.6 6.6 7.5 6.7 6
6.8 6.6 6.7 7.2 6.9 6.8 6.1 5.4
log2(rpm)
8
6
4
Gene AT5G54970
is.expressed_genes TRUE
is.responsive_genes TRUE
is.apcluster E-01
npr3-2 npr4-2
JA
SA
1 0.1 0.01 0.001 0
0 0.001 0.01 0.1 1
7.6 7.7 7.6 7.6 7.5
7.5 7.5 7.6 7.3 7.5
7.2 7.2 7.3 7.2 7.1
6.9 6.7 7 7.2 7
6.8 6.9 7 6.9 7.1
log2(rpm)
7.50
7.25
7.00
6.75
npr3-1 npr4-3
JA
SA
1 0.1 0.01 0.001 0
0 0.001 0.01 0.1 1
7.3 7.4 7.5 7.5 7.2
7.2 7.1 7.1 7.2 7
7.1 7.1 7.2 7.5 7.1
6.7 6.8 7 7 7.1
6.7 6.9 6.6 7.1 6.7
log2(rpm)
7.50
7.25
7.00
6.75

### AT5G55120.1 : VTC5

galactose-1-phosphate guanylyltransferase (GDP)s;GDP-D-glucose phosphorylases;quercetin 4'-O-glucosyltransferases

Encodes a GDP-L-galactose phosphorylase, with similar biochemical properties as VTC2.

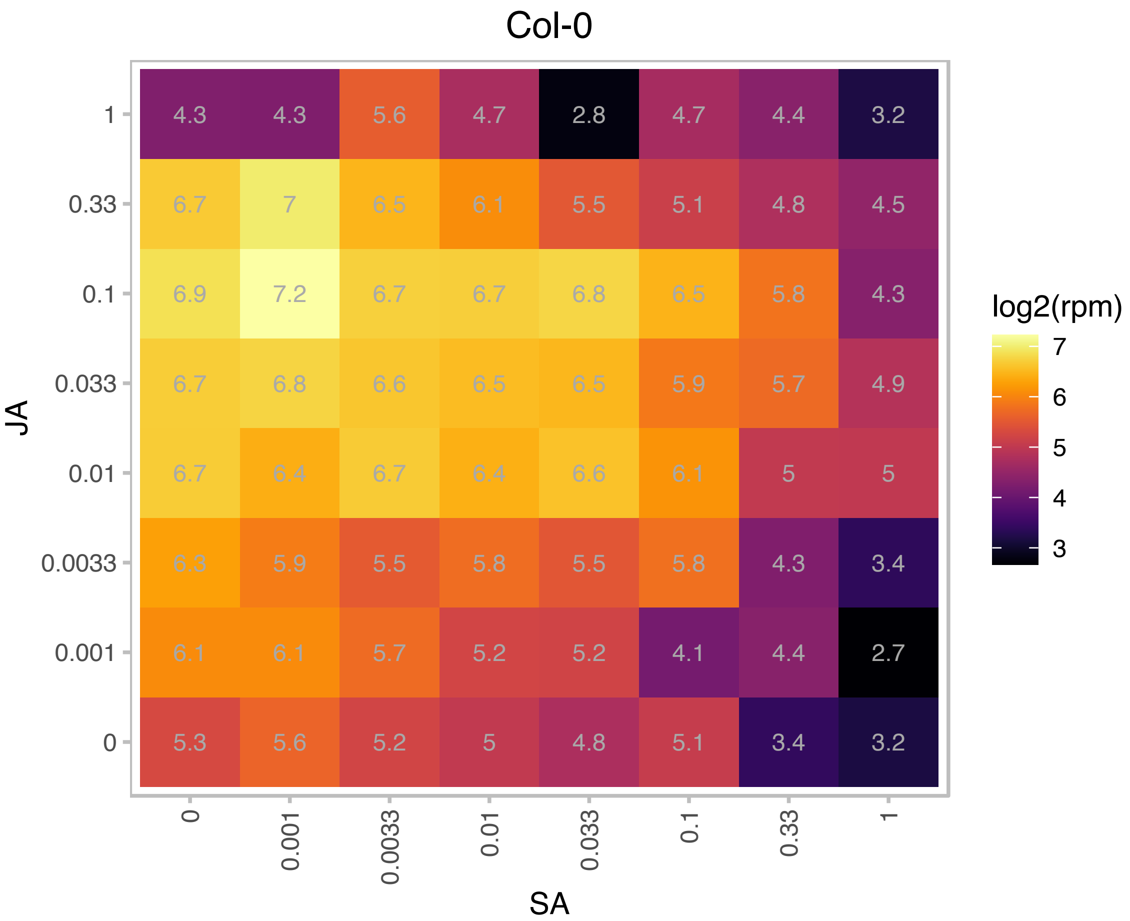


| | |
|---|---|
| Gene | AT5G55120 |
| is.expressed_genes | TRUE |
| is.responsive_genes | TRUE |
| is.apcluster | E-01 |

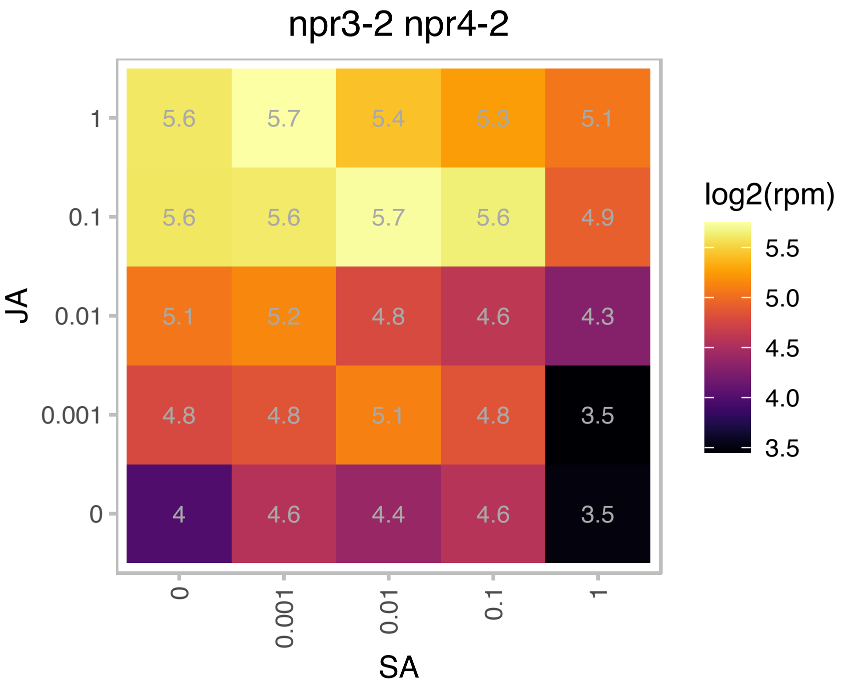


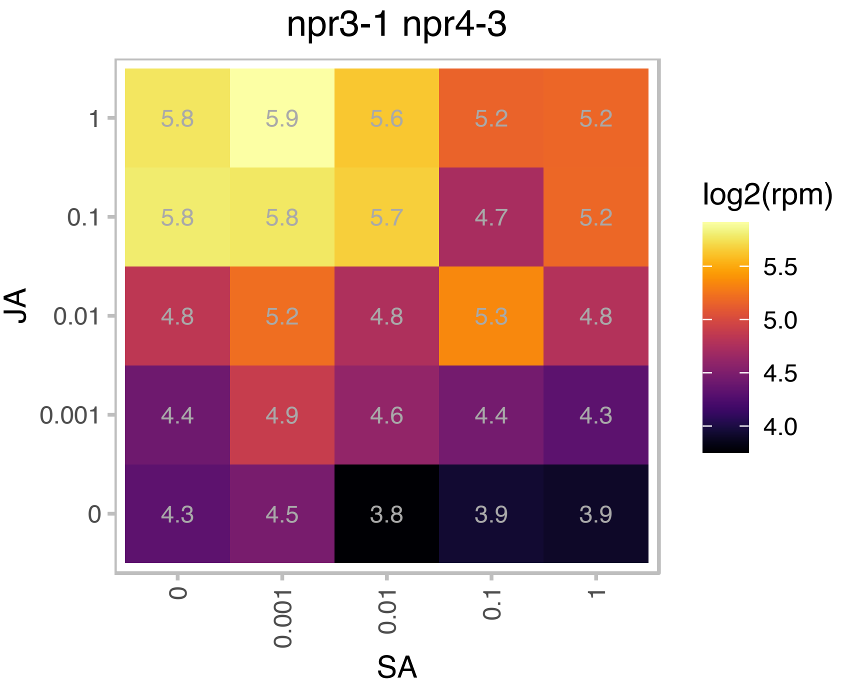

### AT5G57490.1 : ATVDAC4, VDAC4

voltage dependent anion channel 4

Encodes a voltage-dependent anion channel (VDAC: AT3G01280/VDAC1, AT5G67500/VDAC2, AT5G15090/VDAC[illegible] AT5G57490/VDAC4, AT5G15090/VDAC5). VDACs are reported to be porin-type, beta-barrel diffusion pores. They a[illegible] prominently localized in the outer mitochondrial membrane and are involved in metabolite exchange between the organelle and the cytosol.

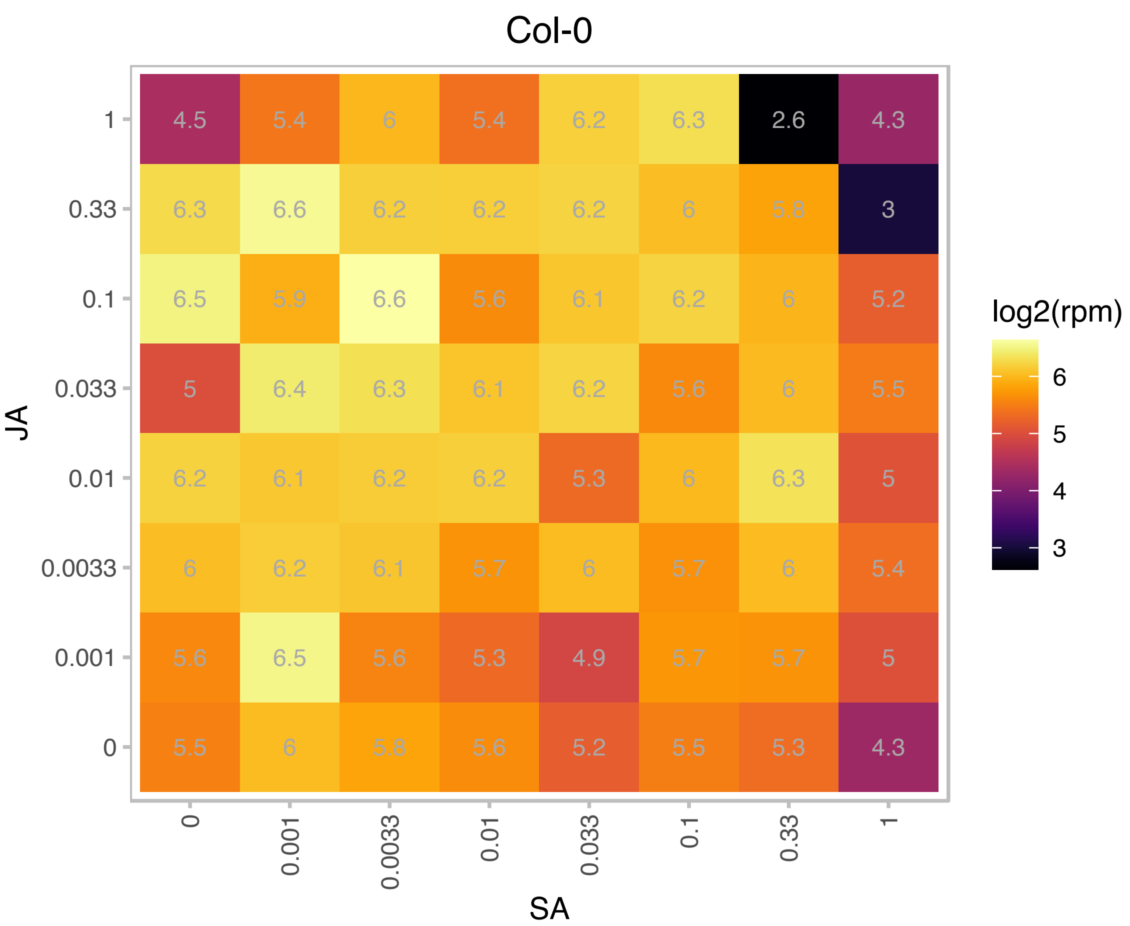


| | |
|---|---|
| Gene | AT5G57490 |
| is.expressed_genes | TRUE |
| is.responsive_genes | TRUE |
| is.apcluster | E-01 |

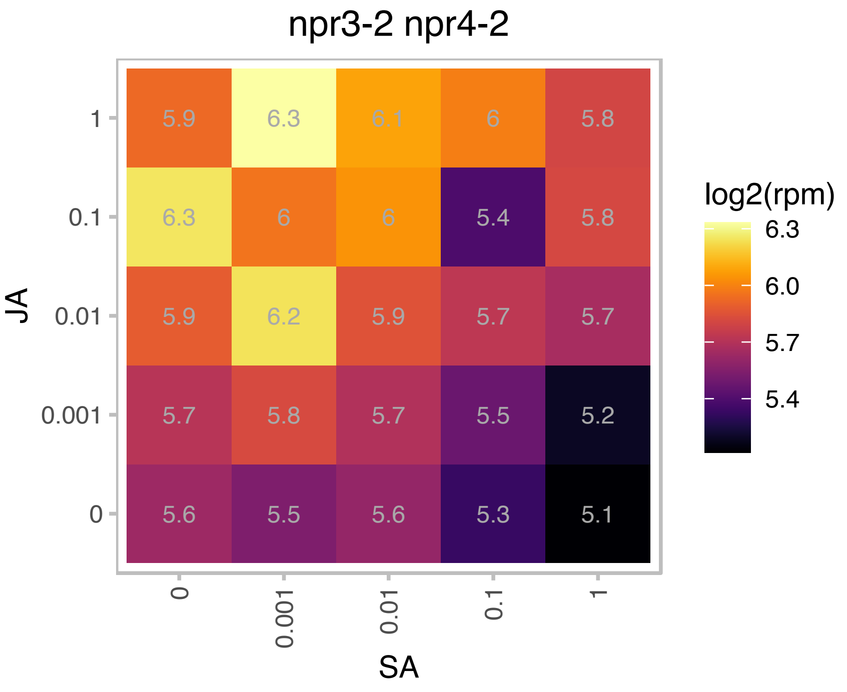


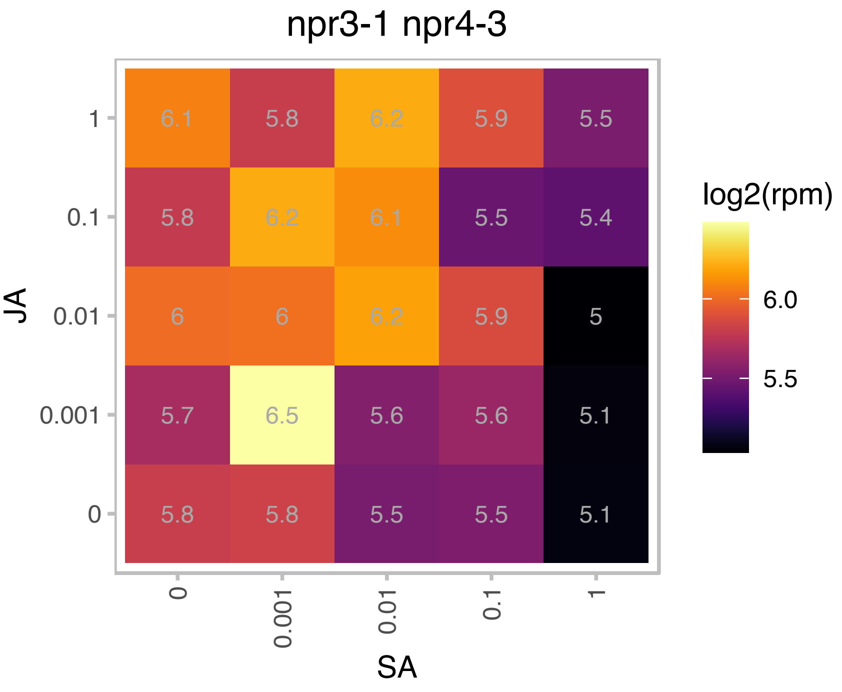

# AT5G62350.1 : NA

Plant invertase/pectin methylesterase inhibitor superfamily protein

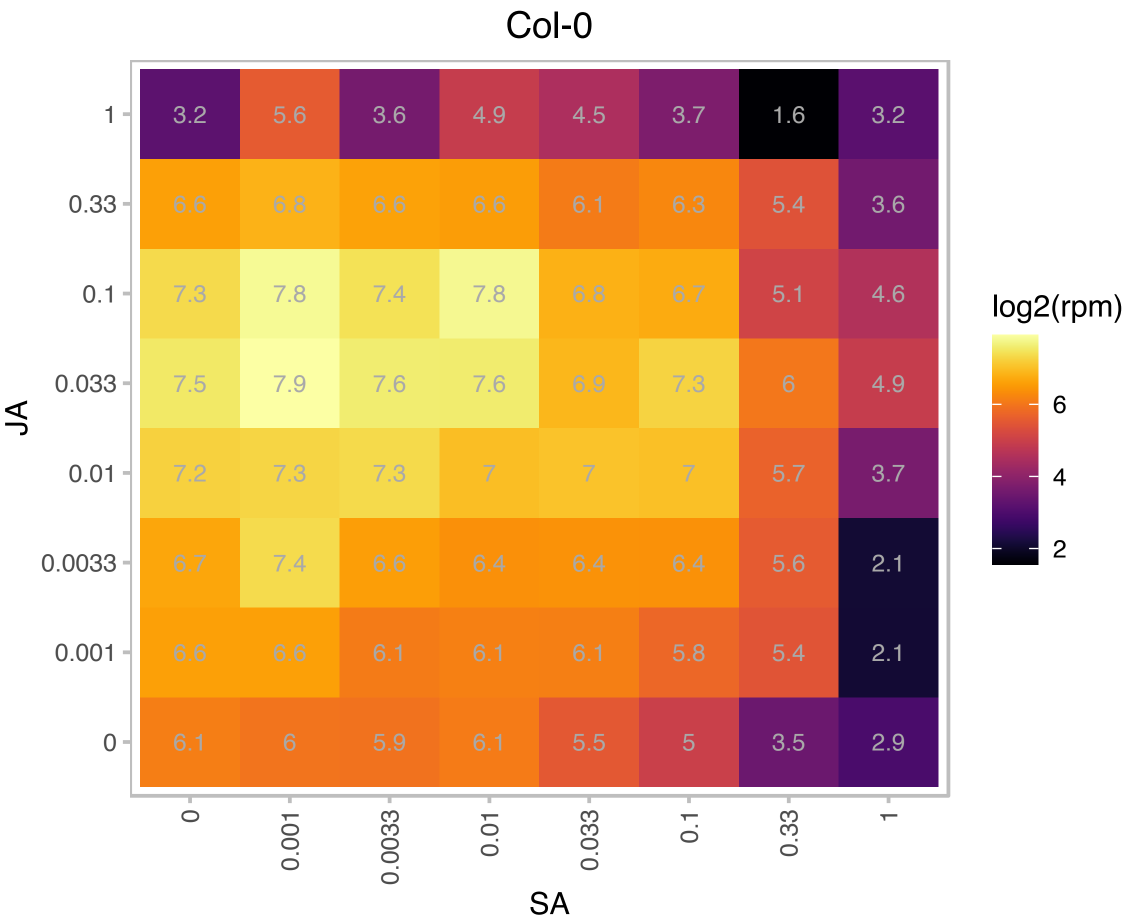


| | |
|---|---|
| Gene | AT5G62350 |
| is.expressed_genes | TRUE |
| is.responsive_genes | TRUE |
| is.apcluster | E-01 |

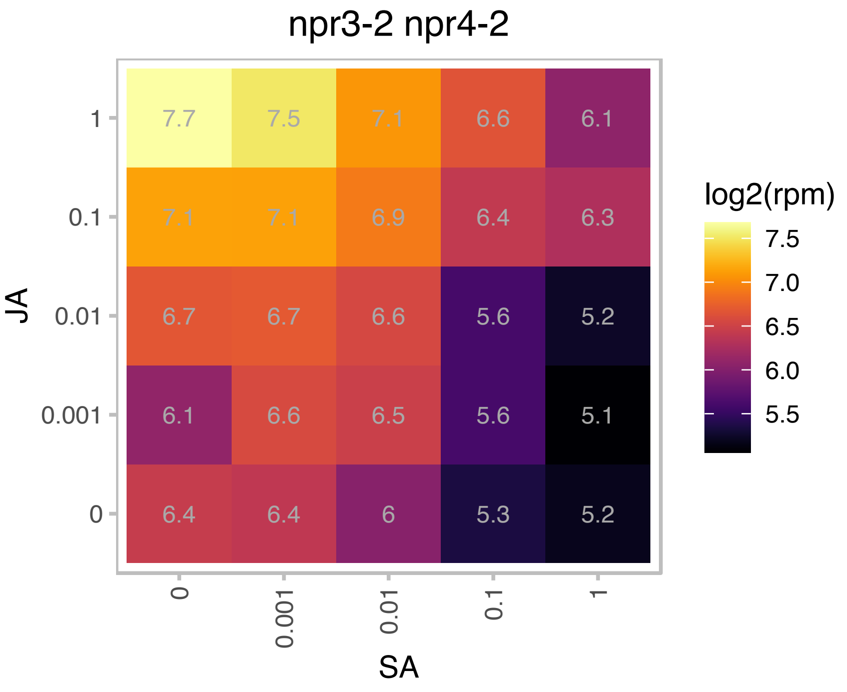


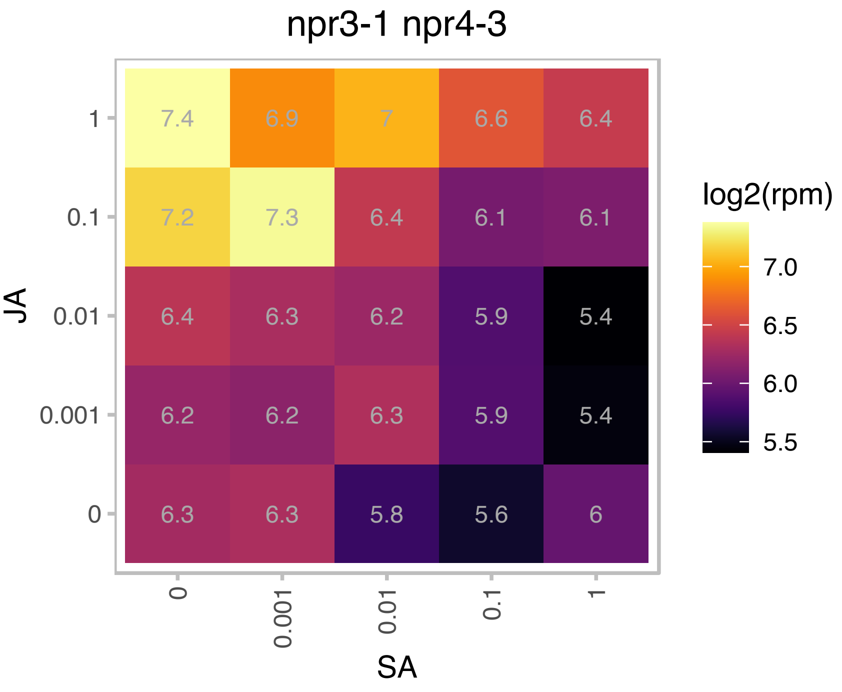

### AT5G62360.1 : PMEI13

Plant invertase/pectin methylesterase inhibitor superfamily protein

Pectin methylesterase inhibitor expressed throughout the plant.

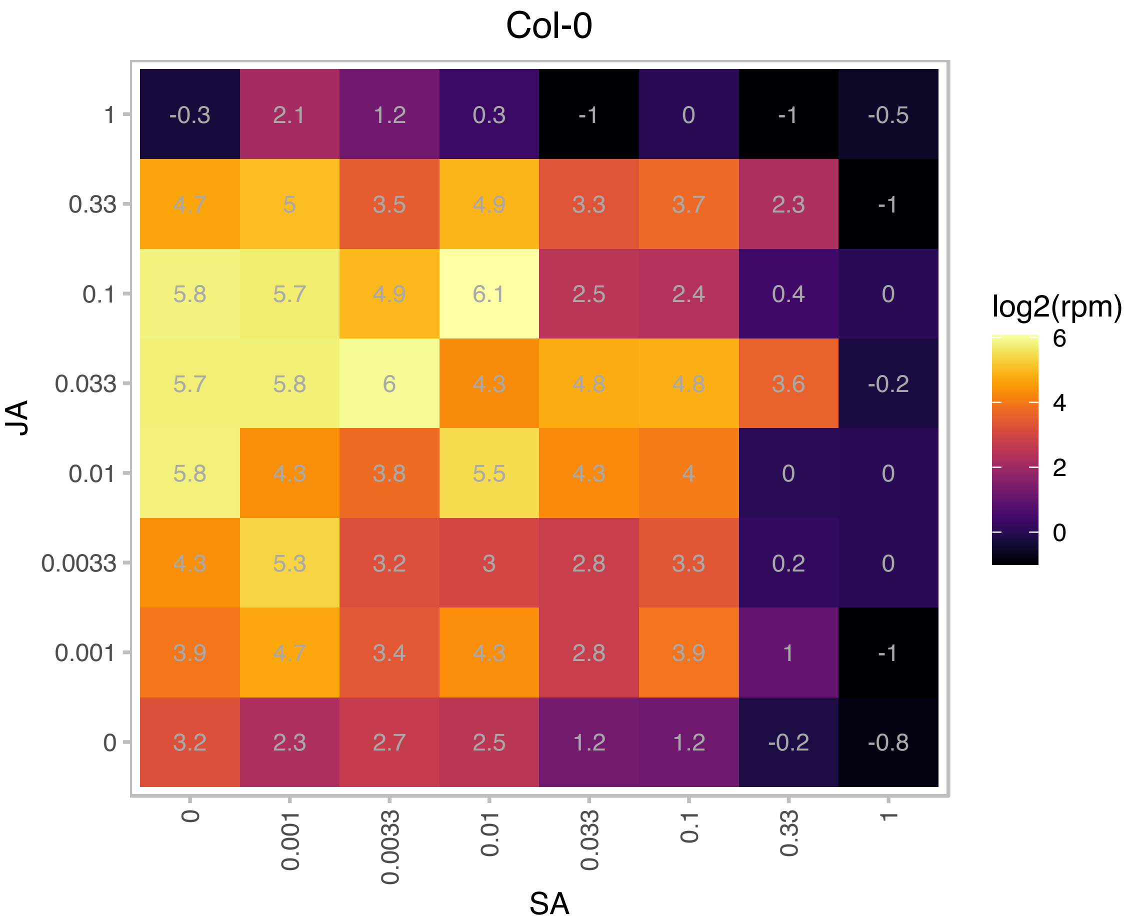


| | |
|---|---|
| Gene | AT5G62360 |
| is.expressed_genes | TRUE |
| is.responsive_genes | TRUE |
| is.apcluster | E-01 |

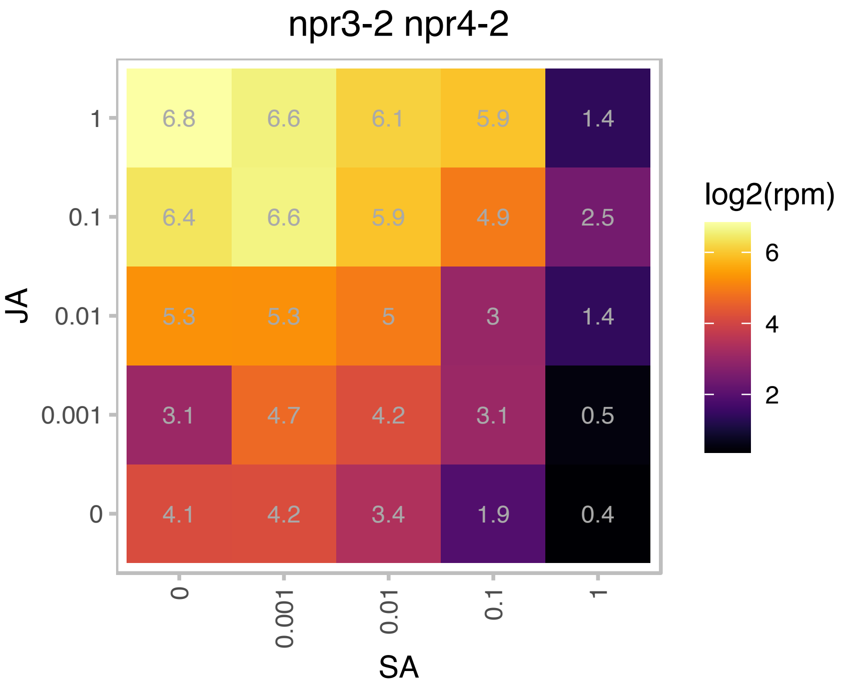


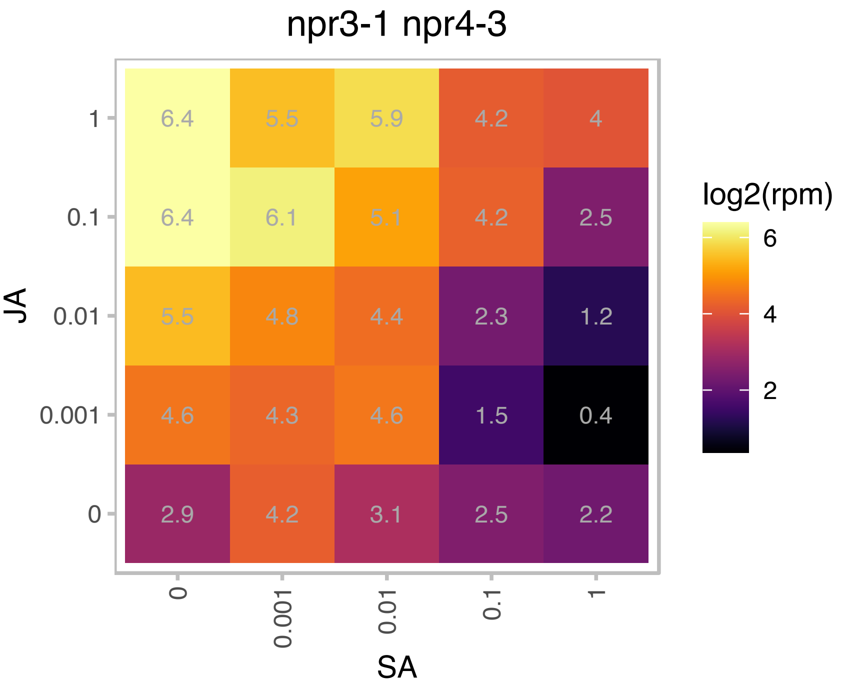

### AT5G62575.1 : SDH7, SDH7B

Encodes subunit 7 of mitochondrial complex II (succinate dehydrogenase complex) and participates in the respiratory chain. It contributes to anchoring succinate dehydrogenase to the inner mitochondrial membrane. The mRNA is cell-to-cell mobile.

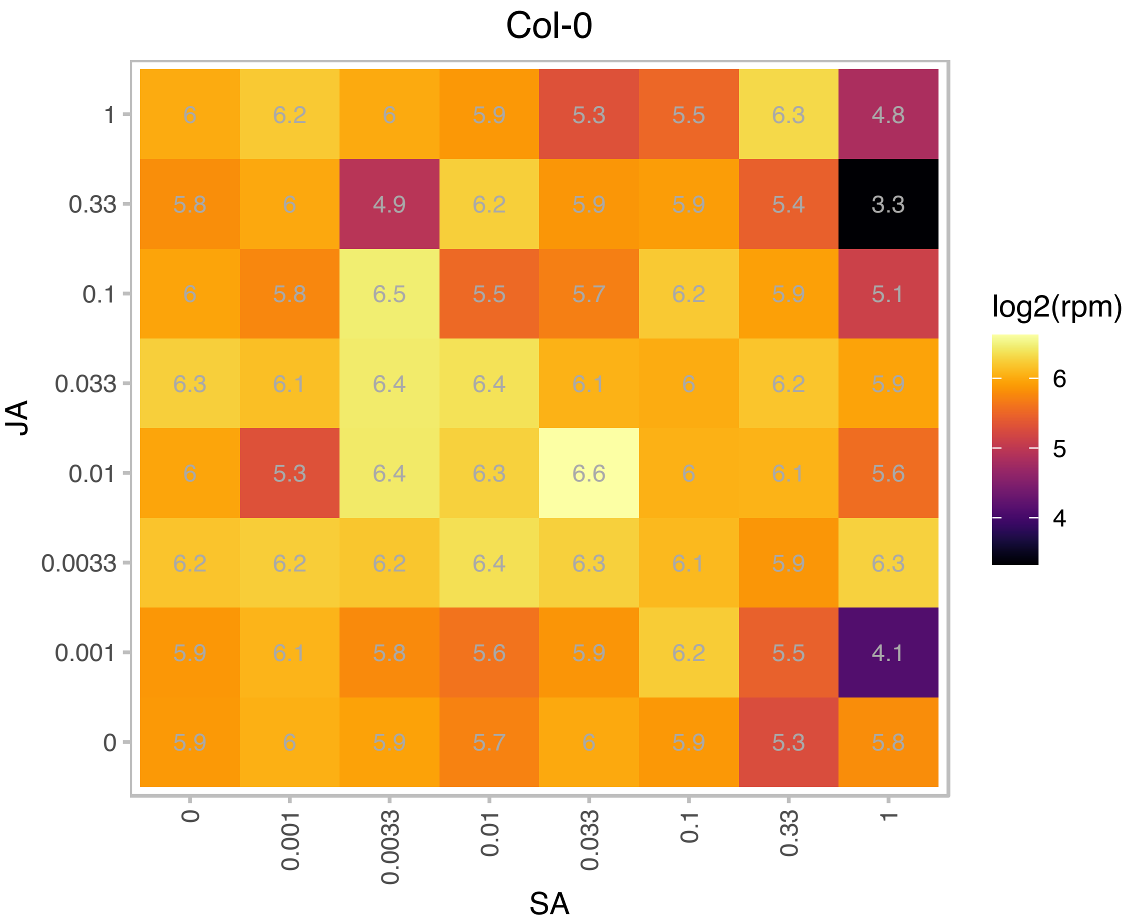


| | |
|---|---|
| Gene | AT5G62575 |
| is.expressed_genes | TRUE |
| is.responsive_genes | TRUE |
| is.apcluster | E-01 |

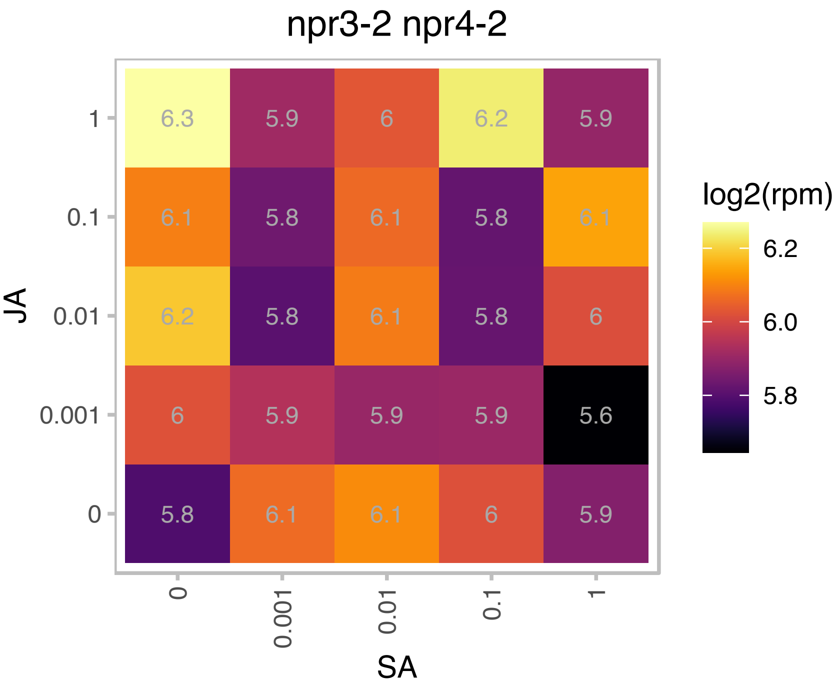


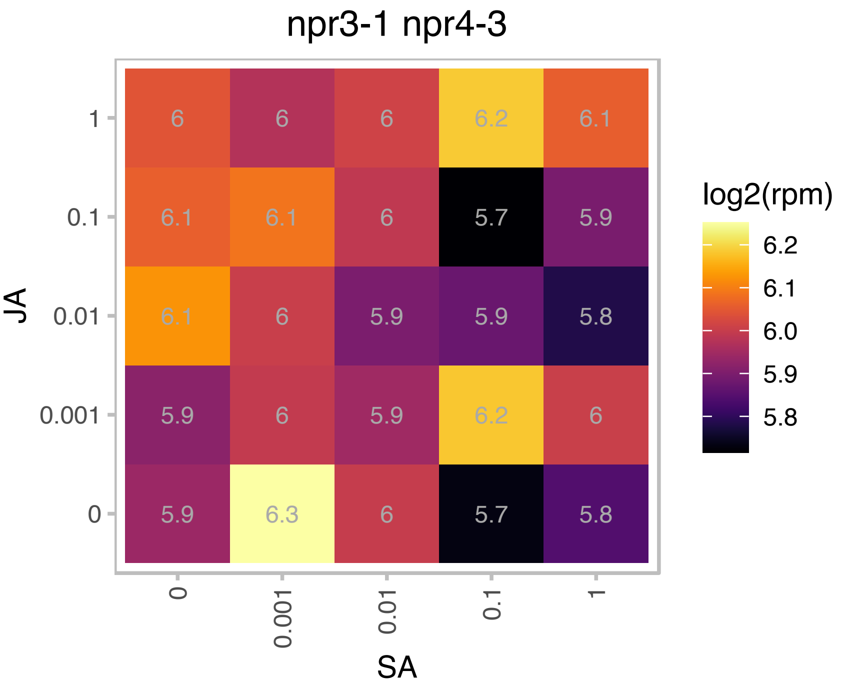

# AT5G67220.1 : NA

FMN-linked oxidoreductases superfamily protein

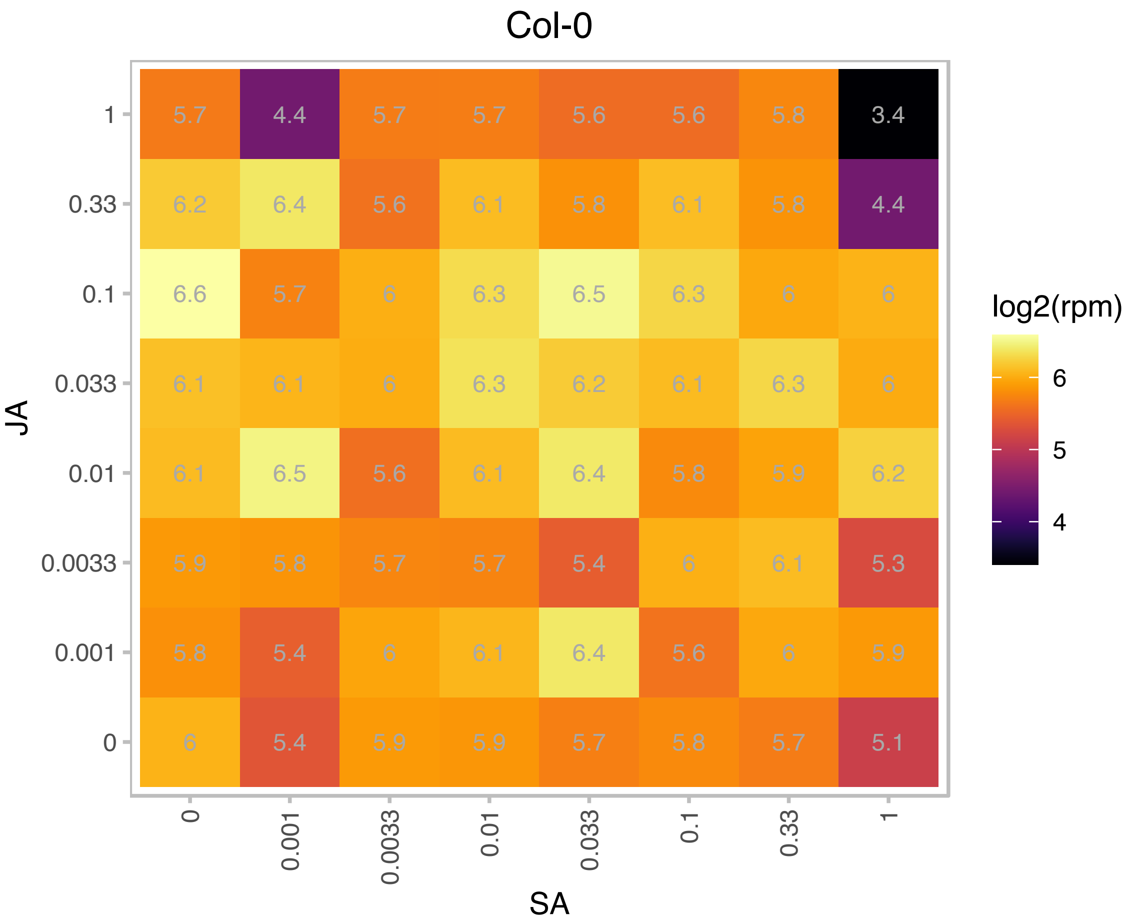


| | |
|---|---|
| Gene | AT5G67220 |
| is.expressed_genes | TRUE |
| is.responsive_genes | TRUE |
| is.apcluster | E-01 |

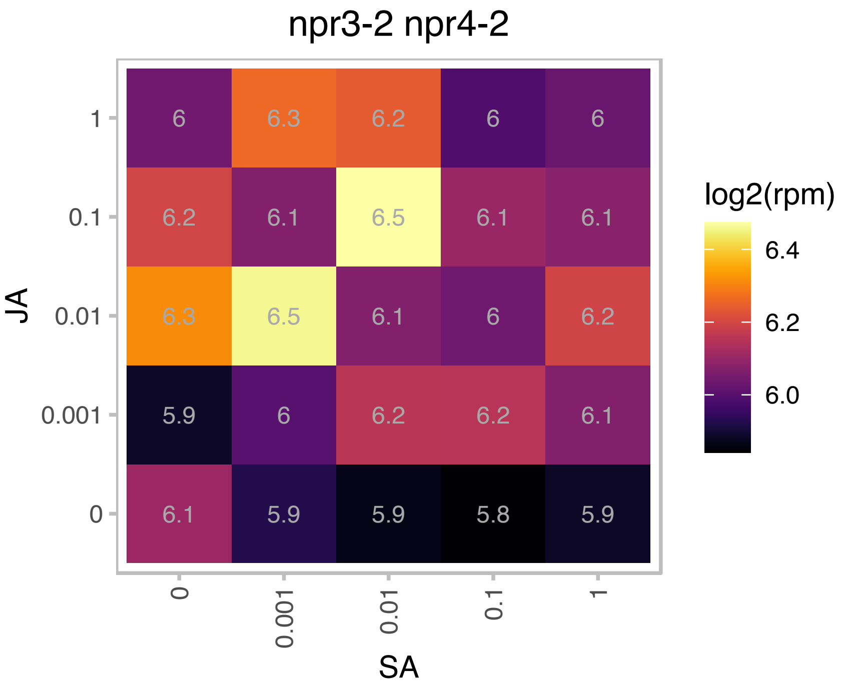


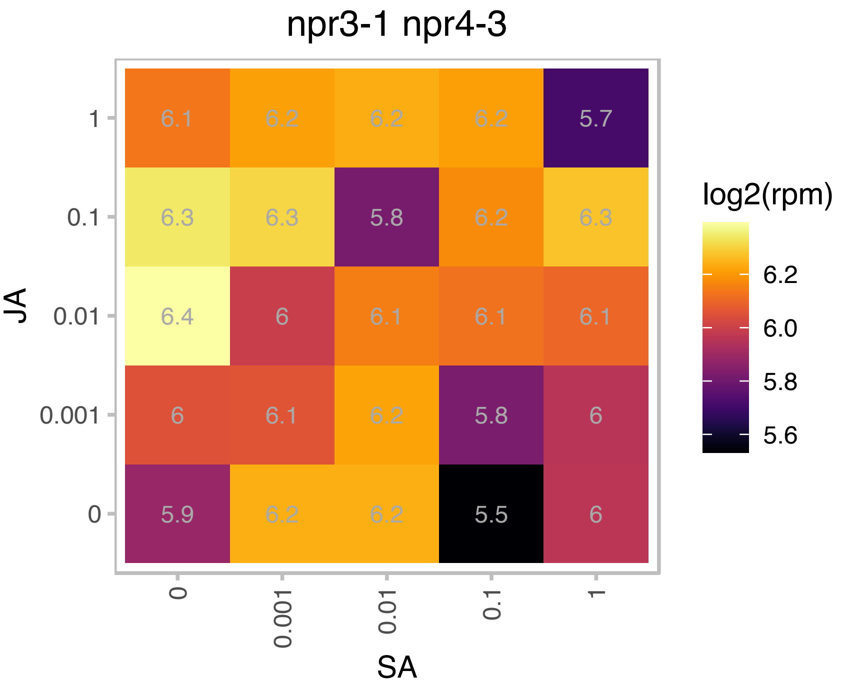

# AT5G67630.1 : NA

P-loop containing nucleoside triphosphate hydrolases superfamily protein

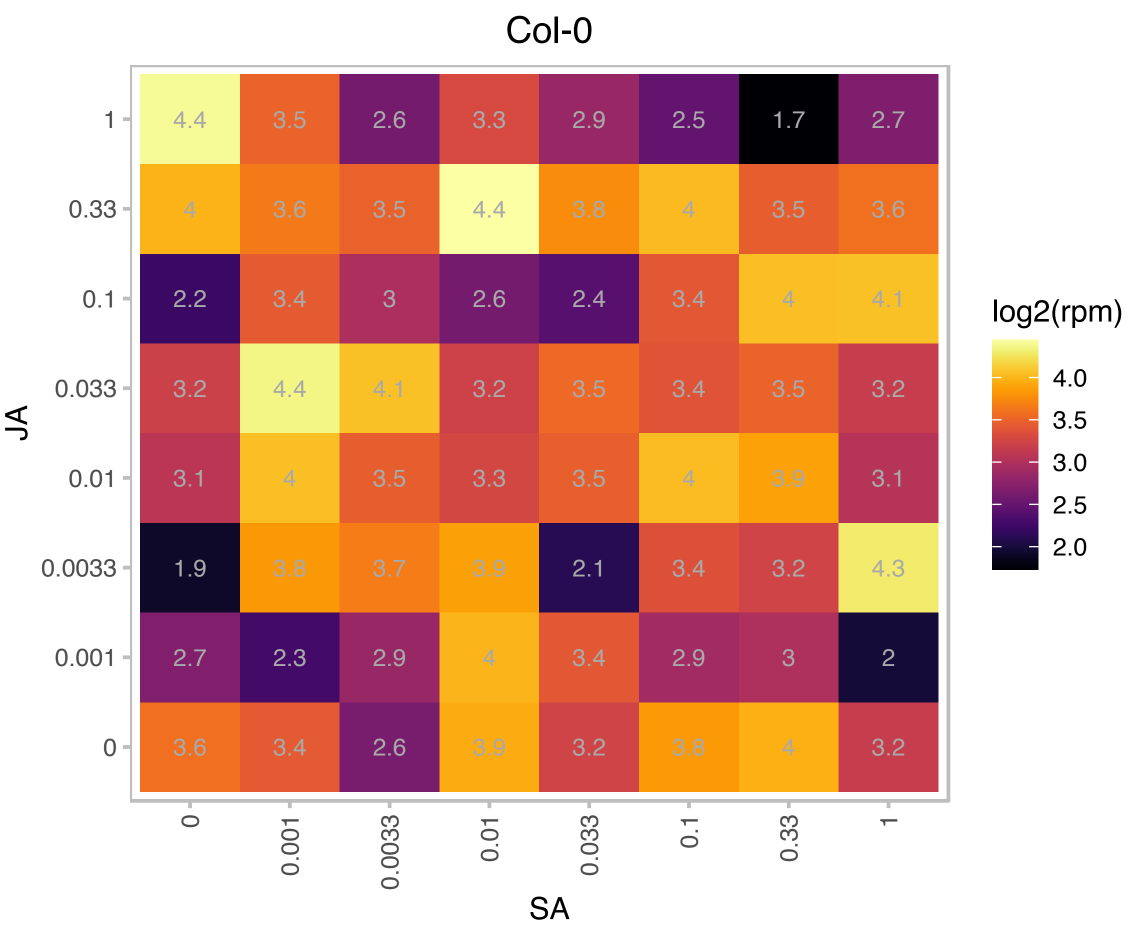


| | |
|---|---|
| Gene | AT5G67630 |
| is.expressed_genes | TRUE |
| is.responsive_genes | TRUE |
| is.apcluster | E-01 |

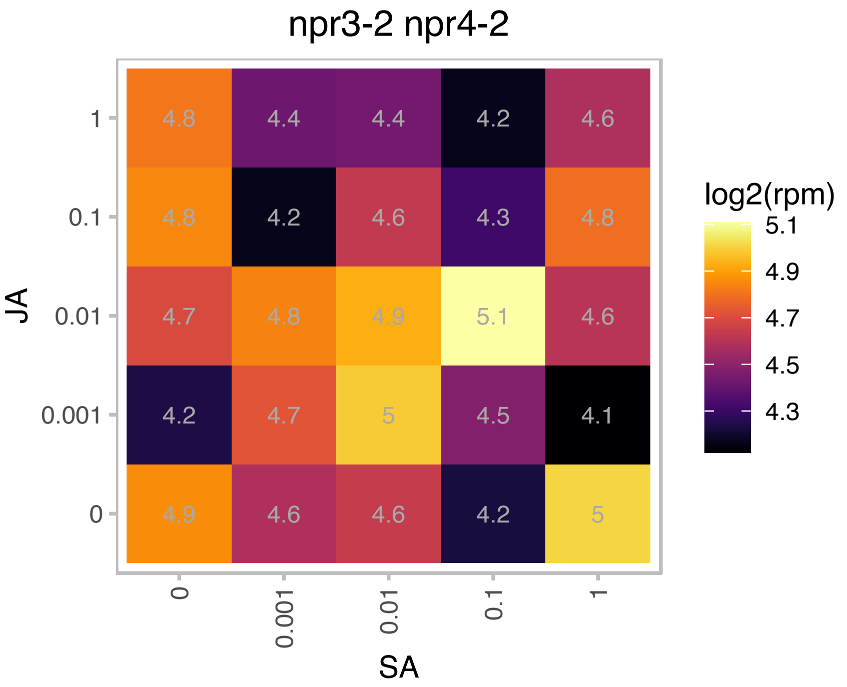


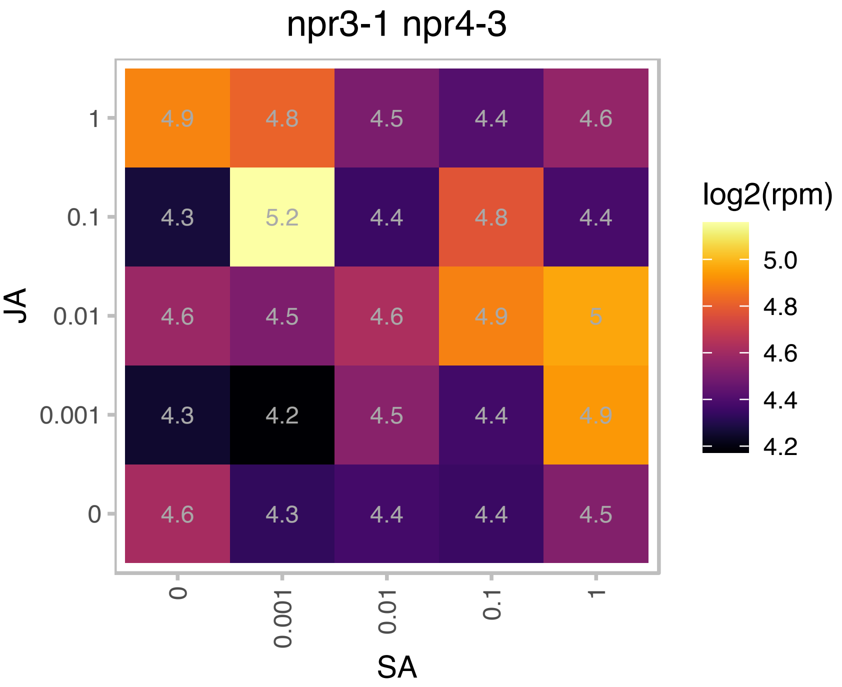

### AT1G06000.1 : UGT89C1

UDP-Glycosyltransferase superfamily protein

encodes a flavonol-7-O-rhamnosyltransferase involved in the formation of rhamnosylated flavonols

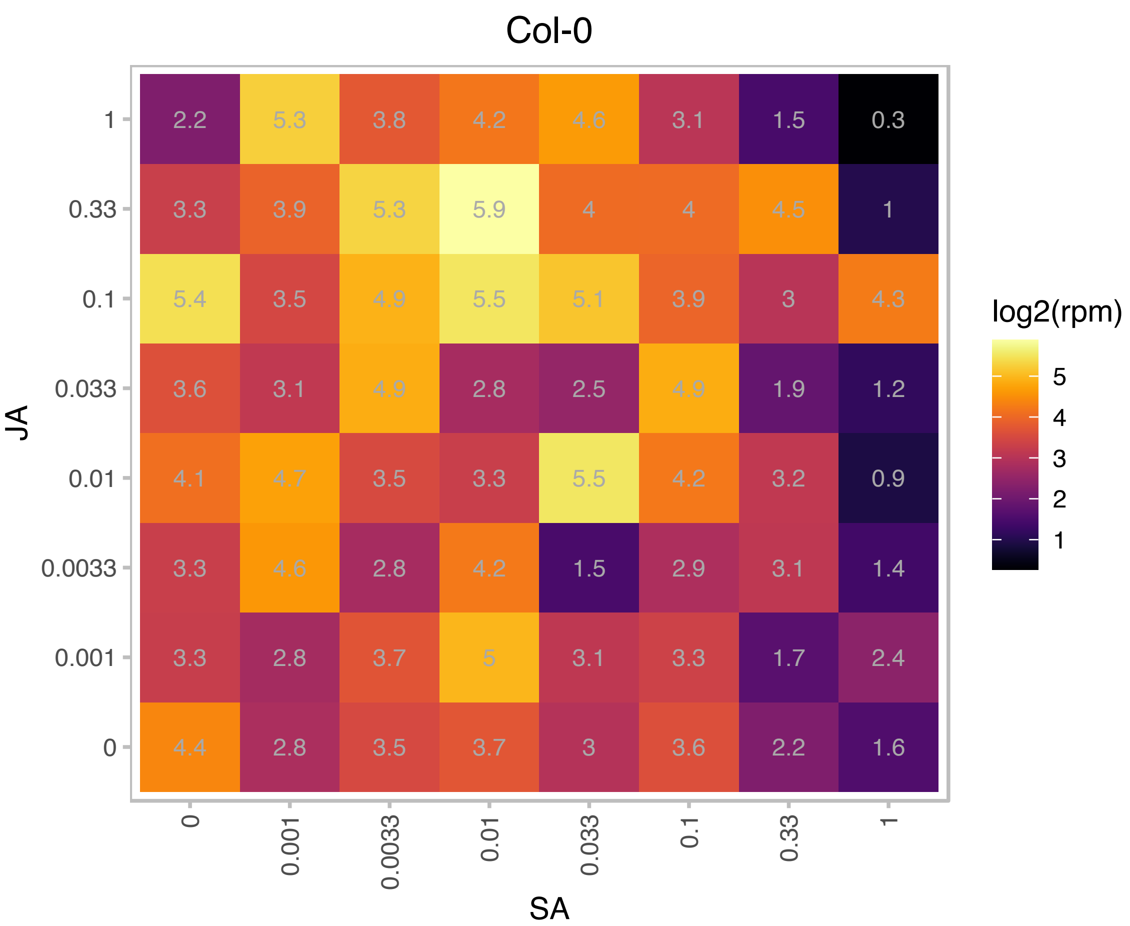


| | |
|---|---|
| Gene | AT1G06000 |
| is.expressed_genes | TRUE |
| is.responsive_genes | TRUE |
| is.apcluster | E-02 |

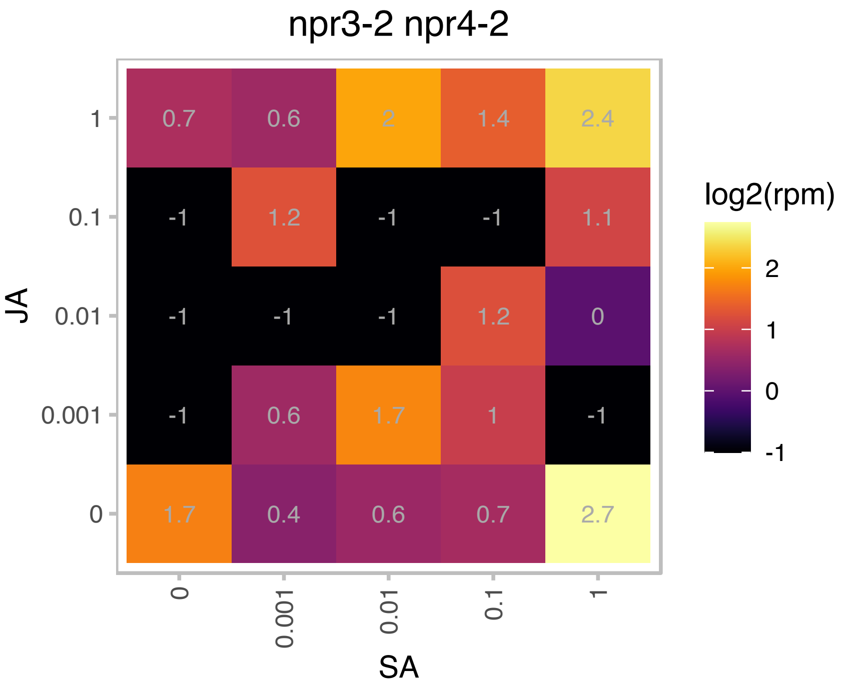


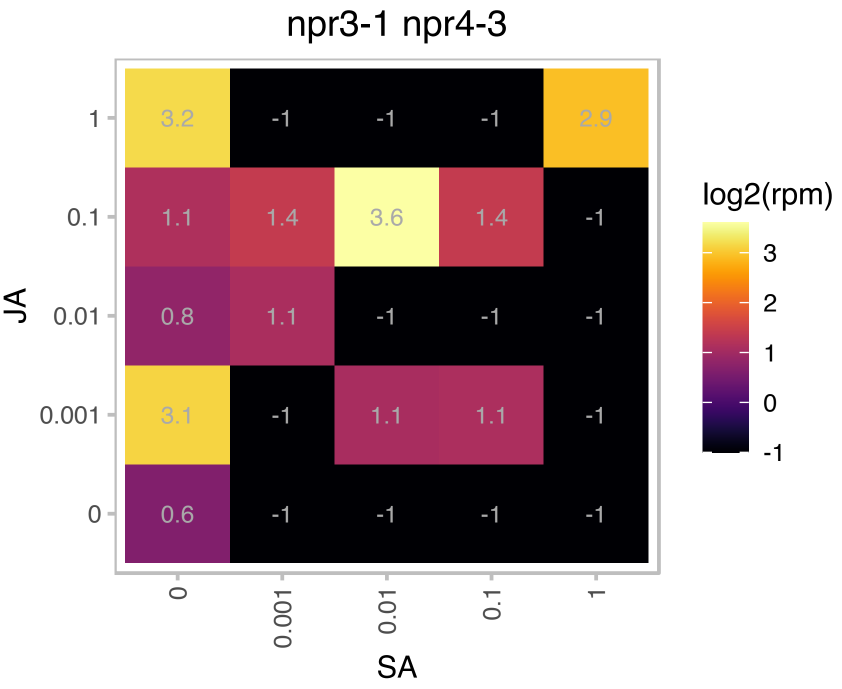

# AT1G06900.1 : NA

Insulinase (Peptidase family M16) family protein

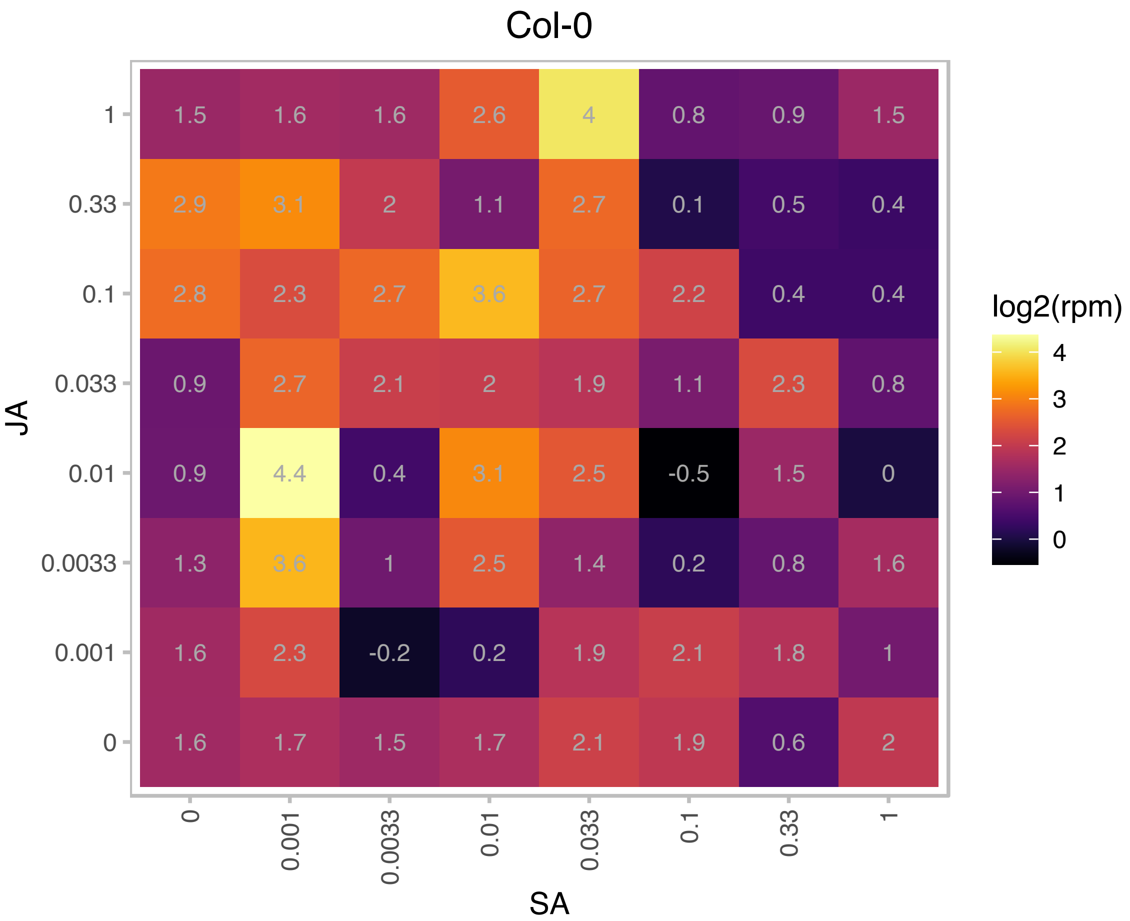


| | |
|---|---|
| Gene | AT1G06900 |
| is.expressed_genes | TRUE |
| is.responsive_genes | TRUE |
| is.apcluster | E-02 |

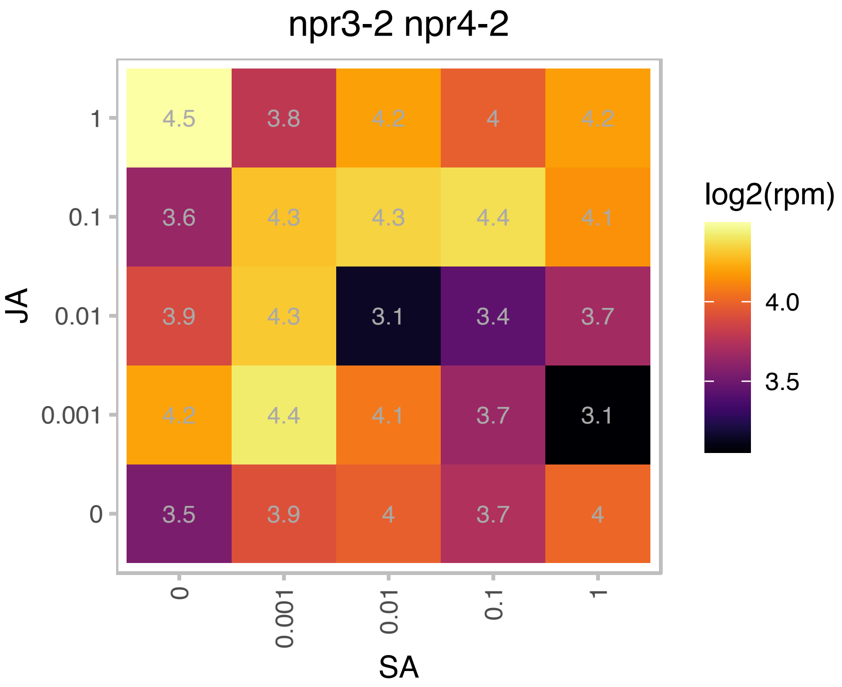


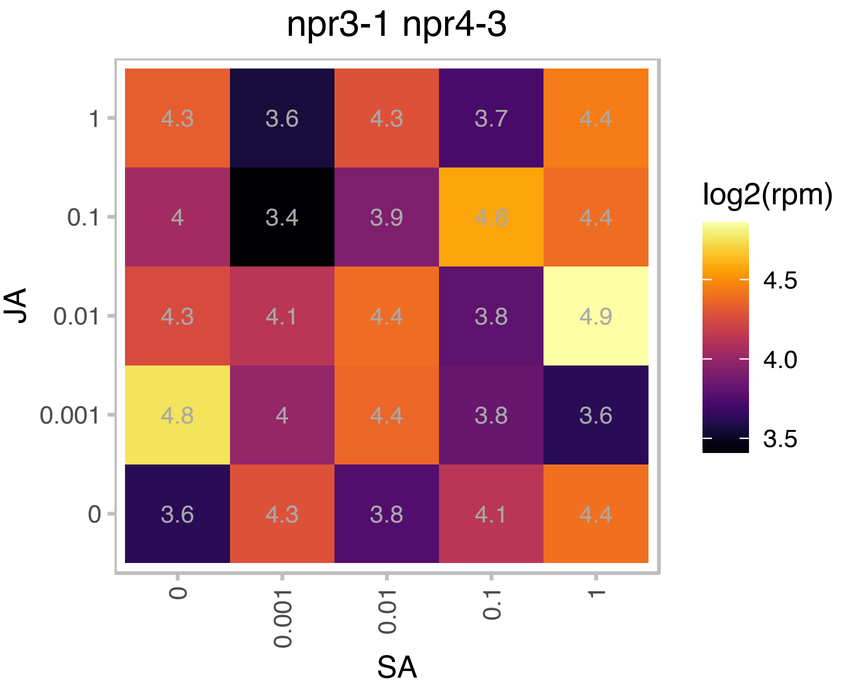

### AT1G17190.1 : ATGSTU26, GSTU26

glutathione S-transferase tau 26

Encodes glutathione transferase belonging to the tau class of GSTs. Naming convention according to Wagner et al. (2002).

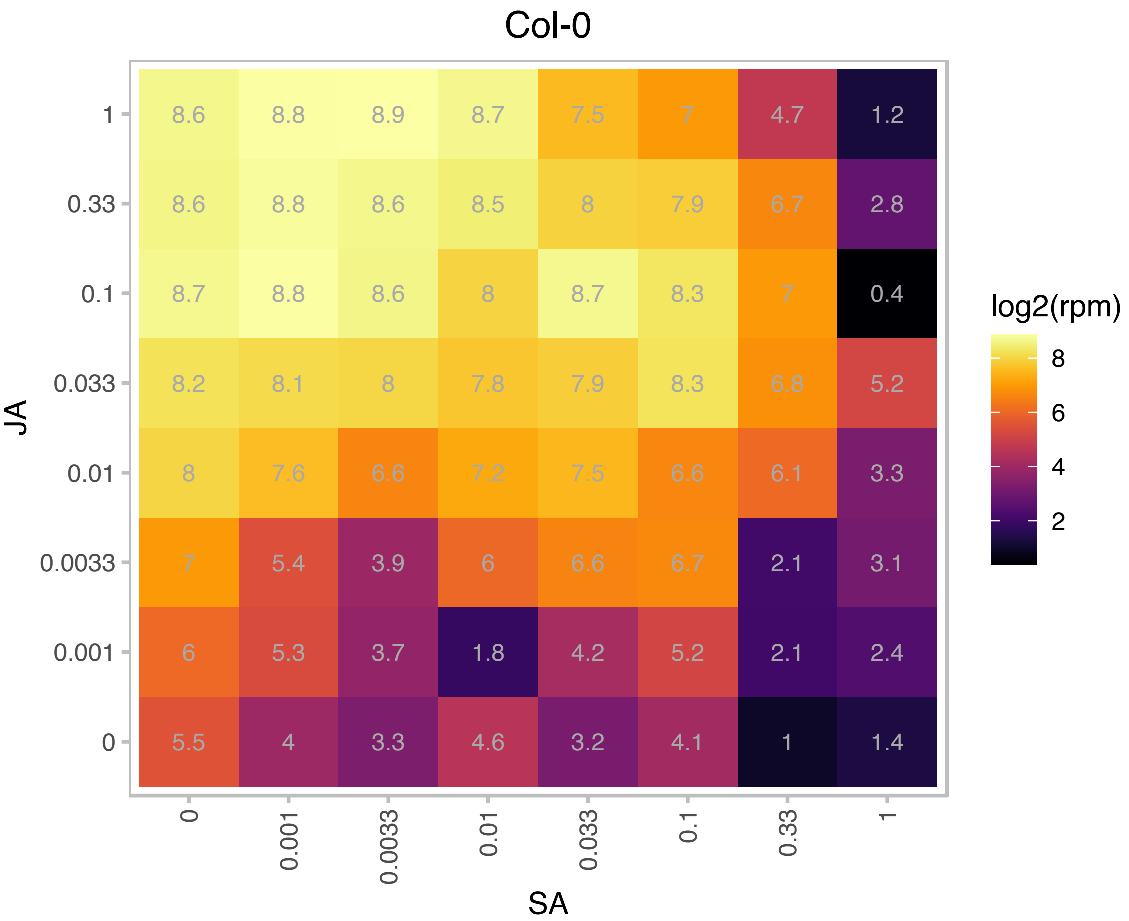


| | |
|---|---|
| Gene | AT1G17190 |
| is.expressed_genes | TRUE |
| is.responsive_genes | TRUE |
| is.apcluster | E-02 |

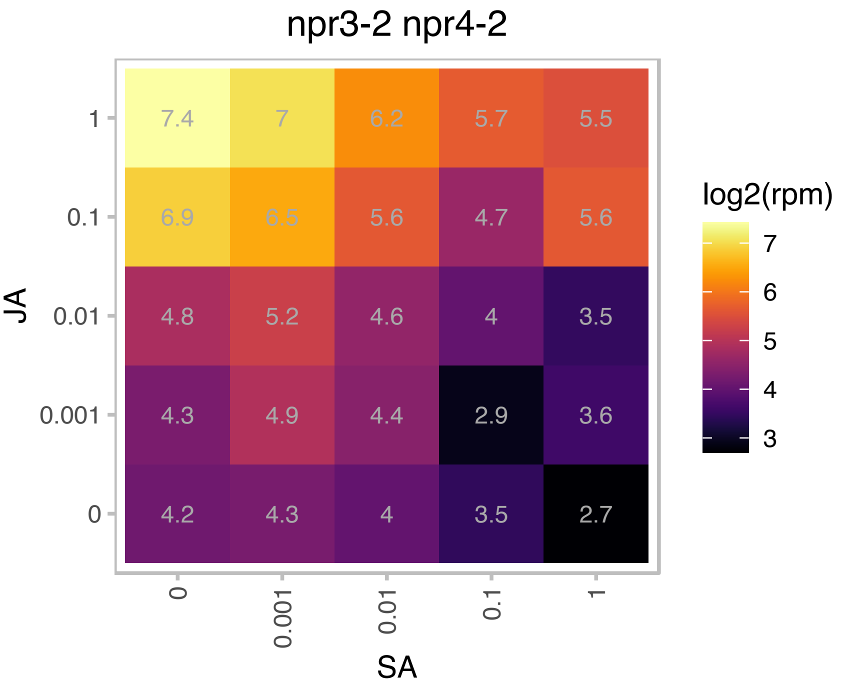


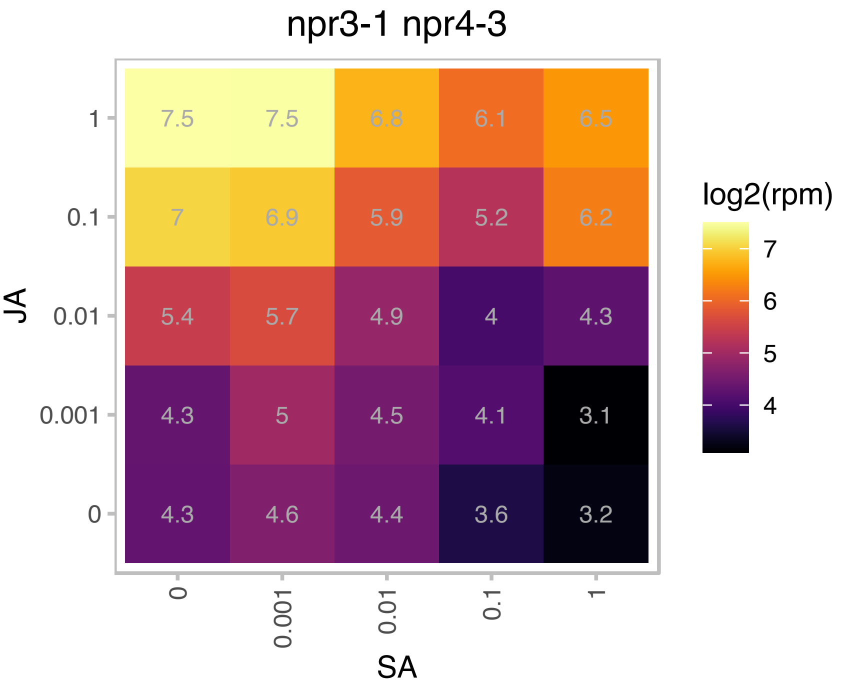

# AT1G31710.1 : NA

Copper amine oxidase family protein

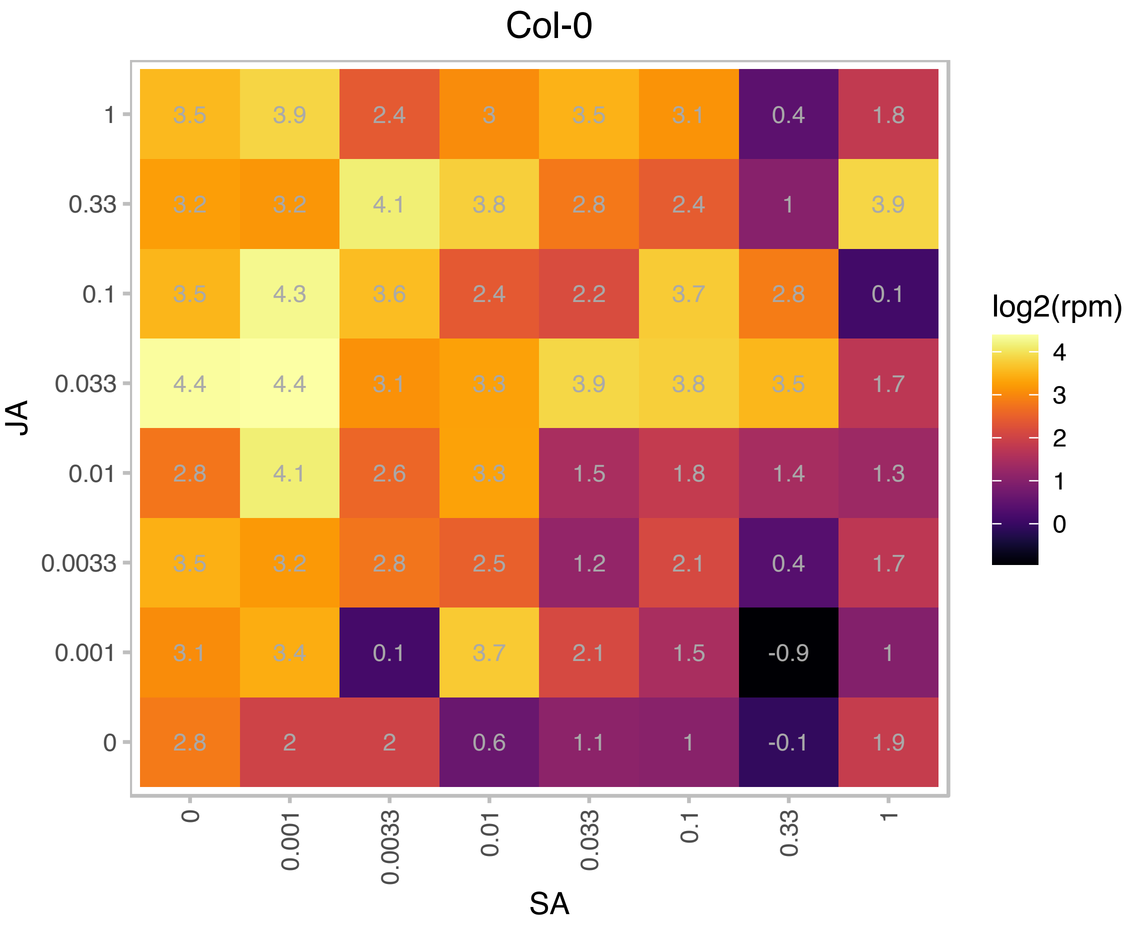


| | |
|---|---|
| Gene | AT1G31710 |
| is.expressed_genes | TRUE |
| is.responsive_genes | TRUE |
| is.apcluster | E-02 |

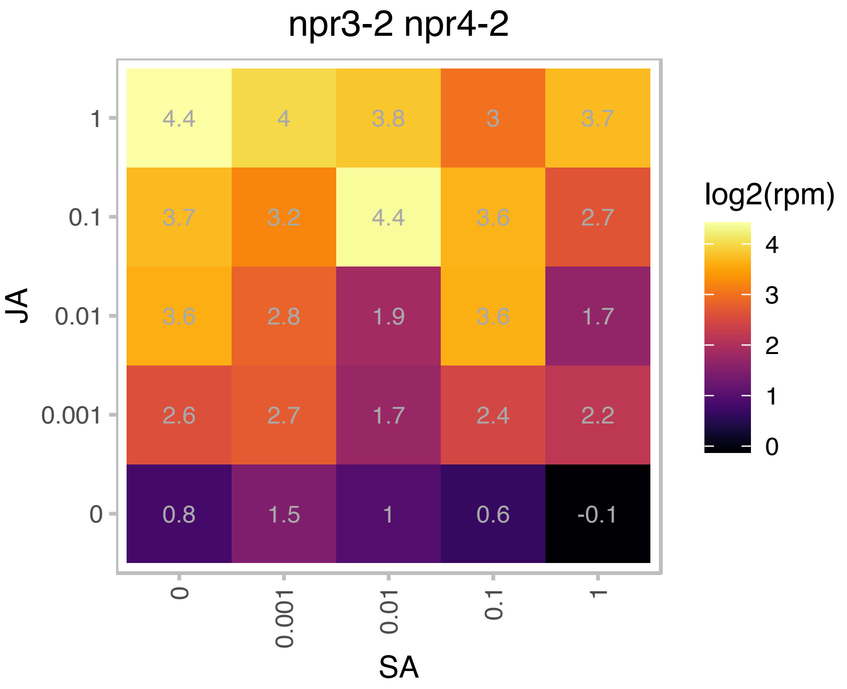


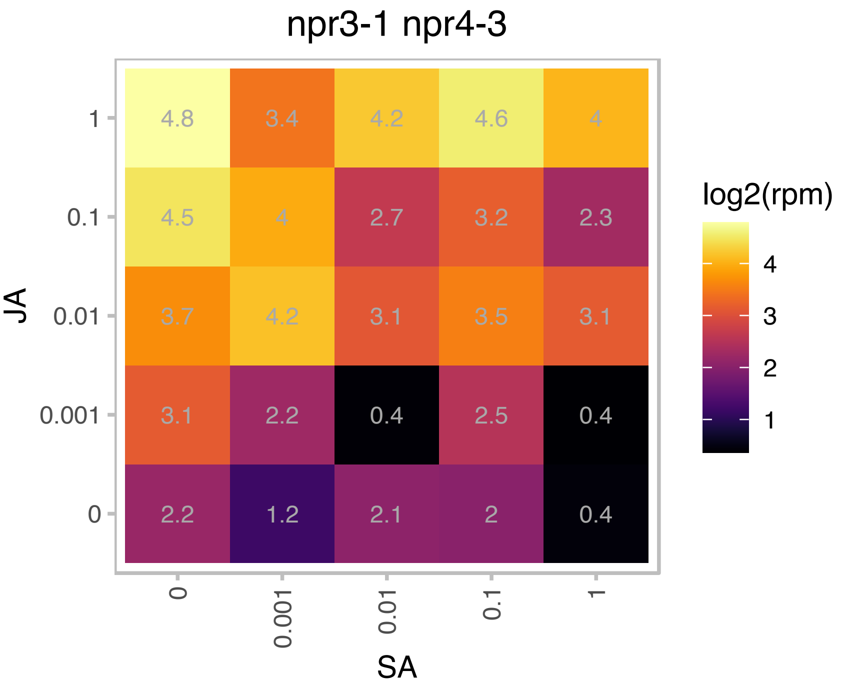

### AT1G35720.1 : ANN1, ANNAT1, AtANN1, ATOXY5, OXY5

annexin 1

Encodes a member of the annexin gene family, a diverse, multigene family of calcium-dependent, membrane-binding proteins. The protein was determined to have peroxidase activity. This activity is thought to be dependent on the presence of post-translational modifications (most likely phosphorylation). The protein was shown to be present as a mixture of monomer and homodimer. The homodimerization seems to be dependent on the presence of Ca2+ or H2O2. The dimerization was prevented by the addition of DTT, β-mercaptoethanol and TCEP. Annat1 mRNA is expressed in flowers, roots,leaves and stems and is most abundant in stems. mRNA levels are increased in response to oxidative stress. Developmental expression patterns suggest a role in Golgi-mediated polysaccharide secretion. It is a Ca 2+-permeable transporter

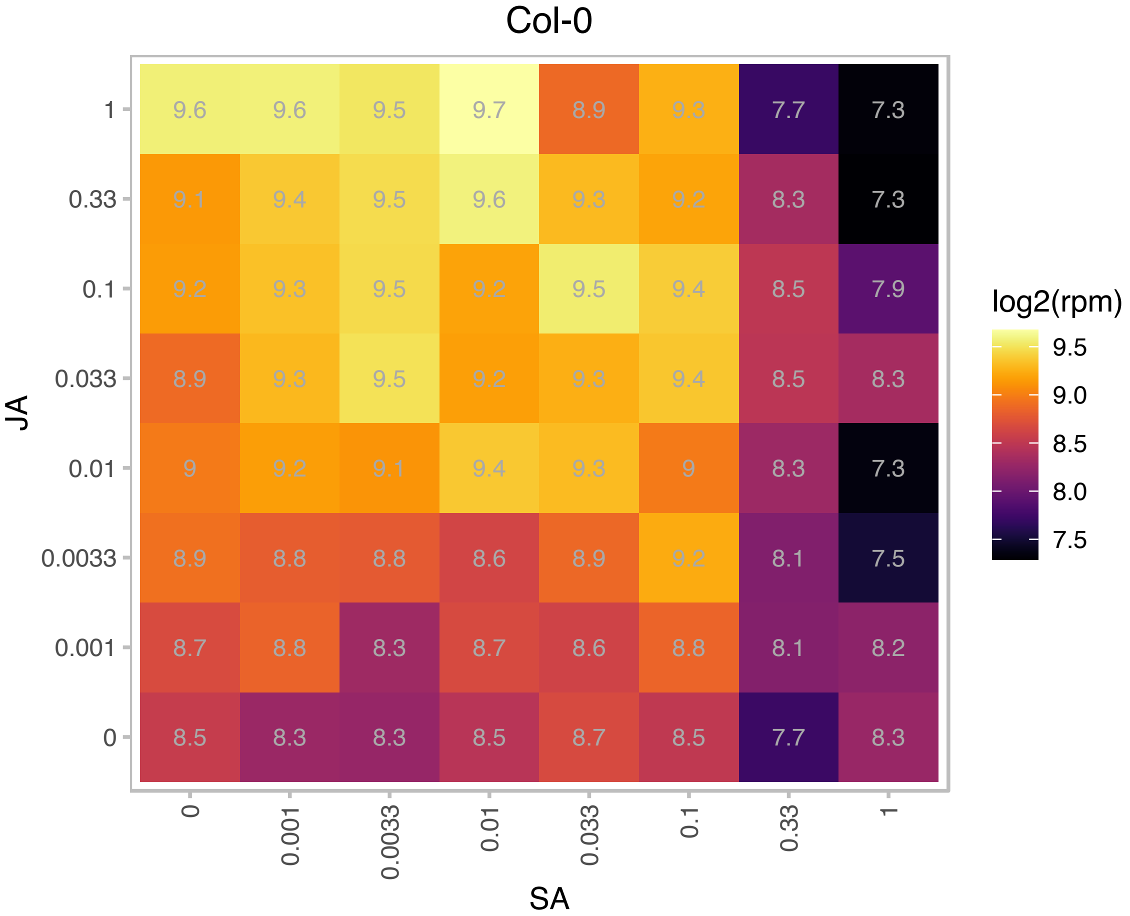


| | |
|---|---|
| Gene | AT1G35720 |
| is.expressed_genes | TRUE |
| is.responsive_genes | TRUE |
| is.apcluster | E-02 |

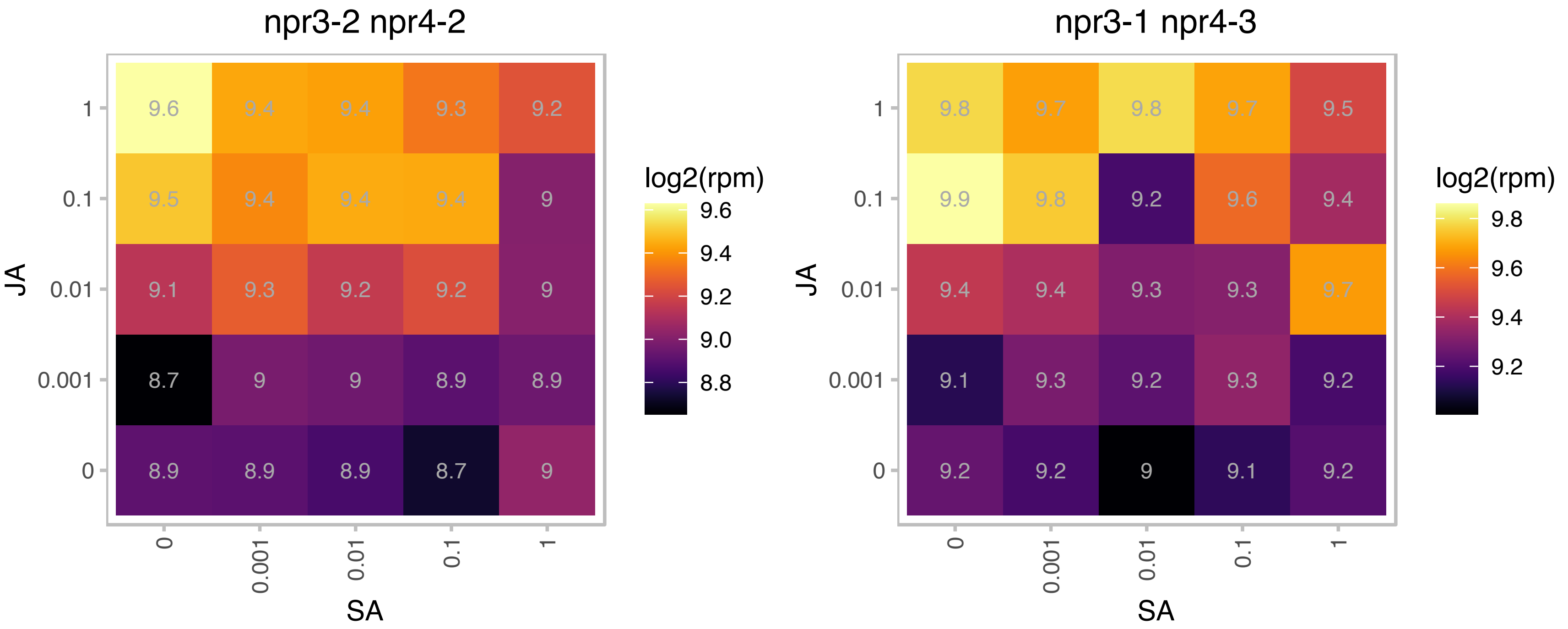

### AT1G45201.1 : ATTLL1, TLL1

triacylglycerol lipase-like 1

Target of AtGRP7 regulation.

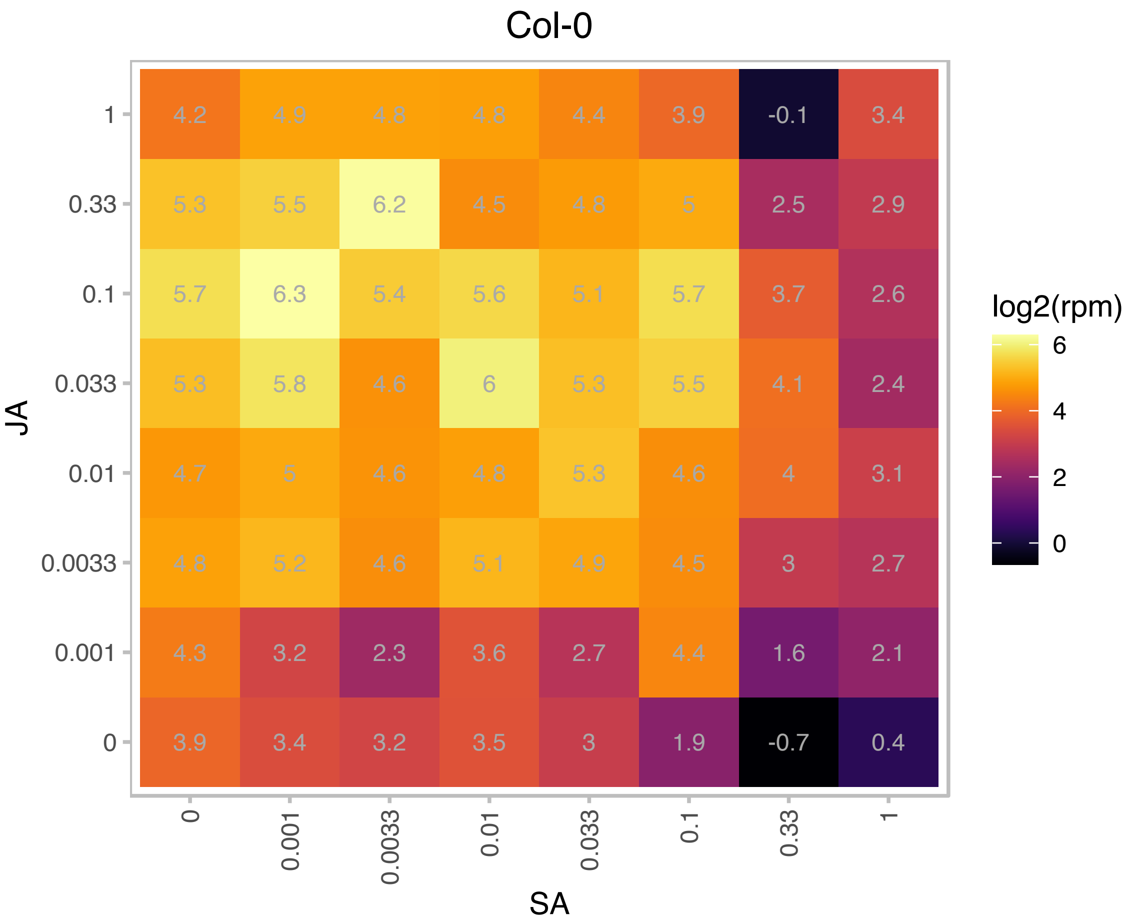


| | |
|---|---|
| Gene | AT1G45201 |
| is.expressed_genes | TRUE |
| is.responsive_genes | TRUE |
| is.apcluster | E-02 |

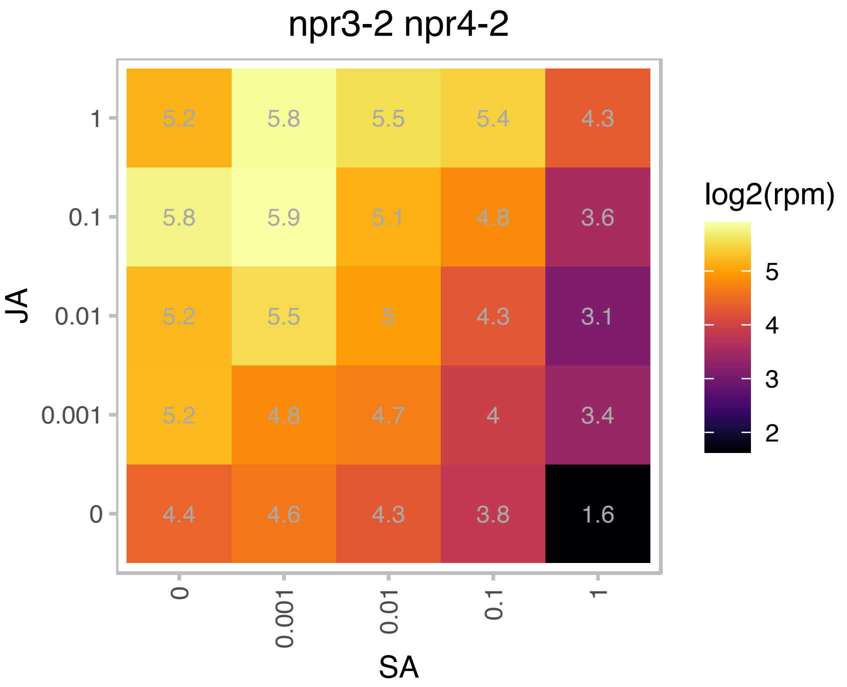


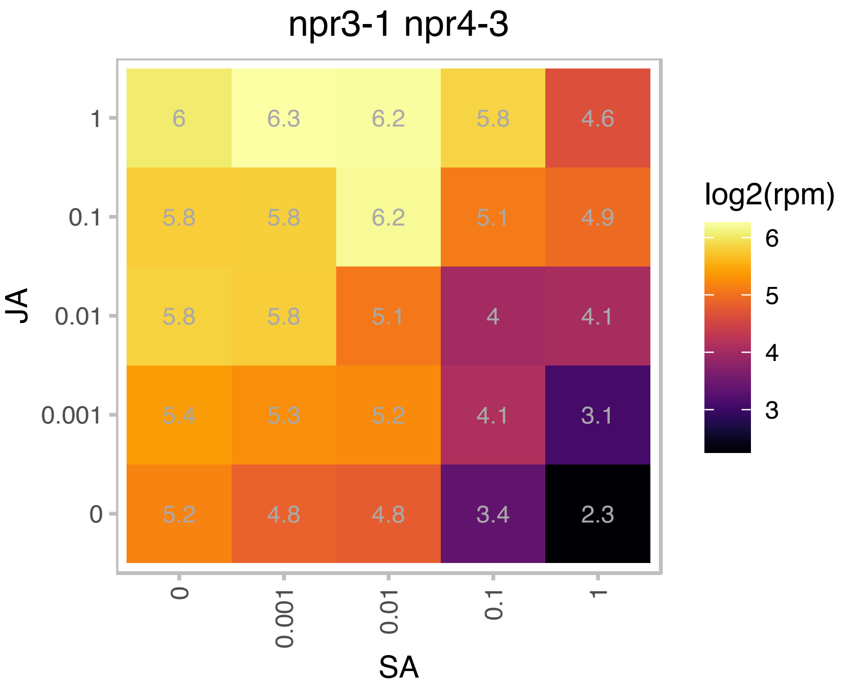

# AT1G49570.1 : NA

Peroxidase superfamily protein

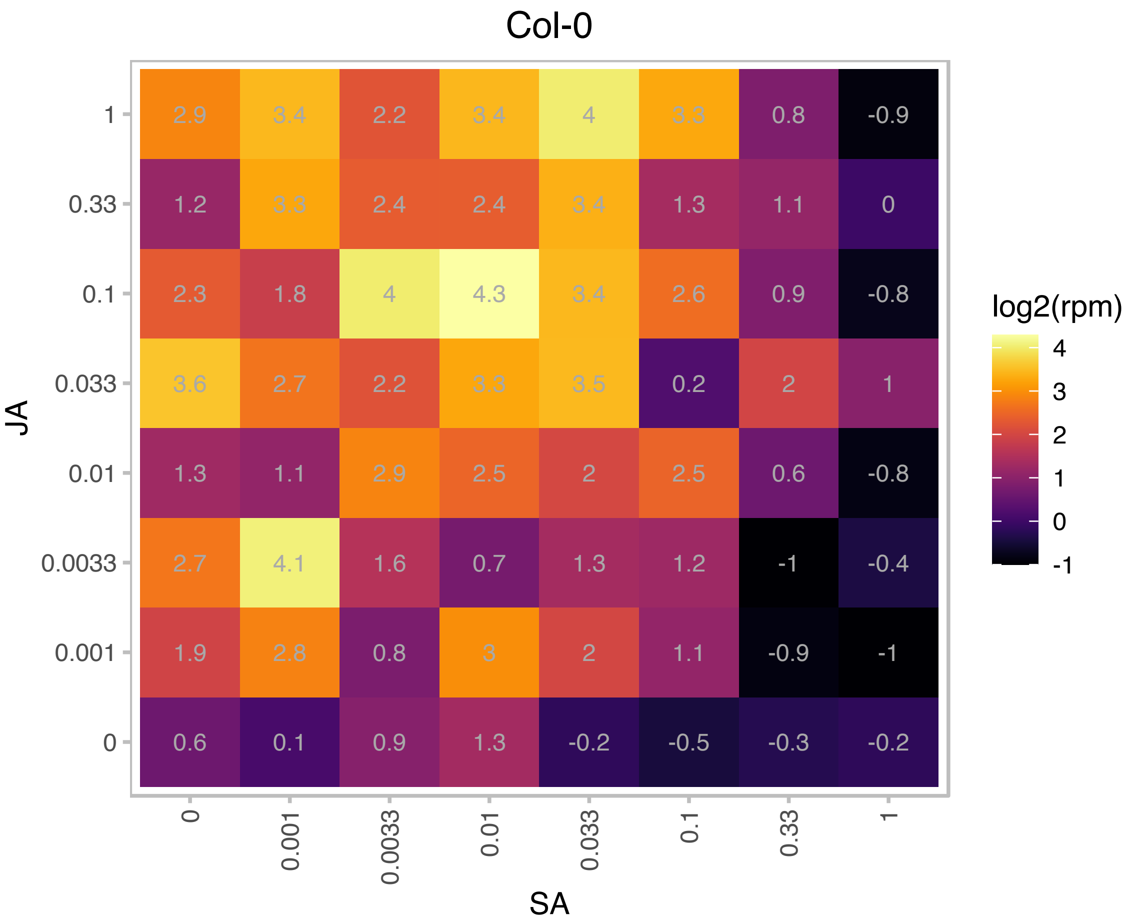


| | |
|---|---|
| Gene | AT1G49570 |
| is.expressed_genes | TRUE |
| is.responsive_genes | TRUE |
| is.apcluster | E-02 |

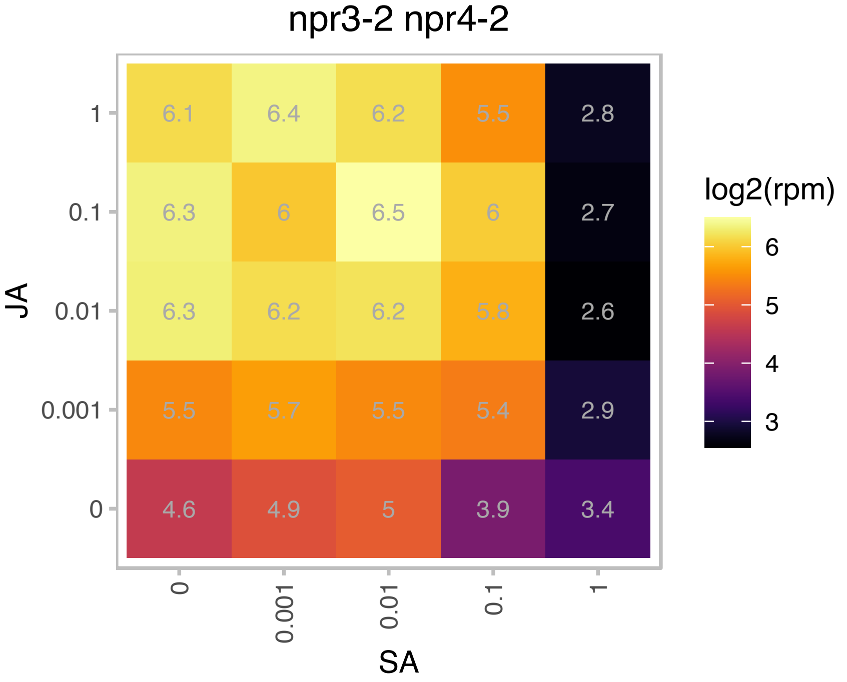


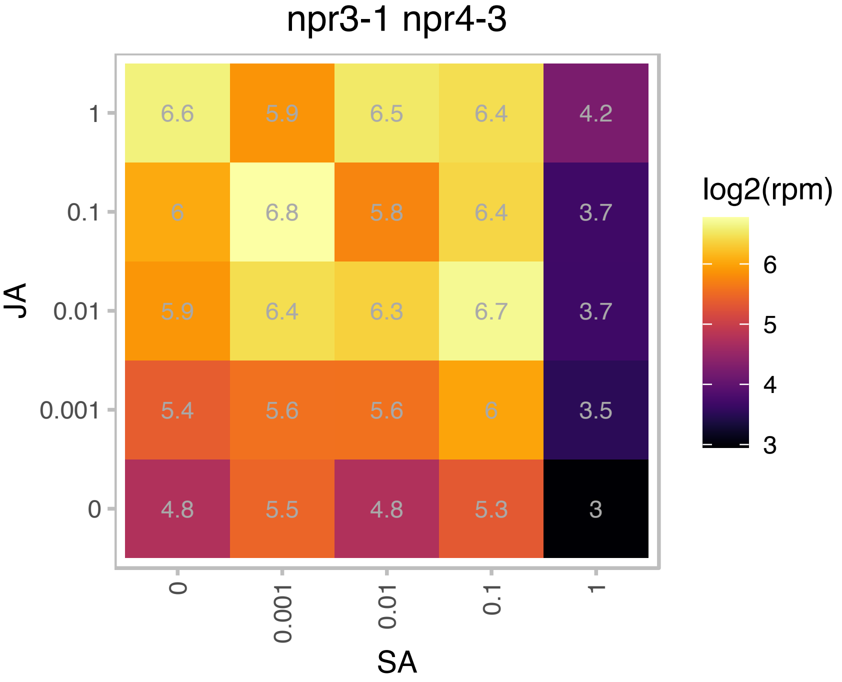

# AT1G51510.1 : Y14

RNA-binding (RRM/RBD/RNP motifs) family protein

This gene is predicted to encode a protein involved in the exon junction complex. Though there is a predicted RNA binding motif, in the Drosophila ortholog (33% identity), this motif mediates interactions with Mago and is not available for RNA binding. The Arabidopsis Y14 protein appears to be predominantly nucleolar, but there is also some evidence for its presence in the cytoplasm.

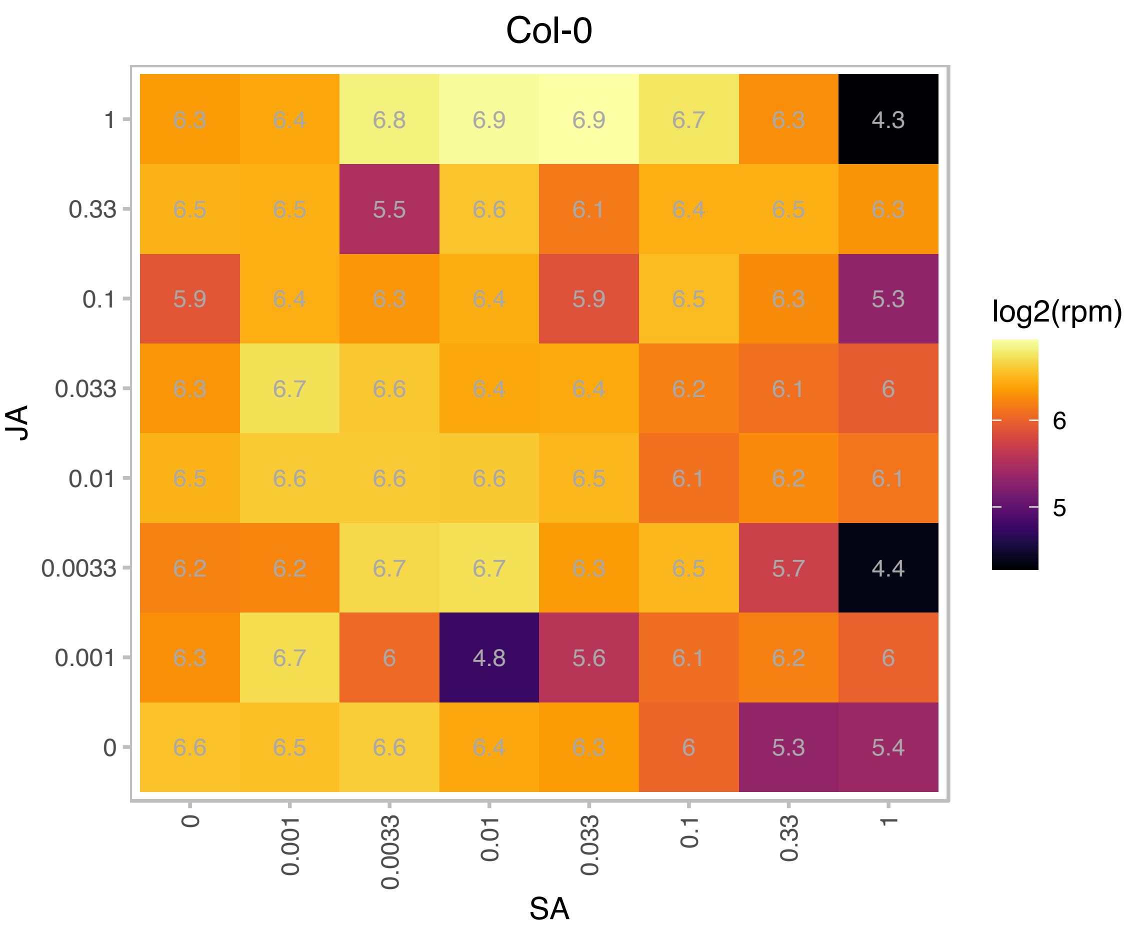


| | |
|---|---|
| Gene | AT1G51510 |
| is.expressed_genes | TRUE |
| is.responsive_genes | TRUE |
| is.apcluster | E-02 |

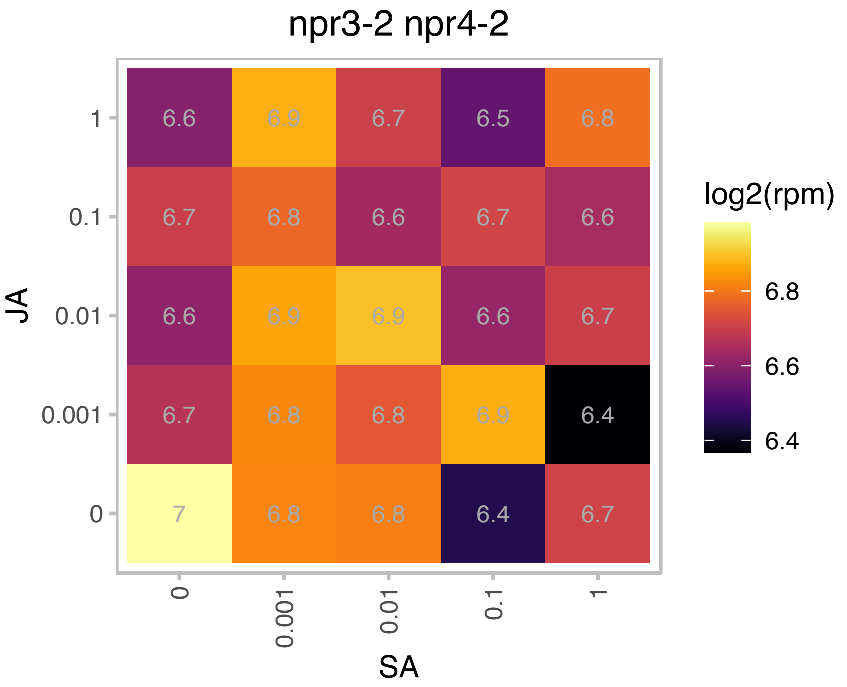


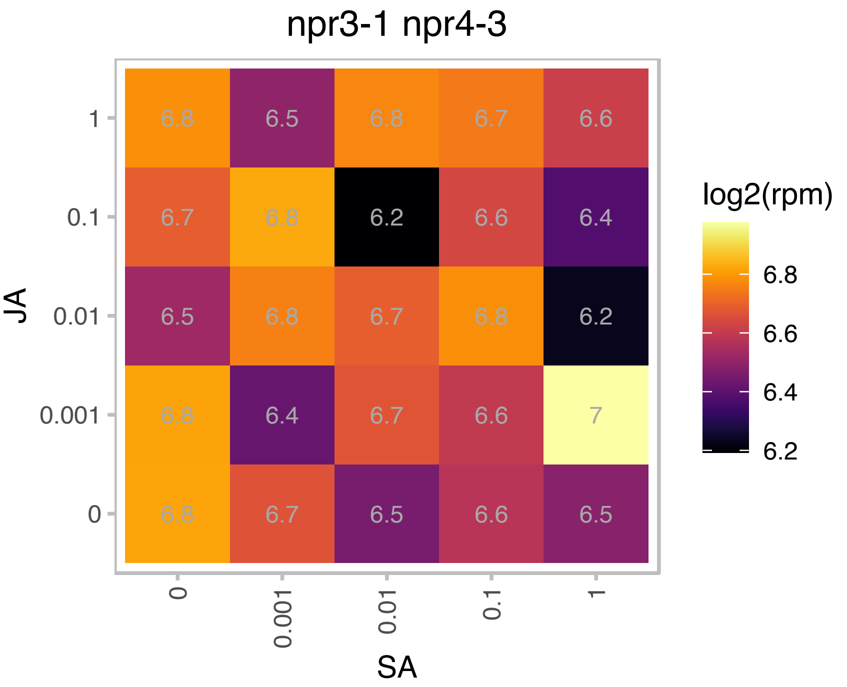

### AT1G52280.1 : AtRABG3d, RABG3d

RAB GTPase homolog G3D

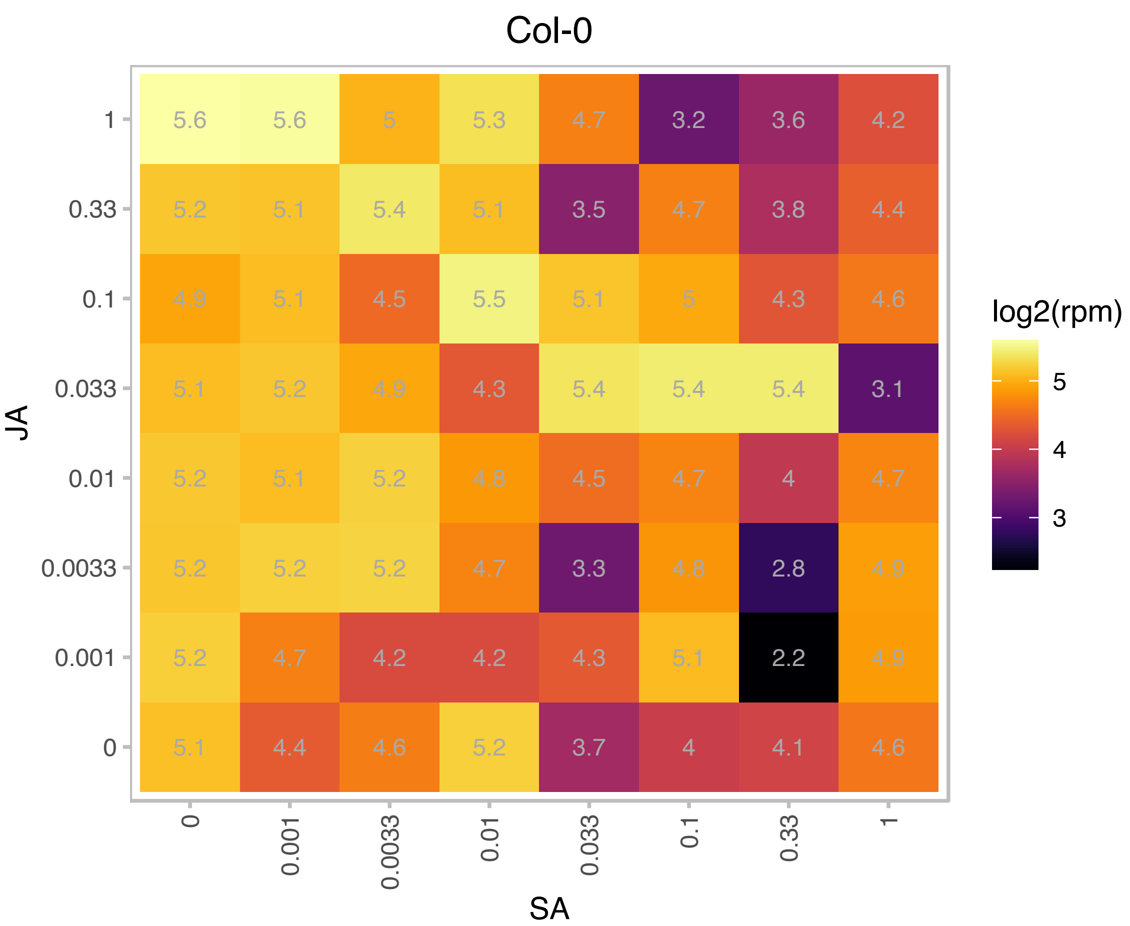


| | |
|---|---|
| Gene | AT1G52280 |
| is.expressed_genes | TRUE |
| is.responsive_genes | TRUE |
| is.apcluster | E-02 |

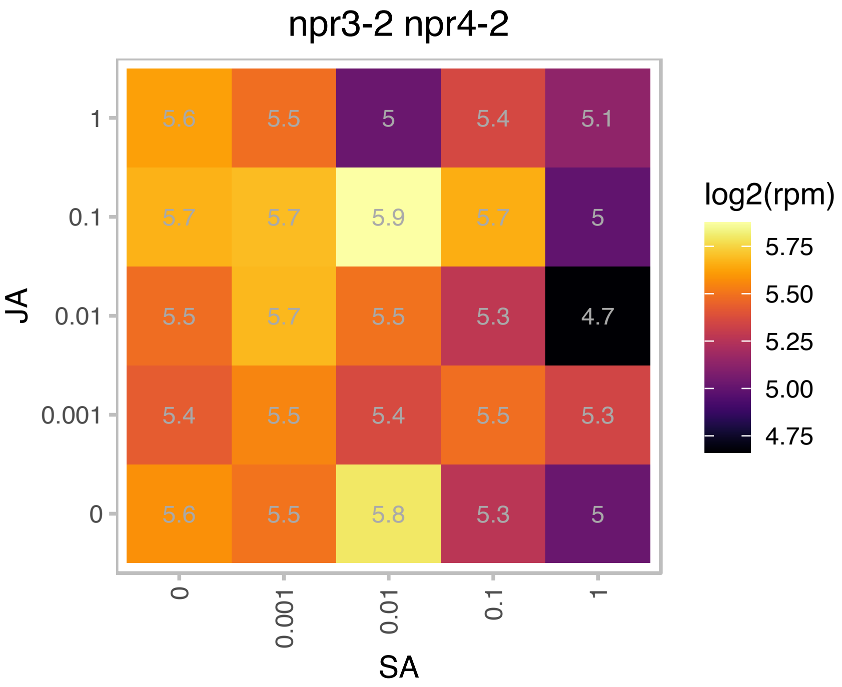


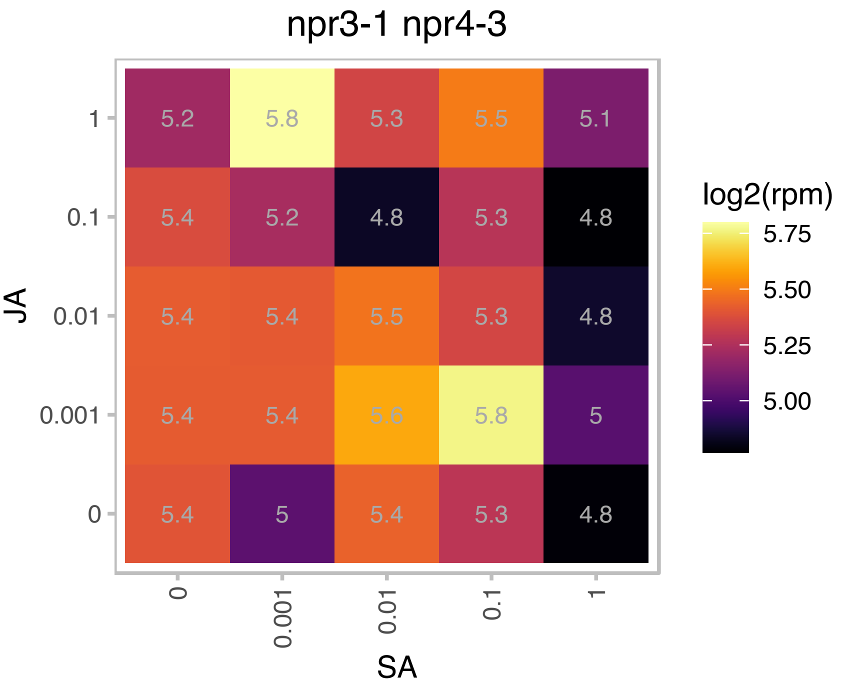

### AT1G54000.1 : GLL22

GDSL-like Lipase/Acylhydrolase superfamily protein

GDSL-motif esterase/acyltransferase/lipase. Enzyme group with broad substrate specificity that may catalyze acyltransfer or hydrolase reactions with lipid and non-lipid substrates.

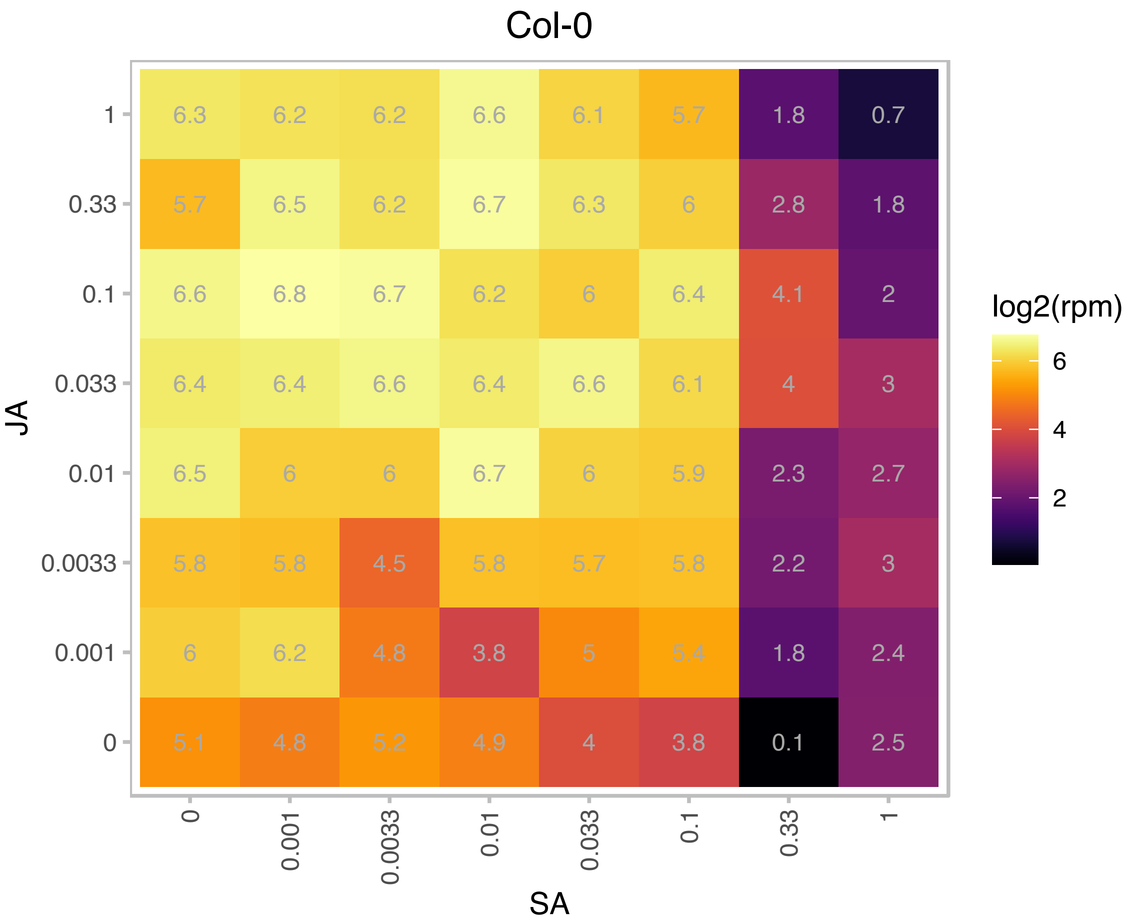


| | |
|---|---|
| Gene | AT1G54000 |
| is.expressed_genes | TRUE |
| is.responsive_genes | TRUE |
| is.apcluster | E-02 |

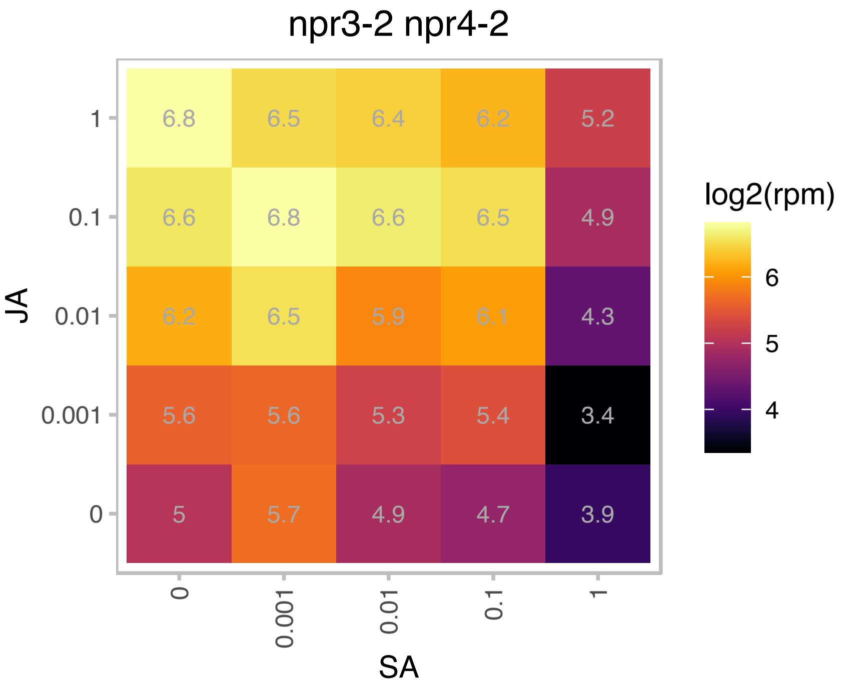


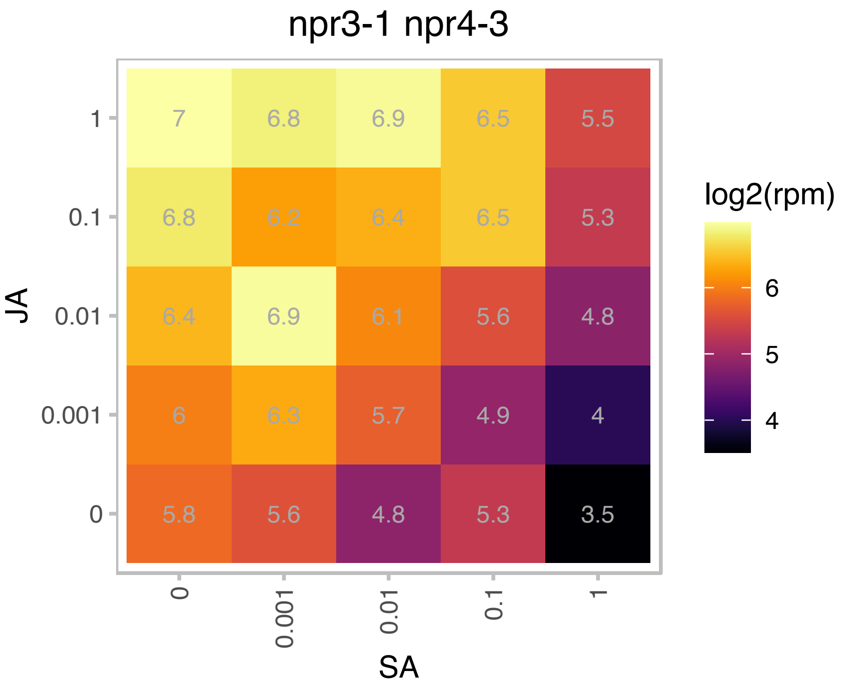

### AT1G54030.1 : ERMO3, GOLD36, MVP1, NUC

GDSL-like Lipase/Acylhydrolase superfamily protein

Encodes a vacuolar protein. Mutation causes organizational defects in the endoplasmic reticulum and aberrant protein trafficking in the plant secretory pathway. The mRNA is cell-to-cell mobile.

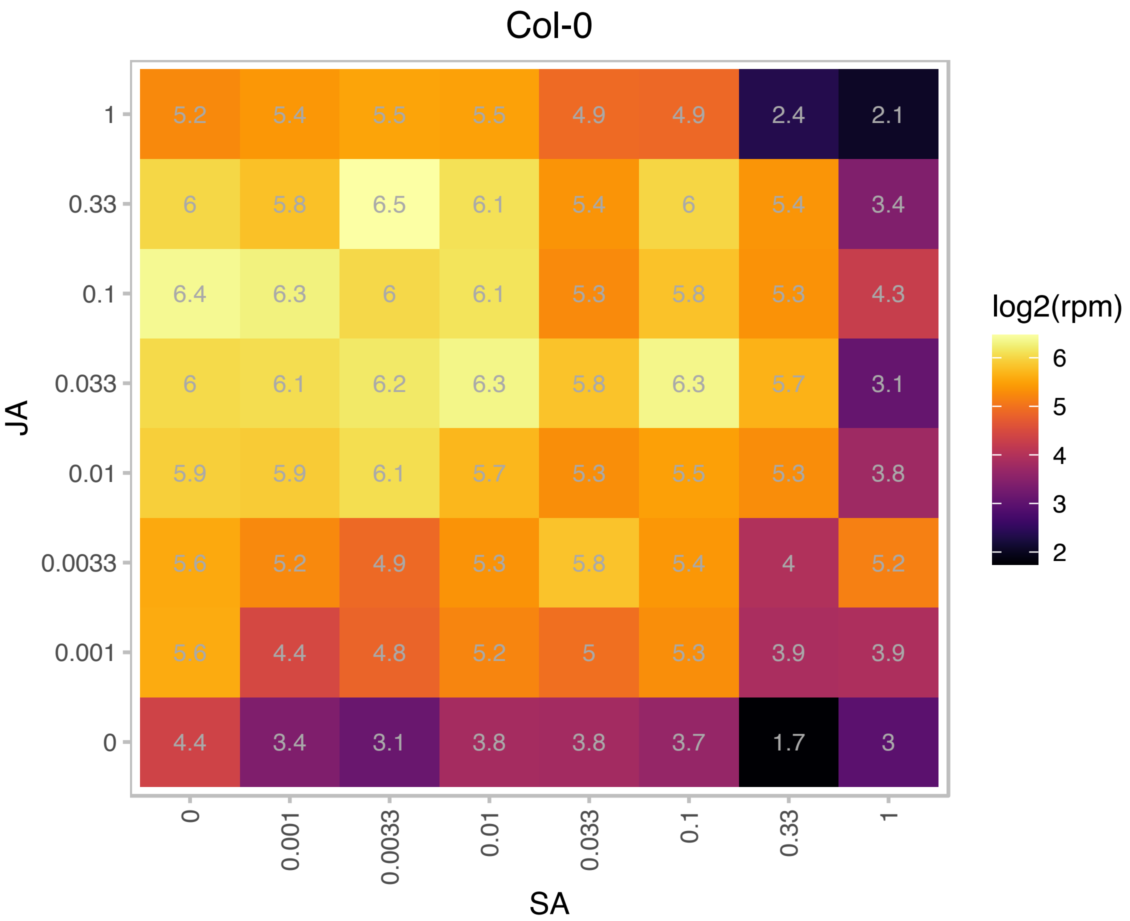


| | |
|---|---|
| Gene | AT1G54030 |
| is.expressed_genes | TRUE |
| is.responsive_genes | TRUE |
| is.apcluster | E-02 |

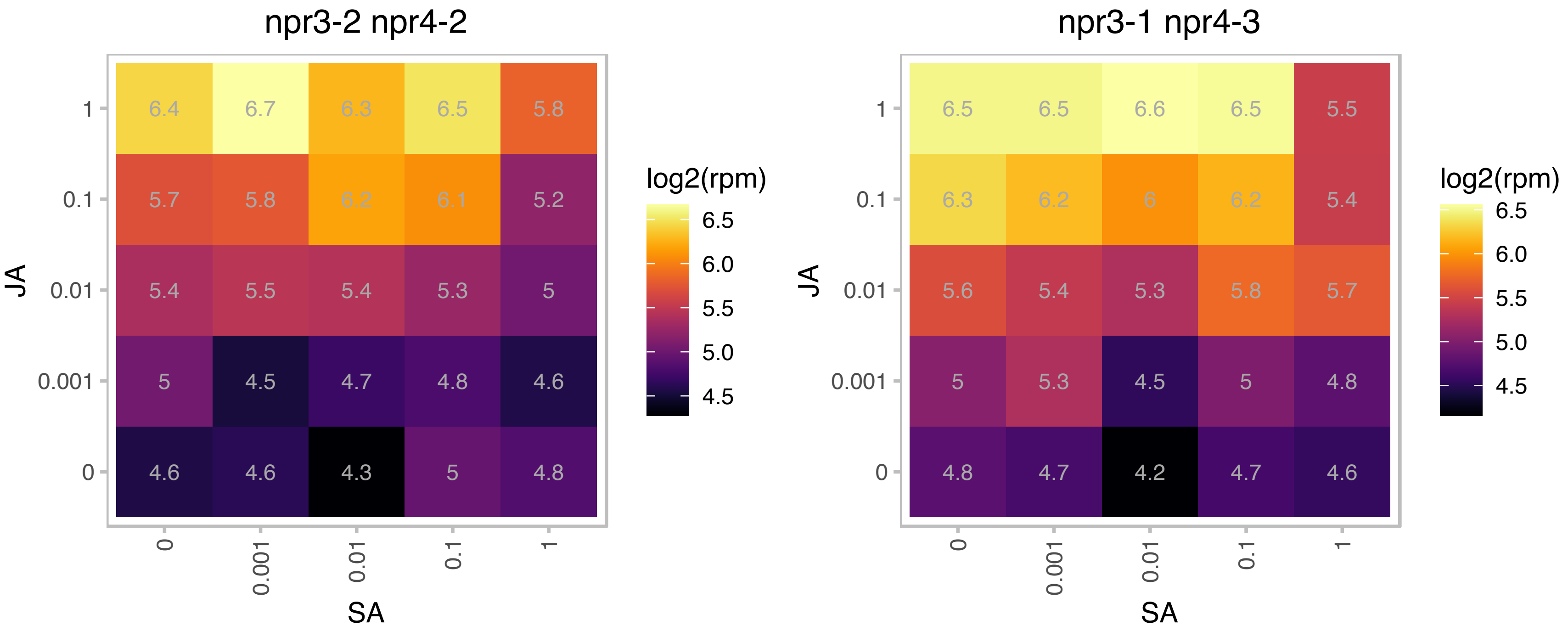

# AT1G62975.1 : NA

basic helix-loop-helix (bHLH) DNA-binding superfamily protein

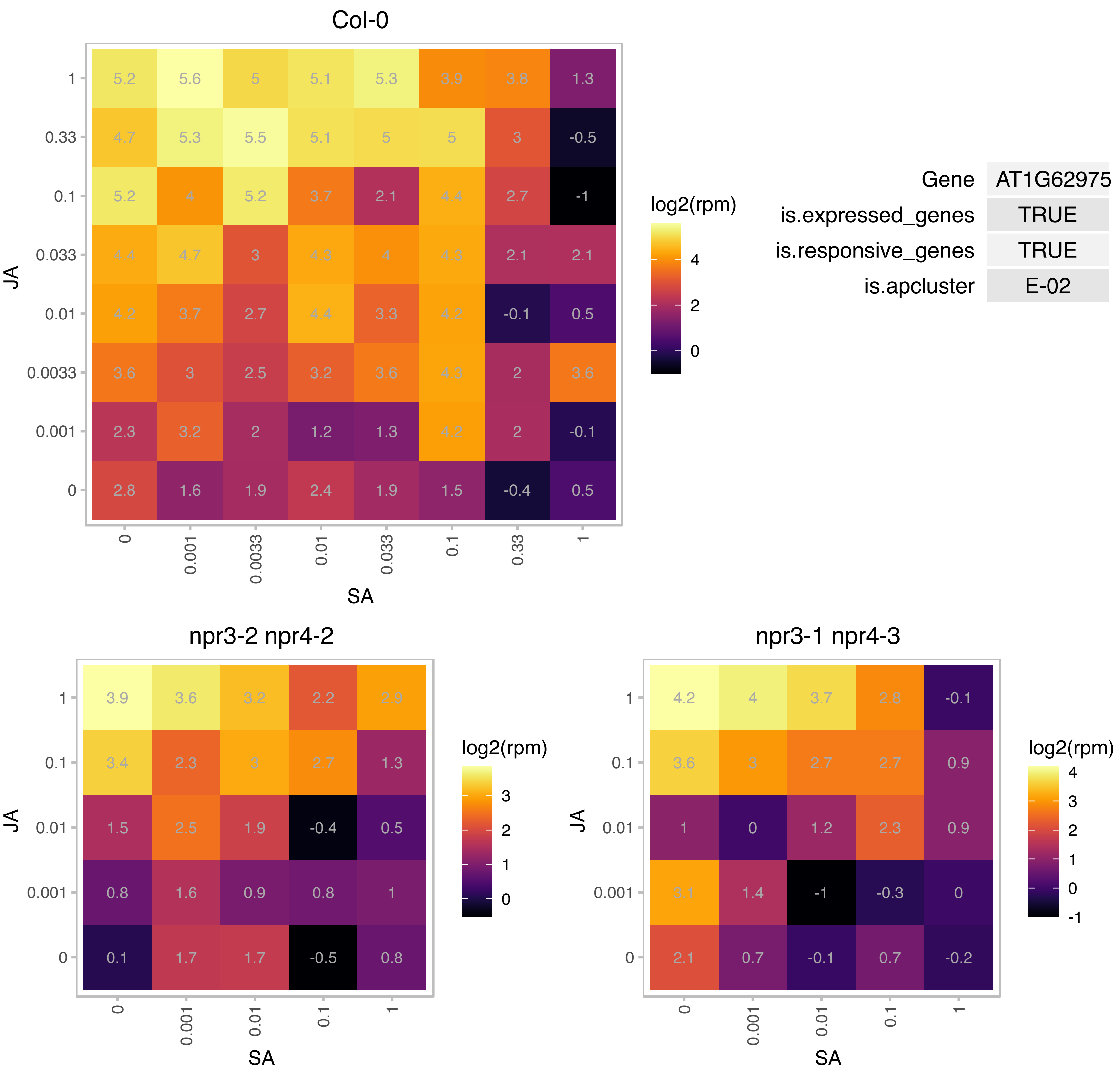

# AT1G69760.1 : NA

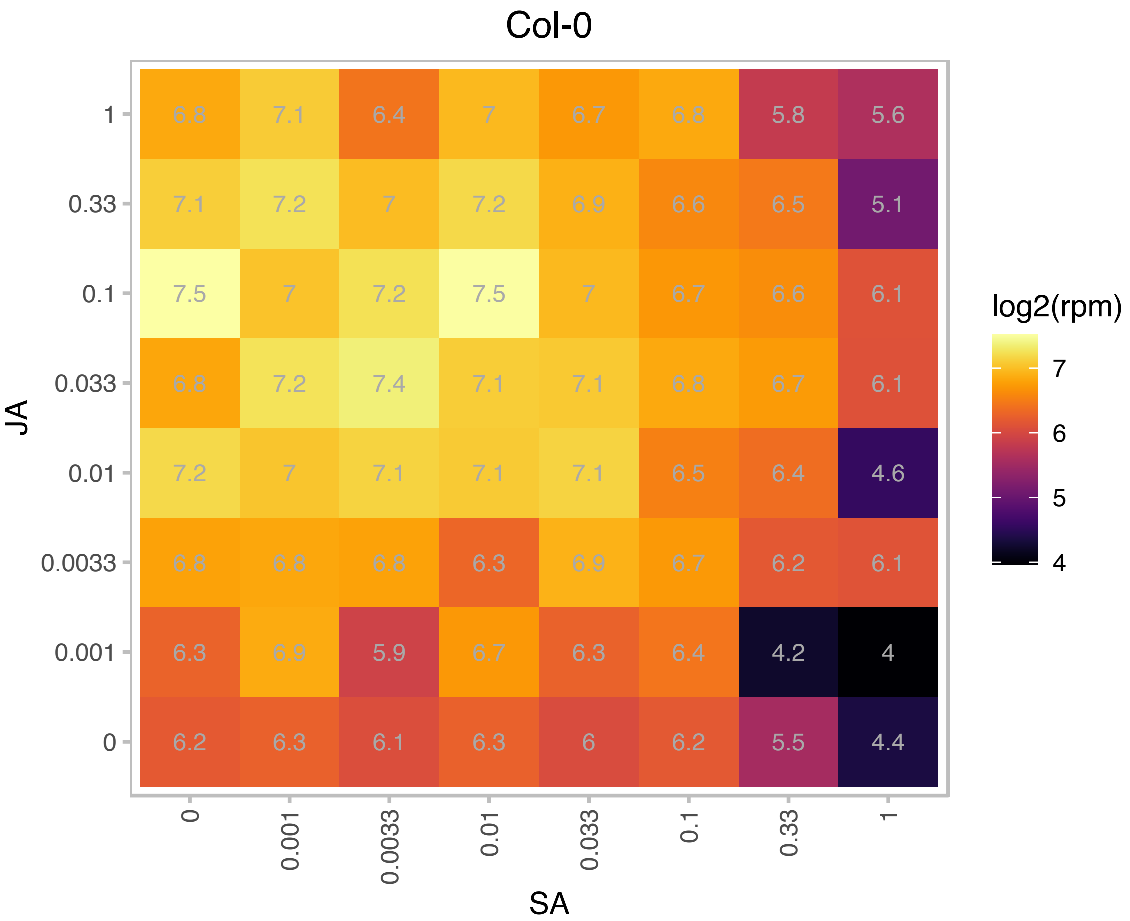


| | |
|---|---|
| Gene | AT1G69760 |
| is.expressed_genes | TRUE |
| is.responsive_genes | TRUE |
| is.apcluster | E-02 |

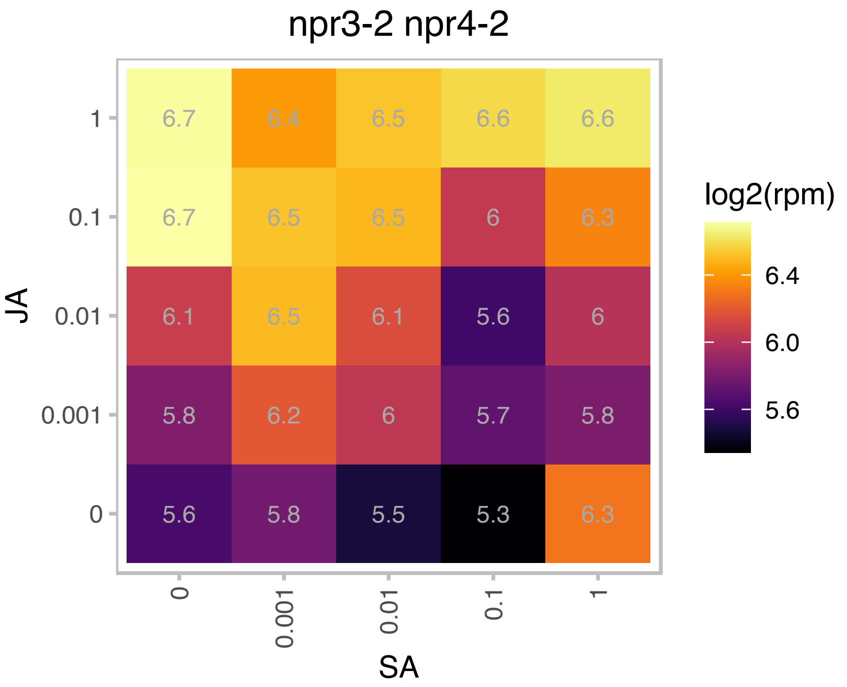


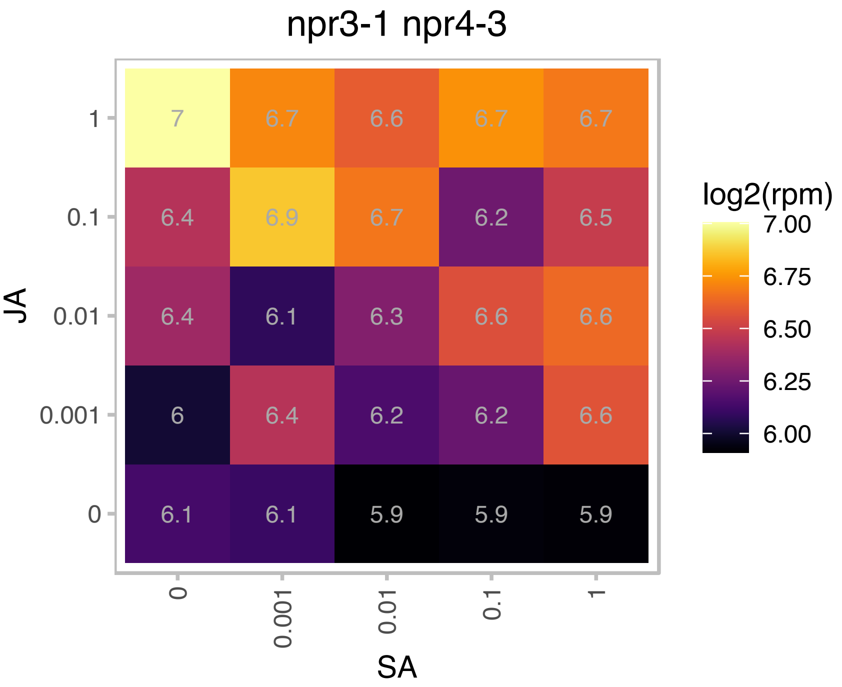

### AT1G76160.1 : sks5

SKU5 similar 5

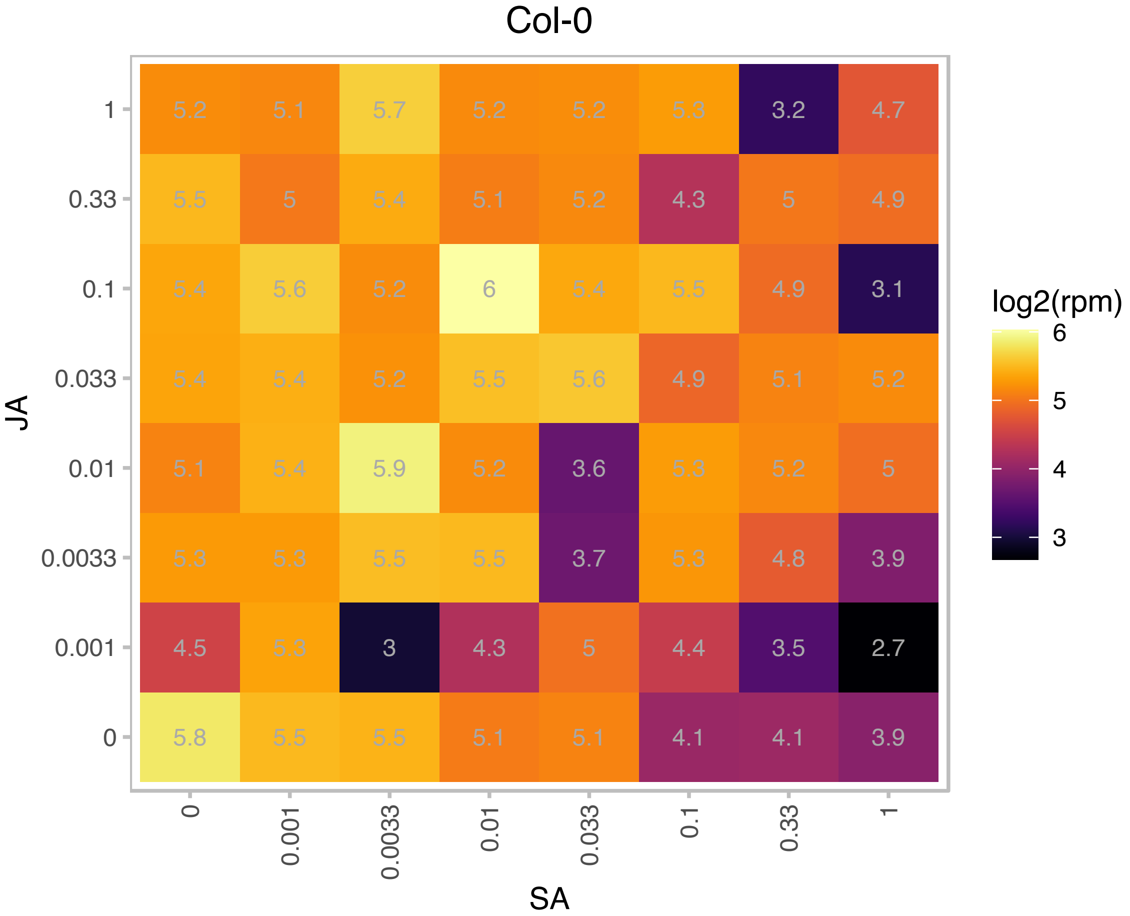


| | |
|---|---|
| Gene | AT1G76160 |
| is.expressed_genes | TRUE |
| is.responsive_genes | TRUE |
| is.apcluster | E-02 |

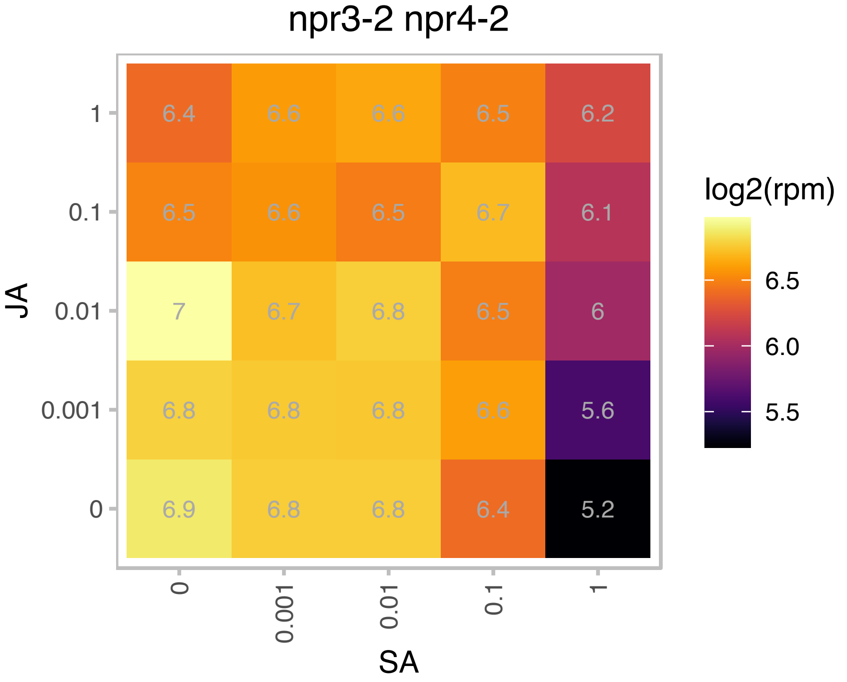


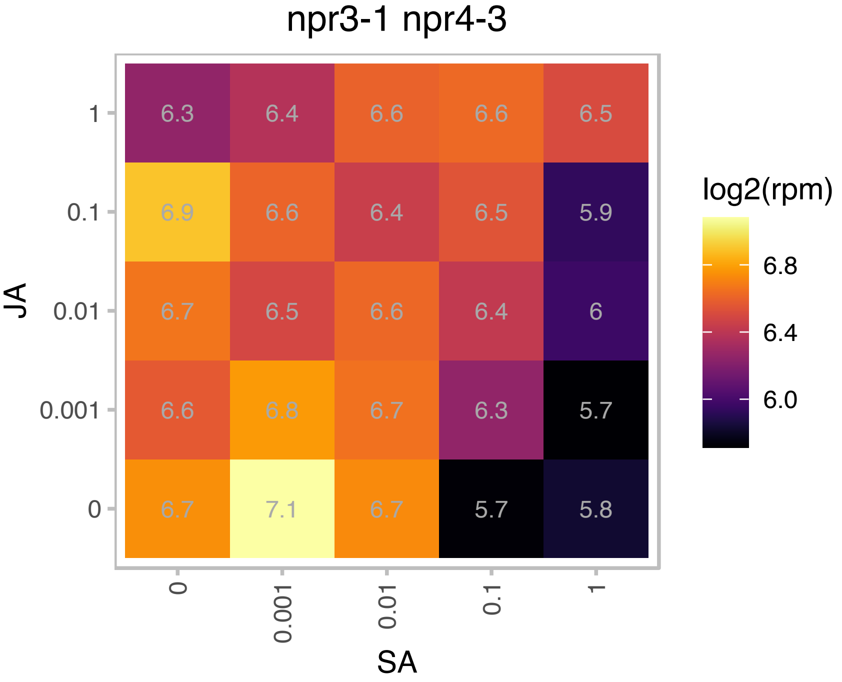

### AT1G76410.1 : ATL8

RING/U-box superfamily protein

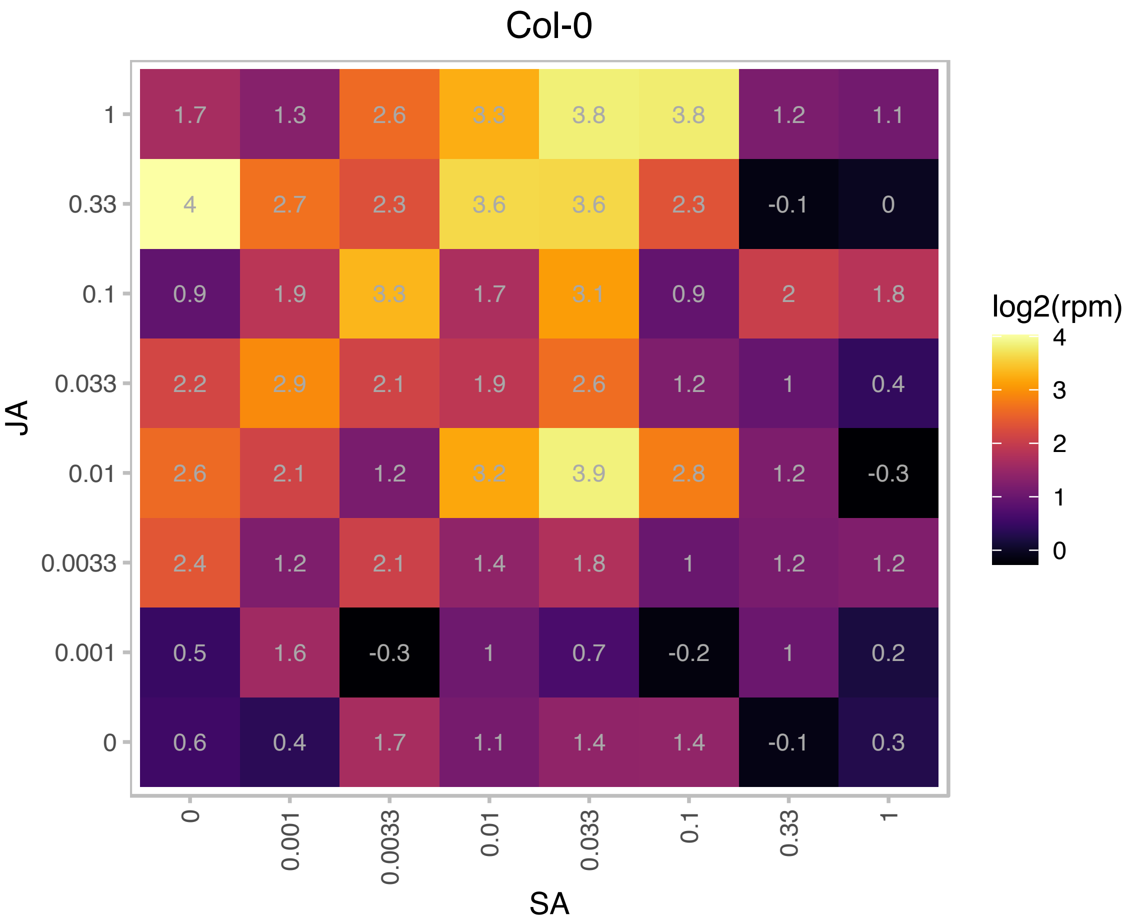


| | |
|---|---|
| Gene | AT1G76410 |
| is.expressed_genes | TRUE |
| is.responsive_genes | TRUE |
| is.apcluster | E-02 |

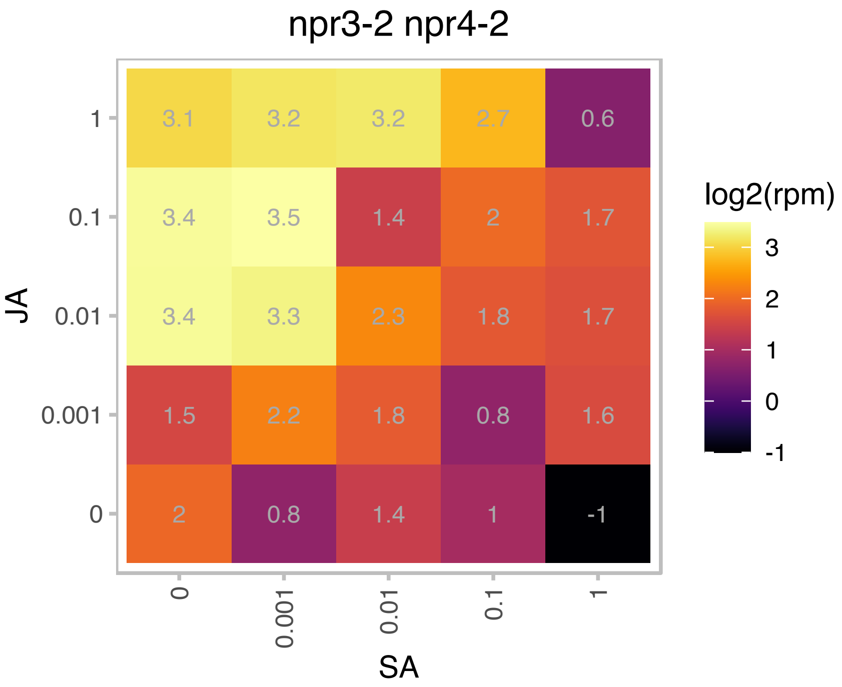


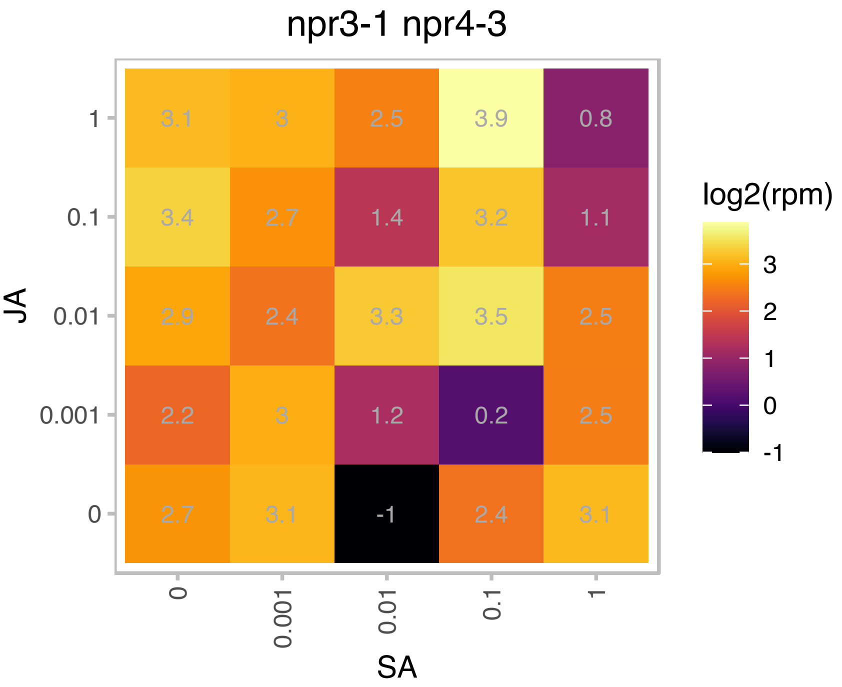

### AT1G76790.1 : IGMT5

O-methyltransferase family protein

Encodes a protein with similarity to N-acetylserotonin O-methyltransferase (ASMT) but it does not have ASMT activity in vitro.

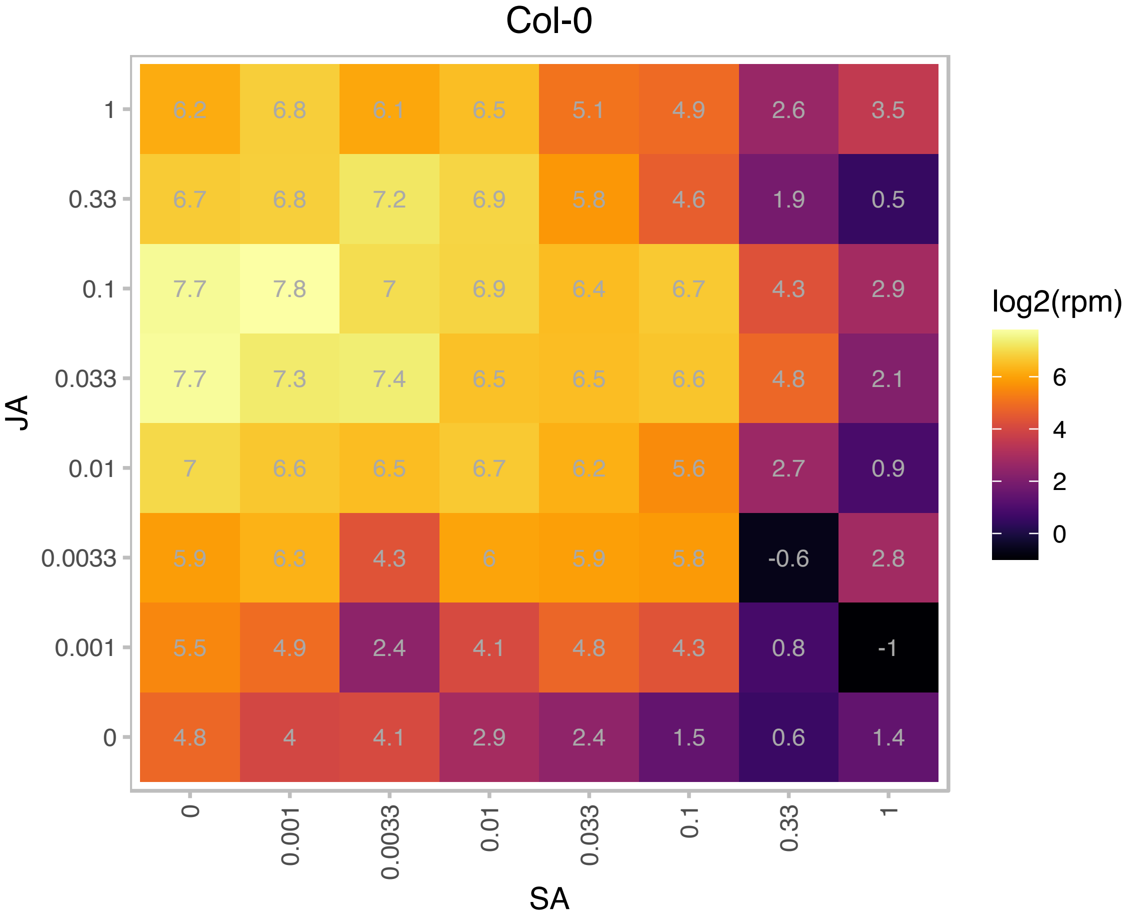


| | |
|---|---|
| Gene | AT1G76790 |
| is.expressed_genes | TRUE |
| is.responsive_genes | TRUE |
| is.apcluster | E-02 |

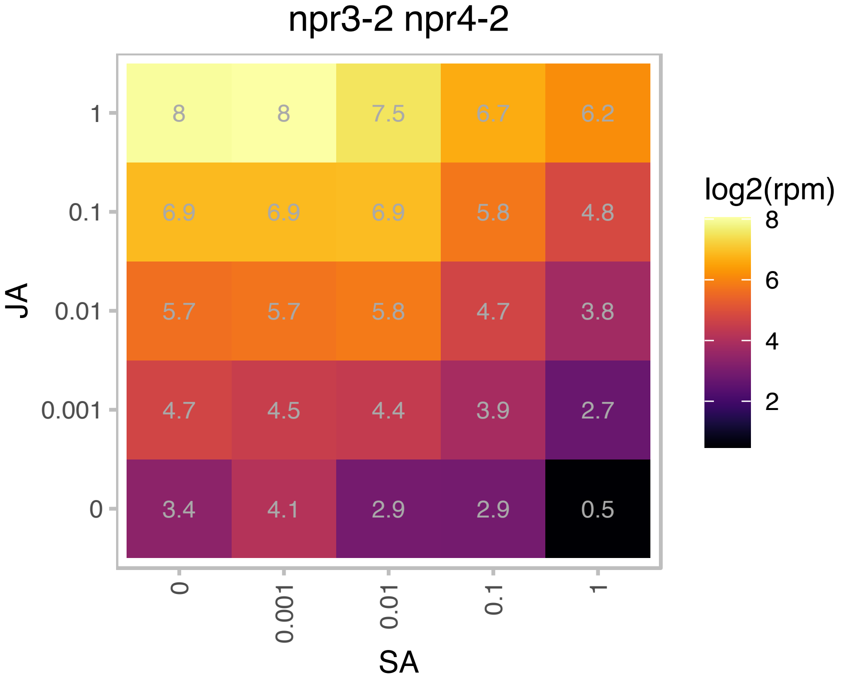


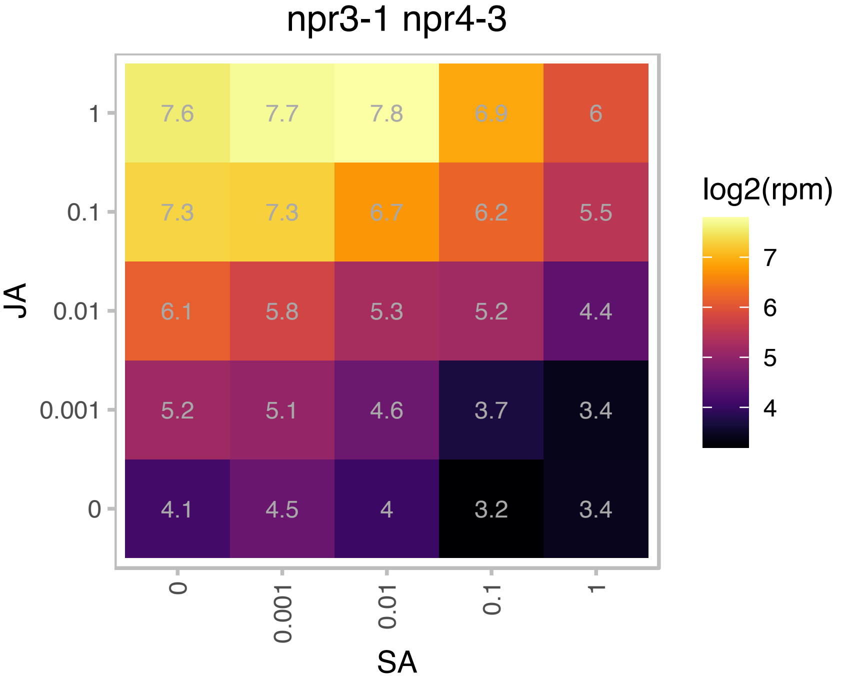

# AT1G79830.1 : GC5

golgin candidate 5

This gene is predicted to encode a protein that functions as a Golgi apparatus structural component known as a golgin in mammals and yeast. A fluorescently-tagged version of GC5 co-localizes with Golgi markers, and this localization appears to be replicated using the C-terminal (139 aa) portion of the protein. The C-terminal portion of the protein can also specifically interact with two members of the Rab family of GTPases (RabH1b and RabH1c).

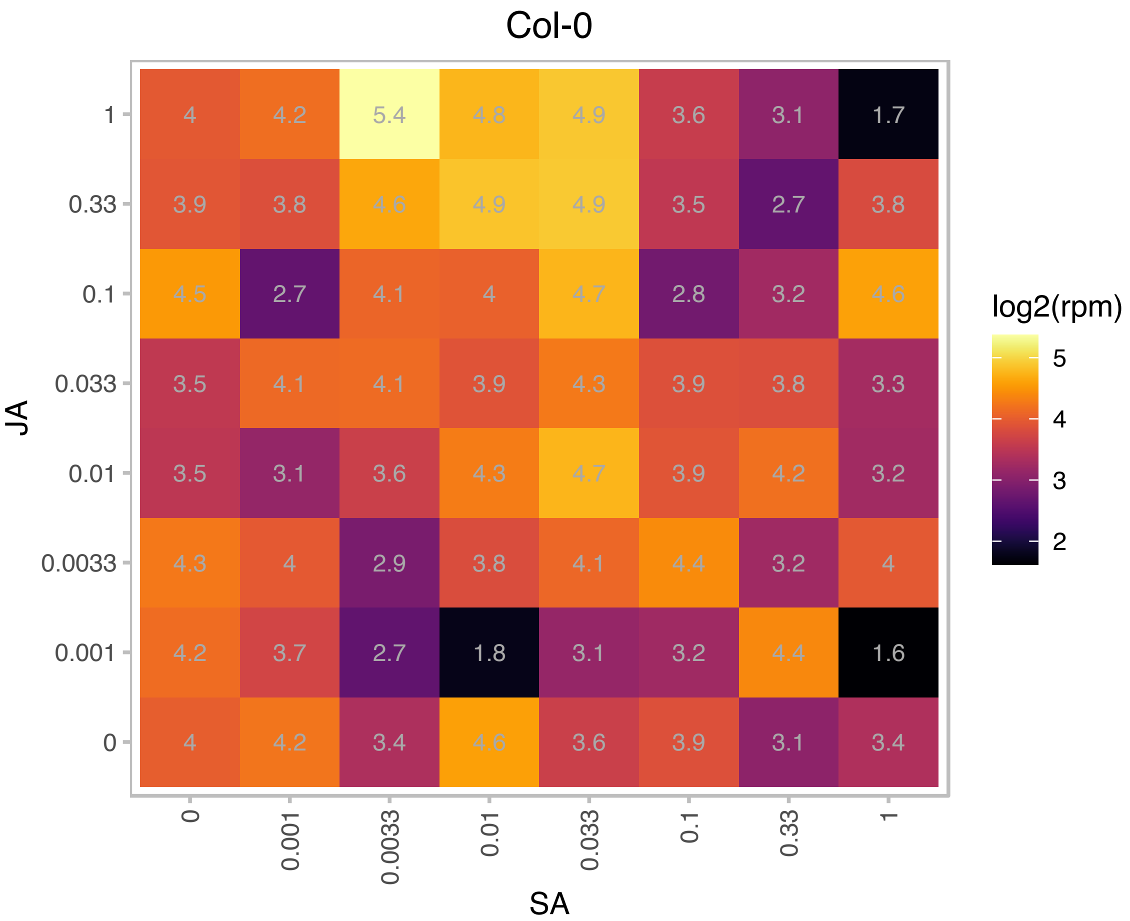


| | |
|---|---|
| Gene | AT1G79830 |
| is.expressed_genes | TRUE |
| is.responsive_genes | TRUE |
| is.apcluster | E-02 |

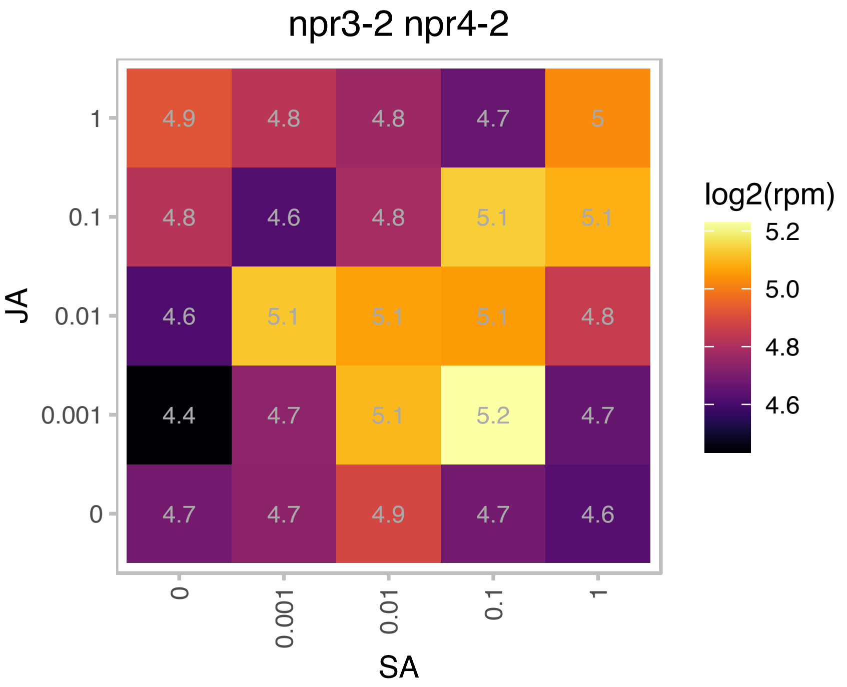


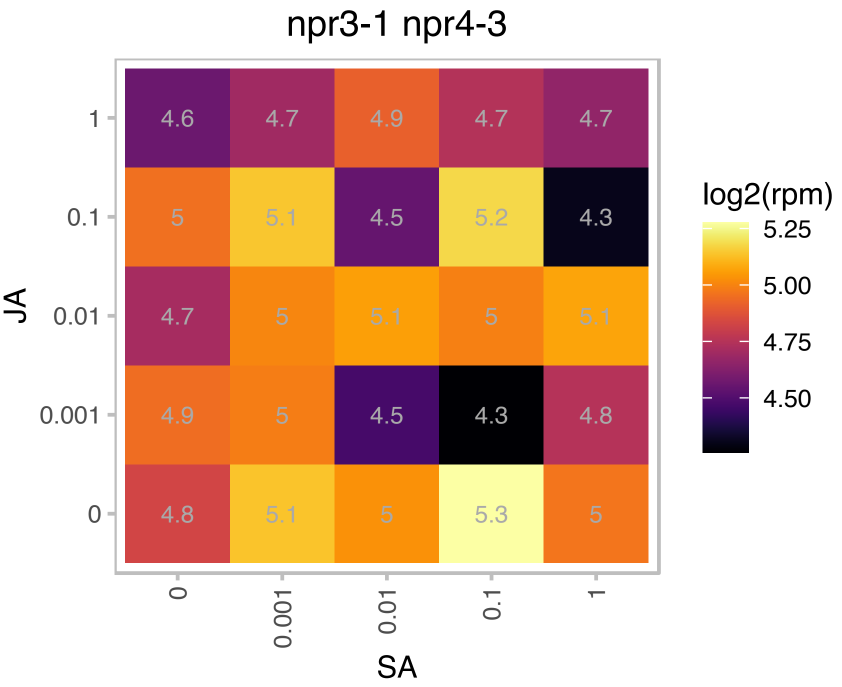

### AT2G01880.1 : ATPAP7, PAP7

purple acid phosphatase 7

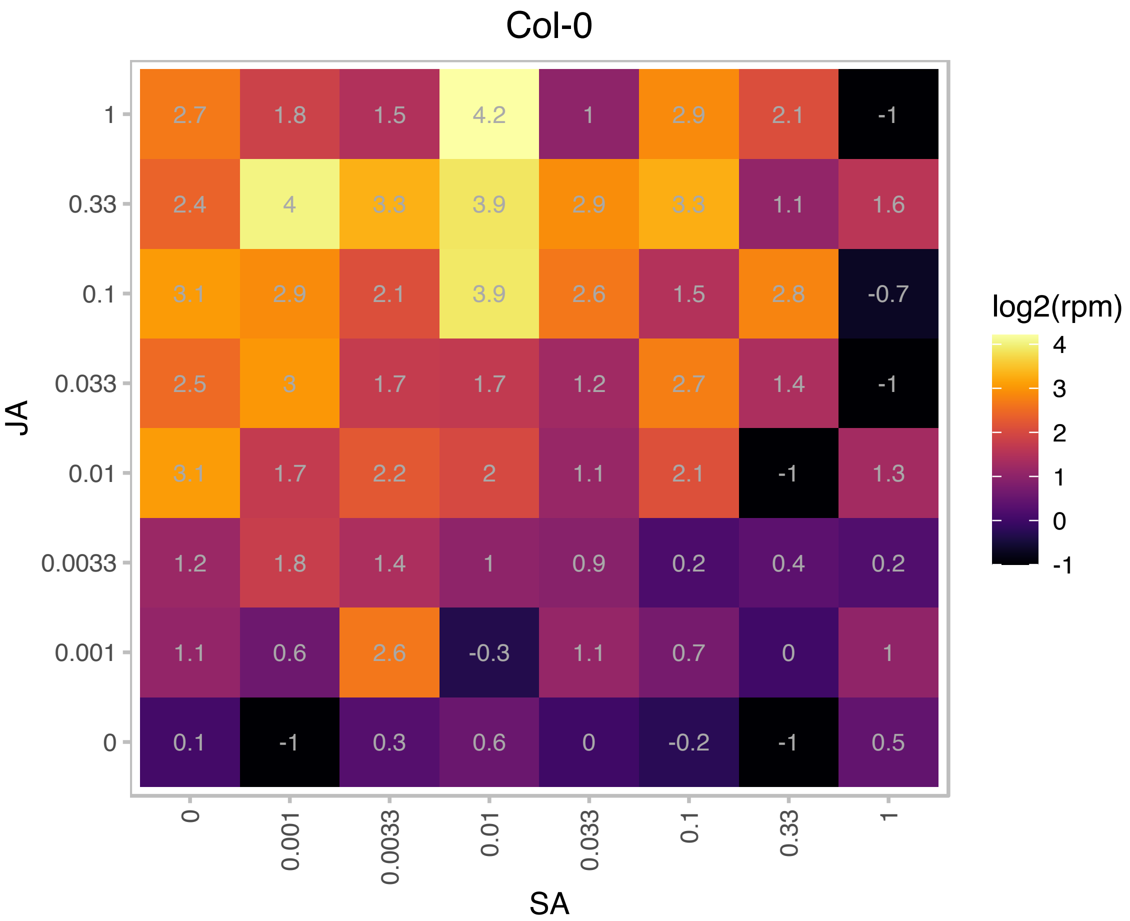


| | |
|---|---|
| Gene | AT2G01880 |
| is.expressed_genes | TRUE |
| is.responsive_genes | TRUE |
| is.apcluster | E-02 |

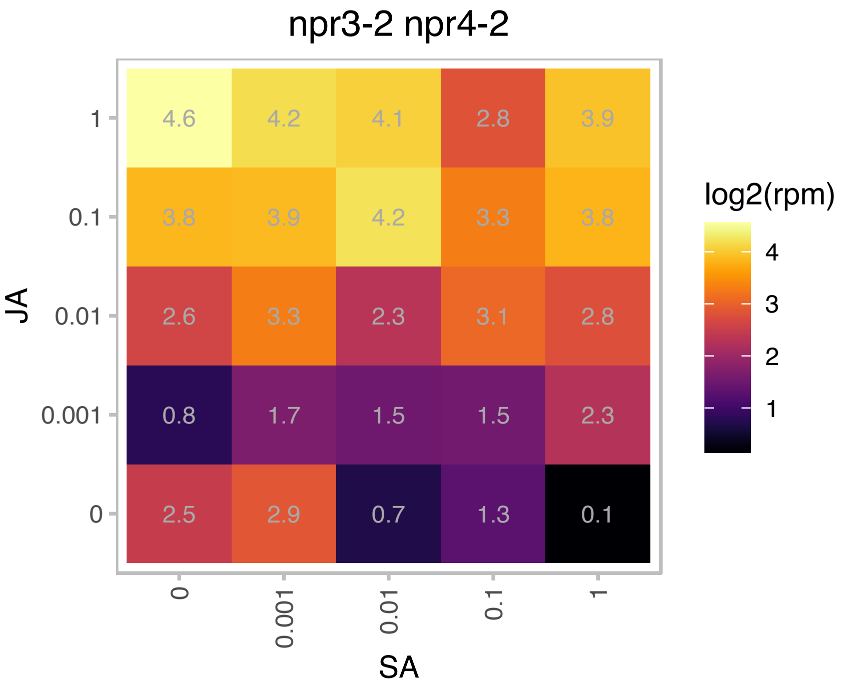


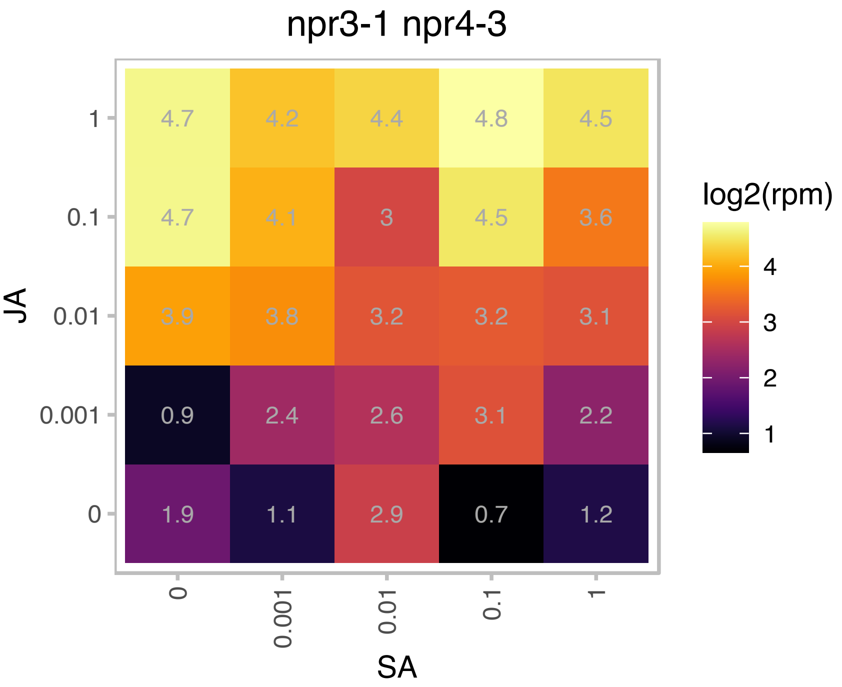

# AT2G08650.1 : NA

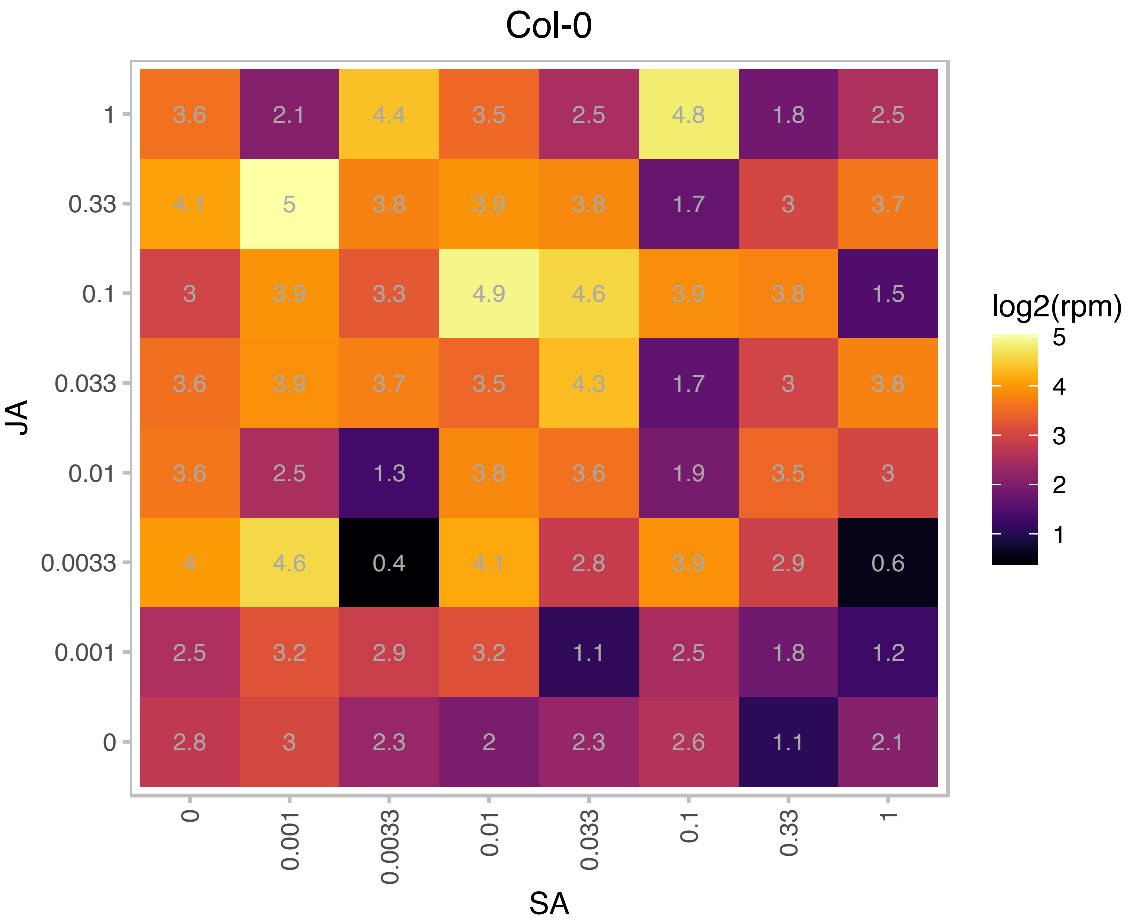


| | |
|---|---|
| Gene | AT2G08650 |
| is.expressed_genes | TRUE |
| is.responsive_genes | TRUE |
| is.apcluster | E-02 |

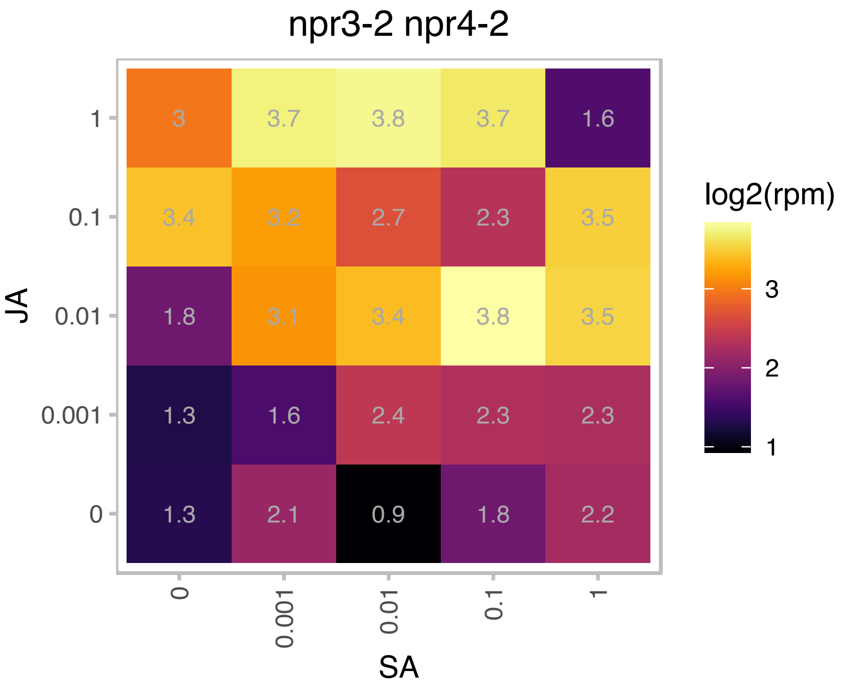


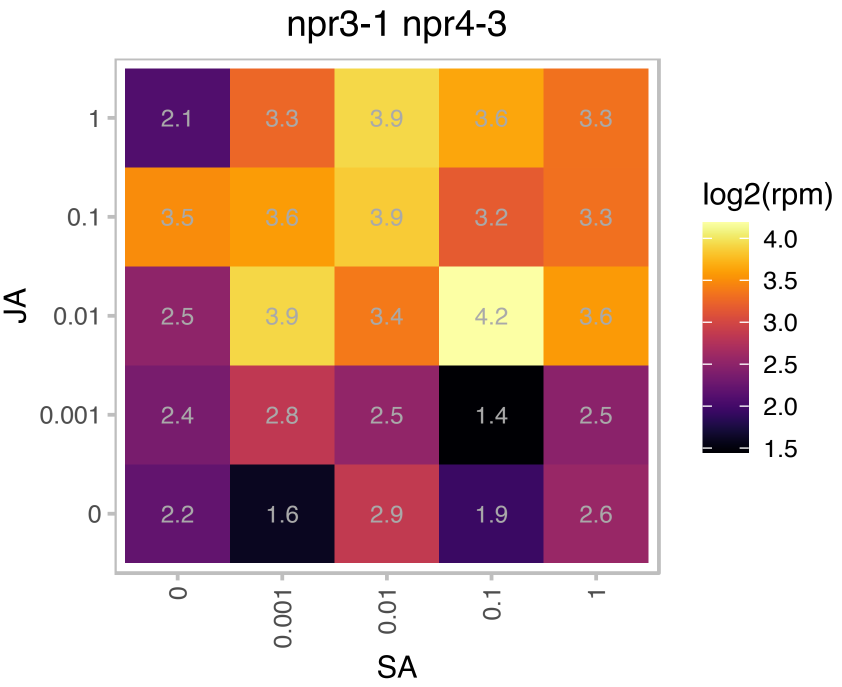

### AT2G22770.1 : NAI1

basic helix-loop-helix (bHLH) DNA-binding superfamily protein

regulates the development of ER bodies. also involves in response to the endophytic fungus Piriformospora indica.

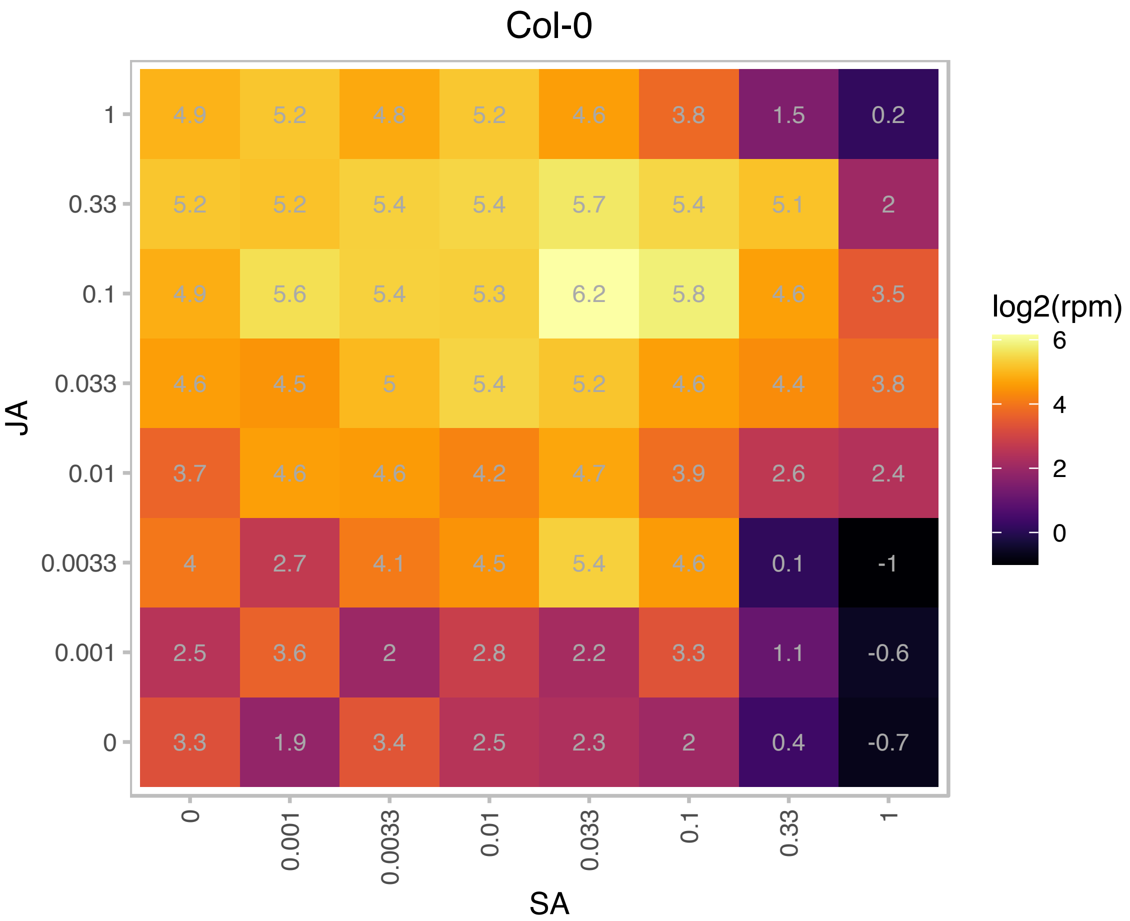


| | |
|---|---|
| Gene | AT2G22770 |
| is.expressed_genes | TRUE |
| is.responsive_genes | TRUE |
| is.apcluster | E-02 |

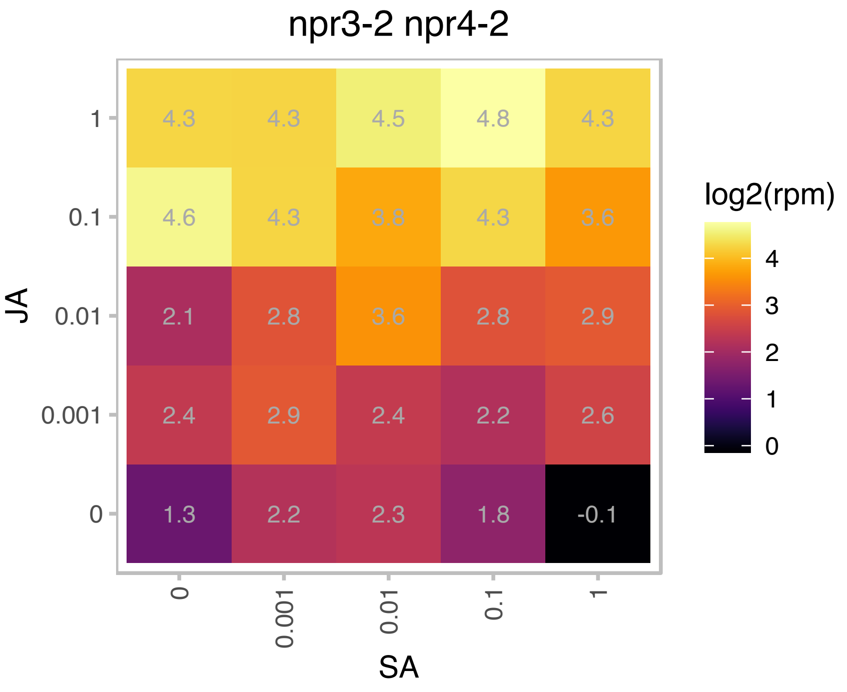


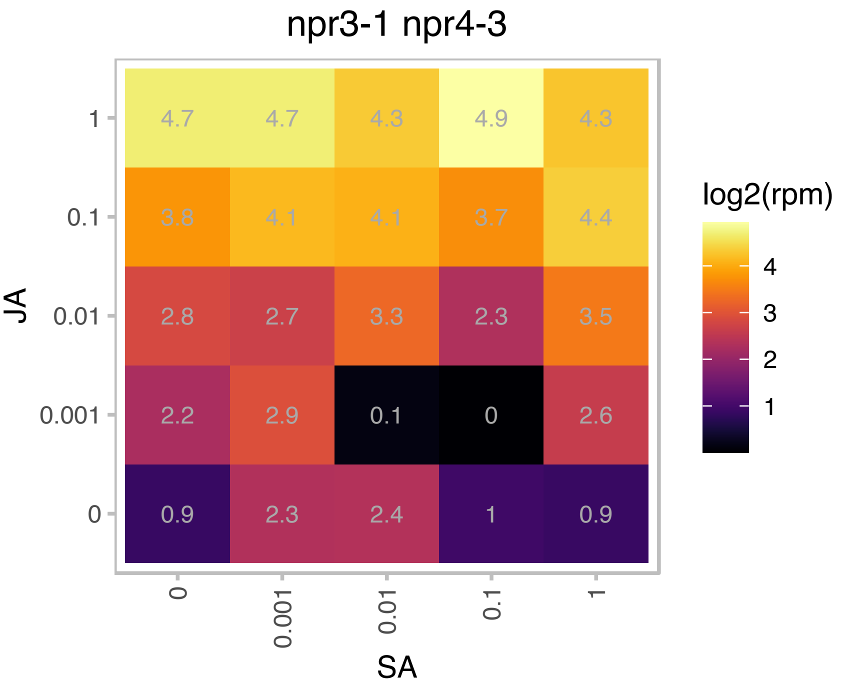

### AT2G23560.1 : ATMES7, MES7

methyl esterase 7

Encodes a protein shown to have carboxylesterase activity, methyl salicylate esterase activity, and methyl IAA esterase activity in vitro. This protein does not act on methyl JA, MeGA4, or MEGA9 in vitro. MES7 appears to be involved in MeSA hydrolysis in planta. Expression of MES7 can restore systemic acquired resistance in SAR-deficient tobacco plants.

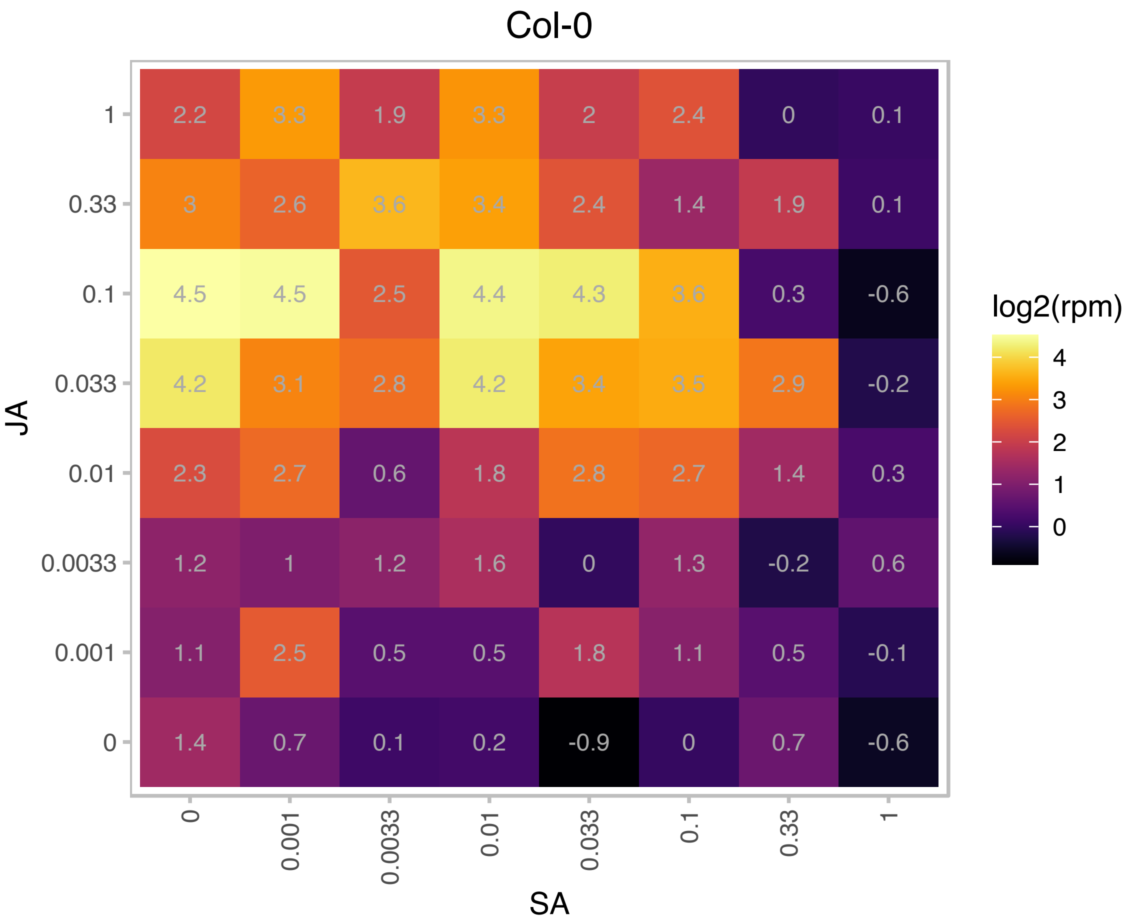


| | |
|---|---|
| Gene | AT2G23560 |
| is.expressed_genes | TRUE |
| is.responsive_genes | TRUE |
| is.apcluster | E-02 |

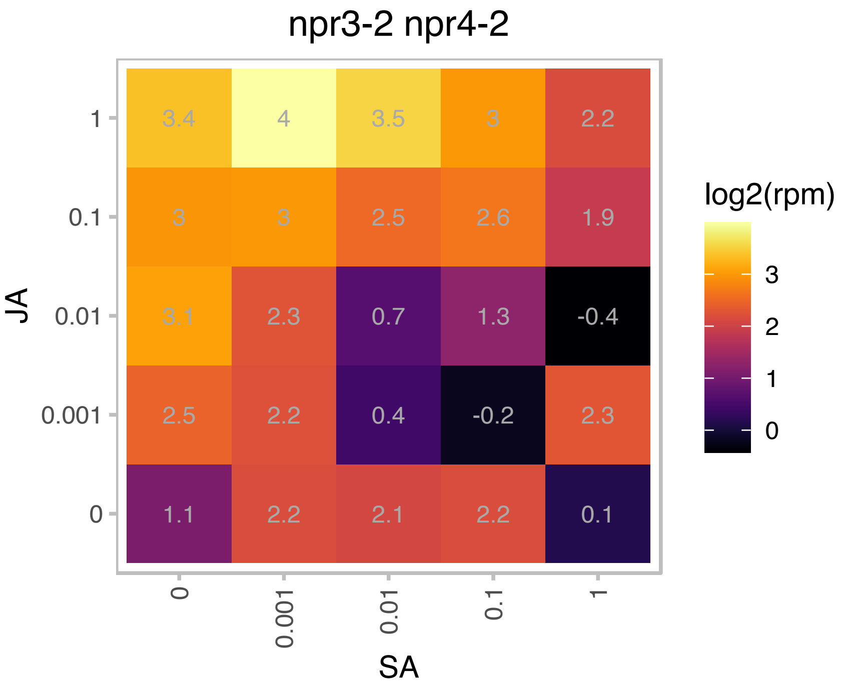


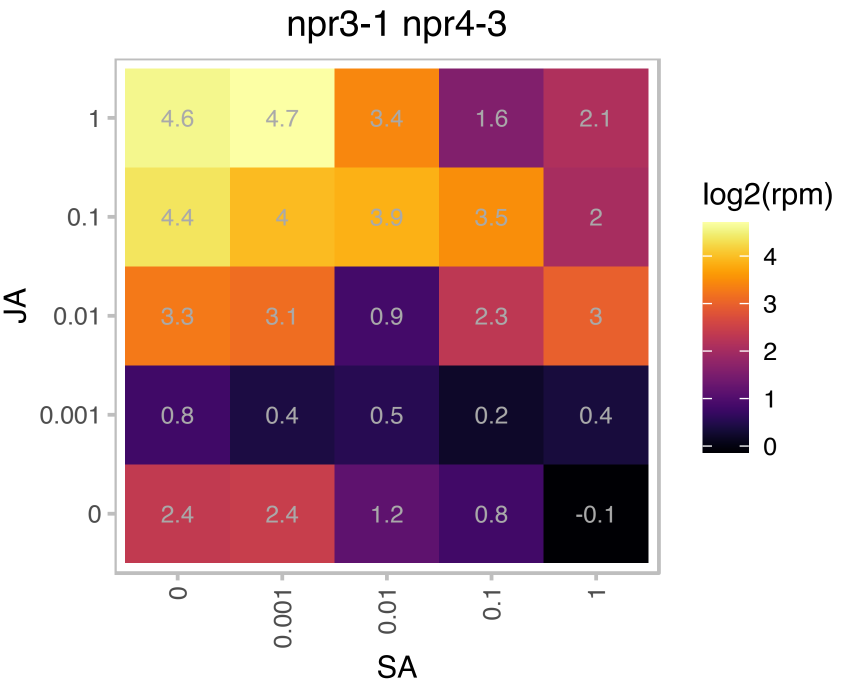

# AT2G25670.1 : NA

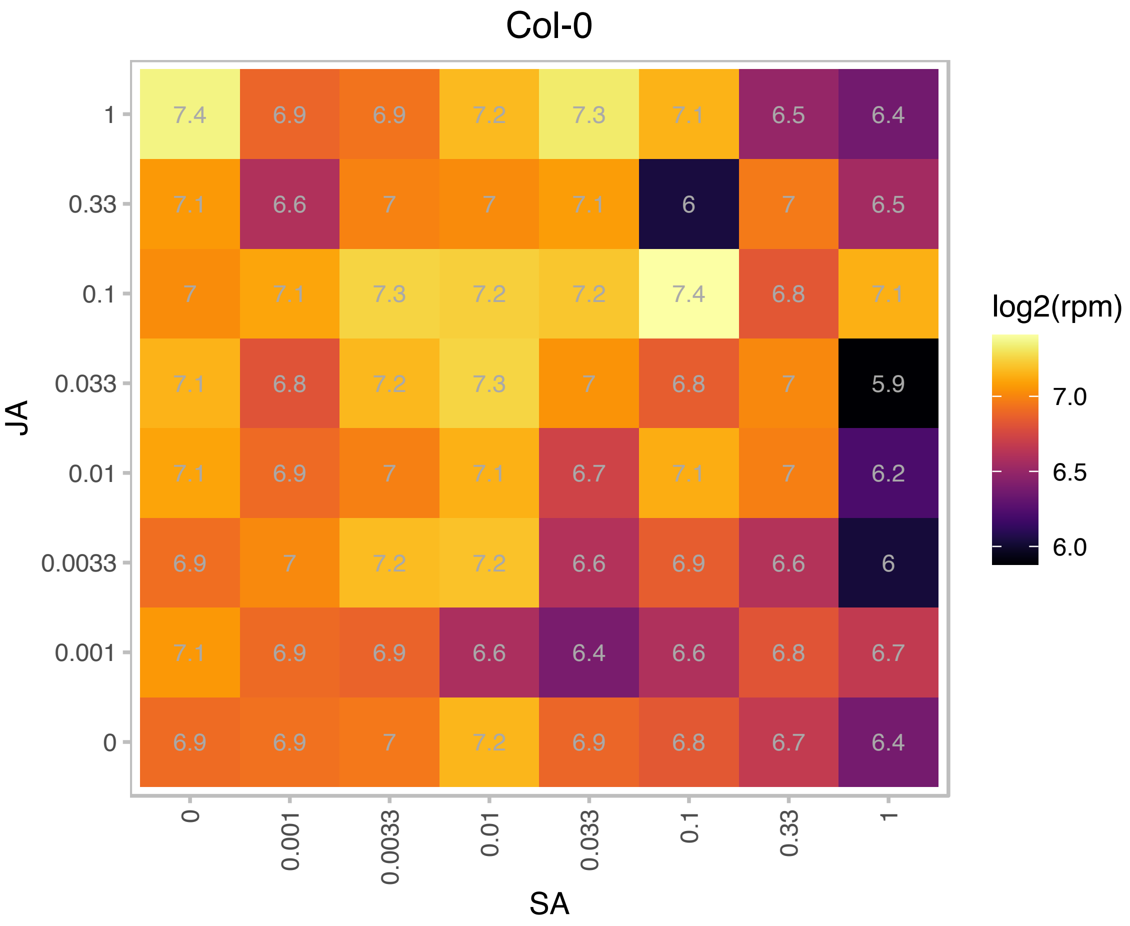


| | |
|---|---|
| Gene | AT2G25670 |
| is.expressed_genes | TRUE |
| is.responsive_genes | TRUE |
| is.apcluster | E-02 |

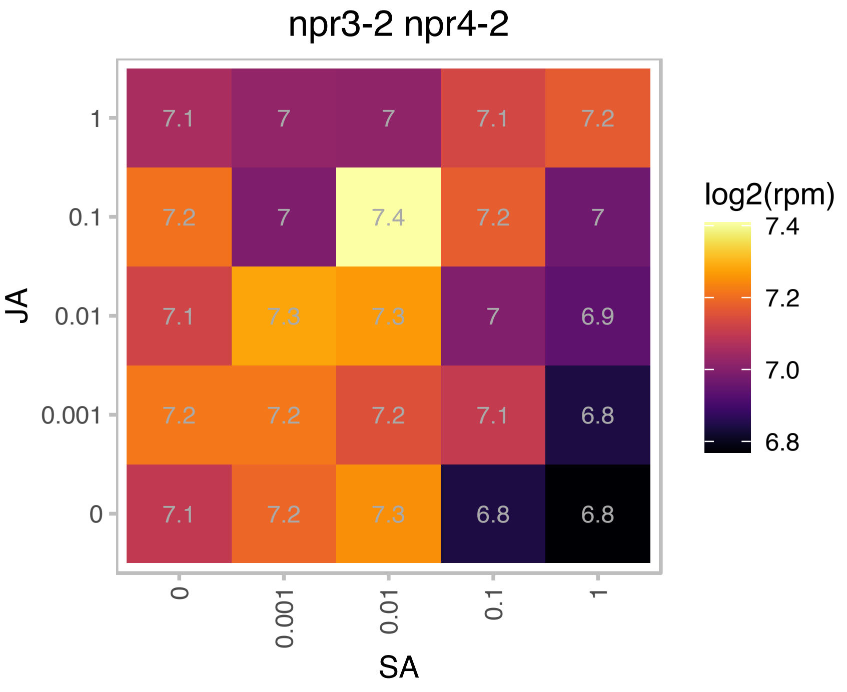


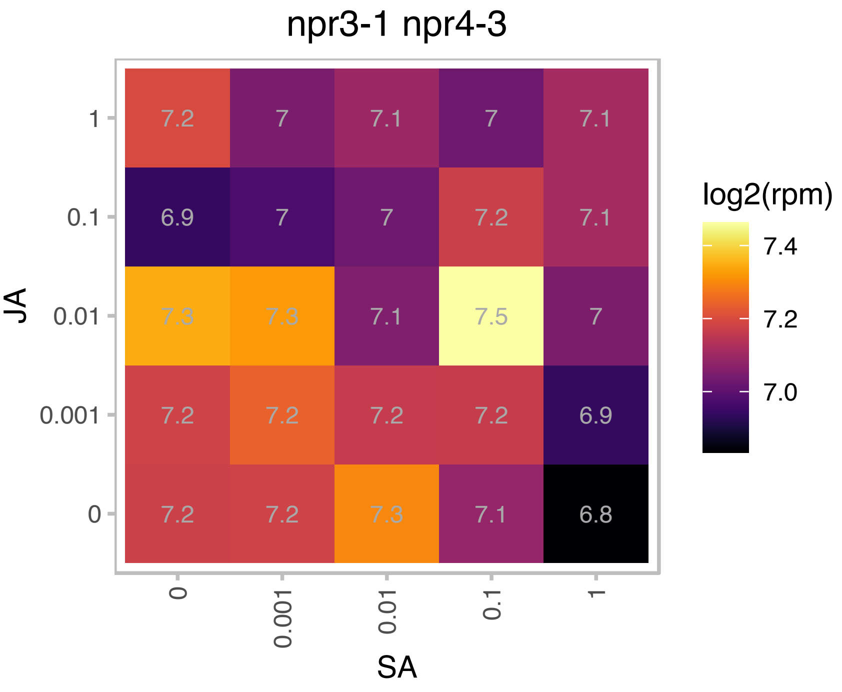

# AT2G26370.1 : NA

MD-2-related lipid recognition domain-containing protein

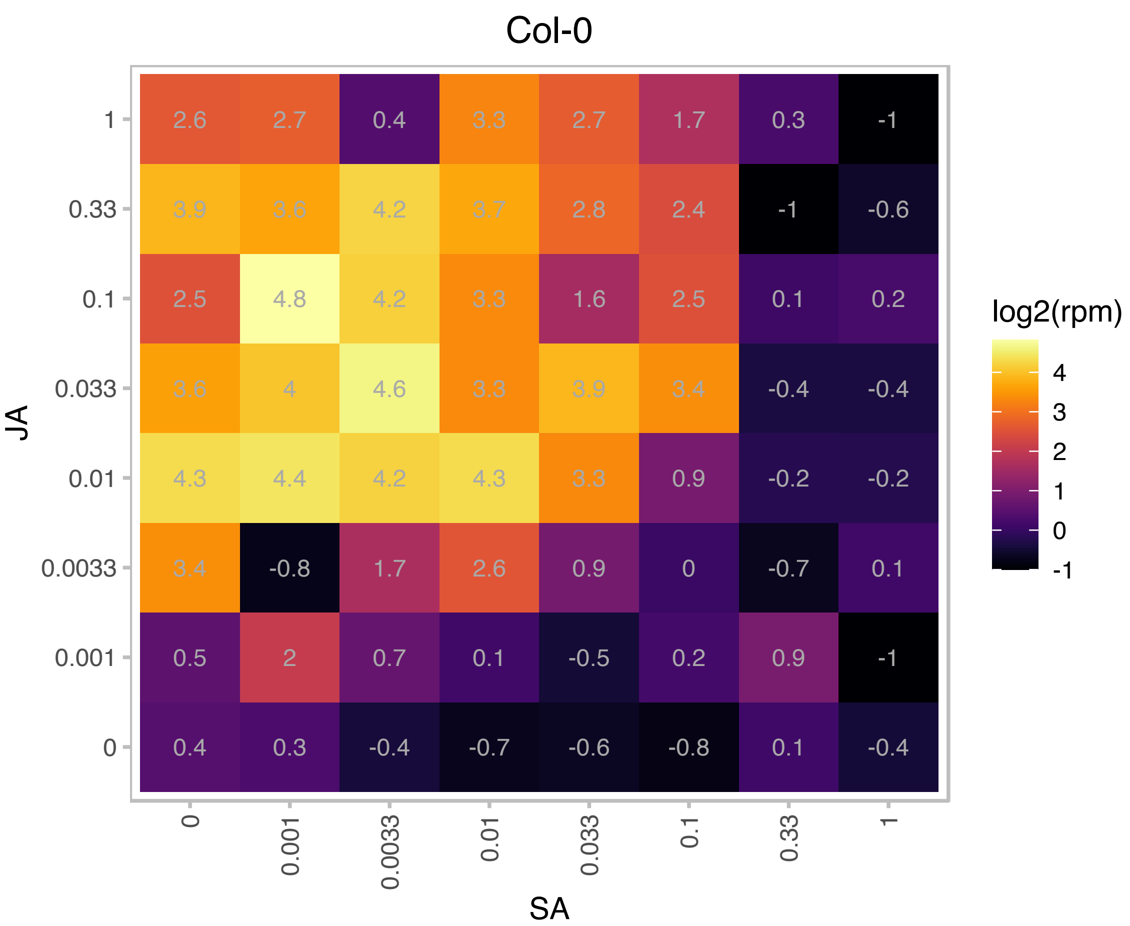


| | |
|---|---|
| Gene | AT2G26370 |
| is.expressed_genes | TRUE |
| is.responsive_genes | TRUE |
| is.apcluster | E-02 |

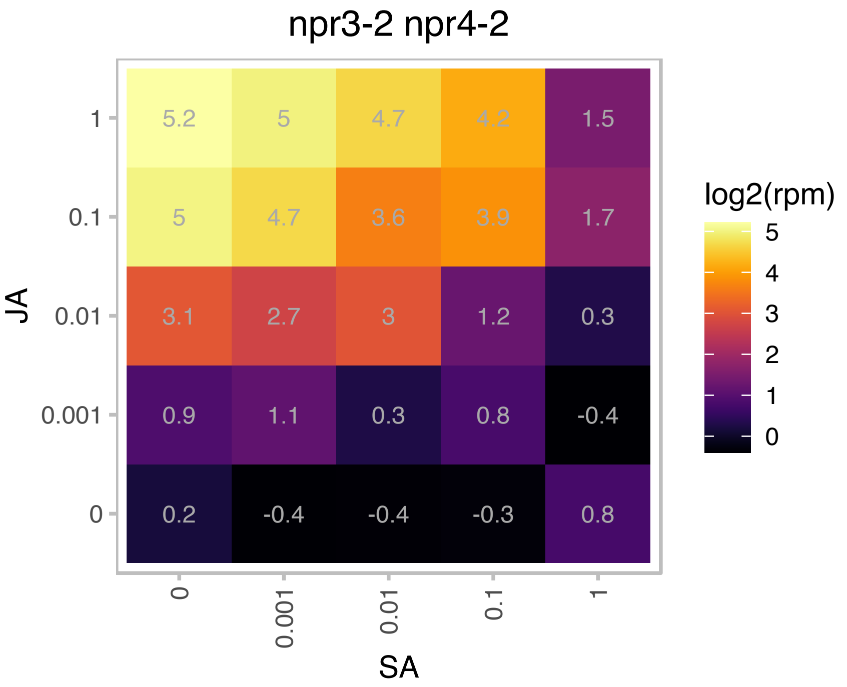


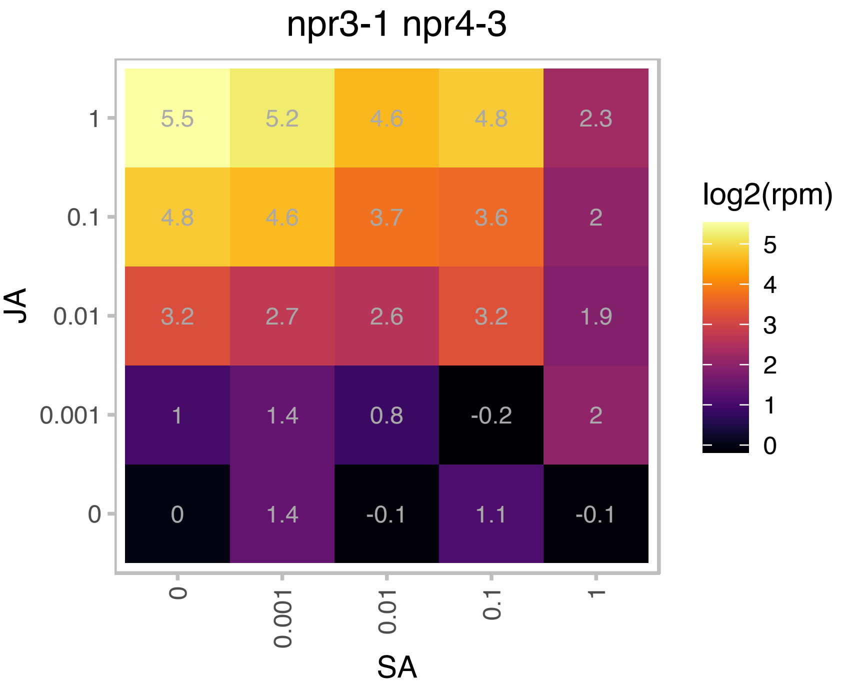

### AT2G30200.1 : EMB3147

catalytics;transferases;[acyl-carrier-protein] S-malonyltransferases;binding

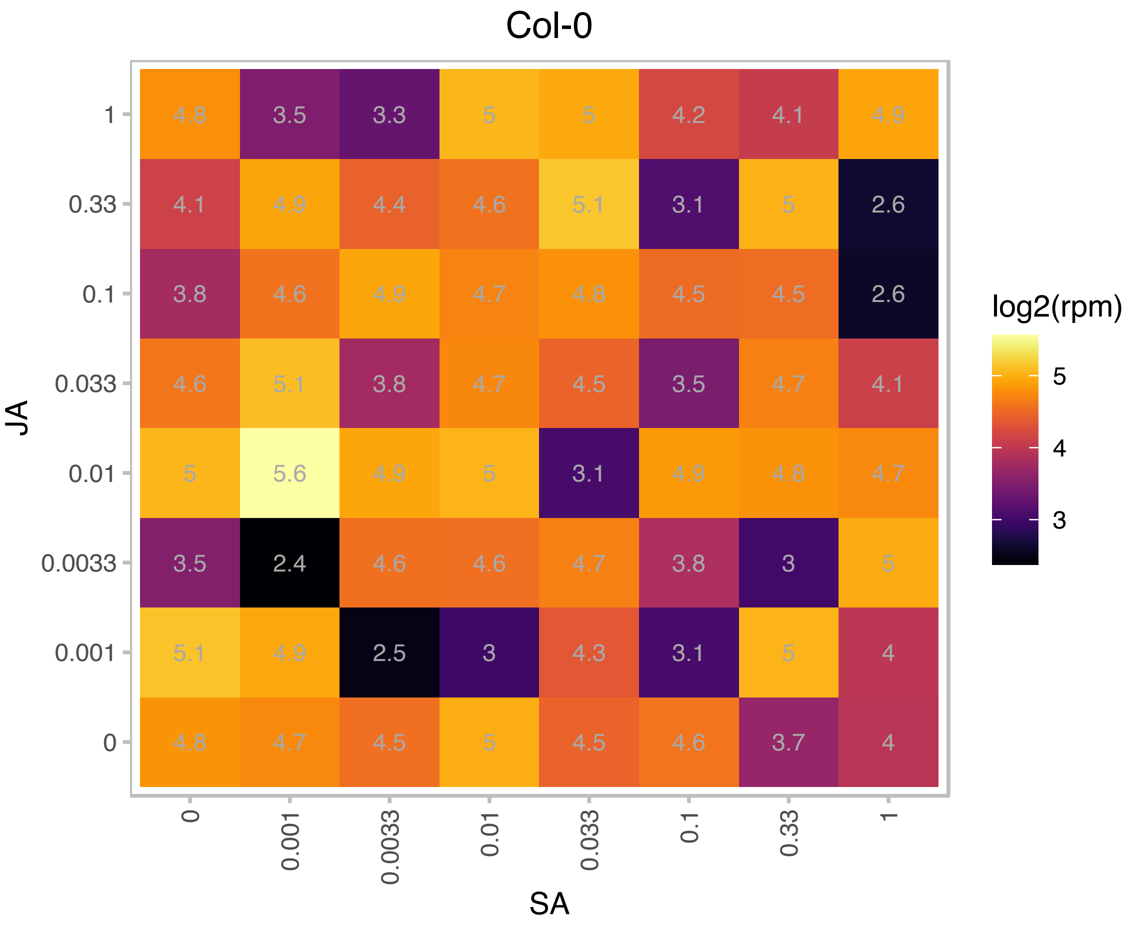


| | |
|---|---|
| Gene | AT2G30200 |
| is.expressed_genes | TRUE |
| is.responsive_genes | TRUE |
| is.apcluster | E-02 |

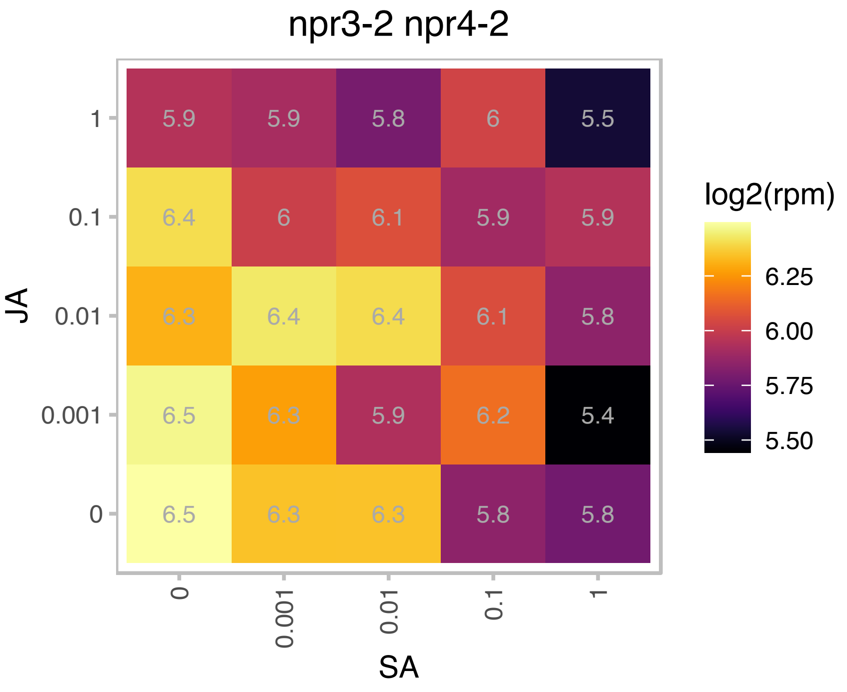


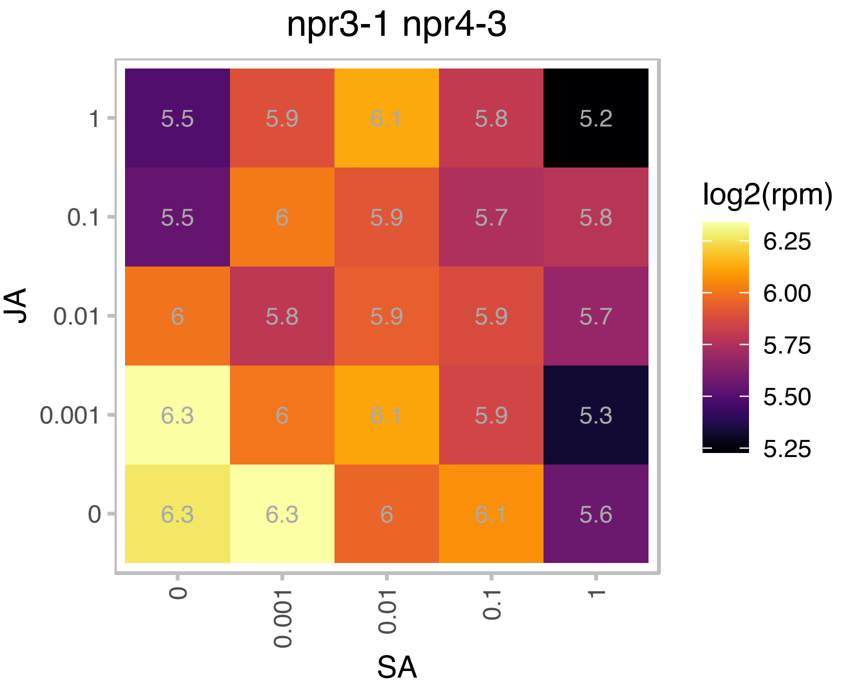

### AT2G36120.1 : DOT1

Glycine-rich protein family
Encodes a glycine rich protein that is involved in leaf vascular patterning. dot1 mutants have an aberrant open-class venation pattern in leaves and cotyledons, as well as several other leaf development defects. The mRNA is cell-to-cell mobile.

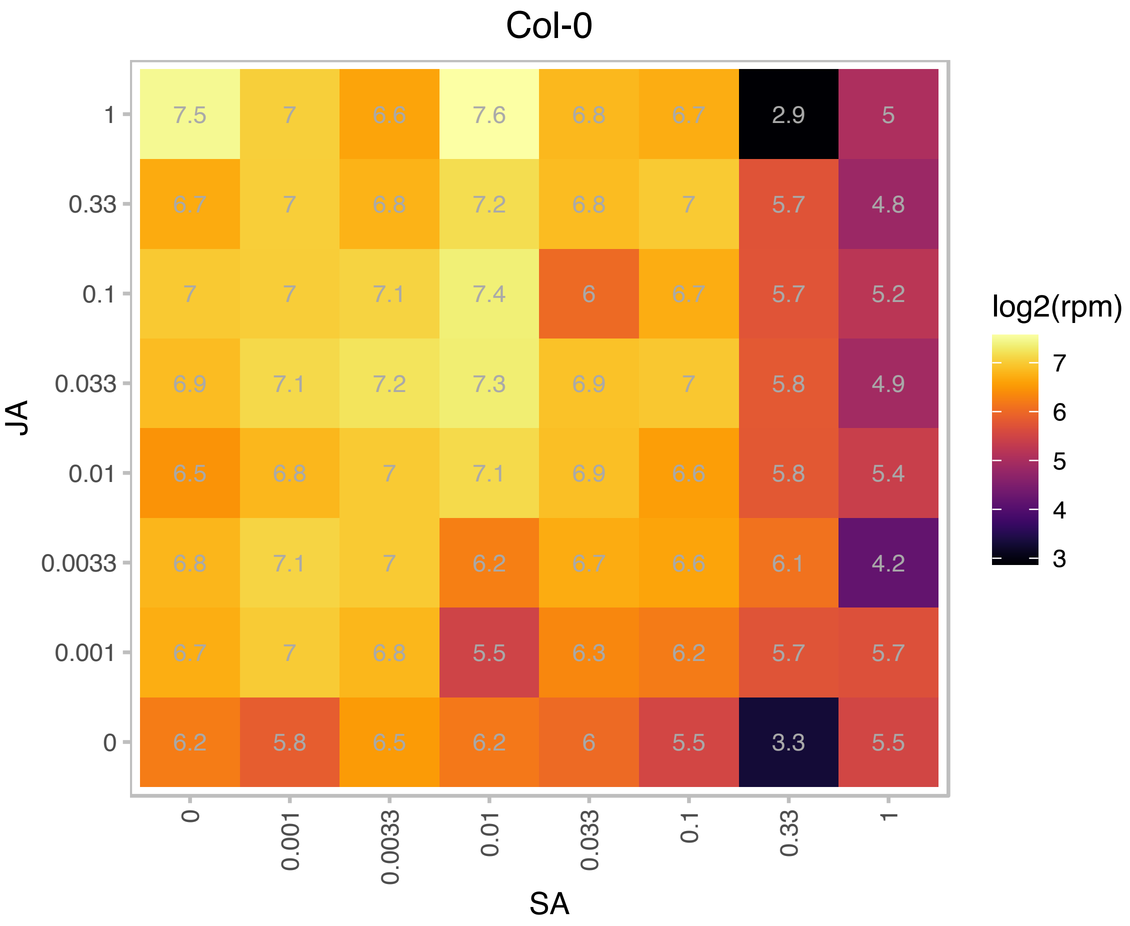


| | |
|---|---|
| Gene | AT2G36120 |
| is.expressed_genes | TRUE |
| is.responsive_genes | TRUE |
| is.apcluster | E-02 |

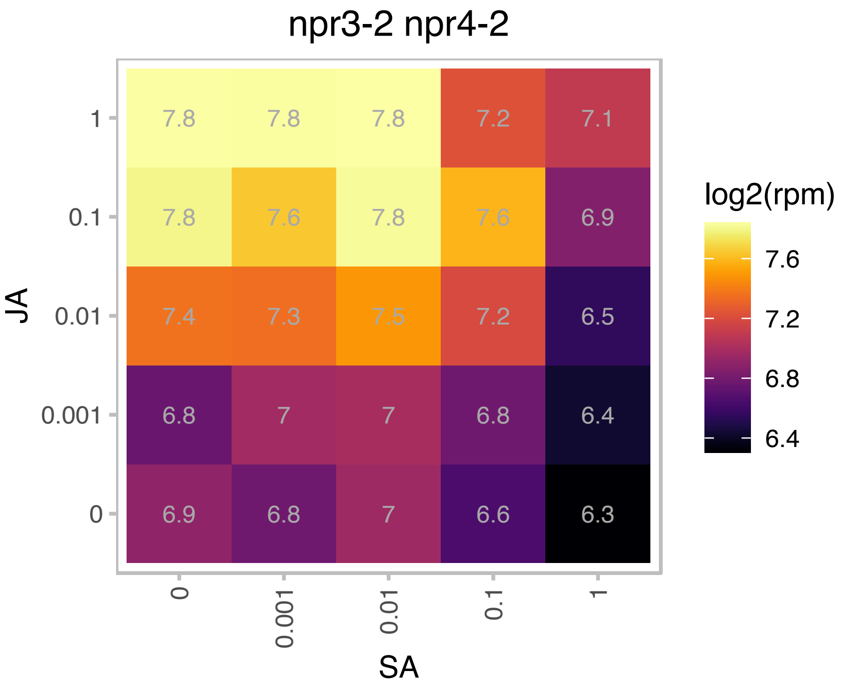


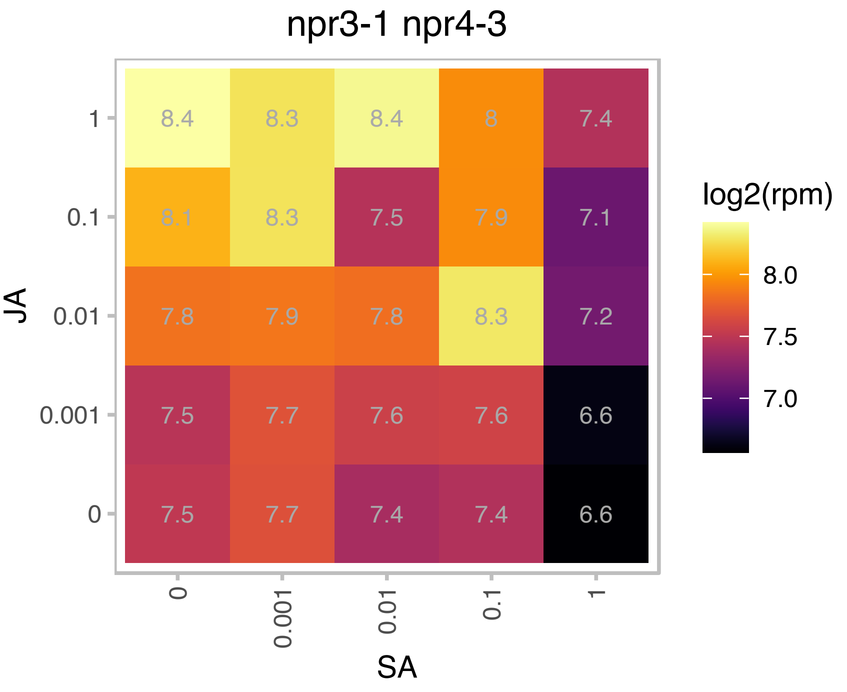

### AT2G40730.1 : CTEXP

Protein kinase family protein with ARM repeat domain

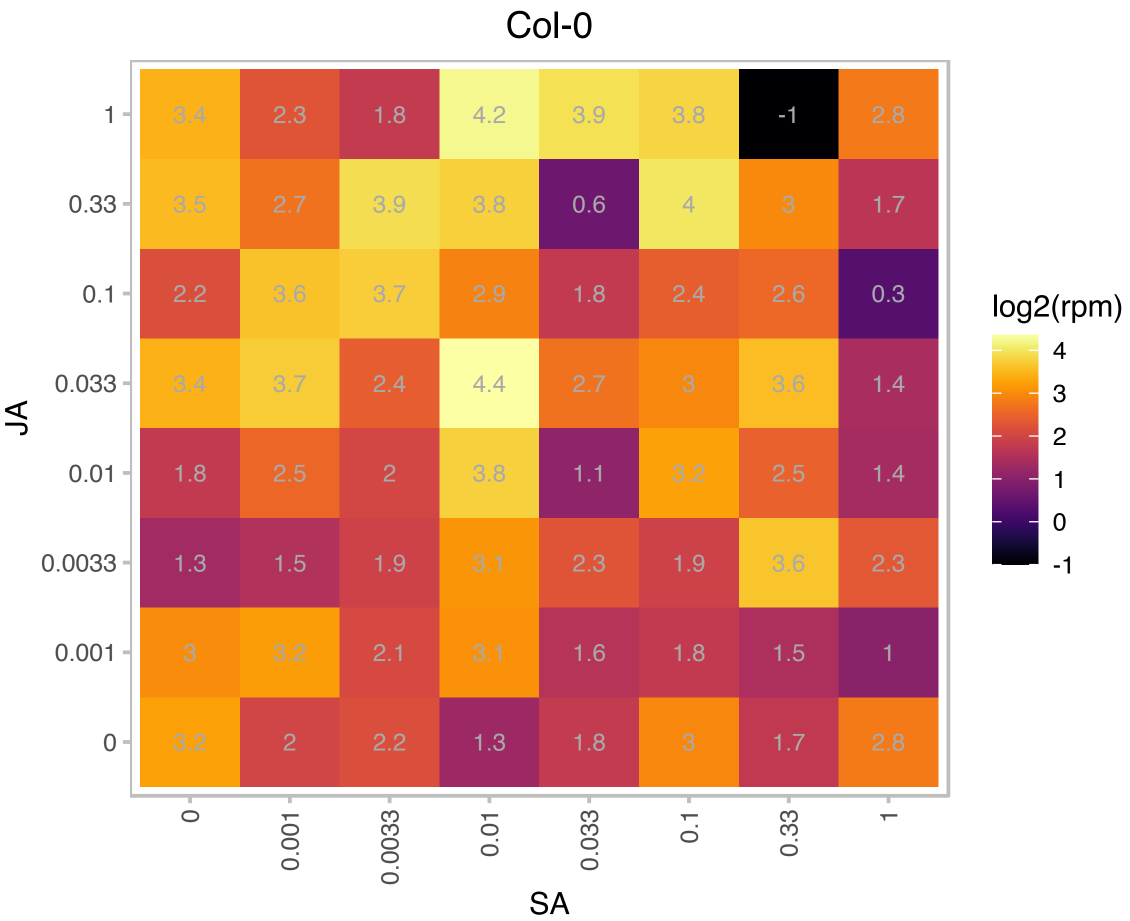


| | |
|---|---|
| Gene | AT2G40730 |
| is.expressed_genes | TRUE |
| is.responsive_genes | TRUE |
| is.apcluster | E-02 |

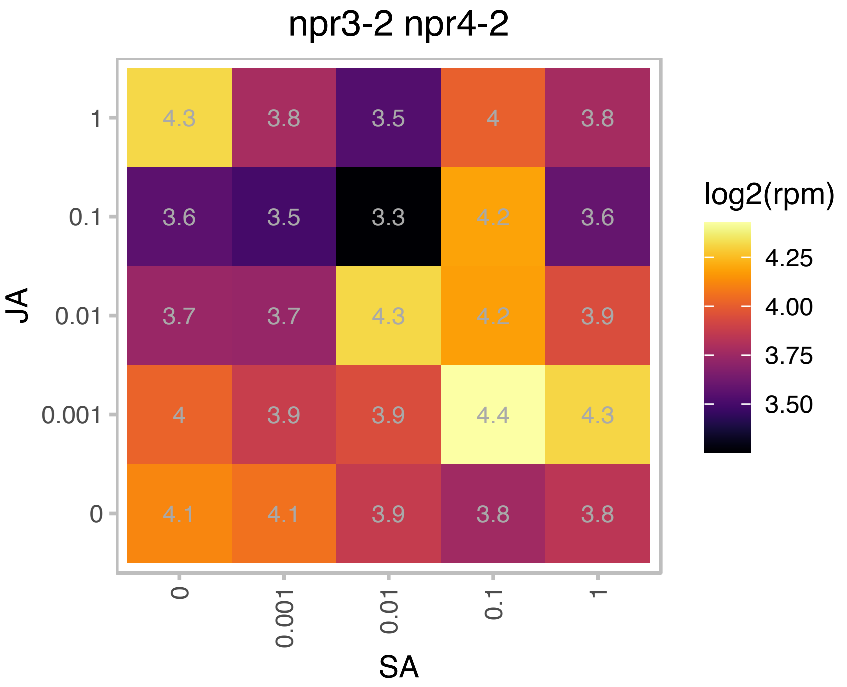


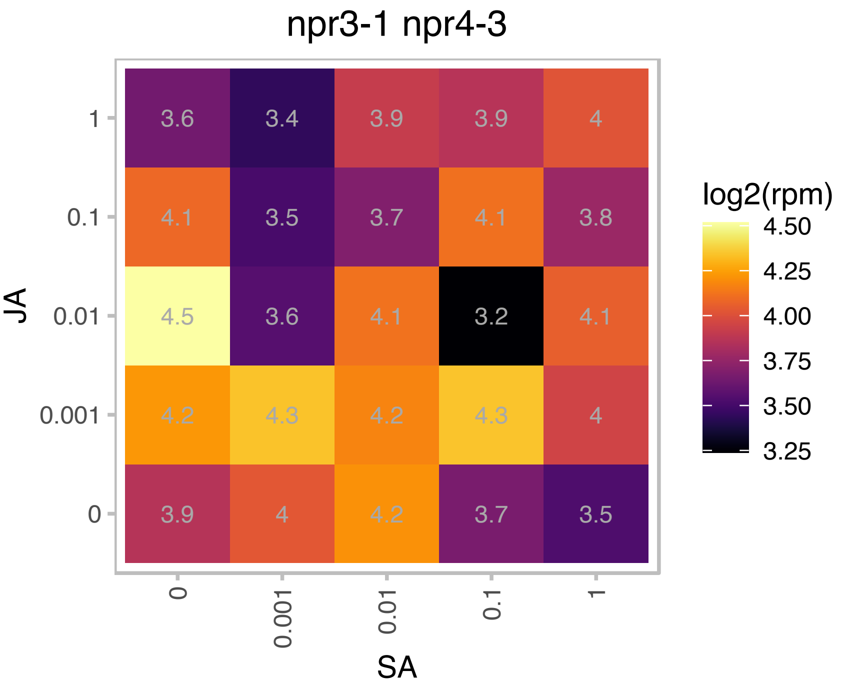

# AT2G48100.1 : NA

Exonuclease family protein

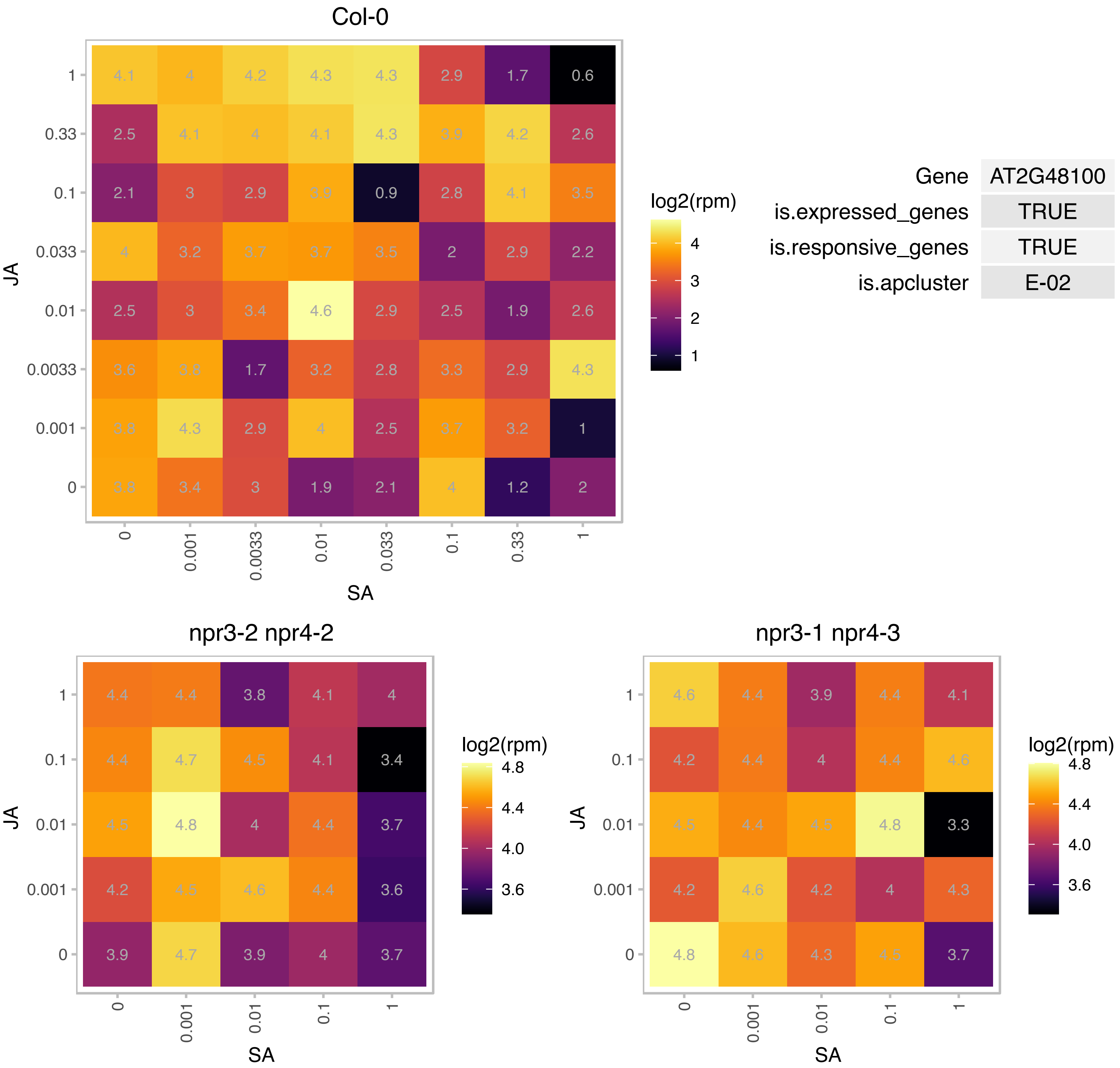


| | |
|---|---|
| Gene | AT2G48100 |
| is.expressed_genes | TRUE |
| is.responsive_genes | TRUE |
| is.apcluster | E-02 |

# AT3G03550.1 : NA

RING/U-box superfamily protein

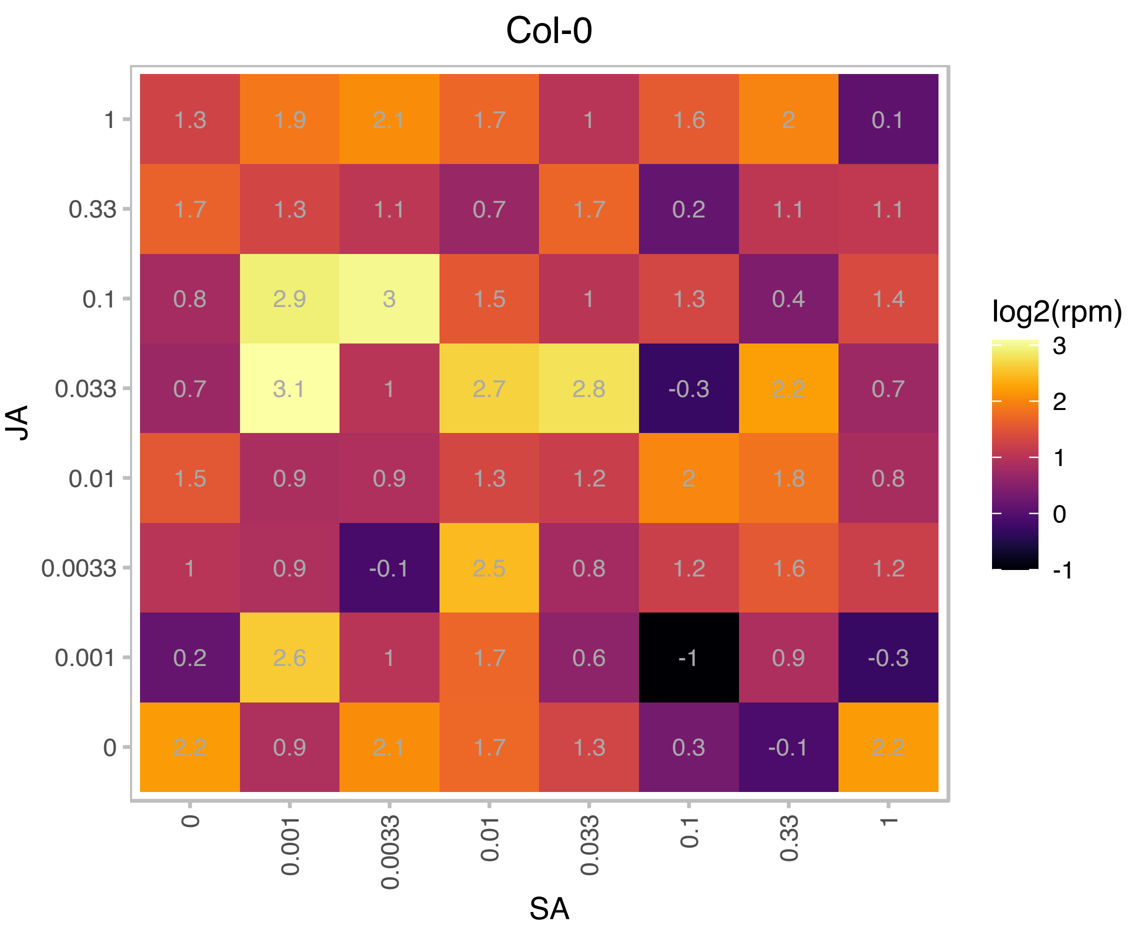


| | |
|---|---|
| Gene | AT3G03550 |
| is.expressed_genes | TRUE |
| is.responsive_genes | TRUE |
| is.apcluster | E-02 |

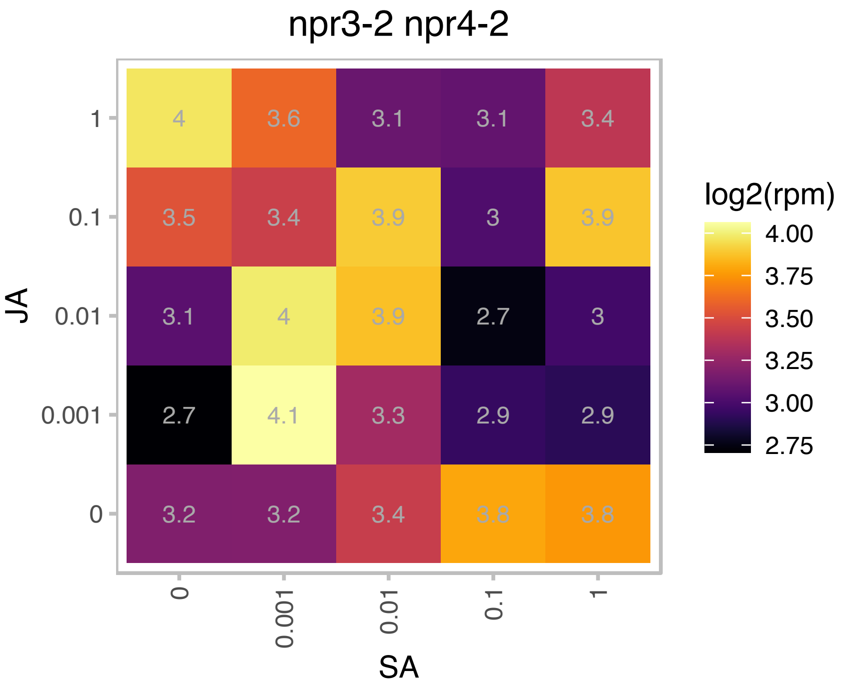


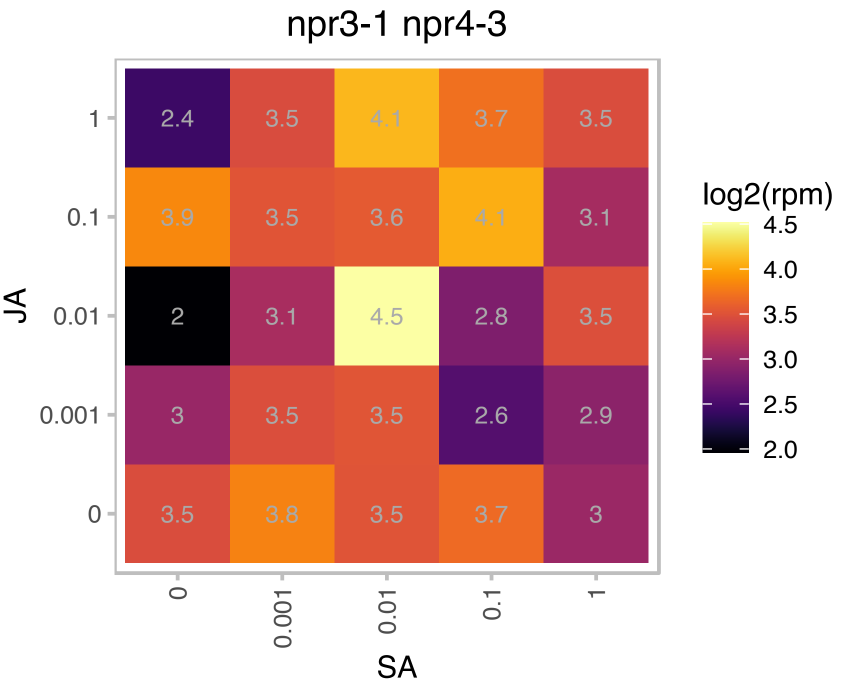

### AT3G04910.1 : ATWNK1, WNK1, ZIK4

with no lysine (K) kinase 1

Serine/threonine protein kinase, whose transcription is regulated by circadian rhythm.

Col-0

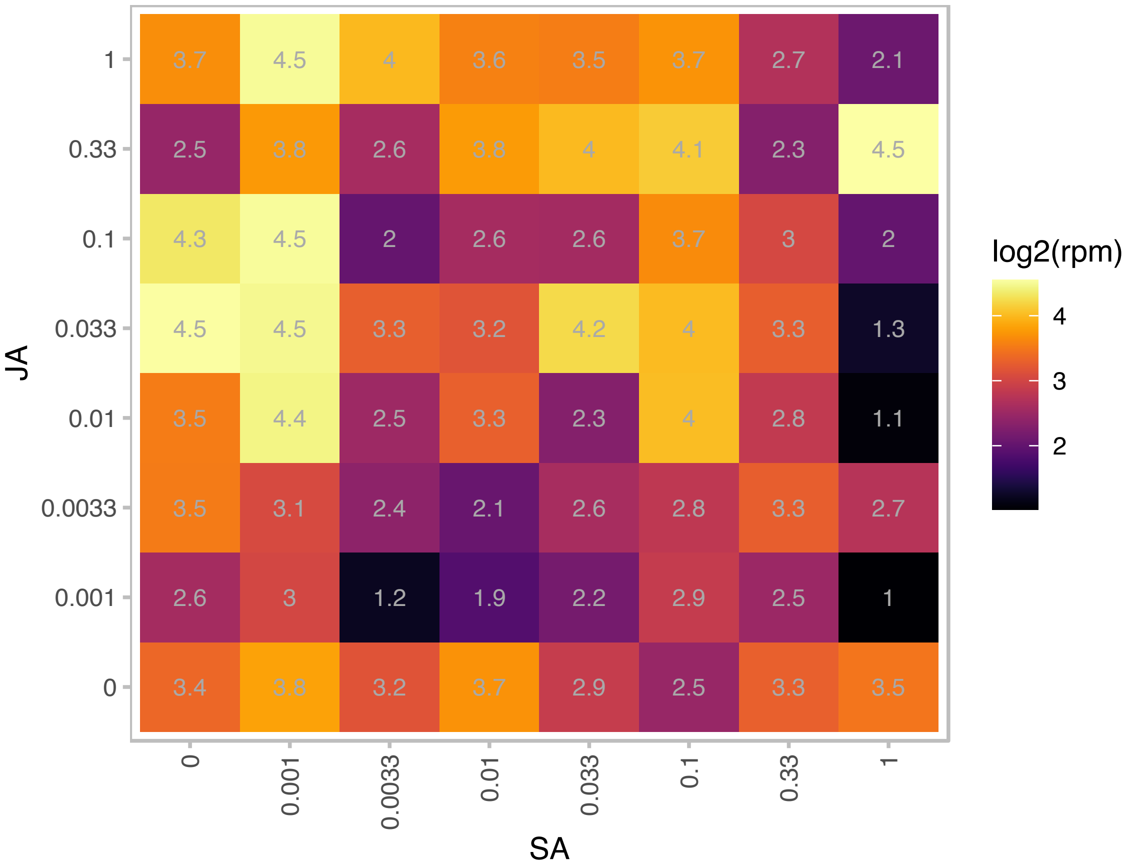


| | |
|---|---|
| Gene | AT3G04910 |
| is.expressed_genes | TRUE |
| is.responsive_genes | TRUE |
| is.apcluster | E-02 |

npr3-2 npr4-2

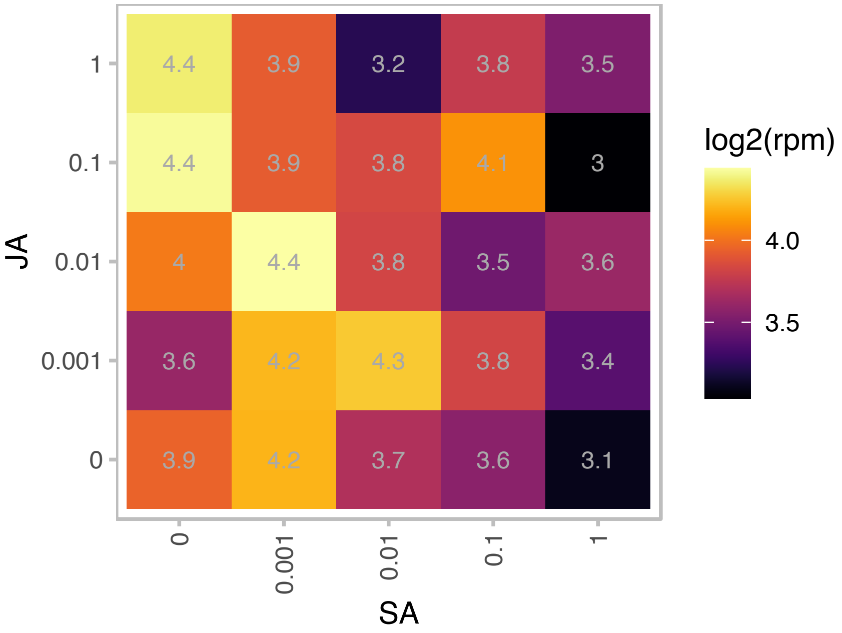


npr3-1 npr4-3

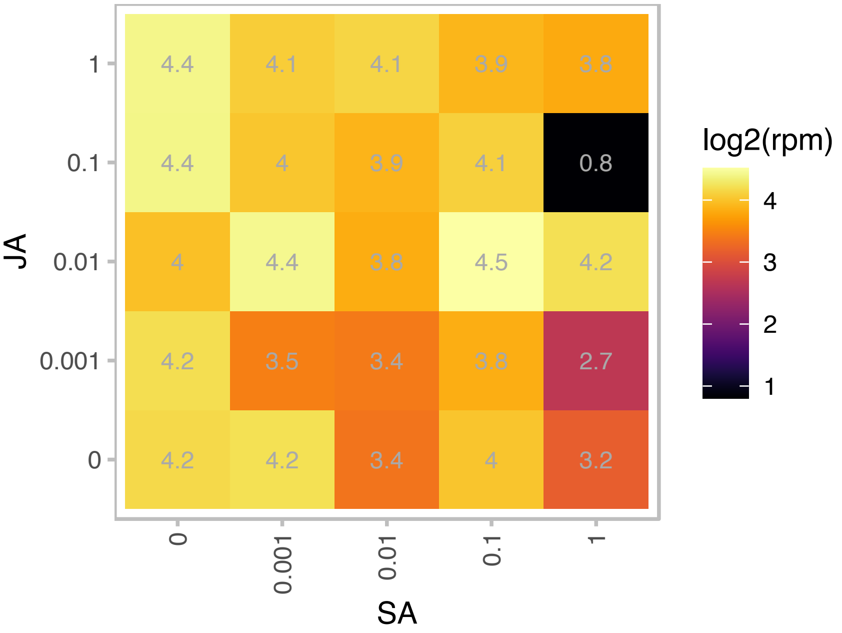

### AT3G09260.1 : BGLU23, LEB, PSR3.1, PYK10

Glycosyl hydrolase superfamily protein

Encodes beta-glucosidase. The major constituent of ER bodies. One of the most abundant proteins in Arabidopsis seedlings. Exist in an soluble (inactive) and non-soluble (active) form, most probably formed in a polymerization process. Involved in the mutualistic interaction between Arabidopsis and the endophytic fungus Piriformospora indica.

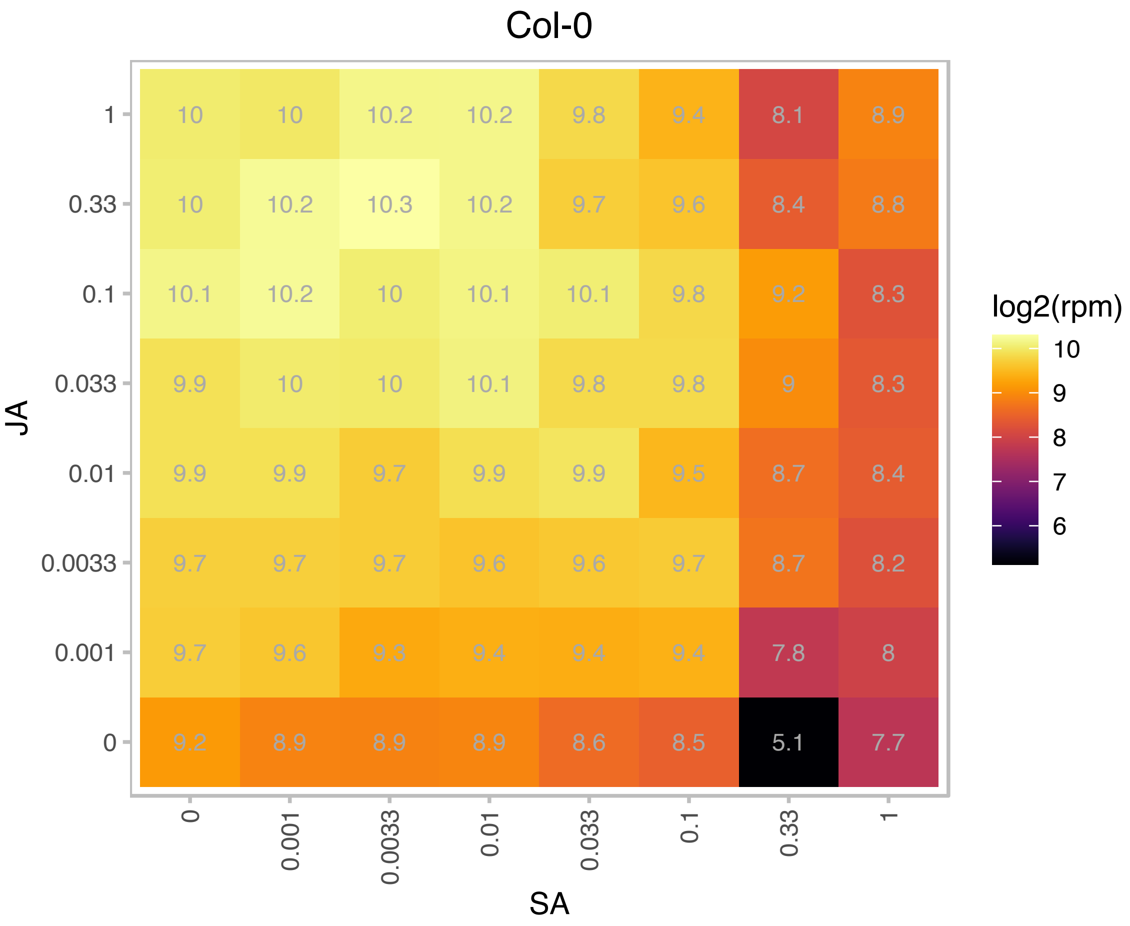


| | |
|---|---|
| Gene | AT3G09260 |
| is.expressed_genes | TRUE |
| is.responsive_genes | TRUE |
| is.apcluster | E-02 |

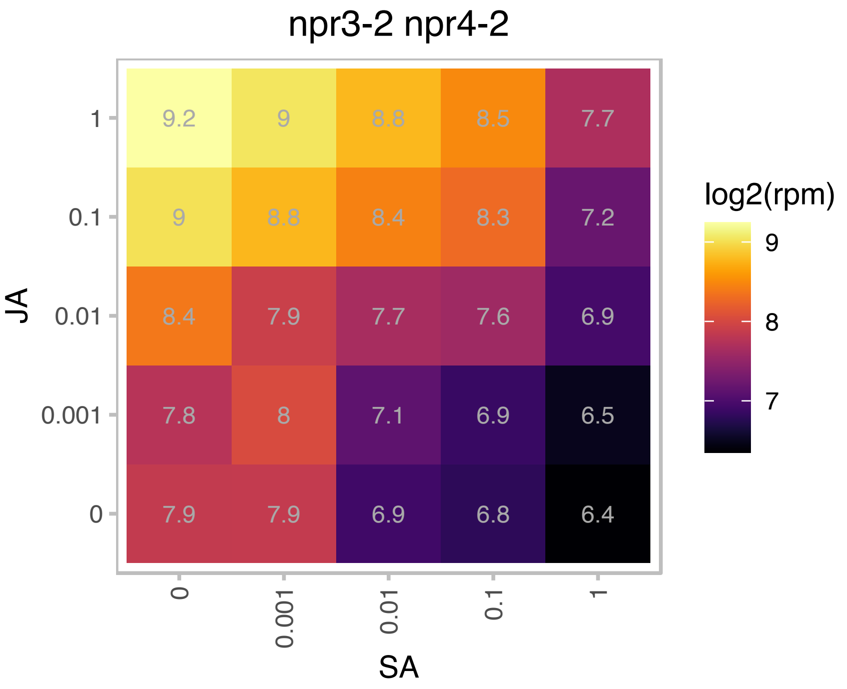


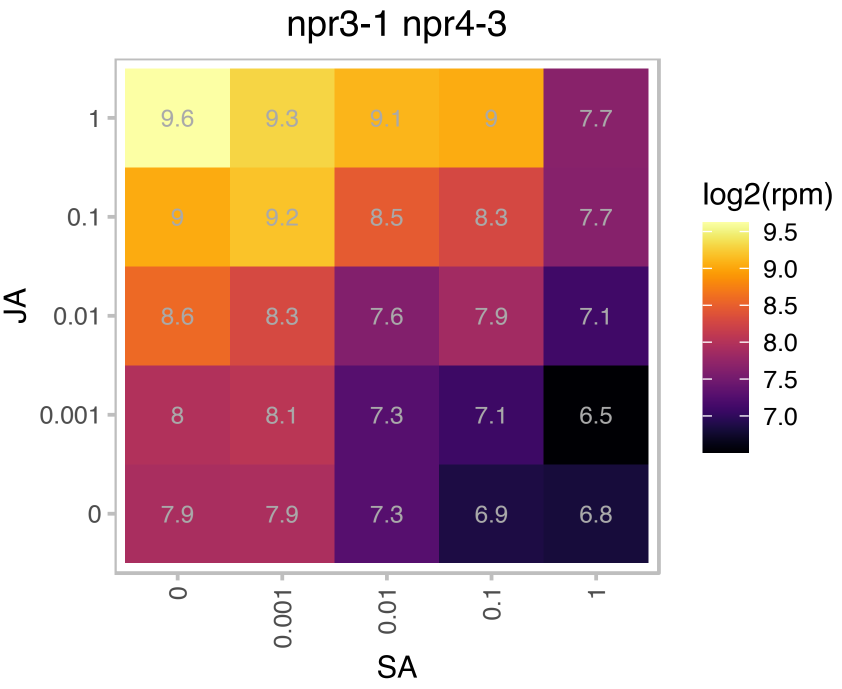

### AT3G15660.1 : ATGRX4, GRX4, GRXS15

glutaredoxin 4

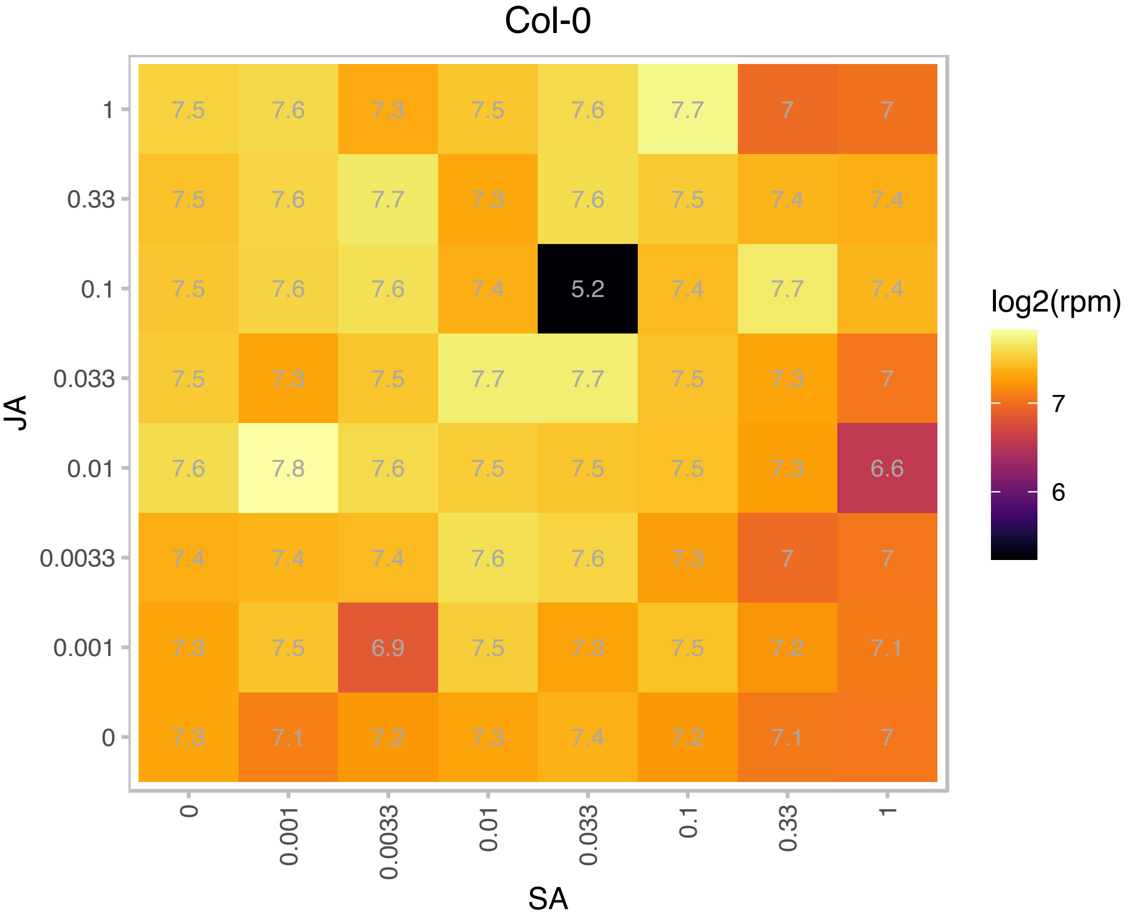


| | |
|---|---|
| Gene | AT3G15660 |
| is.expressed_genes | TRUE |
| is.responsive_genes | TRUE |
| is.apcluster | E-02 |

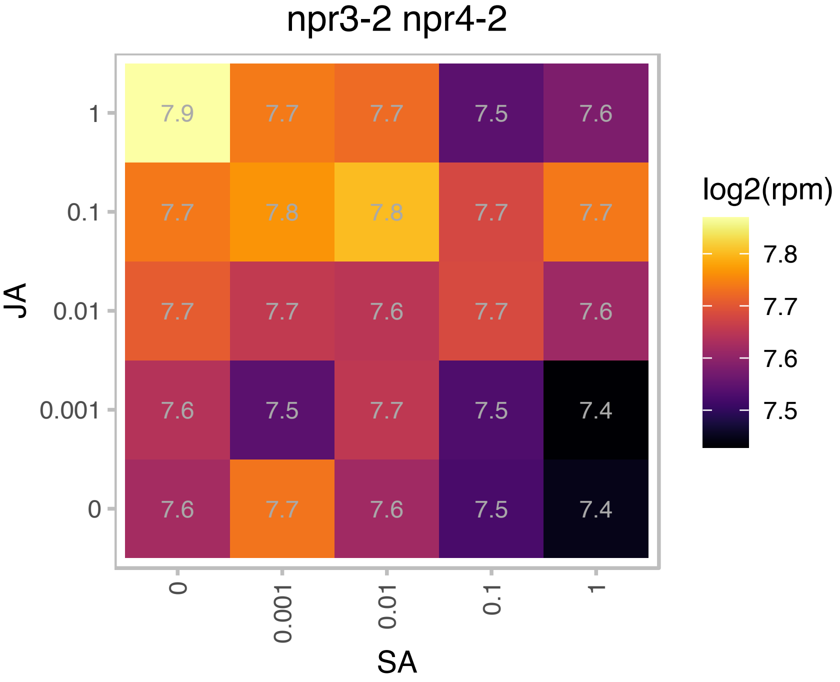


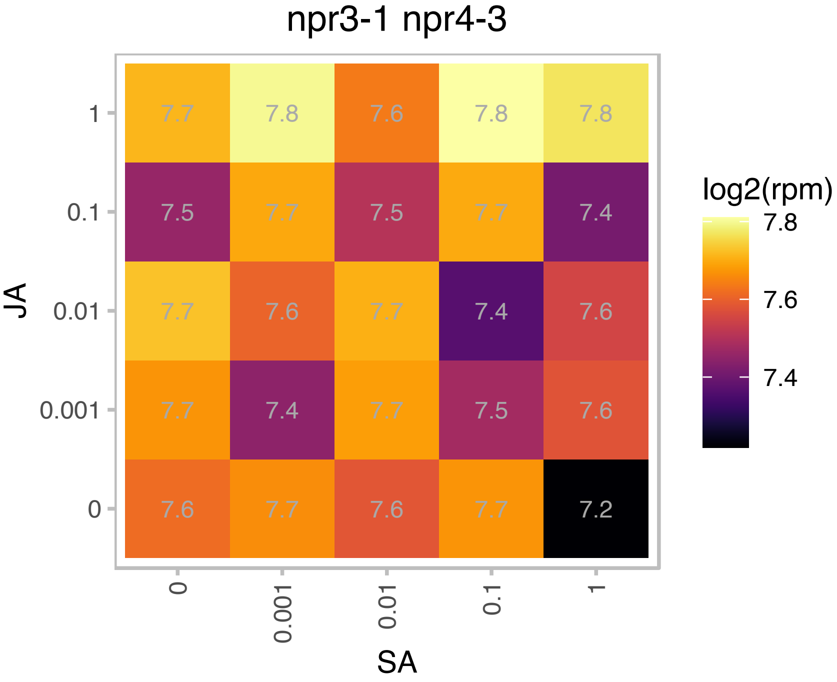

### AT3G15950.1 : NAI2

DNA topoisomerase-related
Similar to TSK-associating protein 1 (TSA1), contains 10 EFE repeats, a novel repeat sequence unique to plants.  Expressed preferentially in the roots.Protein is localized to ER bodies- an endoplasmic reticulum derived structure. Loss of function mutations lack ER bodies.

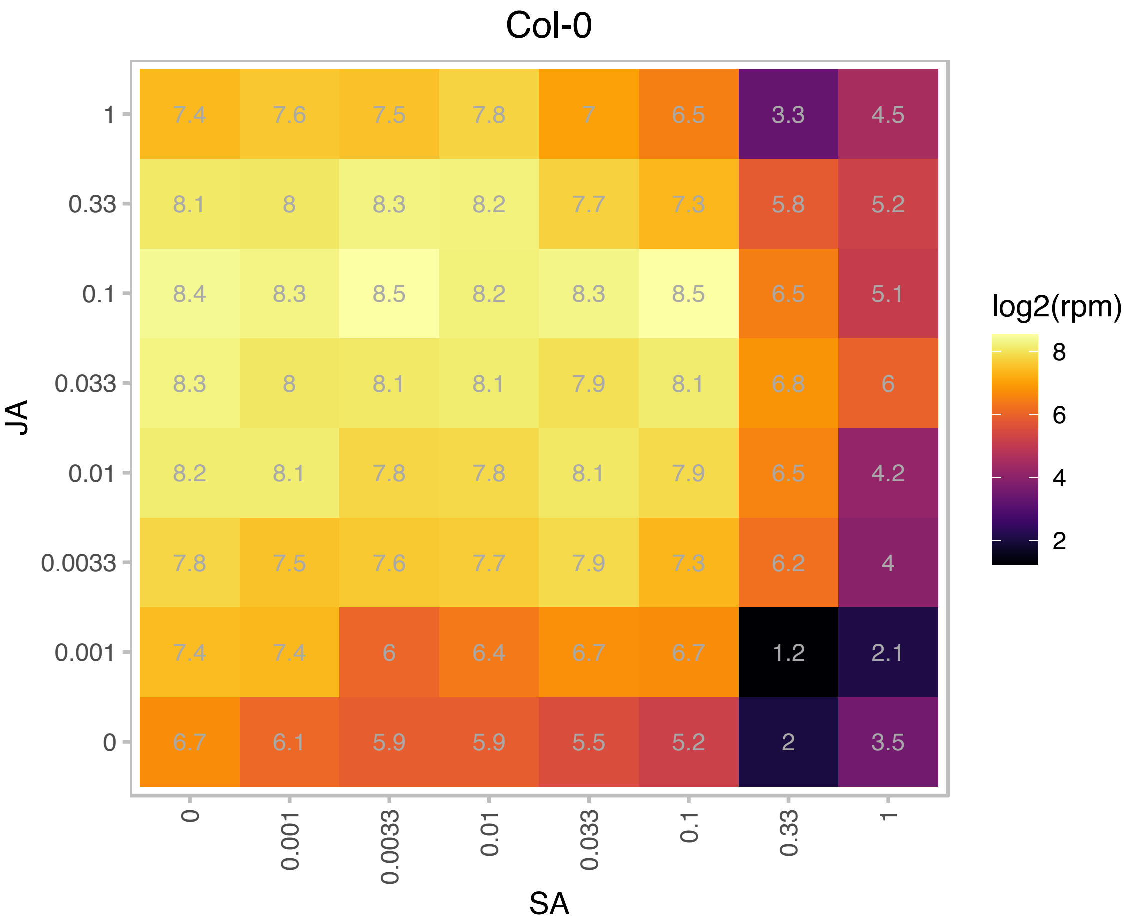


| | |
|---|---|
| Gene | AT3G15950 |
| is.expressed_genes | TRUE |
| is.responsive_genes | TRUE |
| is.apcluster | E-02 |

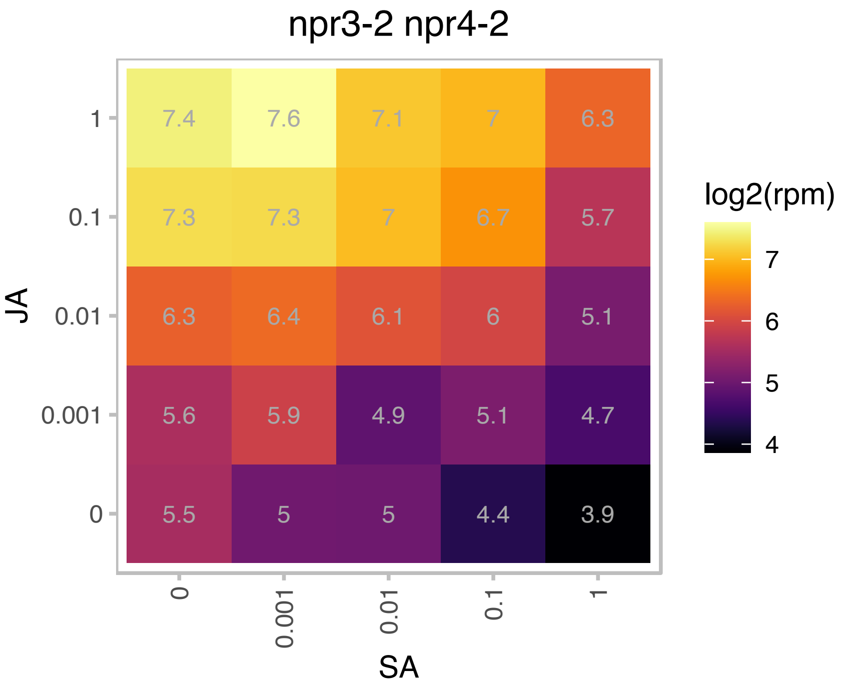


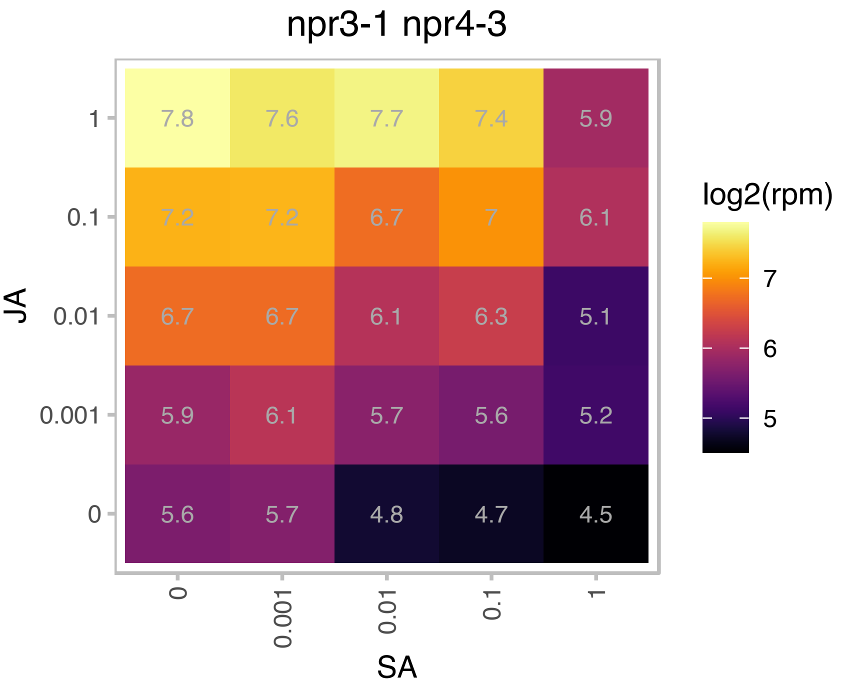

### AT3G16400.1 : ATMLP-470, ATNSP1, NSP1

nitrile specifier protein 1

Encodes a nitrile-specifier protein NSP1 responsible for constitutive and herbivore-induced simple nitrile formation in rosette leaves. NSP1 is one out of five (At3g16400/NSP1, At2g33070/NSP2, At3g16390/NSP3, At3g16410/NSP4 and At5g48180/NSP5) A. thaliana epithiospecifier protein (ESP) homologues that promote simple nitrile, but not epithionitrile or thiocyanate formation. The mRNA is cell-to-cell mobile.

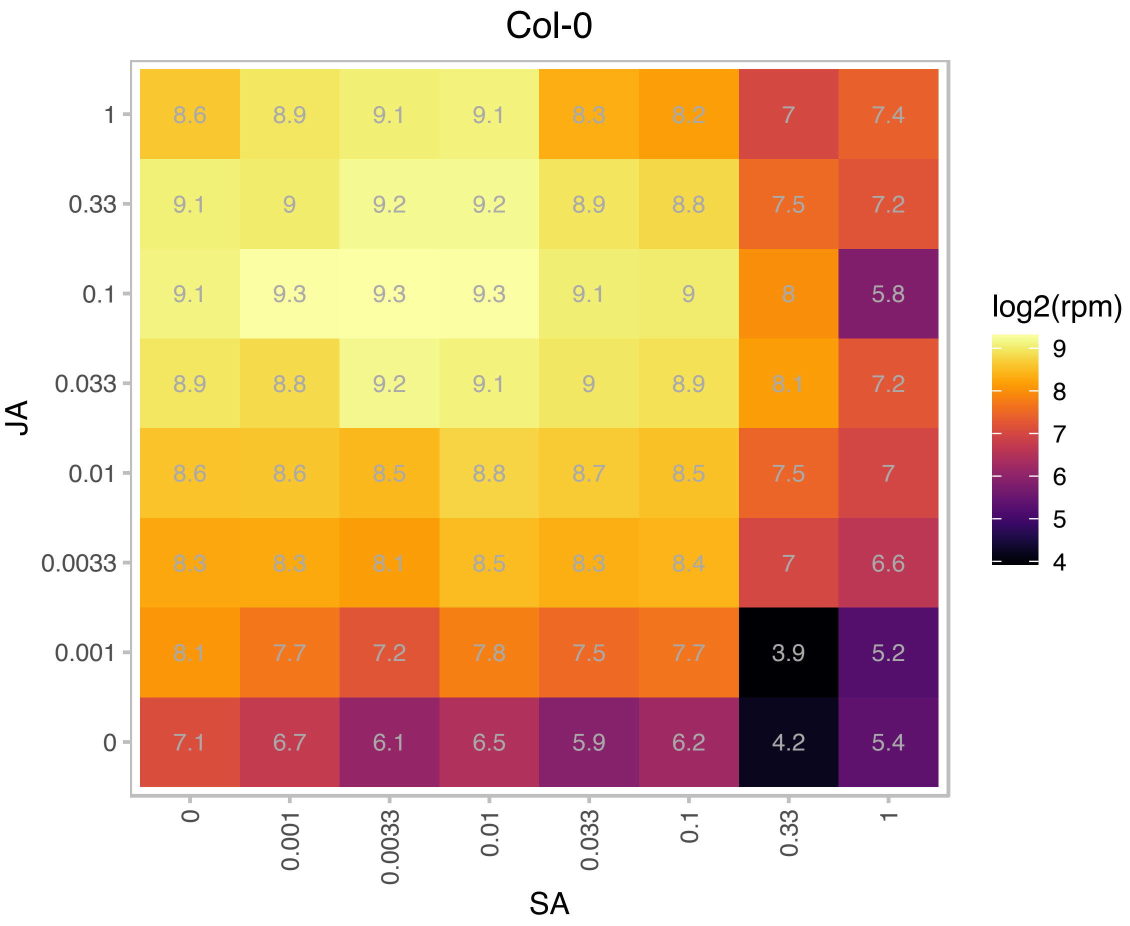


| | |
|---|---|
| Gene | AT3G16400 |
| is.expressed_genes | TRUE |
| is.responsive_genes | TRUE |
| is.apcluster | E-02 |

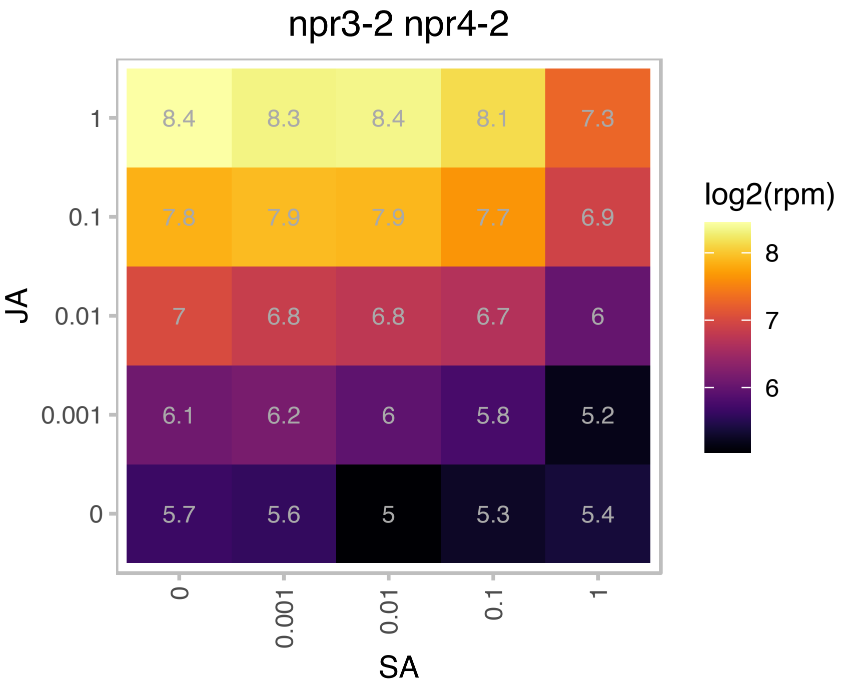


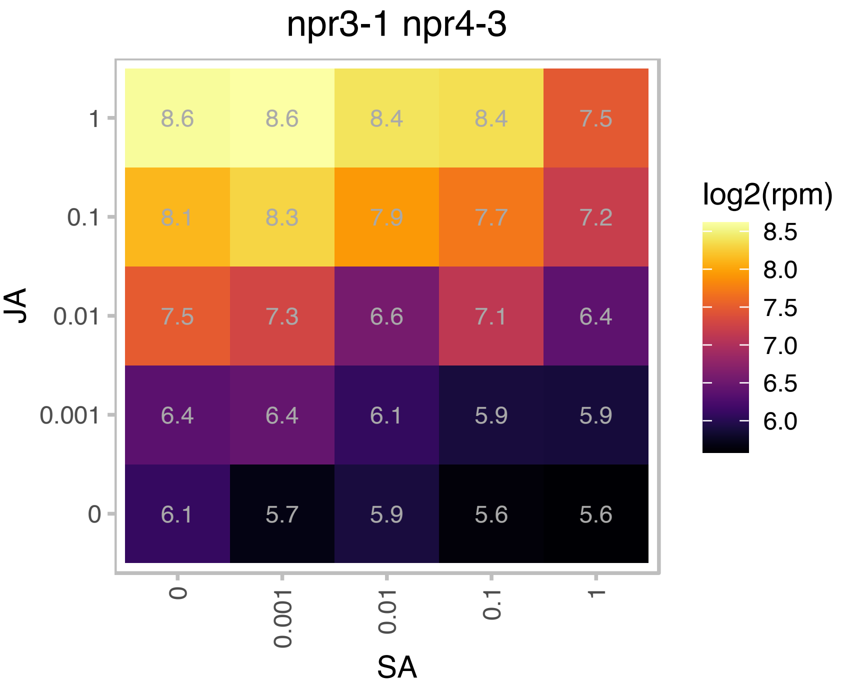

### AT3G16420.1 : JAL30, PBP1, PBPI

PYK10-binding protein 1
The PBP1(PYK10-binding protein 1) assists the PYK10 (beta-glucosidase complex) in its activity and may act like a molecular chaperone that facilitates the correct polymerization of PYK10, when tissues are damaged and subcellular structures are destroyed by pests. The mRNA is cell-to-cell mobile.

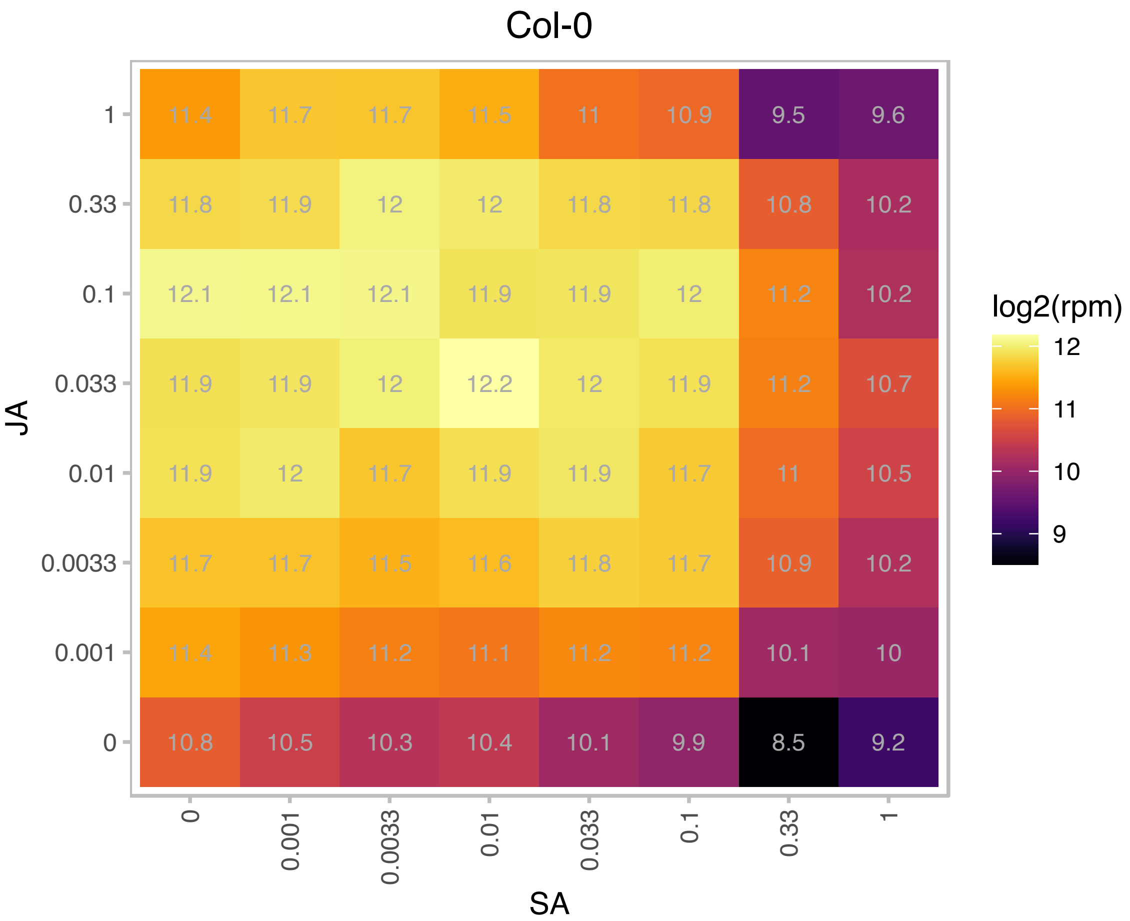


| | |
|---|---|
| Gene | AT3G16420 |
| is.expressed_genes | TRUE |
| is.responsive_genes | TRUE |
| is.apcluster | E-02 |

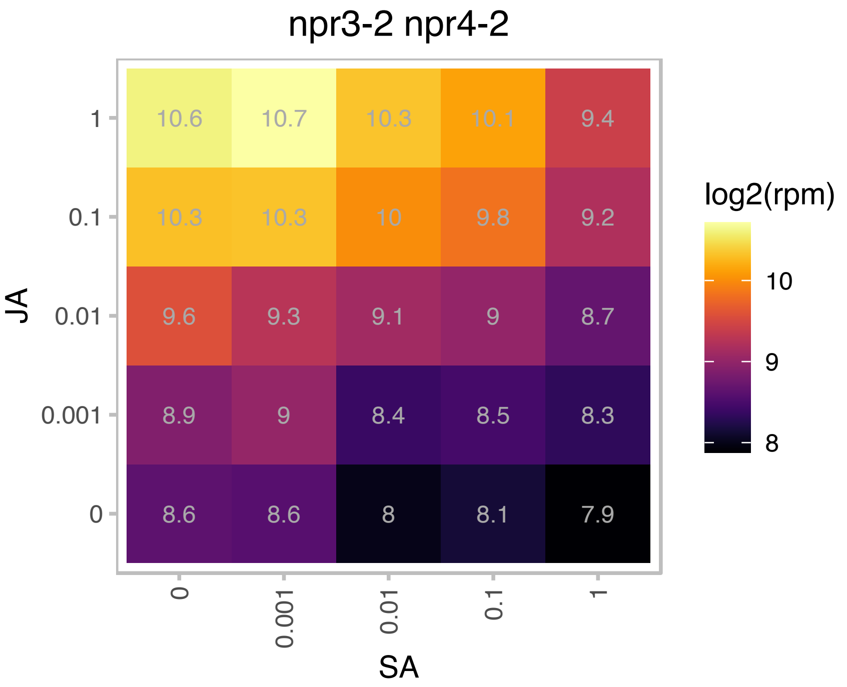


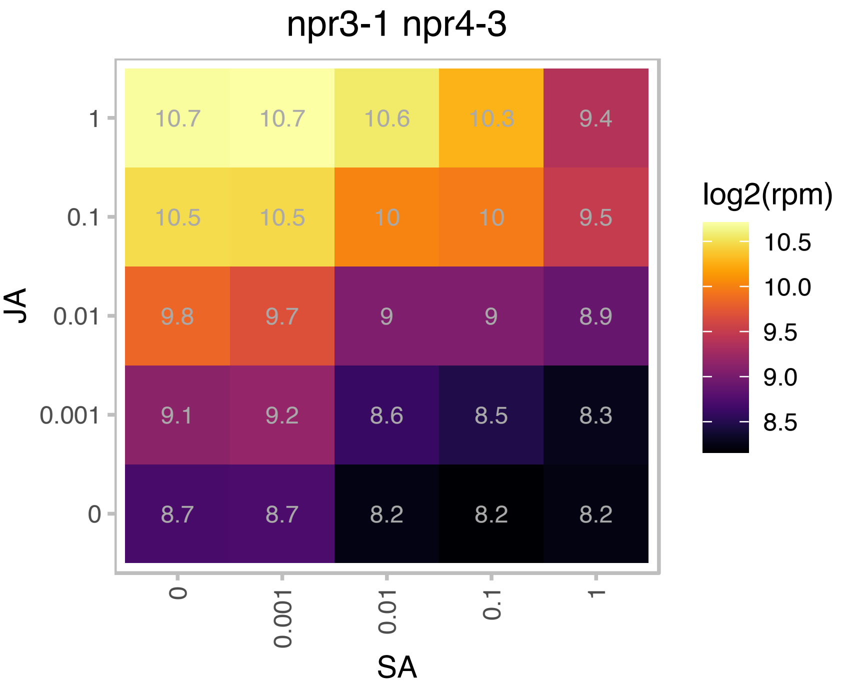

### AT3G16430.1 : JAL31, PBPII

jacalin-related lectin 31

Encodes a protein that increases the beta-glucosidase activities of three scopolin glucosidases in vitro.

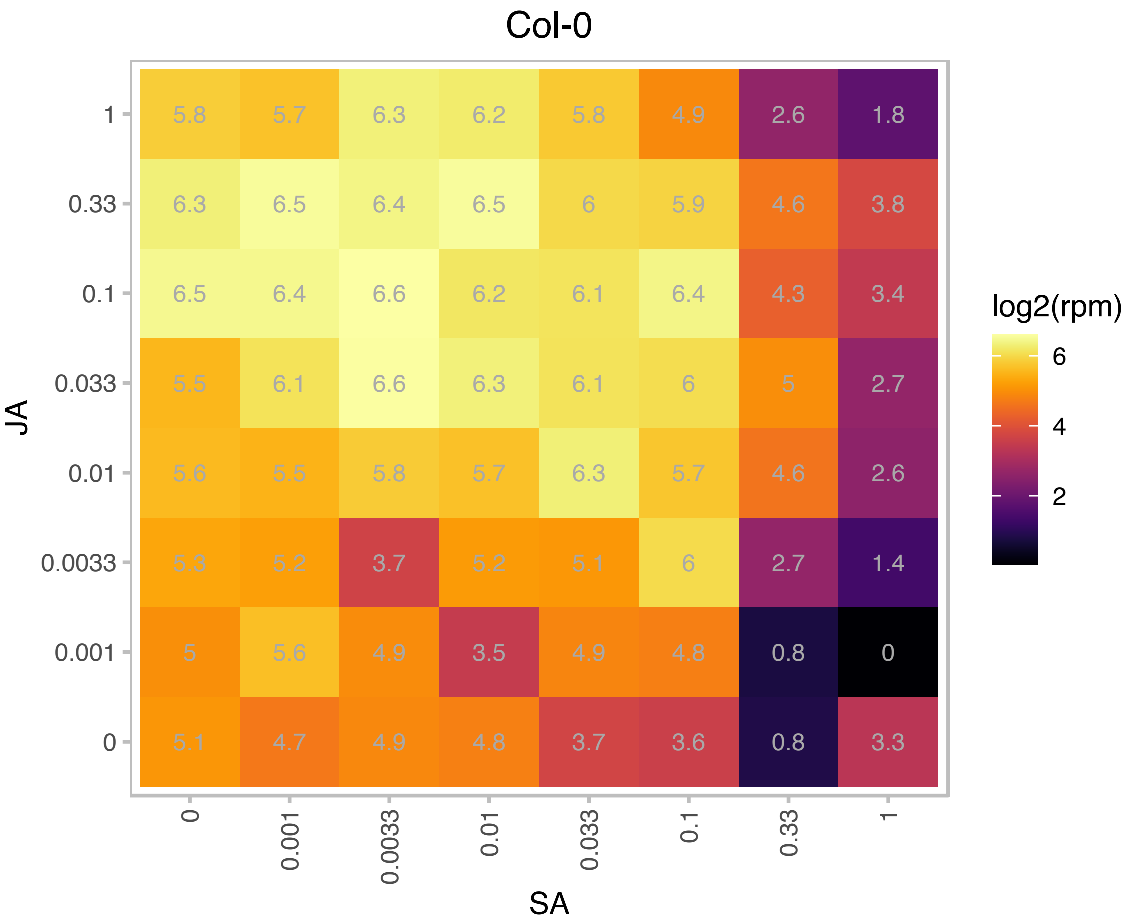


| | |
|---|---|
| Gene | AT3G16430 |
| is.expressed_genes | TRUE |
| is.responsive_genes | TRUE |
| is.apcluster | E-02 |

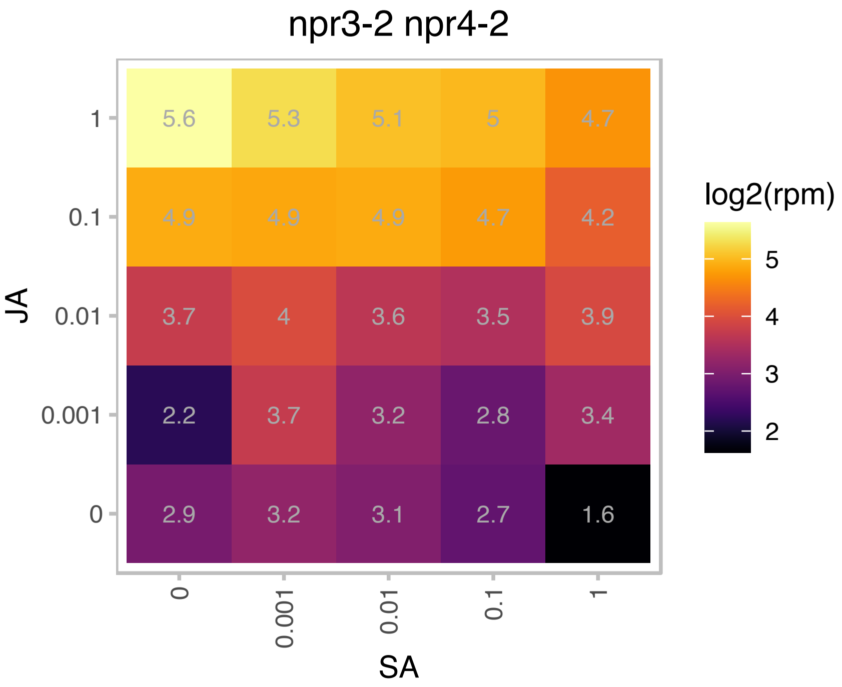


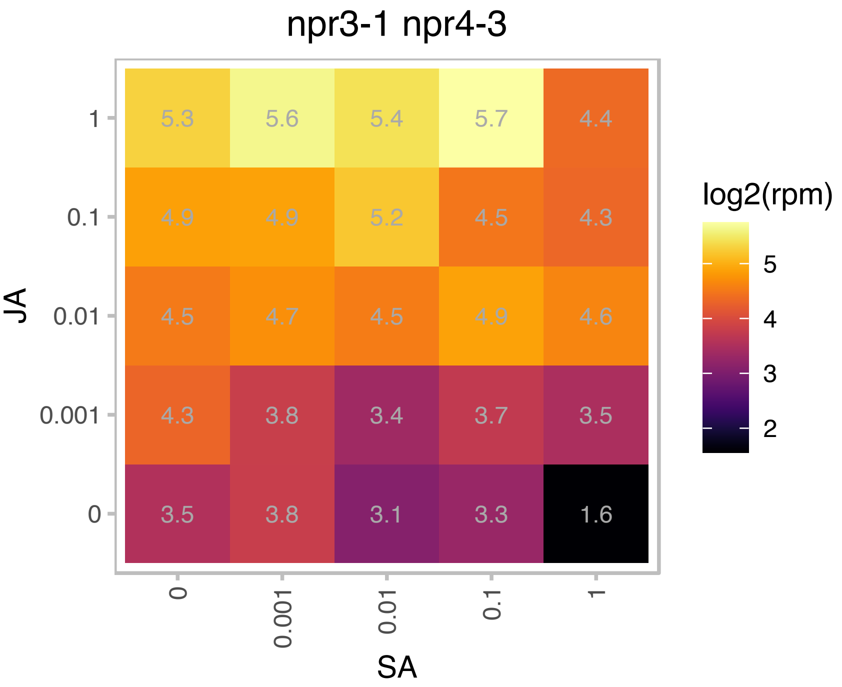

### AT3G16450.1 : JAL33

Mannose-binding lectin superfamily protein

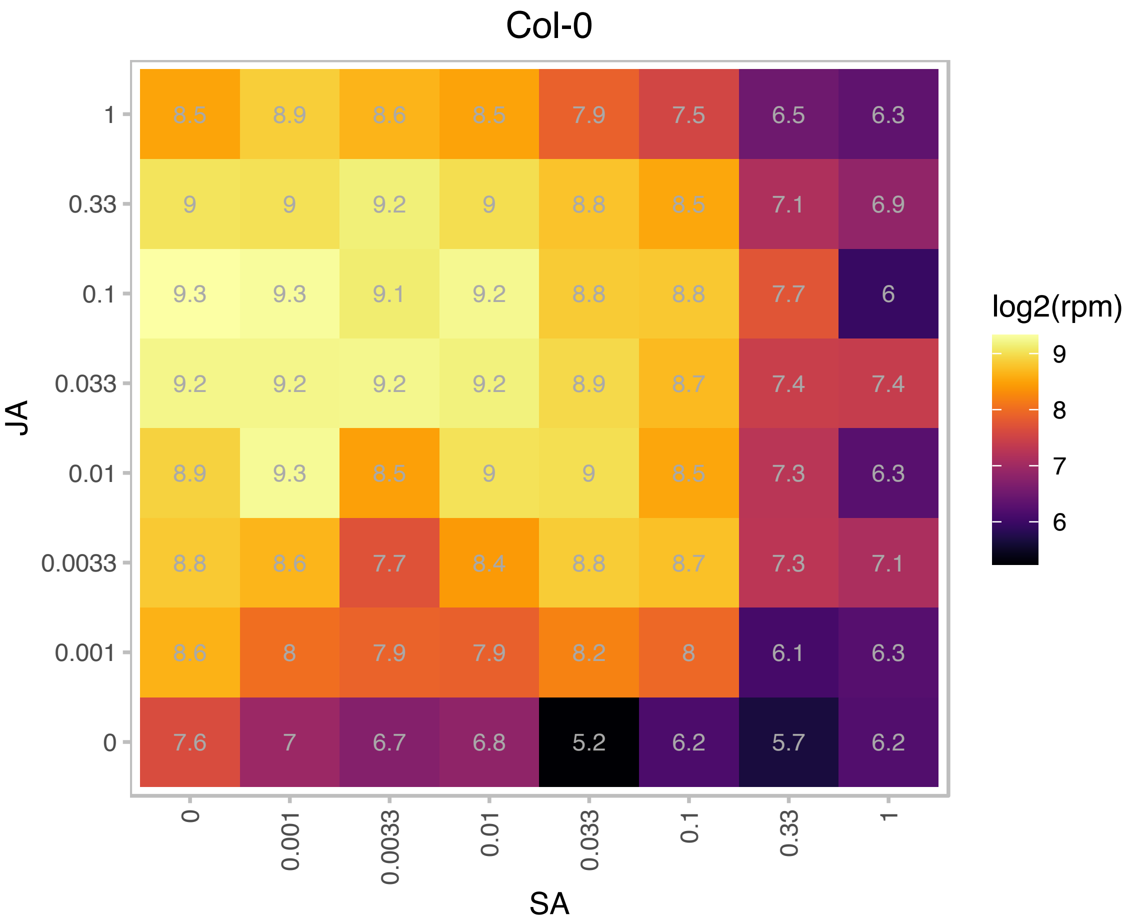


| | |
|---|---|
| Gene | AT3G16450 |
| is.expressed_genes | TRUE |
| is.responsive_genes | TRUE |
| is.apcluster | E-02 |

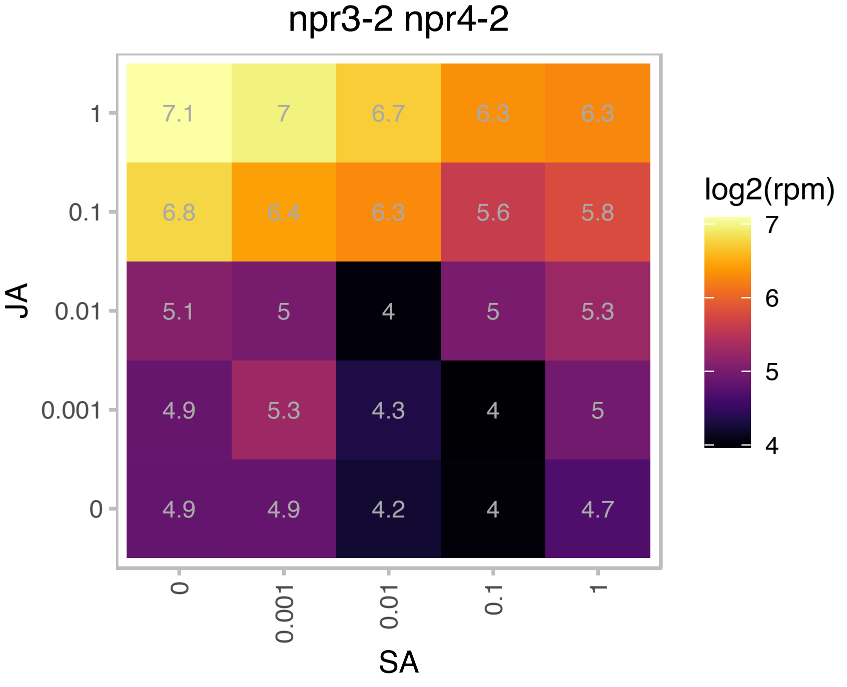


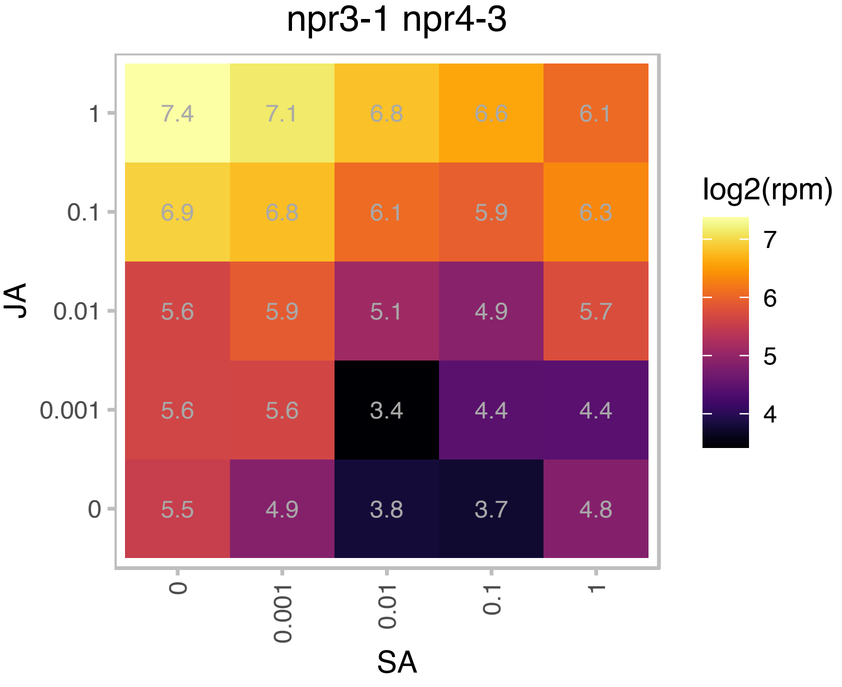

# AT3G21770.1 : NA

Peroxidase superfamily protein

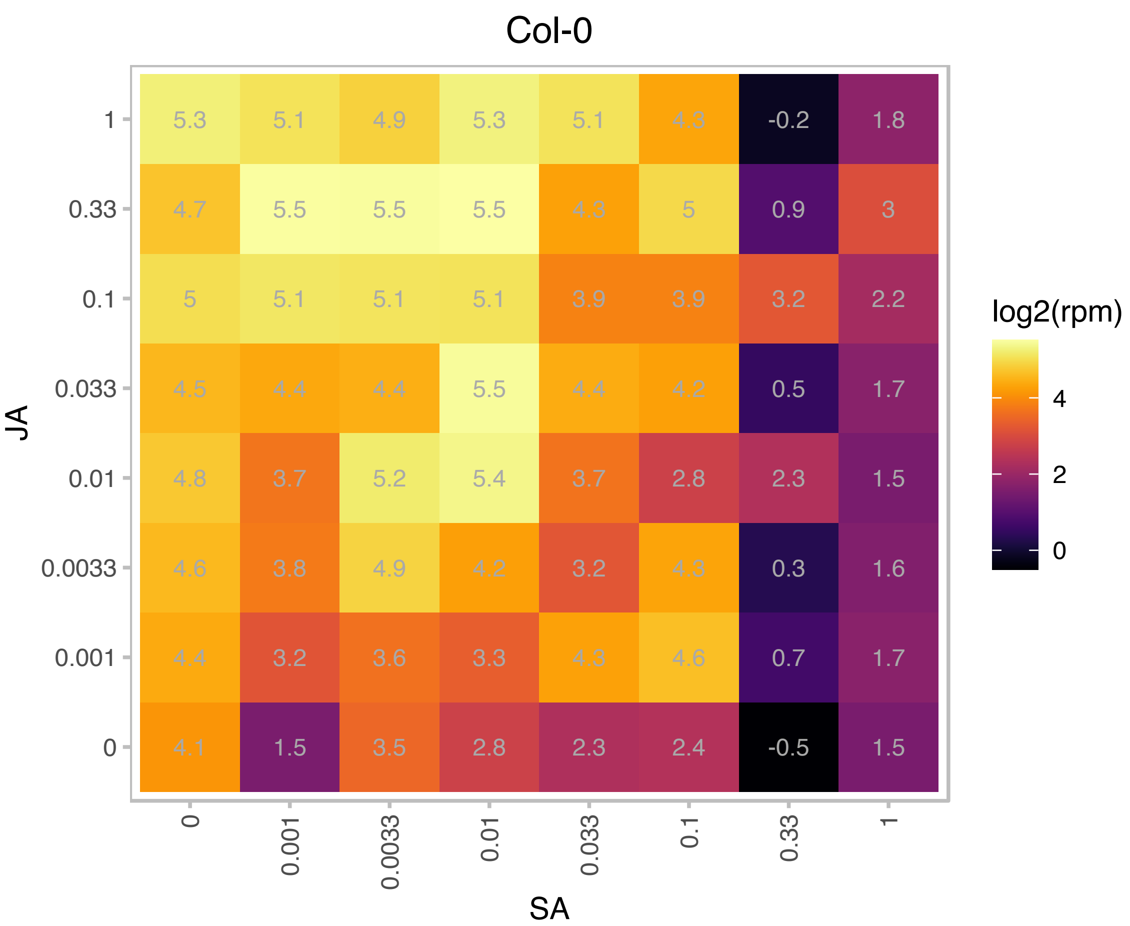


| | |
|---|---|
| Gene | AT3G21770 |
| is.expressed_genes | TRUE |
| is.responsive_genes | TRUE |
| is.apcluster | E-02 |

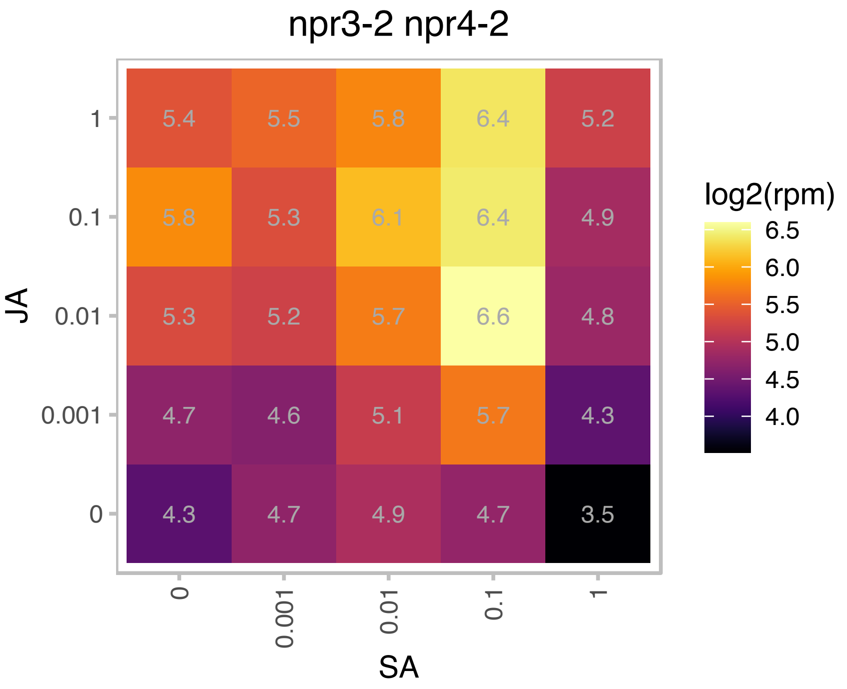


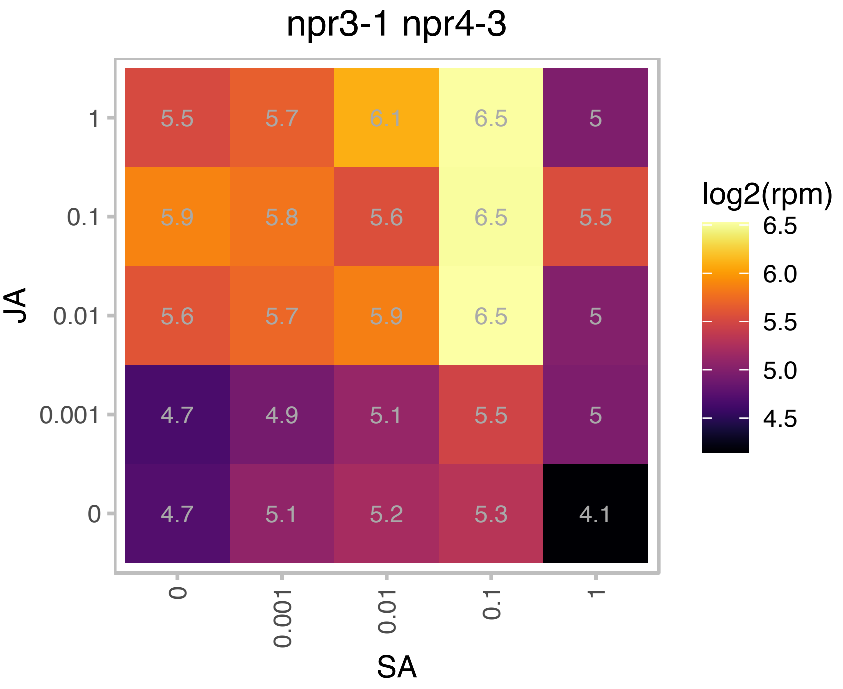

### AT3G22200.1 : GABA-T, HER1, POP2

Genetically redundant with POP3;mediates pollen tube guidance. Double mutants are self sterile; gamma-aminobutyrate transaminase subunit precursor; nuclear gene for mitochondrial product. Encodes gamma-aminobutyrate transaminase that uses pyruvate instead of alpha-ketoglutarate as cosubstrate. Mutations in POP2/HER1 render roots resistant to the inhibitory growth effects of the volatile organic compound E-2-hexenal implicated in plant defense.

Pyridoxal phosphate (PLP)-dependent transferases superfamily protein

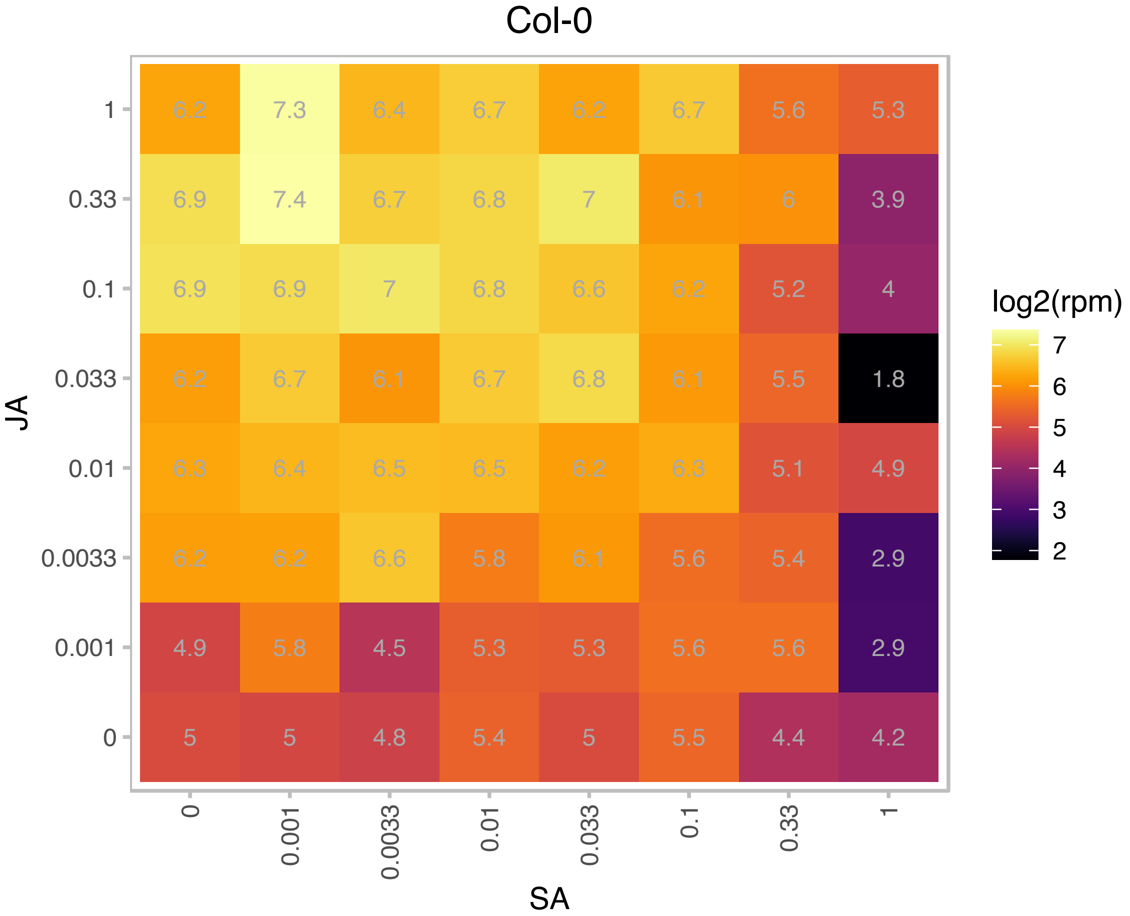


| | |
|---|---|
| Gene | AT3G22200 |
| is.expressed_genes | TRUE |
| is.responsive_genes | TRUE |
| is.apcluster | E-02 |

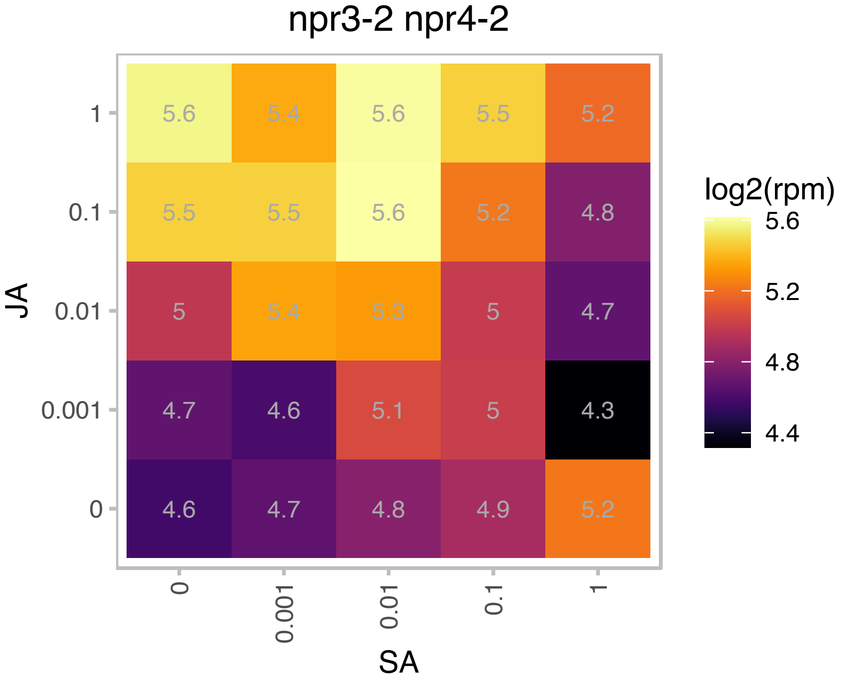


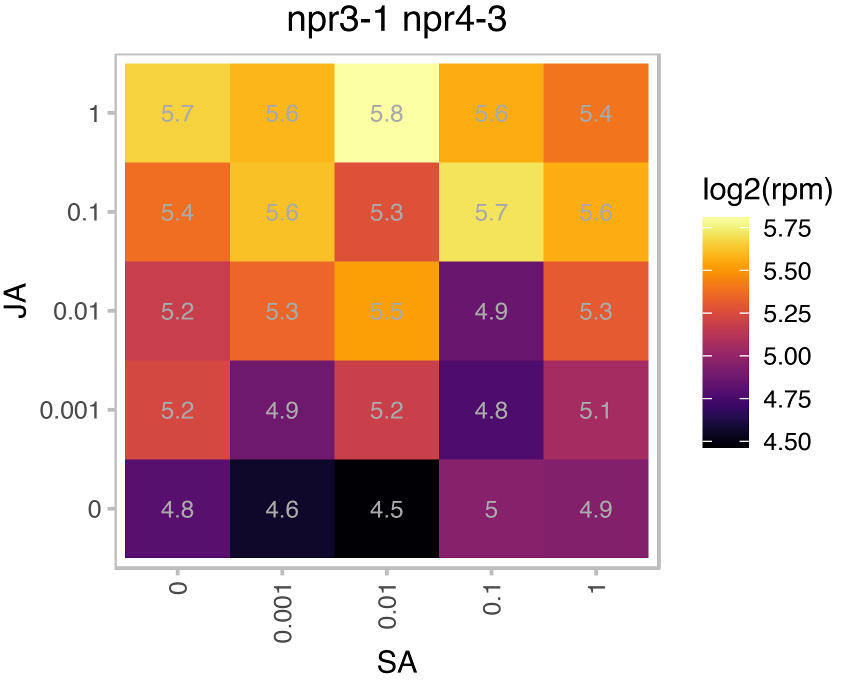

### AT3G23690.1 : CIL2

basic helix-loop-helix (bHLH) DNA-binding superfamily protein

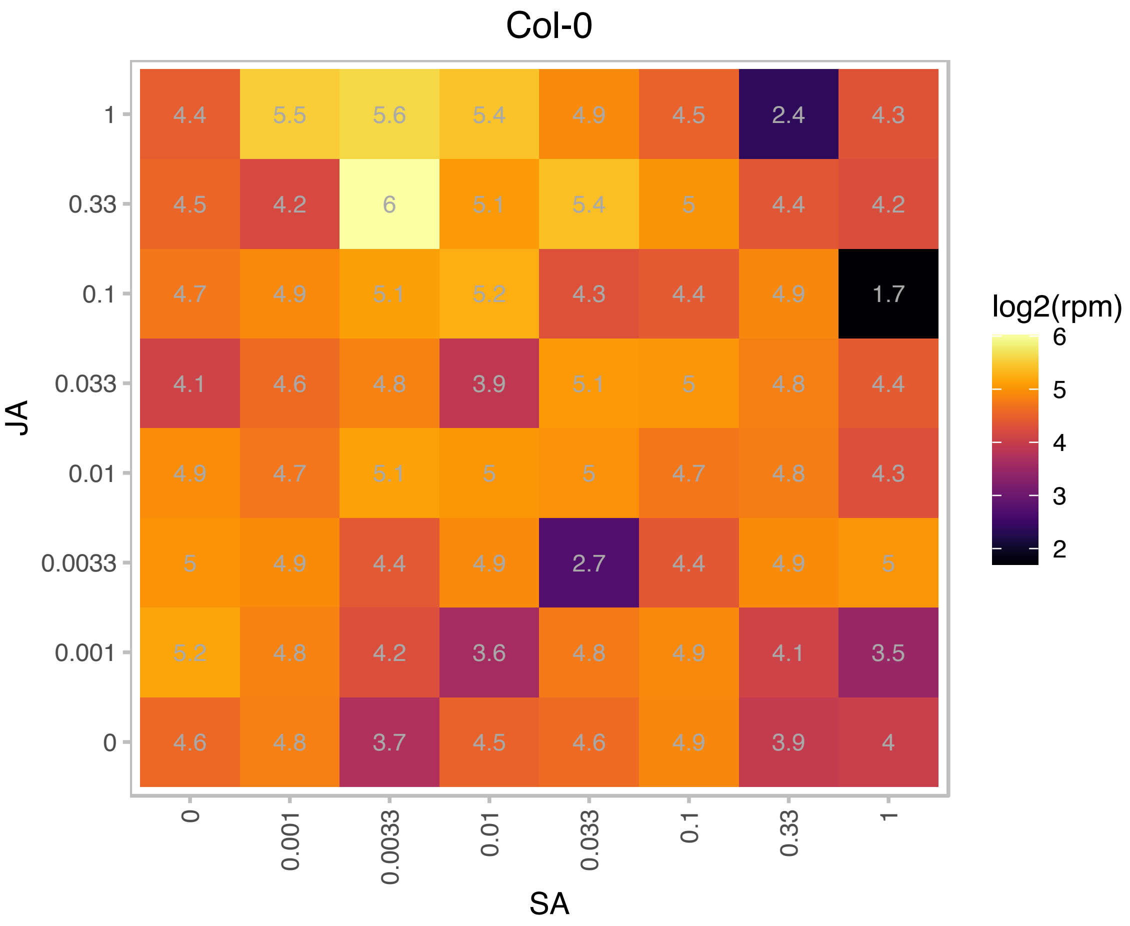


| | |
|---|---|
| Gene | AT3G23690 |
| is.expressed_genes | TRUE |
| is.responsive_genes | TRUE |
| is.apcluster | E-02 |

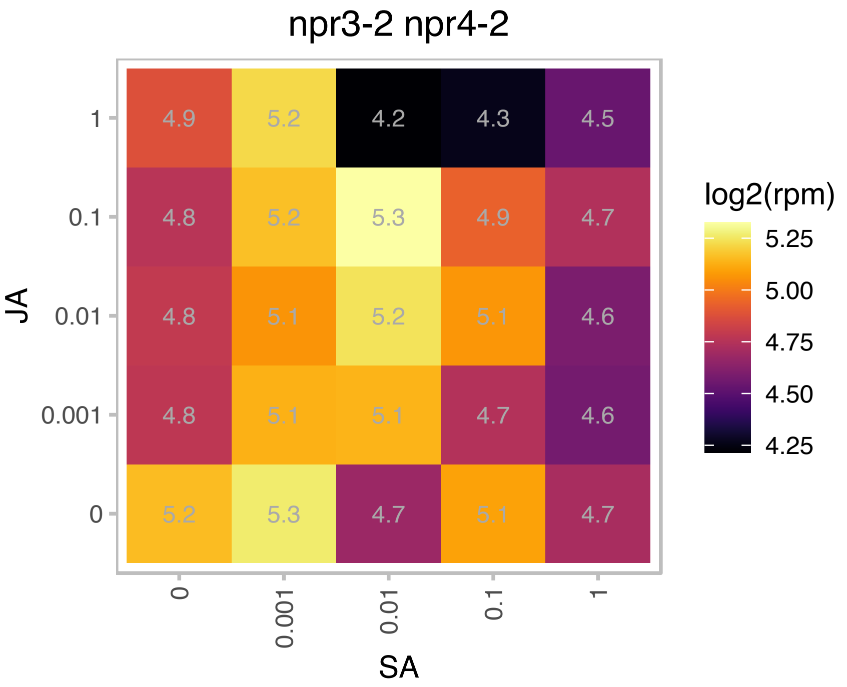


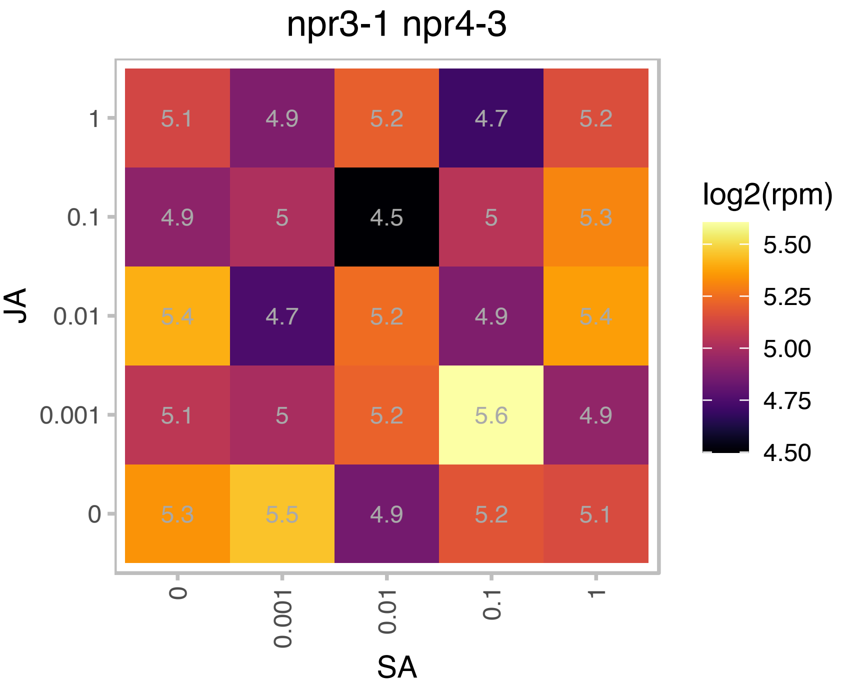

### AT3G25830.1 : ATTPS-CIN, TPS-CIN, TPS-CIN, TPS27

terpene synthase-like sequence-1,8-cineole
Encodes the monoterpene 1,8-cineole synthase, atTPS-Cin. This polypeptide was also shown to synthesize other monoterpenes albeit in minor quantities. The same polypeptide is encoded at two different loci, the result of gene duplication: at3g25820 and at3g25830.

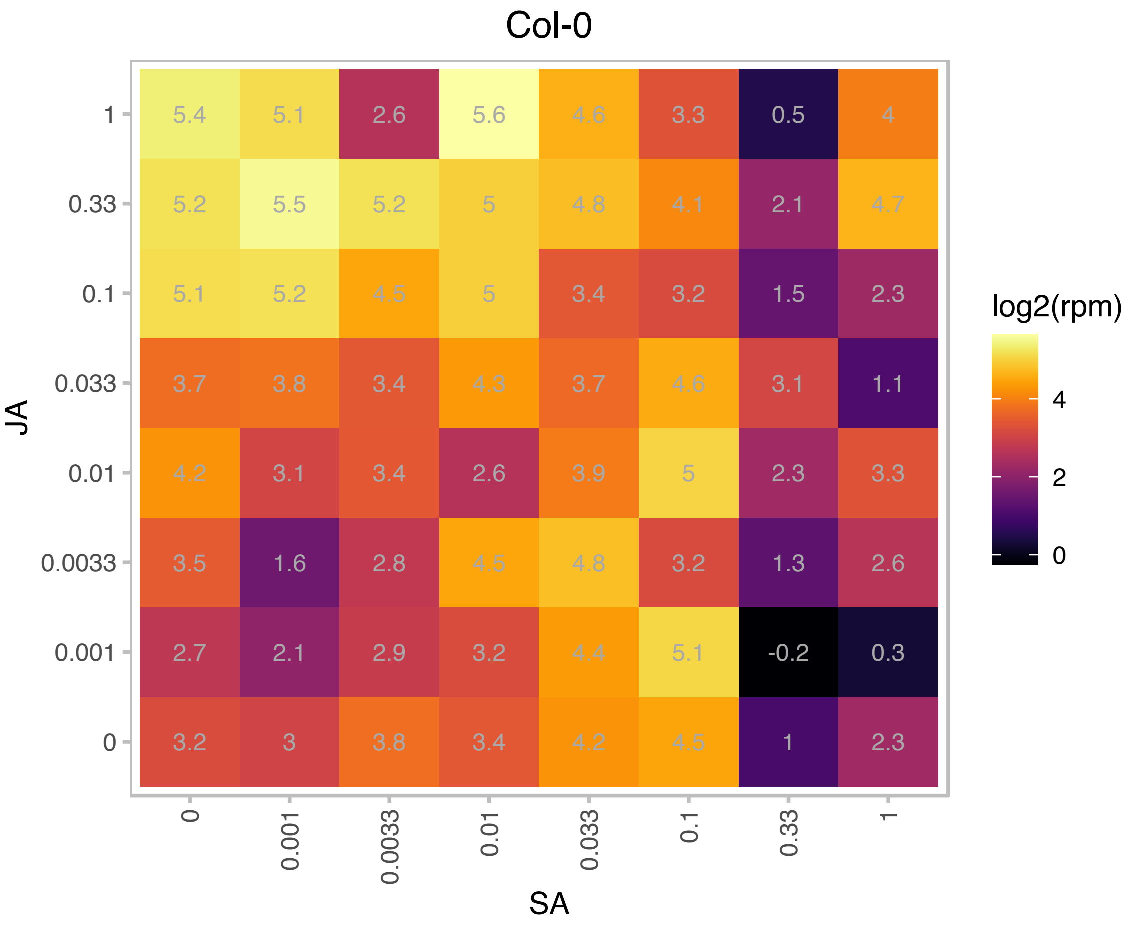


| | |
|---|---|
| Gene | AT3G25830 |
| is.expressed_genes | TRUE |
| is.responsive_genes | TRUE |
| is.apcluster | E-02 |

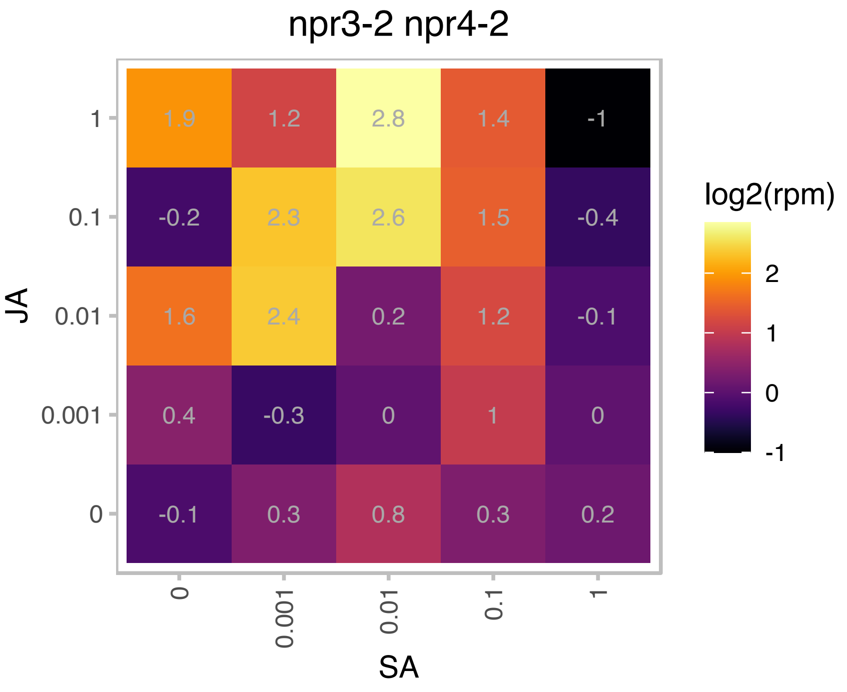


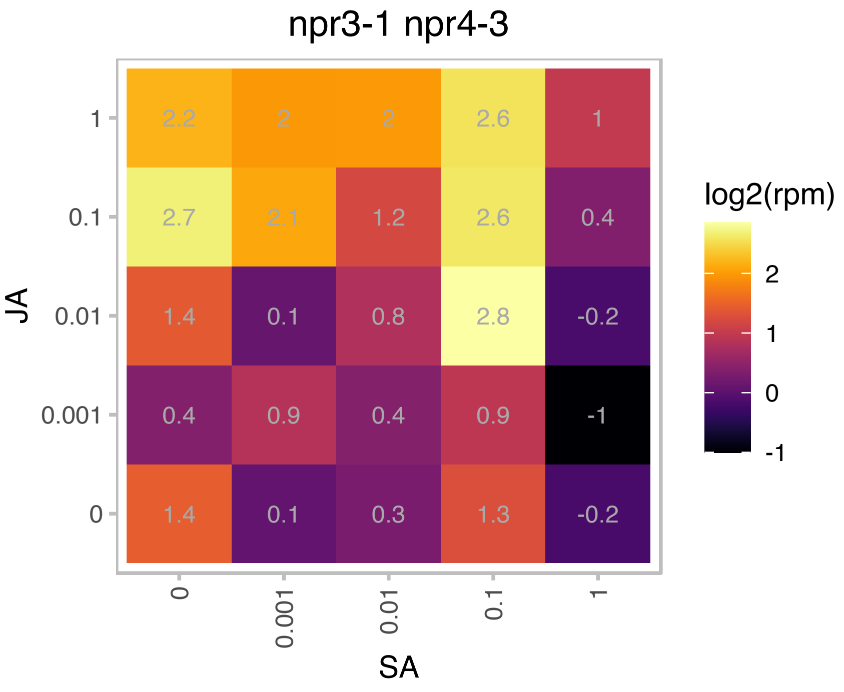

# AT3G26460.1 : NA

Polyketide cyclase/dehydrase and lipid transport superfamily protein

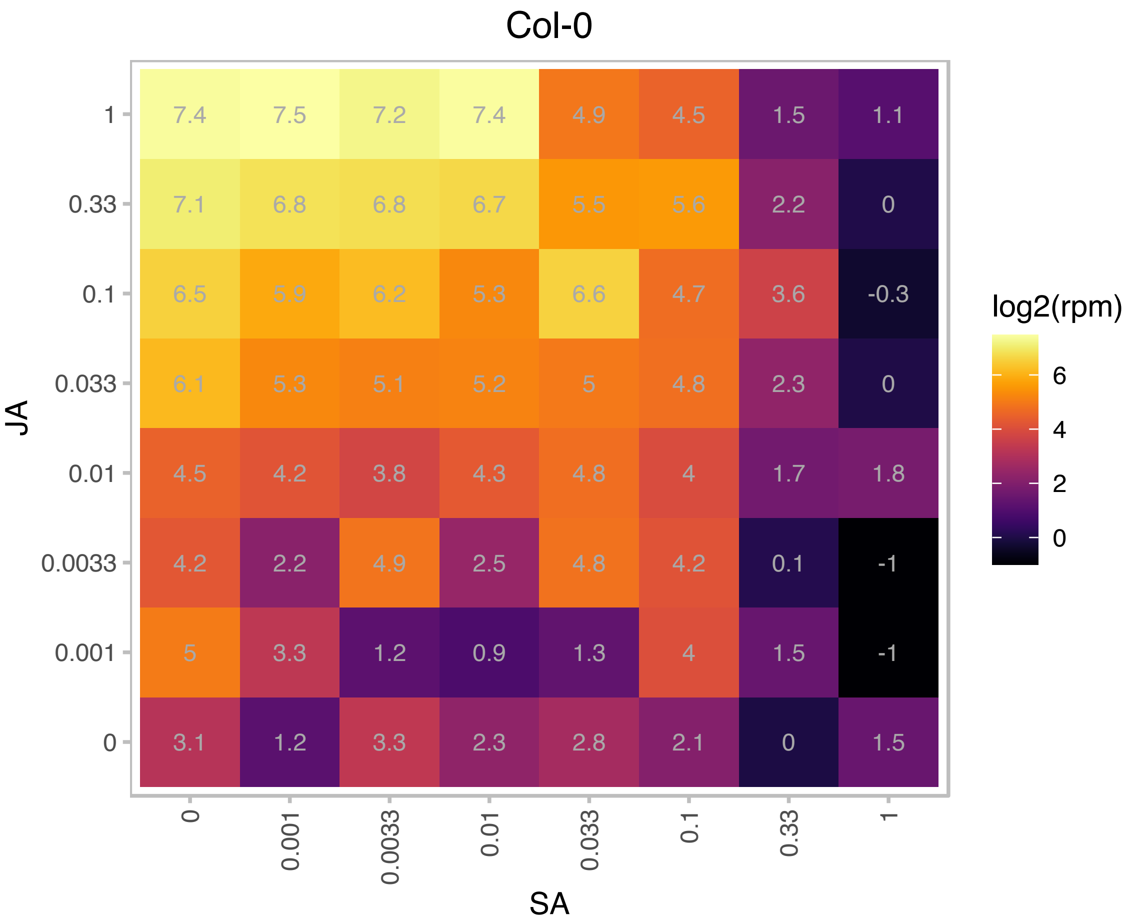


| | |
|---|---|
| Gene | AT3G26460 |
| is.expressed_genes | TRUE |
| is.responsive_genes | TRUE |
| is.apcluster | E-02 |

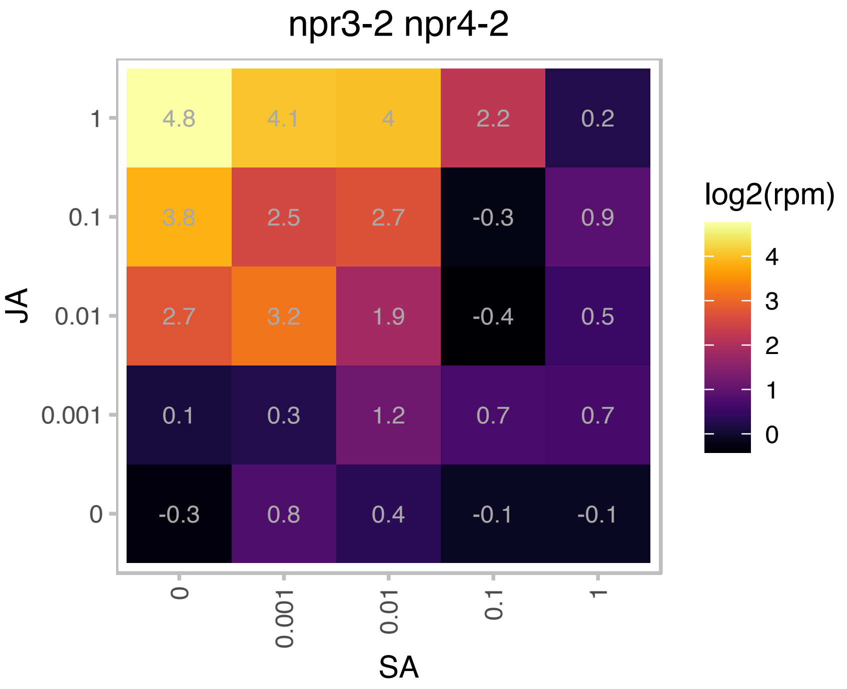


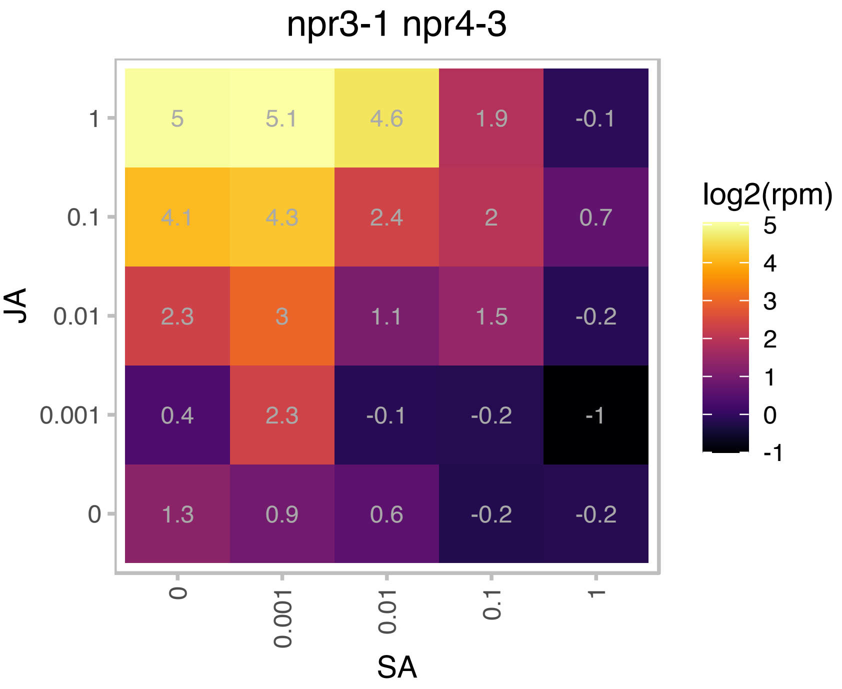

# AT3G46700.1 : NA

UDP-Glycosyltransferase superfamily protein

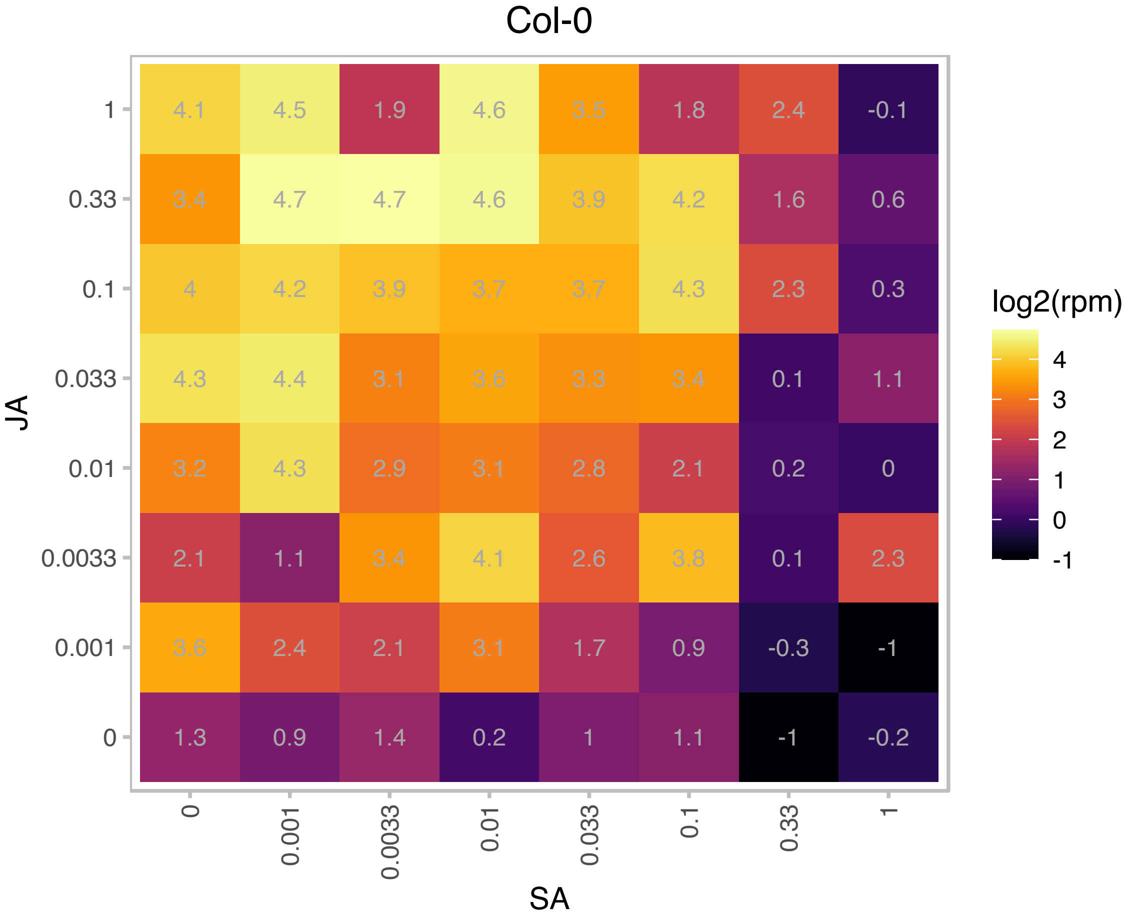


| | |
|---|---|
| Gene | AT3G46700 |
| is.expressed_genes | TRUE |
| is.responsive_genes | TRUE |
| is.apcluster | E-02 |

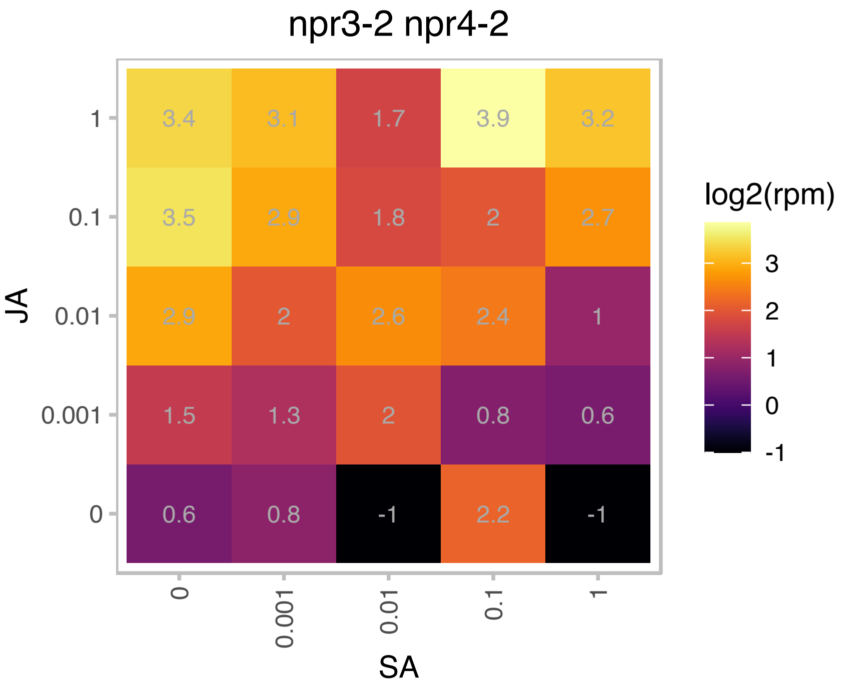


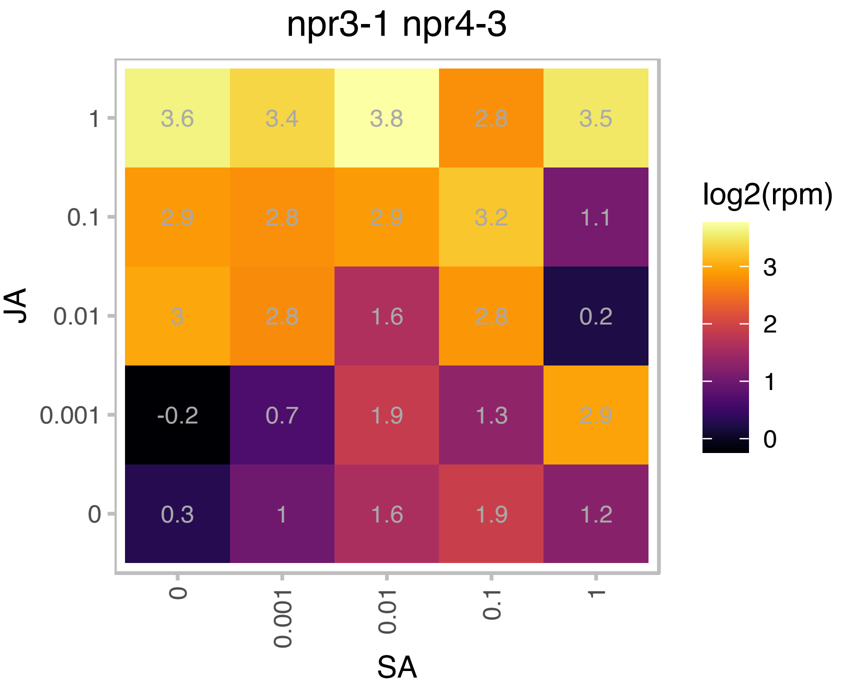

### AT3G58710.1 : ATWRKY69, WRKY69

WRKY DNA-binding protein 69

member of WRKY Transcription Factor; Group II-e

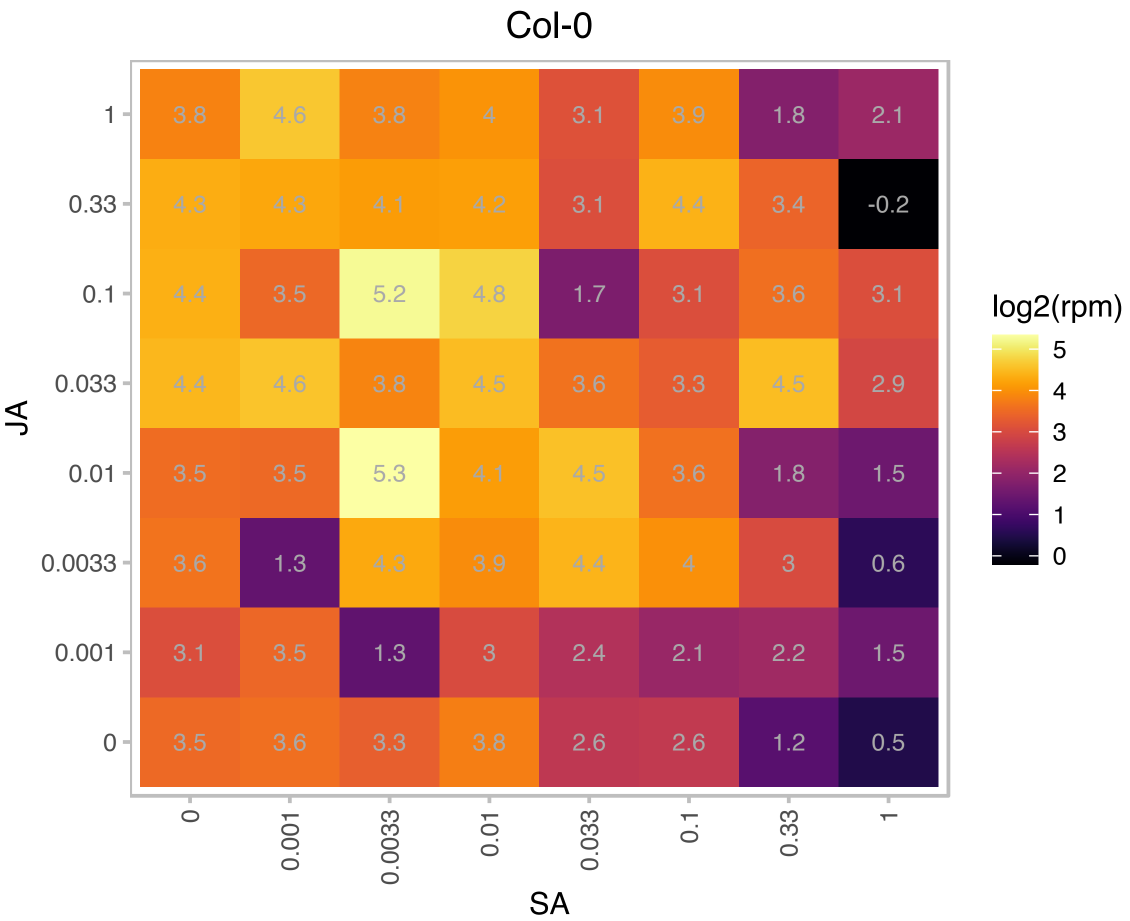


| | |
|---|---|
| Gene | AT3G58710 |
| is.expressed_genes | TRUE |
| is.responsive_genes | TRUE |
| is.apcluster | E-02 |

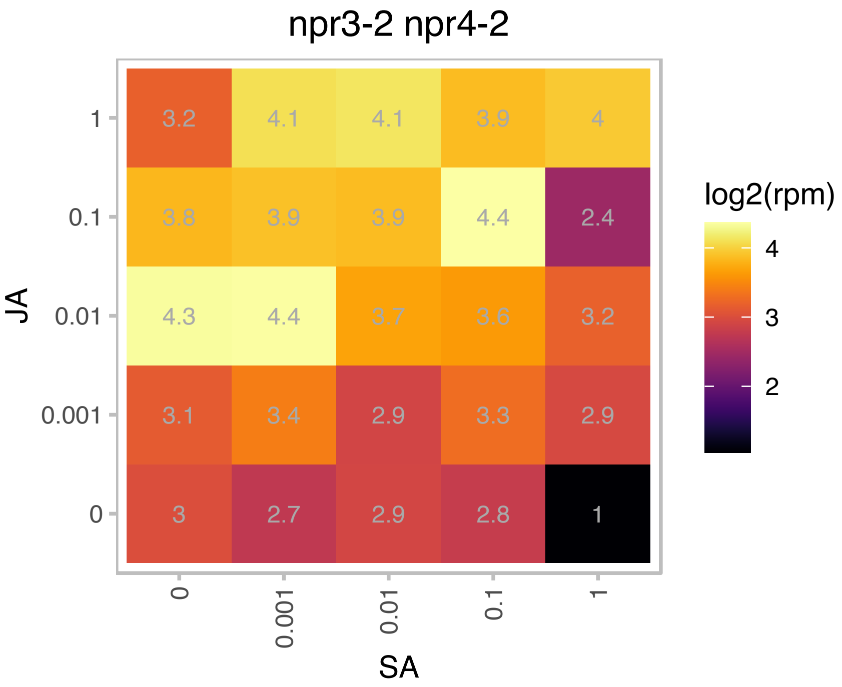


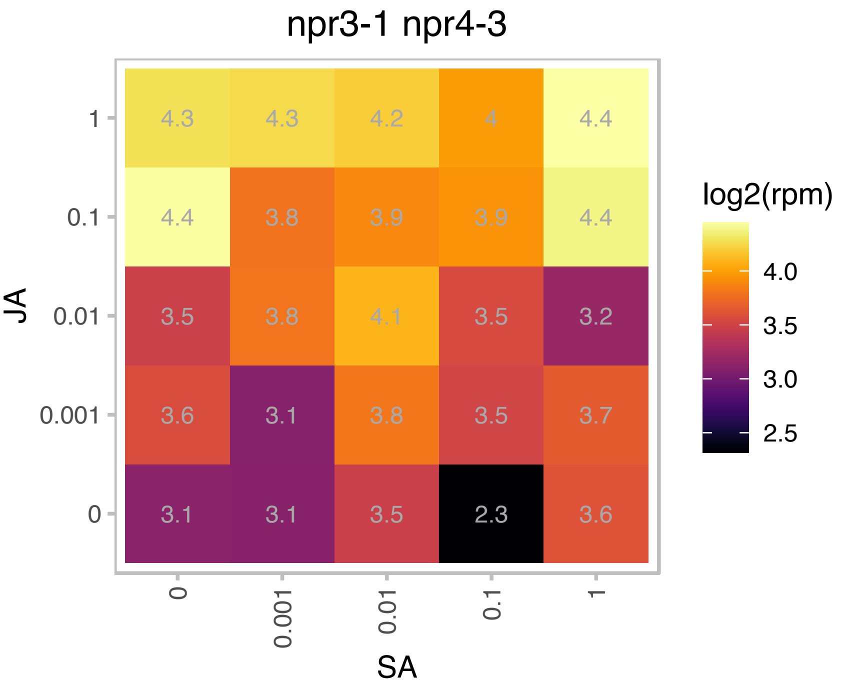

### AT3G59600.1 : NRPB8B, NRPD8B, NRPE8B

RNA polymerase Rpb8

One of two highly similar proteins that can serve as non-catalytic subunits of Nuclear RNA polymerases II, IV and V; homologous to budding yeast RPB8. Probably redundant with At1g54250.

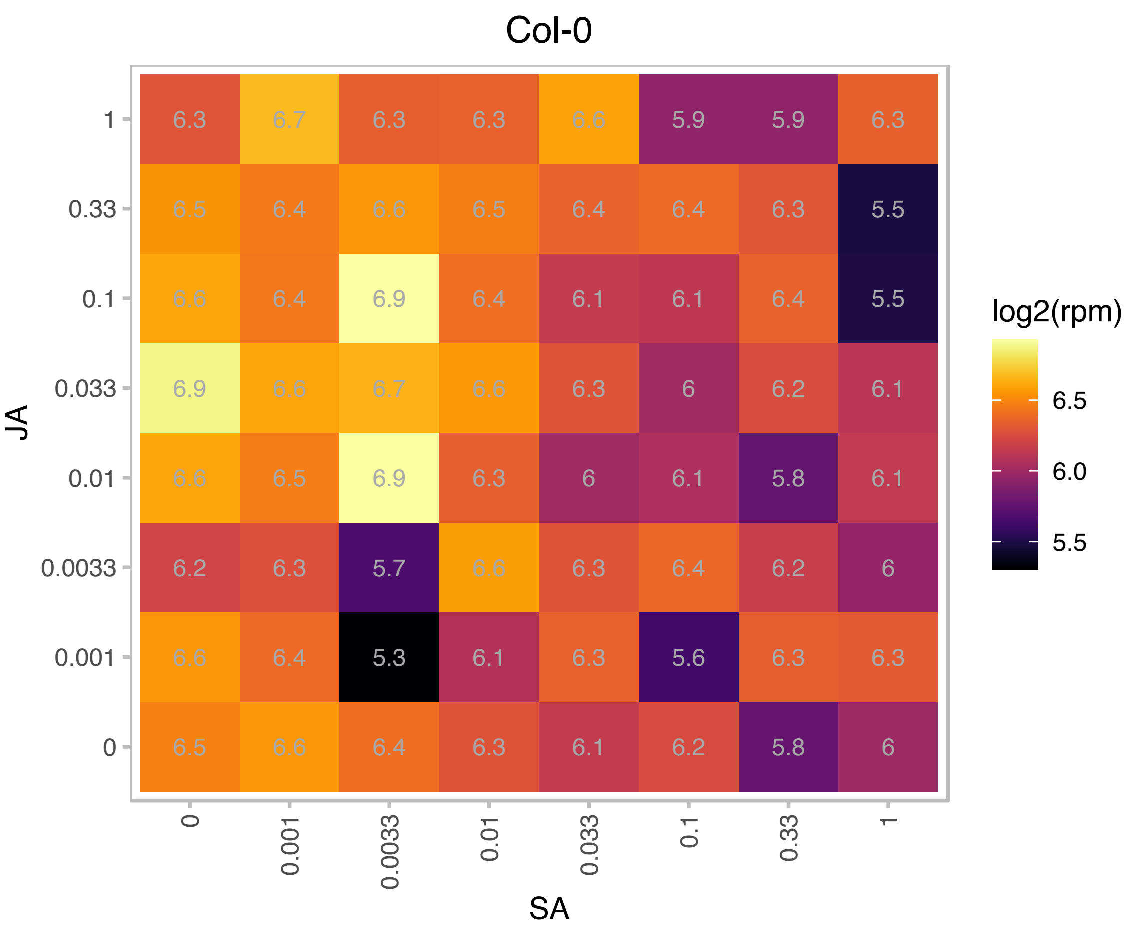


| | |
|---|---|
| Gene | AT3G59600 |
| is.expressed_genes | TRUE |
| is.responsive_genes | TRUE |
| is.apcluster | E-02 |

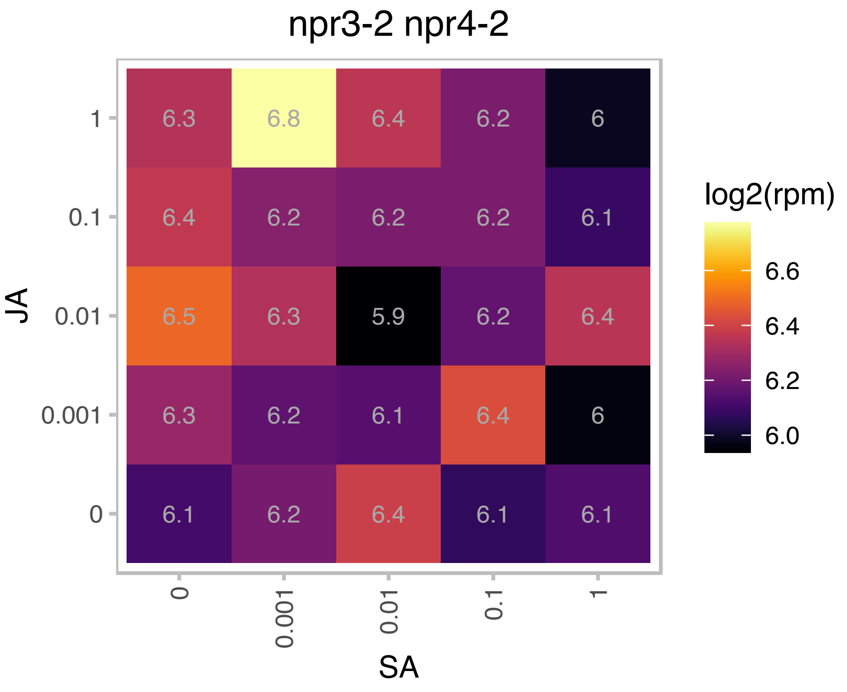


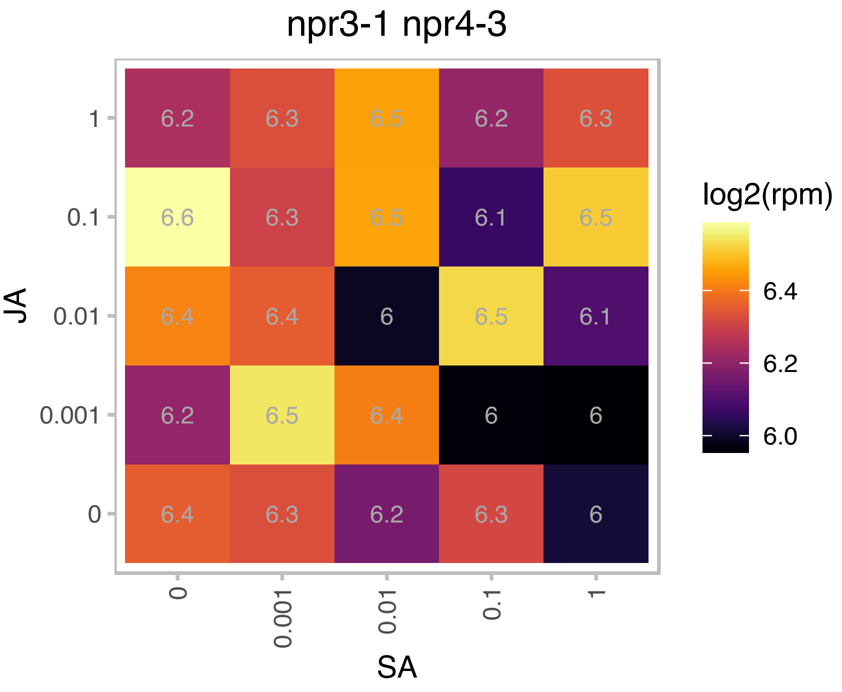

### AT4G01850.1 : AtSAM2, MAT2, SAM-2, SAM2

S-adenosylmethionine synthetase 2

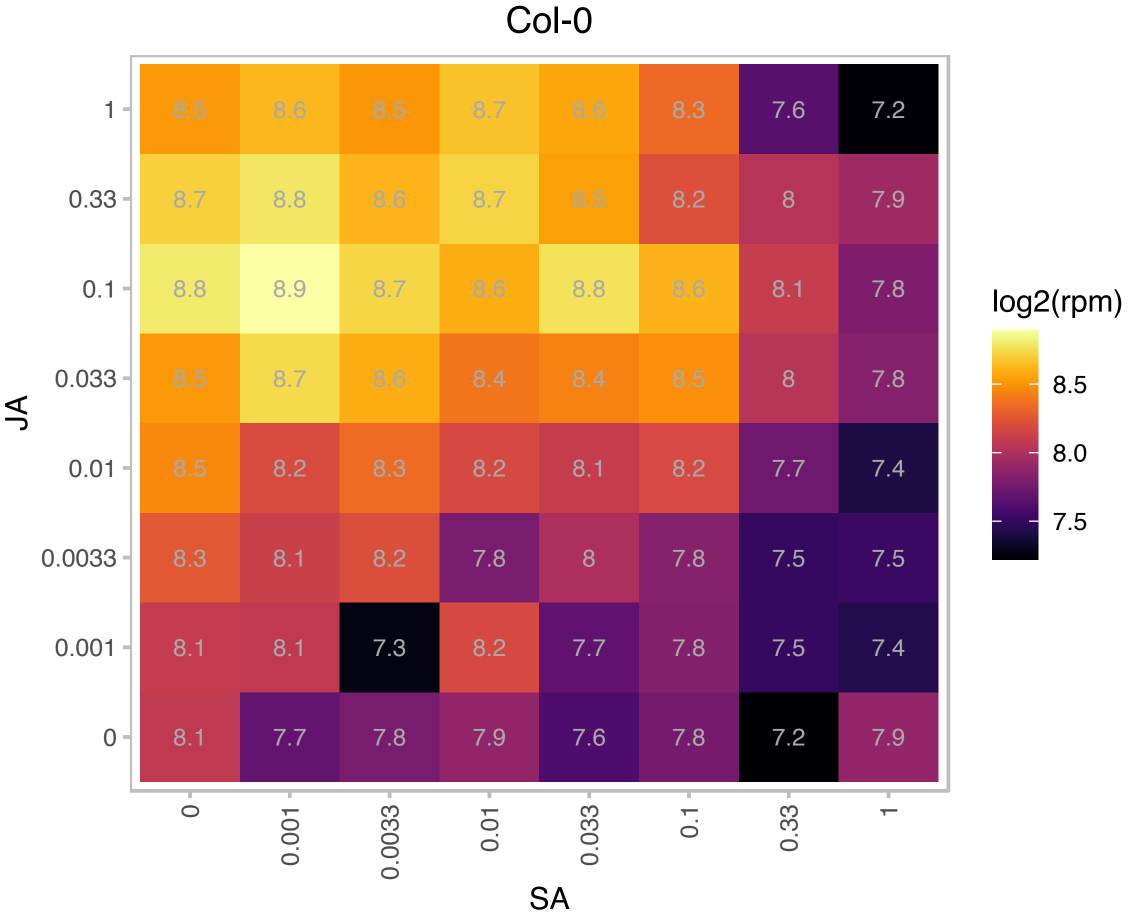


| | |
|---|---|
| Gene | AT4G01850 |
| is.expressed_genes | TRUE |
| is.responsive_genes | TRUE |
| is.apcluster | E-02 |

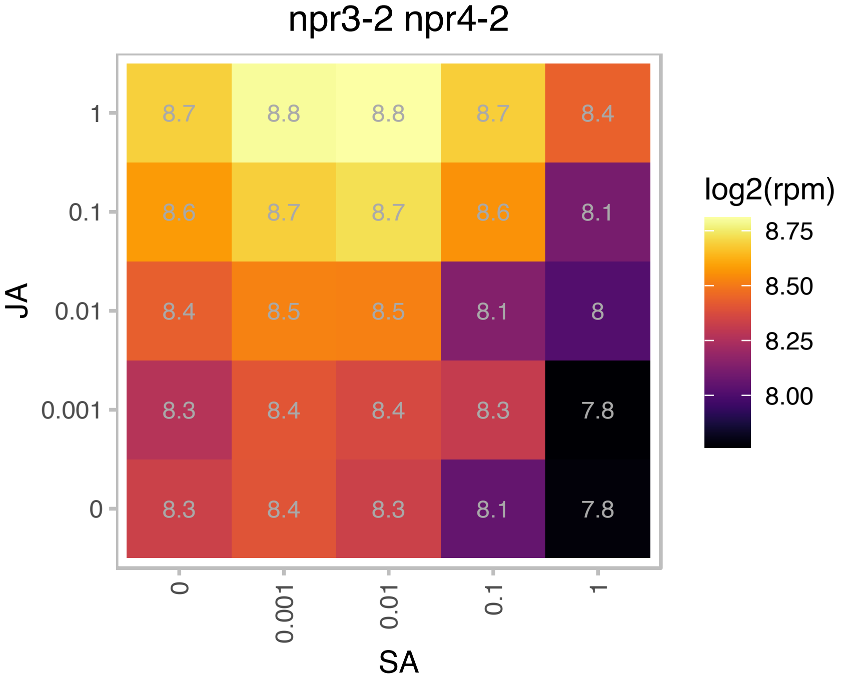


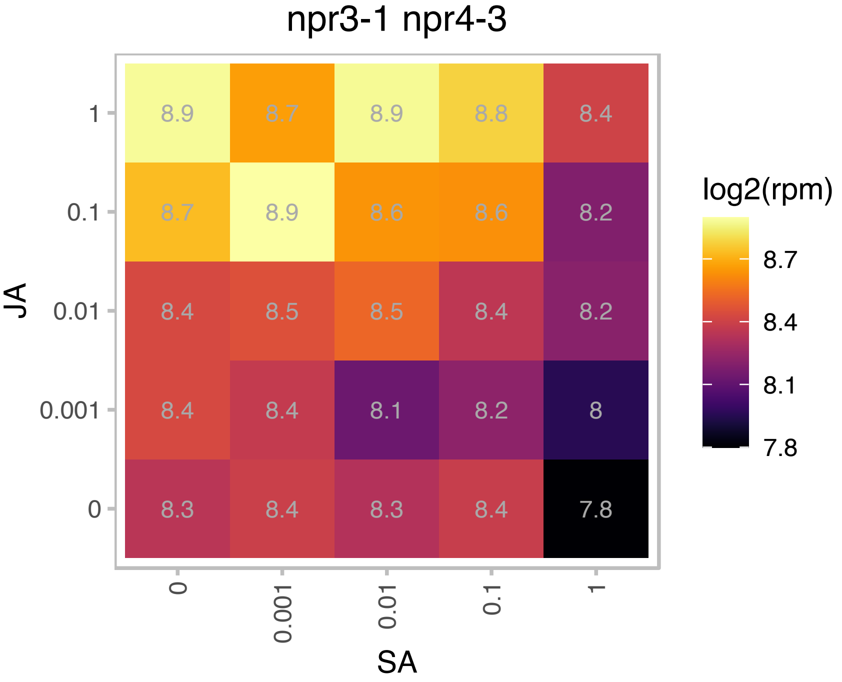

# AT4G11210.1 : NA

Disease resistance-responsive (dirigent-like protein) family protein

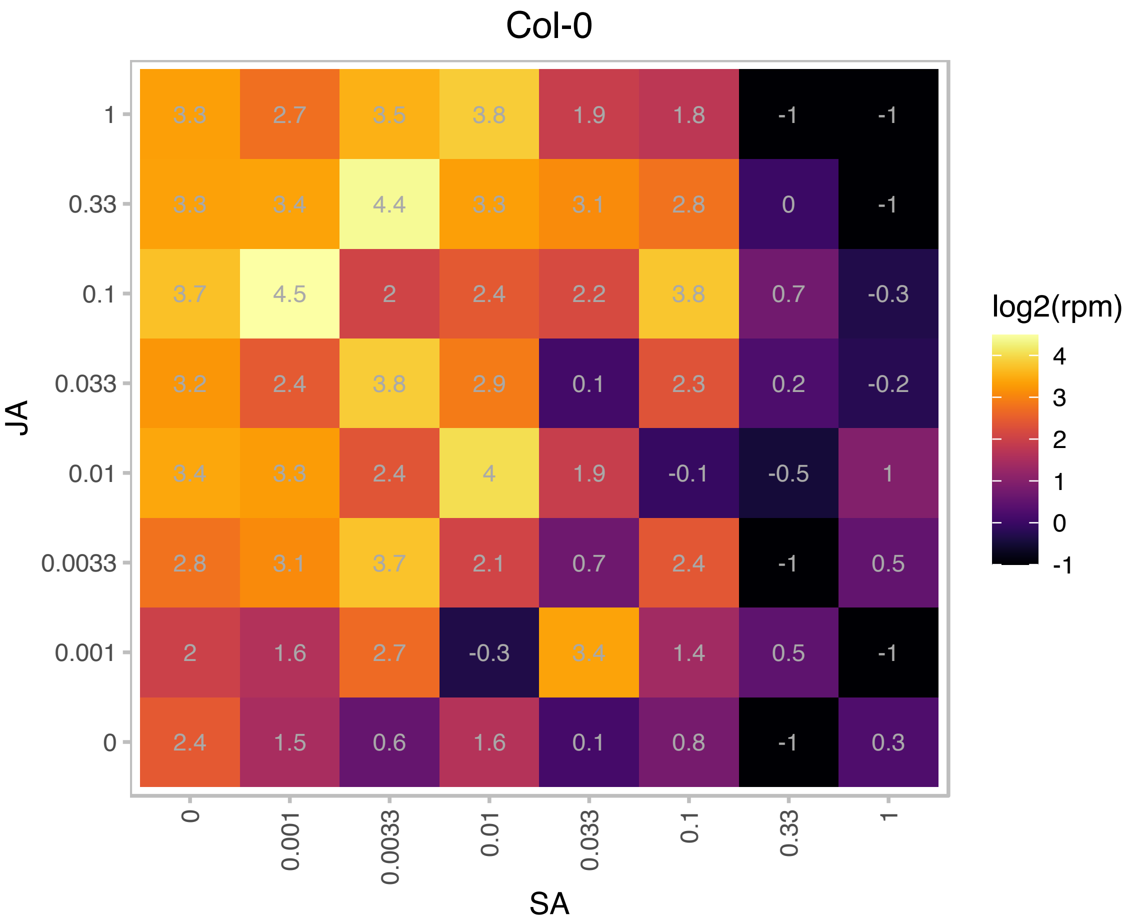


| | |
|---|---|
| Gene | AT4G11210 |
| is.expressed_genes | TRUE |
| is.responsive_genes | TRUE |
| is.apcluster | E-02 |

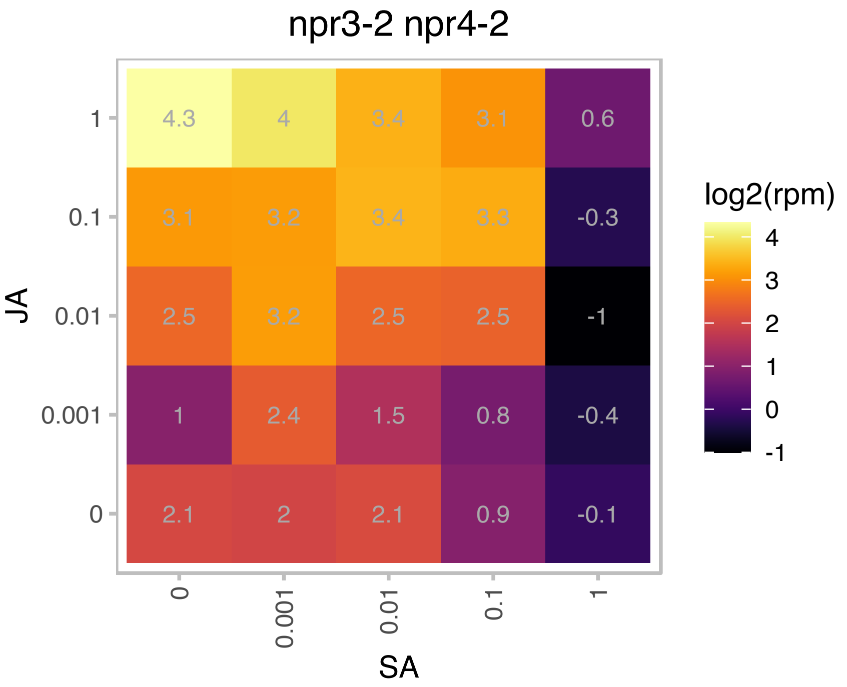


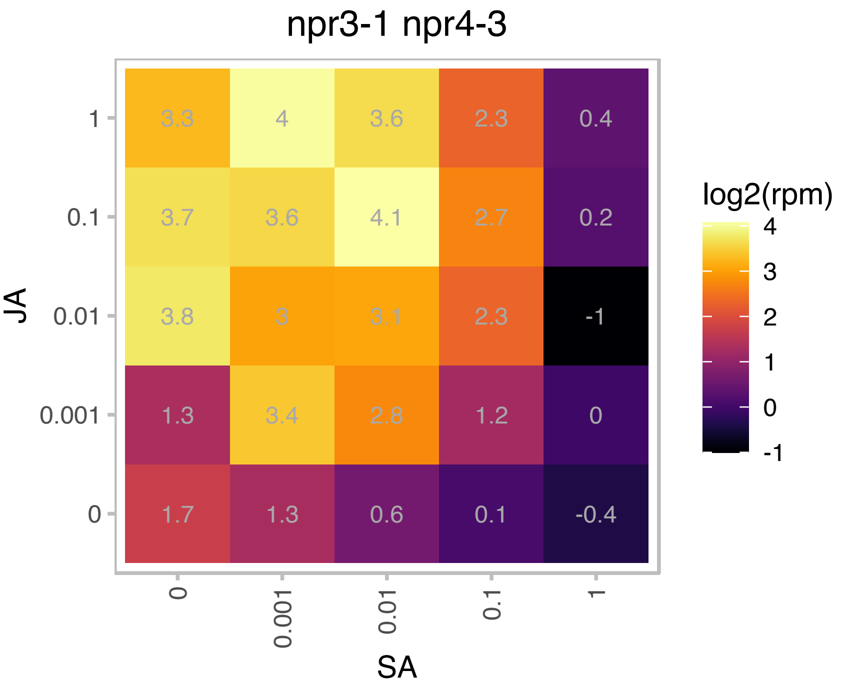

### AT4G13290.1 : CYP71A19

cytochrome P450, family 71, subfamily A, polypeptide 19

putative cytochrome P450

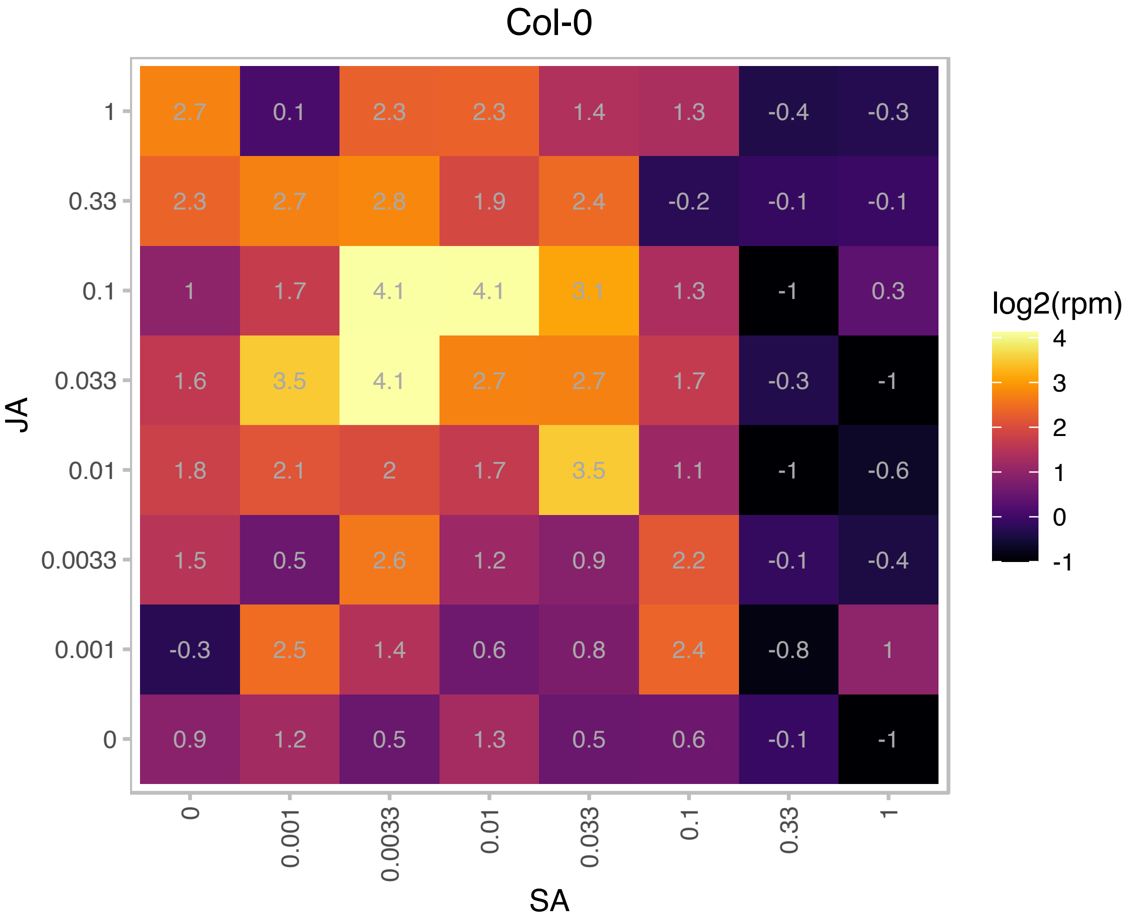


| | |
|---|---|
| Gene | AT4G13290 |
| is.expressed_genes | TRUE |
| is.responsive_genes | TRUE |
| is.apcluster | E-02 |

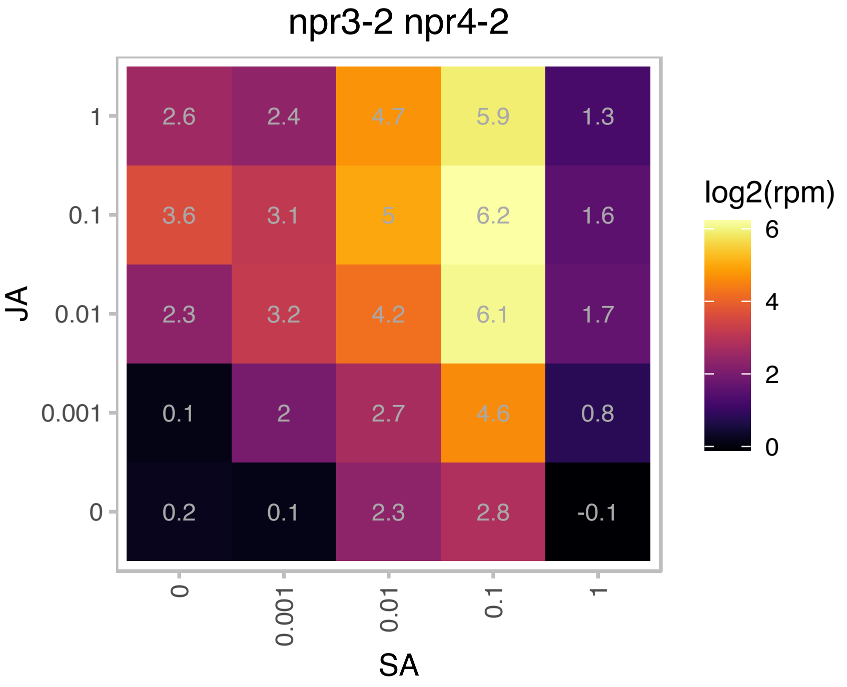


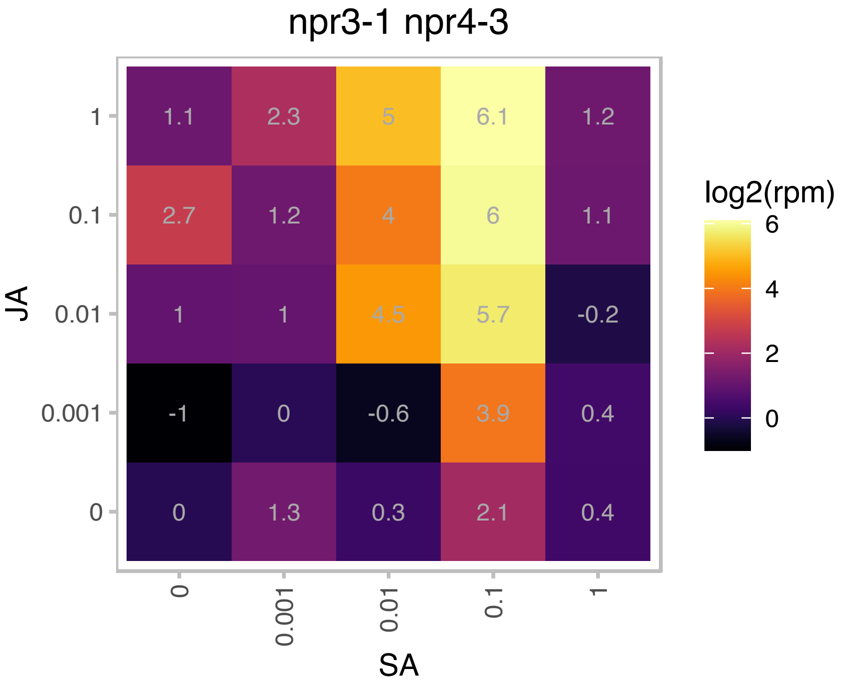

### AT4G13310.1 : CYP71A20

cytochrome P450, family 71, subfamily A, polypeptide 20

putative cytochrome P450

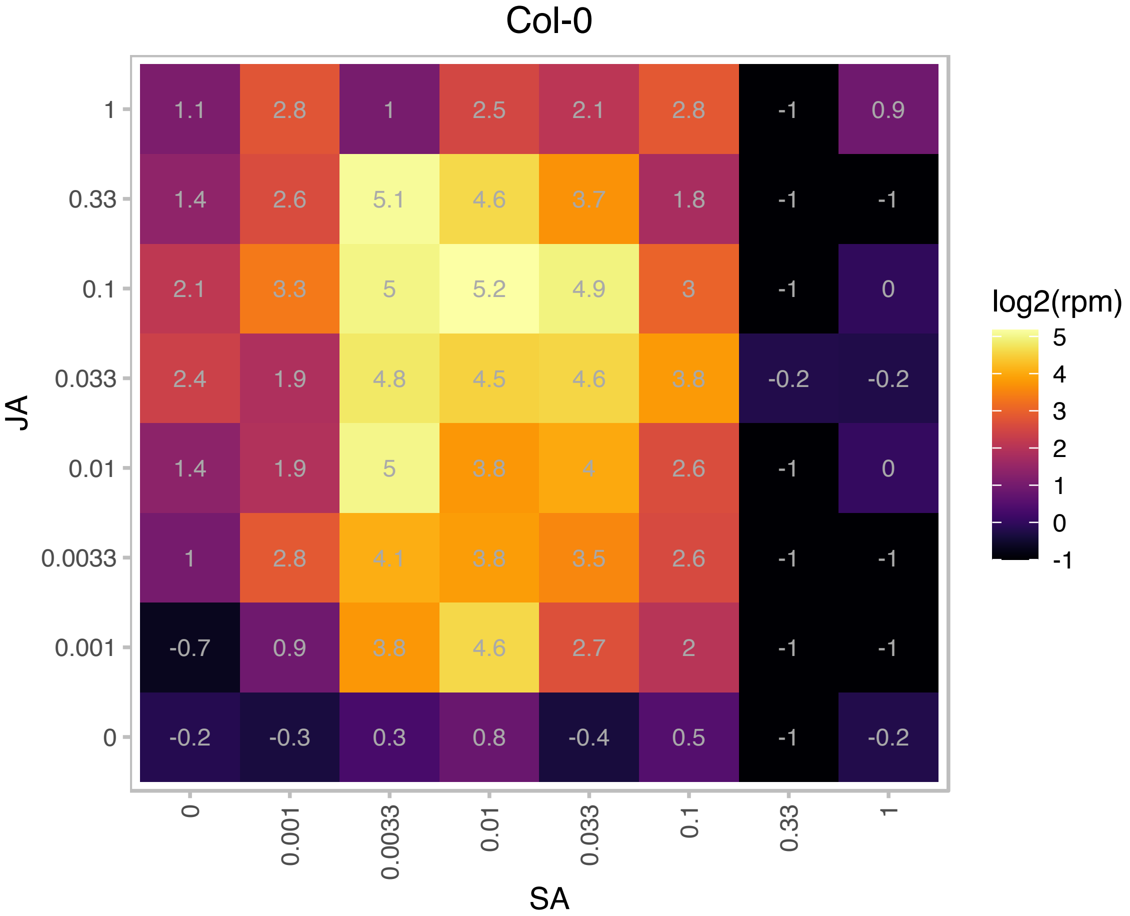


| | |
|---|---|
| Gene | AT4G13310 |
| is.expressed_genes | TRUE |
| is.responsive_genes | TRUE |
| is.apcluster | E-02 |

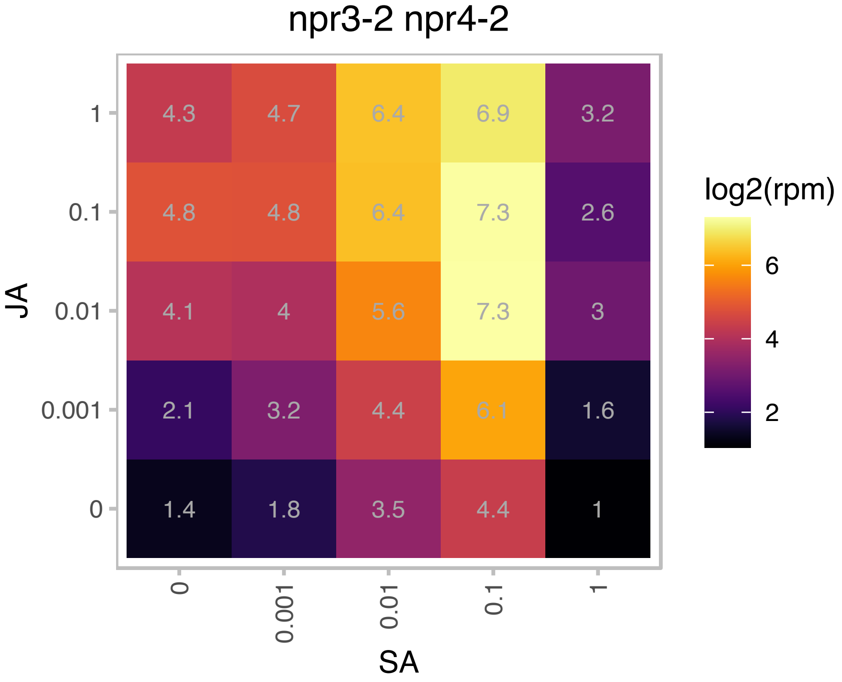


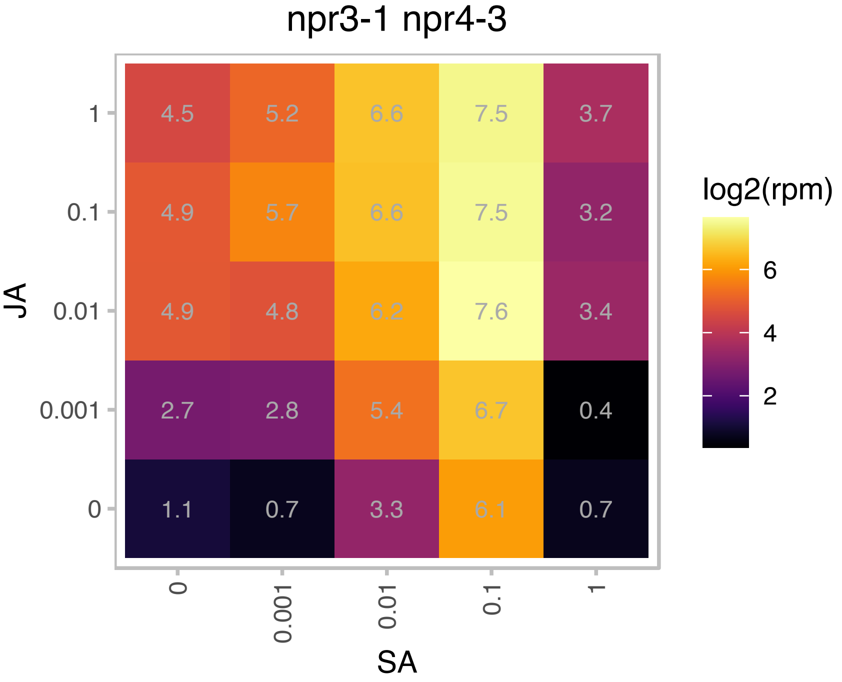

# AT4G13860.1 : NA

RNA-binding (RRM/RBD/RNP motifs) family protein

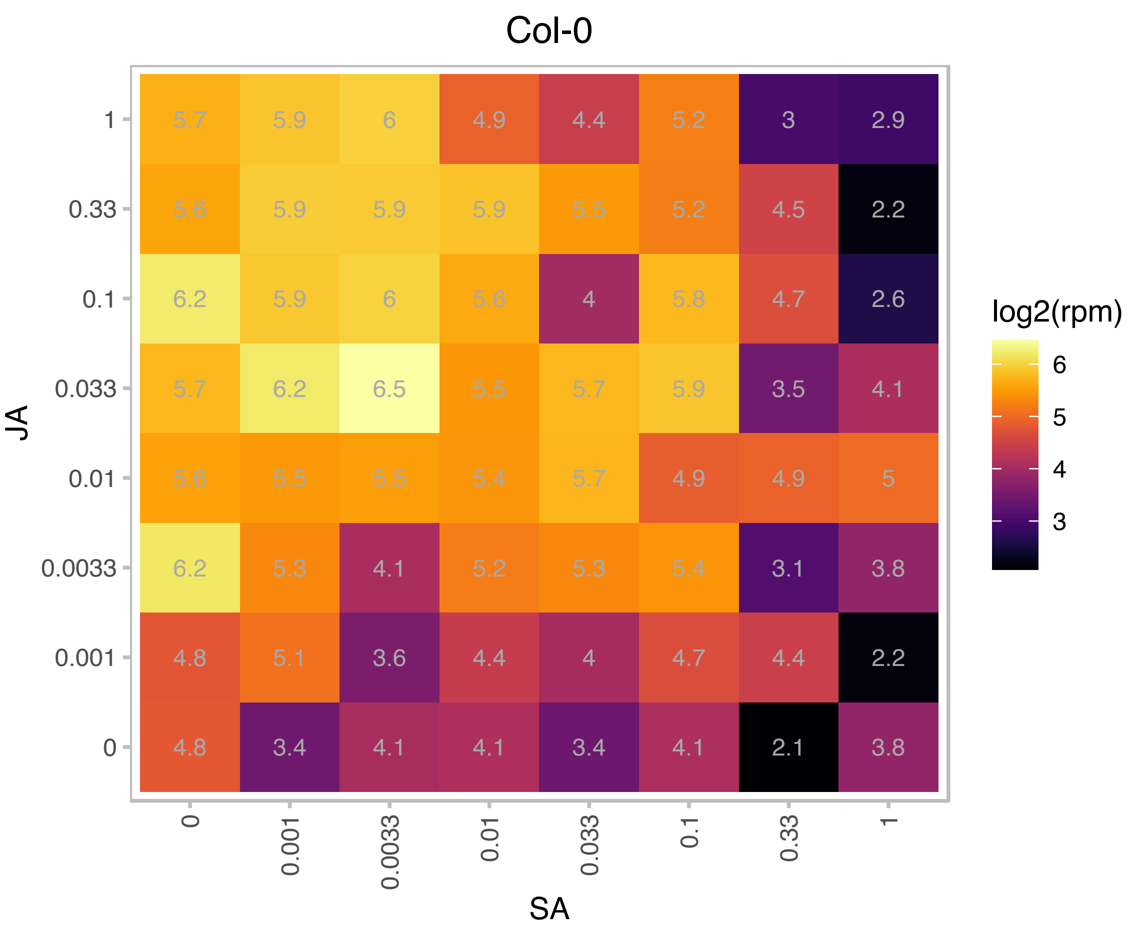


| | |
|---|---|
| Gene | AT4G13860 |
| is.expressed_genes | TRUE |
| is.responsive_genes | TRUE |
| is.apcluster | E-02 |

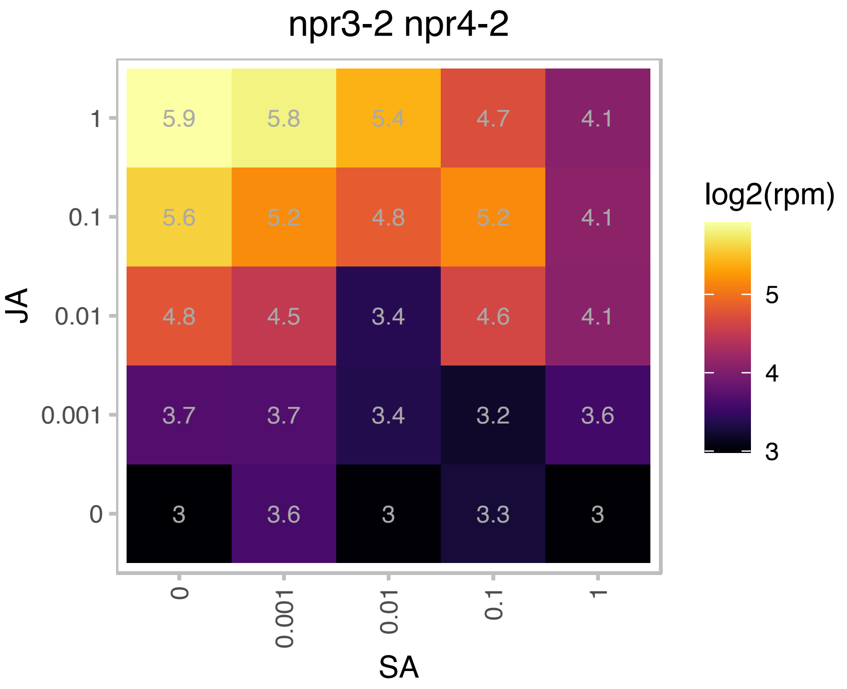


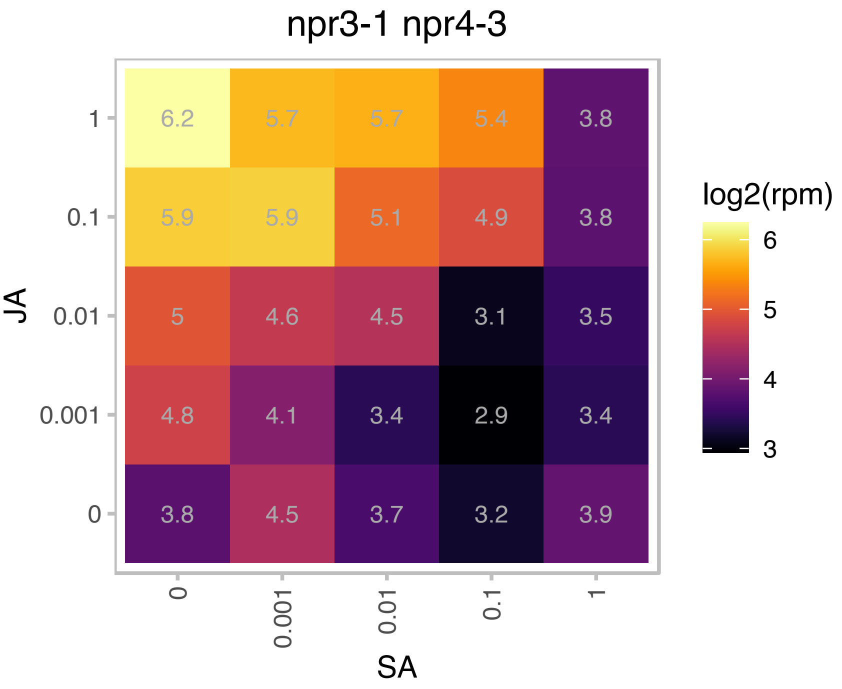

# AT4G16380.1 : NA

Heavy metal transport/detoxification superfamily protein

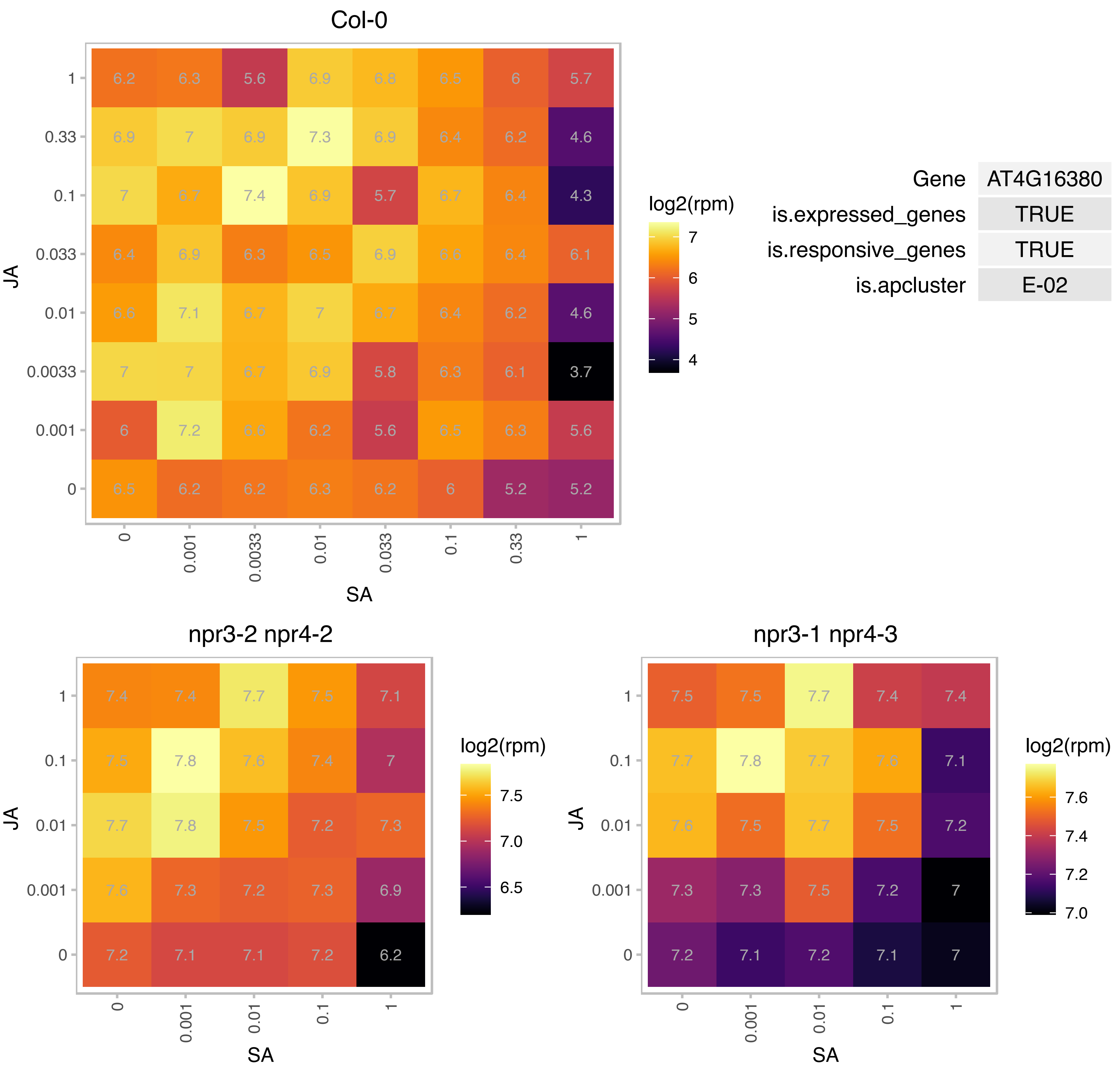


| | |
|---|---|
| Gene | AT4G16380 |
| is.expressed_genes | TRUE |
| is.responsive_genes | TRUE |
| is.apcluster | E-02 |

# AT4G22212.1 : NA

Arabidopsis defensin-like protein

Encodes a defensin-like (DEFL) family protein.

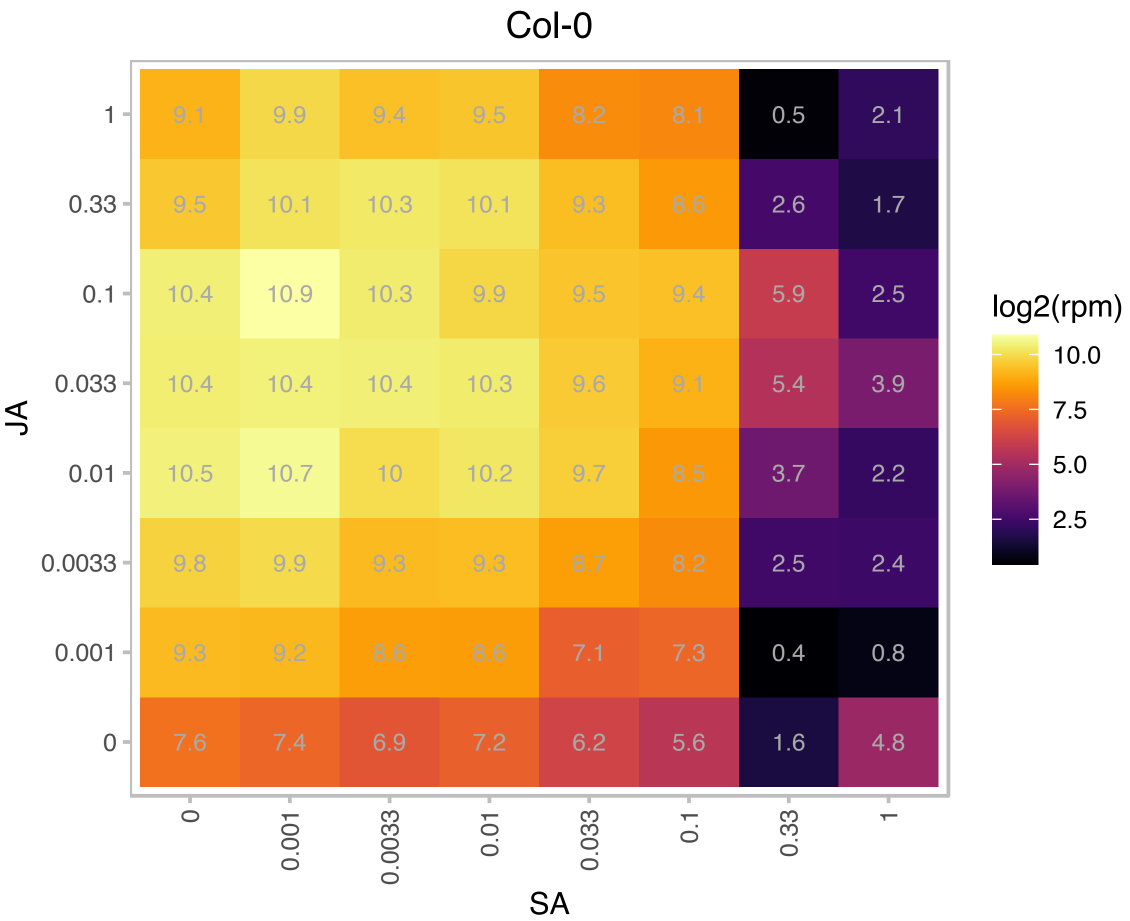


| | |
|---|---|
| Gene | AT4G22212 |
| is.expressed_genes | TRUE |
| is.responsive_genes | TRUE |
| is.apcluster | E-02 |

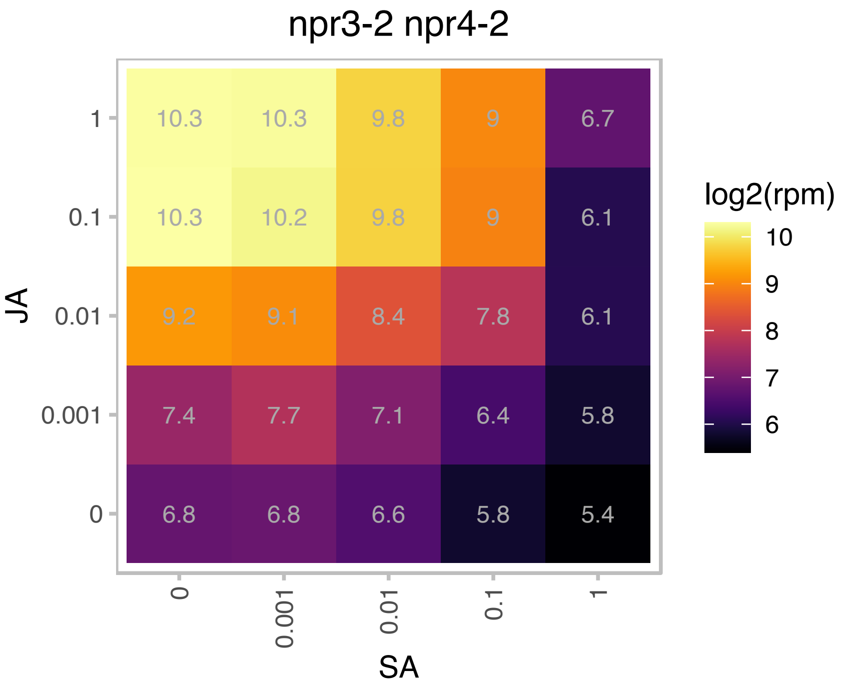


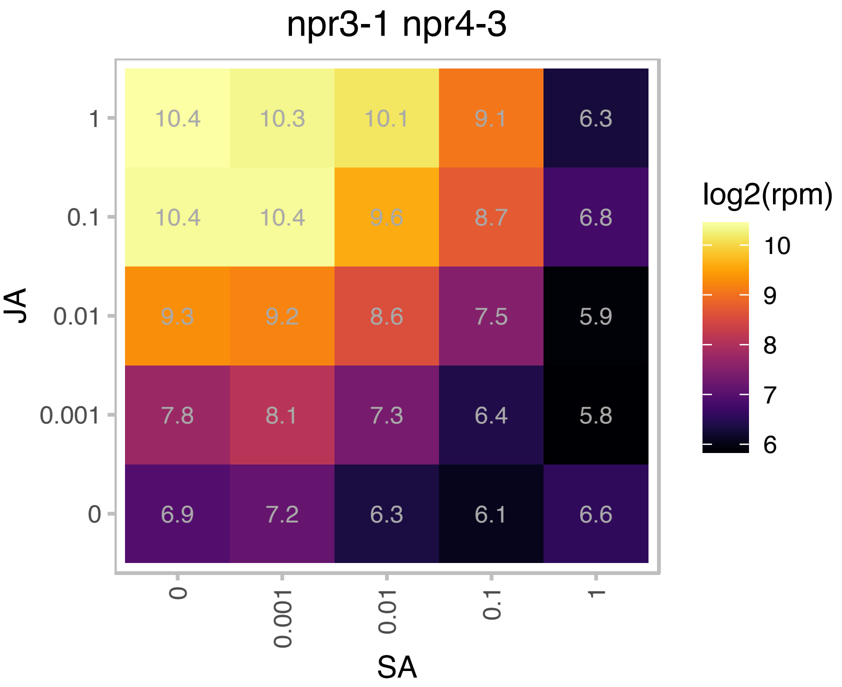

# AT4G22214.1 : NA

Defensin-like (DEFL) family protein

Encodes a defensin-like (DEFL) family protein.

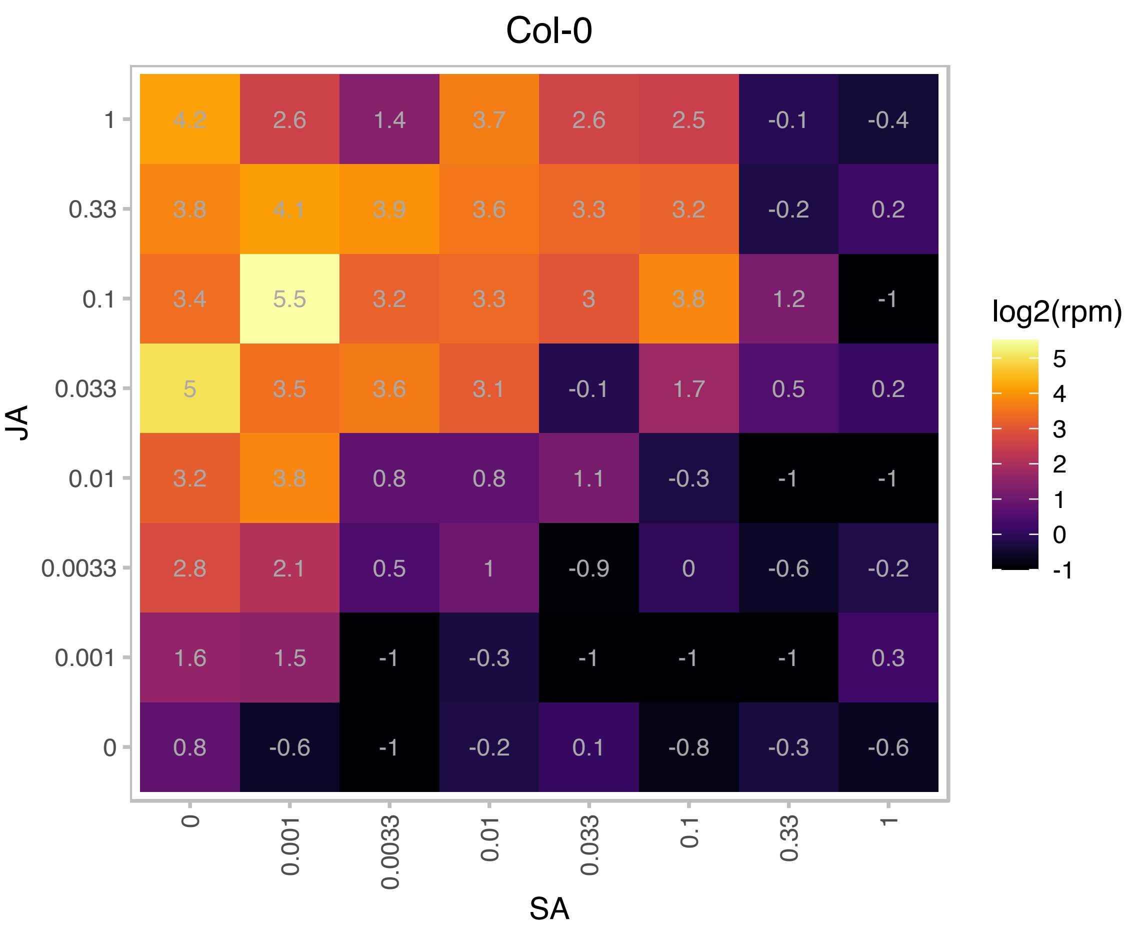


| | |
|---|---|
| Gene | AT4G22214 |
| is.expressed_genes | TRUE |
| is.responsive_genes | TRUE |
| is.apcluster | E-02 |

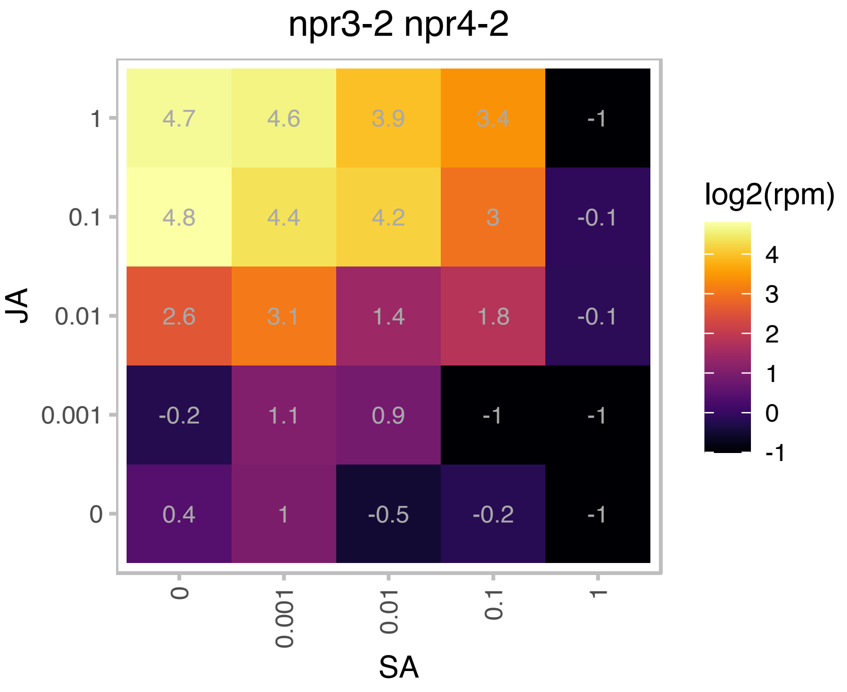


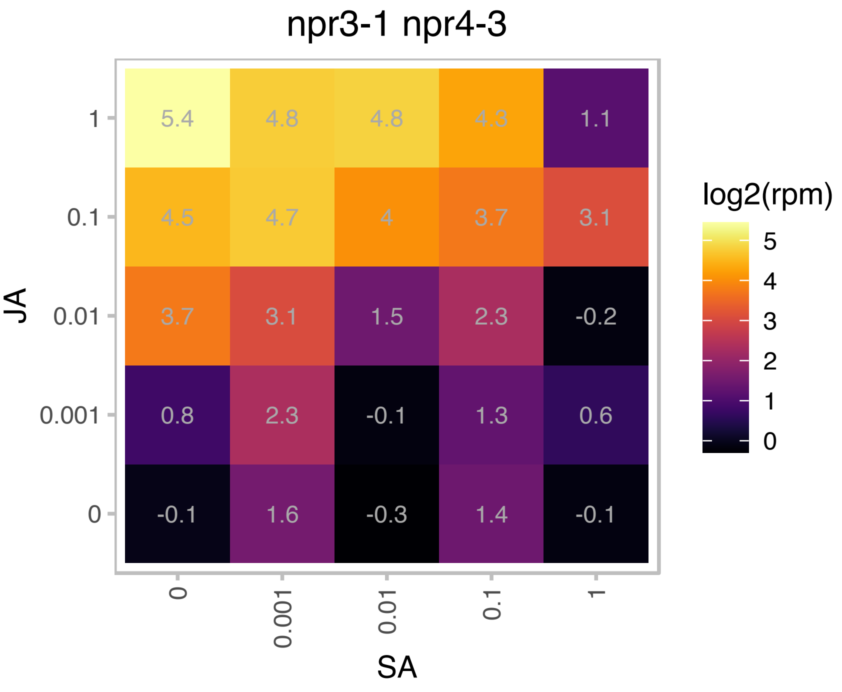

# AT4G22610.1 : NA

Bifunctional inhibitor/lipid-transfer protein/seed storage 2S albumin superfamily protein

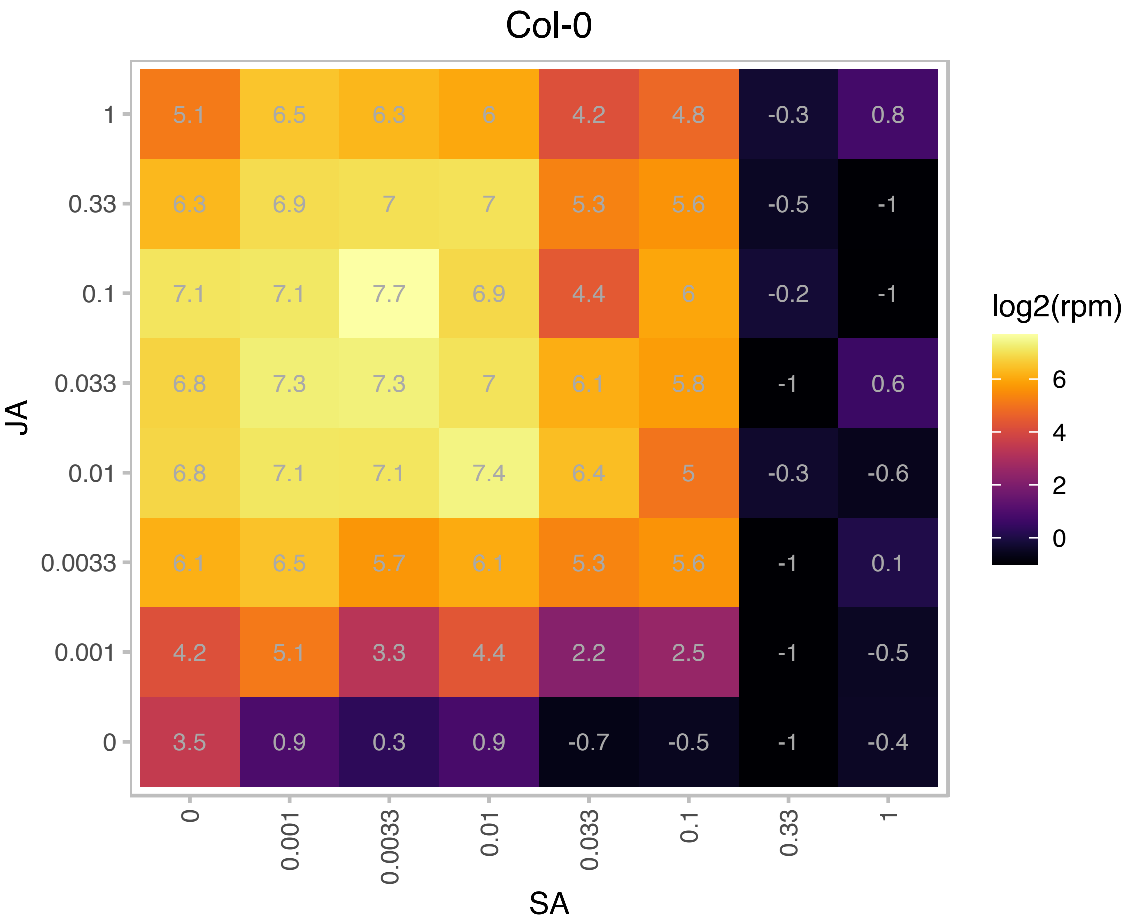


| | |
|---|---|
| Gene | AT4G22610 |
| is.expressed_genes | TRUE |
| is.responsive_genes | TRUE |
| is.apcluster | E-02 |

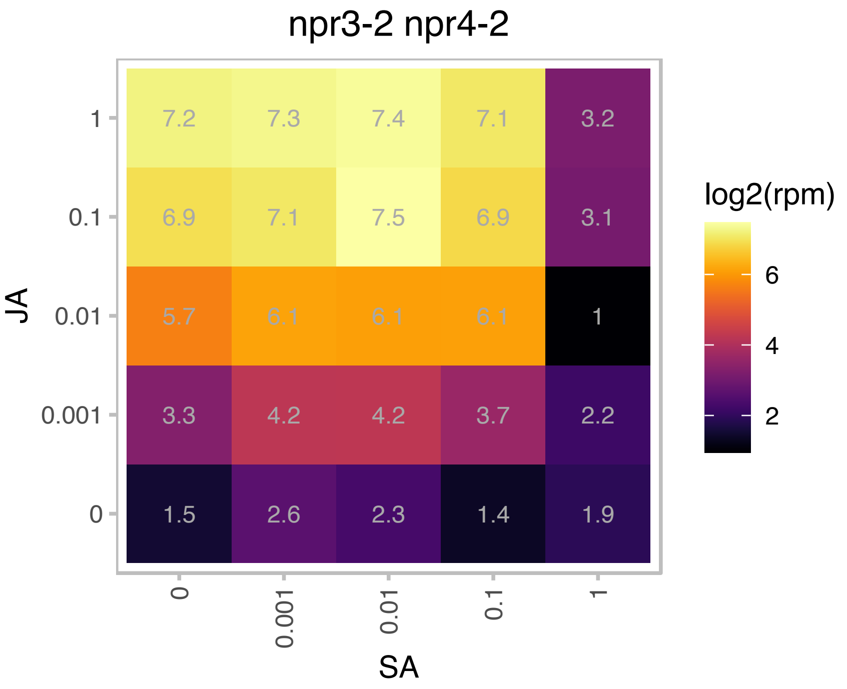


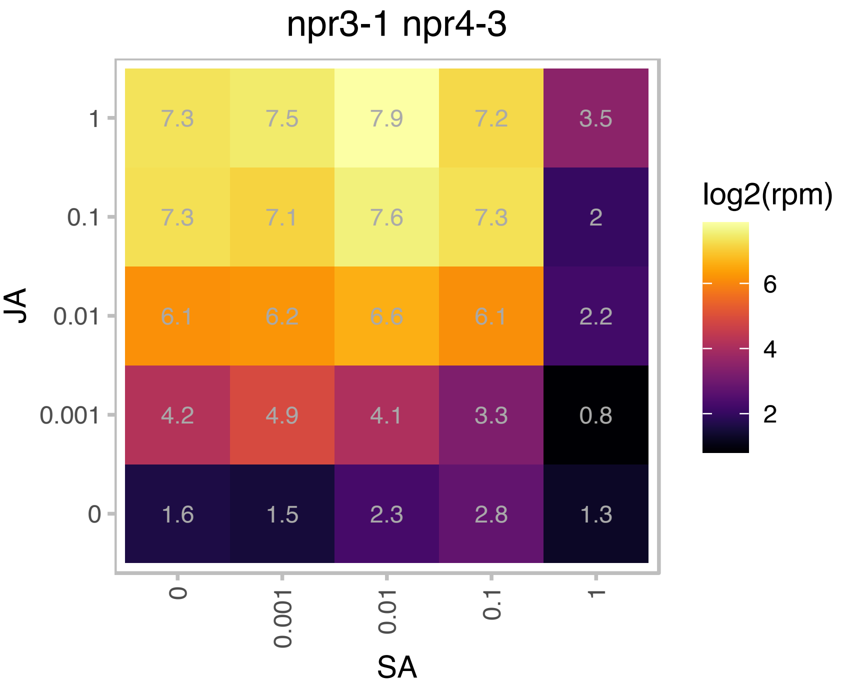

# AT4G23420.1 : NA

NAD(P)-binding Rossmann-fold superfamily protein

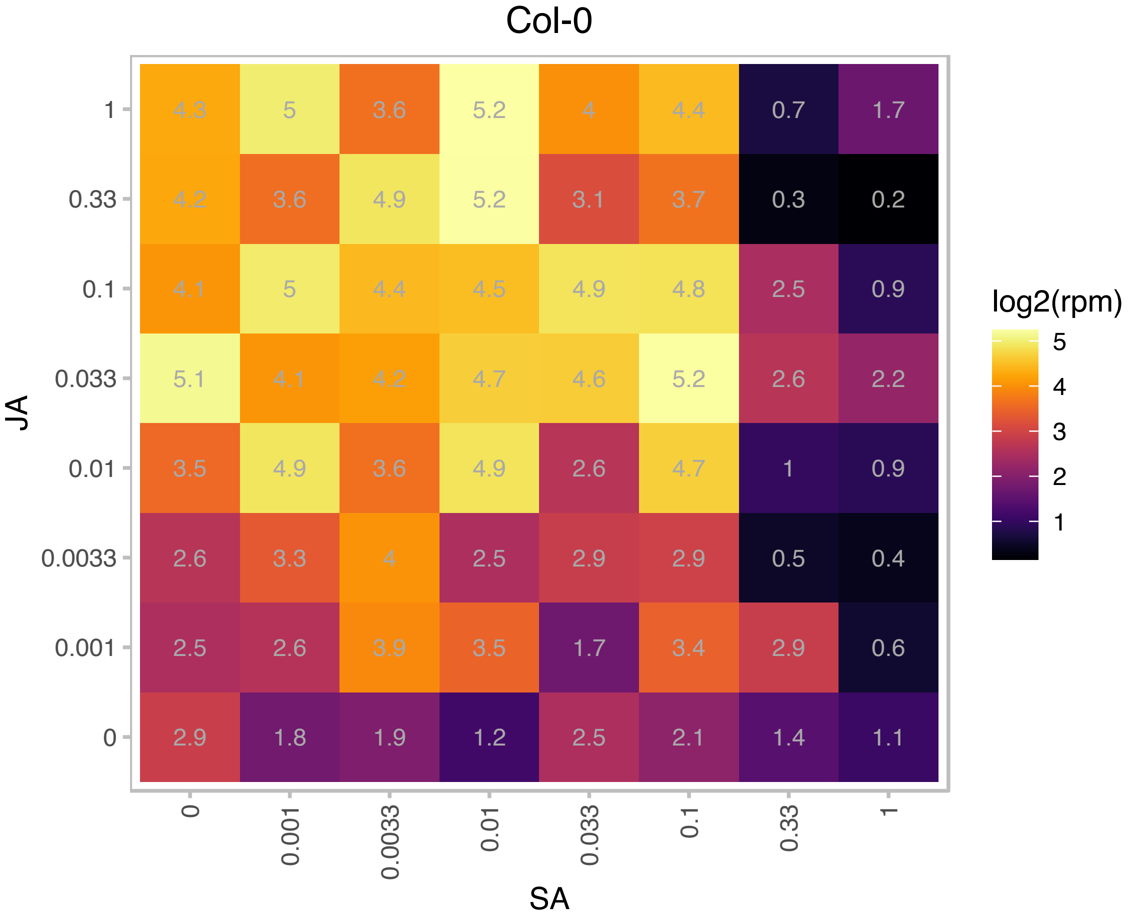


| | |
|---|---|
| Gene | AT4G23420 |
| is.expressed_genes | TRUE |
| is.responsive_genes | TRUE |
| is.apcluster | E-02 |

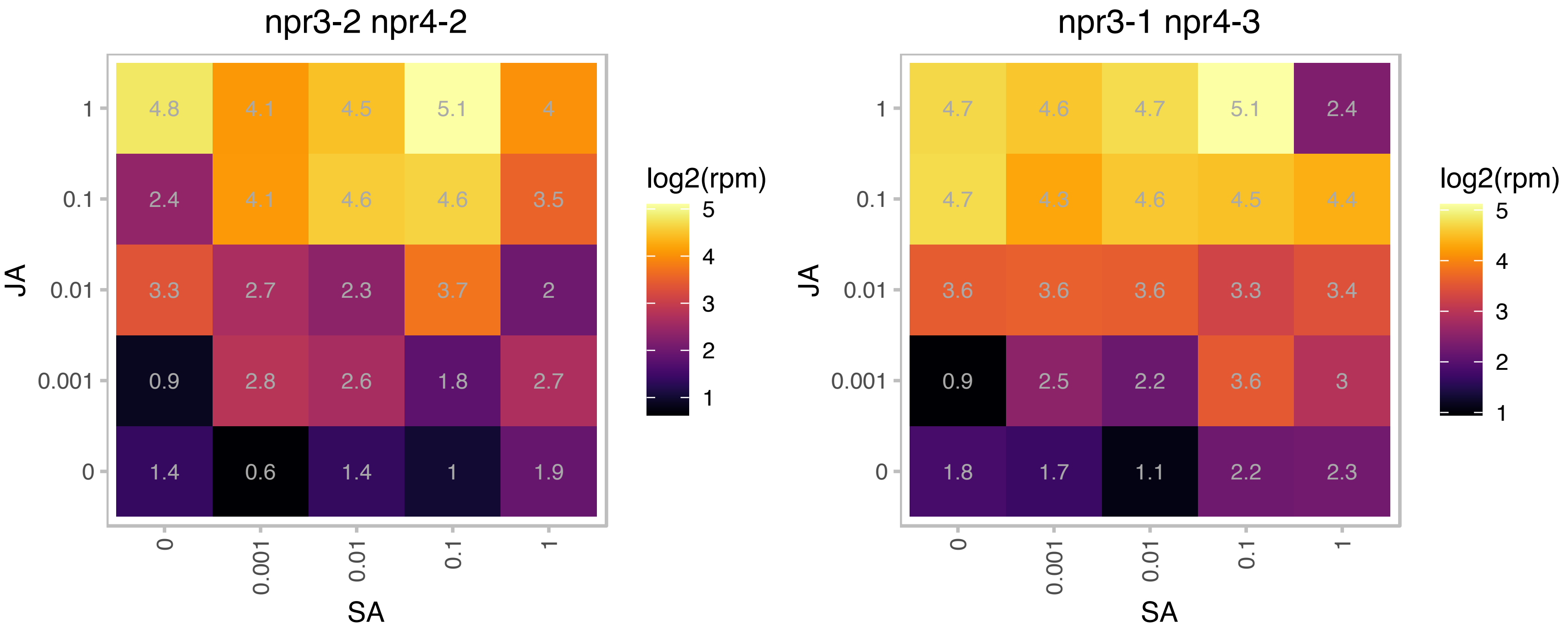

# AT4G23670.1 : NA

Polyketide cyclase/dehydrase and lipid transport superfamily protein

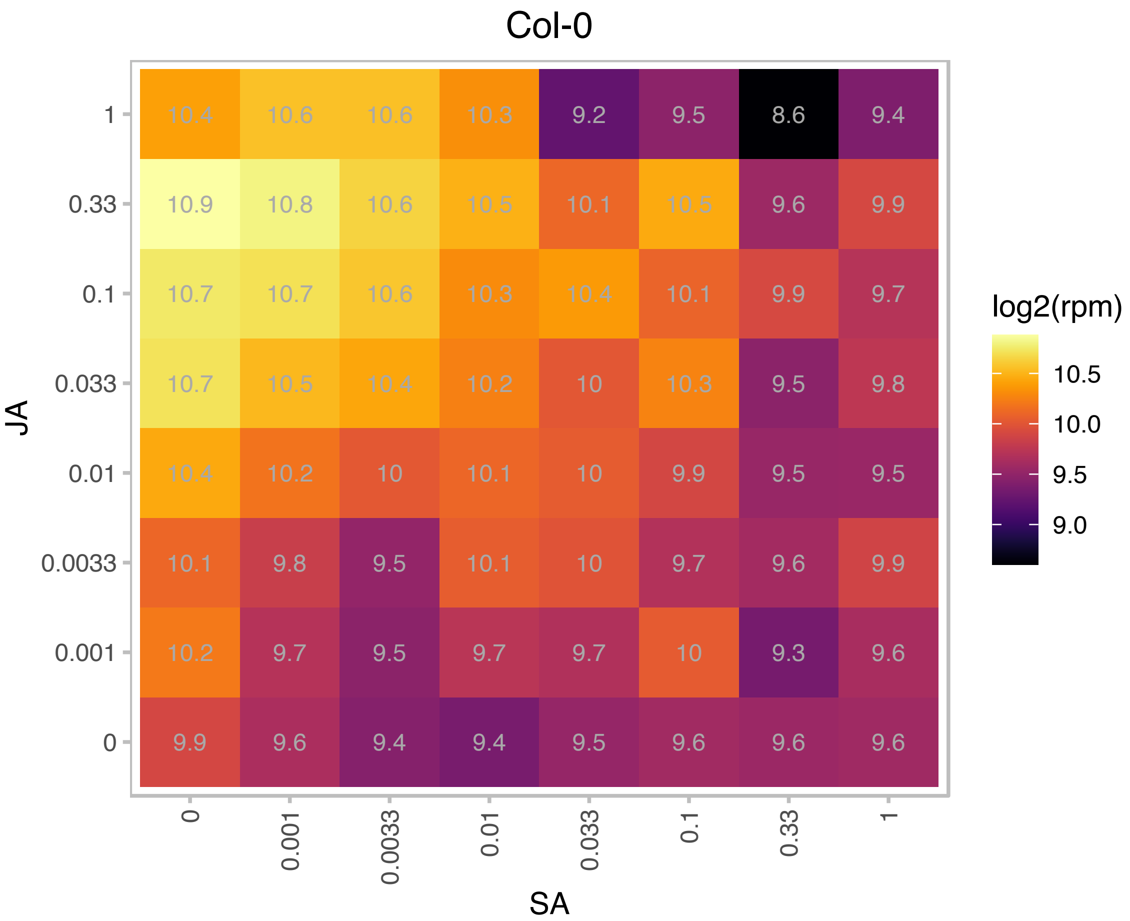


| | |
|---|---|
| Gene | AT4G23670 |
| is.expressed_genes | TRUE |
| is.responsive_genes | TRUE |
| is.apcluster | E-02 |

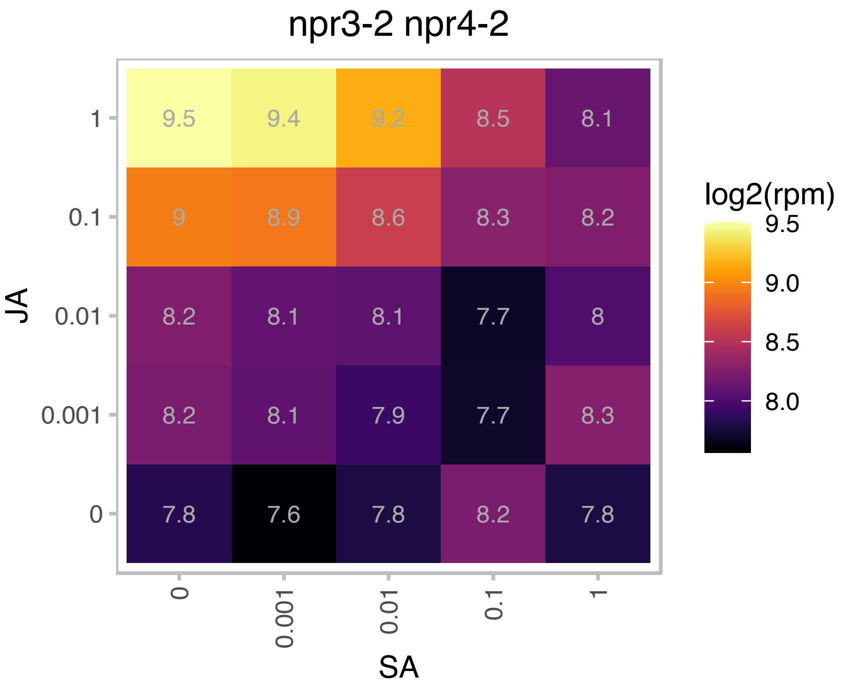


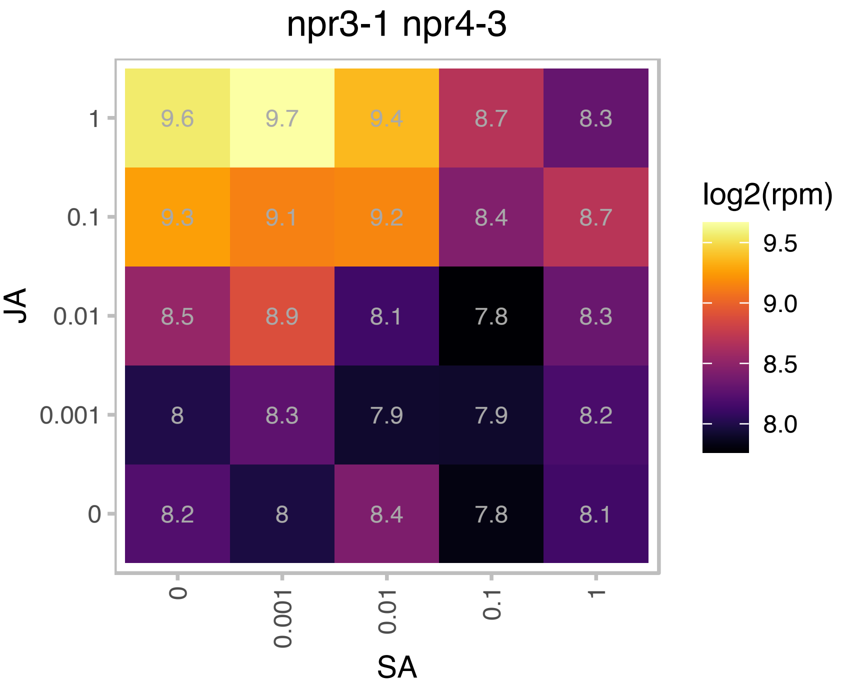

# AT4G24350.1 : NA

Phosphorylase superfamily protein

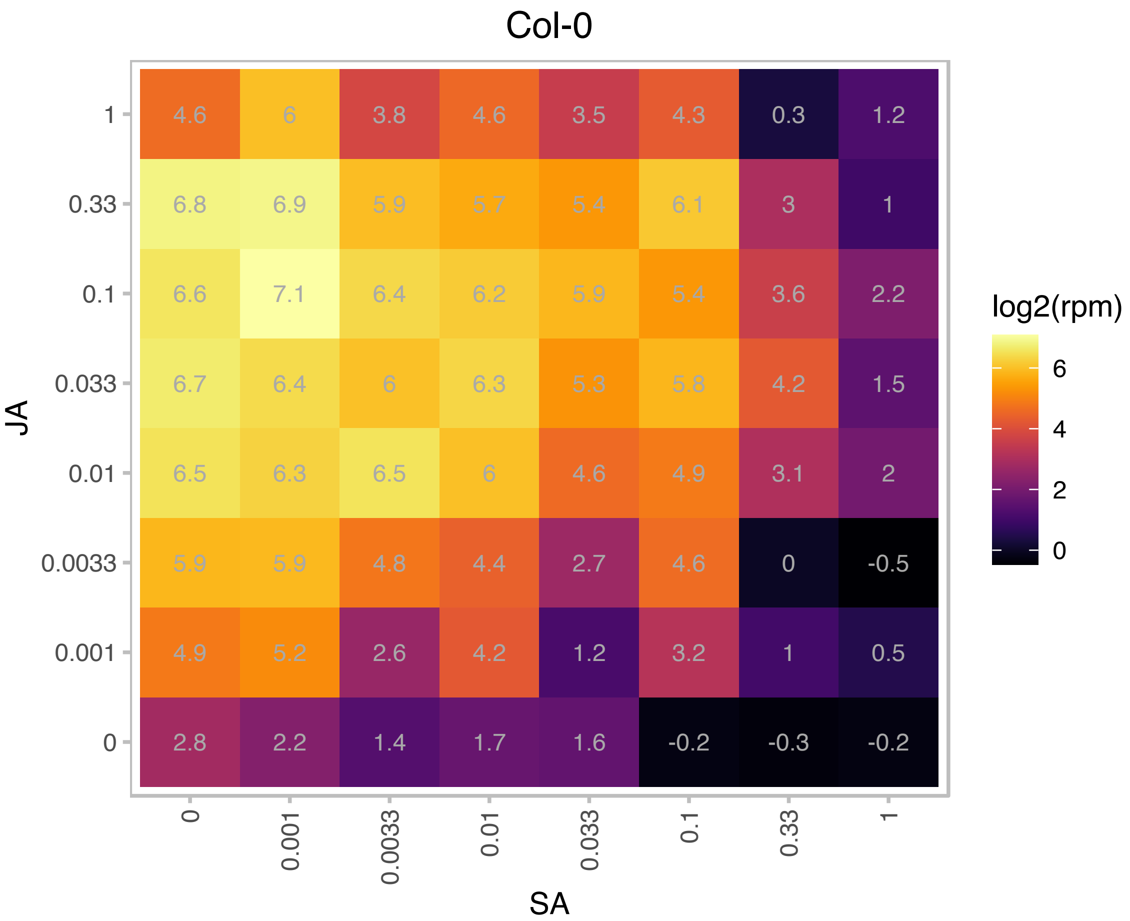


| | |
|---|---|
| Gene | AT4G24350 |
| is.expressed_genes | TRUE |
| is.responsive_genes | TRUE |
| is.apcluster | E-02 |

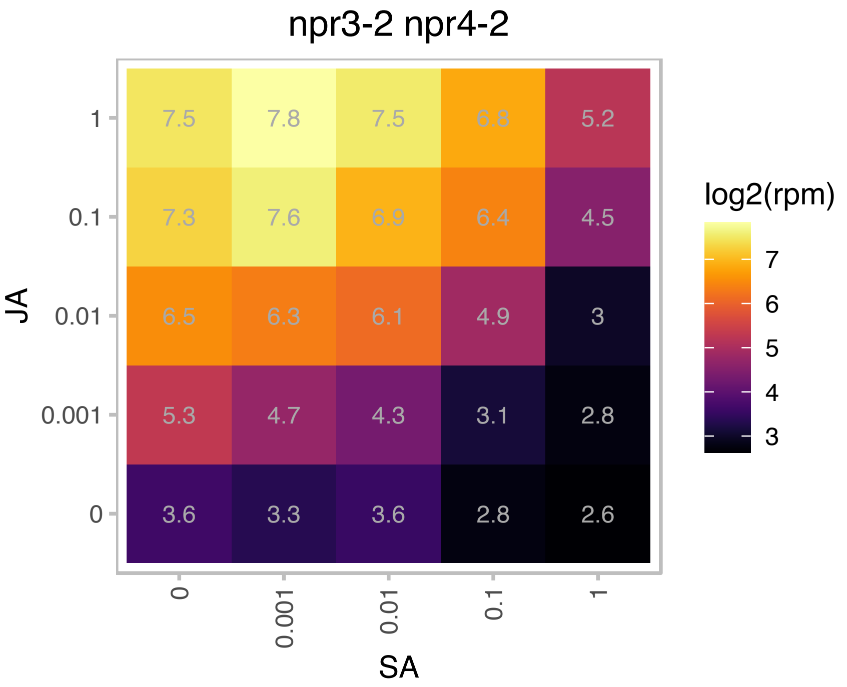


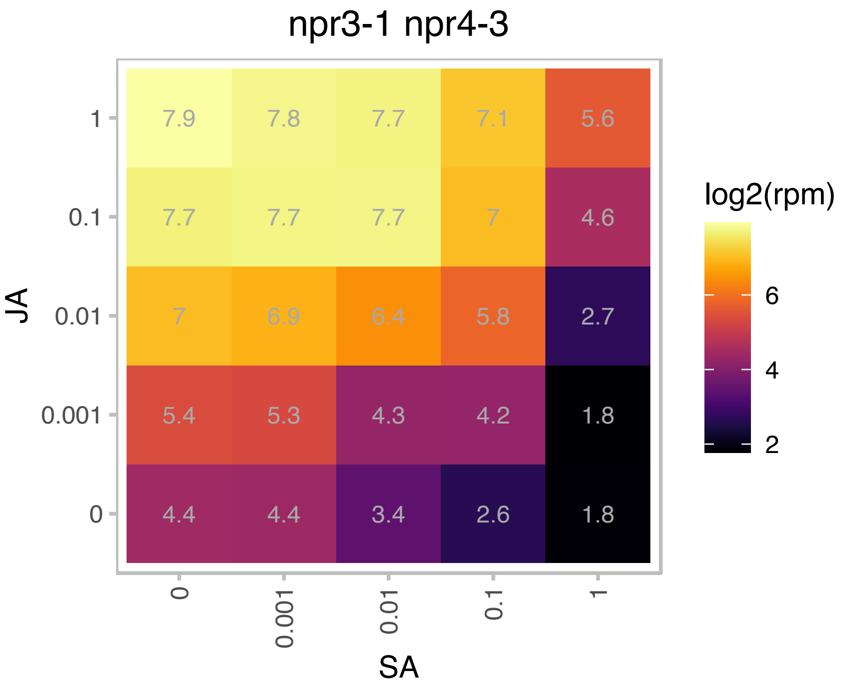

### AT4G30530.1 : GGP1

Class I glutamine amidotransferase-like superfamily protein

Encodes a gamma-glutamyl peptidase, outside the GGT family, that can hydrolyze gamma-glutamyl peptide bonds. The mRNA is cell-to-cell mobile.

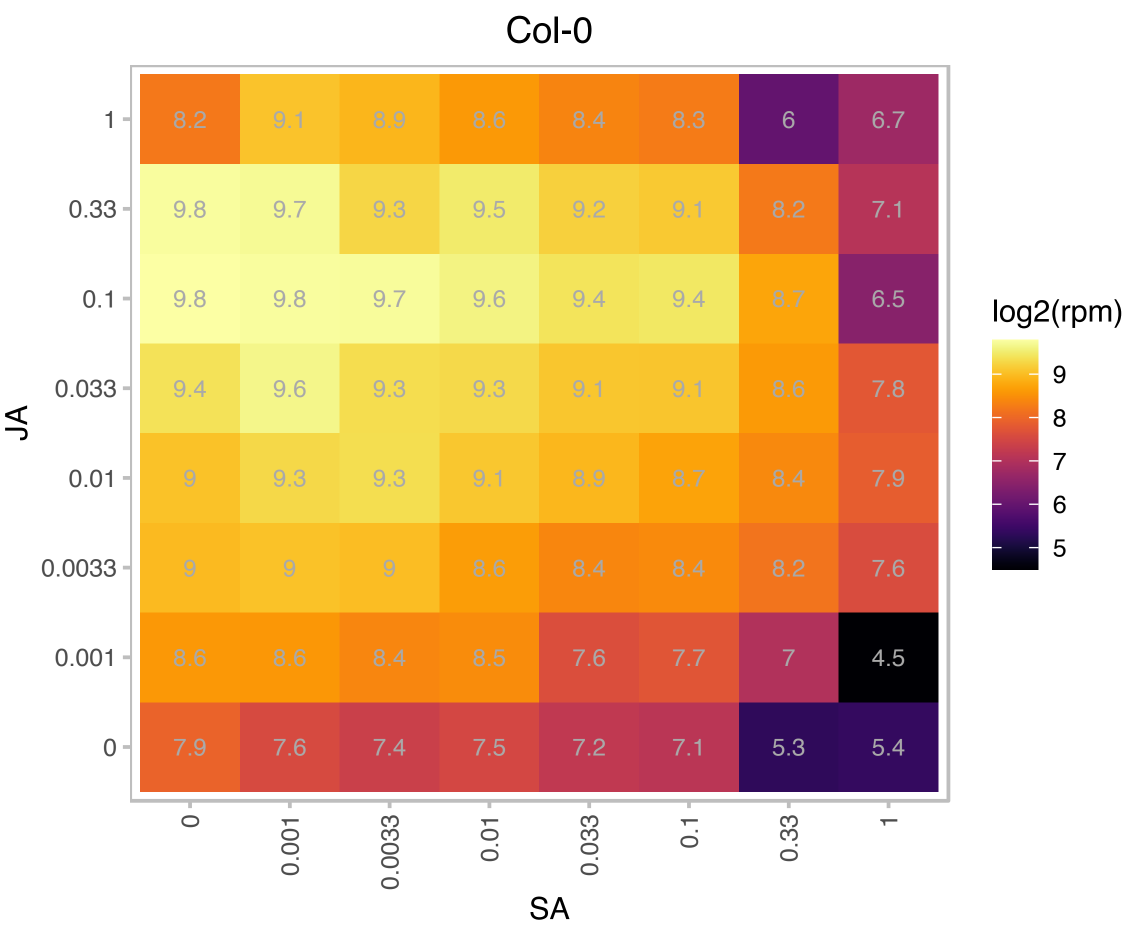


| | |
|---|---|
| Gene | AT4G30530 |
| is.expressed_genes | TRUE |
| is.responsive_genes | TRUE |
| is.apcluster | E-02 |

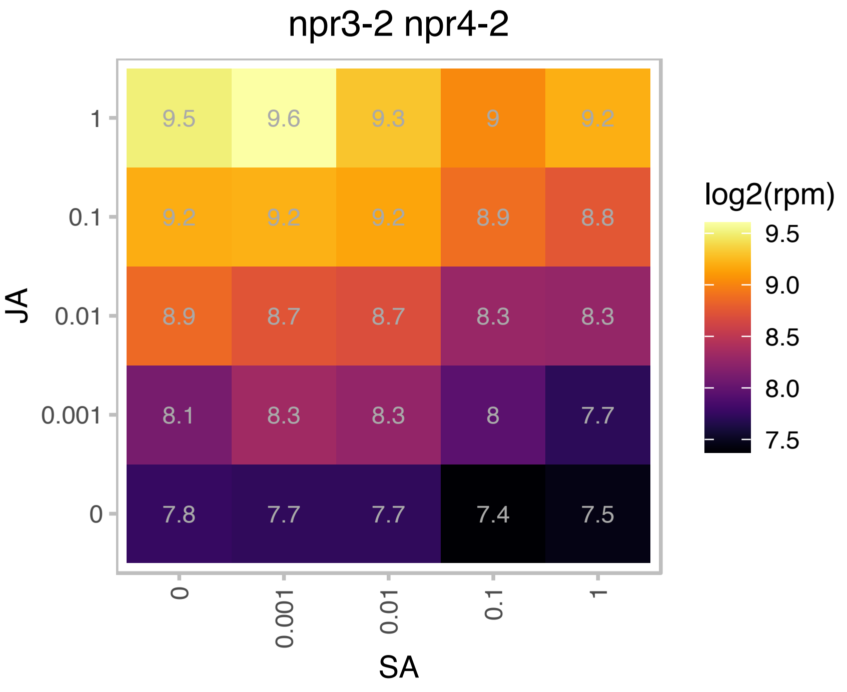


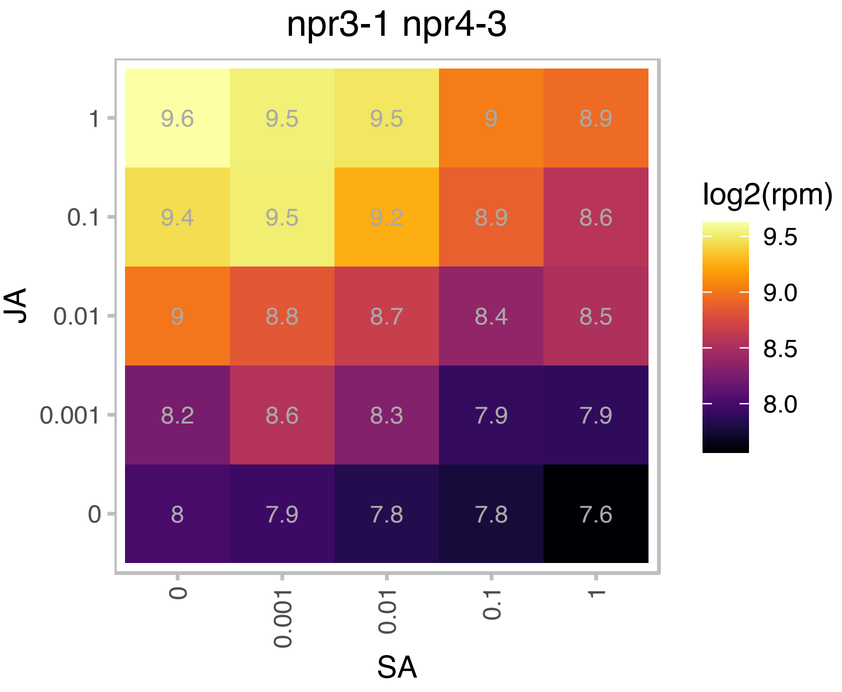

### AT4G37410.1 : CYP81F4

cytochrome P450, family 81, subfamily F, polypeptide 4

member of CYP81F The mRNA is cell-to-cell mobile.

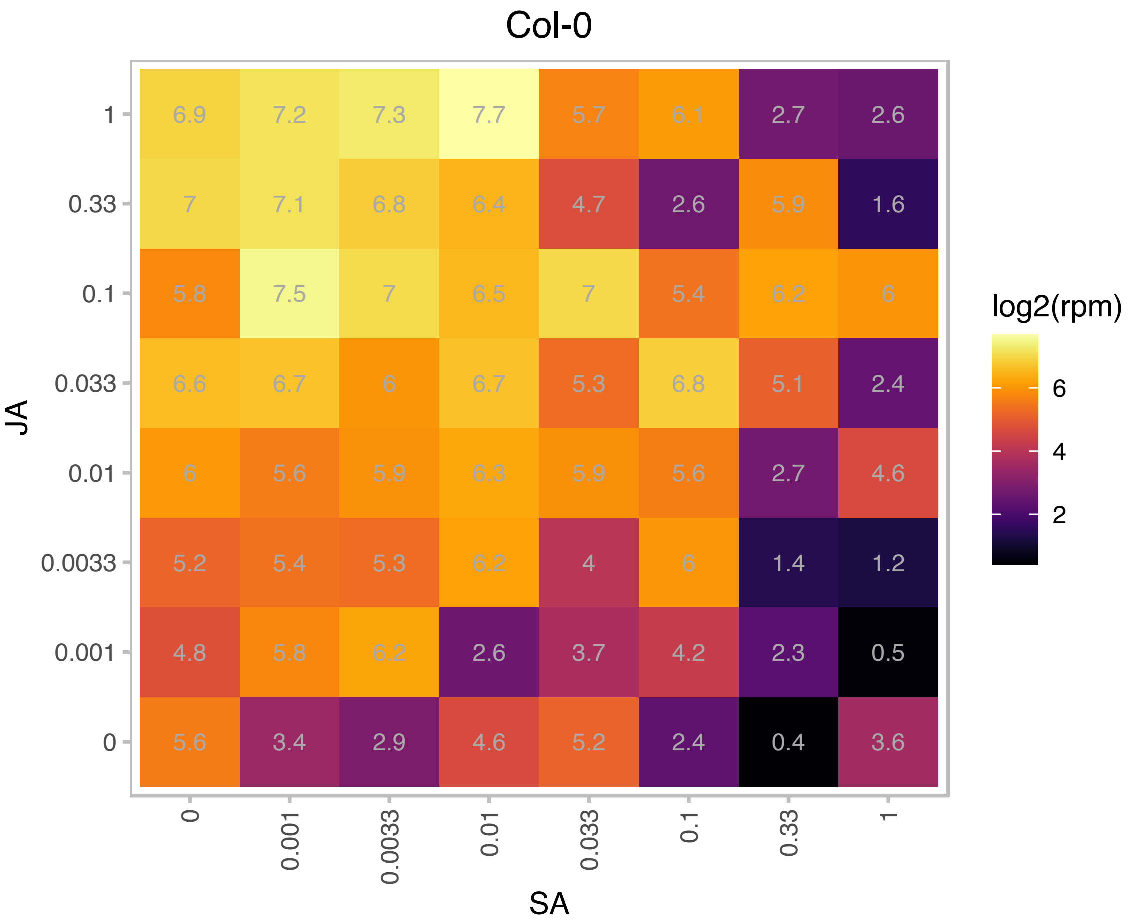


| | |
|---|---|
| Gene | AT4G37410 |
| is.expressed_genes | TRUE |
| is.responsive_genes | TRUE |
| is.apcluster | E-02 |

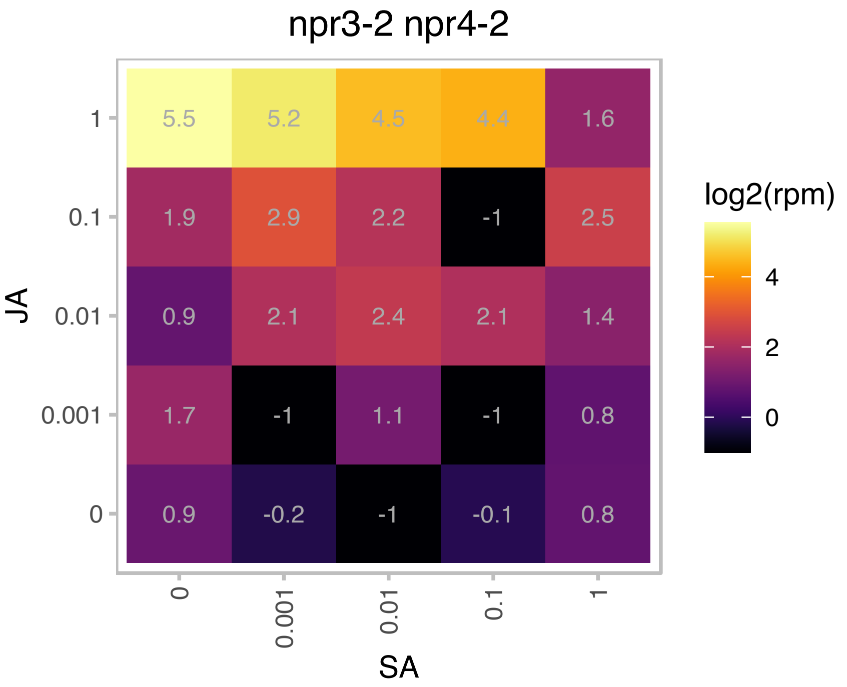


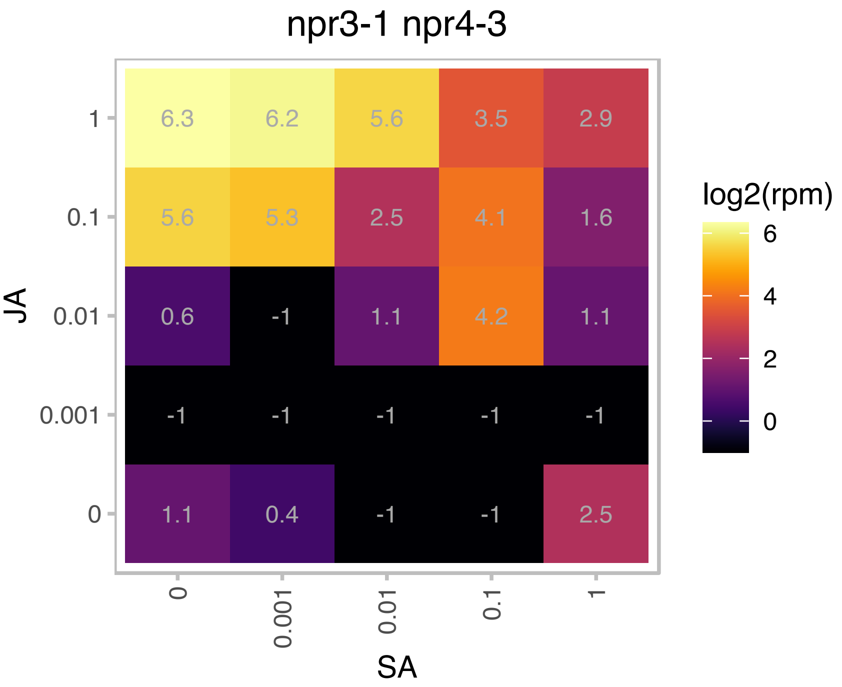

### AT4G39950.1 : CYP79B2

cytochrome P450, family 79, subfamily B, polypeptide 2

Belongs to cytochrome P450 and is involved in tryptophan metabolism. Converts Trp to indo-3-acetaldoxime (IAOx), a precursor to IAA and indole glucosinolates. The mRNA is cell-to-cell mobile.

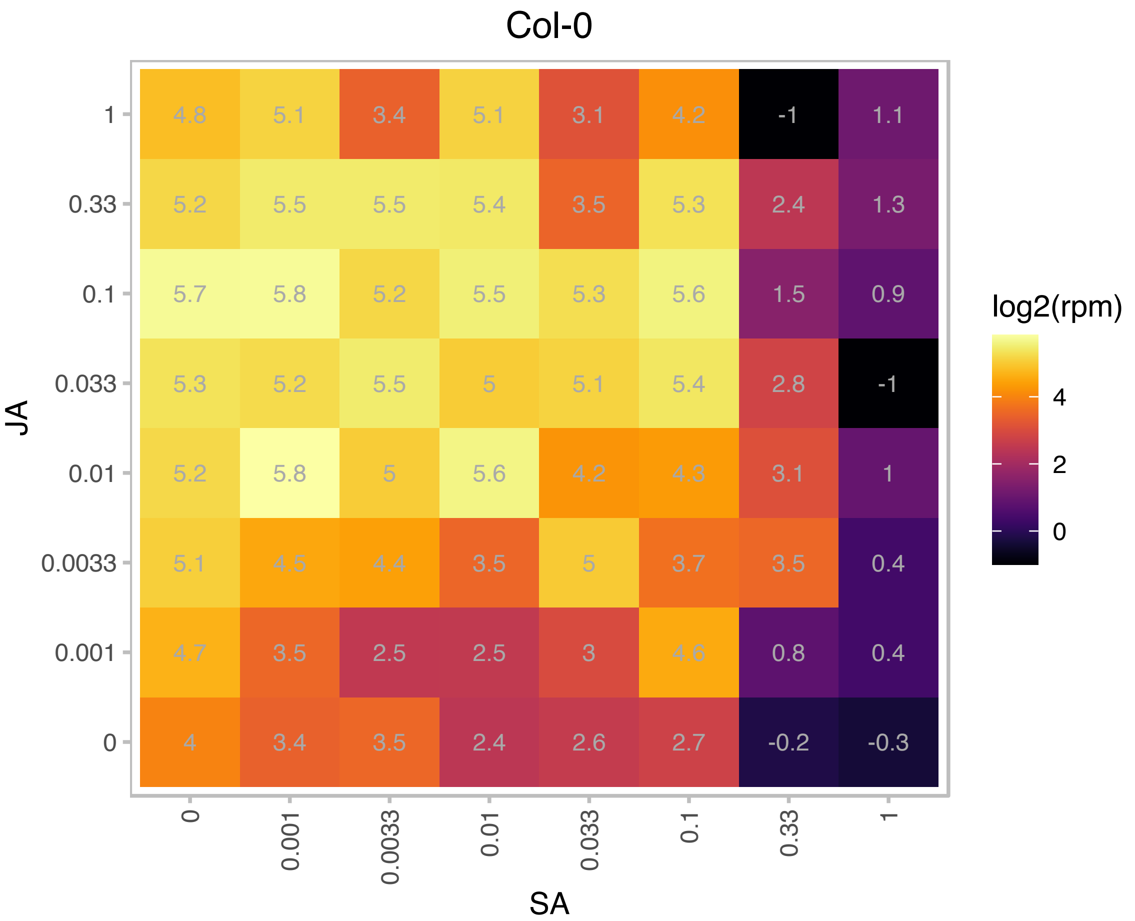


| | |
|---|---|
| Gene | AT4G39950 |
| is.expressed_genes | TRUE |
| is.responsive_genes | TRUE |
| is.apcluster | E-02 |

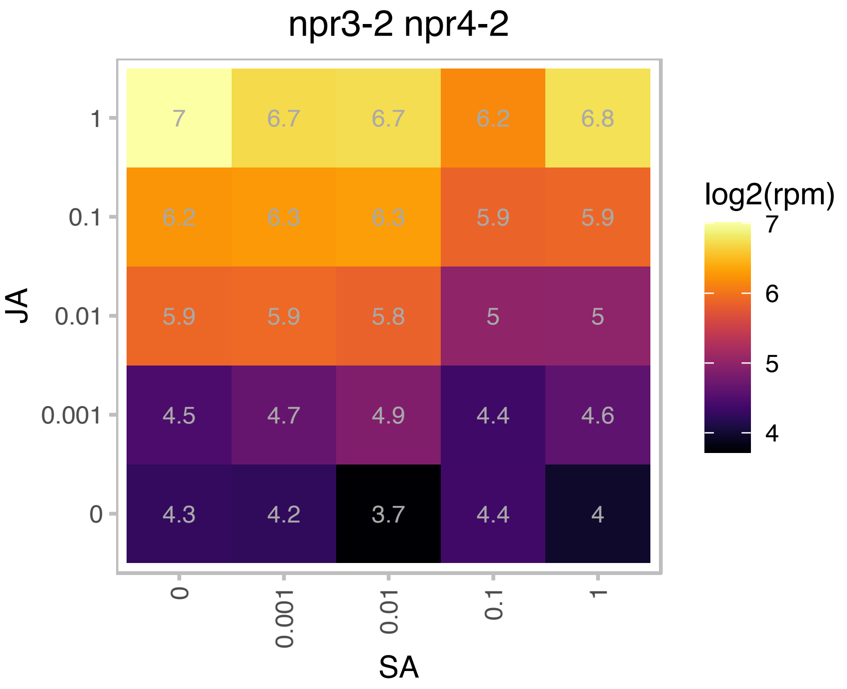


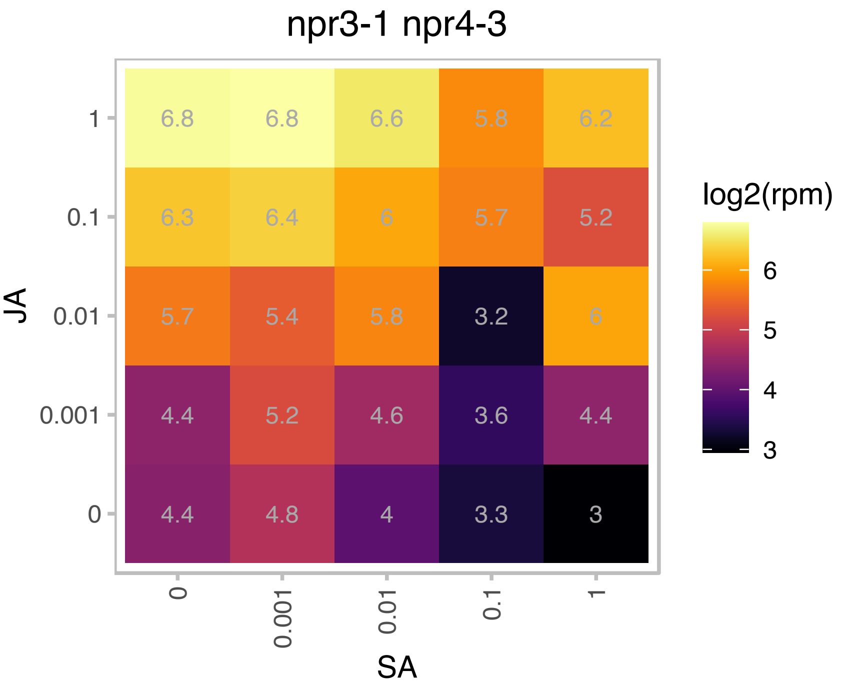

### AT4G39980.1 : AtDAHP1, DAHP1, DHS1

3-deoxy-D-arabino-heptulosonate 7-phosphate synthase 1
Encodes a 2-deoxy-D-arabino-heptulosonate 7-phosphate (DAHP) synthase, which catalyzes the first committed step in aromatic amino acid biosynthesis. Gene expression is induced by wounding and pathogenic bacteria Pseudomonas syringae. The mRNA is cell-to-cell mobile.

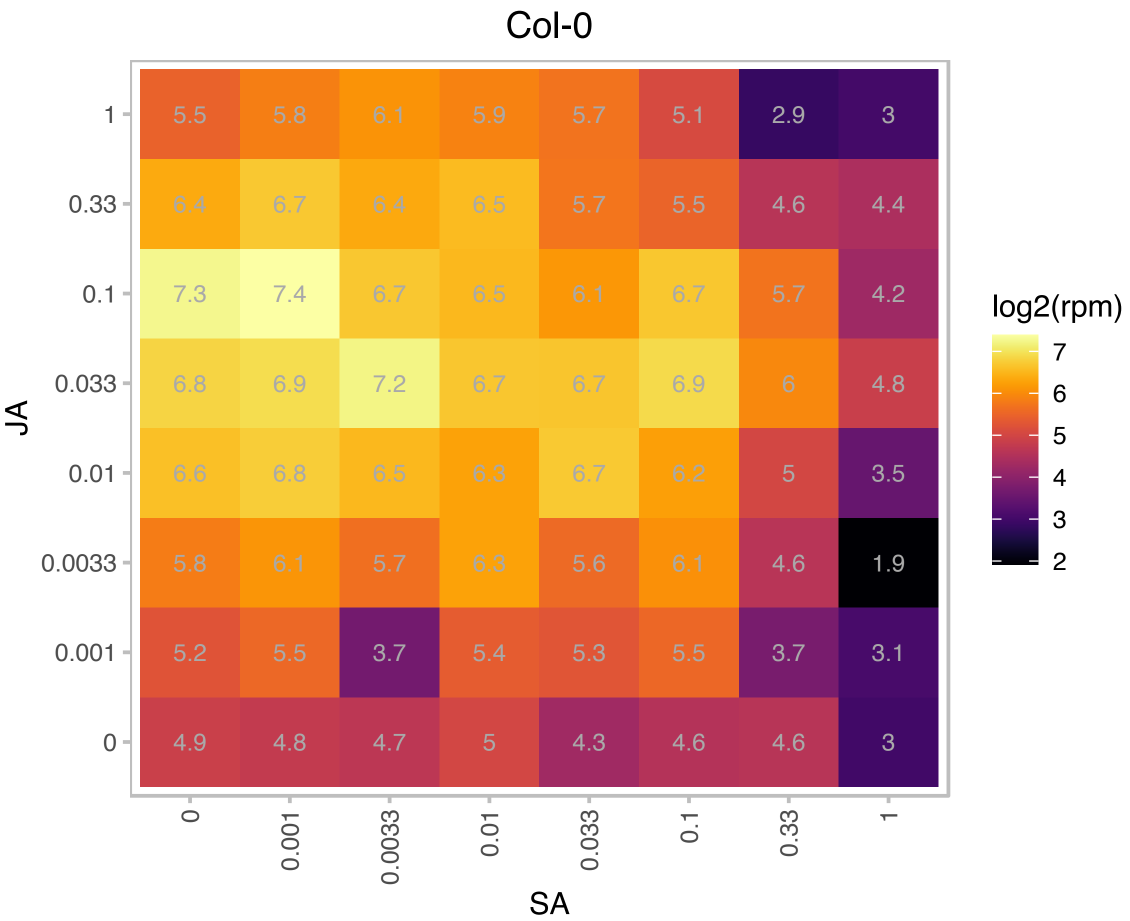


| | |
|---|---|
| Gene | AT4G39980 |
| is.expressed_genes | TRUE |
| is.responsive_genes | TRUE |
| is.apcluster | E-02 |

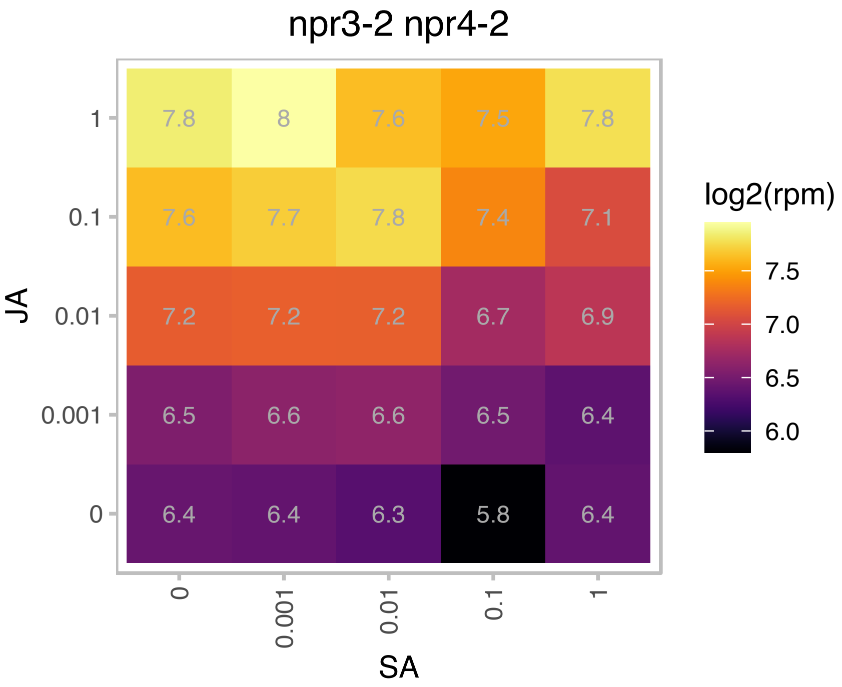


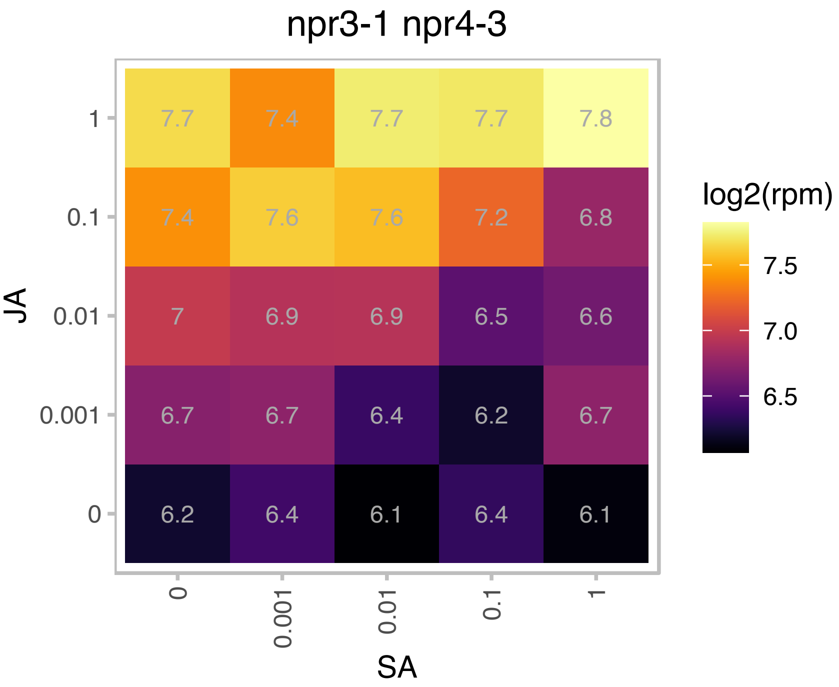

# AT5G08060.1 : NA

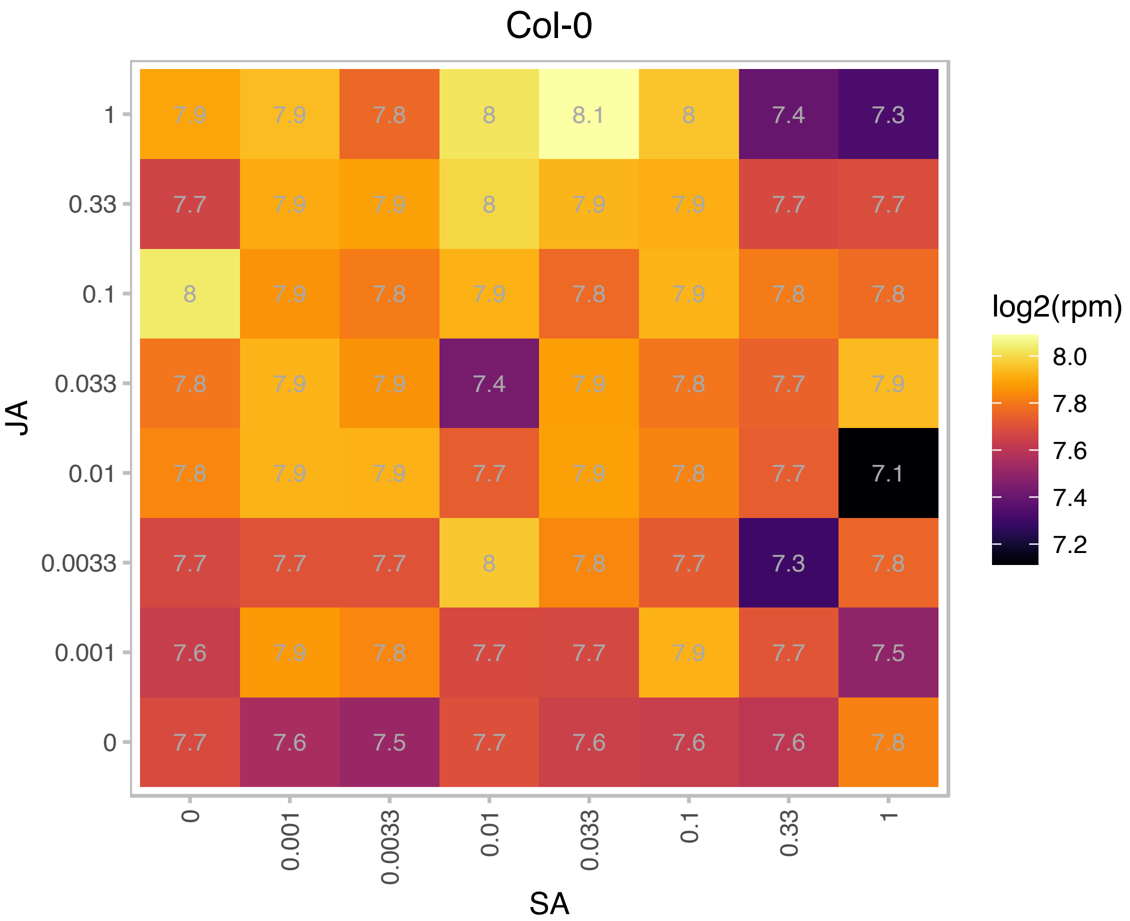


| | |
|---|---|
| Gene | AT5G08060 |
| is.expressed_genes | TRUE |
| is.responsive_genes | TRUE |
| is.apcluster | E-02 |

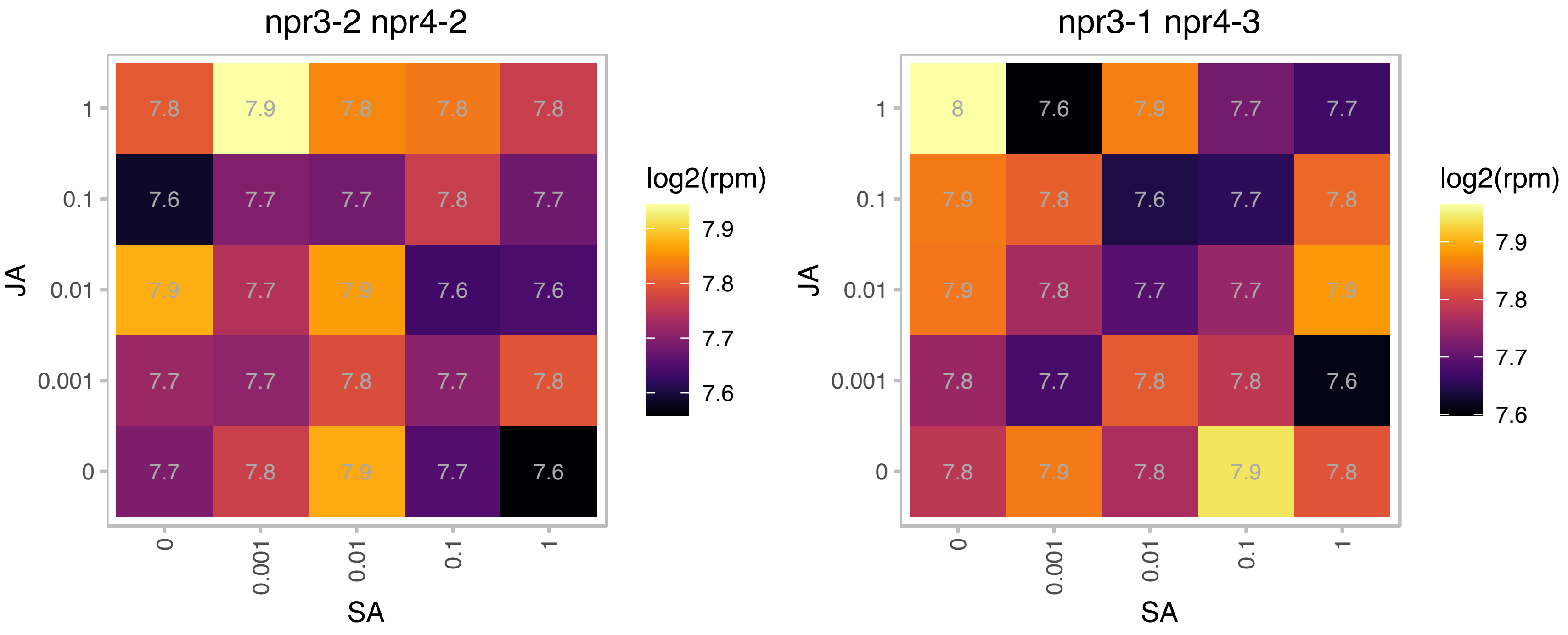

### AT5G15960.1 : KIN1

stress-responsive protein (KIN1) / stress-induced protein (KIN1)

cold and ABA inducible protein kin1, possibly functions as an anti-freeze protein. Transcript level of this gene is induced by cold, ABA, dehydration and osmoticum (mannitol). However, protein activity of GUS fused to the promoter of this gene is inhibited by cold treatment, suggesting an inhibition of the protein by increased transcript level.

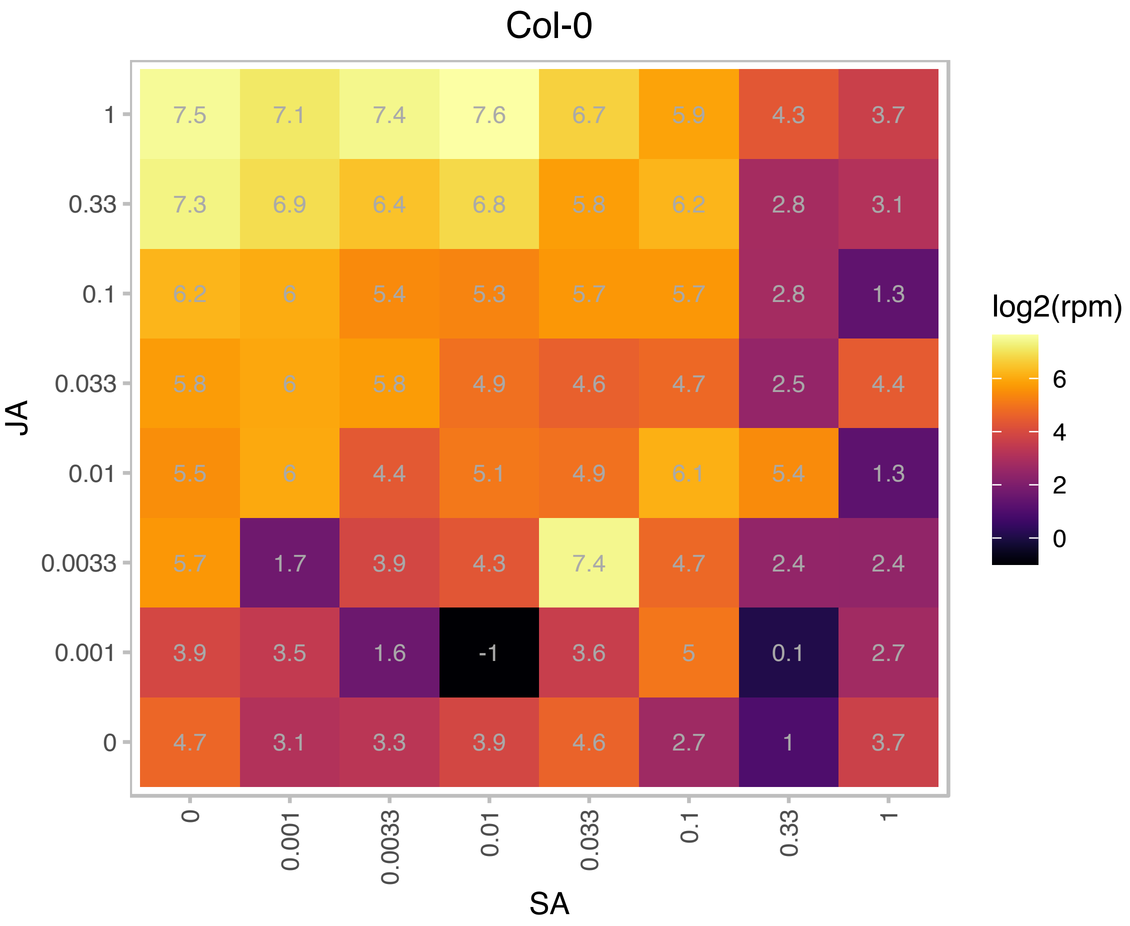


| | |
|---|---|
| Gene | AT5G15960 |
| is.expressed_genes | TRUE |
| is.responsive_genes | TRUE |
| is.apcluster | E-02 |

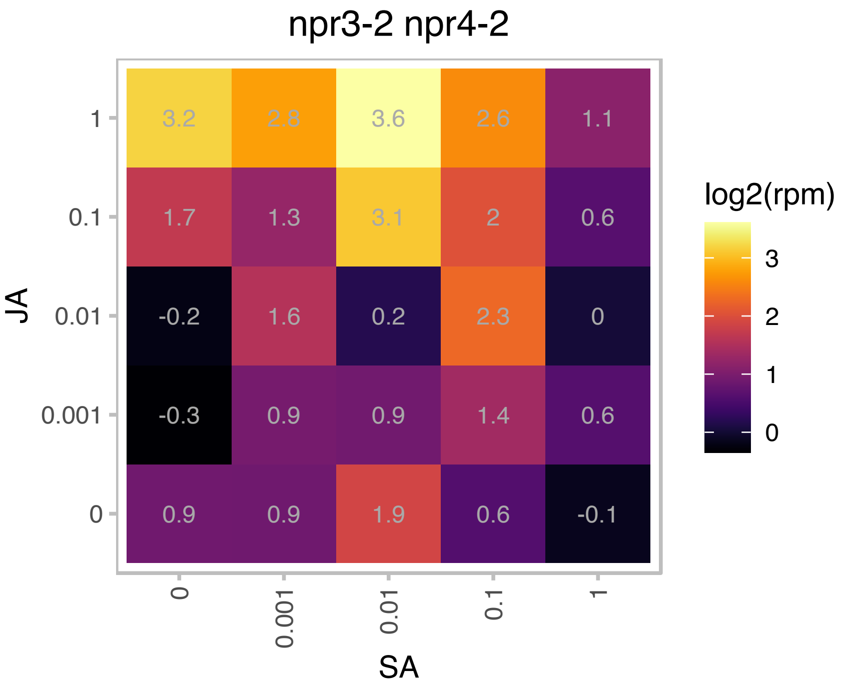


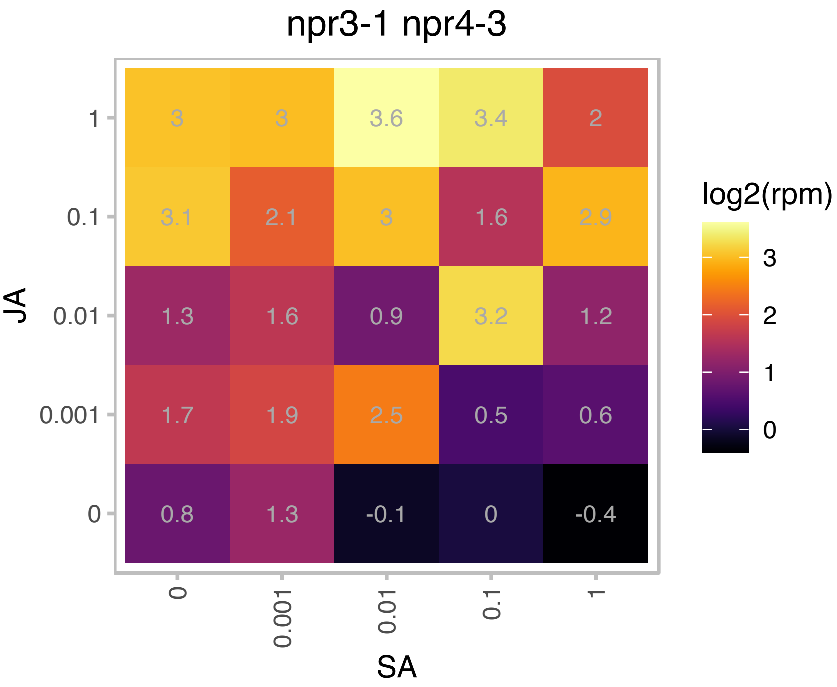

### AT5G19520.1 : ATMSL9, MSL9

mechanosensitive channel of small conductance-like 9

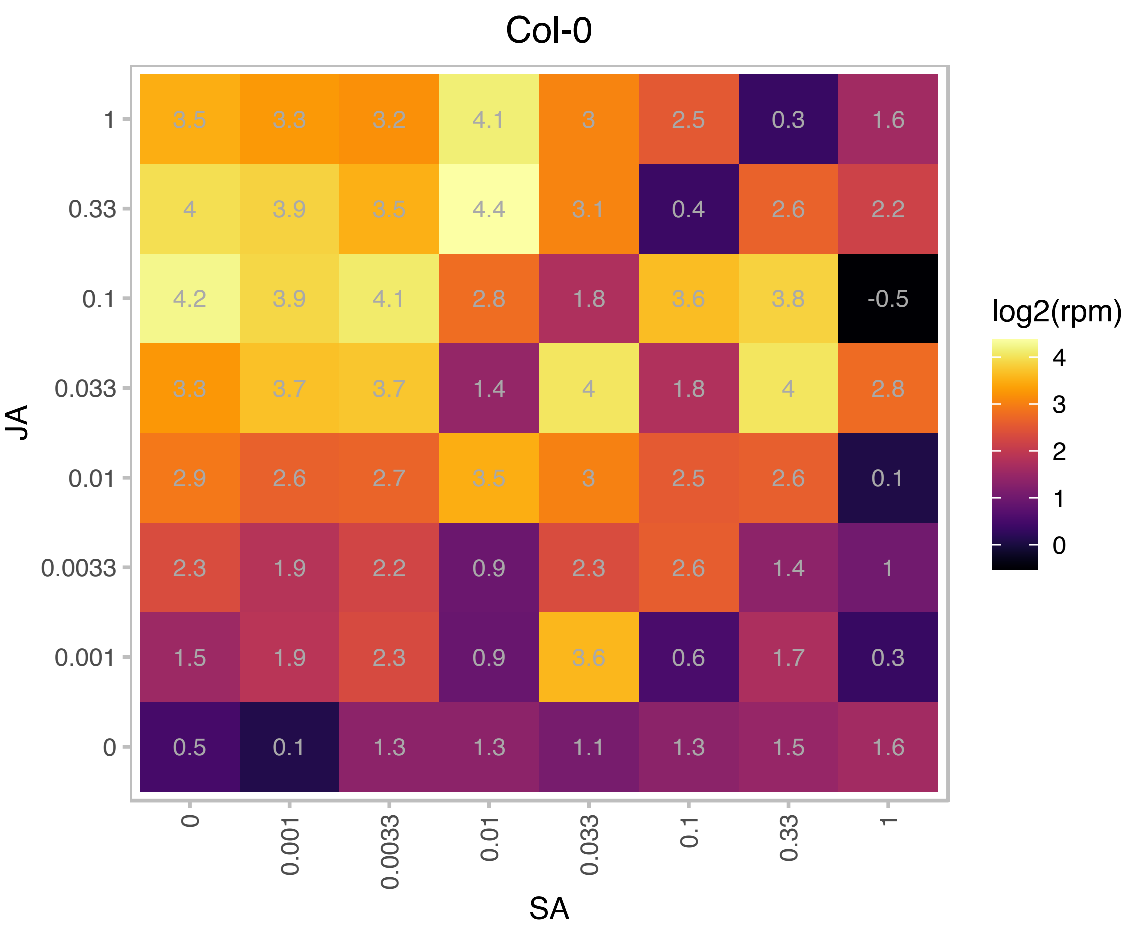


| | |
|---|---|
| Gene | AT5G19520 |
| is.expressed_genes | TRUE |
| is.responsive_genes | TRUE |
| is.apcluster | E-02 |

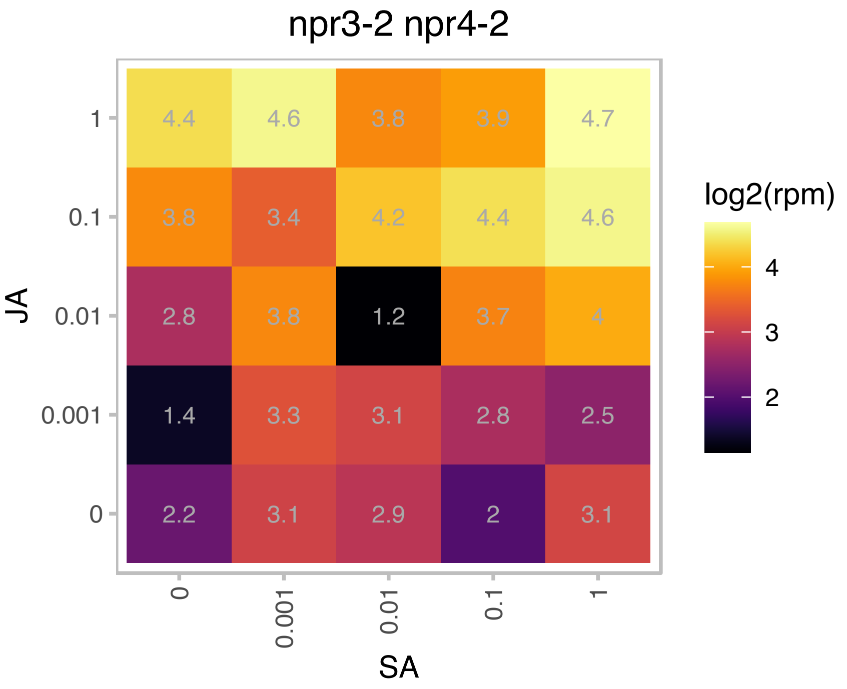


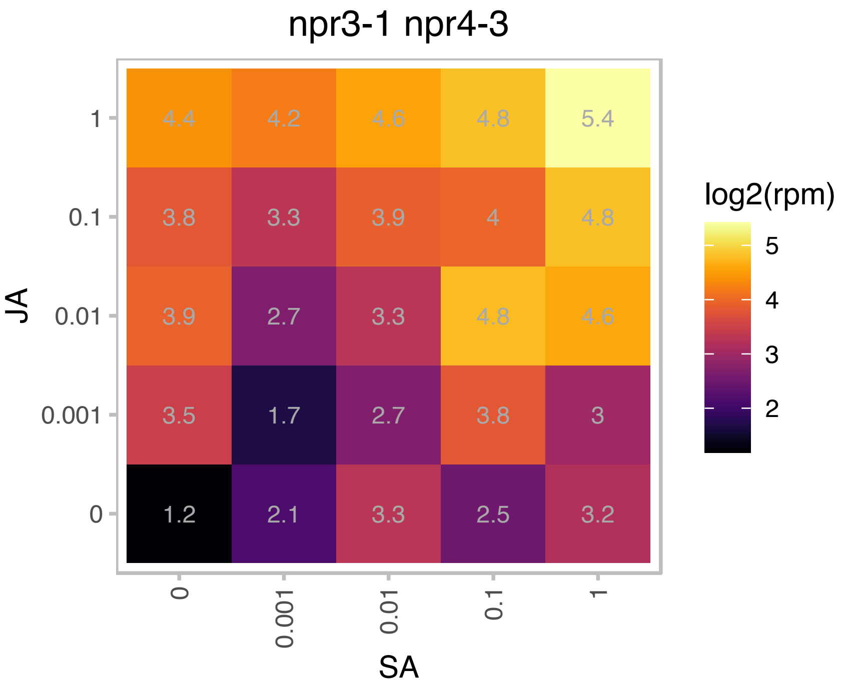

# AT5G20550.1 : NA

2-oxoglutarate (2OG) and Fe(II)-dependent oxygenase superfamily protein

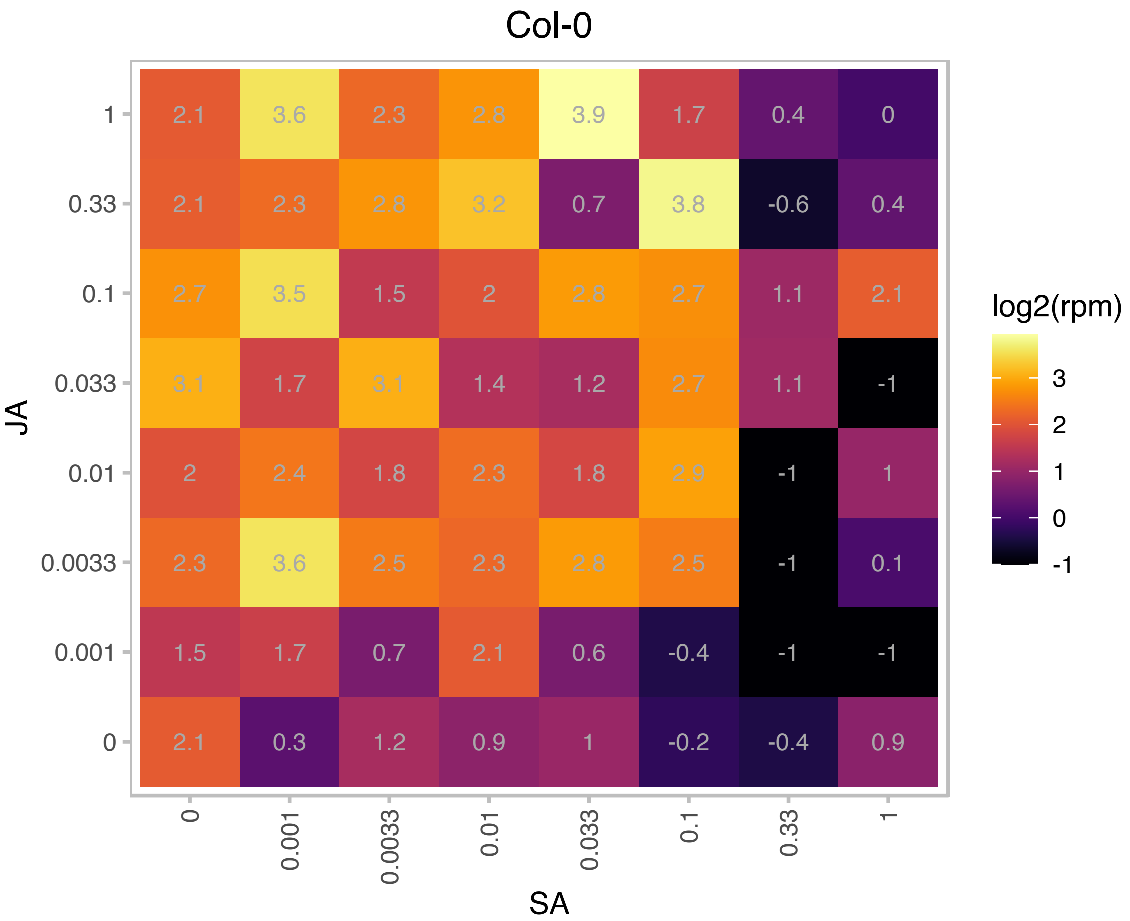


| | |
|---|---|
| Gene | AT5G20550 |
| is.expressed_genes | TRUE |
| is.responsive_genes | TRUE |
| is.apcluster | E-02 |

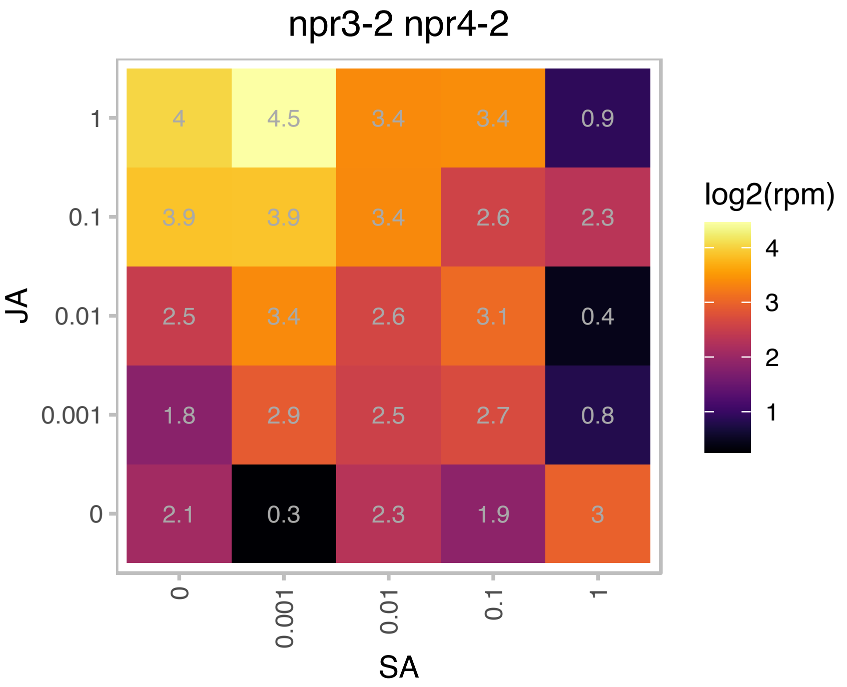


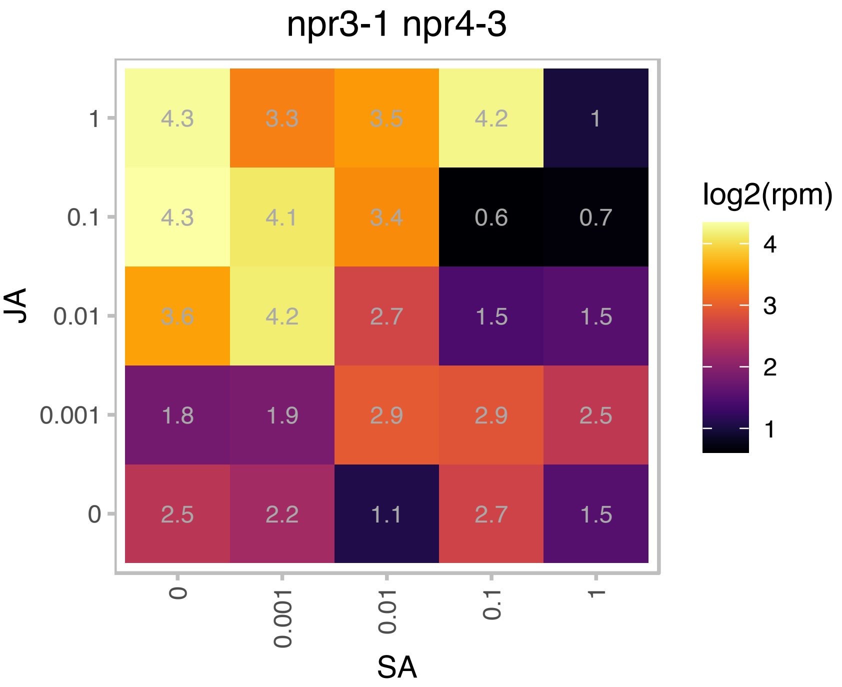

### AT5G23140.1 : CLPP2, NCLPP7

nuclear-encoded CLP protease P7

One of several nuclear-encoded ClpPs (caseinolytic protease). Contains a highly conserved catalytic triad of Ser-type proteases (Ser-His-Asp). The name reflects nomenclature described in Adam et. al (2001).

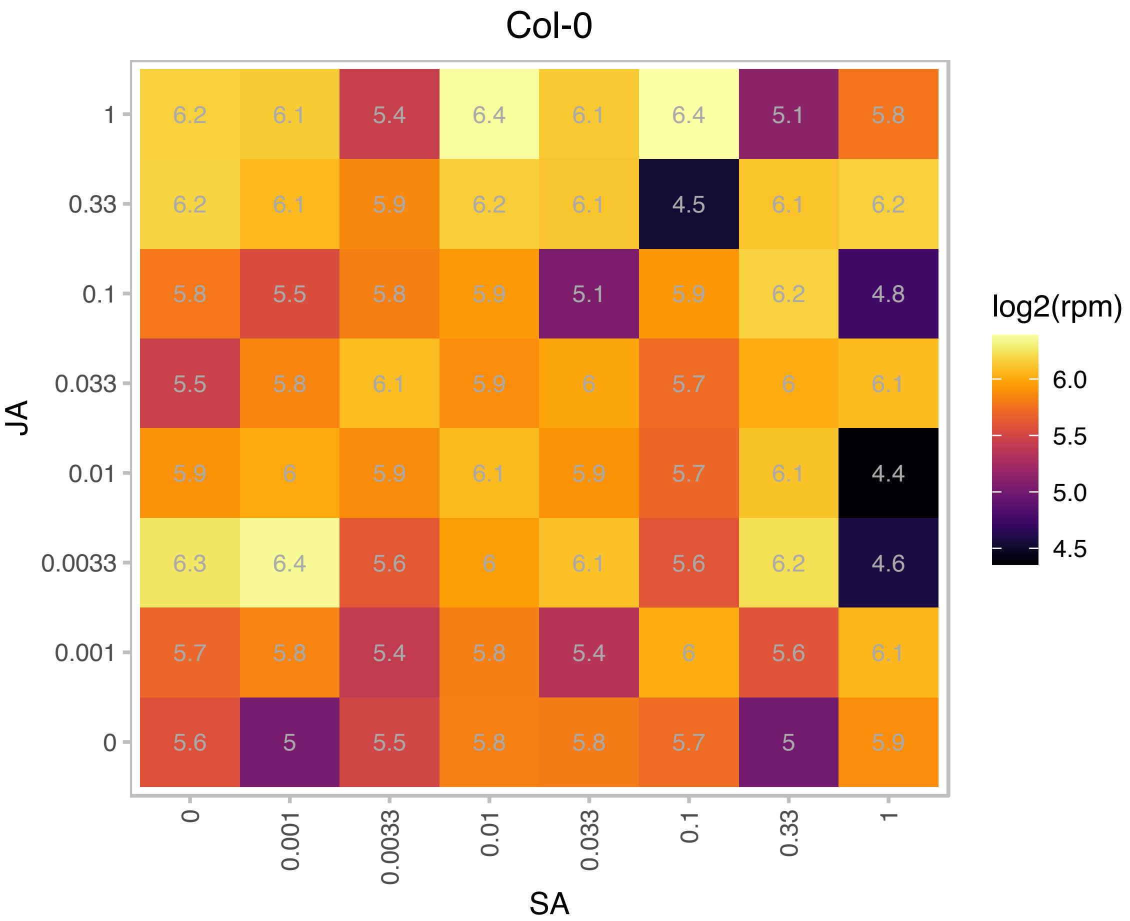


| | |
|---|---|
| Gene | AT5G23140 |
| is.expressed_genes | TRUE |
| is.responsive_genes | TRUE |
| is.apcluster | E-02 |

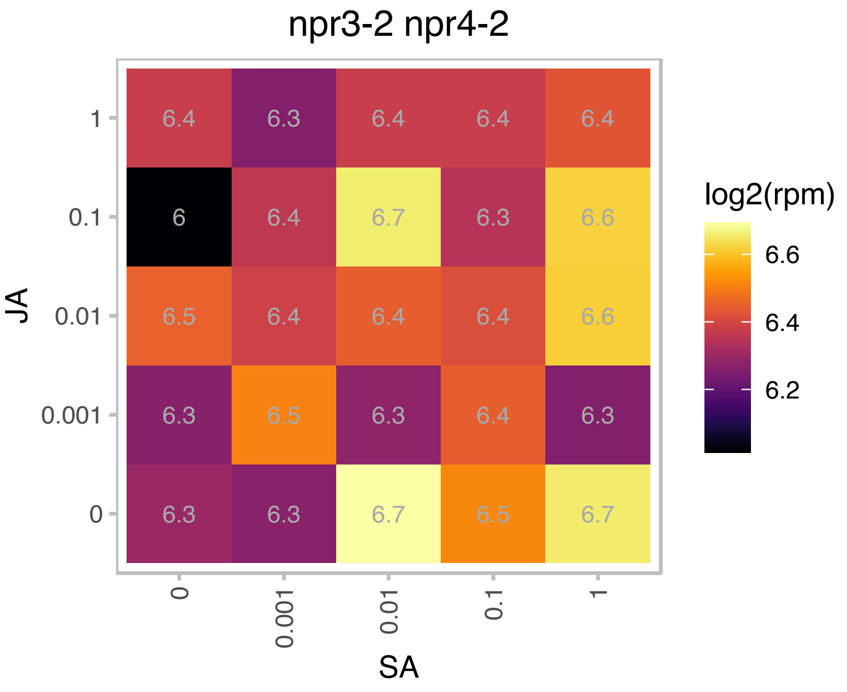


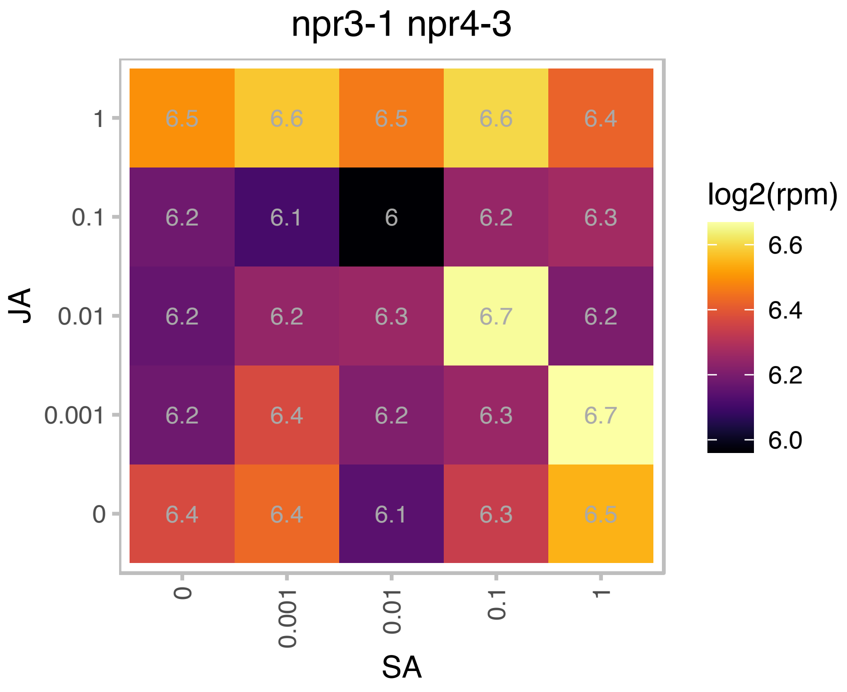

# AT5G23820.1 : ML3

MD-2-related lipid recognition domain-containing protein
ML3 can be modified by NEDD8 and ubiquitin. ML3 expression is regulated by NAI1. ML3 expression is regulated by MeJA, ethylene and wounding. ml3-3 is more susceptible against infections with Alternaria brassicicola and more resistant against infections with Pseudomonas syringae DC3000.

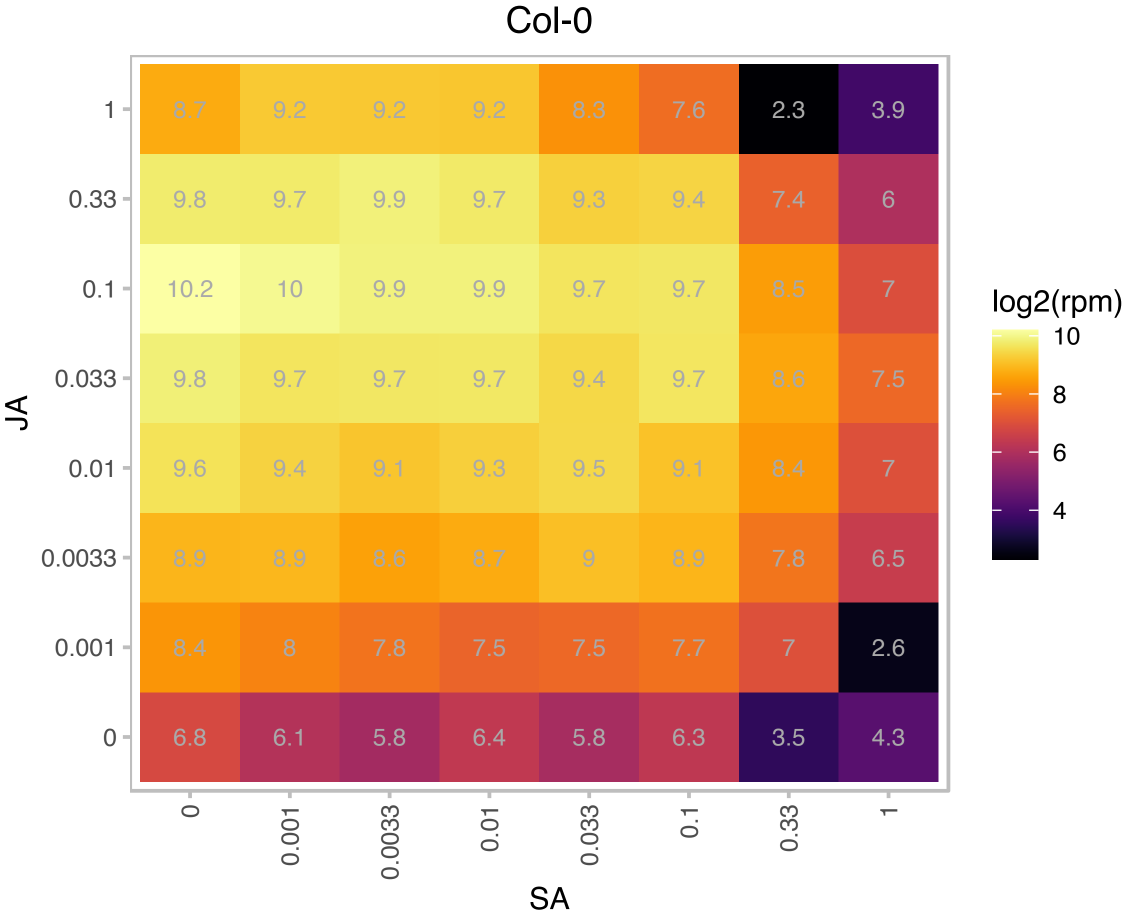


| | |
|---|---|
| Gene | AT5G23820 |
| is.expressed_genes | TRUE |
| is.responsive_genes | TRUE |
| is.apcluster | E-02 |

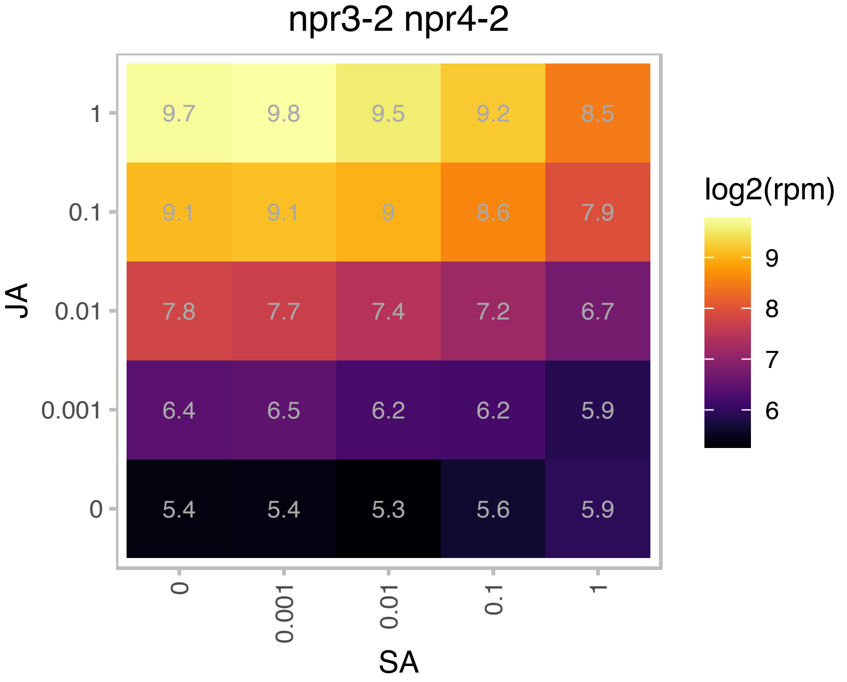


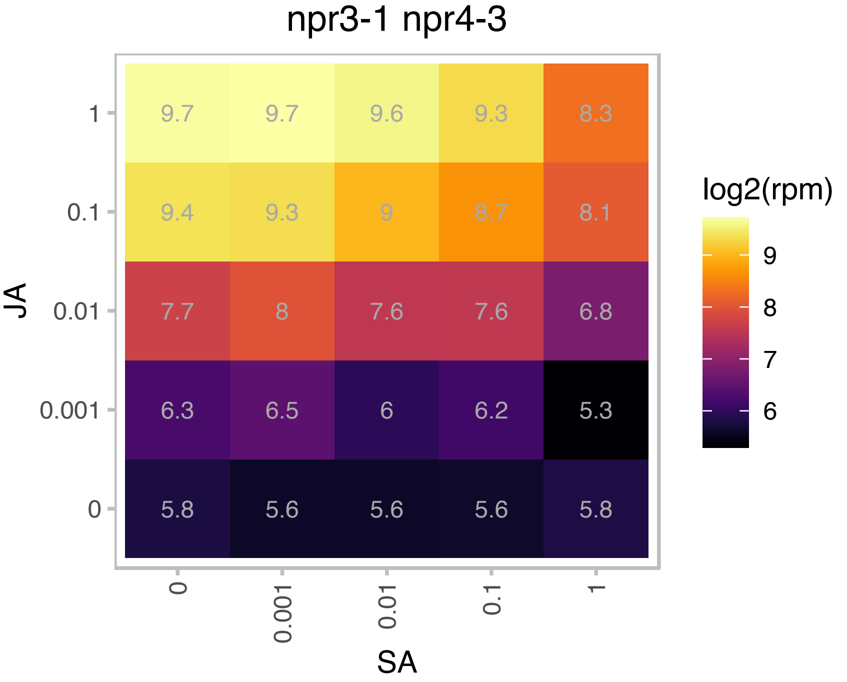

# AT5G23830.1 : NA

MD-2-related lipid recognition domain-containing protein

#### Col-0

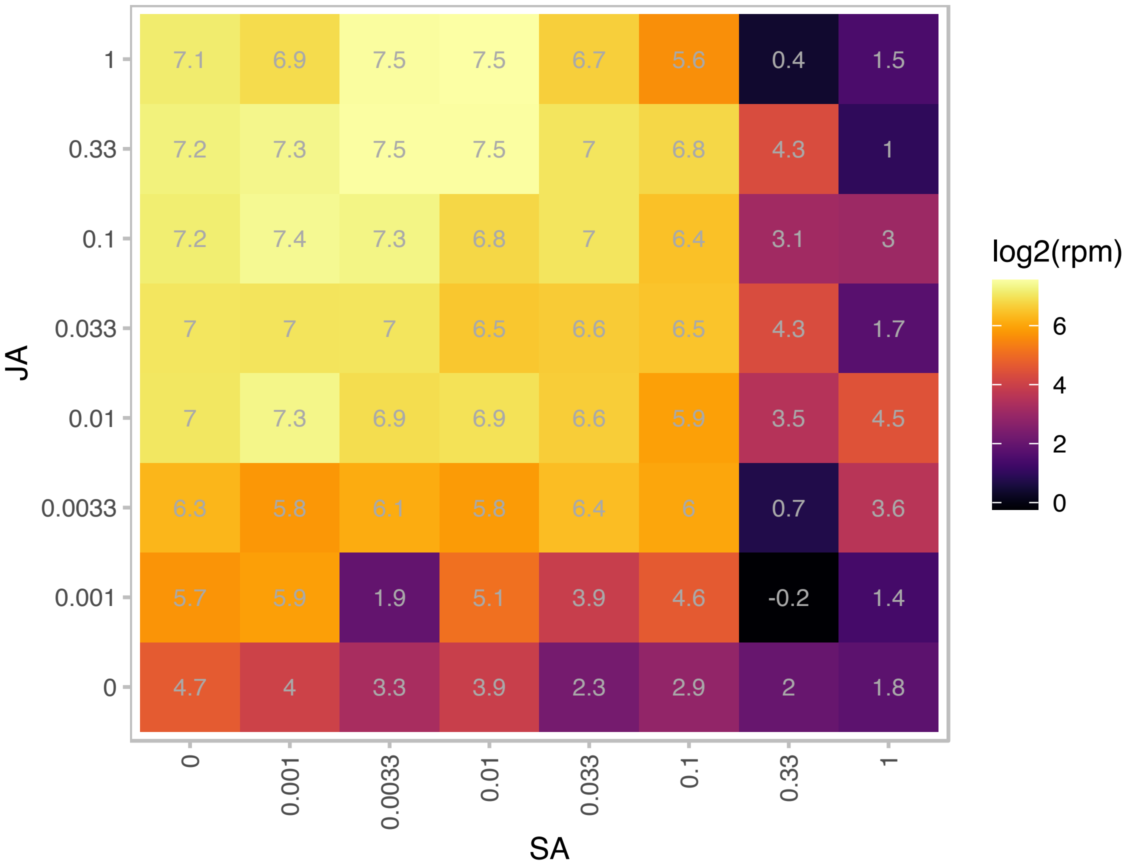


| | |
|---|---|
| Gene | AT5G23830 |
| is.expressed_genes | TRUE |
| is.responsive_genes | TRUE |
| is.apcluster | E-02 |

#### npr3-2 npr4-2

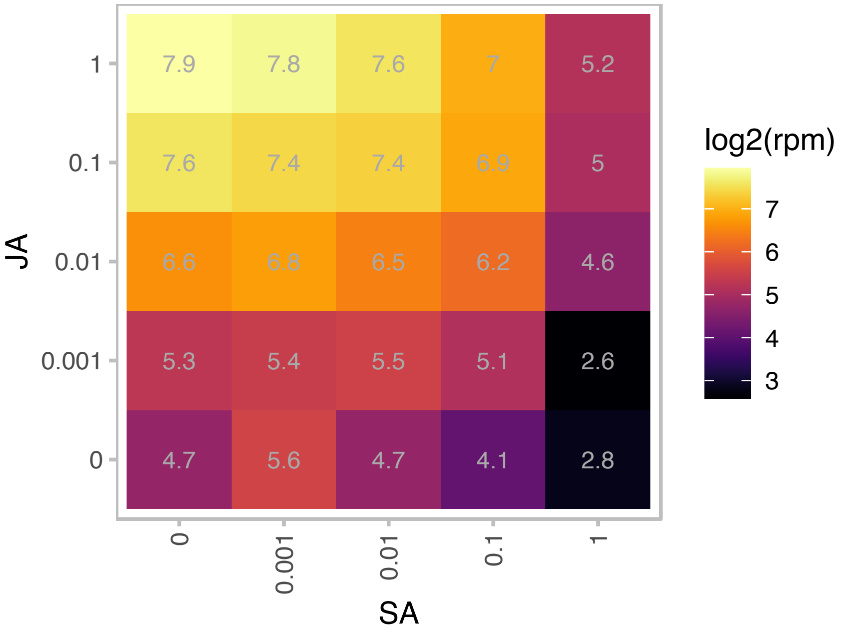


#### npr3-1 npr4-3

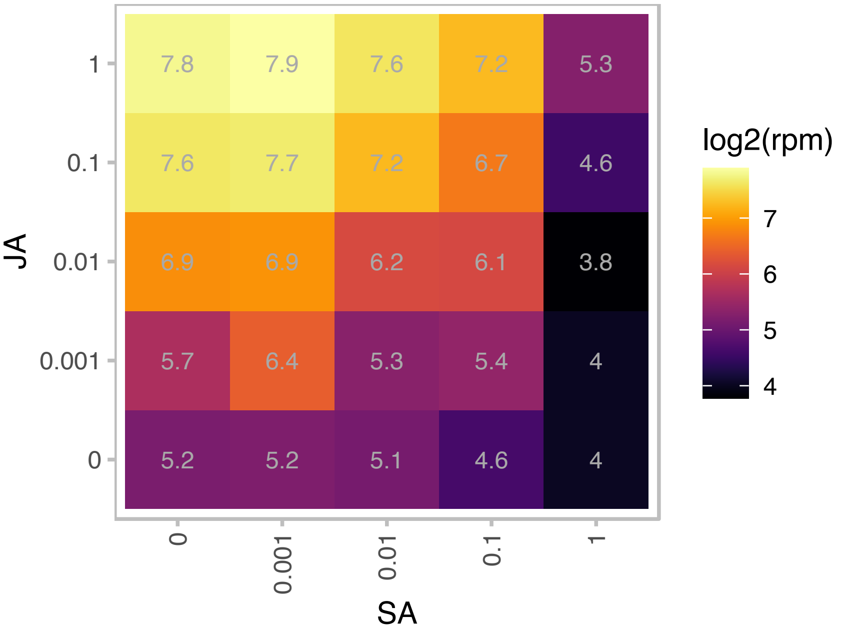

# AT5G47030.1 : NA

ATPase, F1 complex, delta/epsilon subunit

Encodes the mitochondrial ATP synthase subunit delta.

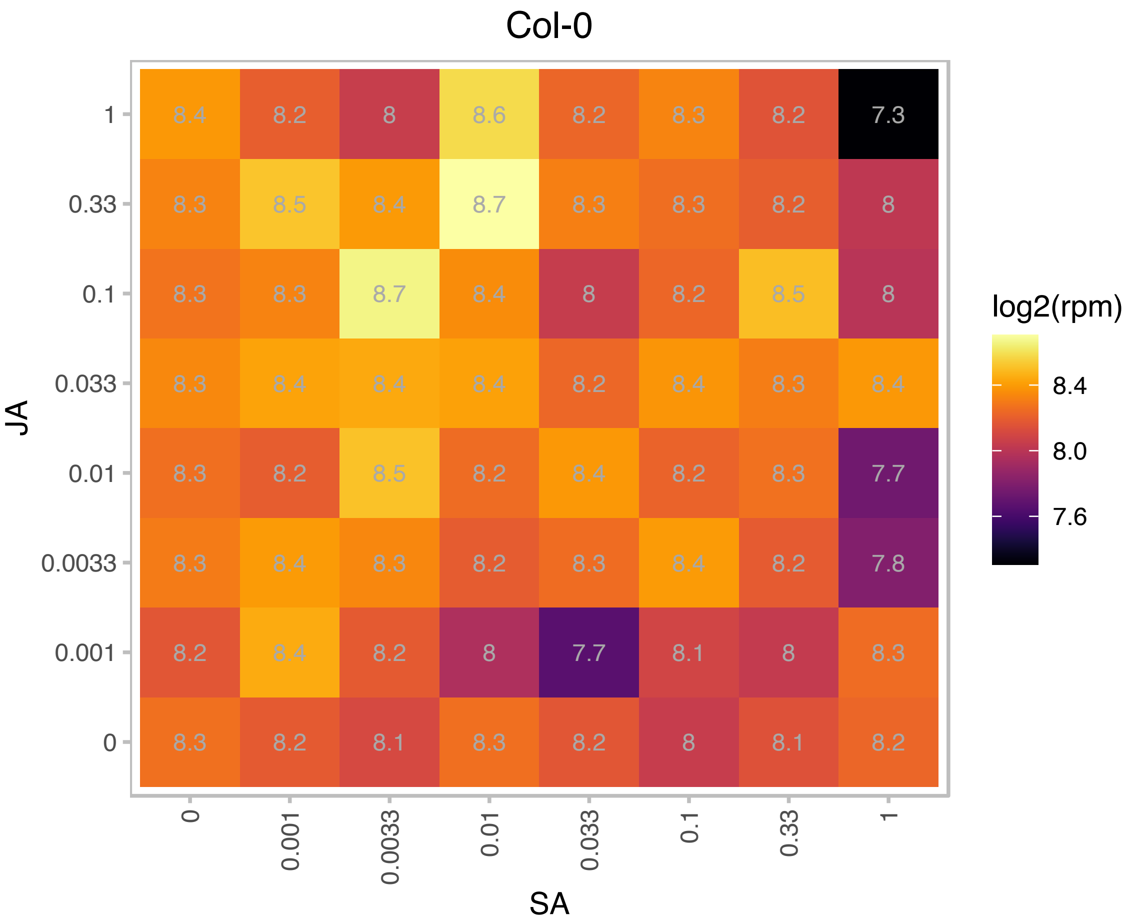


| | |
|---|---|
| Gene | AT5G47030 |
| is.expressed_genes | TRUE |
| is.responsive_genes | TRUE |
| is.apcluster | E-02 |

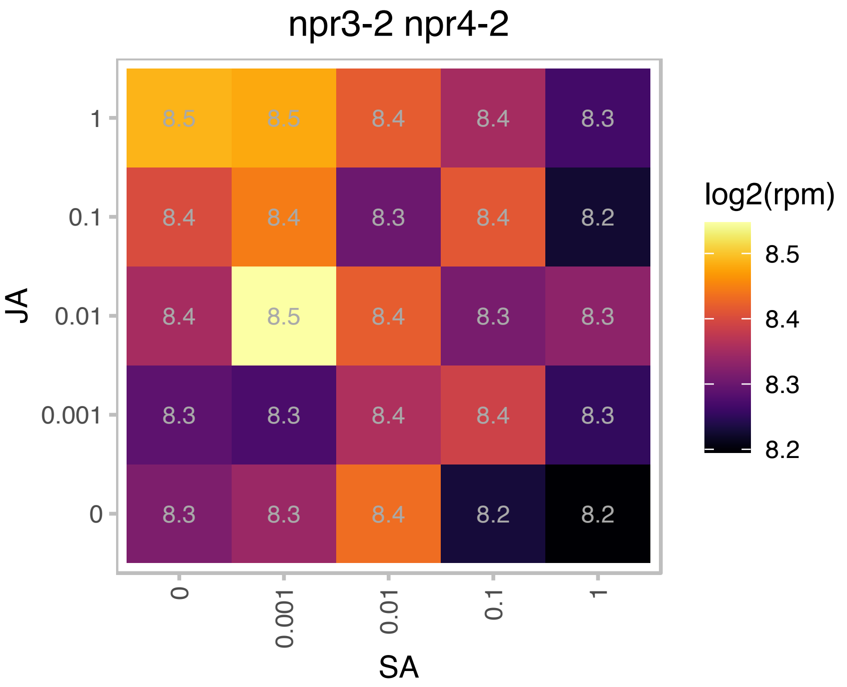


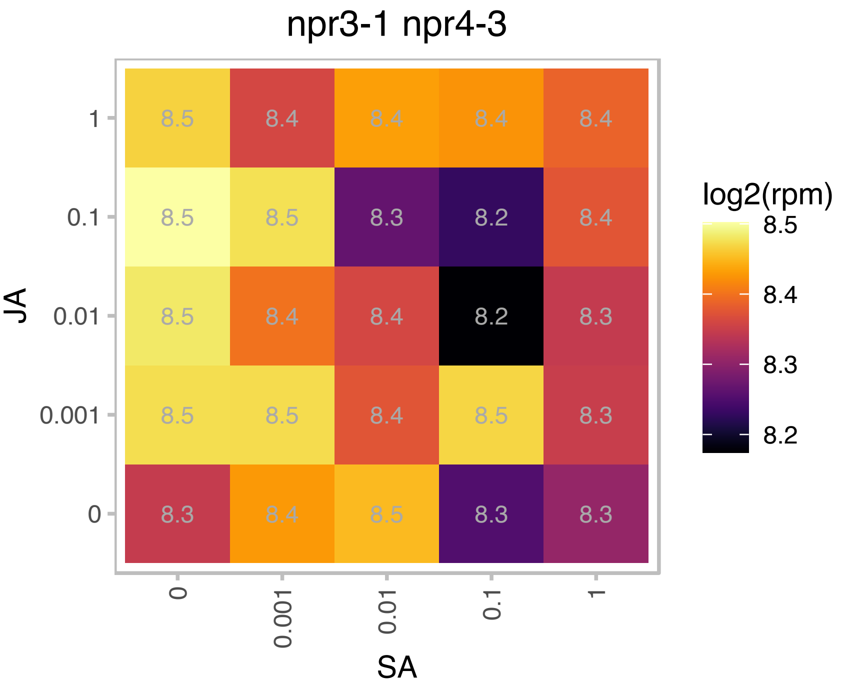

# AT5G47740.1 : NA

Adenine nucleotide alpha hydrolases-like superfamily protein

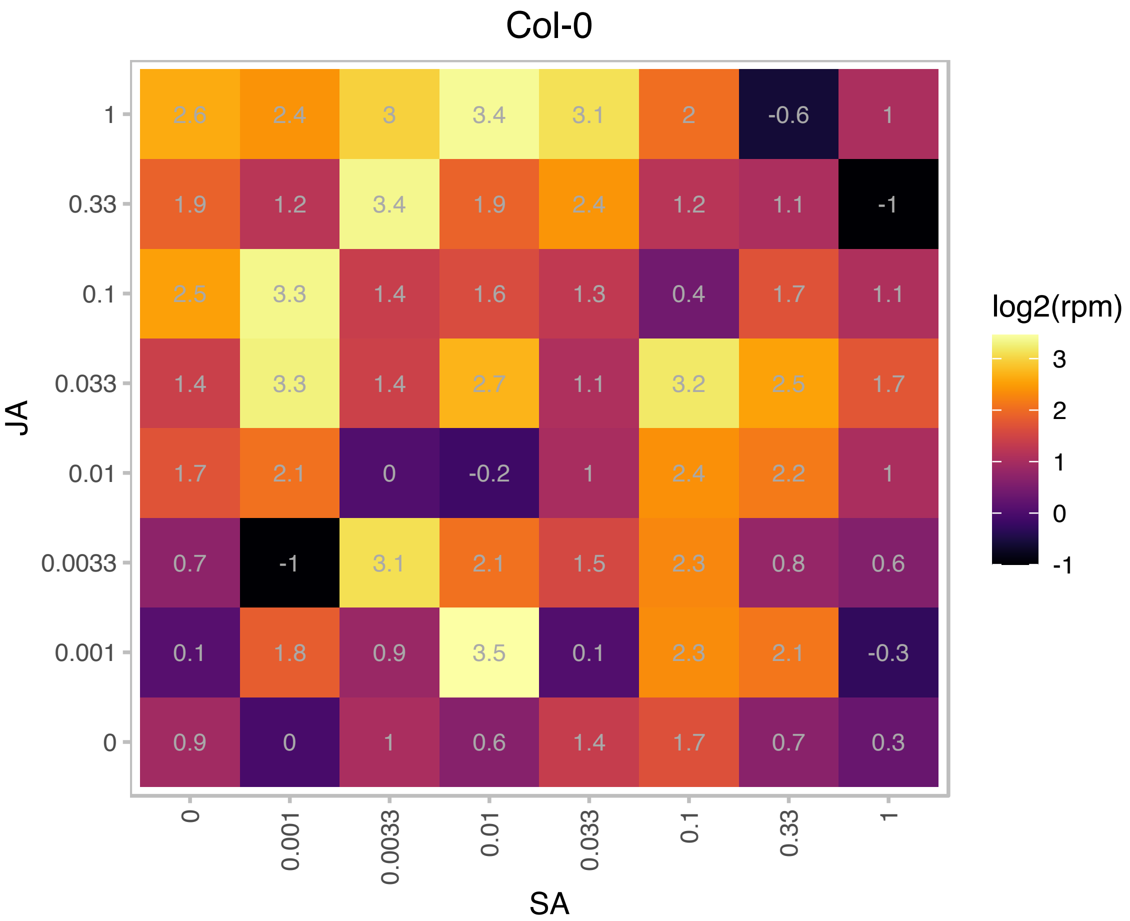


| | |
|---|---|
| Gene | AT5G47740 |
| is.expressed_genes | TRUE |
| is.responsive_genes | TRUE |
| is.apcluster | E-02 |

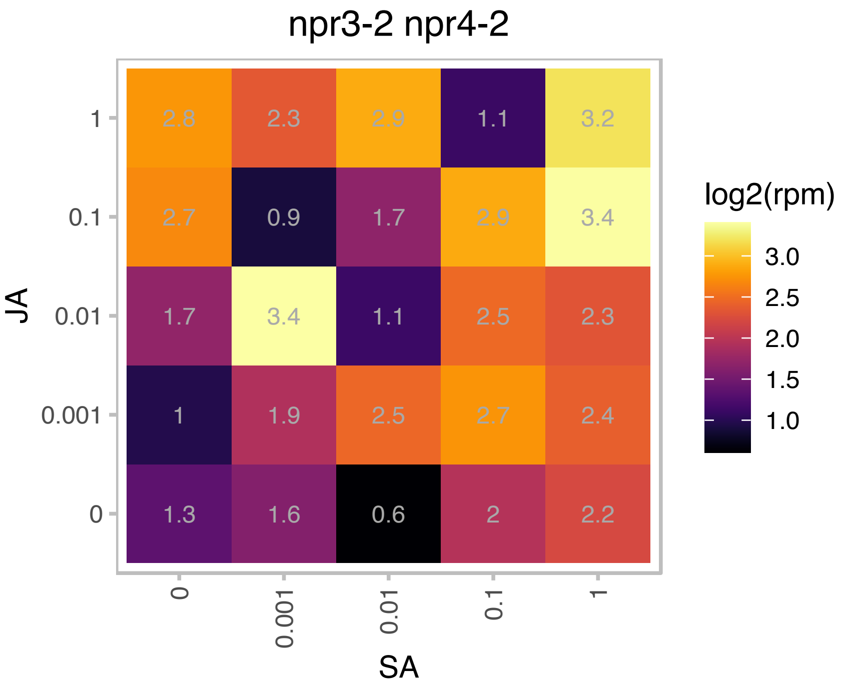


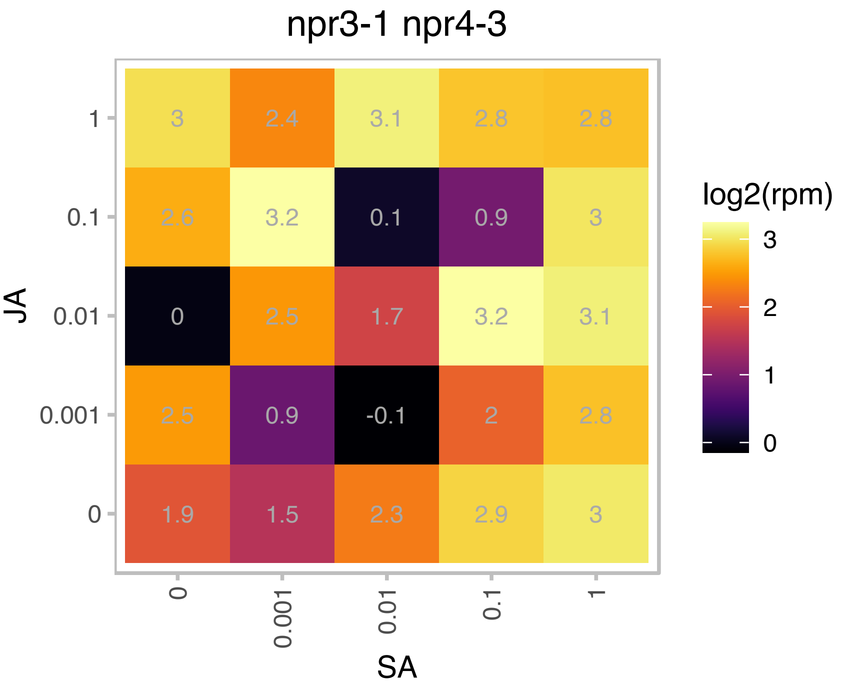

### AT5G47990.1 : CYP705A5, THAD, THAD1

cytochrome P450, family 705, subfamily A, polypeptide 5
Encodes an endomembrane system-expressed member of the CYP705A family of cytochrome P450 enzymes. It ap
to catalyze the addition of a double bond to thalian-diol at carbon 15. Reduced levels of THAD expression lead to a build up of thalian-diol in root extracts. thad1-1 mutants also have longer roots than wild type seedlings and show altered gravitropic responses.

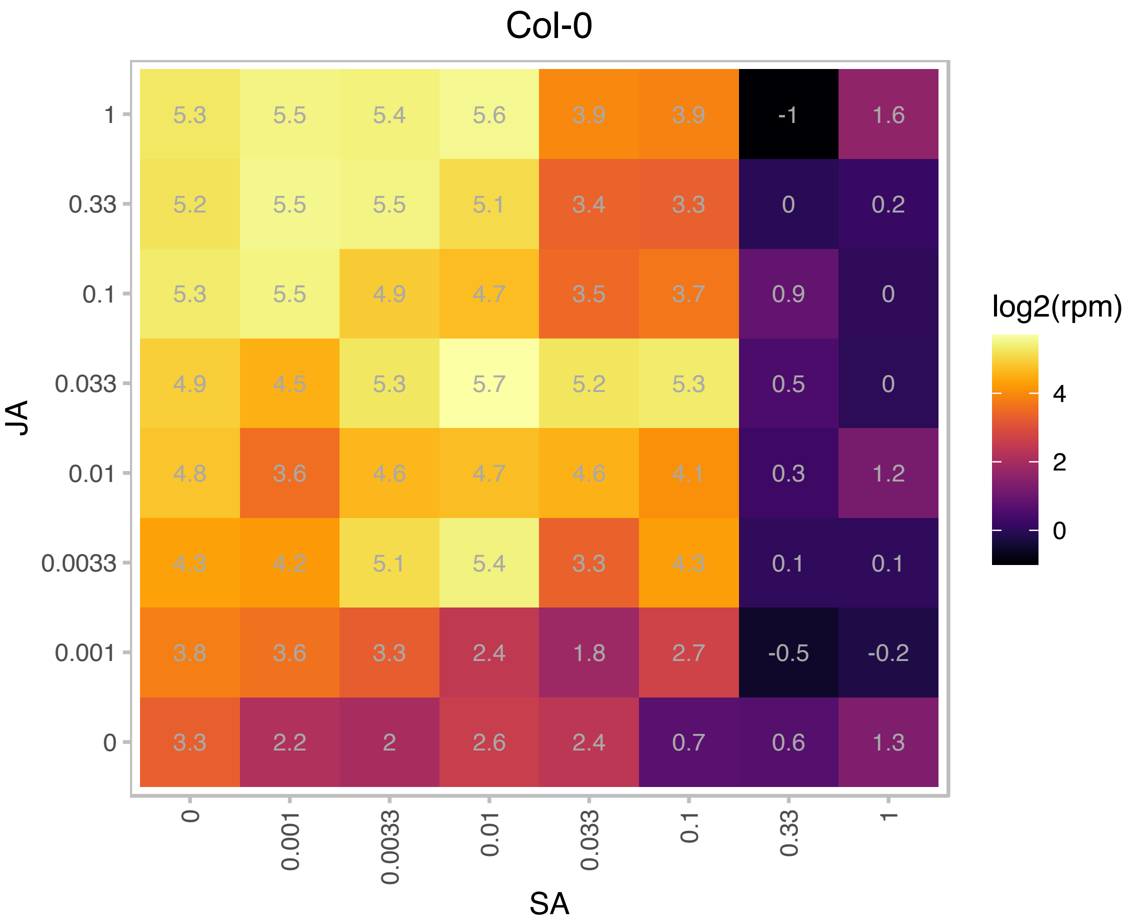


| | |
|---|---|
| Gene | AT5G47990 |
| is.expressed_genes | TRUE |
| is.responsive_genes | TRUE |
| is.apcluster | E-02 |

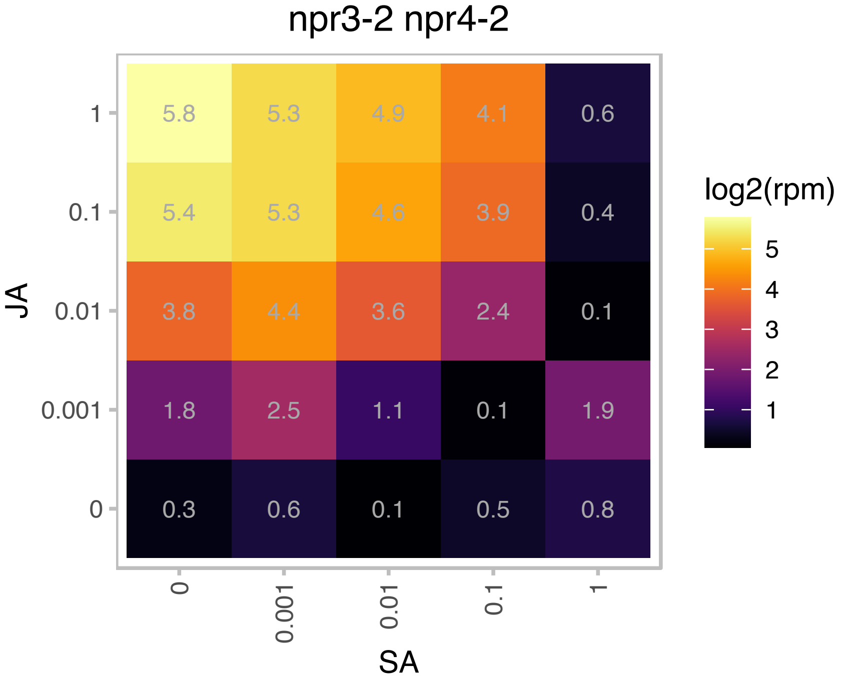


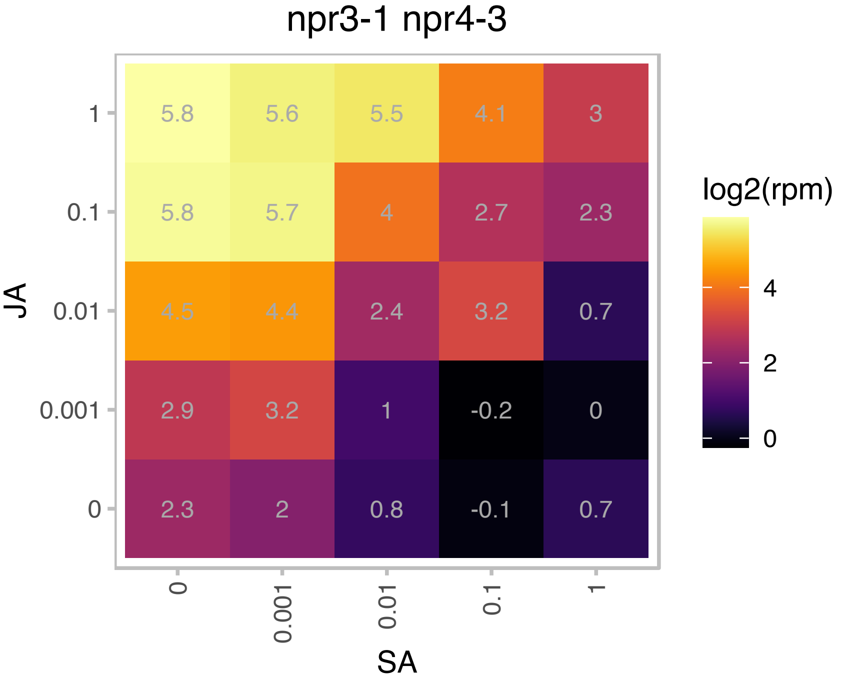

### AT5G48010.1 : AtTHAS1, THAS, THAS1

thalianol synthase 1

Encodes an oxidosqualene cyclase involved in the biosynthesis of thalianol, a tricyclic triterpenoid of unknown function. Overexpression of THAS leads to dwarfing in the aerial tissues of Arabidopsis plants, but increases their root length. THAS is part of a small operon-like cluster of genes (with At5g48000 (THAH) and At5g47990 (THAD)) involved in thalianol metabolism.

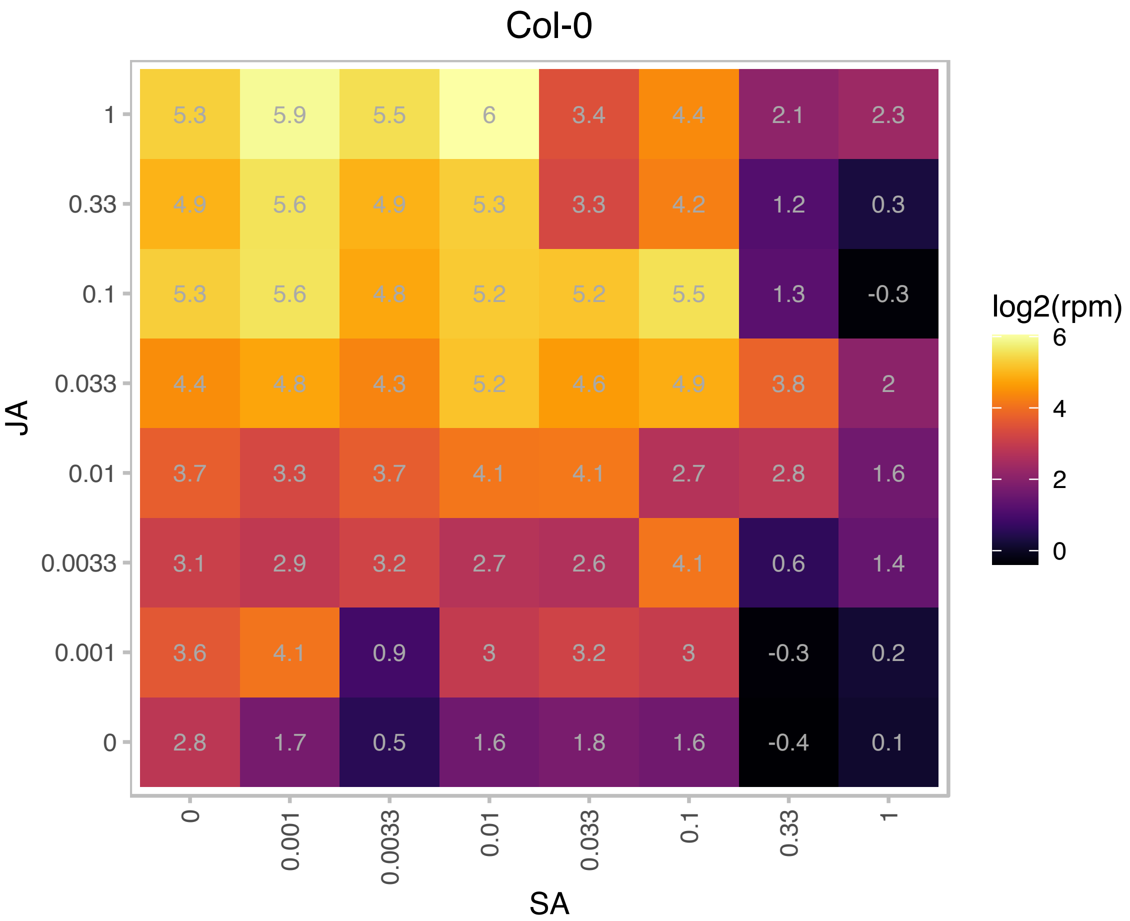


| | |
|---|---|
| Gene | AT5G48010 |
| is.expressed_genes | TRUE |
| is.responsive_genes | TRUE |
| is.apcluster | E-02 |

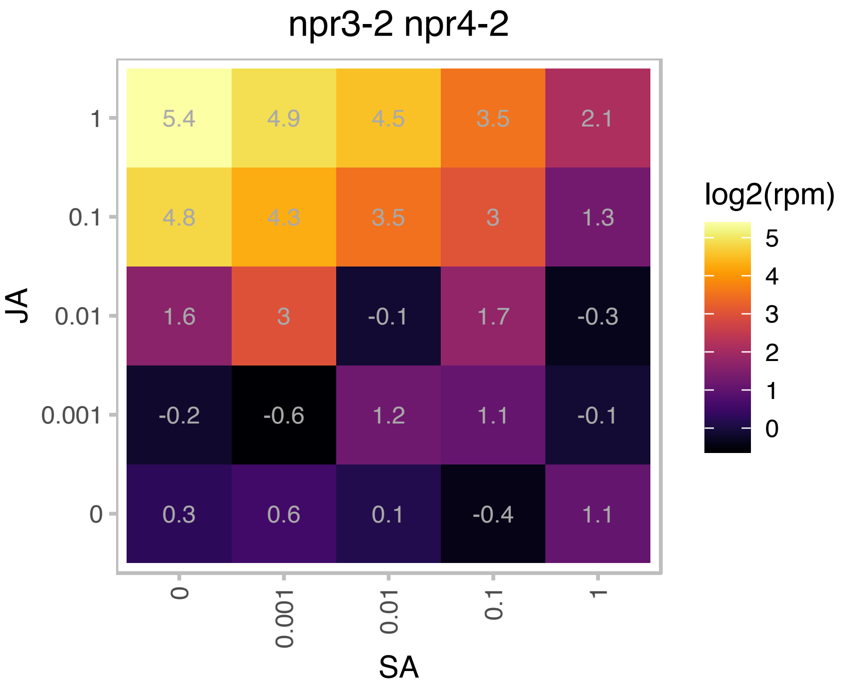


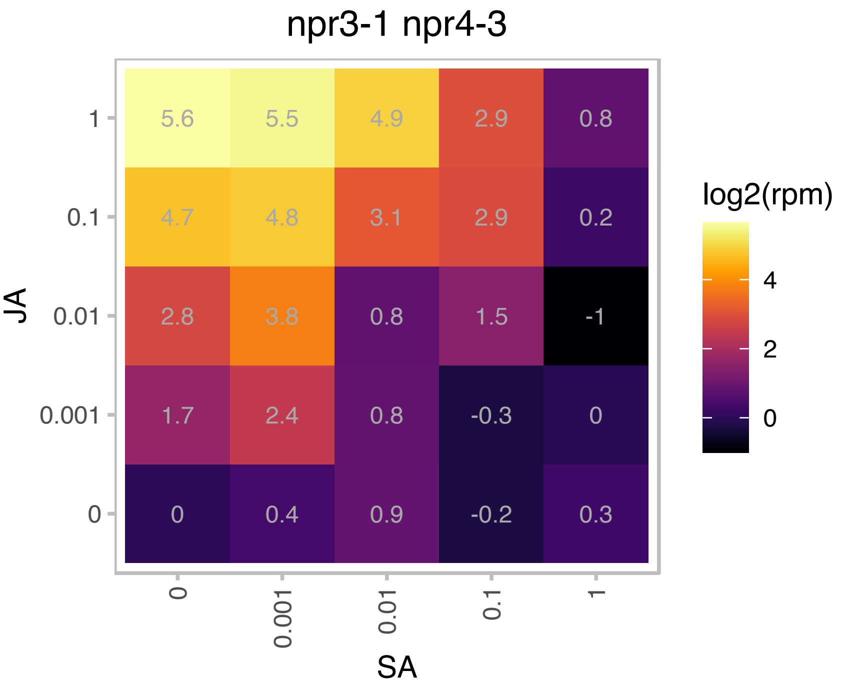

### AT5G49890.1 : ATCLC-C, CLC-C

chloride channel C

member of Anion channel protein family

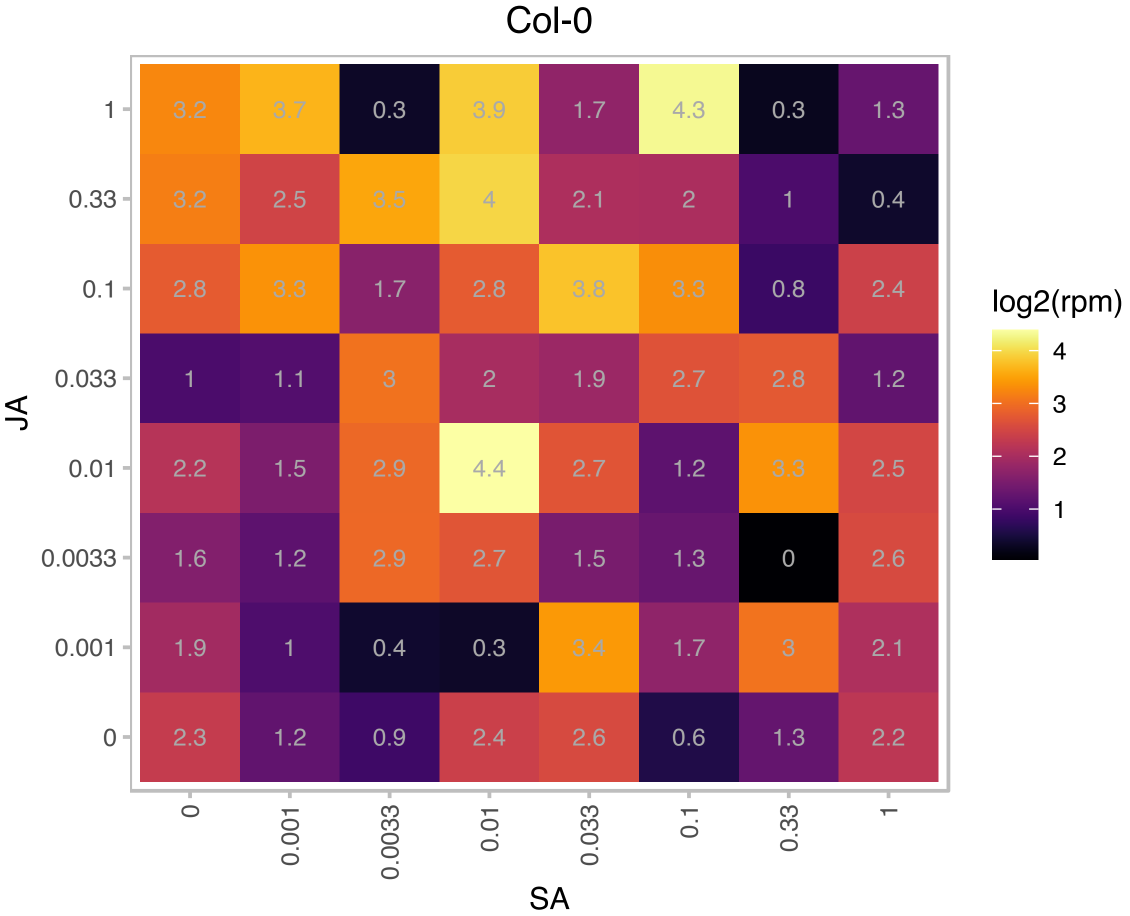


| | |
|---|---|
| Gene | AT5G49890 |
| is.expressed_genes | TRUE |
| is.responsive_genes | TRUE |
| is.apcluster | E-02 |

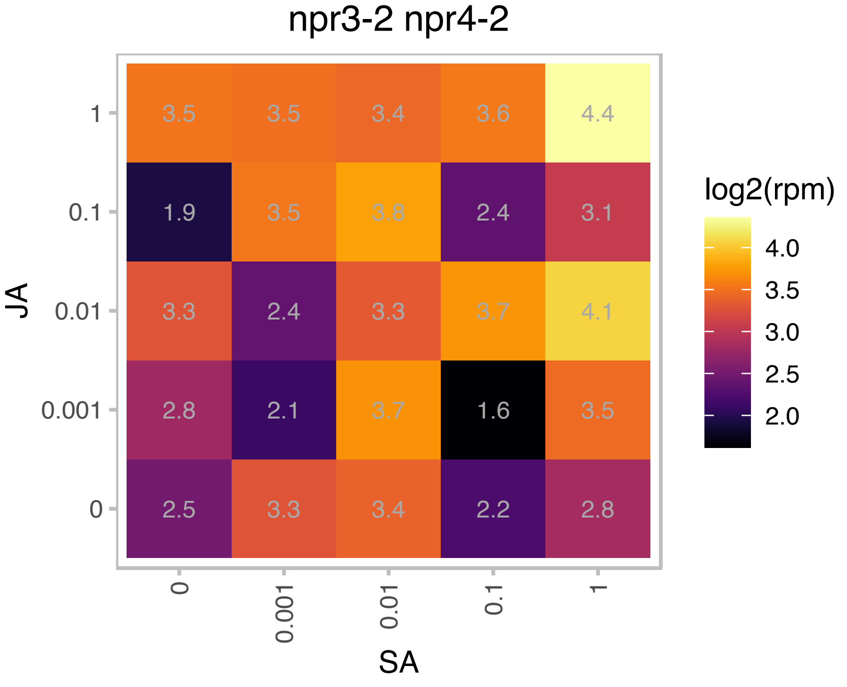


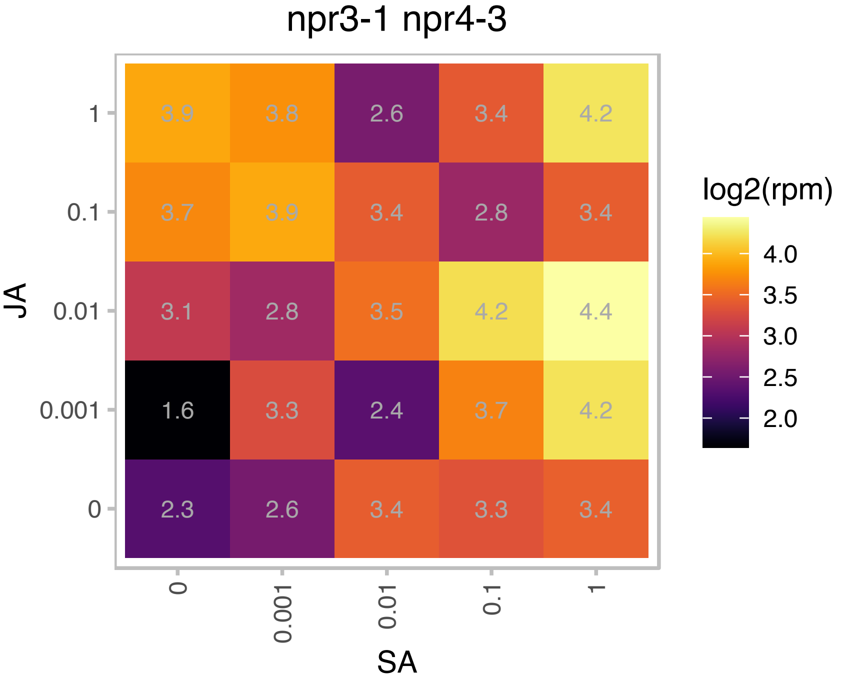

### AT5G54810.1 : ATTSB1, TRP2, TRPB, TSB1

tryptophan synthase beta-subunit 1

A.thaliana tryptophan synthase beta subunit (trpB)

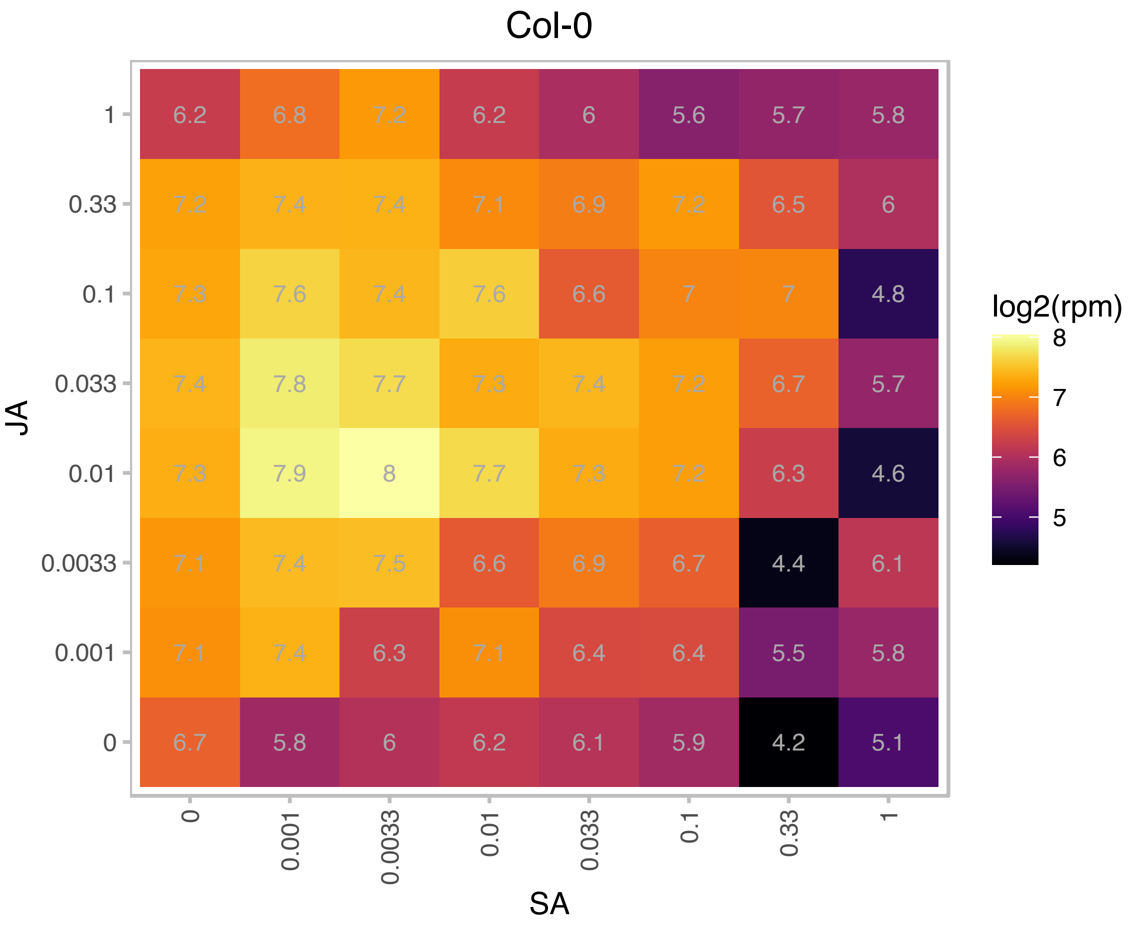


| | |
|---|---|
| Gene | AT5G54810 |
| is.expressed_genes | TRUE |
| is.responsive_genes | TRUE |
| is.apcluster | E-02 |

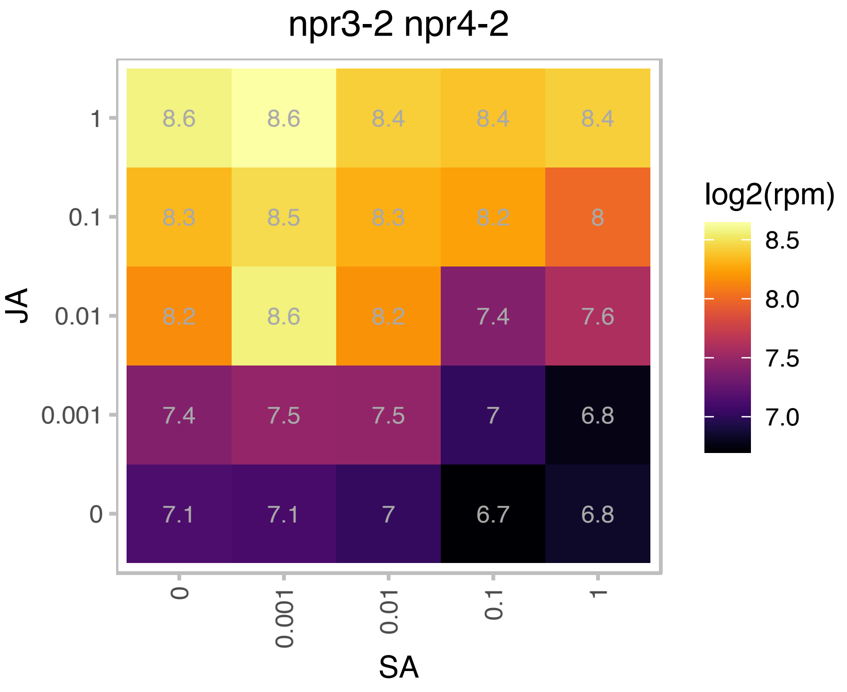


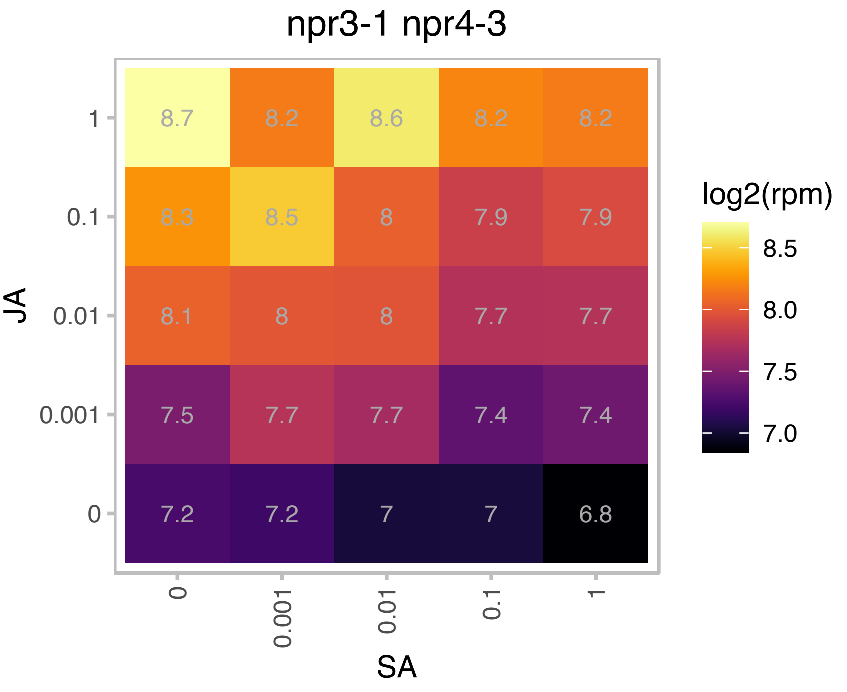

### AT5G59090.1 : ATSBT4.12, SBT4.12

subtilase 4.12

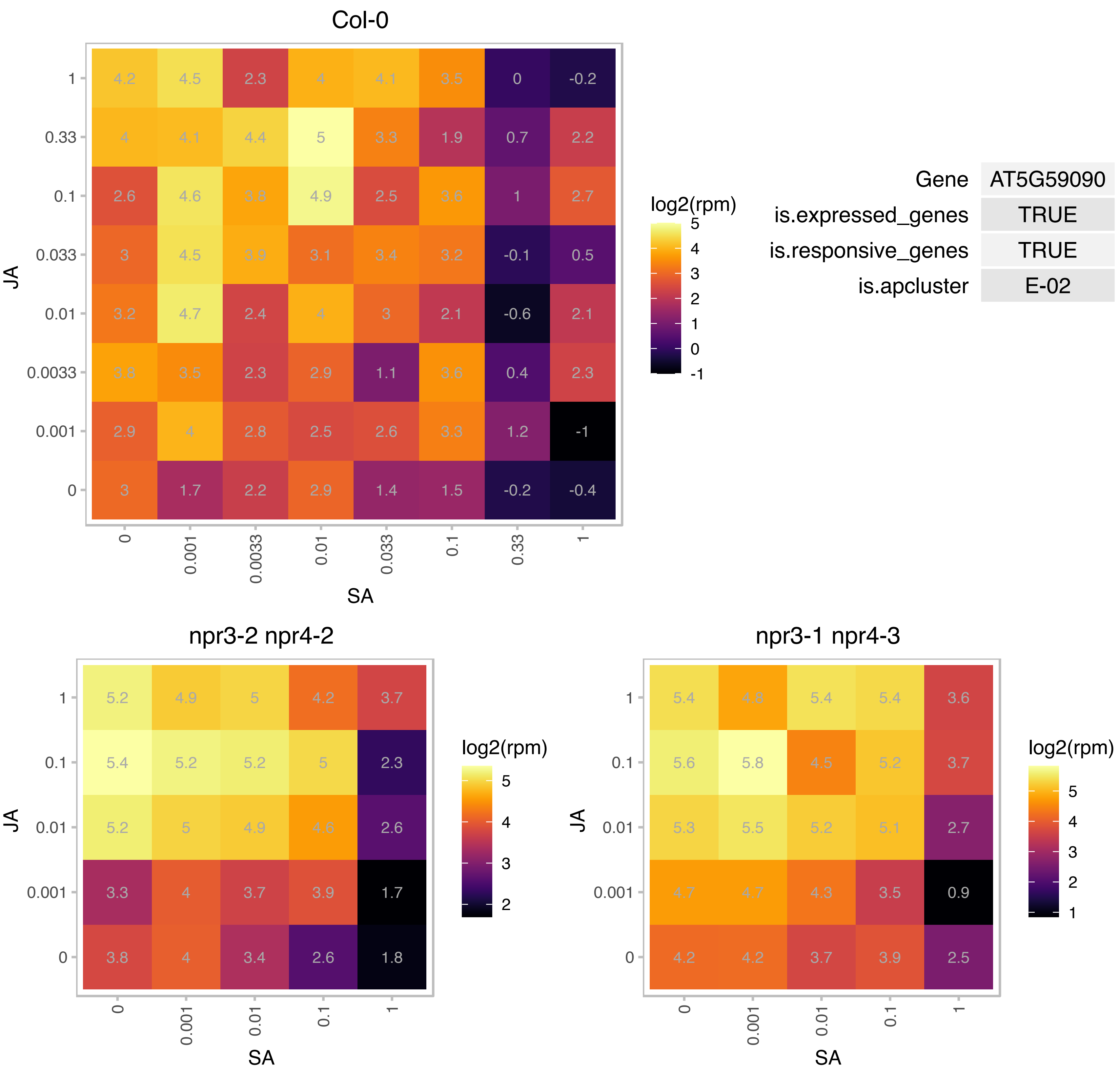

### AT5G60890.1 : AtATR1, ATMYB34, ATR1, MYB34

myb domain protein 34
Myb-like transcription factor that modulates expression of ASA1, a key point of control in the tryptophan pathway; mutant has deregulated expression of ASA1 in dominant allele. Loss of function allele suggests ATR1 also functions at a control point for regulating indole glucosinolate homeostasis.

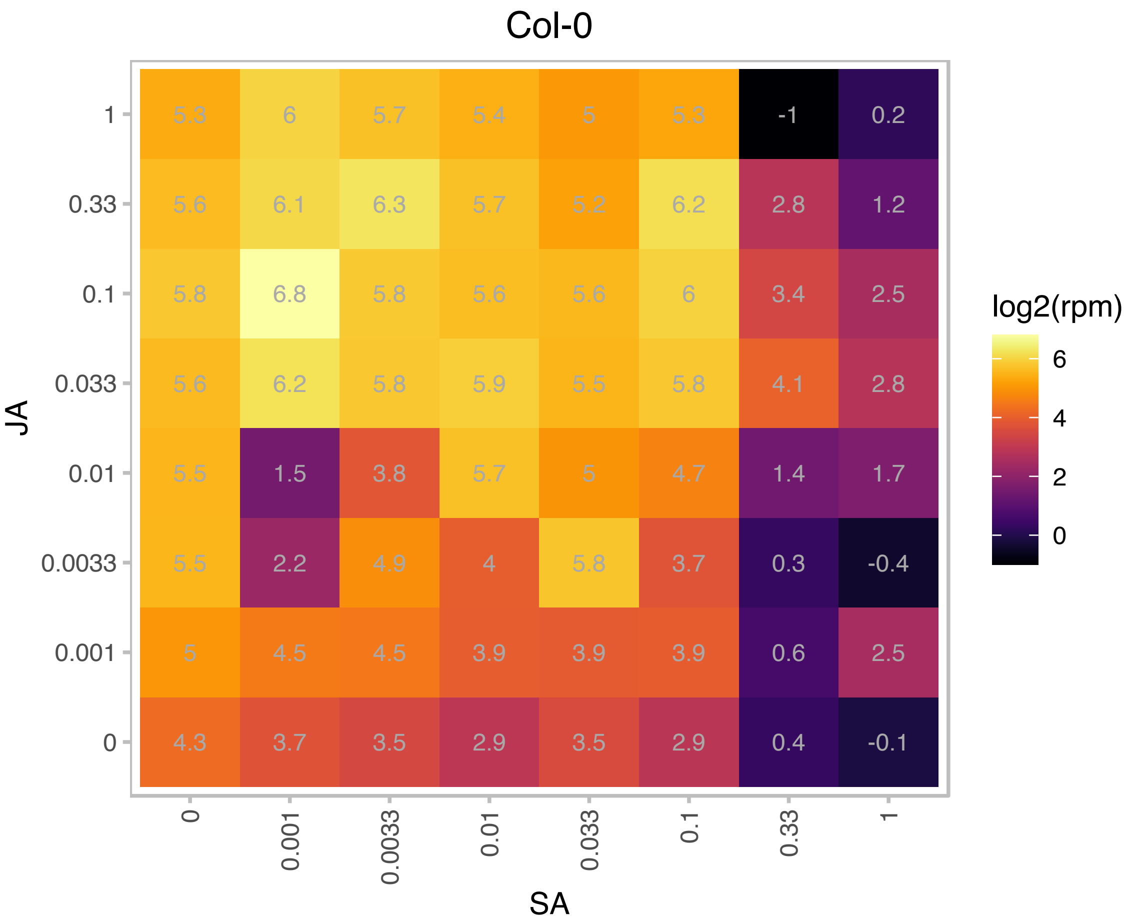


| | |
|---|---|
| Gene | AT5G60890 |
| is.expressed_genes | TRUE |
| is.responsive_genes | TRUE |
| is.apcluster | E-02 |

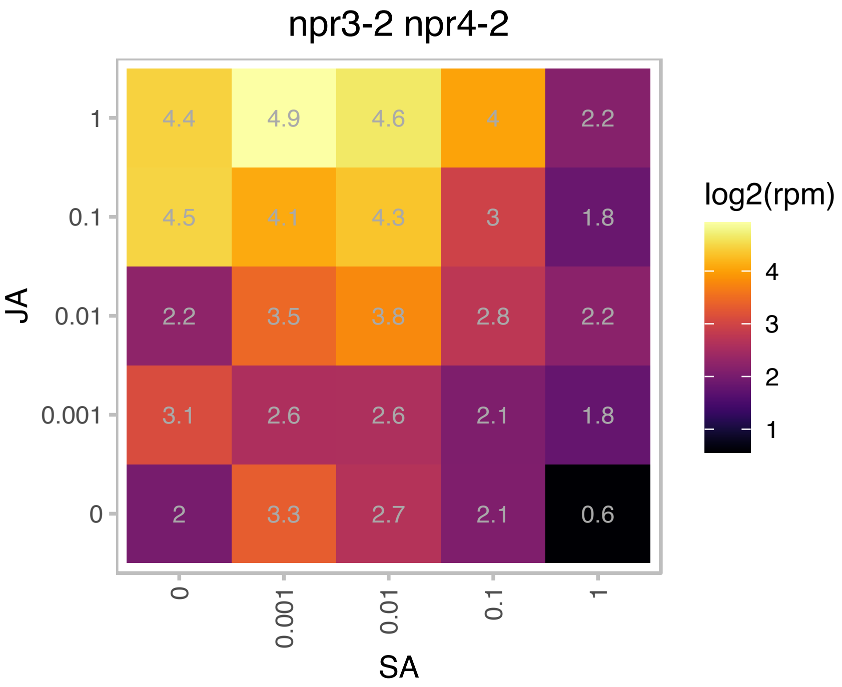


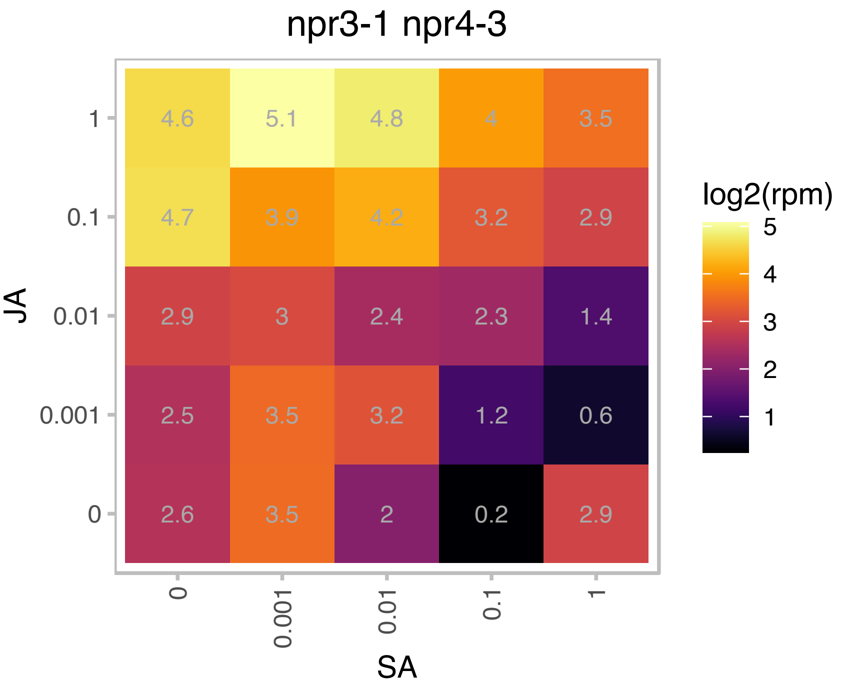

### AT5G63600.1 : ATFLS5, FLS5

flavonol synthase 5

encodes a protein whose sequence is similar to flavonol synthase

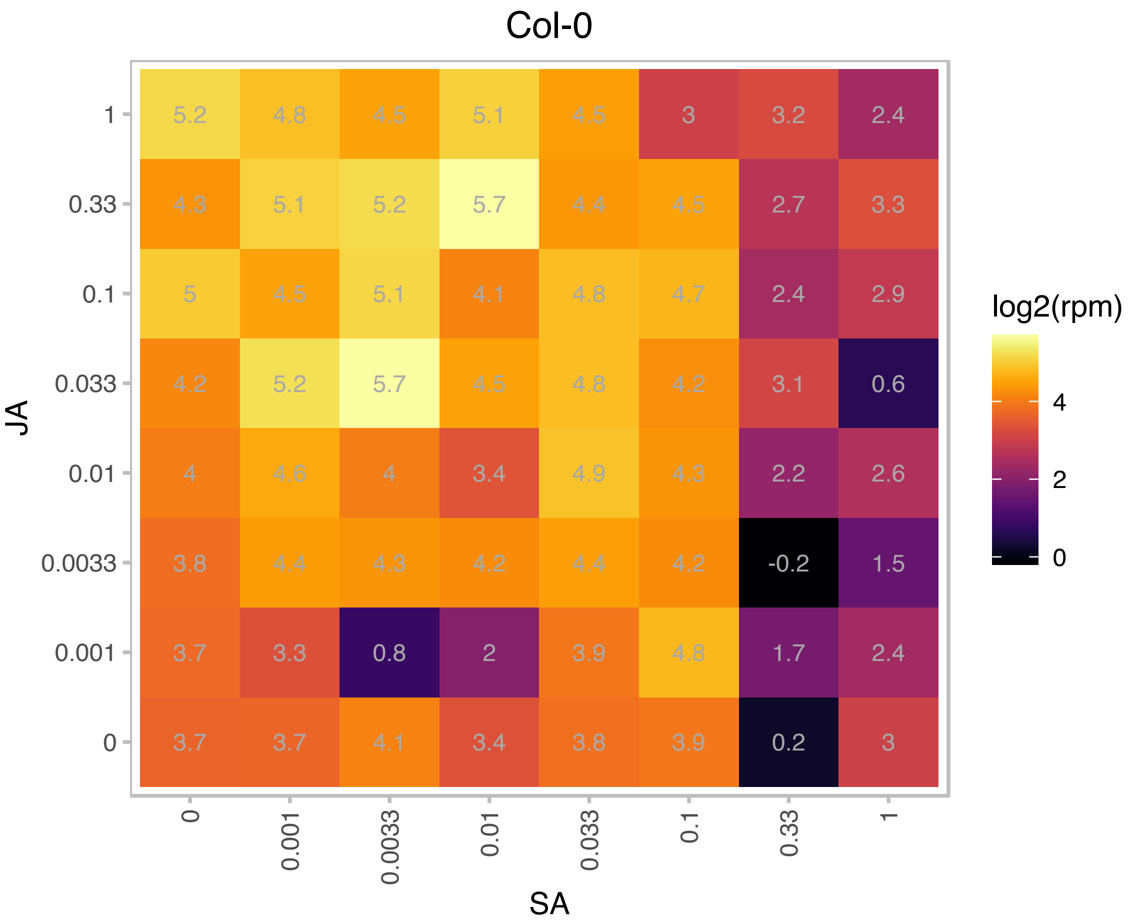


| | |
|---|---|
| Gene | AT5G63600 |
| is.expressed_genes | TRUE |
| is.responsive_genes | TRUE |
| is.apcluster | E-02 |

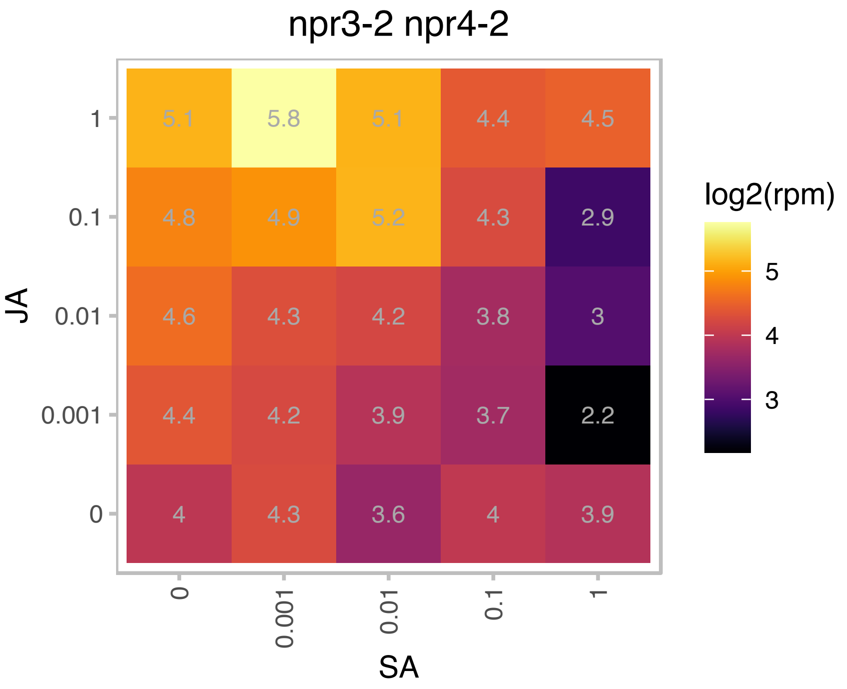


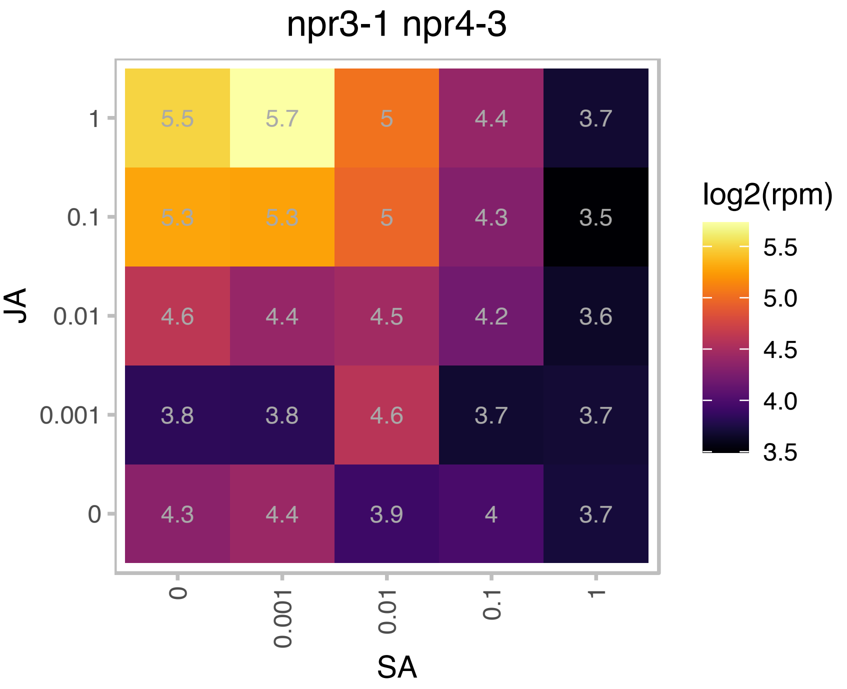

### AT5G63980.1 : ALX8, AtFRY1, ATSAL1, FRY1, HOS2, RO[illegible]

Inositol monophosphatase family protein

Encodes a bifunctional protein that has 3'(2'),5'-bisphosphate nucleotidase and inositol polyphosphate 1-phosphatase activities and rescues sulfur assimilation mutants in yeast. It is involved in the response to cold, drought (negative regulator of drought tolerance), and ABA. Mutants in this gene exhibit enhanced induction of stress genes in response to cold, ABA, salt and dehydration due to higher accumulation of the second messenger, inositol (1,4,5)- triphosphate (IP(3)). Involved in degradation of small mRNAs. Mutants also affect the accumulation of miRNA target cleavage products. Regulates light-dependent repression of hypocotyl elongation and flowering time via its 3'(2'),5'-bisphosphate nucleotidase activity. Its kinase [illegible]

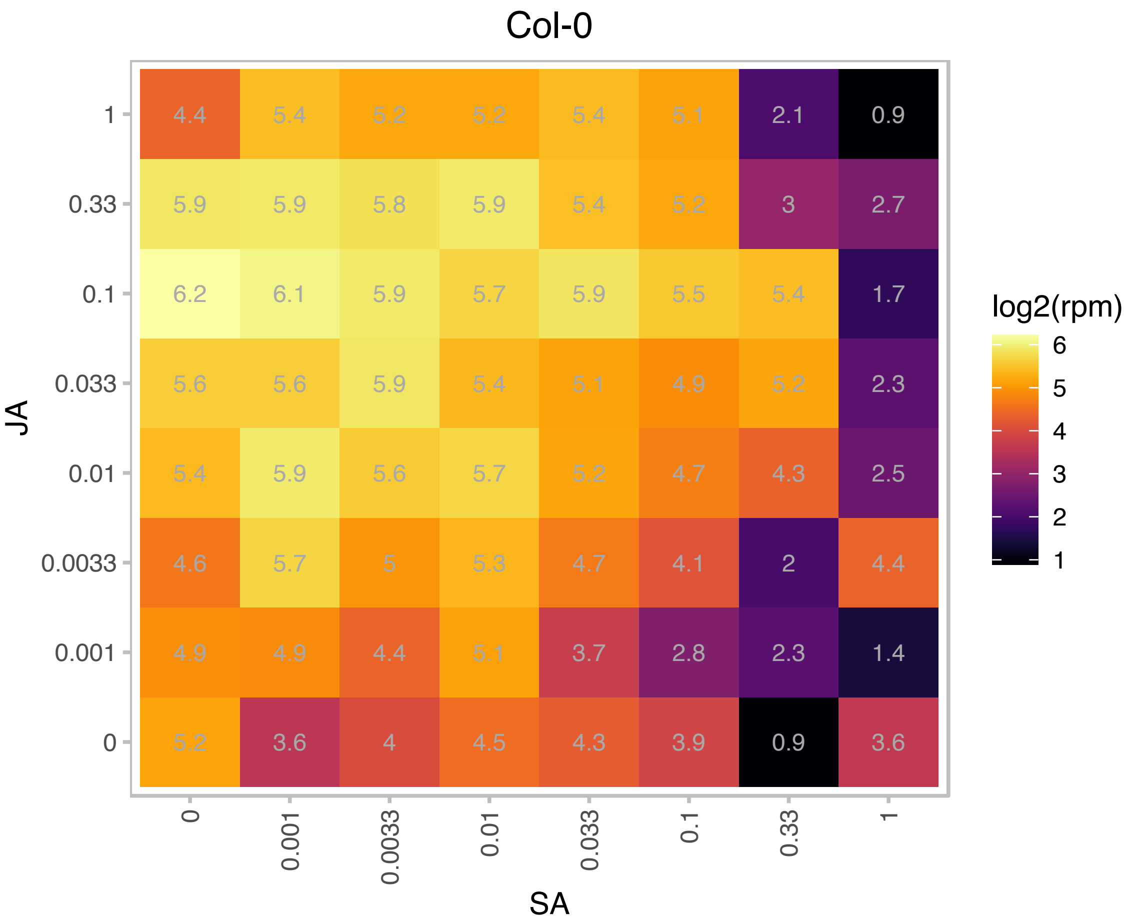


| | |
|---|---|
| Gene | AT5G63980 |
| is.expressed_genes | TRUE |
| is.responsive_genes | TRUE |
| is.apcluster | E-02 |

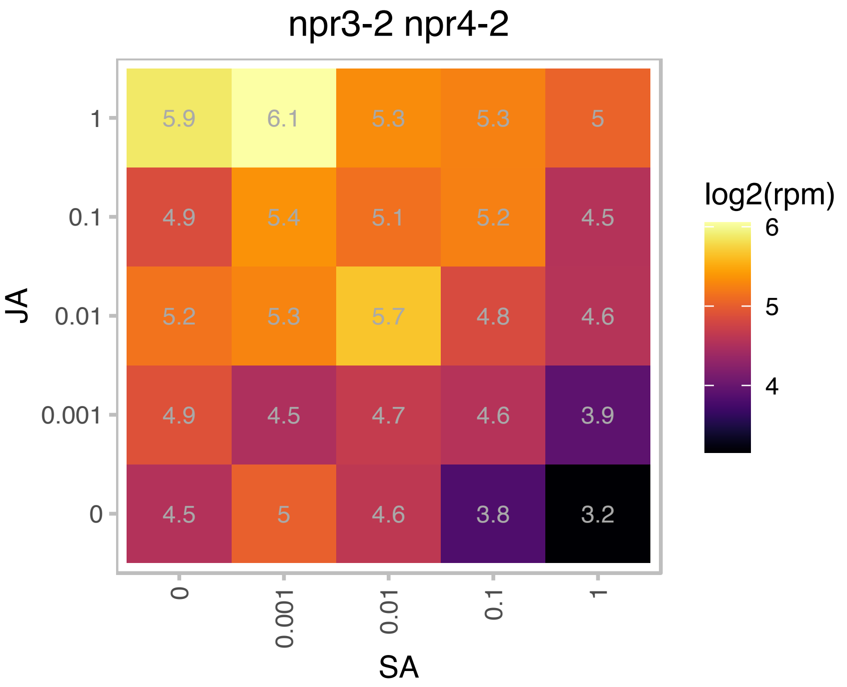


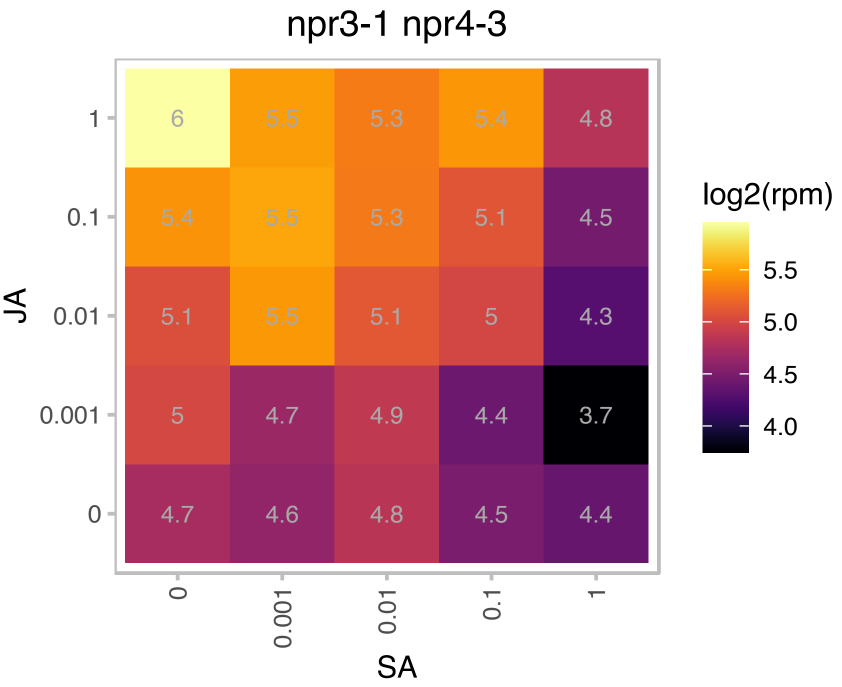

### AT5G65020.1 : ANNAT2, AtANN2

annexin 2
Annexins are calcium binding proteins that are localized in the cytoplasm. When cytosolic Ca2+ increases, they relocate to the plasma membrane.  They may be involved in the Golgi-mediated secretion of polysaccharides.

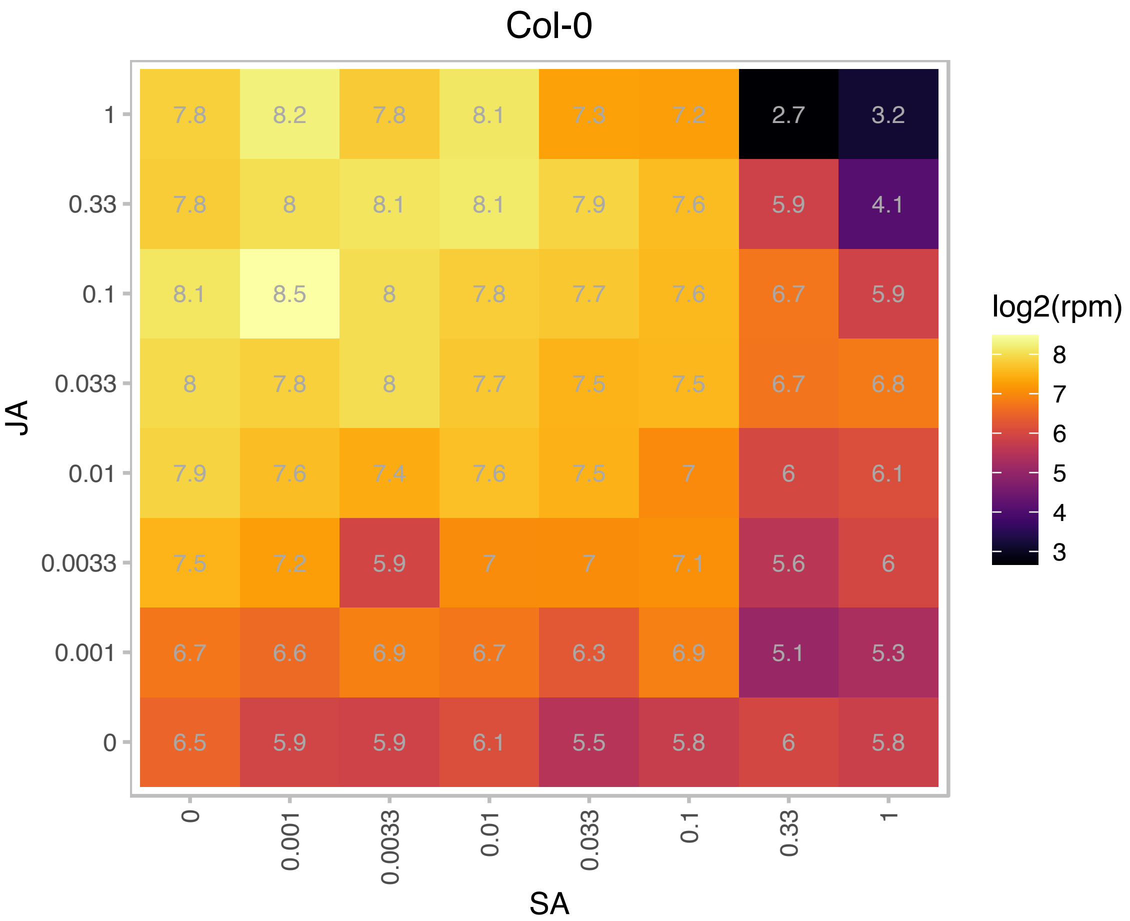


| | |
|---|---|
| Gene | AT5G65020 |
| is.expressed_genes | TRUE |
| is.responsive_genes | TRUE |
| is.apcluster | E-02 |

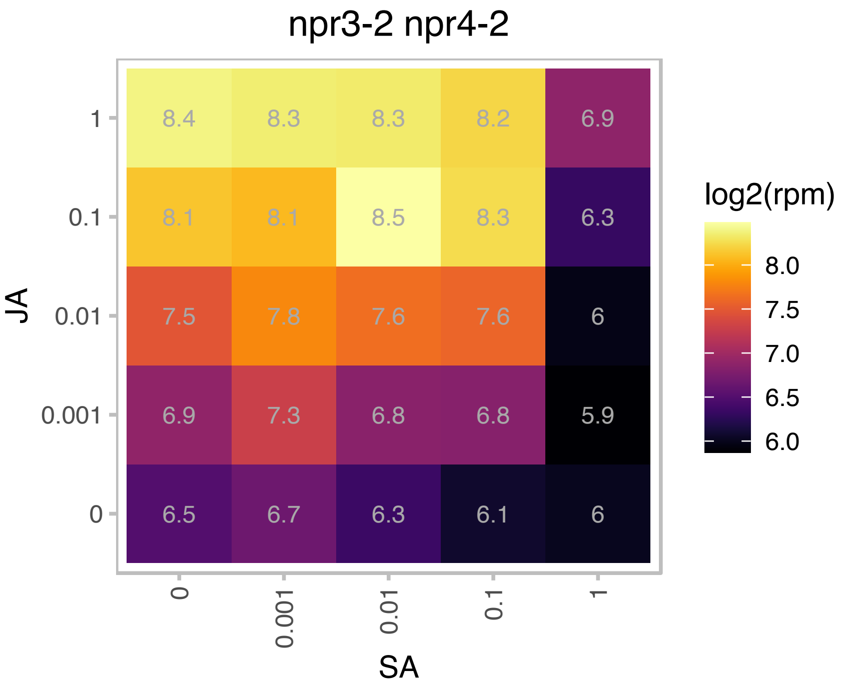


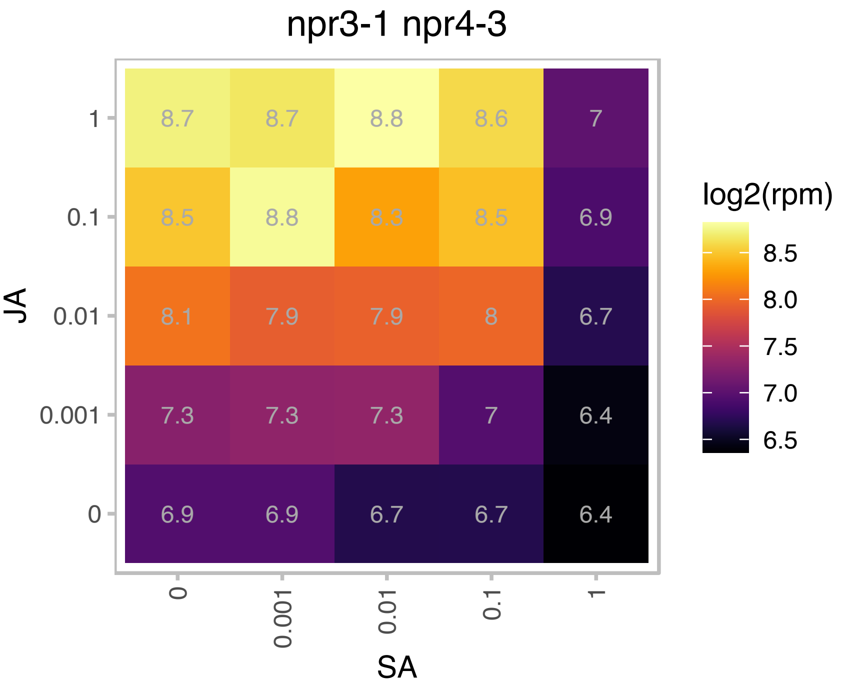

### AT5G67160.1 : EPS1

HXXXD-type acyl-transferase family protein
Encodes a member of the BAHD acyltransferase superfamily. Mutants have enhanced susceptibility to virulent and avirulent pathogens and are defective in pathogen induced SA biosynthesis. EPS1 may act upstream of SA biosynthesis as application of SA can rescue the mutant phenotype.

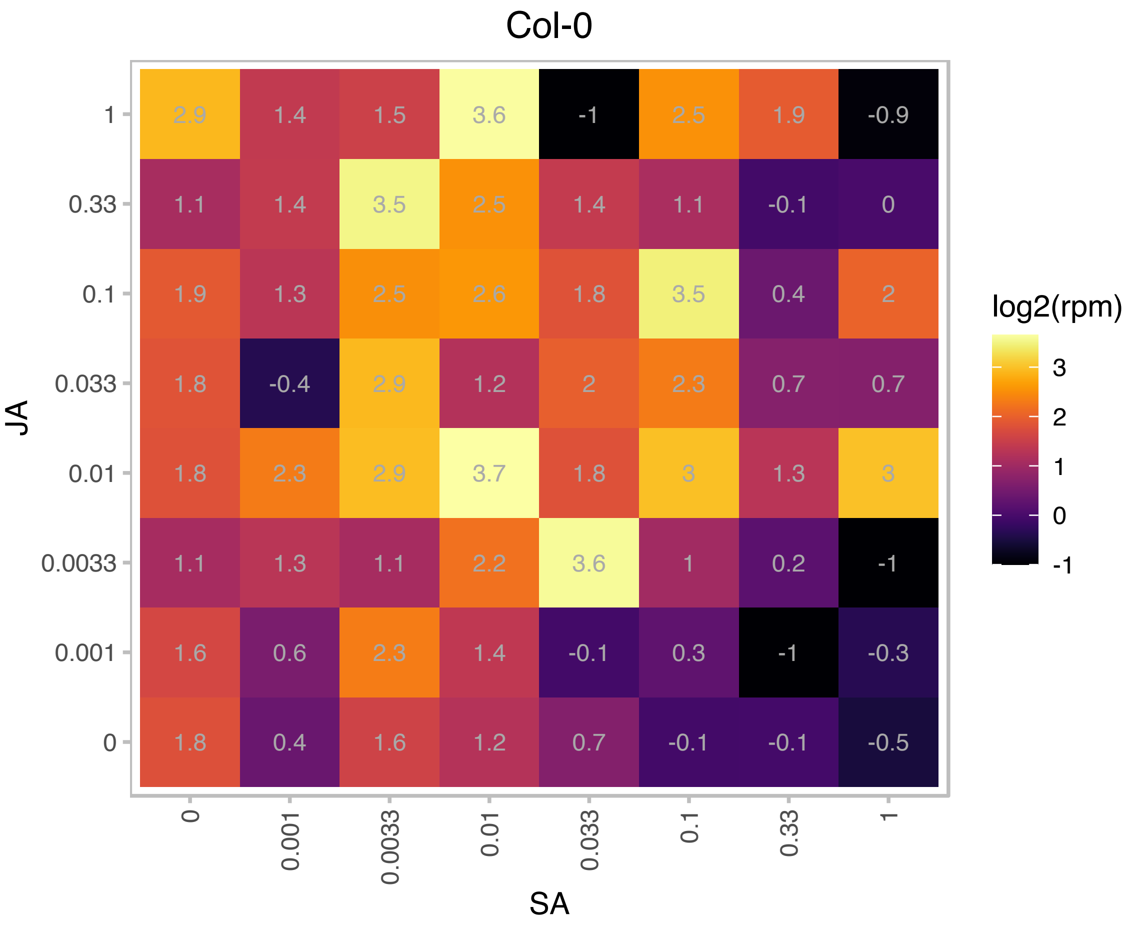


| | |
|---|---|
| Gene | AT5G67160 |
| is.expressed_genes | TRUE |
| is.responsive_genes | TRUE |
| is.apcluster | E-02 |

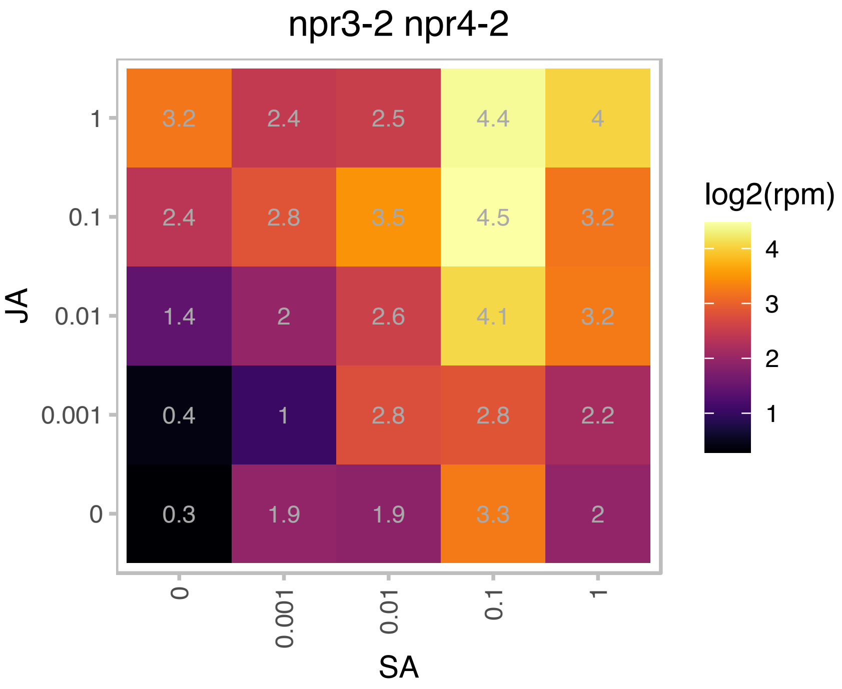


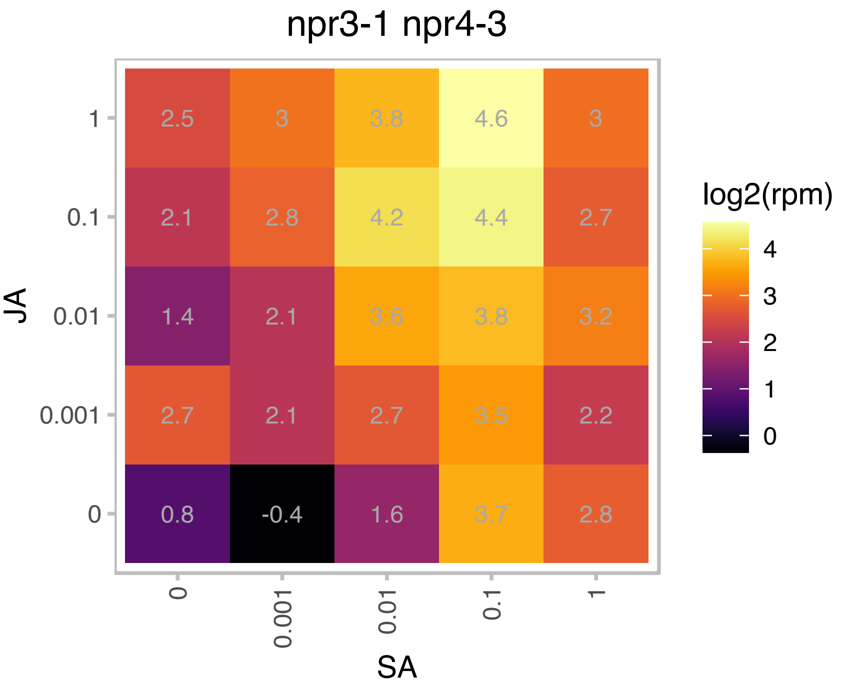

### AT5G67590.1 : FRO1, NDUFS4

NADH-ubiquinone oxidoreductase-related

Mutant leaves have a reduced capacity for cold acclimation, appear water-soaked, leak electrolytes, and accumulate reactive oxygen species constitutively. Encode a protein with high similarity to the 18-kD Fe-S subunit of complex I (NADH dehydrogenase, EC 1.6.5.3) in the mitochondrial electron transfer chain. The mRNA is cell-to-cell mobile.

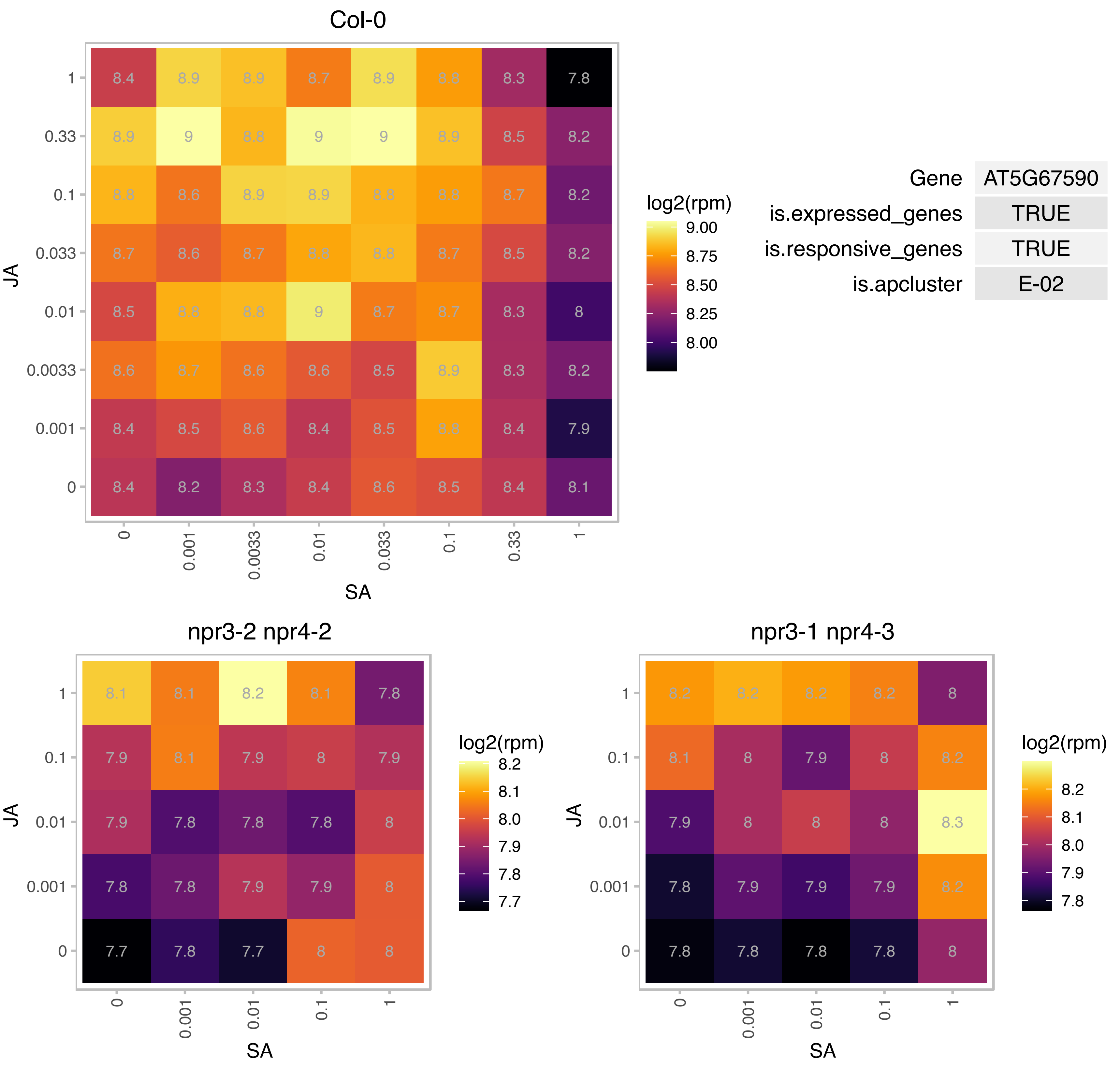


| | |
|---|---|
| Gene | AT5G67590 |
| is.expressed_genes | TRUE |
| is.responsive_genes | TRUE |
| is.apcluster | E-02 |

### ATCG00160.1 : RPS2

ribosomal protein S2

Chloroplast ribosomal protein S2

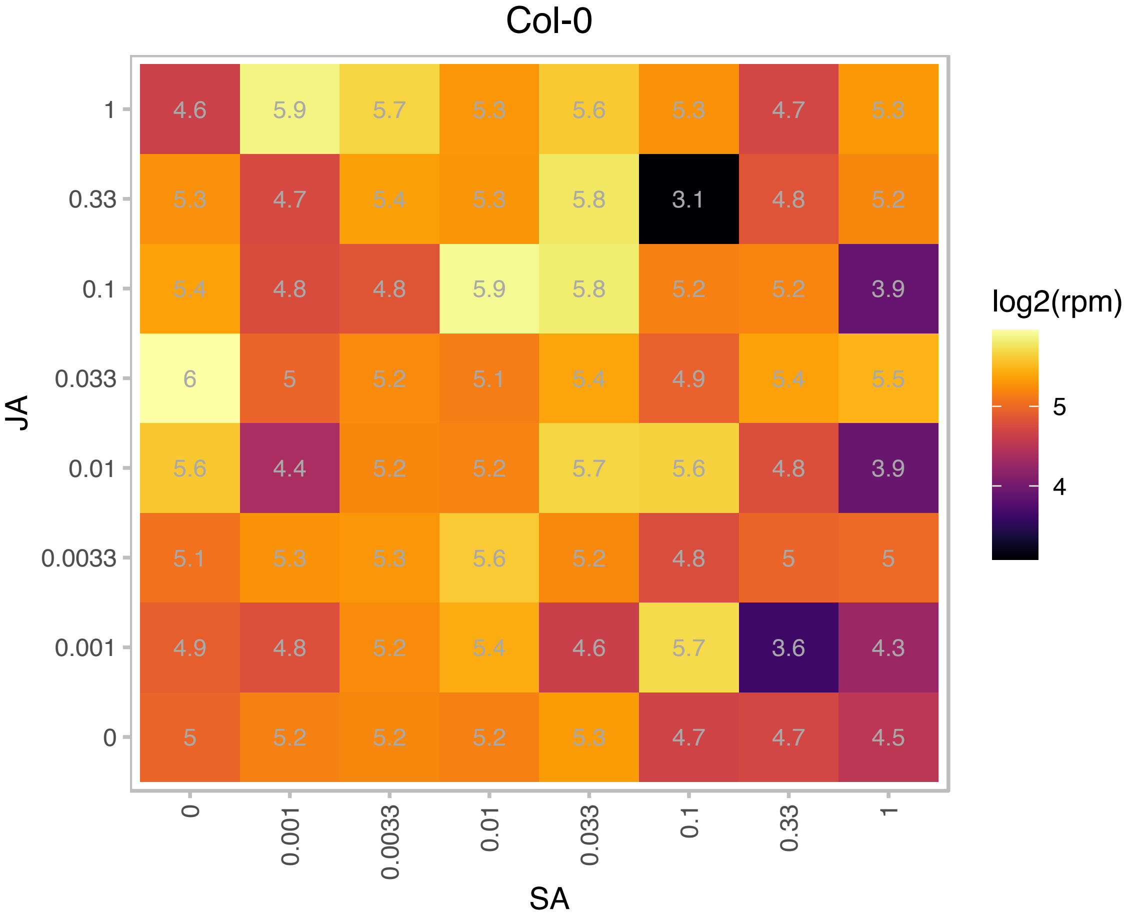


| | |
|---|---|
| Gene | ATCG00160 |
| is.expressed_genes | TRUE |
| is.responsive_genes | TRUE |
| is.apcluster | E-02 |

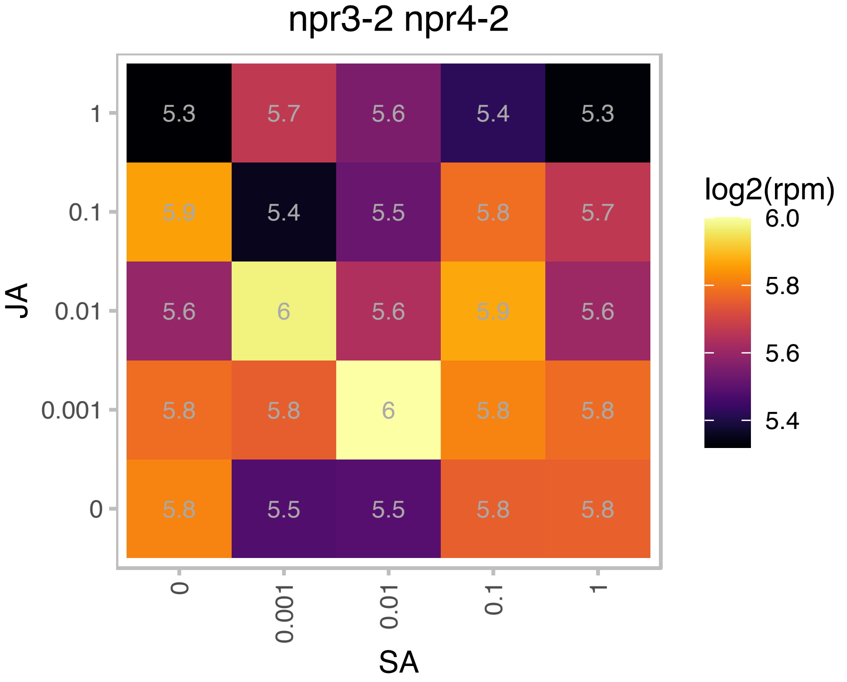


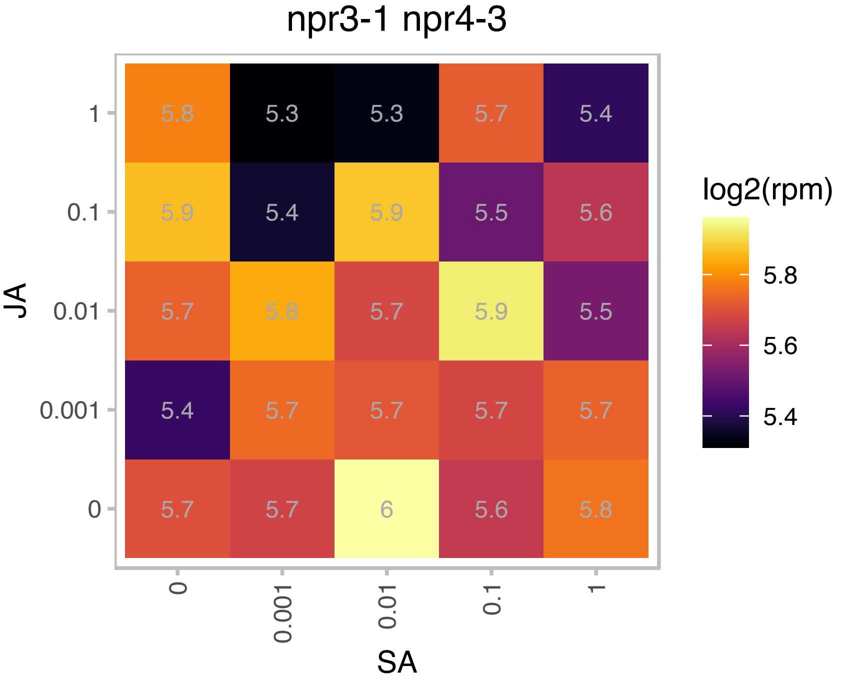

### AT1G07770.1 : RPS15A

ribosomal protein S15A

Encodes cytoplasmic ribosomal protein S15a.

Col-0

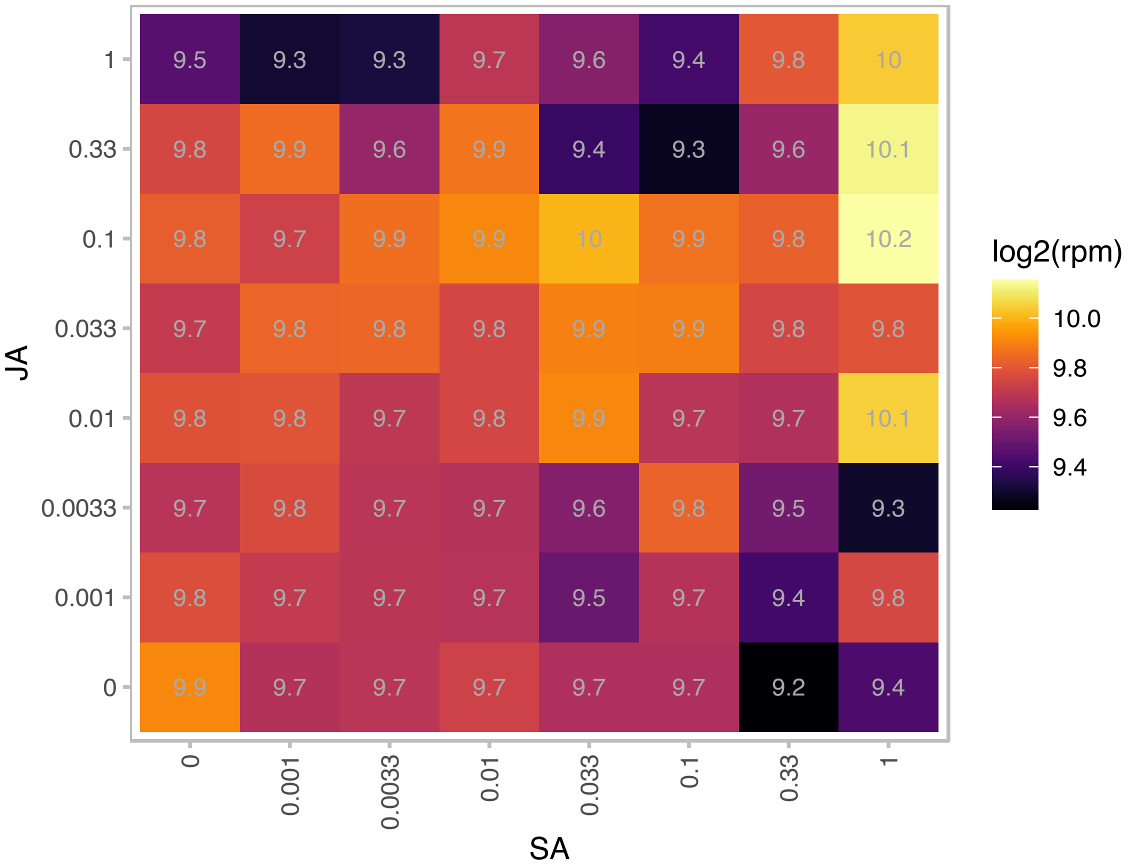


| | |
|---|---|
| Gene | AT1G07770 |
| is.expressed_genes | TRUE |
| is.responsive_genes | TRUE |
| is.apcluster | E-03 |

npr3-2 npr4-2

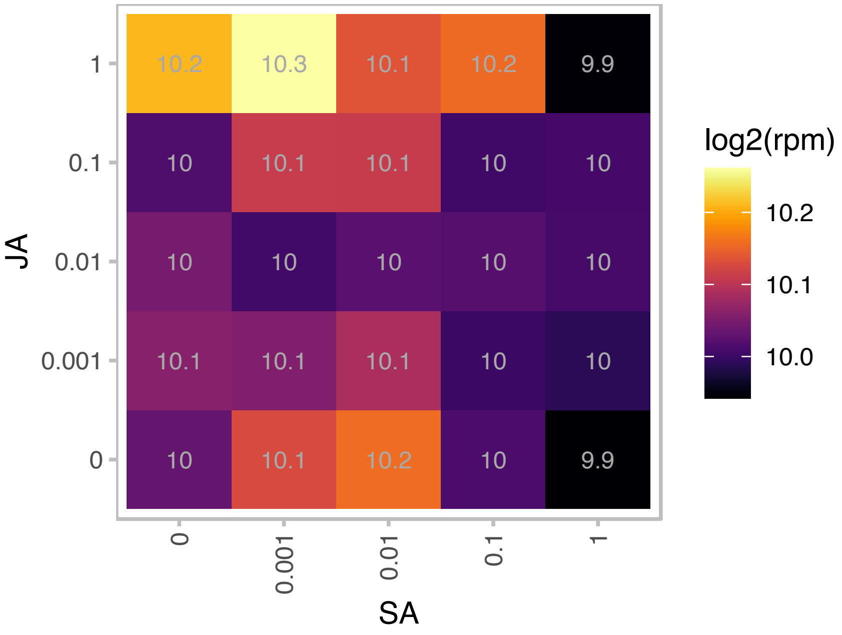


npr3-1 npr4-3

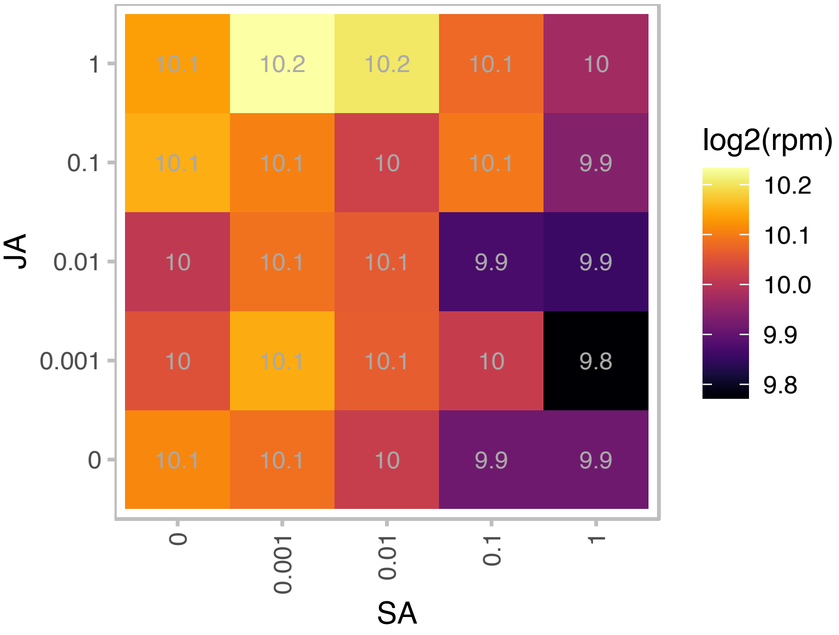

### AT1G10700.1 : PRS3

phosphoribosyl pyrophosphate (PRPP) synthase 3

Encodes a P-independent phosphoribosyl pyrophosphate (PRPP) synthase.

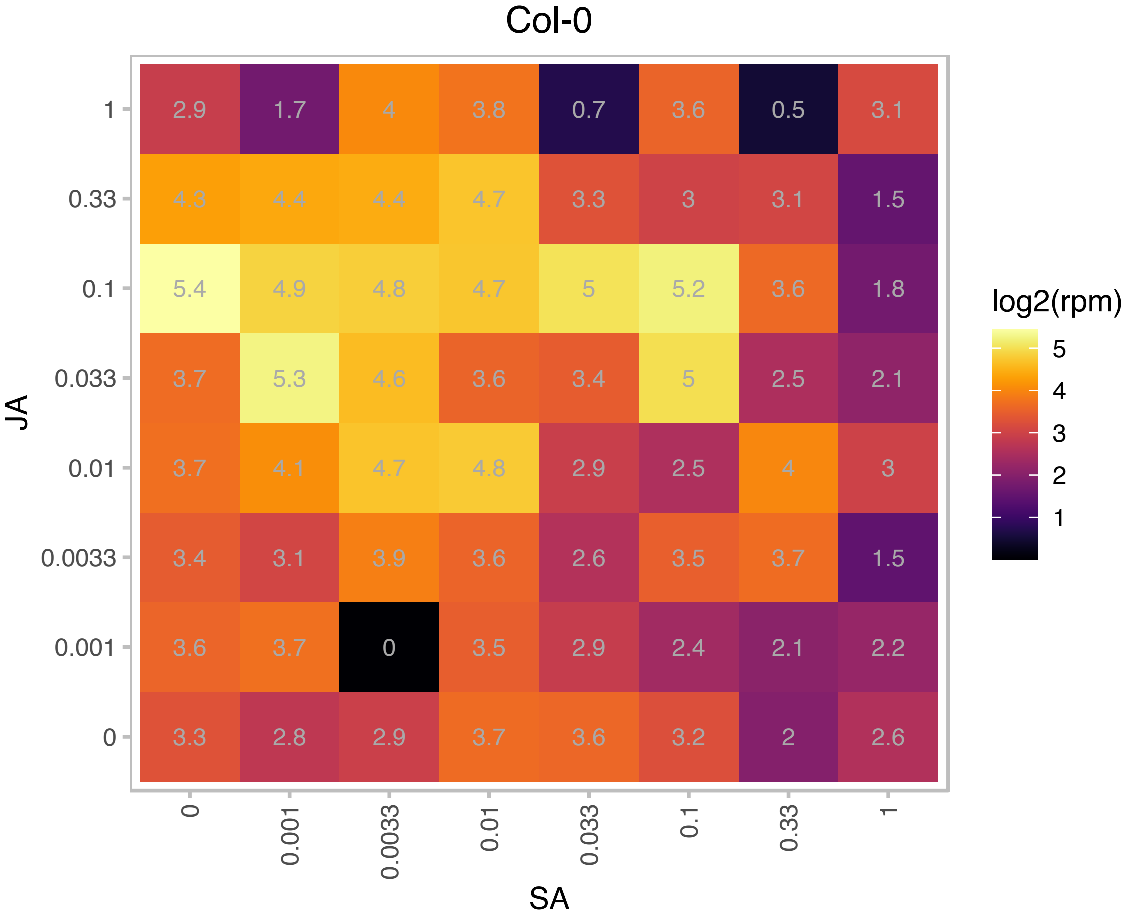


| | |
|---|---|
| Gene | AT1G10700 |
| is.expressed_genes | TRUE |
| is.responsive_genes | TRUE |
| is.apcluster | E-03 |

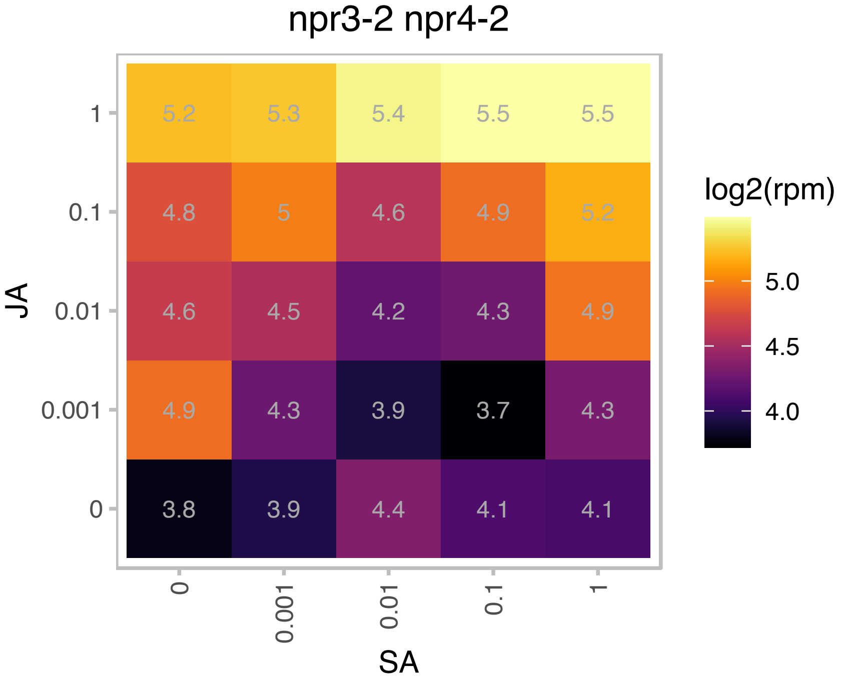


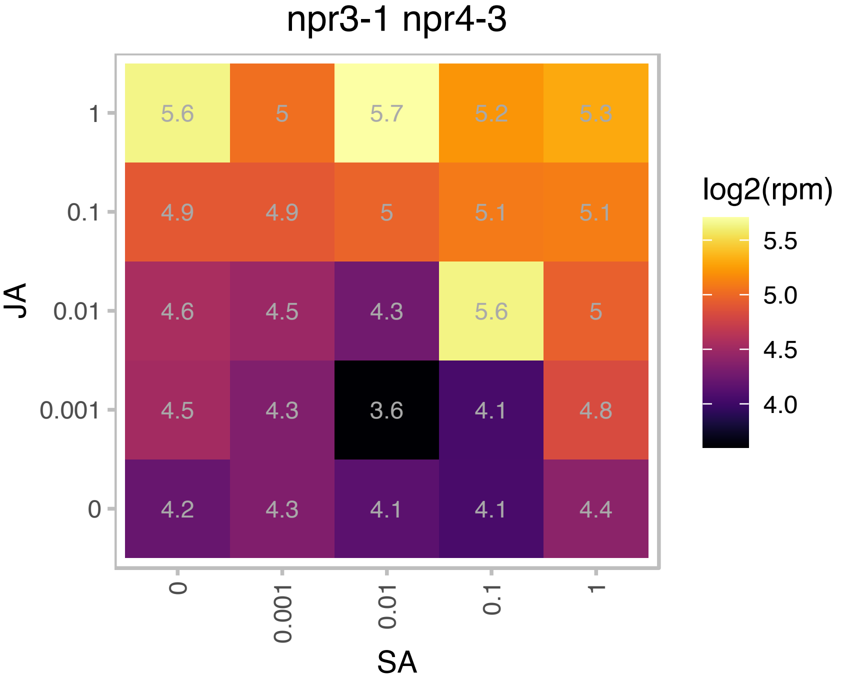

# AT1G14250.1 : NA

GDA1/CD39 nucleoside phosphatase family protein

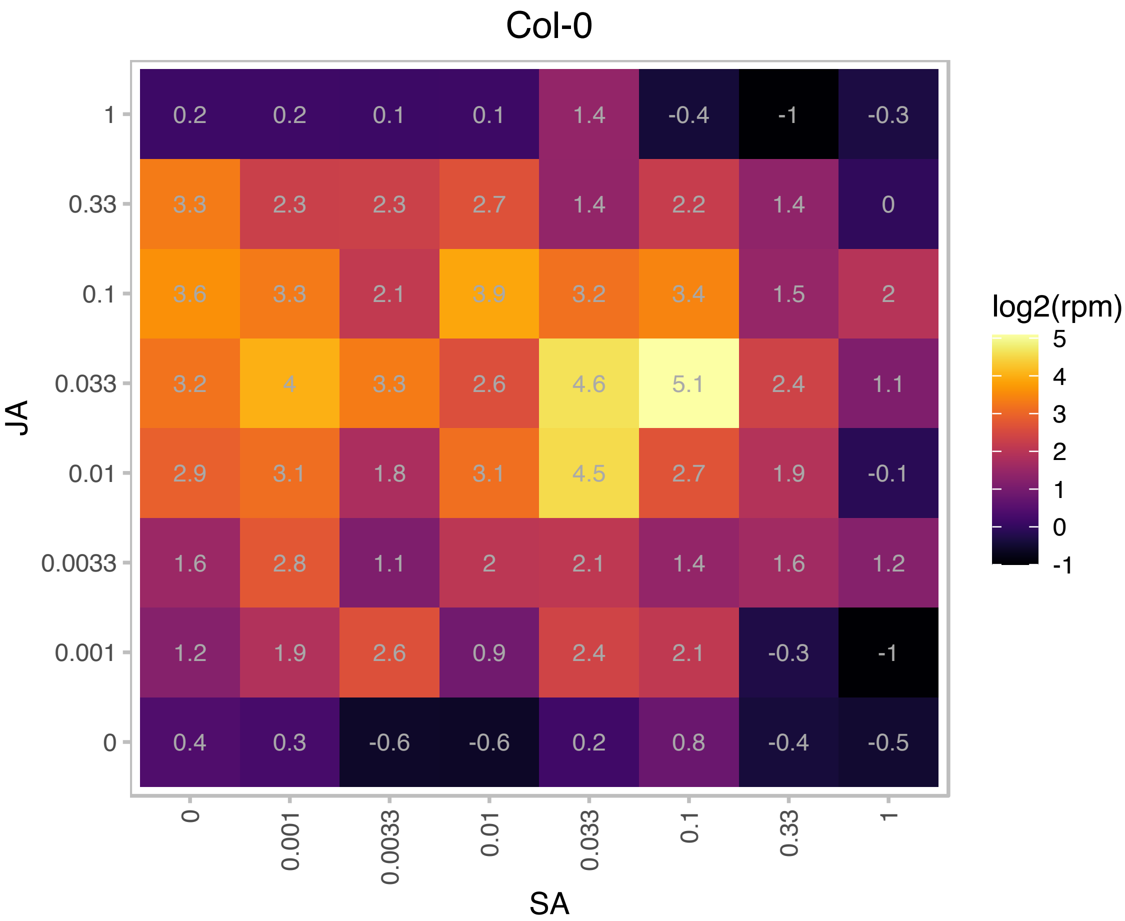


| | |
|---|---|
| Gene | AT1G14250 |
| is.expressed_genes | TRUE |
| is.responsive_genes | TRUE |
| is.apcluster | E-03 |

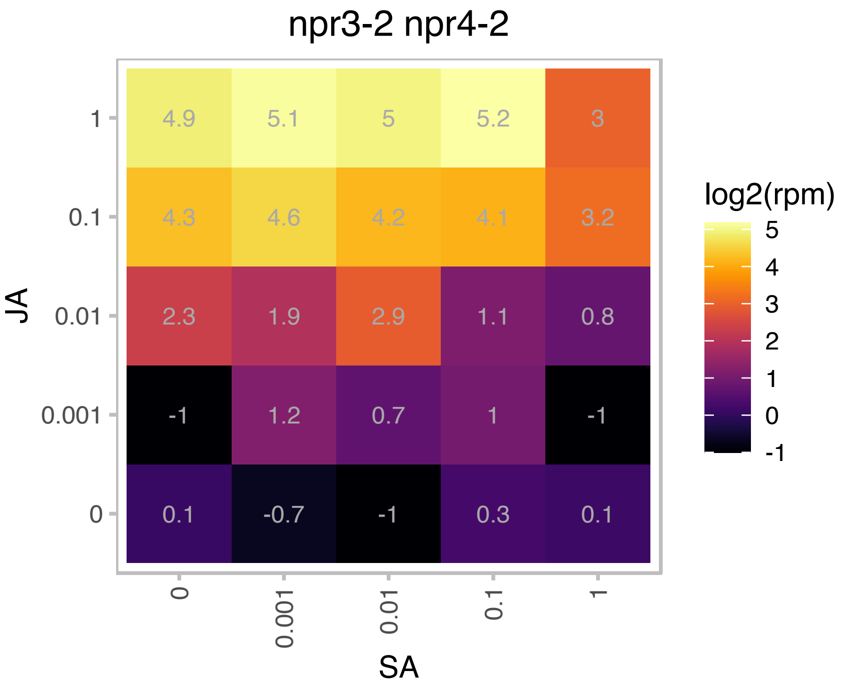


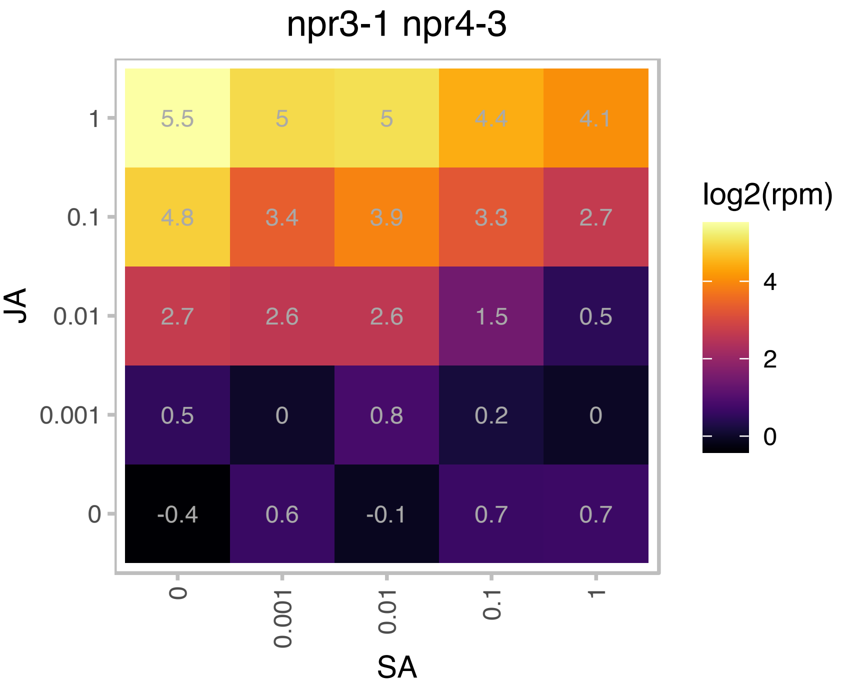

### AT1G19570.1 : ATDHAR1, DHAR1, DHAR5

Encodes a member of the dehydroascorbate reductase gene family. Critical for a mutualistic symbiosis between the host Arabidopsis and the root colonizing fungus Piriformospora indica.Encodes about 50-60% of extractable leaf GSH-dependent DHAR activity, but single knockout mutants show unaltered ascorbate and glutathione status in optimal and oxidative stress conditions (PMID:28381499). Acts redundantly with DHAR2 to oxidize glutathione in response to increased intracelullar hydrogen peroxide (catalase deficiency) . Complementation of a cat2 dhar1 dhar2 dhar3 quadruple mutant with DHAR1 fully restores cat2 phenotype and [illegible]

dehydroascorbate reductase

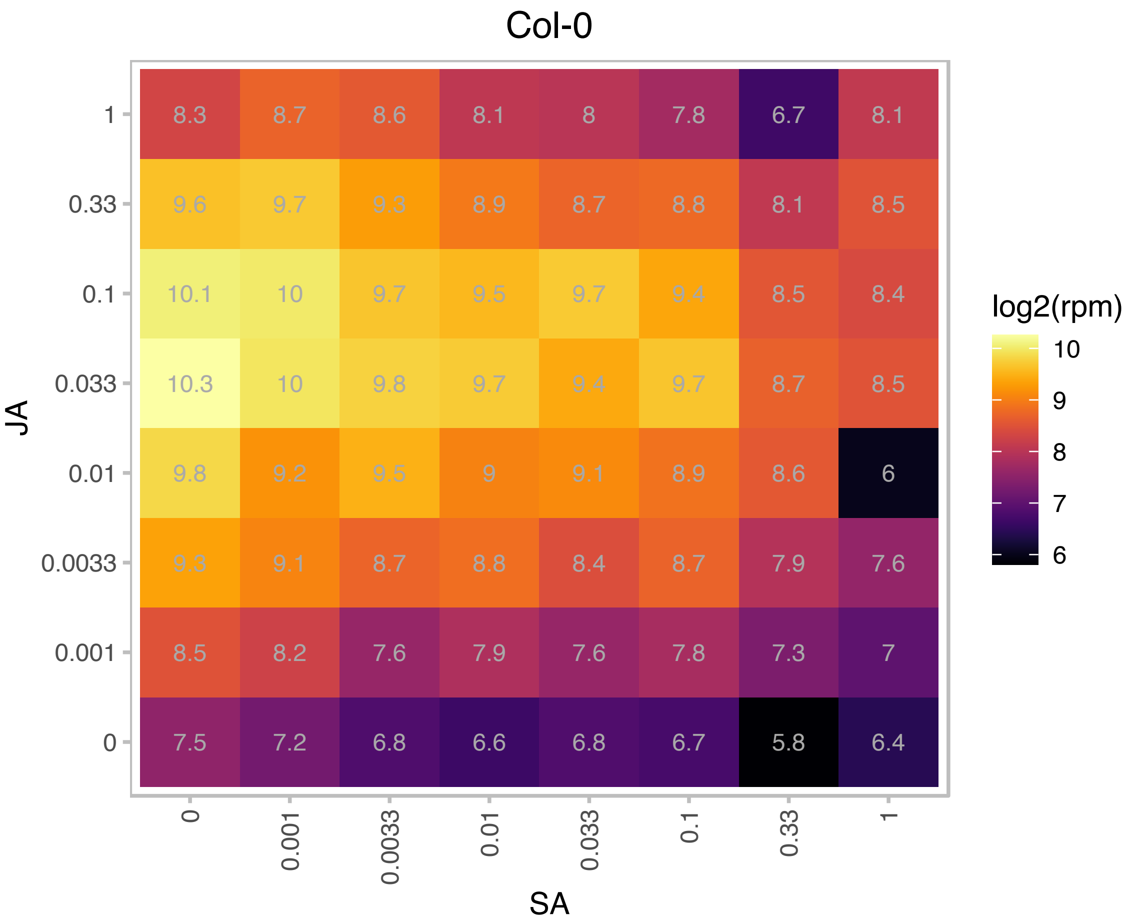


| | |
|---|---|
| Gene | AT1G19570 |
| is.expressed_genes | TRUE |
| is.responsive_genes | TRUE |
| is.apcluster | E-03 |

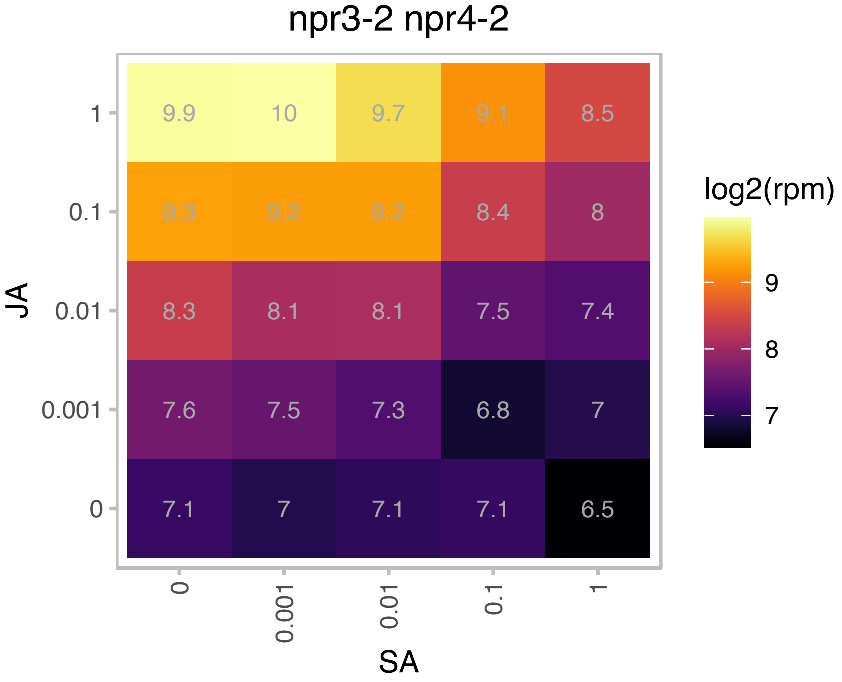


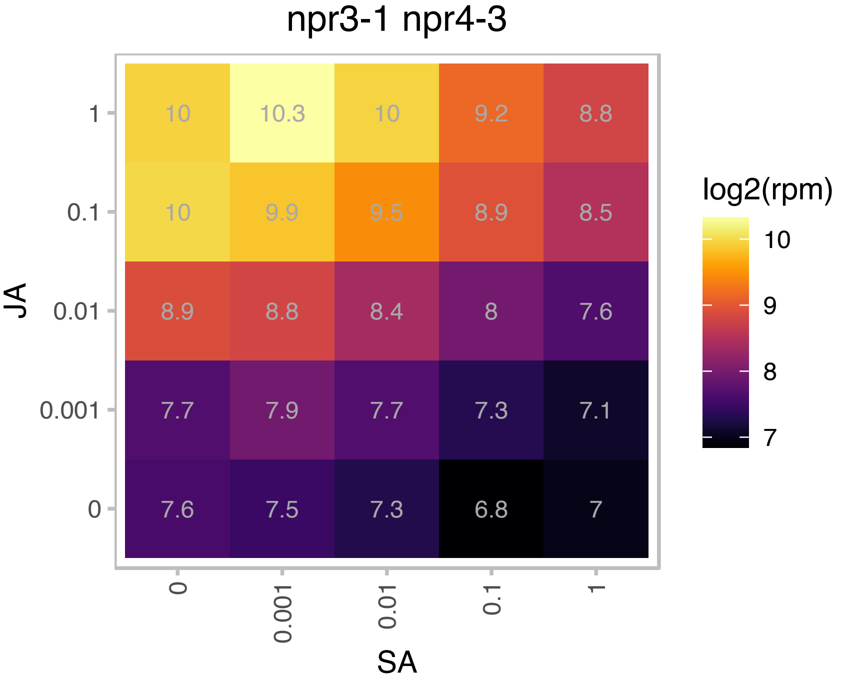

### AT1G49720.1 : ABF1, AtABF1

abscisic acid responsive element-binding factor 1

Identified as a protein that binds to abscisic acid response elements. May mediate transcriptional regulation of ABA responses.

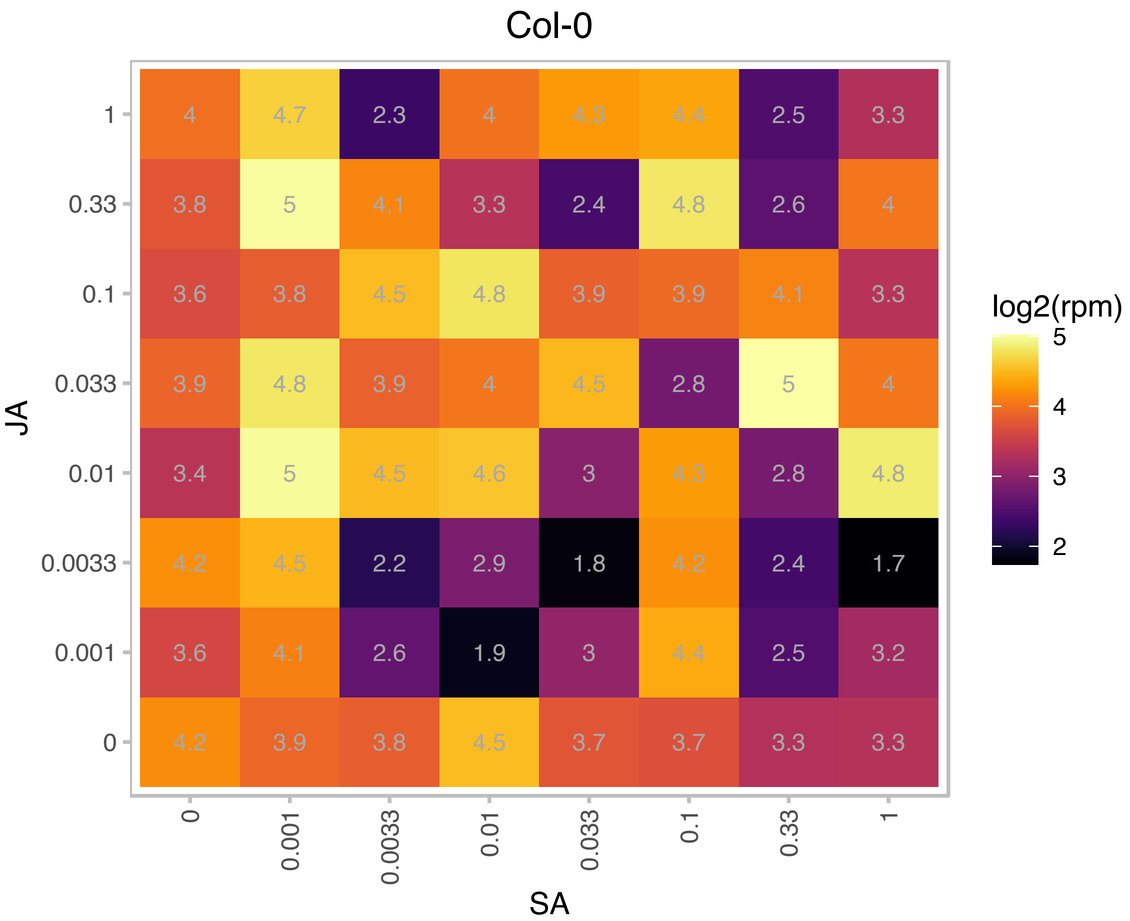


| | |
|---|---|
| Gene | AT1G49720 |
| is.expressed_genes | TRUE |
| is.responsive_genes | TRUE |
| is.apcluster | E-03 |

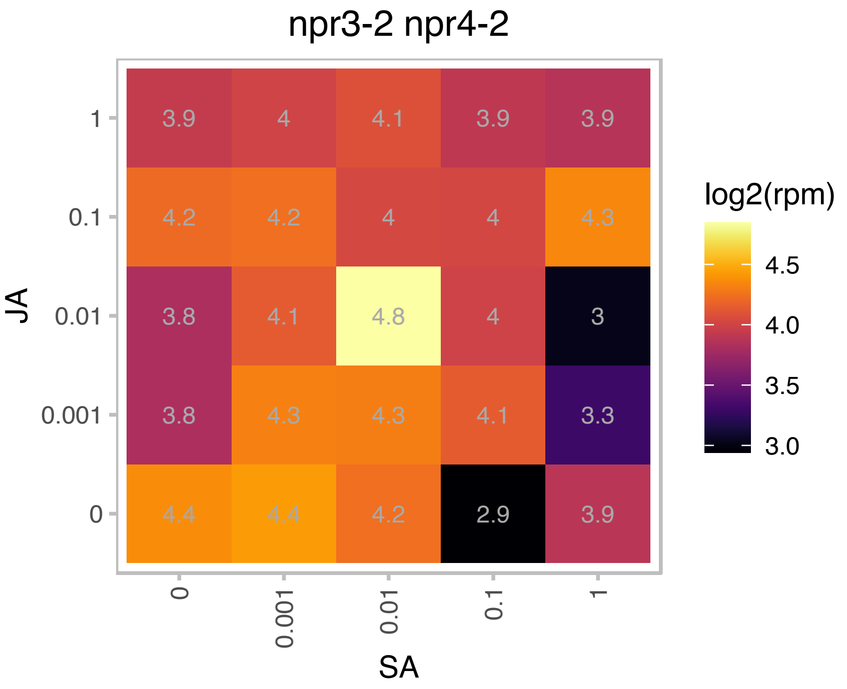


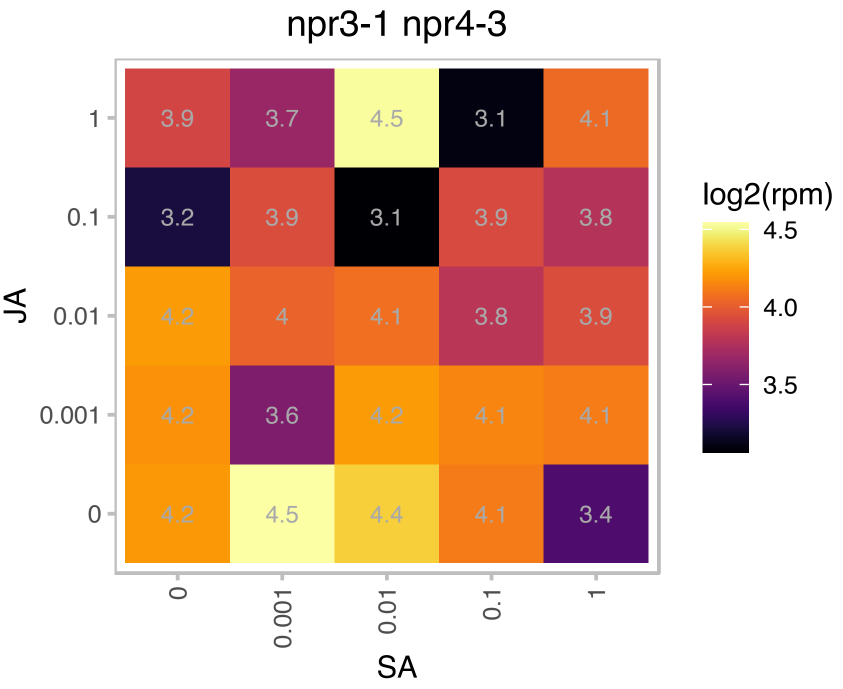

# AT1G52000.1 : NA

Mannose-binding lectin superfamily protein

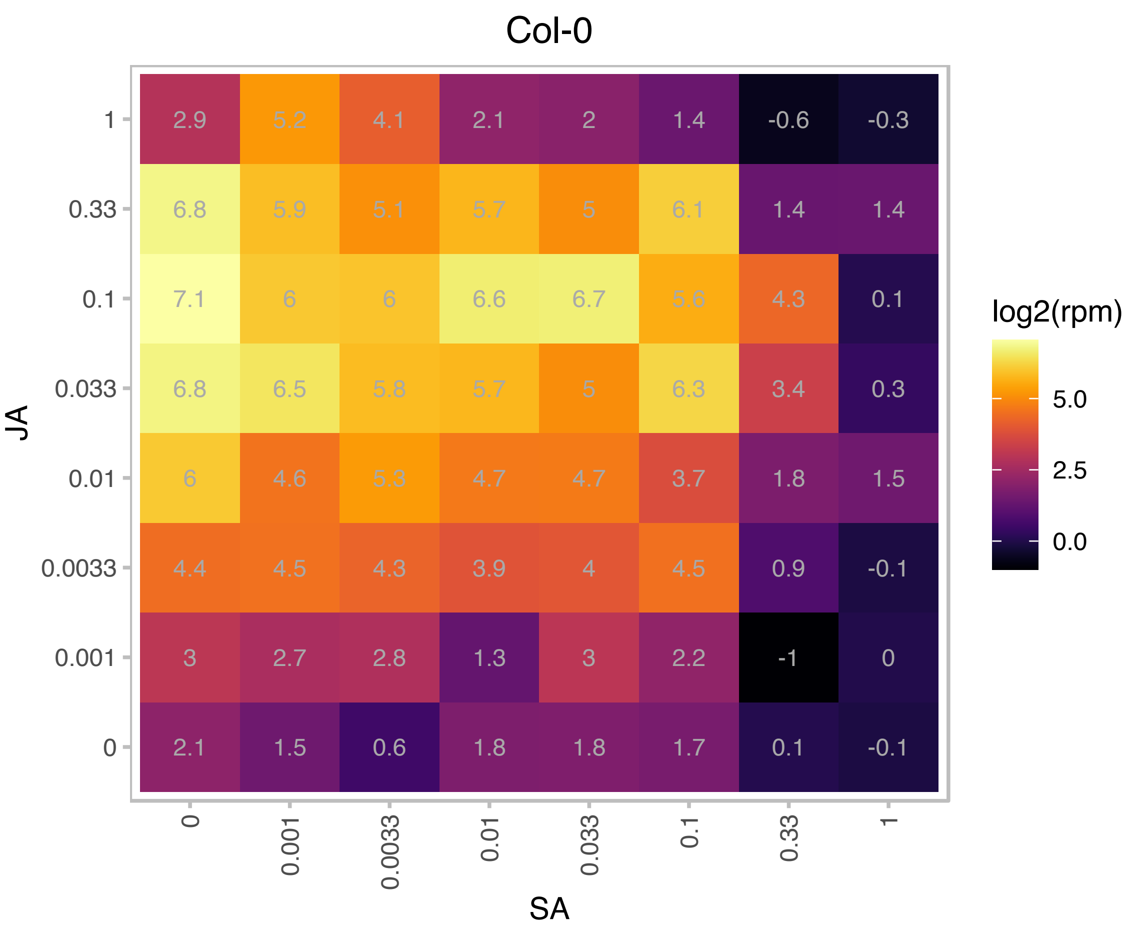


| | |
|---|---|
| Gene | AT1G52000 |
| is.expressed_genes | TRUE |
| is.responsive_genes | TRUE |
| is.apcluster | E-03 |

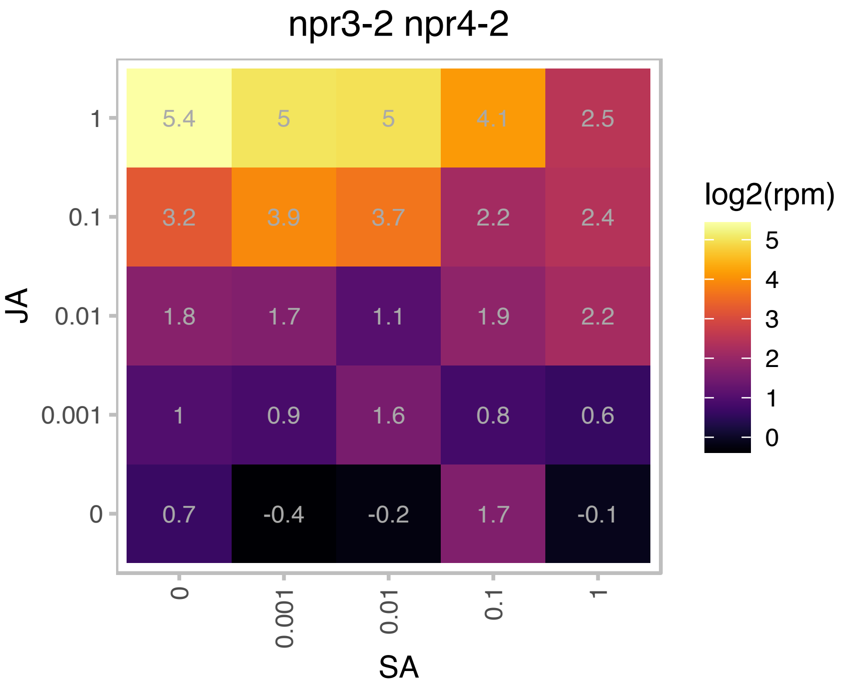


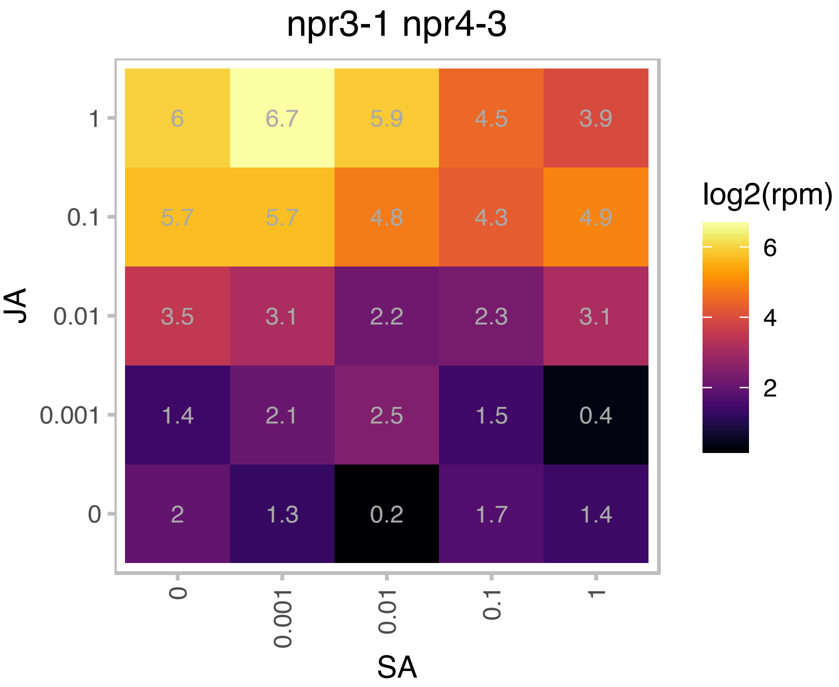

### AT1G52030.1 : F-ATMBP, MBP1.2, MBP2

myrosinase-binding protein 2
Similar to myrosinase binding proteins which may be involved in metabolizing glucosinolates and forming defense compounds to protect against herbivory. Also similar to lectins and other agglutinating factors. Expressed only in flowers.

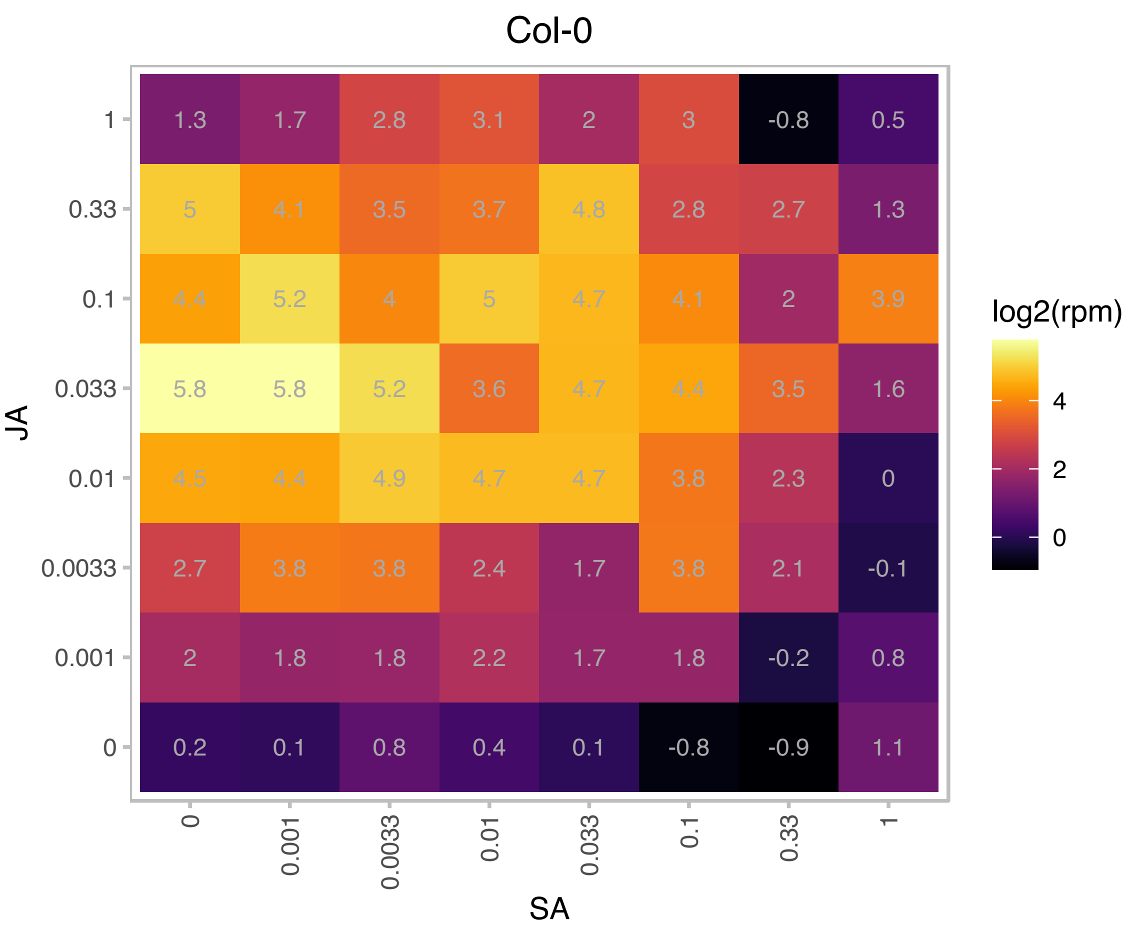


| | |
|---|---|
| Gene | AT1G52030 |
| is.expressed_genes | TRUE |
| is.responsive_genes | TRUE |
| is.apcluster | E-03 |

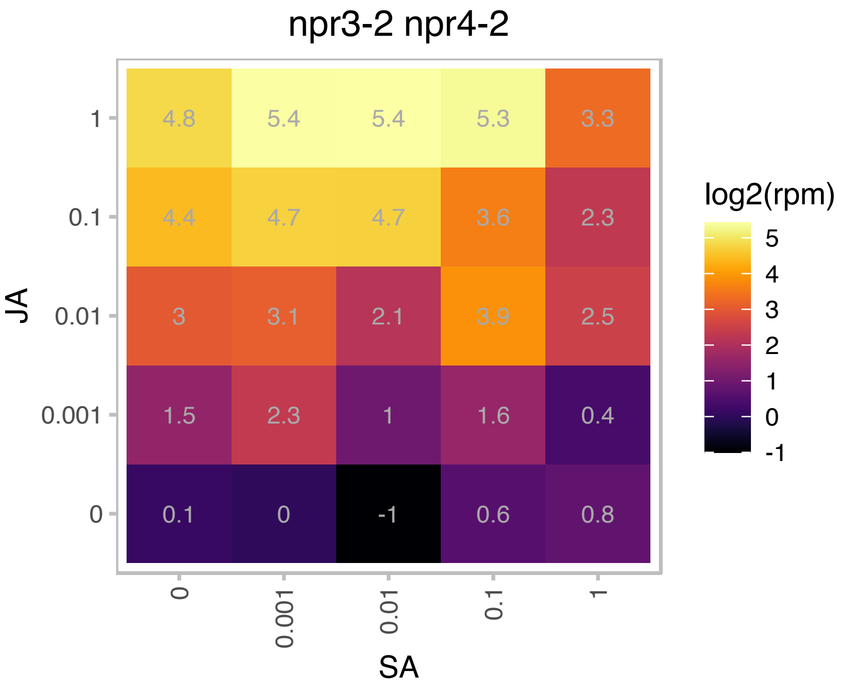


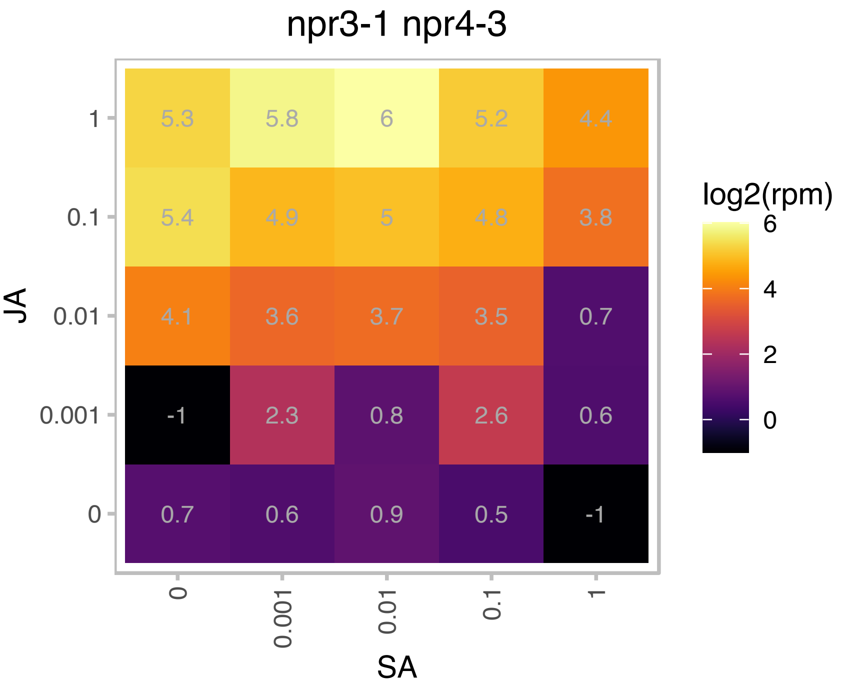

### AT1G52040.1 : ATMBP, MBP1

myrosinase-binding protein 1

Encodes myrosinase-binding protein expressed in flowers.

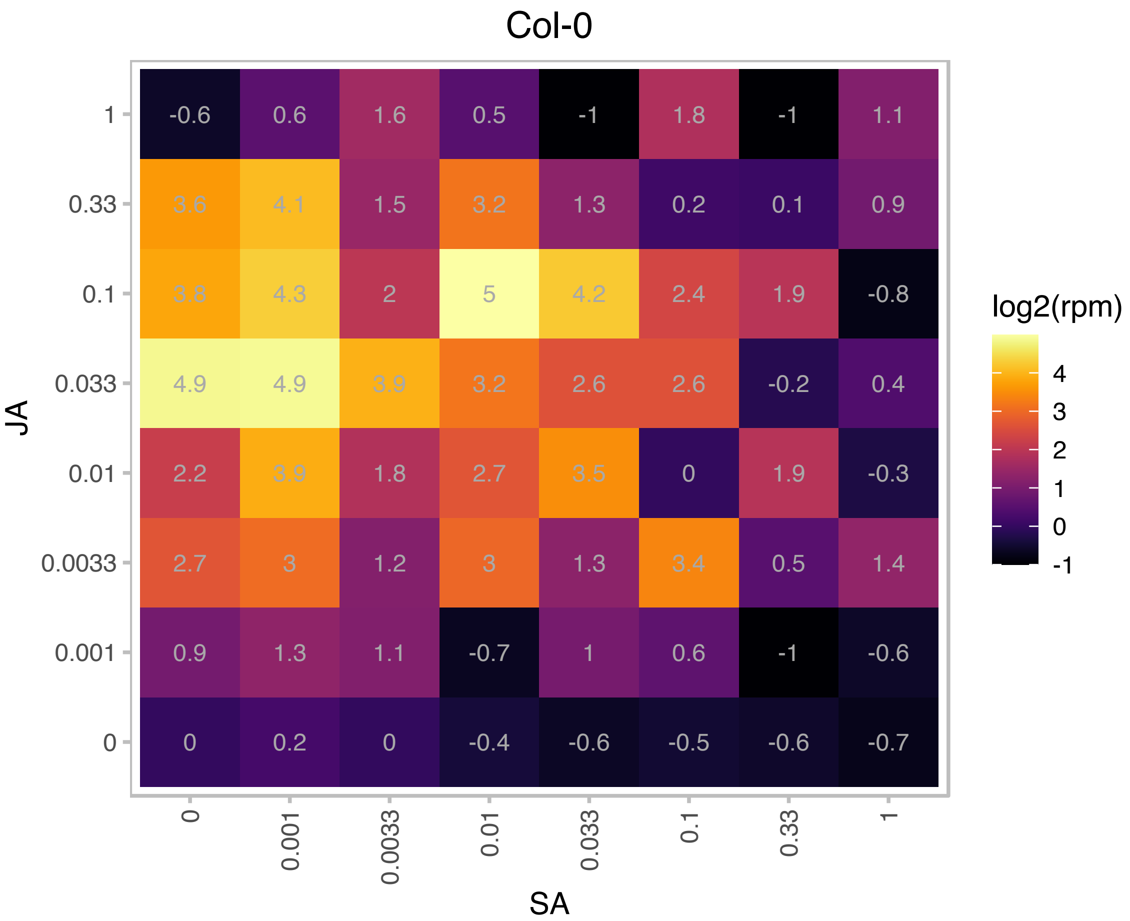


| | |
|---|---|
| Gene | AT1G52040 |
| is.expressed_genes | TRUE |
| is.responsive_genes | TRUE |
| is.apcluster | E-03 |

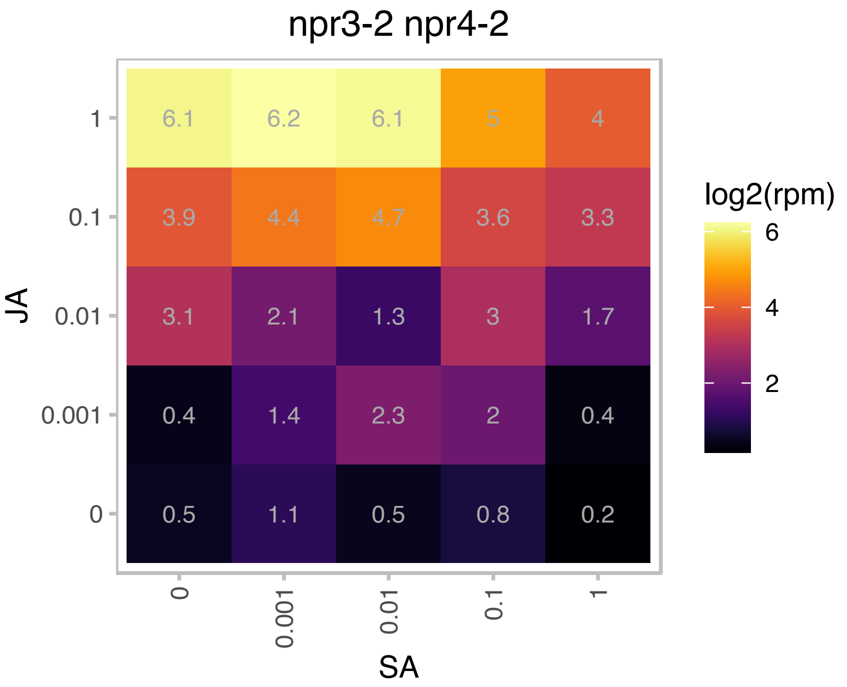


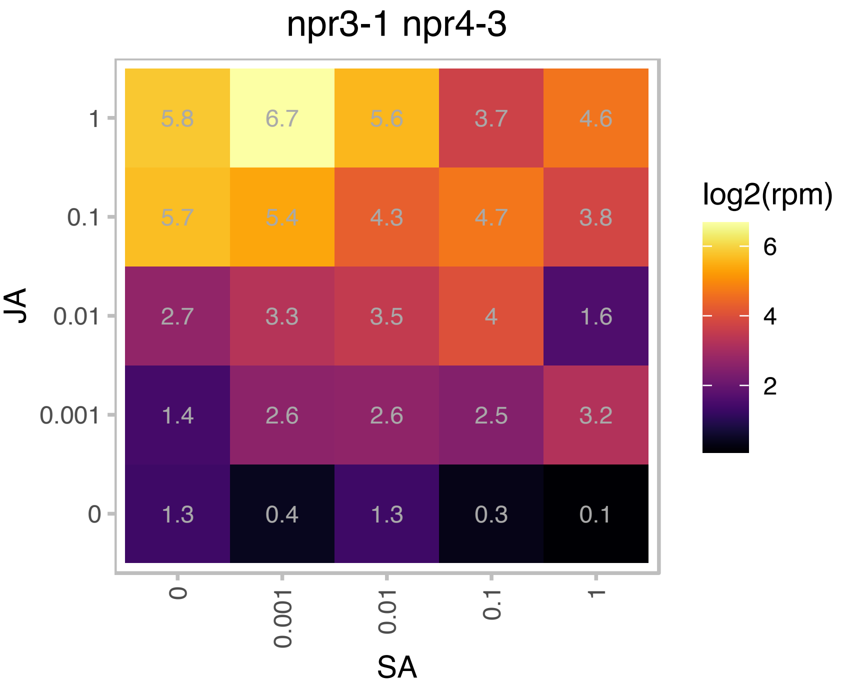

### AT1G52410.1 : AtTSA1, TSA1

TSK-associating protein 1

Contains a novel calcium-binding repeat sequence. Binds TSK in vitro. Localizes to small cytoplasmic vesicles in interphase cells. In cells synchronized for cell division, TSA1 and TSK relocalize to ends of spindle microtubules that are ahead of separating chromatids during metaphase and anaphase of mitosis. May be involved in mitosis together with TSK. Expressed preferentially in the flower and shoot apex. Can form multimers. The mRNA is cell-to-cell mobile.

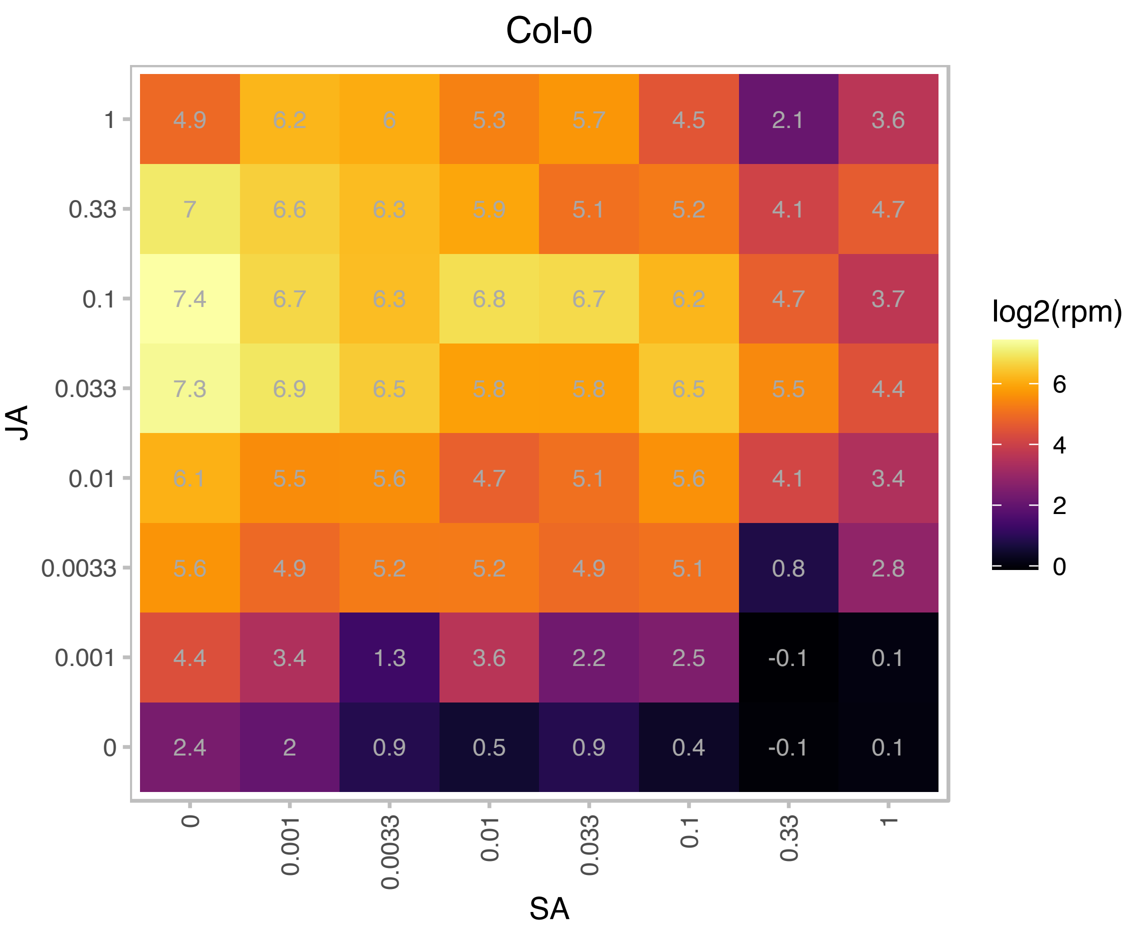


| | |
|---|---|
| Gene | AT1G52410 |
| is.expressed_genes | TRUE |
| is.responsive_genes | TRUE |
| is.apcluster | E-03 |

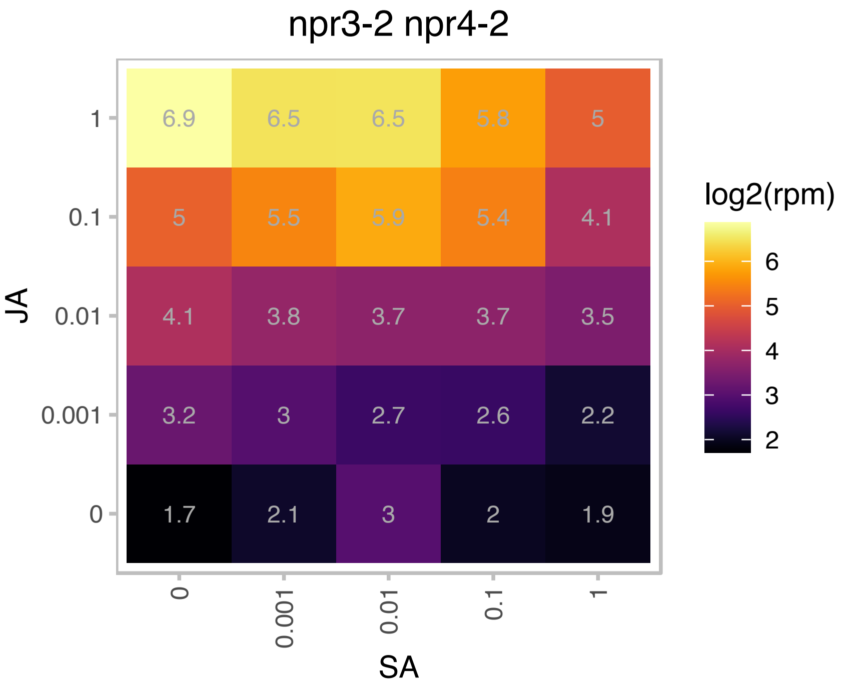


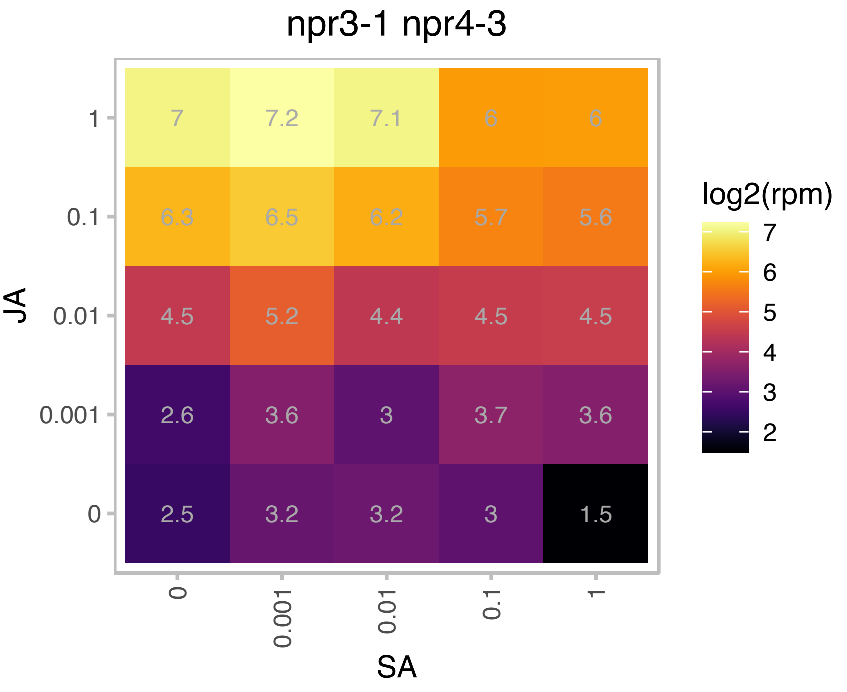

# AT1G53885.1 : NA

Protein of unknown function (DUF581)

| | |
|---|---|
| Gene | AT1G53885 |
| is.expressed_genes | TRUE |
| is.responsive_genes | TRUE |
| is.apcluster | E-03 |

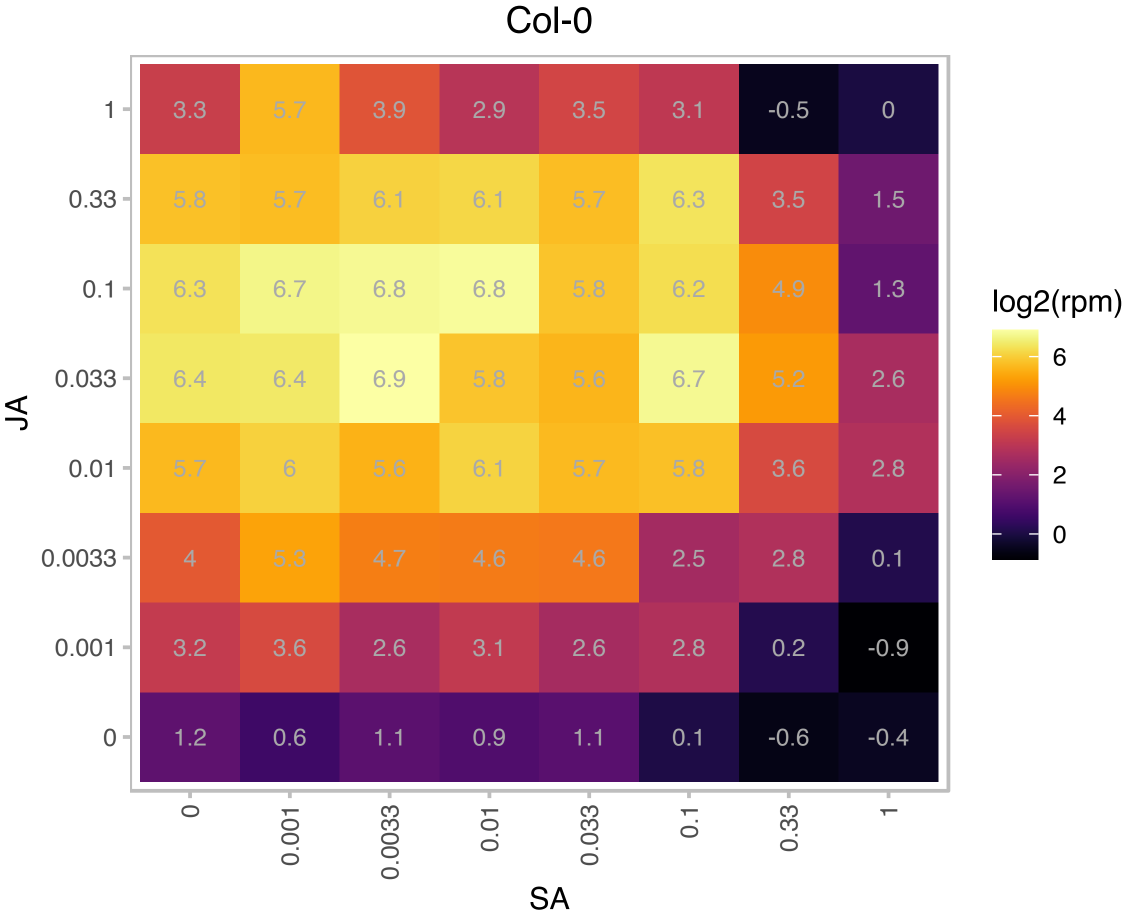


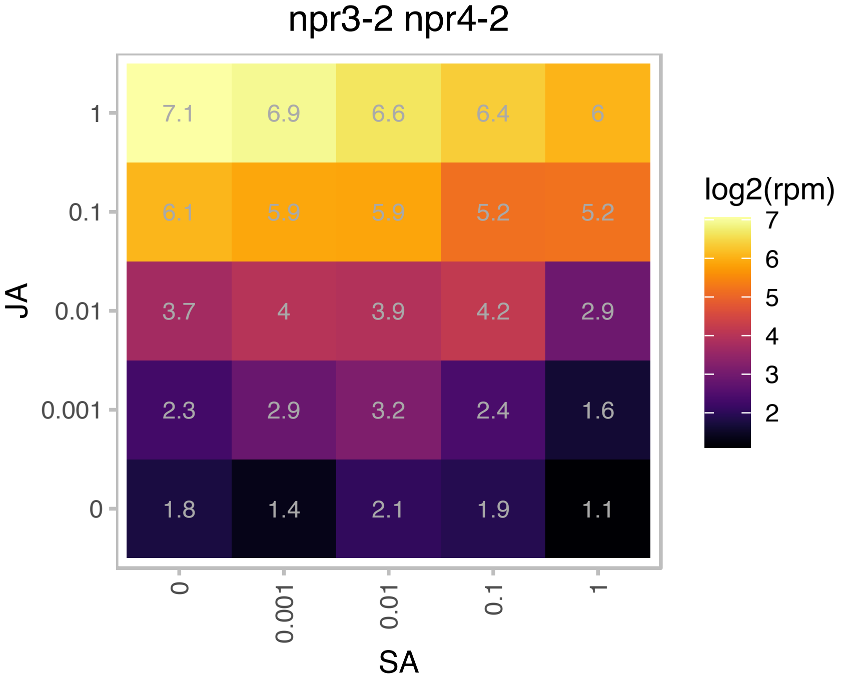


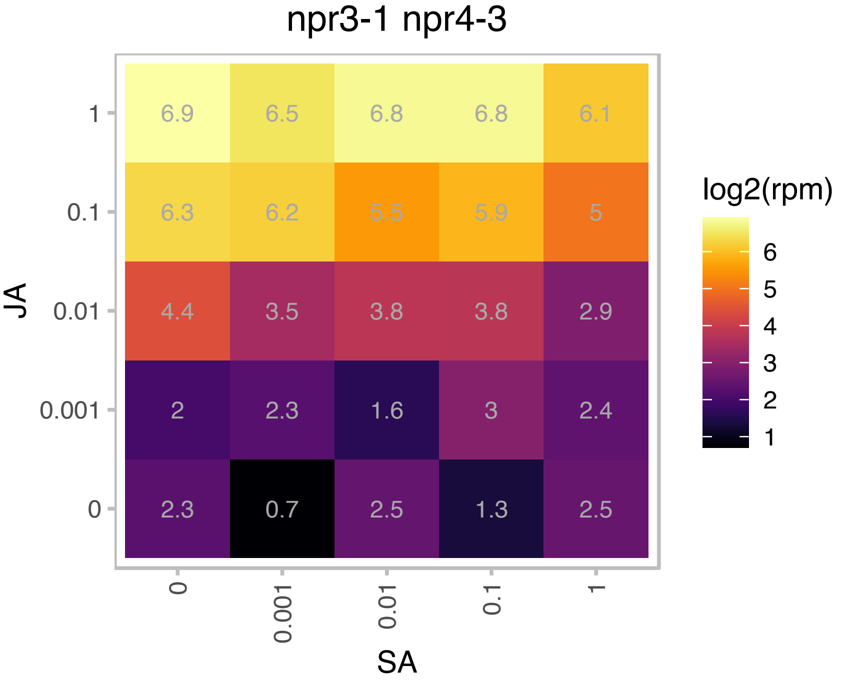

# AT1G53903.1 : NA

Protein of unknown function (DUF581)

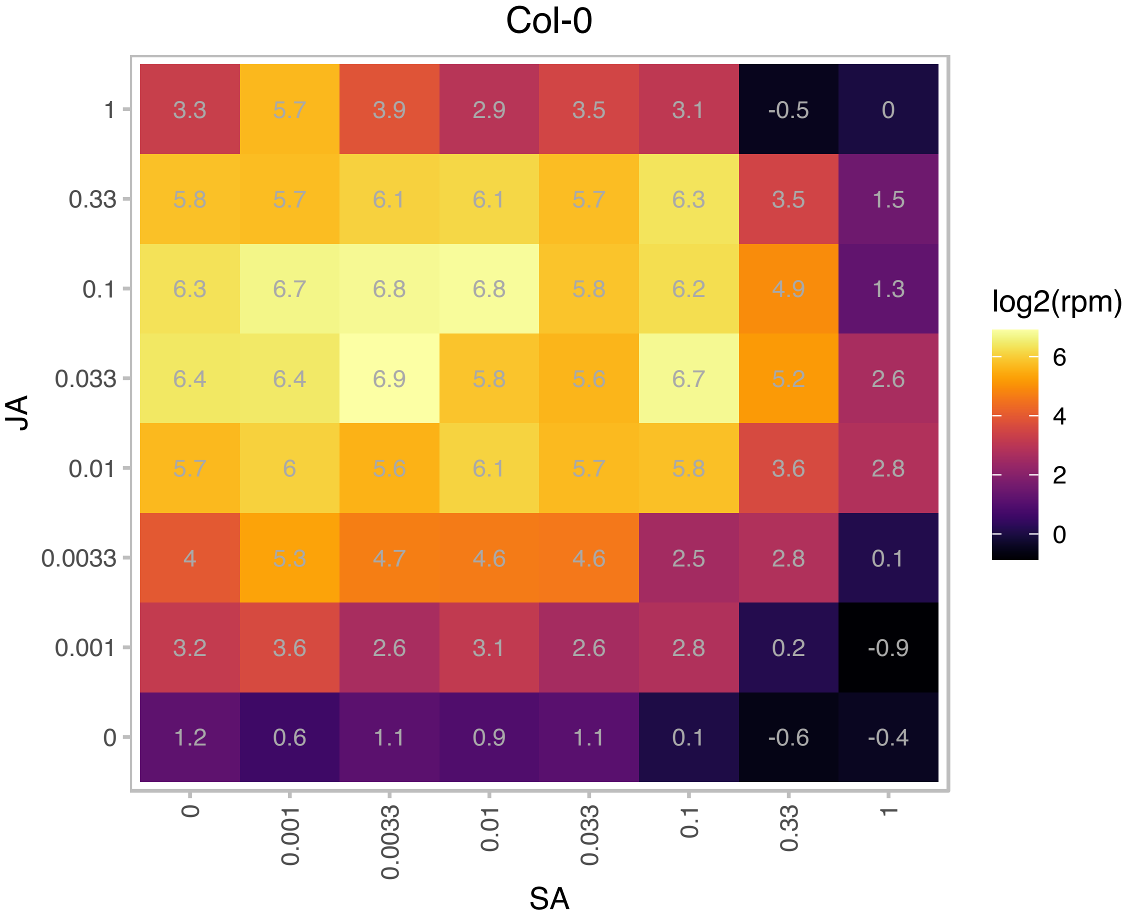


| Gene | AT1G53903 |
|---|---|
| is.expressed_genes | TRUE |
| is.responsive_genes | TRUE |
| is.apcluster | E-03 |

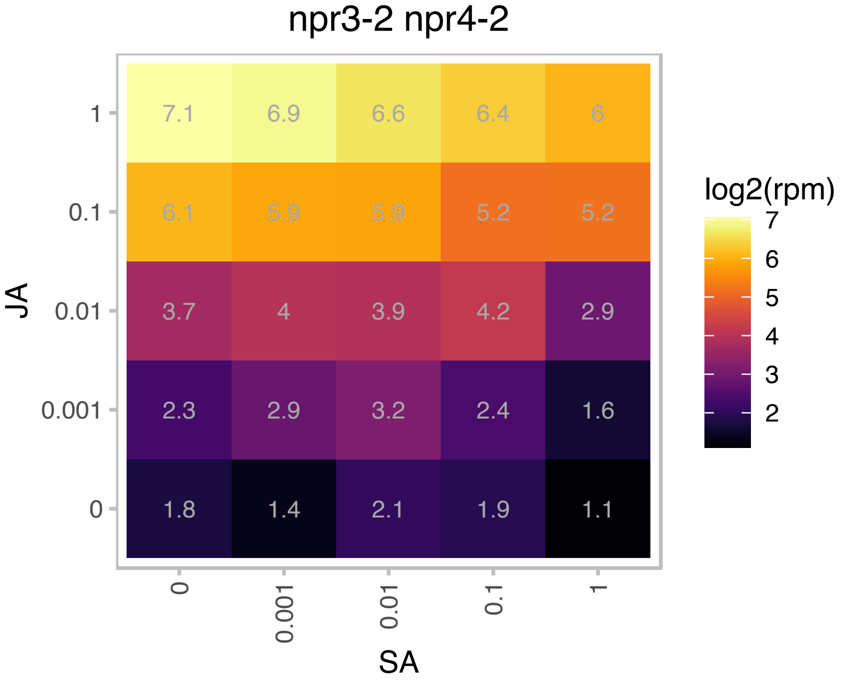


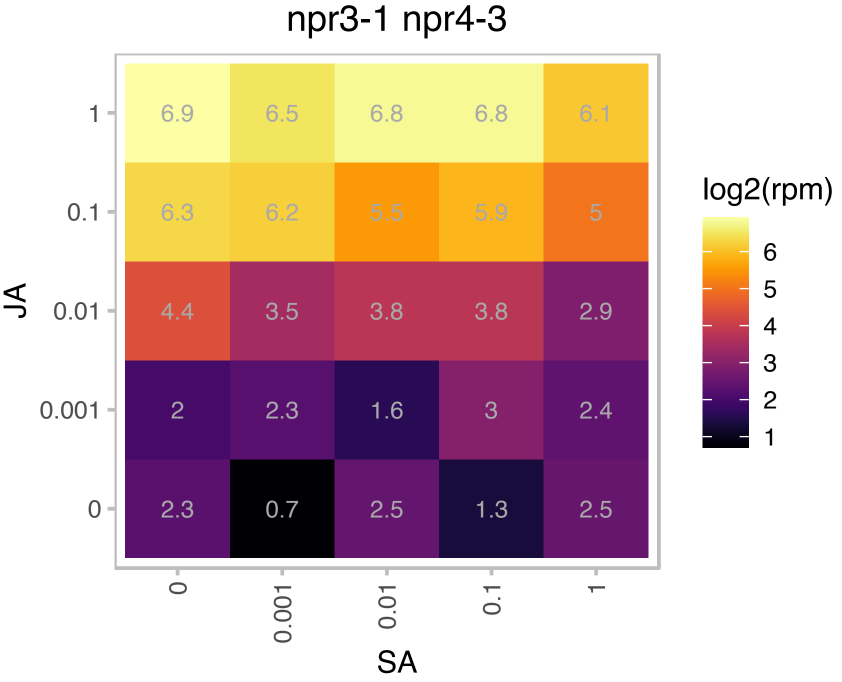

### AT1G60220.1 : OTS1, ULP1D

UB-like protease 1D

Encodes a deSUMOylating enzyme. In vitro it has both peptidase activity and isopeptidase activity: it can cleave C-terminal residues from SUMO to activate it for attachment to a target protein and it can also act on the isopeptide bond between SUMO and another protein. sGFP:OTS1 protein accumulates in the nucleus. Double mutant analysis with ULP1C/OTS2 indicates that these genes are involved in salt stress responses and flowering time regulation. Over-expression of 35S:OTS1 increases salt tolerance and reduces the level of SUMO-conjugated proteins. OTS1 transcript levels do not appear to change in response to salt, but, salt [illegible]

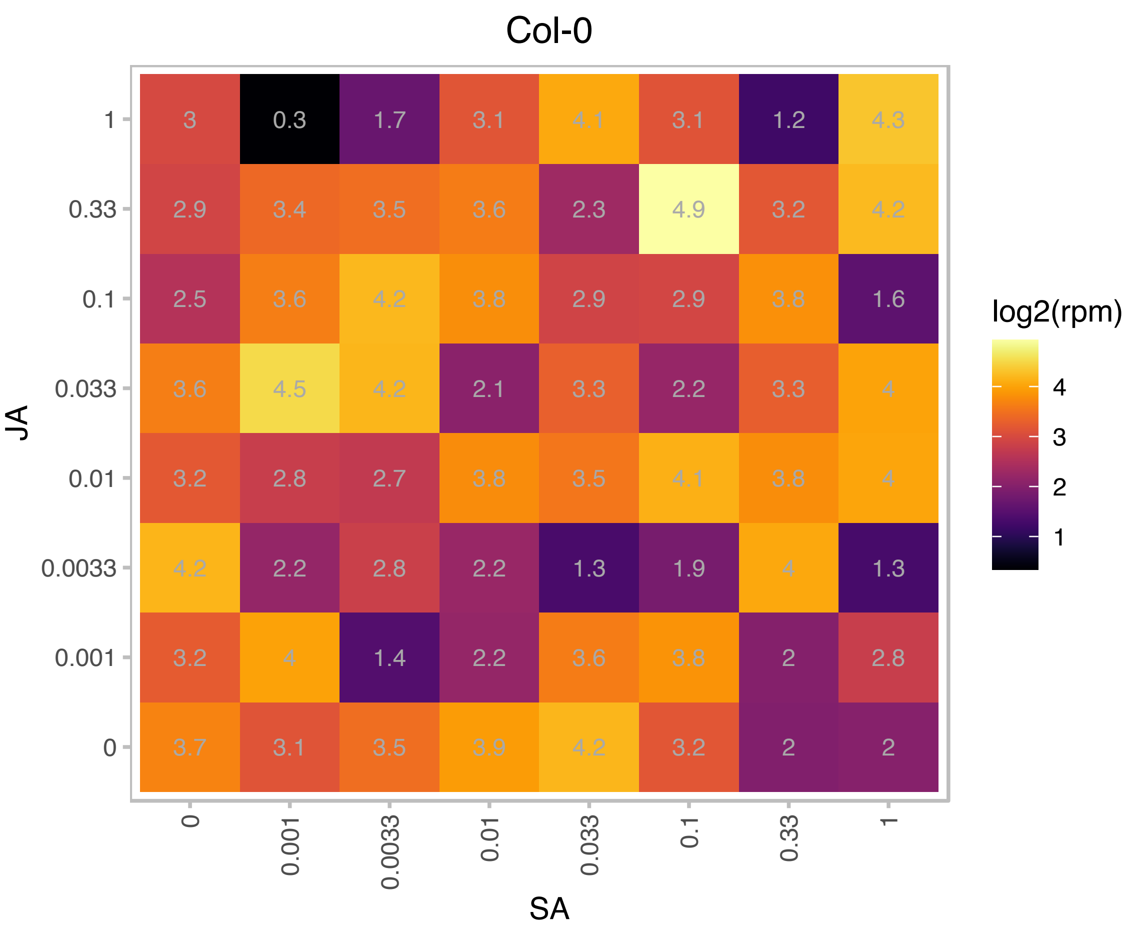


| | |
|---|---|
| Gene | AT1G60220 |
| is.expressed_genes | TRUE |
| is.responsive_genes | TRUE |
| is.apcluster | E-03 |

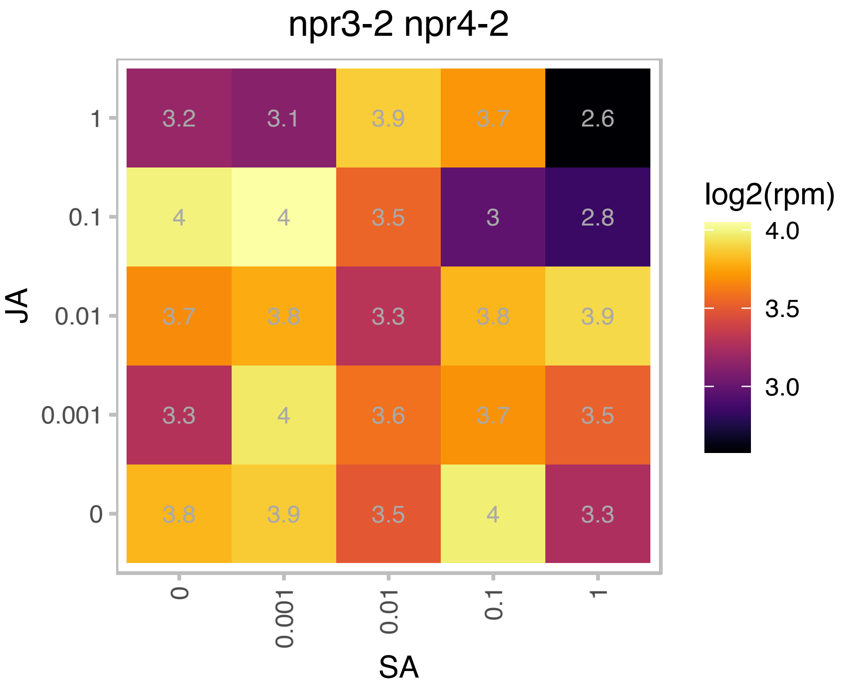


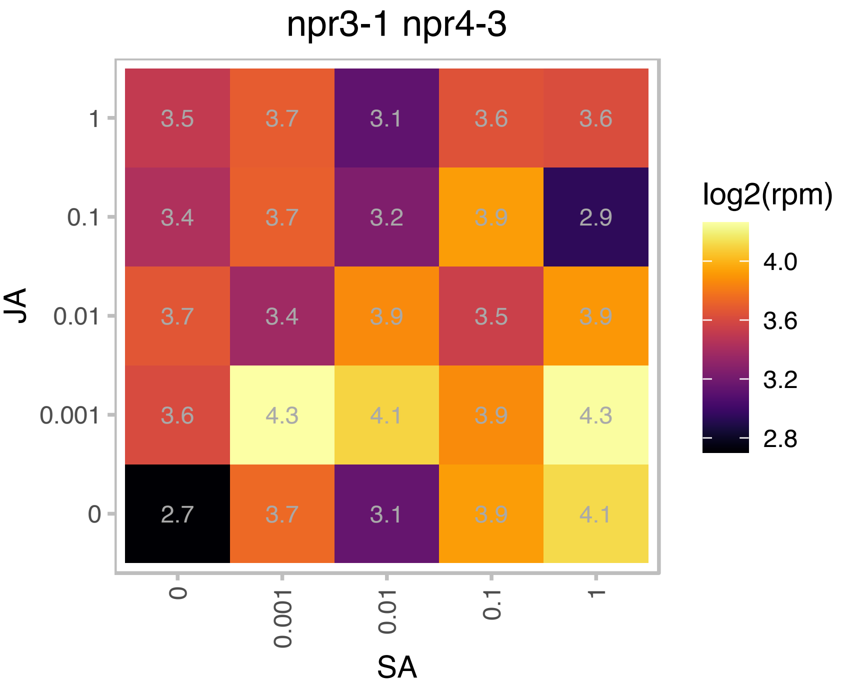

### AT1G60260.1 : BGLU5

beta glucosidase 5

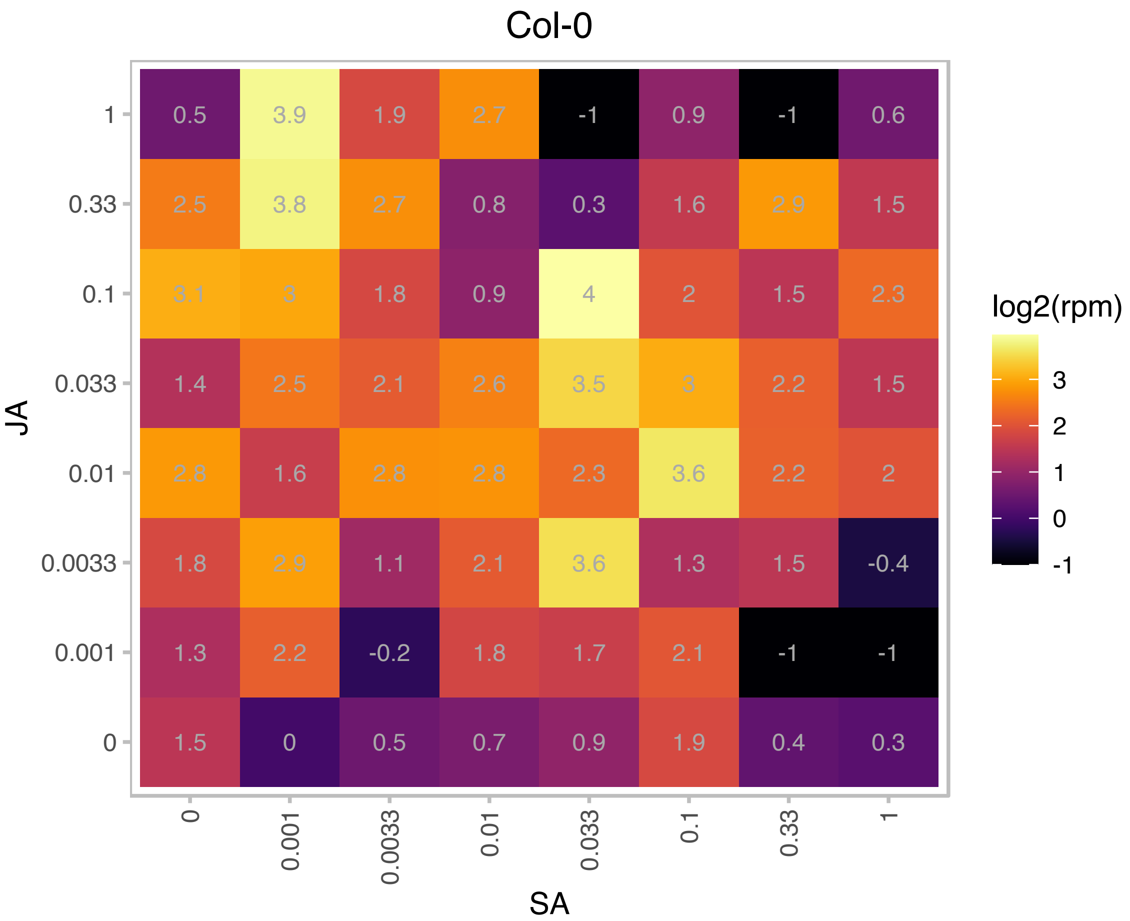


| | |
|---|---|
| Gene | AT1G60260 |
| is.expressed_genes | TRUE |
| is.responsive_genes | TRUE |
| is.apcluster | E-03 |

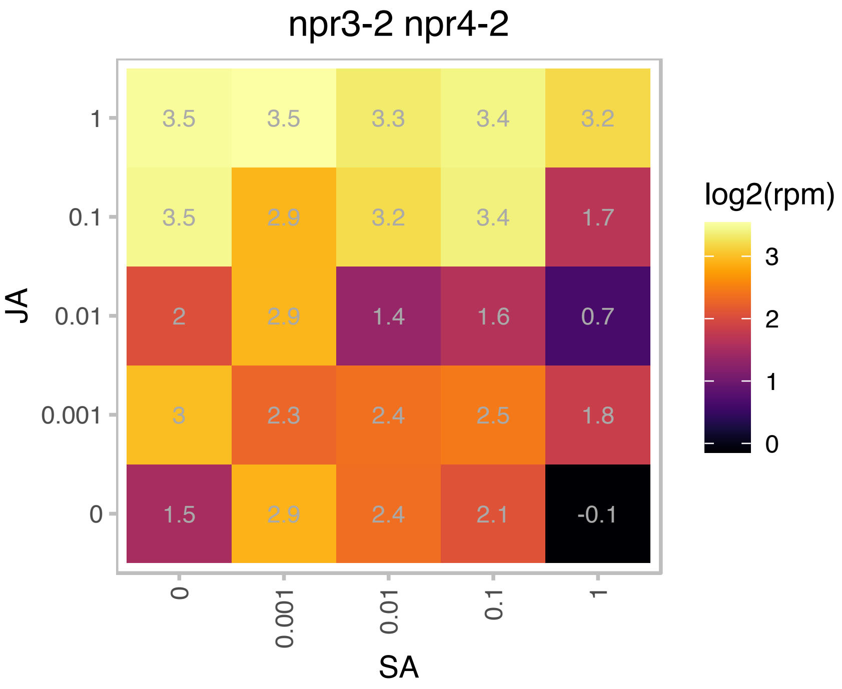


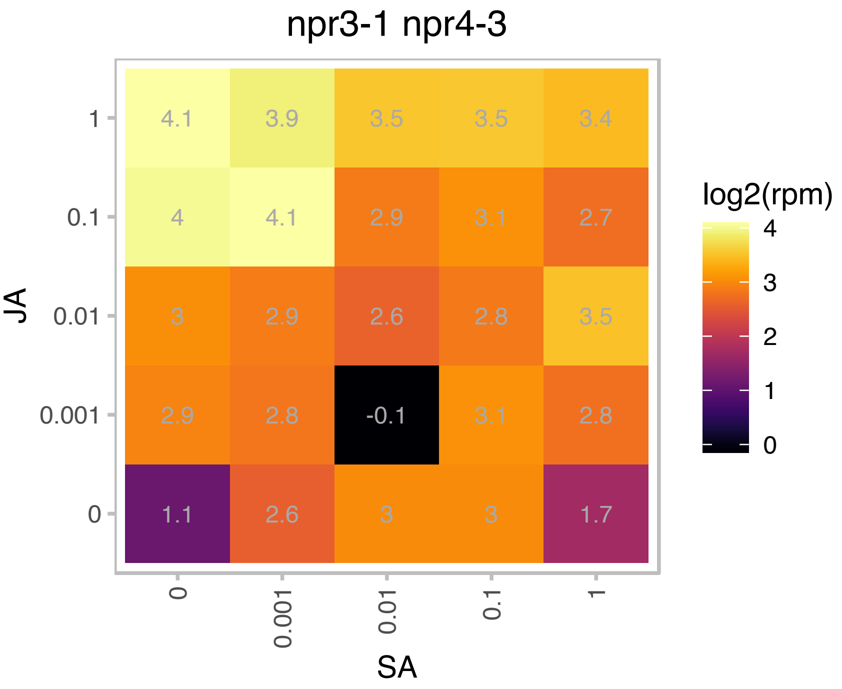

### AT1G64160.1 : AtDIR5, DIR5

Disease resistance-responsive (dirigent-like protein) family protein
Encodes a dirigent protein involved in the synthesis of (-)pinoresinol. Dirigent proteins impart stereoselectivity on the phenoxy radical coupling reaction yielding optically active lignans from two molecules of coniferyl alcohol.

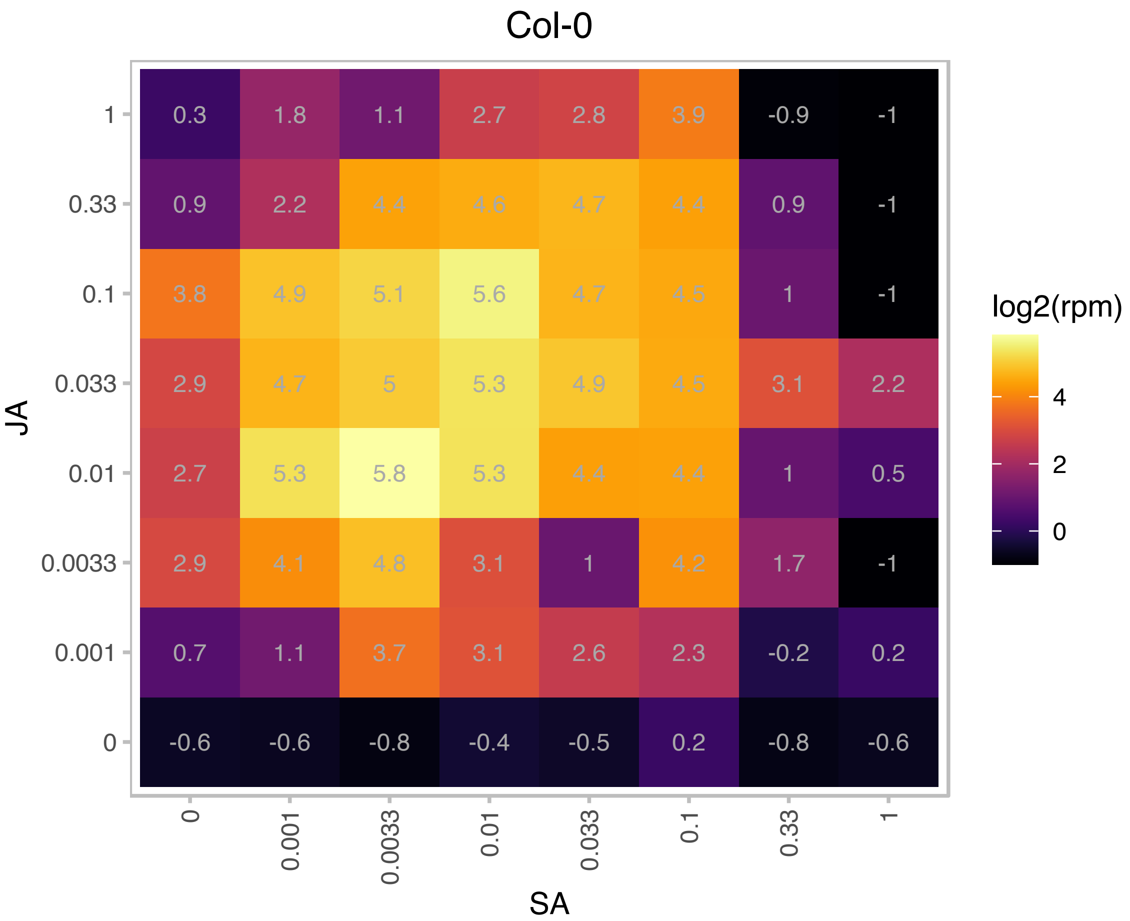


| | |
|---|---|
| Gene | AT1G64160 |
| is.expressed_genes | TRUE |
| is.responsive_genes | TRUE |
| is.apcluster | E-03 |

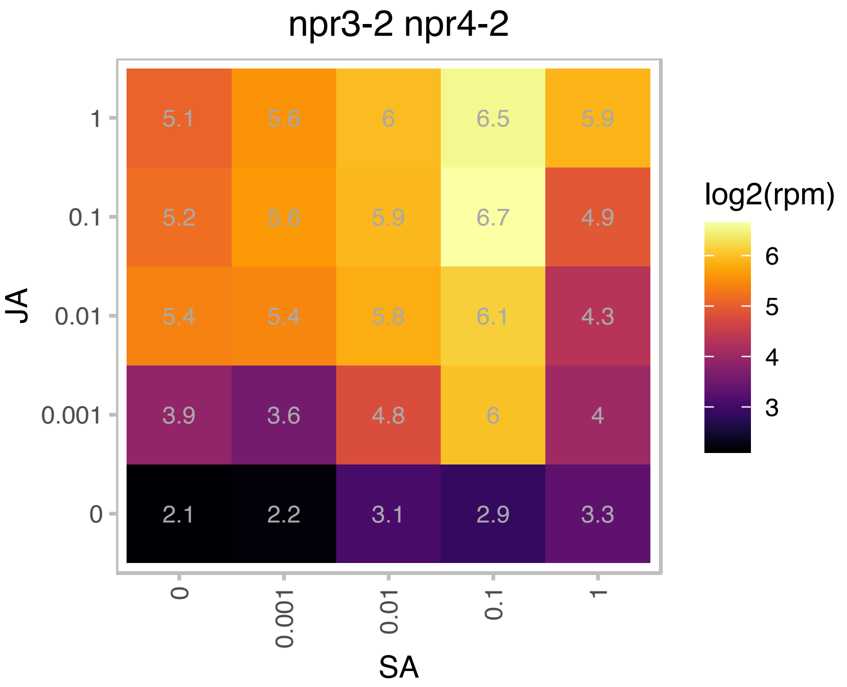


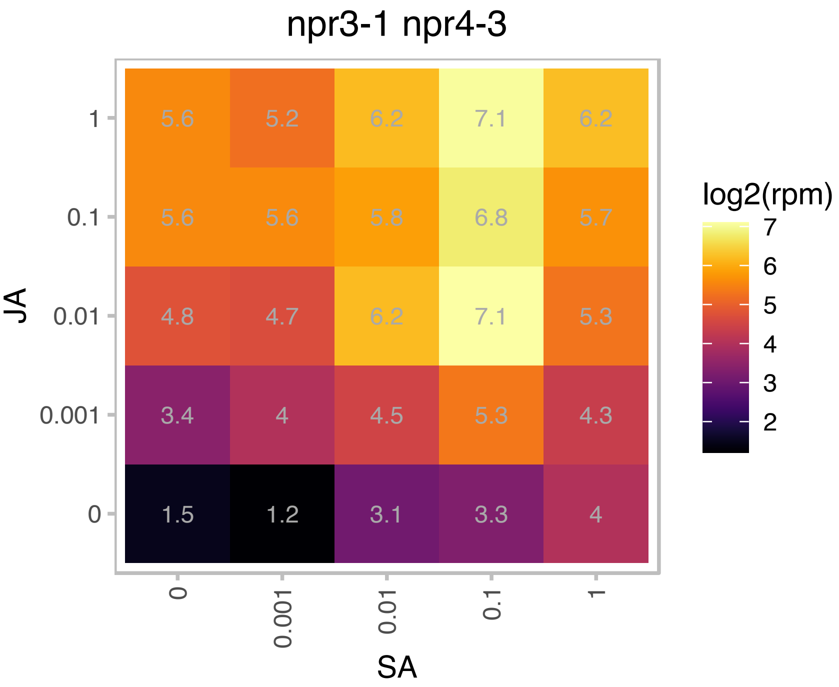

# AT1G69070.1 : NA

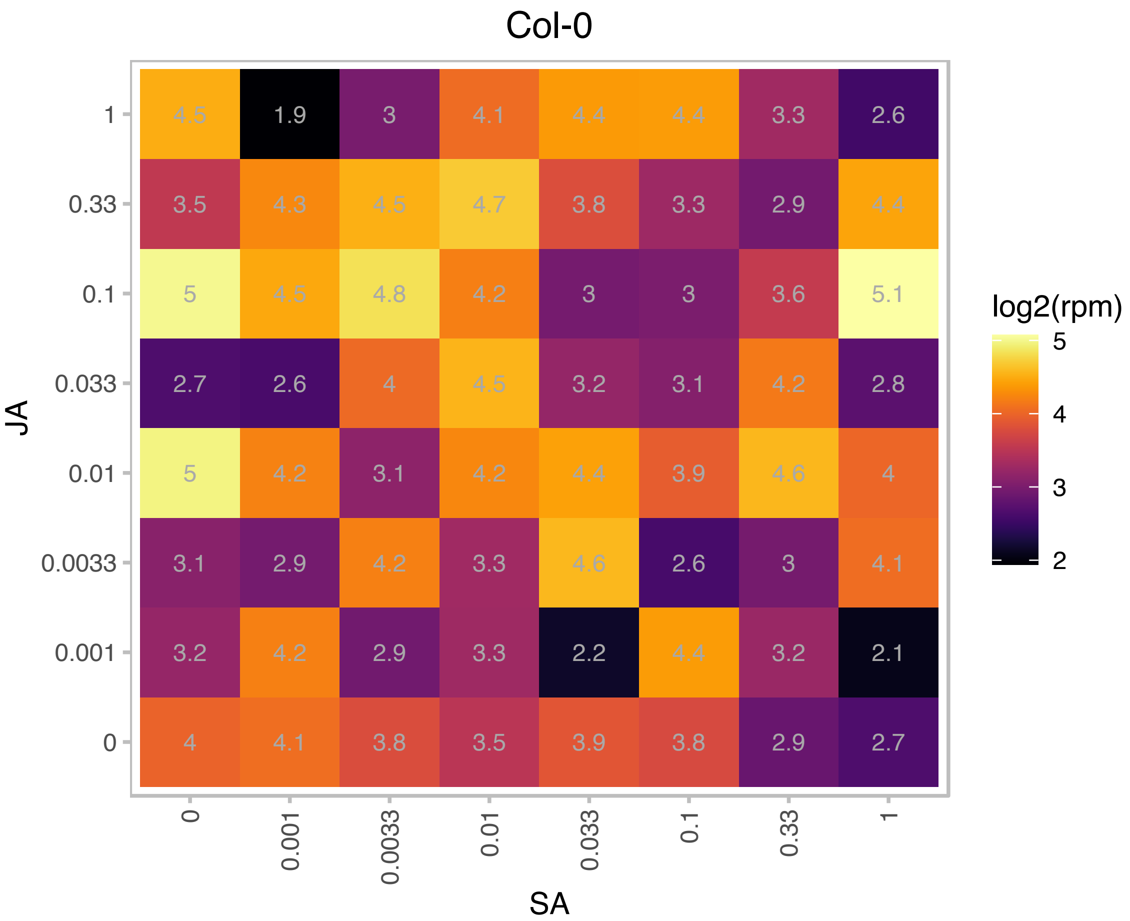


| | |
|---|---|
| Gene | AT1G69070 |
| is.expressed_genes | TRUE |
| is.responsive_genes | TRUE |
| is.apcluster | E-03 |

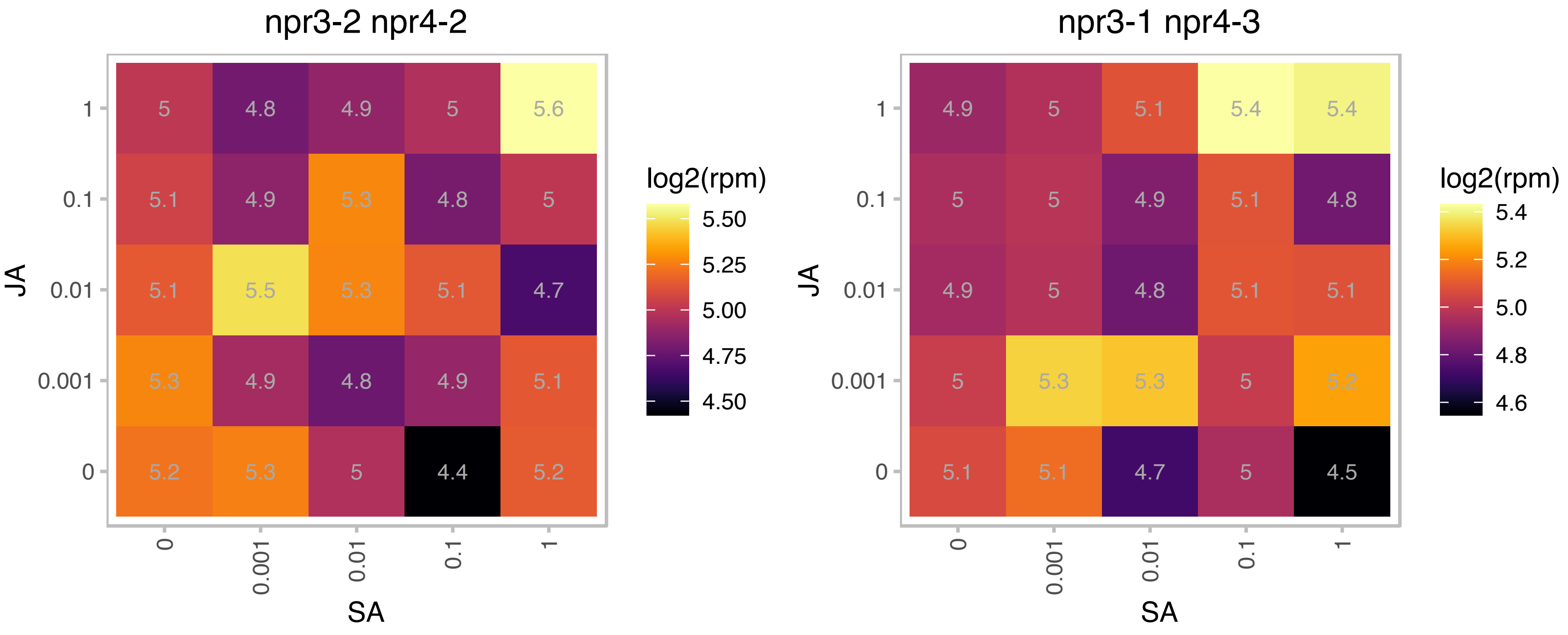

### AT1G69370.1 : AtCM3, cm-3, CM3

chorismate mutase 3

Encodes chorismate mutase 3 (CM3).

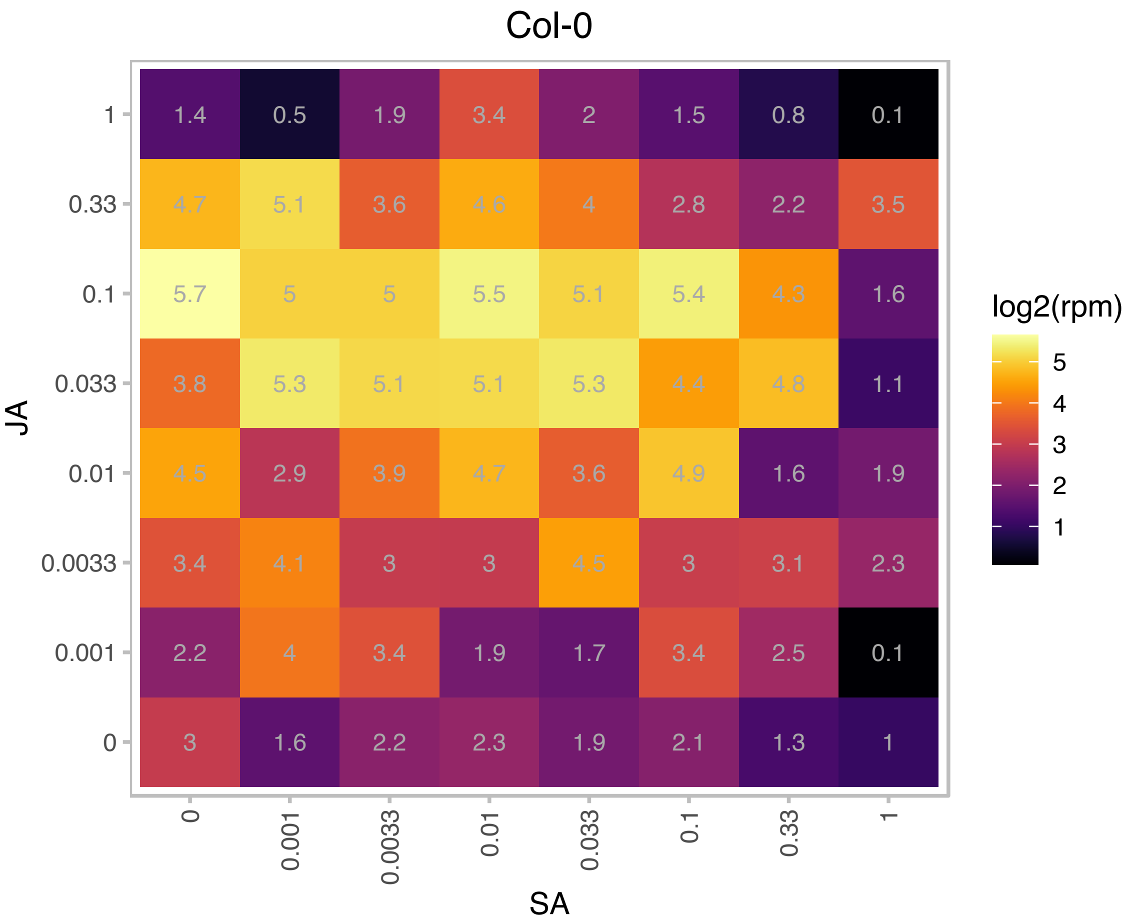


| | |
|---|---|
| Gene | AT1G69370 |
| is.expressed_genes | TRUE |
| is.responsive_genes | TRUE |
| is.apcluster | E-03 |

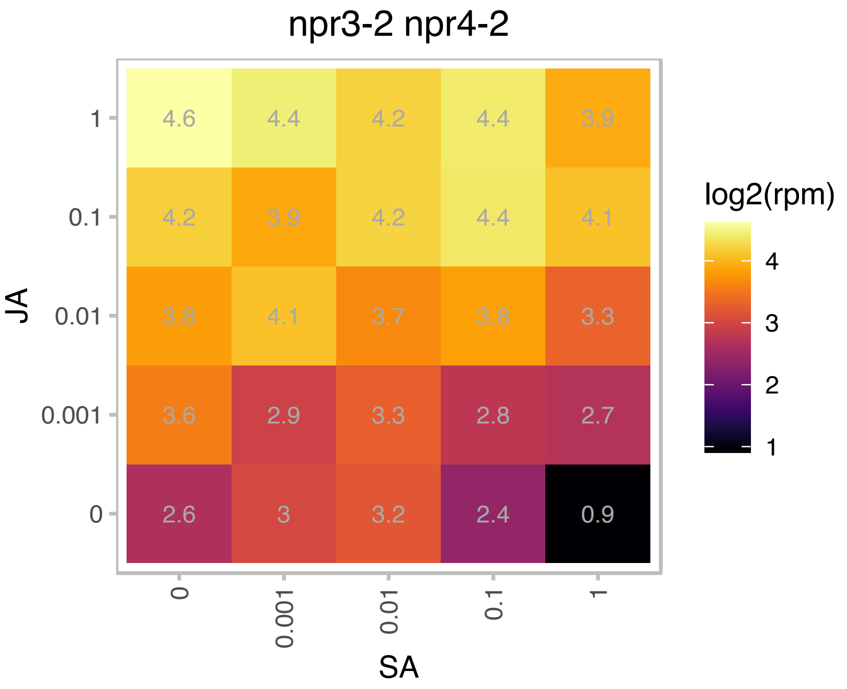


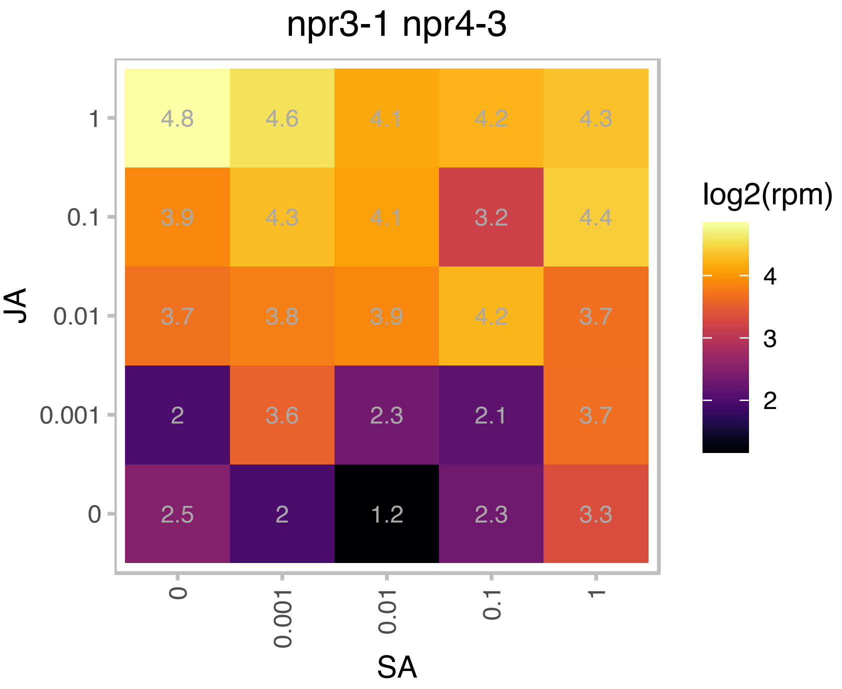

# AT1G70820.1 : NA

phosphoglucomutase, putative / glucose phosphomutase, putative

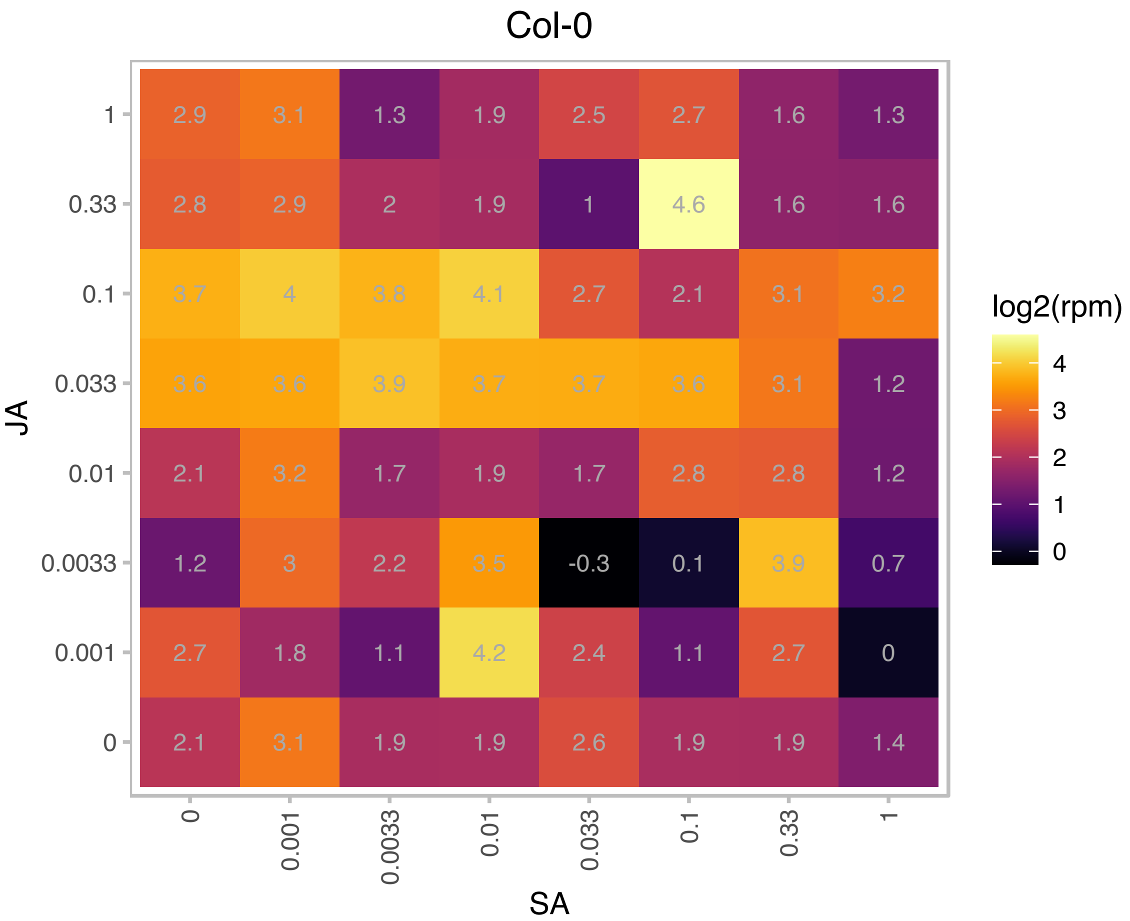


| | |
|---|---|
| Gene | AT1G70820 |
| is.expressed_genes | TRUE |
| is.responsive_genes | TRUE |
| is.apcluster | E-03 |

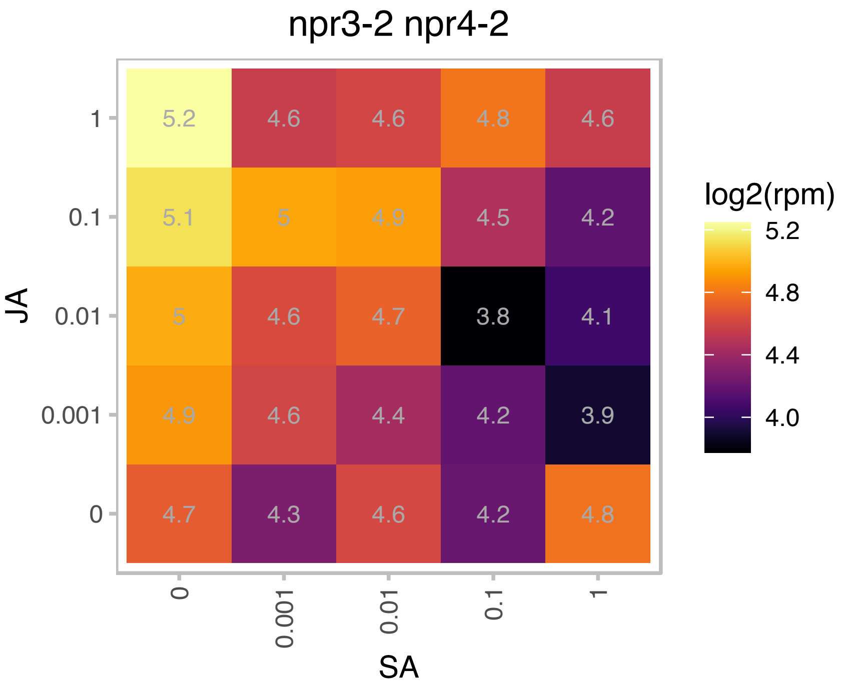


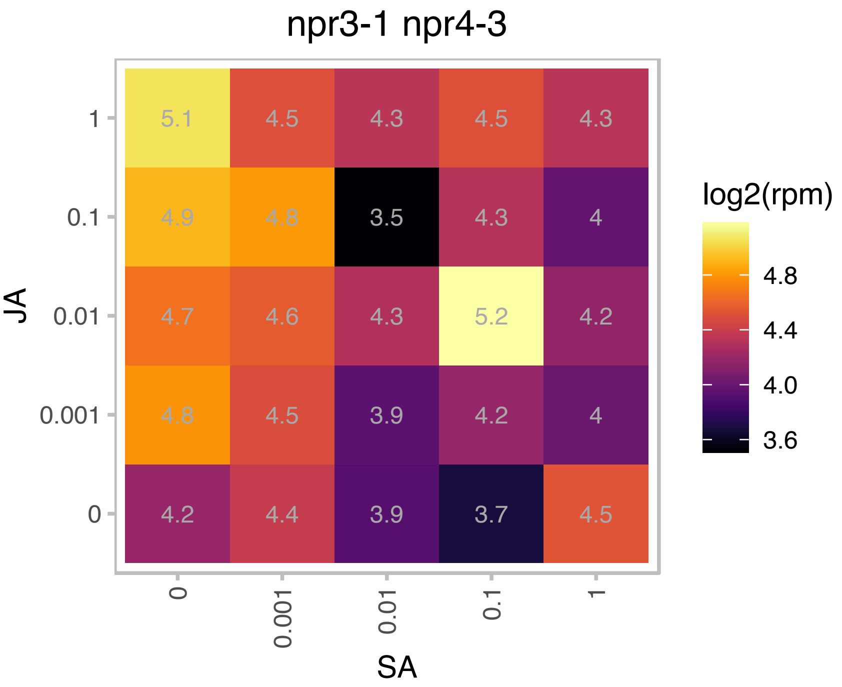

# AT1G73325.1 : NA

Kunitz family trypsin and protease inhibitor protein

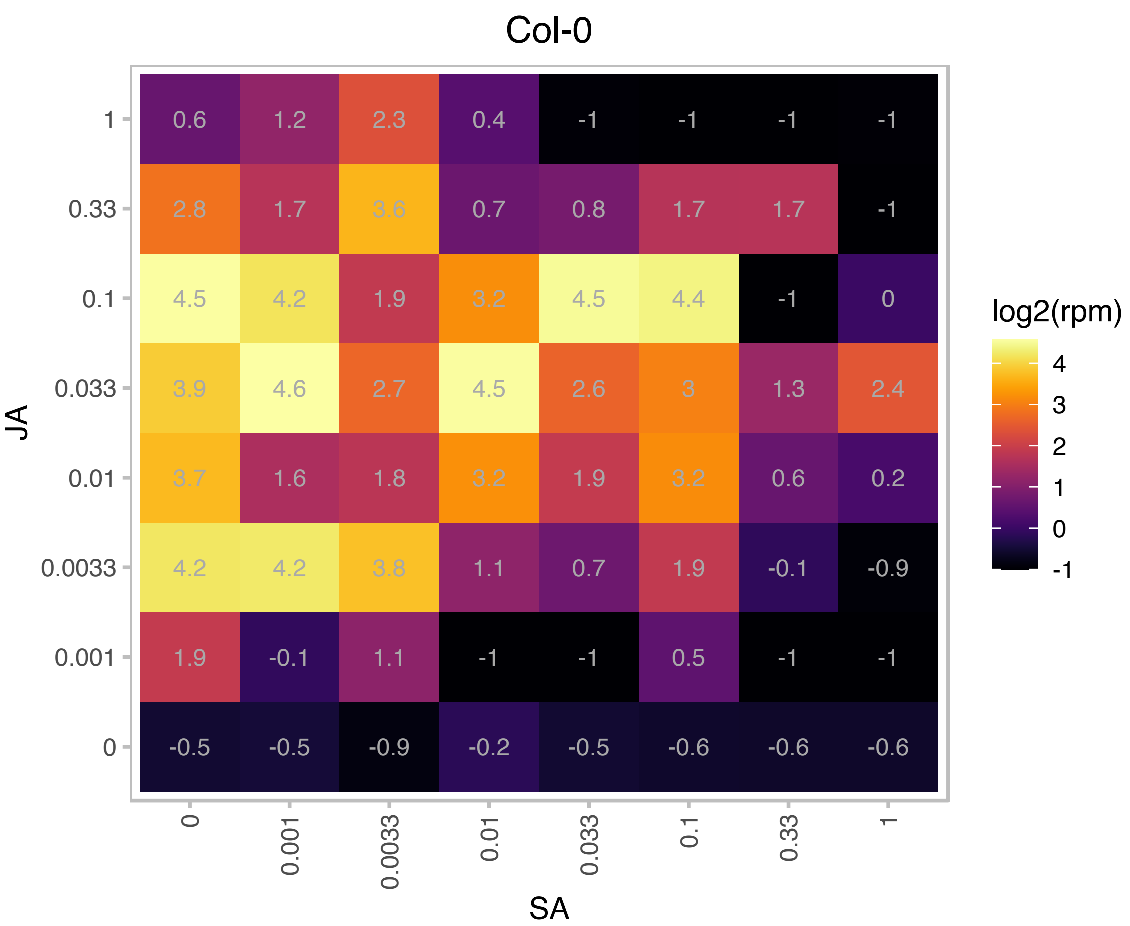


| | |
|---|---|
| Gene | AT1G73325 |
| is.expressed_genes | TRUE |
| is.responsive_genes | TRUE |
| is.apcluster | E-03 |

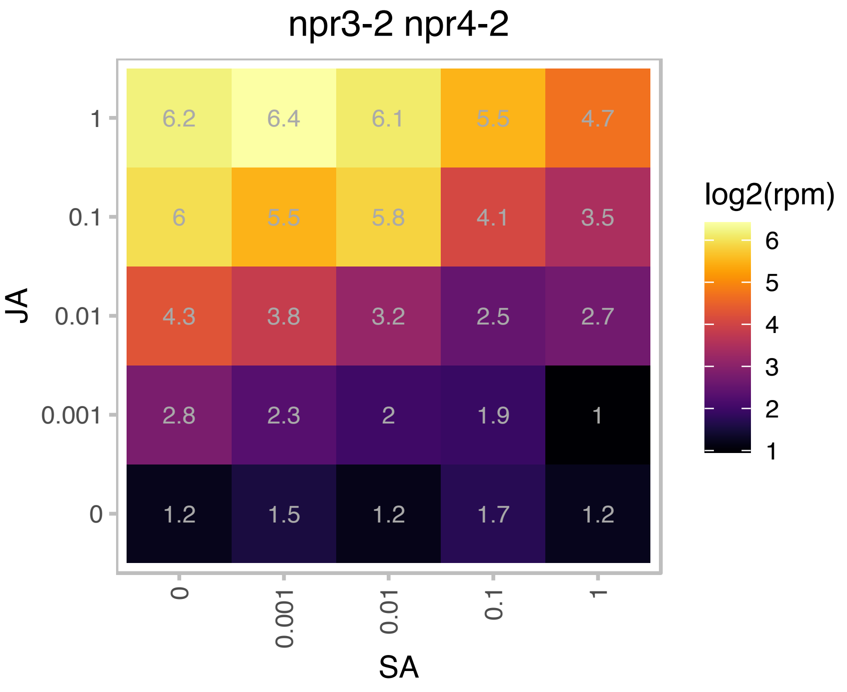


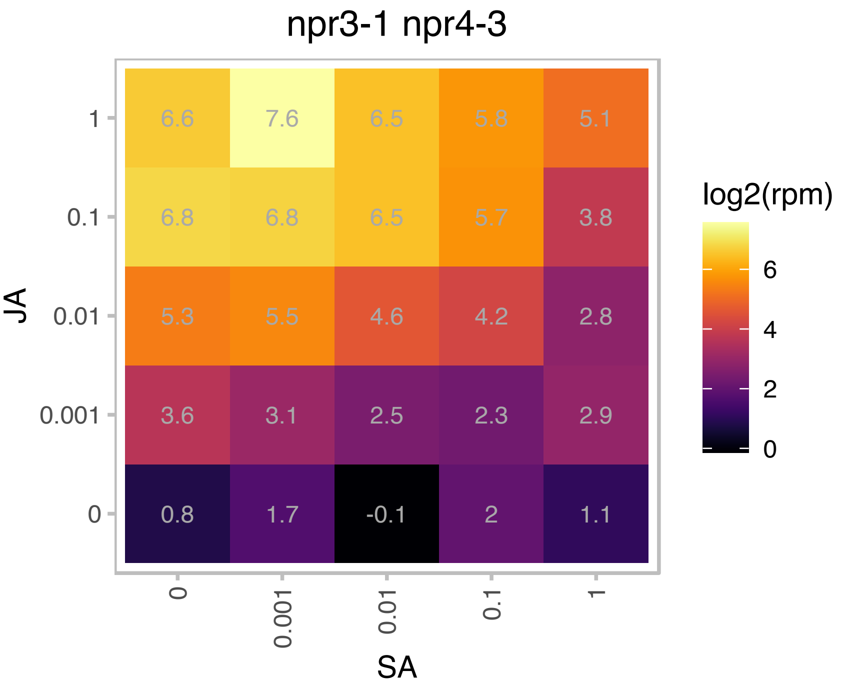

# AT1G76065.1 : NA

LYR family of Fe/S cluster biogenesis protein

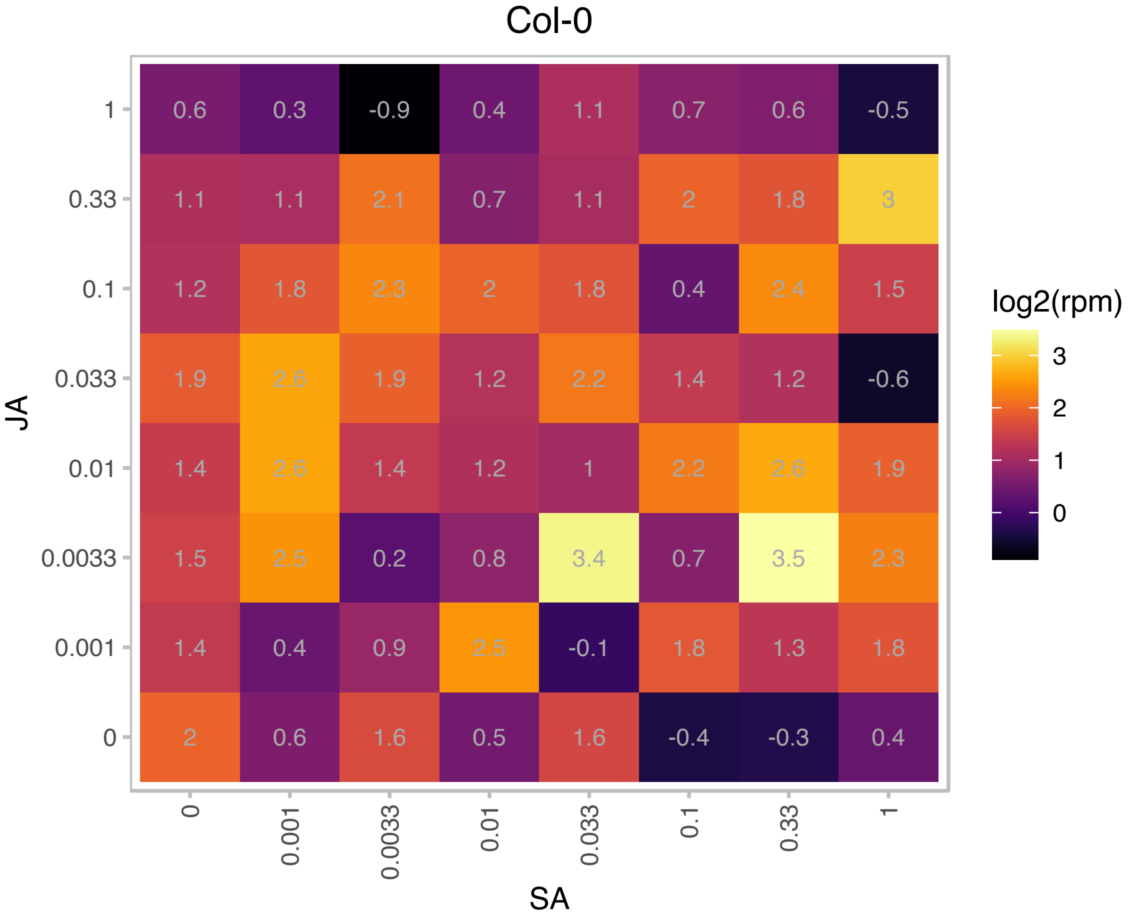


| | |
|---|---|
| Gene | AT1G76065 |
| is.expressed_genes | TRUE |
| is.responsive_genes | TRUE |
| is.apcluster | E-03 |

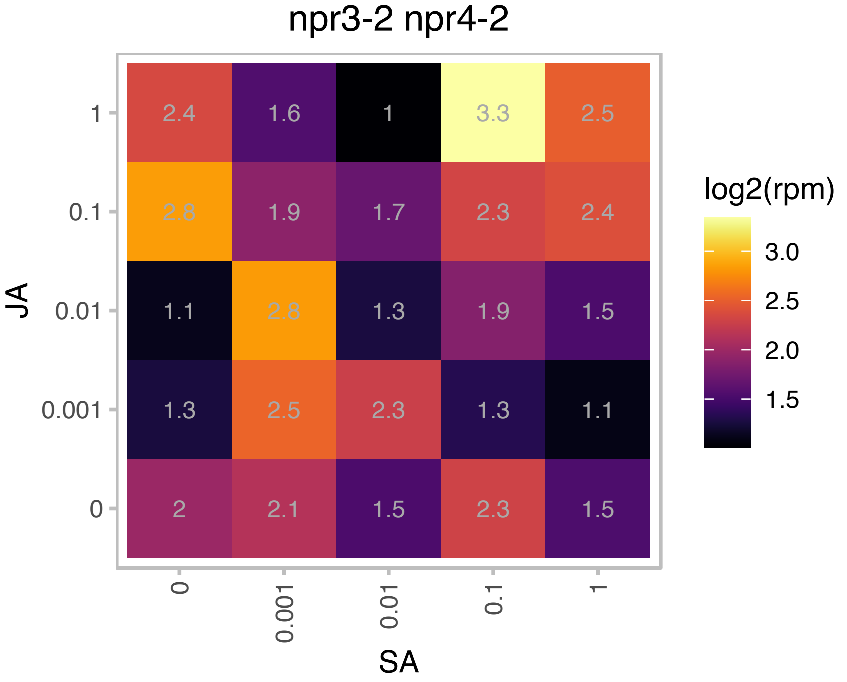


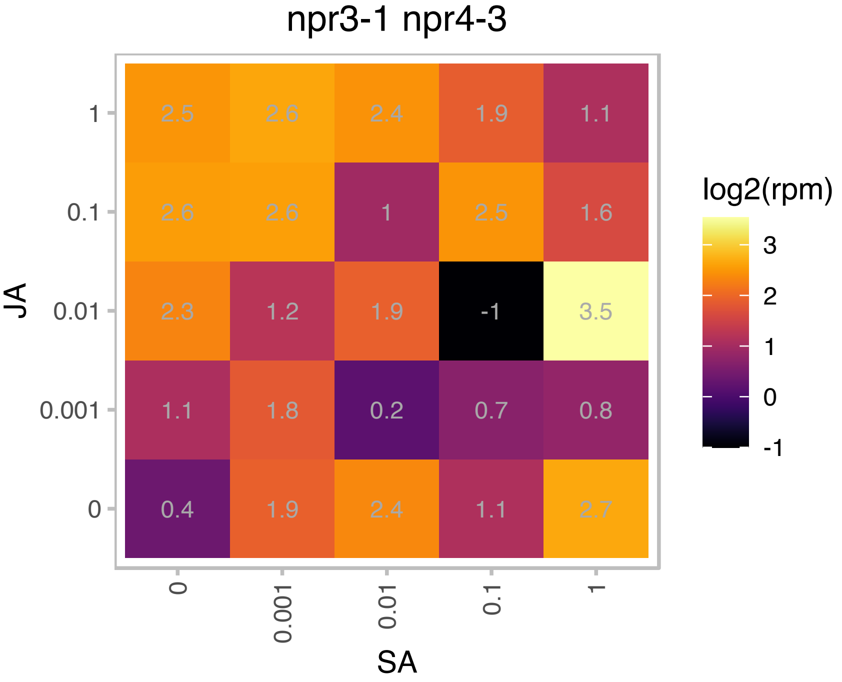

### AT1G76650.1 : CML38

calmodulin-like 38

#### Col-0

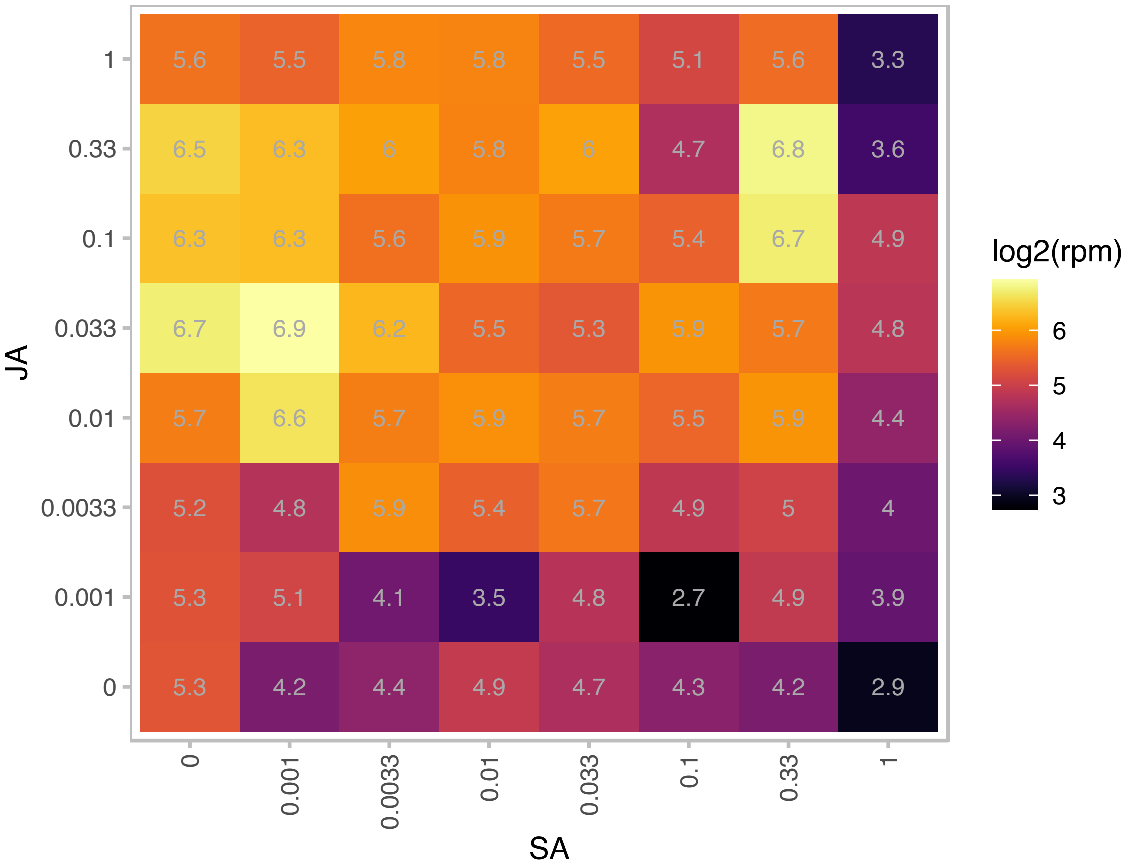


| | |
|---|---|
| Gene | AT1G76650 |
| is.expressed_genes | TRUE |
| is.responsive_genes | TRUE |
| is.apcluster | E-03 |

#### npr3-2 npr4-2

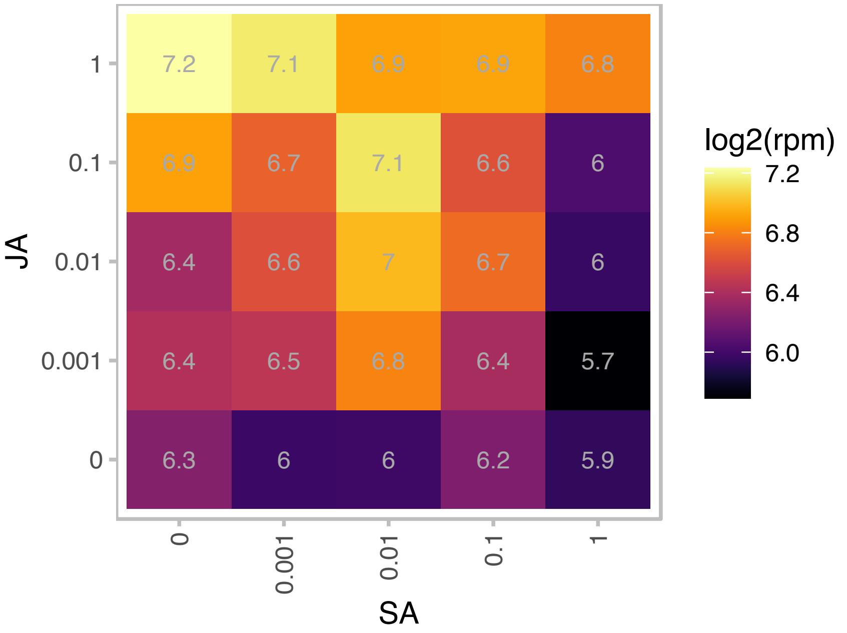


#### npr3-1 npr4-3

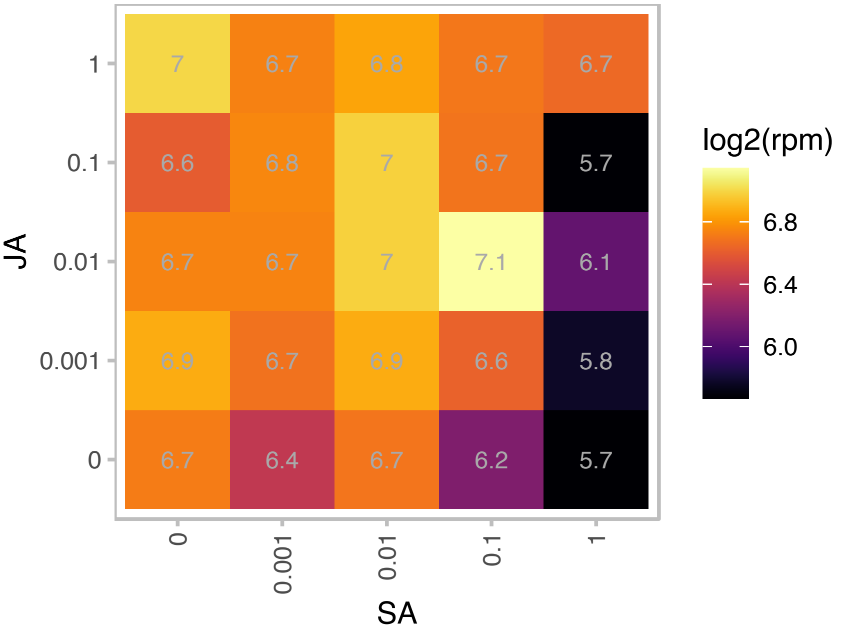

### AT2G14750.1 : AKN1, APK, APK1, ATAKN1

APS kinase

Encodes adenosine-5'-phosphosulfate kinase. Provides activated sulfate for sulfation of secondary metabolites, including the glucosinolates. Essential for pollen viability. The mRNA is cell-to-cell mobile.

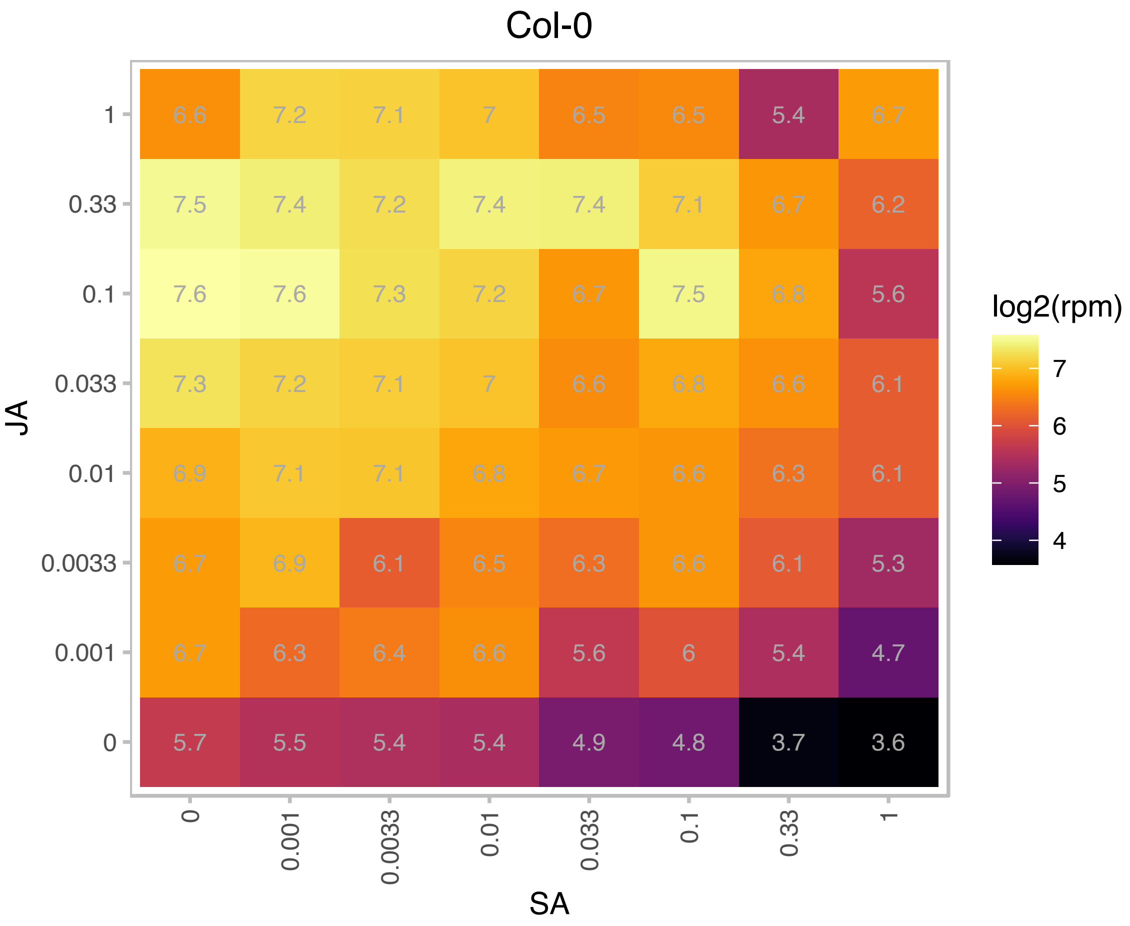


| | |
|---|---|
| Gene | AT2G14750 |
| is.expressed_genes | TRUE |
| is.responsive_genes | TRUE |
| is.apcluster | E-03 |

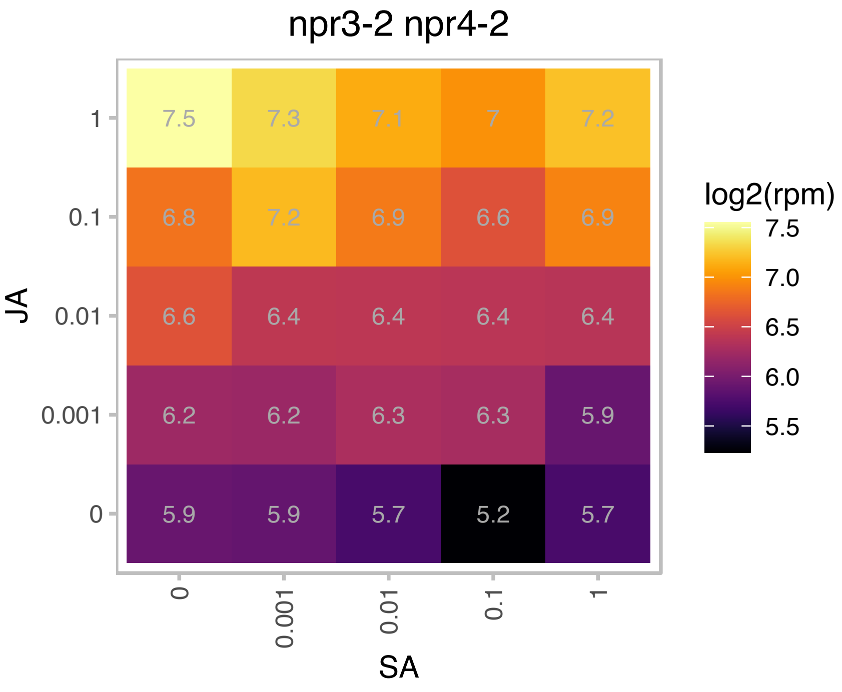


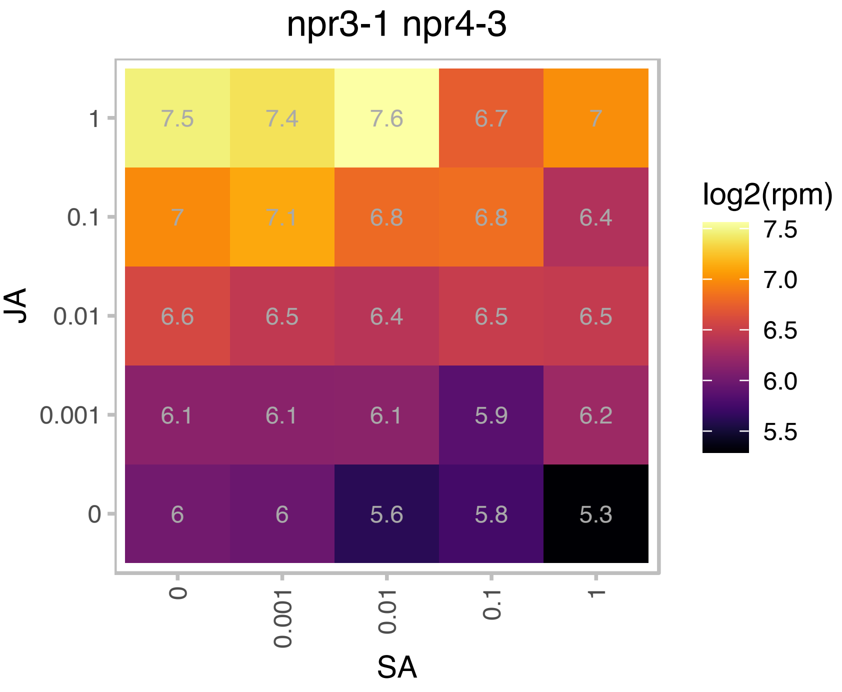

# AT2G15292.1 : NA

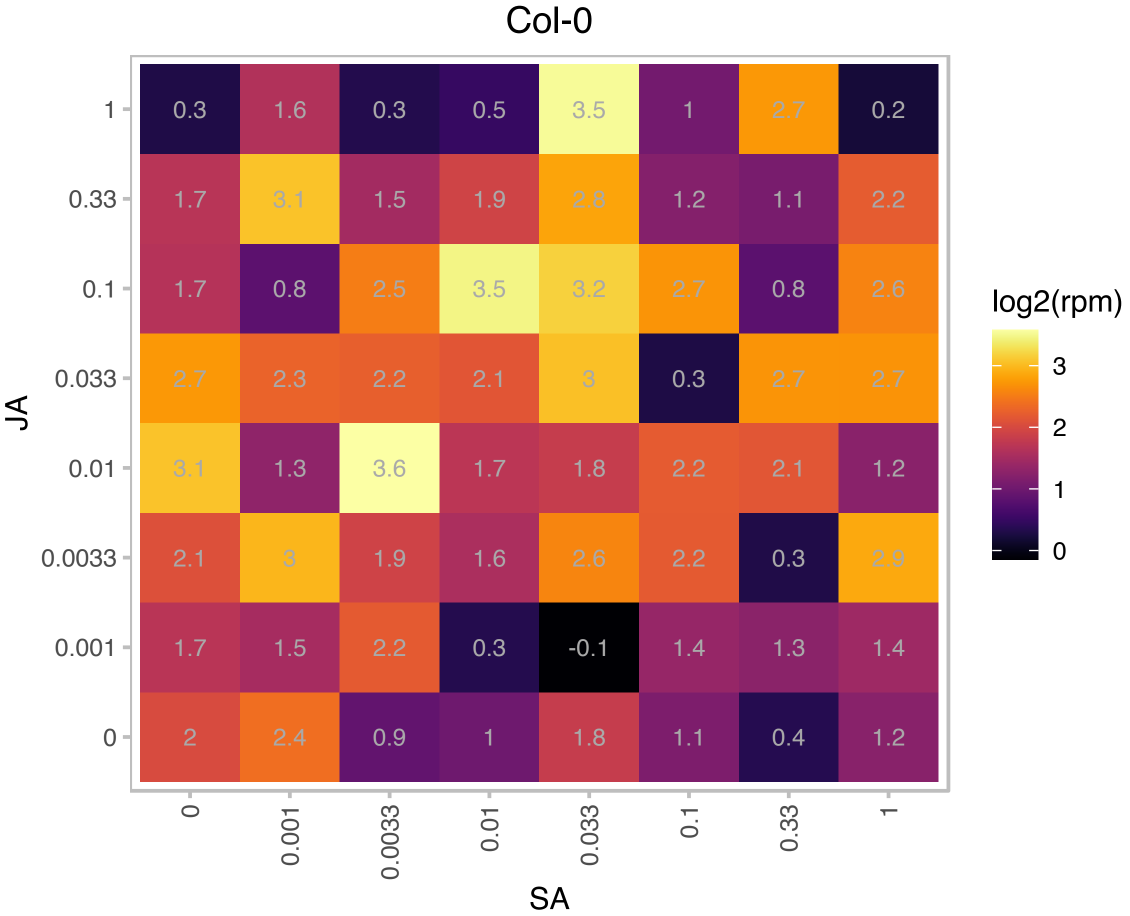


| Gene | AT2G15292 |
|---|---|
| is.expressed_genes | TRUE |
| is.responsive_genes | TRUE |
| is.apcluster | E-03 |

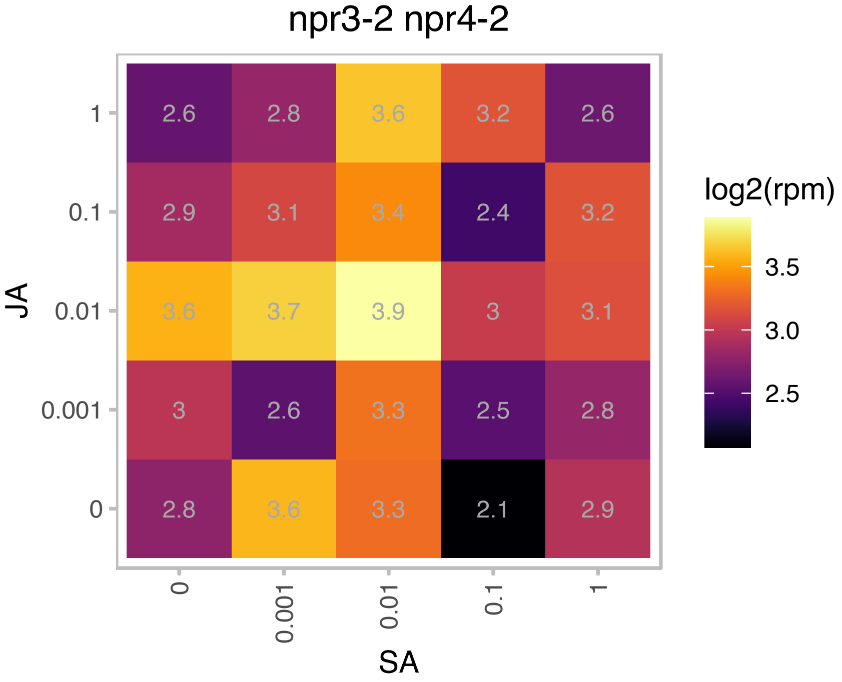


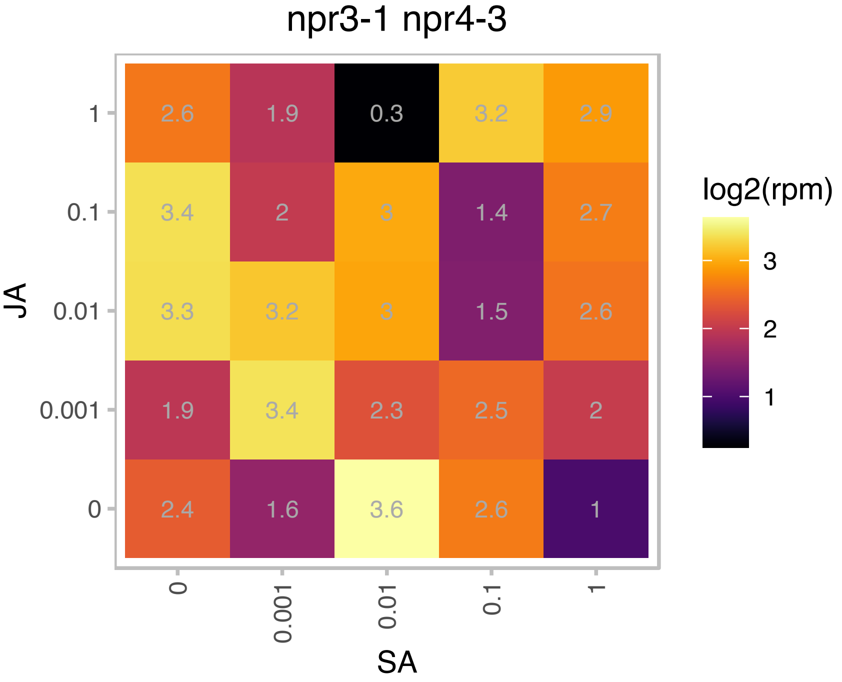

### AT2G39030.1 : NATA1

Acyl-CoA N-acyltransferases (NAT) superfamily protein
Encodes a protein that acts as an ornithine N-delta-acetyltransferase, leading to the formation of N-delta-actetylornithine. This compound is likely used in plant defense and levels of it are increased in Arabidopsis plants in response to MeJA and ABA. The mRNA is cell-to-cell mobile.

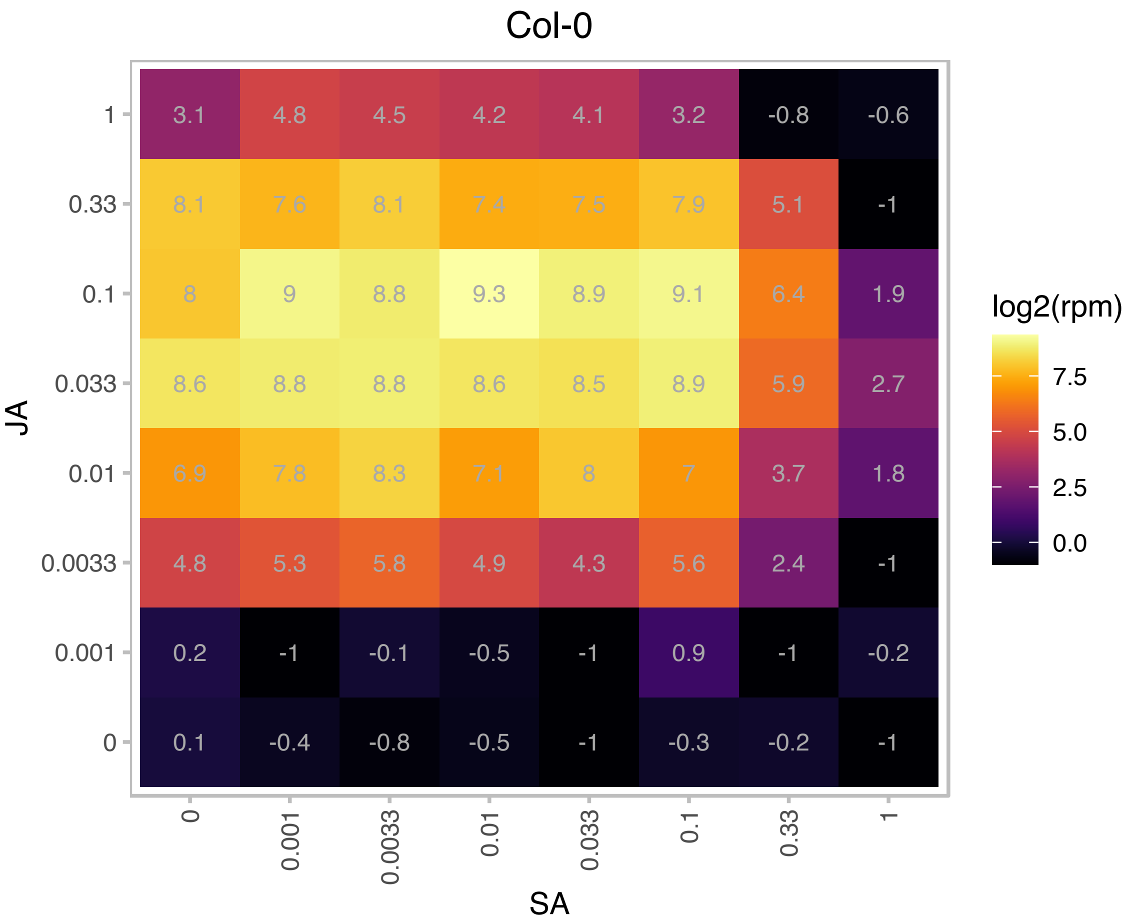


| | |
|---|---|
| Gene | AT2G39030 |
| is.expressed_genes | TRUE |
| is.responsive_genes | TRUE |
| is.apcluster | E-03 |

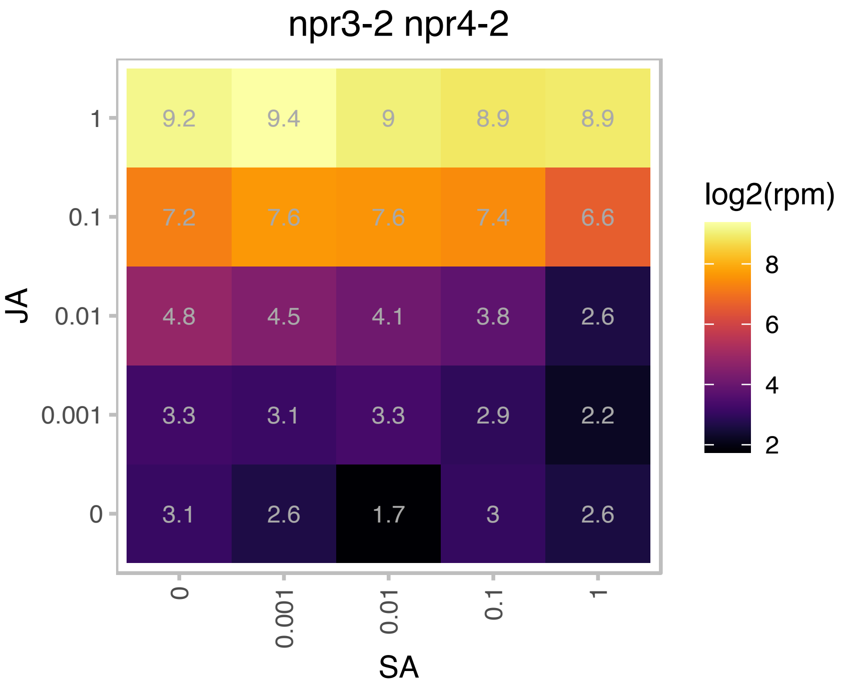


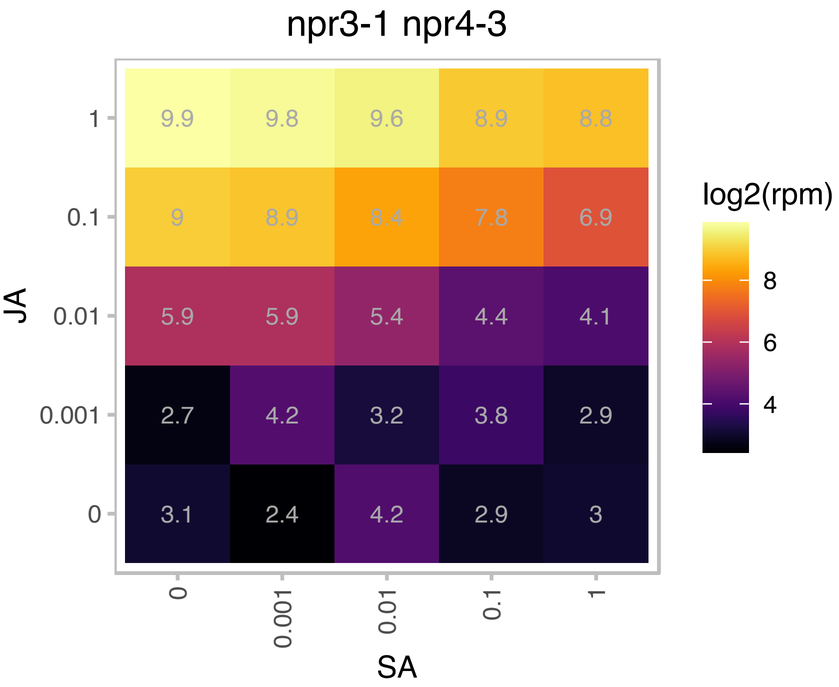

### AT2G39330.1 : JAL23

jacalin-related lectin 23

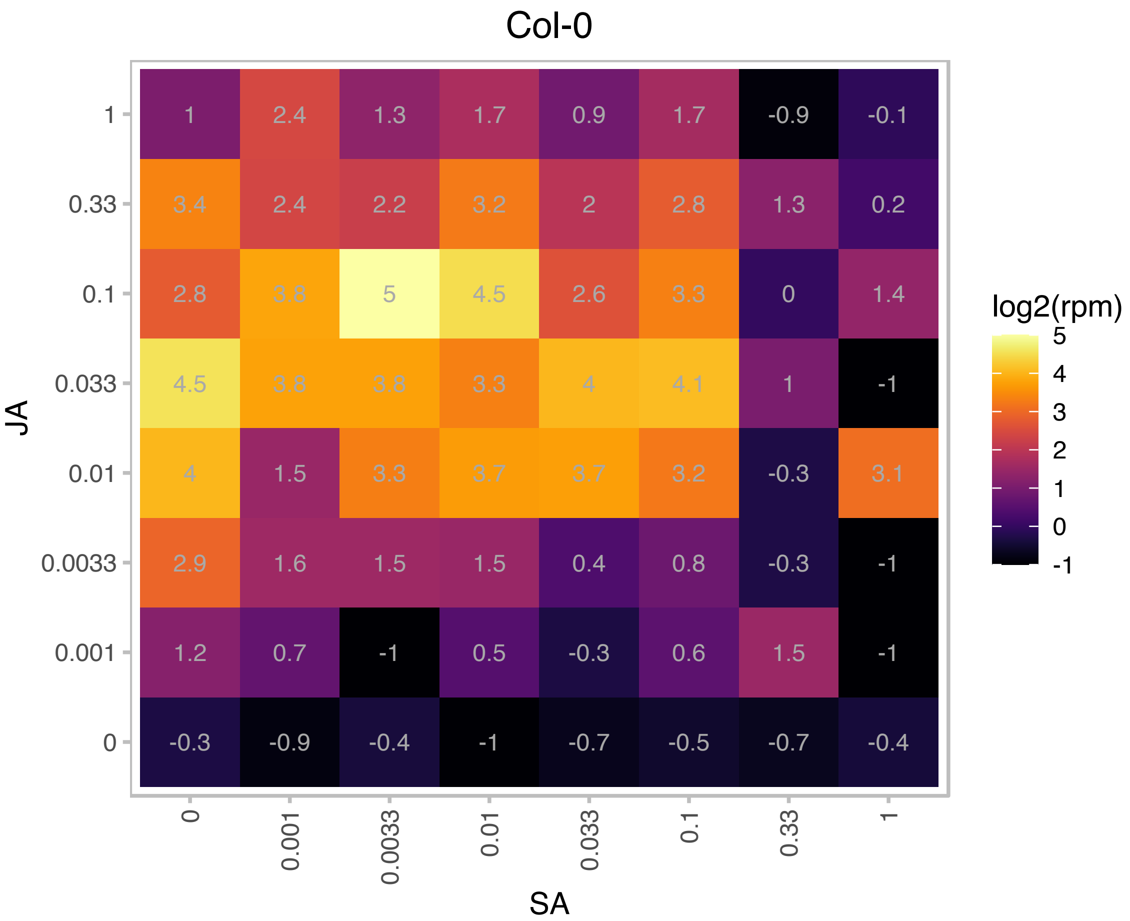


| | |
|---|---|
| Gene | AT2G39330 |
| is.expressed_genes | TRUE |
| is.responsive_genes | TRUE |
| is.apcluster | E-03 |

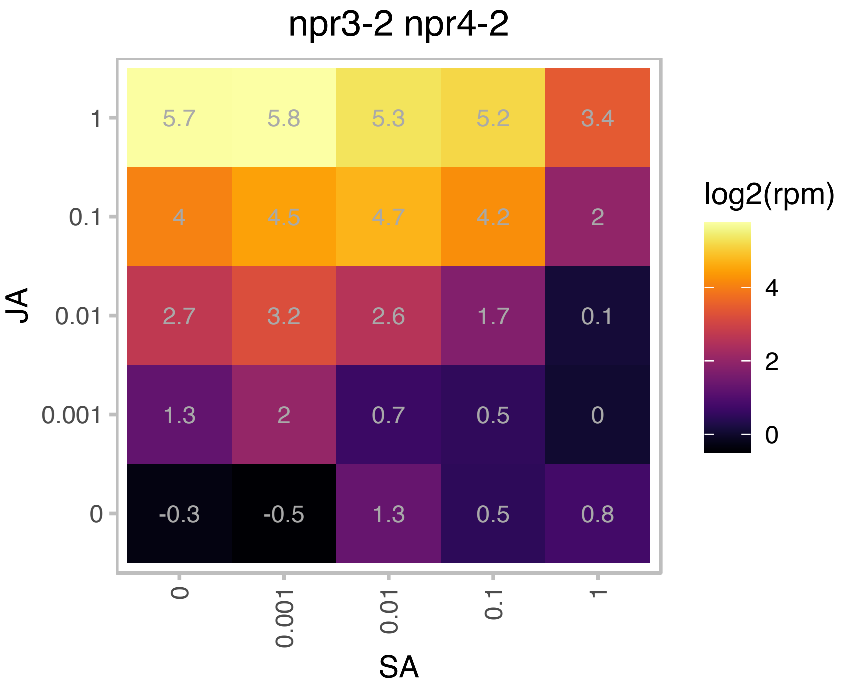


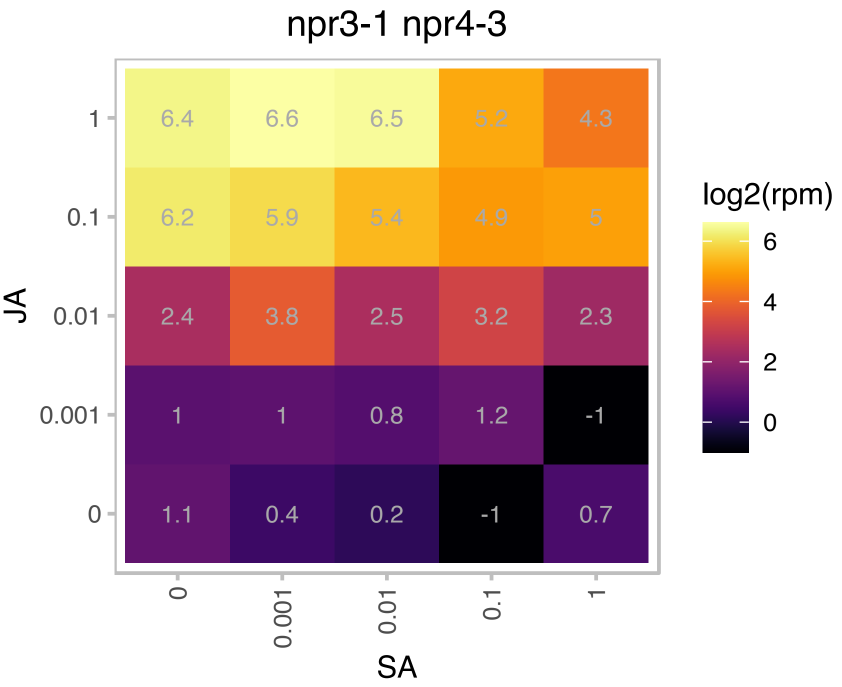

# AT2G42760.1 : NA

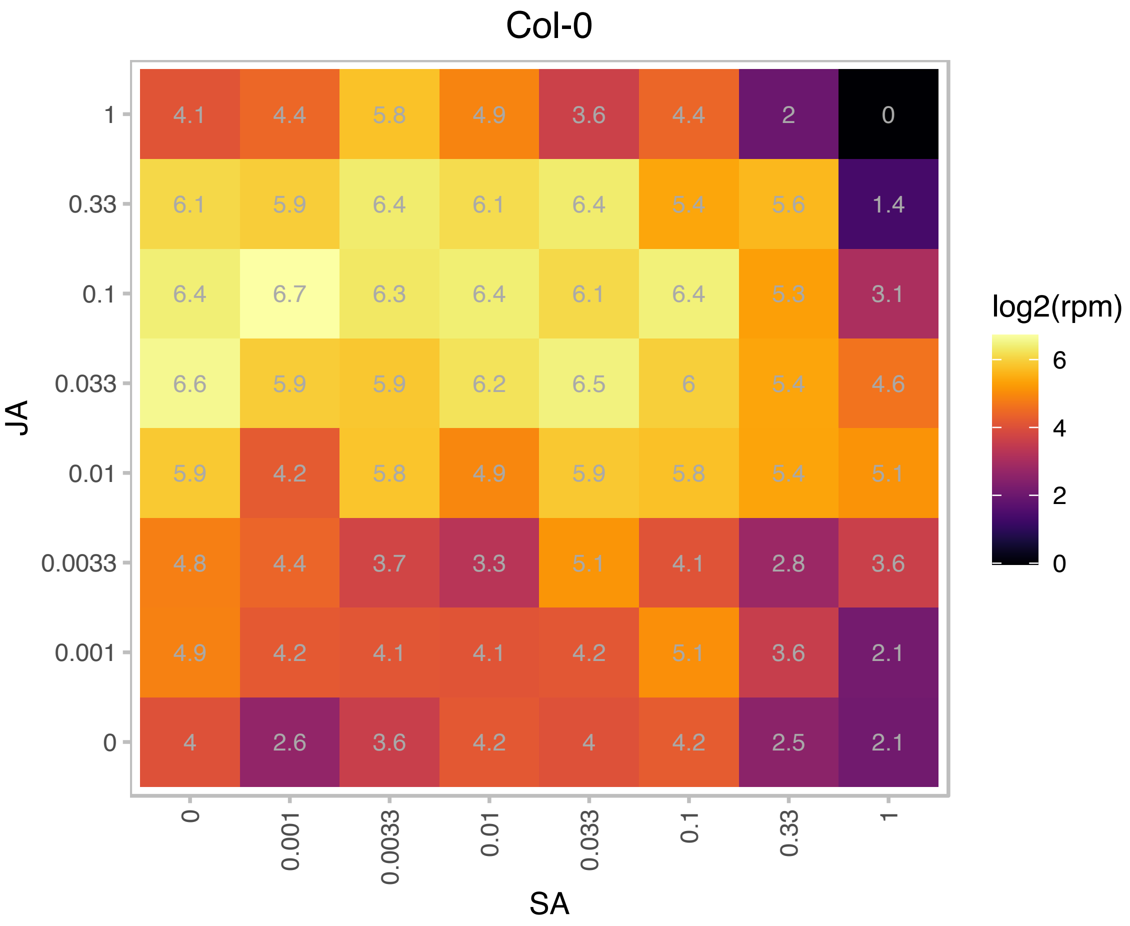


| | |
|---|---|
| Gene | AT2G42760 |
| is.expressed_genes | TRUE |
| is.responsive_genes | TRUE |
| is.apcluster | E-03 |

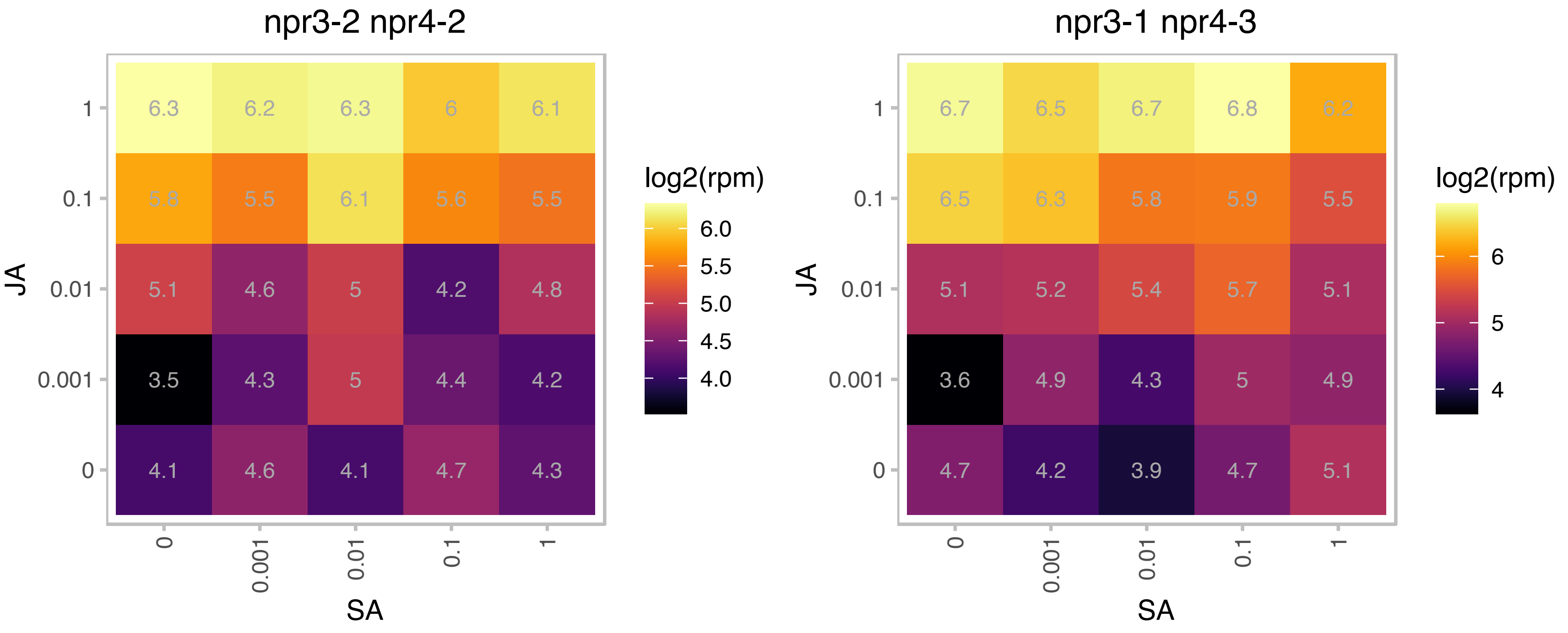

# AT2G43530.1 : NA

Scorpion toxin-like knottin superfamily protein

Encodes a defensin-like (DEFL) family protein. The mRNA is cell-to-cell mobile.

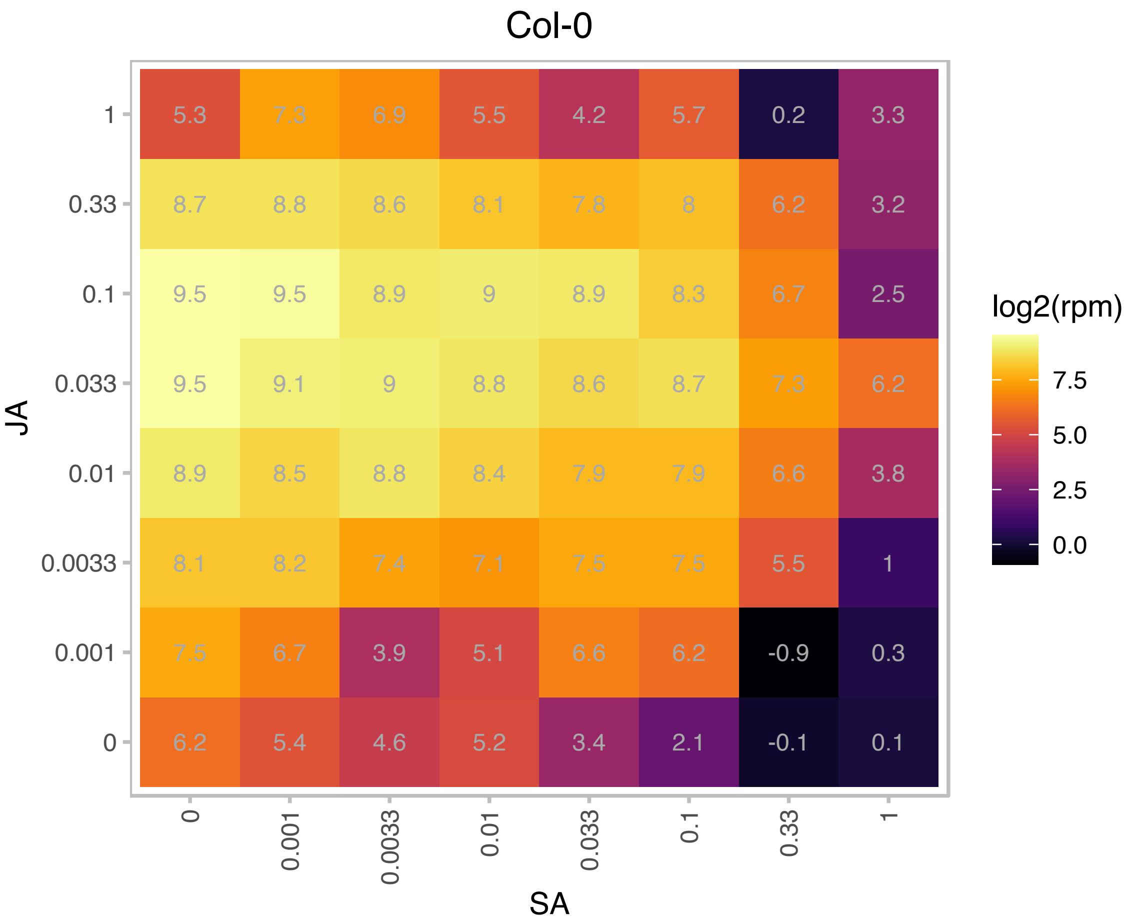


| | |
|---|---|
| Gene | AT2G43530 |
| is.expressed_genes | TRUE |
| is.responsive_genes | TRUE |
| is.apcluster | E-03 |

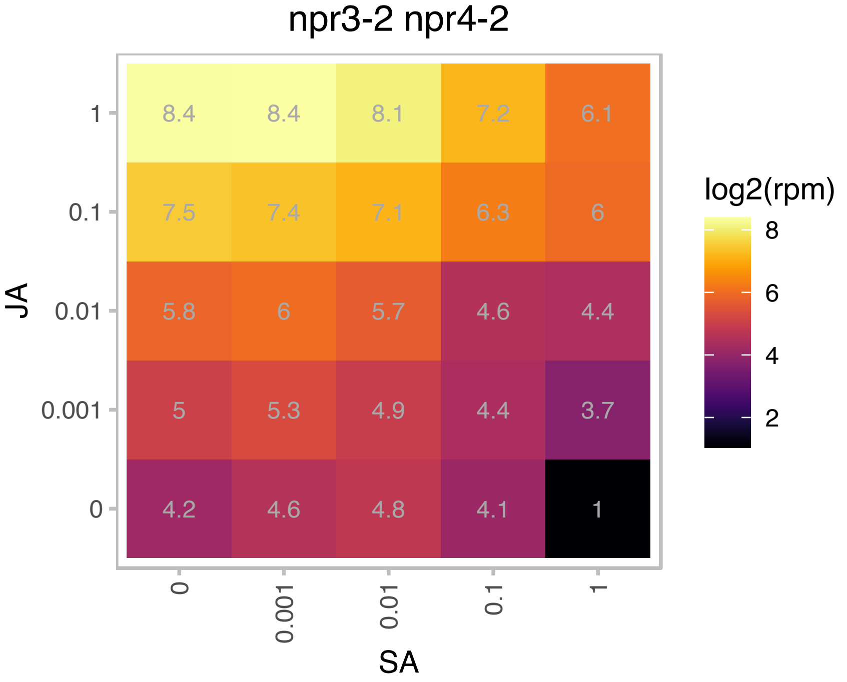


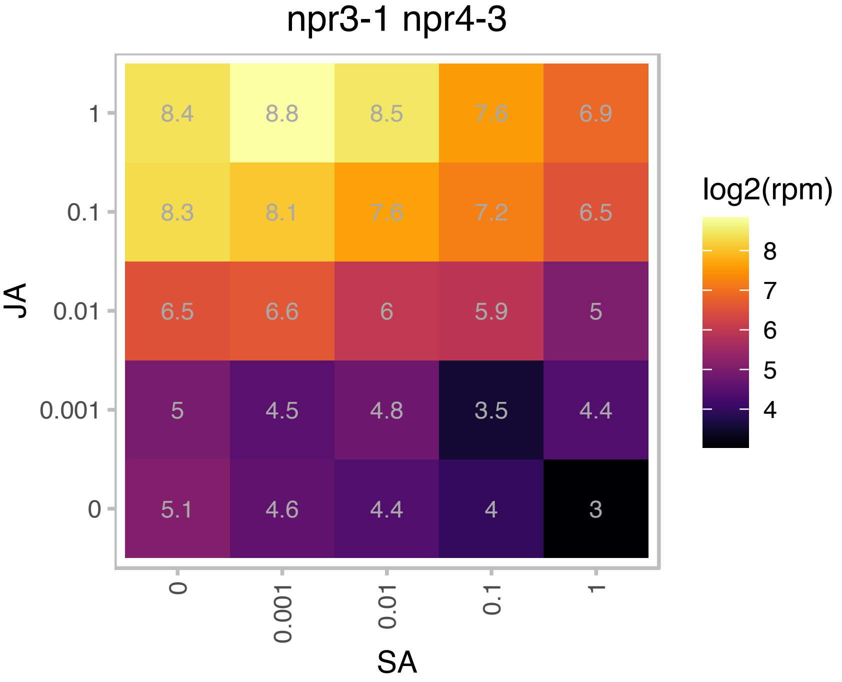

# AT2G43535.1 : NA

Scorpion toxin-like knottin superfamily protein

Encodes a defensin-like (DEFL) family protein.

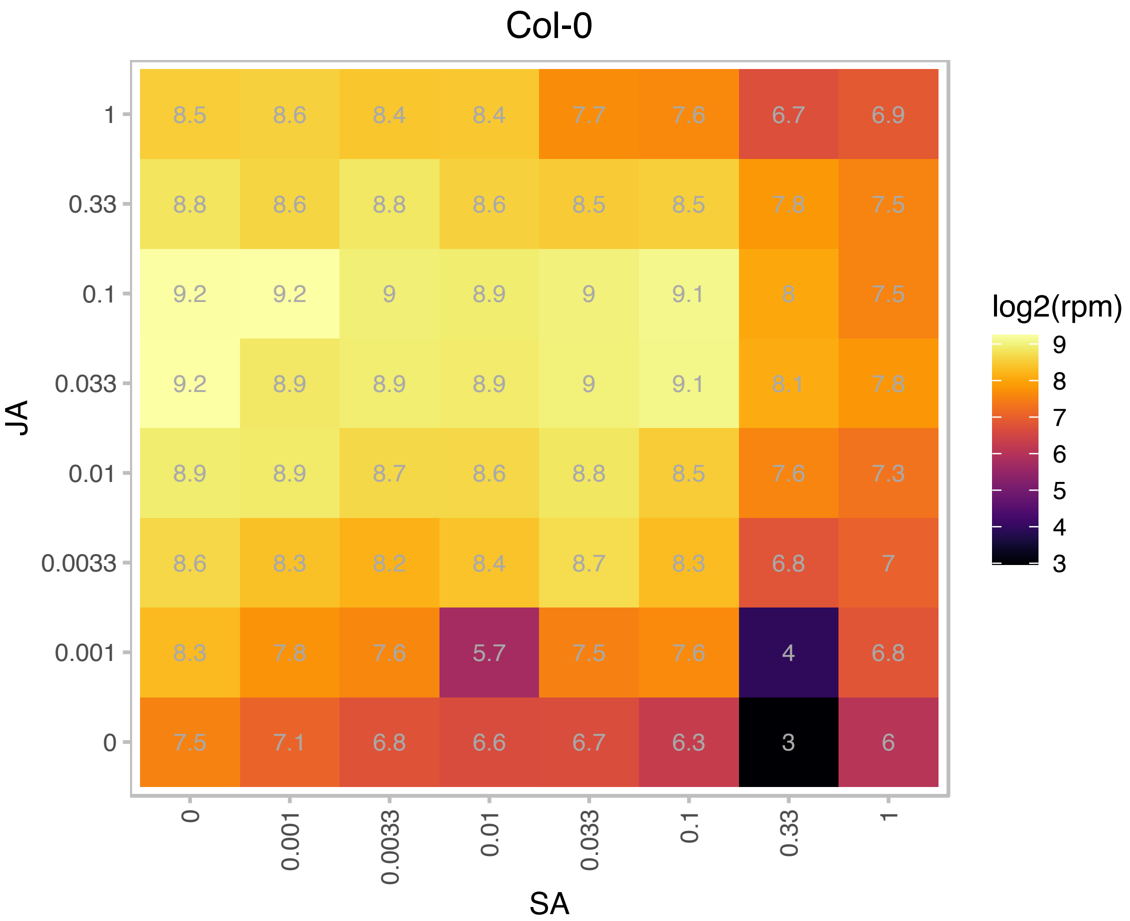


| | |
|---|---|
| Gene | AT2G43535 |
| is.expressed_genes | TRUE |
| is.responsive_genes | TRUE |
| is.apcluster | E-03 |

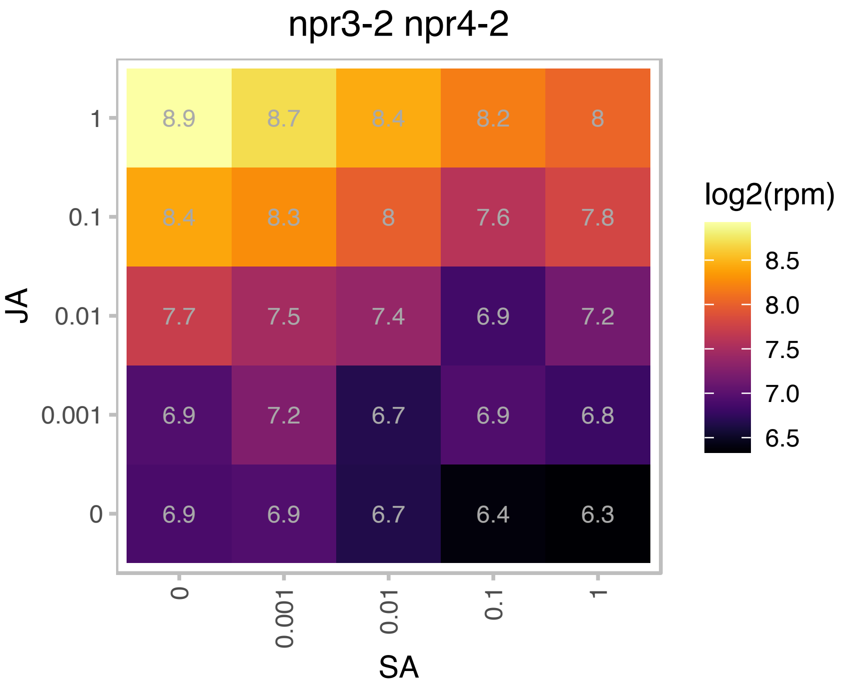


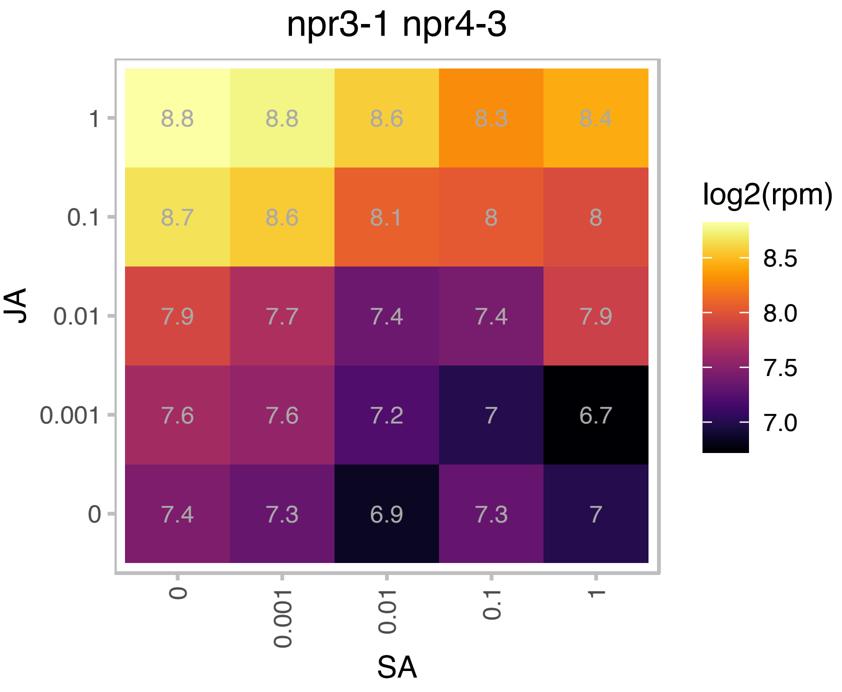

# AT2G43550.1 : NA

Scorpion toxin-like knottin superfamily protein

Encodes a defensin-like (DEFL) family protein.

#### Col-0

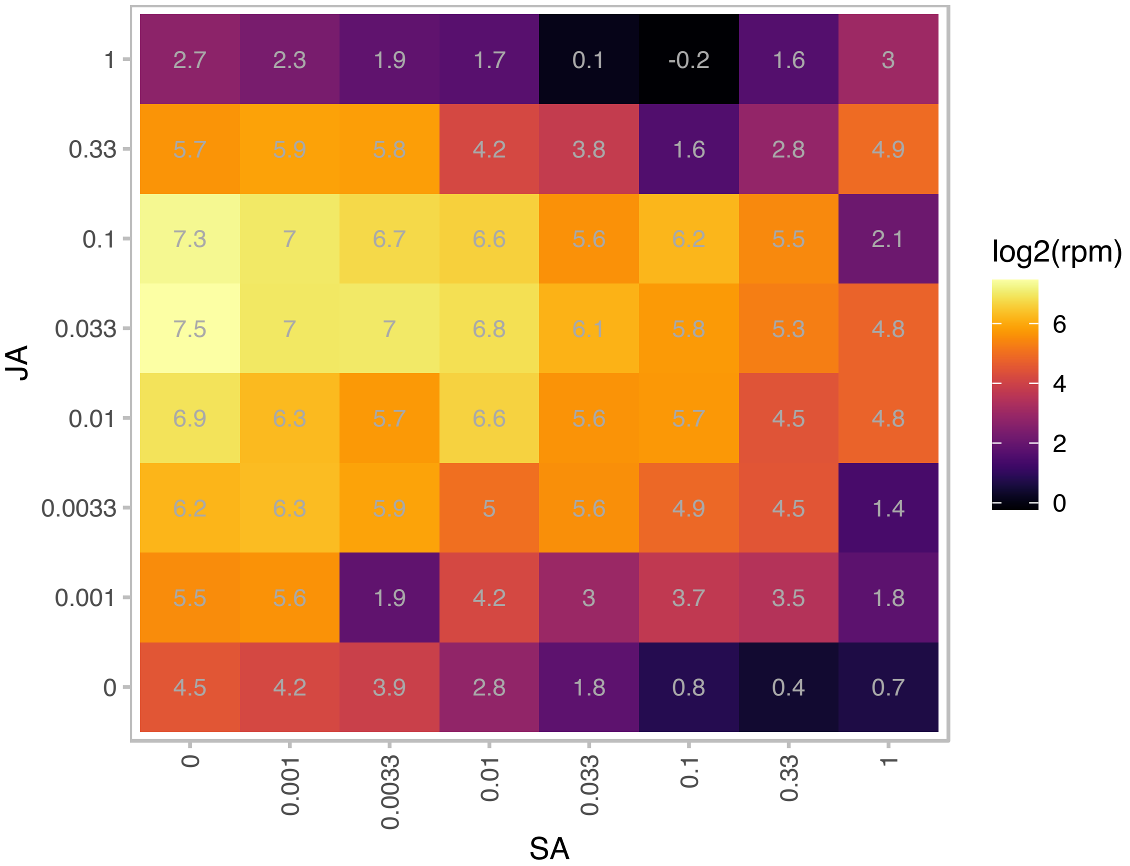


| | |
|---|---|
| Gene | AT2G43550 |
| is.expressed_genes | TRUE |
| is.responsive_genes | TRUE |
| is.apcluster | E-03 |

#### npr3-2 npr4-2

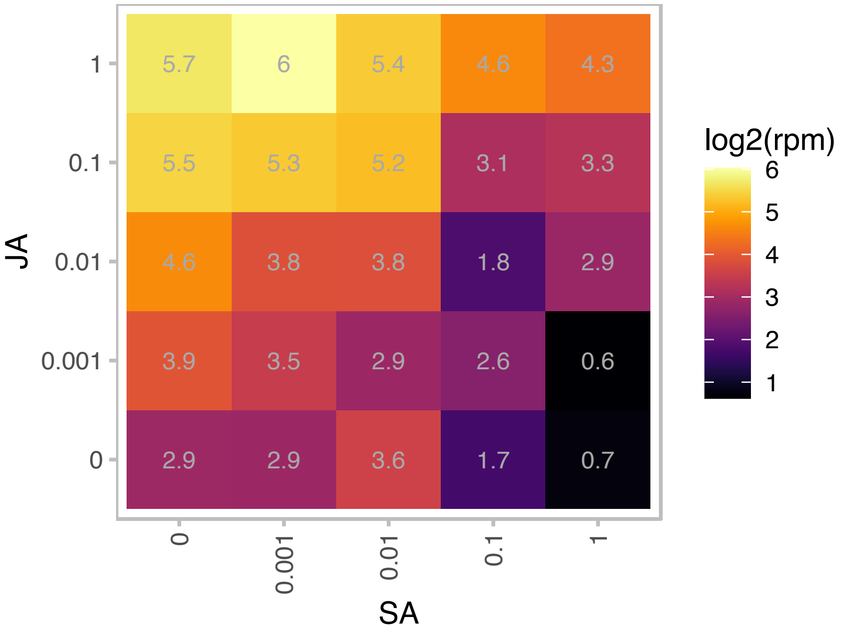


#### npr3-1 npr4-3

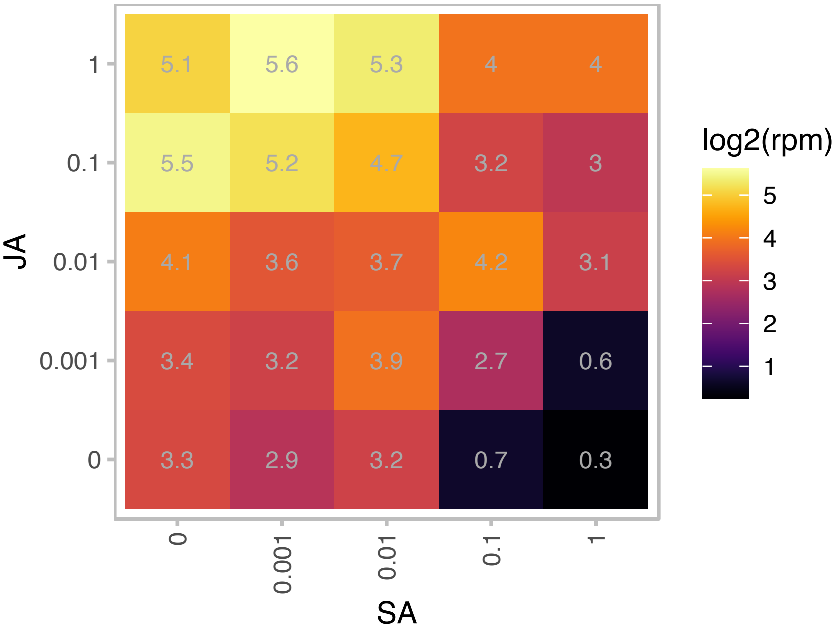

# AT3G05570.1 : NA

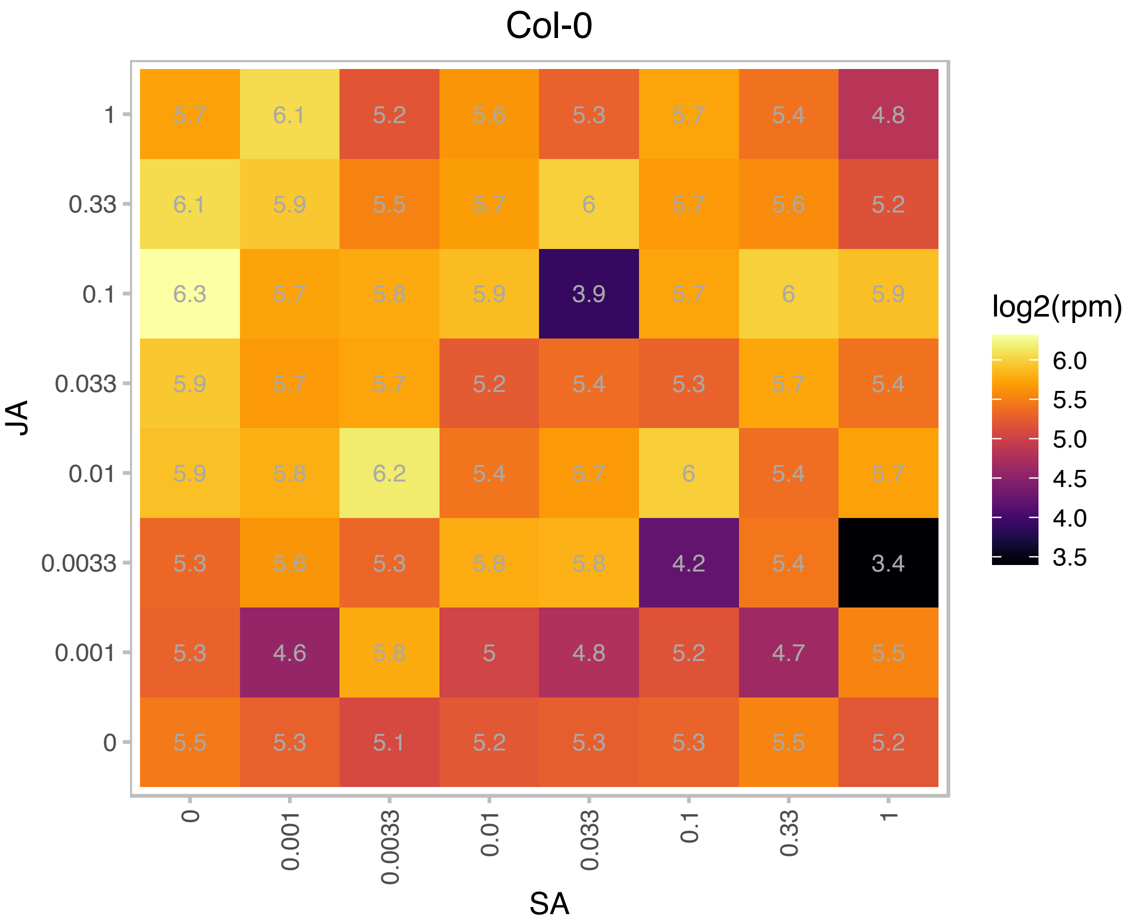


| | |
|---|---|
| Gene | AT3G05570 |
| is.expressed_genes | TRUE |
| is.responsive_genes | TRUE |
| is.apcluster | E-03 |

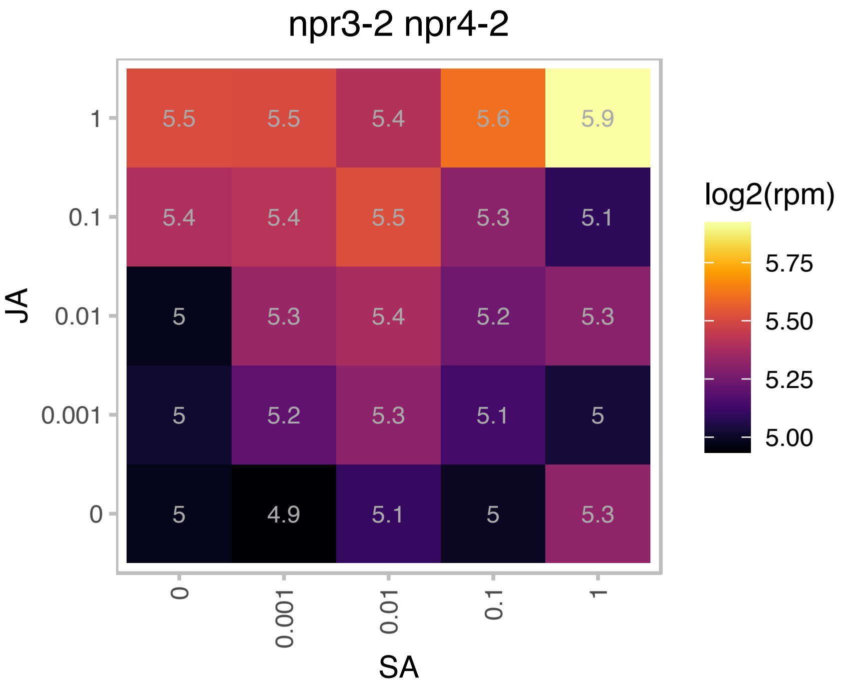


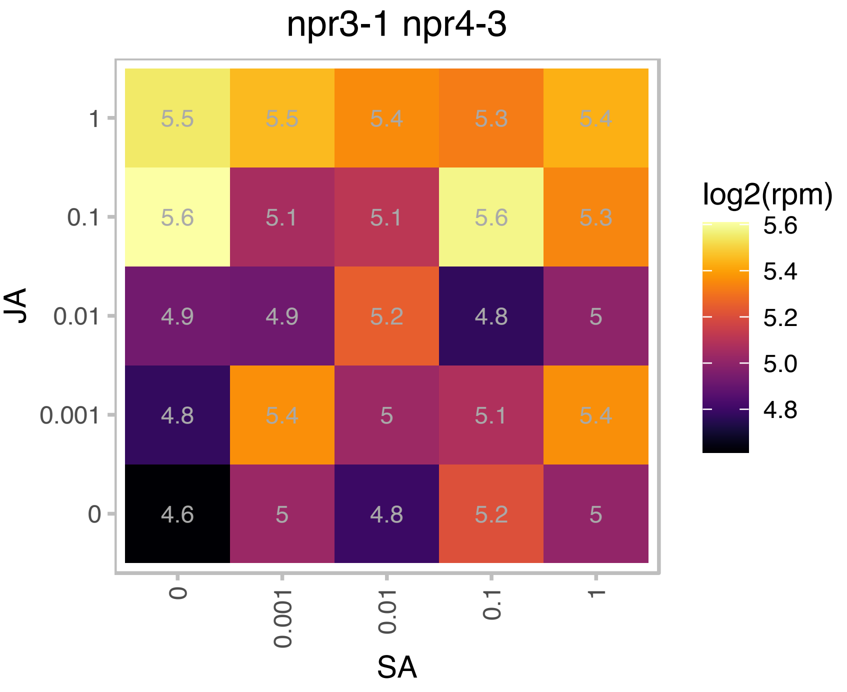

### AT3G06483.1 : ATPDHK, PDK

pyruvate dehydrogenase kinase

Pyruvate dehydrogenase kinase (PDK) specifically phosphorylates the E1α subunit of the pyruvate dehydrogenase complex (PDC) on a Ser residue using ATP as a phosphate donor.  PDK is a unique type of protein kinase having a His-kinase-like sequence but Ser-kinase activity. Site-directed mutagenesis and structural analysis indicate that PDK belongs to the GHKL superfamily.

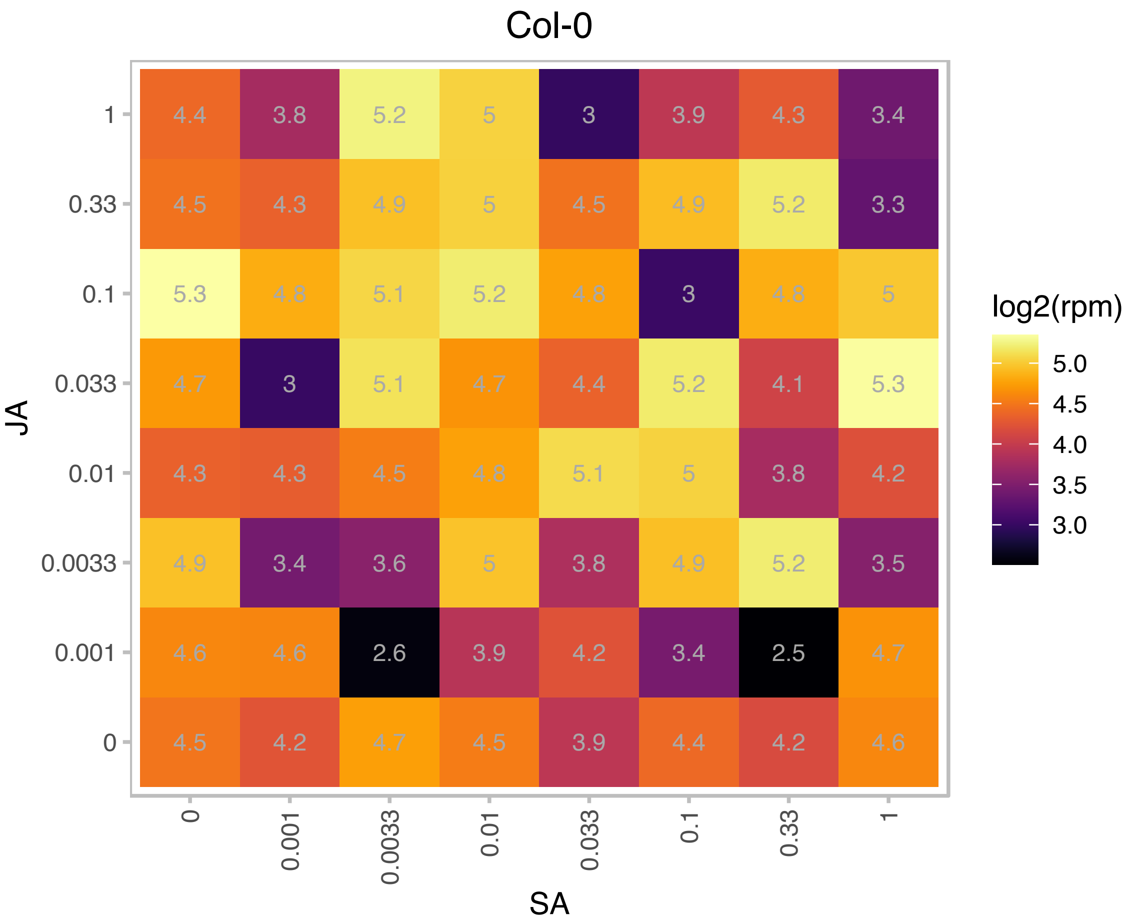


| | |
|---|---|
| Gene | AT3G06483 |
| is.expressed_genes | TRUE |
| is.responsive_genes | TRUE |
| is.apcluster | E-03 |

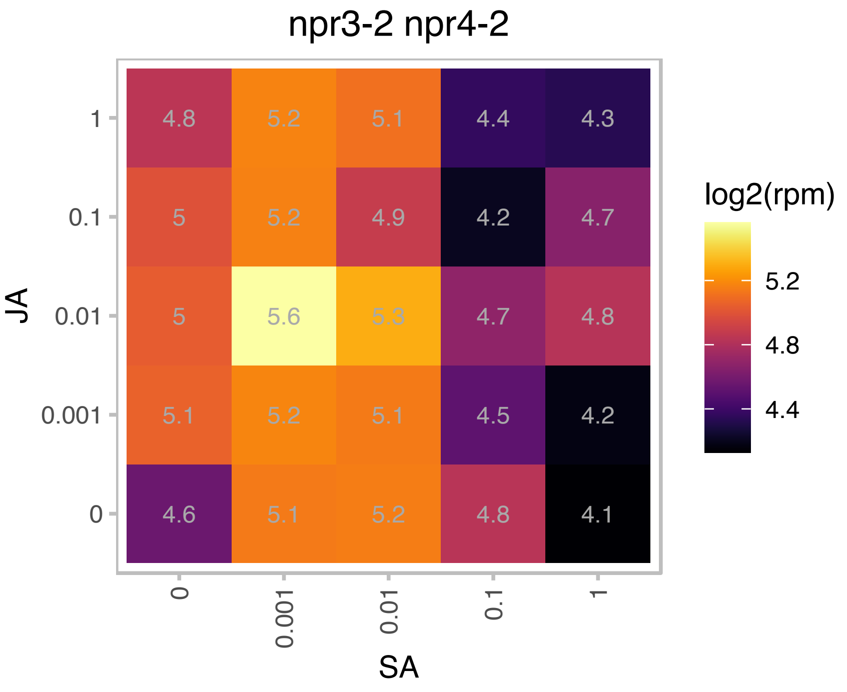


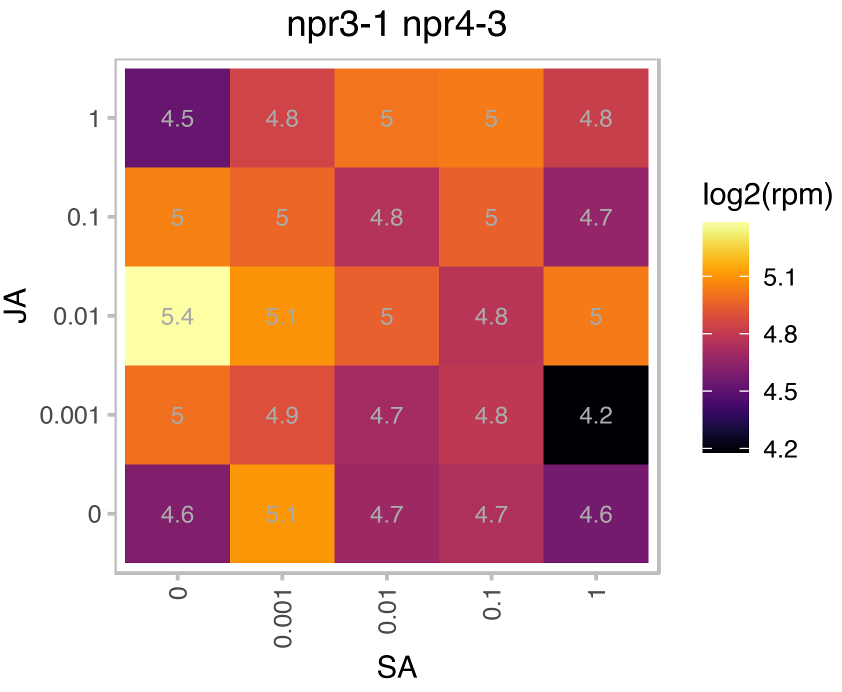

# AT3G07045.1 : NA

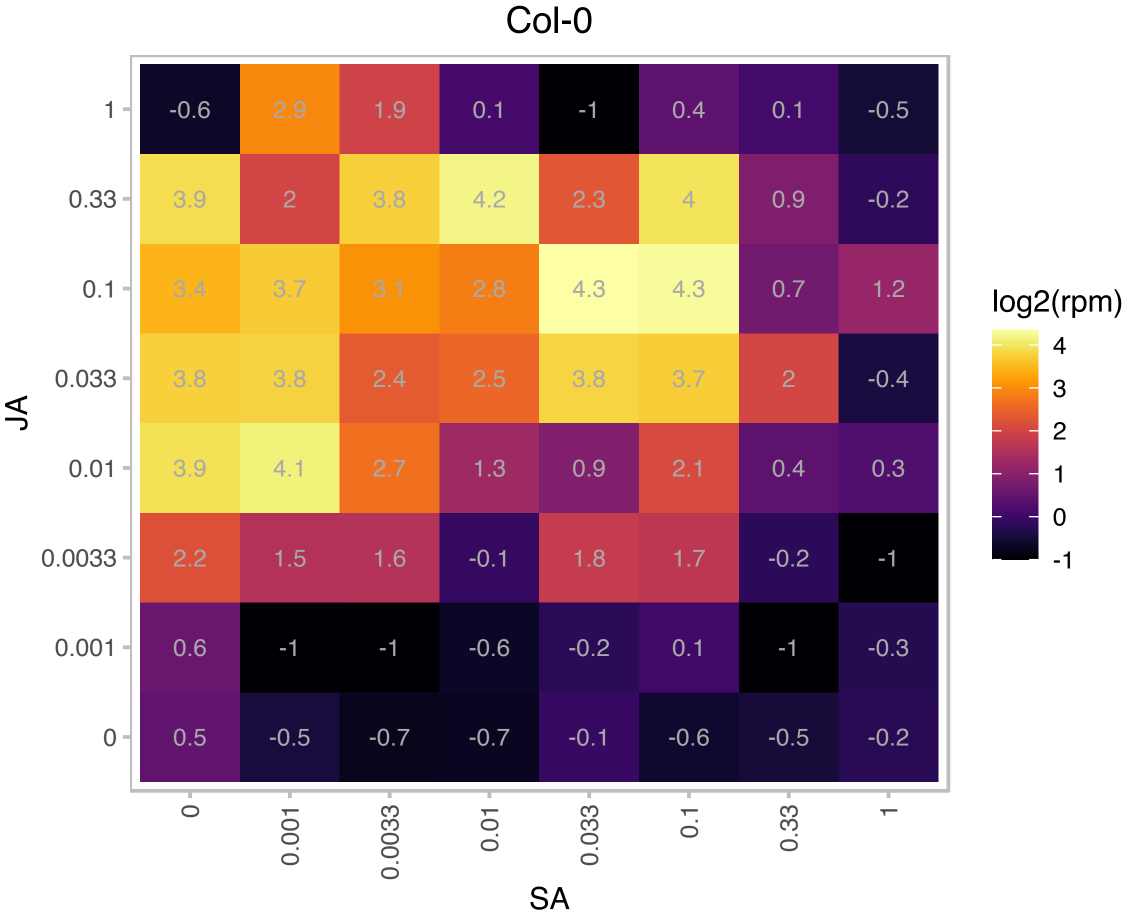


| | |
|---|---|
| Gene | AT3G07045 |
| is.expressed_genes | TRUE |
| is.responsive_genes | TRUE |
| is.apcluster | E-03 |

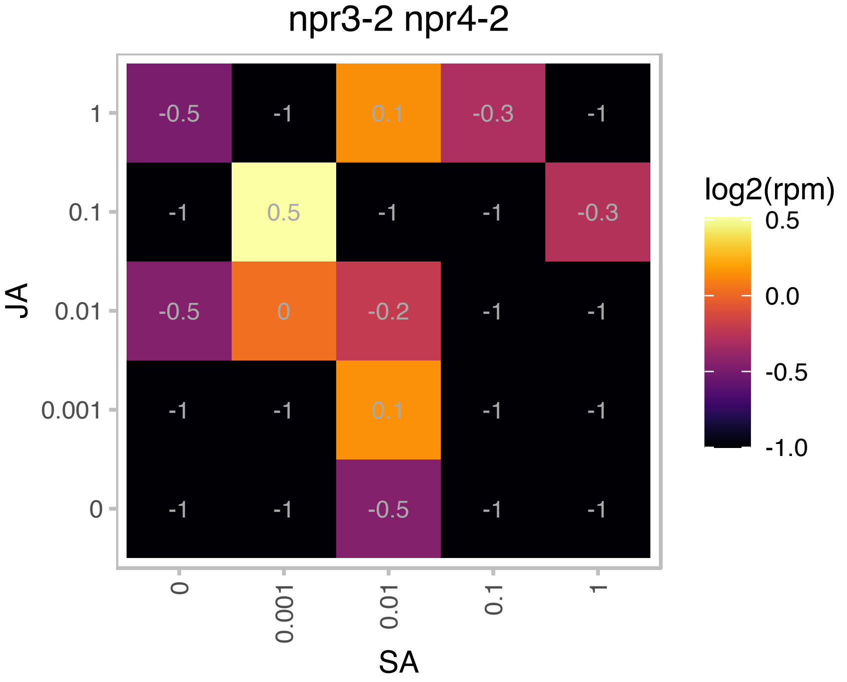


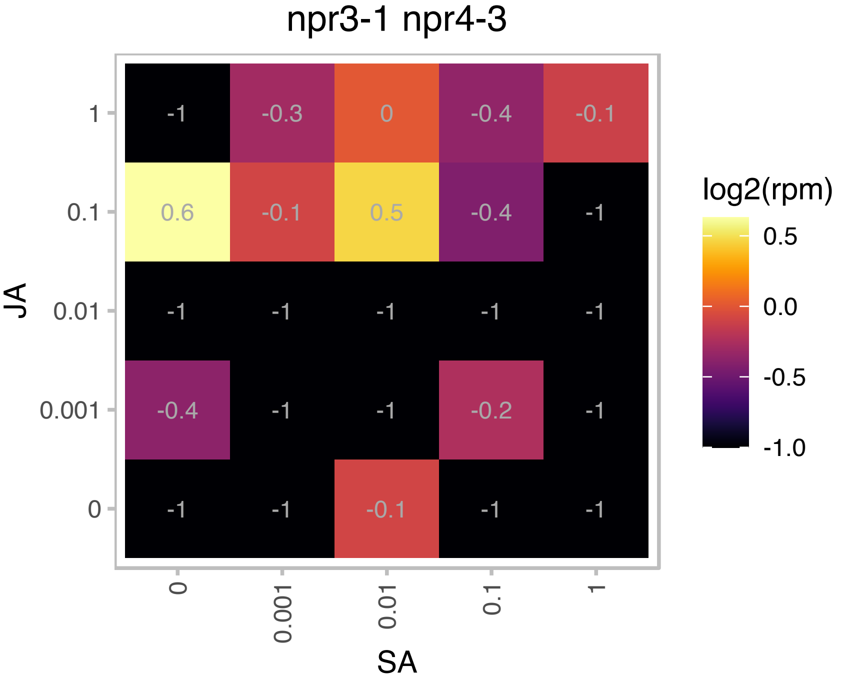

### AT3G16470.1 : AtJAC1, JAL35, JR1

Mannose-binding lectin superfamily protein
Encodes a JA-responsive gene that coordinates with GRP7 in shaping plant development through the regulation of RNA processing in Arabidopsis. AtJAC1 interacts with RNA binding protein GRP7 specifically in the cytoplasm to regulate its nucleocytoplasmic distribution.

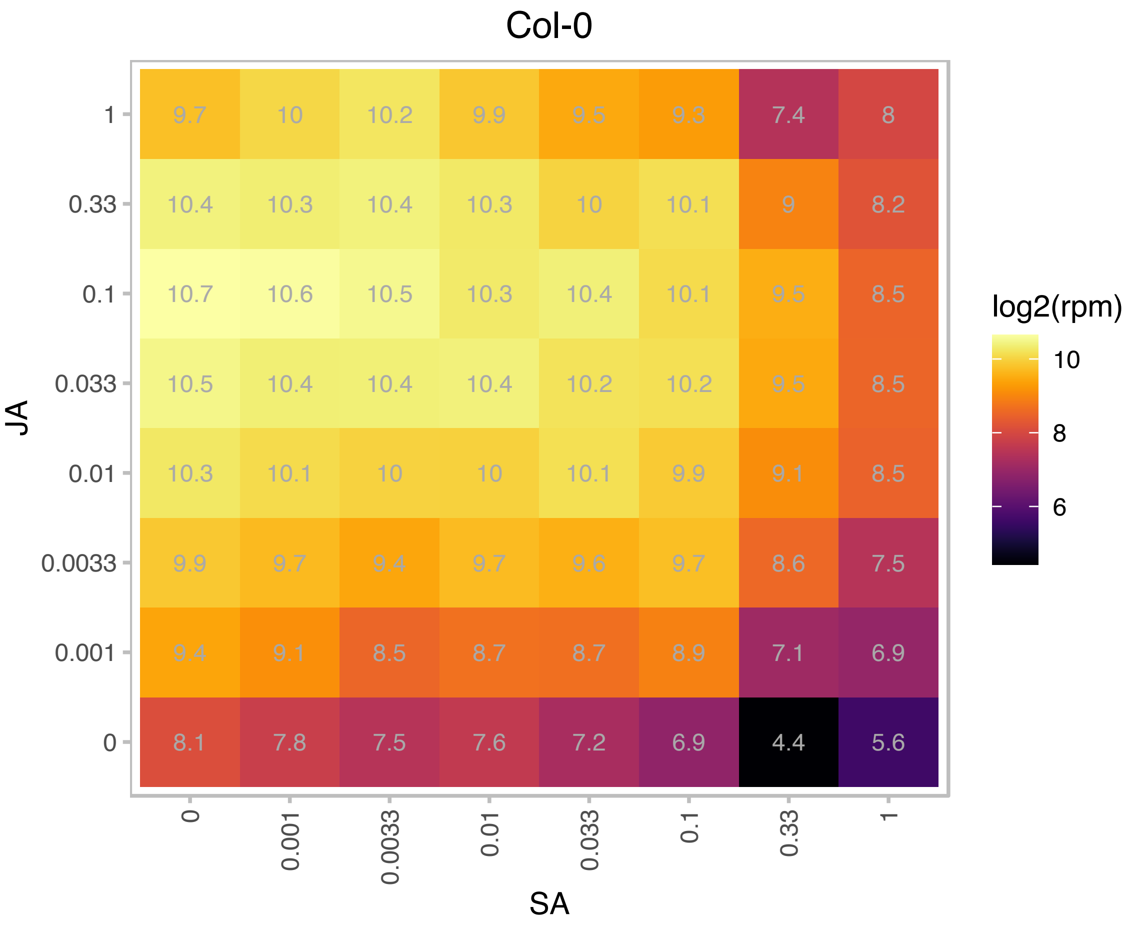


| | |
|---|---|
| Gene | AT3G16470 |
| is.expressed_genes | TRUE |
| is.responsive_genes | TRUE |
| is.apcluster | E-03 |

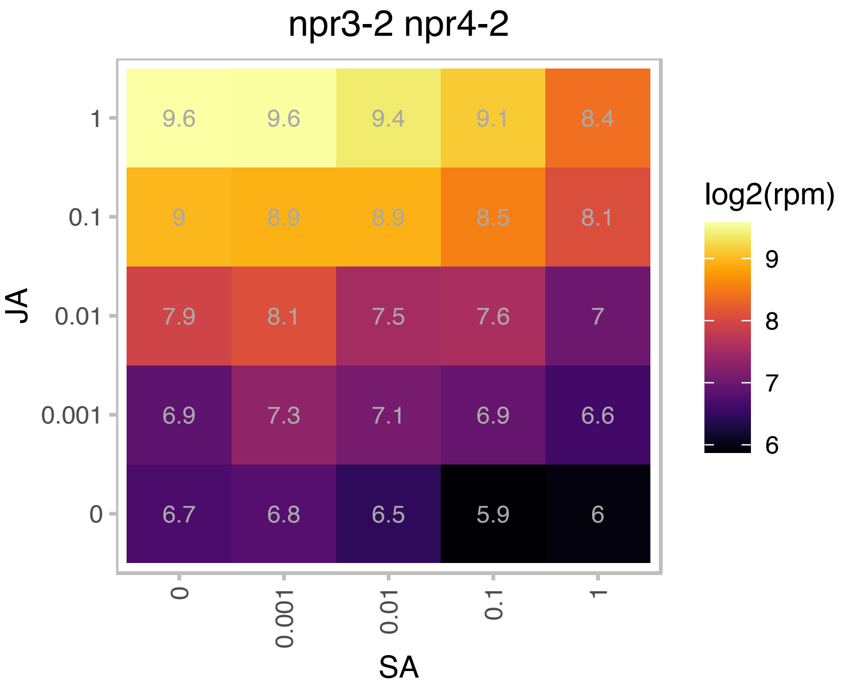


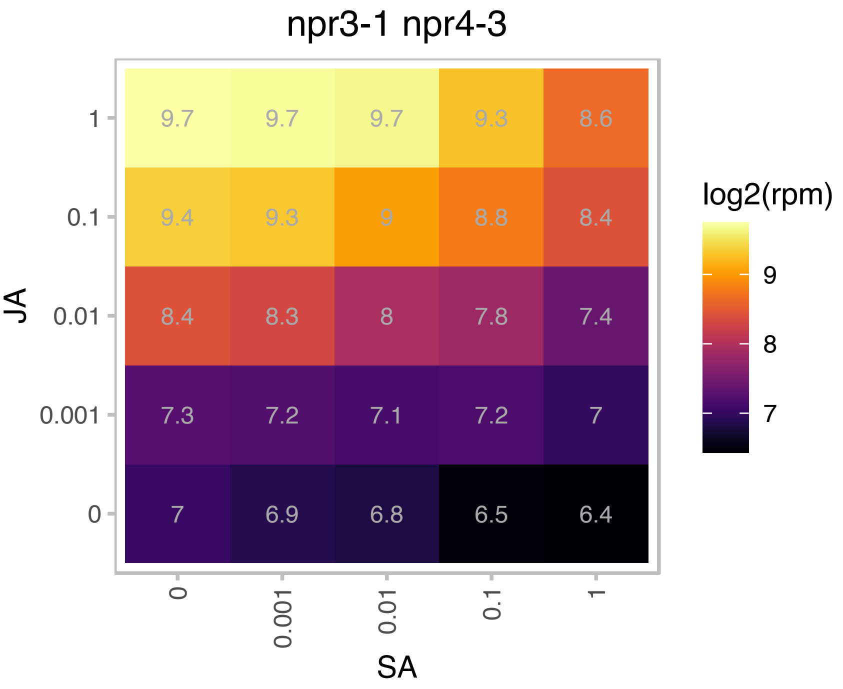

# AT3G28220.1 : NA

TRAF-like family protein

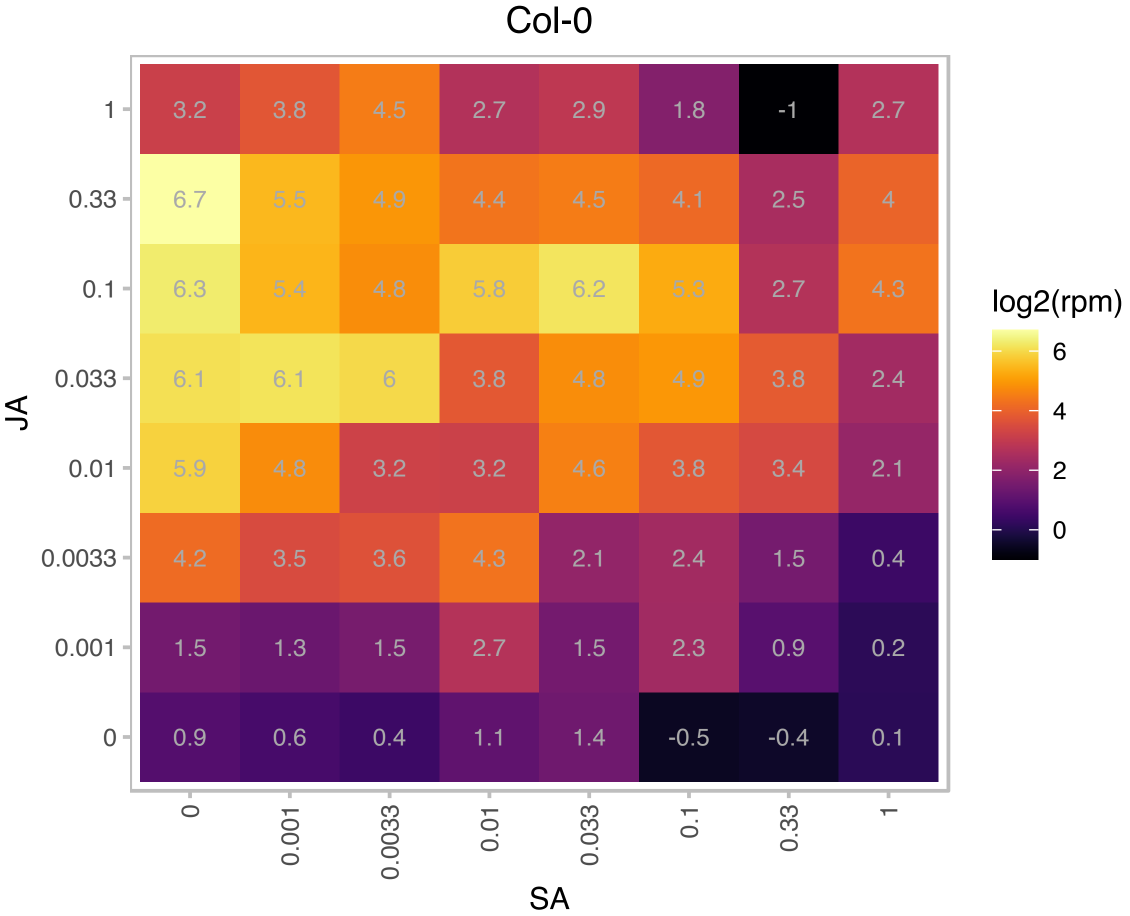


| | |
|---|---|
| Gene | AT3G28220 |
| is.expressed_genes | TRUE |
| is.responsive_genes | TRUE |
| is.apcluster | E-03 |

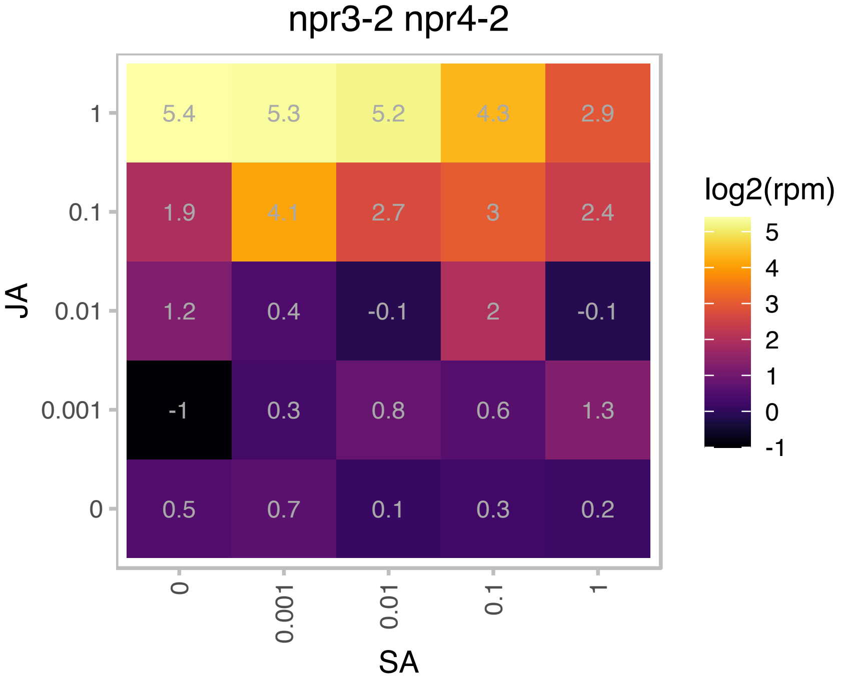


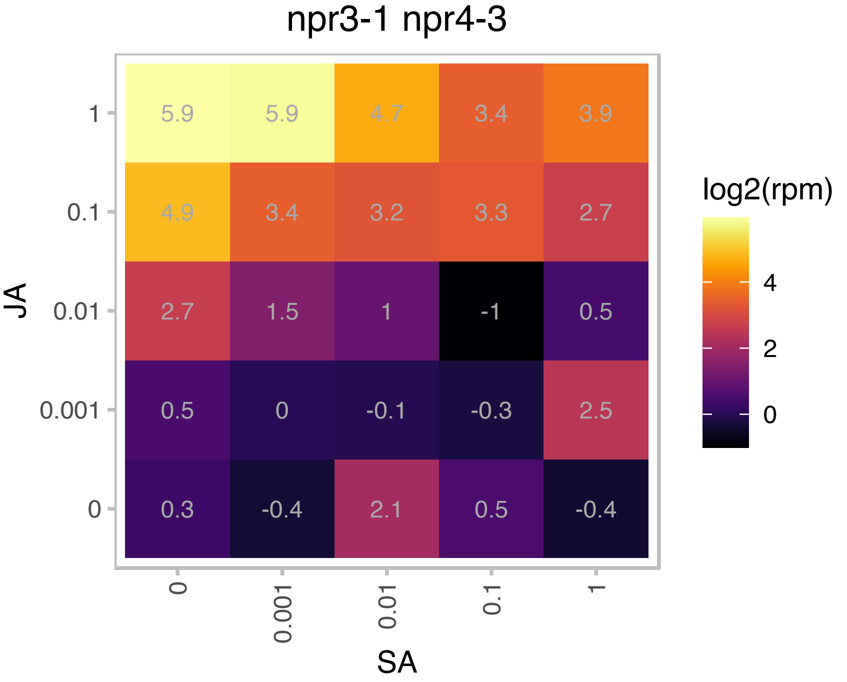

# AT3G28290.1 : AT14A

Protein of unknown function (DUF677)

Encodes a protein with sequence similarity to integrins. Localized to the cytoplasm and plasma membrane. Expressed in all tissues assayed.

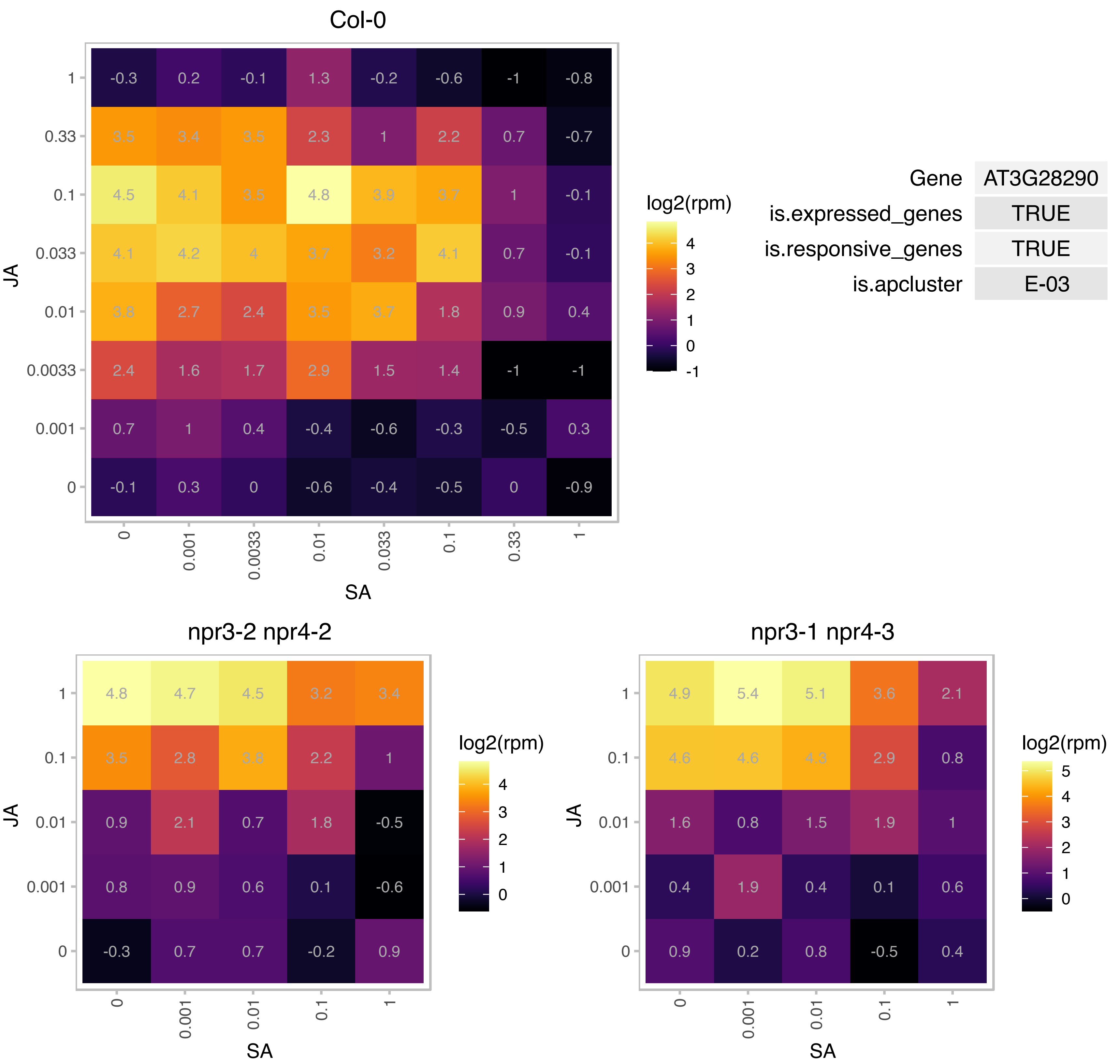


| | |
|---|---|
| Gene | AT3G28290 |
| is.expressed_genes | TRUE |
| is.responsive_genes | TRUE |
| is.apcluster | E-03 |

# AT3G28300.1 : AT14A

Protein of unknown function (DUF677)

Possesses a transmembrane domain and a small region that has sequence similarities to integrins from fungi, insects and humans. Is localized to plasma membrane and cytoplasm.

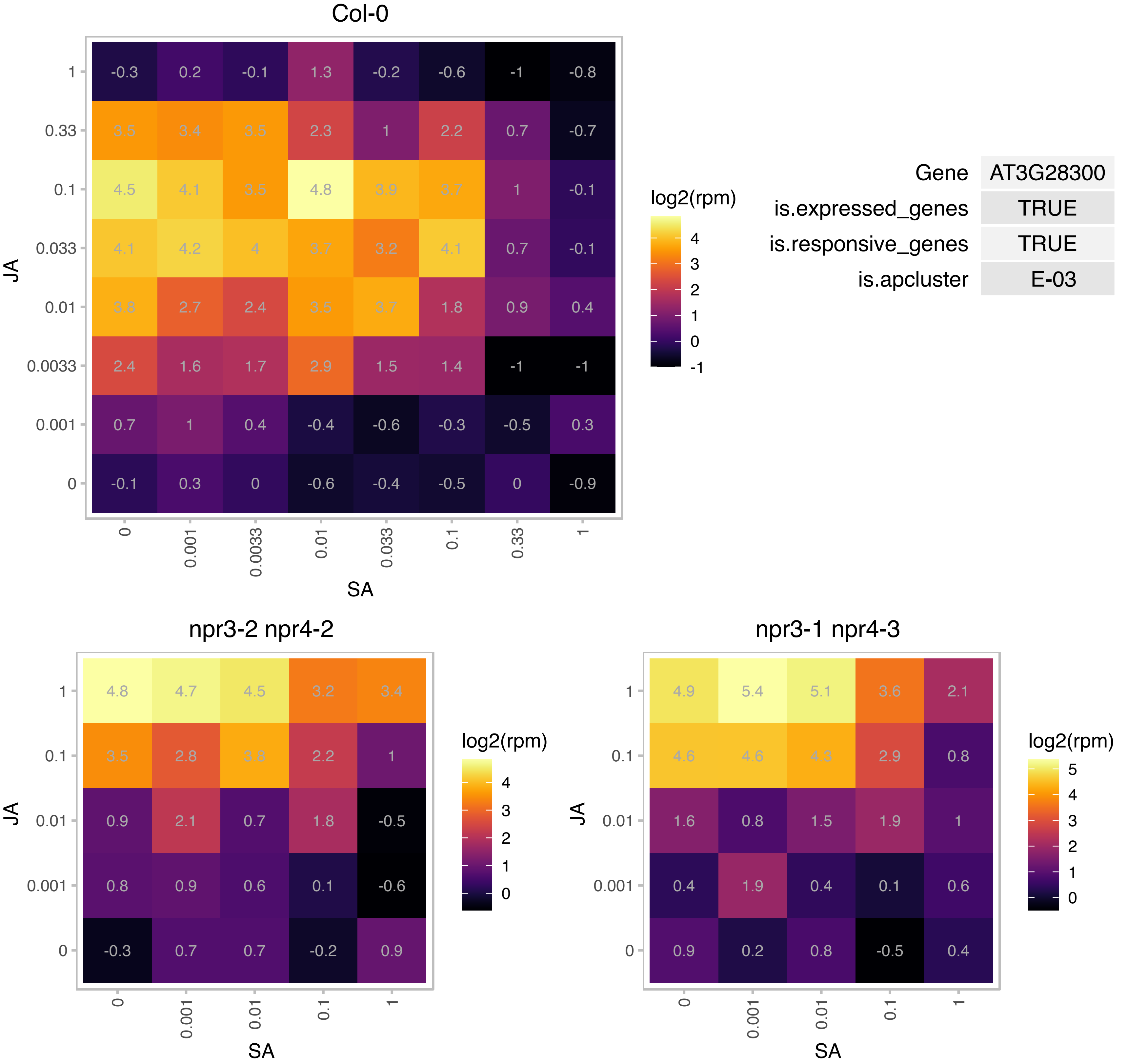


| | |
|---|---|
| Gene | AT3G28300 |
| is.expressed_genes | TRUE |
| is.responsive_genes | TRUE |
| is.apcluster | E-03 |

### AT3G46640.1 : LUX, PCL1

Homeodomain-like superfamily protein

Encodes a myb family transcription factor with a single Myb DNA-binding domain (type SHAQKYF) that is unique to plants and is essential for circadian rhythms, specifically for transcriptional regulation within the circadian clock. LUX is required for normal rhythmic expression of multiple clock outputs in both constant light and darkness. It is coregulated with TOC1 and seems to be repressed by CCA1 and LHY by direct binding of these proteins to the evening element in the LUX promoter. The mRNA is cell-to-cell mobile.

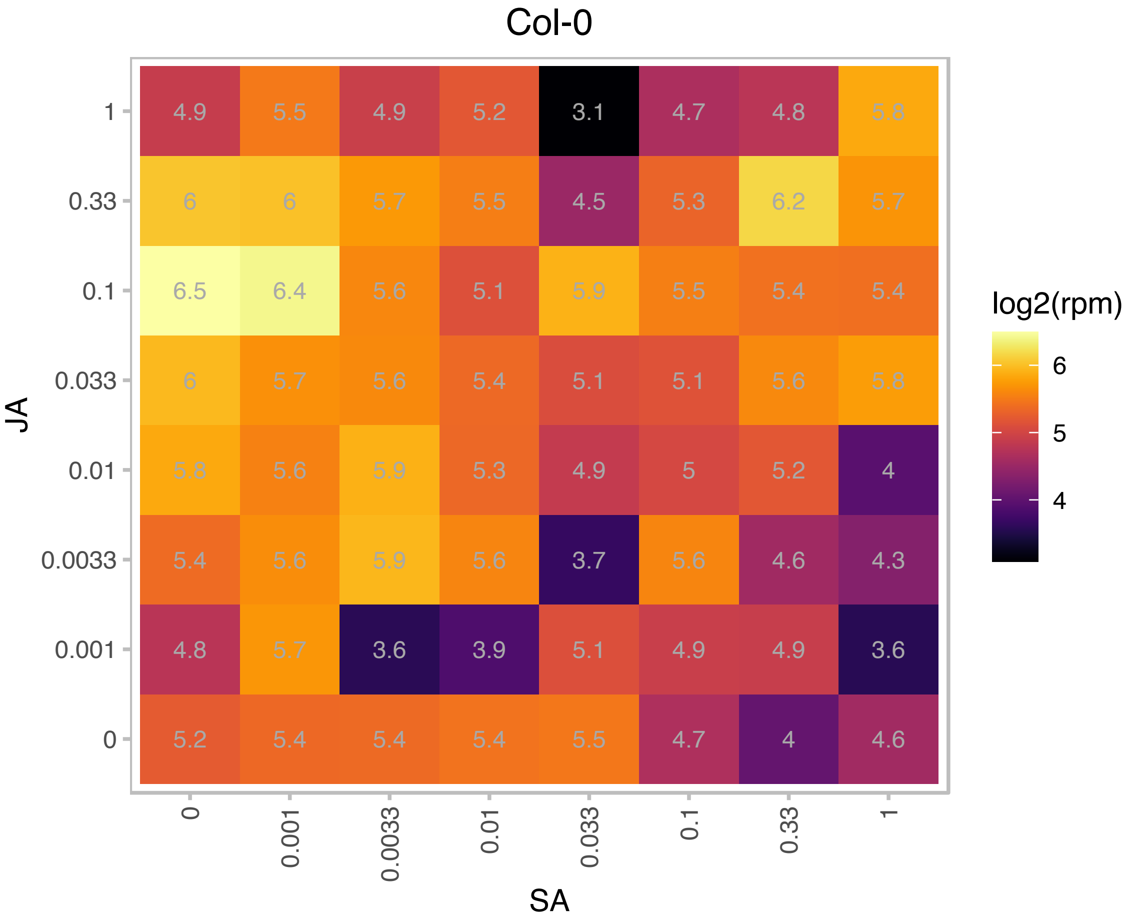


| | |
|---|---|
| Gene | AT3G46640 |
| is.expressed_genes | TRUE |
| is.responsive_genes | TRUE |
| is.apcluster | E-03 |

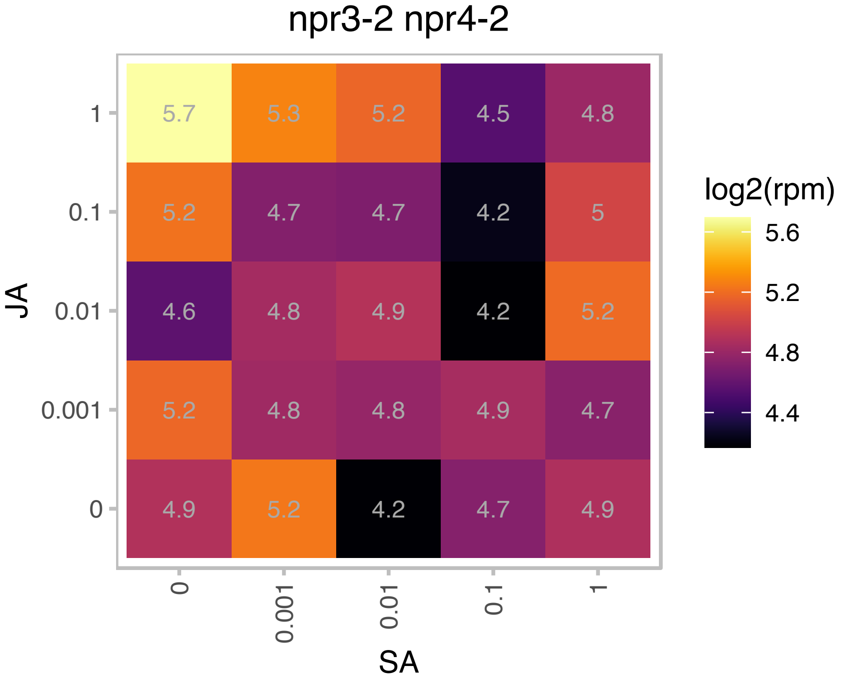


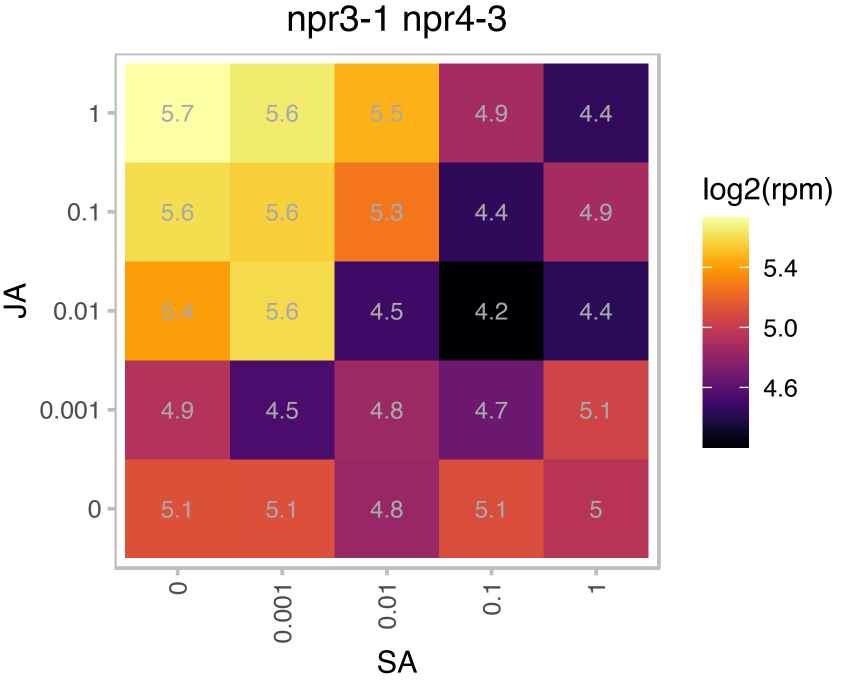

# AT3G46650.1 : NA

UDP-Glycosyltransferase superfamily protein

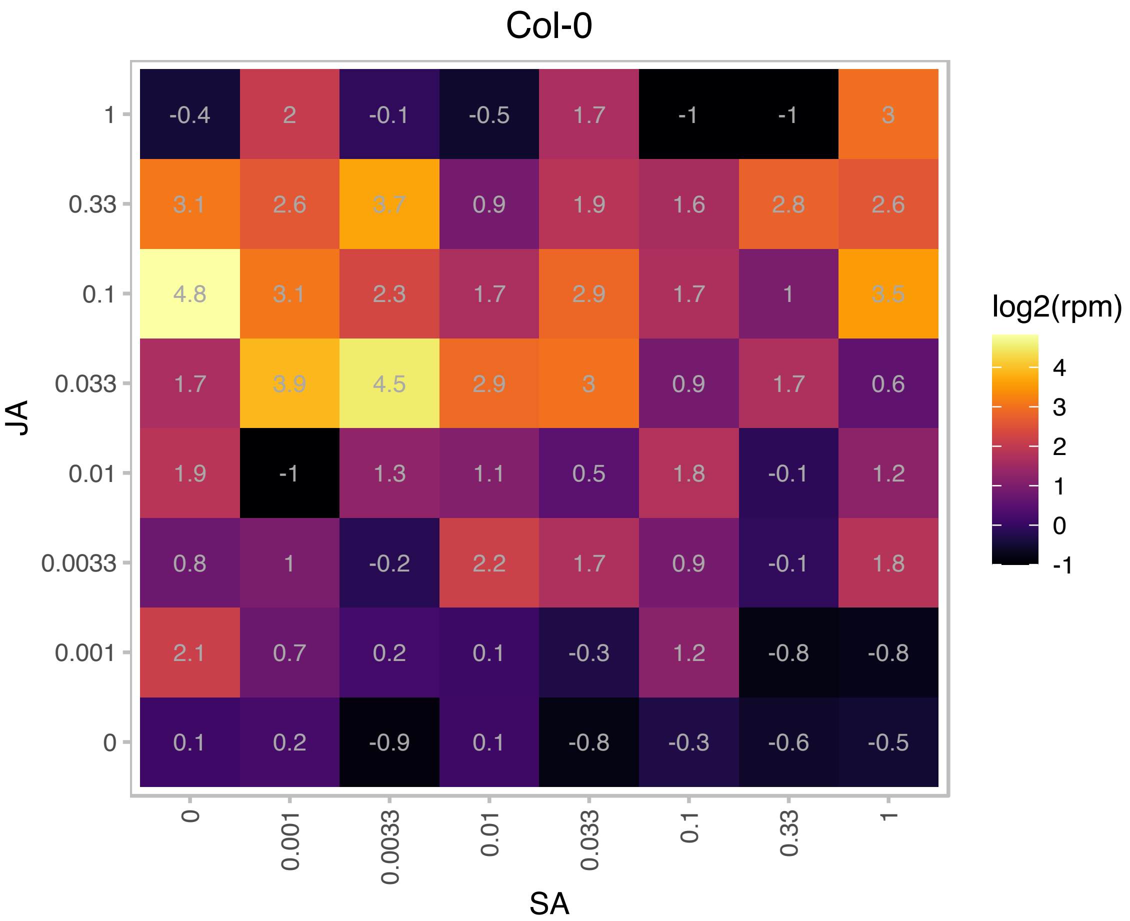


| Gene | AT3G46650 |
| --- | --- |
| is.expressed_genes | TRUE |
| is.responsive_genes | TRUE |
| is.apcluster | E-03 |

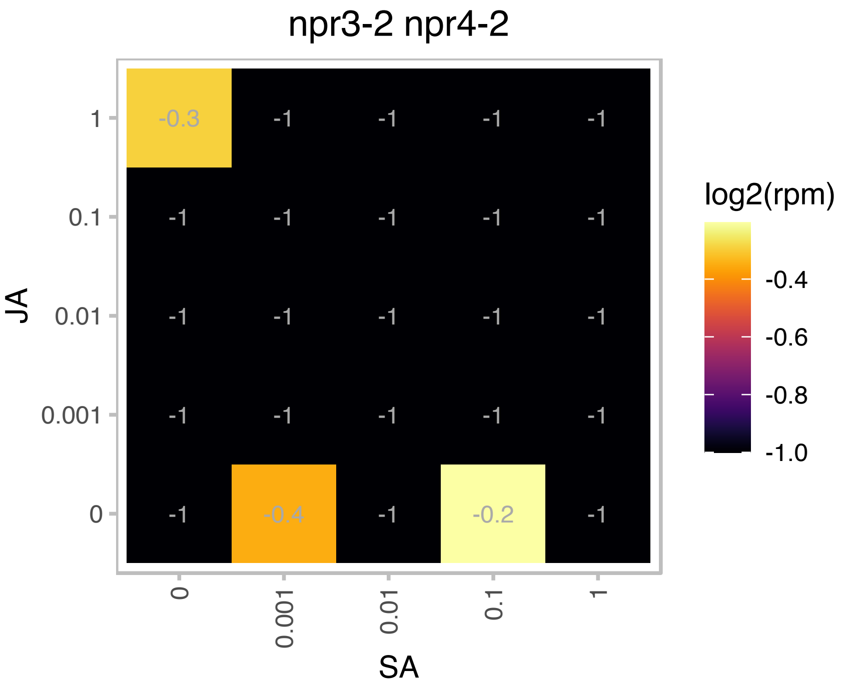


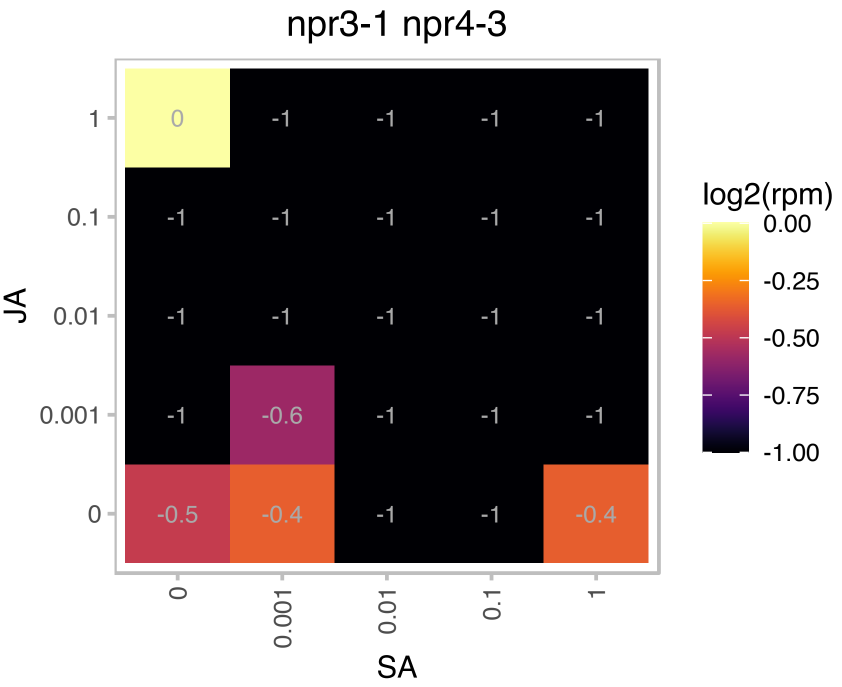

### AT3G49620.1 : DIN11

2-oxoglutarate (2OG) and Fe(II)-dependent oxygenase superfamily protein

encodes a protein similar to 2-oxoacid-dependent dioxygenase. Expression is induced after 24 hours of dark treatment, in senescing leaves and treatment with exogenous photosynthesis inhibitor. Induction of gene expression was suppressed in excised leaves supplied with sugar. The authors suggest that the gene's expression pattern is responding to the level of sugar in the cell.

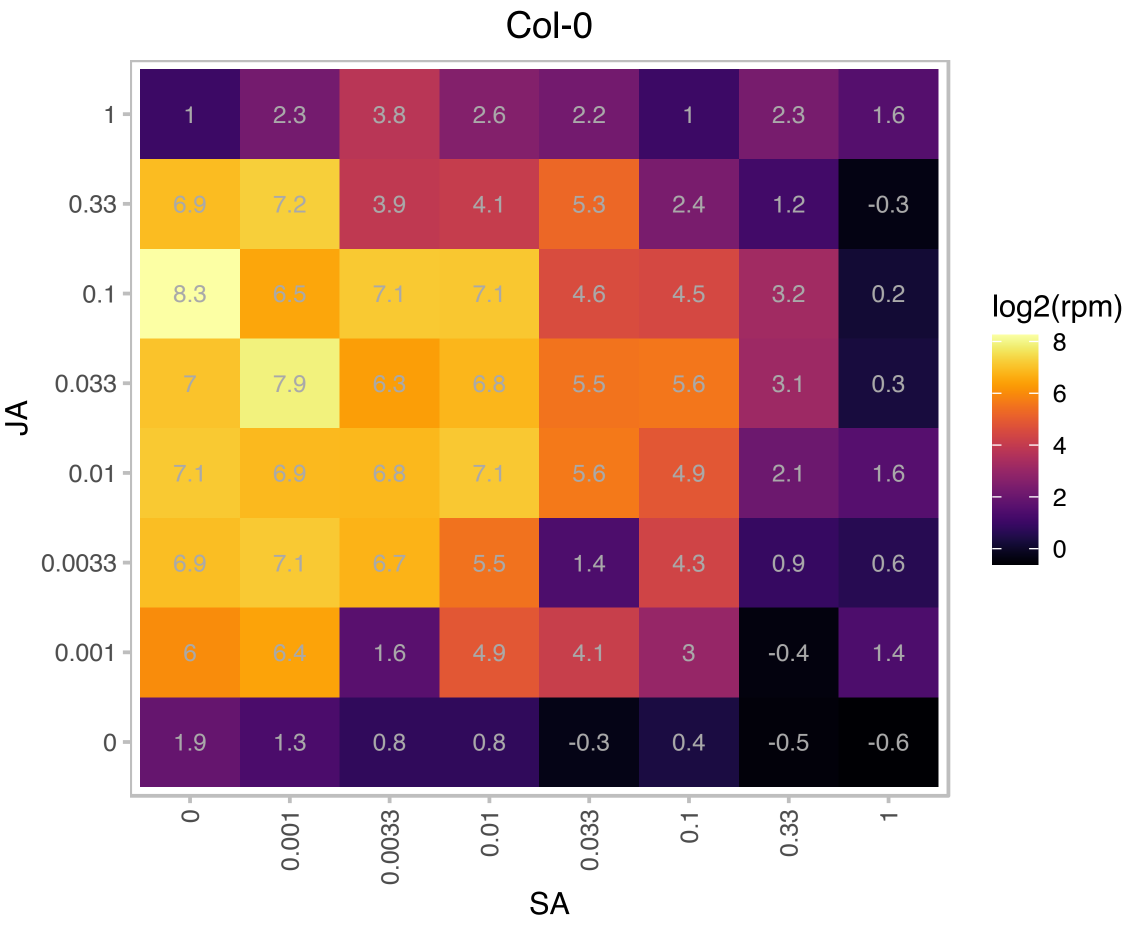


| | |
|---|---|
| Gene | AT3G49620 |
| is.expressed_genes | TRUE |
| is.responsive_genes | TRUE |
| is.apcluster | E-03 |

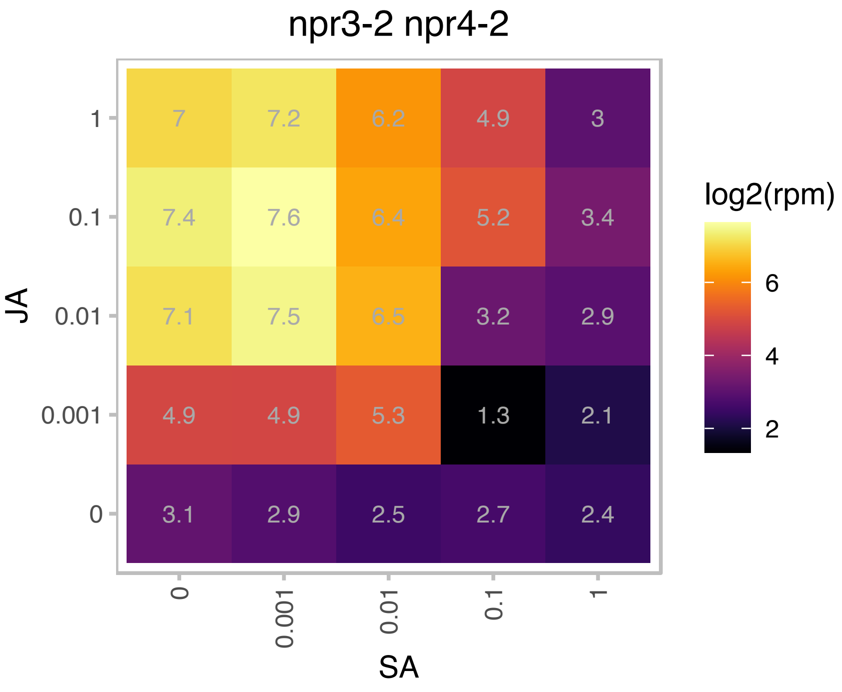


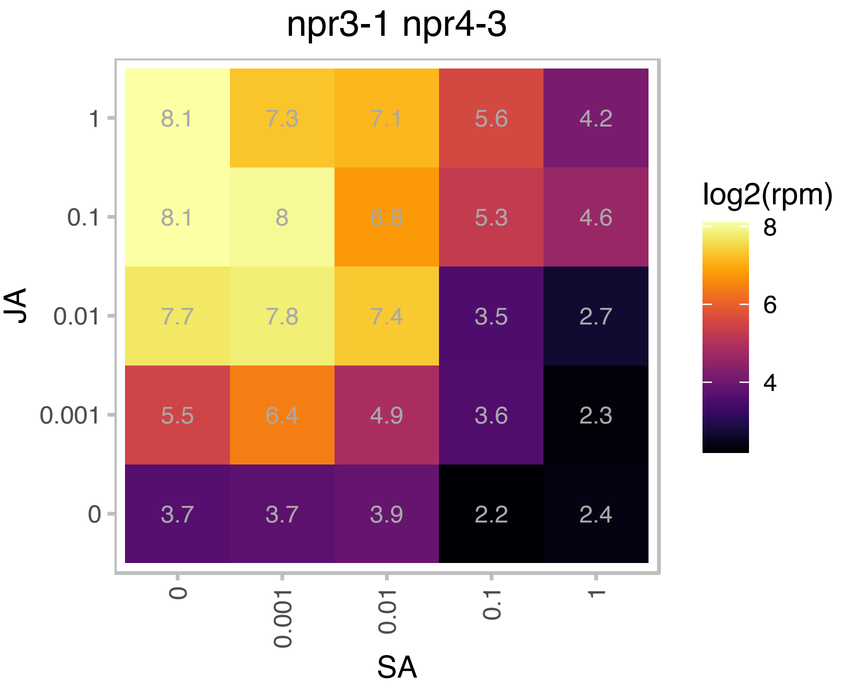

# AT4G02360.1 : NA

Protein of unknown function, DUF538

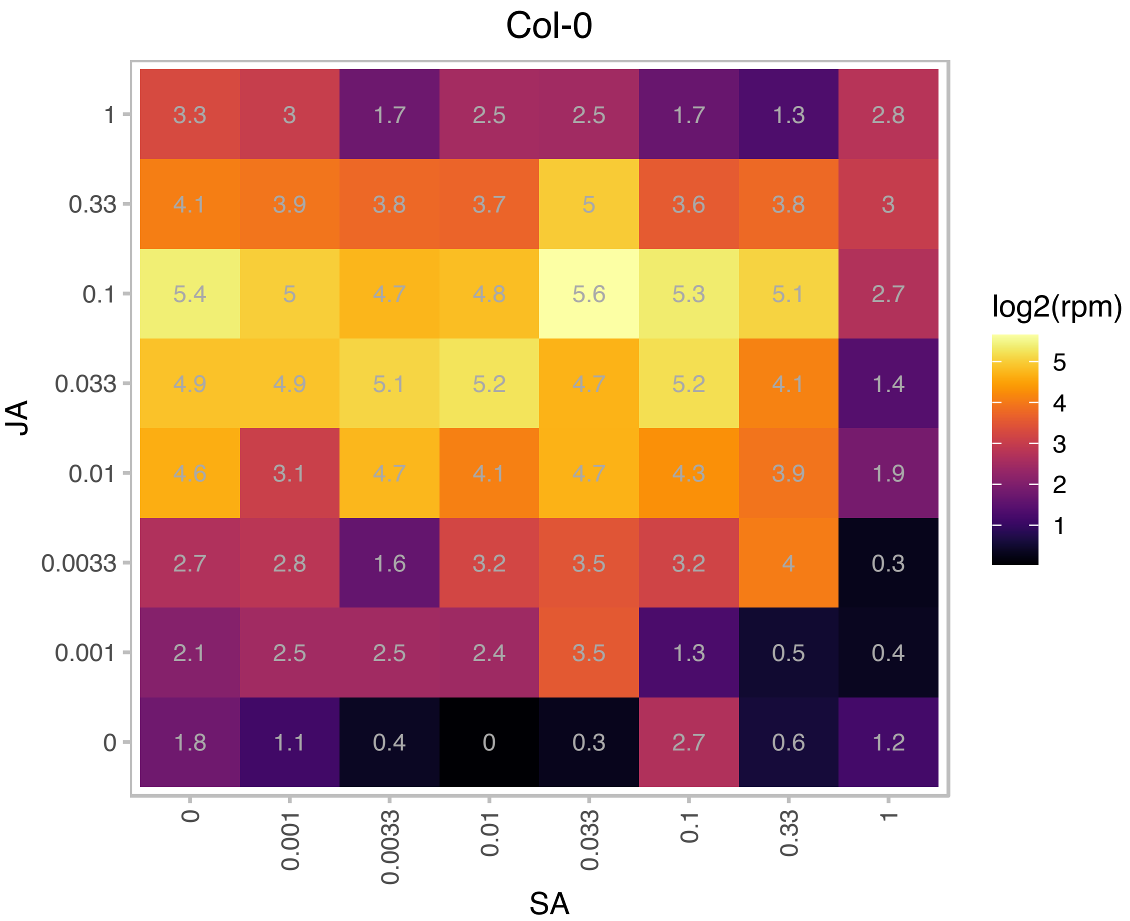


| | |
|---|---|
| Gene | AT4G02360 |
| is.expressed_genes | TRUE |
| is.responsive_genes | TRUE |
| is.apcluster | E-03 |

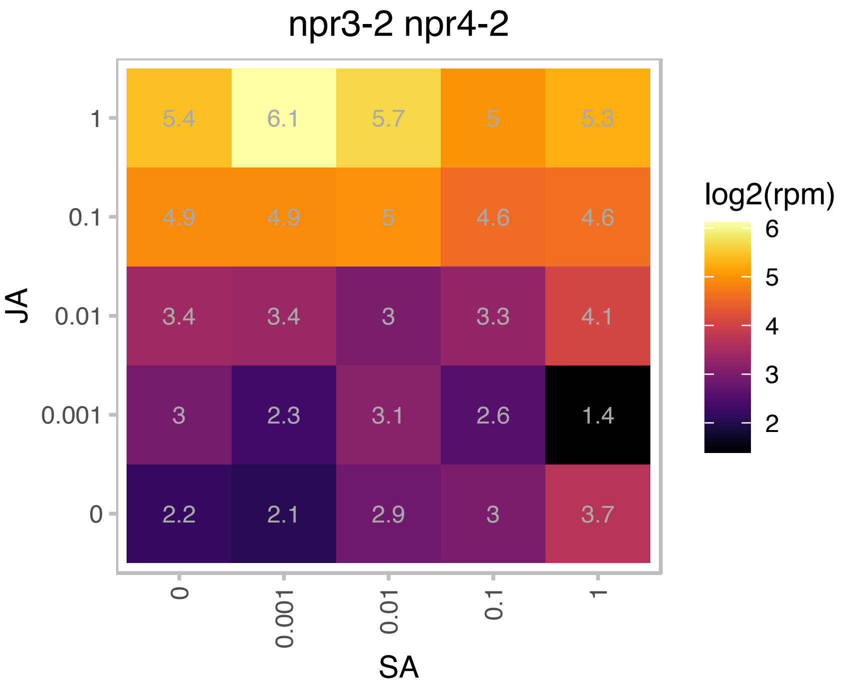


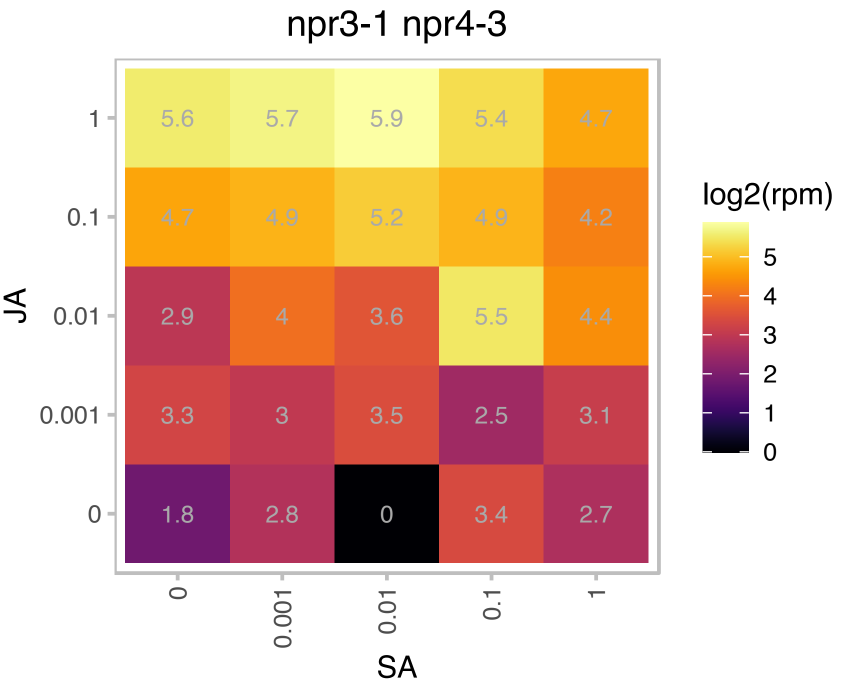

### AT4G08870.1 : ARGAH2

Arginase/deacetylase superfamily protein

Encodes one of the two arginase in the genome. Gene expression is enhanced by methyl jasmonate treatment.  It is involved in the defense response to B.cinerea.

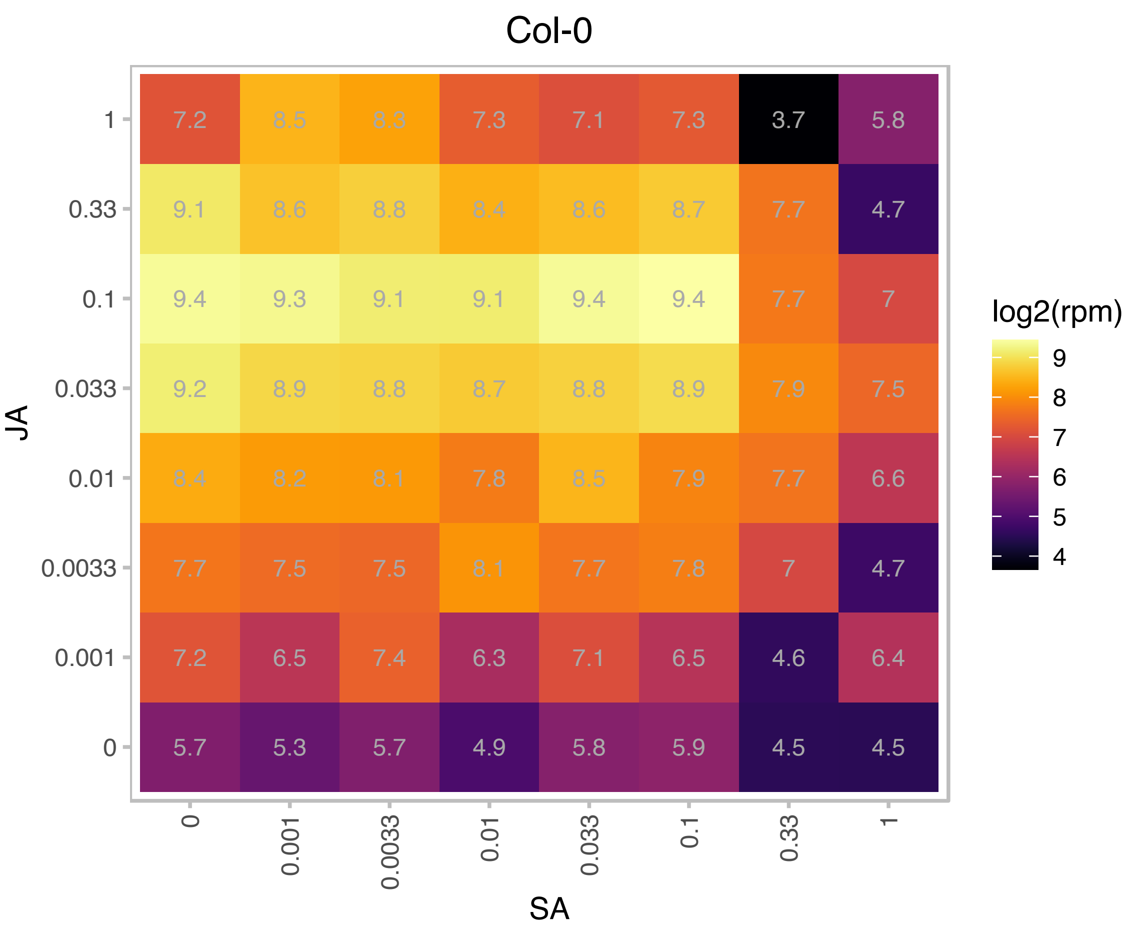


| | |
|---|---|
| Gene | AT4G08870 |
| is.expressed_genes | TRUE |
| is.responsive_genes | TRUE |
| is.apcluster | E-03 |

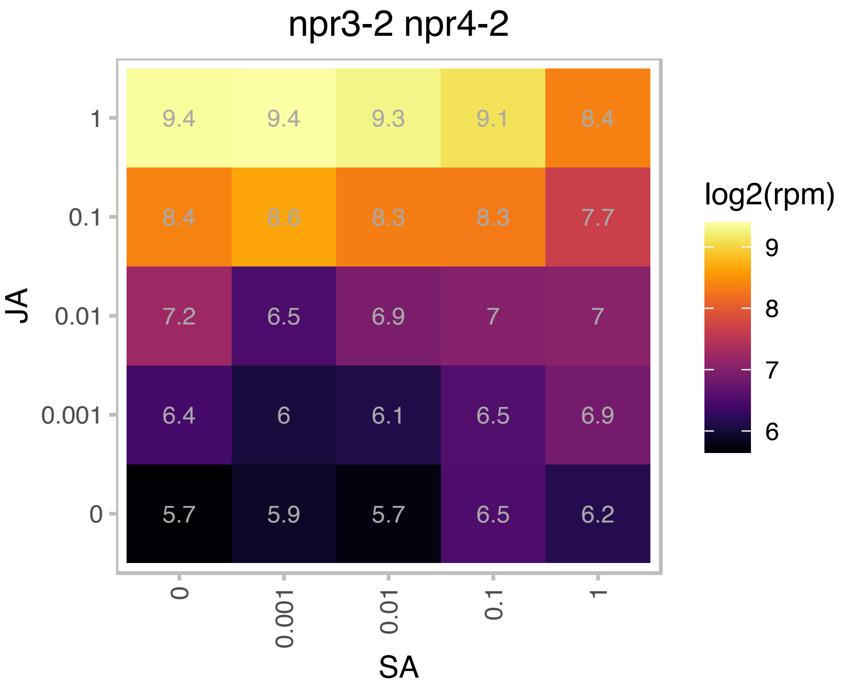


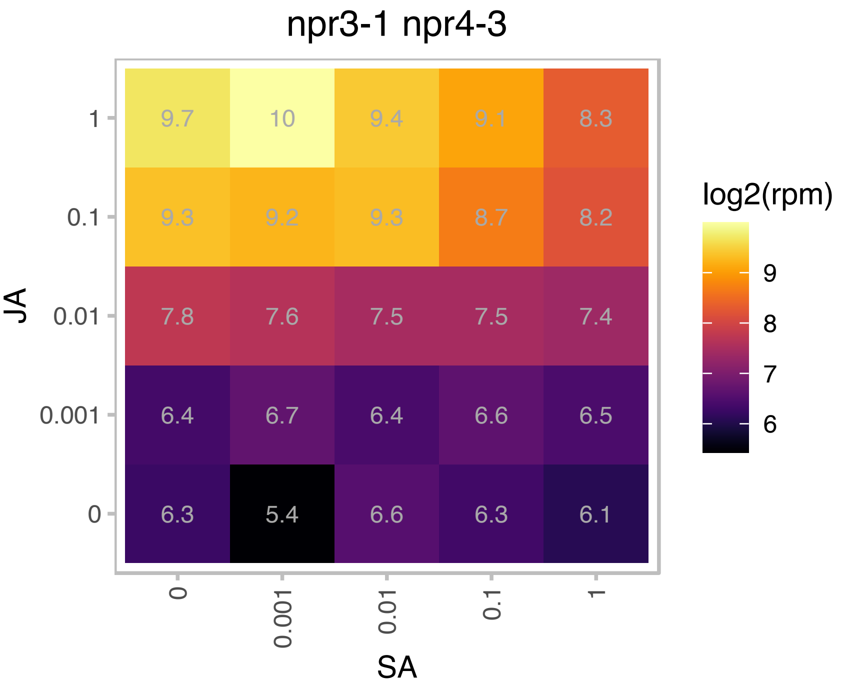

### AT4G15210.1 : AT-BETA-AMY, ATBETA-AMY, BAM5, BMY

beta-amylase 5

cytosolic beta-amylase expressed in rosette leaves and inducible by sugar. RAM1 mutants have reduced beta amylase in leaves and stems.

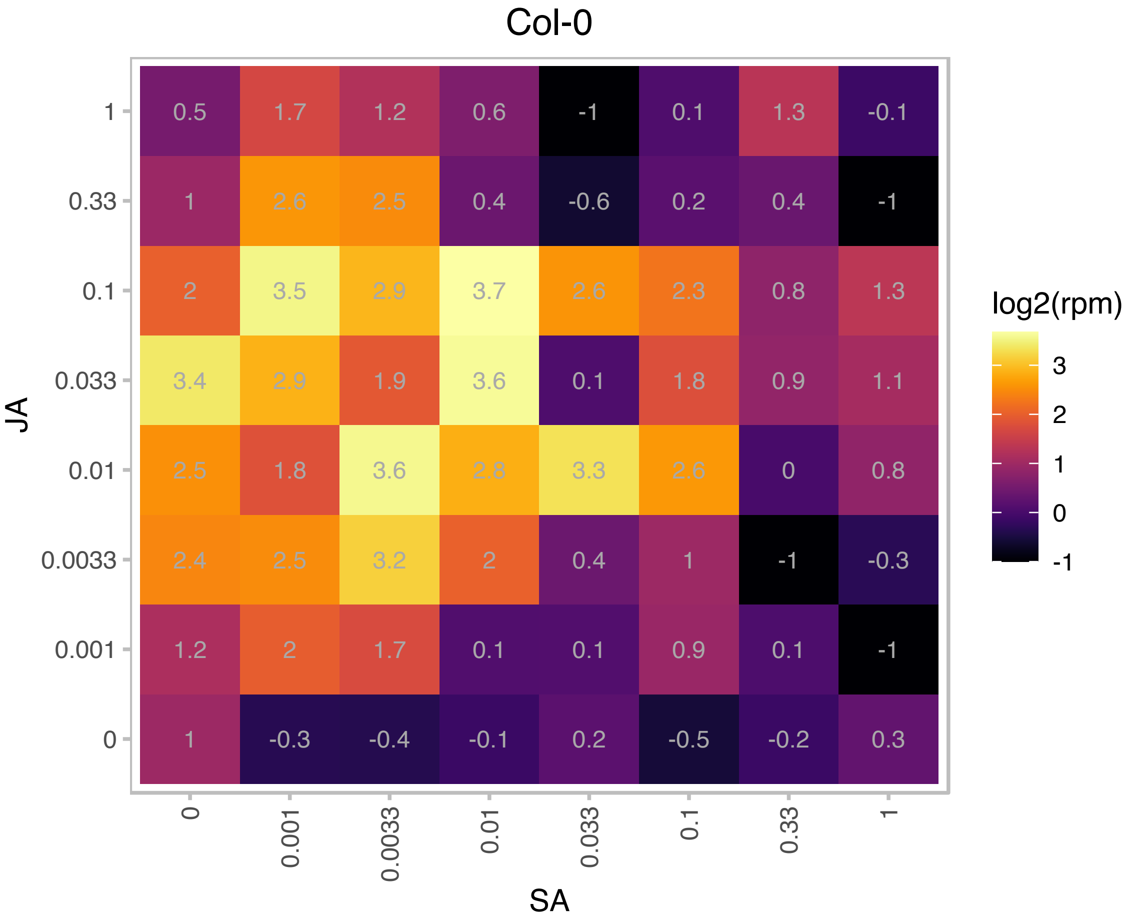


| | |
|---|---|
| Gene | AT4G15210 |
| is.expressed_genes | TRUE |
| is.responsive_genes | TRUE |
| is.apcluster | E-03 |

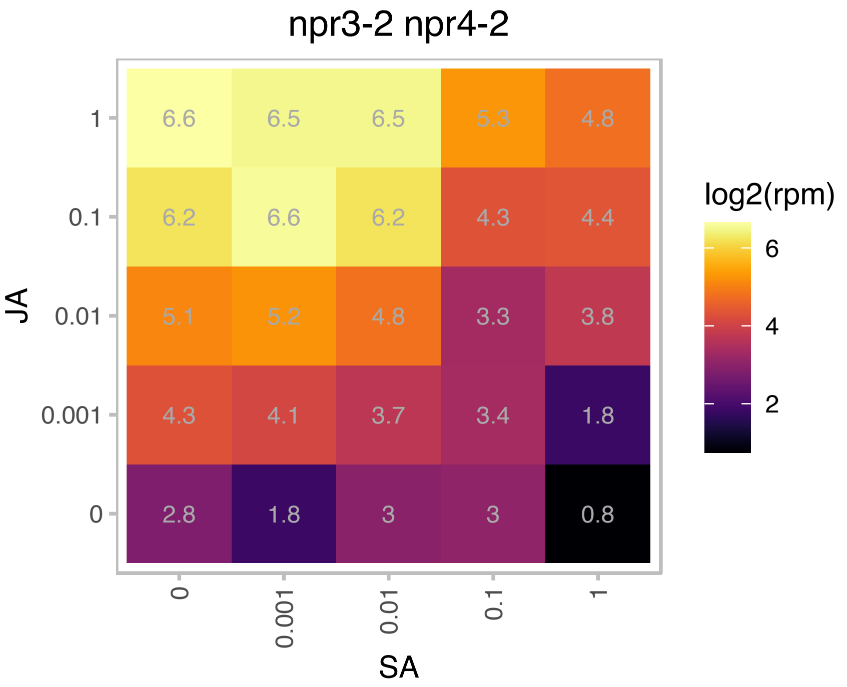


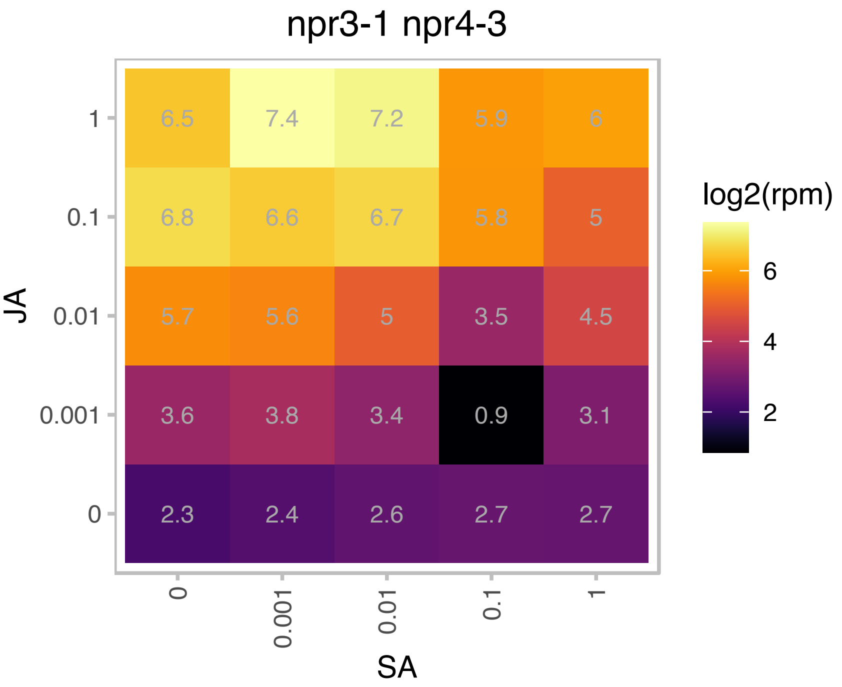

### AT4G15440.1 : CYP74B2, HPL1

hydroperoxide lyase 1
Encodes a hydroperoxide lyase.  Also a member of the CYP74B cytochrome p450 family. In the ecotype Columbia (Col) the gene contains a 10-nucleotide deletion in its first exon that causes it to code for a truncated protein that results in a non-functional hydroperoxide lyase.

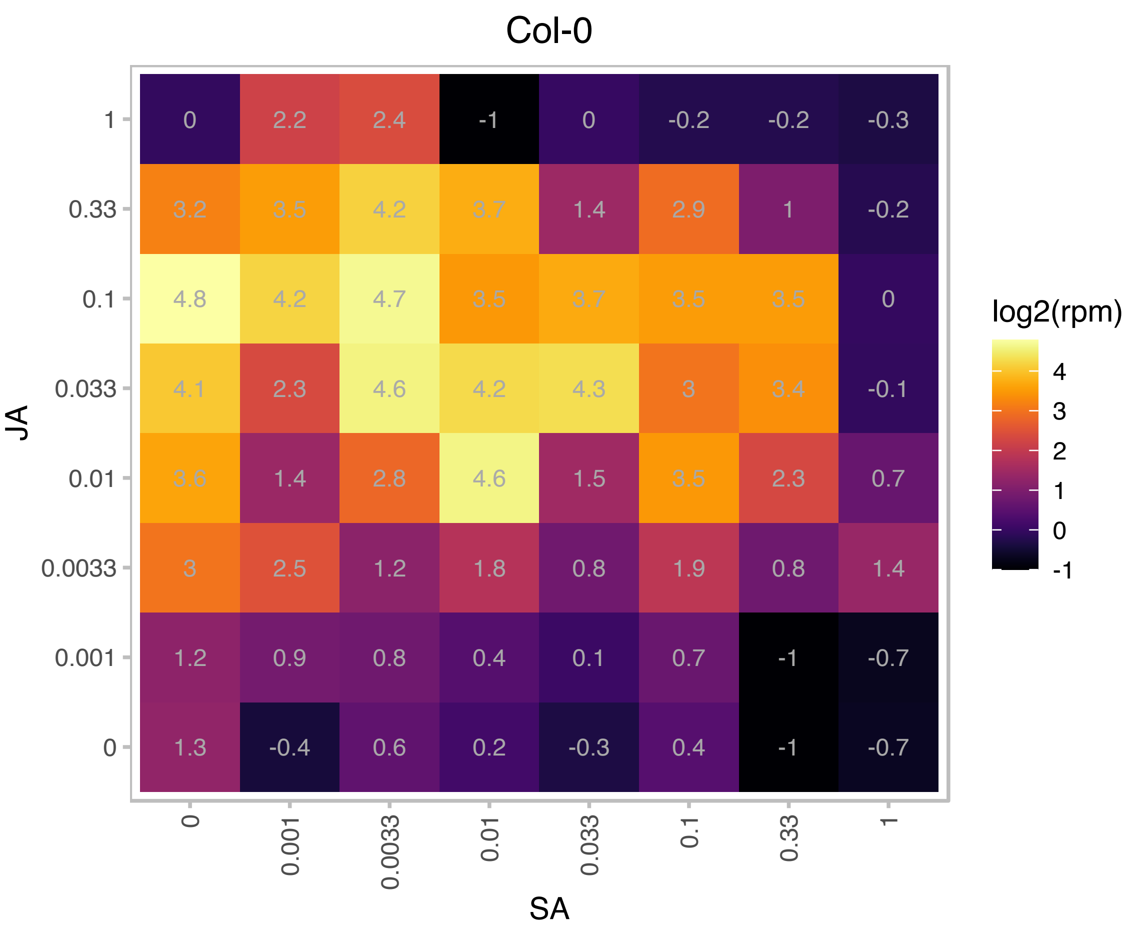


| | |
|---|---|
| Gene | AT4G15440 |
| is.expressed_genes | TRUE |
| is.responsive_genes | TRUE |
| is.apcluster | E-03 |

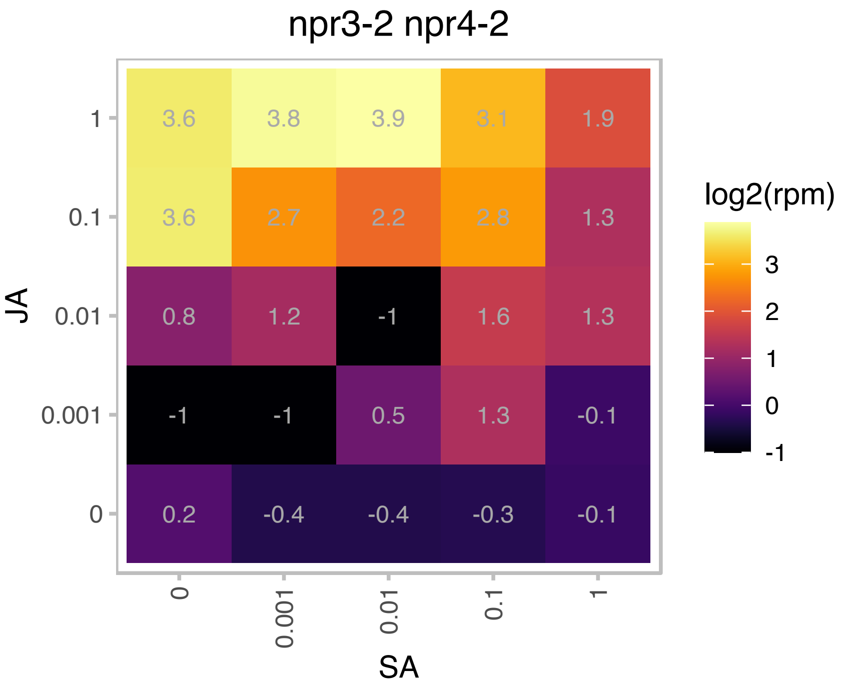


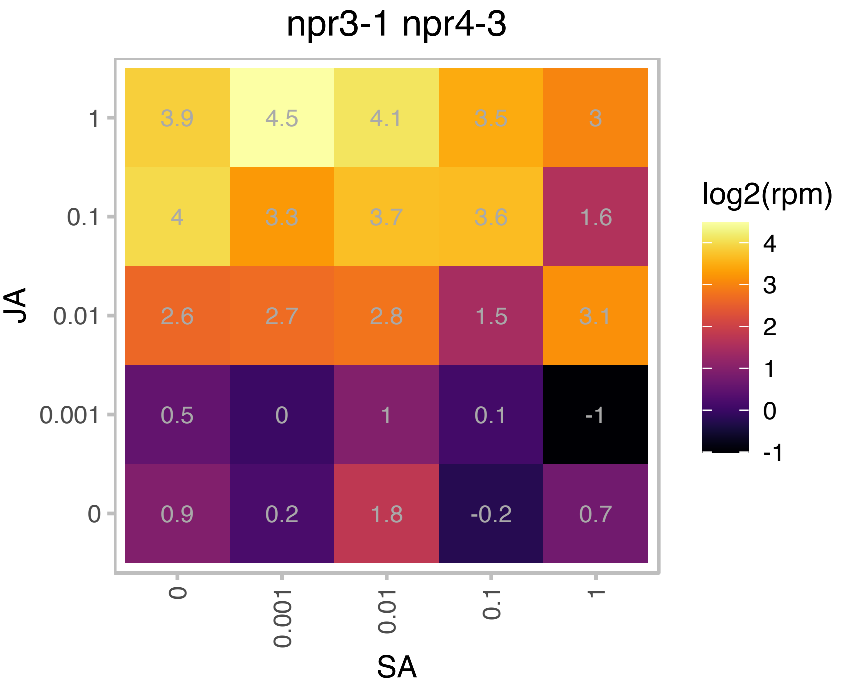

# AT4G17470.1 : NA

alpha/beta-Hydrolases superfamily protein

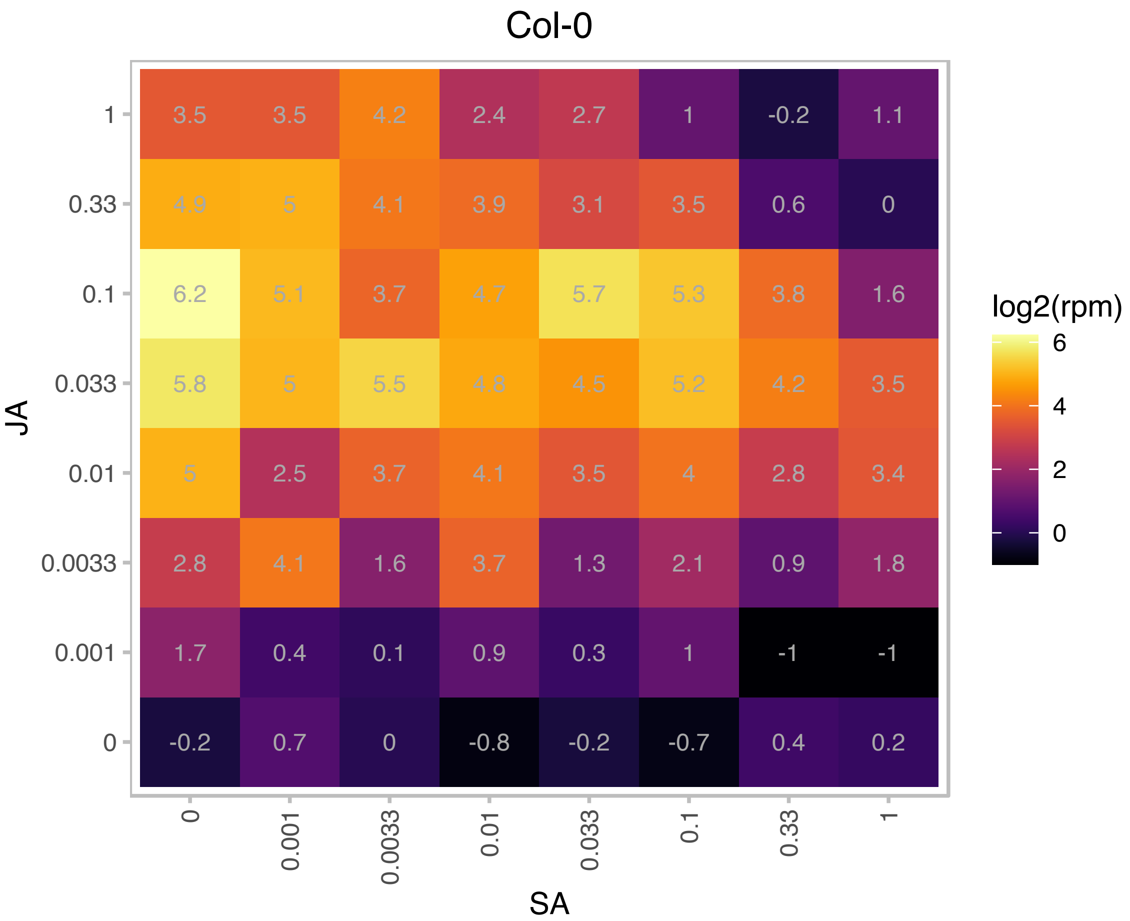


| | |
|---|---|
| Gene | AT4G17470 |
| is.expressed_genes | TRUE |
| is.responsive_genes | TRUE |
| is.apcluster | E-03 |

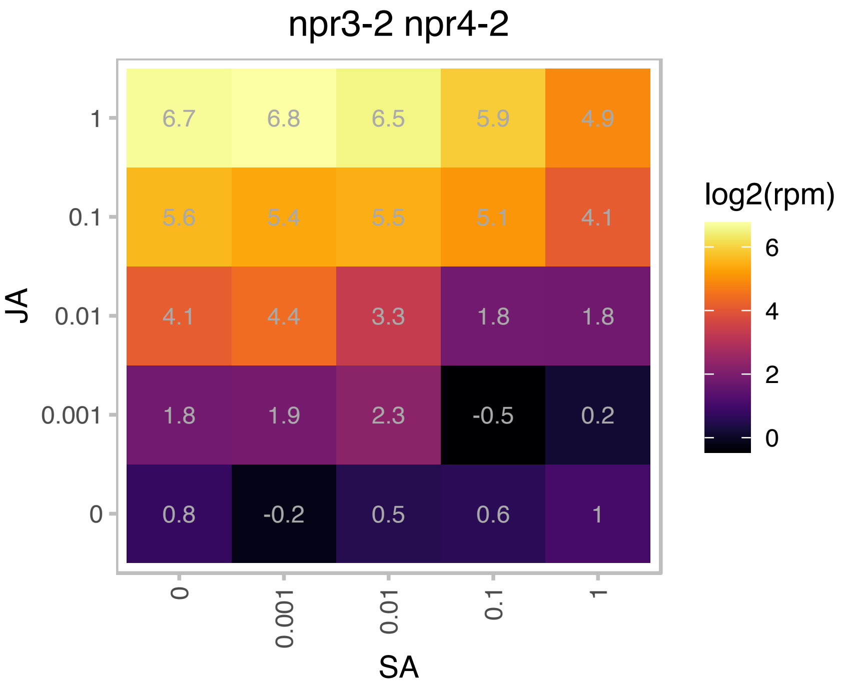


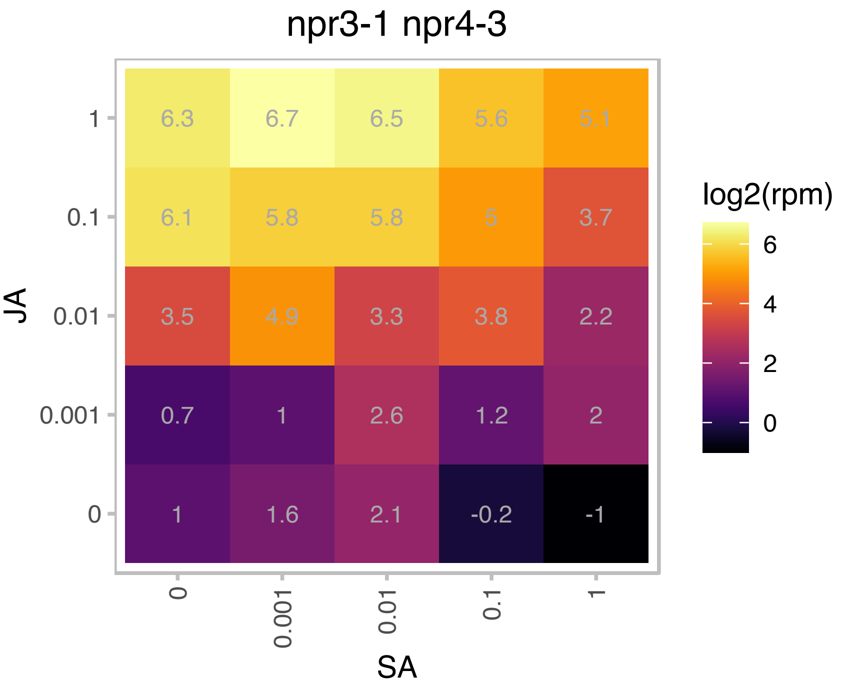

### AT4G21850.1 : ATMSRB9, MSRB9

methionine sulfoxide reductase B9

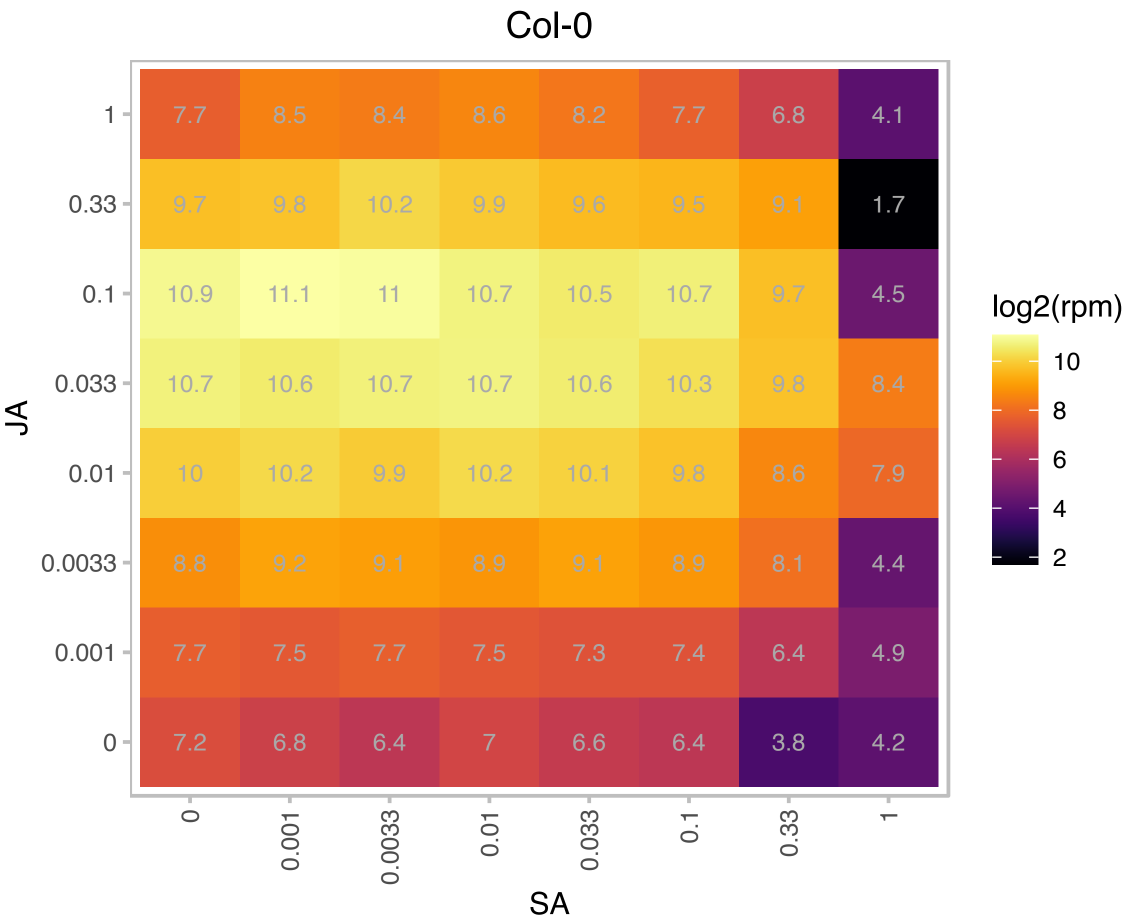


| | |
|---|---|
| Gene | AT4G21850 |
| is.expressed_genes | TRUE |
| is.responsive_genes | TRUE |
| is.apcluster | E-03 |

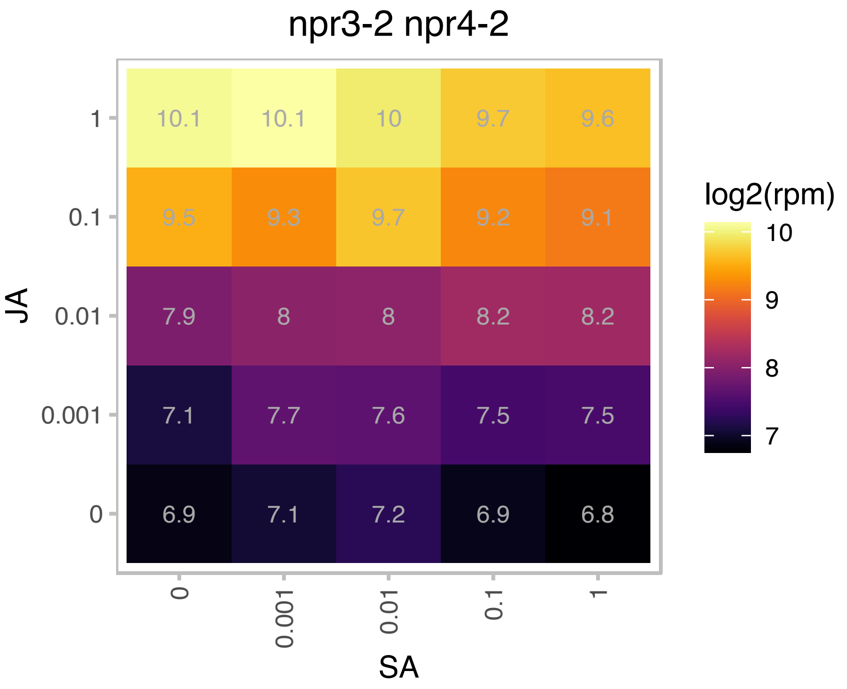


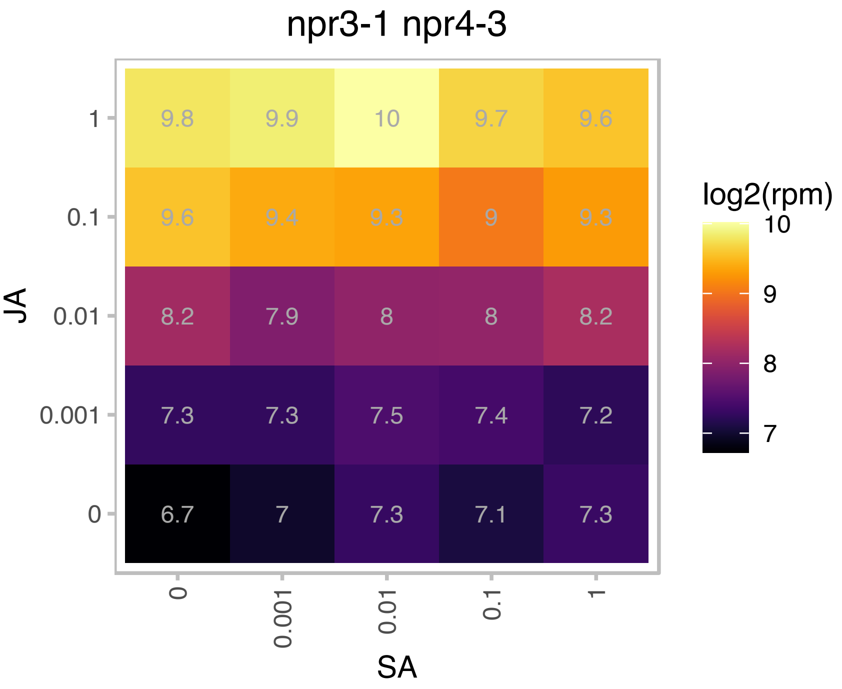

# AT4G24340.1 : NA

Phosphorylase superfamily protein

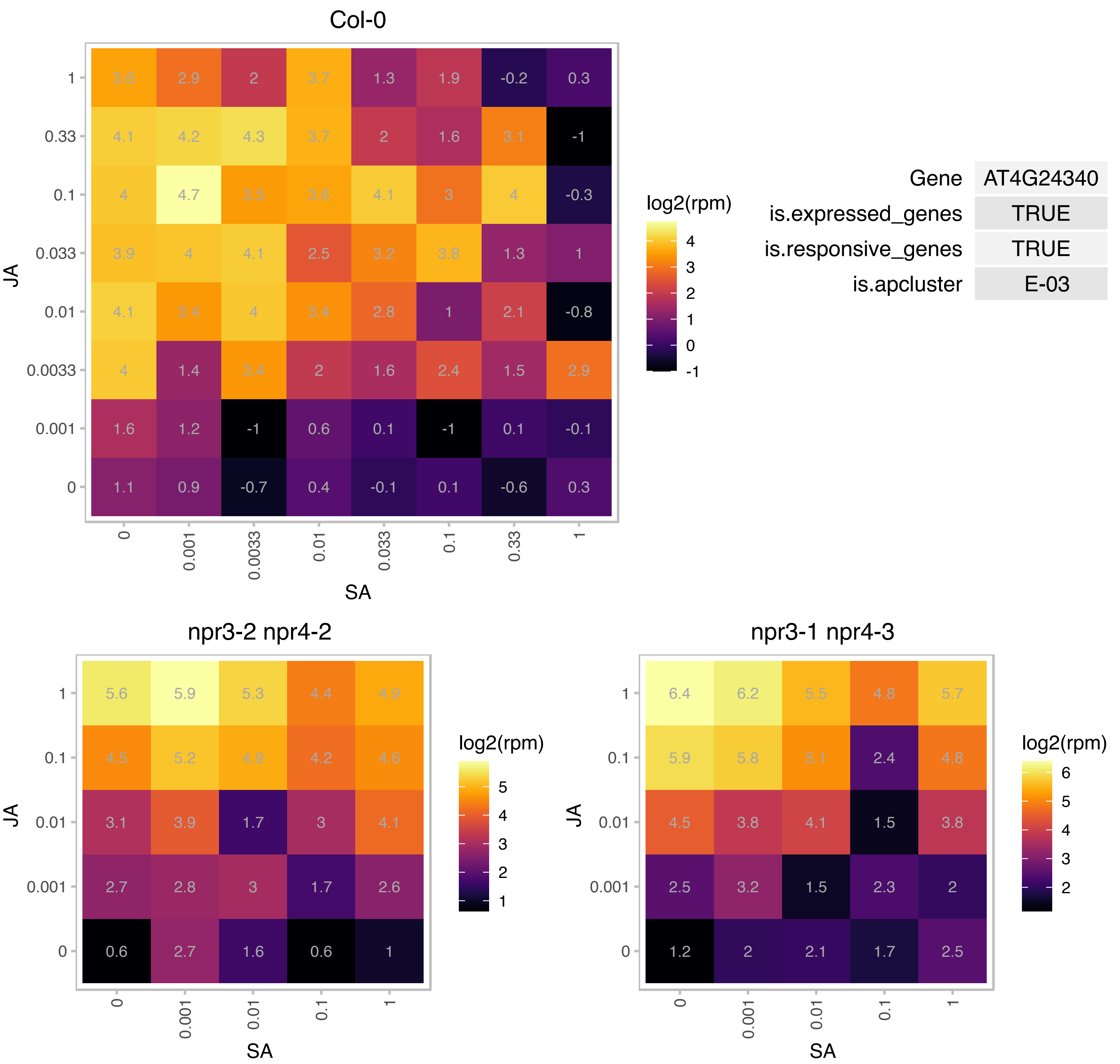


| | |
|---|---|
| Gene | AT4G24340 |
| is.expressed_genes | TRUE |
| is.responsive_genes | TRUE |
| is.apcluster | E-03 |

### AT5G06870.1 : ATPGIP2, PGIP2

polygalacturonase inhibiting protein 2

Encodes a polygalacturonase inhibiting protein involved in plant defense response. PGIPs inhibit the activity of pectin degrading enzymes such as those produced by fungal pathogens. PGIP2 is induced by fungal infection and methyl jasmonate.Suppressed in the proton sensitive stop1-mutant, but the transcription level was recovered by transformation of STOP2. Knockout mutant showed severe damage in the root tip in low Ca and low pH medium.

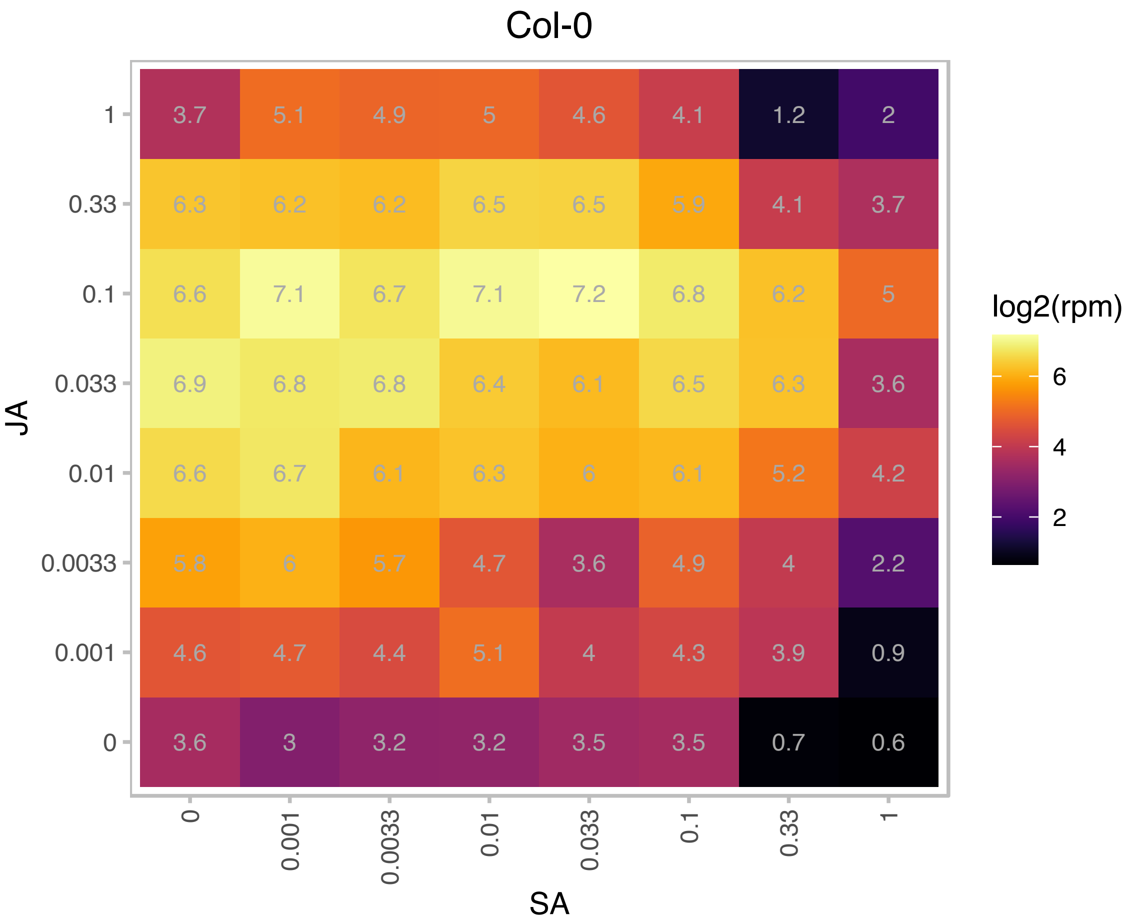


| | |
|---|---|
| Gene | AT5G06870 |
| is.expressed_genes | TRUE |
| is.responsive_genes | TRUE |
| is.apcluster | E-03 |

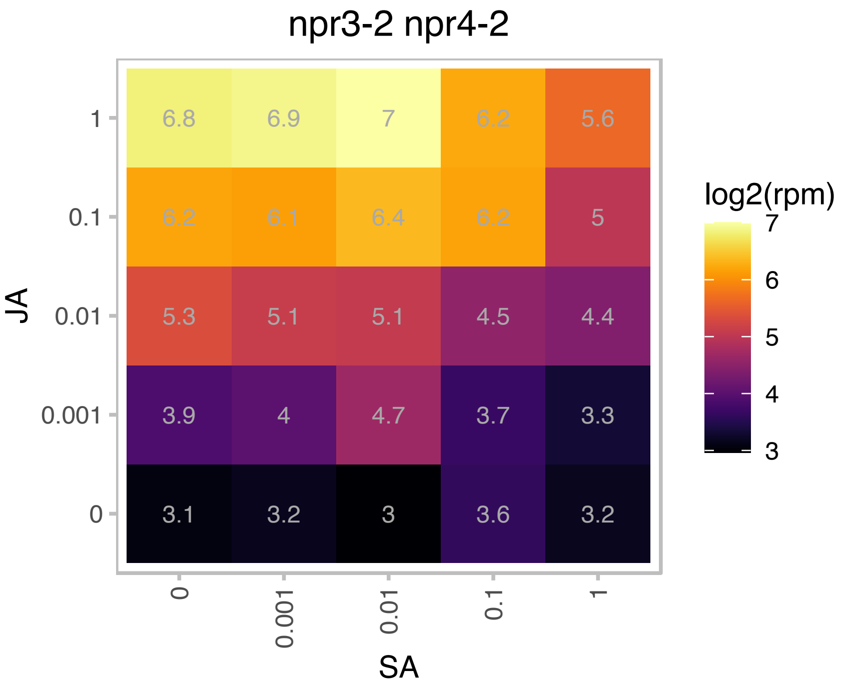


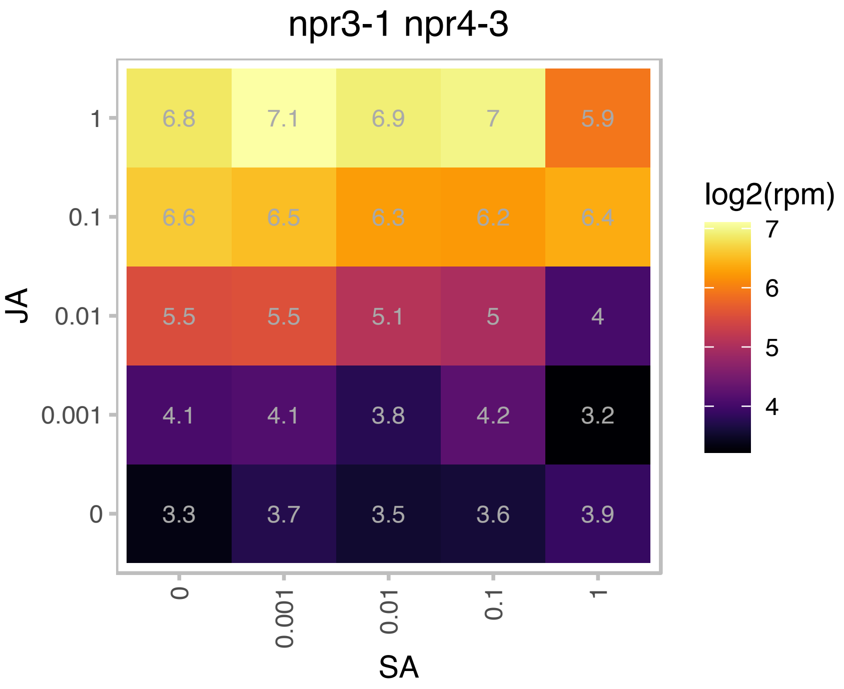

# AT5G14700.1 : NA

NAD(P)-binding Rossmann-fold superfamily protein

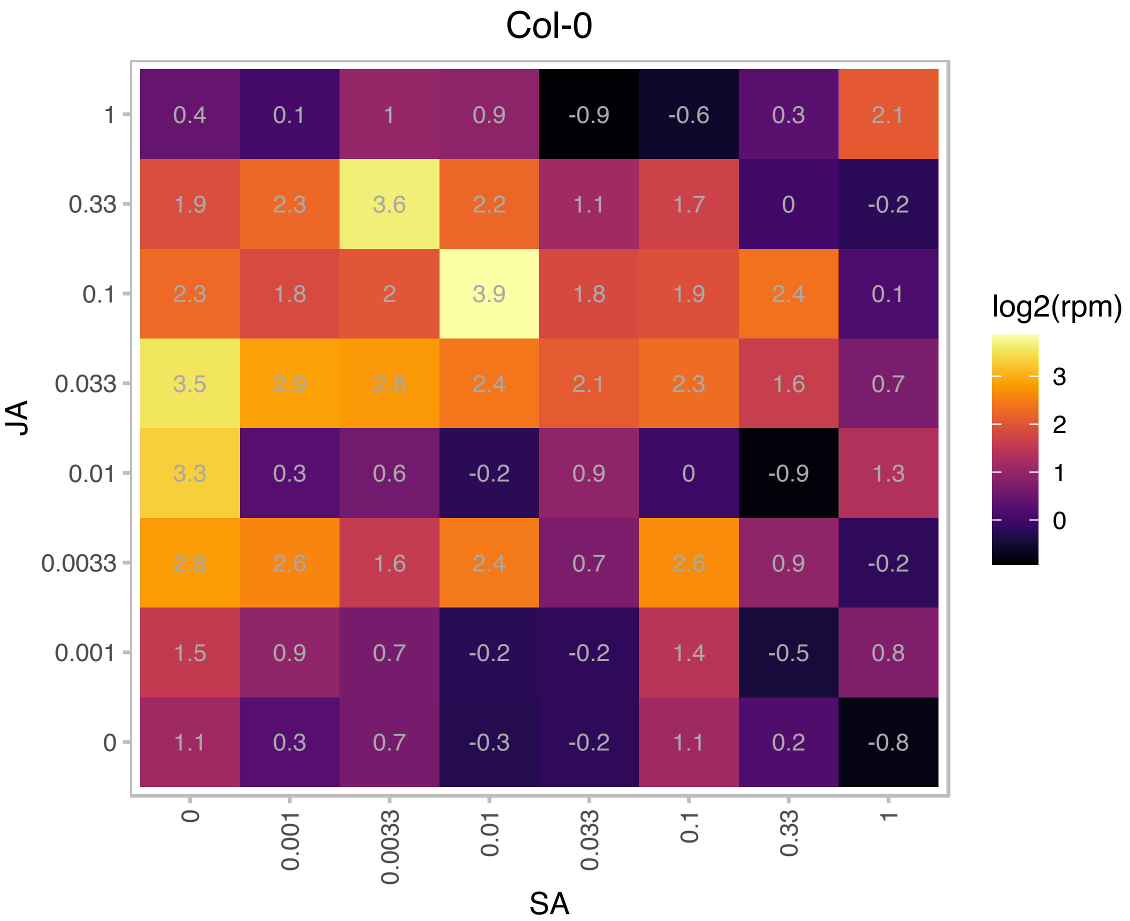


| | |
|---|---|
| Gene | AT5G14700 |
| is.expressed_genes | TRUE |
| is.responsive_genes | TRUE |
| is.apcluster | E-03 |

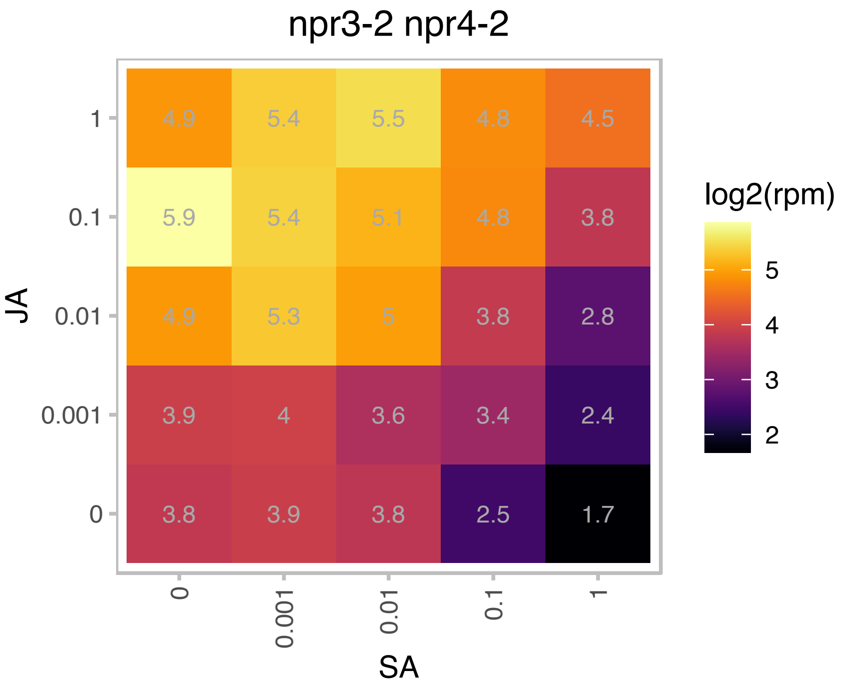


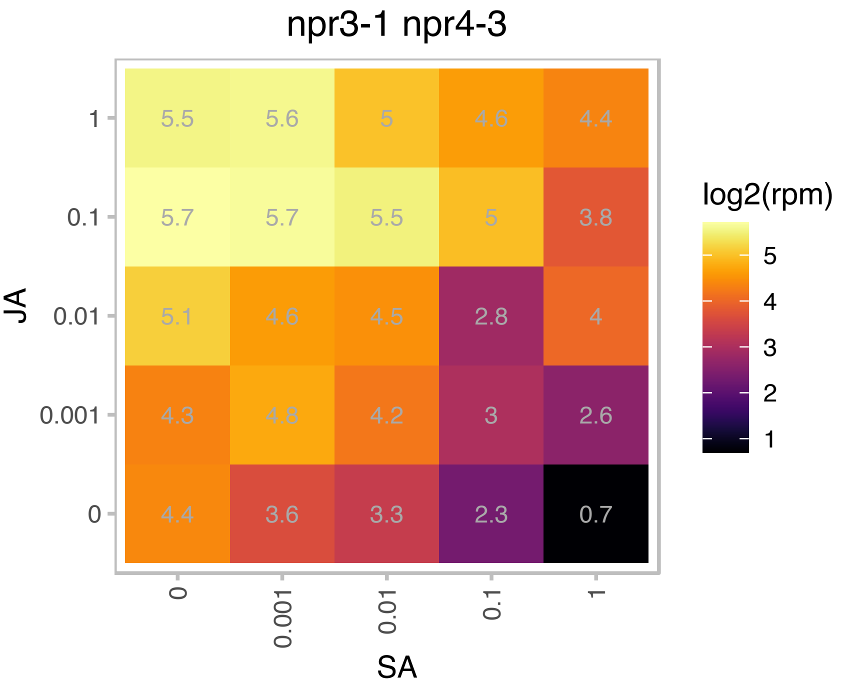

### AT5G24290.1 : MEB2

Vacuolar iron transporter (VIT) family protein

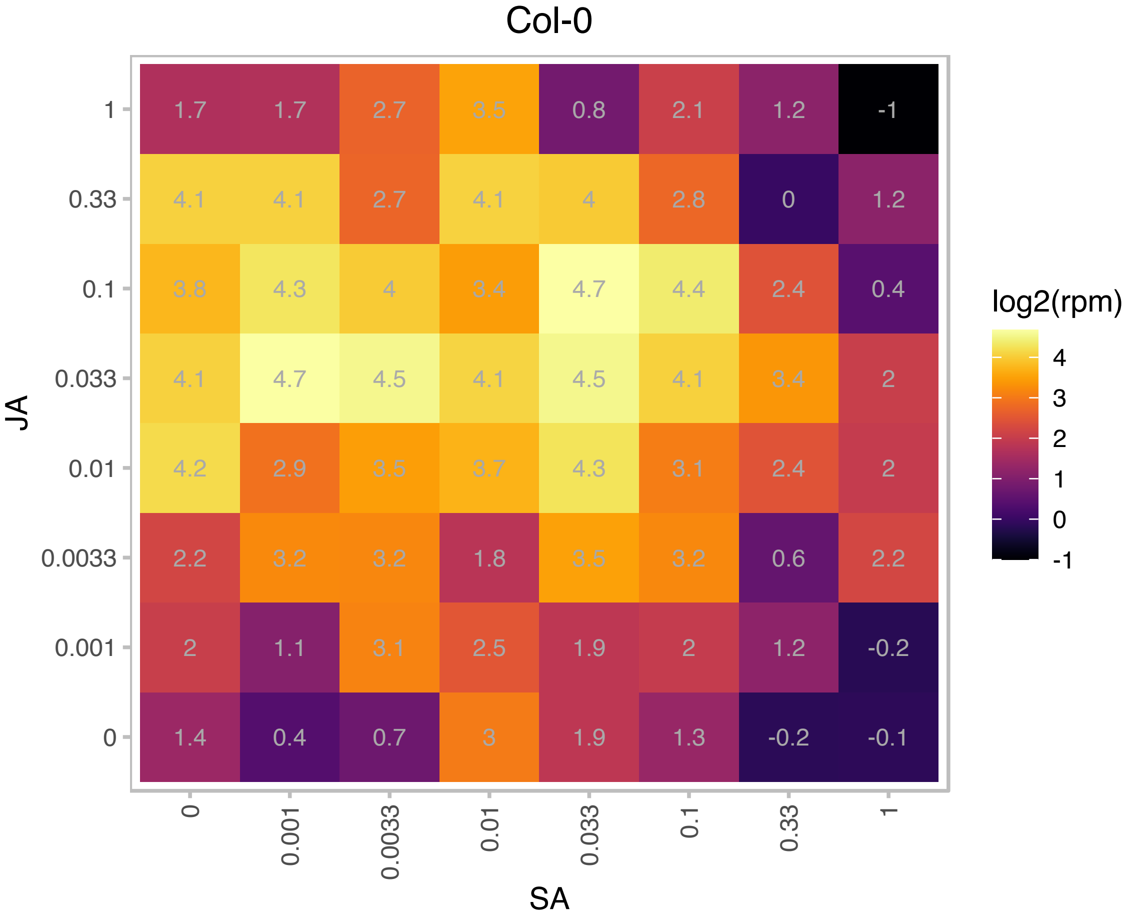


| | |
|---|---|
| Gene | AT5G24290 |
| is.expressed_genes | TRUE |
| is.responsive_genes | TRUE |
| is.apcluster | E-03 |

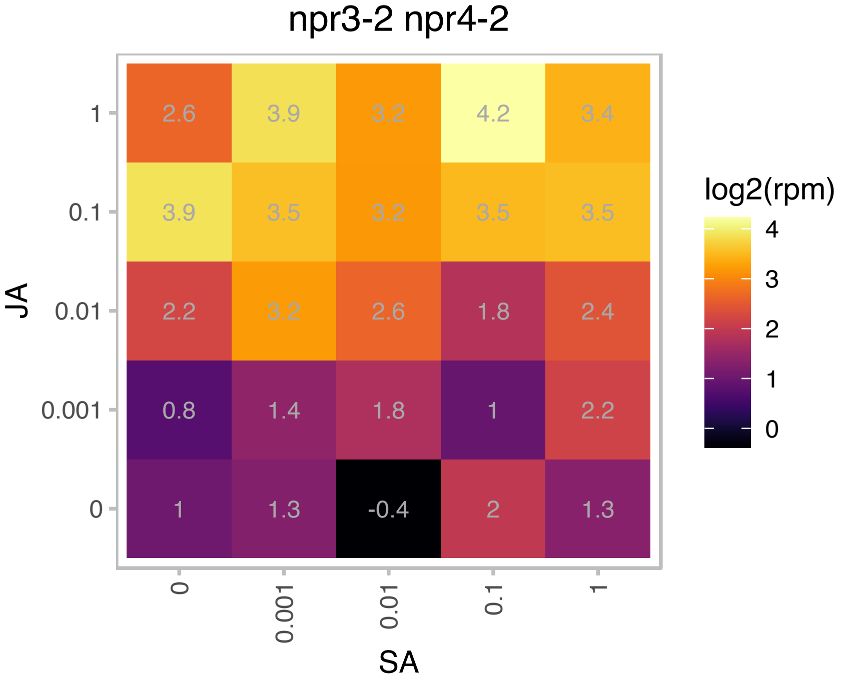


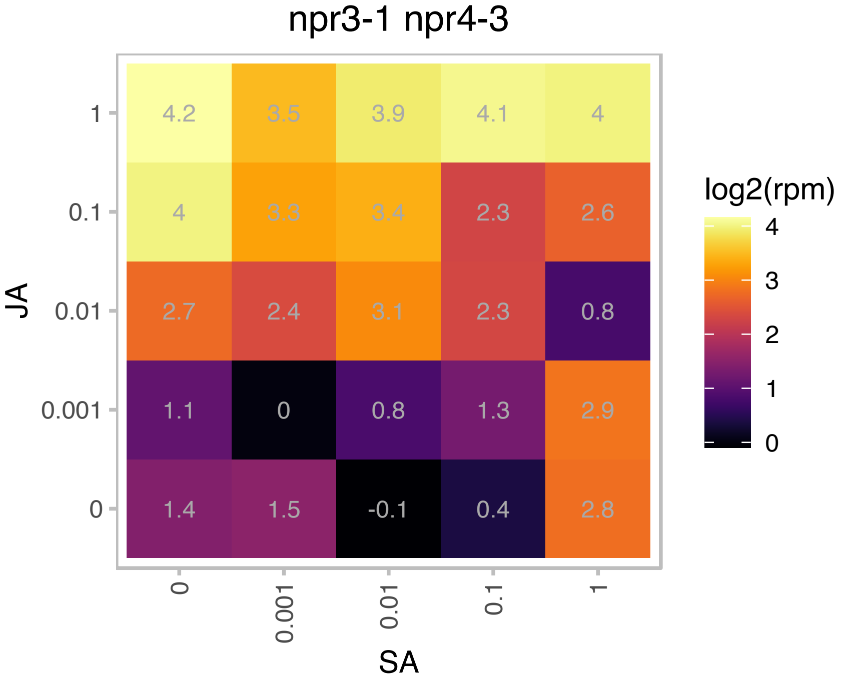

### AT5G24420.1 : PGL5

6-phosphogluconolactonase 5

Encodes a cytosolic 6-phosphogluconolactonase (PGL) thought to be involved in the oxidative pentose-phosphate pathway (OPPP).

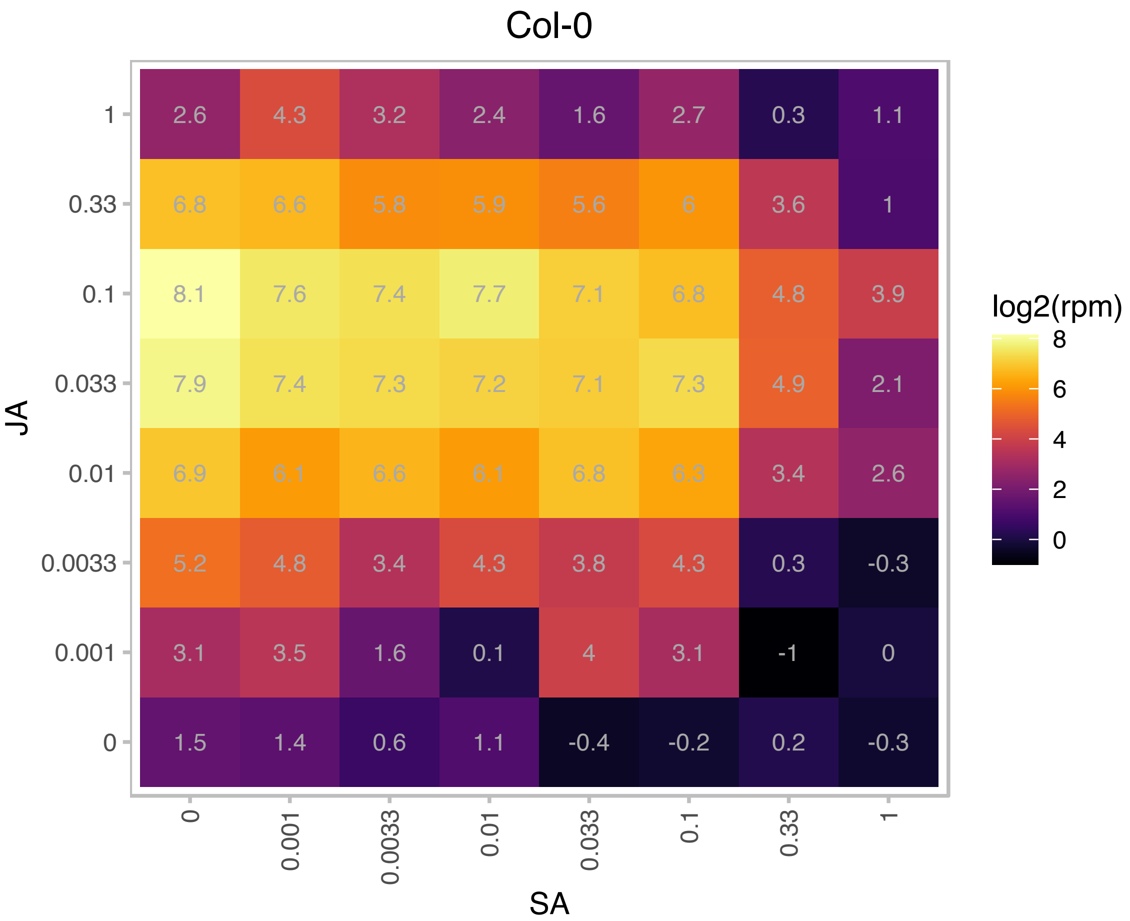


| | |
|---|---|
| Gene | AT5G24420 |
| is.expressed_genes | TRUE |
| is.responsive_genes | TRUE |
| is.apcluster | E-03 |

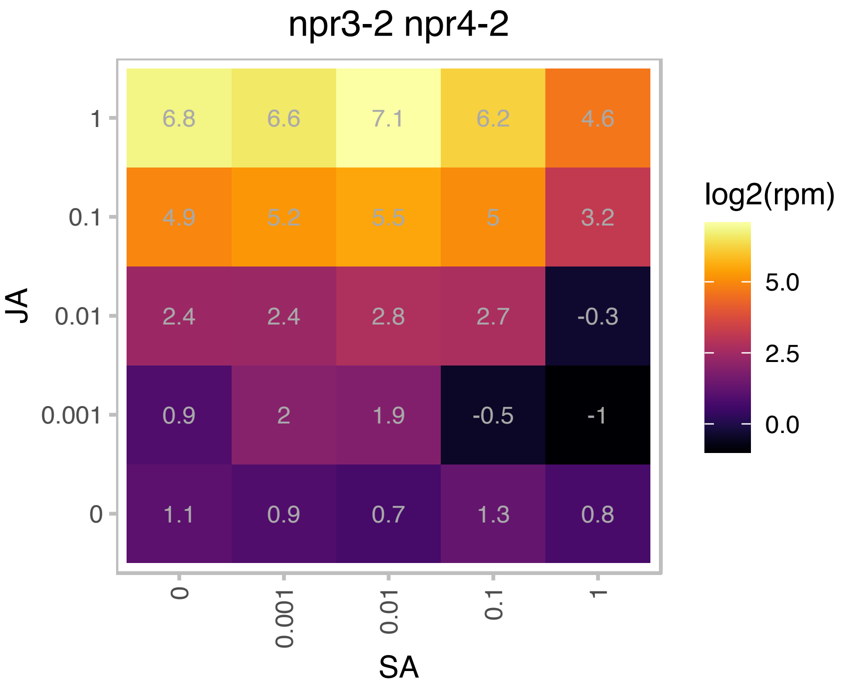


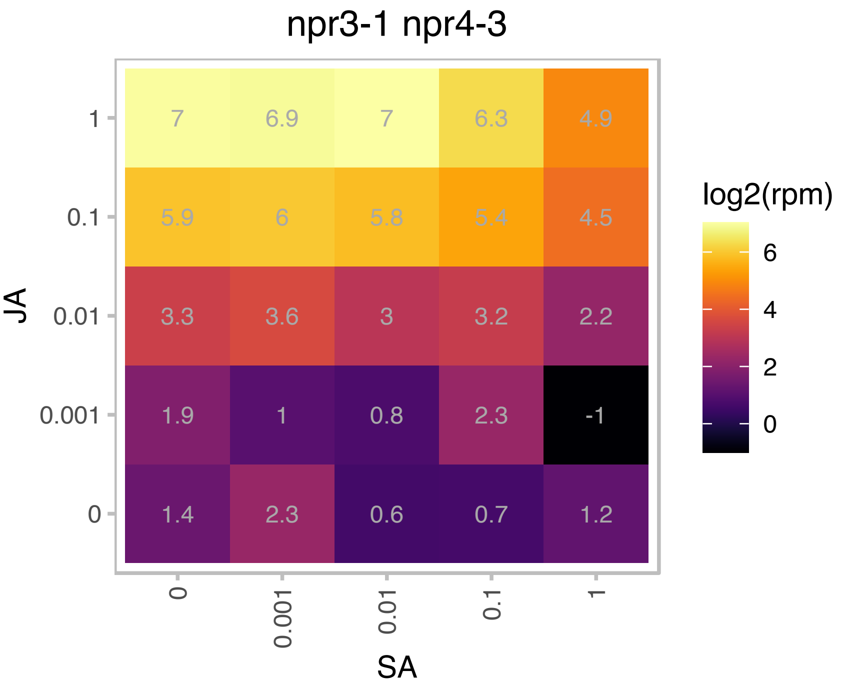

### AT5G24780.1 : ATVSP1, VSP1

vegetative storage protein 1

encodes an acid phosphatase similar to soybean vegetative storage proteins. Gene expression is induced by wounding and jasmonic acid.

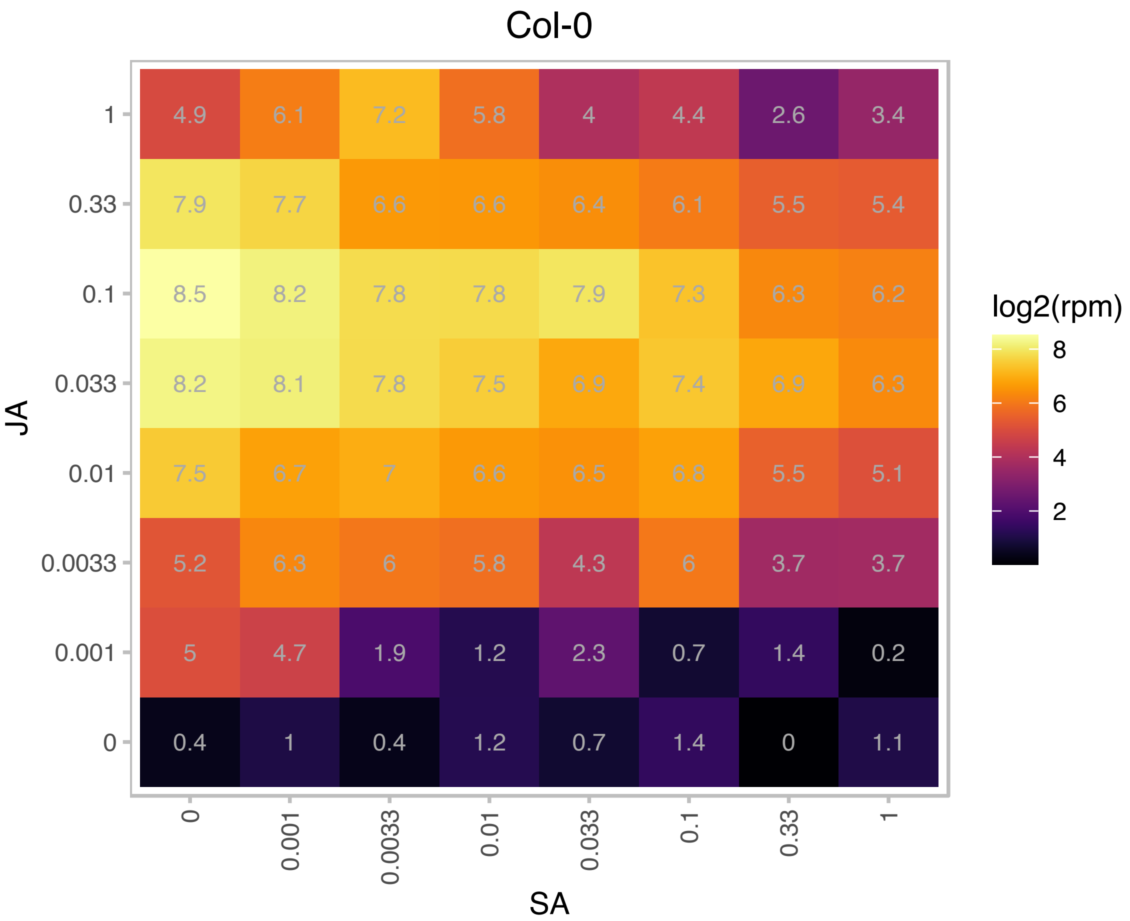


| | |
|---|---|
| Gene | AT5G24780 |
| is.expressed_genes | TRUE |
| is.responsive_genes | TRUE |
| is.apcluster | E-03 |

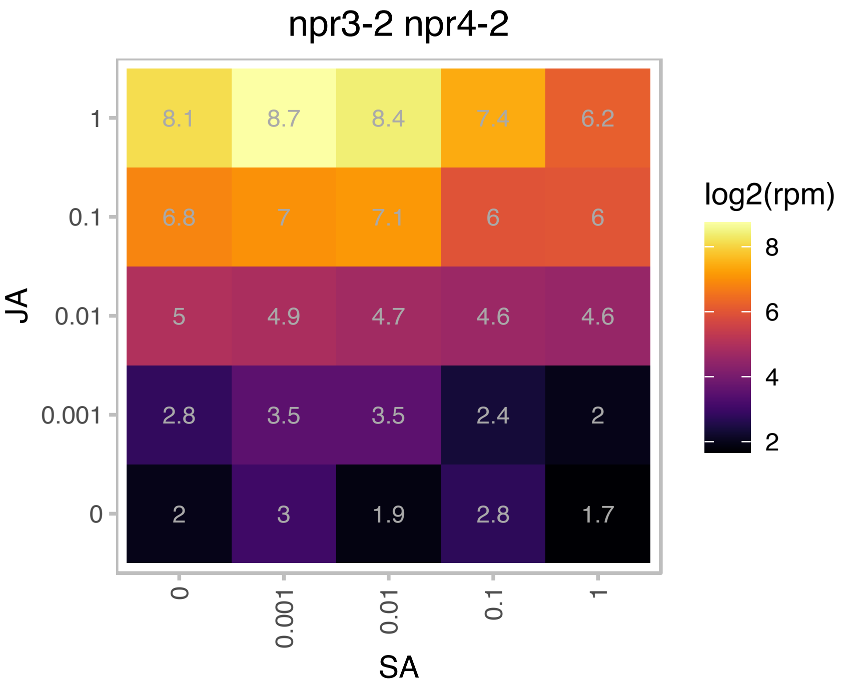


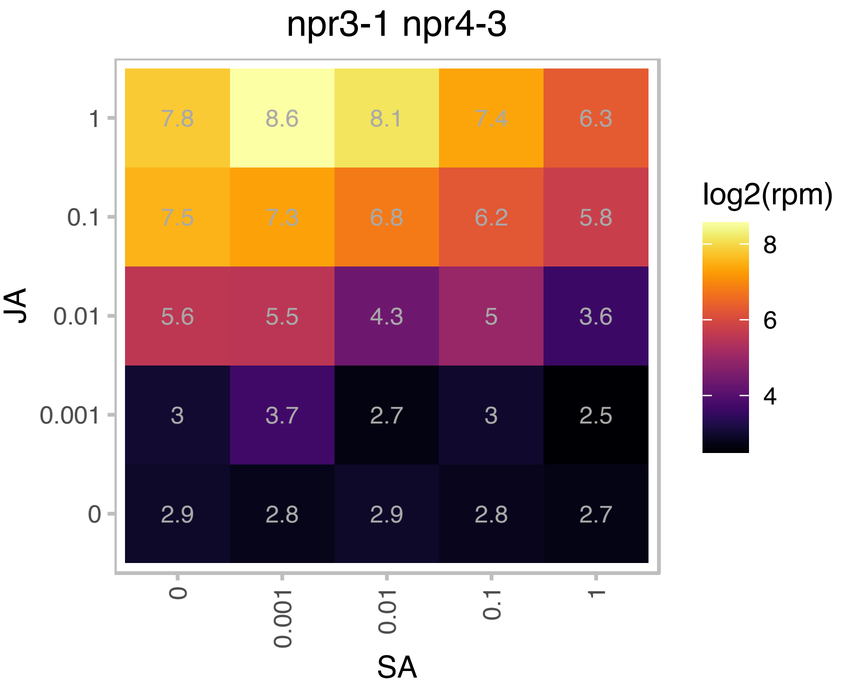

# AT5G41970.1 : NA

Metal-dependent protein hydrolase

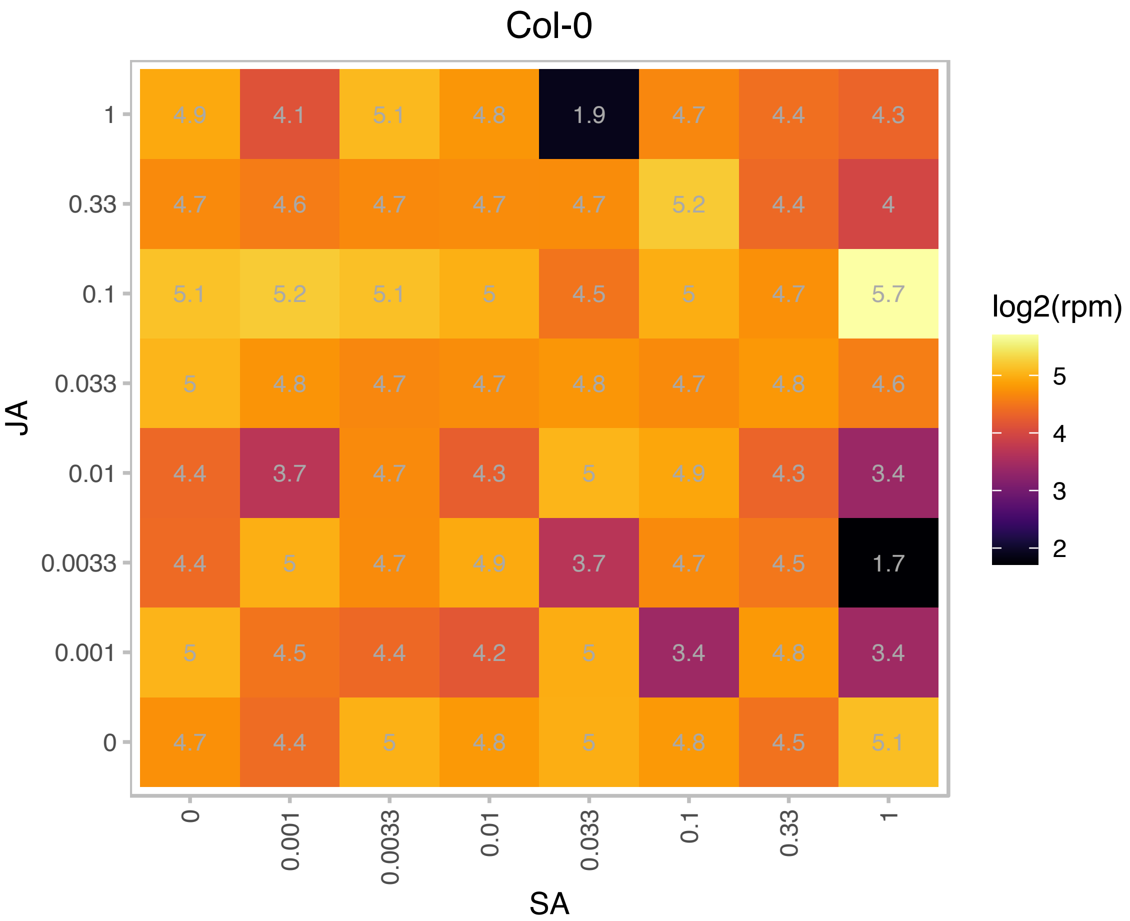


| | |
|---|---|
| Gene | AT5G41970 |
| is.expressed_genes | TRUE |
| is.responsive_genes | TRUE |
| is.apcluster | E-03 |

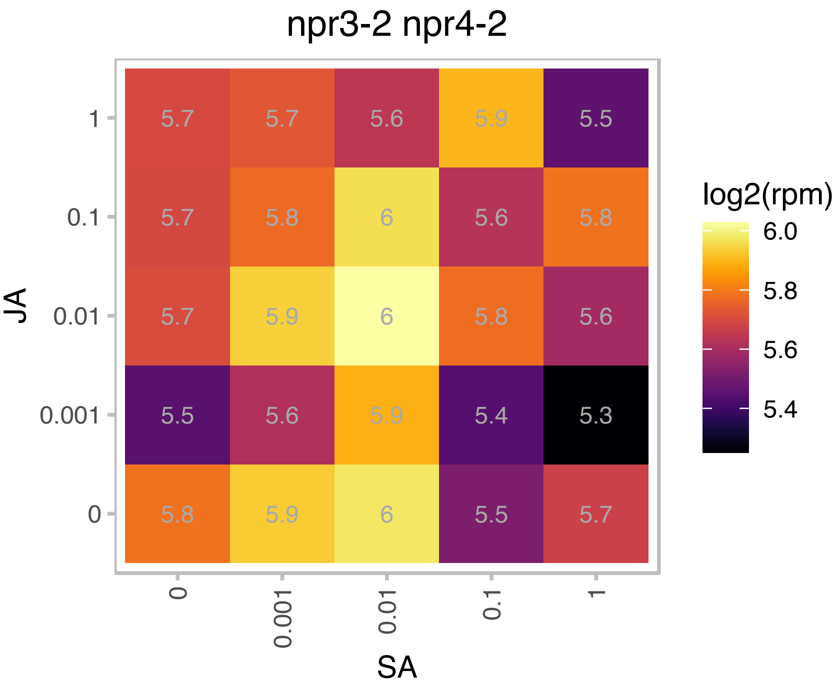


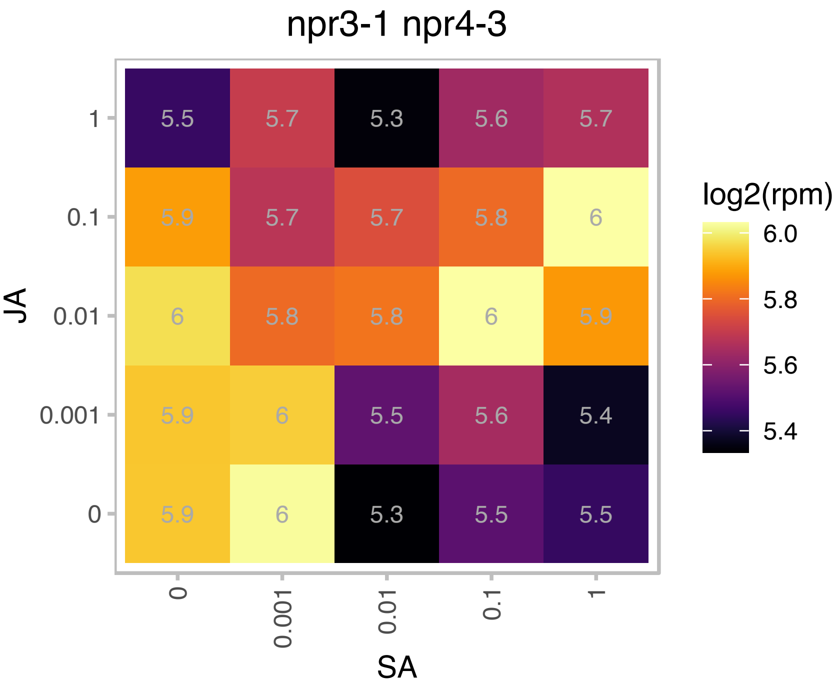

# AT5G44720.1 : NA

Molybdenum cofactor sulfurase family protein

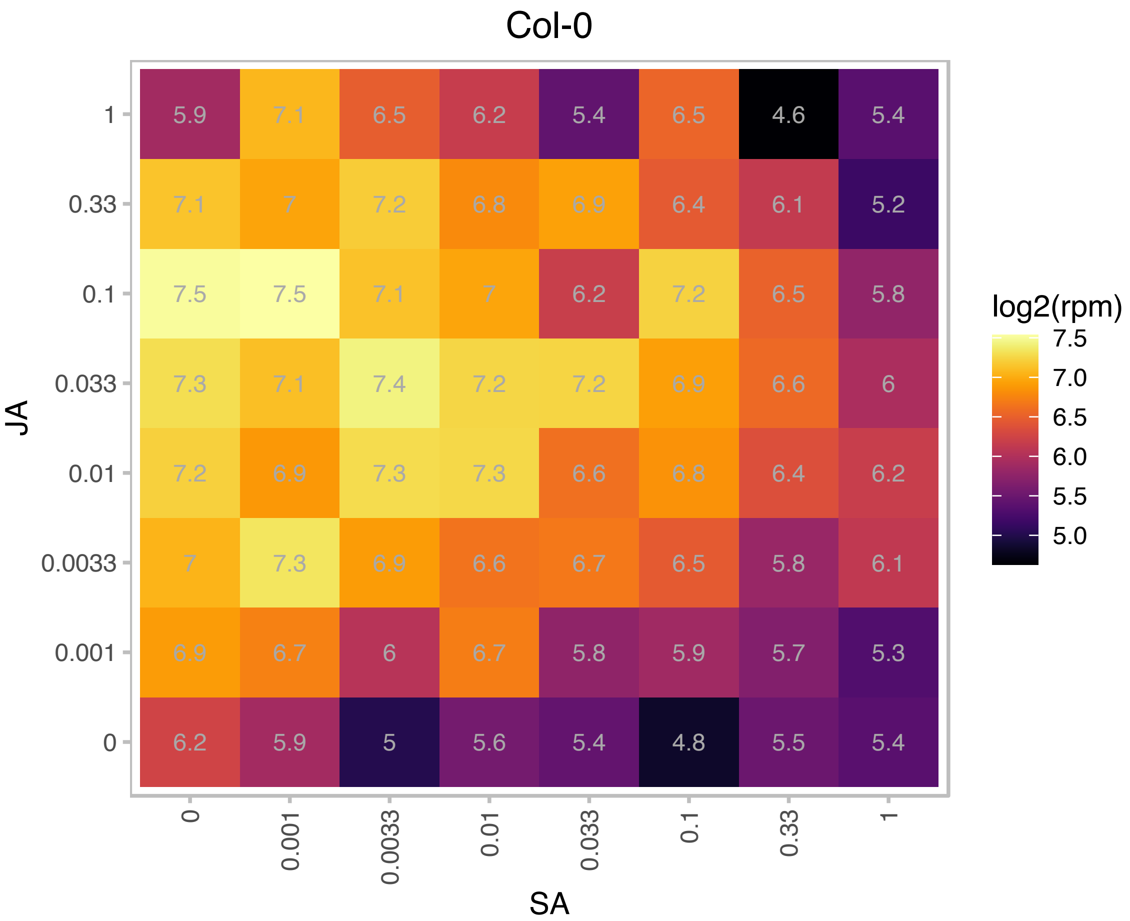


| | |
|---|---|
| Gene | AT5G44720 |
| is.expressed_genes | TRUE |
| is.responsive_genes | TRUE |
| is.apcluster | E-03 |

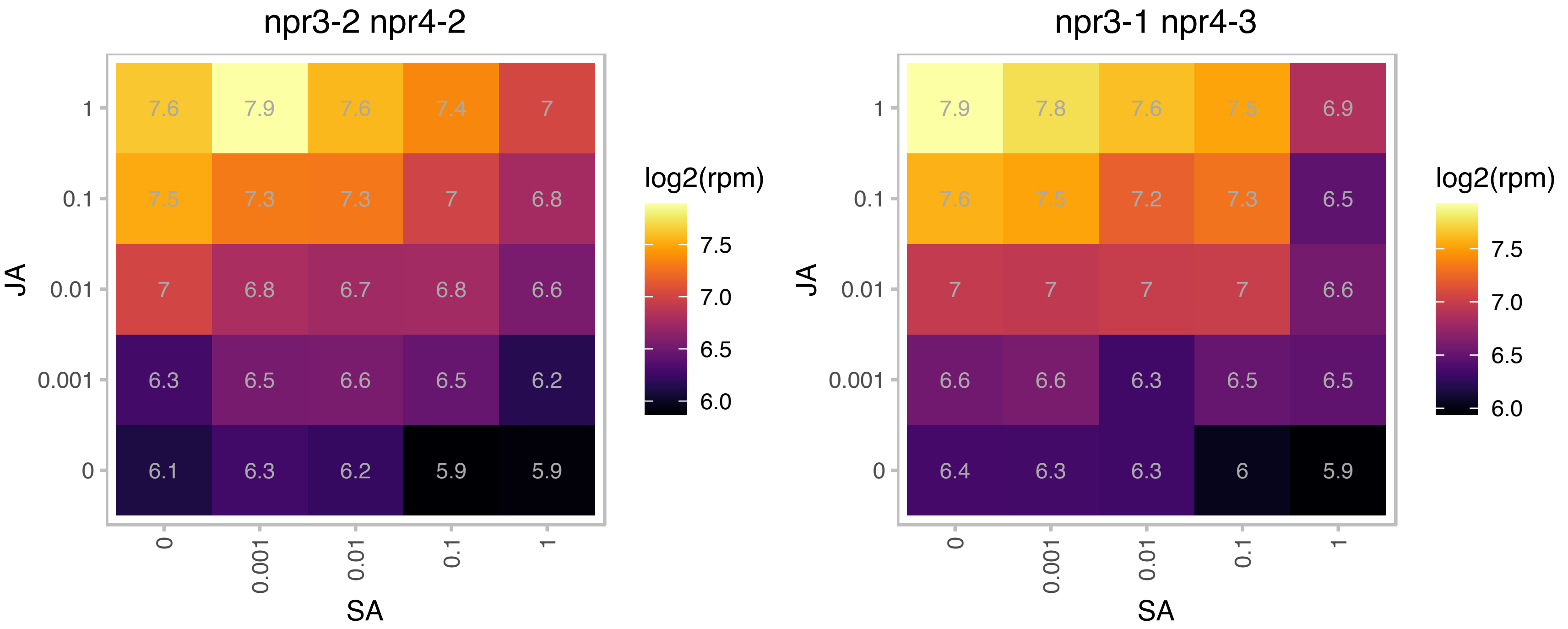

### AT5G60300.1 : DORN1, LecRK-I.9

Concanavalin A-like lectin protein kinase family protein

Lectin Receptor Kinase involved in protein-protein interactions with RGD-containing proteins as potential ligands. Role in cell wall-plasma membrane adhesion.  Involved in Phytophthora resistance.

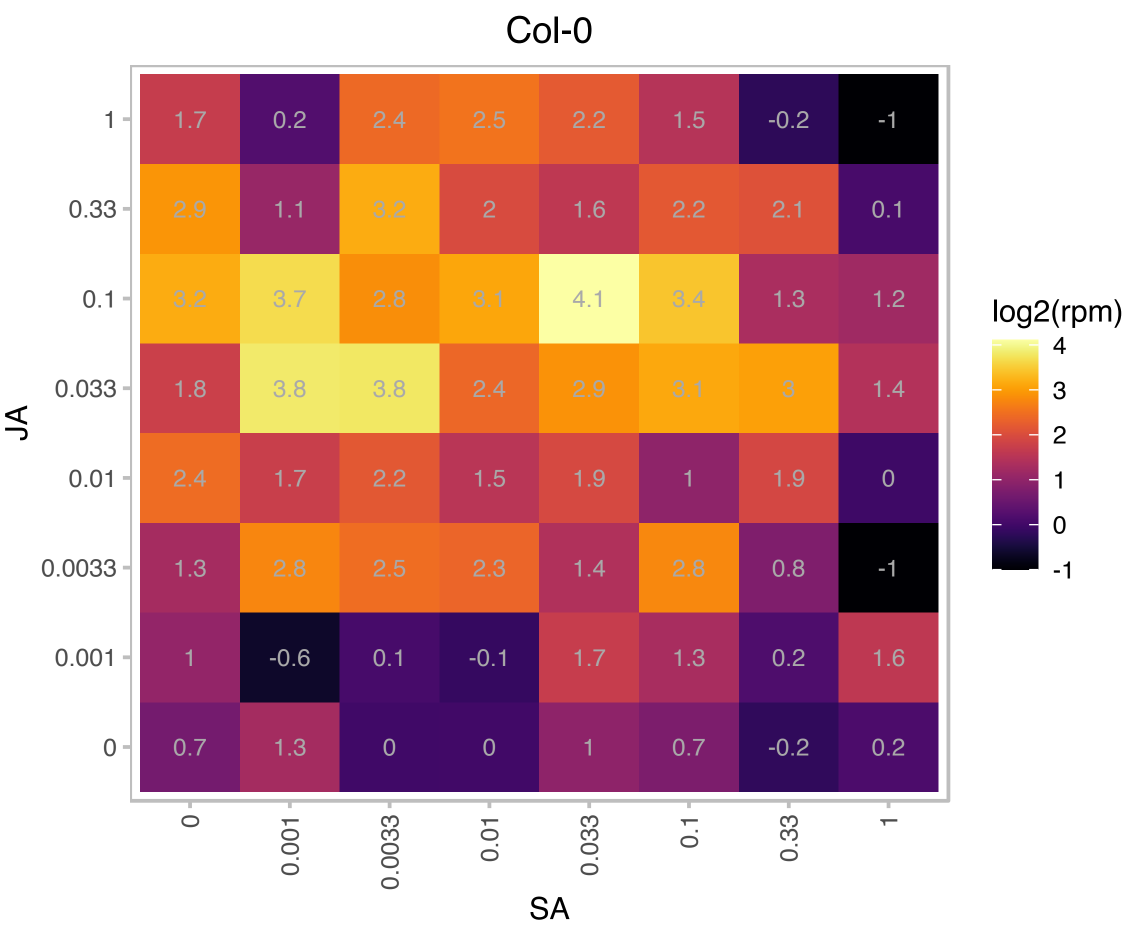


| | |
|---|---|
| Gene | AT5G60300 |
| is.expressed_genes | TRUE |
| is.responsive_genes | TRUE |
| is.apcluster | E-03 |

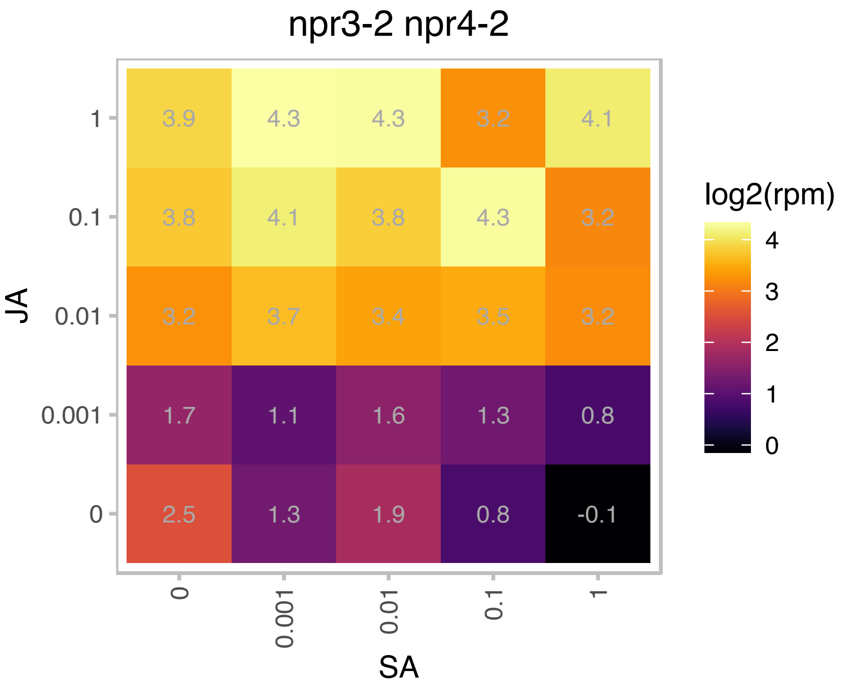


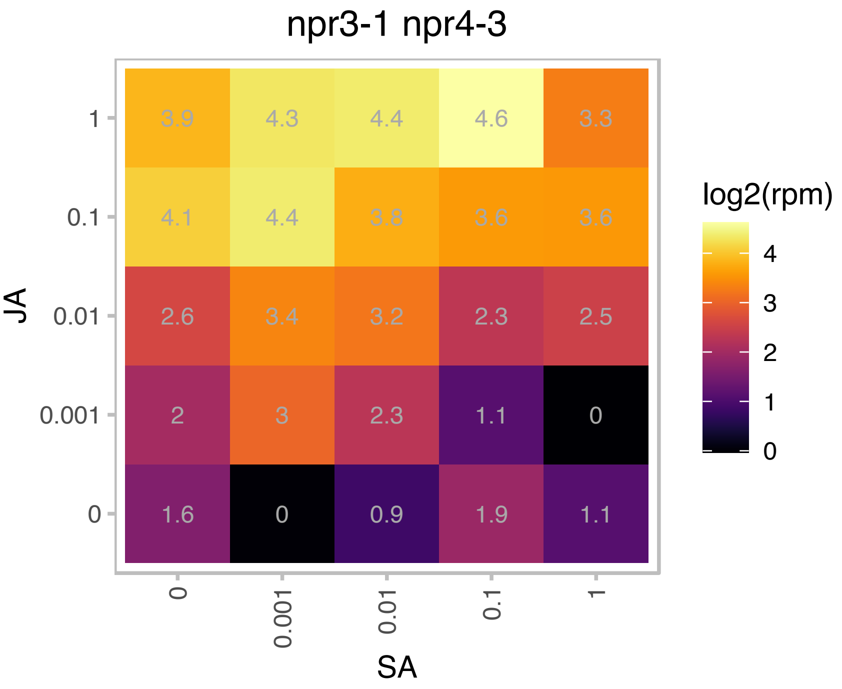

# AT5G63460.1 : NA

SAP domain-containing protein

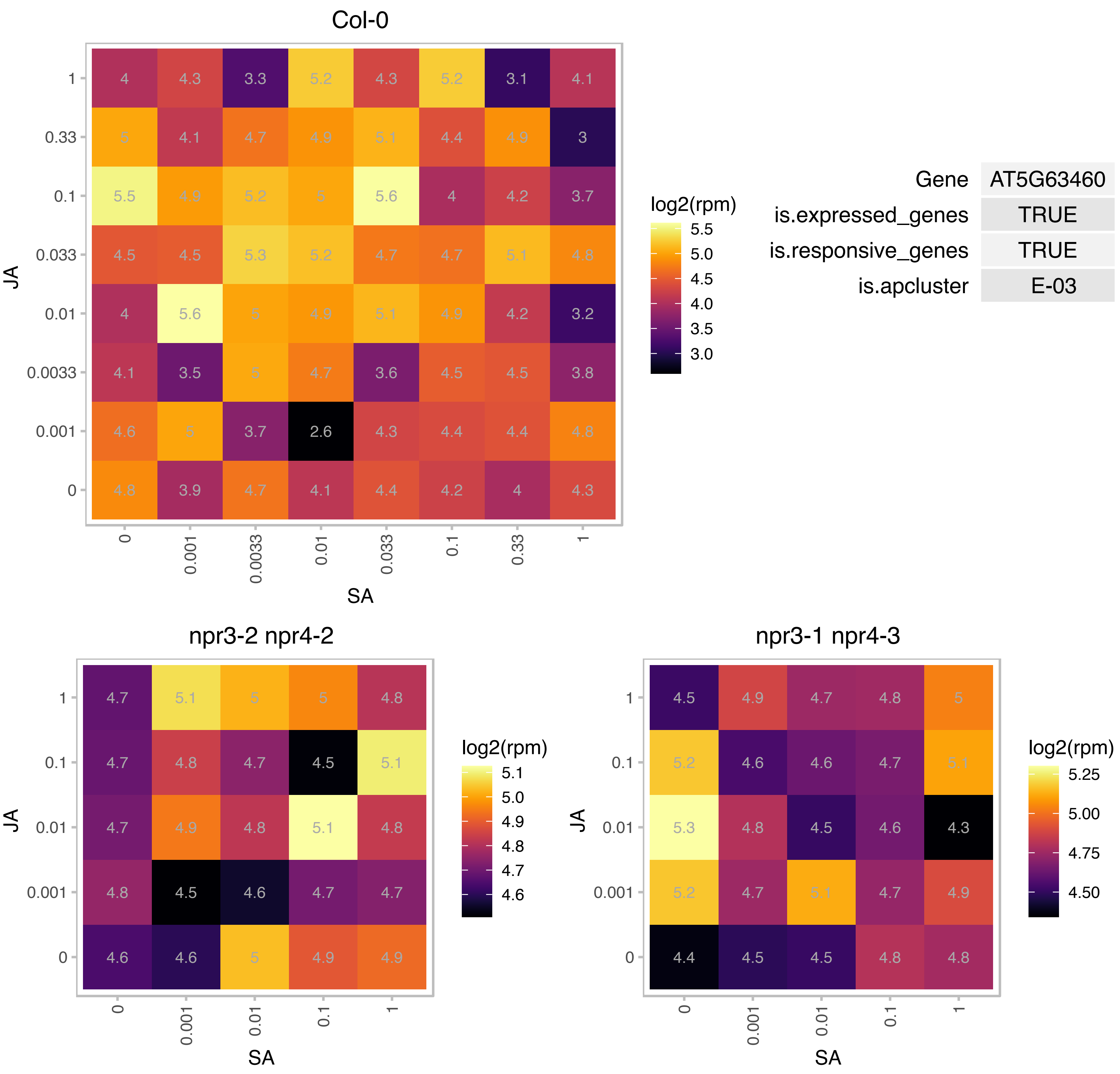

### AT5G67480.1 : ATBT4, BT4

BTB and TAZ domain protein 4

BTB and TAZ domain protein. Located in cytoplasm and expressed in fruit, flower and leaves.

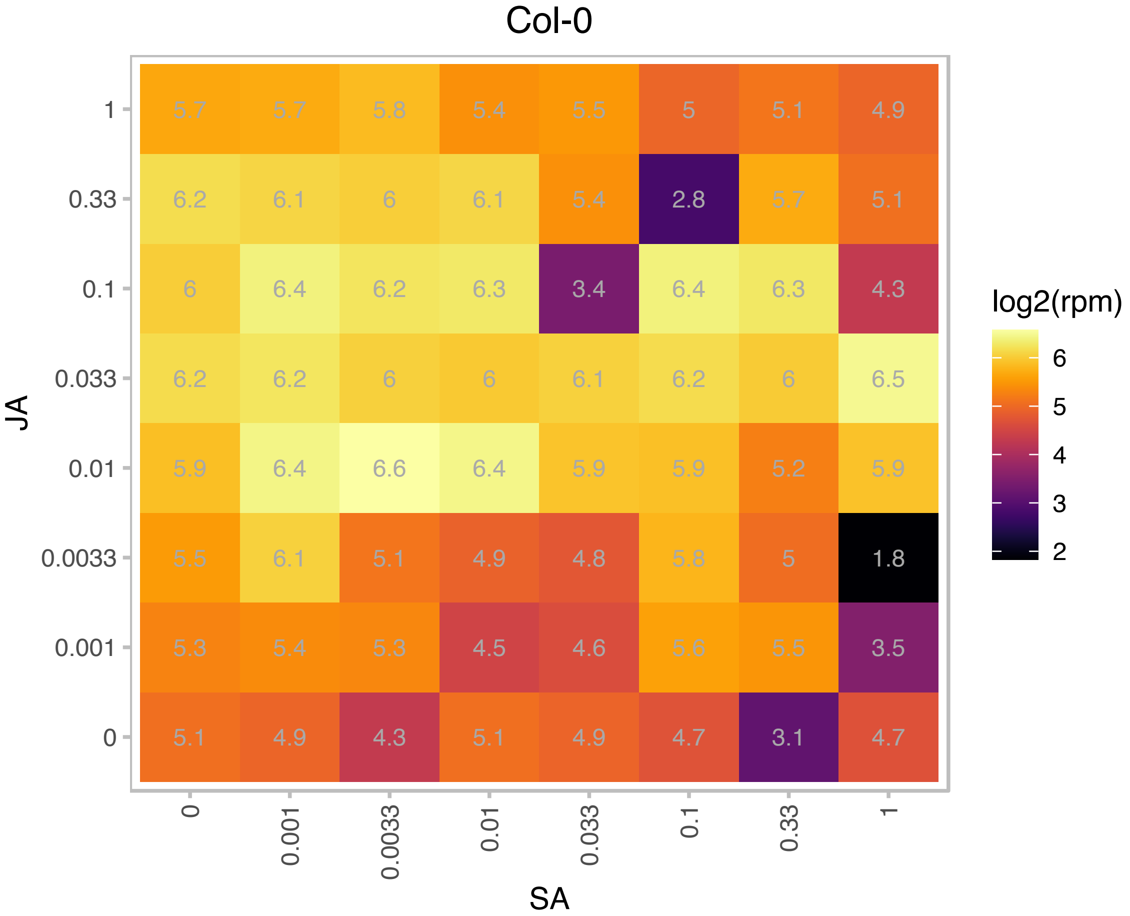


| | |
|---|---|
| Gene | AT5G67480 |
| is.expressed_genes | TRUE |
| is.responsive_genes | TRUE |
| is.apcluster | E-03 |

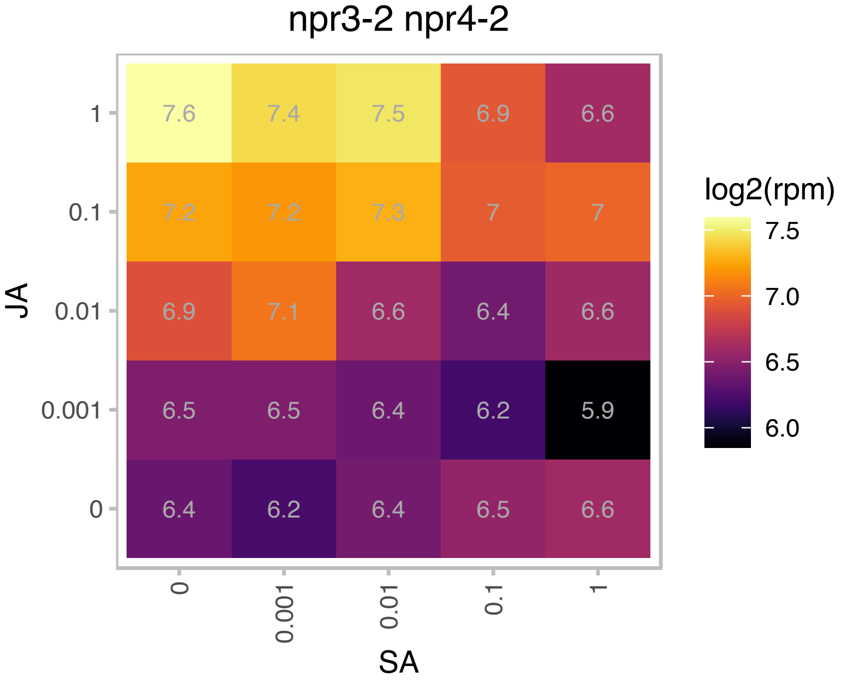


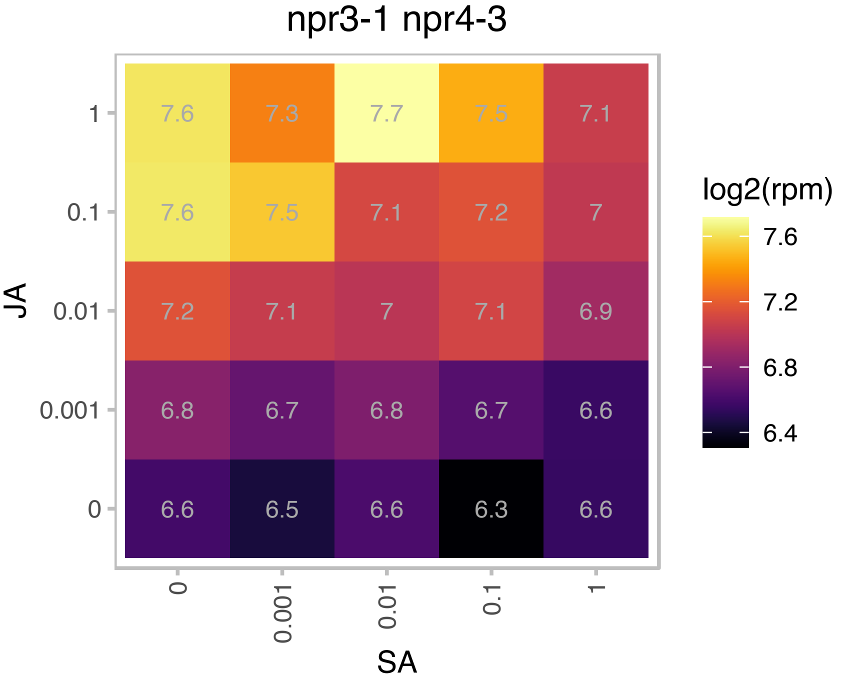

### AT1G15440.1 : ATPWP2, PWP2

periodic tryptophan protein 2

Encodes a nucleolar protein that is a ribosome biogenesis co-factor. Mutants display aberrant RNA processing and female gametophyte development.

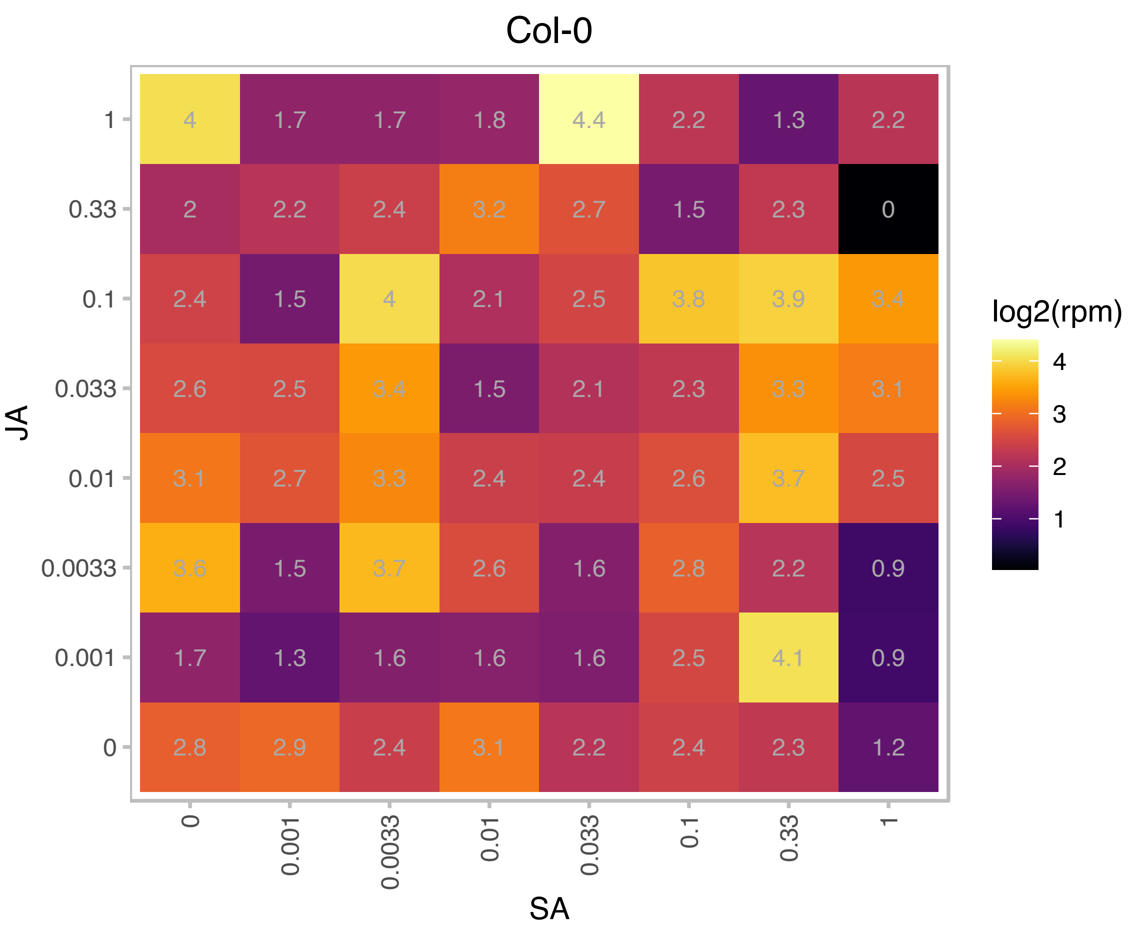


| | |
|---|---|
| Gene | AT1G15440 |
| is.expressed_genes | TRUE |
| is.responsive_genes | TRUE |
| is.apcluster | E-04 |

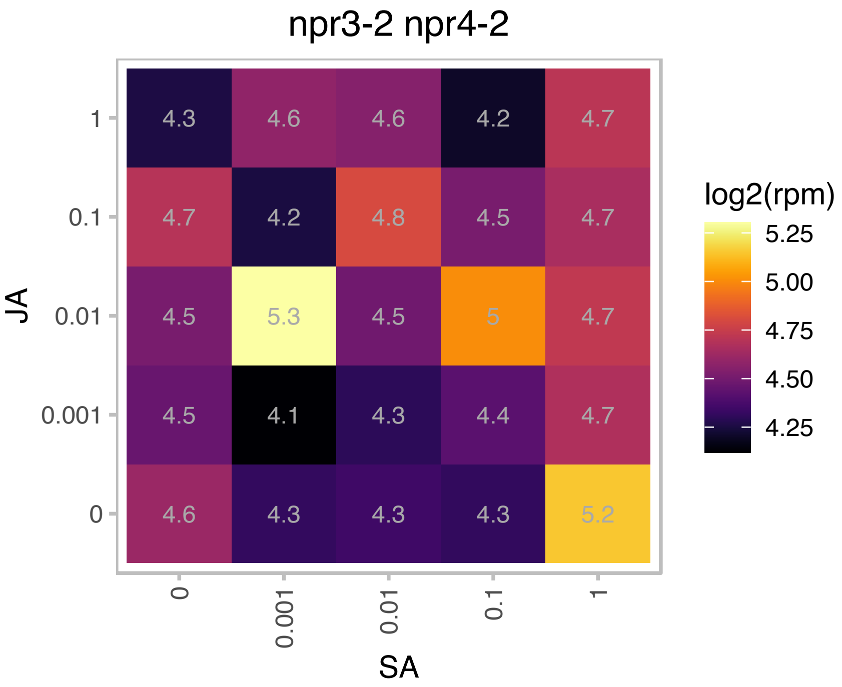


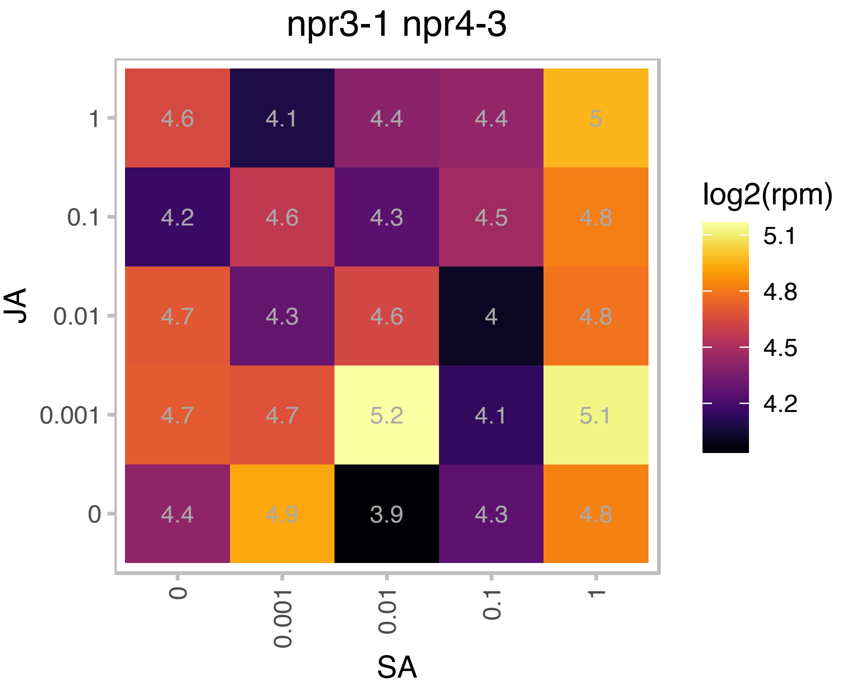

### AT1G16060.1 : ADAP, WRI3

ARIA-interacting double AP2 domain protein

Encodes ADAP, an AP2-domain protein that interacts with ARIA.  ADAP is a positive regulator of the ABA response and is also involved in regulating seedling growth.

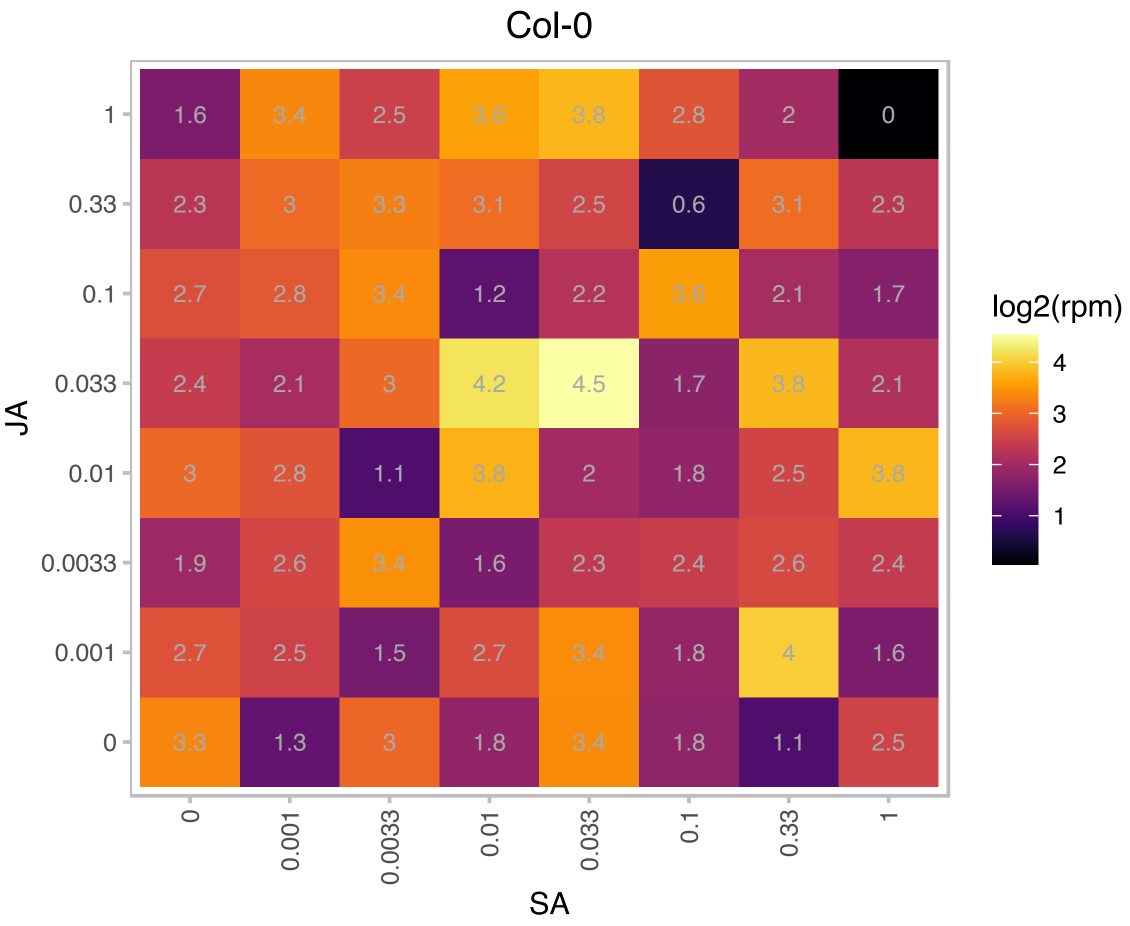


| | |
|---|---|
| Gene | AT1G16060 |
| is.expressed_genes | TRUE |
| is.responsive_genes | TRUE |
| is.apcluster | E-04 |

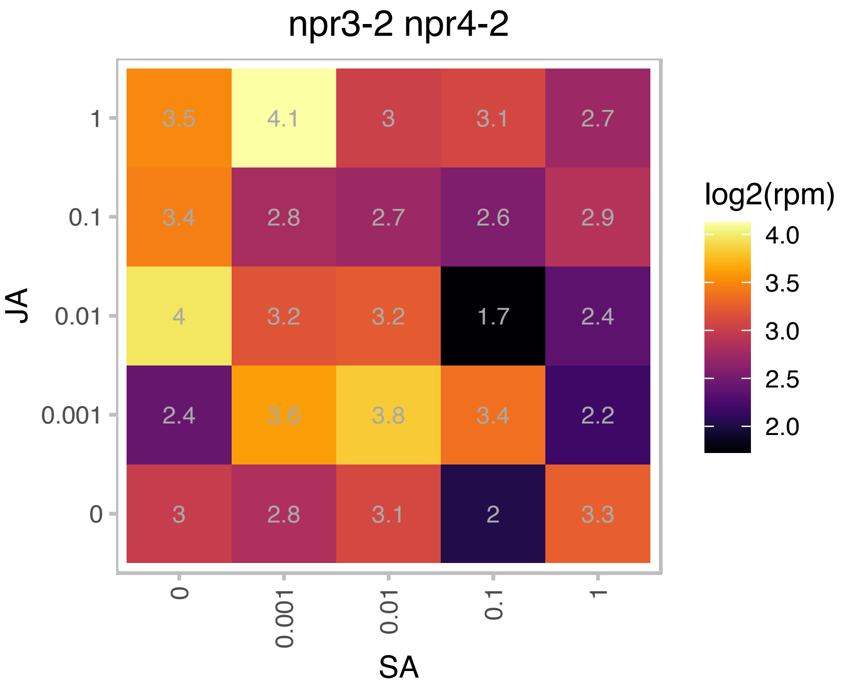


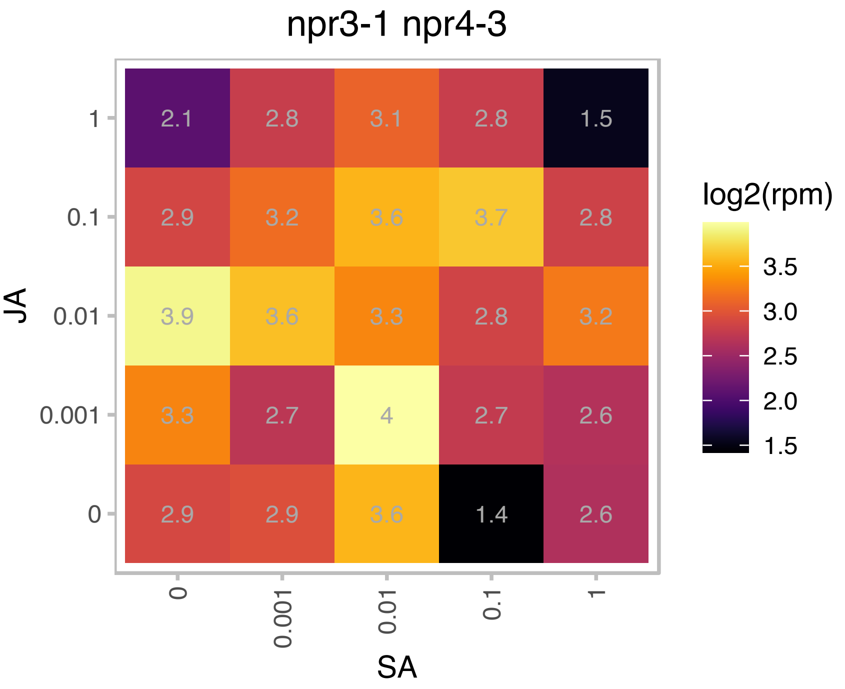

# AT1G16430.1 : NA

Surfeit locus protein 5 subunit 22 of Mediator complex

| | |
|---|---|
| Gene | AT1G16430 |
| is.expressed_genes | TRUE |
| is.responsive_genes | TRUE |
| is.apcluster | E-04 |

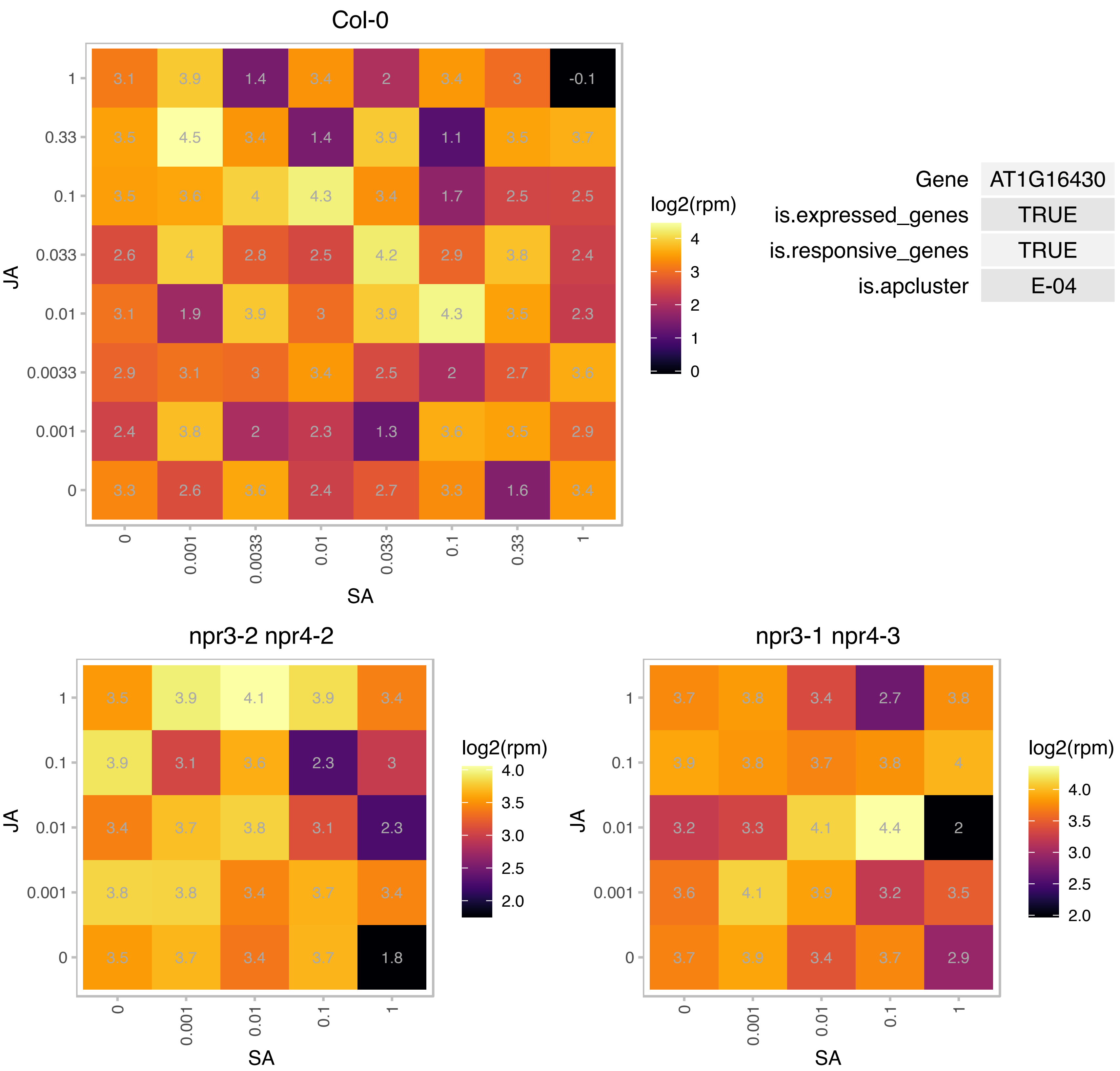

# AT1G16870.1 : NA

mitochondrial 28S ribosomal protein S29-related

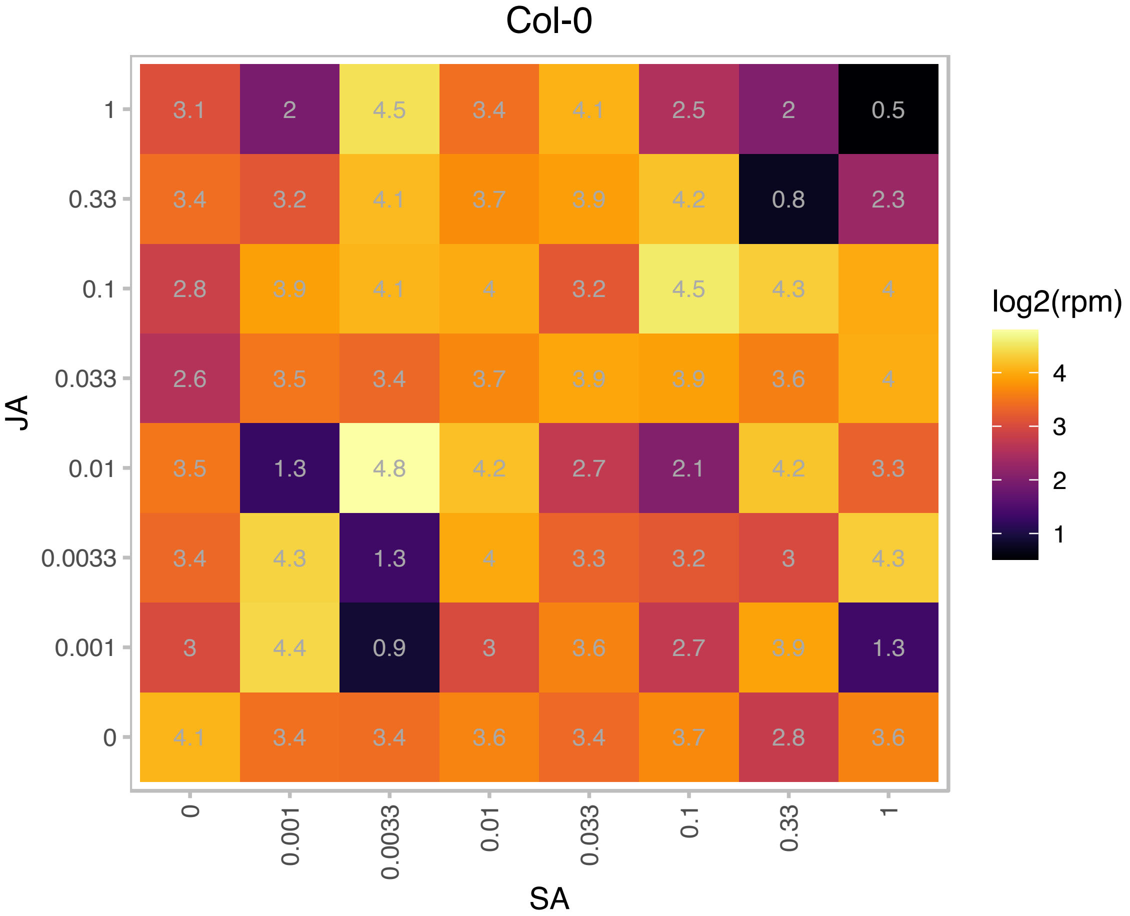


| | |
|---|---|
| Gene | AT1G16870 |
| is.expressed_genes | TRUE |
| is.responsive_genes | TRUE |
| is.apcluster | E-04 |

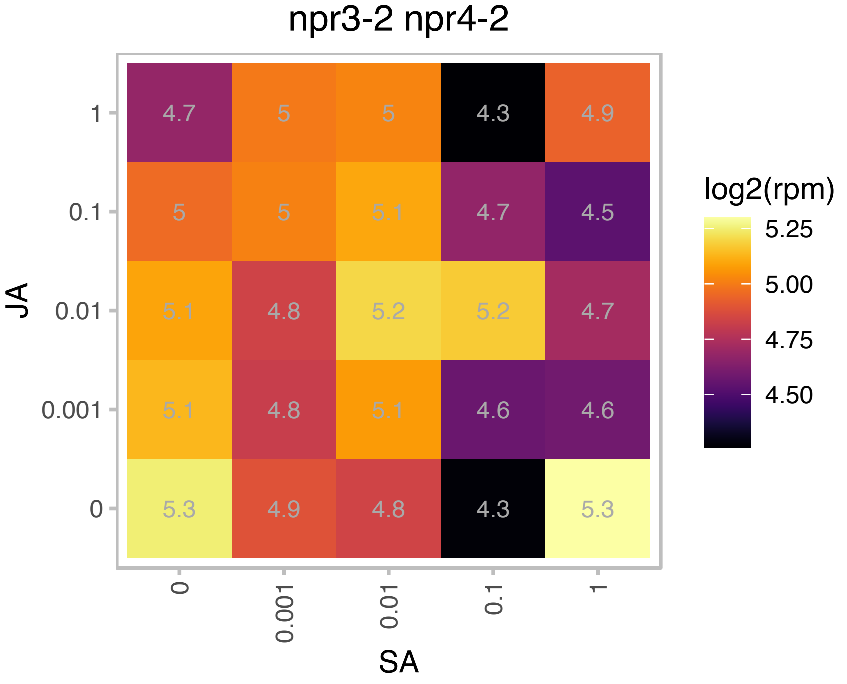


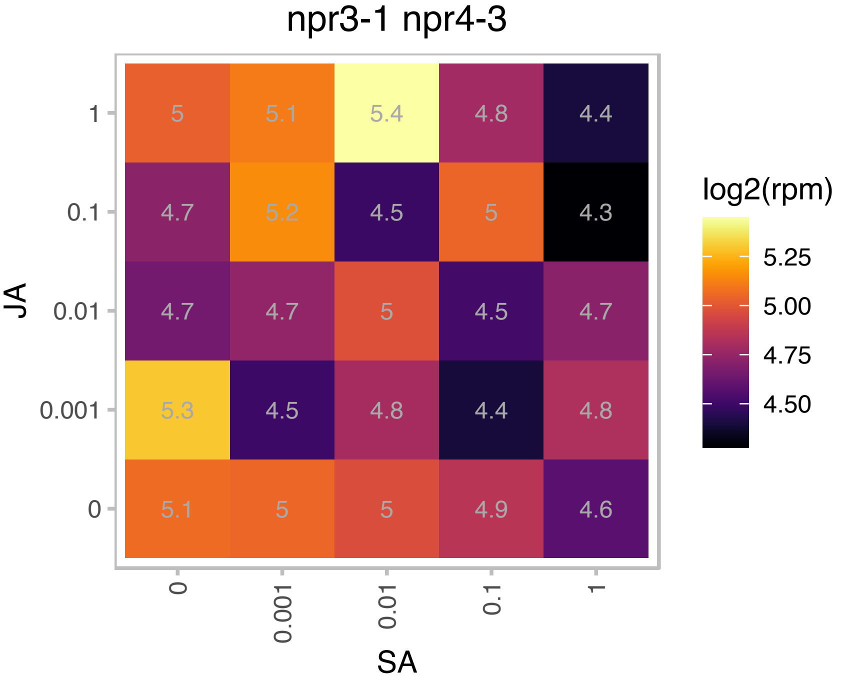

### AT1G26470.1 : SNS1

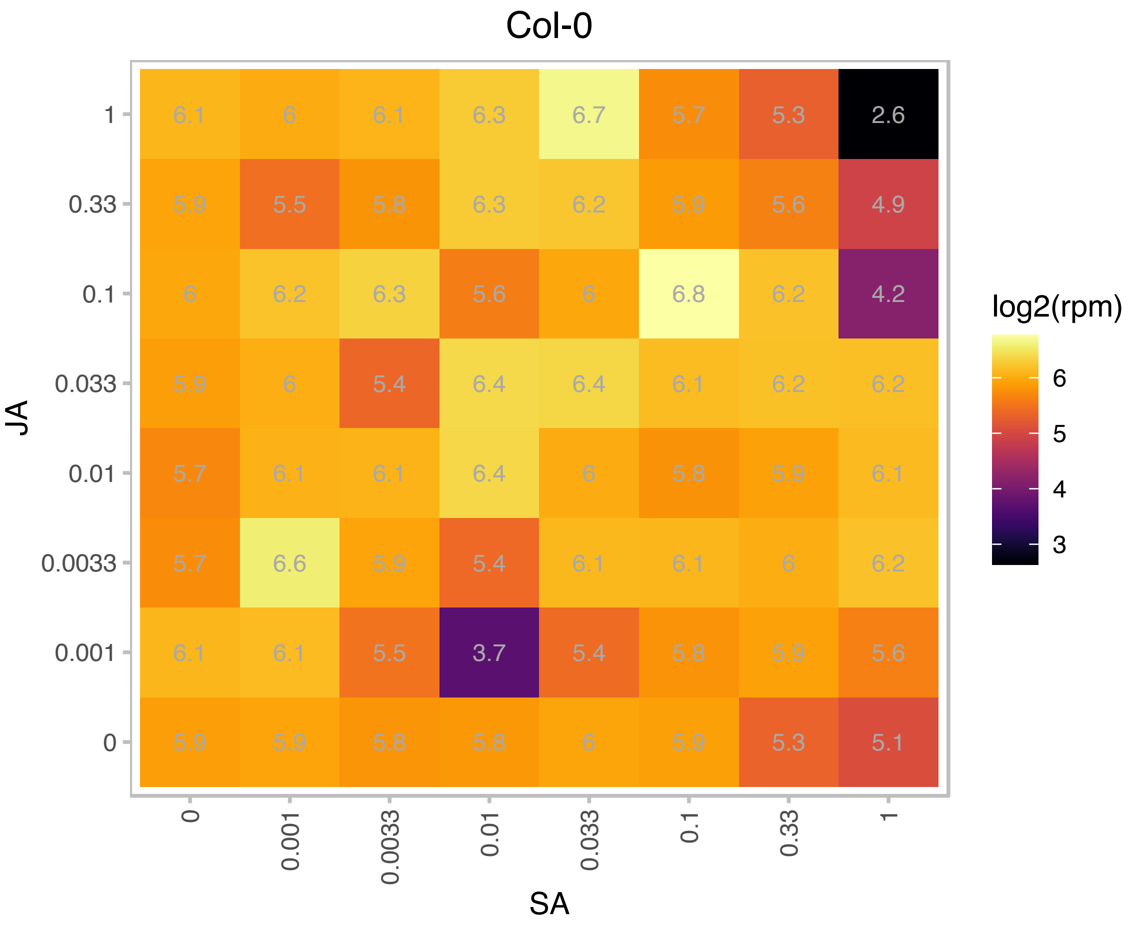


| | |
|---|---|
| Gene | AT1G26470 |
| is.expressed_genes | TRUE |
| is.responsive_genes | TRUE |
| is.apcluster | E-04 |

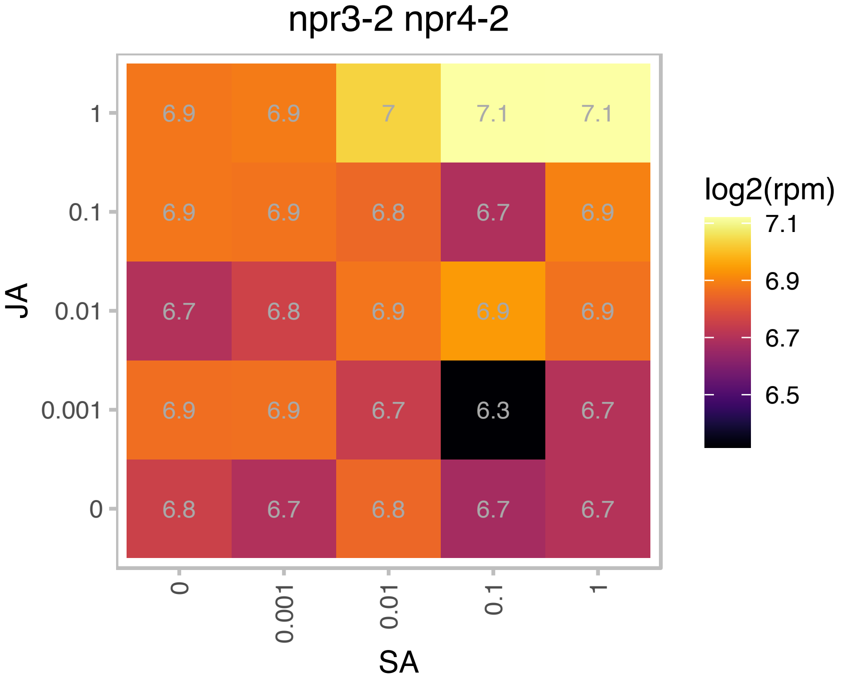


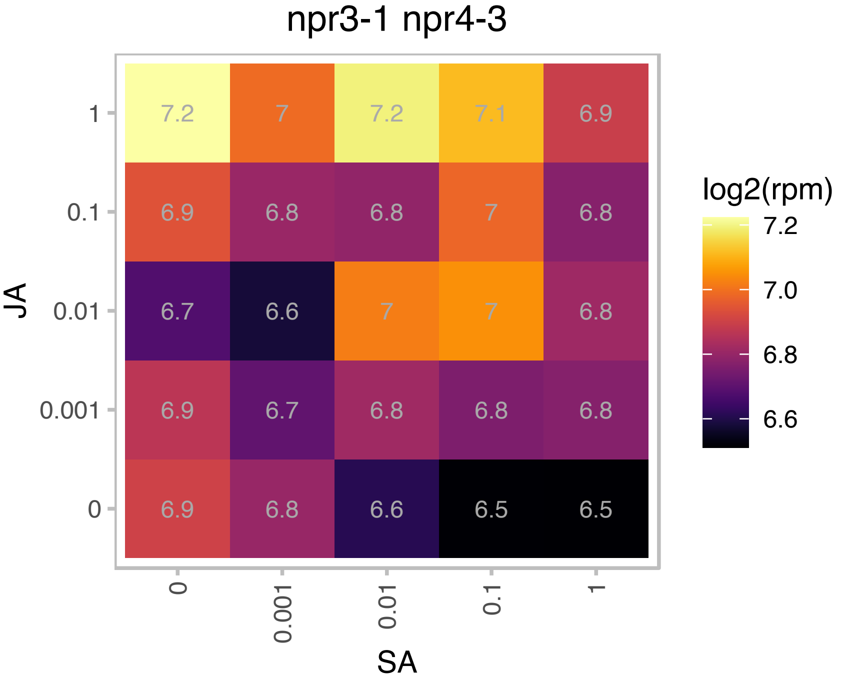

### AT1G30330.1 : ARF6

auxin response factor 6
Encodes a member of the auxin response factor family. Mediates auxin response via expression of auxin regulated genes. Acts redundantly with ARF8 to control stamen elongation and flower maturation. Expression of ARF6 is controlled by miR167.

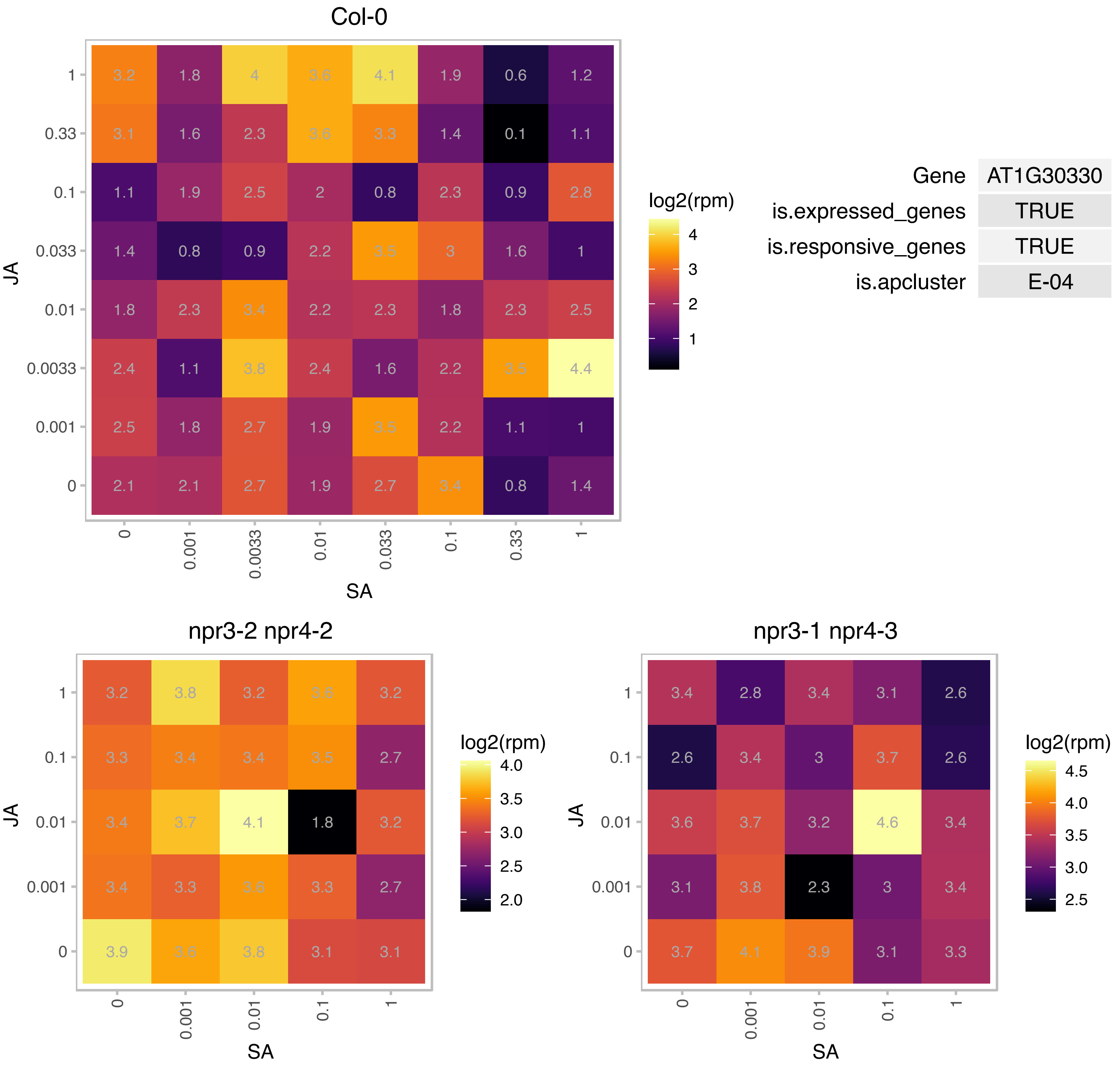


| | |
|---|---|
| Gene | AT1G30330 |
| is.expressed_genes | TRUE |
| is.responsive_genes | TRUE |
| is.apcluster | E-04 |

### AT1G54630.1 : ACP3, AtACP3

acyl carrier protein 3

encodes an acyl carrier protein expressed in leaves, roots, and dry seeds. Gene expression is not regulated by light.

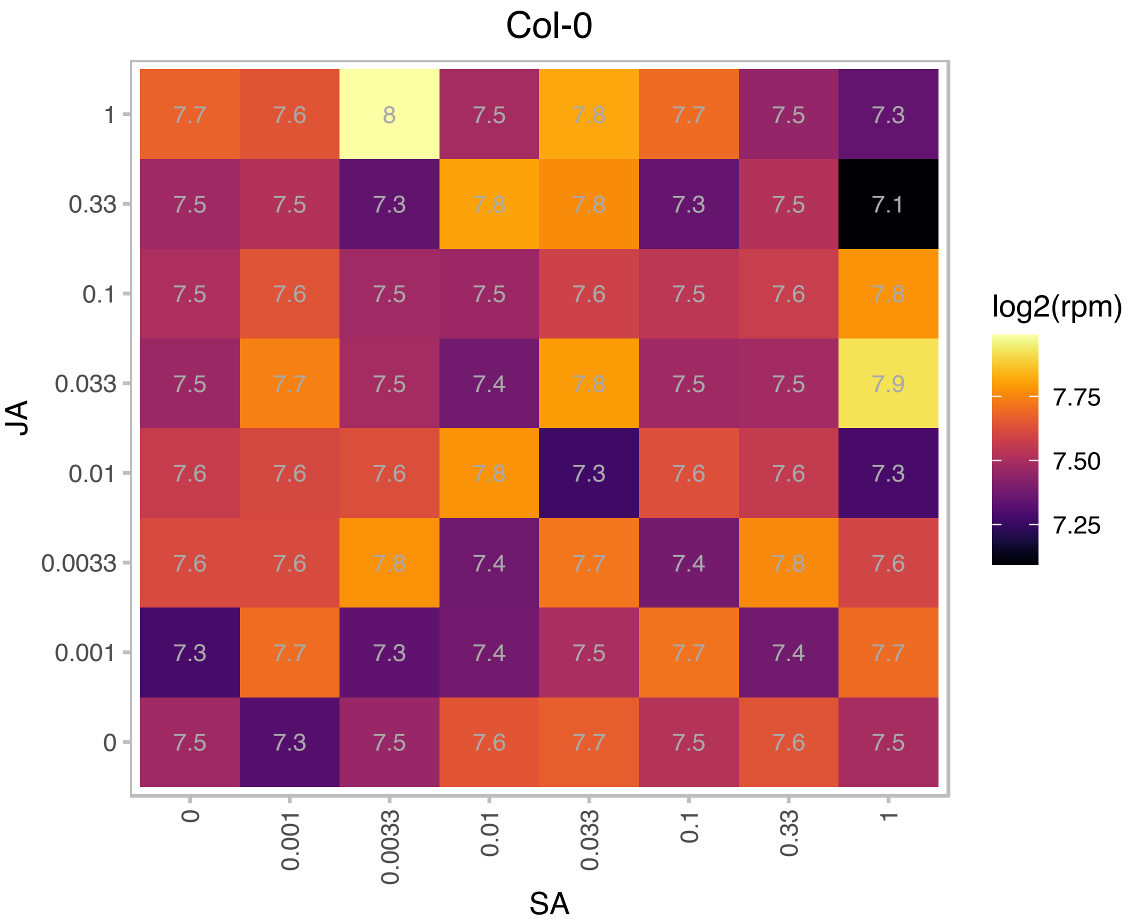


| | |
|---|---|
| Gene | AT1G54630 |
| is.expressed_genes | TRUE |
| is.responsive_genes | TRUE |
| is.apcluster | E-04 |

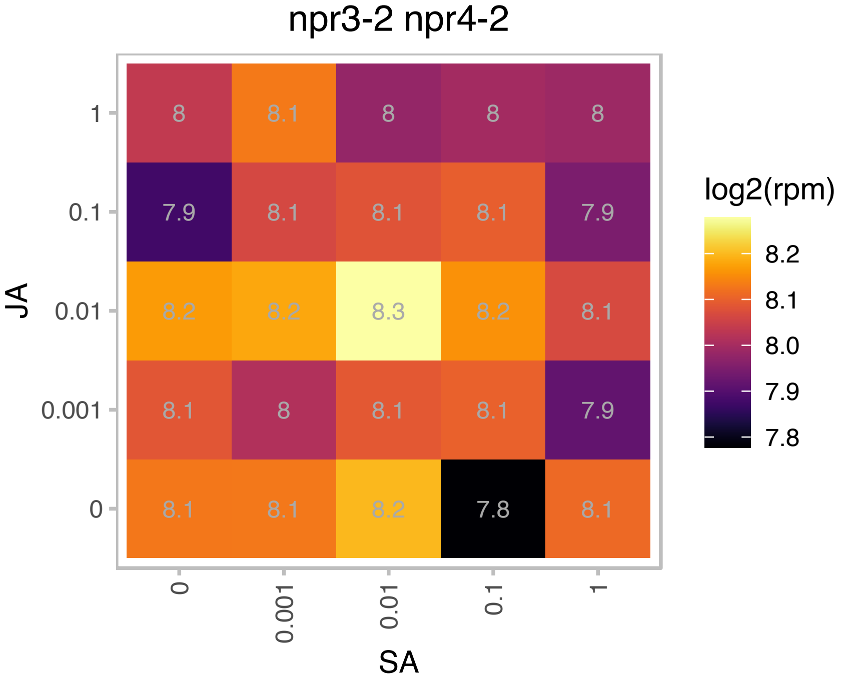


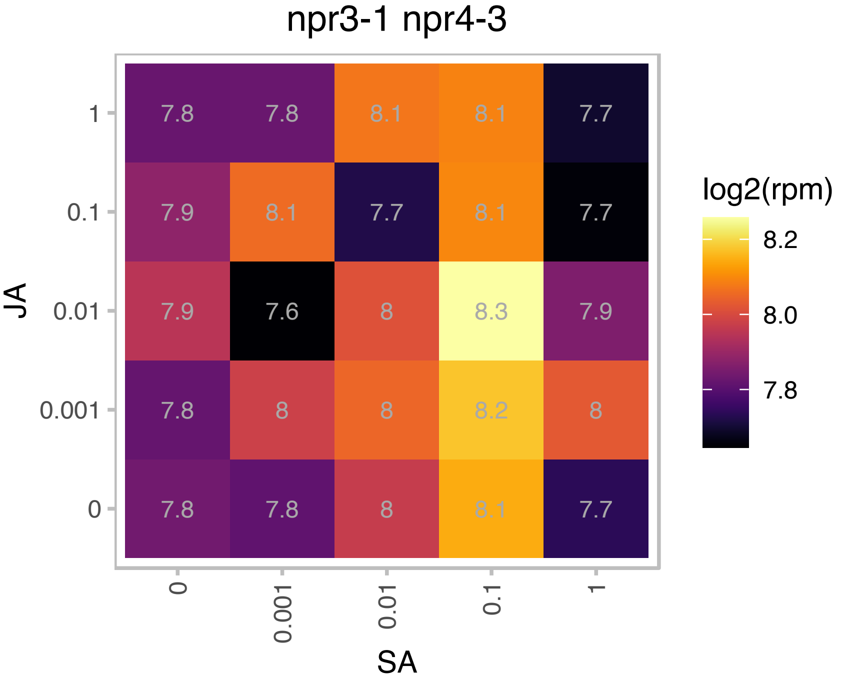

### AT1G66260.1 : ALY3

RNA-binding (RRM/RBD/RNP motifs) family protein

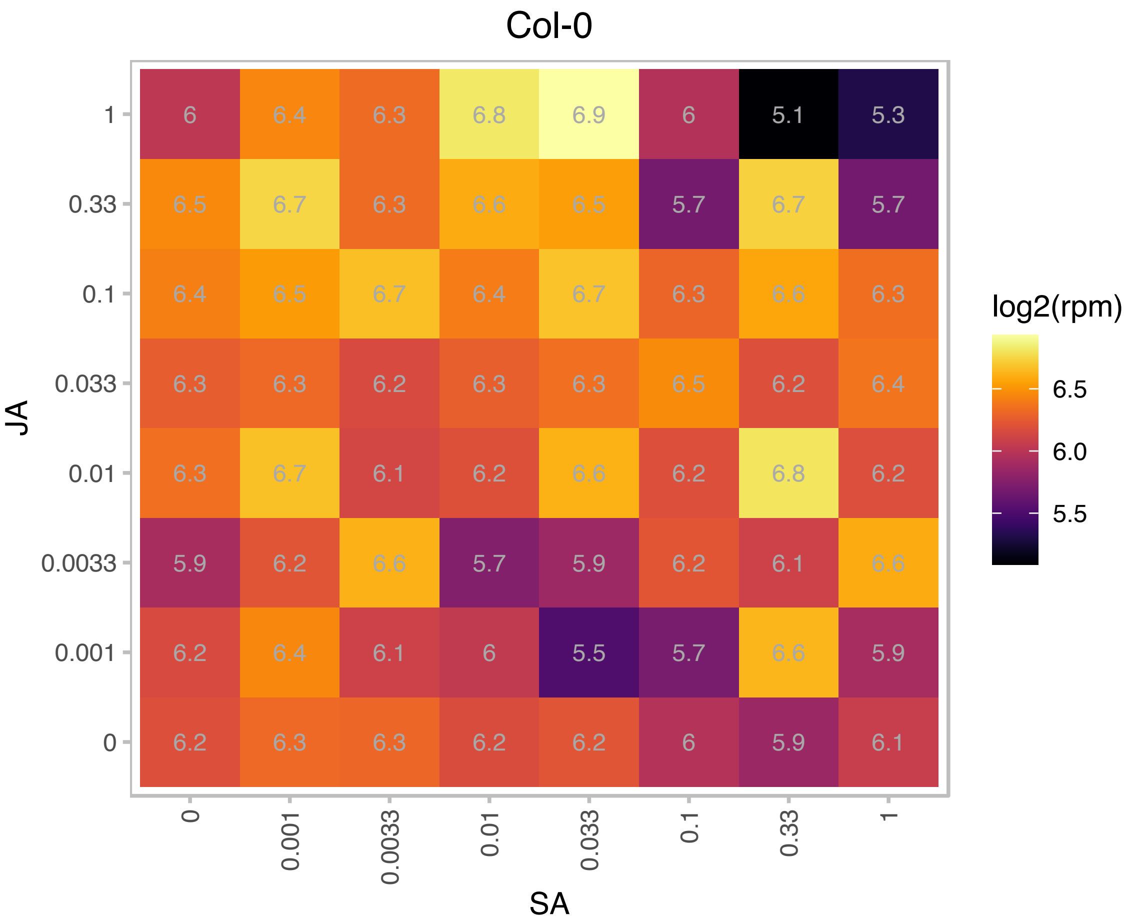


| | |
|---|---|
| Gene | AT1G66260 |
| is.expressed_genes | TRUE |
| is.responsive_genes | TRUE |
| is.apcluster | E-04 |

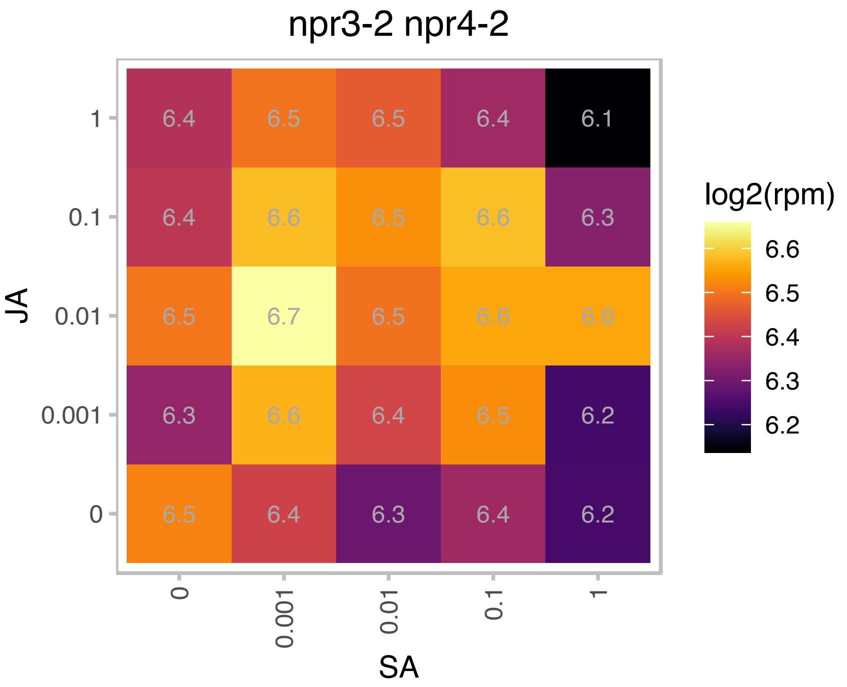


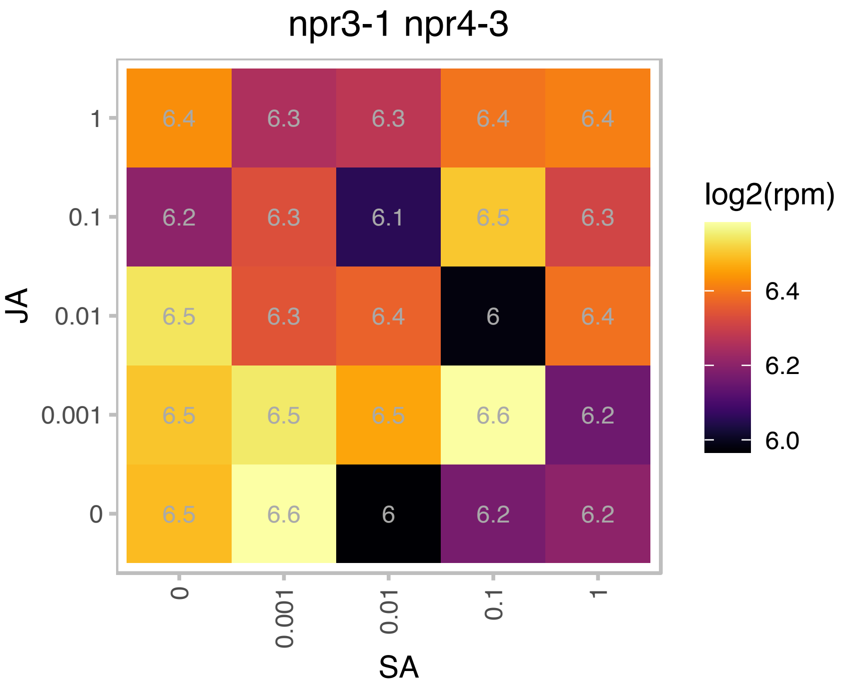

### AT1G74340.1 : DPMS2

dolichol phosphate-mannose biosynthesis regulatory protein-related
Encodes a subunit of the dolichol phosphate mannase synthase (DPMS) complex that may serve as membrane anchors for the catalytic core, DPMS1, or provide catalytic assistance. It is localized in the ER and mediates isoprenyl-linked glycan biogenesis.

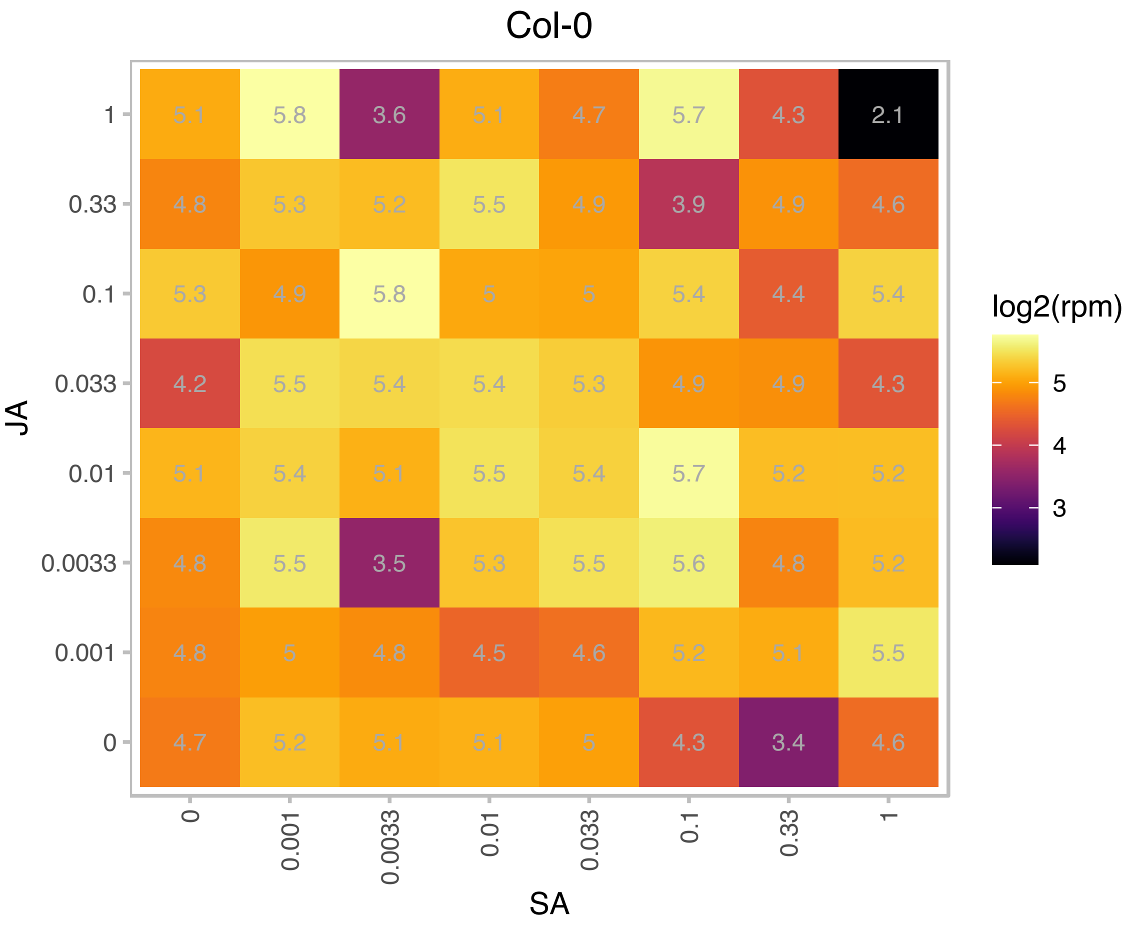


| | |
|---|---|
| Gene | AT1G74340 |
| is.expressed_genes | TRUE |
| is.responsive_genes | TRUE |
| is.apcluster | E-04 |

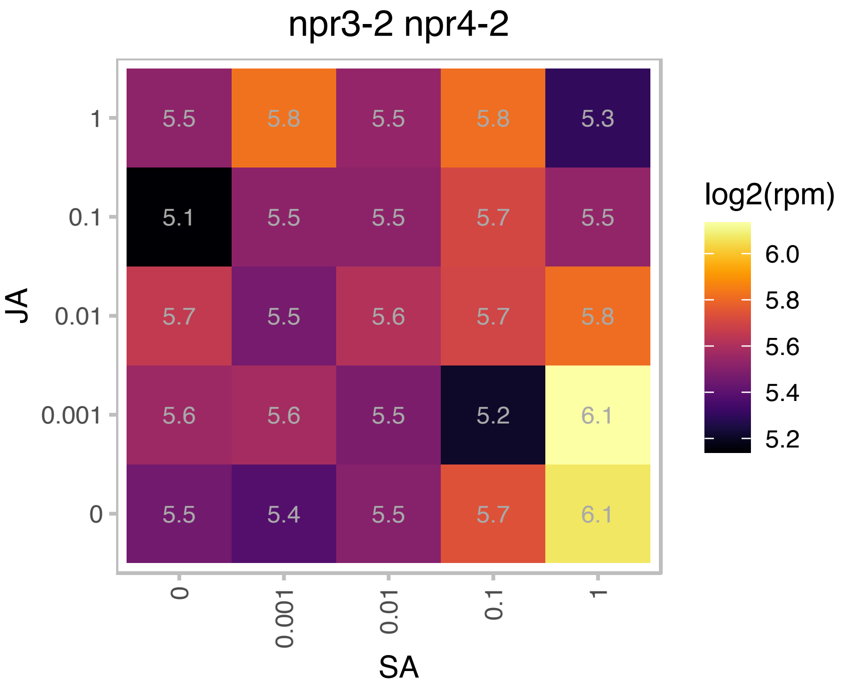


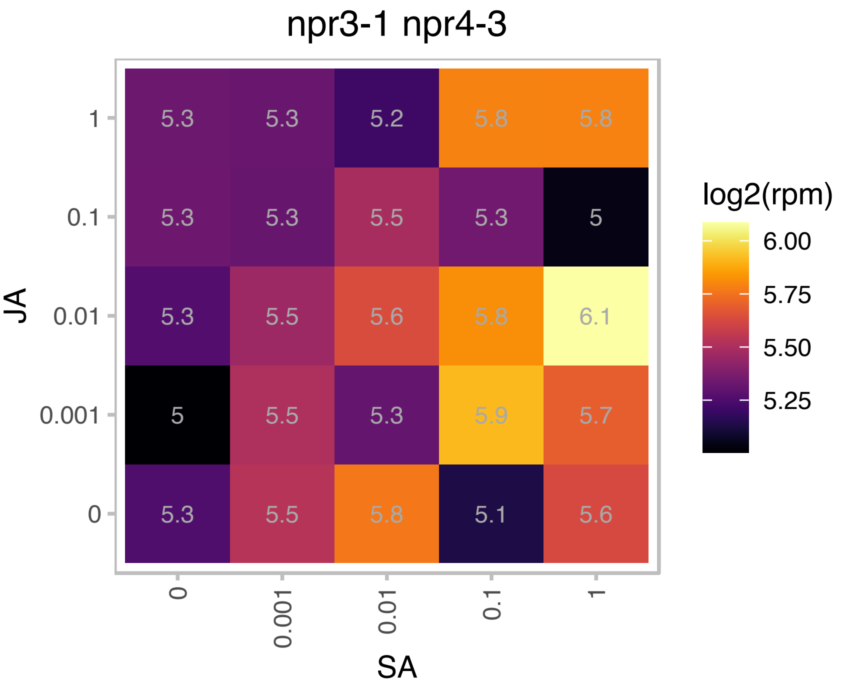

### AT1G74380.1 : XXT5

xyloglucan xylosyltransferase 5

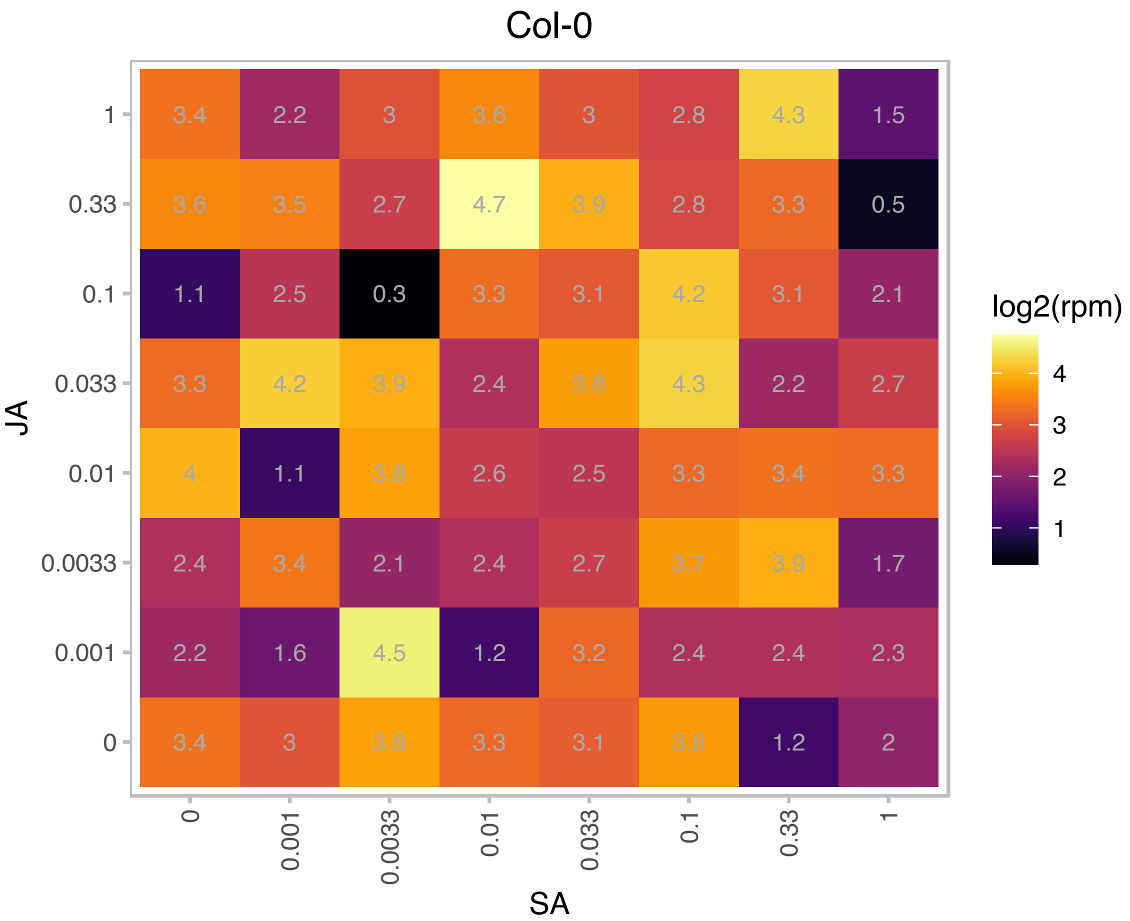


| | |
|---|---|
| Gene | AT1G74380 |
| is.expressed_genes | TRUE |
| is.responsive_genes | TRUE |
| is.apcluster | E-04 |

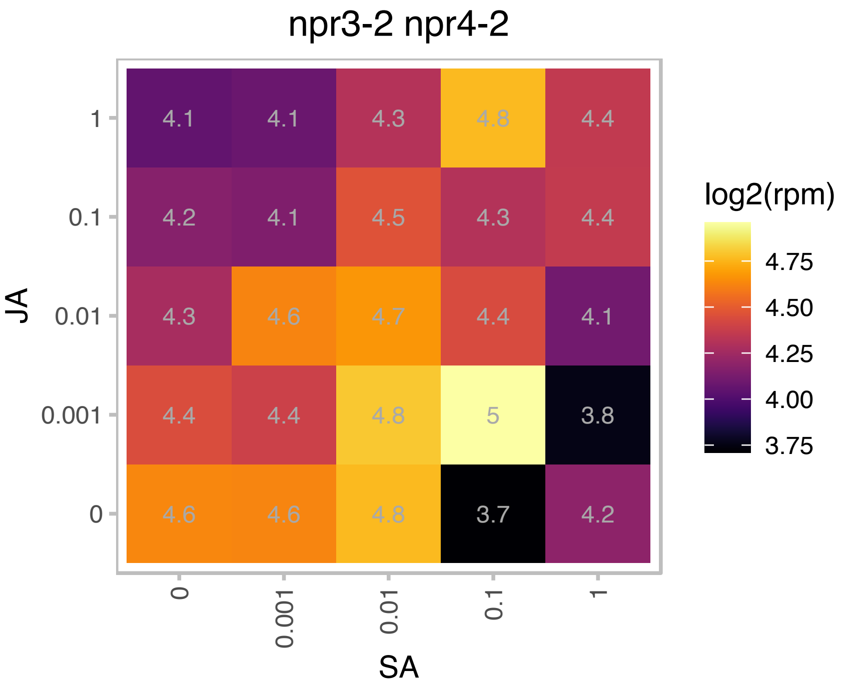


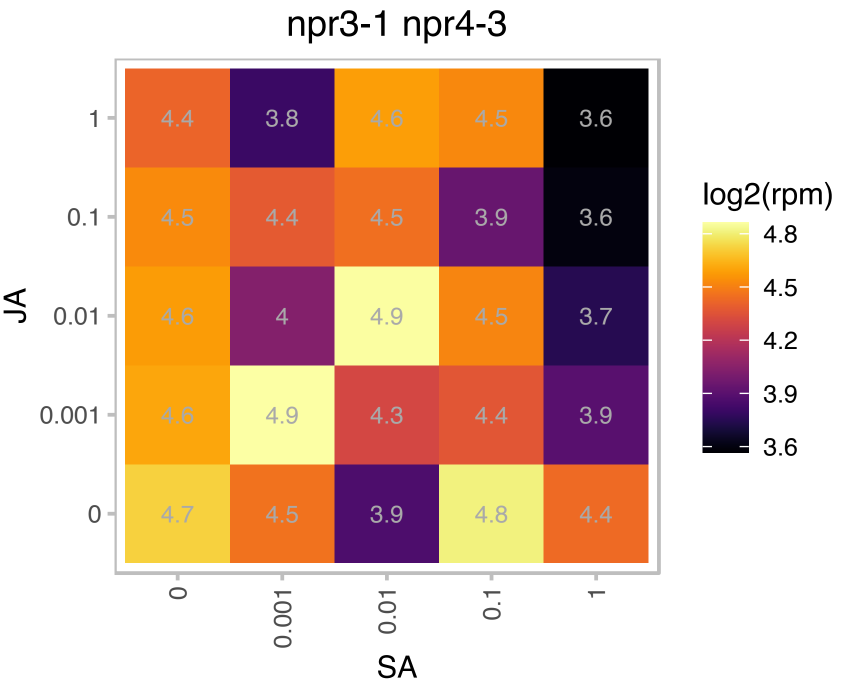

# AT1G80500.1 : NA

SNARE-like superfamily protein

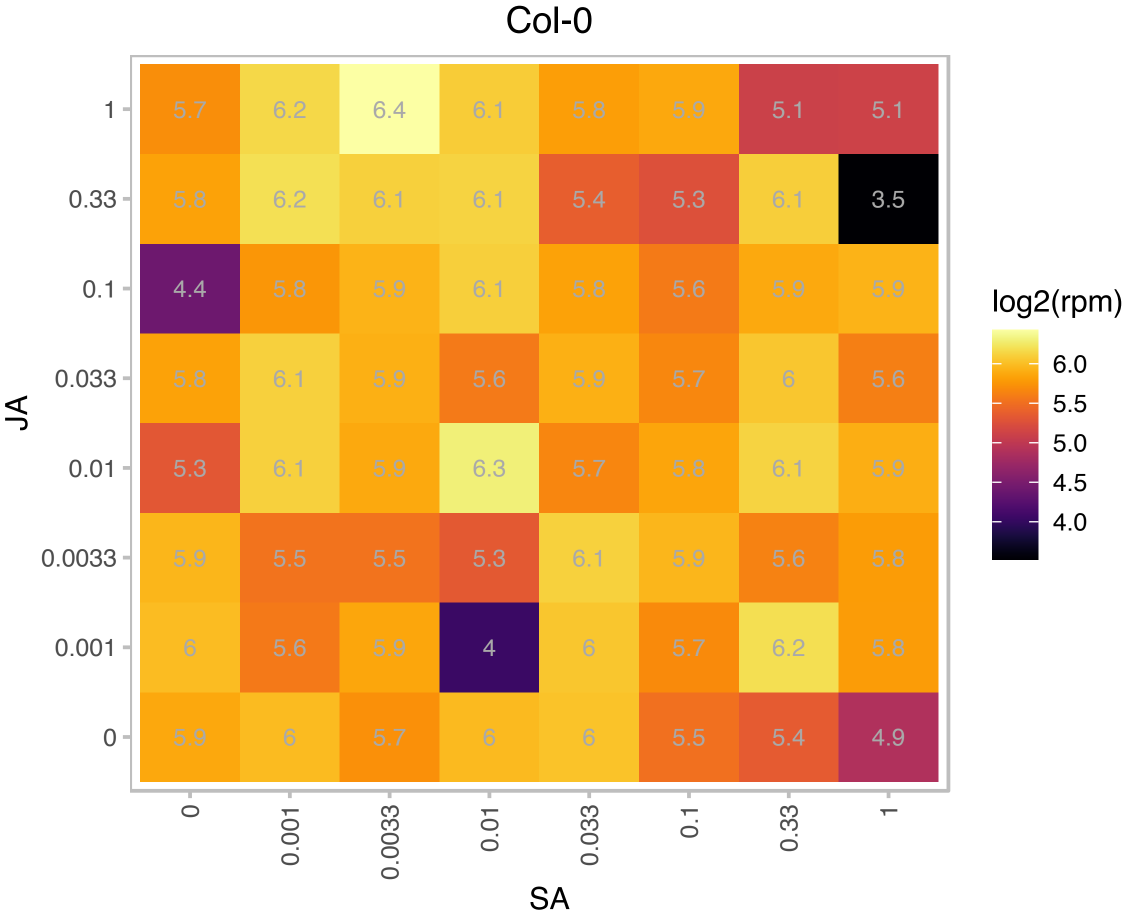


| | |
|---|---|
| Gene | AT1G80500 |
| is.expressed_genes | TRUE |
| is.responsive_genes | TRUE |
| is.apcluster | E-04 |

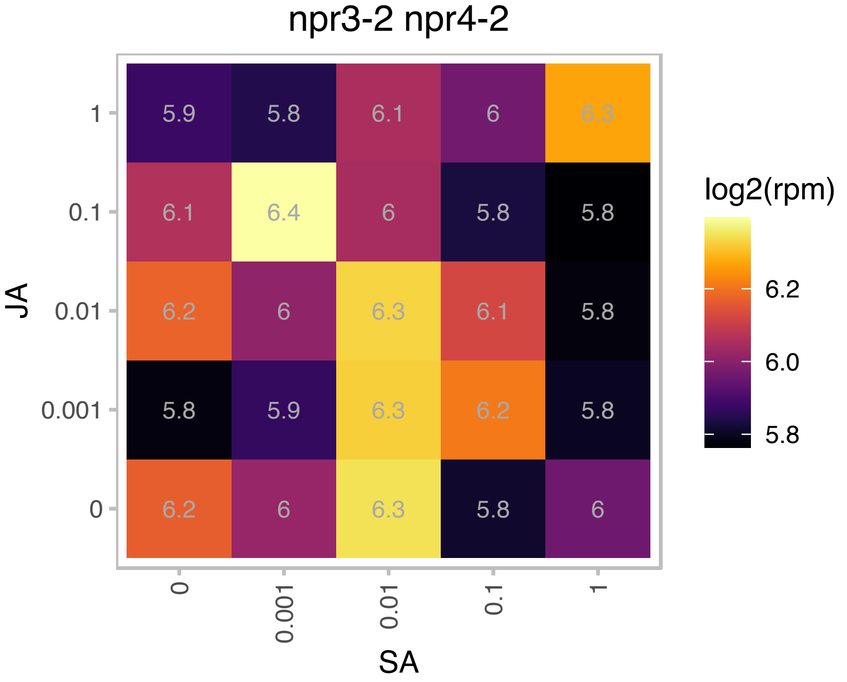


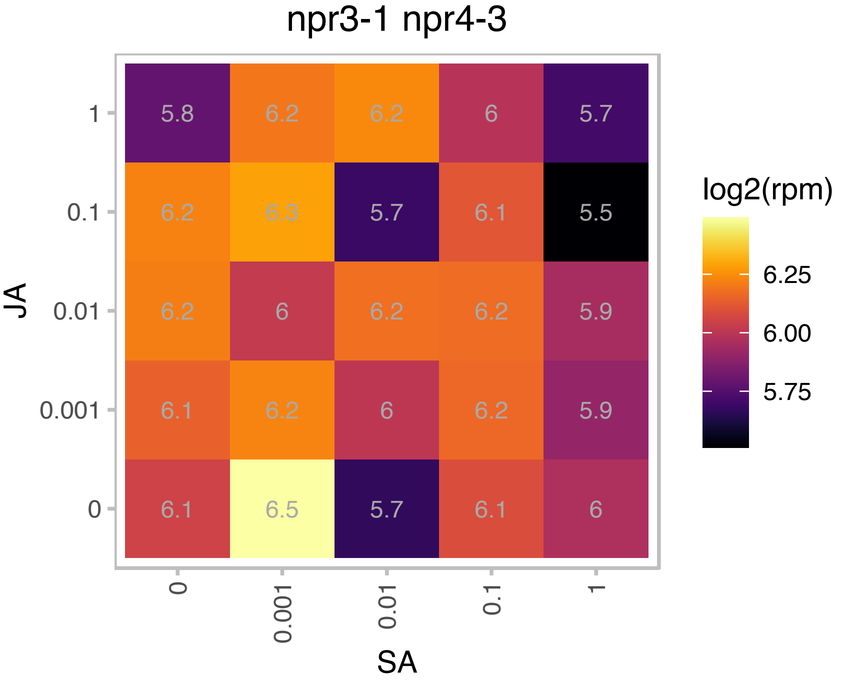

### AT2G16850.1 : PIP2;8, PIP3B

plasma membrane intrinsic protein 2;8

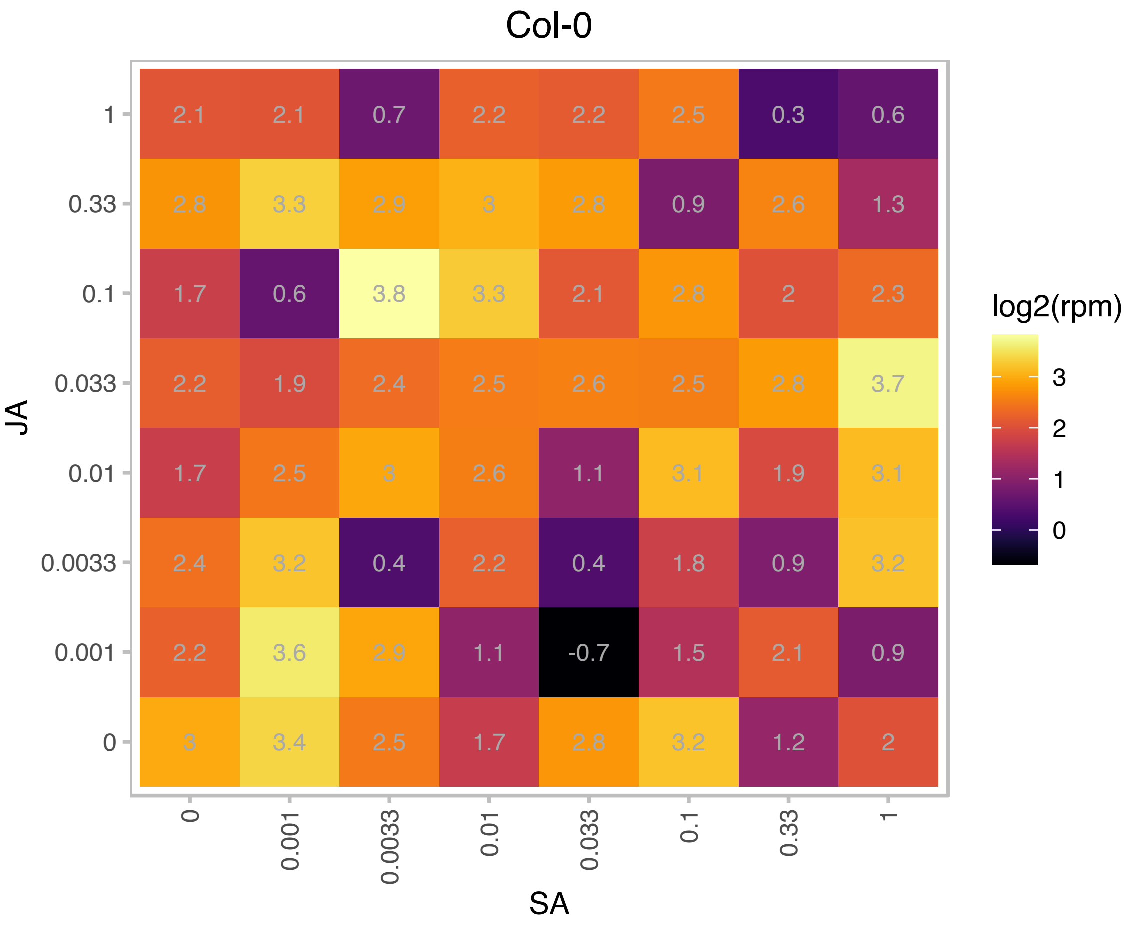


| | |
|---|---|
| Gene | AT2G16850 |
| is.expressed_genes | TRUE |
| is.responsive_genes | TRUE |
| is.apcluster | E-04 |

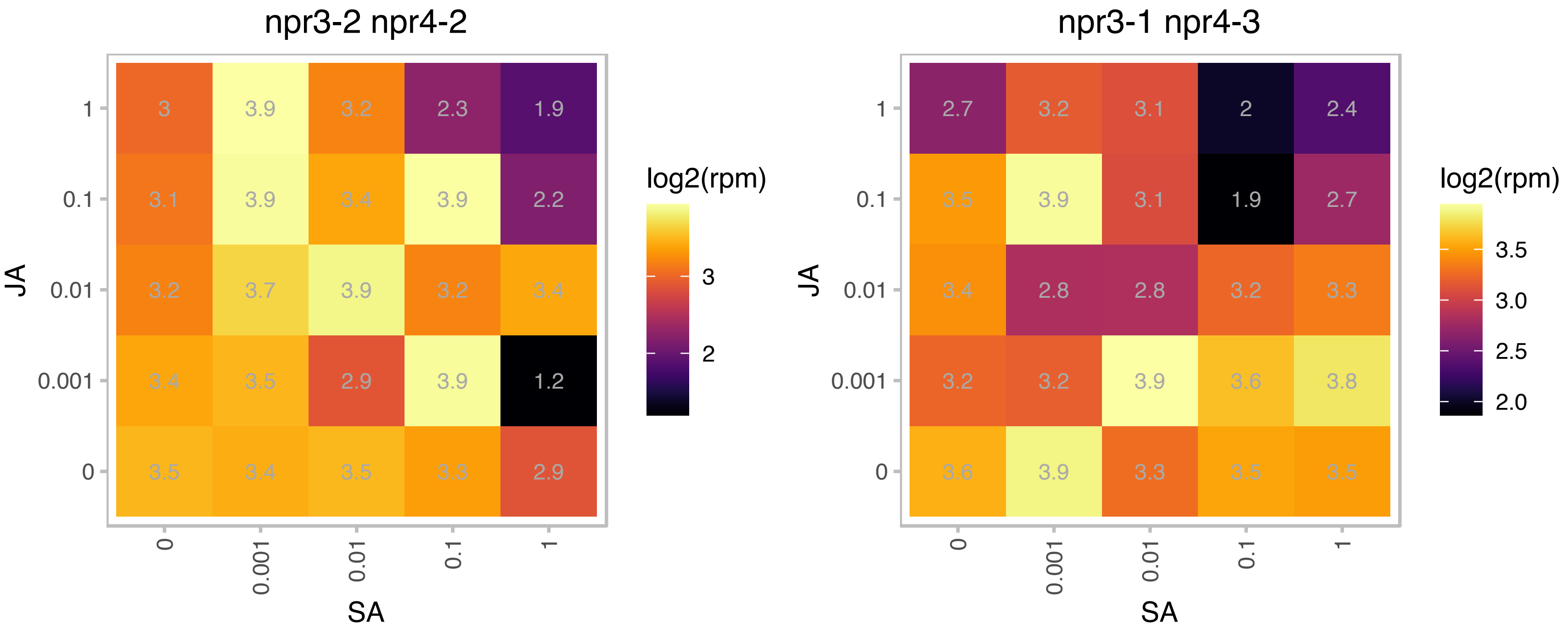

### AT2G28230.1 : MED20, MED20A

TATA-binding related factor (TRF) of subunit 20 of Mediator complex
Encodes one of the subunits of the head domain of the MEDIATOR complex. The complex conveys transcriptional signals by acting as an intermediary between transcription factors and RNA polymerase II. Together with MED18, MED20 is proposed to control the balance of salicylic acid and jasmonate associated defense pathways. It is involved in miRNA biogenesis by recruiting Pol II to promoters of miRNA genes.

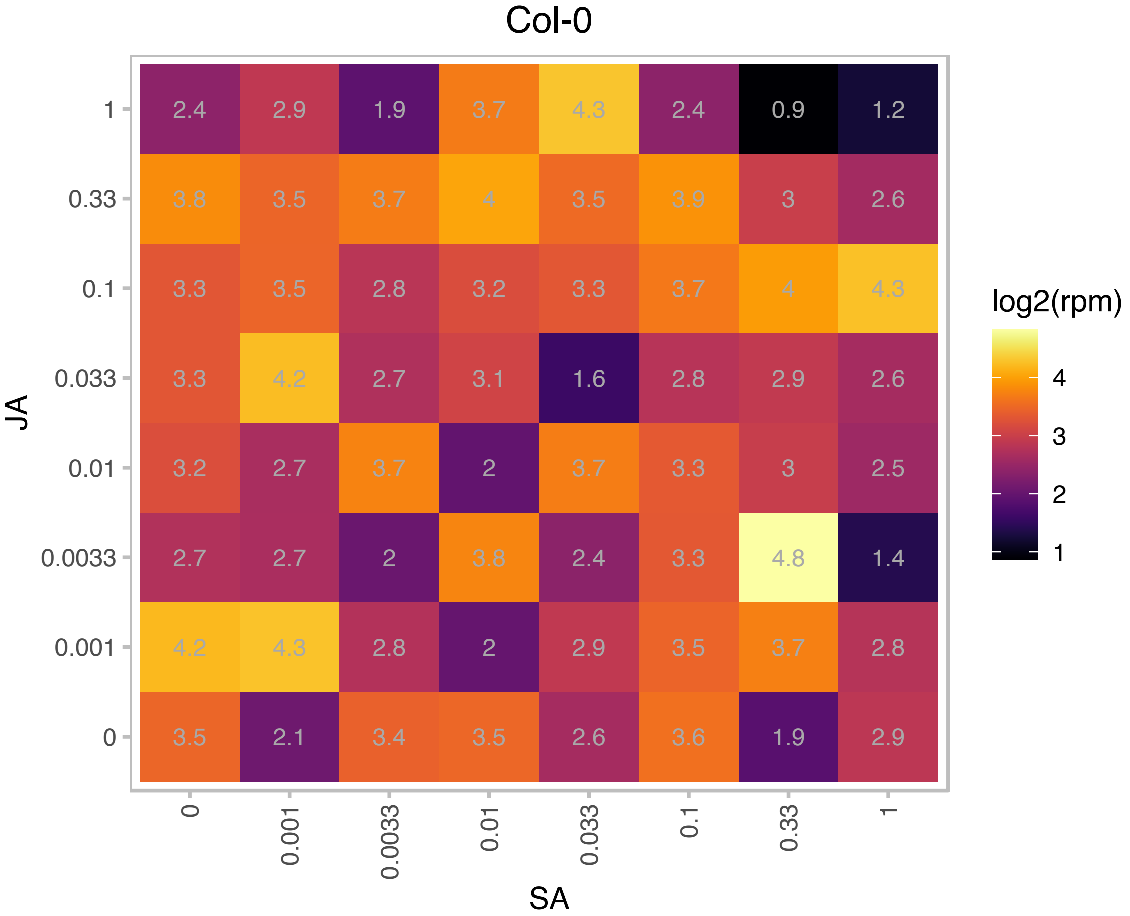


| | |
|---|---|
| Gene | AT2G28230 |
| is.expressed_genes | TRUE |
| is.responsive_genes | TRUE |
| is.apcluster | E-04 |

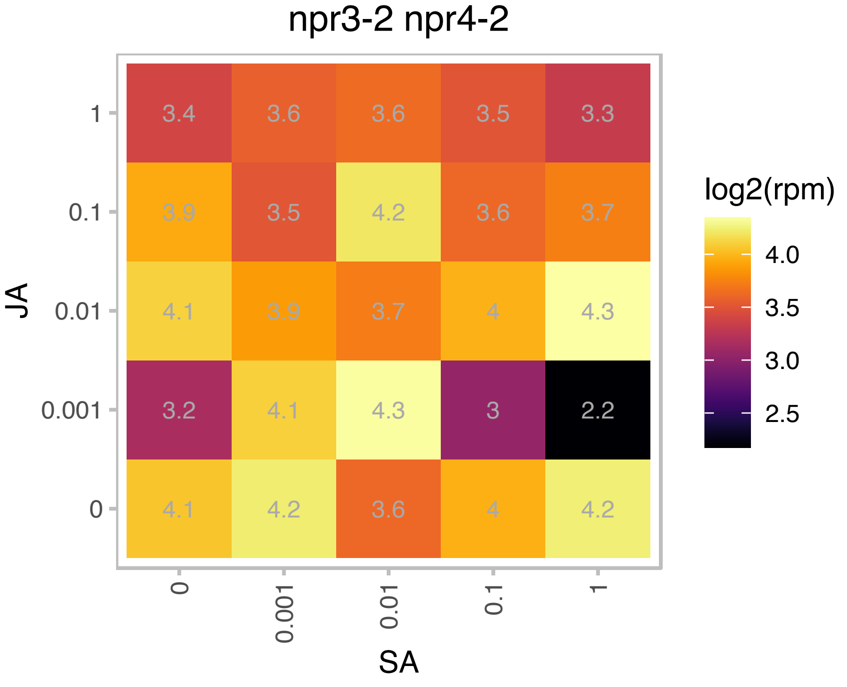


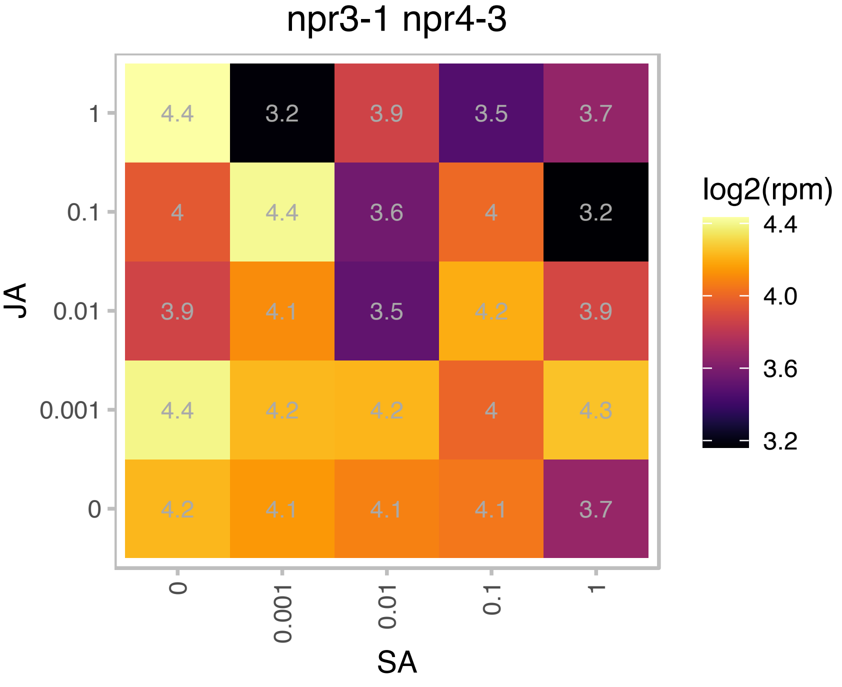

### AT2G29960.1 : ATCYP5, CYP19-4, CYP5

cyclophilin 5
encodes a cyclophilin protein that exhibits peptidylprolyl cis/trans-isomerase and protein refolding activities that were sensitive to cyclosporin A. The protein interacts with GNOM in vitro and is localized to both the cytosolic and membrane fractions. The gene is expressed in the developing embryo.

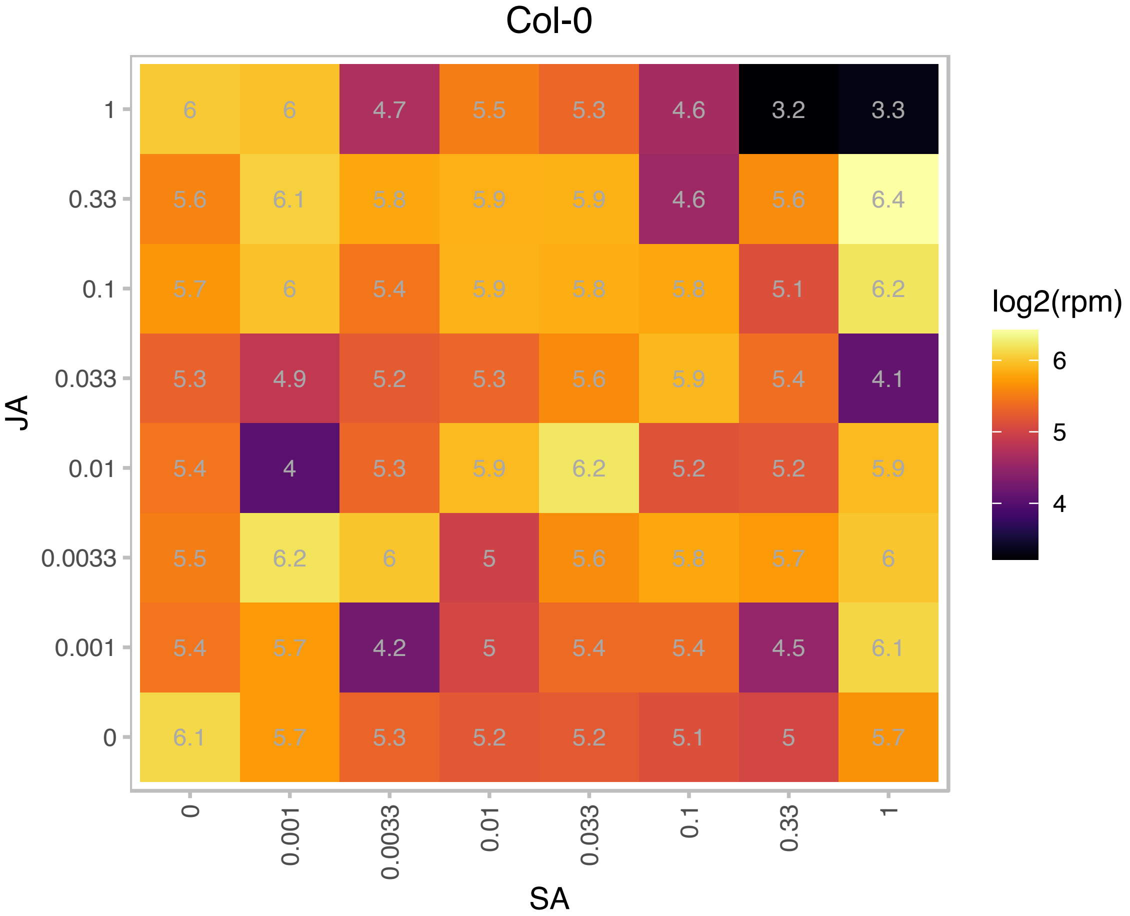


| | |
|---|---|
| Gene | AT2G29960 |
| is.expressed_genes | TRUE |
| is.responsive_genes | TRUE |
| is.apcluster | E-04 |

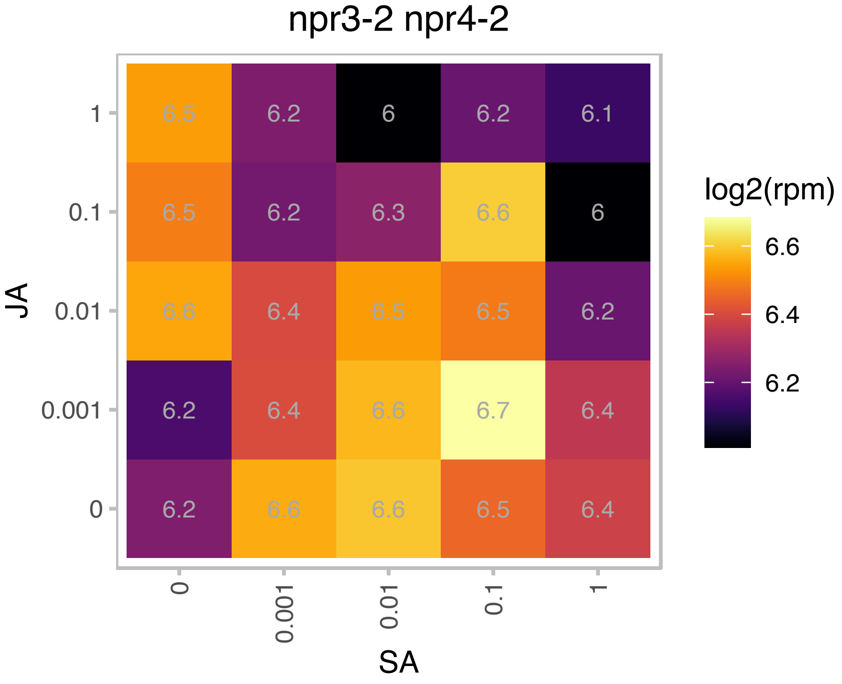


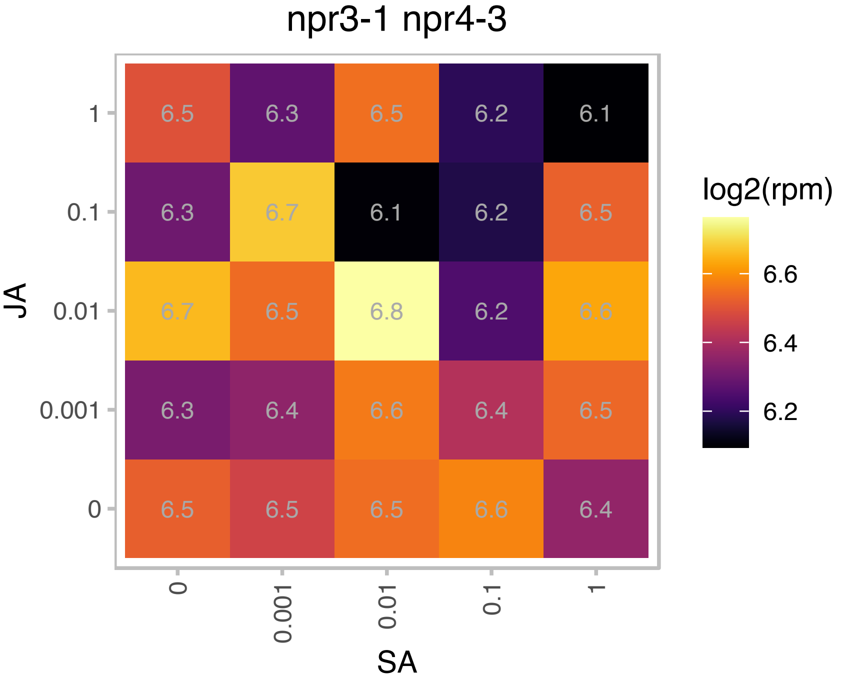

### AT2G29970.1 : SMXL7

Double Clp-N motif-containing P-loop nucleoside triphosphate hydrolases superfamily protein

Encodes a member of an eight-gene family (SMAX1 and SMAX1-like) that has weak similarity to AtHSP101, a ClpB chaperonin required for thermotolerance. The mRNA is cell-to-cell mobile.

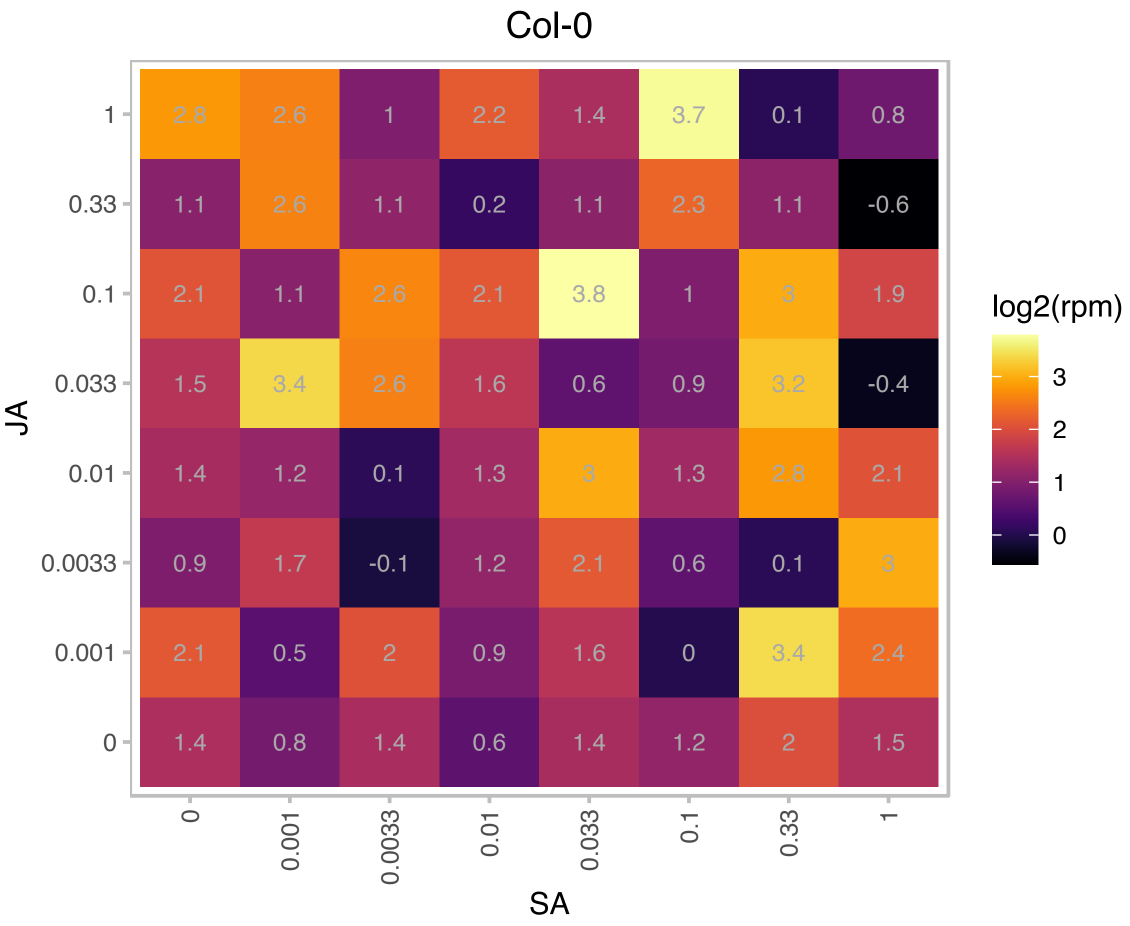


| | |
|---|---|
| Gene | AT2G29970 |
| is.expressed_genes | TRUE |
| is.responsive_genes | TRUE |
| is.apcluster | E-04 |

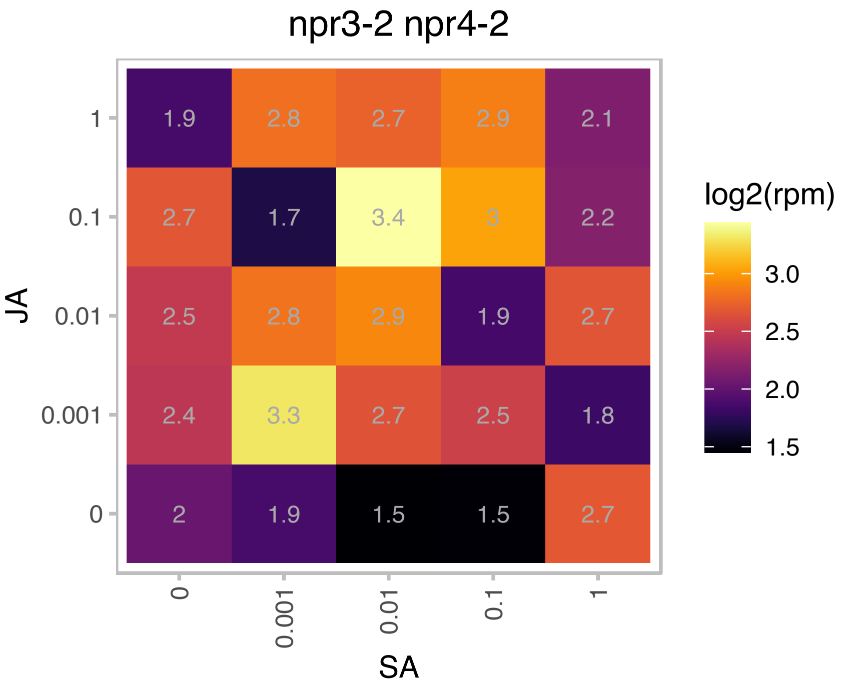


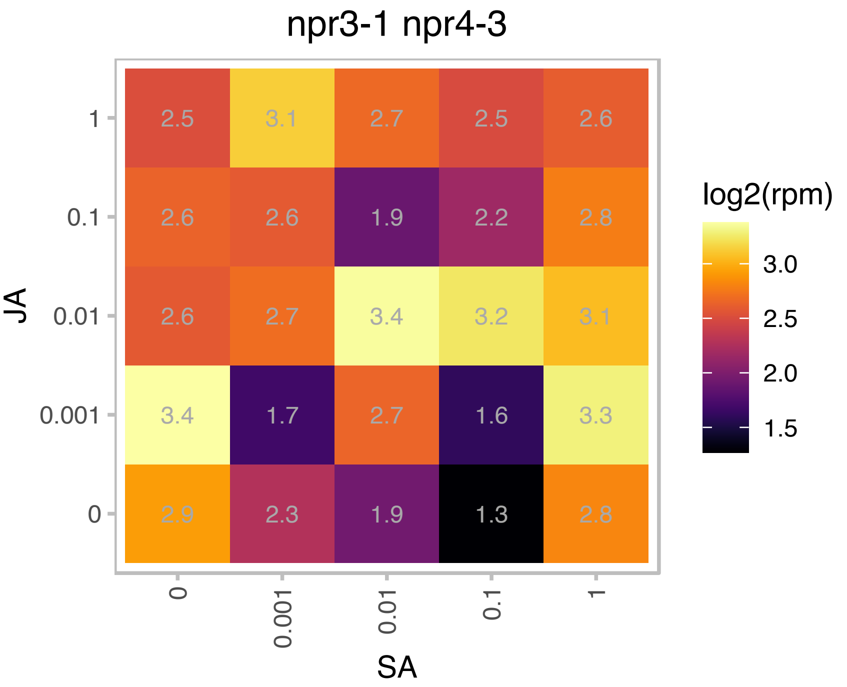

### AT2G30330.1 : BLOS1

GCN5L1 family protein

Putative homolog of mammalian BLOC-1 Subunit 1. Protein - protein interaction with BLOS2 and also with SNX1.Located in endomembrane system and hypothesized to be involved in endomembrane transport.

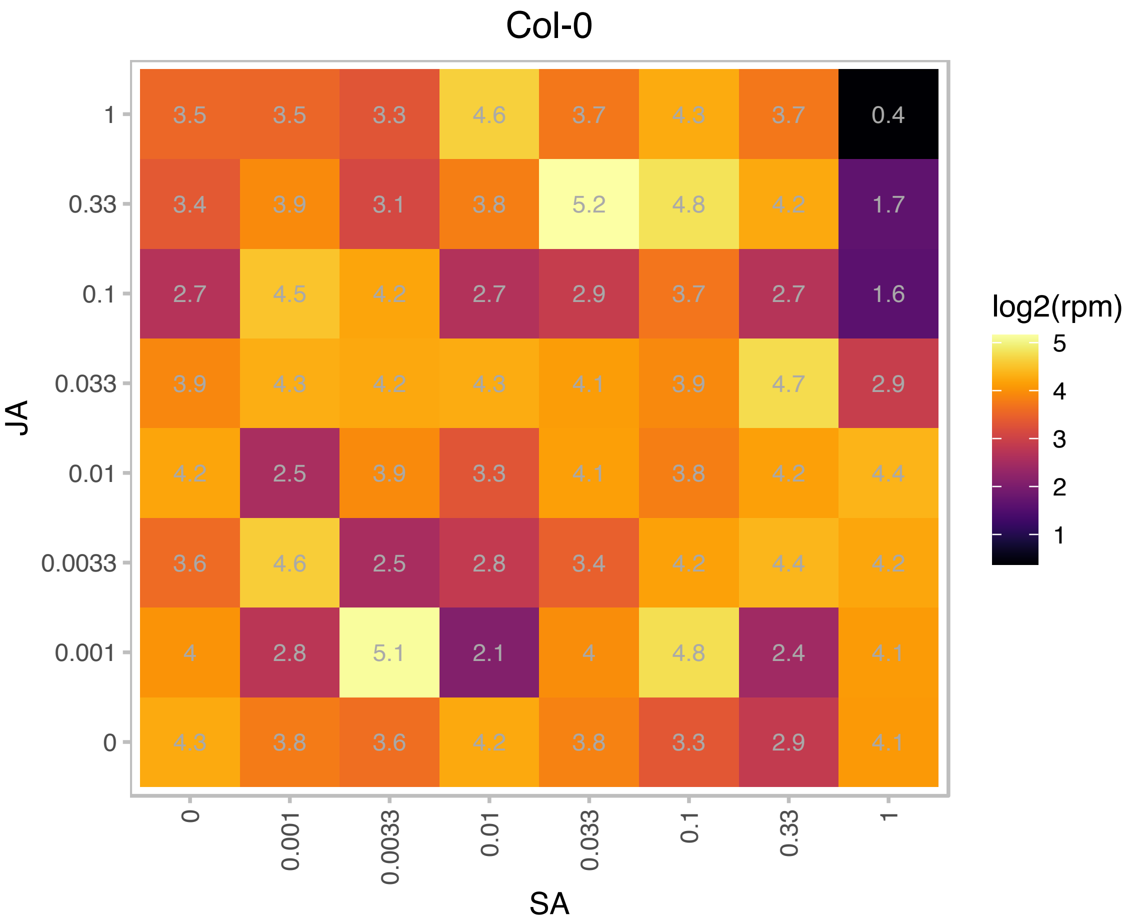


| | |
|---|---|
| Gene | AT2G30330 |
| is.expressed_genes | TRUE |
| is.responsive_genes | TRUE |
| is.apcluster | E-04 |

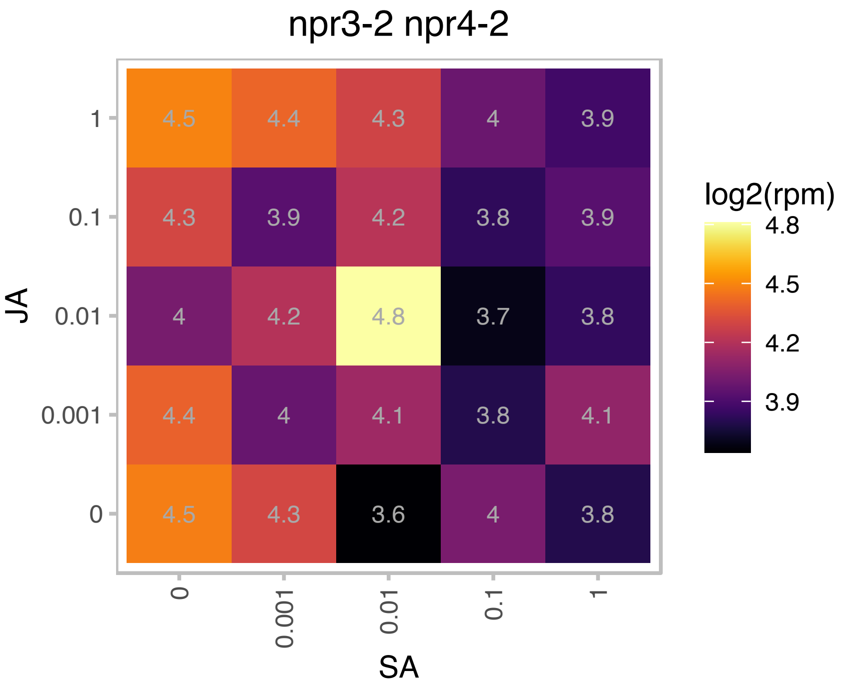


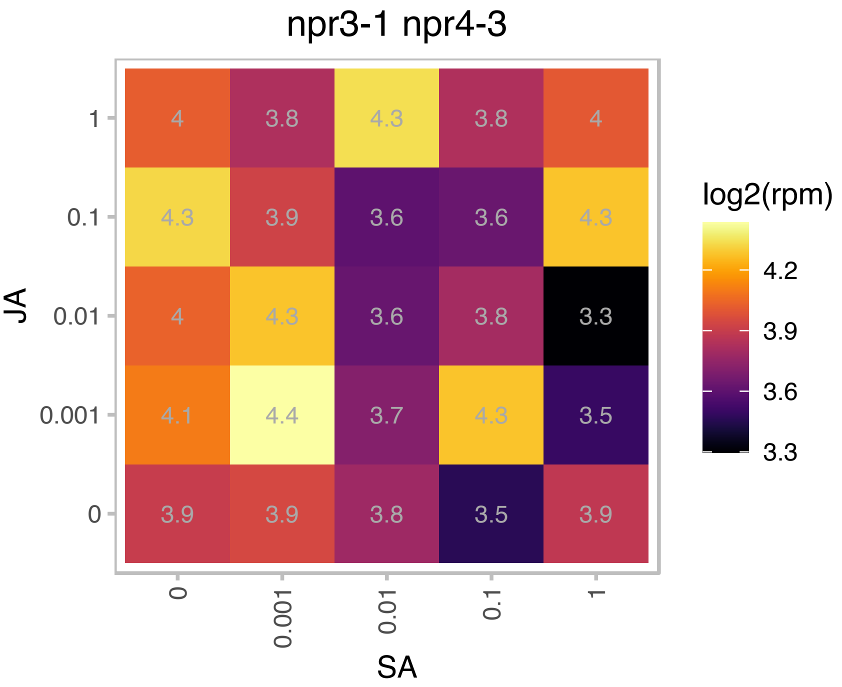

### AT2G44050.1 : COS1, COS1

6,7-dimethyl-8-ribityllumazine synthase / DMRL synthase / lumazine synthase / riboflavin synthase

6,7-dimethyl-8-ribityllumazine synthase / DMRL synthase / lumazine synthase / riboflavin synthase [Arabidopsis thaliana]. Acts in the jasmonic acid signaling pathway. The mRNA is cell-to-cell mobile.

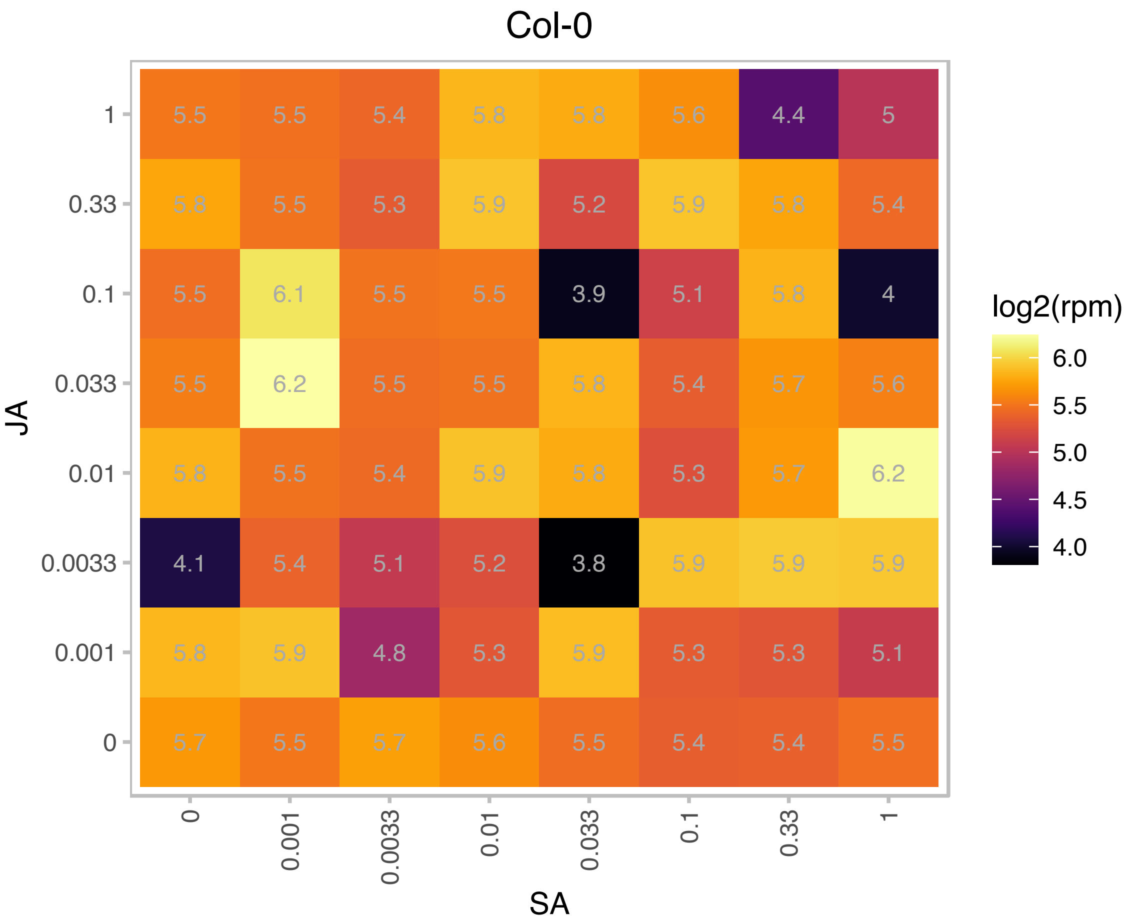


| | |
|---|---|
| Gene | AT2G44050 |
| is.expressed_genes | TRUE |
| is.responsive_genes | TRUE |
| is.apcluster | E-04 |

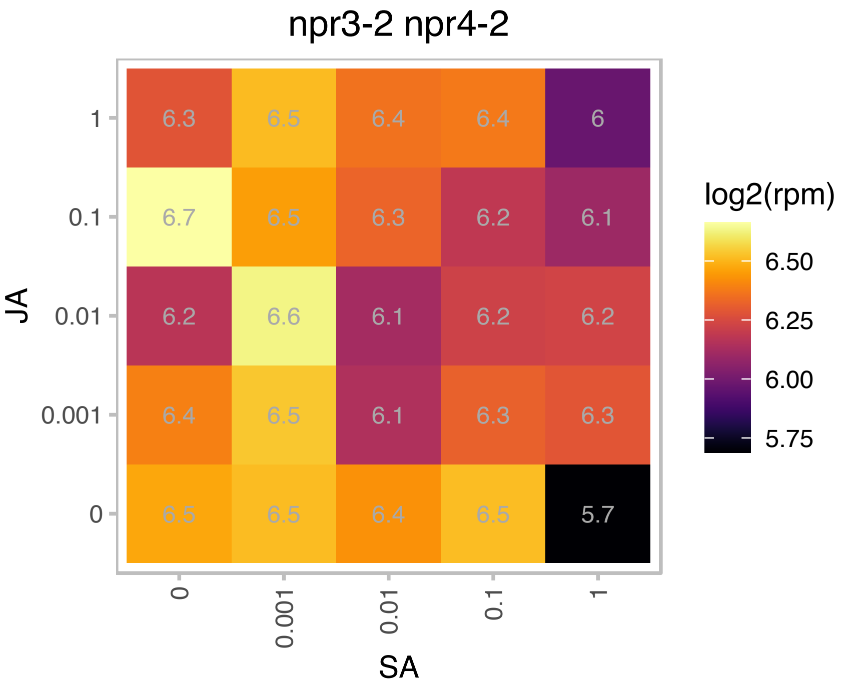


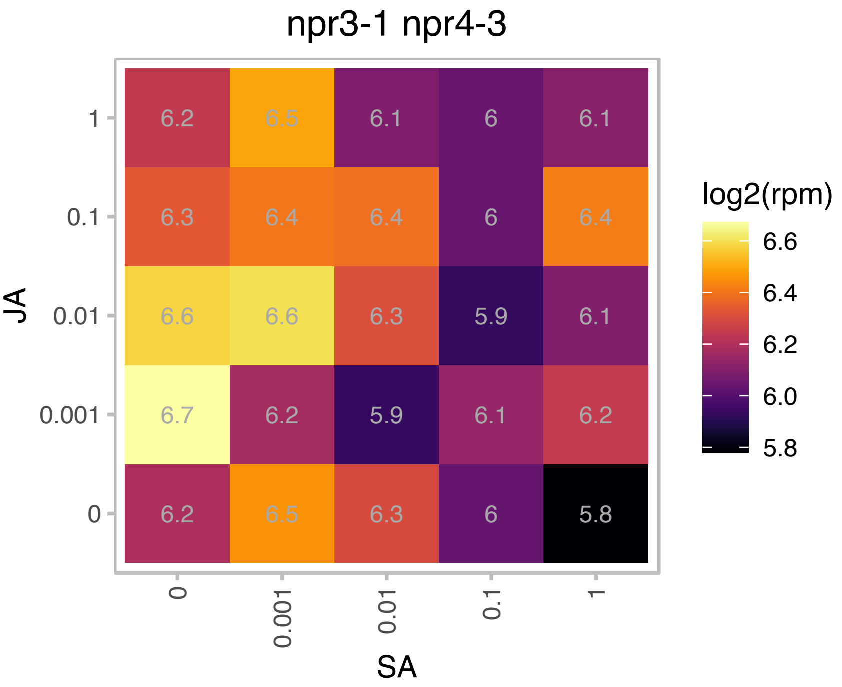

### AT3G03600.1 : RPS2

ribosomal protein S2

Structural component of the mitochondrial ribosome small subunit

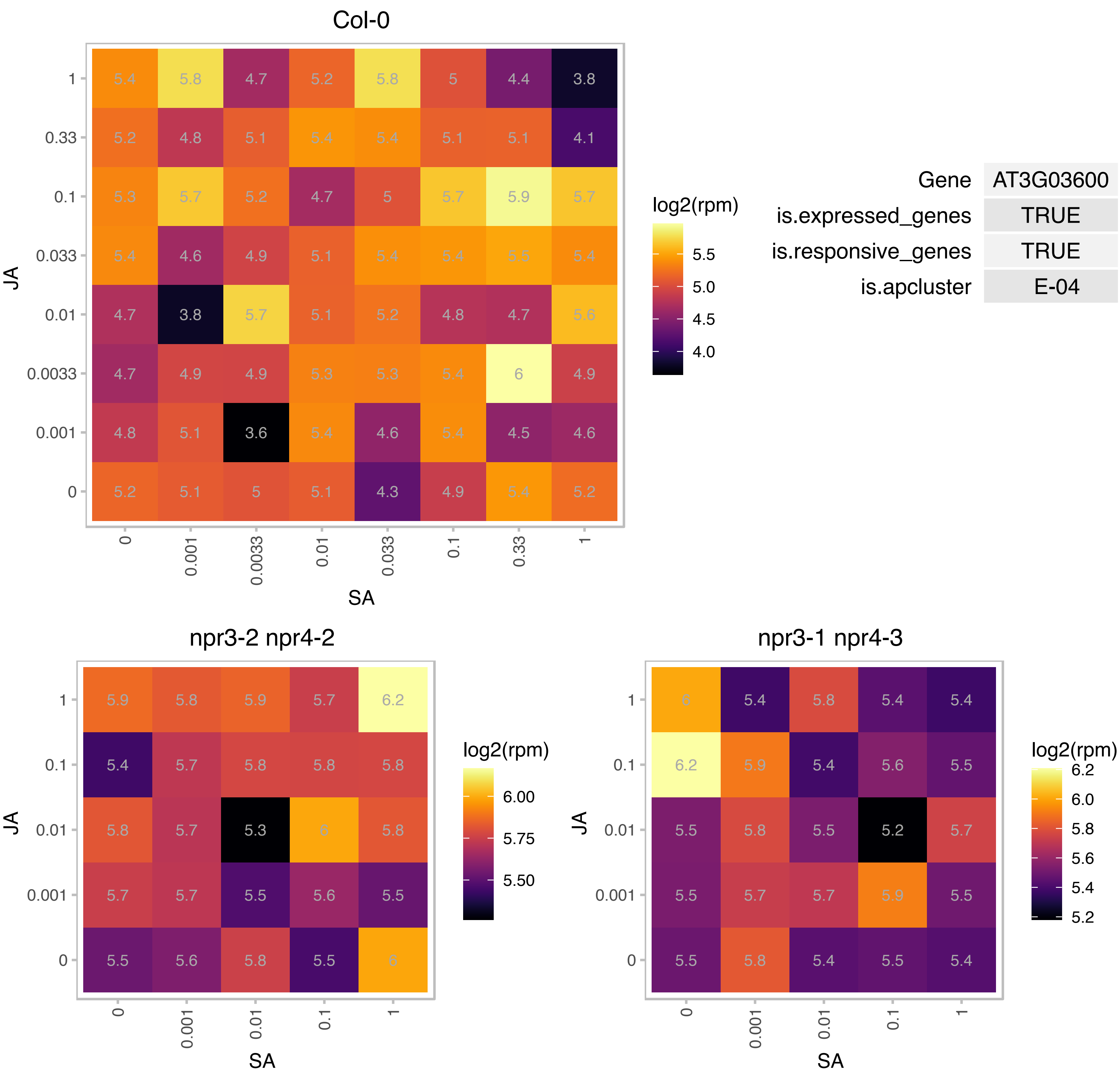


| | |
|---|---|
| Gene | AT3G03600 |
| is.expressed_genes | TRUE |
| is.responsive_genes | TRUE |
| is.apcluster | E-04 |

# AT3G05000.1 : NA

Transport protein particle (TRAPP) component

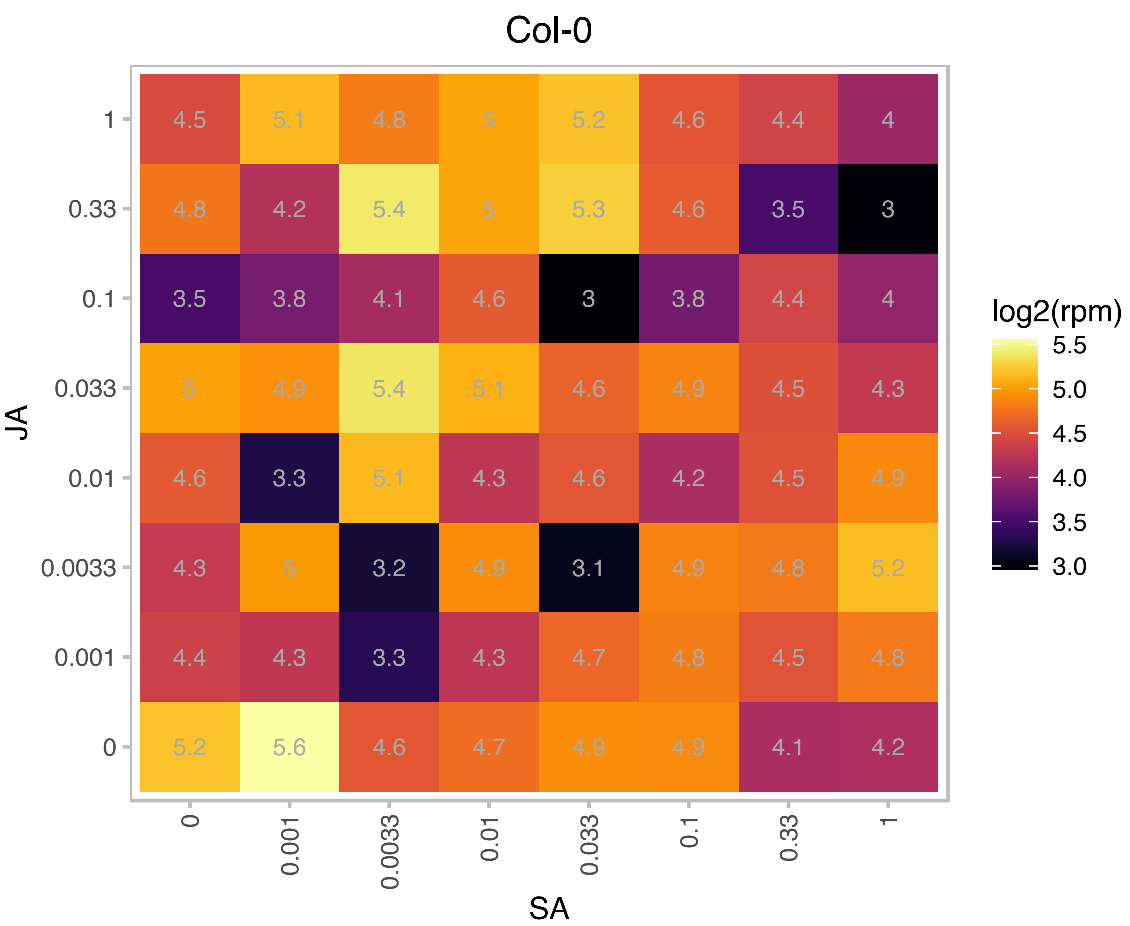


| | |
|---|---|
| Gene | AT3G05000 |
| is.expressed_genes | TRUE |
| is.responsive_genes | TRUE |
| is.apcluster | E-04 |

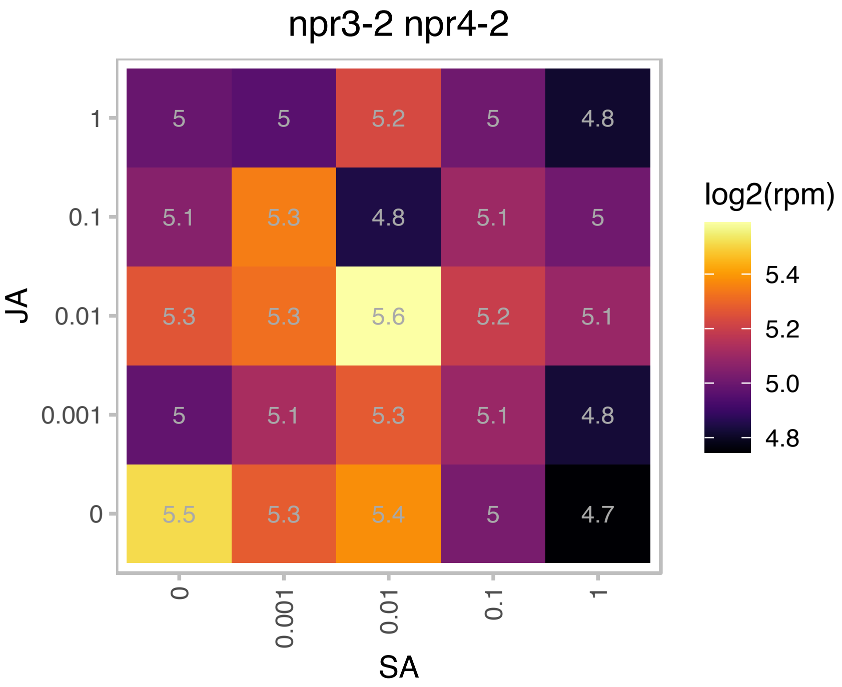


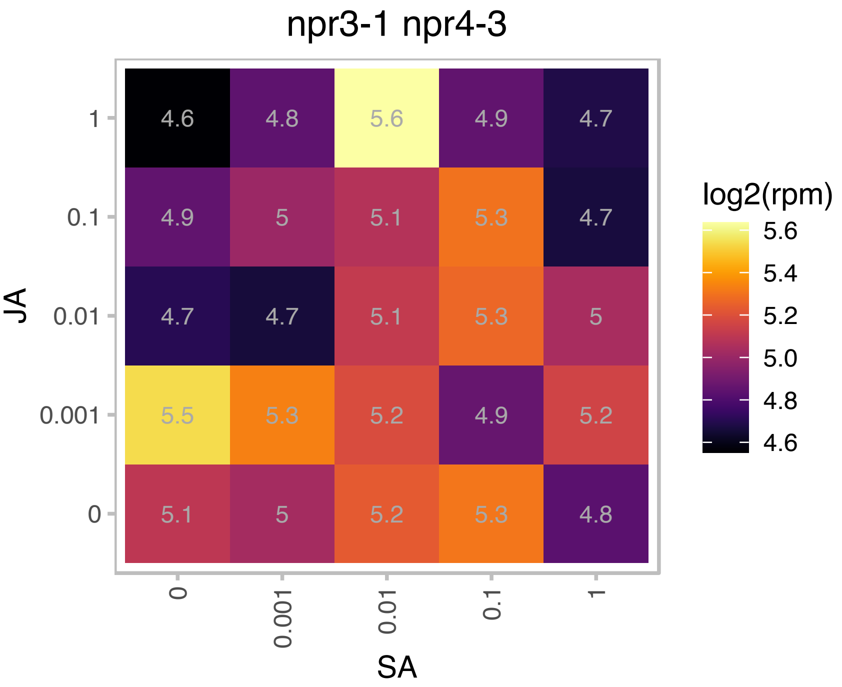

# AT3G18430.1 : NA

Calcium-binding EF-hand family protein

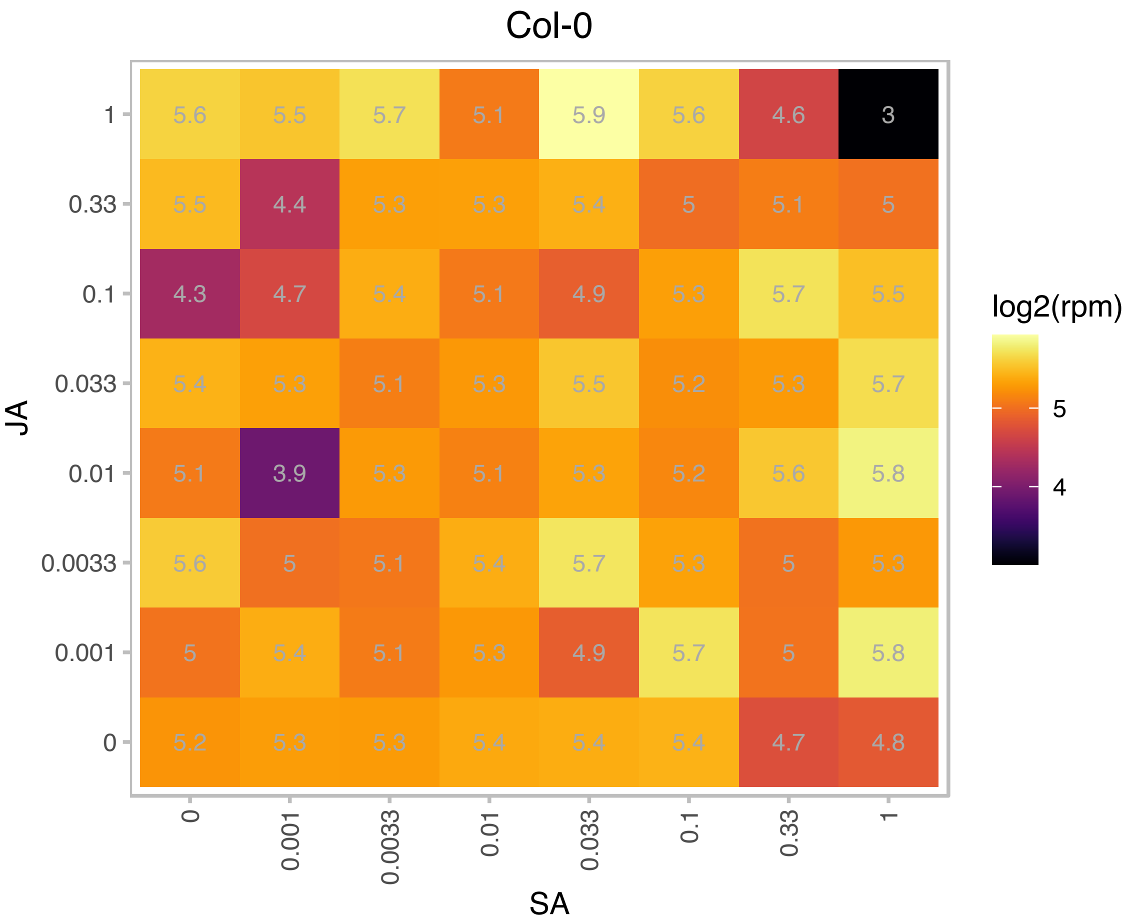


| Gene | AT3G18430 |
|---|---|
| is.expressed_genes | TRUE |
| is.responsive_genes | TRUE |
| is.apcluster | E-04 |

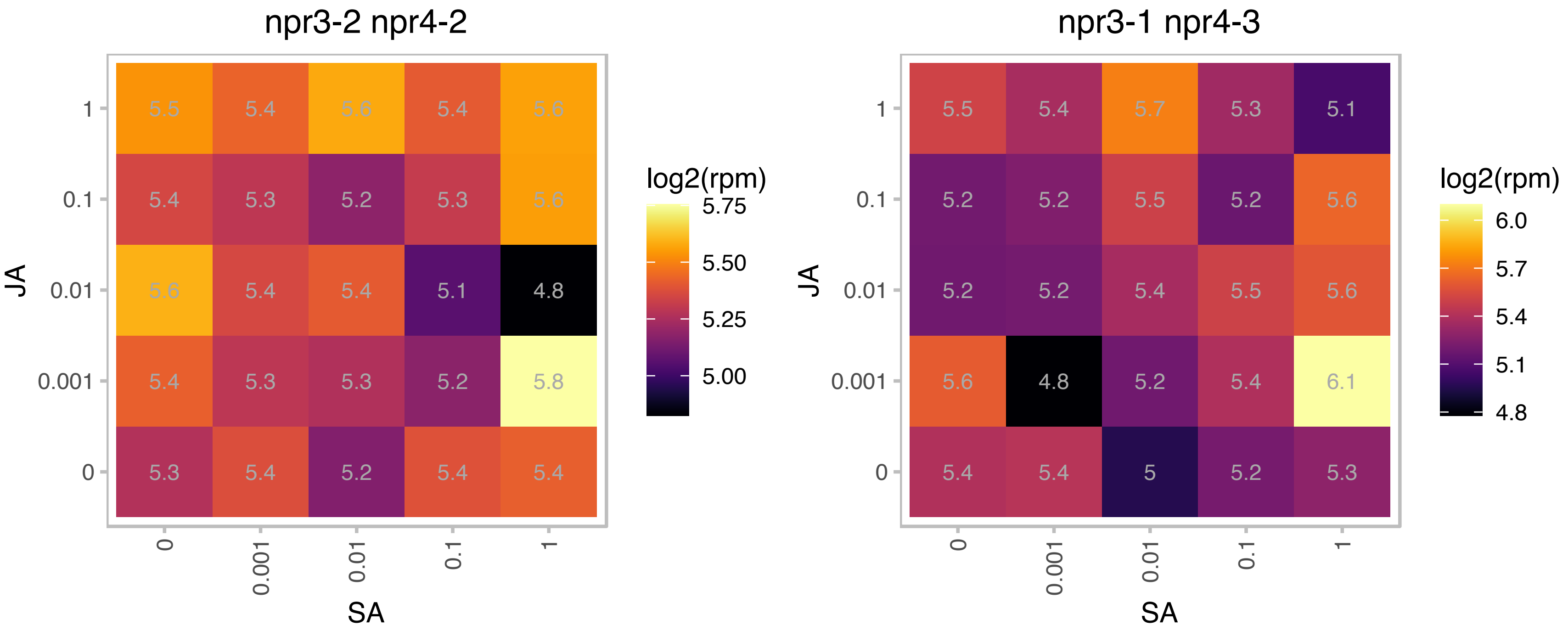

### AT3G20000.1 : TOM40

translocase of the outer mitochondrial membrane 40

Encodes a component of the TOM receptor complex responsible for the recognition and translocation of cytosolically synthesized mitochondrial preproteins. With TOM22, functions as the transit peptide receptor at the surface of the mitochondrial outer membrane and facilitates the movement of preproteins into the translocation pore. The mRNA is cell-to-cell mobile.

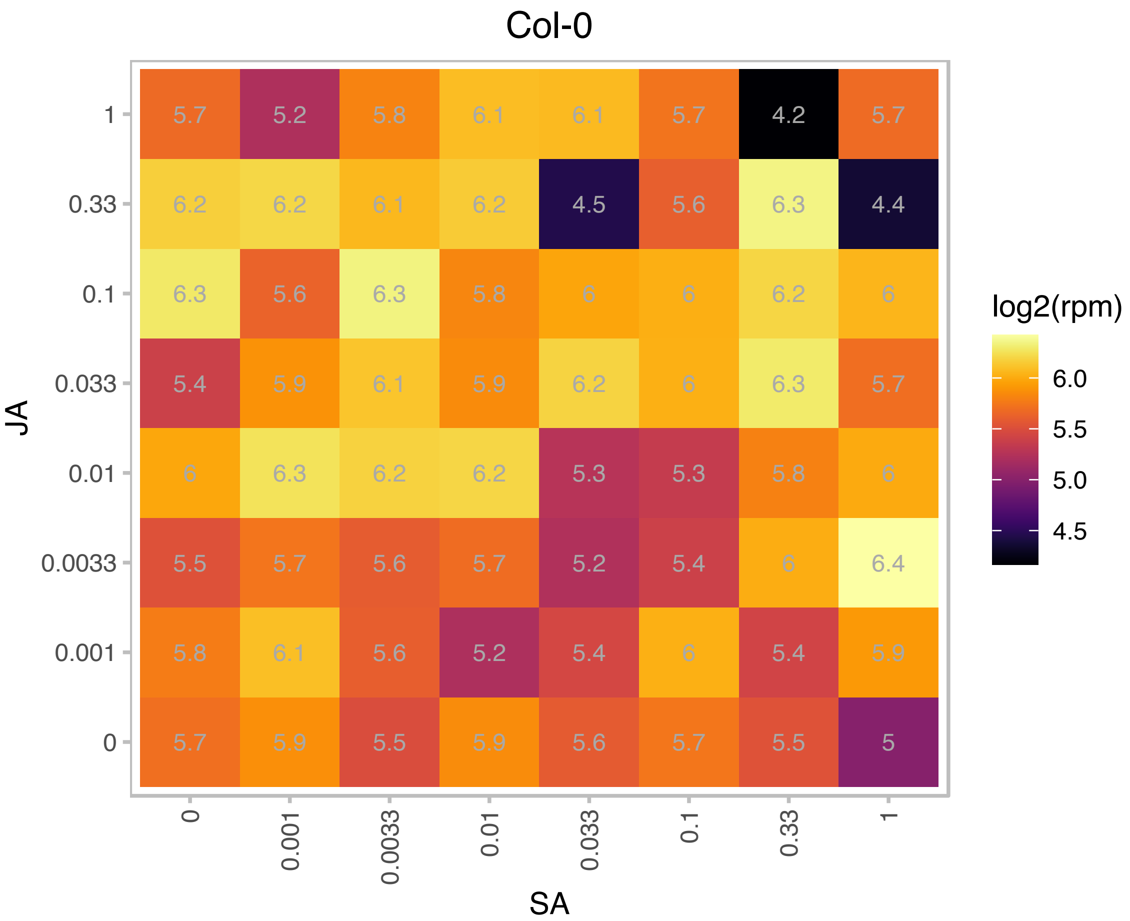


| | |
|---|---|
| Gene | AT3G20000 |
| is.expressed_genes | TRUE |
| is.responsive_genes | TRUE |
| is.apcluster | E-04 |

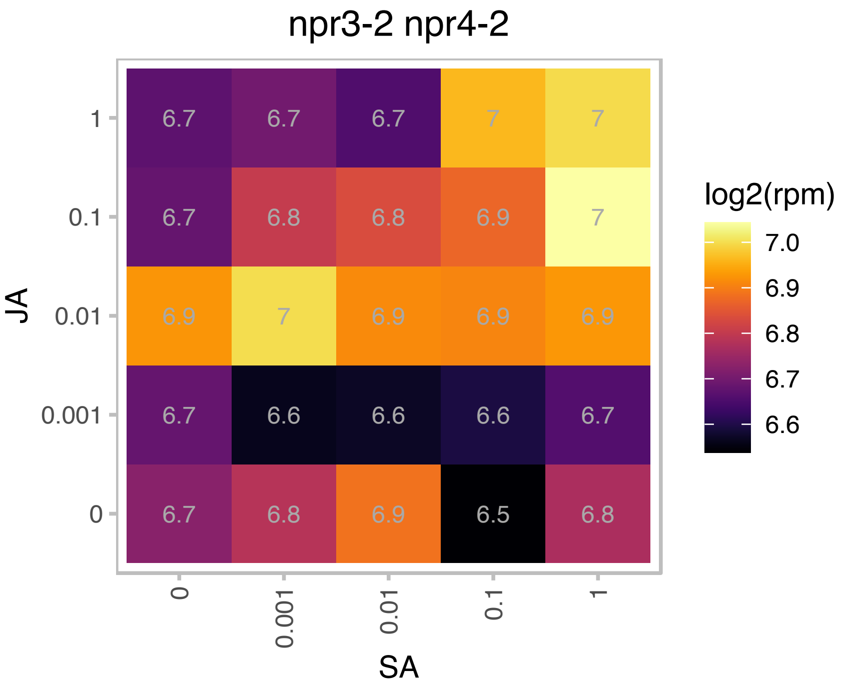


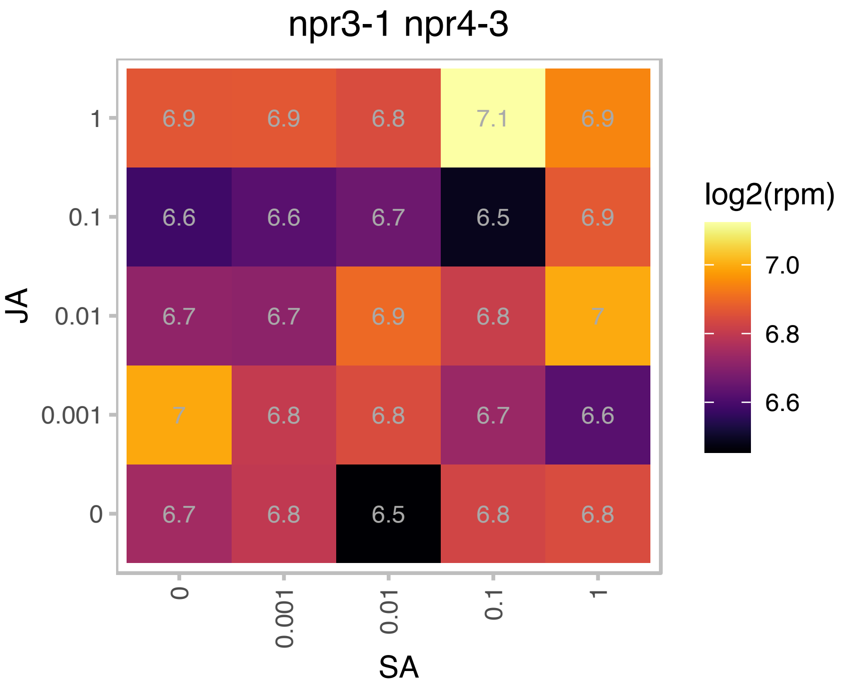

# AT3G27210.1 : NA

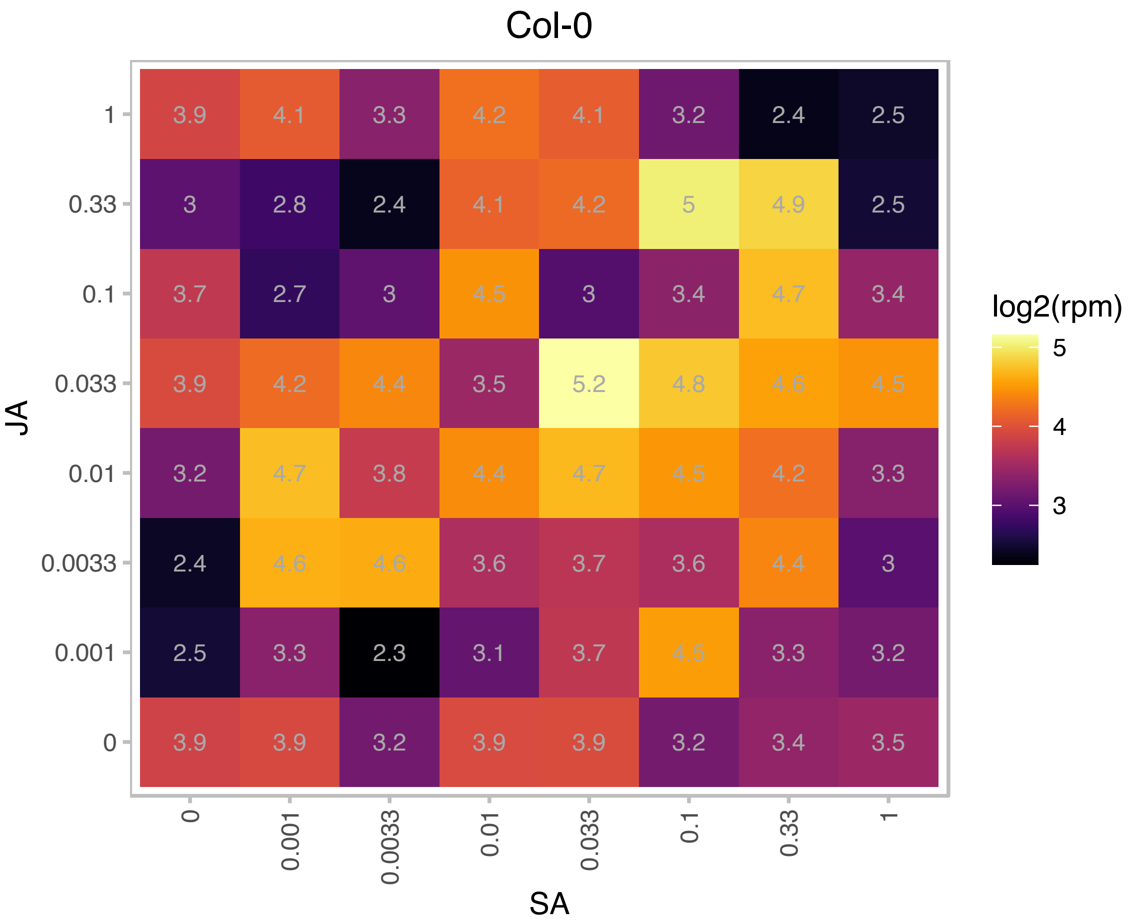


| | |
|---|---|
| Gene | AT3G27210 |
| is.expressed_genes | TRUE |
| is.responsive_genes | TRUE |
| is.apcluster | E-04 |

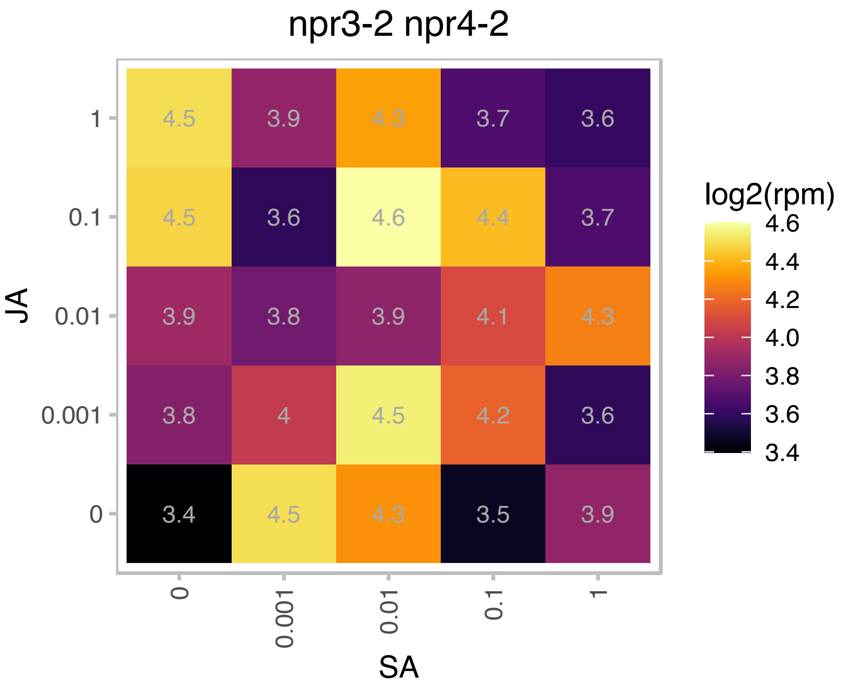


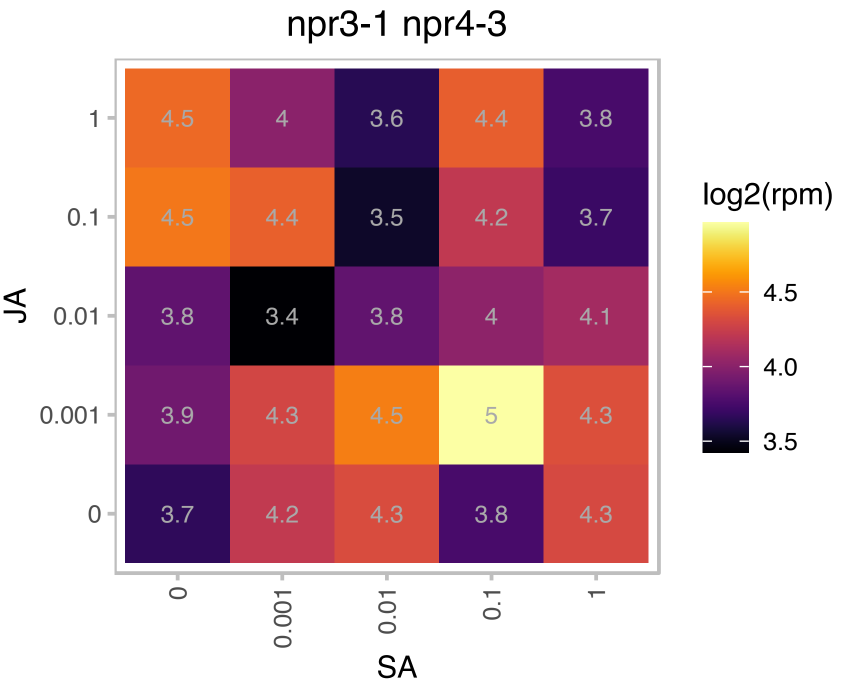

### AT3G56150.1 : ATEIF3C-1, ATTIF3C1, EIF3C, EIF3C-1, TI

eukaryotic translation initiation factor 3C

member of eIF3c - eukaryotic initiation factor 3c

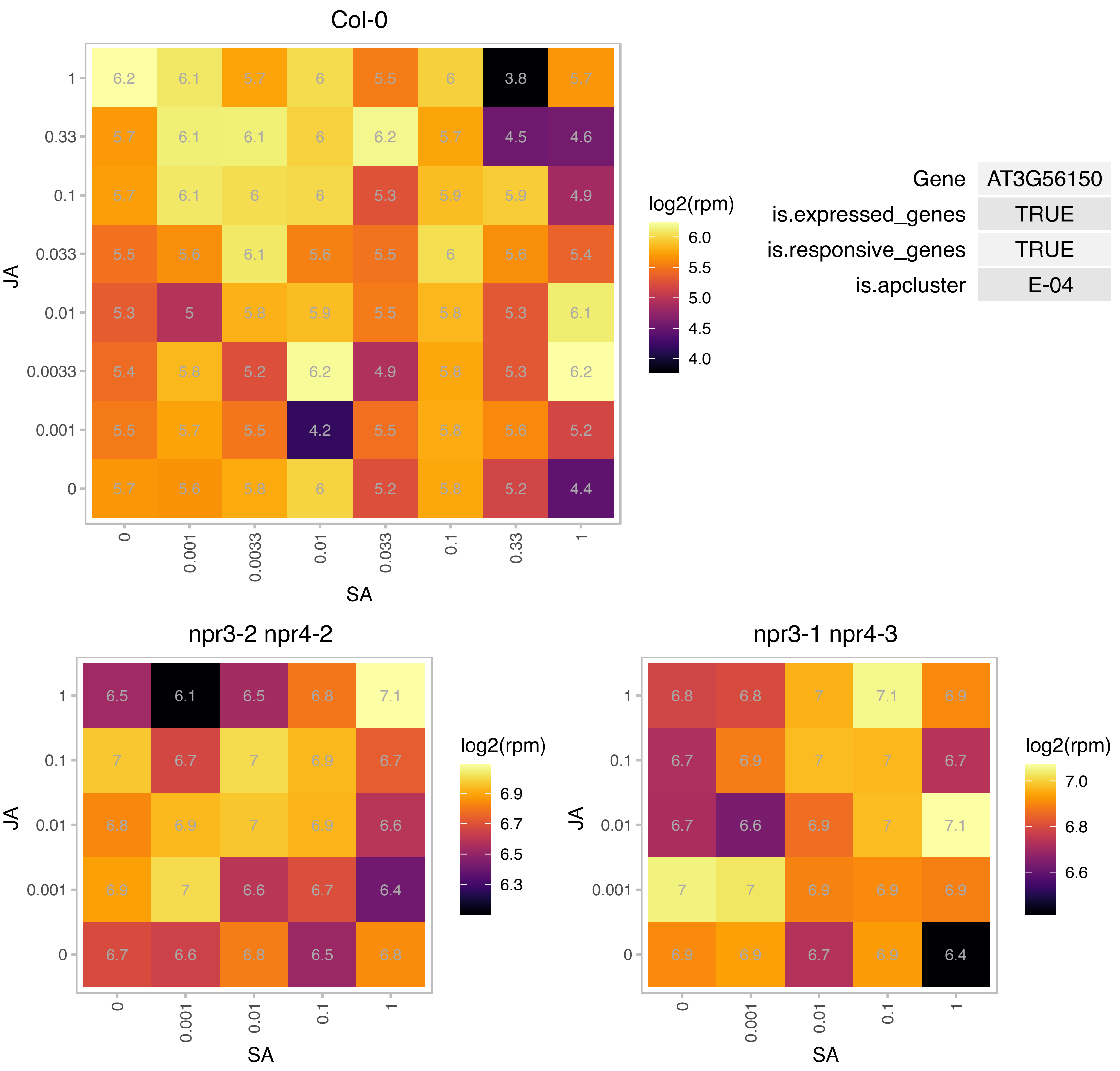

### AT3G60360.1 : EDA14, UTP11

embryo sac development arrest 14

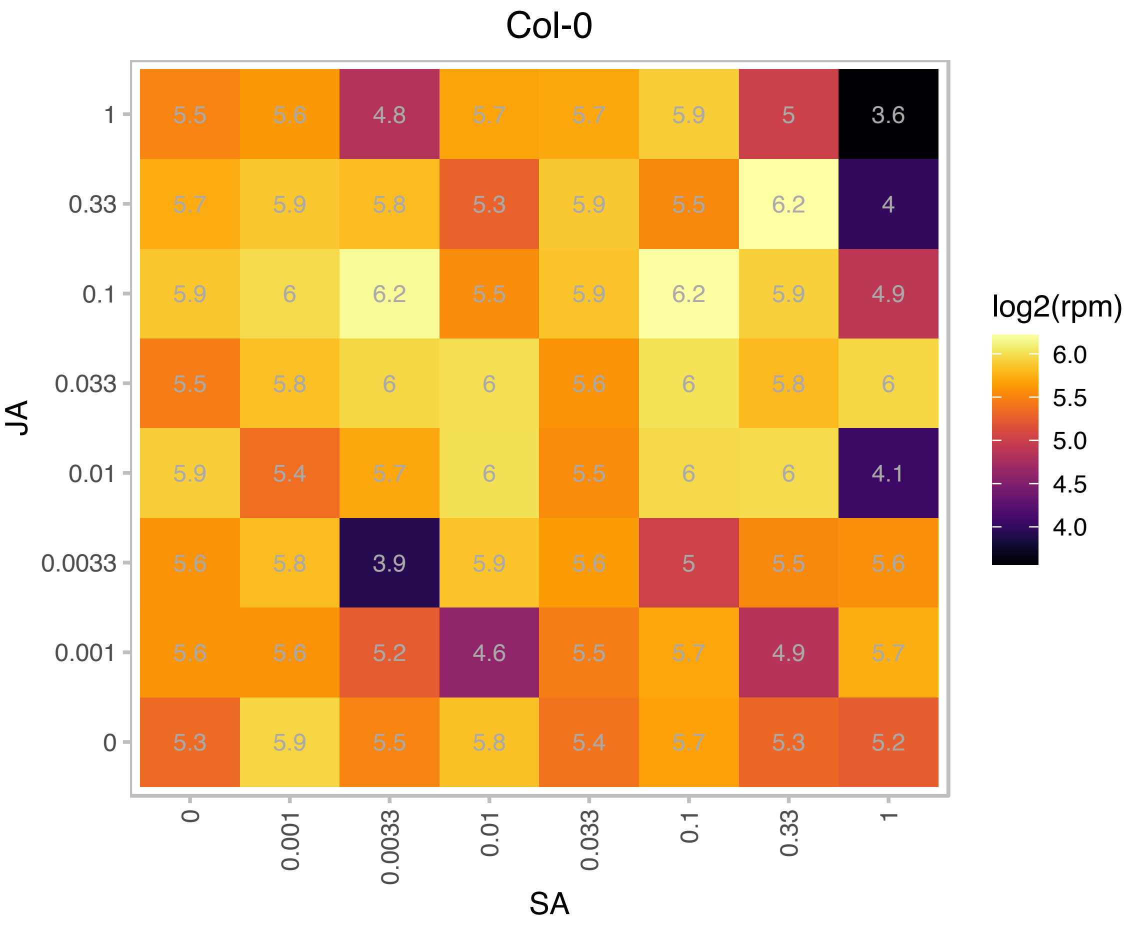


| | |
|---|---|
| Gene | AT3G60360 |
| is.expressed_genes | TRUE |
| is.responsive_genes | TRUE |
| is.apcluster | E-04 |

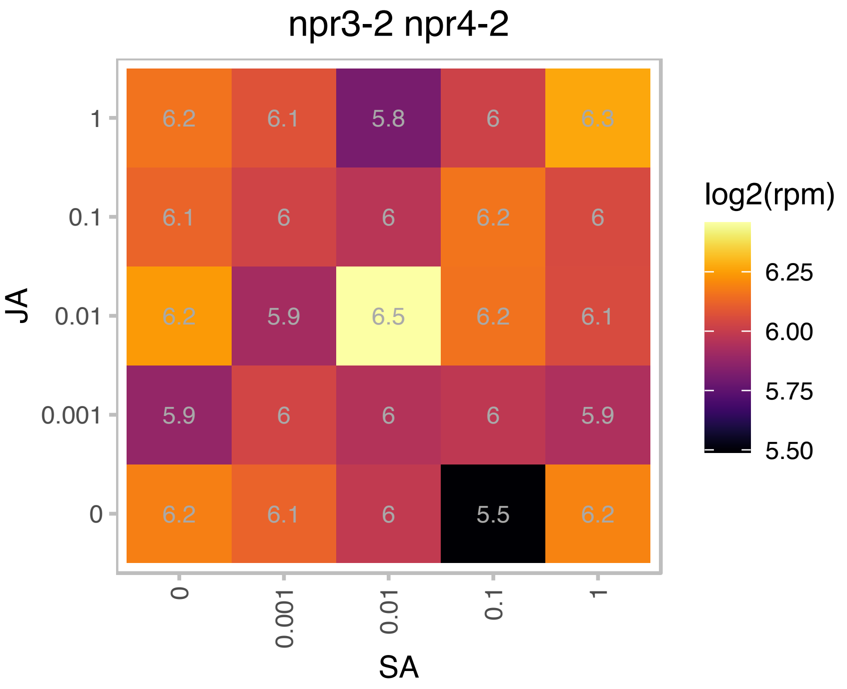


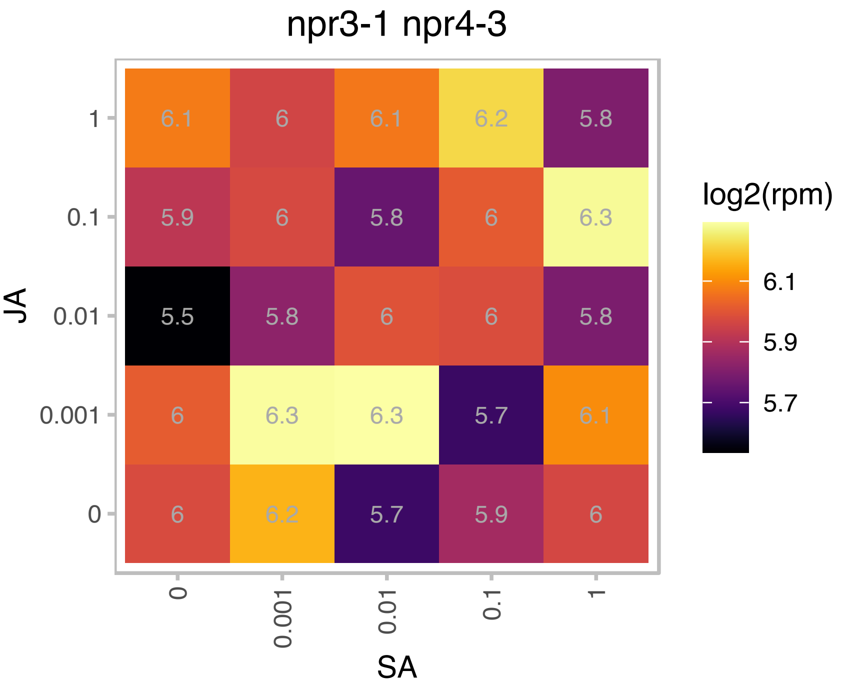

### AT4G01560.1 : MEE49

Ribosomal RNA processing Brix domain protein

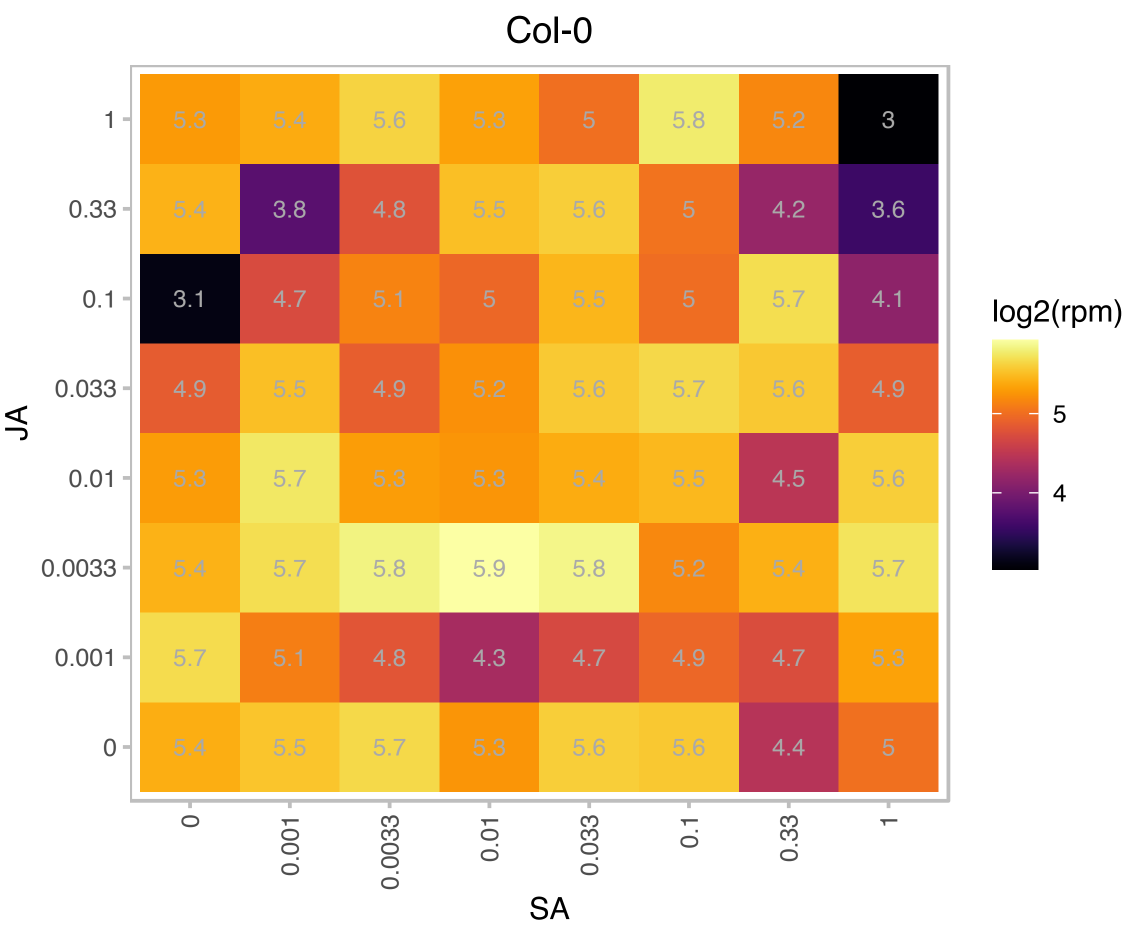


| | |
|---|---|
| Gene | AT4G01560 |
| is.expressed_genes | TRUE |
| is.responsive_genes | TRUE |
| is.apcluster | E-04 |

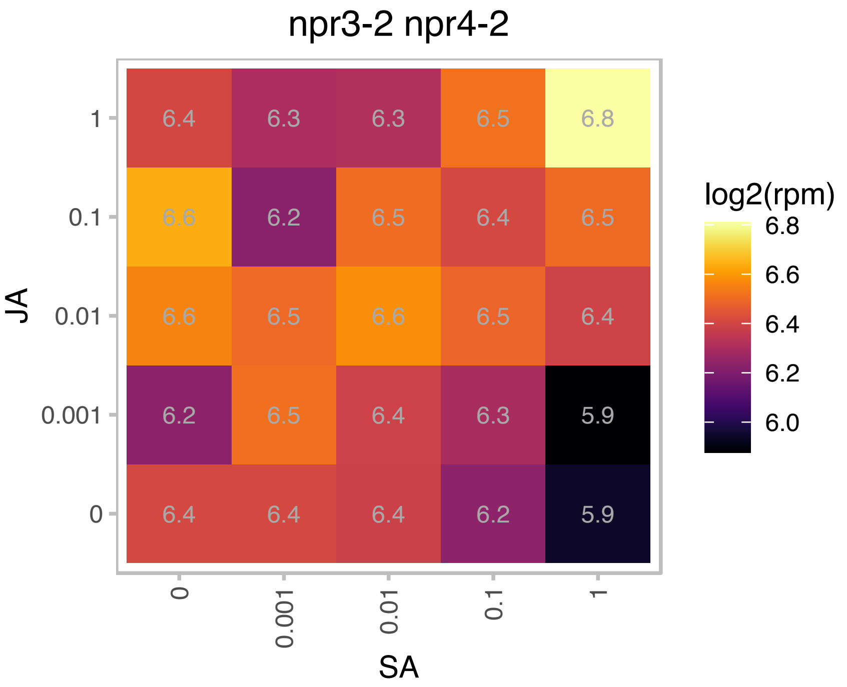


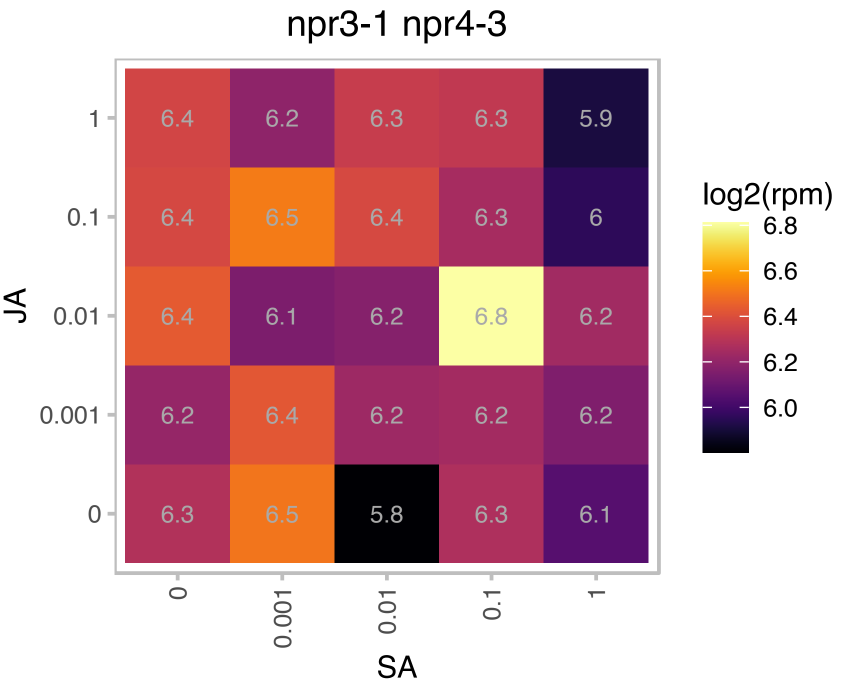

### AT4G29130.1 : ATHXK1, GIN2, HXK1

hexokinase 1
Encodes a hexokinase (HXK1) in the plant glucose-signaling network. Functions as a glucose sensor to interrelate nutrient, light, and hormone signaling networks for controlling growth and development in response to the changing environment.

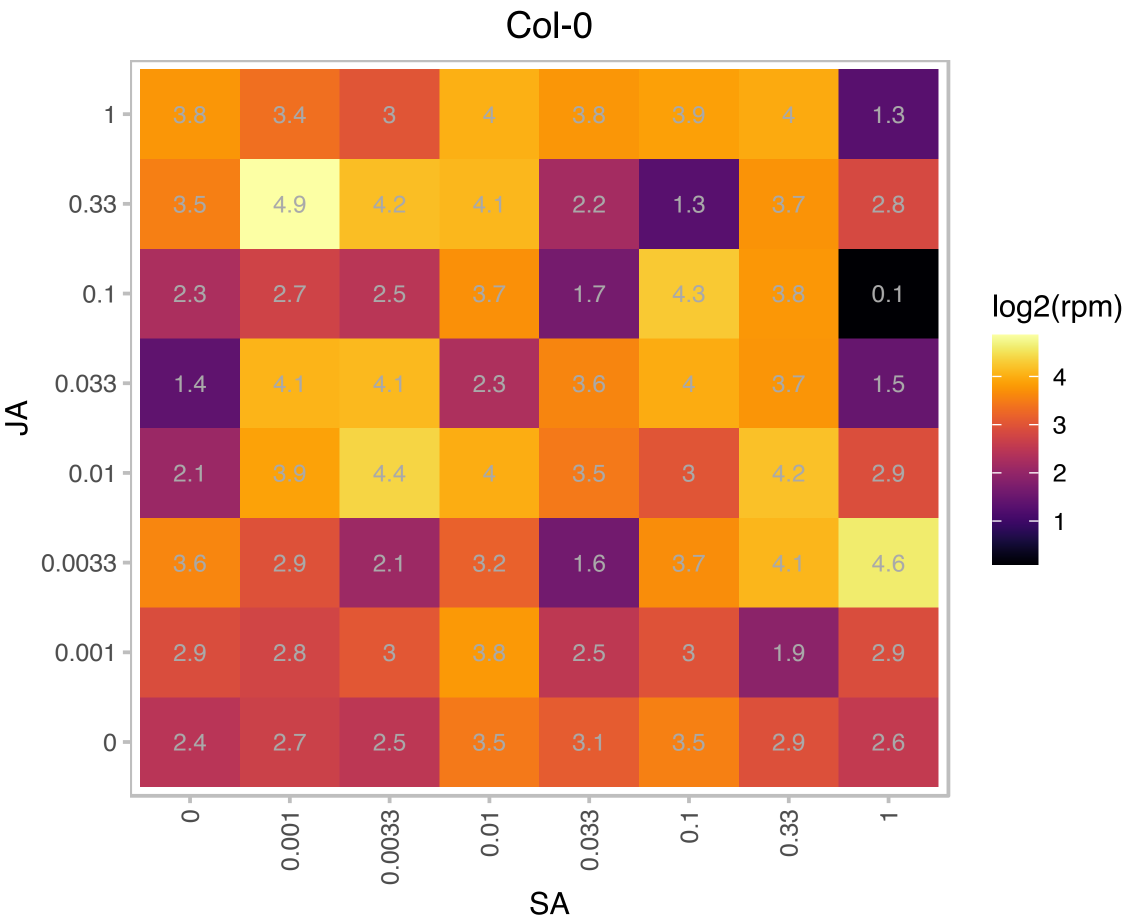


| | |
|---|---|
| Gene | AT4G29130 |
| is.expressed_genes | TRUE |
| is.responsive_genes | TRUE |
| is.apcluster | E-04 |

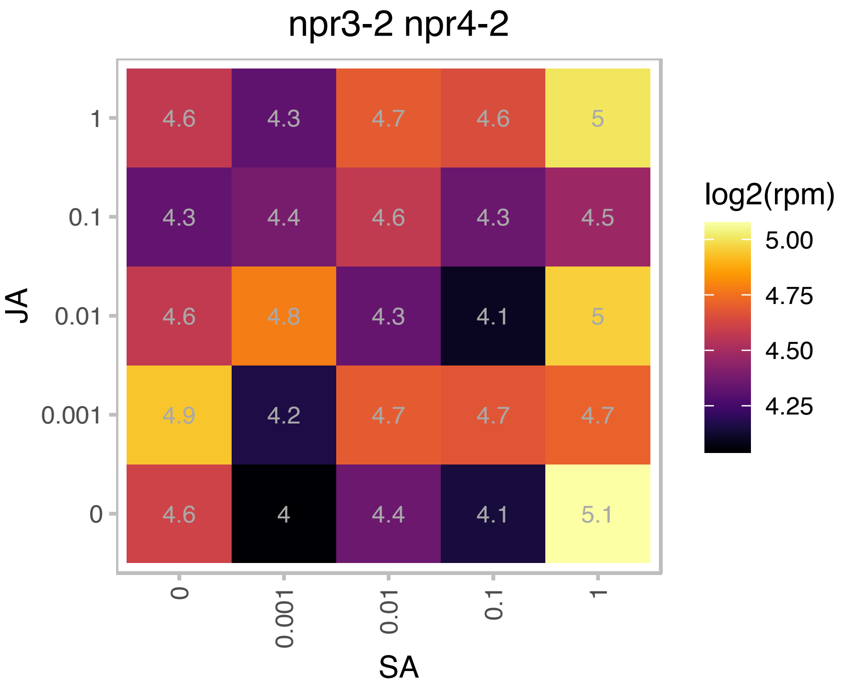


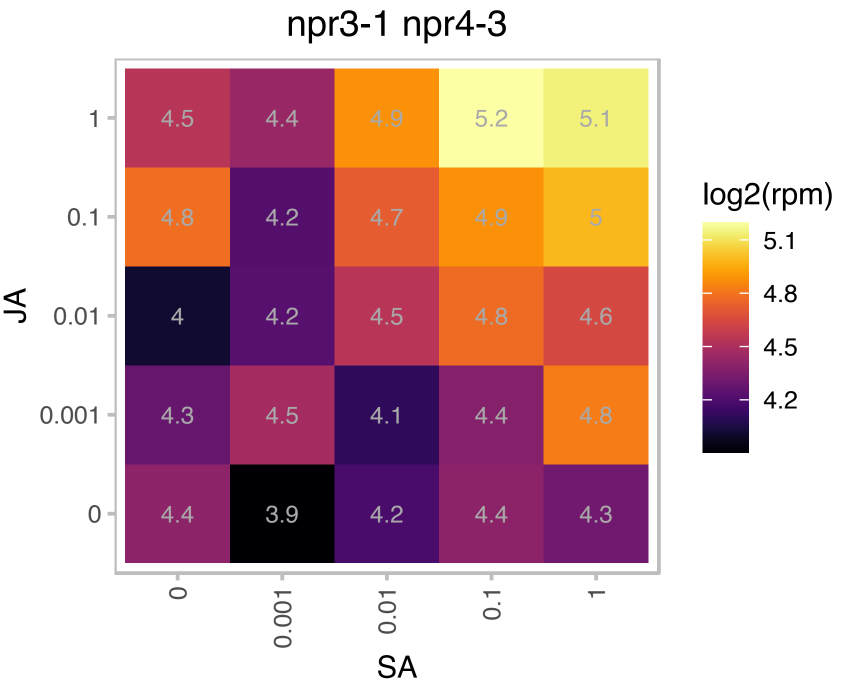

# AT4G30750.1 : NA

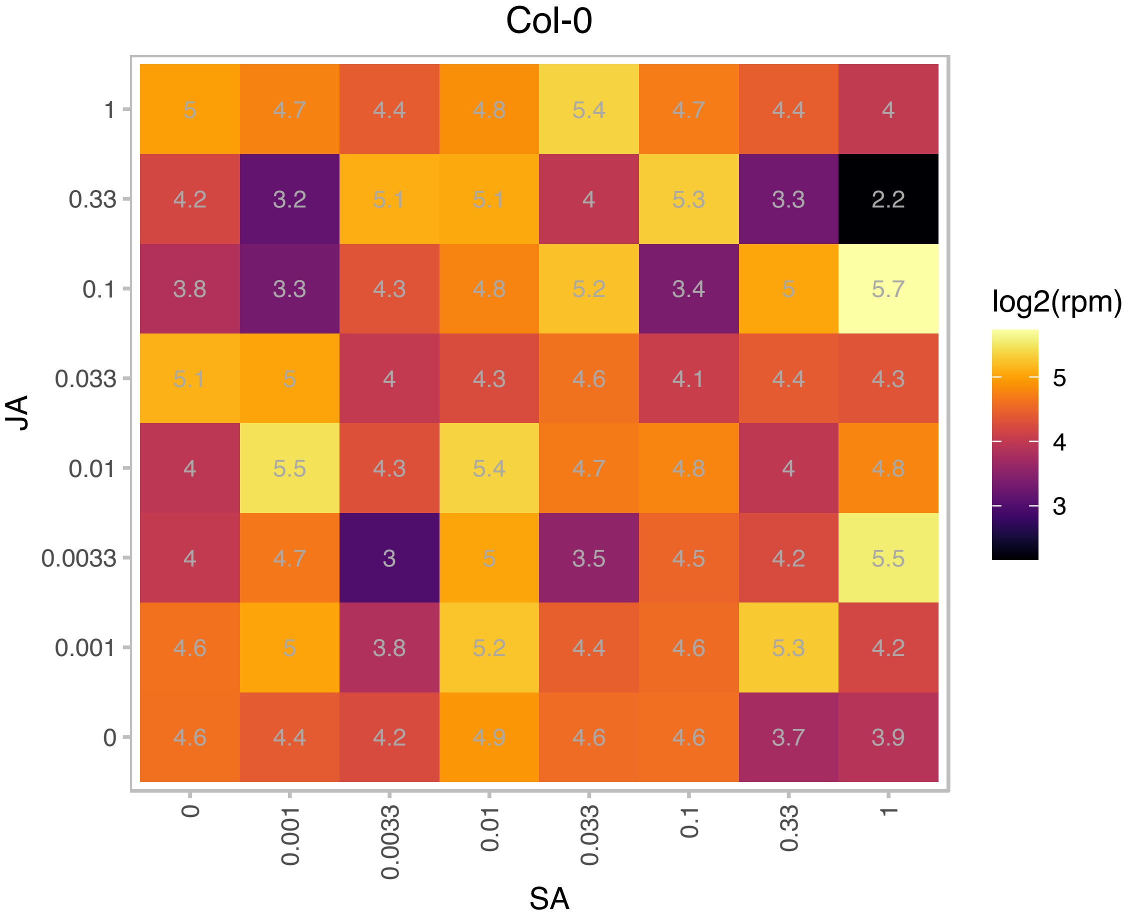


| | |
|---|---|
| Gene | AT4G30750 |
| is.expressed_genes | TRUE |
| is.responsive_genes | TRUE |
| is.apcluster | E-04 |

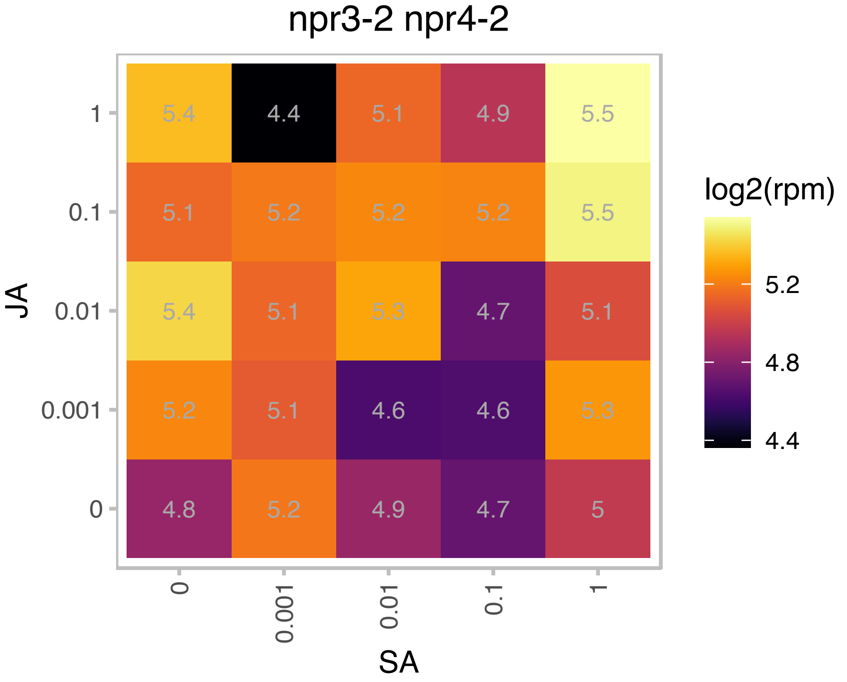


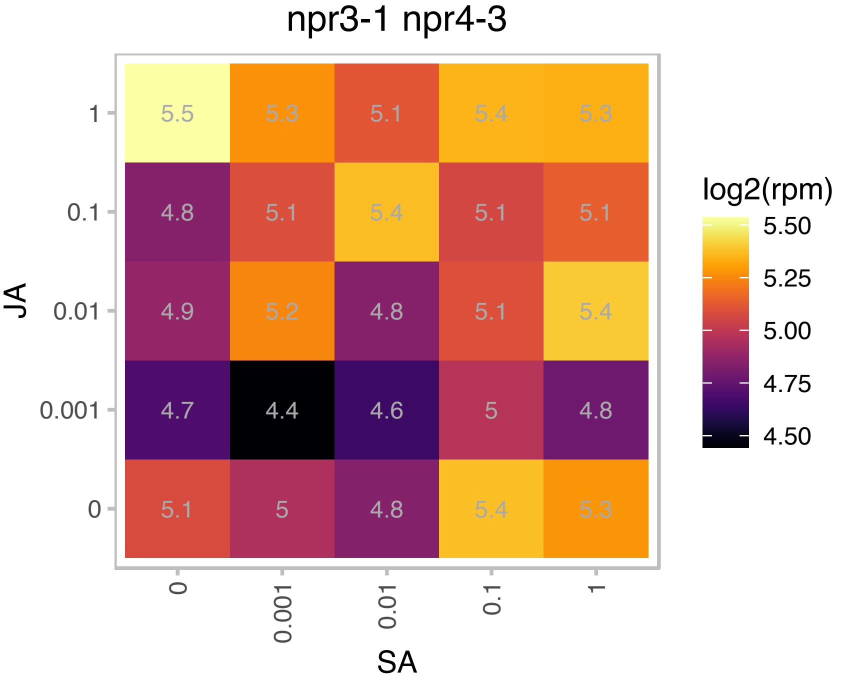

# AT4G30760.1 : NA

Putative endonuclease or glycosyl hydrolase

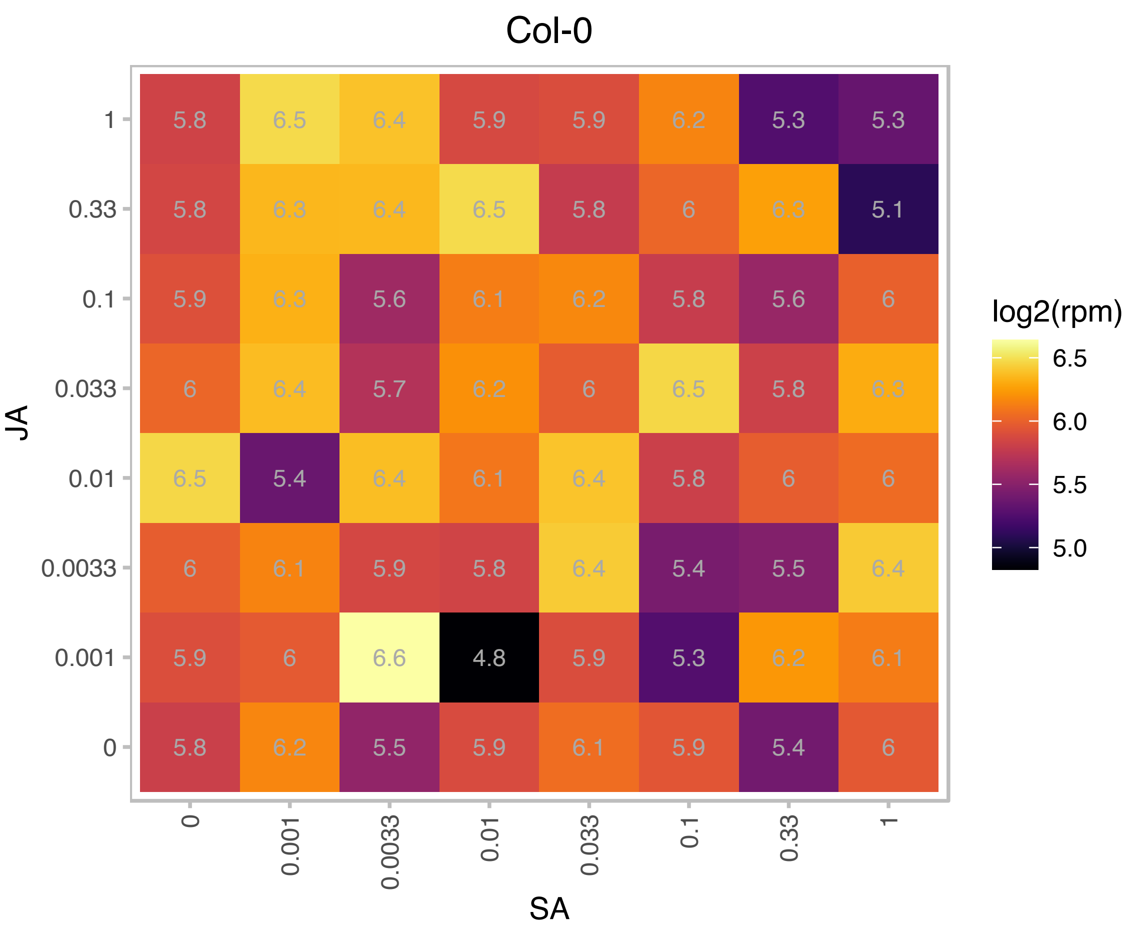


| | |
|---|---|
| Gene | AT4G30760 |
| is.expressed_genes | TRUE |
| is.responsive_genes | TRUE |
| is.apcluster | E-04 |

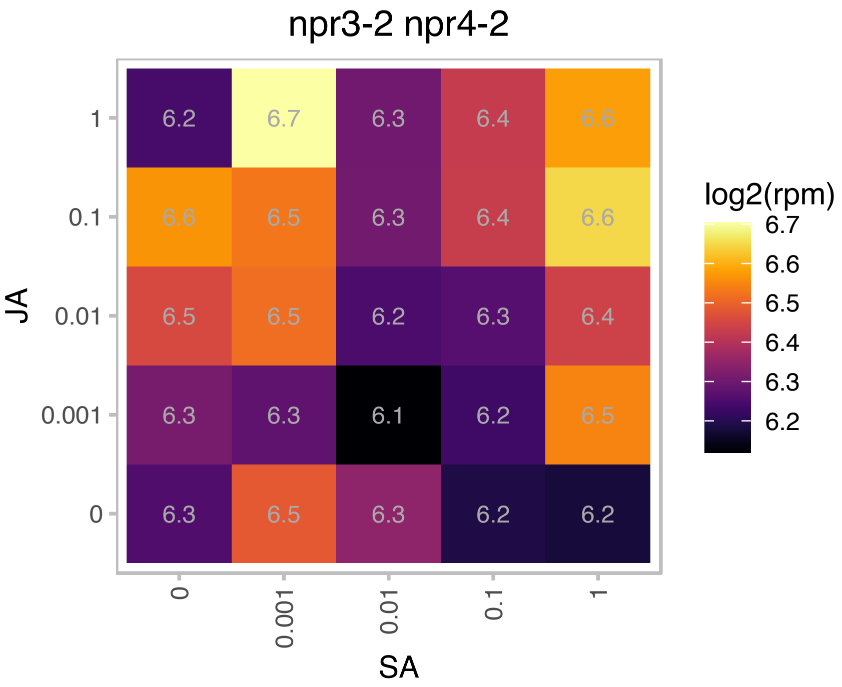


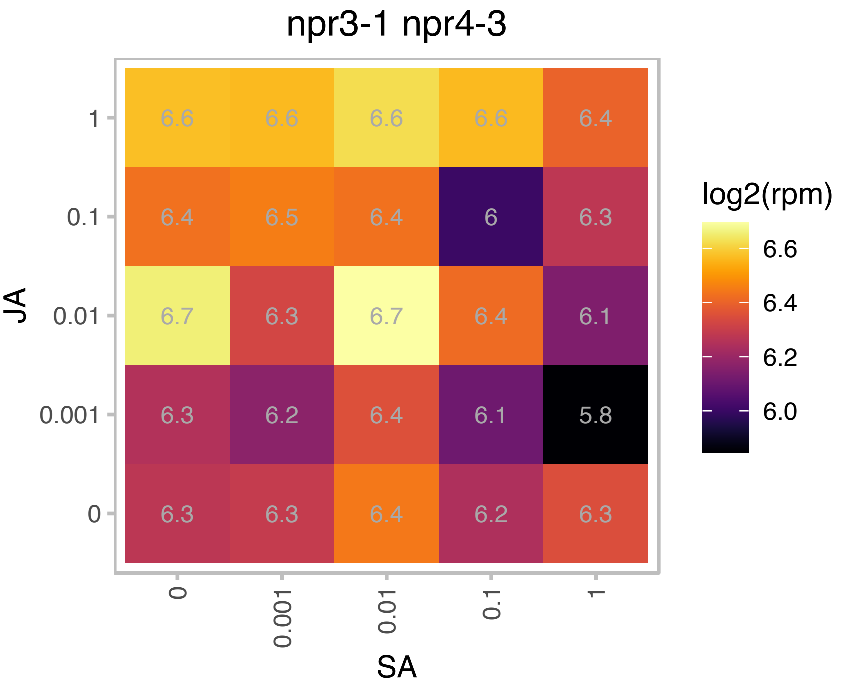

# AT5G02244.1 : NA

Pseudogene of AT3G57870; AHUS5 (SUMO CONJUGATION ENZYME 1); ubiquitin-protein ligase

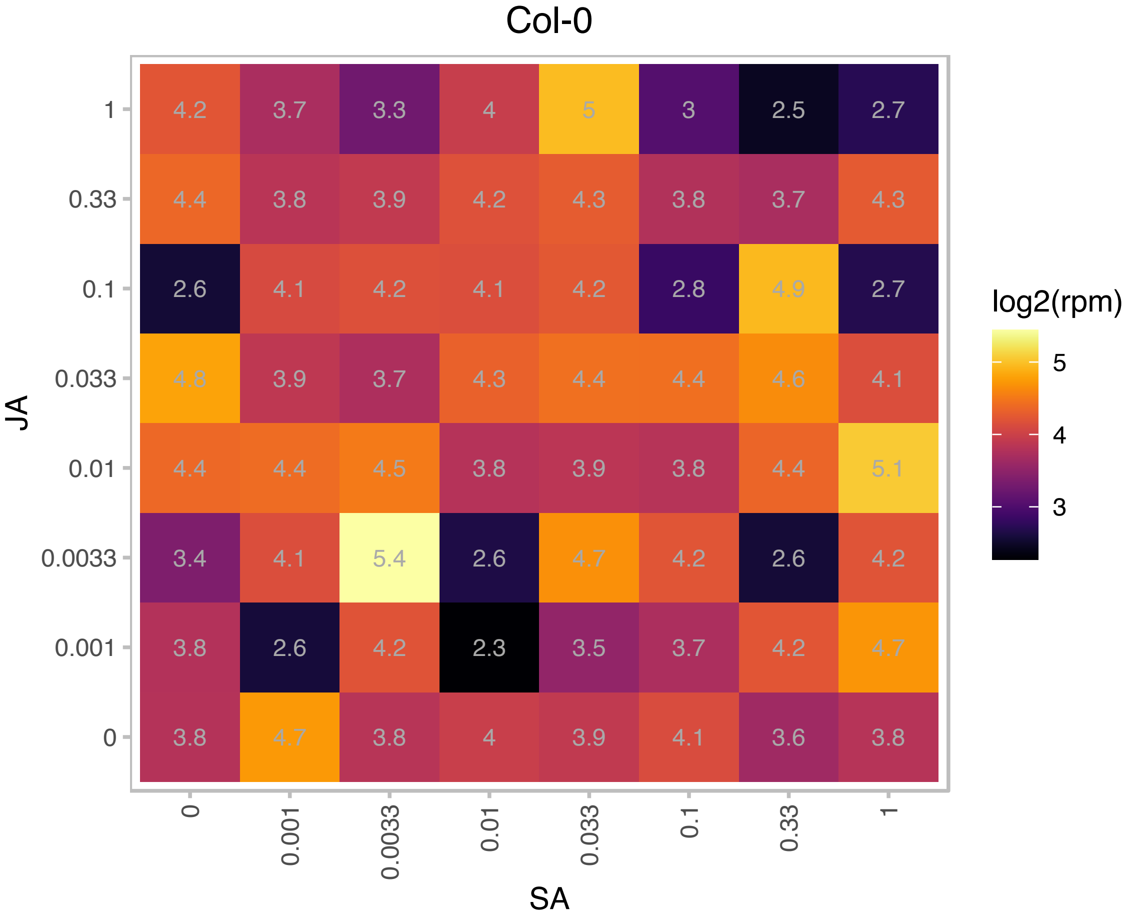


| Gene | AT5G02244 |
| --- | --- |
| is.expressed_genes | TRUE |
| is.responsive_genes | TRUE |
| is.apcluster | E-04 |

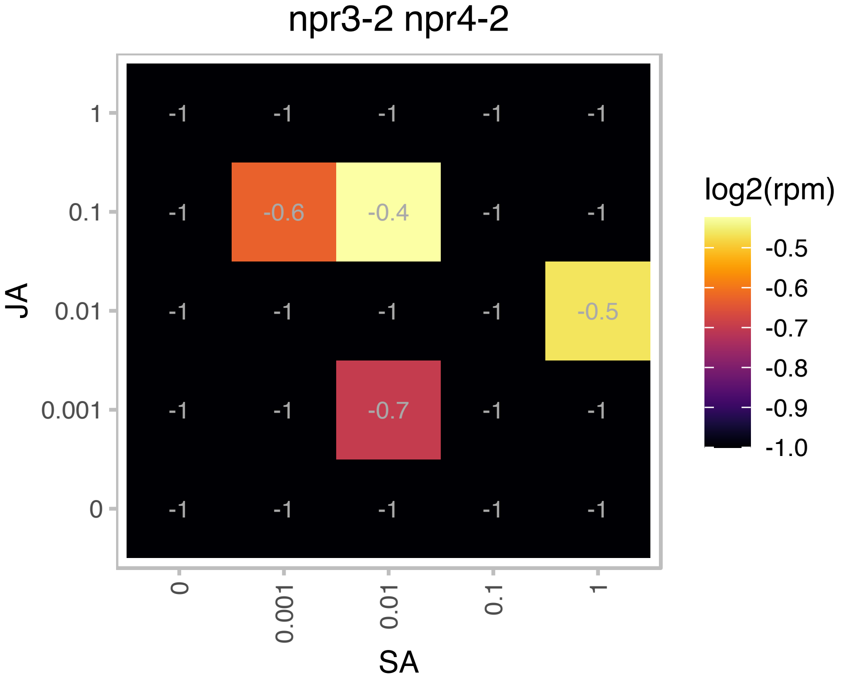


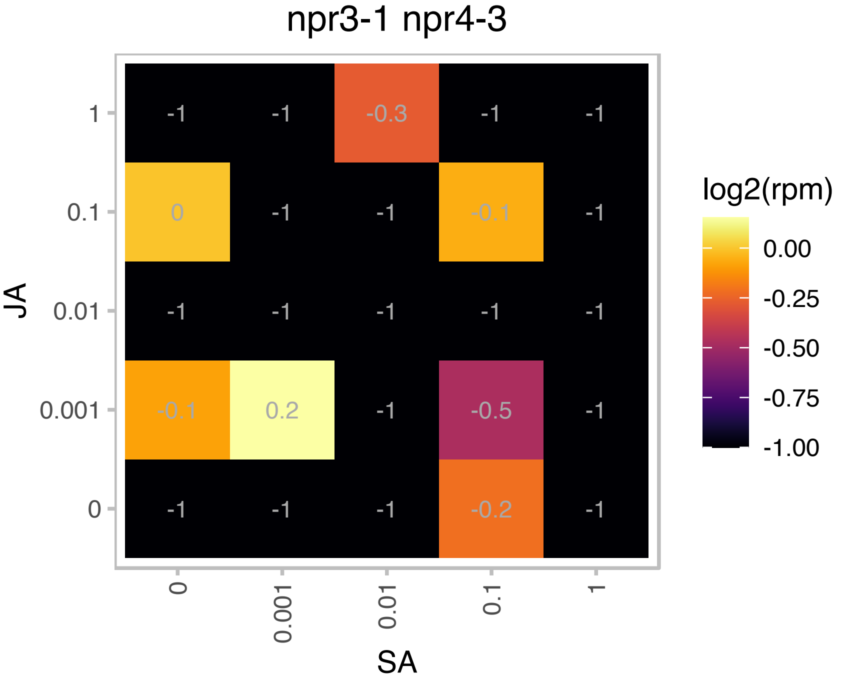

### AT5G07000.1 : ATST2B, ST2B

sulfotransferase 2B
Encodes a member of the sulfotransferase family of proteins. Although it has 85% amino acid identity with ST2A (At5g07010), this protein is not able to transfer a sulfate group to 11- or 12-hydroxyjasmonic acid in vitro. It may be able to act on structurally related jasmonates.

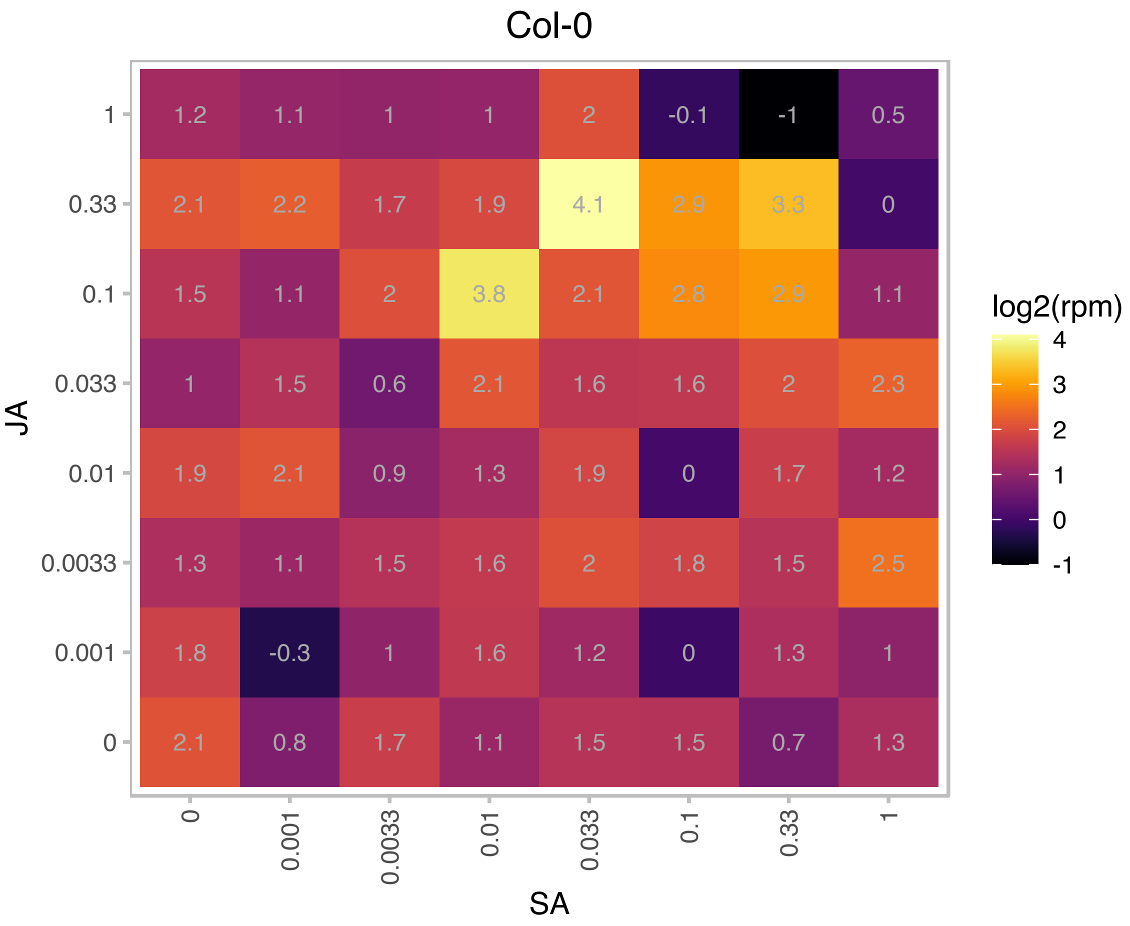


| | |
|---|---|
| Gene | AT5G07000 |
| is.expressed_genes | TRUE |
| is.responsive_genes | TRUE |
| is.apcluster | E-04 |

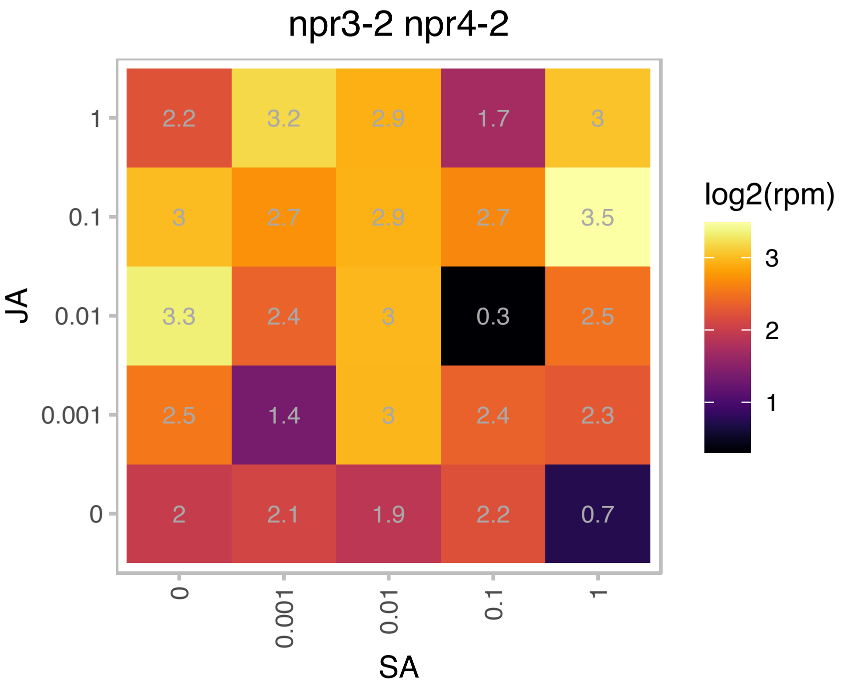


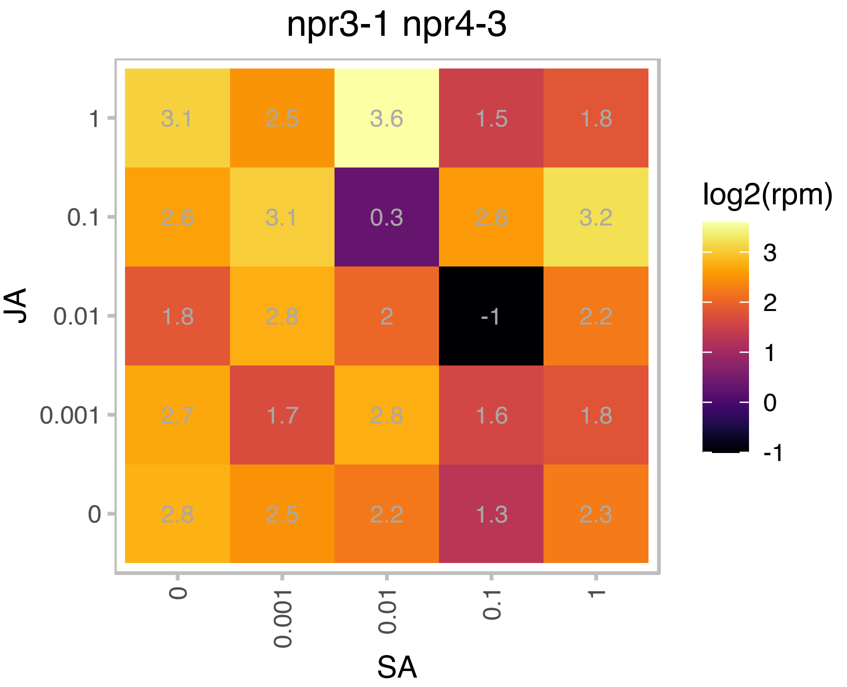

# AT5G16070.1 : NA

TCP-1/cpn60 chaperonin family protein

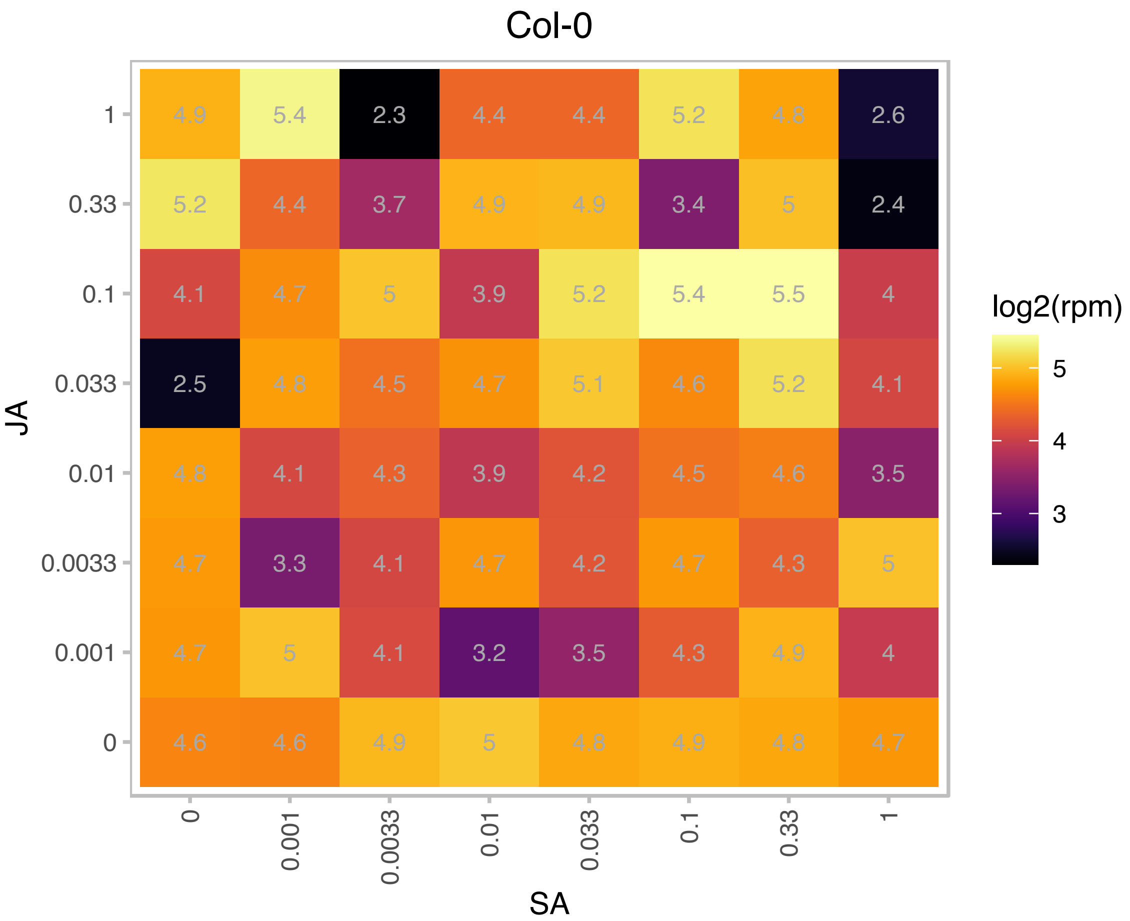


| | |
|---|---|
| Gene | AT5G16070 |
| is.expressed_genes | TRUE |
| is.responsive_genes | TRUE |
| is.apcluster | E-04 |

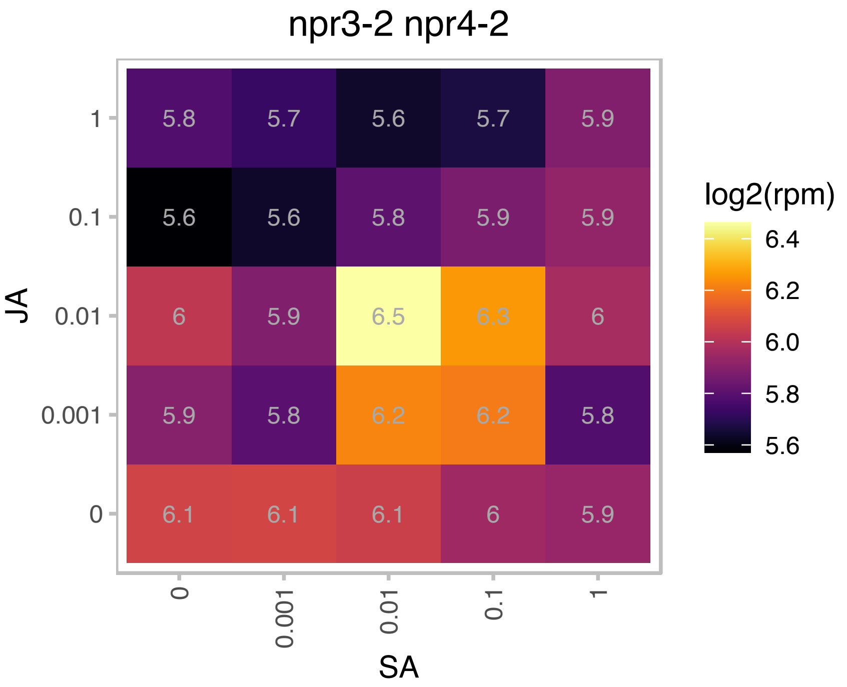


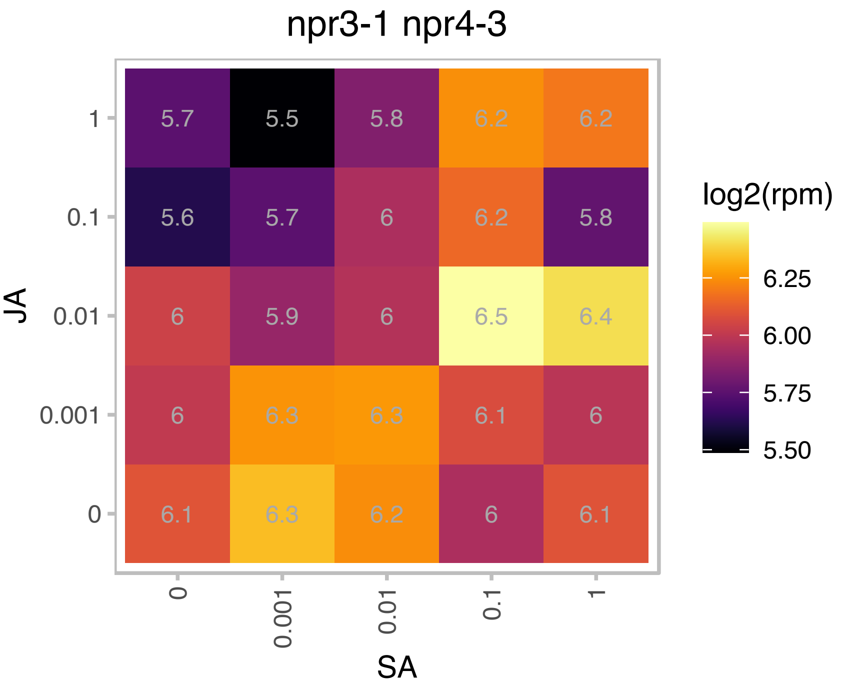

### AT5G60640.1 : ATPDI2, ATPDIL1-4, PDI2, PDIL1-4

PDI-like 1-4

Encodes a protein disulfide isomerase-like (PDIL) protein, a member of a multigene family within the thioredoxin (TRX) superfamily. Unlike several other PDI family members, transcript levels for this gene are not up-regulated in response to three different chemical inducers of ER stress (dithiothreitol, beta-mercaptoethanol, and tunicamycin). However, the level of transcripts for this gene is slightly elevated in atbzip60 mutants.

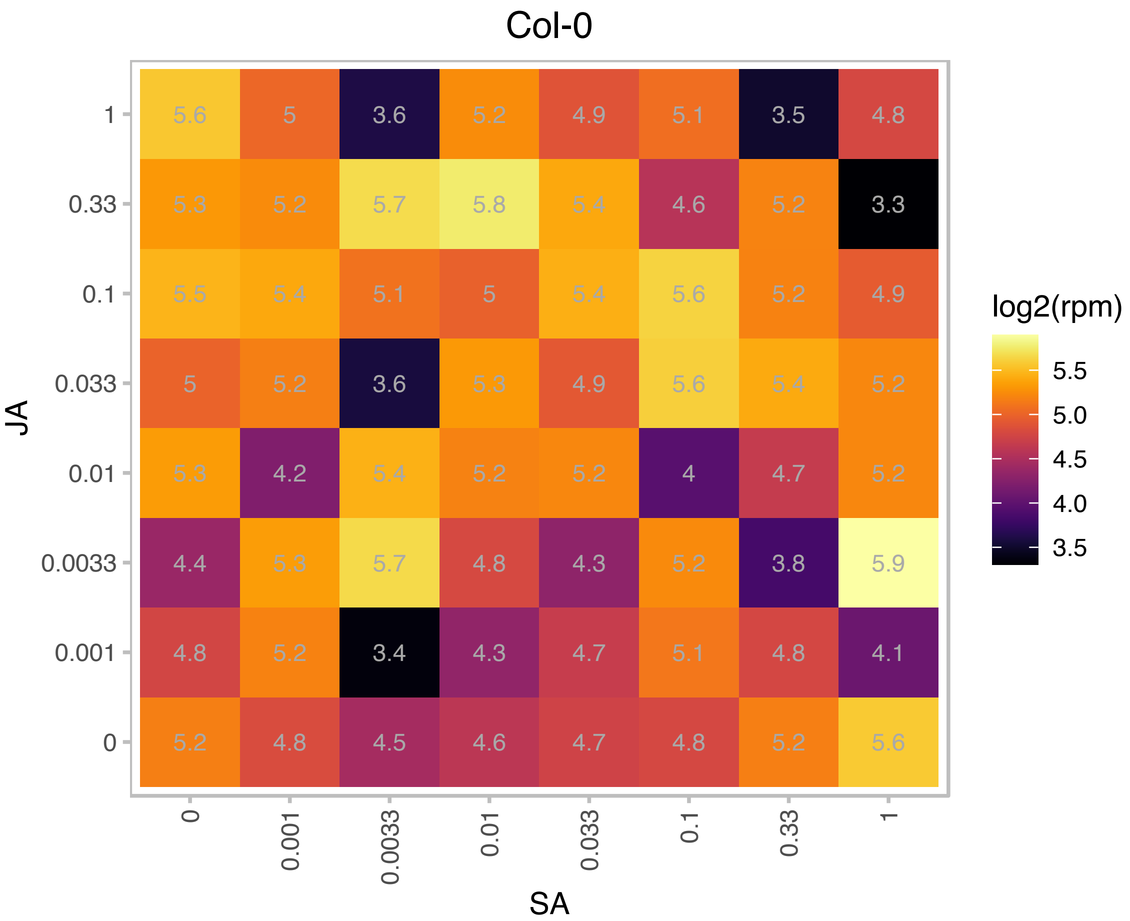


| | |
|---|---|
| Gene | AT5G60640 |
| is.expressed_genes | TRUE |
| is.responsive_genes | TRUE |
| is.apcluster | E-04 |

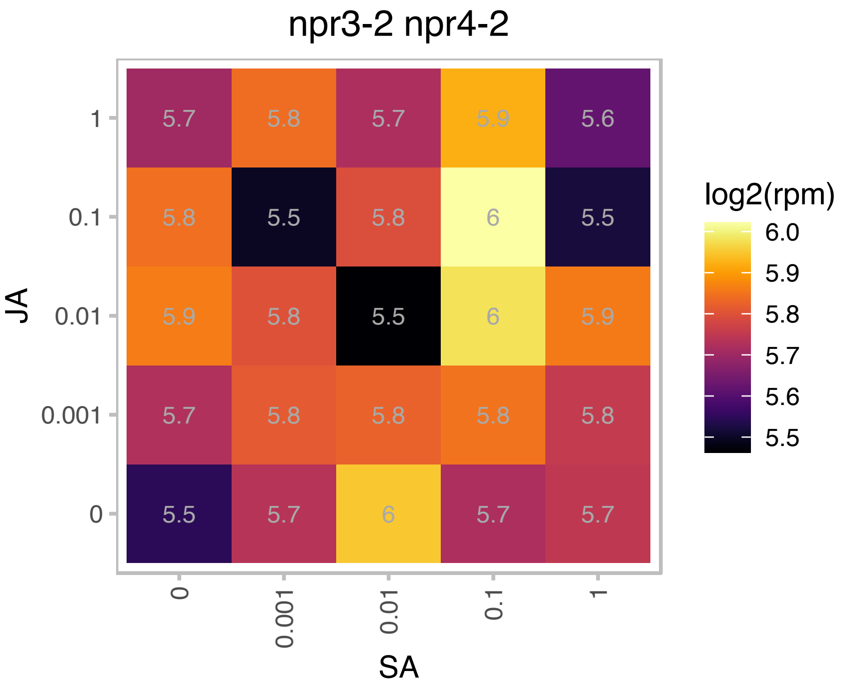


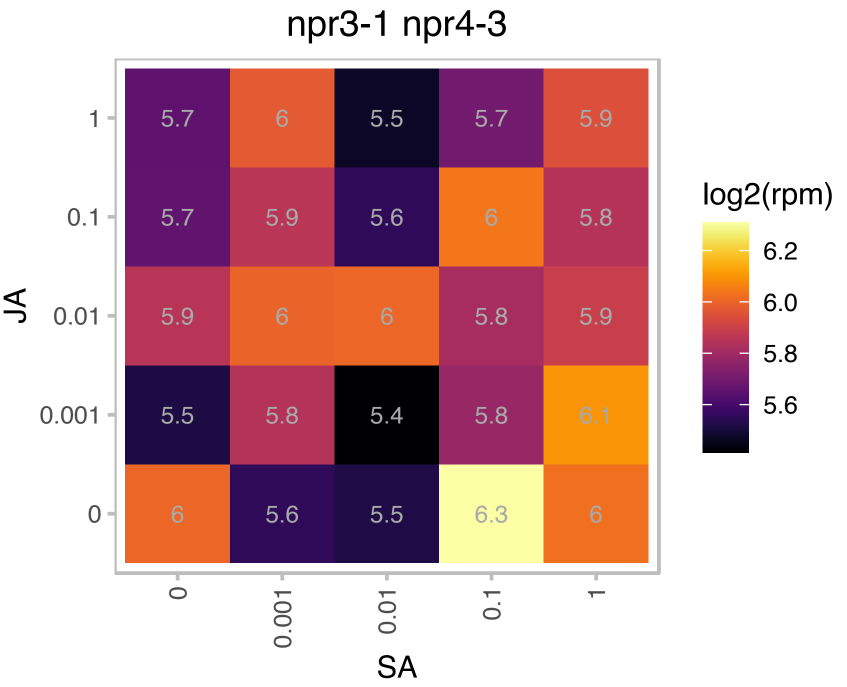

# AT5G63010.1 : NA

Transducin/WD40 repeat-like superfamily protein

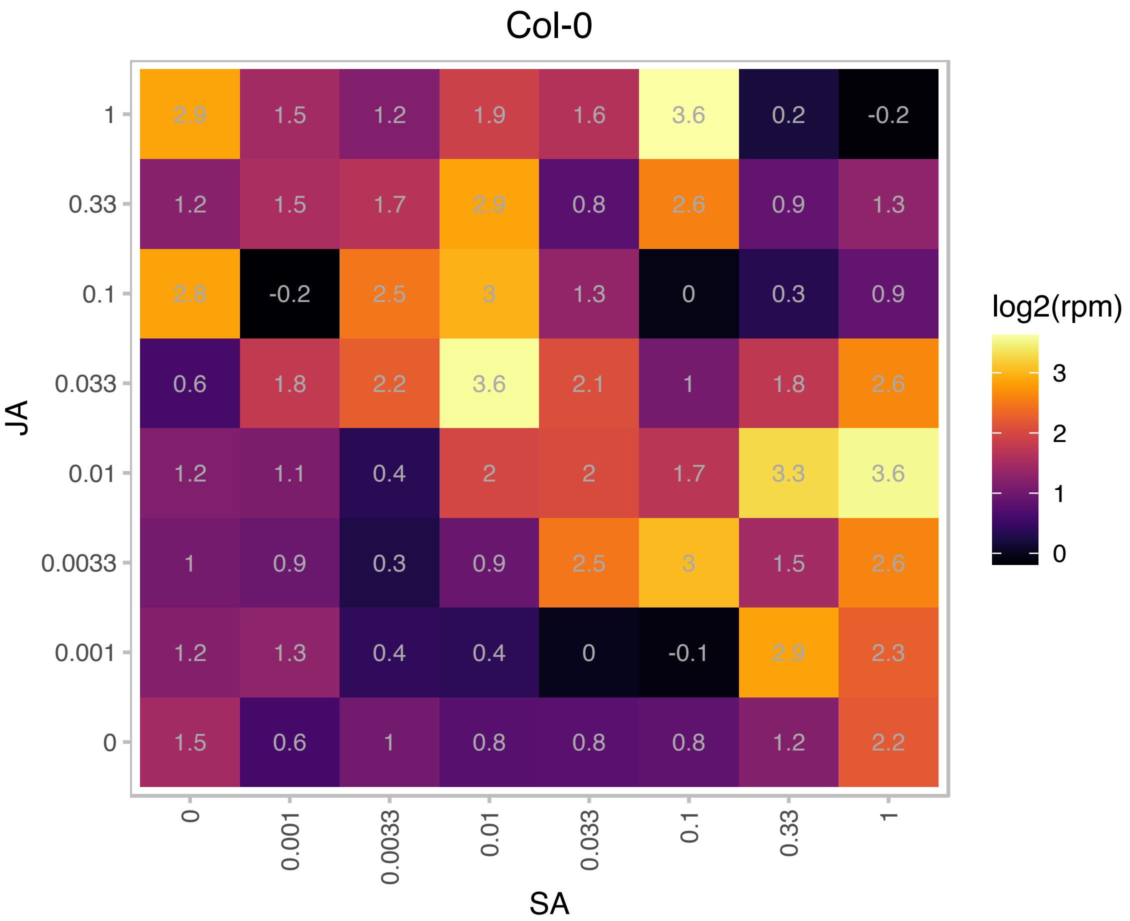


| | |
|---|---|
| Gene | AT5G63010 |
| is.expressed_genes | TRUE |
| is.responsive_genes | TRUE |
| is.apcluster | E-04 |

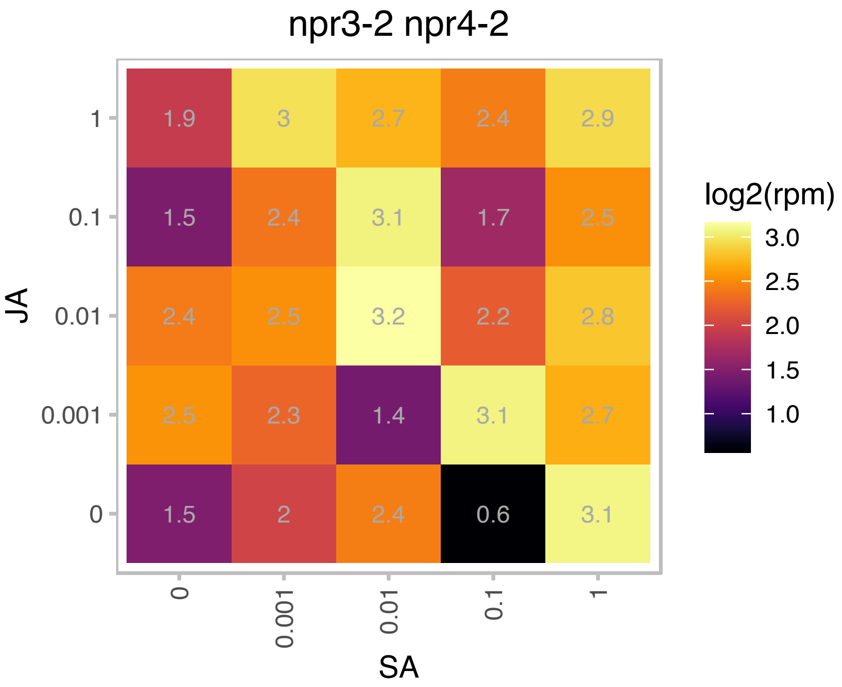


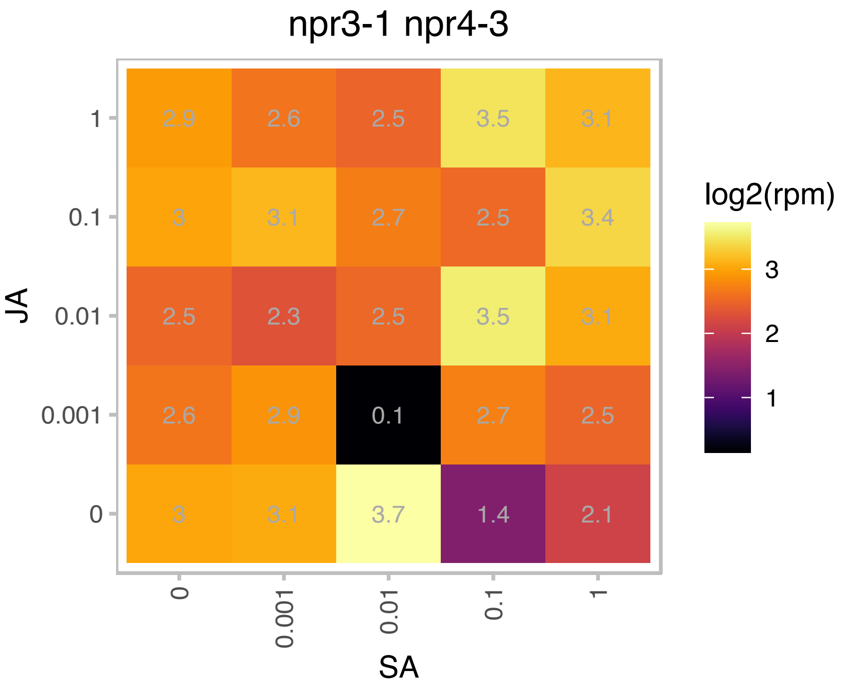

# AT5G65260.1 : NA

RNA-binding (RRM/RBD/RNP motifs) family protein

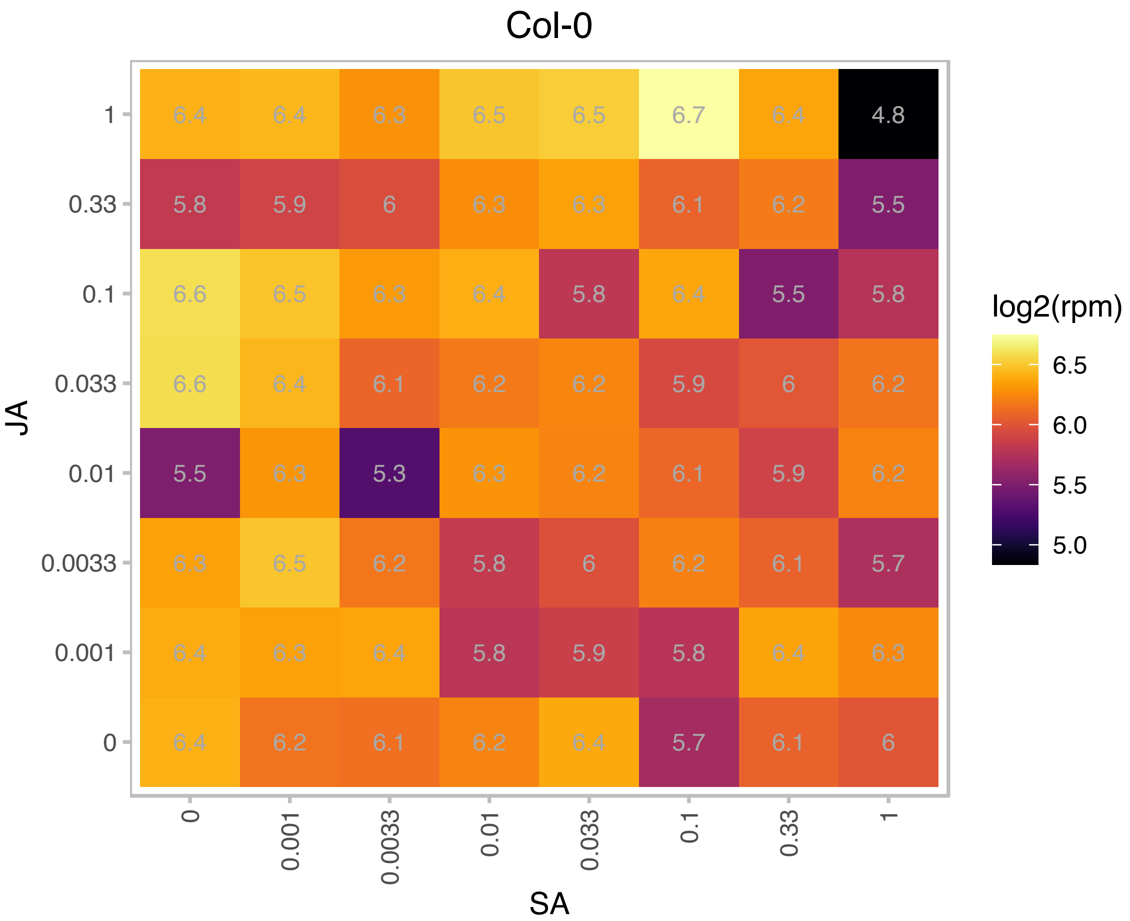


| | |
|---|---|
| Gene | AT5G65260 |
| is.expressed_genes | TRUE |
| is.responsive_genes | TRUE |
| is.apcluster | E-04 |

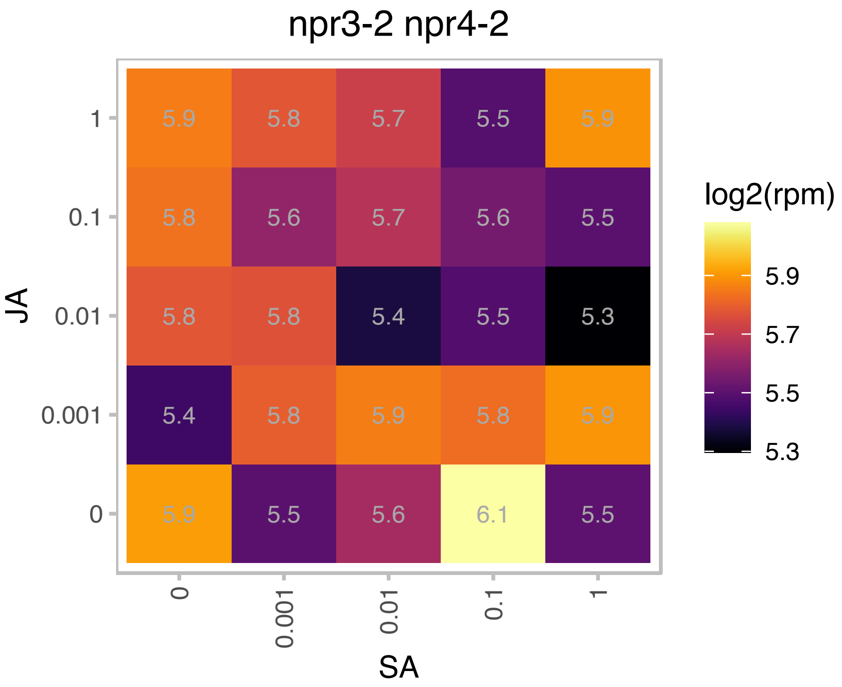


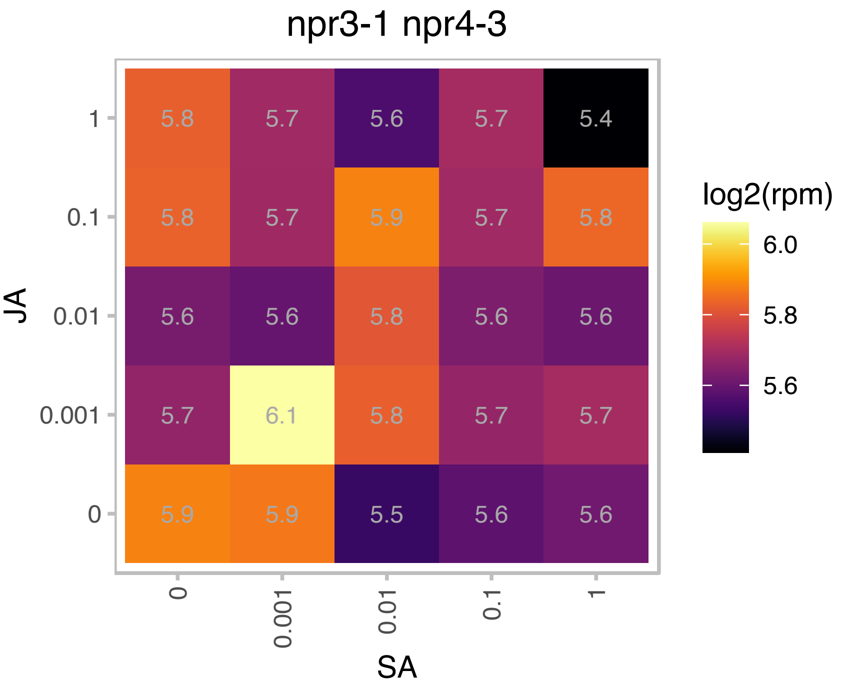

### AT5G66140.1 : PAD2

proteasome alpha subunit D2

Encodes alpha5 subunit of 20S proteosome complex involved in protein degradation.

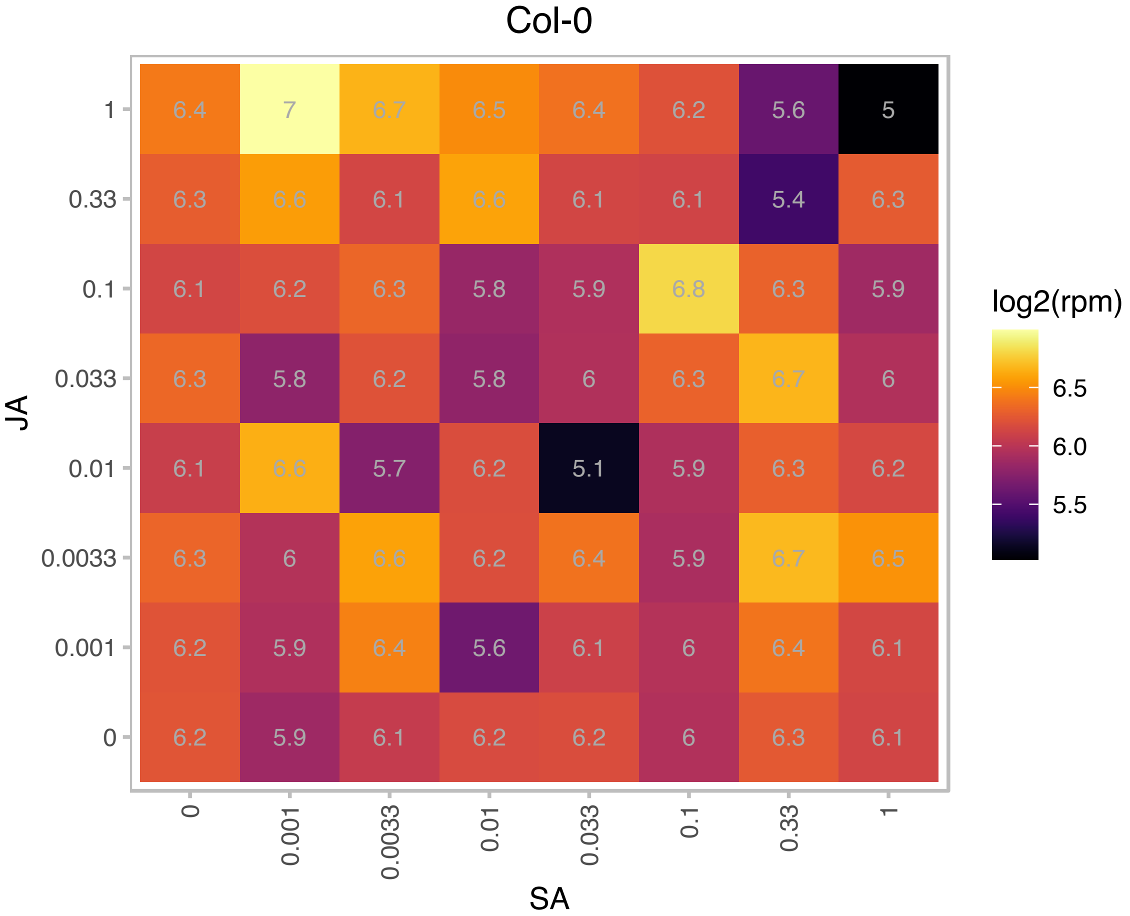


| | |
|---|---|
| Gene | AT5G66140 |
| is.expressed_genes | TRUE |
| is.responsive_genes | TRUE |
| is.apcluster | E-04 |

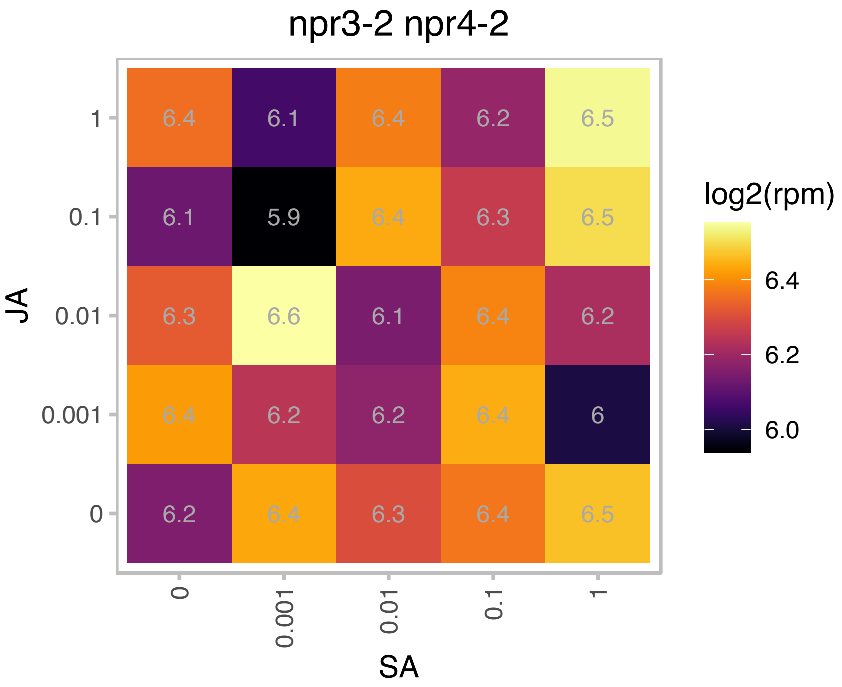


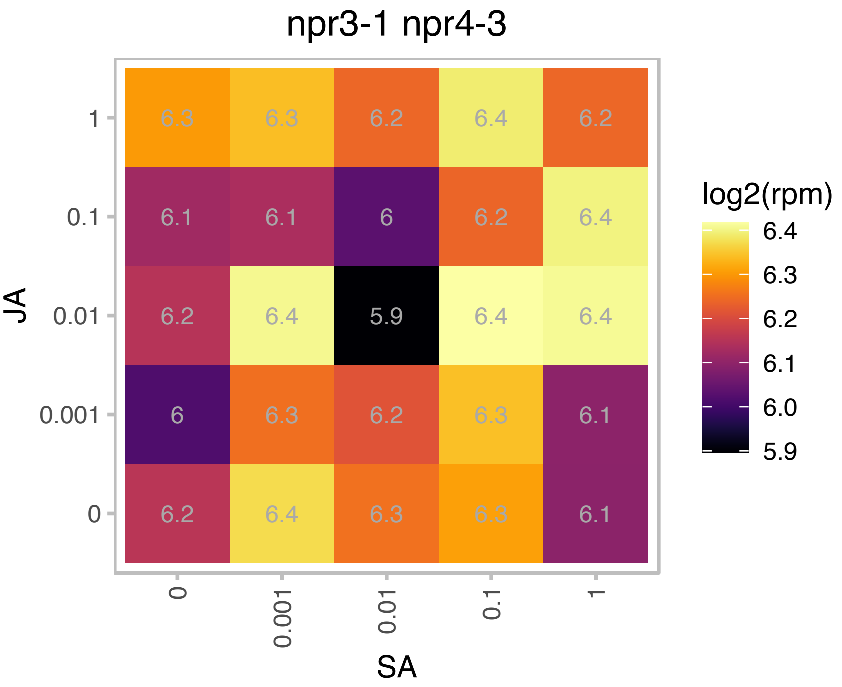

### AT1G06950.1 : ATTIC110, TIC110

translocon at the inner envelope membrane of chloroplasts 110
Encodes a protein thought to be a part of the translocon at the chloroplast inner envelope. Involved in protein import into the chloroplast and chloroplast biogenesis. C-terminal half of Tic110 functions as scaffolds for protein-protein interactions.

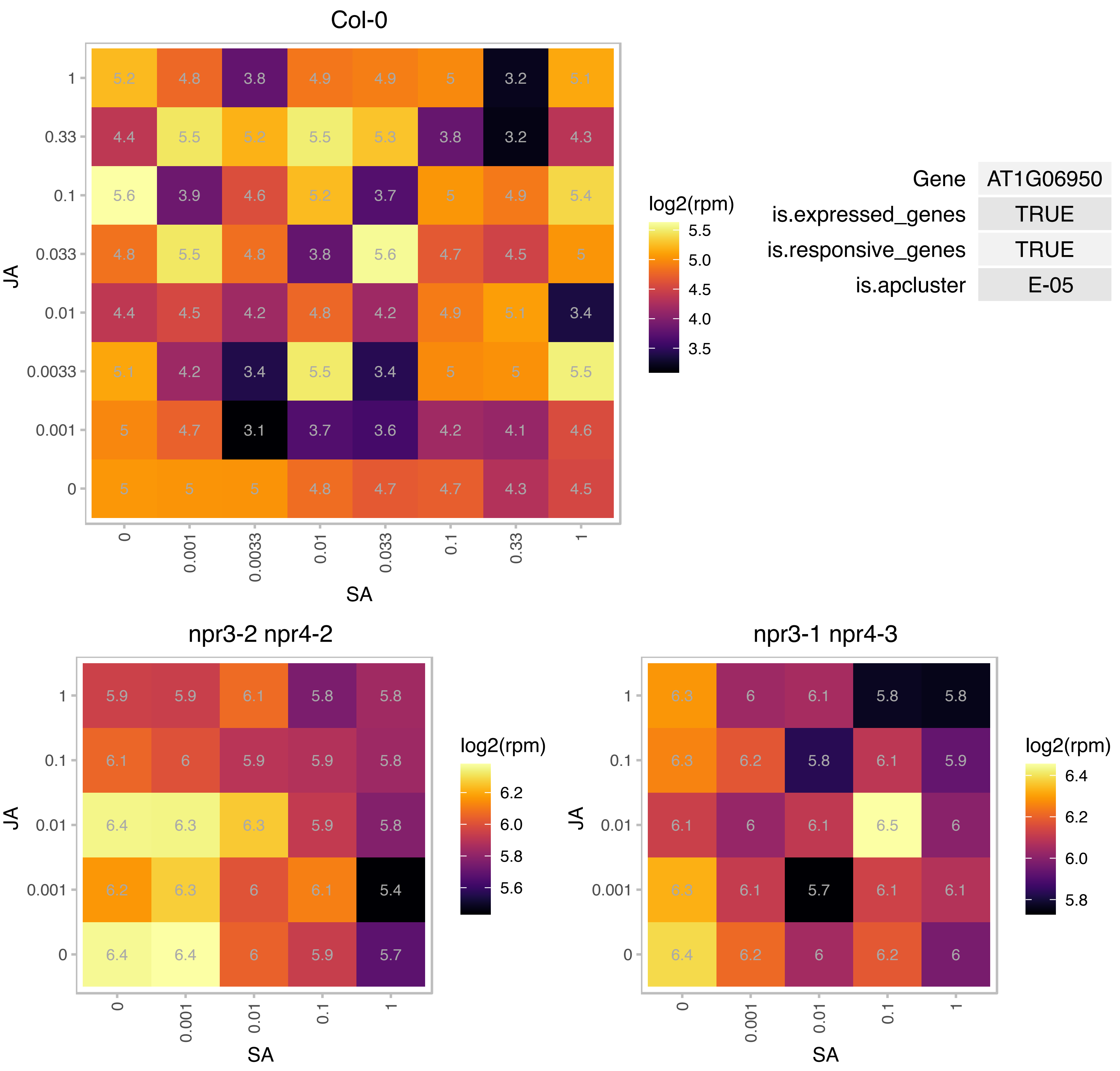


| | |
|---|---|
| Gene | AT1G06950 |
| is.expressed_genes | TRUE |
| is.responsive_genes | TRUE |
| is.apcluster | E-05 |

# AT1G19450.1 : NA

Major facilitator superfamily protein

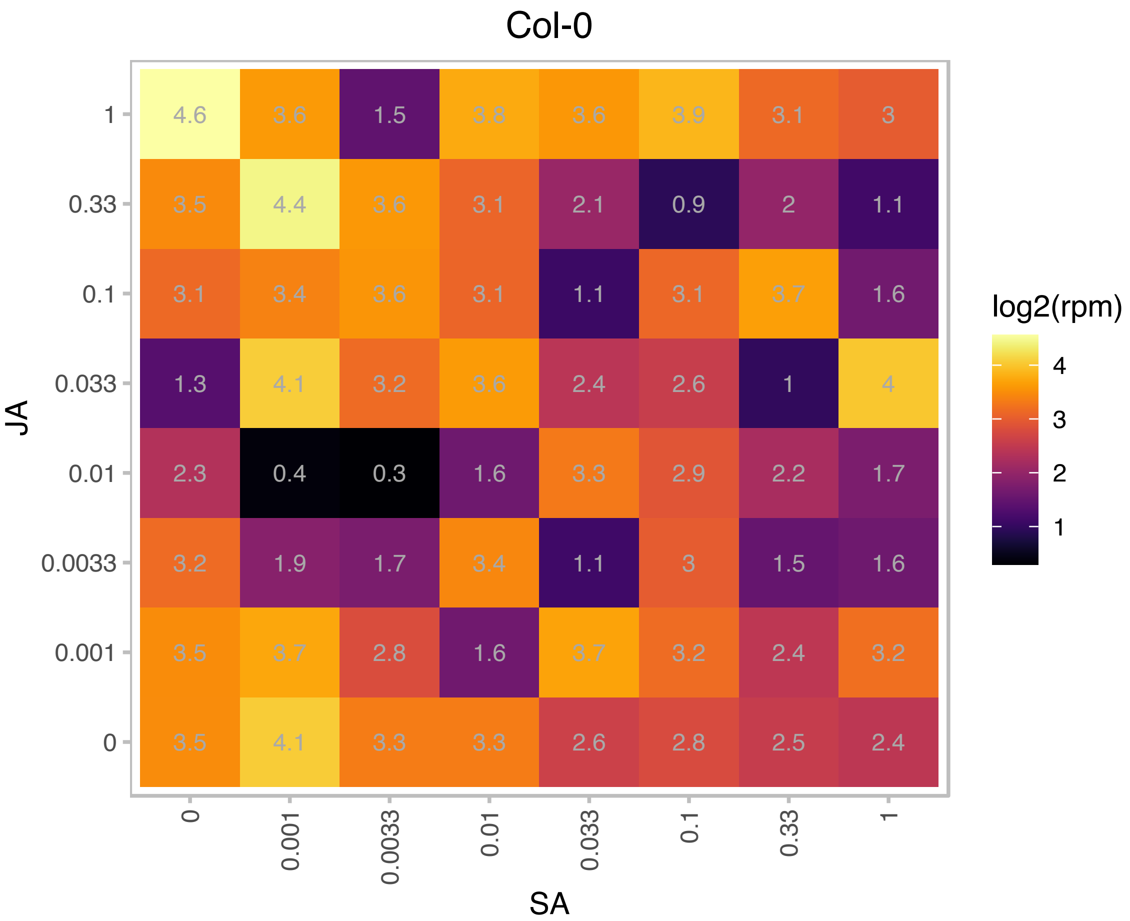


| | |
|---|---|
| Gene | AT1G19450 |
| is.expressed_genes | TRUE |
| is.responsive_genes | TRUE |
| is.apcluster | E-05 |

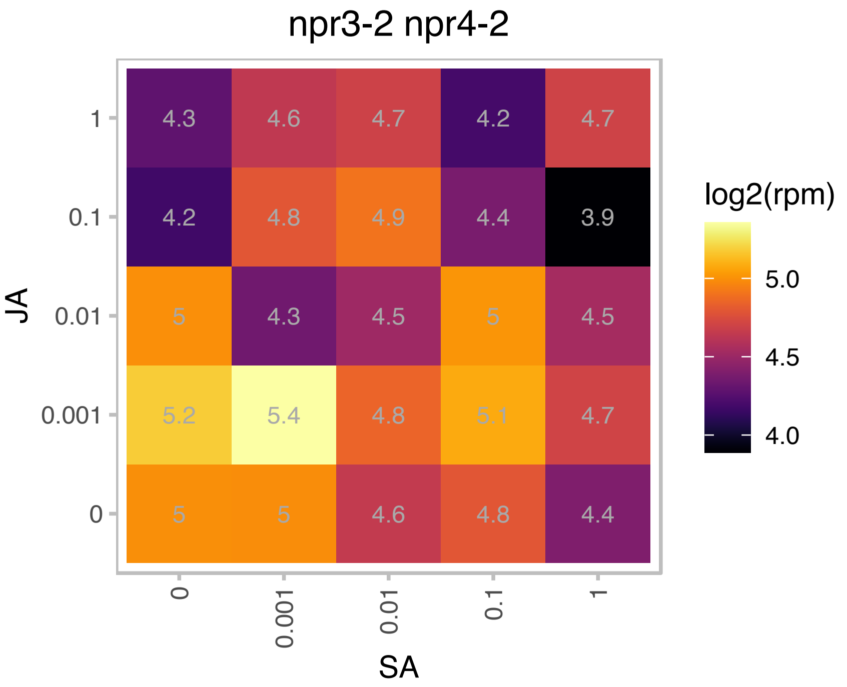


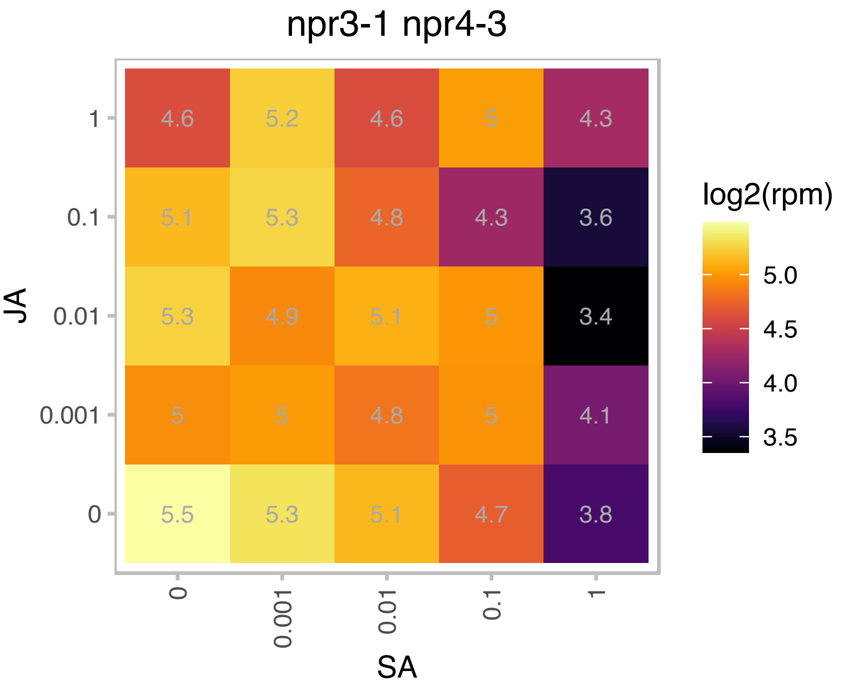

# AT1G21680.1 : NA

DPP6 N-terminal domain-like protein

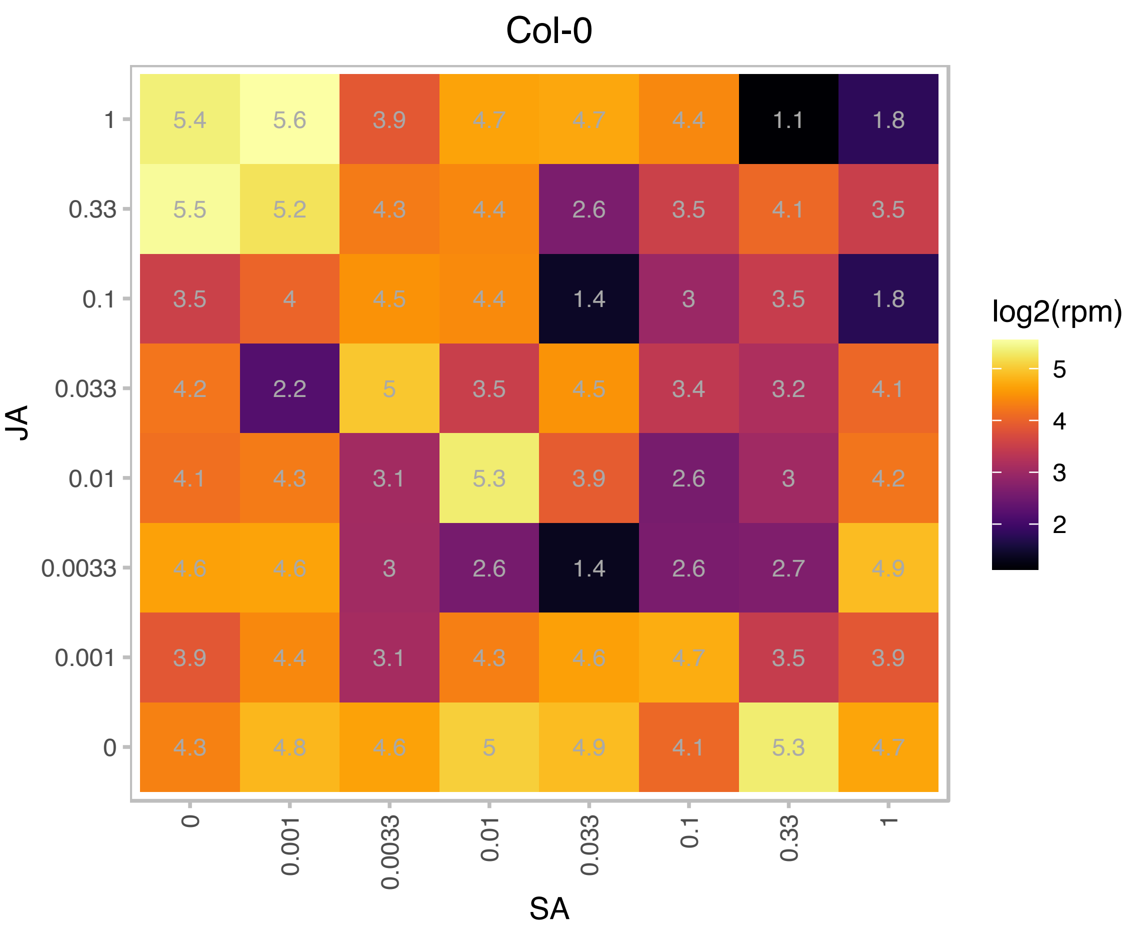


| | |
|---|---|
| Gene | AT1G21680 |
| is.expressed_genes | TRUE |
| is.responsive_genes | TRUE |
| is.apcluster | E-05 |

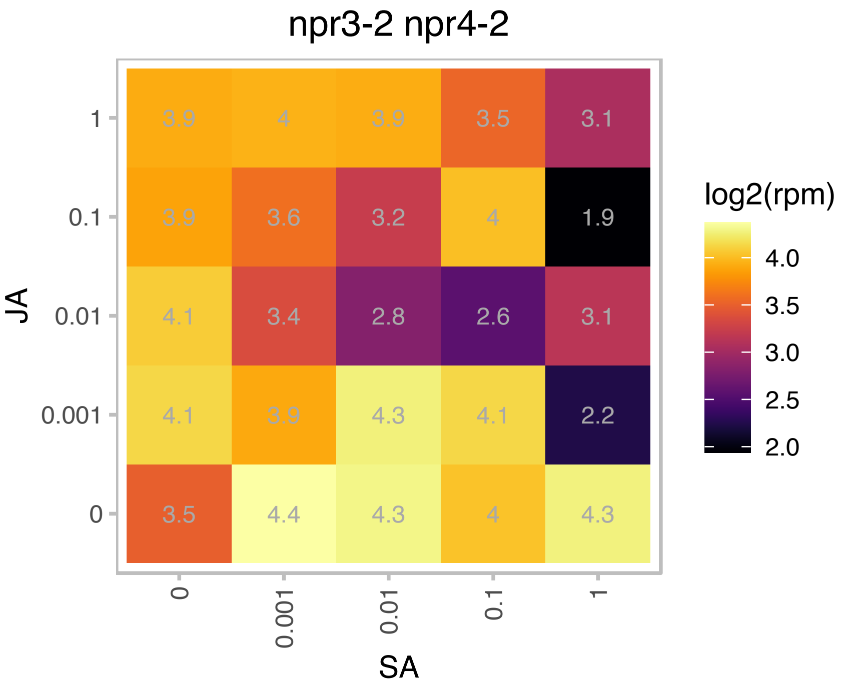


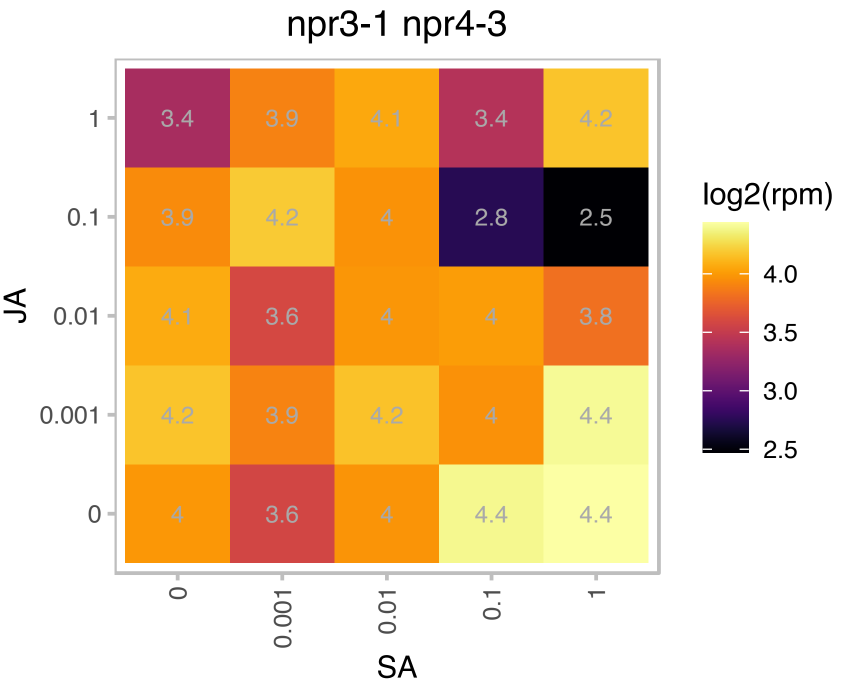

### AT1G31180.1 : ATIMD3, IMD3, IPMDH1

isopropylmalate dehydrogenase 3

The AtIMD3 is one out of 3 genes encoding the enzyme 3-isopropylmalate dehydrogenase involved in leucine biosynthesis in Arabidopsis. Its subcellular location has been targeted to plastids.

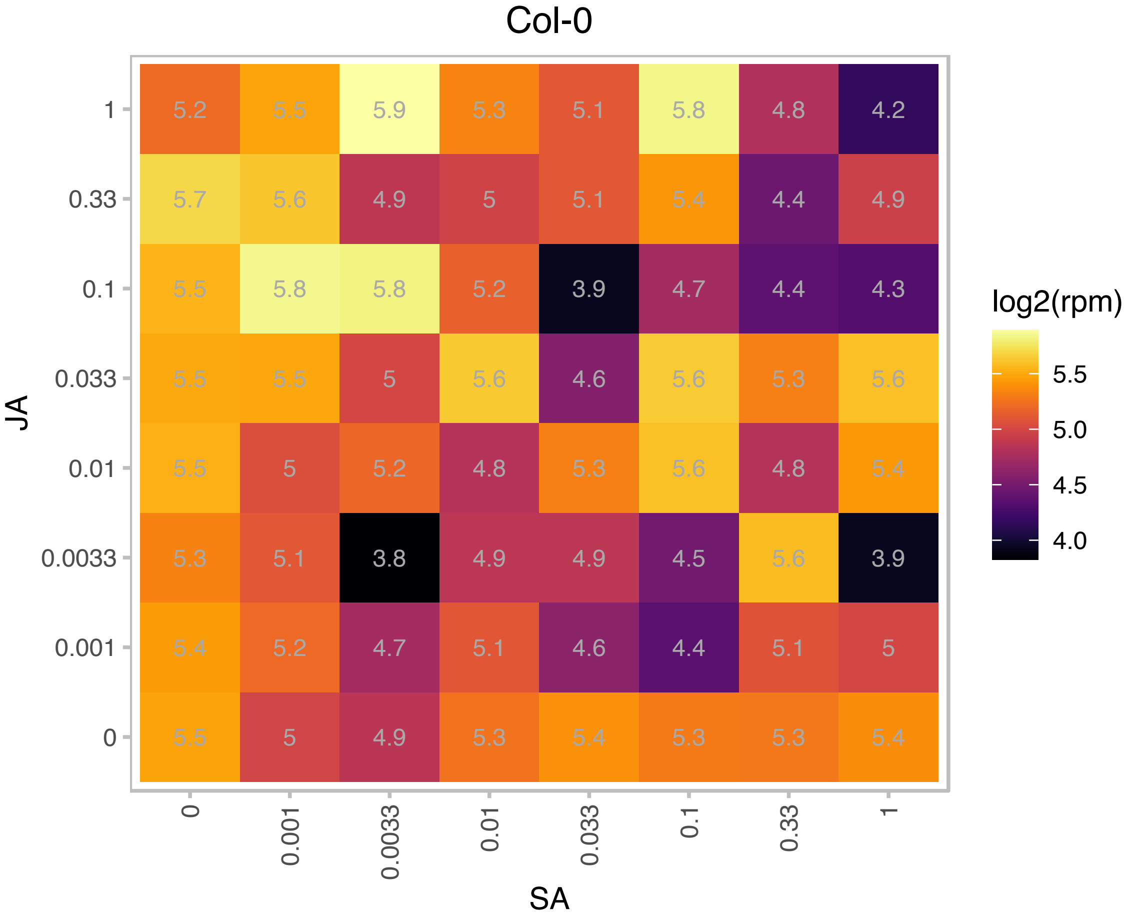


| | |
|---|---|
| Gene | AT1G31180 |
| is.expressed_genes | TRUE |
| is.responsive_genes | TRUE |
| is.apcluster | E-05 |

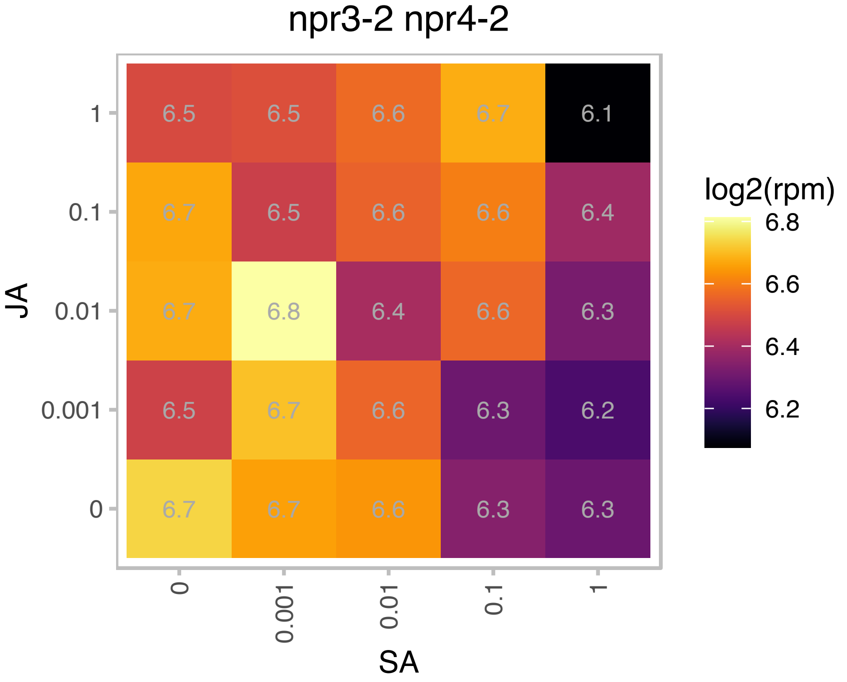


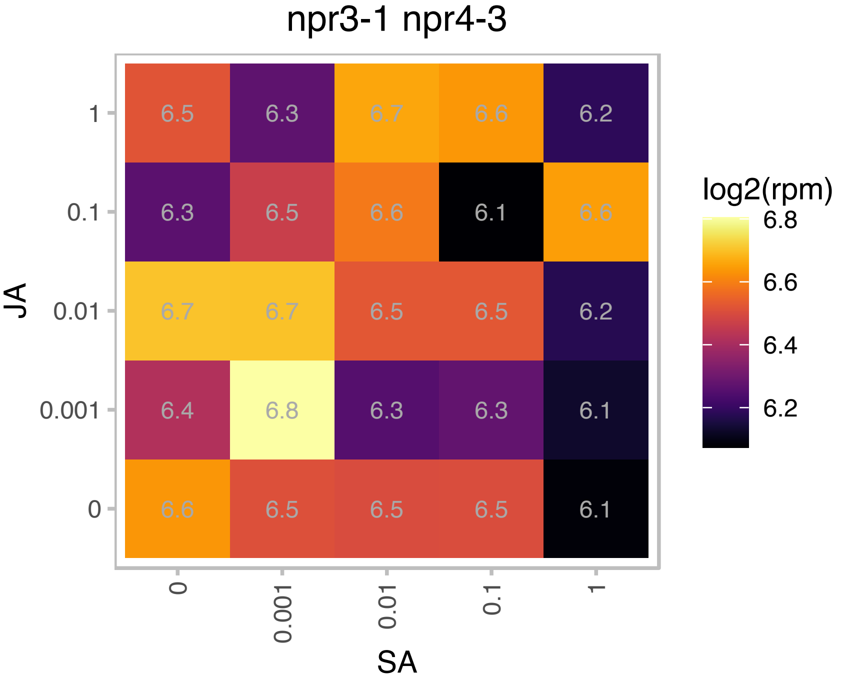

# AT1G54260.1 : NA

winged-helix DNA-binding transcription factor family protein

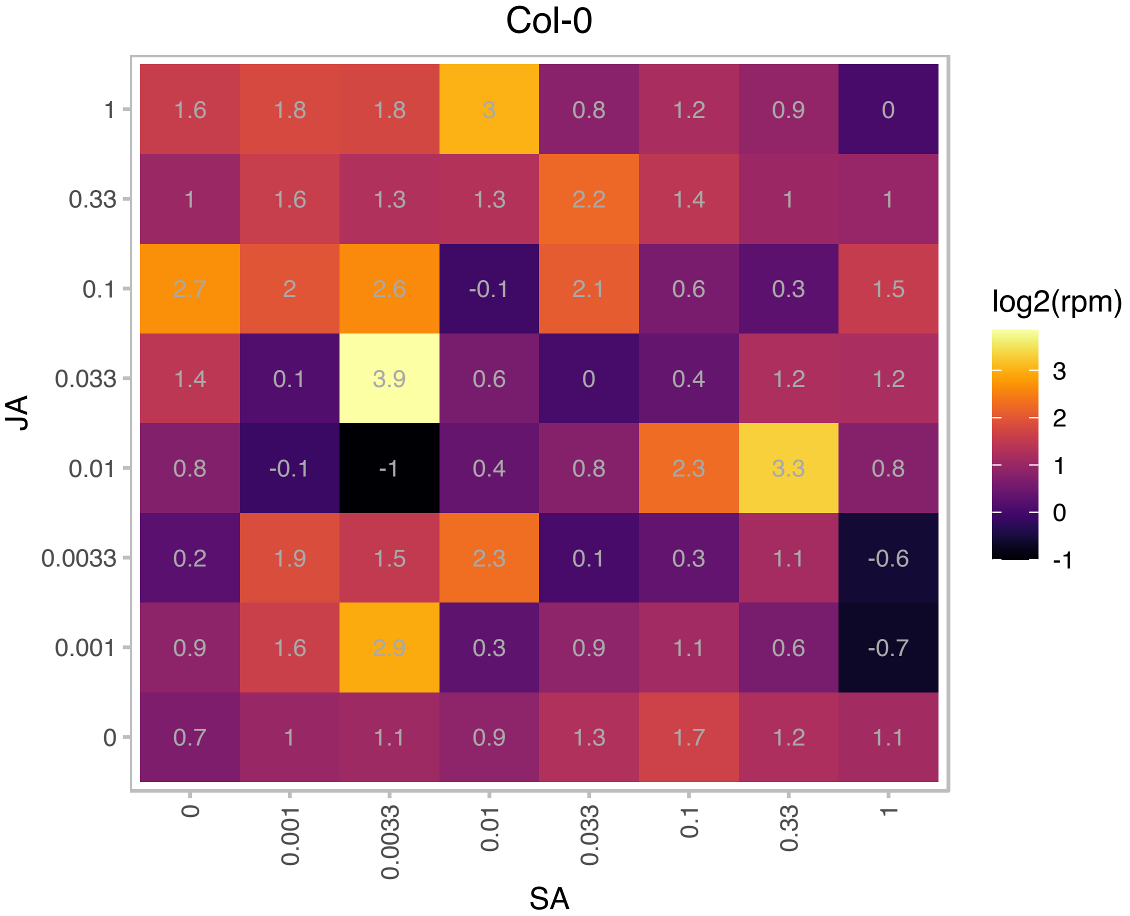


| | |
|---|---|
| Gene | AT1G54260 |
| is.expressed_genes | TRUE |
| is.responsive_genes | TRUE |
| is.apcluster | E-05 |

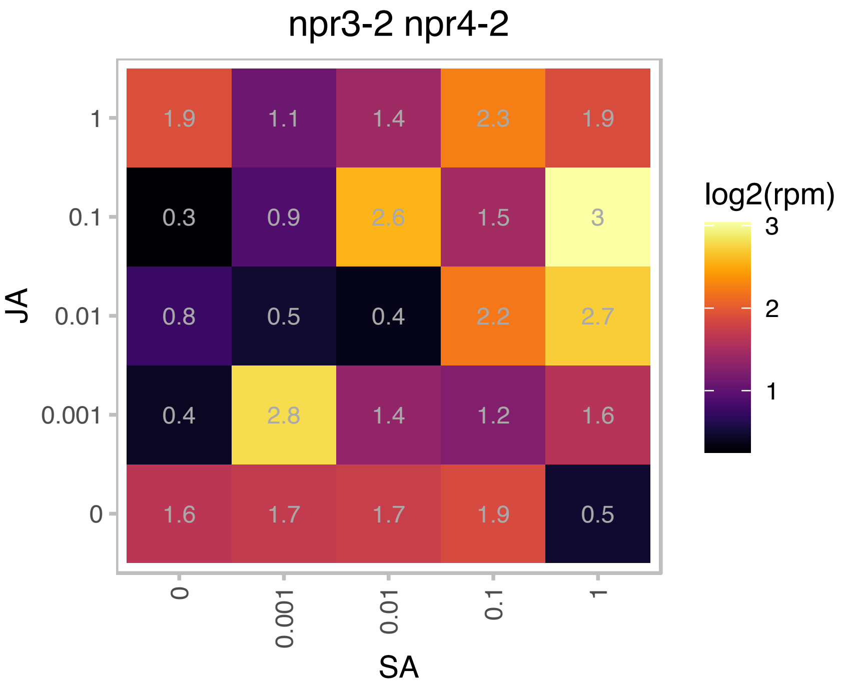


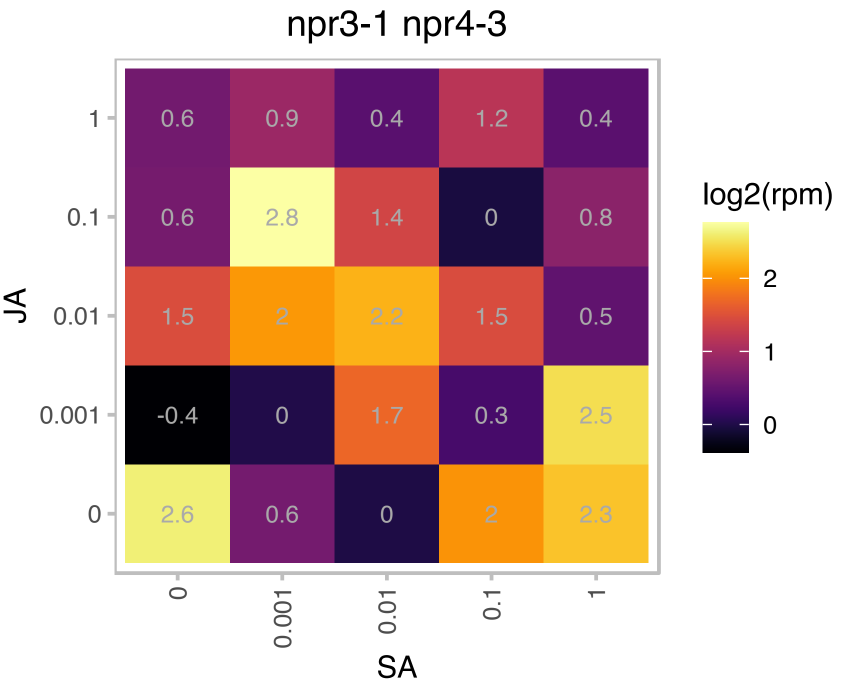

### AT1G70850.1 : MLP34

MLP-like protein 34

| | |
|---|---|
| Gene | AT1G70850 |
| is.expressed_genes | TRUE |
| is.responsive_genes | TRUE |
| is.apcluster | E-05 |

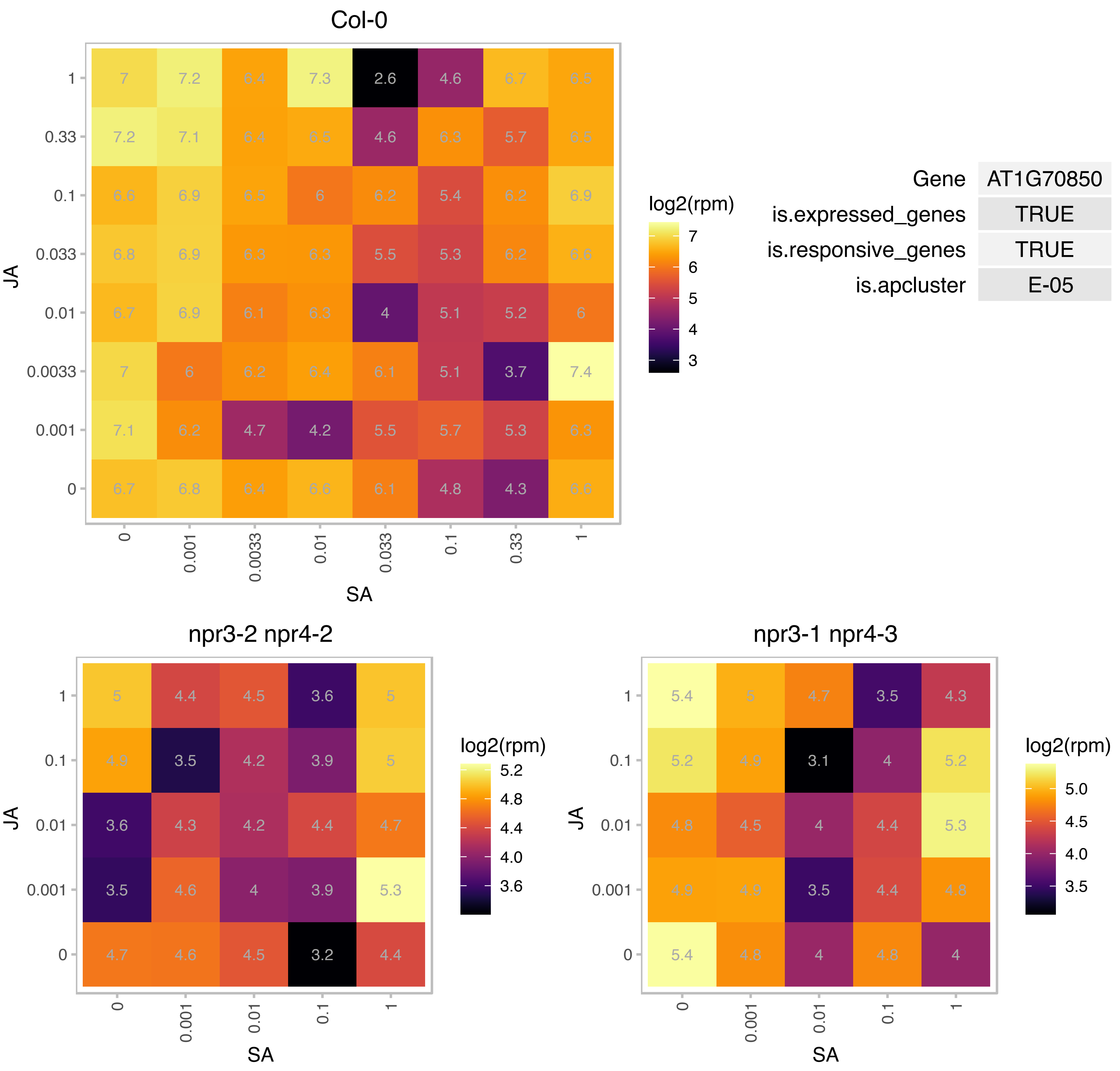

### AT1G73720.1 : SMU1

transducin family protein / WD-40 repeat family protein

Encodes SMU1, a protein involved in RNA splicing.

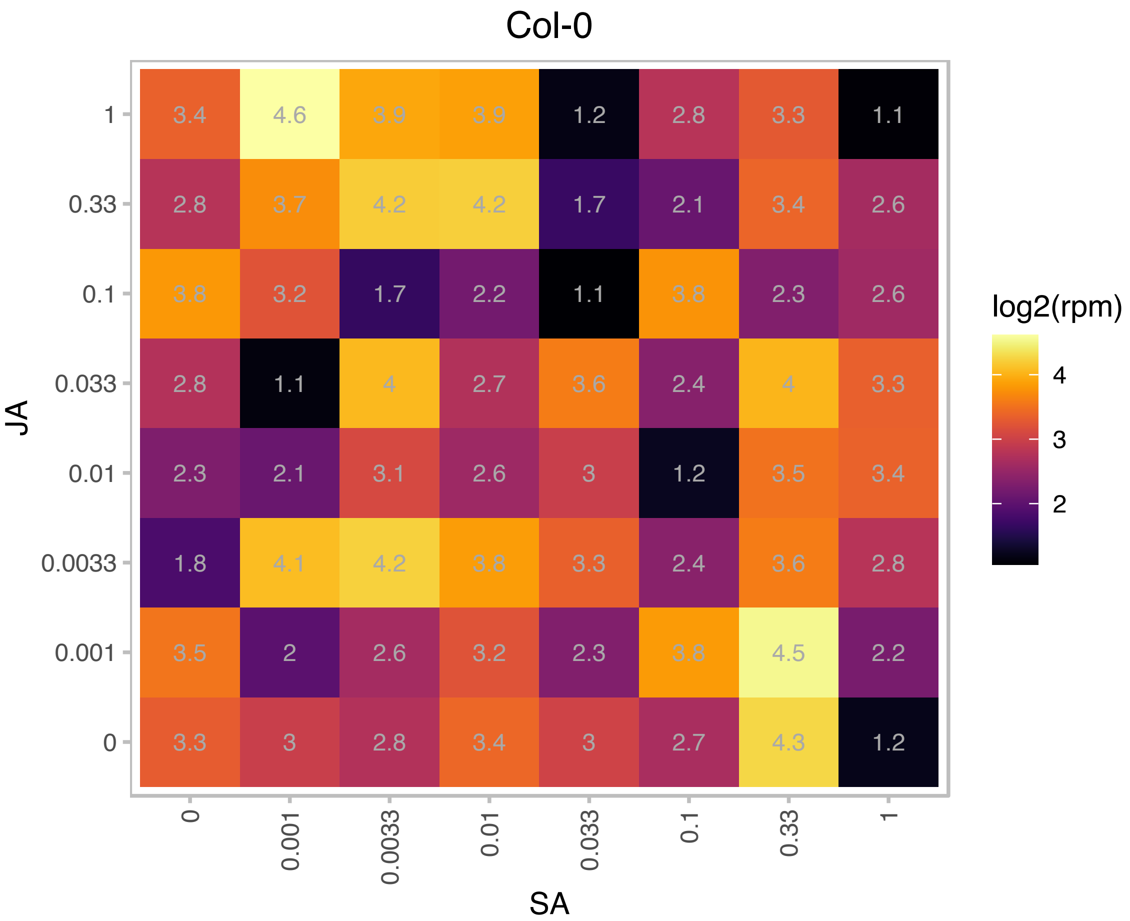


| | |
|---|---|
| Gene | AT1G73720 |
| is.expressed_genes | TRUE |
| is.responsive_genes | TRUE |
| is.apcluster | E-05 |

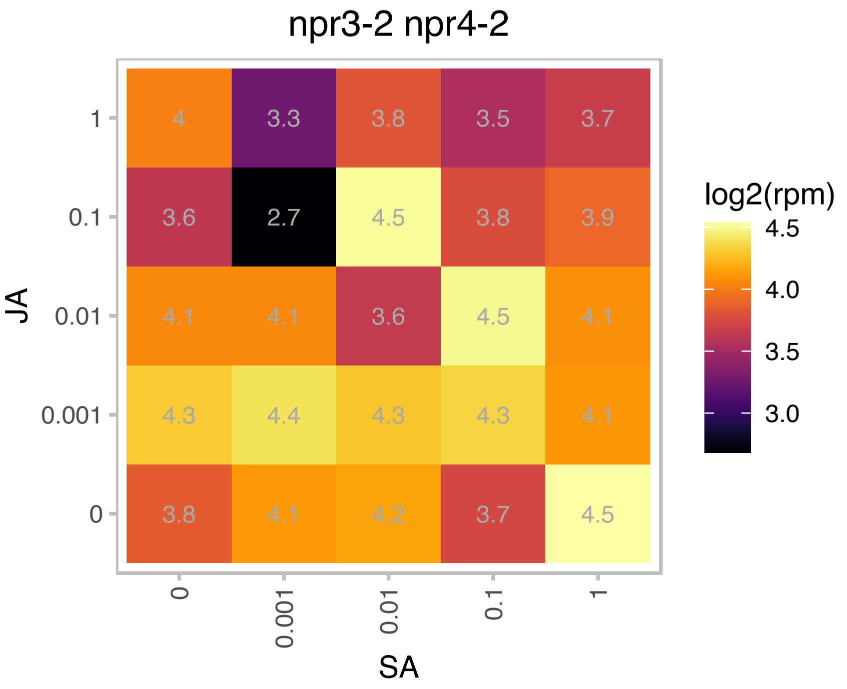


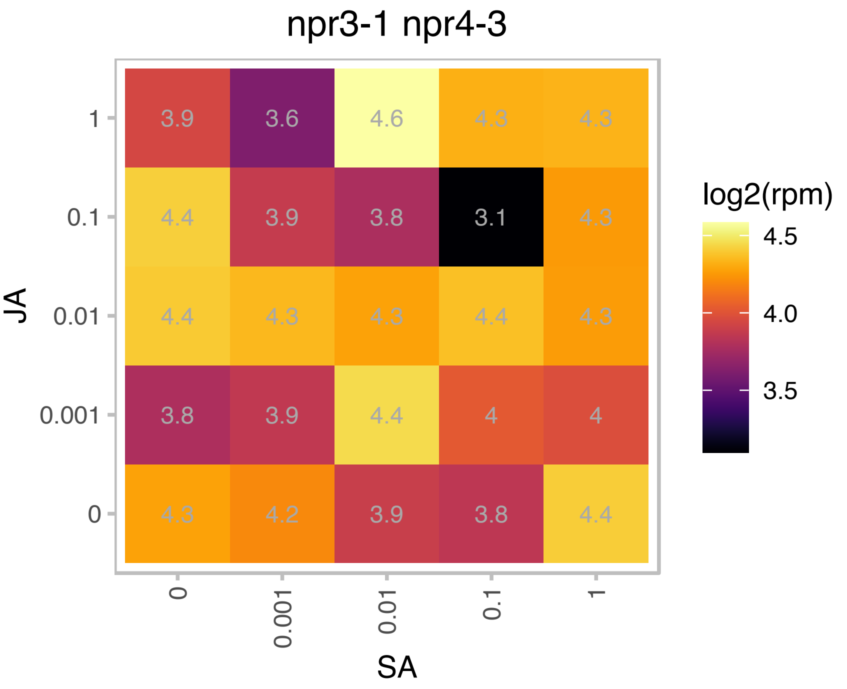

# AT1G79210.1 : NA

N-terminal nucleophile aminohydrolases (Ntn hydrolases) superfamily protein

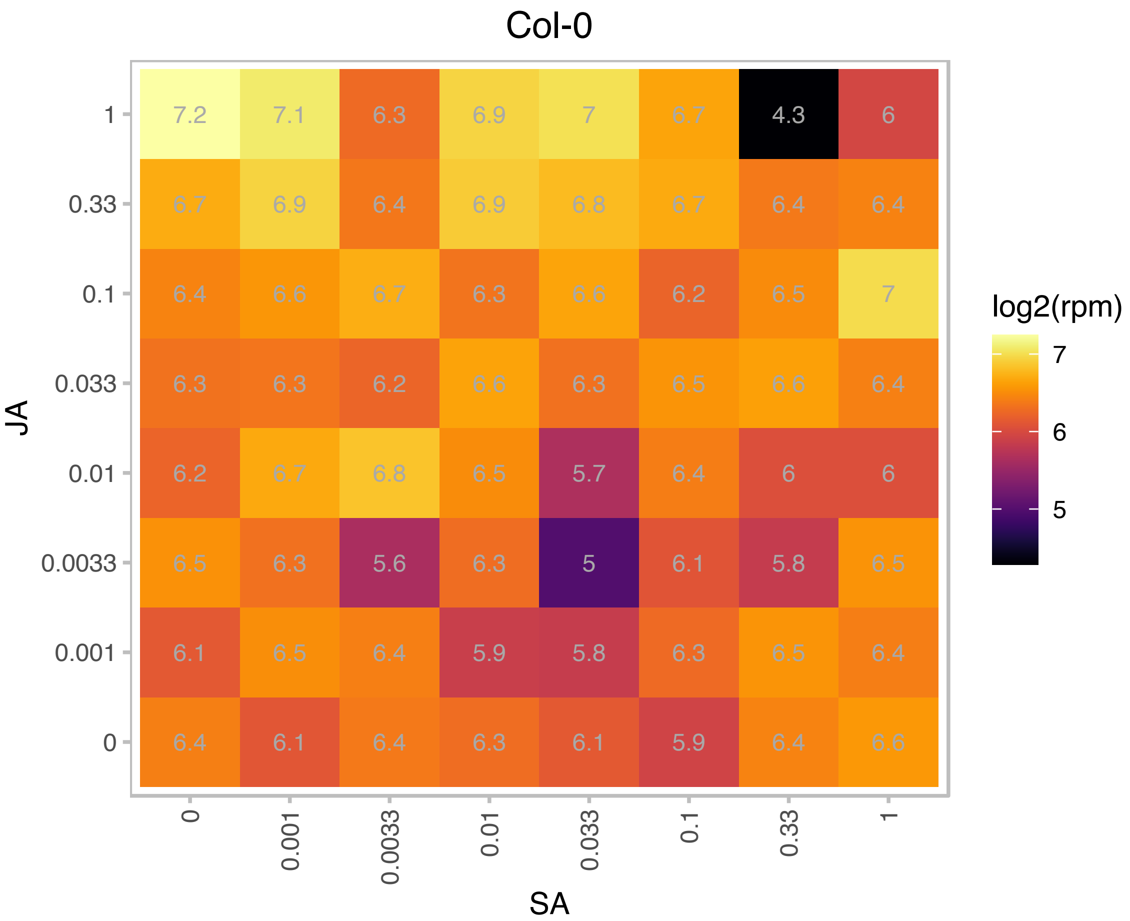


| | |
|---|---|
| Gene | AT1G79210 |
| is.expressed_genes | TRUE |
| is.responsive_genes | TRUE |
| is.apcluster | E-05 |

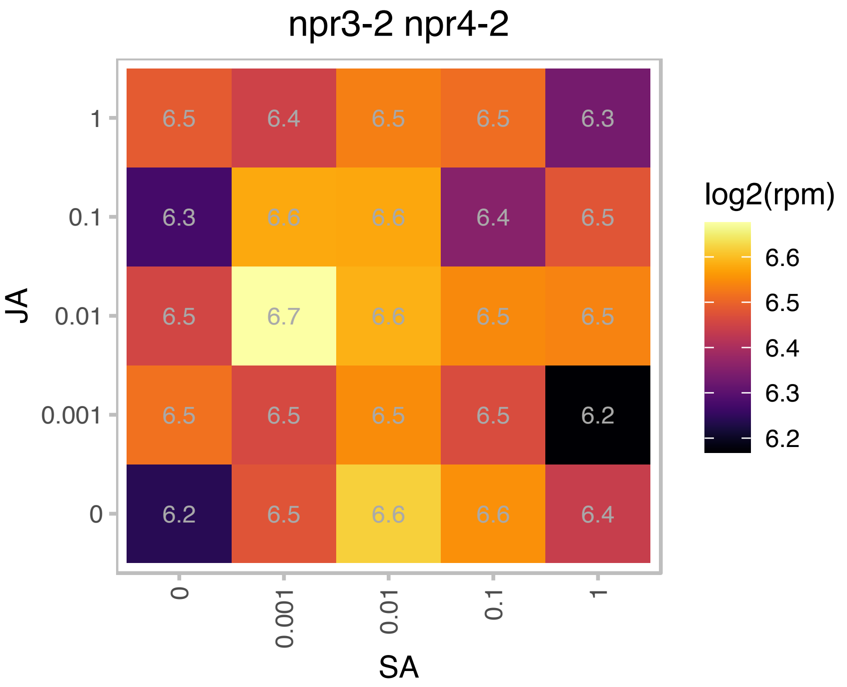


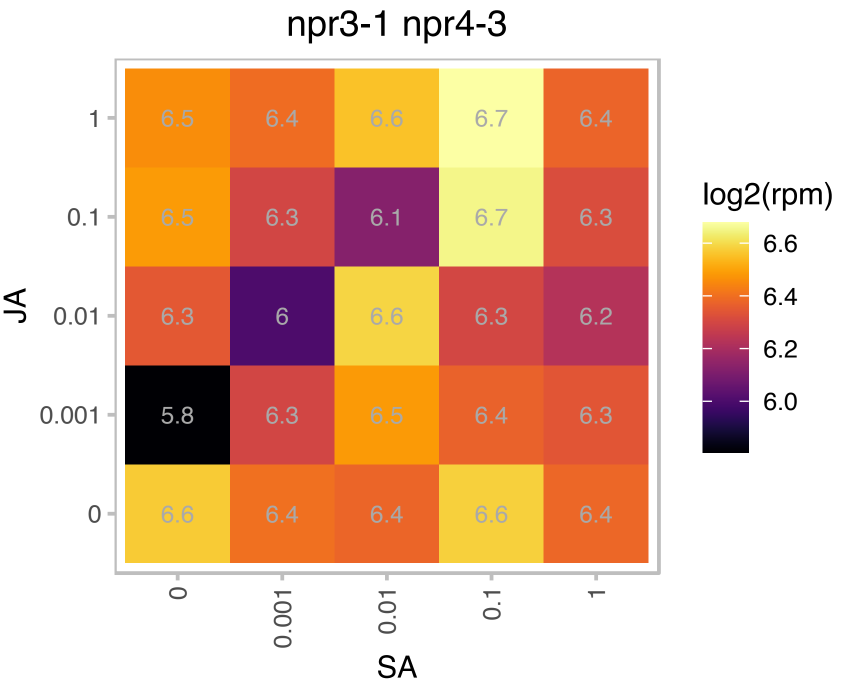

### AT2G01530.1 : MLP329, ZCE2

MLP-like protein 329

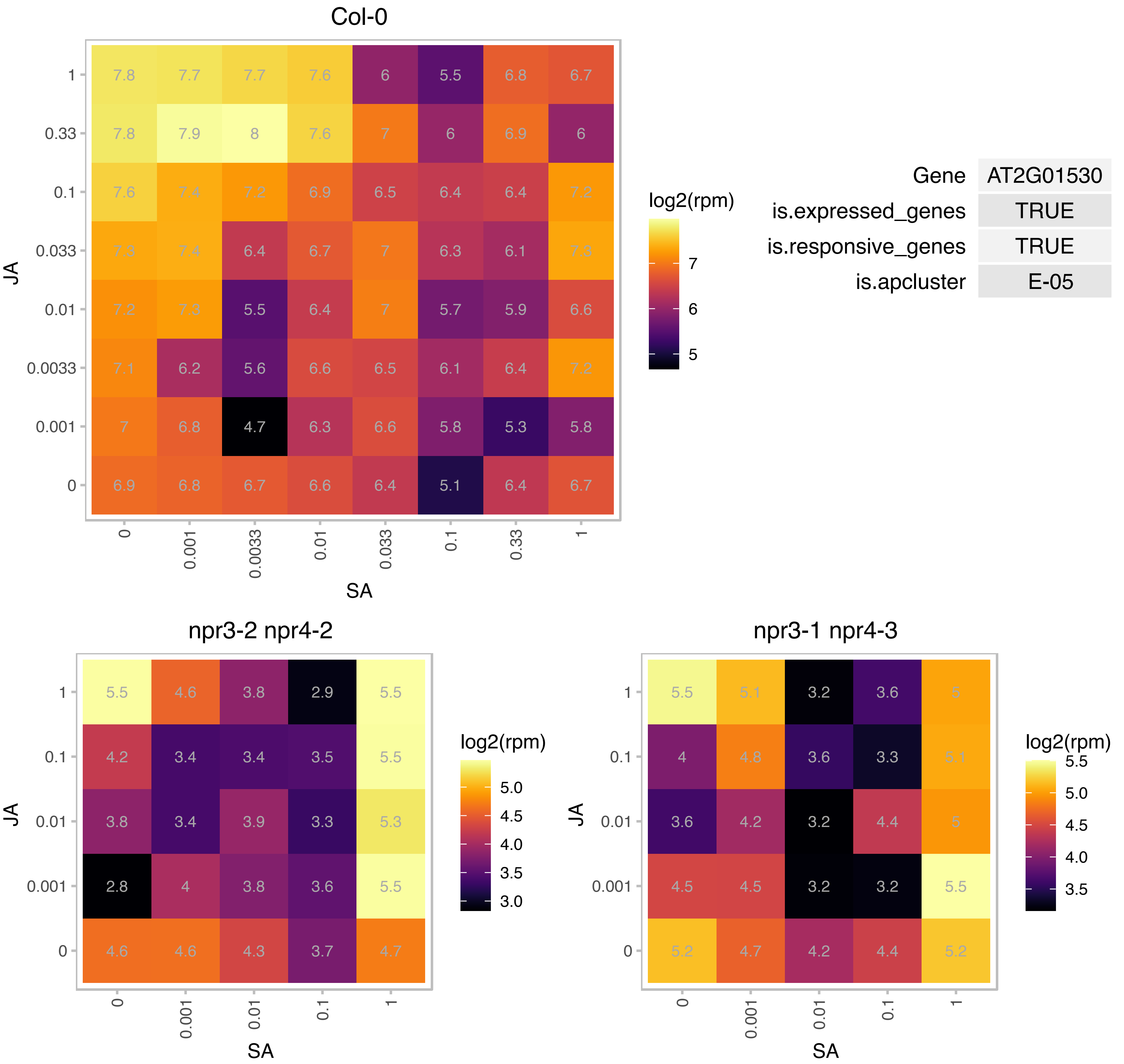


| | |
|---|---|
| Gene | AT2G01530 |
| is.expressed_genes | TRUE |
| is.responsive_genes | TRUE |
| is.apcluster | E-05 |

### AT2G04240.1 : XERICO

RING/U-box superfamily protein
Encodes a small protein with an N-terminal trans-membrane domain and a RING-H2 zinc finger motif located at the C-terminus.  Gene expression is induced by salt and osmotic stress. Transcrips levels are induced by DELLA proteins and repressed by gibberellic acid. Involved in ABA metabolism.

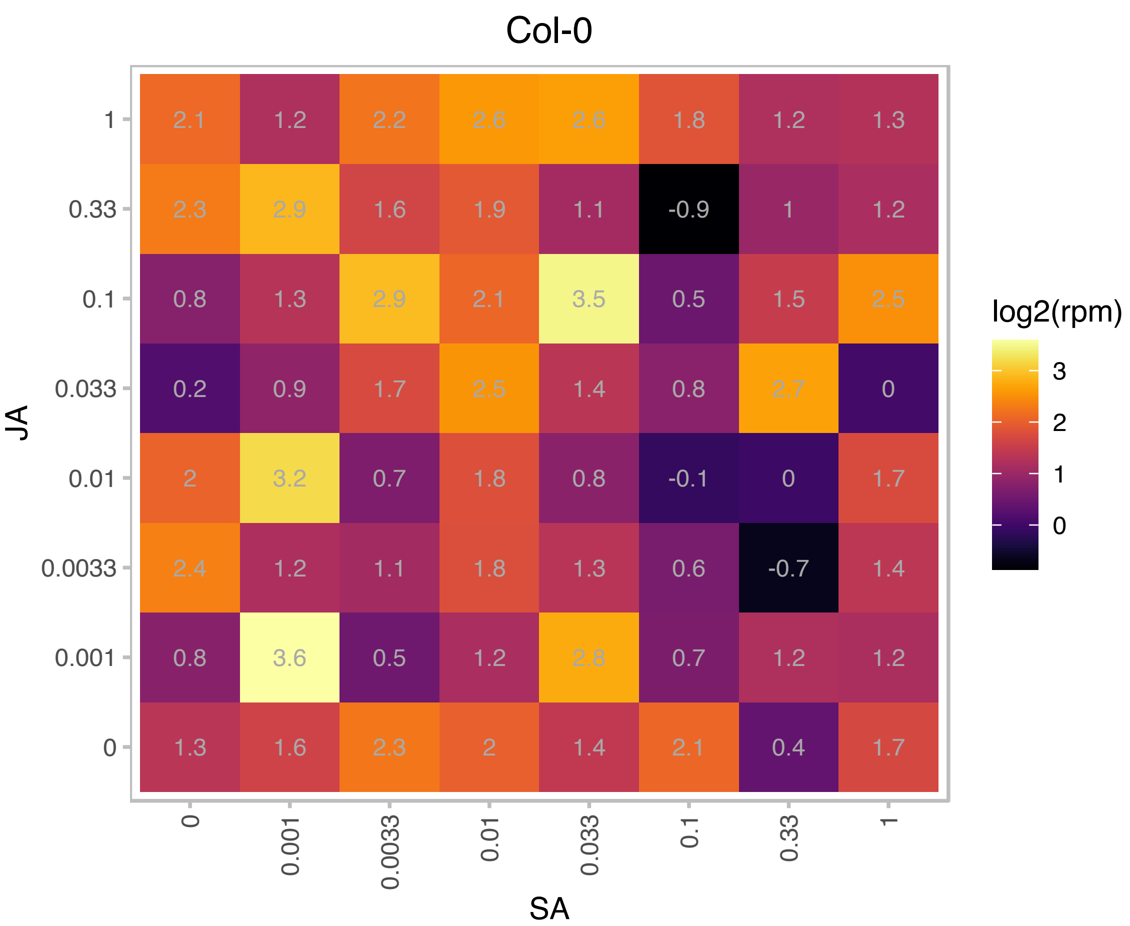


| | |
|---|---|
| Gene | AT2G04240 |
| is.expressed_genes | TRUE |
| is.responsive_genes | TRUE |
| is.apcluster | E-05 |

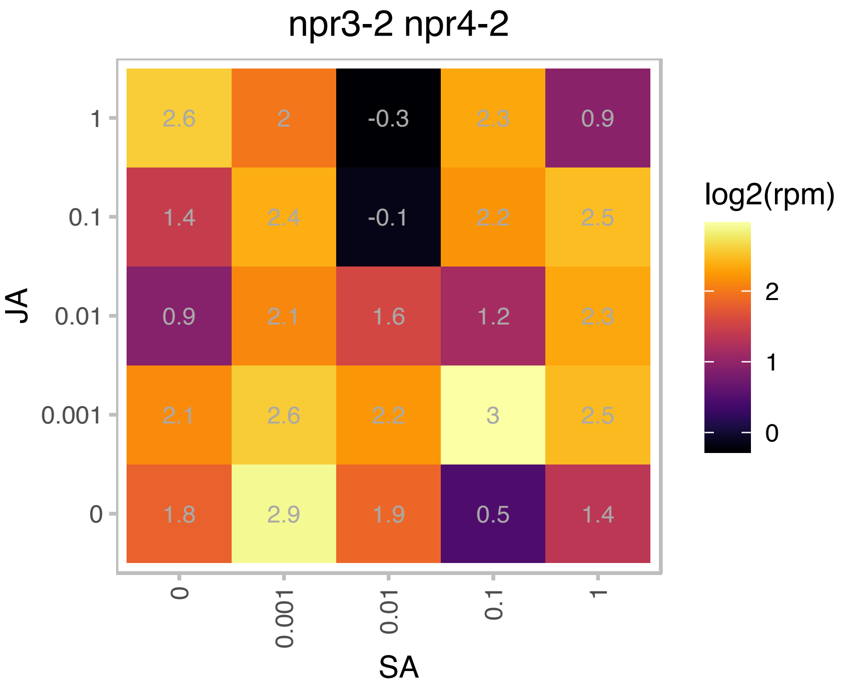


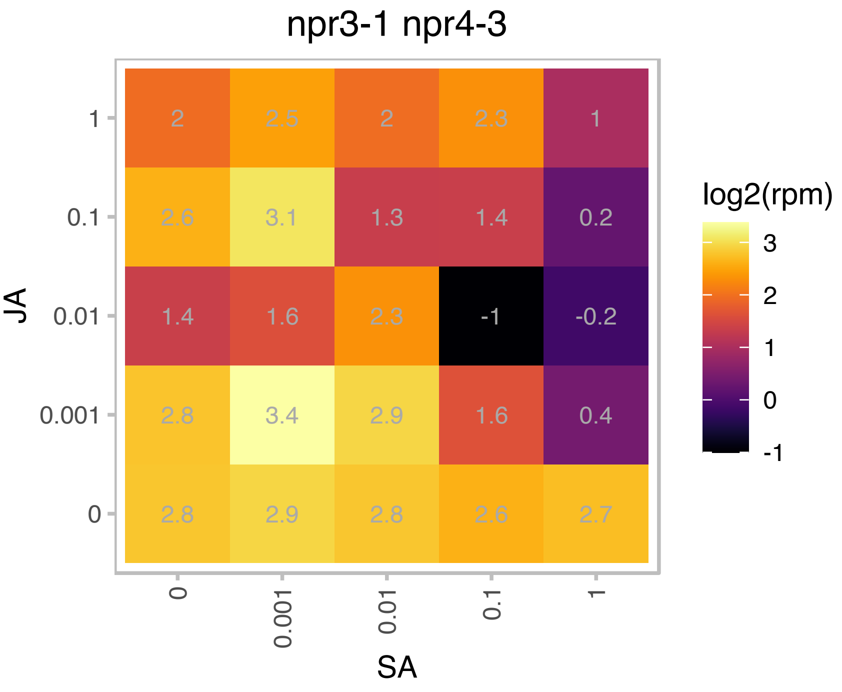

# AT2G30105.1 : NA

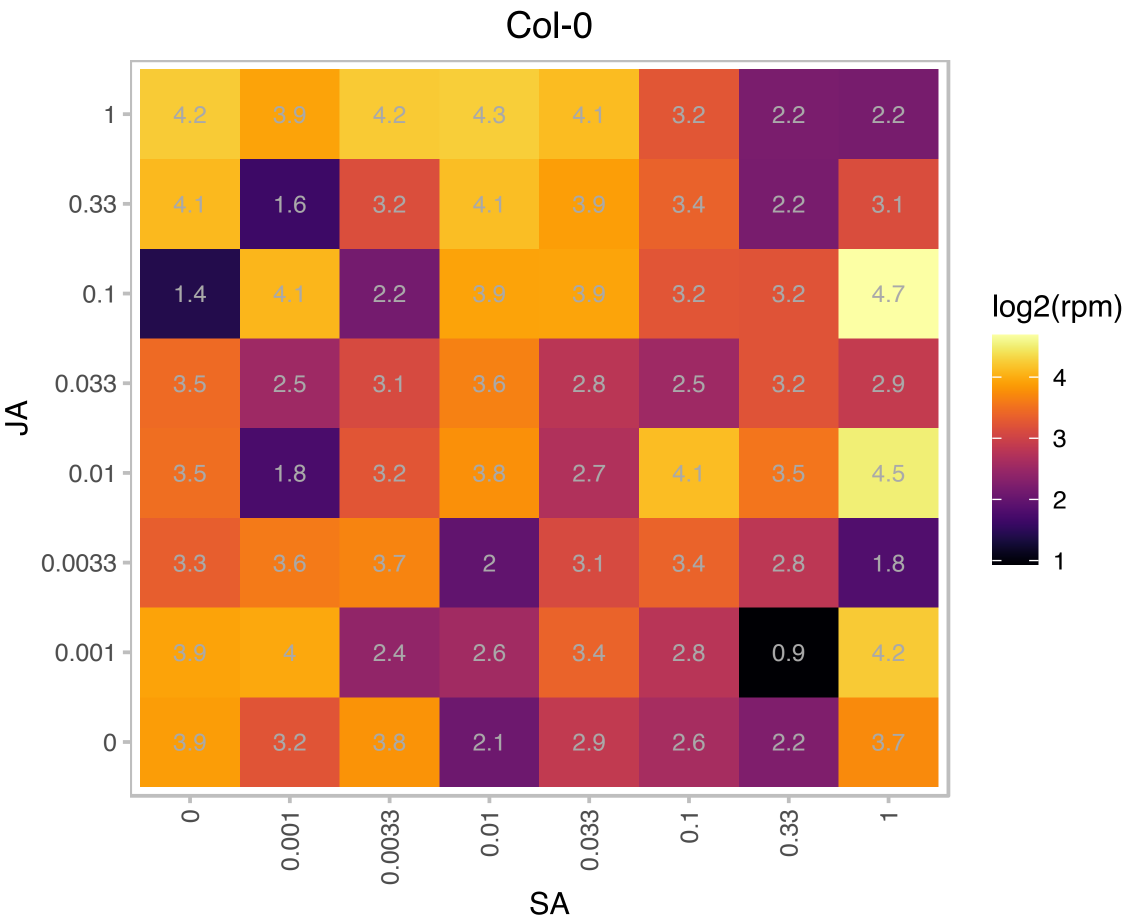


| | |
|---|---|
| Gene | AT2G30105 |
| is.expressed_genes | TRUE |
| is.responsive_genes | TRUE |
| is.apcluster | E-05 |

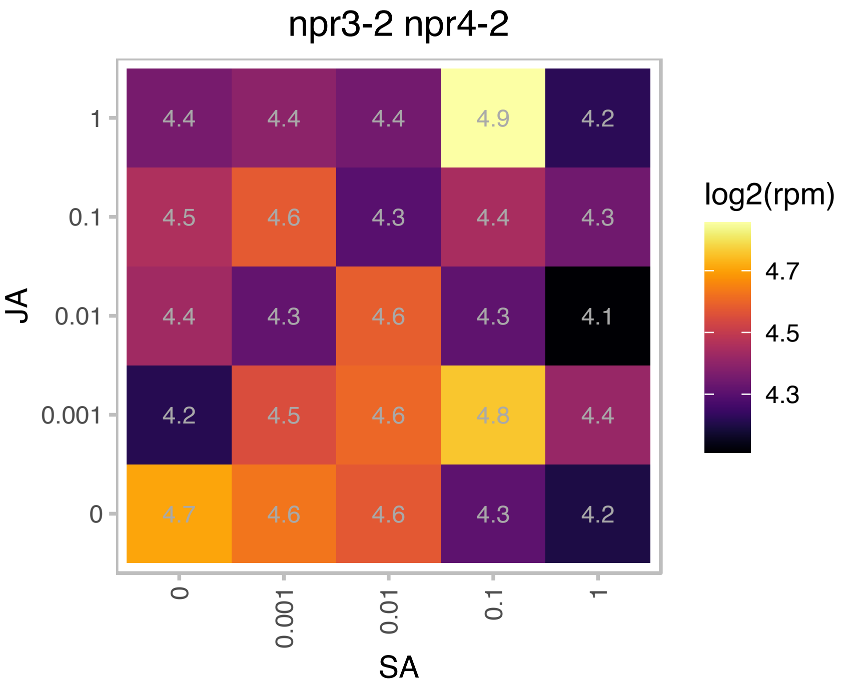


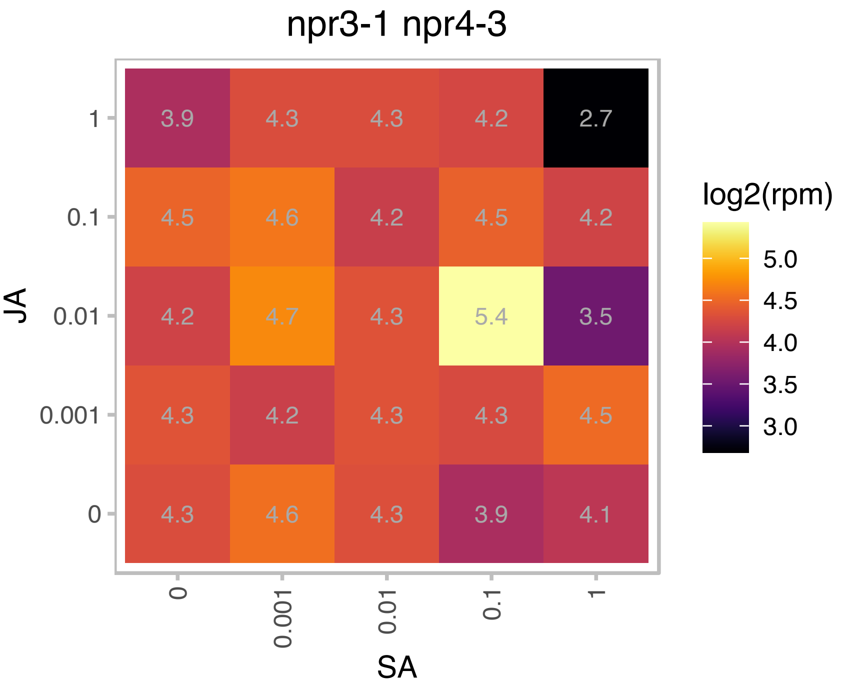

### AT2G36830.1 : GAMMA-TIP, GAMMA-TIP1, TIP1;1

gamma tonoplast intrinsic protein
Encodes a tonoplast intrinsic protein, which functions as water channel. It has also been shown to be able to facilitate the transport of urea and hydrogen peroxide. Highly expressed in vascular tissues of the root, stem, cauline leaves and flowers but not in the apical meristems. The mRNA is cell-to-cell mobile.

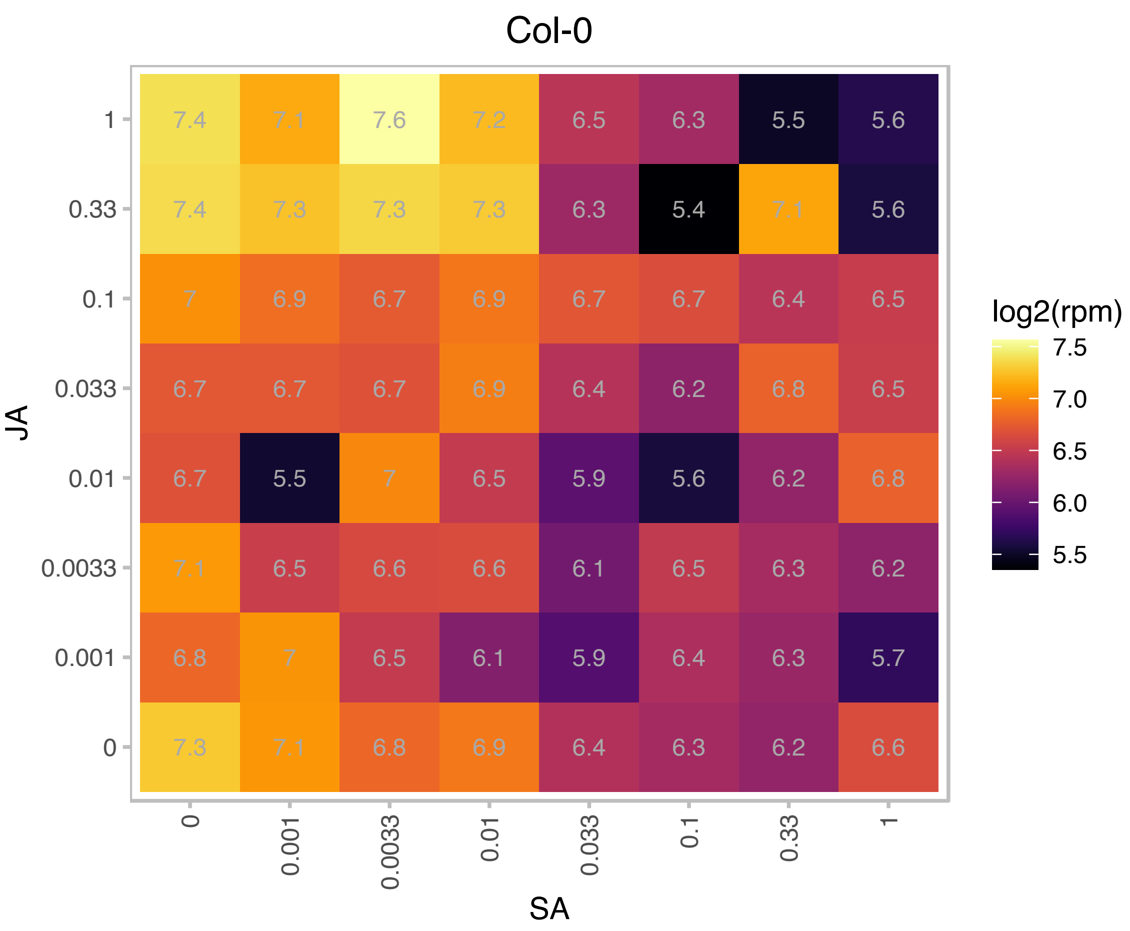


| | |
|---|---|
| Gene | AT2G36830 |
| is.expressed_genes | TRUE |
| is.responsive_genes | TRUE |
| is.apcluster | E-05 |

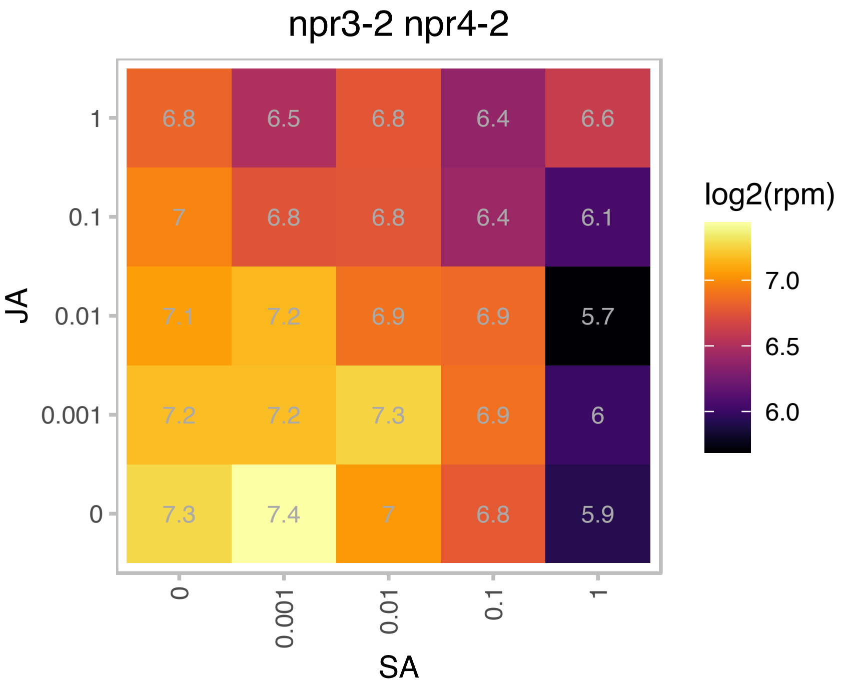


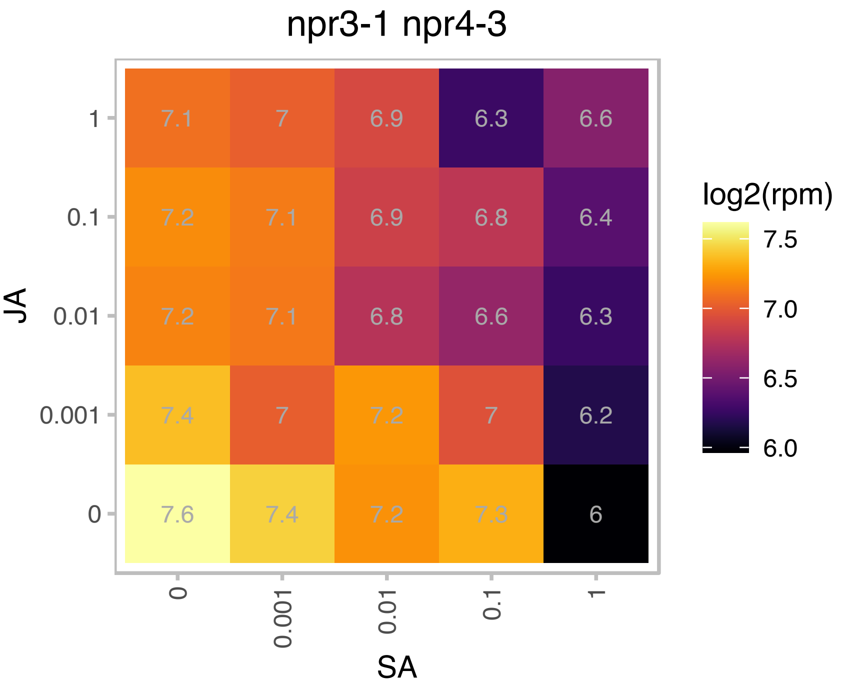

# AT2G42500.1 : PP2A-3

protein phosphatase 2A-3

Encodes one of the isoforms of the catalytic subunit of protein phosphatase 2A: AT1G59830/PP2A-1, AT1G10430/PP2A-2, At2g42500/PP2A-3, At3g58500/PP2A-4 [Plant Molecular Biology (1993) 21:475-485 and (1994 26:523-528; Note that in more recent publications, there is mixed use of gene names for PP2A-3 and PP2A-4 - some refer to At2g42500 as PP2A-3 and some as PP2A-4]. ACR4 phosphorylates the PROTEIN PHOSPHATASE 2 (PP2A-3) catalytic subunit of the PP2A phosphatase holoenzyme and PP2A dephosphorylates ACR4.

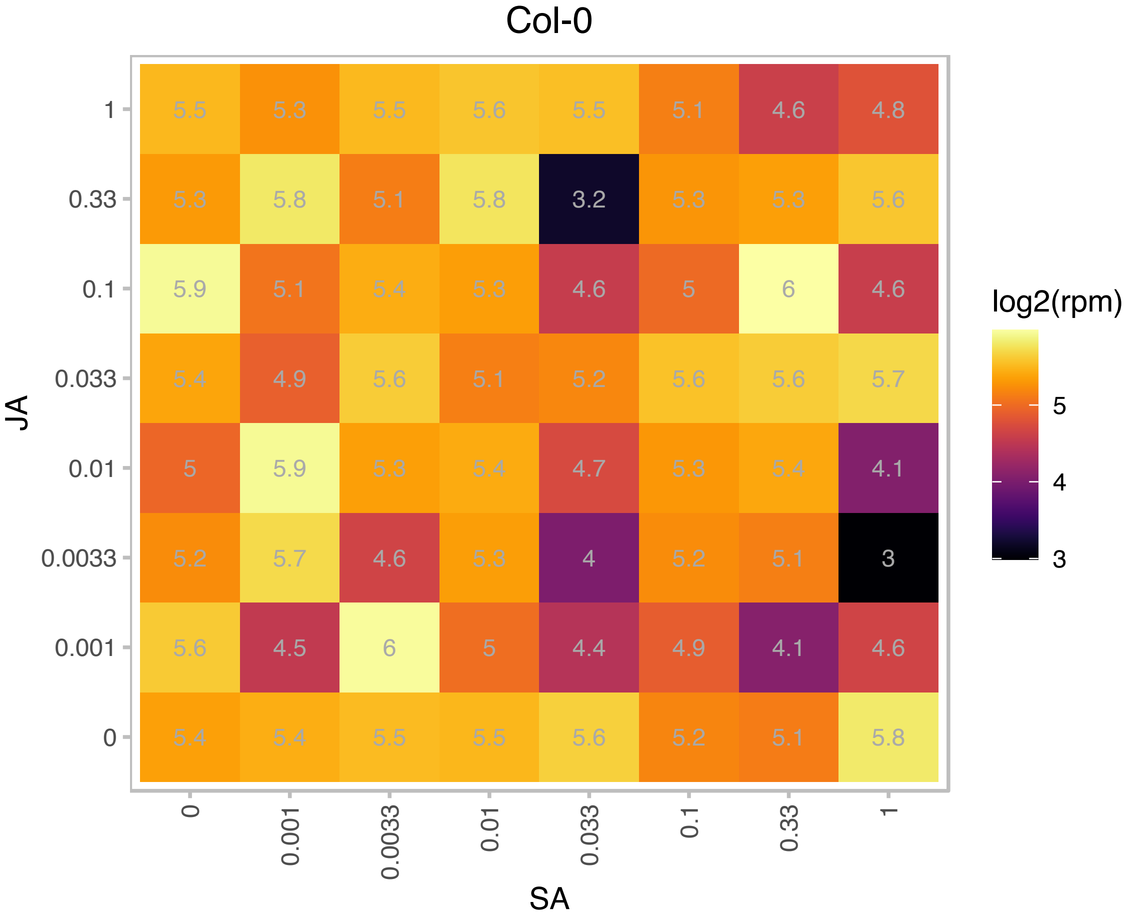


| | |
|---|---|
| Gene | AT2G42500 |
| is.expressed_genes | TRUE |
| is.responsive_genes | TRUE |
| is.apcluster | E-05 |

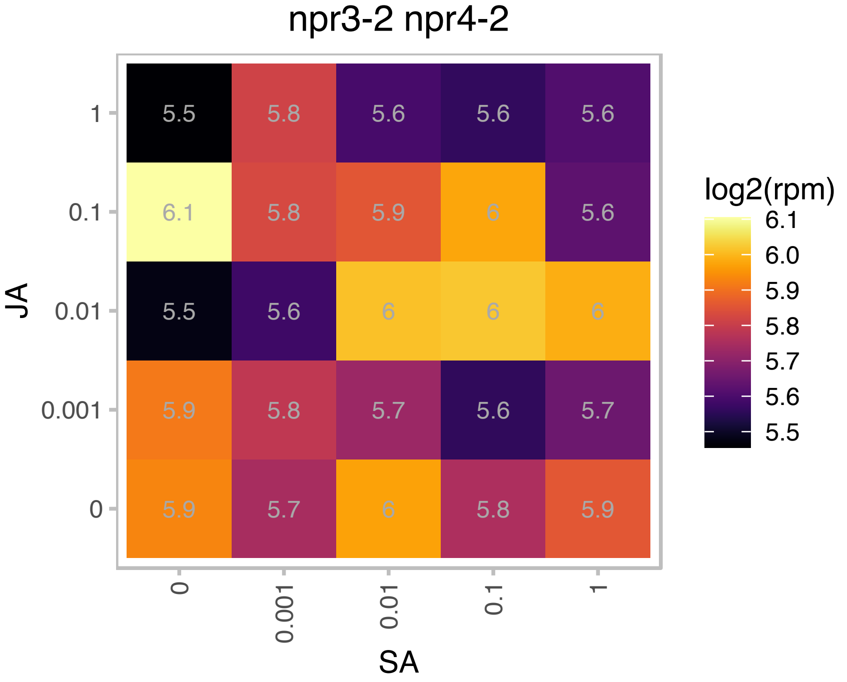


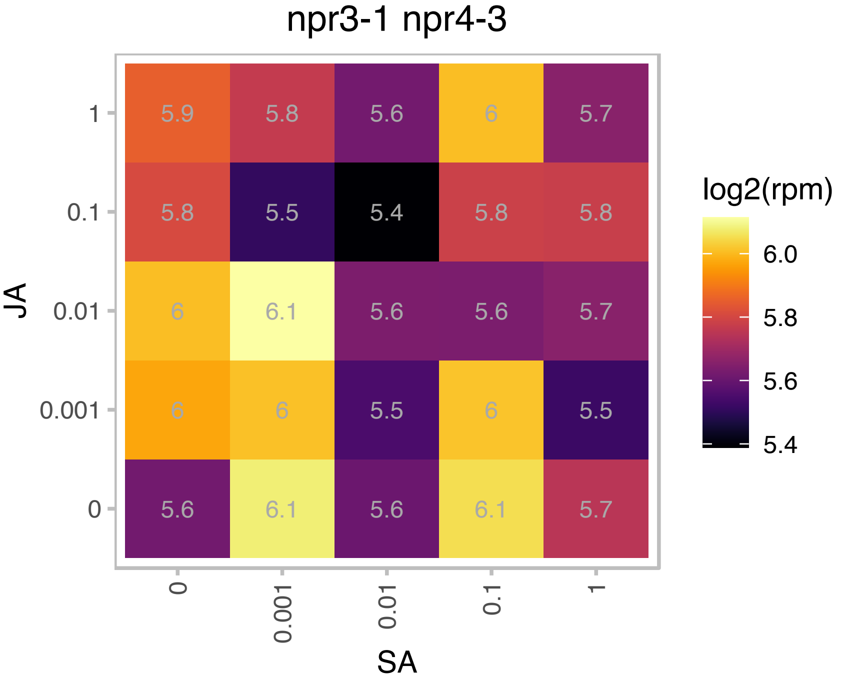

### AT2G44750.1 : TPK2

thiamin pyrophosphokinase 2

Encodes a thiamine pyrophosphokinase capable of producing thiamine pyrophosphate from free thiamine.

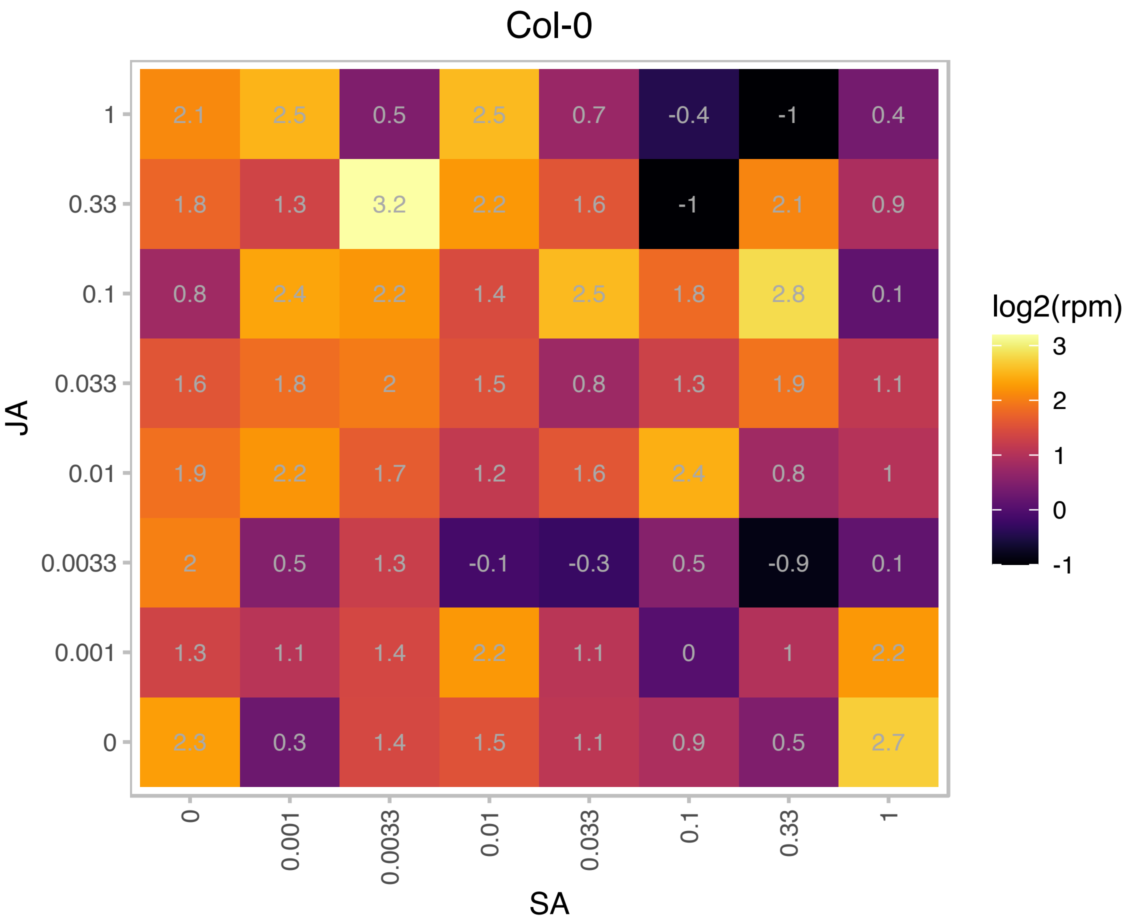


| | |
|---|---|
| Gene | AT2G44750 |
| is.expressed_genes | TRUE |
| is.responsive_genes | TRUE |
| is.apcluster | E-05 |

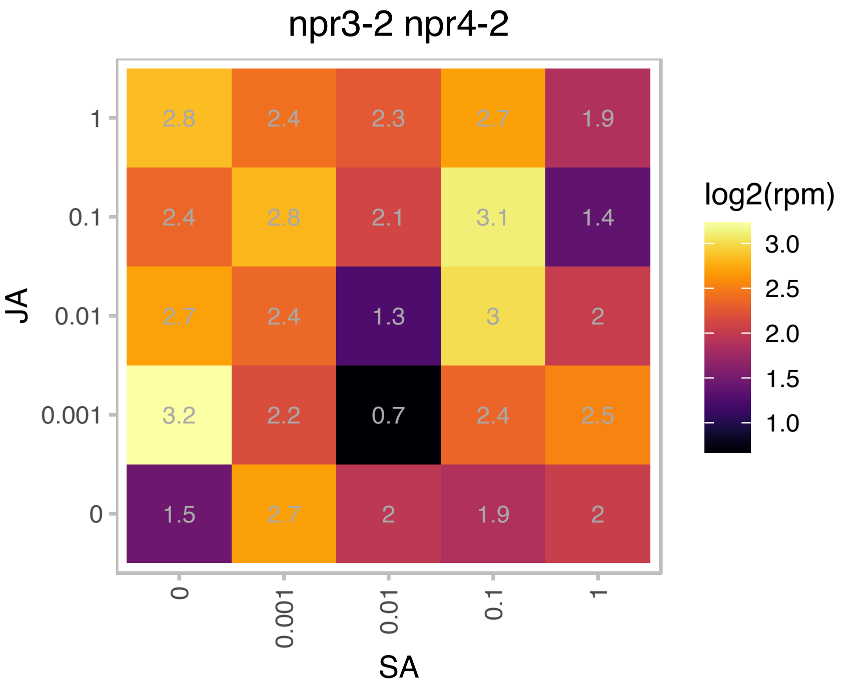


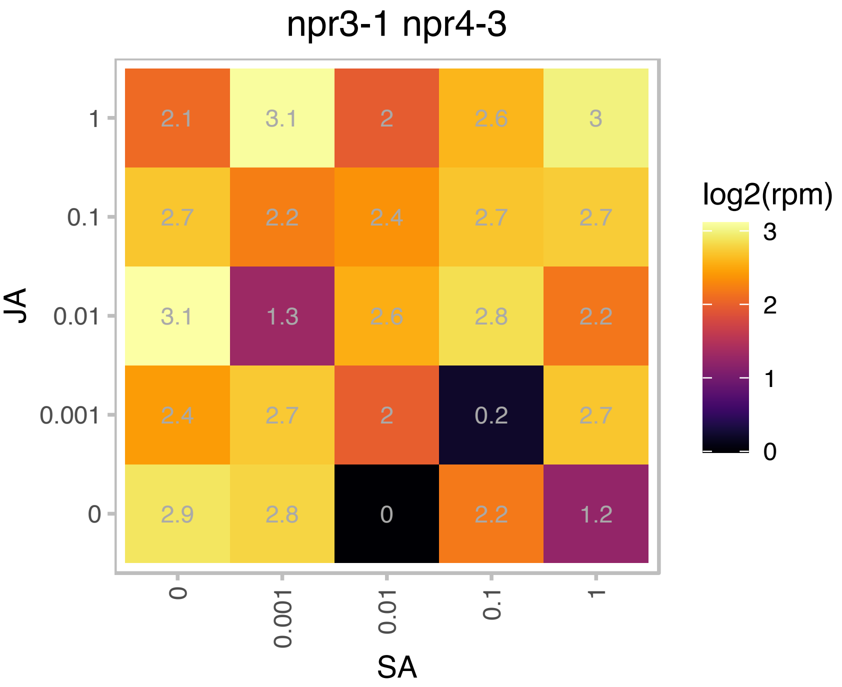

# AT3G08890.1 : NA

Protein of unknown function, DUF538

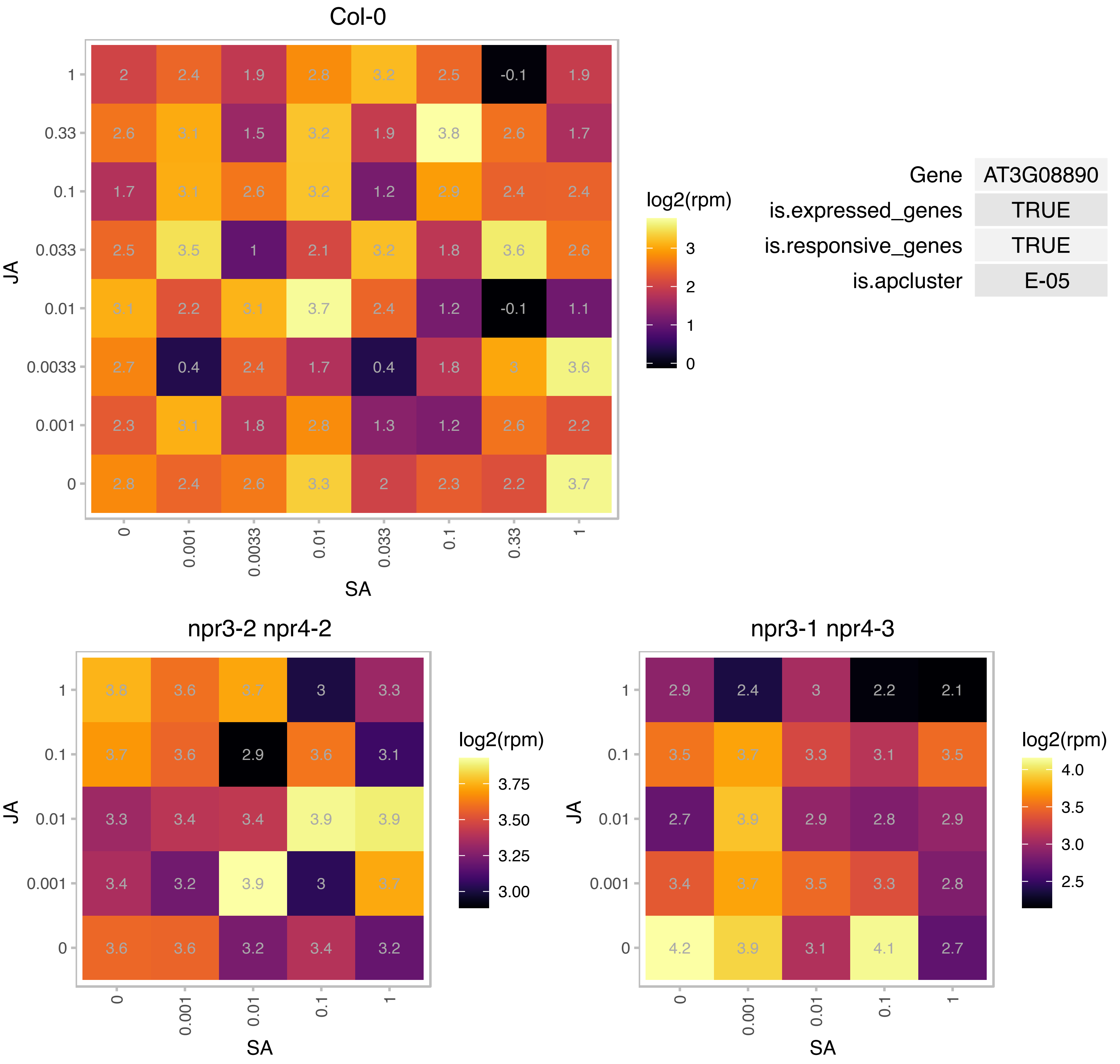


| Gene | AT3G08890 |
| --- | --- |
| is.expressed_genes | TRUE |
| is.responsive_genes | TRUE |
| is.apcluster | E-05 |

# AT3G14390.1 : NA

Pyridoxal-dependent decarboxylase family protein

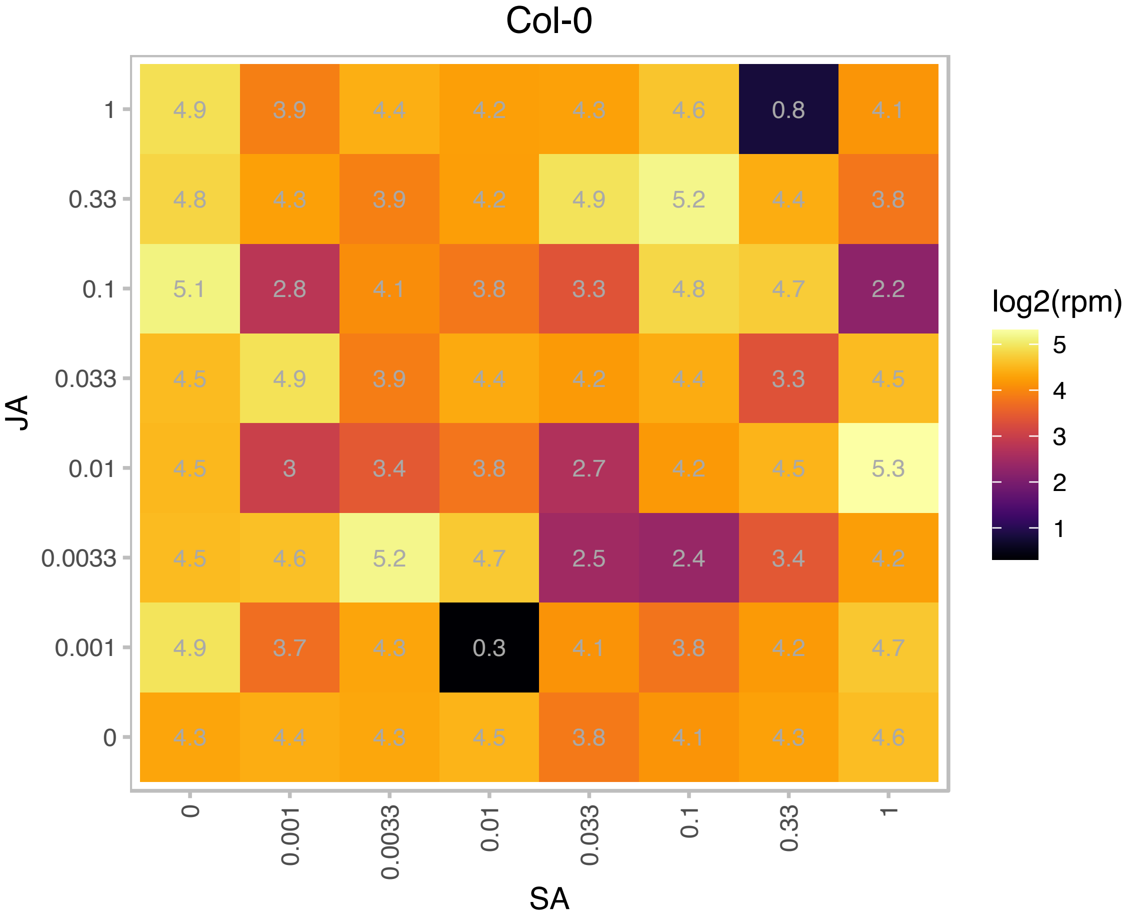


| | |
|---|---|
| Gene | AT3G14390 |
| is.expressed_genes | TRUE |
| is.responsive_genes | TRUE |
| is.apcluster | E-05 |

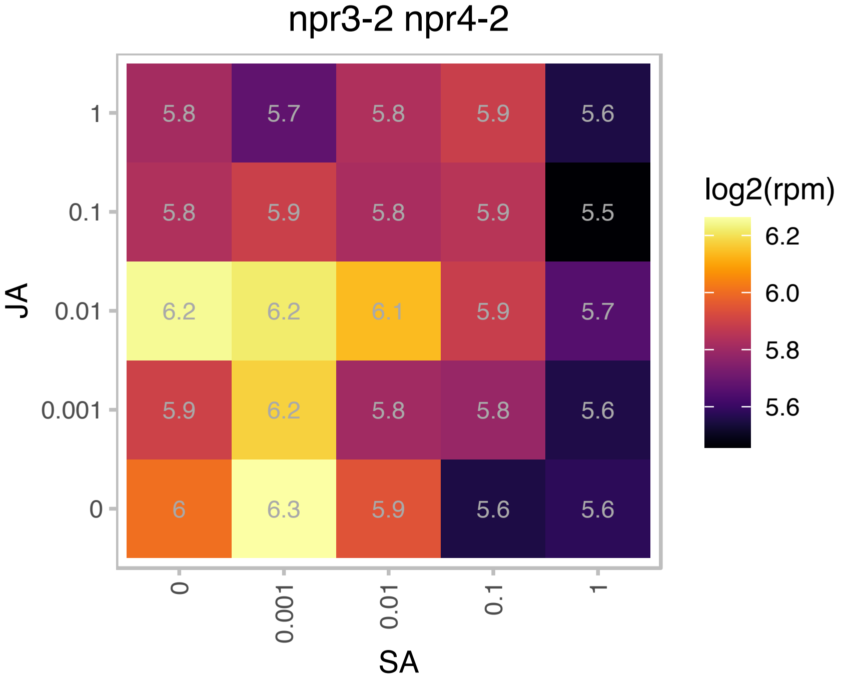


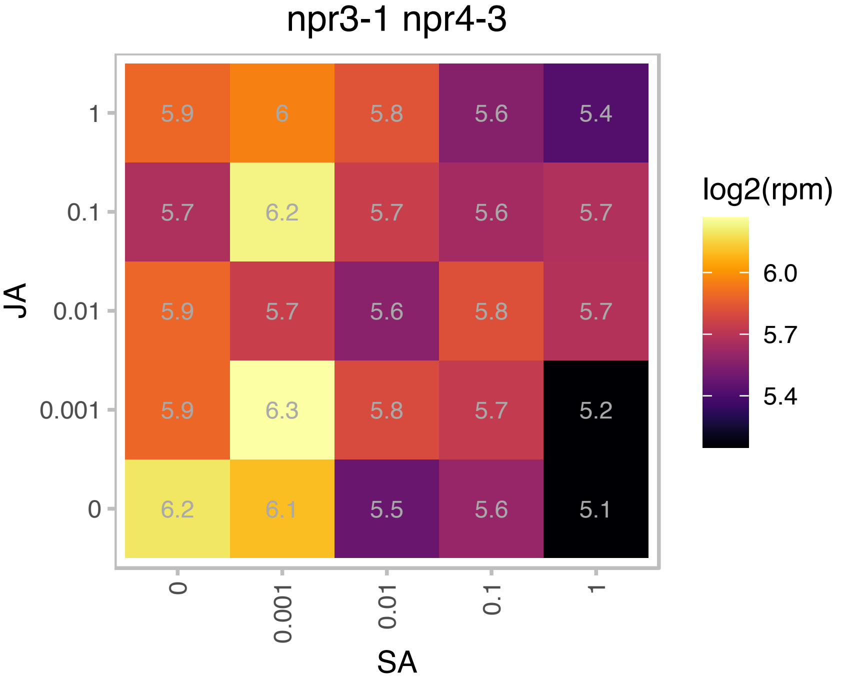

# AT3G24190.1 : NA

Protein kinase superfamily protein

| | |
|---|---|
| Gene | AT3G24190 |
| is.expressed_genes | TRUE |
| is.responsive_genes | TRUE |
| is.apcluster | E-05 |

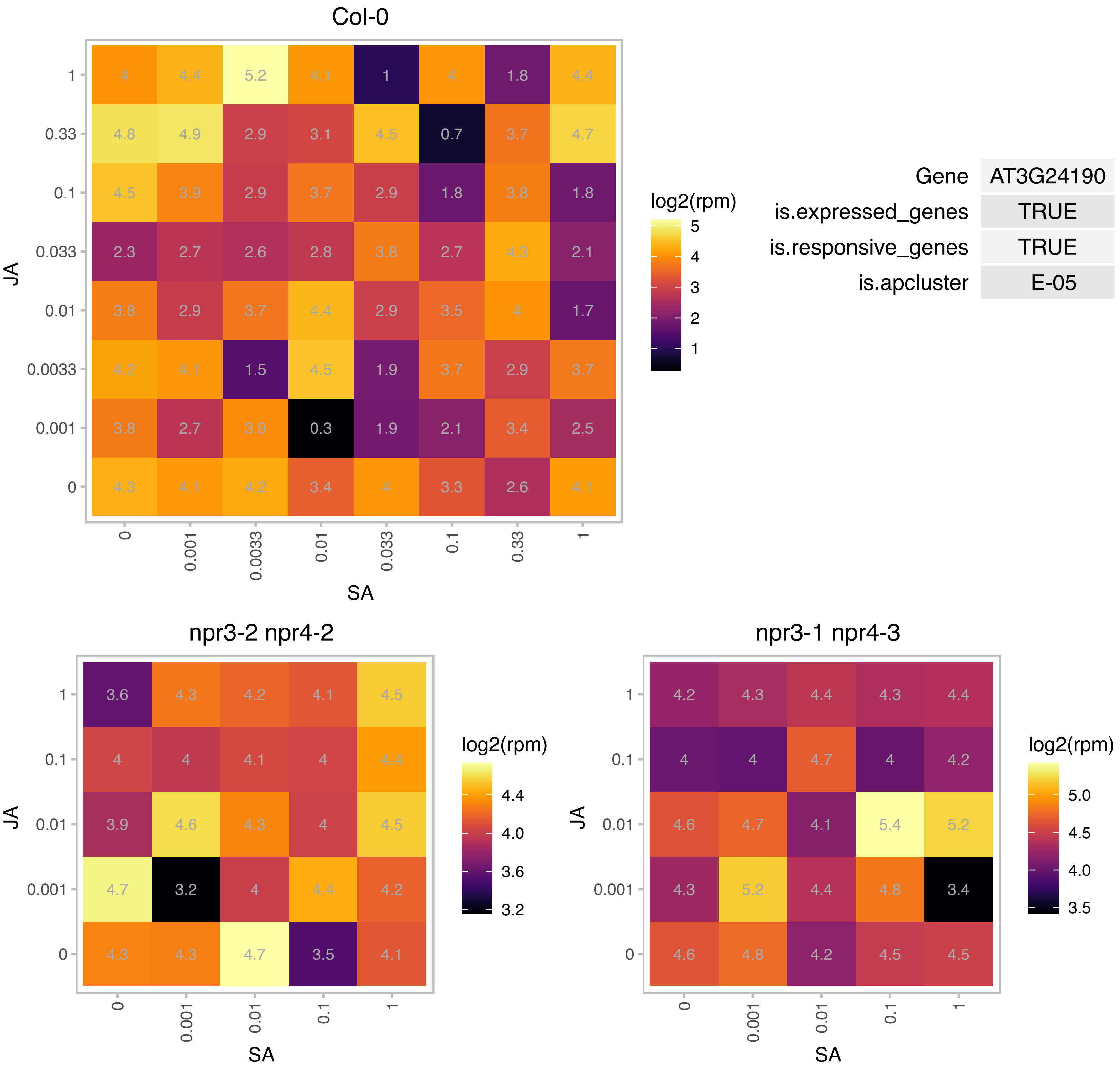

# AT3G25480.1 : NA

Rhodanese/Cell cycle control phosphatase superfamily protein

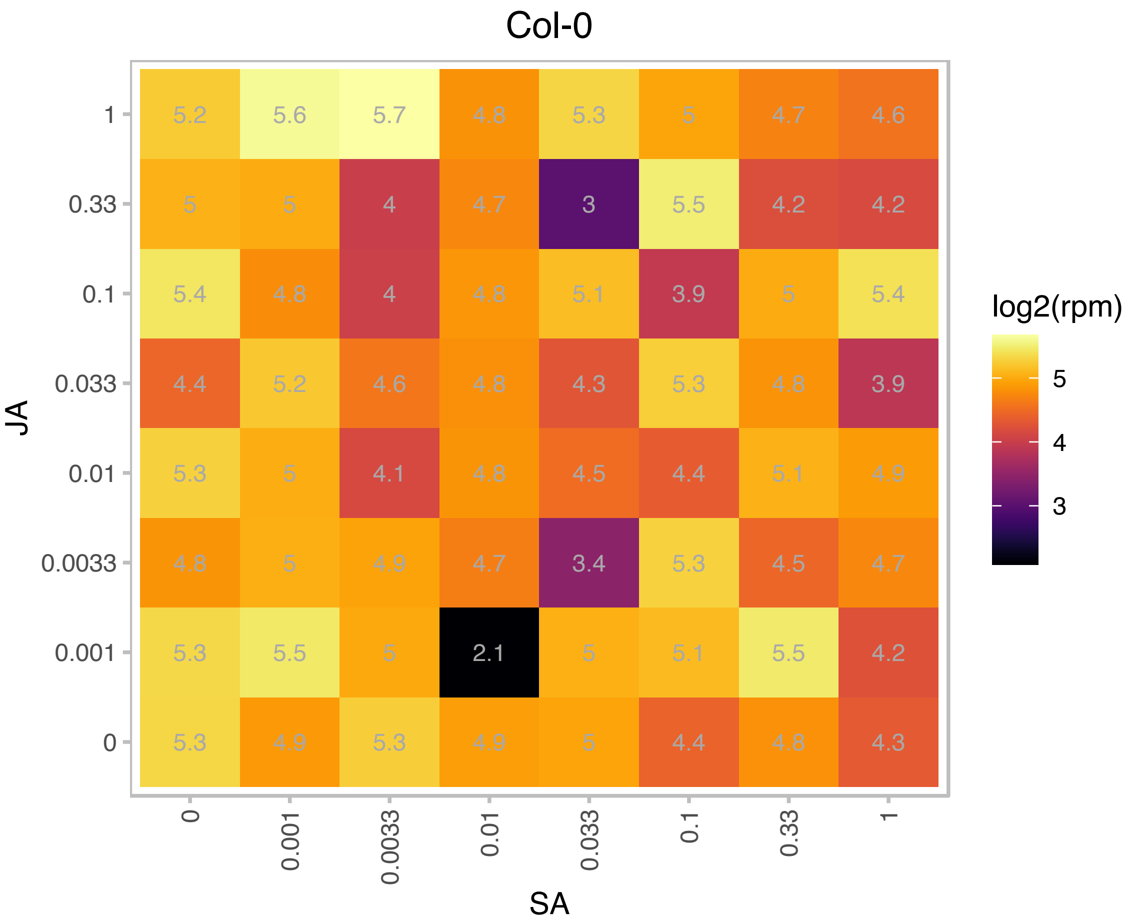


| | |
|---|---|
| Gene | AT3G25480 |
| is.expressed_genes | TRUE |
| is.responsive_genes | TRUE |
| is.apcluster | E-05 |

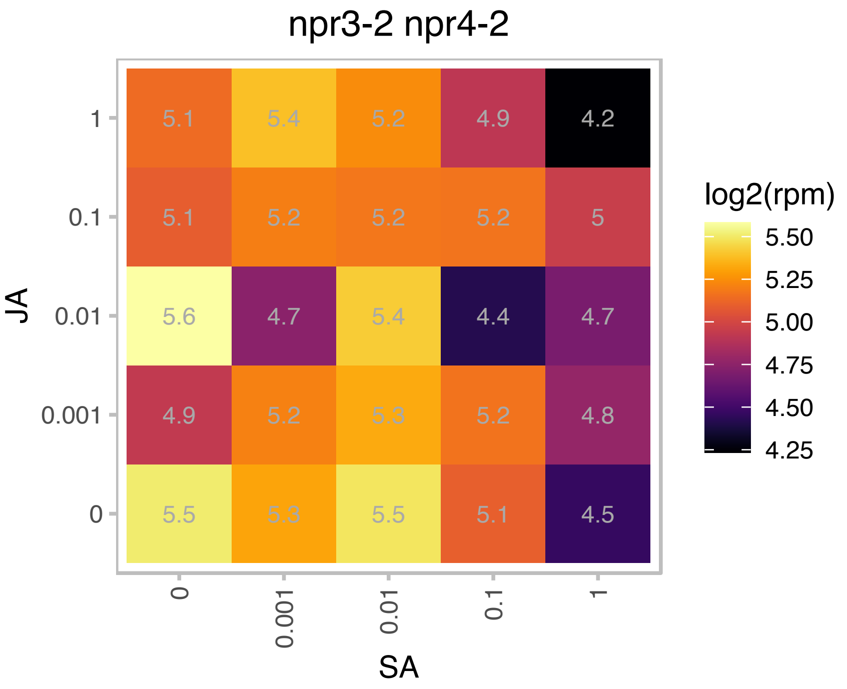


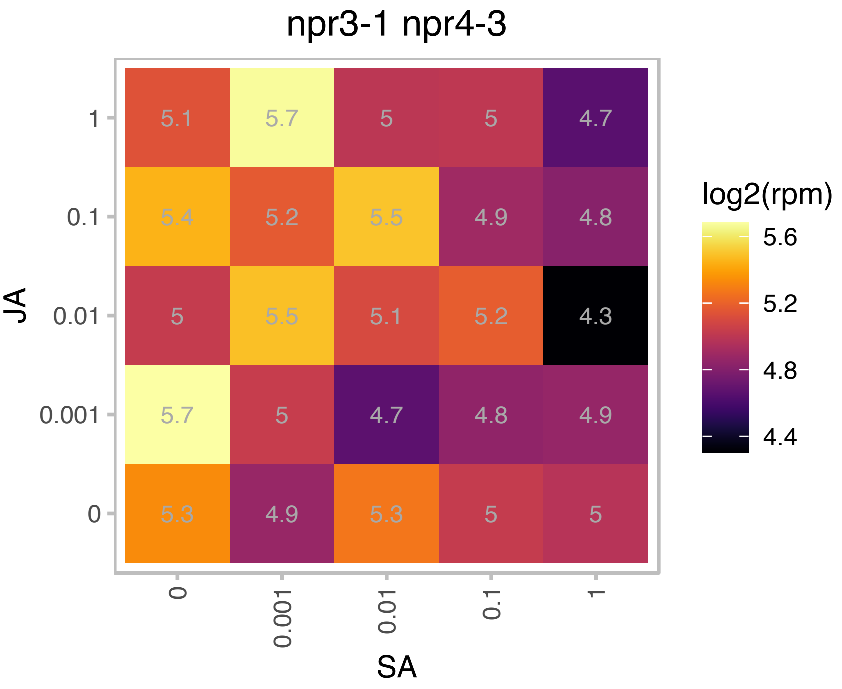

# AT3G26080.1 : NA

plastid-lipid associated protein PAP / fibrillin family protein

| | |
|---|---|
| Gene | AT3G26080 |
| is.expressed_genes | TRUE |
| is.responsive_genes | TRUE |
| is.apcluster | E-05 |

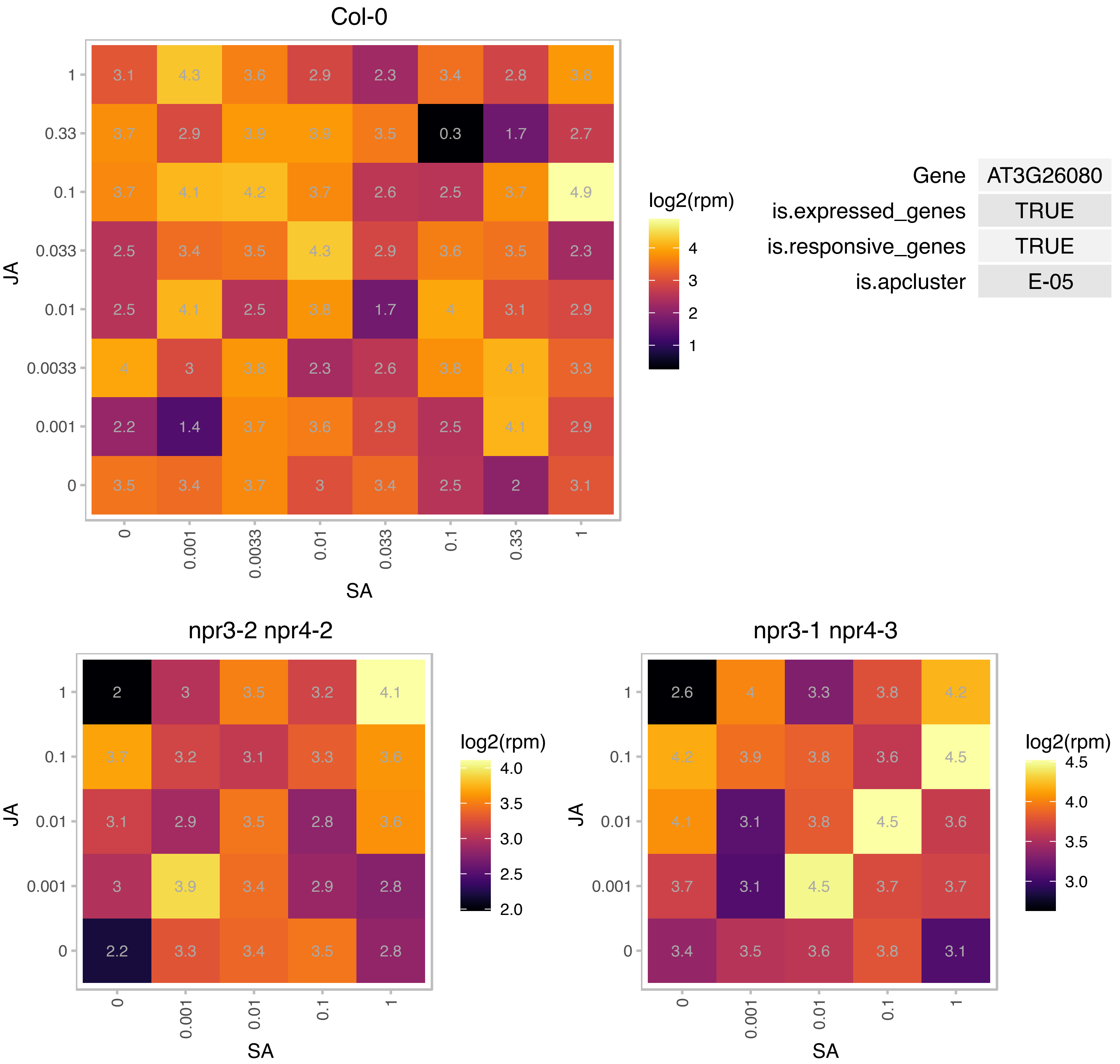

### AT3G60880.1 : DHDPS, DHDPS1

dihydrodipicolinate synthase 1

Encodes a dihydropicolinate synthase involved in lysine biosynthesis. The enzyme is allosterically inhibited by lysine. It is predicted to localize to the cholorplast.

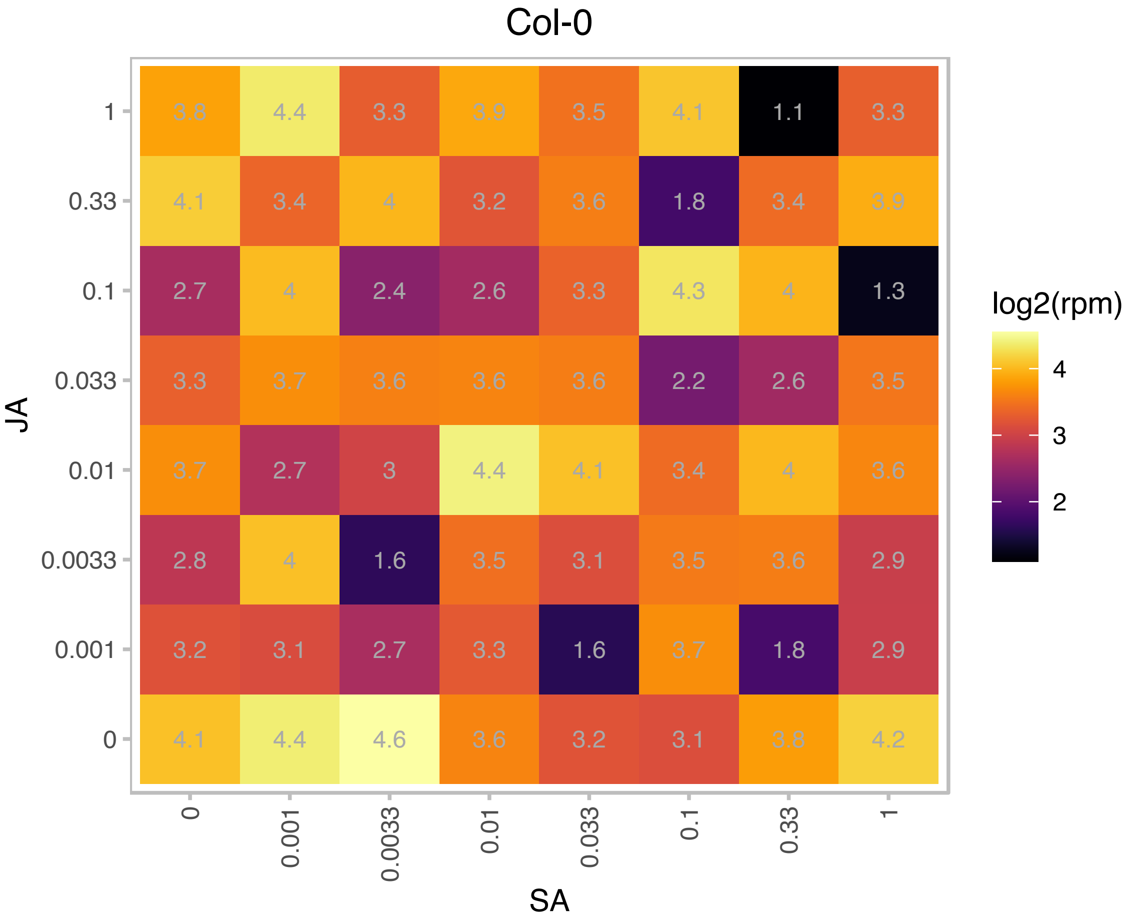


| | |
|---|---|
| Gene | AT3G60880 |
| is.expressed_genes | TRUE |
| is.responsive_genes | TRUE |
| is.apcluster | E-05 |

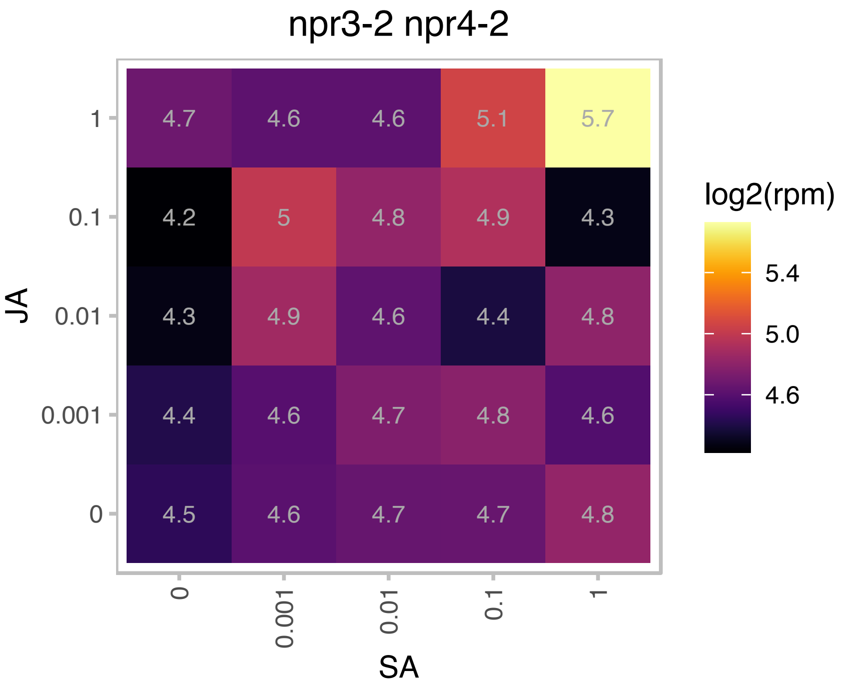


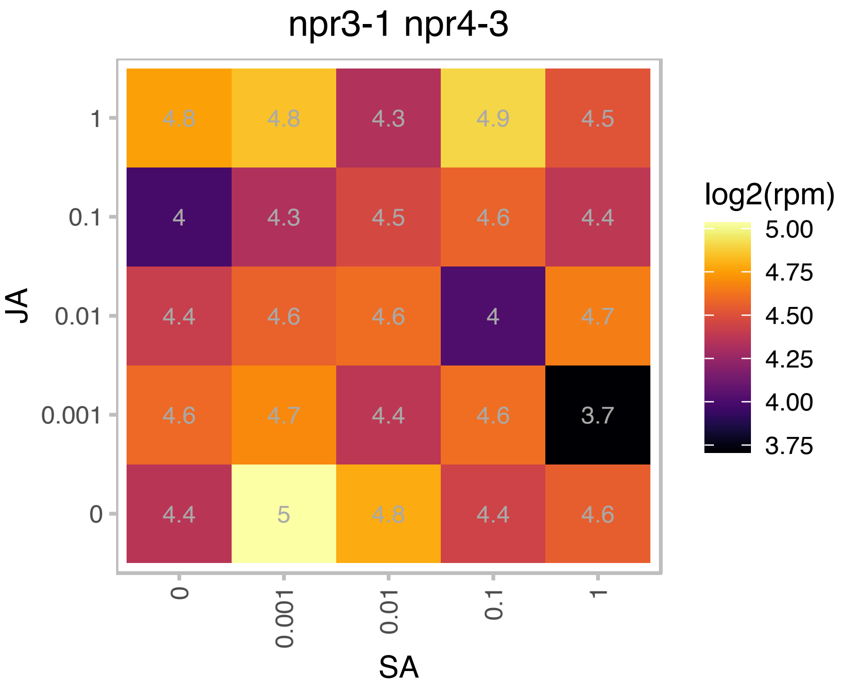

### AT4G10760.1 : EMB1706, MTA

mRNAadenosine methylase

Encodes a member of a core set of mRNA m6A writer proteins and is required for N6-adenosine methylation of mRNA.

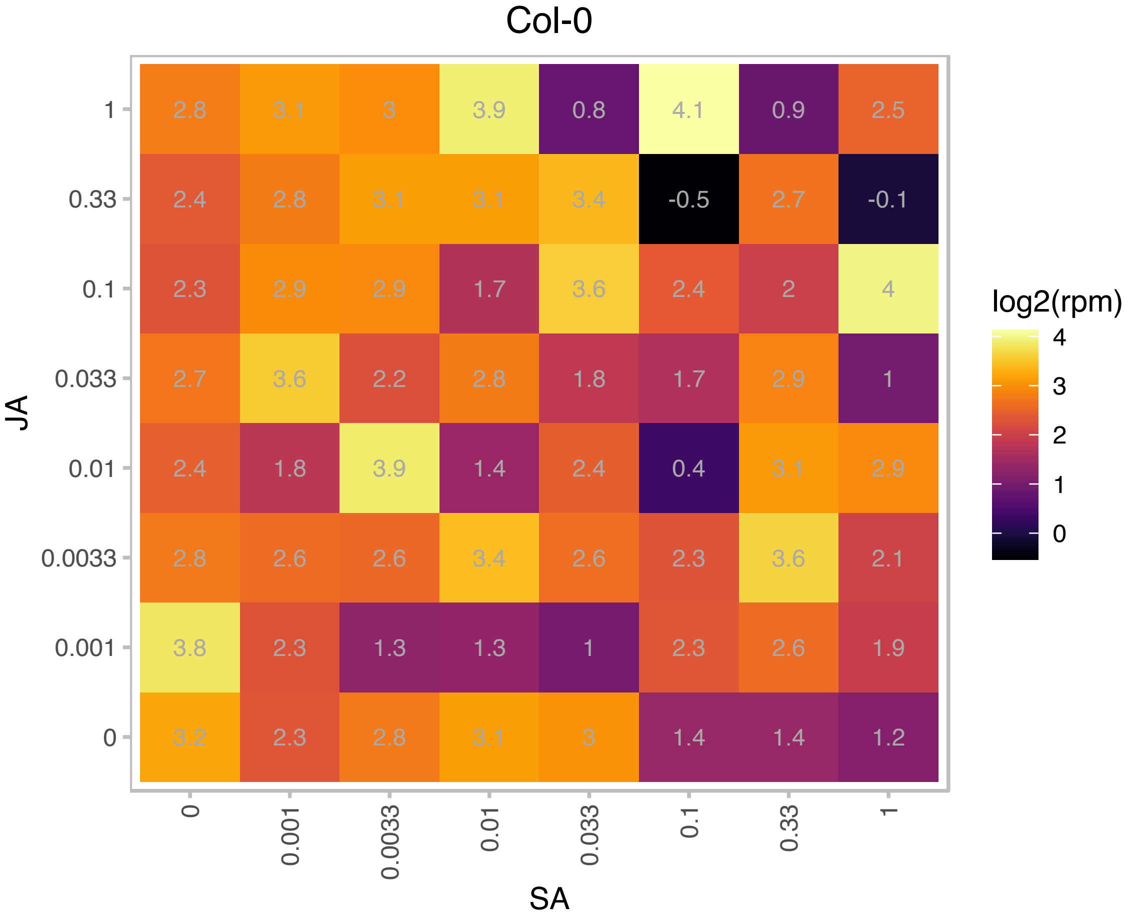


| | |
|---|---|
| Gene | AT4G10760 |
| is.expressed_genes | TRUE |
| is.responsive_genes | TRUE |
| is.apcluster | E-05 |

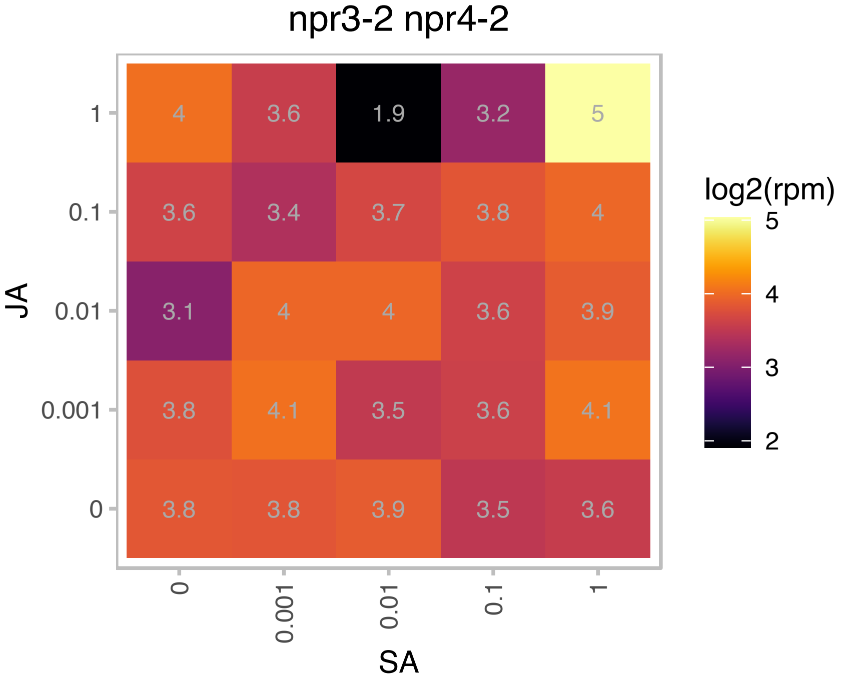


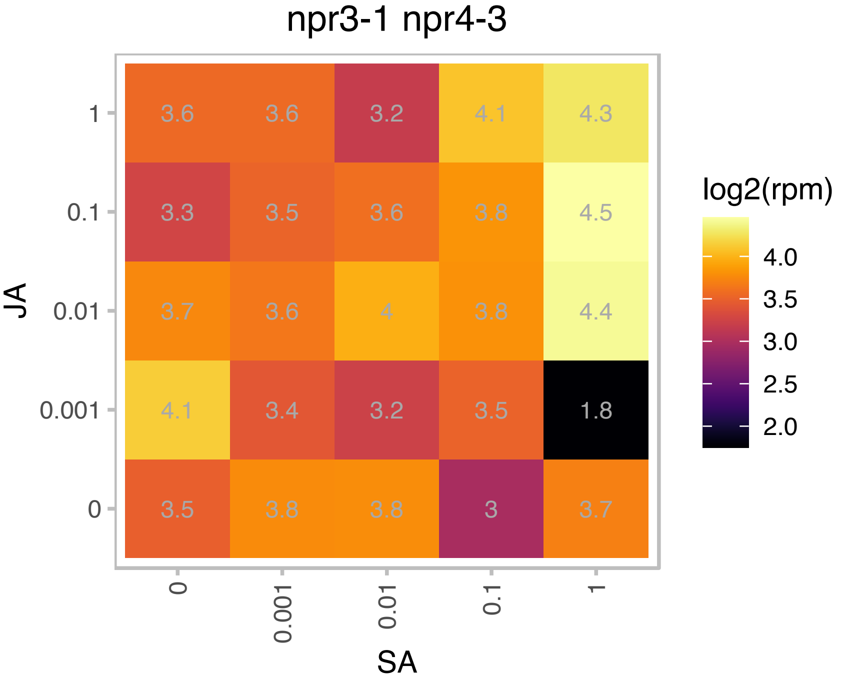

### AT4G11830.1 : PLDGAMMA2

phospholipase D gamma 2

Encodes one of three phospholipase D enzymes of the gamma class.

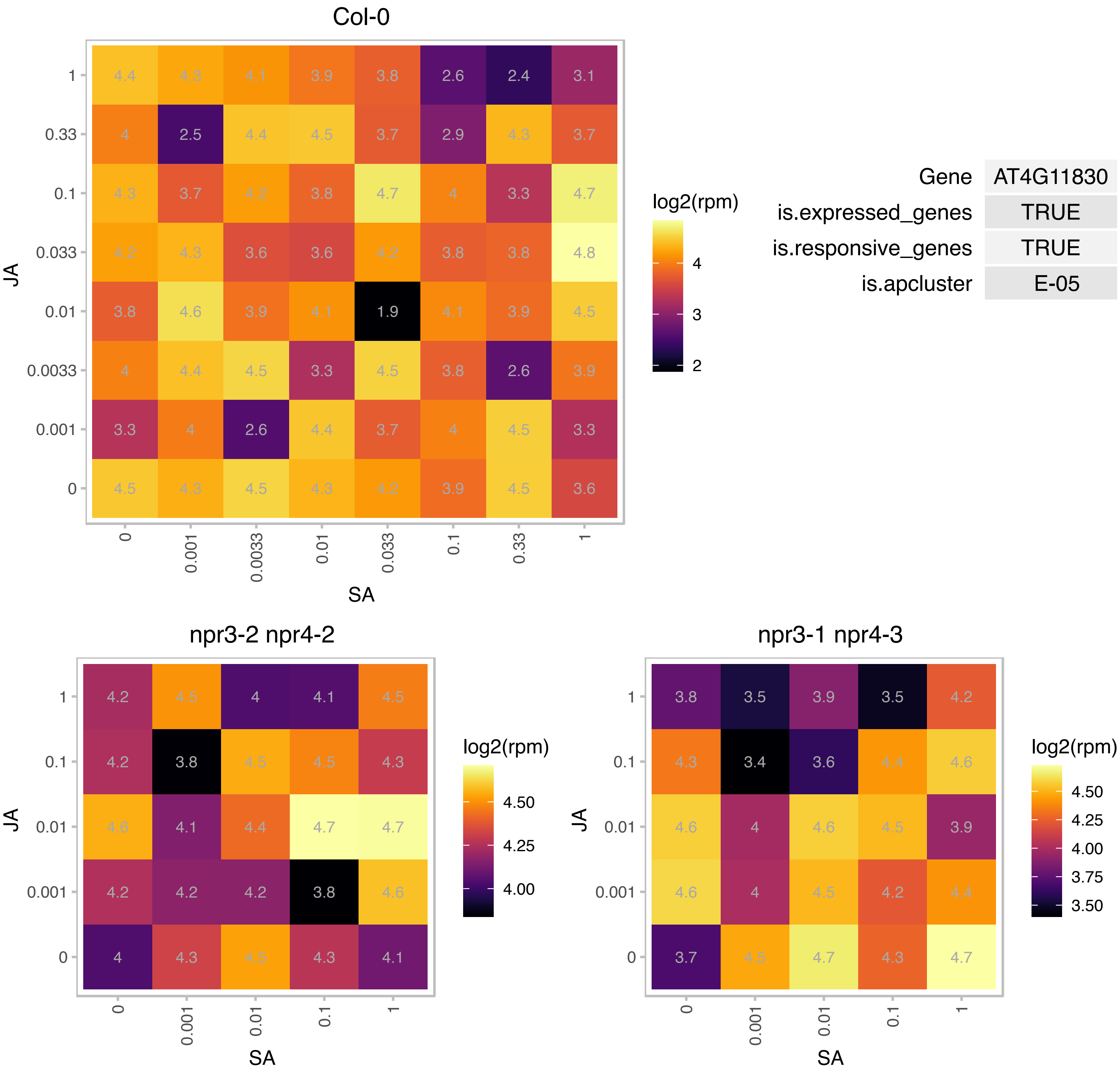


| | |
|---|---|
| Gene | AT4G11830 |
| is.expressed_genes | TRUE |
| is.responsive_genes | TRUE |
| is.apcluster | E-05 |

### AT4G17090.1 : AtBAM3, BAM3, BMY8, CT-BMY

chloroplast beta-amylase

Encodes a beta-amylase targeted to the chloroplast. Transgenic BMY8 RNAi lines fail to accumulate maltose during cold shock suggesting that maltose accumulation coincides with BMY8 expression. Apart from maltose, the sugar content of the RNAi lines were similar to wildtype (glucose and sucrose unaffected).BAM3 activity declines 2 and 4 days after start of cold stress despite an increase in transcript levels. BAM3 activity has a lower temperature optimum than BAM1 (PMID:25293962).

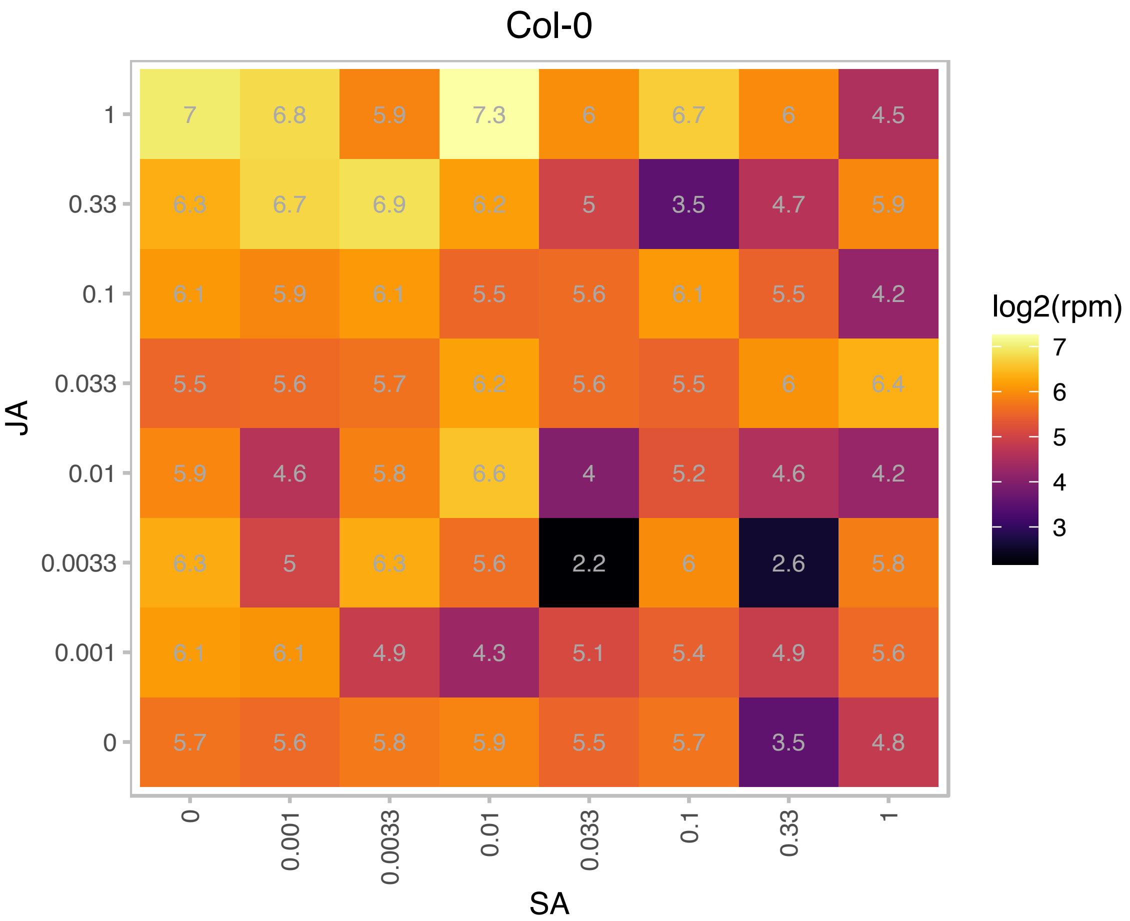


| | |
|---|---|
| Gene | AT4G17090 |
| is.expressed_genes | TRUE |
| is.responsive_genes | TRUE |
| is.apcluster | E-05 |

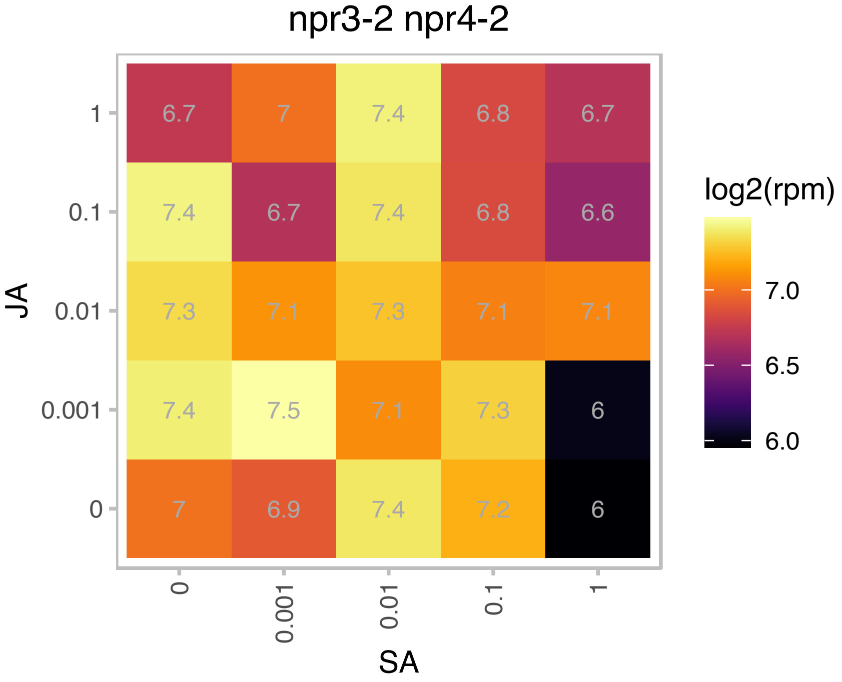


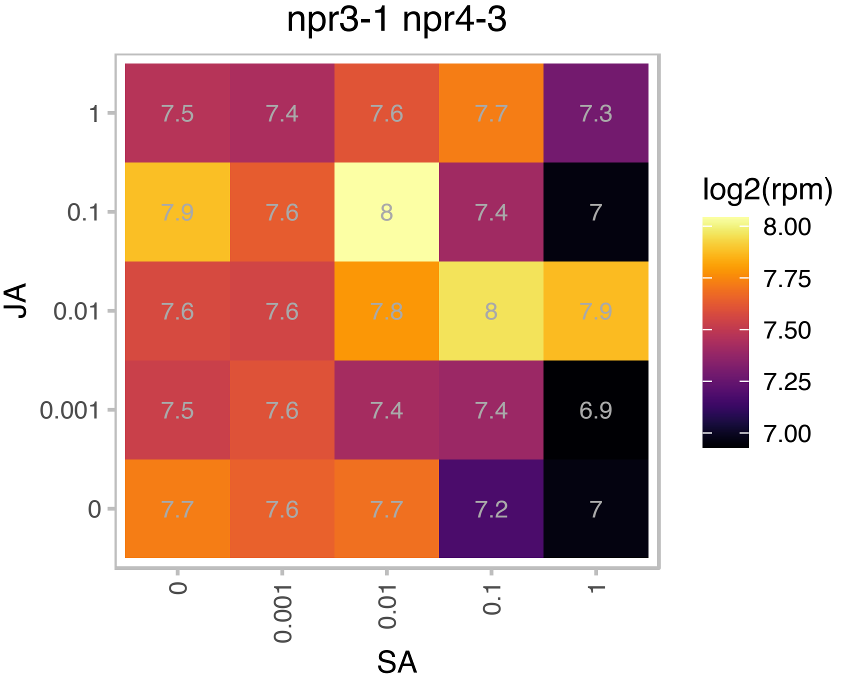

# AT4G17100.1 : NA

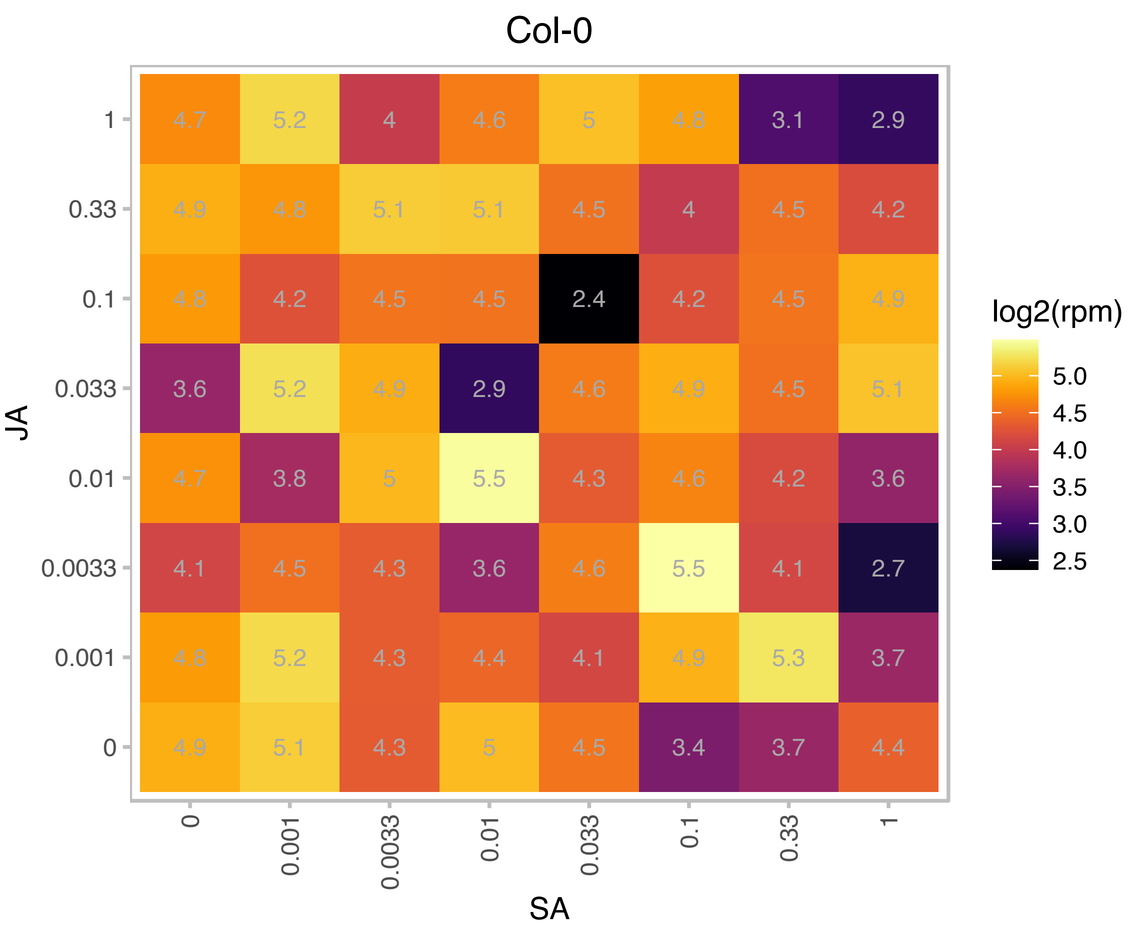


| | |
|---|---|
| Gene | AT4G17100 |
| is.expressed_genes | TRUE |
| is.responsive_genes | TRUE |
| is.apcluster | E-05 |

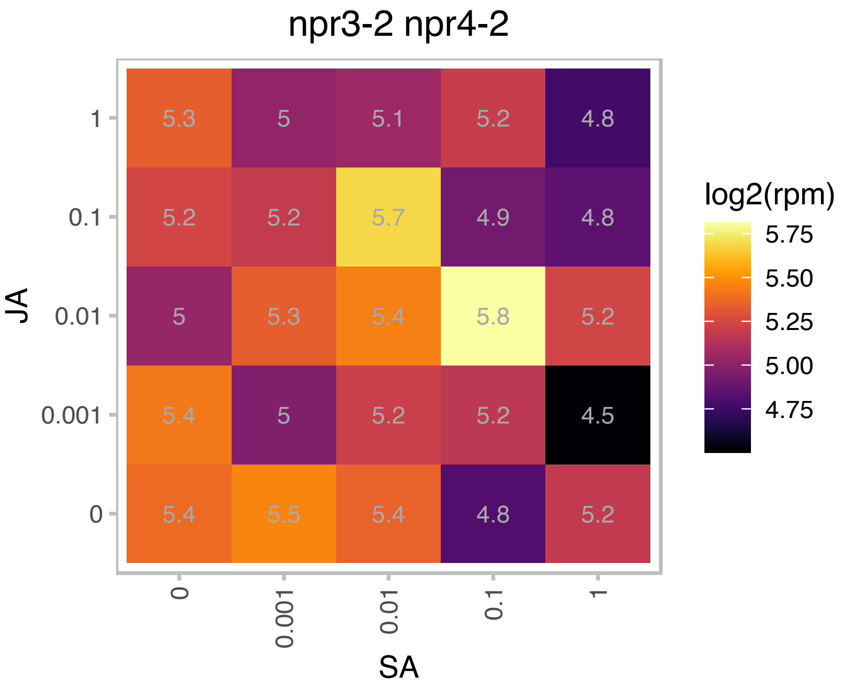


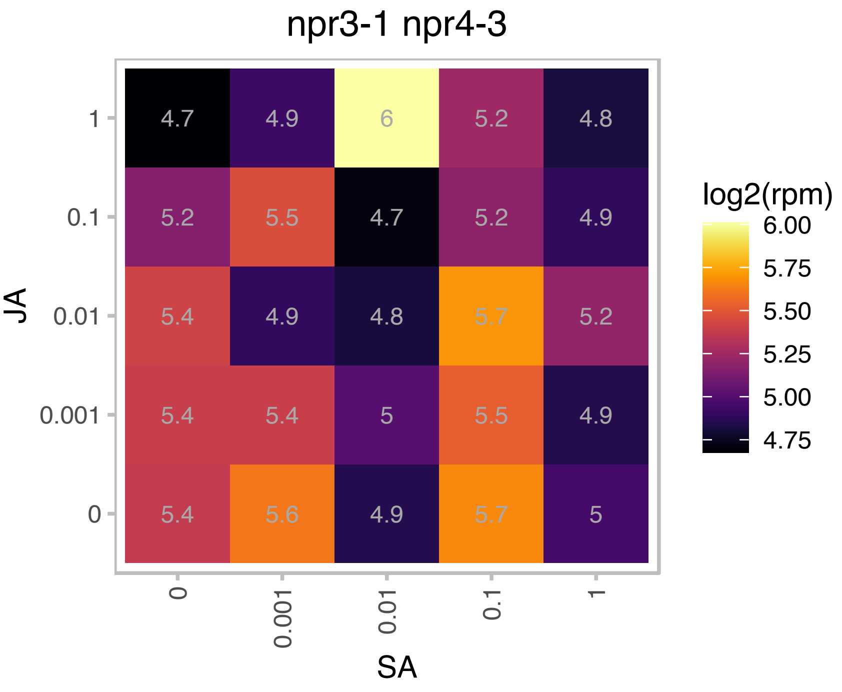

### AT4G19840.1 : AN5, ATPP2-A1, ATPP2A-1, PP2-A1

phloem protein 2-A1

encodes a phloem lectin, similar to phloem lectin in cucumber and celery. Gene is expressed in the phloem, predominantly in the companion cells. The mRNA is cell-to-cell mobile.

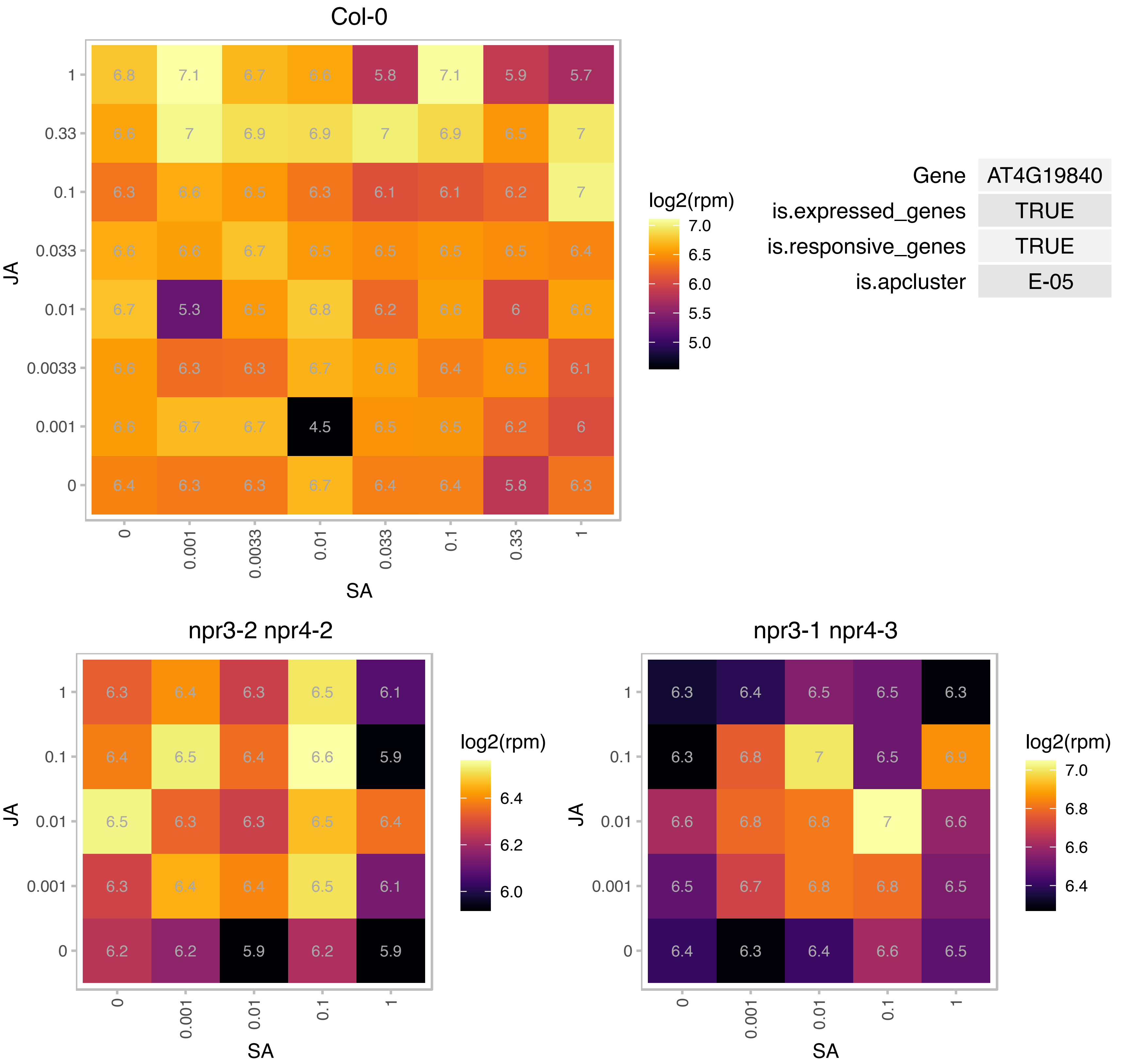


| | |
|---|---|
| Gene | AT4G19840 |
| is.expressed_genes | TRUE |
| is.responsive_genes | TRUE |
| is.apcluster | E-05 |

### AT4G31720.1 : STG1, TAF10, TAFII15

TBP-associated factor II 15

Arabidopsis thaliana putative TBP-associated 15 kDa subunit protein (TAFII15)

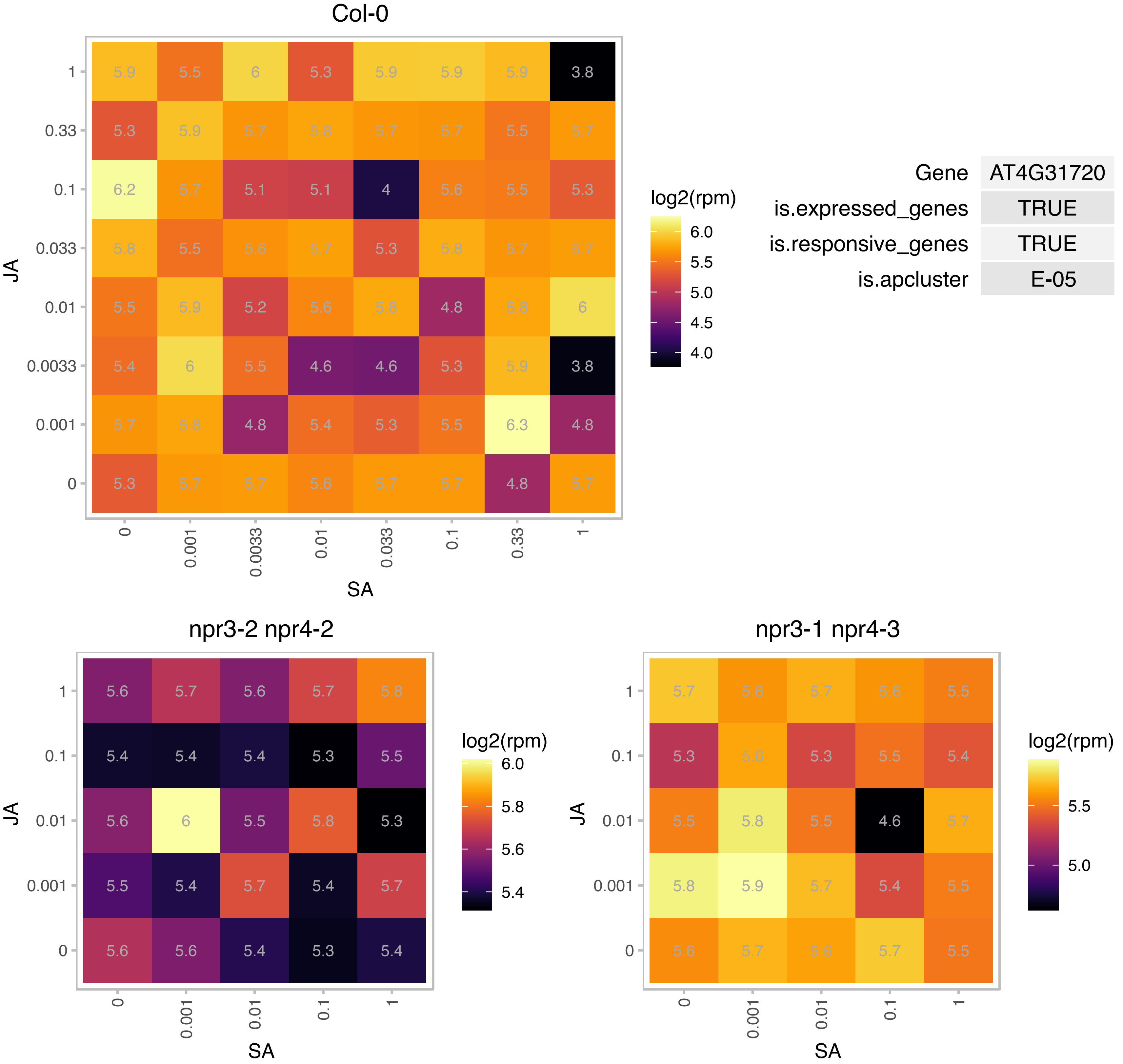


| | |
|---|---|
| Gene | AT4G31720 |
| is.expressed_genes | TRUE |
| is.responsive_genes | TRUE |
| is.apcluster | E-05 |

# AT4G36190.1 : NA

Serine carboxypeptidase S28 family protein

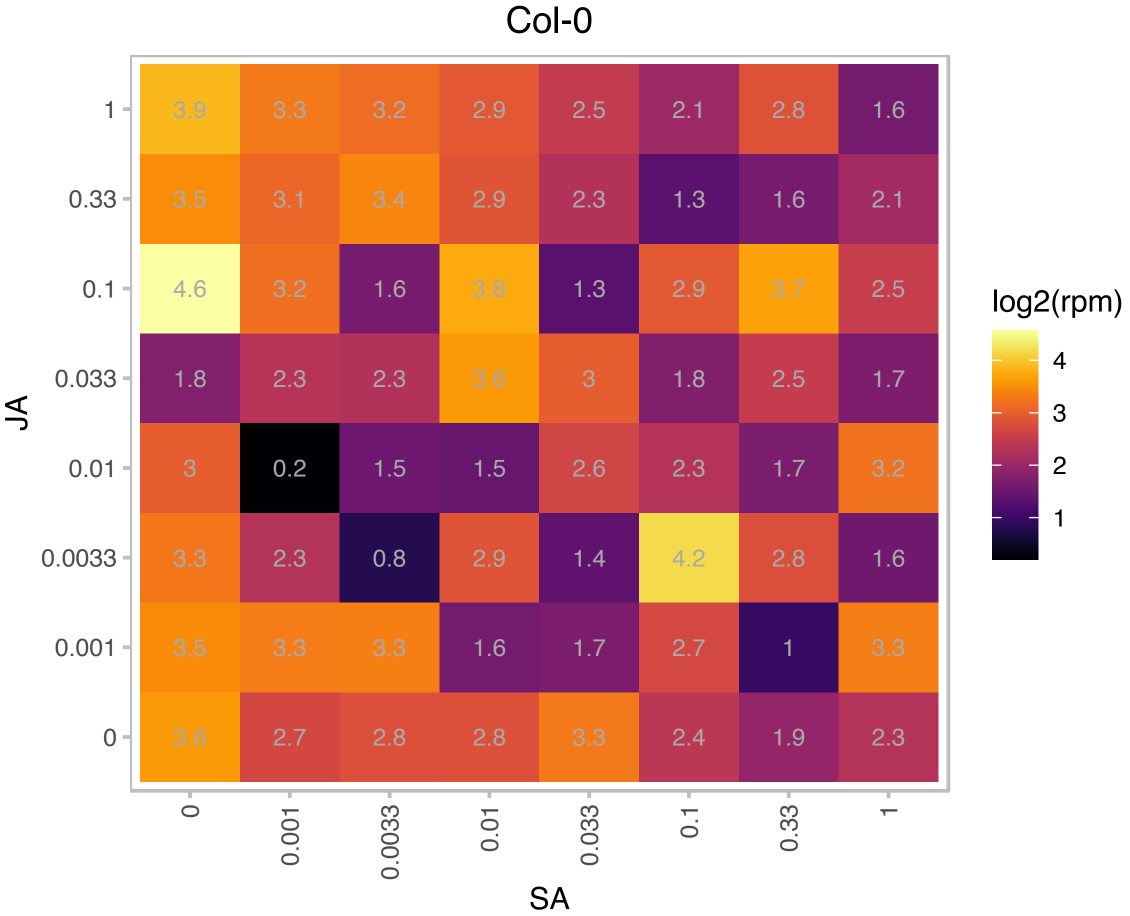


| | |
|---|---|
| Gene | AT4G36190 |
| is.expressed_genes | TRUE |
| is.responsive_genes | TRUE |
| is.apcluster | E-05 |

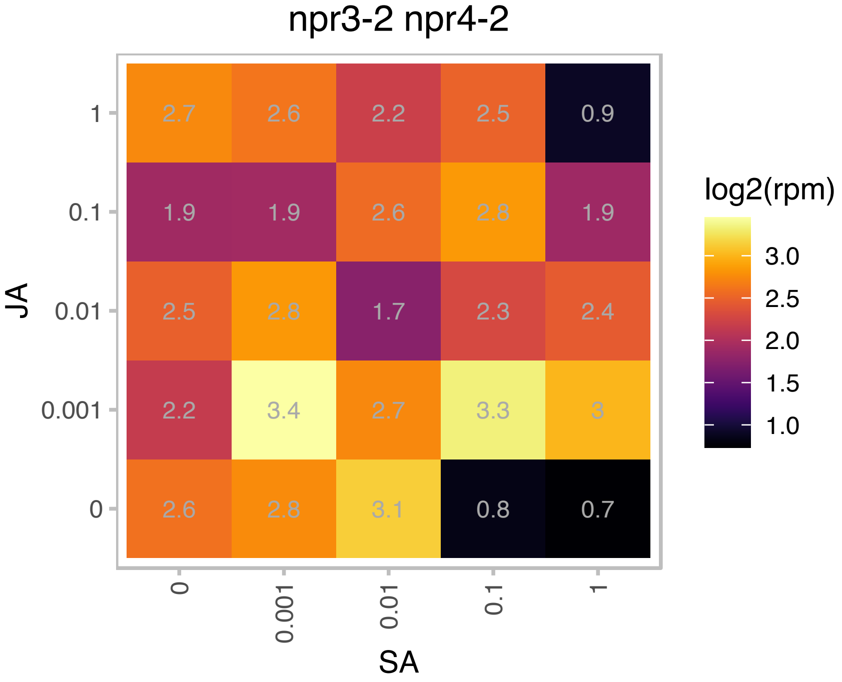


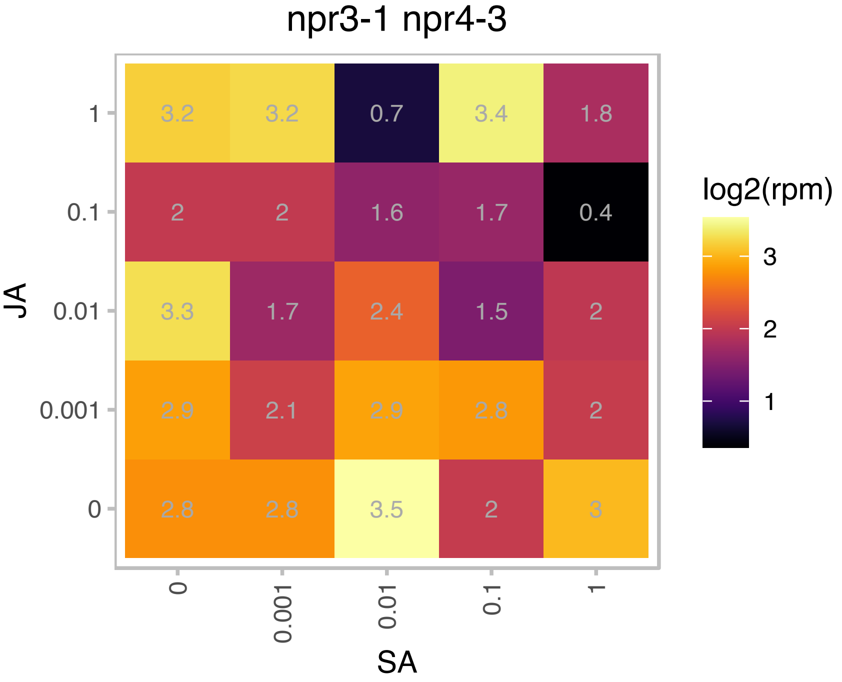

### AT4G37330.1 : CYP81D4

cytochrome P450, family 81, subfamily D, polypeptide 4

member of CYP81D

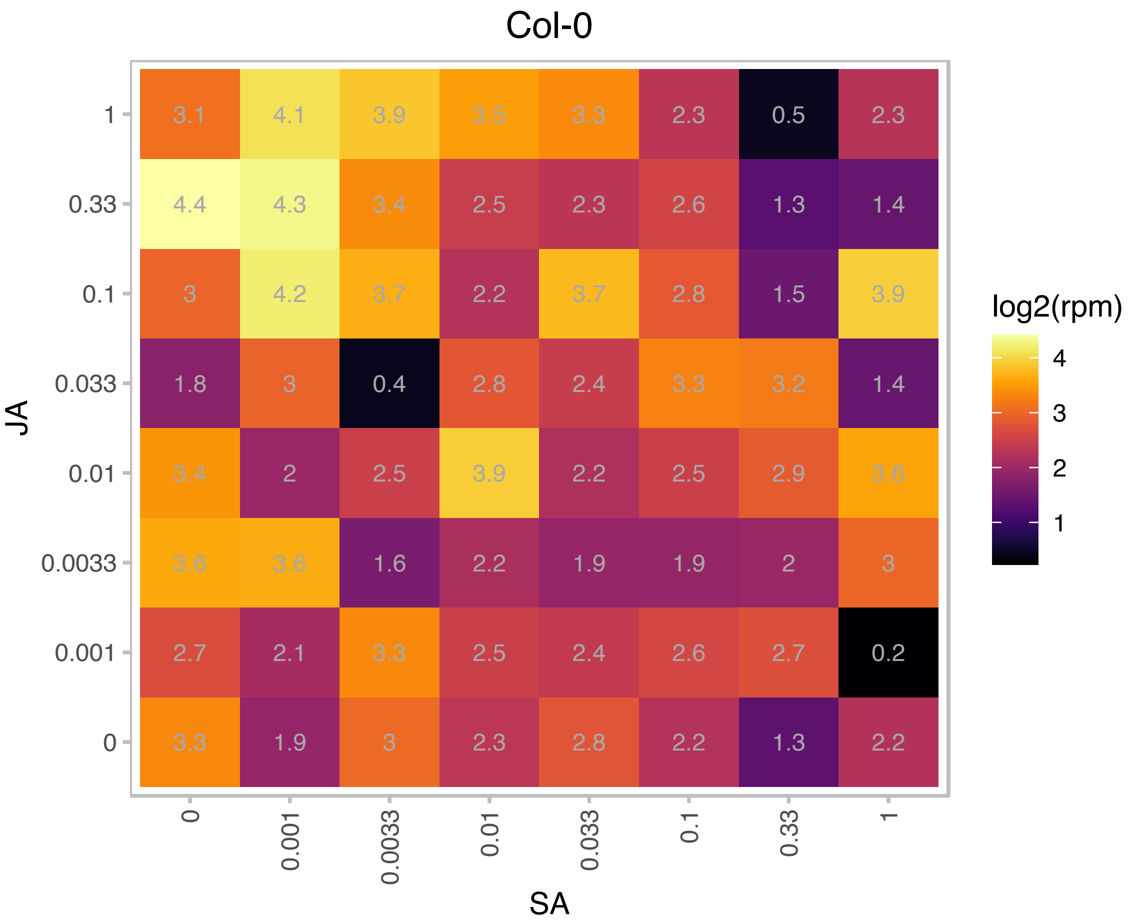


| | |
|---|---|
| Gene | AT4G37330 |
| is.expressed_genes | TRUE |
| is.responsive_genes | TRUE |
| is.apcluster | E-05 |

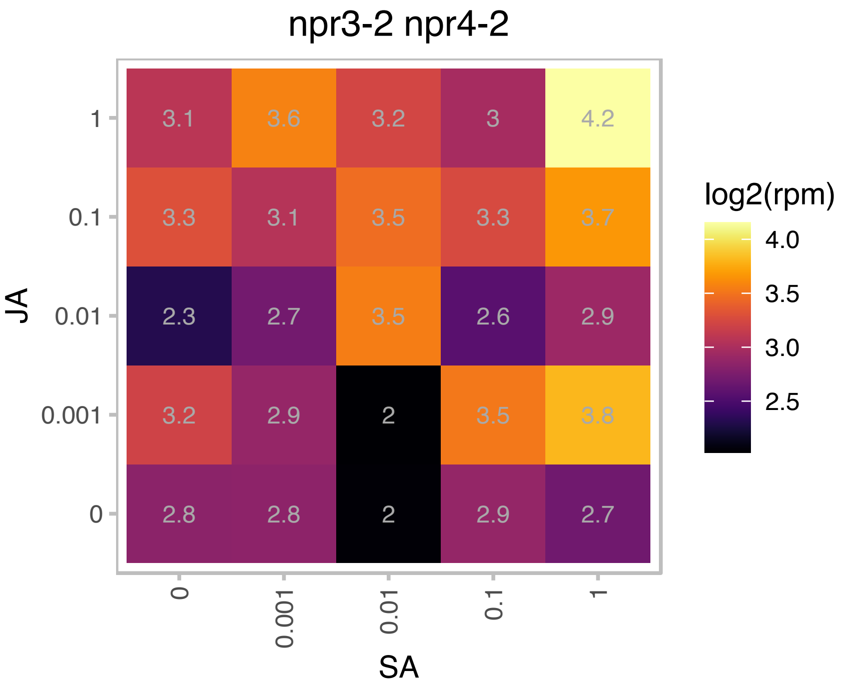


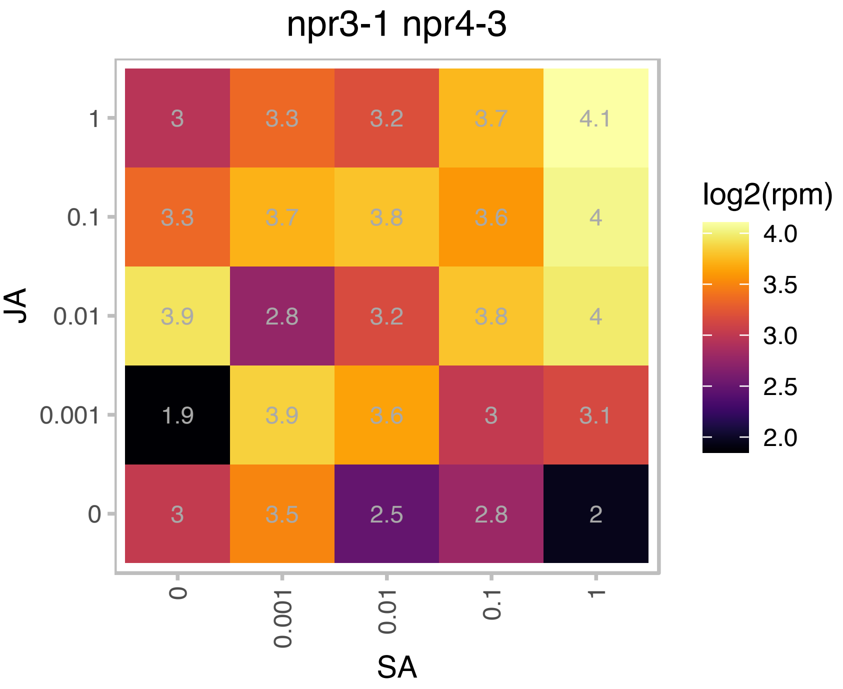

### AT5G01410.1 : ATPDX1, ATPDX1.3, PDX1, PDX1.3, RSR4

Aldolase-type TIM barrel family protein

Encodes a protein predicted to function in tandem with PDX2 to form glutamine amidotransferase complex with involved in vitamin B6 biosynthesis.

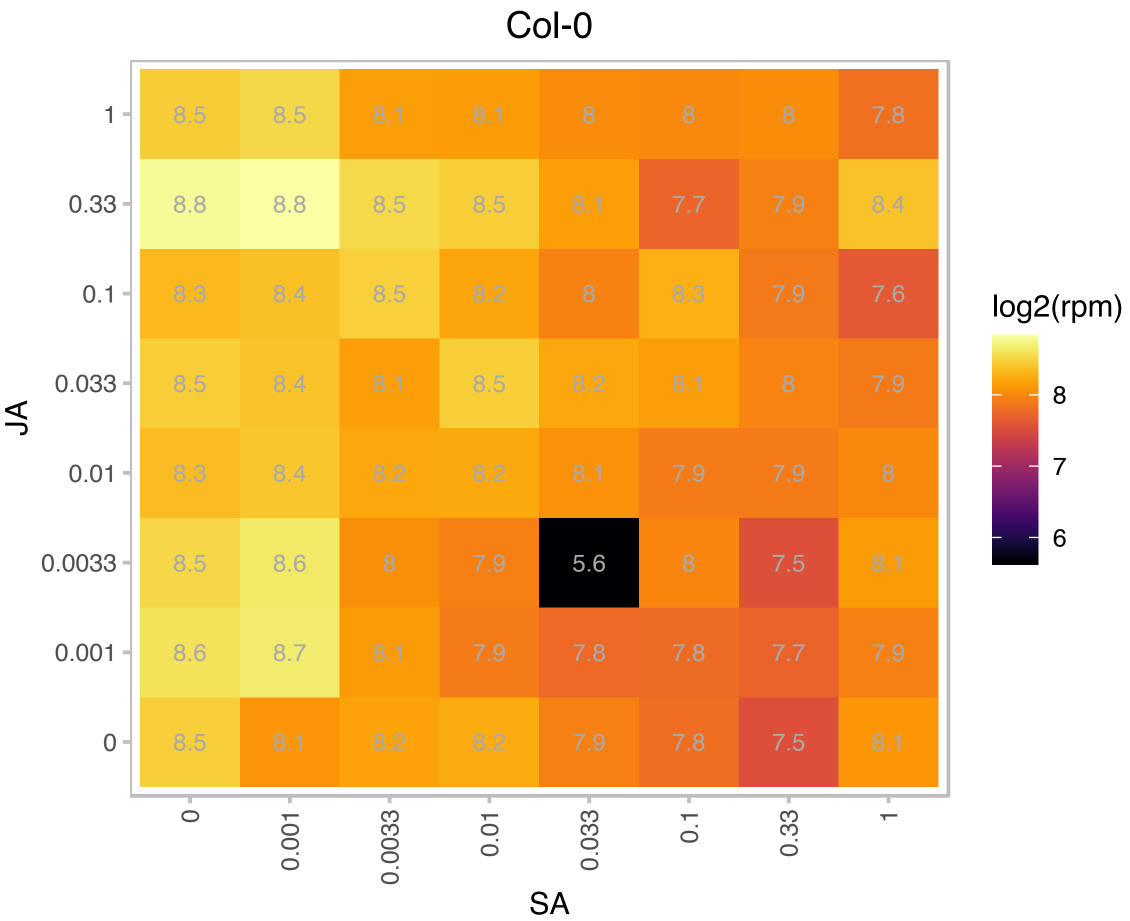


| | |
|---|---|
| Gene | AT5G01410 |
| is.expressed_genes | TRUE |
| is.responsive_genes | TRUE |
| is.apcluster | E-05 |

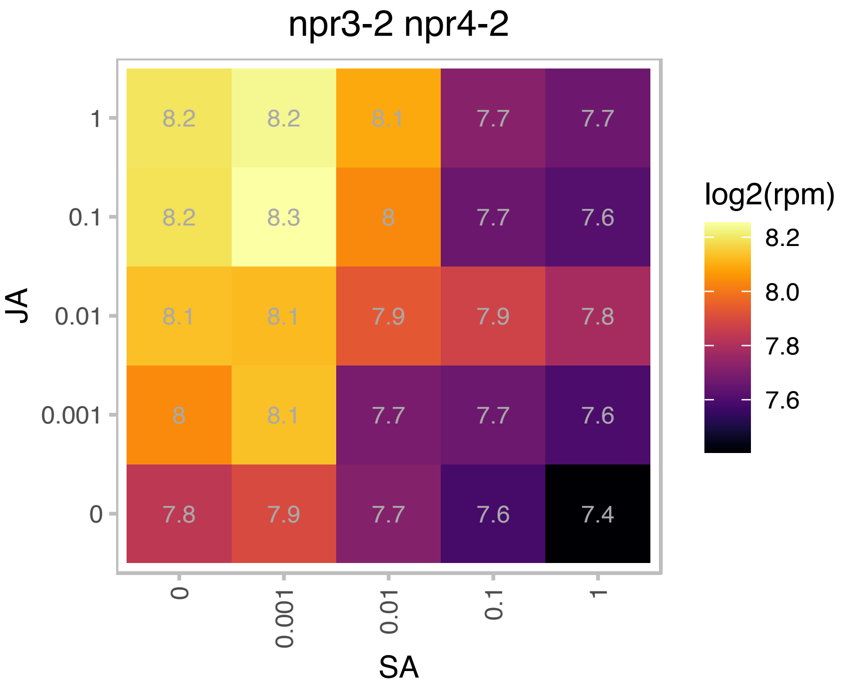


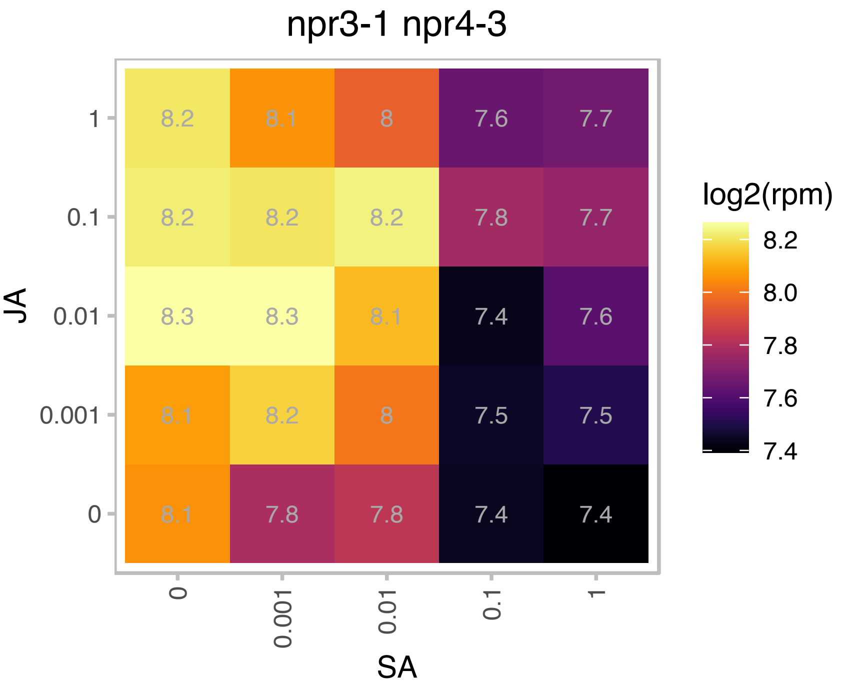

# AT5G03120.1 : NA

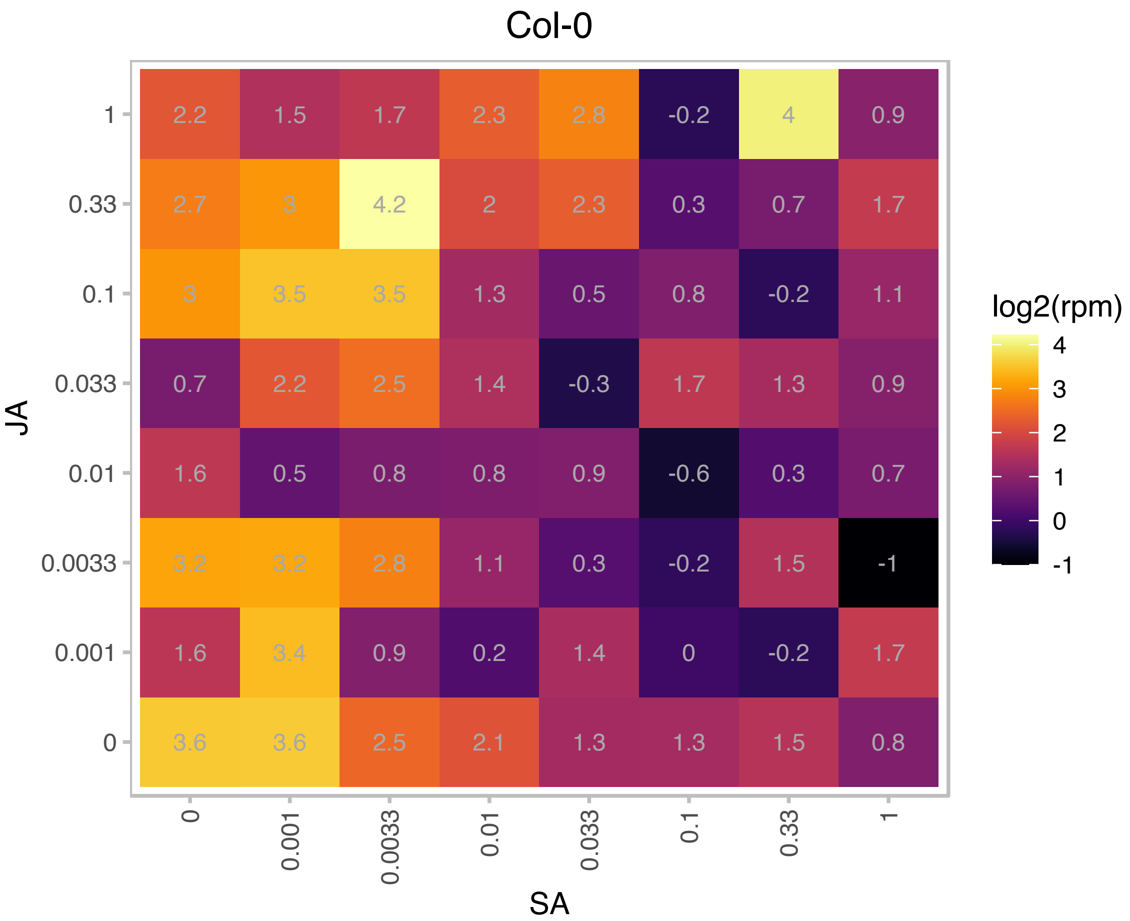


| | |
|---|---|
| Gene | AT5G03120 |
| is.expressed_genes | TRUE |
| is.responsive_genes | TRUE |
| is.apcluster | E-05 |

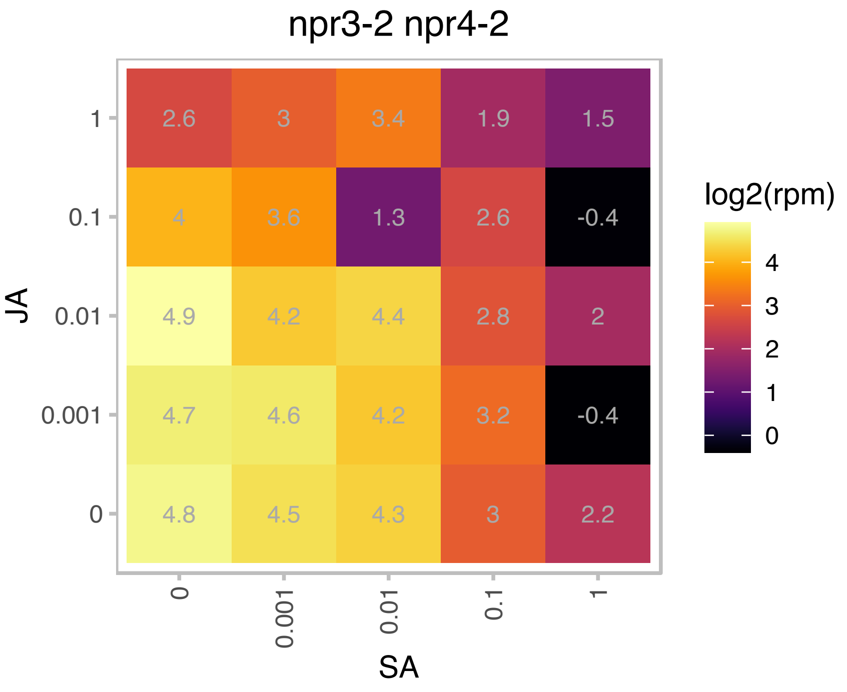


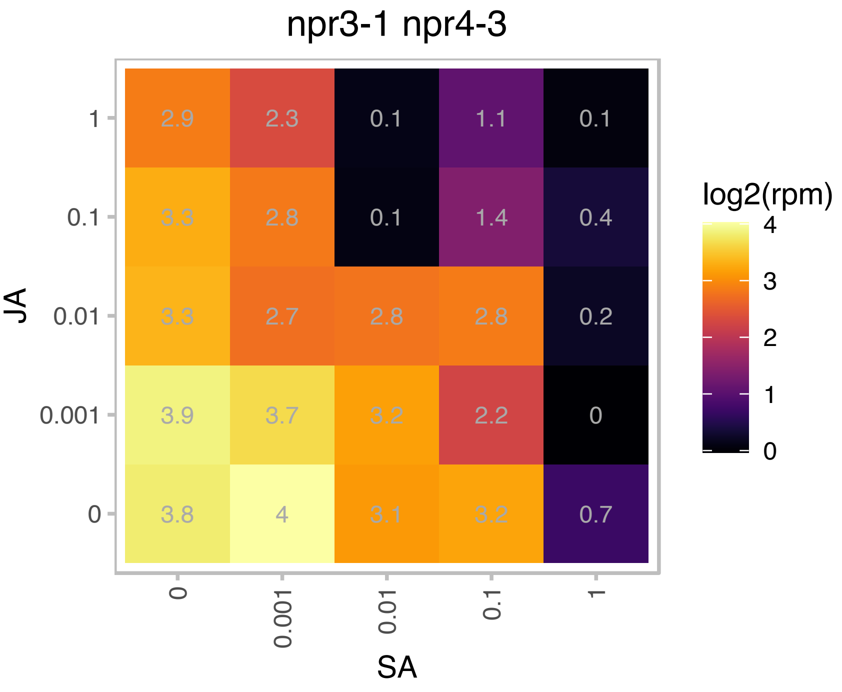

### AT5G26570.1 : ATGWD3, OK1, PWD

catalytics;carbohydrate kinases;phosphoglucan, water dikinases
chloroplastidic phosphoglucan, water dikinase (PWD) which is required for normal degradation of leaf starch in Arabidopsis. NMR analysis of the mutants, suggests that the gene is specifically involved in the phosphorylation of the glucosyl residues of starch at the C3 position.

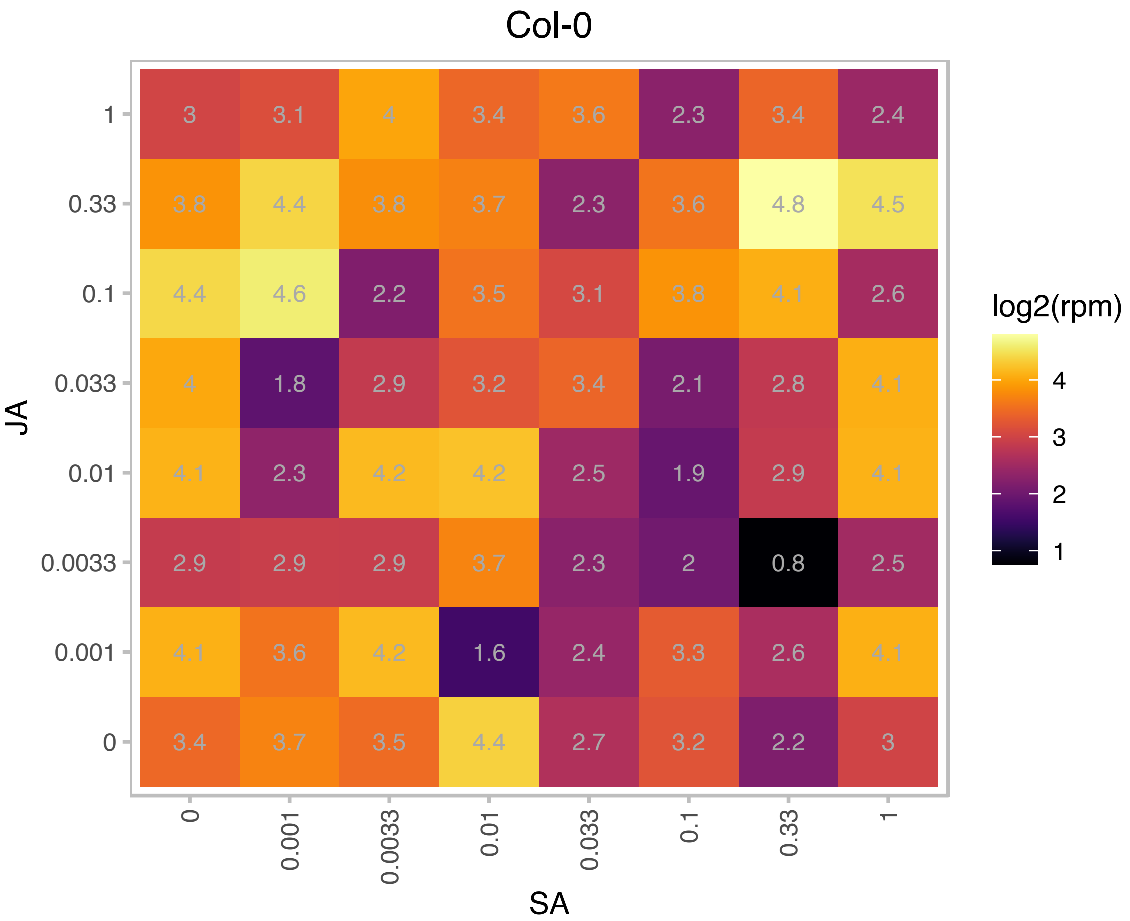


| | |
|---|---|
| Gene | AT5G26570 |
| is.expressed_genes | TRUE |
| is.responsive_genes | TRUE |
| is.apcluster | E-05 |

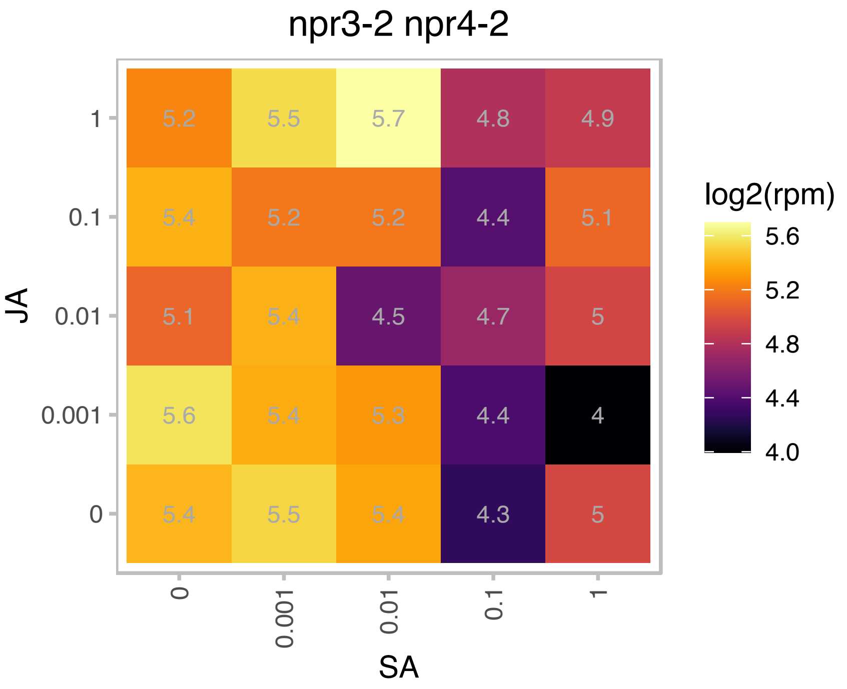


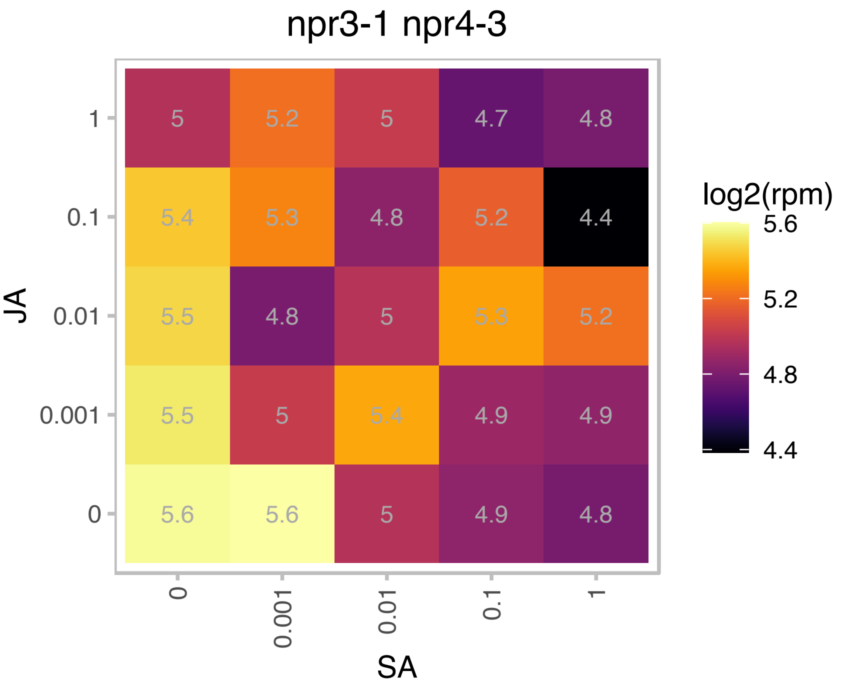

# AT5G58990.1 : NA

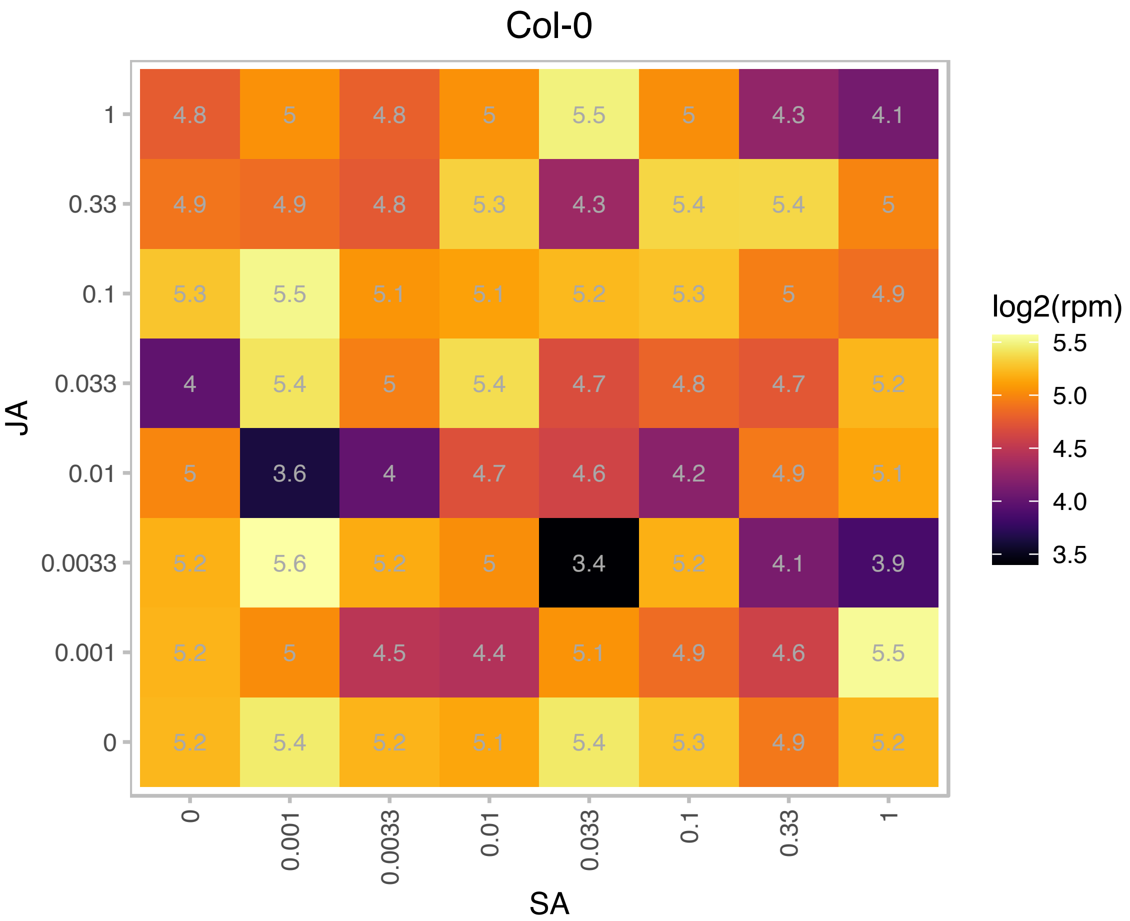


| | |
|---|---|
| Gene | AT5G58990 |
| is.expressed_genes | TRUE |
| is.responsive_genes | TRUE |
| is.apcluster | E-05 |

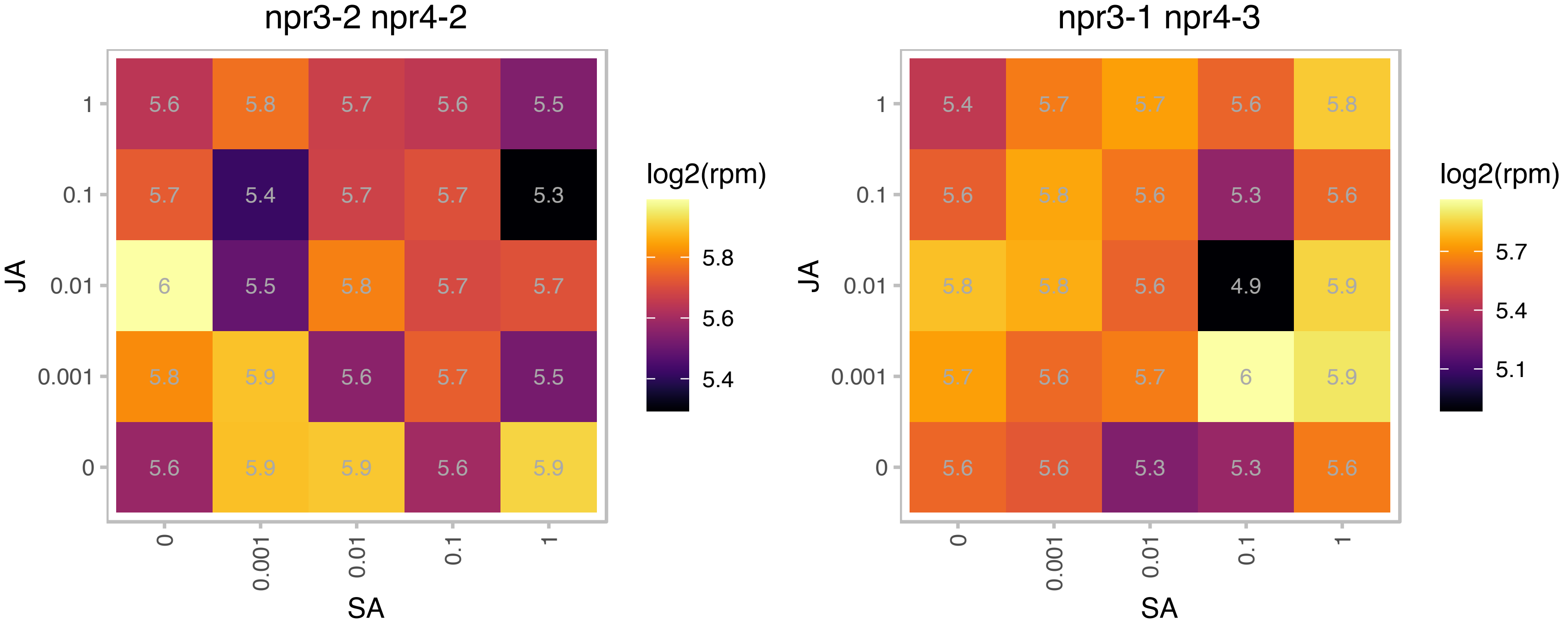

# AT1G01630.1 : NA

Sec14p-like phosphatidylinositol transfer family protein

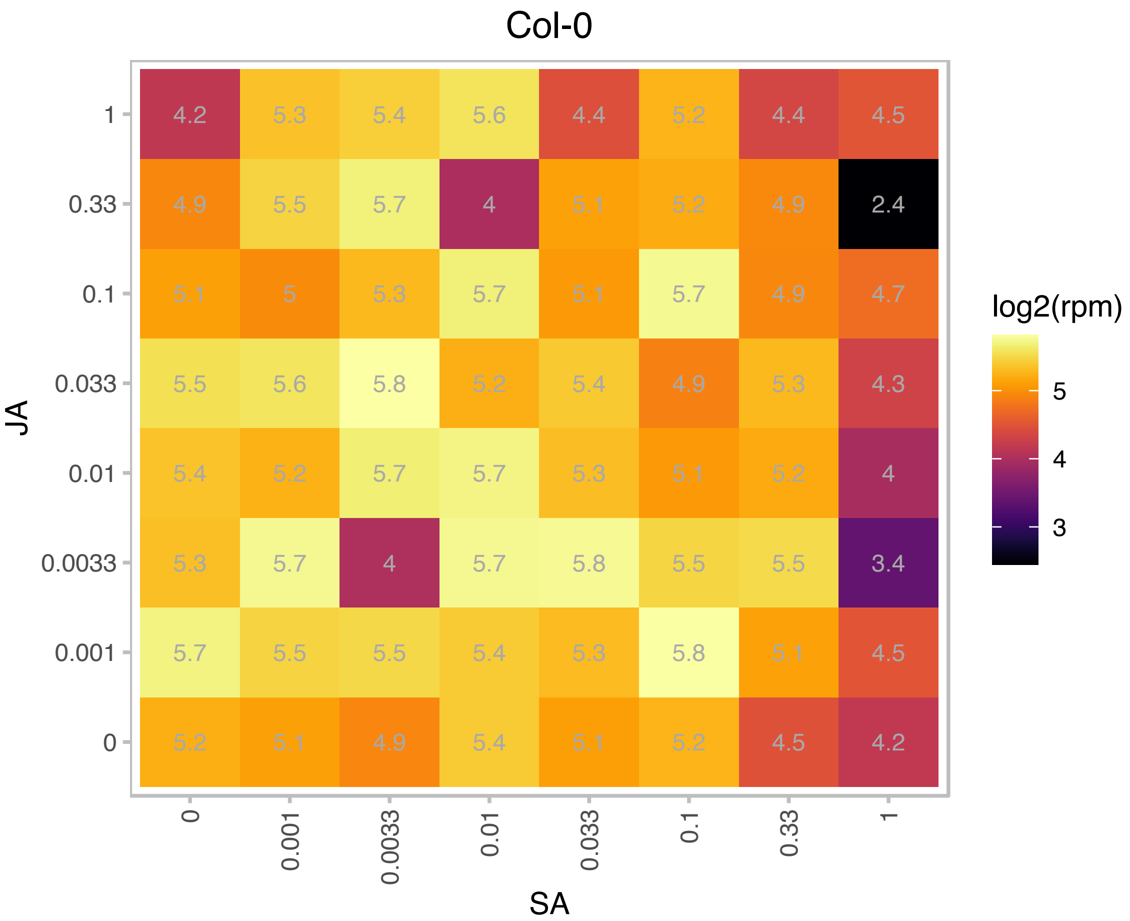


| | |
|---|---|
| Gene | AT1G01630 |
| is.expressed_genes | TRUE |
| is.responsive_genes | TRUE |
| is.apcluster | F-01 |

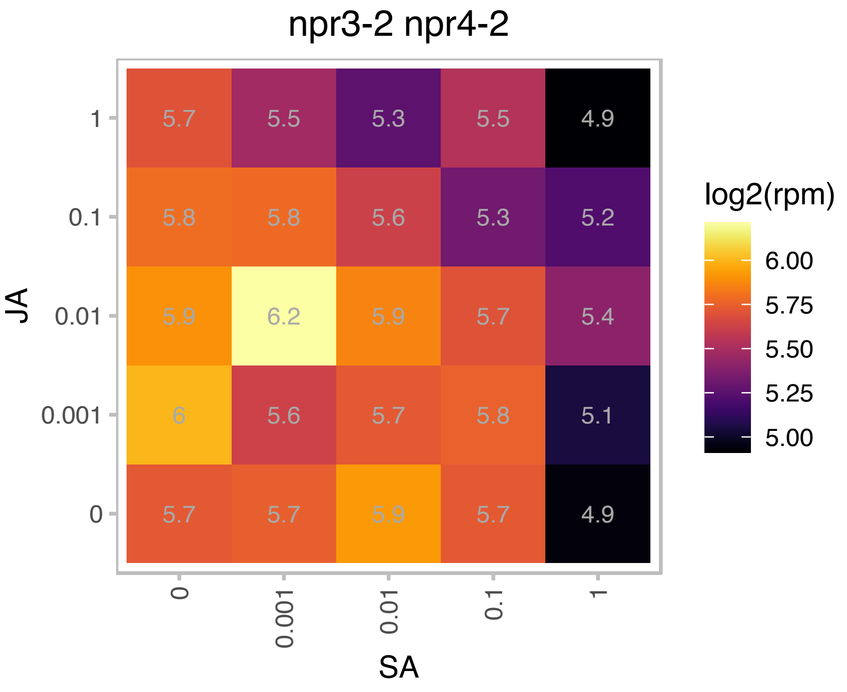


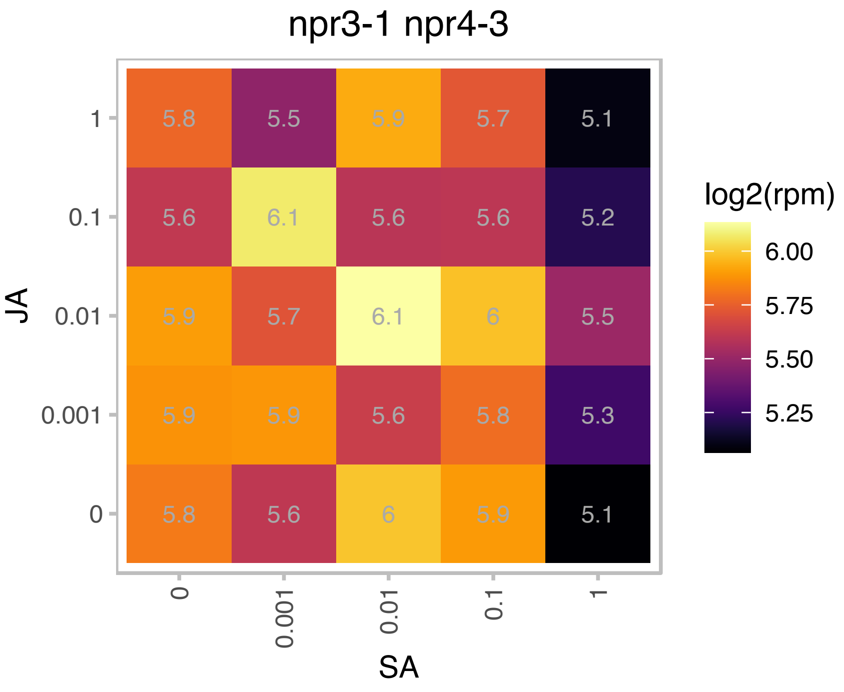

# AT1G01970.1 : NA

Tetratricopeptide repeat (TPR)-like superfamily protein

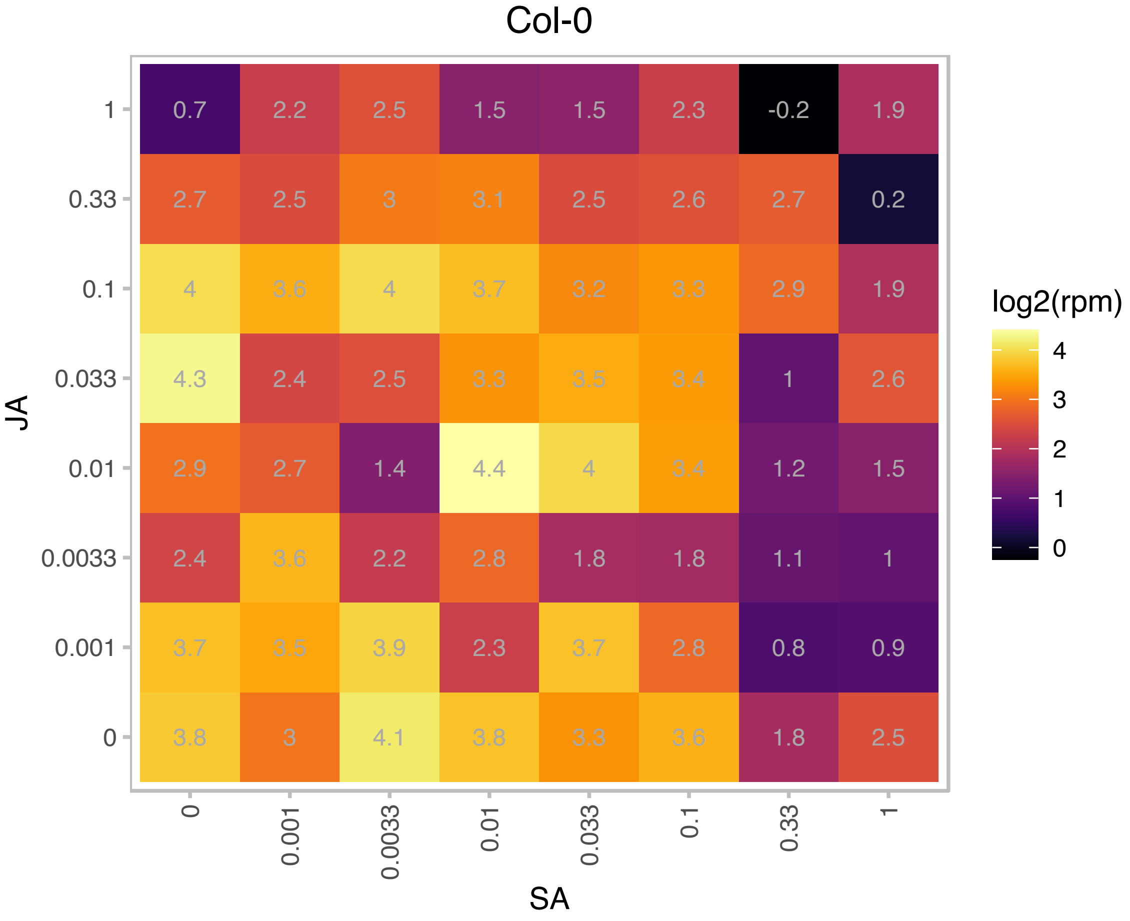


| | |
|---|---|
| Gene | AT1G01970 |
| is.expressed_genes | TRUE |
| is.responsive_genes | TRUE |
| is.apcluster | F-01 |

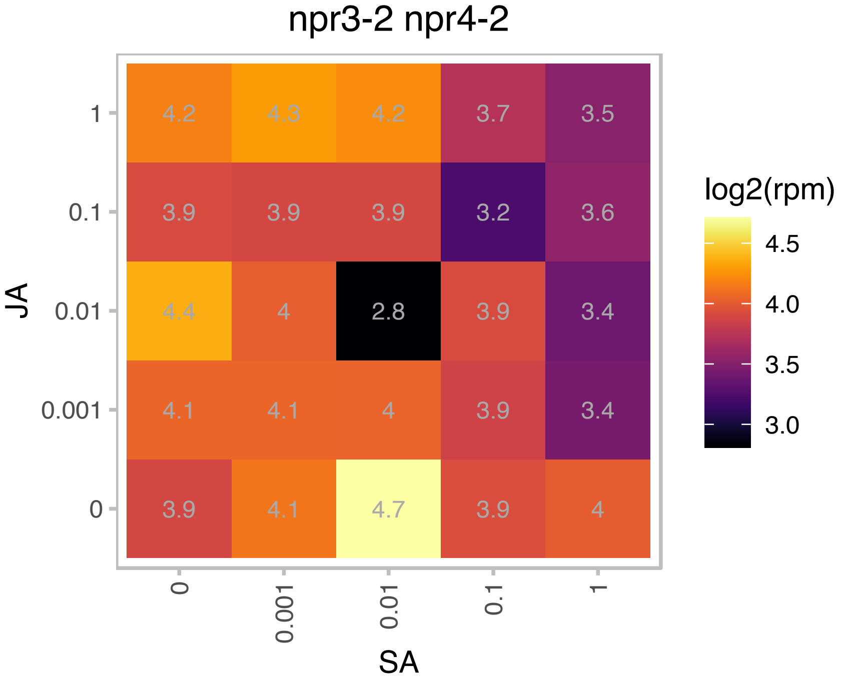


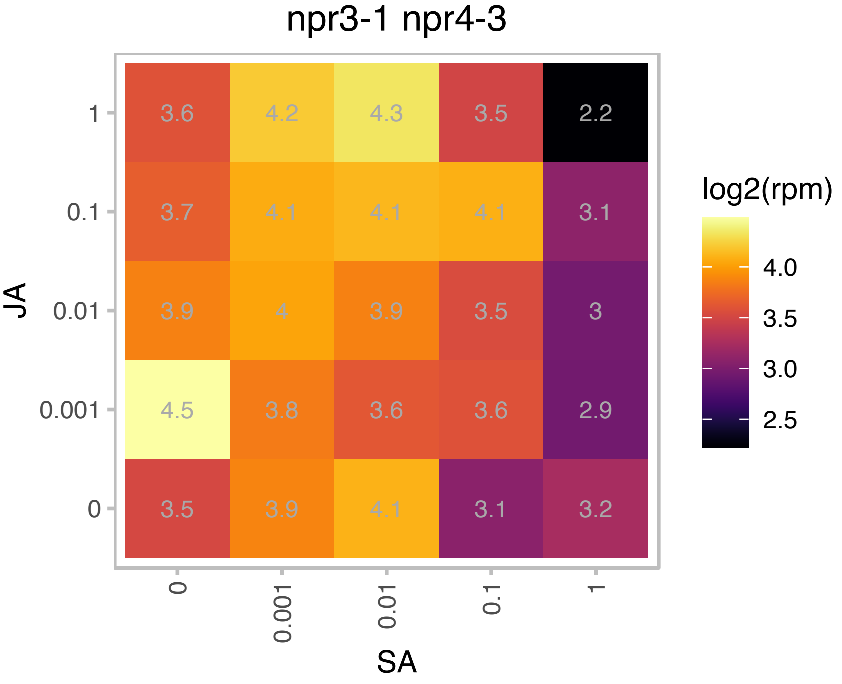

### AT1G03475.1 : ATCPO-I, HEMF1, LIN2

Coproporphyrinogen III oxidase
Encodes coproporphyrinogen III oxidase, a key enzyme in the biosynthetic pathway of chlorophyll and heme, a tetrapyrrole pathway. Mutants express cytological and molecular markers associated with the defense responses, usually activated by pathogen infection.

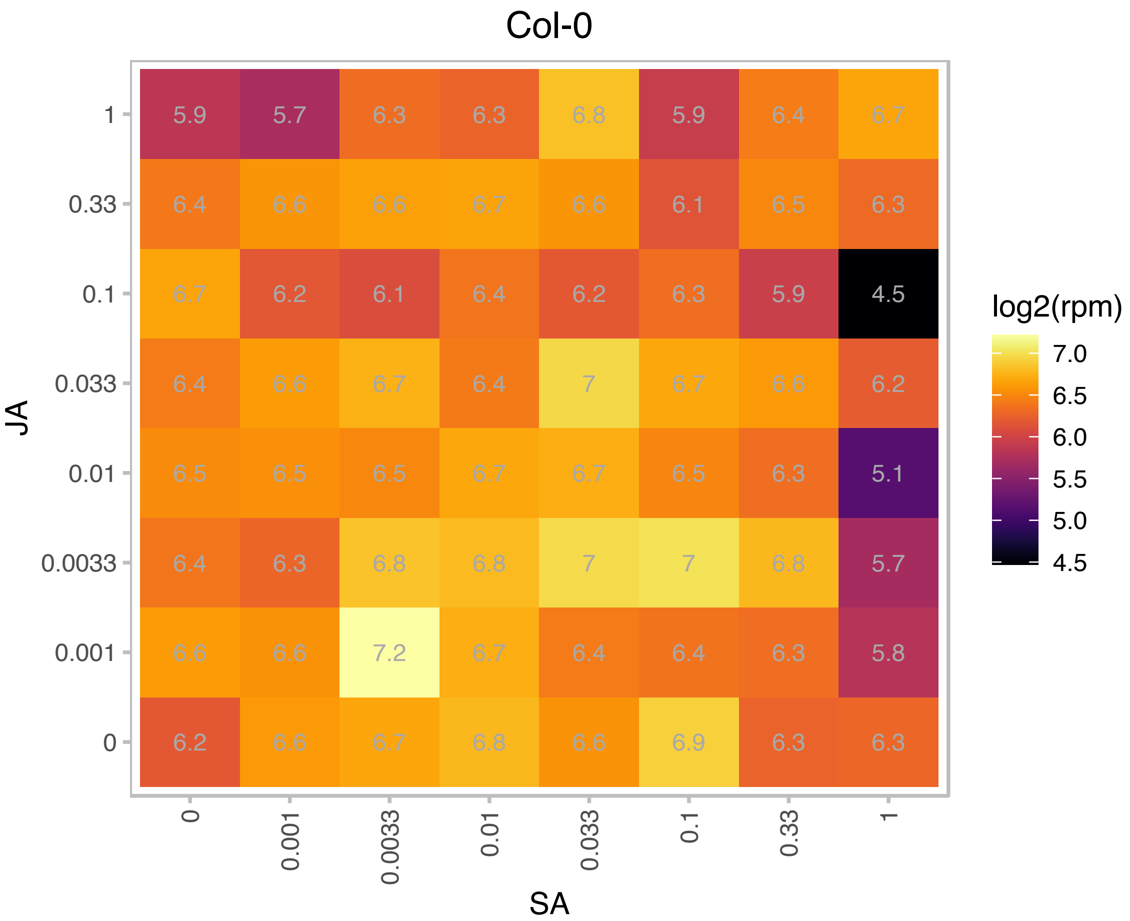


| | |
|---|---|
| Gene | AT1G03475 |
| is.expressed_genes | TRUE |
| is.responsive_genes | TRUE |
| is.apcluster | F-01 |

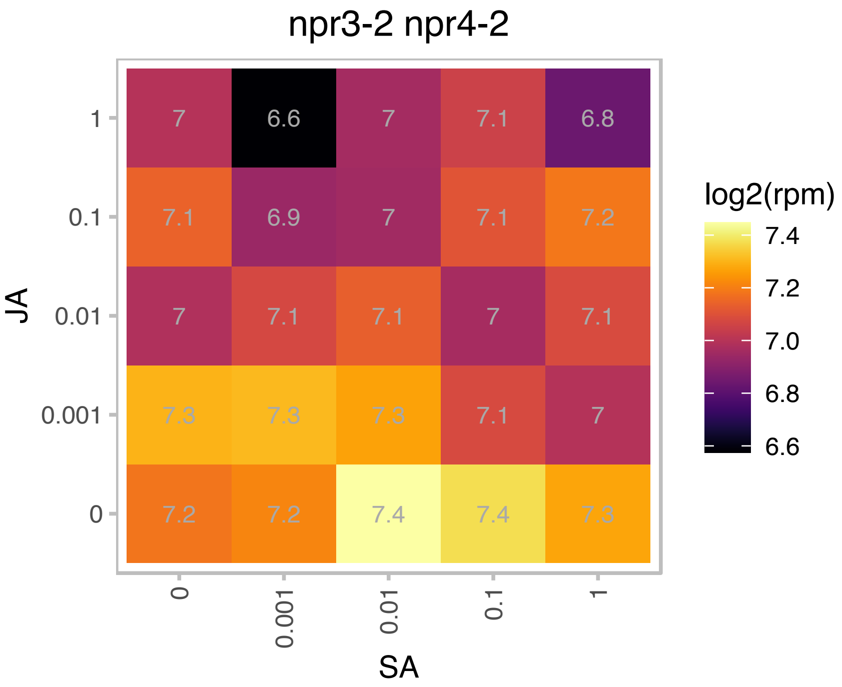


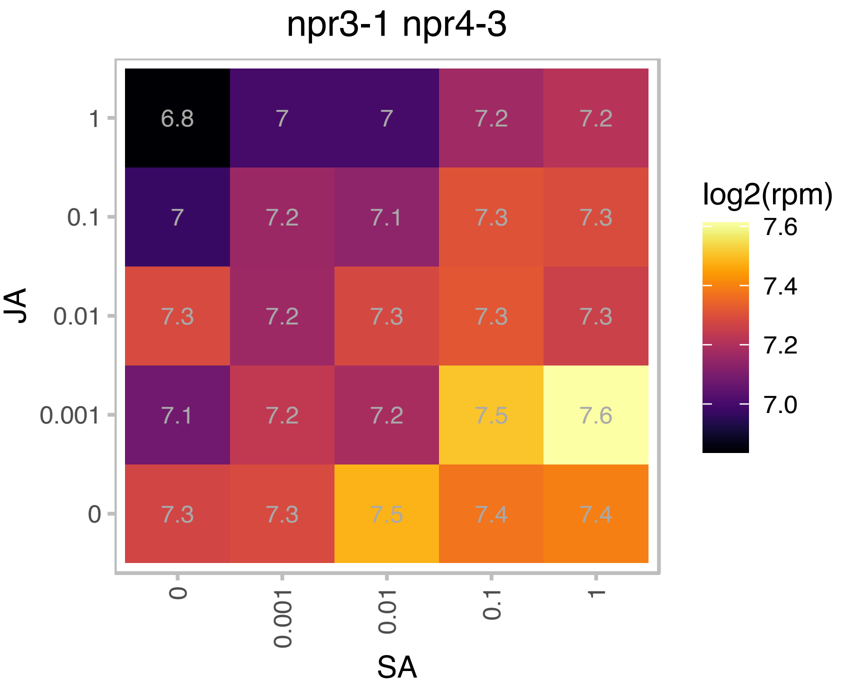

### AT1G04080.1 : PRP39, PRP39a

Tetratricopeptide repeat (TPR)-like superfamily protein

Encodes a U1 small nuclear ribonucleoprotein (snRNP) factor involved in alternative splicing.

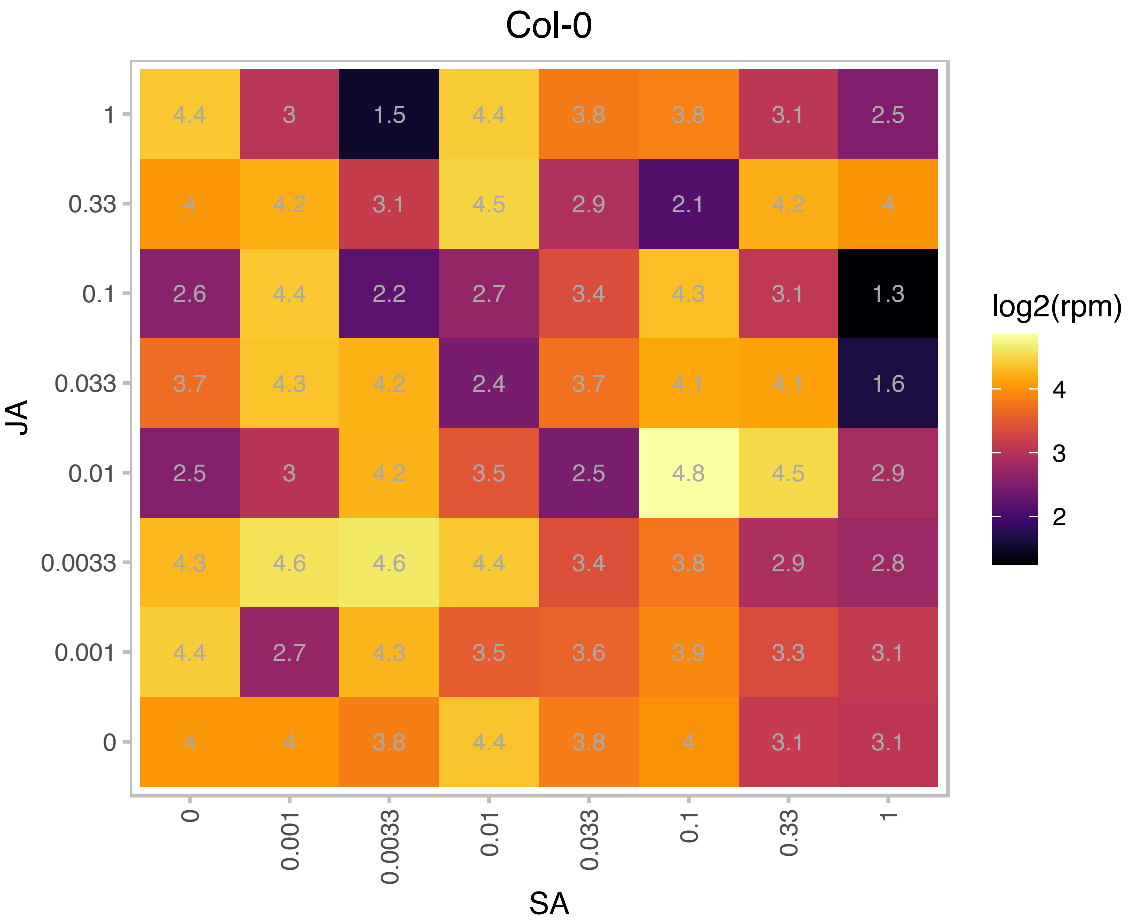


| | |
|---|---|
| Gene | AT1G04080 |
| is.expressed_genes | TRUE |
| is.responsive_genes | TRUE |
| is.apcluster | F-01 |

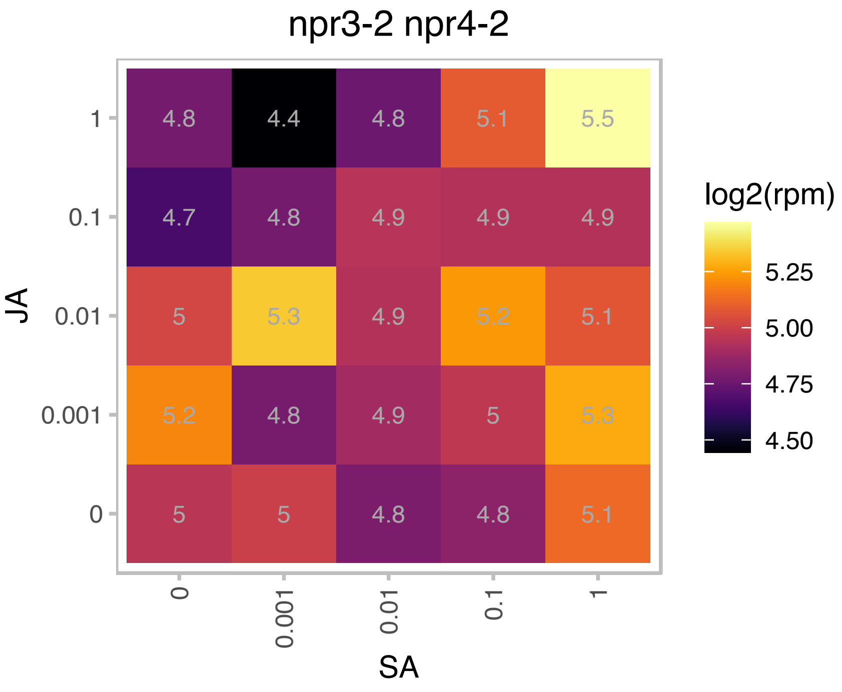


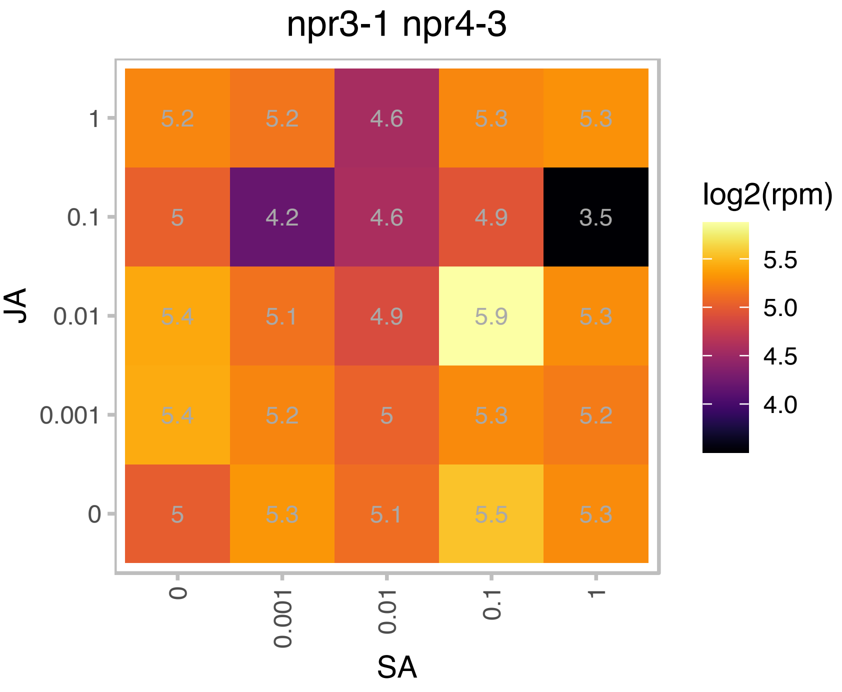

# AT1G04430.1 : NA

S-adenosyl-L-methionine-dependent methyltransferases superfamily protein

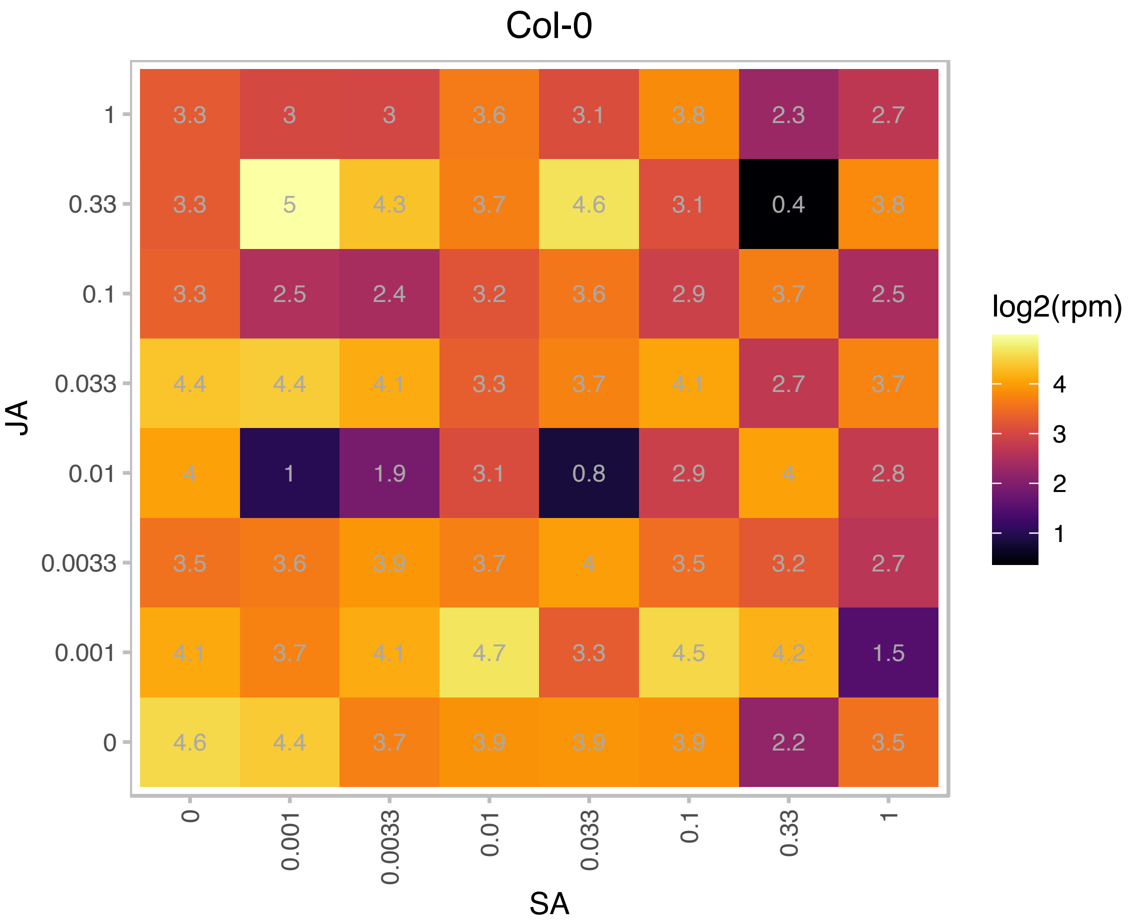


| | |
|---|---|
| Gene | AT1G04430 |
| is.expressed_genes | TRUE |
| is.responsive_genes | TRUE |
| is.apcluster | F-01 |

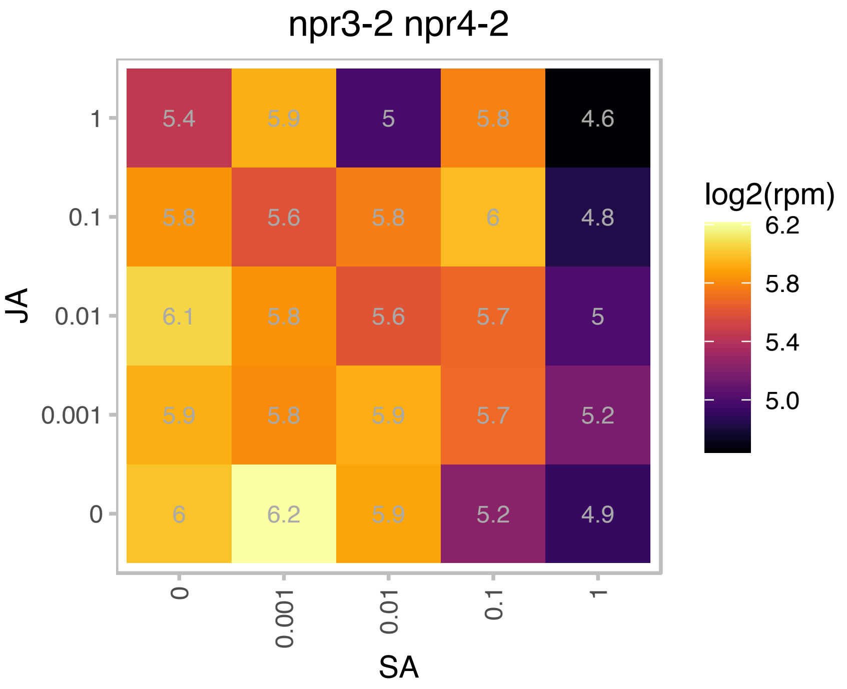


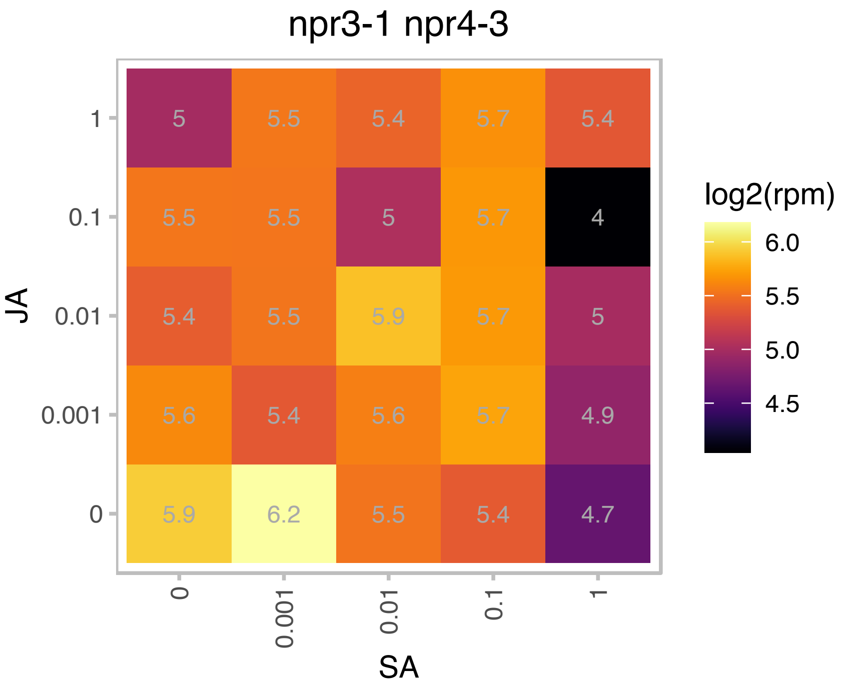

### AT1G04630.1 : MEE4

GRIM-19 protein

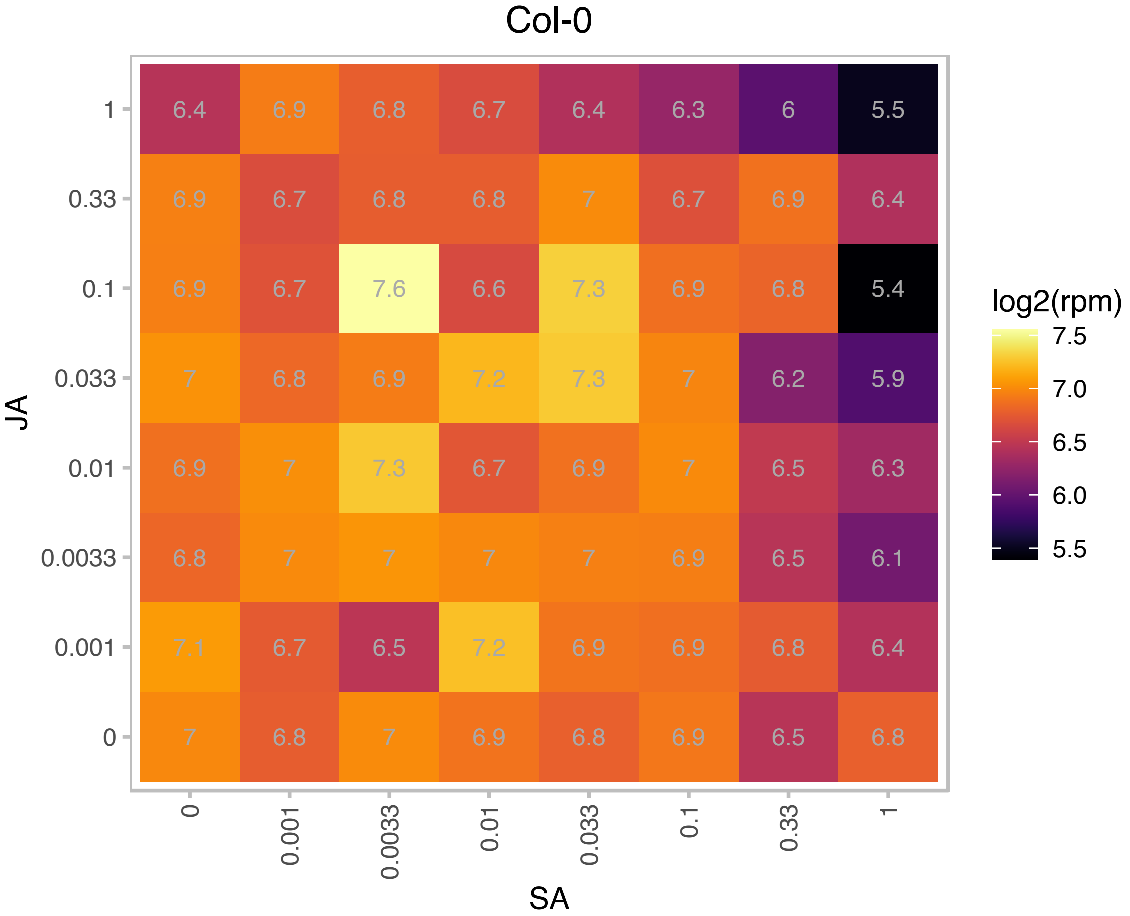


| | |
|---|---|
| Gene | AT1G04630 |
| is.expressed_genes | TRUE |
| is.responsive_genes | TRUE |
| is.apcluster | F-01 |

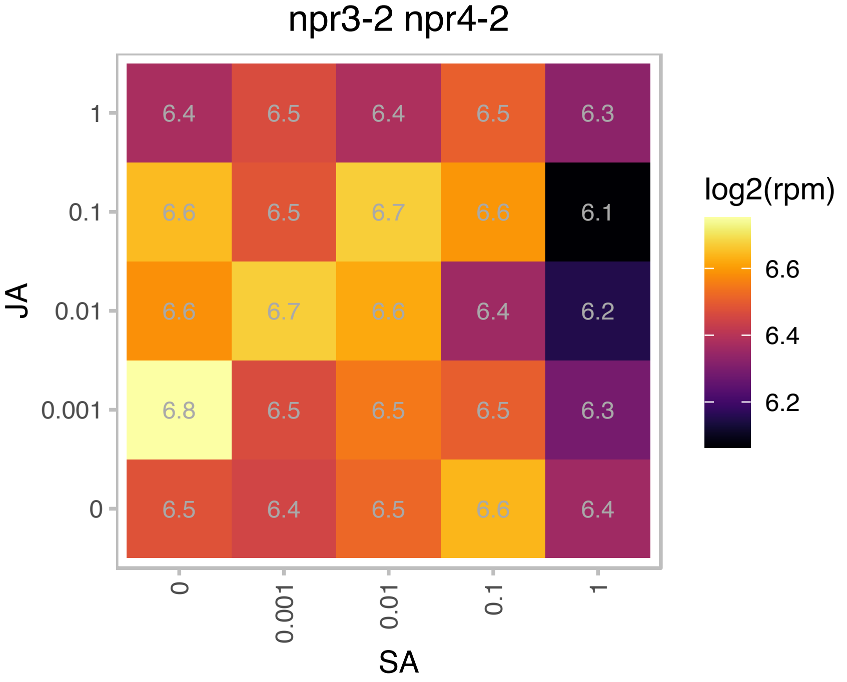


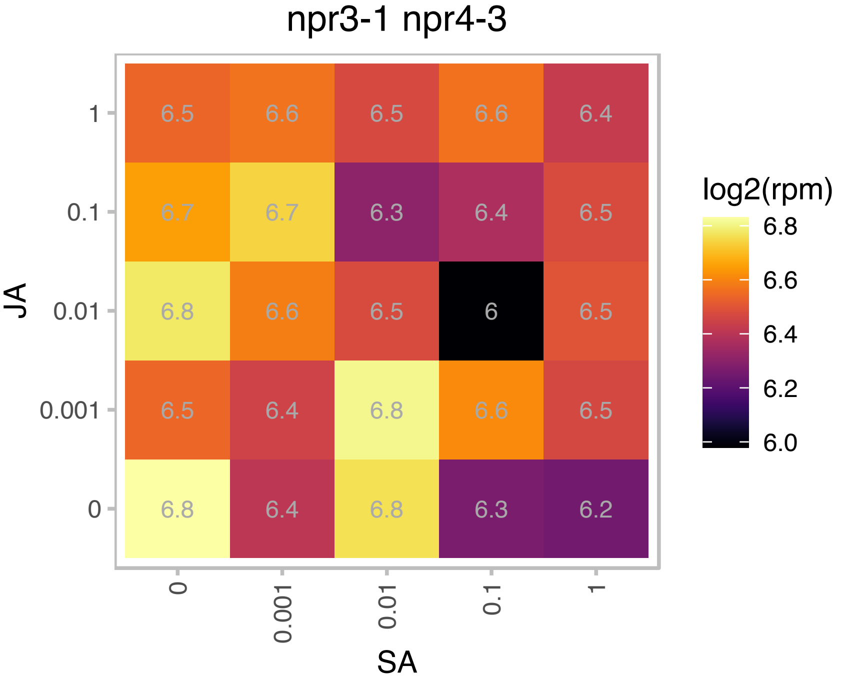

# AT1G05140.1 : NA

Peptidase M50 family protein

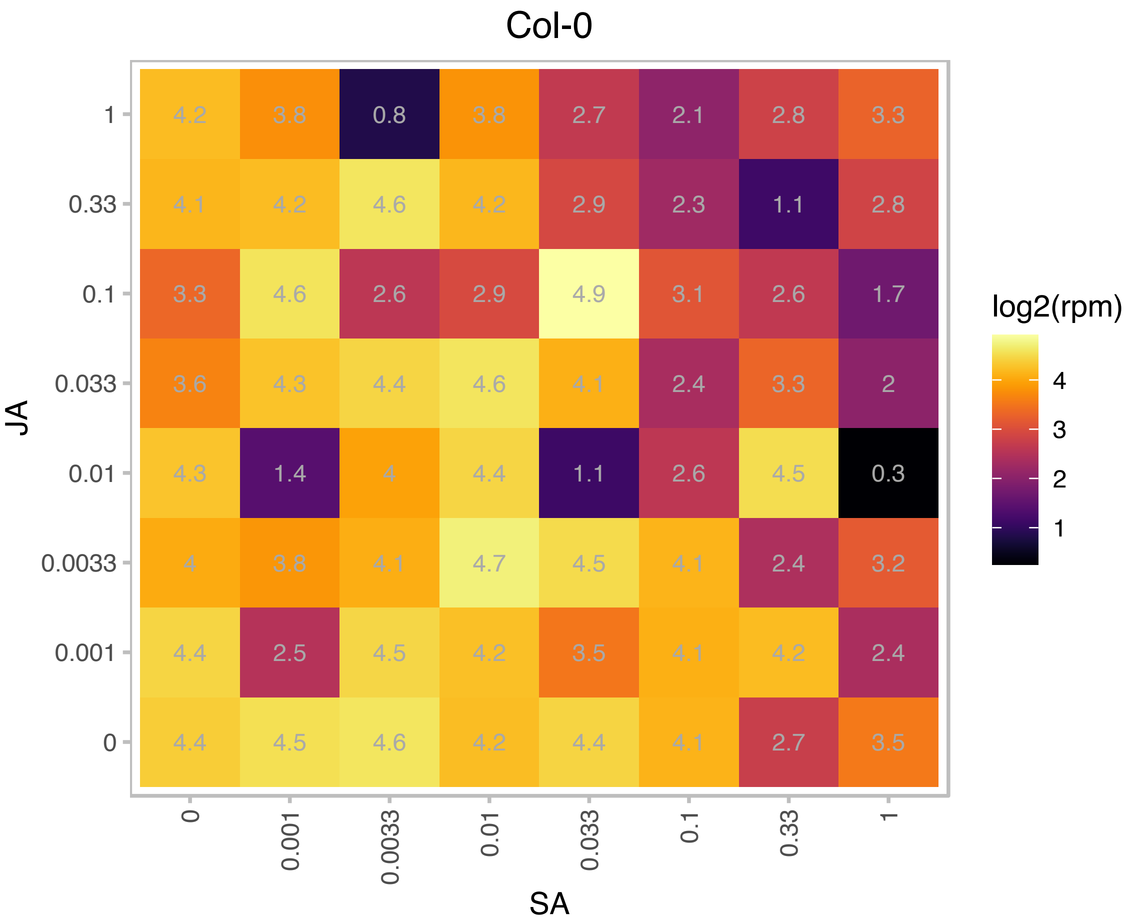


| | |
|---|---|
| Gene | AT1G05140 |
| is.expressed_genes | TRUE |
| is.responsive_genes | TRUE |
| is.apcluster | F-01 |

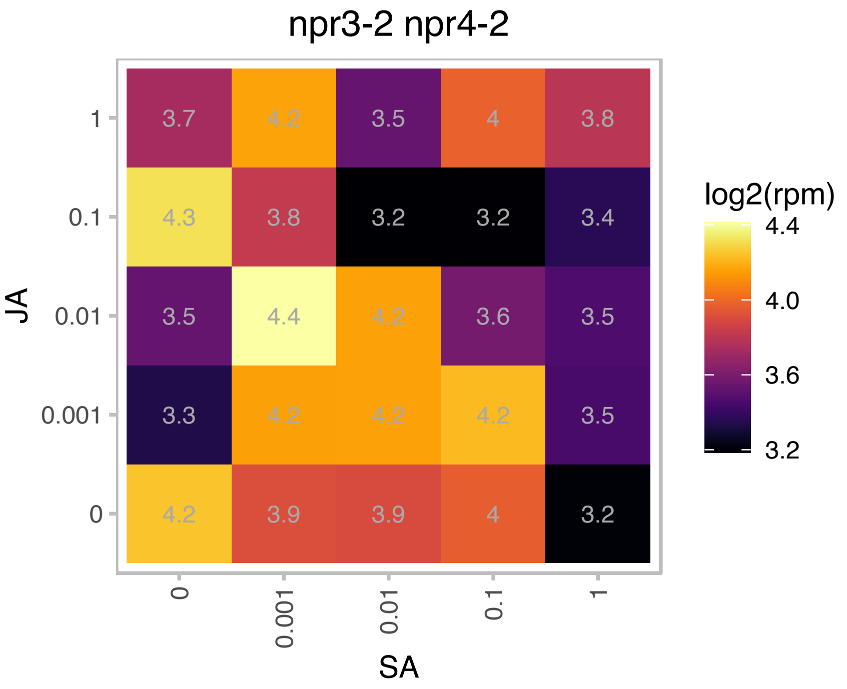


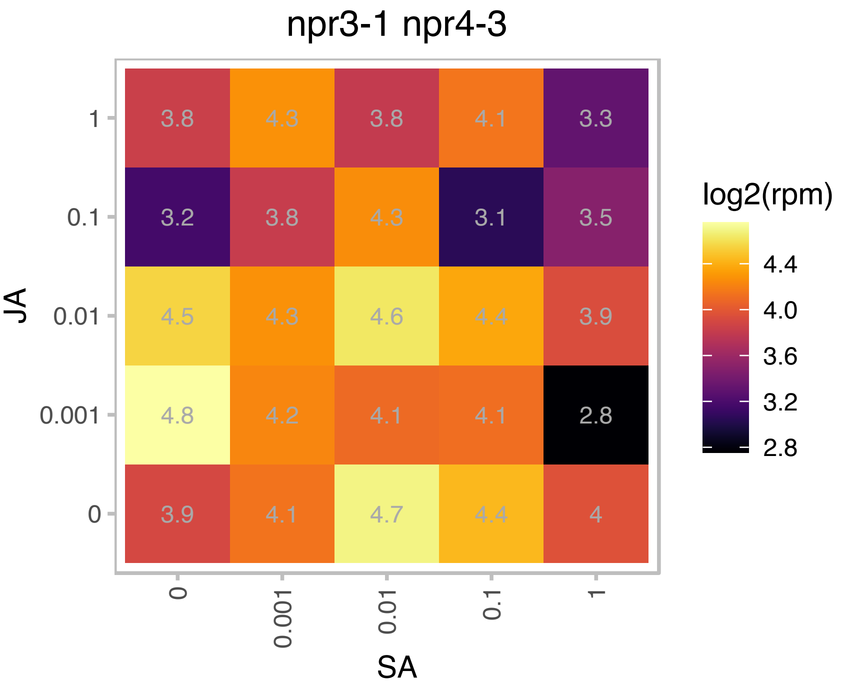

### AT1G05810.1 : ARA, ARA-1, ATRAB11D, ATRABA5E, RAB

RAB GTPase homolog A5E

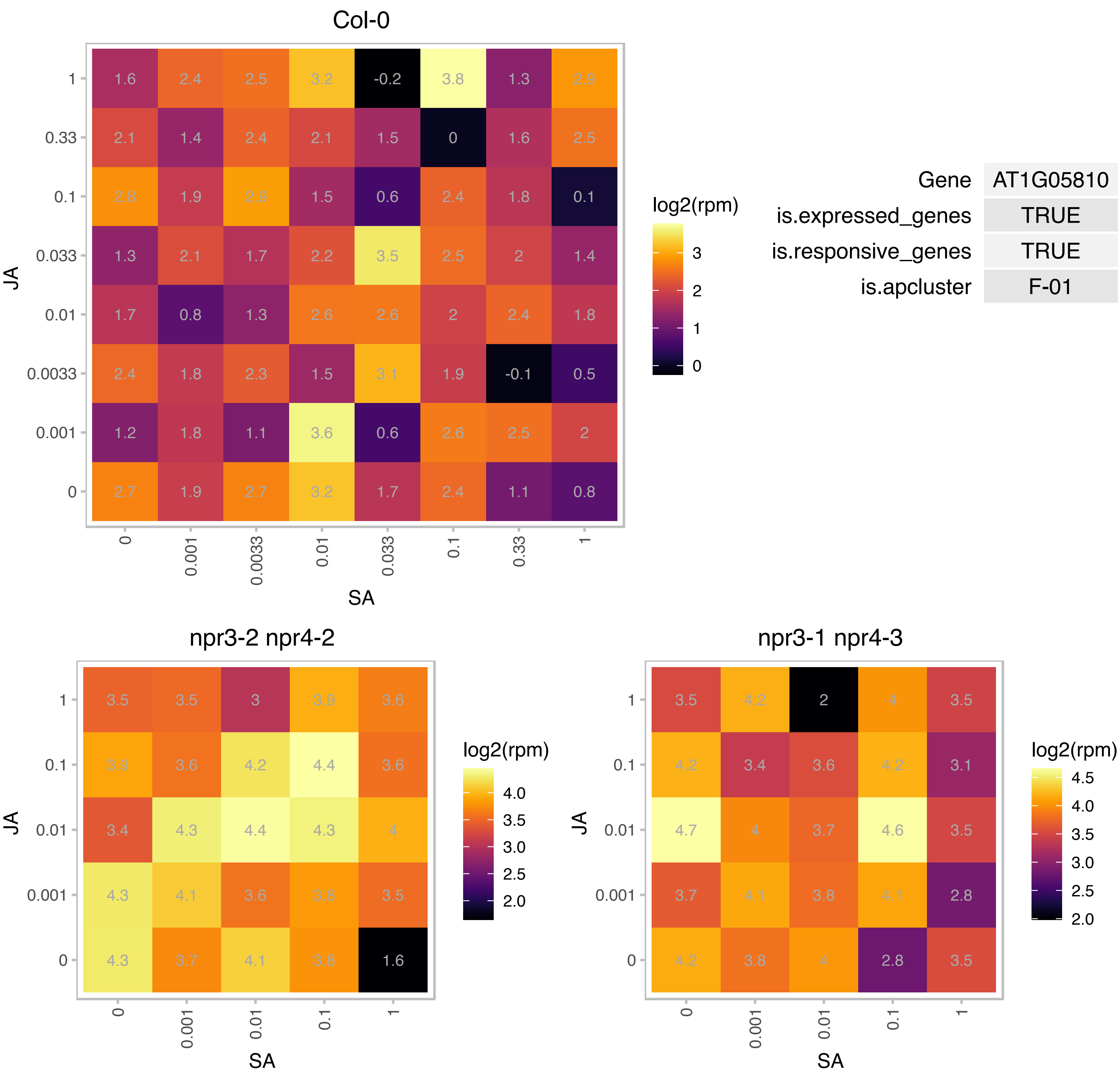


| | |
|---|---|
| Gene | AT1G05810 |
| is.expressed_genes | TRUE |
| is.responsive_genes | TRUE |
| is.apcluster | F-01 |

### AT1G13670.1 : BGL2

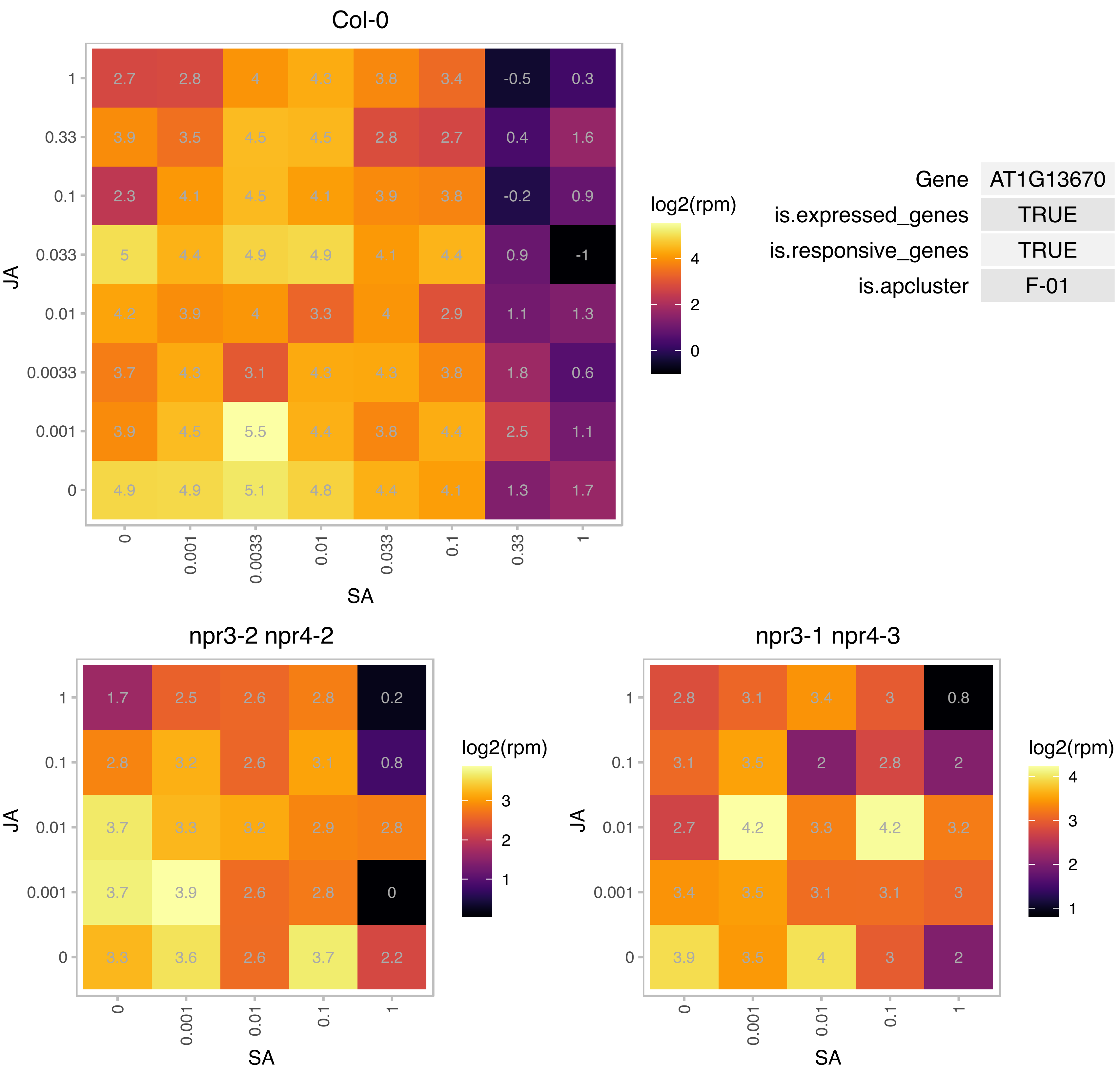

### AT1G17220.1 : FUG1

Translation initiation factor 2, small GTP-binding protein

Encodes a chloroplast localized protein with similarity to translation initiation factor 2. Can complement loss of INFB in E.coli suggesting FUG1 does function as a translation initiation factor in vivo. Identified as a suppressor of the leaf variegation mutant var2-6. Suppression is only seen in hypomorphs as complete loss of function alleles are embryo lethal. The mRNA is cell-to-cell mobile.

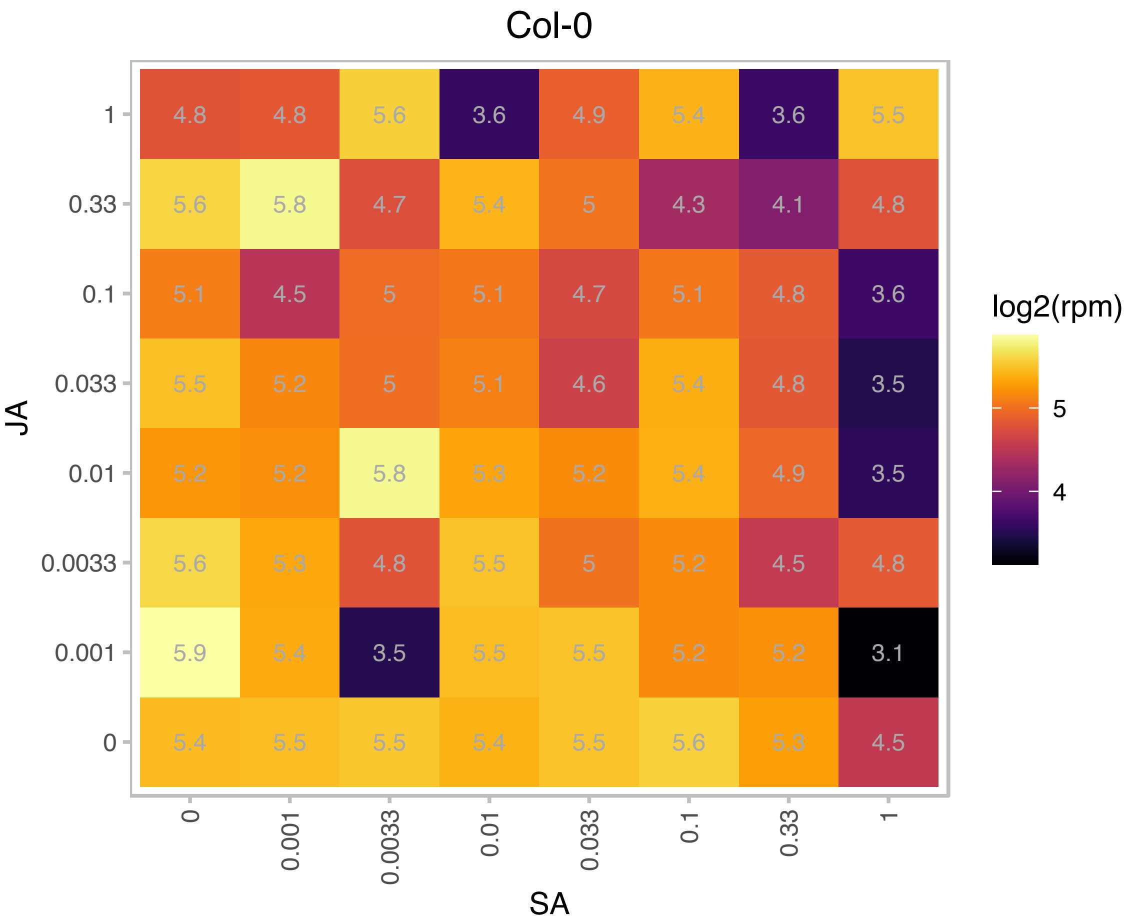


| | |
|---|---|
| Gene | AT1G17220 |
| is.expressed_genes | TRUE |
| is.responsive_genes | TRUE |
| is.apcluster | F-01 |

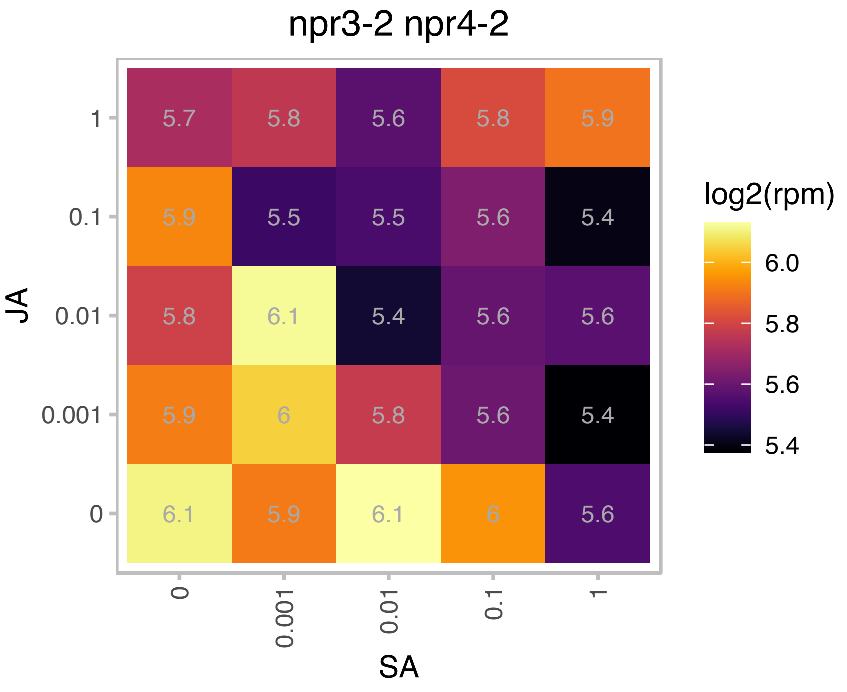


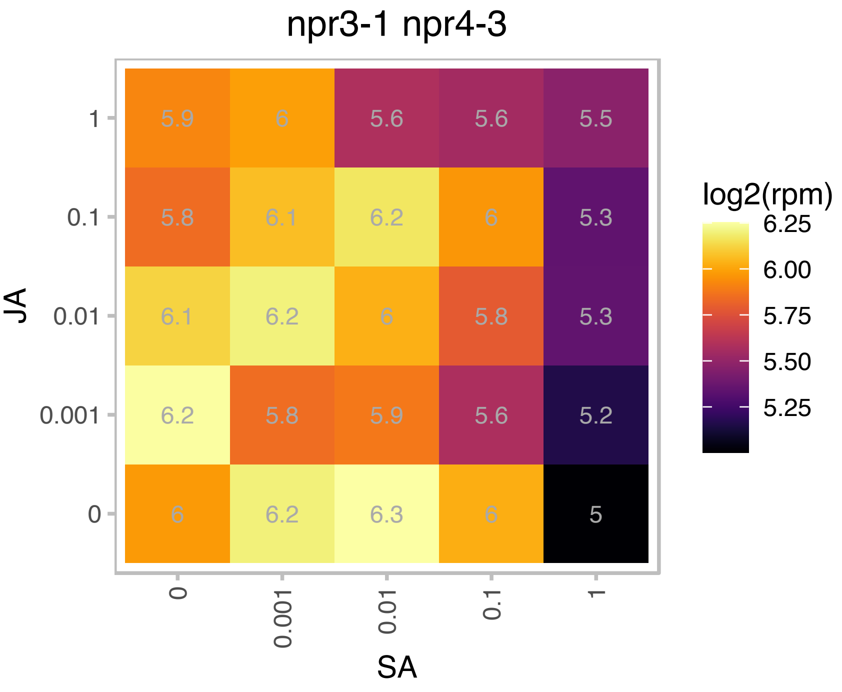

### AT1G20830.1 : MCD1

multiple chloroplast division site 1

Encodes MCD1 (MULTIPLE CHLOROPLAST DIVISION SITE 1). Determines the site of chloroplast division in conc with MinD (AT5G24020).

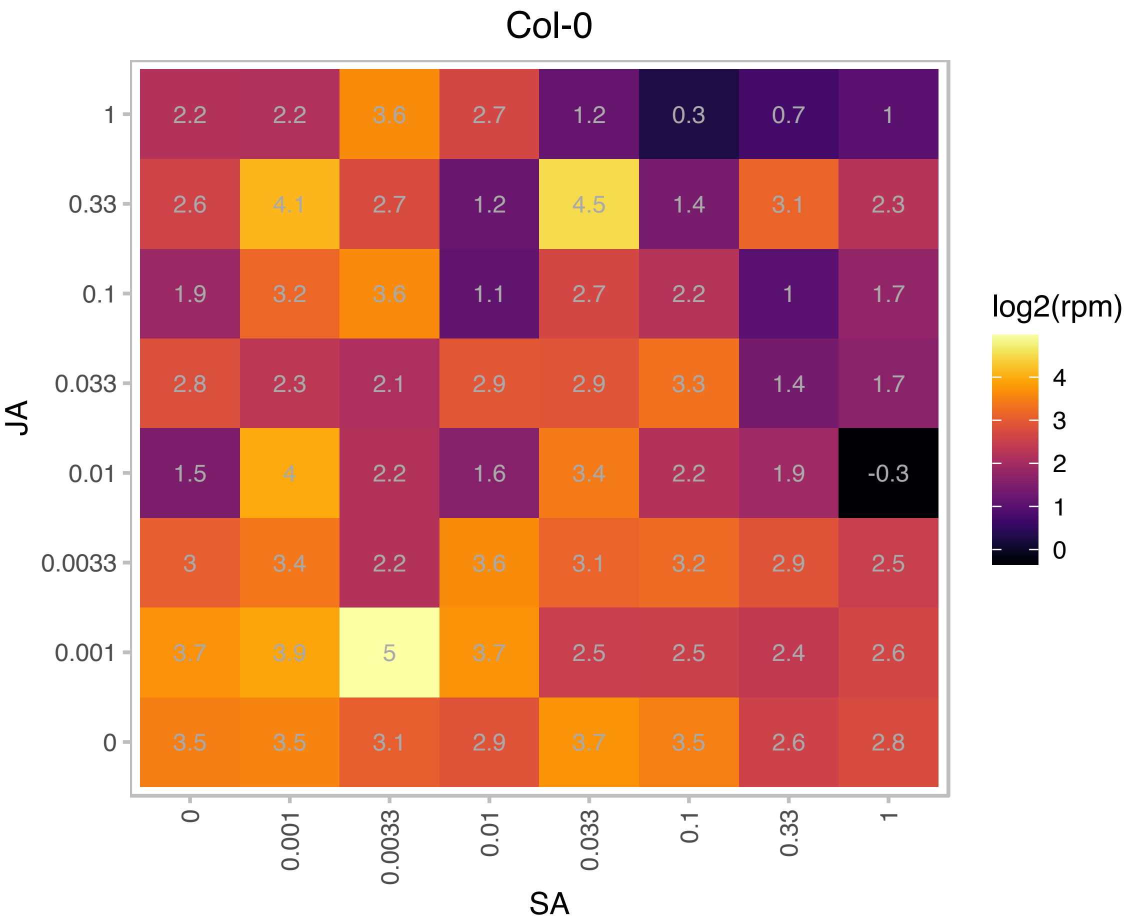


| | |
|---|---|
| Gene | AT1G20830 |
| is.expressed_genes | TRUE |
| is.responsive_genes | TRUE |
| is.apcluster | F-01 |

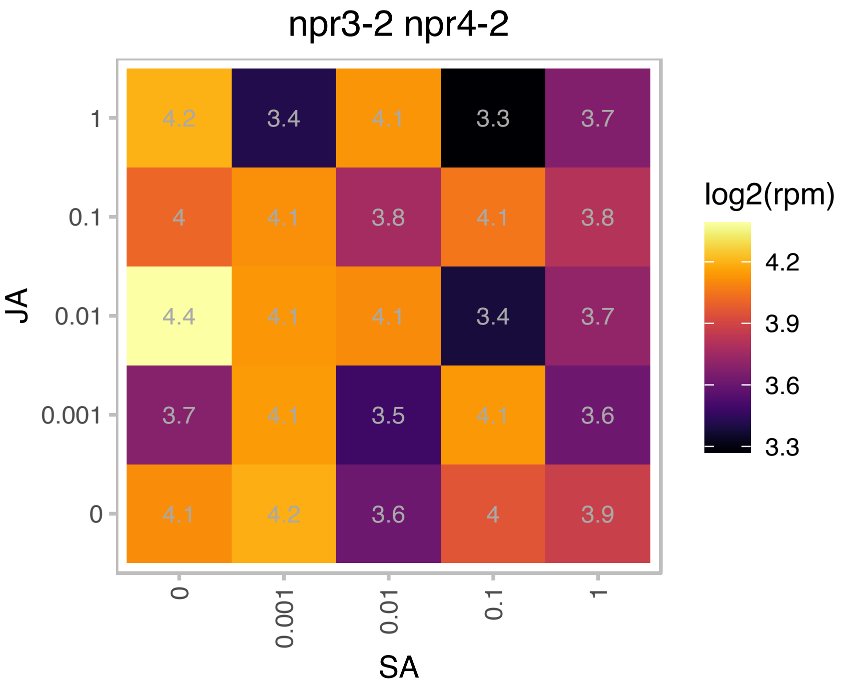


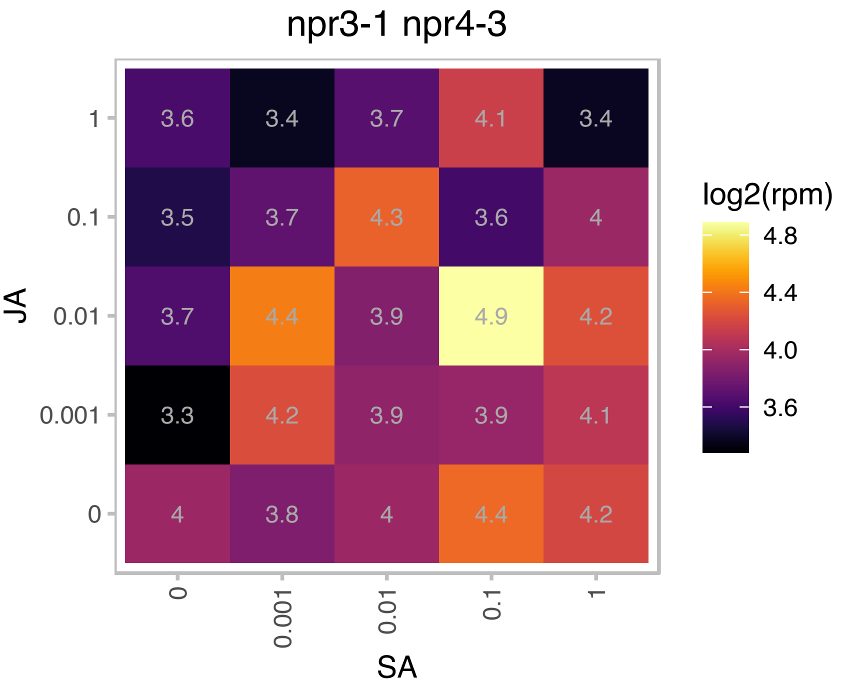

# AT1G22430.1 : NA

GroES-like zinc-binding dehydrogenase family protein

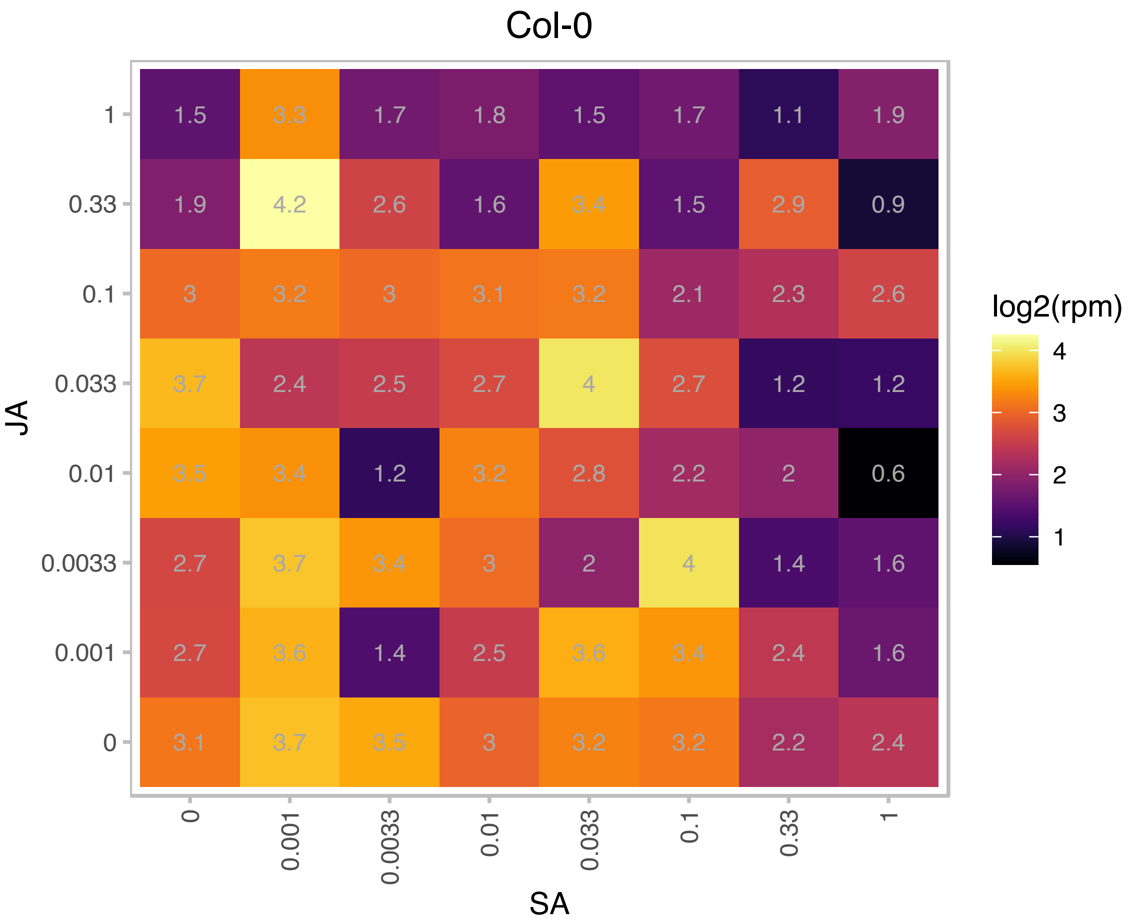


| | |
|---|---|
| Gene | AT1G22430 |
| is.expressed_genes | TRUE |
| is.responsive_genes | TRUE |
| is.apcluster | F-01 |

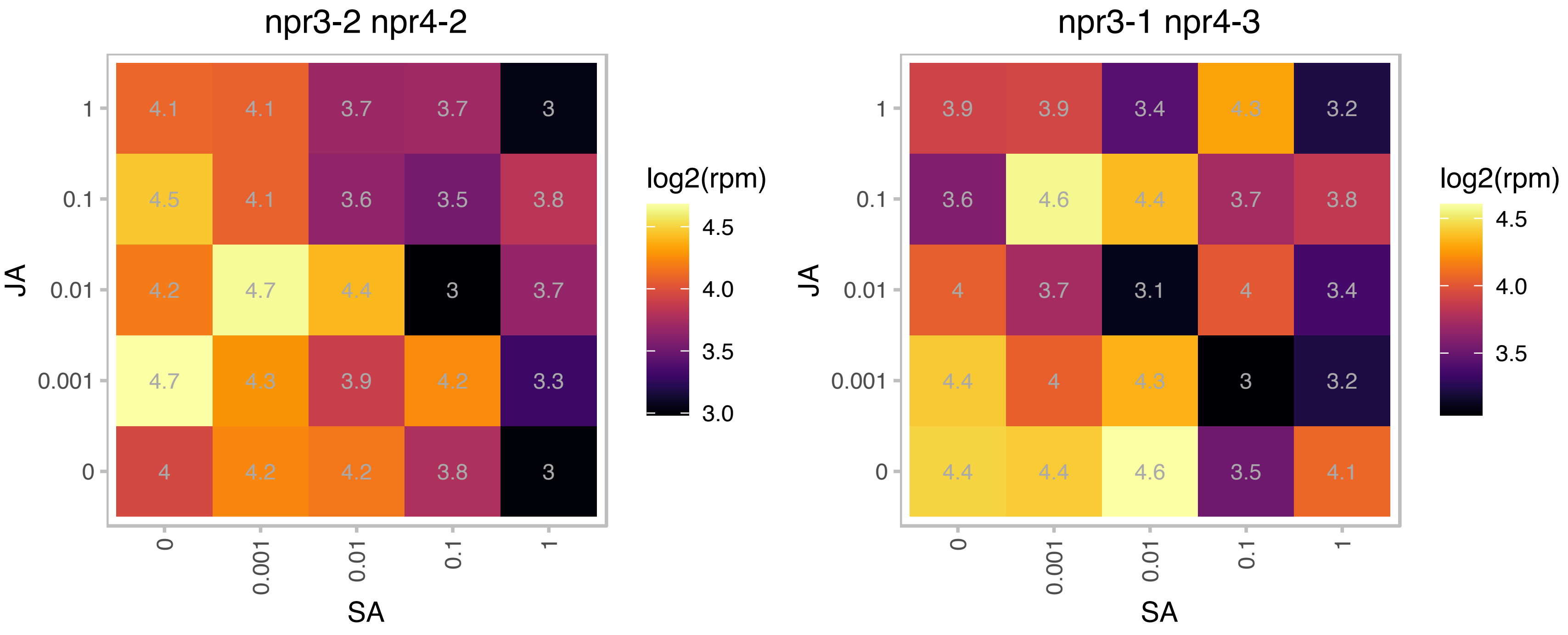

### AT1G24310.1 : NUP54

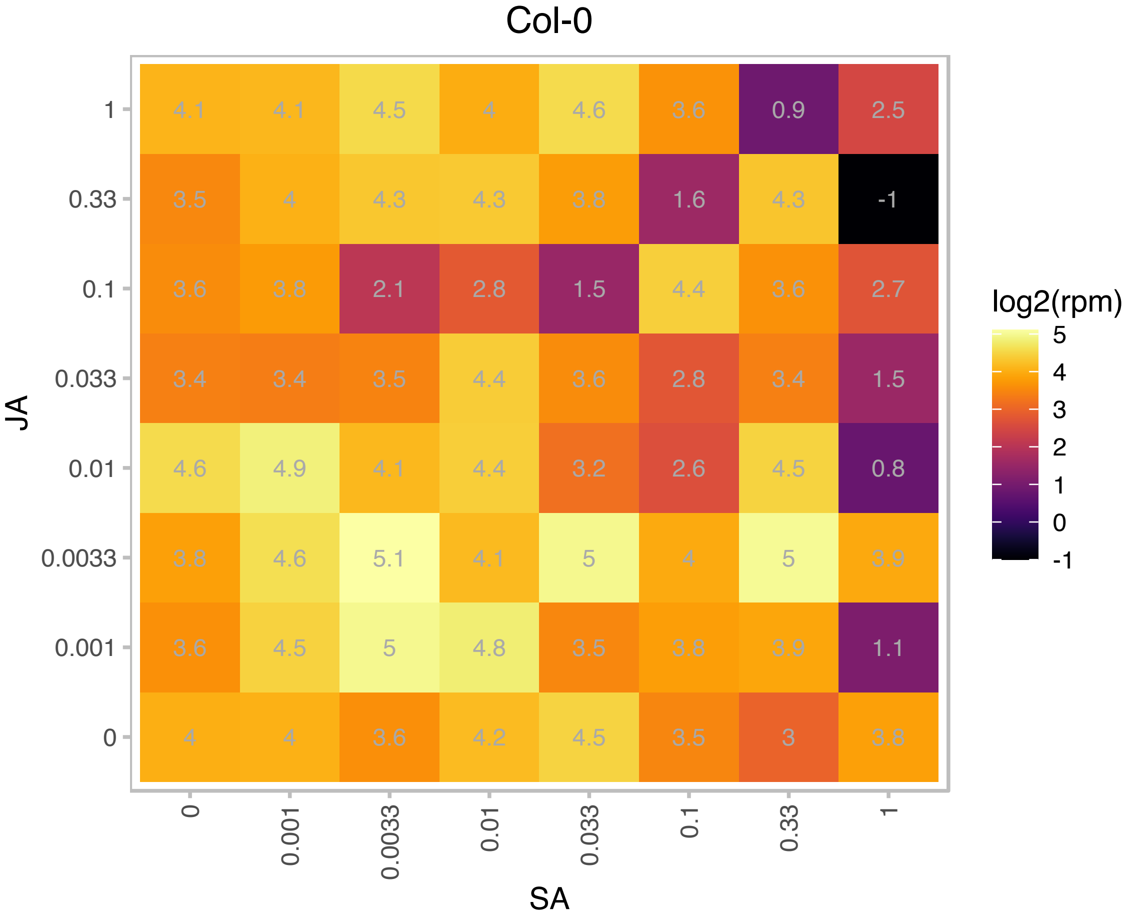


| | |
|---|---|
| Gene | AT1G24310 |
| is.expressed_genes | TRUE |
| is.responsive_genes | TRUE |
| is.apcluster | F-01 |

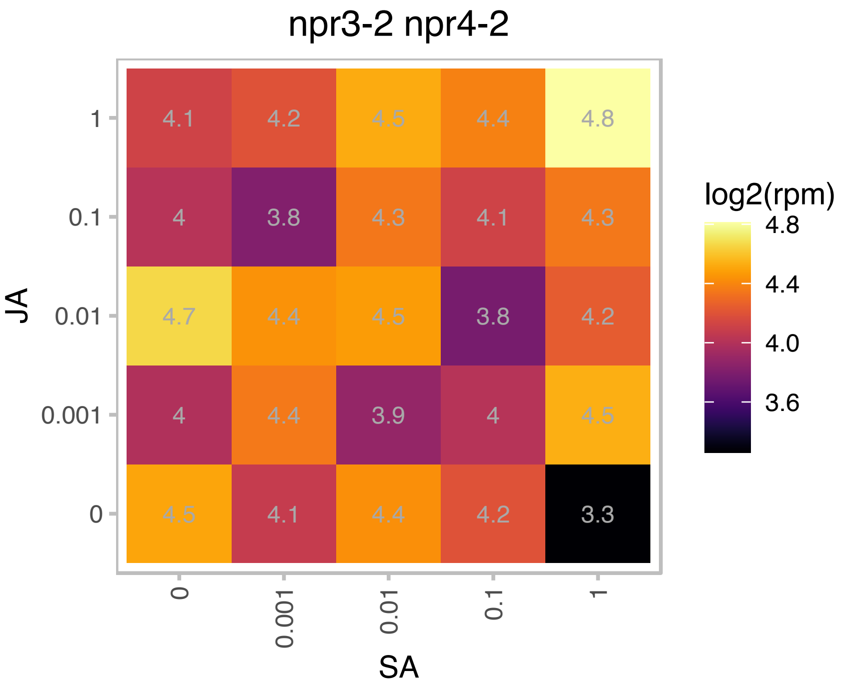


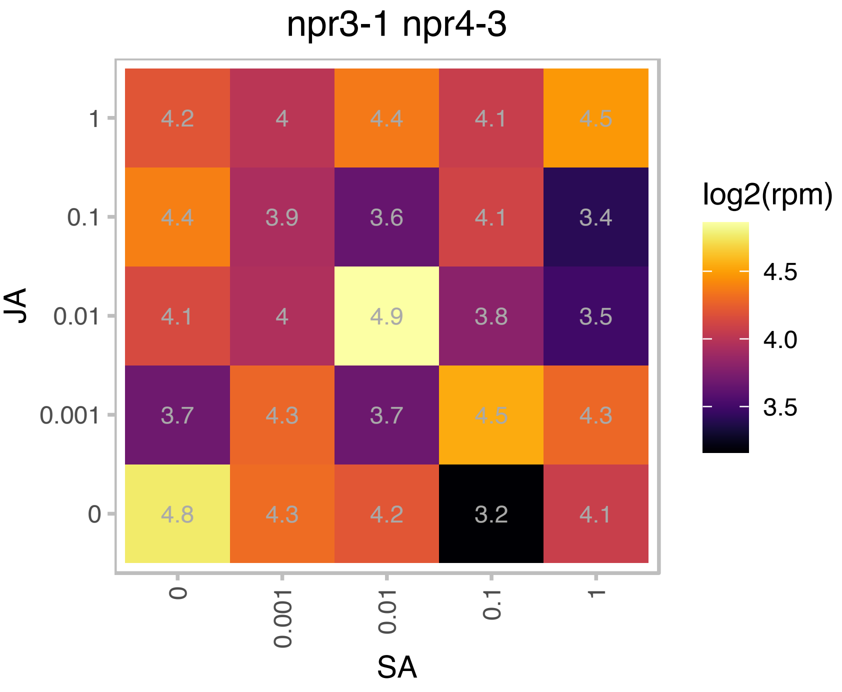

# AT1G28400.1 : NA

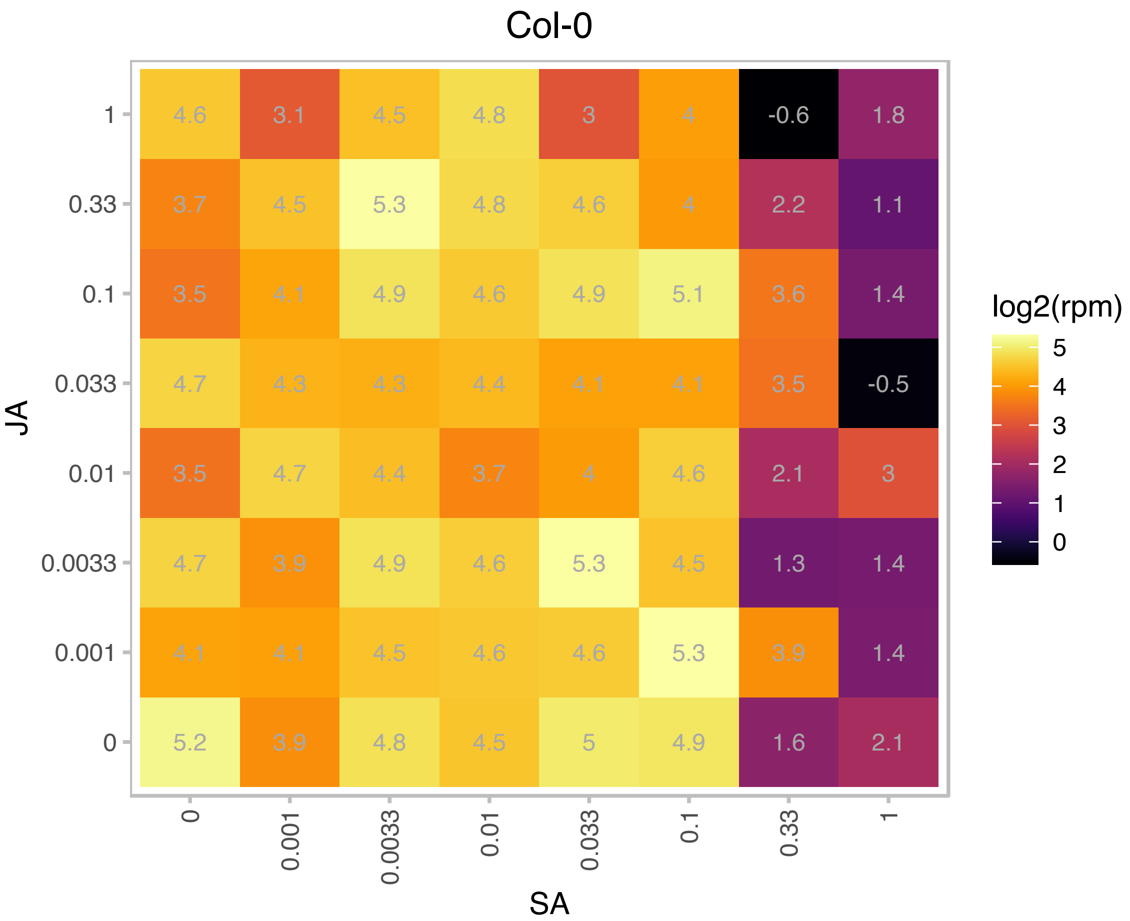


| | |
|---|---|
| Gene | AT1G28400 |
| is.expressed_genes | TRUE |
| is.responsive_genes | TRUE |
| is.apcluster | F-01 |

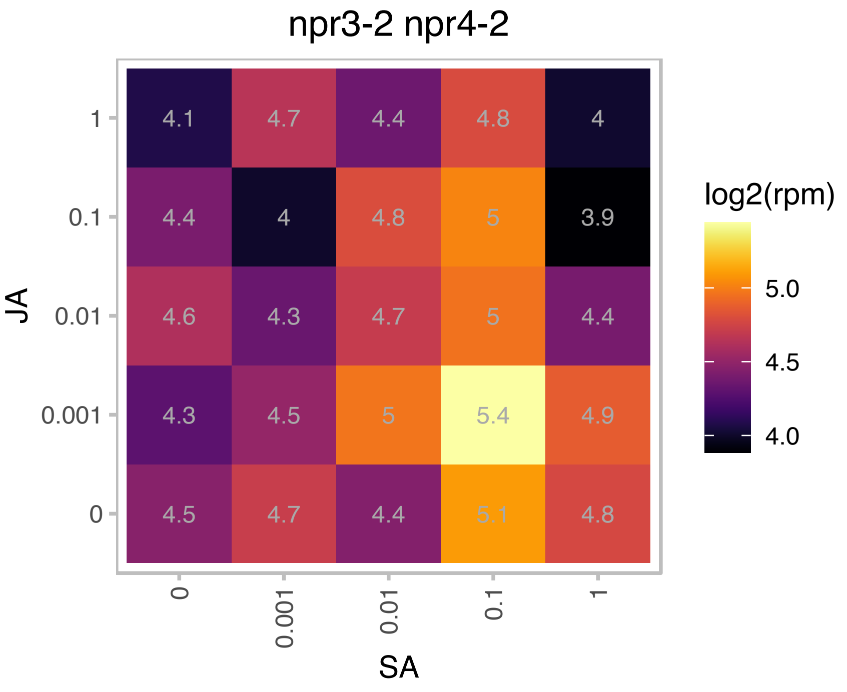


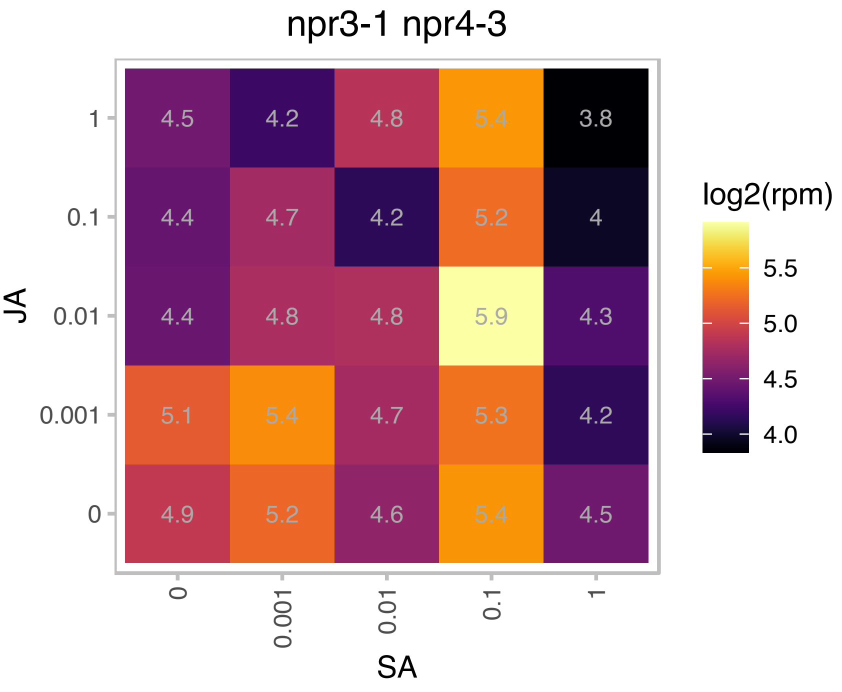

### AT1G44110.1 : CYCA1;1

Cyclin A1;1

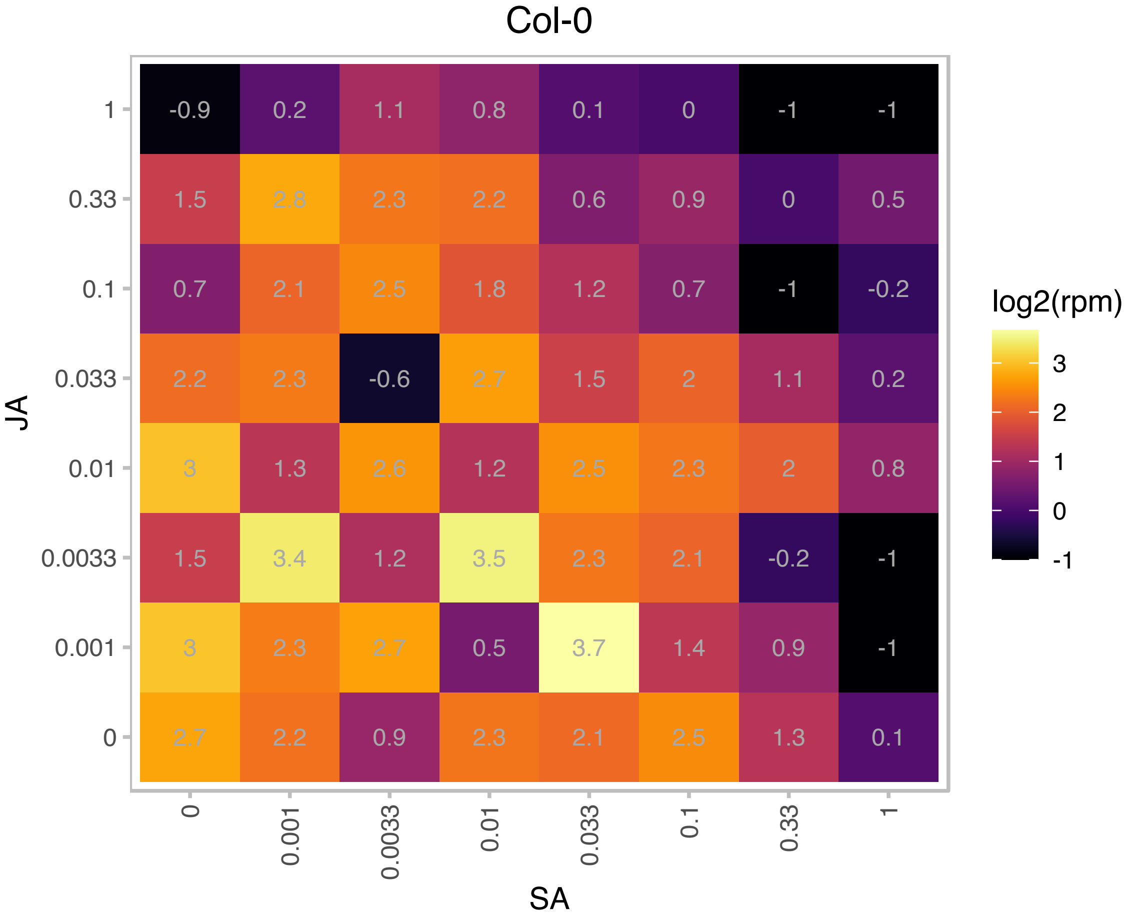


| | |
|---|---|
| Gene | AT1G44110 |
| is.expressed_genes | TRUE |
| is.responsive_genes | TRUE |
| is.apcluster | F-01 |

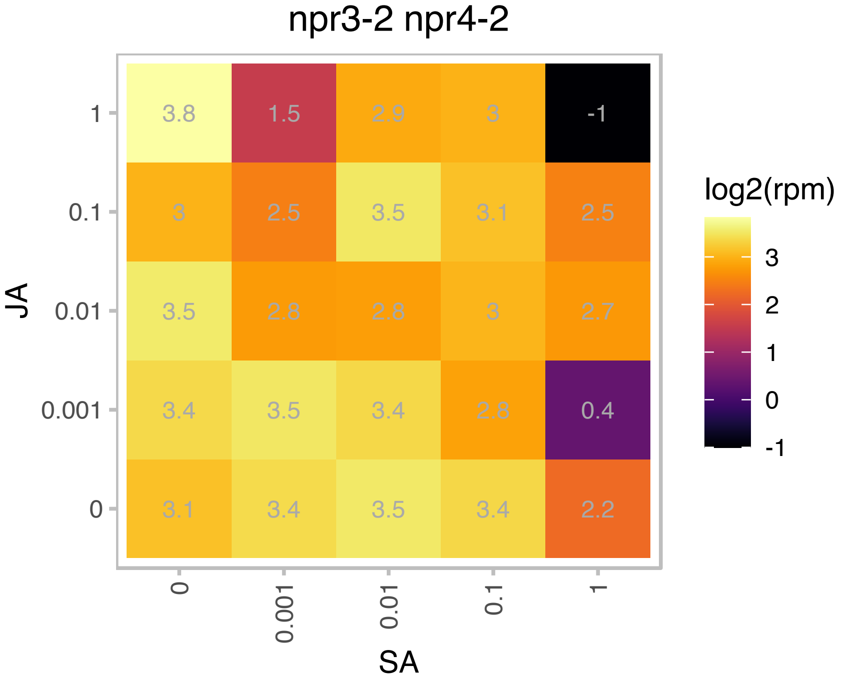


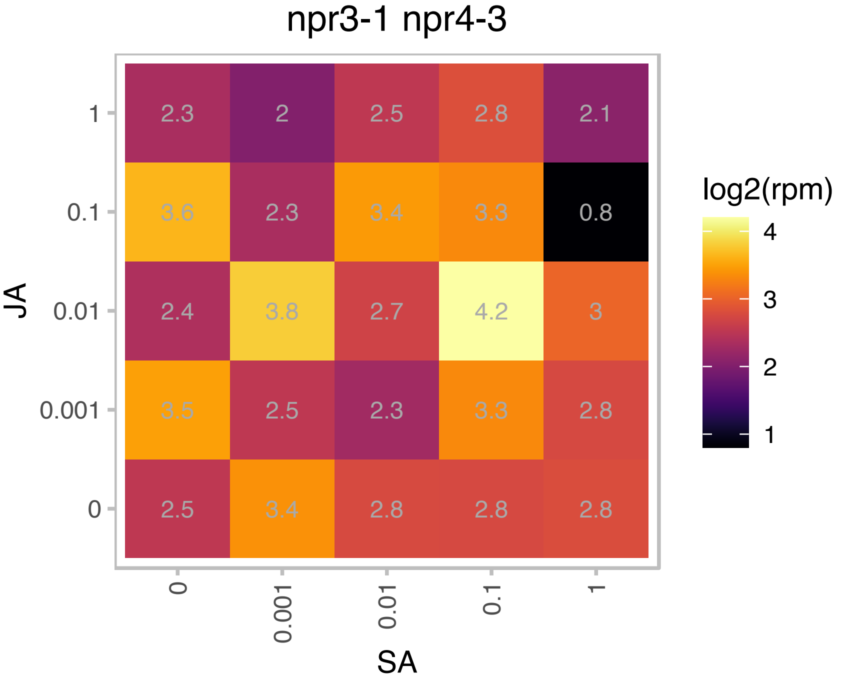

### AT1G49660.1 : AtCXE5, CXE5

carboxyesterase 5

Encodes a protein with carboxylesterase whose activity was tested using pNA.

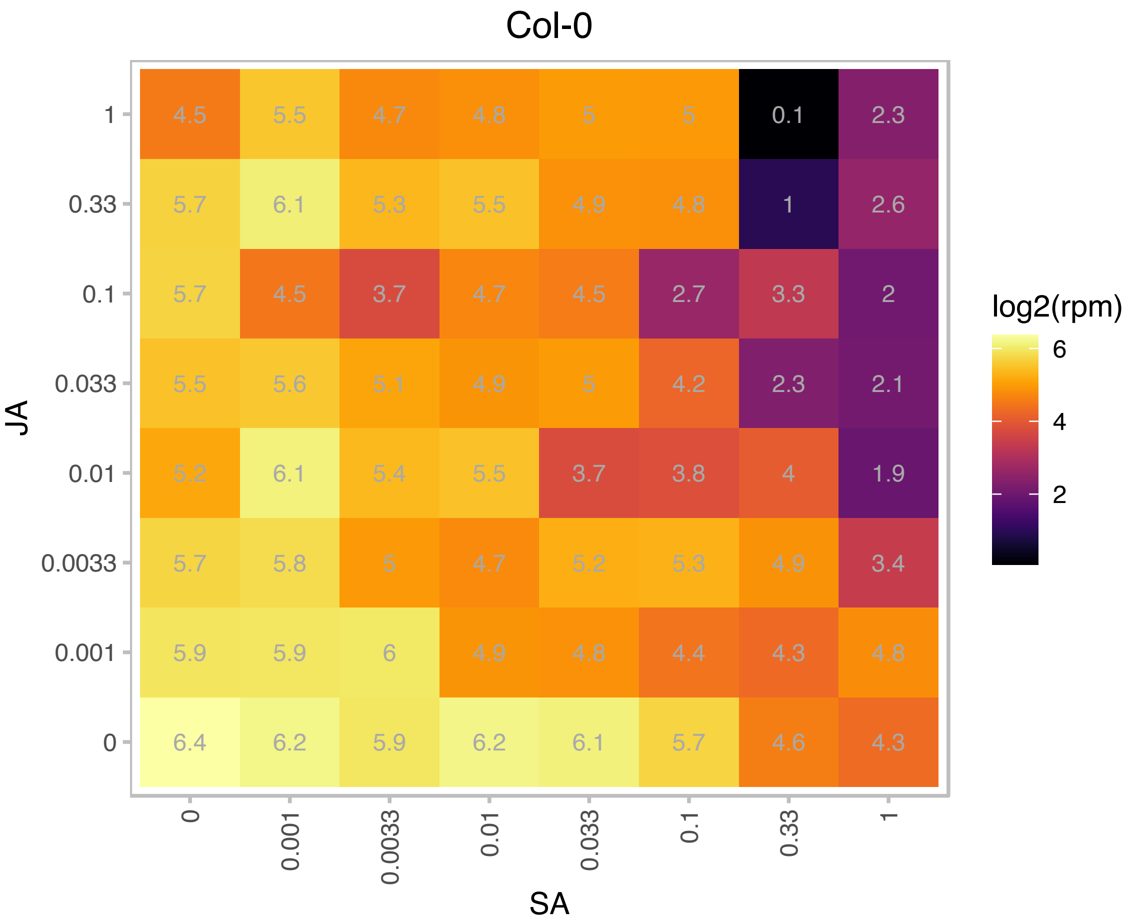


| | |
|---|---|
| Gene | AT1G49660 |
| is.expressed_genes | TRUE |
| is.responsive_genes | TRUE |
| is.apcluster | F-01 |

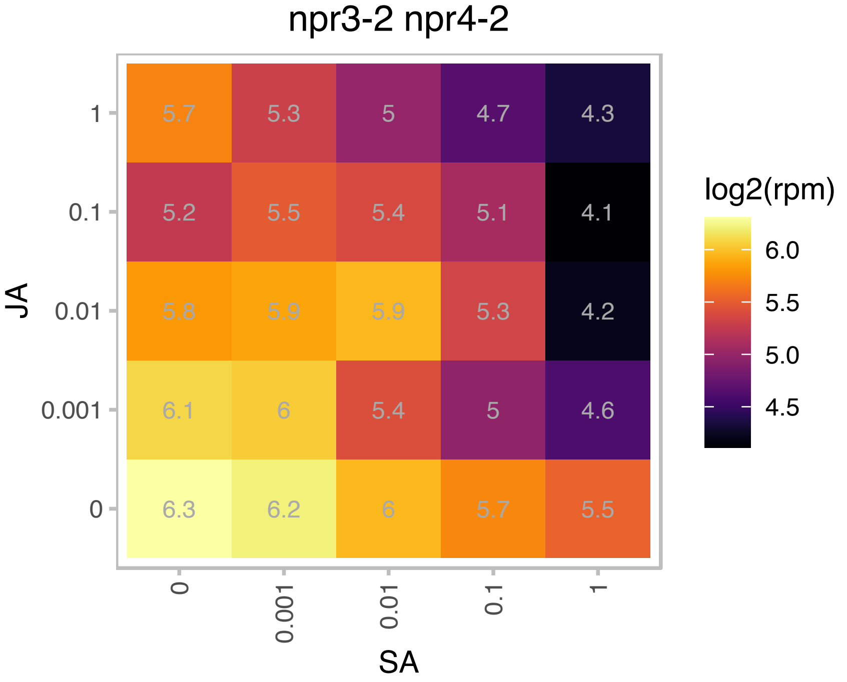


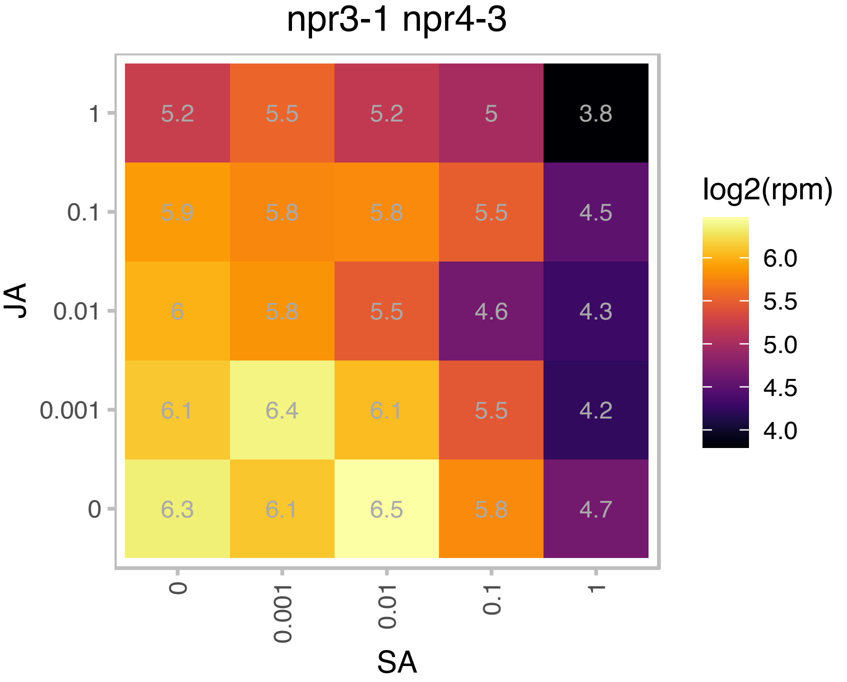

# AT1G53800.1 : NA

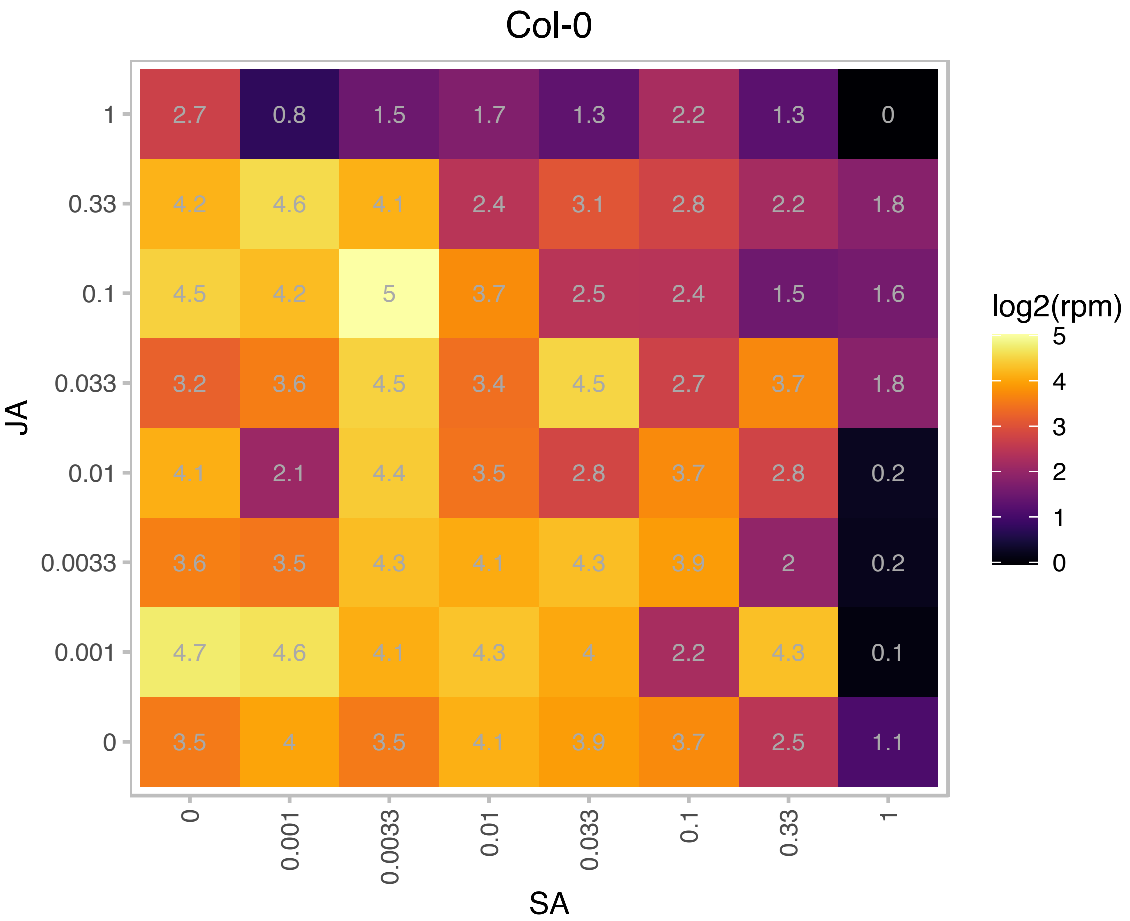


| | |
|---|---|
| Gene | AT1G53800 |
| is.expressed_genes | TRUE |
| is.responsive_genes | TRUE |
| is.apcluster | F-01 |

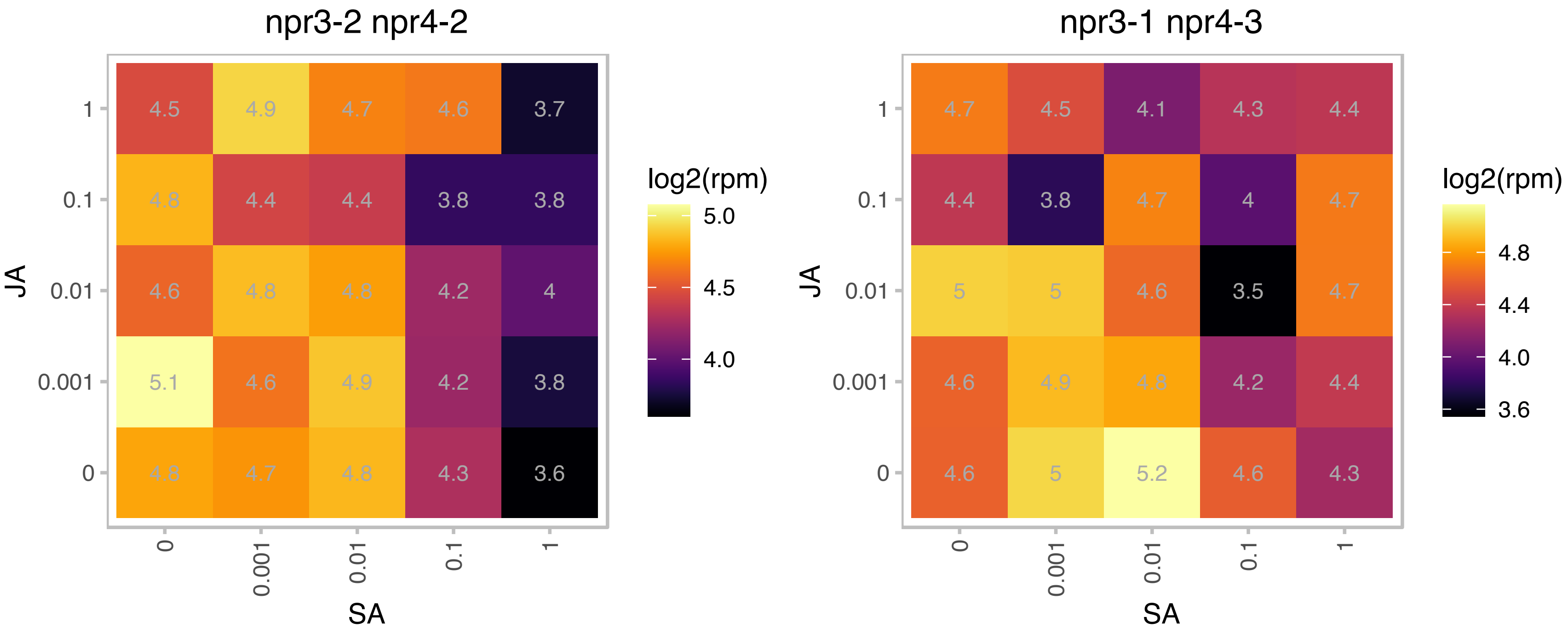

### AT1G63940.1 : MDAR6

monodehydroascorbate reductase 6

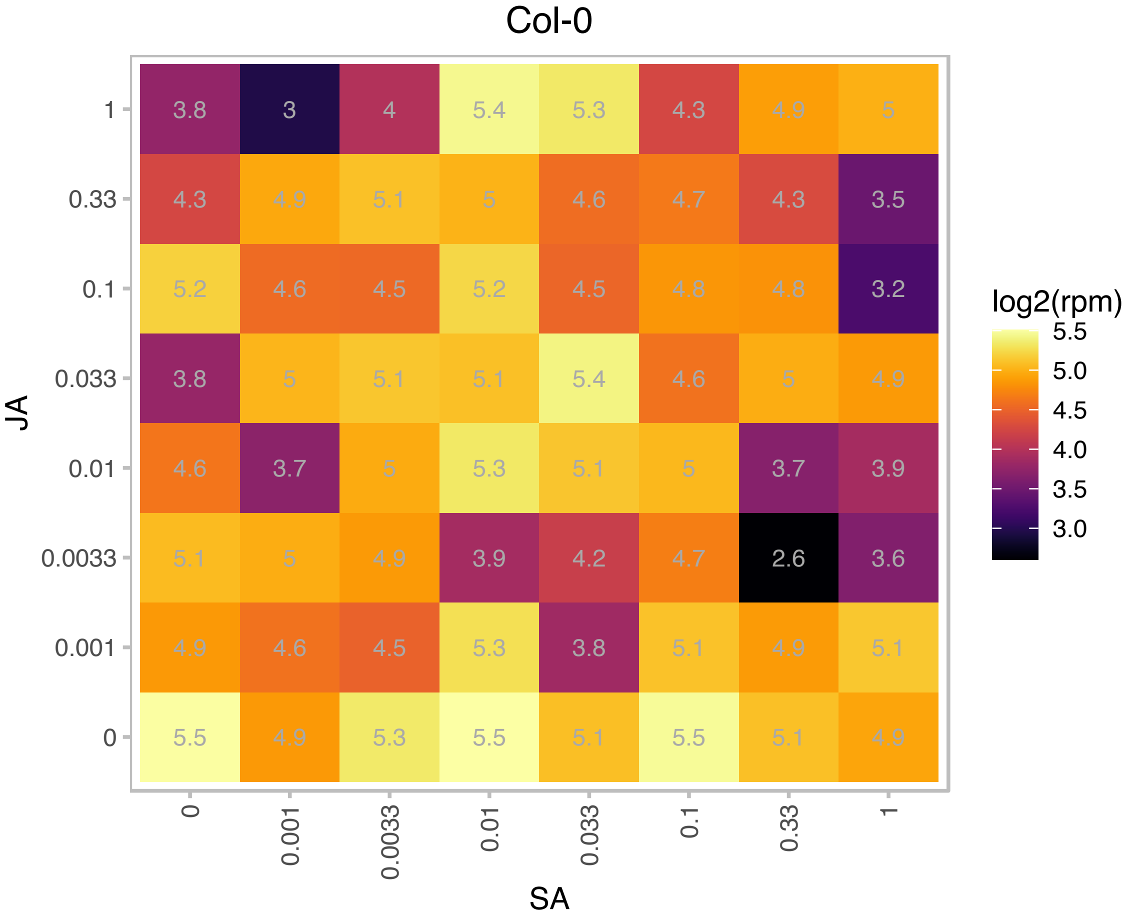


| Gene | AT1G63940 |
|---|---|
| is.expressed_genes | TRUE |
| is.responsive_genes | TRUE |
| is.apcluster | F-01 |

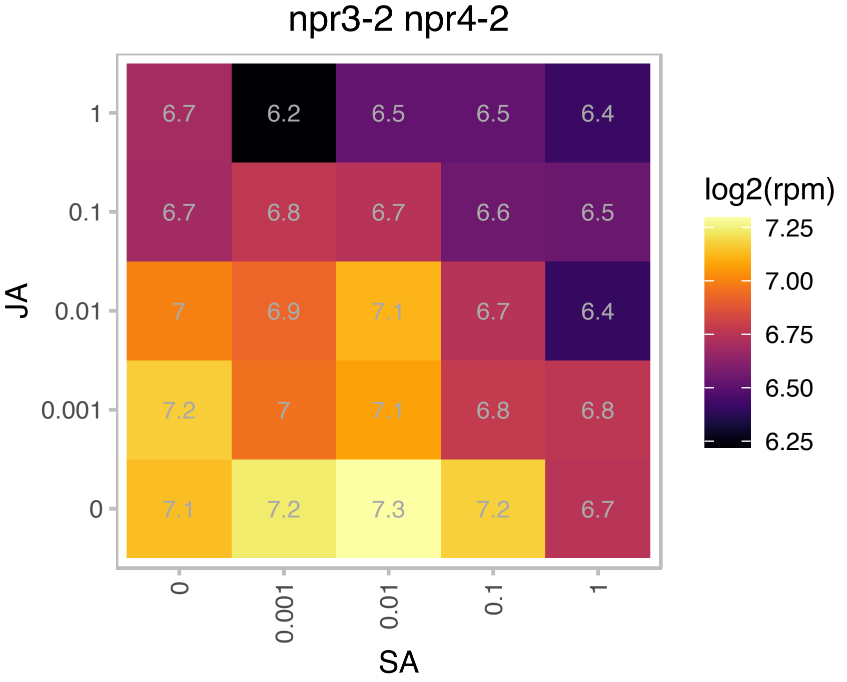


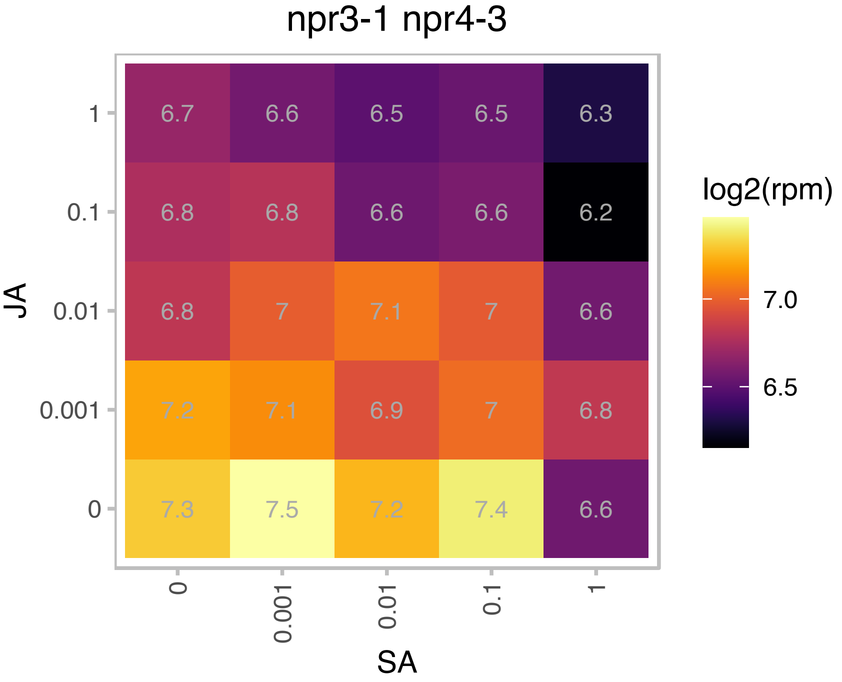

# AT1G65010.1 : NA

Plant protein of unknown function (DUF827).

Encodes a microtubule-associated protein. Putative role in flower development. Comparison of SALK_061426C to Columbia wild type in normal lighting and under low light of 33 micromoles per meter-squared per second resulted in a trend toward earlier bolting in the mutant under low light (P=0.055) (Ann Stapleton and Patrick Pridgen, 2009, personal communication).

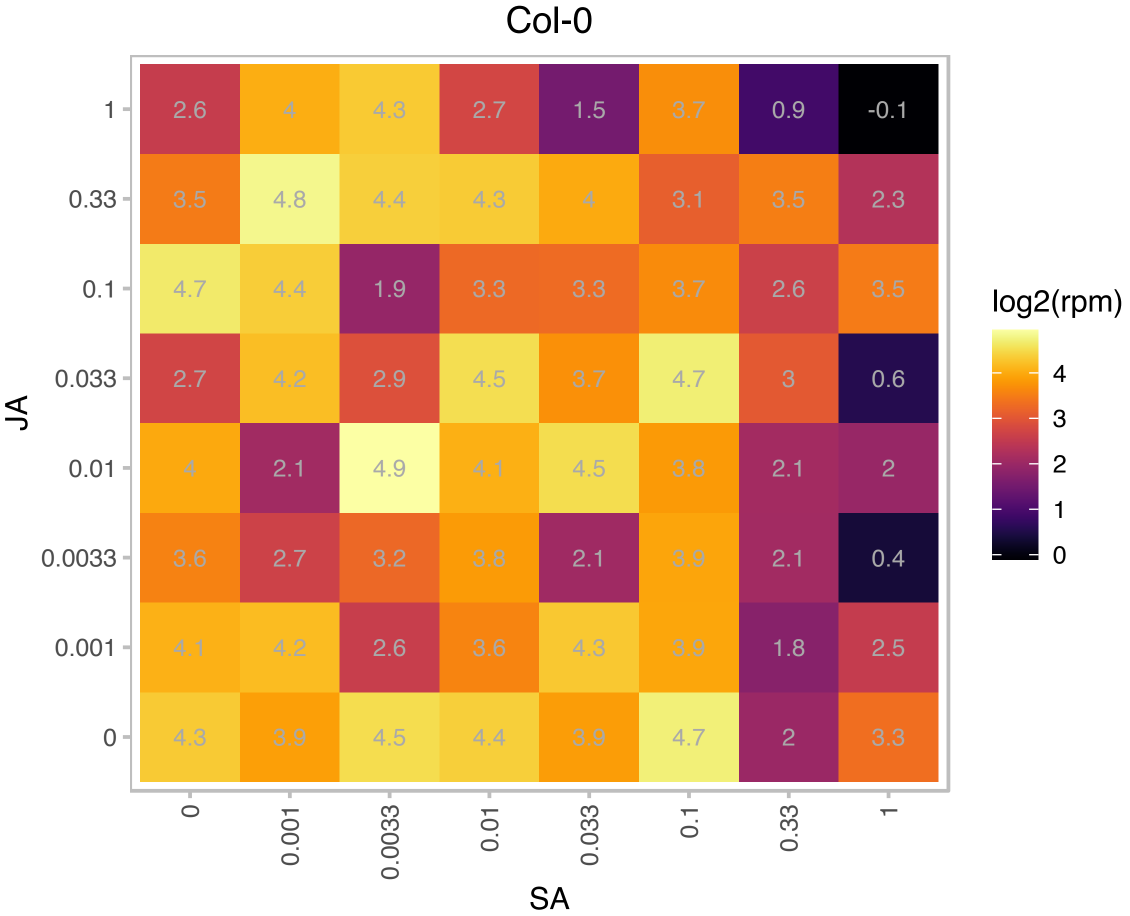


| | |
|---|---|
| Gene | AT1G65010 |
| is.expressed_genes | TRUE |
| is.responsive_genes | TRUE |
| is.apcluster | F-01 |

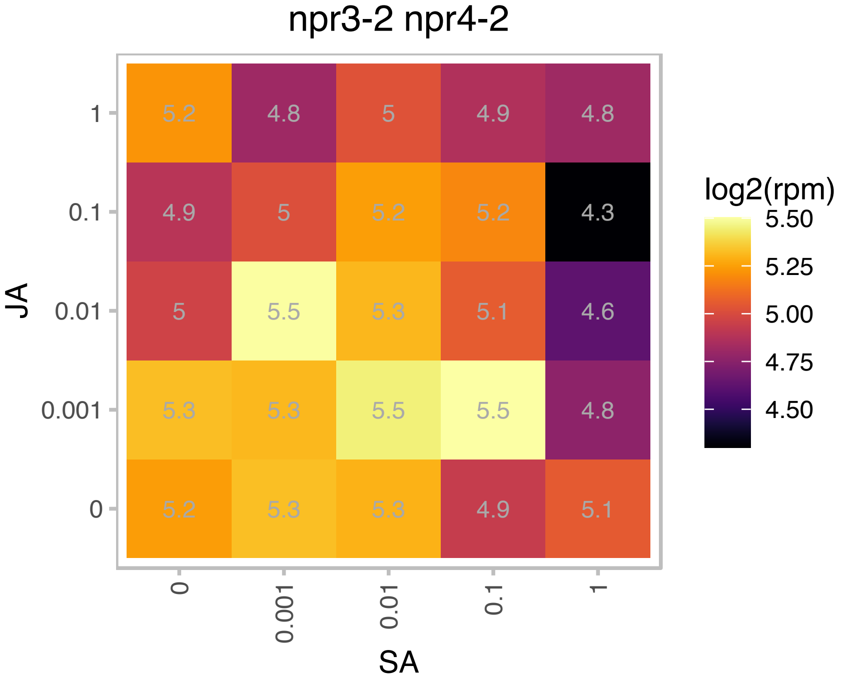


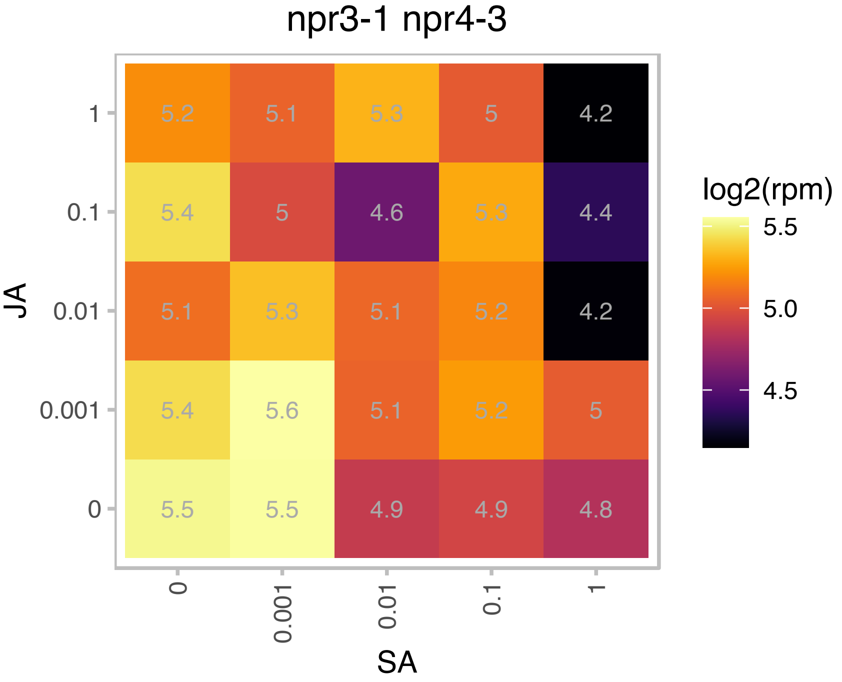

### AT1G65410.1 : ABCI13, ATNAP11, NAP11, TGD3

non-intrinsic ABC protein 11

Encodes a member of NAP subfamily of transporters. Mutations in this gene suppress the low temperature-induced phenotype of Arabidopsis tocopherol-deficient mutant vte2.

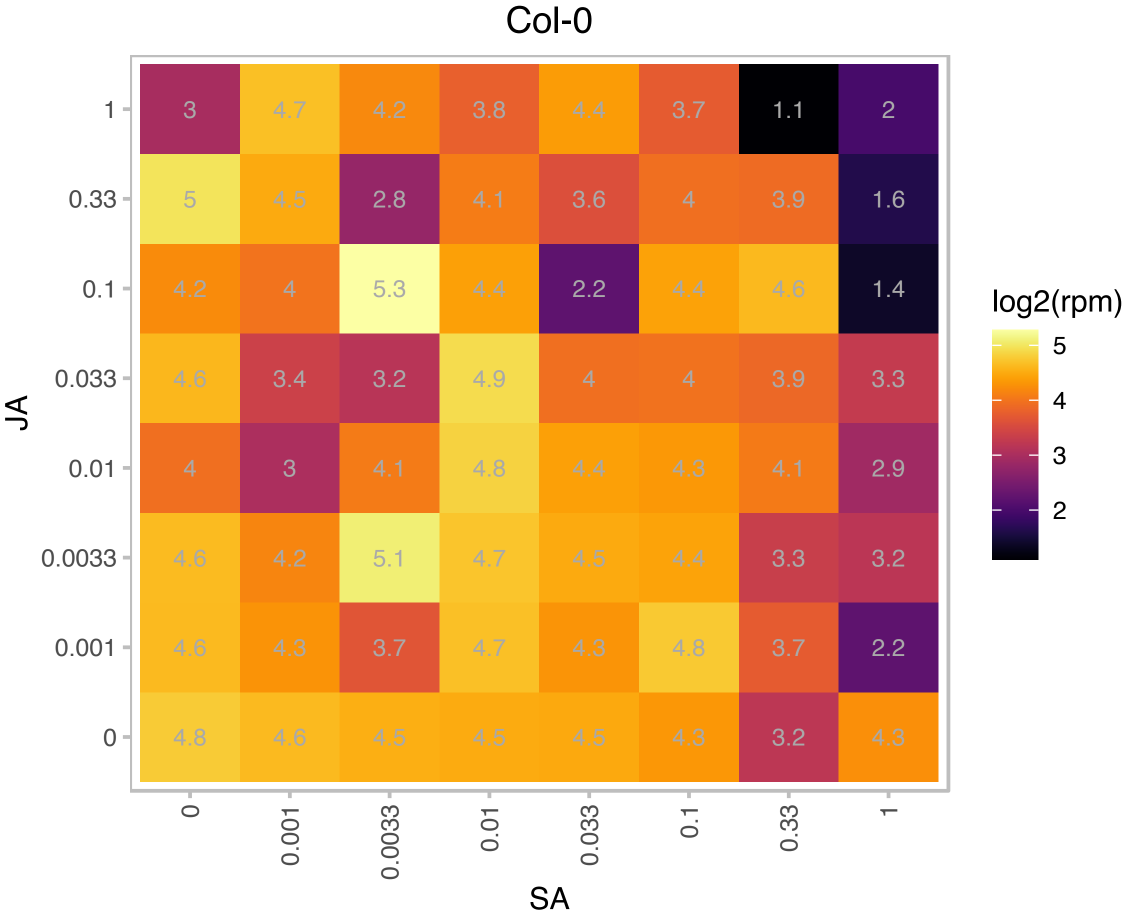


| | |
|---|---|
| Gene | AT1G65410 |
| is.expressed_genes | TRUE |
| is.responsive_genes | TRUE |
| is.apcluster | F-01 |

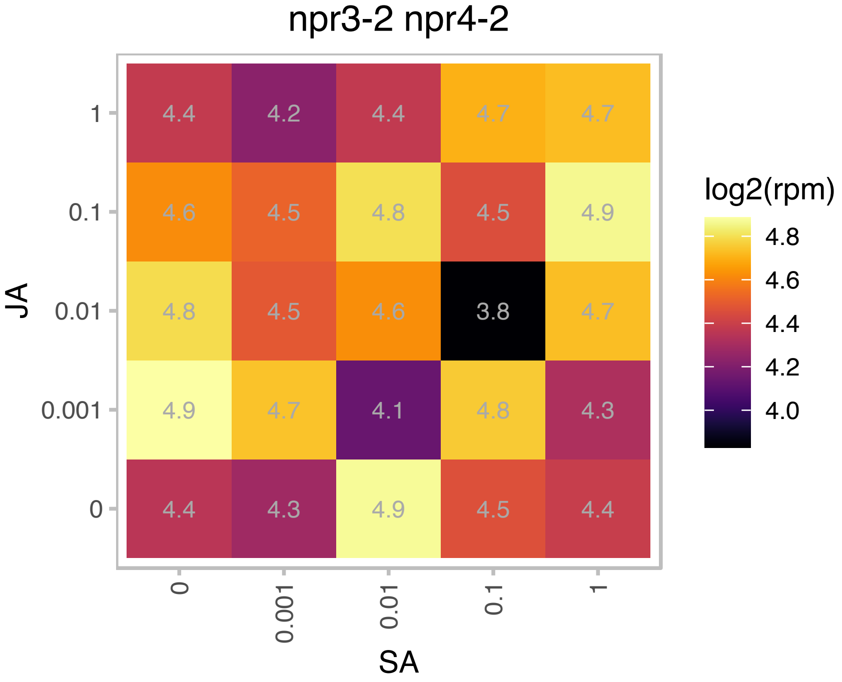


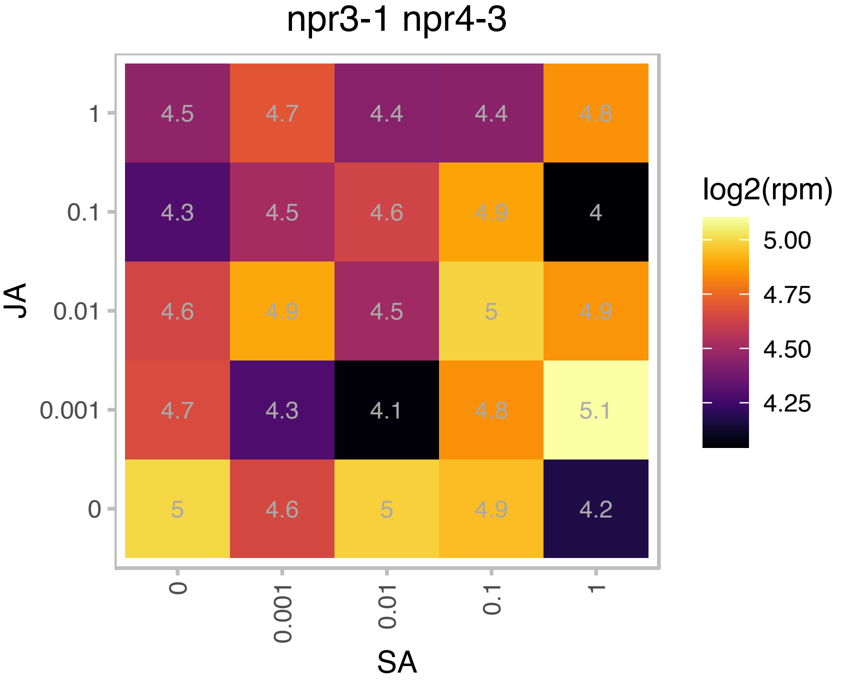

### AT1G68760.1 : ATNUDT1, ATNUDX1, NUDX1, NUDX1

nudix hydrolase 1

Encodes a cytosol-localized nudix hydrolase that hydrolyzes 8-oxo-(d)GTP to its monophosphate form. This protective mechanism prevents the misincorporation of these oxidized nucleotides into DNA and RNA. NUDX1 also has a low level of dihydroneopterin triphosphate pyrophosphatase activity in vitro and may participate in the folate synthesis pathway.

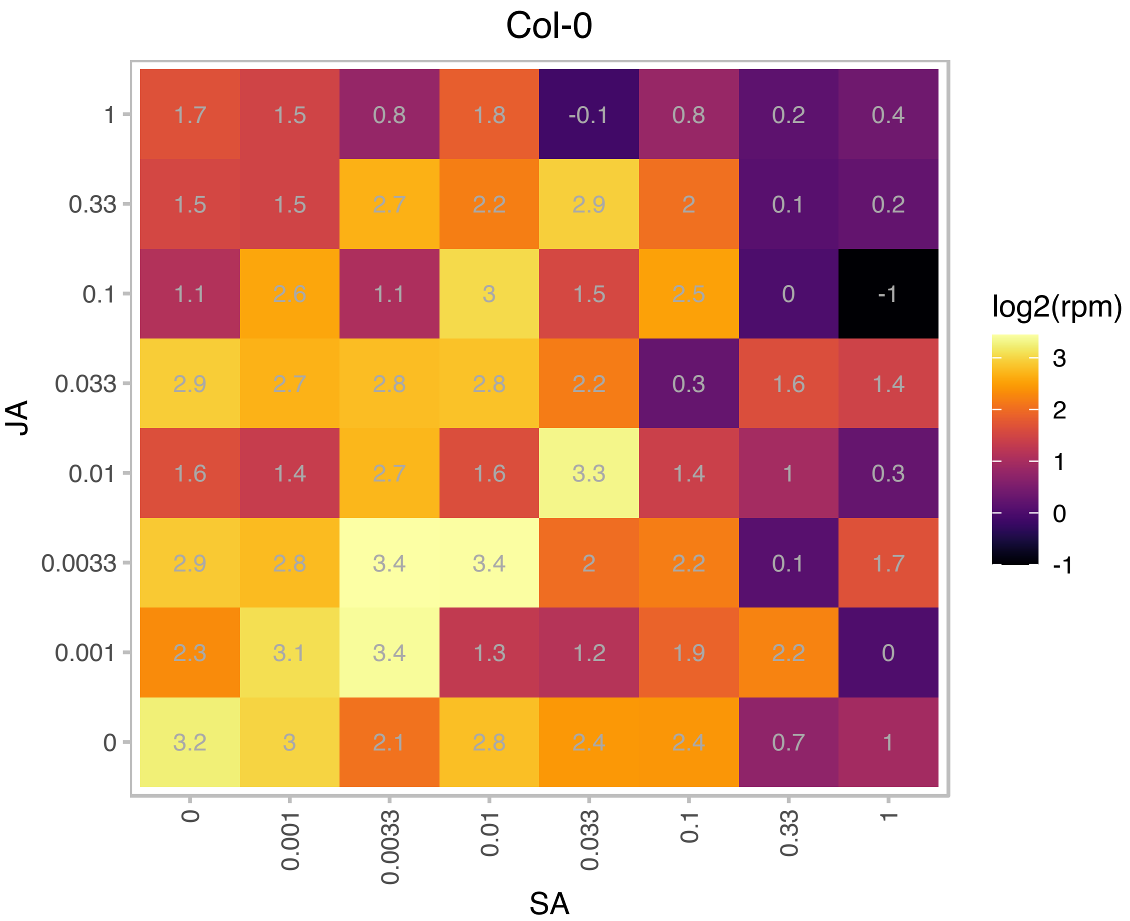


| | |
|---|---|
| Gene | AT1G68760 |
| is.expressed_genes | TRUE |
| is.responsive_genes | TRUE |
| is.apcluster | F-01 |

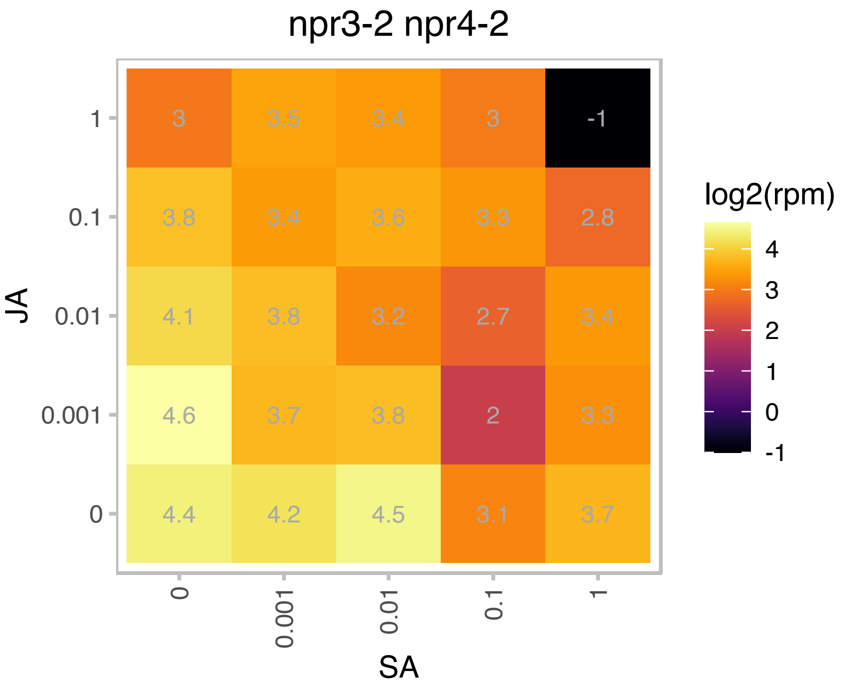


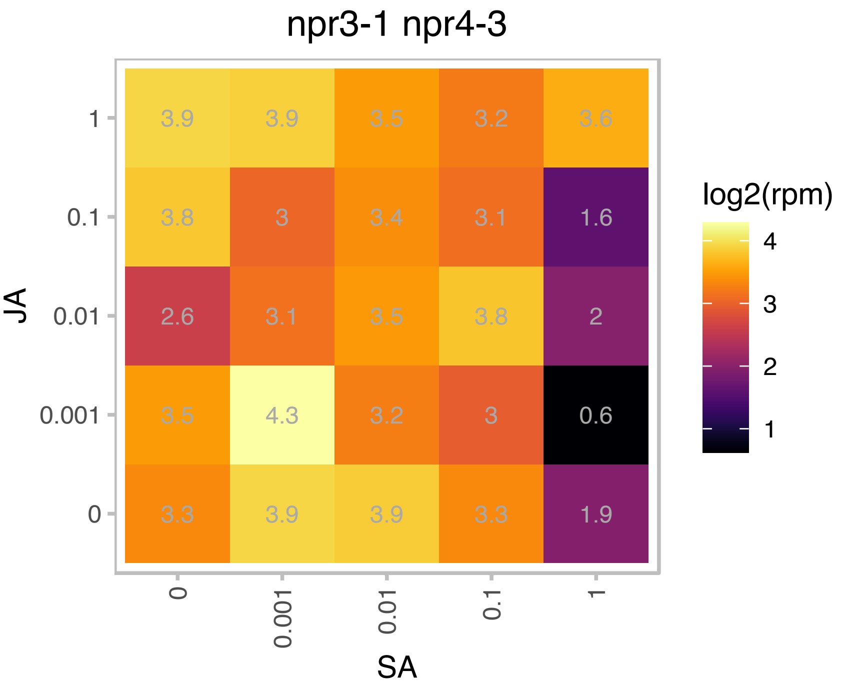

### AT1G70200.1 : HPE1, RBD1

RNA-binding (RRM/RBD/RNP motifs) family protein

Encodes a RNA-Binding Protein RBD1. Promotes chilling tolerance through 23S rRNA processing.

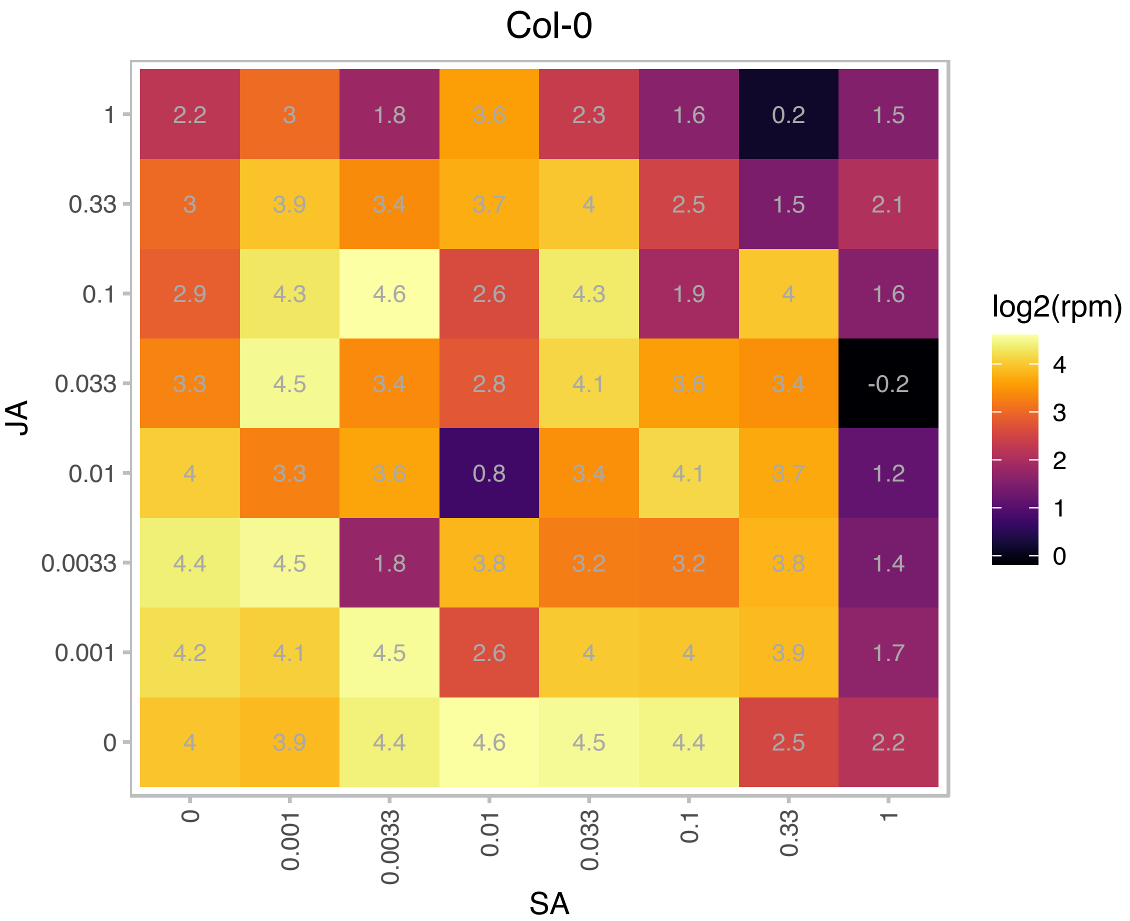


| | |
|---|---|
| Gene | AT1G70200 |
| is.expressed_genes | TRUE |
| is.responsive_genes | TRUE |
| is.apcluster | F-01 |

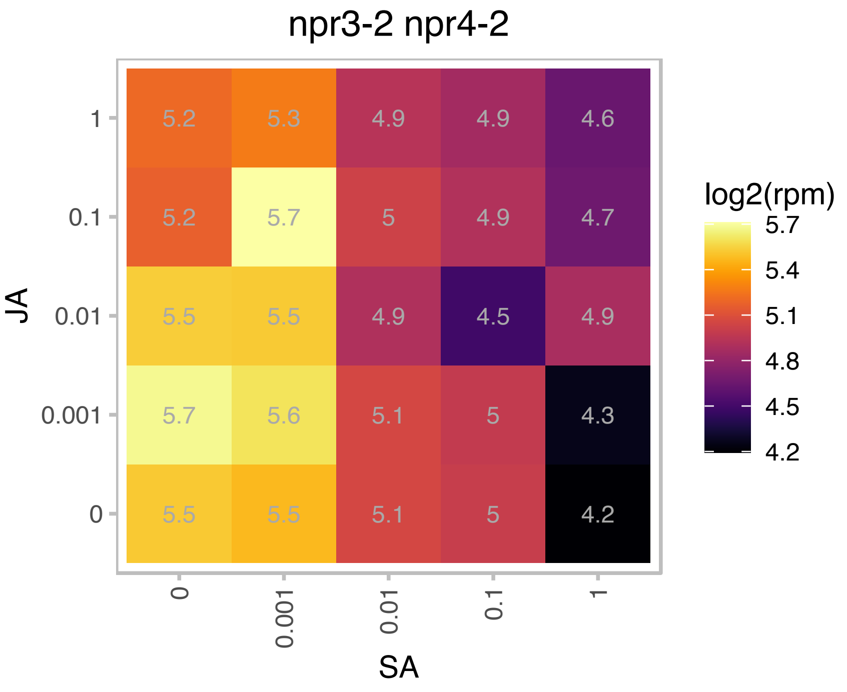


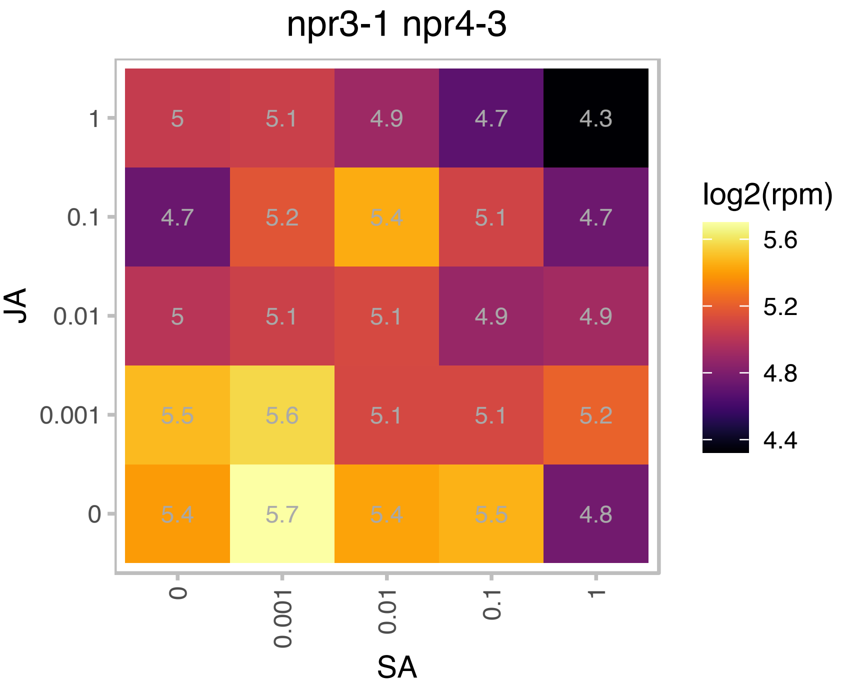

# AT1G75460.1 : NA

ATP-dependent protease La (LON) domain protein

Col-0

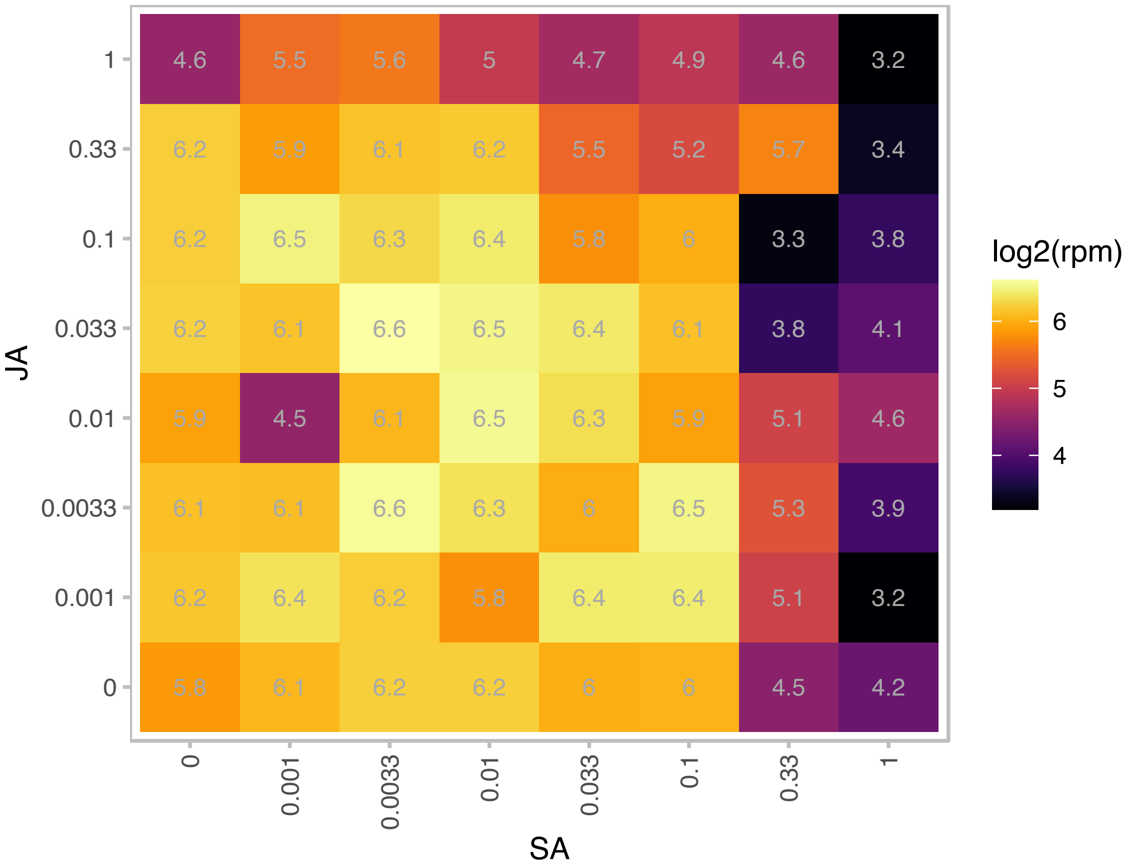


| | |
|---|---|
| Gene | AT1G75460 |
| is.expressed_genes | TRUE |
| is.responsive_genes | TRUE |
| is.apcluster | F-01 |

npr3-2 npr4-2

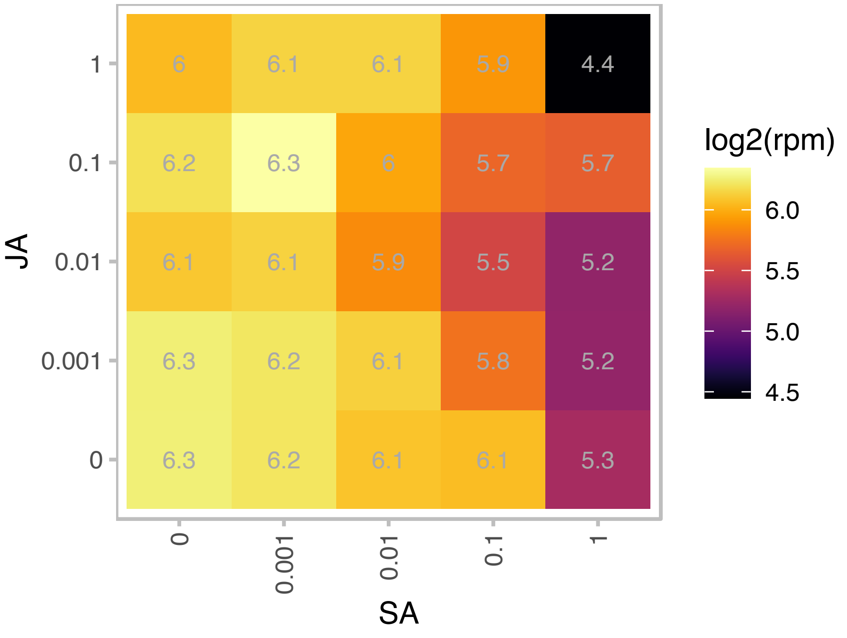


npr3-1 npr4-3

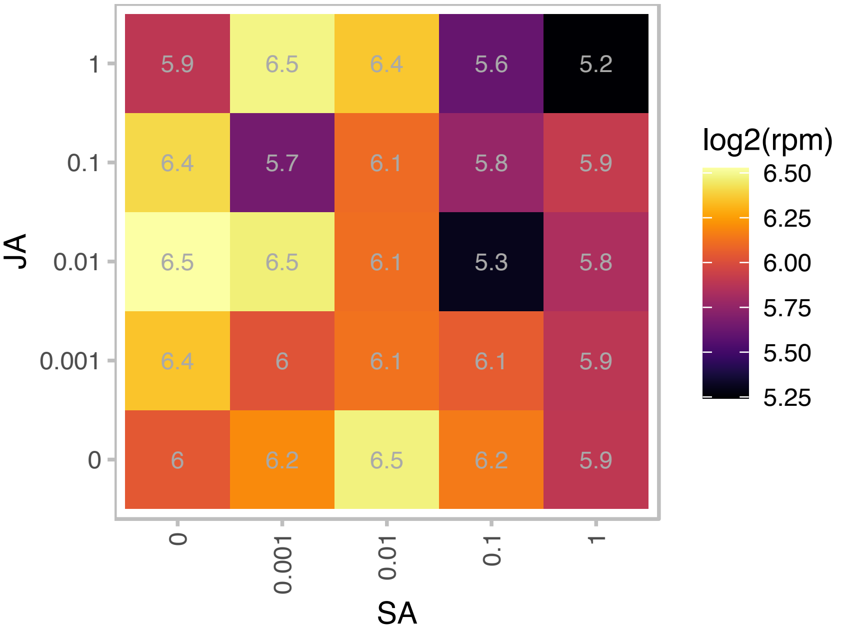

# AT1G78830.1 : NA

Curculin-like (mannose-binding) lectin family protein

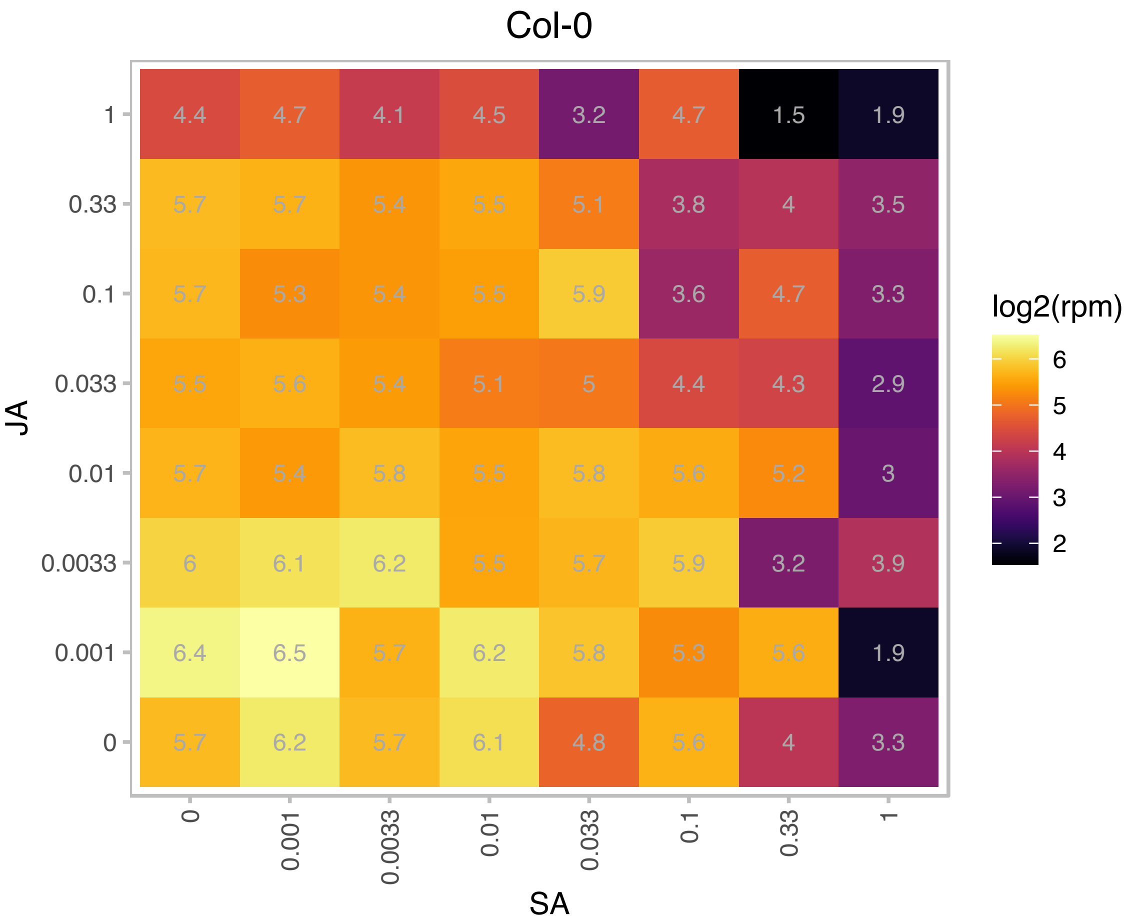


| | |
|---|---|
| Gene | AT1G78830 |
| is.expressed_genes | TRUE |
| is.responsive_genes | TRUE |
| is.apcluster | F-01 |

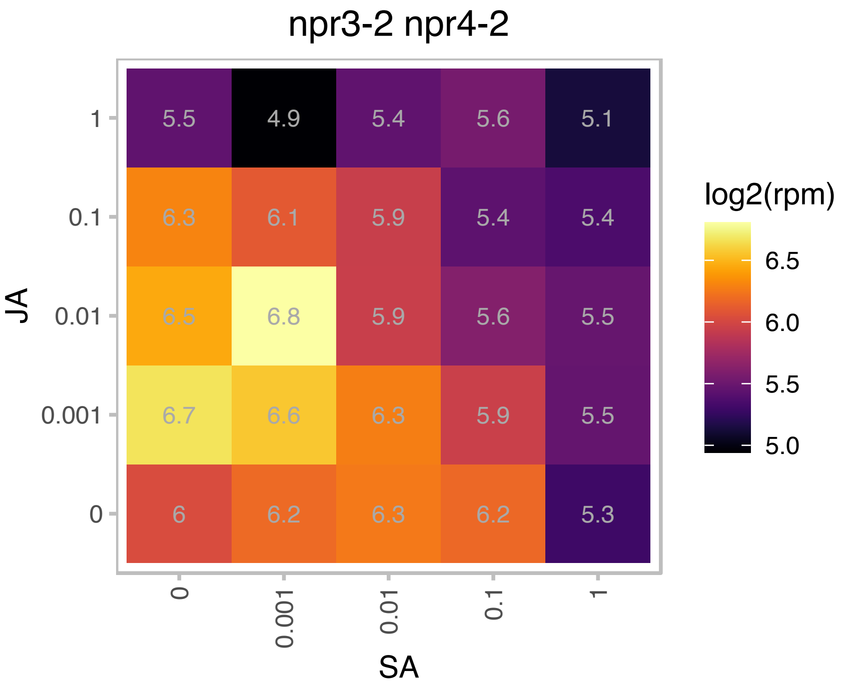


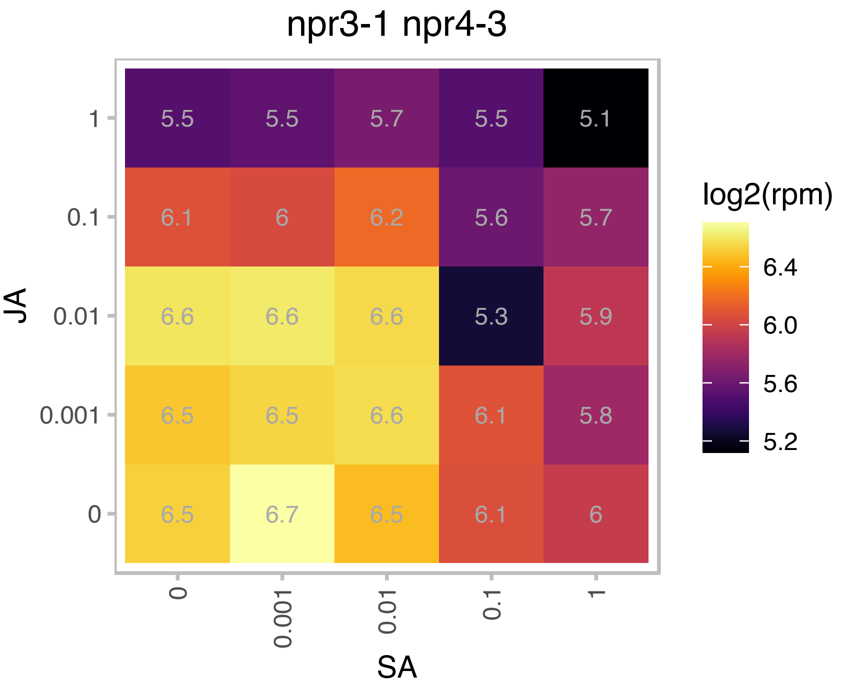

### AT1G79850.1 : CS17, PDE347, PRPS17, RPS17

ribosomal protein S17

nuclear-encoded 30S chloroplast ribosomal protein S17

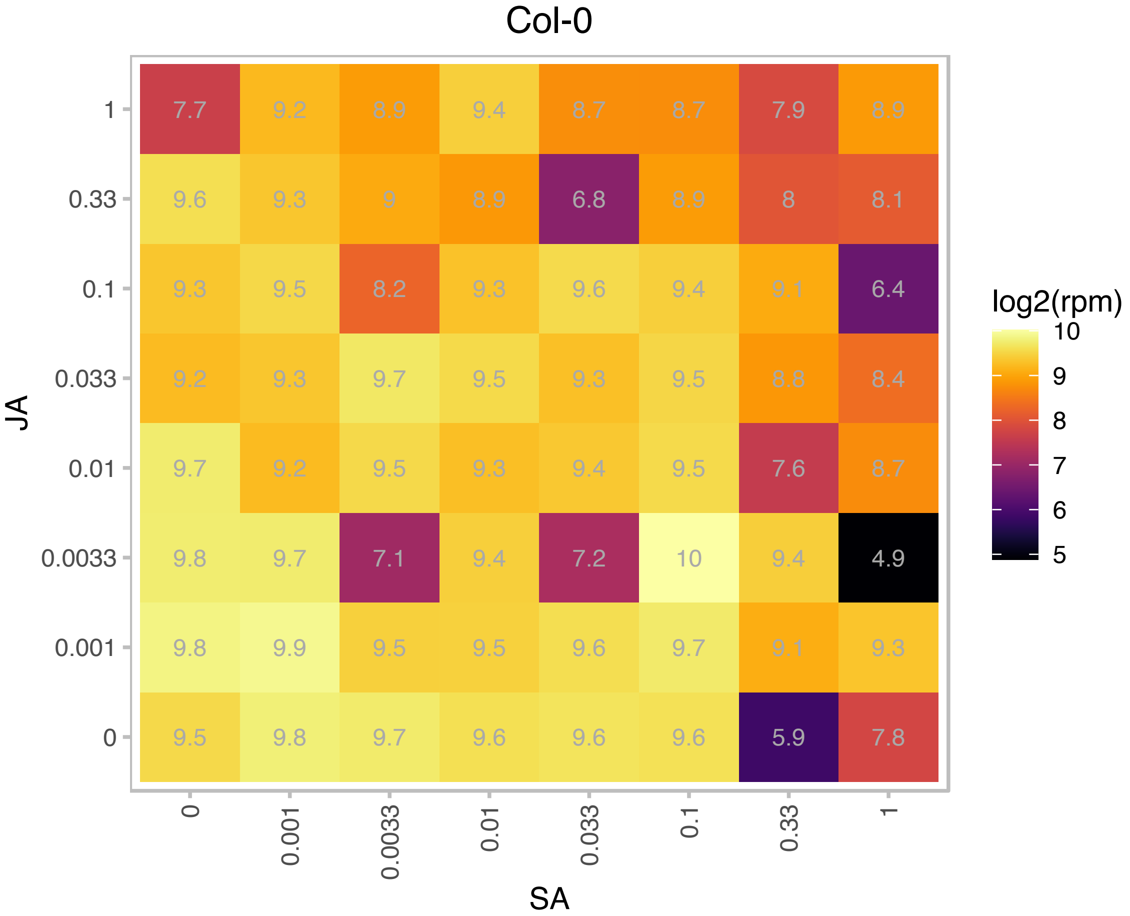


| | |
|---|---|
| Gene | AT1G79850 |
| is.expressed_genes | TRUE |
| is.responsive_genes | TRUE |
| is.apcluster | F-01 |

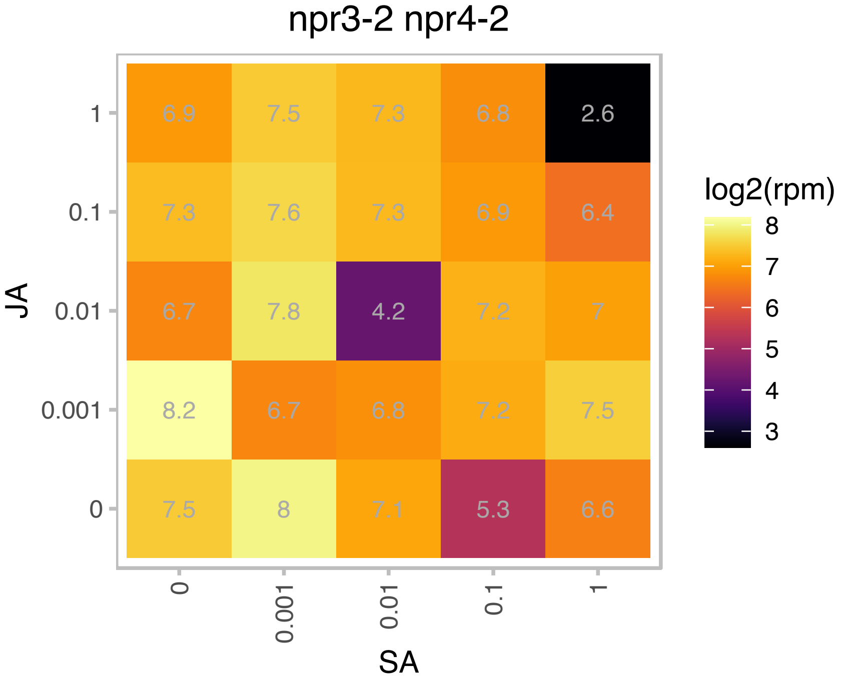


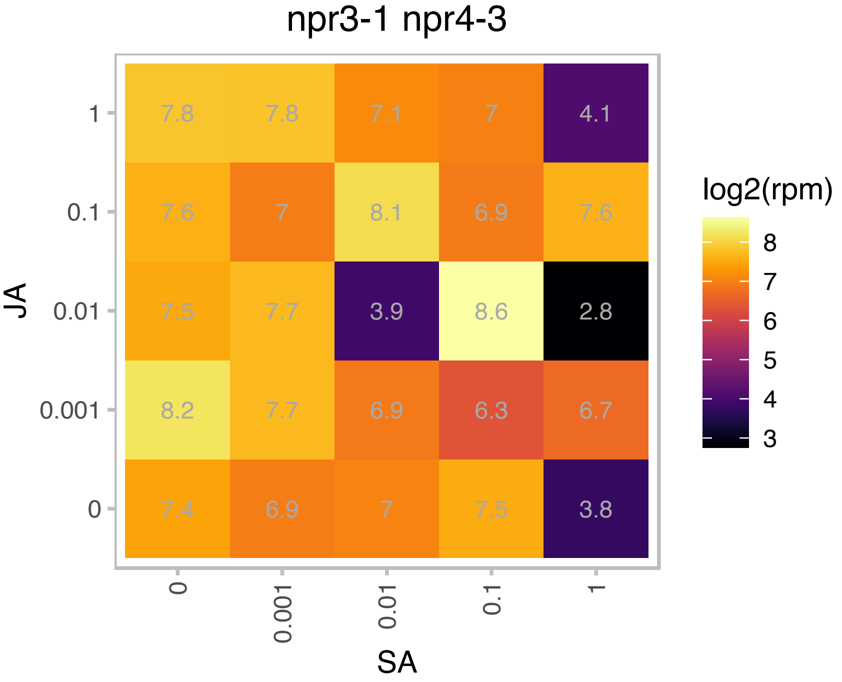

# AT2G01870.1 : NA

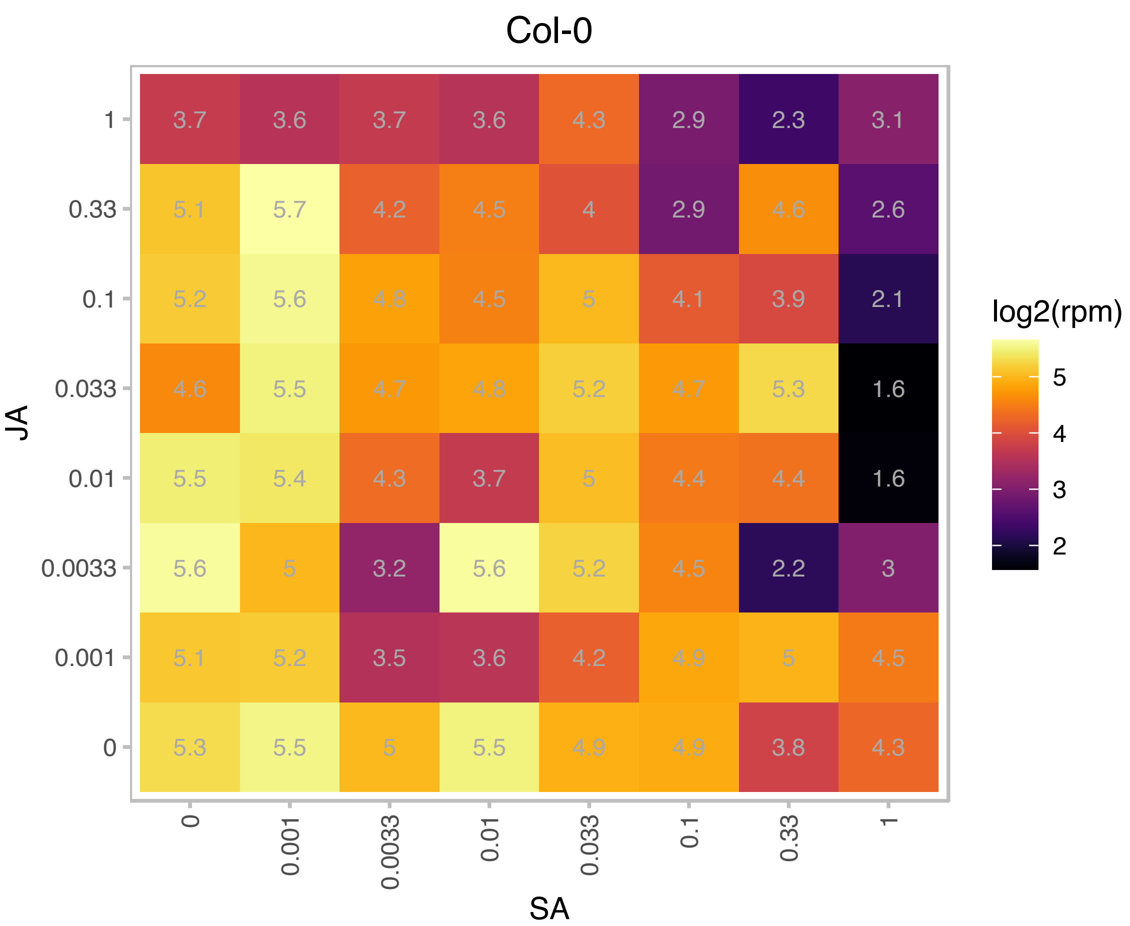


| | |
|---|---|
| Gene | AT2G01870 |
| is.expressed_genes | TRUE |
| is.responsive_genes | TRUE |
| is.apcluster | F-01 |

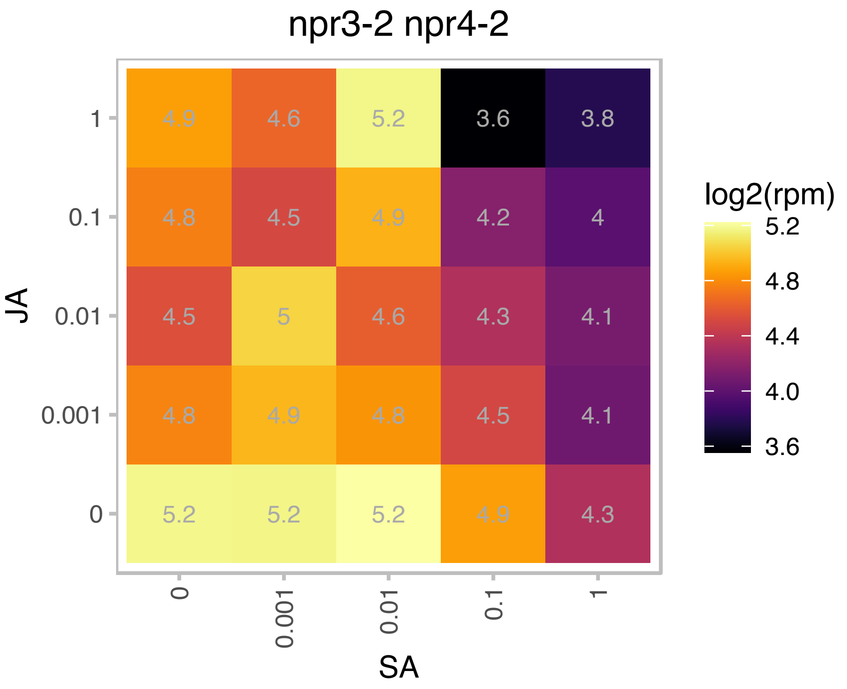


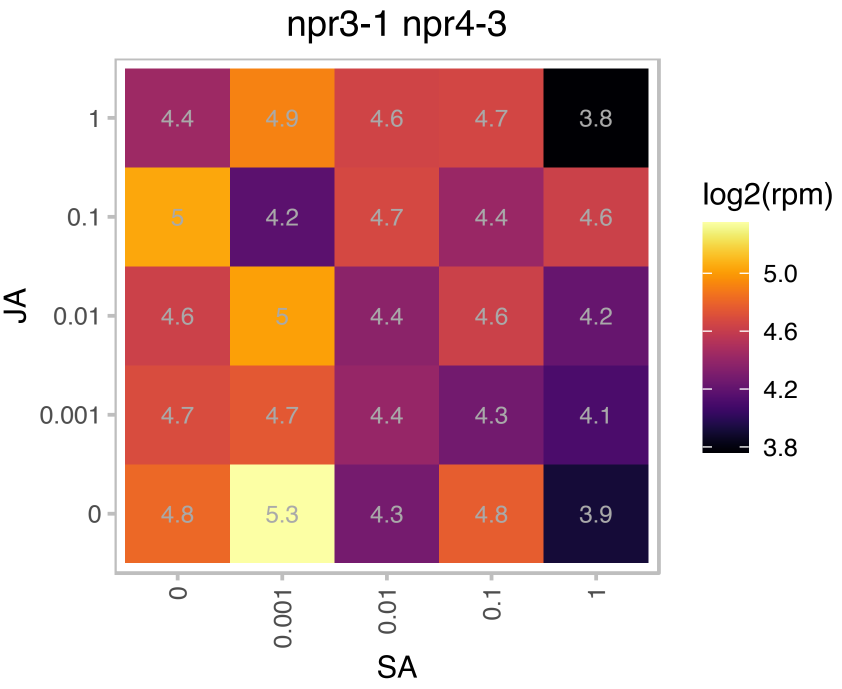

### AT2G16400.1 : BLH7

BEL1-like homeodomain 7

Col-0

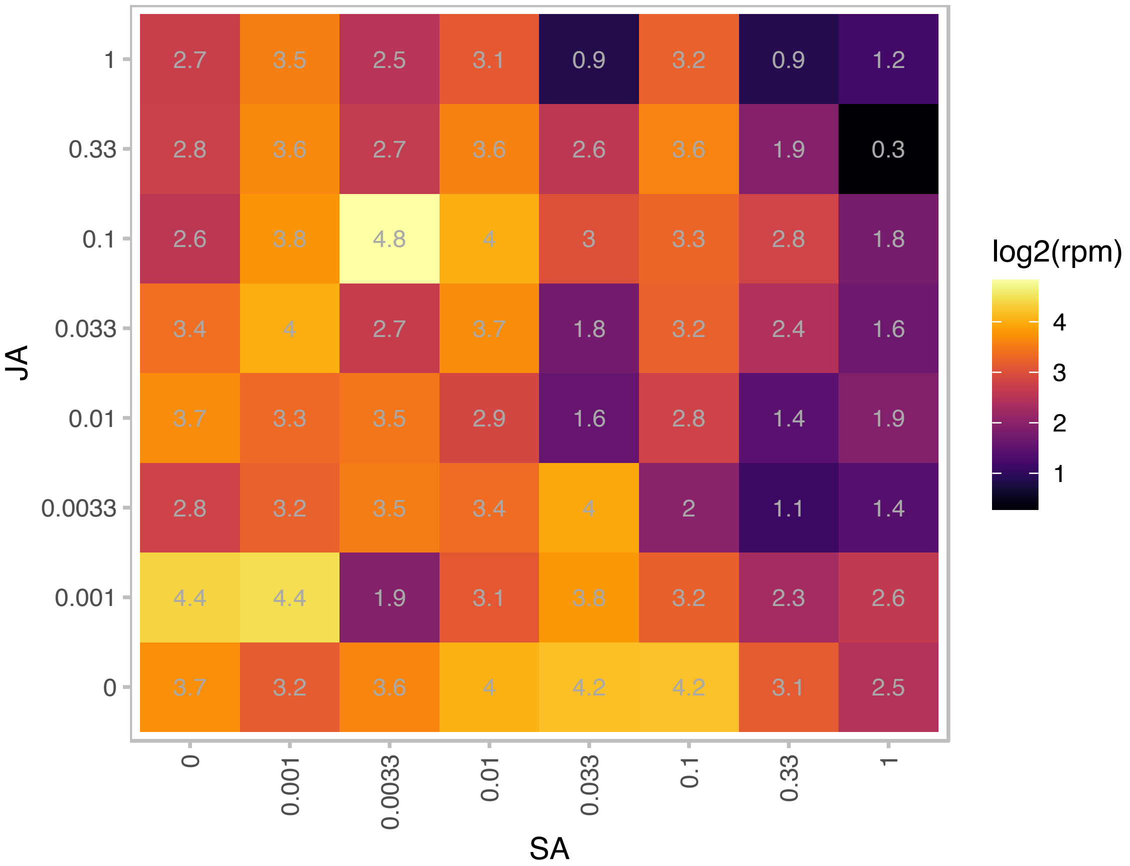


| | |
|---|---|
| Gene | AT2G16400 |
| is.expressed_genes | TRUE |
| is.responsive_genes | TRUE |
| is.apcluster | F-01 |

npr3-2 npr4-2

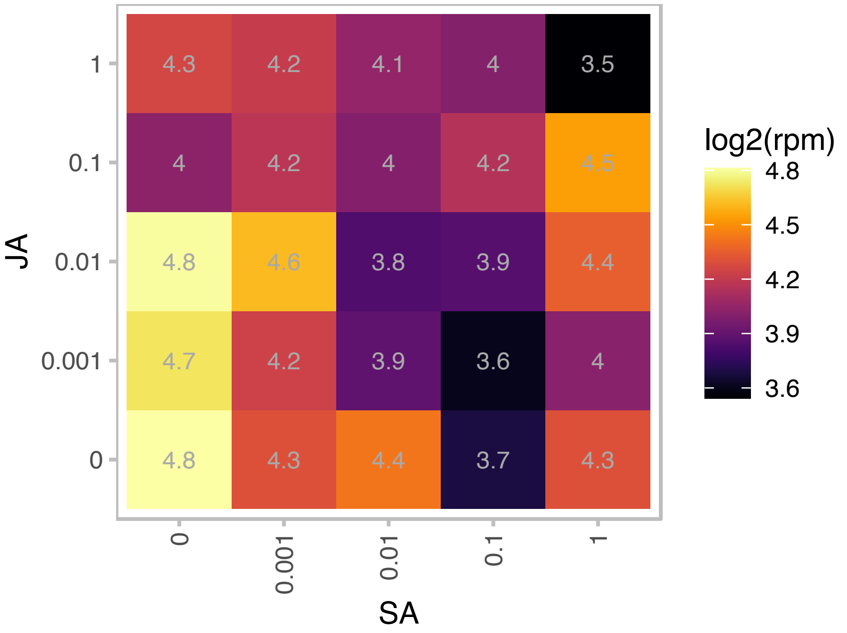


npr3-1 npr4-3

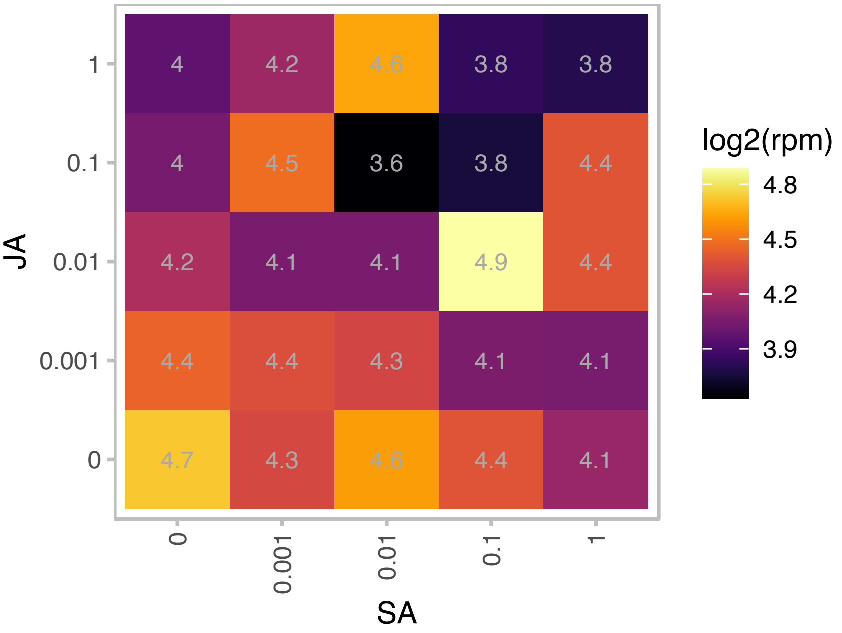

# AT2G20515.1 : NA

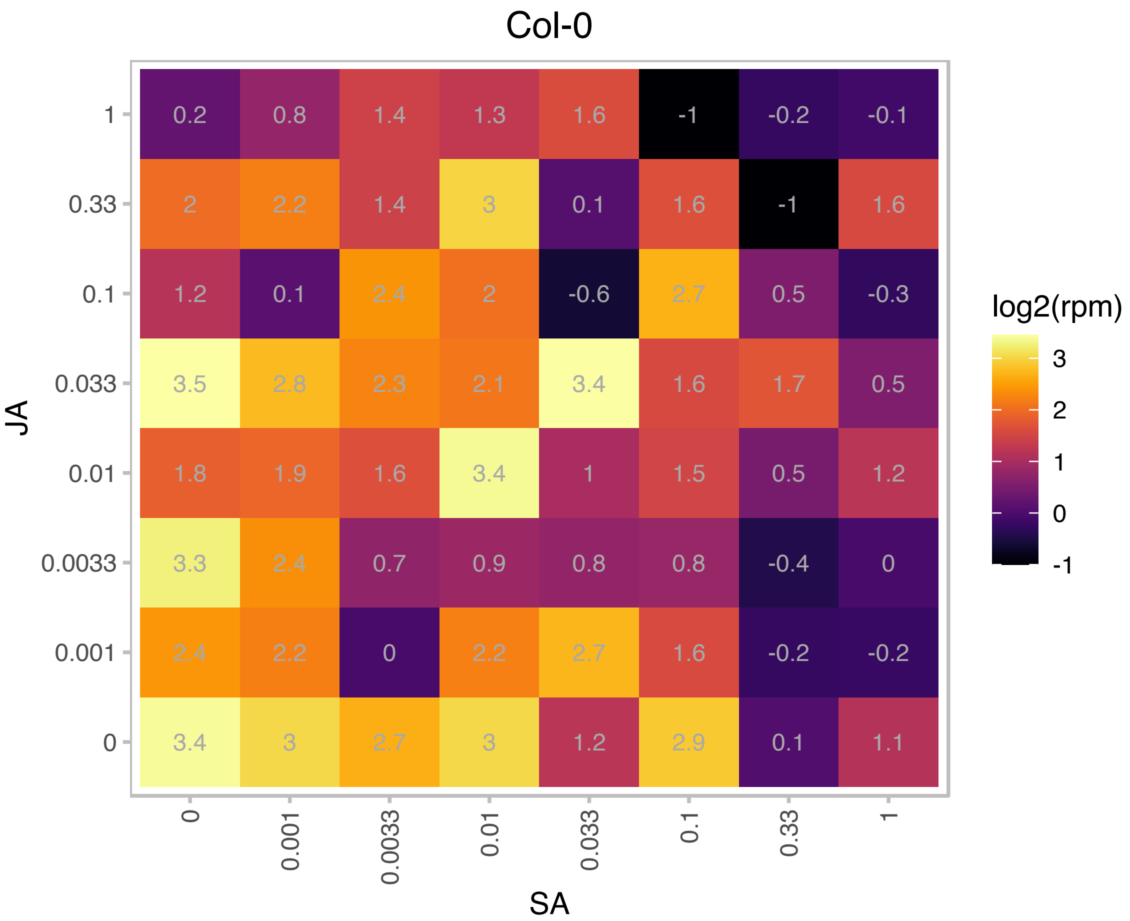


| | |
|---|---|
| Gene | AT2G20515 |
| is.expressed_genes | TRUE |
| is.responsive_genes | TRUE |
| is.apcluster | F-01 |

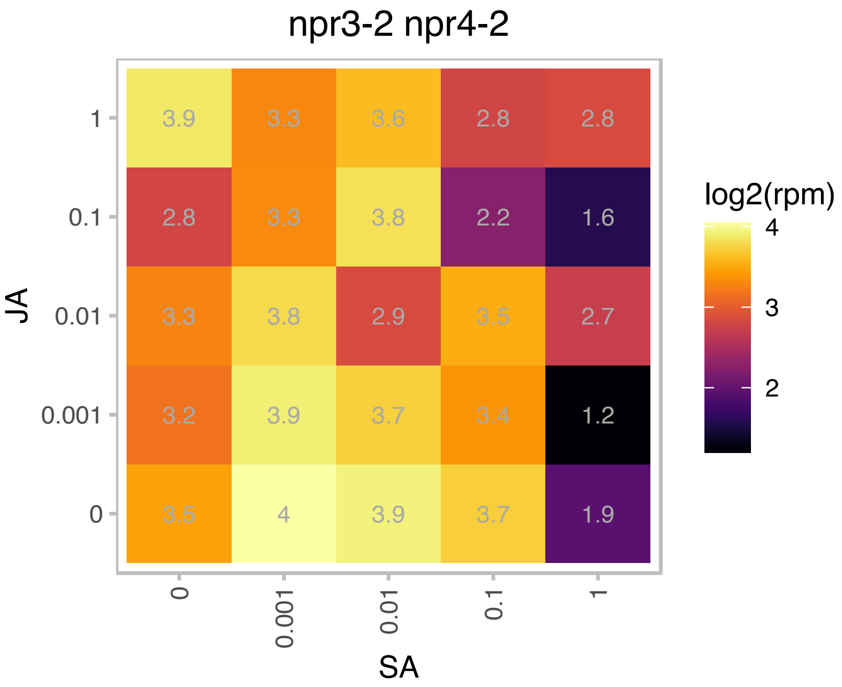


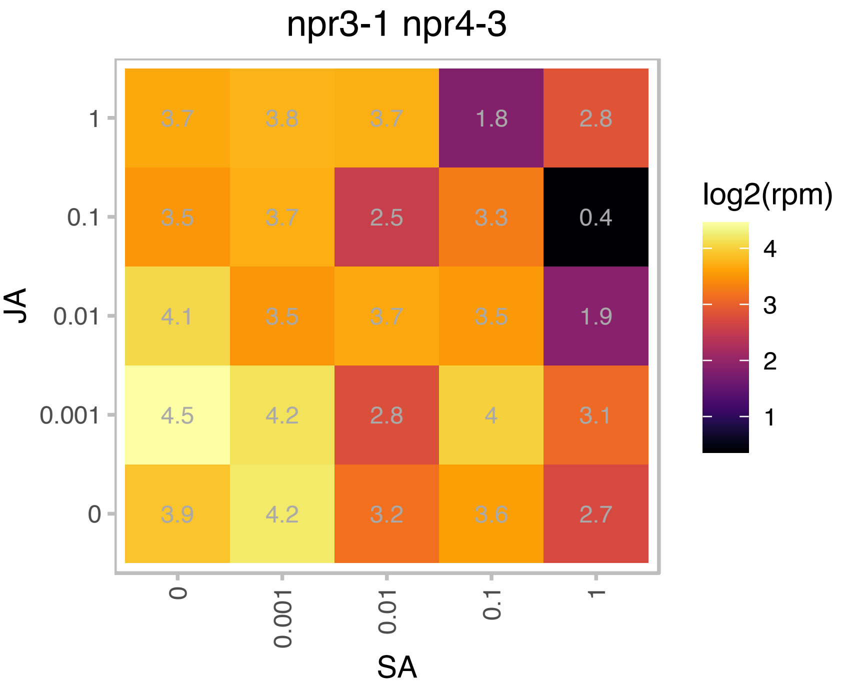

# AT2G24395.1 : NA

chaperone protein dnaJ-related

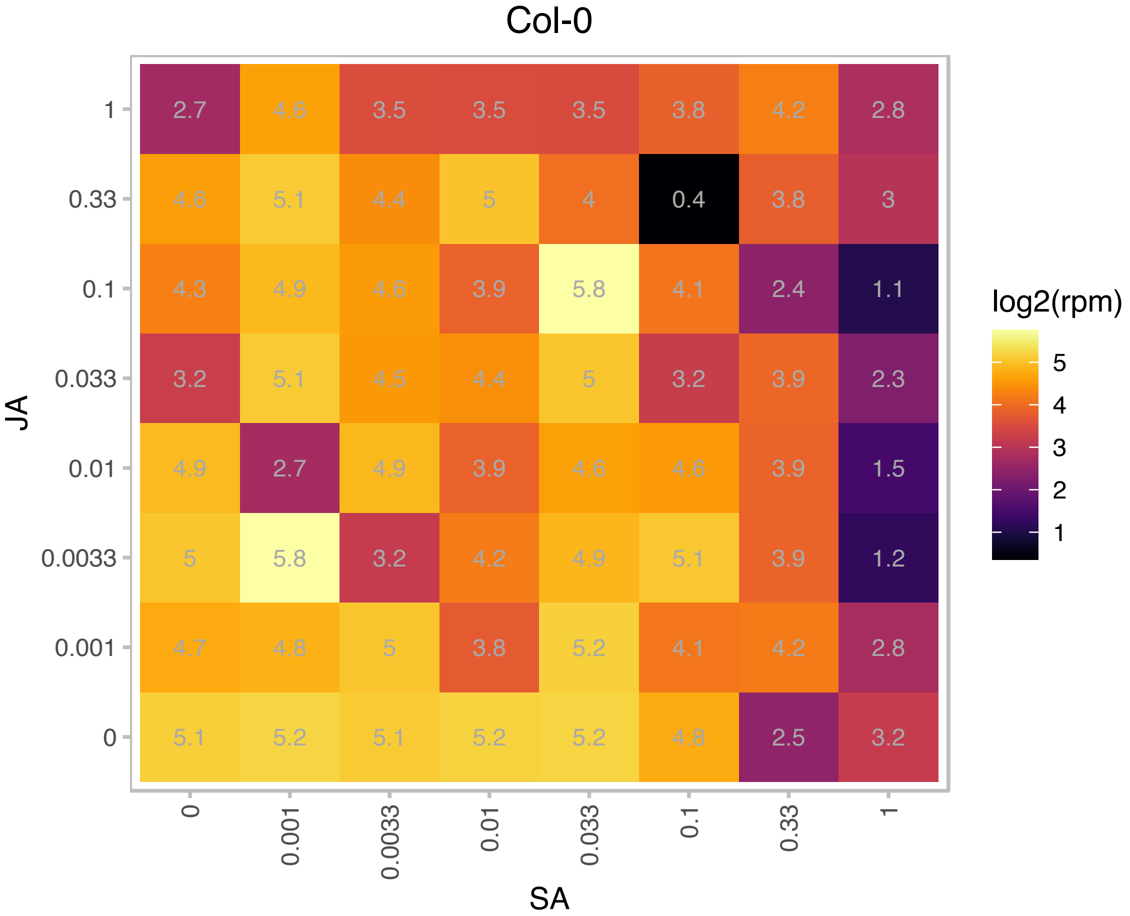


| | |
|---|---|
| Gene | AT2G24395 |
| is.expressed_genes | TRUE |
| is.responsive_genes | TRUE |
| is.apcluster | F-01 |

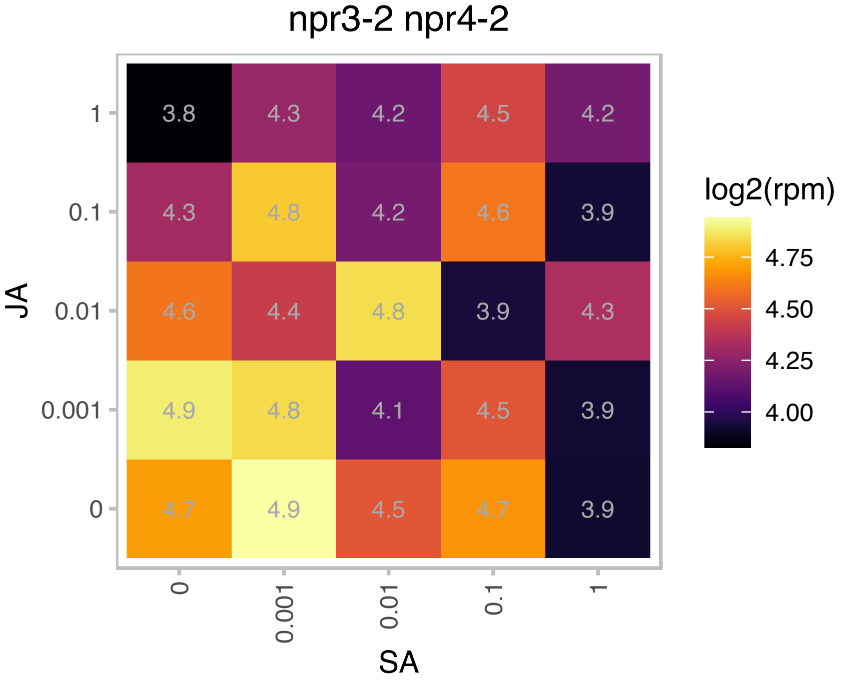


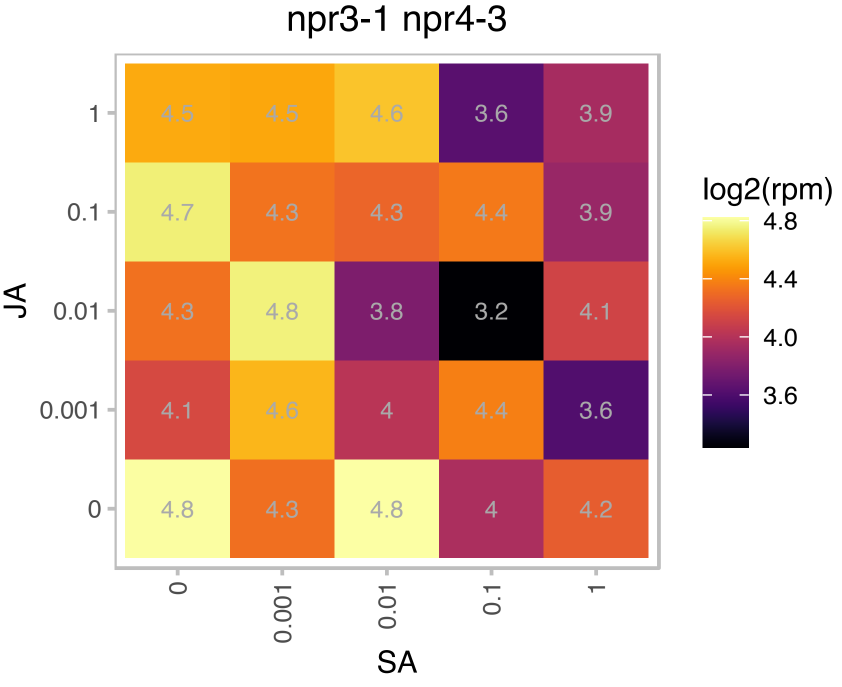

### AT2G27970.1 : CKS2

CDK-subunit 2

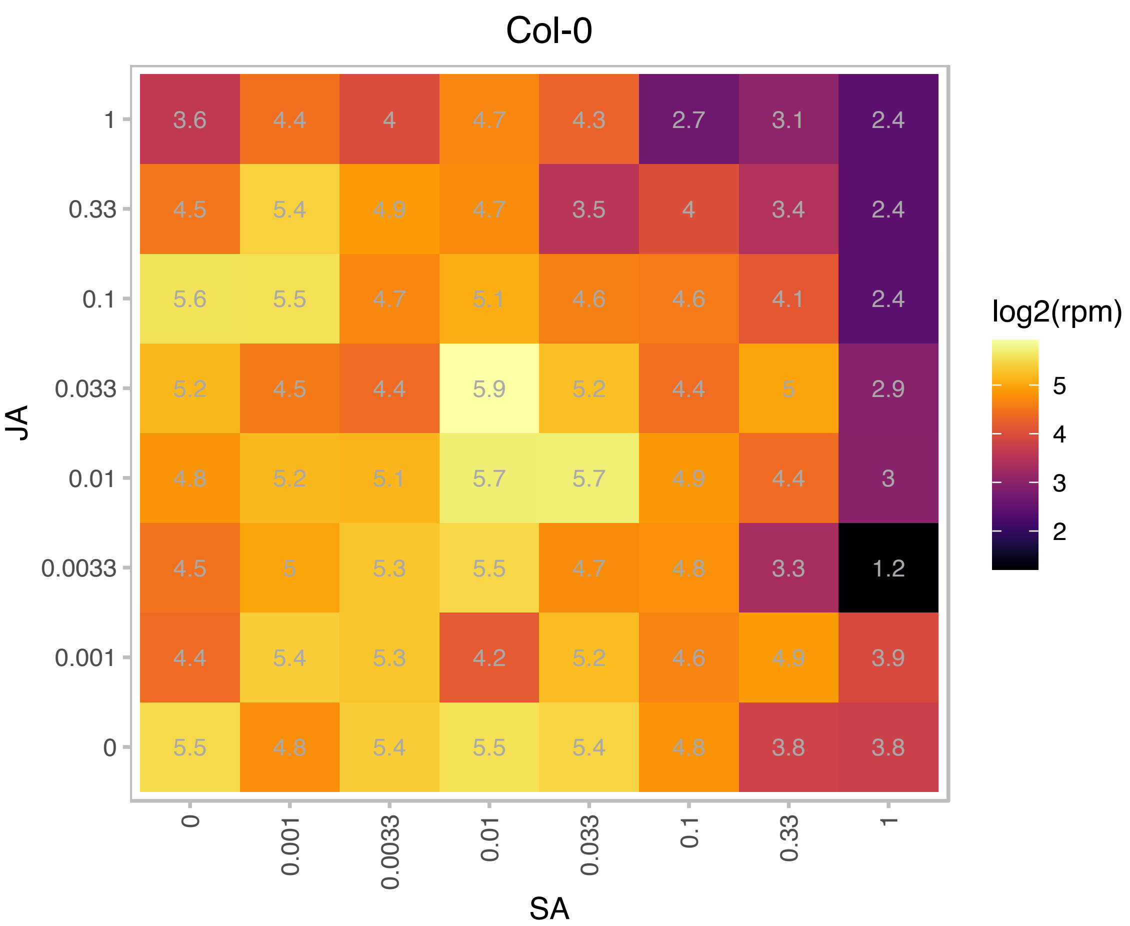


| | |
|---|---|
| Gene | AT2G27970 |
| is.expressed_genes | TRUE |
| is.responsive_genes | TRUE |
| is.apcluster | F-01 |

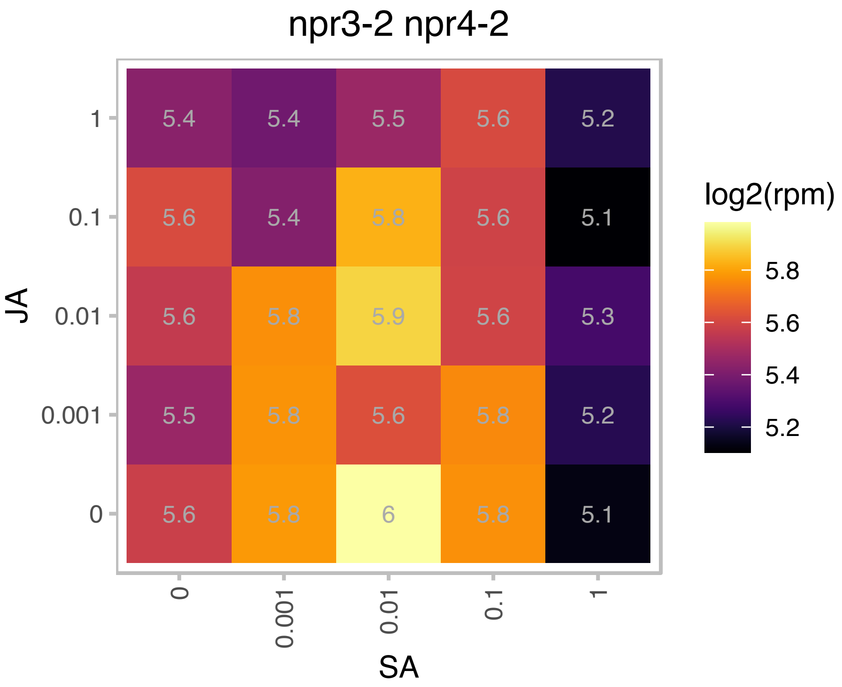


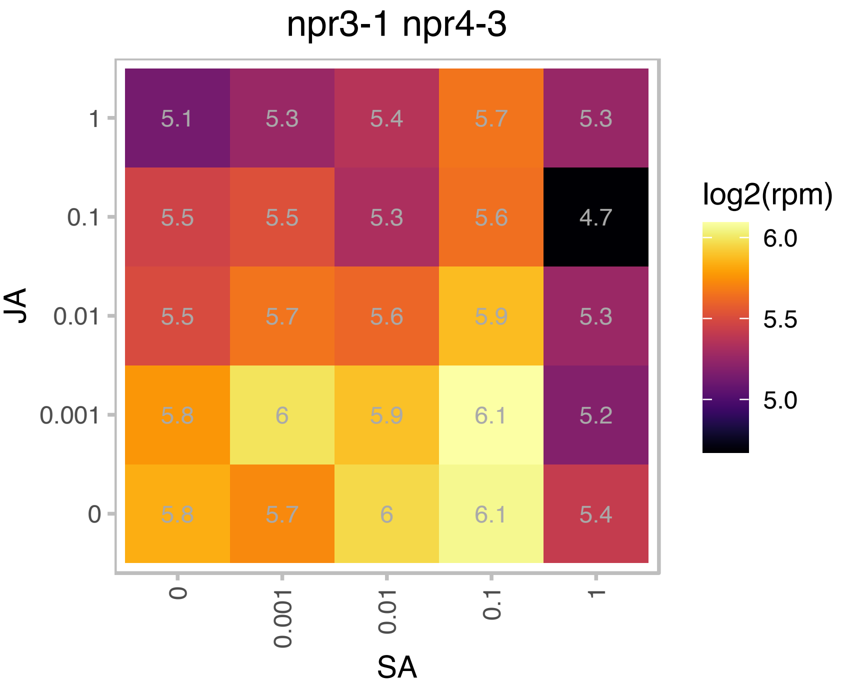

# AT2G33220.1 : NA

GRIM-19 protein

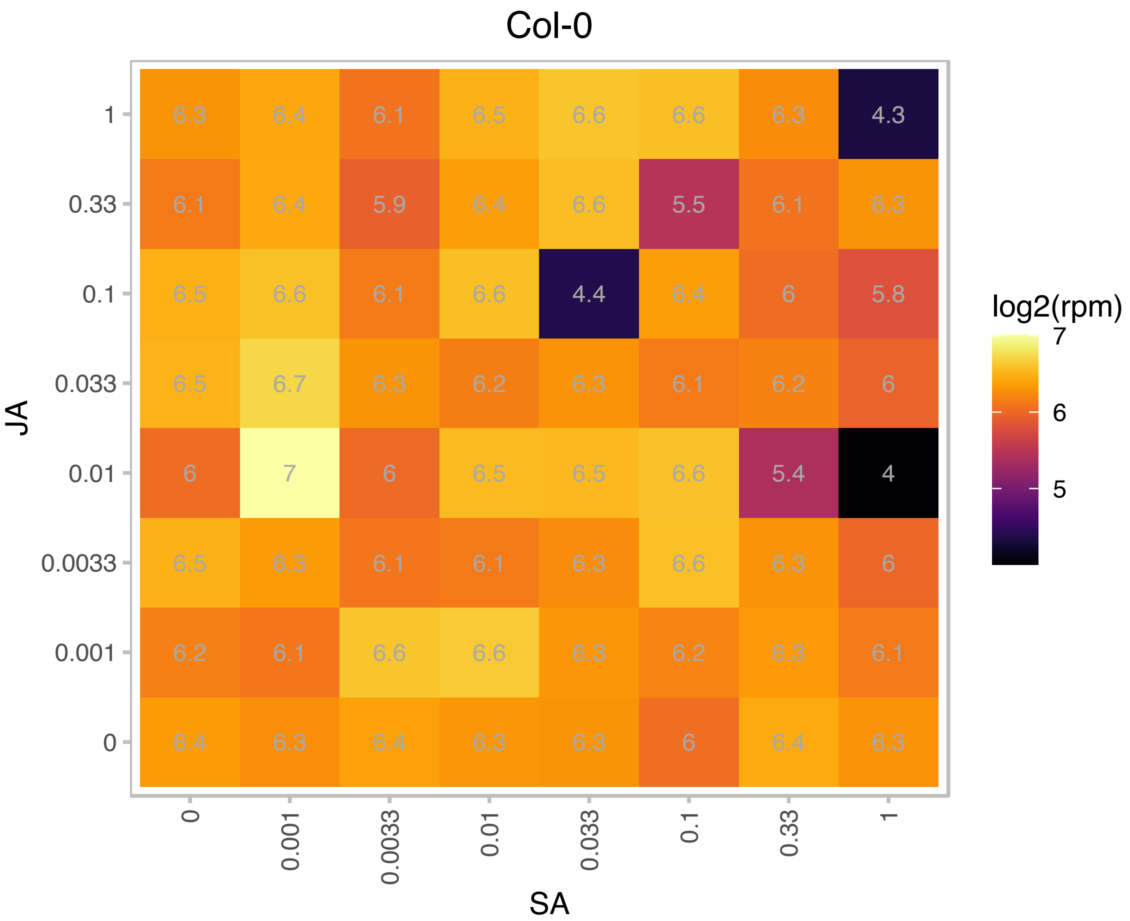


| | |
|---|---|
| Gene | AT2G33220 |
| is.expressed_genes | TRUE |
| is.responsive_genes | TRUE |
| is.apcluster | F-01 |

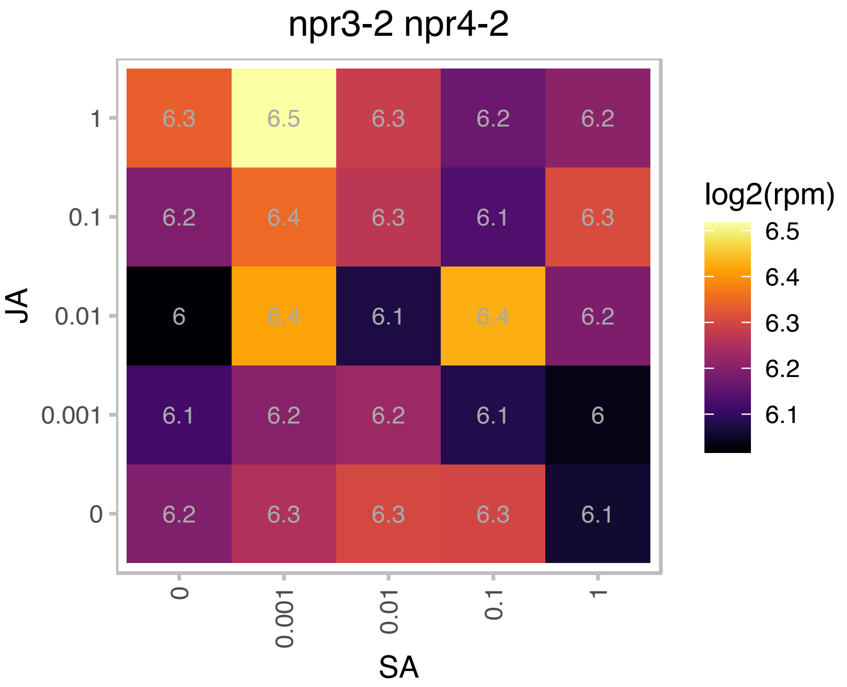


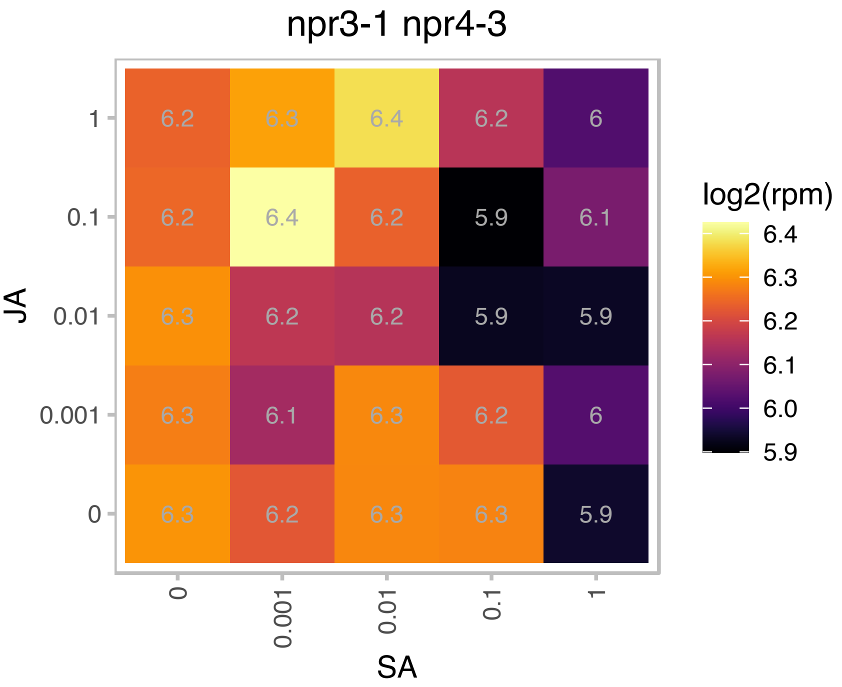

### AT2G34620.1 : mTERF10

Mitochondrial transcription termination factor family protein

Mitochondrial transcription termination factor family member.

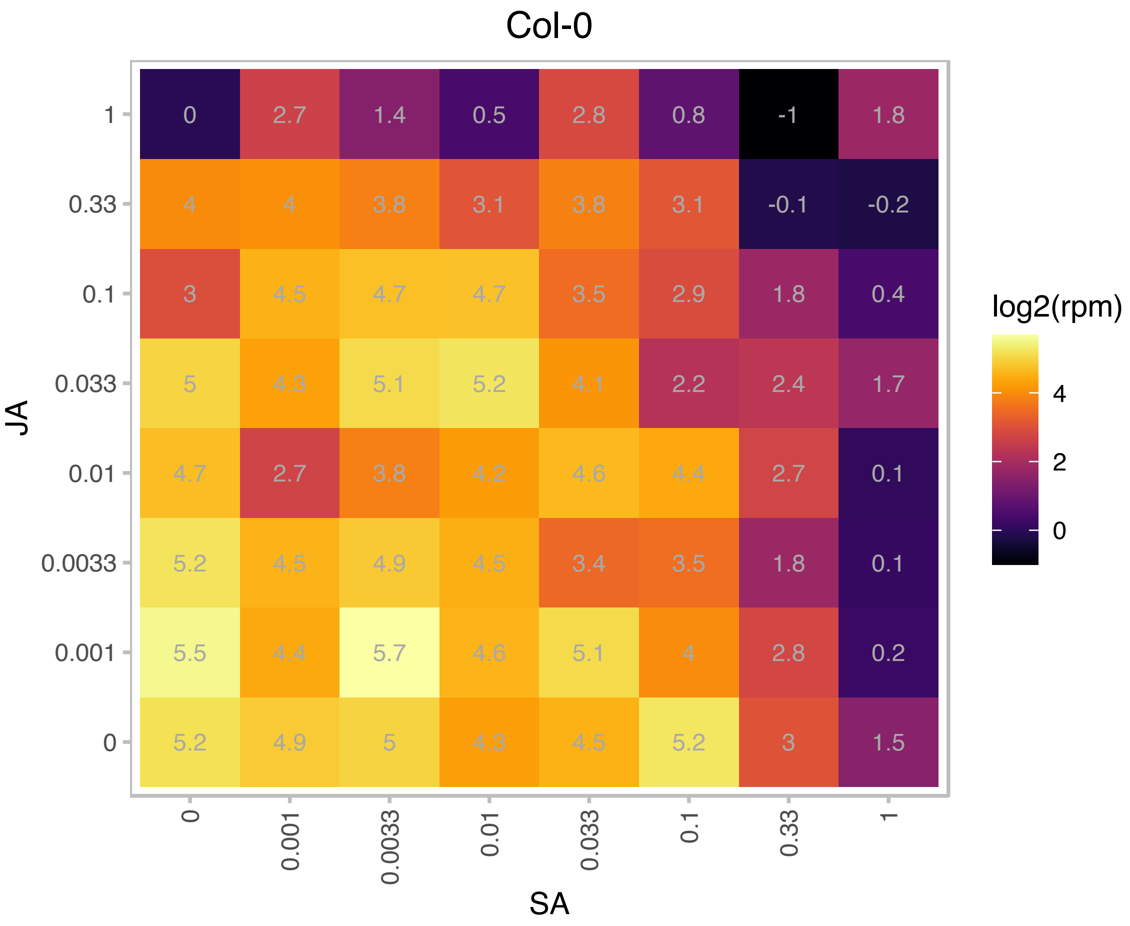


| | |
|---|---|
| Gene | AT2G34620 |
| is.expressed_genes | TRUE |
| is.responsive_genes | TRUE |
| is.apcluster | F-01 |

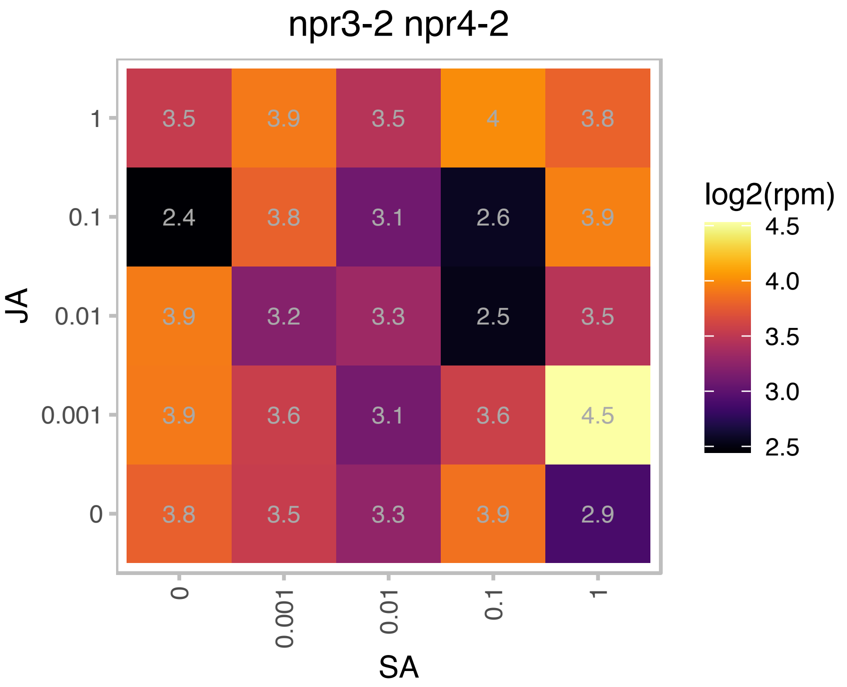


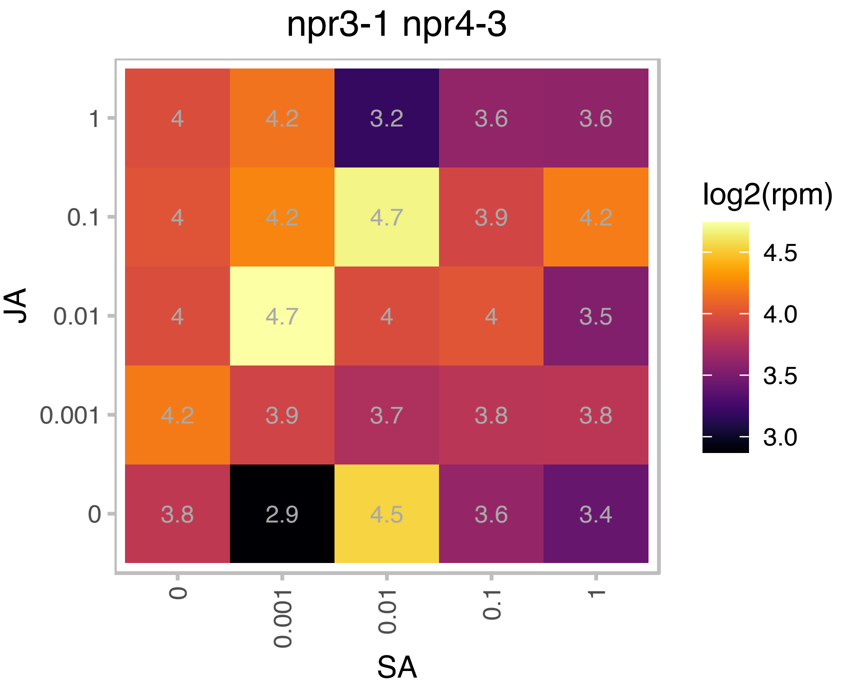

# AT2G36240.1 : NA

pentatricopeptide (PPR) repeat-containing protein

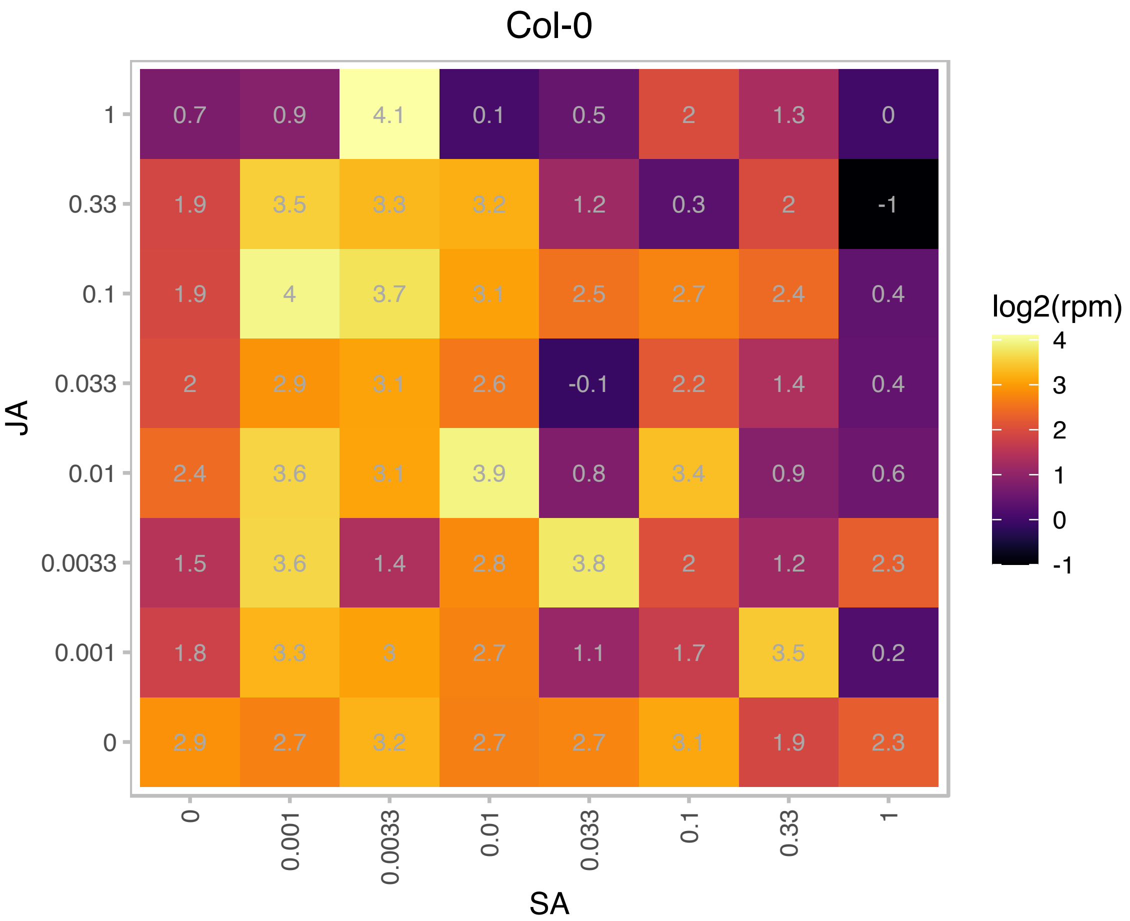


| | |
|---|---|
| Gene | AT2G36240 |
| is.expressed_genes | TRUE |
| is.responsive_genes | TRUE |
| is.apcluster | F-01 |

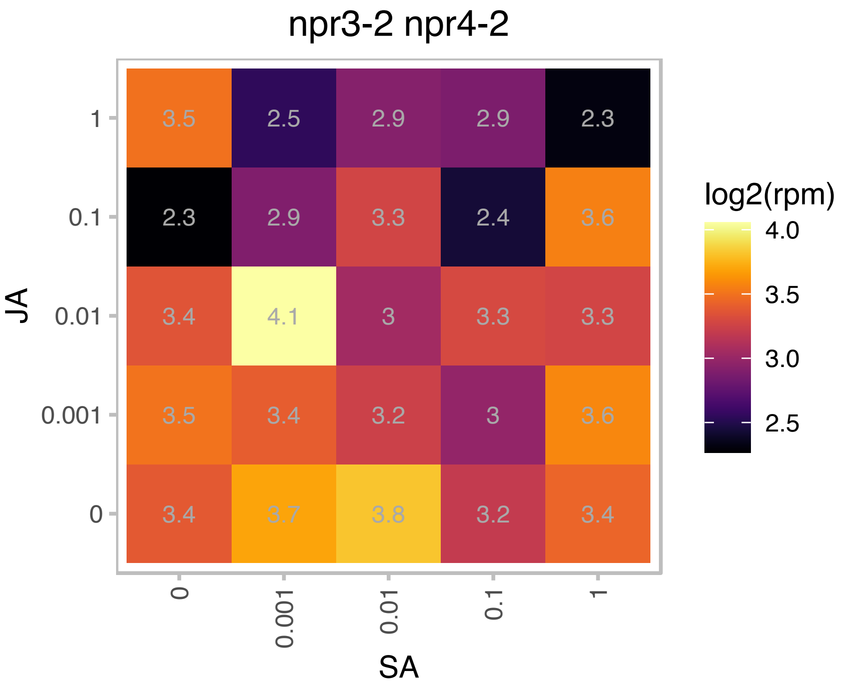


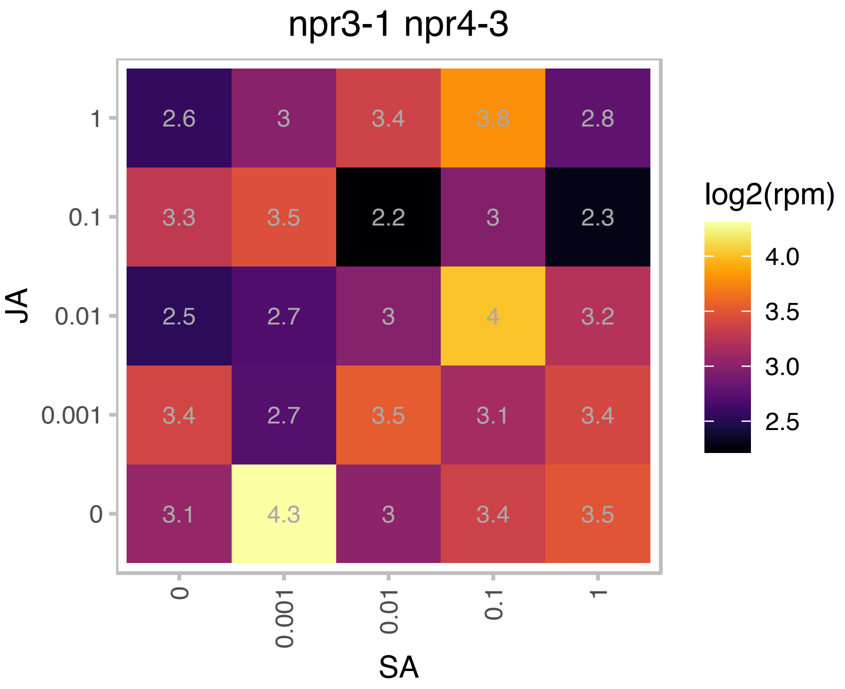

# AT2G38450.1 : NA

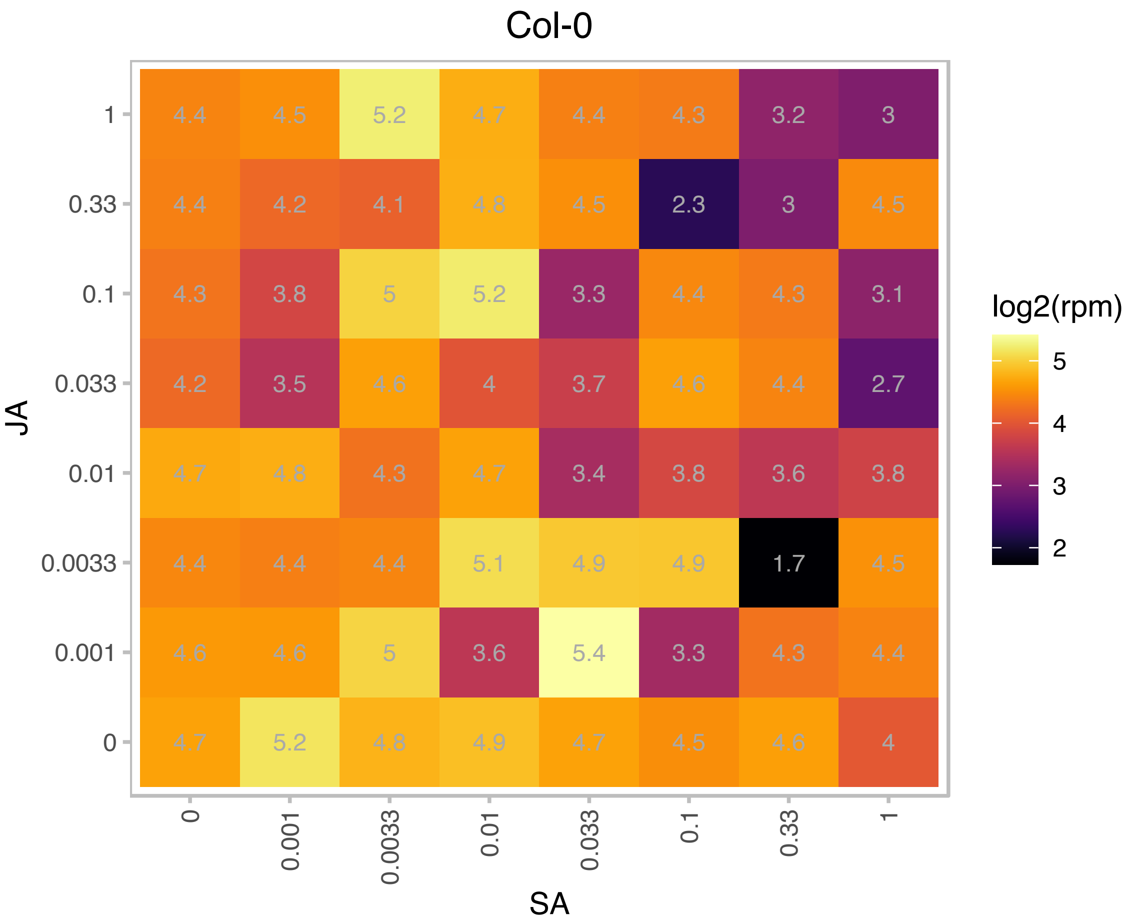


| | |
|---|---|
| Gene | AT2G38450 |
| is.expressed_genes | TRUE |
| is.responsive_genes | TRUE |
| is.apcluster | F-01 |

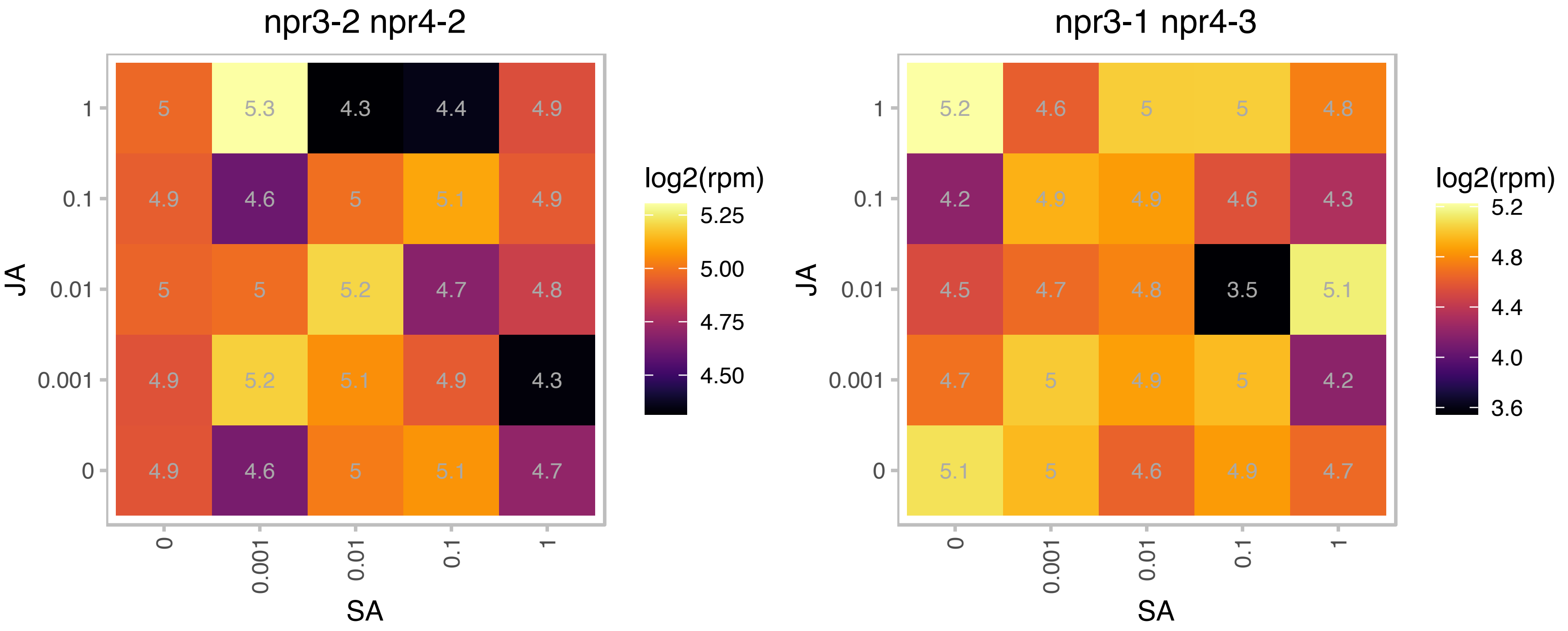

### AT2G39900.1 : WLIM2a

GATA type zinc finger transcription factor family protein
Encodes a member of the Arabidopsis LIM proteins: a family of actin bundlers with distinct expression patterns. WLIM1, WLIM2a, and WLIM2b are widely expressed, whereas PLIM2a, PLIM2b, and PLIM2c are predominantly expressed in pollen. Regulates actin cytoskeleton organization.

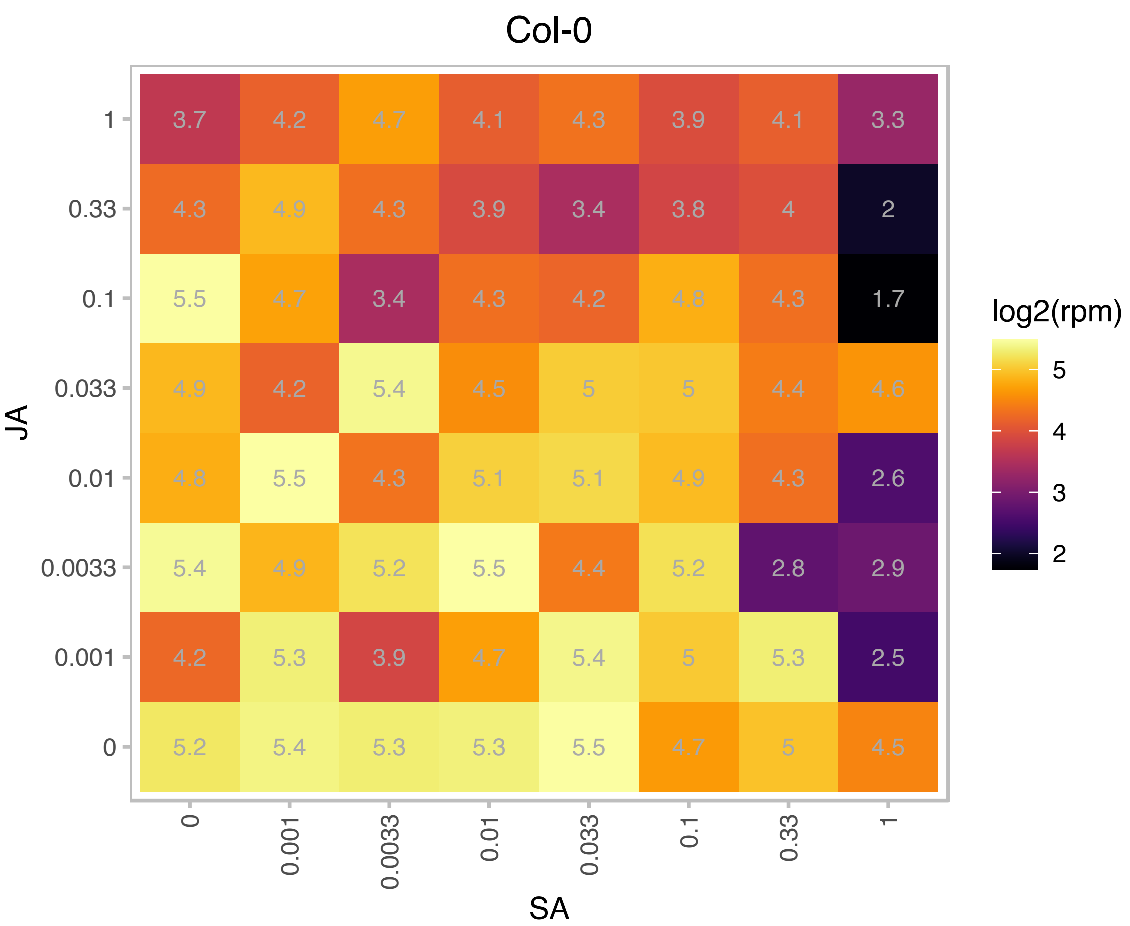


| | |
|---|---|
| Gene | AT2G39900 |
| is.expressed_genes | TRUE |
| is.responsive_genes | TRUE |
| is.apcluster | F-01 |

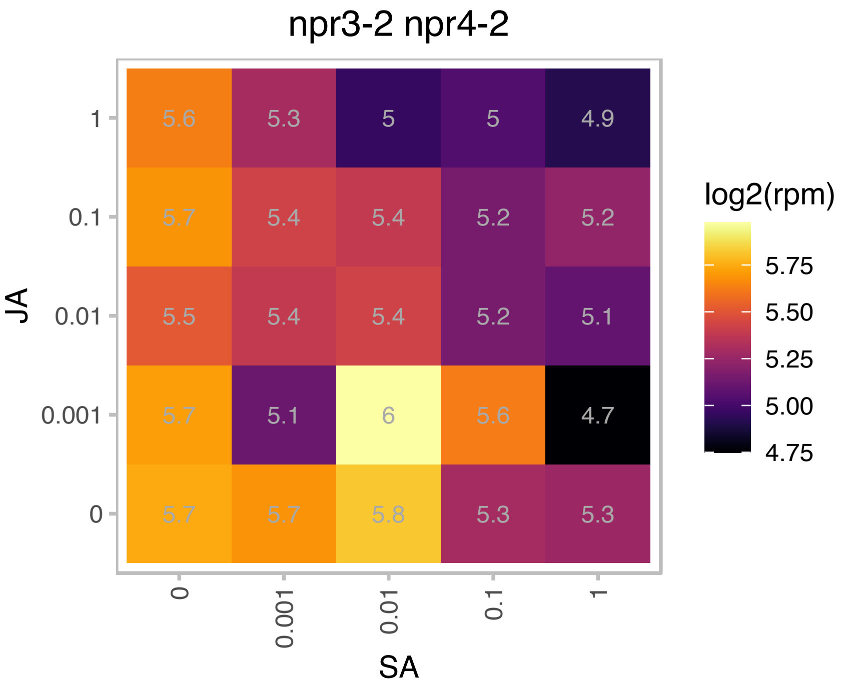


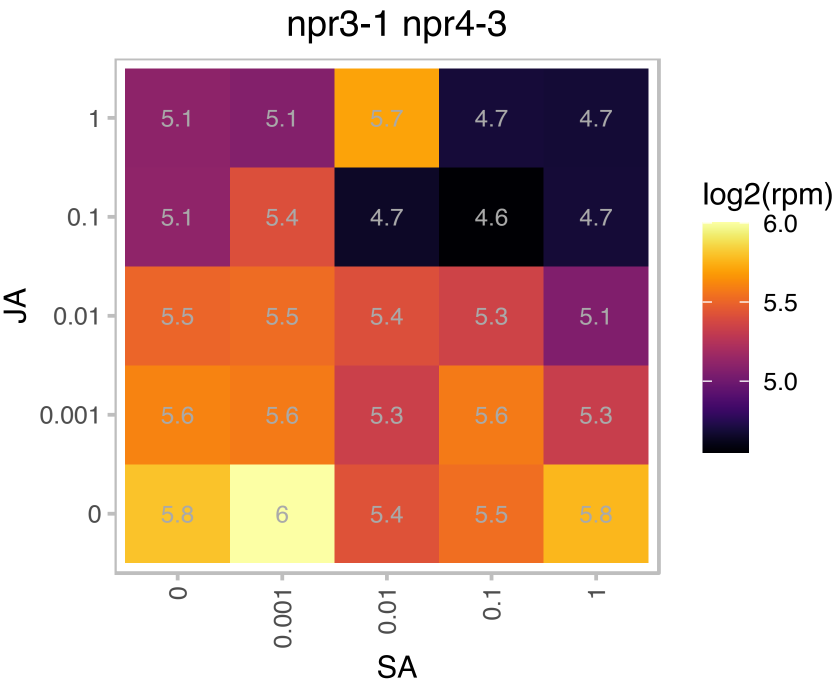

# AT2G40660.1 : NA

Nucleic acid-binding, OB-fold-like protein

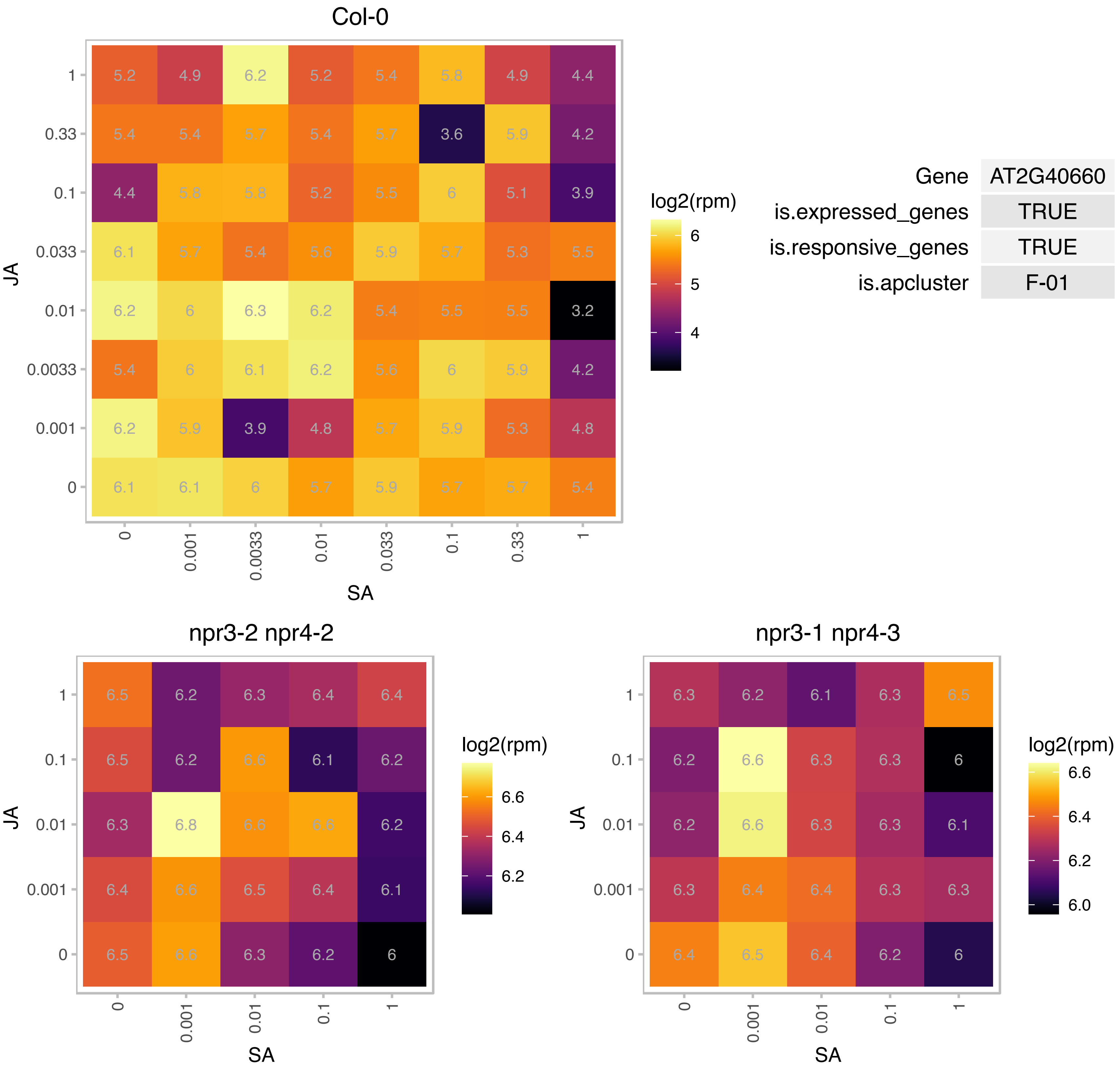


| | |
|---|---|
| Gene | AT2G40660 |
| is.expressed_genes | TRUE |
| is.responsive_genes | TRUE |
| is.apcluster | F-01 |

### AT2G41010.1 : ATCAMBP25, CAMBP25

calmodulin (CAM)-binding protein of 25 kDa

Encodes a novel calmodulin binding protein whose gene expression is induced by dehydration and ionic (salt) and non-ionic (mannitol) osmotic stress. Lines over-expressing this gene are more sensitive and anti-sense lines are more tolerant to osmotic stress, suggesting this gene may be a negative regulator of response to osmotic stress.

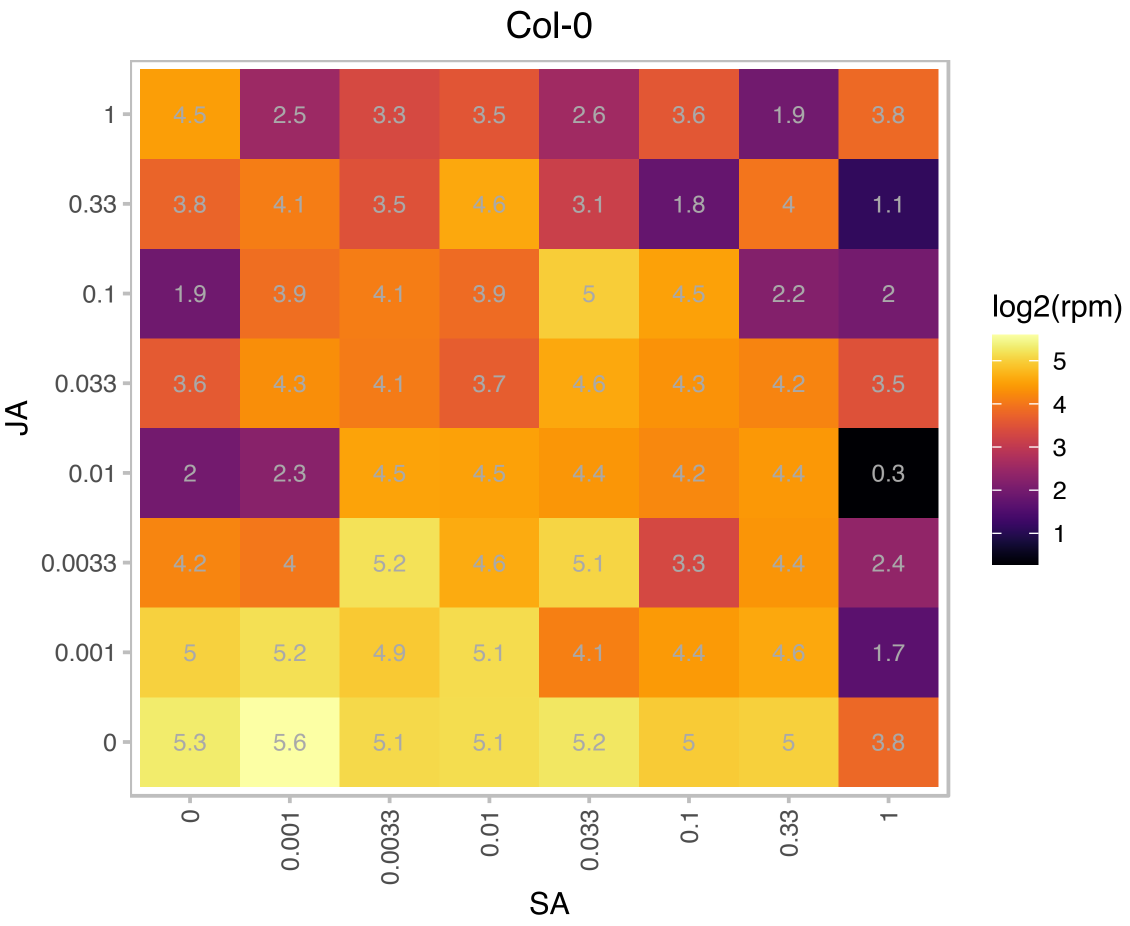


| | |
|---|---|
| Gene | AT2G41010 |
| is.expressed_genes | TRUE |
| is.responsive_genes | TRUE |
| is.apcluster | F-01 |

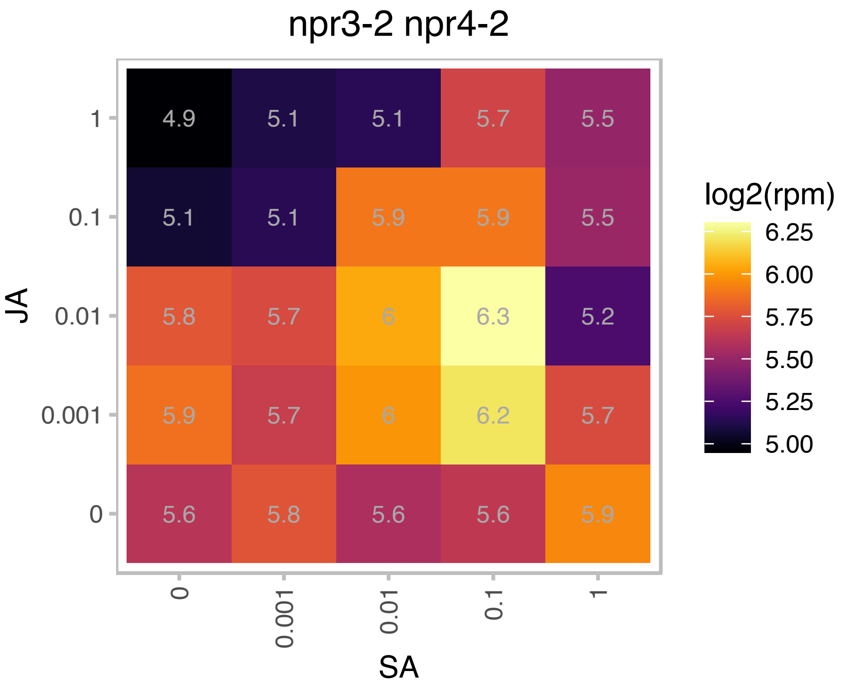


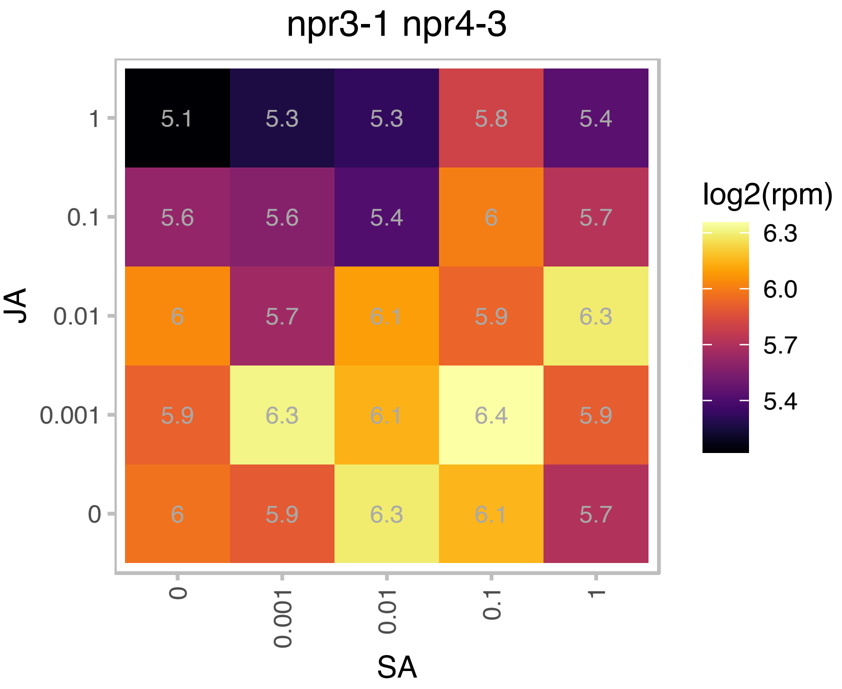

# AT2G44310.1 : NA

Calcium-binding EF-hand family protein

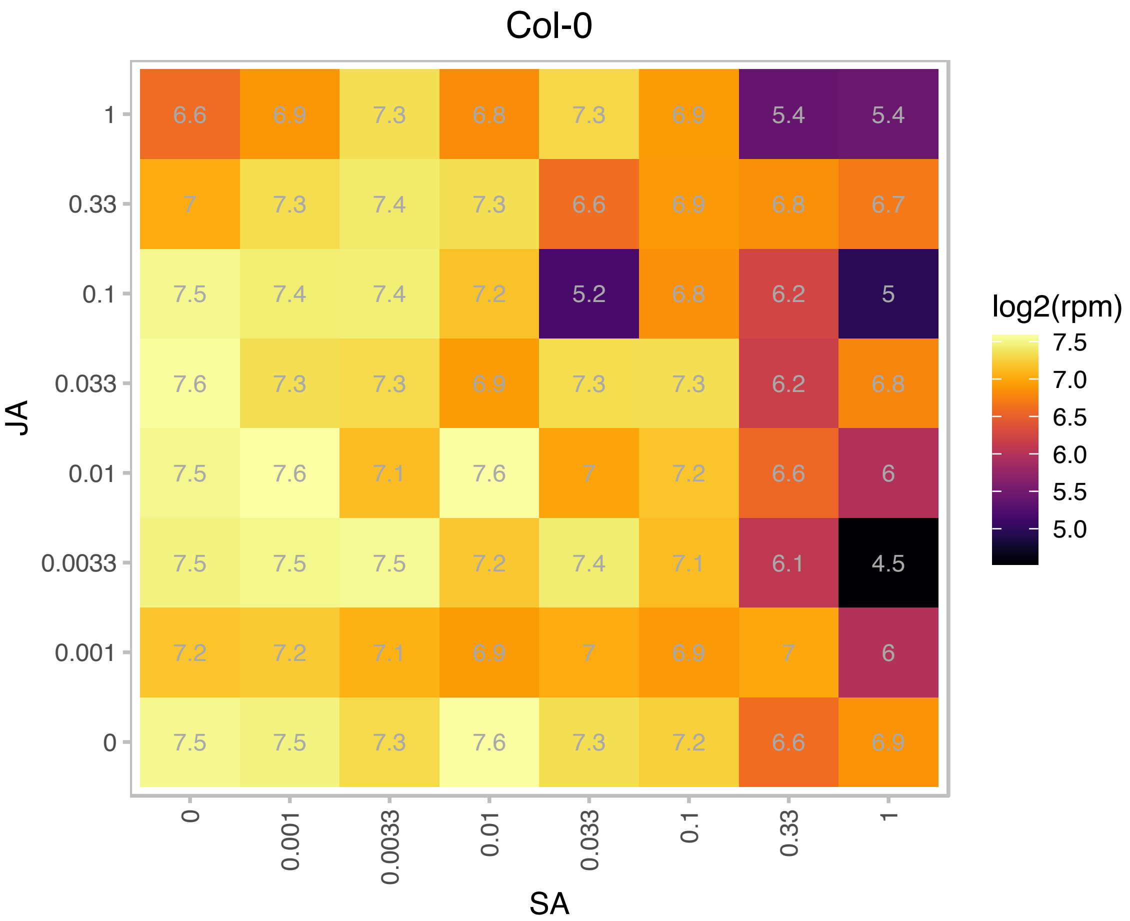


| | |
|---|---|
| Gene | AT2G44310 |
| is.expressed_genes | TRUE |
| is.responsive_genes | TRUE |
| is.apcluster | F-01 |

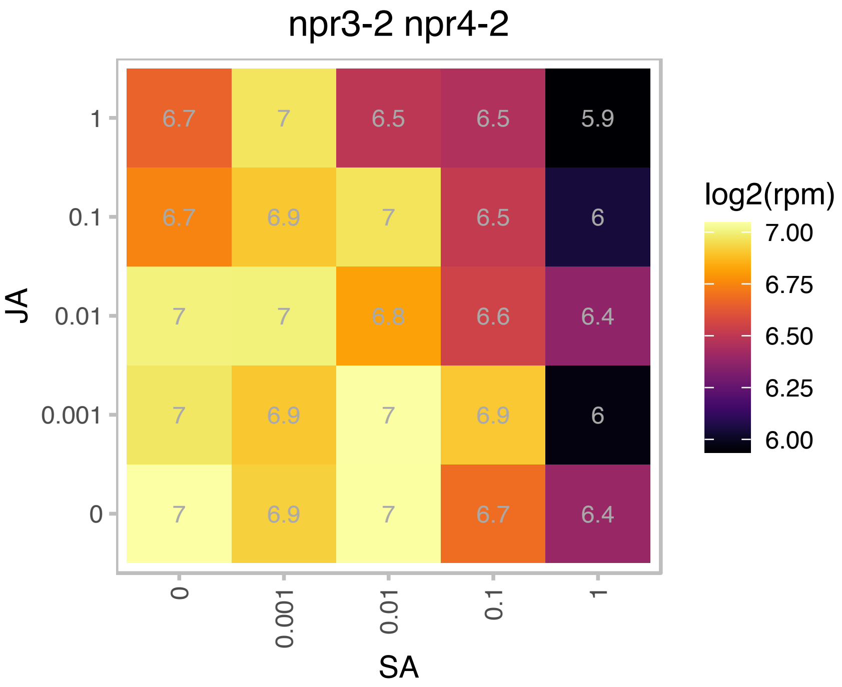


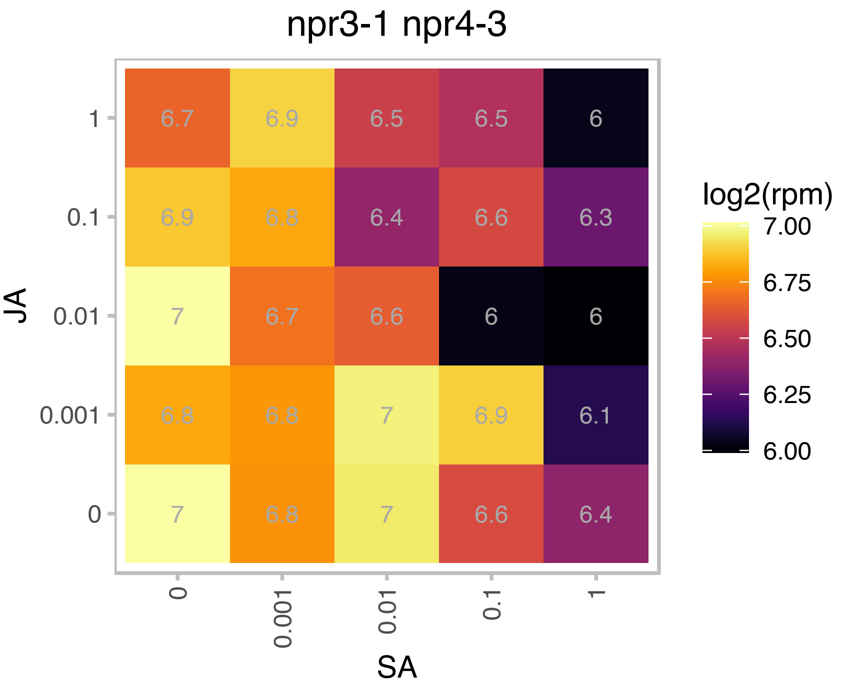

### AT2G47270.1 : UPB1

sequence-specific DNA binding transcription factors;transcription regulators

Encodes UPBEAT1 (UPB1), a transcription factor with a bHLH domain. Regulates the expression of a set of peroxidases that modulate the balance of reactive oxygen species (ROS) between the zones of cell proliferation and the zone of cell elongation where differentiation begins. Disruption of UPB1 activity alters this ROS balance, leading to a delay in the onset of differentiation.

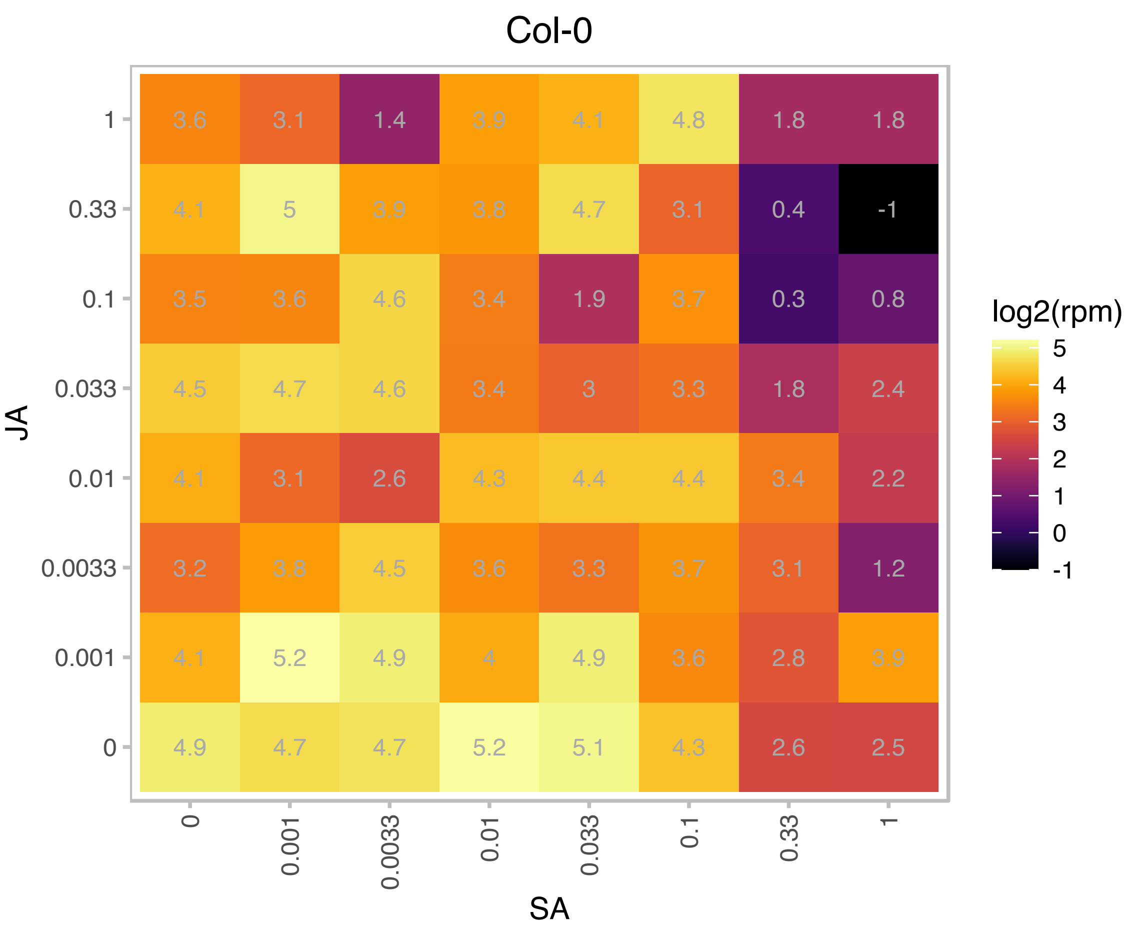


| | |
|---|---|
| Gene | AT2G47270 |
| is.expressed_genes | TRUE |
| is.responsive_genes | TRUE |
| is.apcluster | F-01 |

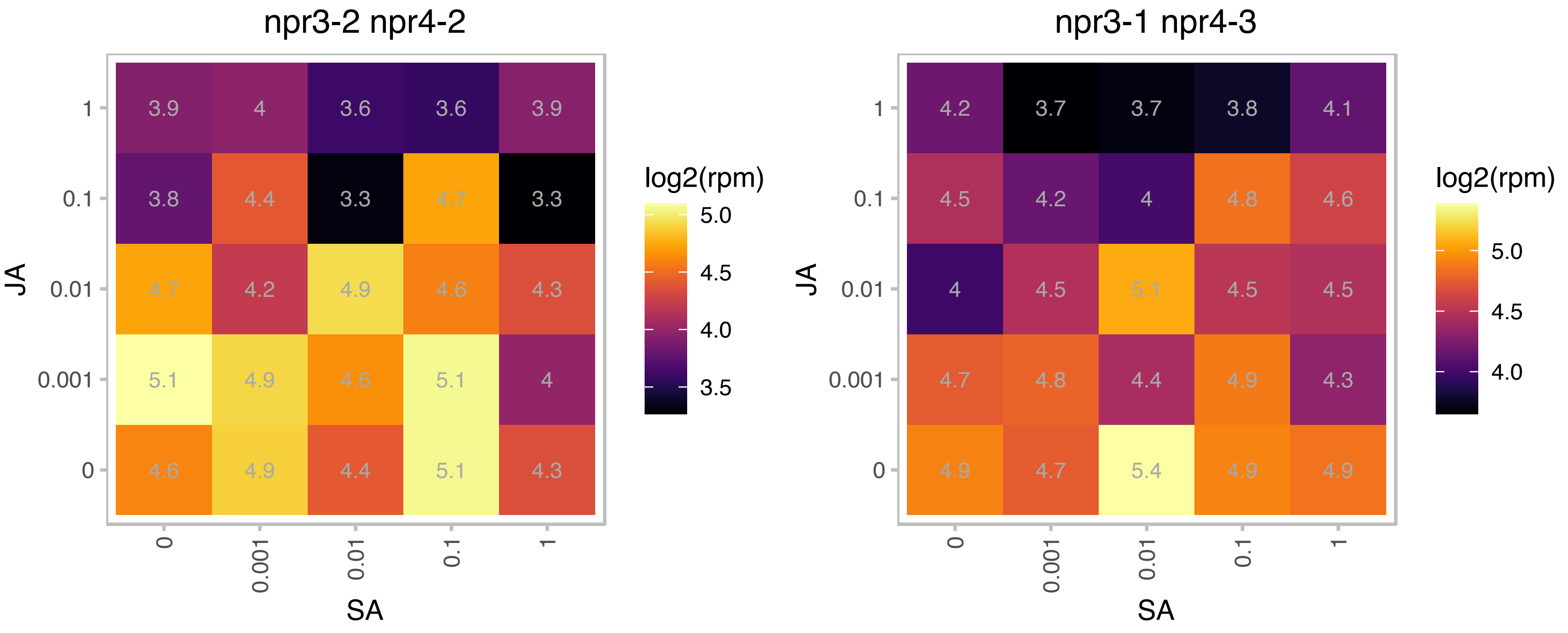

### AT3G01370.1 : ATCFM2, CFM2

CRM family member 2

Encodes a protein containing a CRM domain that is involved in group I and group II intron splicing.

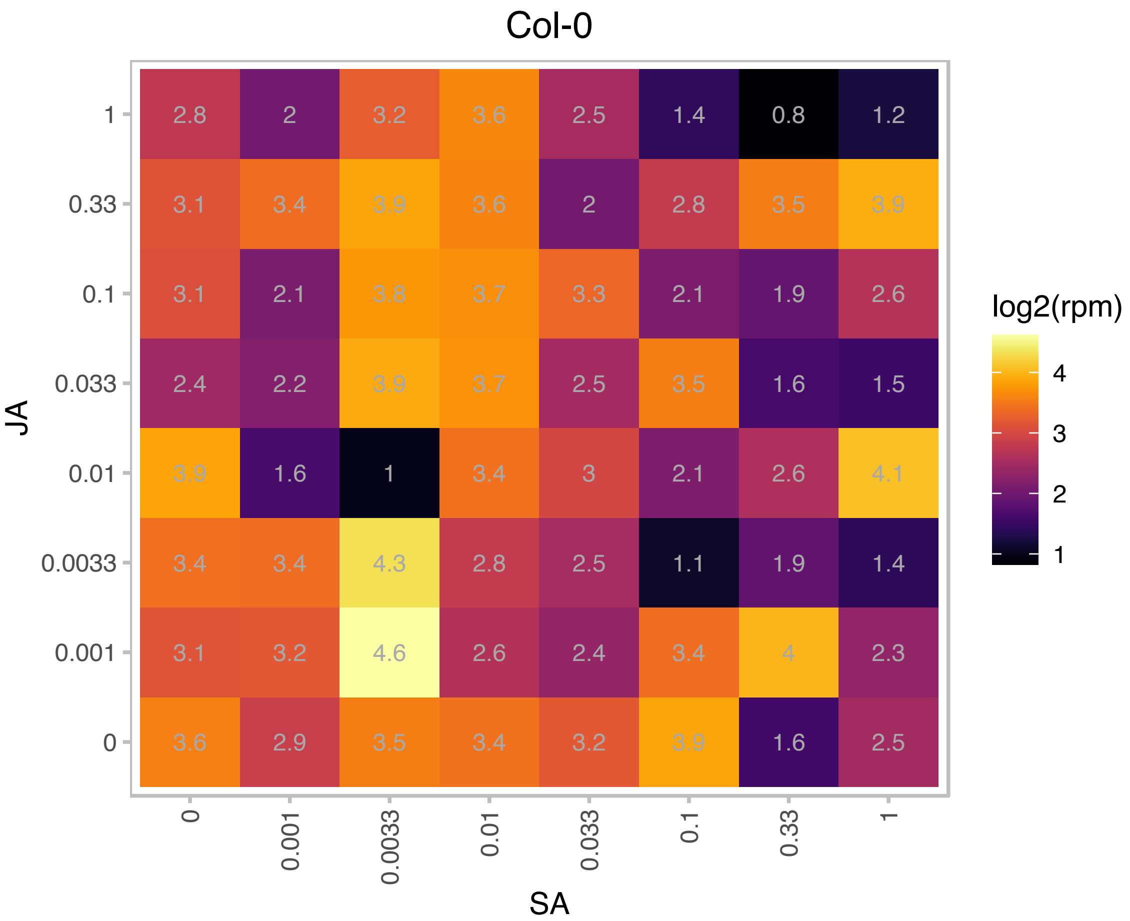


| | |
|---|---|
| Gene | AT3G01370 |
| is.expressed_genes | TRUE |
| is.responsive_genes | TRUE |
| is.apcluster | F-01 |

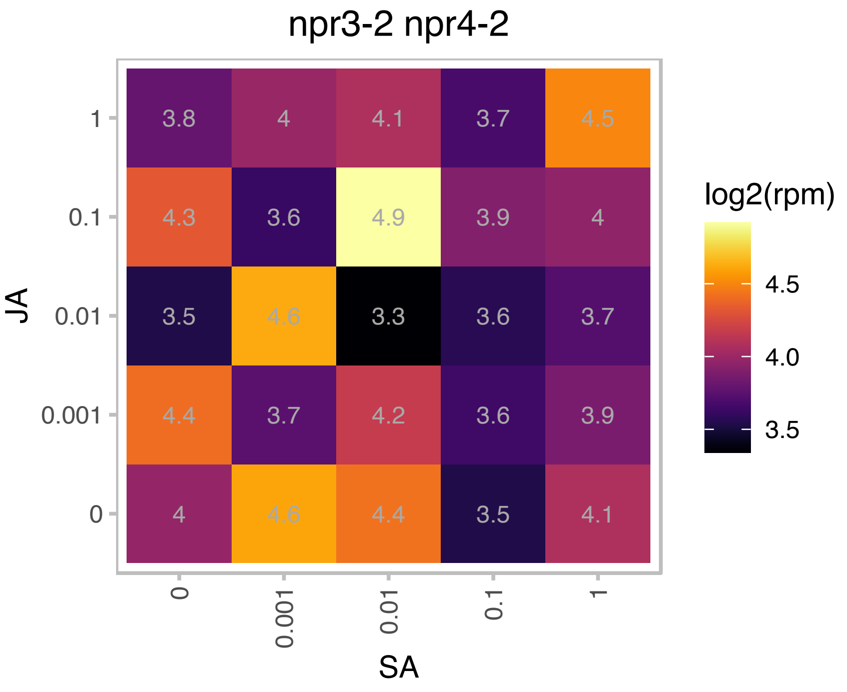


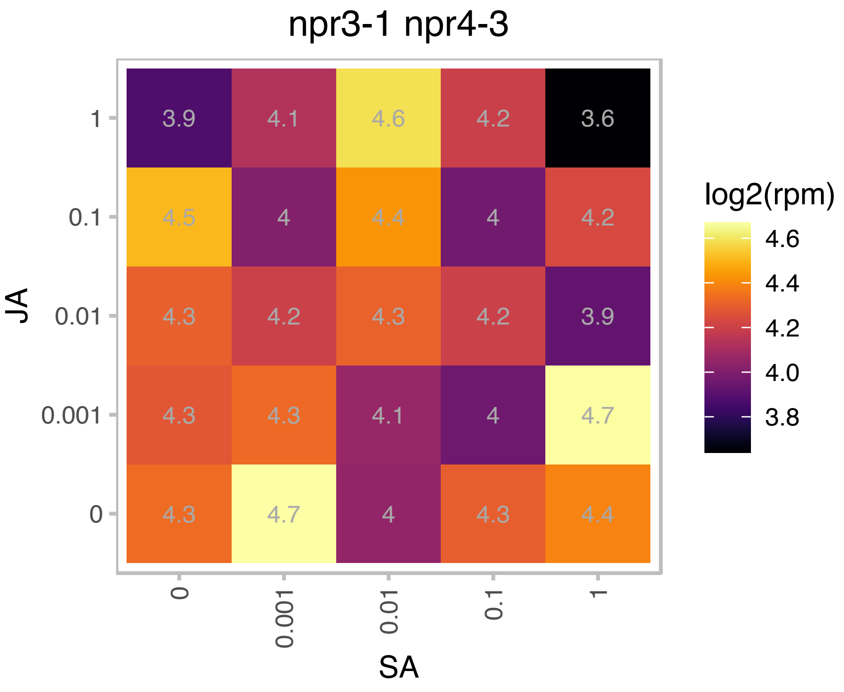

# AT3G07310.1 : NA

Protein of unknown function (DUF760)

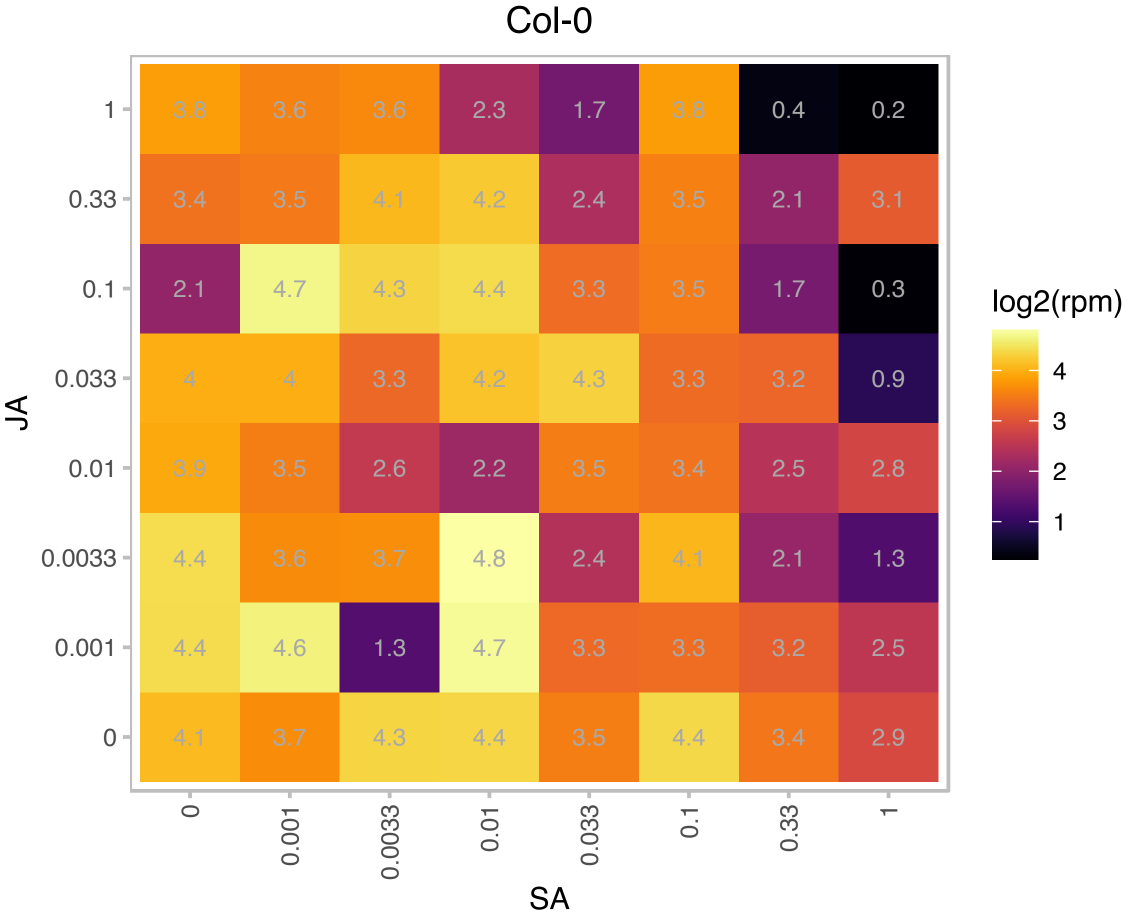


| | |
|---|---|
| Gene | AT3G07310 |
| is.expressed_genes | TRUE |
| is.responsive_genes | TRUE |
| is.apcluster | F-01 |

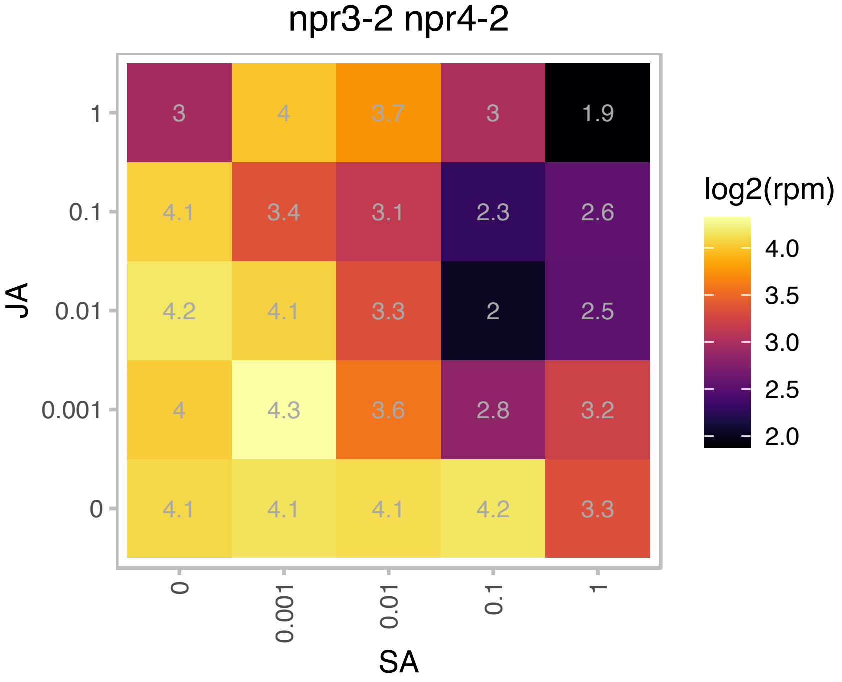


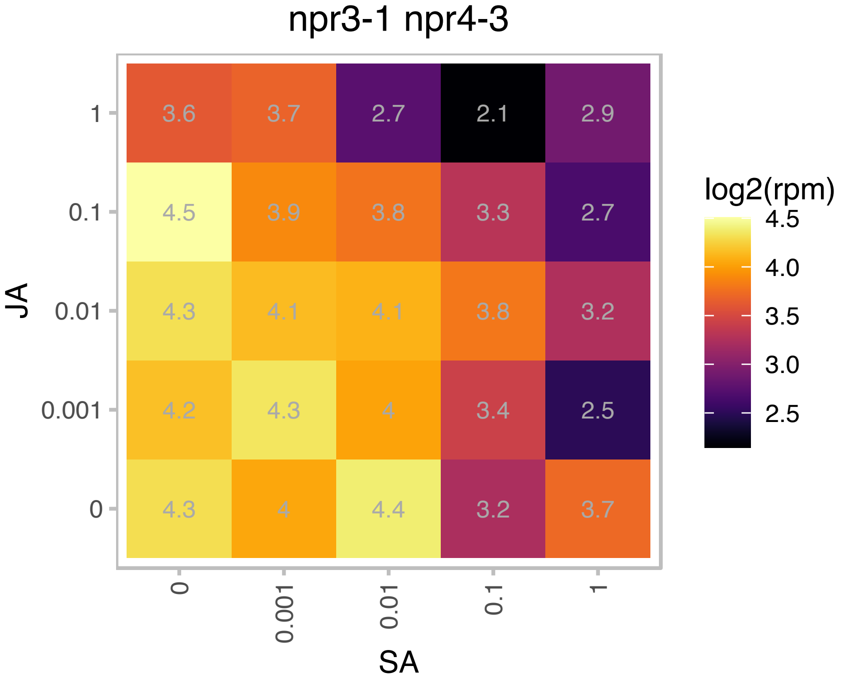

### AT3G14930.1 : HEME1

Uroporphyrinogen decarboxylase

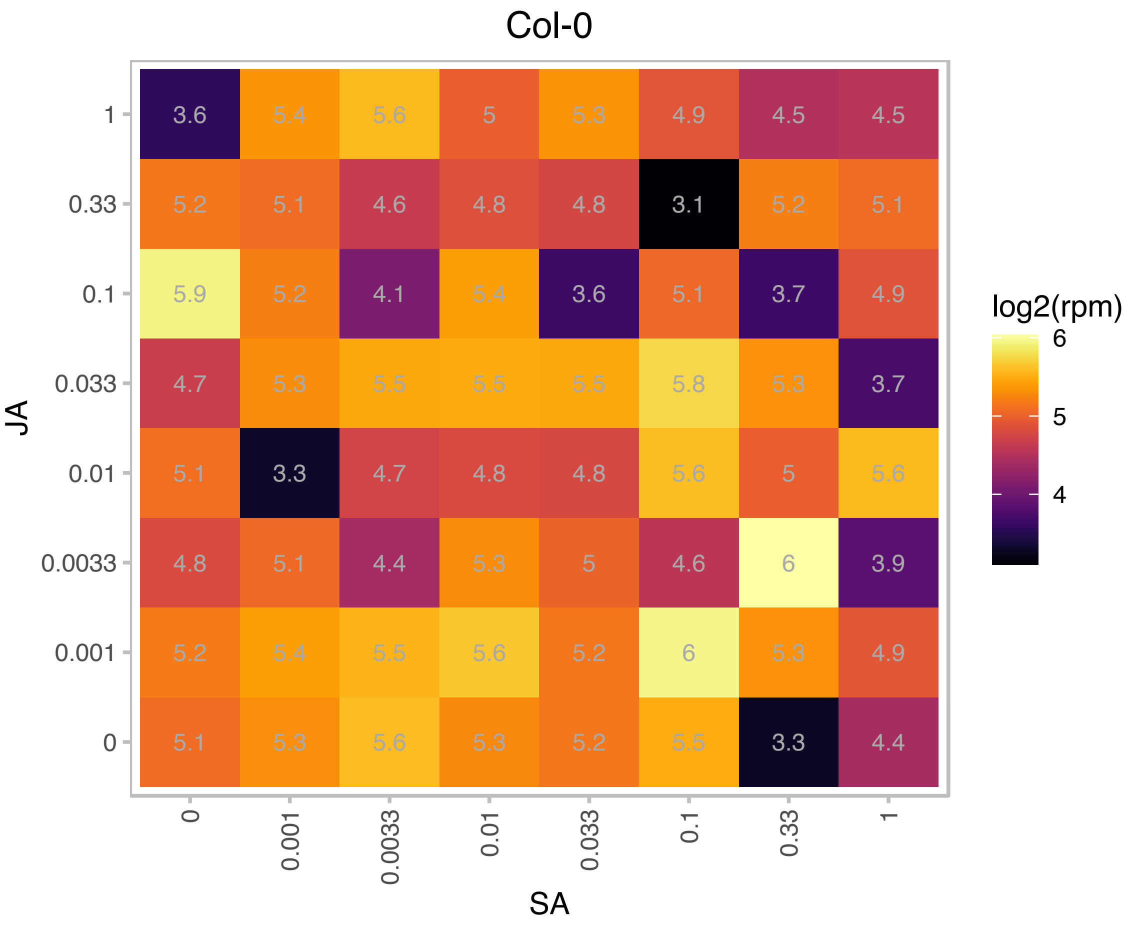


| | |
|---|---|
| Gene | AT3G14930 |
| is.expressed_genes | TRUE |
| is.responsive_genes | TRUE |
| is.apcluster | F-01 |

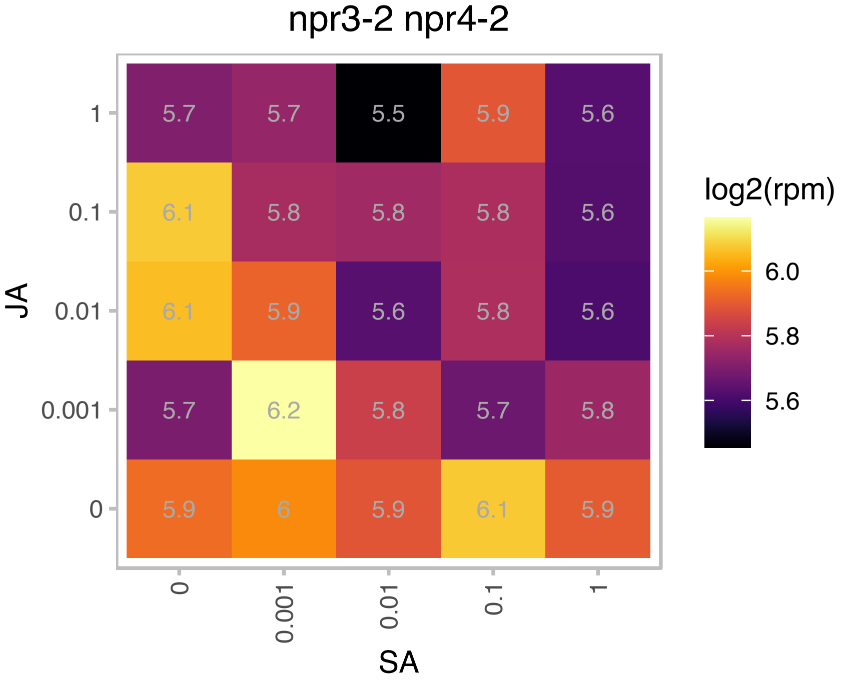


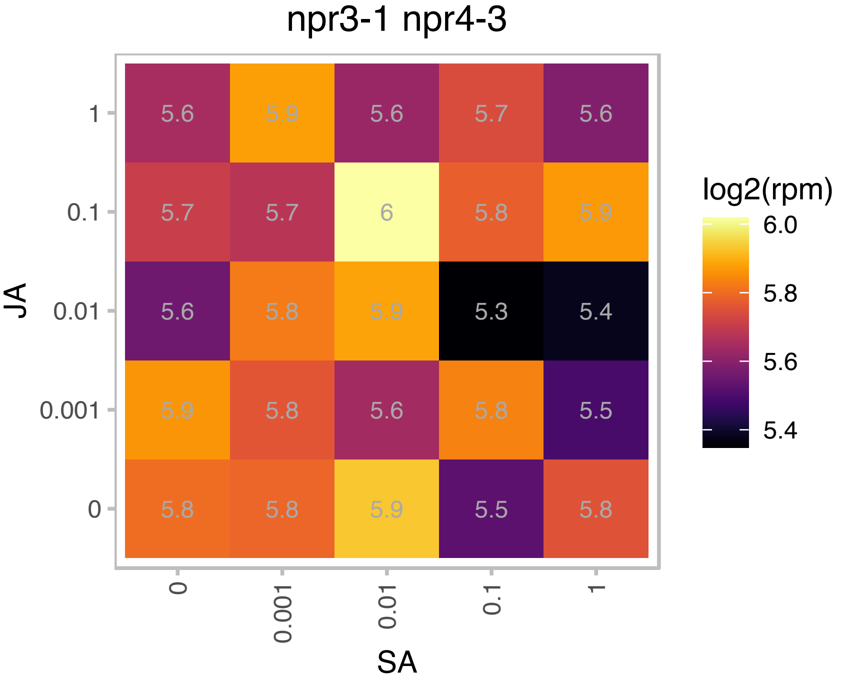

# AT3G16660.1 : NA

Pollen Ole e 1 allergen and extensin family protein

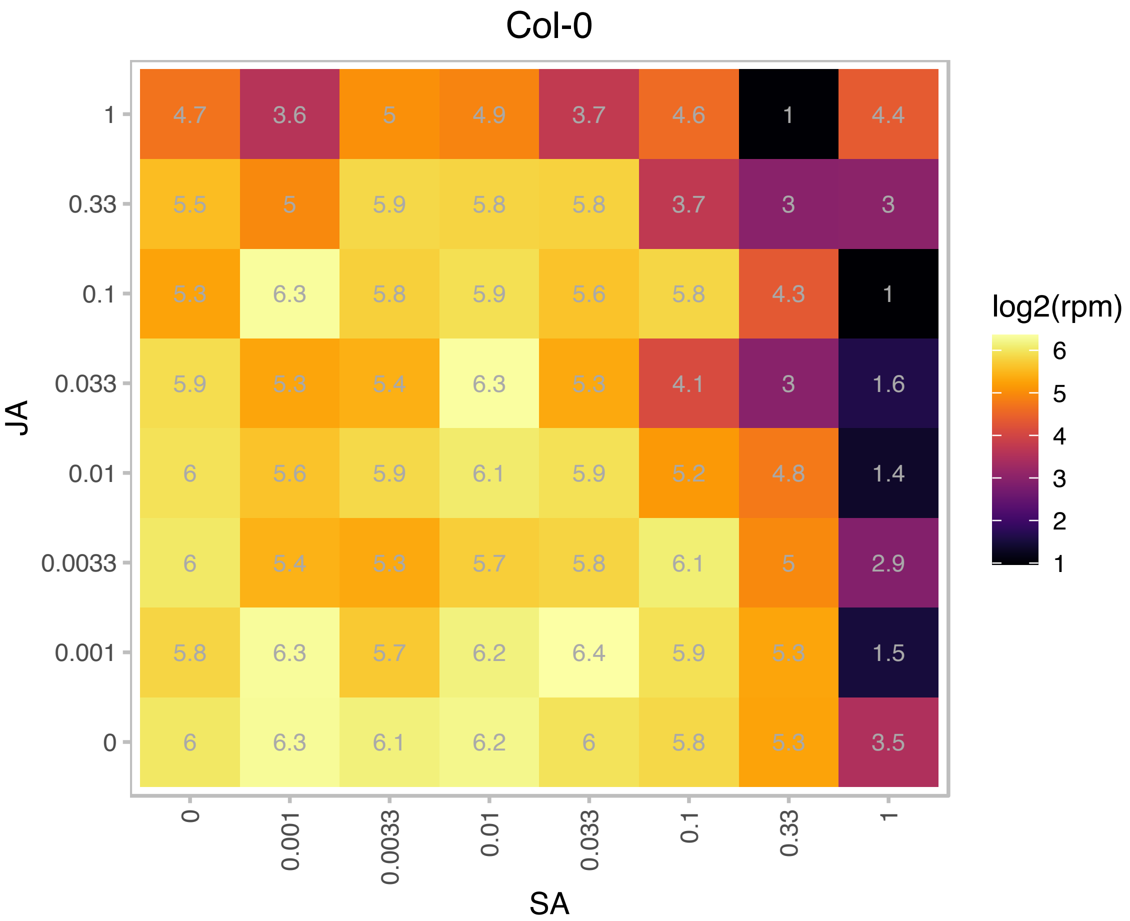


| | |
|---|---|
| Gene | AT3G16660 |
| is.expressed_genes | TRUE |
| is.responsive_genes | TRUE |
| is.apcluster | F-01 |

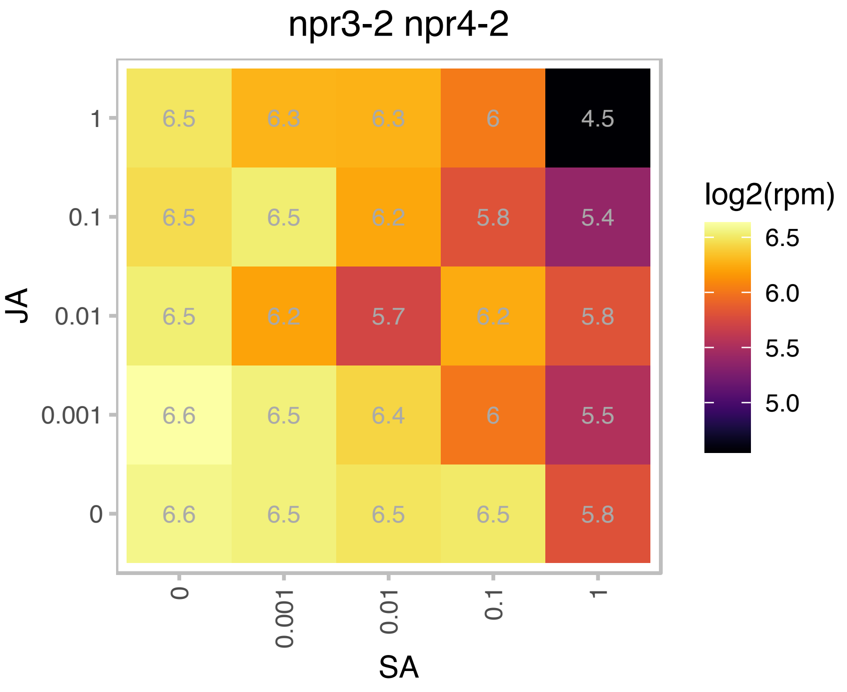


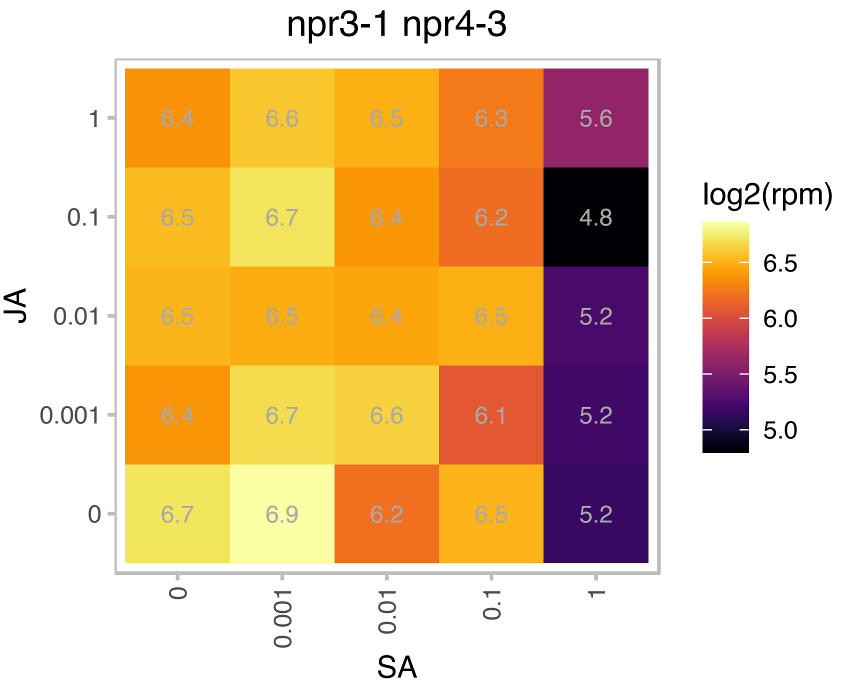

# AT3G16670.1 : NA

Pollen Ole e 1 allergen and extensin family protein

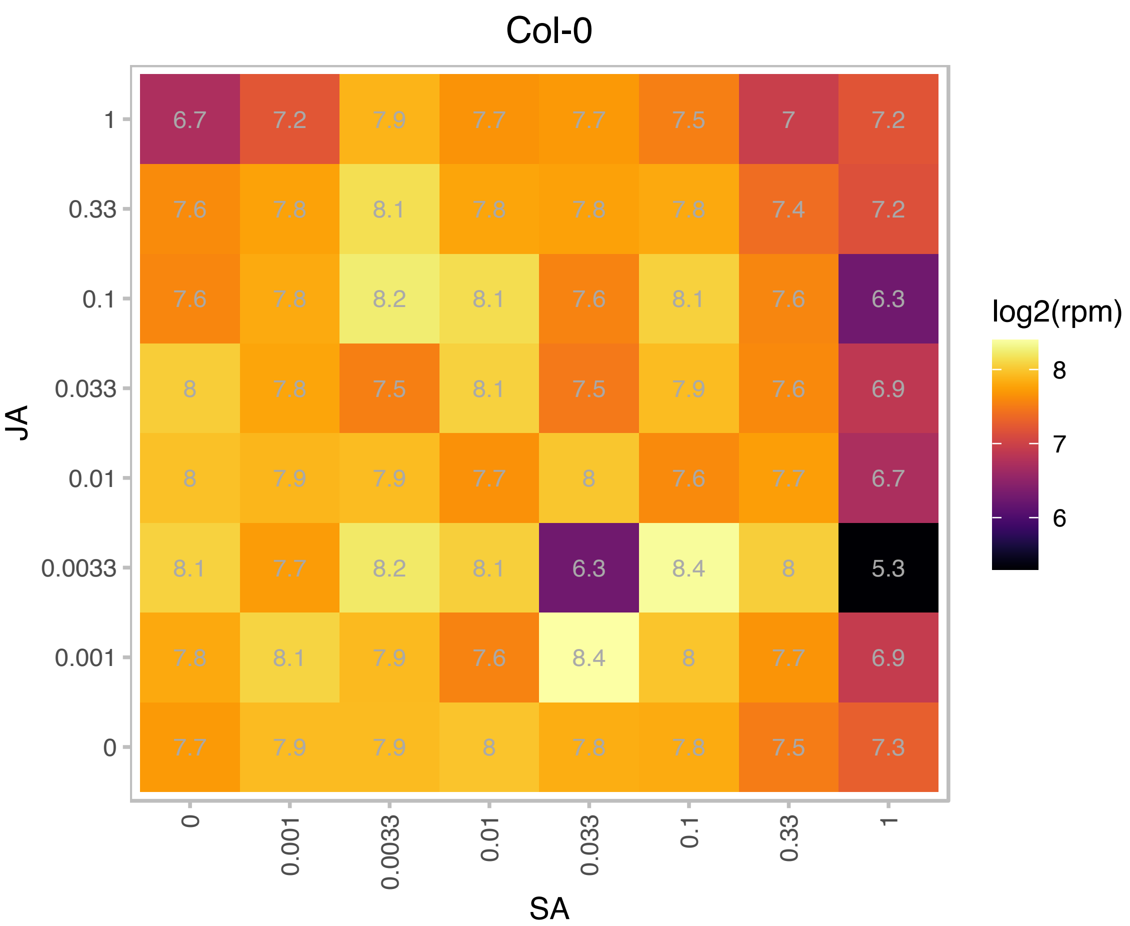


| | |
|---|---|
| Gene | AT3G16670 |
| is.expressed_genes | TRUE |
| is.responsive_genes | TRUE |
| is.apcluster | F-01 |

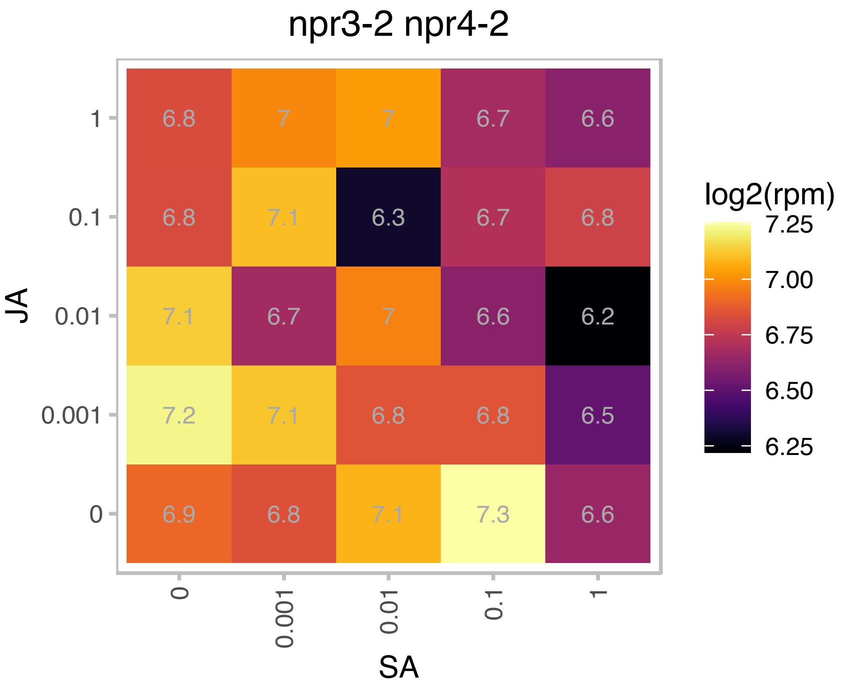


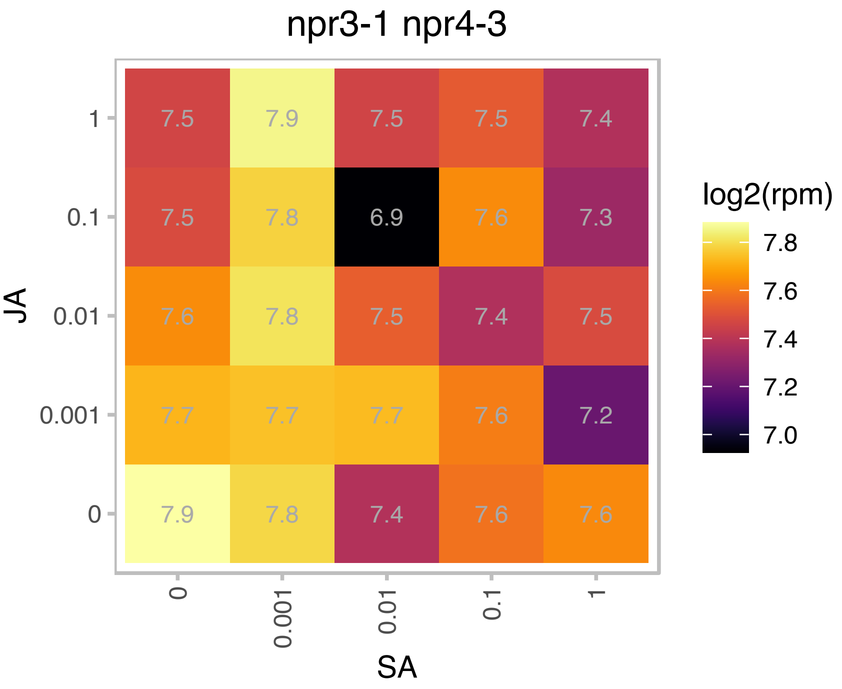

### AT3G18080.1 : BGLU44

B-S glucosidase 44

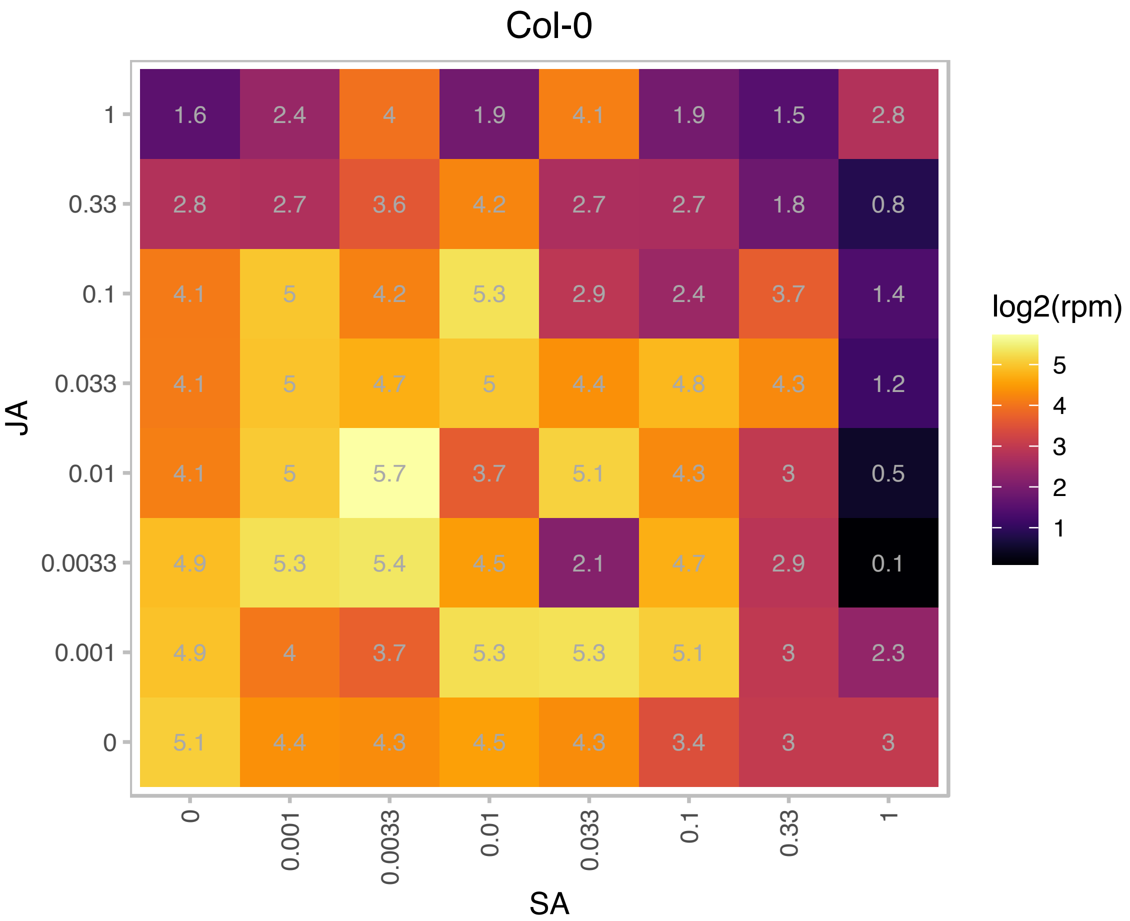


| | |
|---|---|
| Gene | AT3G18080 |
| is.expressed_genes | TRUE |
| is.responsive_genes | TRUE |
| is.apcluster | F-01 |

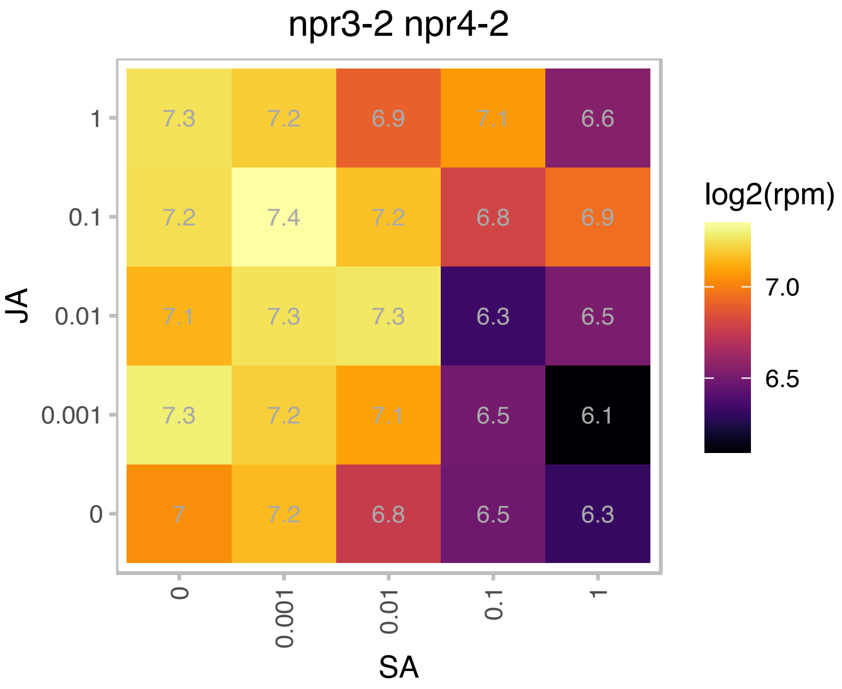


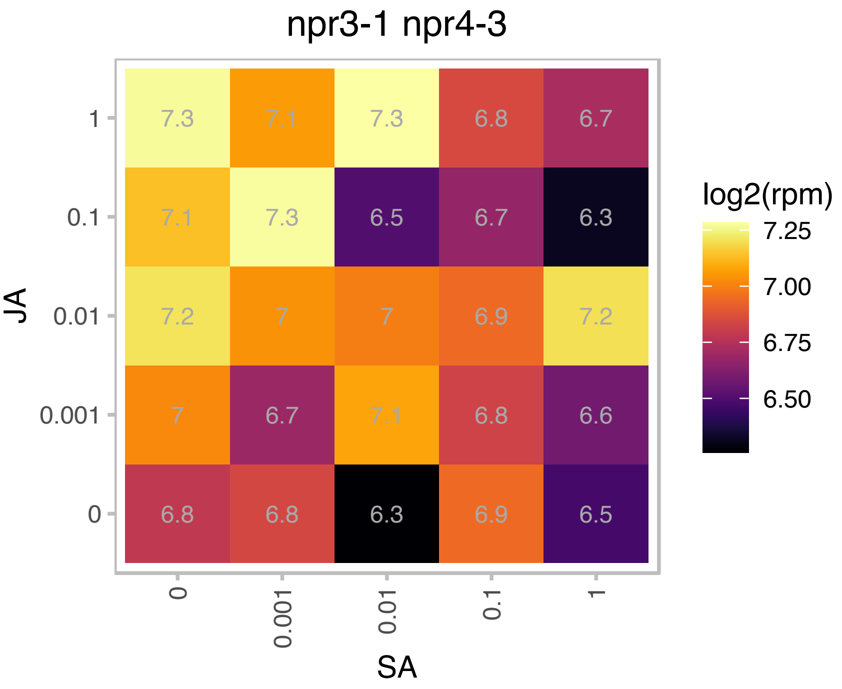

### AT3G19800.1 : DUF177B

Protein of unknown function (DUF177)

Encodes the DUF177B version of the two DUF177 proteins in Arabidopsis.  This version differs from DUF177A in containing a 23 aa insertion compared to the DUF177A sequence.

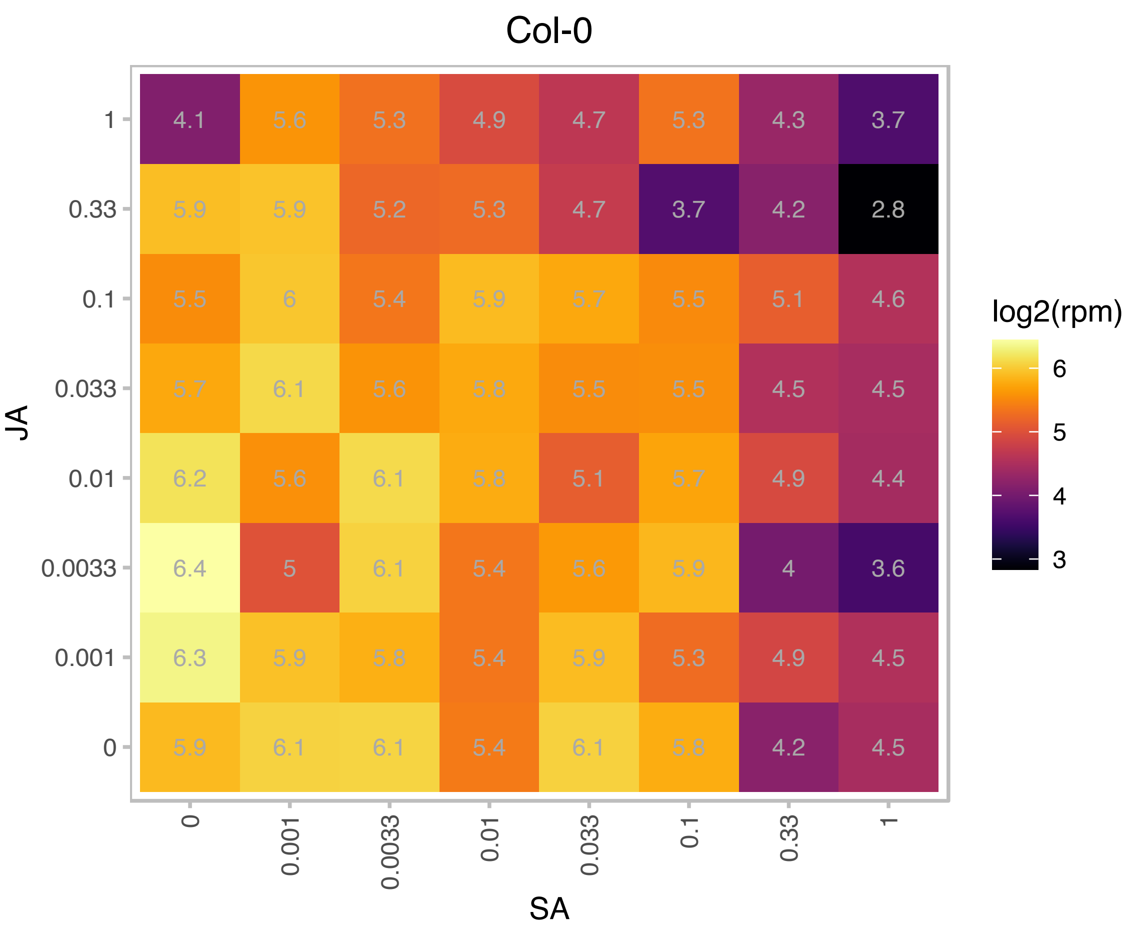


| | |
|---|---|
| Gene | AT3G19800 |
| is.expressed_genes | TRUE |
| is.responsive_genes | TRUE |
| is.apcluster | F-01 |

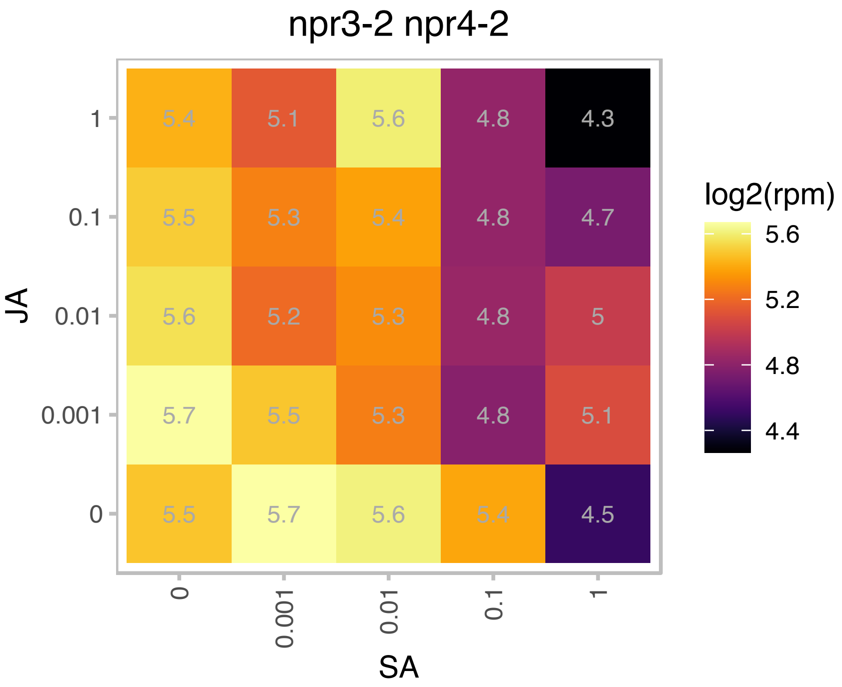


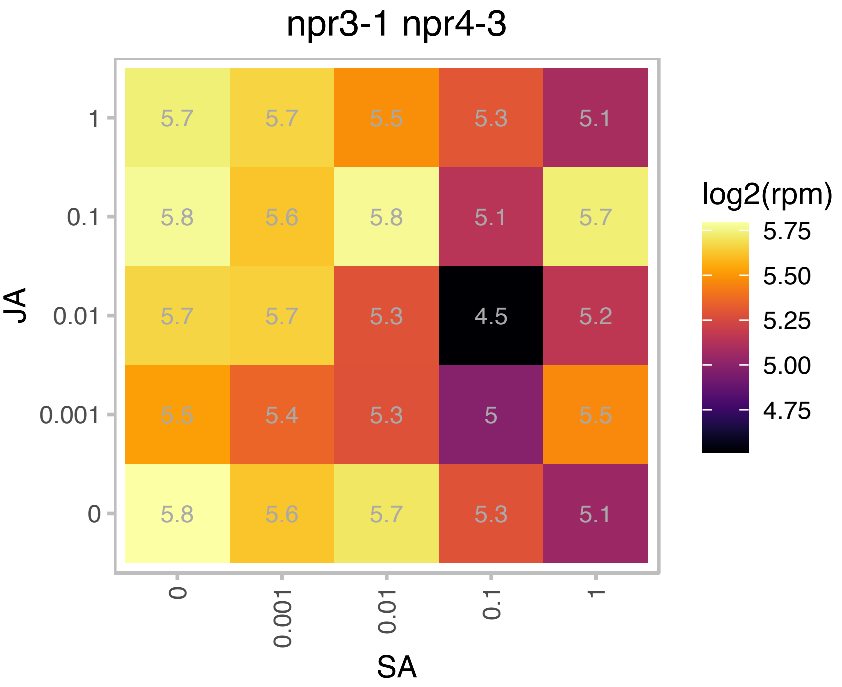

# AT3G22210.1 : NA

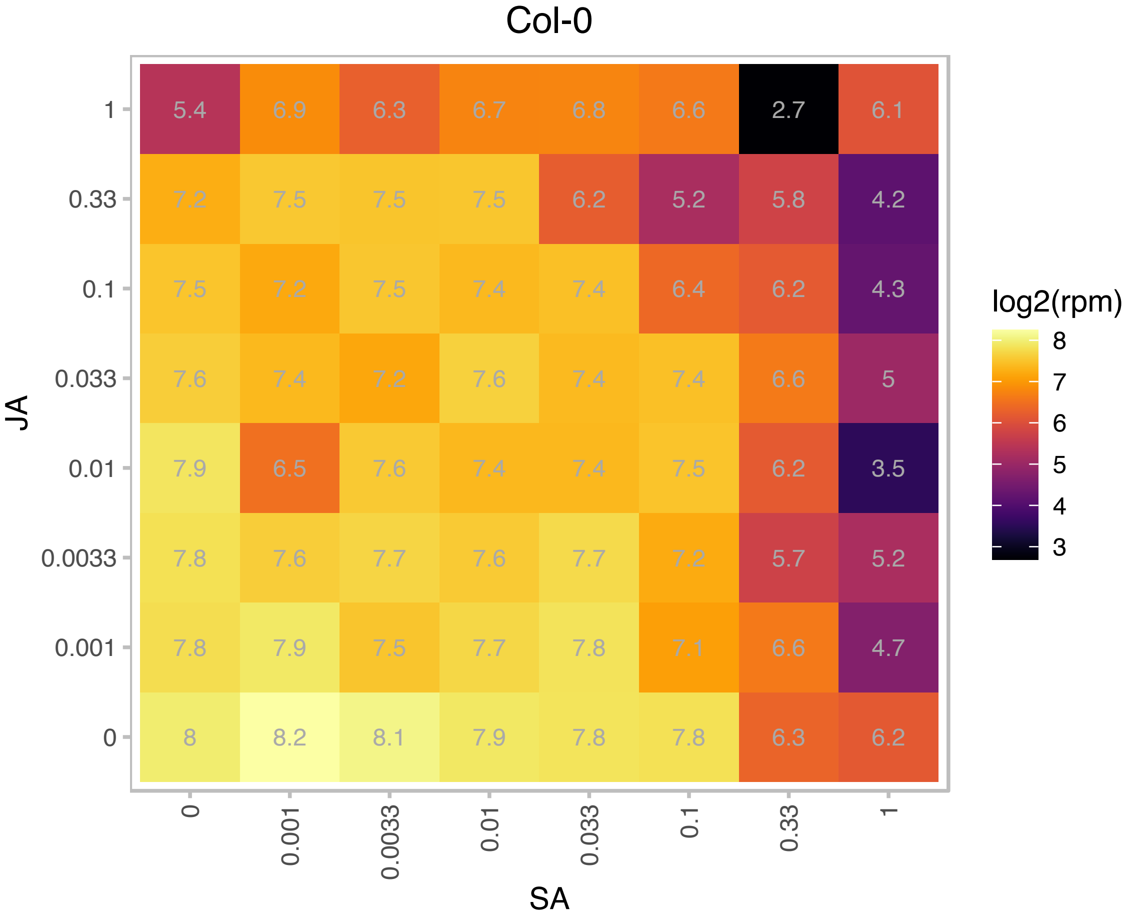


| | |
|---|---|
| Gene | AT3G22210 |
| is.expressed_genes | TRUE |
| is.responsive_genes | TRUE |
| is.apcluster | F-01 |

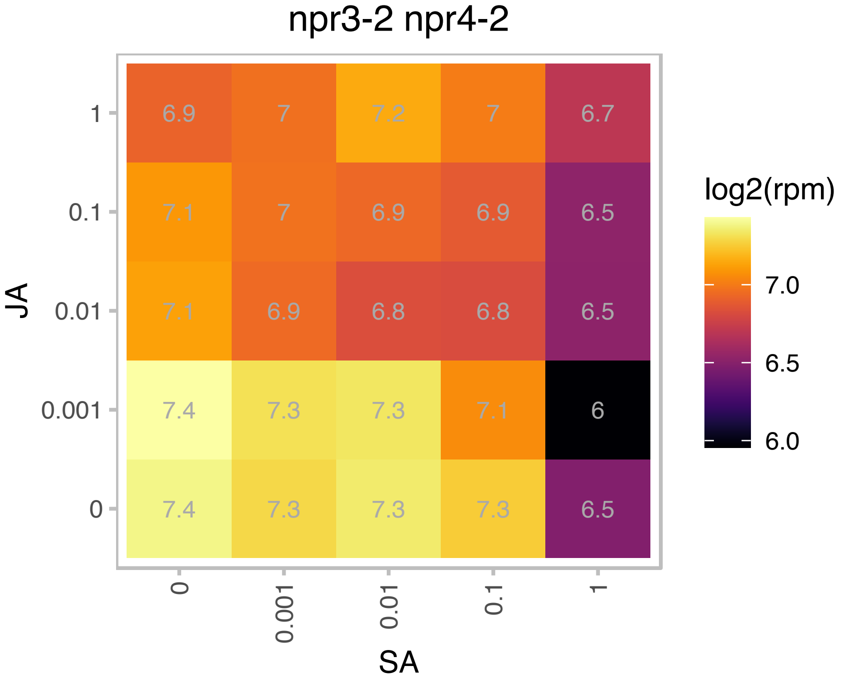


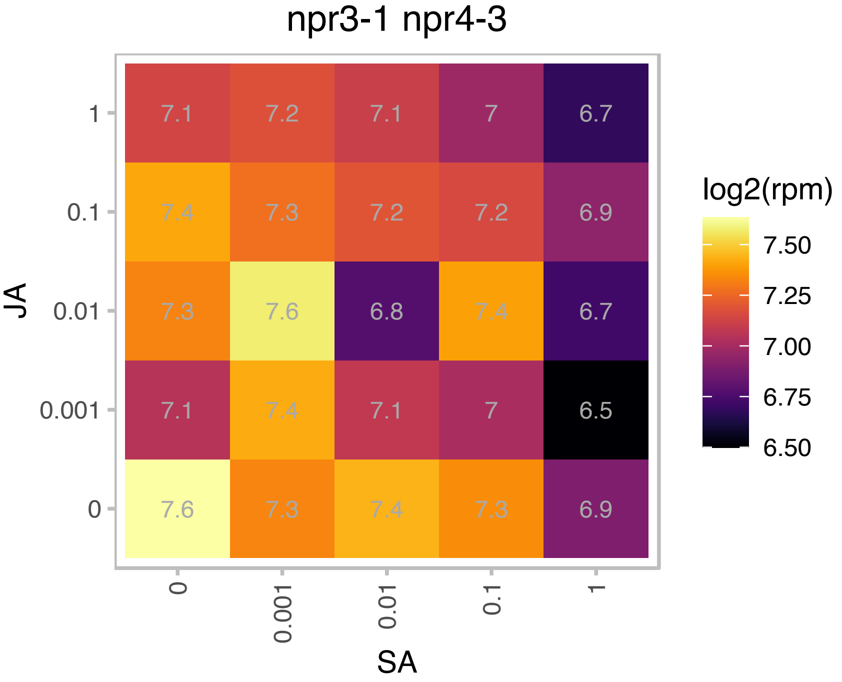

### AT3G46580.1 : ATMBD5, MBD05, MBD5

methyl-CPG-binding domain protein 5

Protein containing a putative methyl-CpG-binding domain.Has sequence similarity to human MBD proteins.

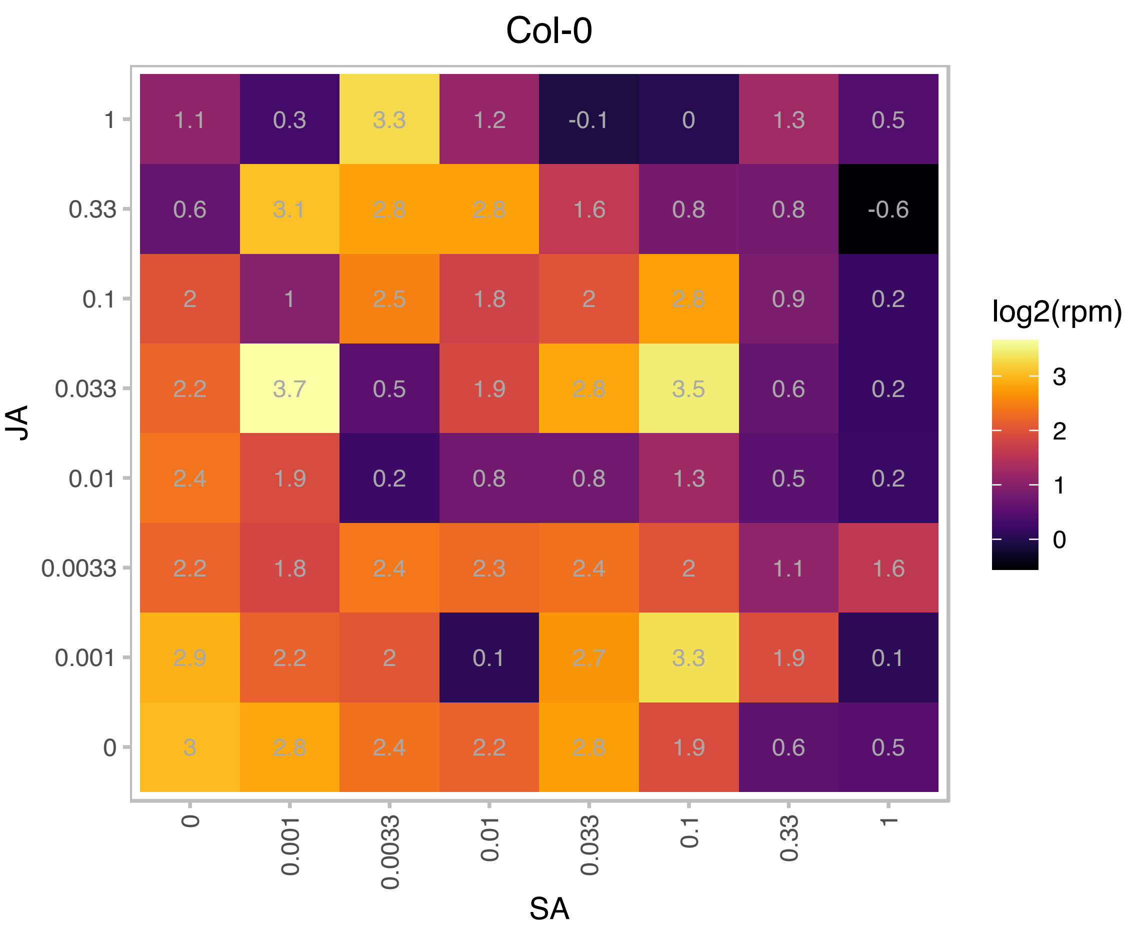


| | |
|---|---|
| Gene | AT3G46580 |
| is.expressed_genes | TRUE |
| is.responsive_genes | TRUE |
| is.apcluster | F-01 |

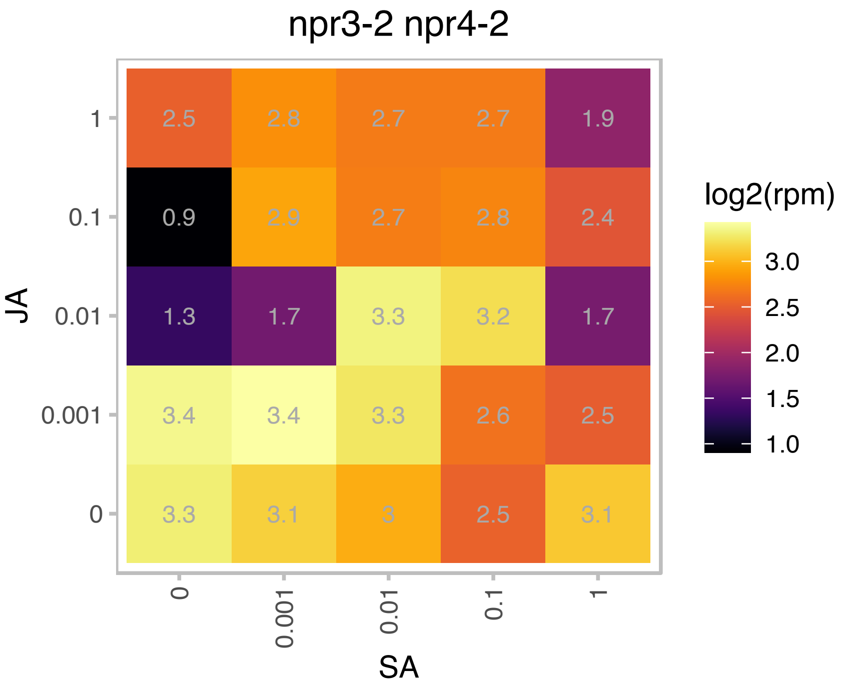


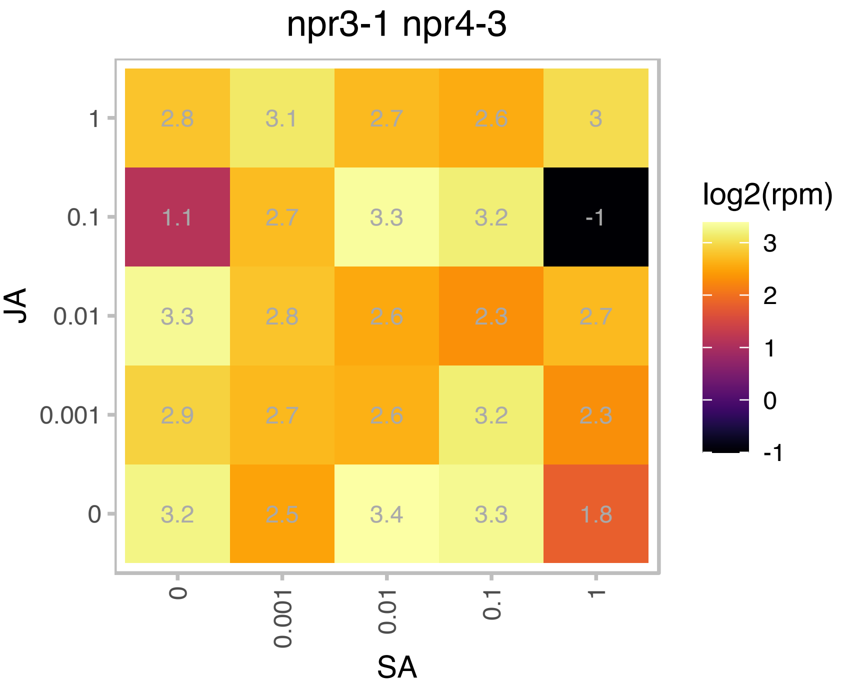

### AT3G51610.1 : NPU

Encodes a membrane protein NO PRIMEXINE AND PLASMA MEMBRANE UNDULATION (NPU). Involved in prime
deposition and plasma membrane undulation during early pollen wall development.

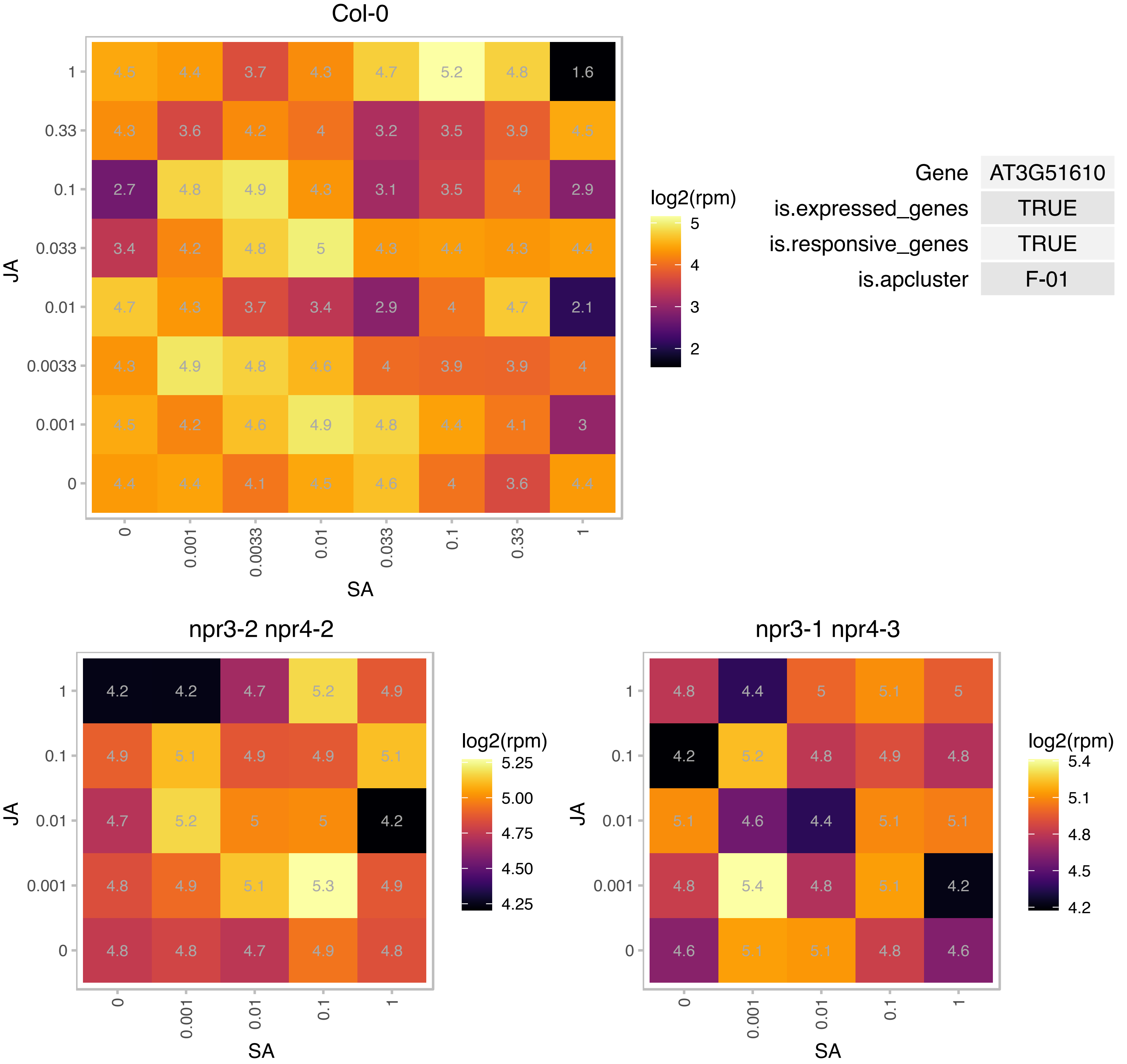


| | |
|---|---|
| Gene | AT3G51610 |
| is.expressed_genes | TRUE |
| is.responsive_genes | TRUE |
| is.apcluster | F-01 |

# AT3G54040.1 : NA

PAR1 protein

| Gene | AT3G54040 |
|---|---|
| is.expressed_genes | TRUE |
| is.responsive_genes | TRUE |
| is.apcluster | F-01 |

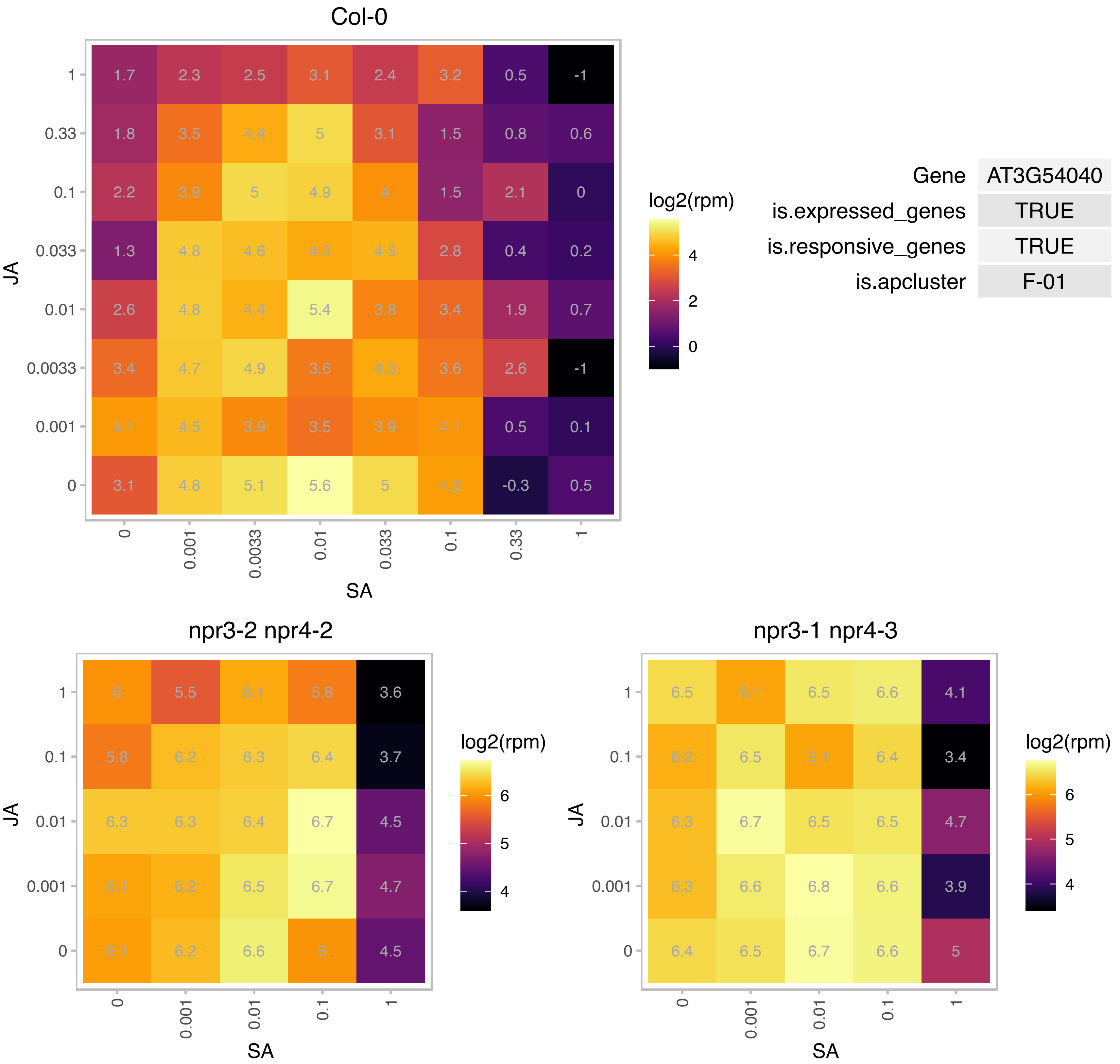

# AT3G55170.1 : NA

Ribosomal L29 family protein

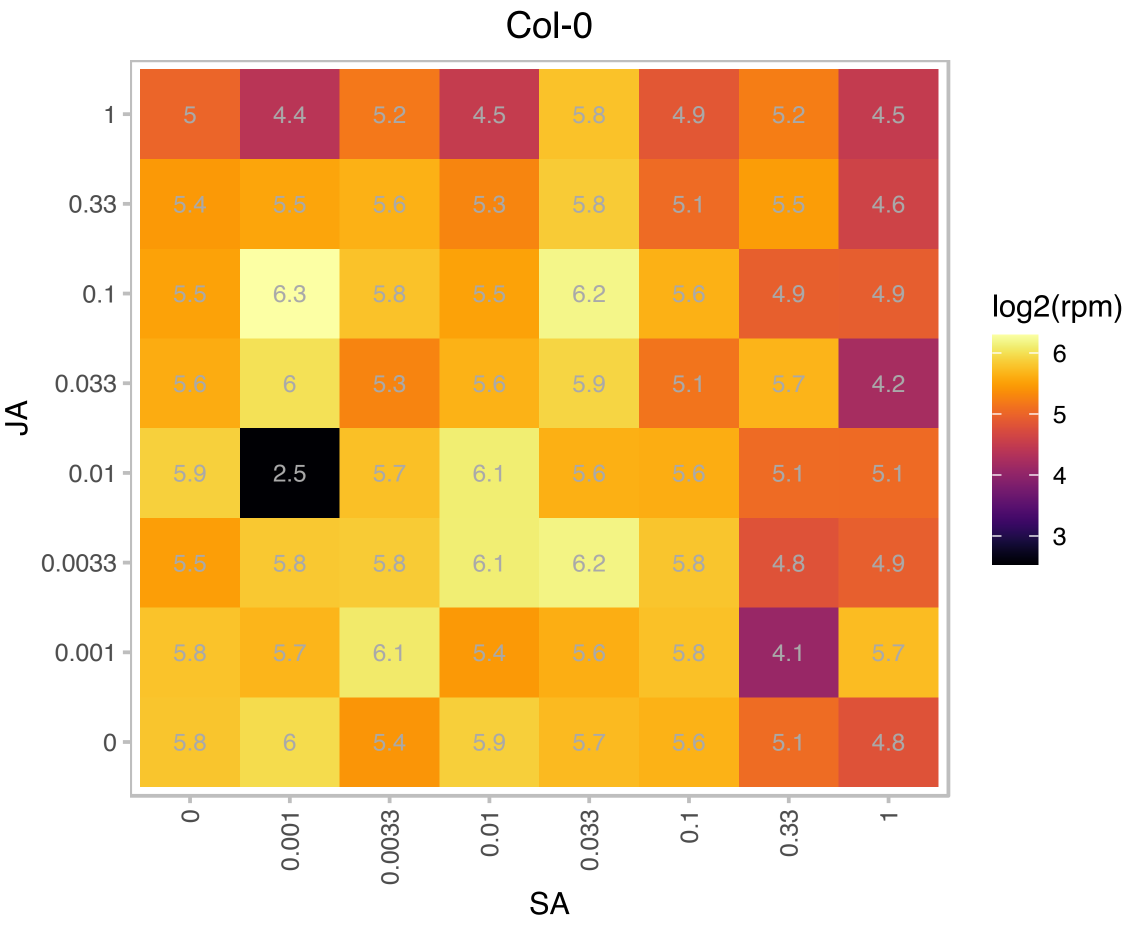


| | |
|---|---|
| Gene | AT3G55170 |
| is.expressed_genes | TRUE |
| is.responsive_genes | TRUE |
| is.apcluster | F-01 |

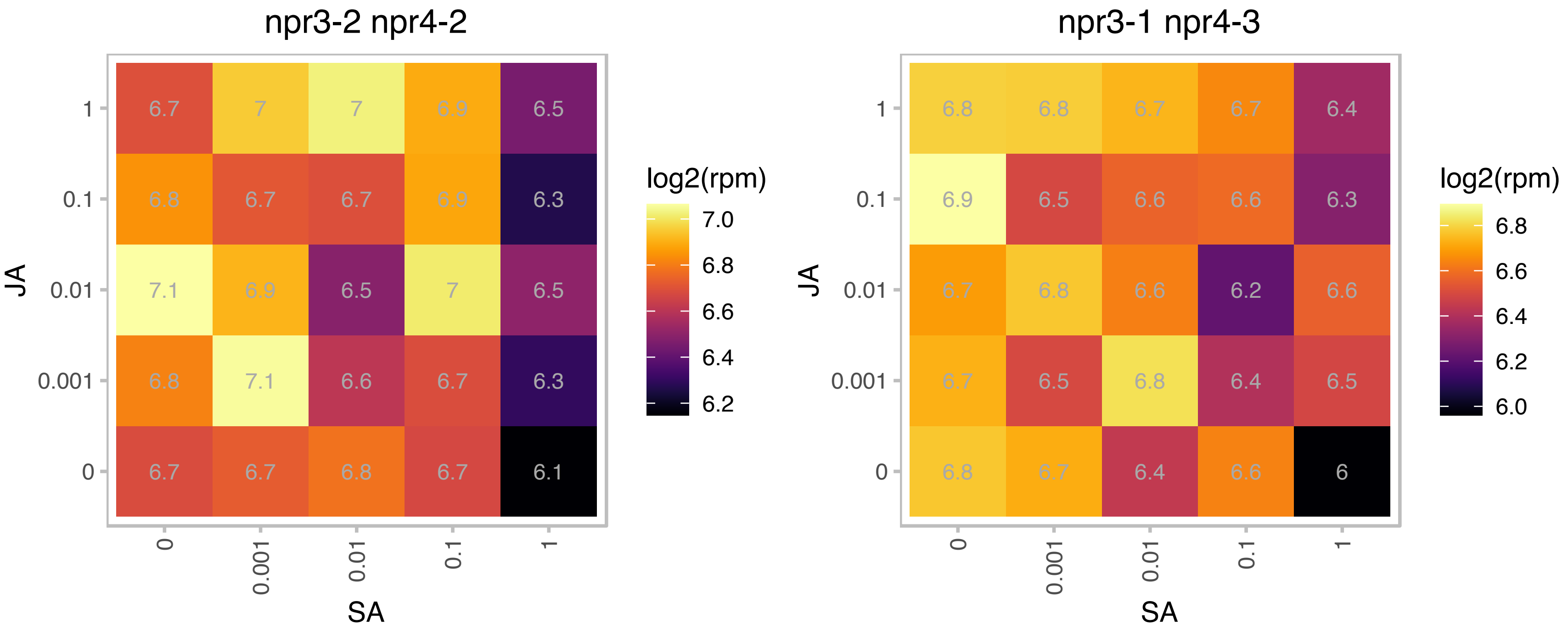

# AT3G57930.1 : NA

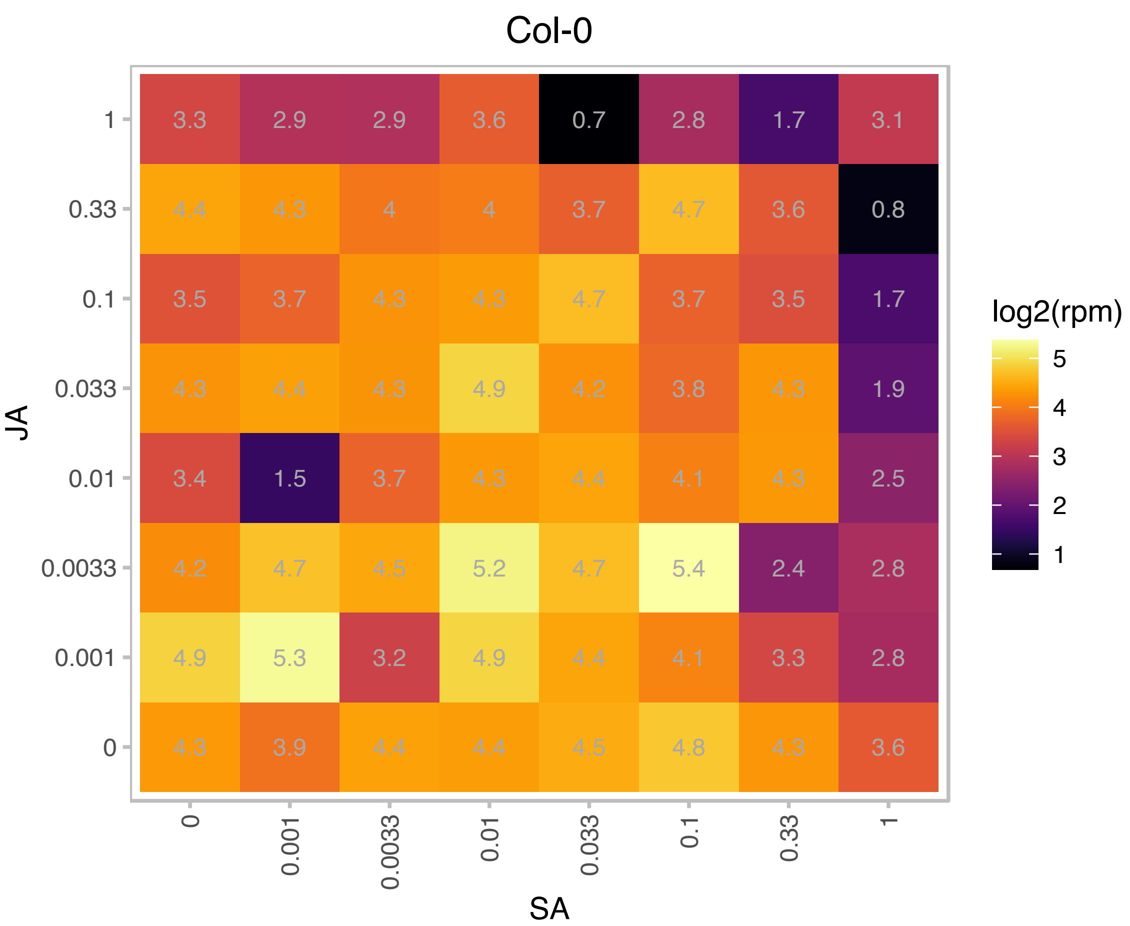


| | |
|---|---|
| Gene | AT3G57930 |
| is.expressed_genes | TRUE |
| is.responsive_genes | TRUE |
| is.apcluster | F-01 |

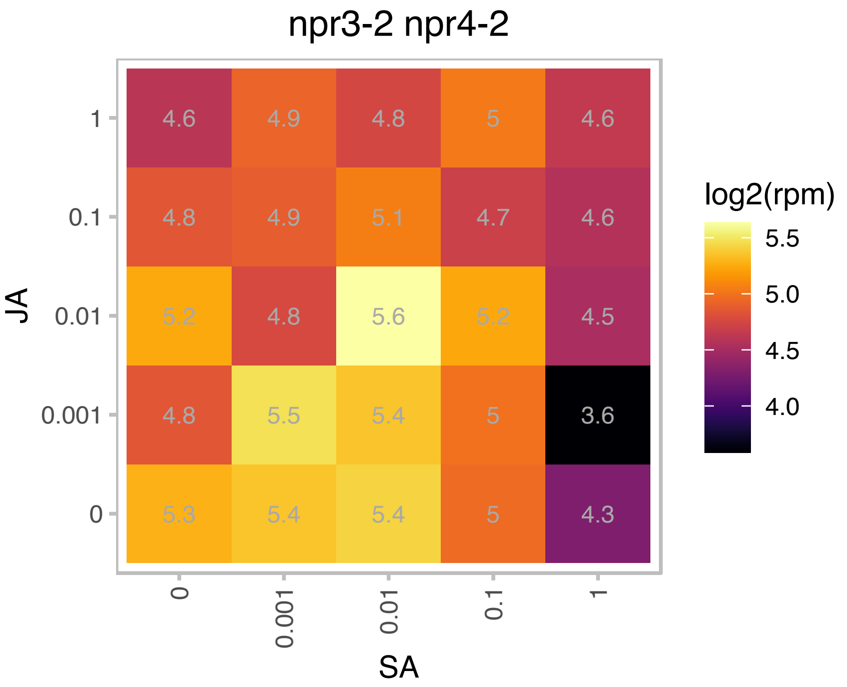


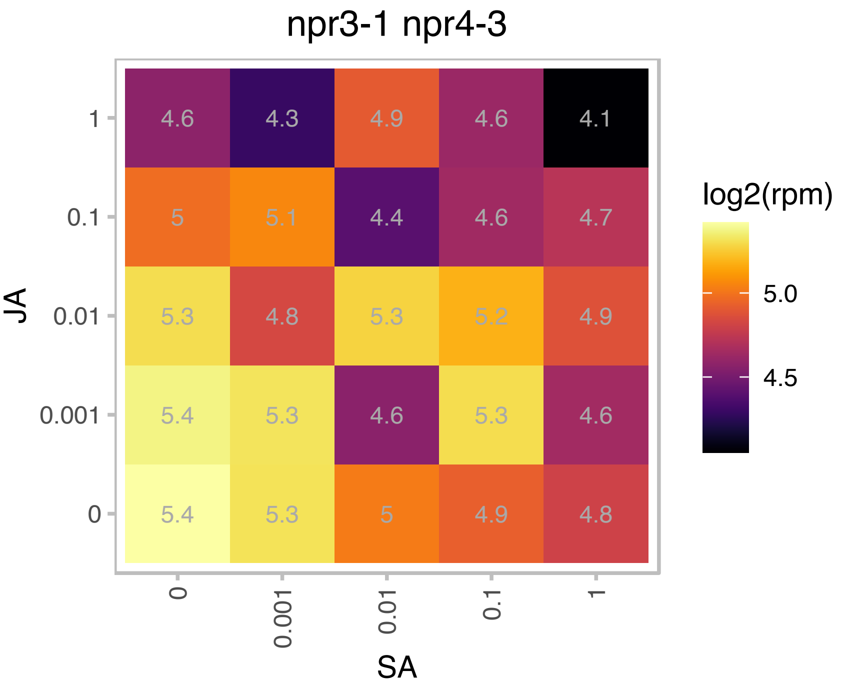

# AT3G60370.1 : NA

Encodes an immunophilin, FKBP20-2, that belongs to the FK-506 binding protein (FKBP) subfamily functioning as peptidyl-prolyl isomerases (PPIases) in protein folding. FKBP20-2 has a unique pair of cysteines at the C
FKBP-like peptidyl-prolyl cis-trans isomerase family protein
terminus and was found to be reduced by thioredoxin (Trx) (itself reduced by NADPH by means of NADP-Trx reductase). The FKBP20-2 protein, which contains only two of the five amino acids required for catalysis, showed a low level of PPIase activity that was unaffected on reduction by Trx.  Genetic disruption of the FKBP20-2 gene provide evidence that FKBP20-2 participates specifically in the accumulation of the PSII [illegible]

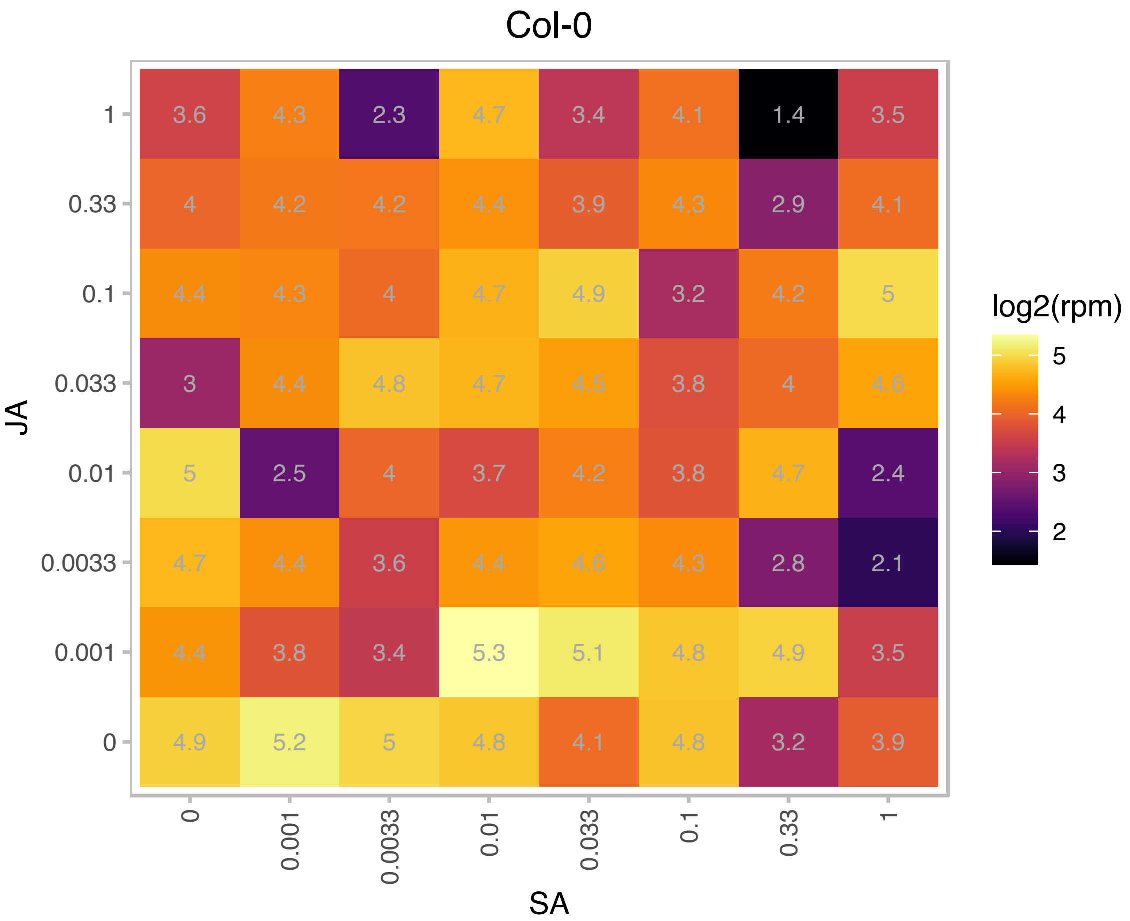


| | |
|---|---|
| Gene | AT3G60370 |
| is.expressed_genes | TRUE |
| is.responsive_genes | TRUE |
| is.apcluster | F-01 |

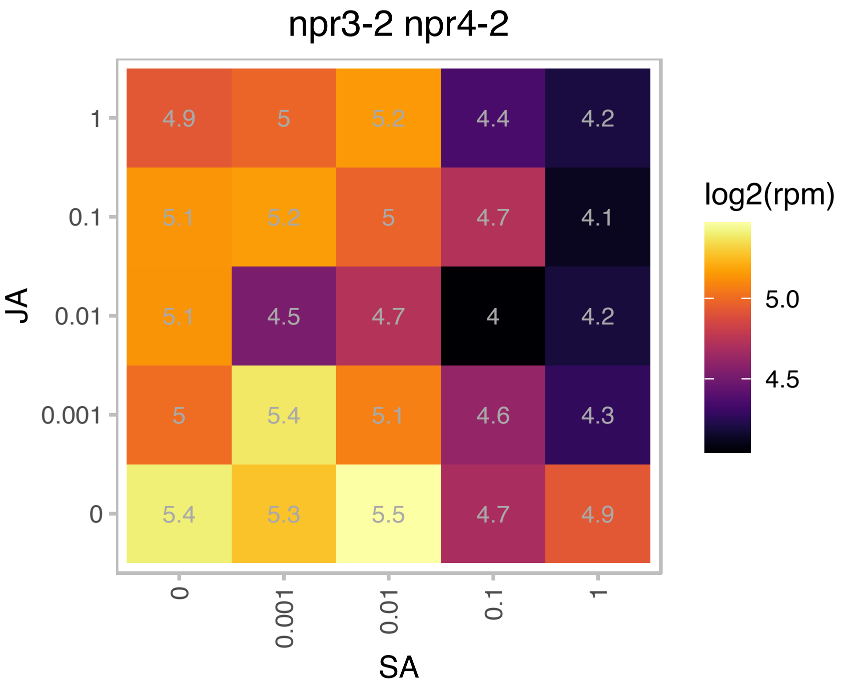


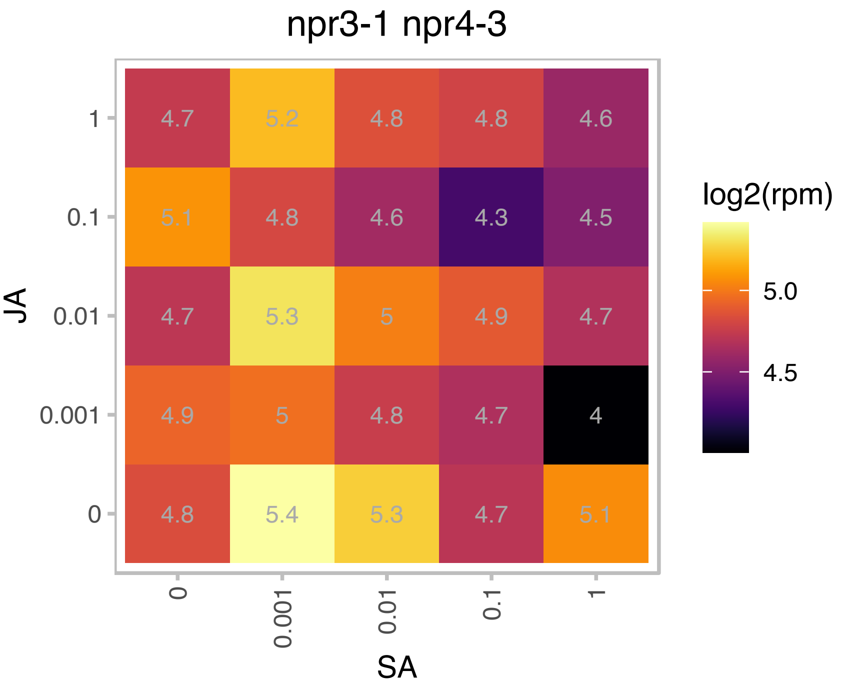

### AT4G00355.1 : ATI2

Encodes an Atg8-interacting protein that is partially associated with the ER during favorable growth conditions and becomes mainly associated with a spherical compartment that dynamically moves along the ER network.

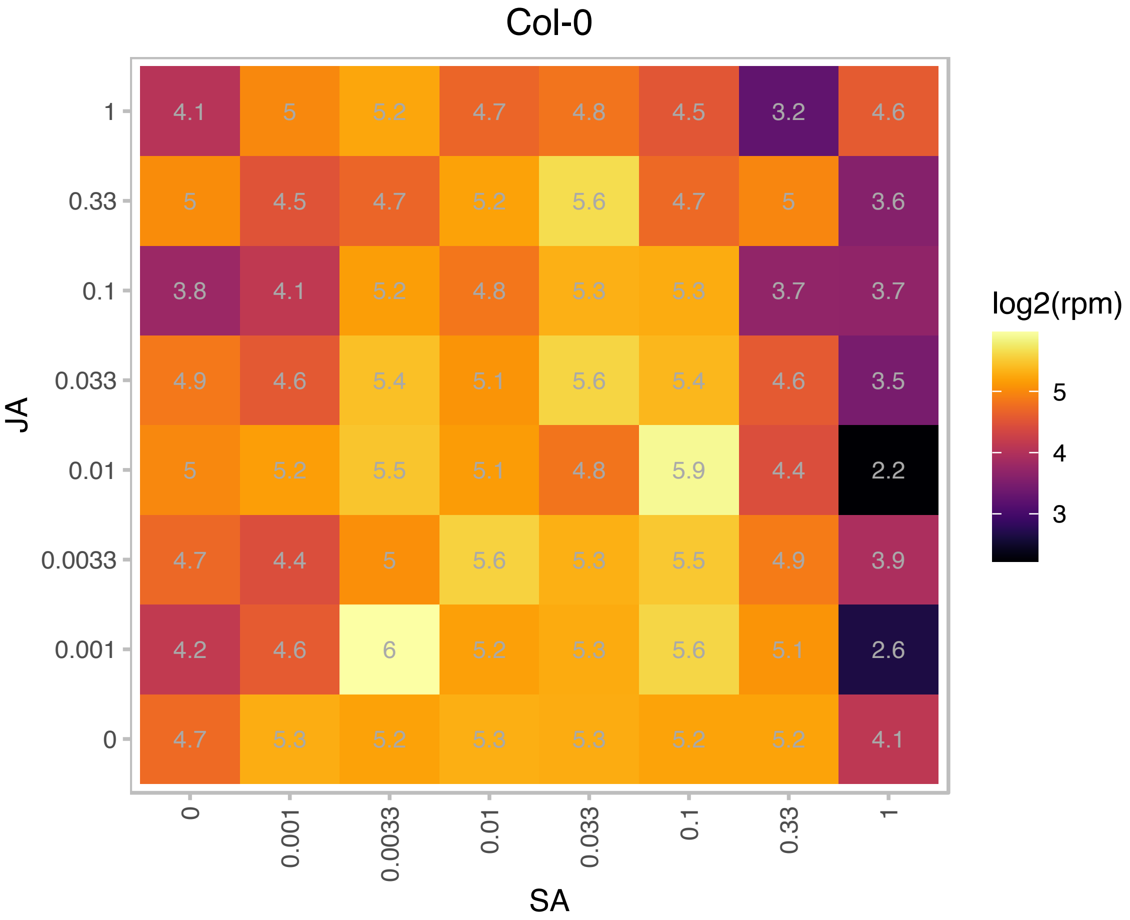


| | |
|---|---|
| Gene | AT4G00355 |
| is.expressed_genes | TRUE |
| is.responsive_genes | TRUE |
| is.apcluster | F-01 |

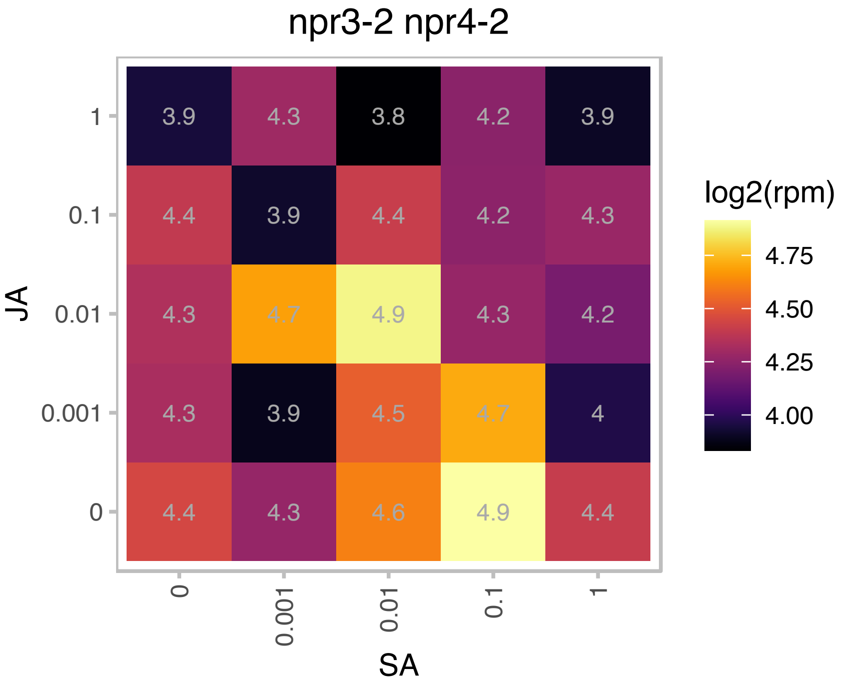


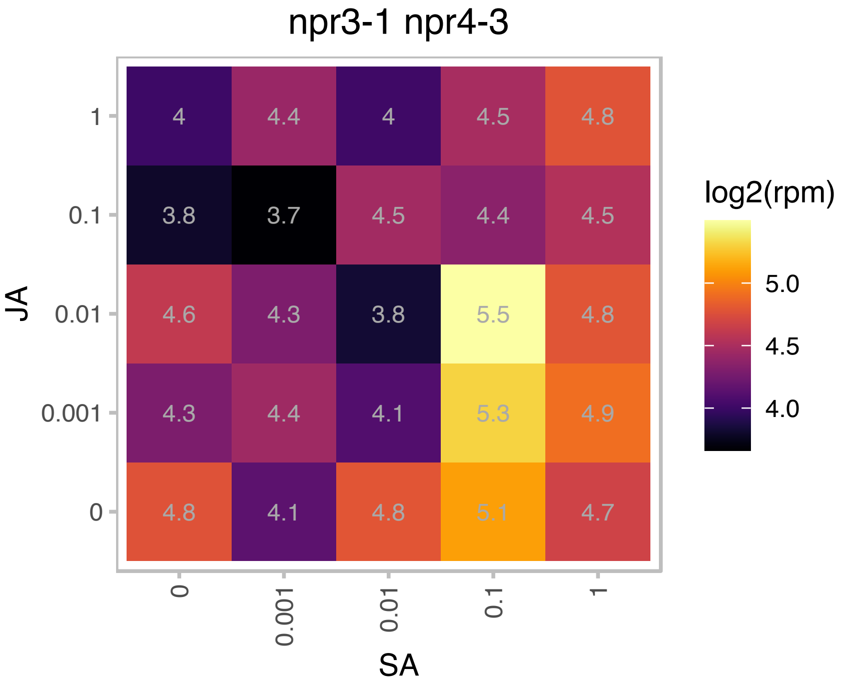

# AT4G05195.1 : NA

| | |
|---|---|
| Gene | AT4G05195 |
| is.expressed_genes | TRUE |
| is.responsive_genes | TRUE |
| is.apcluster | F-01 |

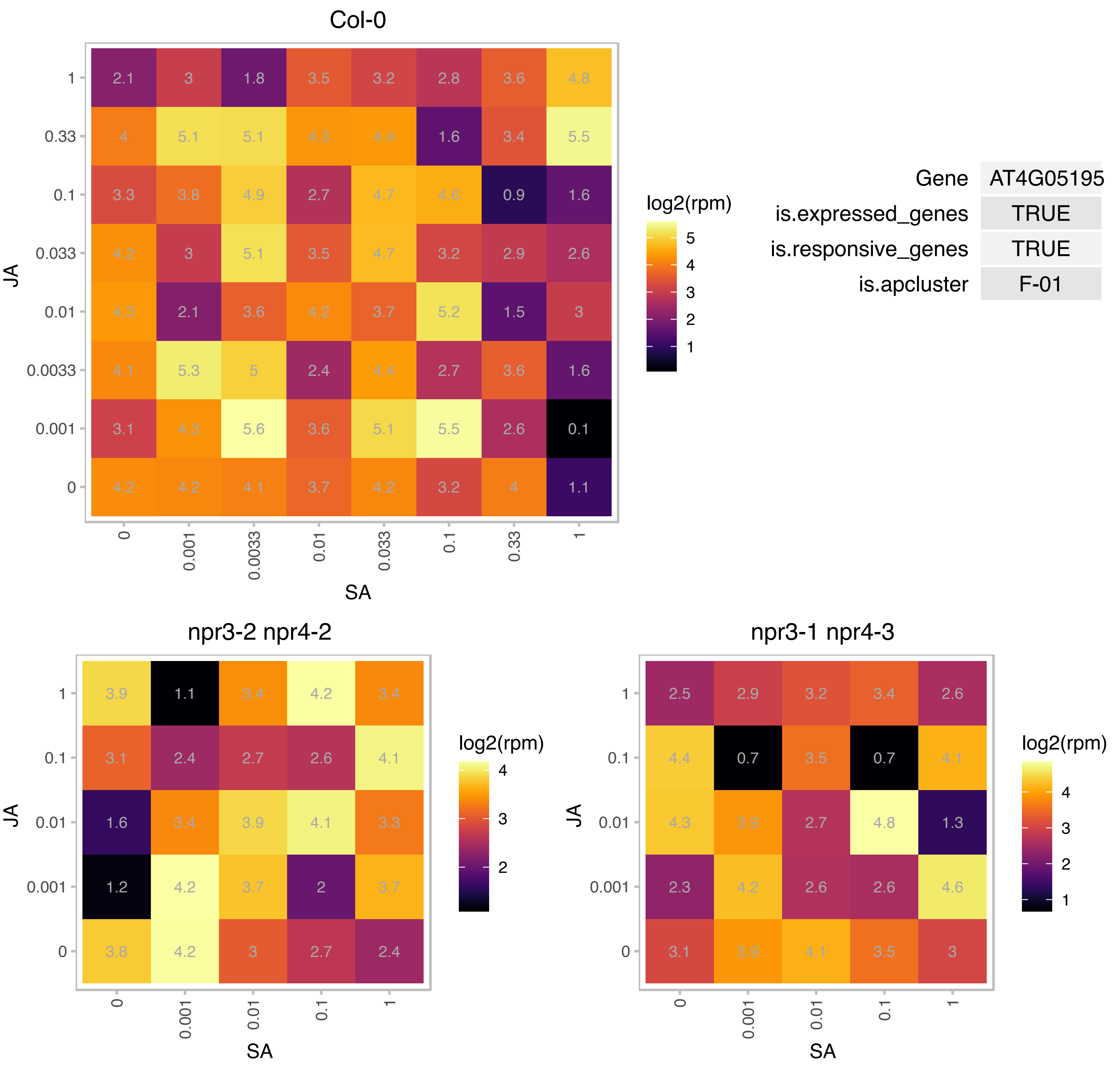

### AT4G09620.1 : mTERF12

Mitochondrial transcription termination factor family protein

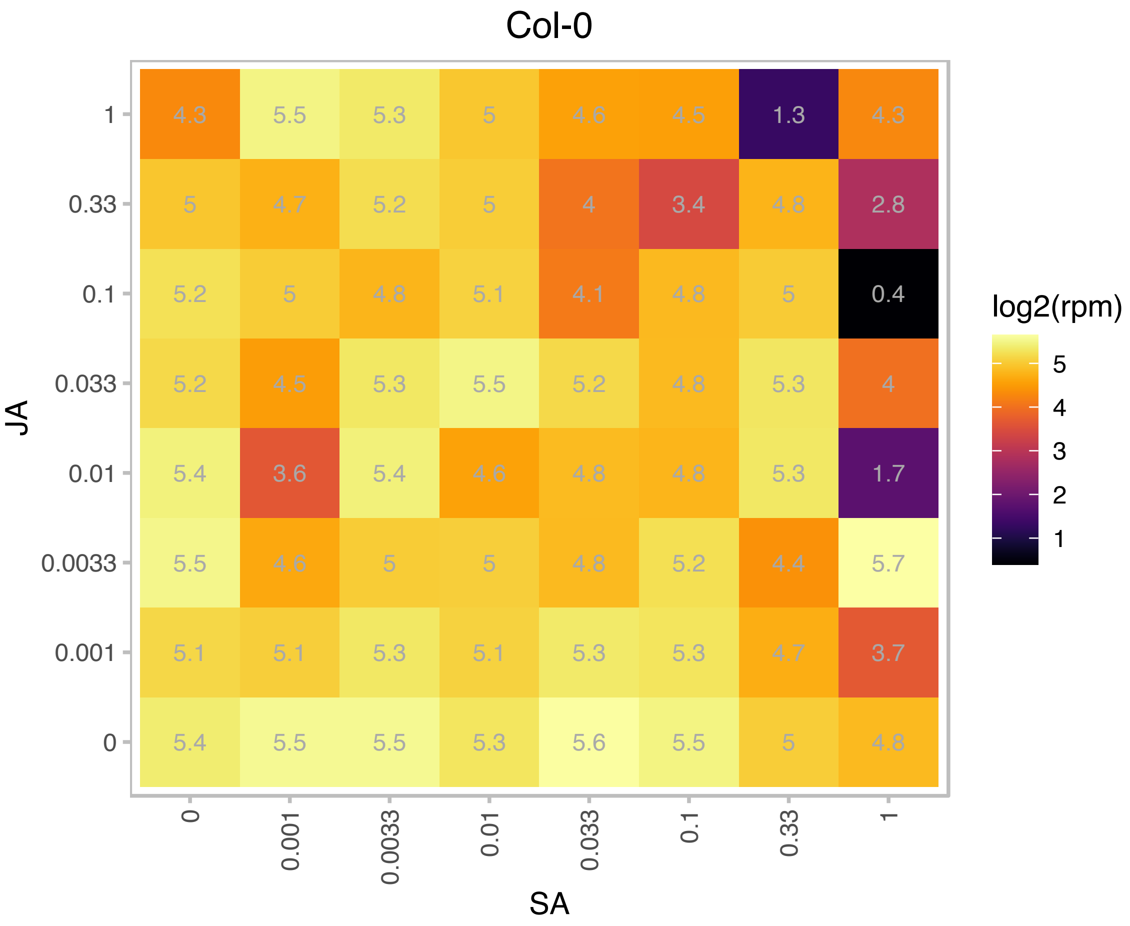


| | |
|---|---|
| Gene | AT4G09620 |
| is.expressed_genes | TRUE |
| is.responsive_genes | TRUE |
| is.apcluster | F-01 |

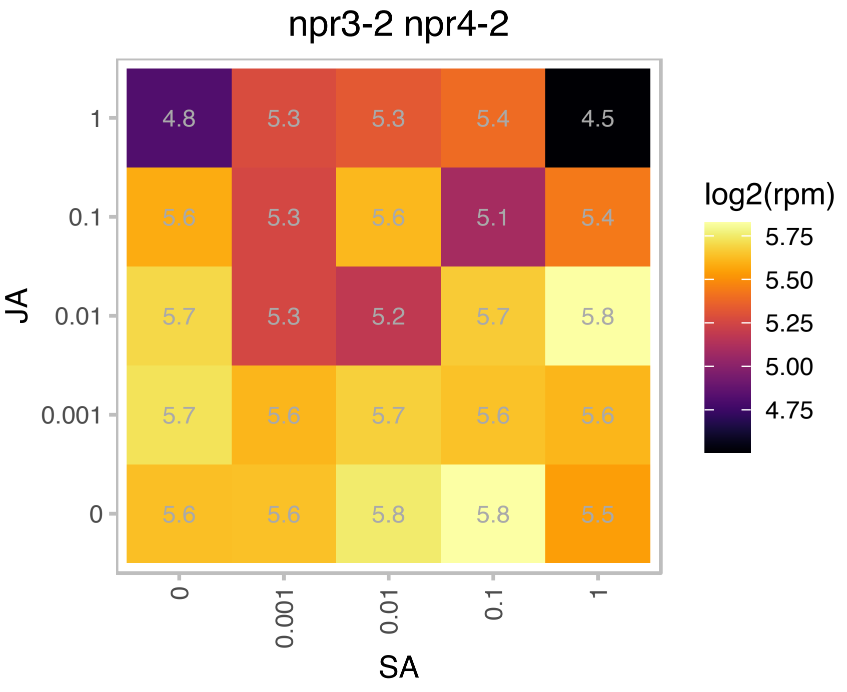


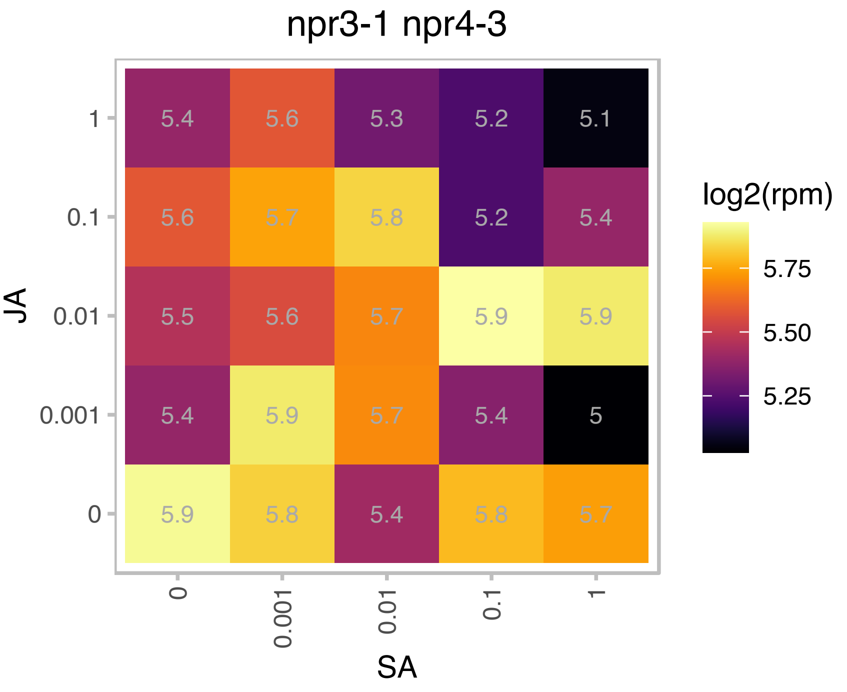

# AT4G13170.1 : NA

Ribosomal protein L13 family protein

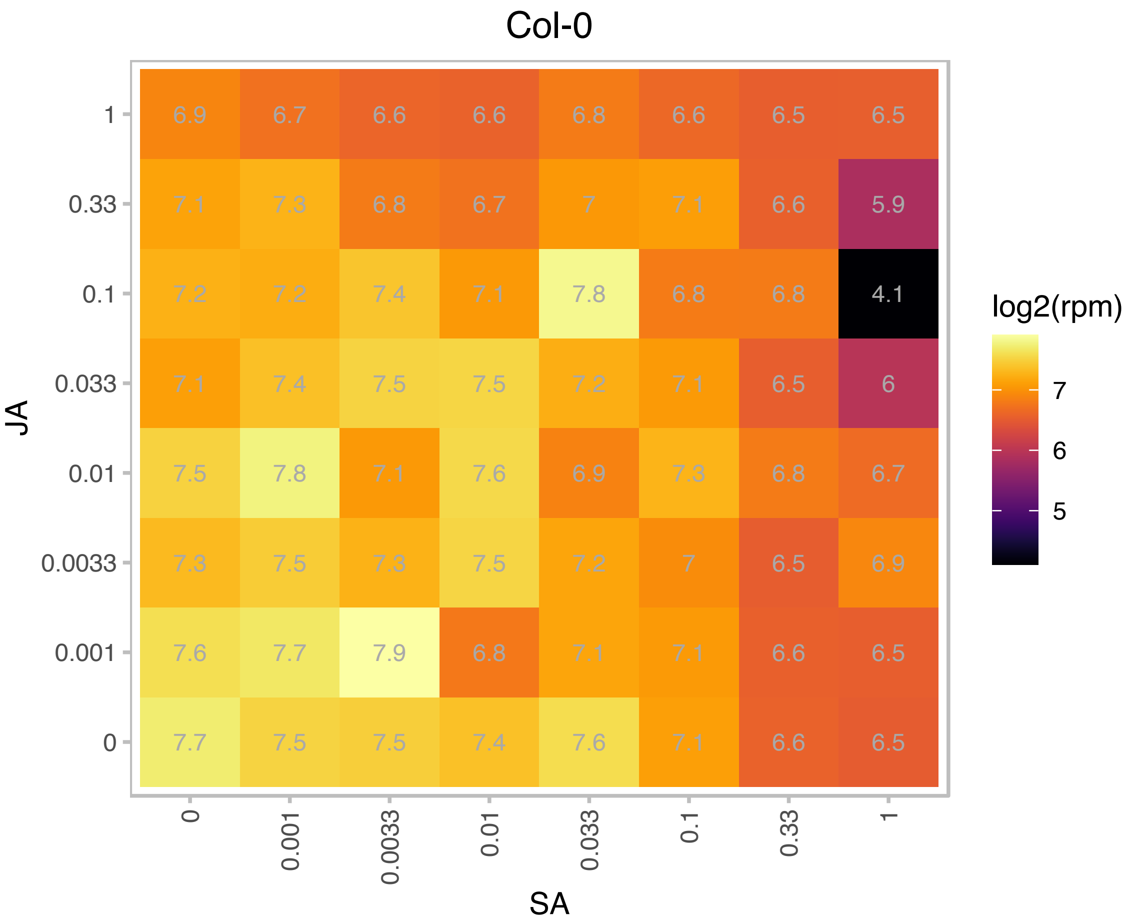


| | |
|---|---|
| Gene | AT4G13170 |
| is.expressed_genes | TRUE |
| is.responsive_genes | TRUE |
| is.apcluster | F-01 |

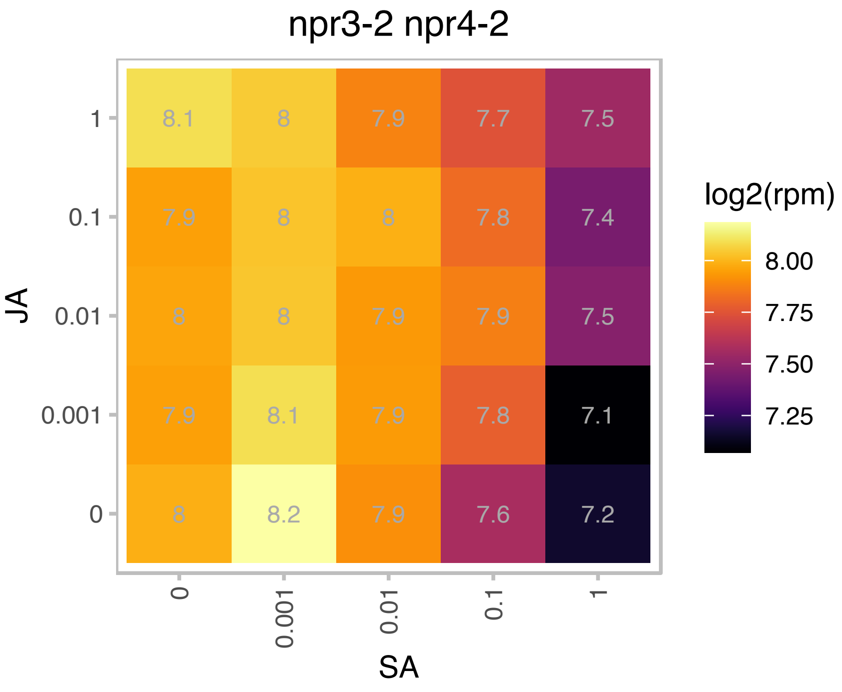


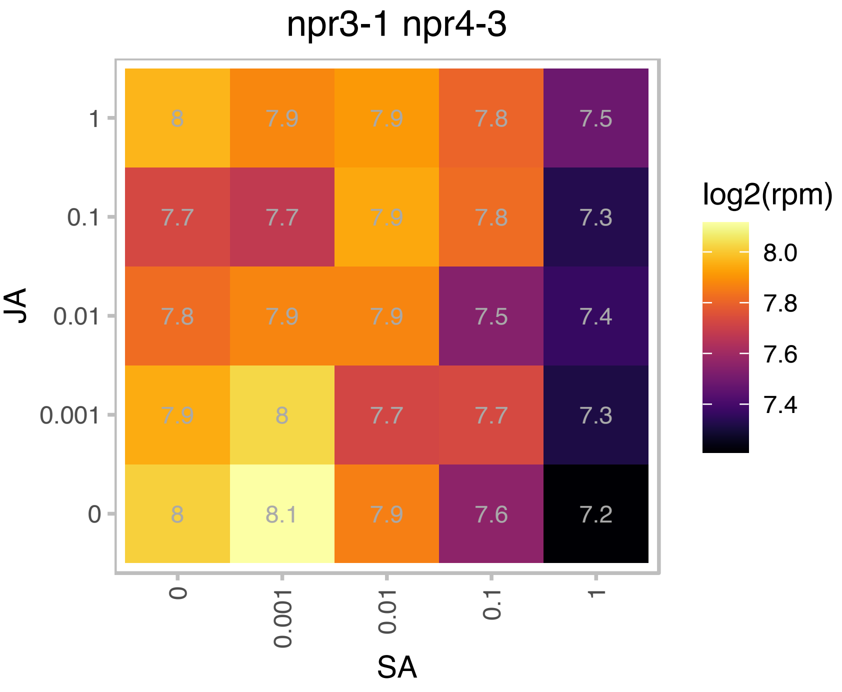

### AT4G15690.1 : GRXS5, ROXY12

Thioredoxin superfamily protein

Encodes a member of the CC-type glutaredoxin (ROXY) family that has been shown to interact with the transcription factor TGA2 and suppress ORA59 promoter activity.

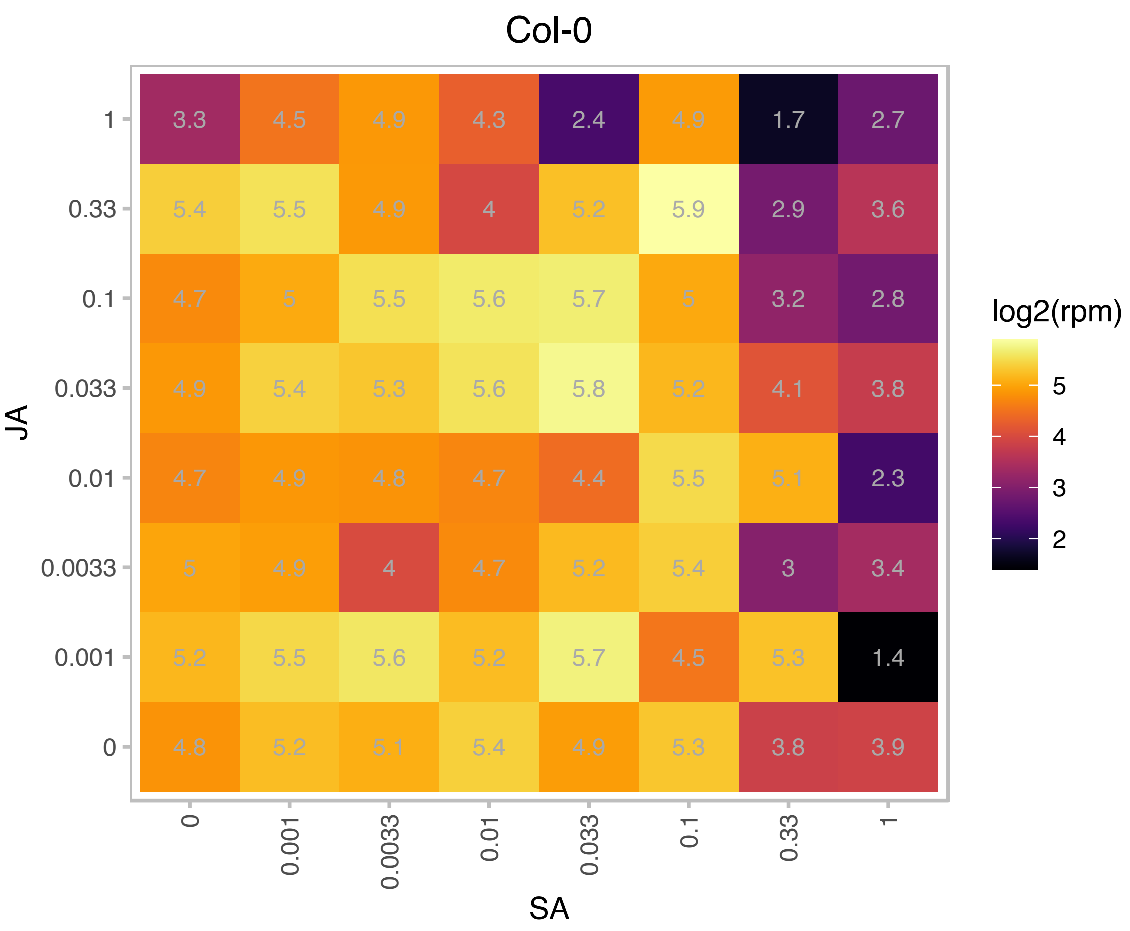


| | |
|---|---|
| Gene | AT4G15690 |
| is.expressed_genes | TRUE |
| is.responsive_genes | TRUE |
| is.apcluster | F-01 |

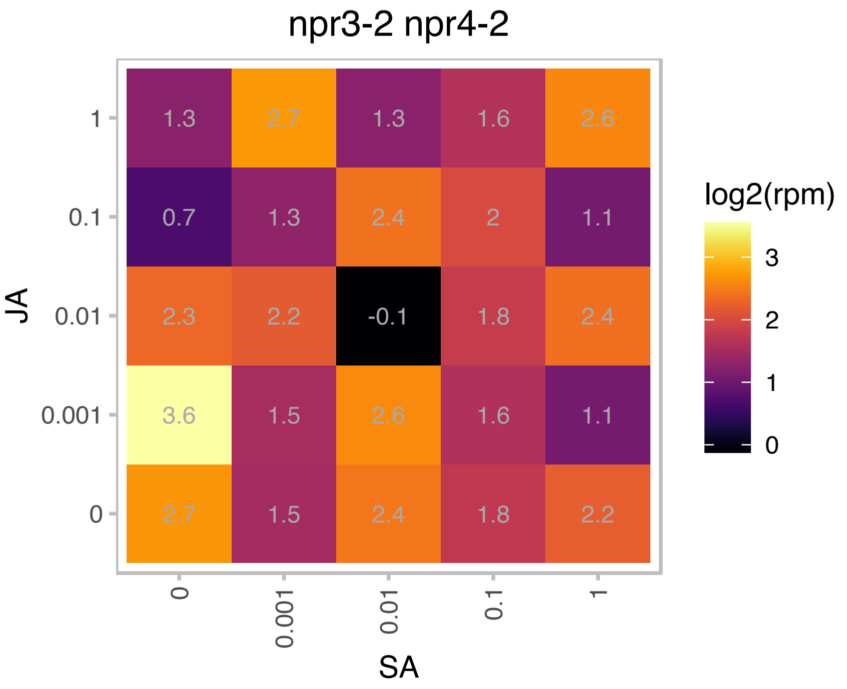


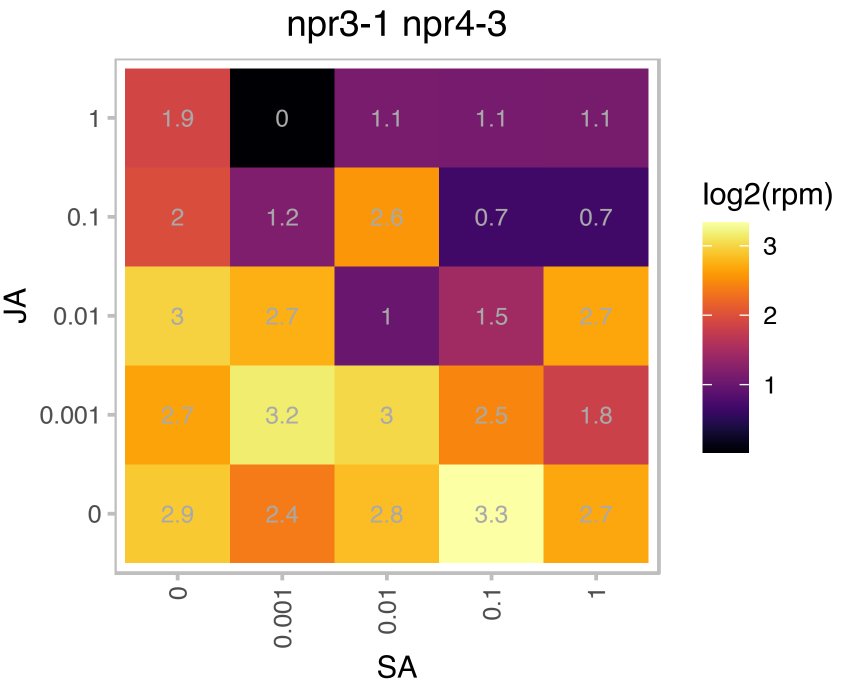

### AT4G20780.1 : CML42

calmodulin like 42

Calcium sensor involved in trichome branching.

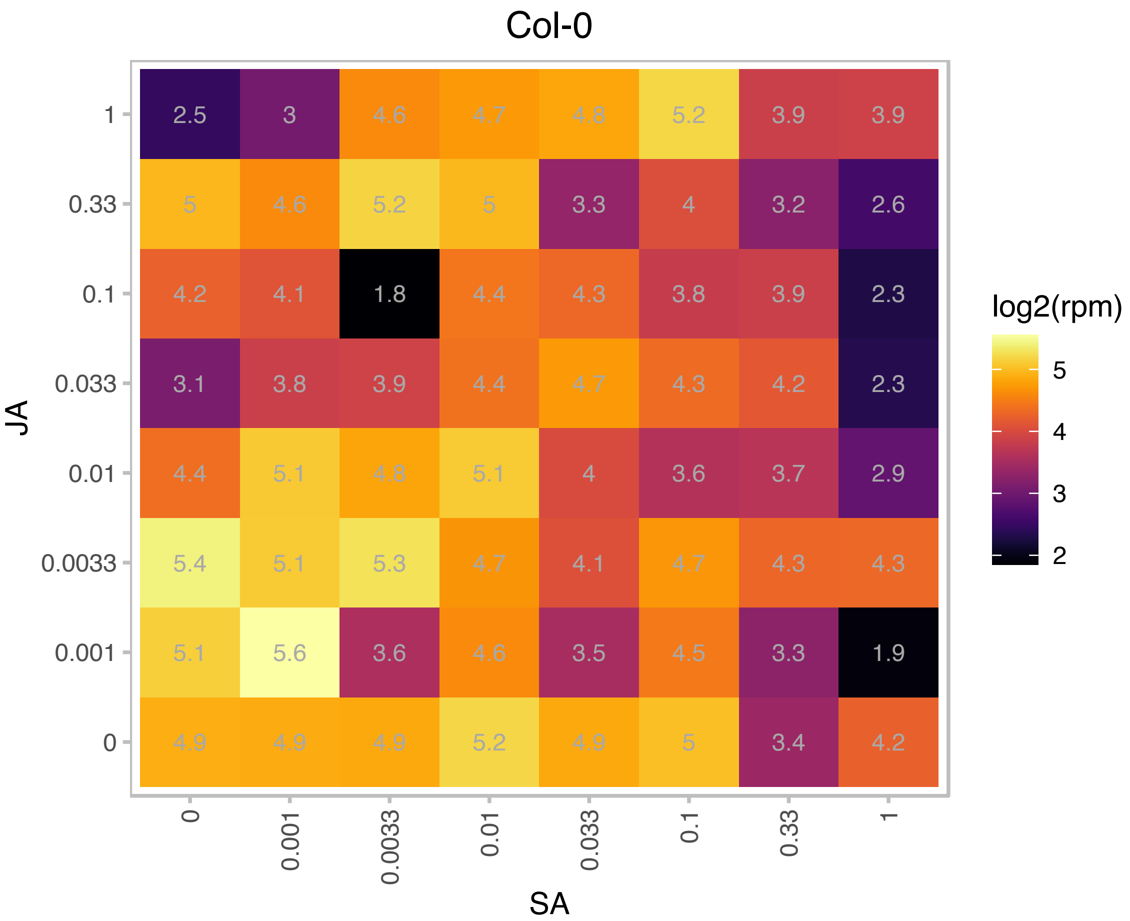


| | |
|---|---|
| Gene | AT4G20780 |
| is.expressed_genes | TRUE |
| is.responsive_genes | TRUE |
| is.apcluster | F-01 |

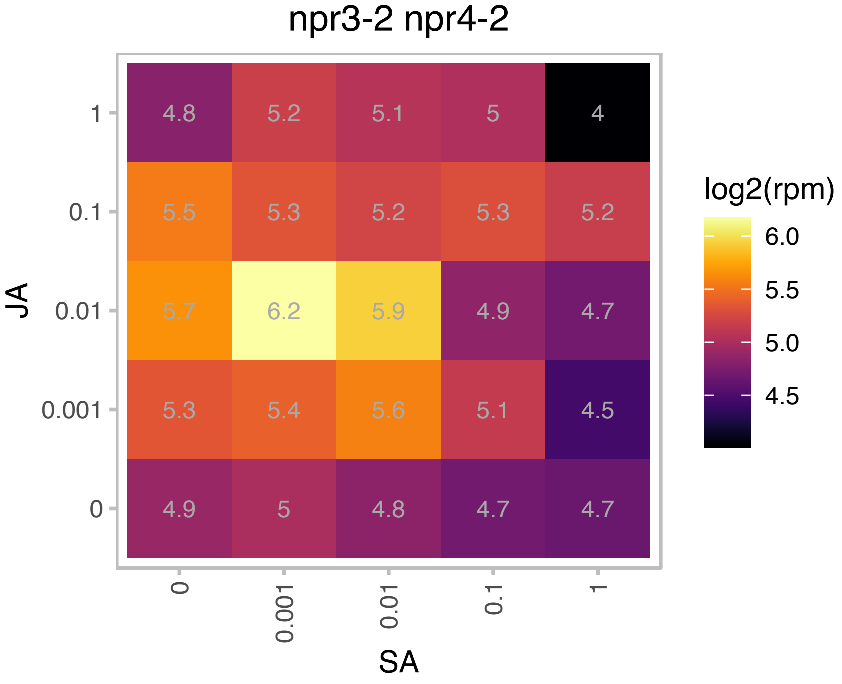


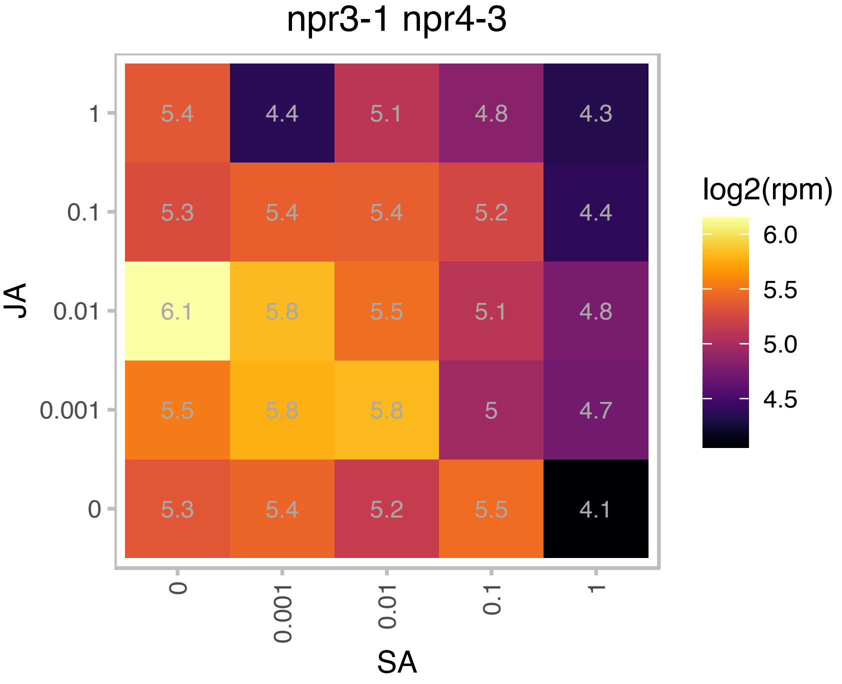

# AT4G26288.1 : NA

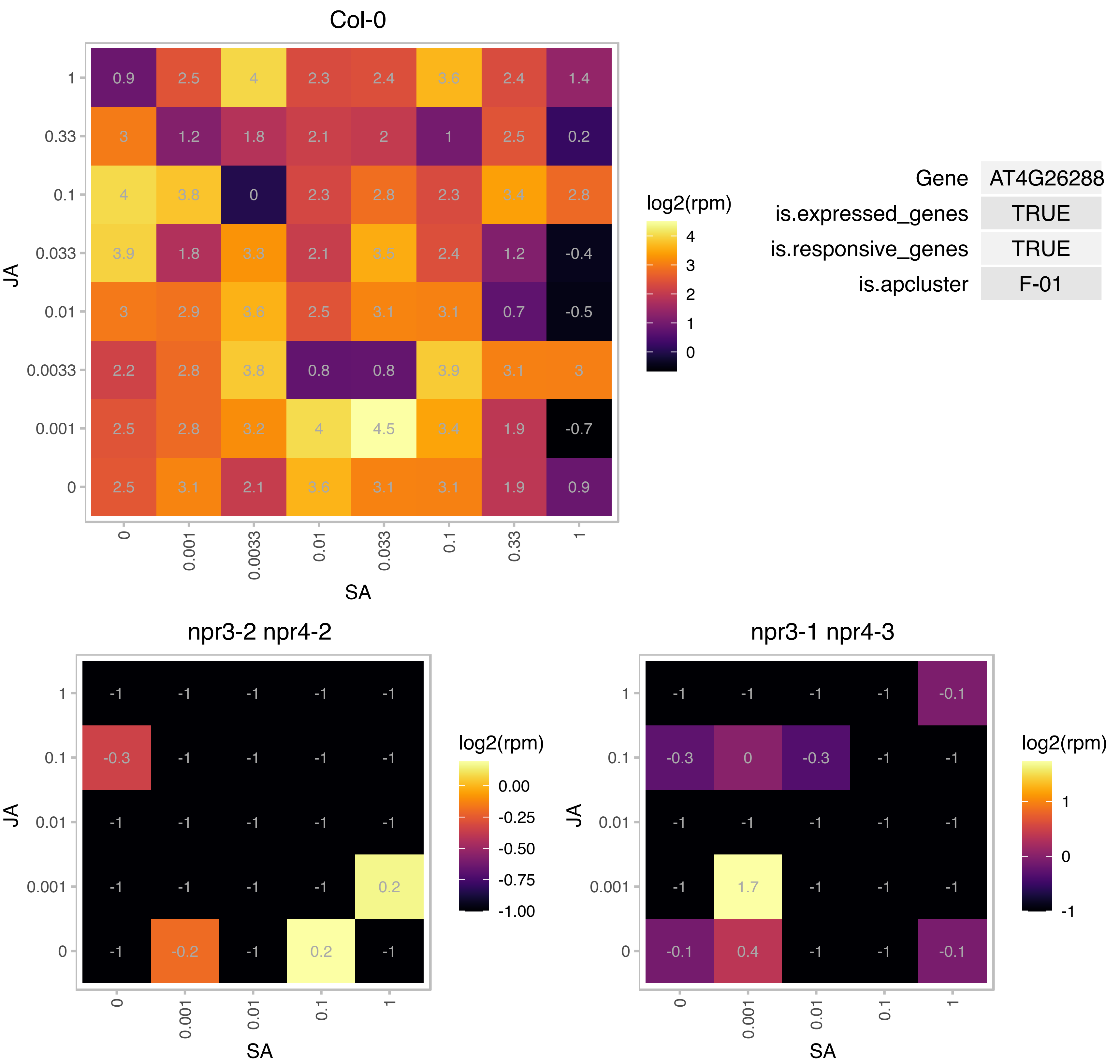


| | |
|---|---|
| Gene | AT4G26288 |
| is.expressed_genes | TRUE |
| is.responsive_genes | TRUE |
| is.apcluster | F-01 |

### AT4G28080.1 : REC2

Tetratricopeptide repeat (TPR)-like superfamily protein

Encodes REDUCED CHLOROPLAST COVERAGE 2 (REC2). Along with REC1 and REC3 it contributes to establis size of the chloroplast compartment.

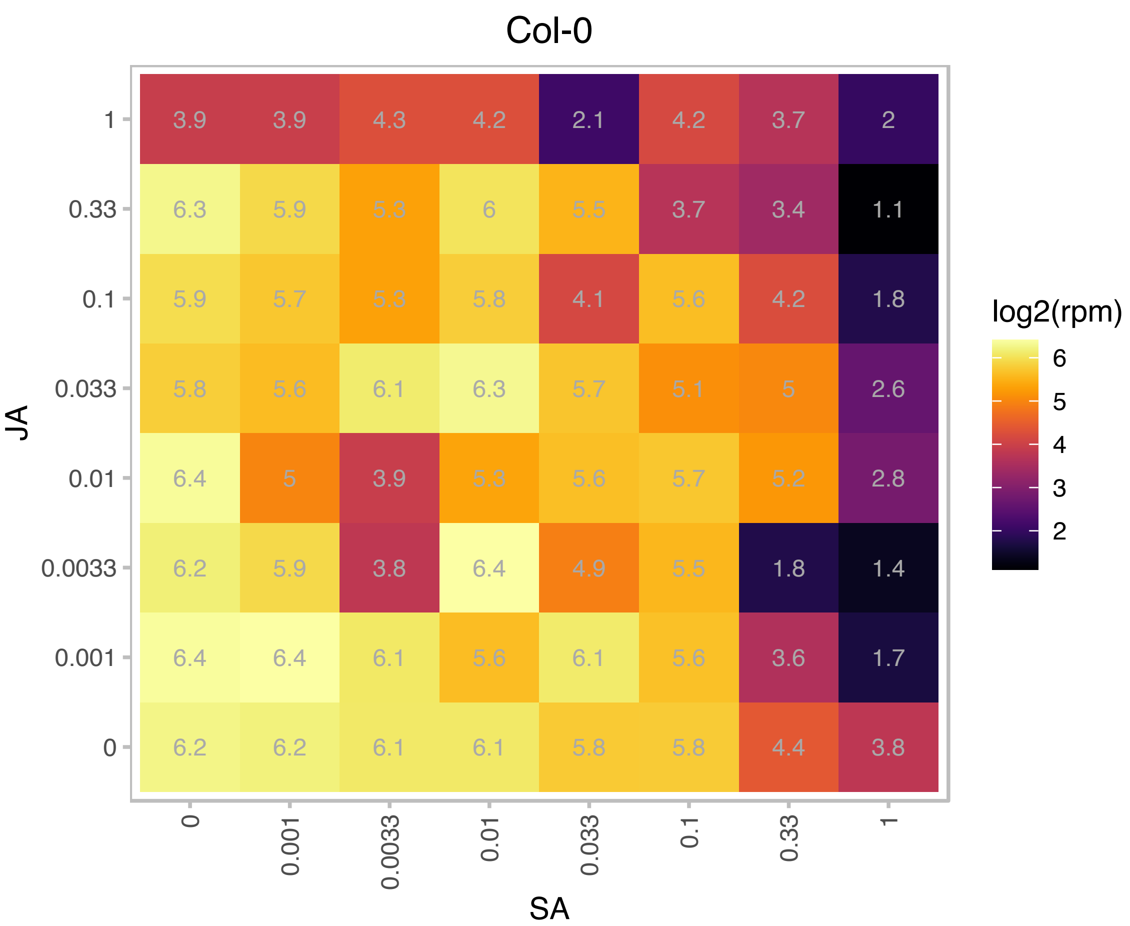


| | |
|---|---|
| Gene | AT4G28080 |
| is.expressed_genes | TRUE |
| is.responsive_genes | TRUE |
| is.apcluster | F-01 |

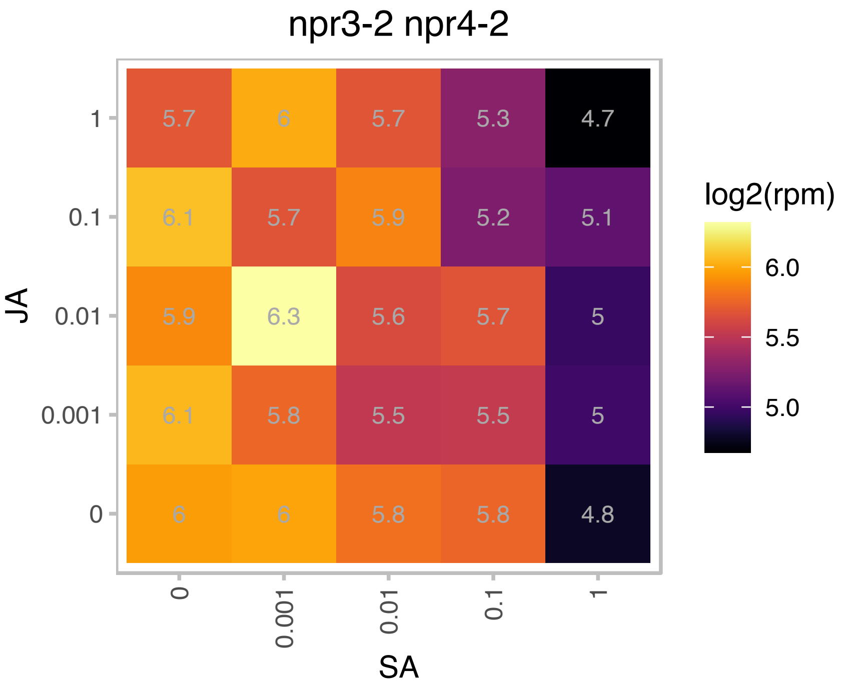


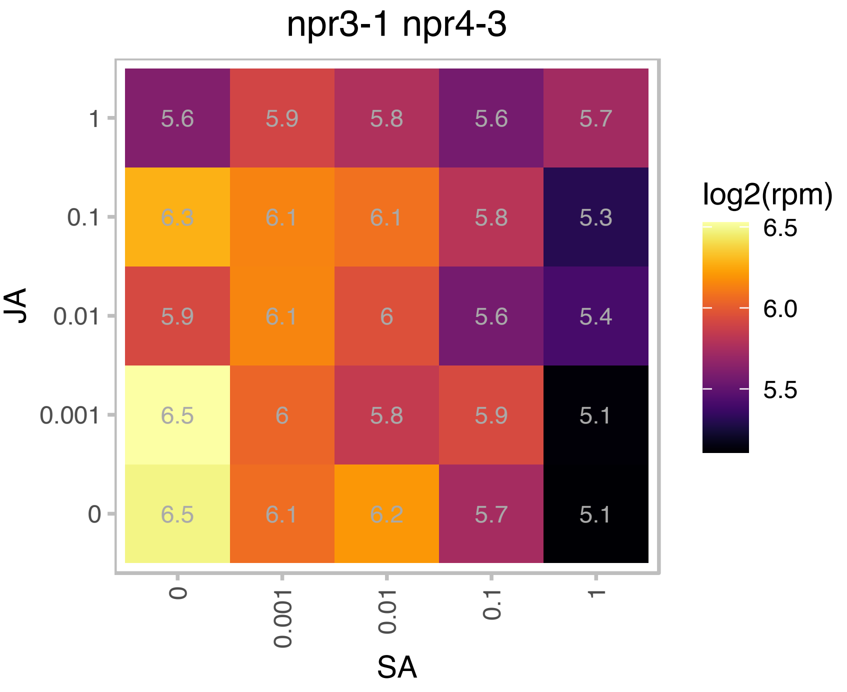

# AT4G28240.1 : NA

Wound-responsive family protein

| | |
|---|---|
| Gene | AT4G28240 |
| is.expressed_genes | TRUE |
| is.responsive_genes | TRUE |
| is.apcluster | F-01 |

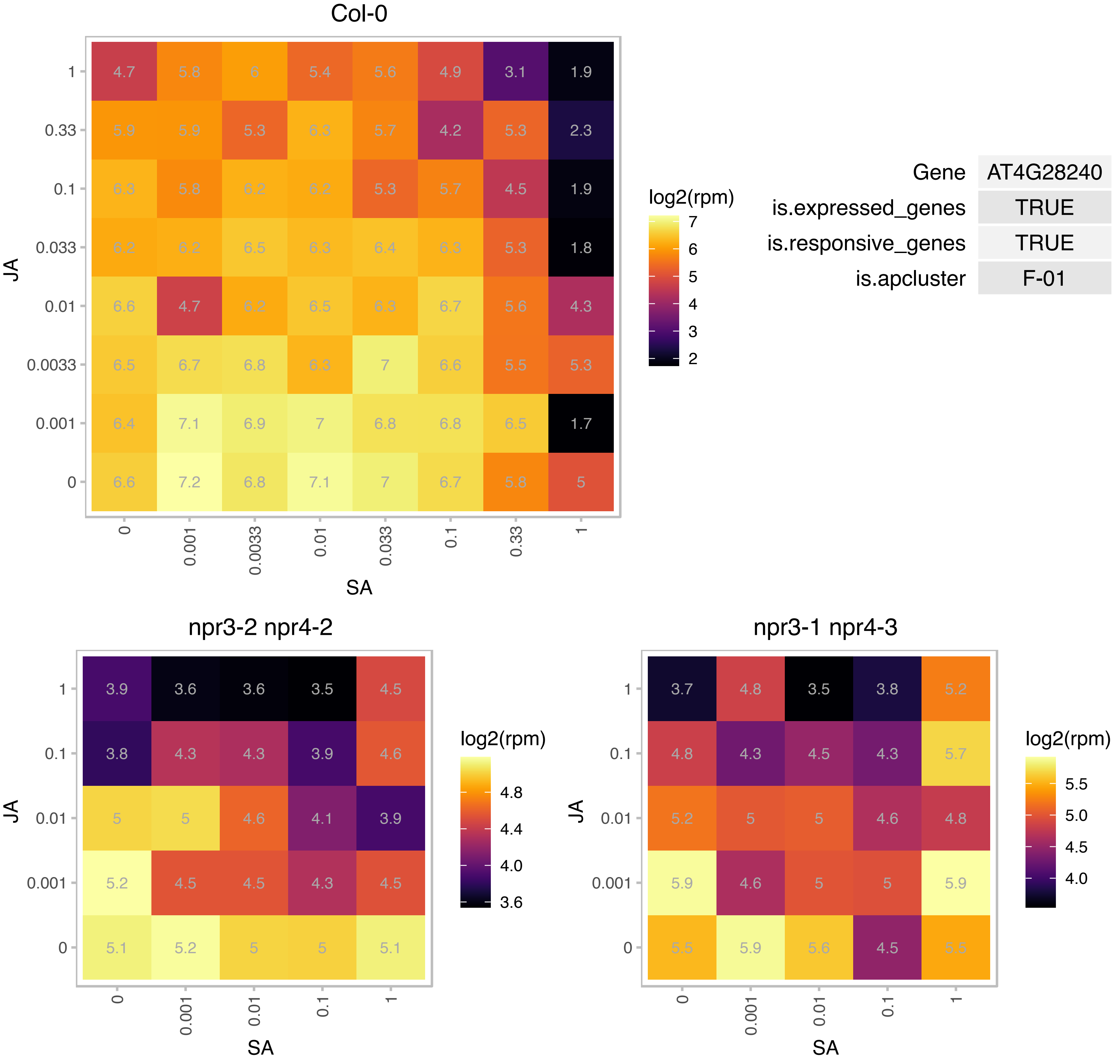

# AT4G28740.1 : NA

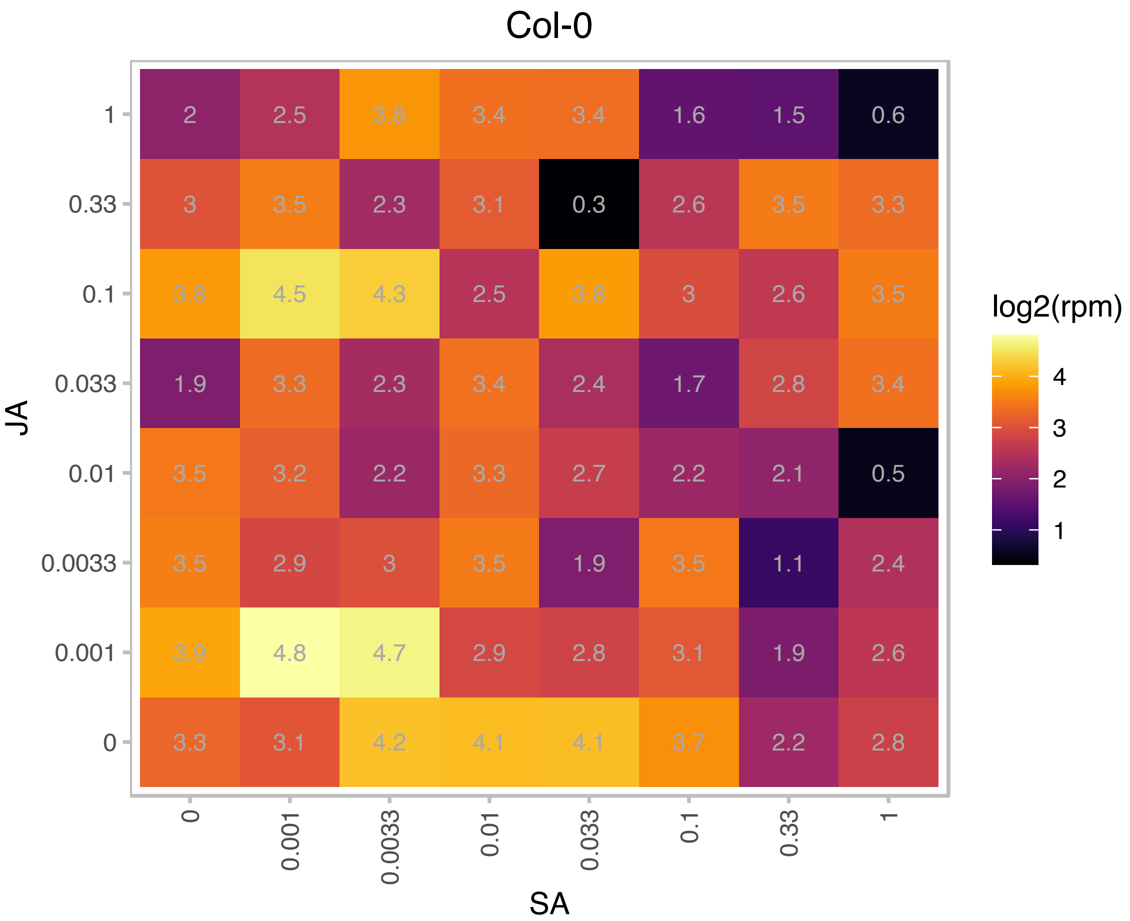


| | |
|---|---|
| Gene | AT4G28740 |
| is.expressed_genes | TRUE |
| is.responsive_genes | TRUE |
| is.apcluster | F-01 |

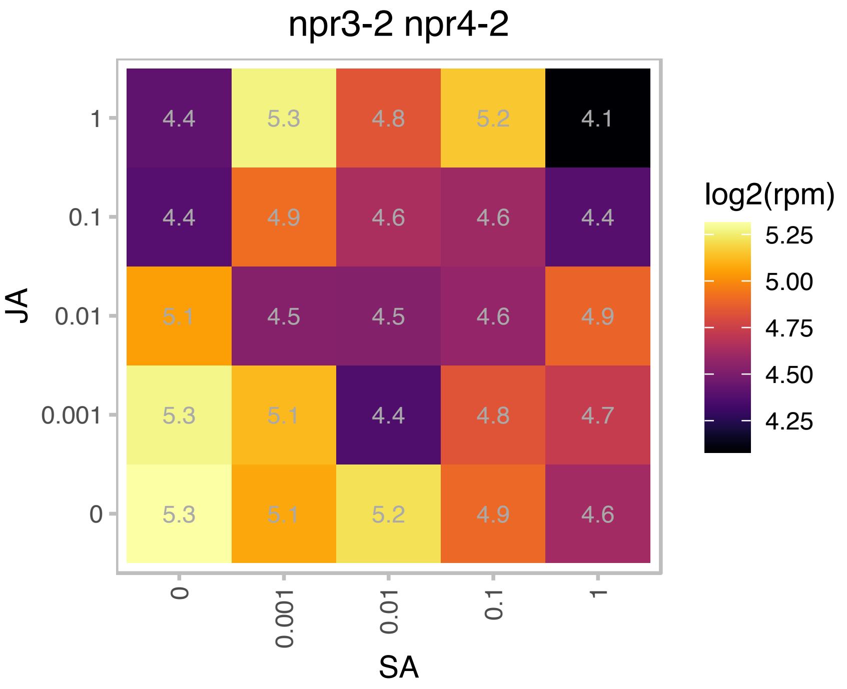


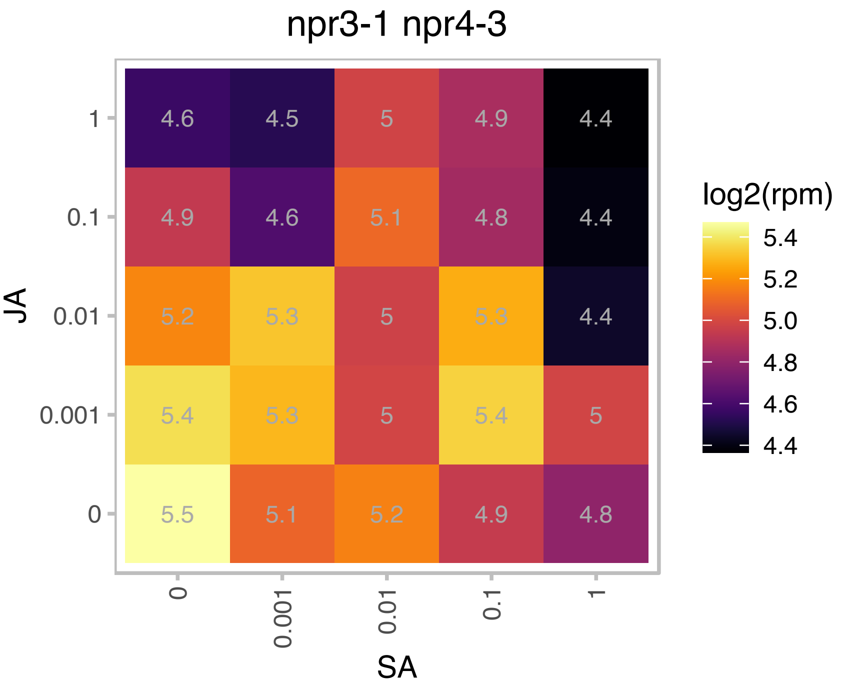

### AT4G29060.1 : emb2726

elongation factor Ts family protein

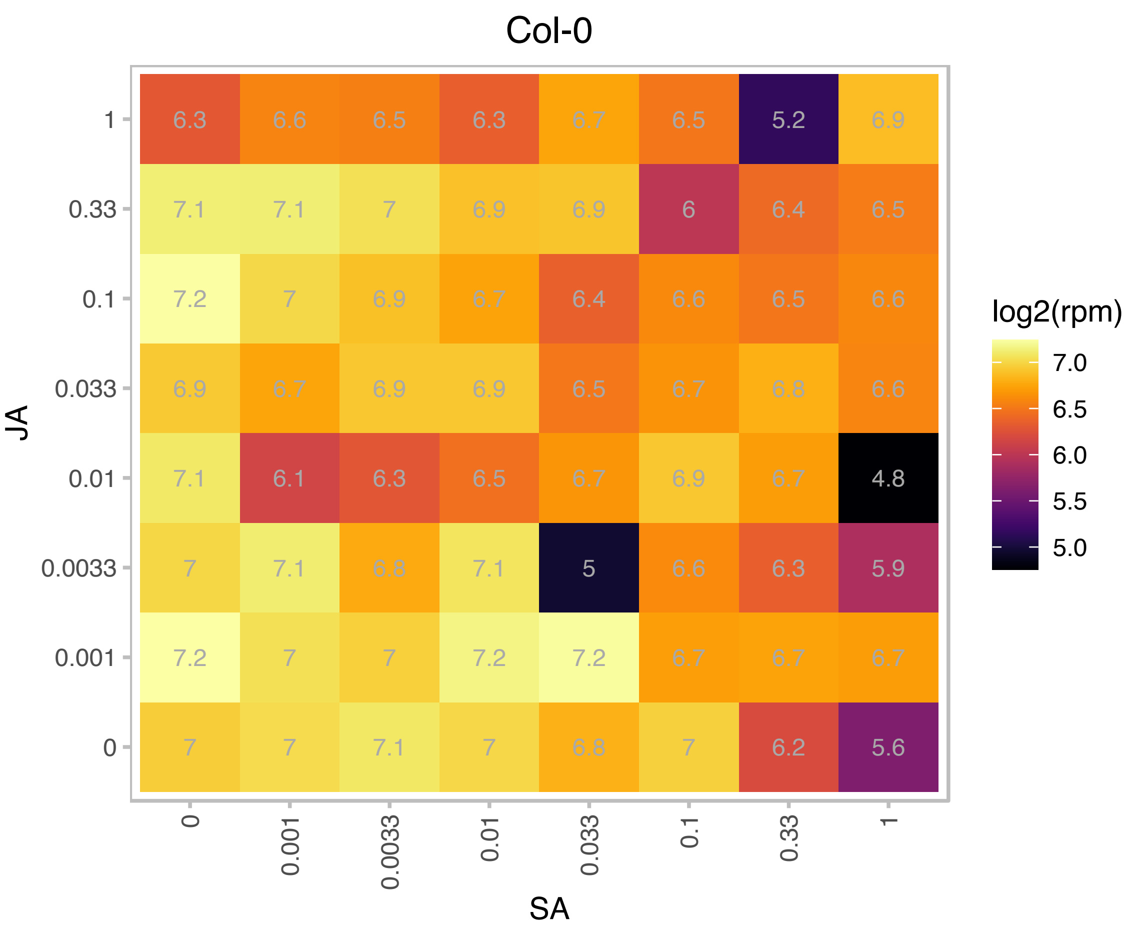


| | |
|---|---|
| Gene | AT4G29060 |
| is.expressed_genes | TRUE |
| is.responsive_genes | TRUE |
| is.apcluster | F-01 |

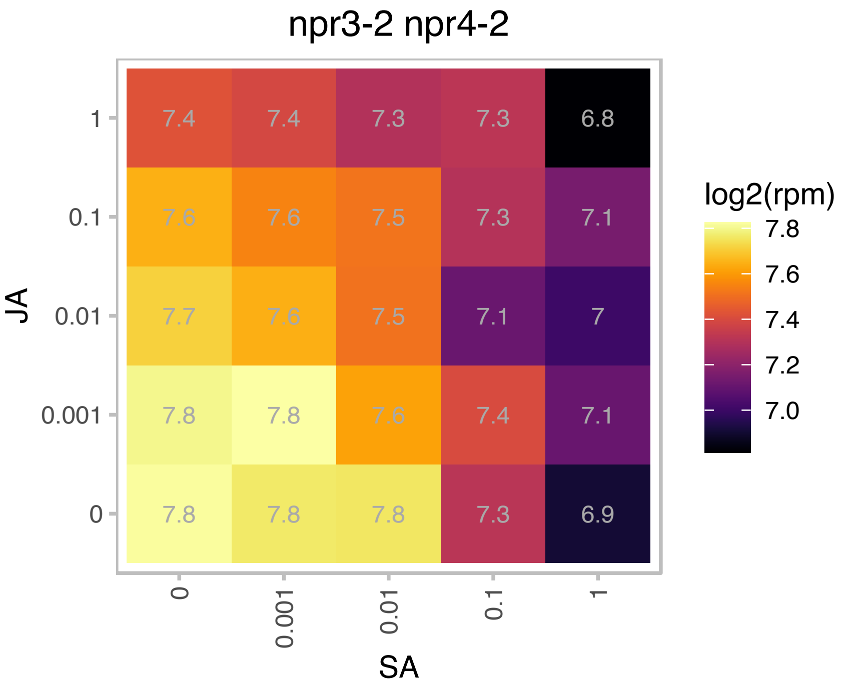


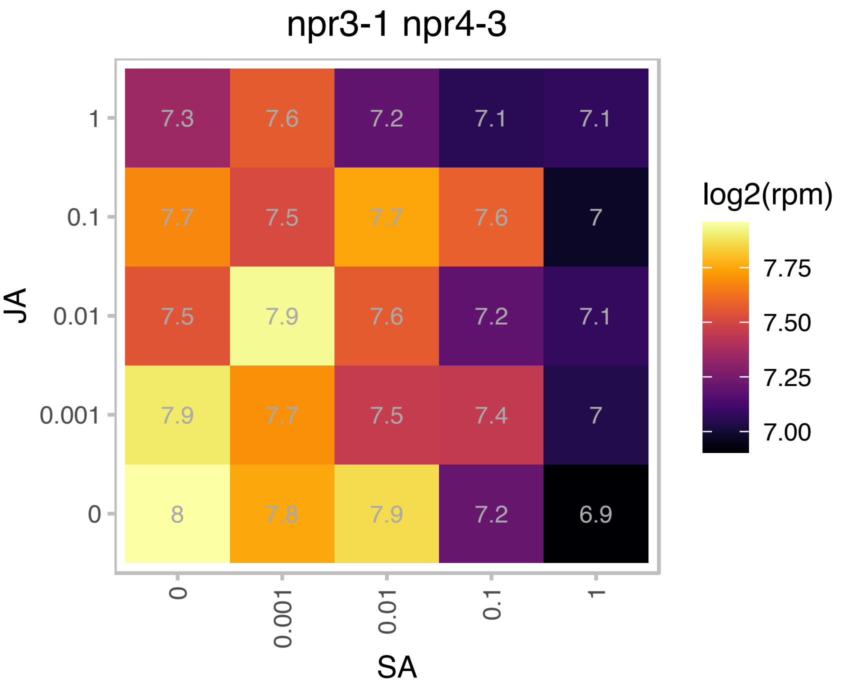

# AT4G29110.1 : NA

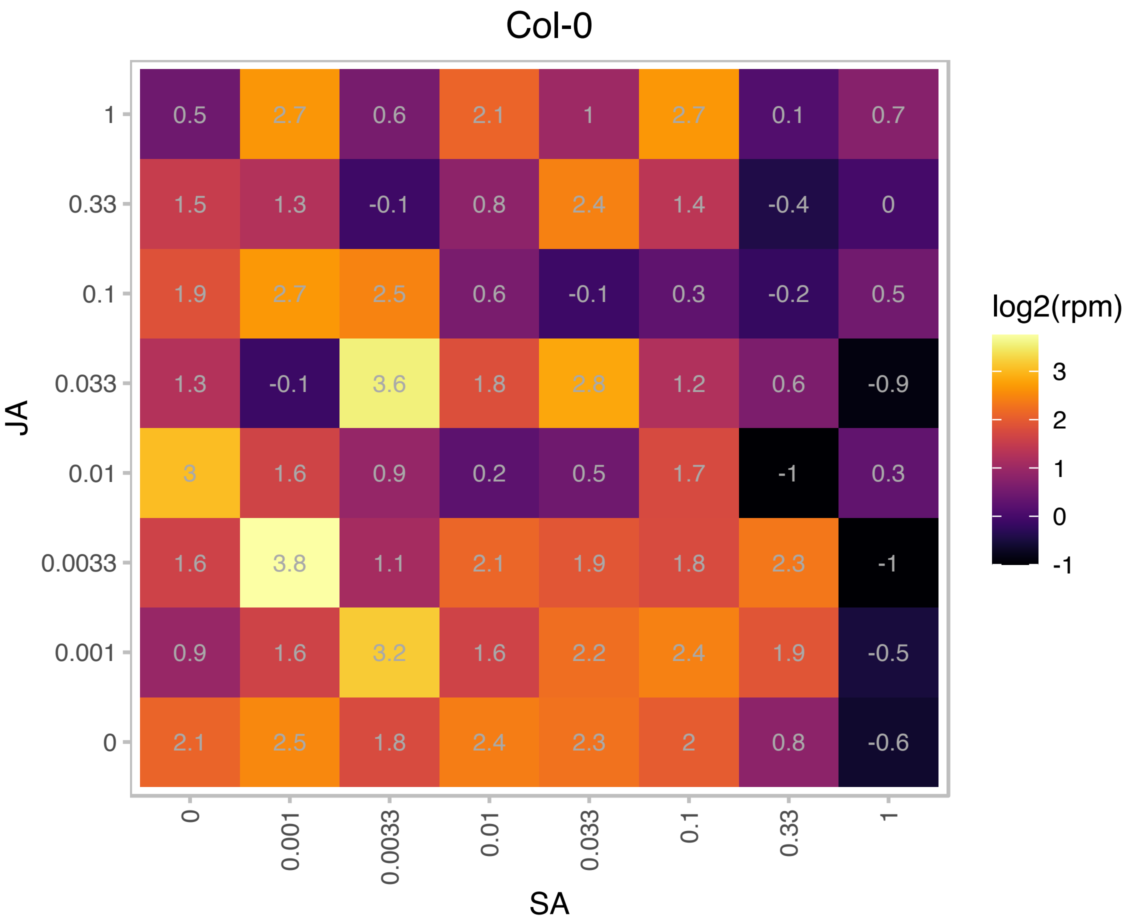


| | |
|---|---|
| Gene | AT4G29110 |
| is.expressed_genes | TRUE |
| is.responsive_genes | TRUE |
| is.apcluster | F-01 |

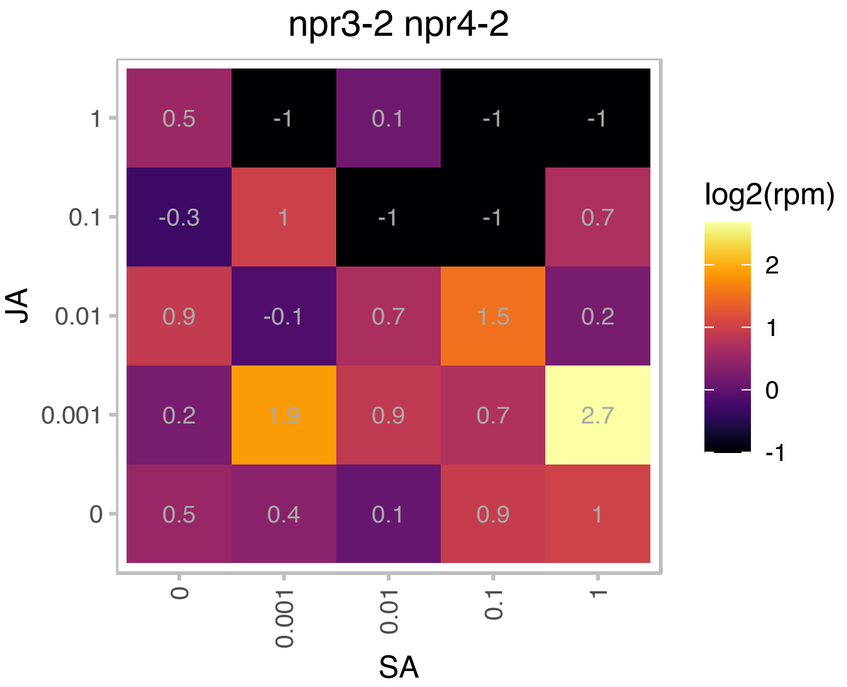


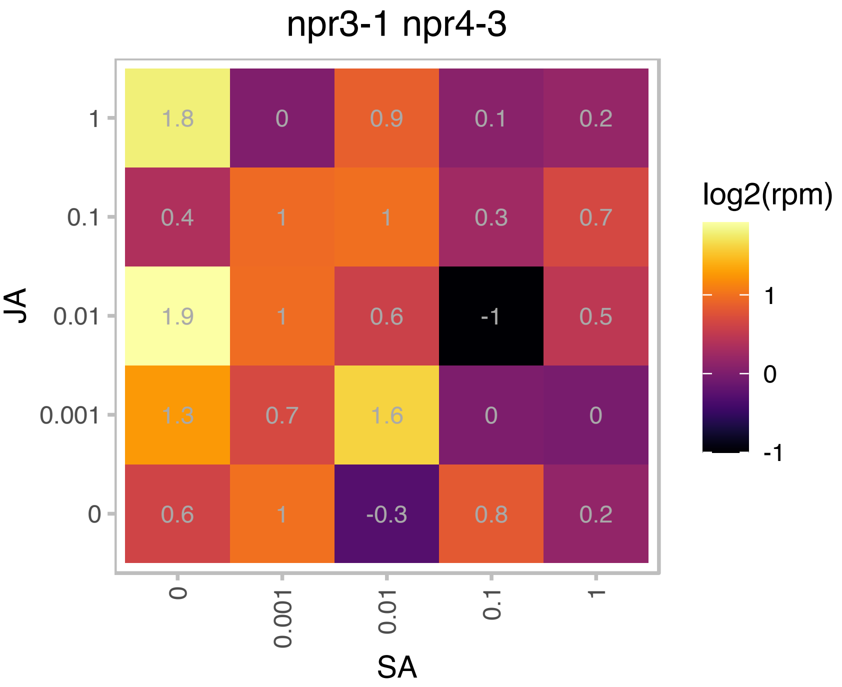

# AT4G33960.1 : NA

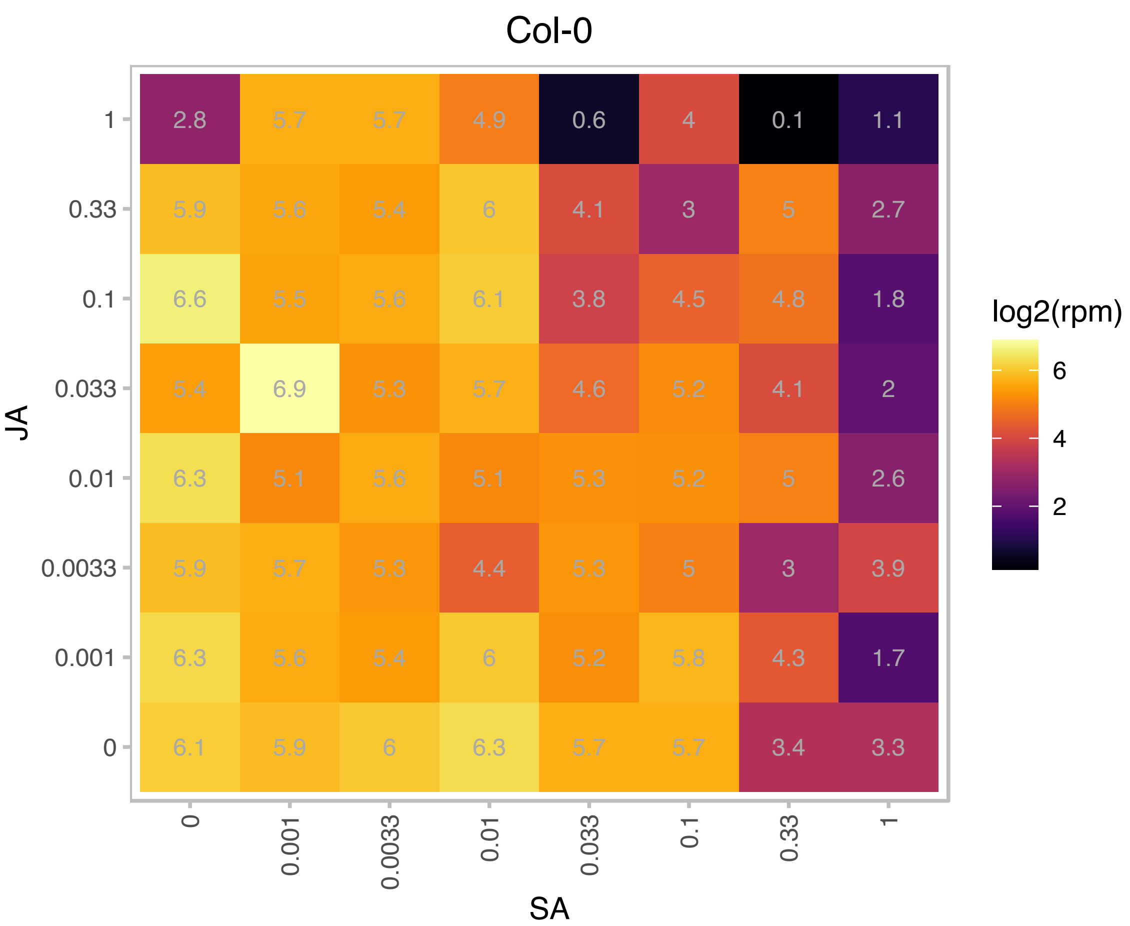


| | |
|---|---|
| Gene | AT4G33960 |
| is.expressed_genes | TRUE |
| is.responsive_genes | TRUE |
| is.apcluster | F-01 |

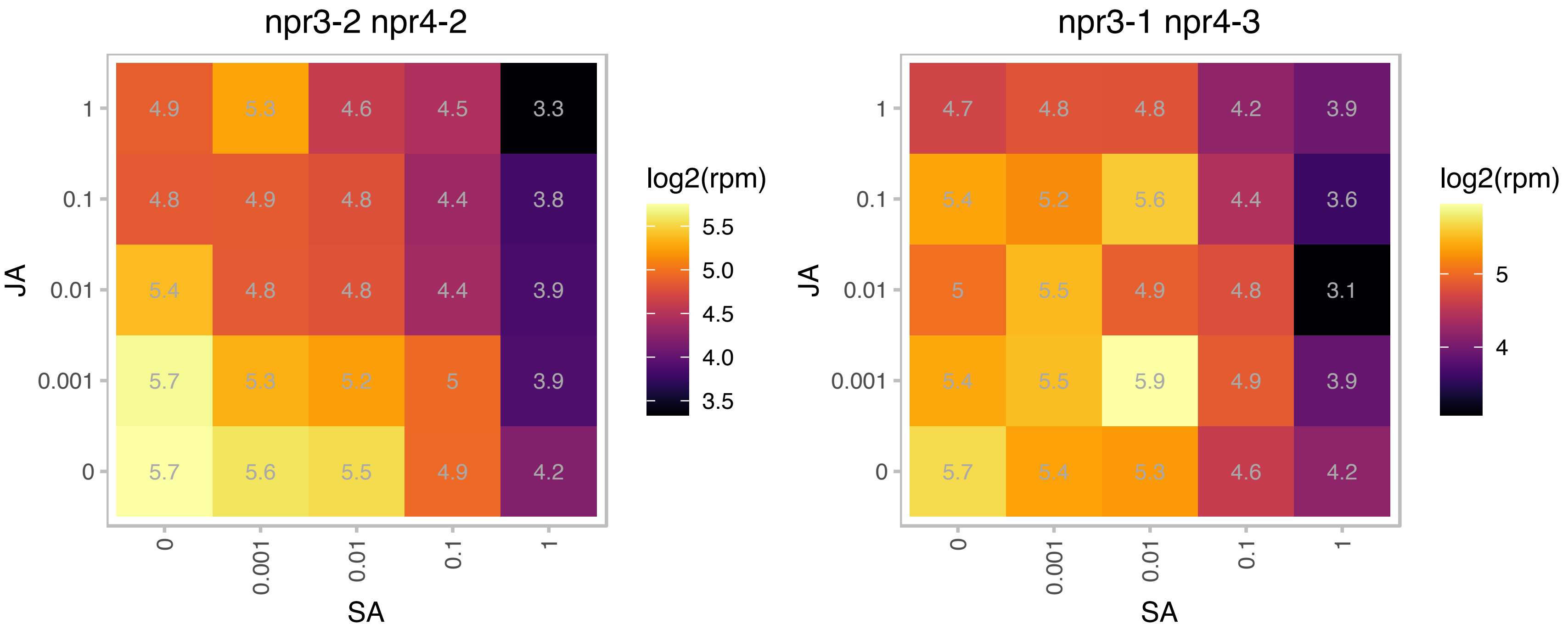

### AT4G36410.1 : UBC17

ubiquitin-conjugating enzyme 17

ubiquitin-conjugating enzyme

| | |
|---|---|
| Gene | AT4G36410 |
| is.expressed_genes | TRUE |
| is.responsive_genes | TRUE |
| is.apcluster | F-01 |

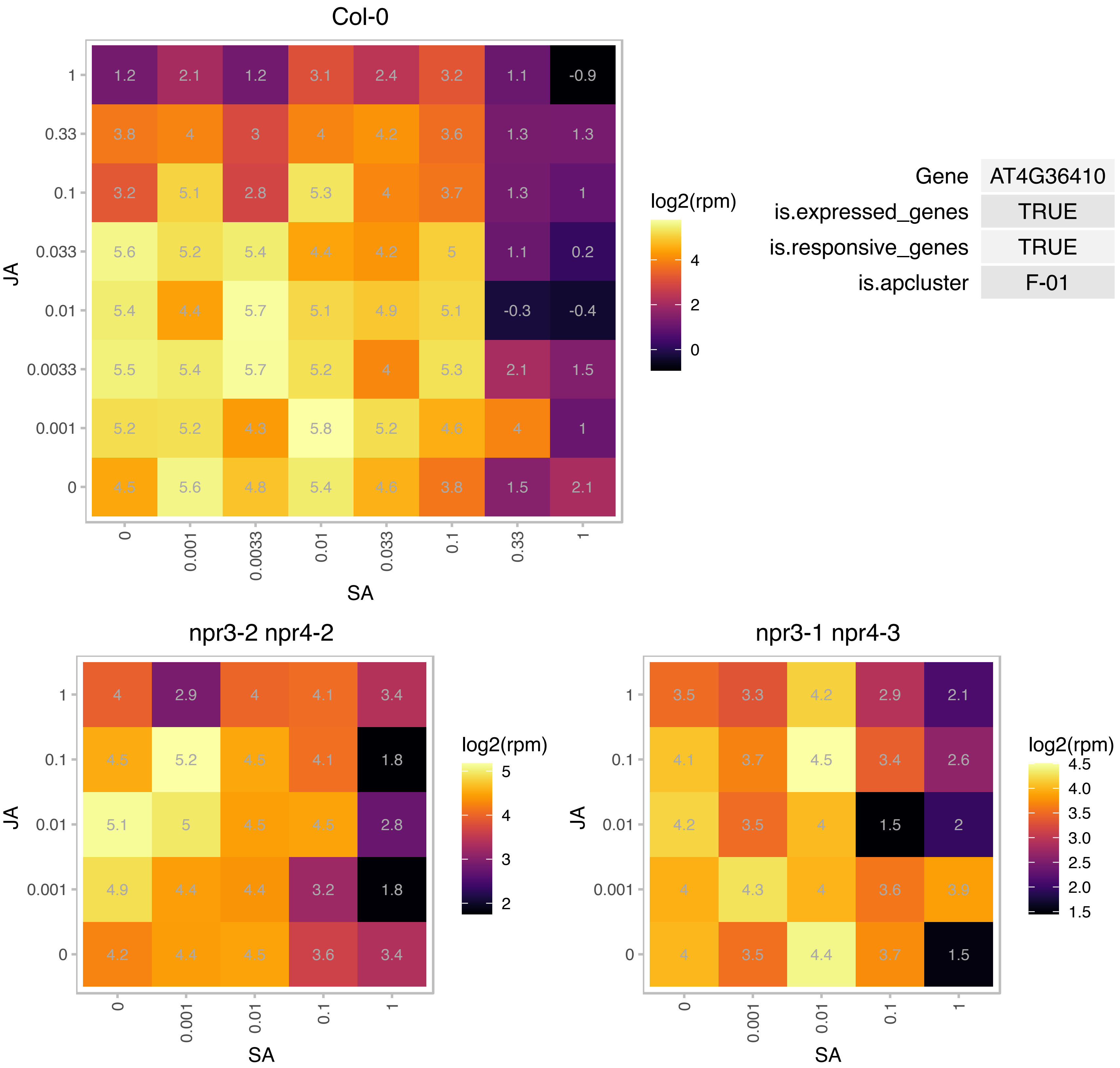

### AT4G40060.1 : ATHB-16, ATHB16, HB16

homeobox protein 16

Encodes a homeodomain leucine zipper class I (HD-Zip I) protein.

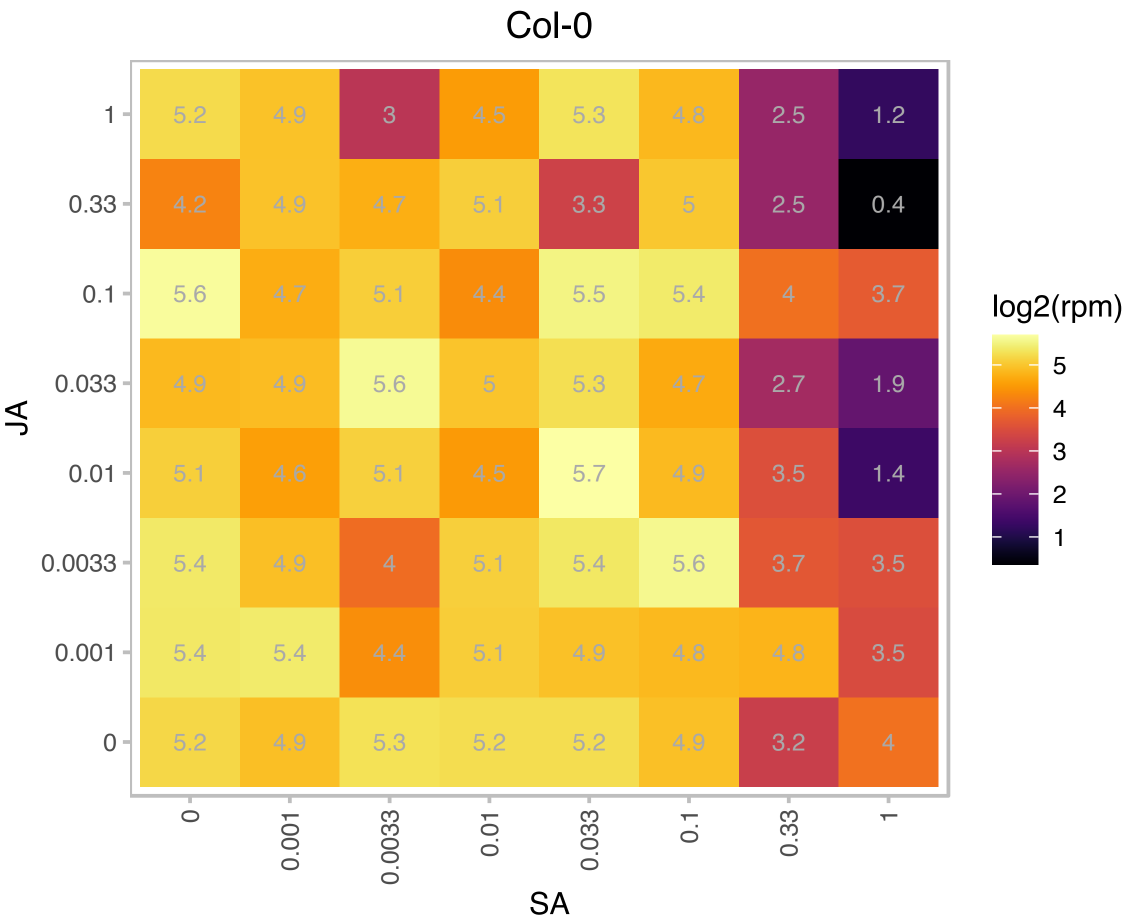


| | |
|---|---|
| Gene | AT4G40060 |
| is.expressed_genes | TRUE |
| is.responsive_genes | TRUE |
| is.apcluster | F-01 |

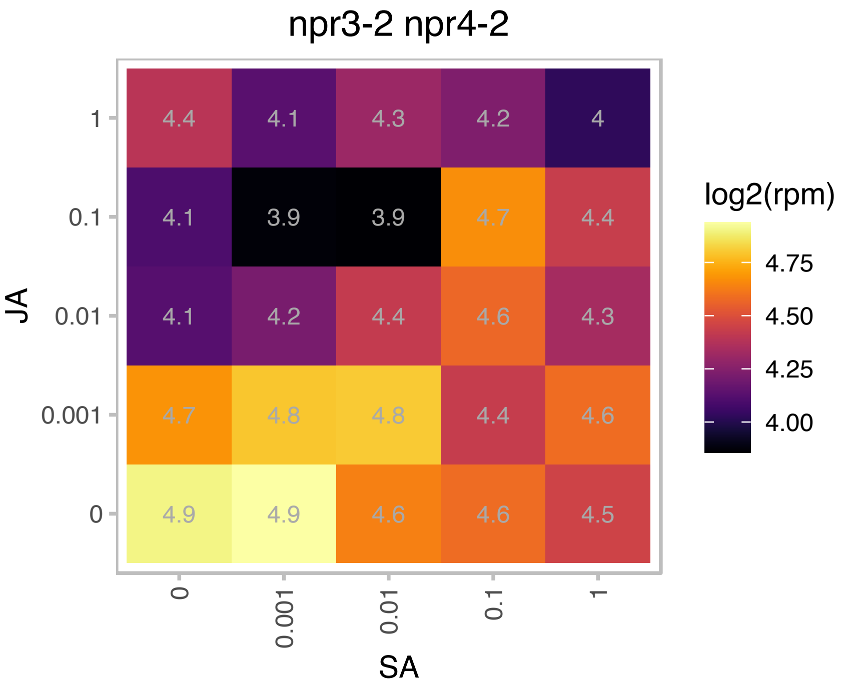


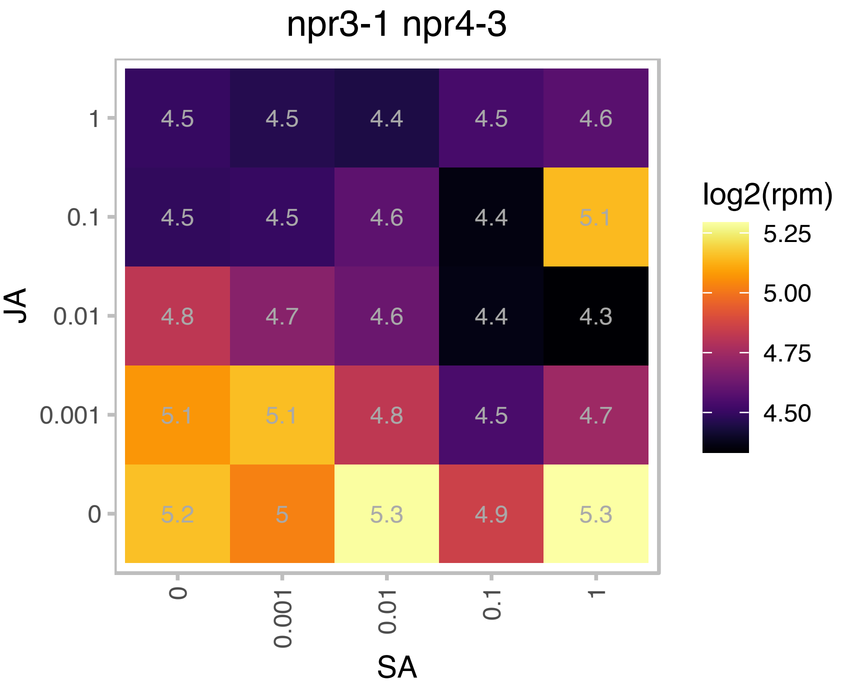

### AT5G05000.1 : ATTOC34, OEP34, TOC34

translocon at the outer envelope membrane of chloroplasts 34

Outer membrane GTPase protein that may function in import of nuclear encoded proteins into the chloroplast. Phosphorylation of the G-domains regulate translocon assembly.

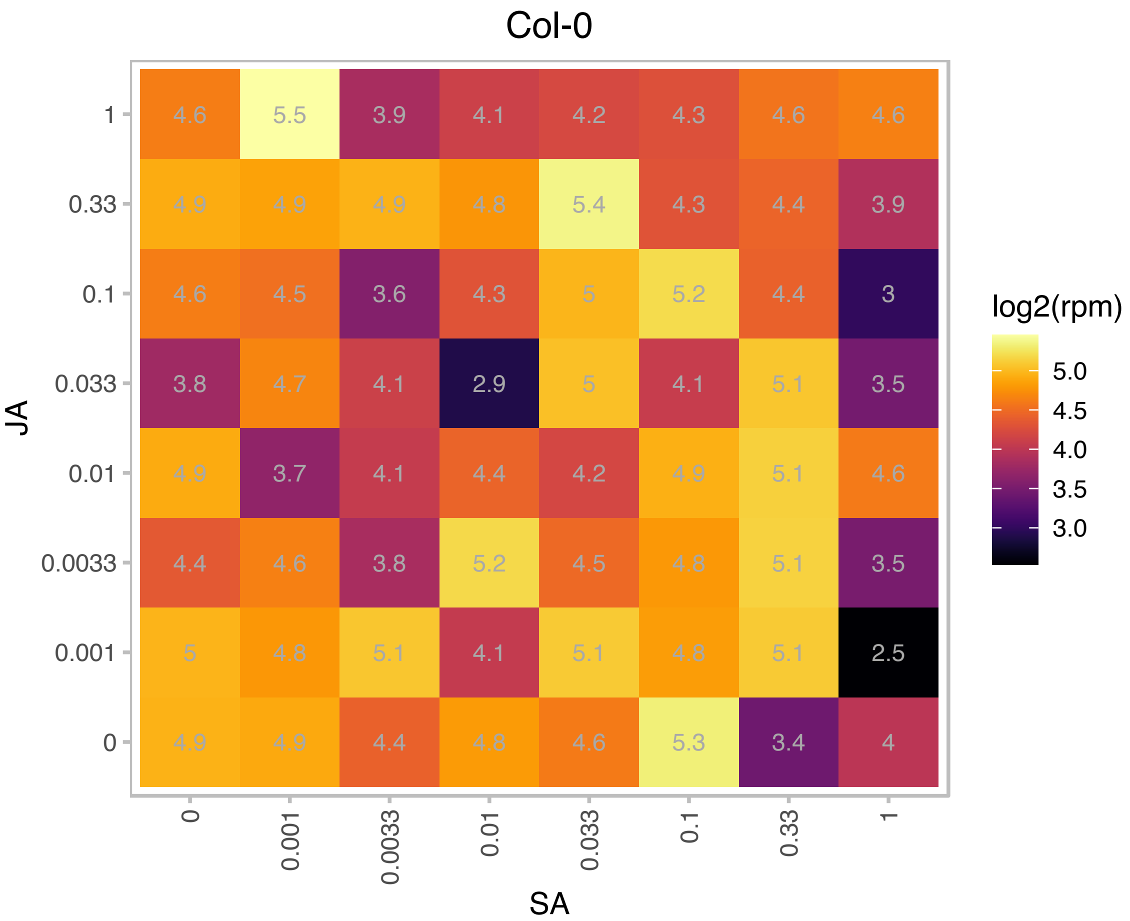


| | |
|---|---|
| Gene | AT5G05000 |
| is.expressed_genes | TRUE |
| is.responsive_genes | TRUE |
| is.apcluster | F-01 |

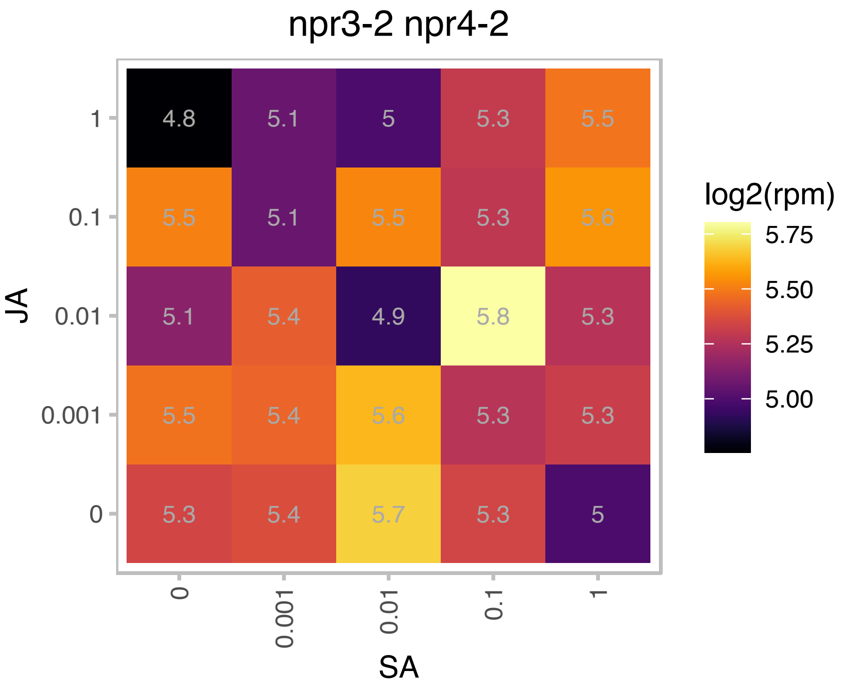


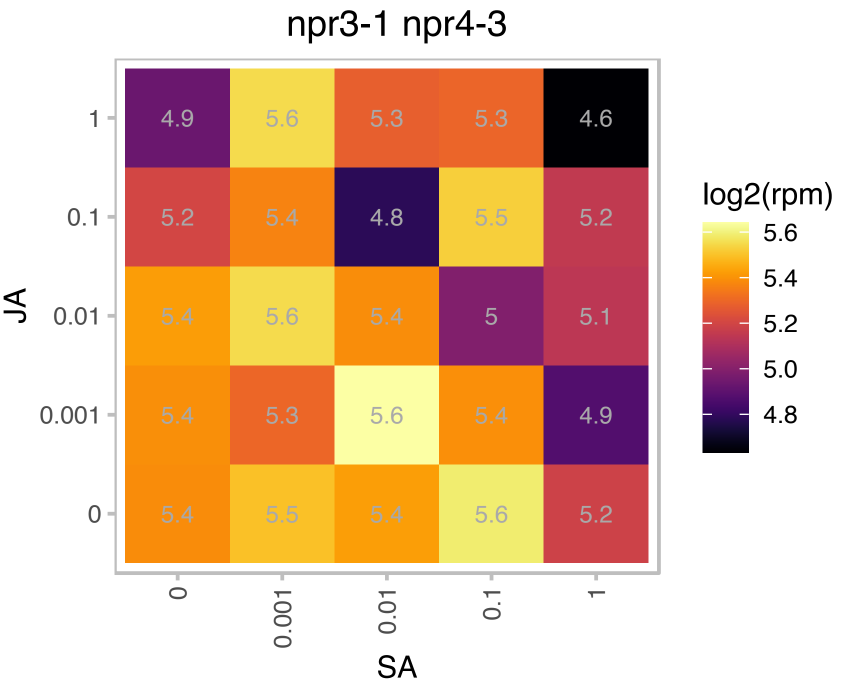

### AT5G08050.1 : RIQ1

Protein of unknown function (DUF1118)

Encodes a grana core localized protein. Mutant plants have reduced NPQ, affected organization of light-havesting complex II and an enhanced grana stacking.

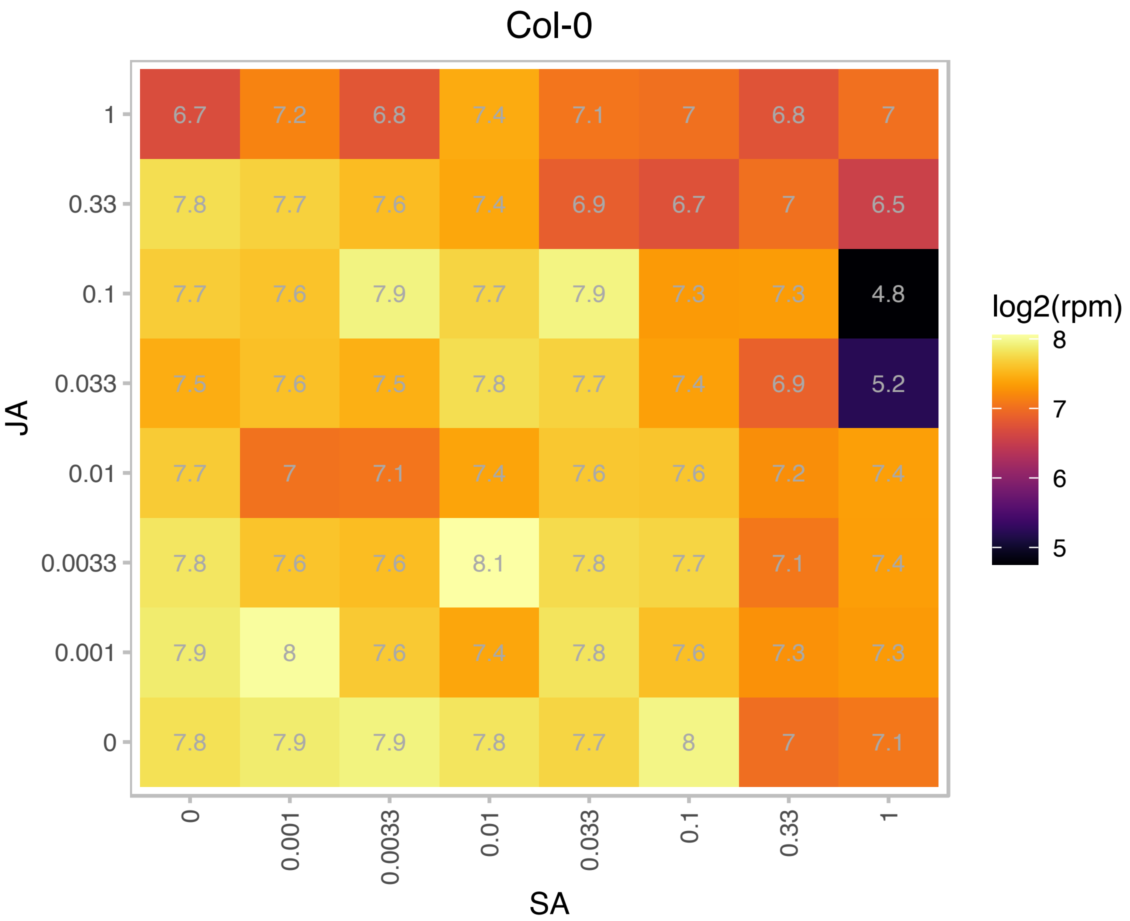


| | |
|---|---|
| Gene | AT5G08050 |
| is.expressed_genes | TRUE |
| is.responsive_genes | TRUE |
| is.apcluster | F-01 |

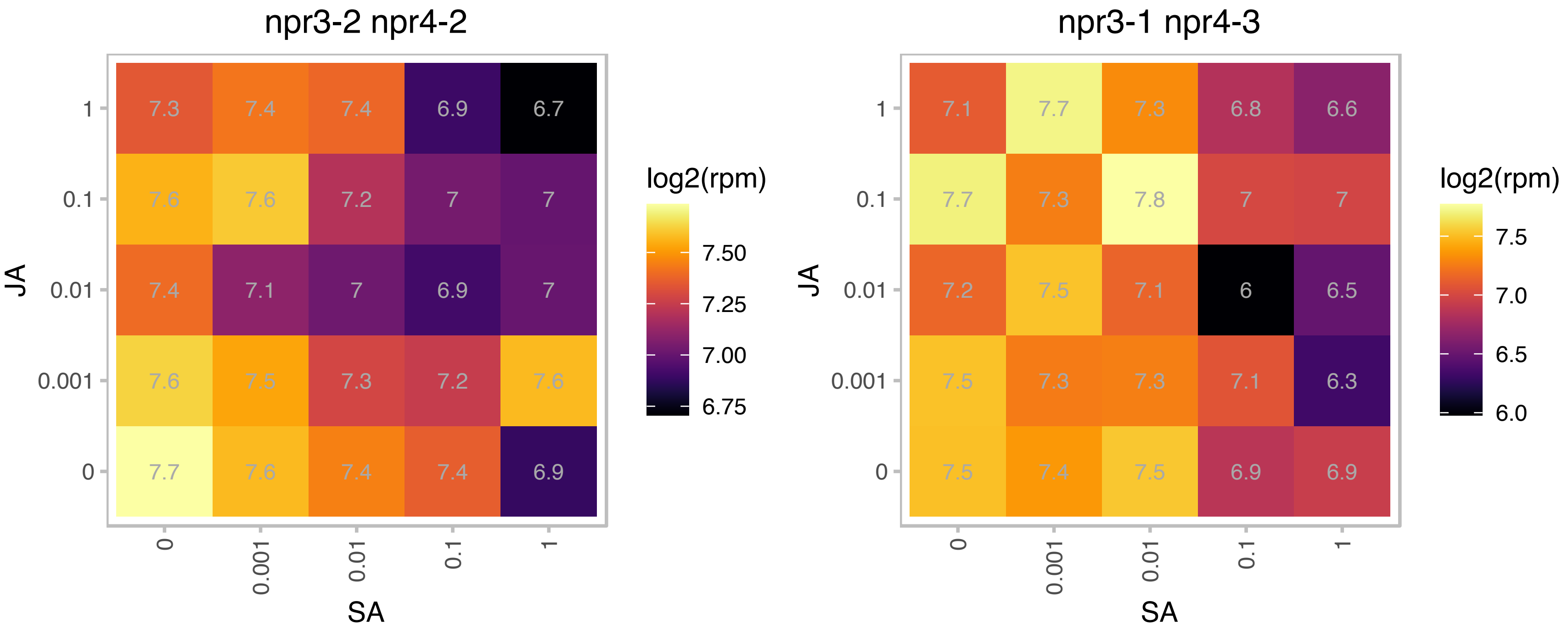

# AT5G08400.1 : NA

Protein of unknown function (DUF3531)

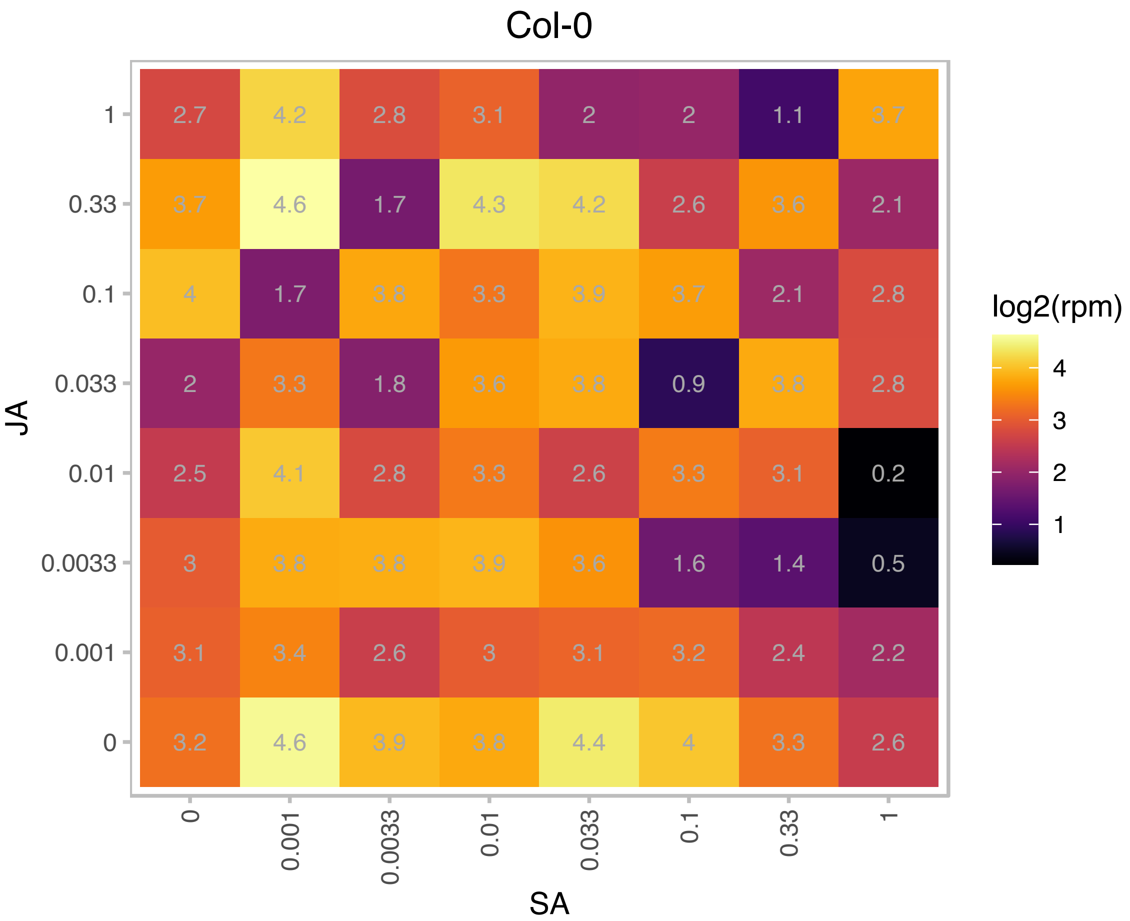


| | |
|---|---|
| Gene | AT5G08400 |
| is.expressed_genes | TRUE |
| is.responsive_genes | TRUE |
| is.apcluster | F-01 |

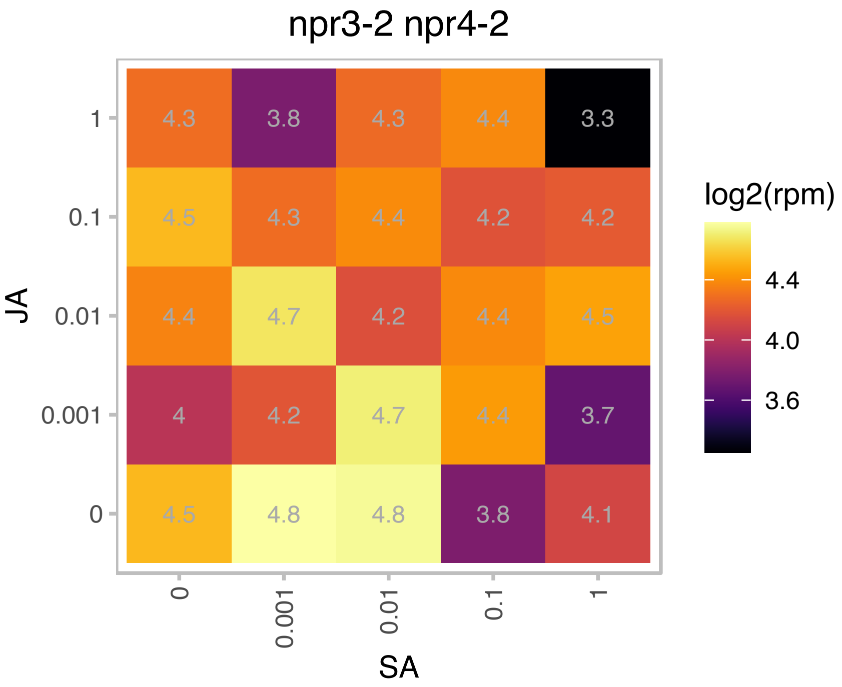


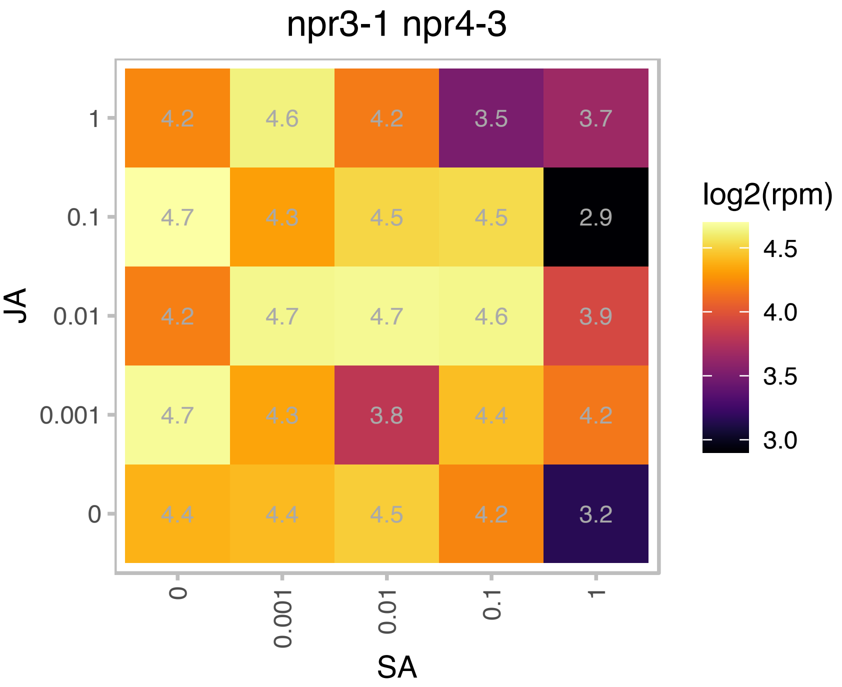

### AT5G08640.1 : ATFLS1, FLS, FLS1

flavonol synthase 1

Encodes a flavonol synthase that catalyzes formation of flavonols from dihydroflavonols. Co-expressed with CHI and CHS (qRT-PCR).

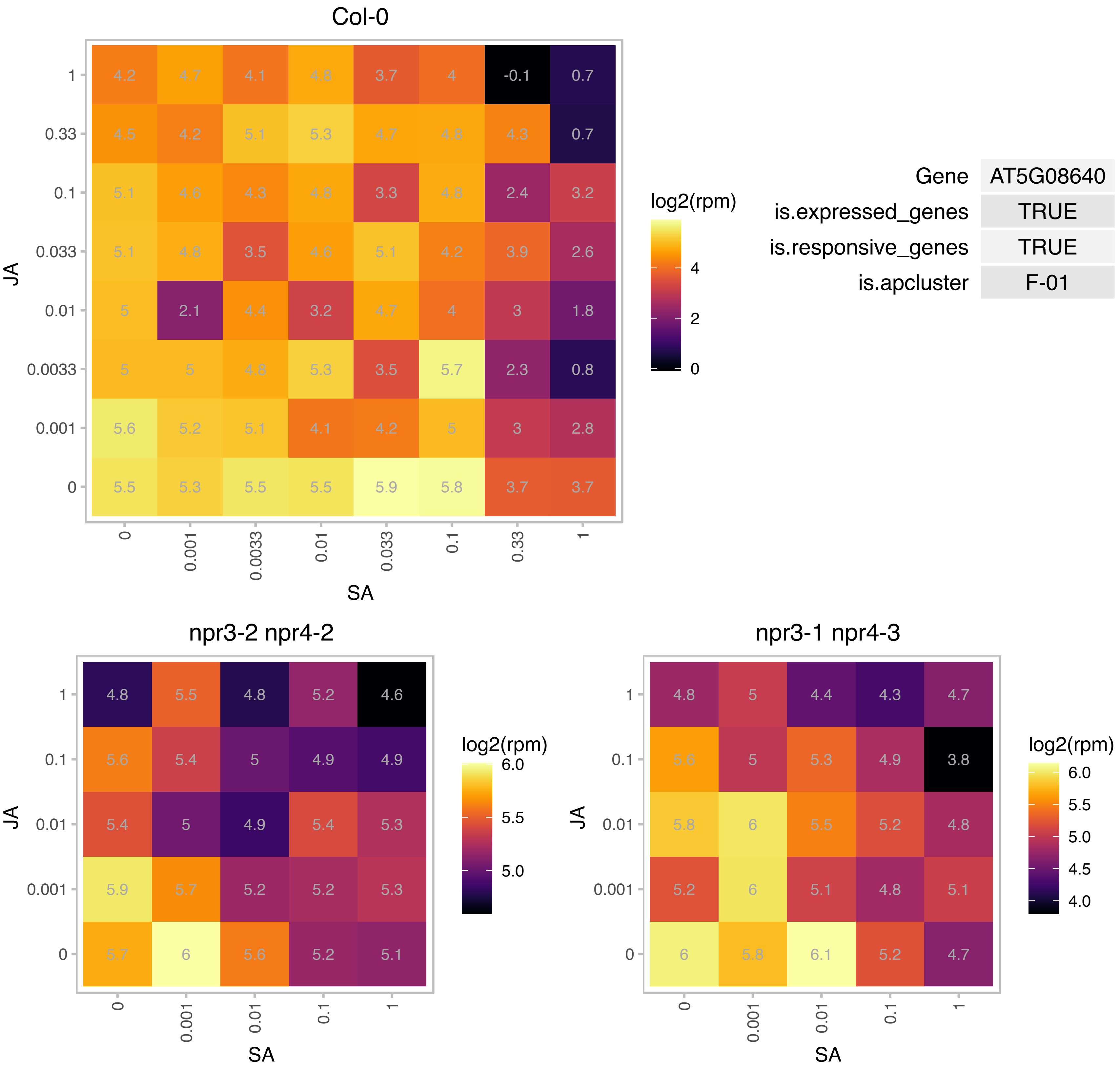

# AT5G09240.1 : NA

ssDNA-binding transcriptional regulator

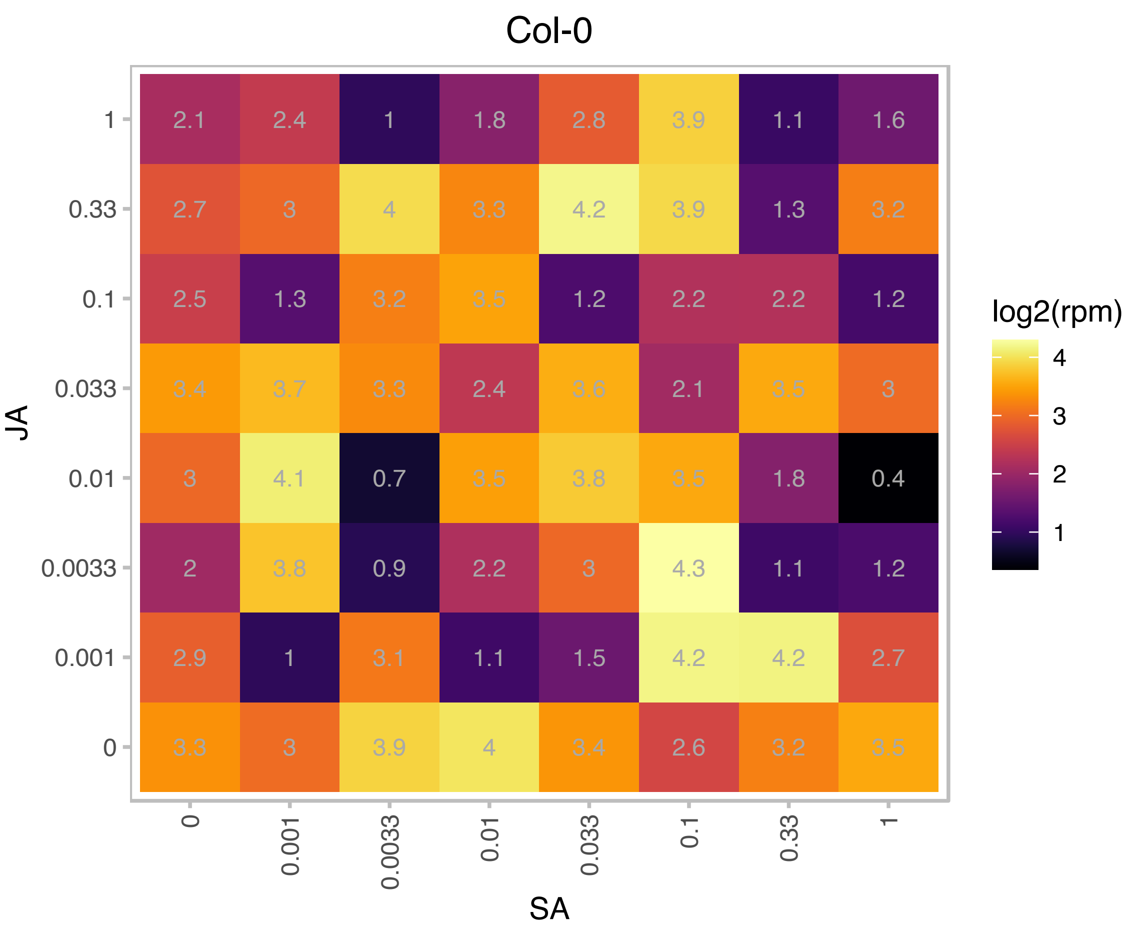


| | |
|---|---|
| Gene | AT5G09240 |
| is.expressed_genes | TRUE |
| is.responsive_genes | TRUE |
| is.apcluster | F-01 |

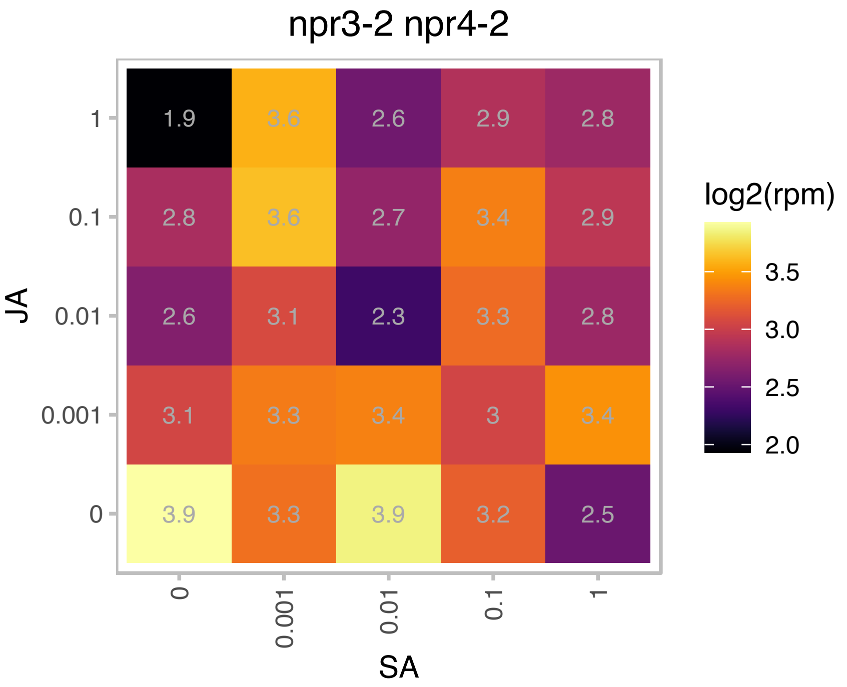


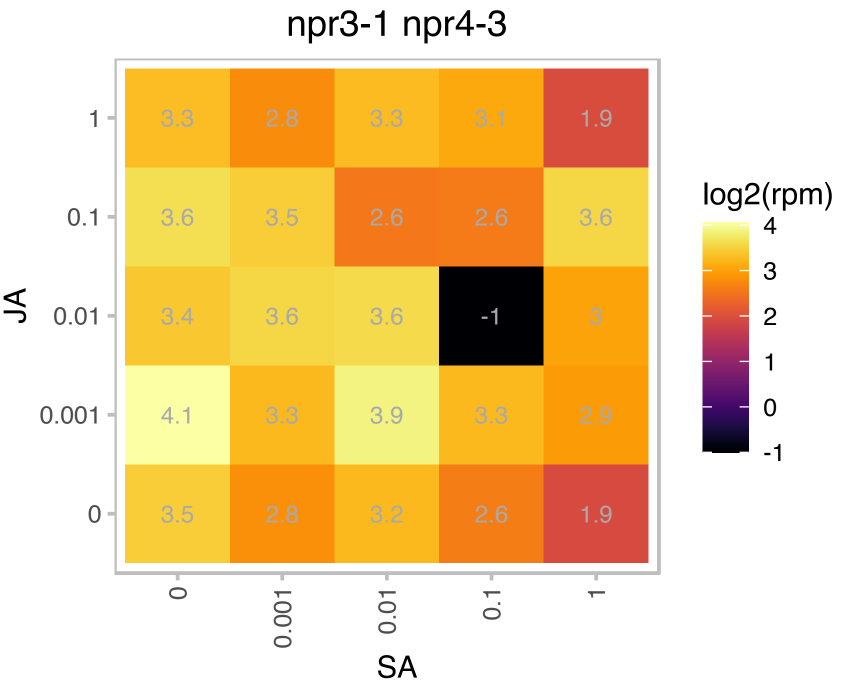

### AT5G10430.1 : AGP4, ATAGP4, JAGGER

arabinogalactan protein 4
Encodes arabinogalactan-protein (AGP4) that is expressed in female reproductive tissues. It is involved in promoting degeneration of the persistent synergid after fertilization. In mutant ovules, the persistent synergid does not degrade resulting in polytuby.

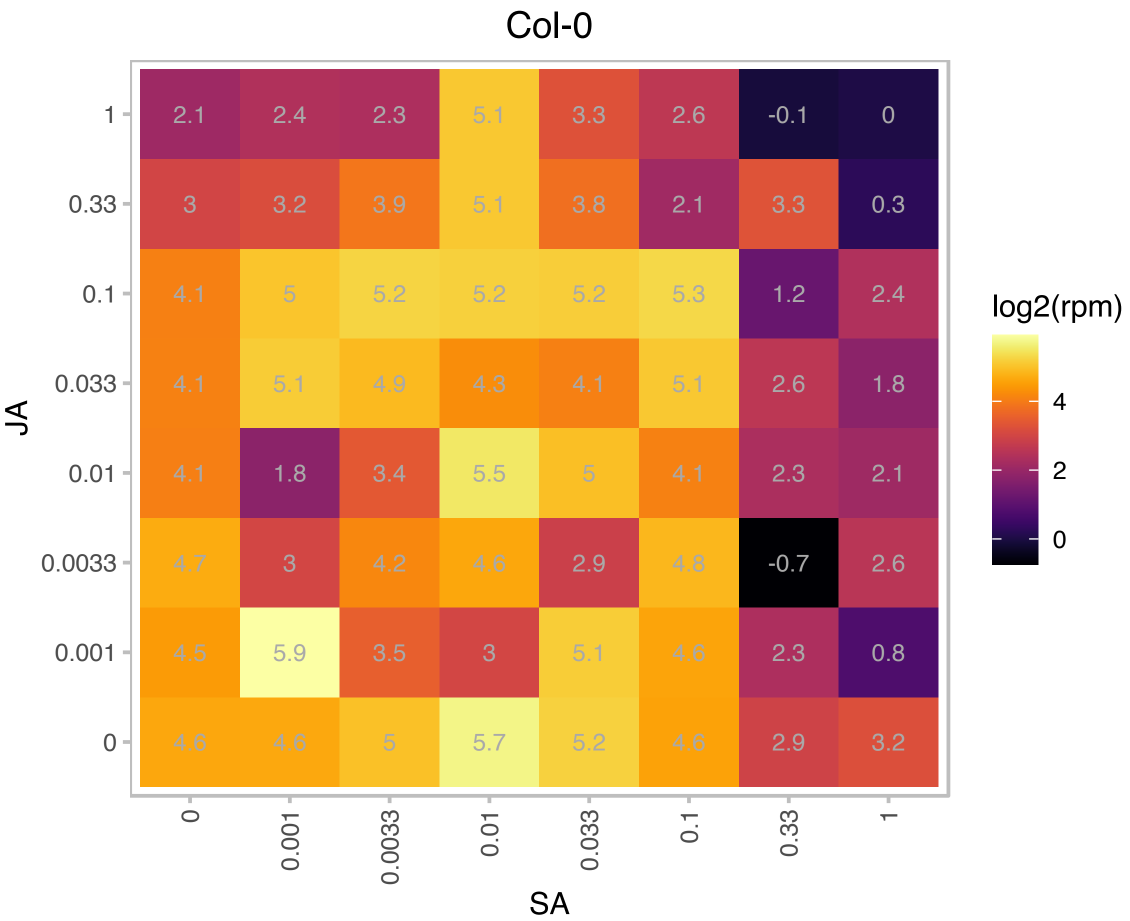


| | |
|---|---|
| Gene | AT5G10430 |
| is.expressed_genes | TRUE |
| is.responsive_genes | TRUE |
| is.apcluster | F-01 |

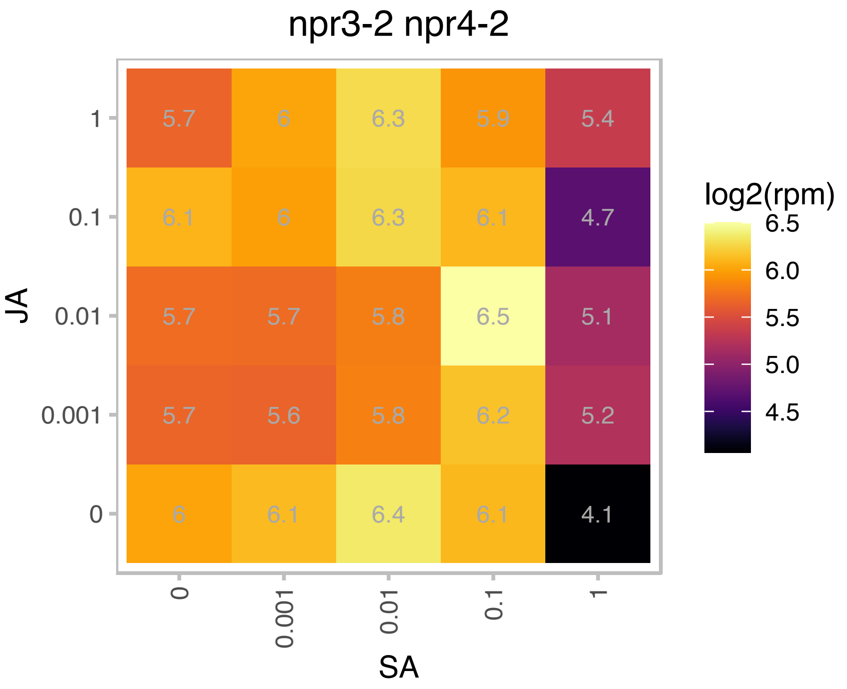


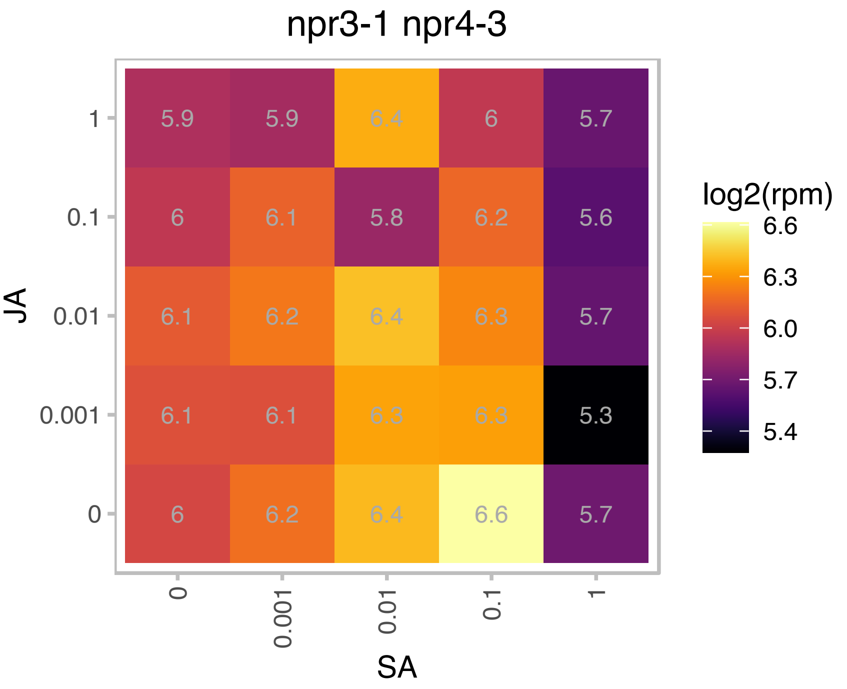

# AT5G11090.1 : NA

serine-rich protein-related

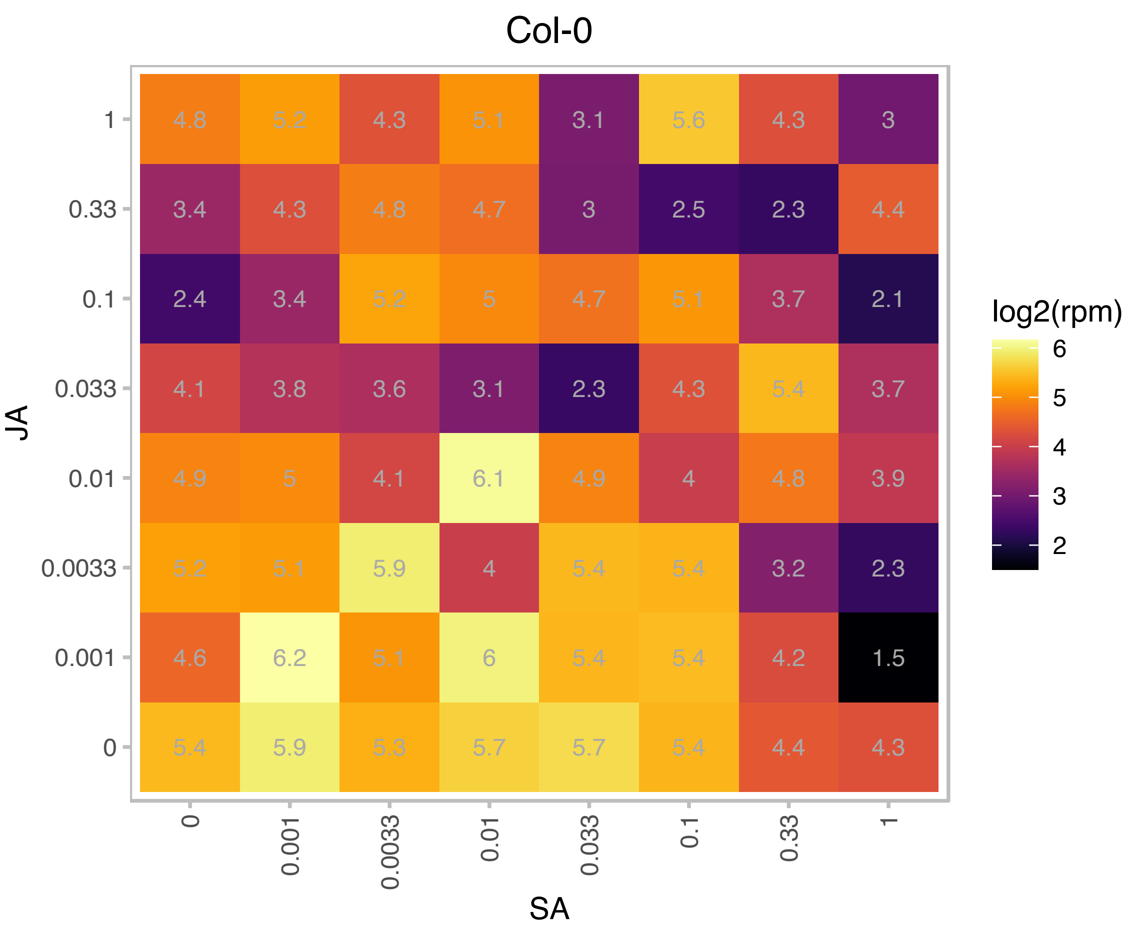


| | |
|---|---|
| Gene | AT5G11090 |
| is.expressed_genes | TRUE |
| is.responsive_genes | TRUE |
| is.apcluster | F-01 |

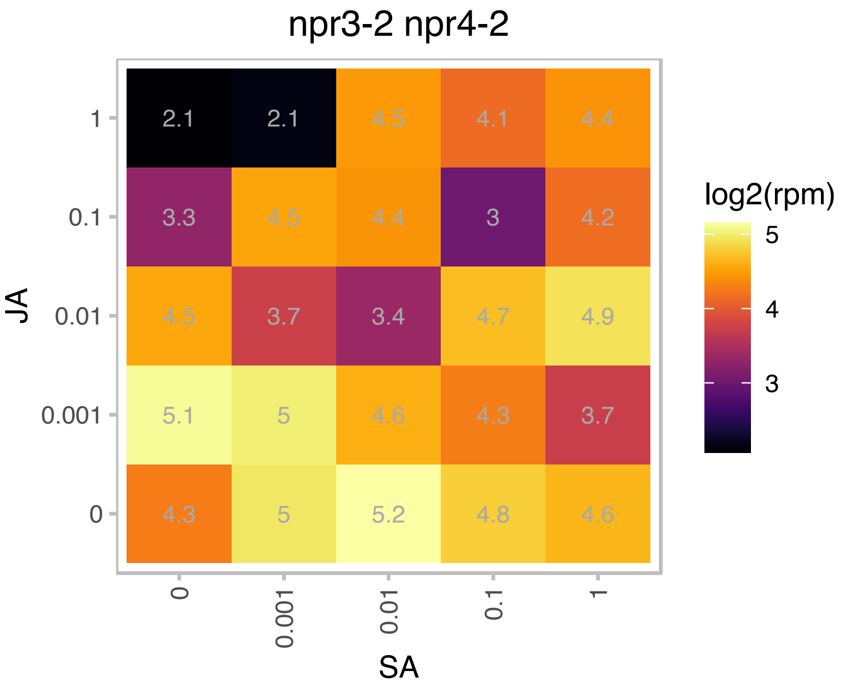


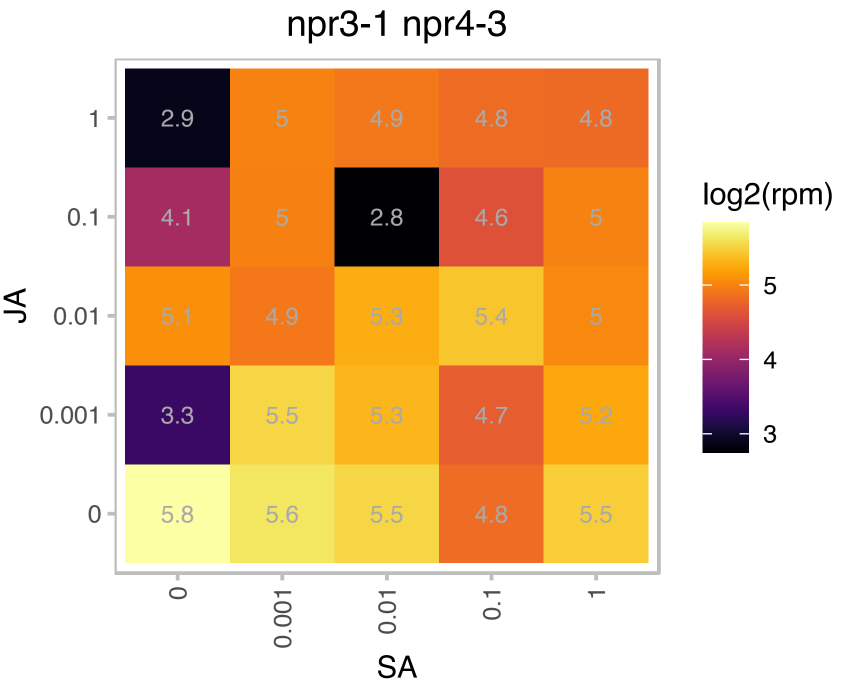

# AT5G12110.1 : NA

Glutathione S-transferase, C-terminal-like;Translation elongation factor EF1B/ribosomal protein S6

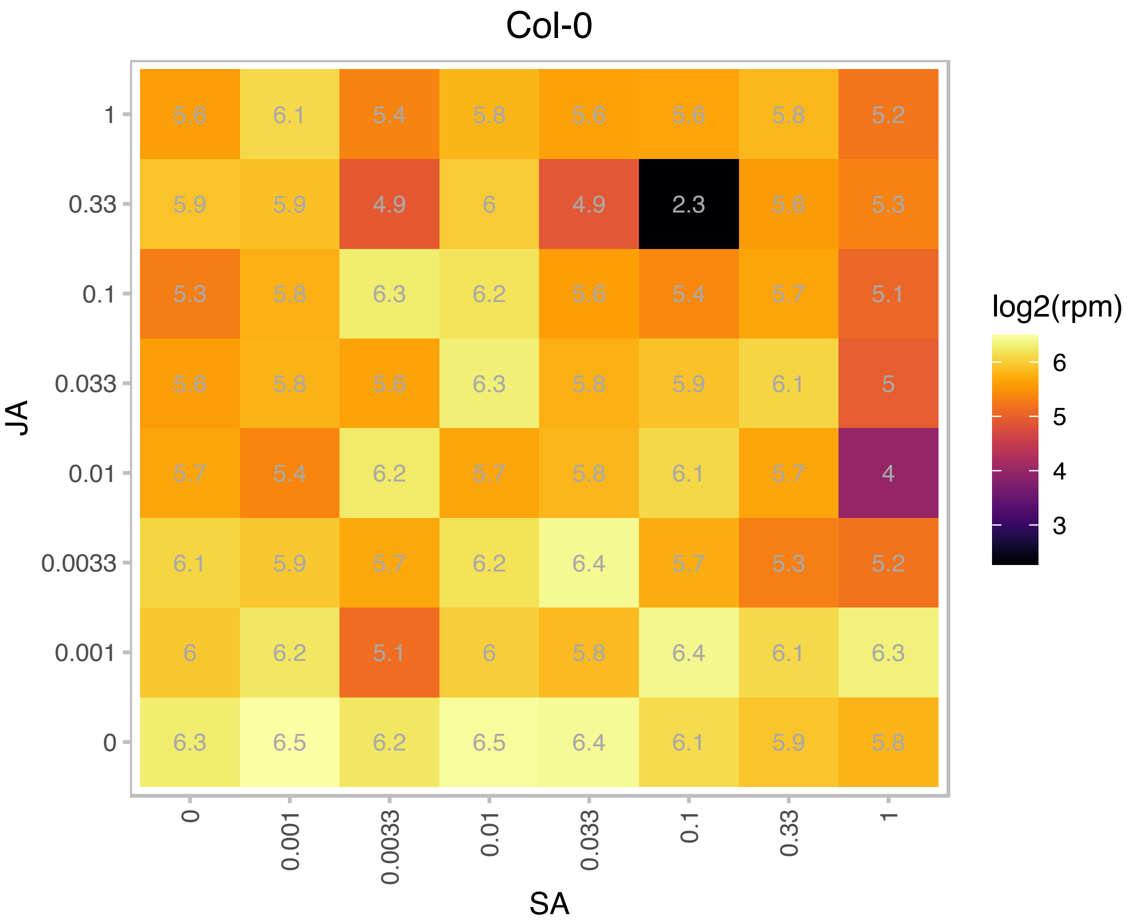


| | |
|---|---|
| Gene | AT5G12110 |
| is.expressed_genes | TRUE |
| is.responsive_genes | TRUE |
| is.apcluster | F-01 |

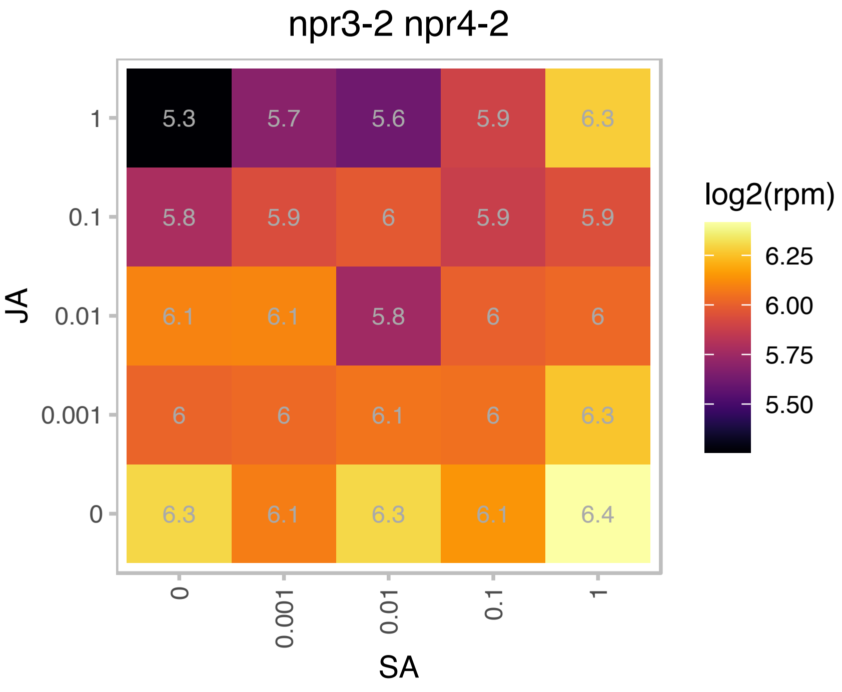


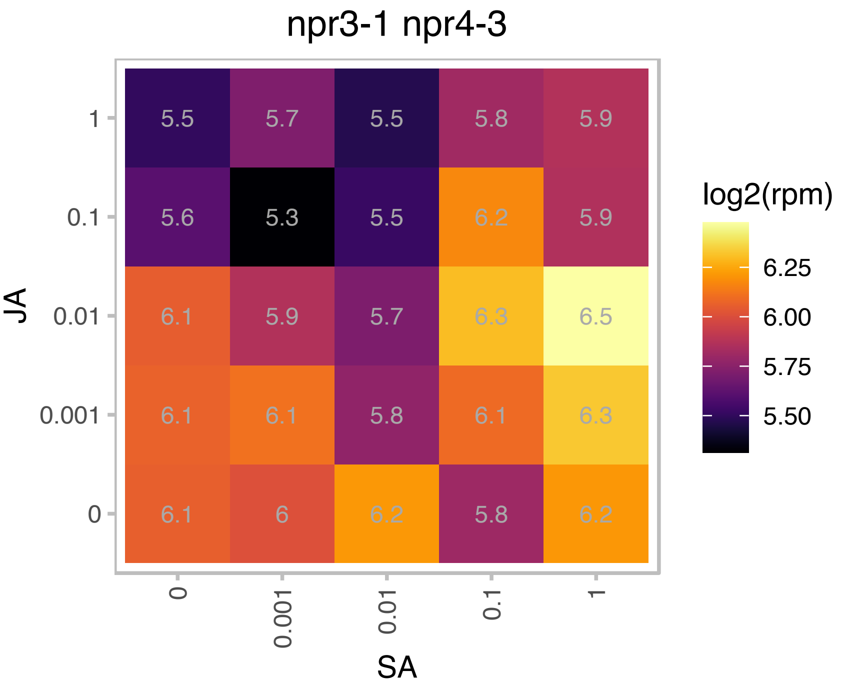

### AT5G14920.1 : GASA14

Gibberellin-regulated family protein
Encodes a GASA domain containing protein that regulates increases in plant growth through GA-induced and DELLA-dependent signal transduction and that can increase abiotic stress resistance by reducing ROS accumulation.

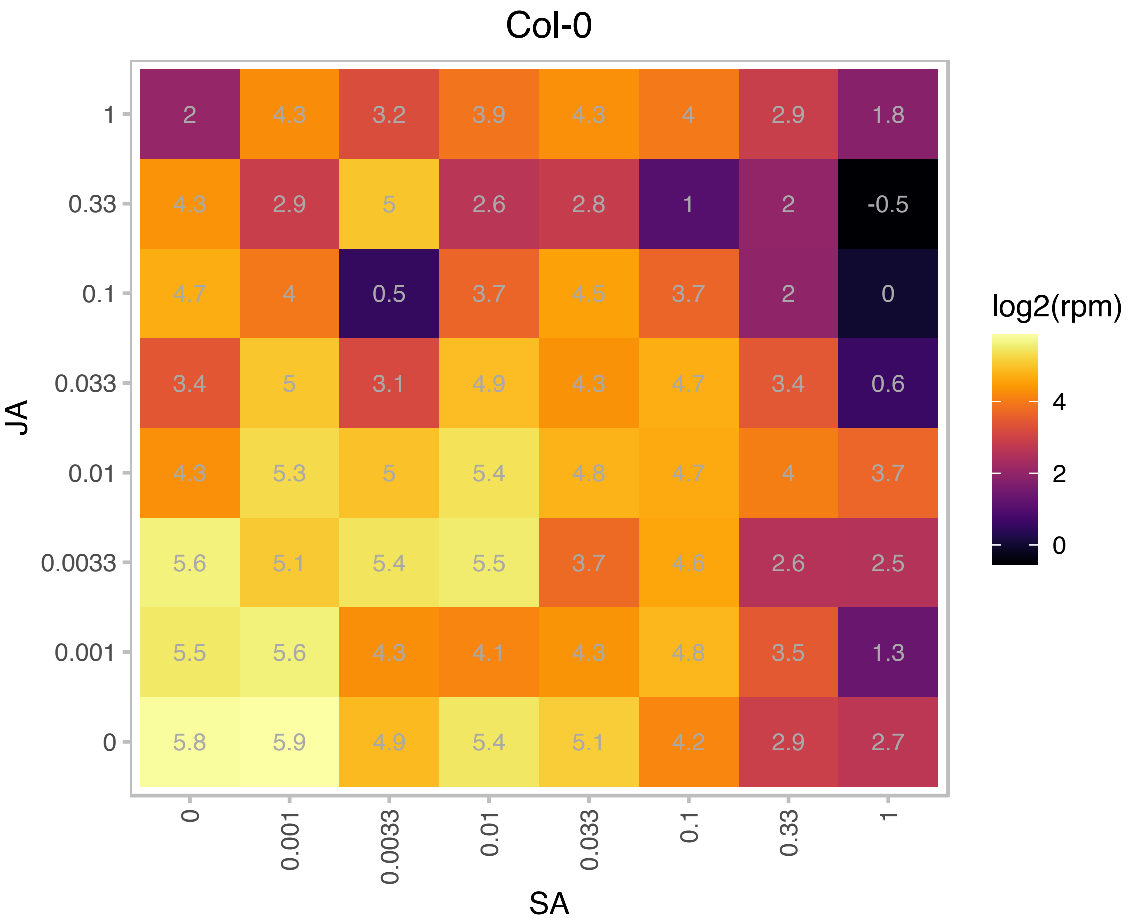


| | |
|---|---|
| Gene | AT5G14920 |
| is.expressed_genes | TRUE |
| is.responsive_genes | TRUE |
| is.apcluster | F-01 |

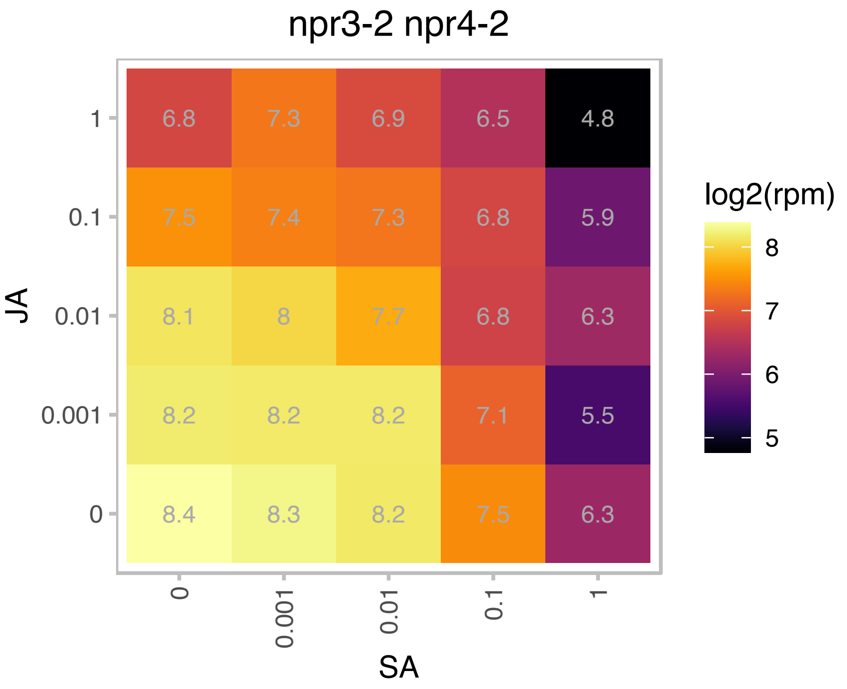


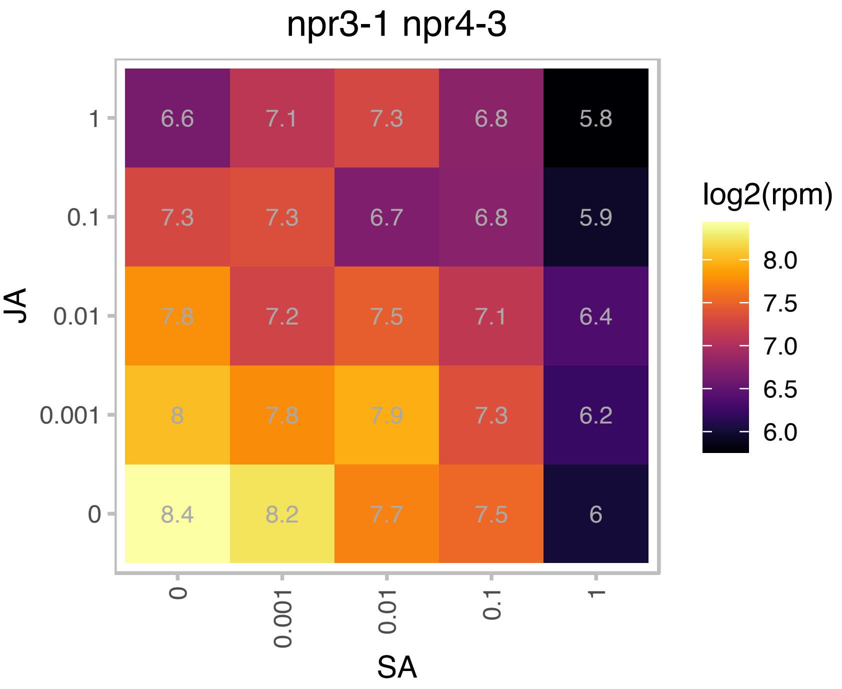

### AT5G15802.1 : HCF222

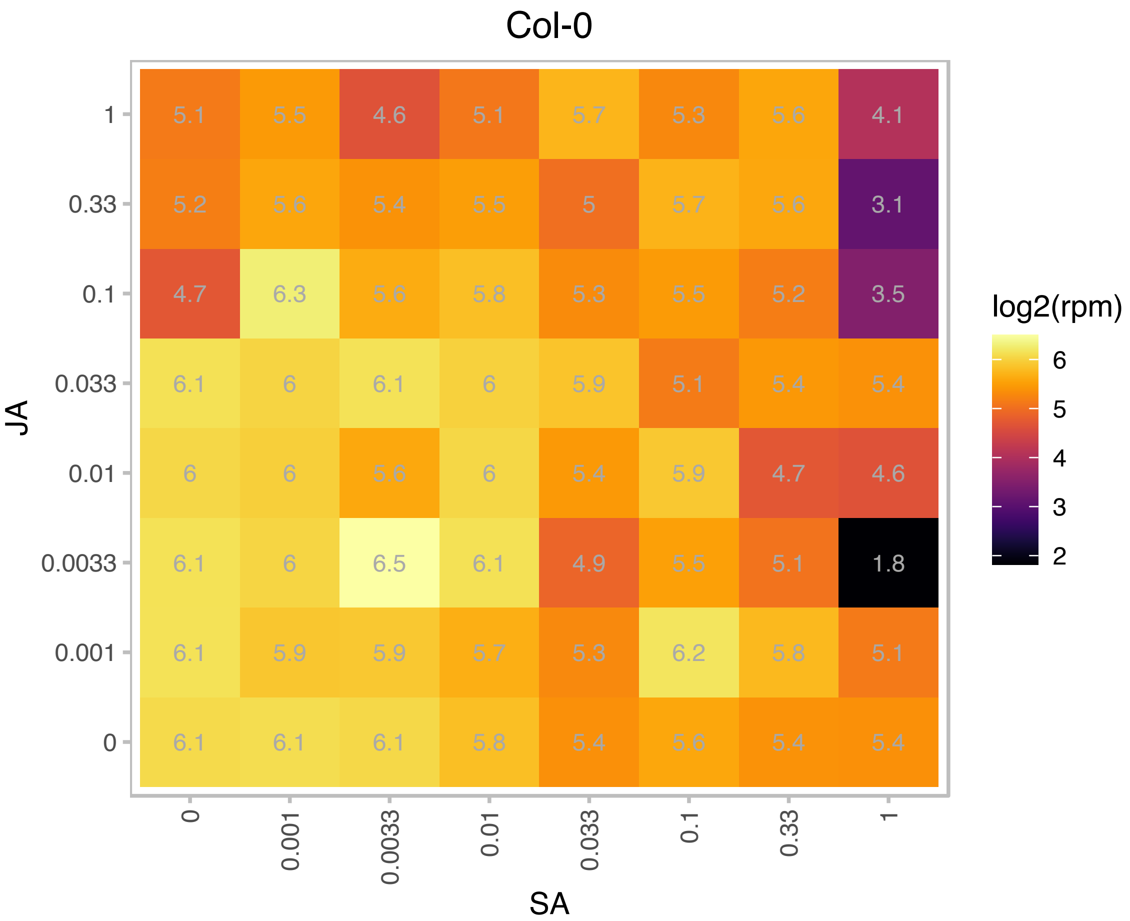


| | |
|---|---|
| Gene | AT5G15802 |
| is.expressed_genes | TRUE |
| is.responsive_genes | TRUE |
| is.apcluster | F-01 |

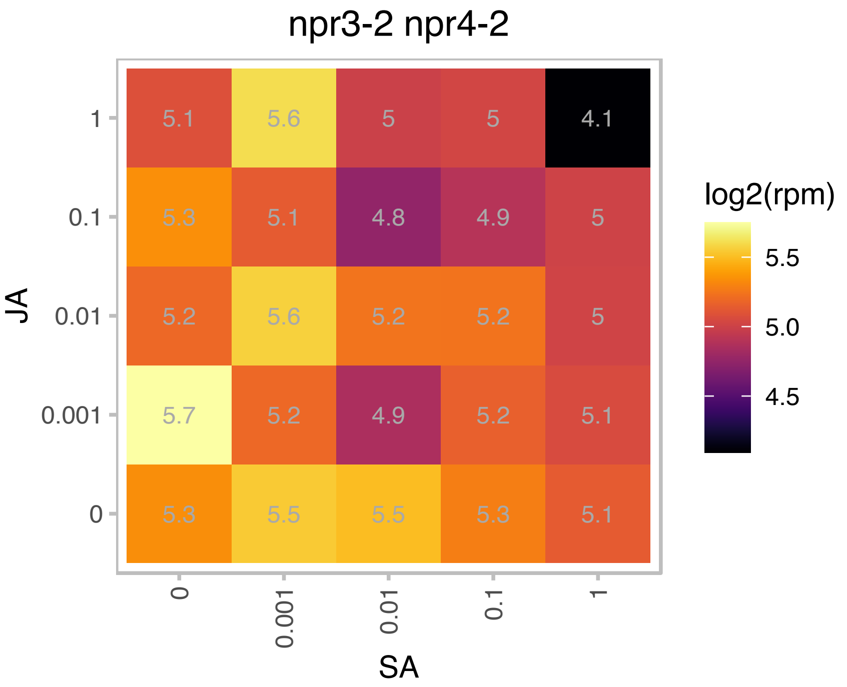


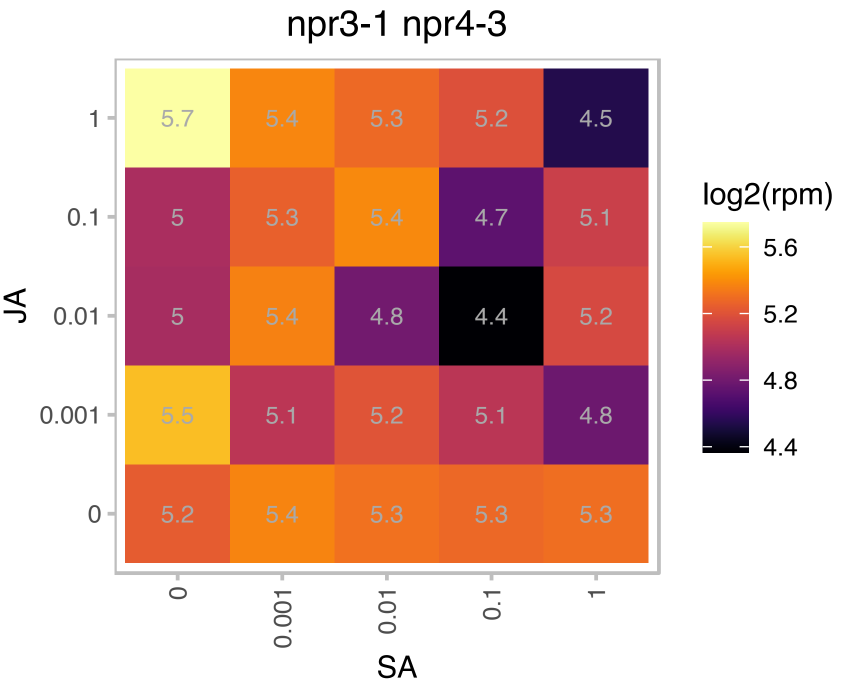

### AT5G22640.1 : emb1211, TIC100

MORN (Membrane Occupation and Recognition Nexus) repeat-containing protein

EMB1211 is a MORN (multiple membrane occupation and recognition nexus) motif containing protein involved in embryo development and chloroplast biogenesis. The mRNA is cell-to-cell mobile.

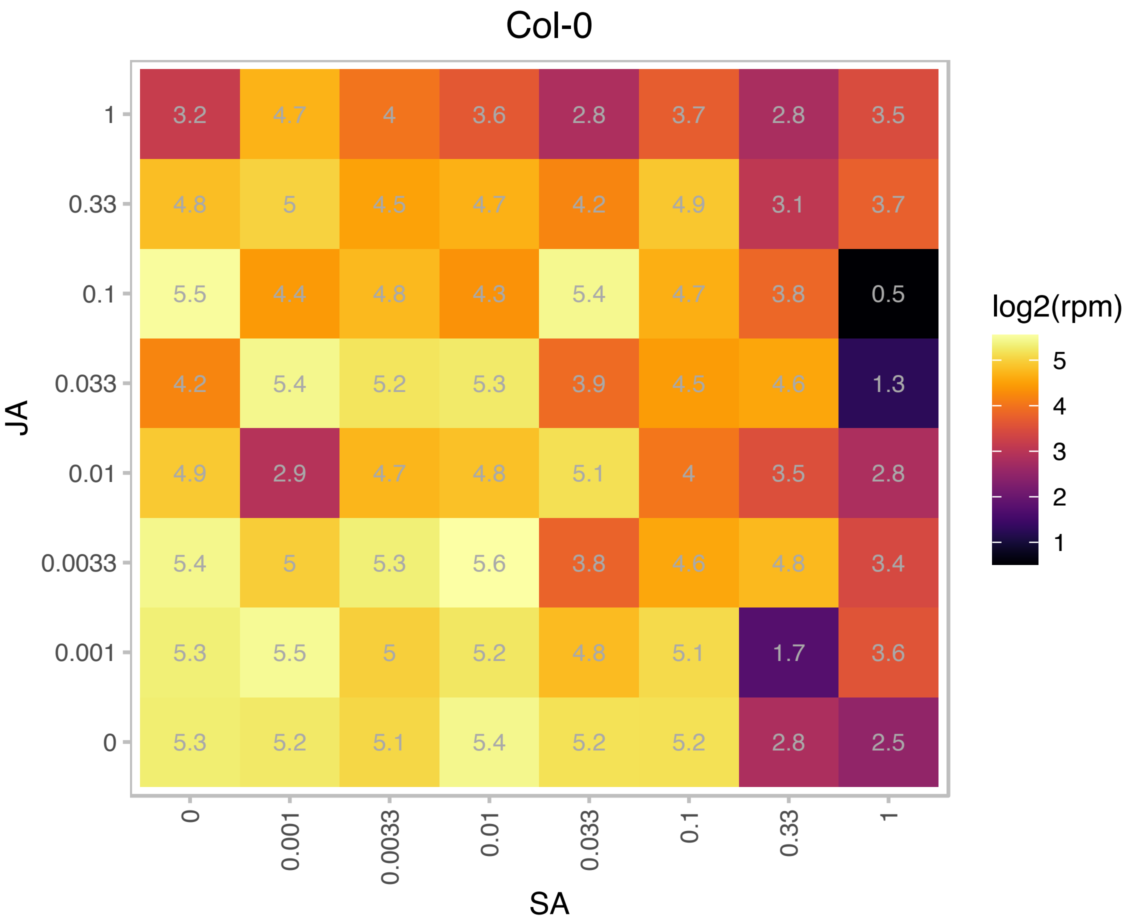


| | |
|---|---|
| Gene | AT5G22640 |
| is.expressed_genes | TRUE |
| is.responsive_genes | TRUE |
| is.apcluster | F-01 |

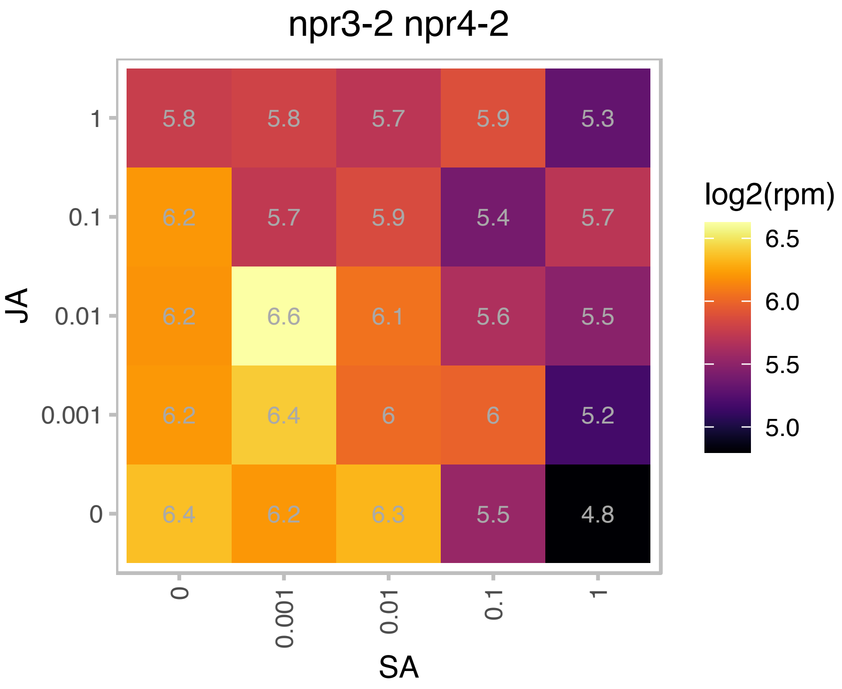


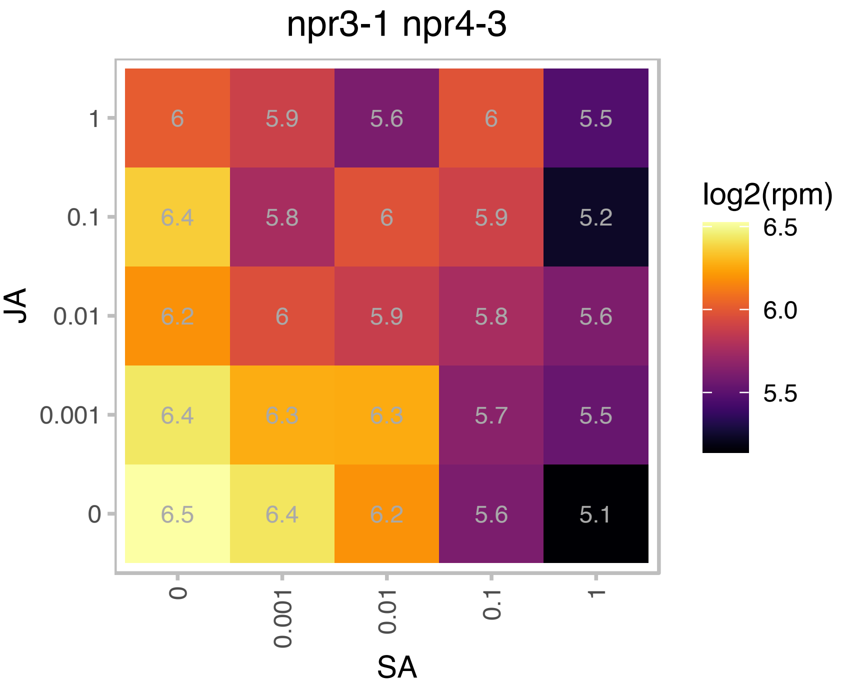

# AT5G23920.1 : NA

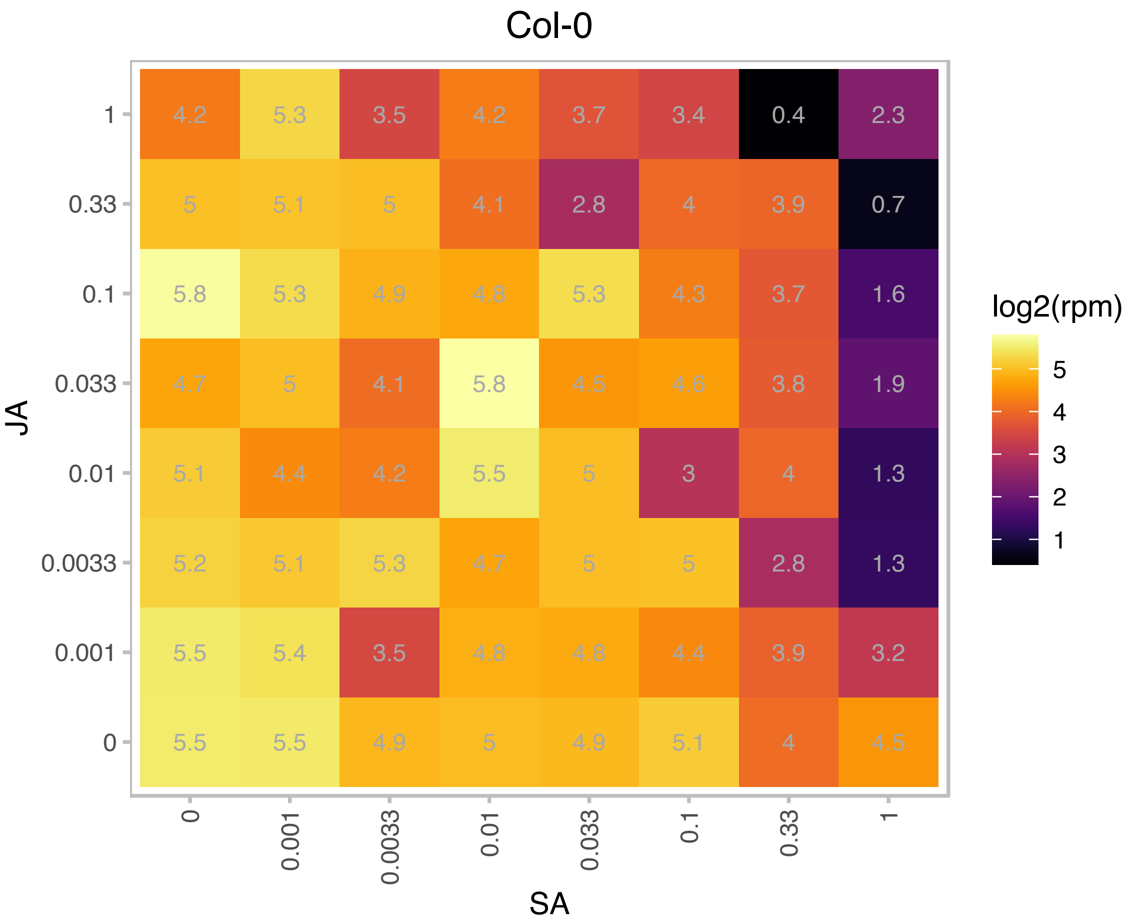


| | |
|---|---|
| Gene | AT5G23920 |
| is.expressed_genes | TRUE |
| is.responsive_genes | TRUE |
| is.apcluster | F-01 |

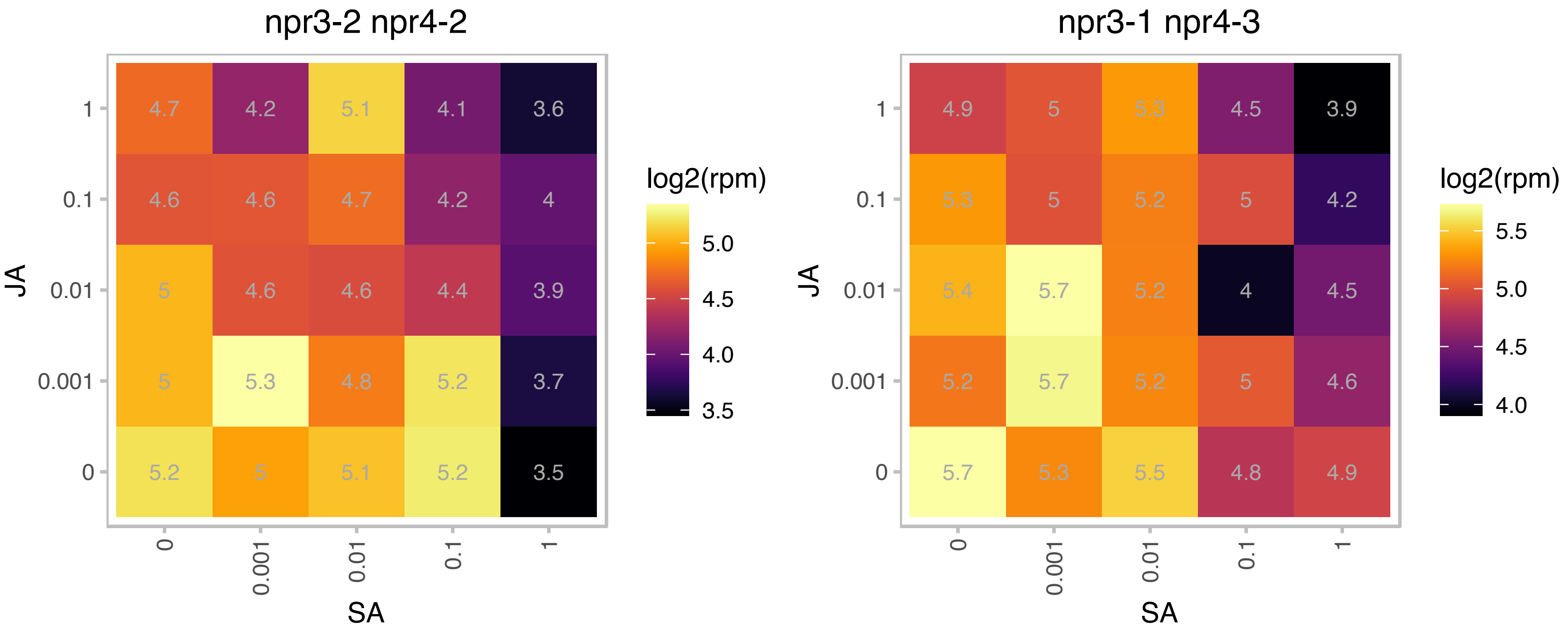

# AT5G28500.1 : NA

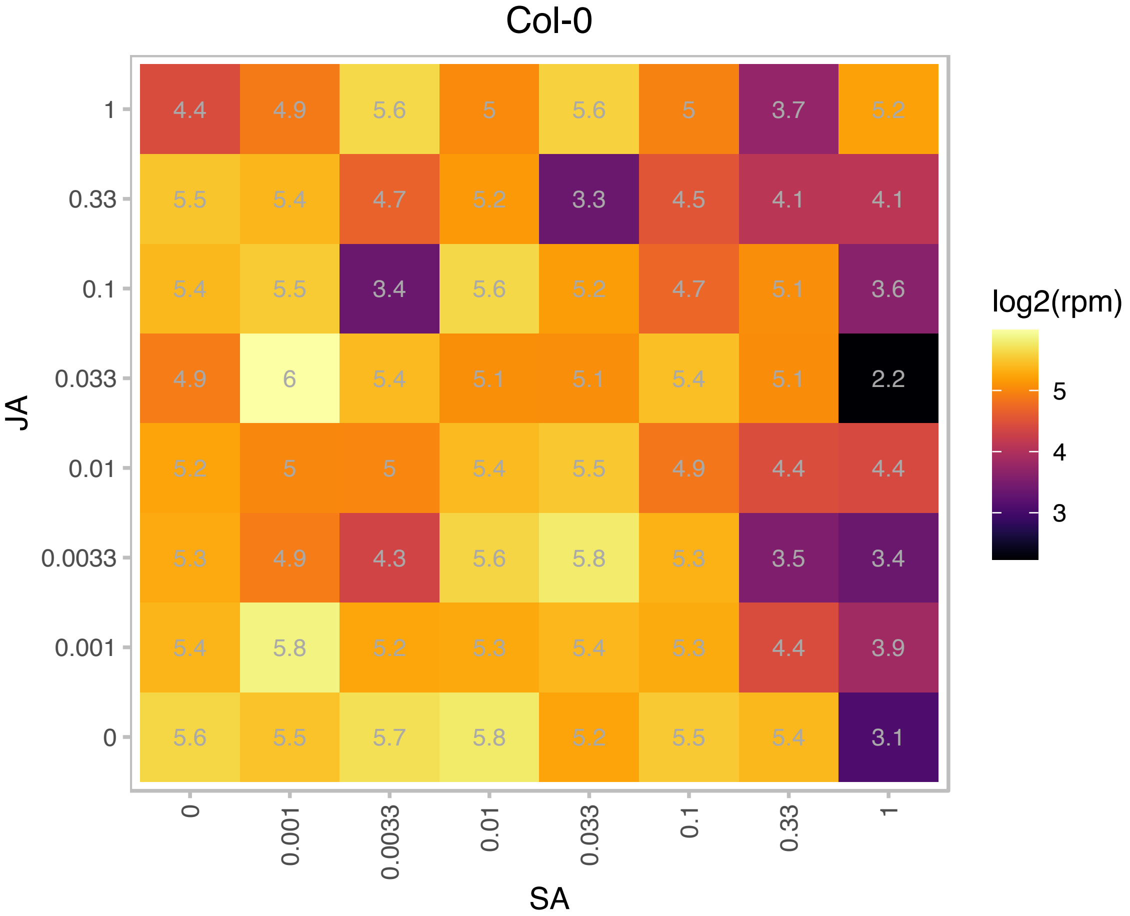


| | |
|---|---|
| Gene | AT5G28500 |
| is.expressed_genes | TRUE |
| is.responsive_genes | TRUE |
| is.apcluster | F-01 |

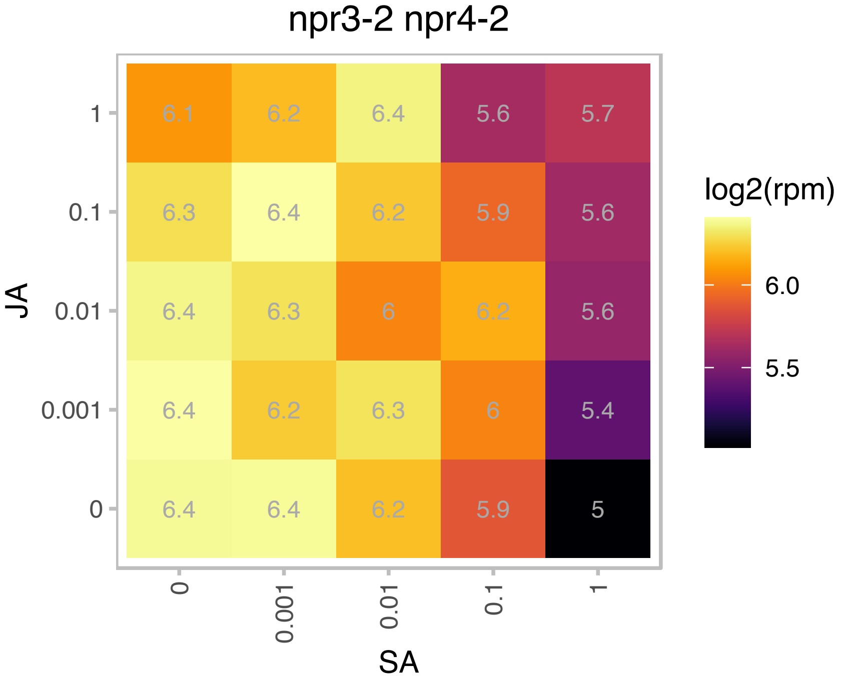


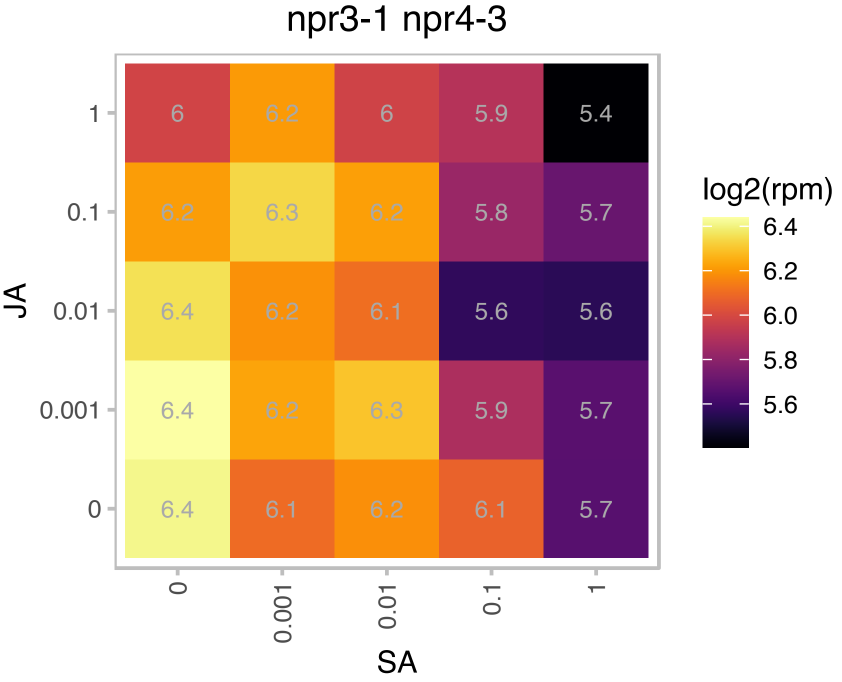

# AT5G37475.1 : NA

Translation initiation factor eIF3 subunit

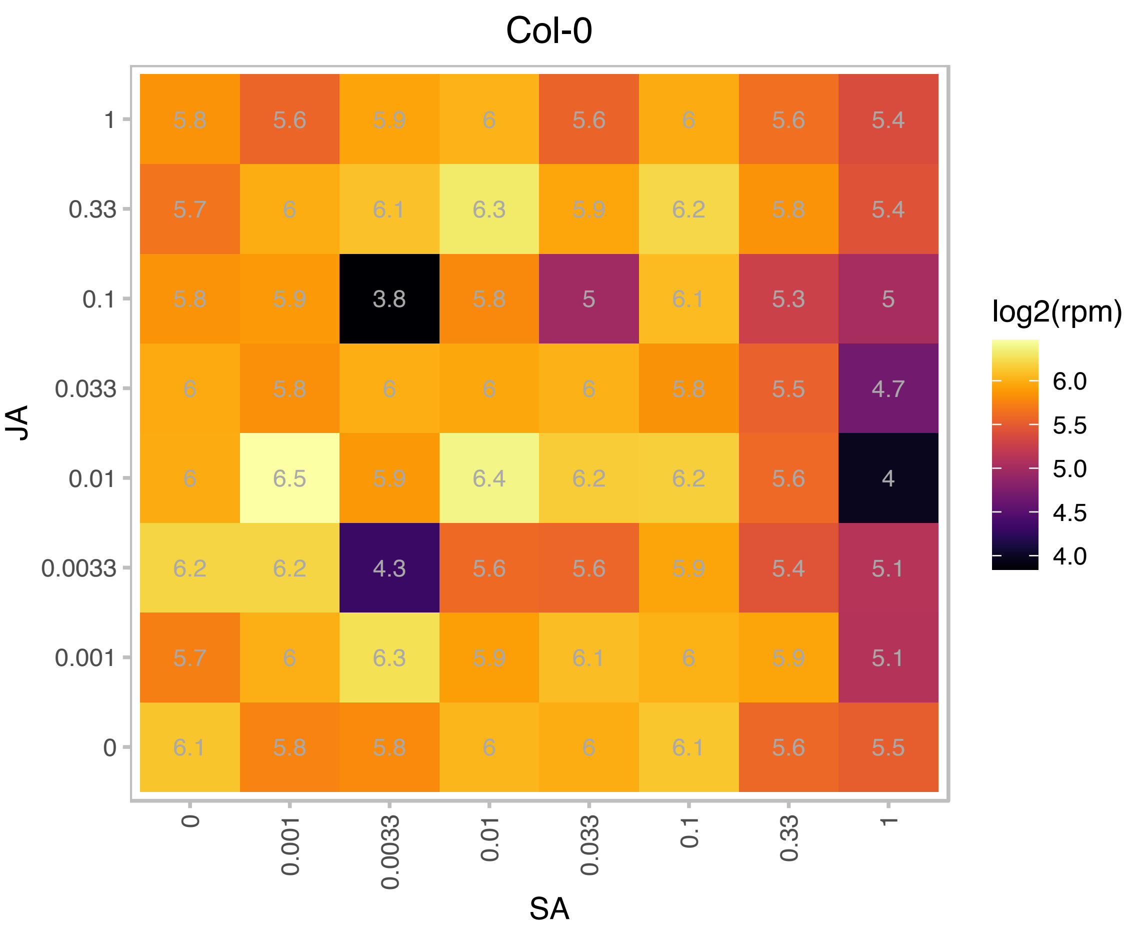


| | |
|---|---|
| Gene | AT5G37475 |
| is.expressed_genes | TRUE |
| is.responsive_genes | TRUE |
| is.apcluster | F-01 |

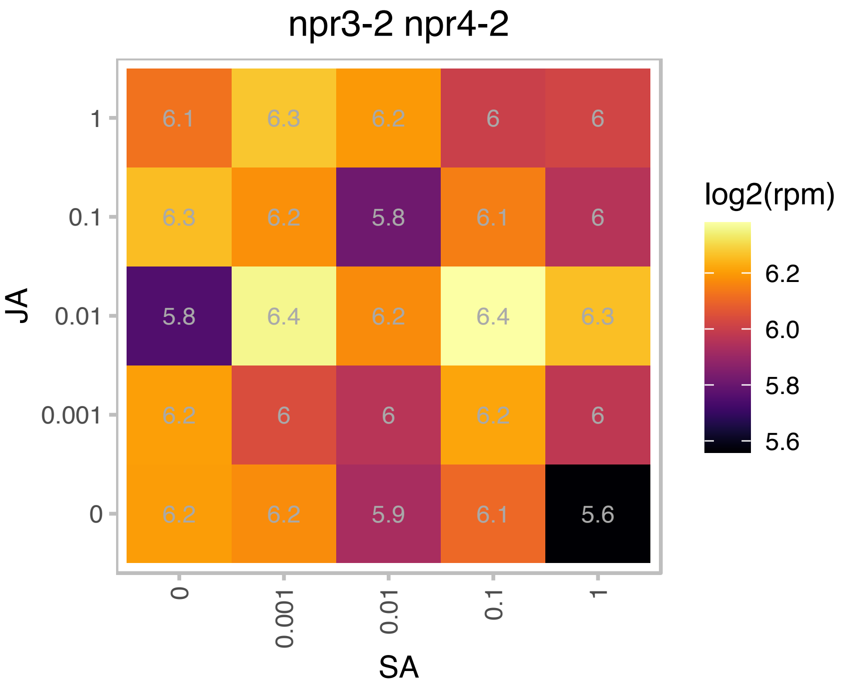


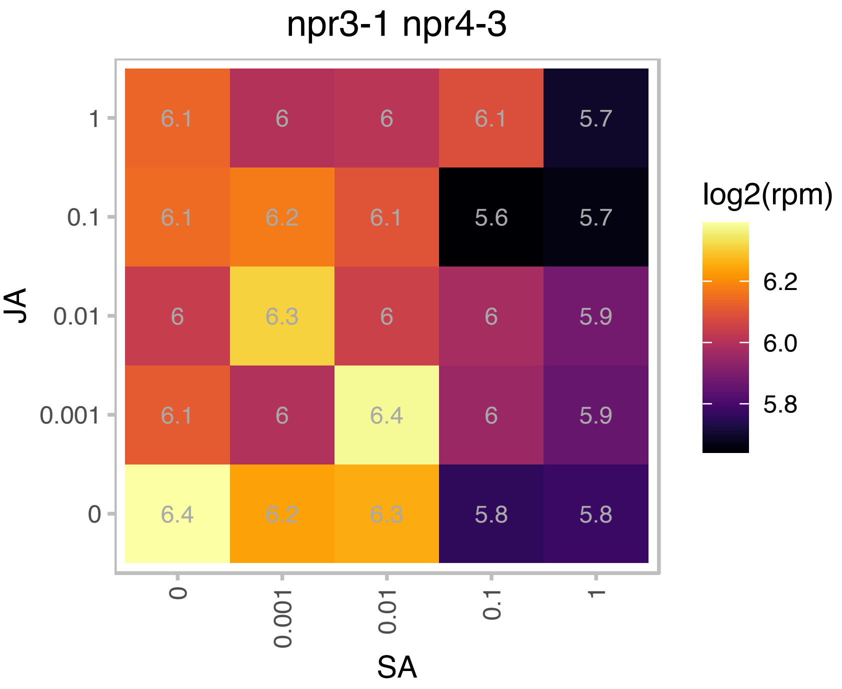

### AT5G42750.1 : BKI1

Encodes a plasma-membrane associated phosphoprotein that interacts directly with the kinase domain of BRI1 through the evolutionarily conserved C-terminal BIM motif binding to the C-lobe of the BRI1 kinase domain. It interferes with the interaction between BRI1 with its signalling partner, the plasma membrane localised LRR-receptor kinase BAK1 by inhibiting the transphosphorylation to keep BRI1 at a basal level of activity. It is phosphorylated by BRI1 at Ser270 & Ser274 and at tyrosine site Tyr211 and dissociates from plasma membrane to end up in the cytosol after phosphorylation. Its loss-of-function mutant shows higher sensitivity to BR

BRI1 kinase inhibitor 1

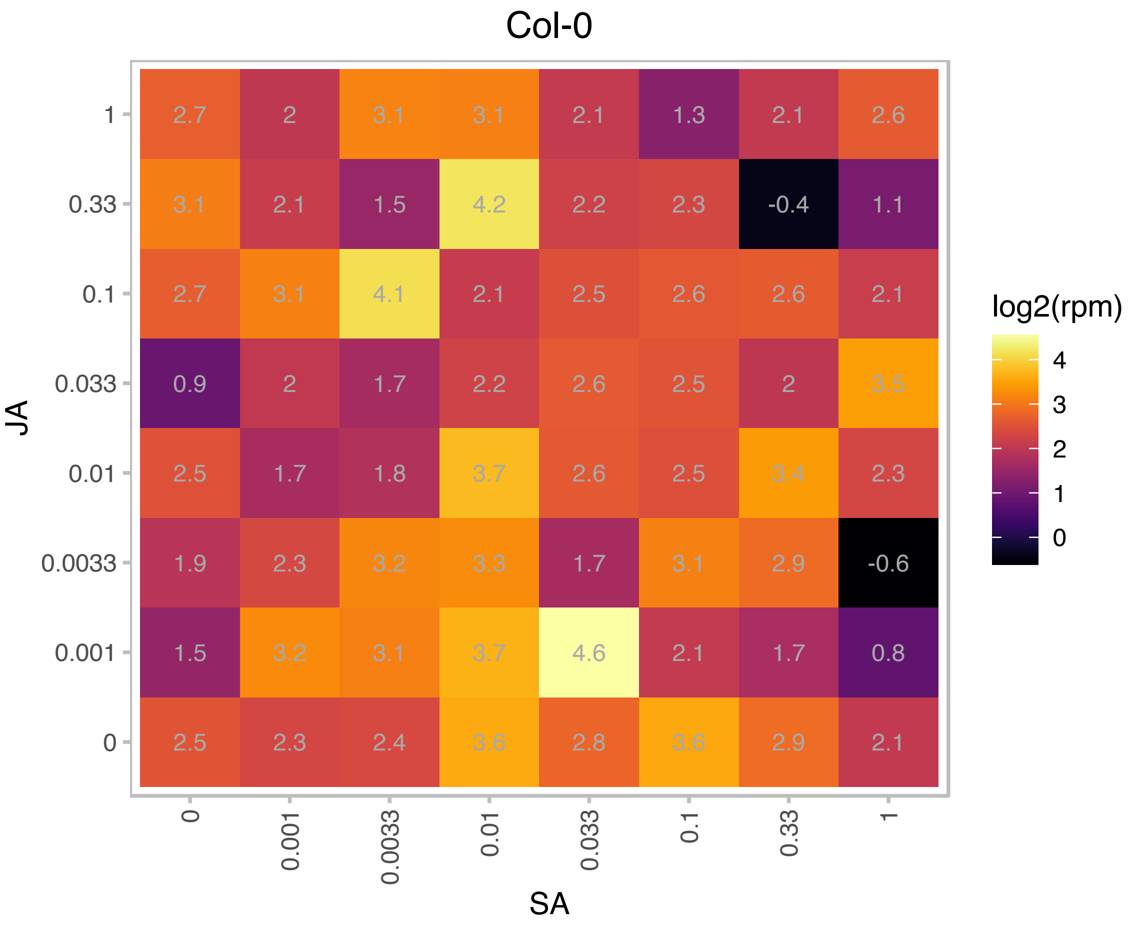


| | |
|---|---|
| Gene | AT5G42750 |
| is.expressed_genes | TRUE |
| is.responsive_genes | TRUE |
| is.apcluster | F-01 |

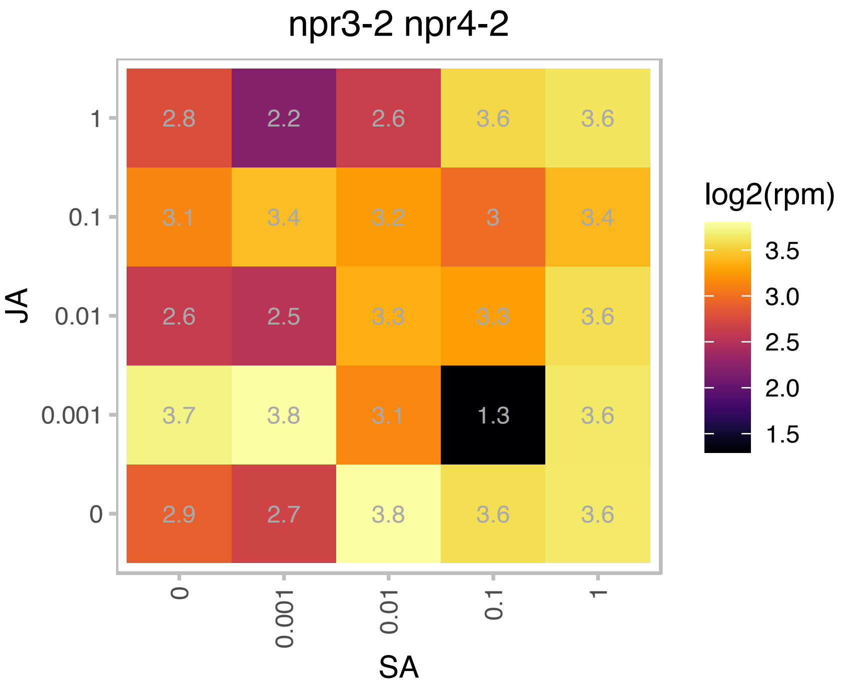


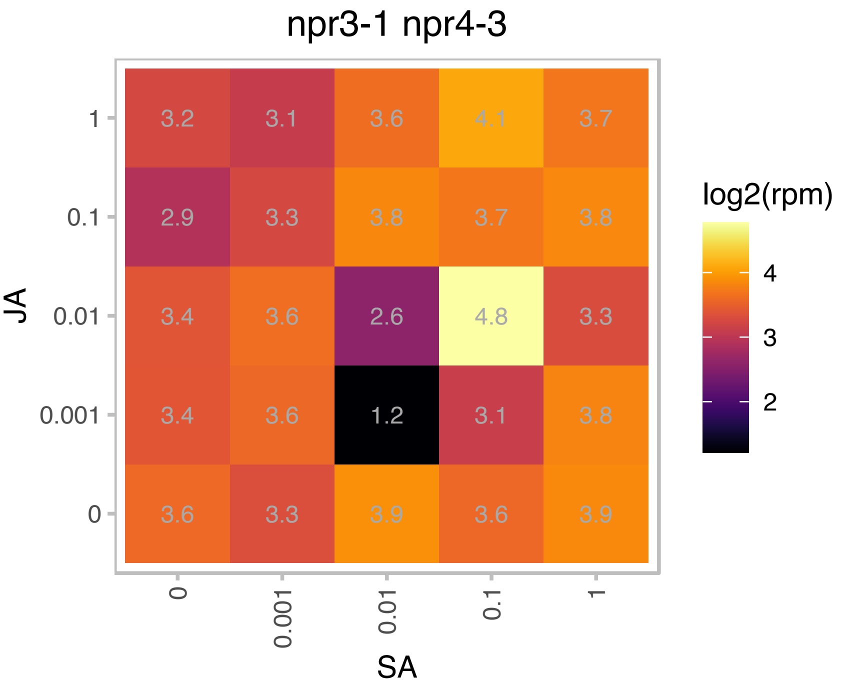

# AT5G44060.1 : NA

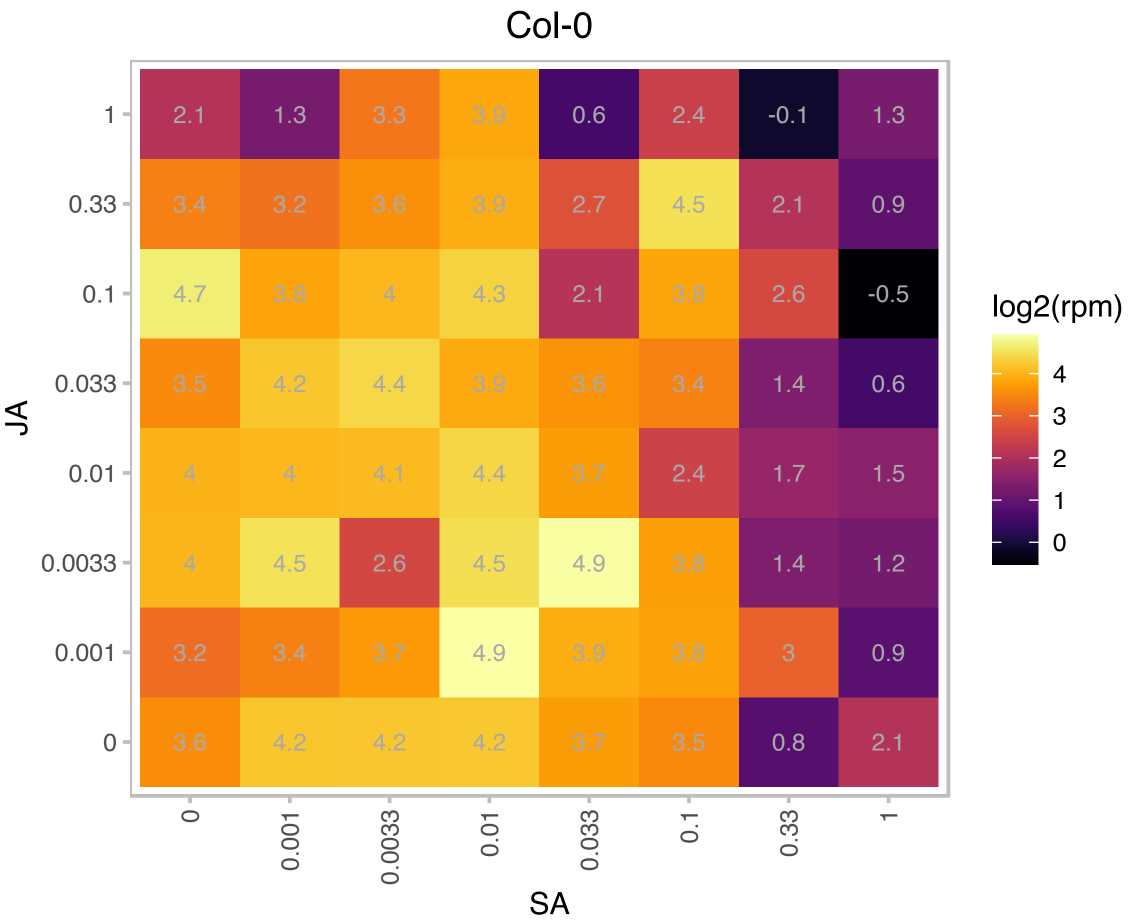


| | |
|---|---|
| Gene | AT5G44060 |
| is.expressed_genes | TRUE |
| is.responsive_genes | TRUE |
| is.apcluster | F-01 |

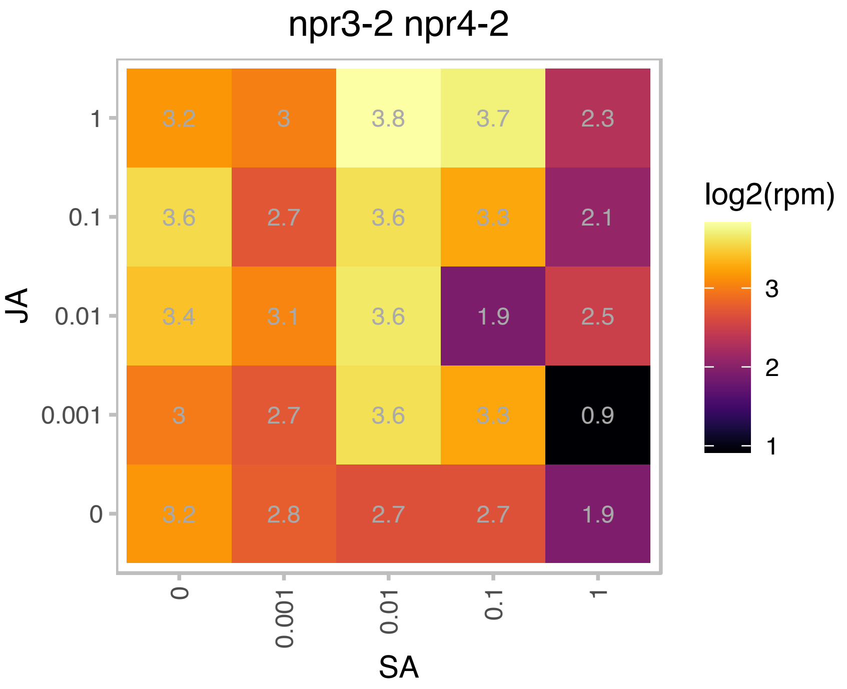


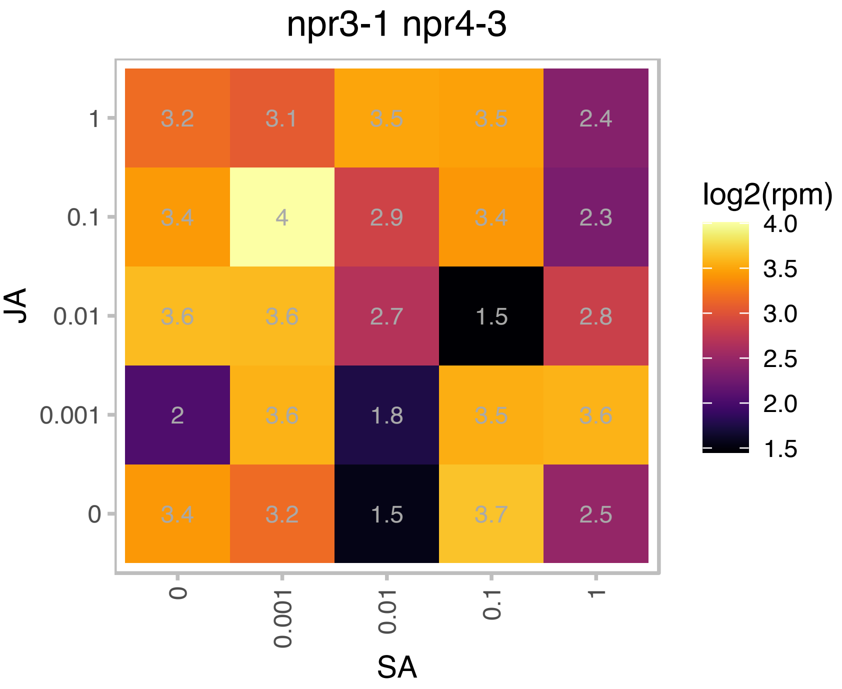

# AT5G45670.1 : NA

GDSL-like Lipase/Acylhydrolase superfamily protein

GDSL-motif esterase/acyltransferase/lipase. Enzyme group with broad substrate specificity that may catalyze acyltransfer or hydrolase reactions with lipid and non-lipid substrates.

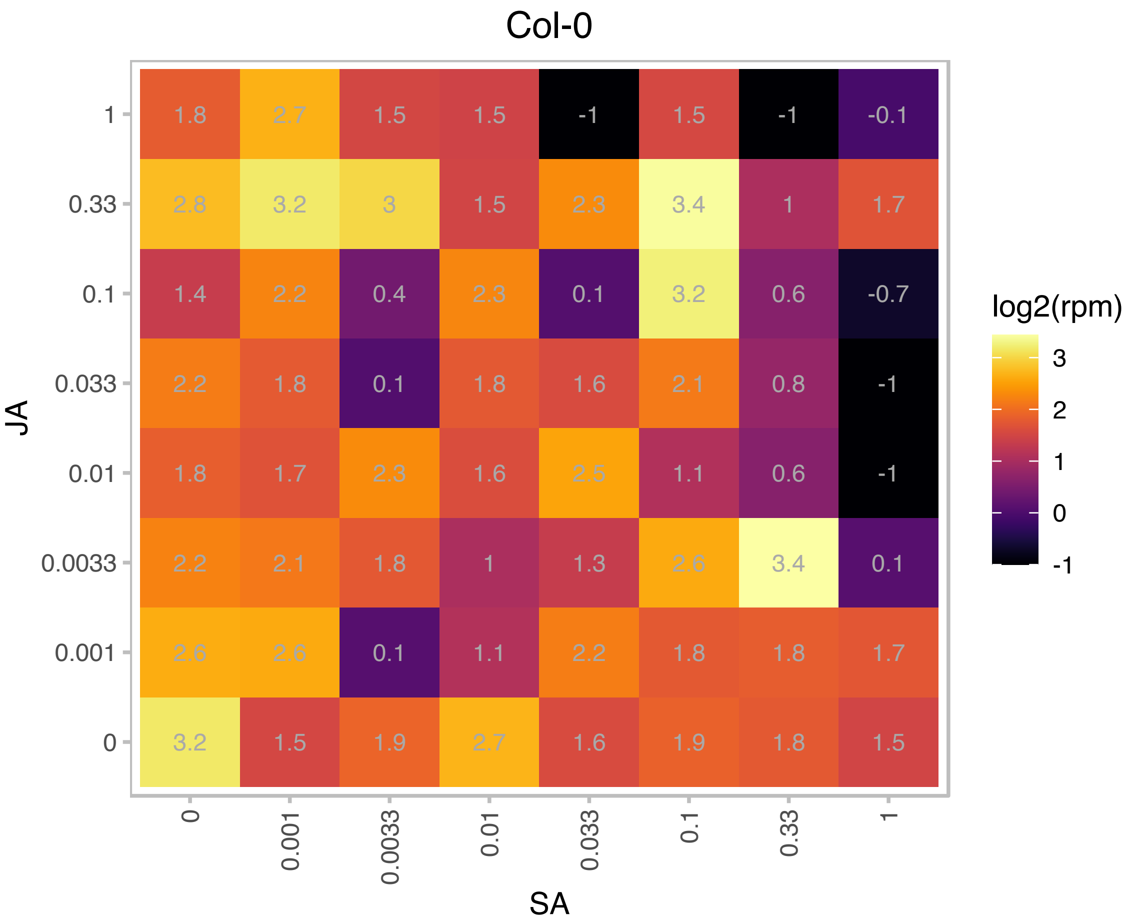


| | |
|---|---|
| Gene | AT5G45670 |
| is.expressed_genes | TRUE |
| is.responsive_genes | TRUE |
| is.apcluster | F-01 |

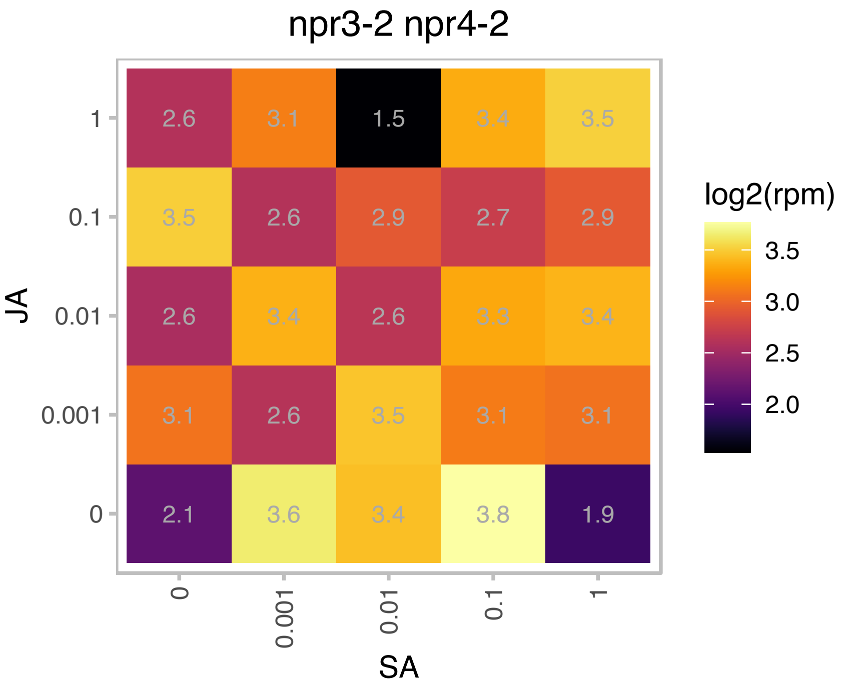


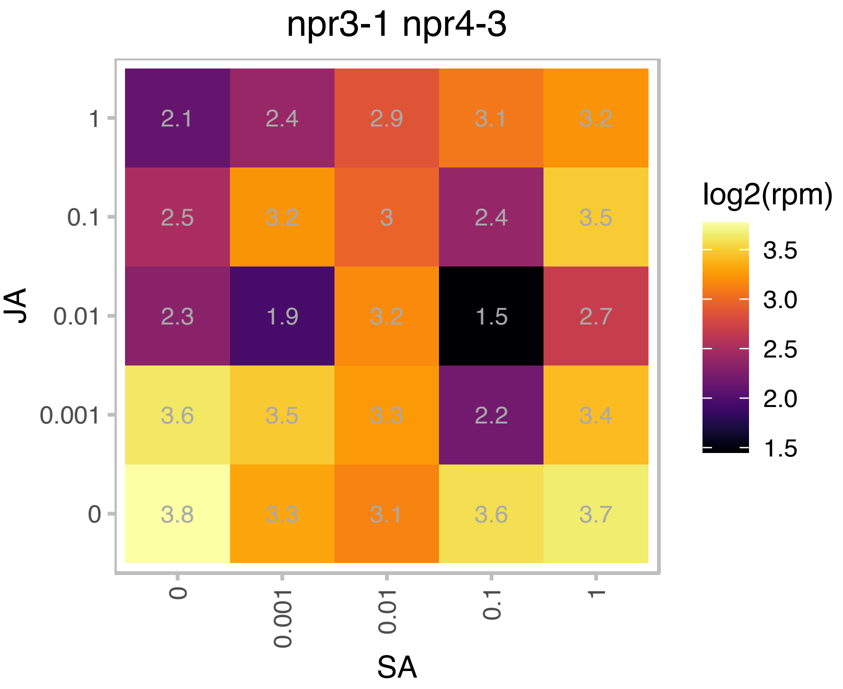

### AT5G47390.1 : KUA1, MYBH

myb-like transcription factor family protein

Encodes a circadian-regulated transcription factor which specifically controls cell expansion during leaf development by controlling ROS homeostasis. The mRNA is cell-to-cell mobile.

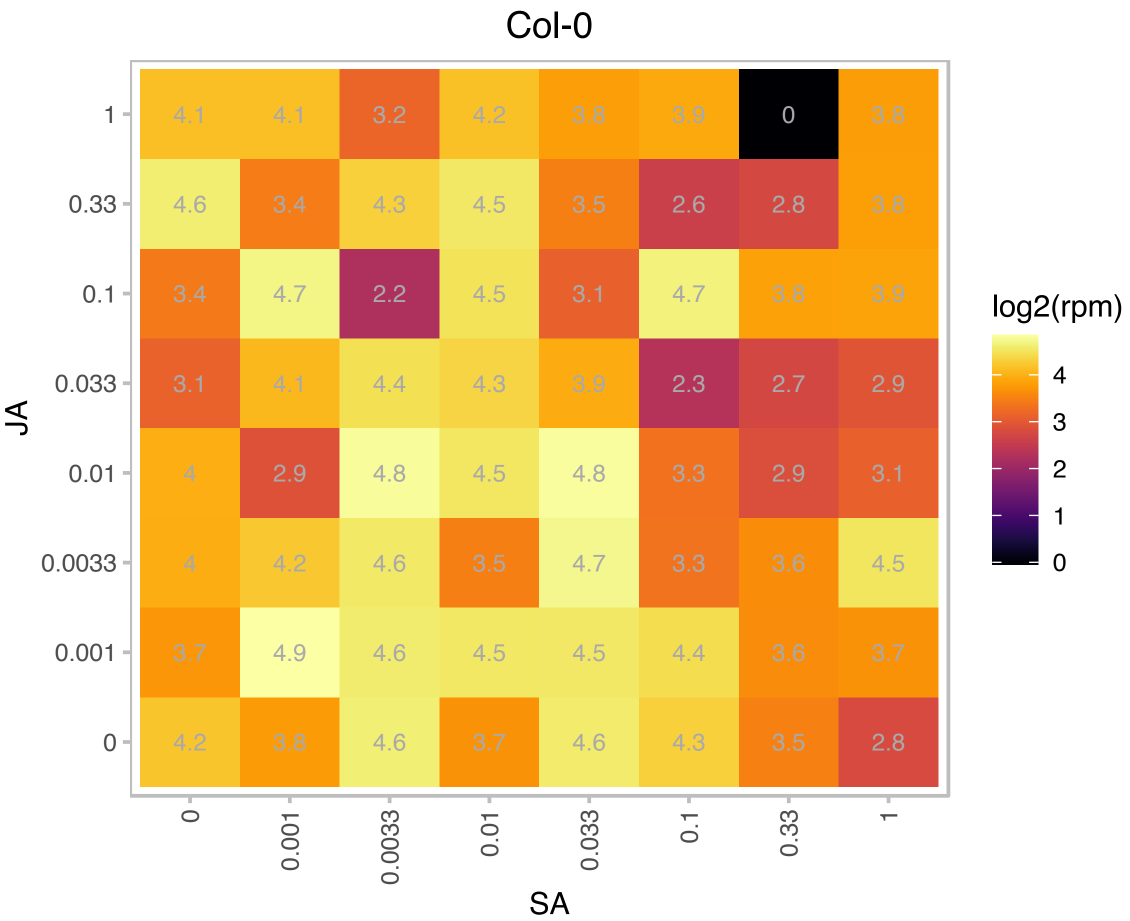


| | |
|---|---|
| Gene | AT5G47390 |
| is.expressed_genes | TRUE |
| is.responsive_genes | TRUE |
| is.apcluster | F-01 |

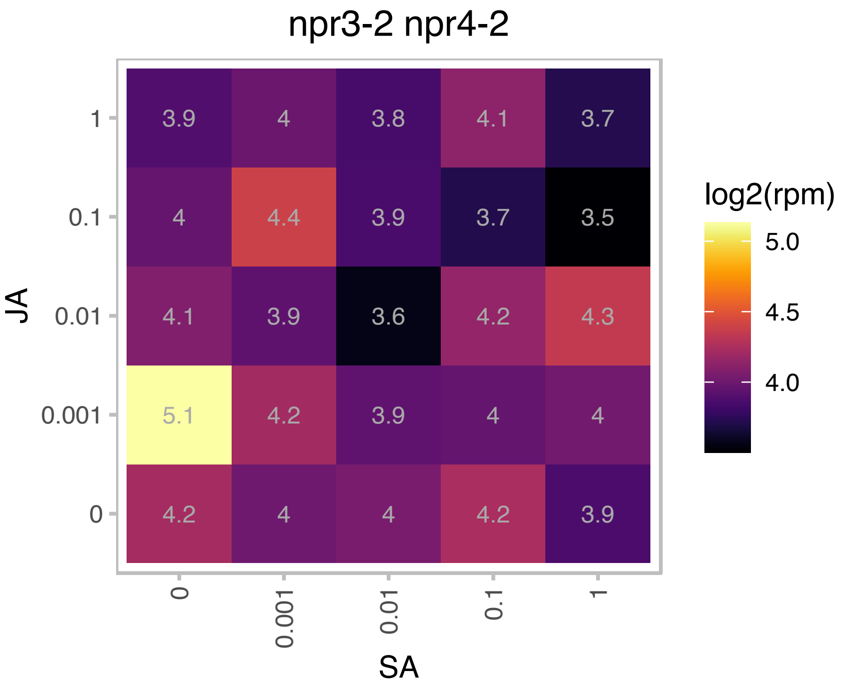


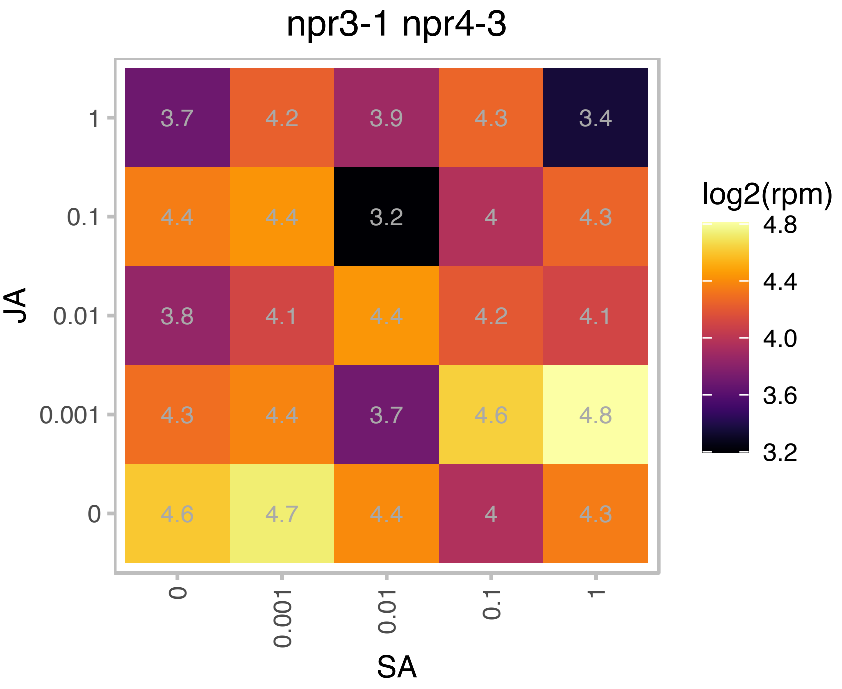

### AT5G47780.1 : GAUT4

galacturonosyltransferase 4

Encodes a protein with putative galacturonosyltransferase activity. The mRNA is cell-to-cell mobile.

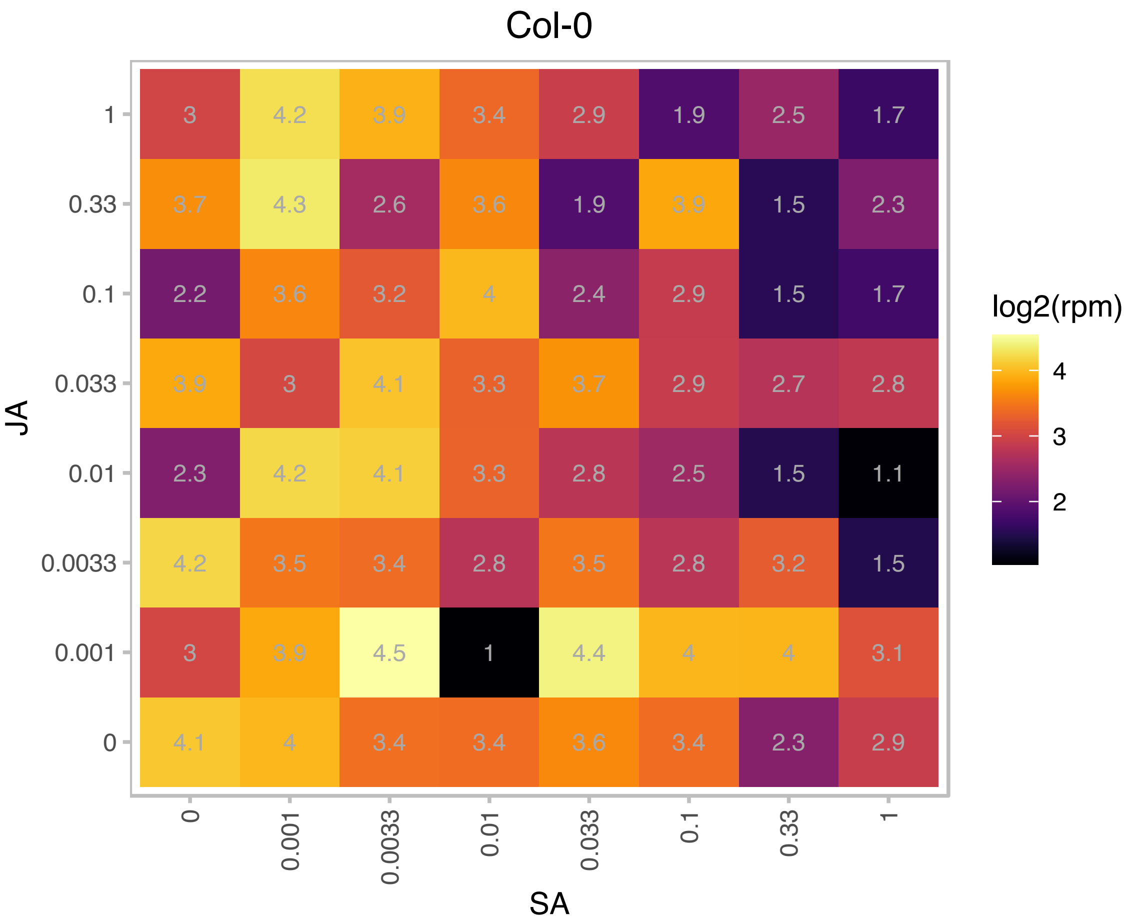


| | |
|---|---|
| Gene | AT5G47780 |
| is.expressed_genes | TRUE |
| is.responsive_genes | TRUE |
| is.apcluster | F-01 |

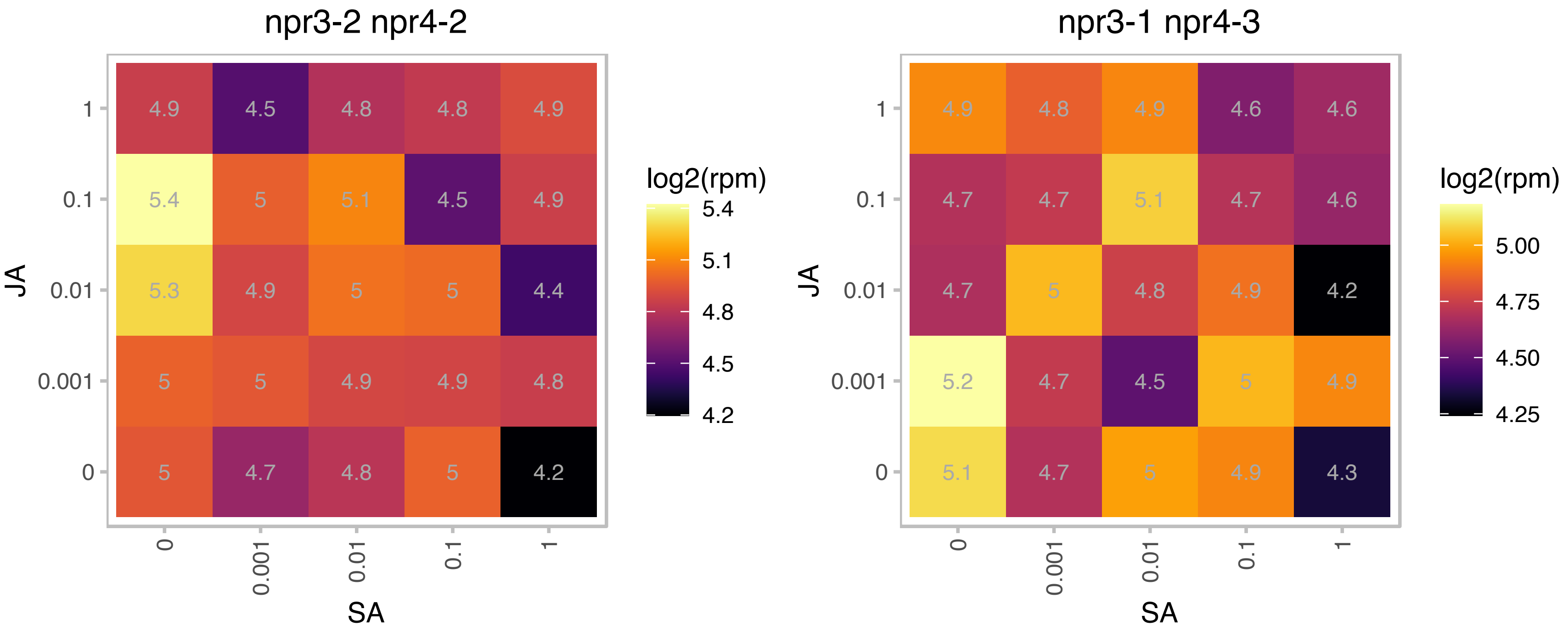

### AT5G49360.1 : ATBXL1, BXL1

beta-xylosidase 1

Encodes a bifunctional {beta}-D-xylosidase/{alpha}-L-arabinofuranosidase required for pectic arabinan modification.  Located in the extracellular matrix. Gene is expressed specifically in tissues undergoing secondary wall thickening. This is a member of glycosyl hydrolase family 3 and has six other closely related members.

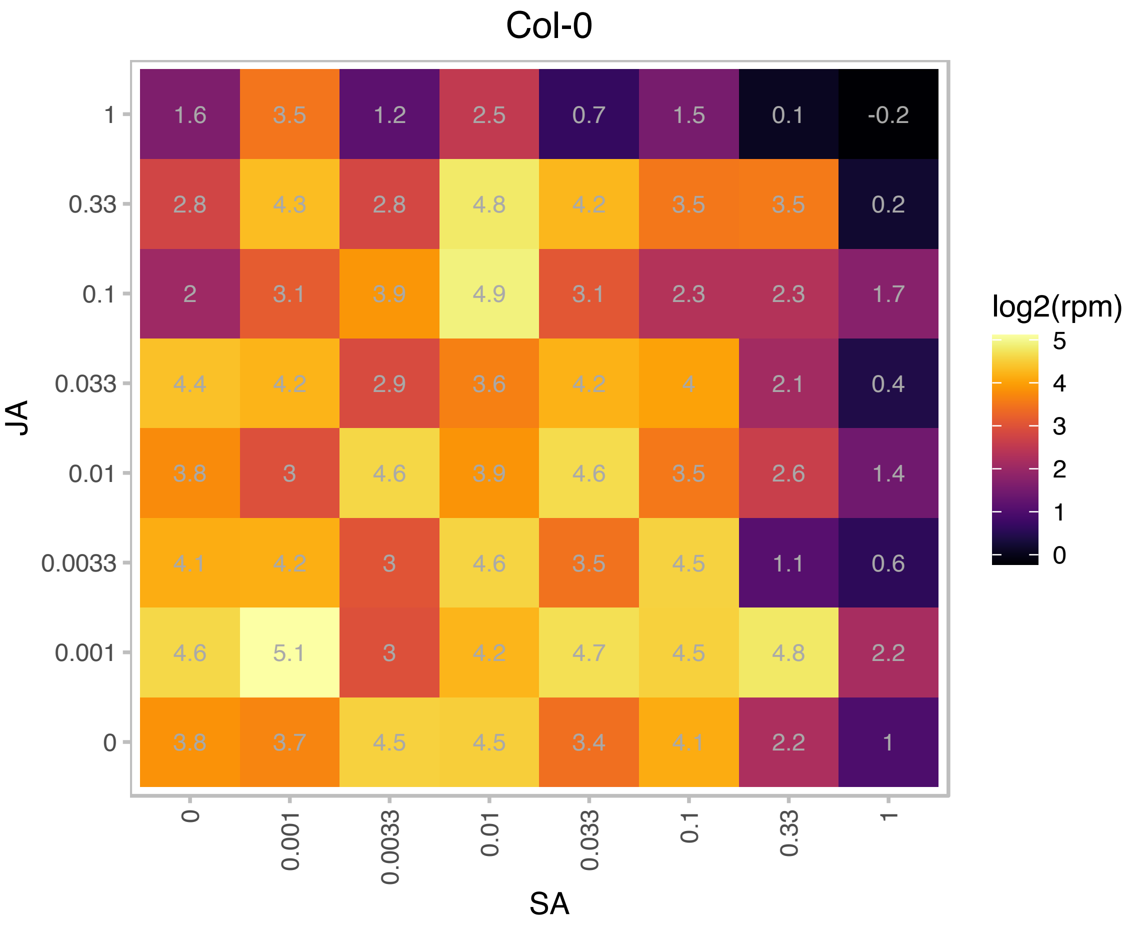


| | |
|---|---|
| Gene | AT5G49360 |
| is.expressed_genes | TRUE |
| is.responsive_genes | TRUE |
| is.apcluster | F-01 |

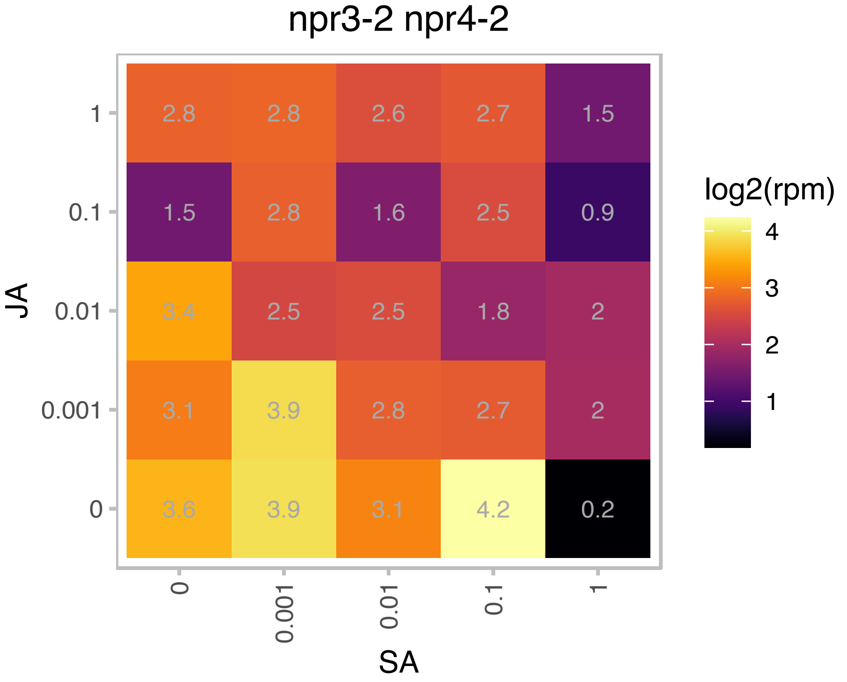


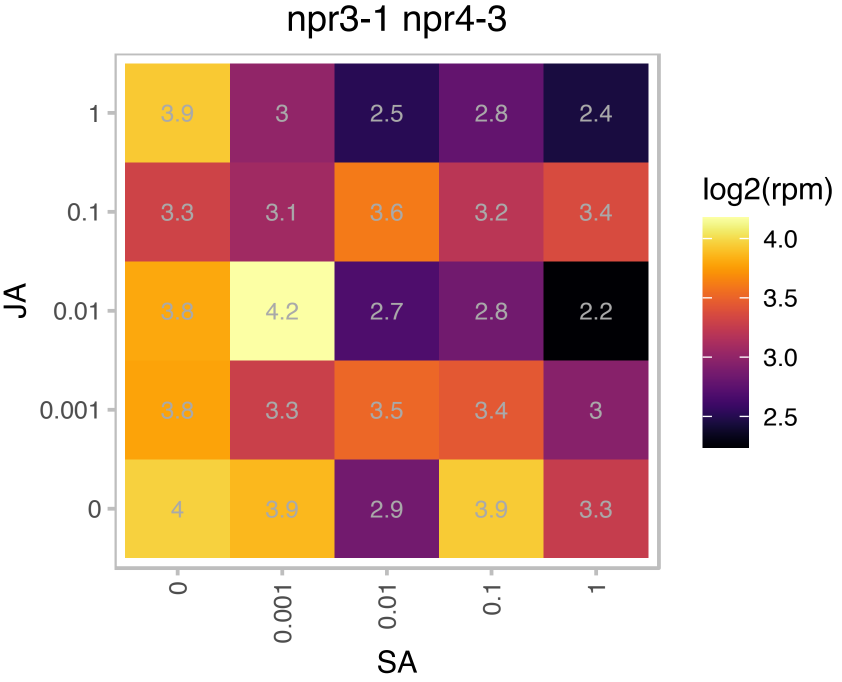

### AT5G49910.1 : cpHsc70-2, HSC70-7

chloroplast heat shock protein 70-2

Stromal heat shock protein involved in protein import into chloroplast. The mRNA is cell-to-cell mobile.

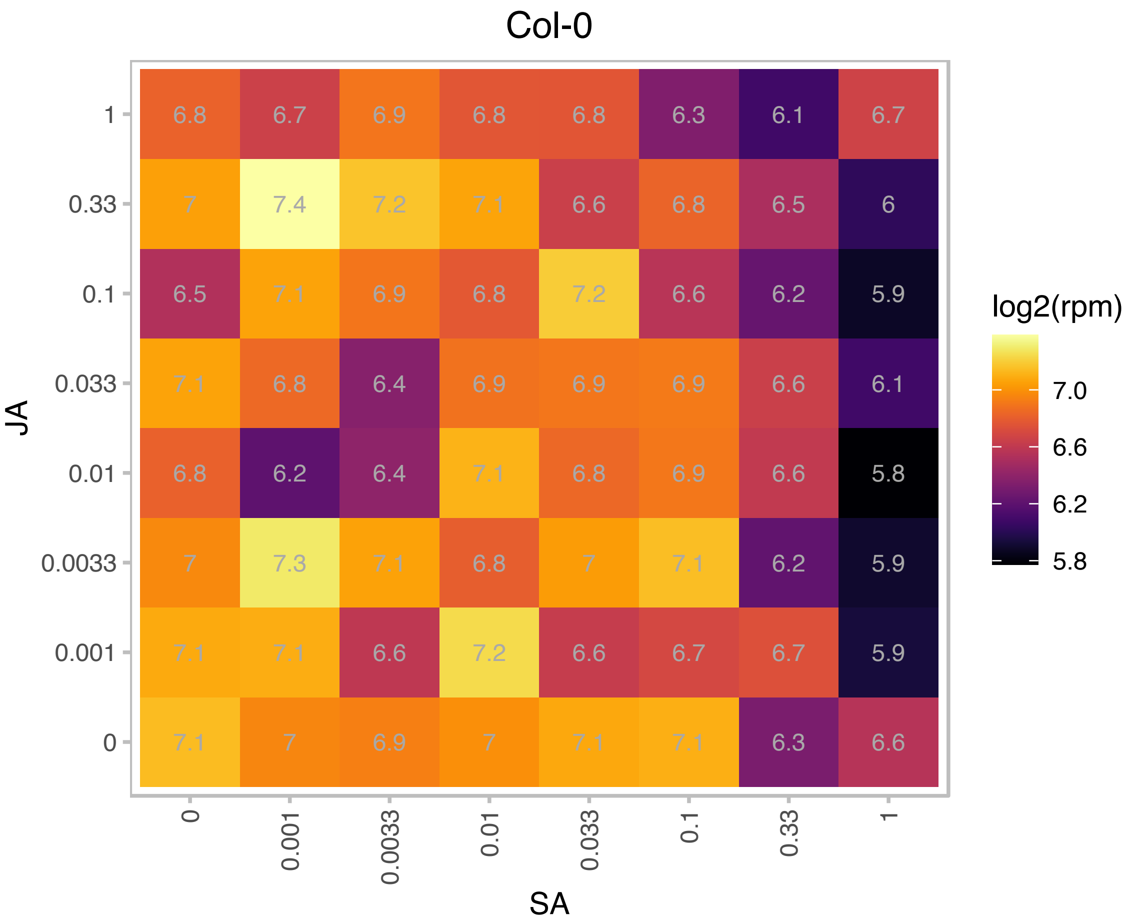


| | |
|---|---|
| Gene | AT5G49910 |
| is.expressed_genes | TRUE |
| is.responsive_genes | TRUE |
| is.apcluster | F-01 |

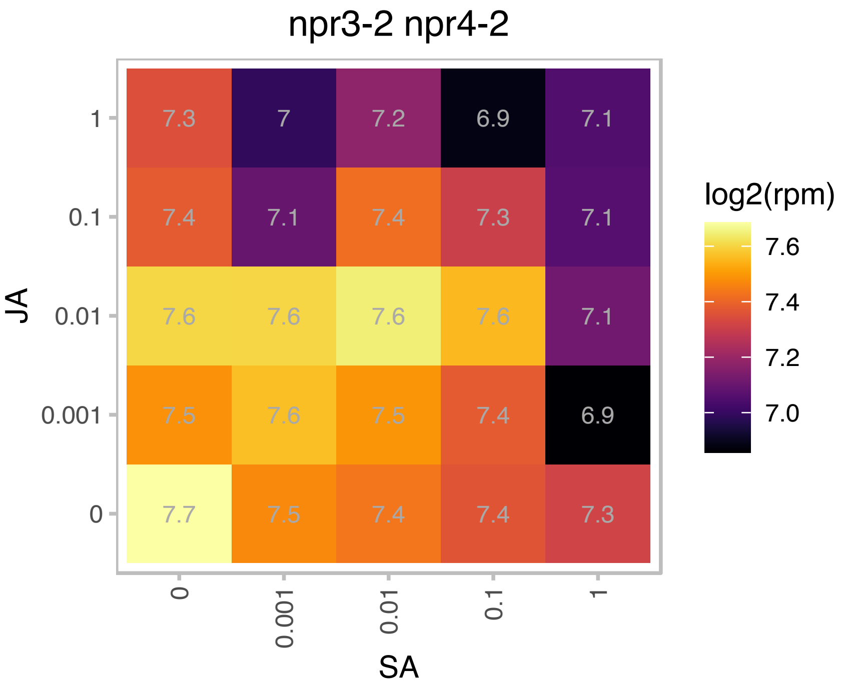


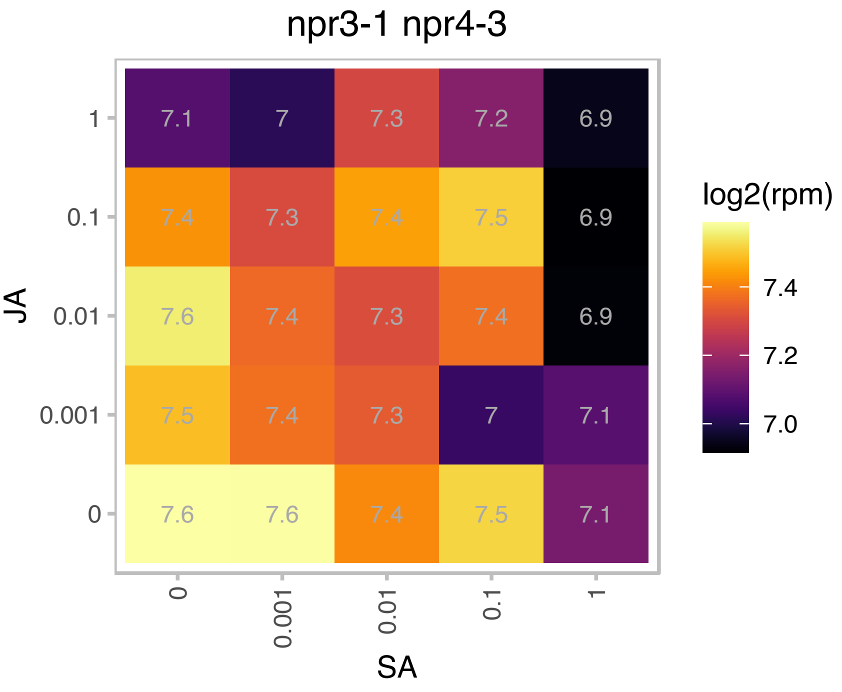

# AT5G50335.1 : NA

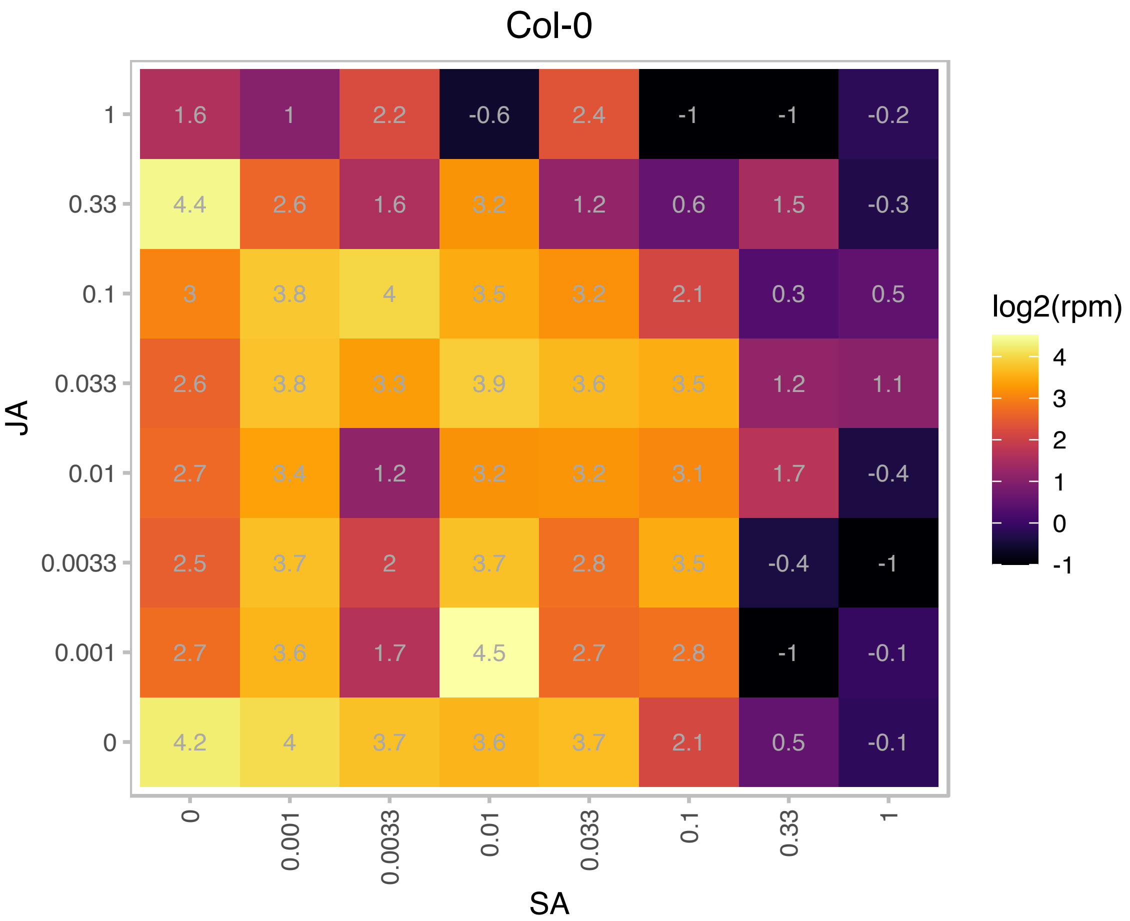


| | |
|---|---|
| Gene | AT5G50335 |
| is.expressed_genes | TRUE |
| is.responsive_genes | TRUE |
| is.apcluster | F-01 |

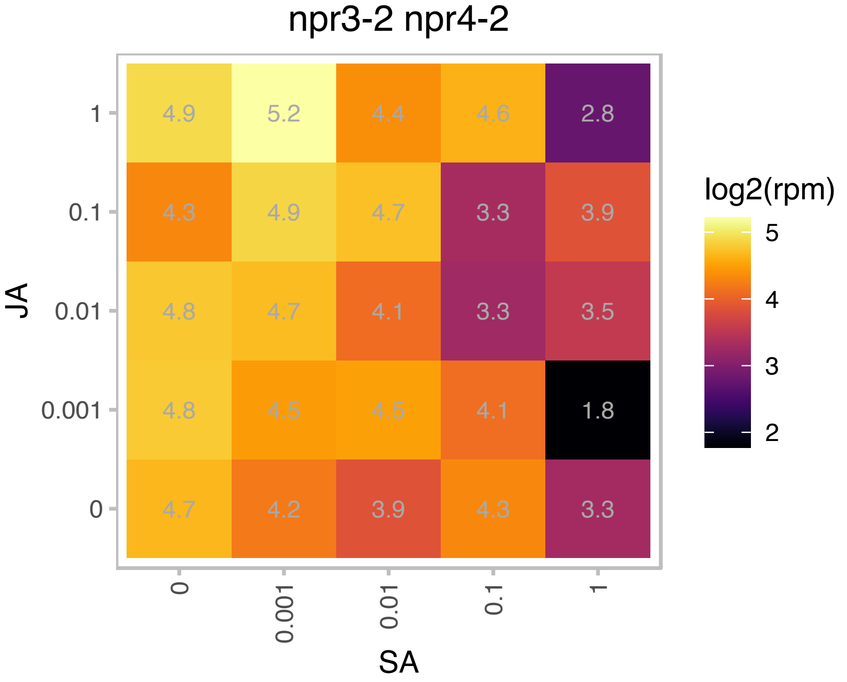


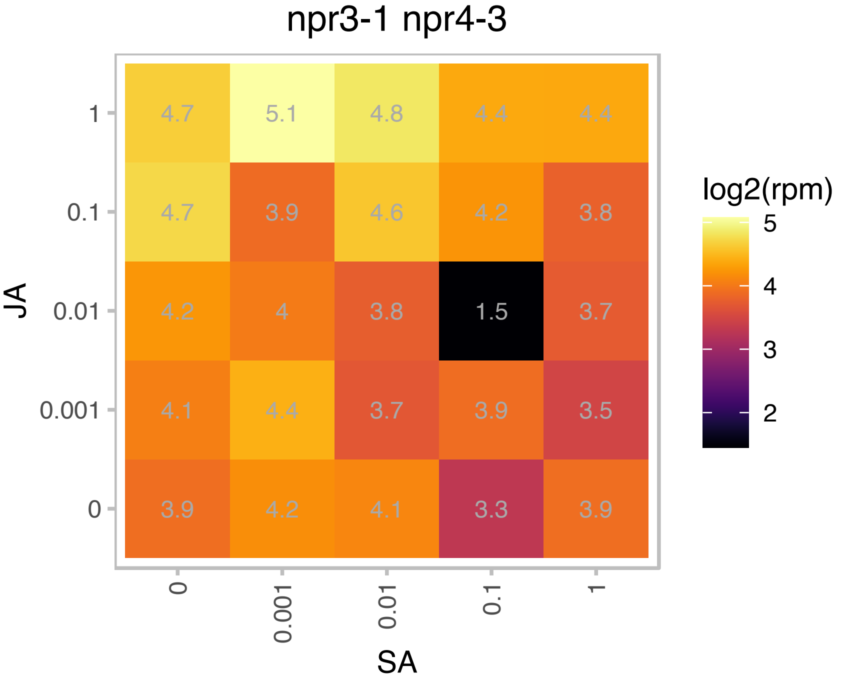

# AT5G57170.1 : NA

Tautomerase/MIF superfamily protein

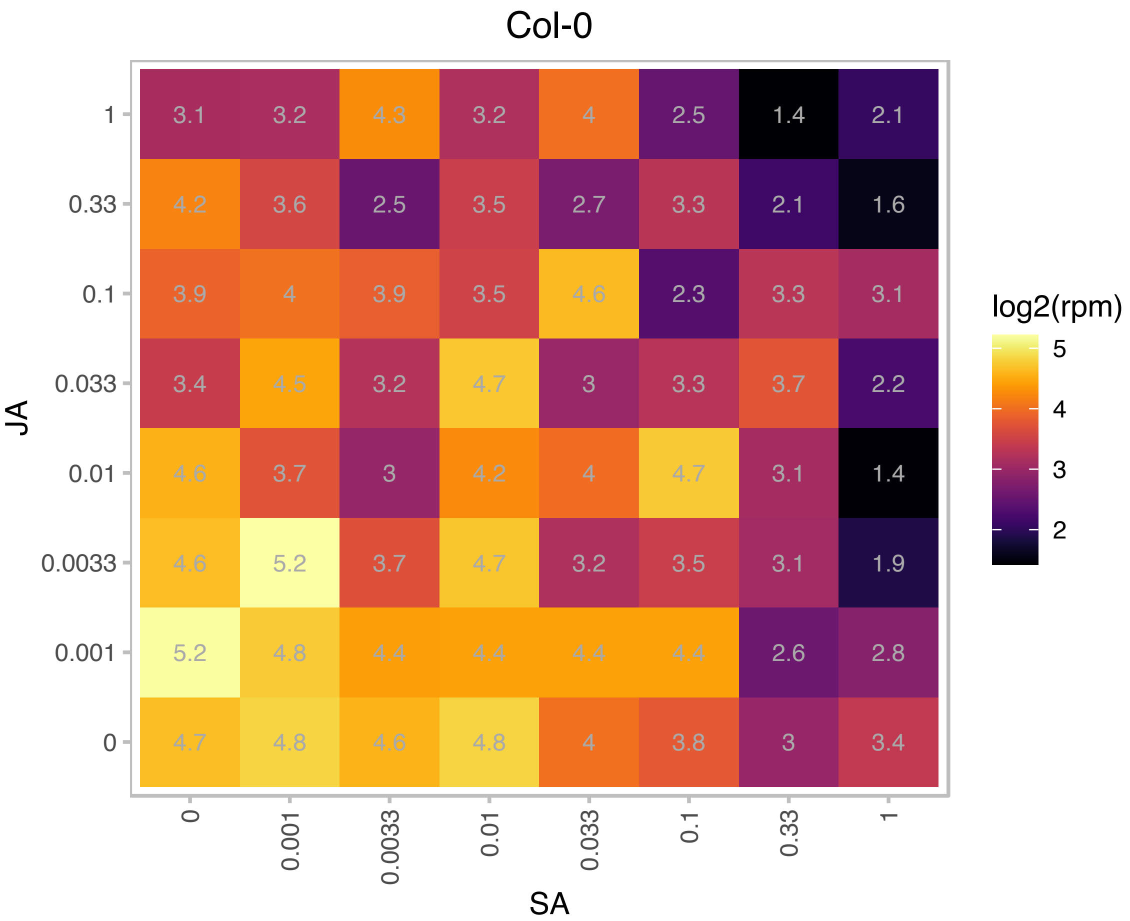


| | |
|---|---|
| Gene | AT5G57170 |
| is.expressed_genes | TRUE |
| is.responsive_genes | TRUE |
| is.apcluster | F-01 |

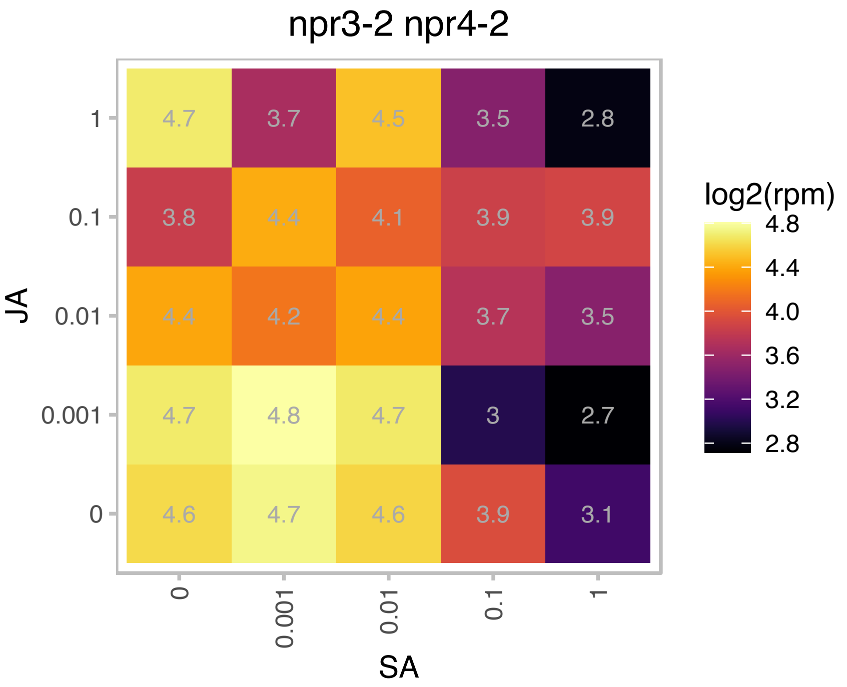


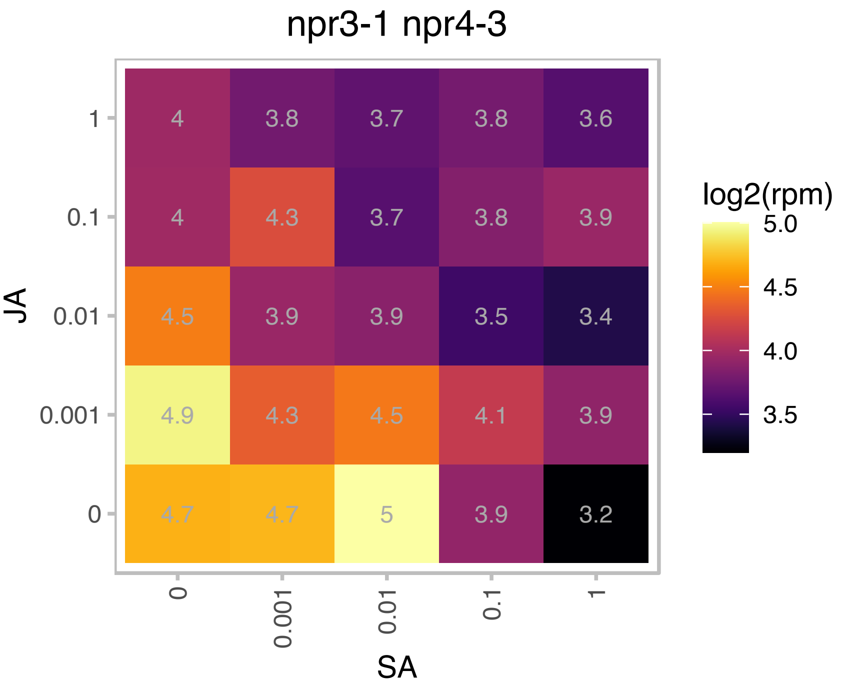

# AT5G59770.1 : NA

Protein-tyrosine phosphatase-like, PTPLA

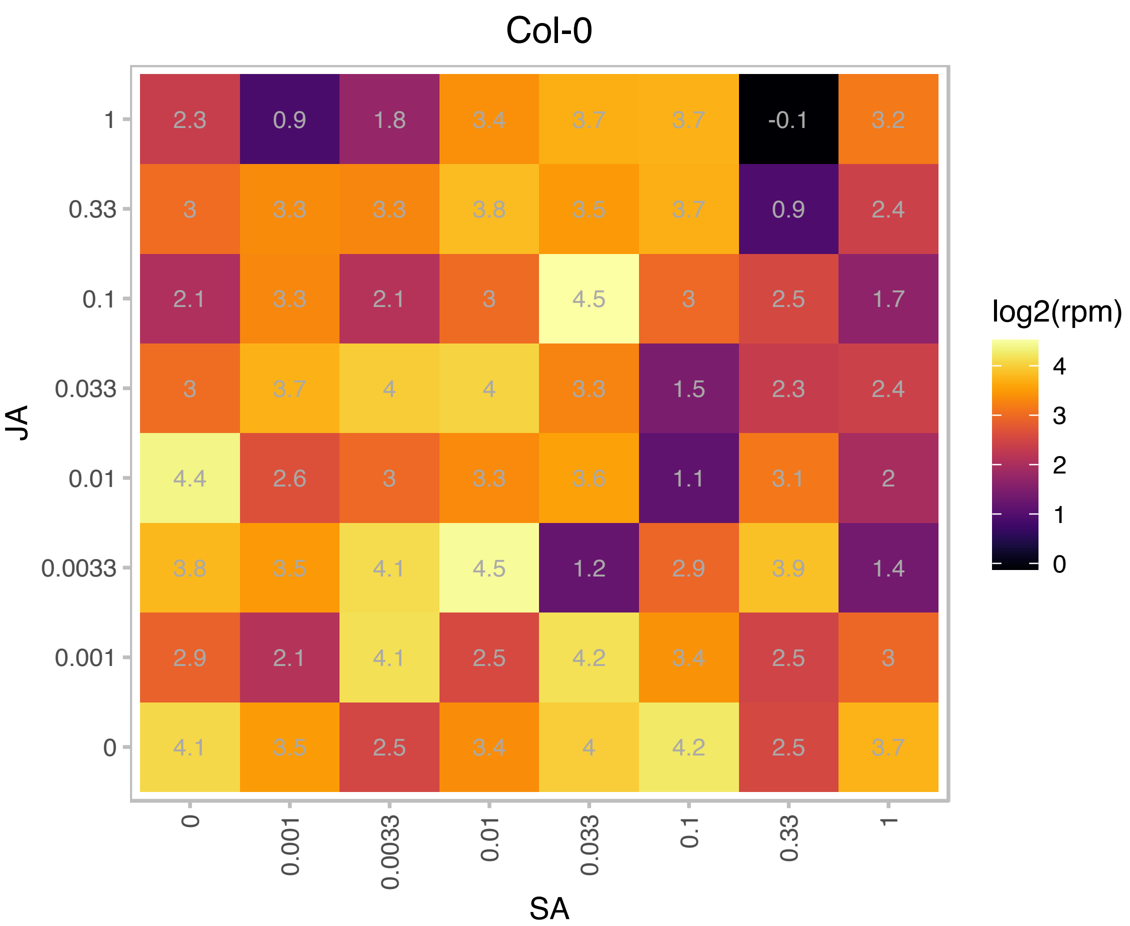


| | |
|---|---|
| Gene | AT5G59770 |
| is.expressed_genes | TRUE |
| is.responsive_genes | TRUE |
| is.apcluster | F-01 |

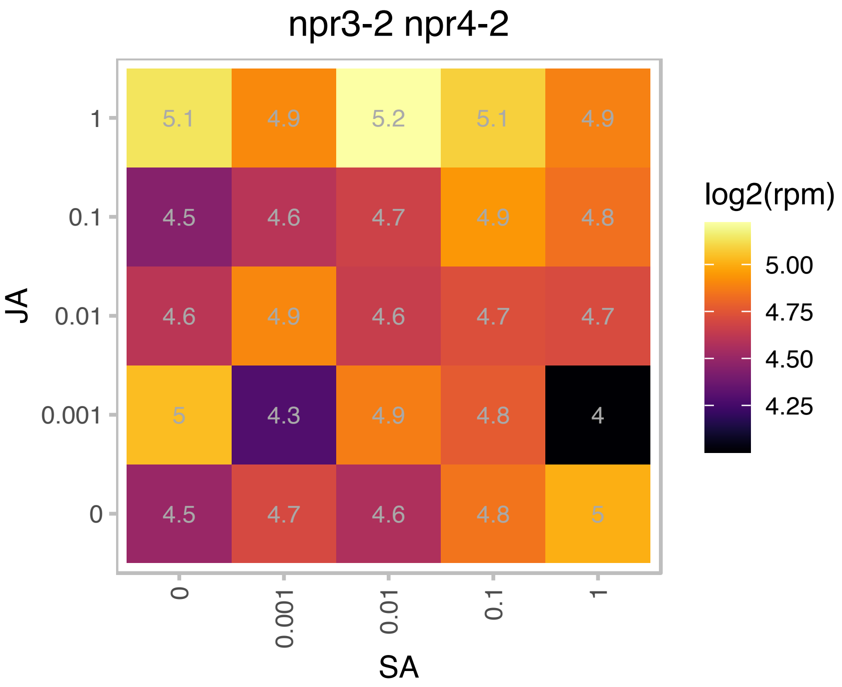


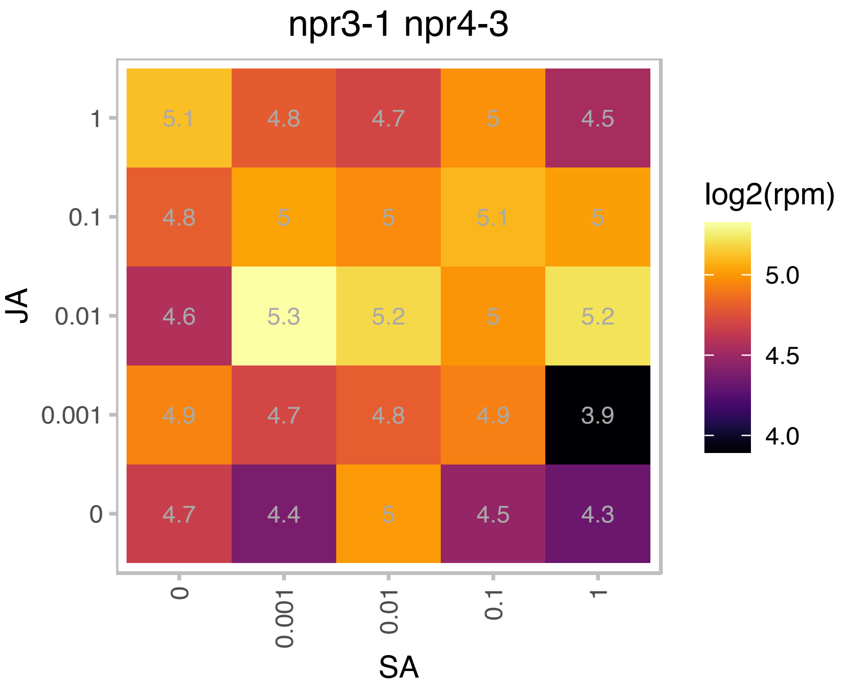

# AT5G60680.1 : NA

Protein of unknown function, DUF584

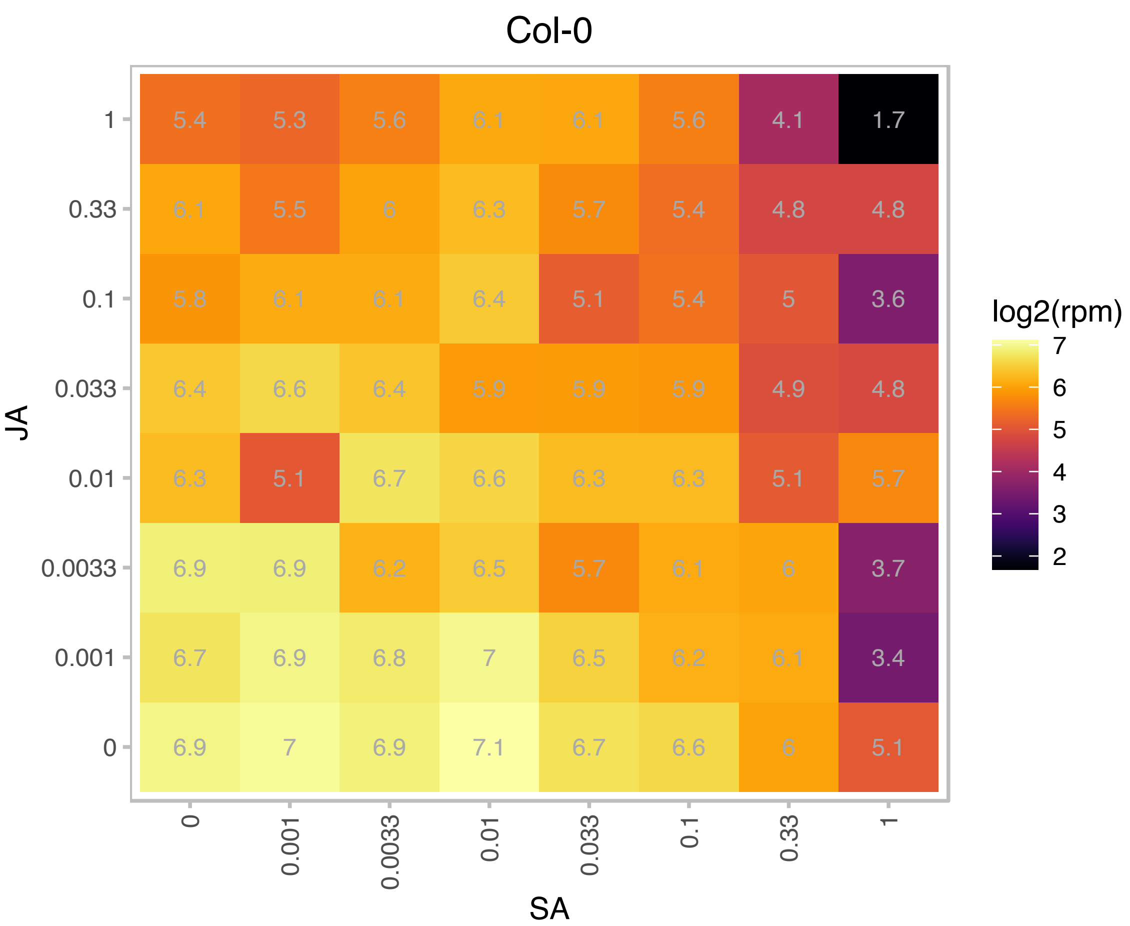


| | |
|---|---|
| Gene | AT5G60680 |
| is.expressed_genes | TRUE |
| is.responsive_genes | TRUE |
| is.apcluster | F-01 |

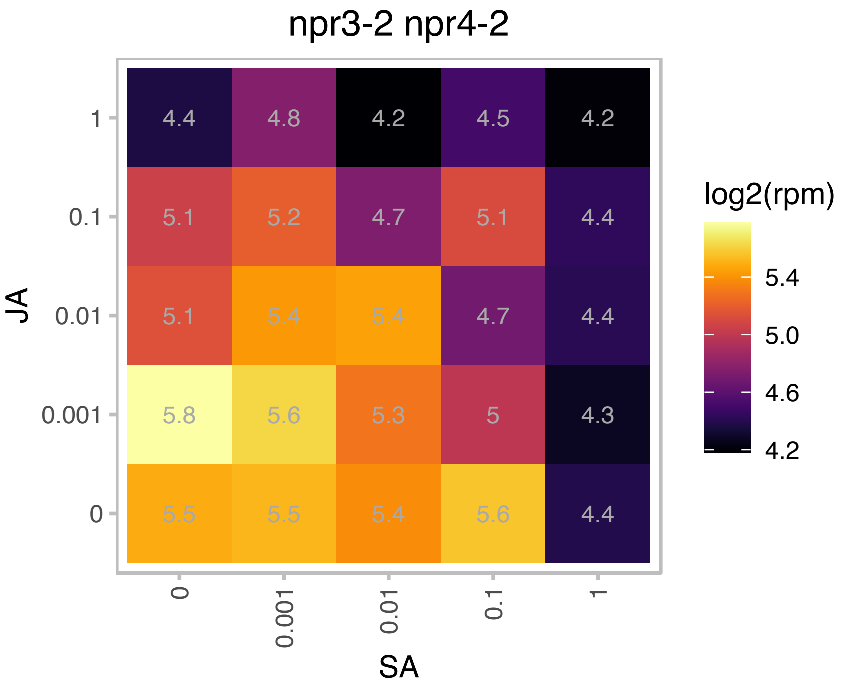


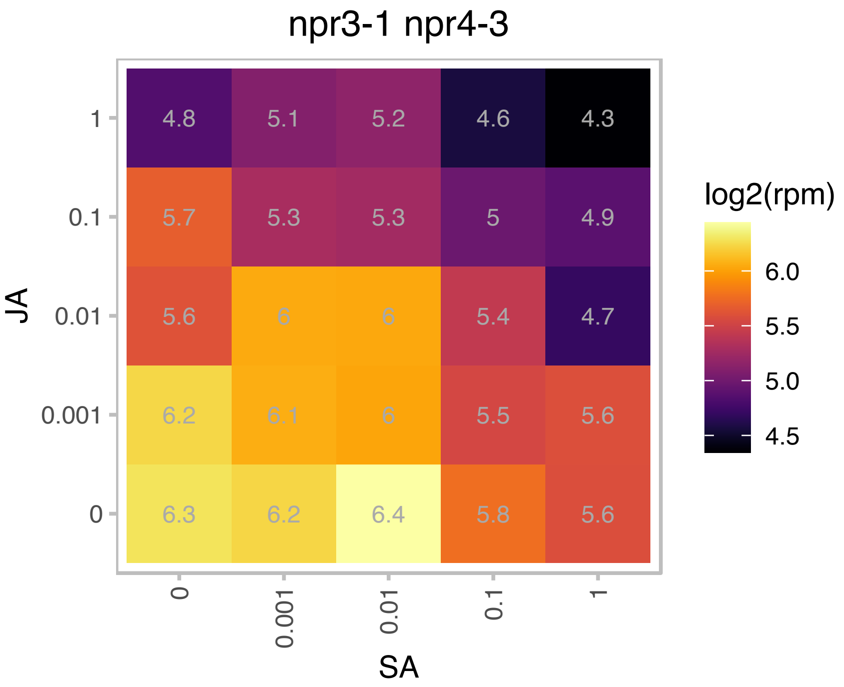

### AT5G61130.1 : AtPDCB1, PDCB1

plasmodesmata callose-binding protein 1

Encodes a plasmodesmatal localized protein with callose binding activity. The function of PDCB1 is unknown but overexpression leads to the hyperaccumulation of callose.

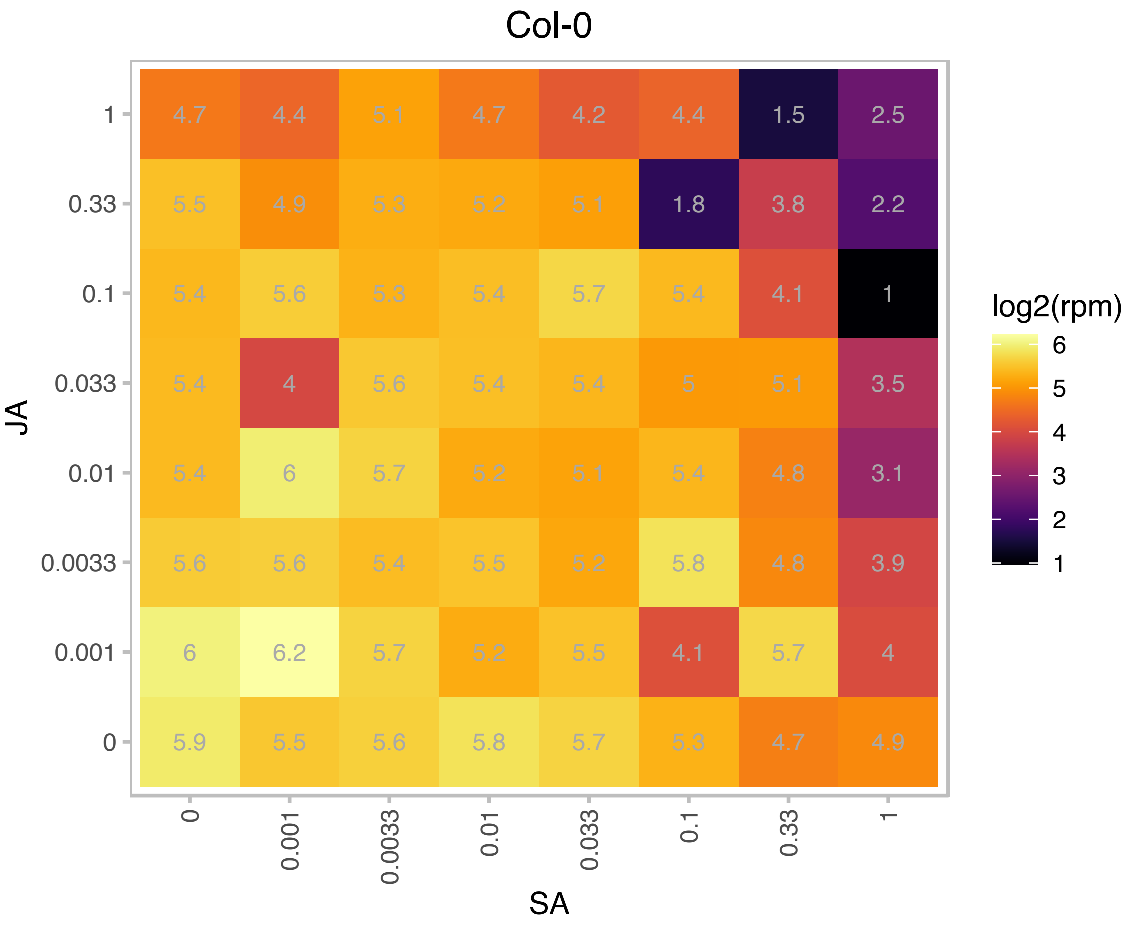


| | |
|---|---|
| Gene | AT5G61130 |
| is.expressed_genes | TRUE |
| is.responsive_genes | TRUE |
| is.apcluster | F-01 |

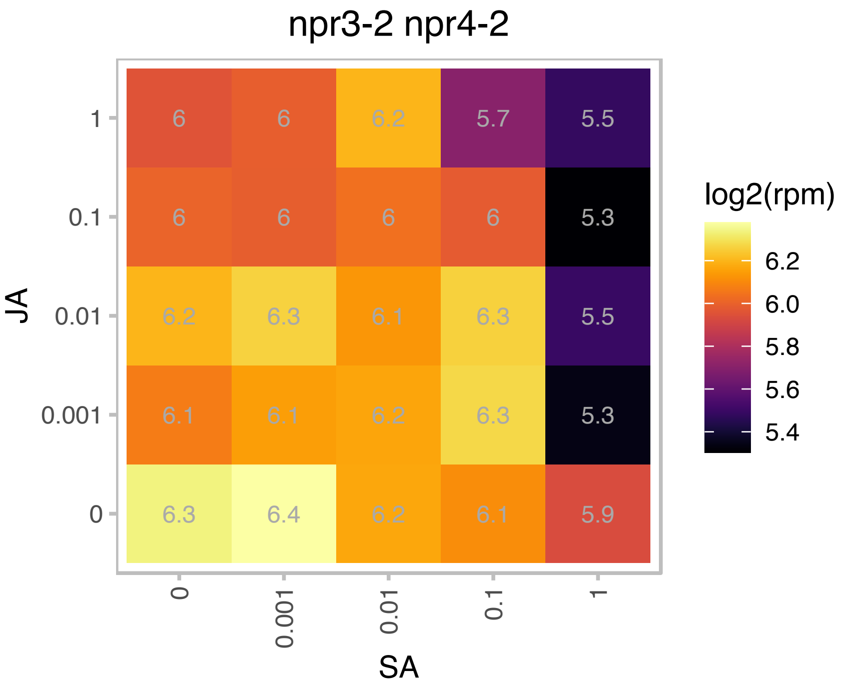


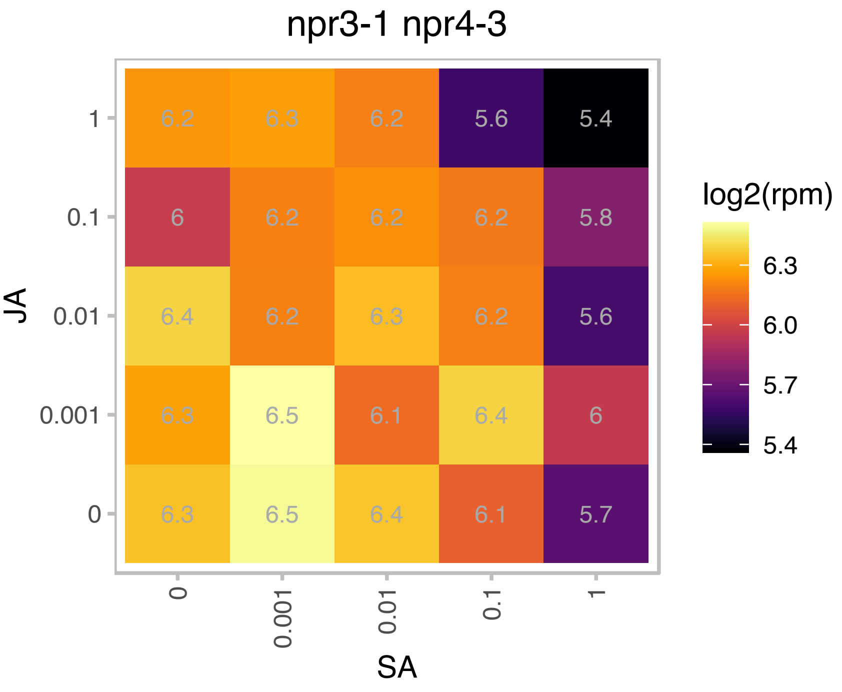

### AT5G61590.1 : DEWAX, ERF107

Integrase-type DNA-binding superfamily protein

Encodes an AP2/ERF-type transcription factor that is preferentially expressed in the epidermis and induced by darkness and negatively regulates cuticular wax biosynthesis.

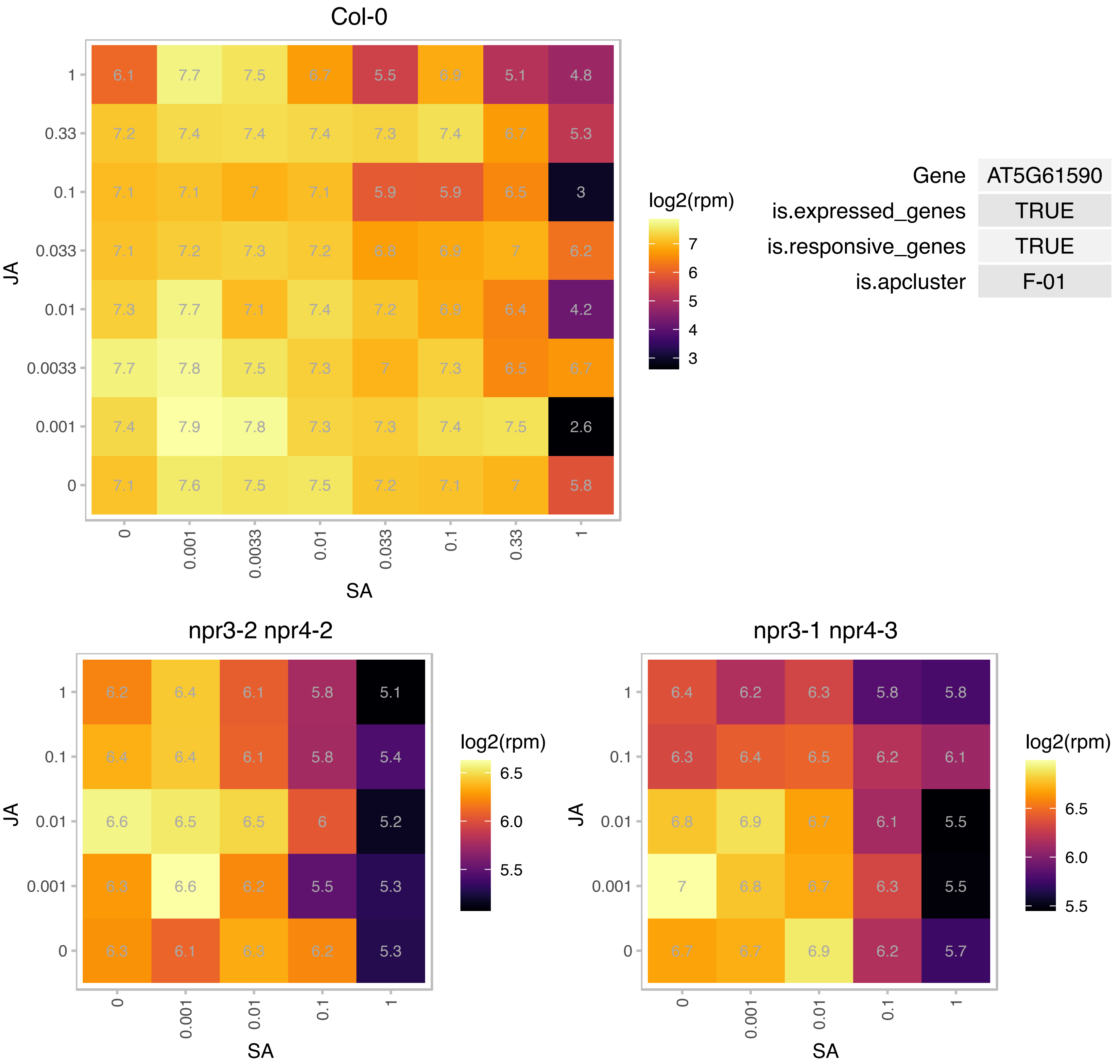


| | |
|---|---|
| Gene | AT5G61590 |
| is.expressed_genes | TRUE |
| is.responsive_genes | TRUE |
| is.apcluster | F-01 |

### AT5G65640.1 : bHLH093, NFL

bHLH093/NFL encodes a bHLH transcription factor involved in GA mediated control of flowering time. Mutants are non-flowering in short days and phenotype can be reversed with GA application. Based on the expression of GA biosynthetic genes in the mutant, it likely acts through regulation of GA metabolism. Its expression shows developmental stage and tissue specificity. In short days it is expressed mainly in root tips and SAM, with weak expression in cotyledons throughout development. In LD GUS activity was observed in the hypocotyl and in root tips and SAM throughout the developmental stages.

beta HLH protein 93

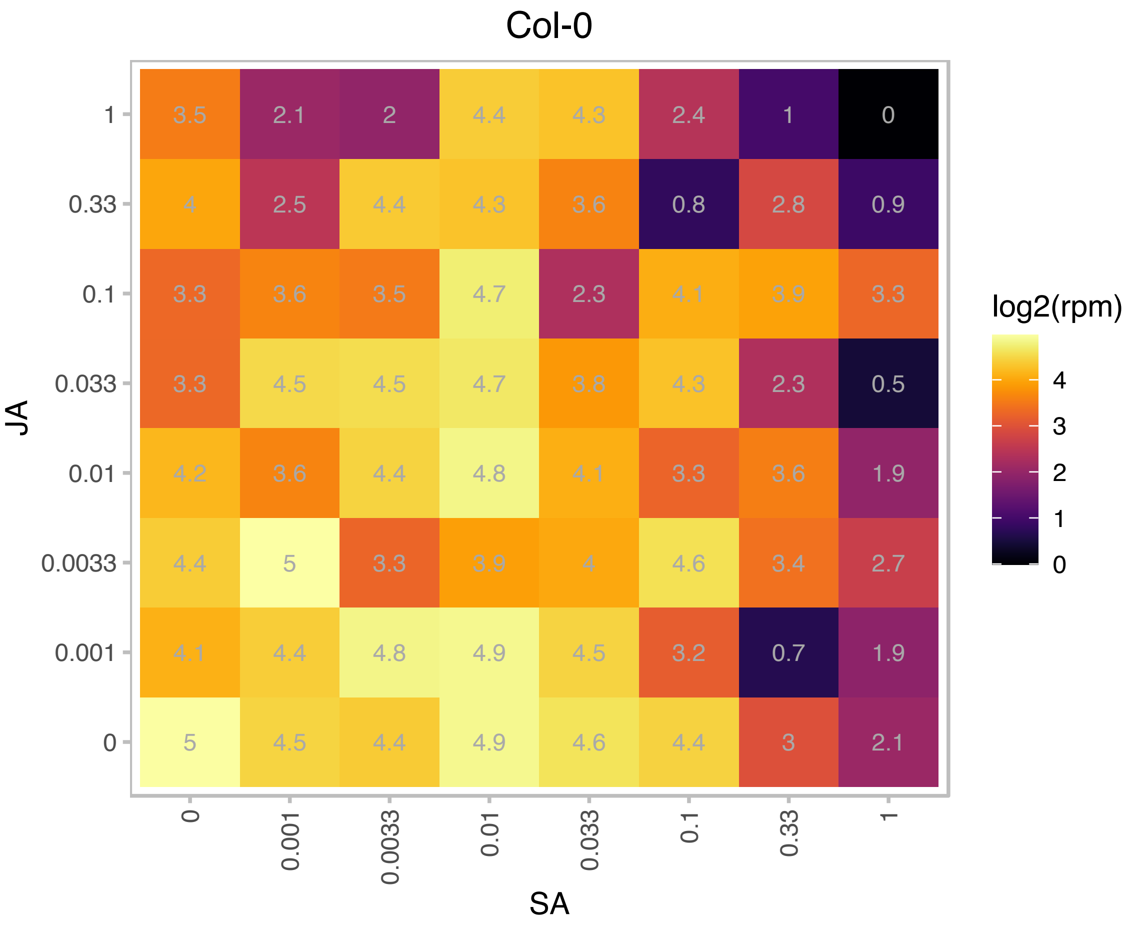


| | |
|---|---|
| Gene | AT5G65640 |
| is.expressed_genes | TRUE |
| is.responsive_genes | TRUE |
| is.apcluster | F-01 |

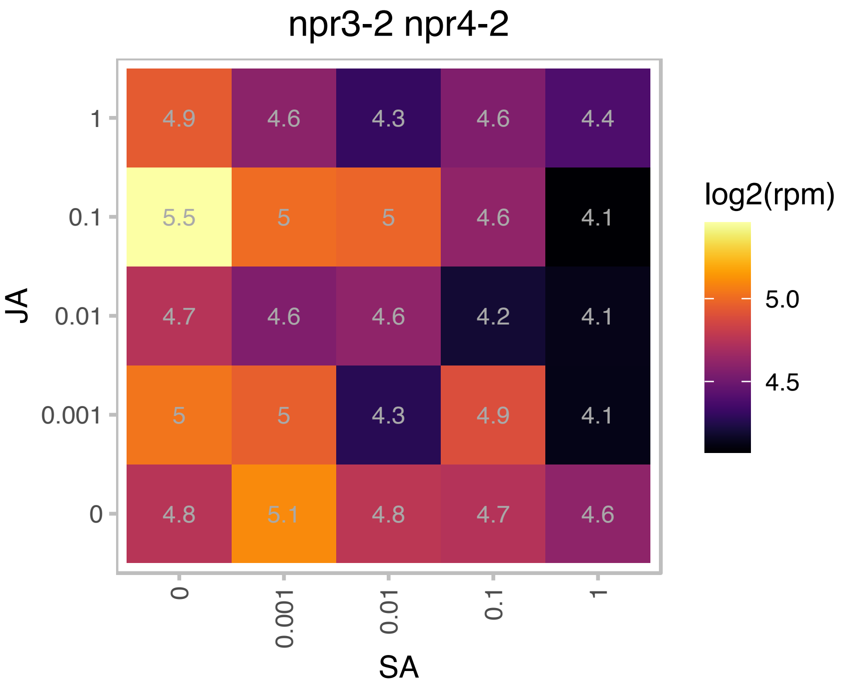


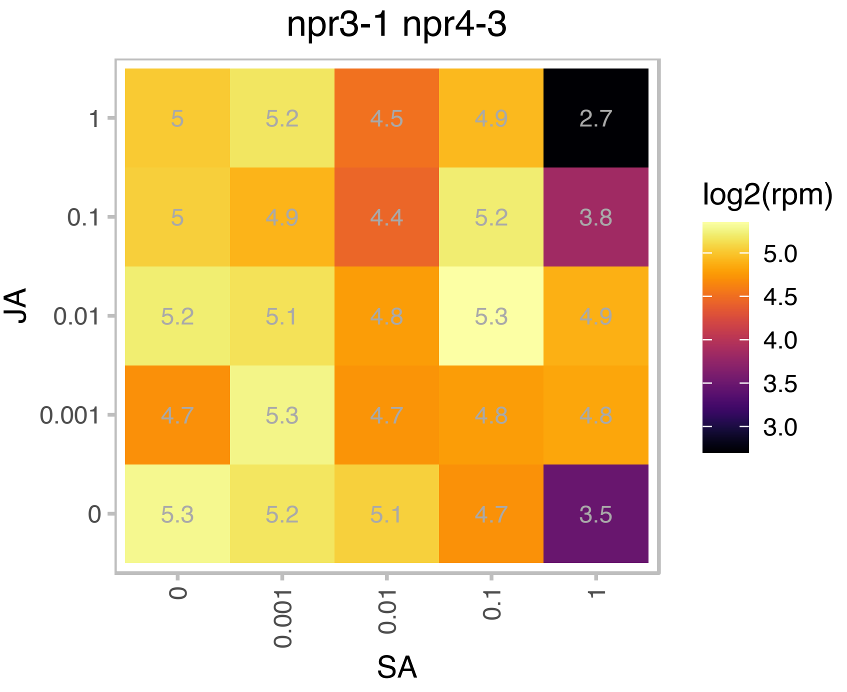

# AT5G65860.1 : NA

ankyrin repeat family protein

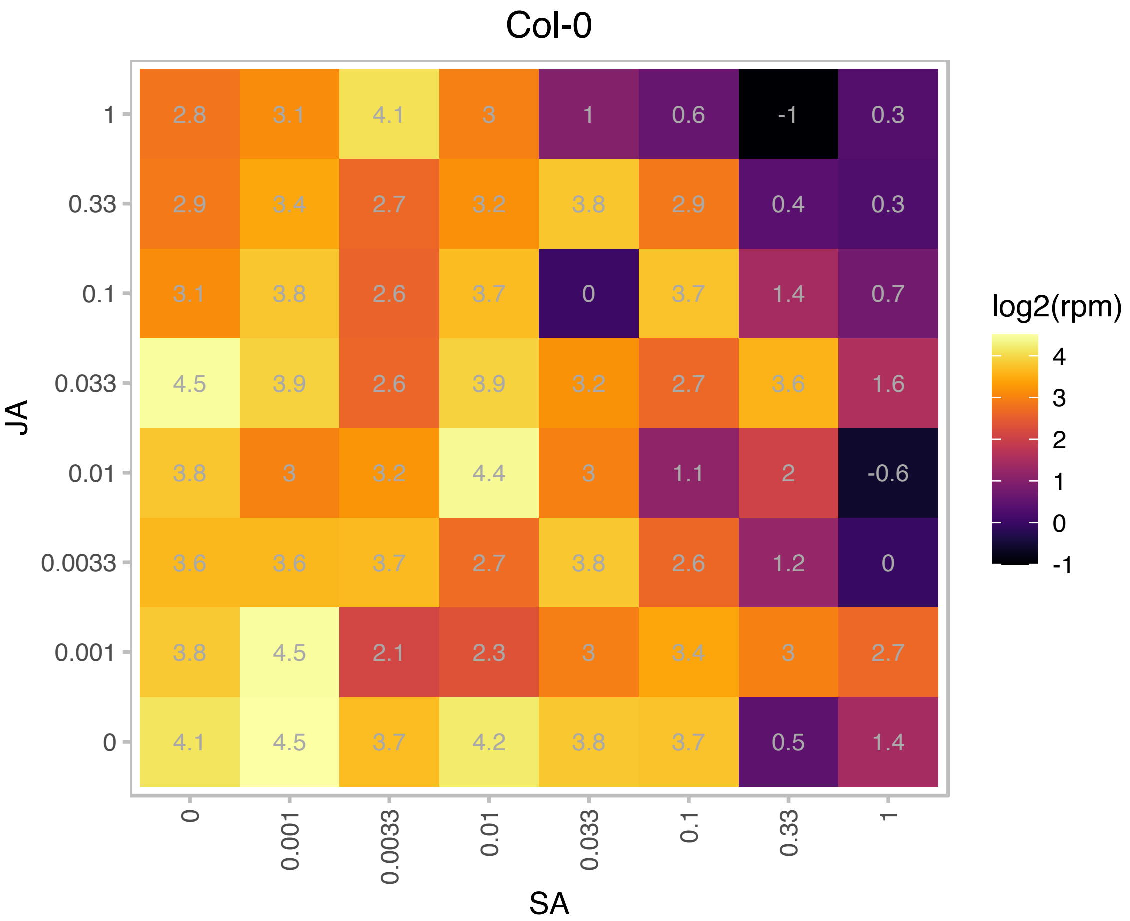

| | |
|---|---|
| Gene | AT5G65860 |
| is.expressed_genes | TRUE |
| is.responsive_genes | TRUE |
| is.apcluster | F-01 |

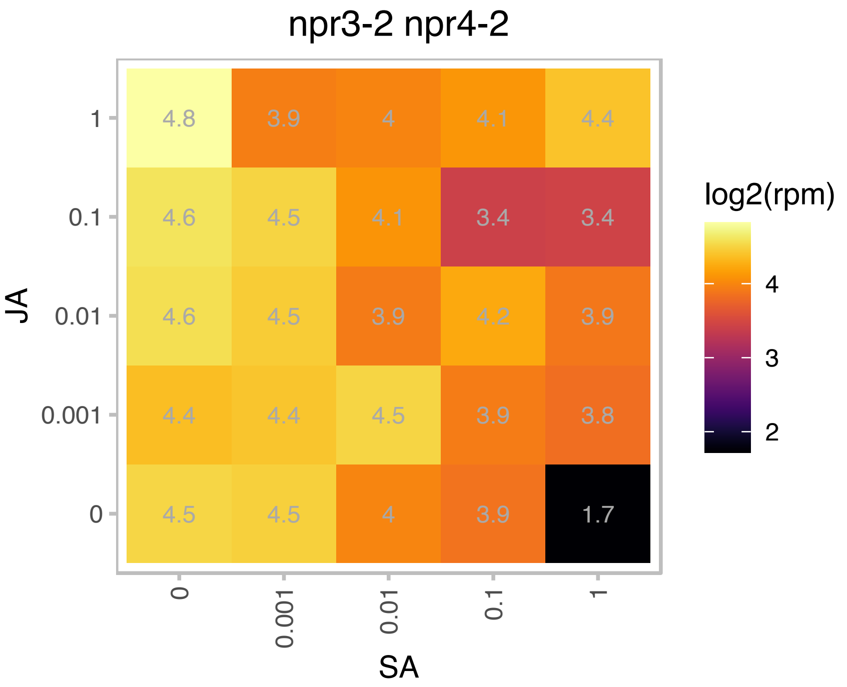

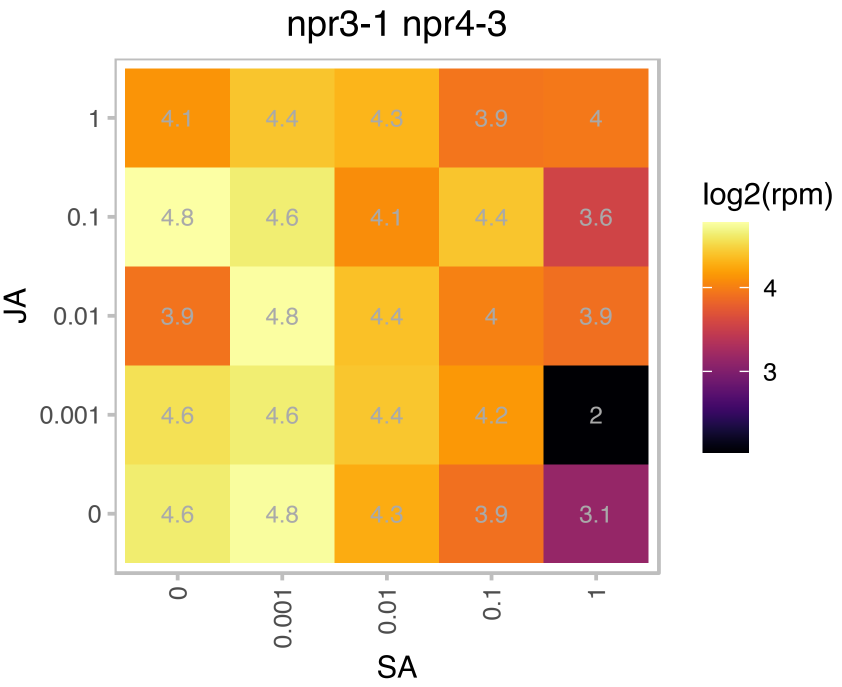

# AT5G66090.1 : NA

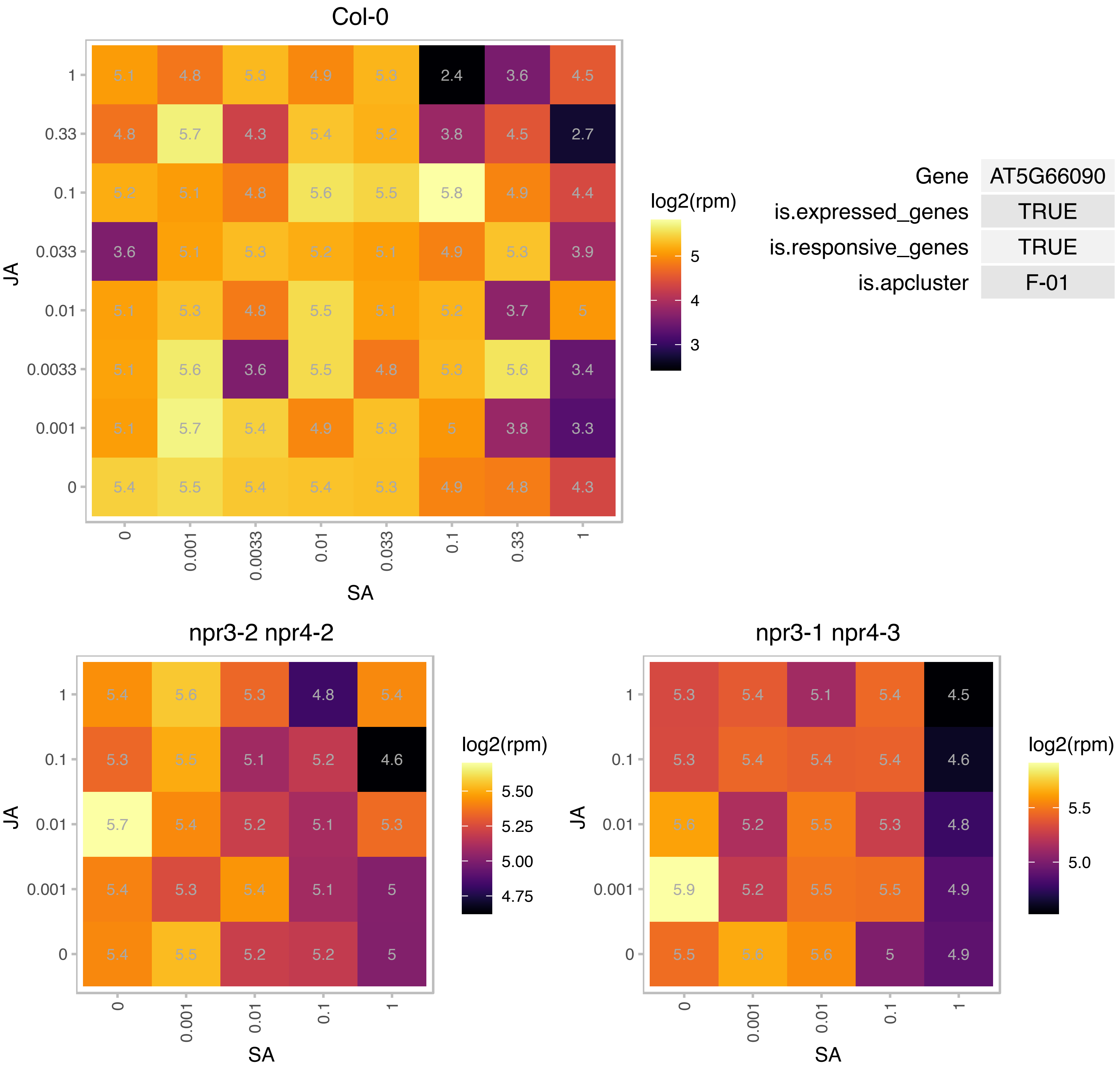

### ATCG00380.1 : RPS4

chloroplast ribosomal protein S4

Chloroplast encoded ribosomal protein S4

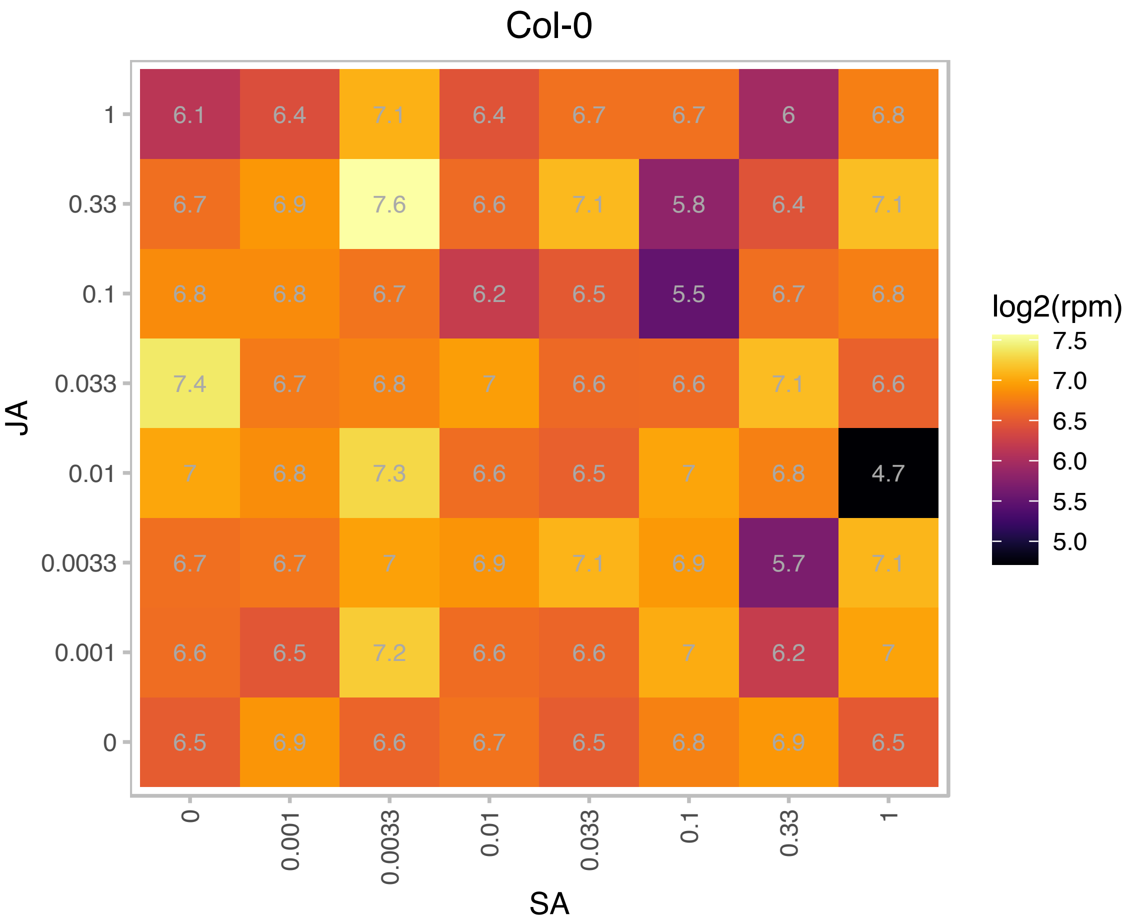


| | |
|---|---|
| Gene | ATCG00380 |
| is.expressed_genes | TRUE |
| is.responsive_genes | TRUE |
| is.apcluster | F-01 |

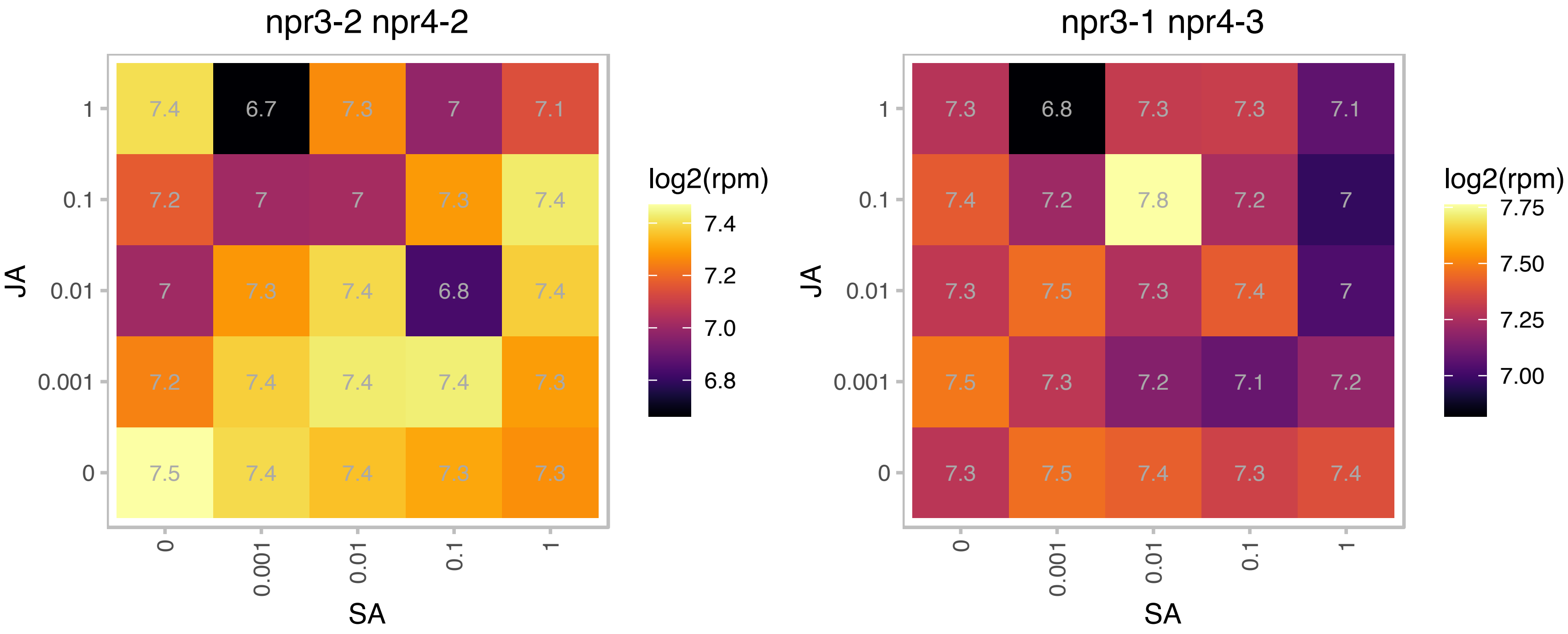

### AT1G01160.1 : GIF2

GRF1-interacting factor 2

Arabidopsis thaliana GRF1-interacting factor 2 (GIF2) mRNA

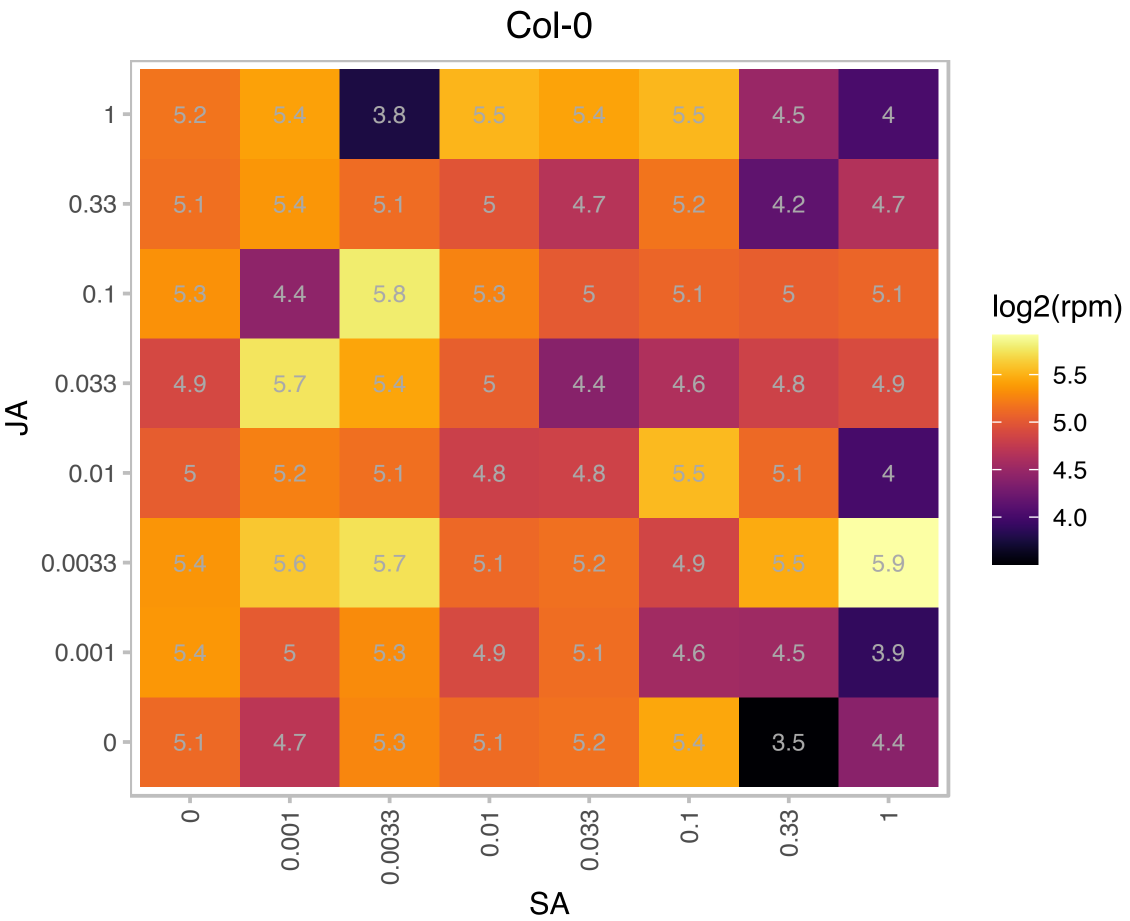


| | |
|---|---|
| Gene | AT1G01160 |
| is.expressed_genes | TRUE |
| is.responsive_genes | TRUE |
| is.apcluster | F-02 |

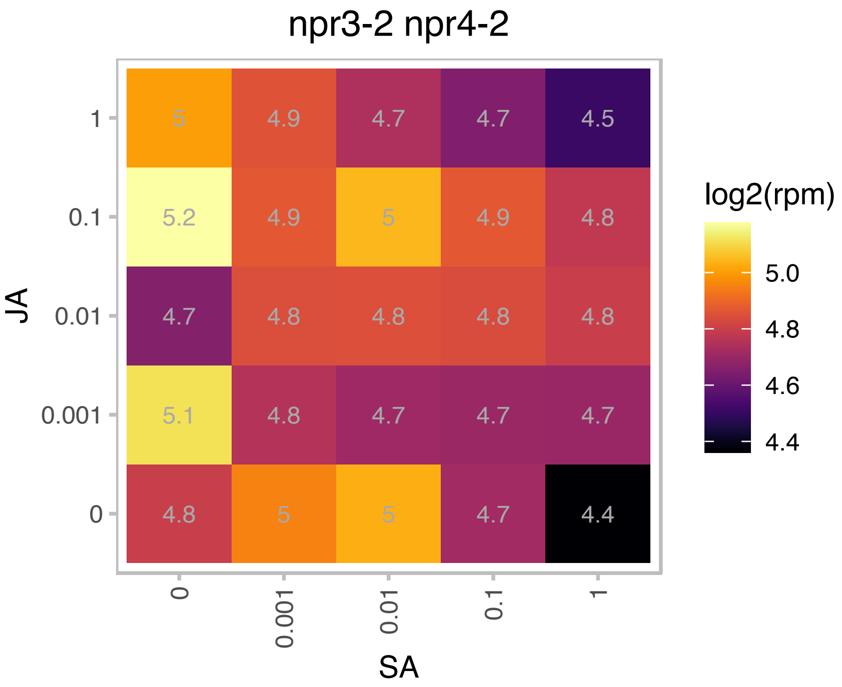


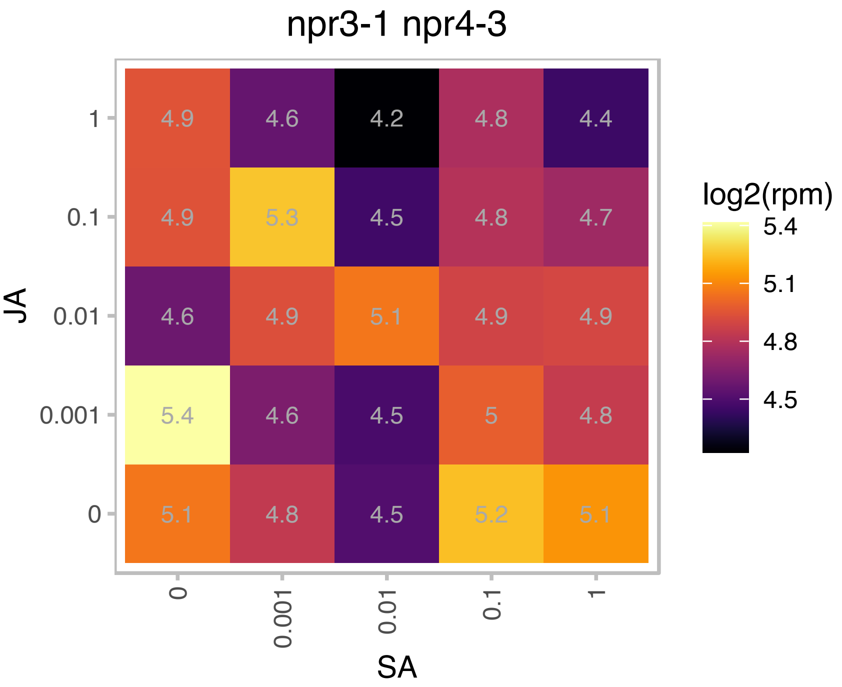

### AT1G01750.1 : ADF11

actin depolymerizing factor 11

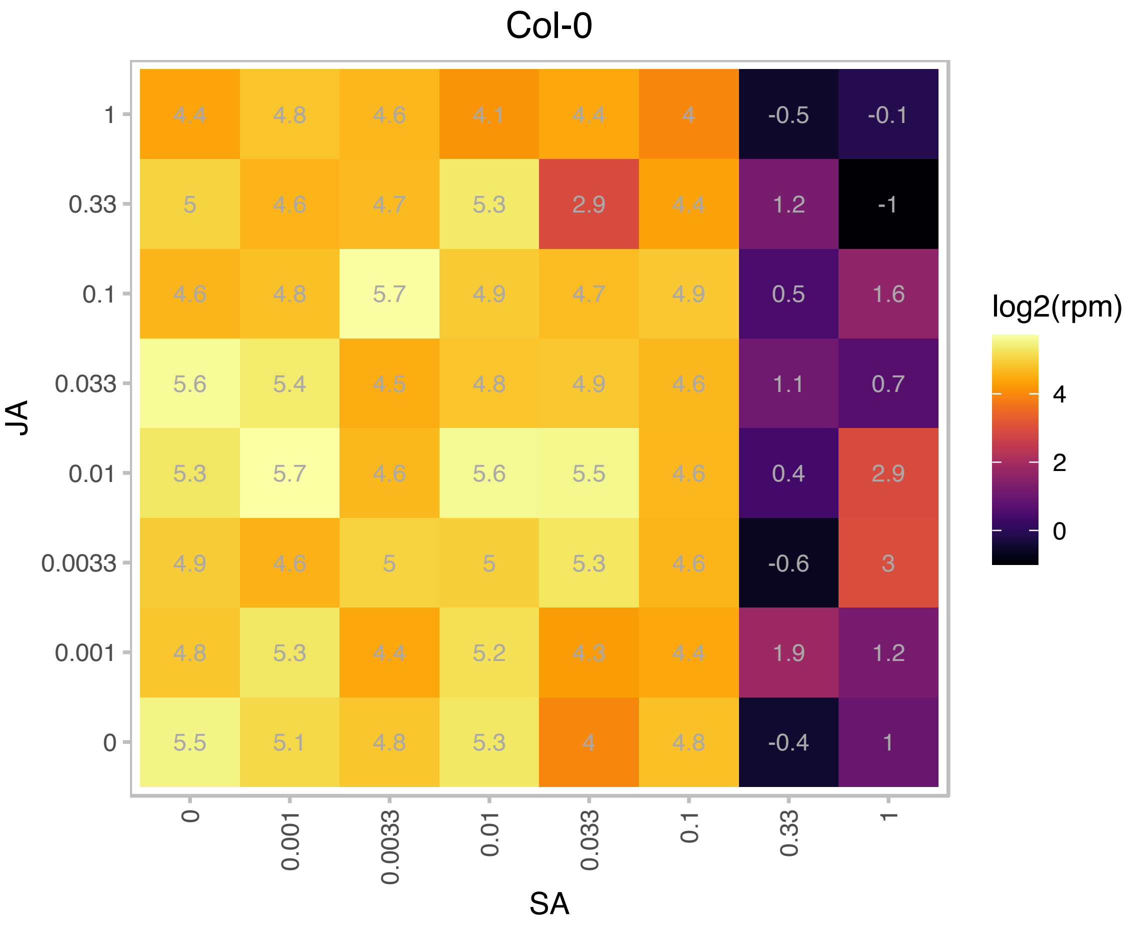


| | |
|---|---|
| Gene | AT1G01750 |
| is.expressed_genes | TRUE |
| is.responsive_genes | TRUE |
| is.apcluster | F-02 |

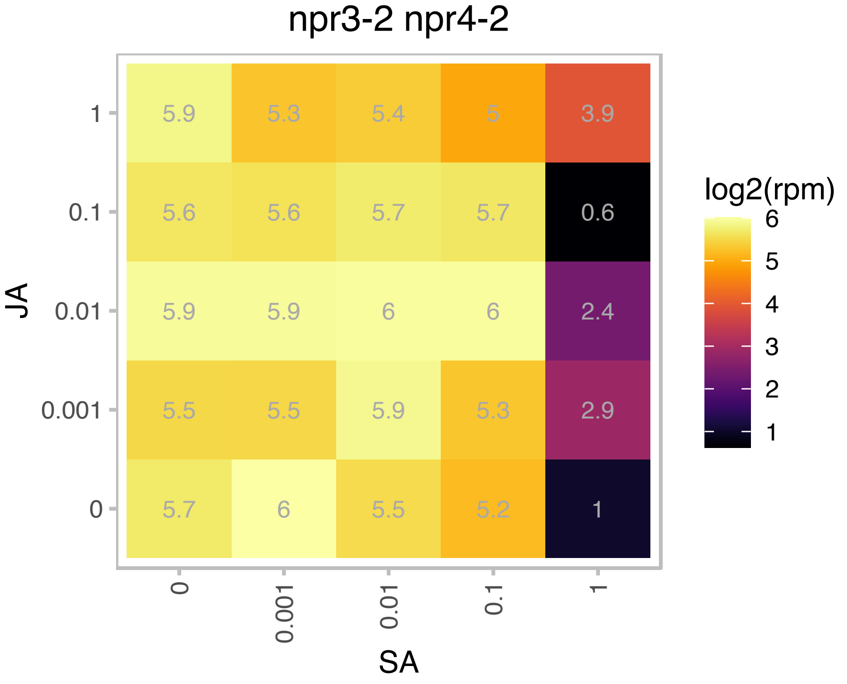


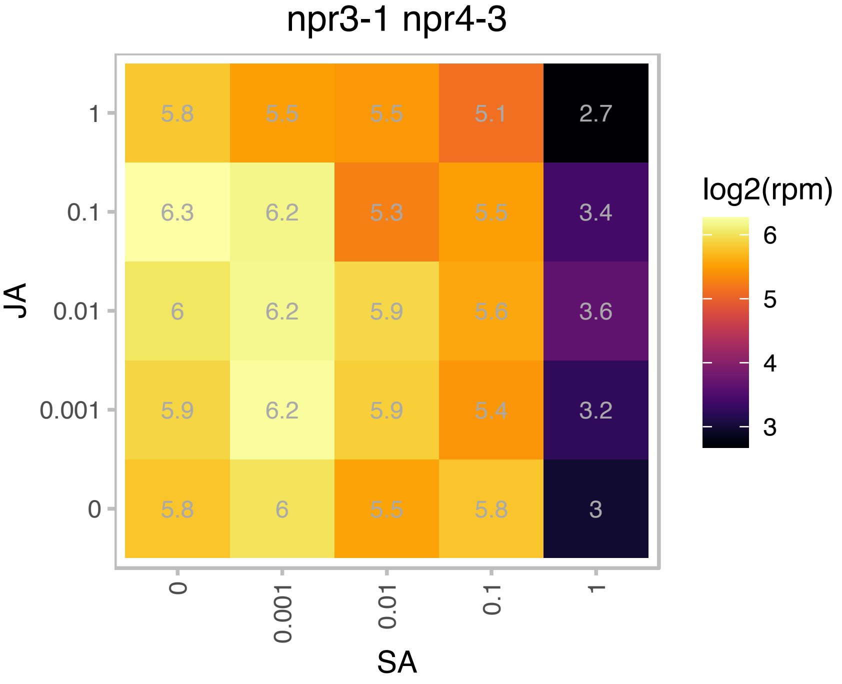

### AT1G02330.1 : CSU2

Encodes a nuclear coiled-coil domain-containing protein that interacts with and negatively regulates COP1's E3 ubiquitin ligase activity, and represses COP1 mediated HY5 degradation in cell-free extracts.

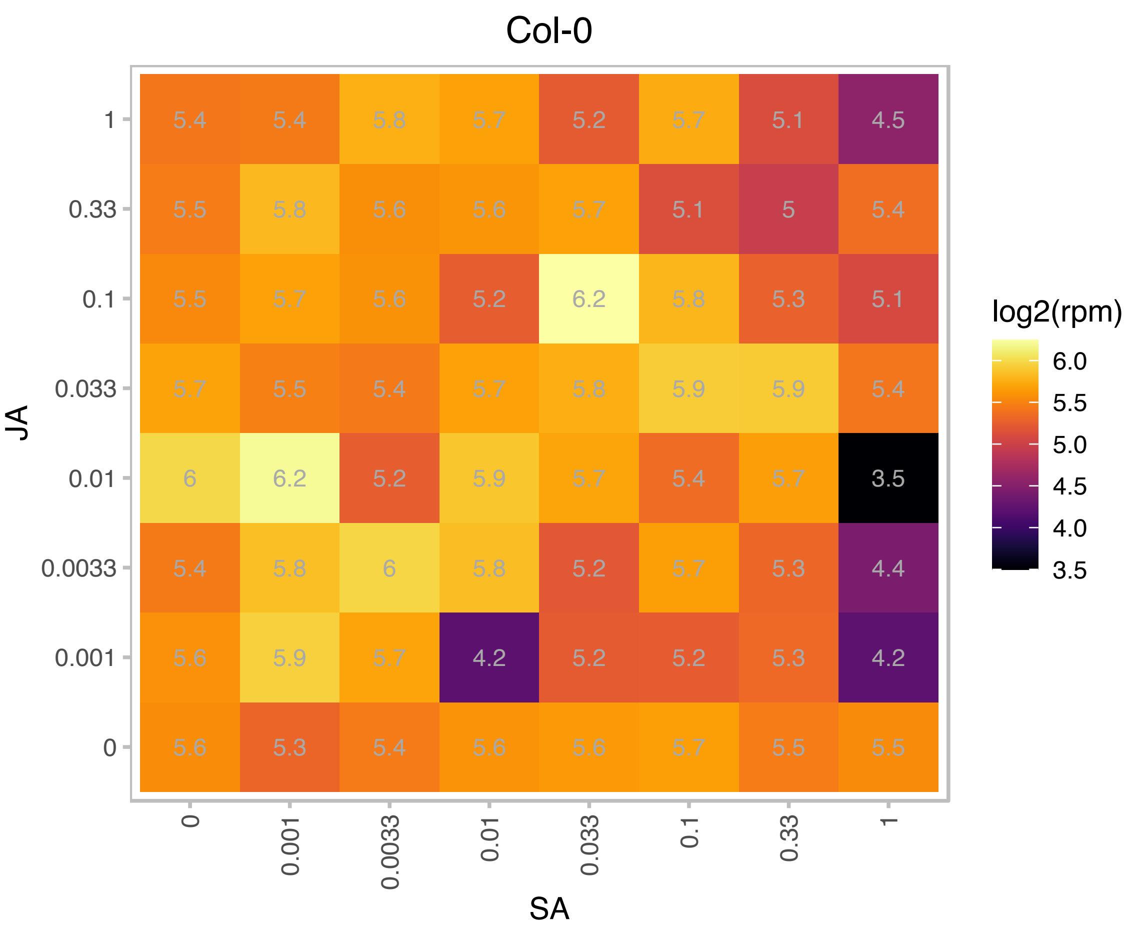


| | |
|---|---|
| Gene | AT1G02330 |
| is.expressed_genes | TRUE |
| is.responsive_genes | TRUE |
| is.apcluster | F-02 |

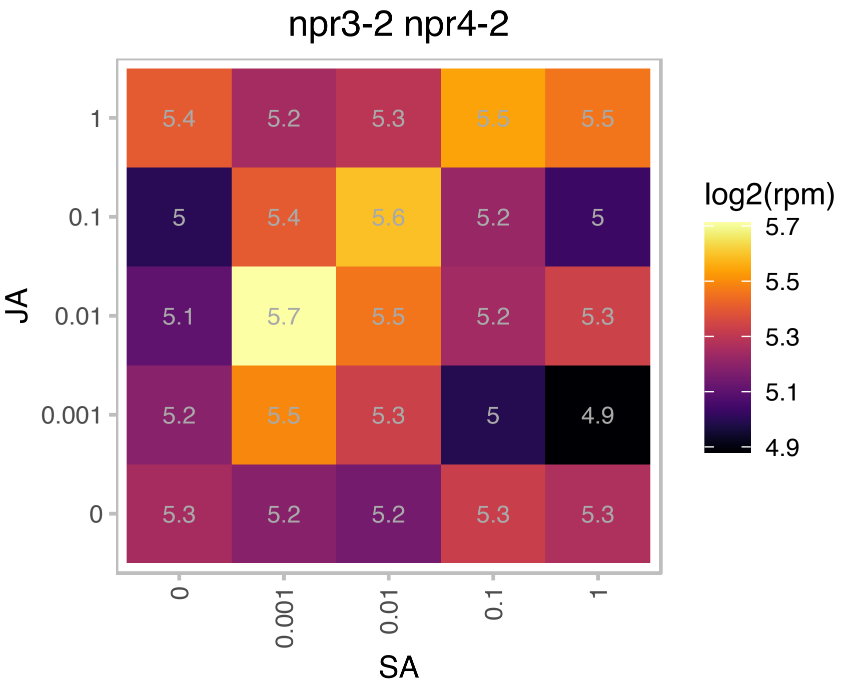


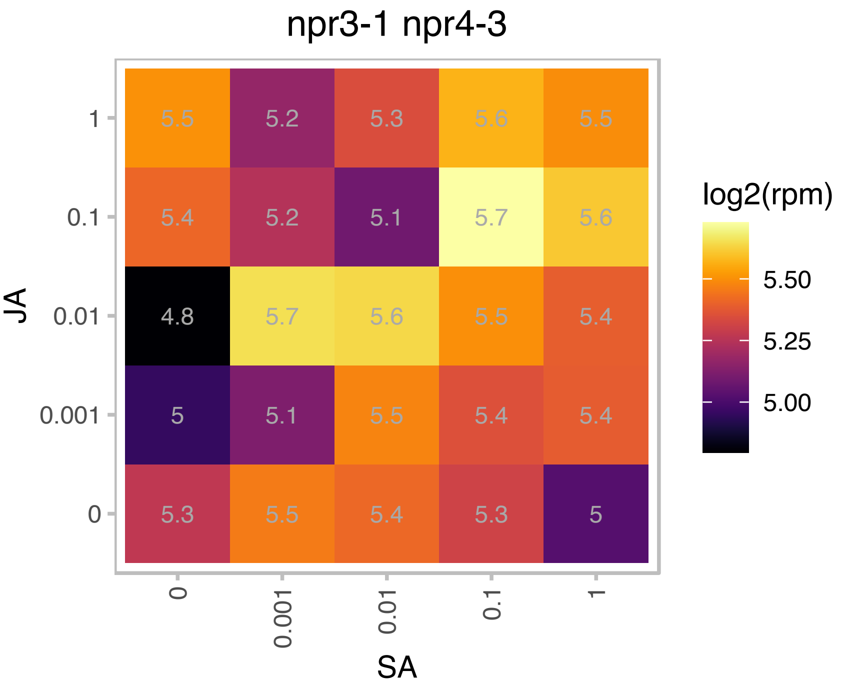

### AT1G05250.1 : AtPRX1, PRX2

Peroxidase superfamily protein

Encodes a cationic cell-wall-bound peroxidase homolog that is involved in the lignification of cell walls.

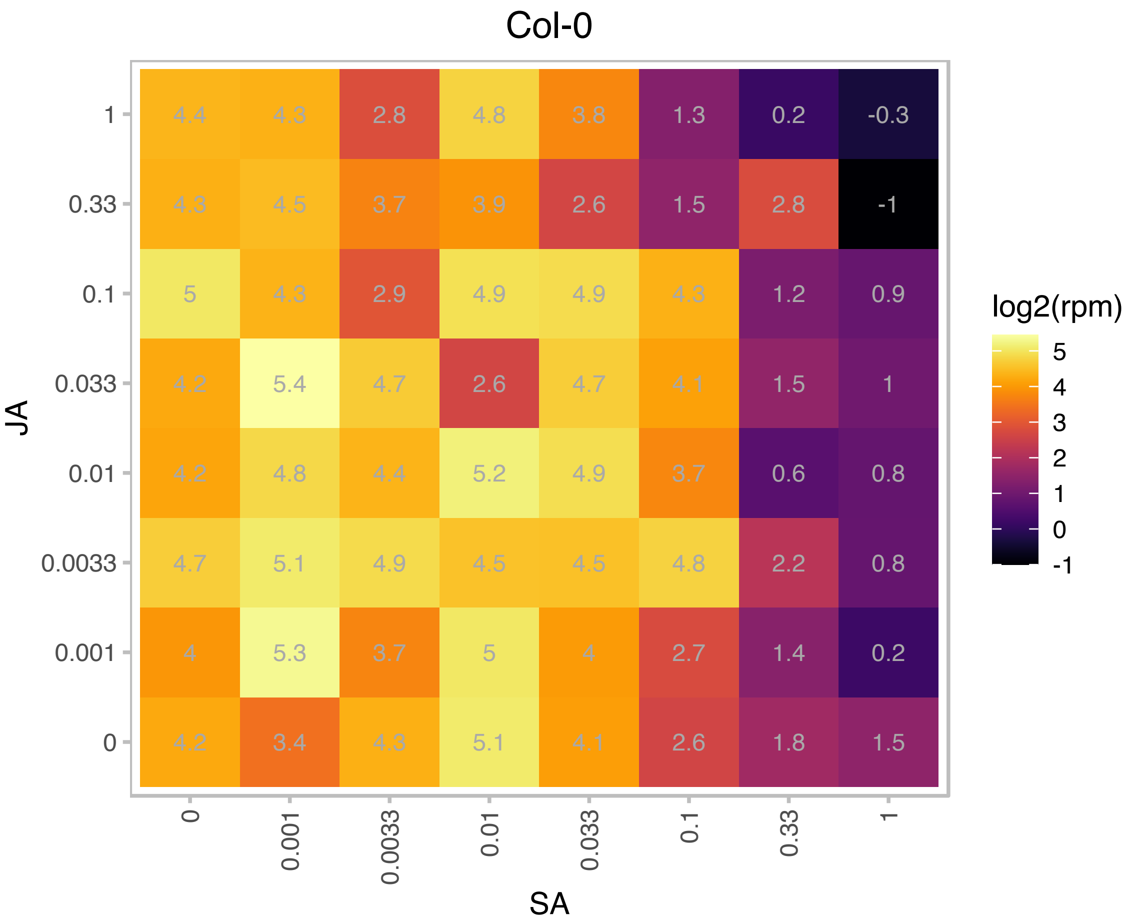


| | |
|---|---|
| Gene | AT1G05250 |
| is.expressed_genes | TRUE |
| is.responsive_genes | TRUE |
| is.apcluster | F-02 |

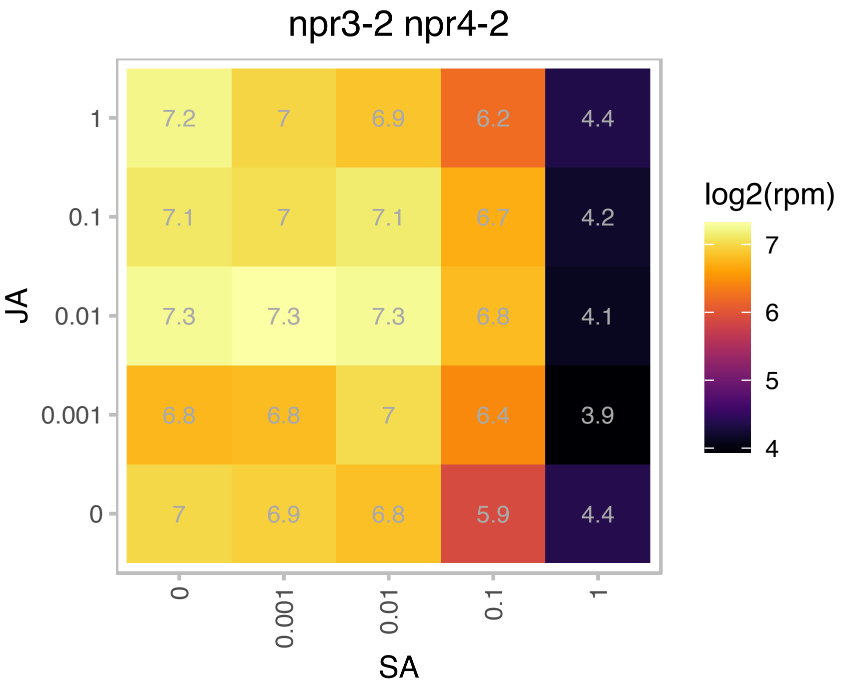


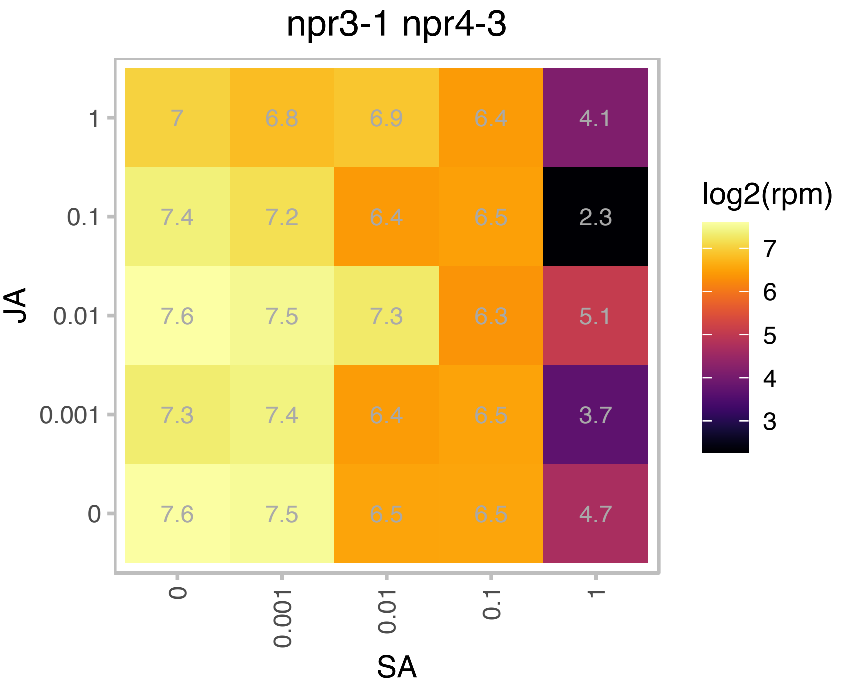

### AT1G07890.1 : APX1, ATAPX01, ATAPX1, CS1, MEE6

ascorbate peroxidase 1

Encodes a cytosolic ascorbate peroxidase APX1. Ascorbate peroxidases are enzymes that scavenge hydrogen peroxide in plant cells. Eight types of APX have been described for Arabidopsis: three cytosolic (APX1, APX2, APX6), two chloroplastic types (stromal sAPX, thylakoid tAPX), and three microsomal (APX3, APX4, APX5) isoforms. At least part of the induction of heat shock proteins during light stress in Arabidopsis is mediated by H2O2 that is scavenged by APX1. Expression of the gene is downregulated in the presence of paraquat, an inducer of photoxidative stress. The mRNA is cell-to-cell mobile.

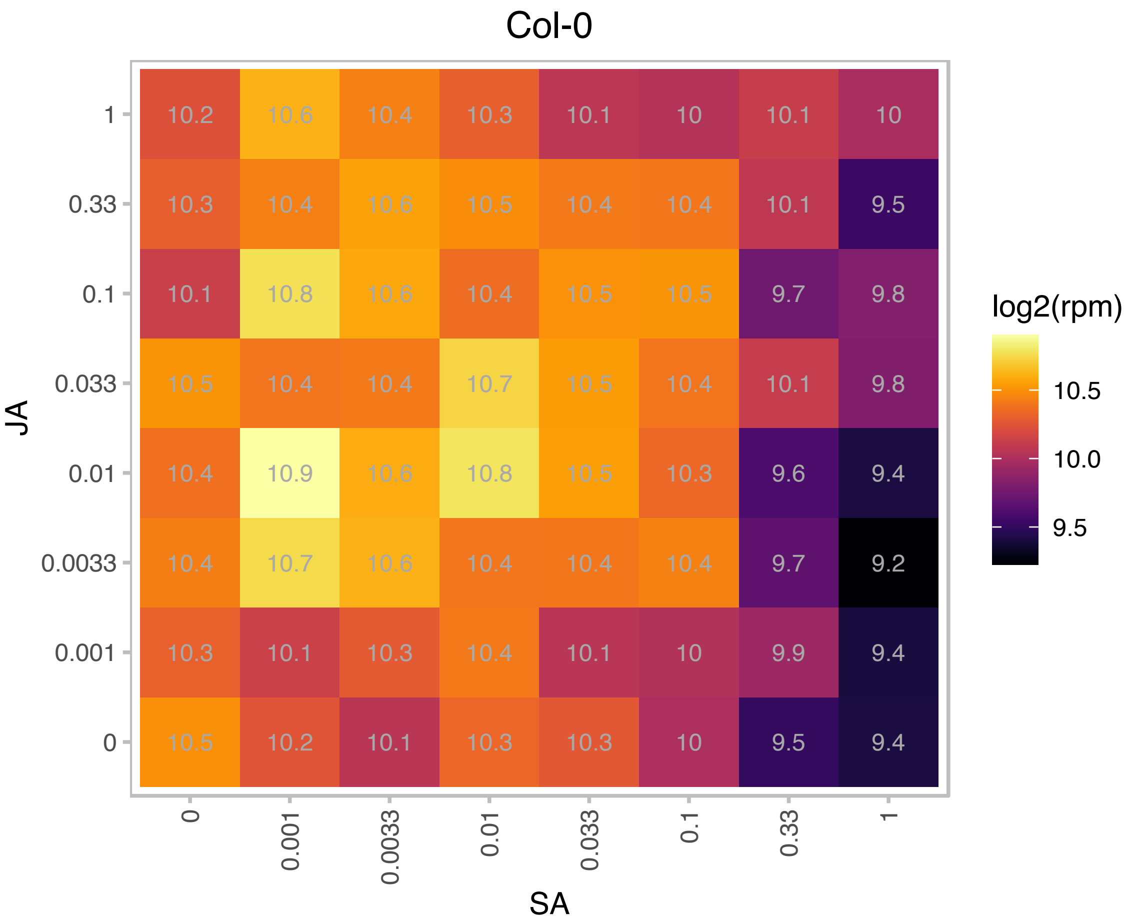


| | |
|---|---|
| Gene | AT1G07890 |
| is.expressed_genes | TRUE |
| is.responsive_genes | TRUE |
| is.apcluster | F-02 |

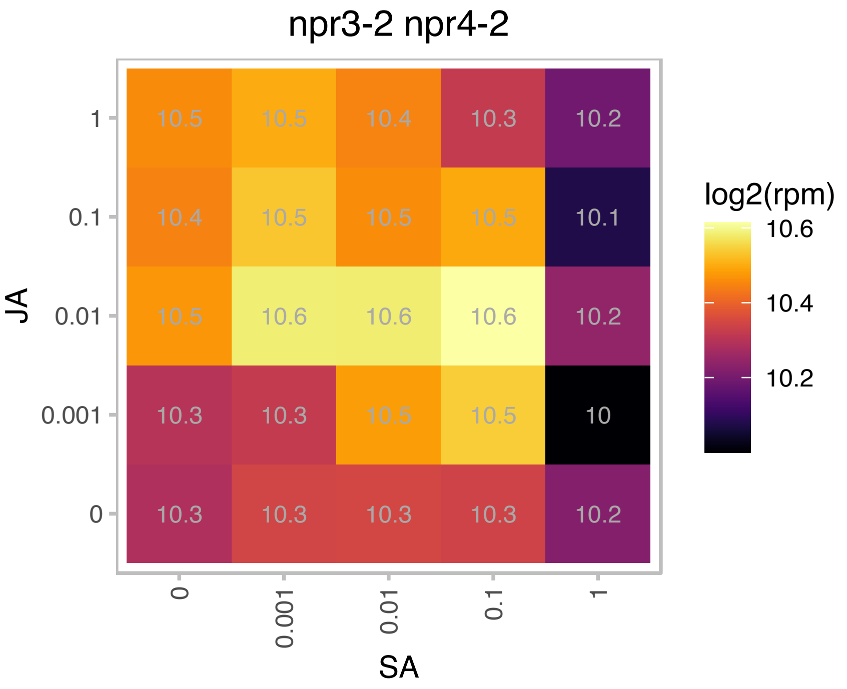


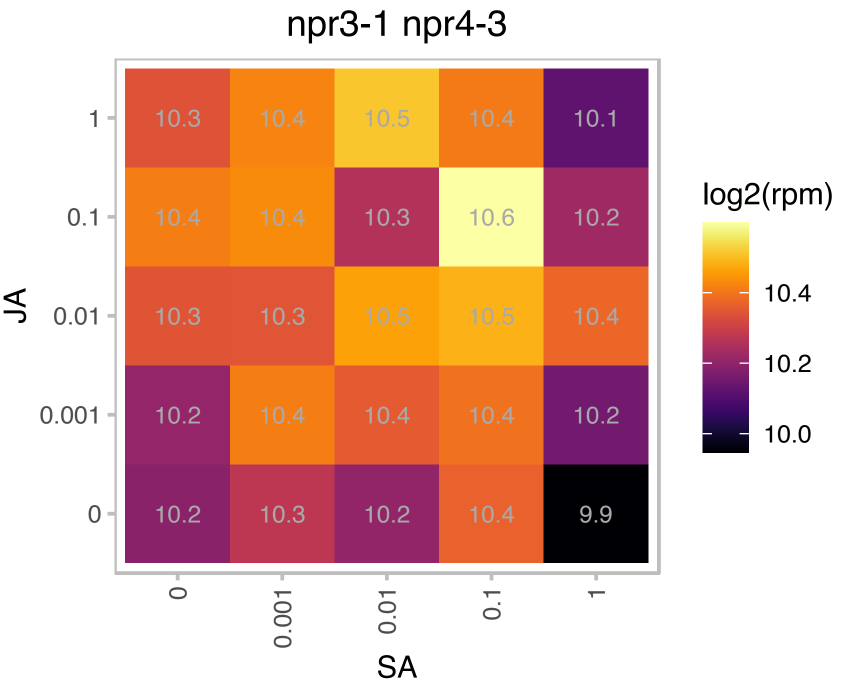

### AT1G11430.1 : MORF9, RIP9

plastid developmental protein DAG, putative

Encodes a protein involved in RNA editing in chloroplasts. The mRNA is cell-to-cell mobile.

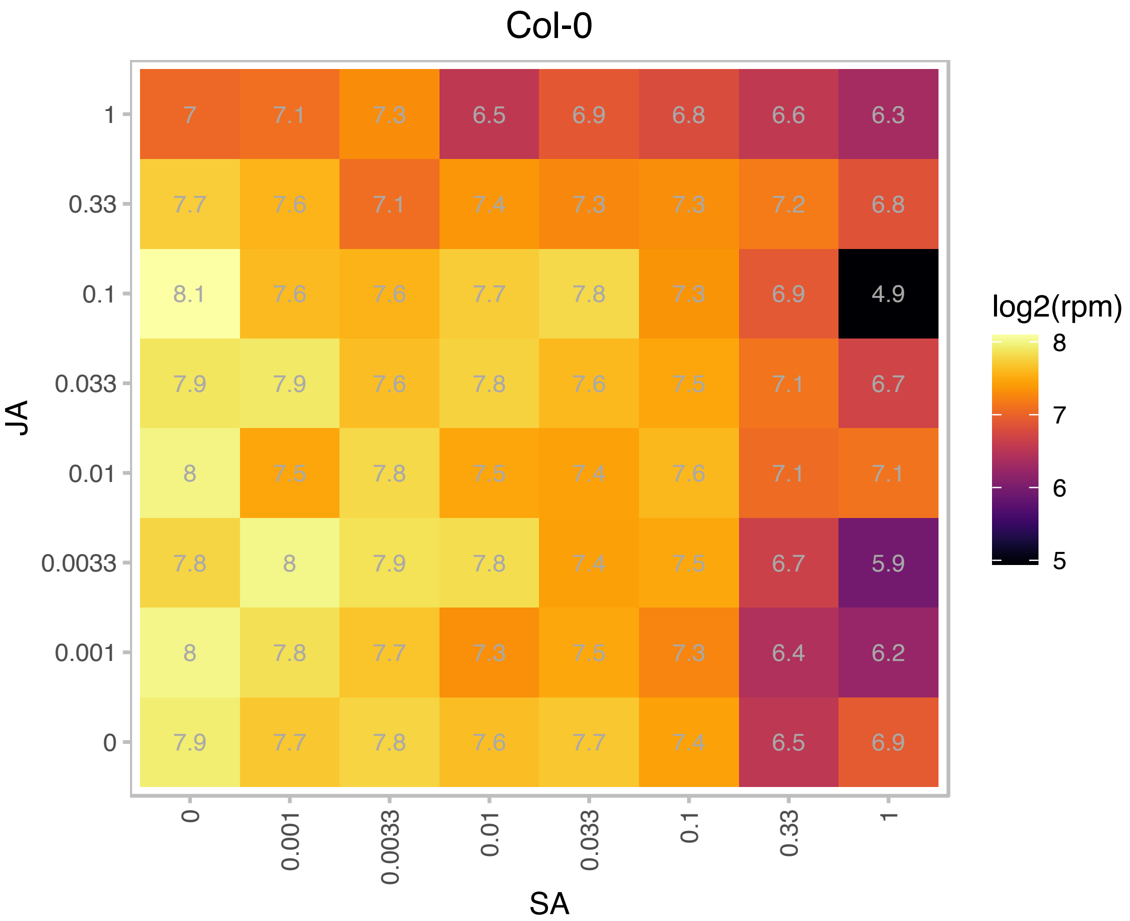


| | |
|---|---|
| Gene | AT1G11430 |
| is.expressed_genes | TRUE |
| is.responsive_genes | TRUE |
| is.apcluster | F-02 |

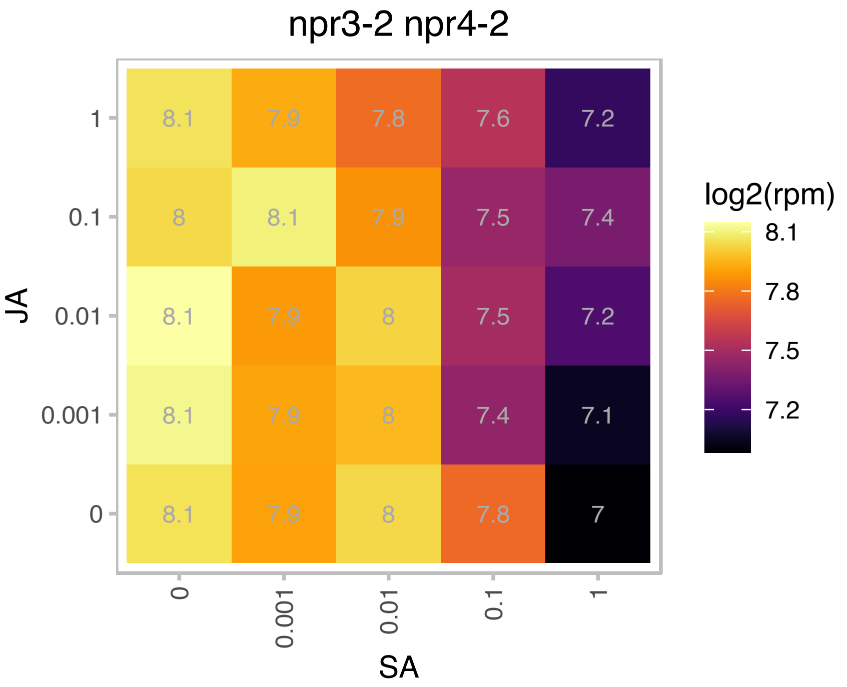


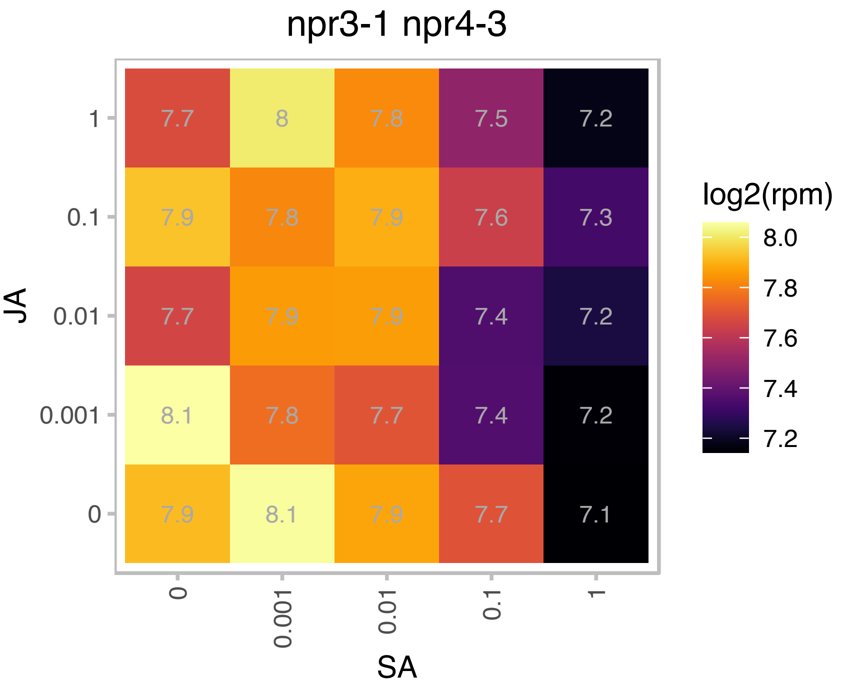

# AT1G11700.1 : NA

Protein of unknown function, DUF584

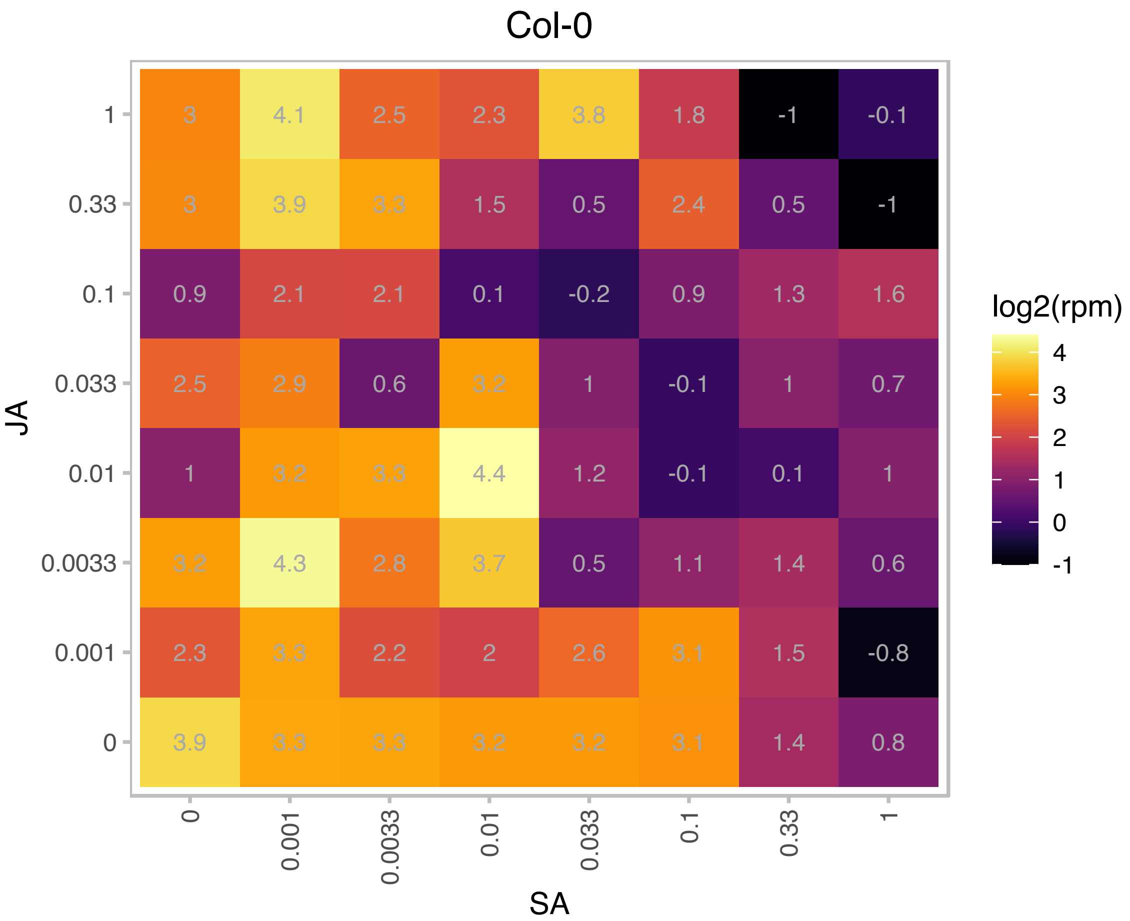


| | |
|---|---|
| Gene | AT1G11700 |
| is.expressed_genes | TRUE |
| is.responsive_genes | TRUE |
| is.apcluster | F-02 |

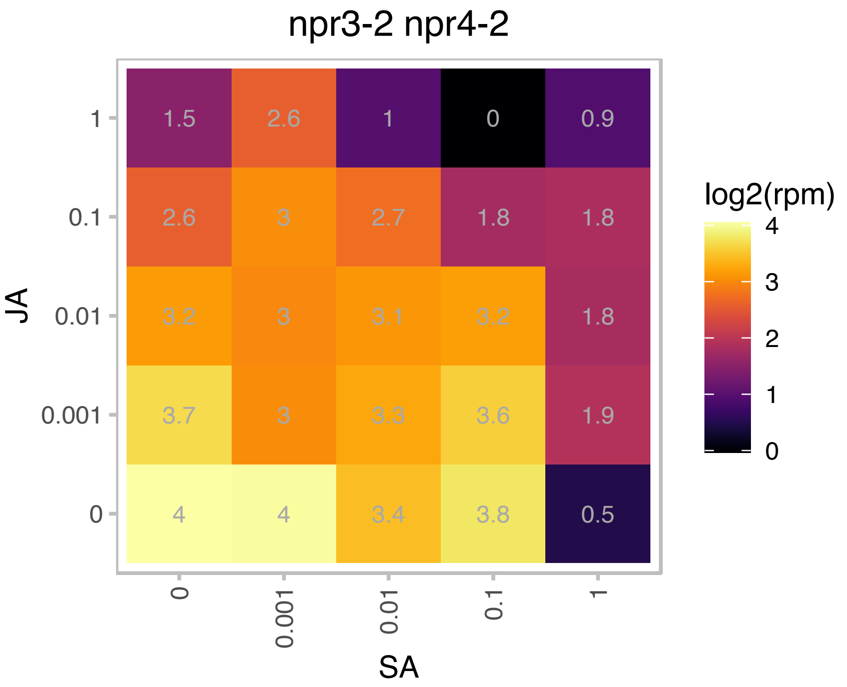


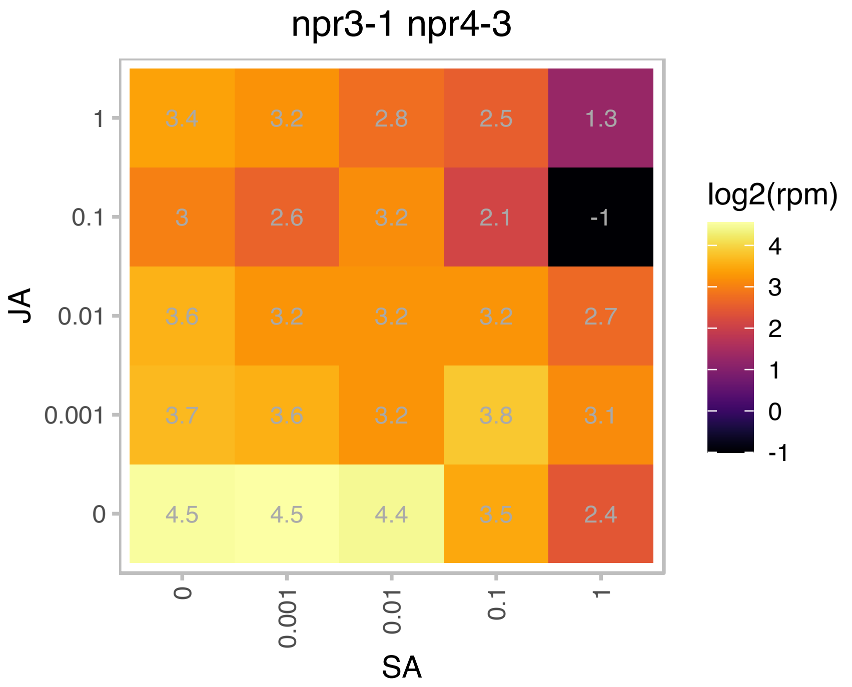

### AT1G12090.1 : ELP

extensin-like protein

extensin-like protein (ELP)

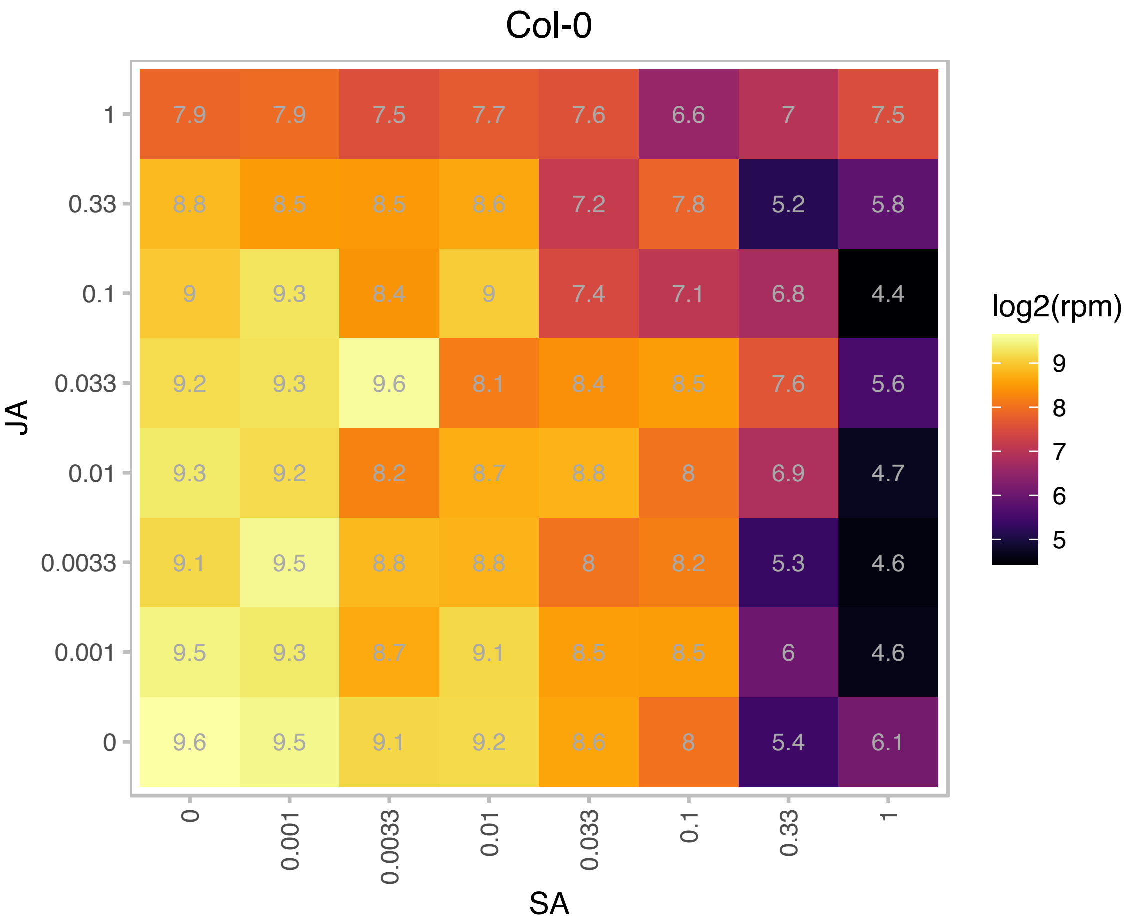


| | |
|---|---|
| Gene | AT1G12090 |
| is.expressed_genes | TRUE |
| is.responsive_genes | TRUE |
| is.apcluster | F-02 |

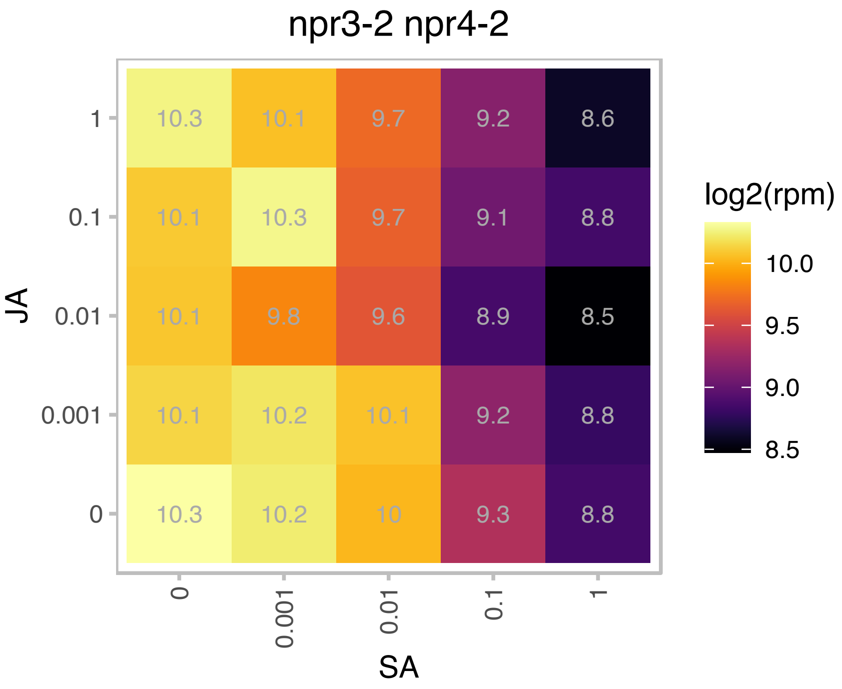


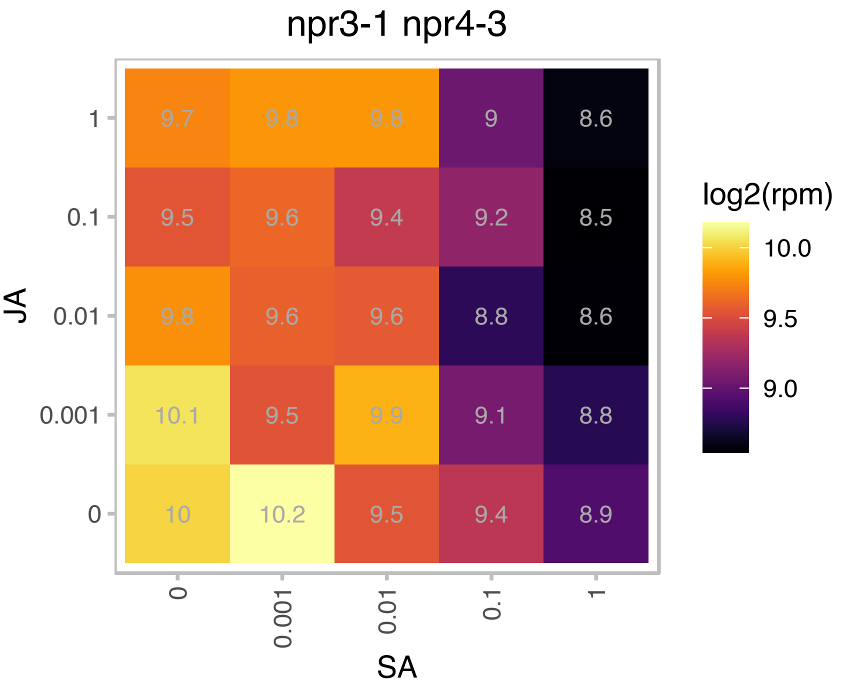

### AT1G12800.1 : SDP

Nucleic acid-binding, OB-fold-like protein
SDP is a chloroplast localized RNA binding protein that is required for plastid rRNA processing. Plants harboring a mutation in SDP have numerous defects including reduced chlorophyll content, poor growth, yellow leaves and abnormal chloroplasts.

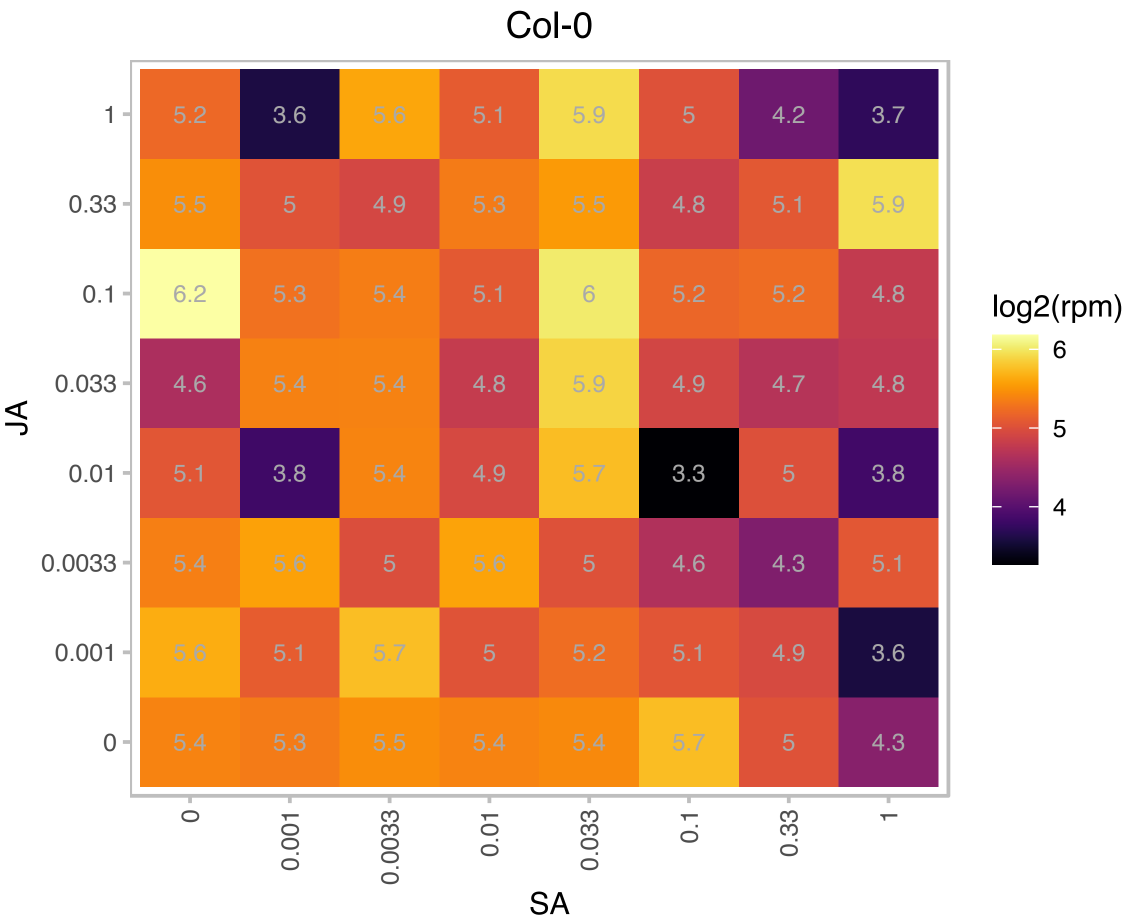


| | |
|---|---|
| Gene | AT1G12800 |
| is.expressed_genes | TRUE |
| is.responsive_genes | TRUE |
| is.apcluster | F-02 |

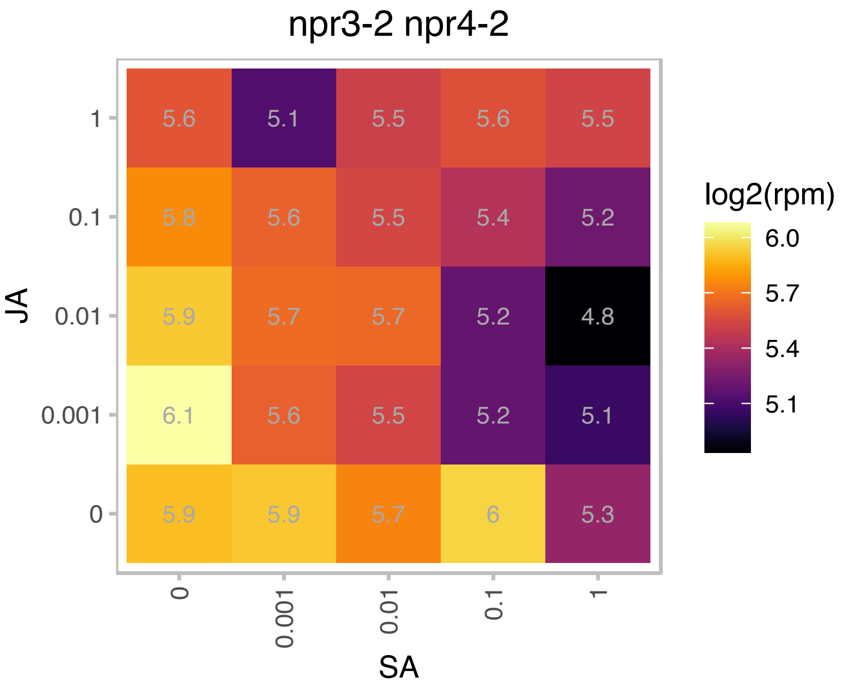


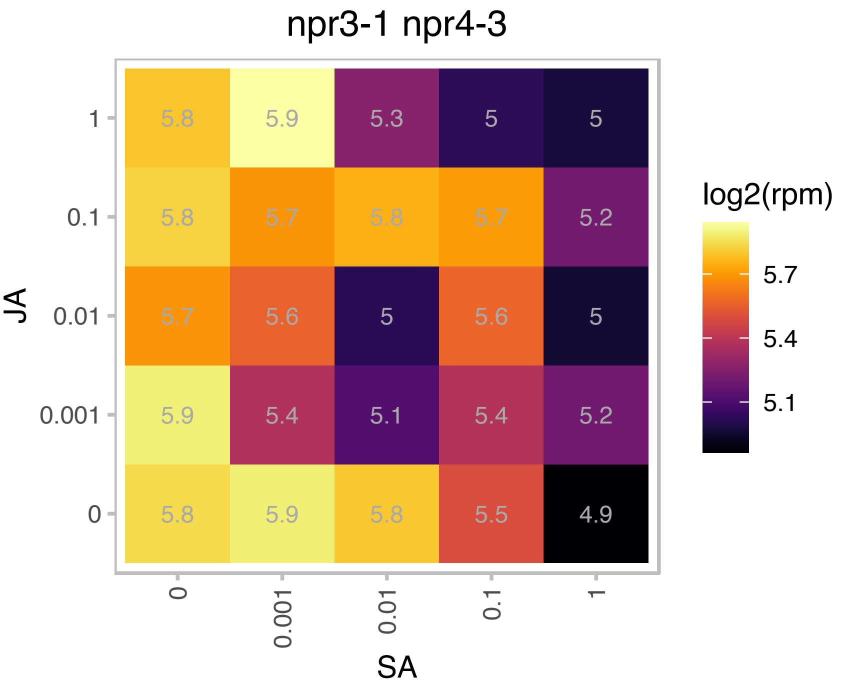

### AT1G13590.1 : ATPSK1, PSK1

phytosulfokine 1 precursor

Encodes a phytosulfokine-alpha (PSK) precursor, a unique plant peptide growth factor first described in Asparagus.

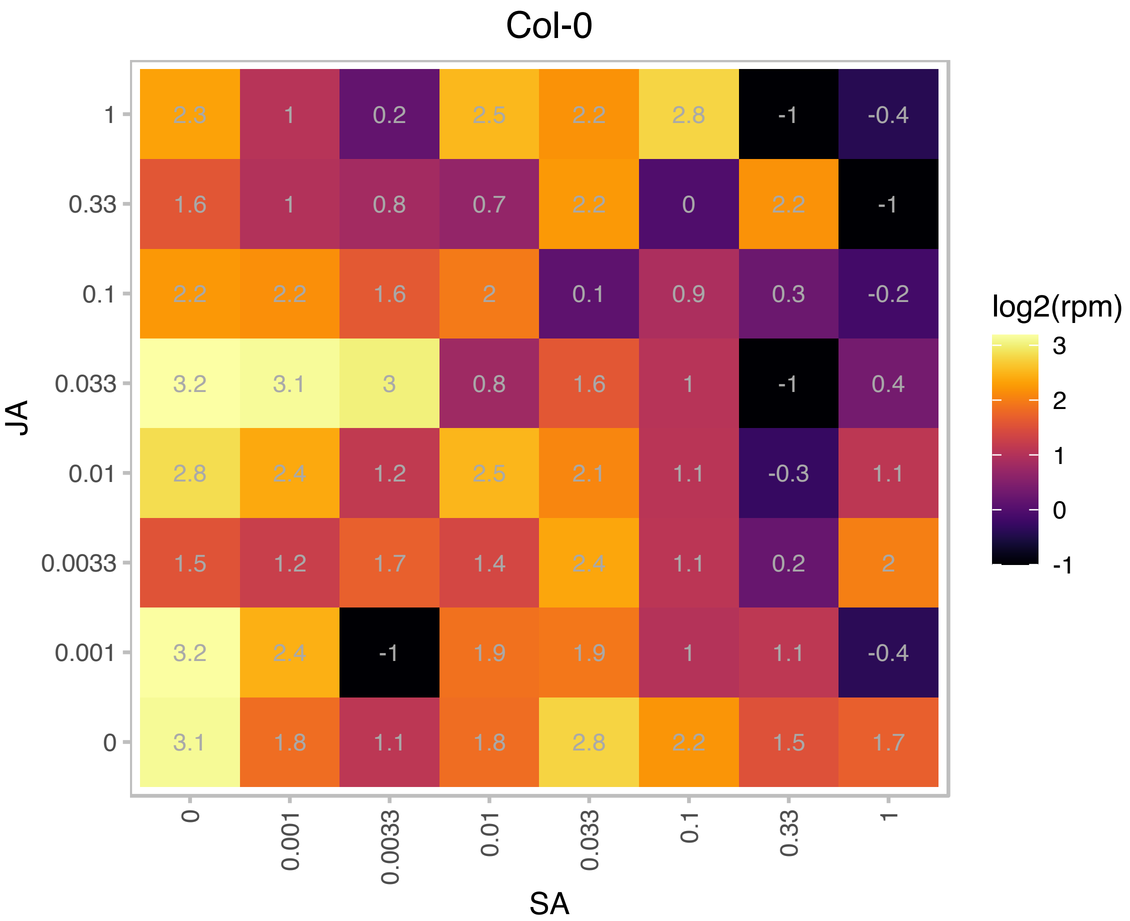


| | |
|---|---|
| Gene | AT1G13590 |
| is.expressed_genes | TRUE |
| is.responsive_genes | TRUE |
| is.apcluster | F-02 |

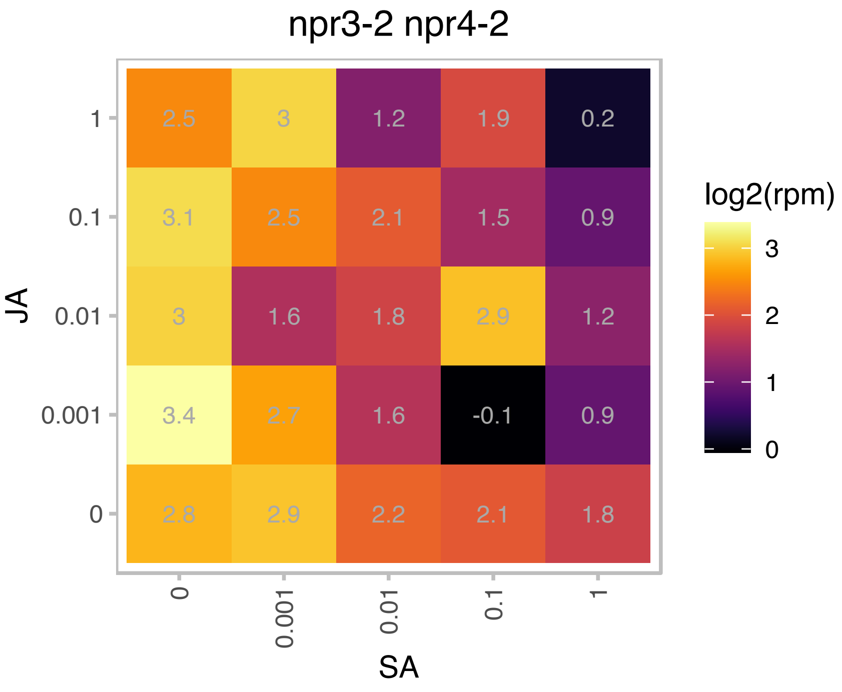


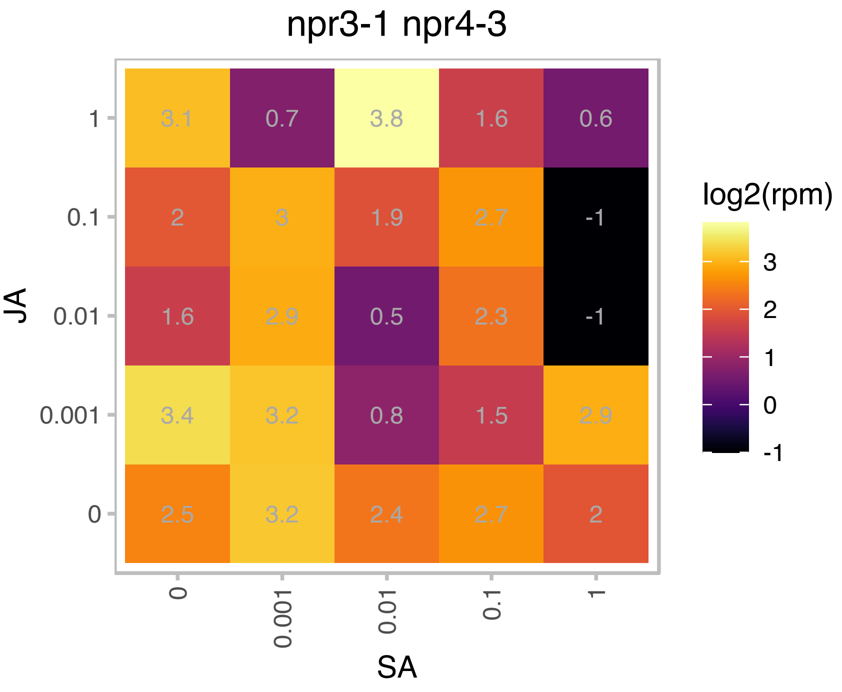

### AT1G15690.1 : AtAVP1, ATAVP3, AtVHP1;1, AVP-3, AVP1,

Encodes a H(+)-translocating (pyrophosphate-energized) inorganic pyrophosphatase (H(+)-PPase; EC 3.6.1.1) located in the vacuolar membrane. Expression is found in all tissues examined, including meristems and floral organ primordium. Expression is particularly enhanced in pollen, and is repressed by light. Over expression and loss of function phenotypes suggest AVP1 is involved in regulation of apoplastic pH and auxin transport. The effect on auxin transport likely involves effects of extracellular pH on subcellular localization of auxin efflux carriers such as PIN1. The mRNA is cell-to-cell mobile.

Inorganic H pyrophosphatase family protein

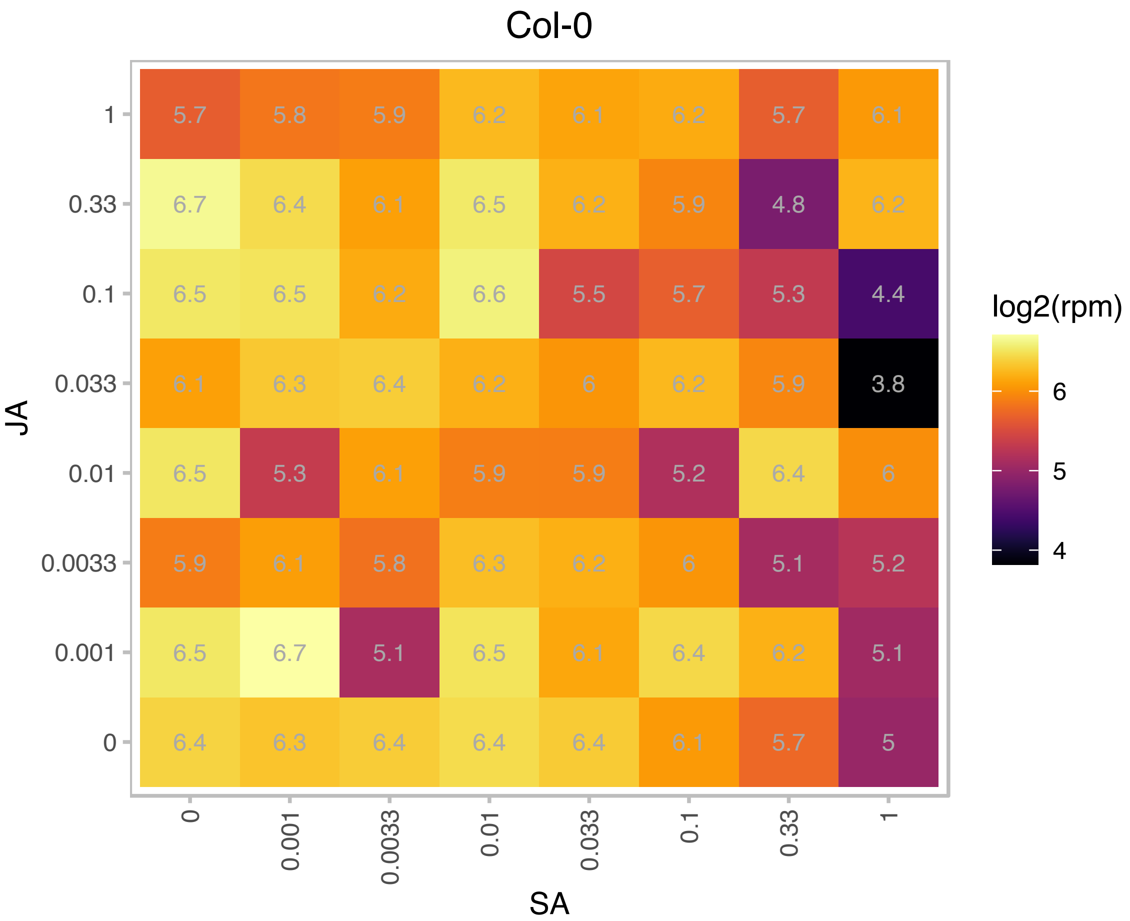


| | |
|---|---|
| Gene | AT1G15690 |
| is.expressed_genes | TRUE |
| is.responsive_genes | TRUE |
| is.apcluster | F-02 |

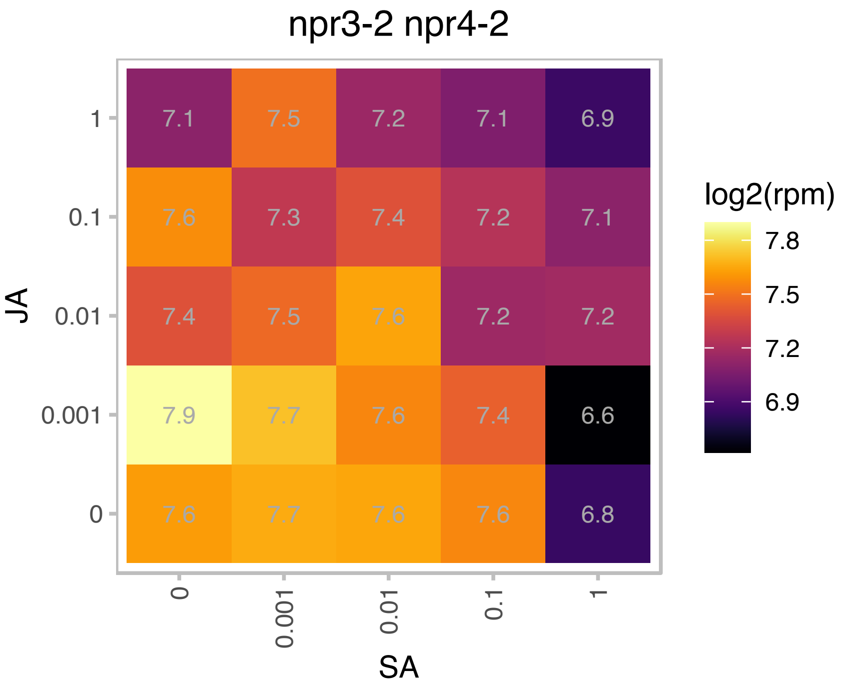


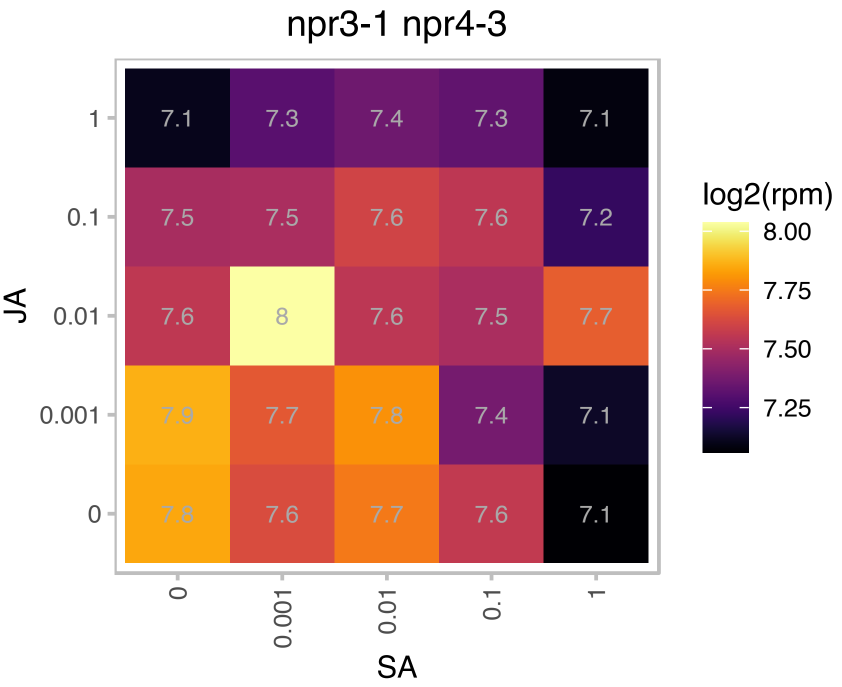

### AT1G16400.1 : CYP79F2

cytochrome P450, family 79, subfamily F, polypeptide 2

Encodes cytochrome P450 CYP79F2.

Col-0

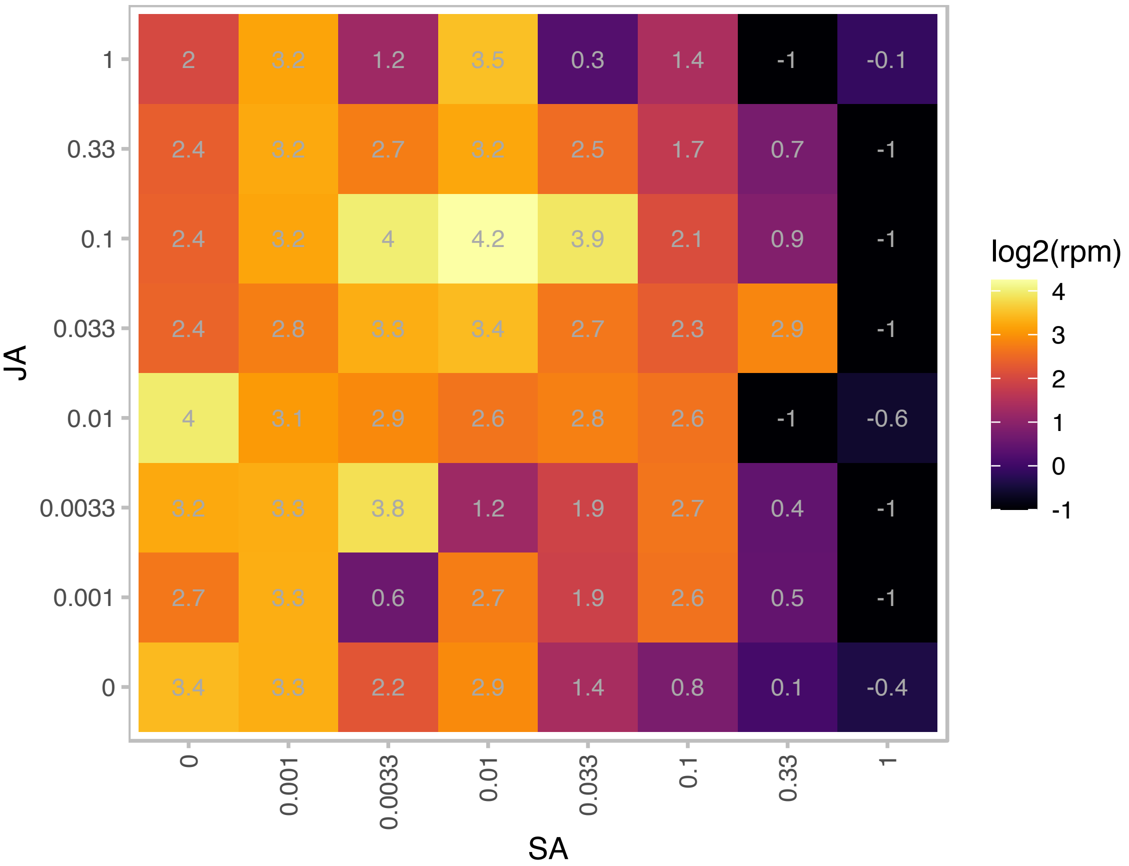


| | |
|---|---|
| Gene | AT1G16400 |
| is.expressed_genes | TRUE |
| is.responsive_genes | TRUE |
| is.apcluster | F-02 |

npr3-2 npr4-2

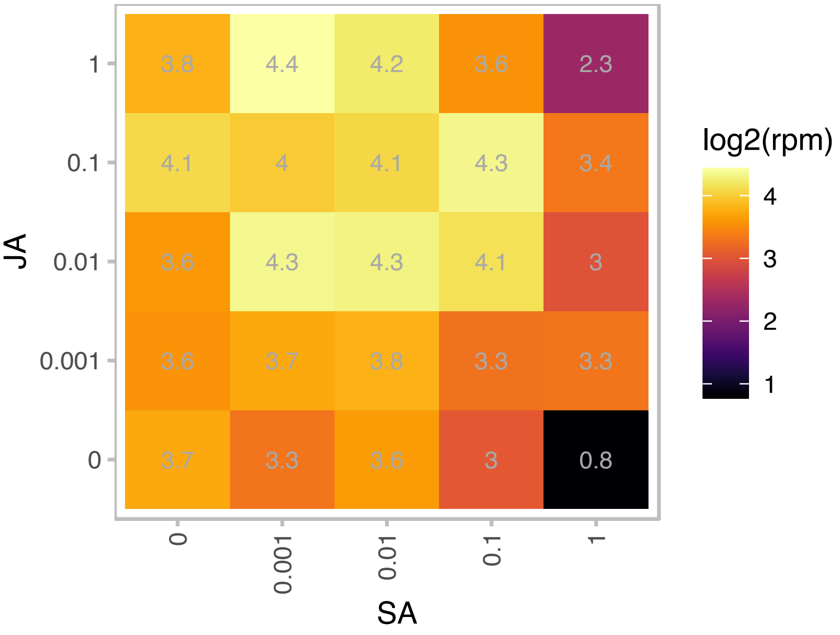


npr3-1 npr4-3

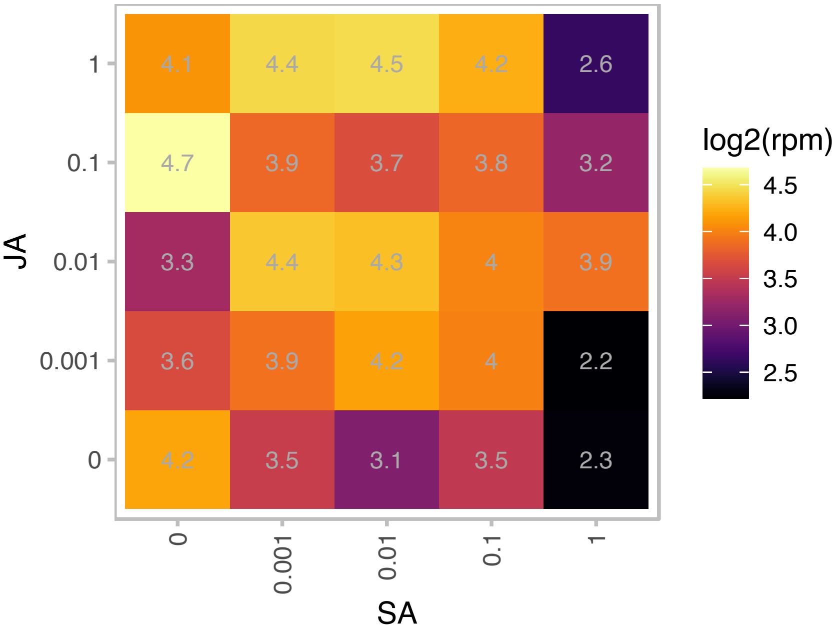

# AT1G19740.1 : NA

ATP-dependent protease La (LON) domain protein

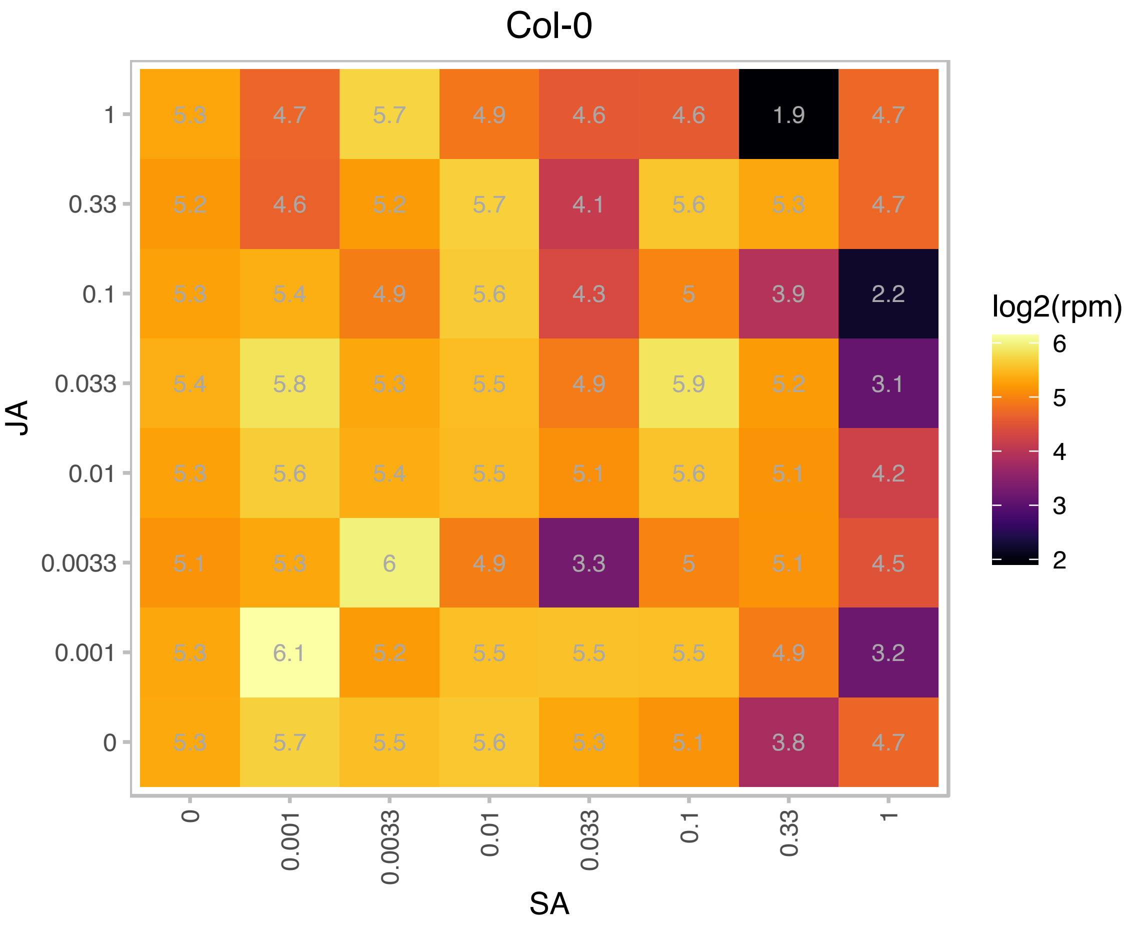


| | |
|---|---|
| Gene | AT1G19740 |
| is.expressed_genes | TRUE |
| is.responsive_genes | TRUE |
| is.apcluster | F-02 |

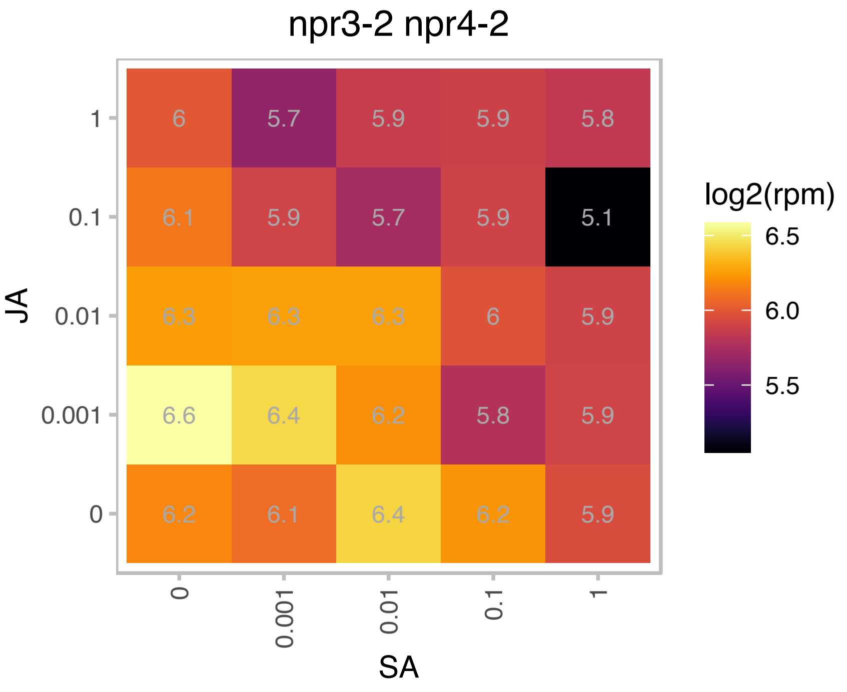


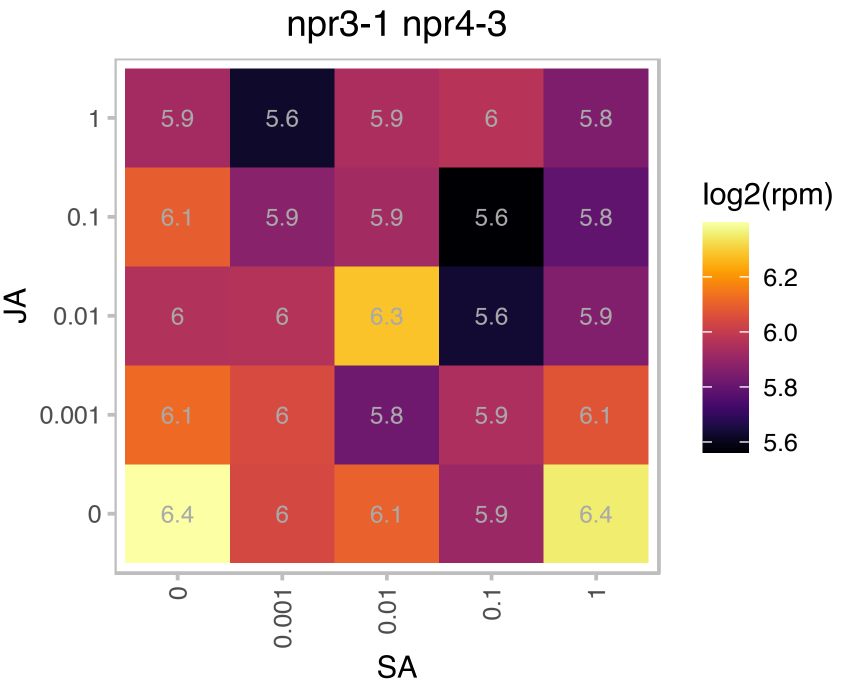

### AT1G20620.1 : ATCAT3, CAT3, SEN2

catalase 3

Catalase, catalyzes the breakdown of hydrogen peroxide (H2O2) into water and oxygen. The mRNA is cell-to-cell mobile.

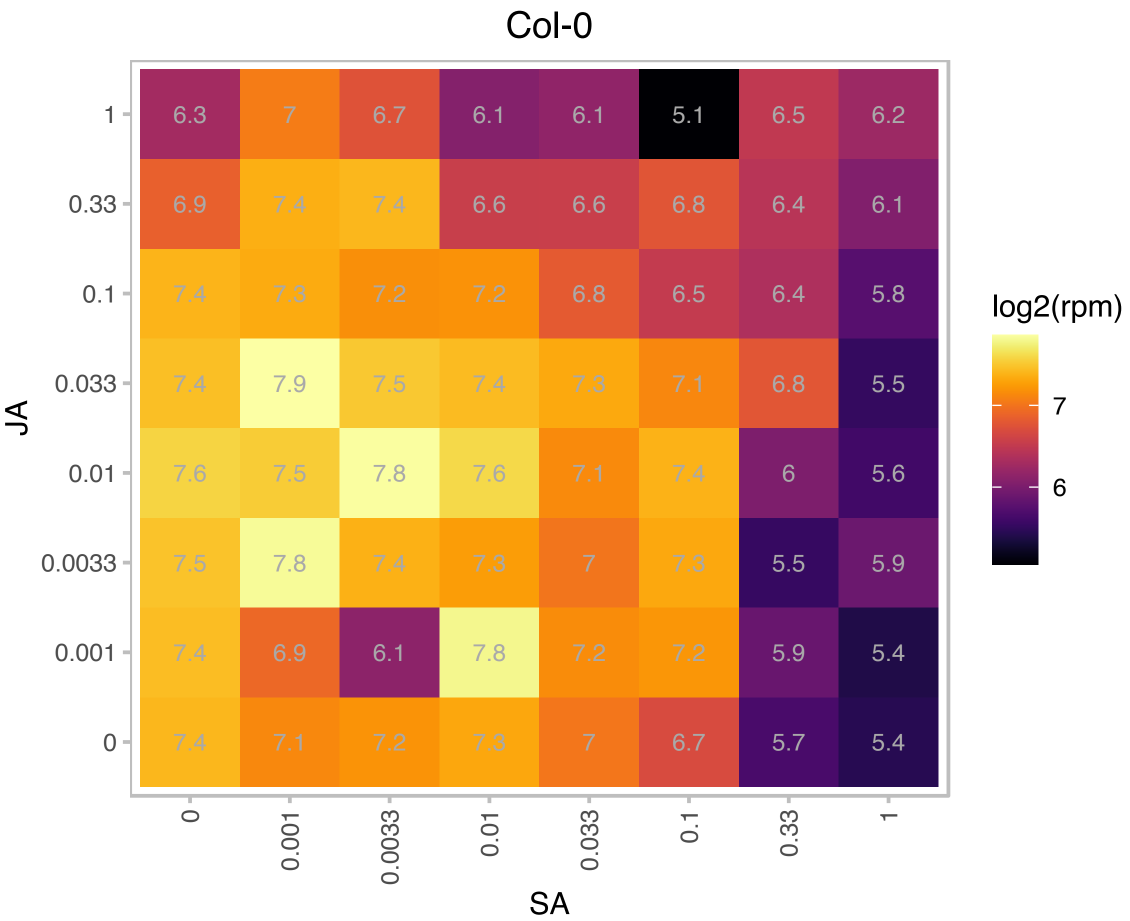


| | |
|---|---|
| Gene | AT1G20620 |
| is.expressed_genes | TRUE |
| is.responsive_genes | TRUE |
| is.apcluster | F-02 |

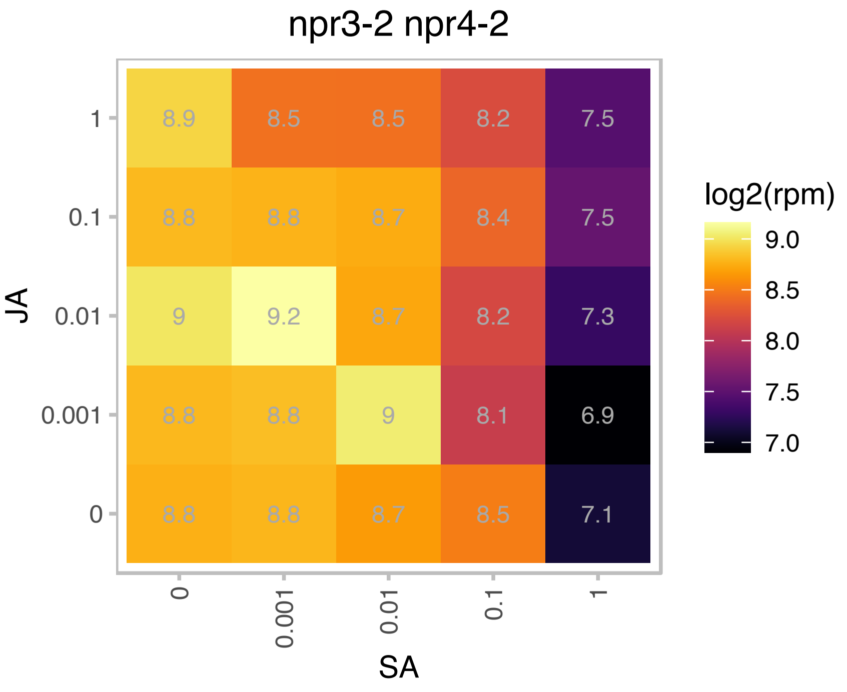


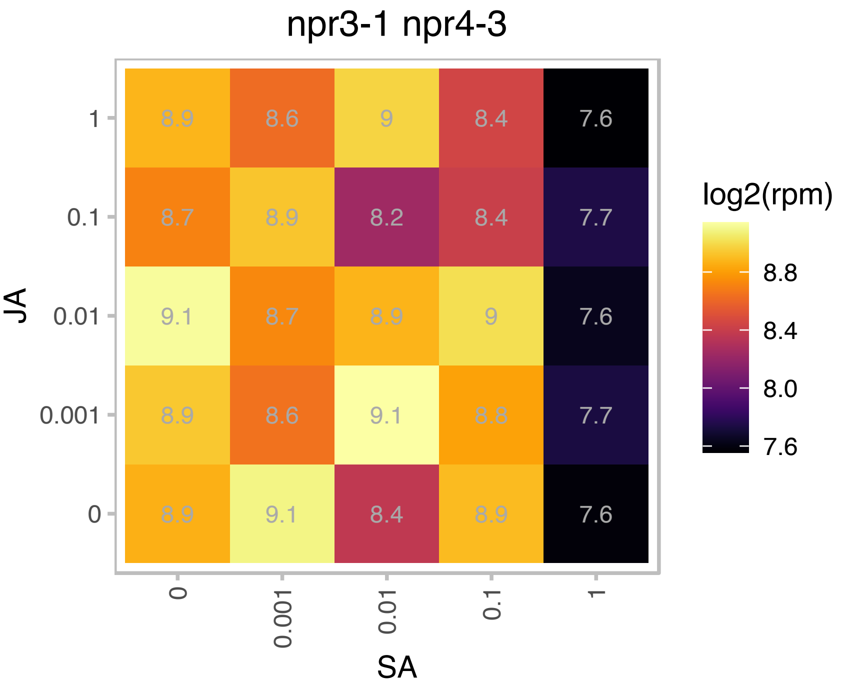

# AT1G24510.1 : NA

TCP-1/cpn60 chaperonin family protein

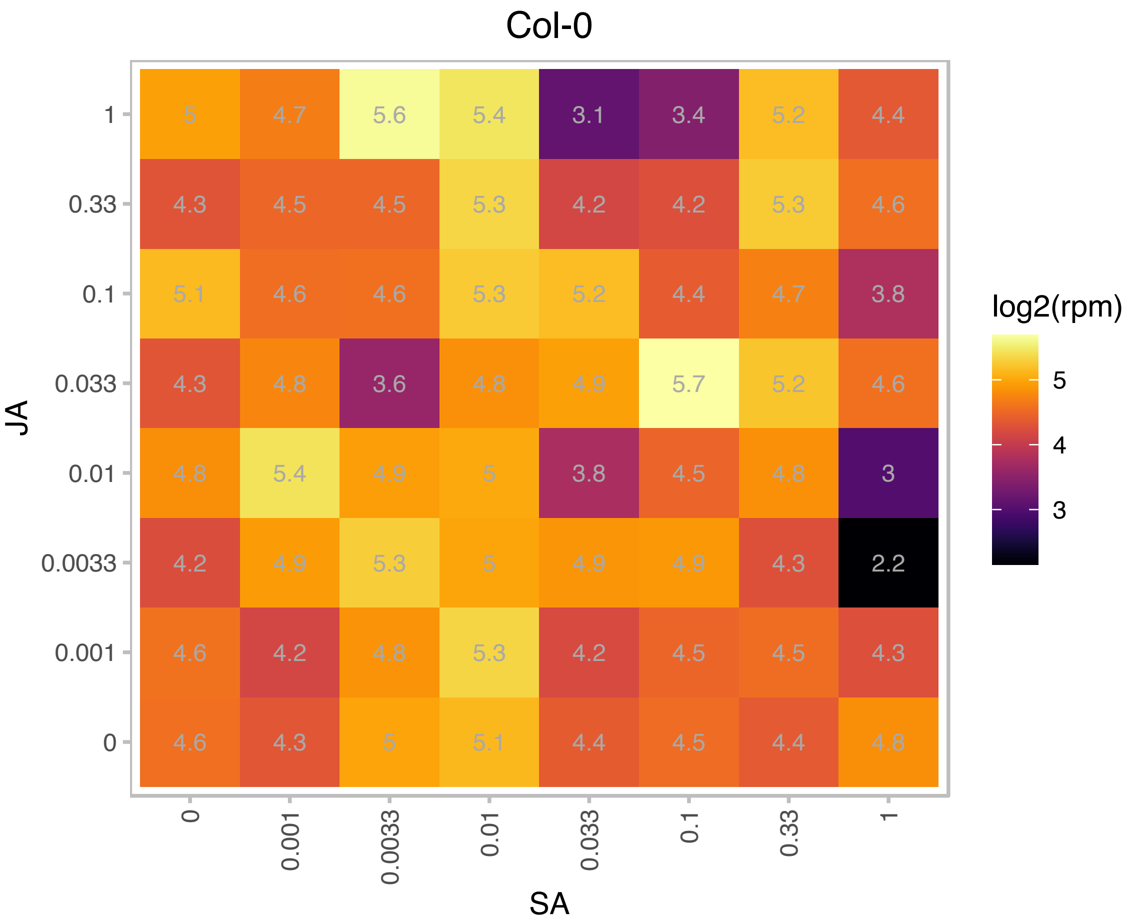


| | |
|---|---|
| Gene | AT1G24510 |
| is.expressed_genes | TRUE |
| is.responsive_genes | TRUE |
| is.apcluster | F-02 |

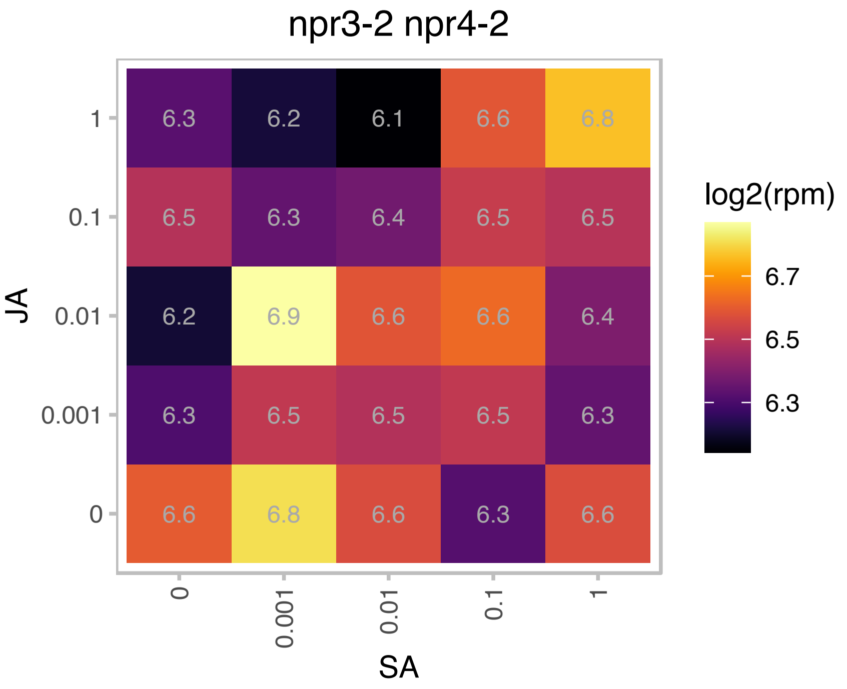


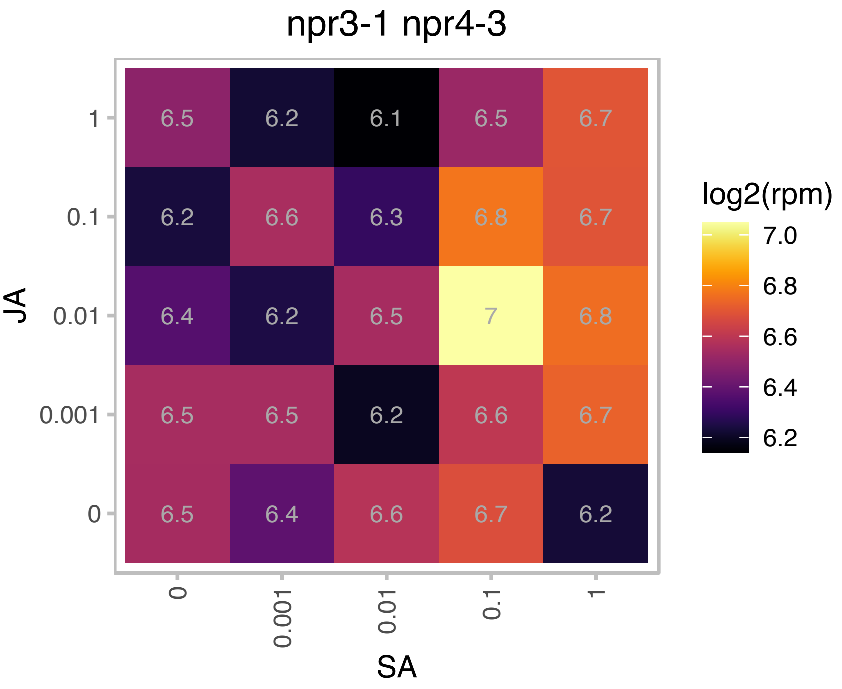

### AT1G26630.1 : ATELF5A-2, ELF5A-2, FBR12

Eukaryotic translation initiation factor 5A-1 (eIF-5A 1) protein
Eukaryotic translation initiation factor 5A-2. Involved in programmed cell death triggered as a response to pseudomonas syringae infection. Loss of function mutants are more resistant to infection. The mRNA is cell-to-cell mobile.

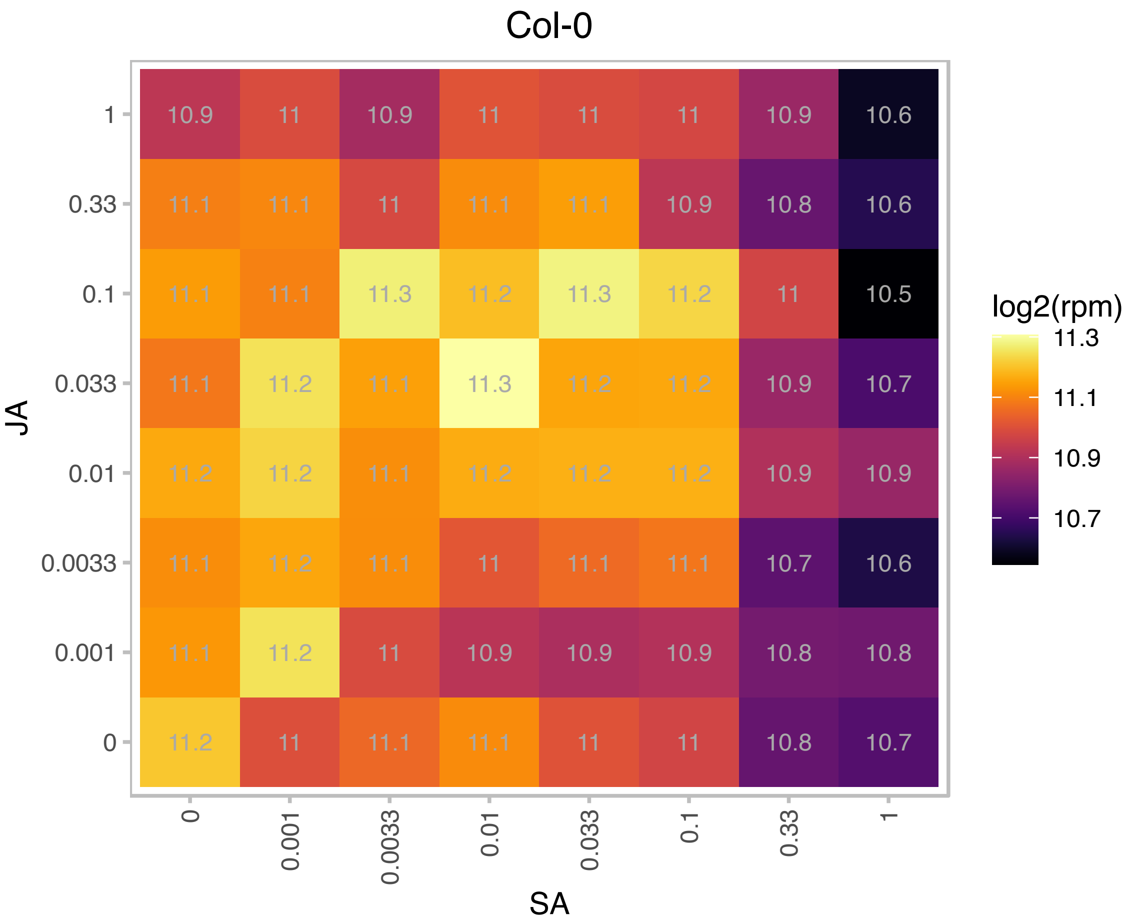


| | |
|---|---|
| Gene | AT1G26630 |
| is.expressed_genes | TRUE |
| is.responsive_genes | TRUE |
| is.apcluster | F-02 |

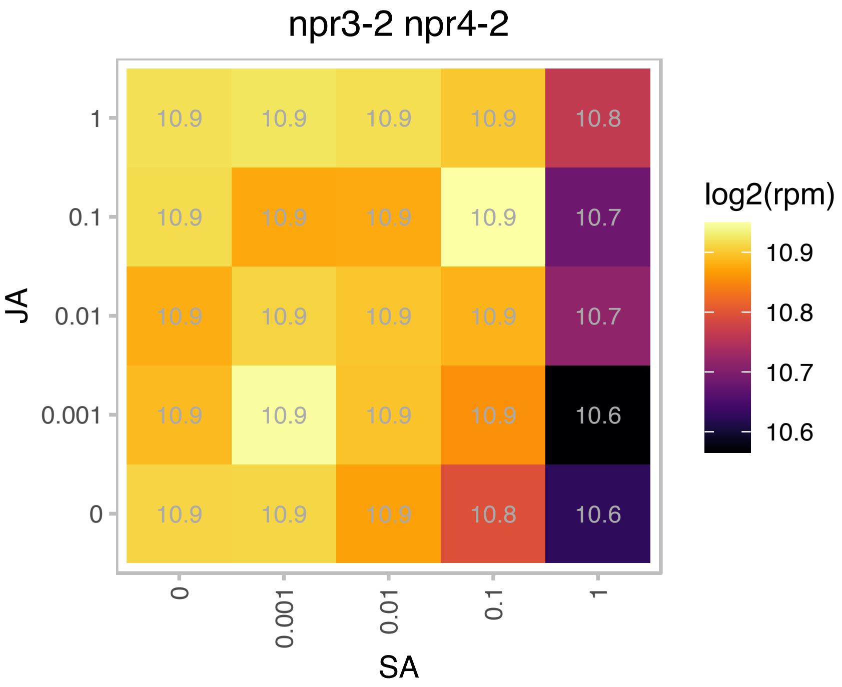


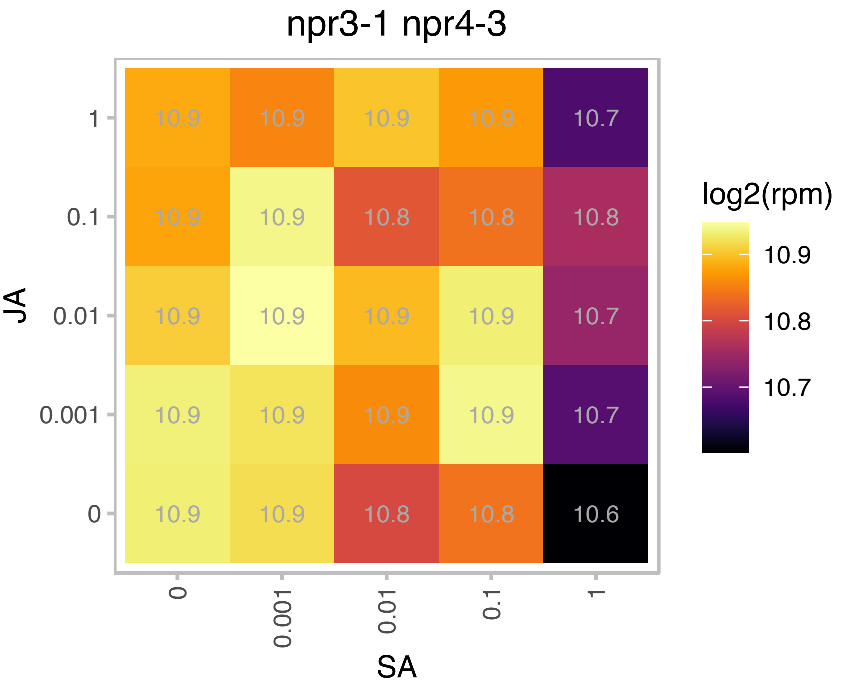

# AT1G29965.1 : NA

Ribosomal protein L18ae/LX family protein

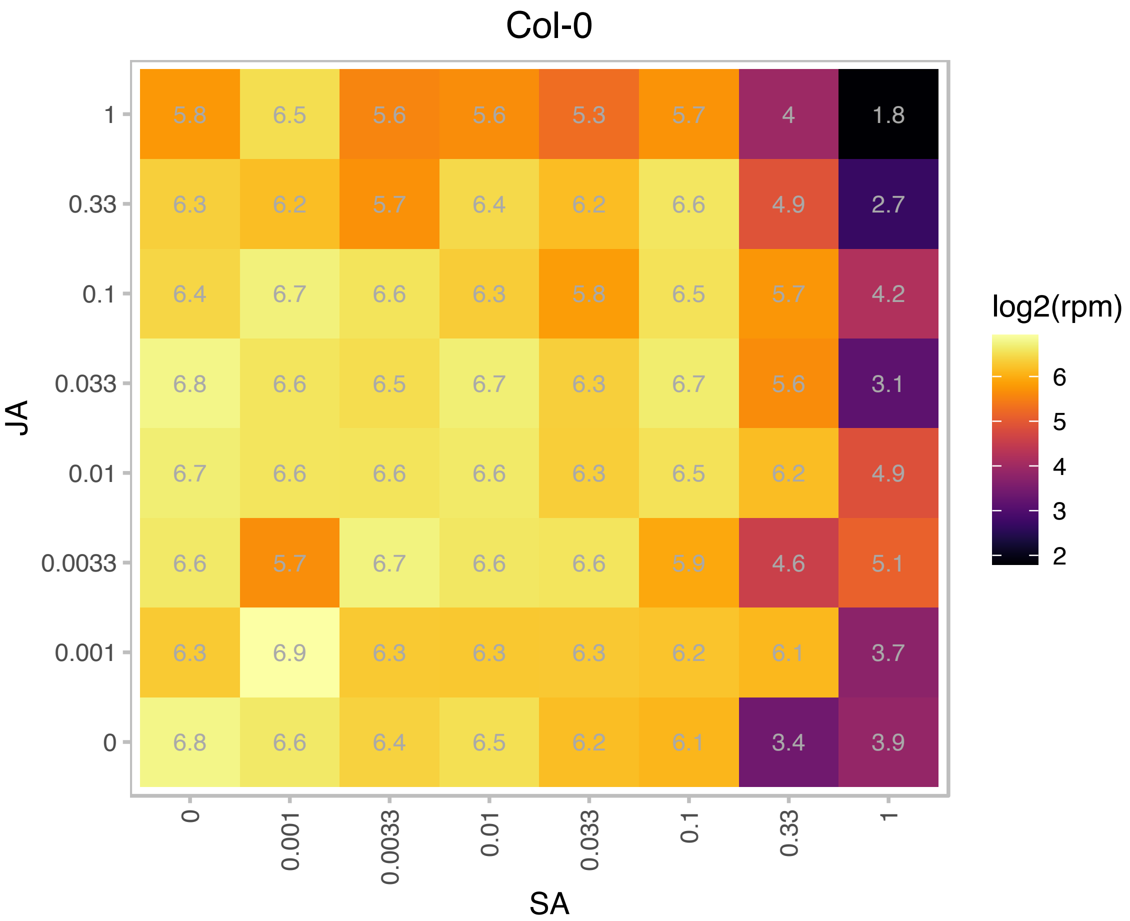


| | |
|---|---|
| Gene | AT1G29965 |
| is.expressed_genes | TRUE |
| is.responsive_genes | TRUE |
| is.apcluster | F-02 |

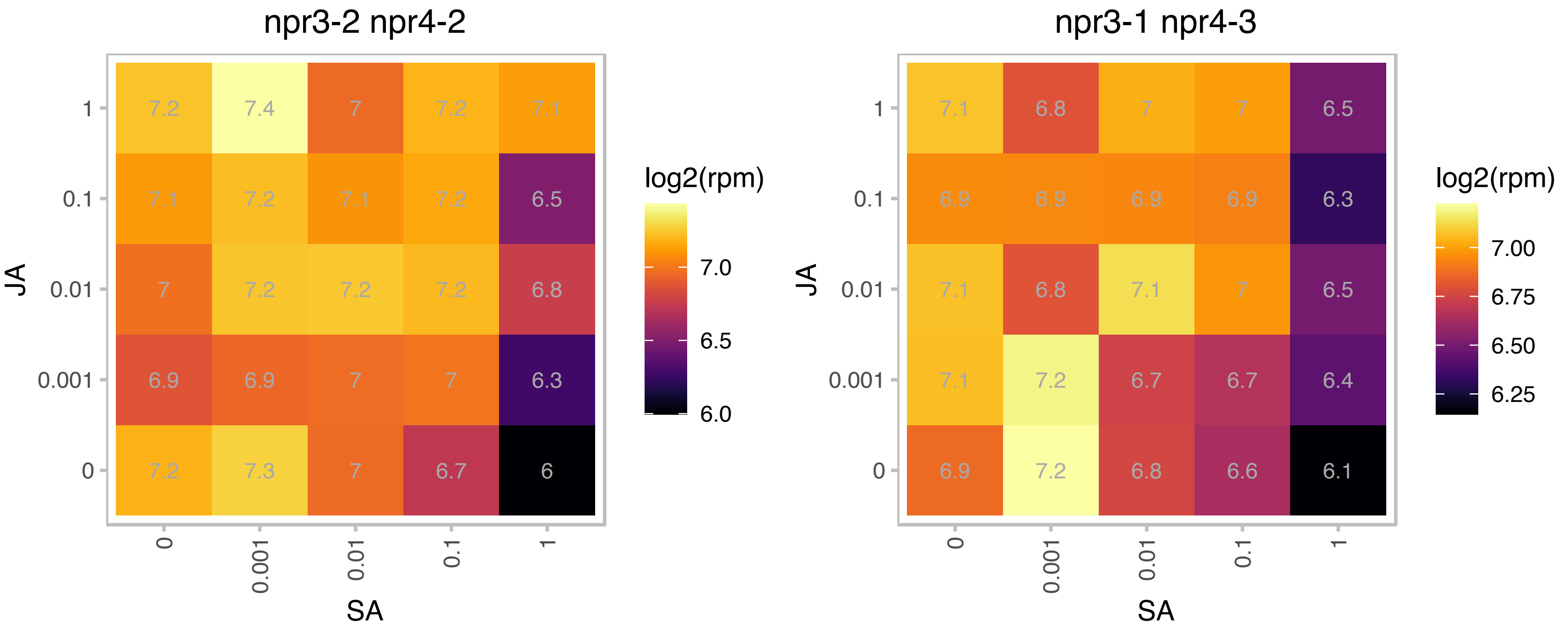

### AT1G30690.1 : PATL4

Sec14p-like phosphatidylinositol transfer family protein

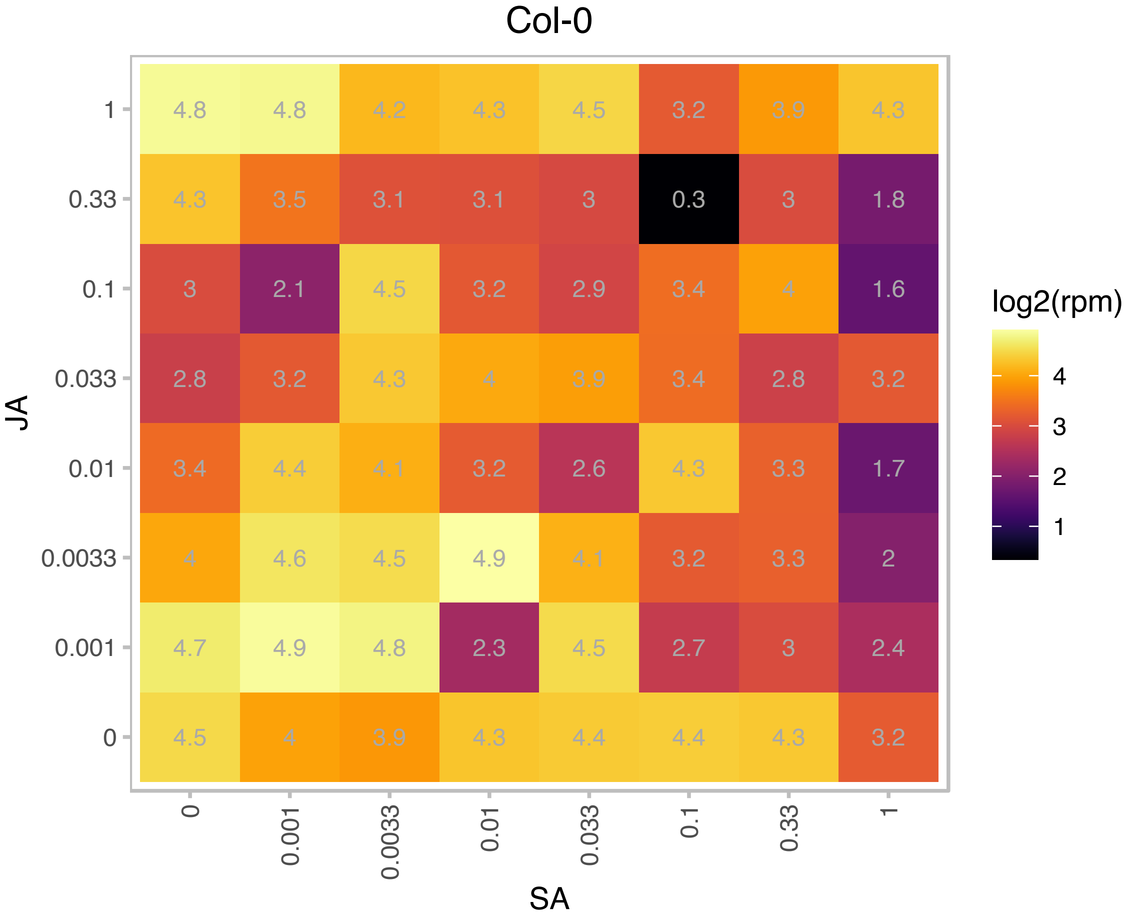


| | |
|---|---|
| Gene | AT1G30690 |
| is.expressed_genes | TRUE |
| is.responsive_genes | TRUE |
| is.apcluster | F-02 |

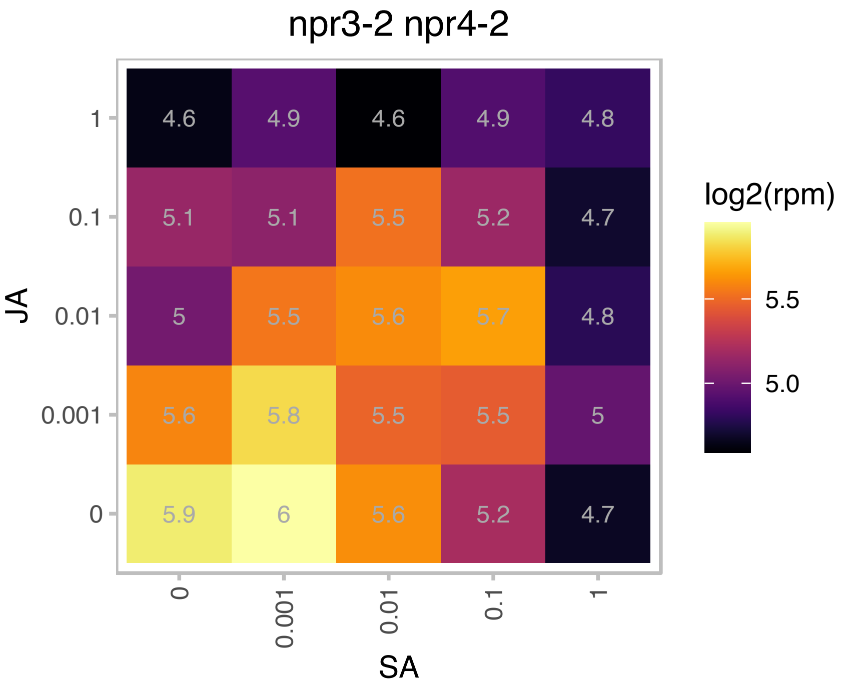


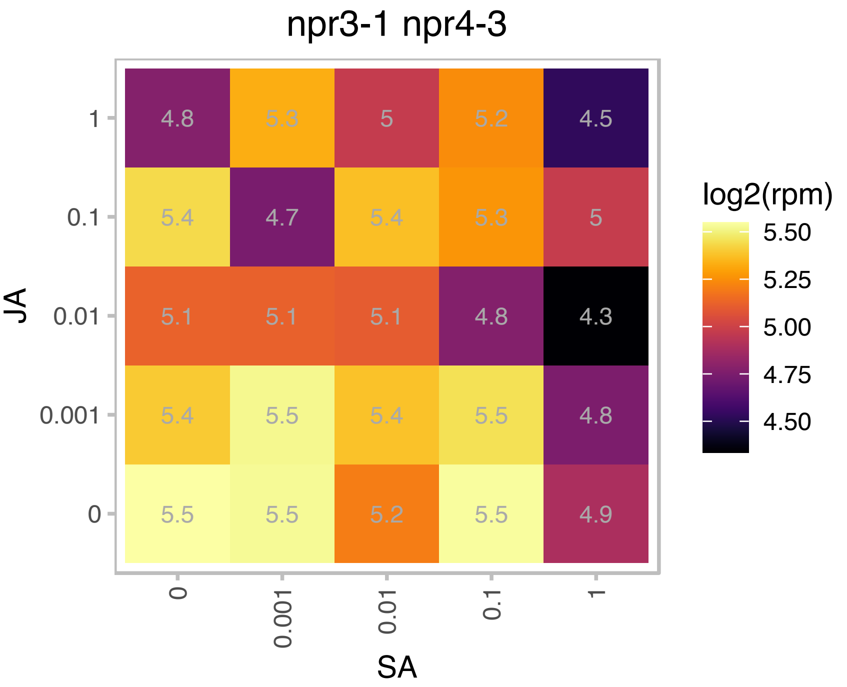

### AT1G31230.1 : AK-HSDH, AK-HSDH I

aspartate kinase-homoserine dehydrogenase i

Encodes a bifunctional aspartate kinase/homoserine dehydrogenase.  These two activities catalyze the first and the third steps toward the synthesis of the essential amino acids threonine, isoleucine and methionine.

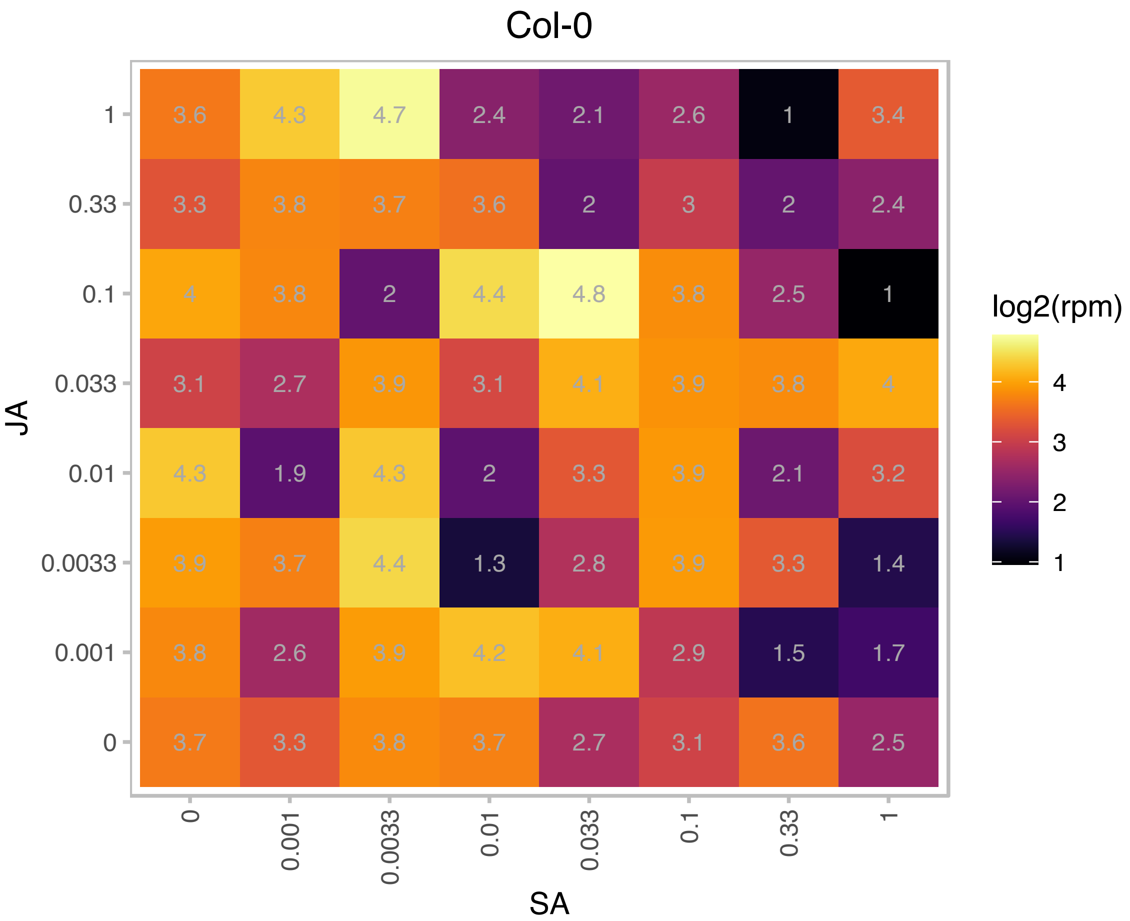


| | |
|---|---|
| Gene | AT1G31230 |
| is.expressed_genes | TRUE |
| is.responsive_genes | TRUE |
| is.apcluster | F-02 |

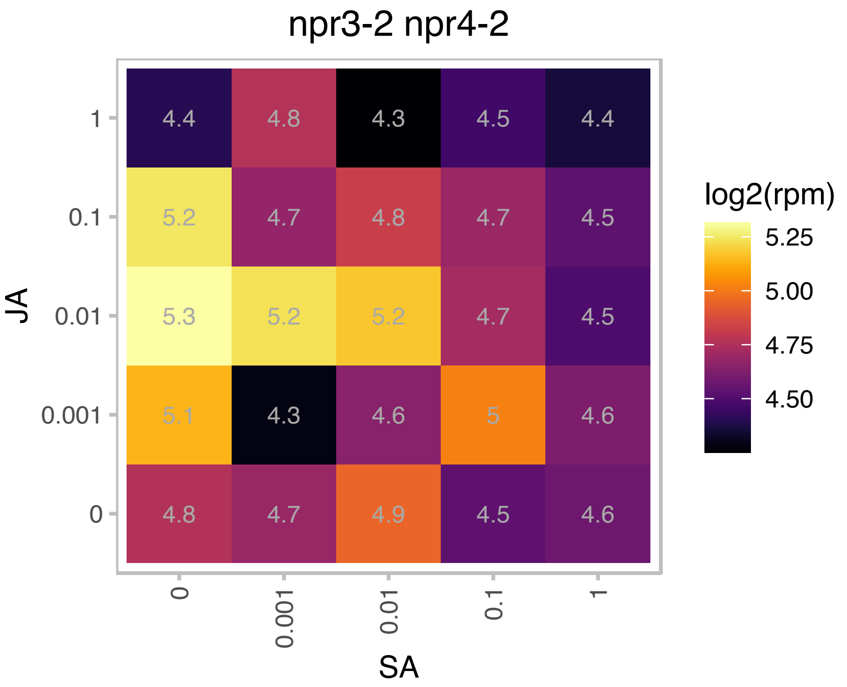


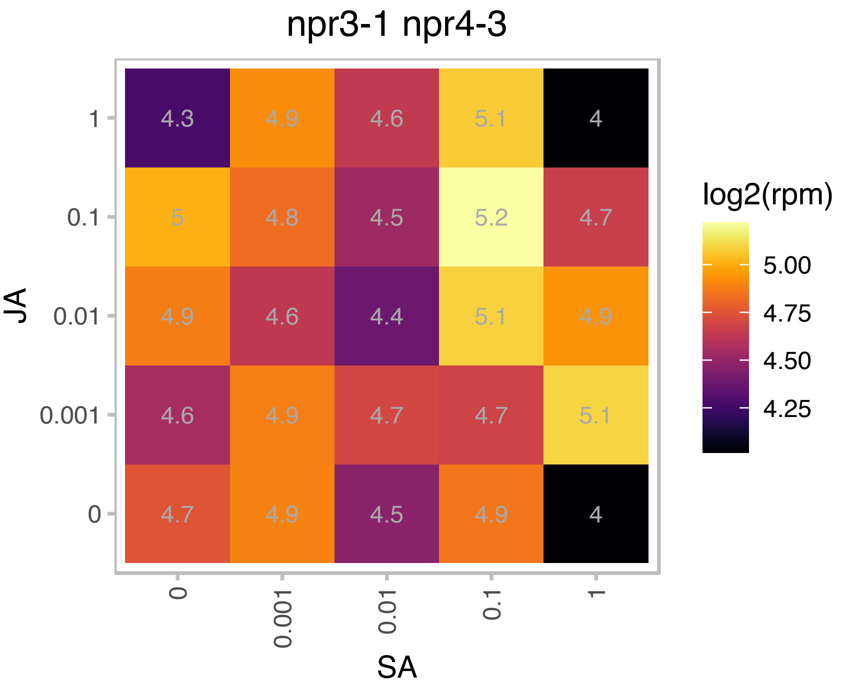

### AT1G31860.1 : AT-IE, HISN2

histidine biosynthesis bifunctional protein (HISIE)

encodes a bifunctional protein that has phosphoribosyl-ATP pyrophosphohydrolase (PRA-PH) and phosphoribosyl-AMP cyclohydrolase (PRA-CH) activities.

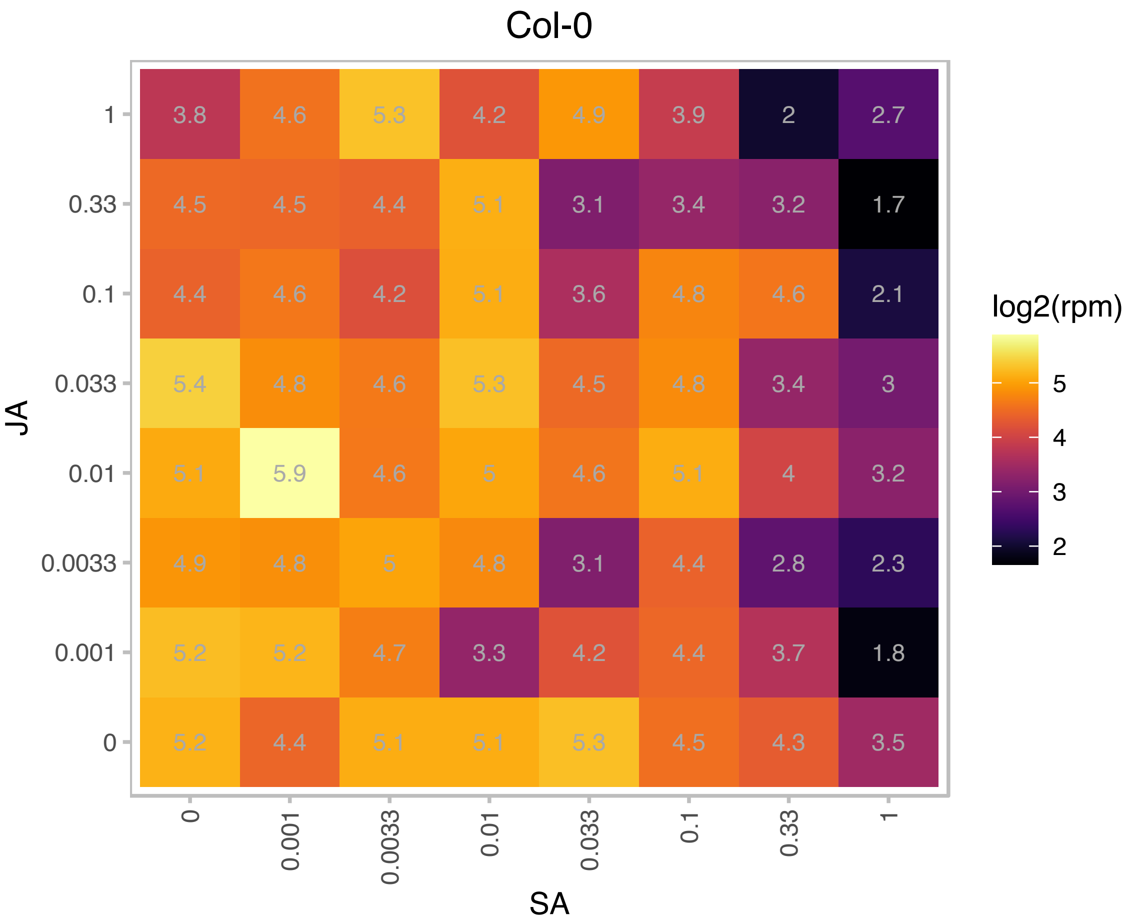


| | |
|---|---|
| Gene | AT1G31860 |
| is.expressed_genes | TRUE |
| is.responsive_genes | TRUE |
| is.apcluster | F-02 |

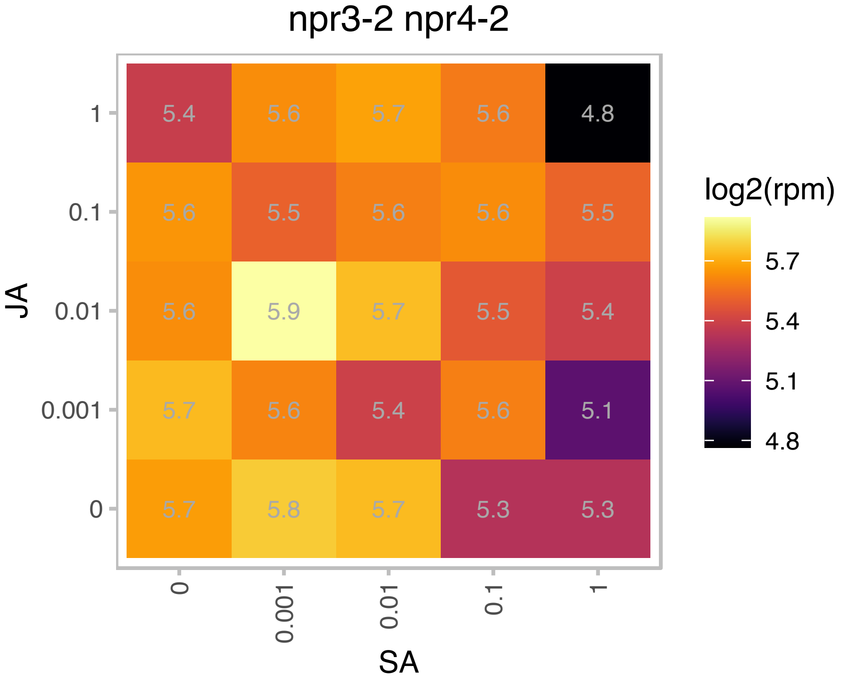


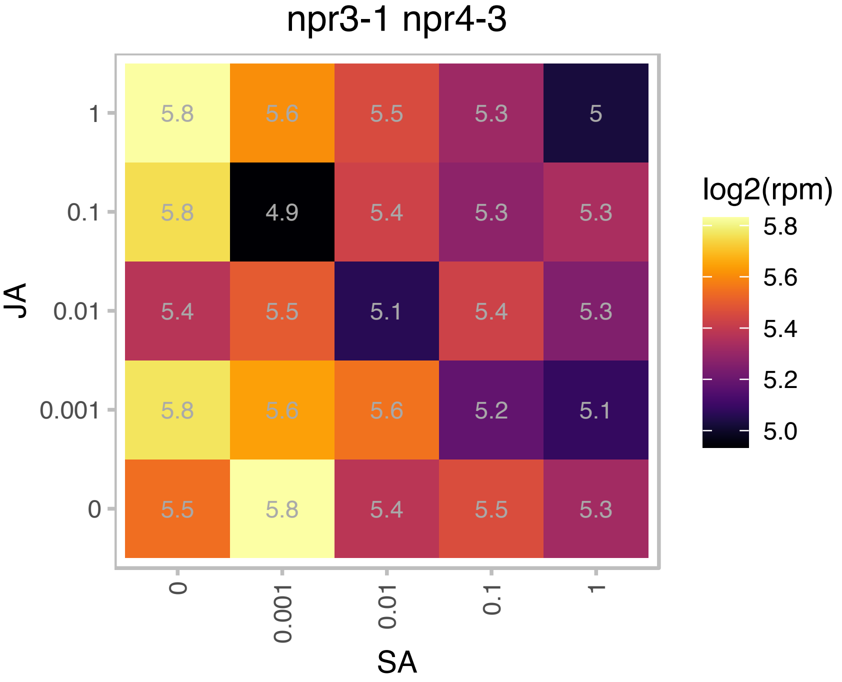

### AT1G31935.1 : sORF12

other RNA

Has been identified as a translated small open reading frame by ribosome profiling.

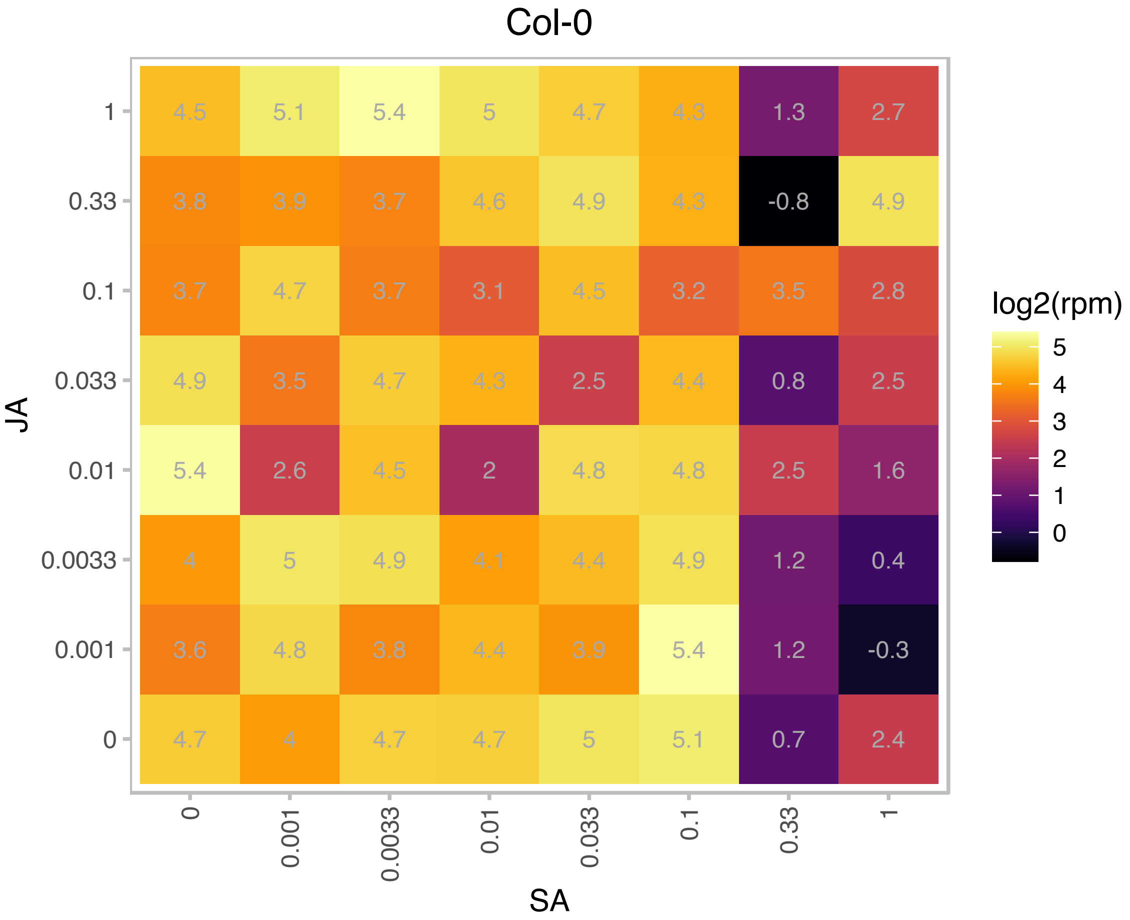


| | |
|---|---|
| Gene | AT1G31935 |
| is.expressed_genes | TRUE |
| is.responsive_genes | TRUE |
| is.apcluster | F-02 |

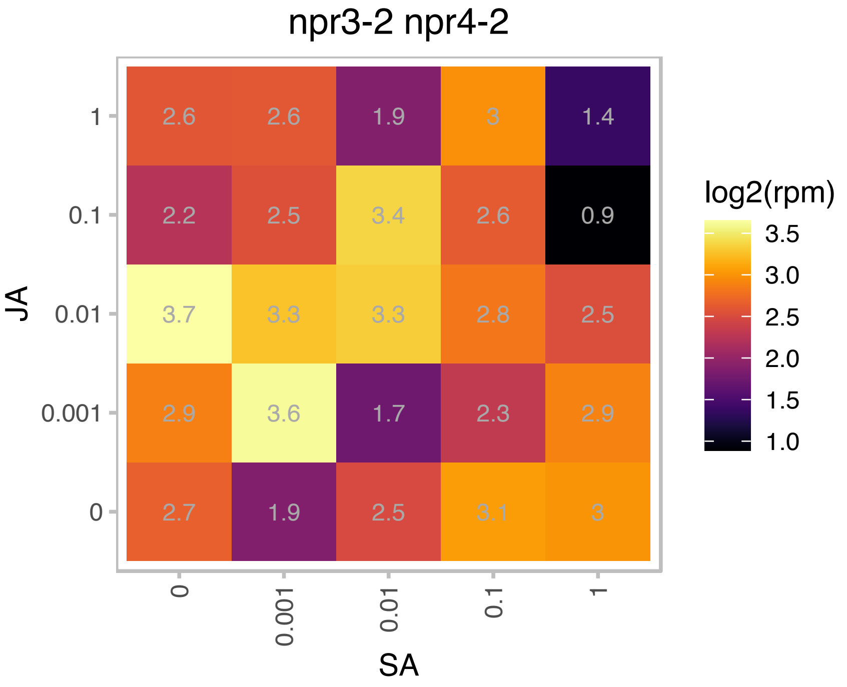


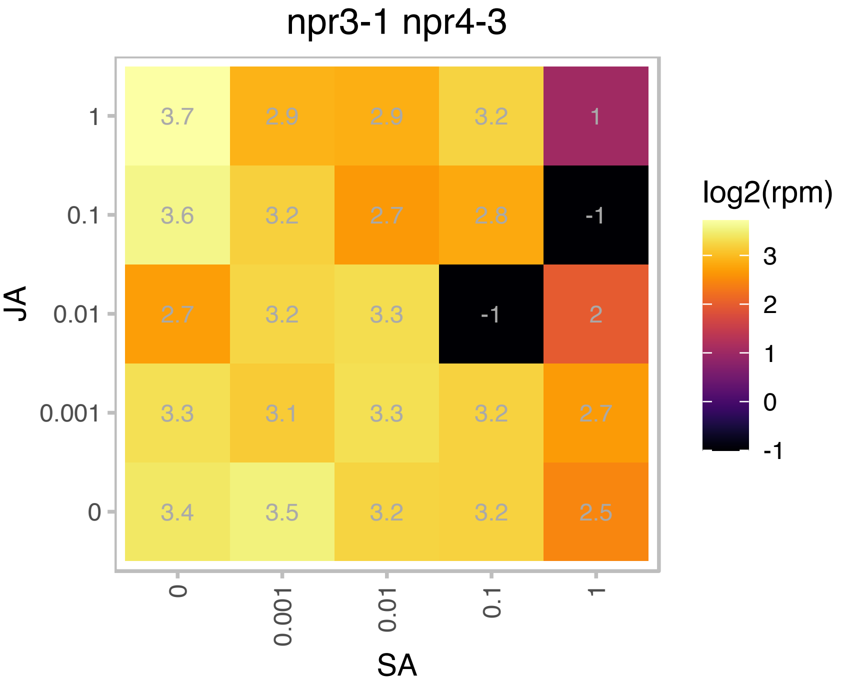

# AT1G32520.1 : NA

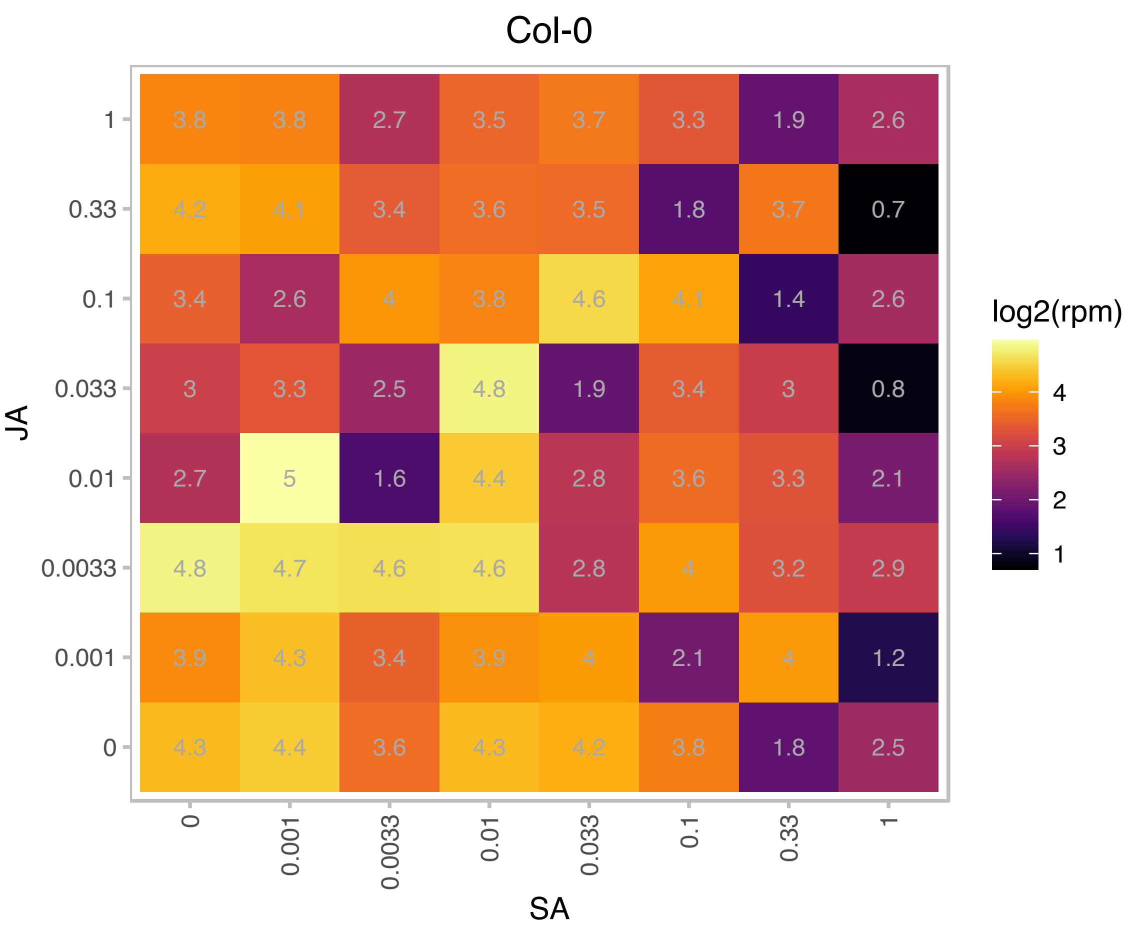


| | |
|---|---|
| Gene | AT1G32520 |
| is.expressed_genes | TRUE |
| is.responsive_genes | TRUE |
| is.apcluster | F-02 |

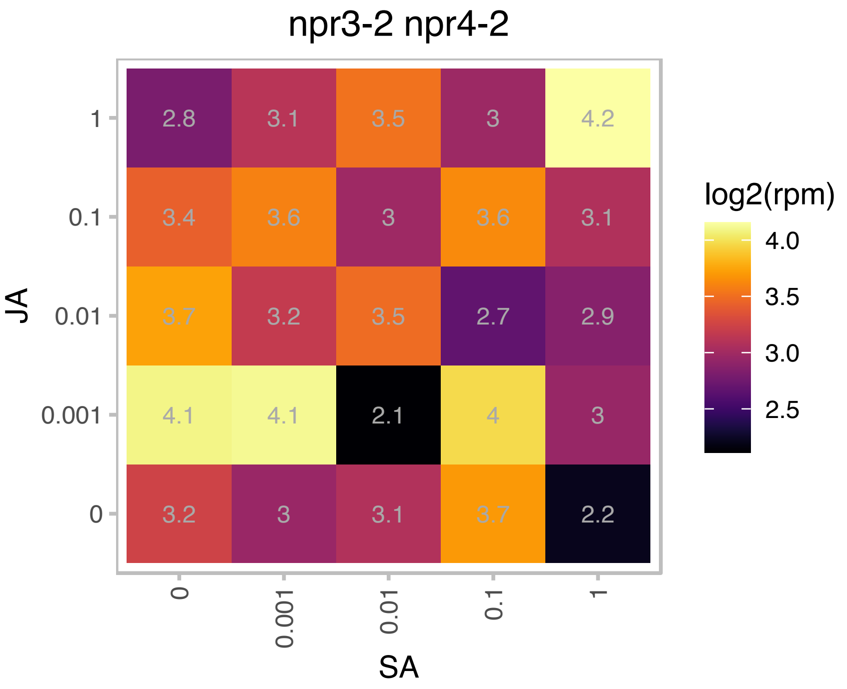


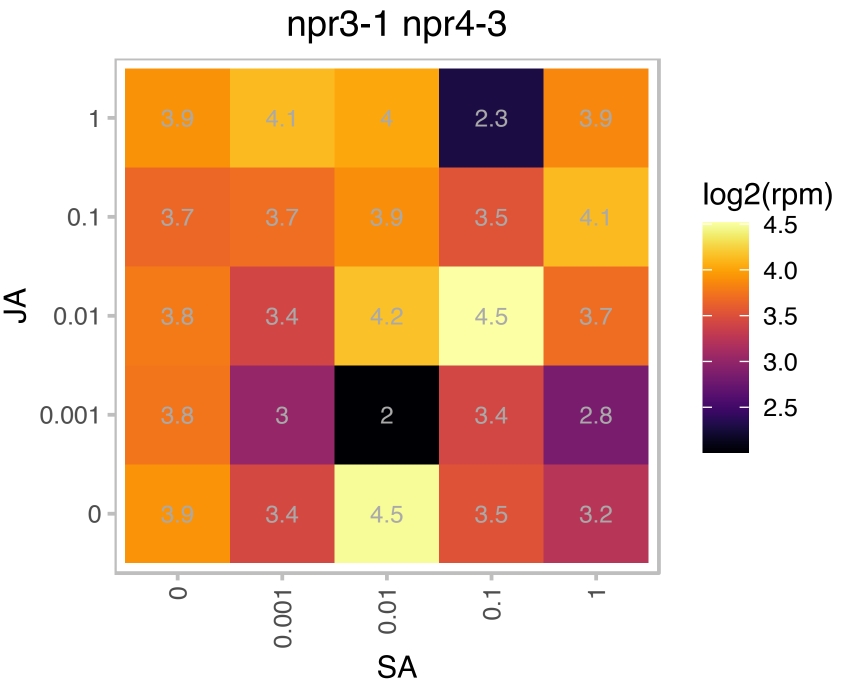

### AT1G35460.1 : CFLAP2, FBH1

basic helix-loop-helix (bHLH) DNA-binding superfamily protein

Encodes a bHLH transcription factor involved in CFL1-mediated regulation of cuticle development. Overexpression leads to abnormal cuticle development.

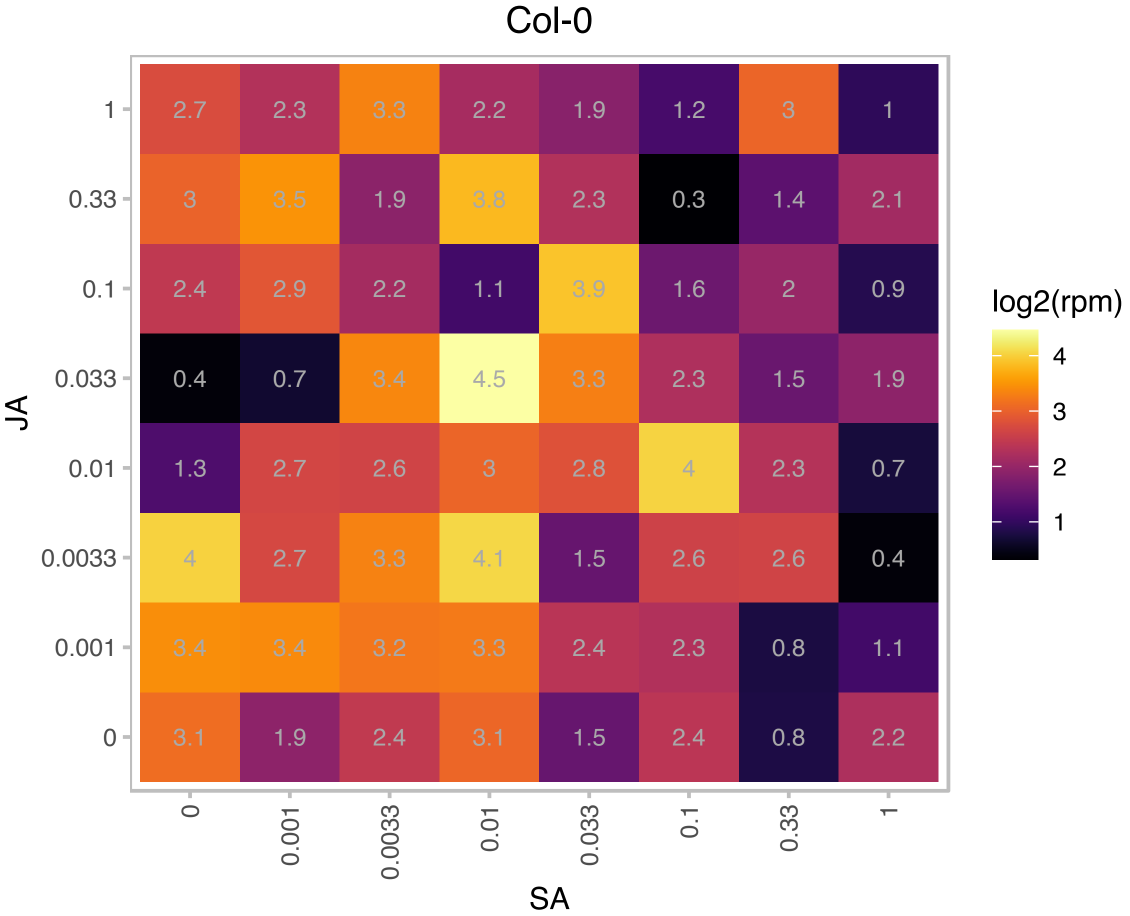


| | |
|---|---|
| Gene | AT1G35460 |
| is.expressed_genes | TRUE |
| is.responsive_genes | TRUE |
| is.apcluster | F-02 |

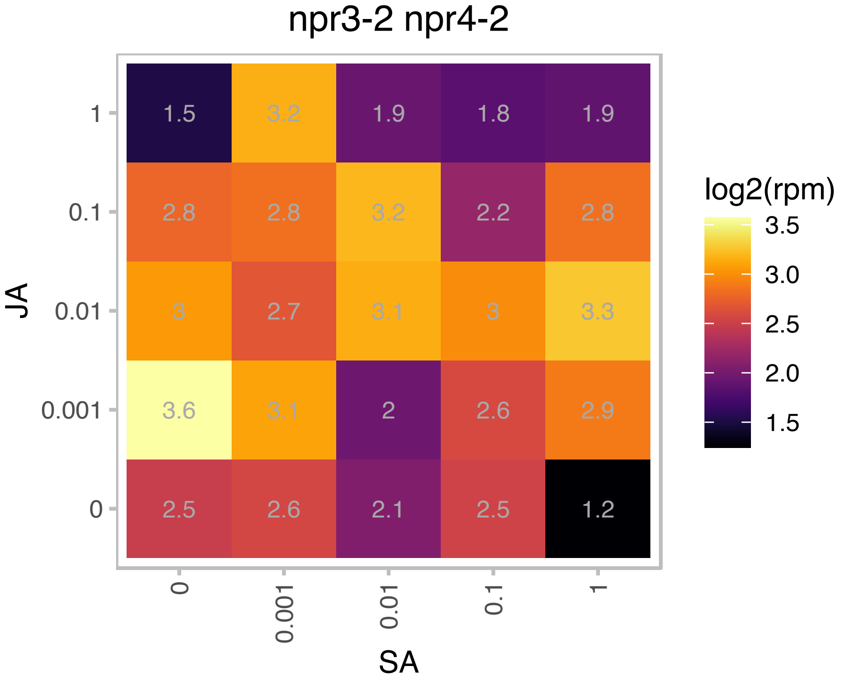


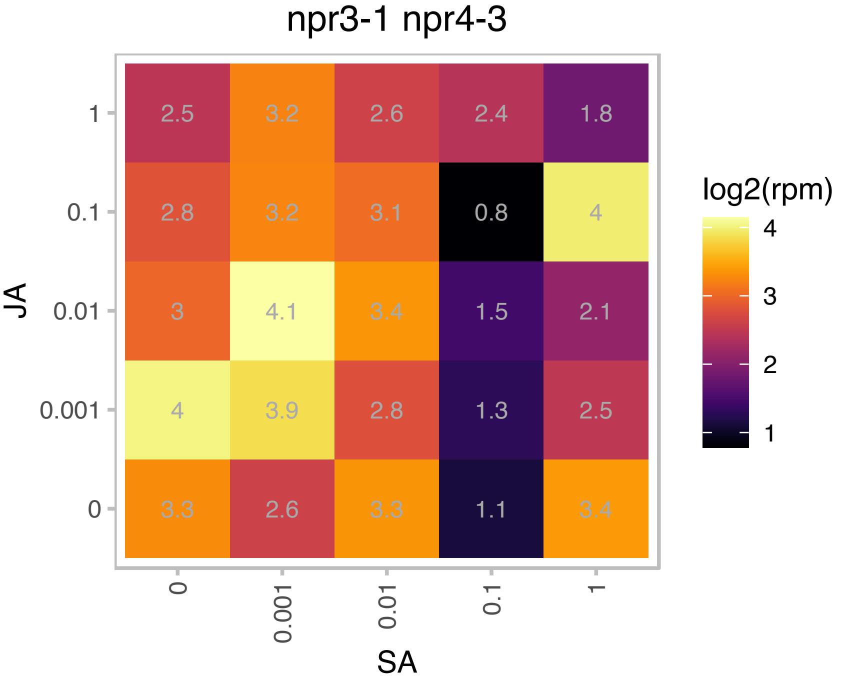

### AT1G43170.1 : ARP1, emb2207, RP1, RPL3A

ribosomal protein 1

Encodes a cytoplasmic ribosomal protein. The mRNA is cell-to-cell mobile.

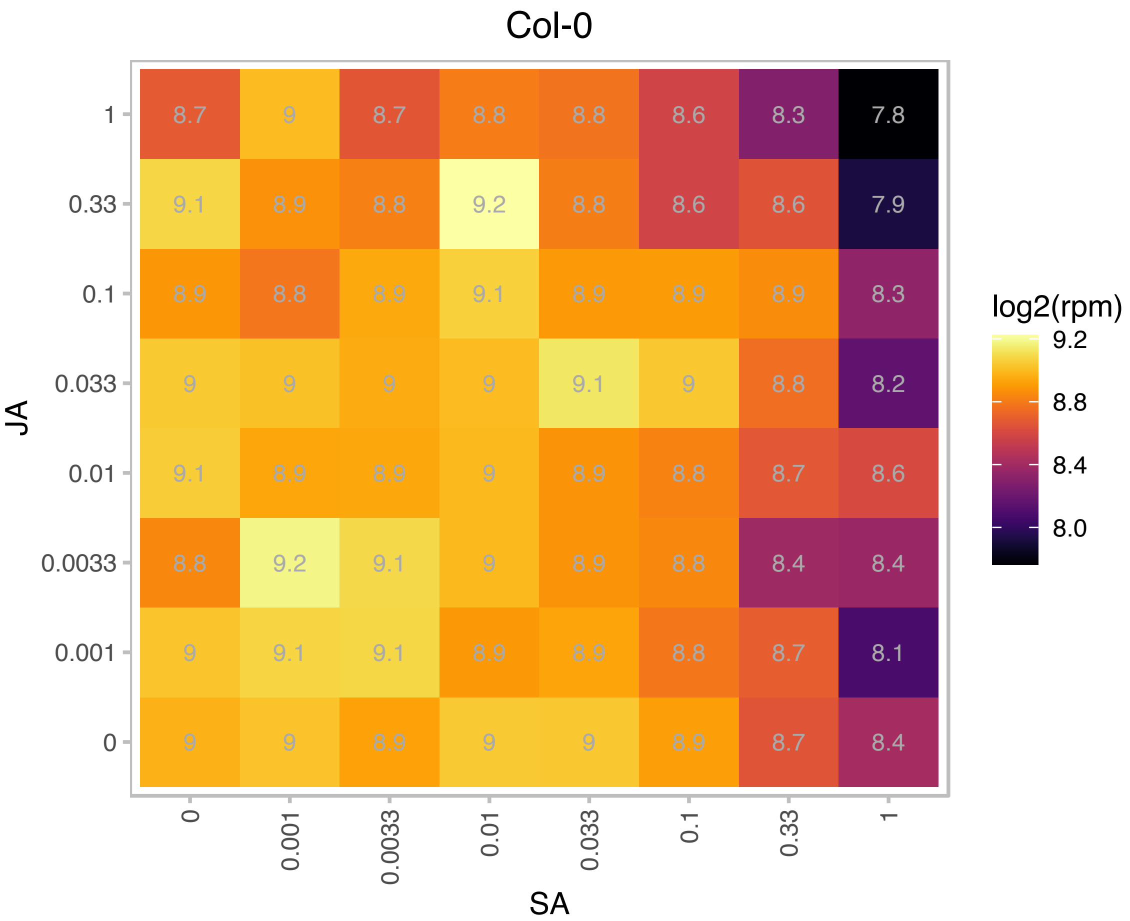


| | |
|---|---|
| Gene | AT1G43170 |
| is.expressed_genes | TRUE |
| is.responsive_genes | TRUE |
| is.apcluster | F-02 |

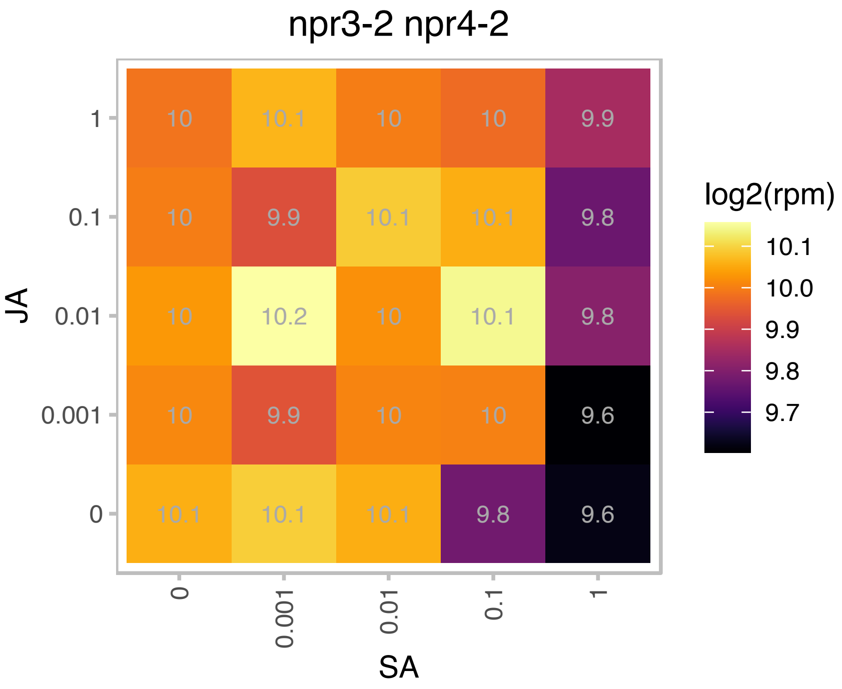


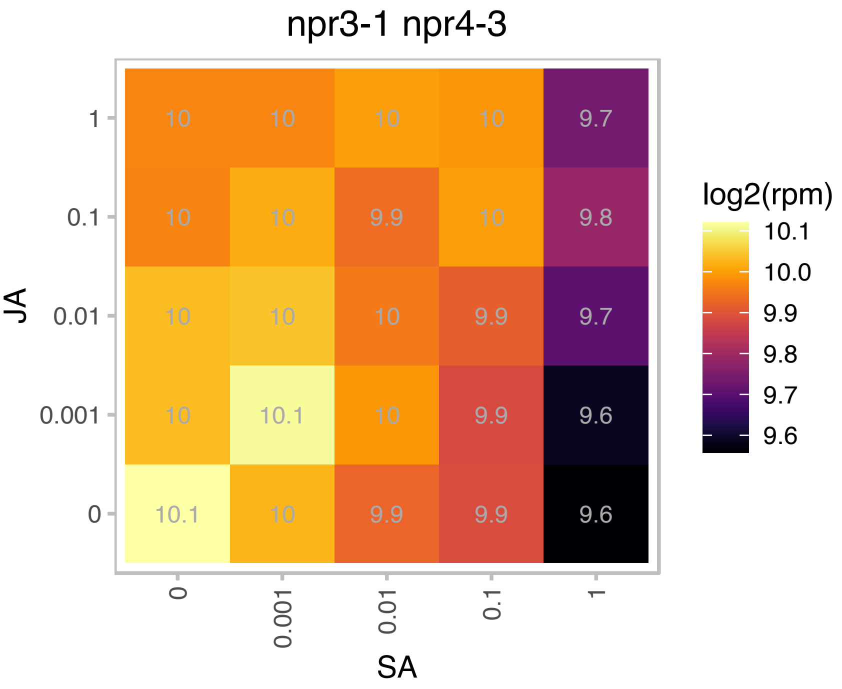

### AT1G44100.1 : AAP5

amino acid permease 5

amino acid permease 5

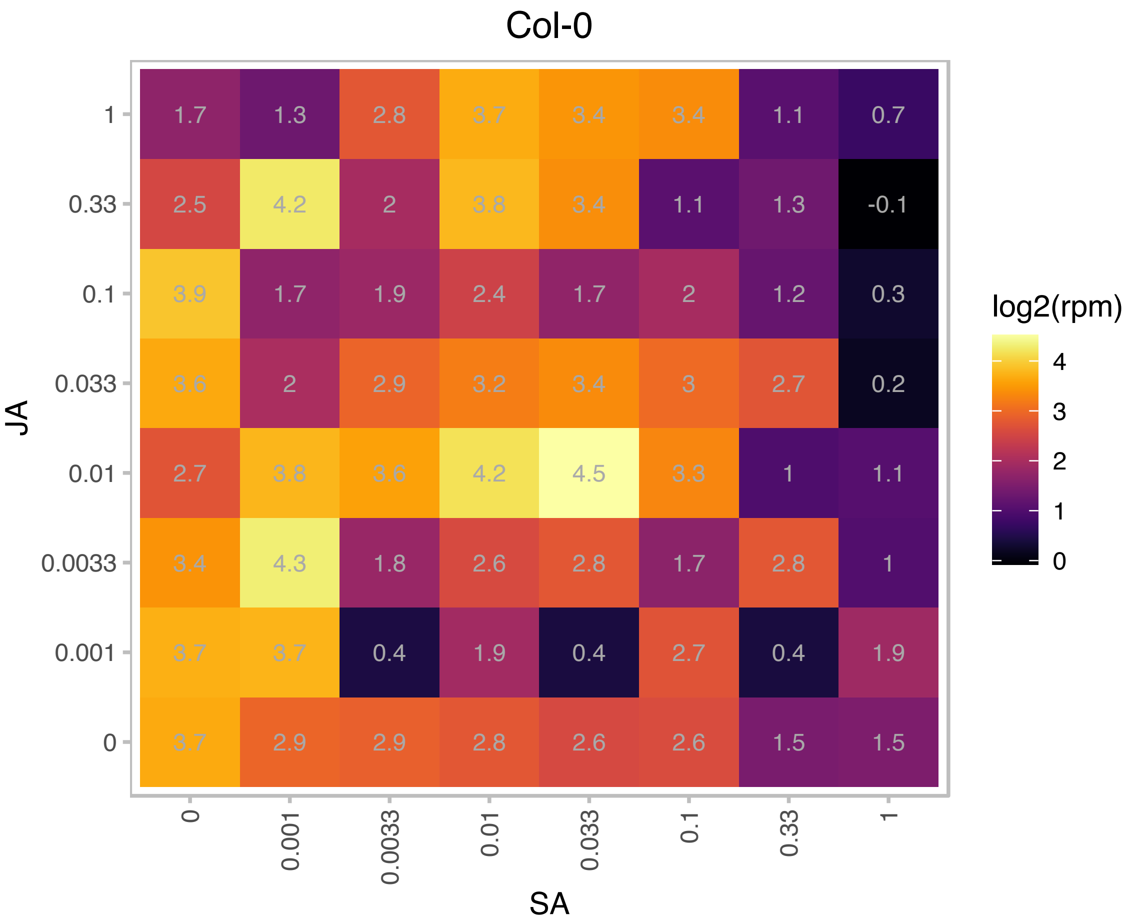


| | |
|---|---|
| Gene | AT1G44100 |
| is.expressed_genes | TRUE |
| is.responsive_genes | TRUE |
| is.apcluster | F-02 |

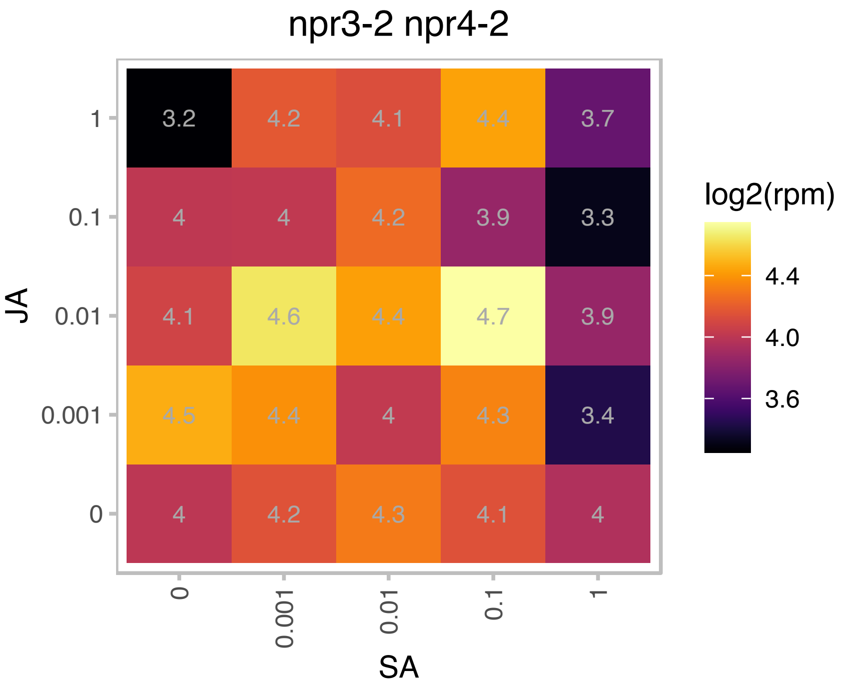


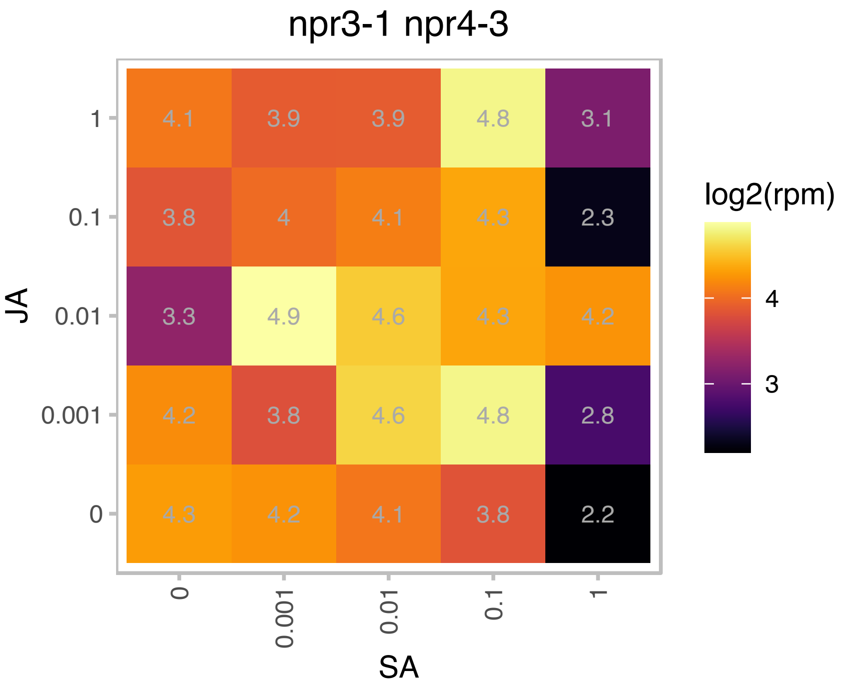

# AT1G44835.1 : NA

YbaK/aminoacyl-tRNA synthetase-associated domain

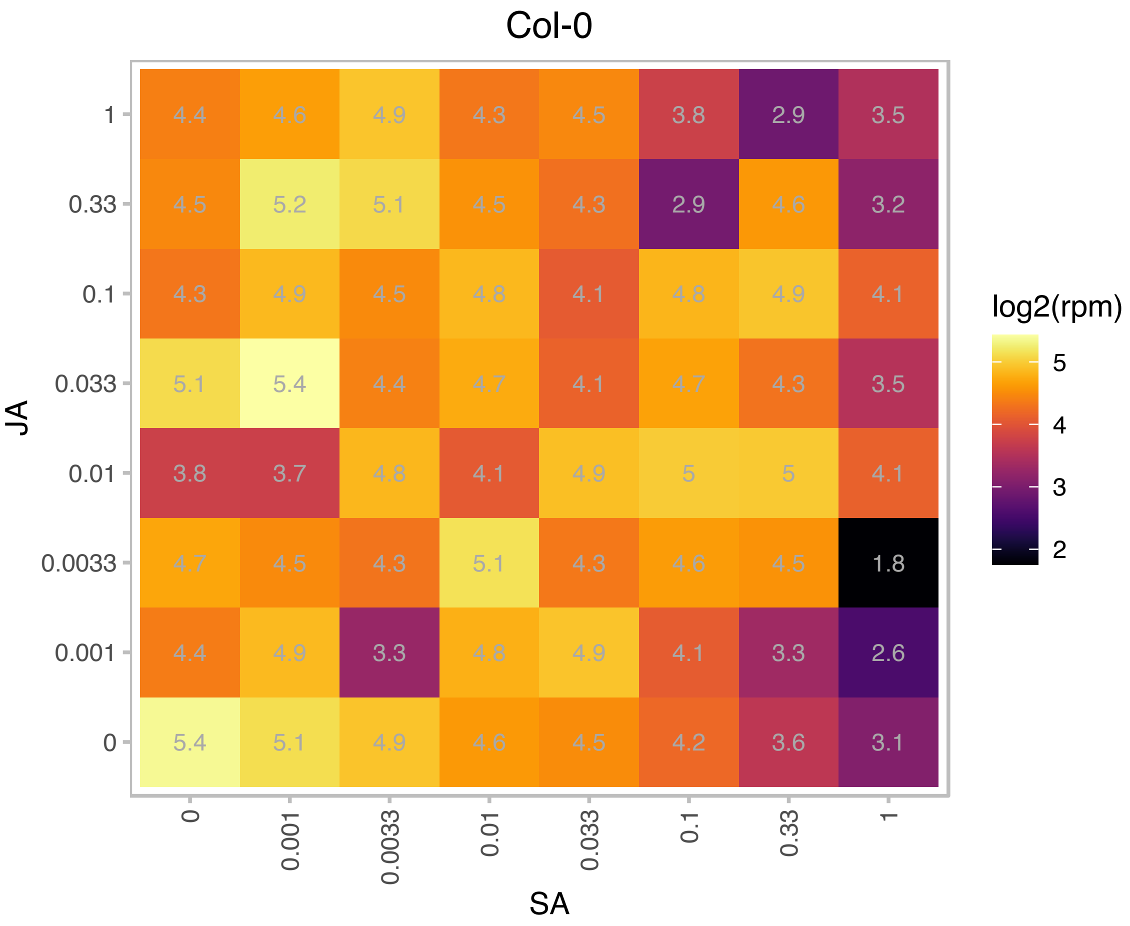


| | |
|---|---|
| Gene | AT1G44835 |
| is.expressed_genes | TRUE |
| is.responsive_genes | TRUE |
| is.apcluster | F-02 |

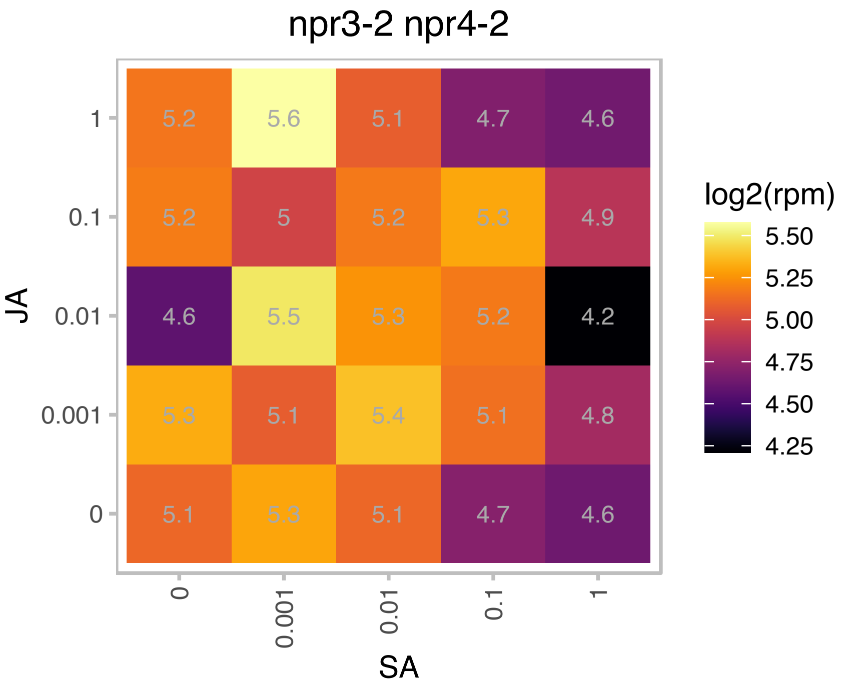


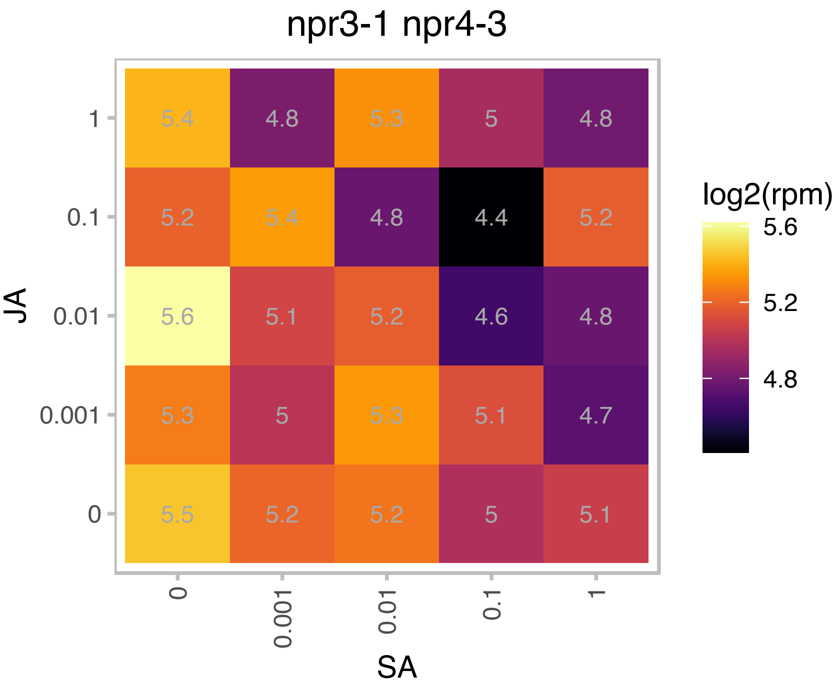

### AT1G49760.1 : PAB8, PABP8

poly(A) binding protein 8

polyadenylate-binding protein, putative / PABP, putative, similar to poly(A)-binding protein GB:AAF66825 GI:7673359 from (Nicotiana tabacum). Highly and ubiquitously expressed. Member of the class II PABP family.

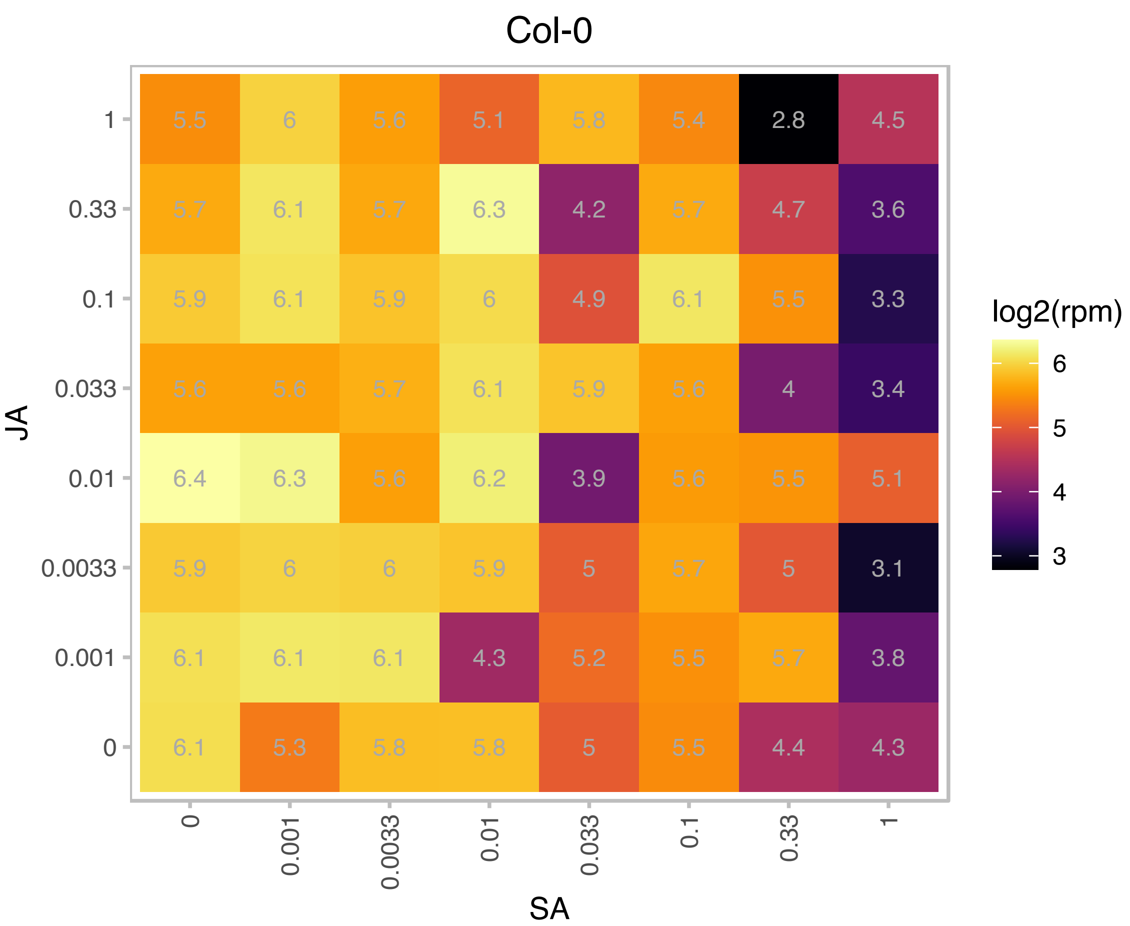


| | |
|---|---|
| Gene | AT1G49760 |
| is.expressed_genes | TRUE |
| is.responsive_genes | TRUE |
| is.apcluster | F-02 |

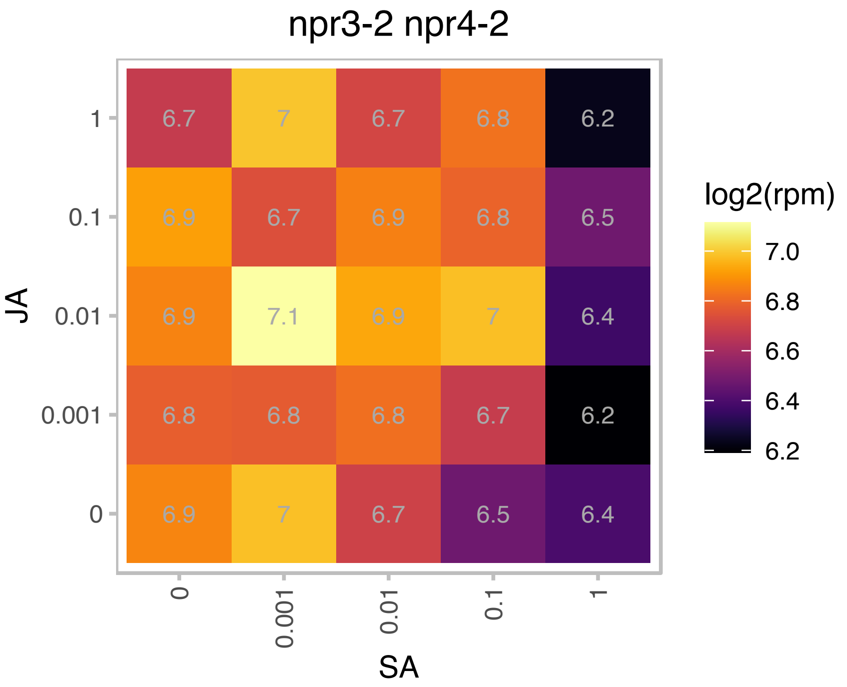


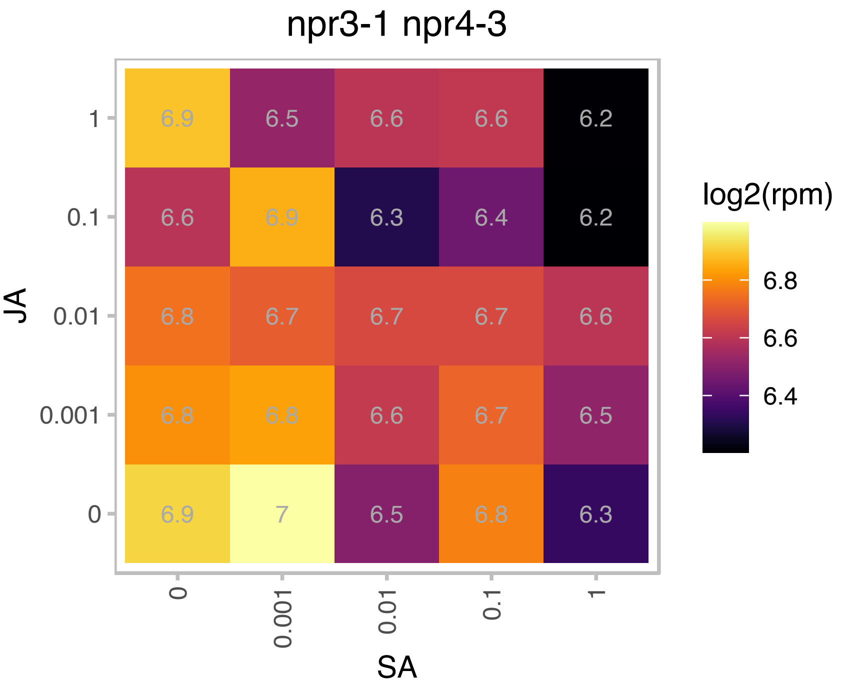

### AT1G54390.1 : ING2

PHD finger protein-related

ING2 encodes a member of the Inhibitor of Growth family of nuclear-localized PhD domain containing homeodomain proteins. Binds to H3K4 di or trimethylated DNA.

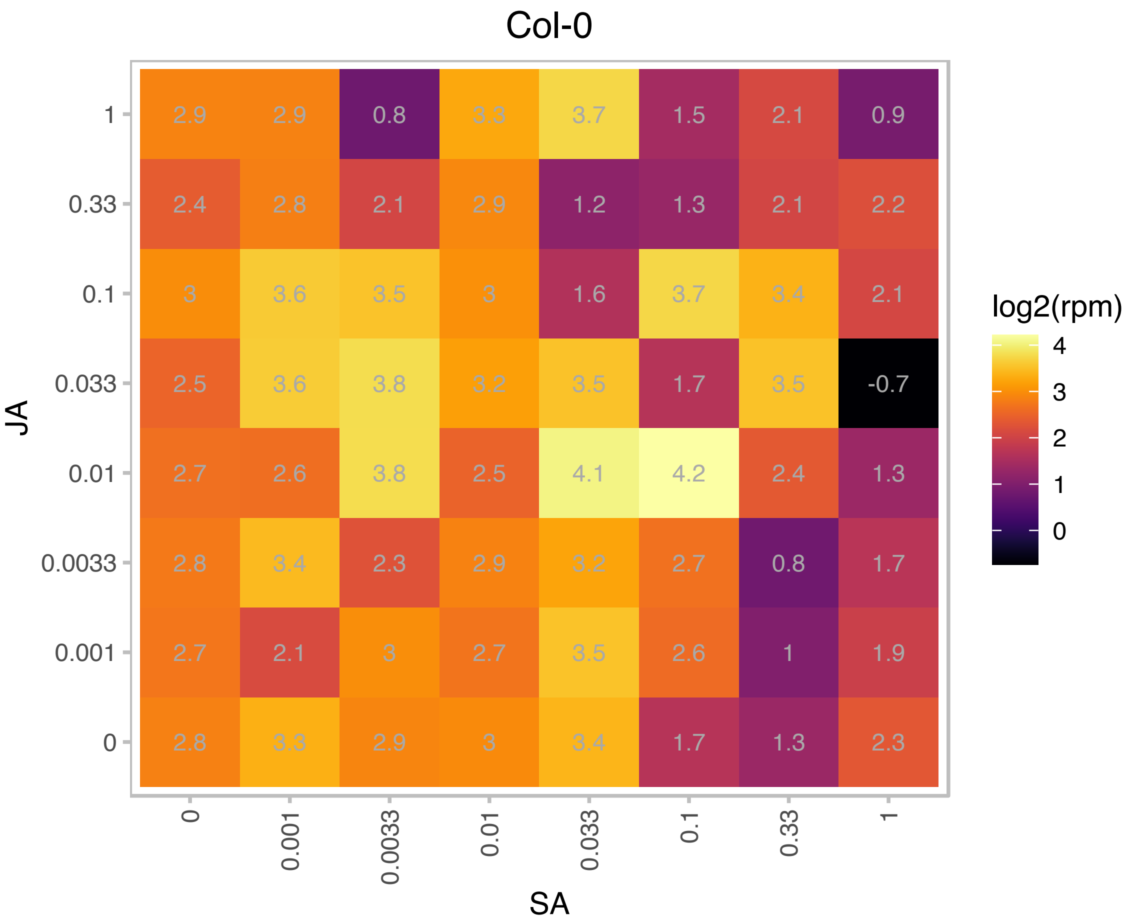


| | |
|---|---|
| Gene | AT1G54390 |
| is.expressed_genes | TRUE |
| is.responsive_genes | TRUE |
| is.apcluster | F-02 |

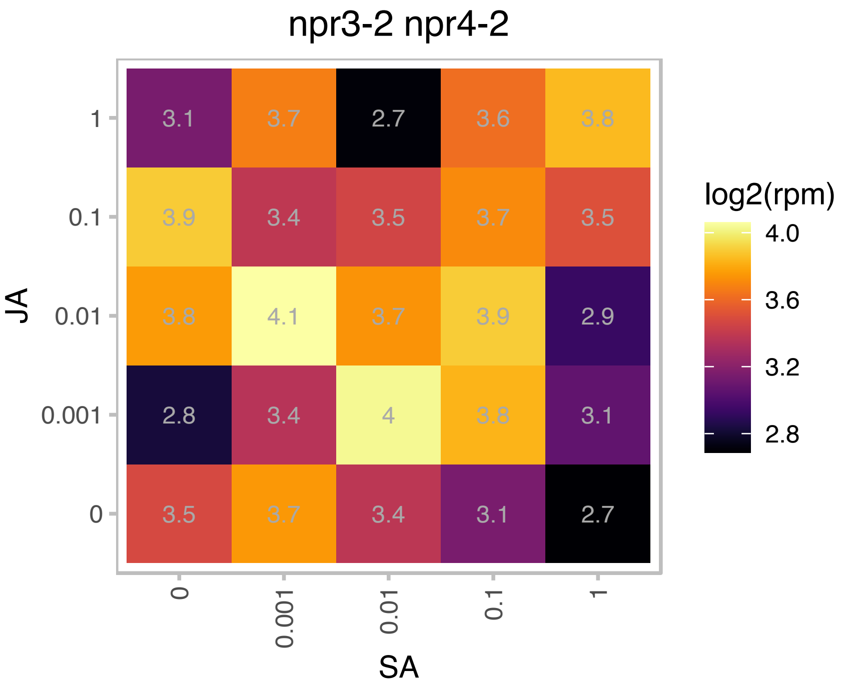


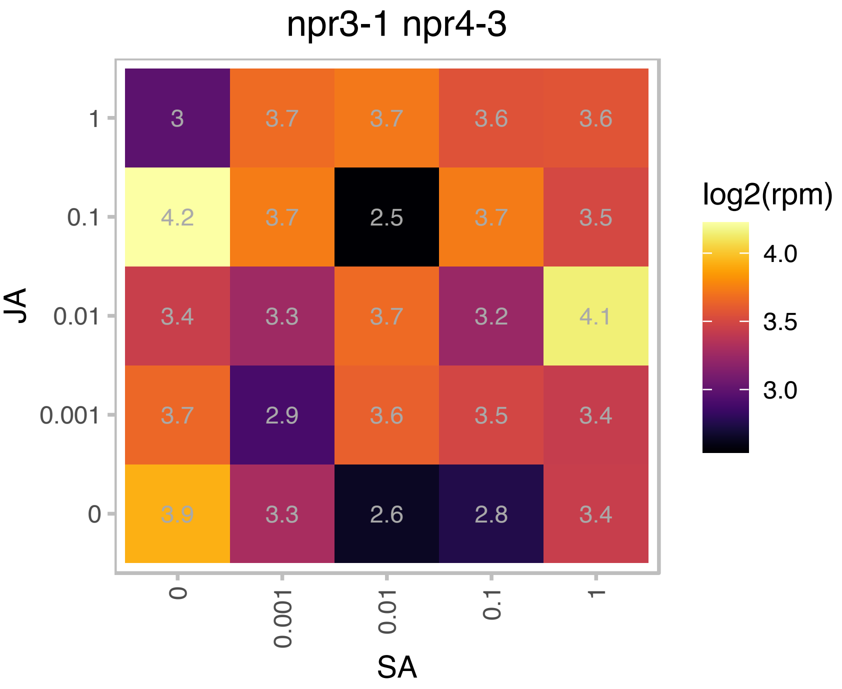

### AT1G54970.1 : ATPRP1, PRP1, RHS7

proline-rich protein 1

encodes a proline-rich protein that is specifically expressed in the root. The mRNA is cell-to-cell mobile.

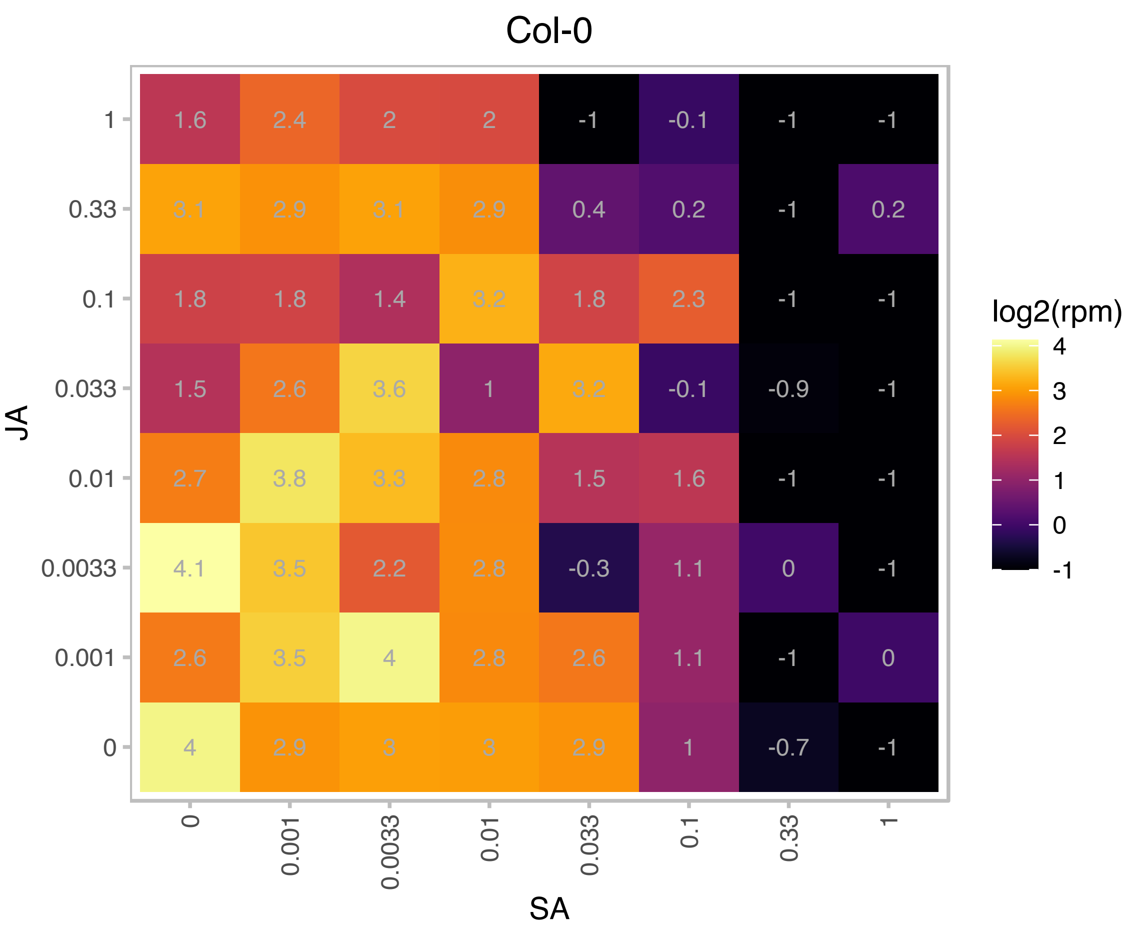


| | |
|---|---|
| Gene | AT1G54970 |
| is.expressed_genes | TRUE |
| is.responsive_genes | TRUE |
| is.apcluster | F-02 |

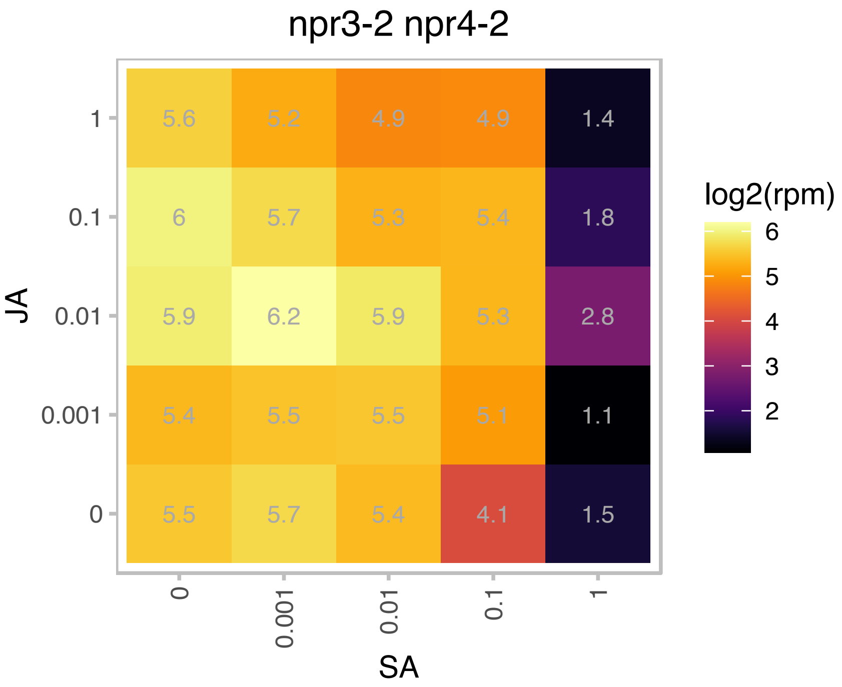


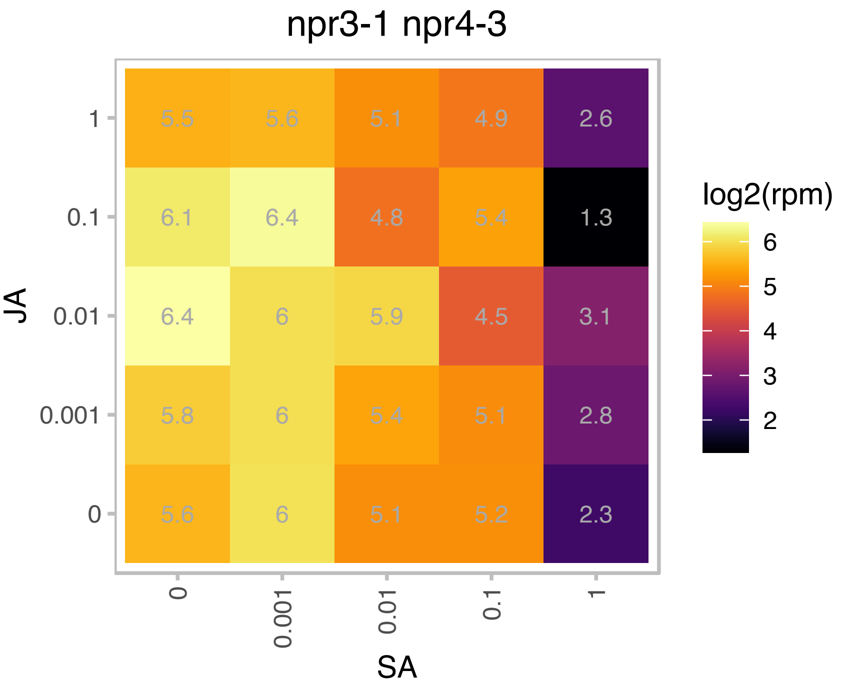

### AT1G55350.1 : ATDEK1, DEK1, EMB1275, EMB80

calpain-type cysteine protease family.

Similar to maize DEK1, a gene encoding a membrane protein of the calpain gene superfamily required for aleurone cell development in the endosperm of maize grains. A key component of the embryonic L1 cell-layer specification pathway. It localizes to membranes and undergoes intramolecular autolytic cleavage events that release the calpain domain into the cytoplasm.

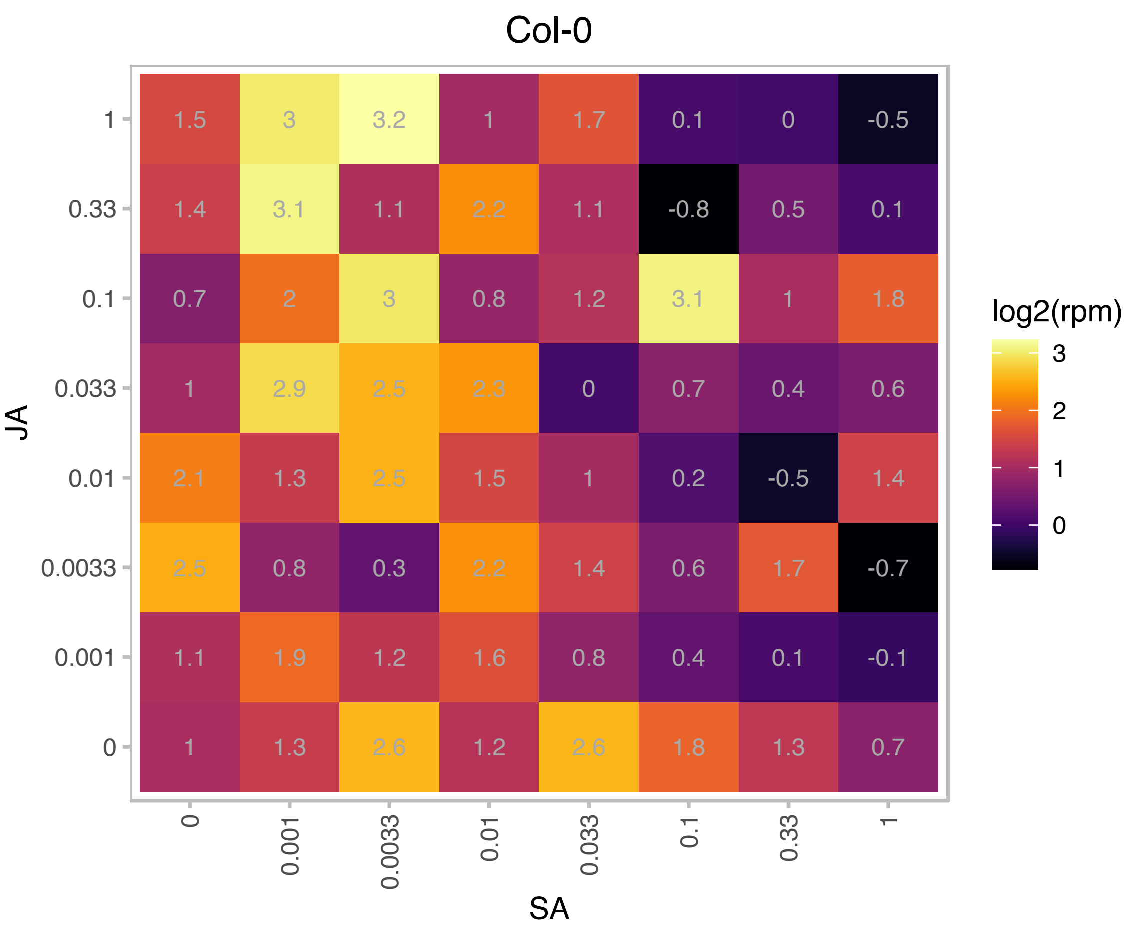


| | |
|---|---|
| Gene | AT1G55350 |
| is.expressed_genes | TRUE |
| is.responsive_genes | TRUE |
| is.apcluster | F-02 |

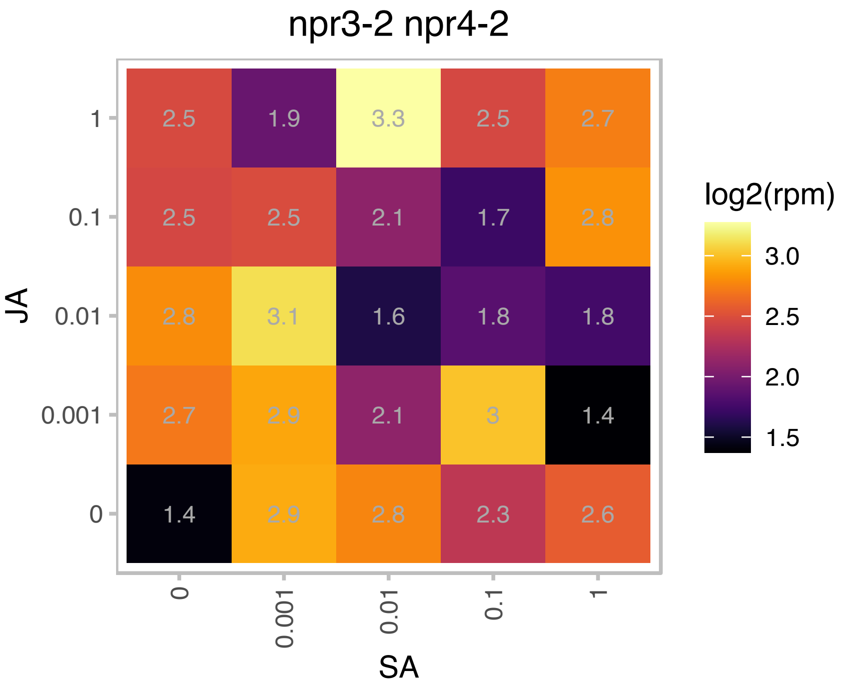


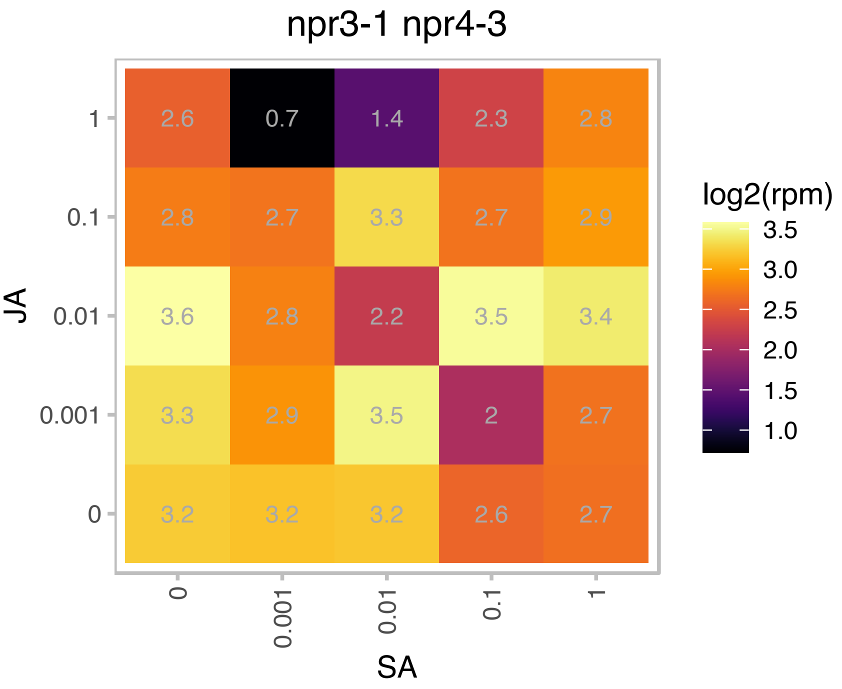

### AT1G72360.1 : AtERF73, ERF73, HRE1

Integrase-type DNA-binding superfamily protein

Encodes a member of the ERF (ethylene response factor) subfamily B-2 of ERF/AP2 transcription factor family. The protein contains one AP2 domain. There are 5 members in this subfamily including RAP2.2 AND RAP2.12.

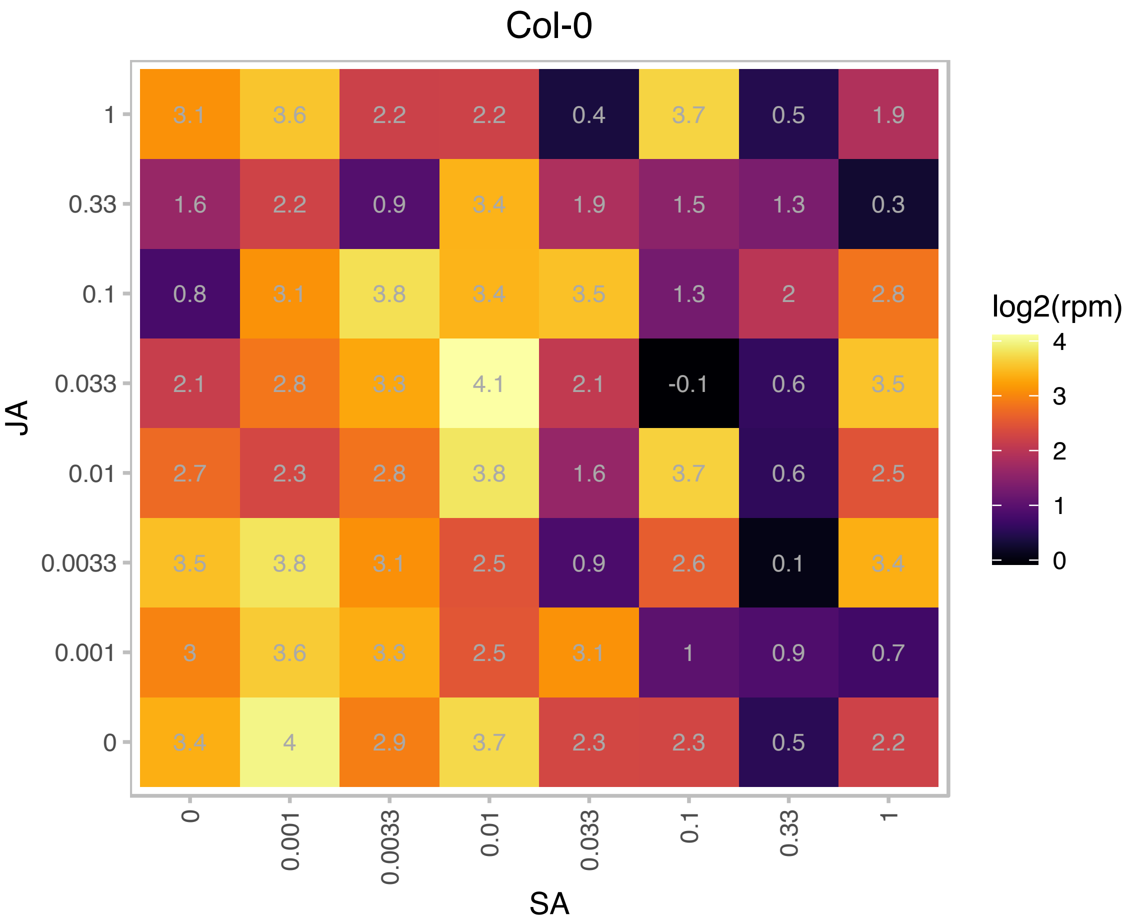


| | |
|---|---|
| Gene | AT1G72360 |
| is.expressed_genes | TRUE |
| is.responsive_genes | TRUE |
| is.apcluster | F-02 |

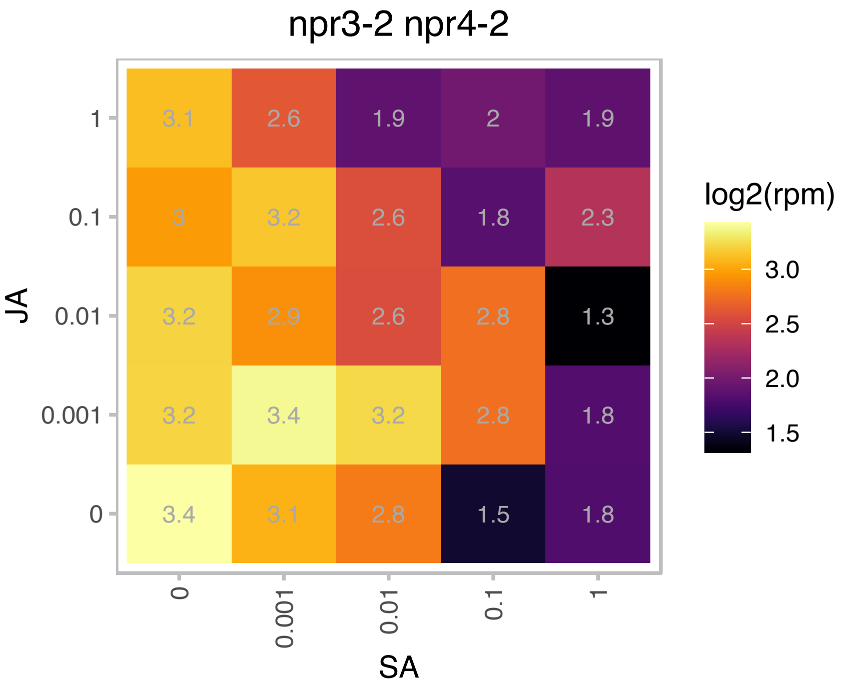


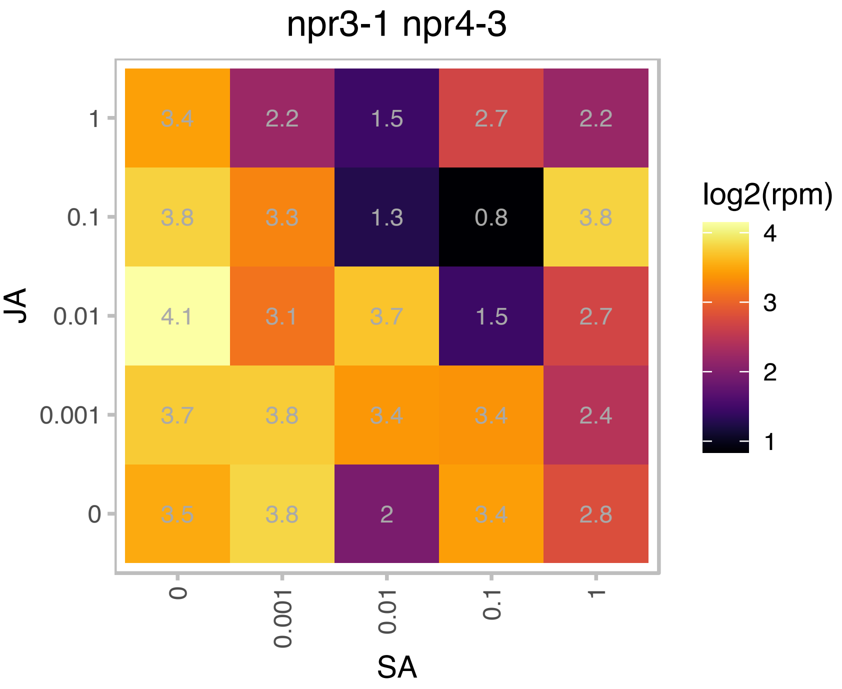

### AT1G72430.1 : SAUR78

SAUR-like auxin-responsive protein family

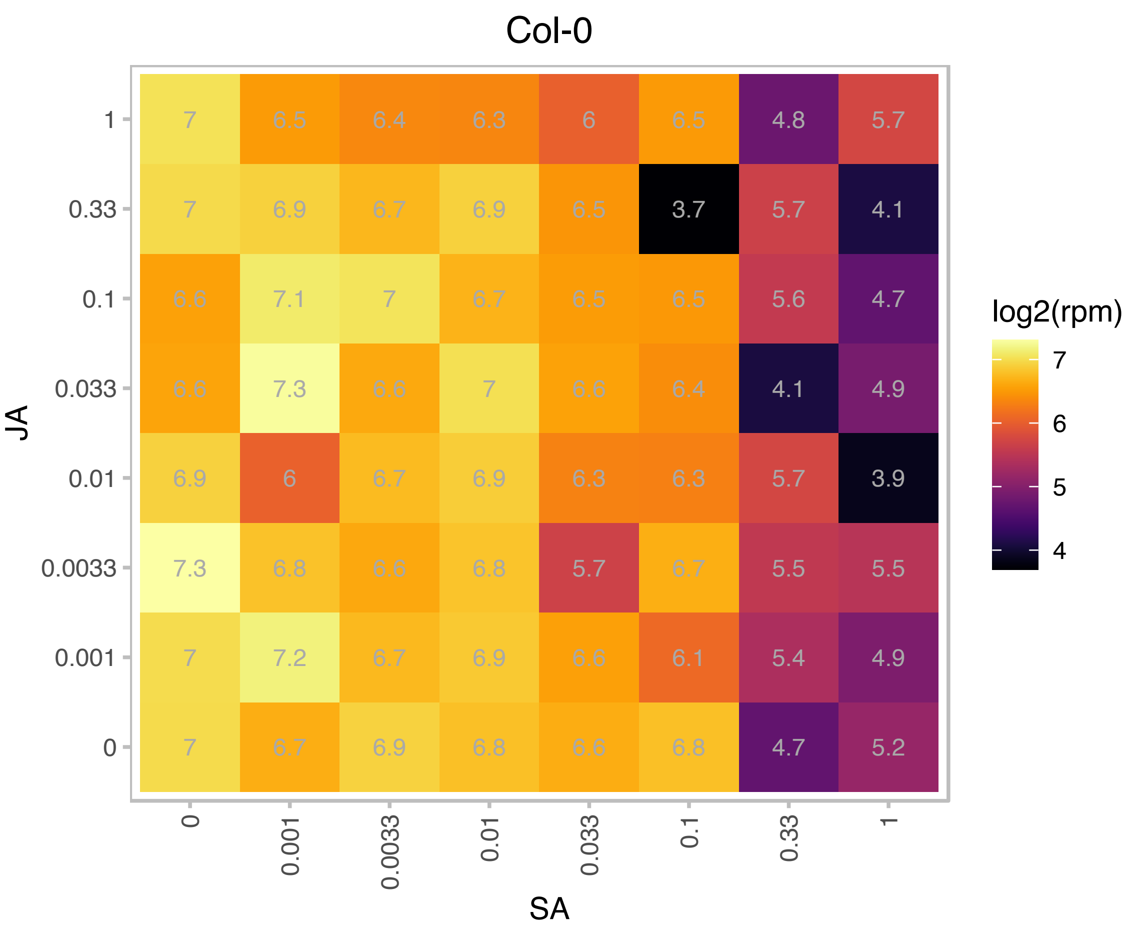


| | |
|---|---|
| Gene | AT1G72430 |
| is.expressed_genes | TRUE |
| is.responsive_genes | TRUE |
| is.apcluster | F-02 |

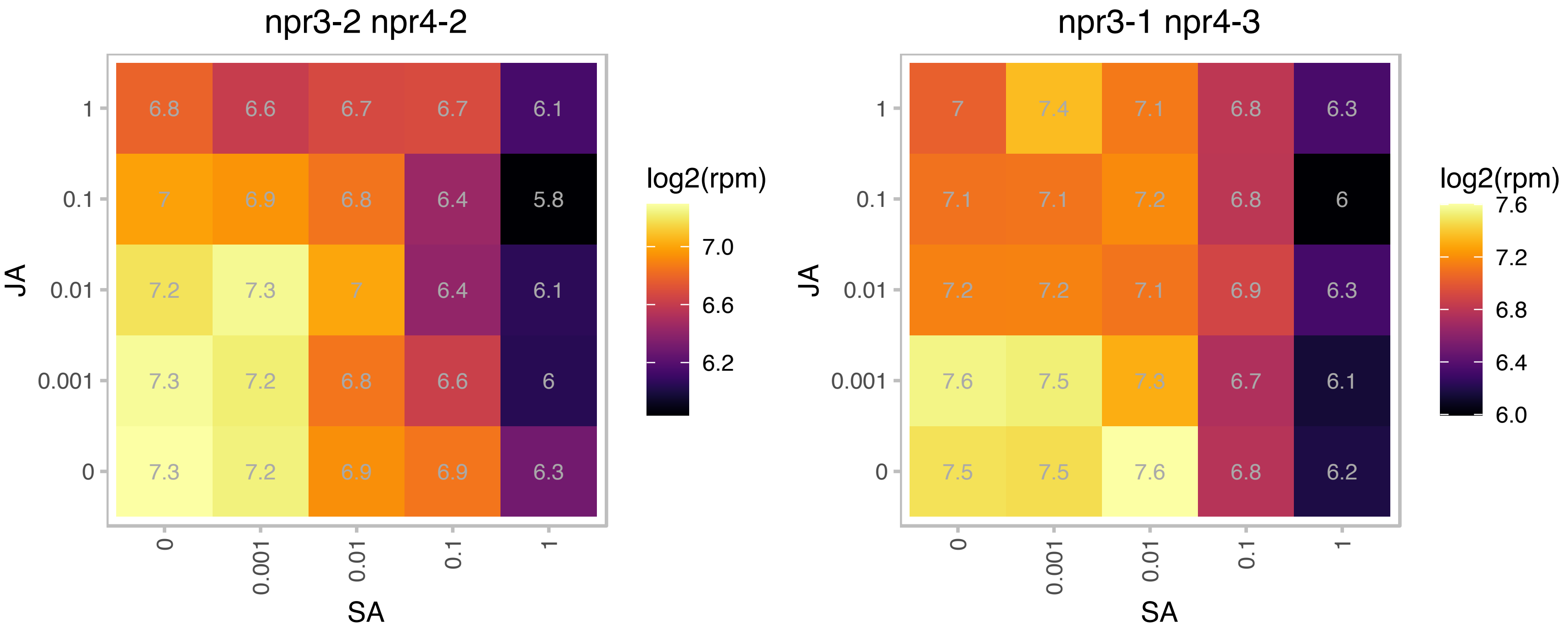

### AT1G74090.1 : ATSOT18, ATST5B, SOT18

desulfo-glucosinolate sulfotransferase 18
encodes a desulfoglucosinolate sulfotransferase, involved in the final step of glucosinolate core structure biosynthesis. Has a broad-substrate specificity with preference with methionine-derived desulfoglucosinolates.

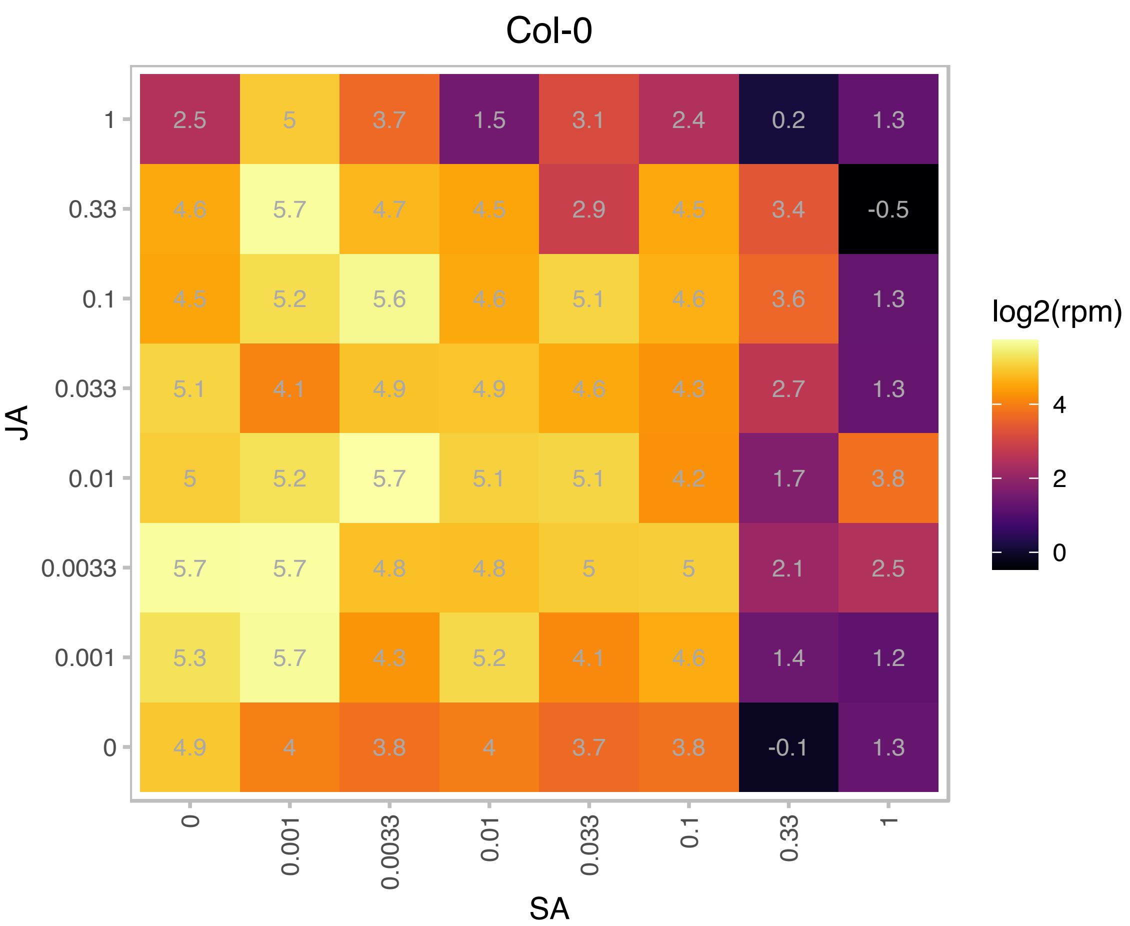


| | |
|---|---|
| Gene | AT1G74090 |
| is.expressed_genes | TRUE |
| is.responsive_genes | TRUE |
| is.apcluster | F-02 |

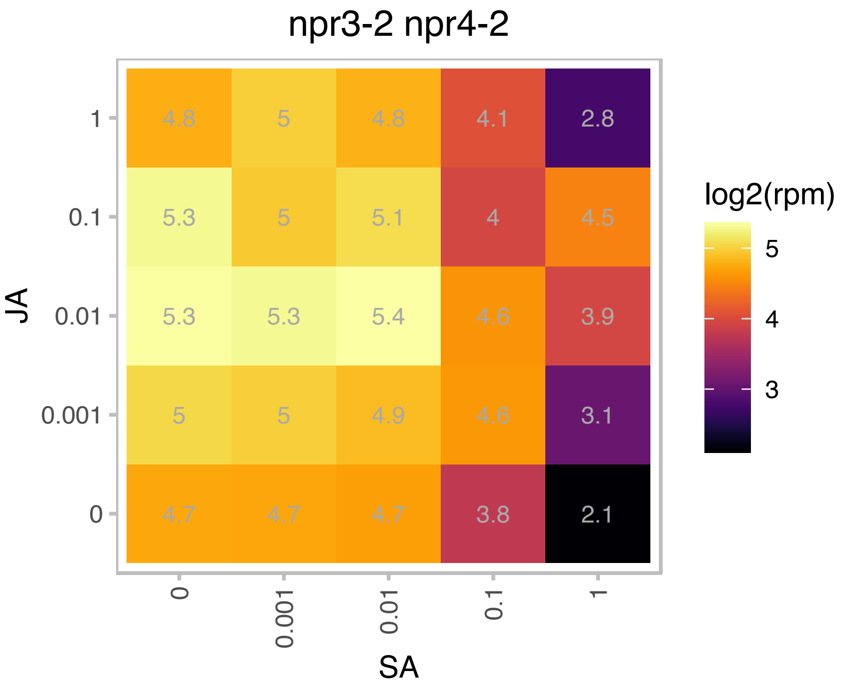


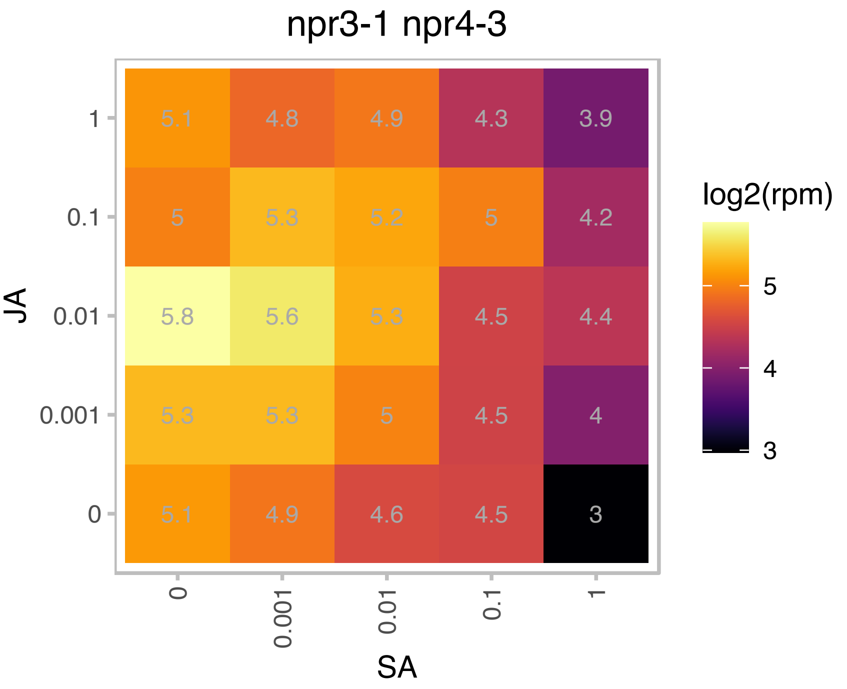

### AT1G74500.1 : ATBS1, bHLH135, BS1, PRE3, TMO7

activation-tagged BRI1(brassinosteroid-insensitive 1)-suppressor 1
Encodes a basic helix/loop/helix transcription factor that acts downstream of MP in root initiation. TMO7 protein moves to the hypophysis and to vascular cells, contributing to MP-dependent root formation. Promotes the correct definition of the hypophysis cell division plane.

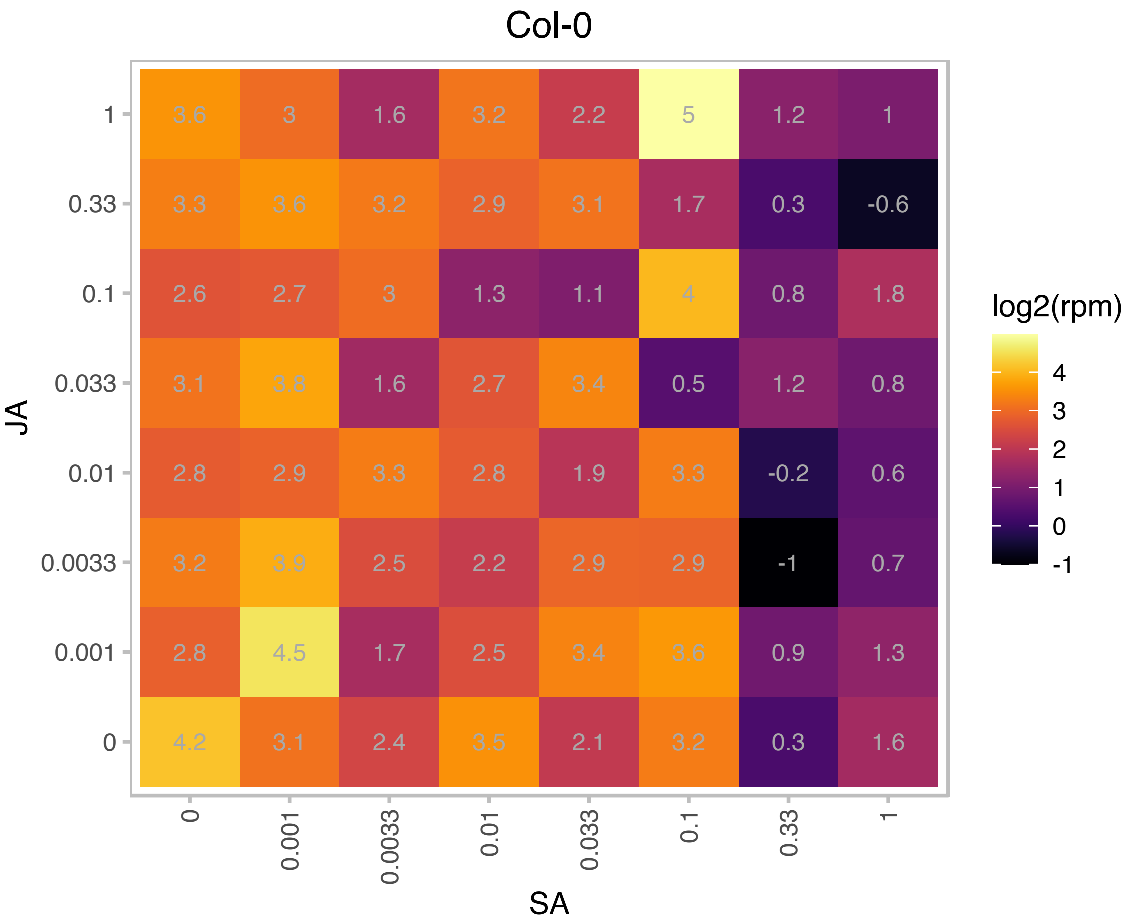


| | |
|---|---|
| Gene | AT1G74500 |
| is.expressed_genes | TRUE |
| is.responsive_genes | TRUE |
| is.apcluster | F-02 |

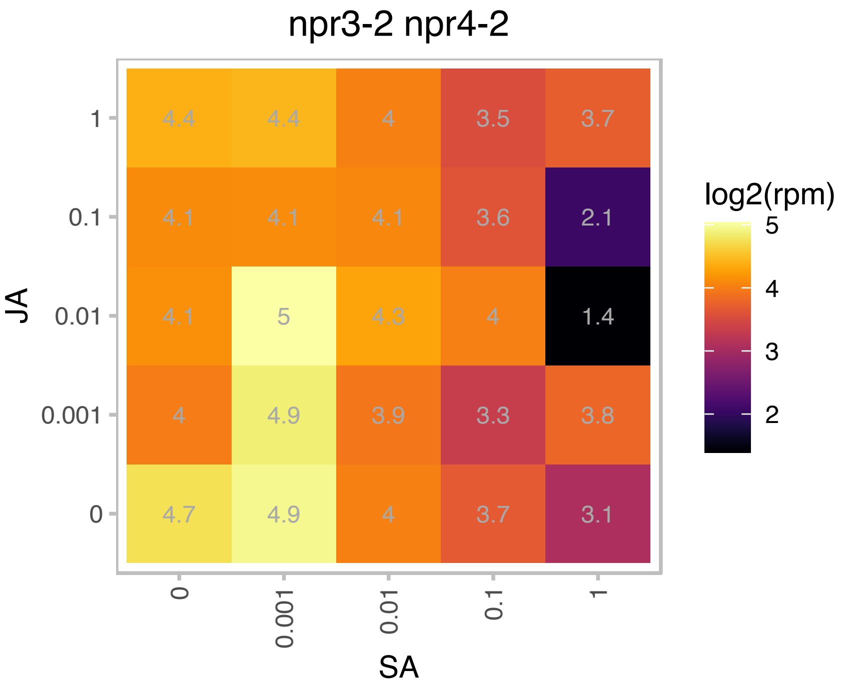


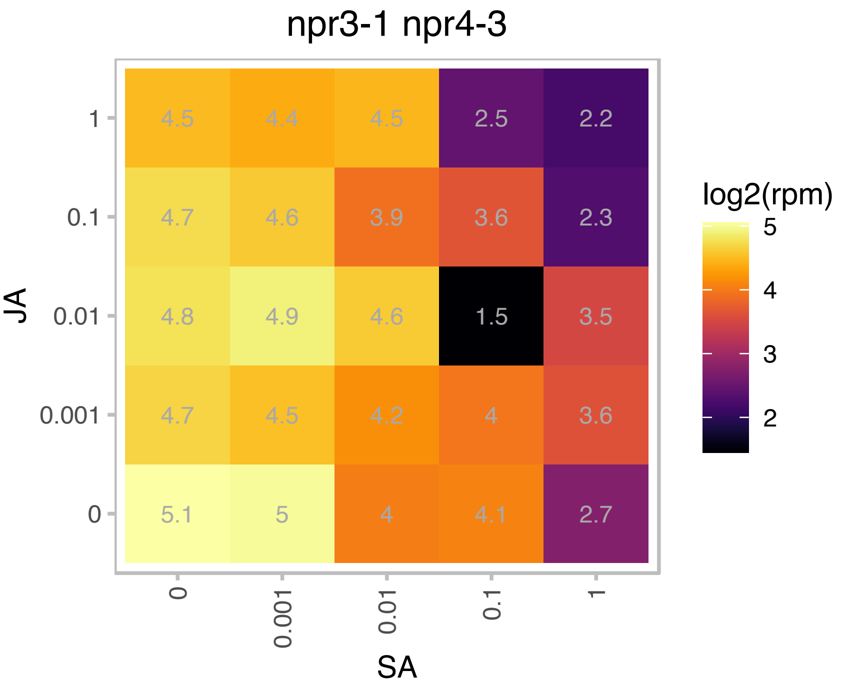

# AT1G74840.1 : NA

Homeodomain-like superfamily protein

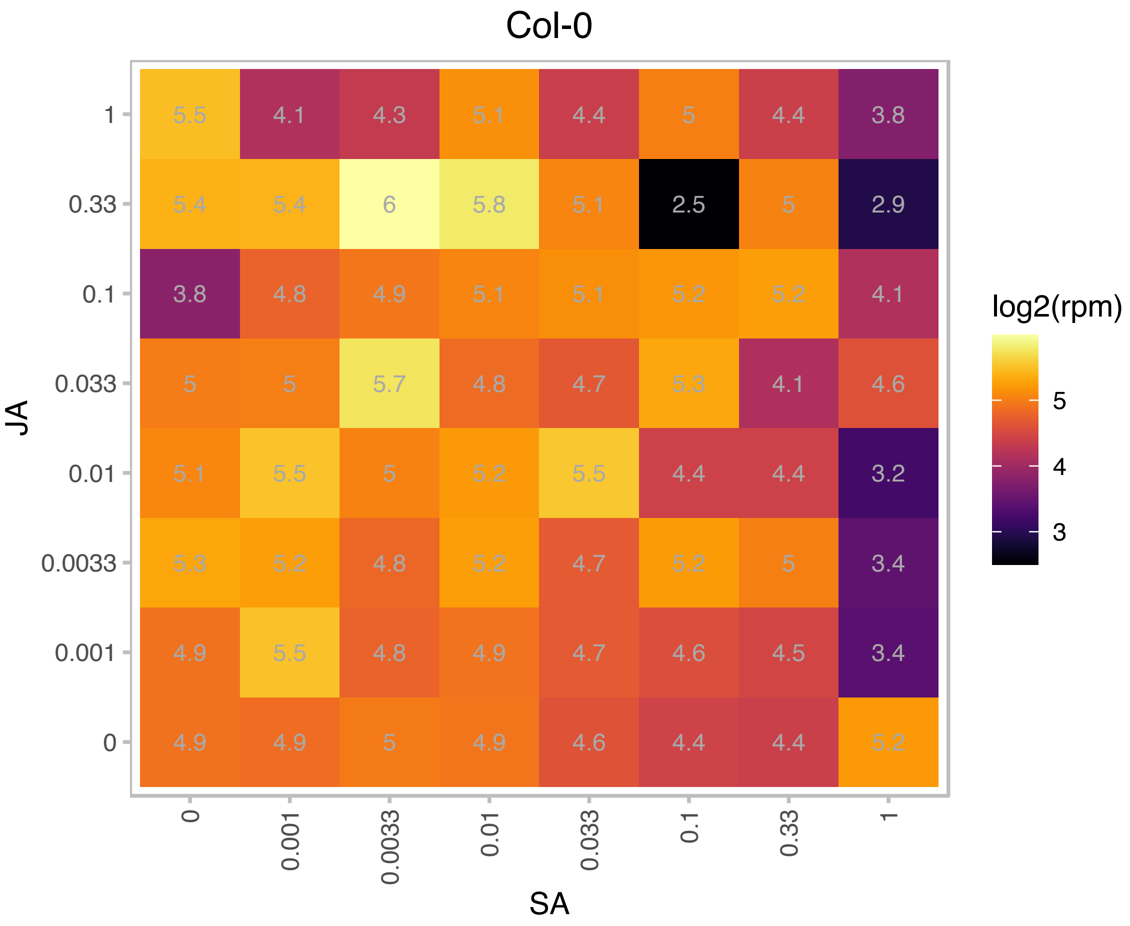


| | |
|---|---|
| Gene | AT1G74840 |
| is.expressed_genes | TRUE |
| is.responsive_genes | TRUE |
| is.apcluster | F-02 |

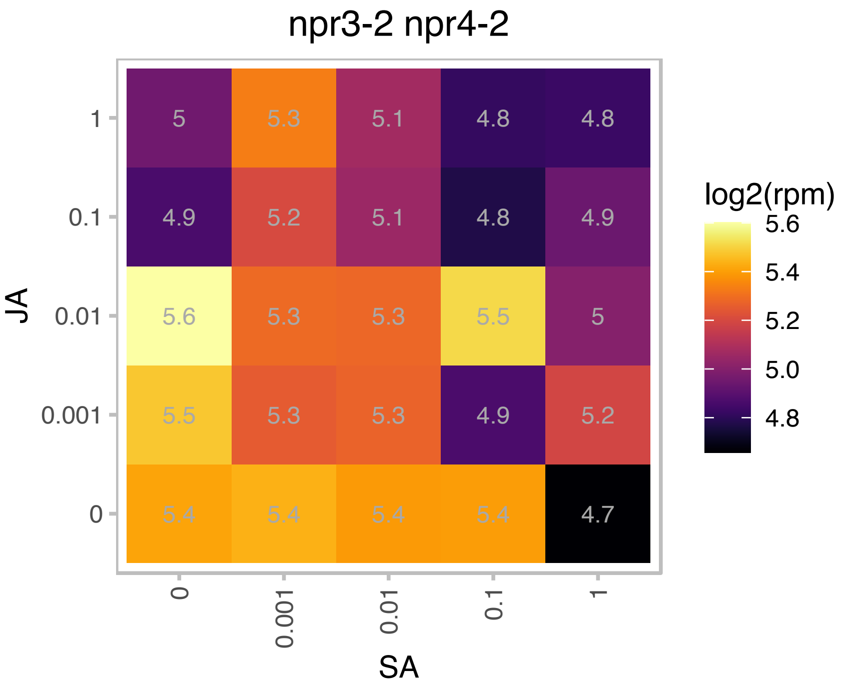


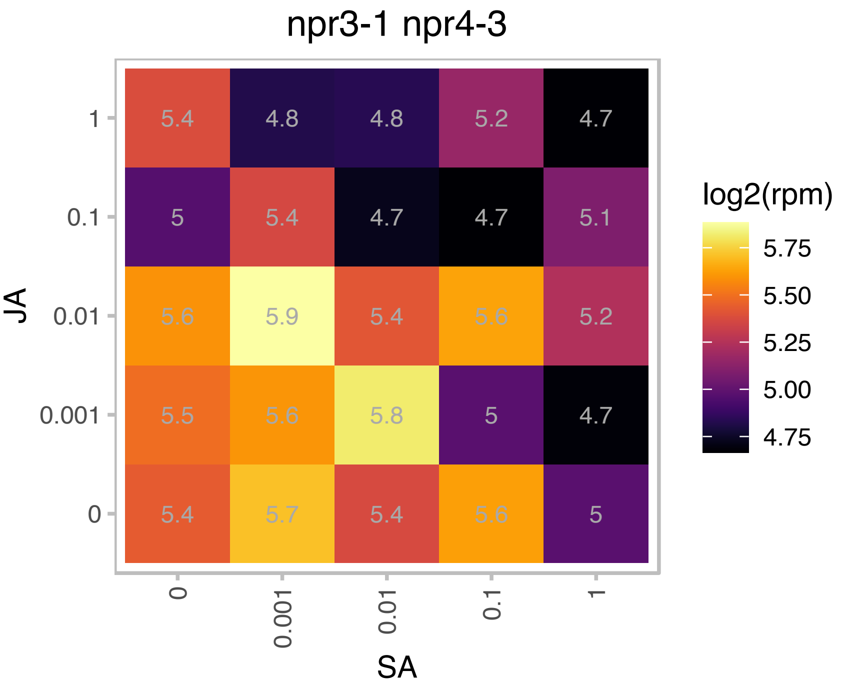

### AT1G77330.1 : ACO5

2-oxoglutarate (2OG) and Fe(II)-dependent oxygenase superfamily protein

similar to 1-aminocyclopropane-1-carboxylate oxidase GI:3386565 from (Sorghum bicolor)

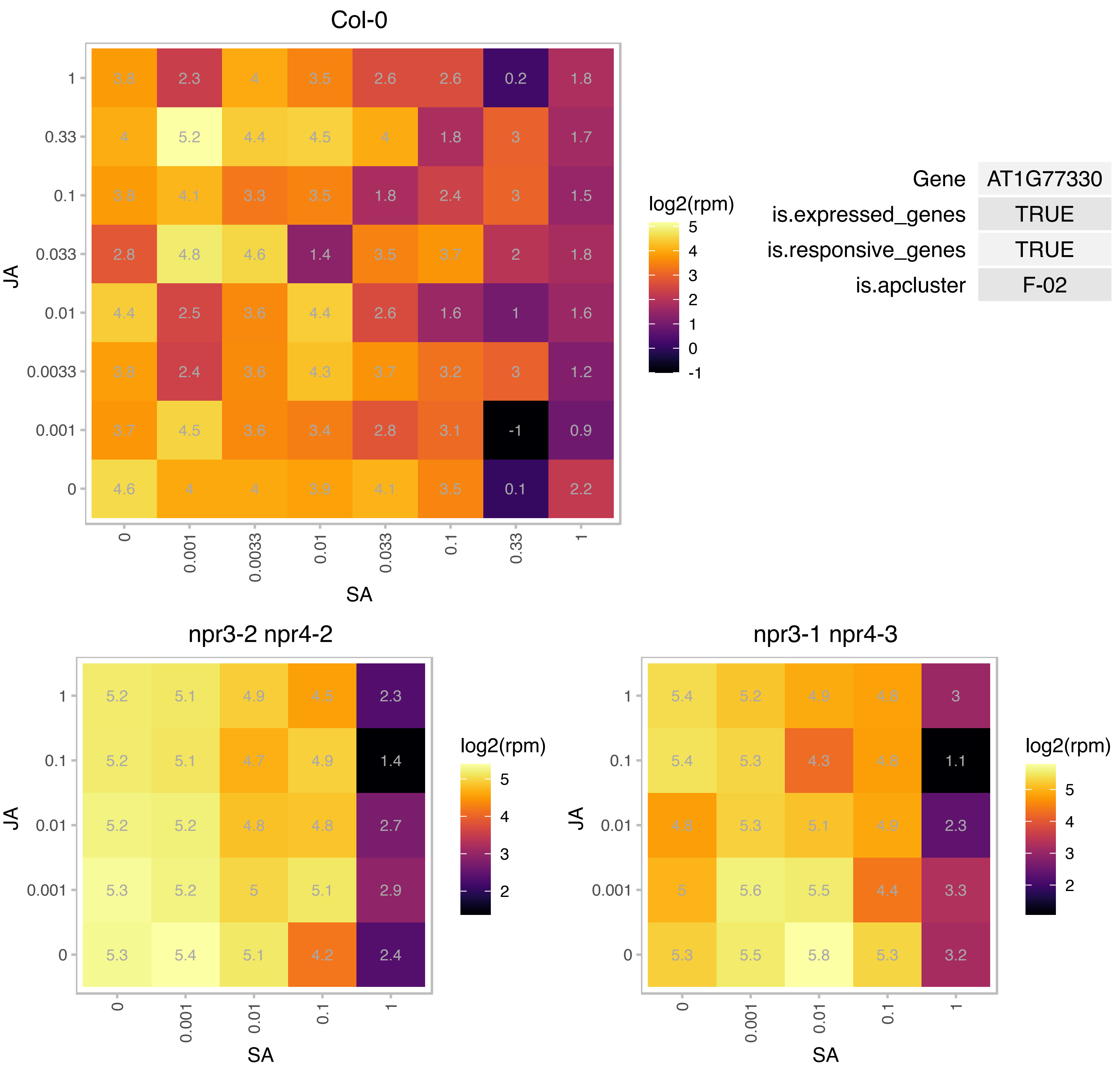


| | |
|---|---|
| Gene | AT1G77330 |
| is.expressed_genes | TRUE |
| is.responsive_genes | TRUE |
| is.apcluster | F-02 |

### AT1G78370.1 : ATGSTU20, GSTU20

glutathione S-transferase TAU 20

Encodes glutathione transferase belonging to the tau class of GSTs. Naming convention according to Wagner et al. (2002).

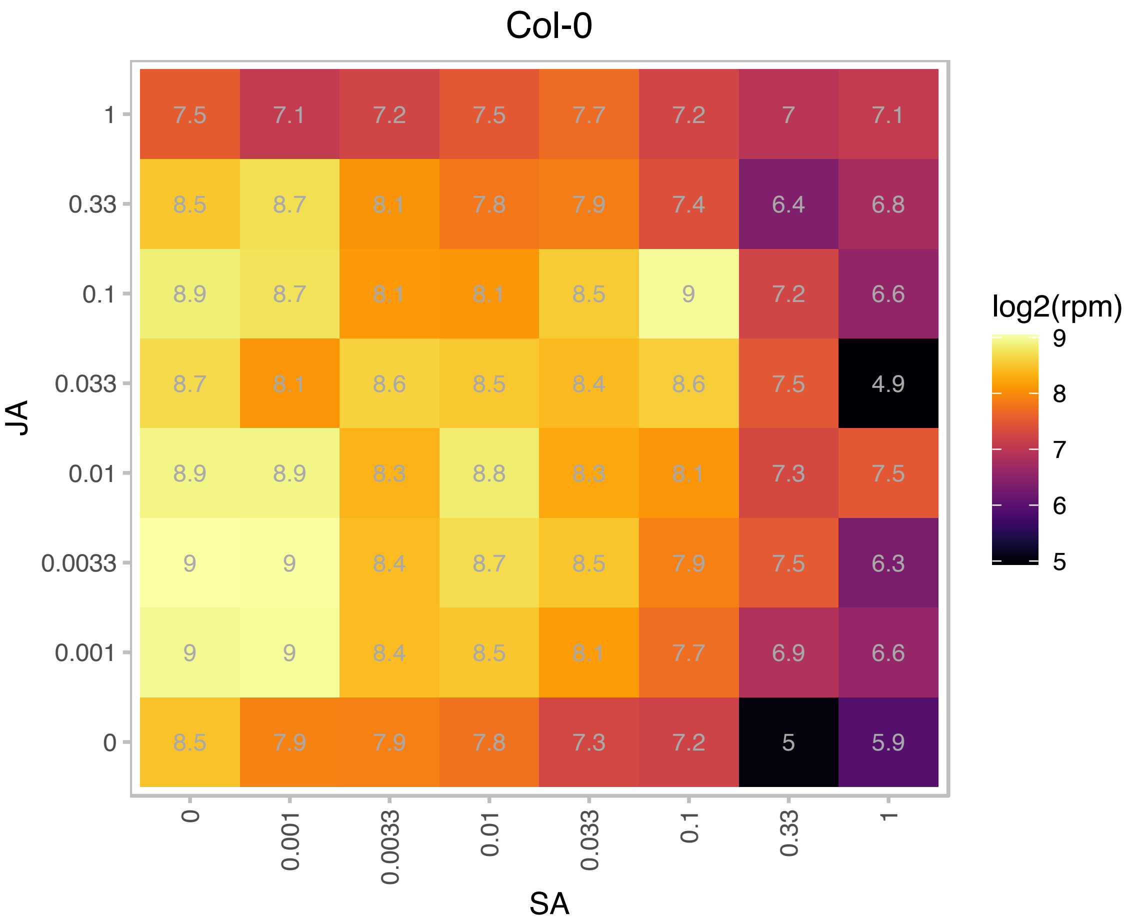


| | |
|---|---|
| Gene | AT1G78370 |
| is.expressed_genes | TRUE |
| is.responsive_genes | TRUE |
| is.apcluster | F-02 |

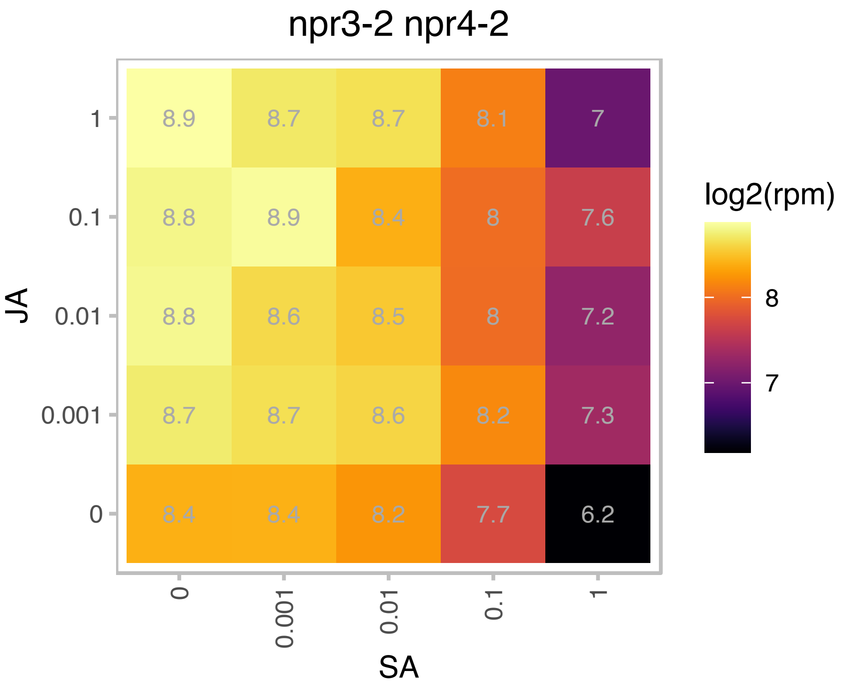


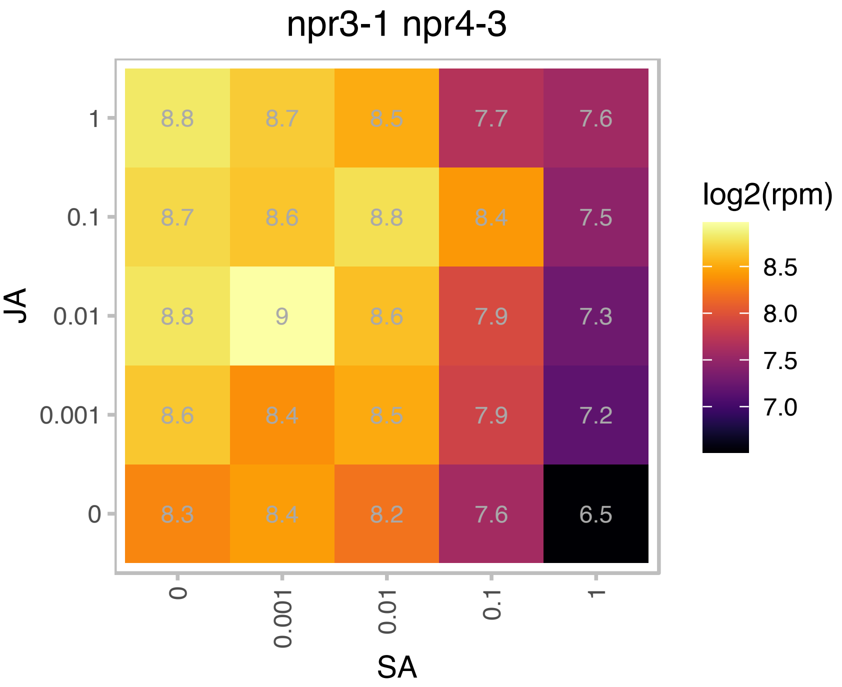

### AT2G05830.1 : MTI1, MTI1

NagB/RpiA/CoA transferase-like superfamily protein

Encodes a 5-methylthioribose-1-phosphate isomerase.

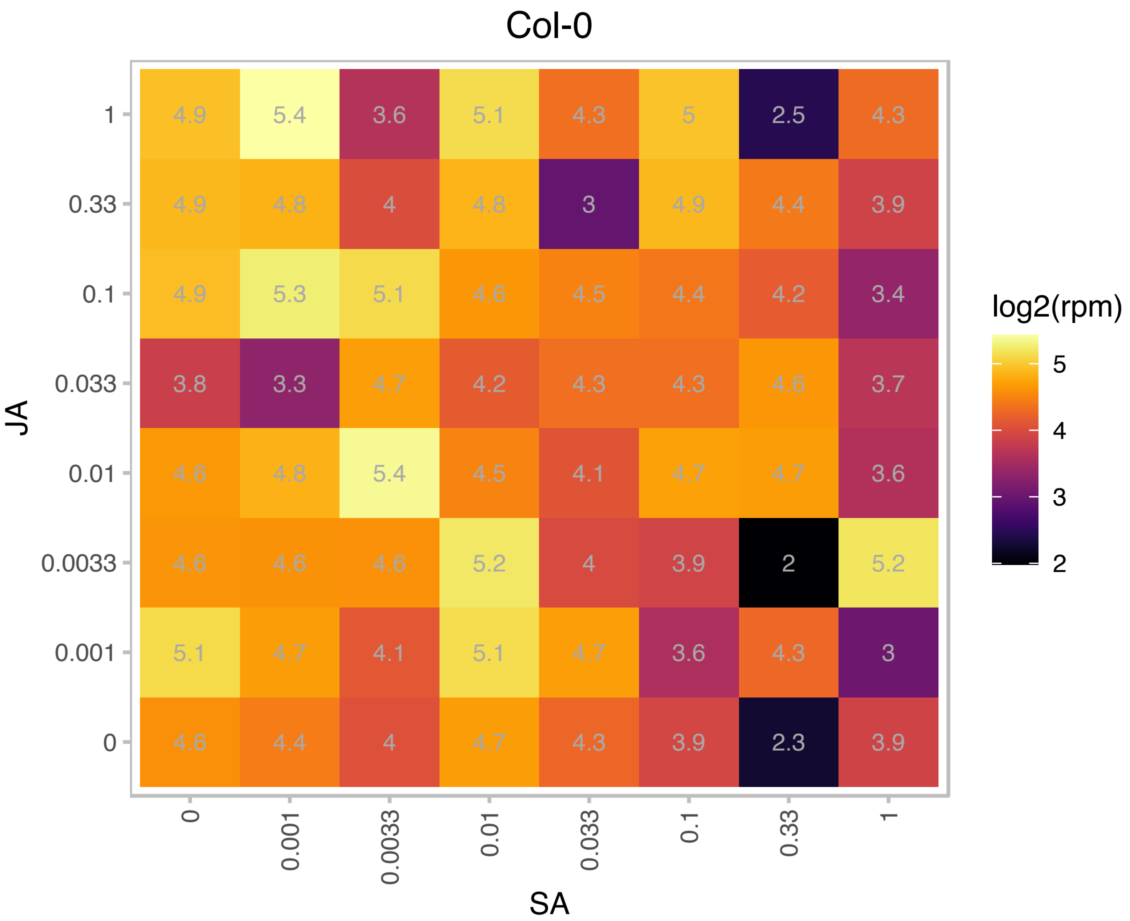


| | |
|---|---|
| Gene | AT2G05830 |
| is.expressed_genes | TRUE |
| is.responsive_genes | TRUE |
| is.apcluster | F-02 |

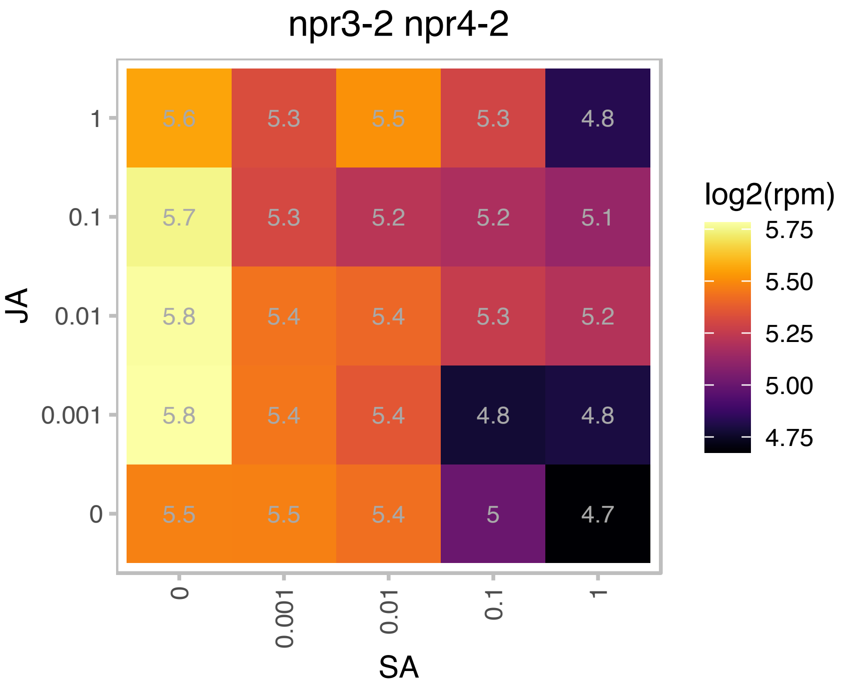


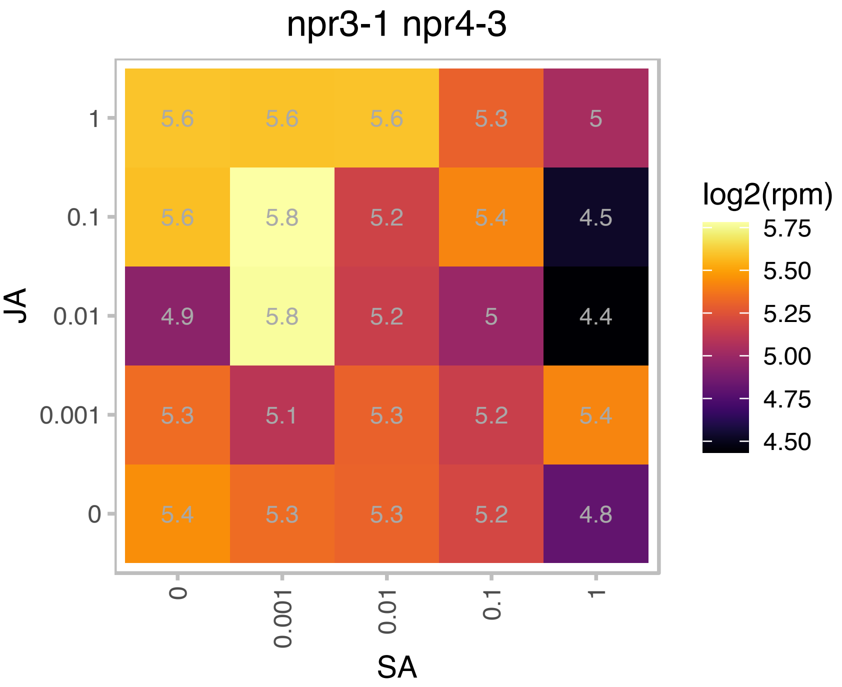

### AT2G05920.1 : SBT1.8

Subtilase family protein

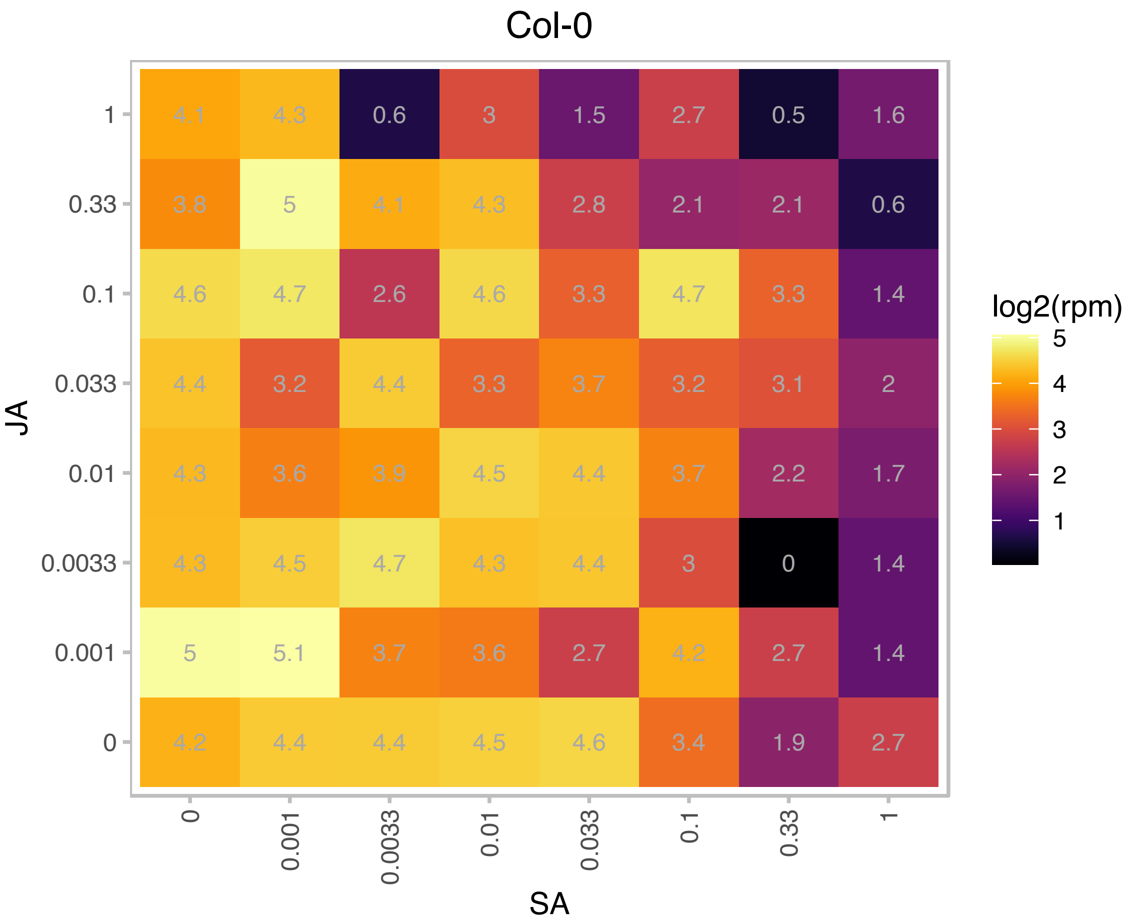


| | |
|---|---|
| Gene | AT2G05920 |
| is.expressed_genes | TRUE |
| is.responsive_genes | TRUE |
| is.apcluster | F-02 |

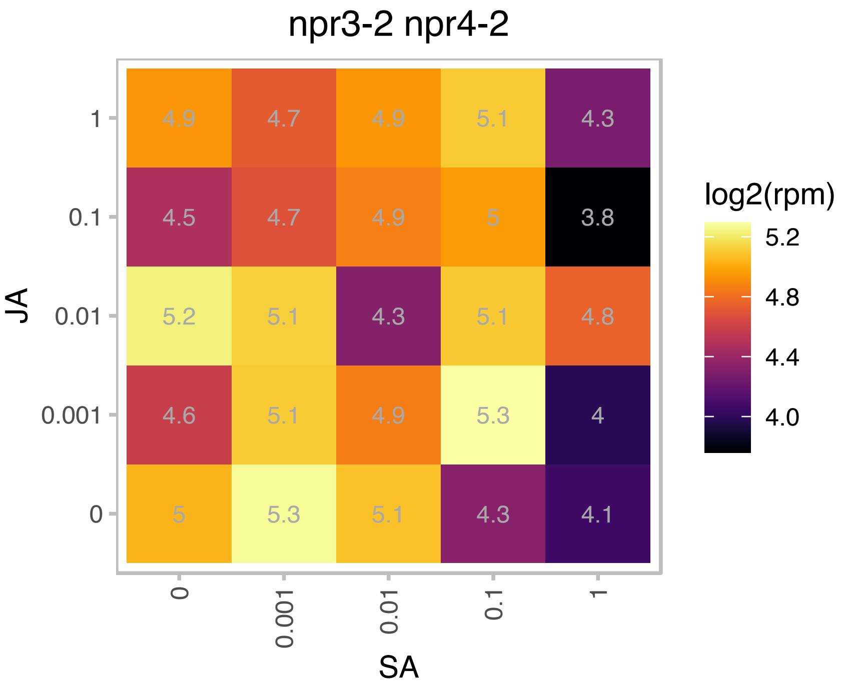


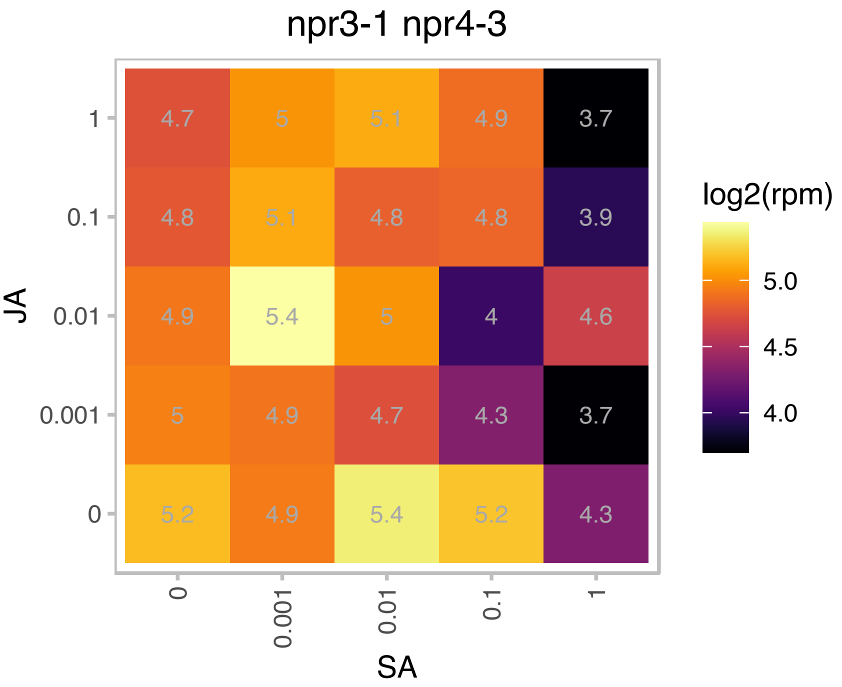

# AT2G10940.1 : NA

Bifunctional inhibitor/lipid-transfer protein/seed storage 2S albumin superfamily protein

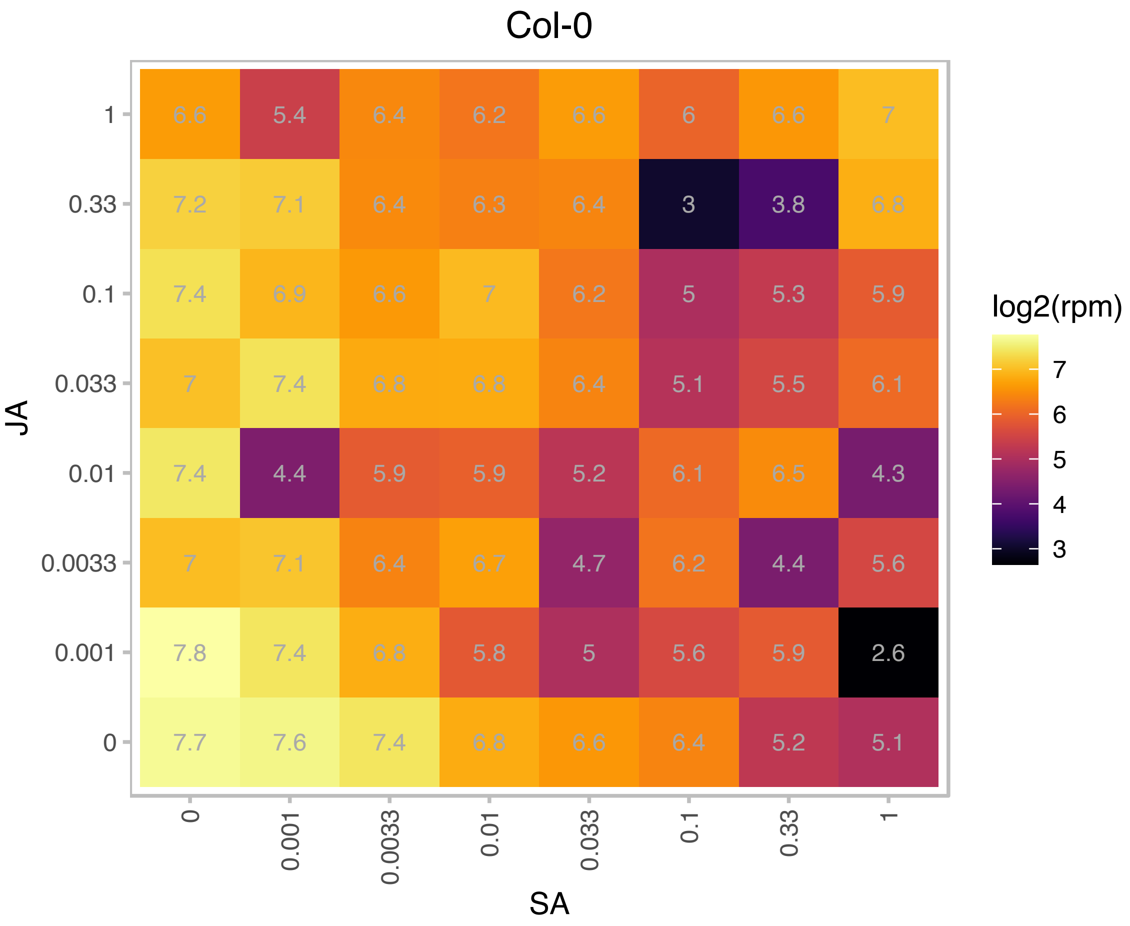


| | |
|---|---|
| Gene | AT2G10940 |
| is.expressed_genes | TRUE |
| is.responsive_genes | TRUE |
| is.apcluster | F-02 |

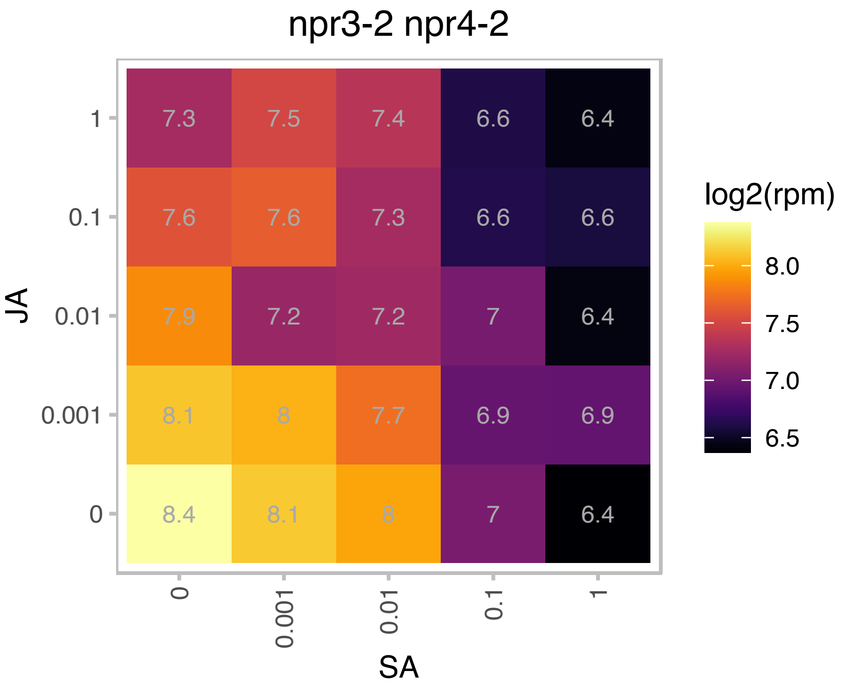


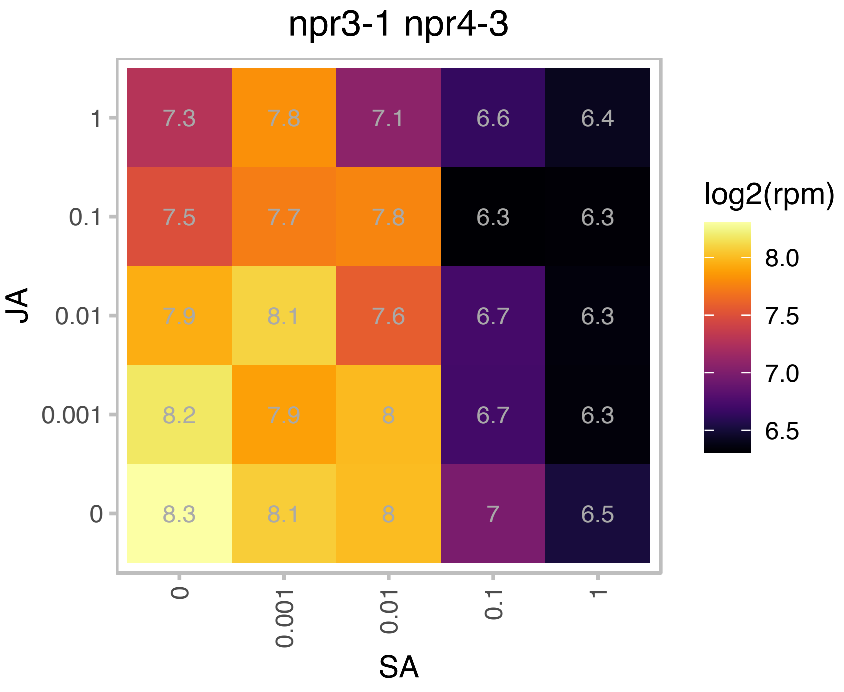

### AT2G17560.1 : HMGB4, NFD04, NFD4

high mobility group B4

Encodes a protein belonging to the subgroup of HMGB (high mobility group B) proteins that have a distinctive DNA-binding motif, the HMG-box domain.  The motif confers non-sequence specific interaction with linear DNA and structure-specific binding to distorted DNA sites.  The HMGB proteins are involved in the assembly of nucleoprotein complexes. Can be phosphorylated by CK2alpha.

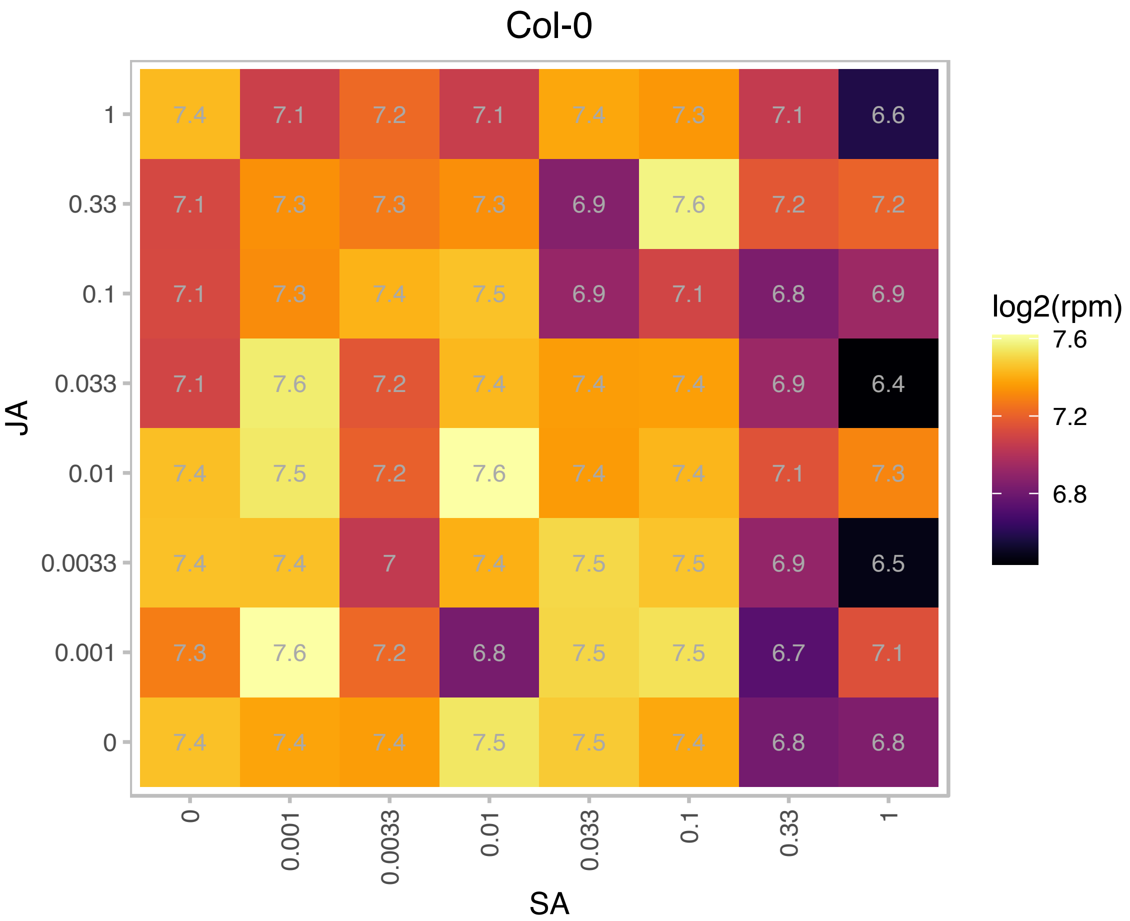


| | |
|---|---|
| Gene | AT2G17560 |
| is.expressed_genes | TRUE |
| is.responsive_genes | TRUE |
| is.apcluster | F-02 |

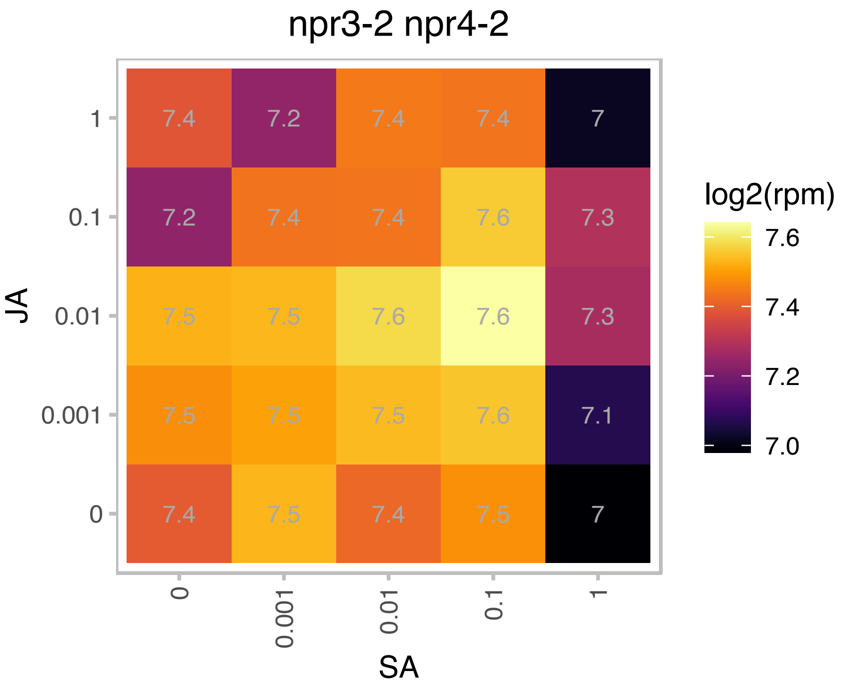


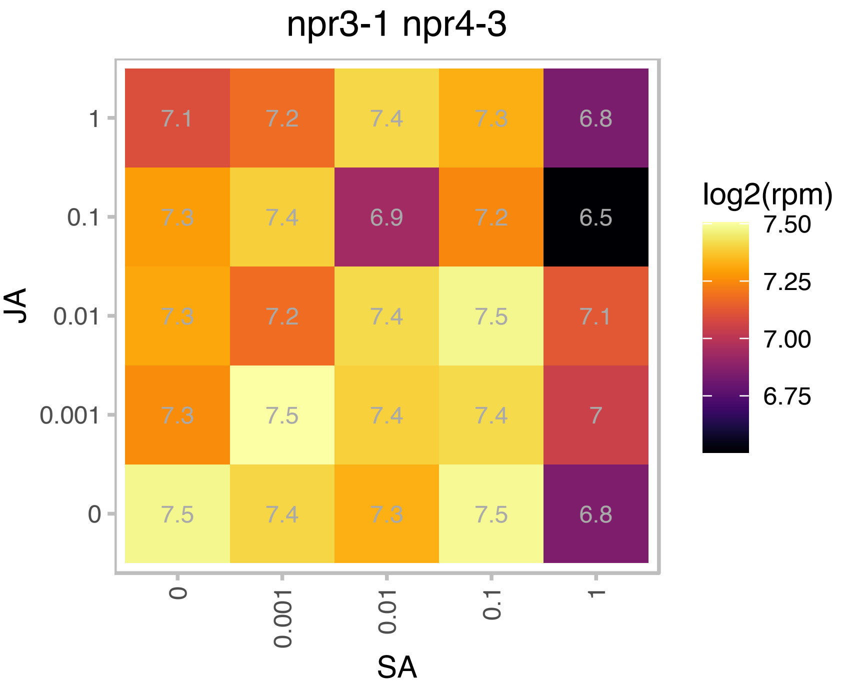

### AT2G18990.1 : TXND9

thioredoxin domain-containing protein 9 homolog

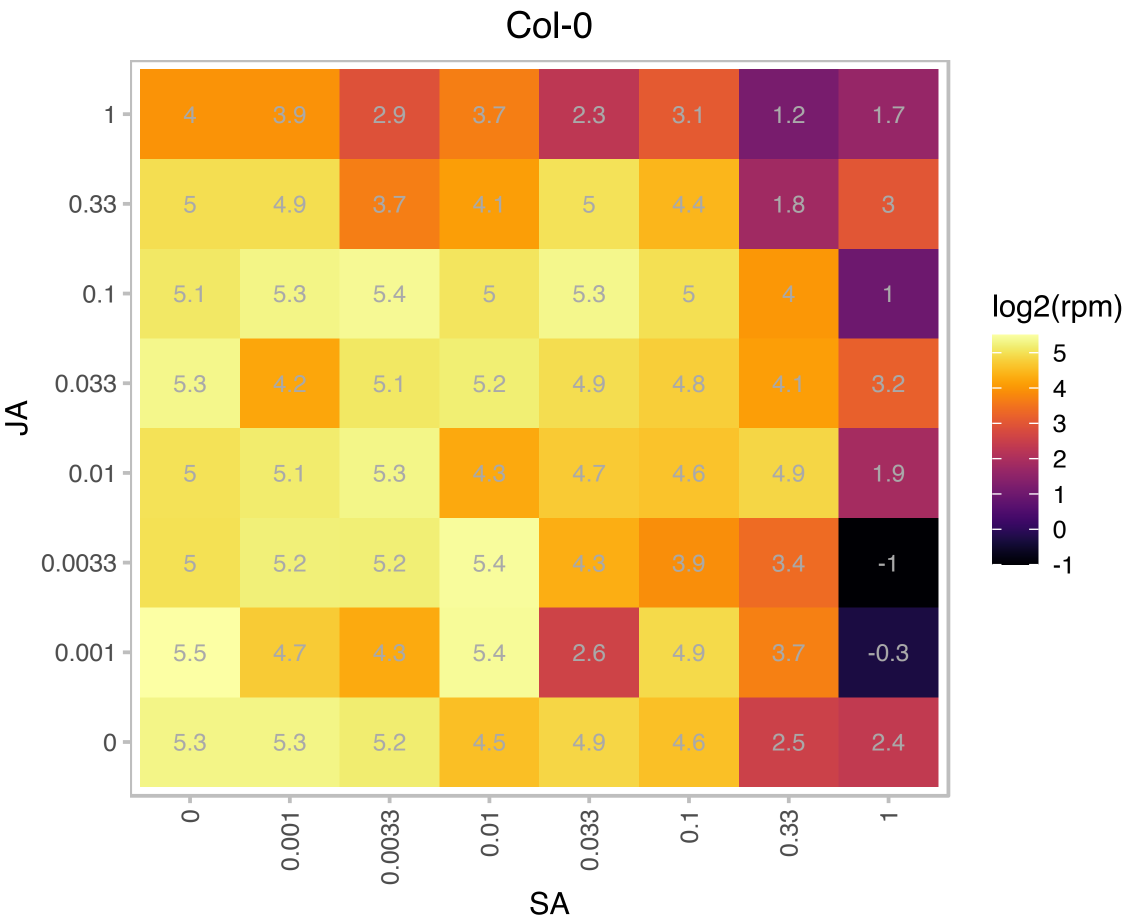


| | |
|---|---|
| Gene | AT2G18990 |
| is.expressed_genes | TRUE |
| is.responsive_genes | TRUE |
| is.apcluster | F-02 |

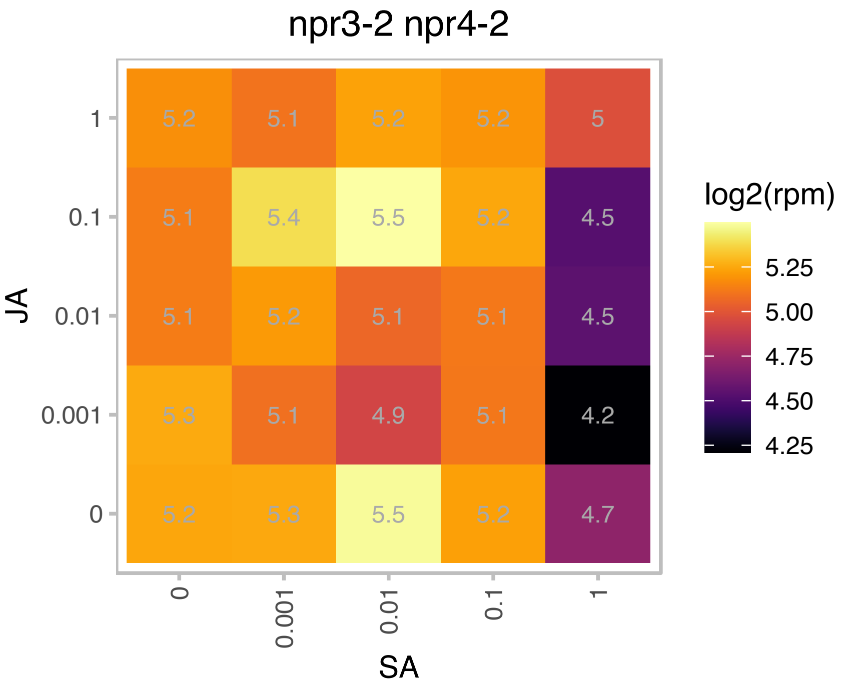


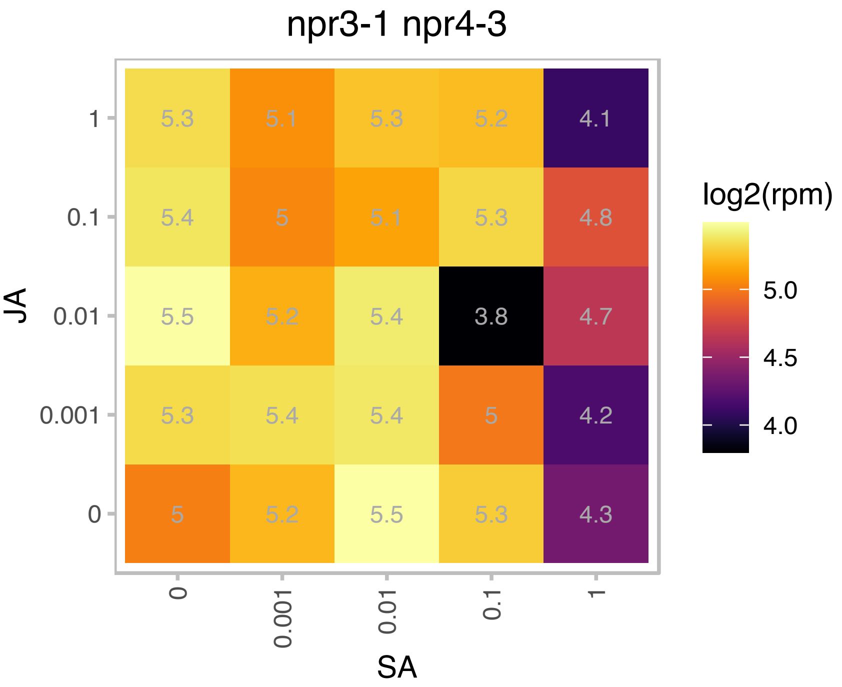

# AT2G22230.1 : NA

Thioesterase superfamily protein

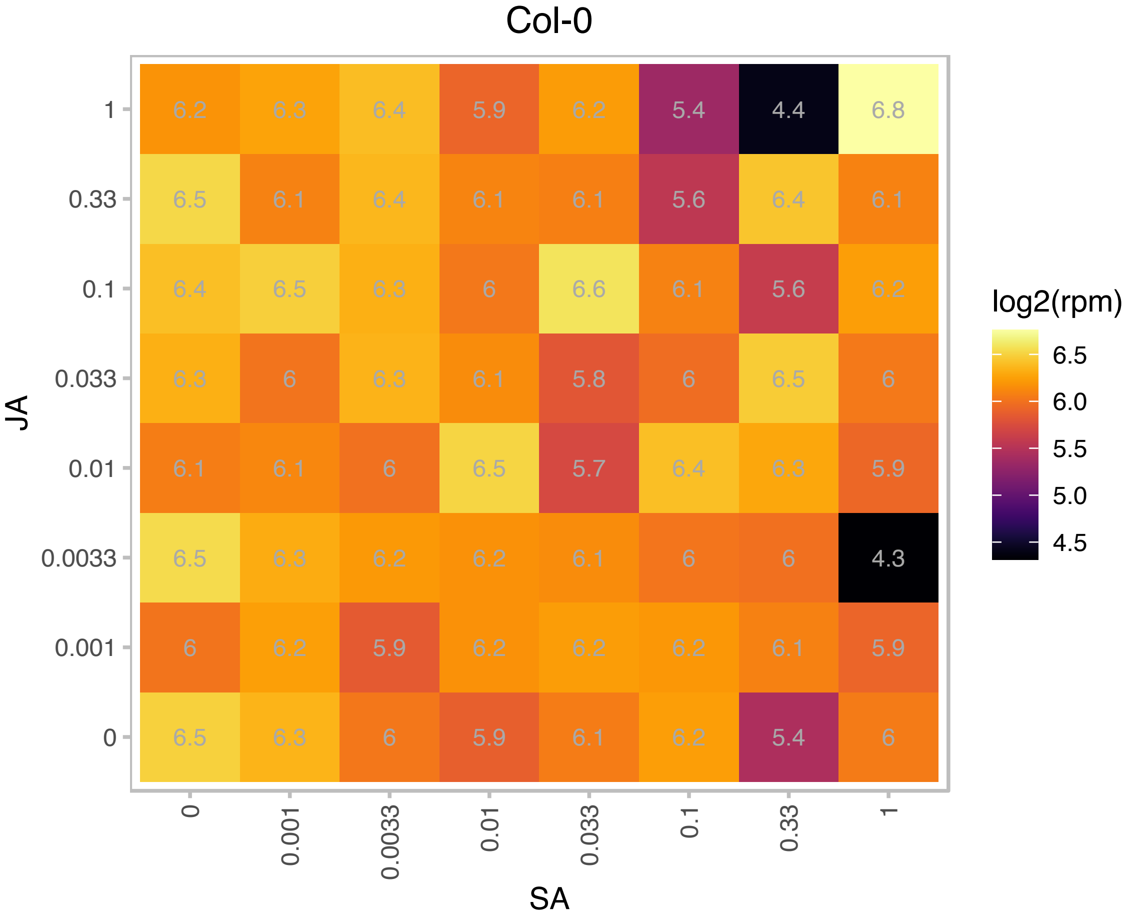


| | |
|---|---|
| Gene | AT2G22230 |
| is.expressed_genes | TRUE |
| is.responsive_genes | TRUE |
| is.apcluster | F-02 |

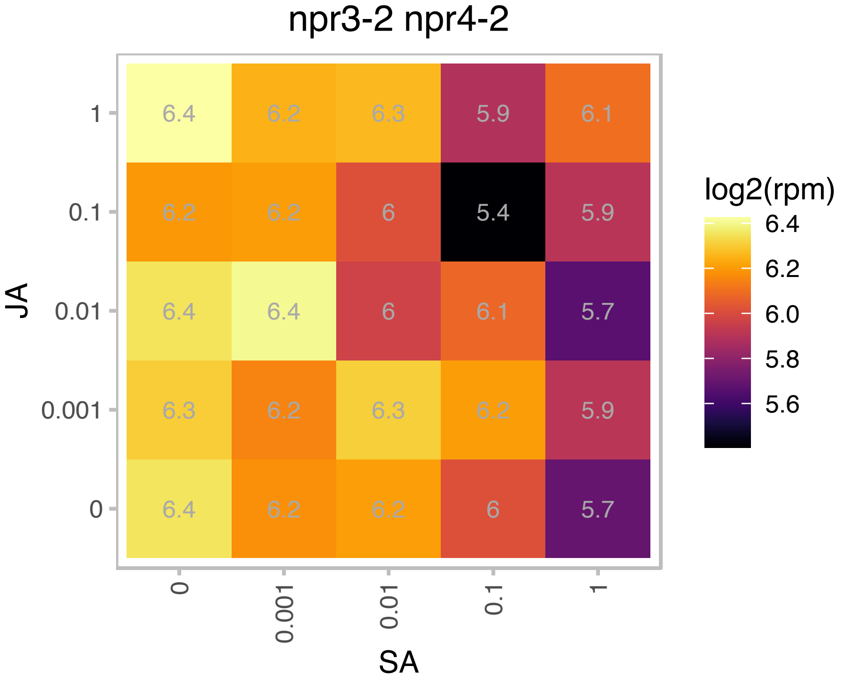


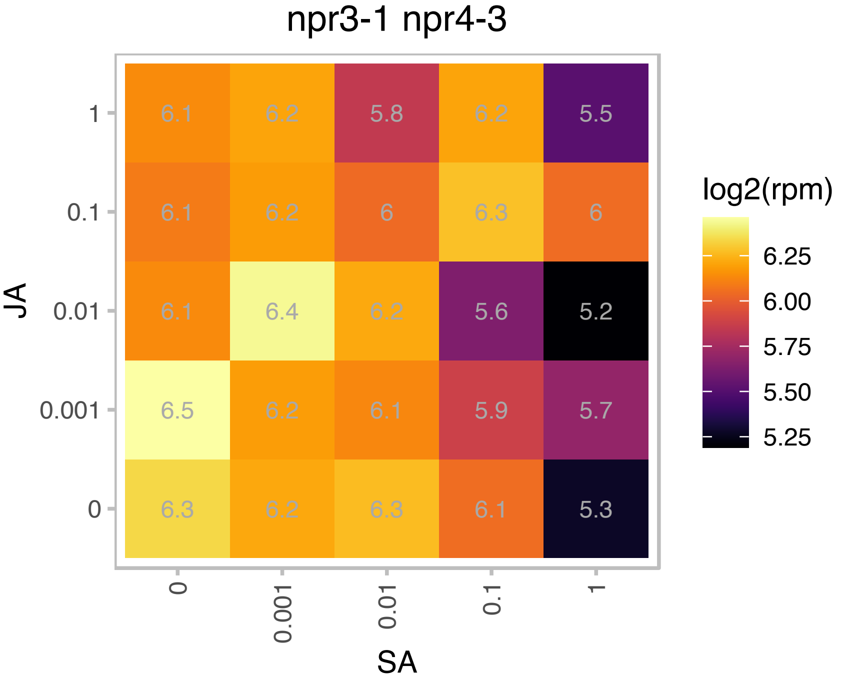

### AT2G24980.1 : EXT6

Proline-rich extensin-like family protein

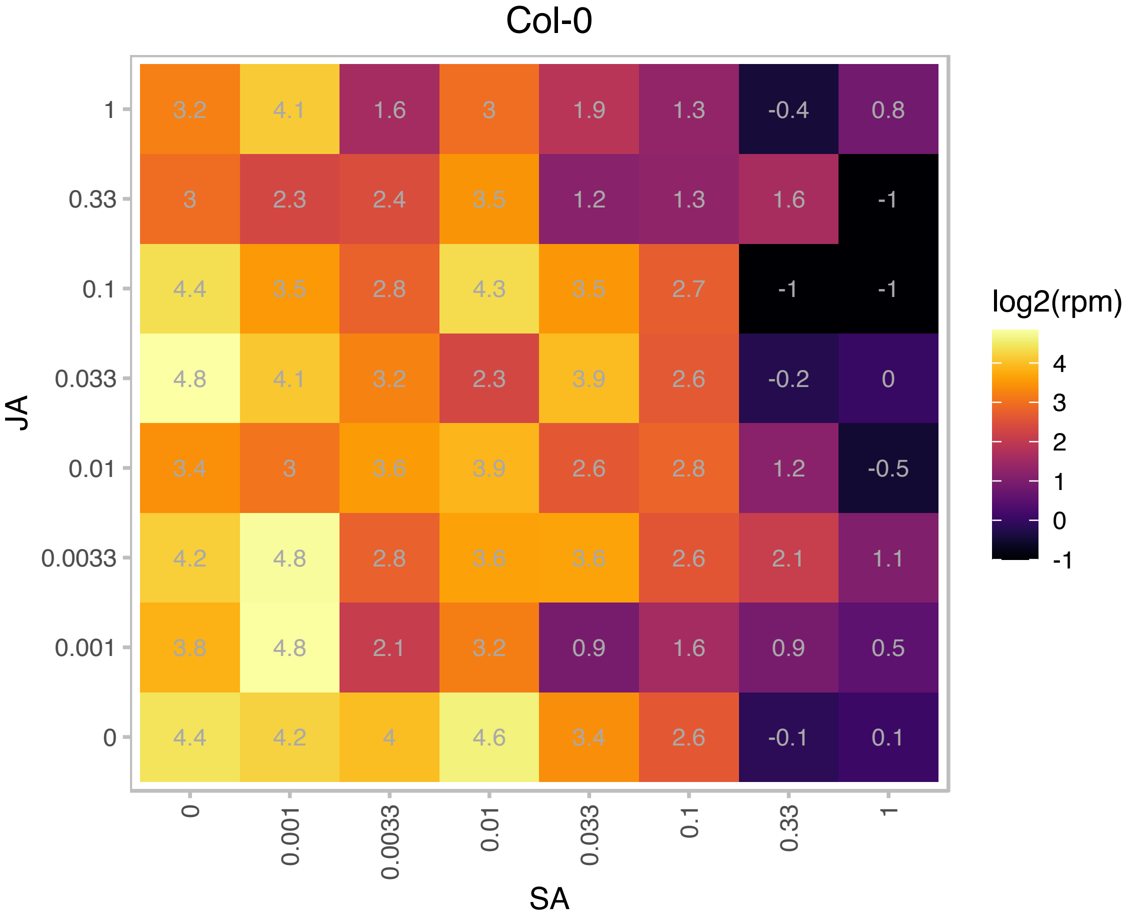


| | |
|---|---|
| Gene | AT2G24980 |
| is.expressed_genes | TRUE |
| is.responsive_genes | TRUE |
| is.apcluster | F-02 |

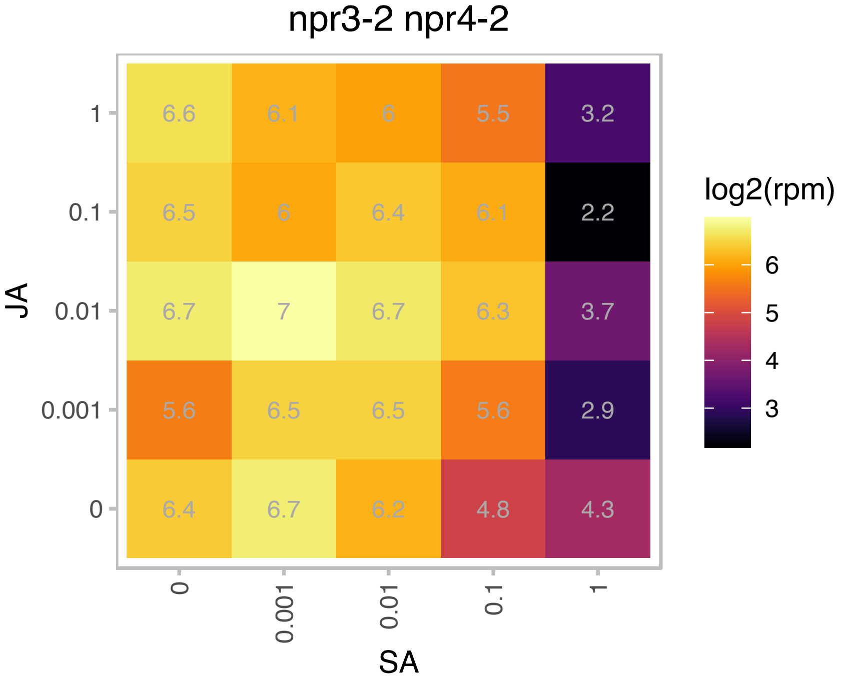


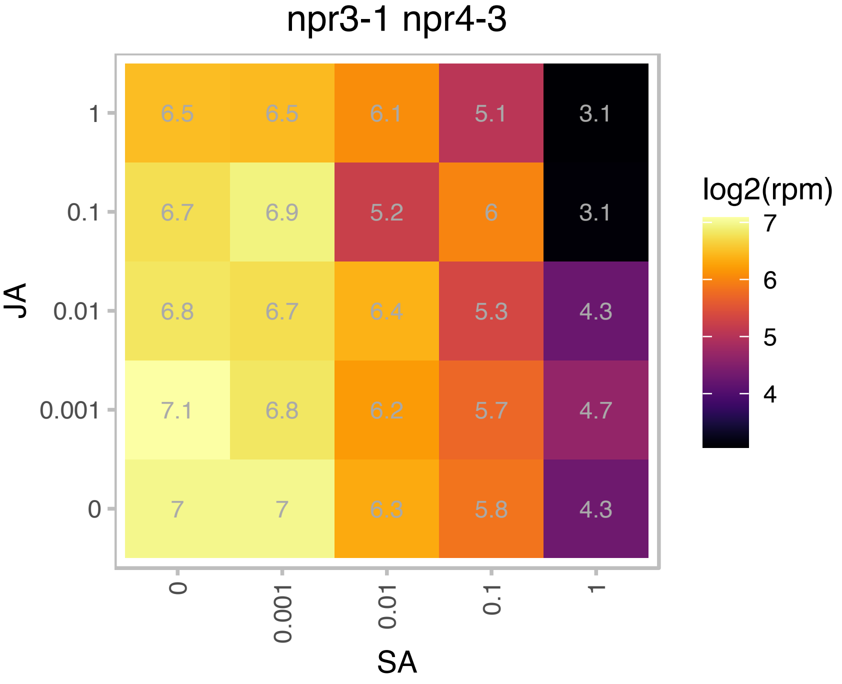

# AT2G28600.1 : NA

P-loop containing nucleoside triphosphate hydrolases superfamily protein

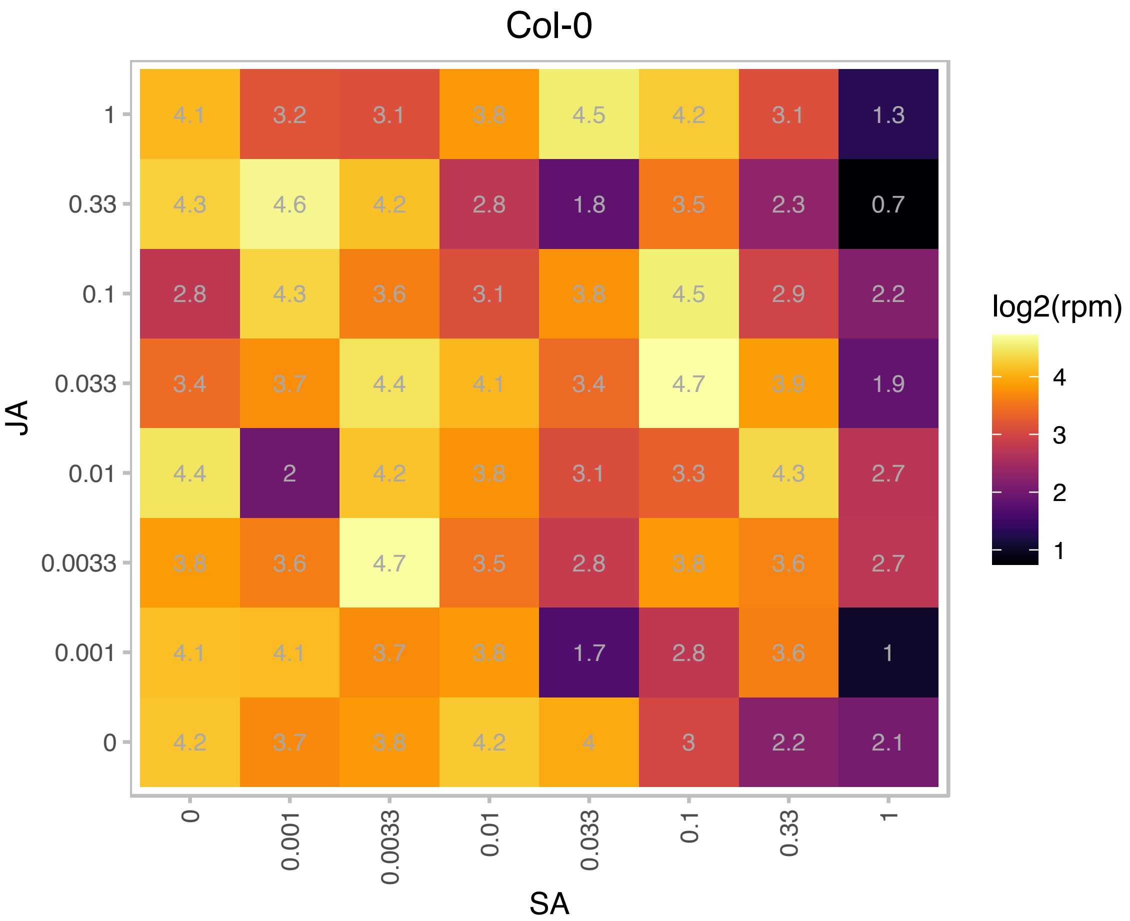


| | |
|---|---|
| Gene | AT2G28600 |
| is.expressed_genes | TRUE |
| is.responsive_genes | TRUE |
| is.apcluster | F-02 |

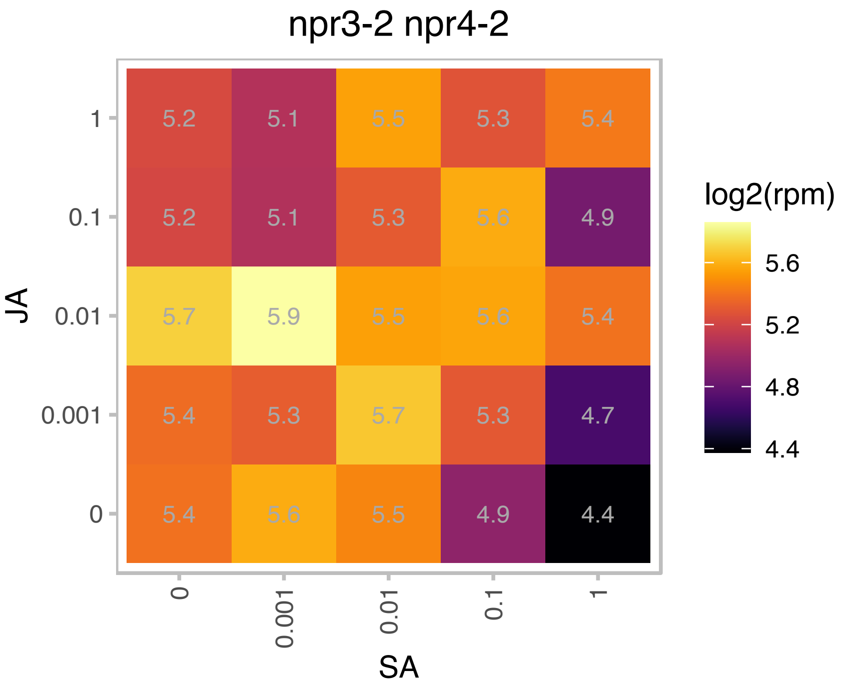


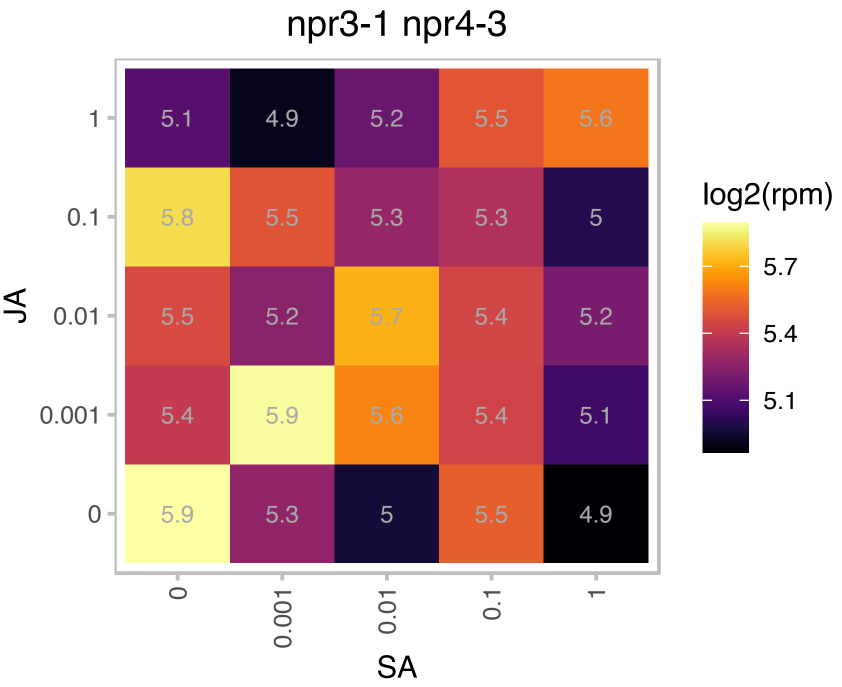

# AT2G29170.1 : NA

NAD(P)-binding Rossmann-fold superfamily protein

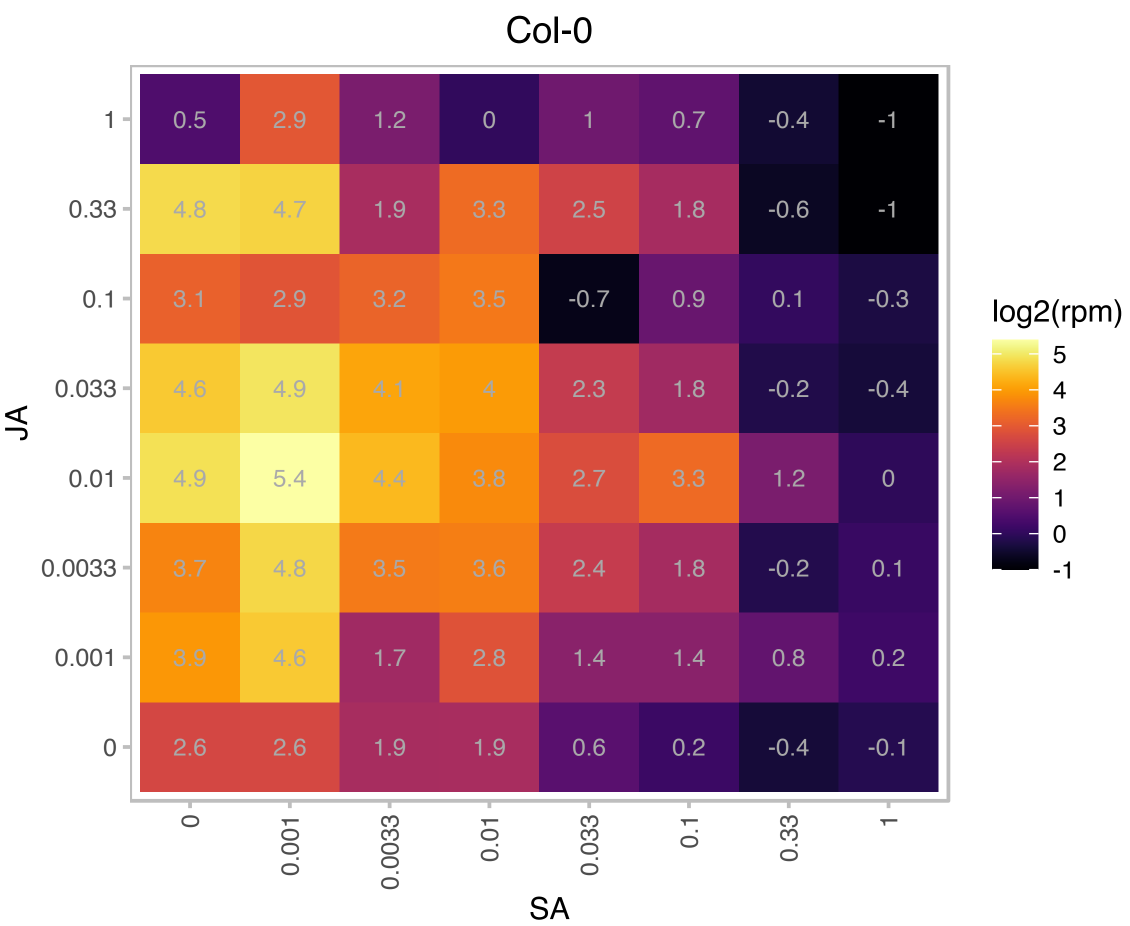


| | |
|---|---|
| Gene | AT2G29170 |
| is.expressed_genes | TRUE |
| is.responsive_genes | TRUE |
| is.apcluster | F-02 |

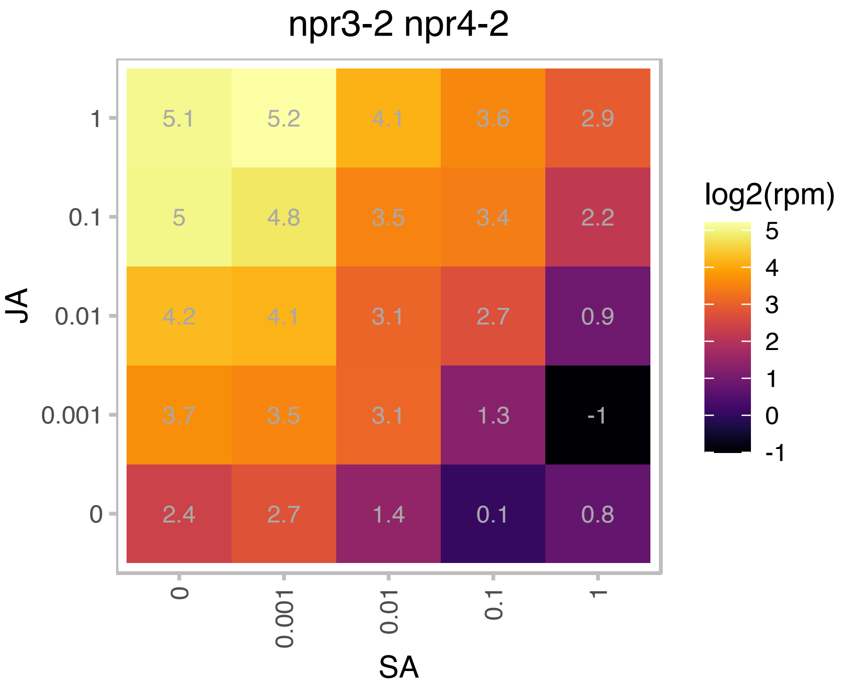


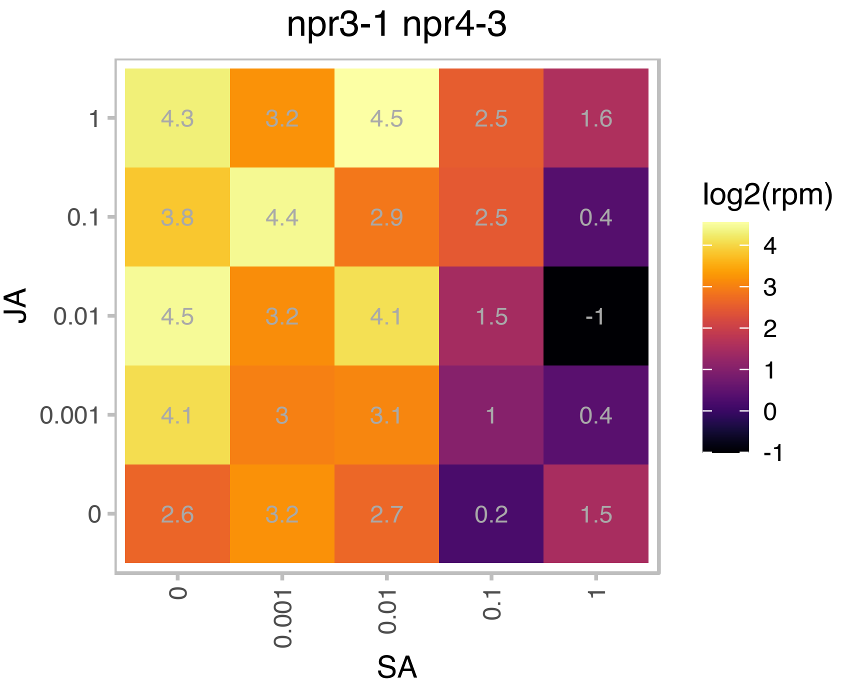

# AT2G31560.1 : NA

Protein of unknown function (DUF1685)

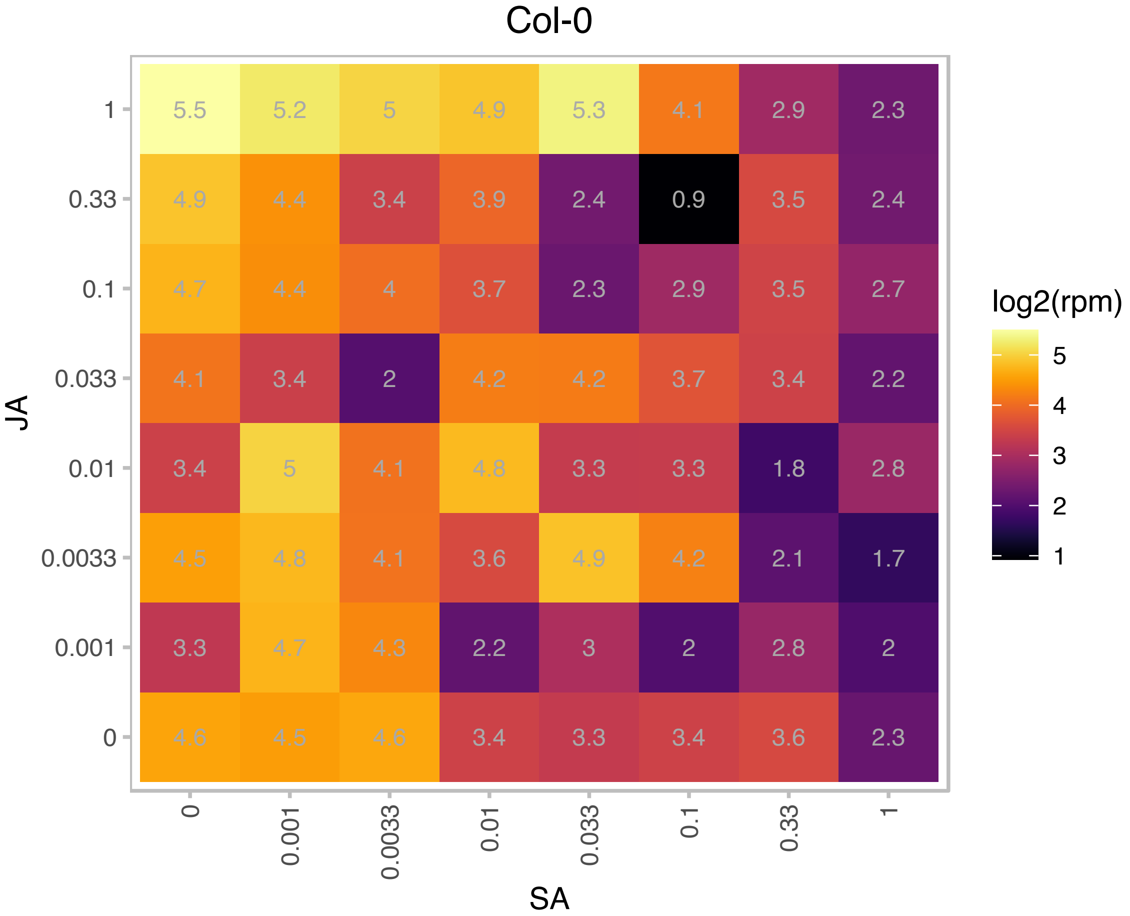


| | |
|---|---|
| Gene | AT2G31560 |
| is.expressed_genes | TRUE |
| is.responsive_genes | TRUE |
| is.apcluster | F-02 |

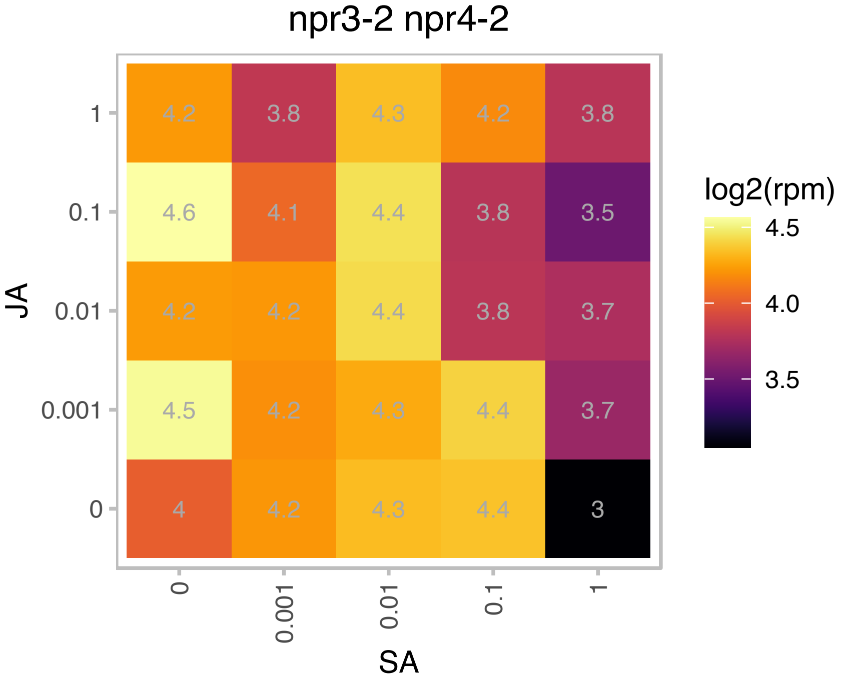


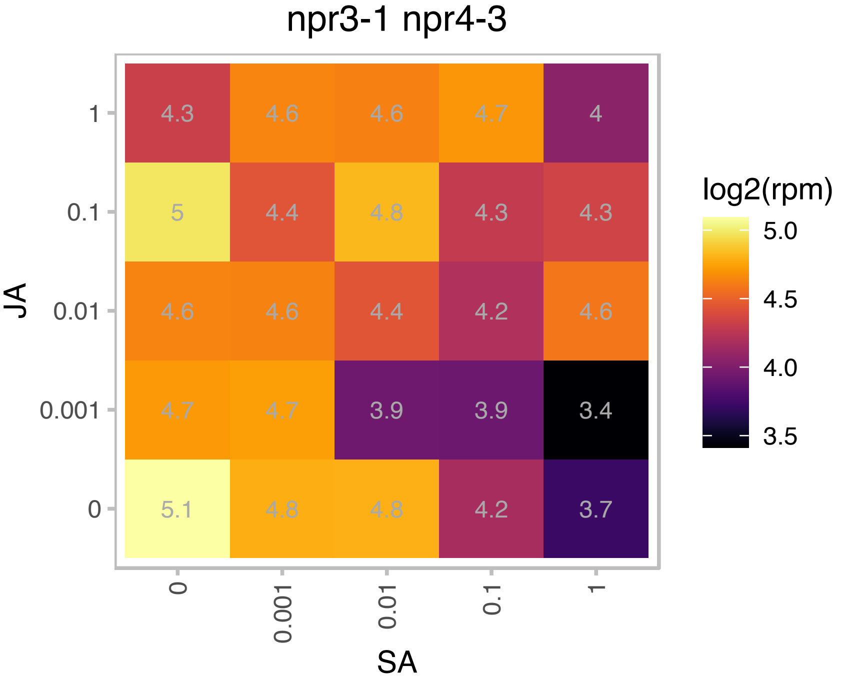

### AT2G37170.1 : PIP2;2, PIP2B

plasma membrane intrinsic protein 2
a member of the plasma membrane intrinsic protein subfamily PIP2. localizes to the plasma membrane and exhibits water transport activity in Xenopus oocyte. expressed specifically in the vascular bundles and protein level increases slightly during leaf dev

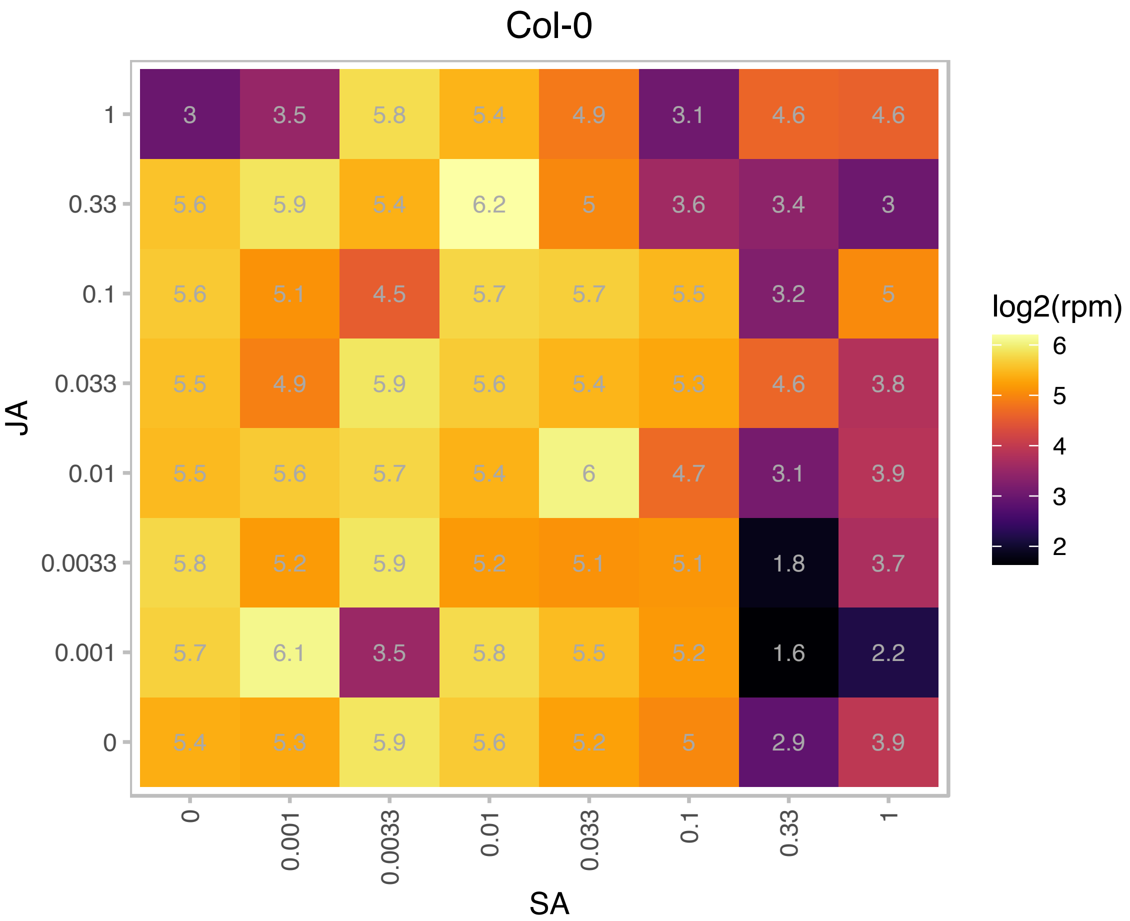


| | |
|---|---|
| Gene | AT2G37170 |
| is.expressed_genes | TRUE |
| is.responsive_genes | TRUE |
| is.apcluster | F-02 |

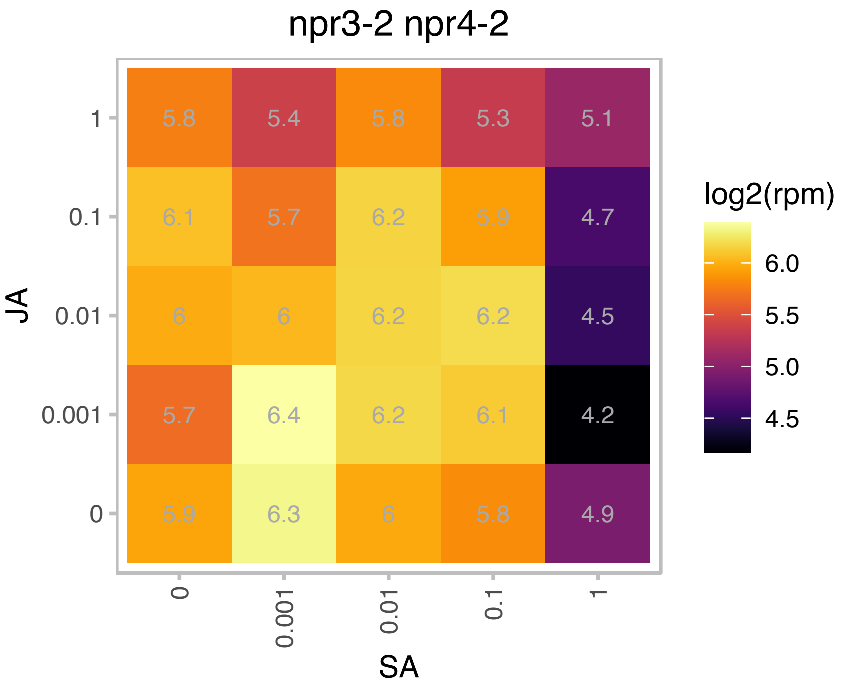


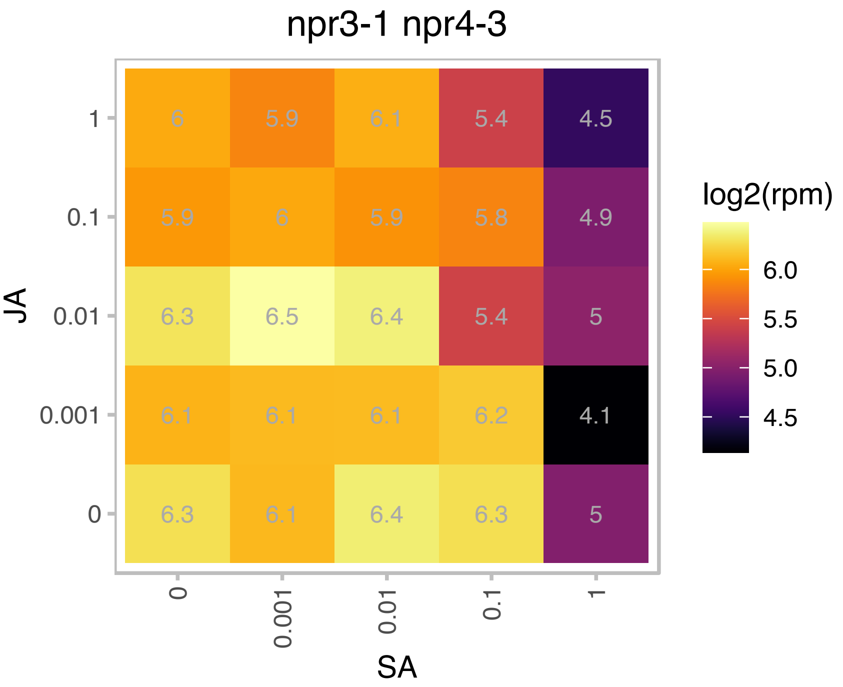

### AT2G37790.1 : AKR4C10

NAD(P)-linked oxidoreductase superfamily protein

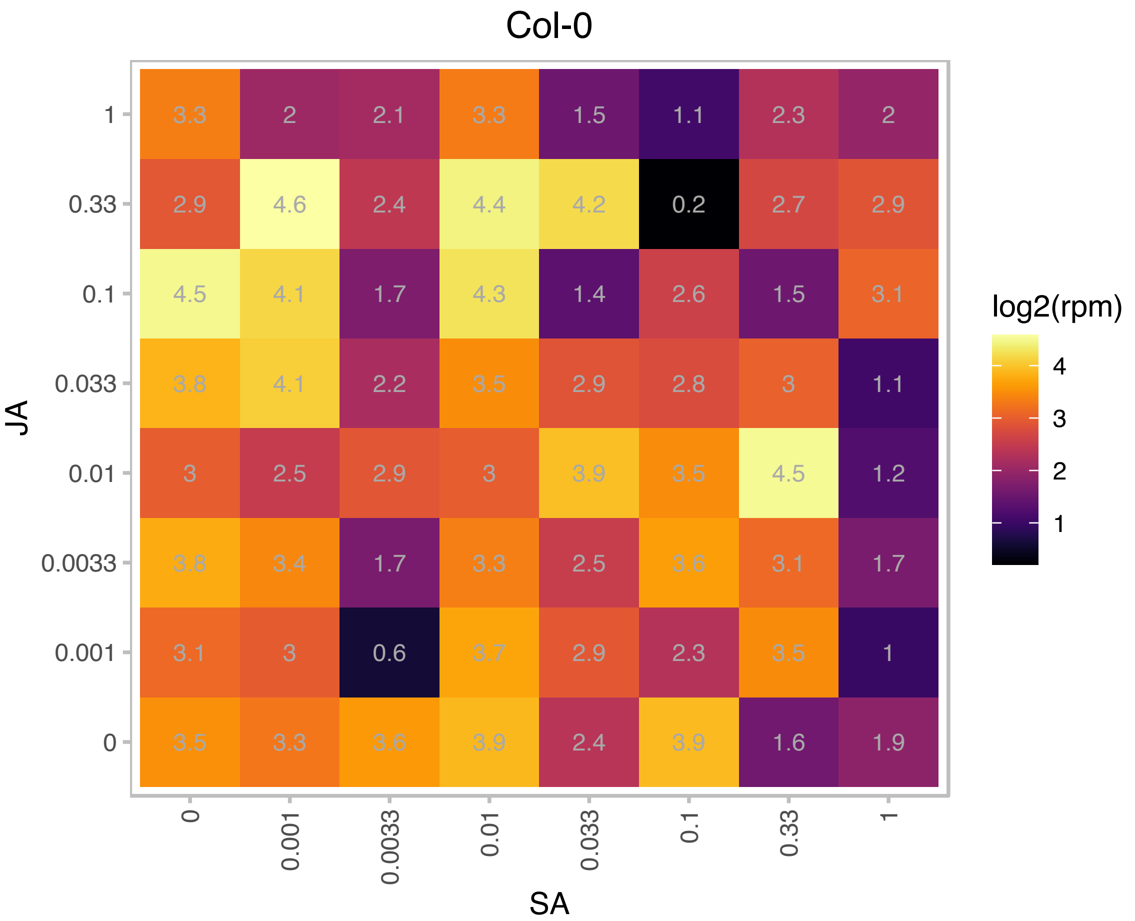


| | |
|---|---|
| Gene | AT2G37790 |
| is.expressed_genes | TRUE |
| is.responsive_genes | TRUE |
| is.apcluster | F-02 |

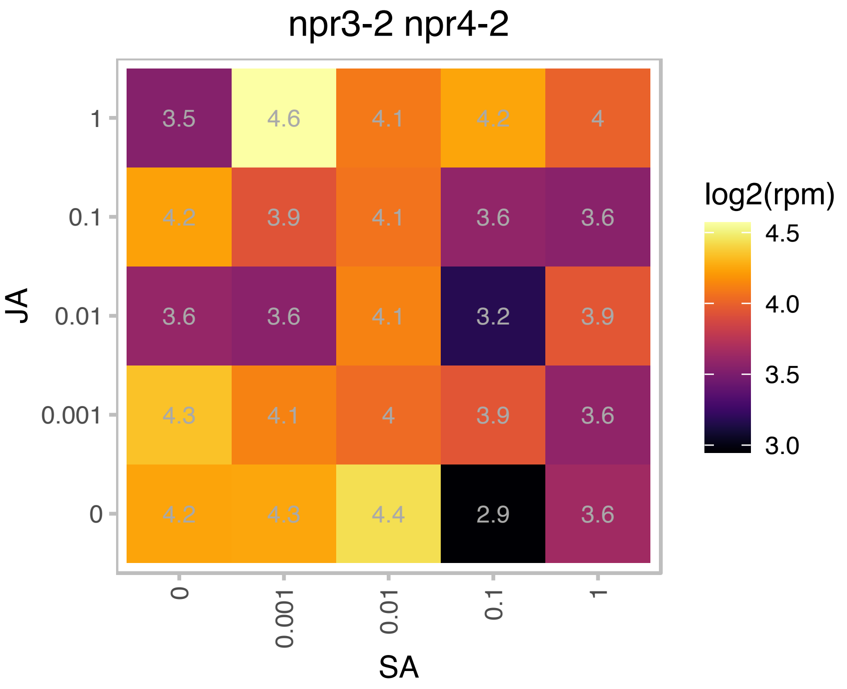


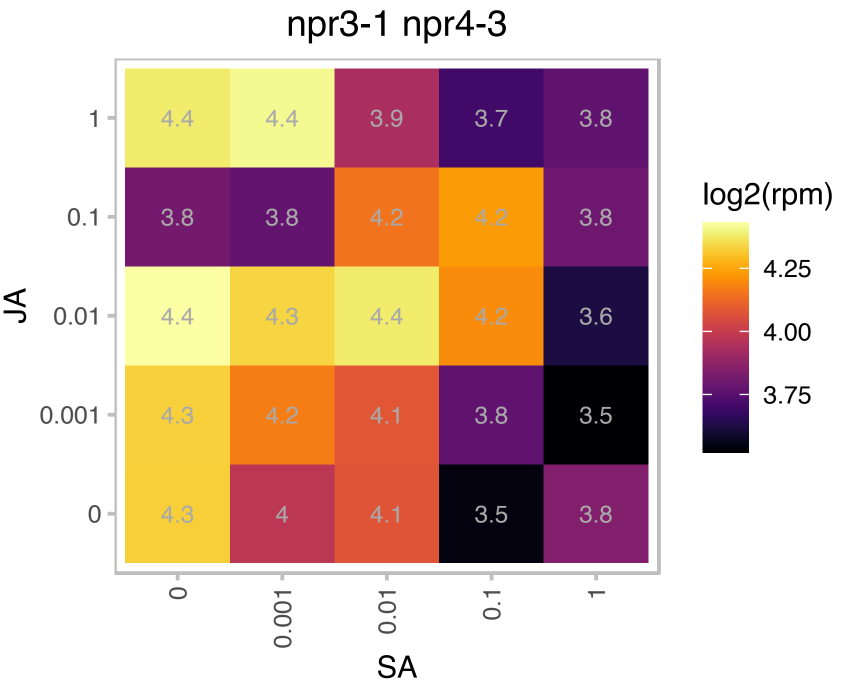

# AT2G37990.1 : NA

ribosome biogenesis regulatory protein (RRS1) family protein

| | |
|---|---|
| Gene | AT2G37990 |
| is.expressed_genes | TRUE |
| is.responsive_genes | TRUE |
| is.apcluster | F-02 |

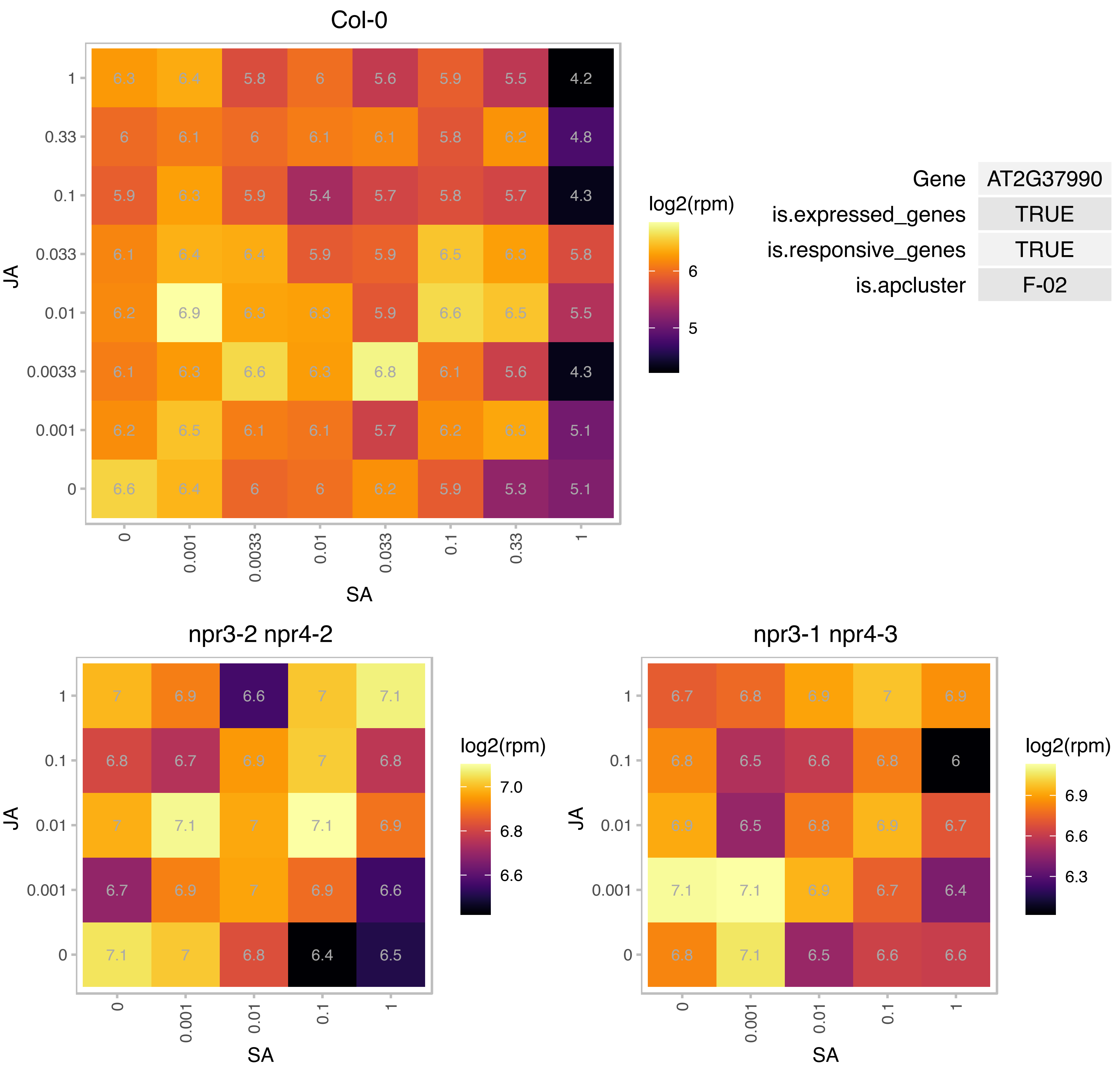

### AT2G39990.1 : AteIF3f, EIF2, eIF3F

eukaryotic translation initiation factor 2

translation initiation factor eIF2 p47 subunit homolog

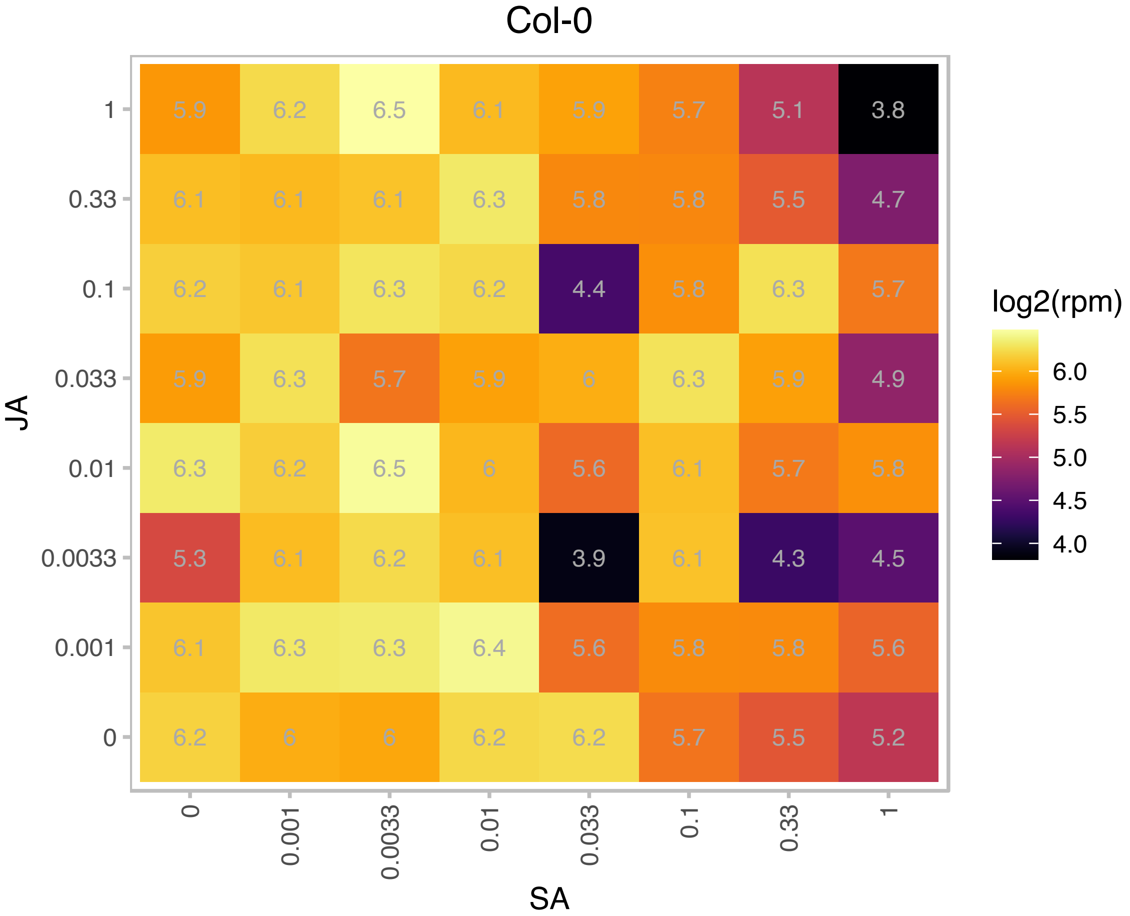


| | |
|---|---|
| Gene | AT2G39990 |
| is.expressed_genes | TRUE |
| is.responsive_genes | TRUE |
| is.apcluster | F-02 |

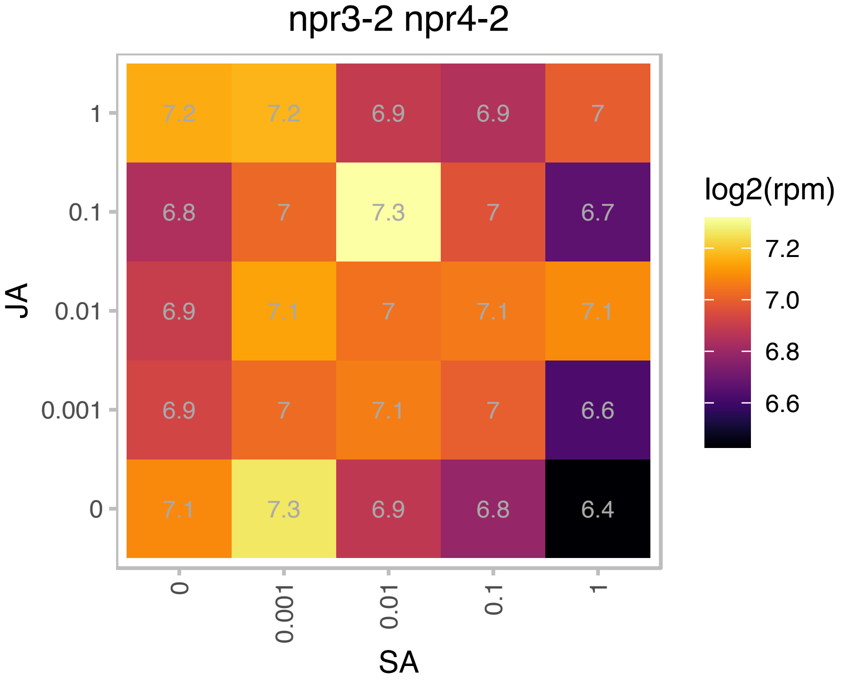


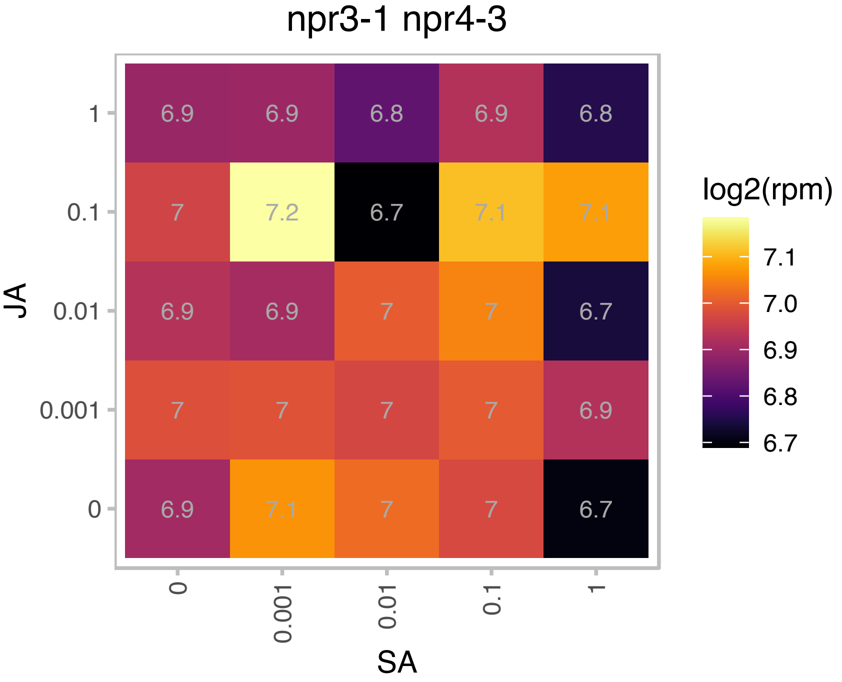

### AT2G40400.1 : BPG3, RER5

Protein of unknown function (DUF399 and DUF3411)

Encodes a chloroplast localized protein of unknown function that is involved in regulation of chloroplast development.

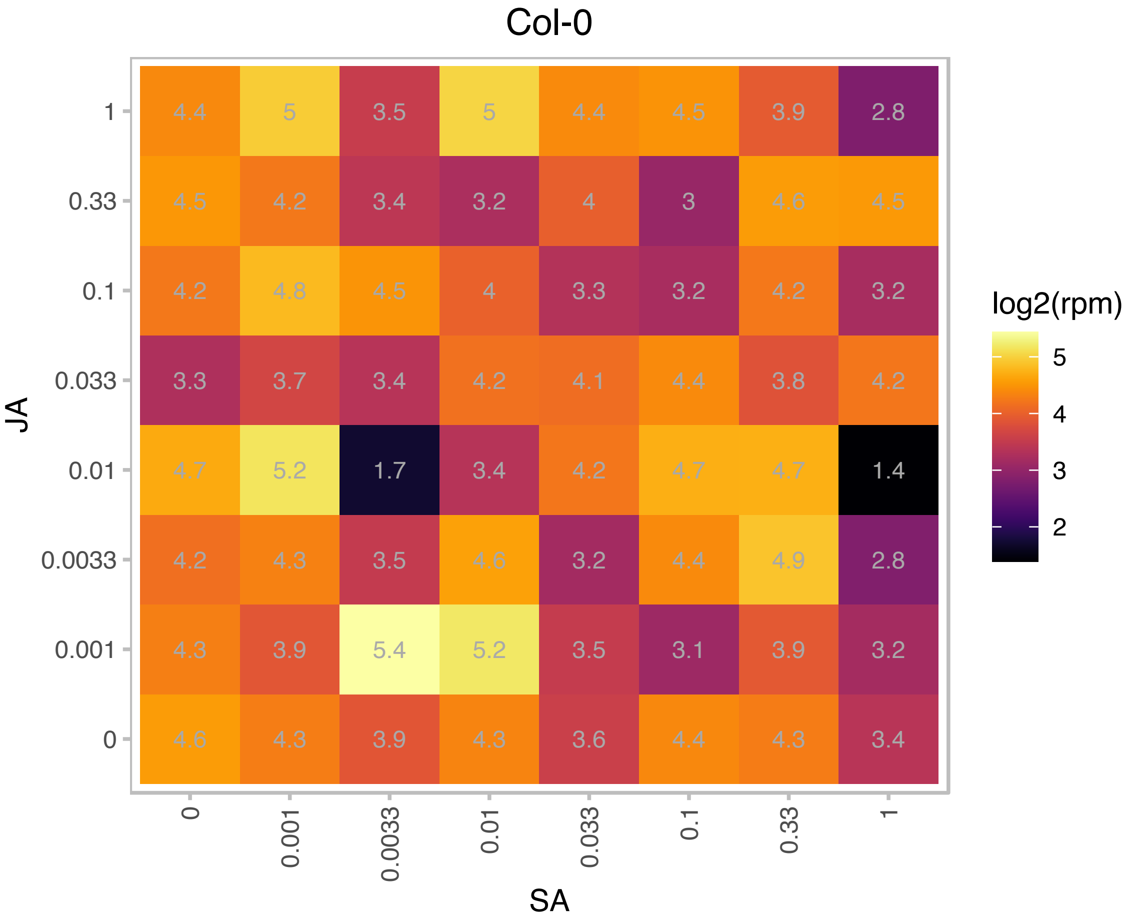


| | |
|---|---|
| Gene | AT2G40400 |
| is.expressed_genes | TRUE |
| is.responsive_genes | TRUE |
| is.apcluster | F-02 |

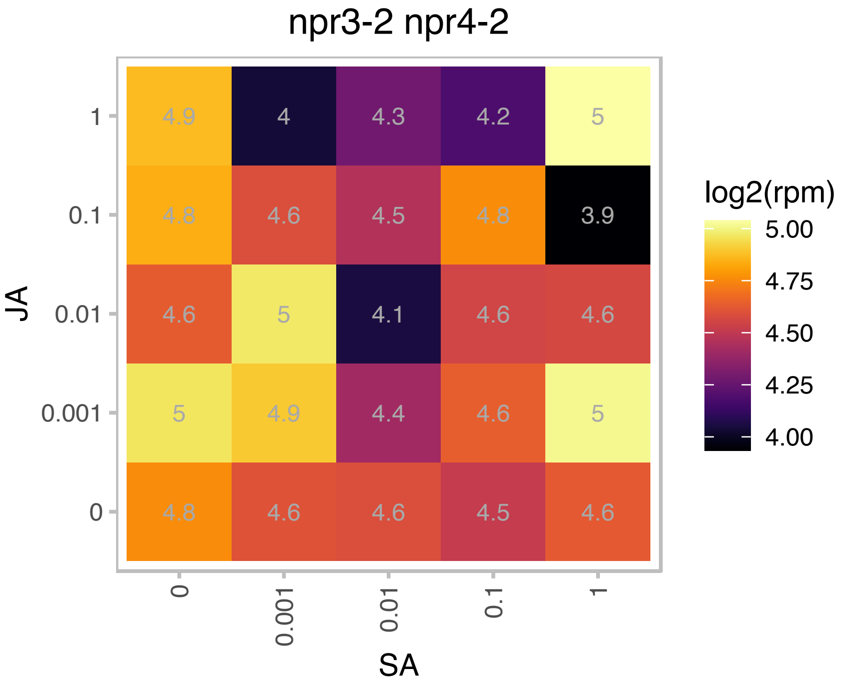


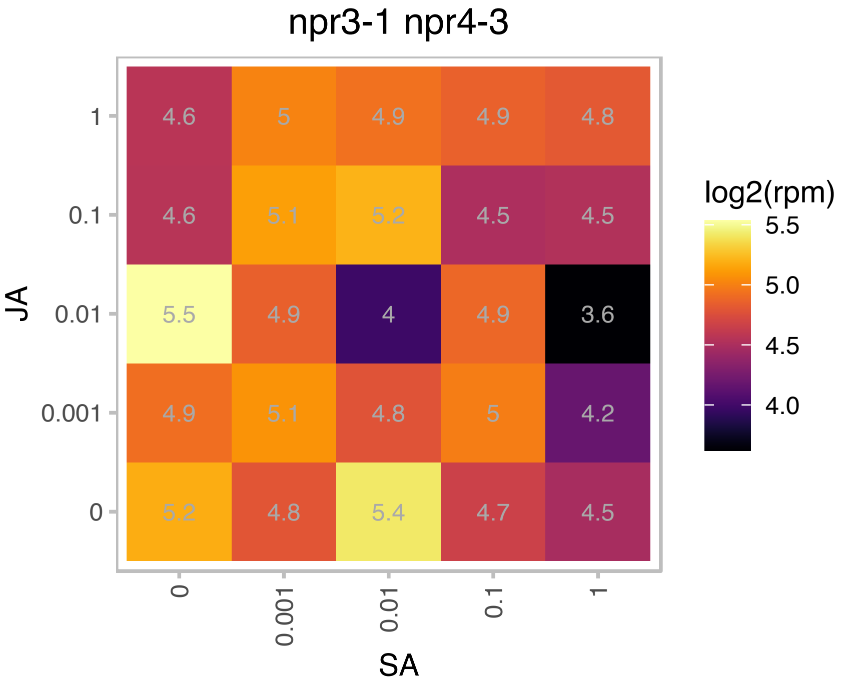

### AT2G40430.1 : SMO4

Encodes a homolog of yeast NOP53 that is an important regulator of cell division during organ growth.

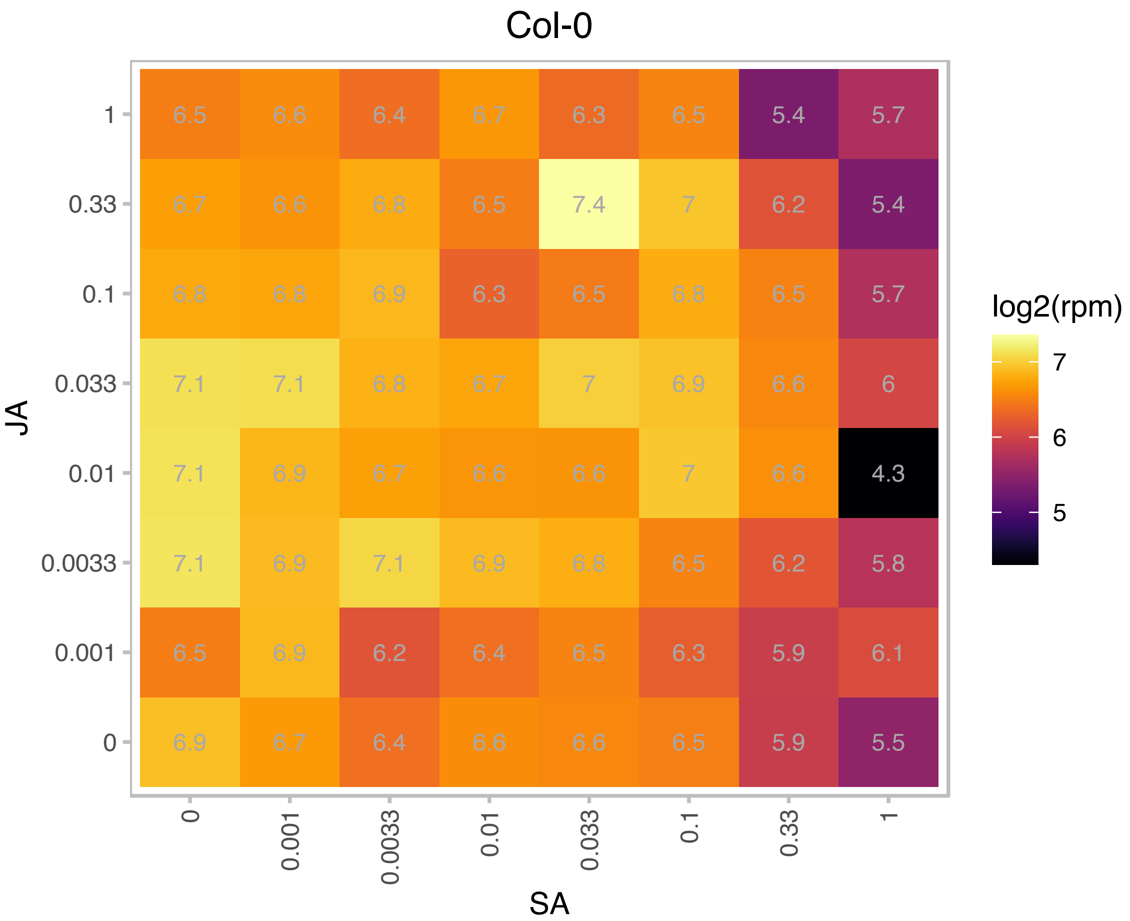


| | |
|---|---|
| Gene | AT2G40430 |
| is.expressed_genes | TRUE |
| is.responsive_genes | TRUE |
| is.apcluster | F-02 |

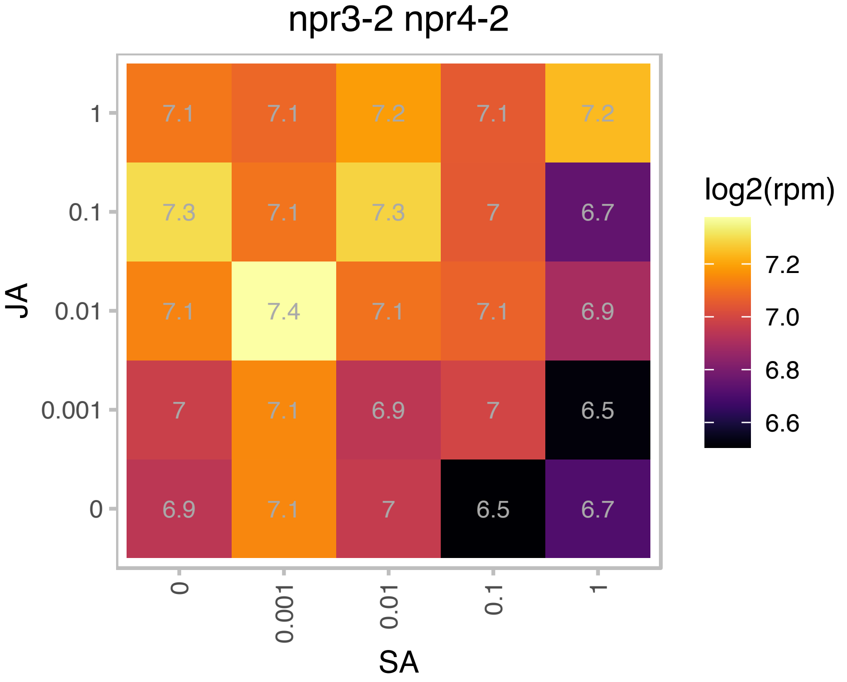


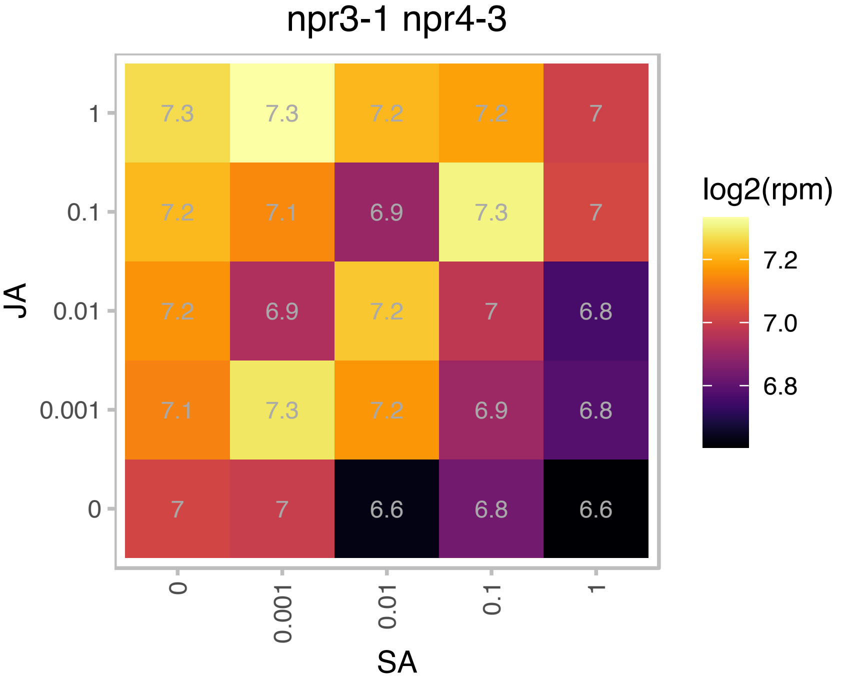

### AT2G43100.1 : ATLEUD1, IPMI2

isopropylmalate isomerase 2

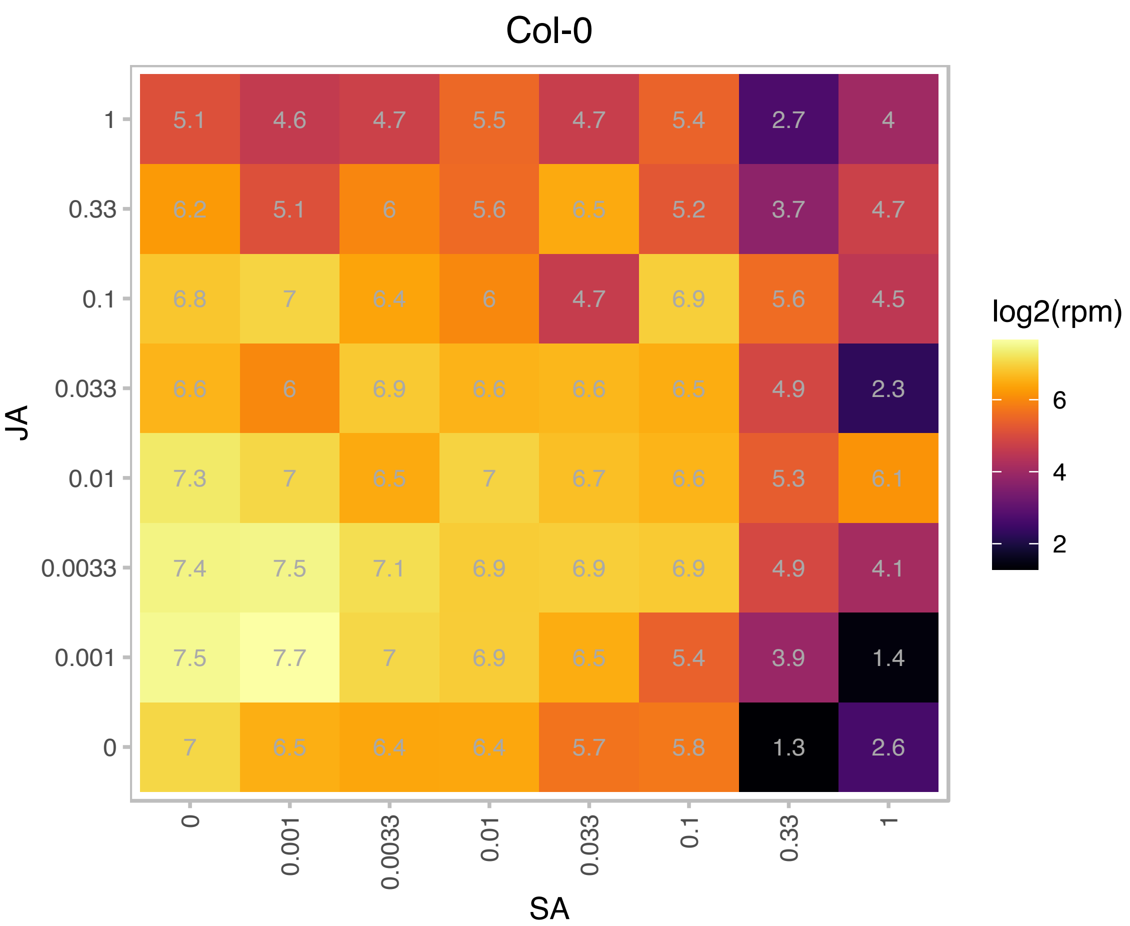


| | |
|---|---|
| Gene | AT2G43100 |
| is.expressed_genes | TRUE |
| is.responsive_genes | TRUE |
| is.apcluster | F-02 |

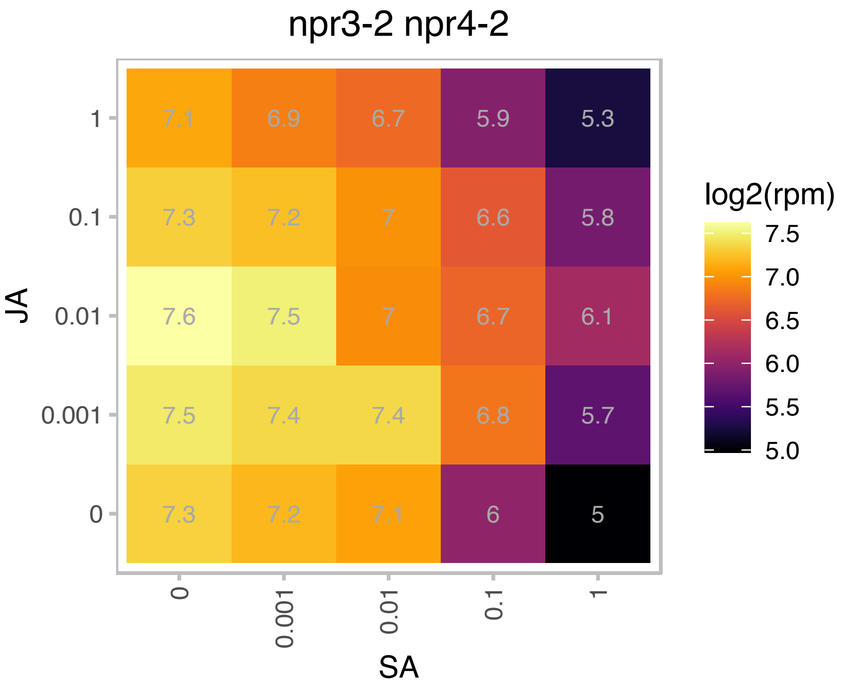


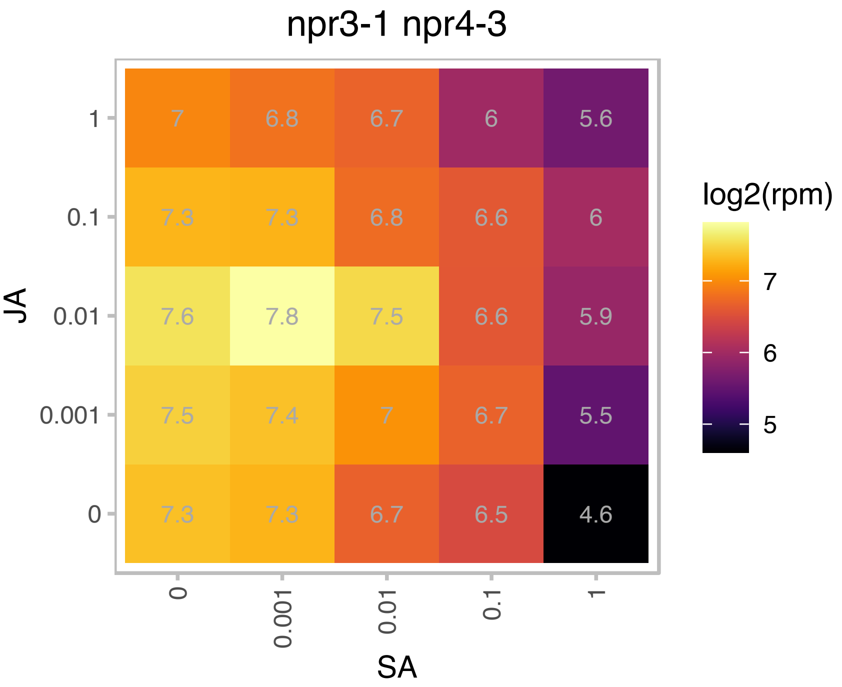

# AT2G43150.1 : NA

Proline-rich extensin-like family protein

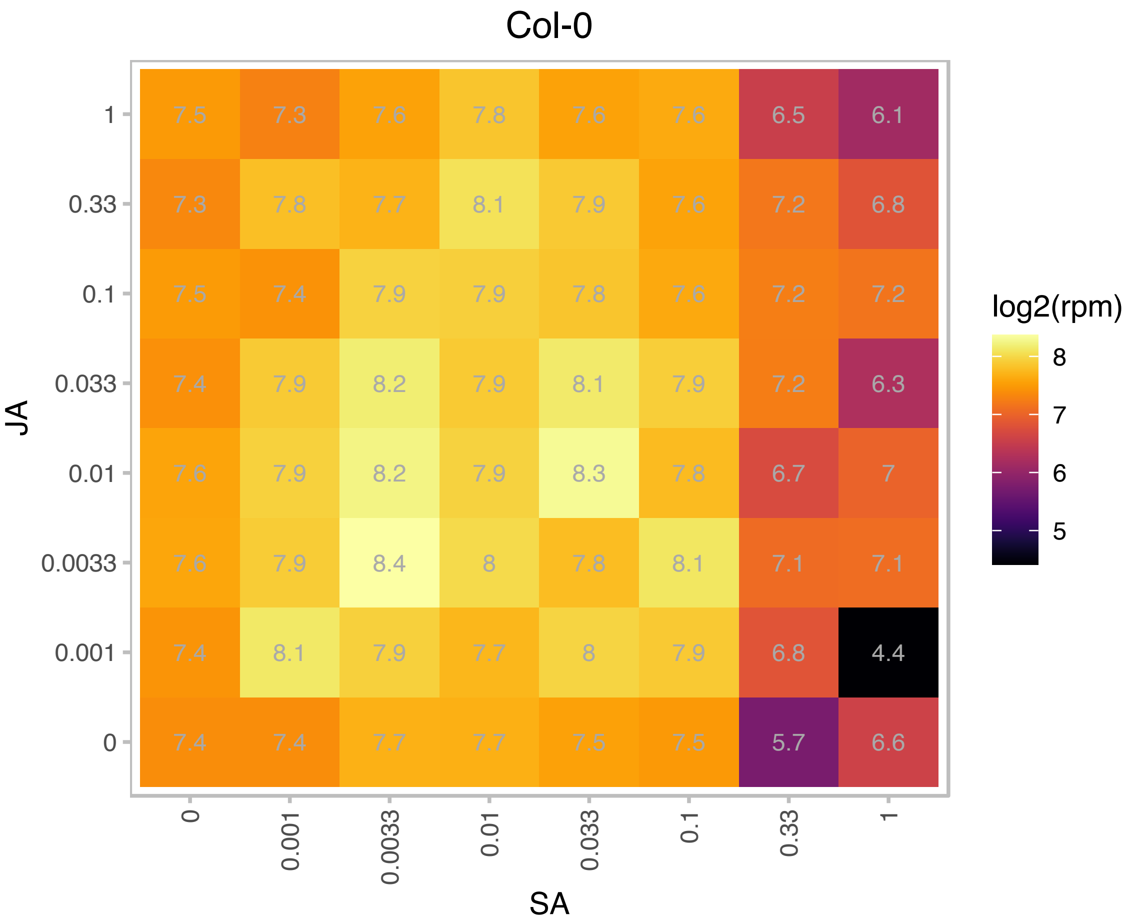


| | |
|---|---|
| Gene | AT2G43150 |
| is.expressed_genes | TRUE |
| is.responsive_genes | TRUE |
| is.apcluster | F-02 |

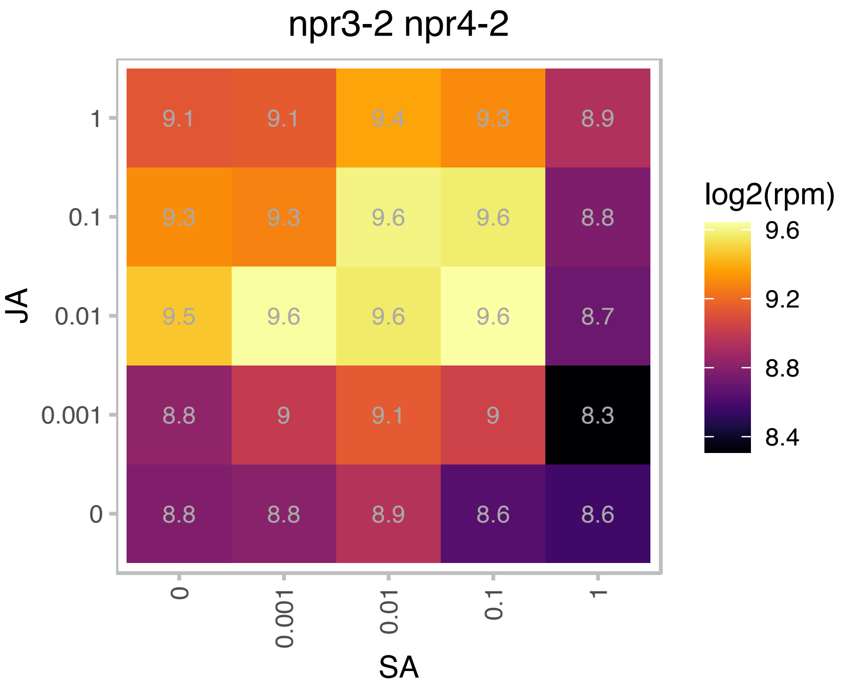


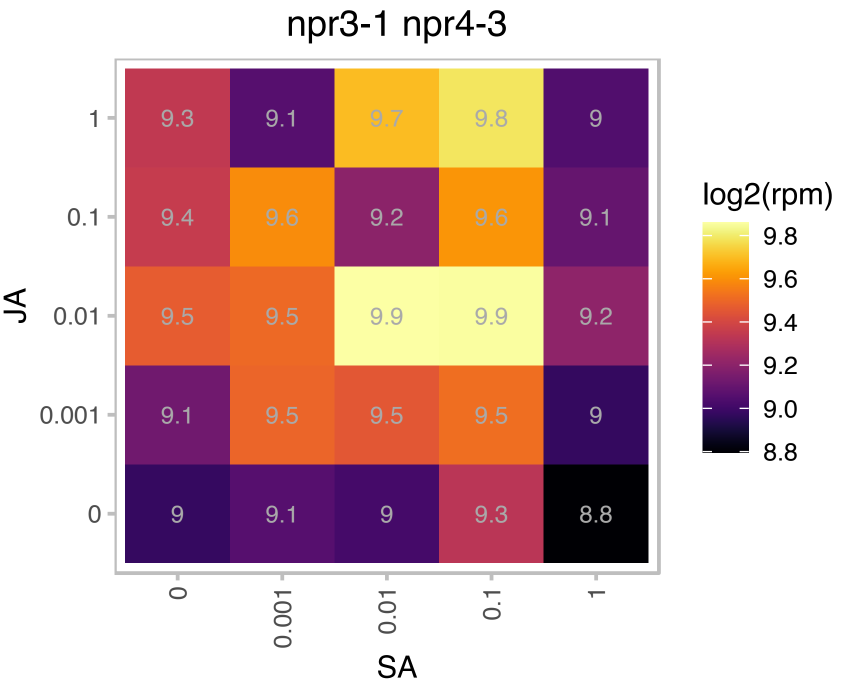

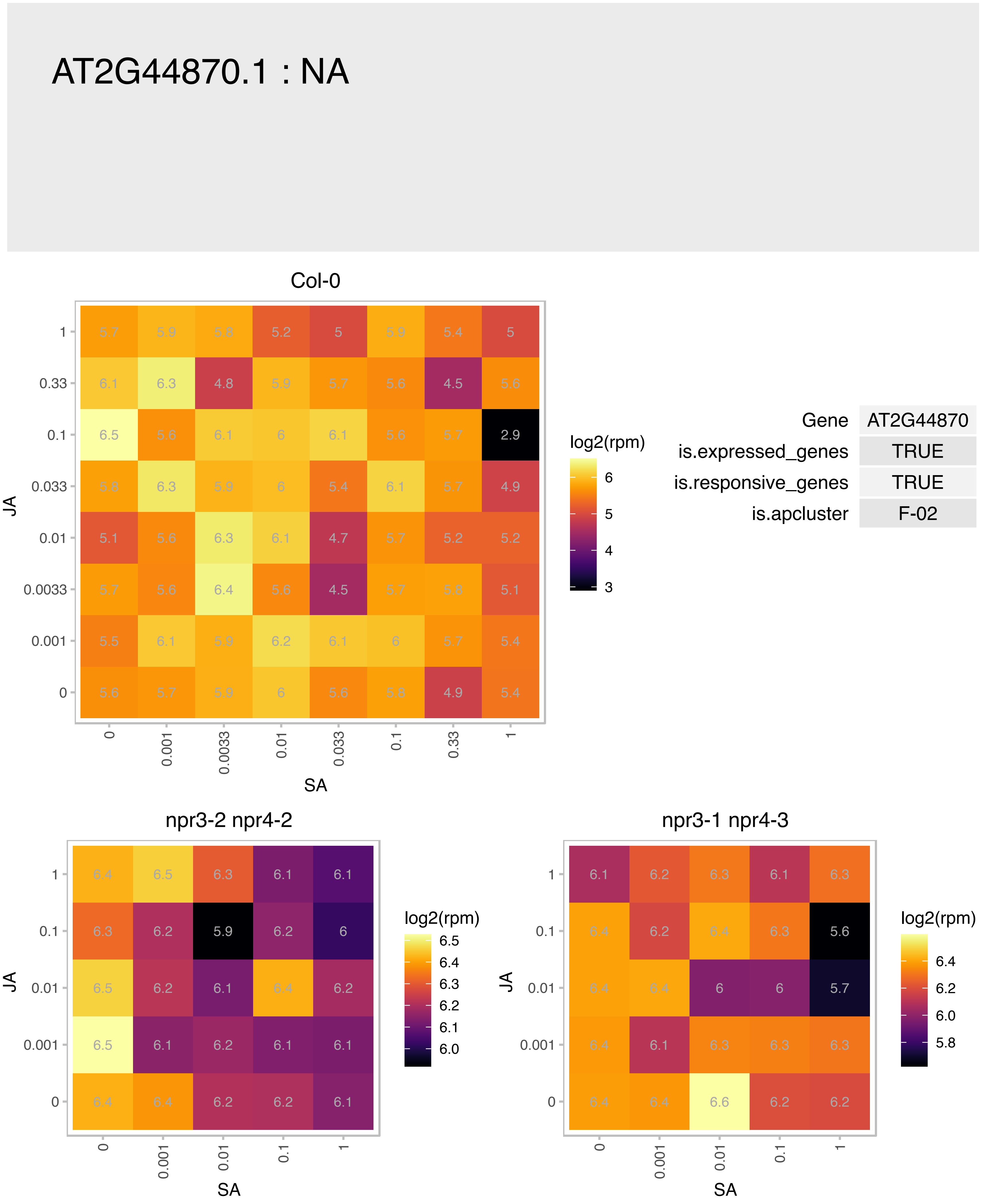
AT2G44870.1 : NA
Col-0
JA
SA
log2(rpm)
Gene AT2G44870
is.expressed_genes TRUE
is.responsive_genes TRUE
is.apcluster F-02
npr3-2 npr4-2
npr3-1 npr4-3

# AT3G02020.1 : AK3

aspartate kinase 3

encodes a monofunctional aspartate kinase

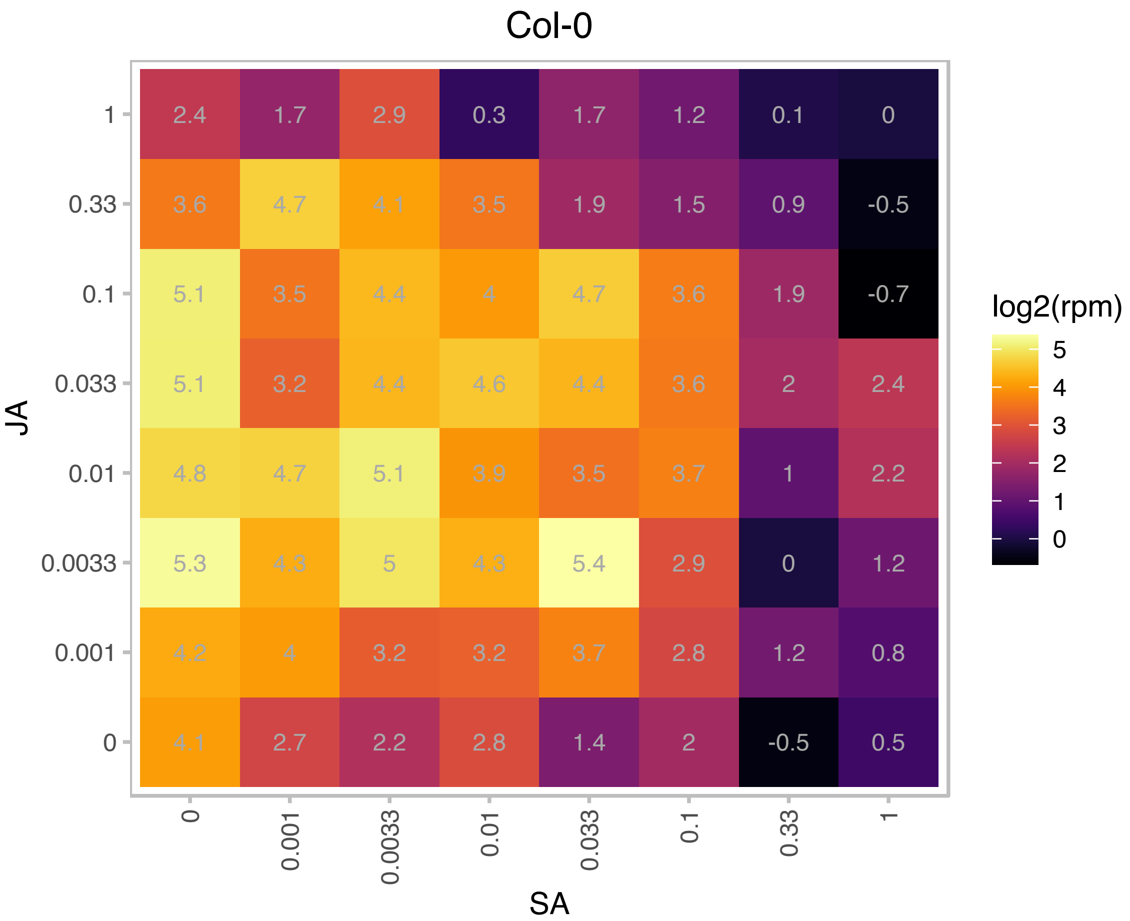


| | |
|---|---|
| Gene | AT3G02020 |
| is.expressed_genes | TRUE |
| is.responsive_genes | TRUE |
| is.apcluster | F-02 |

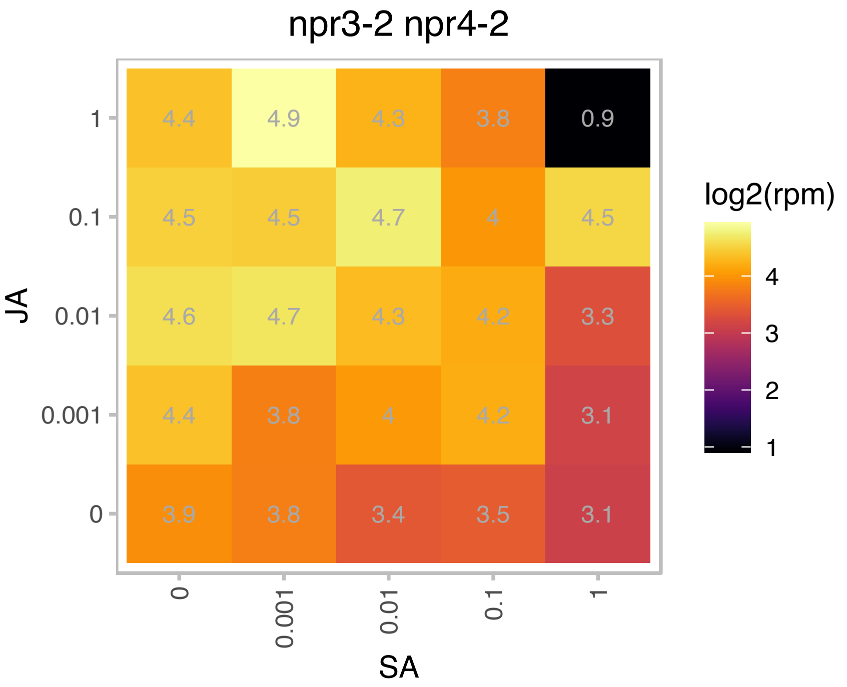


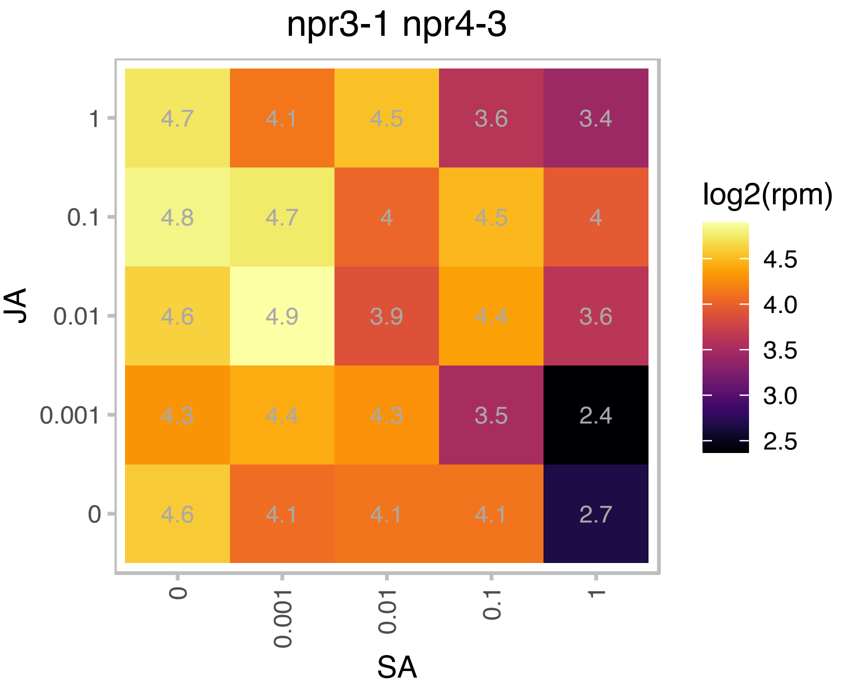

### AT3G03190.1 : ATGSTF11, ATGSTF6, GSTF11

glutathione S-transferase F11

Encodes glutathione transferase belonging to the phi class of GSTs. Naming convention according to Wagner et al. (2002).

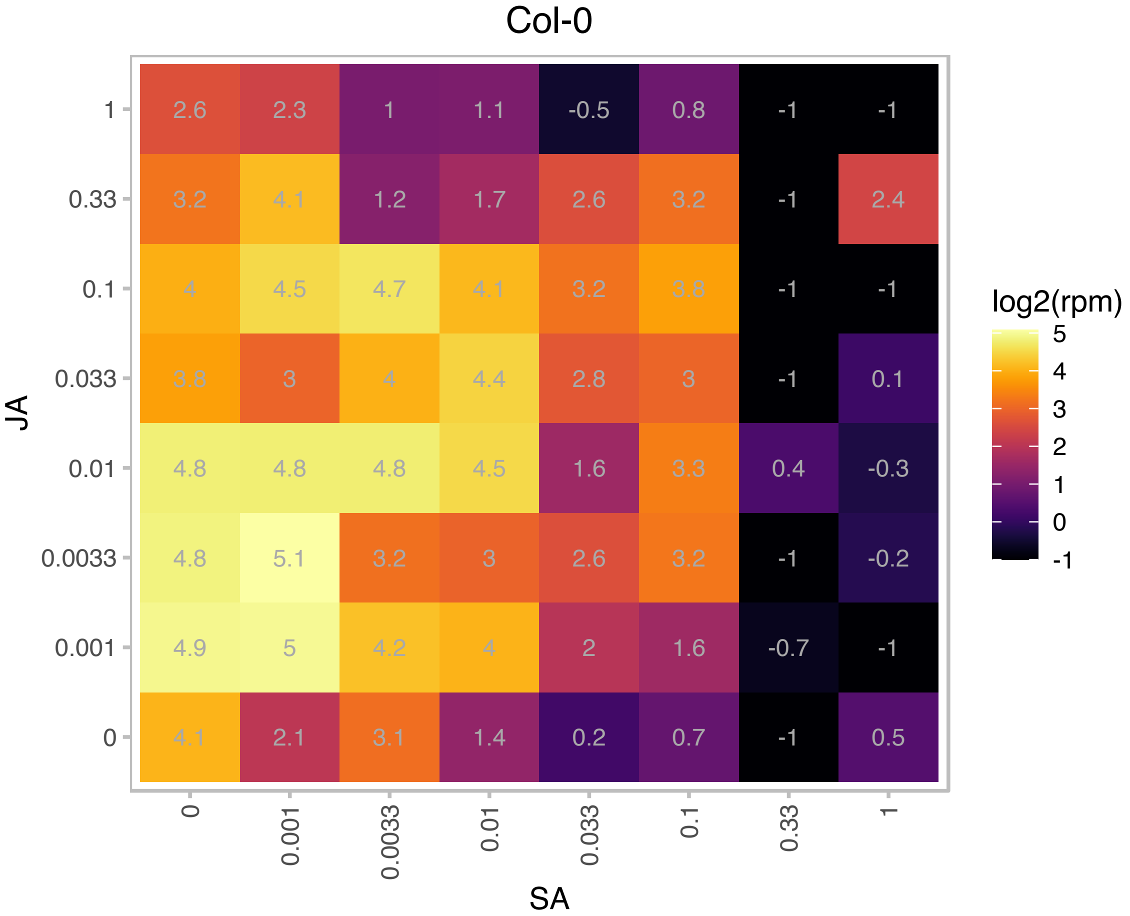


| | |
|---|---|
| Gene | AT3G03190 |
| is.expressed_genes | TRUE |
| is.responsive_genes | TRUE |
| is.apcluster | F-02 |

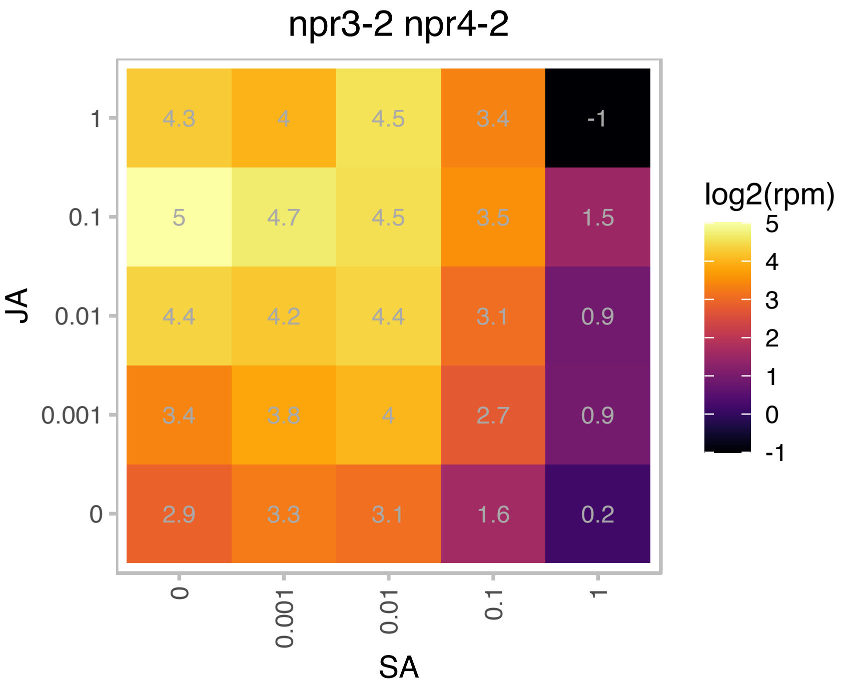


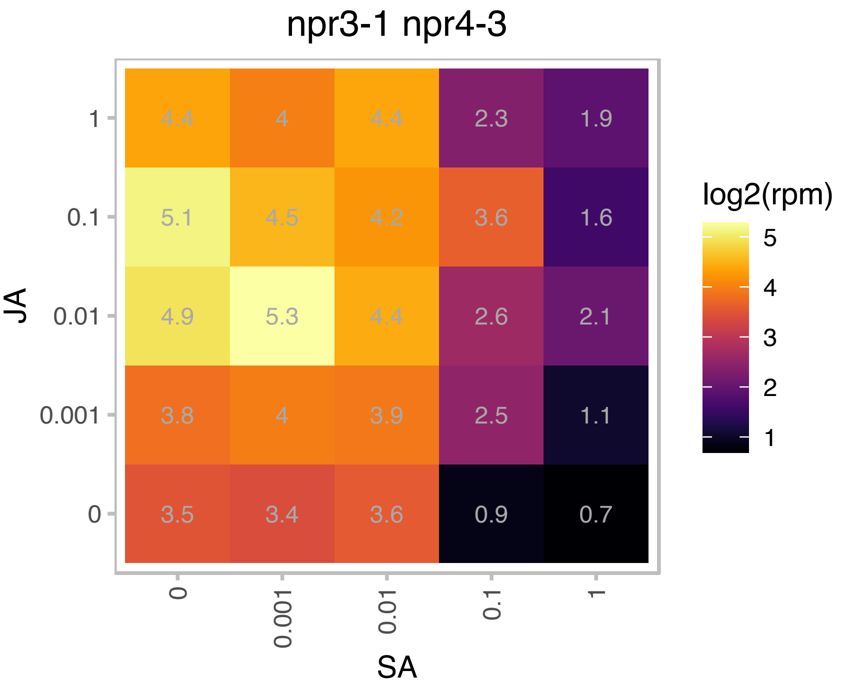

### AT3G04550.1 : RAF1

Encodes an ancillary chaperone protein that functions in Rubisco biogenesis. RAF1 dimers function in the assembly of the large subunit of Rubisco. Co-expression of RAF1 and rbcL in tobacco cells results in increased photosynthesis and plant growth. The mRNA is cell-to-cell mobile.

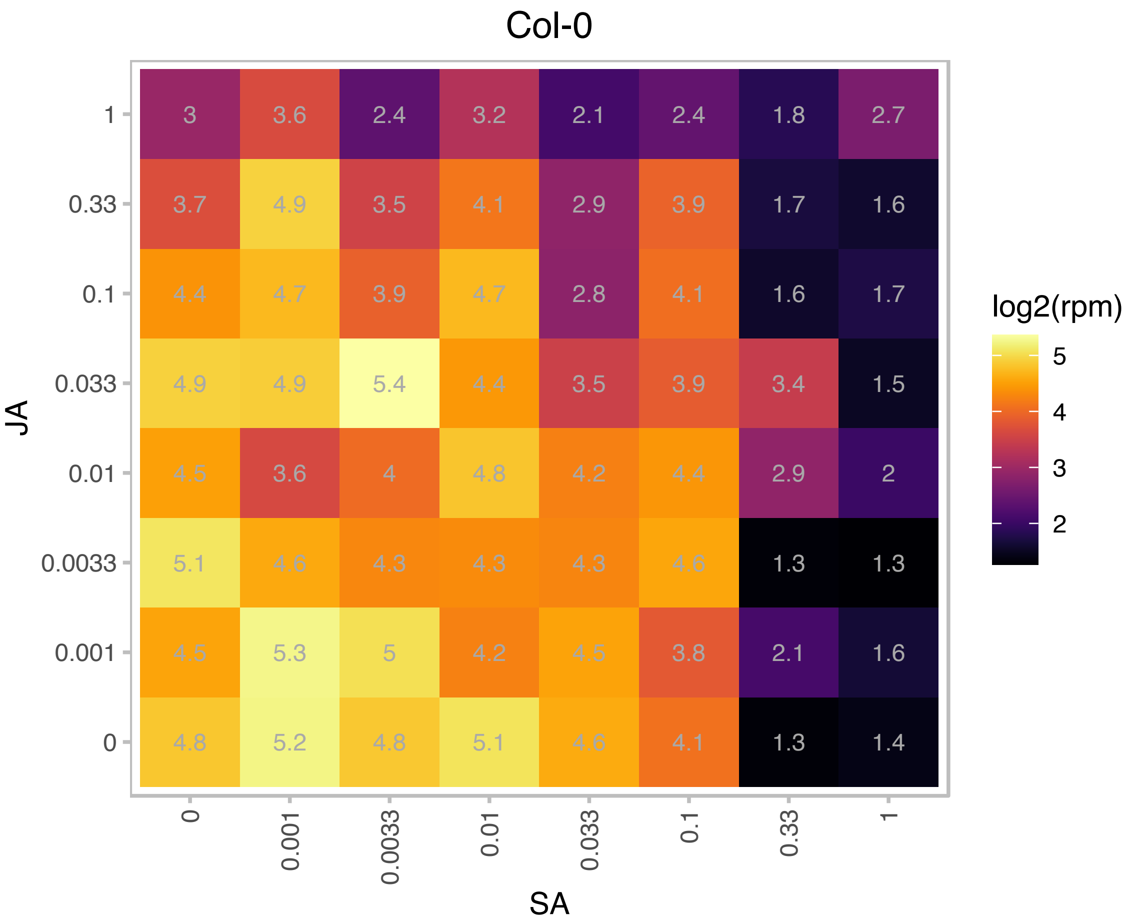


| | |
|---|---|
| Gene | AT3G04550 |
| is.expressed_genes | TRUE |
| is.responsive_genes | TRUE |
| is.apcluster | F-02 |

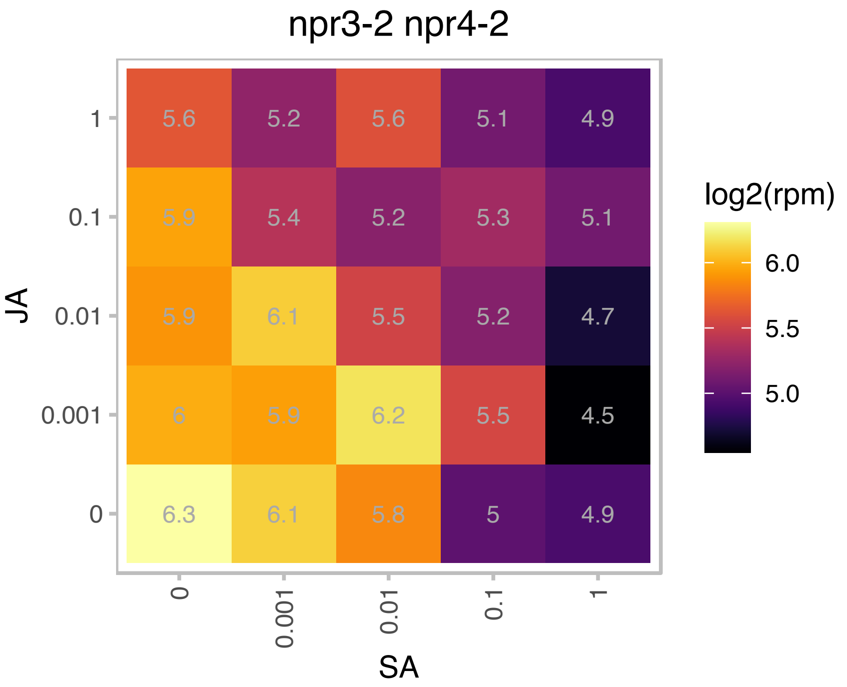


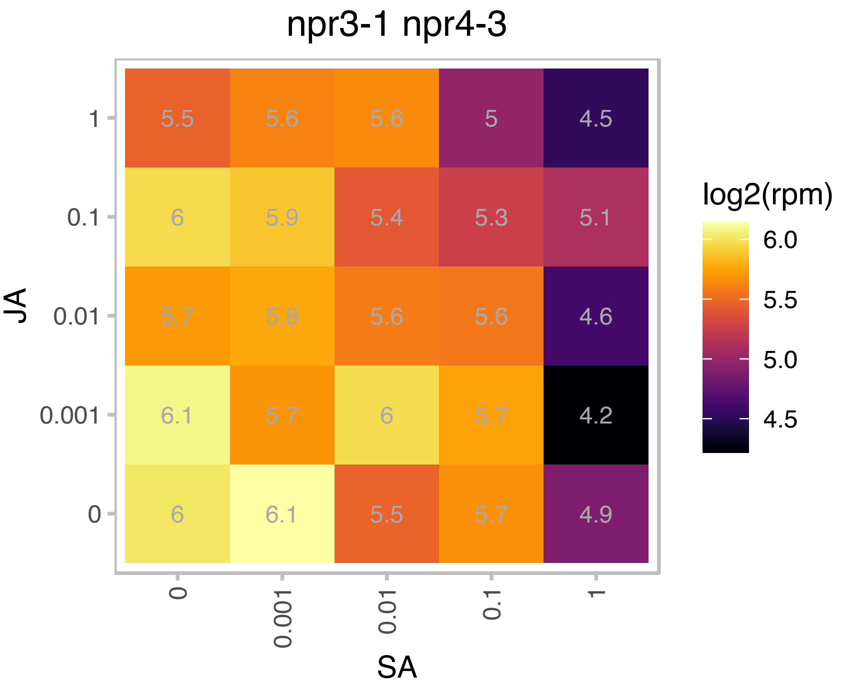

### AT3G04720.1 : AtPR4, HEL, PR-4, PR4

pathogenesis-related 4

Encodes a protein similar to the antifungal chitin-binding protein hevein from rubber tree latex. mRNA levels increase in response to ethylene and turnip crinkle virus infection. The mRNA is cell-to-cell mobile.

Col-0

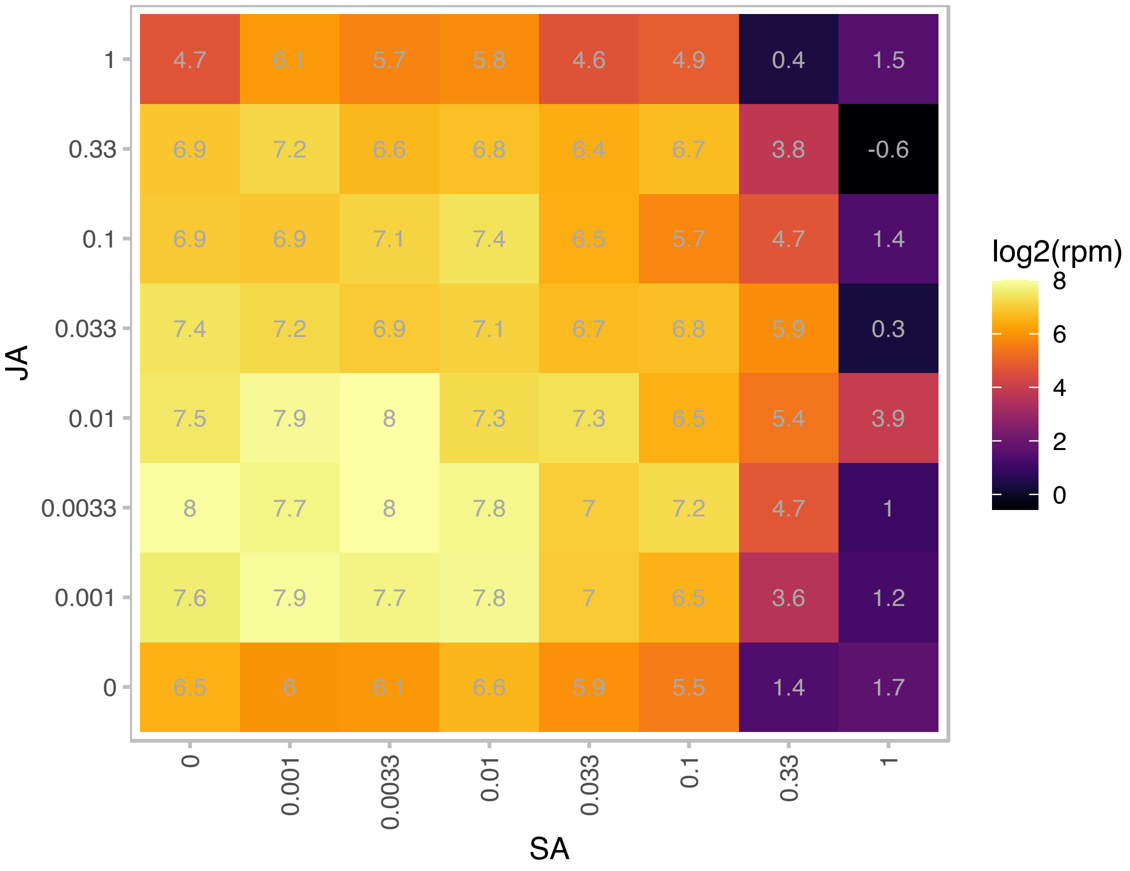


| | |
|---|---|
| Gene | AT3G04720 |
| is.expressed_genes | TRUE |
| is.responsive_genes | TRUE |
| is.apcluster | F-02 |

npr3-2 npr4-2

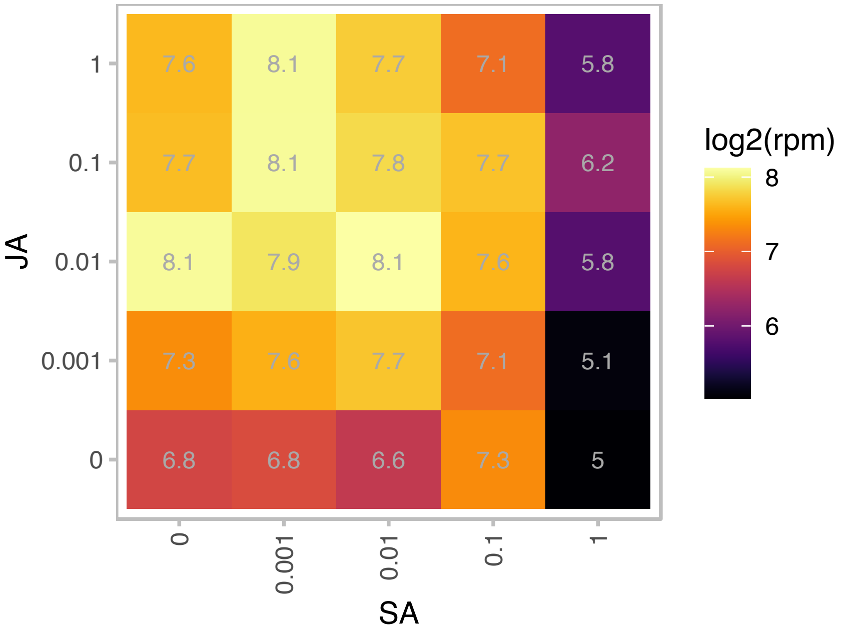


npr3-1 npr4-3

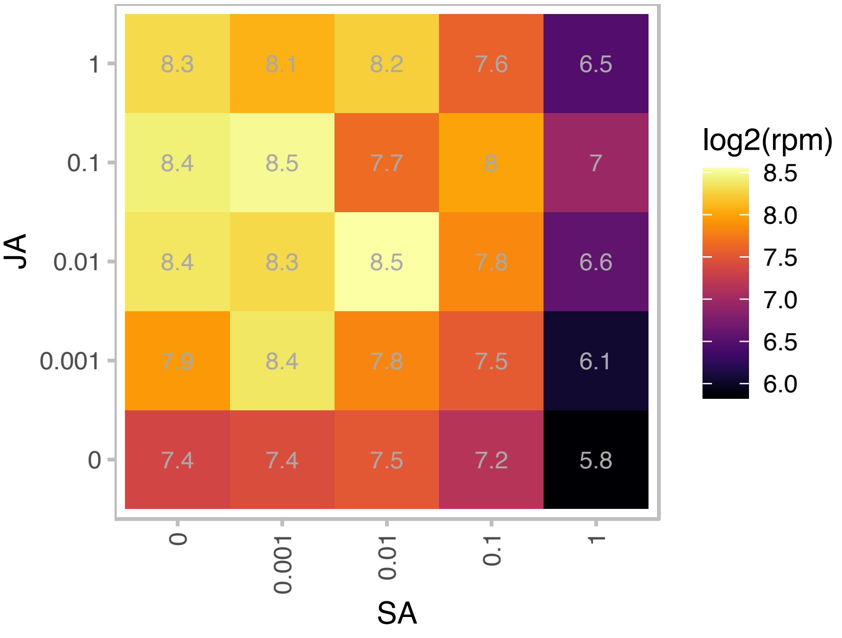

### AT3G09630.1 : SAC56

Ribosomal protein L4/L1 family

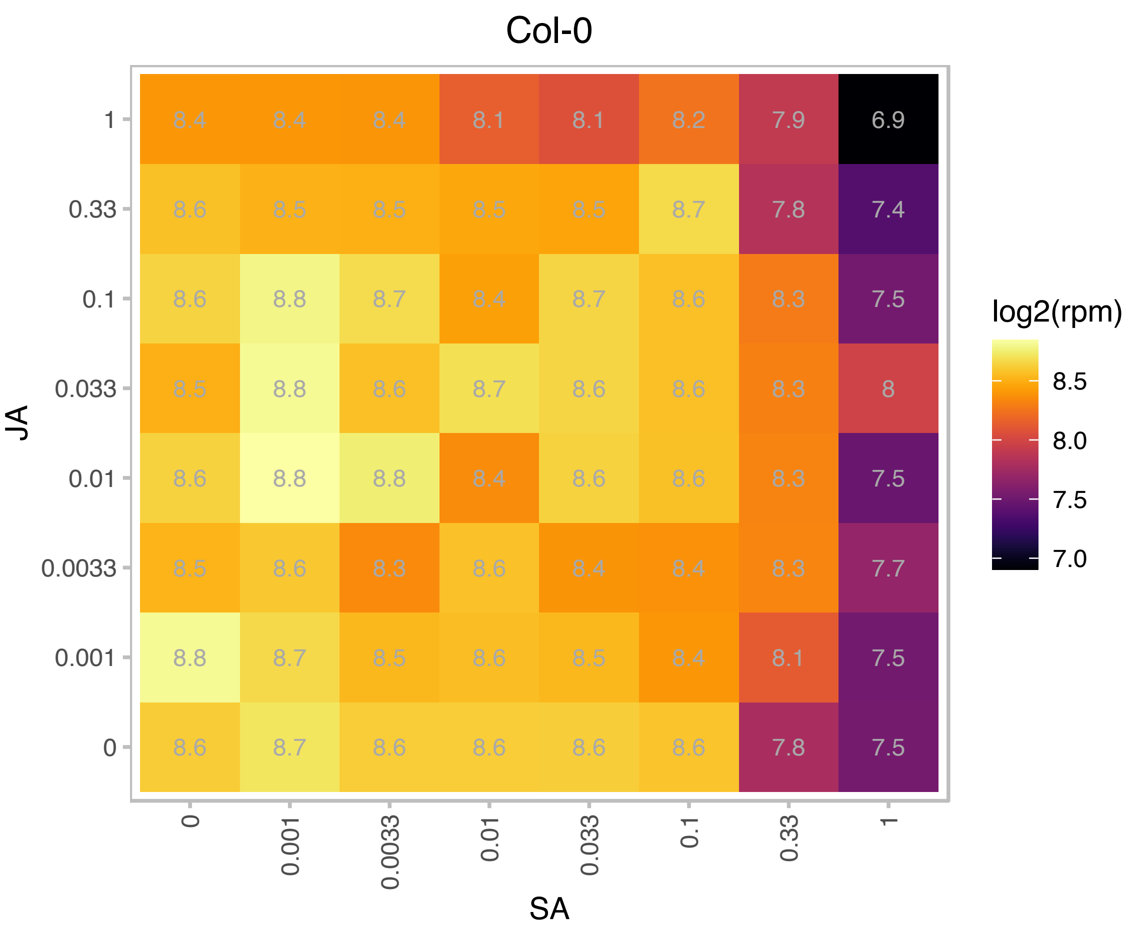


| | |
|---|---|
| Gene | AT3G09630 |
| is.expressed_genes | TRUE |
| is.responsive_genes | TRUE |
| is.apcluster | F-02 |

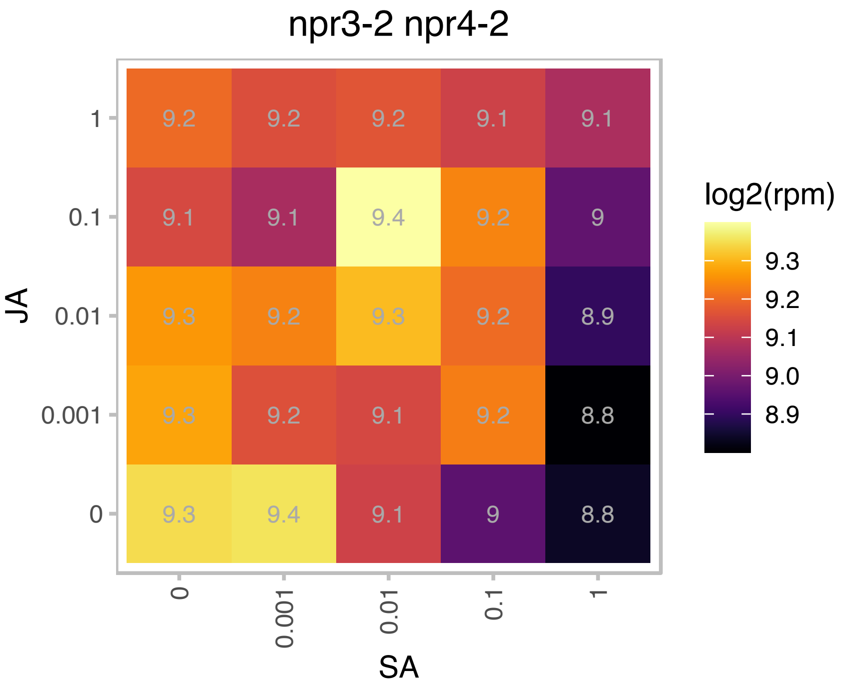


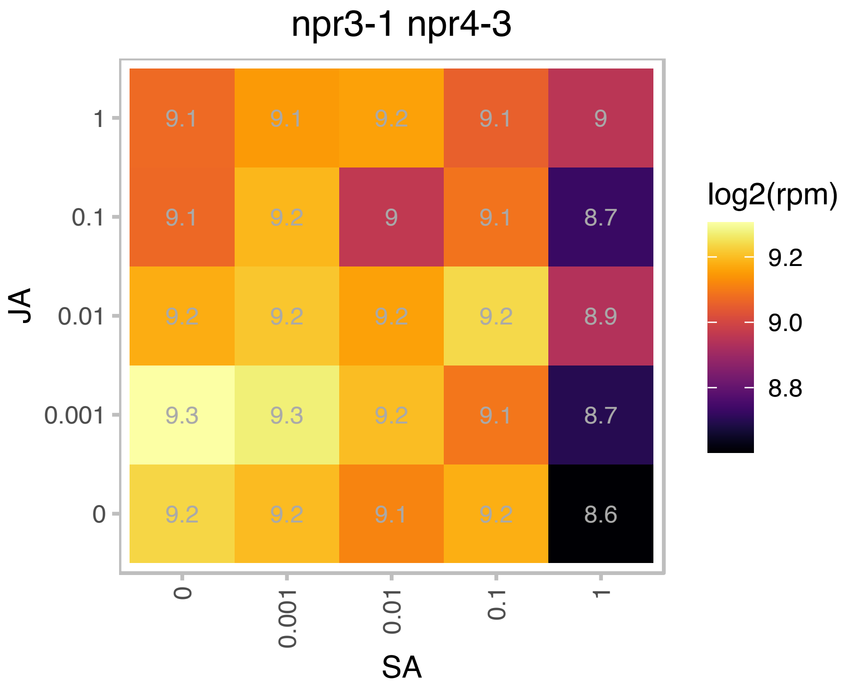

### AT3G11400.1 : ATEIF3G1, EIF3G1

eukaryotic translation initiation factor 3G1

One of the 2 genes that code for the G subunit of eukaryotic initiation factor 3 (EIF3). The mRNA is cell-to-cell mobile.

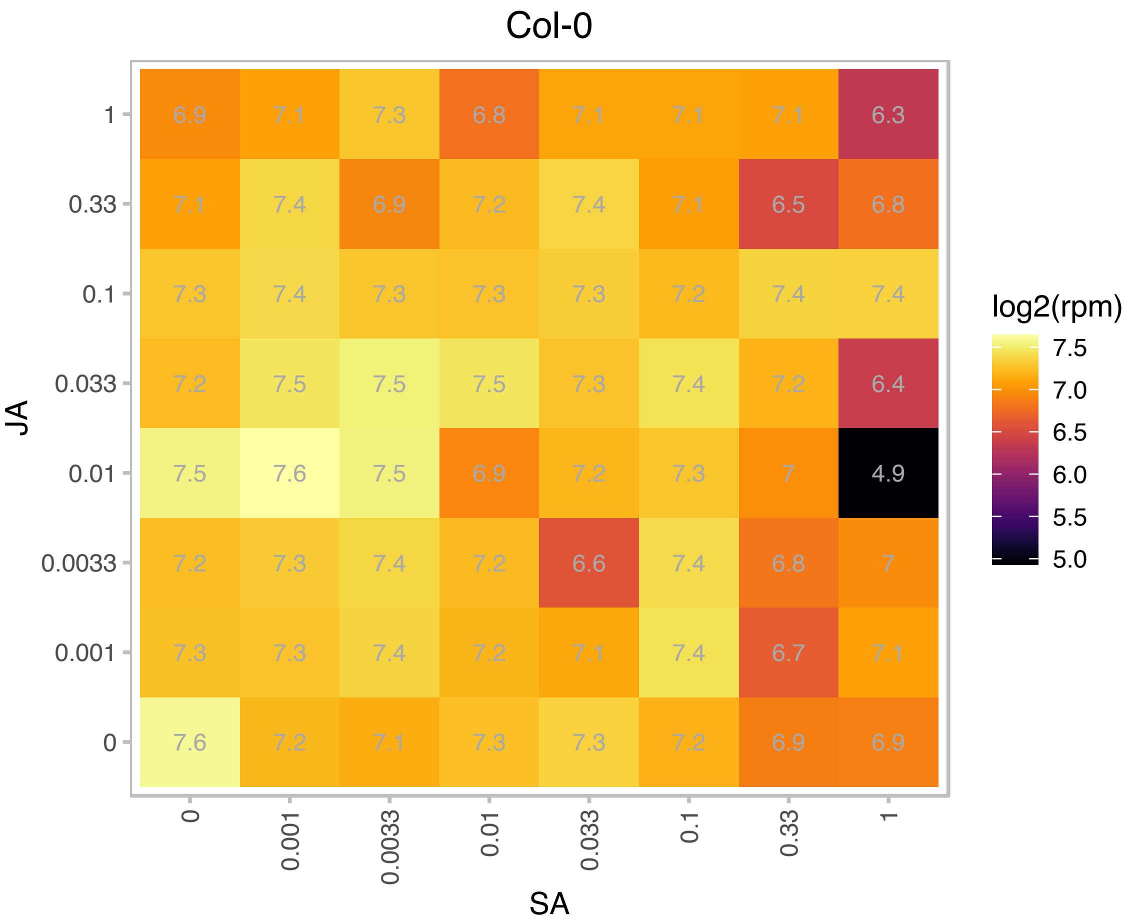


| | |
|---|---|
| Gene | AT3G11400 |
| is.expressed_genes | TRUE |
| is.responsive_genes | TRUE |
| is.apcluster | F-02 |

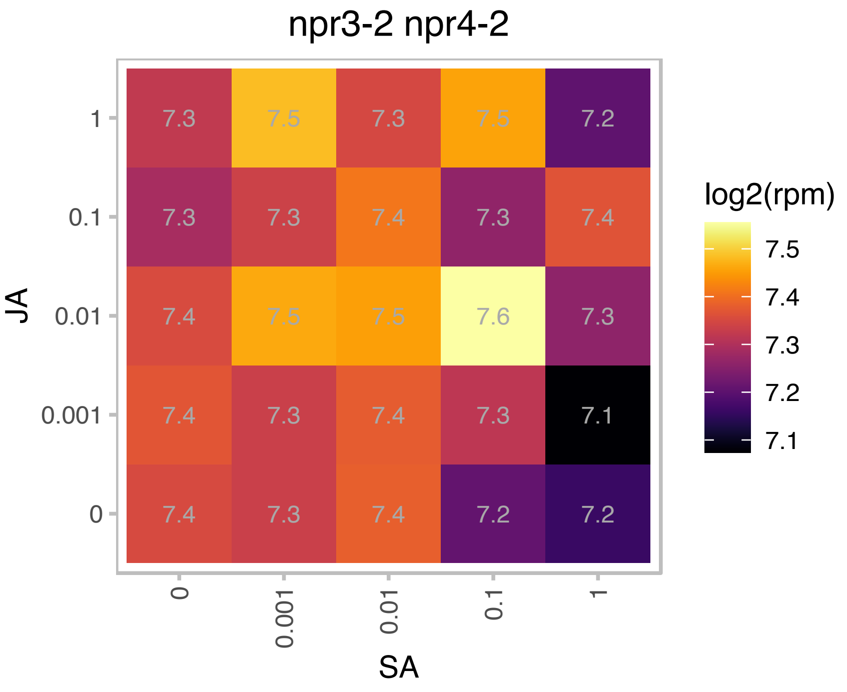


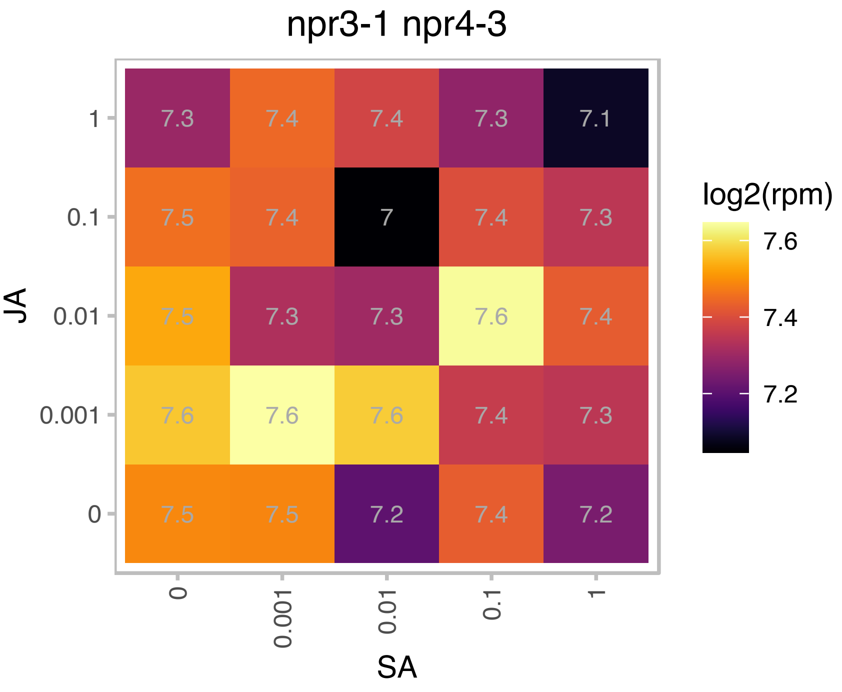

# AT3G12370.1 : NA

Ribosomal protein L10 family protein

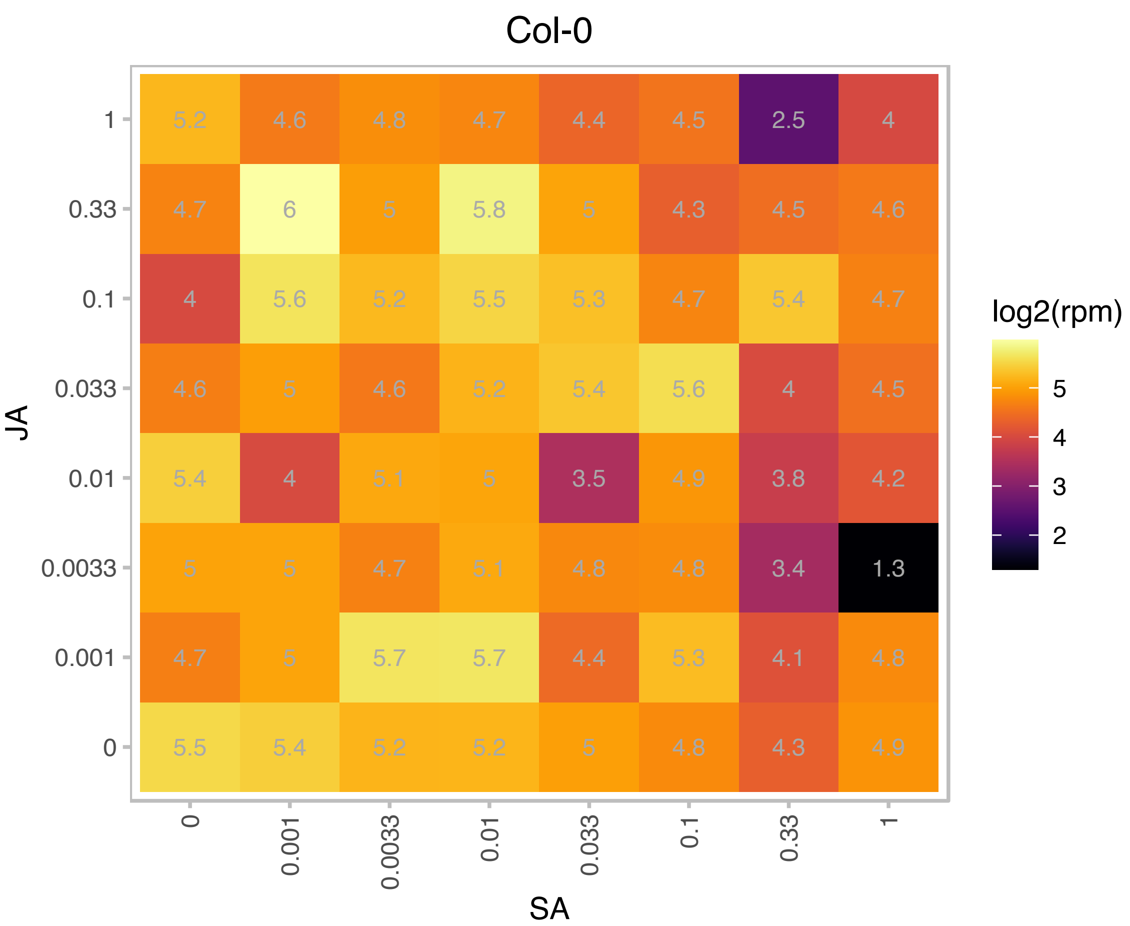


| | |
|---|---|
| Gene | AT3G12370 |
| is.expressed_genes | TRUE |
| is.responsive_genes | TRUE |
| is.apcluster | F-02 |

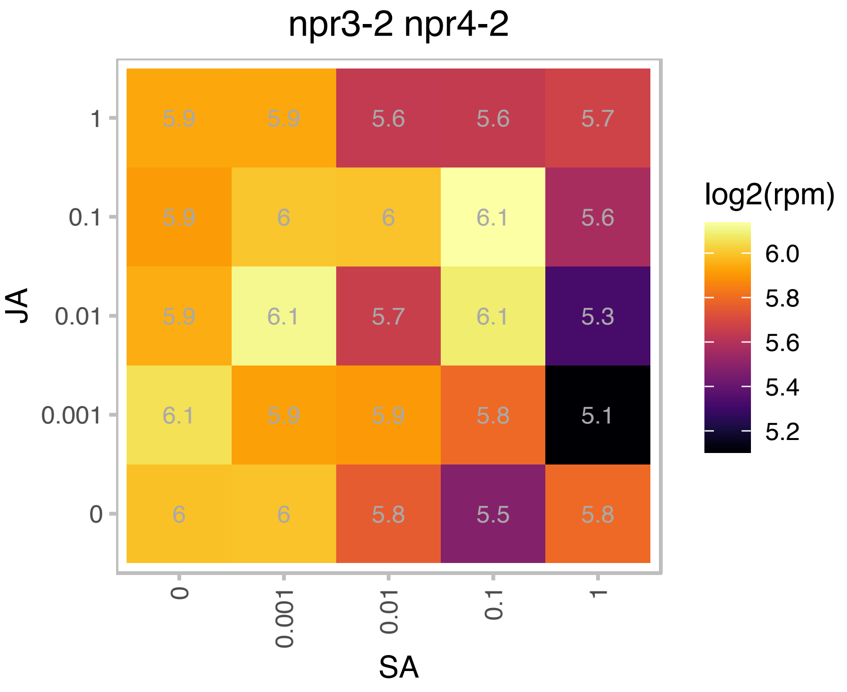


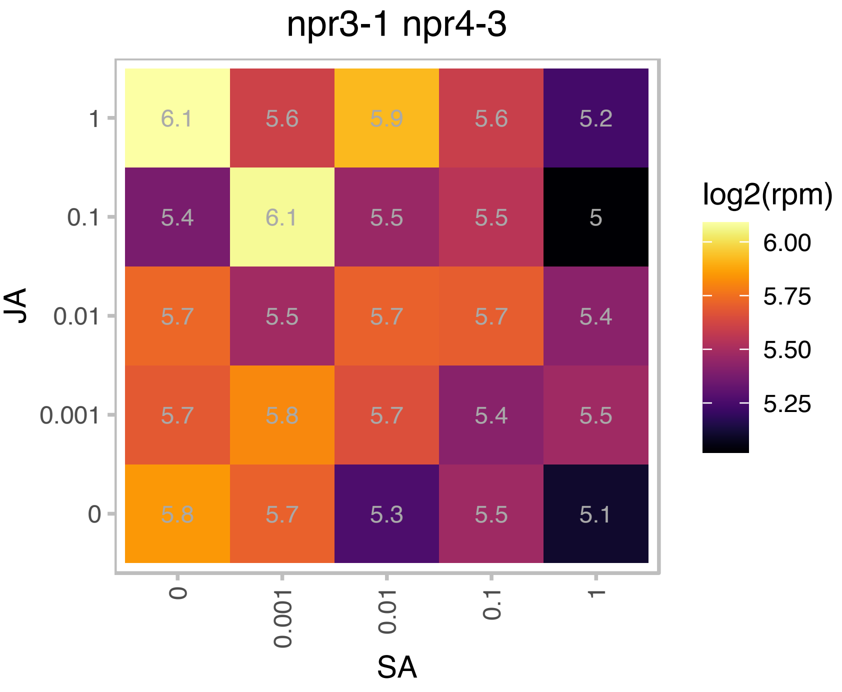

### AT3G14310.1 : ATPME3, OZS2, PME3

pectin methylesterase 3

encodes a pectin methylesterase, targeted by a cellulose binding protein (CBP) from the parasitic nematode Heterodera schachtii during parasitism.

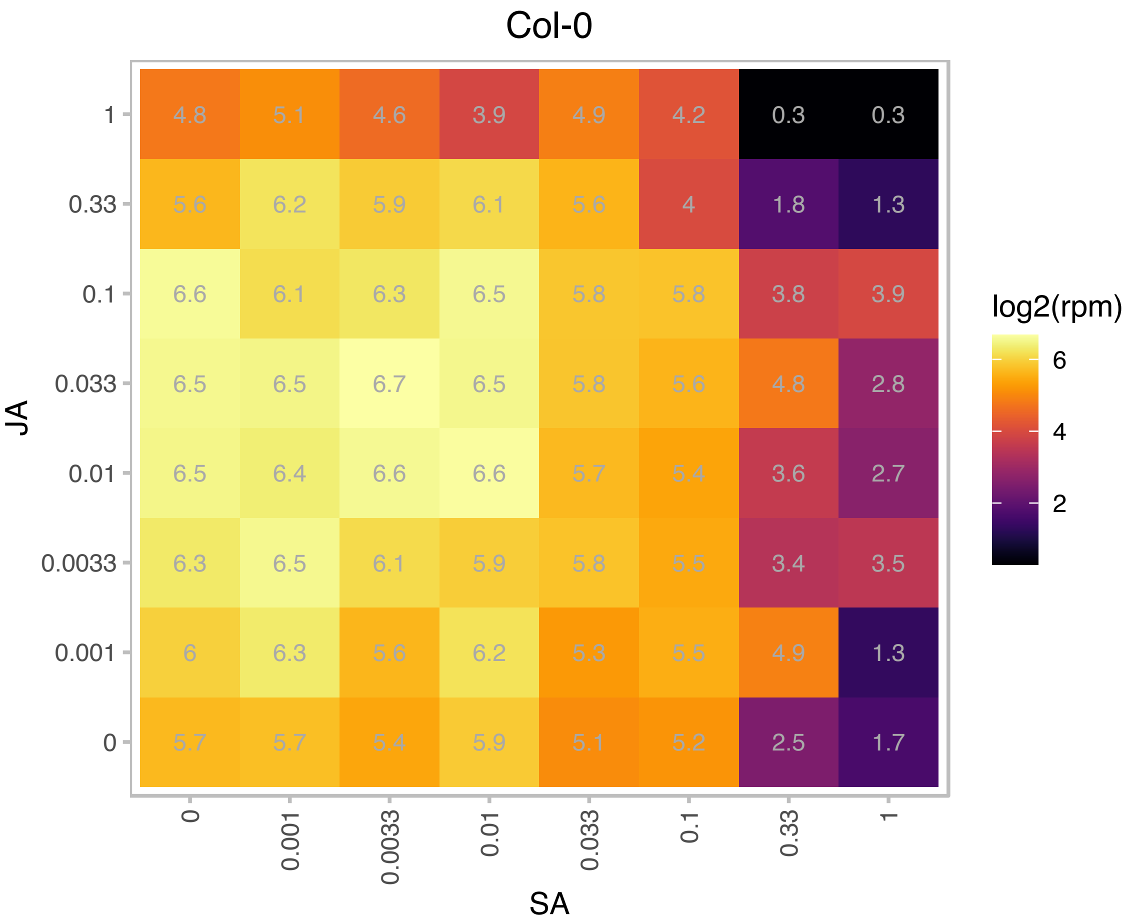


| | |
|---|---|
| Gene | AT3G14310 |
| is.expressed_genes | TRUE |
| is.responsive_genes | TRUE |
| is.apcluster | F-02 |

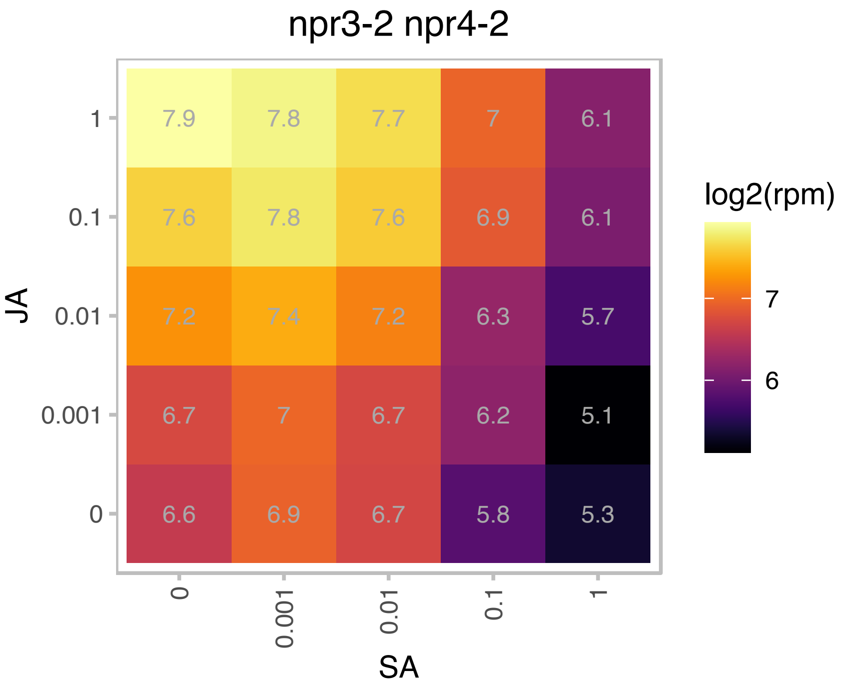


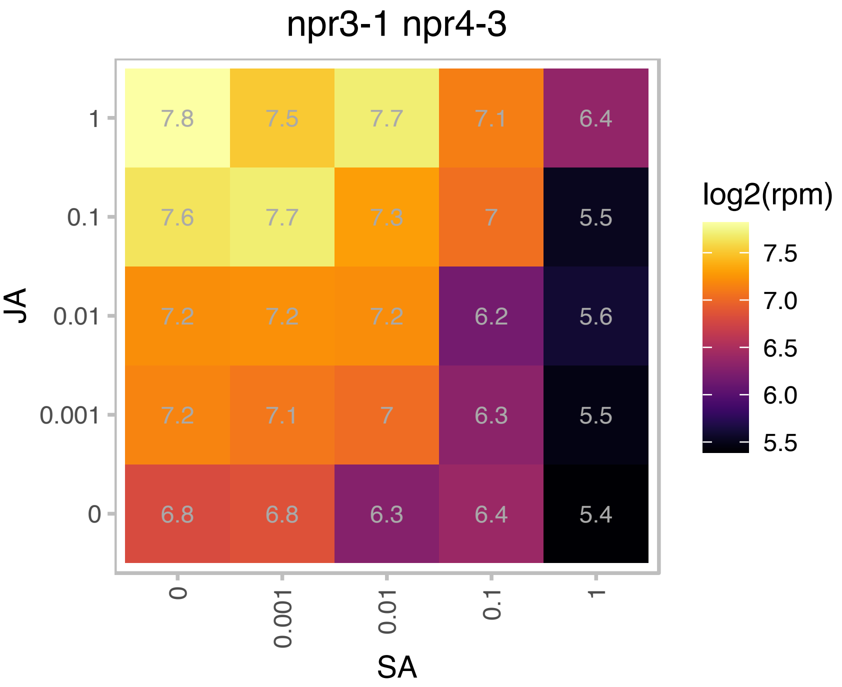

# AT3G15356.1 : NA

Legume lectin family protein

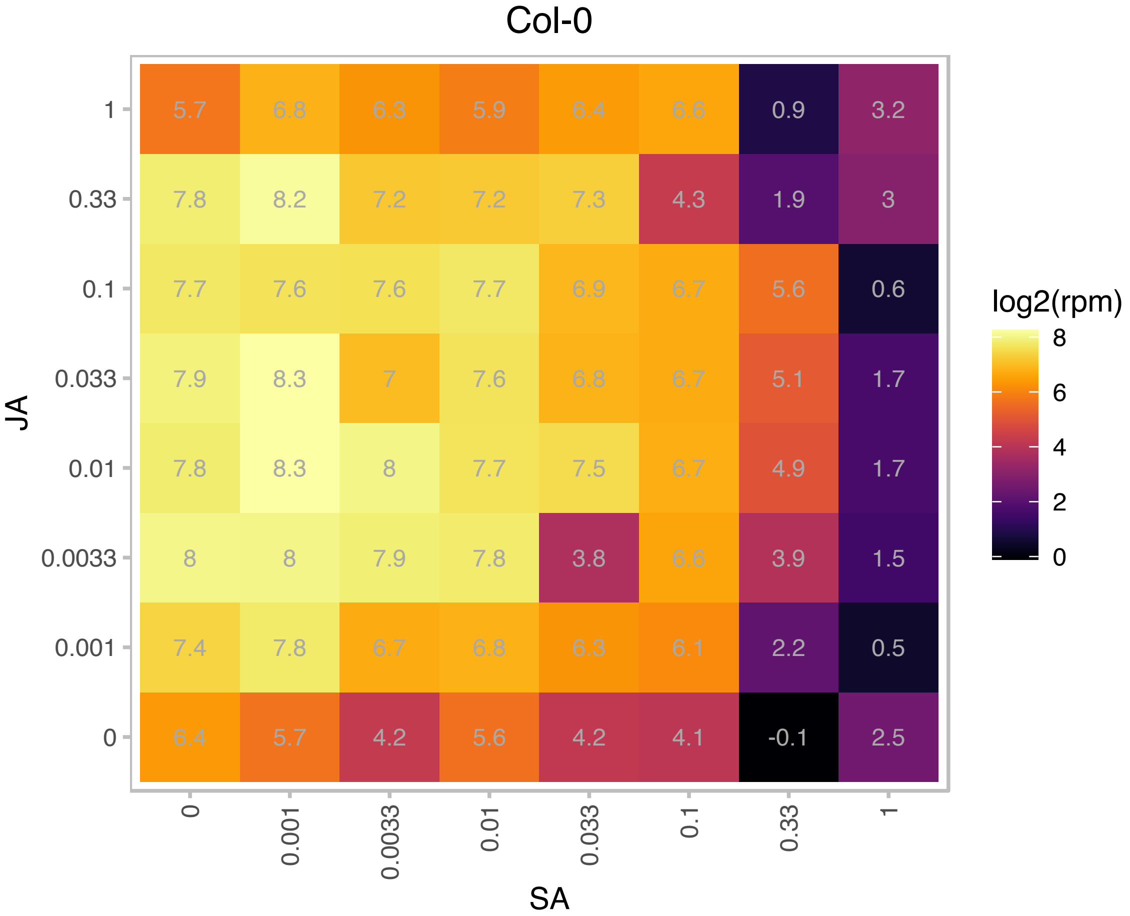


| Gene | AT3G15356 |
|---|---|
| is.expressed_genes | TRUE |
| is.responsive_genes | TRUE |
| is.apcluster | F-02 |

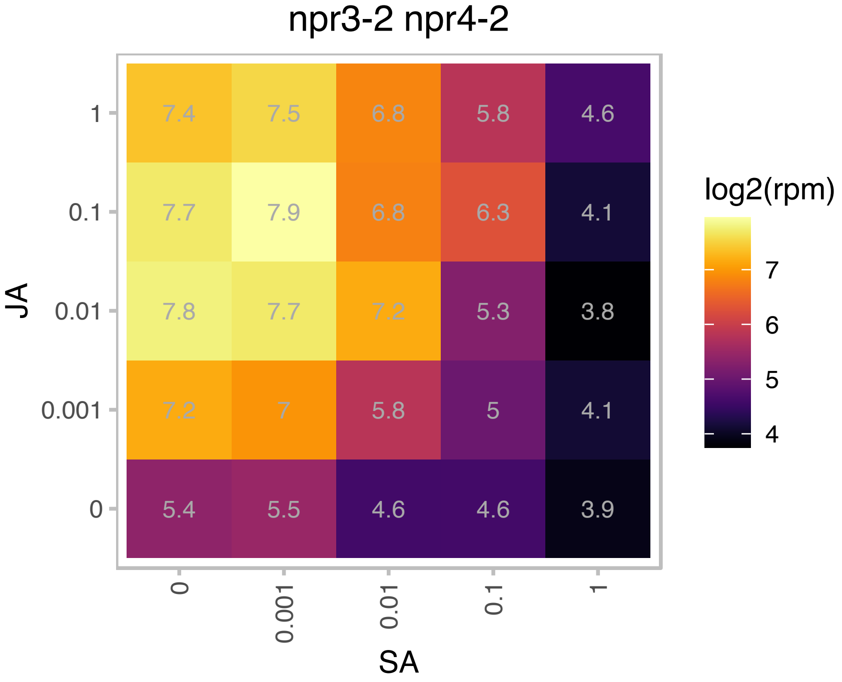


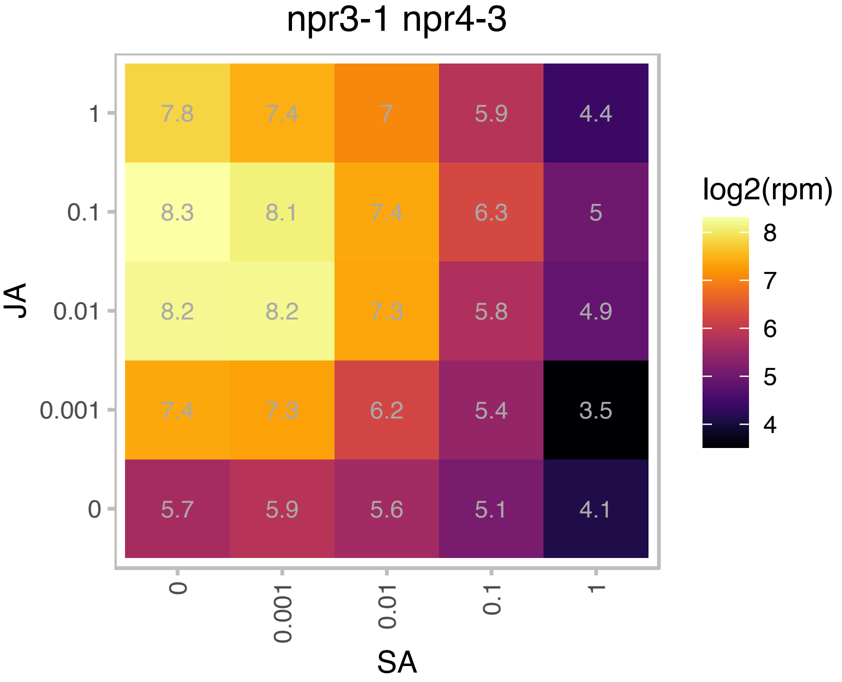

# AT3G16350.1 : NA

Homeodomain-like superfamily protein

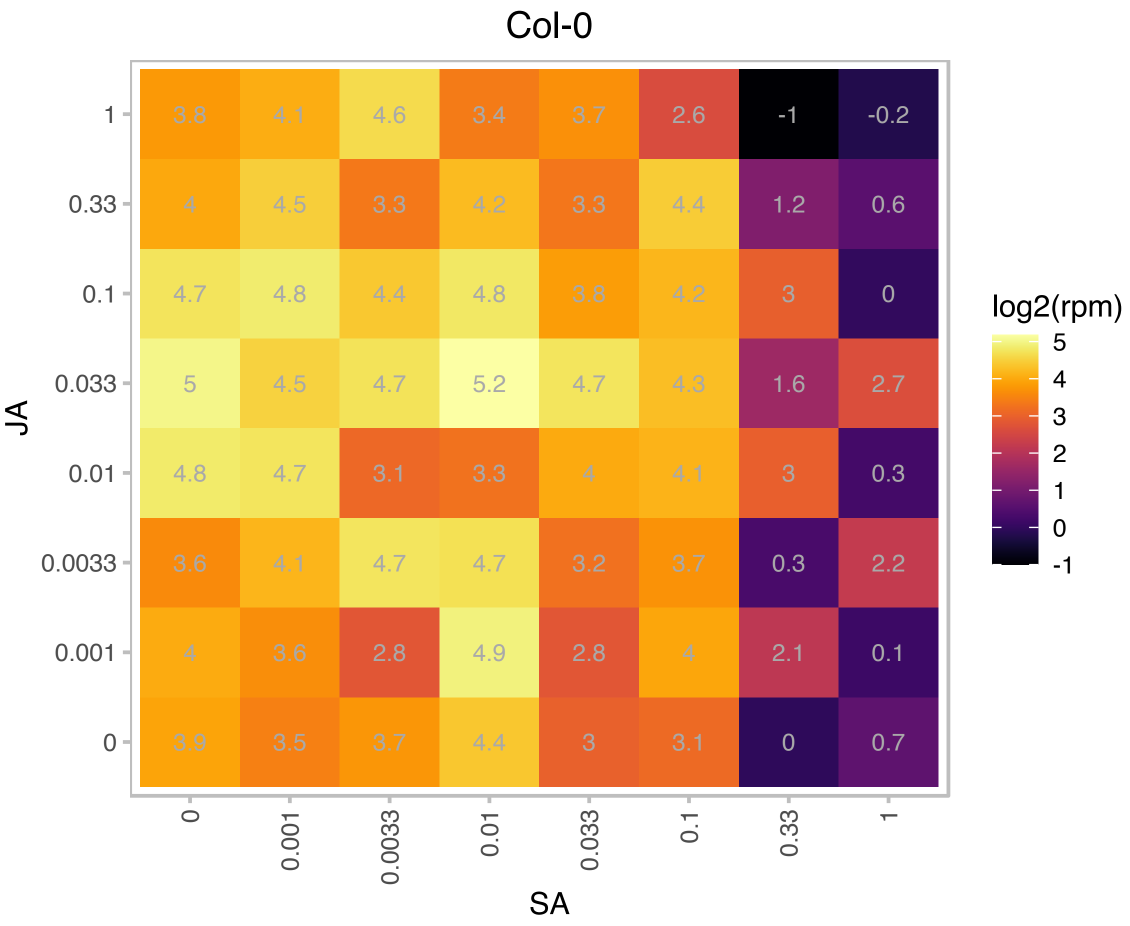


| | |
|---|---|
| Gene | AT3G16350 |
| is.expressed_genes | TRUE |
| is.responsive_genes | TRUE |
| is.apcluster | F-02 |

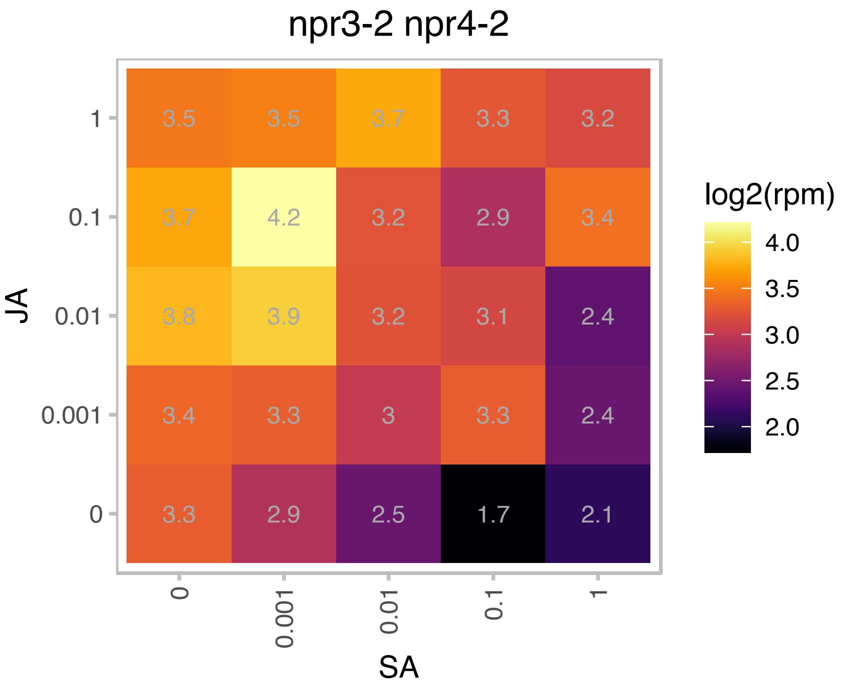


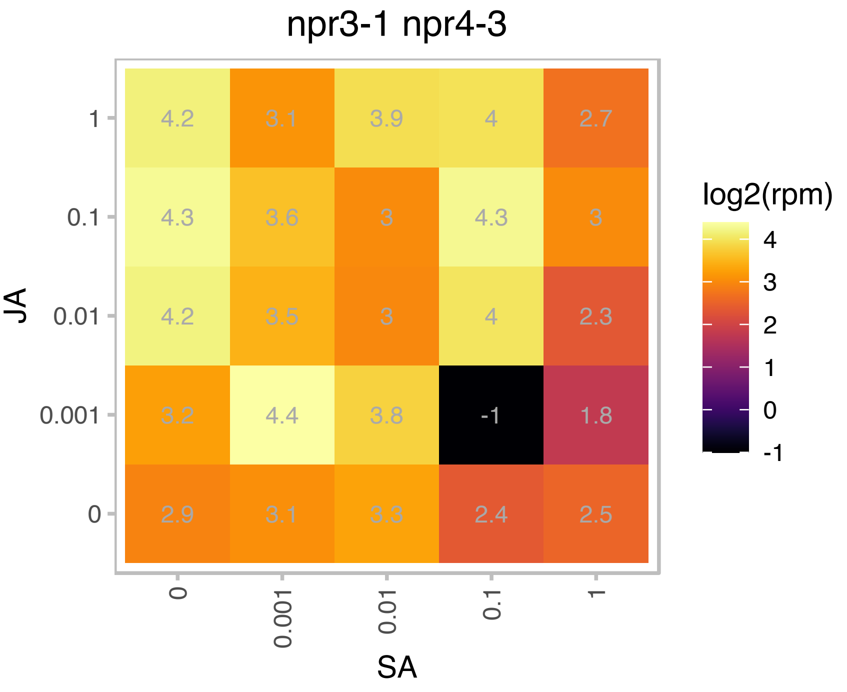

# AT3G17120.1 : NA

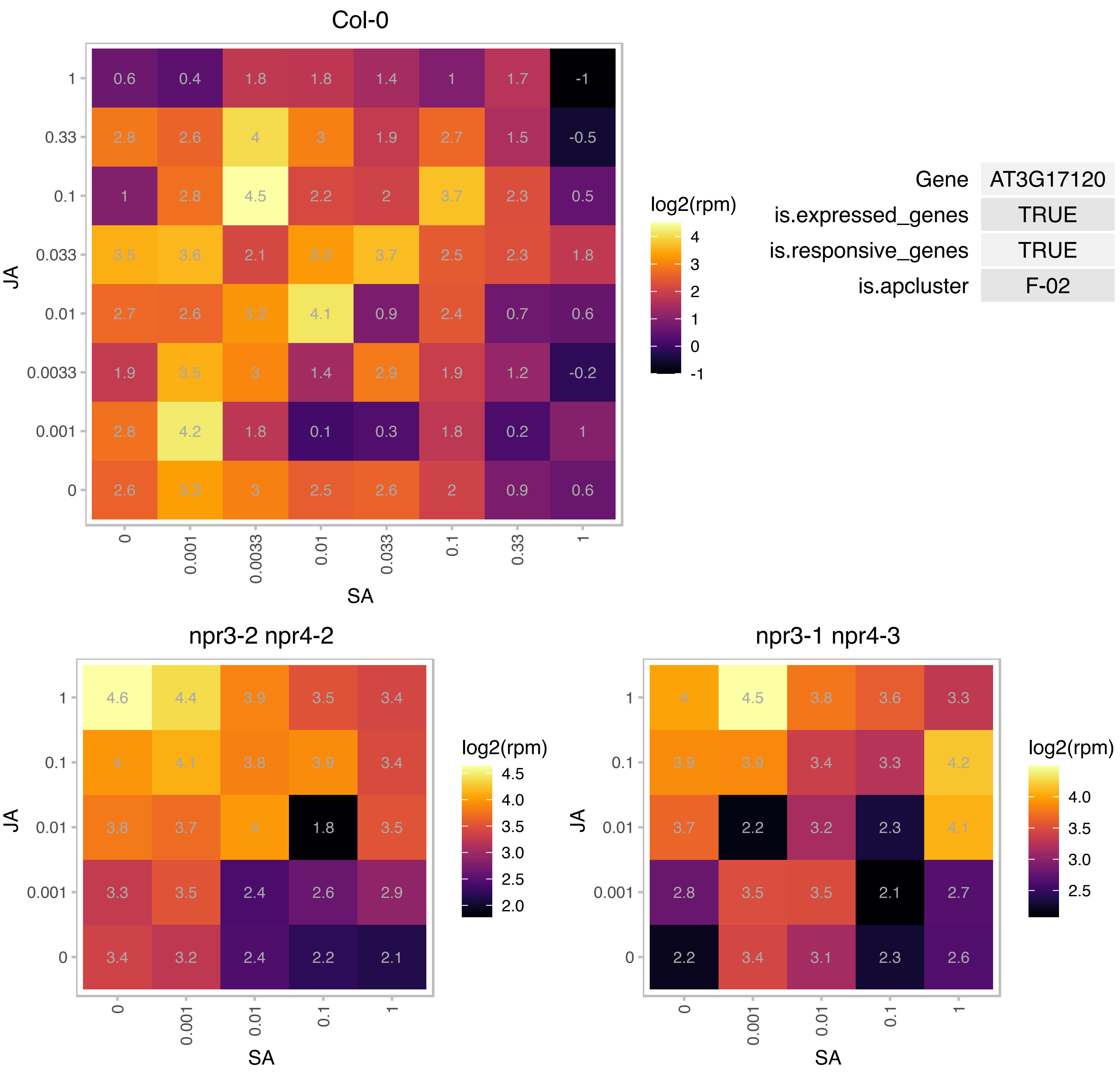

# AT3G17670.1 : NA

tetratricopeptide repeat (TPR)-containing protein

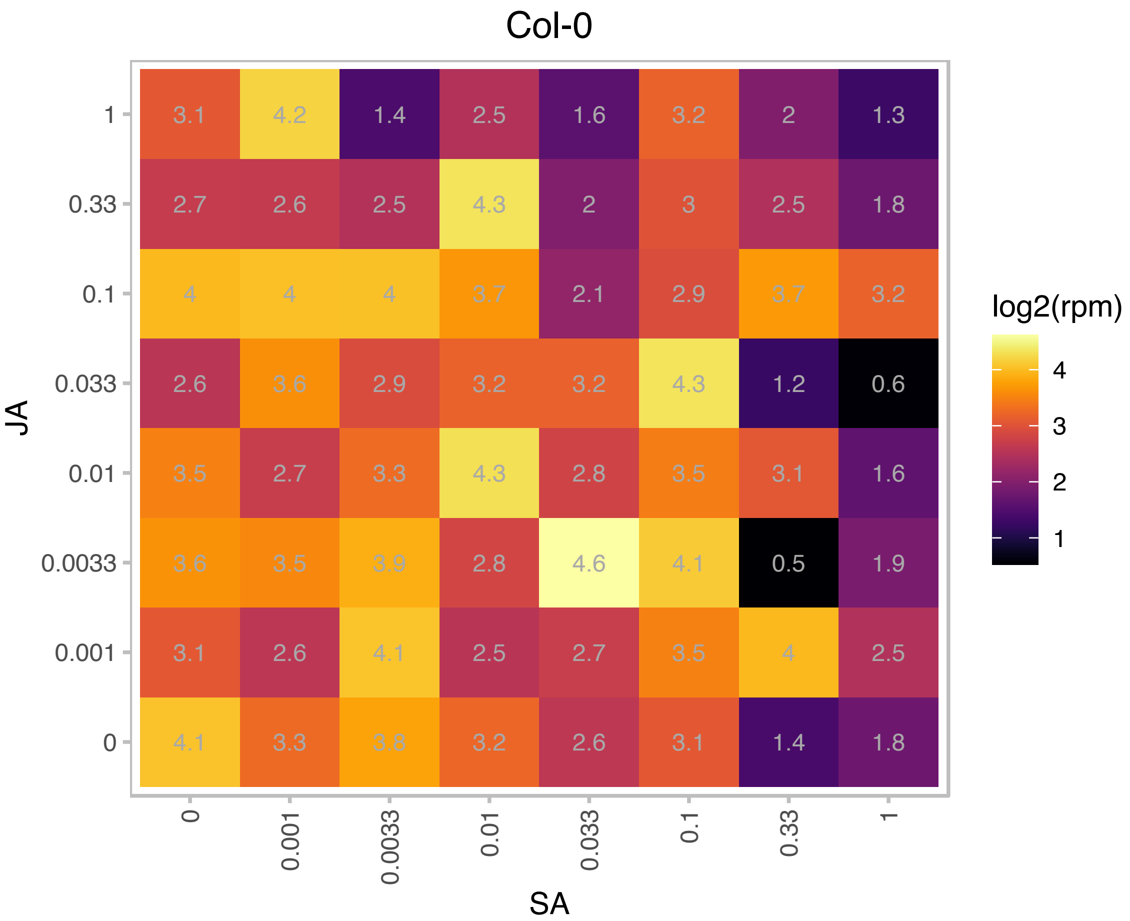


| | |
|---|---|
| Gene | AT3G17670 |
| is.expressed_genes | TRUE |
| is.responsive_genes | TRUE |
| is.apcluster | F-02 |

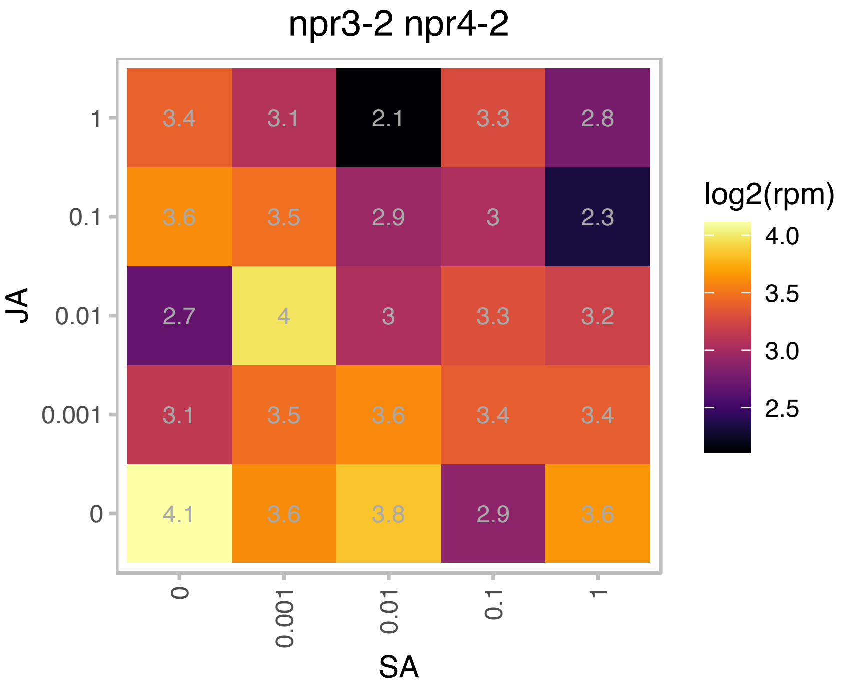


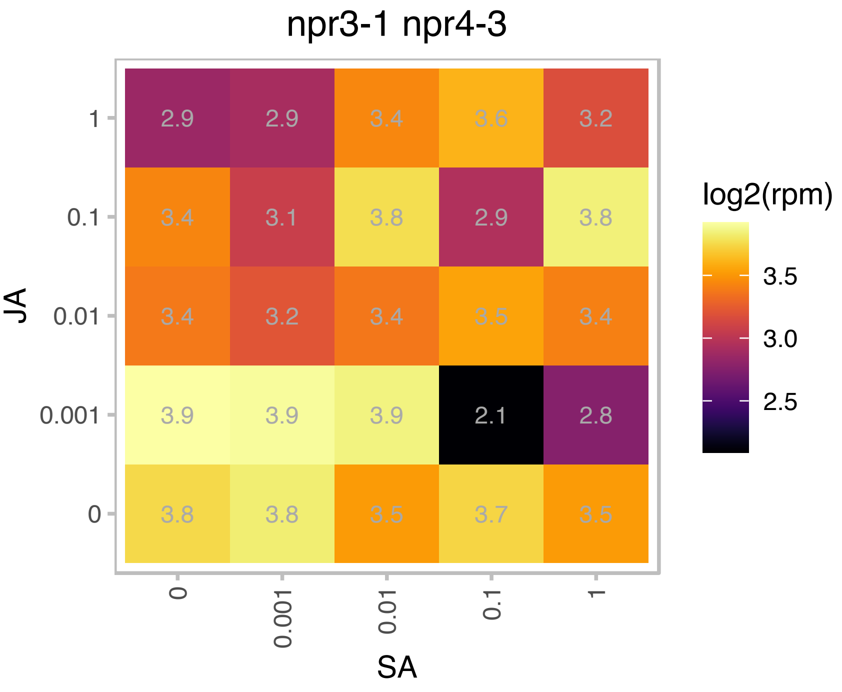

### AT3G19710.1 : BCAT4

branched-chain aminotransferase4
Belongs to the branched-chain amino acid aminotransferase gene family. Encodes a methionine-oxo-acid transaminase. Involved in the methionine chain elongation pathway that leads to the ultimate biosynthesis of methionine-derived glucosinolates.

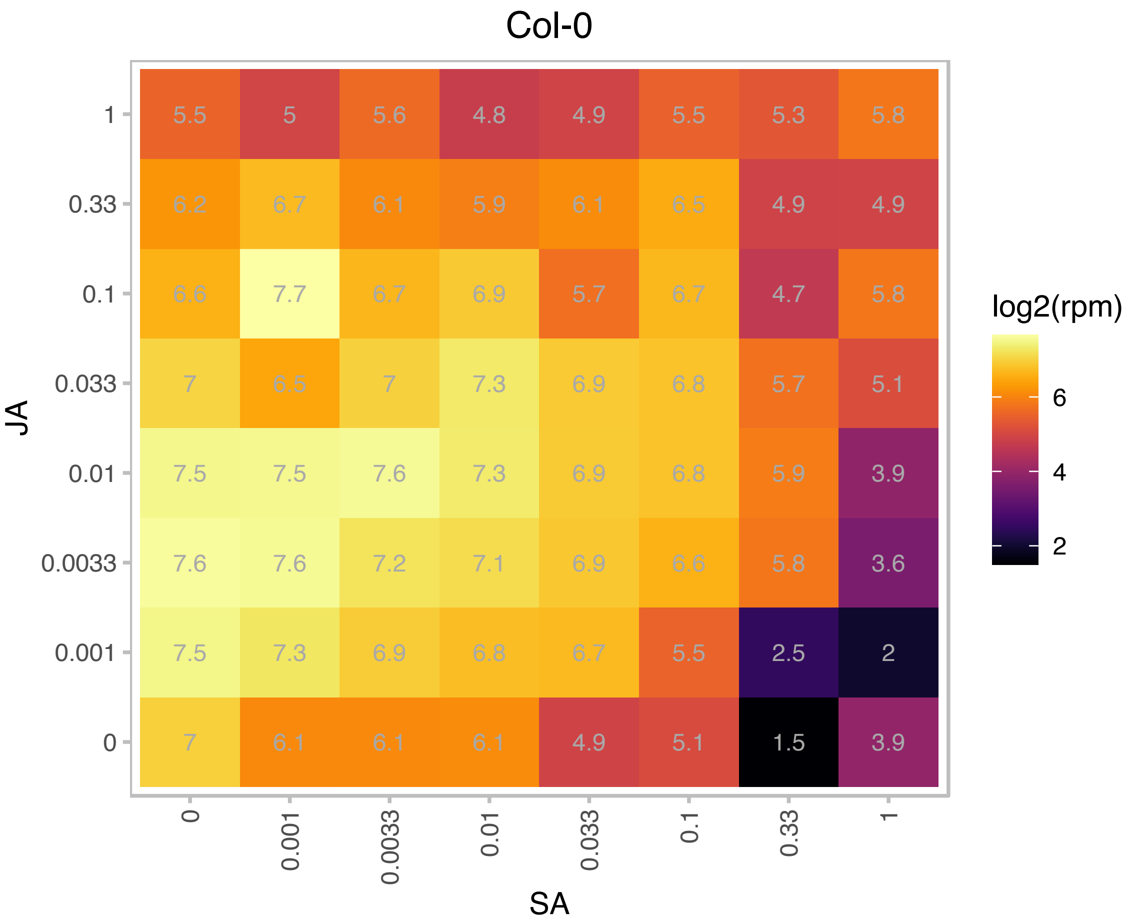


| | |
|---|---|
| Gene | AT3G19710 |
| is.expressed_genes | TRUE |
| is.responsive_genes | TRUE |
| is.apcluster | F-02 |

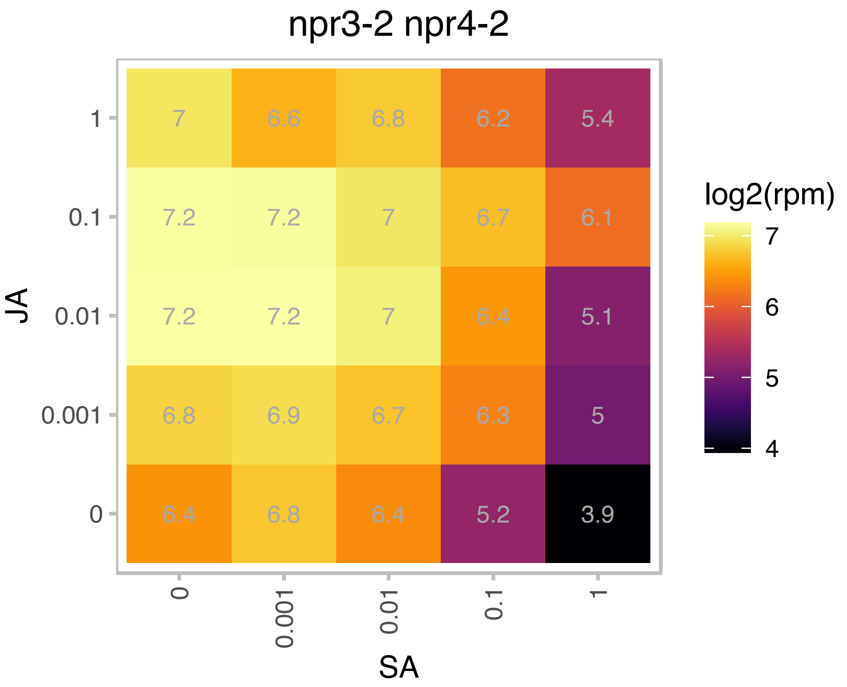


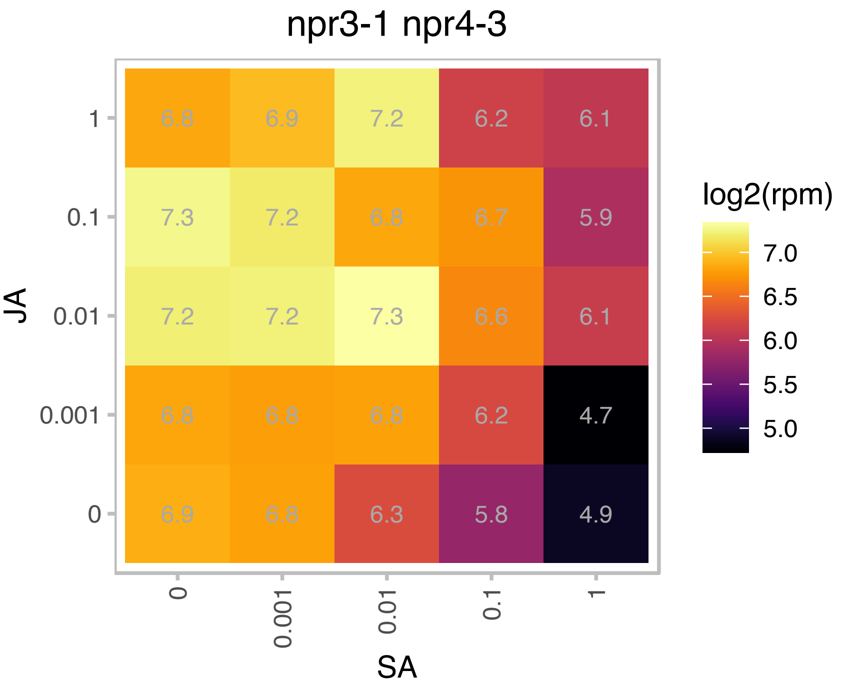

### AT3G21510.1 : AHP1

histidine-containing phosphotransmitter 1
Encodes AHP1, one of the six Arabidopsis thaliana histidine phosphotransfer proteins (AHPs). AHPs function as redundant positive regulators of cytokinin signaling. Members of the AHP gene family include: AT3G21510 (AHP1), AT3G29350 (AHP2), AT5G39340 (AHP3), AT3G16360 (AHP4), AT1G03430 (AHP5) and AT1G80100 (AHP6

| | |
|---|---|
| Gene | AT3G21510 |
| is.expressed_genes | TRUE |
| is.responsive_genes | TRUE |
| is.apcluster | F-02 |

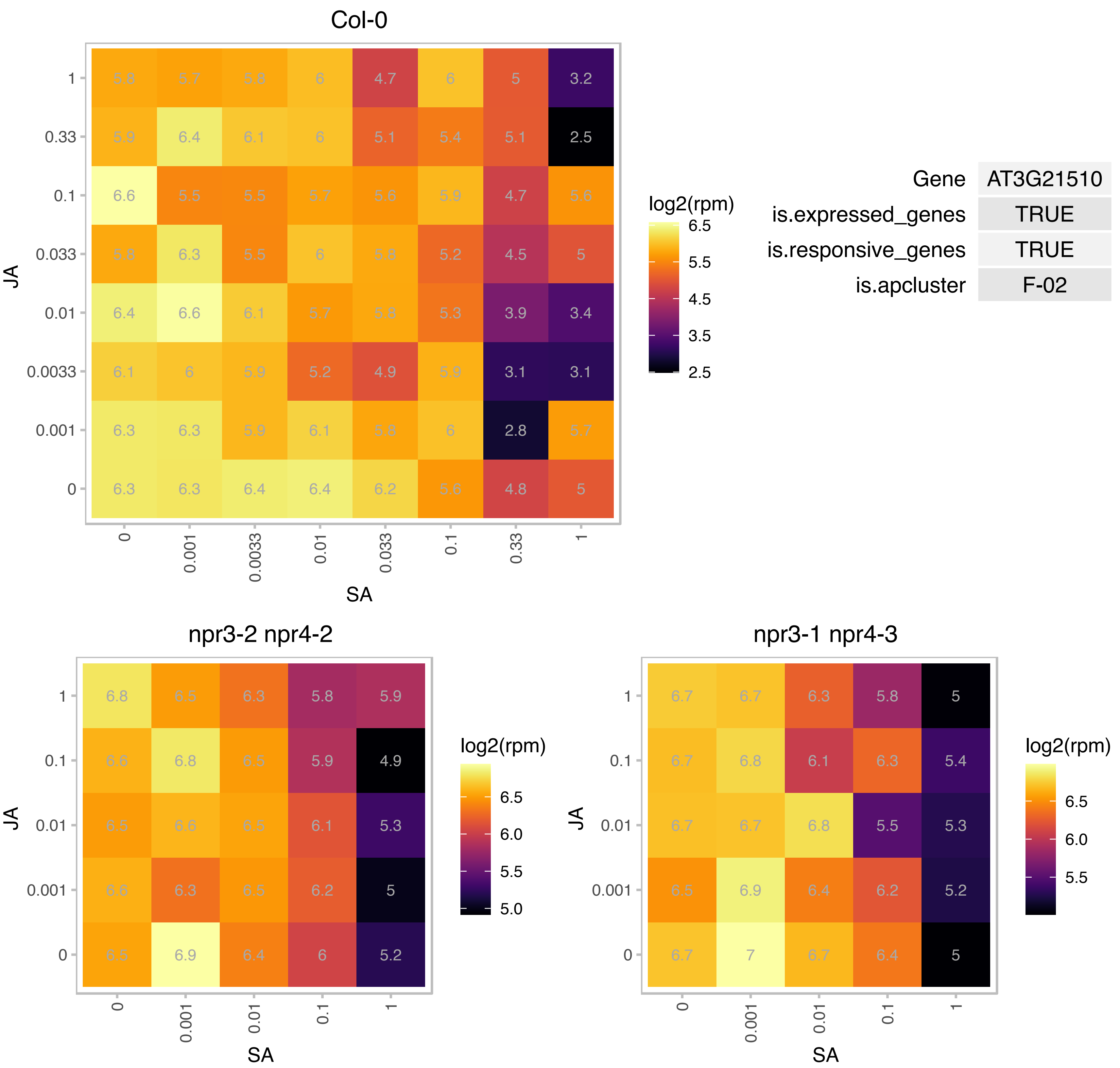

# AT3G24506.1 : NA

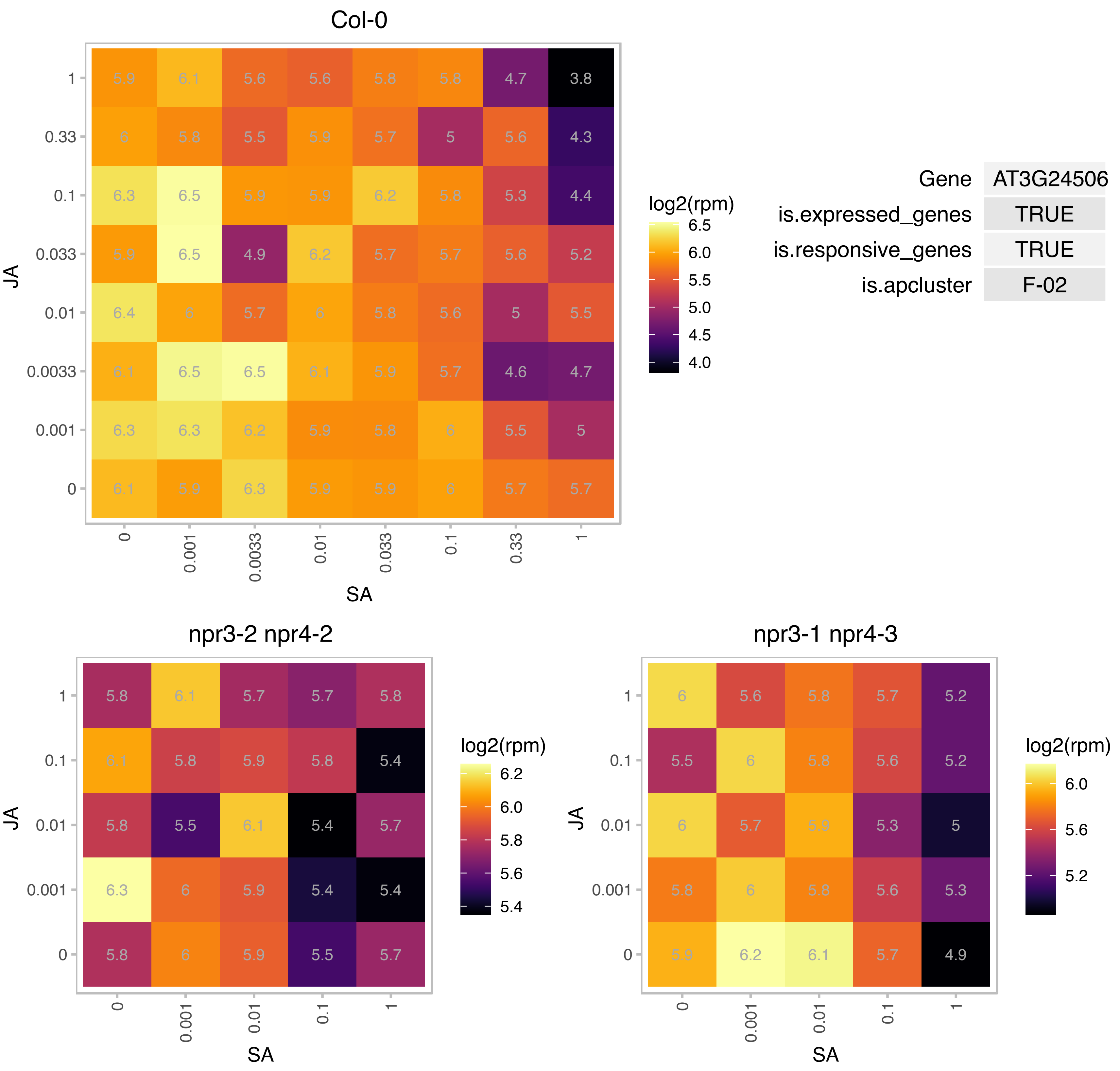


| | |
|---|---|
| Gene | AT3G24506 |
| is.expressed_genes | TRUE |
| is.responsive_genes | TRUE |
| is.apcluster | F-02 |

### AT3G27080.1 : TOM20-3

translocase of outer membrane 20 kDa subunit 3

Component of the TOM complex involved in transport of nuclear-encoded mitochondrial proteins

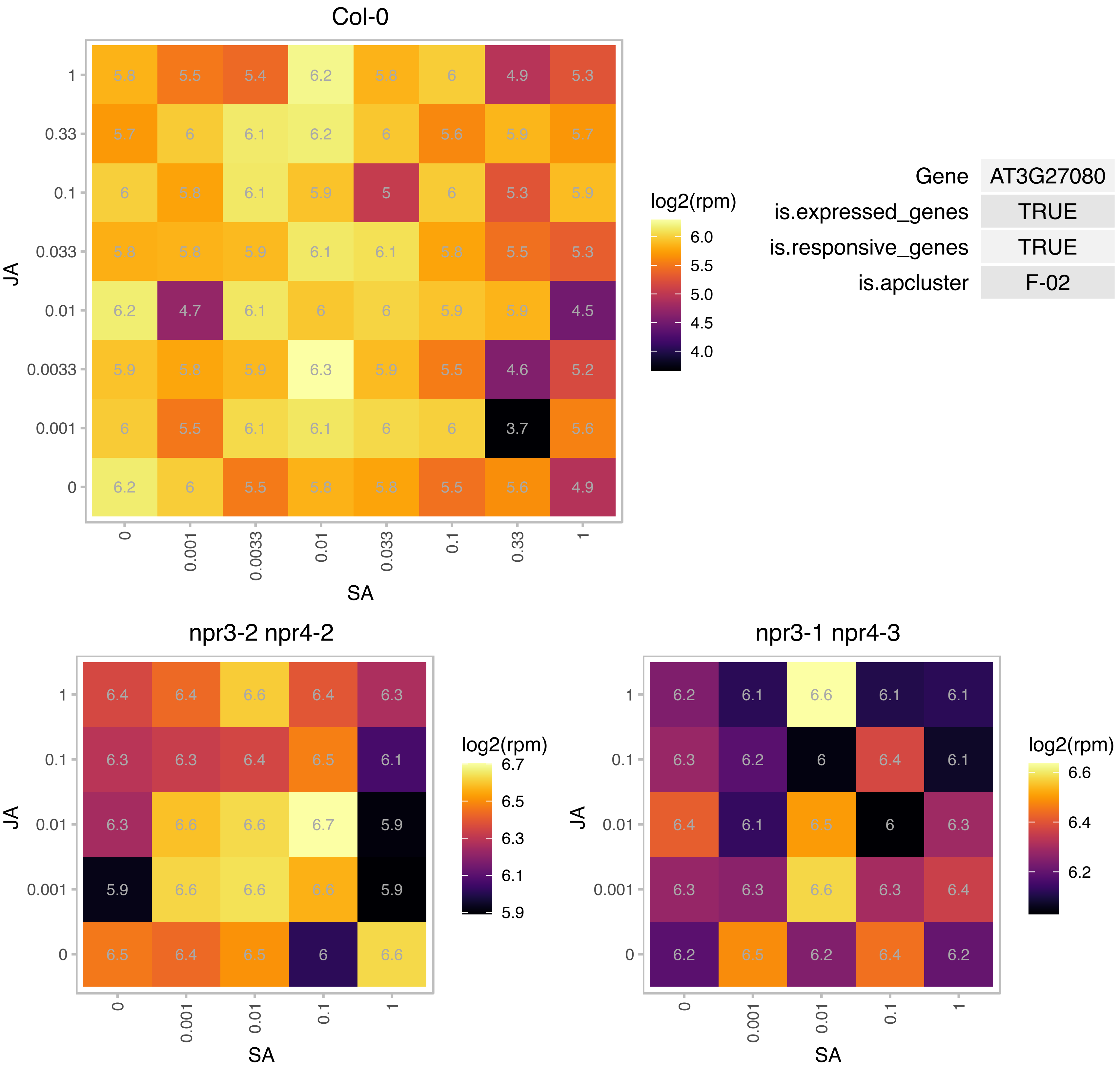


| | |
|---|---|
| Gene | AT3G27080 |
| is.expressed_genes | TRUE |
| is.responsive_genes | TRUE |
| is.apcluster | F-02 |

# AT3G28550.1 : NA

Proline-rich extensin-like family protein

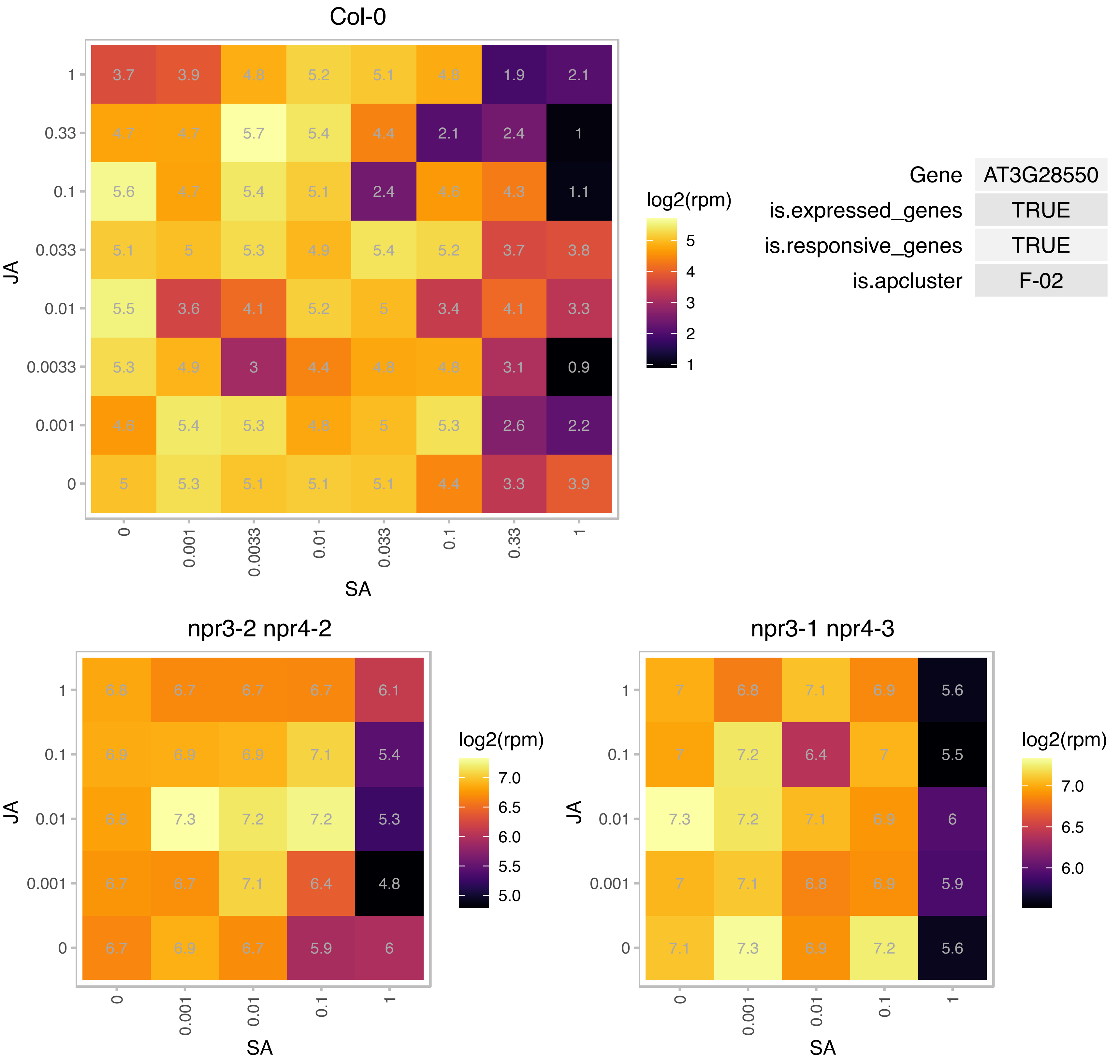


| | |
|---|---|
| Gene | AT3G28550 |
| is.expressed_genes | TRUE |
| is.responsive_genes | TRUE |
| is.apcluster | F-02 |

# AT3G47510.1 : NA

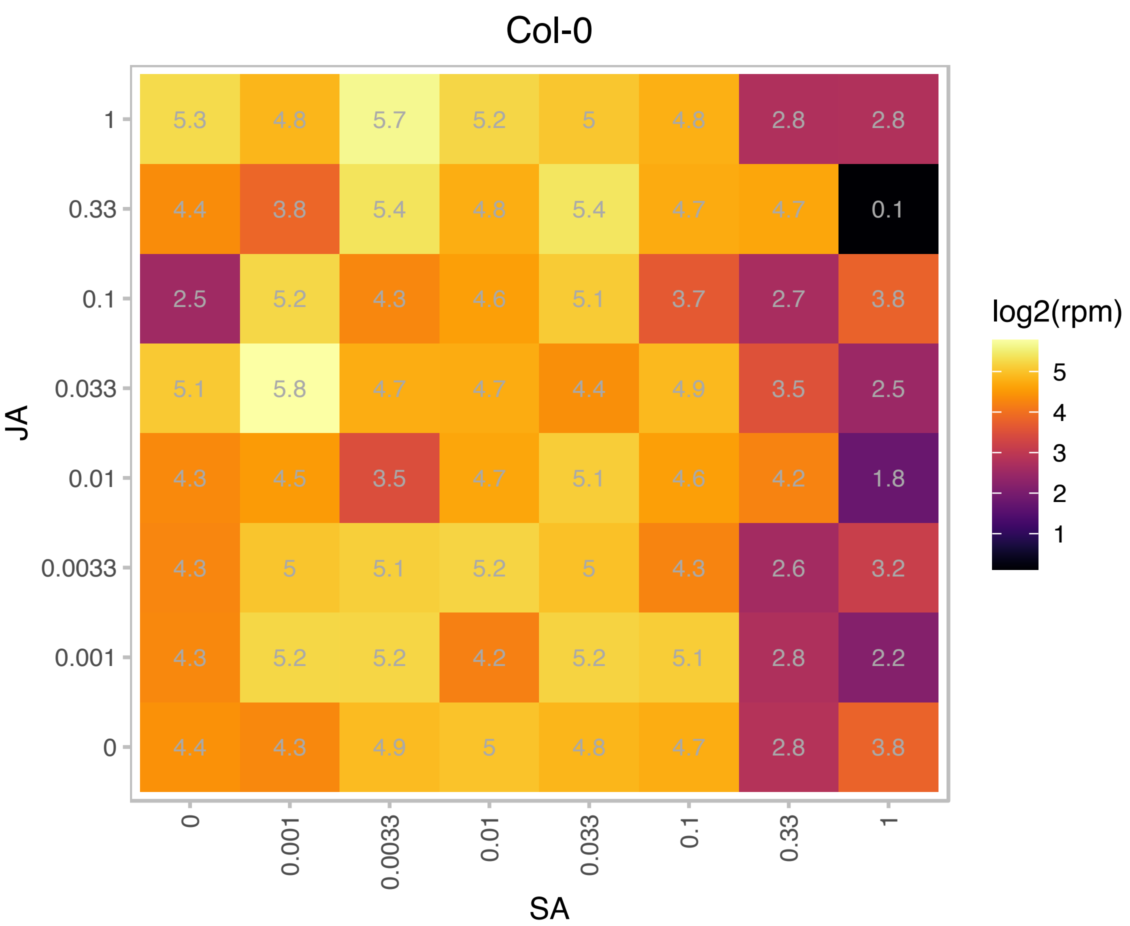


| | |
|---|---|
| Gene | AT3G47510 |
| is.expressed_genes | TRUE |
| is.responsive_genes | TRUE |
| is.apcluster | F-02 |

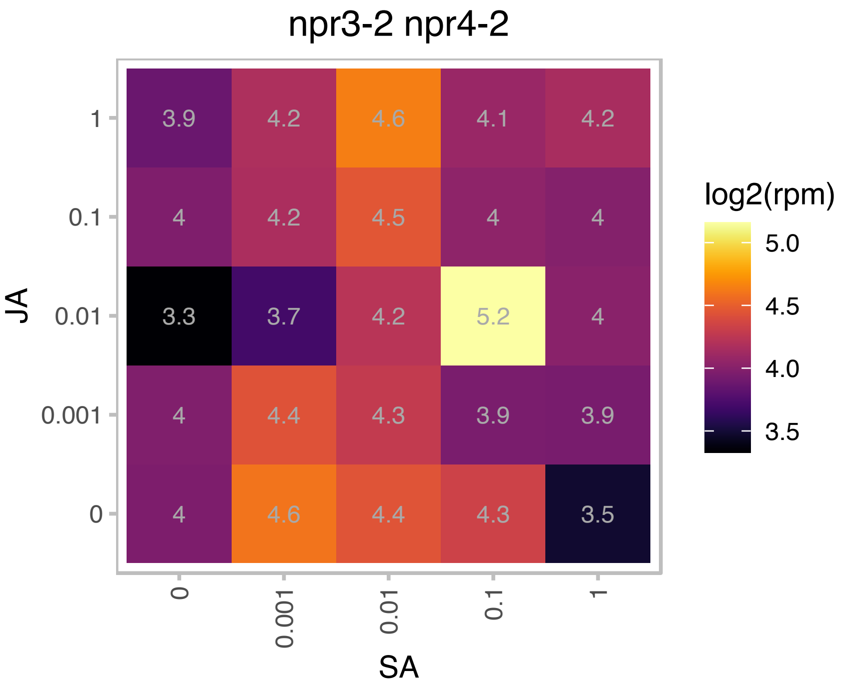


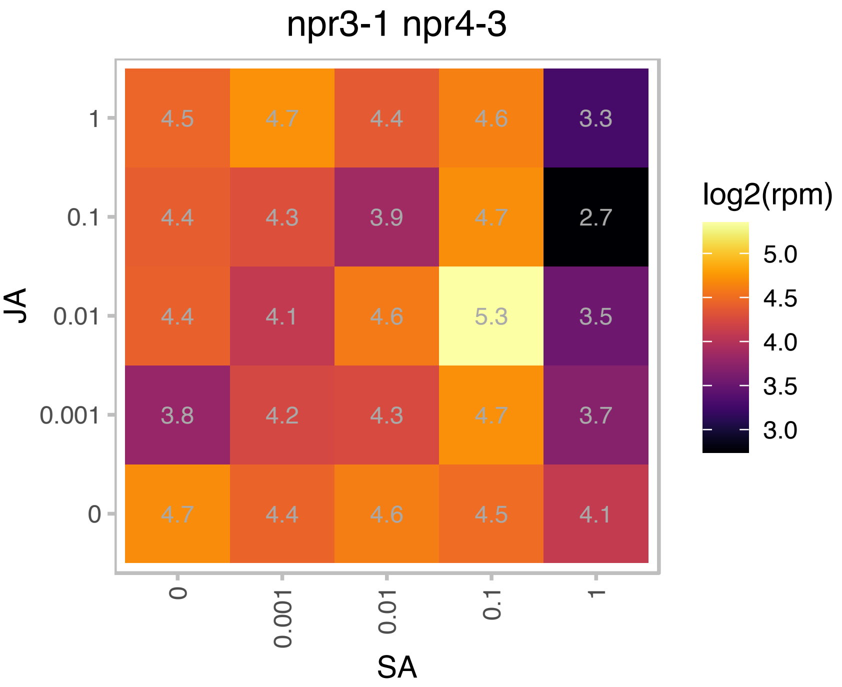

### AT3G48590.1 : ATHAP5A, HAP5A, NF-YC1

nuclear factor Y, subunit C1

Encodes a protein with similarity to a subunit of the CCAAT promoter motif binding complex of yeast.One of two members of this class (HAP5A) and expressed in vegetative and reproductive tissues.

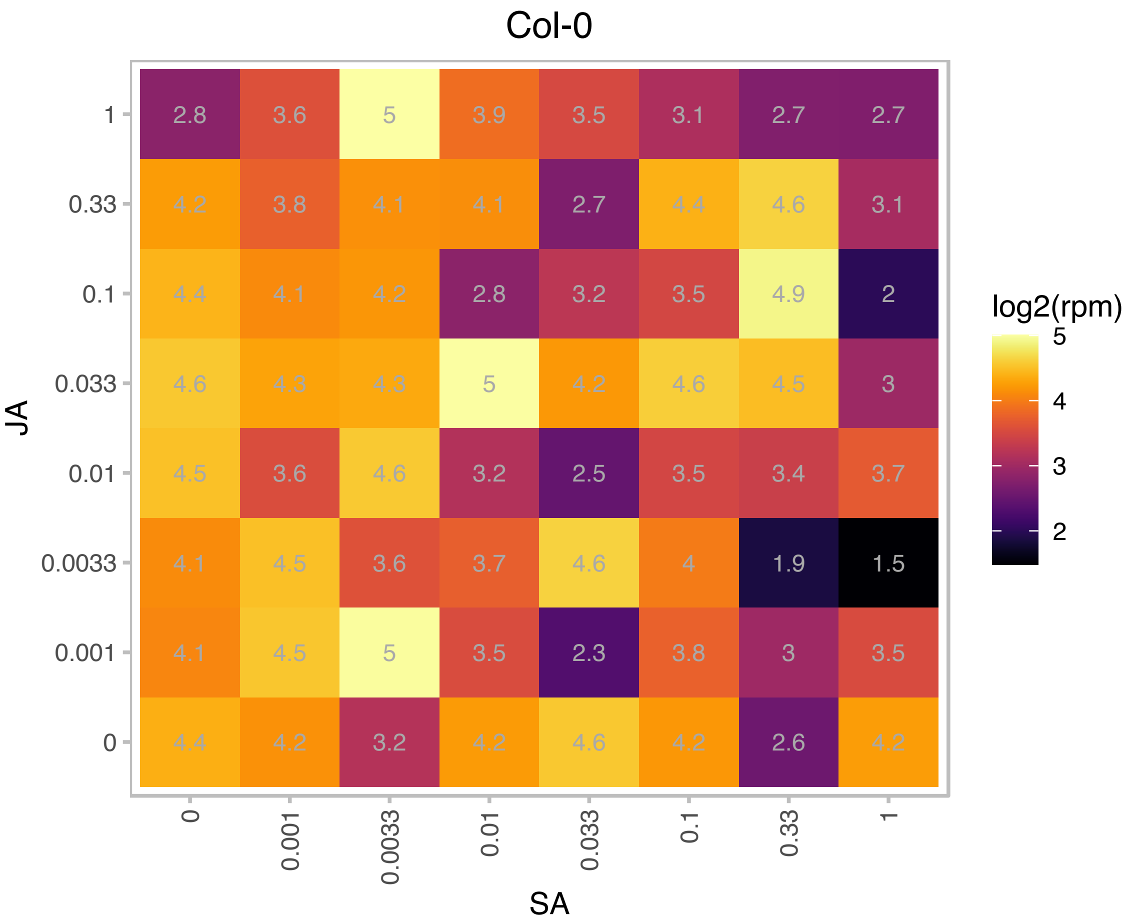


| | |
|---|---|
| Gene | AT3G48590 |
| is.expressed_genes | TRUE |
| is.responsive_genes | TRUE |
| is.apcluster | F-02 |

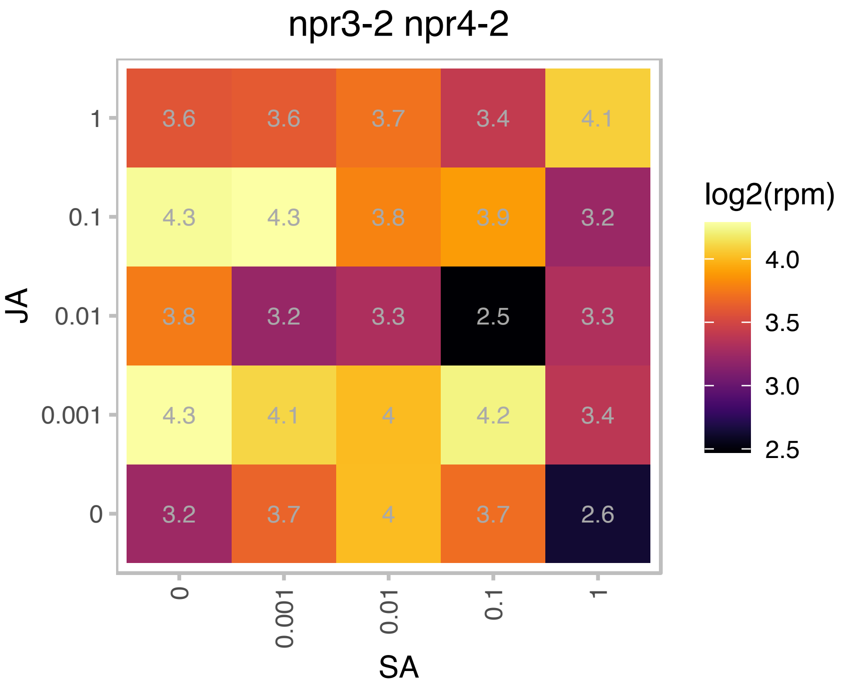


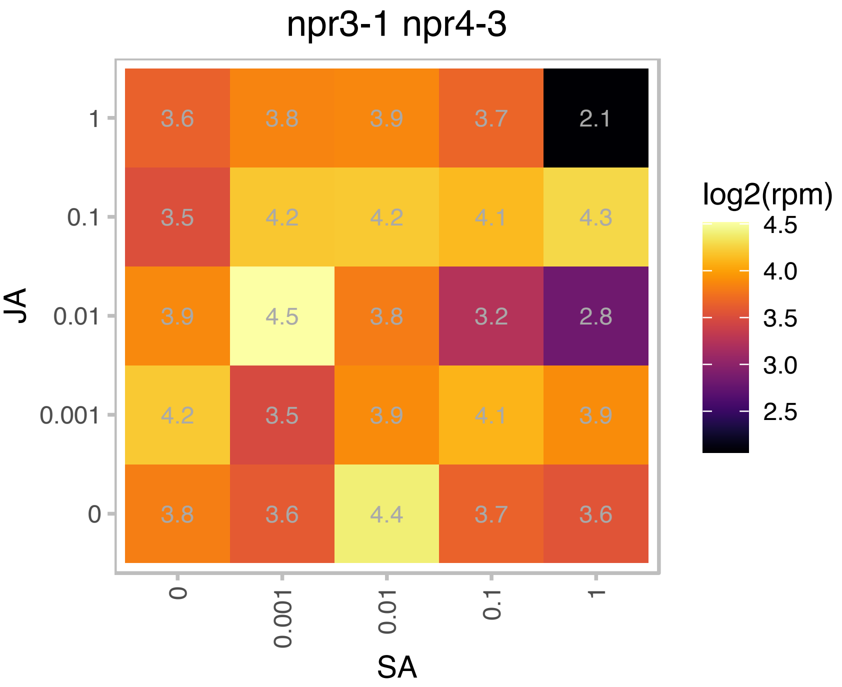

### AT3G49560.1 : HP30, Tric1

Mitochondrial import inner membrane translocase subunit Tim17/Tim22/Tim23 family protein

HP30/Tric1 is a component of the mitochondrial protein translocation complex and is involved in tRNA transport along with HP30-2/Tric2.

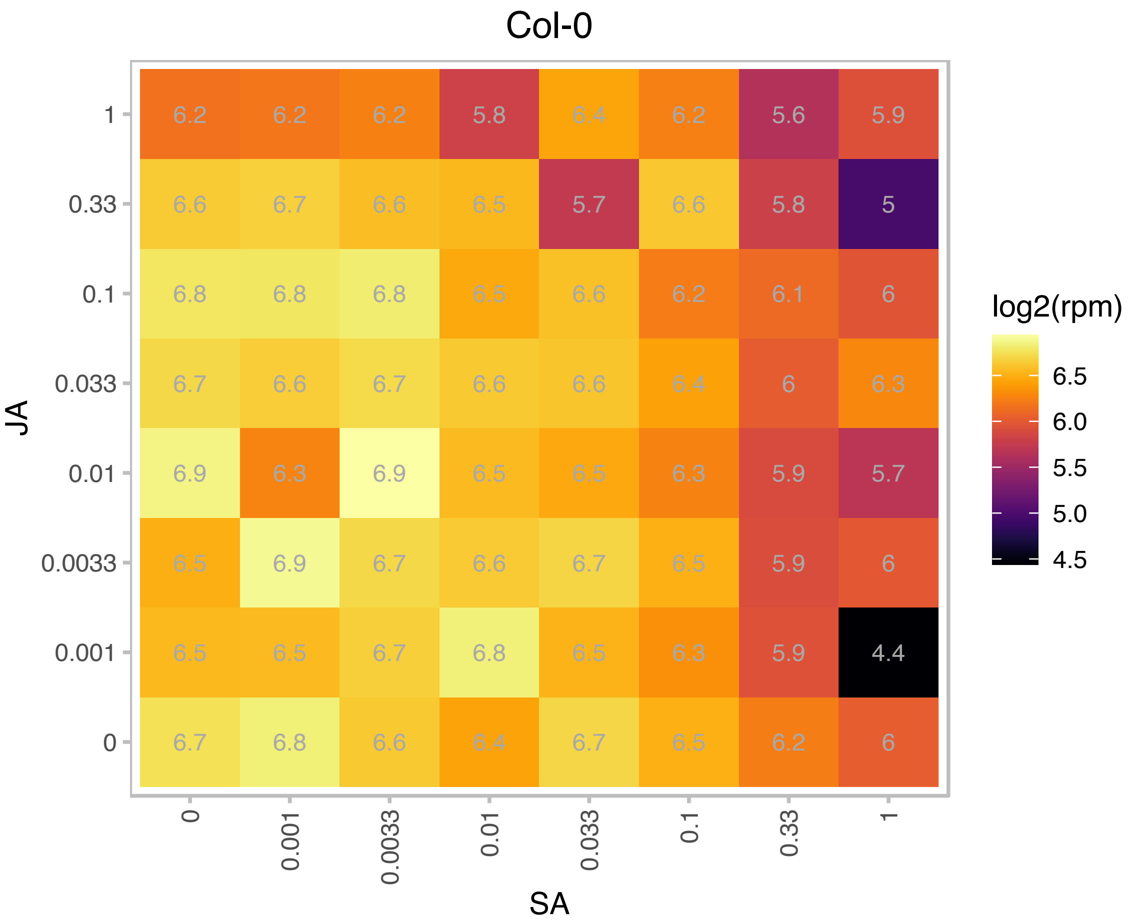


| | |
|---|---|
| Gene | AT3G49560 |
| is.expressed_genes | TRUE |
| is.responsive_genes | TRUE |
| is.apcluster | F-02 |

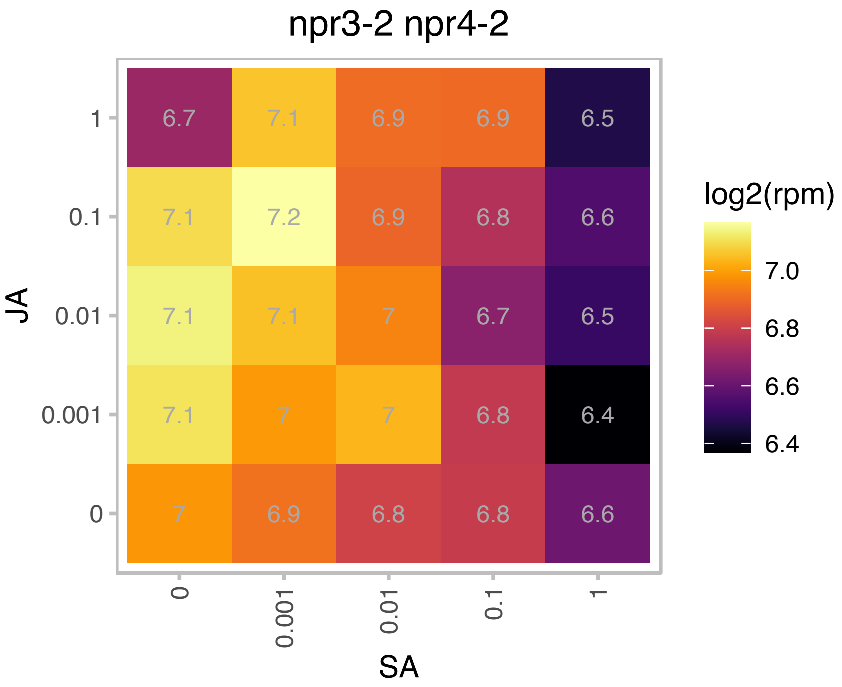


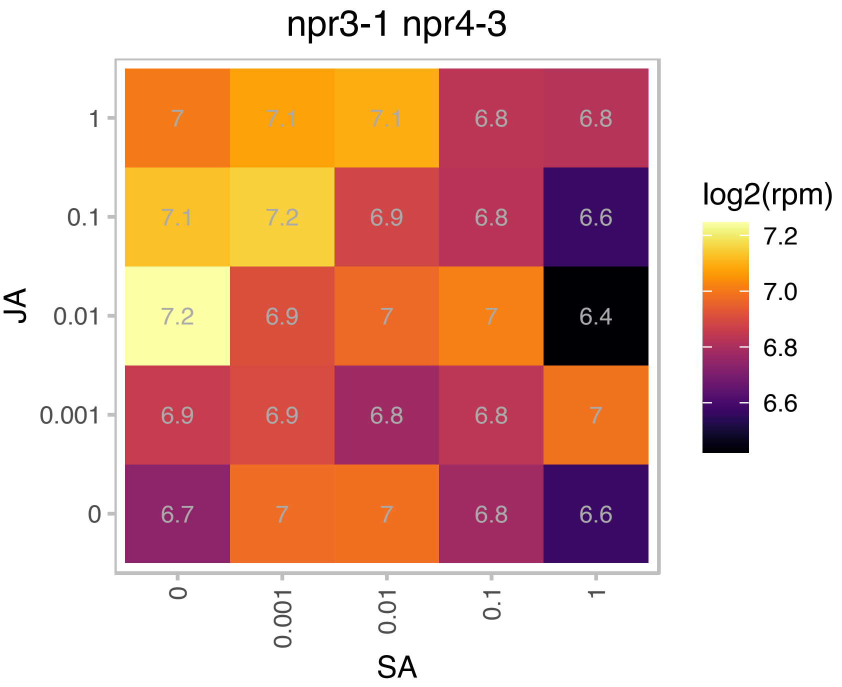

### AT3G49720.1 : CGR2

Encodes CGR2 (cotton Golgi-related 2). CGR2 and a close homologue CGR3 have overlapping roles in plant growt
and pectin methylesterification in the Golgi apparatus.

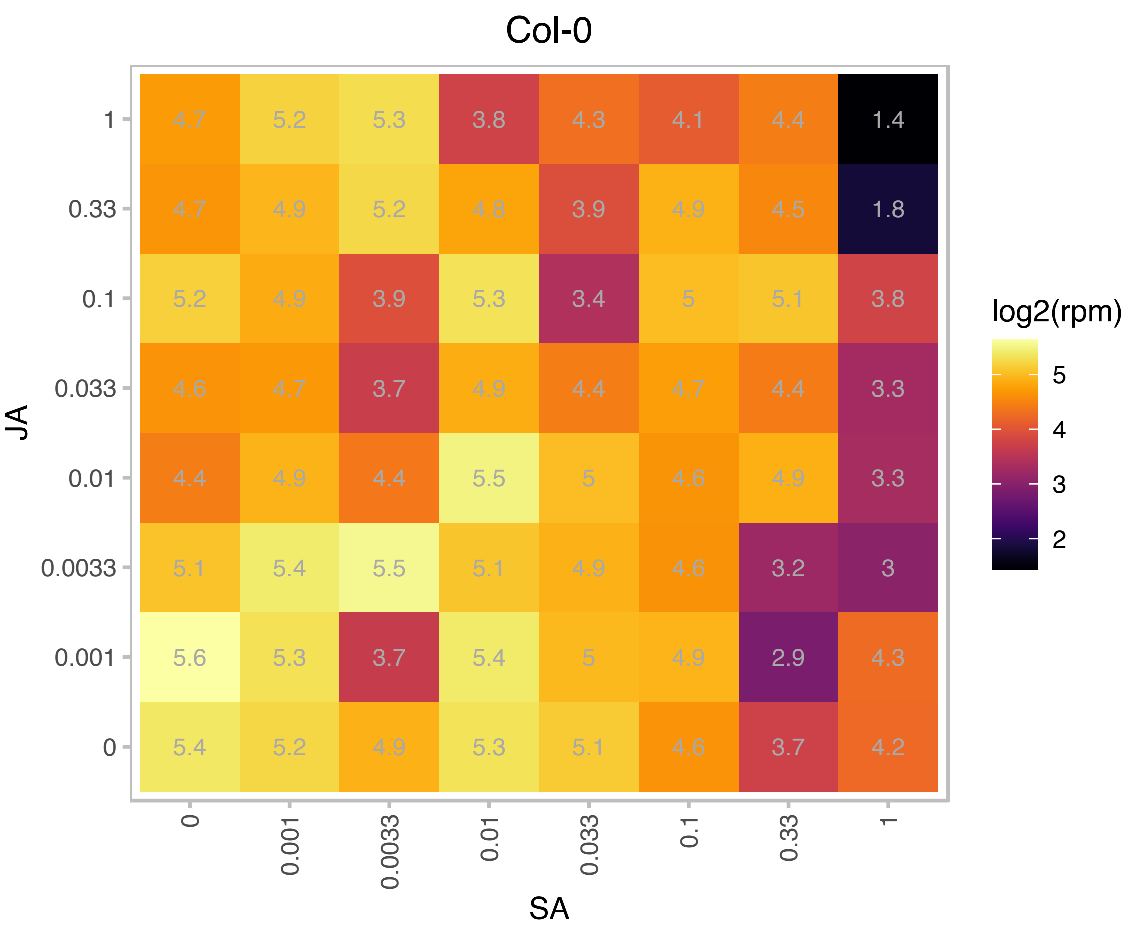


| | |
|---|---|
| Gene | AT3G49720 |
| is.expressed_genes | TRUE |
| is.responsive_genes | TRUE |
| is.apcluster | F-02 |

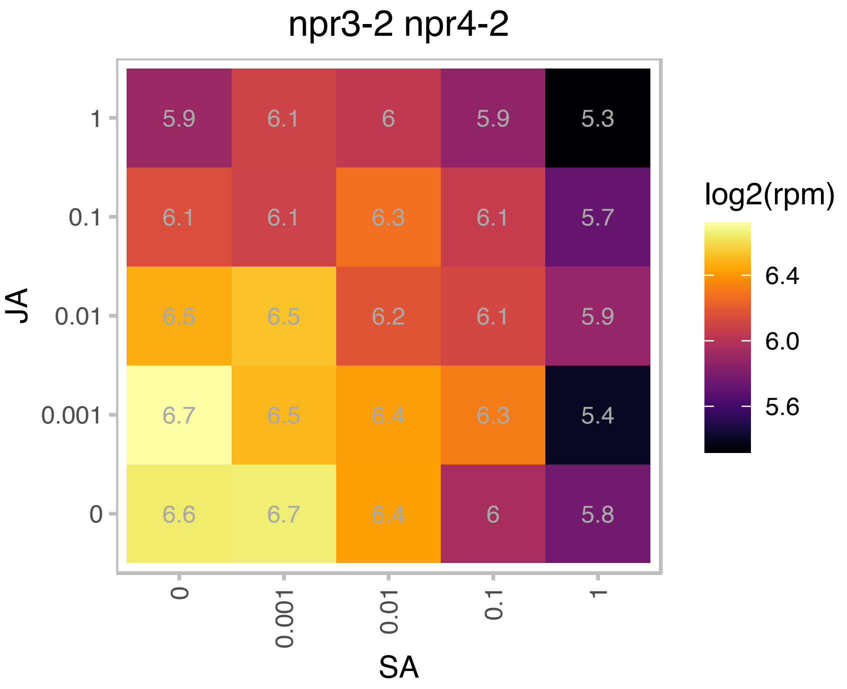


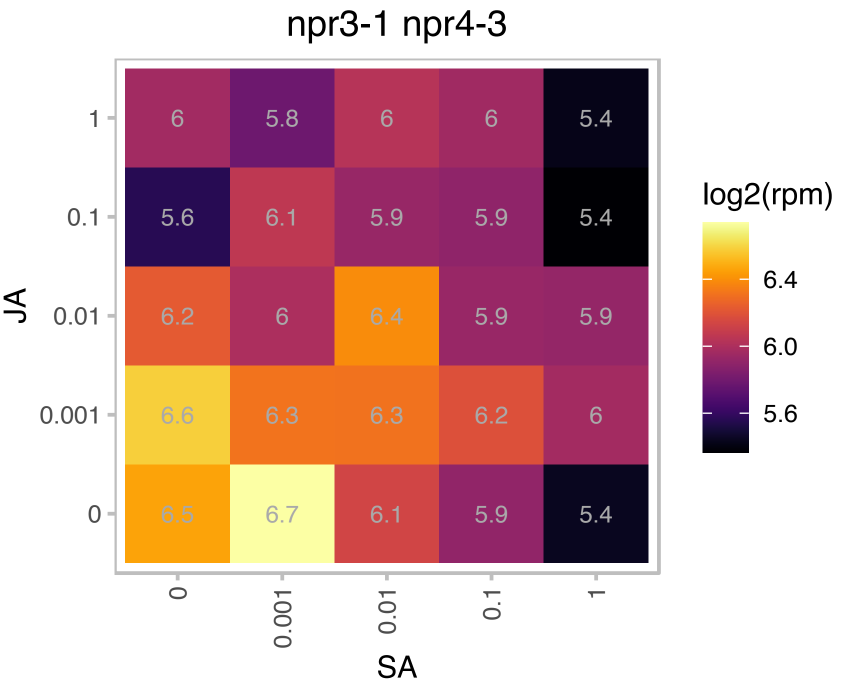

### AT3G49870.1 : ARLA1C, ATARLA1C

ADP-ribosylation factor-like A1C

A member of ARF-like GTPase family. A thaliana has 21 members, in two subfamilies, ARF and ARF-like (ARL) GTPases.

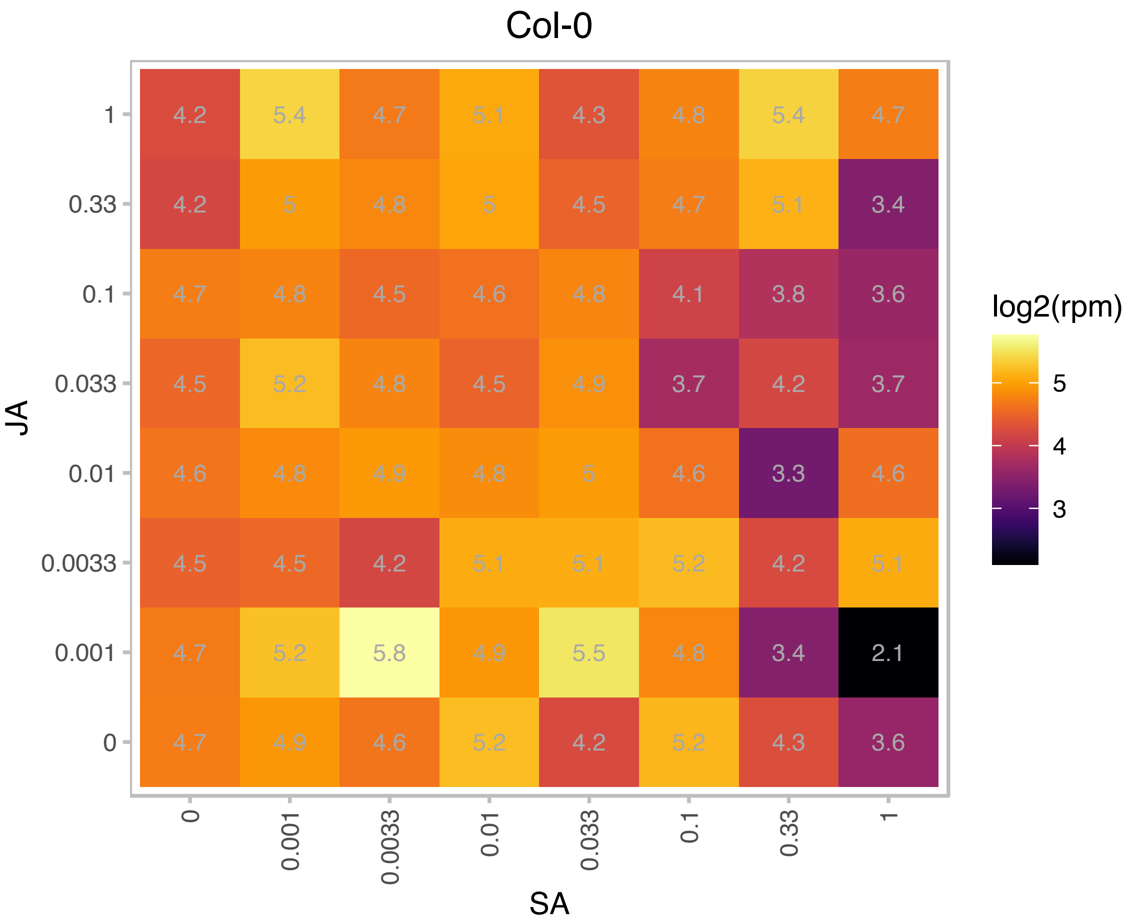


| Gene | AT3G49870 |
|---|---|
| is.expressed_genes | TRUE |
| is.responsive_genes | TRUE |
| is.apcluster | F-02 |

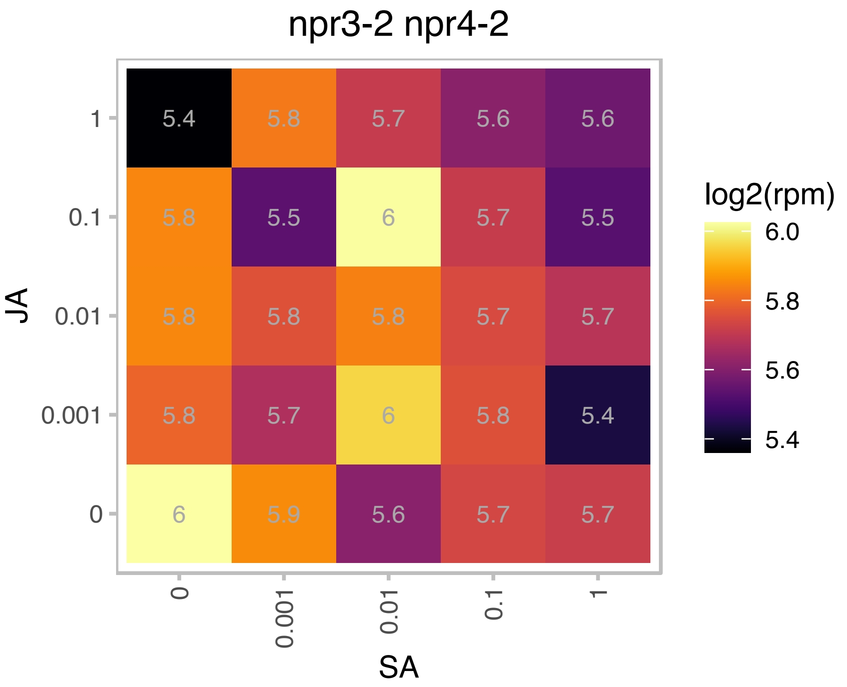


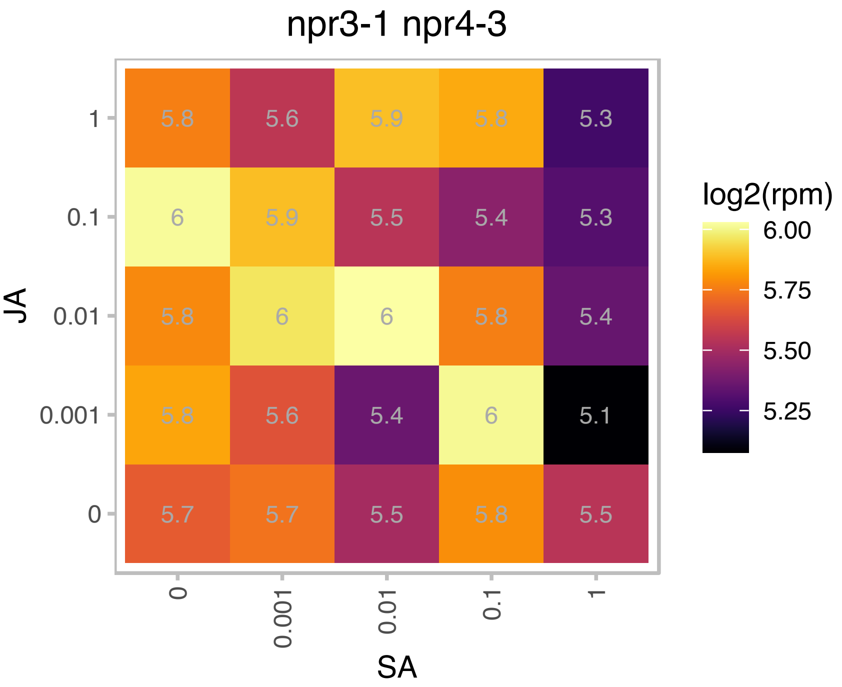

# AT3G51240.1 : F3'H, F3H, TT6

flavanone 3-hydroxylase

Encodes flavanone 3-hydroxylase that is coordinately expressed with chalcone synthase and chalcone isomerases. Regulates flavonoid biosynthesis. Not responsive to auxin or ethylene stimulus (qRT-PCR).

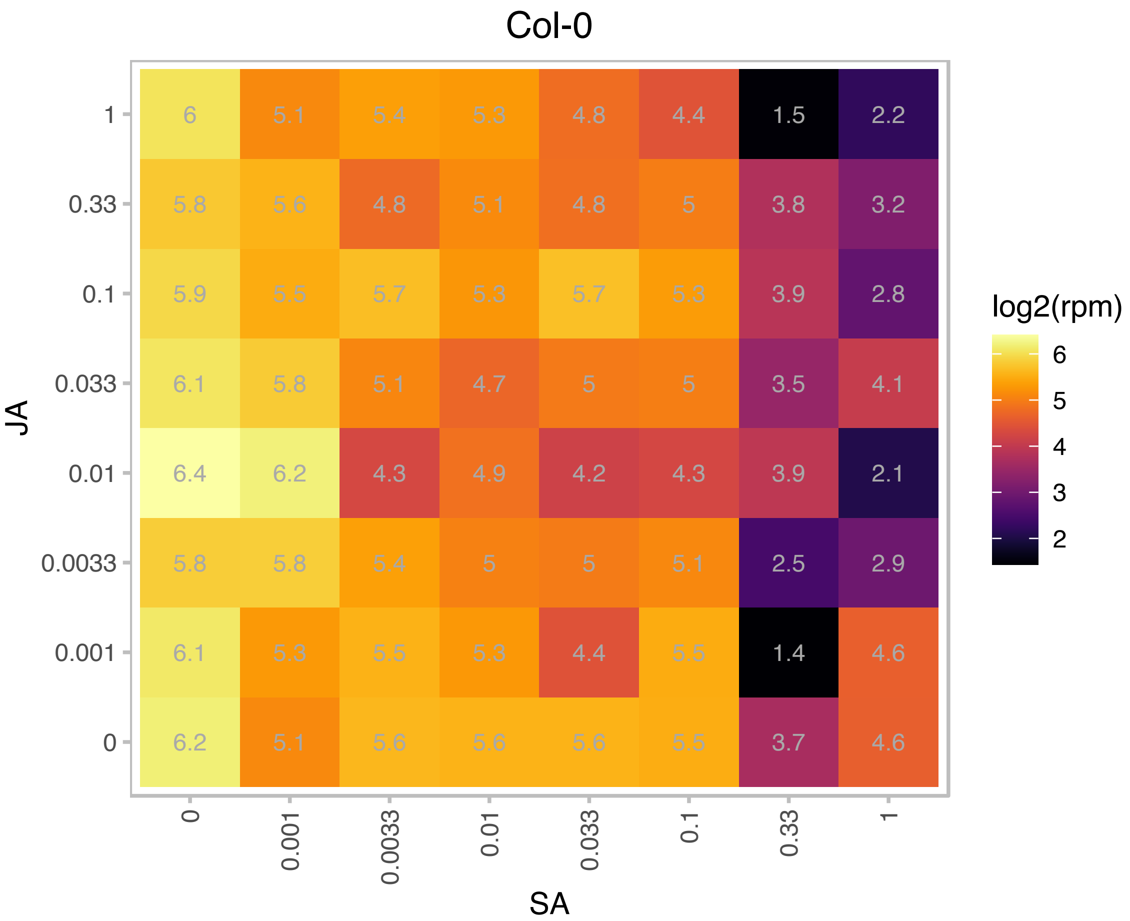


| | |
|---|---|
| Gene | AT3G51240 |
| is.expressed_genes | TRUE |
| is.responsive_genes | TRUE |
| is.apcluster | F-02 |

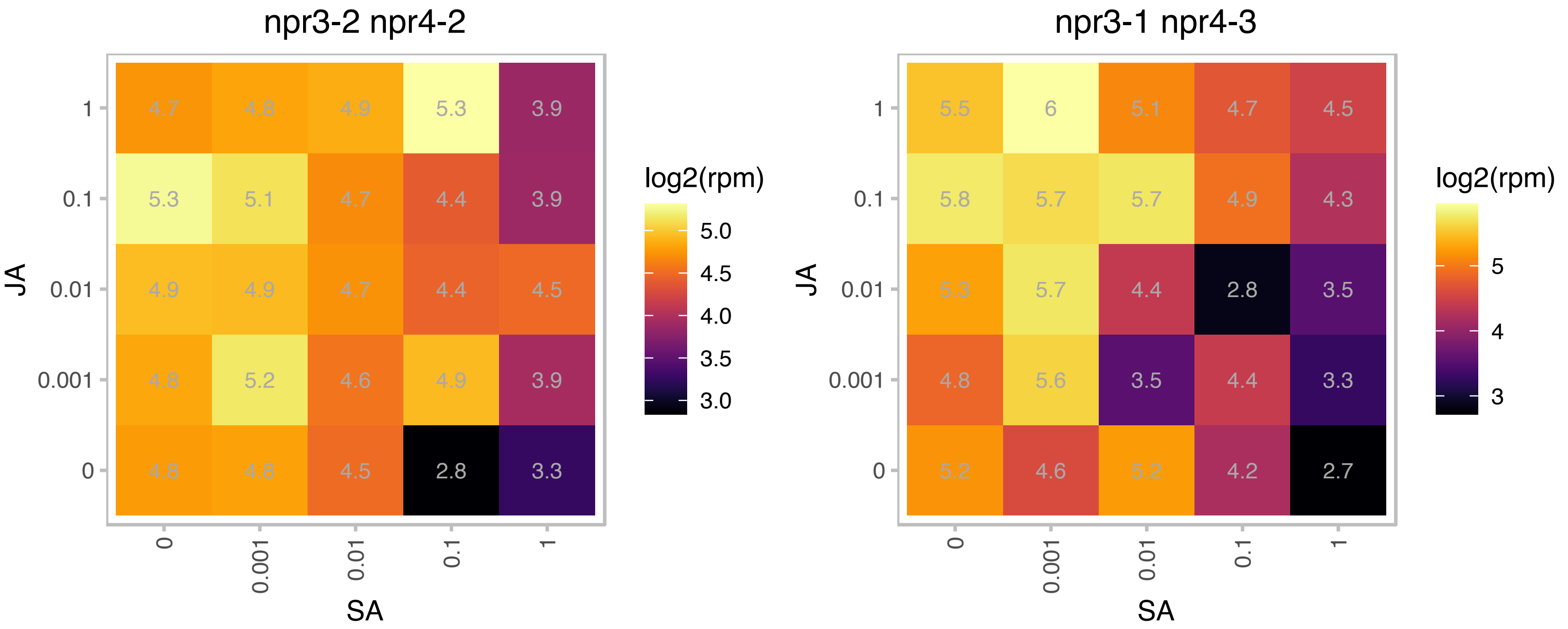

### AT3G51800.1 : ATEBP1, ATG2, EBP1, G2p

metallopeptidase M24 family protein

putative nuclear DNA-binding protein G2p (AtG2) mRNA,

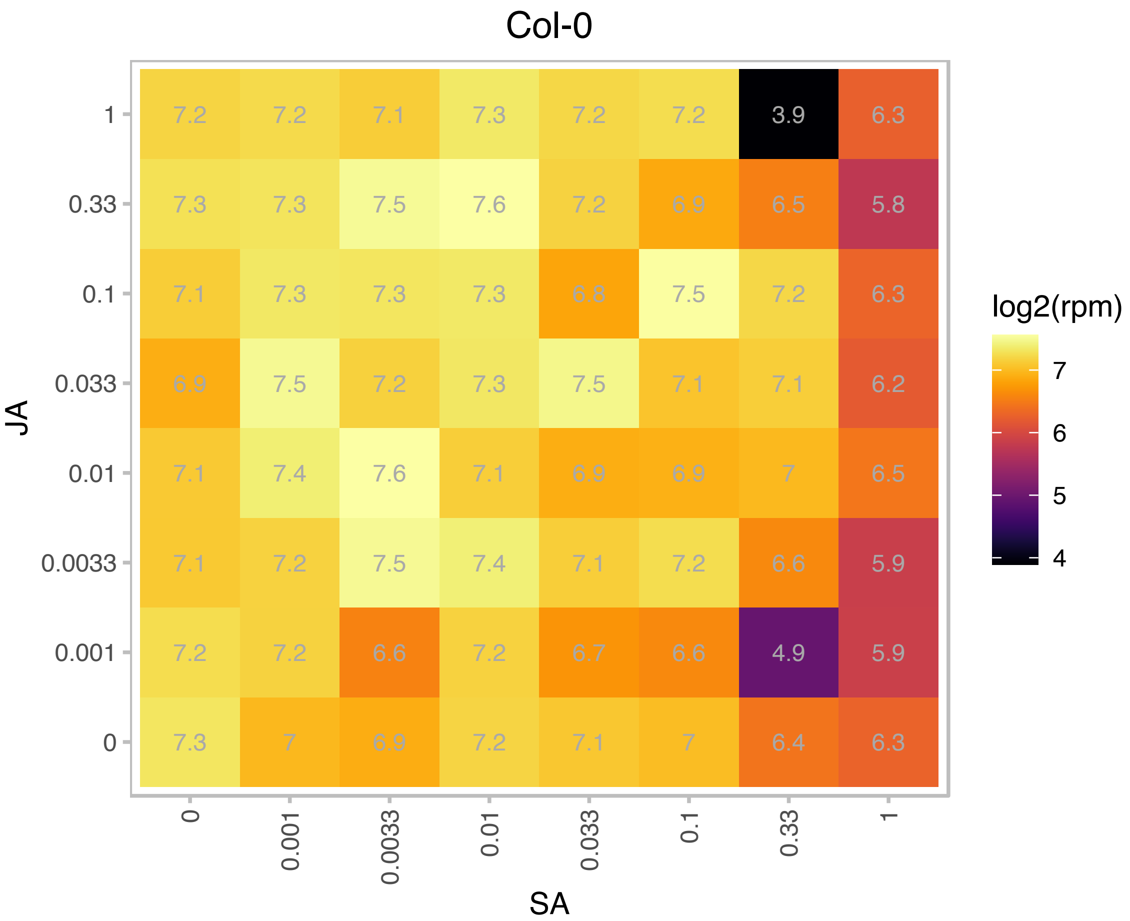


| | |
|---|---|
| Gene | AT3G51800 |
| is.expressed_genes | TRUE |
| is.responsive_genes | TRUE |
| is.apcluster | F-02 |

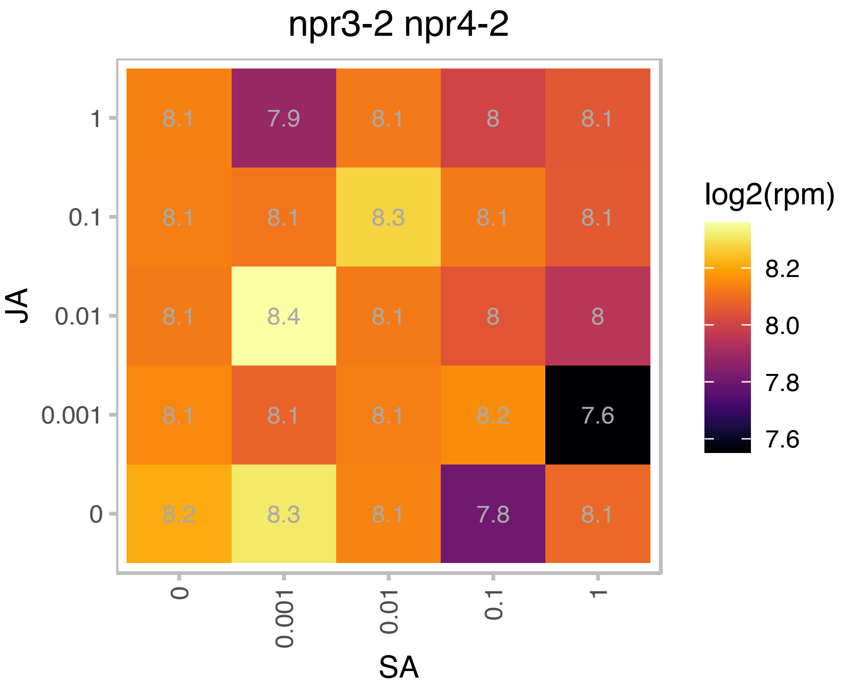


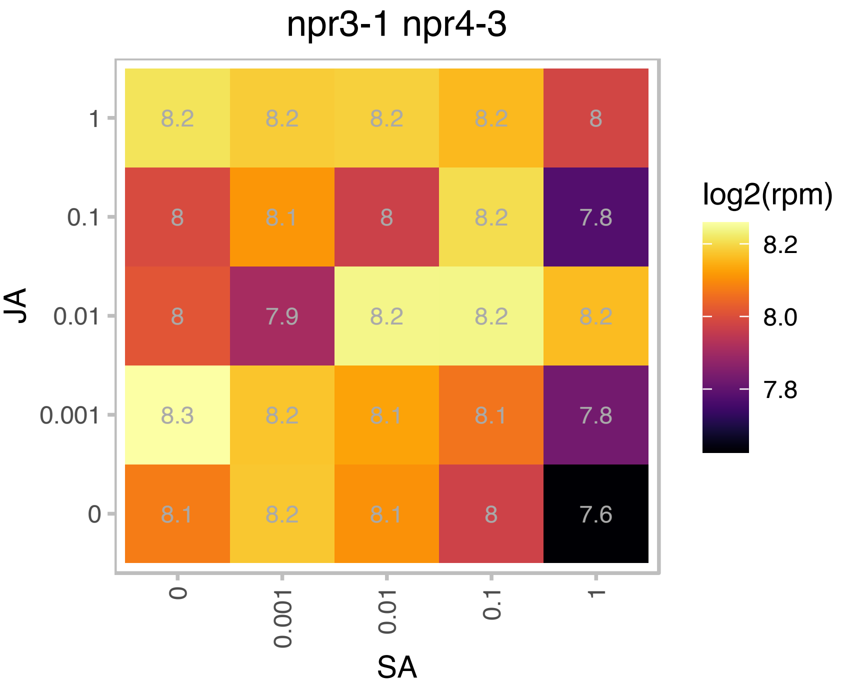

# AT3G52230.1 : NA

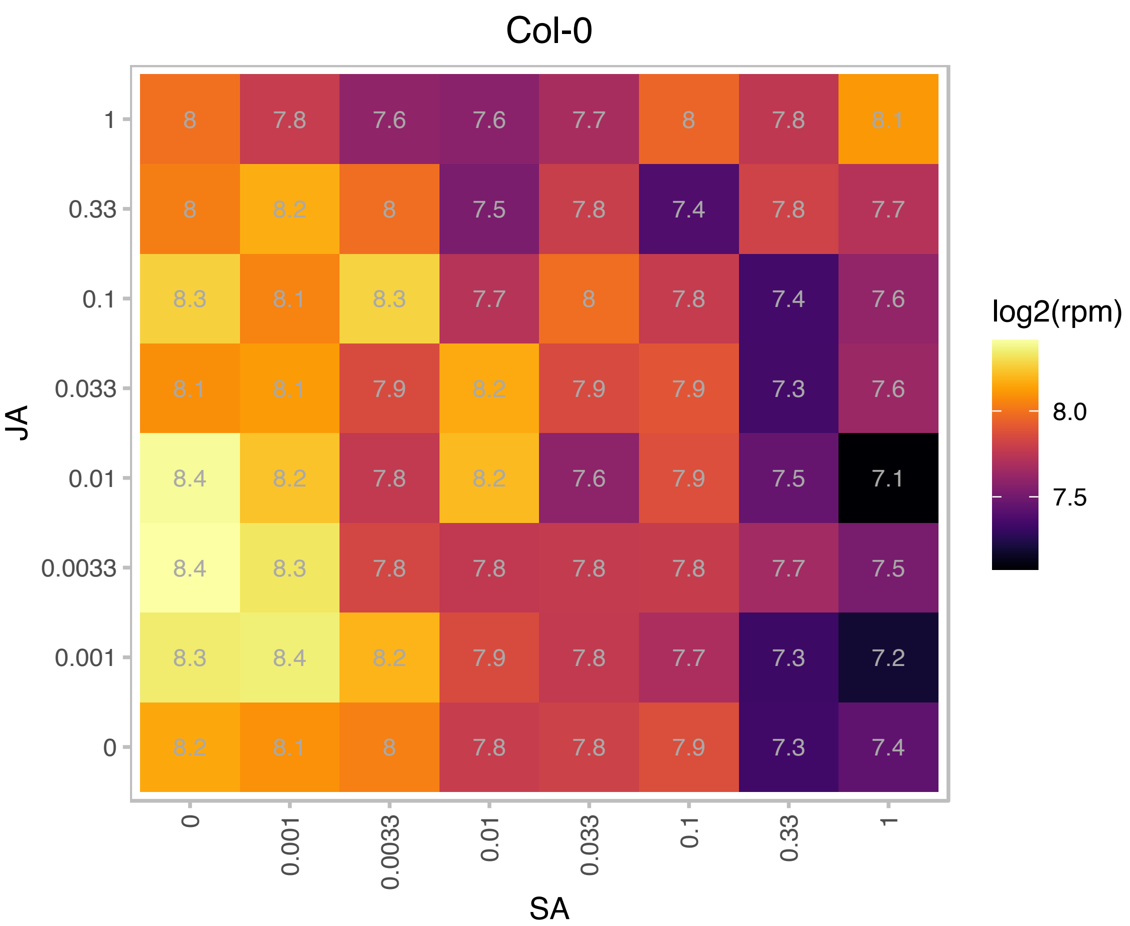


| | |
|---|---|
| Gene | AT3G52230 |
| is.expressed_genes | TRUE |
| is.responsive_genes | TRUE |
| is.apcluster | F-02 |

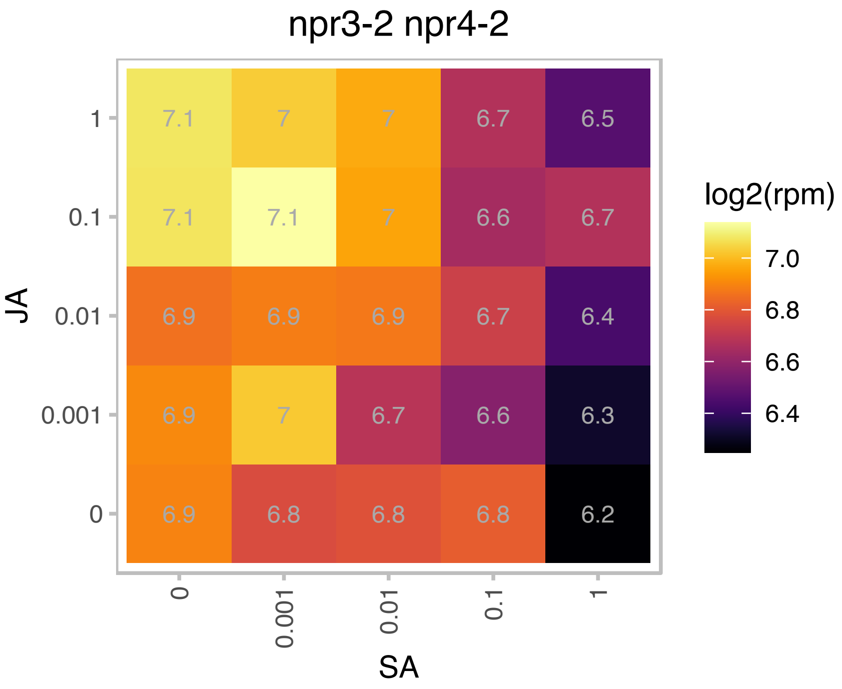


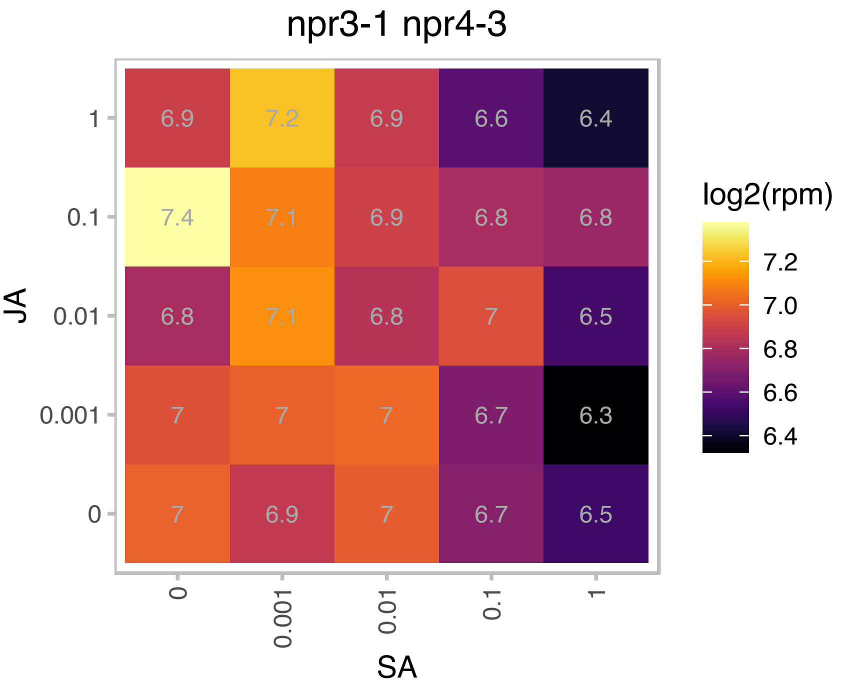

### AT3G53750.1 : ACT3

actin 3

Member of the Actin gene family. Expressed in mature pollen.

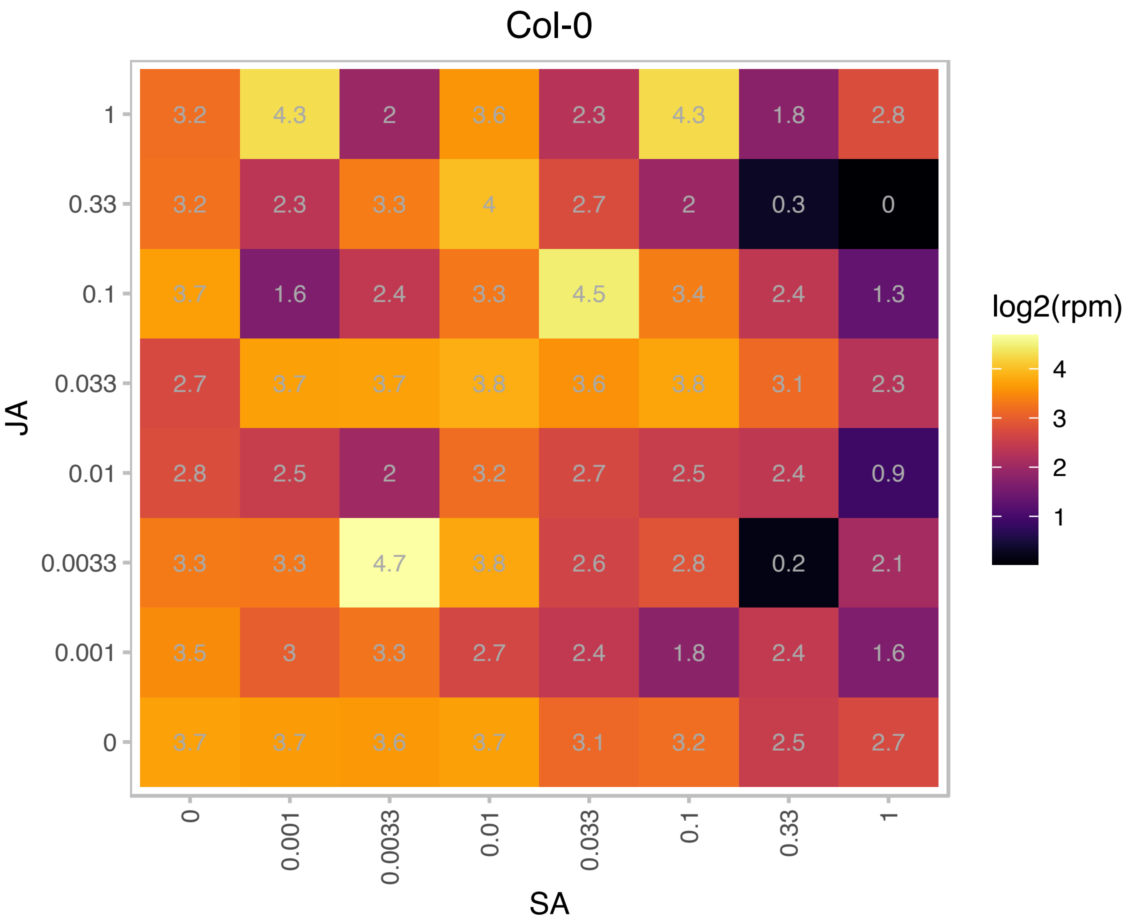


| | |
|---|---|
| Gene | AT3G53750 |
| is.expressed_genes | TRUE |
| is.responsive_genes | TRUE |
| is.apcluster | F-02 |

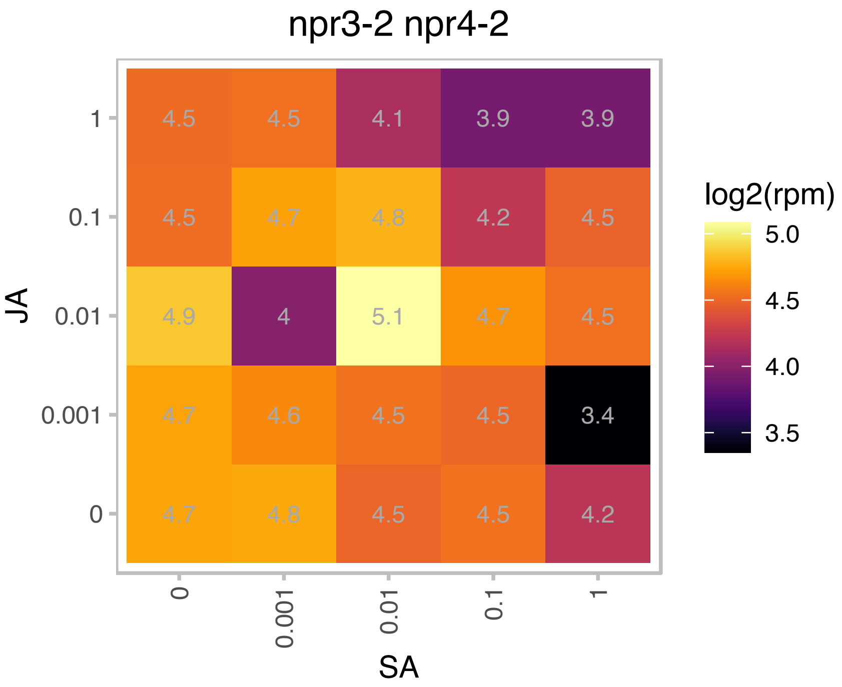


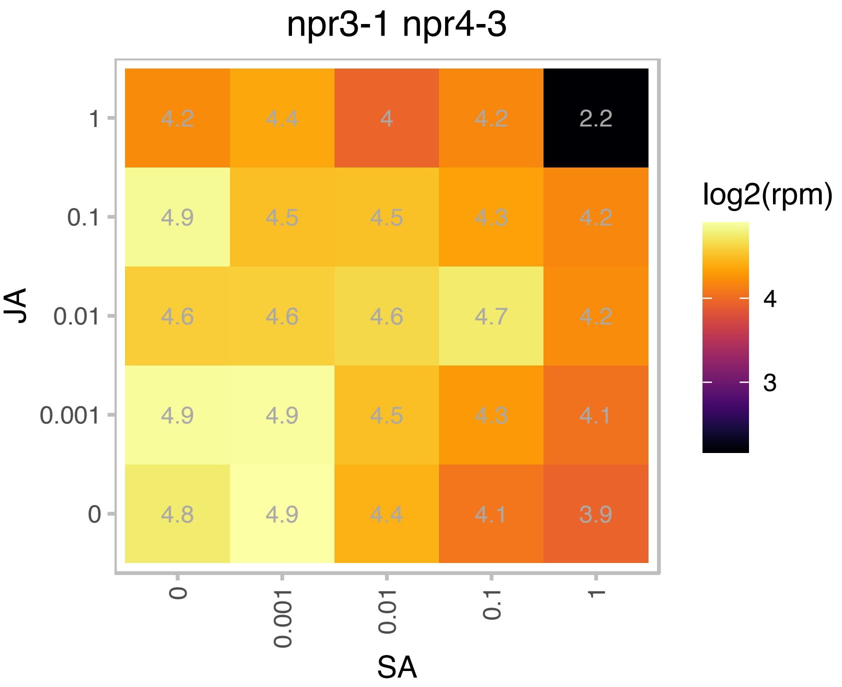

# AT3G54580.1 : NA

Proline-rich extensin-like family protein

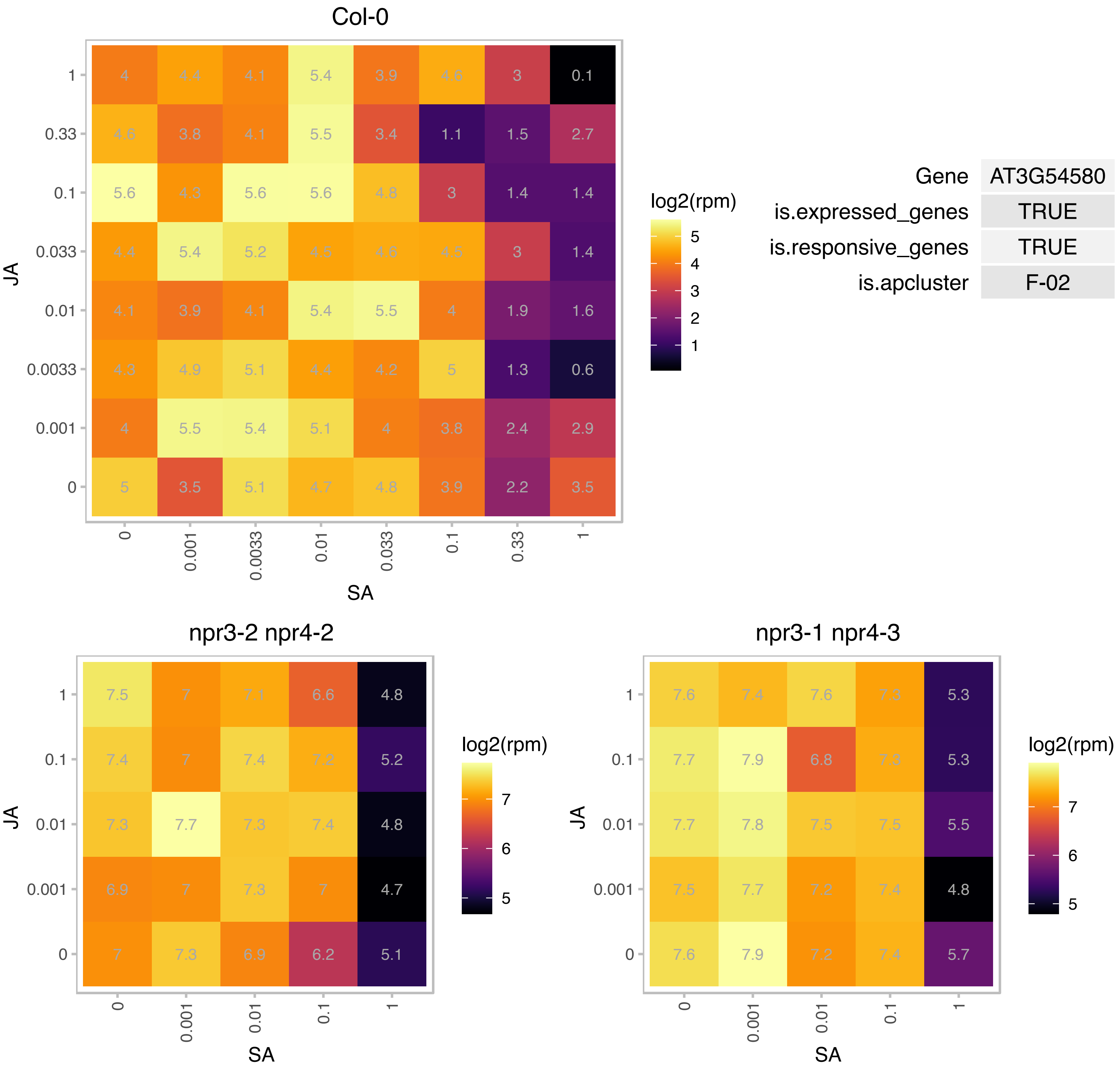


| | |
|---|---|
| Gene | AT3G54580 |
| is.expressed_genes | TRUE |
| is.responsive_genes | TRUE |
| is.apcluster | F-02 |

### AT3G54590.1 : ATHRGP1, HRGP1

hydroxyproline-rich glycoprotein

Encodes a hydroxyproline-rich glycoprotein. The mRNA is cell-to-cell mobile.

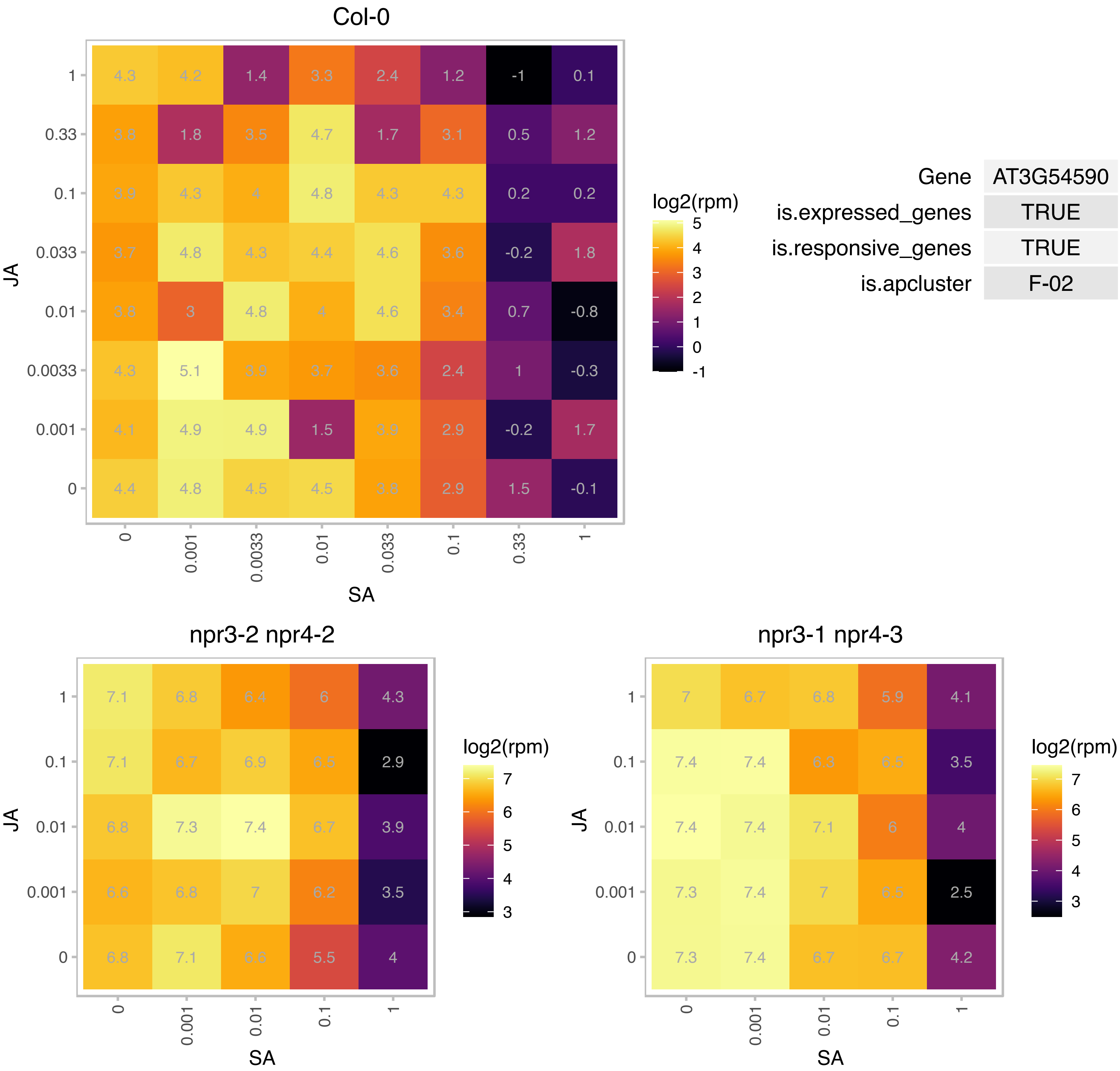


| | |
|---|---|
| Gene | AT3G54590 |
| is.expressed_genes | TRUE |
| is.responsive_genes | TRUE |
| is.apcluster | F-02 |

# AT3G54600.1 : DJ-1f, DJ1F

Class I glutamine amidotransferase-like superfamily protein

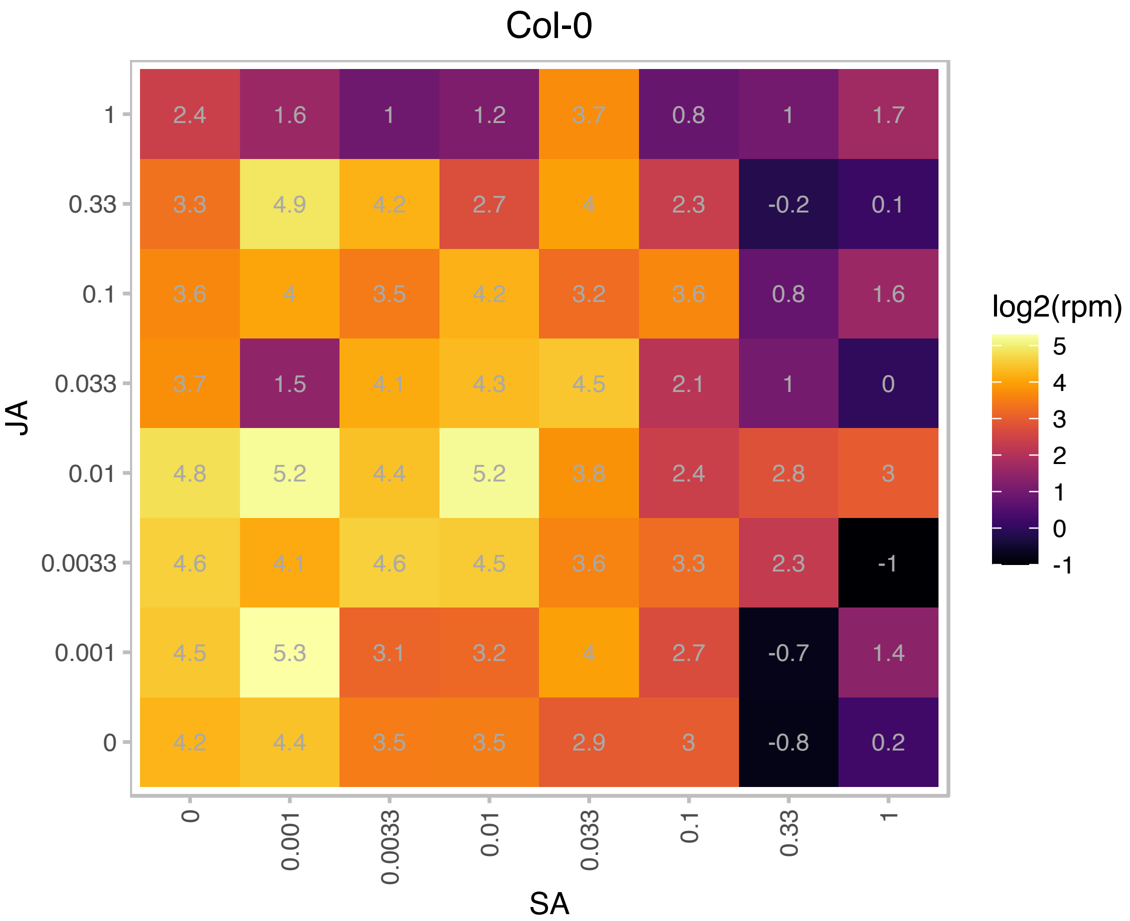


| | |
|---|---|
| Gene | AT3G54600 |
| is.expressed_genes | TRUE |
| is.responsive_genes | TRUE |
| is.apcluster | F-02 |

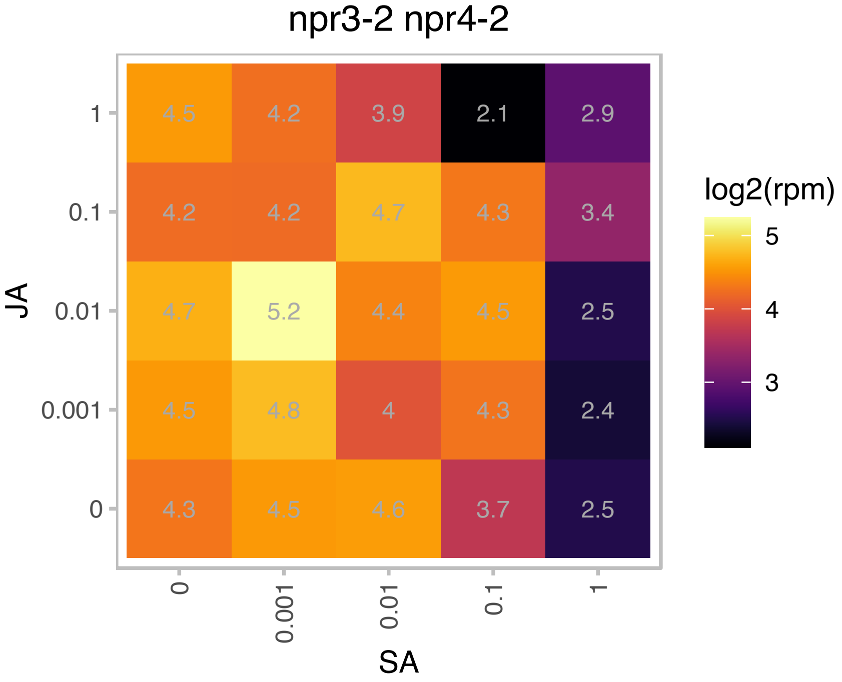


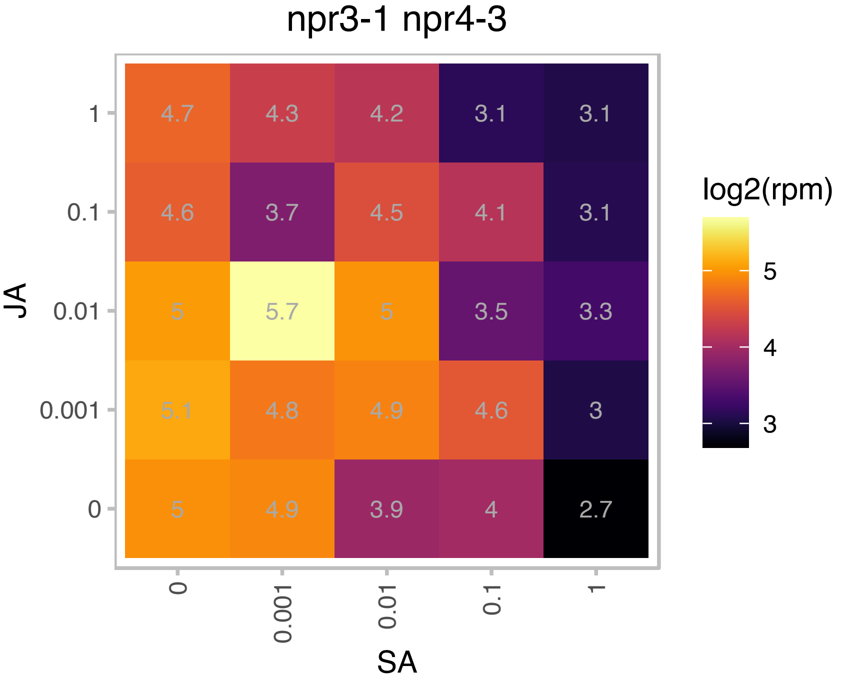

### AT3G56090.1 : ATFER3, FER3

ferritin 3
Encodes FERRITIN 3, AtFER3.  Ferritins are a class of 24-mer multi-meric proteins found in all kingdoms of life.  Function as the main iron store in mammals.  Evidence suggests that Arabidopsis ferritins are essential to protect cells against oxidative damage, but they do not constitute the major iron pool.

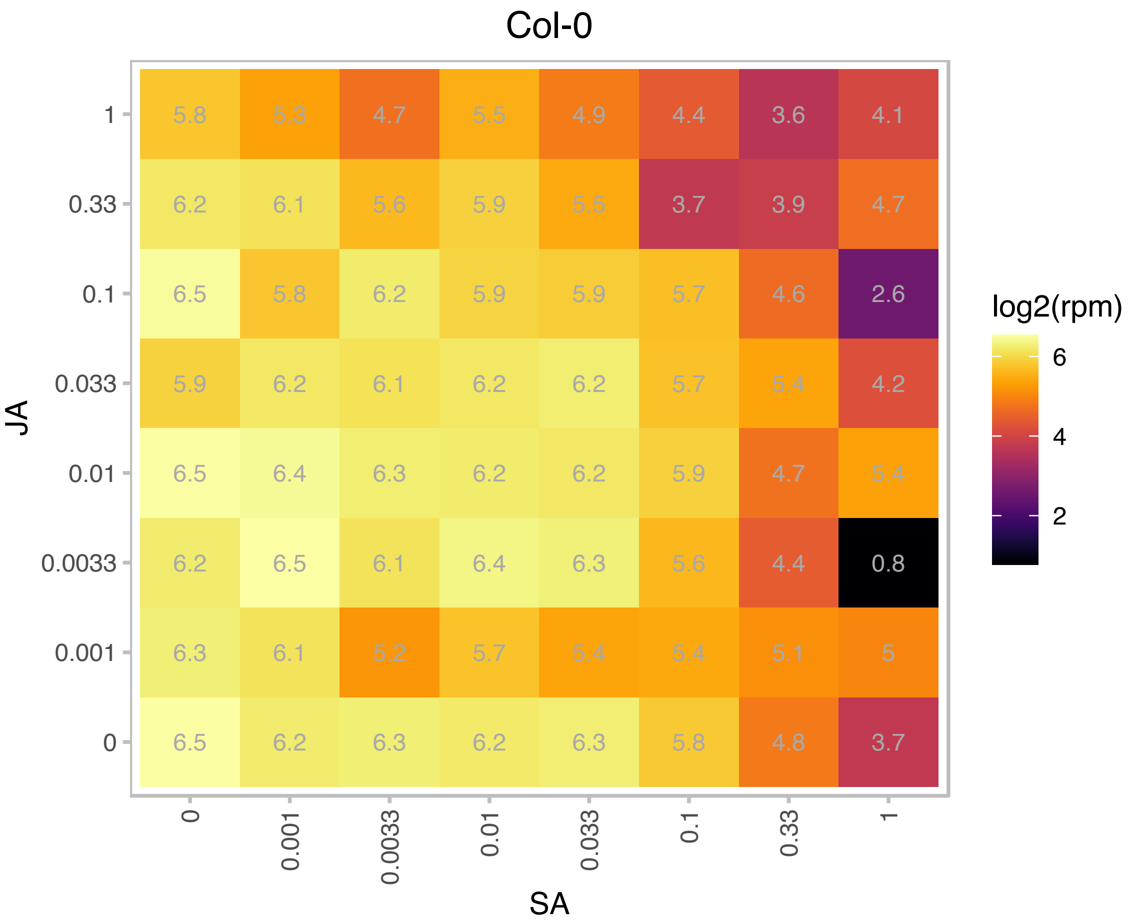


| | |
|---|---|
| Gene | AT3G56090 |
| is.expressed_genes | TRUE |
| is.responsive_genes | TRUE |
| is.apcluster | F-02 |

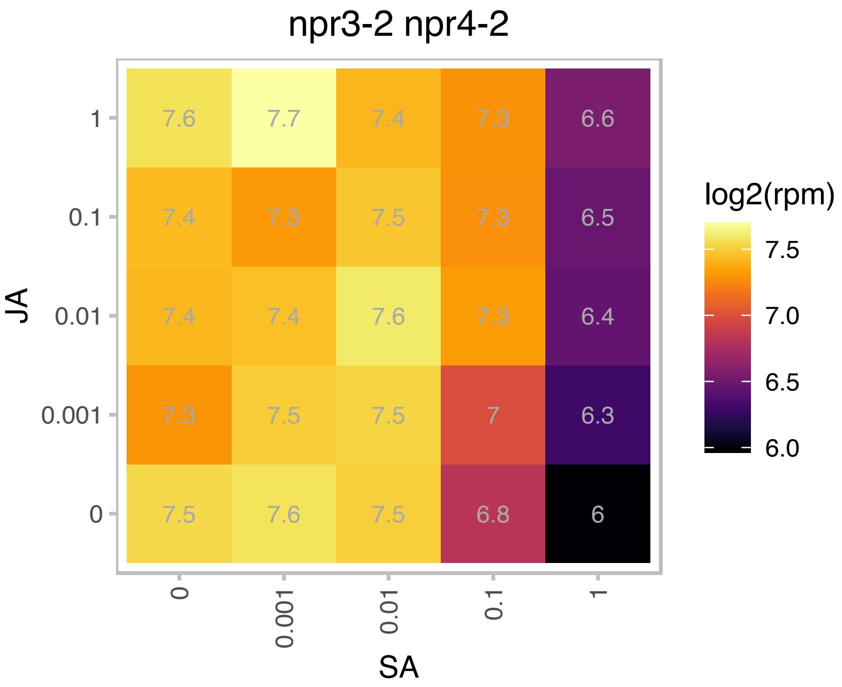


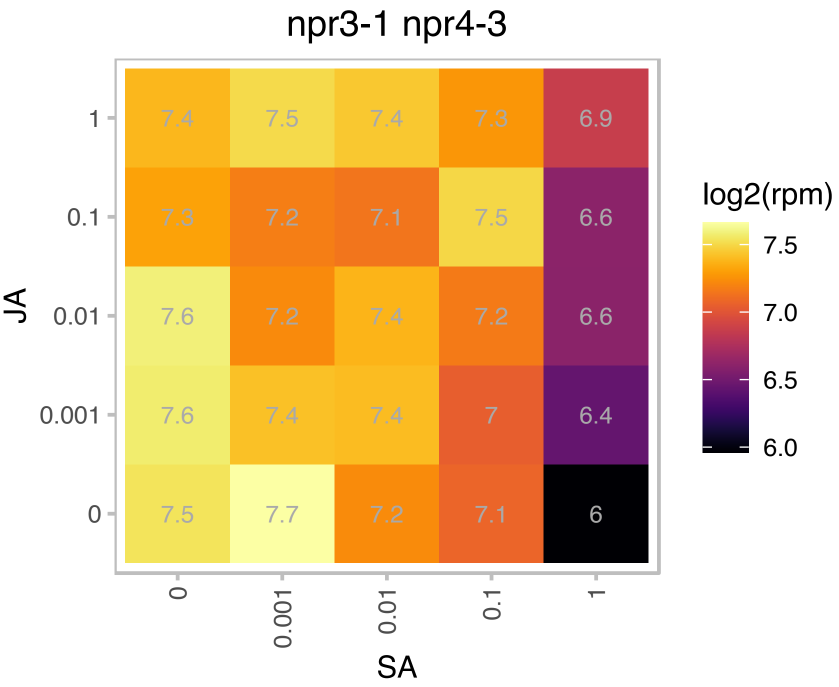

# AT3G57000.1 : NA

nucleolar essential protein-related

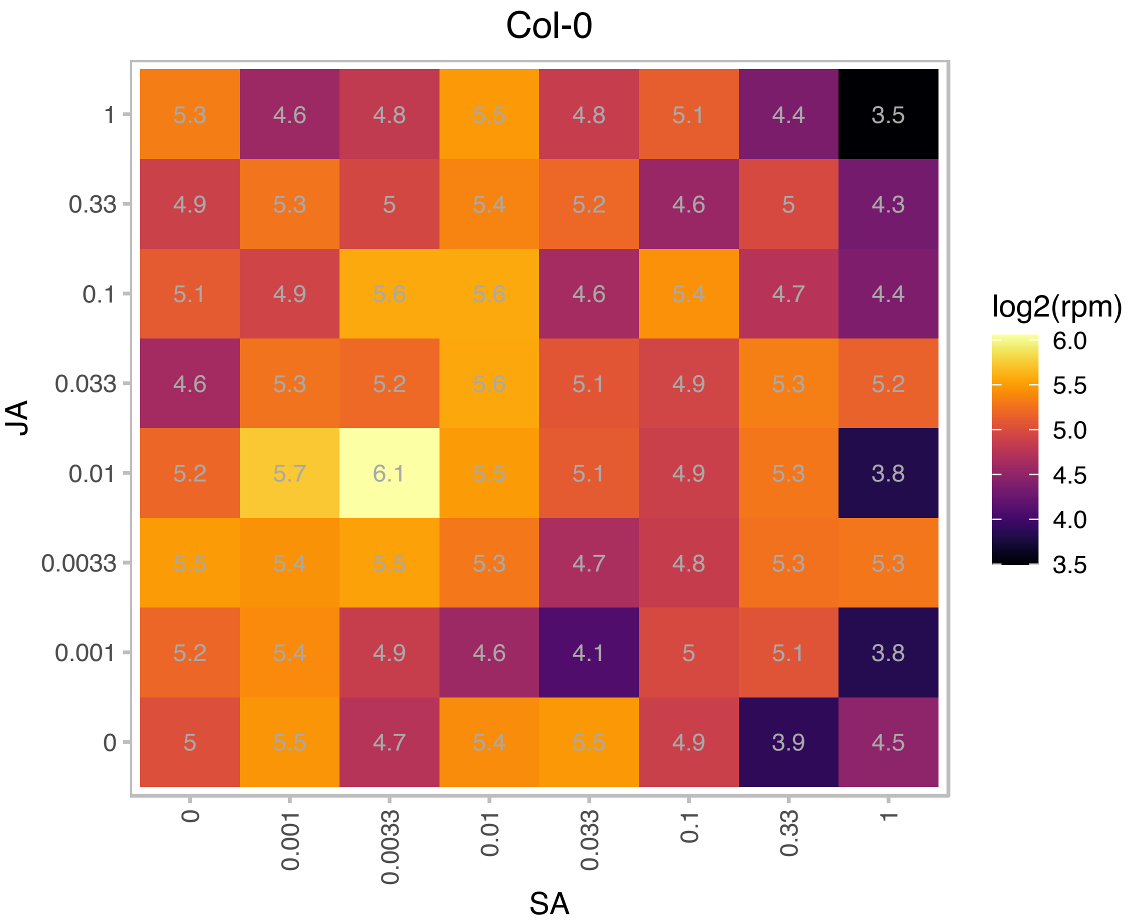


| | |
|---|---|
| Gene | AT3G57000 |
| is.expressed_genes | TRUE |
| is.responsive_genes | TRUE |
| is.apcluster | F-02 |

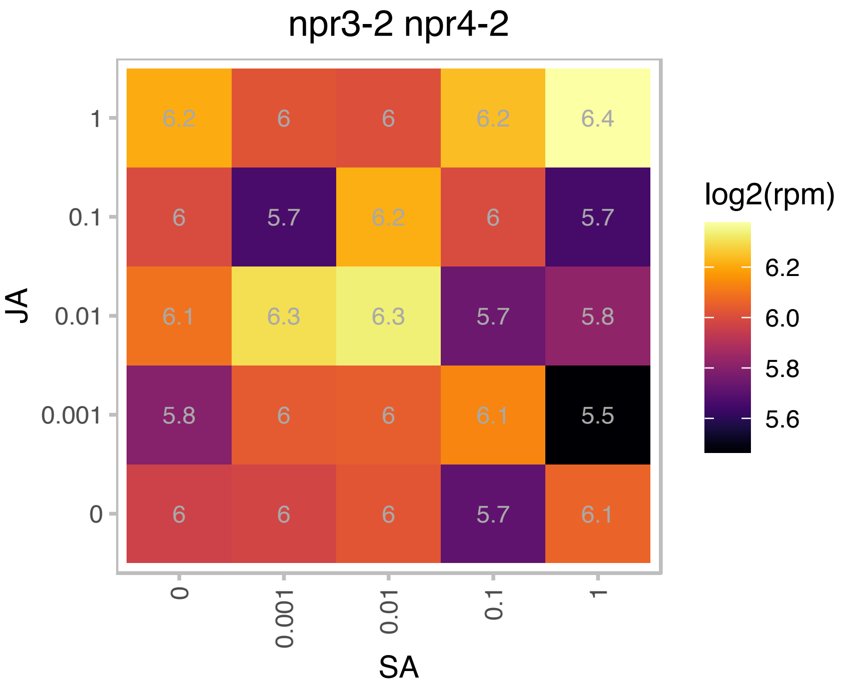


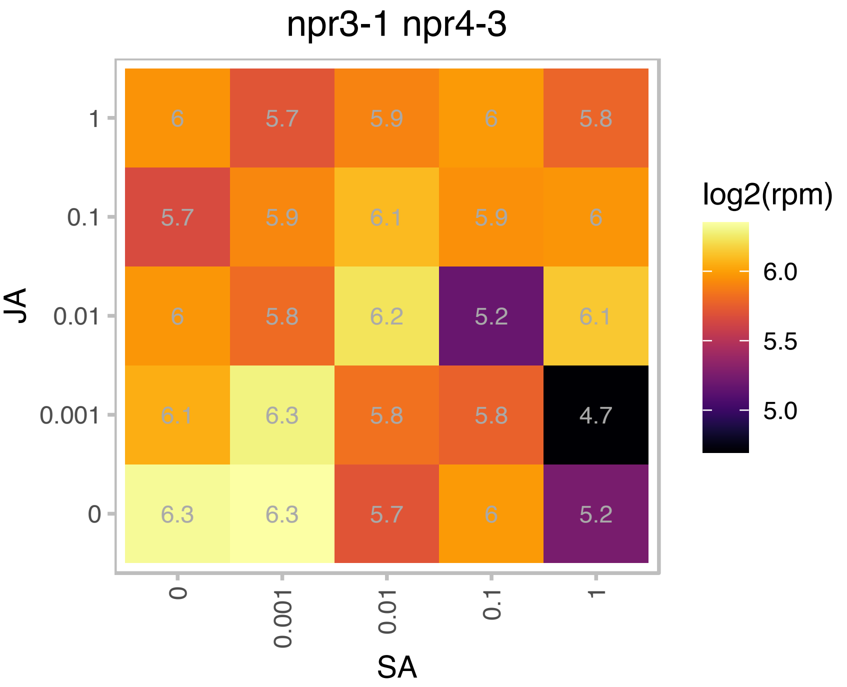

### AT3G57280.1 : AtFAX1, FAX1

Transmembrane proteins 14C
Encodes a chloroplast inner envelope localized member of the Tmemb_14 gene family. FAX1 is involved in fatty acid and lipid homeostasis and likely functions as a fatty acid transporter that exports fatty acids from the plastid. The mRNA is cell-to-cell mobile.

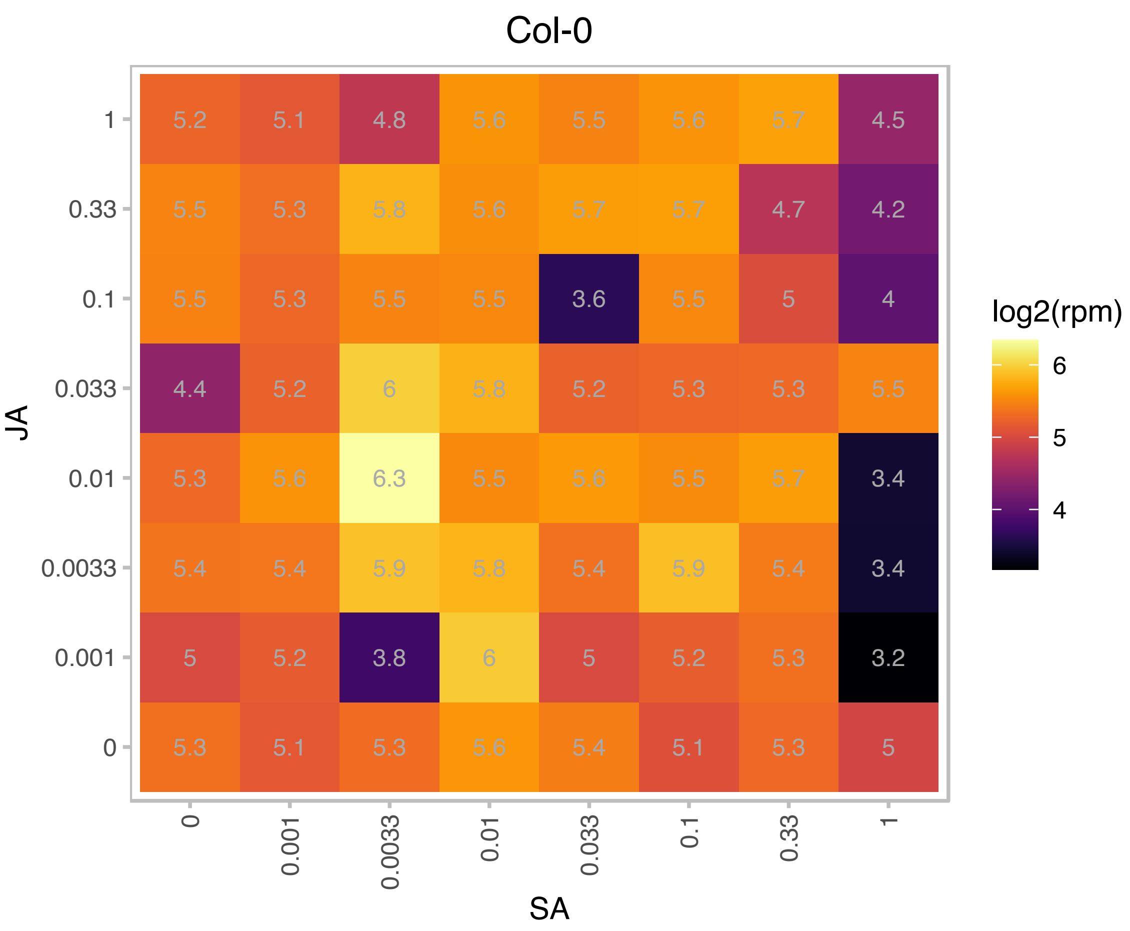


| | |
|---|---|
| Gene | AT3G57280 |
| is.expressed_genes | TRUE |
| is.responsive_genes | TRUE |
| is.apcluster | F-02 |

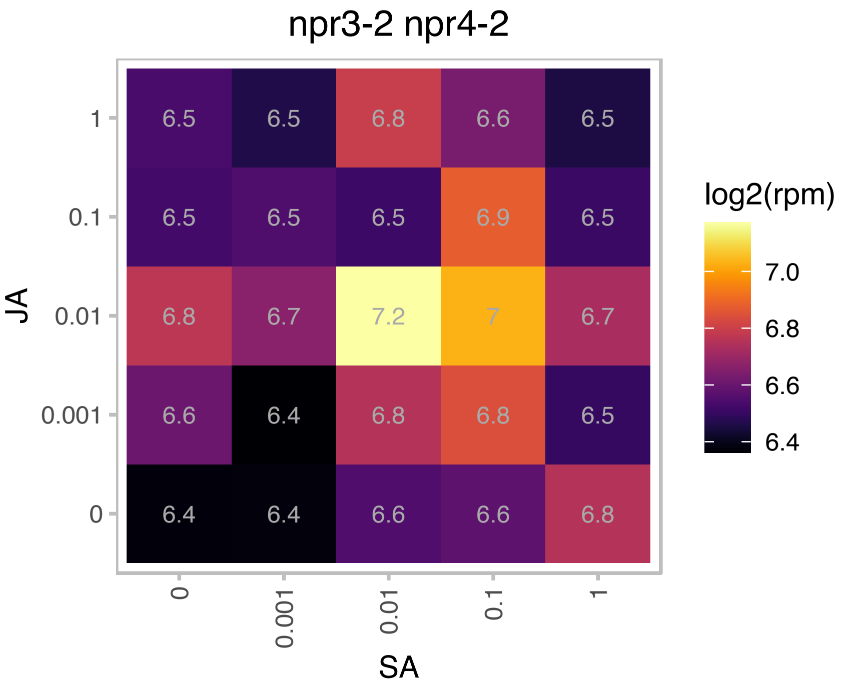


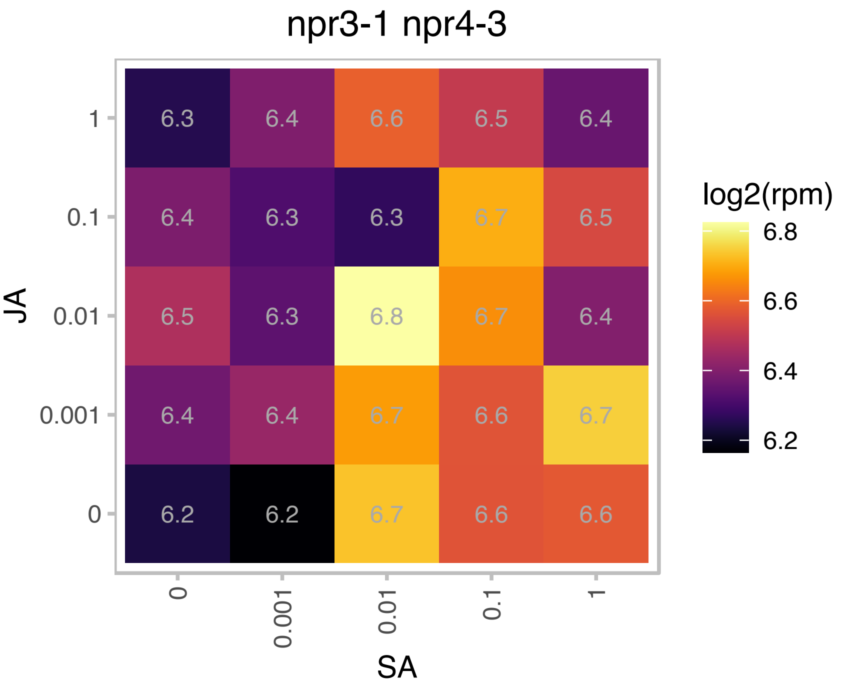

# AT3G61370.1 : NA

Protein of unknown function (DUF3245)

| | |
|---|---|
| Gene | AT3G61370 |
| is.expressed_genes | TRUE |
| is.responsive_genes | TRUE |
| is.apcluster | F-02 |

Col-0

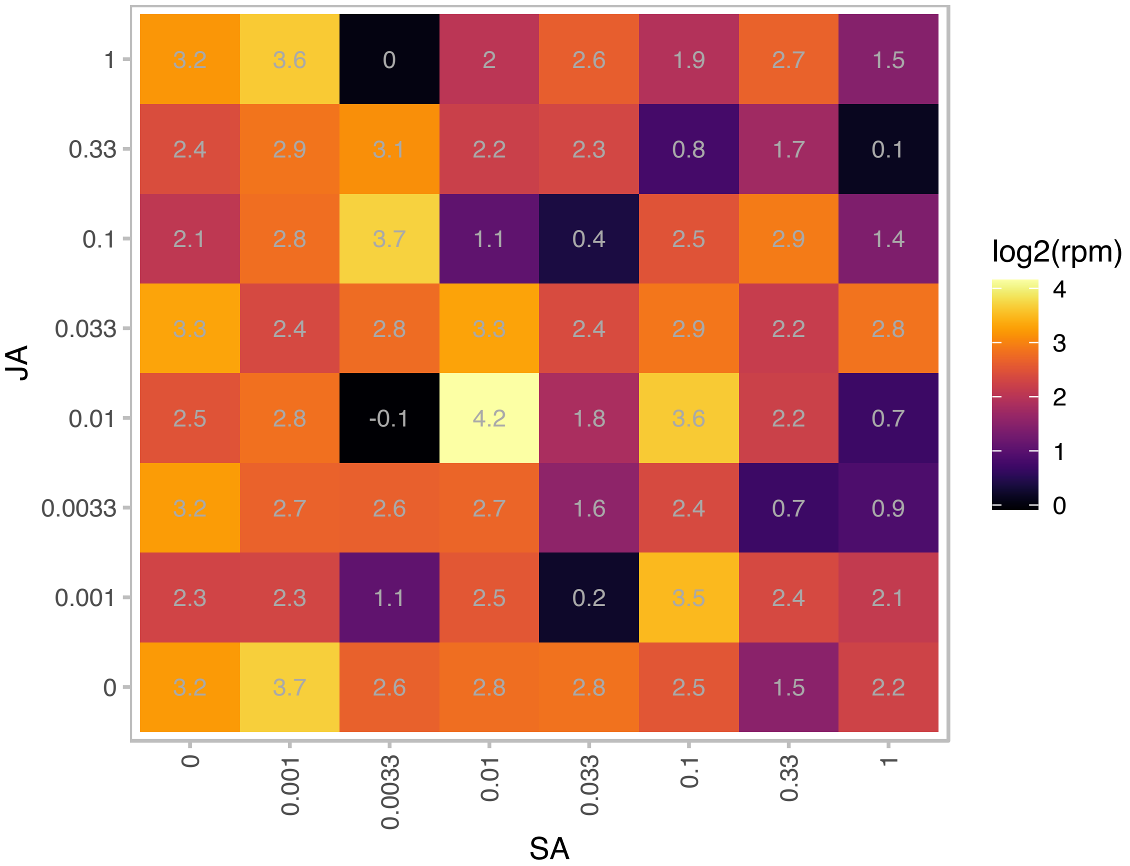


npr3-2 npr4-2

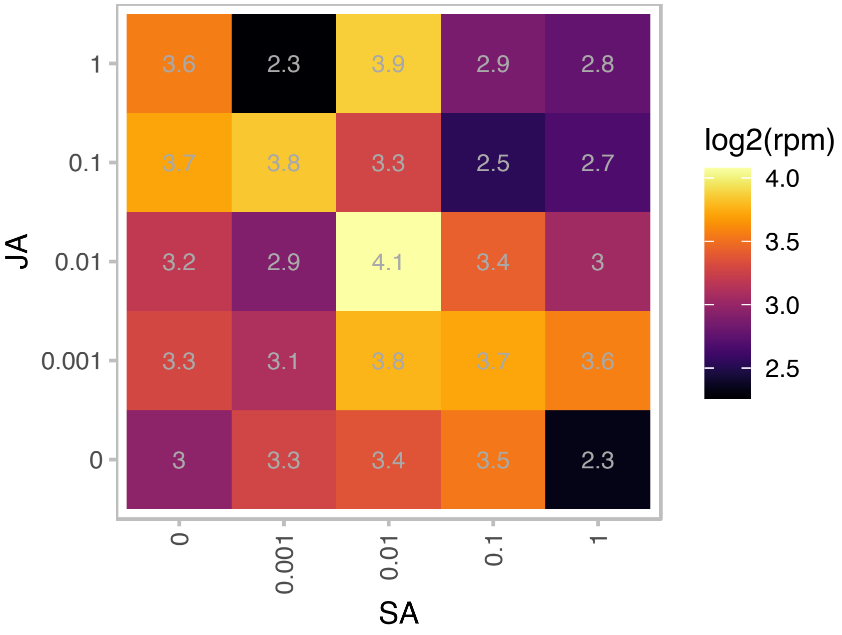


npr3-1 npr4-3

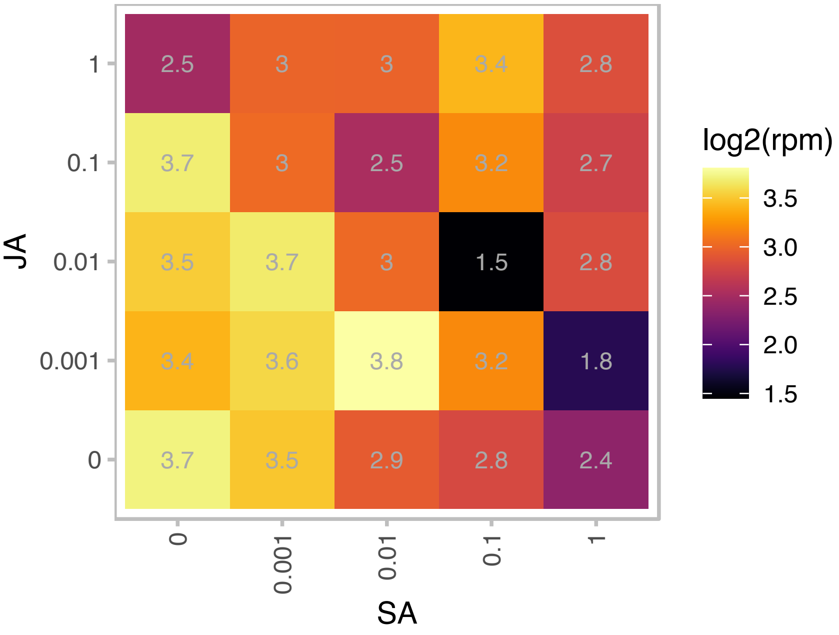

# AT3G62400.1 : NA

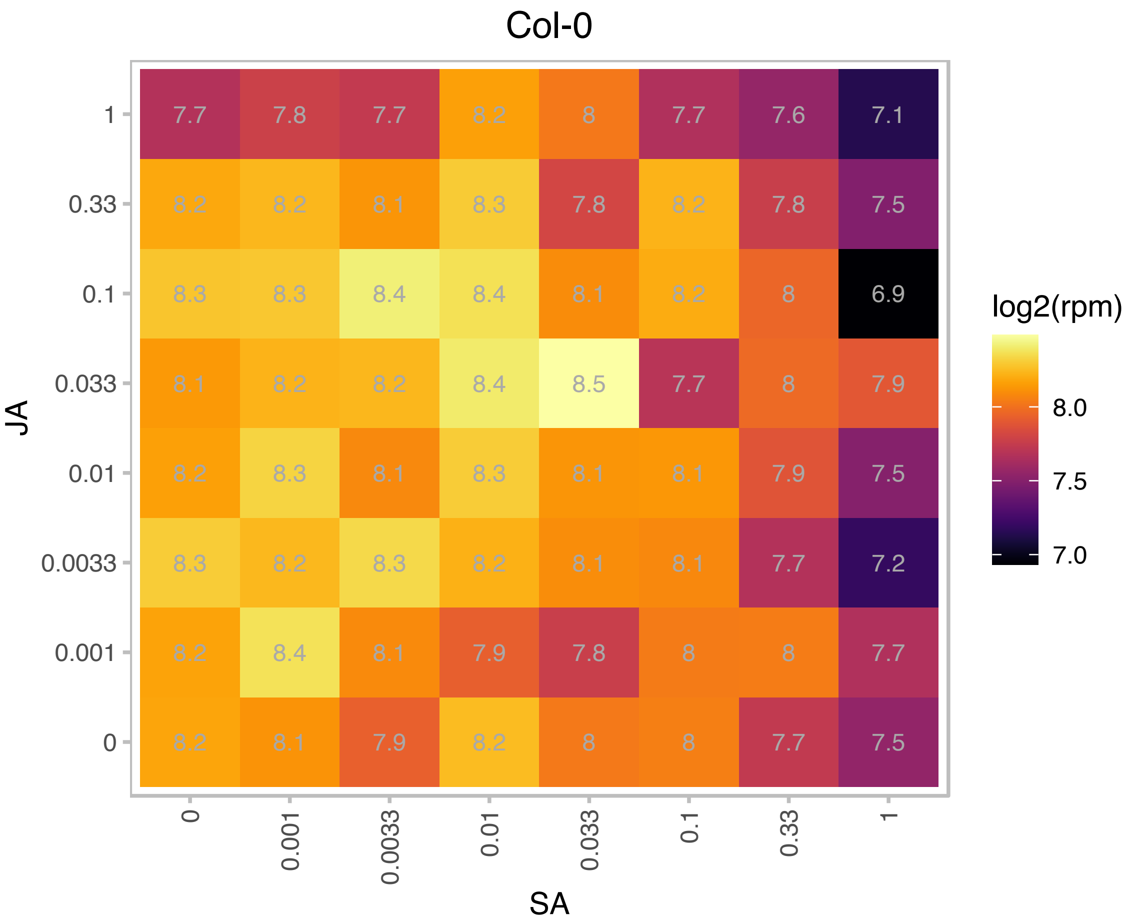


| | |
|---|---|
| Gene | AT3G62400 |
| is.expressed_genes | TRUE |
| is.responsive_genes | TRUE |
| is.apcluster | F-02 |

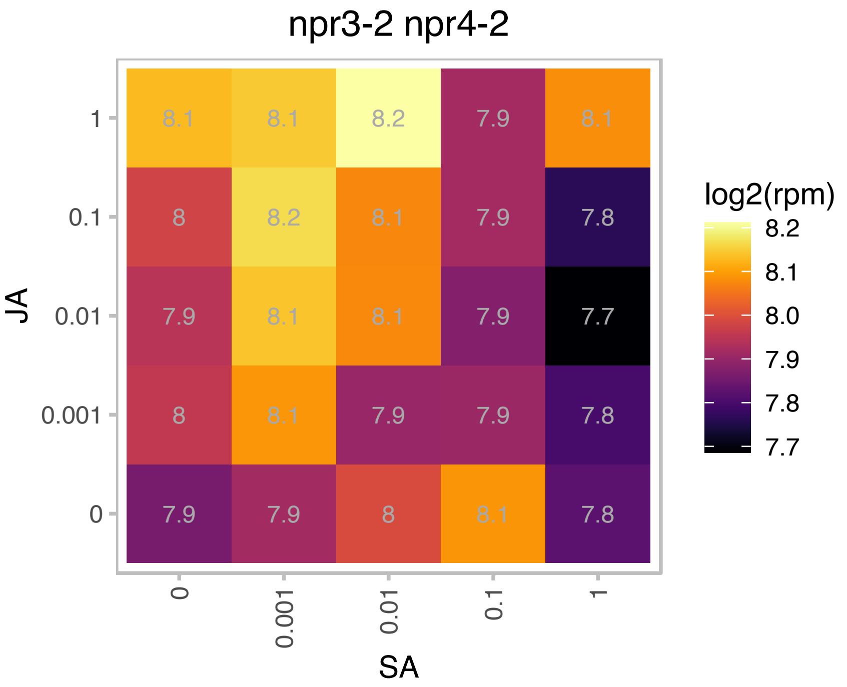


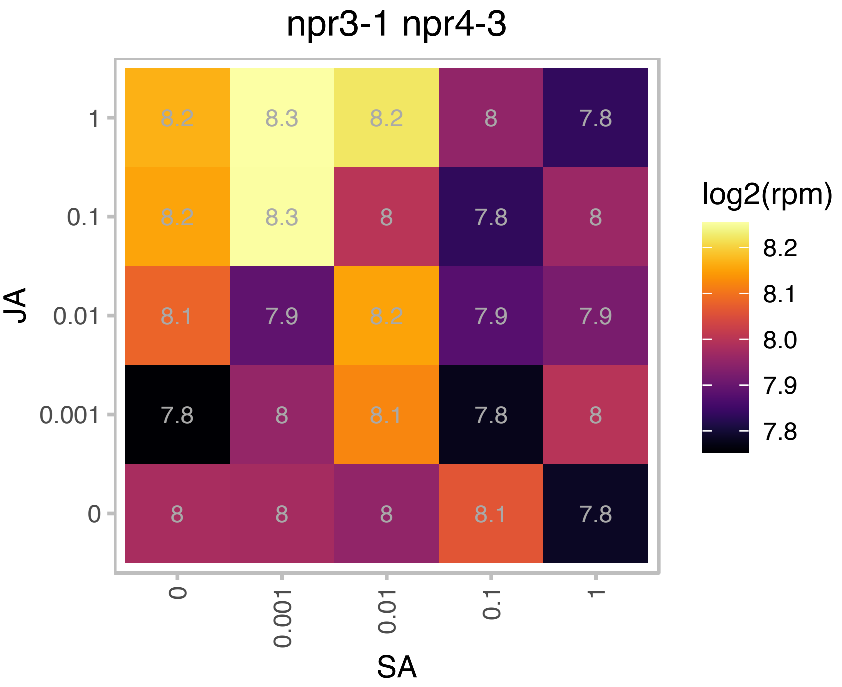

### AT3G62680.1 : ATPRP3, PRP3

proline-rich protein 3

Proline-rich protein The mRNA is cell-to-cell mobile.

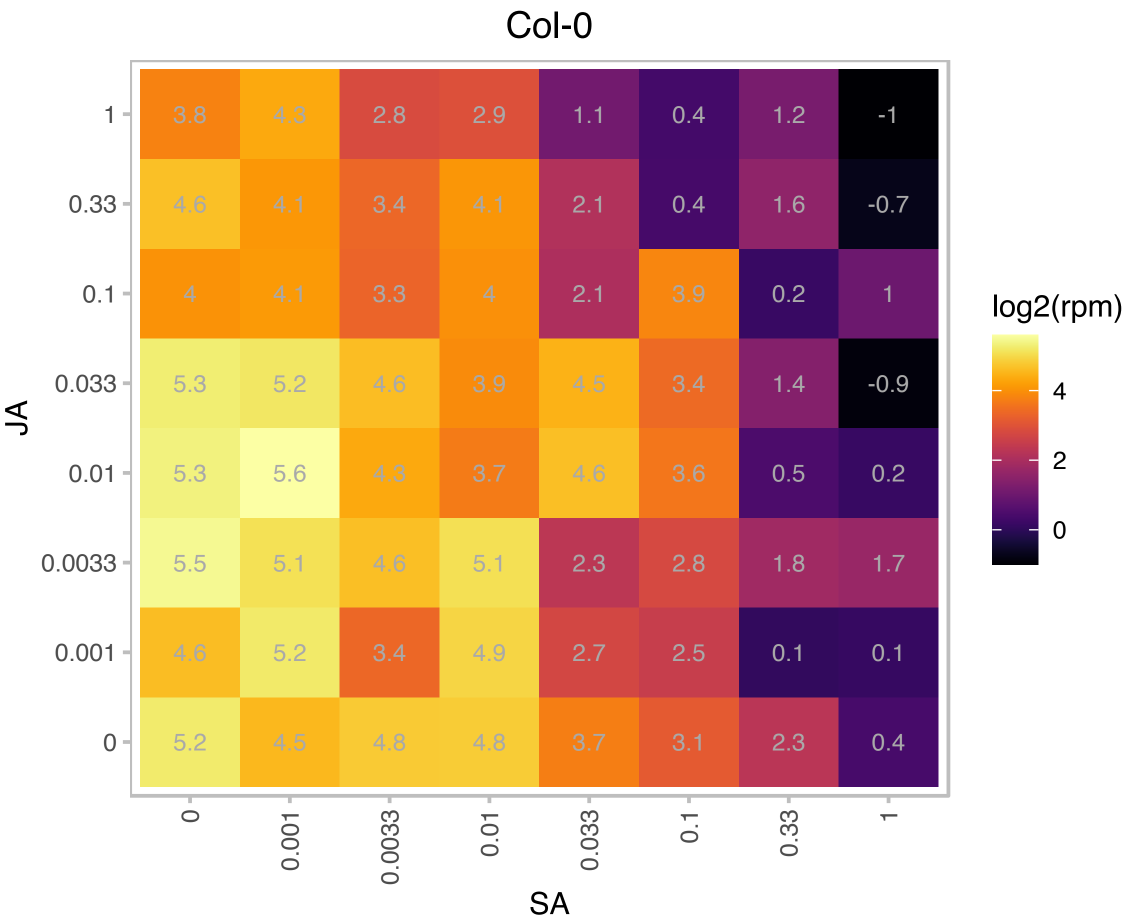


| | |
|---|---|
| Gene | AT3G62680 |
| is.expressed_genes | TRUE |
| is.responsive_genes | TRUE |
| is.apcluster | F-02 |

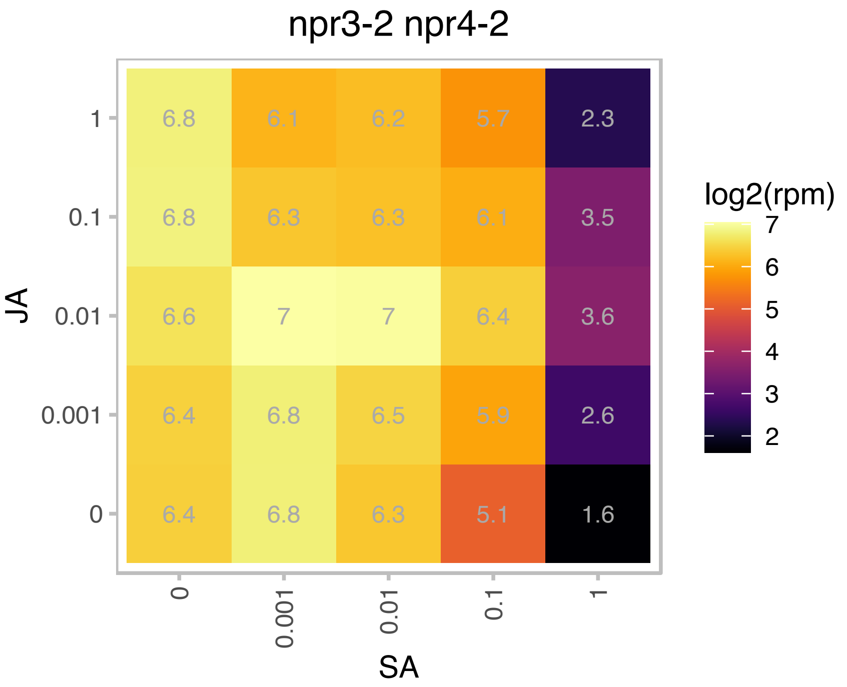


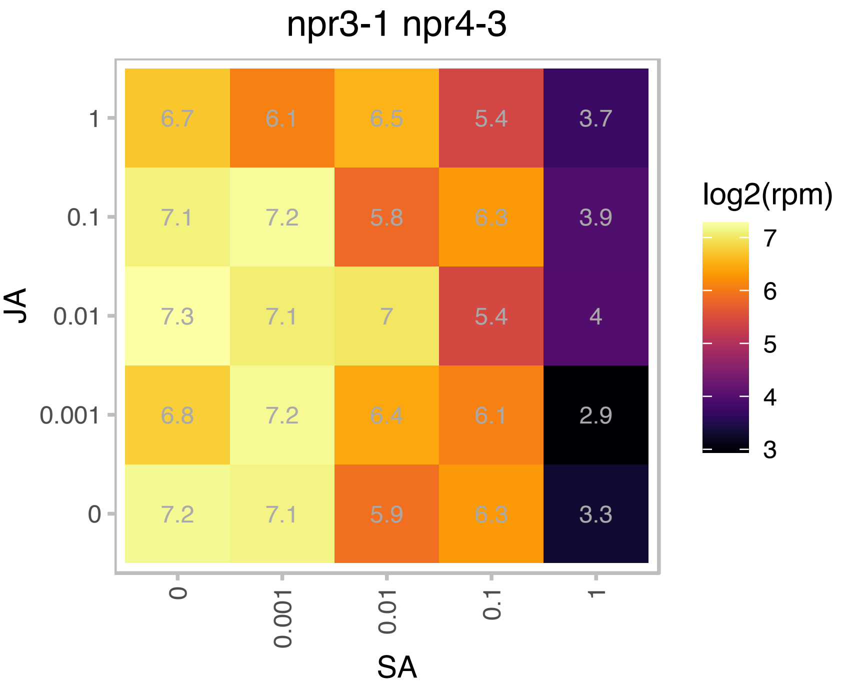

### AT3G63130.1 : ATRANGAP1, RANGAP1

RAN GTPase activating protein 1

Encodes a RAN GTPase activating protein involved in nuclear import, cell plate formation and mitotic spindle formation.

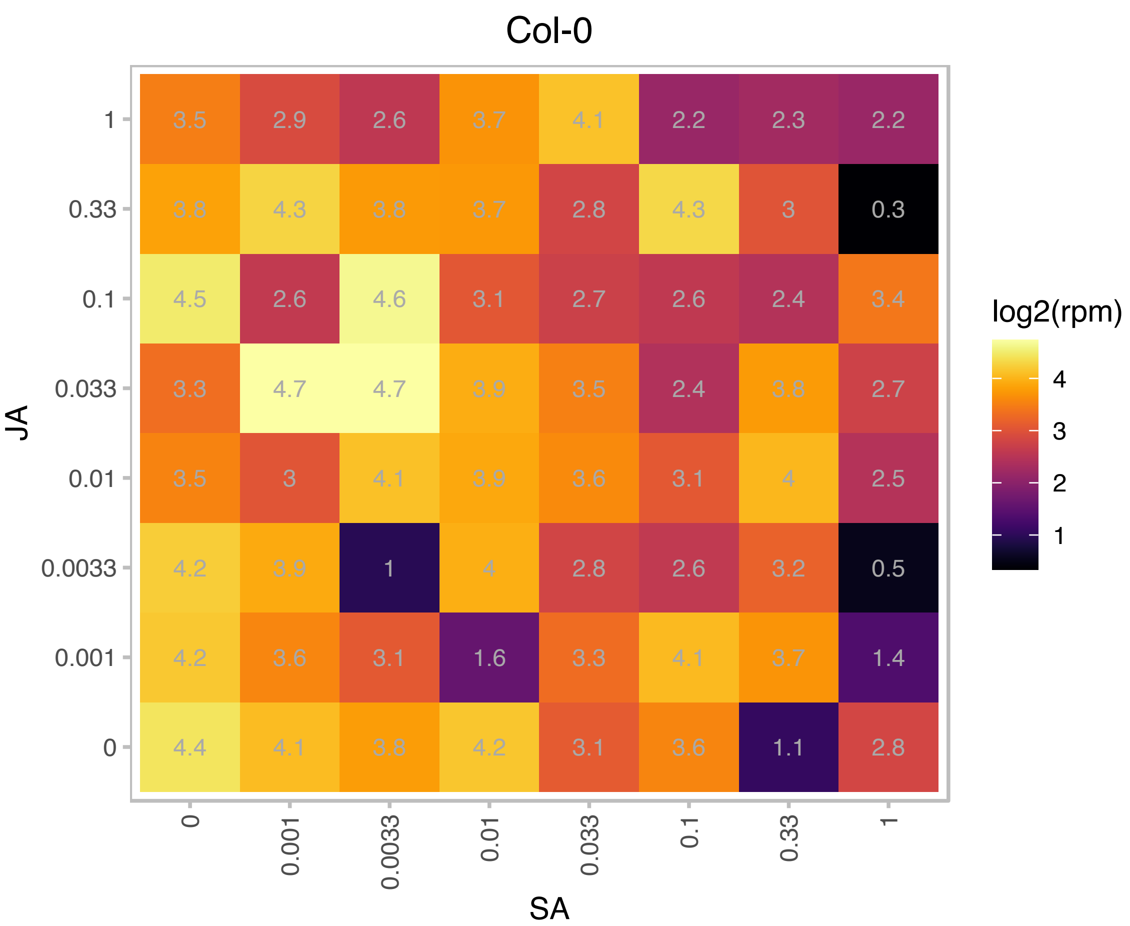


| | |
|---|---|
| Gene | AT3G63130 |
| is.expressed_genes | TRUE |
| is.responsive_genes | TRUE |
| is.apcluster | F-02 |

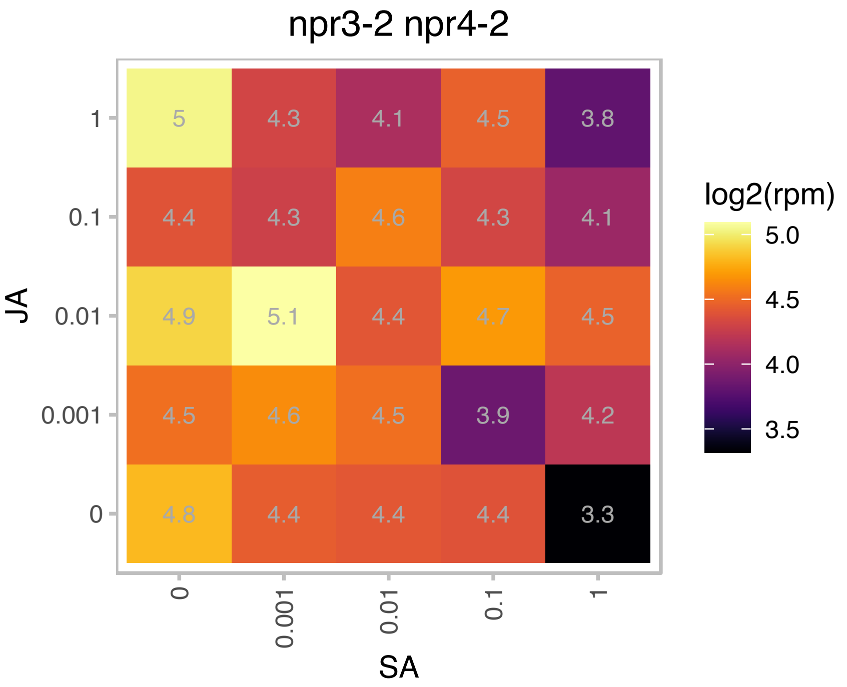


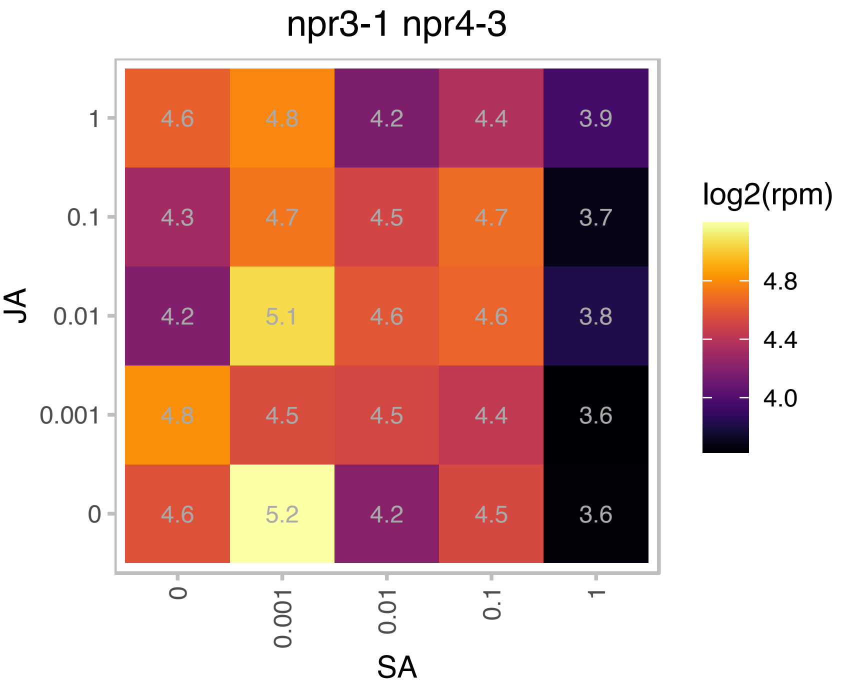

### AT3G63200.1 : PLA IIIB, PLP9

PATATIN-like protein 9

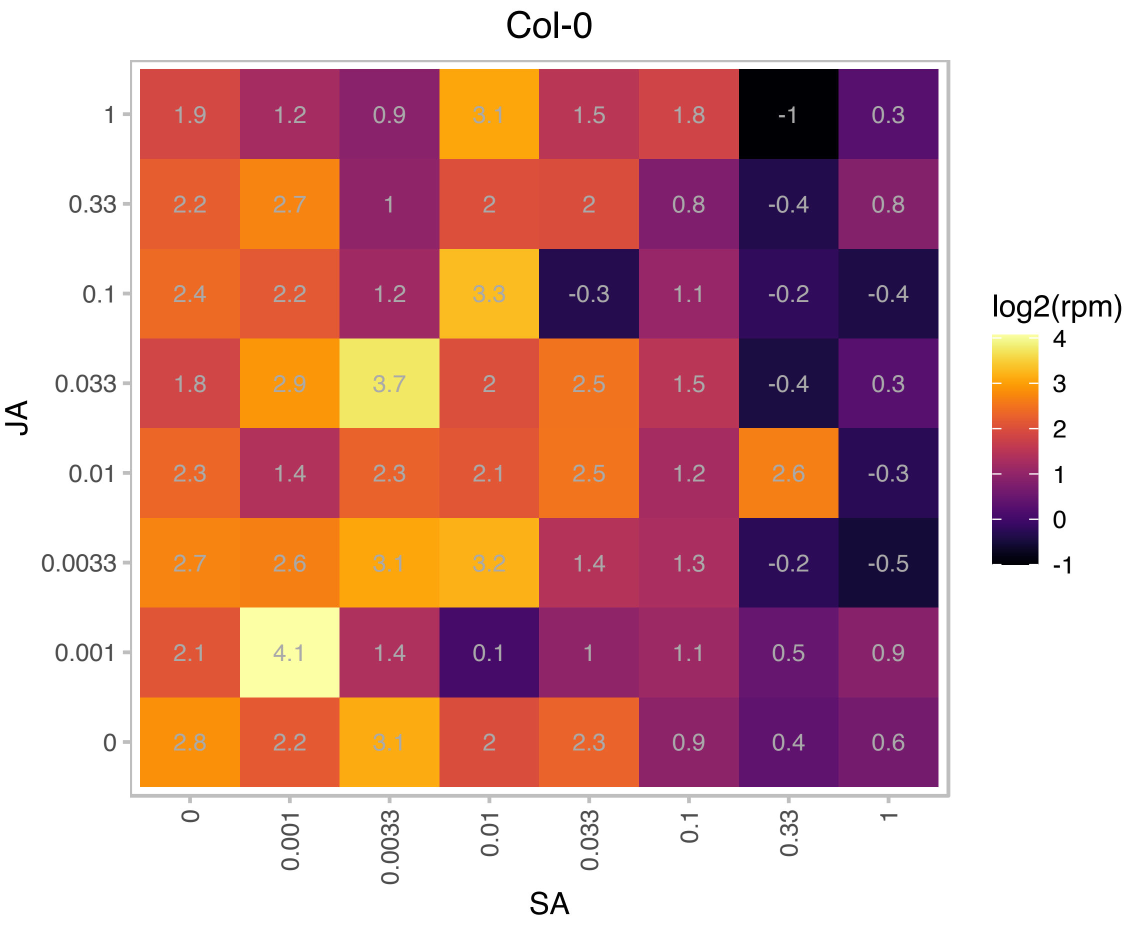


| | |
|---|---|
| Gene | AT3G63200 |
| is.expressed_genes | TRUE |
| is.responsive_genes | TRUE |
| is.apcluster | F-02 |

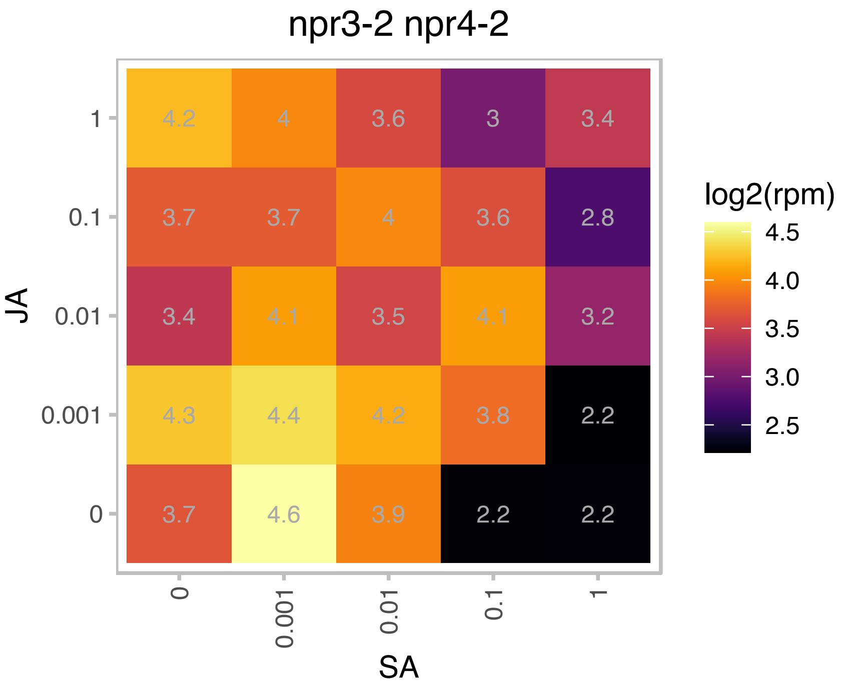


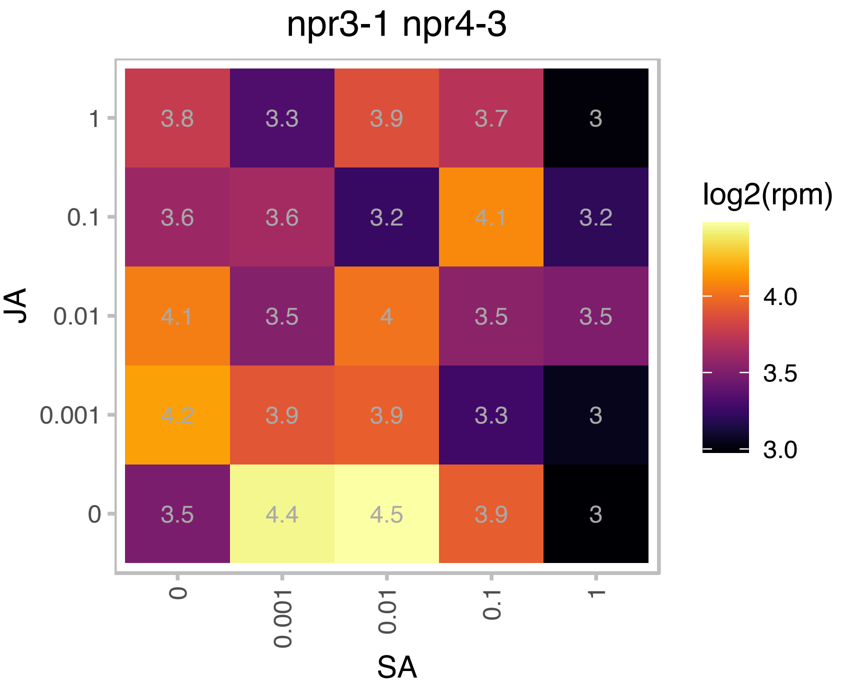

### AT4G02270.1 : RHS13, SRPP

root hair specific 13

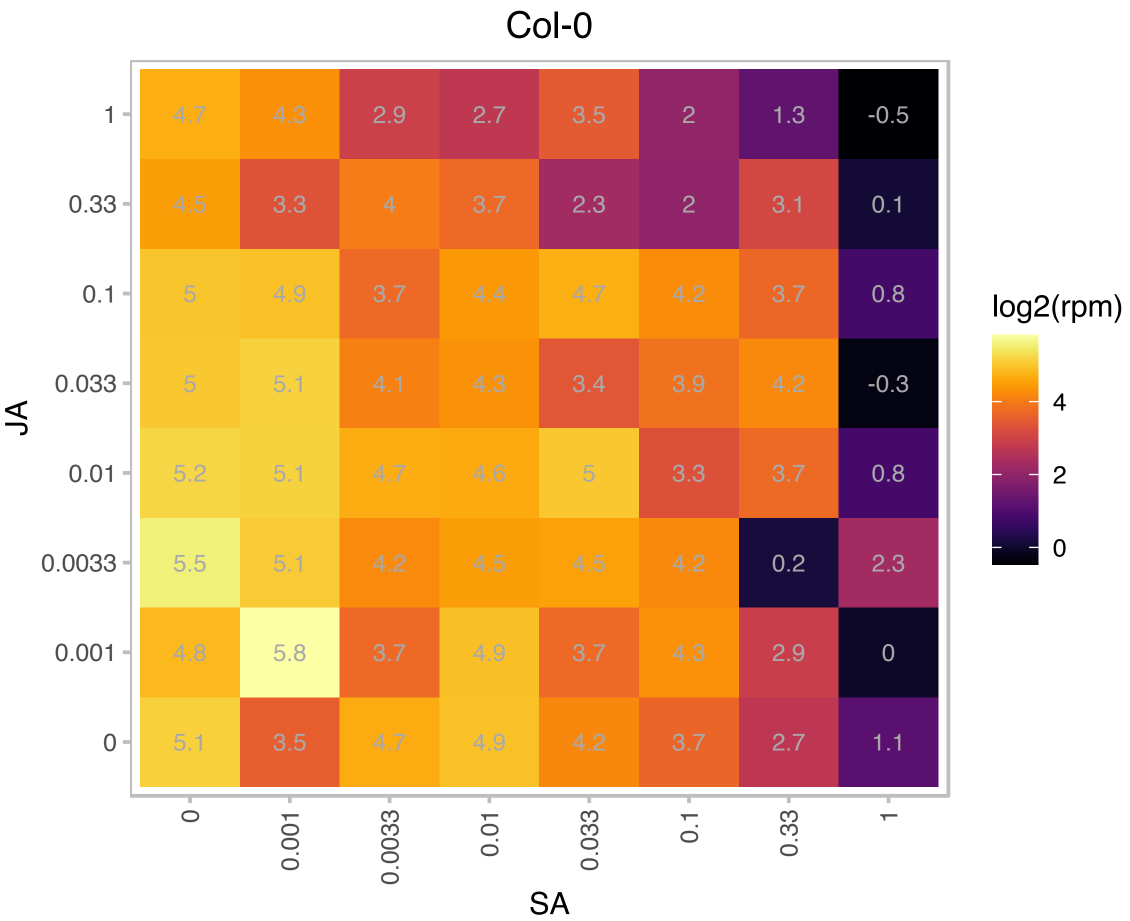


| | |
|---|---|
| Gene | AT4G02270 |
| is.expressed_genes | TRUE |
| is.responsive_genes | TRUE |
| is.apcluster | F-02 |

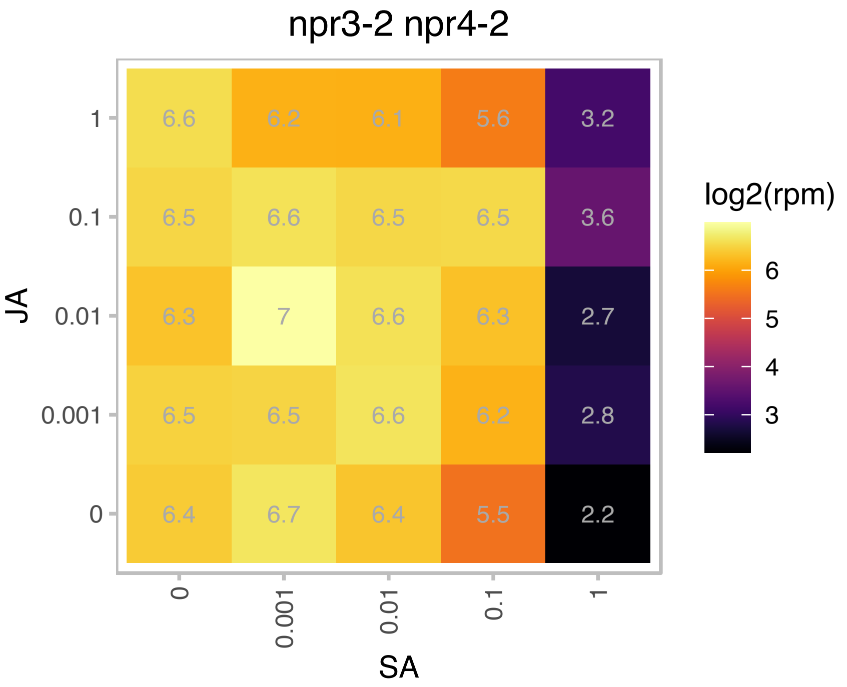


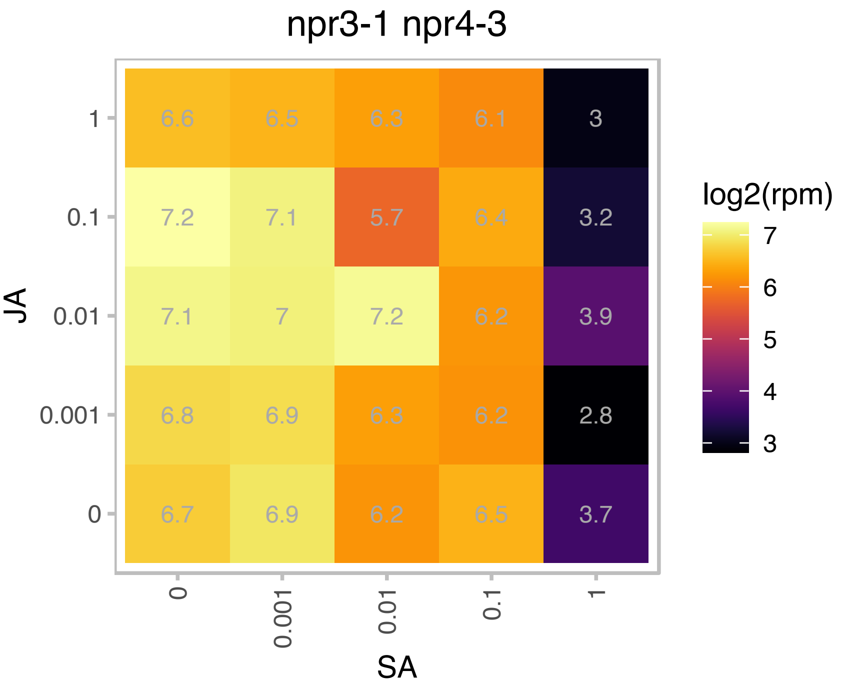

### AT4G09000.1 : GF14 CHI, GRF1

Encodes a 14-3-3 gene, designated GRF1 chi (for general regulatory factor1-G-box factor 14-3-3 homolog
general regulatory factor 1
isoform chi). The major native forms of 14-3-3s are homo- and hetero-dimers, the biological functions of which are to interact physically with specific client proteins and thereby effect a change in the client. As a result, 14-3-3s are involved in a vast array of processes such as the response to stress, cell-cycle control, and apoptosis, serving as adapters, activators, and repressors. There are currently 133 full-length sequences available.

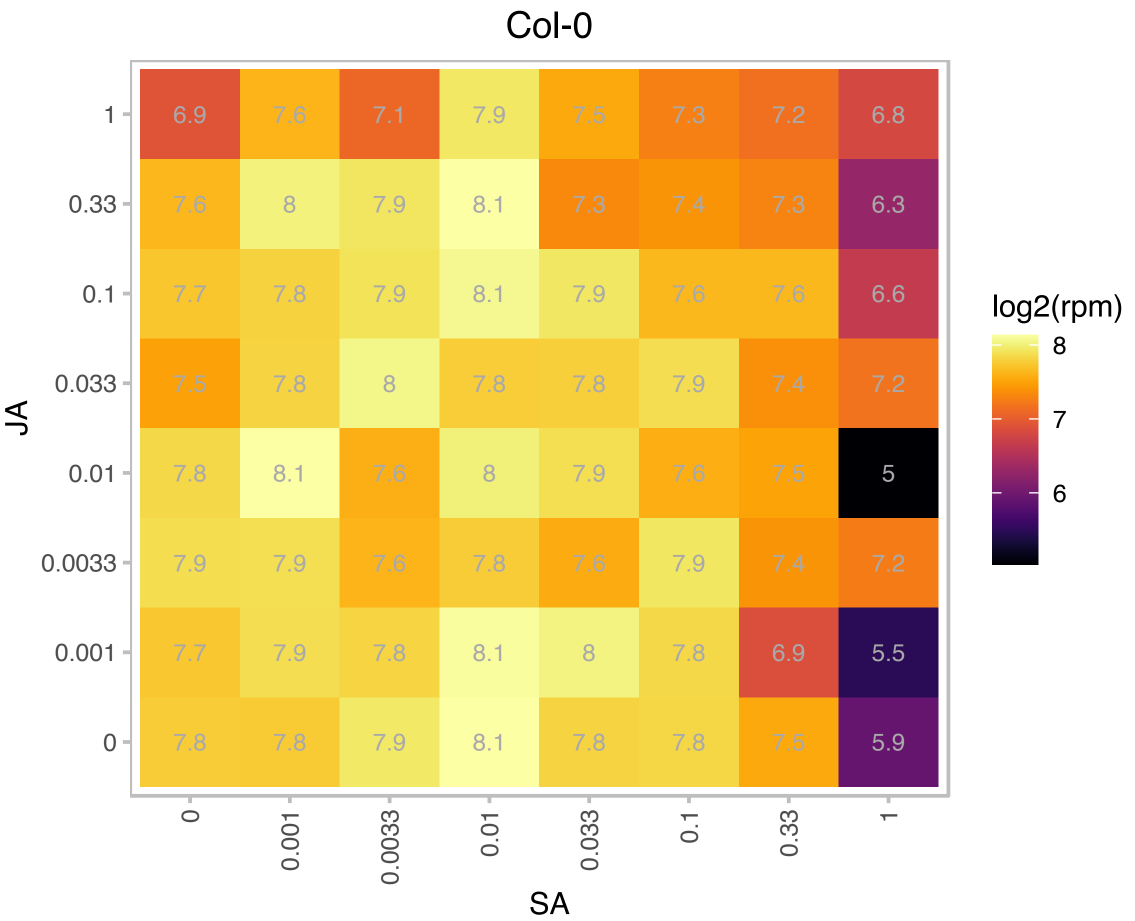


| | |
|---|---|
| Gene | AT4G09000 |
| is.expressed_genes | TRUE |
| is.responsive_genes | TRUE |
| is.apcluster | F-02 |

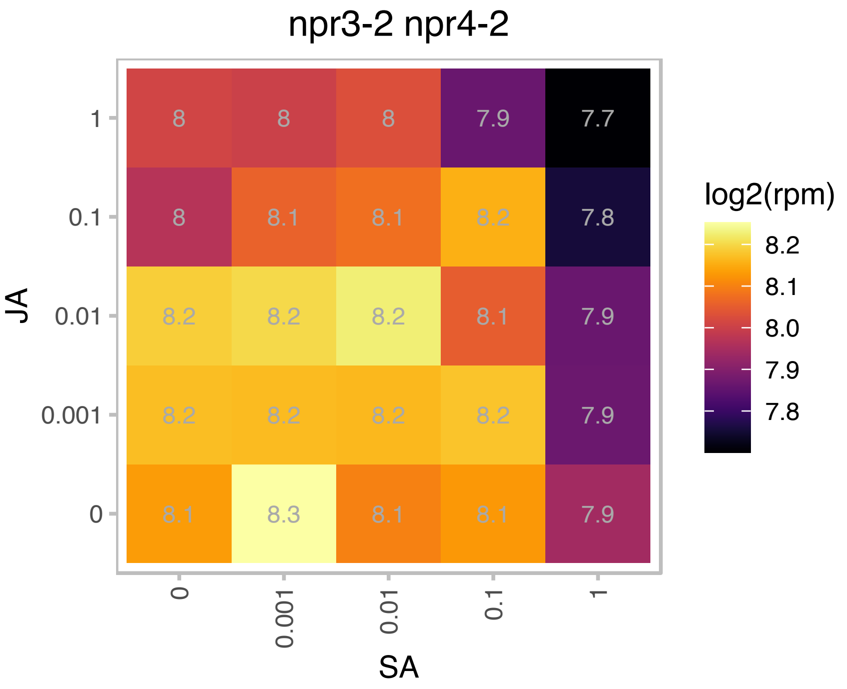


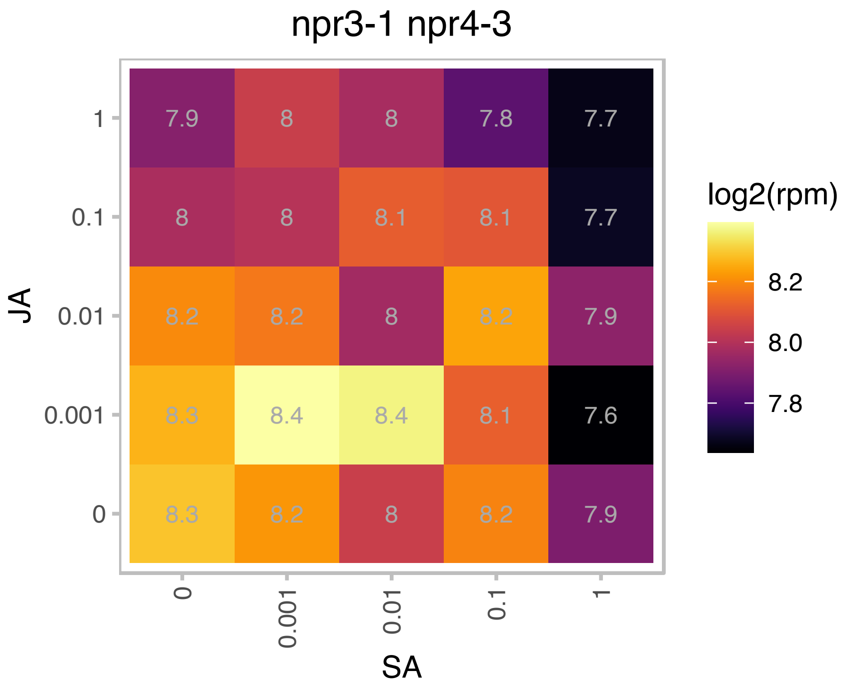

# AT4G09670.1 : NA

Oxidoreductase family protein

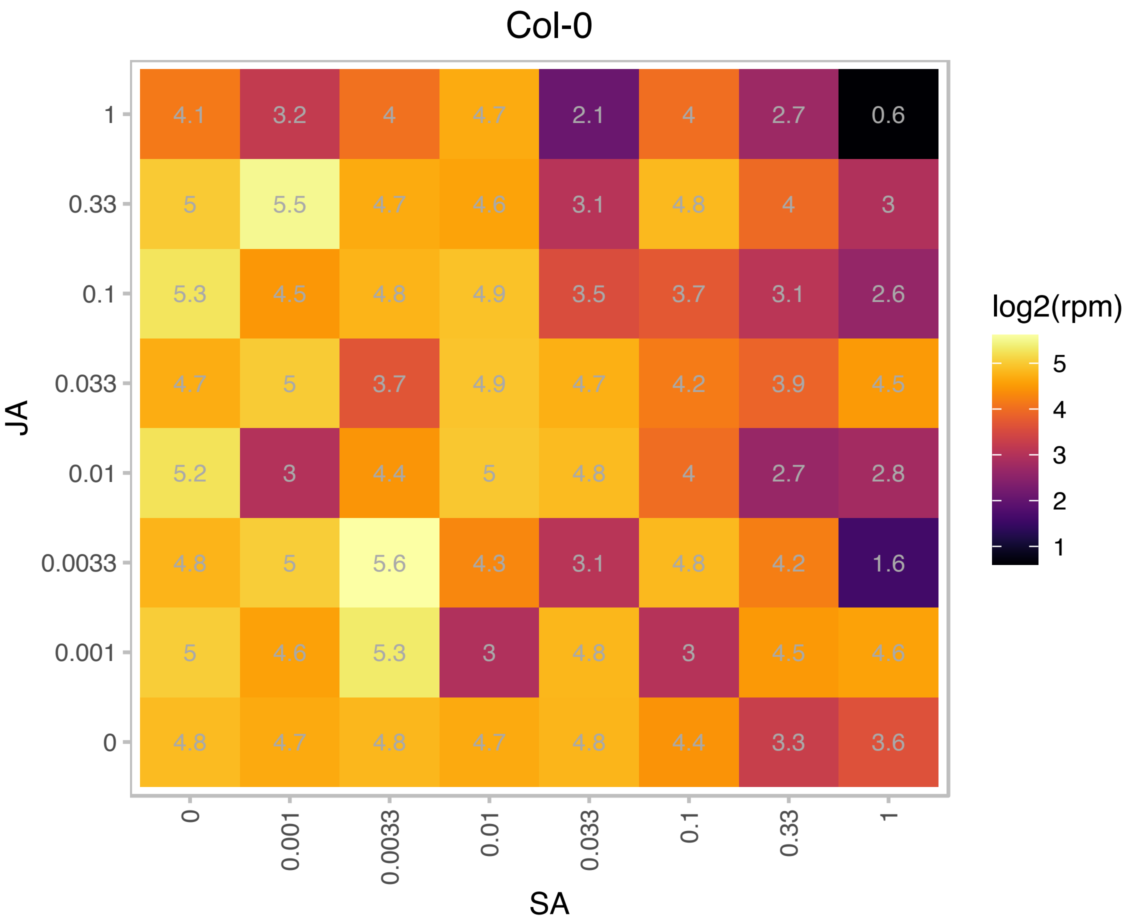


| | |
|---|---|
| Gene | AT4G09670 |
| is.expressed_genes | TRUE |
| is.responsive_genes | TRUE |
| is.apcluster | F-02 |

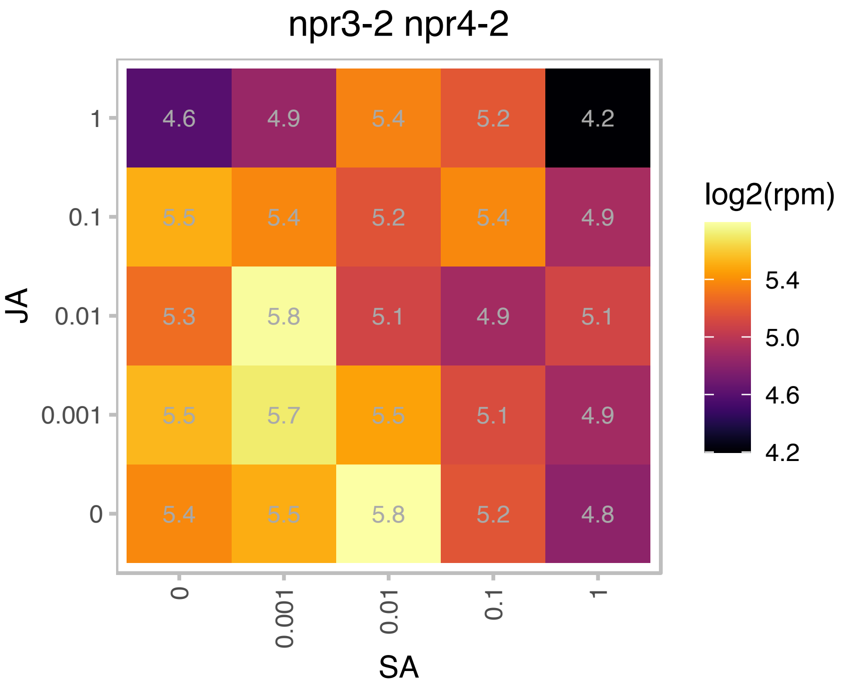


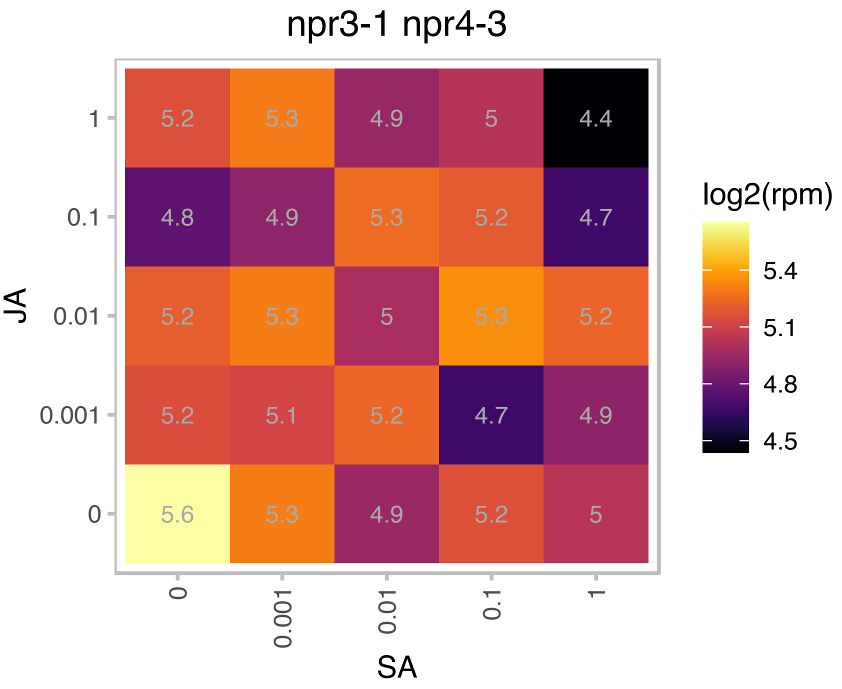

# AT4G11190.1 : NA

Disease resistance-responsive (dirigent-like protein) family protein

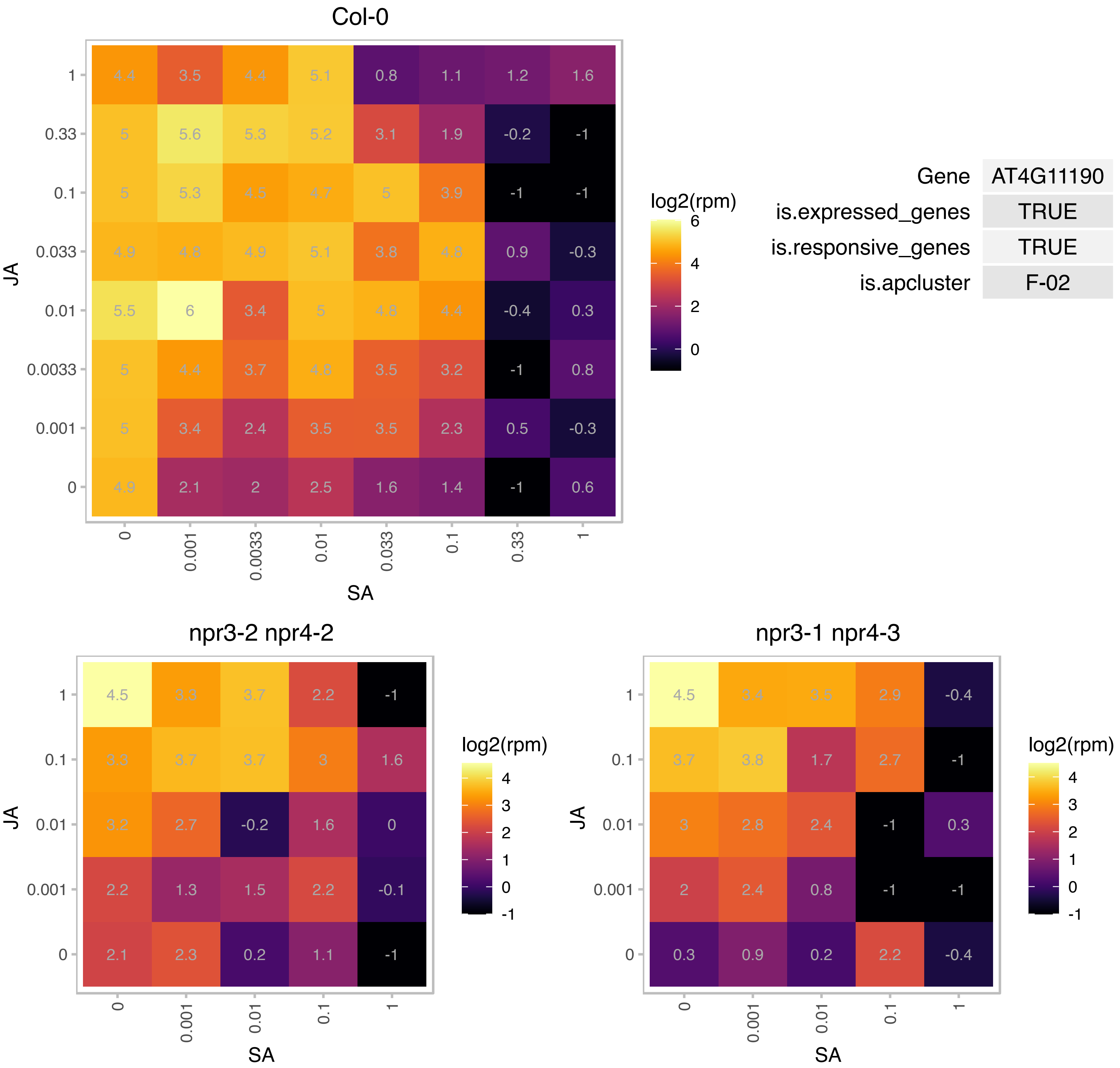


| | |
|---|---|
| Gene | AT4G11190 |
| is.expressed_genes | TRUE |
| is.responsive_genes | TRUE |
| is.apcluster | F-02 |

### AT4G11320.1 : AtCP2, CP2

Papain family cysteine protease

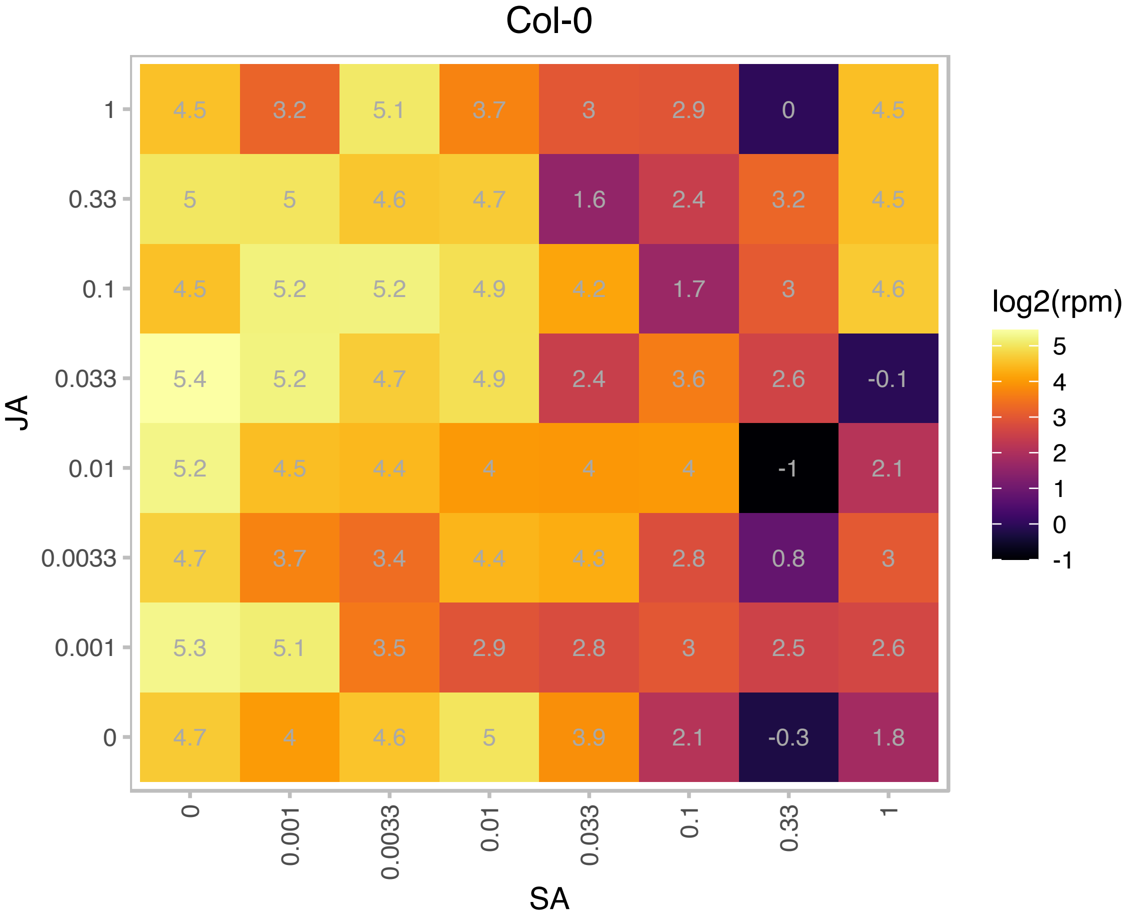


| | |
|---|---|
| Gene | AT4G11320 |
| is.expressed_genes | TRUE |
| is.responsive_genes | TRUE |
| is.apcluster | F-02 |

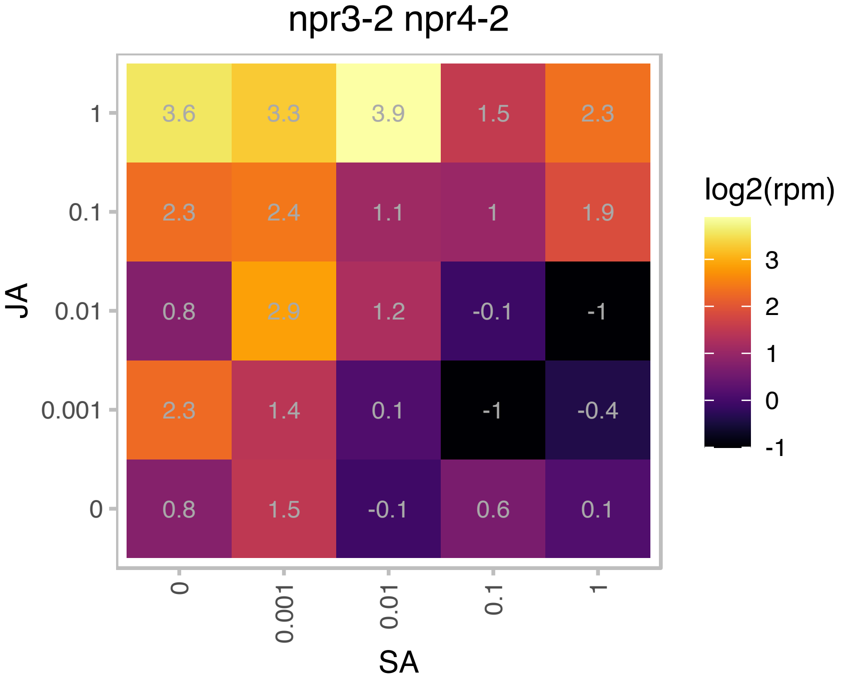


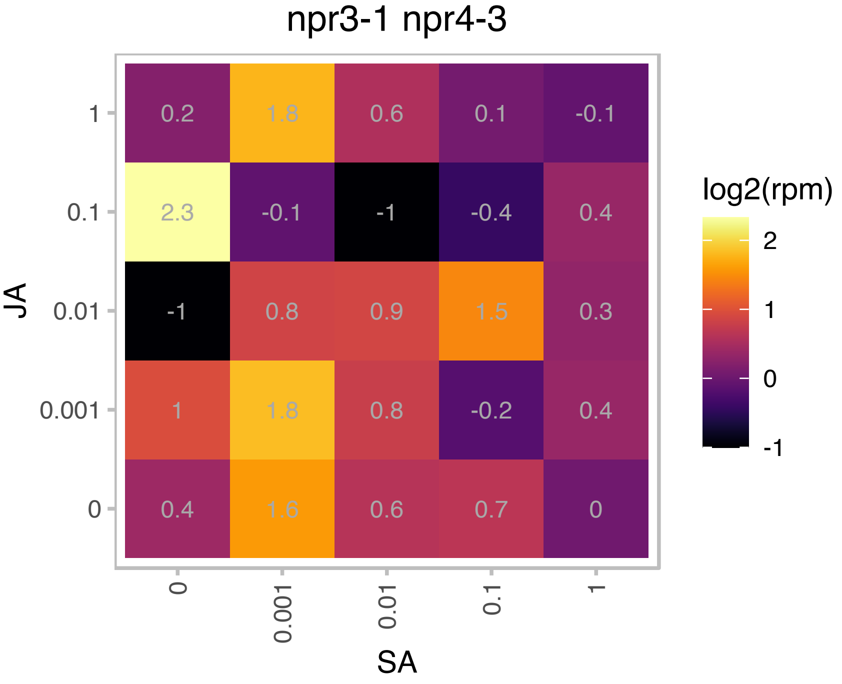

# AT4G12520.1 : NA

Bifunctional inhibitor/lipid-transfer protein/seed storage 2S albumin superfamily protein

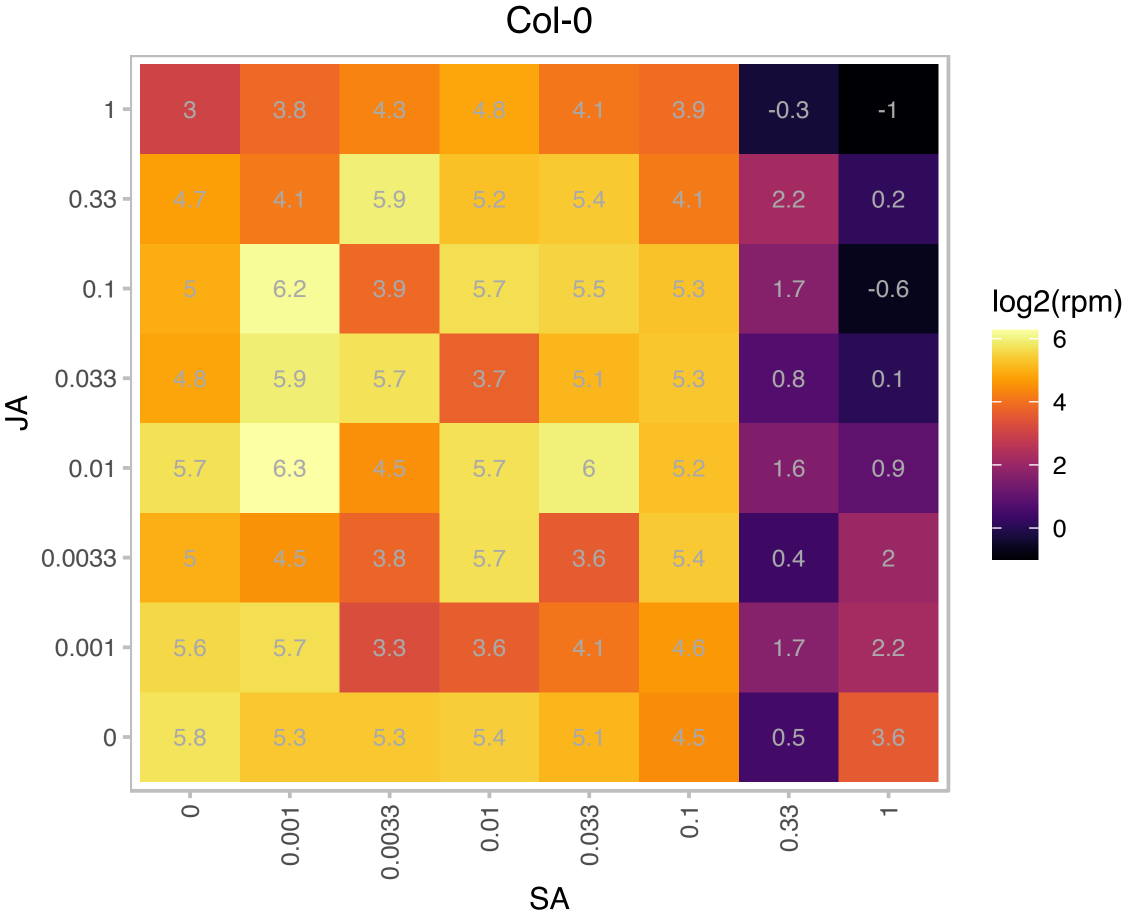


| | |
|---|---|
| Gene | AT4G12520 |
| is.expressed_genes | TRUE |
| is.responsive_genes | TRUE |
| is.apcluster | F-02 |

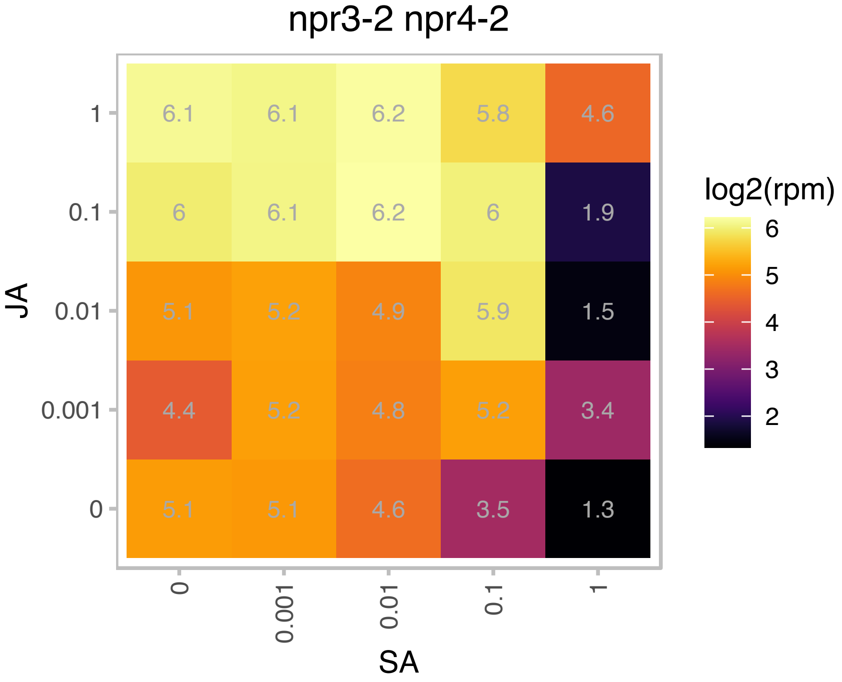


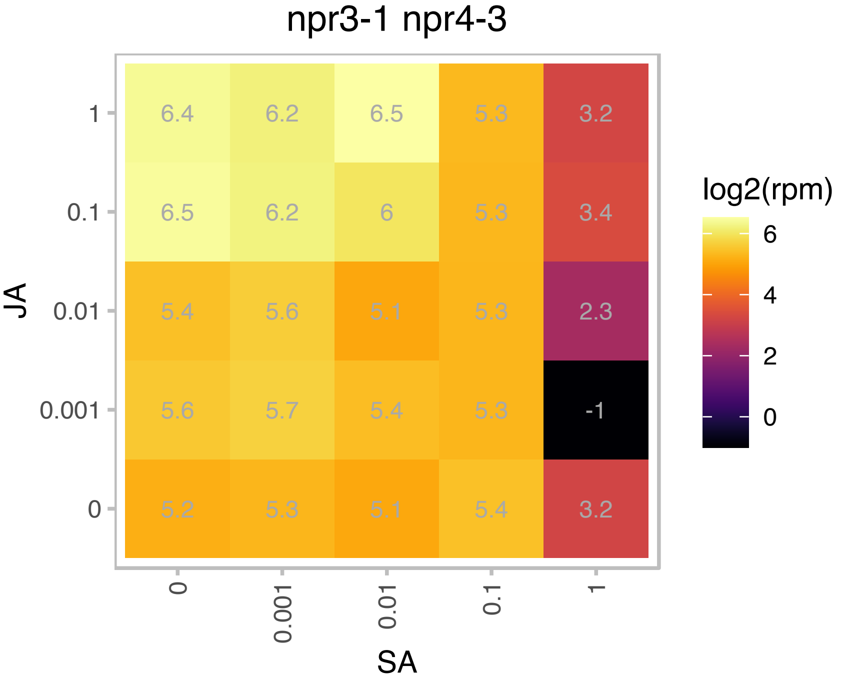

# AT4G12545.1 : NA

Bifunctional inhibitor/lipid-transfer protein/seed storage 2S albumin superfamily protein

| | |
|---|---|
| Gene | AT4G12545 |
| is.expressed_genes | TRUE |
| is.responsive_genes | TRUE |
| is.apcluster | F-02 |

Col-0

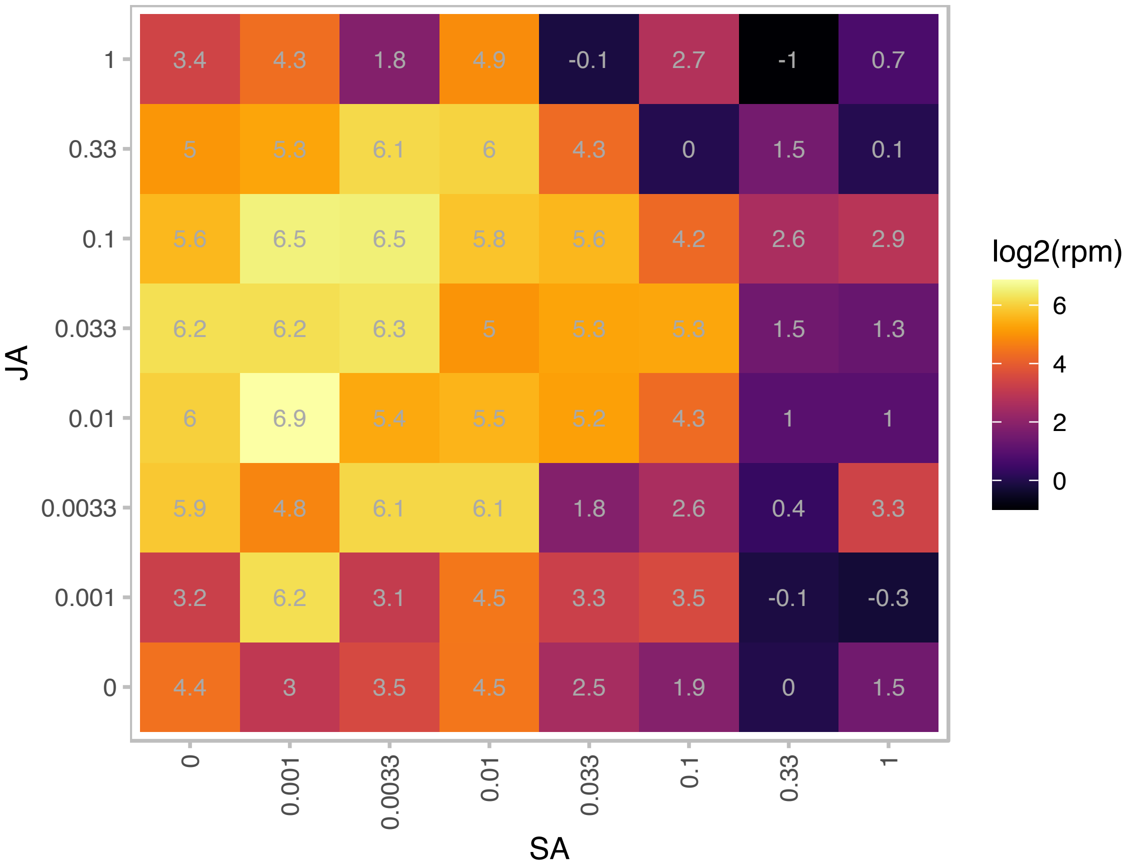


npr3-2 npr4-2

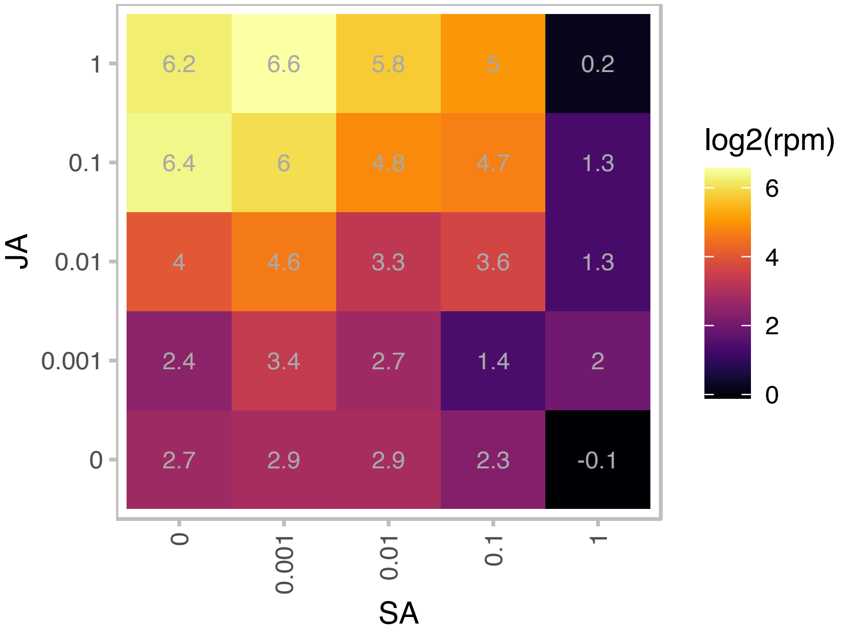


npr3-1 npr4-3

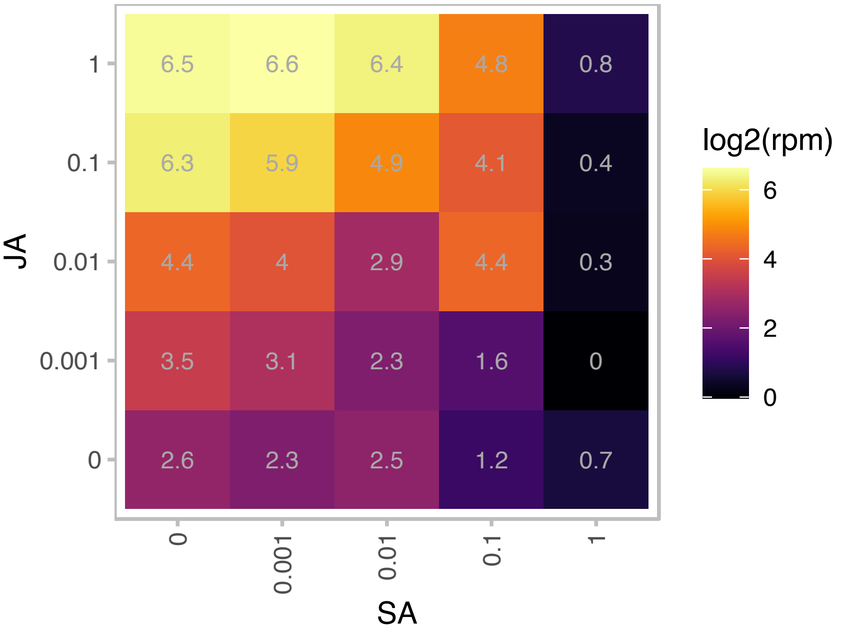

### AT4G13390.1 : EXT12

Proline-rich extensin-like family protein

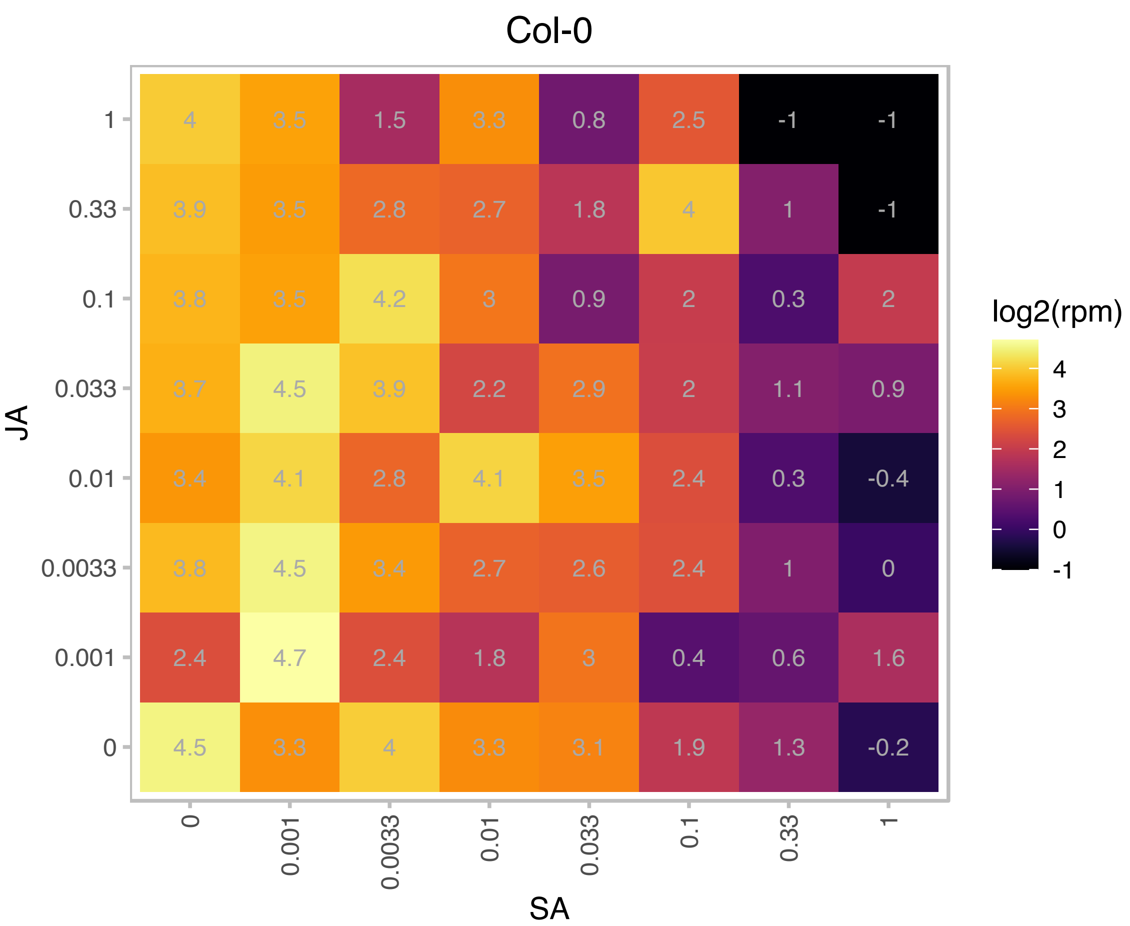


| Gene | AT4G13390 |
| --- | --- |
| is.expressed_genes | TRUE |
| is.responsive_genes | TRUE |
| is.apcluster | F-02 |

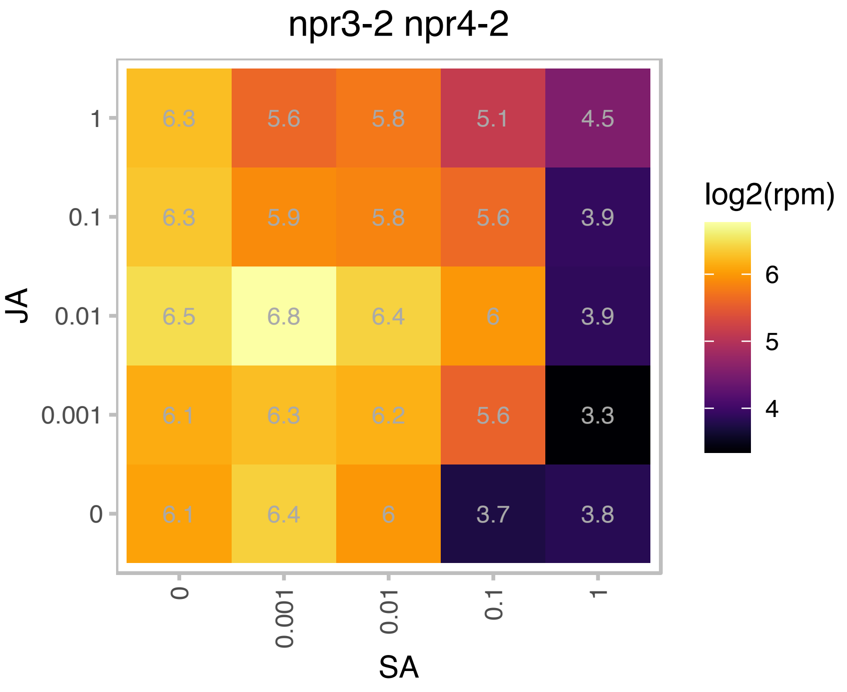


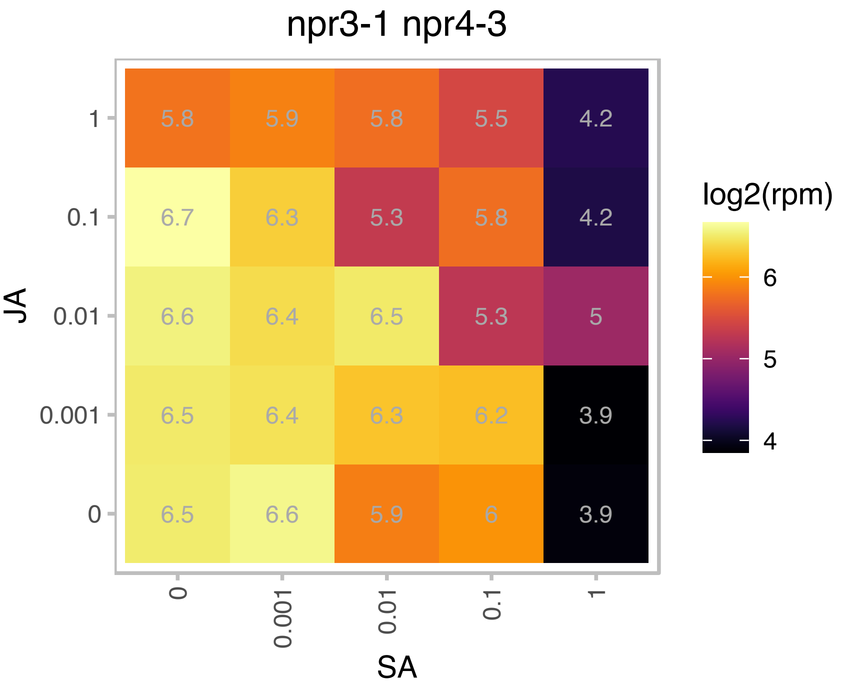

### AT4G13950.1 : RALFL31

ralf-like 31
Member of a diversely expressed predicted peptide family showing sequence similarity to tobacco Rapid Alkalinization Factor (RALF), and is believed to play an essential role in the physiology of Arabidopsis. Consists of a single exon and is characterized by a conserved C-terminal motif and N-terminal signal peptide.

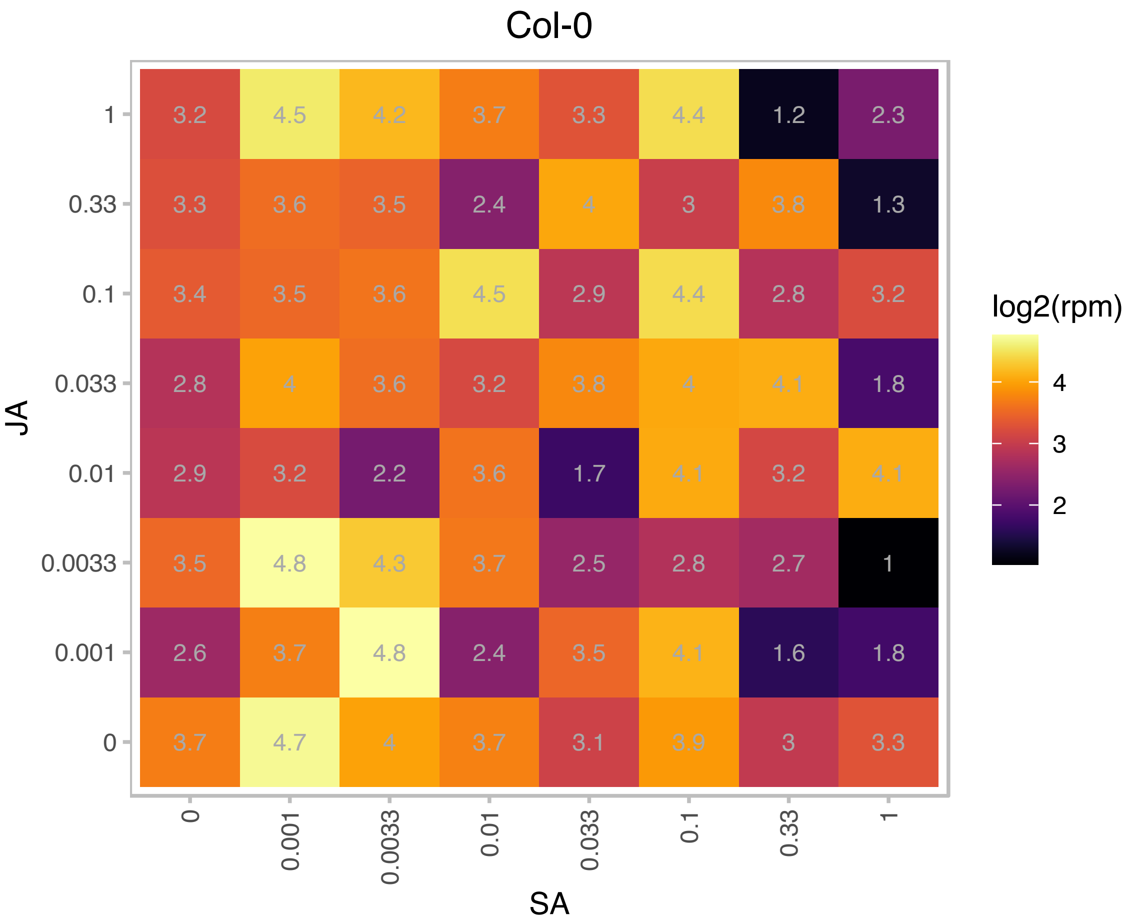


| | |
|---|---|
| Gene | AT4G13950 |
| is.expressed_genes | TRUE |
| is.responsive_genes | TRUE |
| is.apcluster | F-02 |

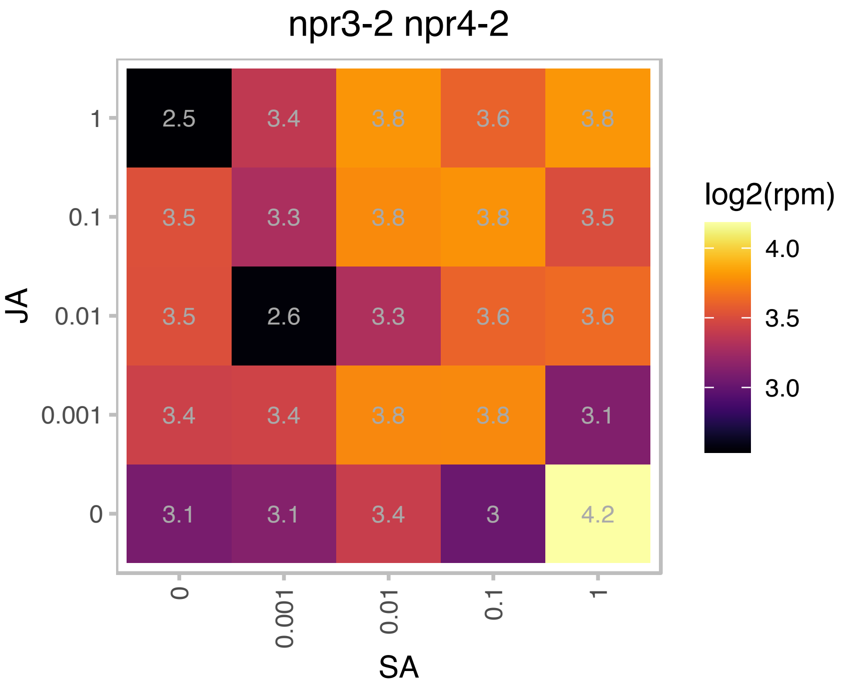


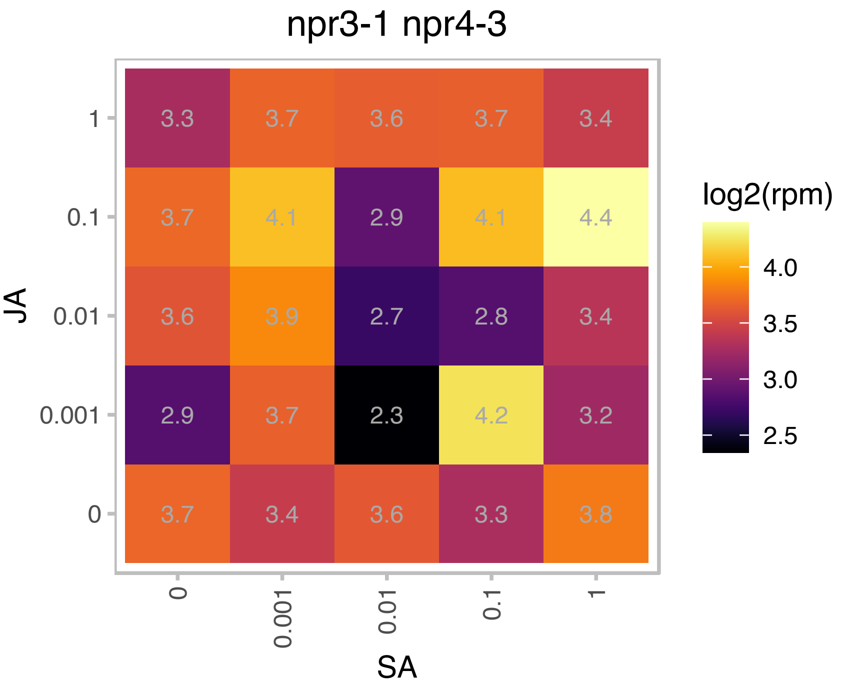

### AT4G14300.1 : AtRNP1, RBGD4

RNA-binding (RRM/RBD/RNP motifs) family protein

Belongs to a member of the RNA-binding glycine-rich (RBG) gene superfamily.

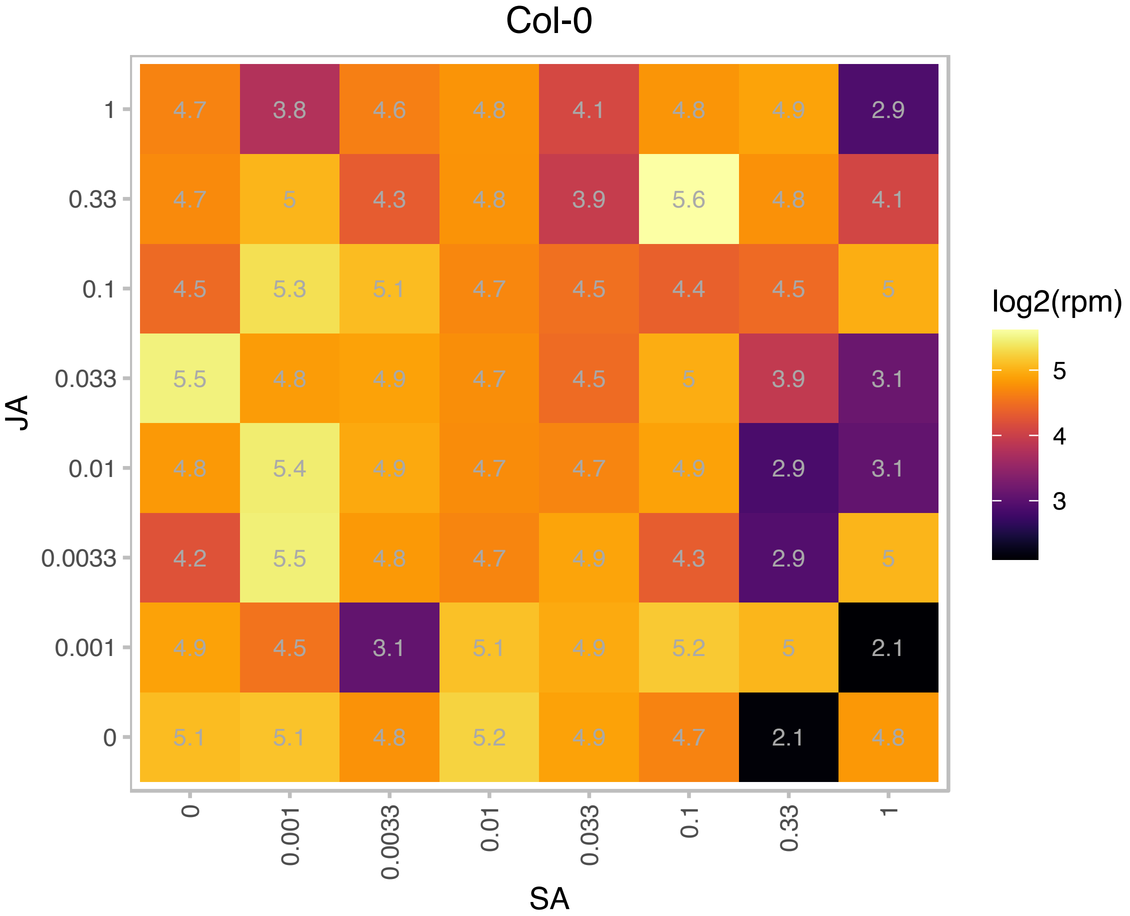


| | |
|---|---|
| Gene | AT4G14300 |
| is.expressed_genes | TRUE |
| is.responsive_genes | TRUE |
| is.apcluster | F-02 |

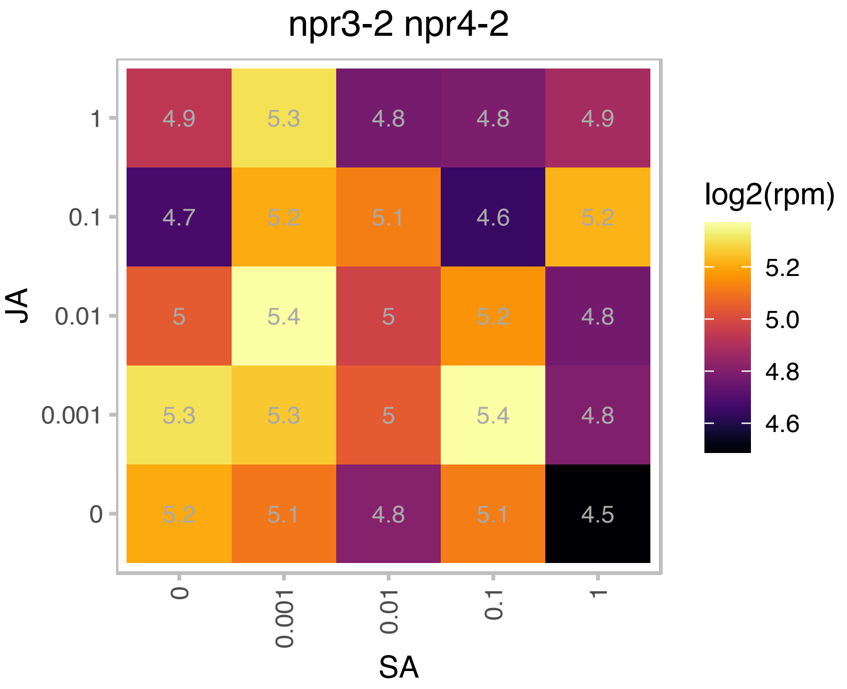


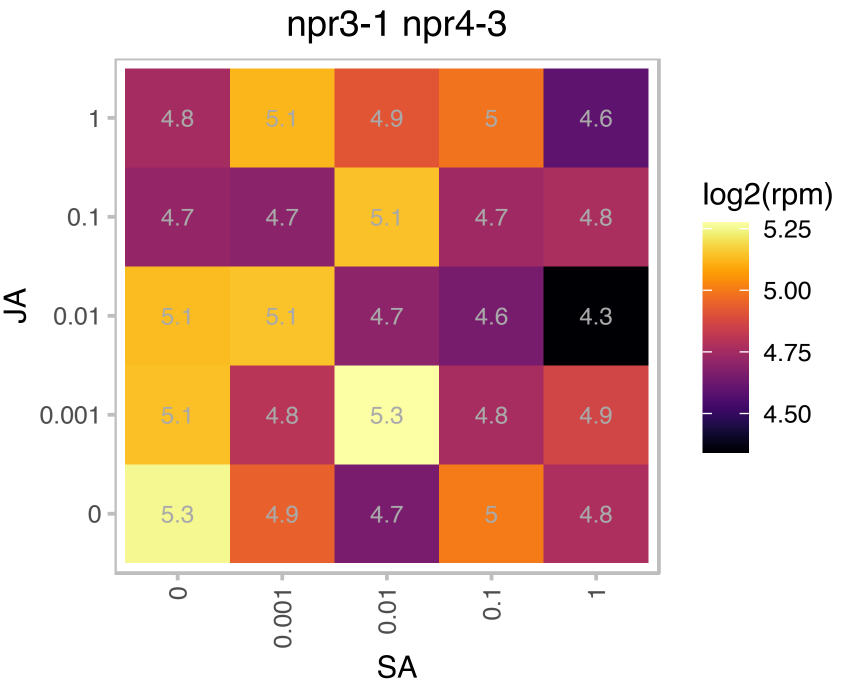

### AT4G14960.1 : TUA6

Tubulin/FtsZ family protein

Encodes an alpha-tubulin isoform required for right handed helical growth.

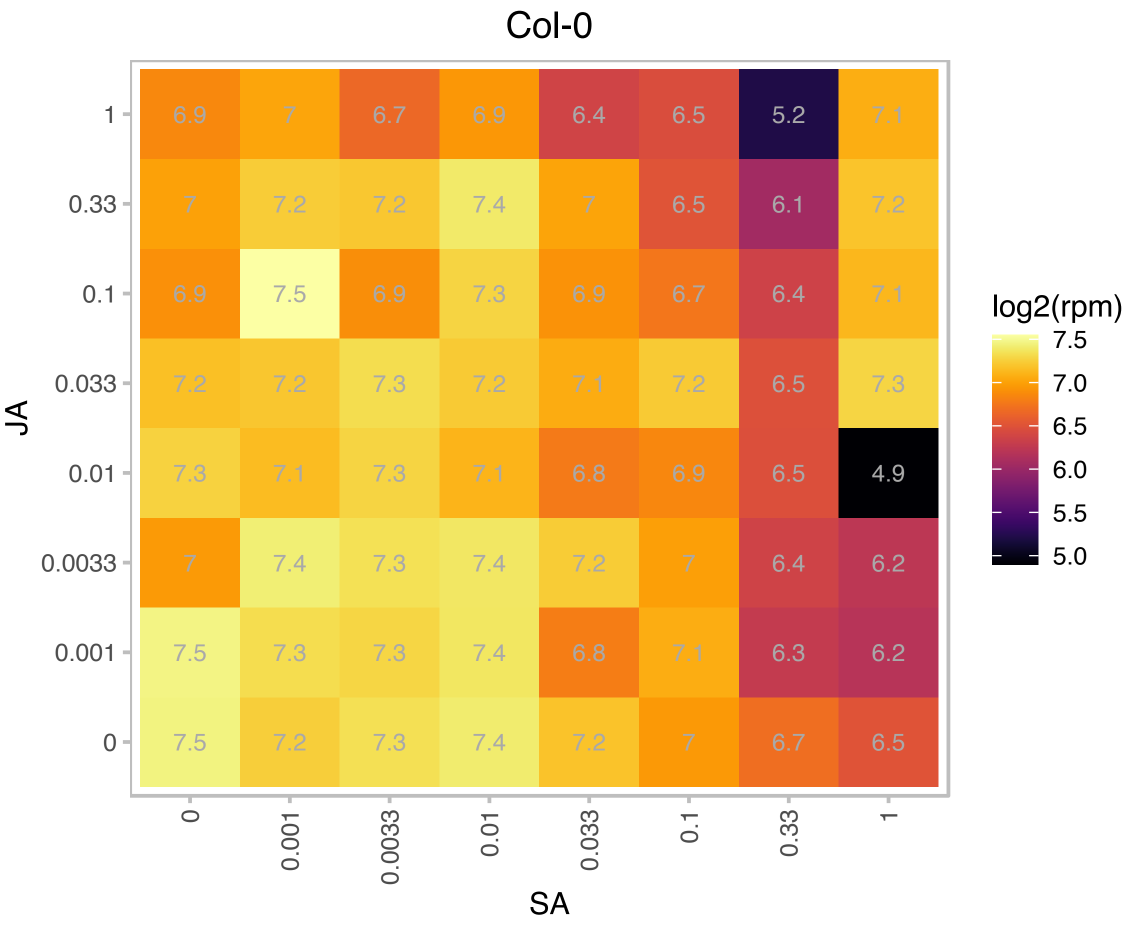


| | |
|---|---|
| Gene | AT4G14960 |
| is.expressed_genes | TRUE |
| is.responsive_genes | TRUE |
| is.apcluster | F-02 |

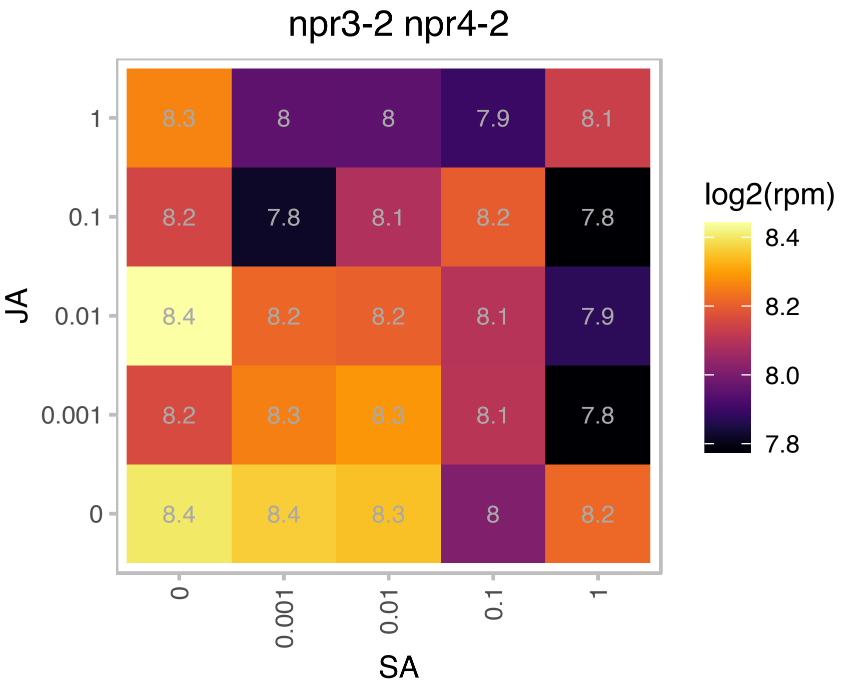


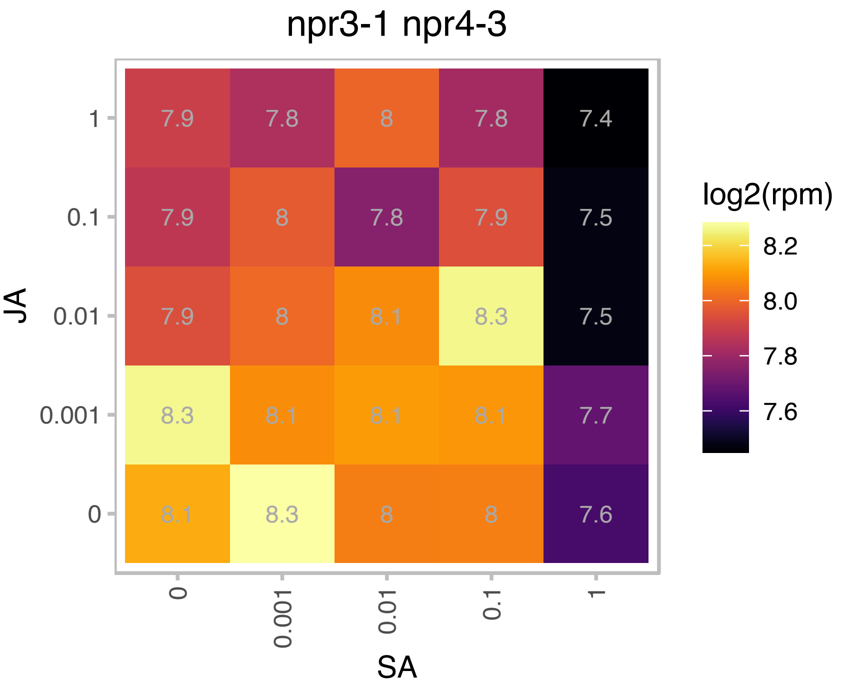

# AT4G15390.1 : NA

HXXXD-type acyl-transferase family protein

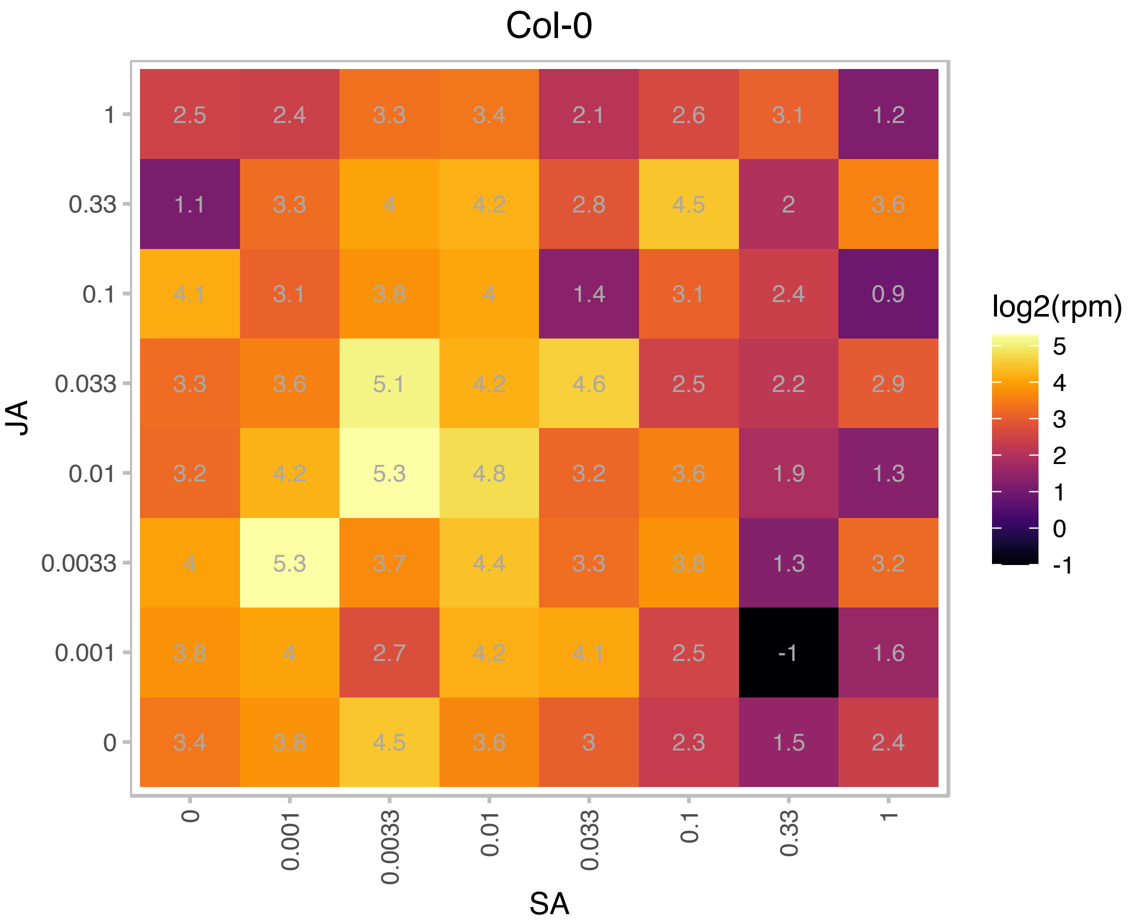


| | |
|---|---|
| Gene | AT4G15390 |
| is.expressed_genes | TRUE |
| is.responsive_genes | TRUE |
| is.apcluster | F-02 |

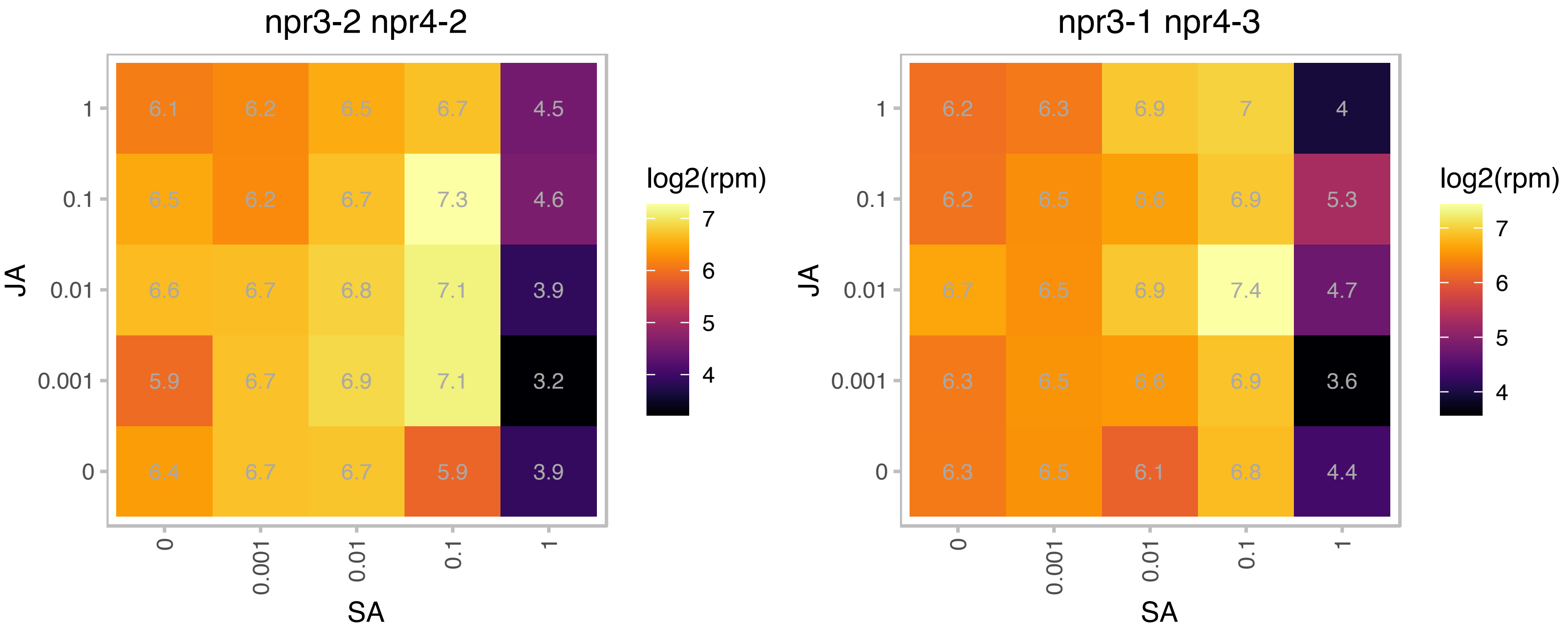

### AT4G15680.1 : GRXS4, ROXY13

Thioredoxin superfamily protein

Encodes a member of the CC-type glutaredoxin (ROXY) family.

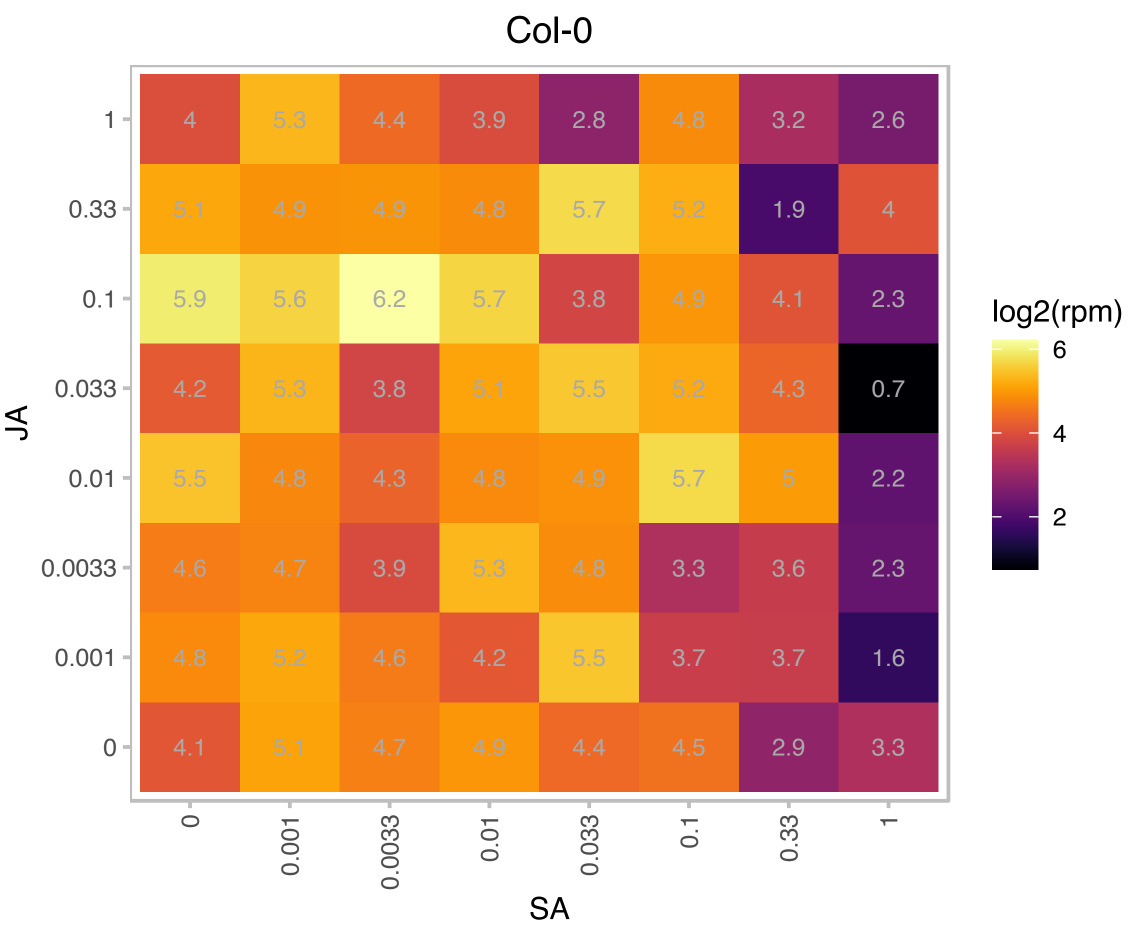


| | |
|---|---|
| Gene | AT4G15680 |
| is.expressed_genes | TRUE |
| is.responsive_genes | TRUE |
| is.apcluster | F-02 |

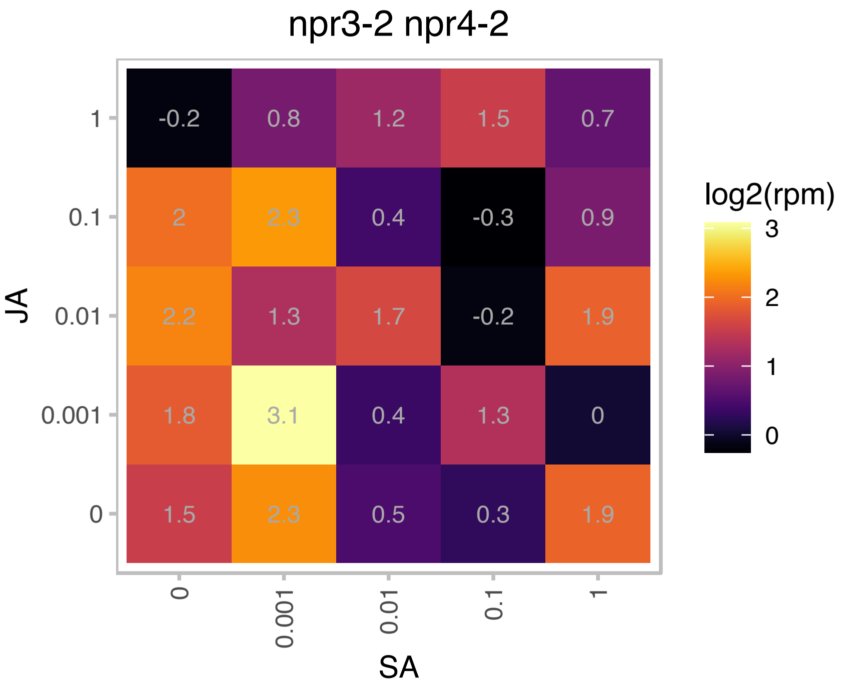


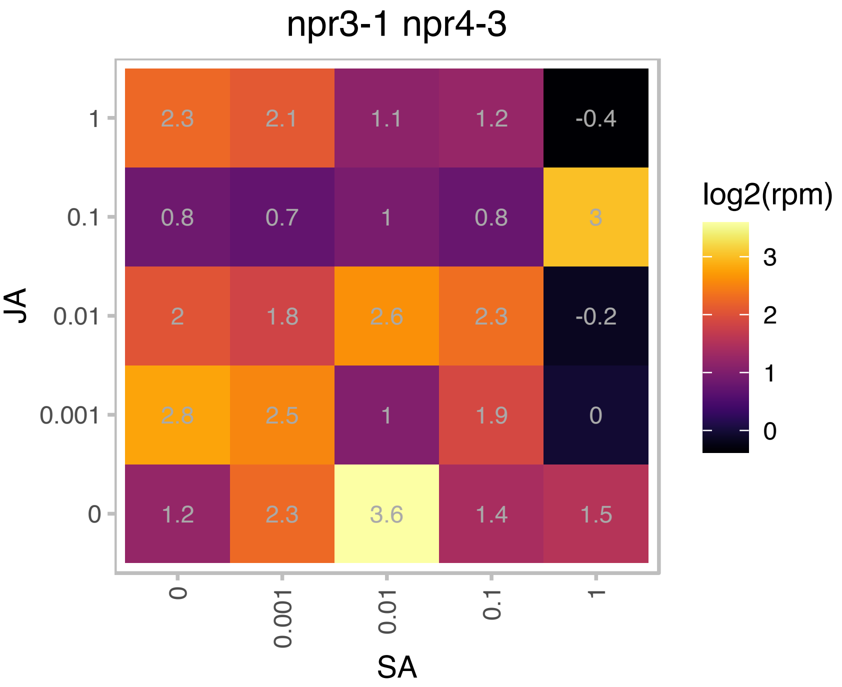

# AT4G15990.1 : NA

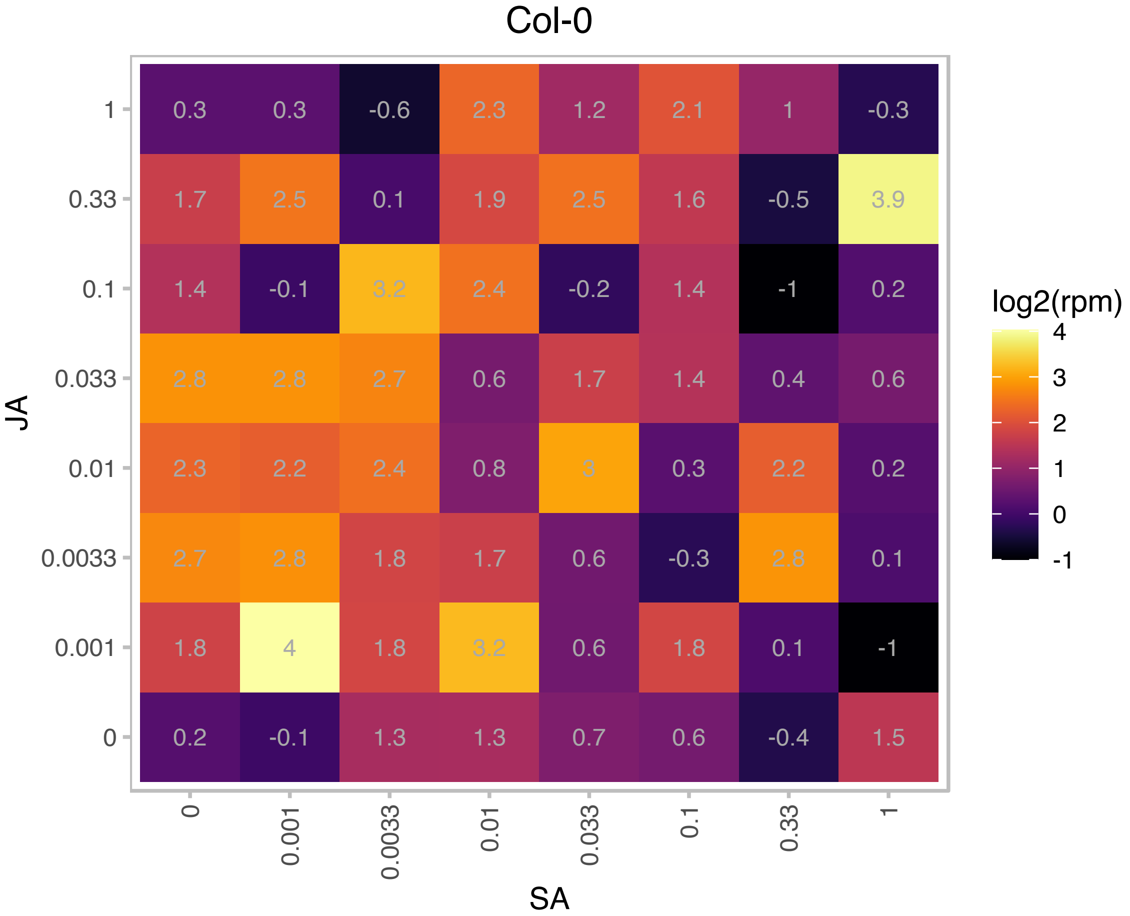


| | |
|---|---|
| Gene | AT4G15990 |
| is.expressed_genes | TRUE |
| is.responsive_genes | TRUE |
| is.apcluster | F-02 |

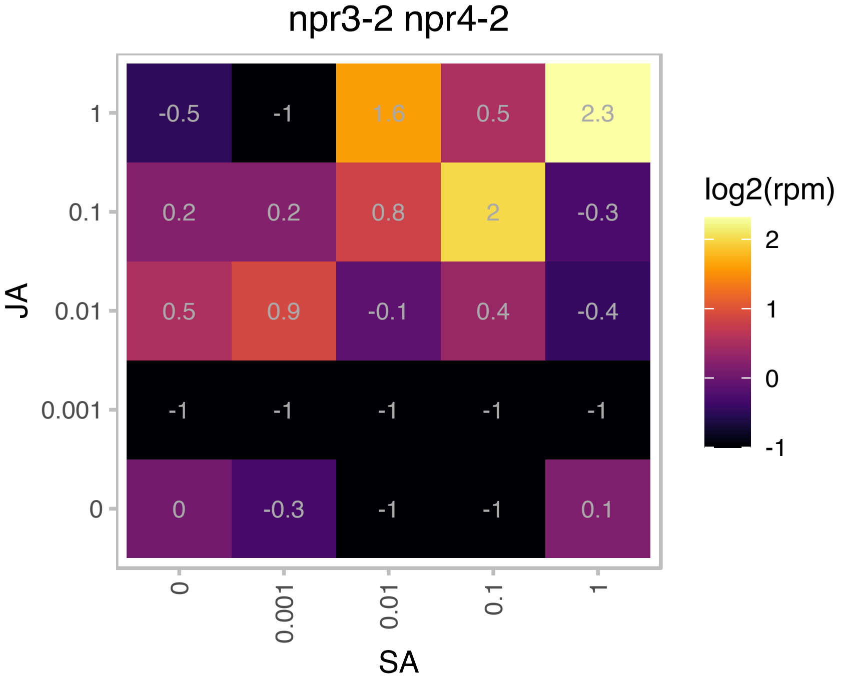


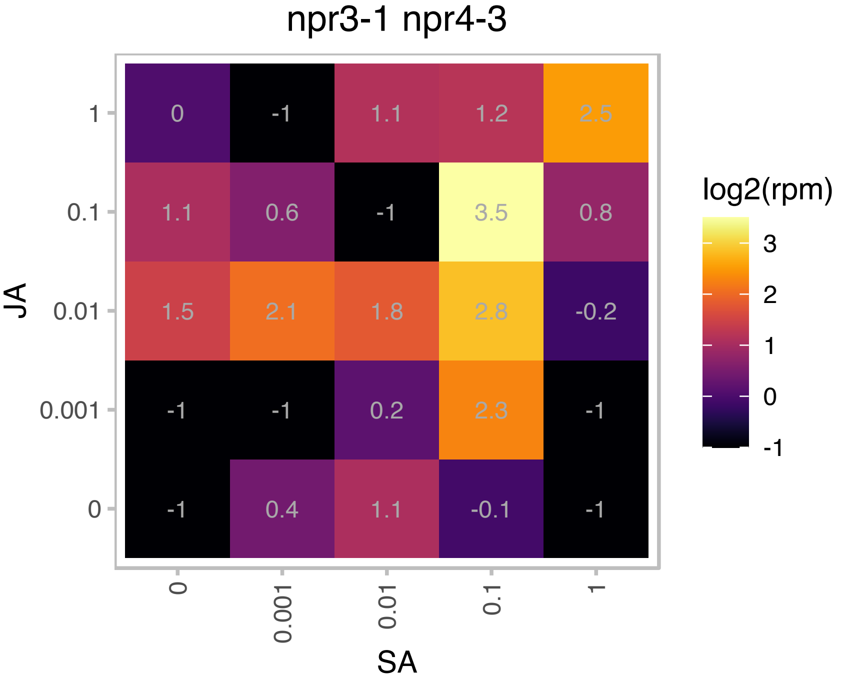

# AT4G17520.1 : NA

Hyaluronan / mRNA binding family

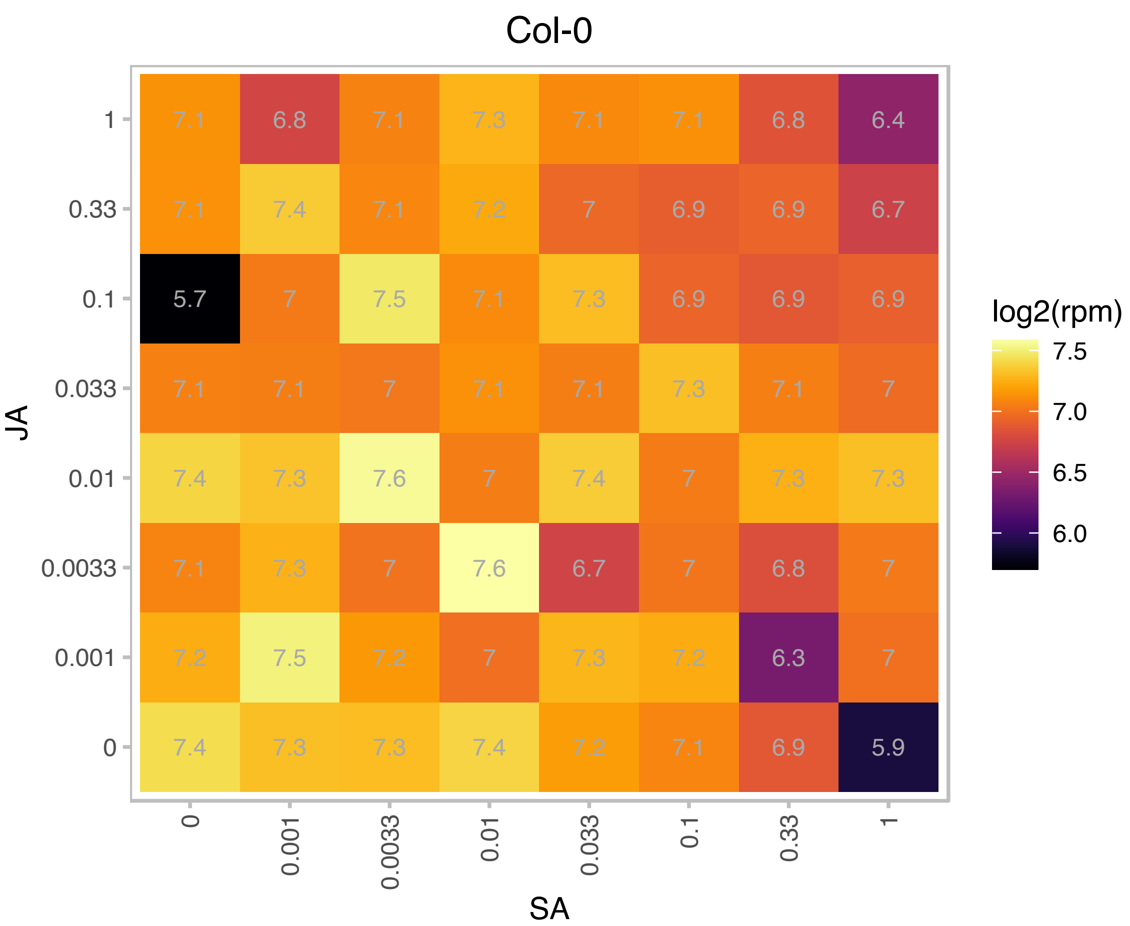


| | |
|---|---|
| Gene | AT4G17520 |
| is.expressed_genes | TRUE |
| is.responsive_genes | TRUE |
| is.apcluster | F-02 |

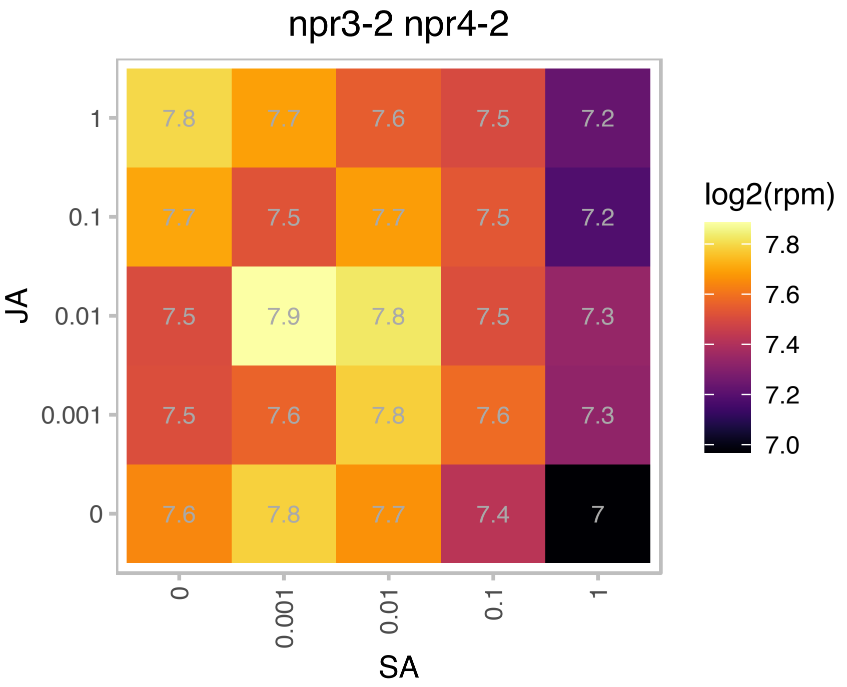


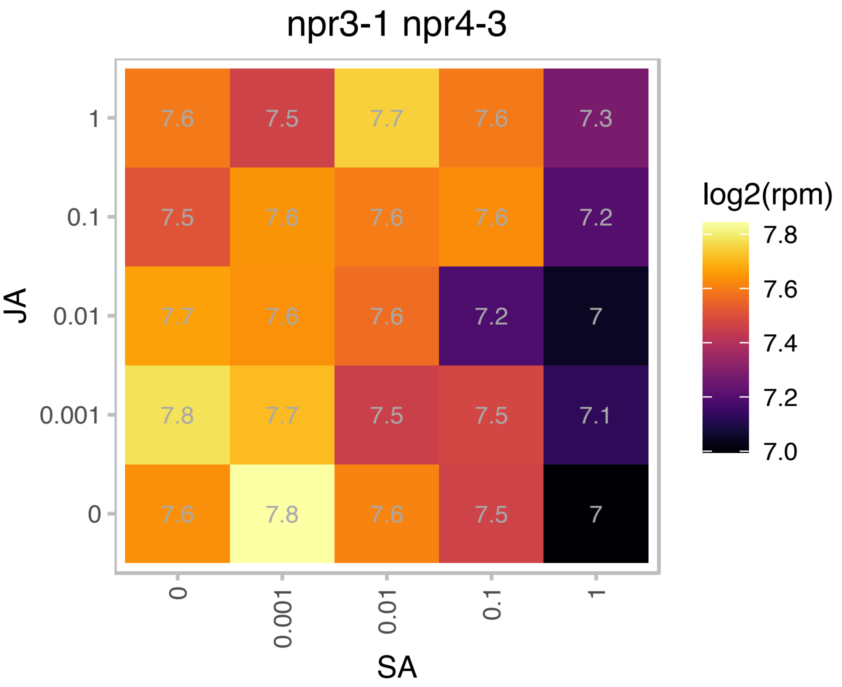

### AT4G18040.1 : AT.EIF4E1, CUM1, EIF4E, eIF4E1

eukaryotic translation initiation factor 4E

eIF4E protein. The cum1 mutation affects the local spreading of CMV within the inoculated leaf, delaying accumulation of cucumber mosaic virus coat protein.

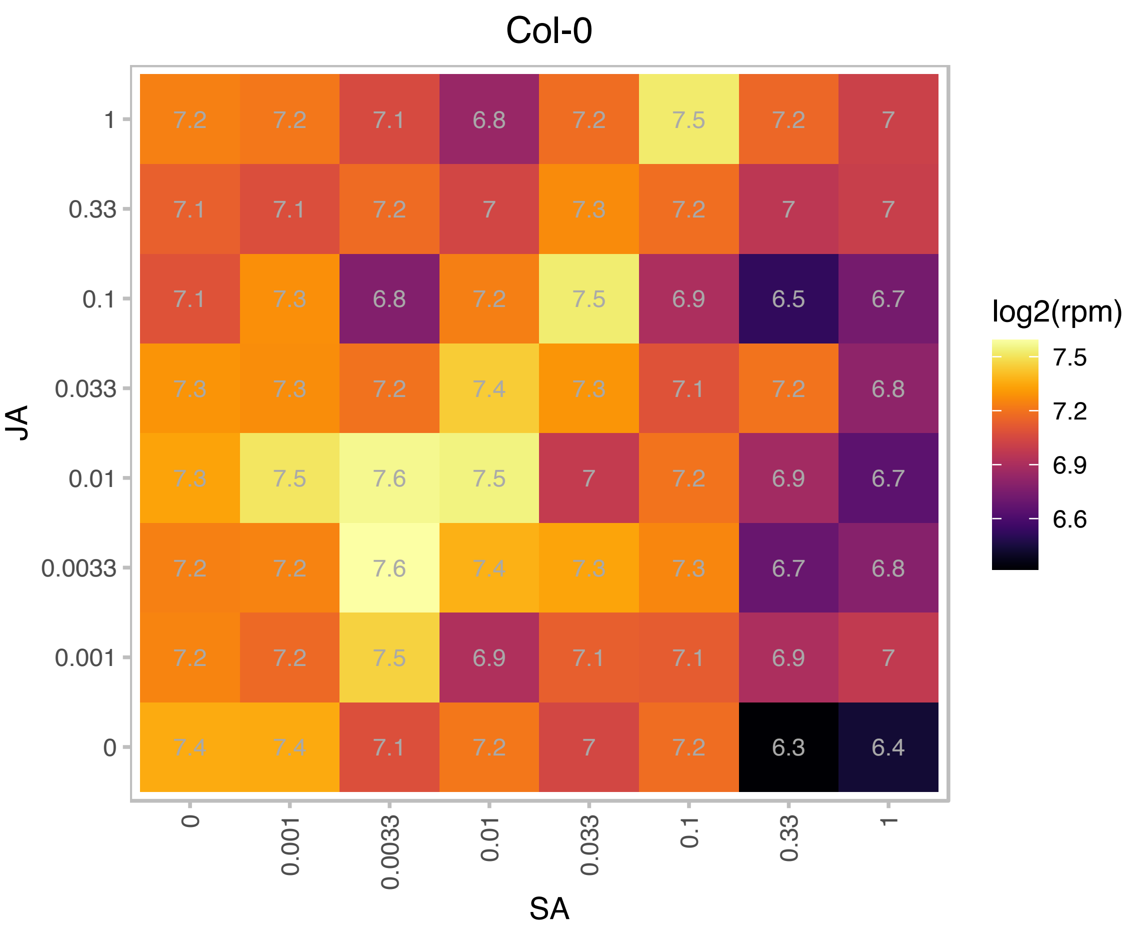


| | |
|---|---|
| Gene | AT4G18040 |
| is.expressed_genes | TRUE |
| is.responsive_genes | TRUE |
| is.apcluster | F-02 |

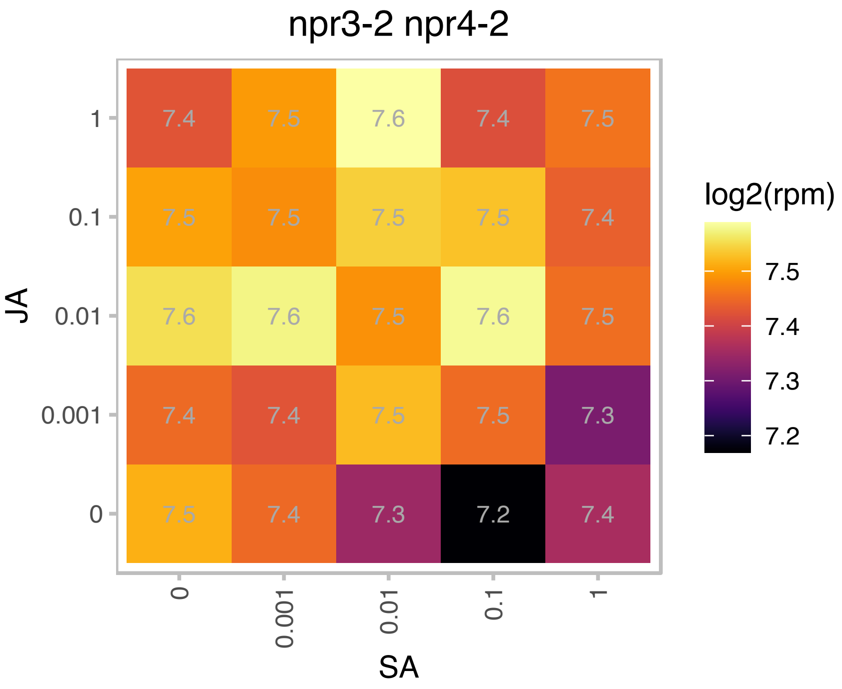


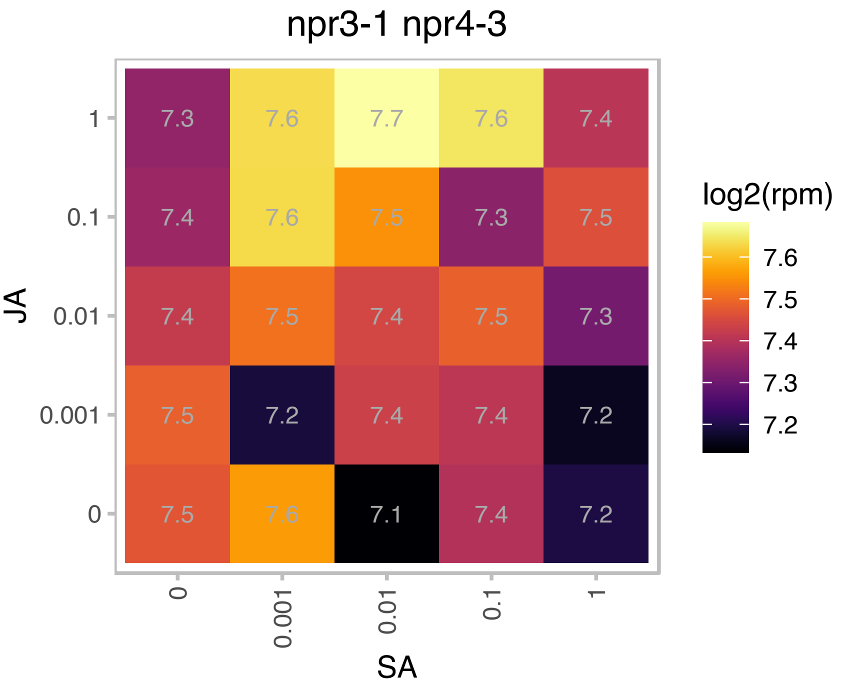

# AT4G19610.1 : NA

nucleotide binding;nucleic acid binding;RNA binding

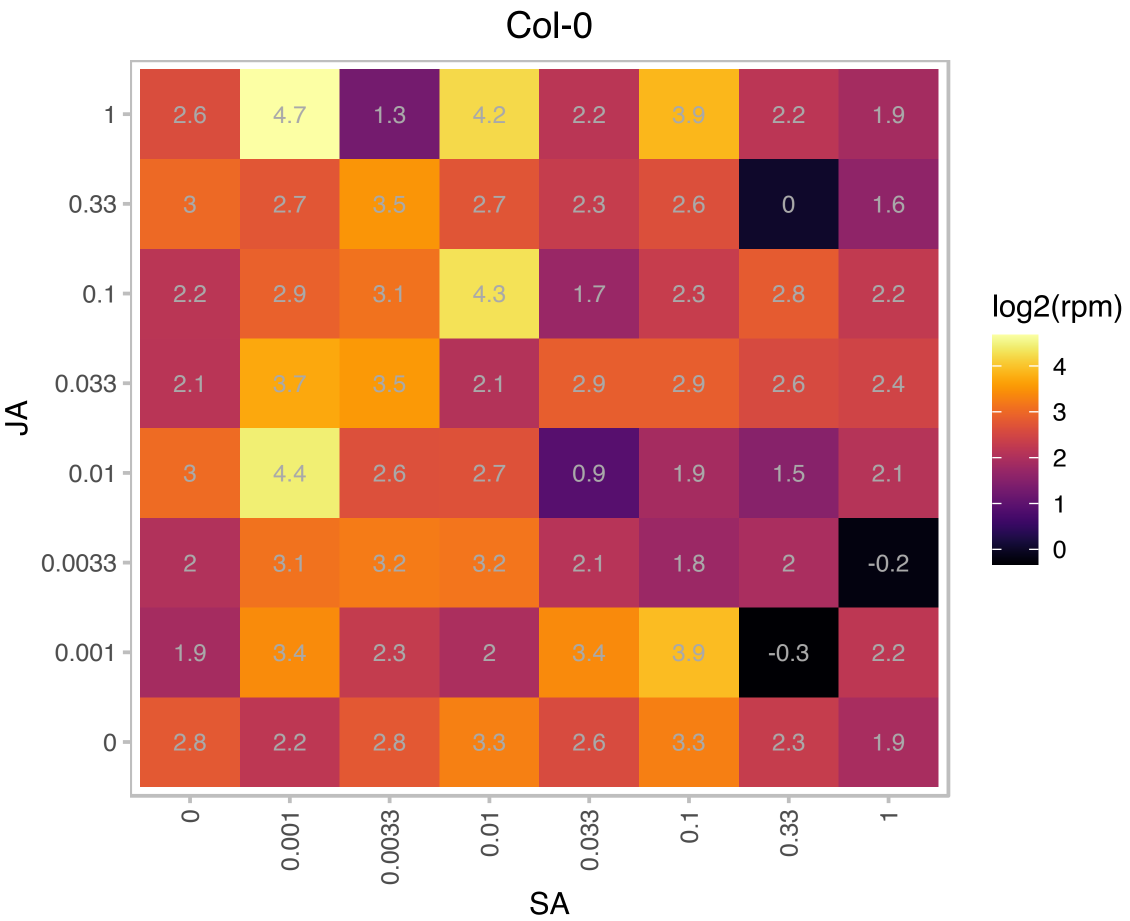


| | |
|---|---|
| Gene | AT4G19610 |
| is.expressed_genes | TRUE |
| is.responsive_genes | TRUE |
| is.apcluster | F-02 |

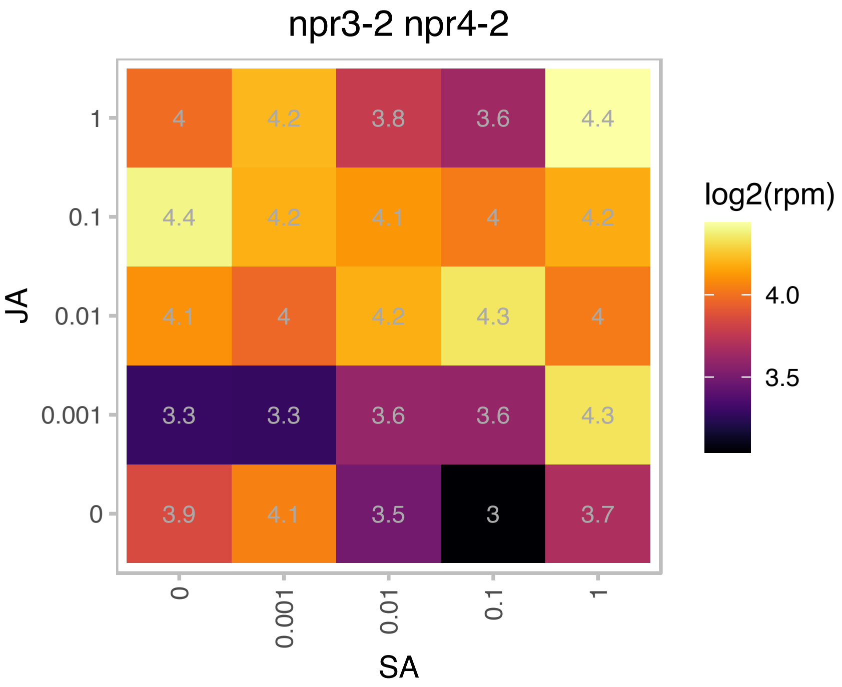


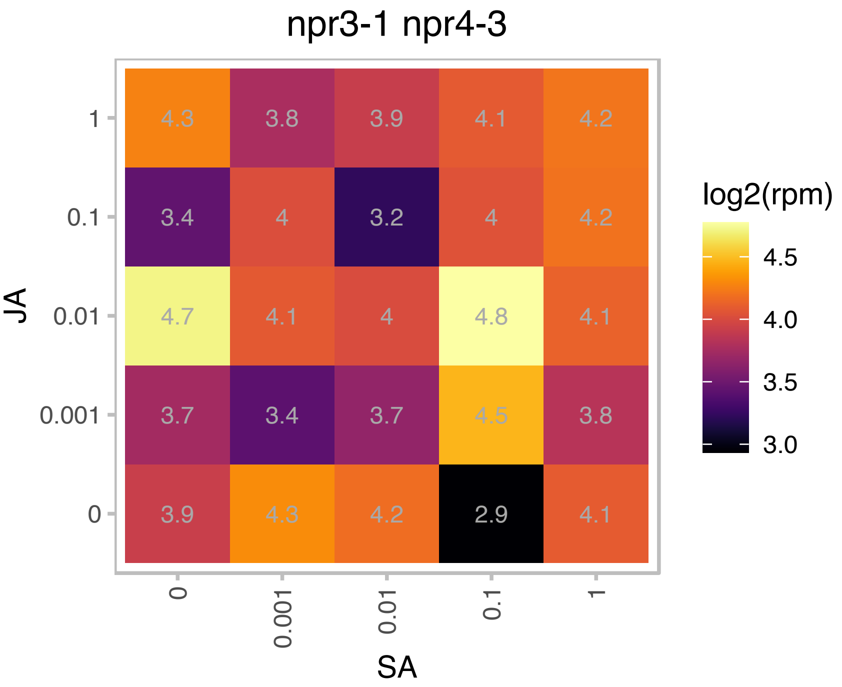

# AT4G21192.1 : NA

Cytochrome c oxidase biogenesis protein Cmc1-like

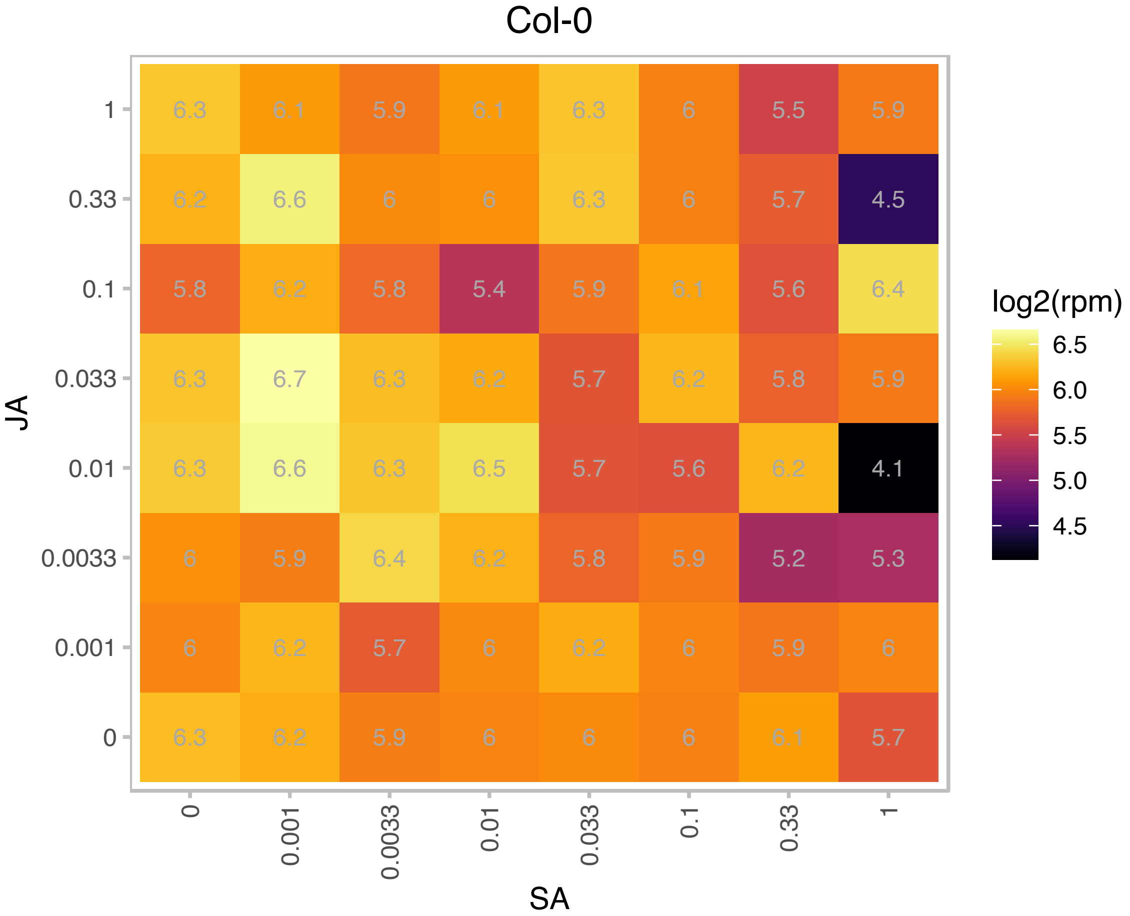


| | |
|---|---|
| Gene | AT4G21192 |
| is.expressed_genes | TRUE |
| is.responsive_genes | TRUE |
| is.apcluster | F-02 |

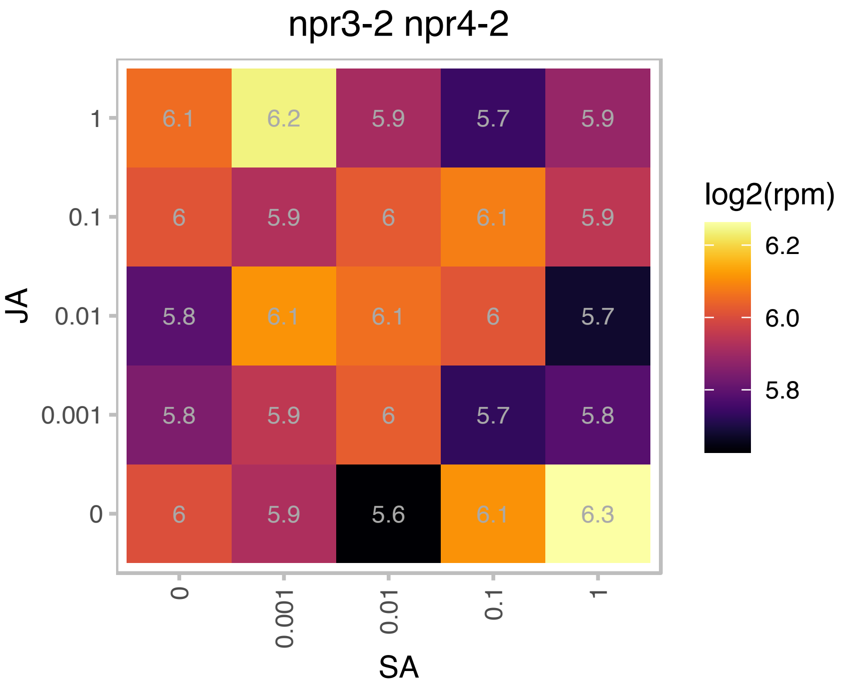


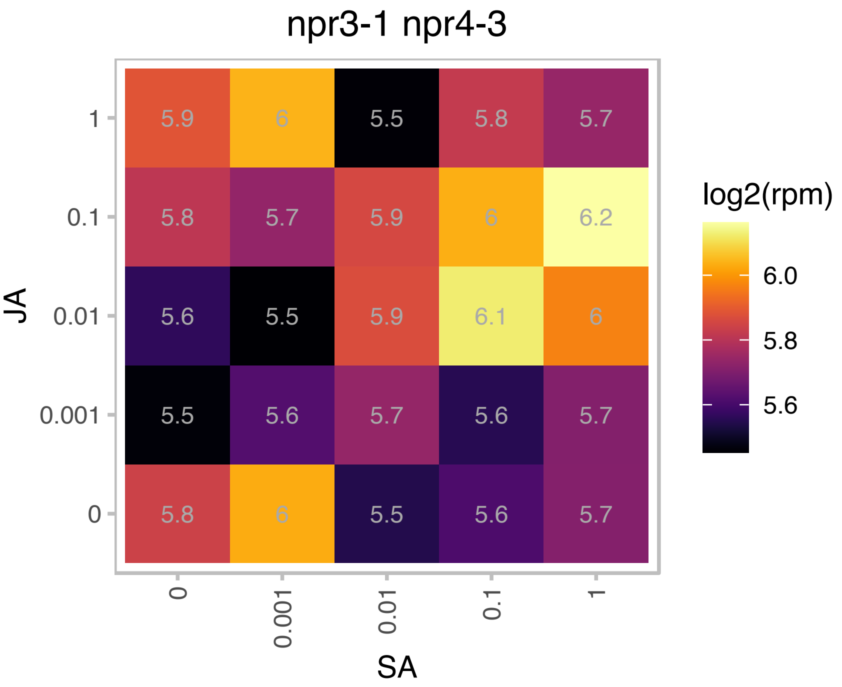

### AT4G22690.1 : CYP706A1

cytochrome P450, family 706, subfamily A, polypeptide 1

member of CYP706A The mRNA is cell-to-cell mobile.

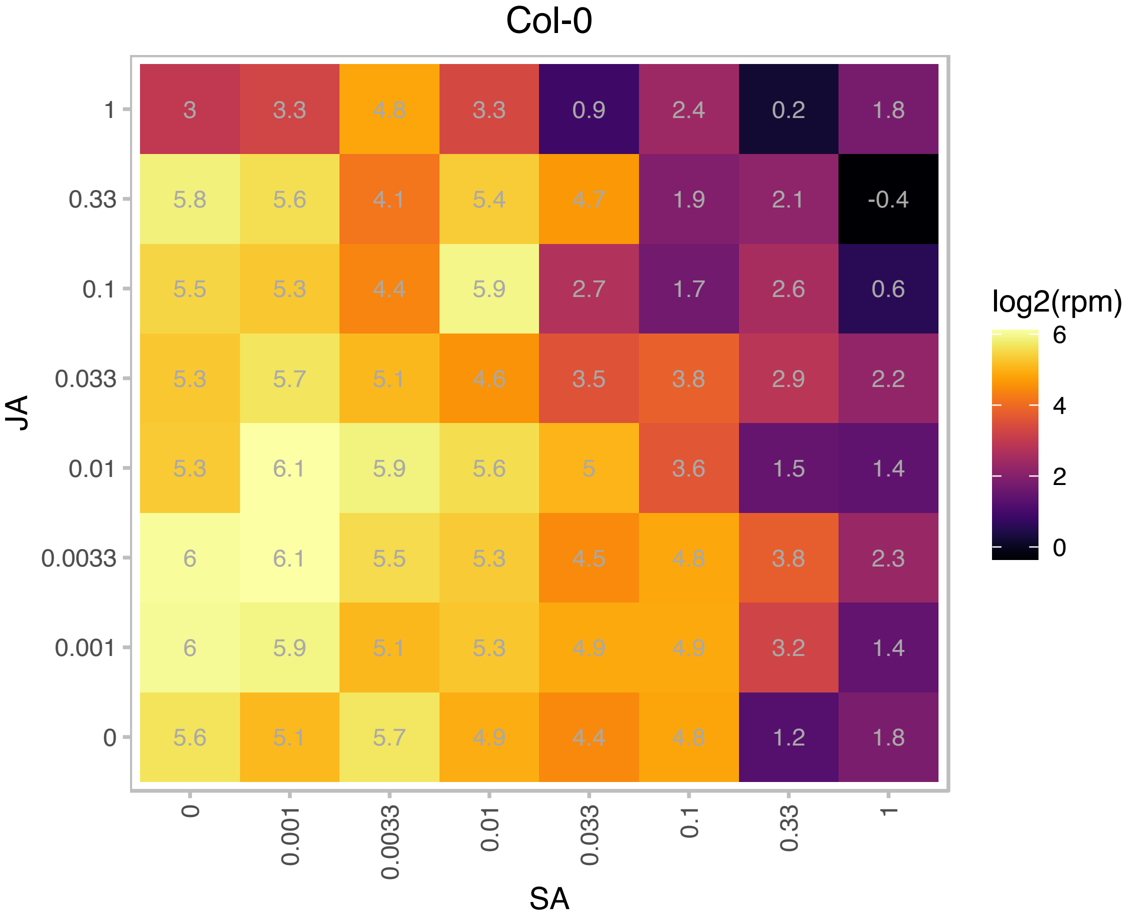


| | |
|---|---|
| Gene | AT4G22690 |
| is.expressed_genes | TRUE |
| is.responsive_genes | TRUE |
| is.apcluster | F-02 |

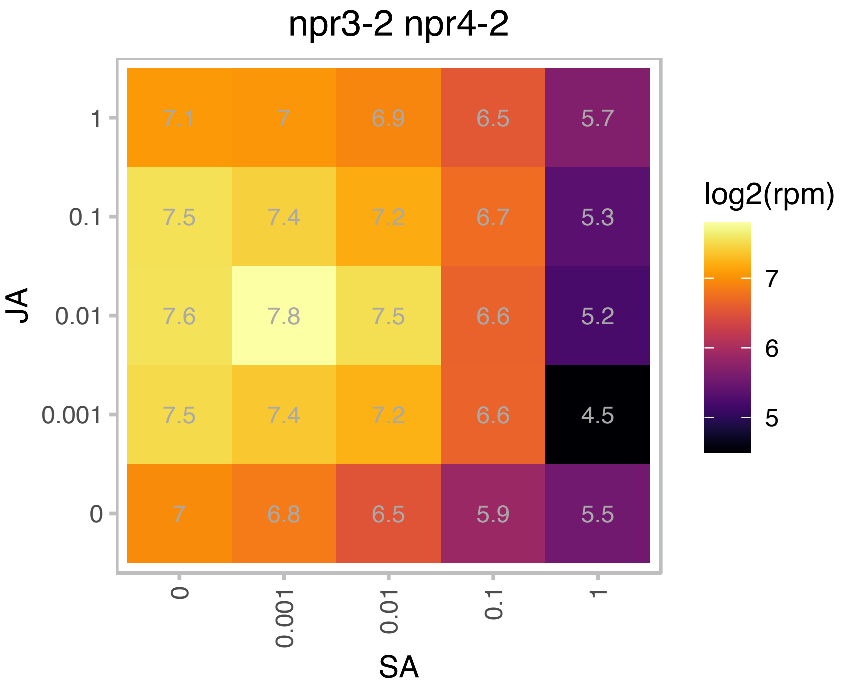


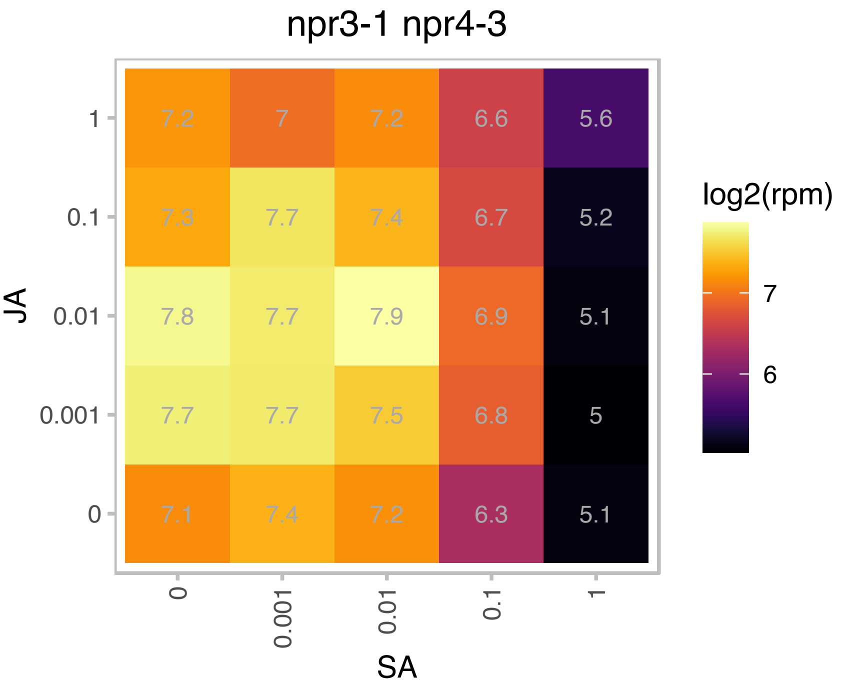

### AT4G23940.1 : ARC1, FtsHi1

FtsH extracellular protease family

Encodes FtsHi1. Localizes to the chloroplast envelope membrane. Functions in chloroplast biogenesis and division.

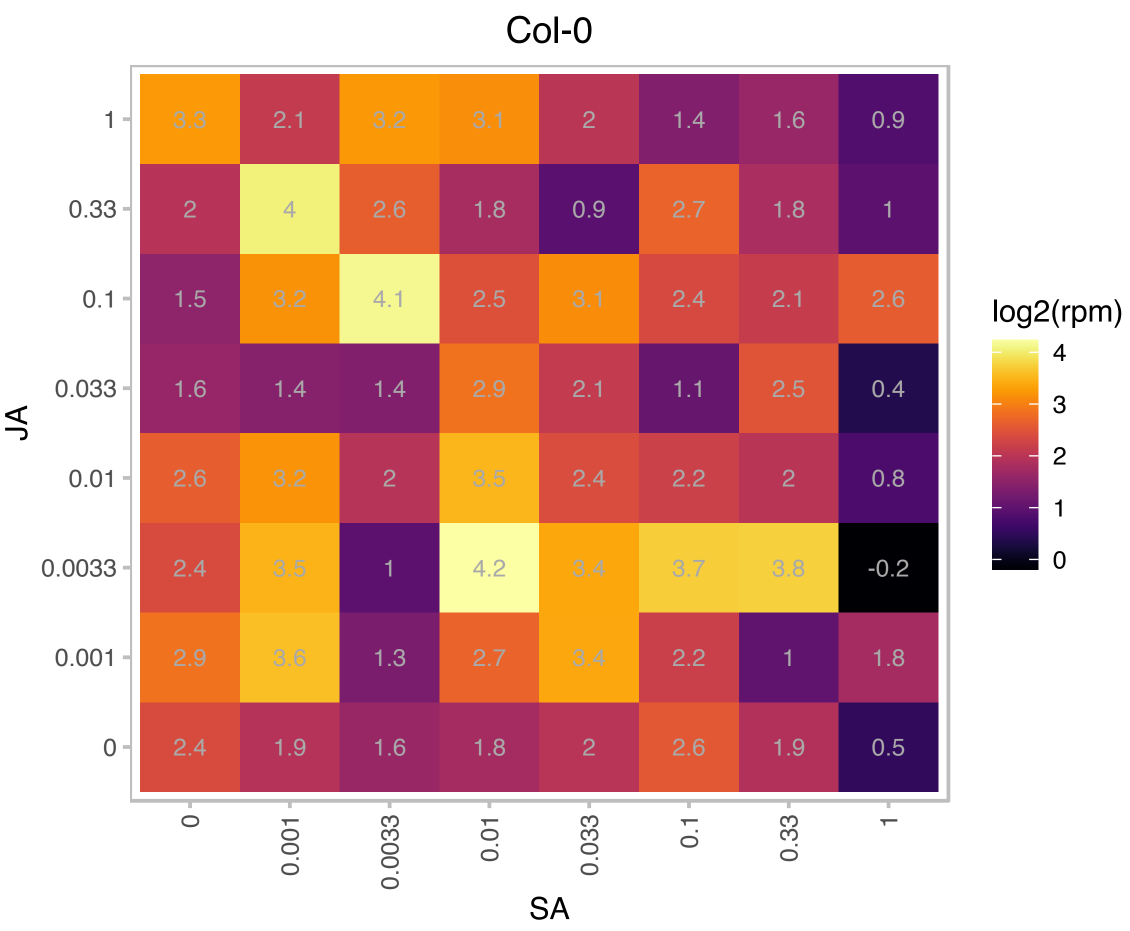


| | |
|---|---|
| Gene | AT4G23940 |
| is.expressed_genes | TRUE |
| is.responsive_genes | TRUE |
| is.apcluster | F-02 |

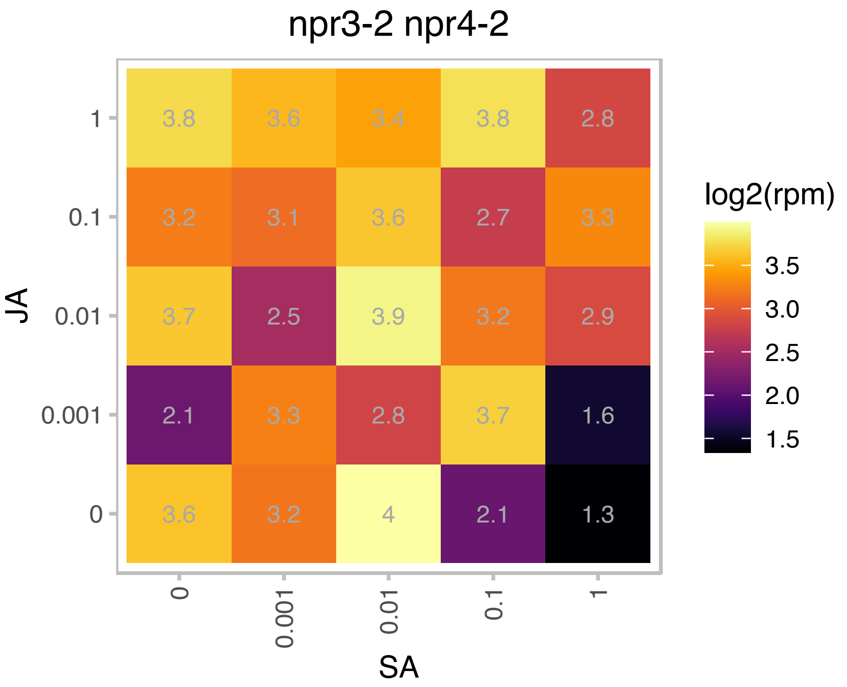


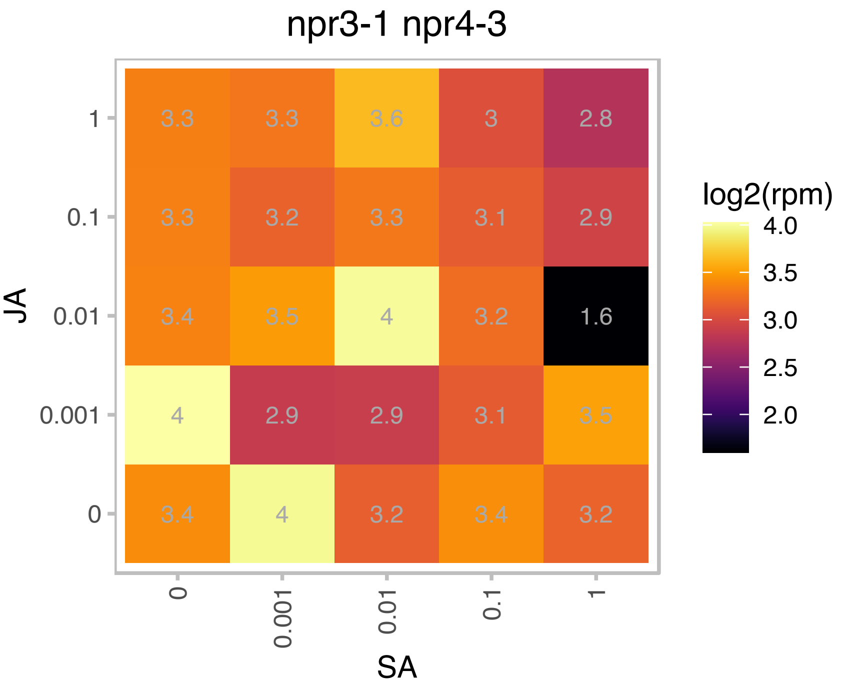

### AT4G24660.1 : ATHB22, HB22, MEE68, ZHD2

homeobox protein 22

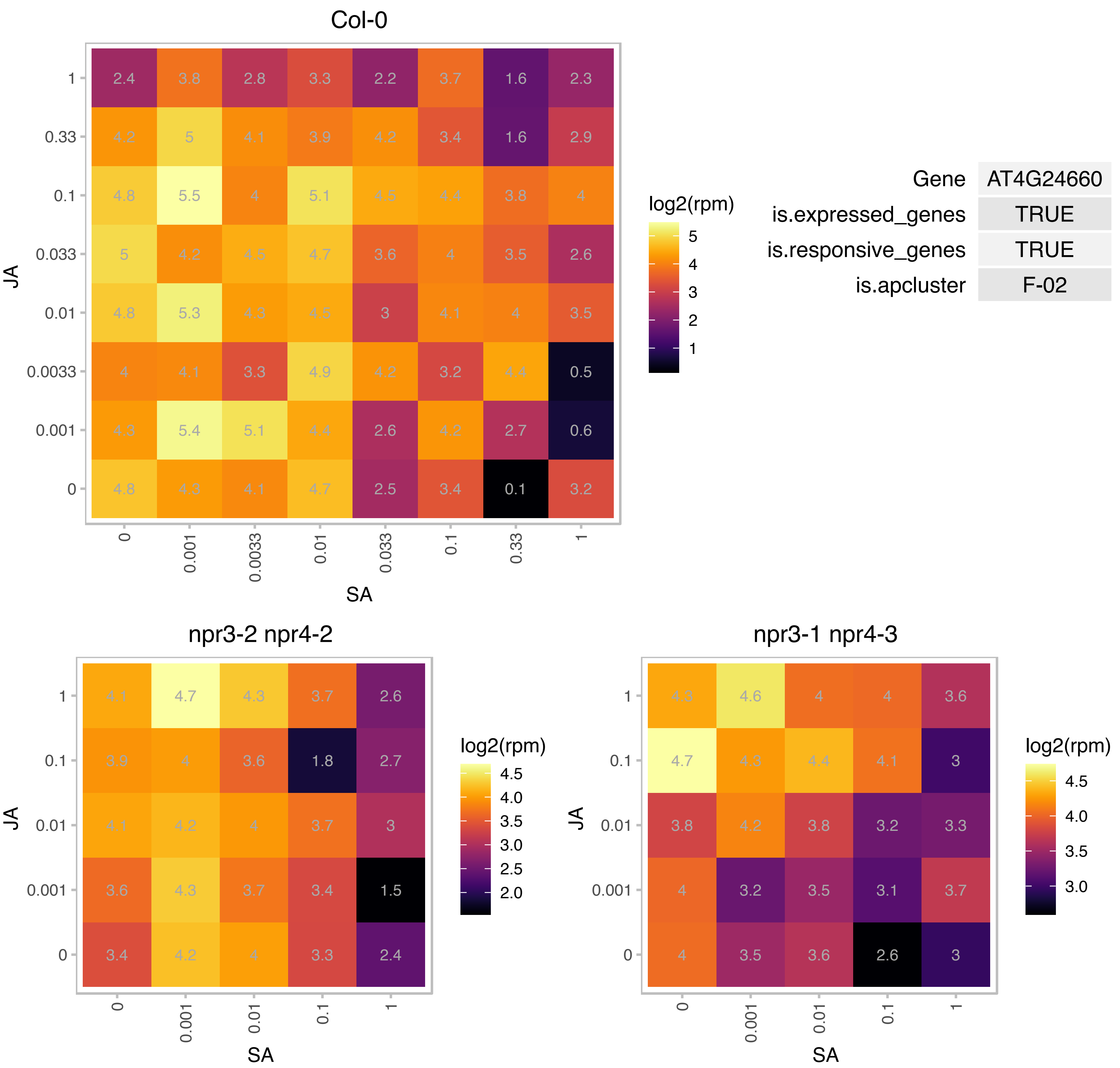


| | |
|---|---|
| Gene | AT4G24660 |
| is.expressed_genes | TRUE |
| is.responsive_genes | TRUE |
| is.apcluster | F-02 |

# AT4G29590.1 : NA

S-adenosyl-L-methionine-dependent methyltransferases superfamily protein

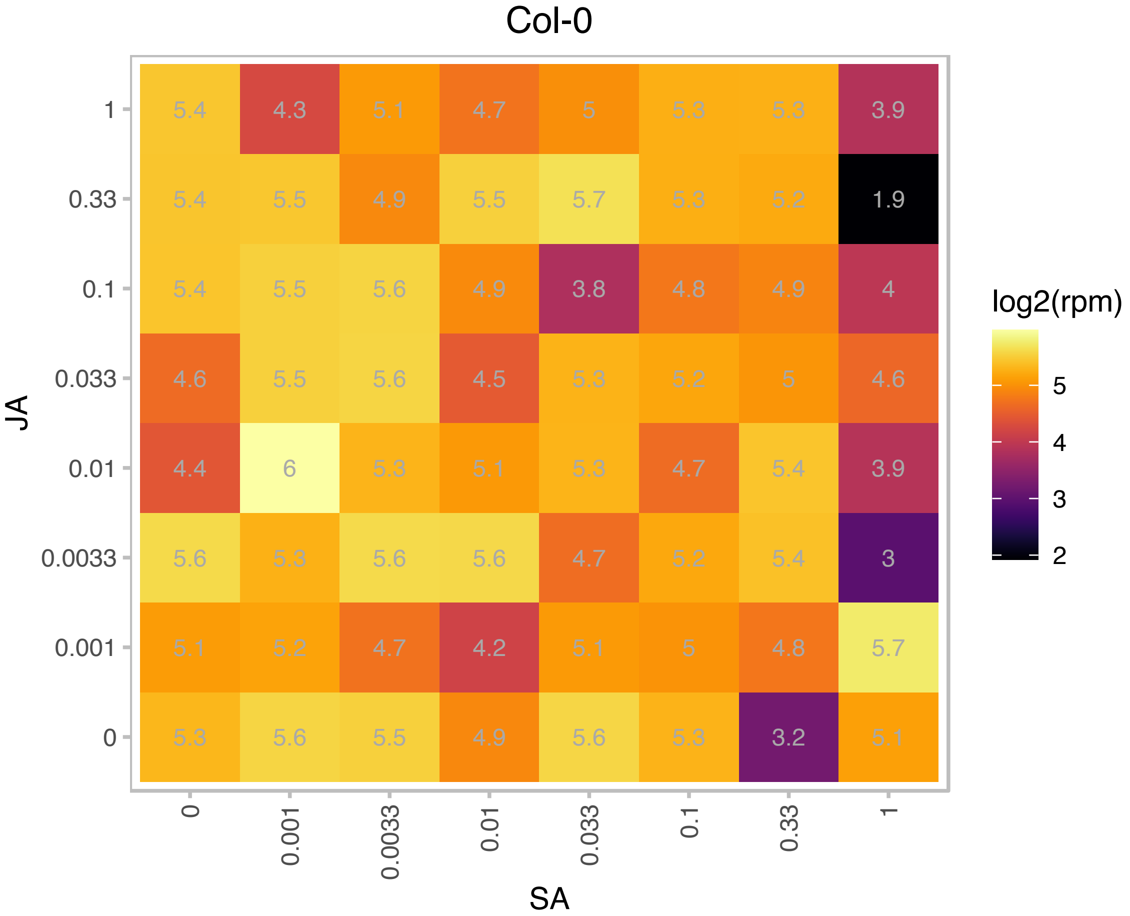


| | |
|---|---|
| Gene | AT4G29590 |
| is.expressed_genes | TRUE |
| is.responsive_genes | TRUE |
| is.apcluster | F-02 |

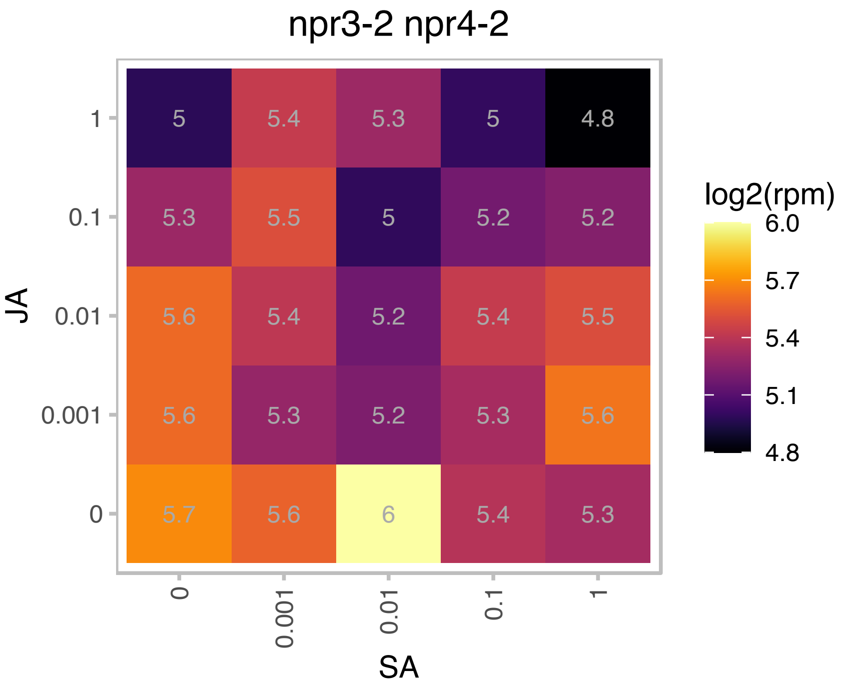


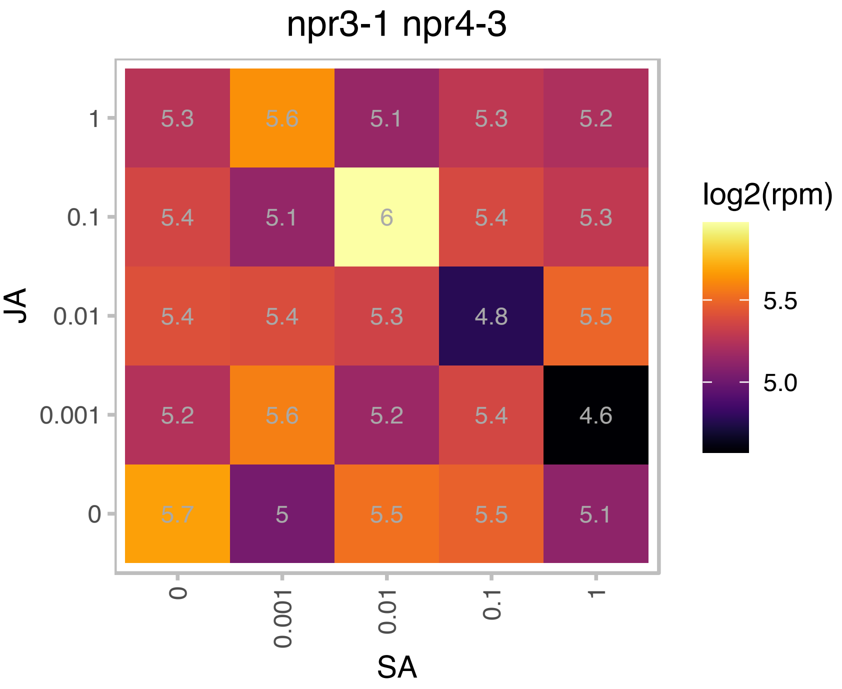

# AT4G30170.1 : NA

Peroxidase family protein

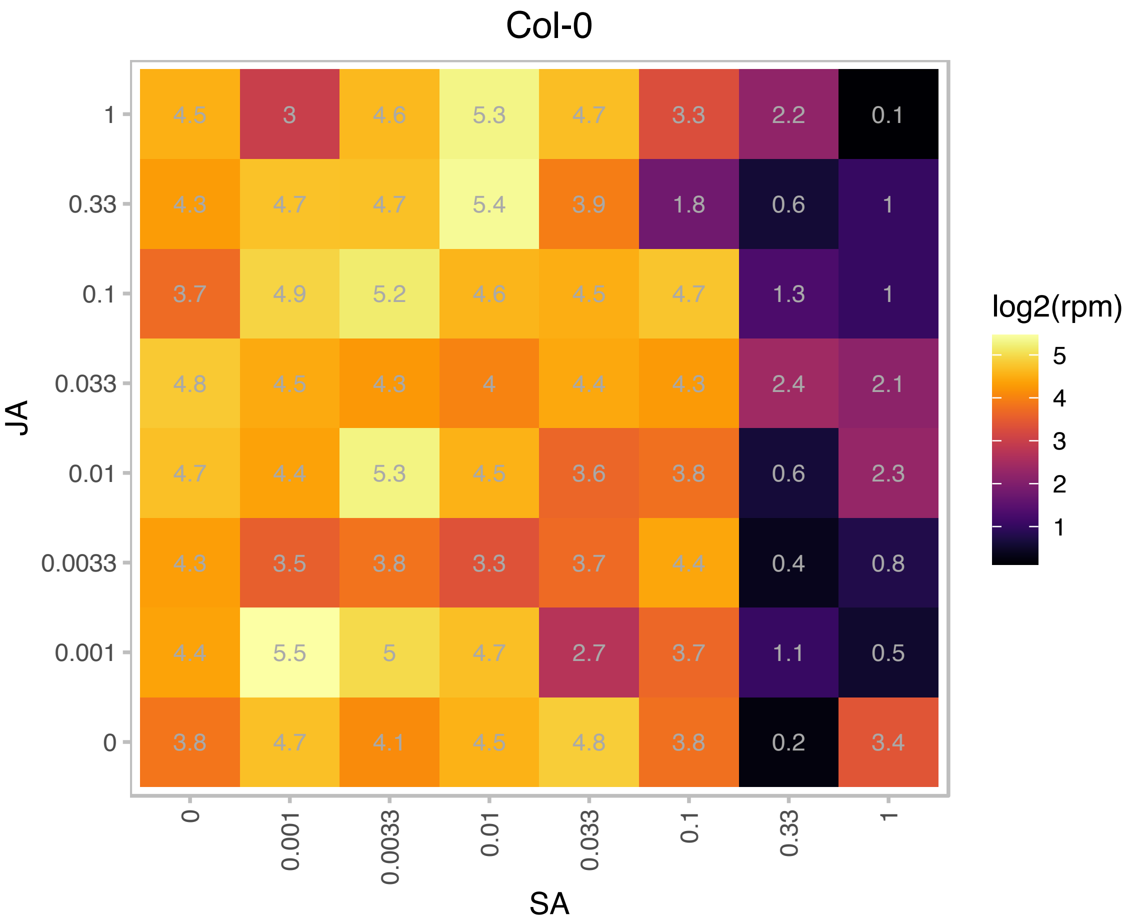


| | |
|---|---|
| Gene | AT4G30170 |
| is.expressed_genes | TRUE |
| is.responsive_genes | TRUE |
| is.apcluster | F-02 |

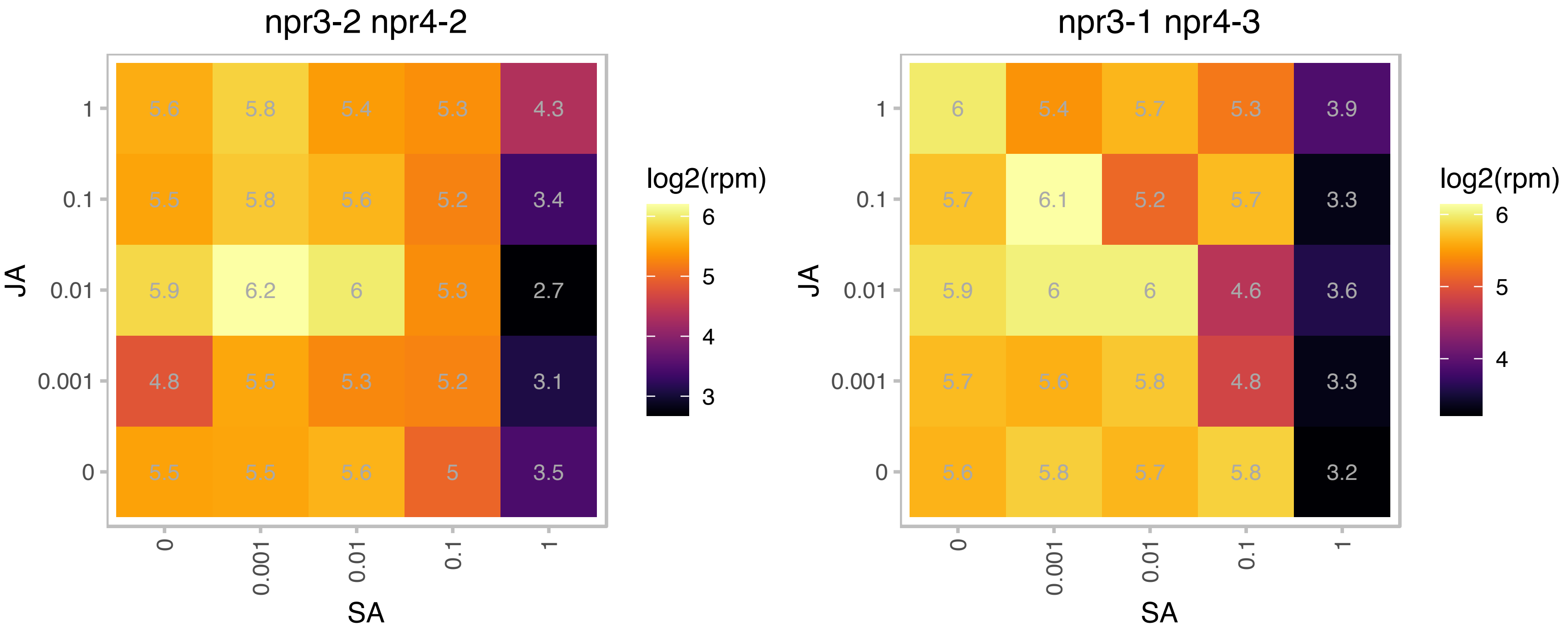

# AT4G36130.1 : NA

Ribosomal protein L2 family

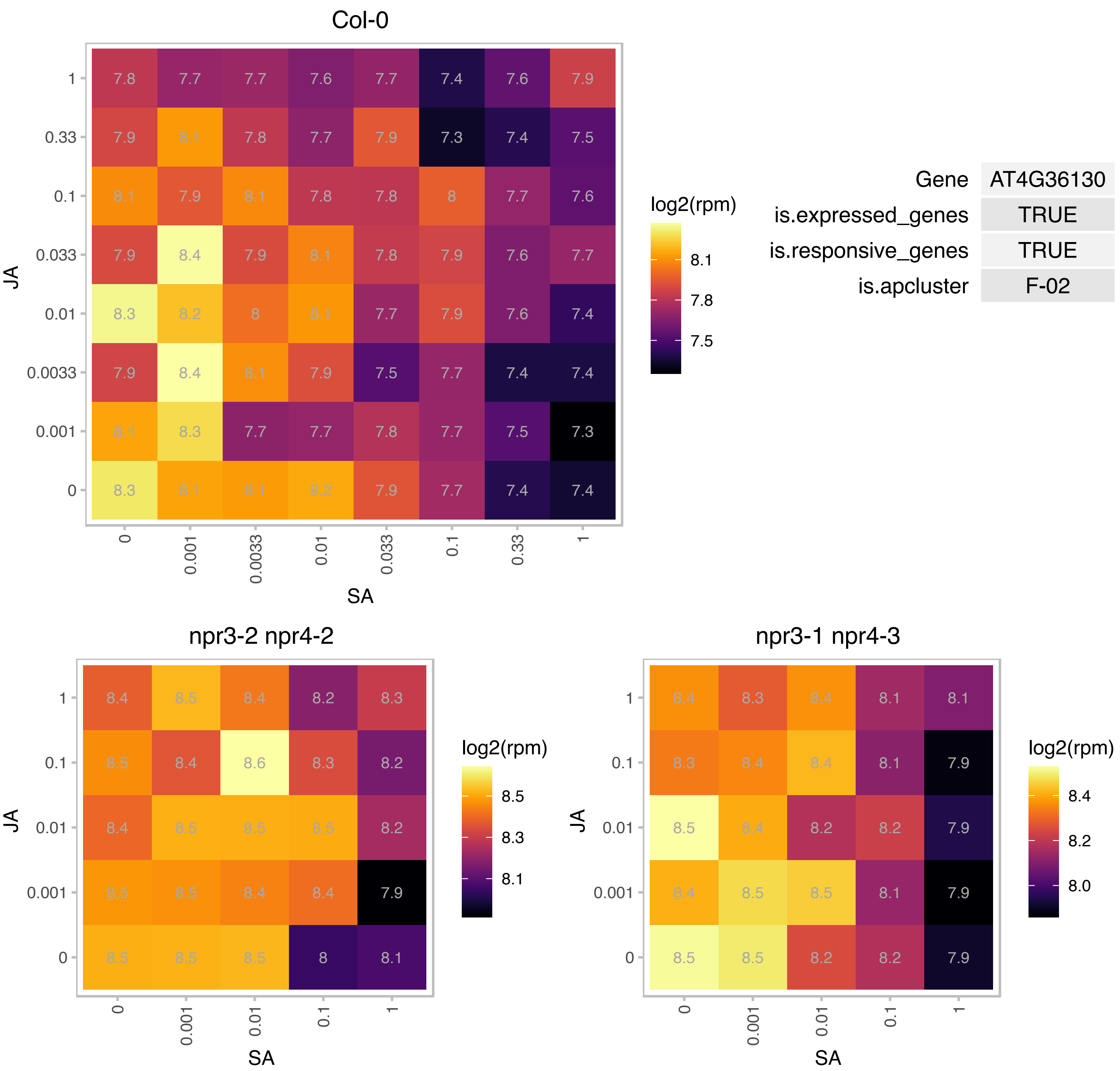


| | |
|---|---|
| Gene | AT4G36130 |
| is.expressed_genes | TRUE |
| is.responsive_genes | TRUE |
| is.apcluster | F-02 |

### AT4G37260.1 : ATMYB73, MYB73

myb domain protein 73

Member of the R2R3 factor gene family. The mRNA is cell-to-cell mobile.

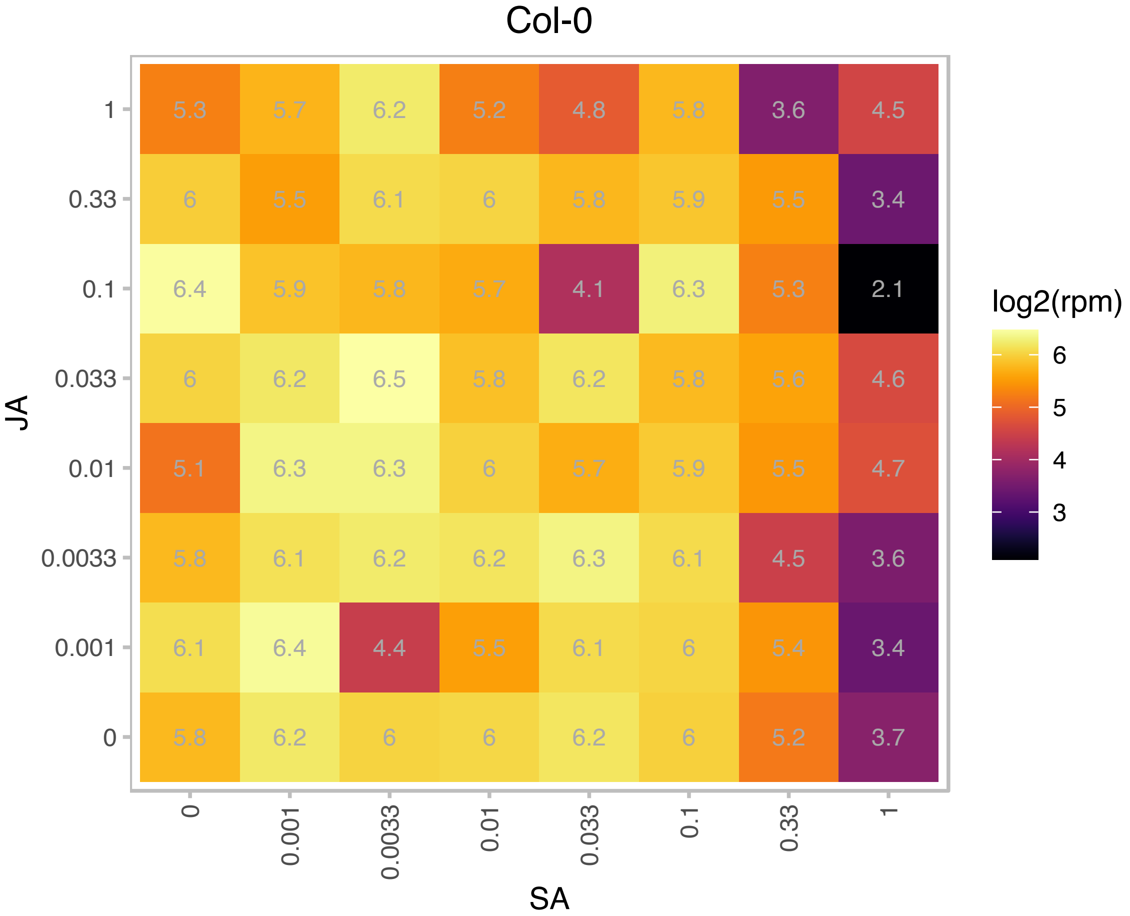


| | |
|---|---|
| Gene | AT4G37260 |
| is.expressed_genes | TRUE |
| is.responsive_genes | TRUE |
| is.apcluster | F-02 |

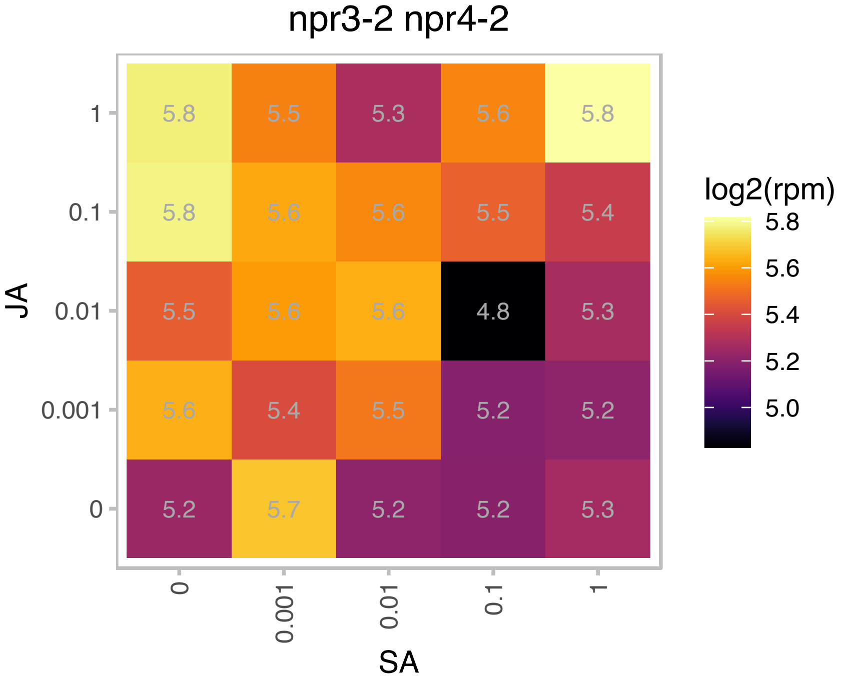


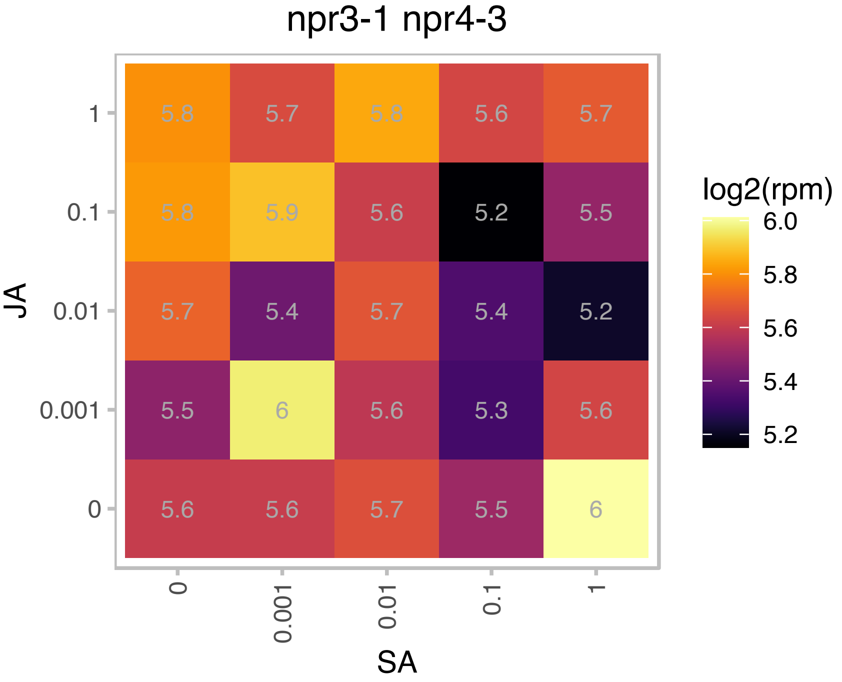

### AT4G37540.1 : LBD39

LOB domain-containing protein 39

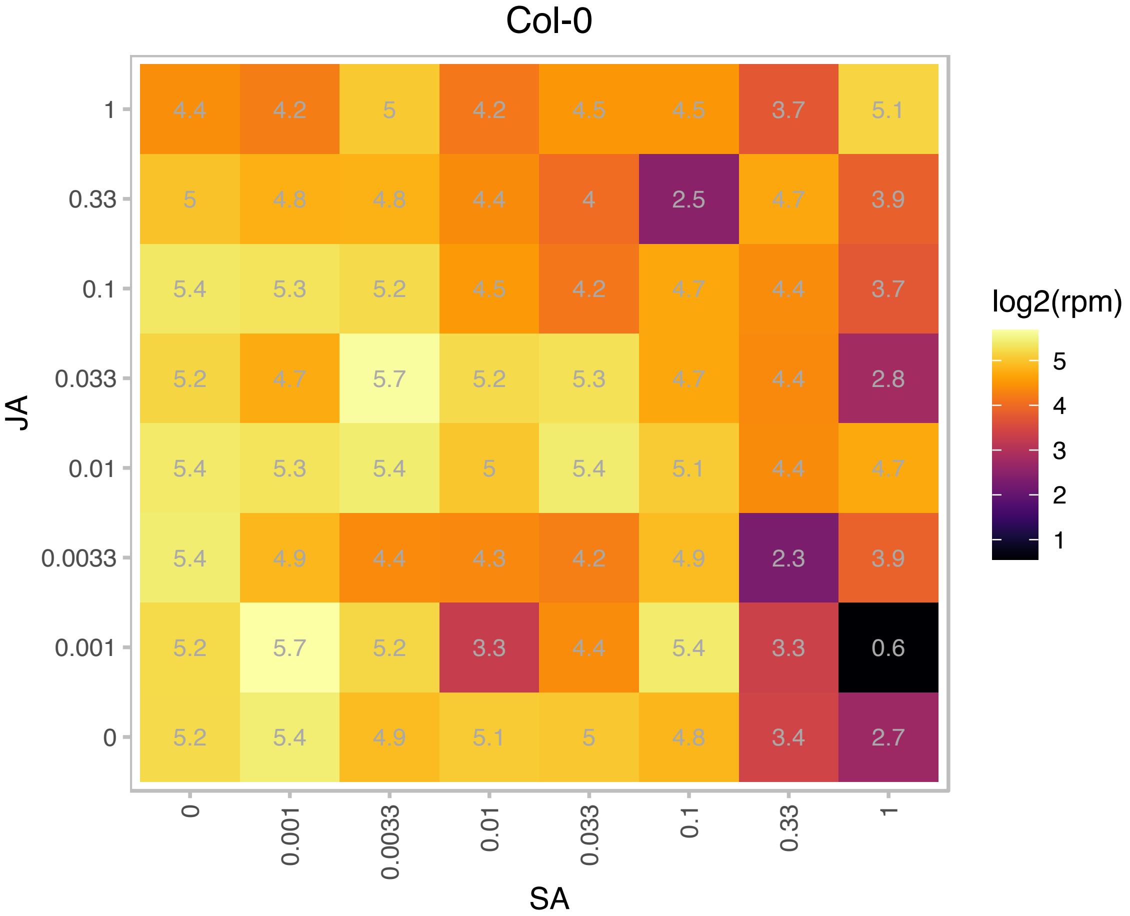


| | |
|---|---|
| Gene | AT4G37540 |
| is.expressed_genes | TRUE |
| is.responsive_genes | TRUE |
| is.apcluster | F-02 |

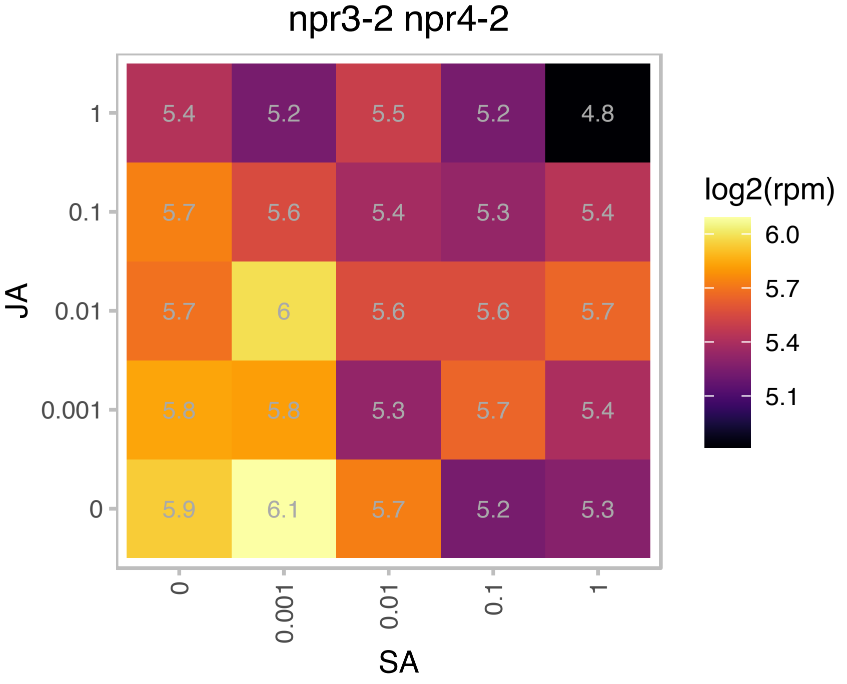


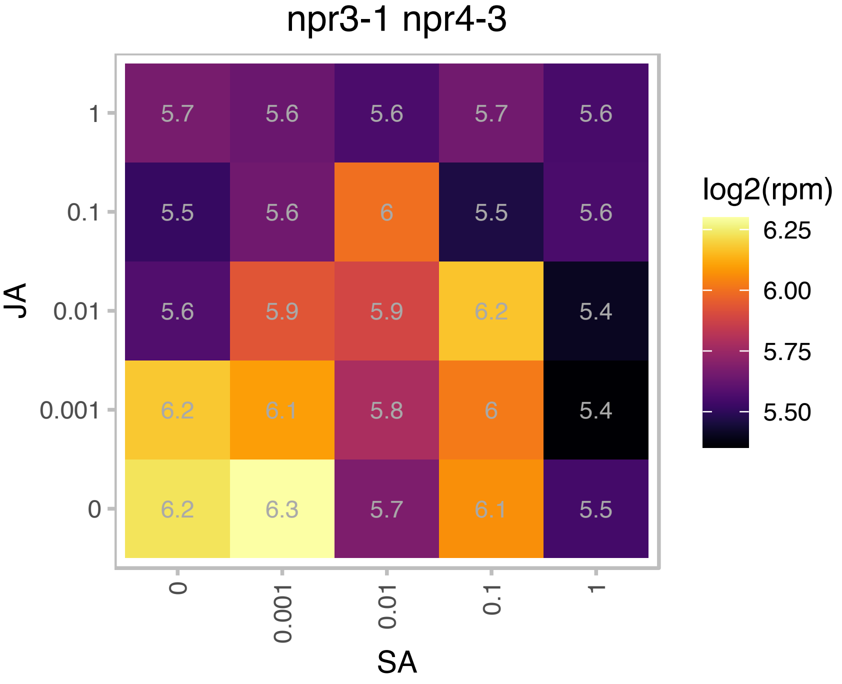

# AT4G38225.1 : NA

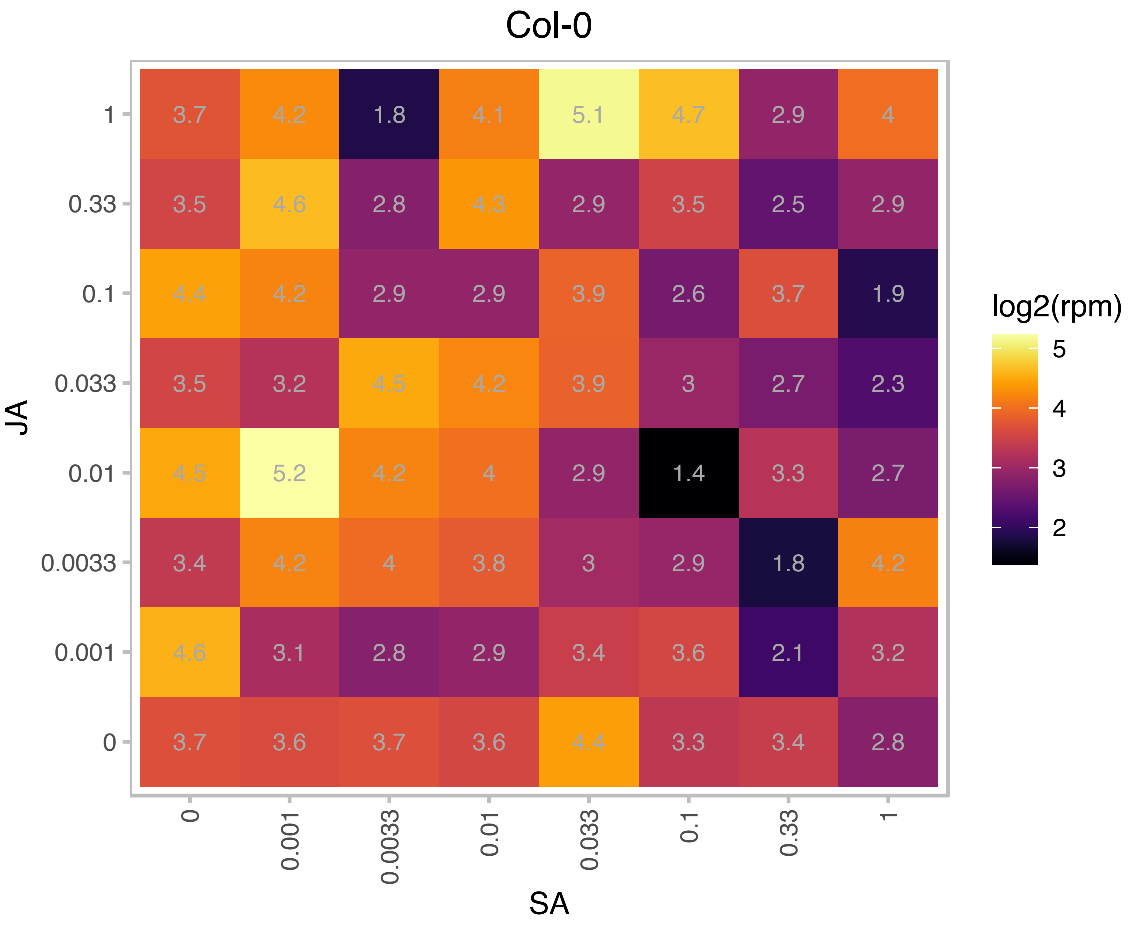


| | |
|---|---|
| Gene | AT4G38225 |
| is.expressed_genes | TRUE |
| is.responsive_genes | TRUE |
| is.apcluster | F-02 |

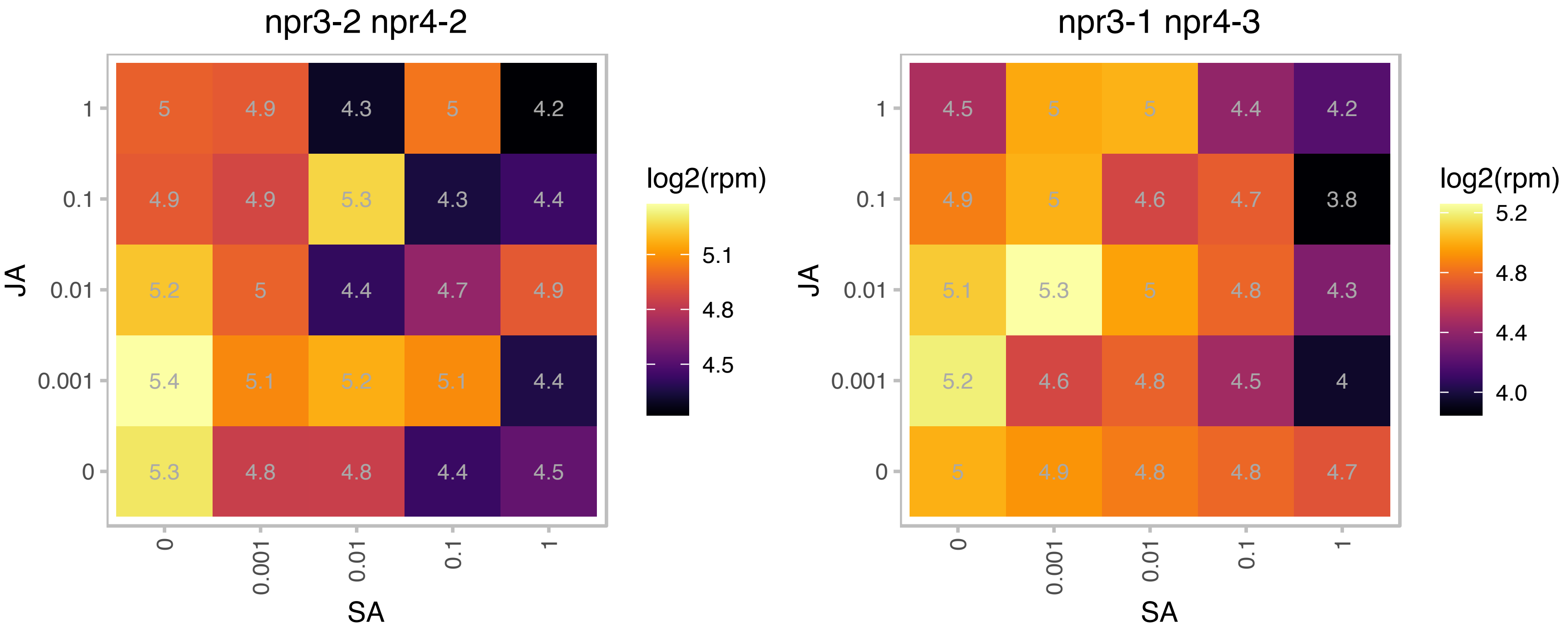

### AT4G39350.1 : ATCESA2, ATH-A, CESA2

cellulose synthase A2
Encodes a cellulose synthase isomer, related to CESA6. As inferred from the null role of secondary wall-type CesAs, included in a set of five primary wall-type CesAs that may support trichome cell wall thickening. The mRNA is cell-to-cell mobile.

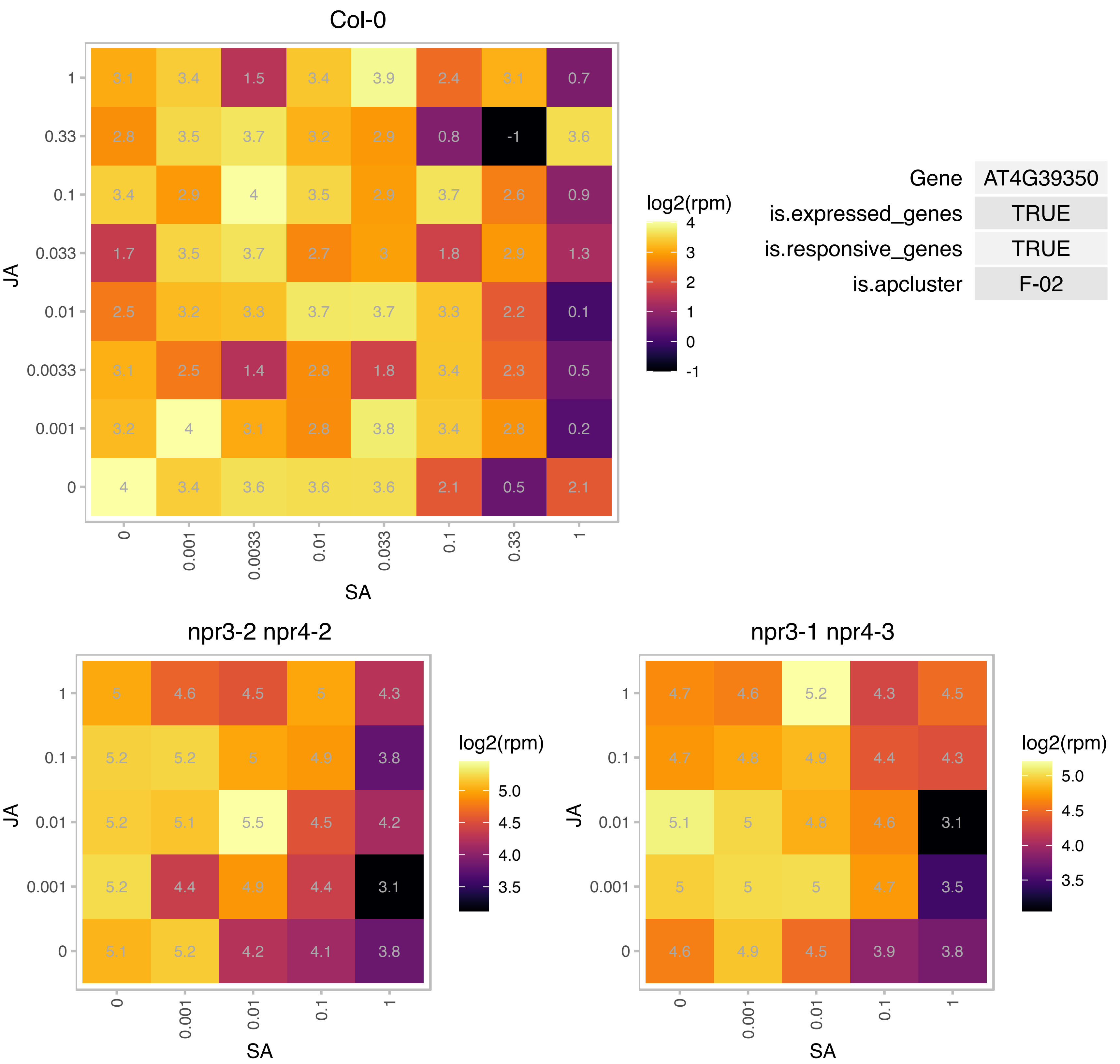


| | |
|---|---|
| Gene | AT4G39350 |
| is.expressed_genes | TRUE |
| is.responsive_genes | TRUE |
| is.apcluster | F-02 |

### AT4G40090.1 : AGP3

arabinogalactan protein 3

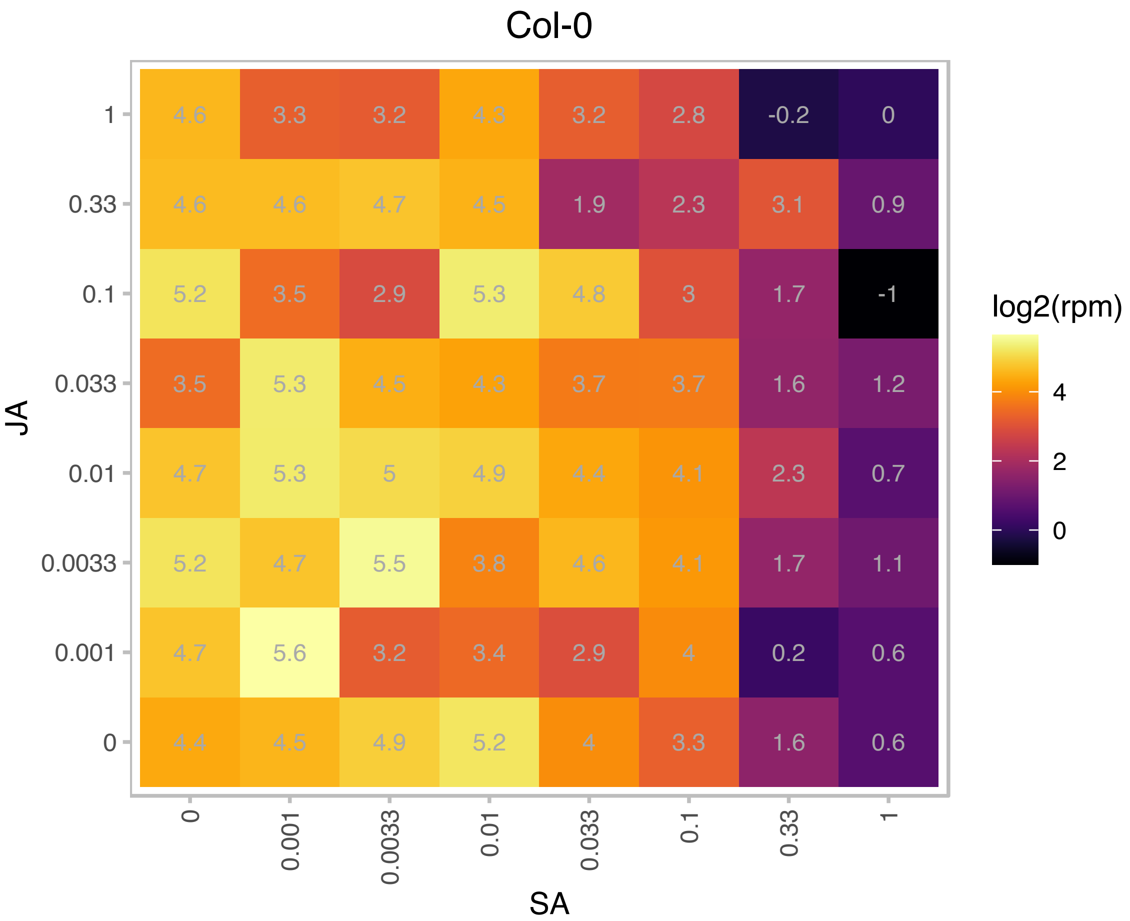


| | |
|---|---|
| Gene | AT4G40090 |
| is.expressed_genes | TRUE |
| is.responsive_genes | TRUE |
| is.apcluster | F-02 |

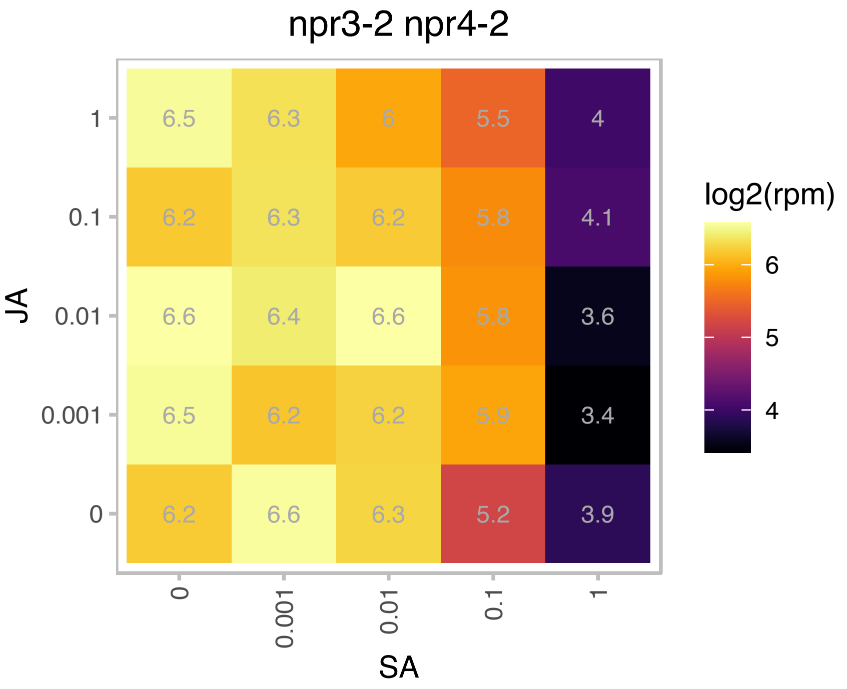


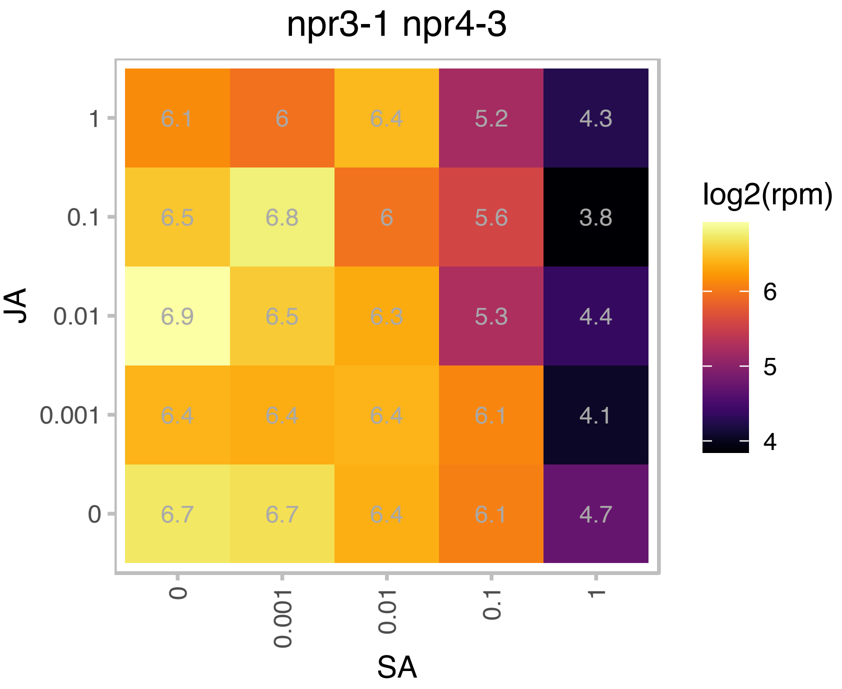

### AT5G01600.1 : ATFER1, FER1

ferretin 1
Encodes a ferretin protein that is targeted to the chloroplast. Member of a Ferritin gene family. Gene expression is induced in response to iron overload and by nitric oxide. Expression of the gene is downregulated in the presence of paraquat, an inducer of photoxidative stress.

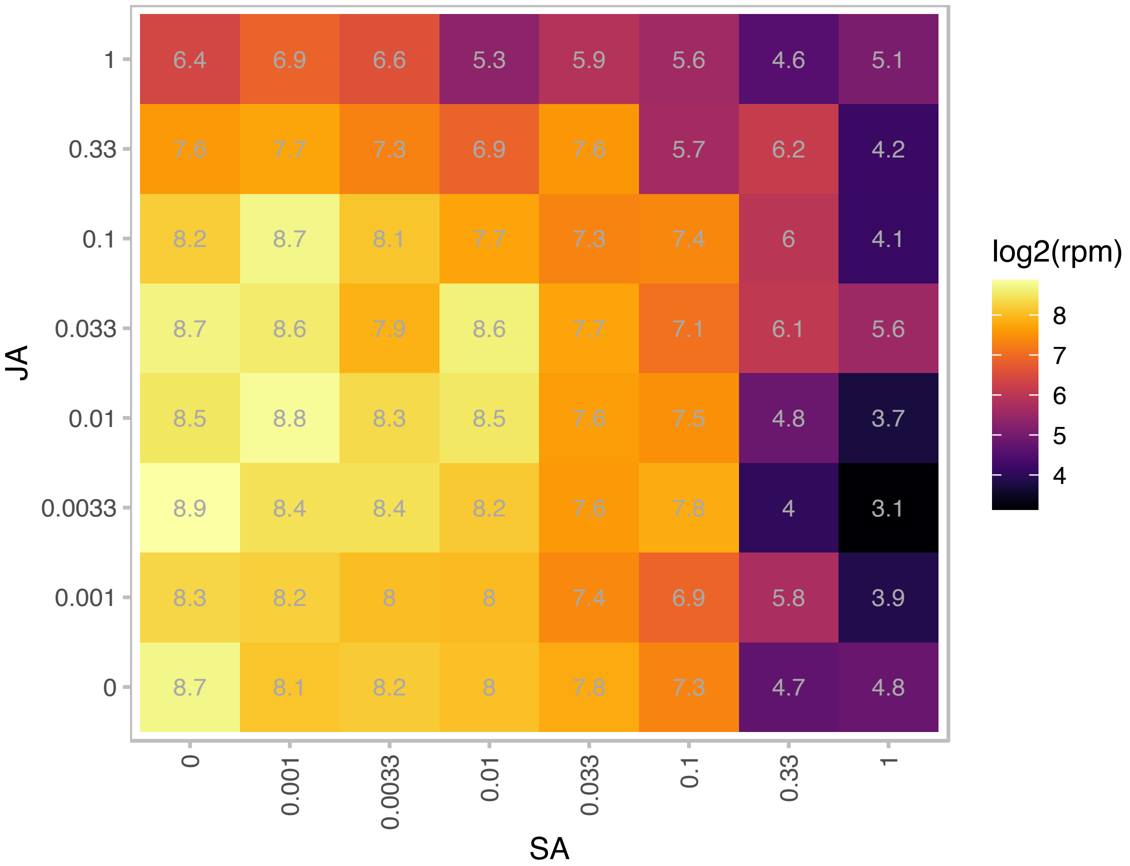


| | |
|---|---|
| Gene | AT5G01600 |
| is.expressed_genes | TRUE |
| is.responsive_genes | TRUE |
| is.apcluster | F-02 |

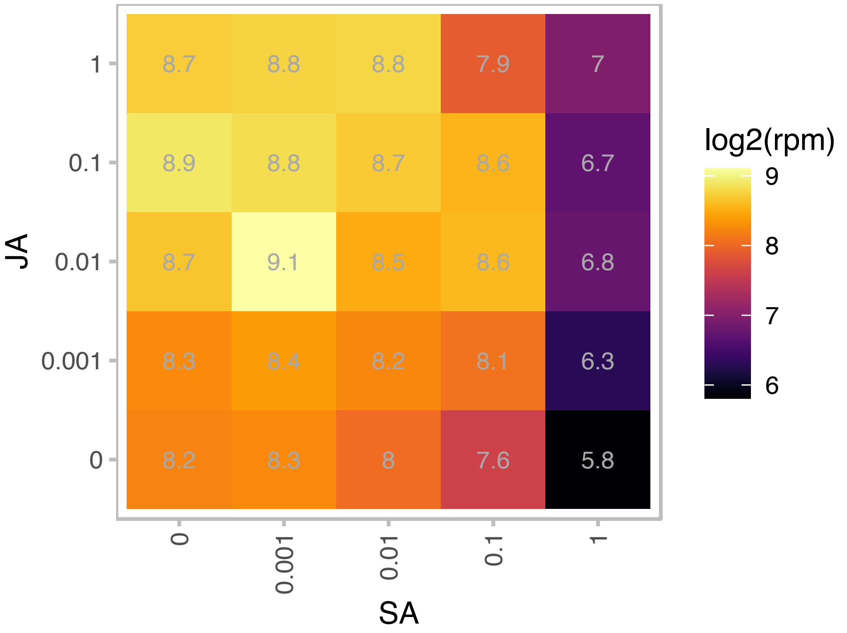


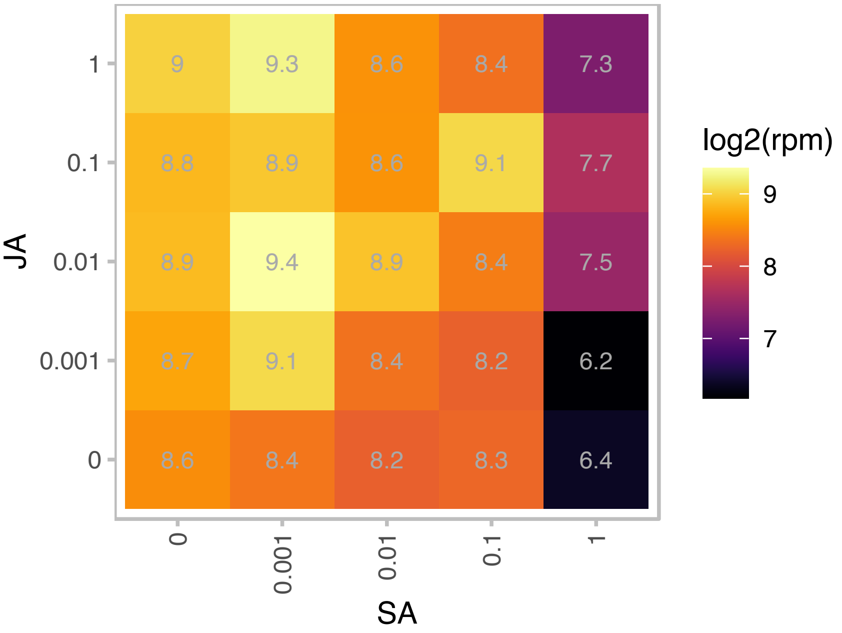

# AT5G03230.1 : NA

Protein of unknown function, DUF584

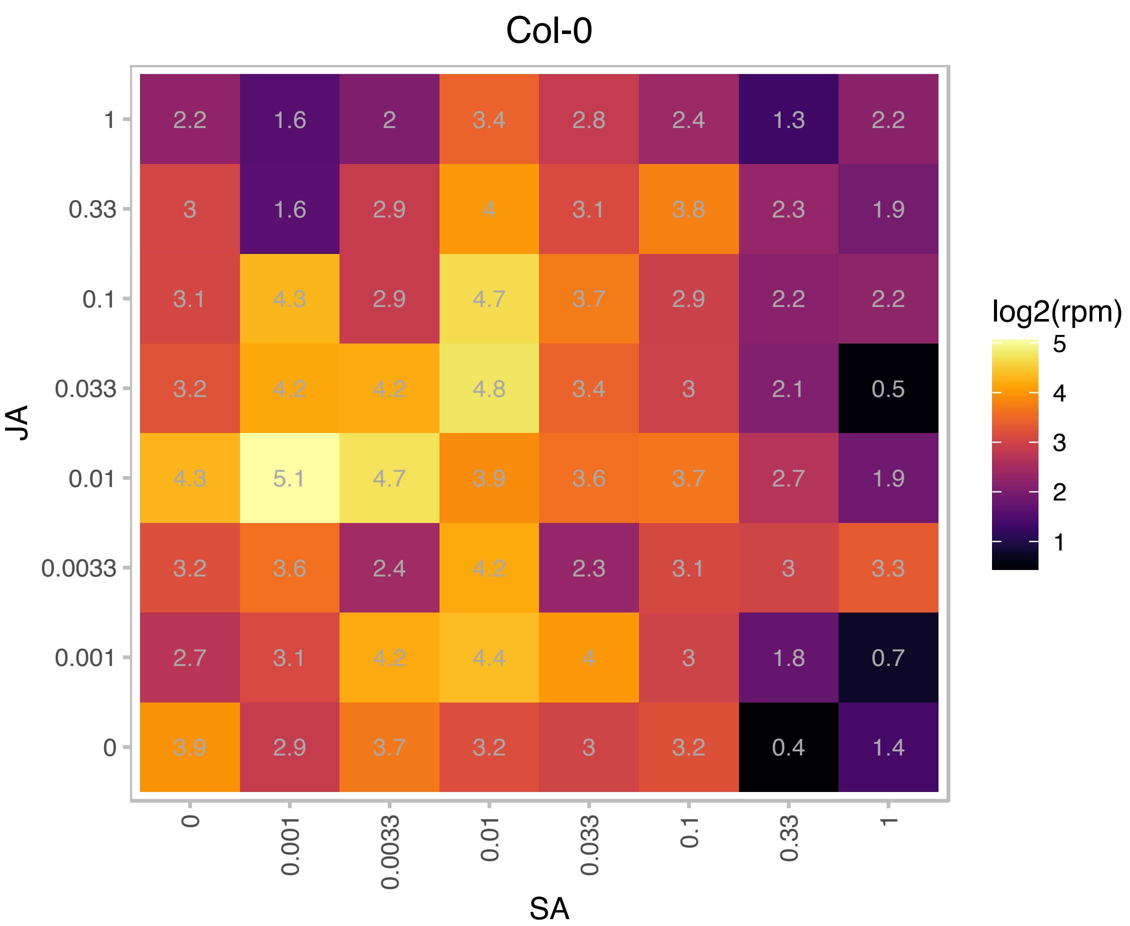


| | |
|---|---|
| Gene | AT5G03230 |
| is.expressed_genes | TRUE |
| is.responsive_genes | TRUE |
| is.apcluster | F-02 |

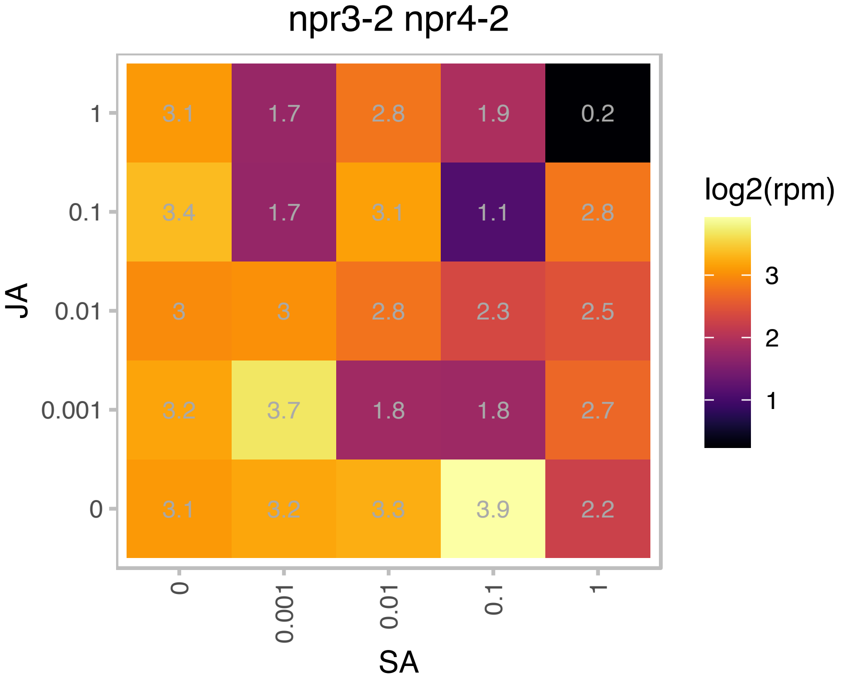


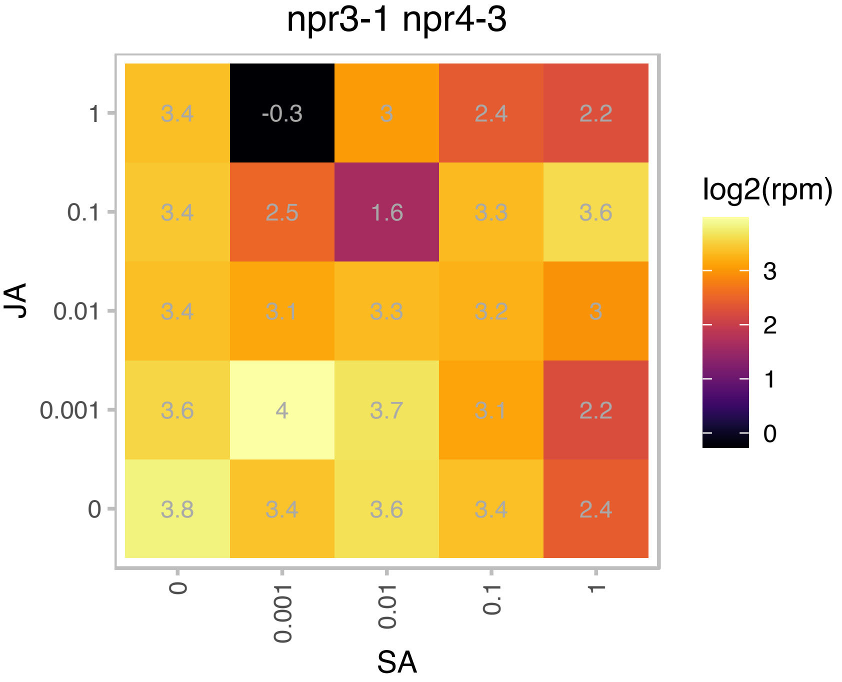

# AT5G05210.1 : NA

Surfeit locus protein 6

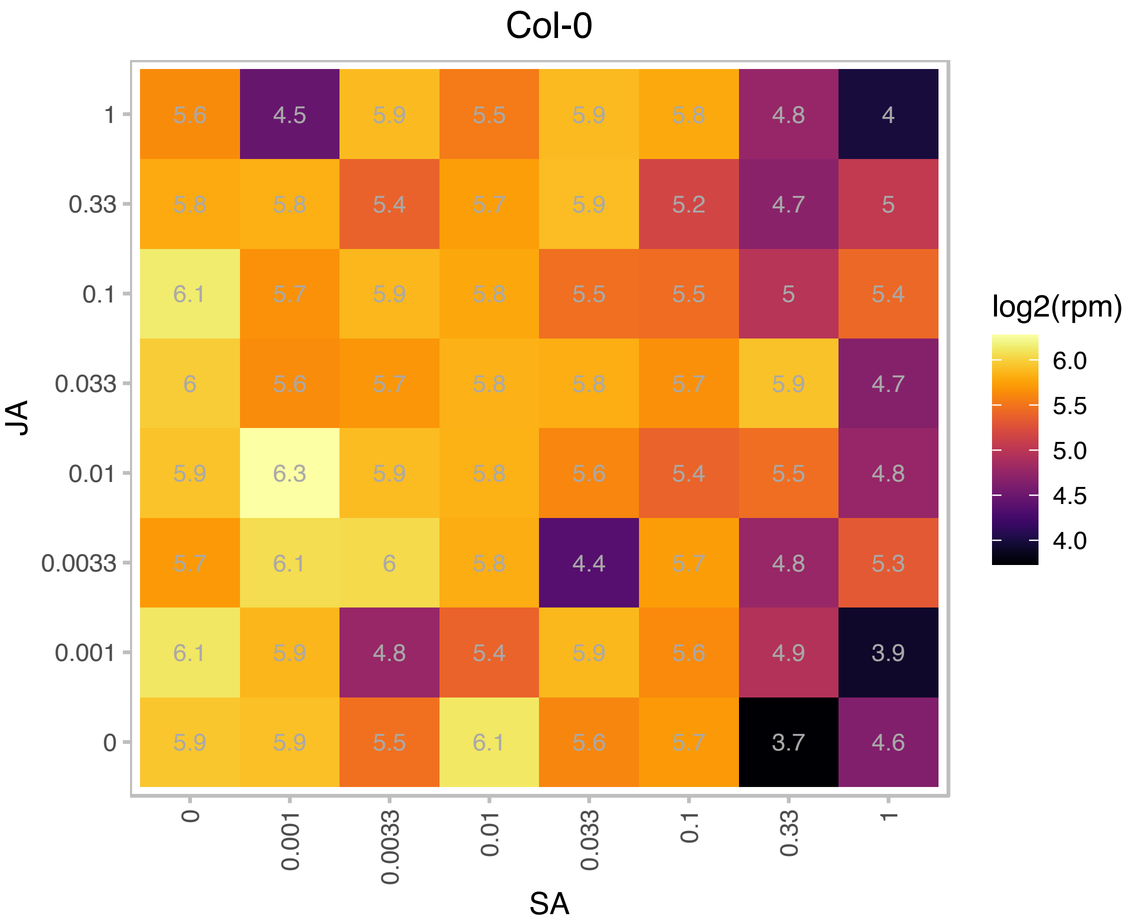


| | |
|---|---|
| Gene | AT5G05210 |
| is.expressed_genes | TRUE |
| is.responsive_genes | TRUE |
| is.apcluster | F-02 |

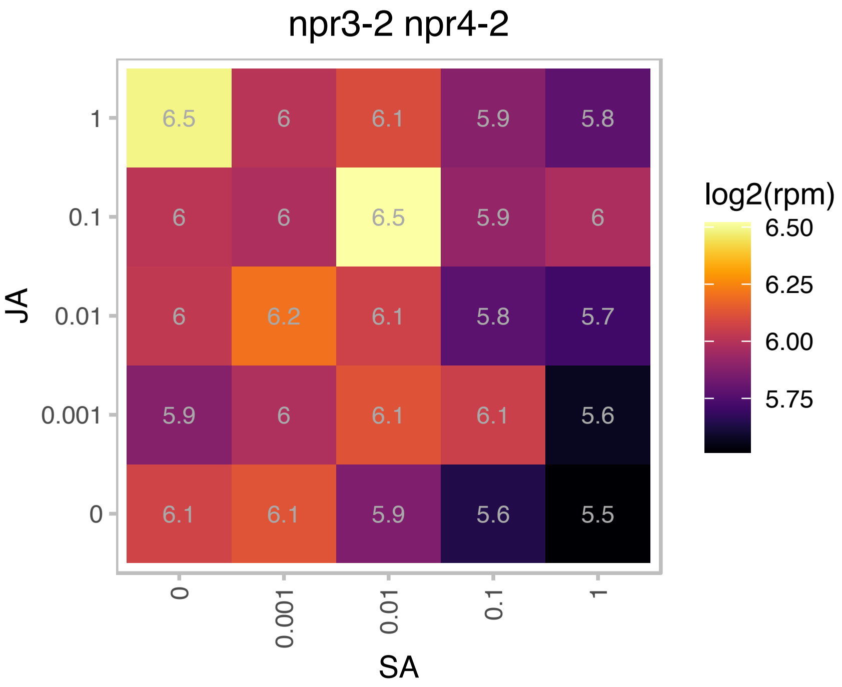


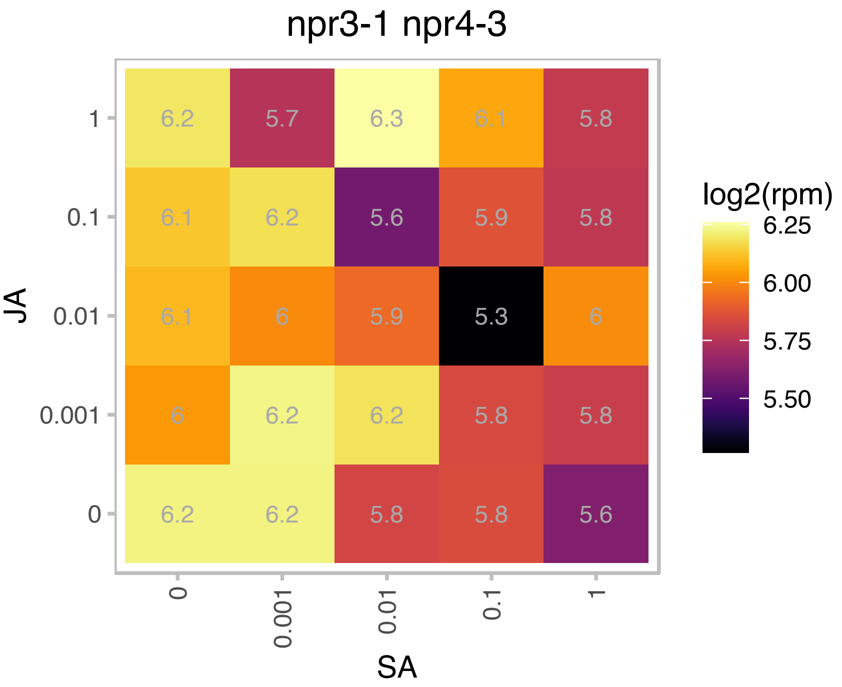

### AT5G05500.1 : AtPRPL1, MOP10, PRPL1

Pollen Ole e 1 allergen and extensin family protein

Encodes Proline-rich protein-like PRPL1, controls elongation of root hairs.

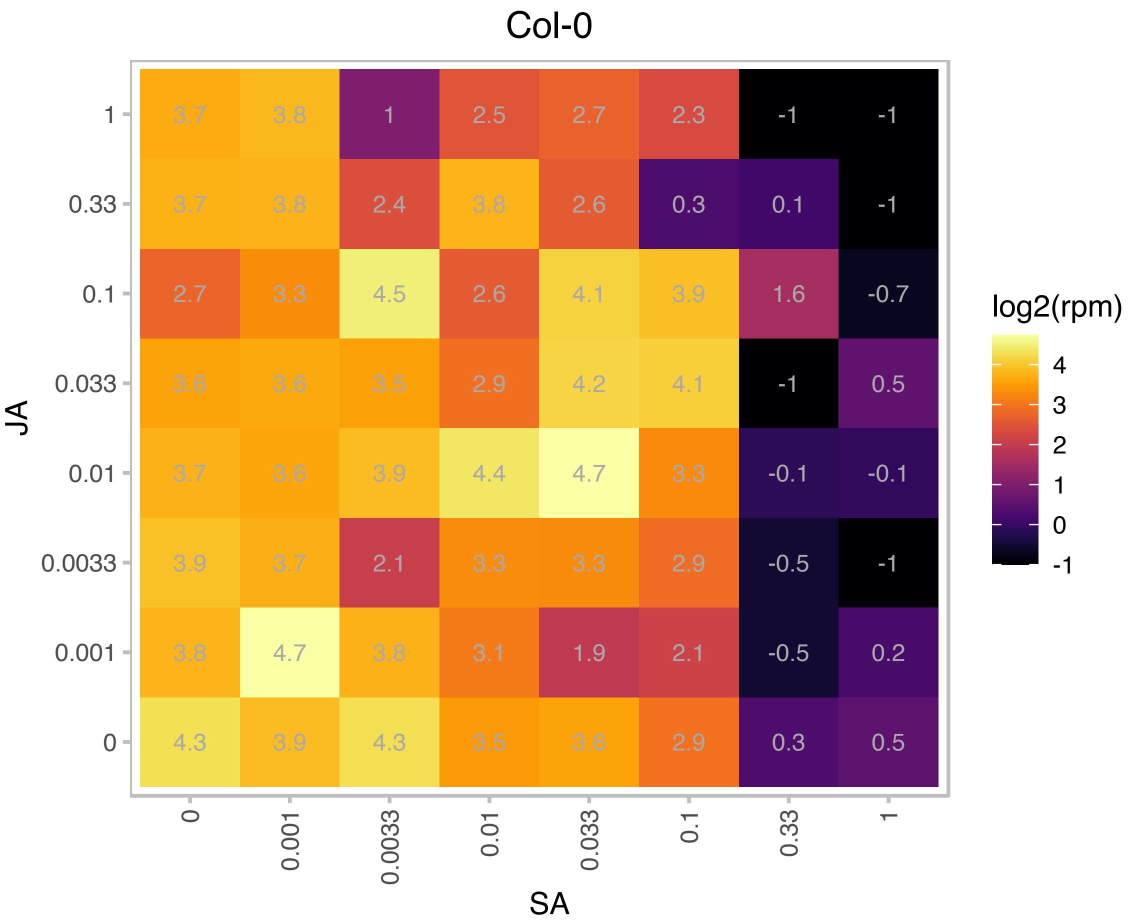


| | |
|---|---|
| Gene | AT5G05500 |
| is.expressed_genes | TRUE |
| is.responsive_genes | TRUE |
| is.apcluster | F-02 |

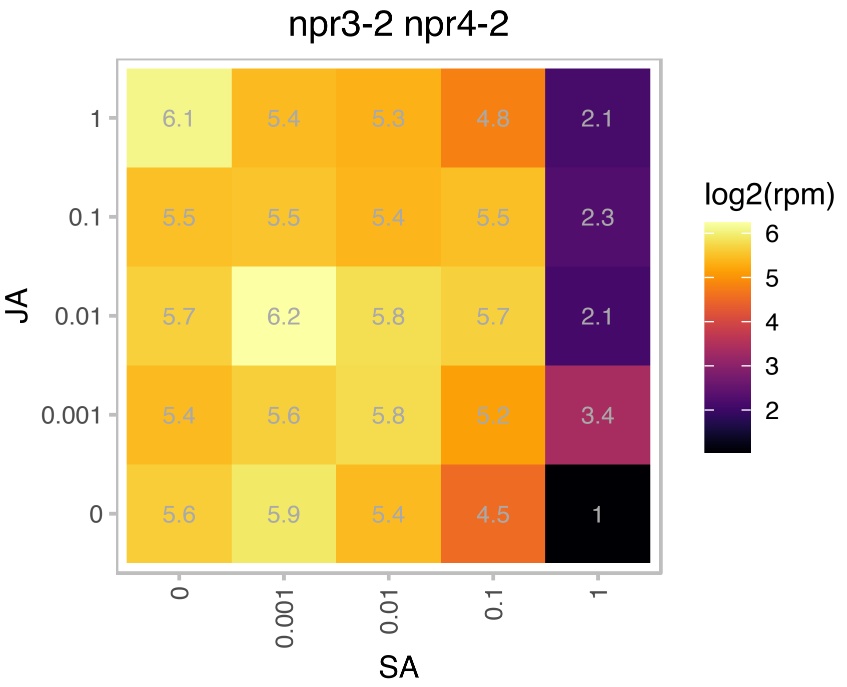


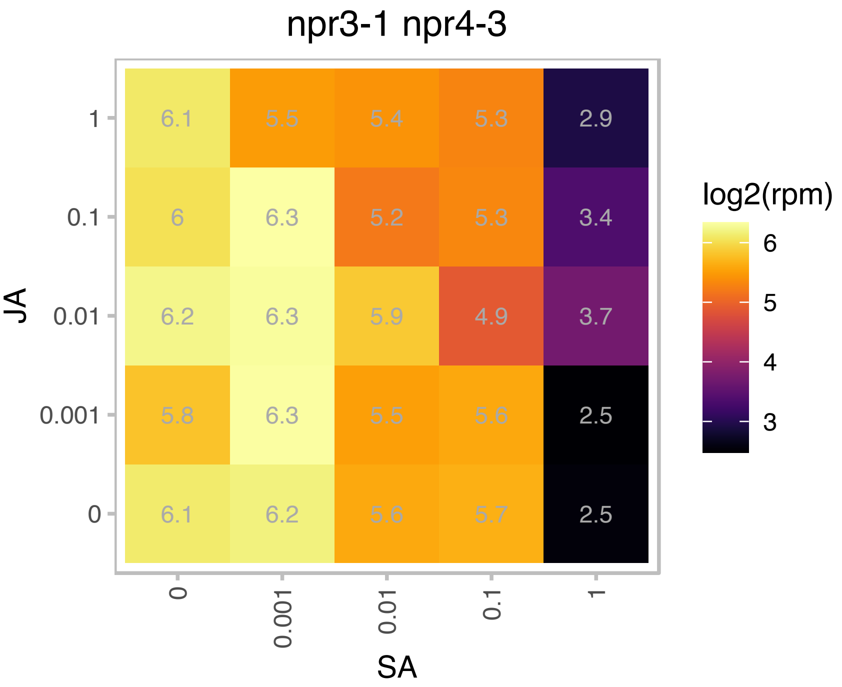

# AT5G06630.1 : NA

proline-rich extensin-like family protein

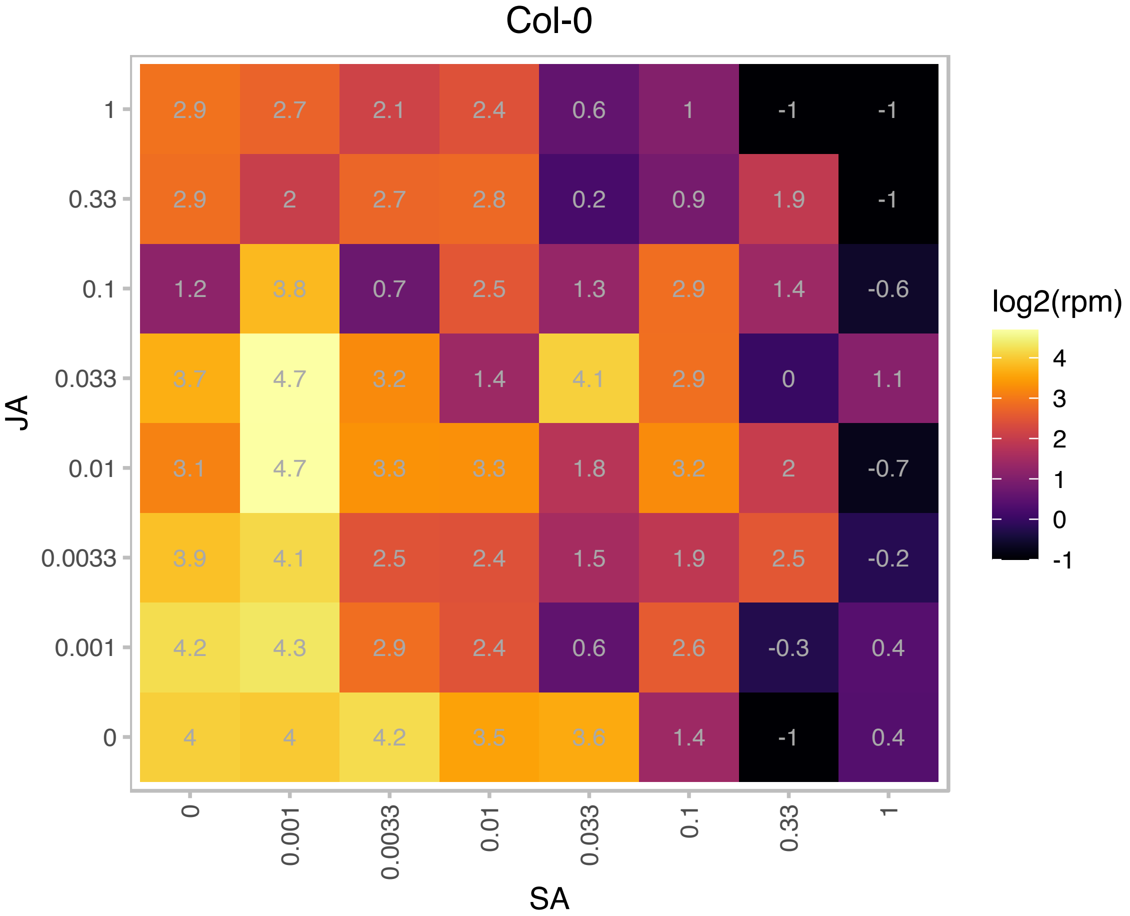


| | |
|---|---|
| Gene | AT5G06630 |
| is.expressed_genes | TRUE |
| is.responsive_genes | TRUE |
| is.apcluster | F-02 |

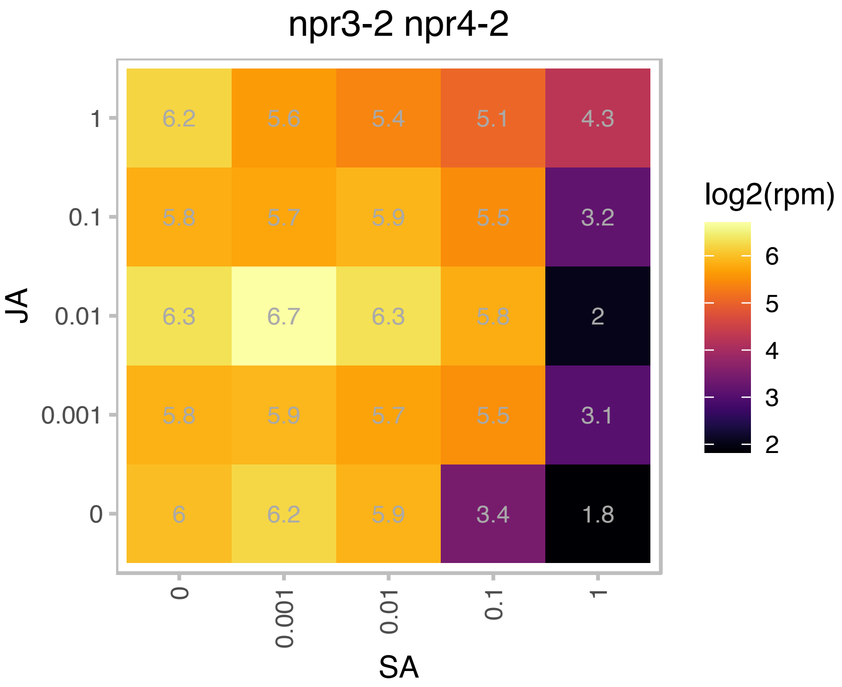


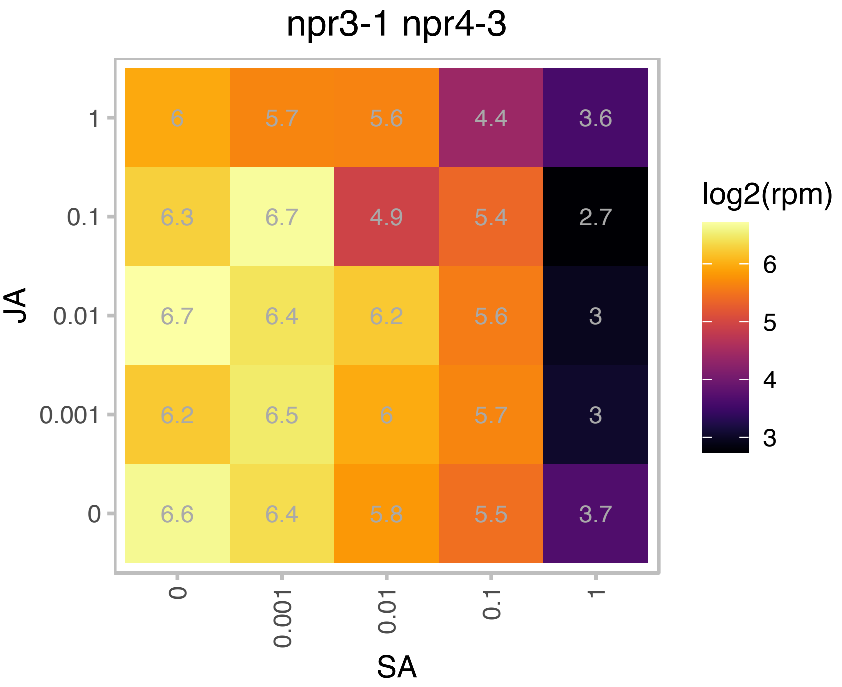

### AT5G06640.1 : EXT10

Proline-rich extensin-like family protein

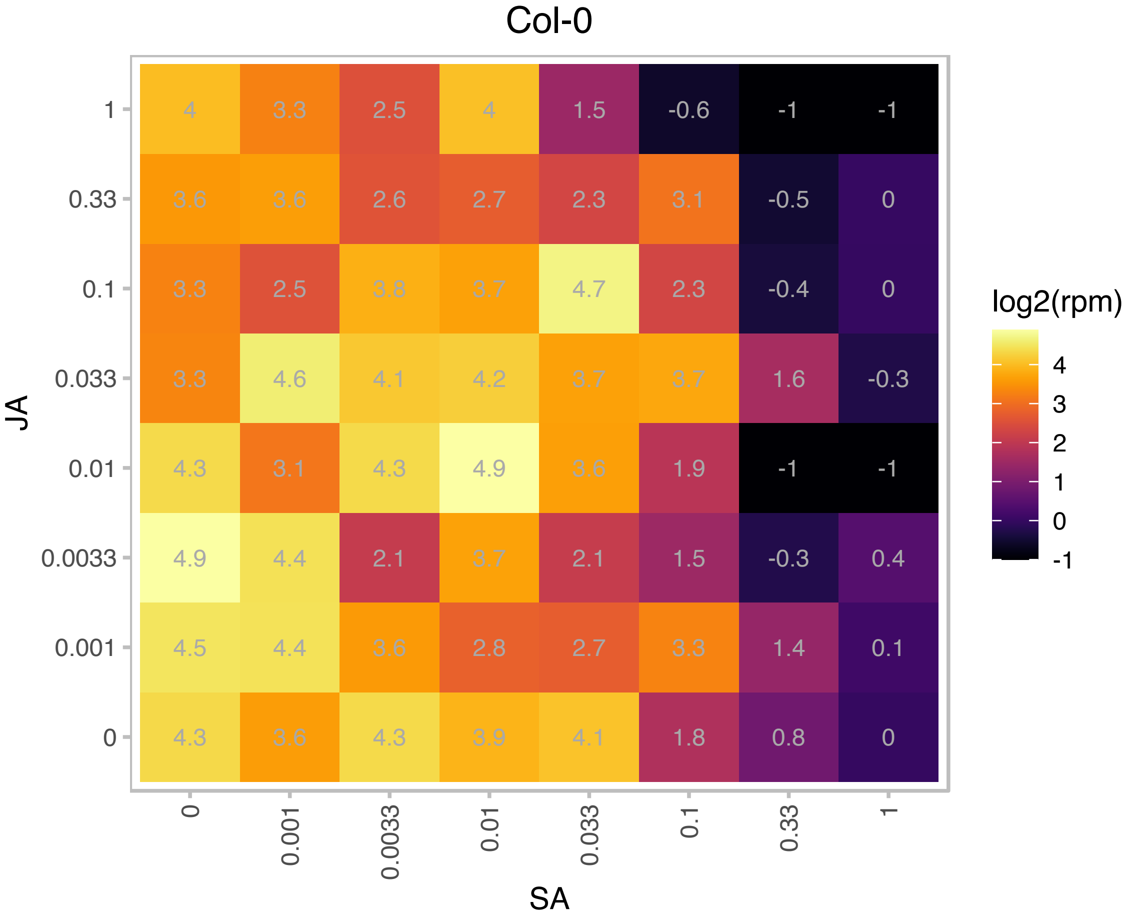


| | |
|---|---|
| Gene | AT5G06640 |
| is.expressed_genes | TRUE |
| is.responsive_genes | TRUE |
| is.apcluster | F-02 |

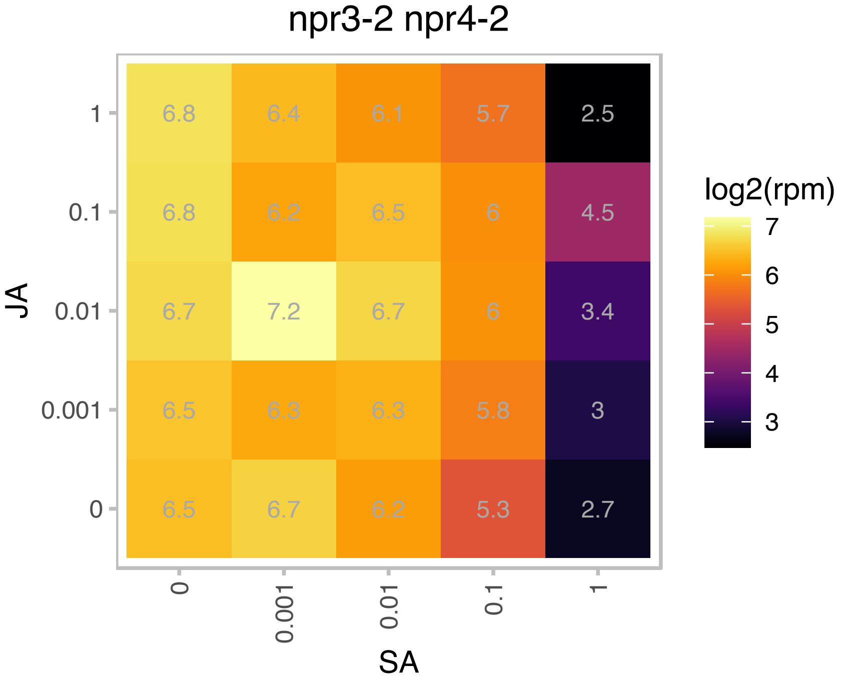


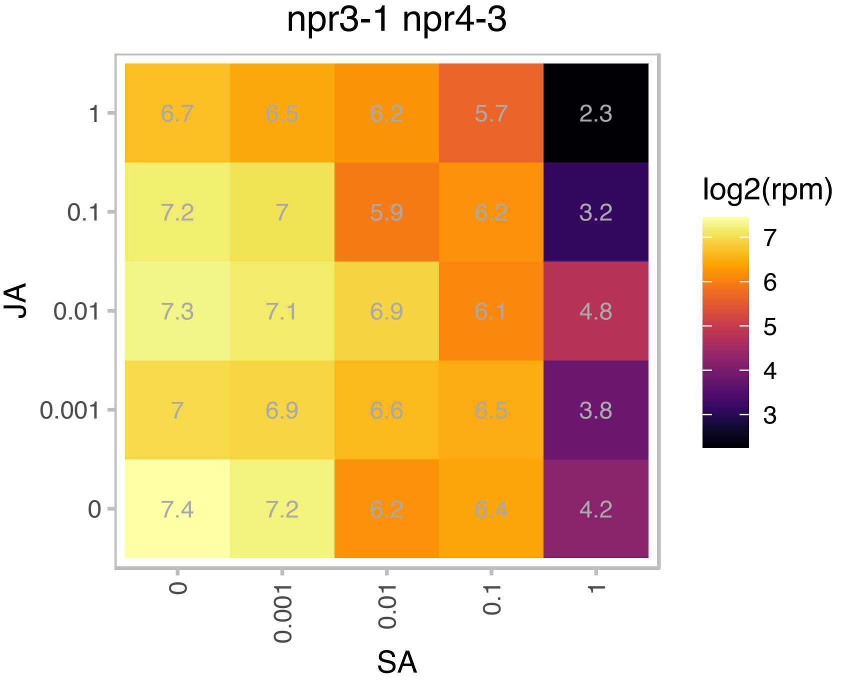

### AT5G07100.1 : WRKY26

WRKY DNA-binding protein 26

Encodes WRKY DNA-binding protein 26 (WRKY26).

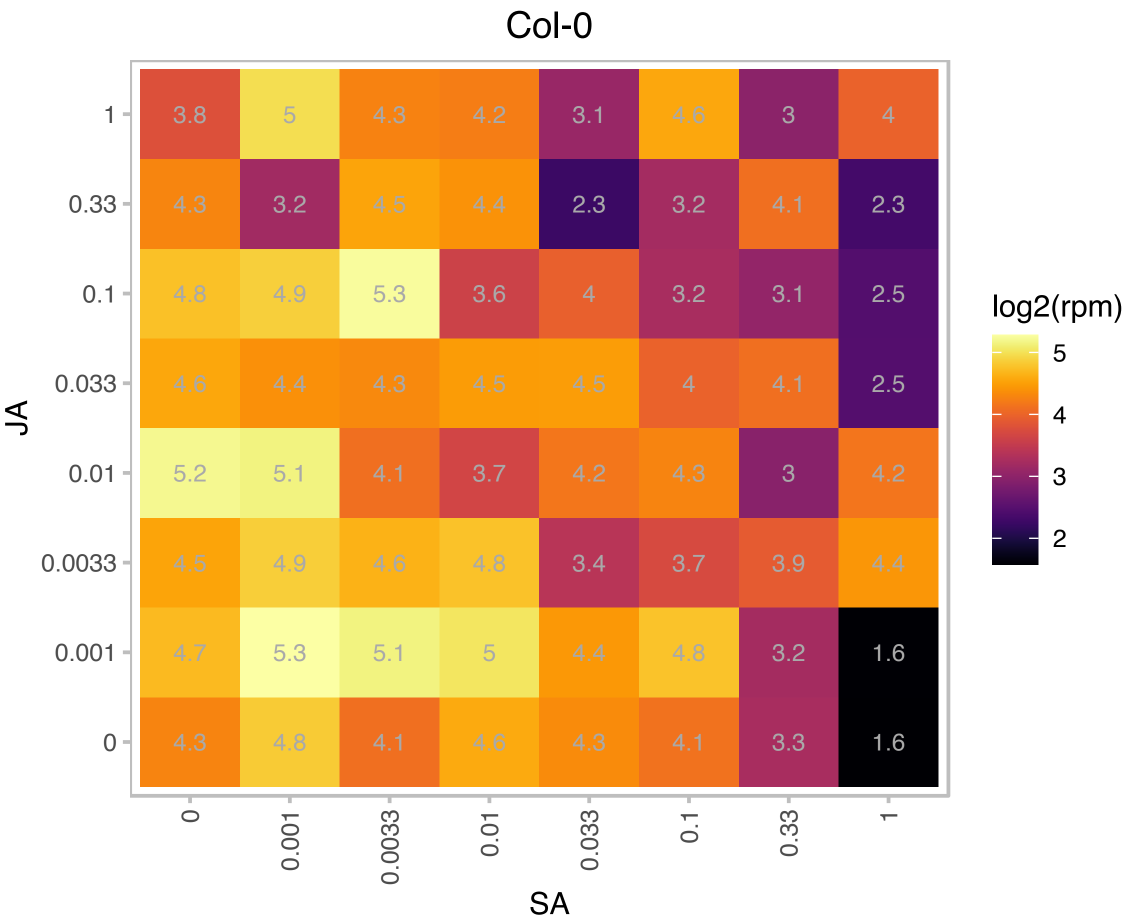


| | |
|---|---|
| Gene | AT5G07100 |
| is.expressed_genes | TRUE |
| is.responsive_genes | TRUE |
| is.apcluster | F-02 |

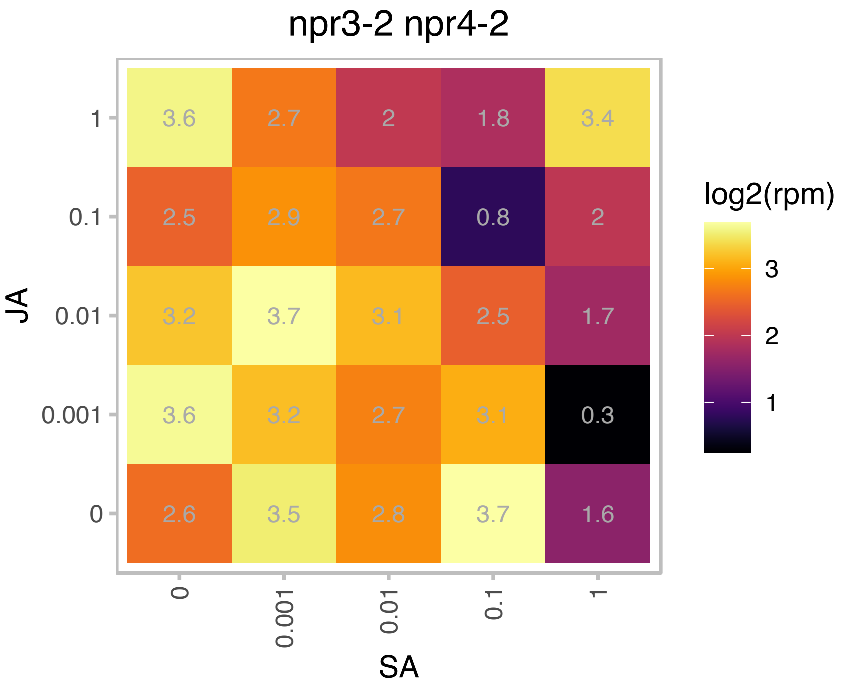


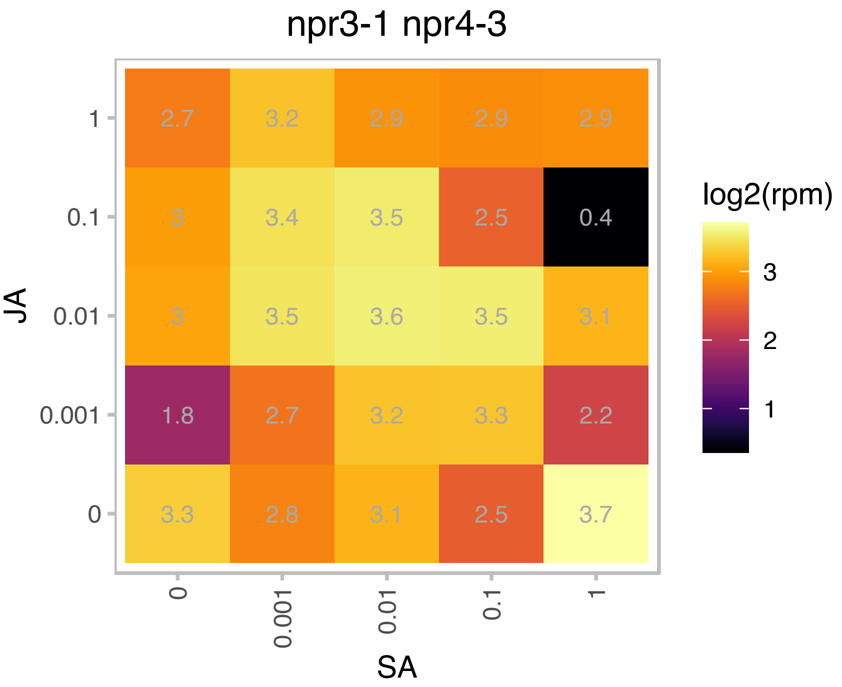

# AT5G11430.1 : NA

SPOC domain / Transcription elongation factor S-II protein

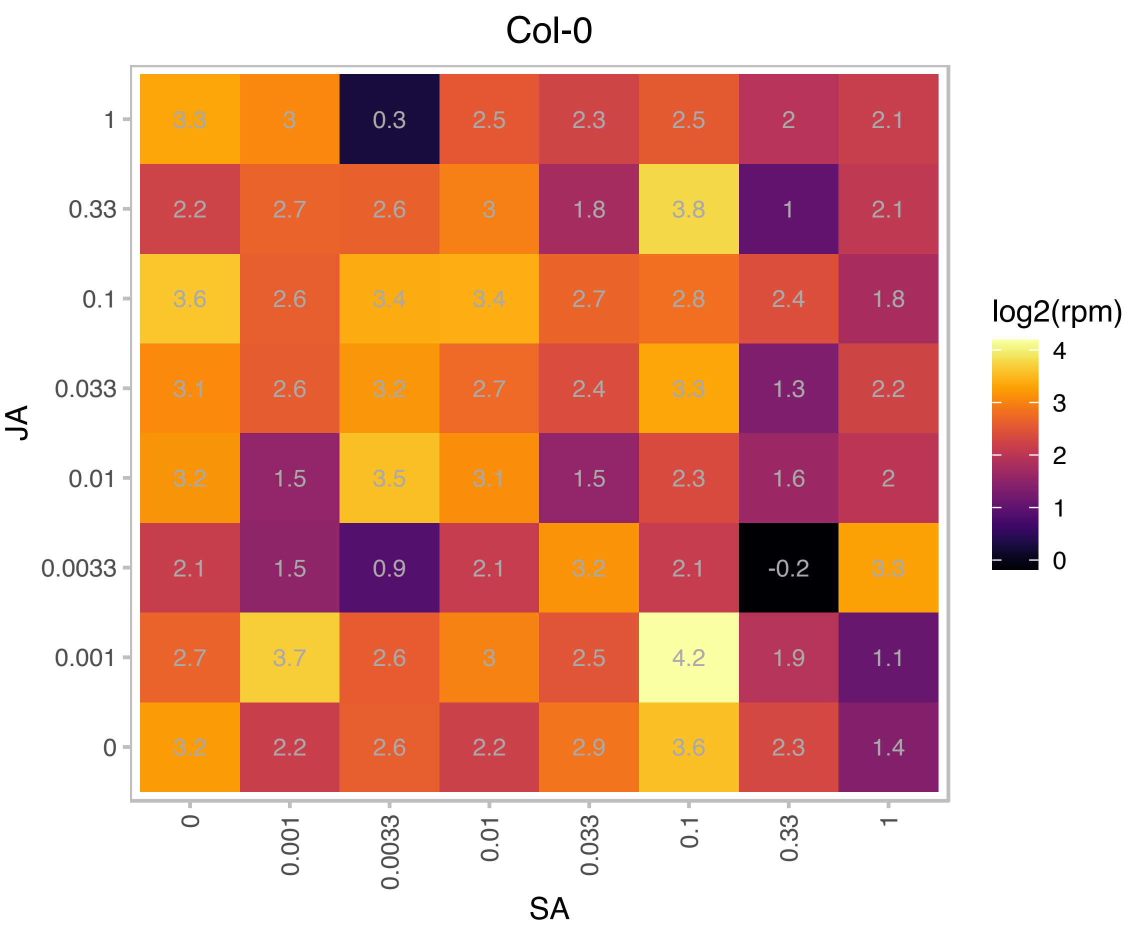


| | |
|---|---|
| Gene | AT5G11430 |
| is.expressed_genes | TRUE |
| is.responsive_genes | TRUE |
| is.apcluster | F-02 |

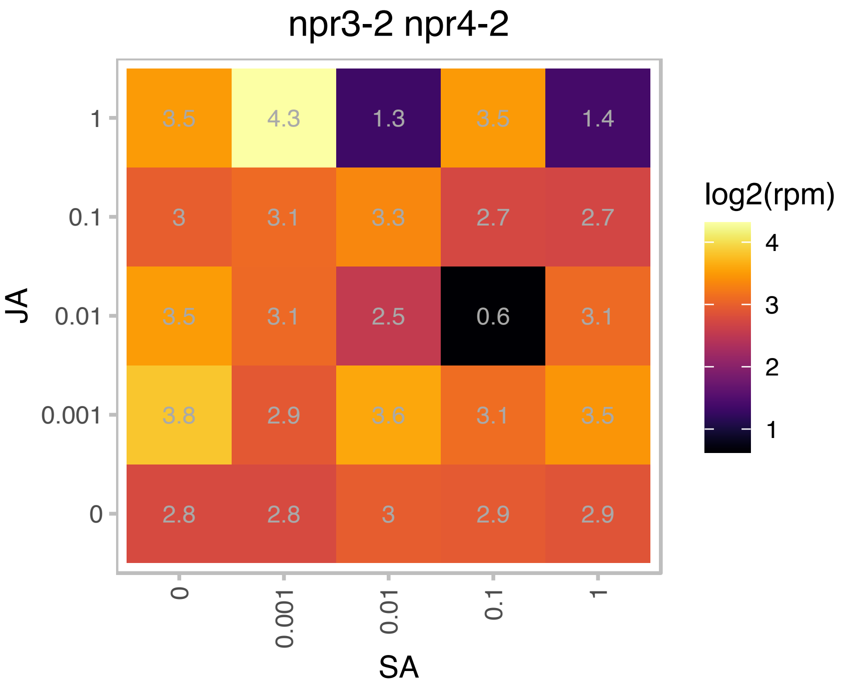


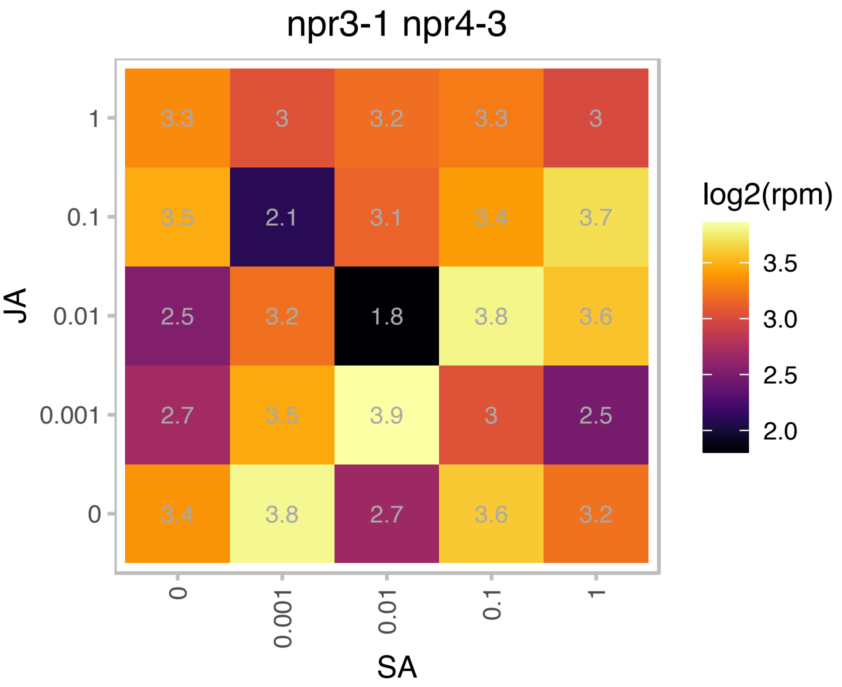

### AT5G14200.1 : ATIMD1, IMD1

isopropylmalate dehydrogenase 1

The AtIMD1 is one out of 3 genes encoding the enzyme 3-isopropylmalate dehydrogenase involved in leucine biosynthesis in Arabidopsis. Its subcellular location has been targeted to plastids. Encodes methylthioalkylmalate dehydrogenase. Involved in glucosinolate biosynthesis, in methionine chain elongation. The mRNA is cell-to-cell mobile.

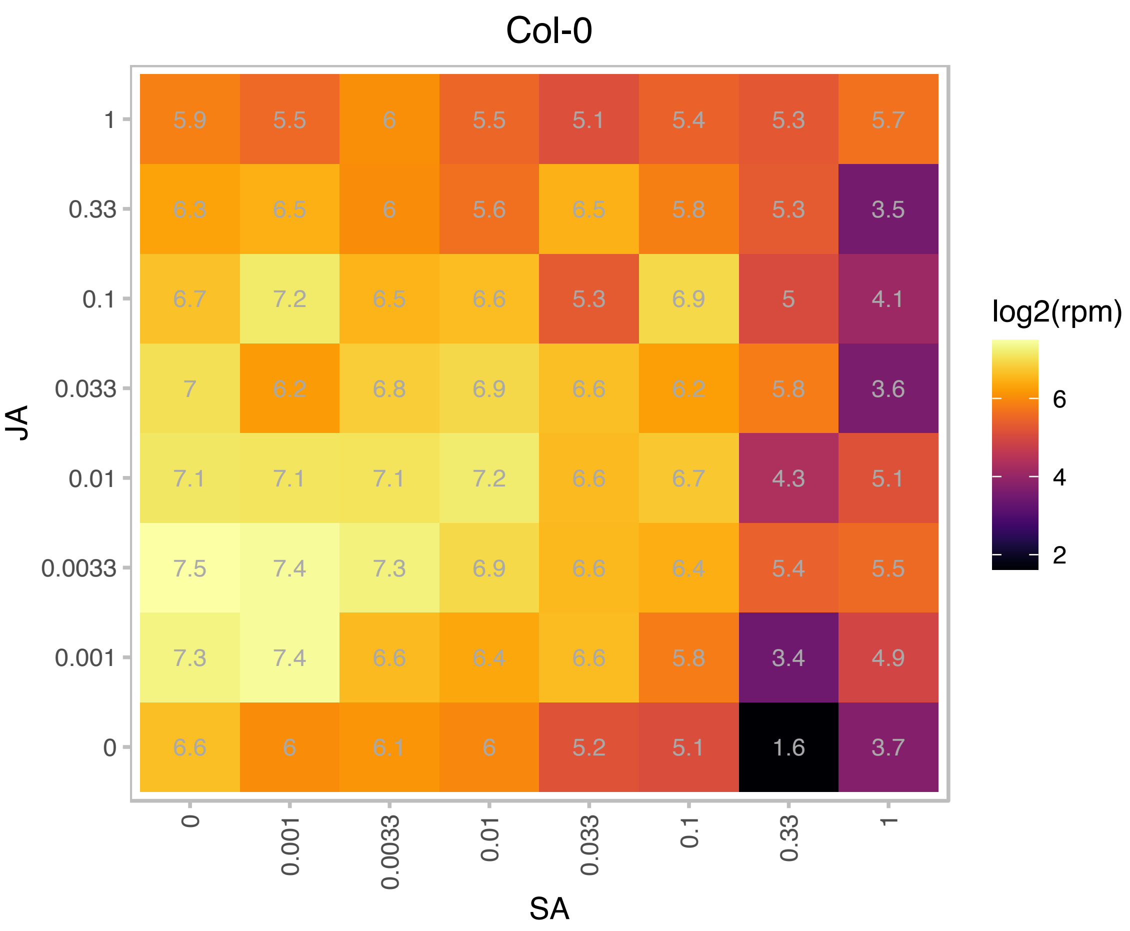


| | |
|---|---|
| Gene | AT5G14200 |
| is.expressed_genes | TRUE |
| is.responsive_genes | TRUE |
| is.apcluster | F-02 |

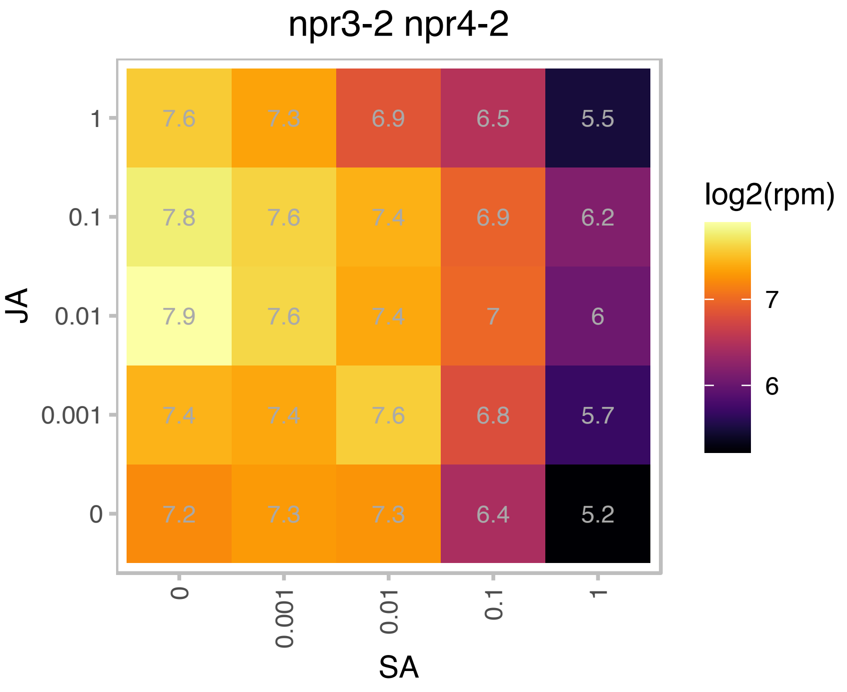


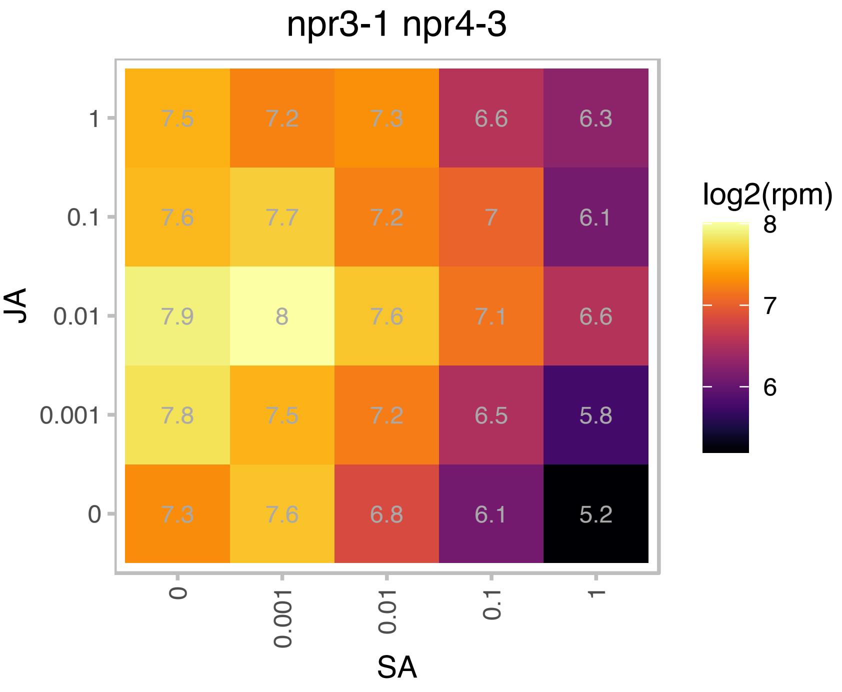

# AT5G14460.1 : NA

Pseudouridine synthase family protein

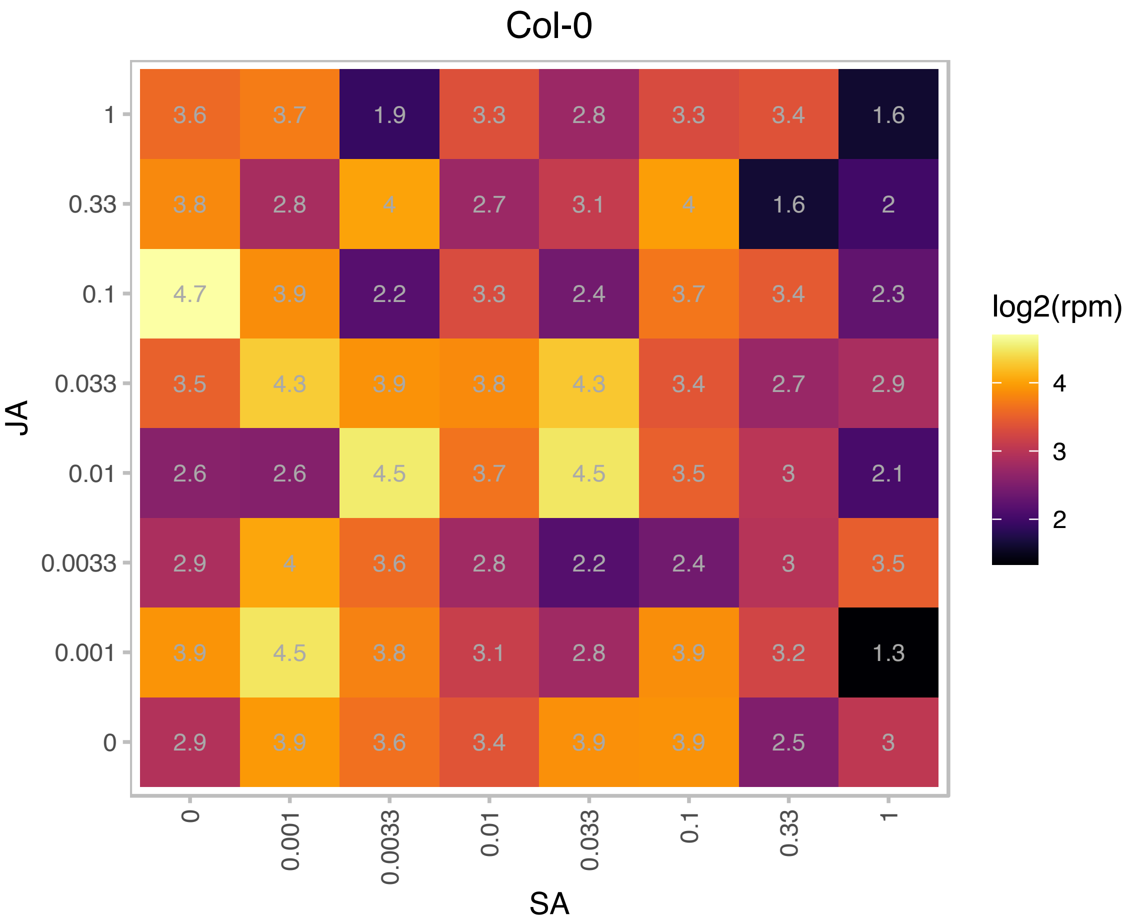


| | |
|---|---|
| Gene | AT5G14460 |
| is.expressed_genes | TRUE |
| is.responsive_genes | TRUE |
| is.apcluster | F-02 |

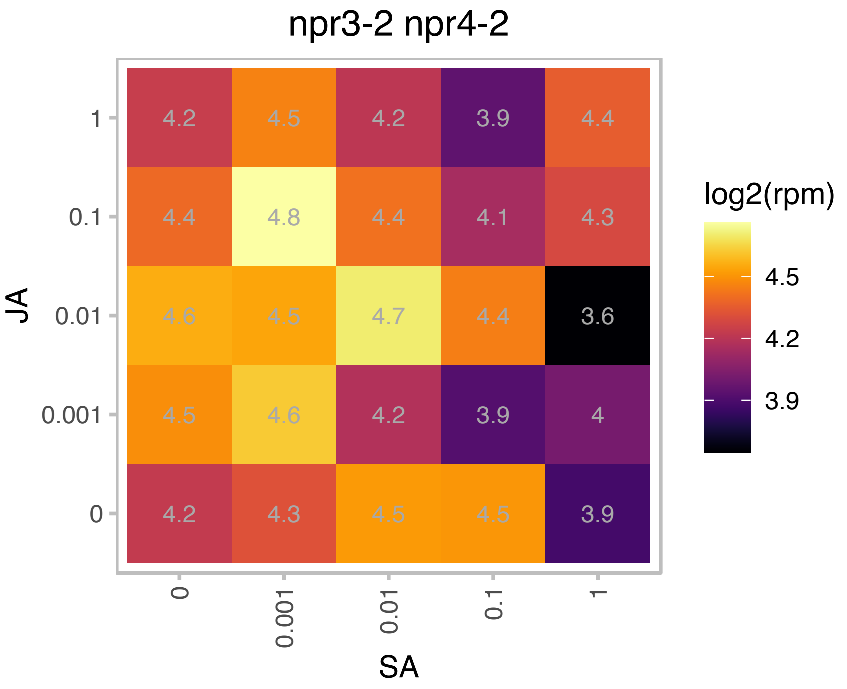


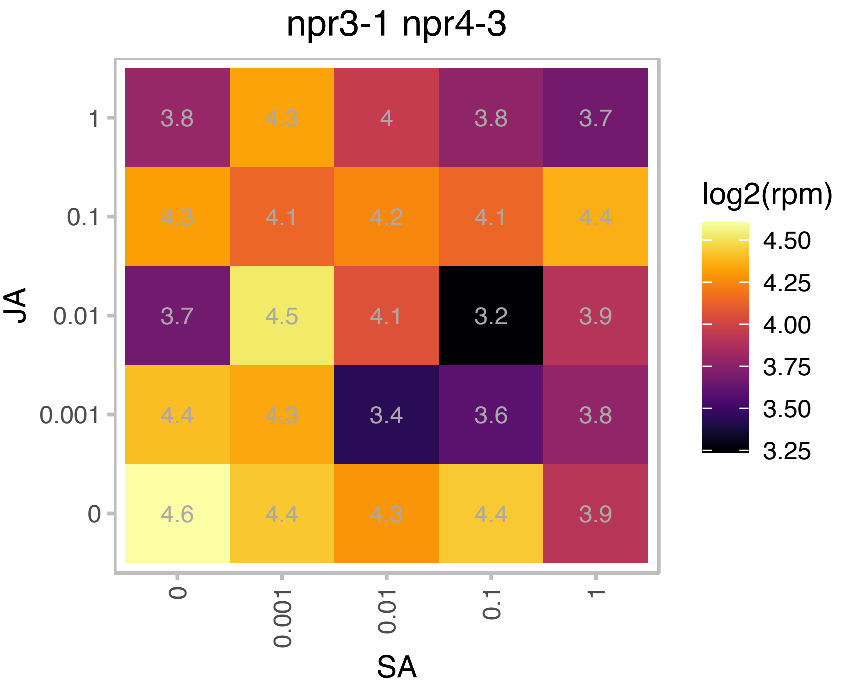

### AT5G16390.1 : BCCP, BCCP-1, BCCP1, CAC1, CAC1-A, C

chloroplastic acetylcoenzyme A carboxylase 1

Encodes for the biotin carboxyl-carrier subunit of the multi-enzyme plastidial acetyl-coenzyme A carboxylase complex.

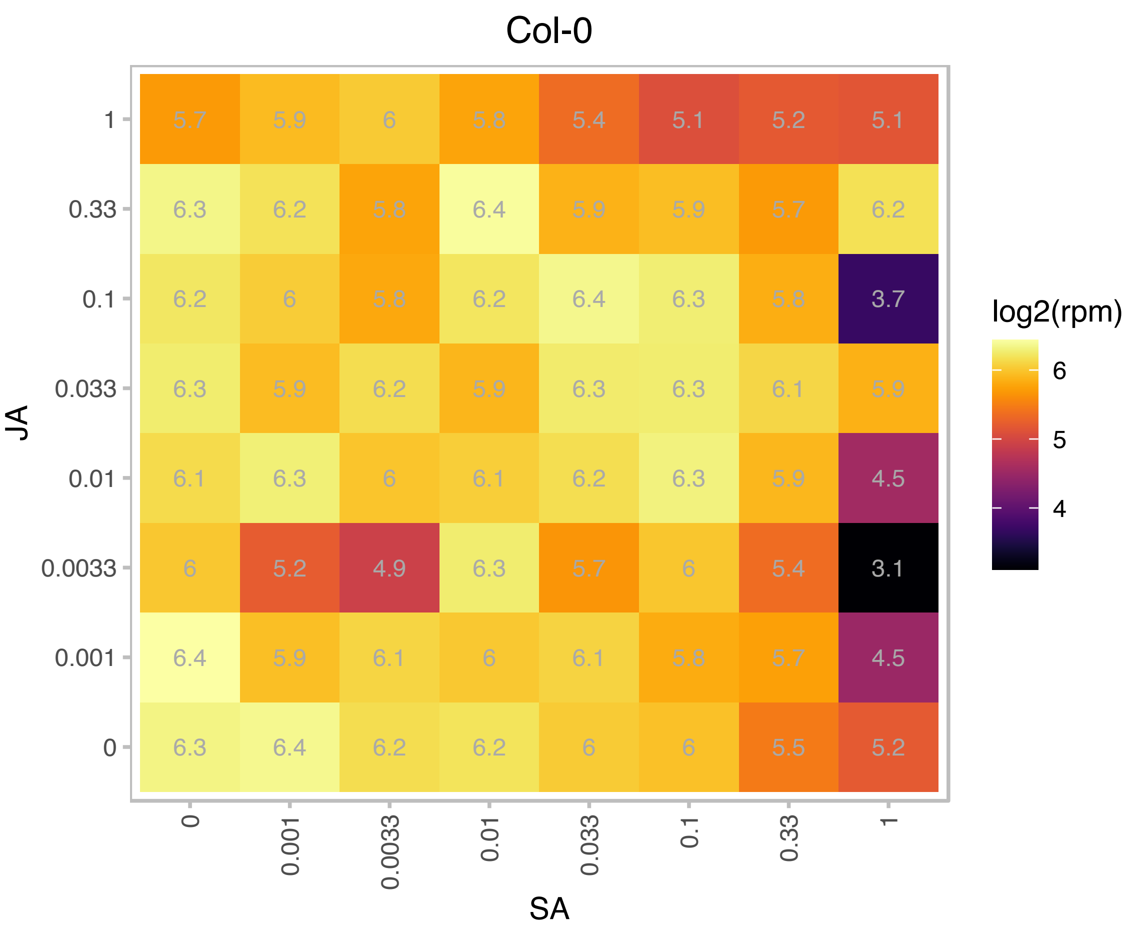


| | |
|---|---|
| Gene | AT5G16390 |
| is.expressed_genes | TRUE |
| is.responsive_genes | TRUE |
| is.apcluster | F-02 |

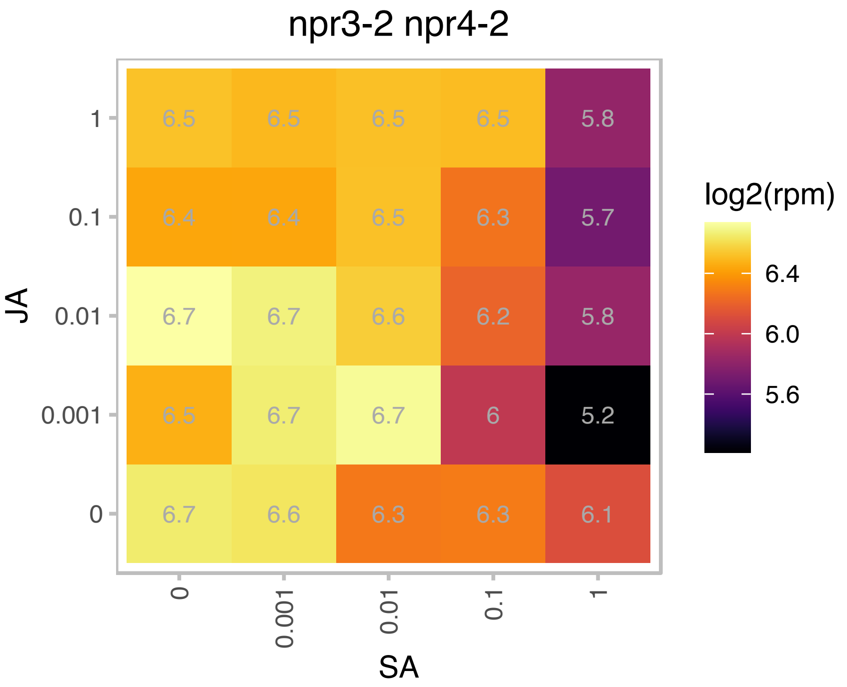


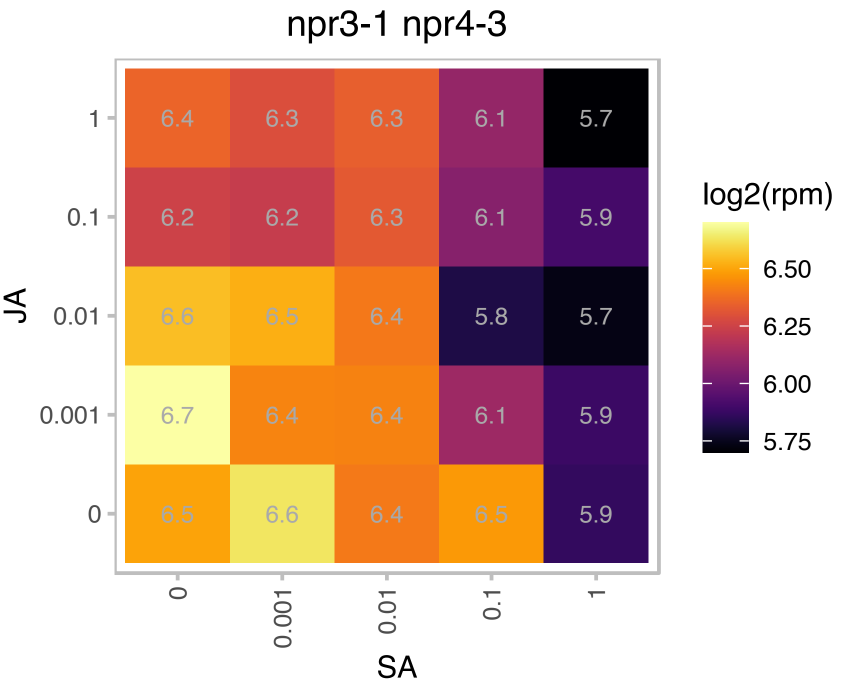

### AT5G23020.1 : IMS2, MAM-L, MAM3

2-isopropylmalate synthase 2

methylthioalkymalate synthase-like. Also known as 2-isopropylmalate synthase (IMS2). encodes a methylthioalkylmalate synthase involved in the biosynthesis of aliphatic glucosinolates which accepts all the omega-methylthio-2-oxoalkanoic acids needed to form the known C3 to C8 glucosinolates in Arabidopsis. The mRNA is cell-to-cell mobile.

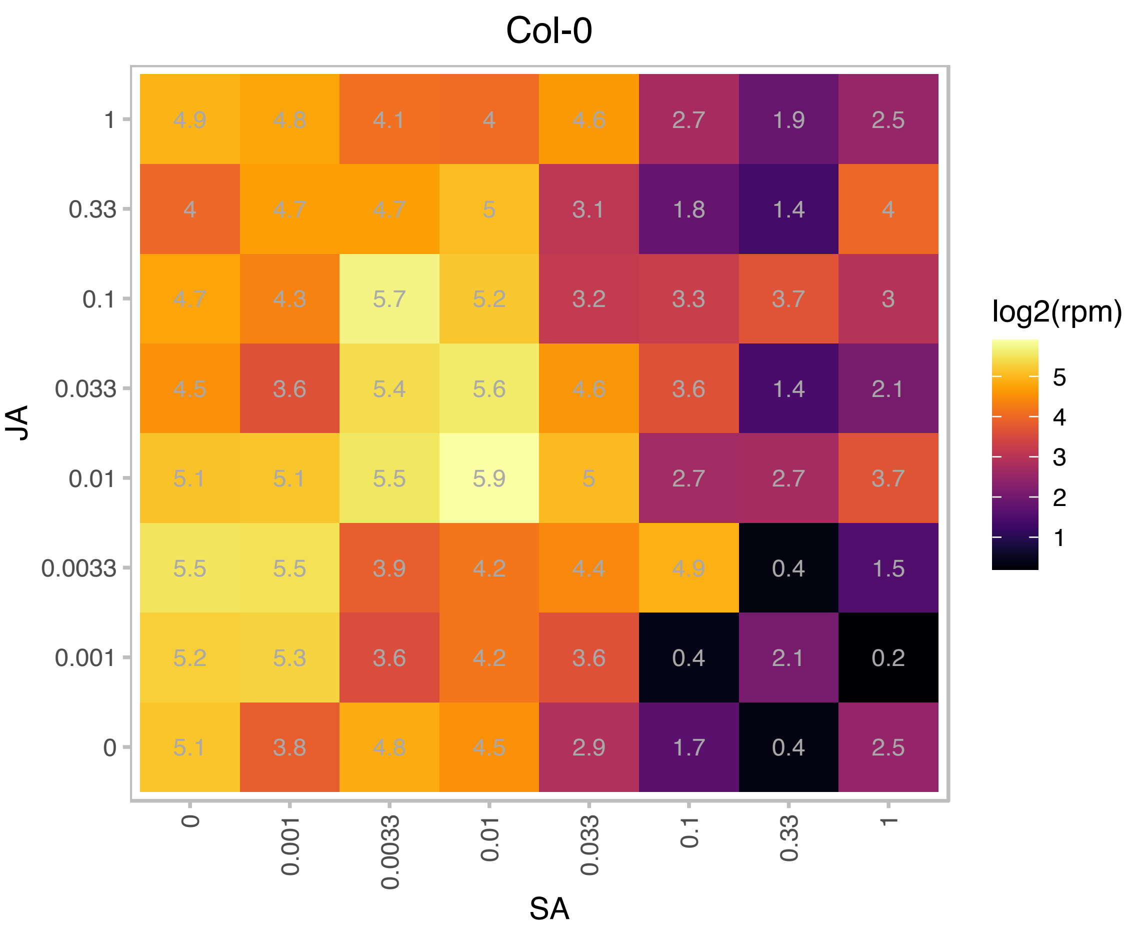


| | |
|---|---|
| Gene | AT5G23020 |
| is.expressed_genes | TRUE |
| is.responsive_genes | TRUE |
| is.apcluster | F-02 |

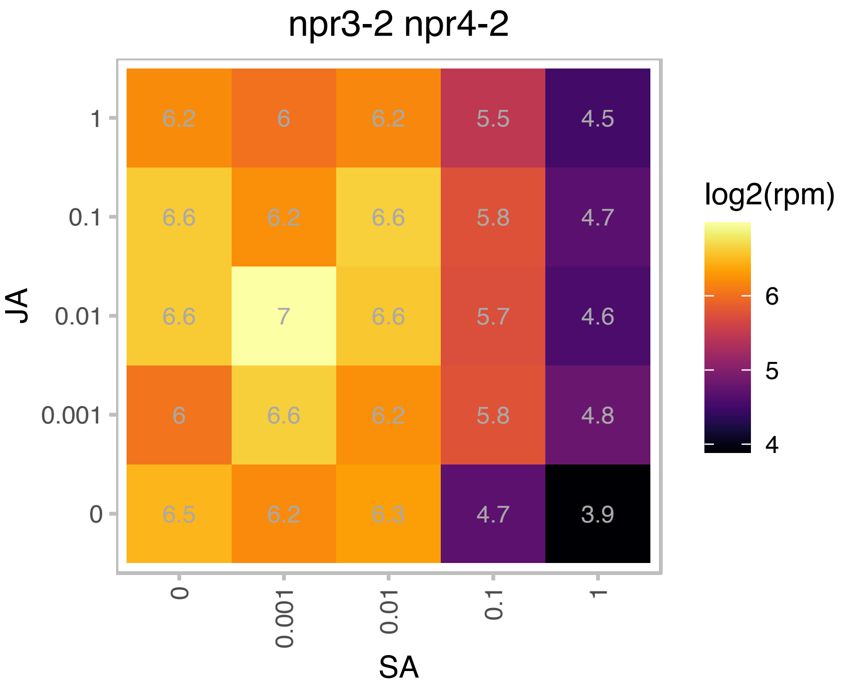


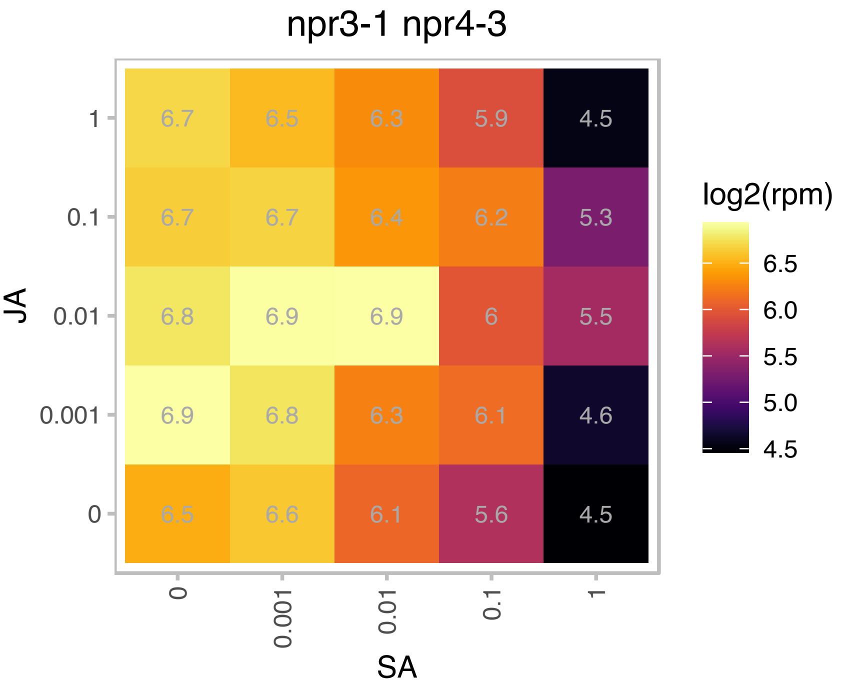

# AT5G27120.1 : NA

NOP56-like pre RNA processing ribonucleoprotein
SAR DNA-binding protein, putative, strong similarity to SAR DNA-binding protein-1 (Pisum sativum) GI:3132696; contains Pfam profile PF01798: Putative snoRNA binding domain; has similarity to MAR binding NOP58 protein
The mRNA is cell-to-cell mobile.

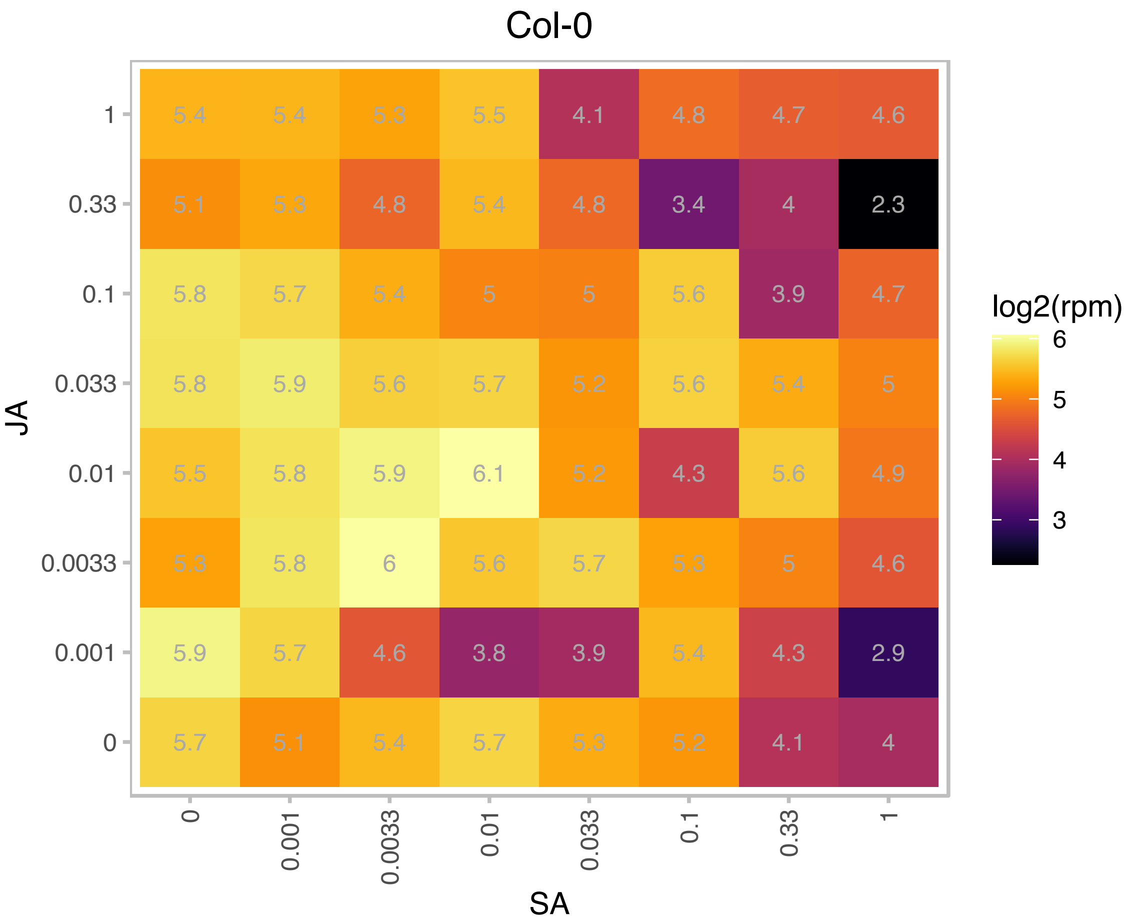


| | |
|---|---|
| Gene | AT5G27120 |
| is.expressed_genes | TRUE |
| is.responsive_genes | TRUE |
| is.apcluster | F-02 |

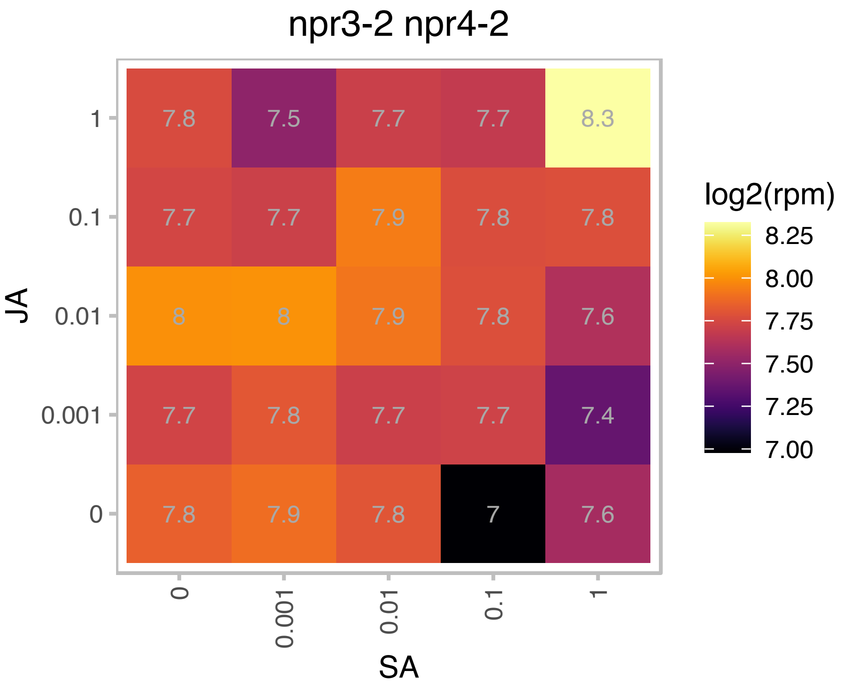


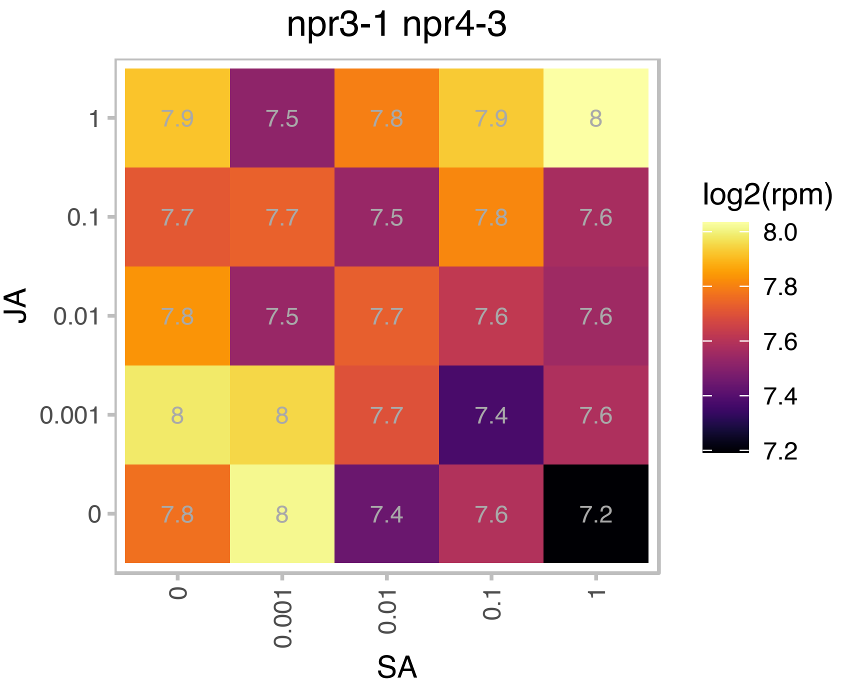

### AT5G40850.1 : AtUPM1, UPM1

urophorphyrin methylase 1
Encodes a urophorphyrin III methylase that catalyzes S-adenosyl-L-methionine-dependent transmethylation in a multistep process involving the formation of a covalently linked complex with S-adenosyl-L-methionine. The mRNA is cell-to-cell mobile.

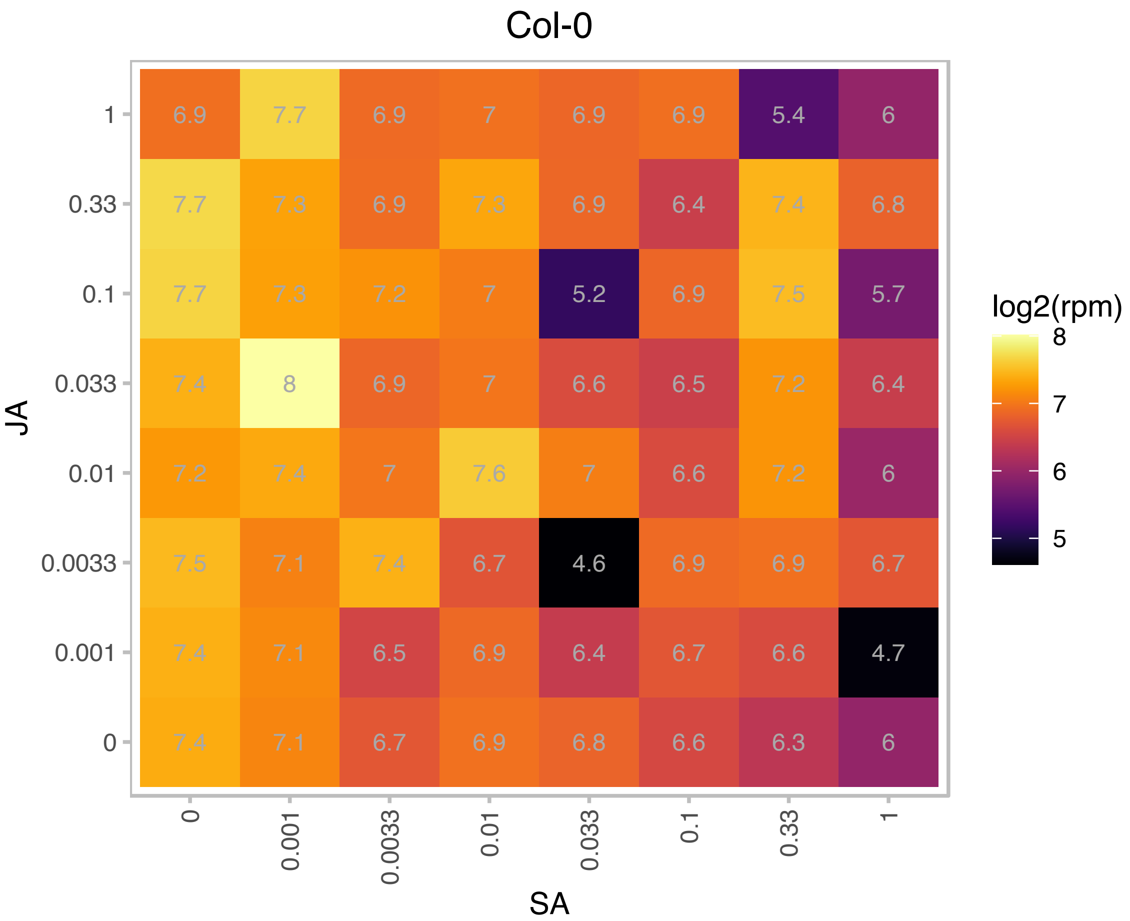


| | |
|---|---|
| Gene | AT5G40850 |
| is.expressed_genes | TRUE |
| is.responsive_genes | TRUE |
| is.apcluster | F-02 |

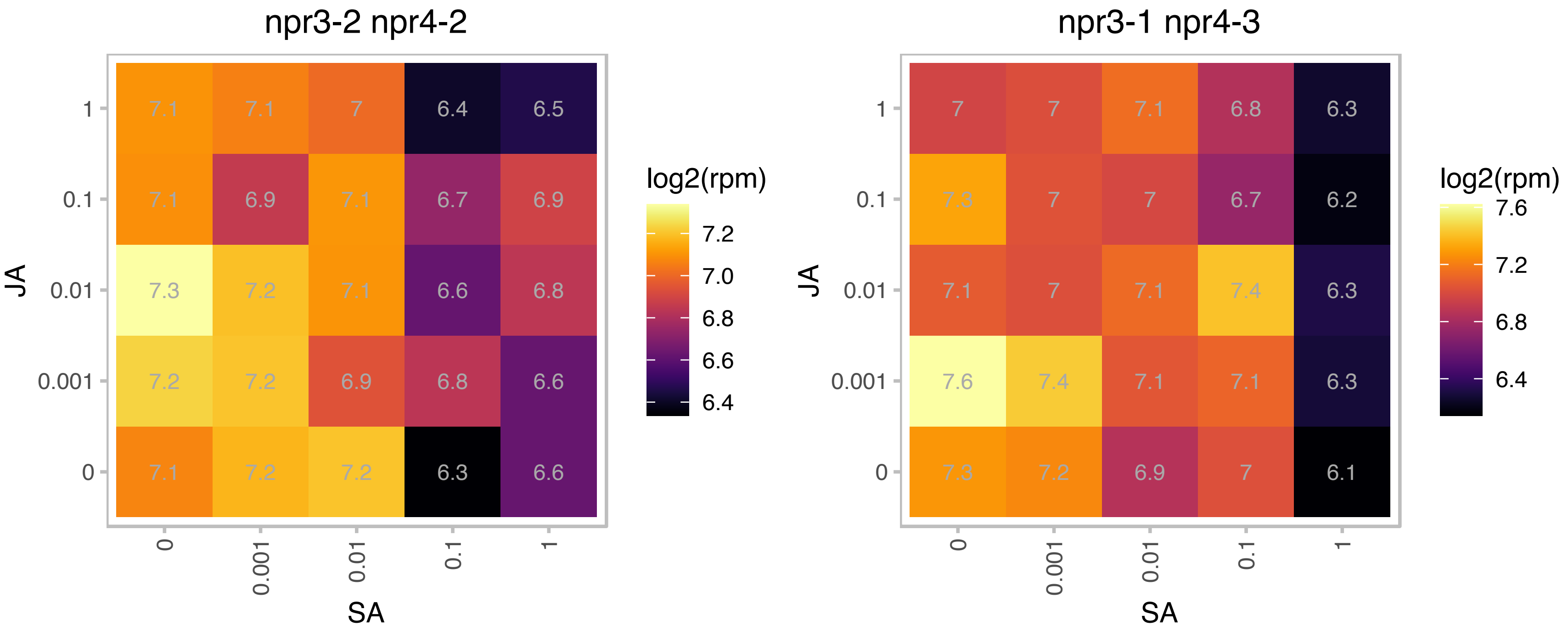

# AT5G43720.1 : NA

Protein of unknown function (DUF2361)

Col-0

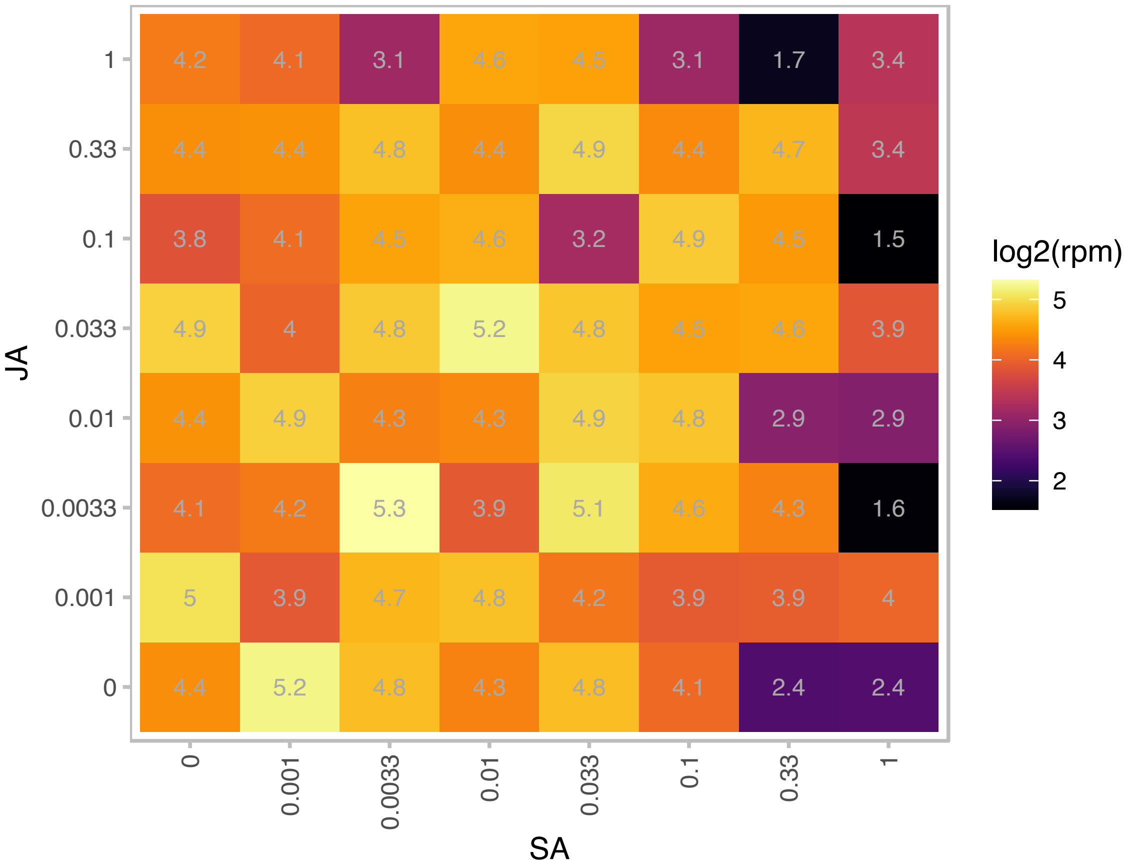


| | |
|---|---|
| Gene | AT5G43720 |
| is.expressed_genes | TRUE |
| is.responsive_genes | TRUE |
| is.apcluster | F-02 |

npr3-2 npr4-2

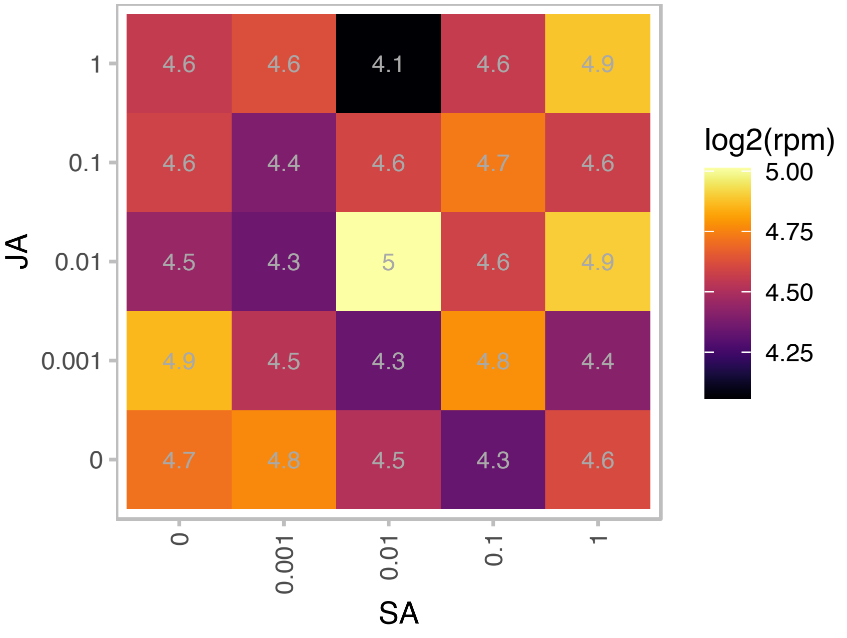


npr3-1 npr4-3

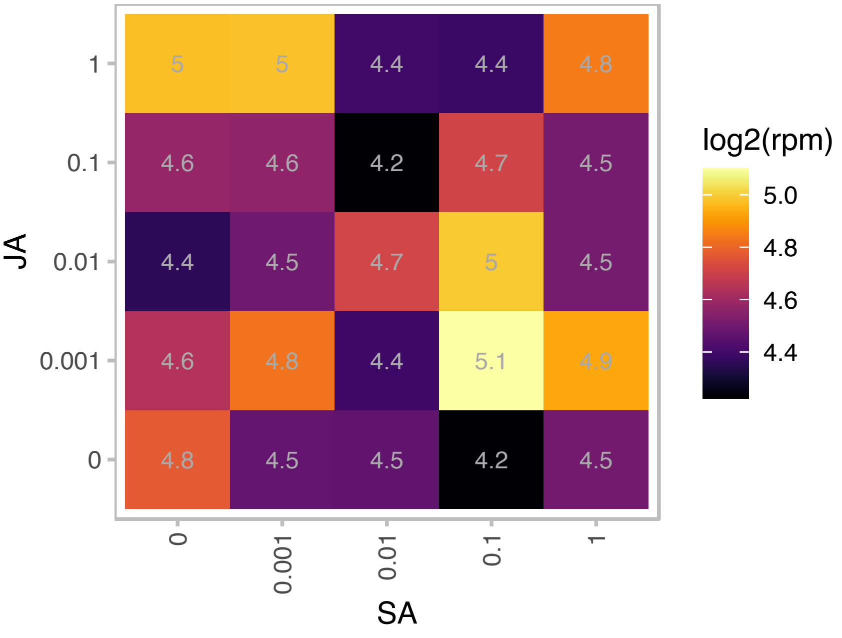

### AT5G47910.1 : ATRBOHD, RBOHD

respiratory burst oxidase homologue D
NADPH/respiratory burst oxidase protein D (RbohD).Interacts with AtrbohF gene to fine tune the spatial control of ROI production and hypersensitive response to cell in and around infection site. The mRNA is cell-to-cell mobile.

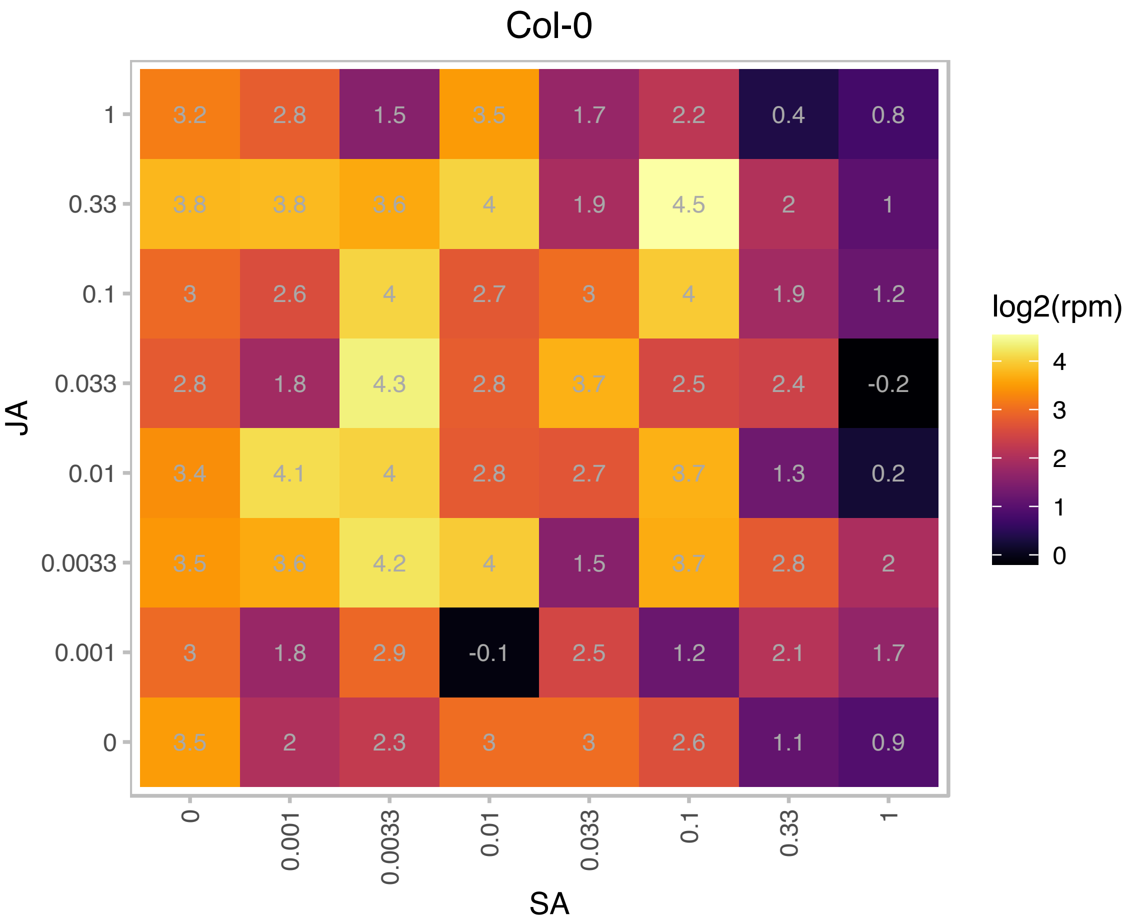


| | |
|---|---|
| Gene | AT5G47910 |
| is.expressed_genes | TRUE |
| is.responsive_genes | TRUE |
| is.apcluster | F-02 |

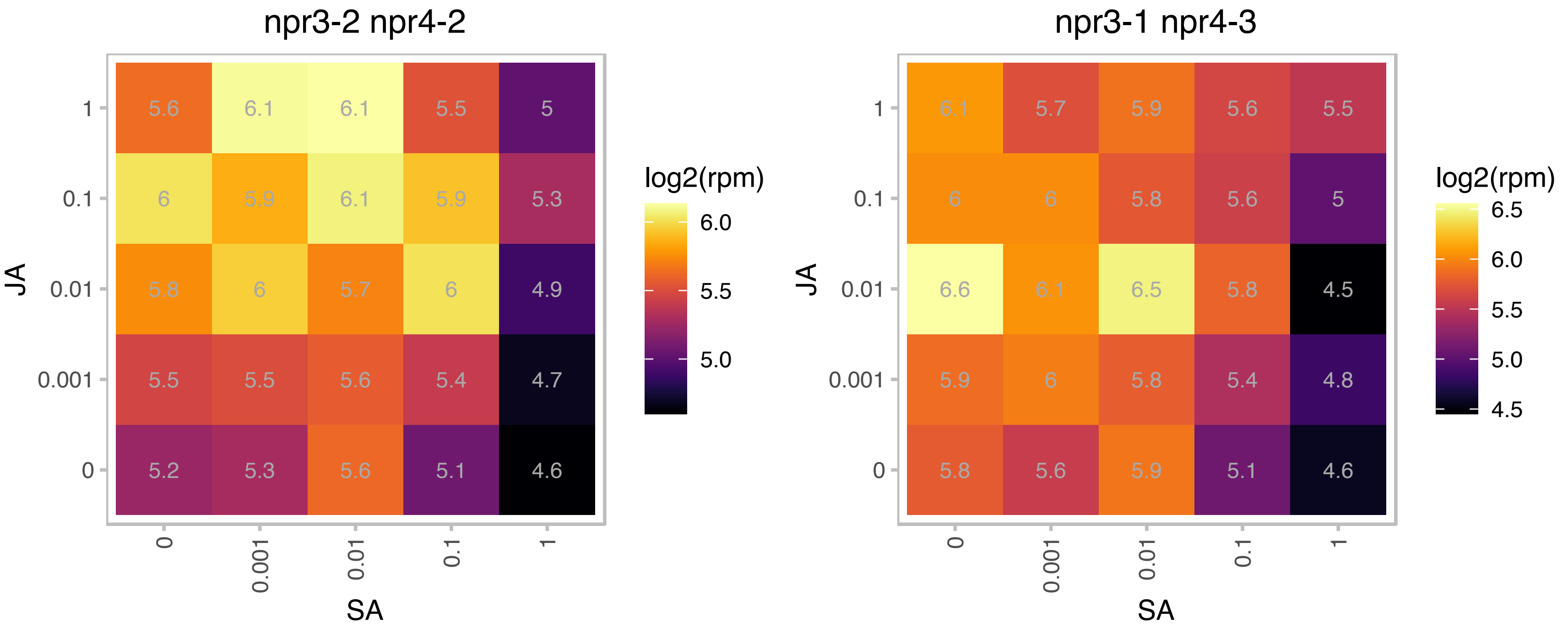

# AT5G51840.1 : NA

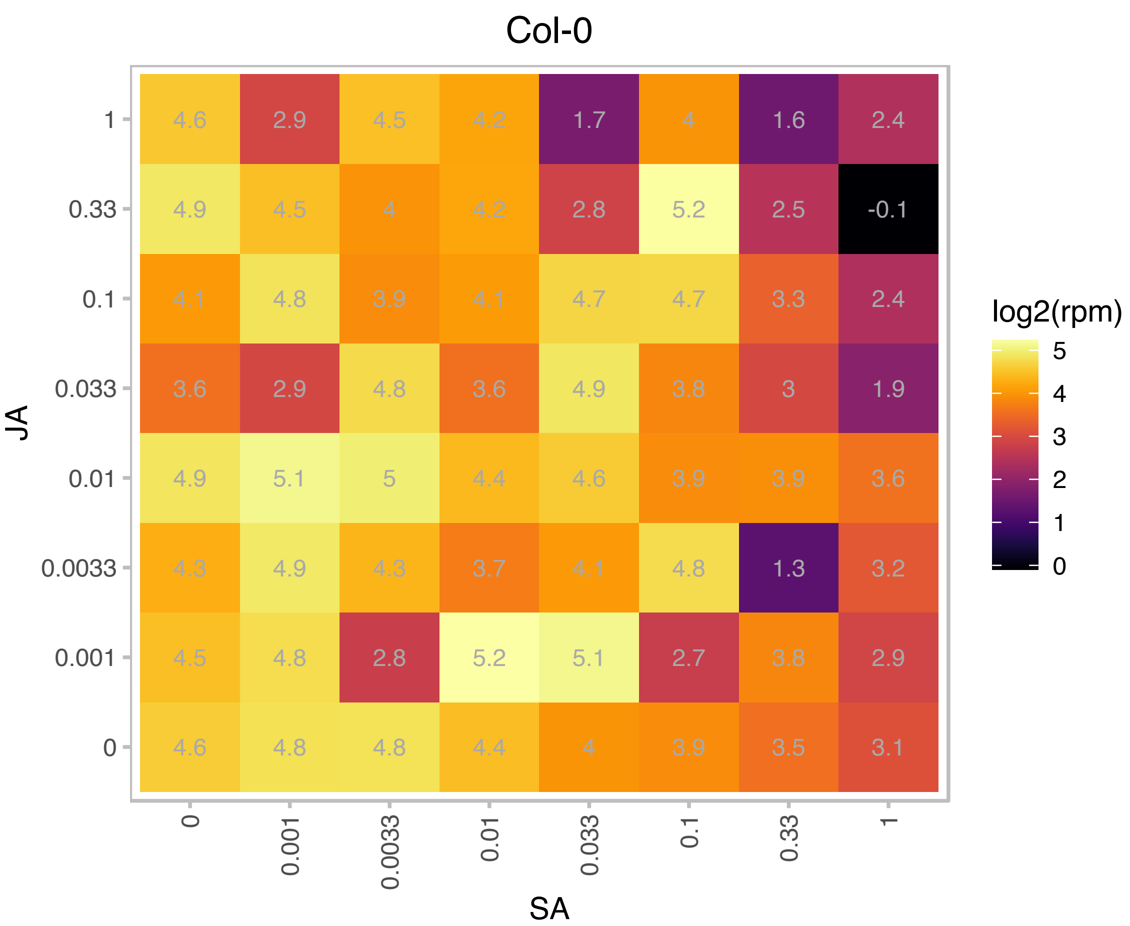


| | |
|---|---|
| Gene | AT5G51840 |
| is.expressed_genes | TRUE |
| is.responsive_genes | TRUE |
| is.apcluster | F-02 |

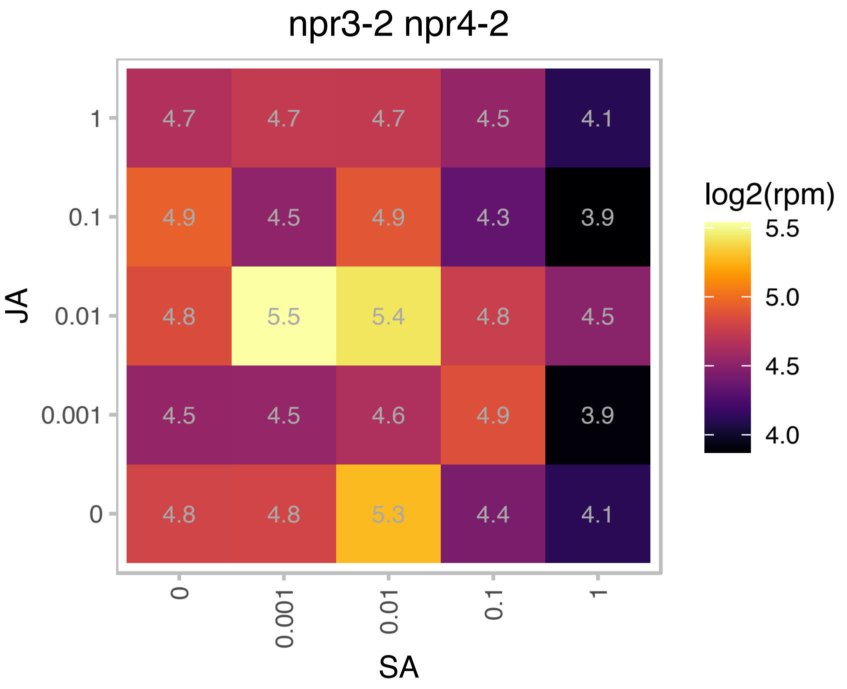


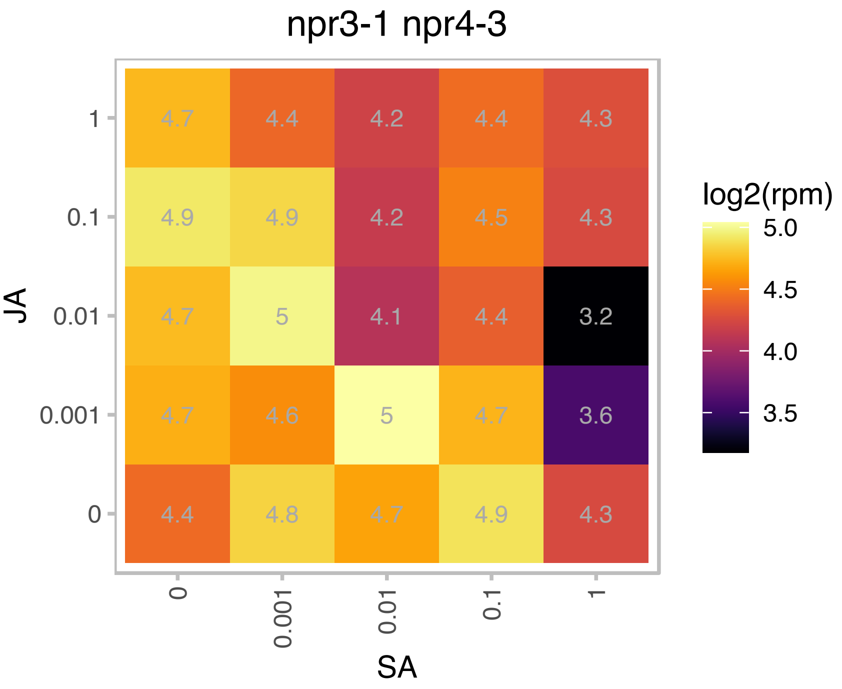

### AT5G57625.1 : AtCAPE5

CAP (Cysteine-rich secretory proteins, Antigen 5, and Pathogenesis-related 1 protein) superfamily protein

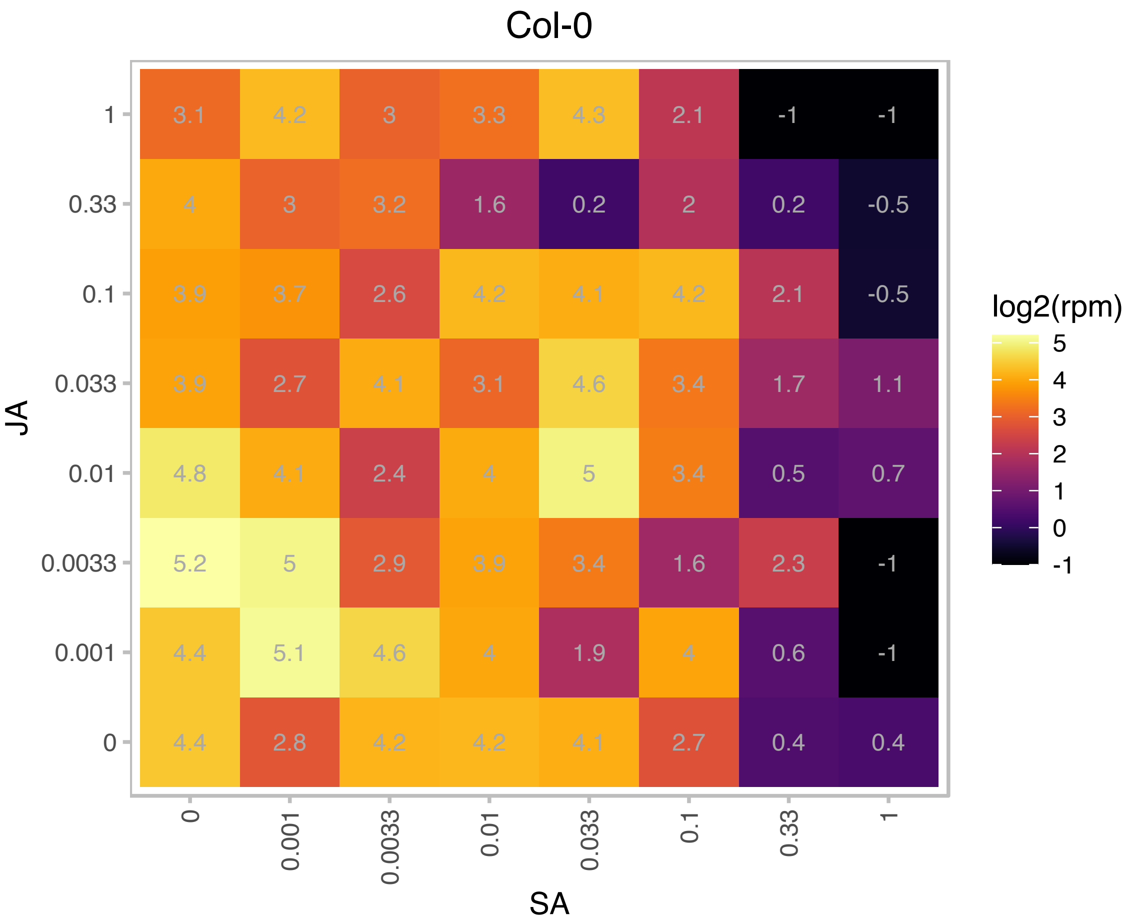


| | |
|---|---|
| Gene | AT5G57625 |
| is.expressed_genes | TRUE |
| is.responsive_genes | TRUE |
| is.apcluster | F-02 |

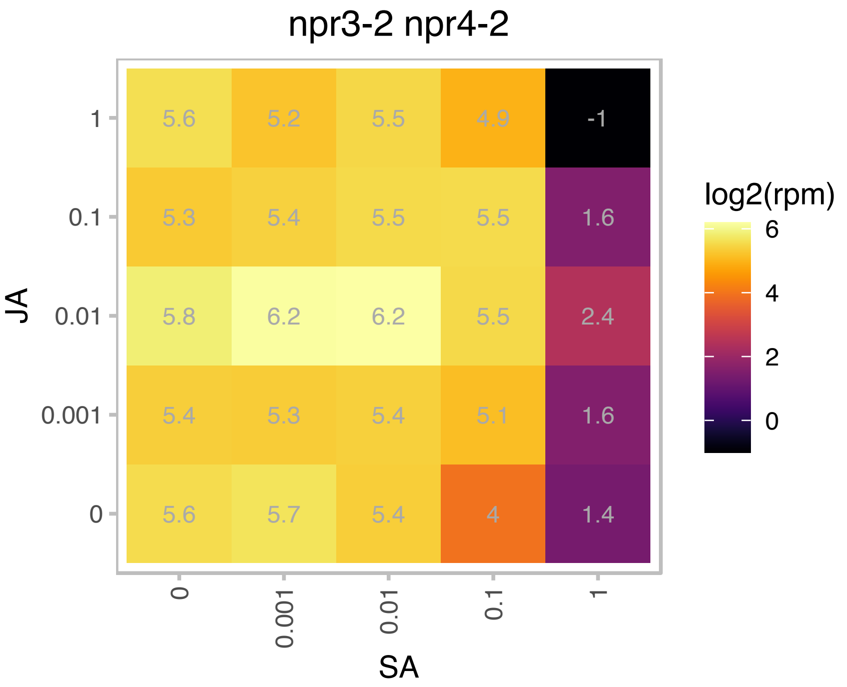


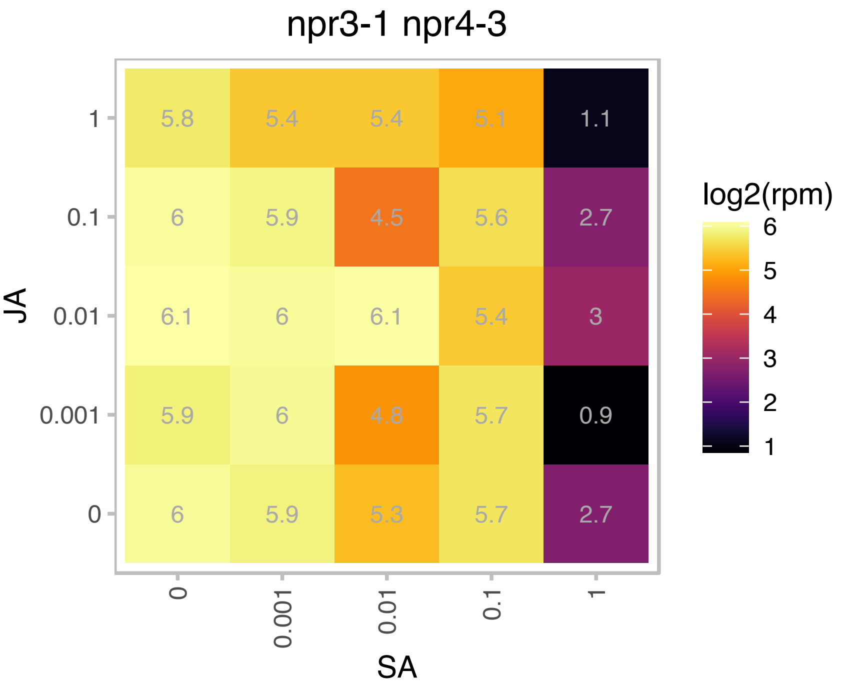

# AT5G58420.1 : NA

Ribosomal protein S4 (RPS4A) family protein

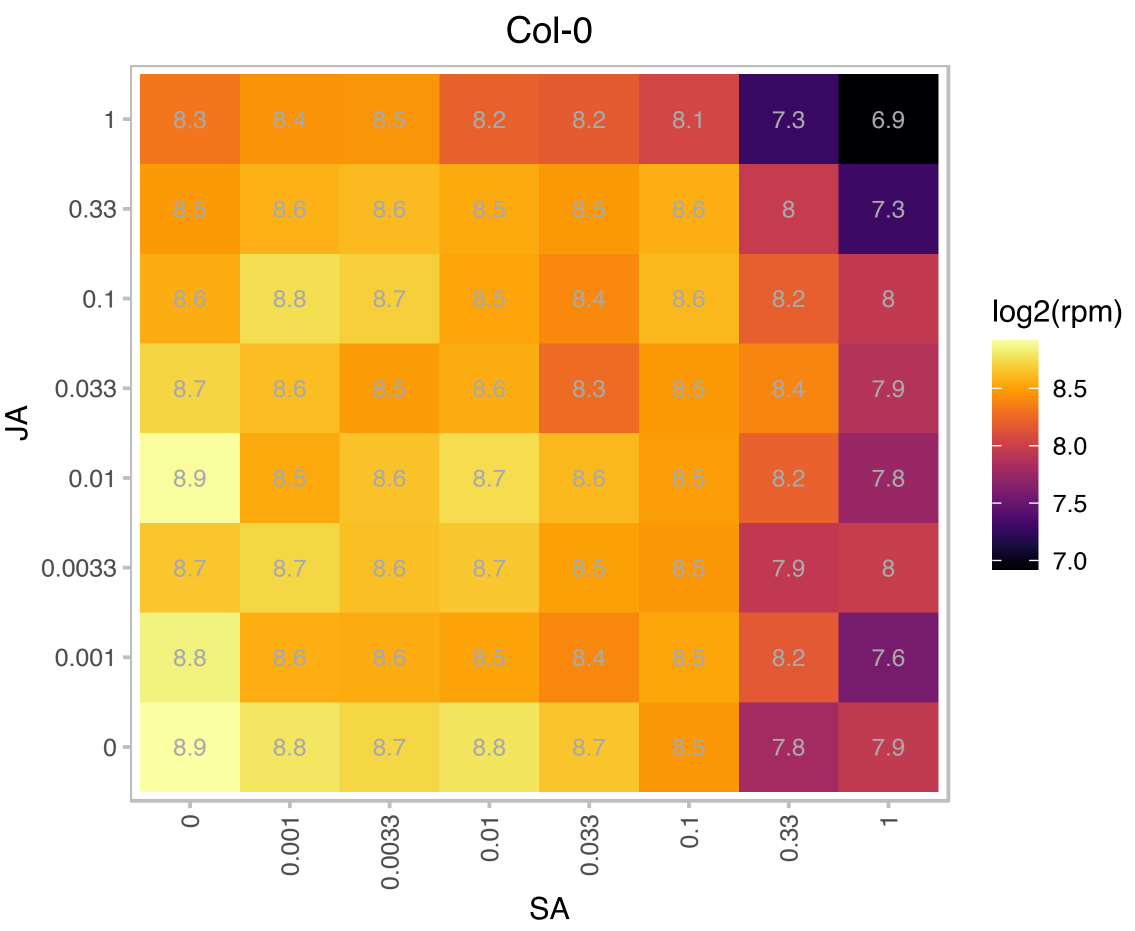


| | |
|---|---|
| Gene | AT5G58420 |
| is.expressed_genes | TRUE |
| is.responsive_genes | TRUE |
| is.apcluster | F-02 |

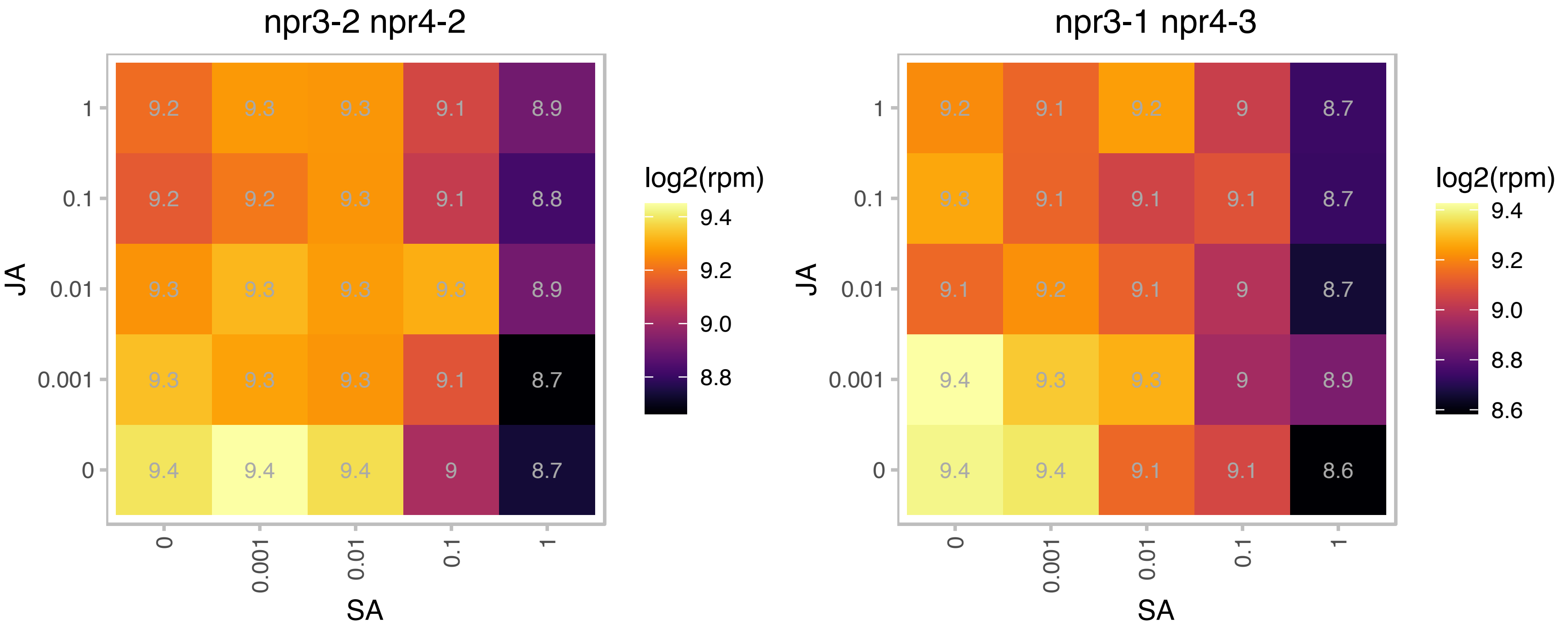

# AT5G58900.1 : NA

Homeodomain-like transcriptional regulator

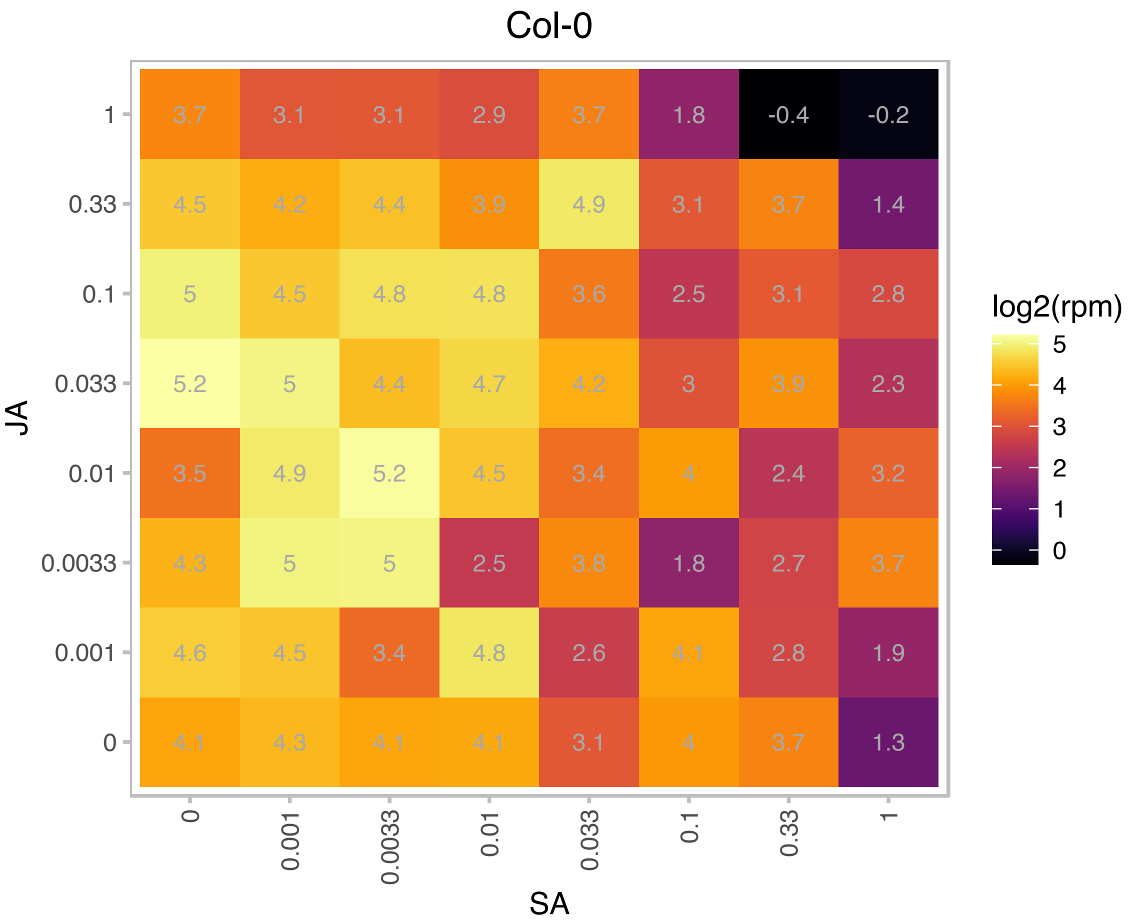


| | |
|---|---|
| Gene | AT5G58900 |
| is.expressed_genes | TRUE |
| is.responsive_genes | TRUE |
| is.apcluster | F-02 |

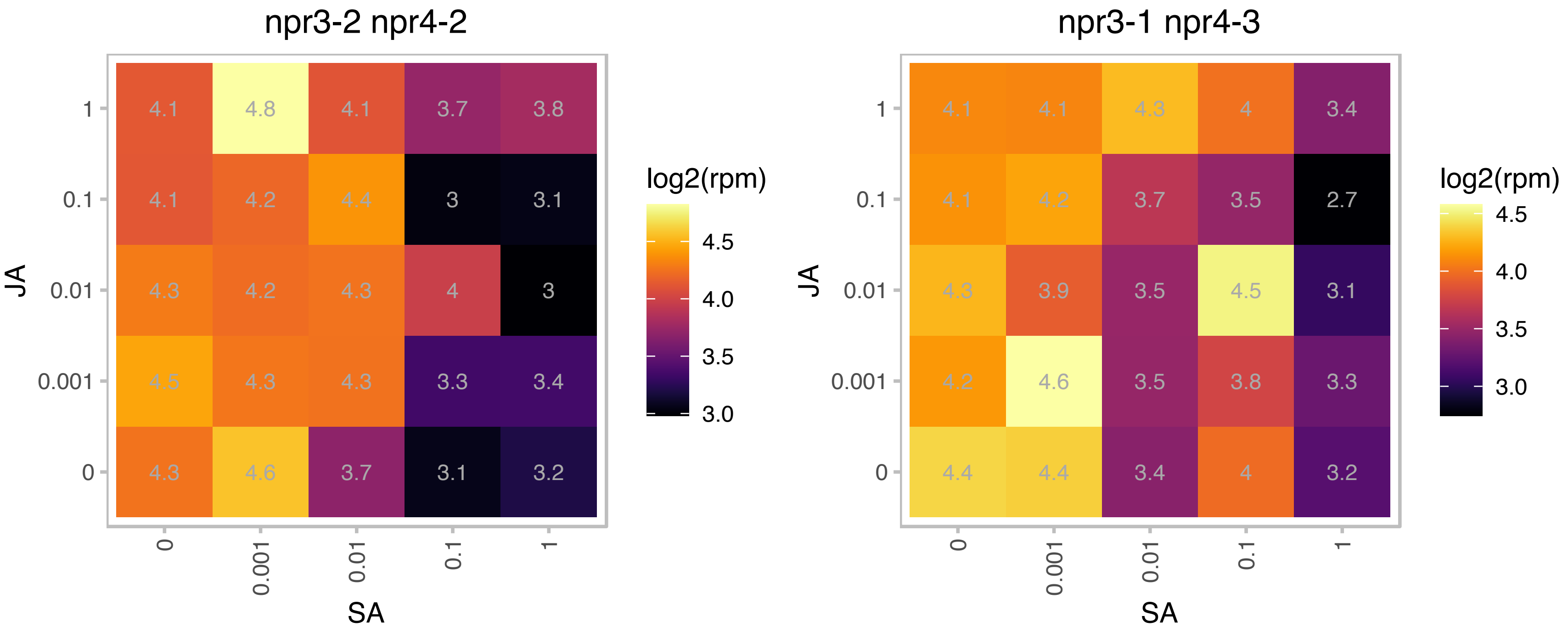

# AT5G60030.1 : NA

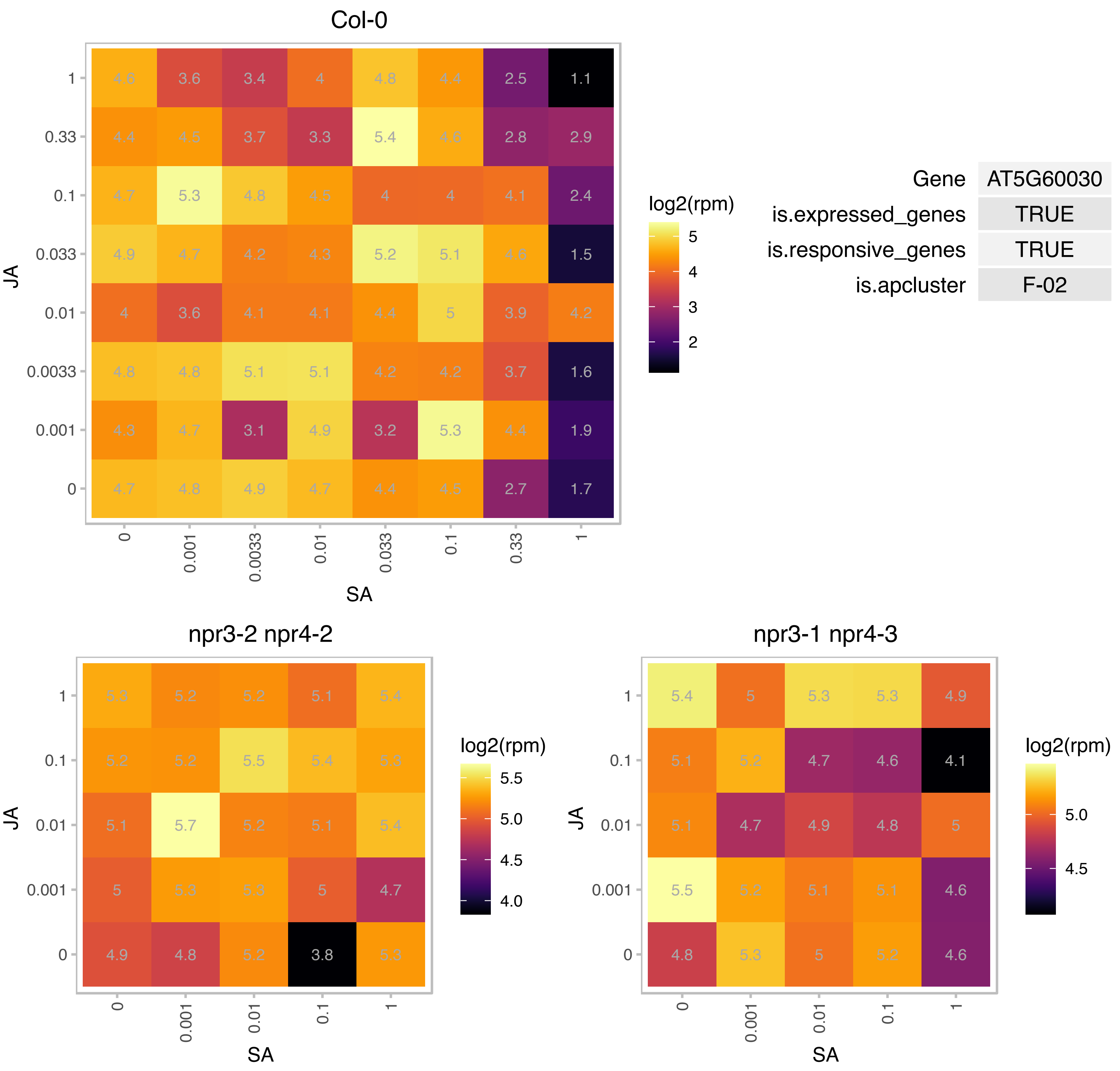


| Gene | AT5G60030 |
|---|---|
| is.expressed_genes | TRUE |
| is.responsive_genes | TRUE |
| is.apcluster | F-02 |

# AT5G61310.1 : NA

Cytochrome c oxidase subunit Vc family protein

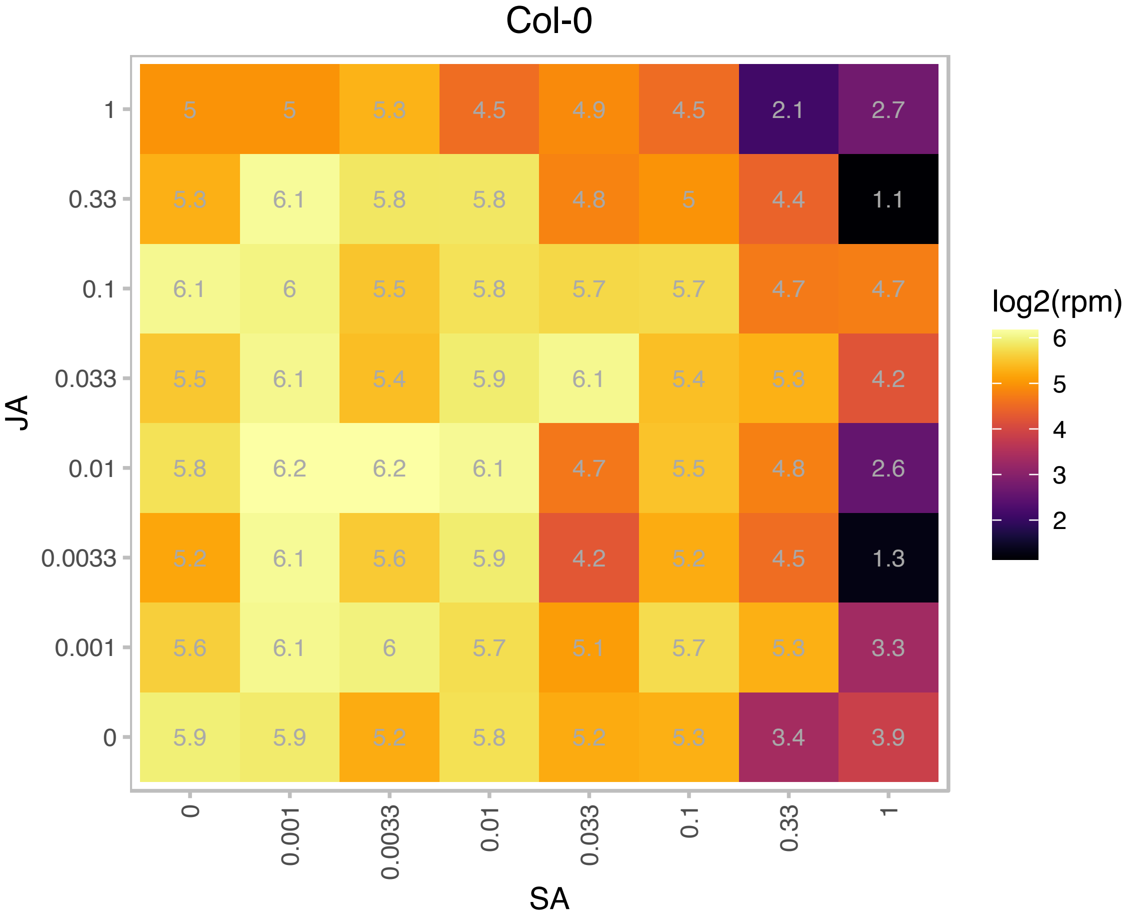


| | |
|---|---|
| Gene | AT5G61310 |
| is.expressed_genes | TRUE |
| is.responsive_genes | TRUE |
| is.apcluster | F-02 |

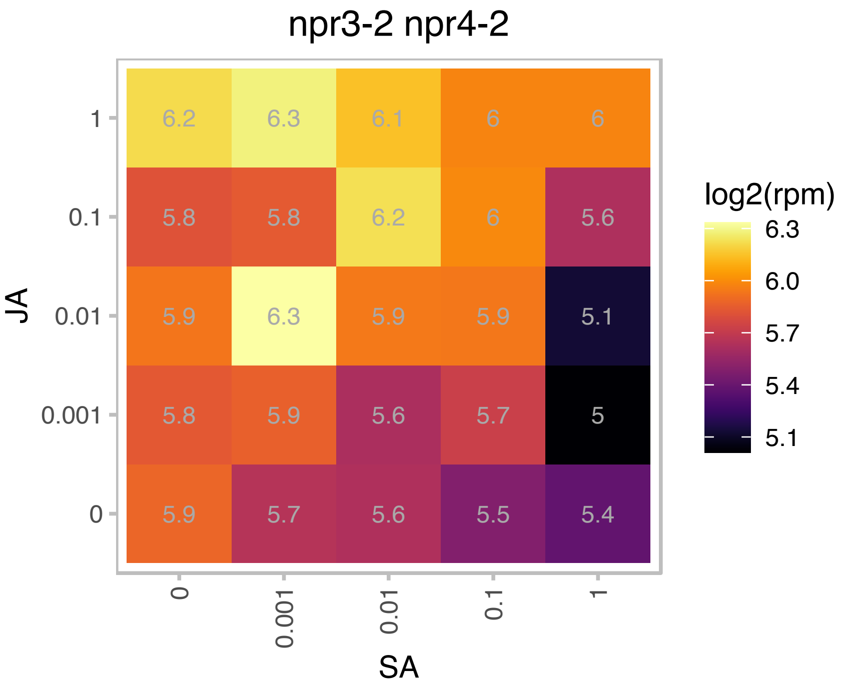


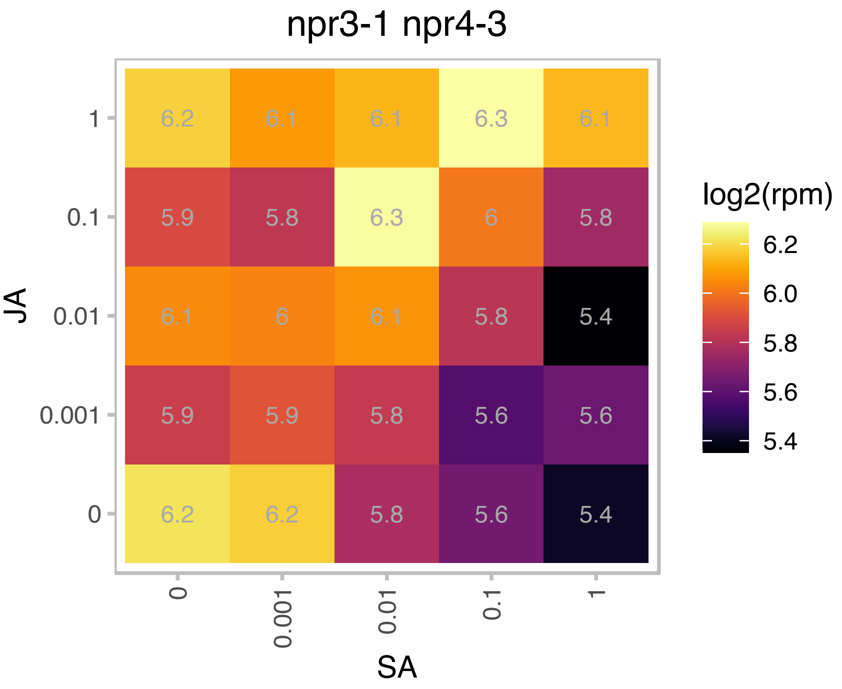

# AT5G61330.1 : NA

rRNA processing protein-related

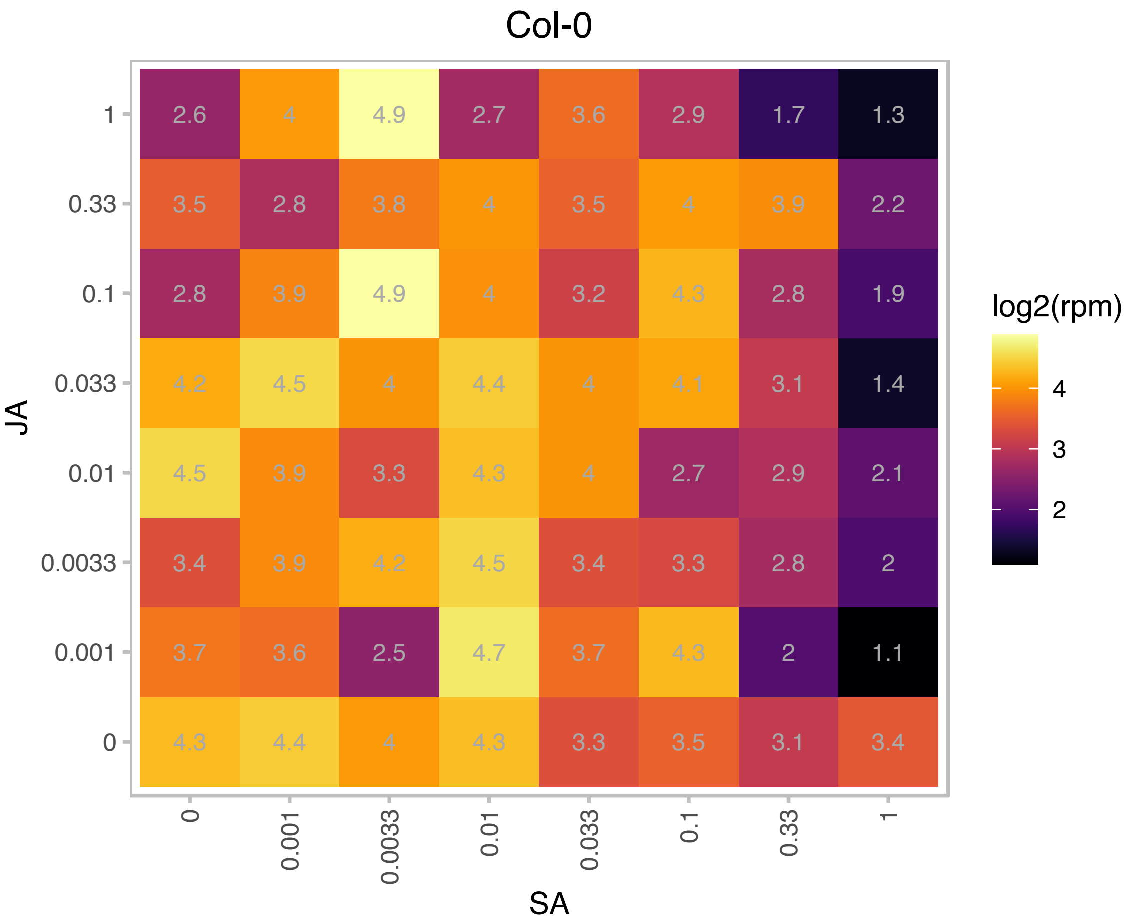


| | |
|---|---|
| Gene | AT5G61330 |
| is.expressed_genes | TRUE |
| is.responsive_genes | TRUE |
| is.apcluster | F-02 |

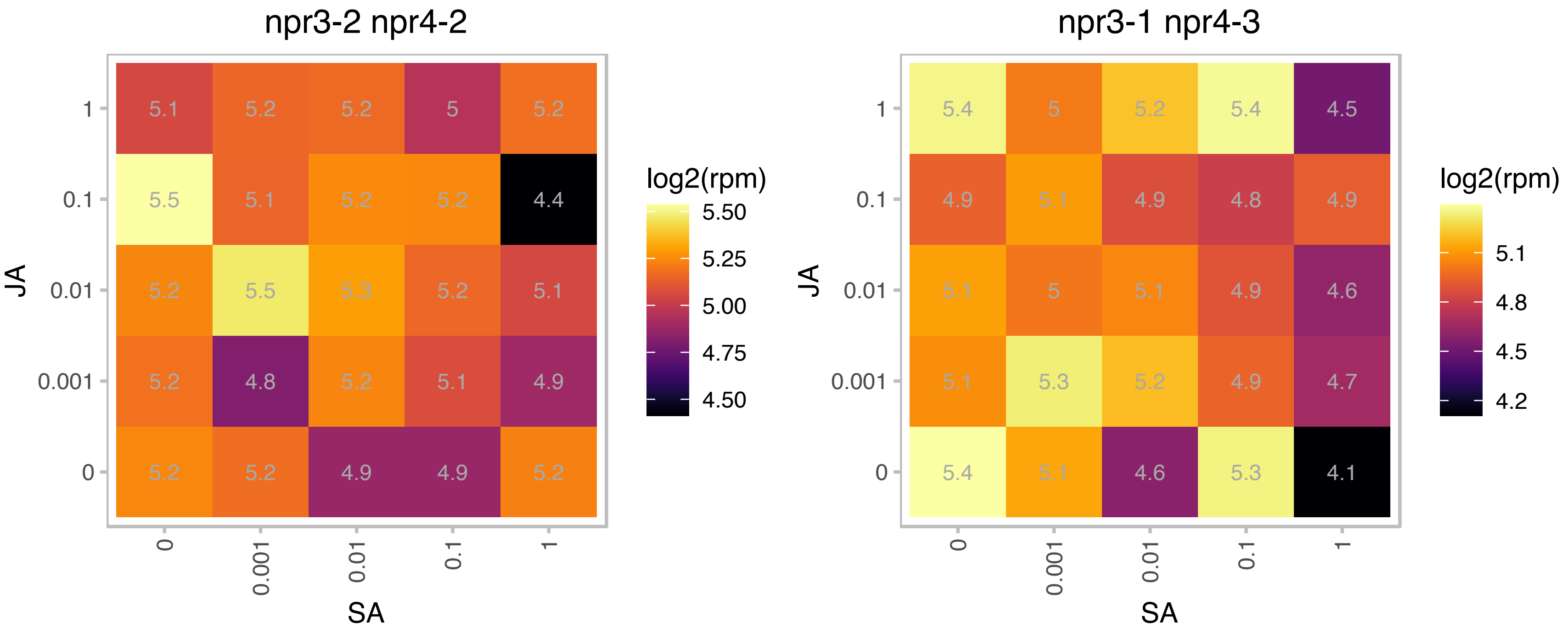

### AT5G63660.1 : LCR74, PDF2.5

Scorpion toxin-like knottin superfamily protein

Predicted to encode a PR (pathogenesis-related) protein. Belongs to the plant defensin (PDF) family with the following members: At1g75830/PDF1.1, At5g44420/PDF1.2a, At2g26020/PDF1.2b, At5g44430/PDF1.2c, At2g26010/PDF1.3, At1g19610/PDF1.4, At1g55010/PDF1.5, At2g02120/PDF2.1, At2g02100/PDF2.2, At2g02130/PD[illegible] At1g61070/PDF2.4, At5g63660/PDF2.5, At2g02140/PDF2.6, At5g38330/PDF3.1 and At4g30070/PDF3.2.

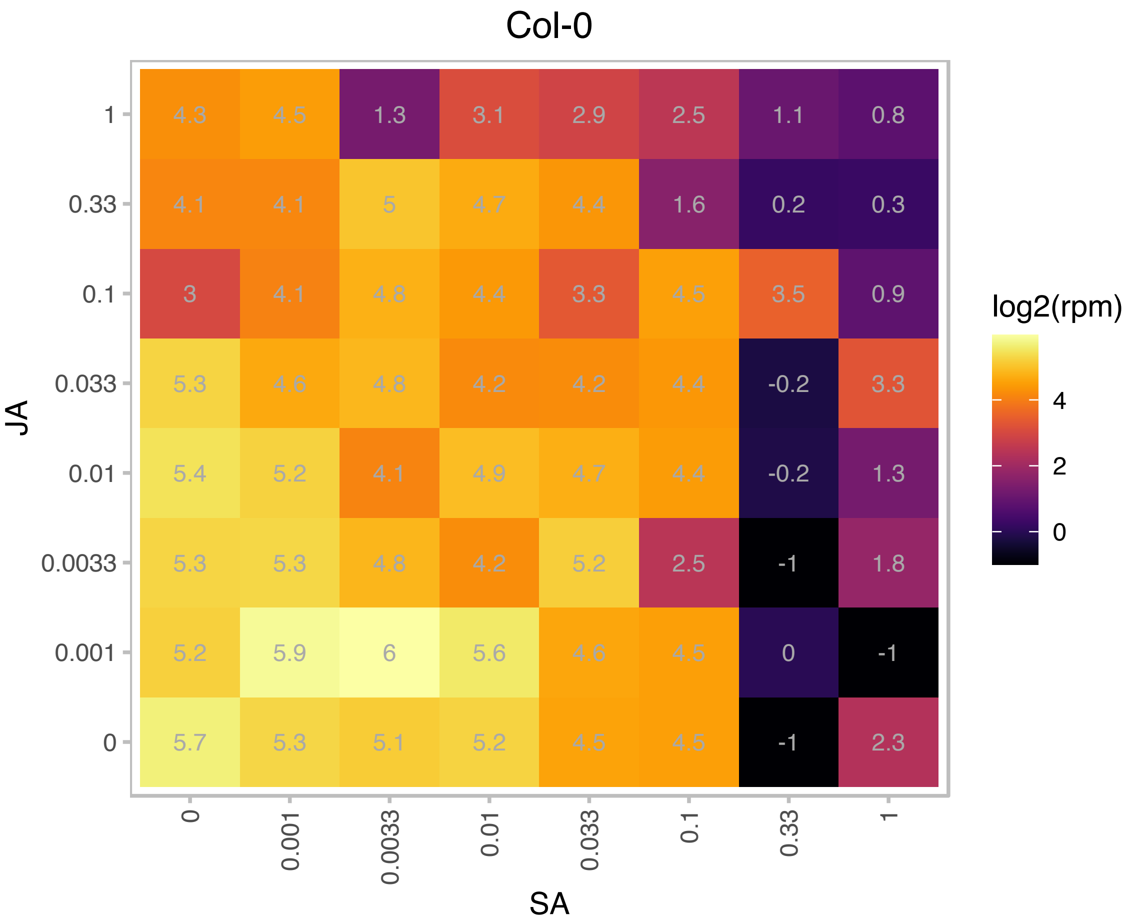


| | |
|---|---|
| Gene | AT5G63660 |
| is.expressed_genes | TRUE |
| is.responsive_genes | TRUE |
| is.apcluster | F-02 |

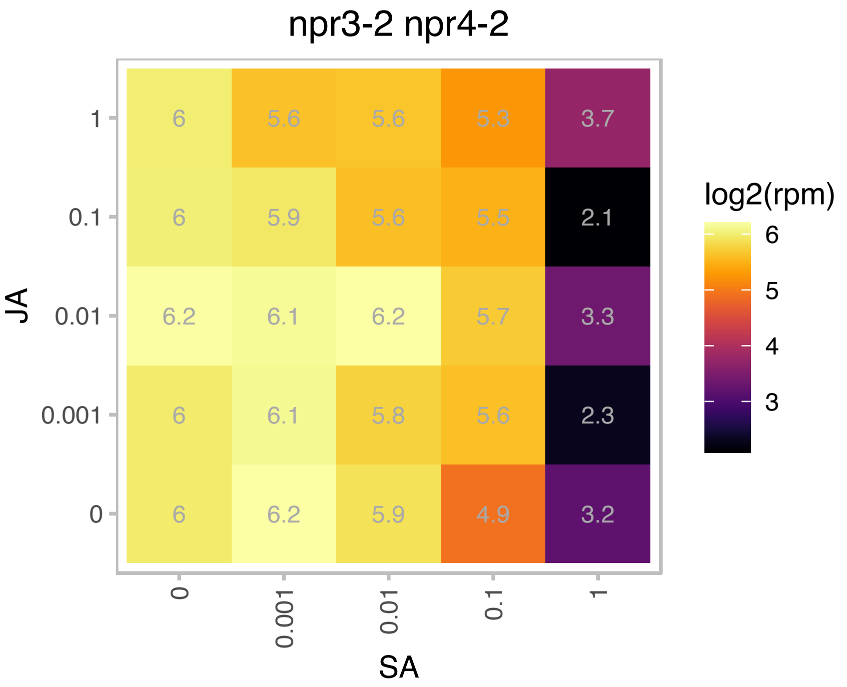


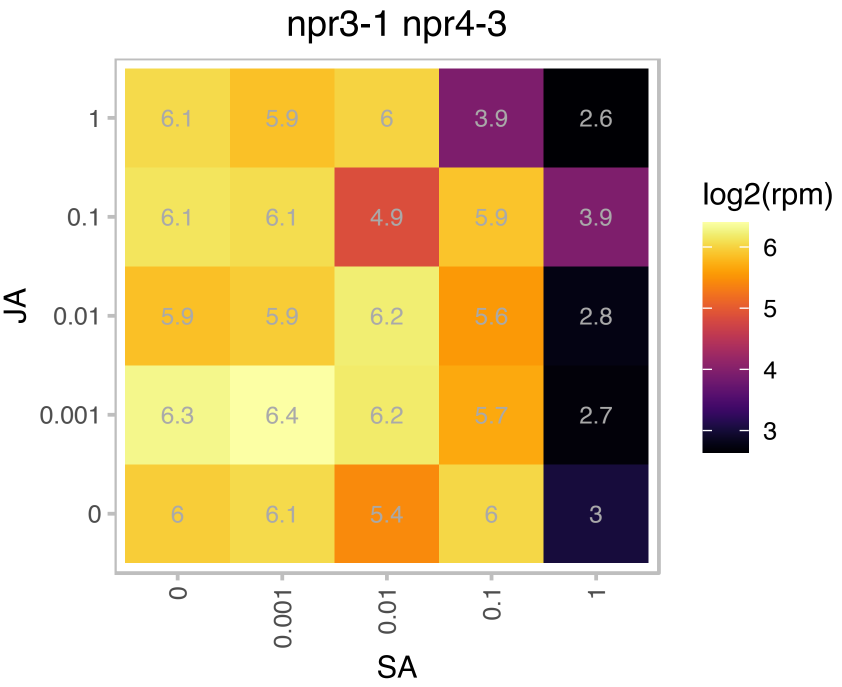

### AT5G65310.1 : ATHB-5, ATHB5, HB5

homeobox protein 5

Encodes a class I HDZip (homeodomain-leucine zipper) protein that is a positive regulator of ABA-responsiveness, mediating the inhibitory effect of ABA on growth during seedling establishment.

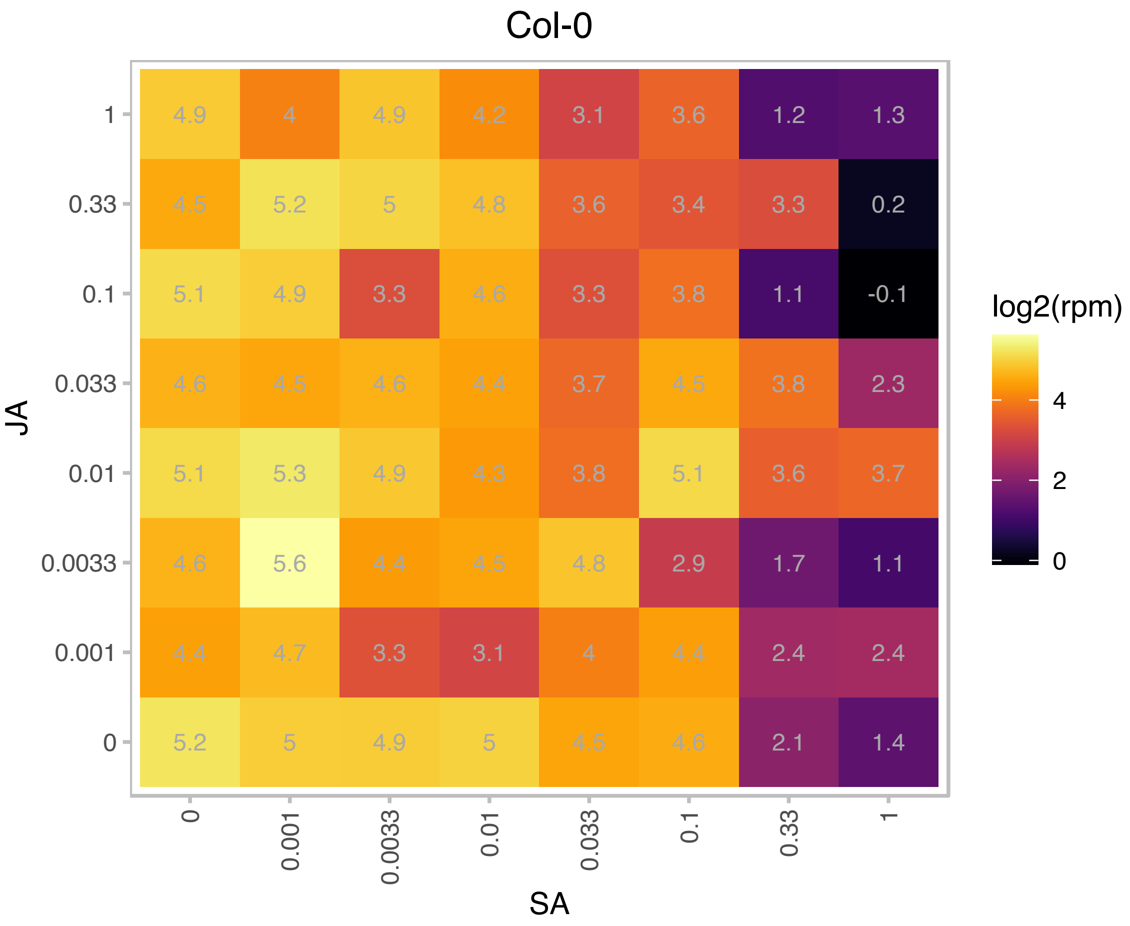


| | |
|---|---|
| Gene | AT5G65310 |
| is.expressed_genes | TRUE |
| is.responsive_genes | TRUE |
| is.apcluster | F-02 |

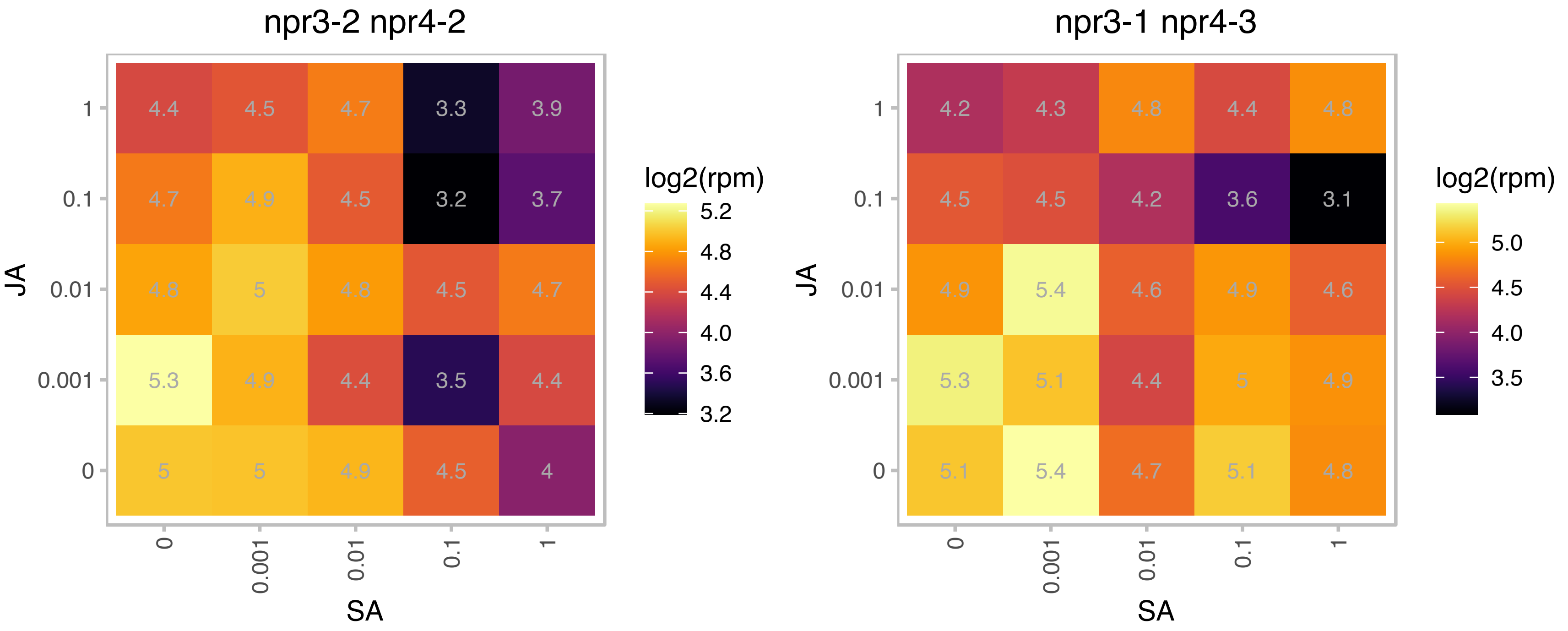

### AT5G67400.1 : RHS19

root hair specific 19

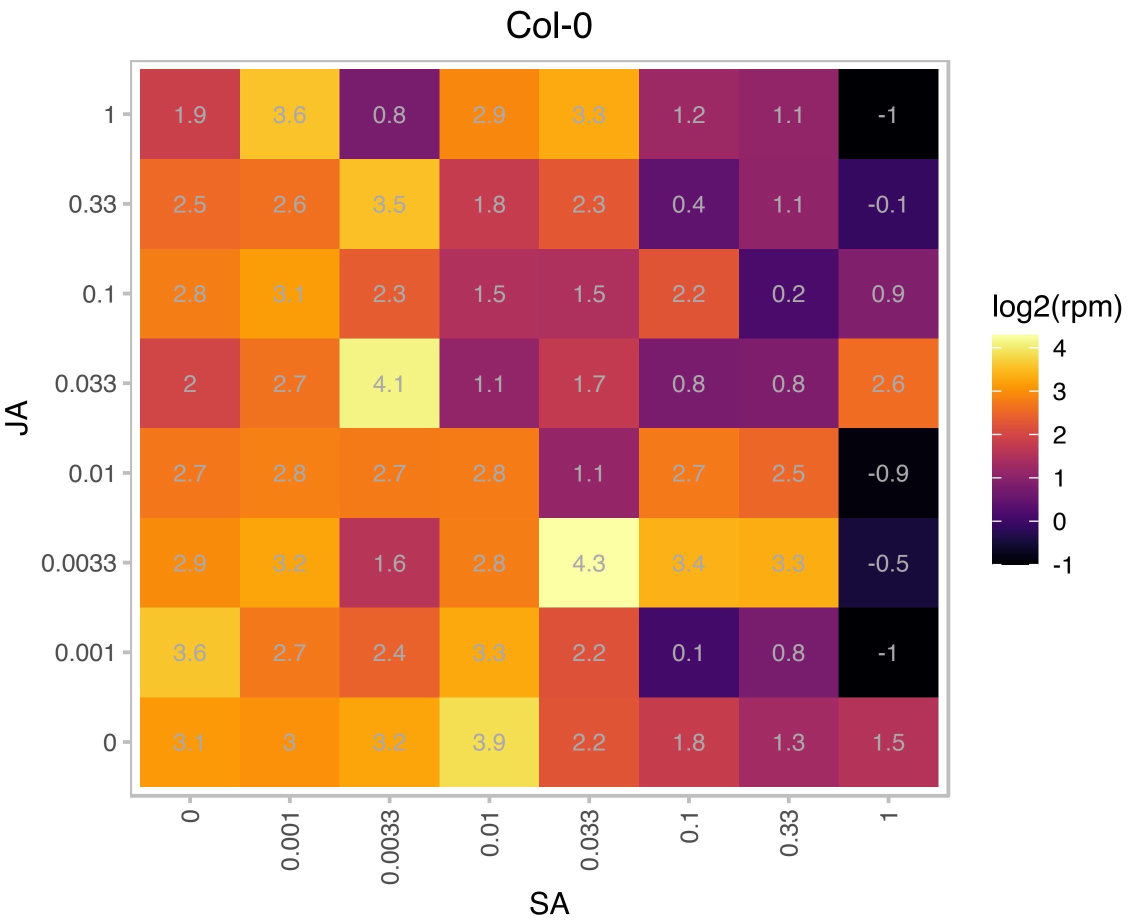


| | |
|---|---|
| Gene | AT5G67400 |
| is.expressed_genes | TRUE |
| is.responsive_genes | TRUE |
| is.apcluster | F-02 |

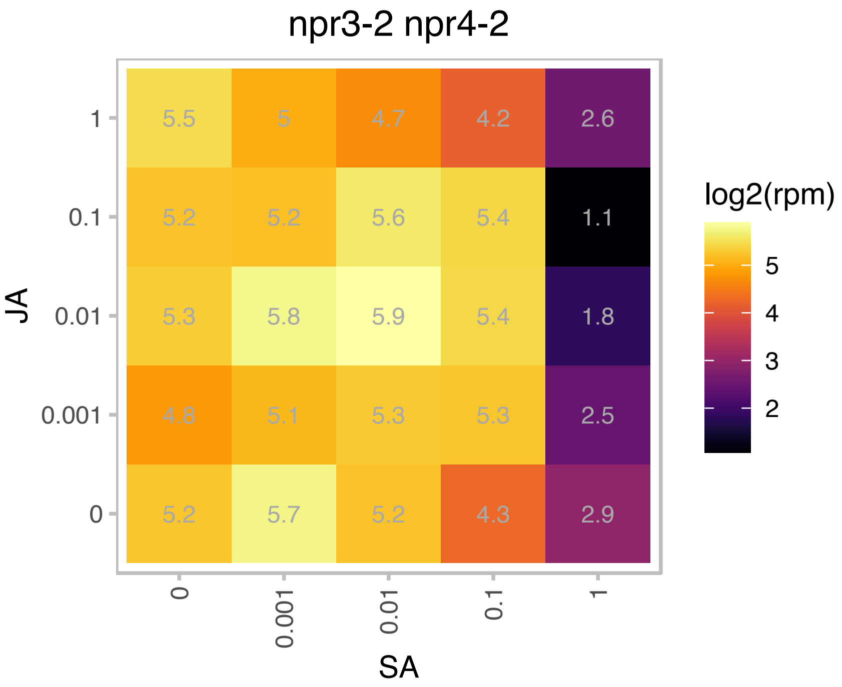


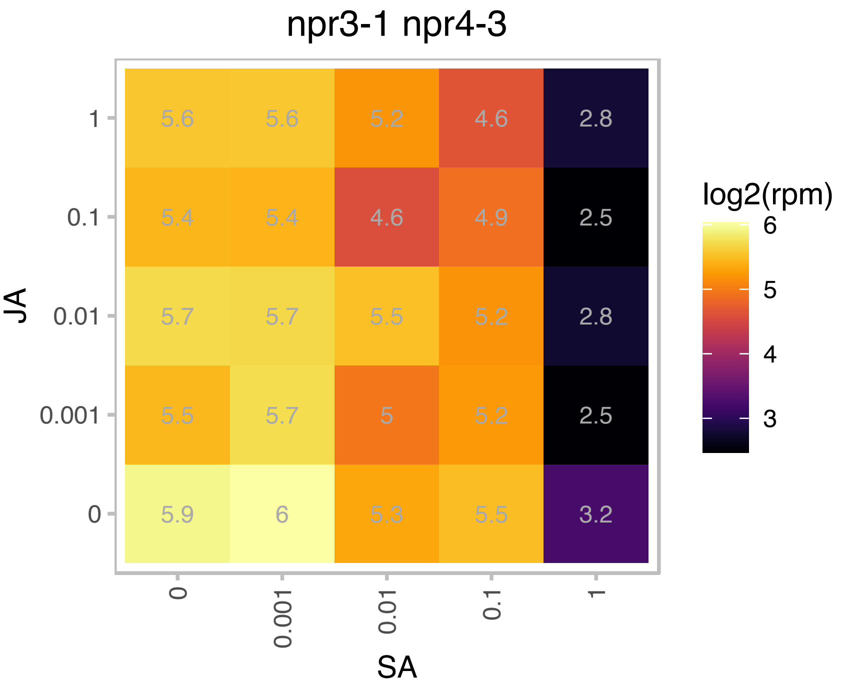

### AT1G03870.1 : FLA9

FASCICLIN-like arabinoogalactan 9

fasciclin-like arabinogalactan-protein 9 (Fla9)

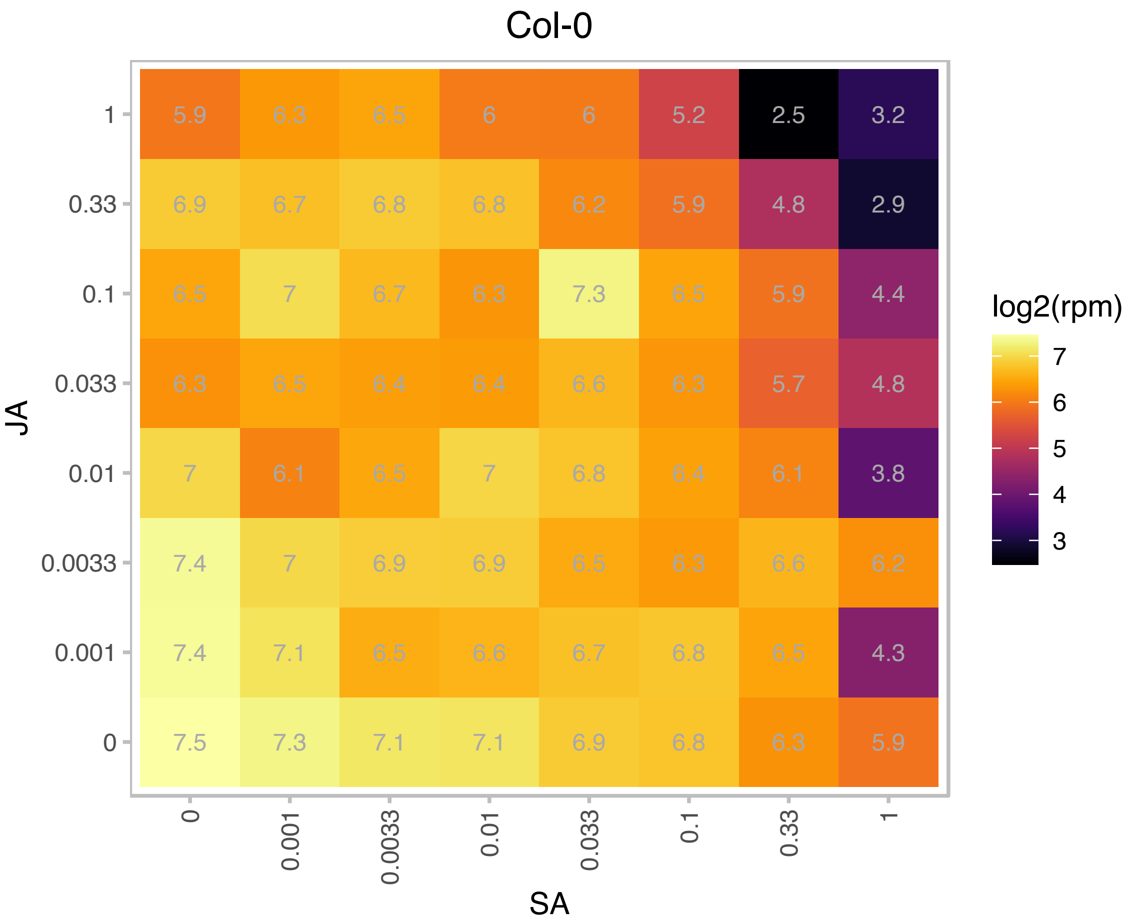


| | |
|---|---|
| Gene | AT1G03870 |
| is.expressed_genes | TRUE |
| is.responsive_genes | TRUE |
| is.apcluster | F-03 |

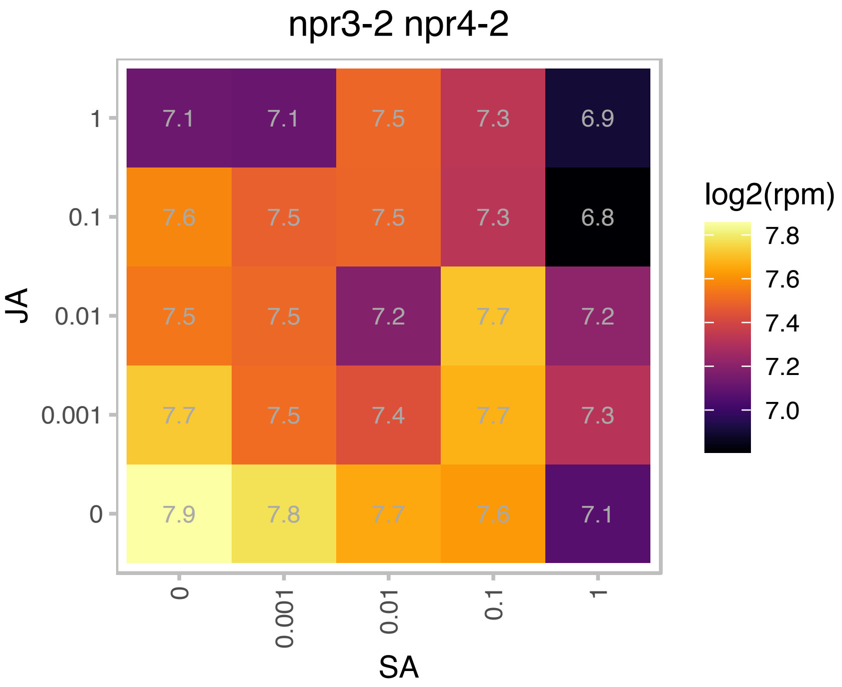


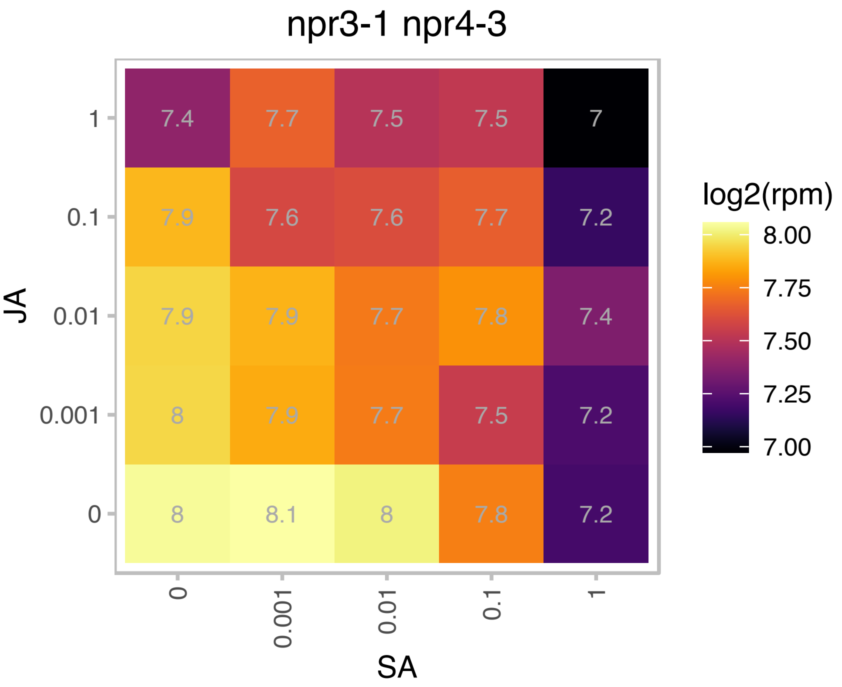

# AT1G07210.1 : NA

Ribosomal protein S18

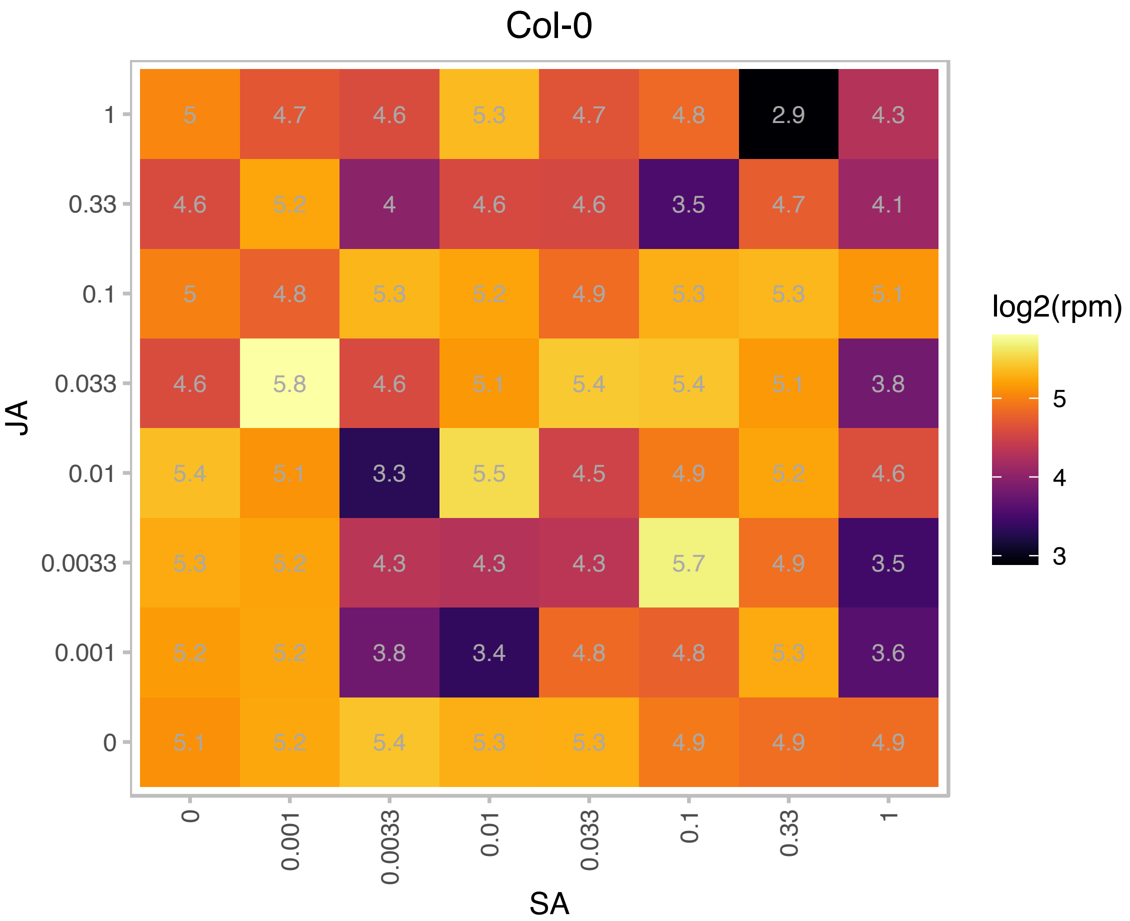


| | |
|---|---|
| Gene | AT1G07210 |
| is.expressed_genes | TRUE |
| is.responsive_genes | TRUE |
| is.apcluster | F-03 |

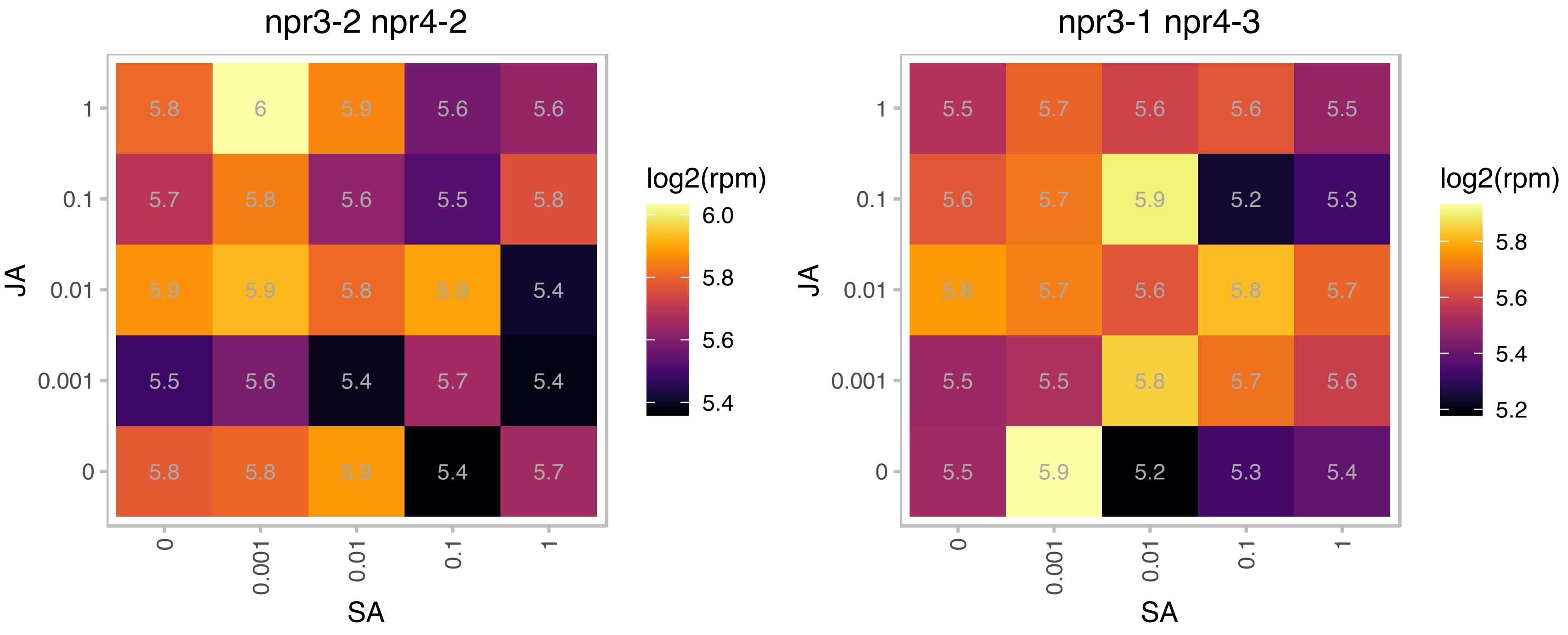

### AT1G12770.1 : EMB1586, ISE1

P-loop containing nucleoside triphosphate hydrolases superfamily protein

Encodes a DEAD-box RNA helicase that localizes to mitochondria and is essential for regulating cell-to-cell transport via plasmodesmata. The mRNA is cell-to-cell mobile.

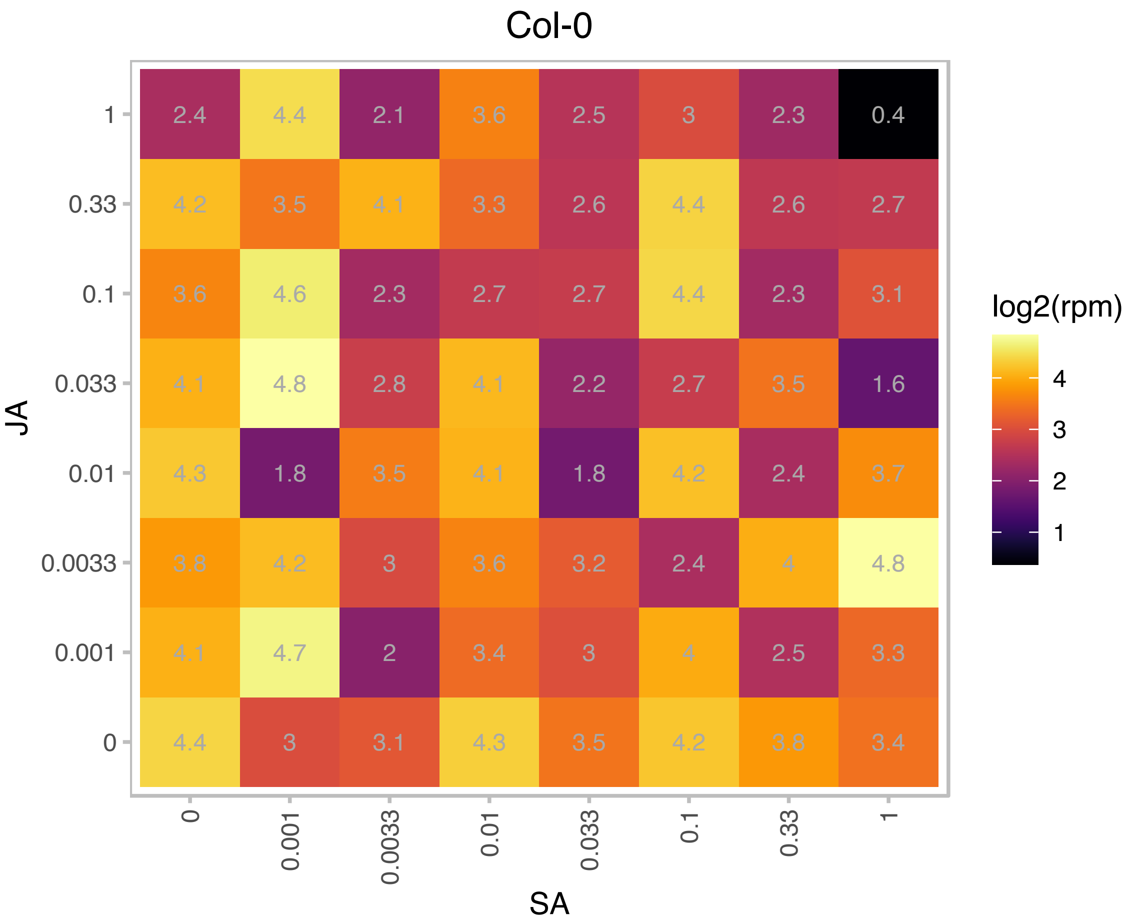


| | |
|---|---|
| Gene | AT1G12770 |
| is.expressed_genes | TRUE |
| is.responsive_genes | TRUE |
| is.apcluster | F-03 |

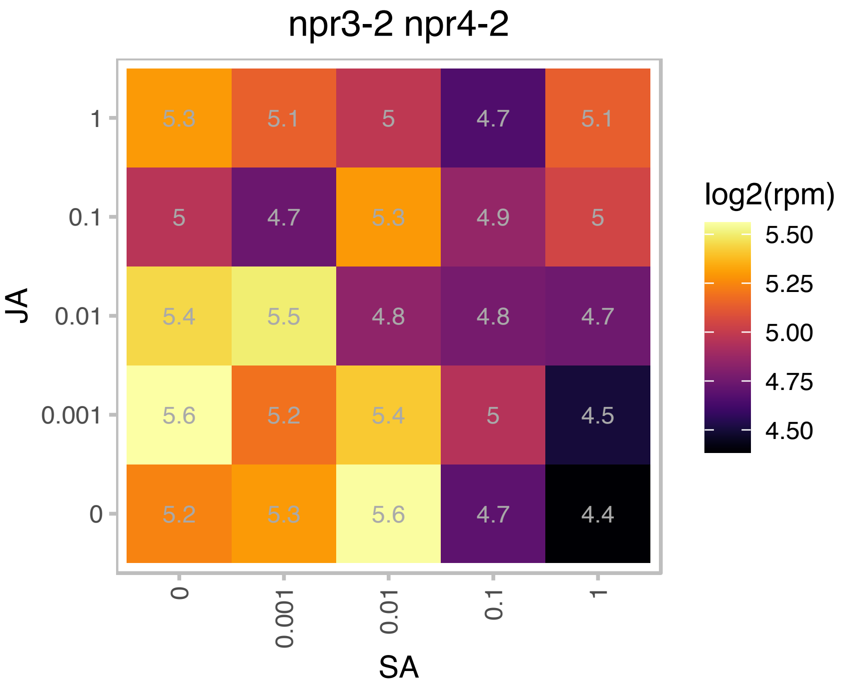


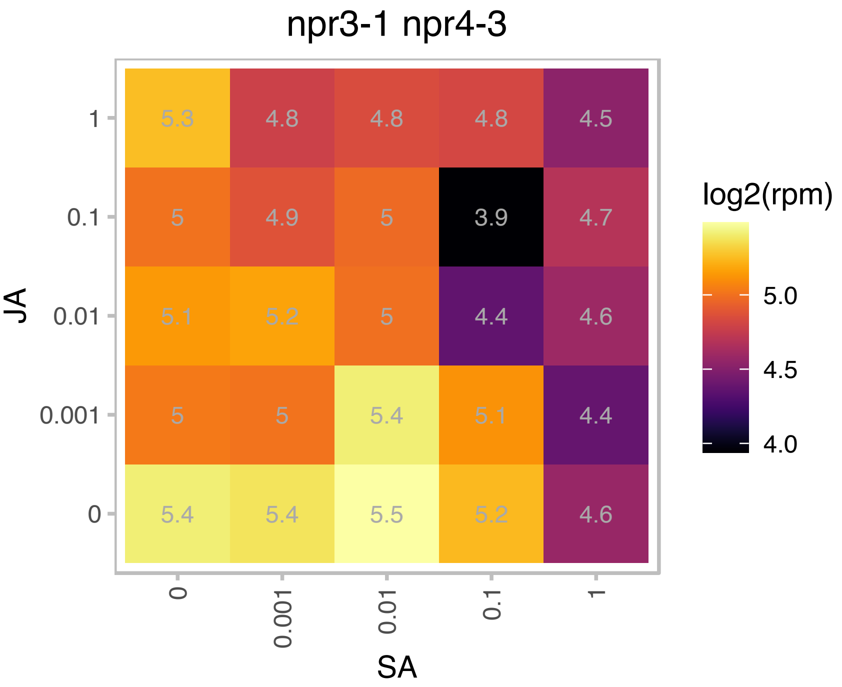

# AT1G14710.1 : NA

hydroxyproline-rich glycoprotein family protein

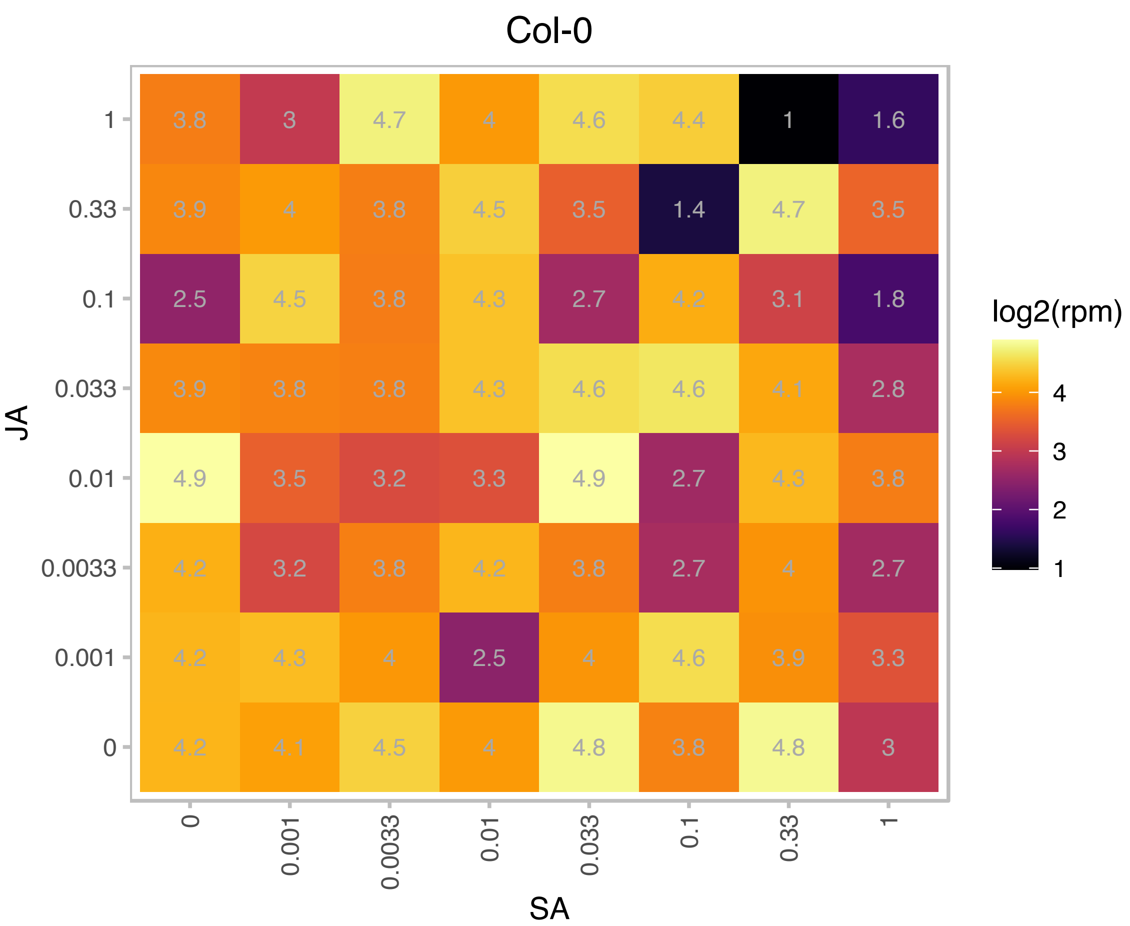


| | |
|---|---|
| Gene | AT1G14710 |
| is.expressed_genes | TRUE |
| is.responsive_genes | TRUE |
| is.apcluster | F-03 |

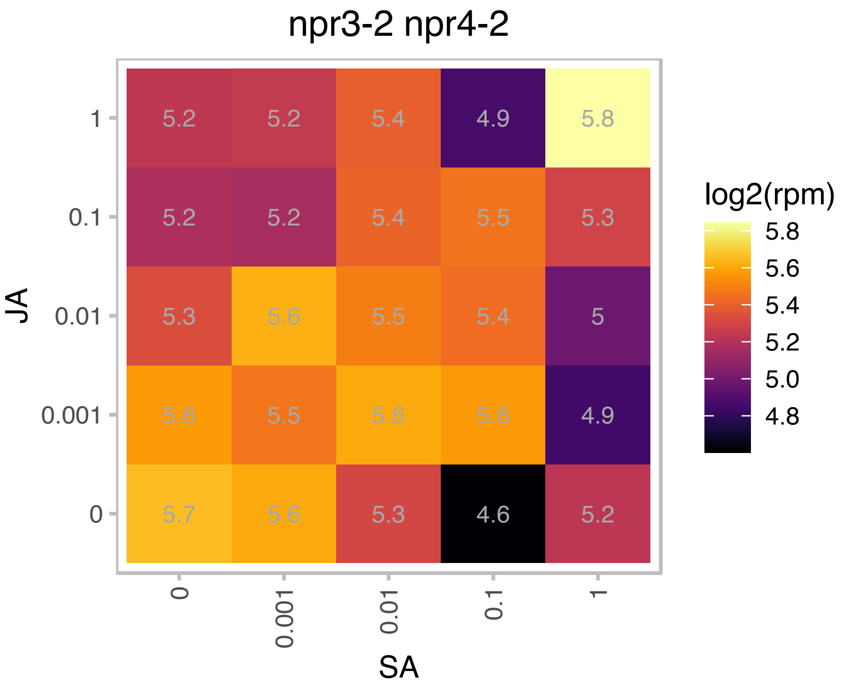


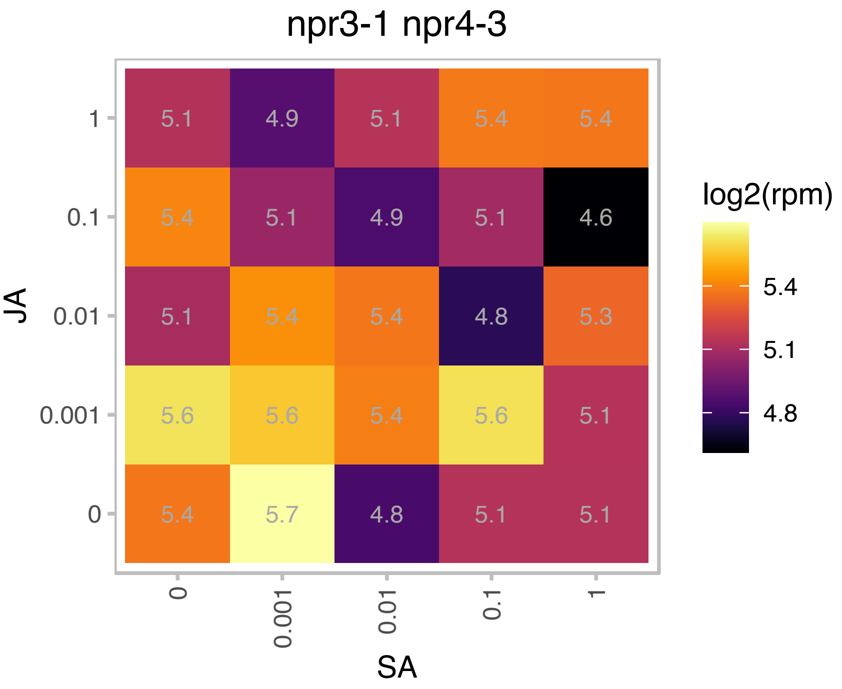

### AT1G14720.1 : ATXTH28, EXGT-A2, XTH28, XTR2

xyloglucan endotransglucosylase/hydrolase 28

member of Glycoside Hydrolase Family 16

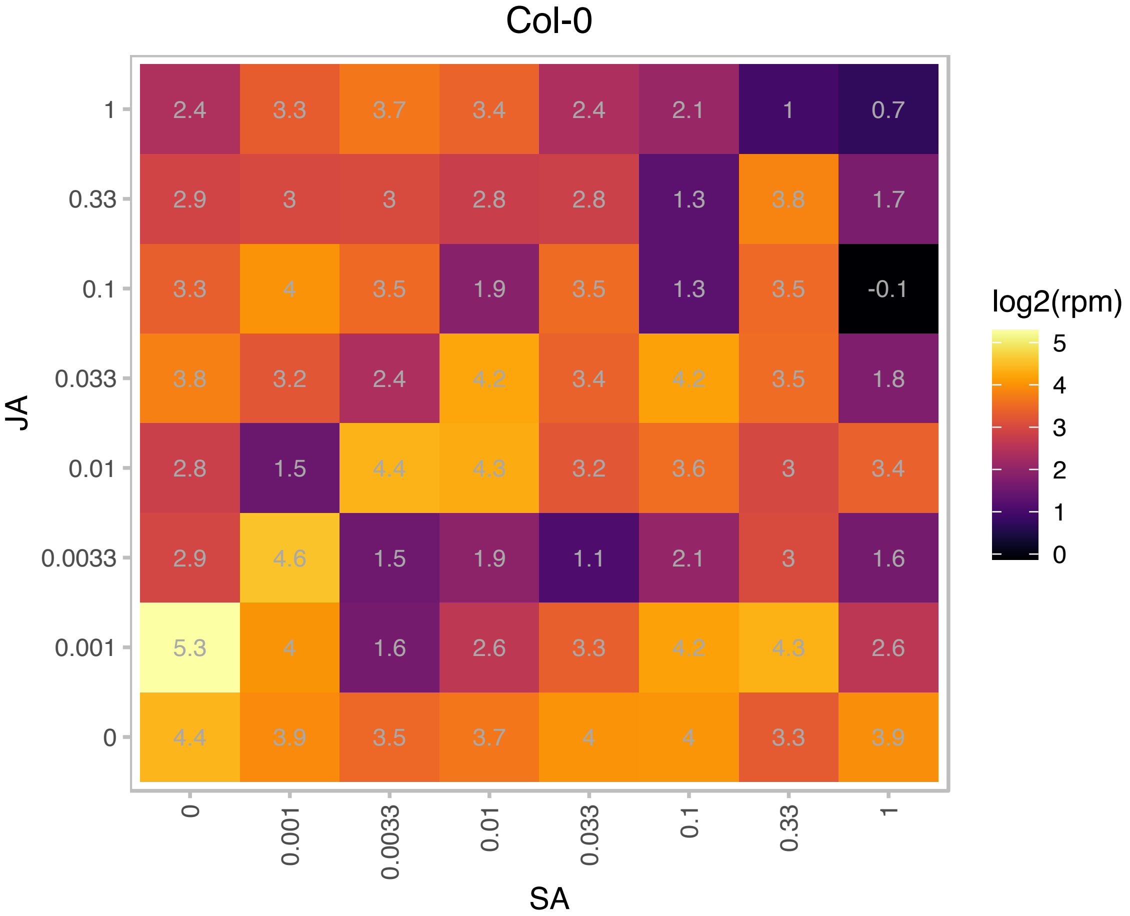


| | |
|---|---|
| Gene | AT1G14720 |
| is.expressed_genes | TRUE |
| is.responsive_genes | TRUE |
| is.apcluster | F-03 |

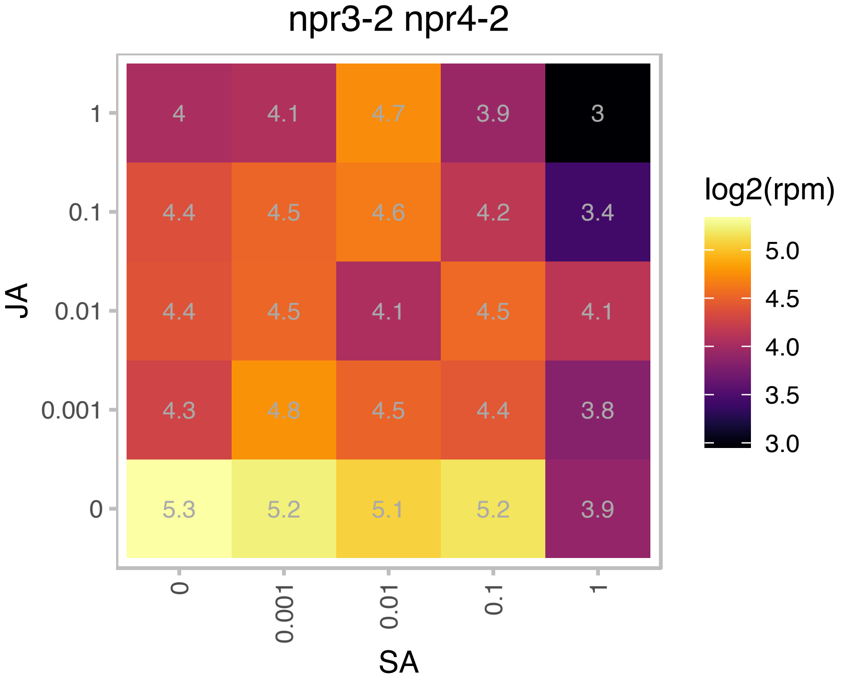


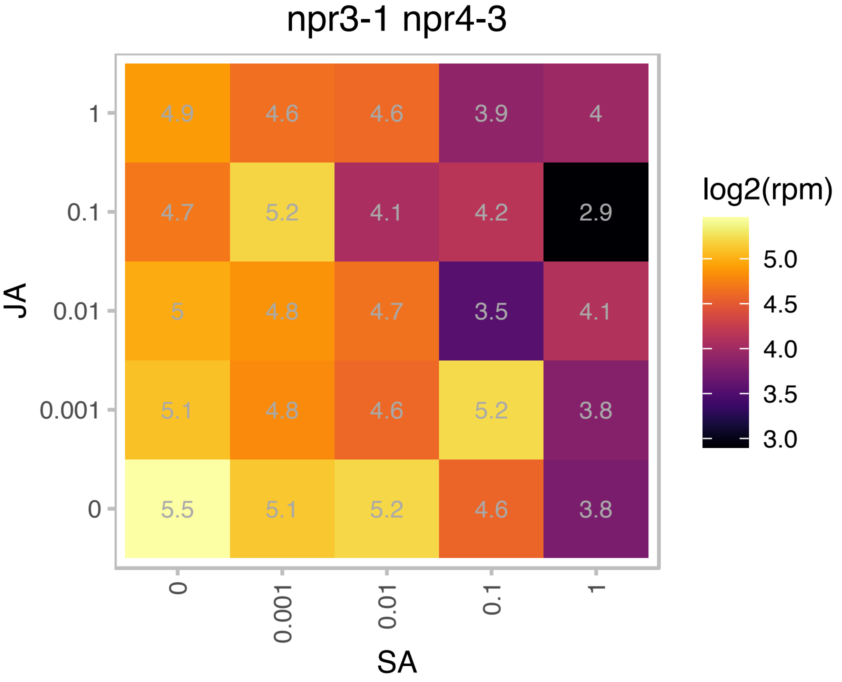

# AT1G17510.1 : NA

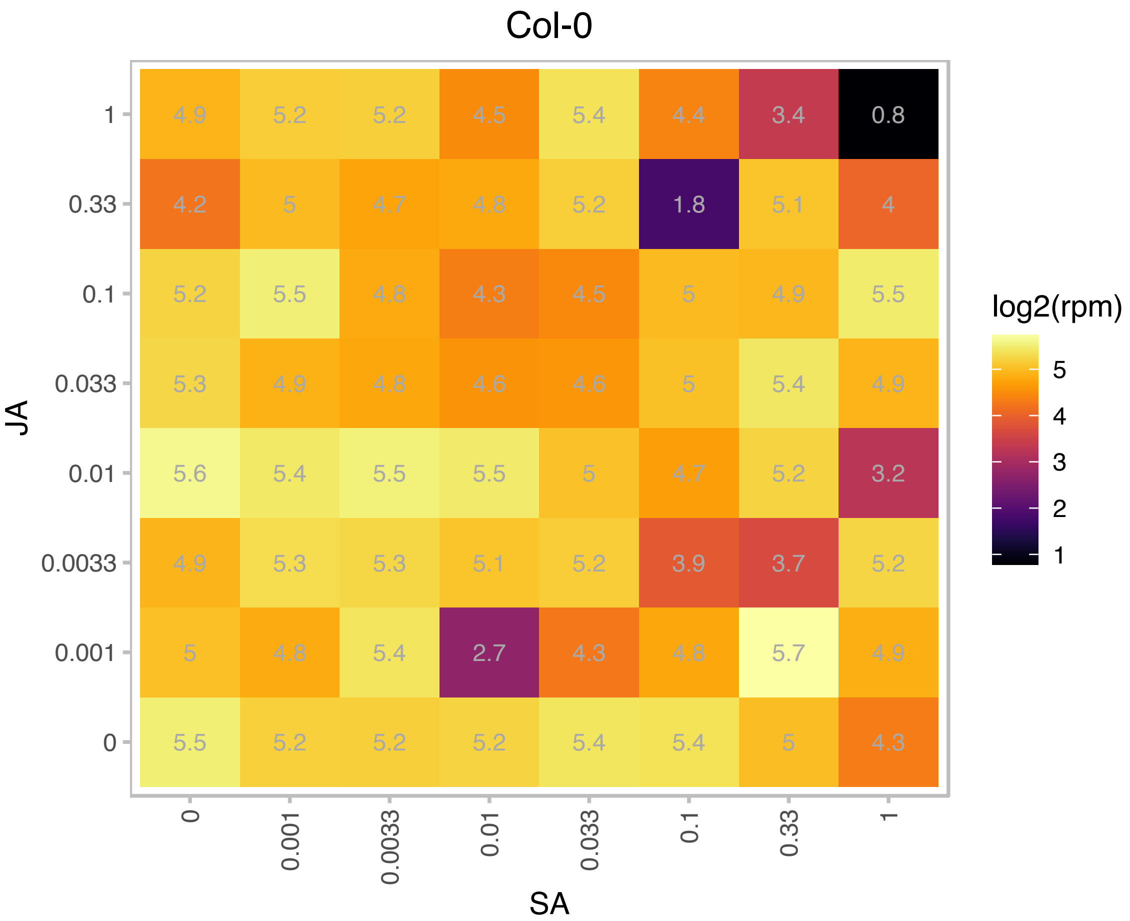


| | |
|---|---|
| Gene | AT1G17510 |
| is.expressed_genes | TRUE |
| is.responsive_genes | TRUE |
| is.apcluster | F-03 |

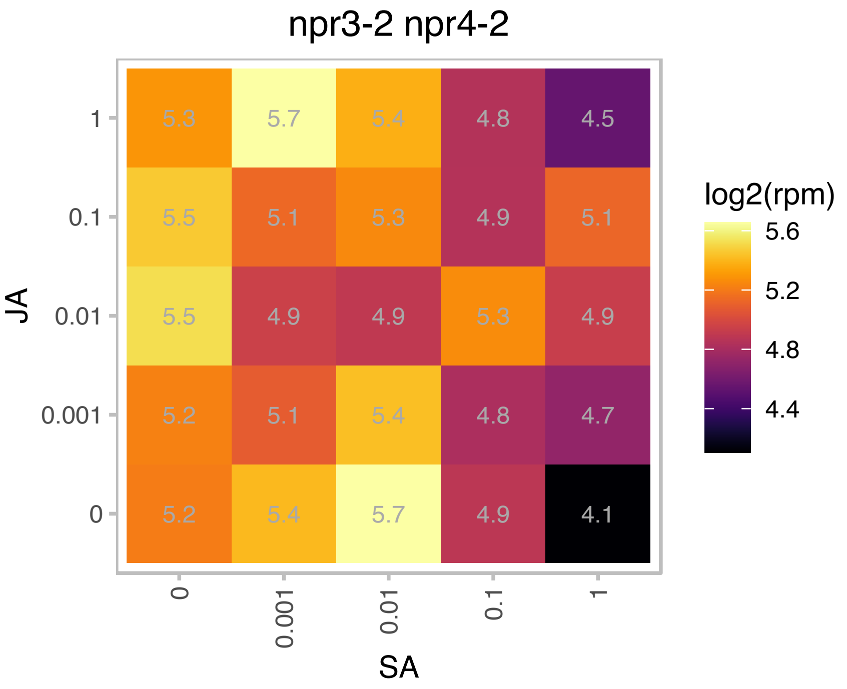


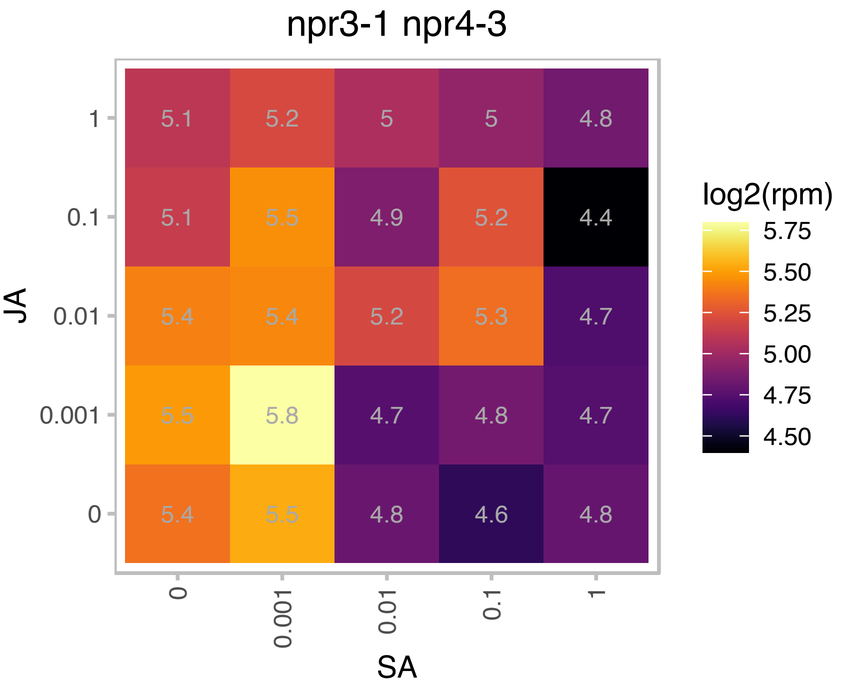

### AT1G18620.1 : LNG4, TRM3

Member of a small gene family in Arabidopsis. Quadruple mutants in this family display defects in cell elongation.

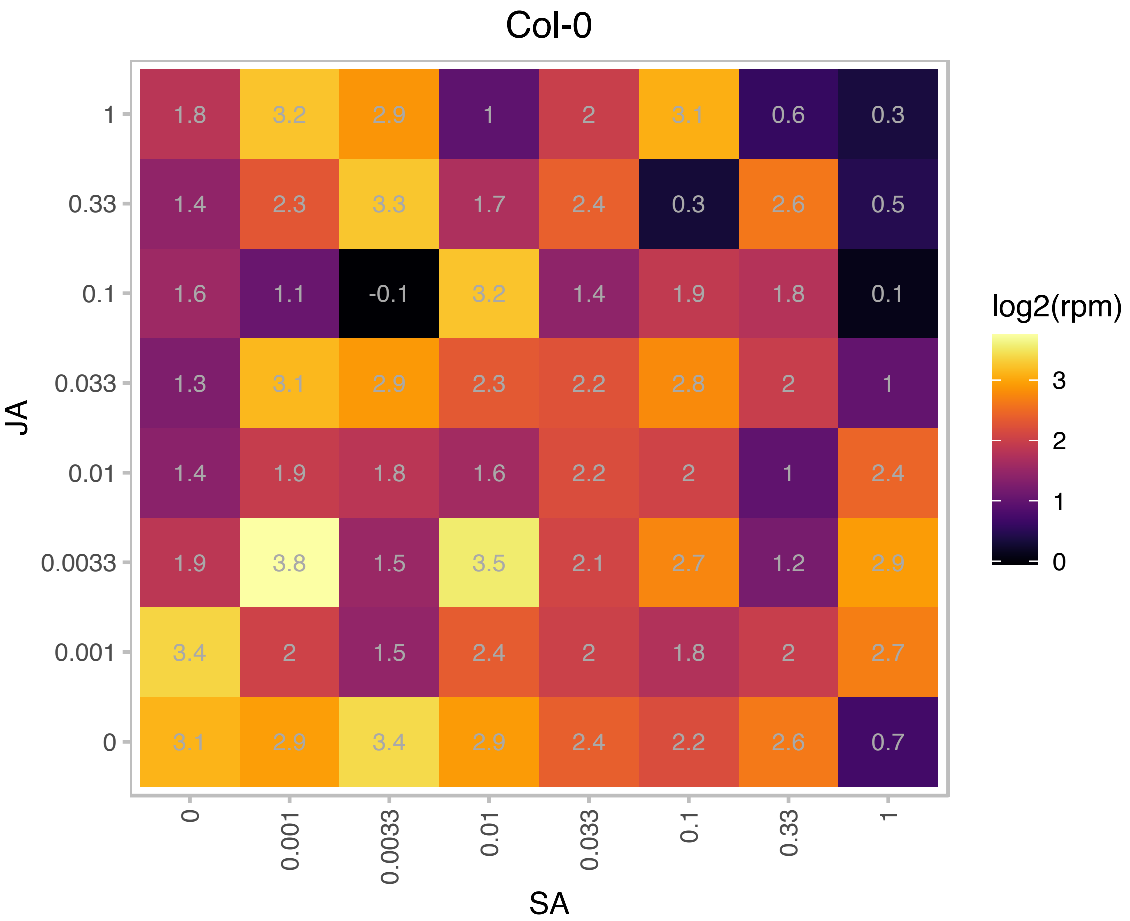


| | |
|---|---|
| Gene | AT1G18620 |
| is.expressed_genes | TRUE |
| is.responsive_genes | TRUE |
| is.apcluster | F-03 |

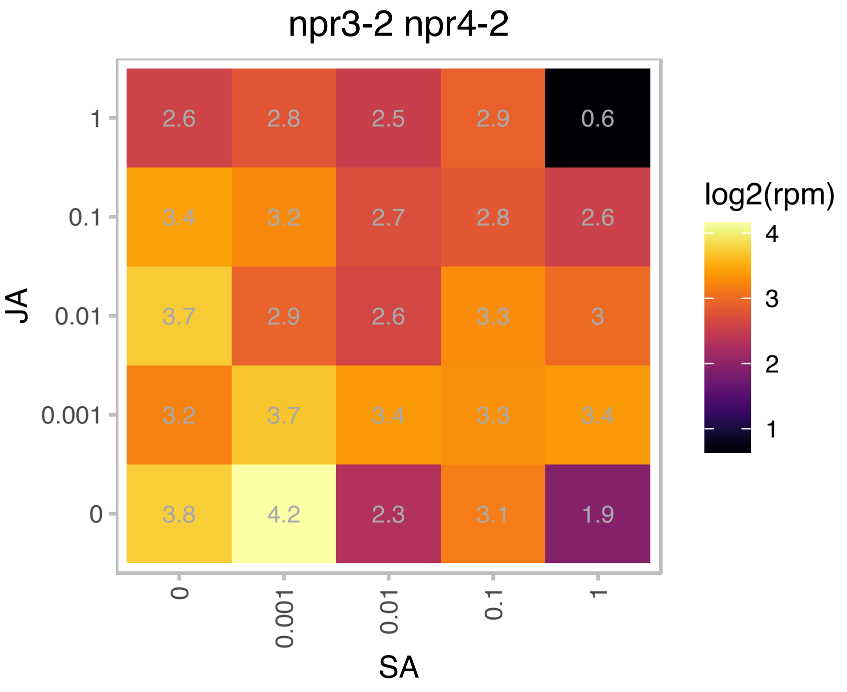


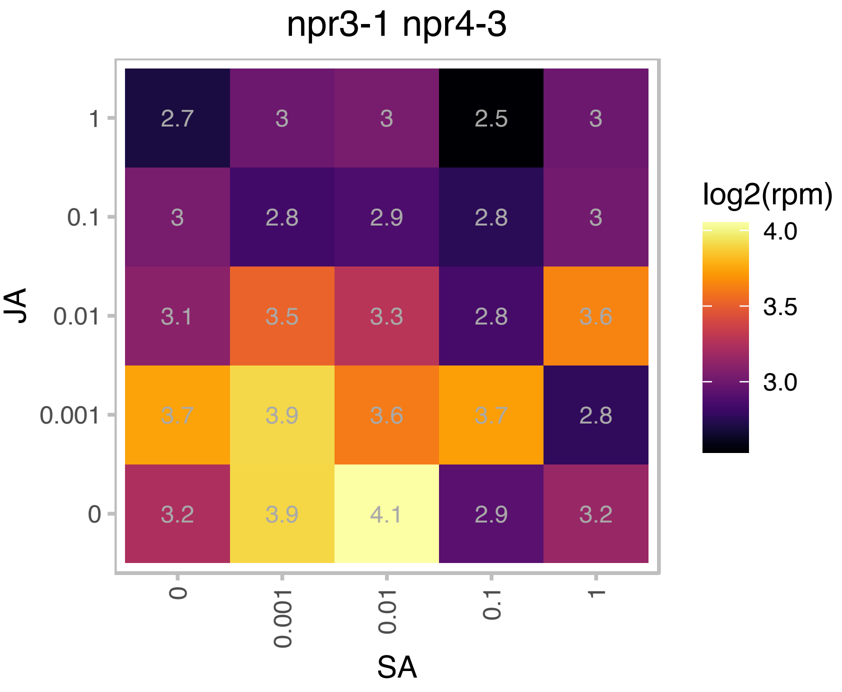

# AT1G19240.1 : NA

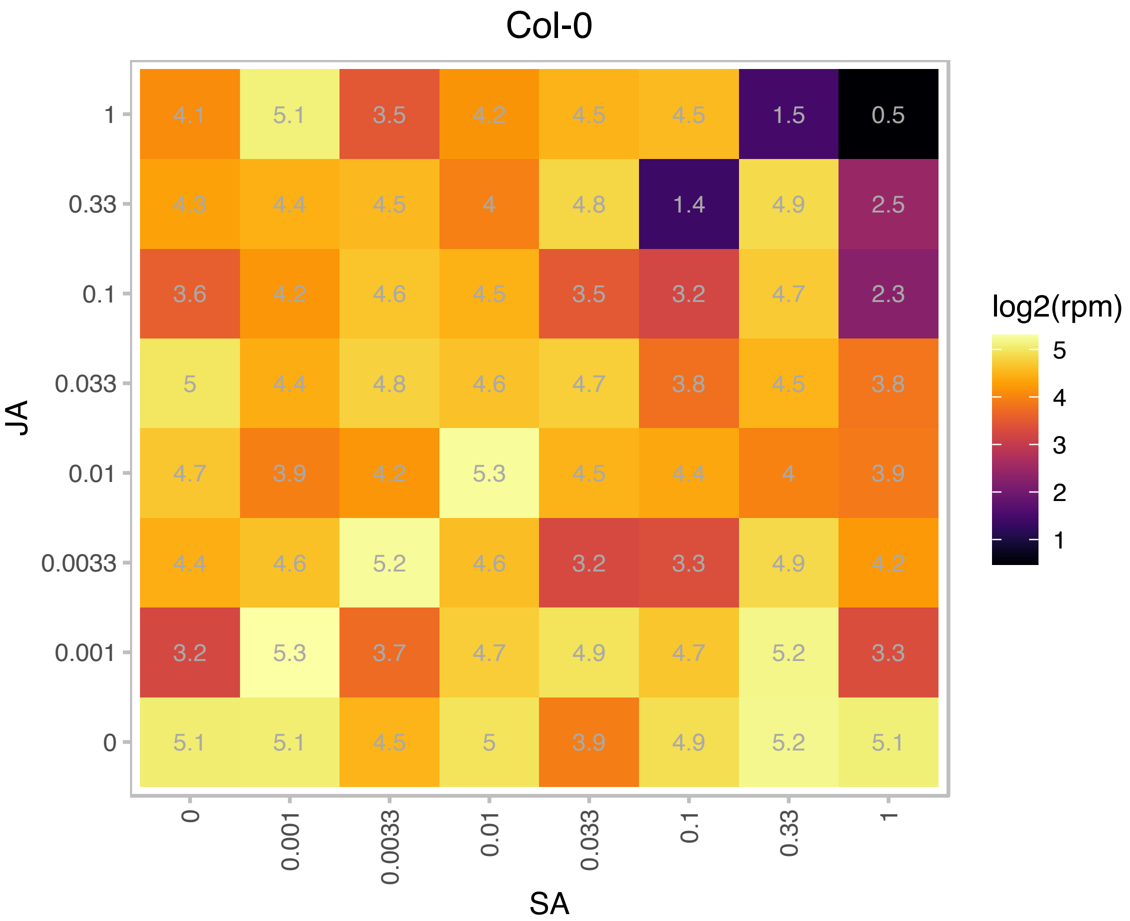


| | |
|---|---|
| Gene | AT1G19240 |
| is.expressed_genes | TRUE |
| is.responsive_genes | TRUE |
| is.apcluster | F-03 |

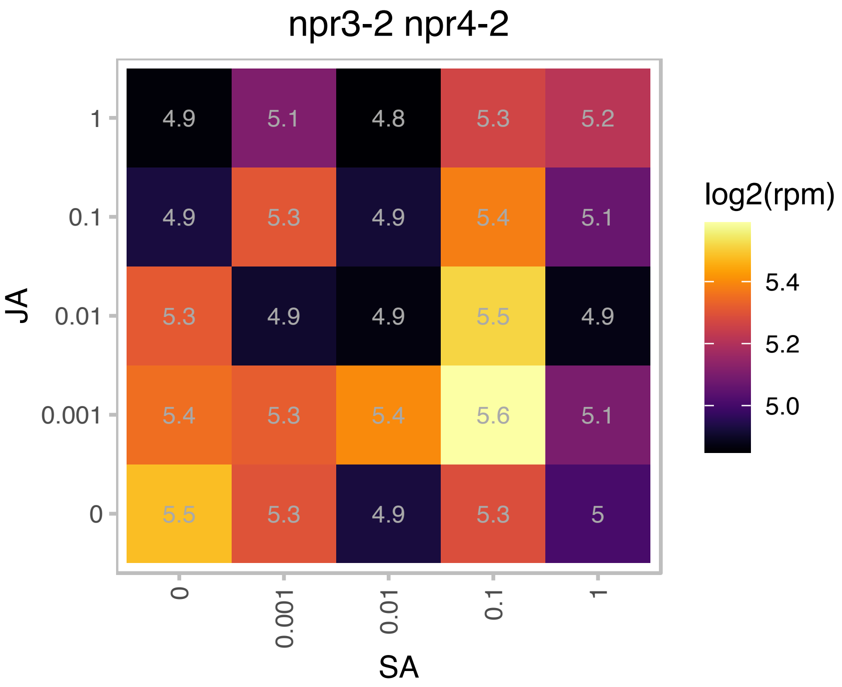


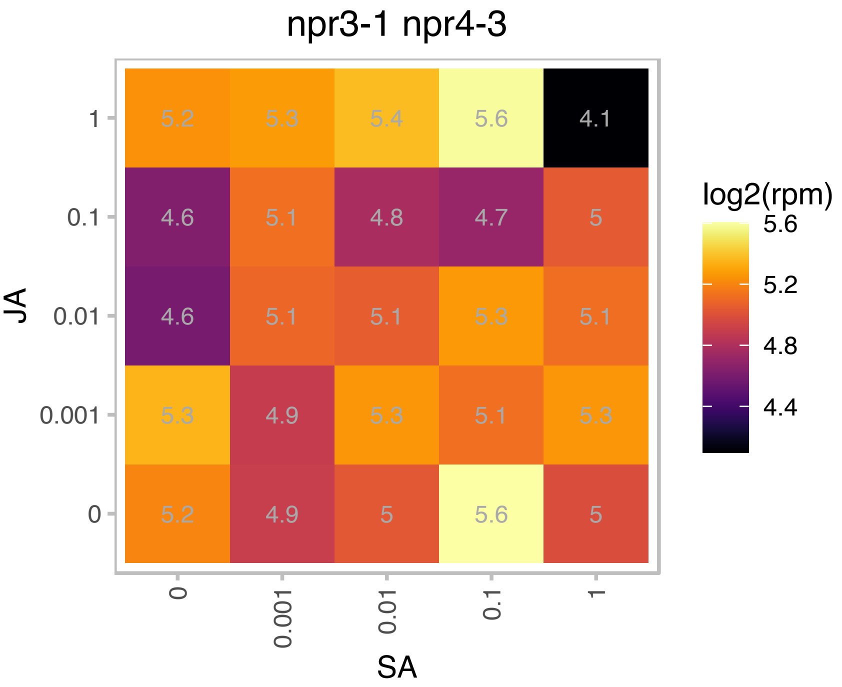

# AT1G20950.1 : NA

Phosphofructokinase family protein

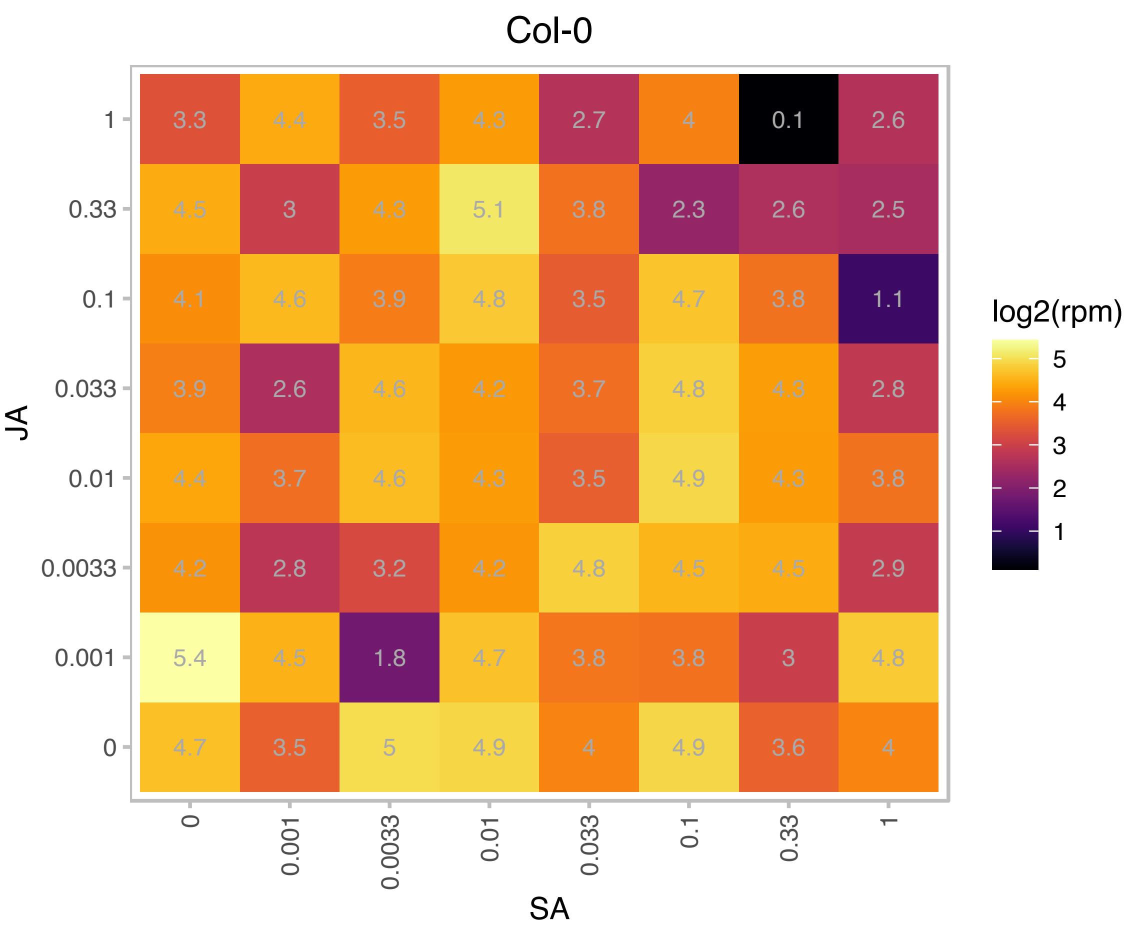


| | |
|---|---|
| Gene | AT1G20950 |
| is.expressed_genes | TRUE |
| is.responsive_genes | TRUE |
| is.apcluster | F-03 |

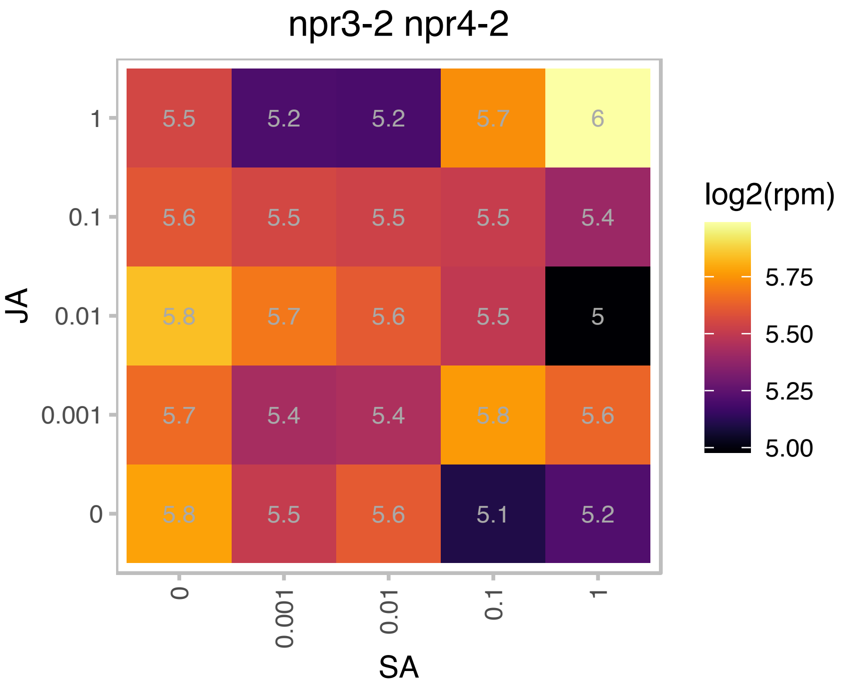


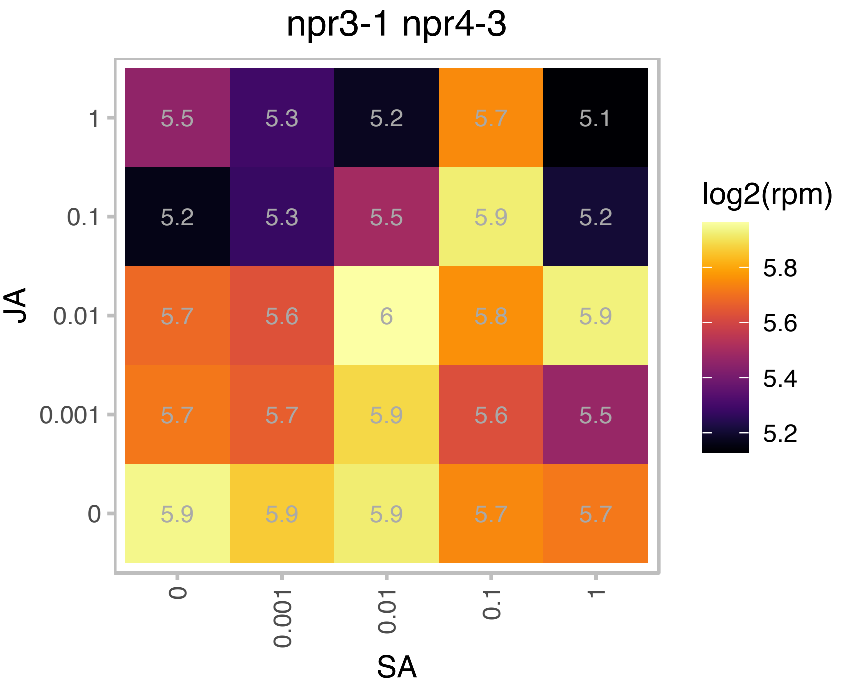

# AT1G24050.1 : NA

RNA-processing, Lsm domain

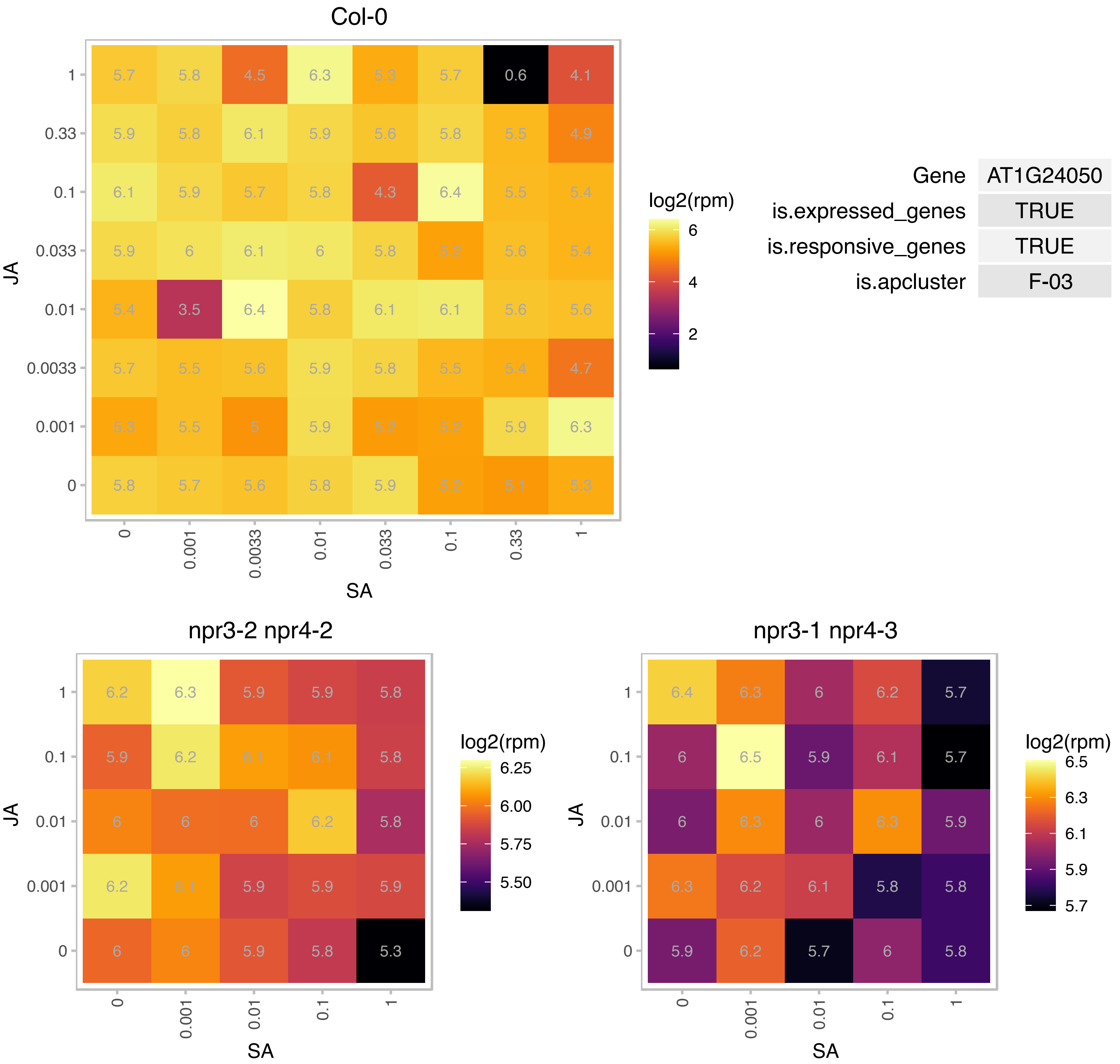


| | |
|---|---|
| Gene | AT1G24050 |
| is.expressed_genes | TRUE |
| is.responsive_genes | TRUE |
| is.apcluster | F-03 |

# AT1G24160.1 : NA

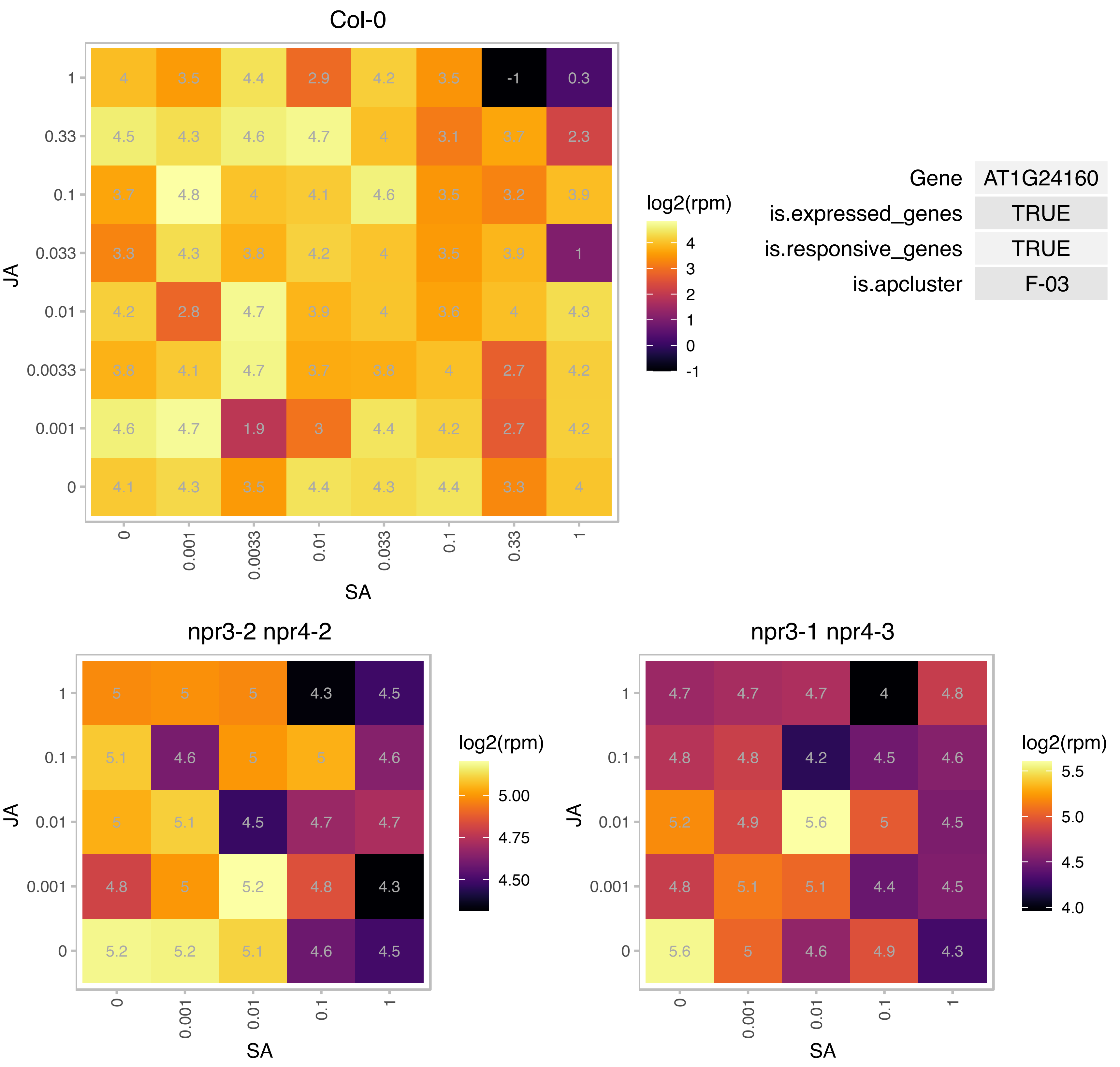


| | |
|---|---|
| Gene | AT1G24160 |
| is.expressed_genes | TRUE |
| is.responsive_genes | TRUE |
| is.apcluster | F-03 |

### AT1G29900.1 : CARB, VEN3

carbamoyl phosphate synthetase B
Encodes carbamoyl phosphate synthetase (CPS) large subunit (CARB), also named as VEN3. Heterologous expression of the Arabidopsis VEN3 and VEN6 genes in a CPS-deficient Escherichia coli strain fully restored bacterial growth in minimal medium, demonstrating the enzymatic activity of the VEN3 and VEN6 proteins.

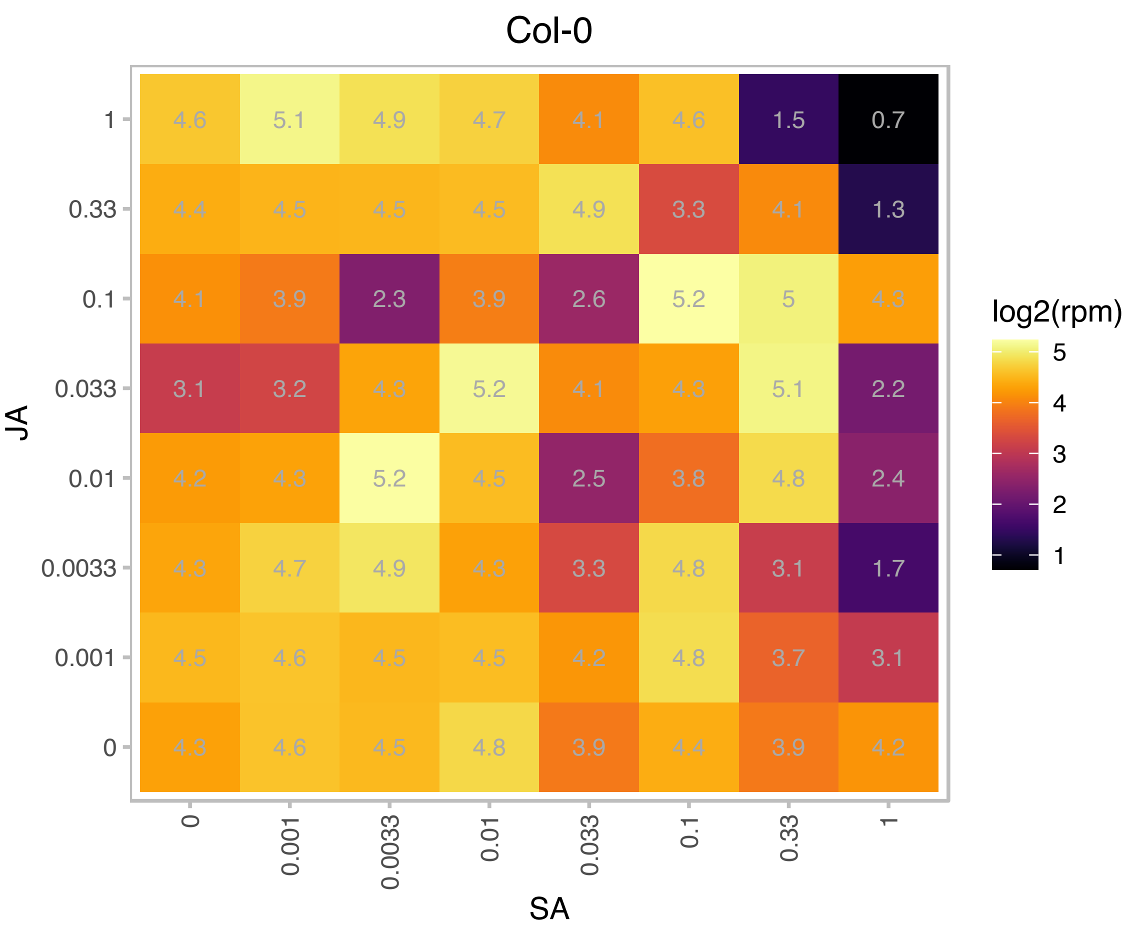


| | |
|---|---|
| Gene | AT1G29900 |
| is.expressed_genes | TRUE |
| is.responsive_genes | TRUE |
| is.apcluster | F-03 |

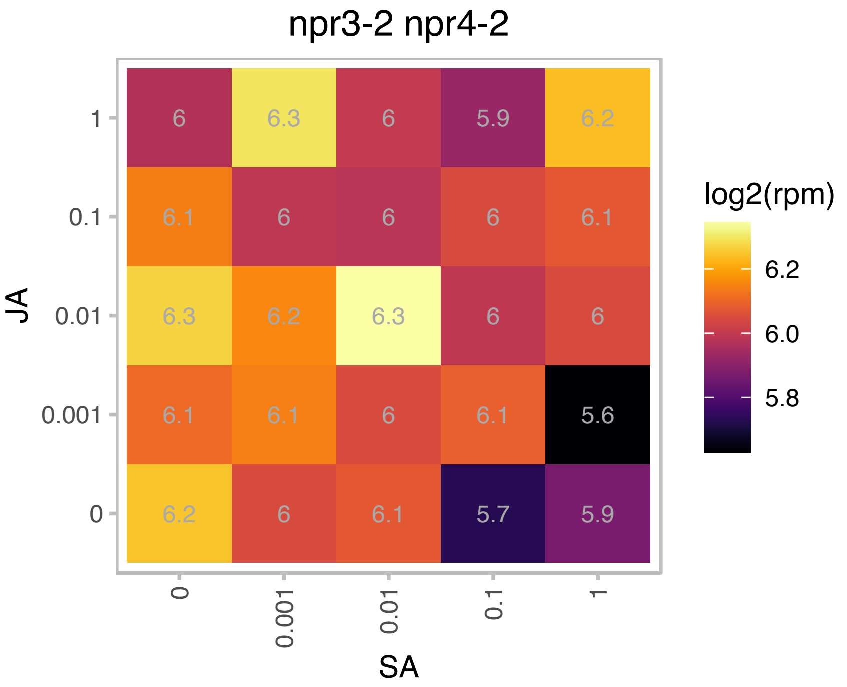


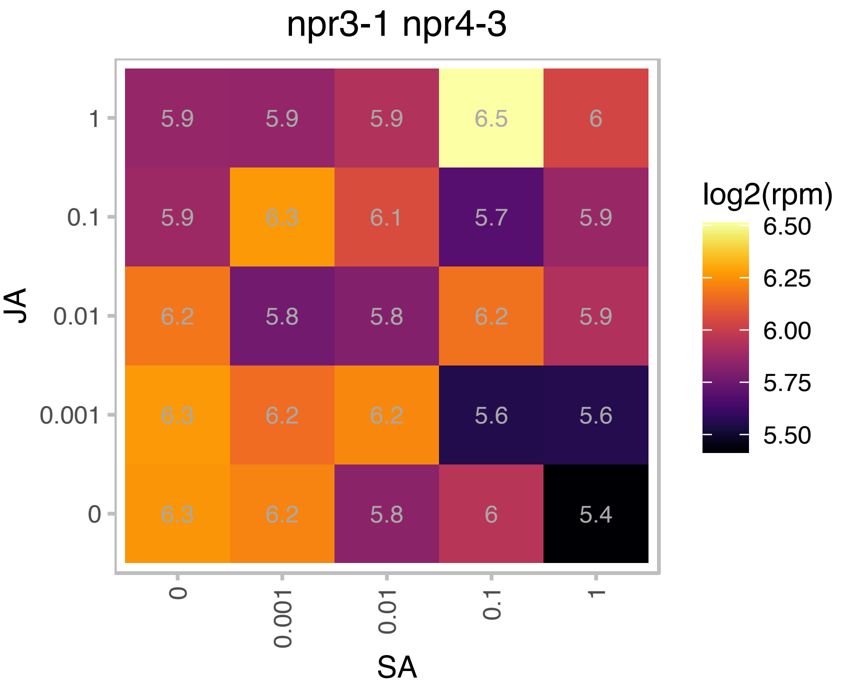

### AT1G31817.1 : NFD3

Ribosomal L18p/L5e family protein

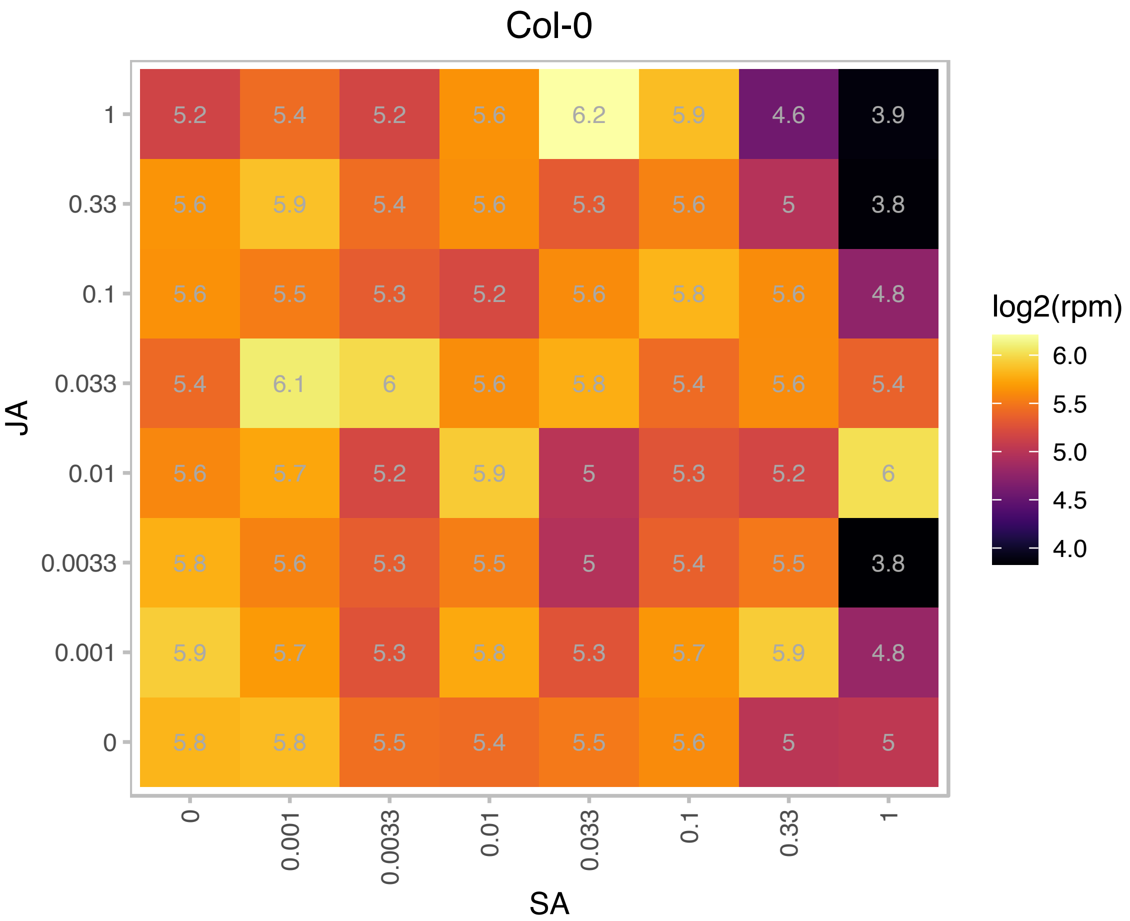


| | |
|---|---|
| Gene | AT1G31817 |
| is.expressed_genes | TRUE |
| is.responsive_genes | TRUE |
| is.apcluster | F-03 |

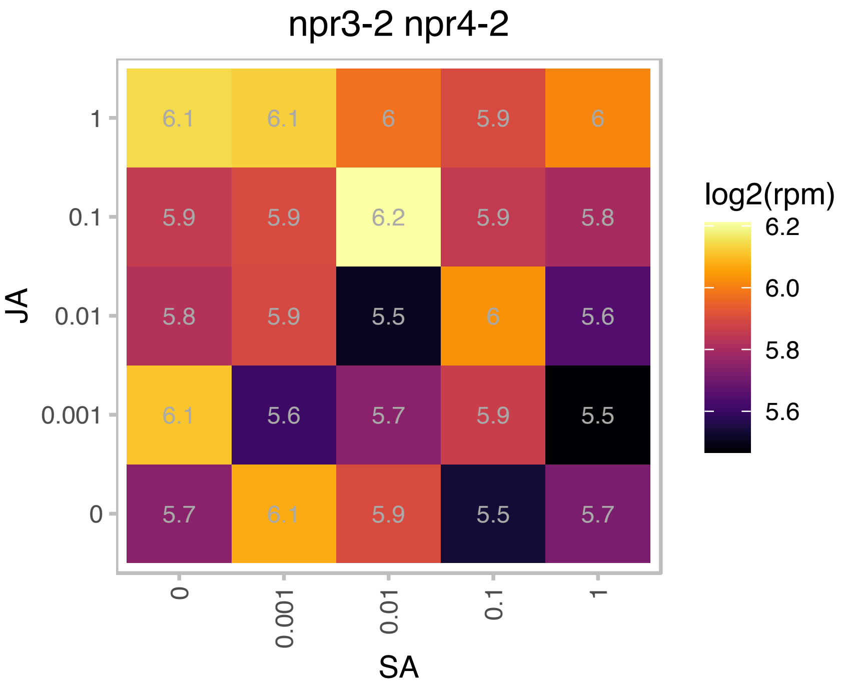


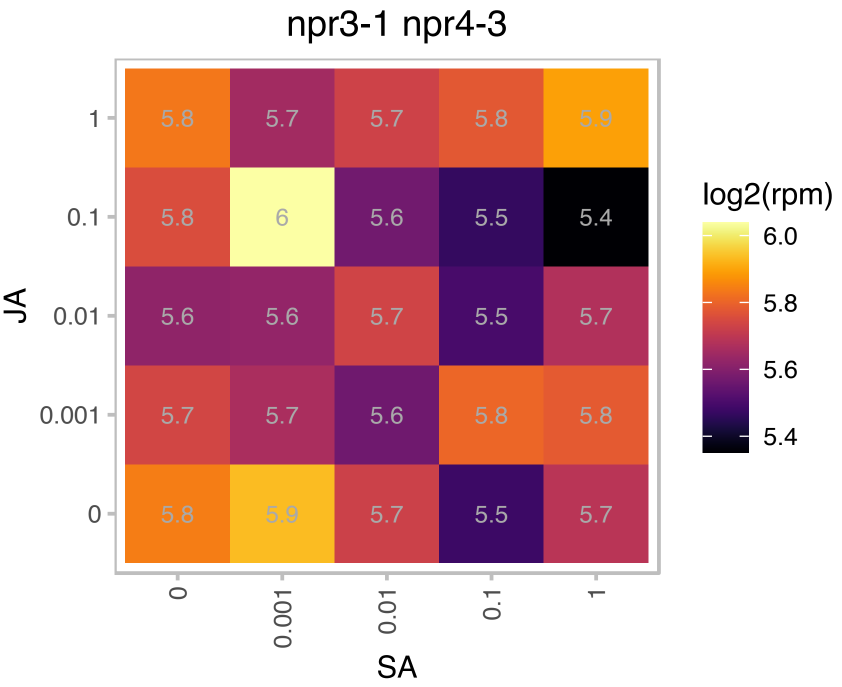

# AT1G36390.1 : NA

Co-chaperone GrpE family protein

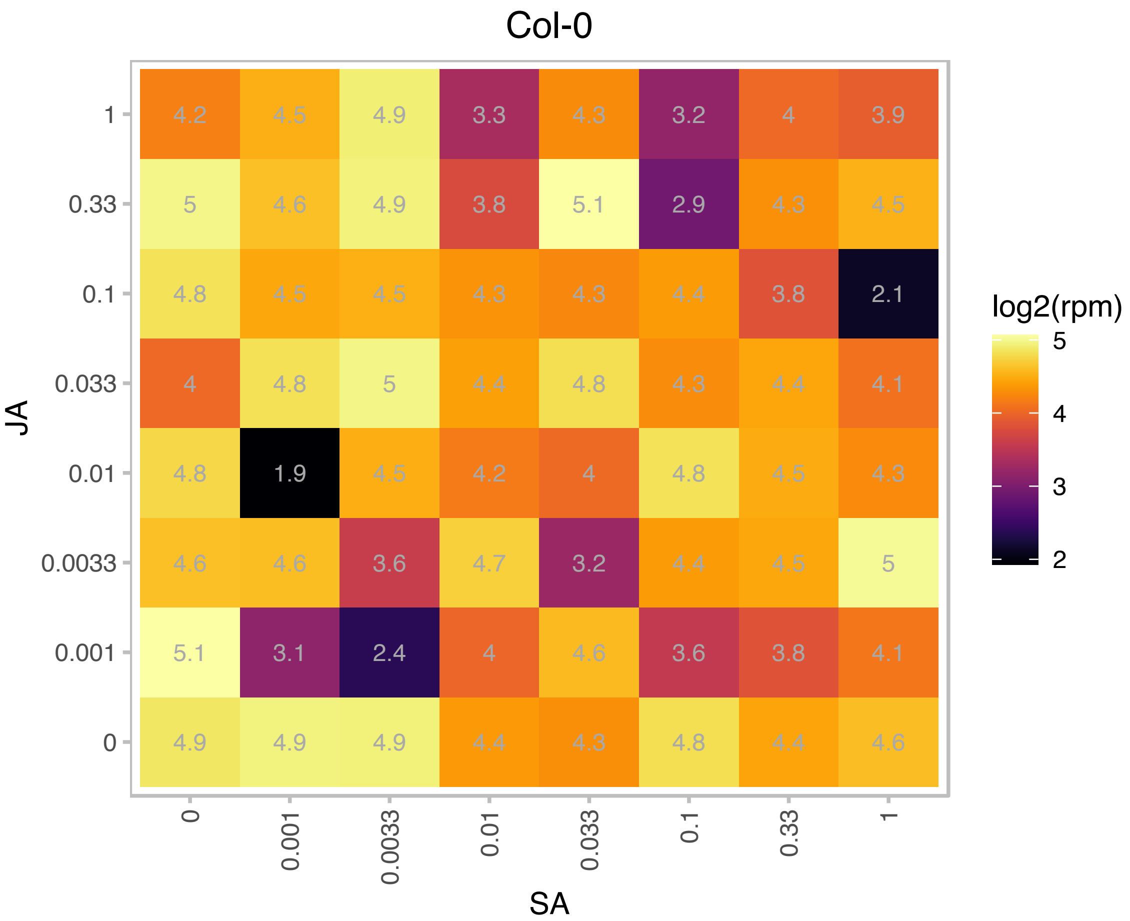


| | |
|---|---|
| Gene | AT1G36390 |
| is.expressed_genes | TRUE |
| is.responsive_genes | TRUE |
| is.apcluster | F-03 |

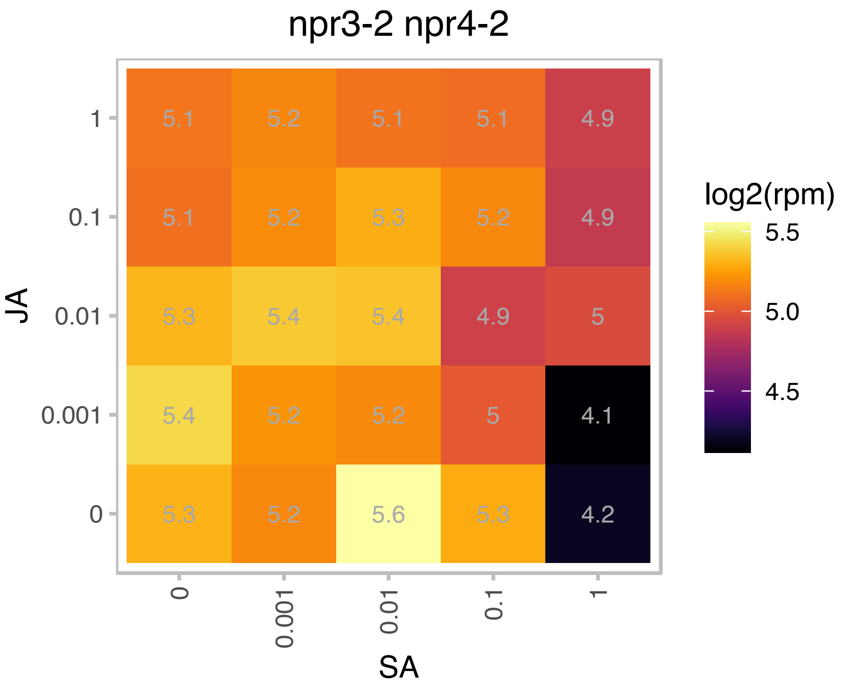


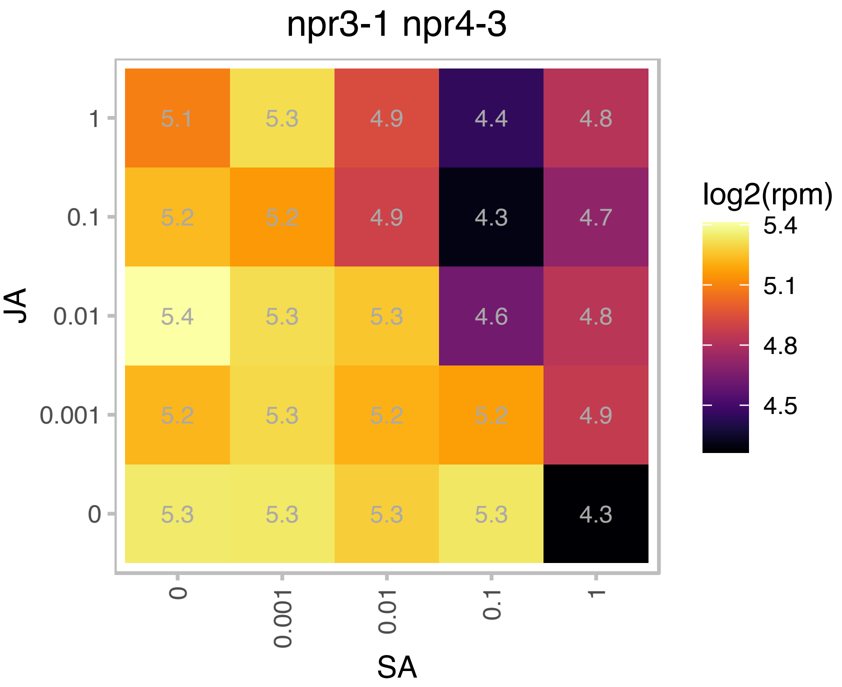

### AT1G48920.1 : ATNUC-L1, NUC-L1, NUC1, PARL1

Encodes ATNUC-L1 (NUCLEOLIN LIKE 1), the predominant form of the two nucleolin proteins found in Arabidopsis.
nucleolin like 1
This protein is involved in rRNA processing, ribosome biosynthesis, and vascular pattern formation. PARL1 localizes to the nucleolus and parl1 mutants accumulate elevated levels of the unspliced 35S pre-rRNA. parl1 mutants also have defects in cotyledon, leaf, sepal, and petal vein patterning and have reduced stature, reduced fertility, increased bushiness, and reduced root length. The sugar-induced expression of ribosome proteins is also reduced in parl1 mutants. The mRNA is cell-to-cell mobile.

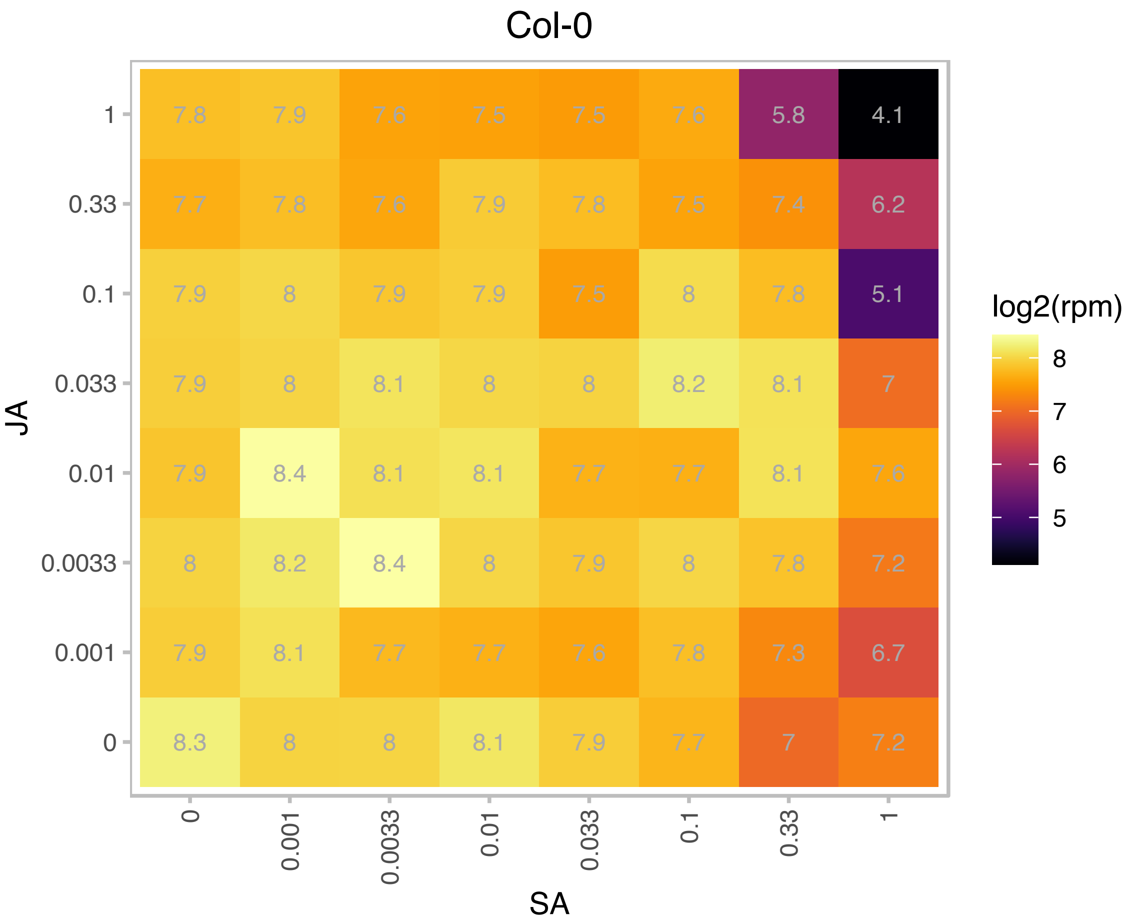


| | |
|---|---|
| Gene | AT1G48920 |
| is.expressed_genes | TRUE |
| is.responsive_genes | TRUE |
| is.apcluster | F-03 |

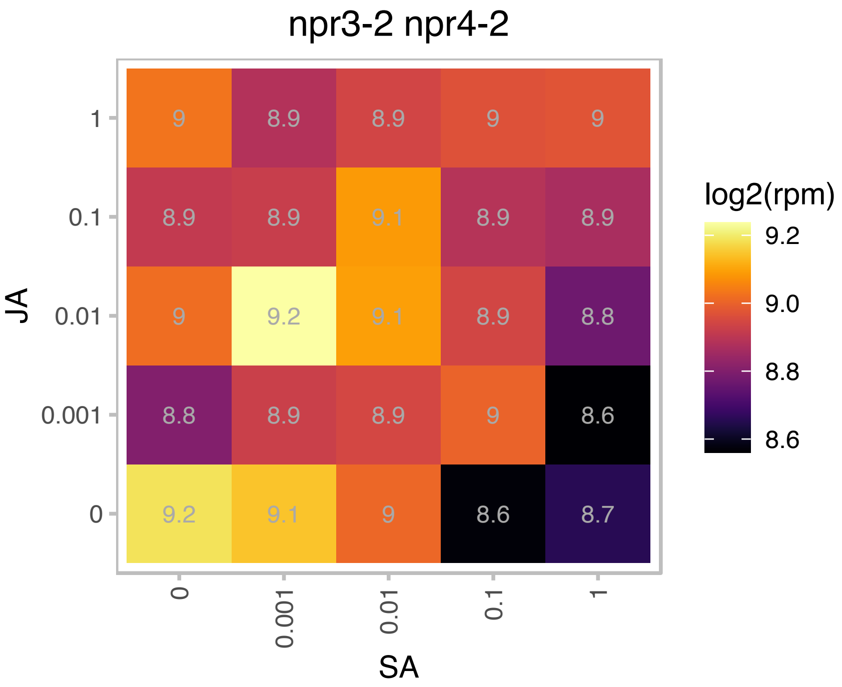


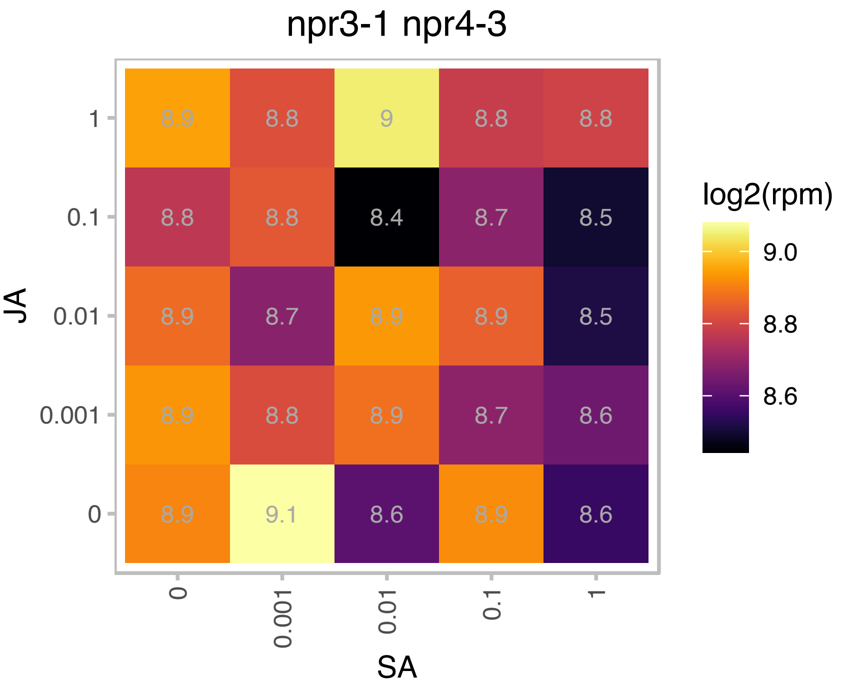

### AT1G54060.1 : ASIL1

6B-interacting protein 1-like 1

Member of the trihelix DNA binding protein family. Nuclear localized. Involved in repressing seed maturation genes during seed germination and seedling development.

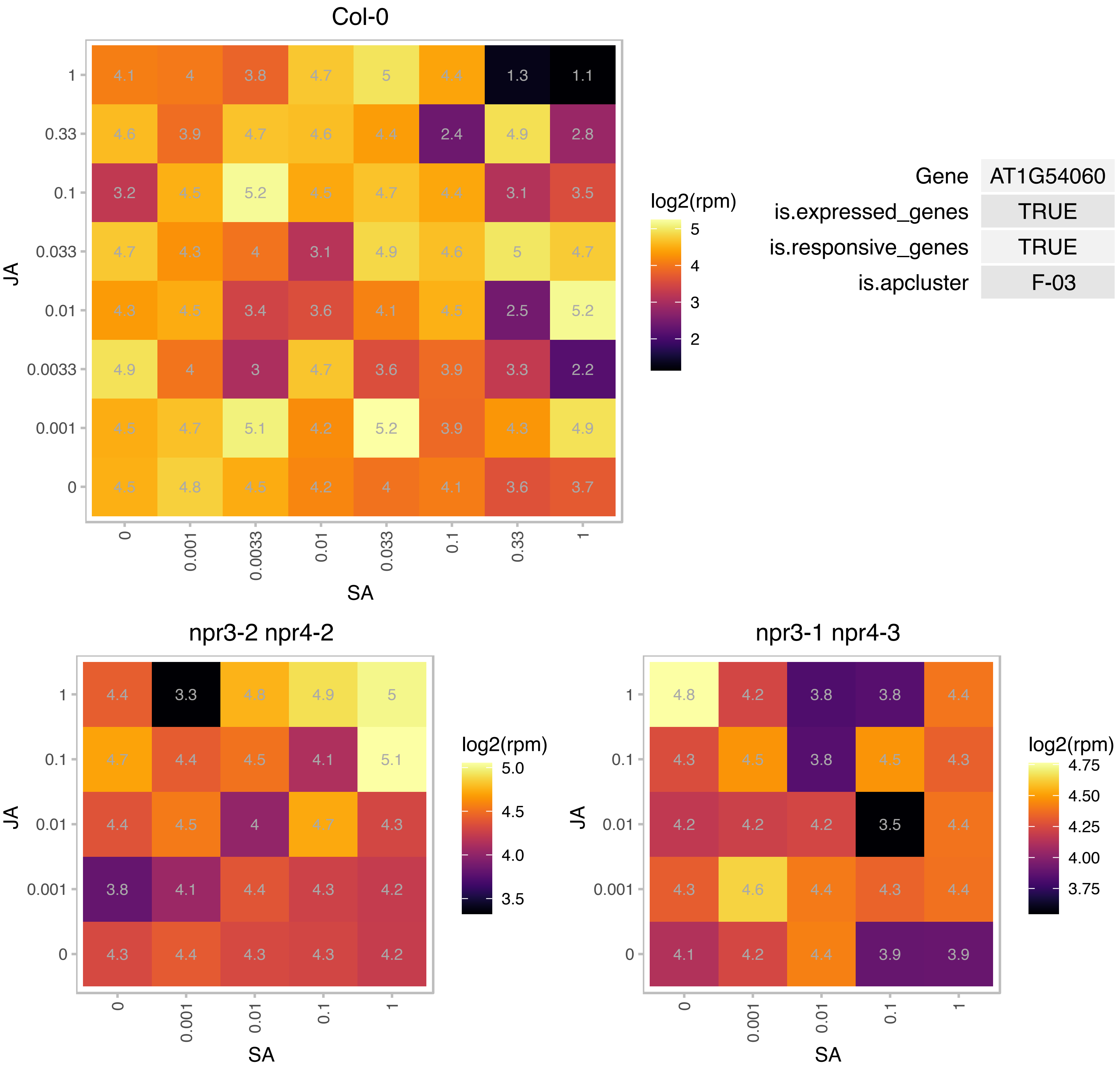


| | |
|---|---|
| Gene | AT1G54060 |
| is.expressed_genes | TRUE |
| is.responsive_genes | TRUE |
| is.apcluster | F-03 |

### AT1G54350.1 : ABCD2

ABC transporter family protein

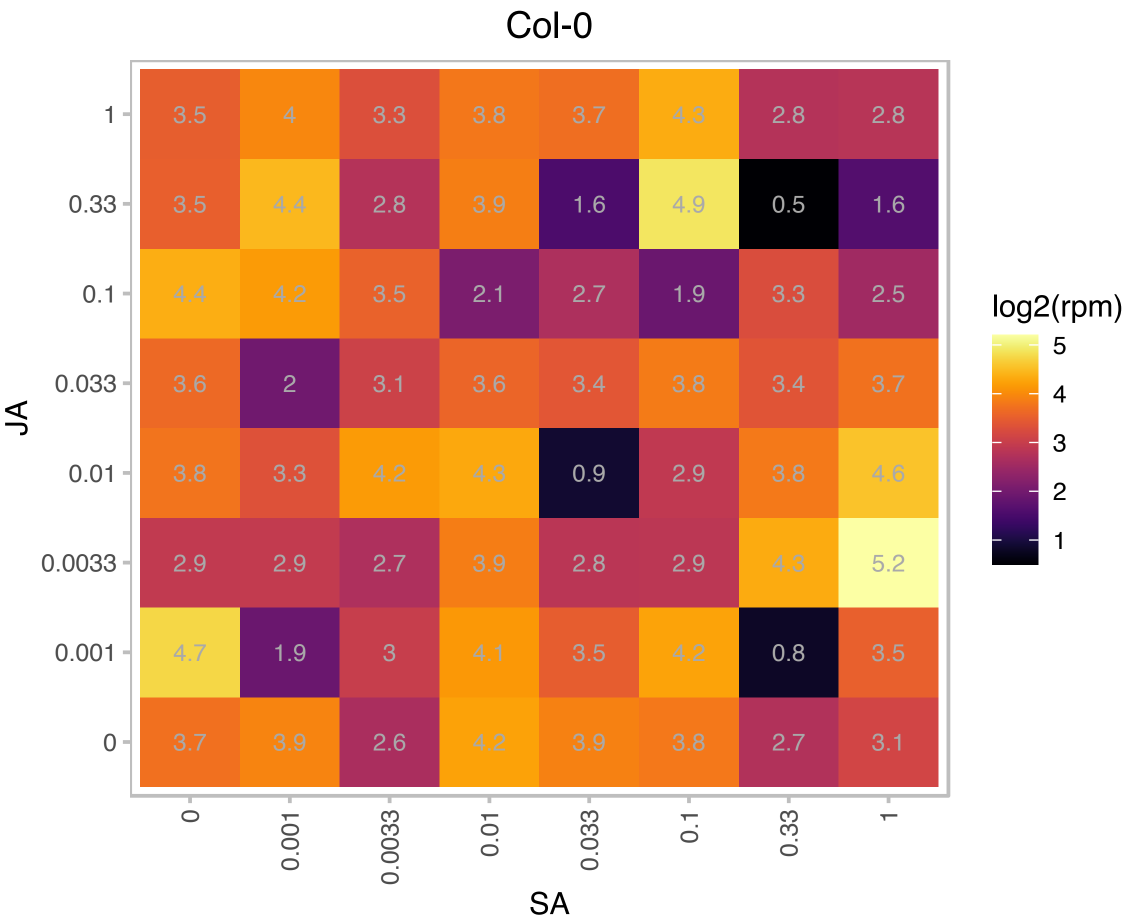


| | |
|---|---|
| Gene | AT1G54350 |
| is.expressed_genes | TRUE |
| is.responsive_genes | TRUE |
| is.apcluster | F-03 |

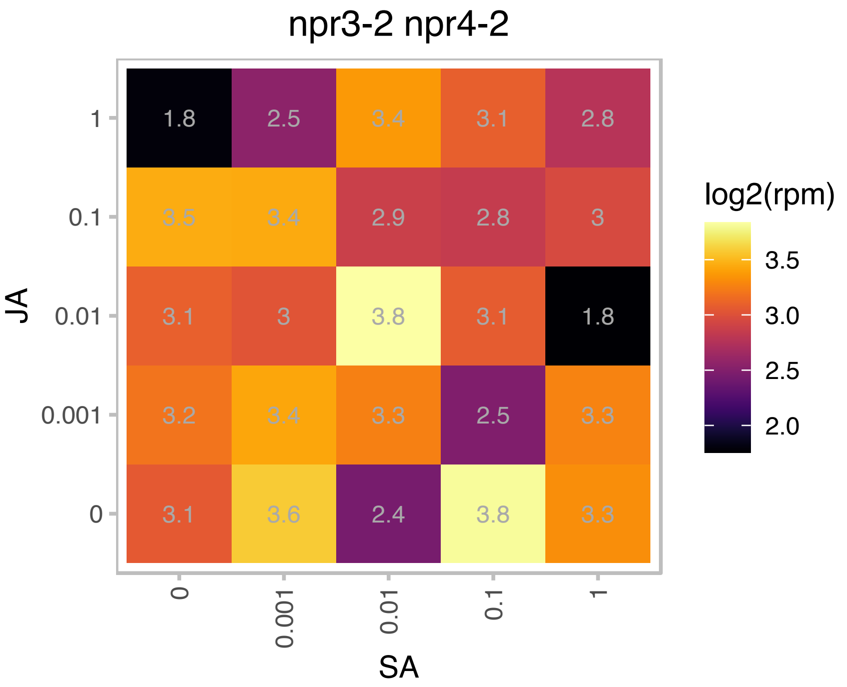


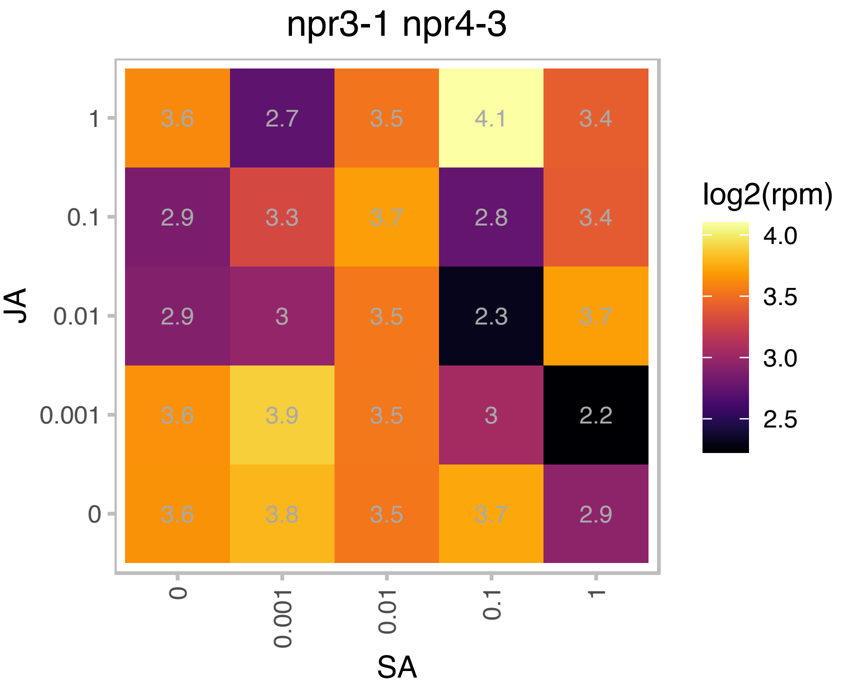

### AT1G54580.1 : ACP2, AtACP2

acyl carrier protein 2
encodes an acyl carrier protein expressed in leaves, roots, and dry seeds. Gene expression is not regulated by light but downregulated by starvation and upregulated by increased level of exogenous sucrose. The mRNA is cell-to-cell mobile.

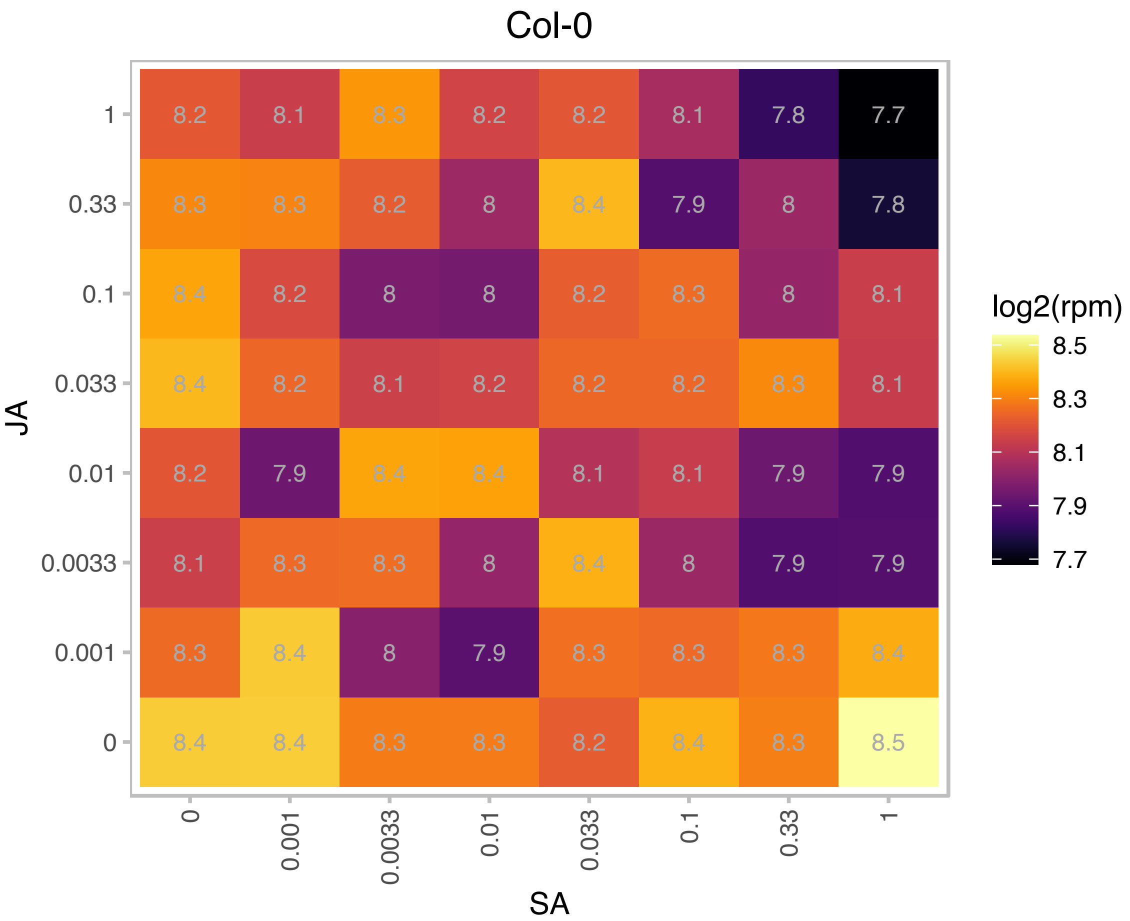


| | |
|---|---|
| Gene | AT1G54580 |
| is.expressed_genes | TRUE |
| is.responsive_genes | TRUE |
| is.apcluster | F-03 |

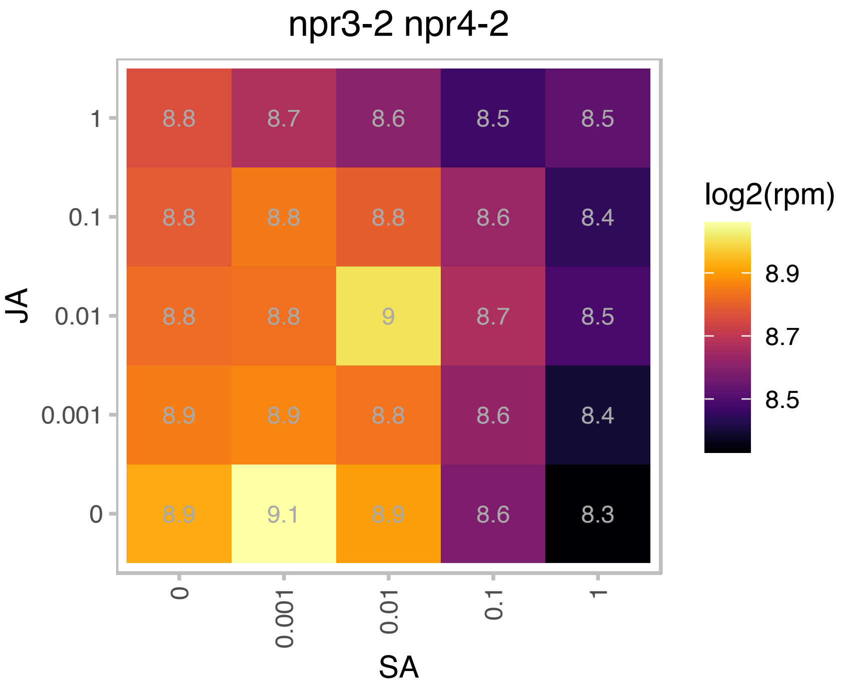


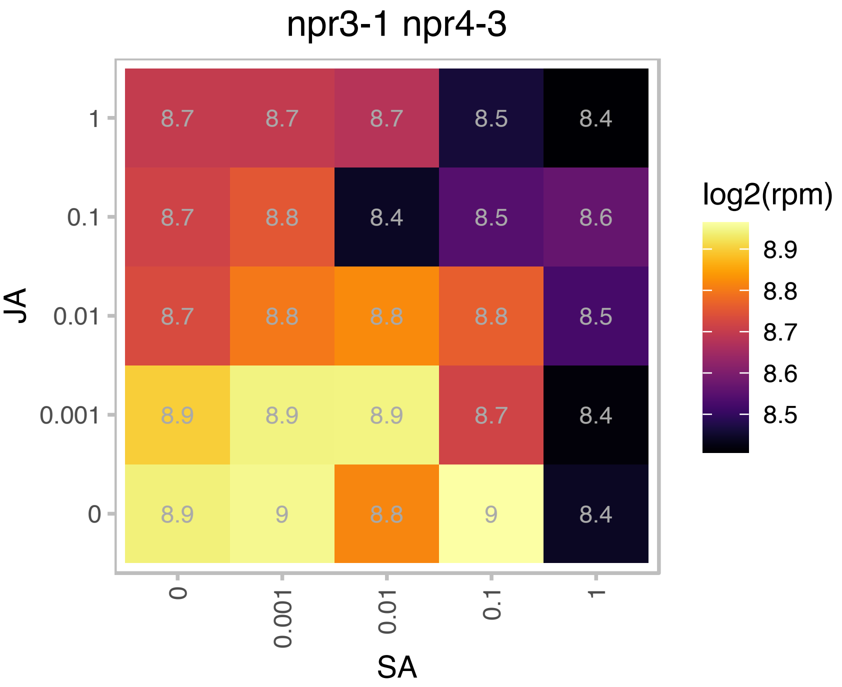

### AT1G55805.1 : BolA1, BolA2

BolA-like family protein

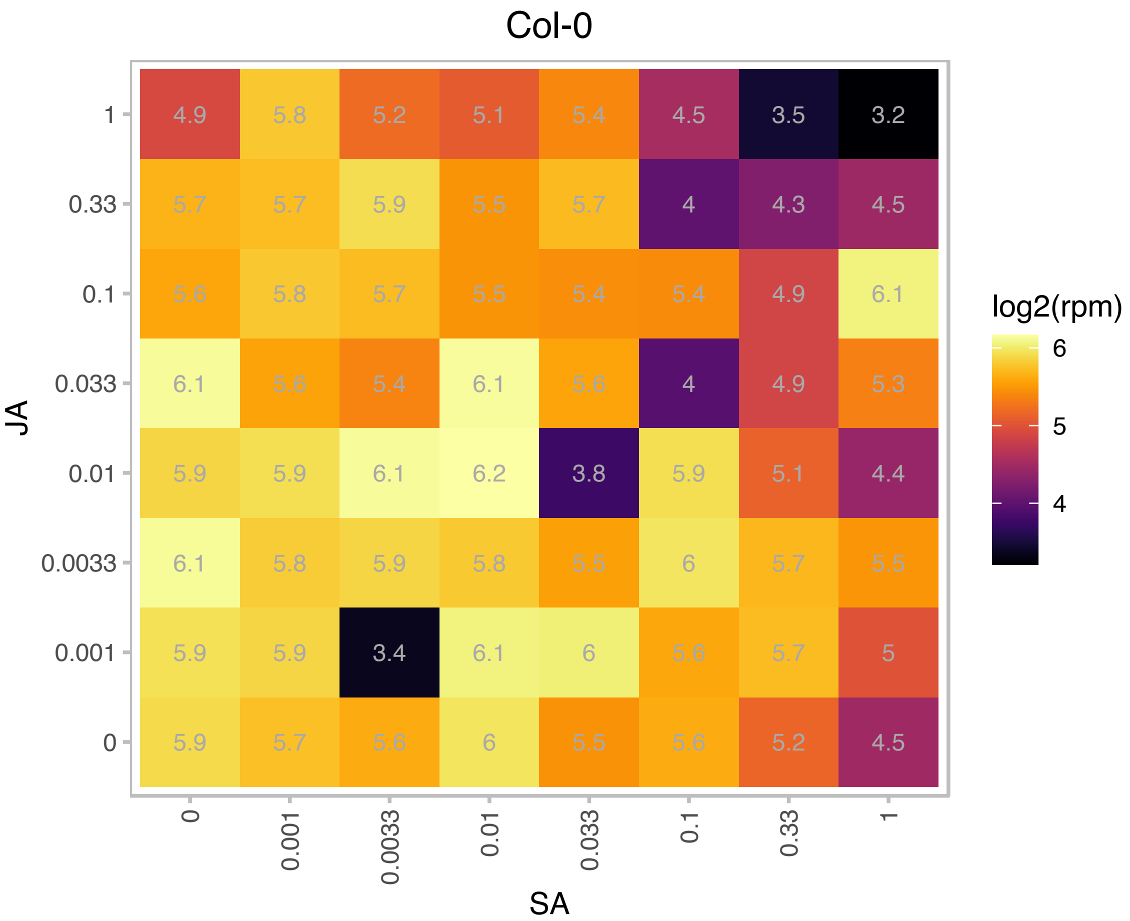


| | |
|---|---|
| Gene | AT1G55805 |
| is.expressed_genes | TRUE |
| is.responsive_genes | TRUE |
| is.apcluster | F-03 |

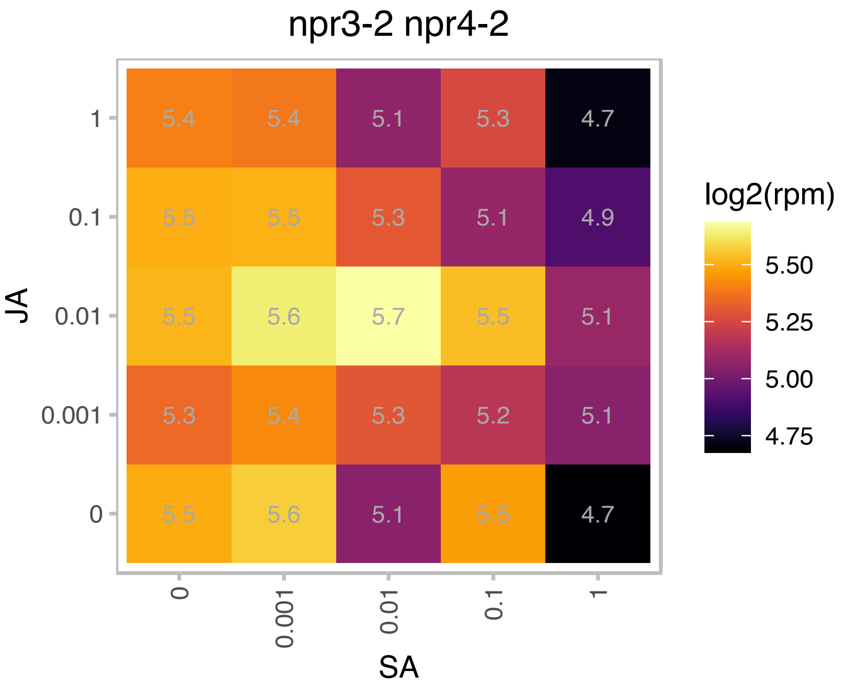


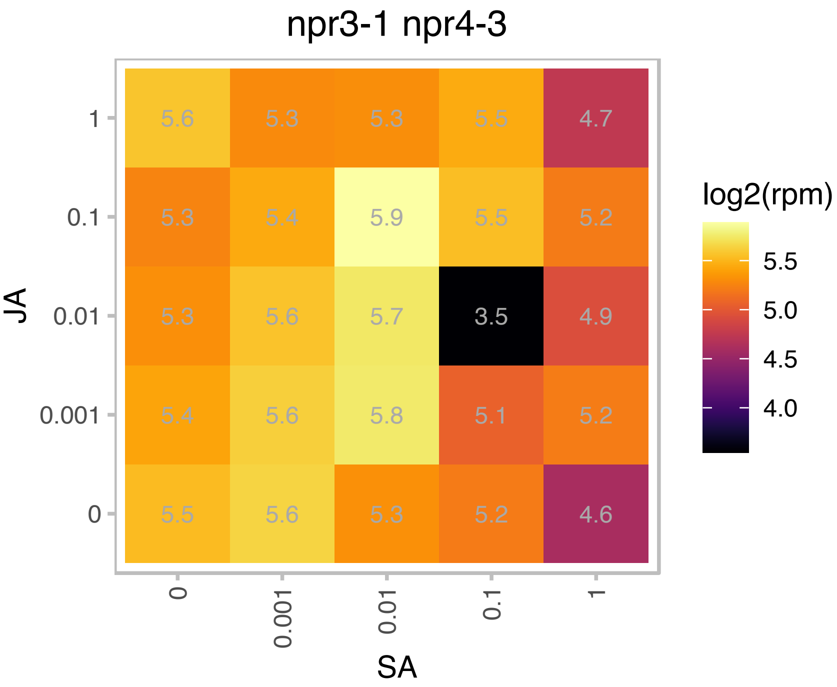

### AT1G68060.1 : ATMAP70-1, MAP70-1

microtubule-associated proteins 70-1

Encodes a microtubule associated protein (MAP70-1). Expressed in all tissues.

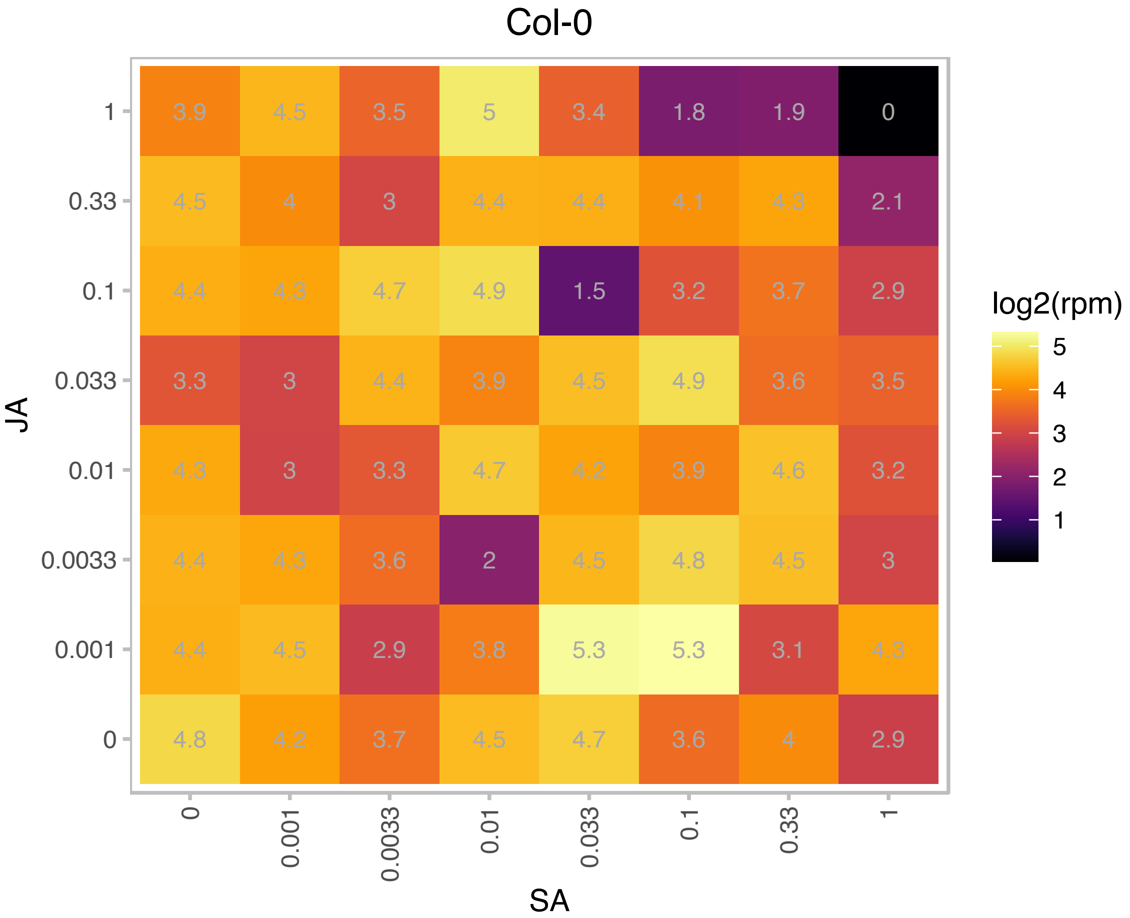


| | |
|---|---|
| Gene | AT1G68060 |
| is.expressed_genes | TRUE |
| is.responsive_genes | TRUE |
| is.apcluster | F-03 |

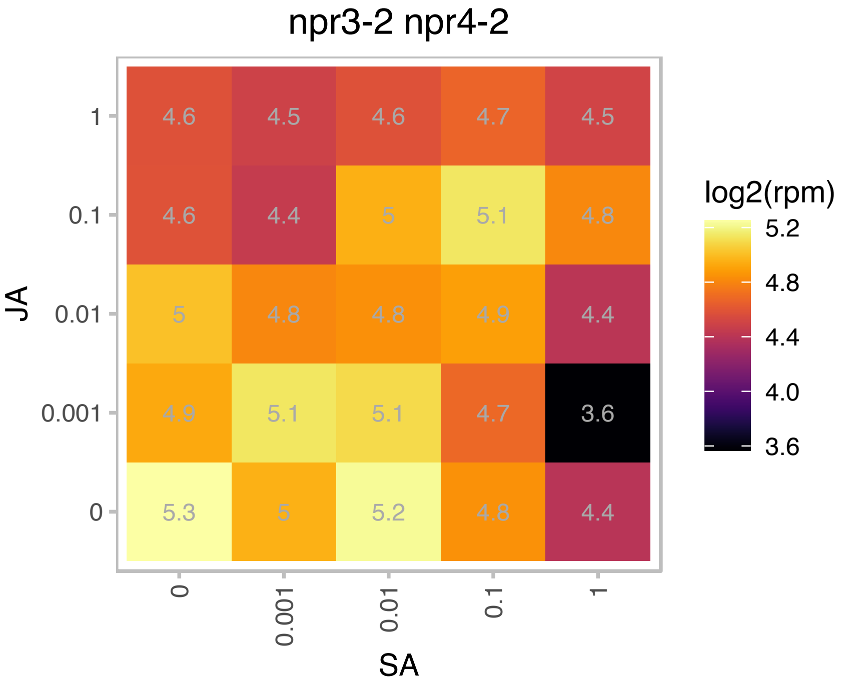


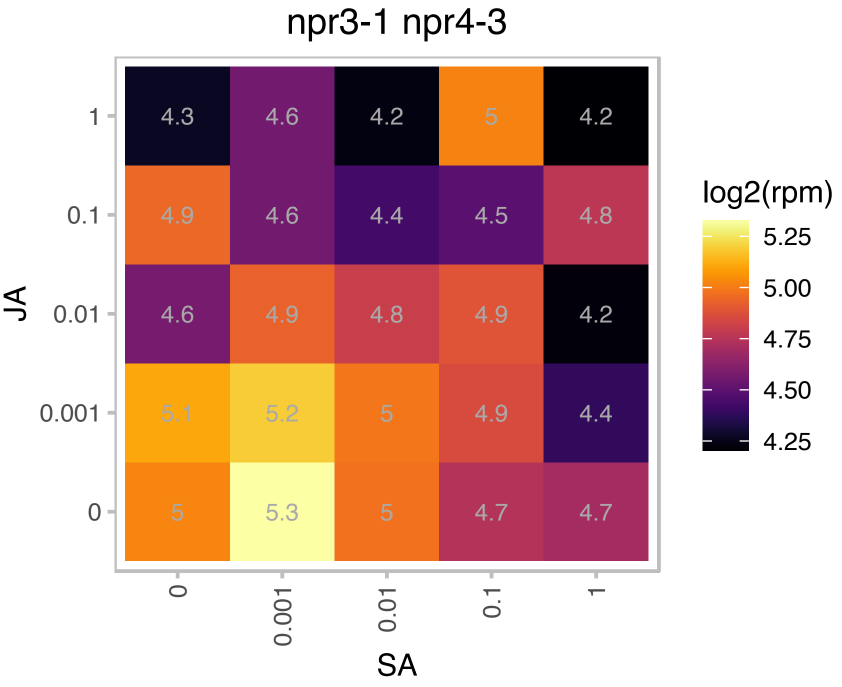

# AT1G71430.1 : NA

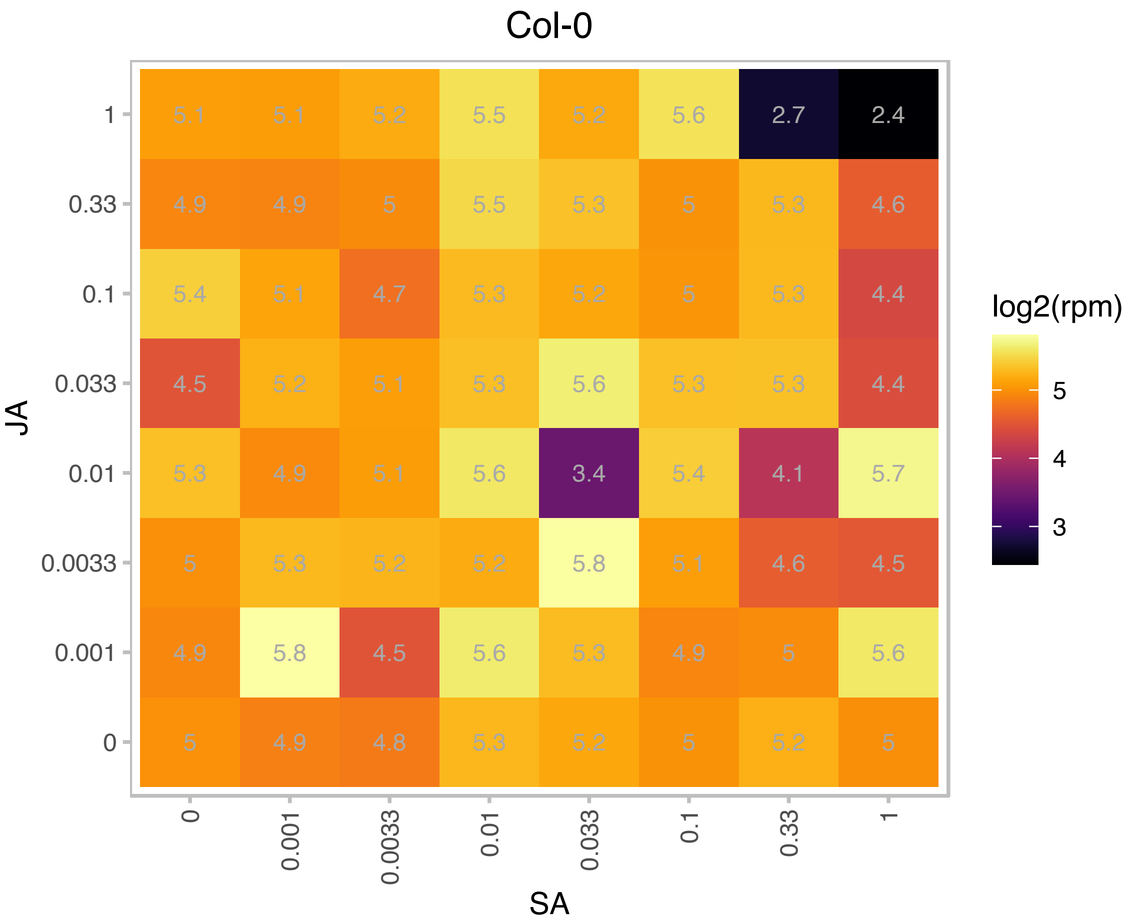


| | |
|---|---|
| Gene | AT1G71430 |
| is.expressed_genes | TRUE |
| is.responsive_genes | TRUE |
| is.apcluster | F-03 |

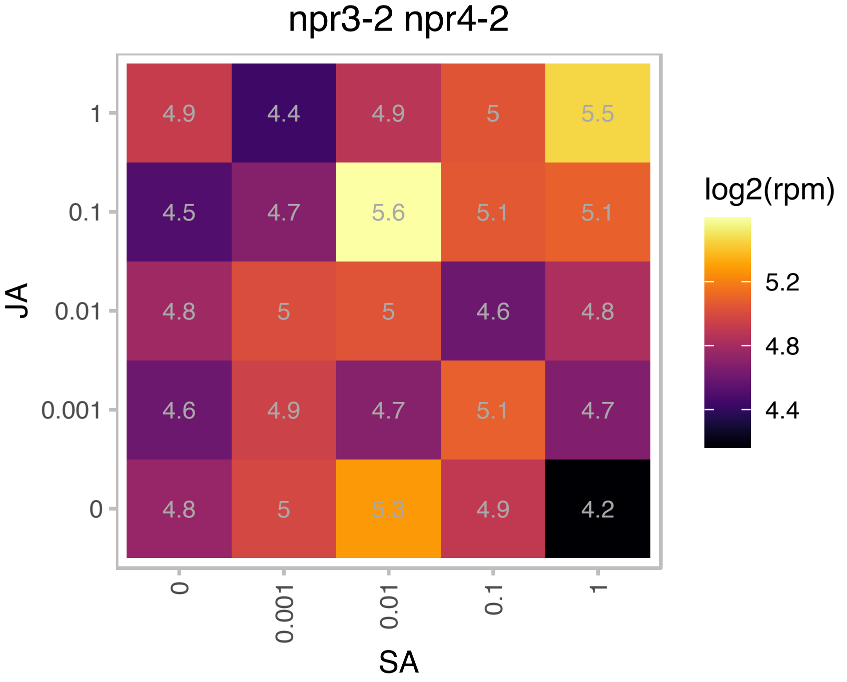


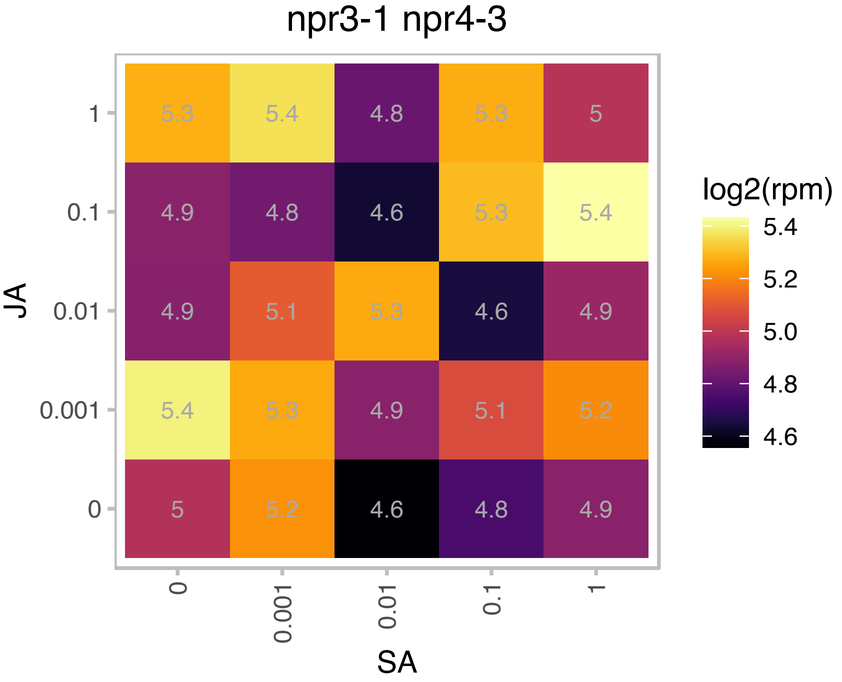

# AT1G73940.1 : NA

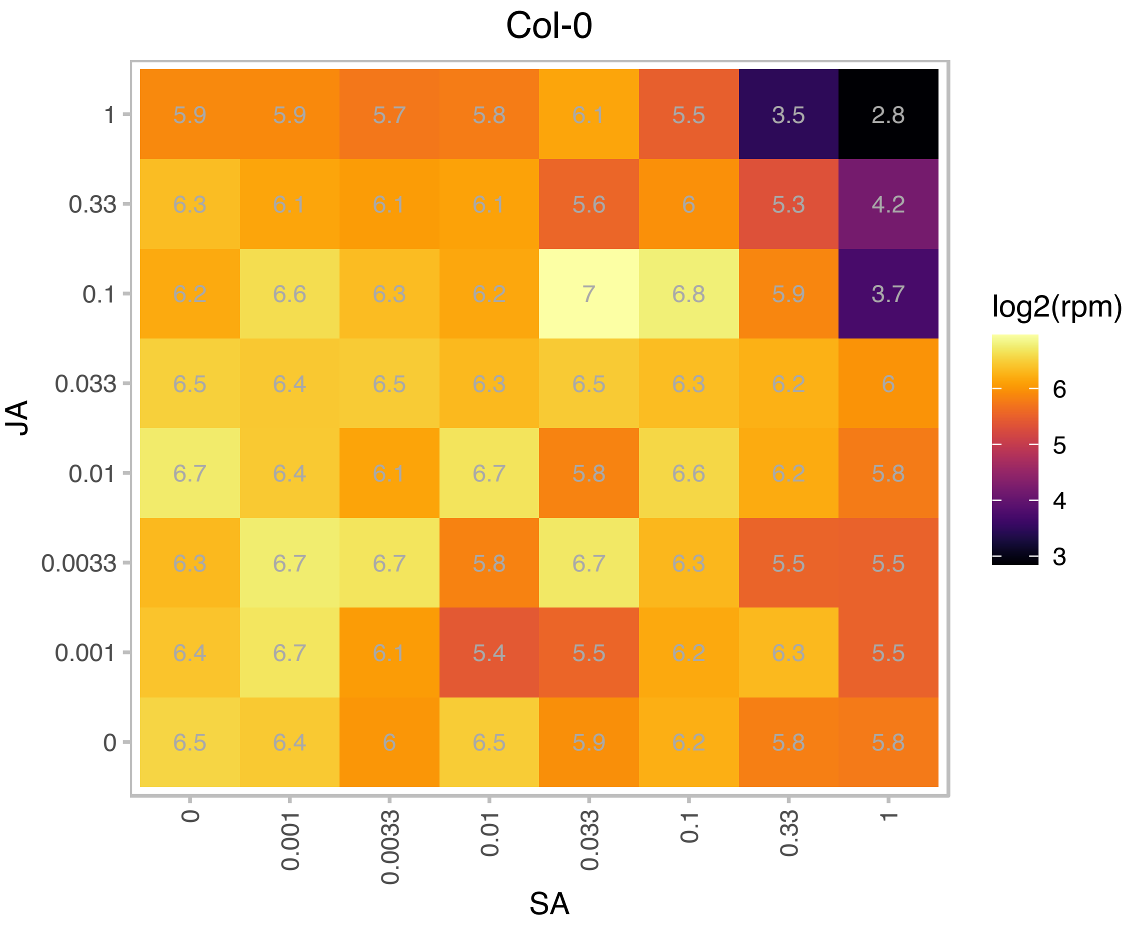


| | |
|---|---|
| Gene | AT1G73940 |
| is.expressed_genes | TRUE |
| is.responsive_genes | TRUE |
| is.apcluster | F-03 |

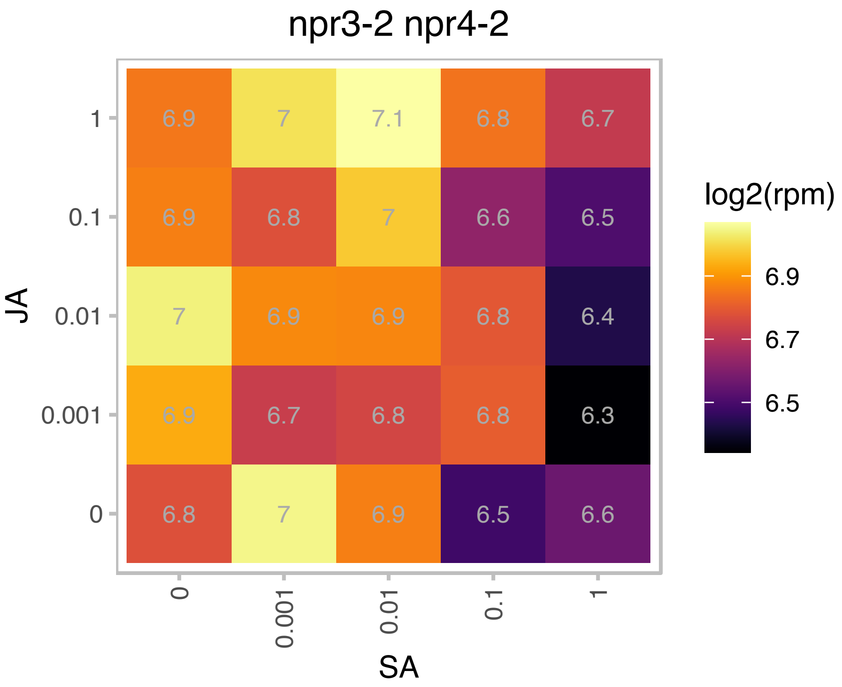


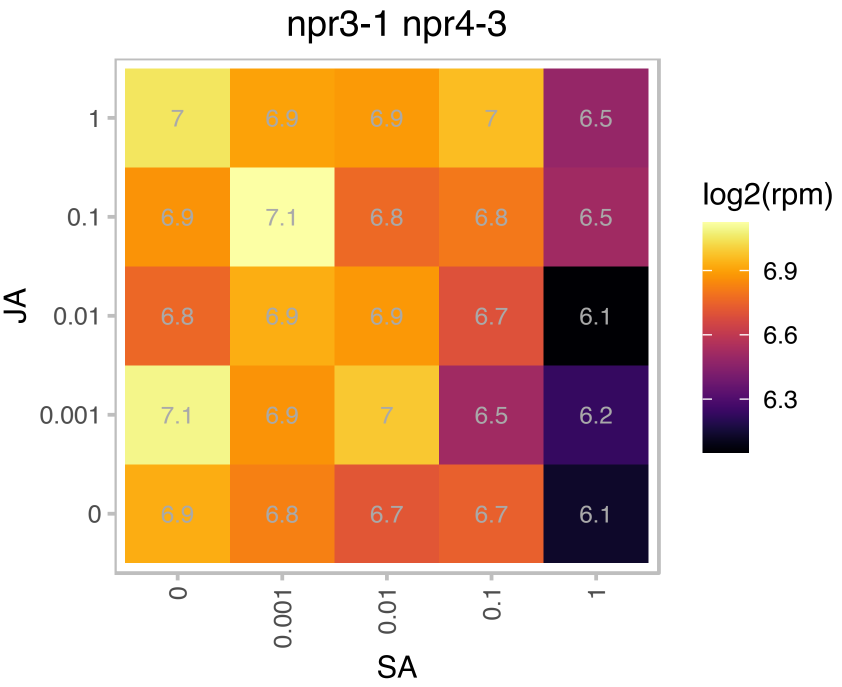

### AT2G02130.1 : LCR68, PDF2.3

Predicted to encode a PR (pathogenesis-related) protein. Belongs to the plant defensin (PDF) family with the following members: At1g75830/PDF1.1, At5g44420/PDF1.2a, At2g26020/PDF1.2b, At5g44430/PDF1.2c, At2g26010/PDF1.3, At1g19610/PDF1.4, At1g55010/PDF1.5, At2g02120/PDF2.1, At2g02100/PDF2.2, At2g02130/PD At1g61070/PDF2.4, At5g63660/PDF2.5, At2g02140/PDF2.6, At5g38330/PDF3.1 and At4g30070/PDF3.2. The mRN cell-to-cell mobile.

low-molecular-weight cysteine-rich 68

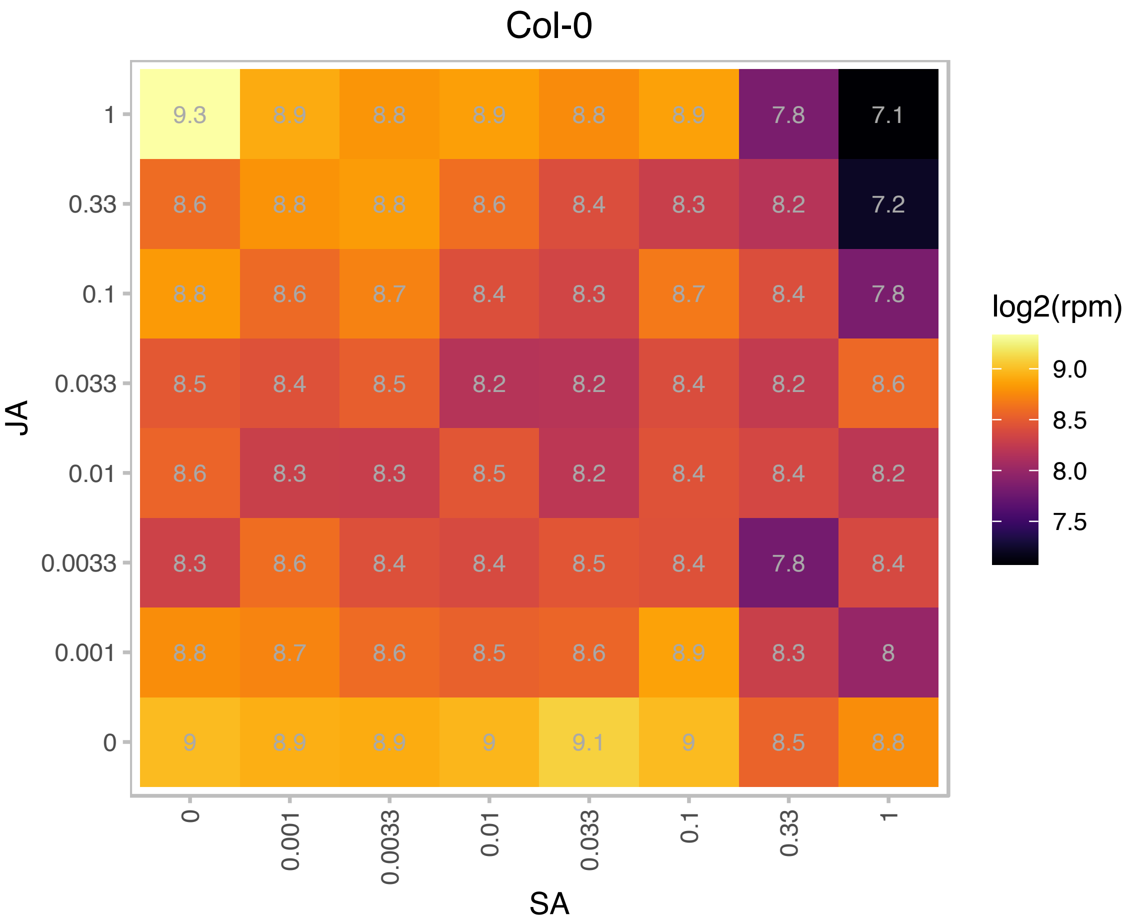


| | |
|---|---|
| Gene | AT2G02130 |
| is.expressed_genes | TRUE |
| is.responsive_genes | TRUE |
| is.apcluster | F-03 |

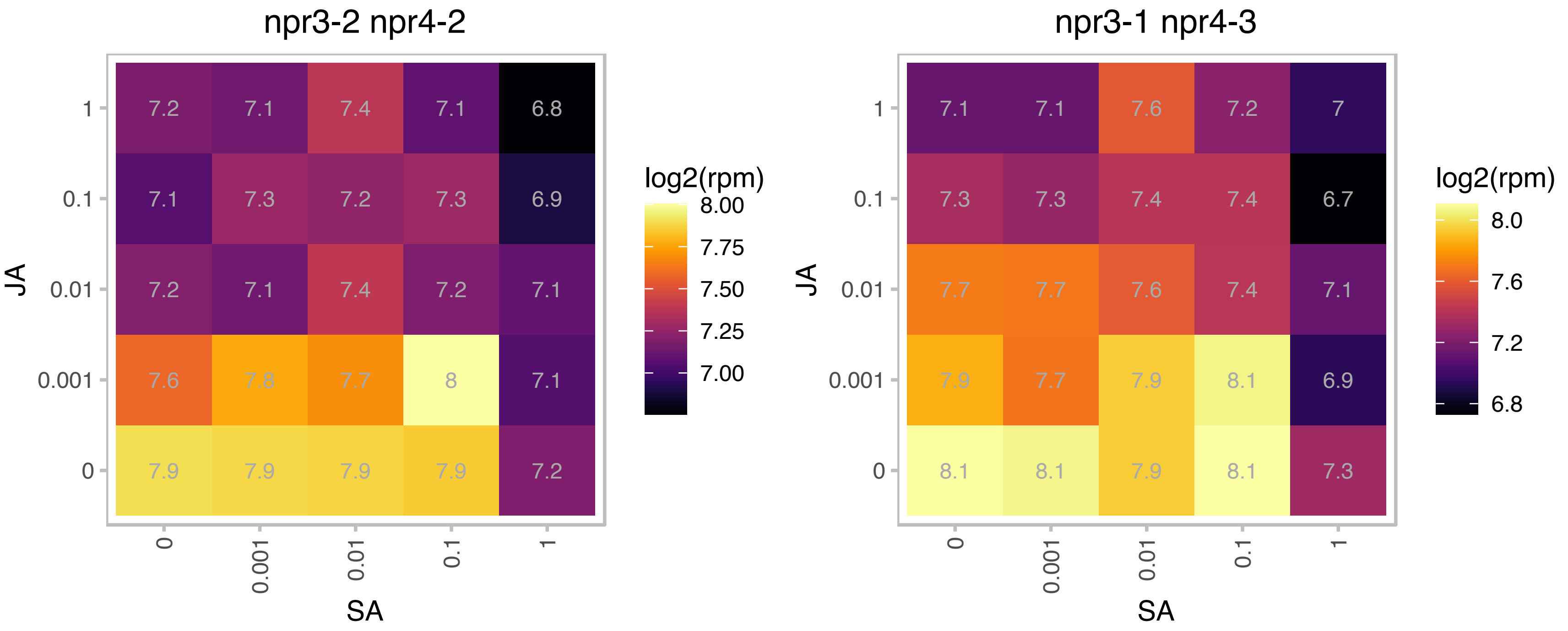

# AT2G19680.1 : NA

Mitochondrial ATP synthase subunit G protein

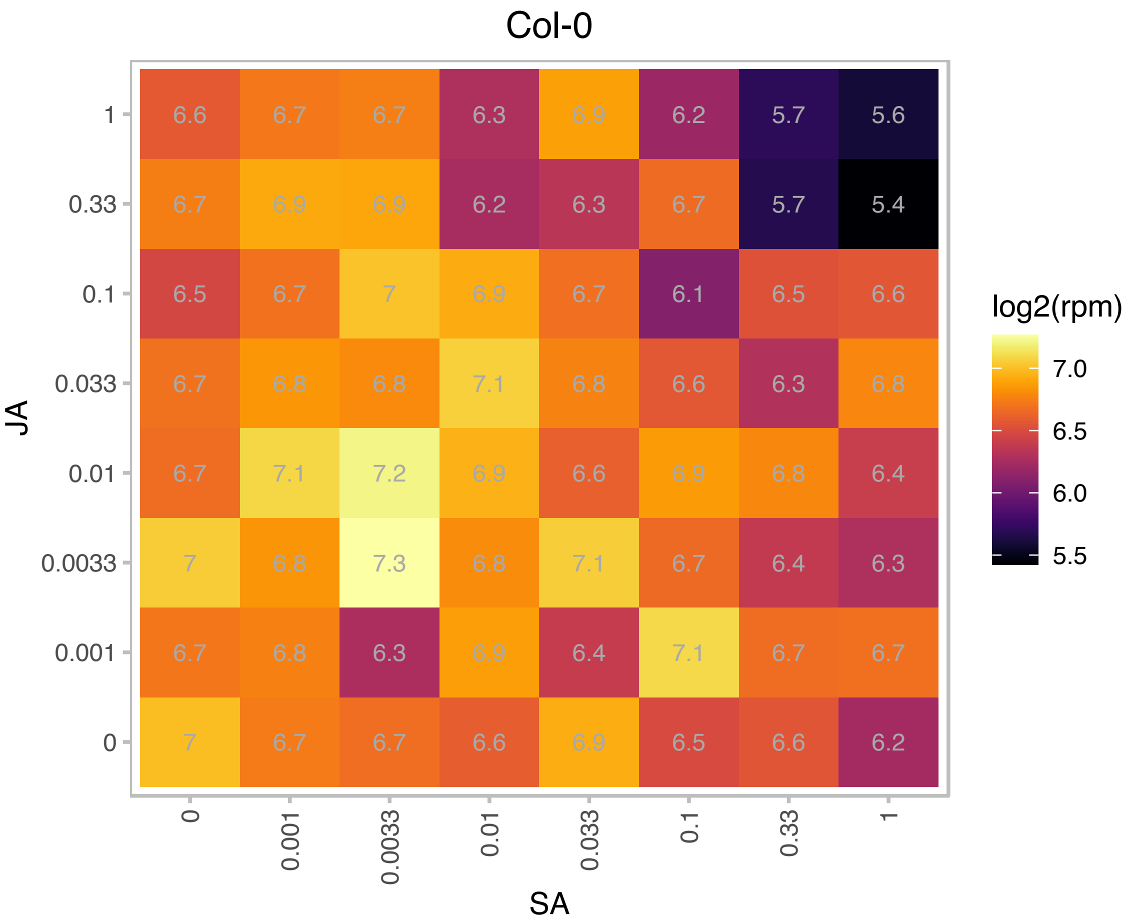


| | |
|---|---|
| Gene | AT2G19680 |
| is.expressed_genes | TRUE |
| is.responsive_genes | TRUE |
| is.apcluster | F-03 |

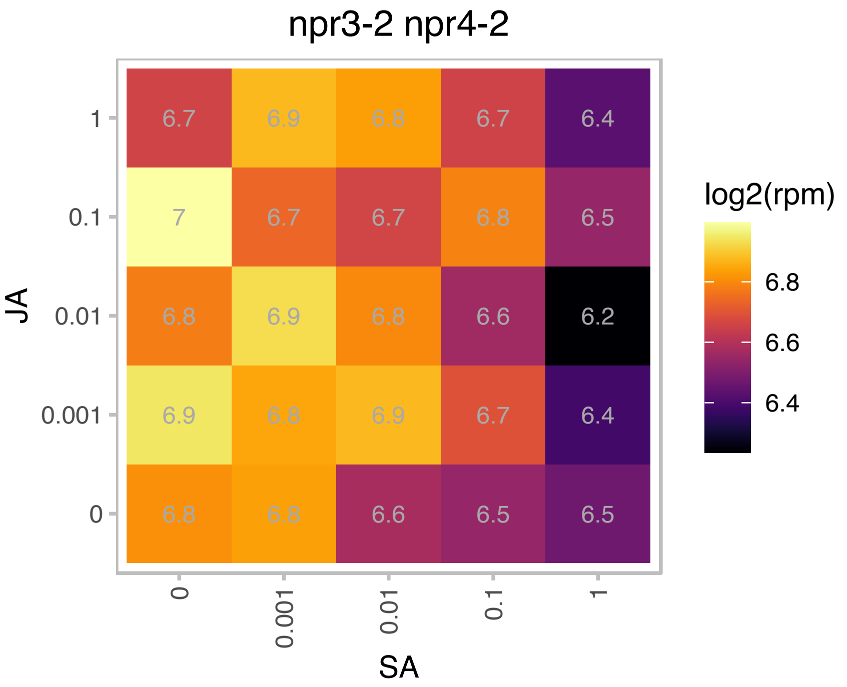


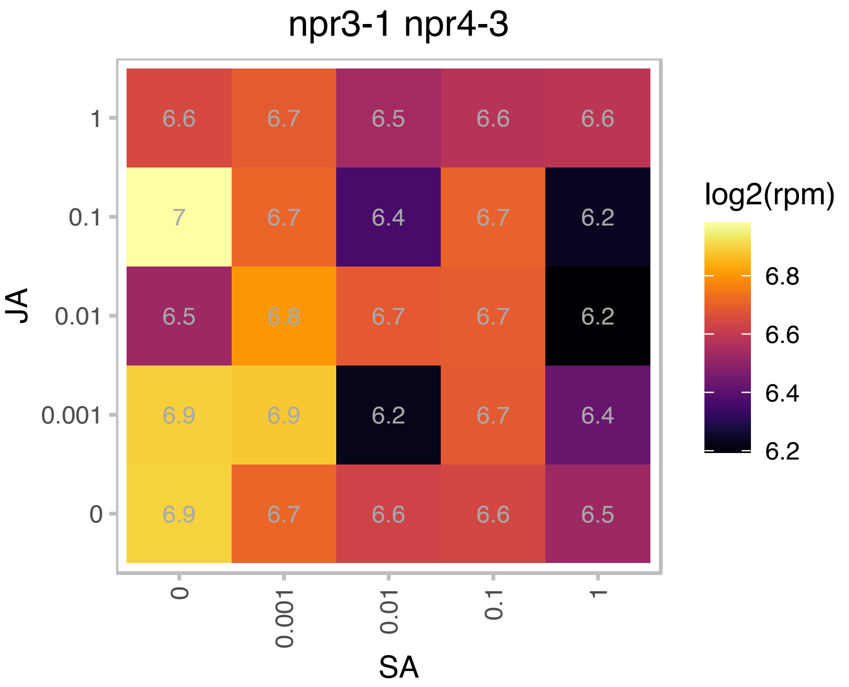

### AT2G21045.1 : HAC1

Rhodanese/Cell cycle control phosphatase superfamily protein

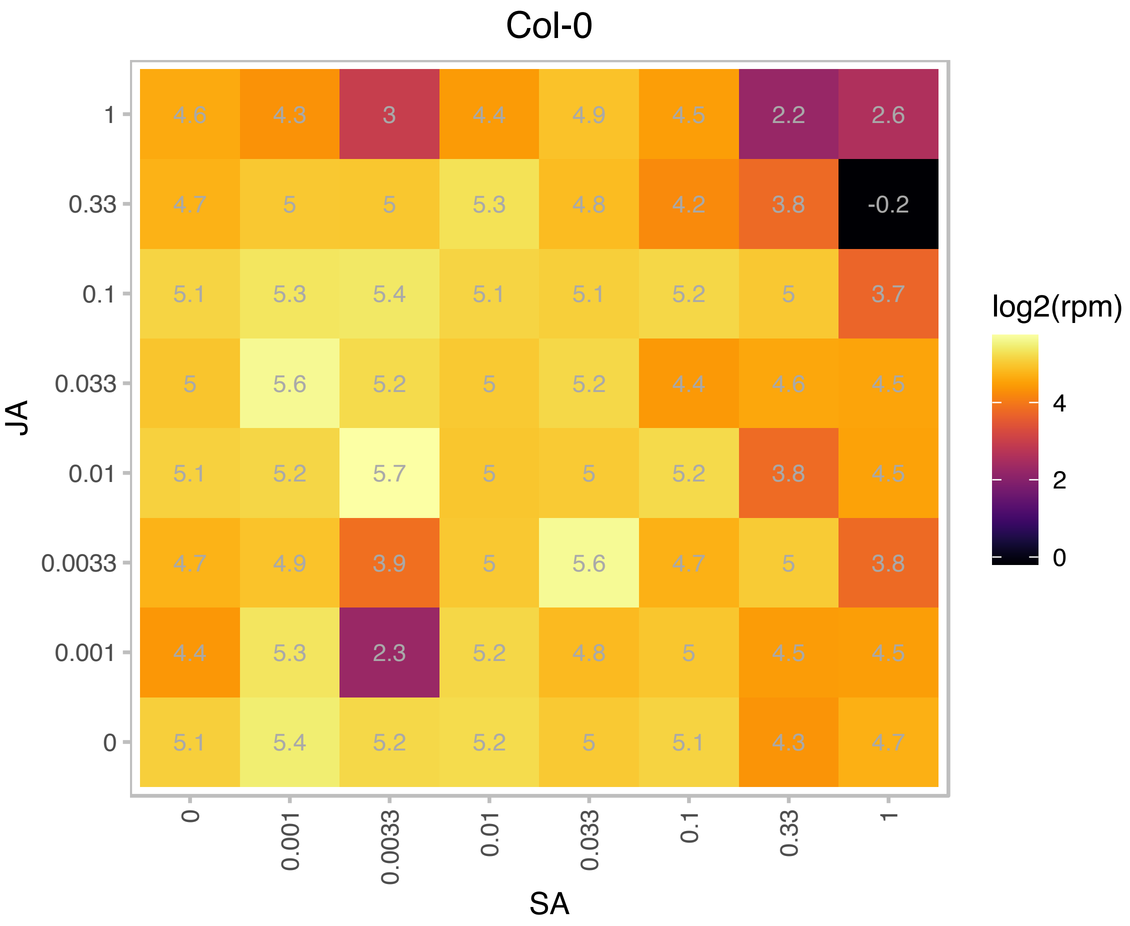


| | |
|---|---|
| Gene | AT2G21045 |
| is.expressed_genes | TRUE |
| is.responsive_genes | TRUE |
| is.apcluster | F-03 |

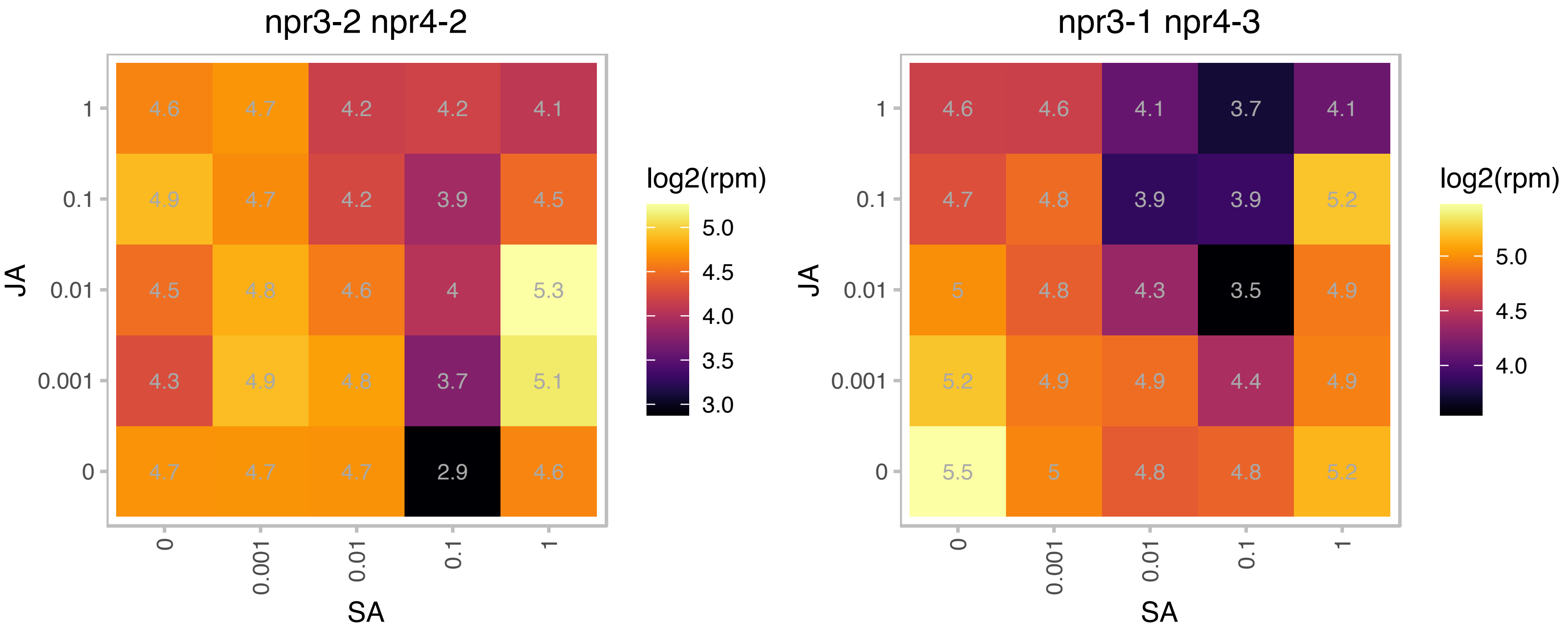

### AT2G24790.1 : ATCOL3, BBX4, COL3

CONSTANS-like 3
Positive regulator of photomorphogenesis that acts downstream of COP1 but can promote lateral root development independently of COP1 and also function as a daylength-sensitive regulator of shoot branching. The mRNA is cell-to-cell mobile.

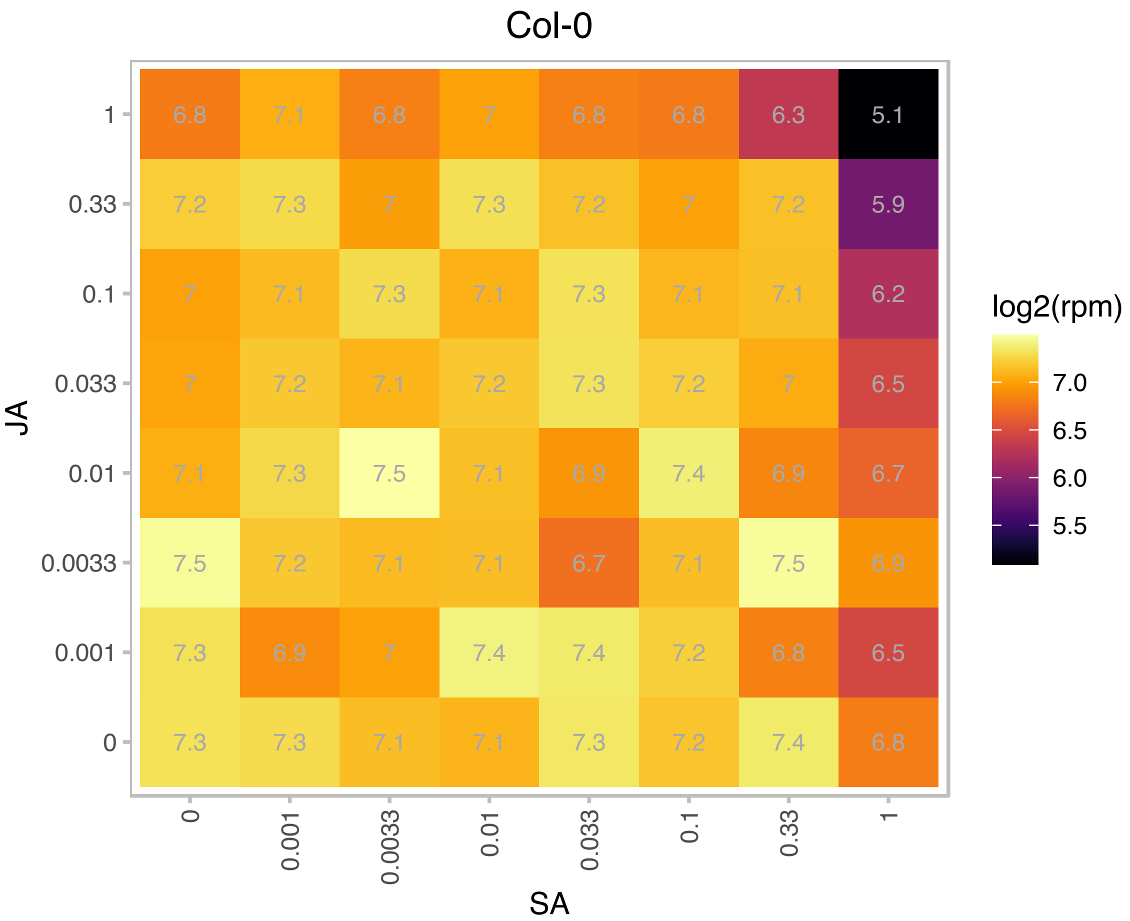


| | |
|---|---|
| Gene | AT2G24790 |
| is.expressed_genes | TRUE |
| is.responsive_genes | TRUE |
| is.apcluster | F-03 |

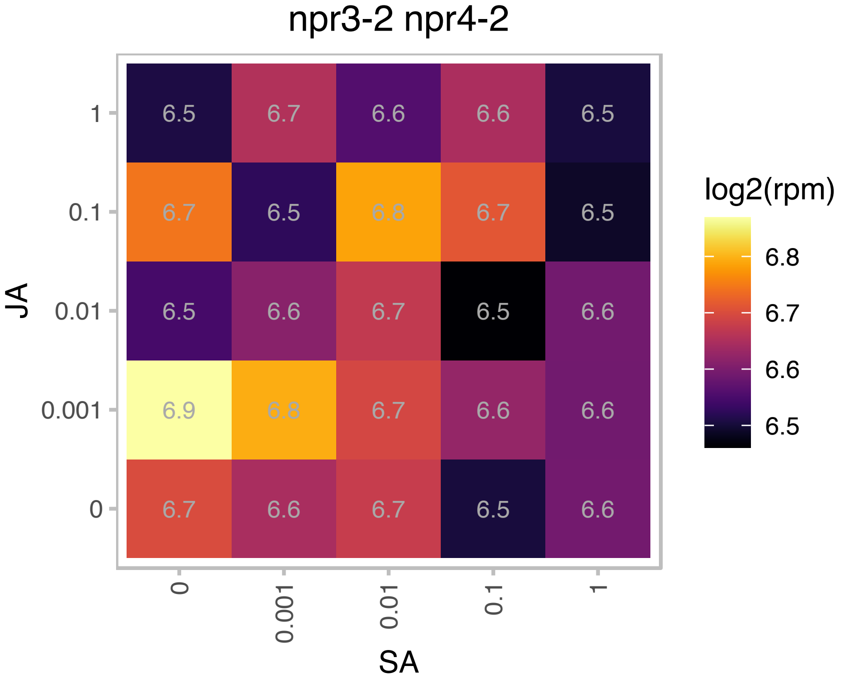


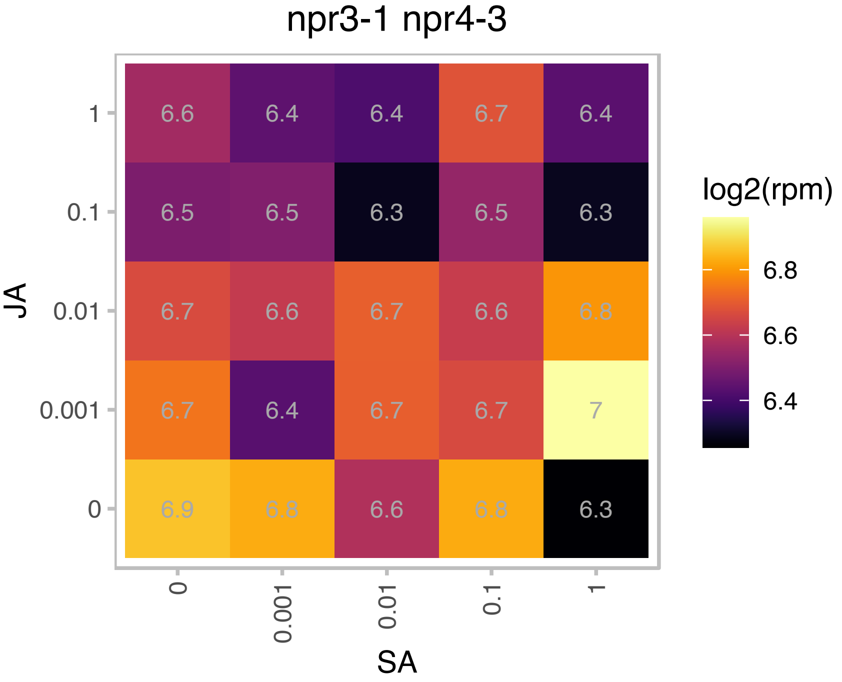

### AT2G30280.1 : DMS4, RDM4

RNA-directed DNA methylation 4

Encodes RDM4, a transcriptional regulator functioning in RNA-directed DNA methylation and plant development.

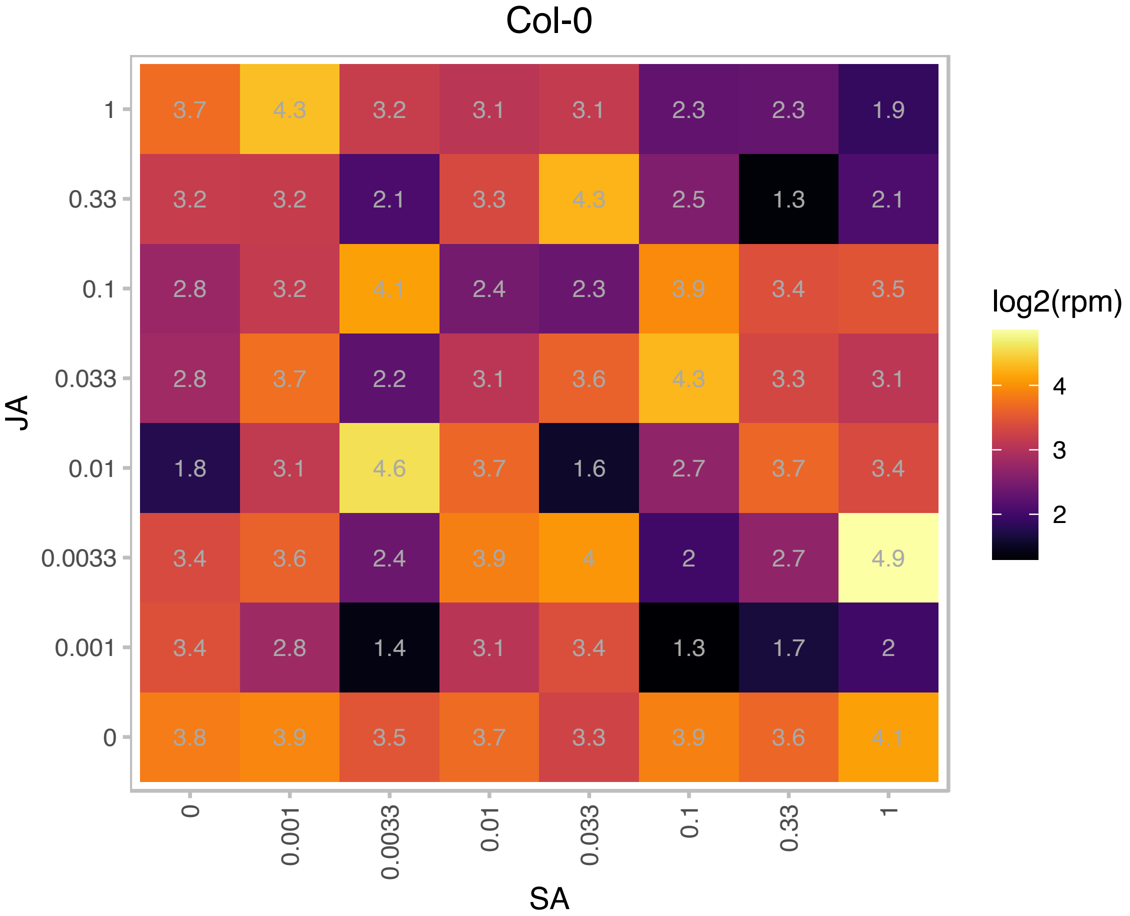


| | |
|---|---|
| Gene | AT2G30280 |
| is.expressed_genes | TRUE |
| is.responsive_genes | TRUE |
| is.apcluster | F-03 |

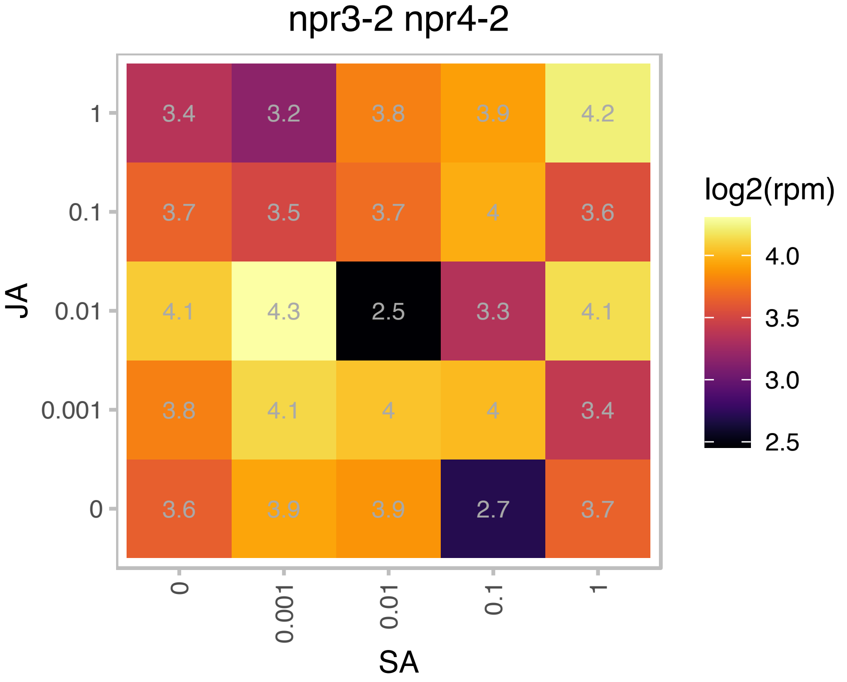


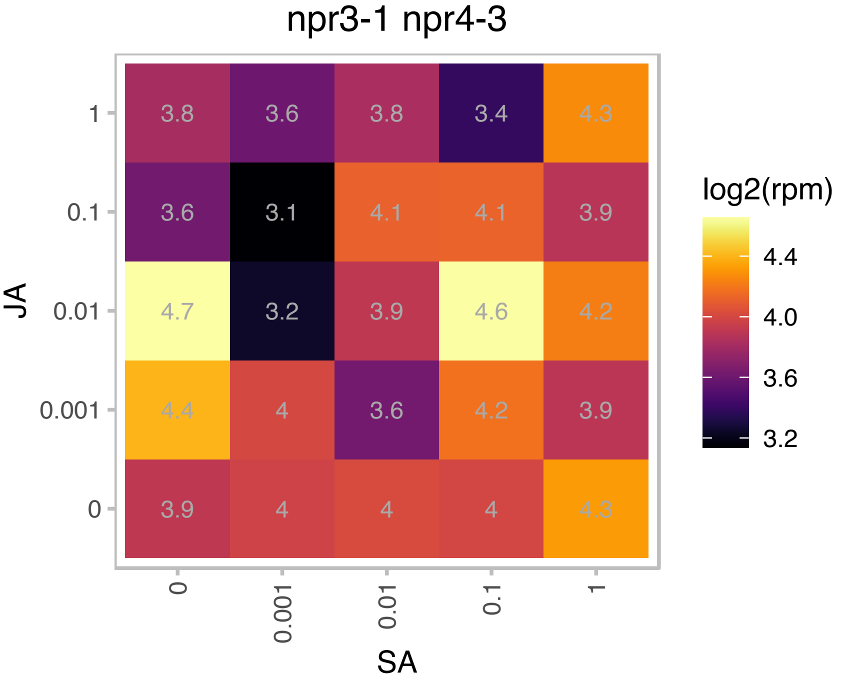

### AT2G31140.1 : AtIMP2

Peptidase S24/S26A/S26B/S26C family protein

| | |
|---|---|
| Gene | AT2G31140 |
| is.expressed_genes | TRUE |
| is.responsive_genes | TRUE |
| is.apcluster | F-03 |

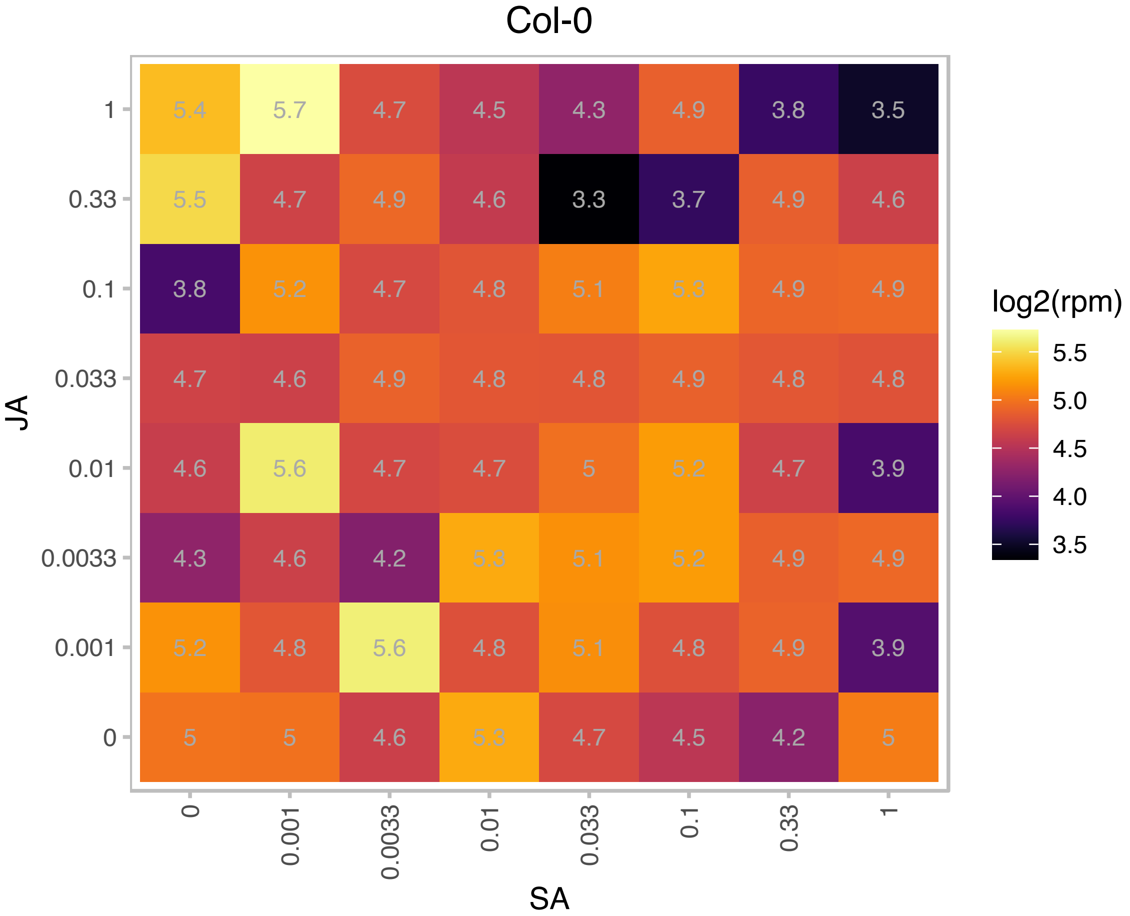


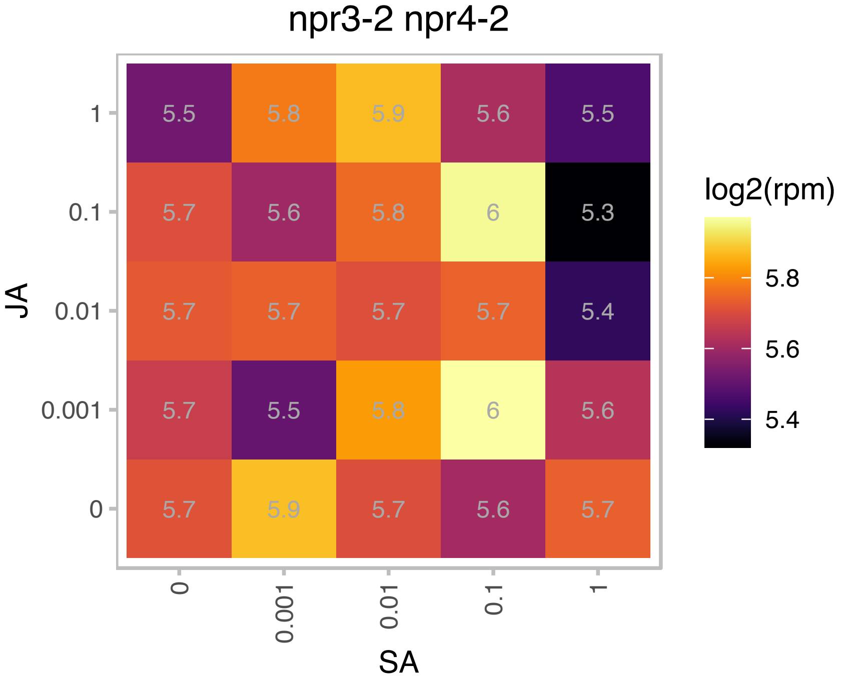


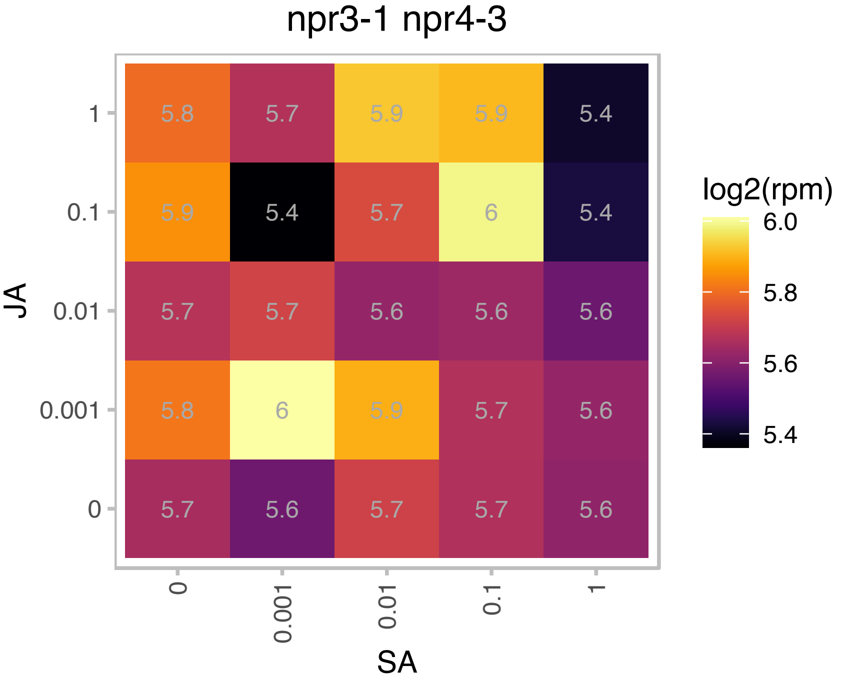

# AT2G31610.1 : NA

Ribosomal protein S3 family protein

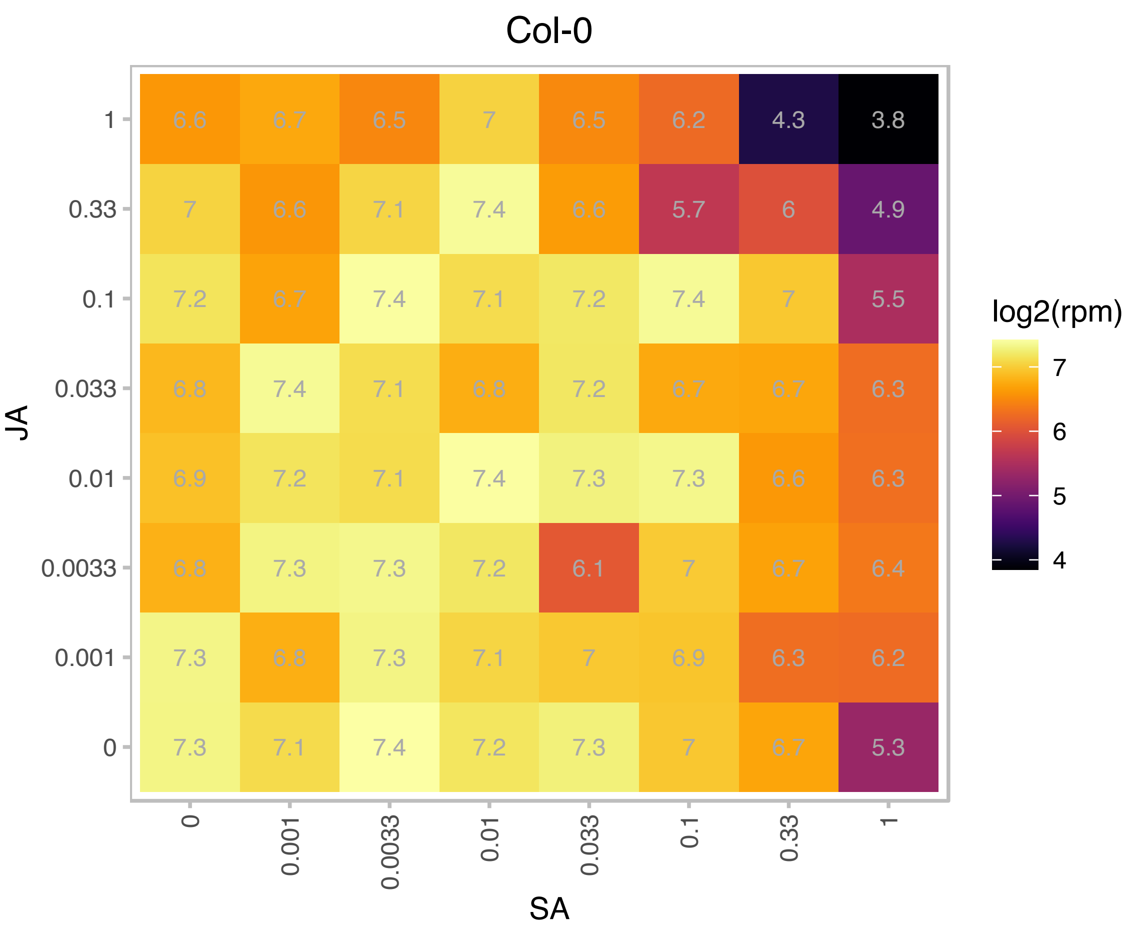


| | |
|---|---|
| Gene | AT2G31610 |
| is.expressed_genes | TRUE |
| is.responsive_genes | TRUE |
| is.apcluster | F-03 |

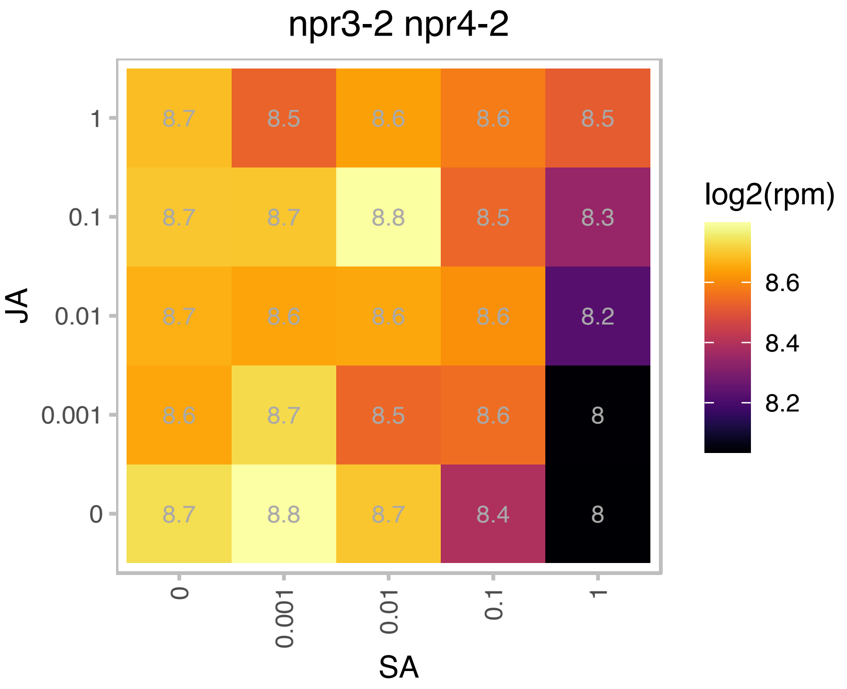


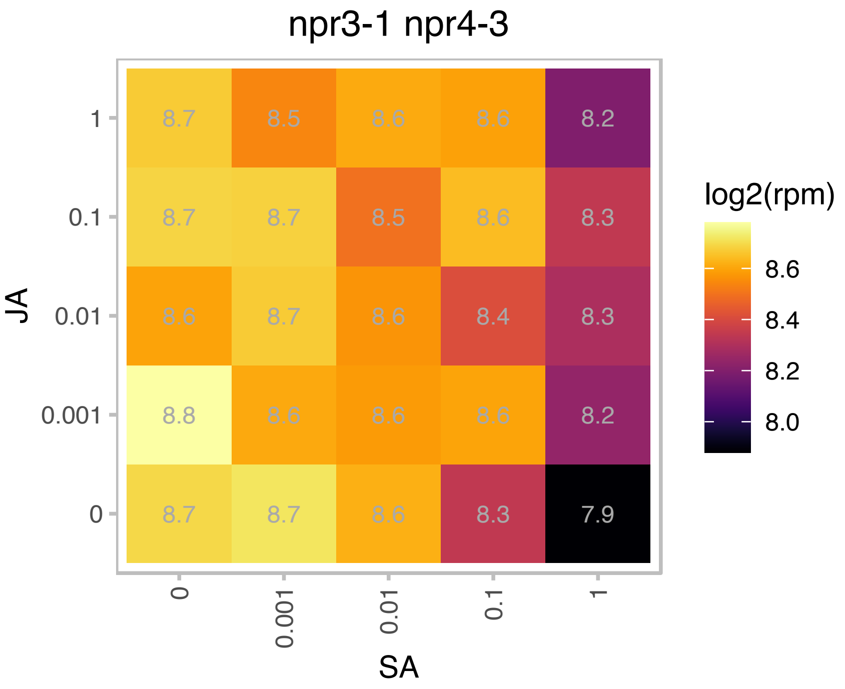

# AT2G35830.1 : NA

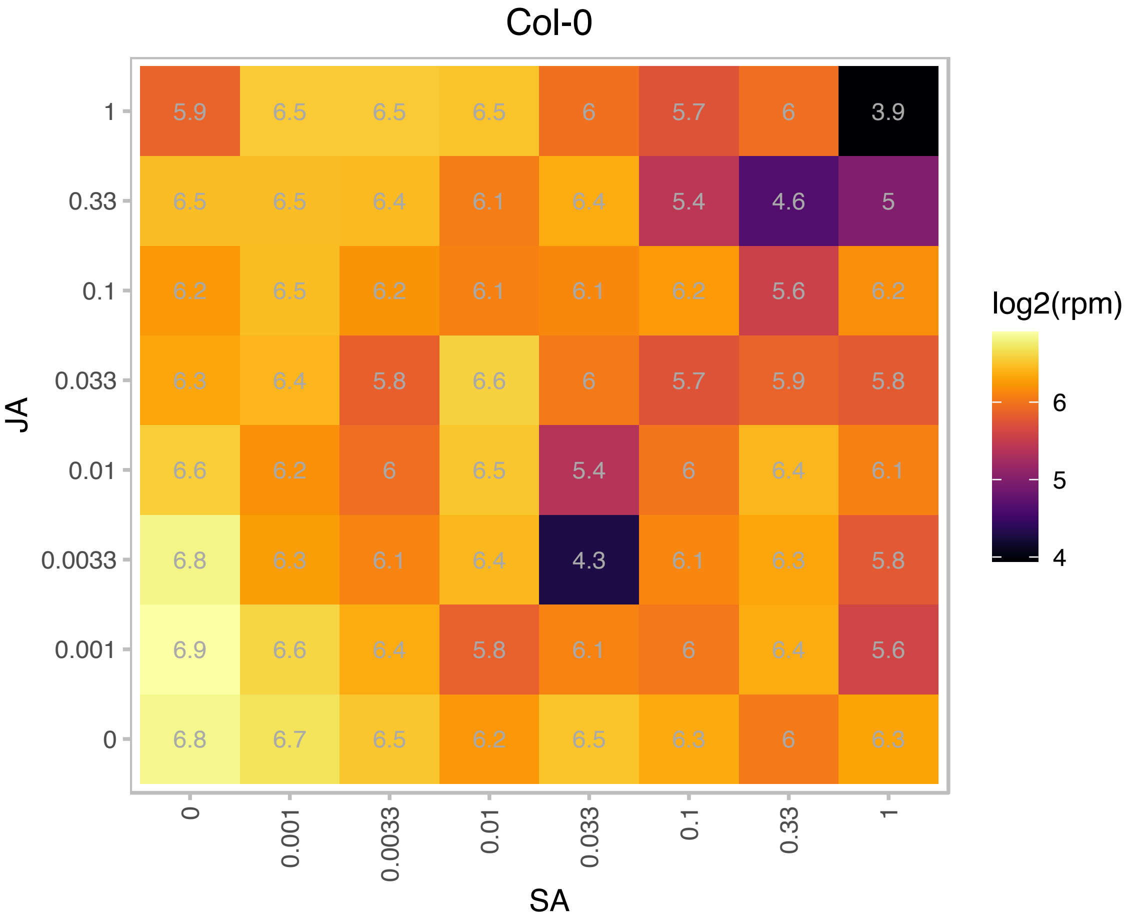


| | |
|---|---|
| Gene | AT2G35830 |
| is.expressed_genes | TRUE |
| is.responsive_genes | TRUE |
| is.apcluster | F-03 |

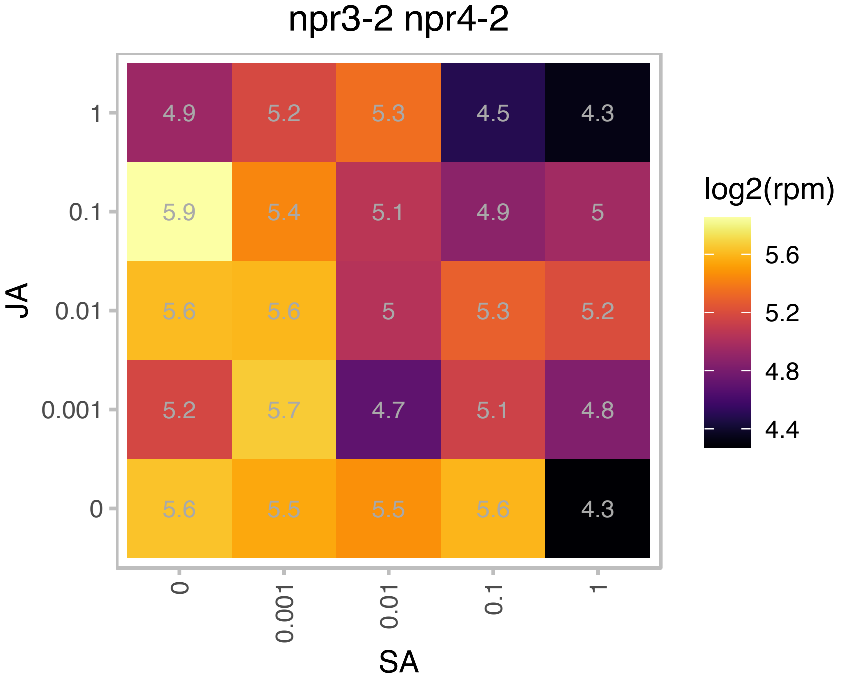


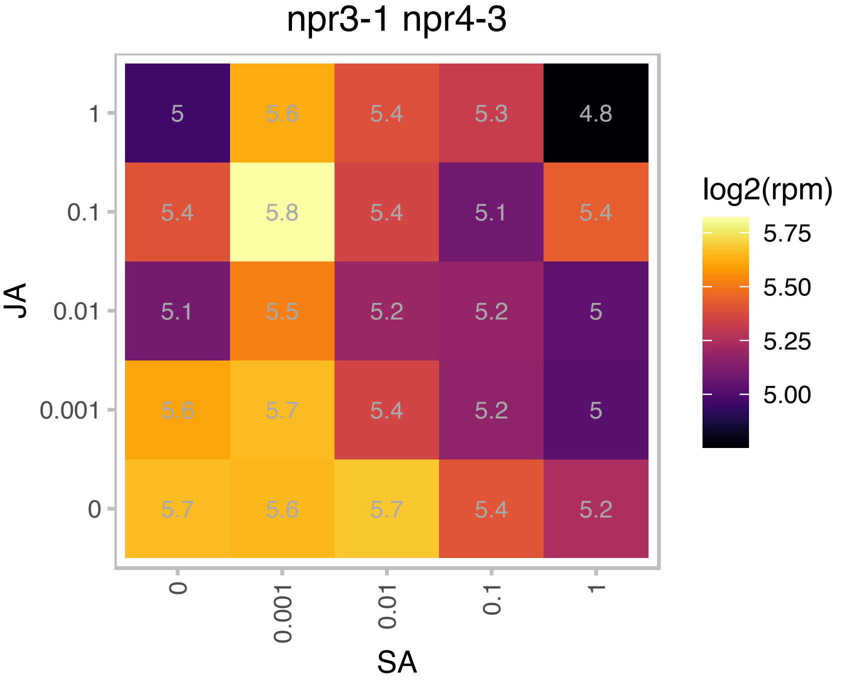

# AT2G37975.1 : NA

Yos1-like protein

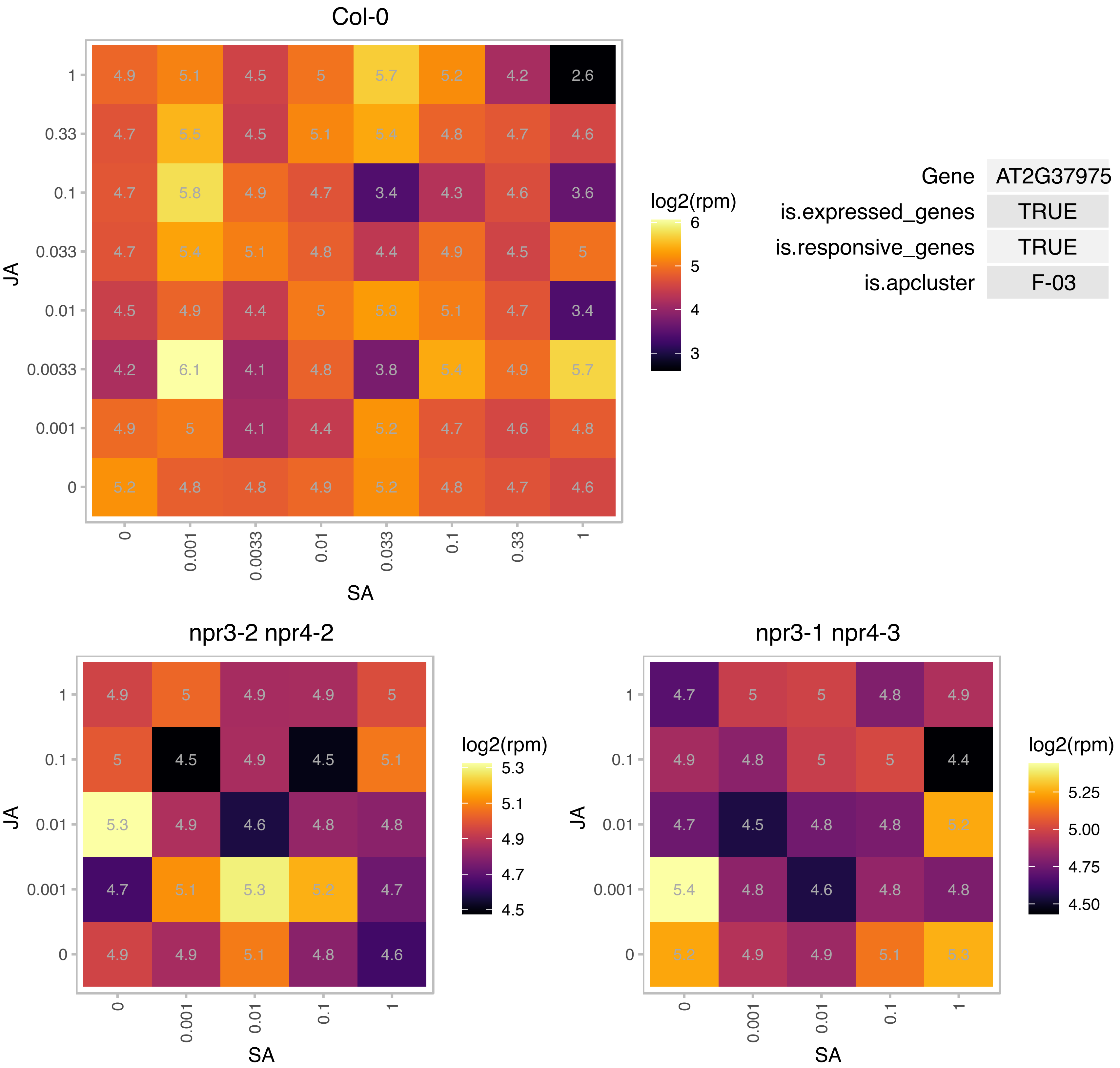


| | |
|---|---|
| Gene | AT2G37975 |
| is.expressed_genes | TRUE |
| is.responsive_genes | TRUE |
| is.apcluster | F-03 |

### AT2G39010.1 : PIP2;6, PIP2E

plasma membrane intrinsic protein 2E

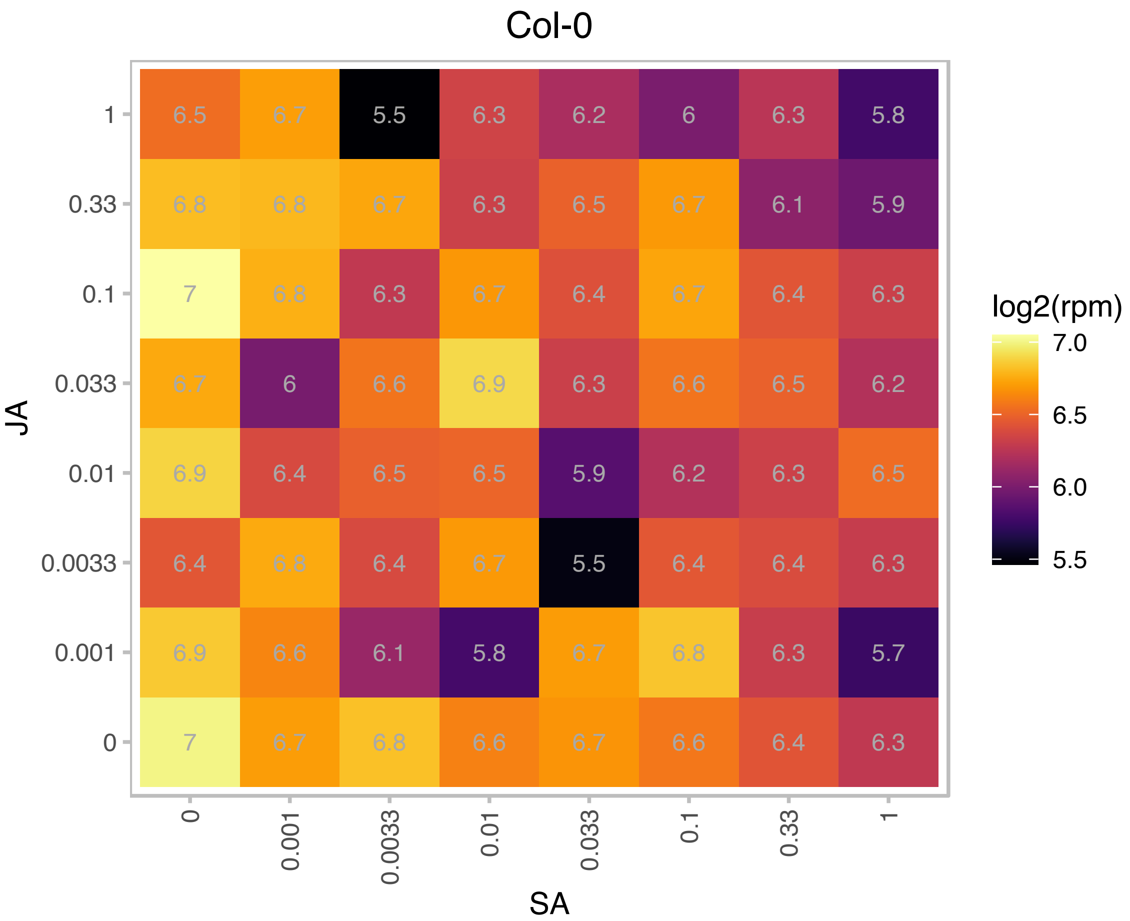


| | |
|---|---|
| Gene | AT2G39010 |
| is.expressed_genes | TRUE |
| is.responsive_genes | TRUE |
| is.apcluster | F-03 |

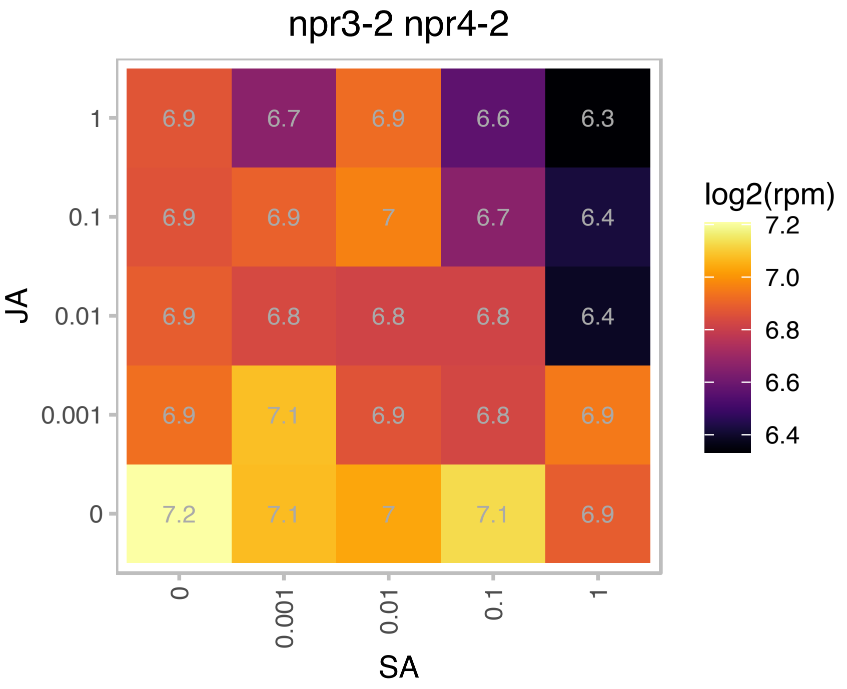


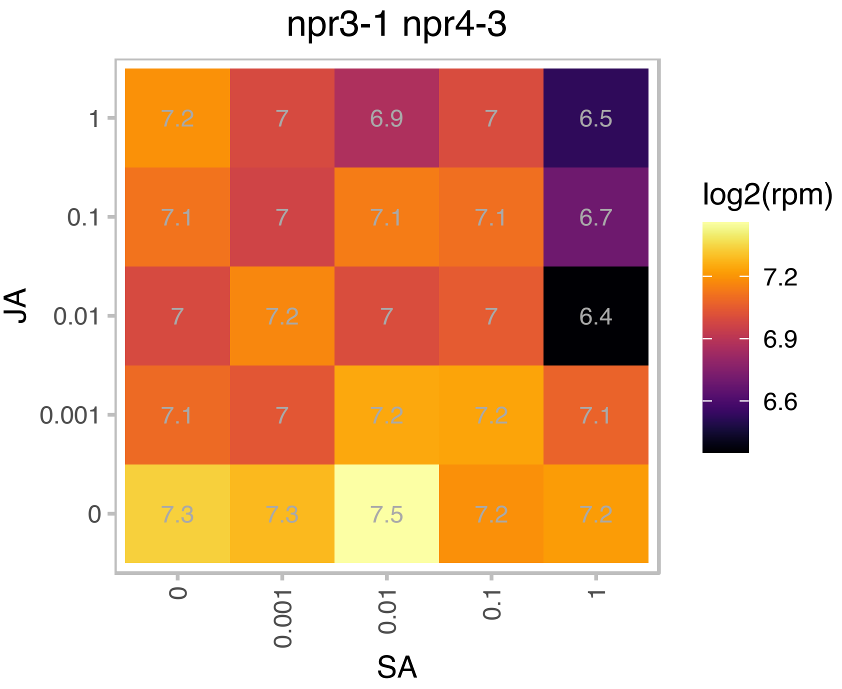

# AT2G41650.1 : NA

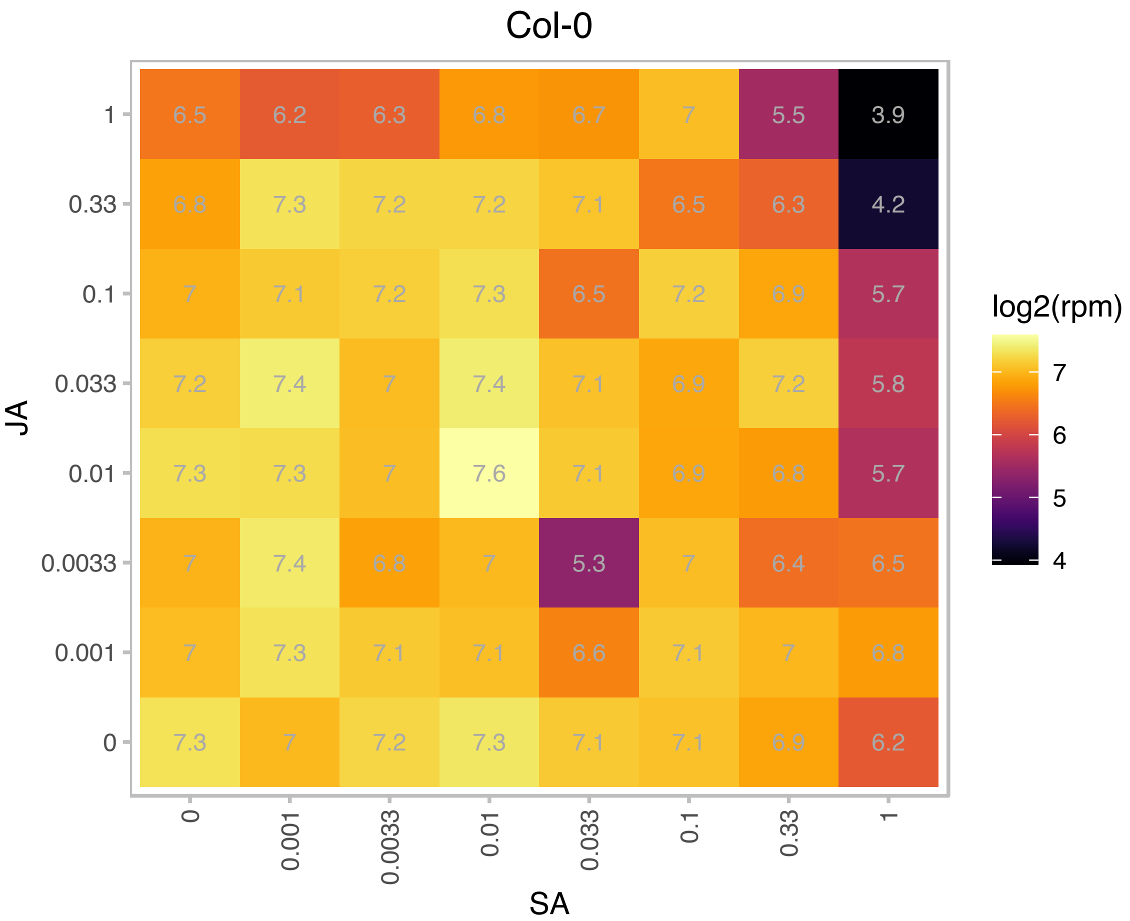


| | |
|---|---|
| Gene | AT2G41650 |
| is.expressed_genes | TRUE |
| is.responsive_genes | TRUE |
| is.apcluster | F-03 |

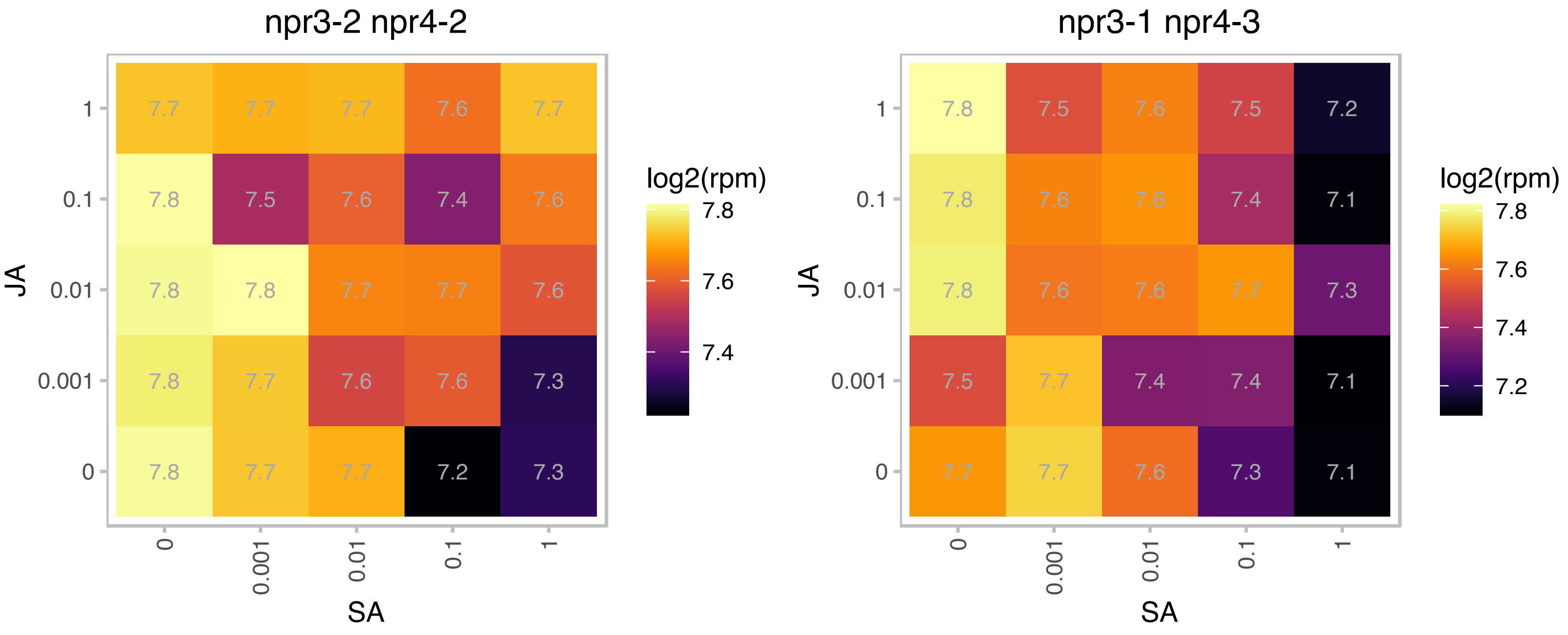

### AT2G42840.1 : PDF1

protodermal factor 1

Encodes a putative extracellular proline-rich protein is exclusively expressed in the L1 layer of vegetative, inflorescence and floral meristems and the protoderm of organ primordia.

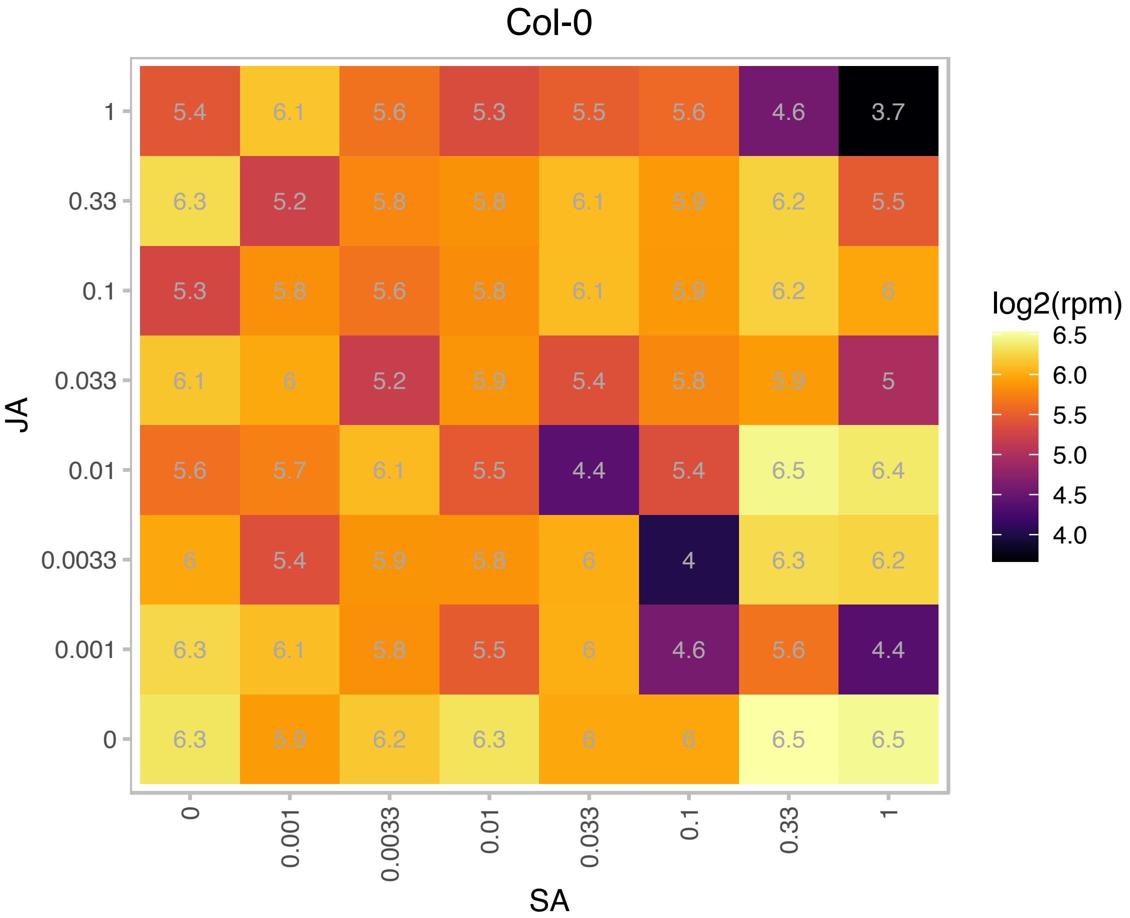


| | |
|---|---|
| Gene | AT2G42840 |
| is.expressed_genes | TRUE |
| is.responsive_genes | TRUE |
| is.apcluster | F-03 |

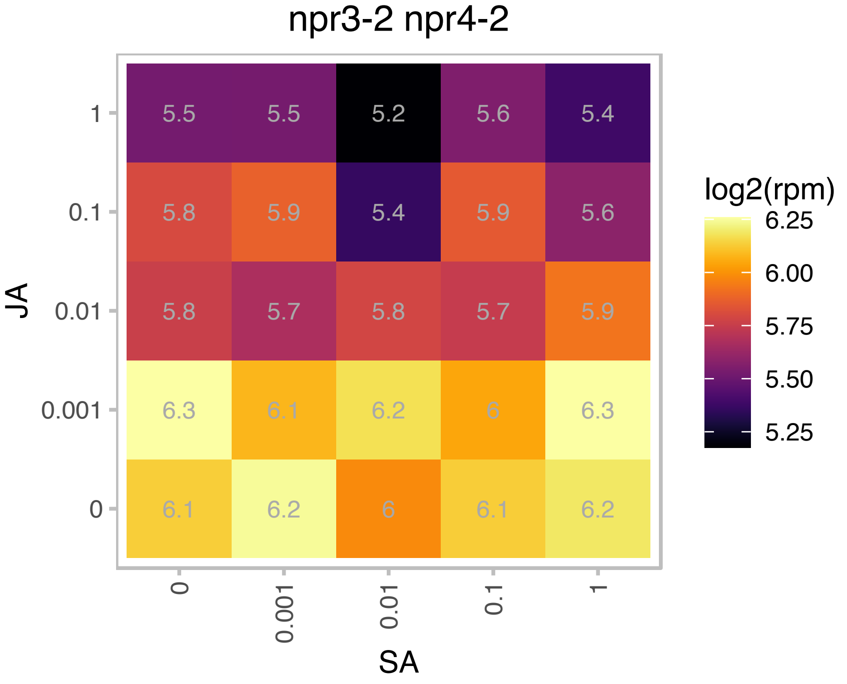


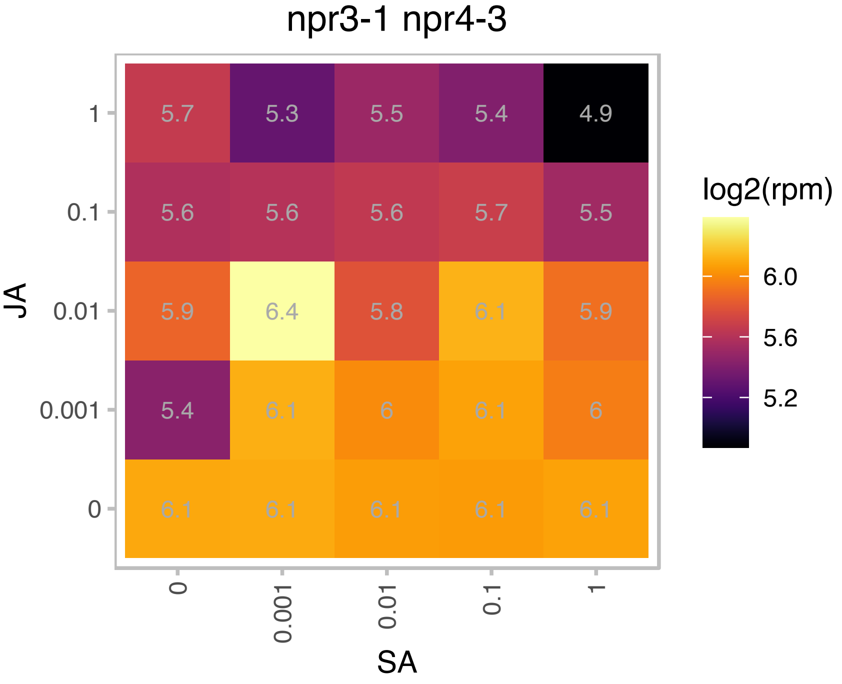

# AT2G43590.1 : NA

Chitinase family protein

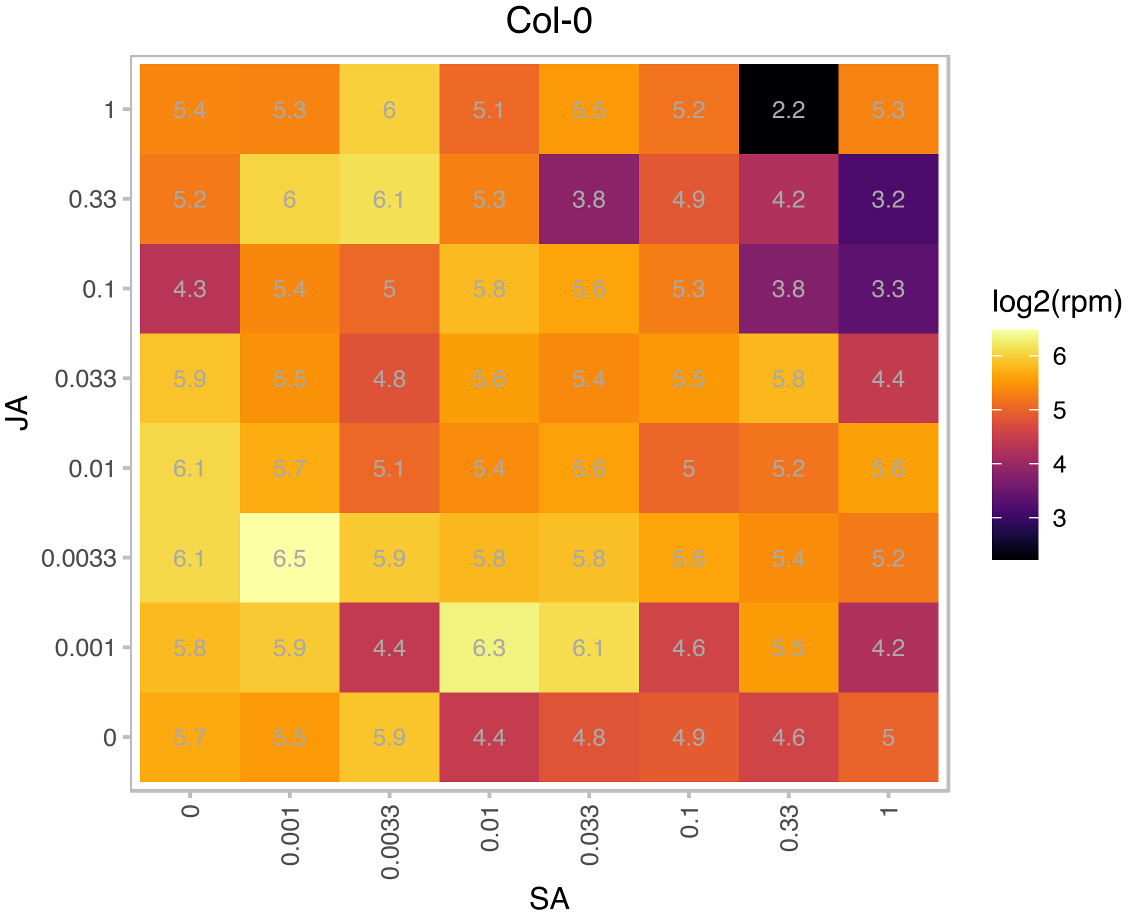


| | |
|---|---|
| Gene | AT2G43590 |
| is.expressed_genes | TRUE |
| is.responsive_genes | TRUE |
| is.apcluster | F-03 |

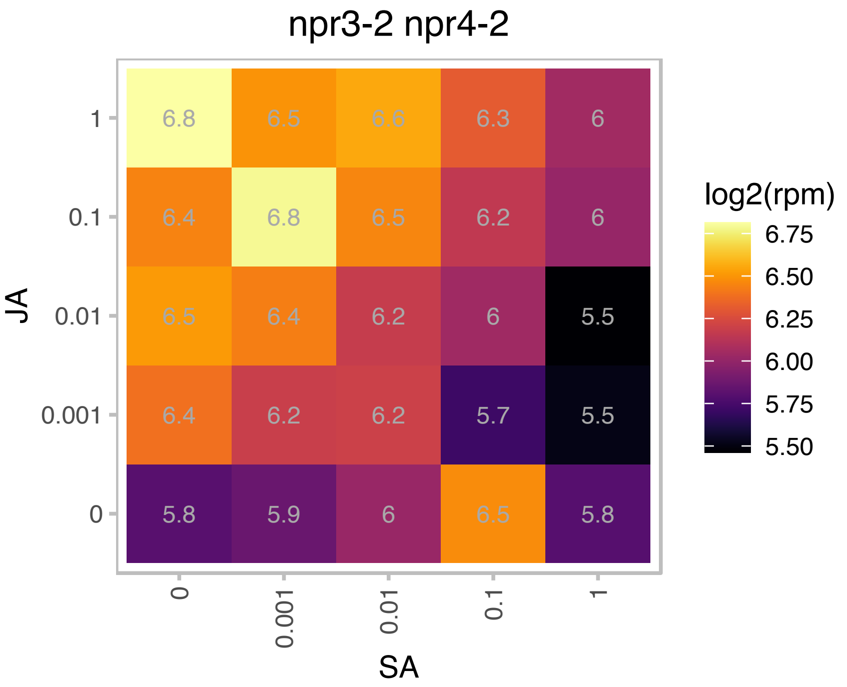


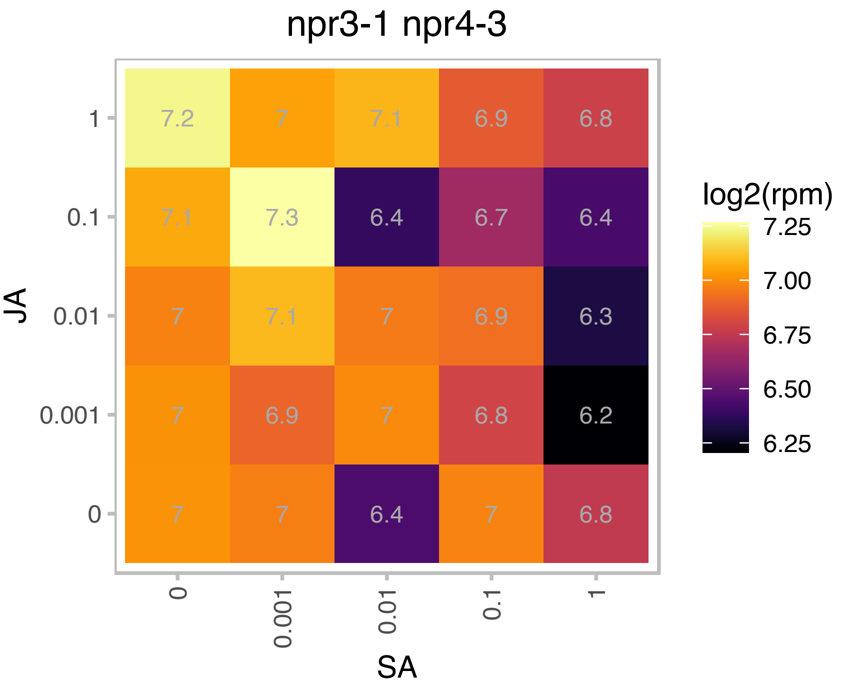

### AT2G43760.1 : CNX6

molybdopterin biosynthesis MoaE family protein

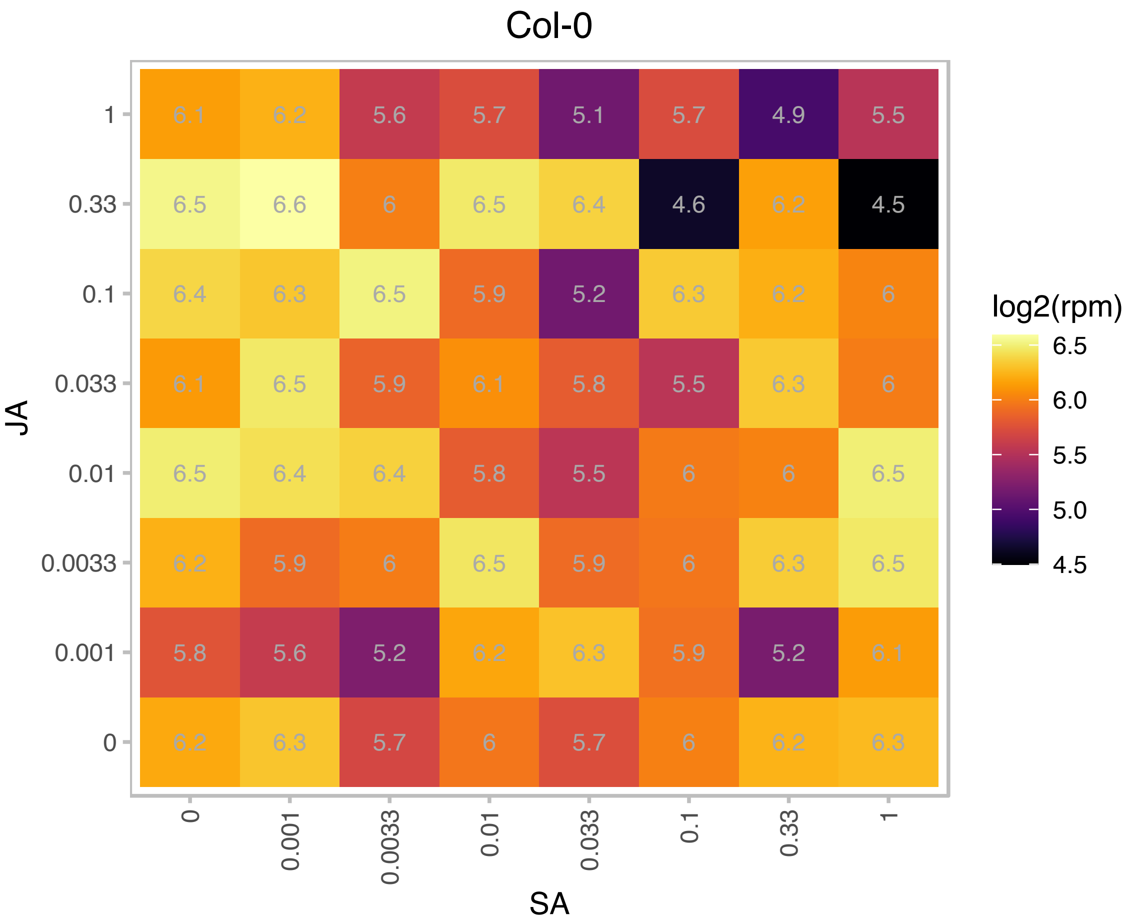


| | |
|---|---|
| Gene | AT2G43760 |
| is.expressed_genes | TRUE |
| is.responsive_genes | TRUE |
| is.apcluster | F-03 |

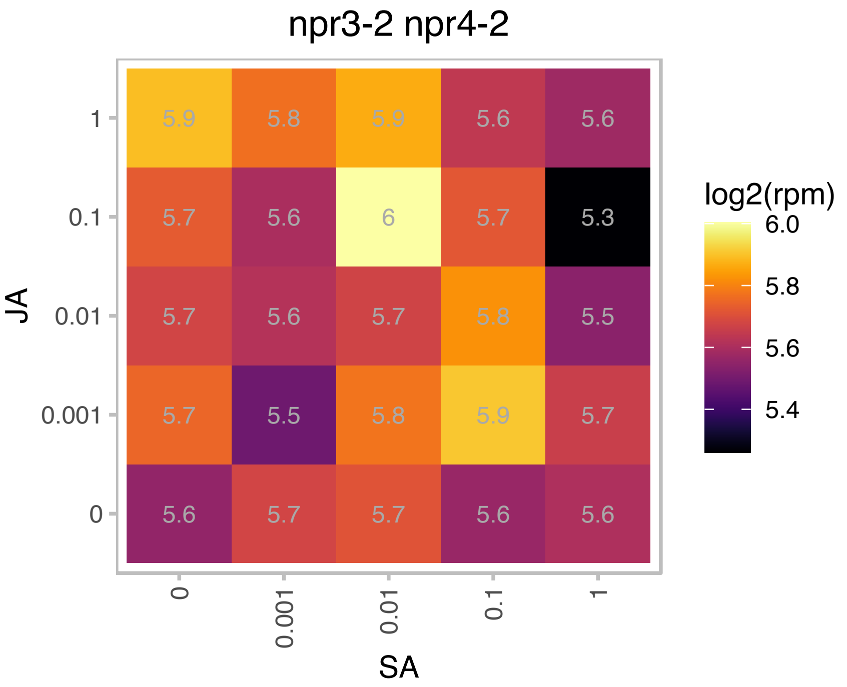


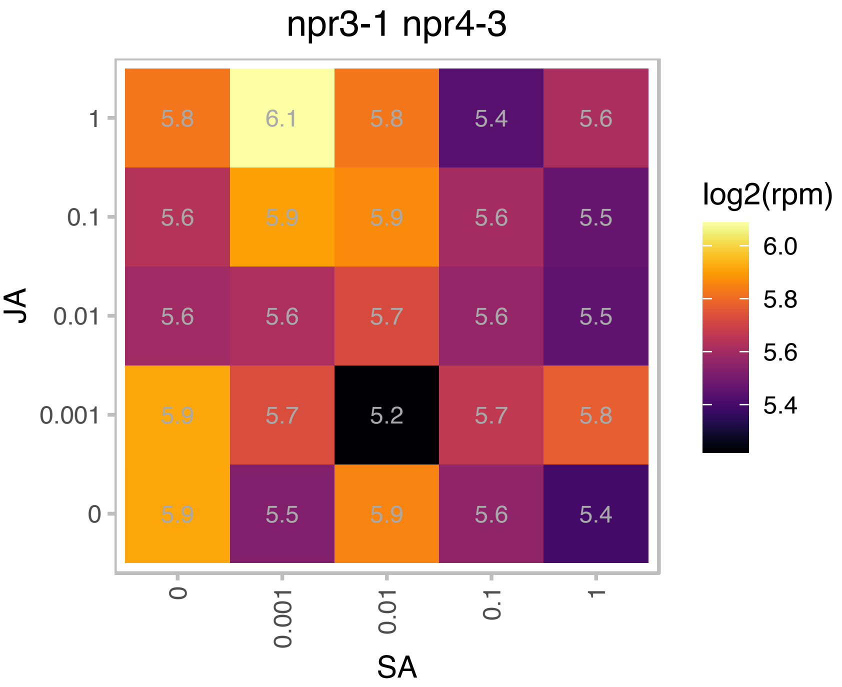

# AT2G45530.1 : NA

RING/U-box superfamily protein

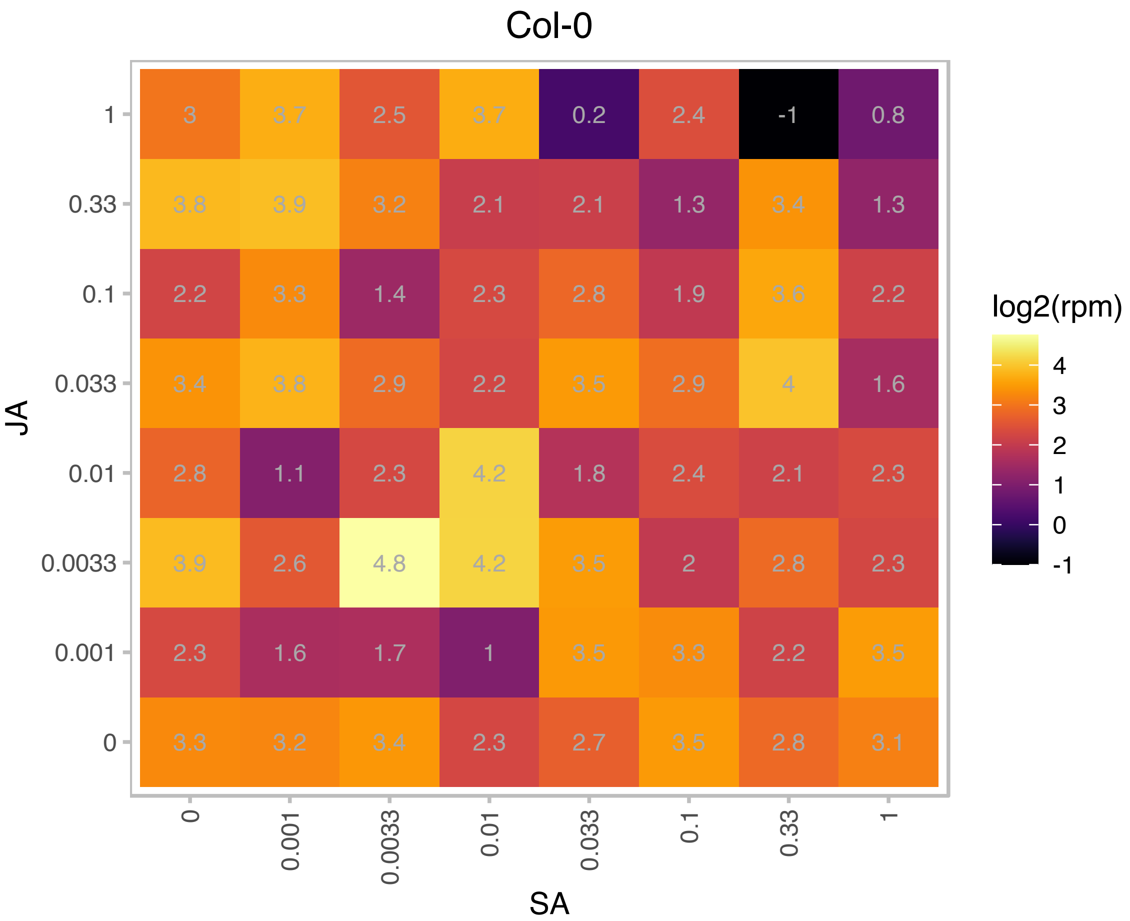


| | |
|---|---|
| Gene | AT2G45530 |
| is.expressed_genes | TRUE |
| is.responsive_genes | TRUE |
| is.apcluster | F-03 |

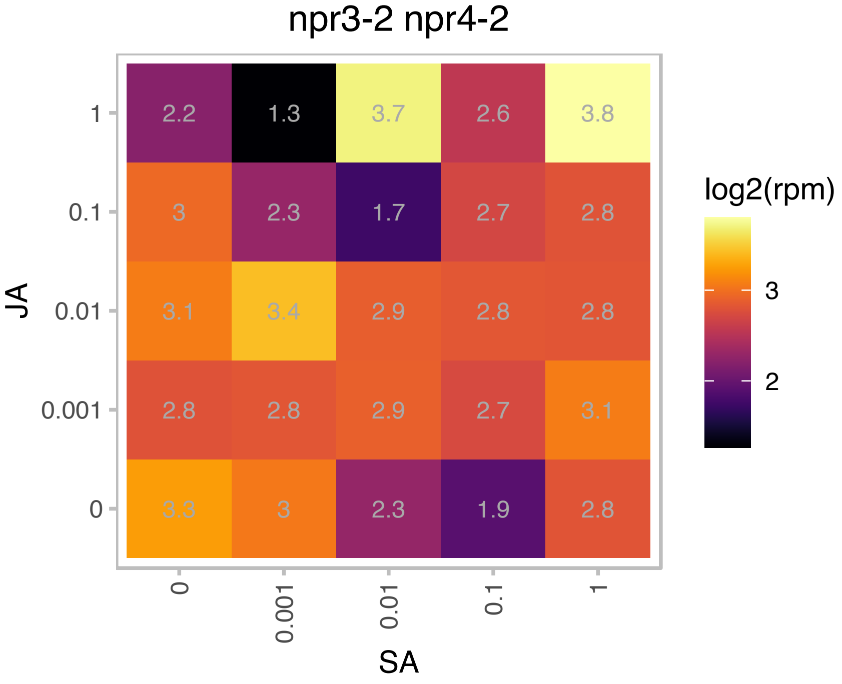


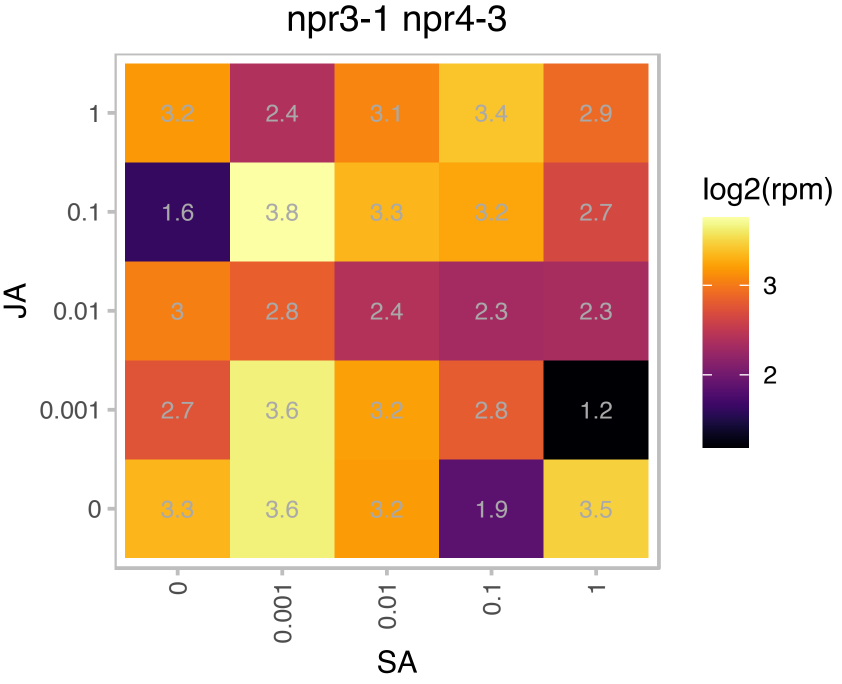

### AT2G45960.1 : ATHH2, PIP1;2, PIP1B, TMP-A

plasma membrane intrinsic protein 1B
a member of the plasma membrane intrinsic protein subfamily PIP1. localizes to the plasma membrane and exhibits water transport activity in Xenopus oocyte. expressed ubiquitously and protein level decreases slightly during leaf development.

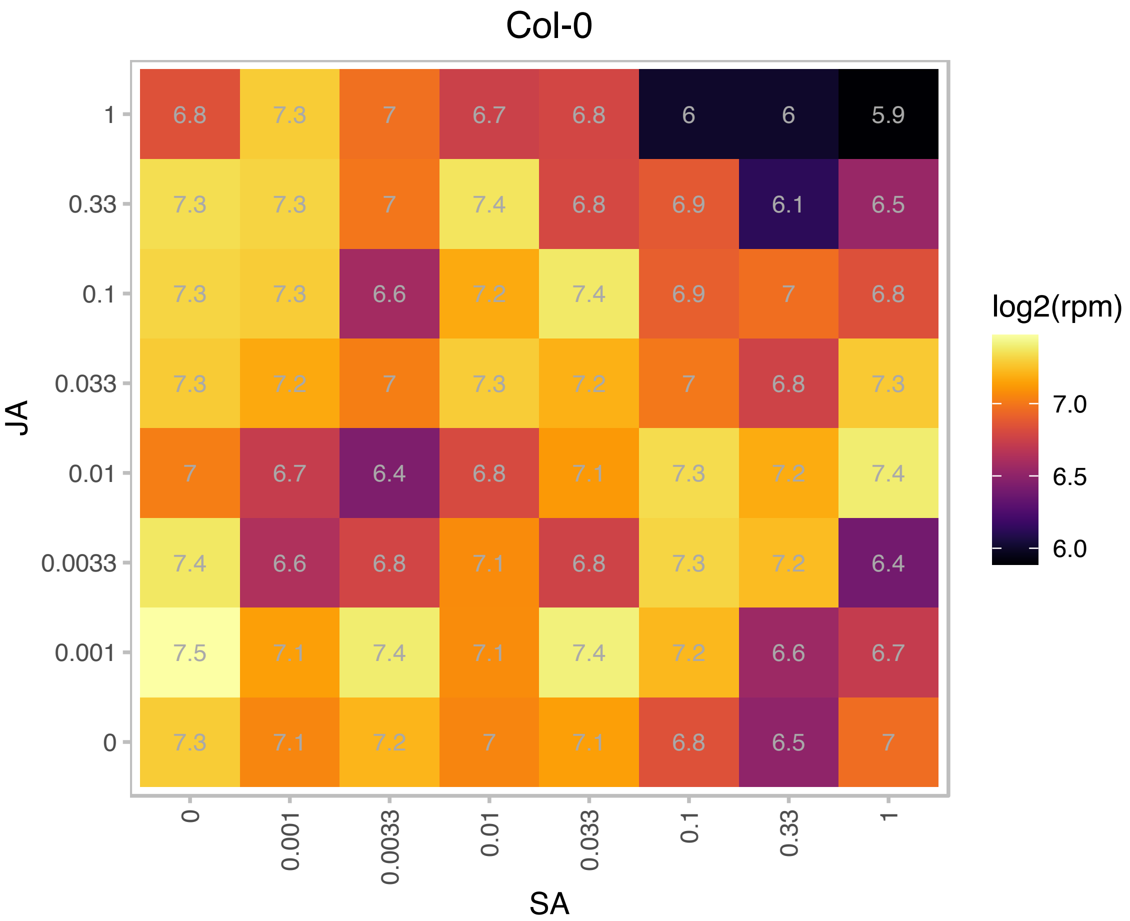


| | |
|---|---|
| Gene | AT2G45960 |
| is.expressed_genes | TRUE |
| is.responsive_genes | TRUE |
| is.apcluster | F-03 |

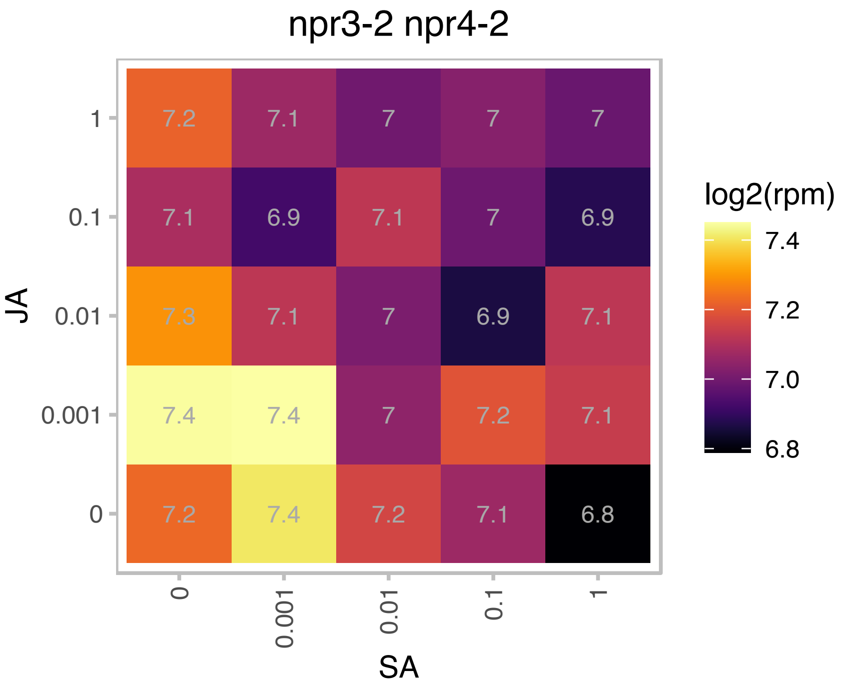


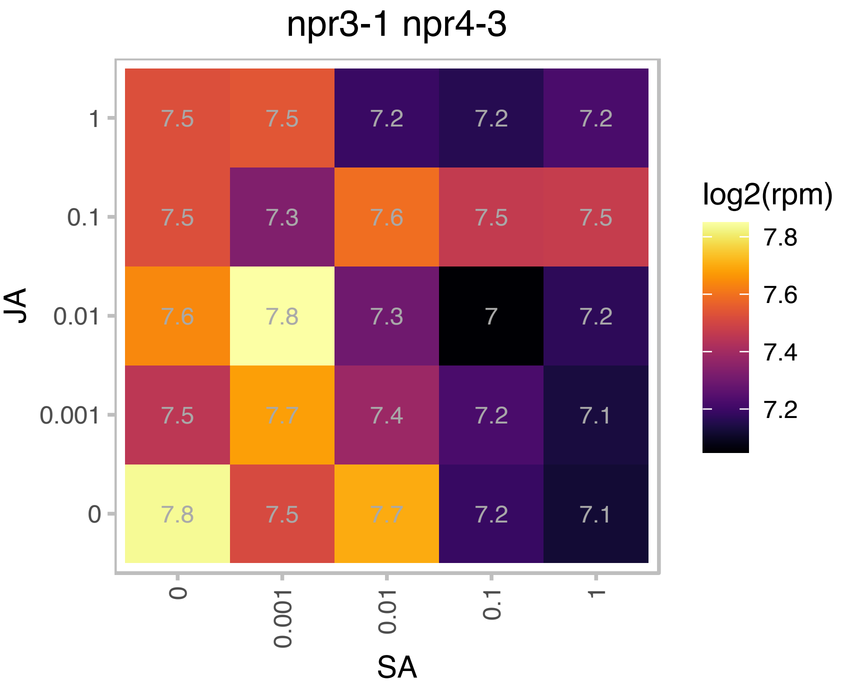

### AT2G47240.1 : CER8, LACS1

AMP-dependent synthetase and ligase family protein

Encodes an acyl-CoA synthetase that acts on long-chain and very-long-chain fatty acids, involved in cuticular wax and cutin biosynthesis The mRNA is cell-to-cell mobile.

Col-0

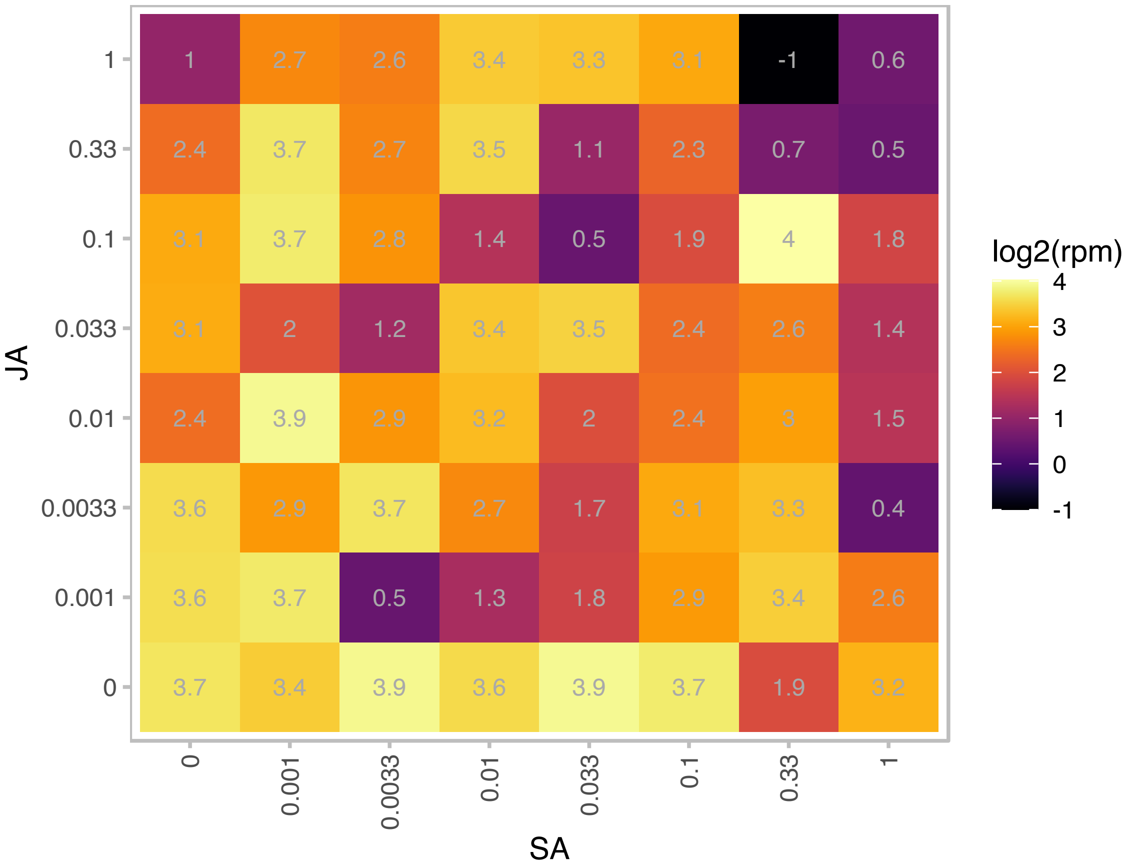


| | |
|---|---|
| Gene | AT2G47240 |
| is.expressed_genes | TRUE |
| is.responsive_genes | TRUE |
| is.apcluster | F-03 |

npr3-2 npr4-2

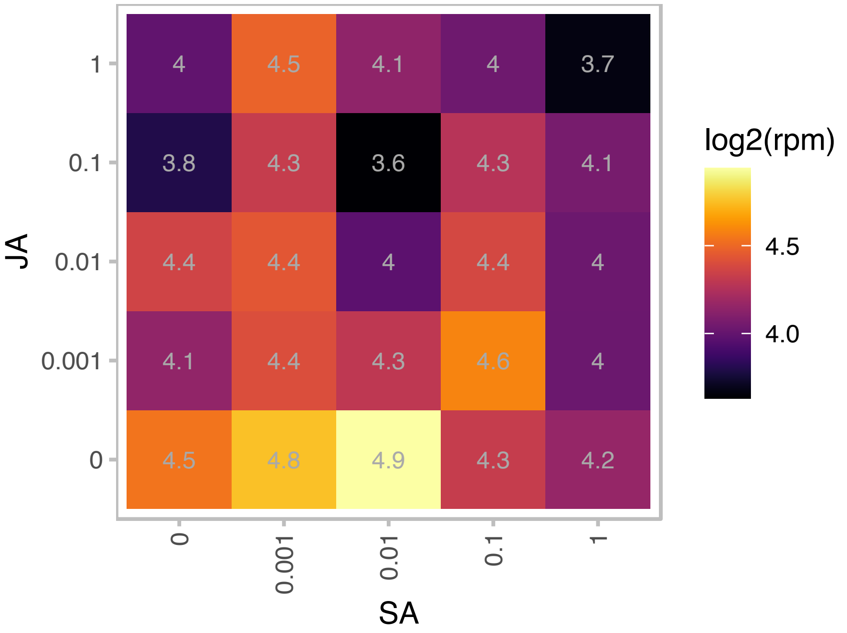


npr3-1 npr4-3

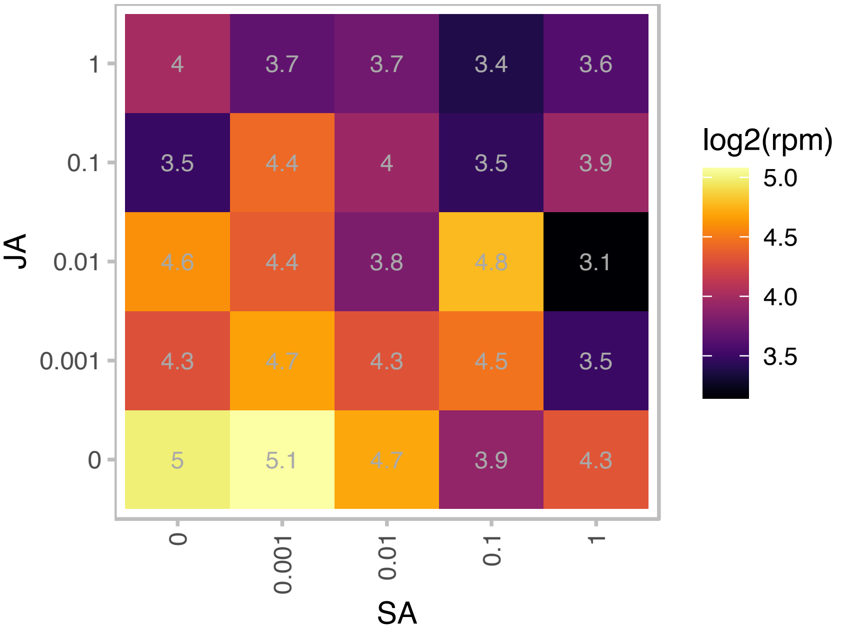

# AT3G00590.1 : NA

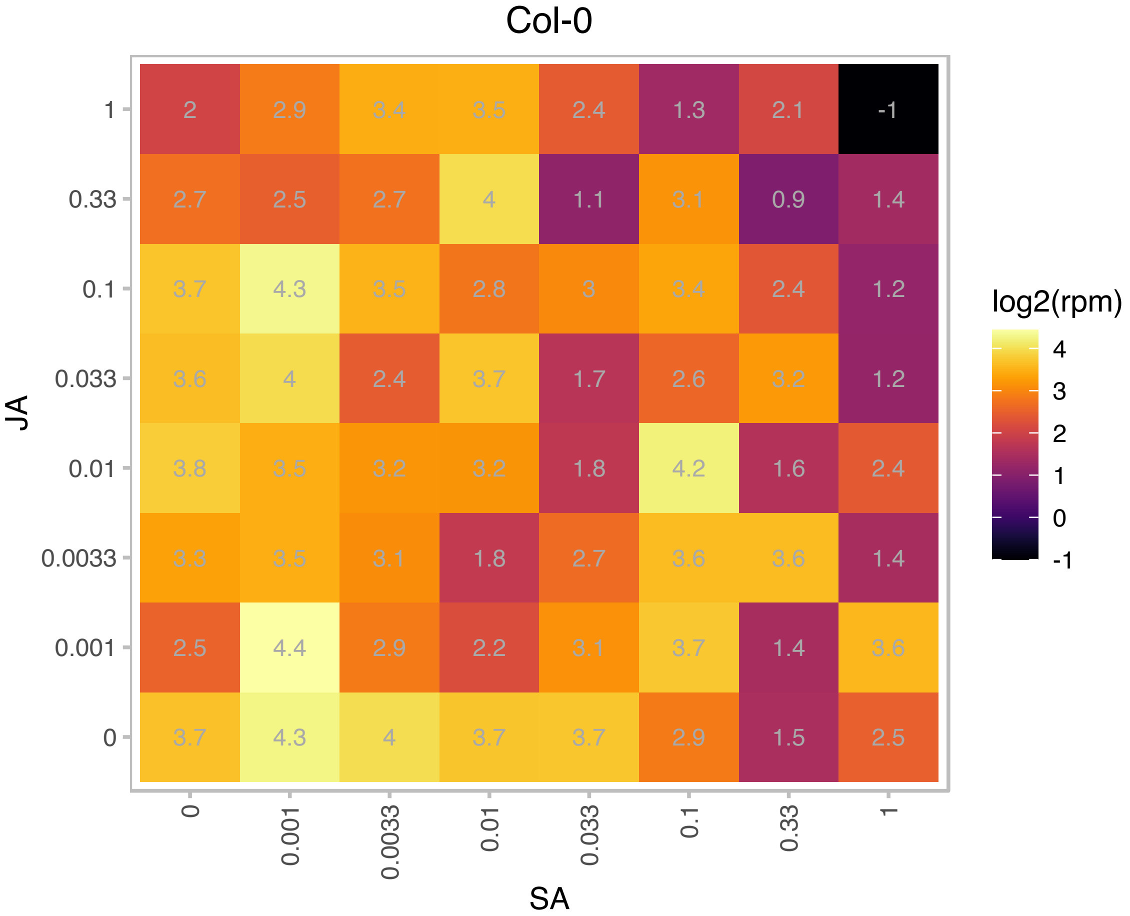


| | |
|---|---|
| Gene | AT3G00590 |
| is.expressed_genes | TRUE |
| is.responsive_genes | TRUE |
| is.apcluster | F-03 |

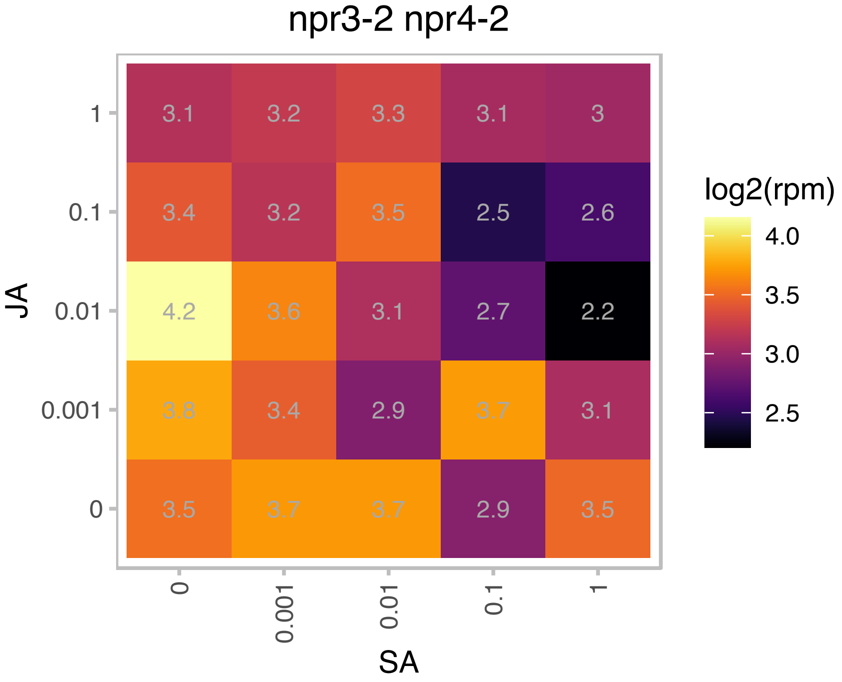


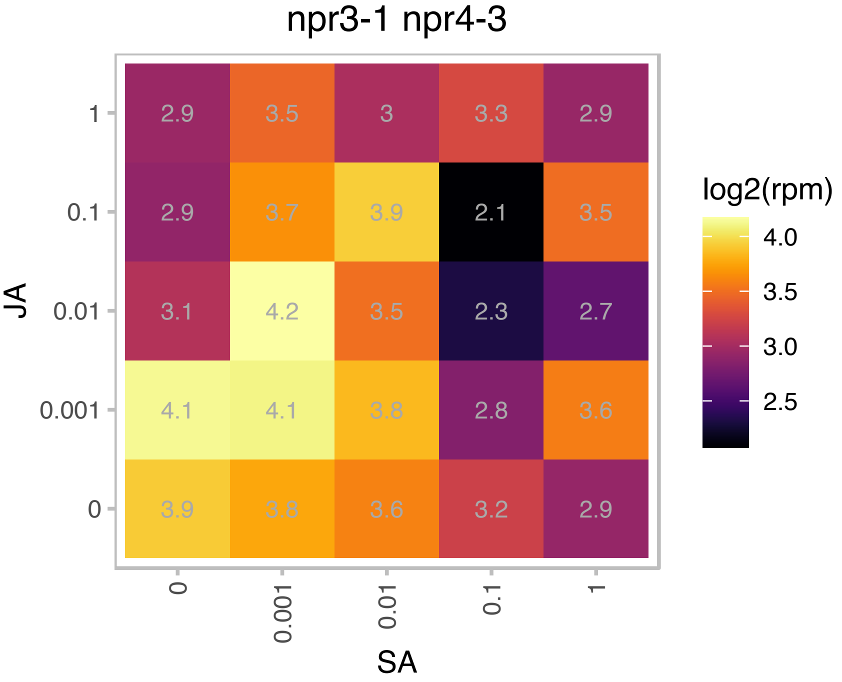

# AT3G03960.1 : NA

TCP-1/cpn60 chaperonin family protein

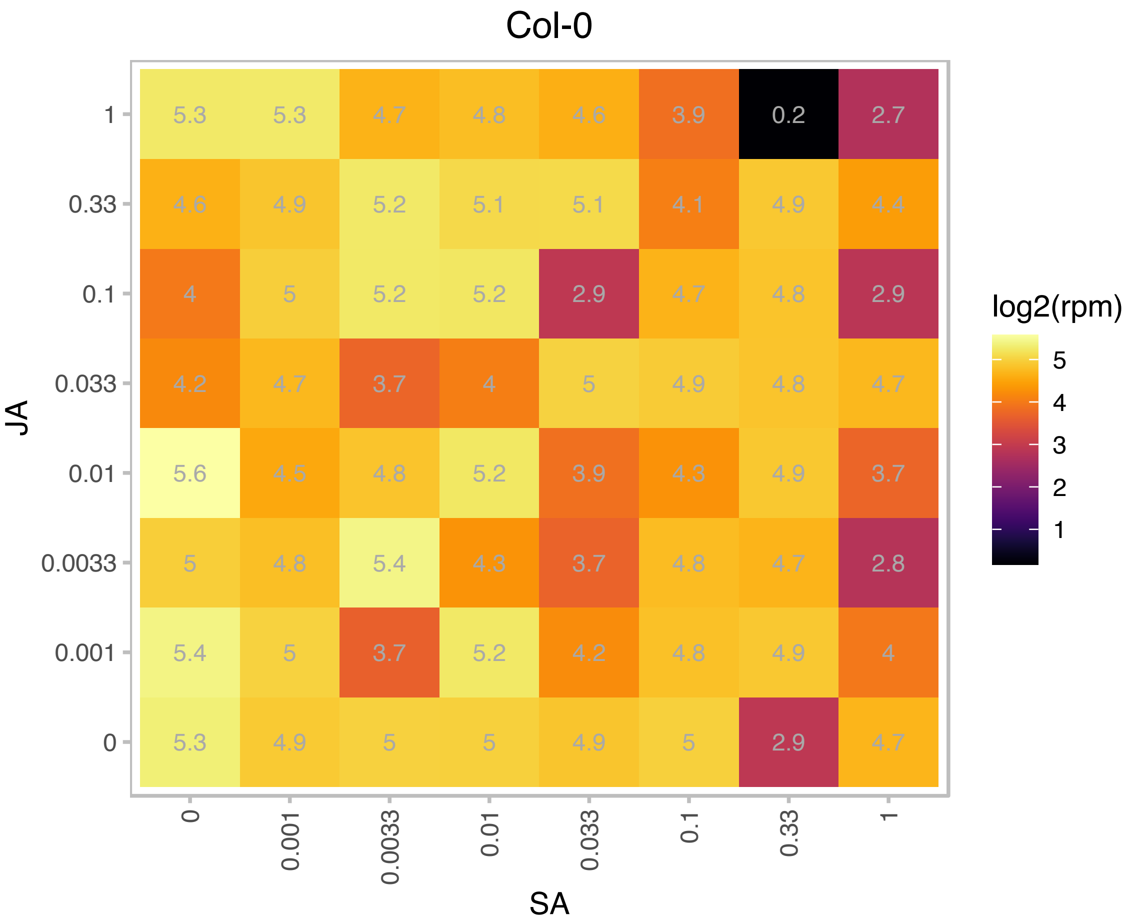


| | |
|---|---|
| Gene | AT3G03960 |
| is.expressed_genes | TRUE |
| is.responsive_genes | TRUE |
| is.apcluster | F-03 |

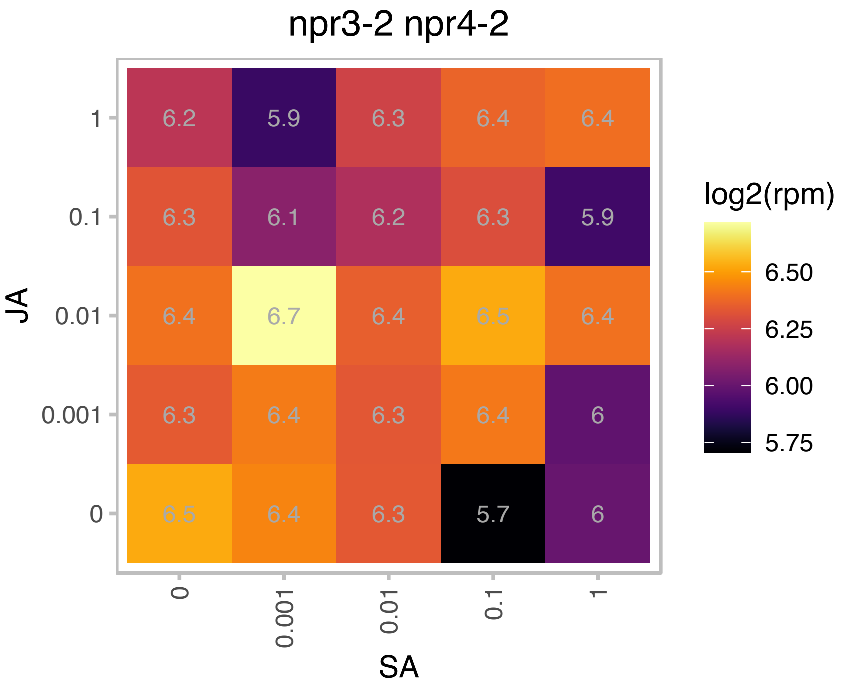


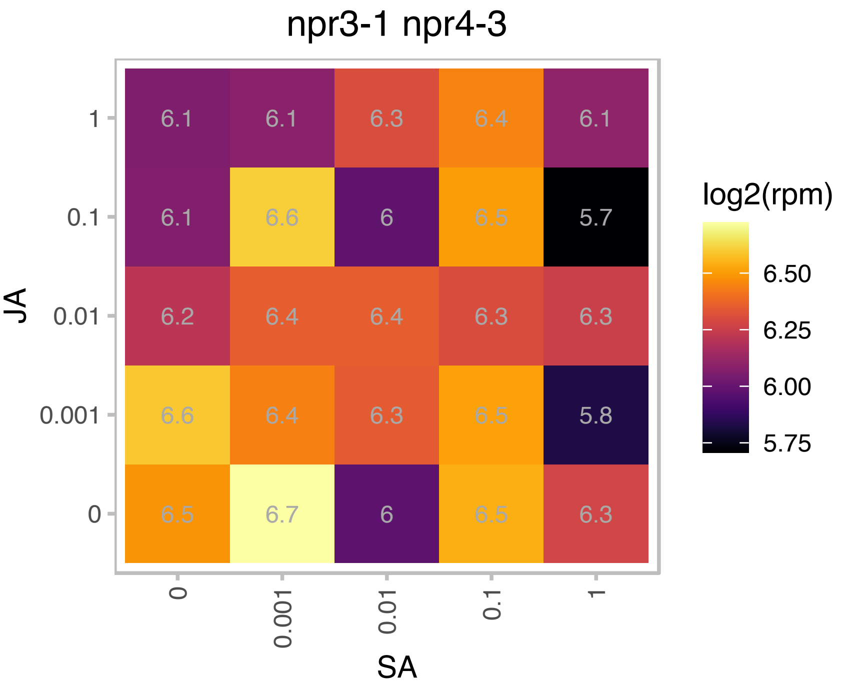

# AT3G05060.1 : NA

NOP56-like pre RNA processing ribonucleoprotein

SAR DNA-binding protein, putative, strong similarity to SAR DNA-binding protein-1 (Pisum sativum) GI:3132696; contains Pfam profile PF01798: Putative snoRNA binding domain; encodes NOP58-like protein

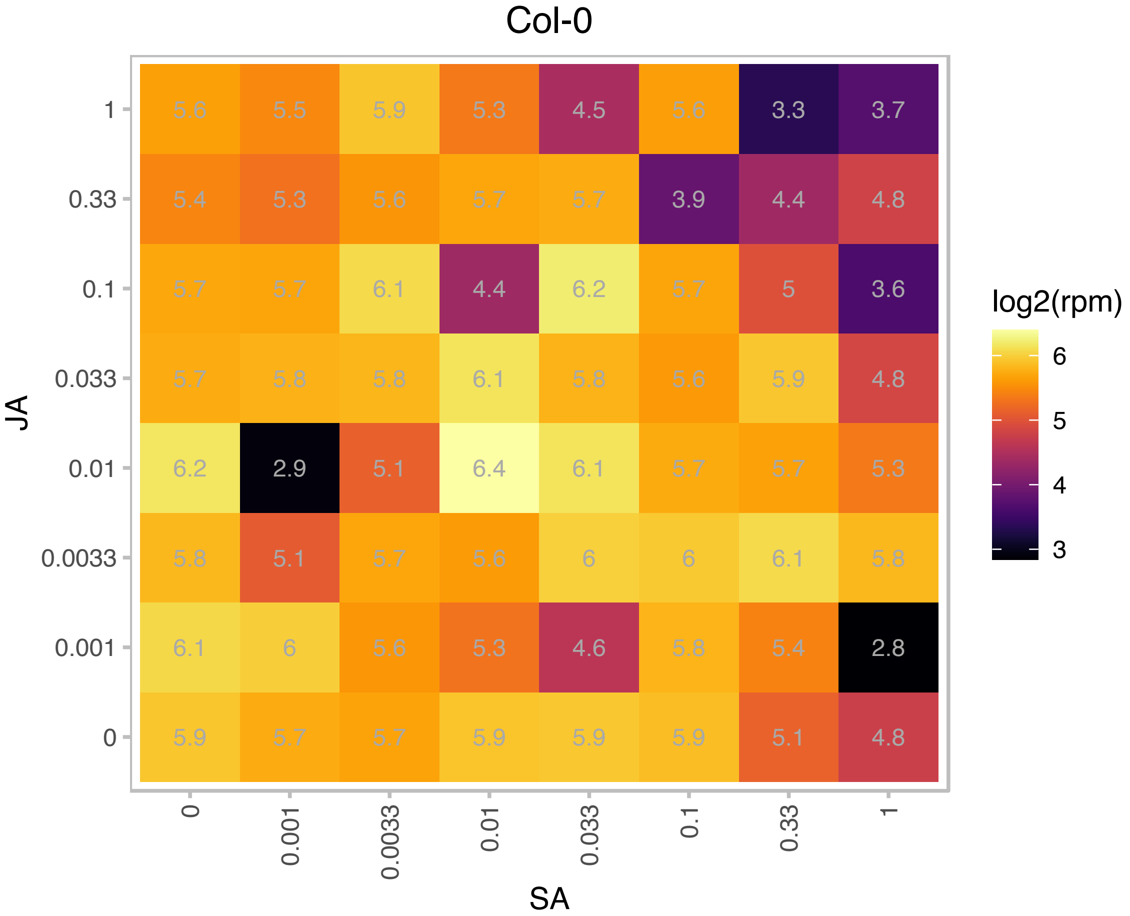


| | |
|---|---|
| Gene | AT3G05060 |
| is.expressed_genes | TRUE |
| is.responsive_genes | TRUE |
| is.apcluster | F-03 |

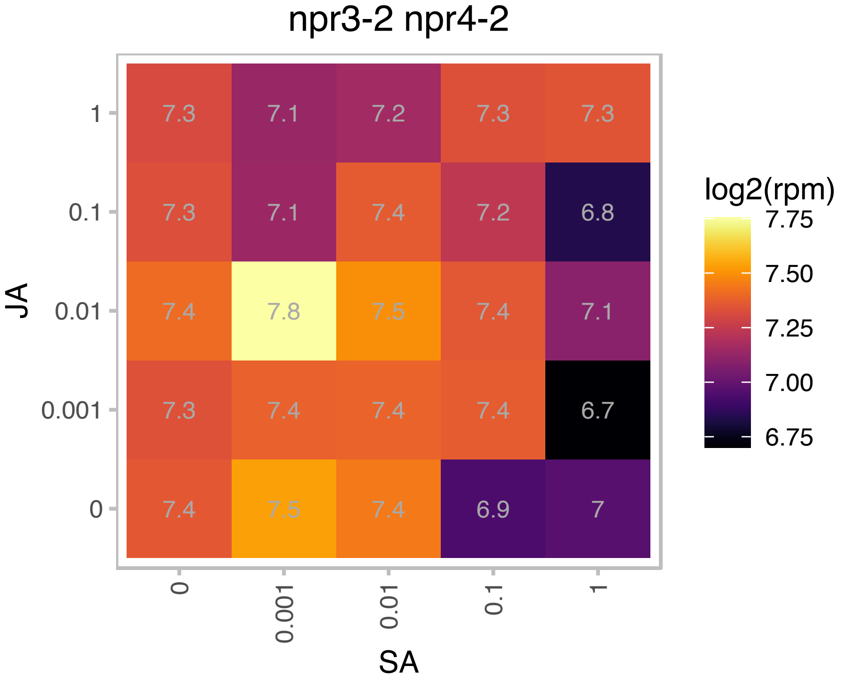


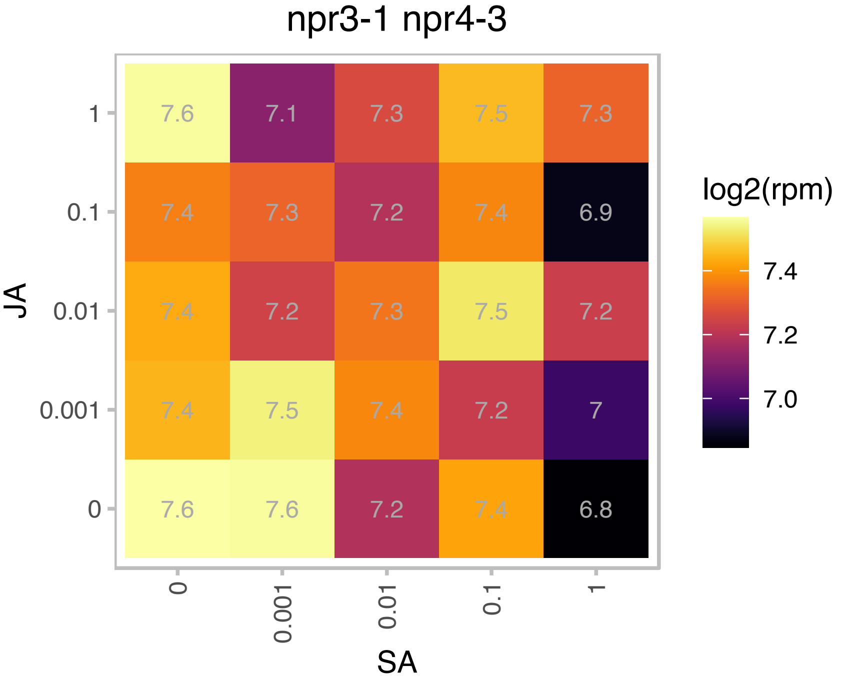

### AT3G05890.1 : LTI6b, RCI2B

Low temperature and salt responsive protein family

Col-0

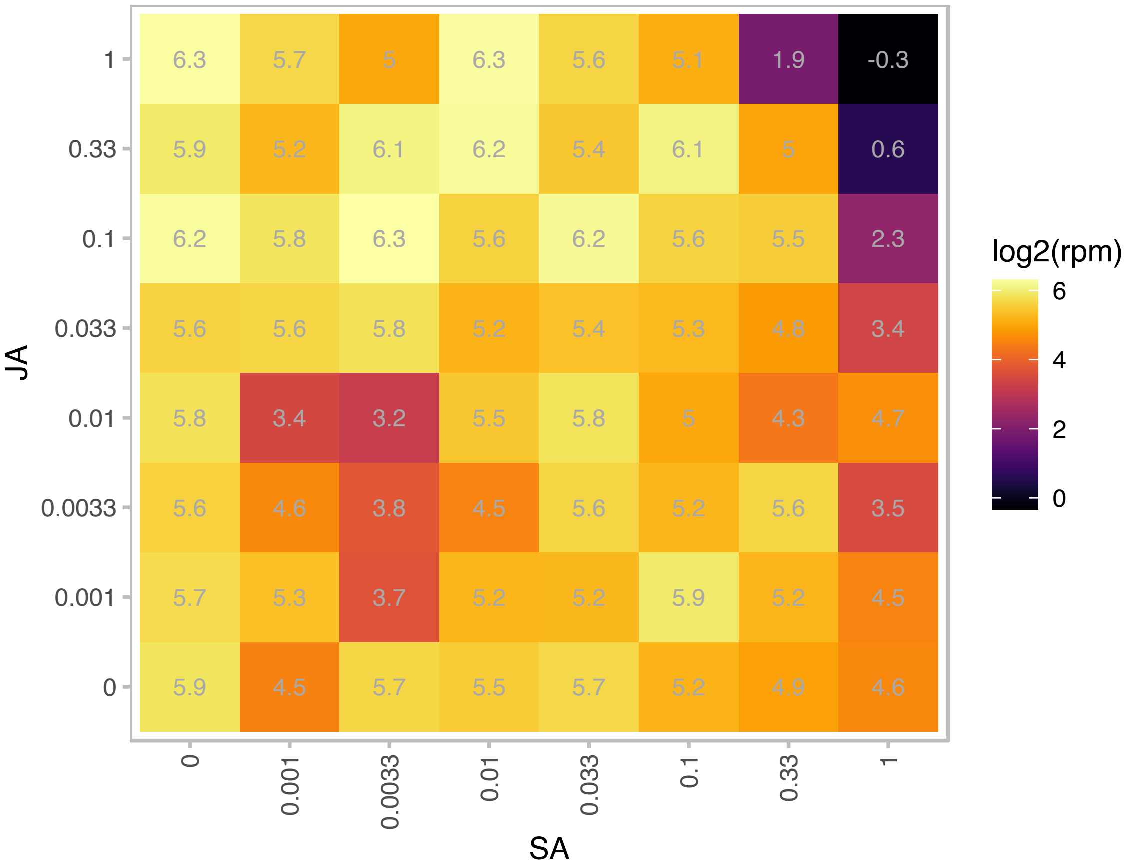


| | |
|---|---|
| Gene | AT3G05890 |
| is.expressed_genes | TRUE |
| is.responsive_genes | TRUE |
| is.apcluster | F-03 |

npr3-2 npr4-2

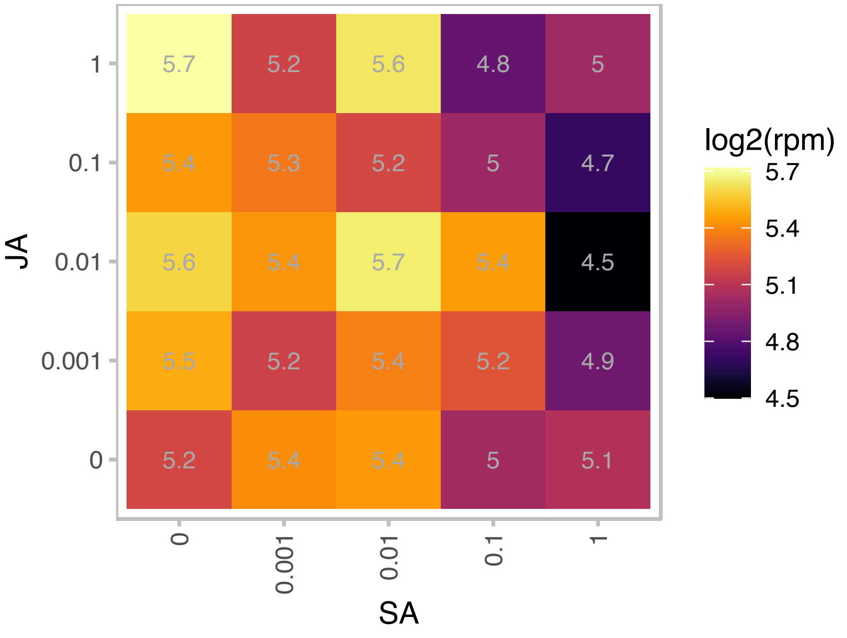


npr3-1 npr4-3

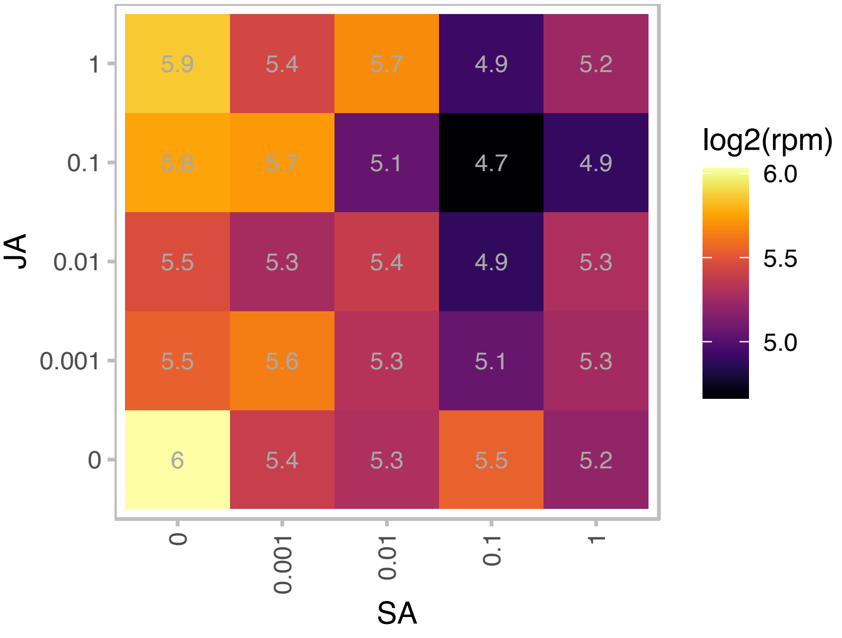

# AT3G07230.1 : NA

wound-responsive protein-related

| | |
|---|---|
| Gene | AT3G07230 |
| is.expressed_genes | TRUE |
| is.responsive_genes | TRUE |
| is.apcluster | F-03 |

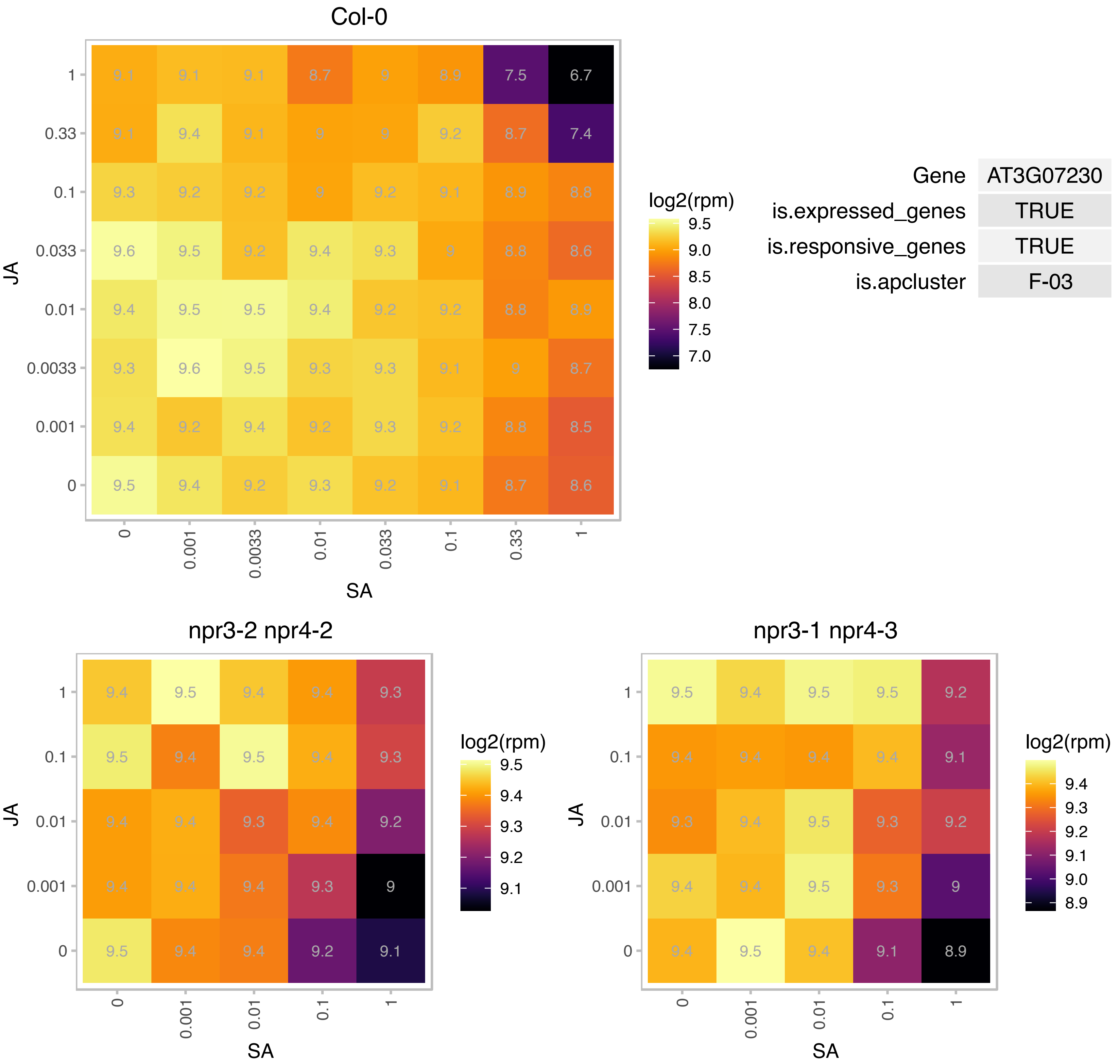

### AT3G16520.1 : UGT88A1

UDP-glucosyl transferase 88A1

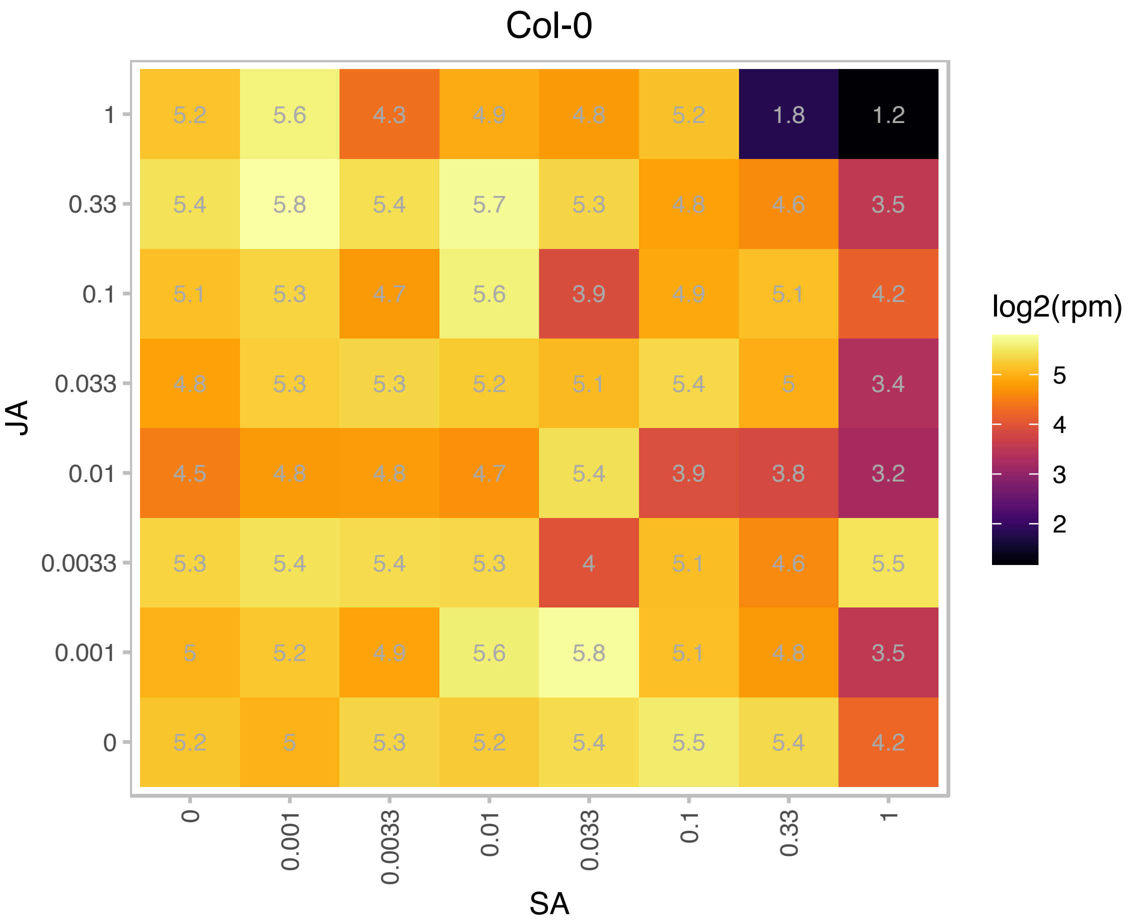


| | |
|---|---|
| Gene | AT3G16520 |
| is.expressed_genes | TRUE |
| is.responsive_genes | TRUE |
| is.apcluster | F-03 |

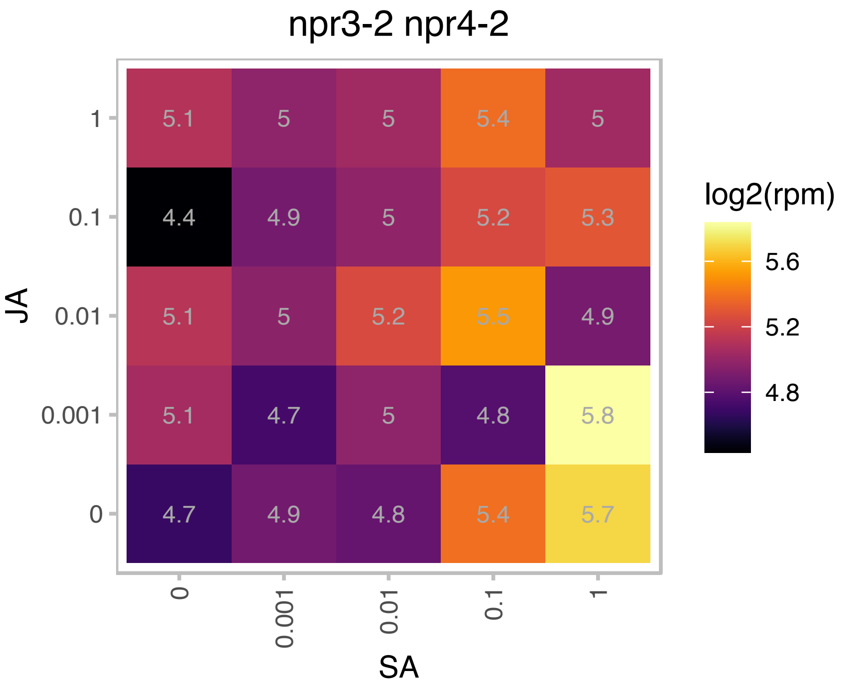


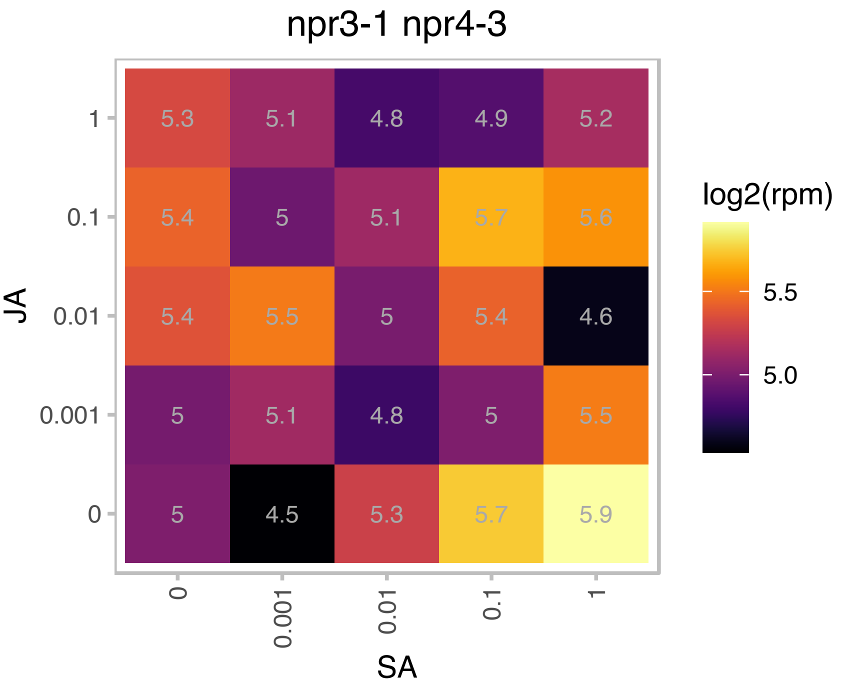

# AT3G18420.1 : SG1

Protein prenylyltransferase superfamily protein

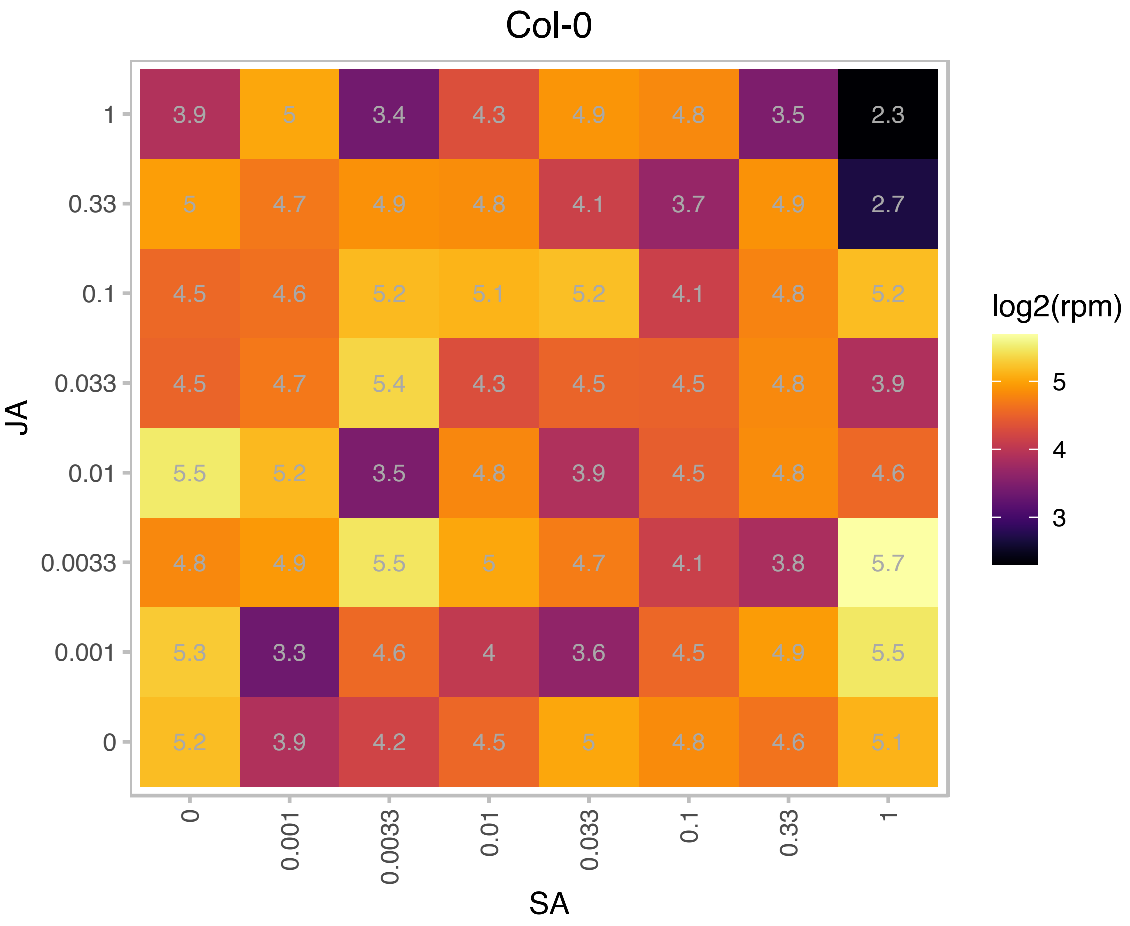

| | |
|---|---|
| Gene | AT3G18420 |
| is.expressed_genes | TRUE |
| is.responsive_genes | TRUE |
| is.apcluster | F-03 |

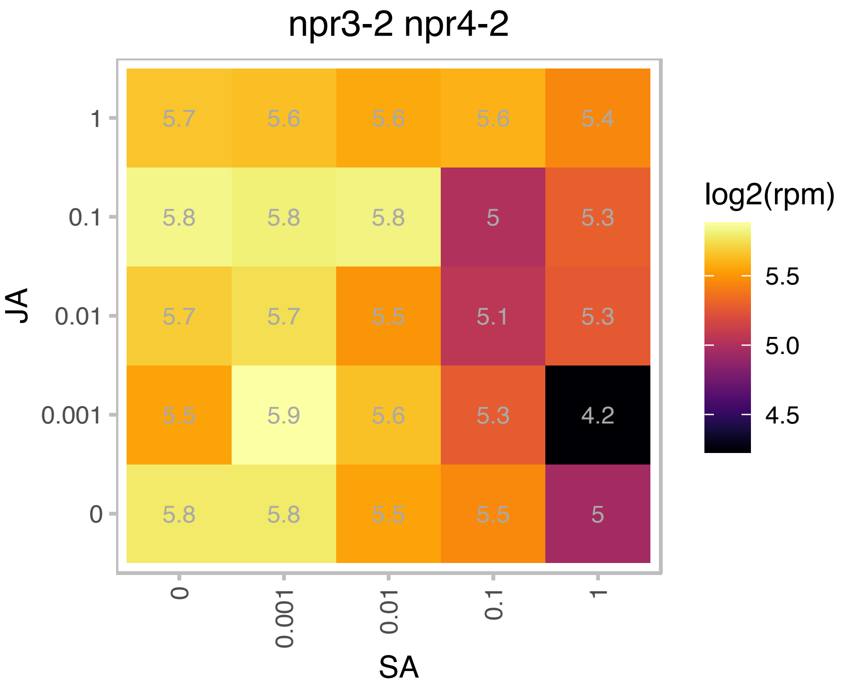

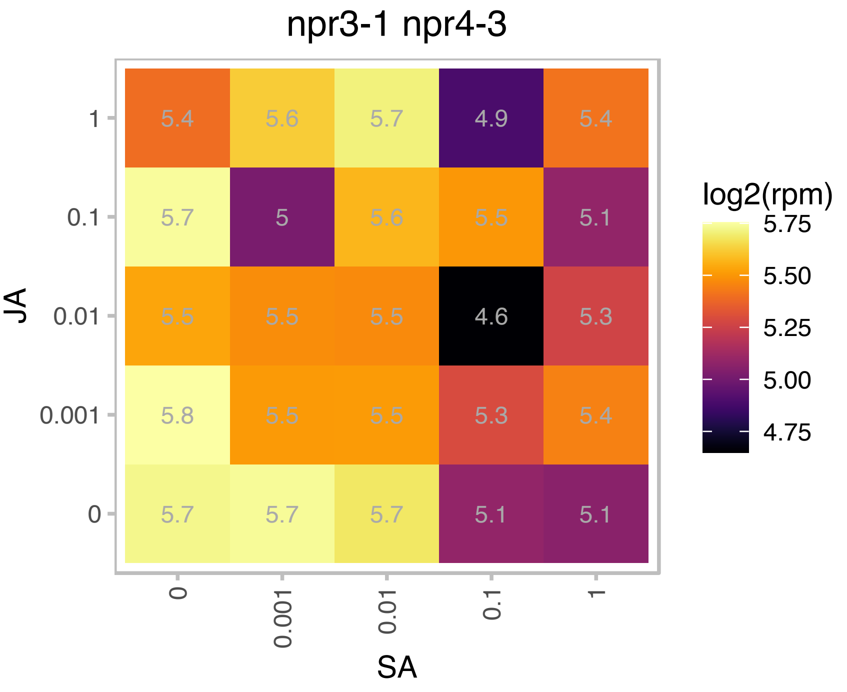

### AT3G22480.1 : PDF2

prefoldin 2

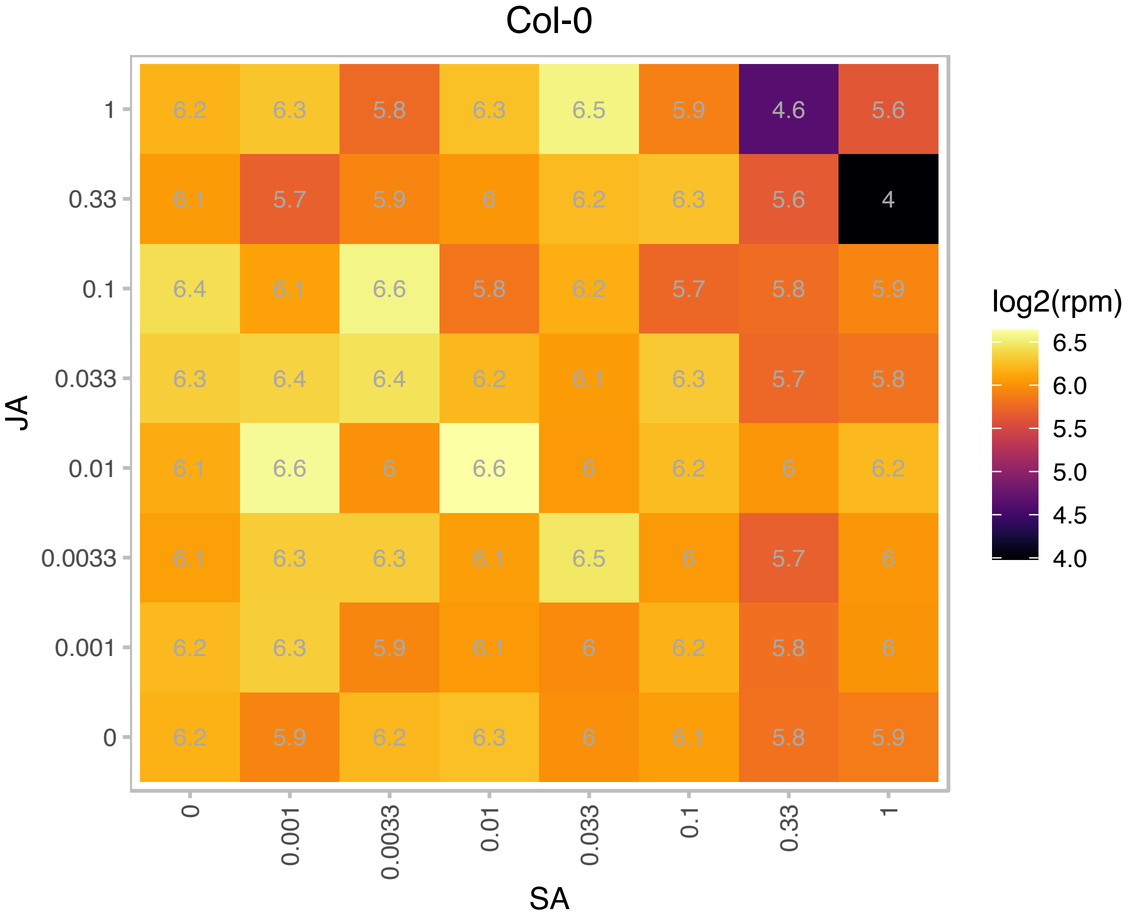


| | |
|---|---|
| Gene | AT3G22480 |
| is.expressed_genes | TRUE |
| is.responsive_genes | TRUE |
| is.apcluster | F-03 |

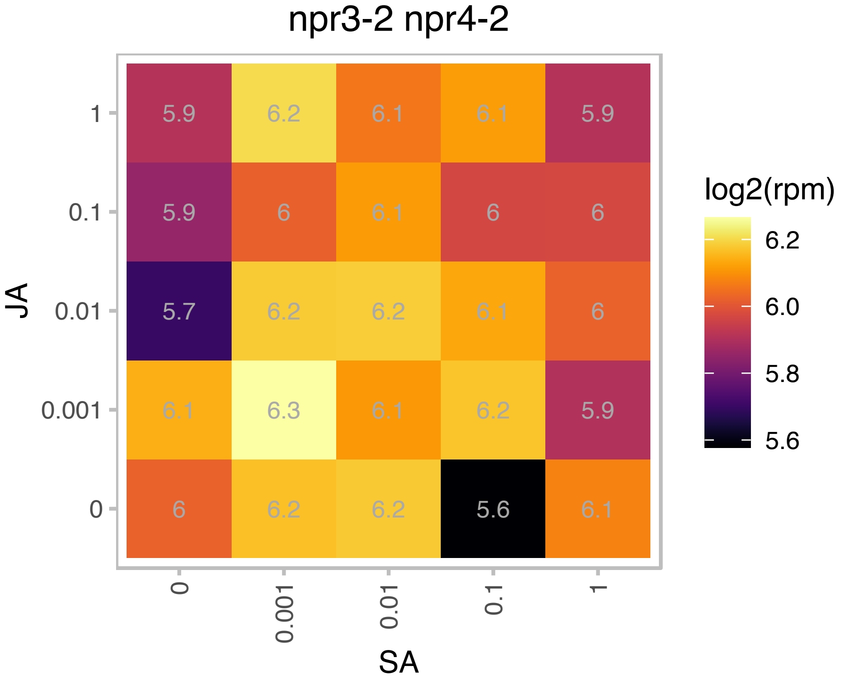


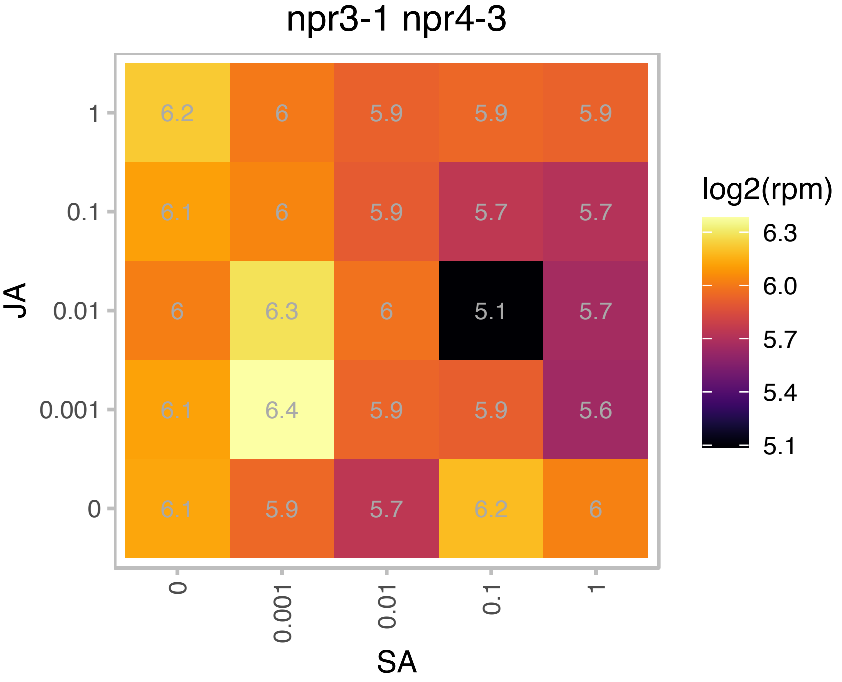

### AT3G23810.1 : ATSAHH2, SAHH2

S-adenosyl-l-homocysteine (SAH) hydrolase 2

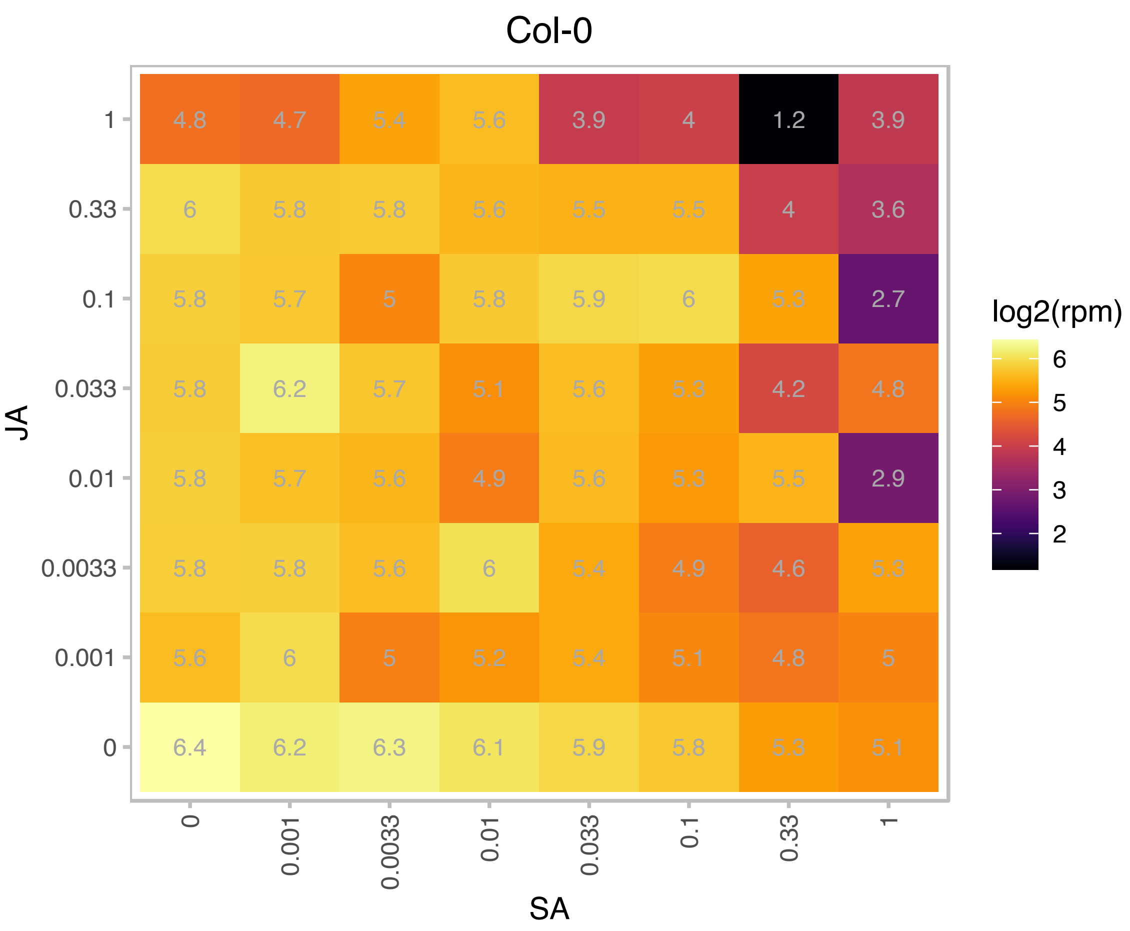


| | |
|---|---|
| Gene | AT3G23810 |
| is.expressed_genes | TRUE |
| is.responsive_genes | TRUE |
| is.apcluster | F-03 |

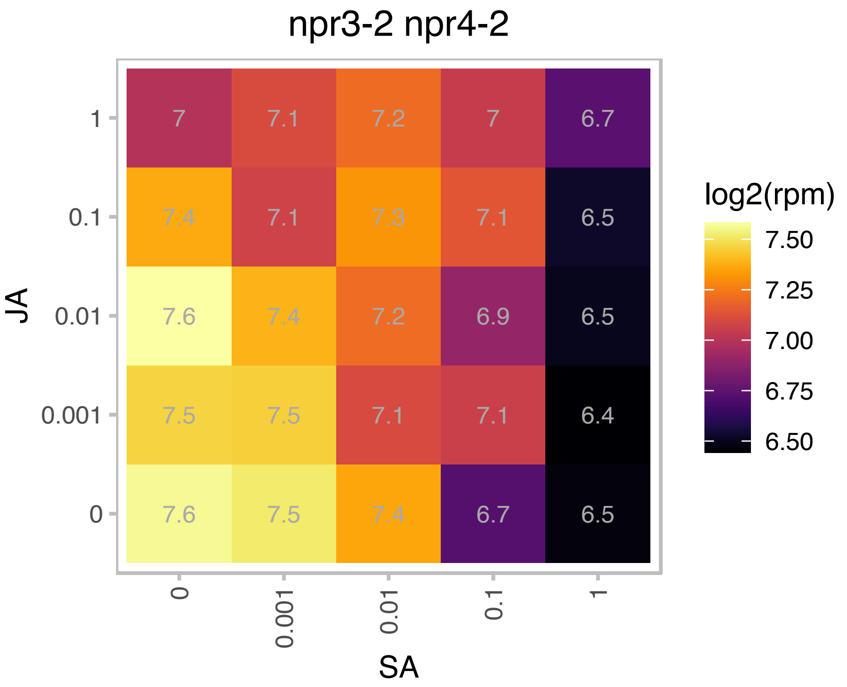


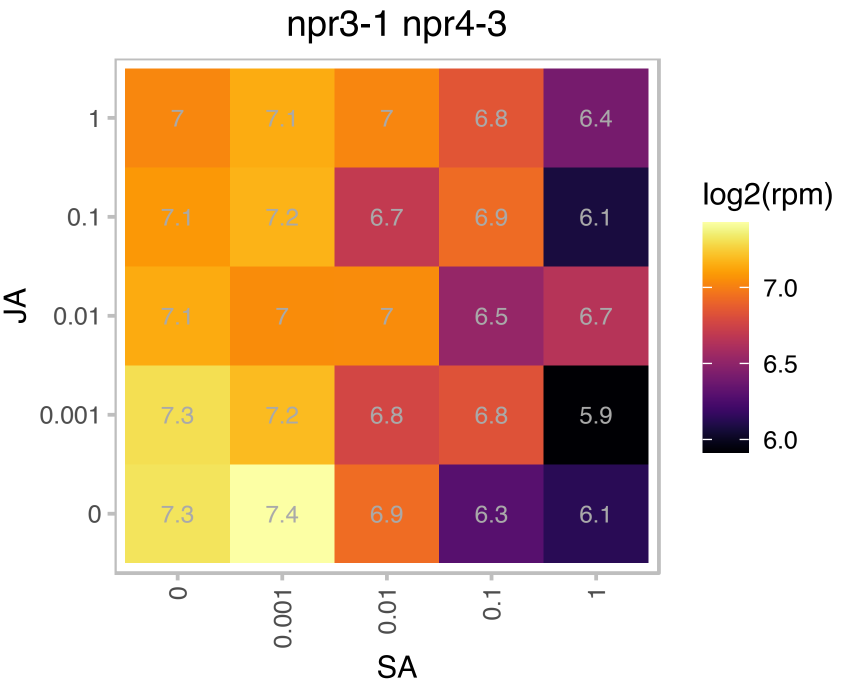

### AT3G25040.1 : ERD2B

endoplasmic reticulum retention defective 2B

Encodes ERD2b. a homolog of the yeast endoplasmic reticulum retention receptor ERD2.  Mutations in ERD2b compromise EFR but not FLS2 signaling. The mRNA is cell-to-cell mobile.

Col-0

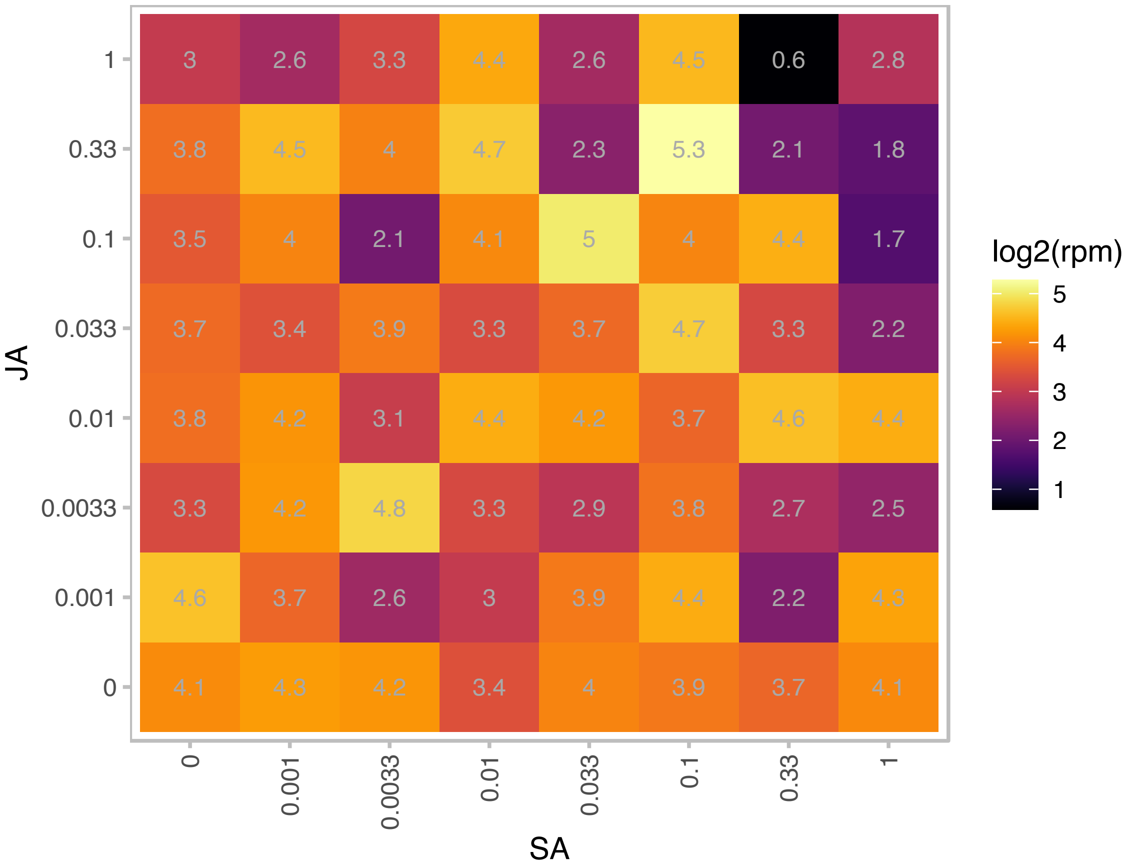


| | |
|---|---|
| Gene | AT3G25040 |
| is.expressed_genes | TRUE |
| is.responsive_genes | TRUE |
| is.apcluster | F-03 |

npr3-2 npr4-2

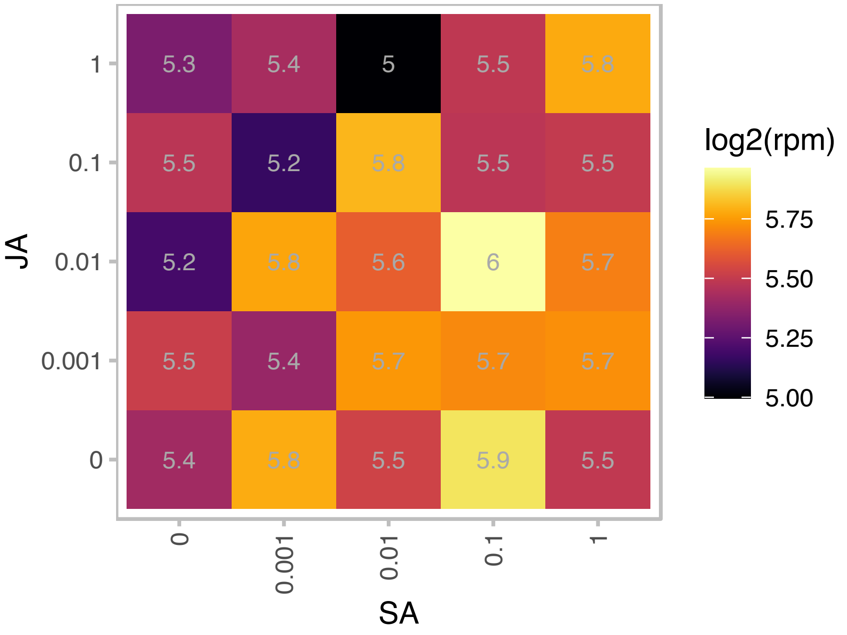


npr3-1 npr4-3

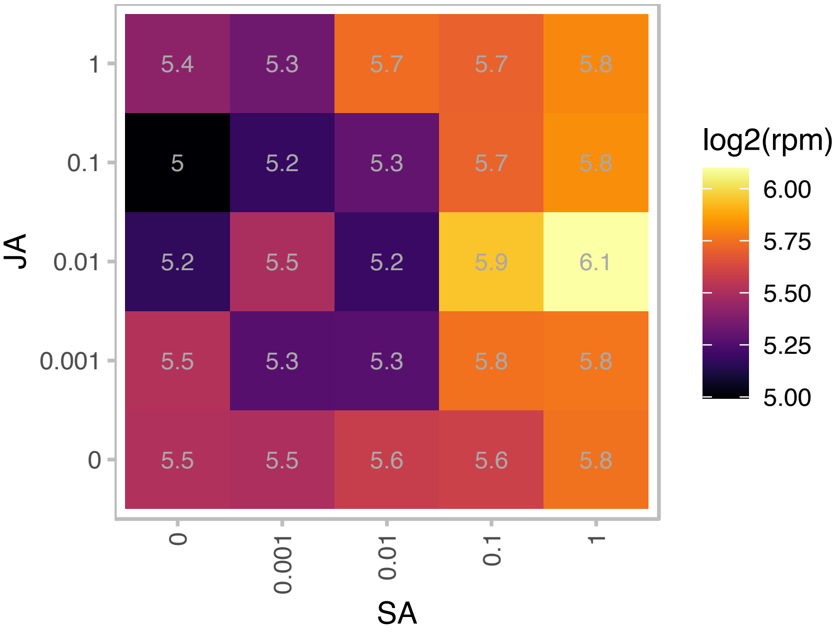

### AT3G26570.1 : ORF02, PHT2;1

phosphate transporter 2;1

low affinity phosphate transporter

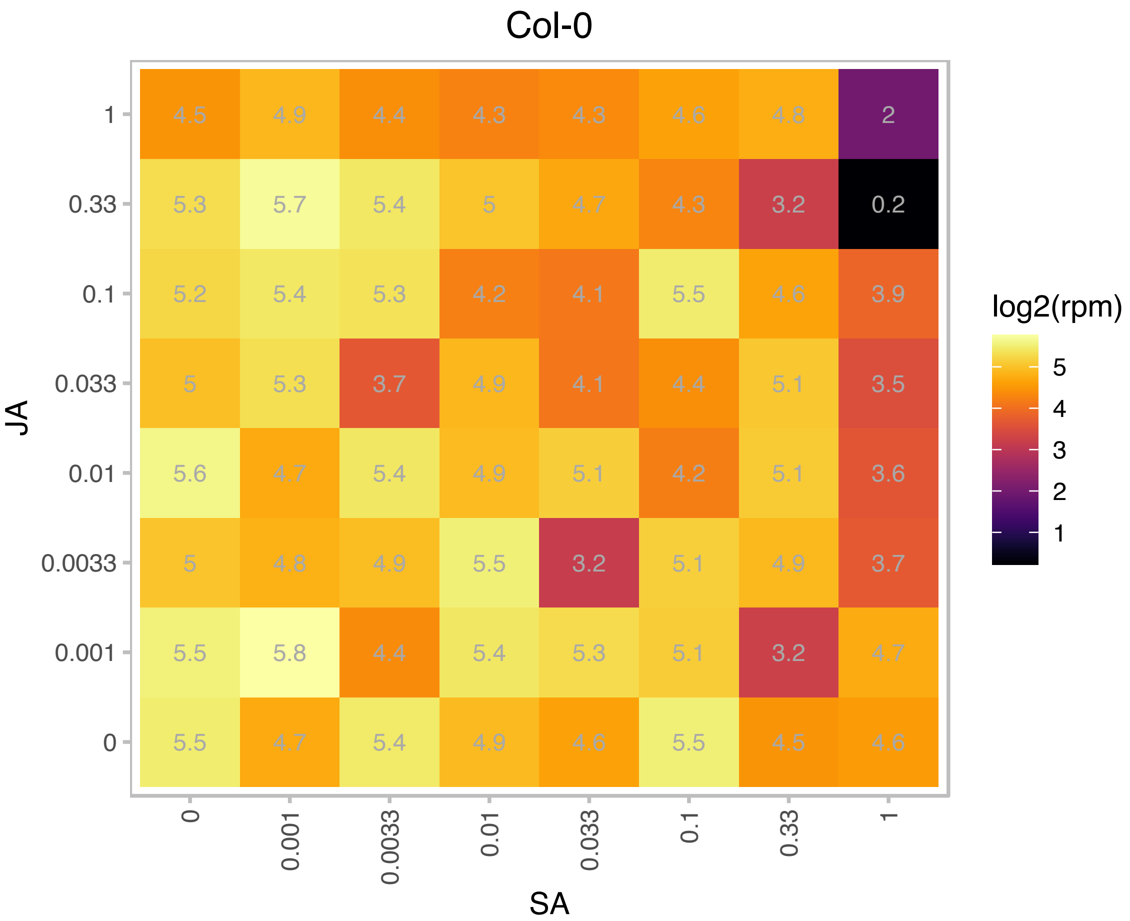


| | |
|---|---|
| Gene | AT3G26570 |
| is.expressed_genes | TRUE |
| is.responsive_genes | TRUE |
| is.apcluster | F-03 |

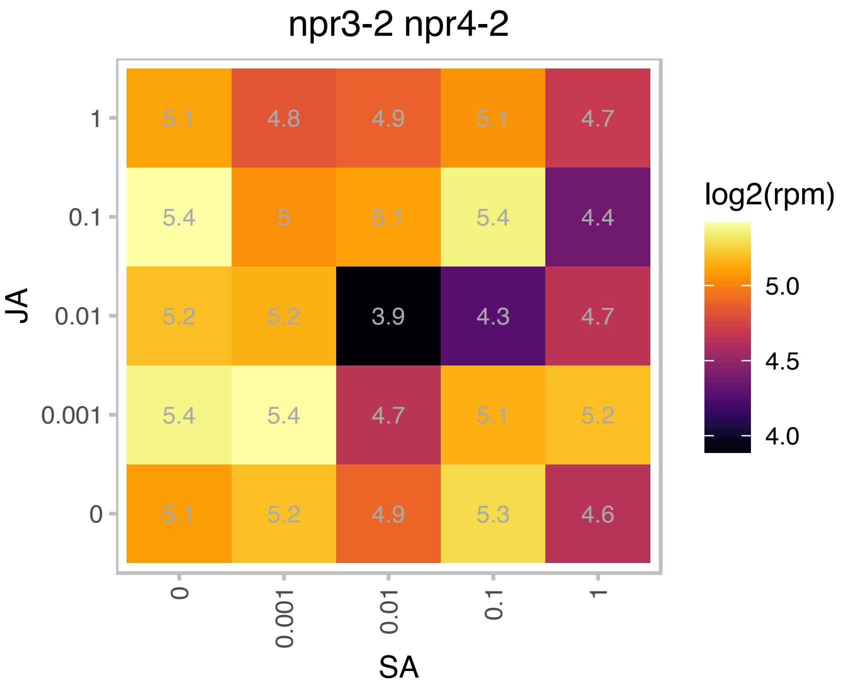


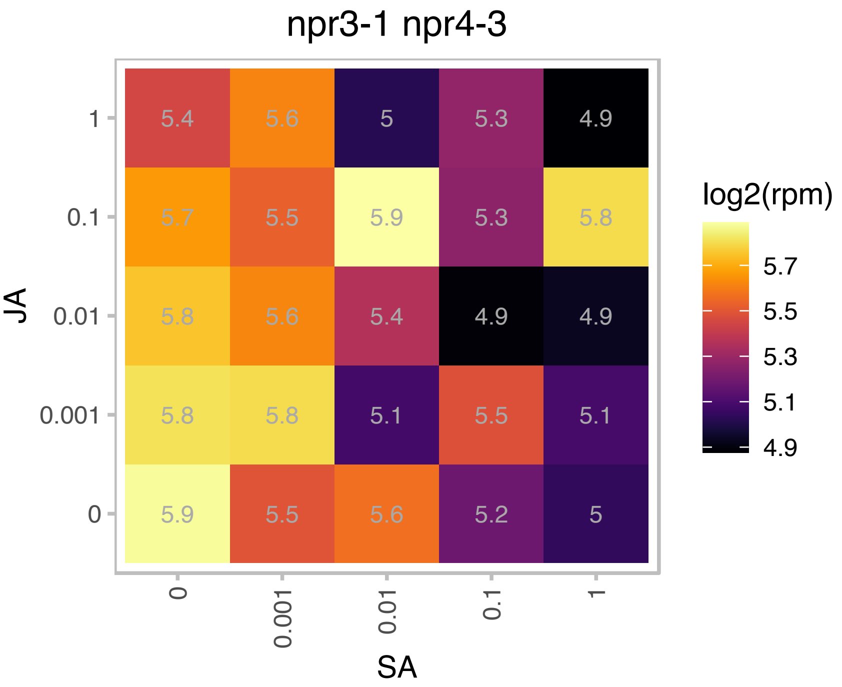

# AT3G28956.1 : NA

RNA polymerase II, Rpb4, core protein

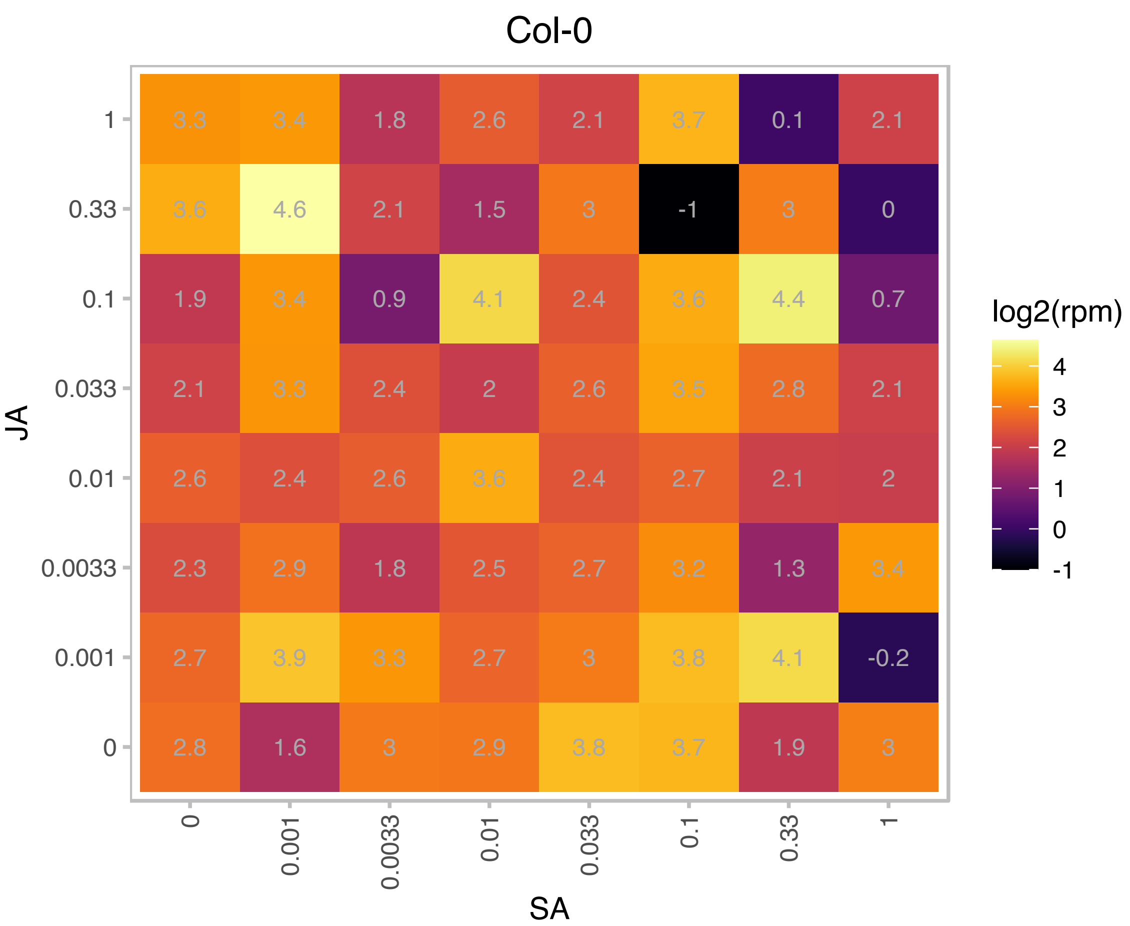


| | |
|---|---|
| Gene | AT3G28956 |
| is.expressed_genes | TRUE |
| is.responsive_genes | TRUE |
| is.apcluster | F-03 |

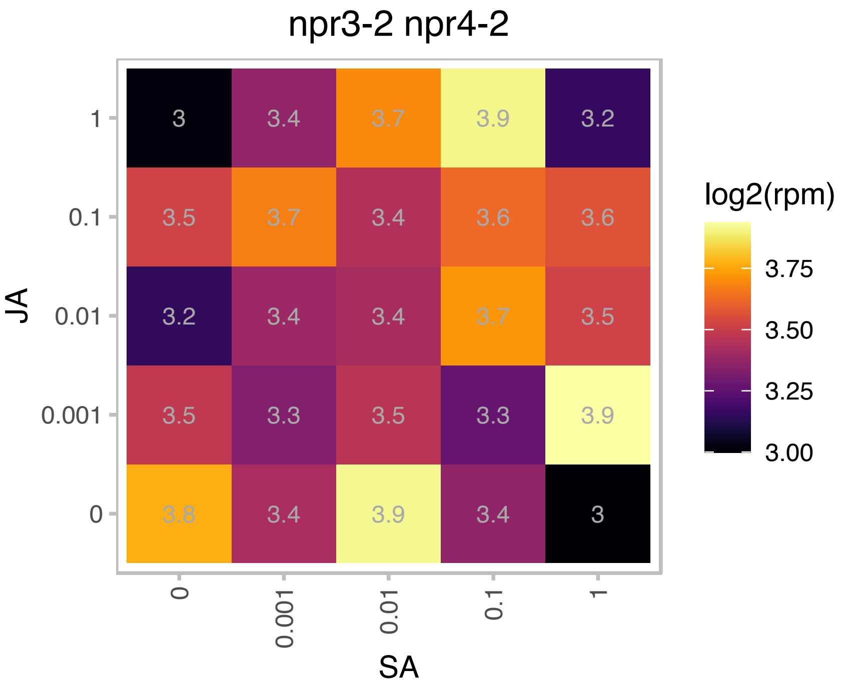


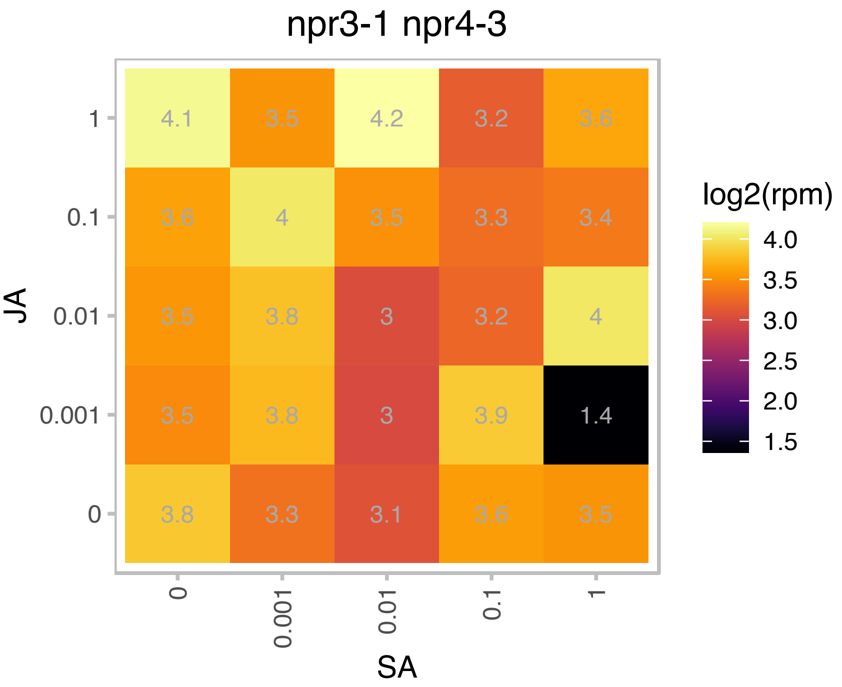

# AT3G49990.1 : NA

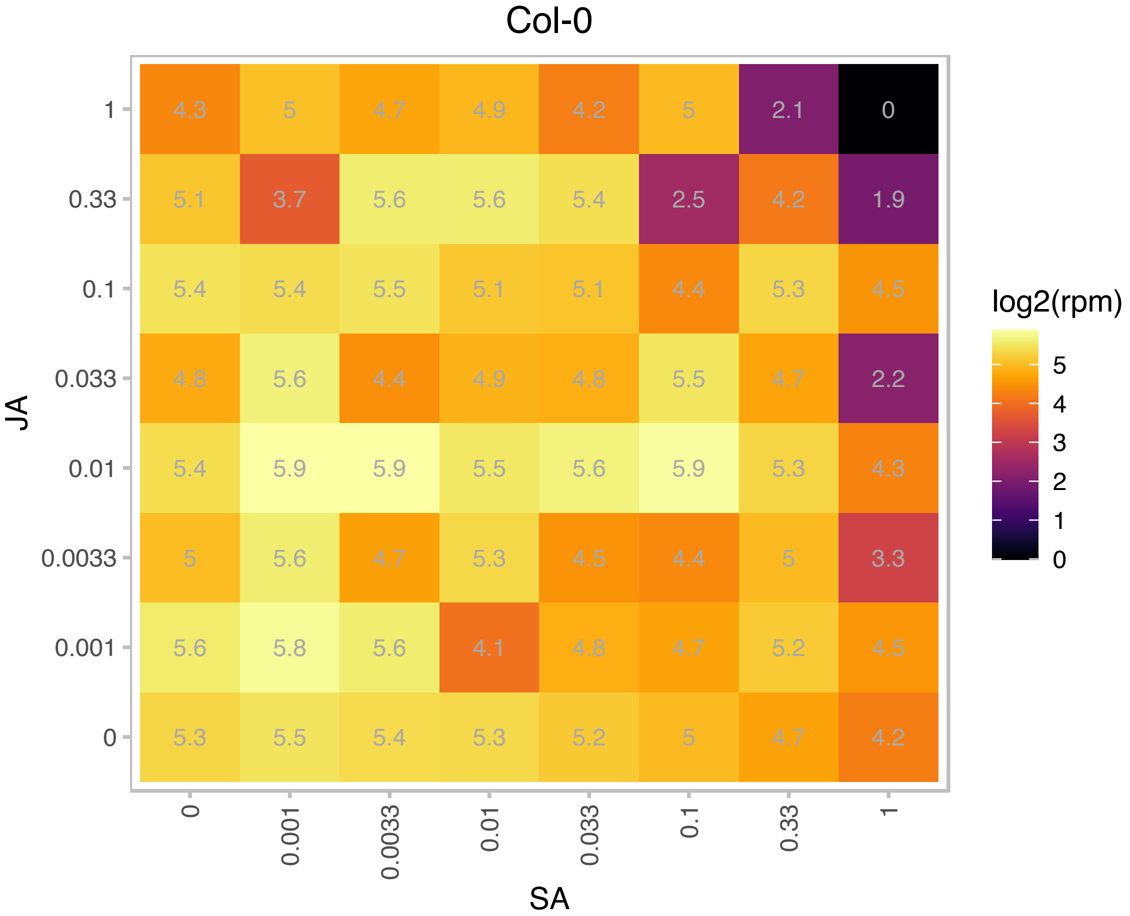


| | |
|---|---|
| Gene | AT3G49990 |
| is.expressed_genes | TRUE |
| is.responsive_genes | TRUE |
| is.apcluster | F-03 |

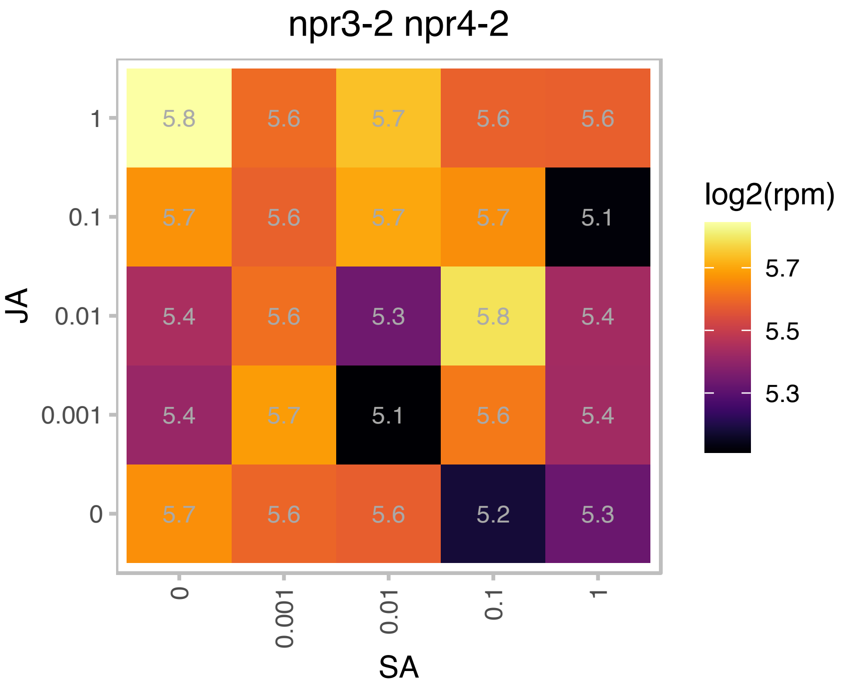


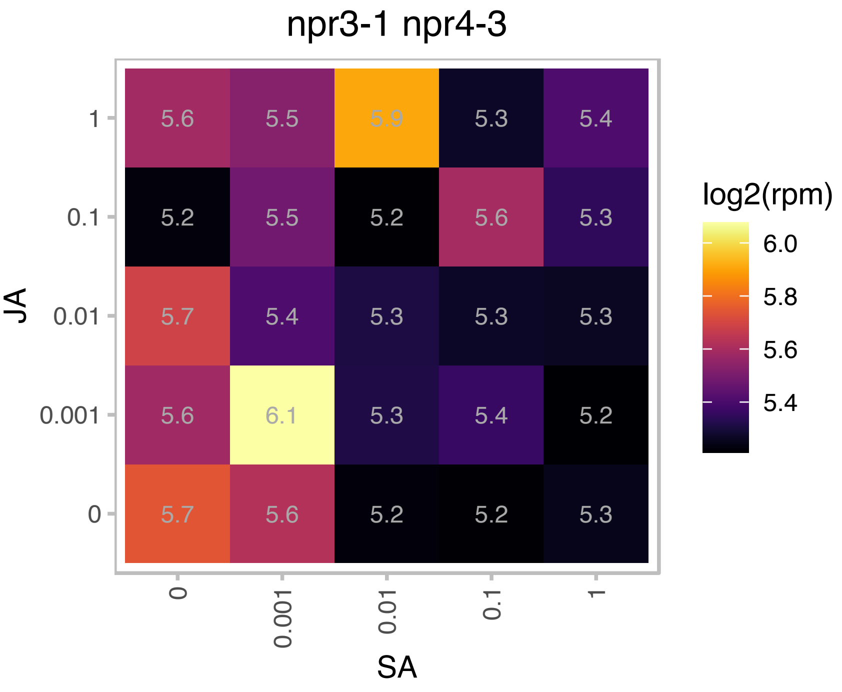

### AT3G50070.1 : CYCD3;3

CYCLIN D3;3

Encode CYCD3;3, a CYCD3 D-type cyclin.  Important for determining cell number in developing lateral organs. Mediating cytokinin effects in apical growth and development.

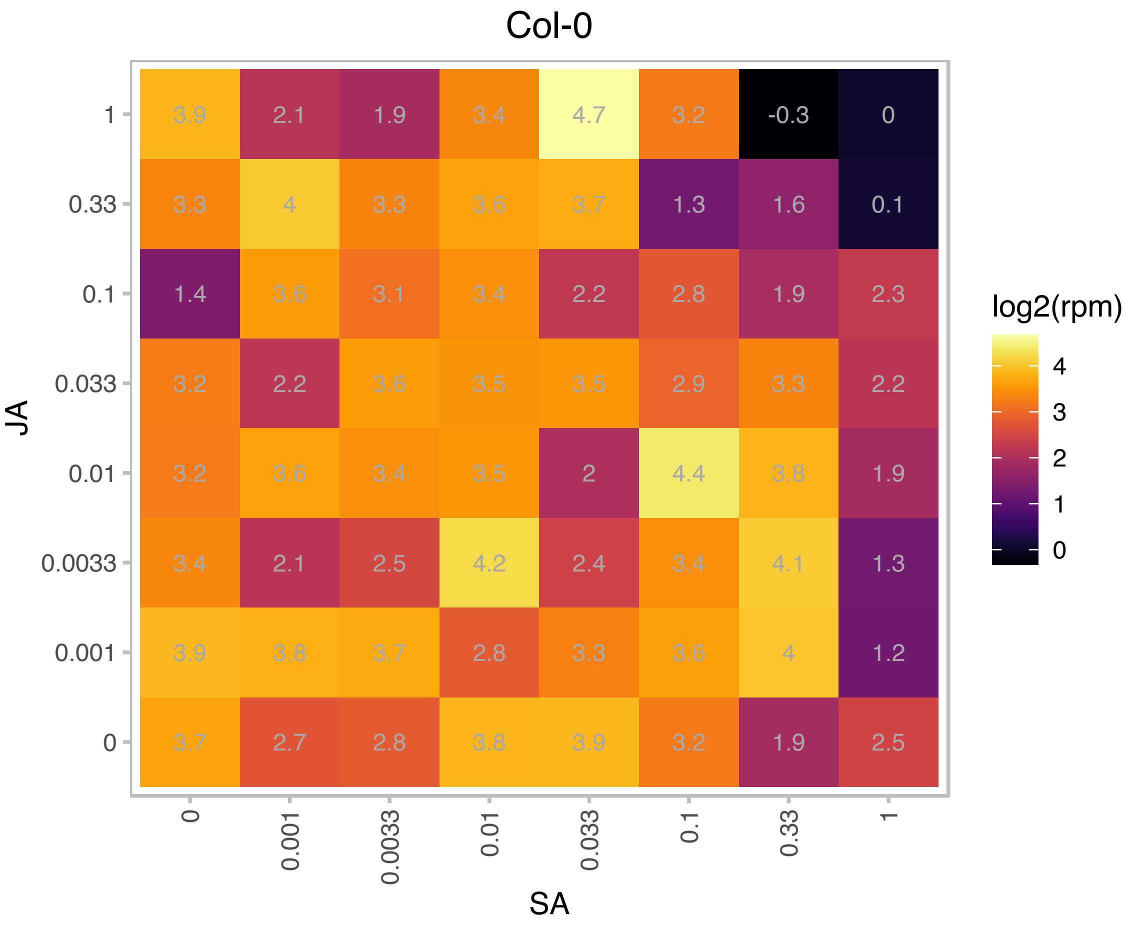


| | |
|---|---|
| Gene | AT3G50070 |
| is.expressed_genes | TRUE |
| is.responsive_genes | TRUE |
| is.apcluster | F-03 |

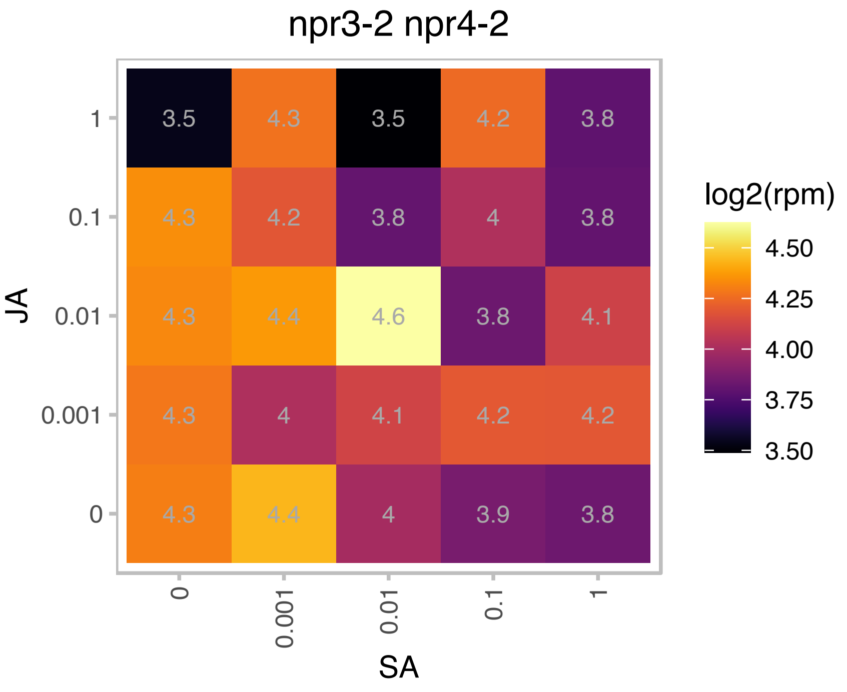


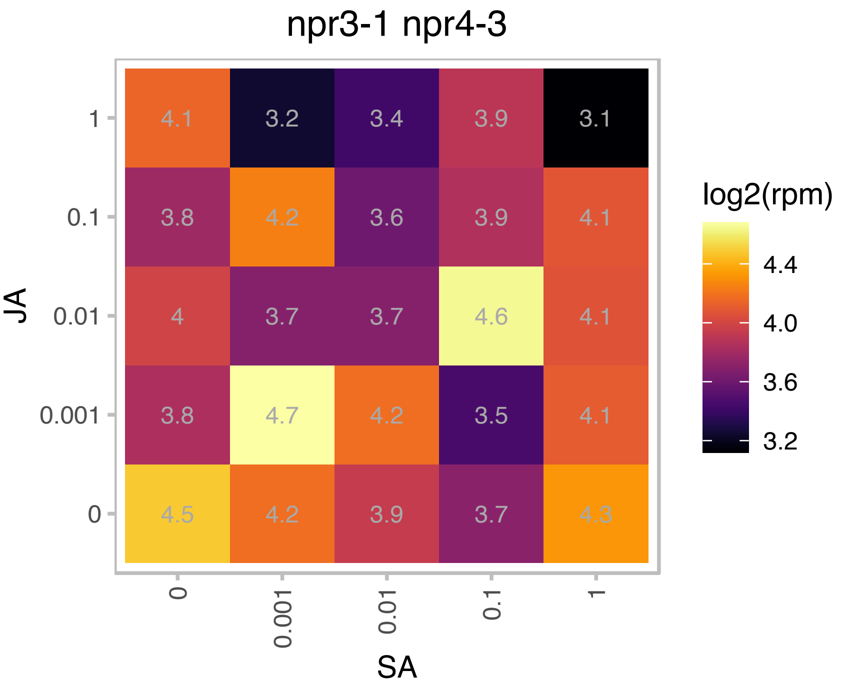

# AT3G54400.1 : NA

Eukaryotic aspartyl protease family protein

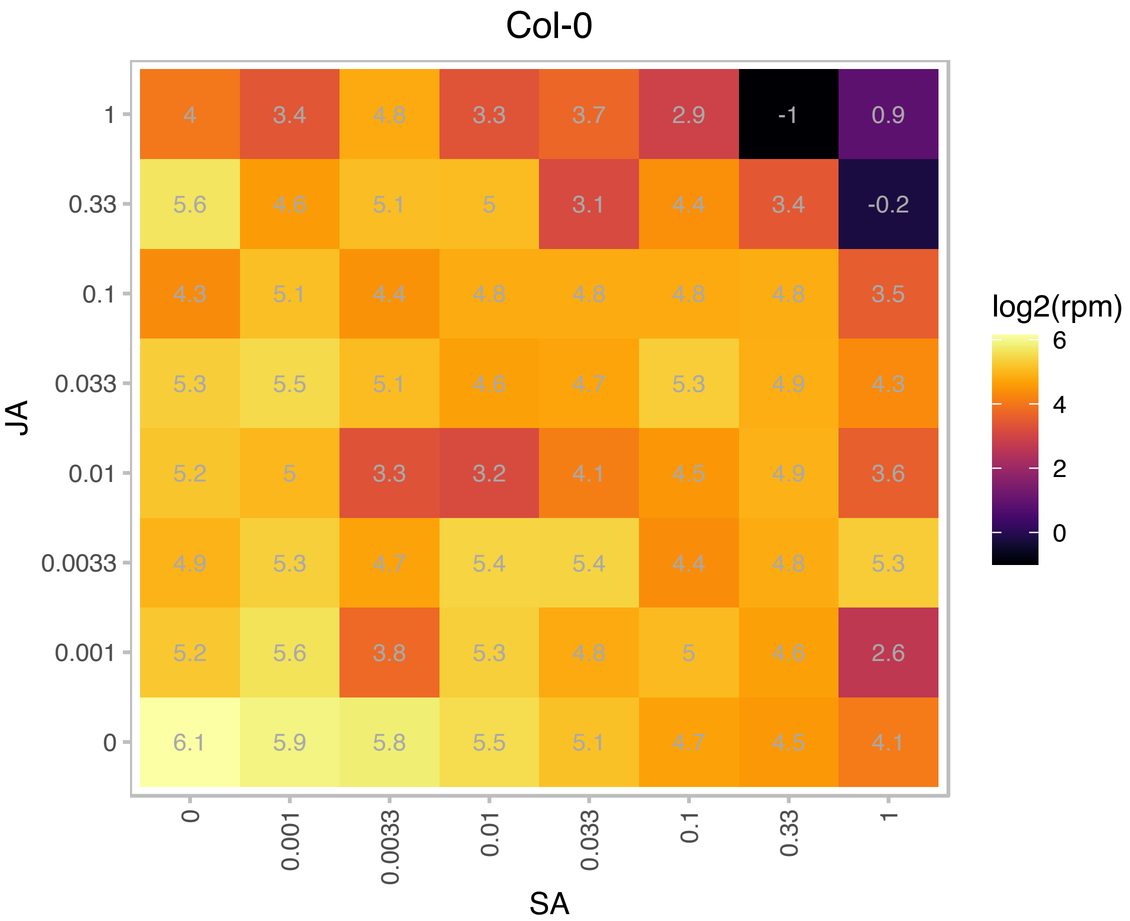


| | |
|---|---|
| Gene | AT3G54400 |
| is.expressed_genes | TRUE |
| is.responsive_genes | TRUE |
| is.apcluster | F-03 |

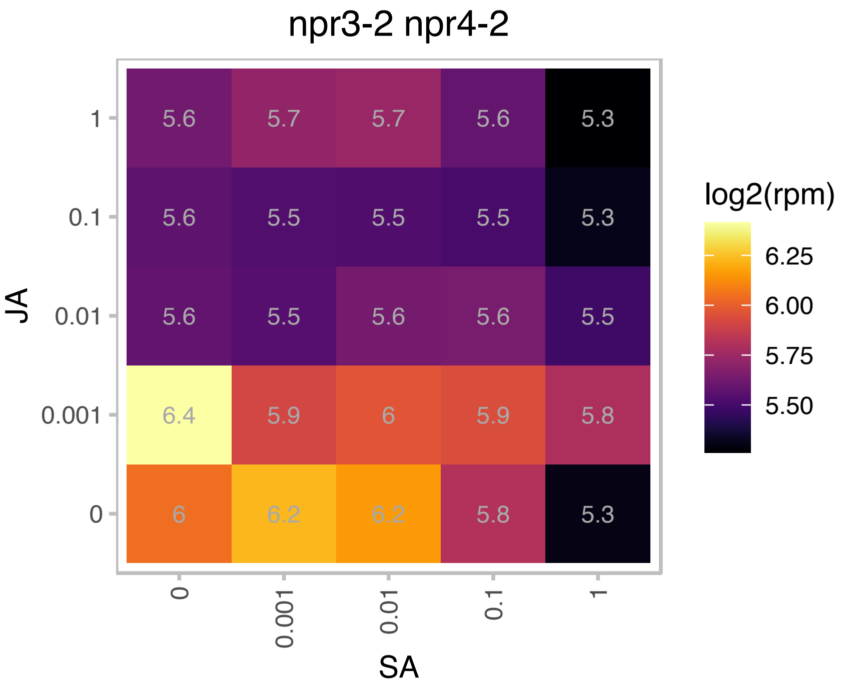


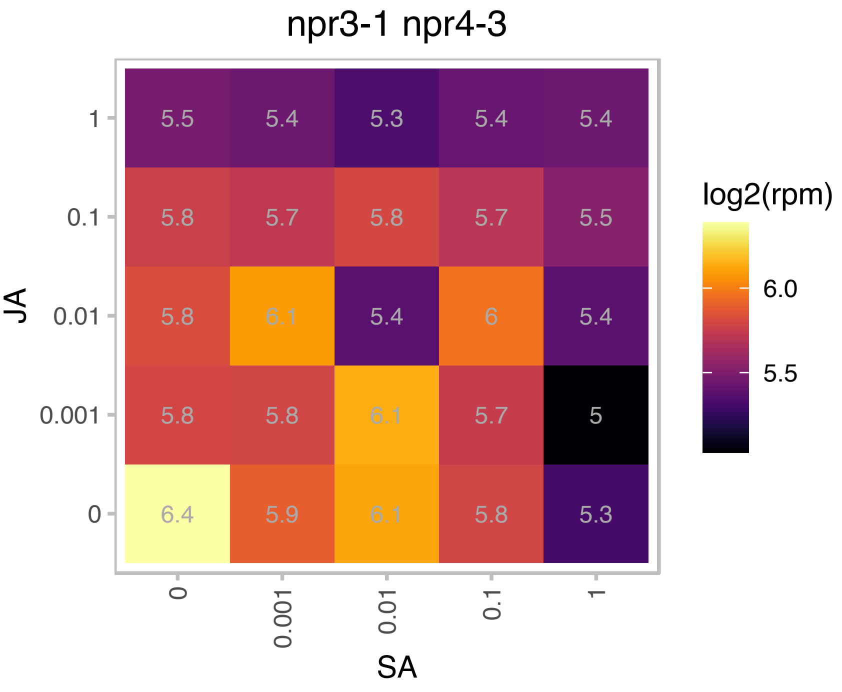

# AT3G59890.1 : NA

Dihydrodipicolinate reductase, bacterial/plant

| | |
|---|---|
| Gene | AT3G59890 |
| is.expressed_genes | TRUE |
| is.responsive_genes | TRUE |
| is.apcluster | F-03 |

#### Col-0

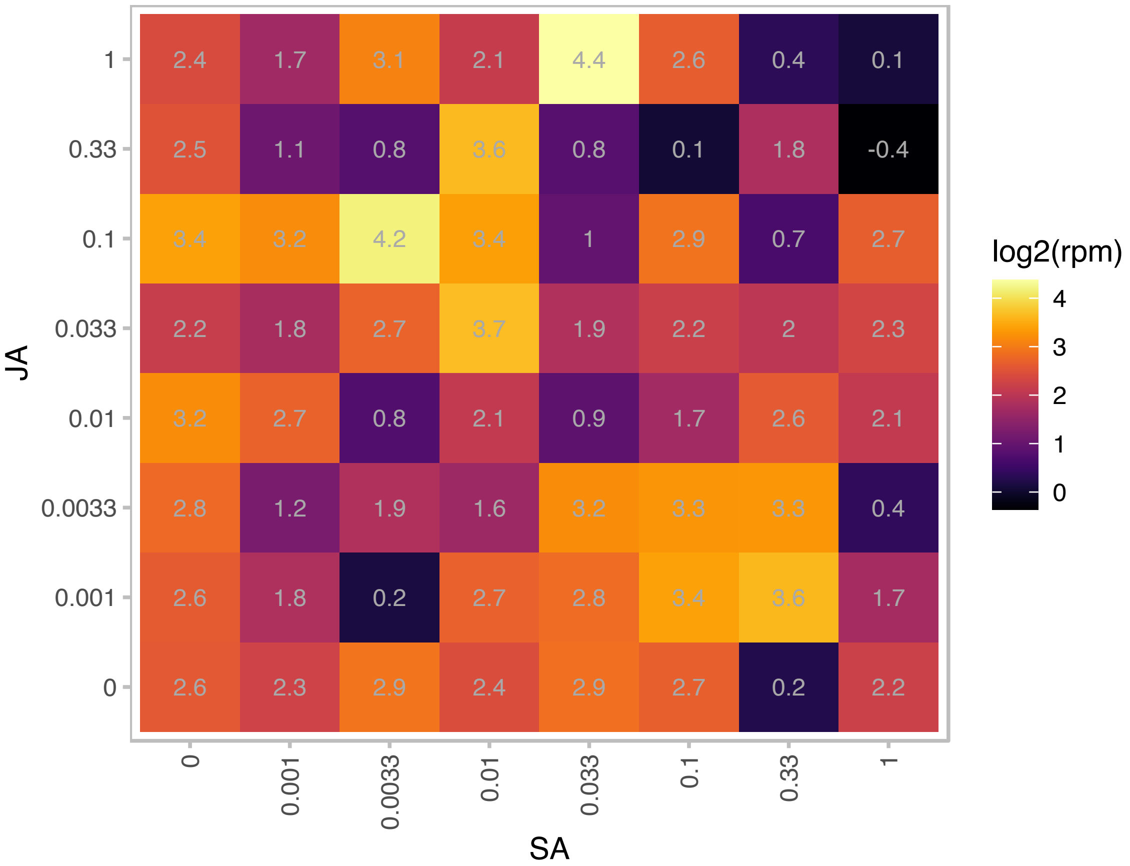


#### npr3-2 npr4-2

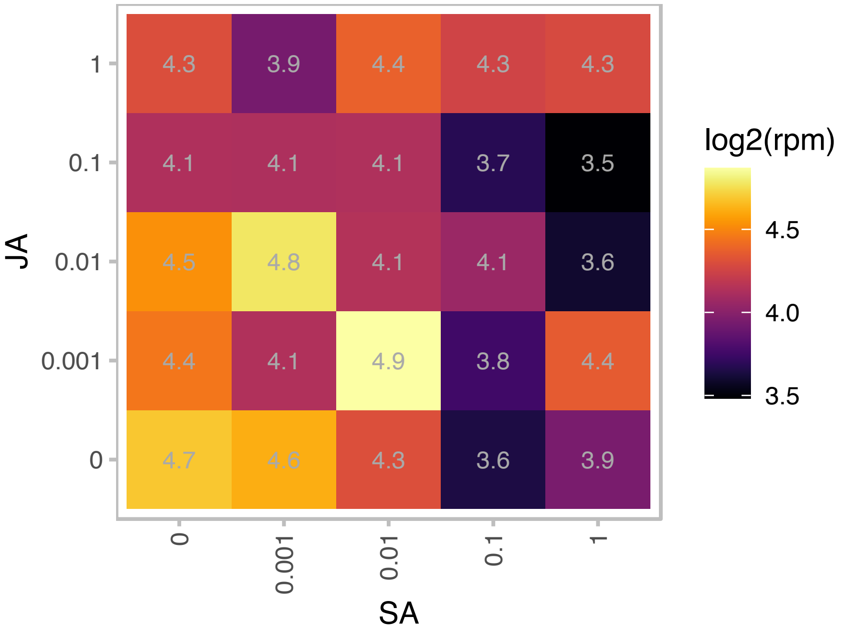


#### npr3-1 npr4-3

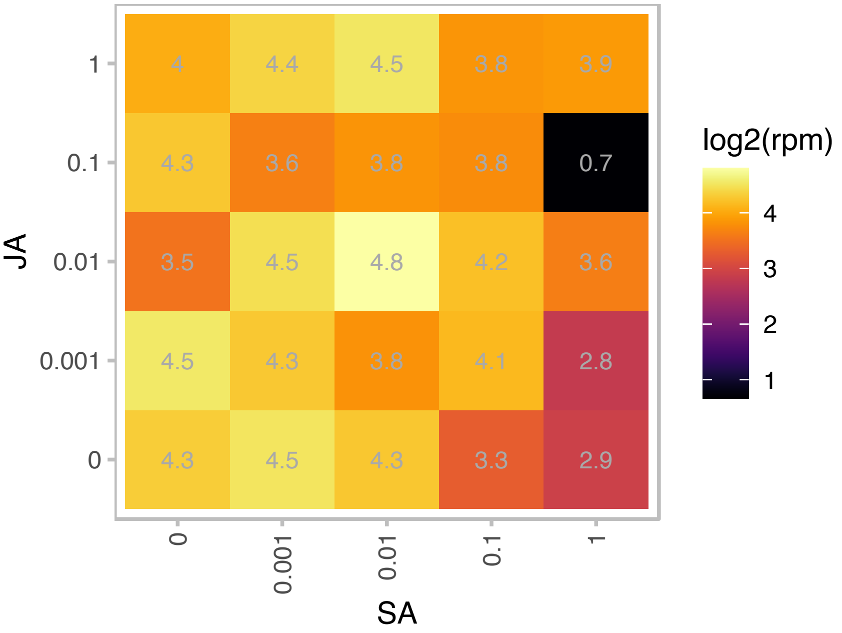

### AT3G63170.1 : AtFAP1, FAP1

Chalcone-flavanone isomerase family protein

Encodes a plastid stroma localized fatty acid binding protein involved in fatty acid metabolism.

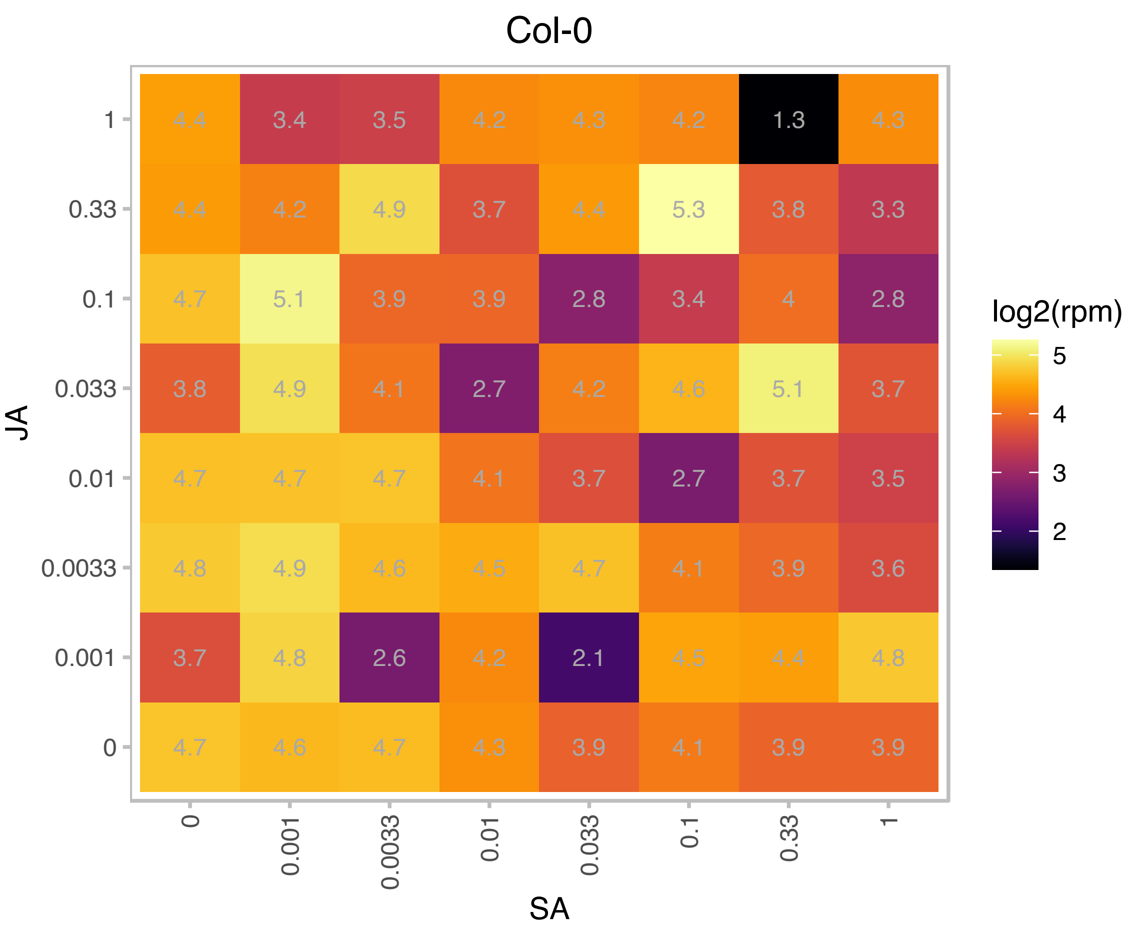


| | |
|---|---|
| Gene | AT3G63170 |
| is.expressed_genes | TRUE |
| is.responsive_genes | TRUE |
| is.apcluster | F-03 |

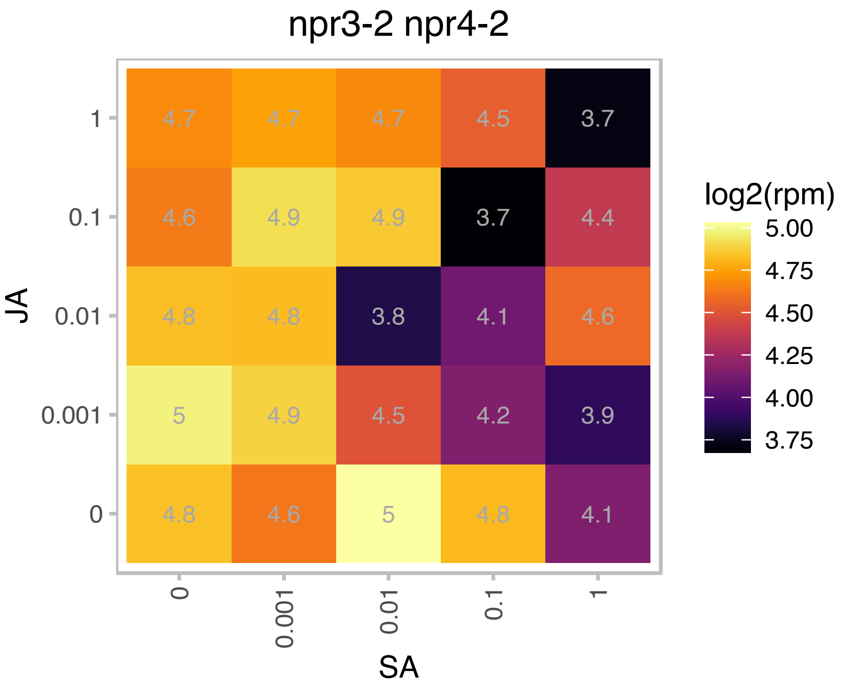


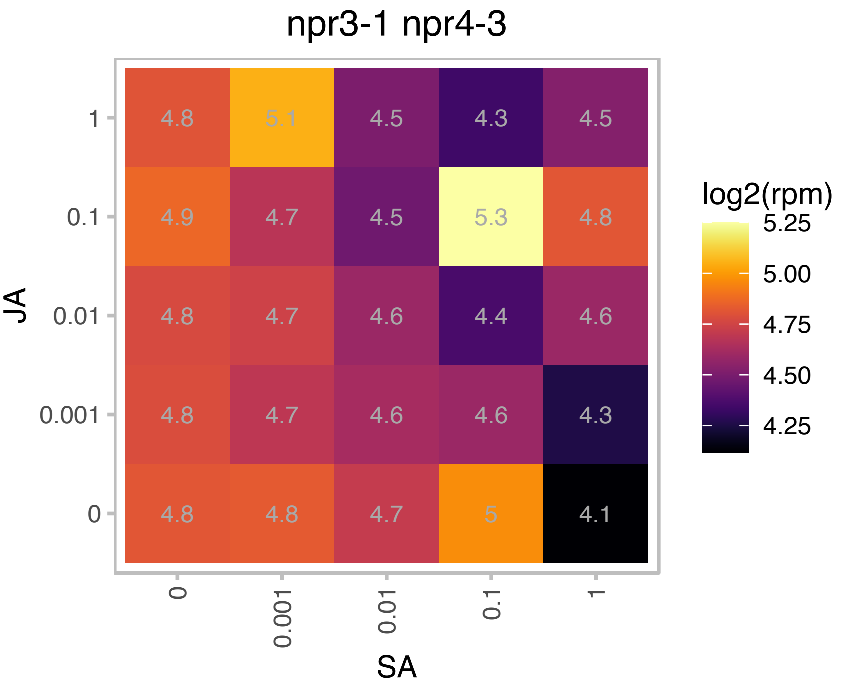

# AT4G03180.1 : NA

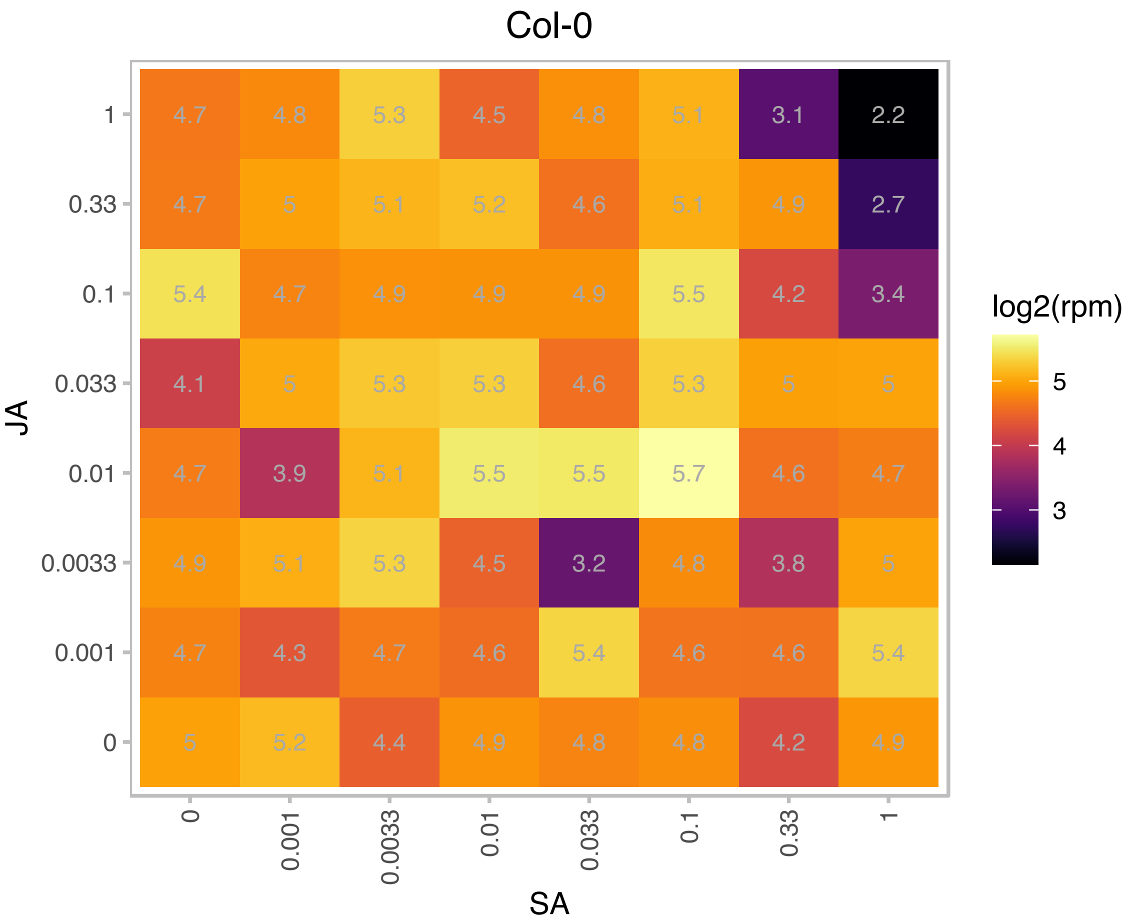


| | |
|---|---|
| Gene | AT4G03180 |
| is.expressed_genes | TRUE |
| is.responsive_genes | TRUE |
| is.apcluster | F-03 |

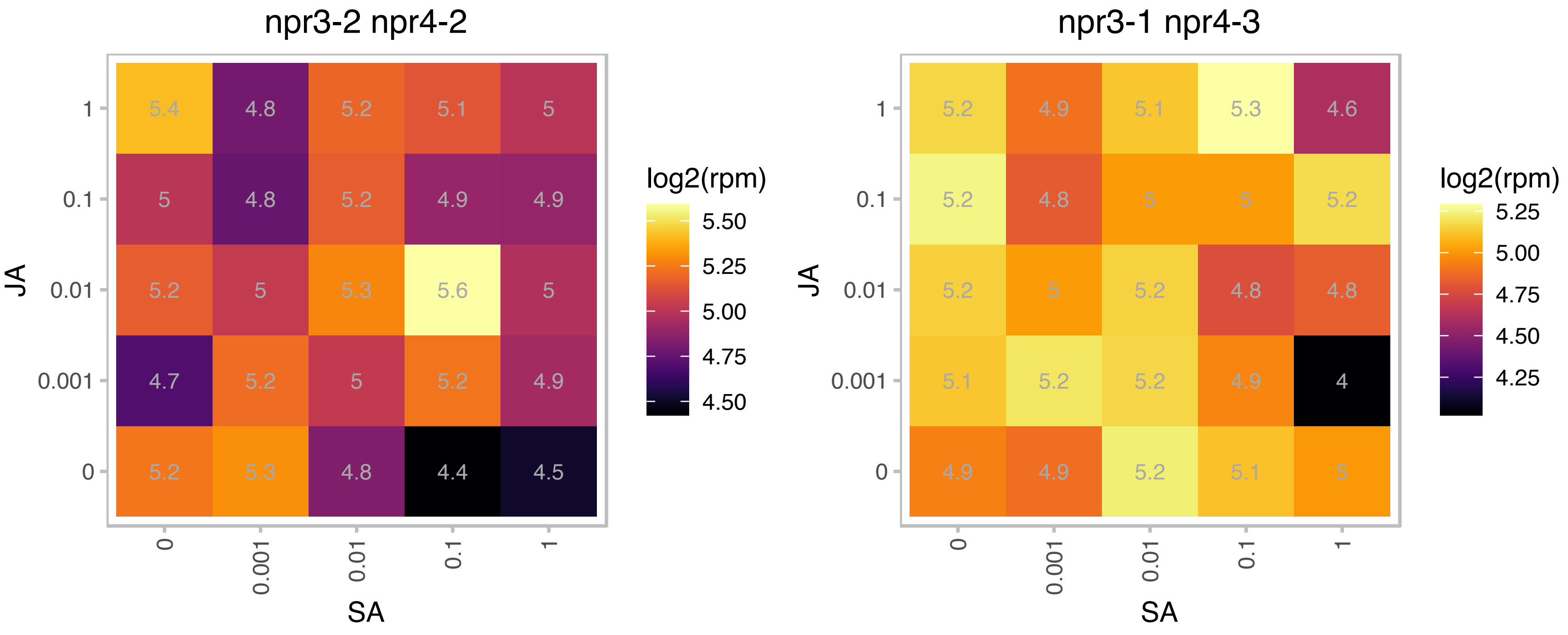

### AT4G11280.1 : ACS6, ATACS6

1-aminocyclopropane-1-carboxylic acid (acc) synthase 6

encodes a a member of the 1-aminocyclopropane-1-carboxylate (ACC) synthase (S-adenosyl-L-methionine methylthioadenosine-lyase, EC 4.4.1.14) gene family The mRNA is cell-to-cell mobile.

Col-0

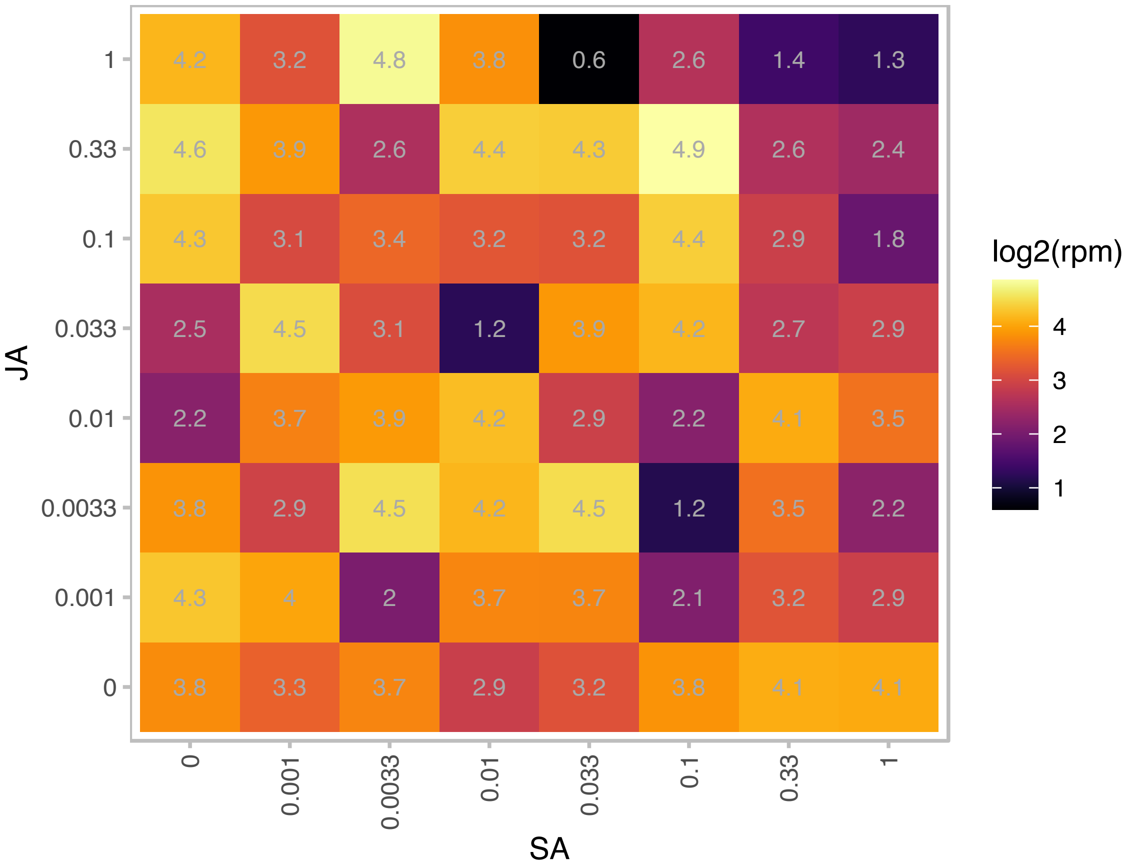


| | |
|---|---|
| Gene | AT4G11280 |
| is.expressed_genes | TRUE |
| is.responsive_genes | TRUE |
| is.apcluster | F-03 |

npr3-2 npr4-2

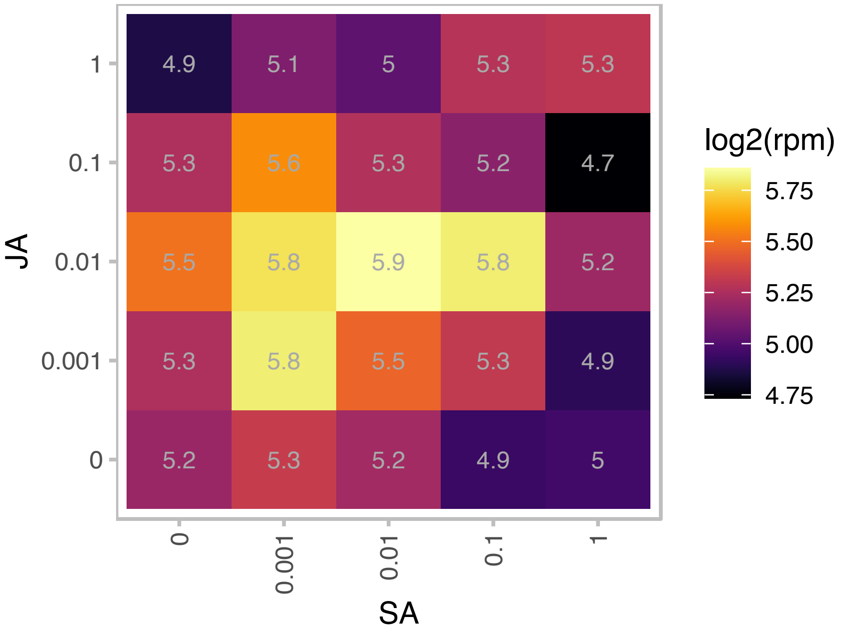


npr3-1 npr4-3

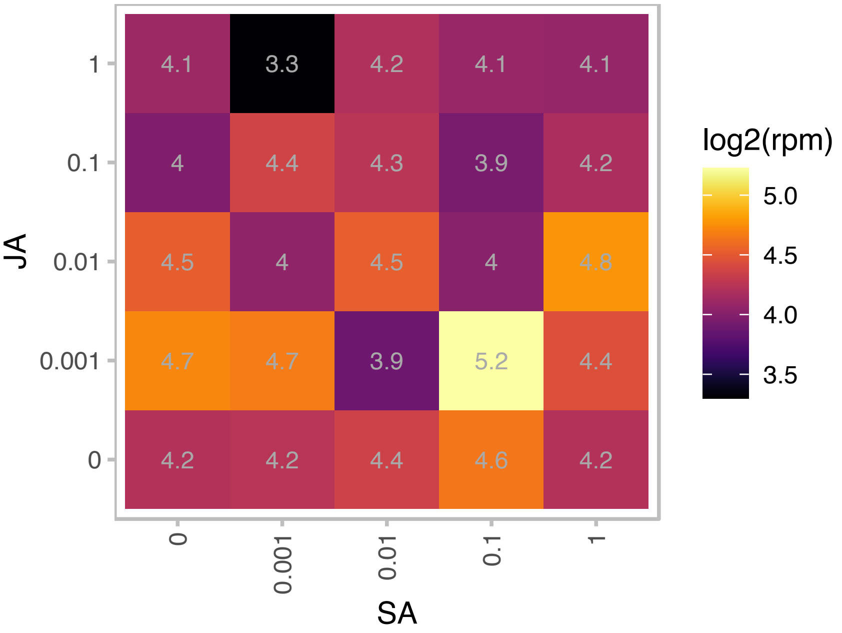

### AT4G14770.1 : ATTCX2, TCX2

TESMIN/TSO1-like CXC 2

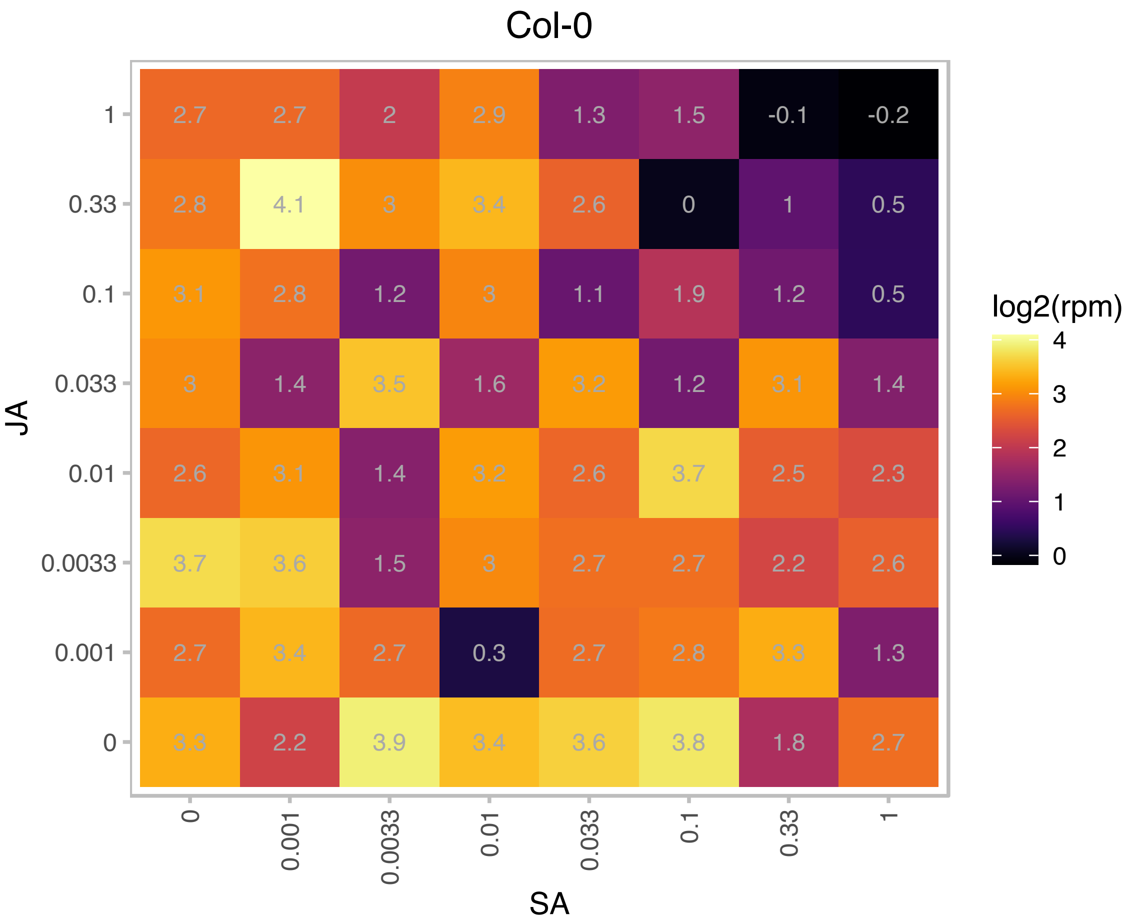


| | |
|---|---|
| Gene | AT4G14770 |
| is.expressed_genes | TRUE |
| is.responsive_genes | TRUE |
| is.apcluster | F-03 |

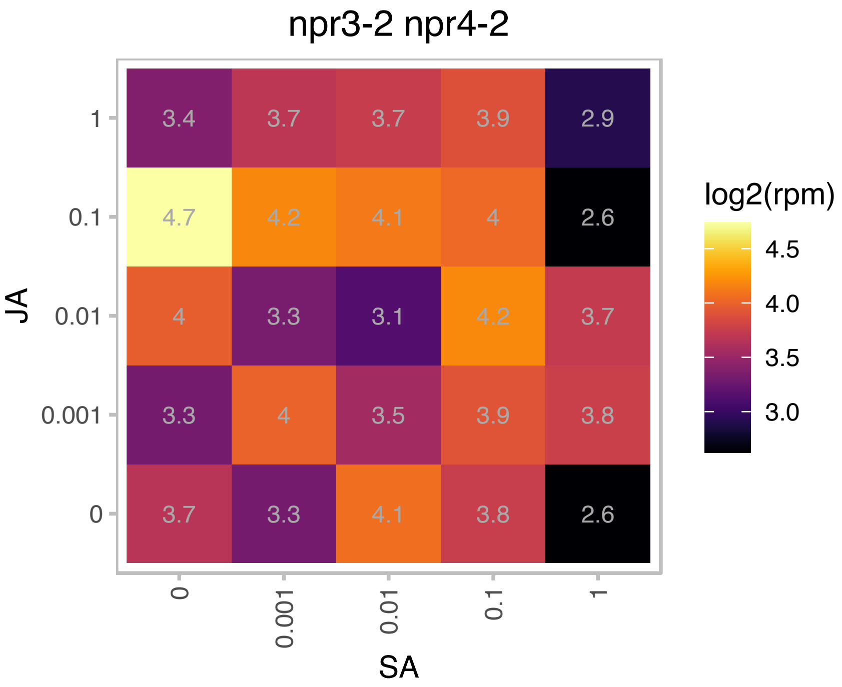


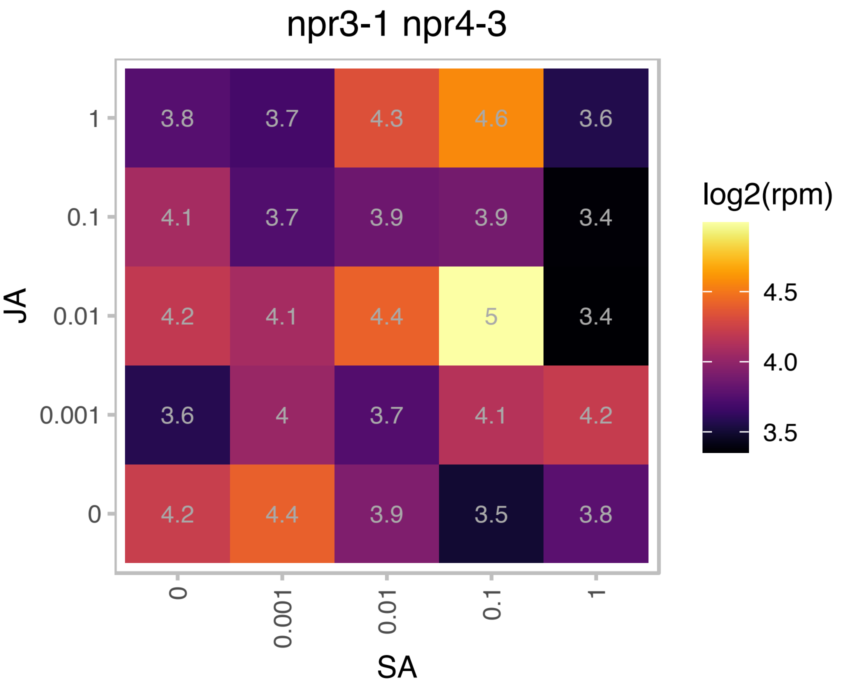

# AT4G15160.1 : NA

Bifunctional inhibitor/lipid-transfer protein/seed storage 2S albumin superfamily protein

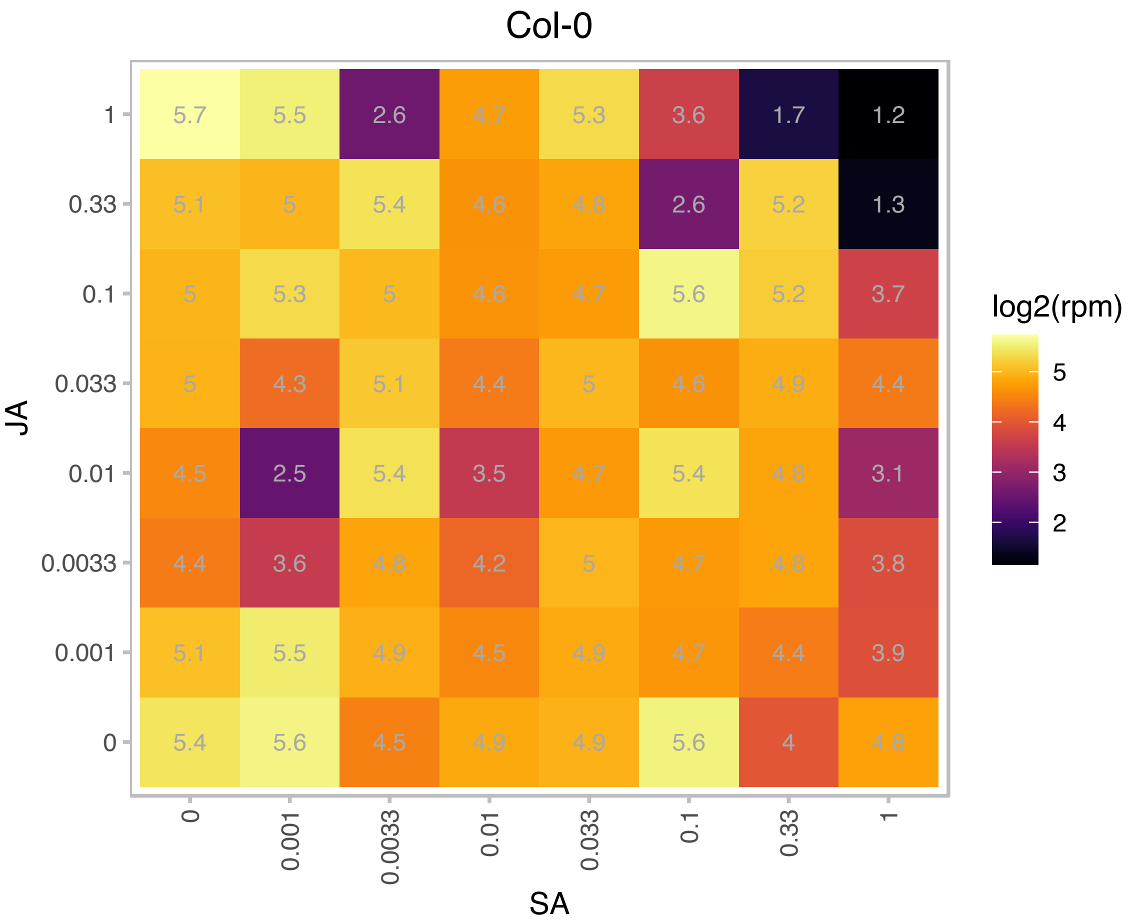


| | |
|---|---|
| Gene | AT4G15160 |
| is.expressed_genes | TRUE |
| is.responsive_genes | TRUE |
| is.apcluster | F-03 |

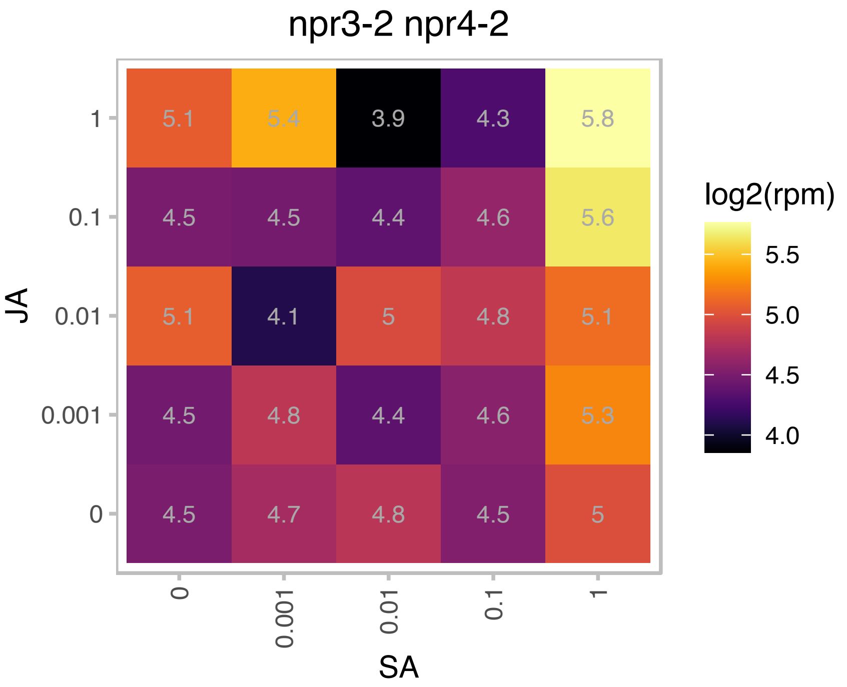


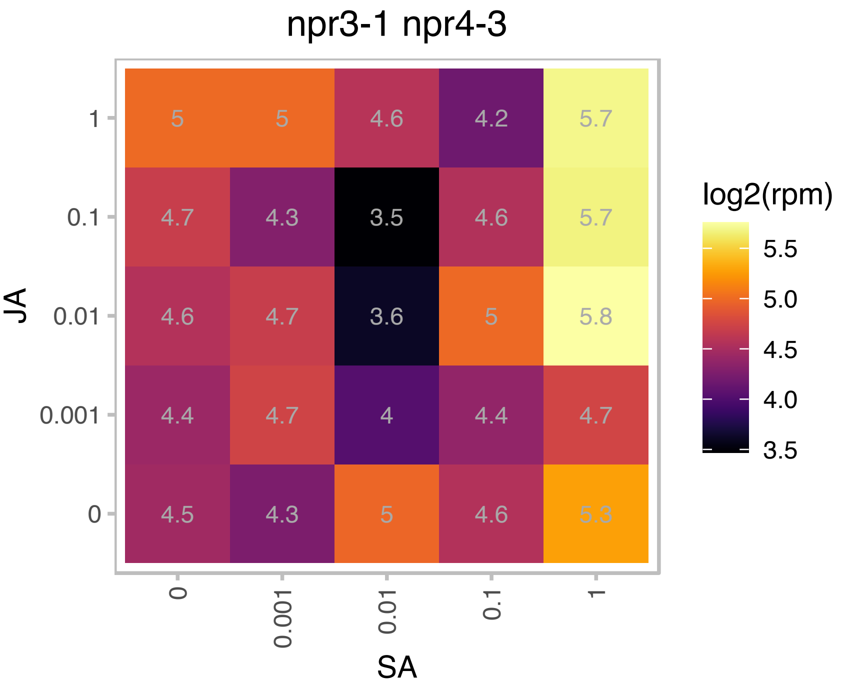

# AT4G19150.1 : NA

Ankyrin repeat family protein

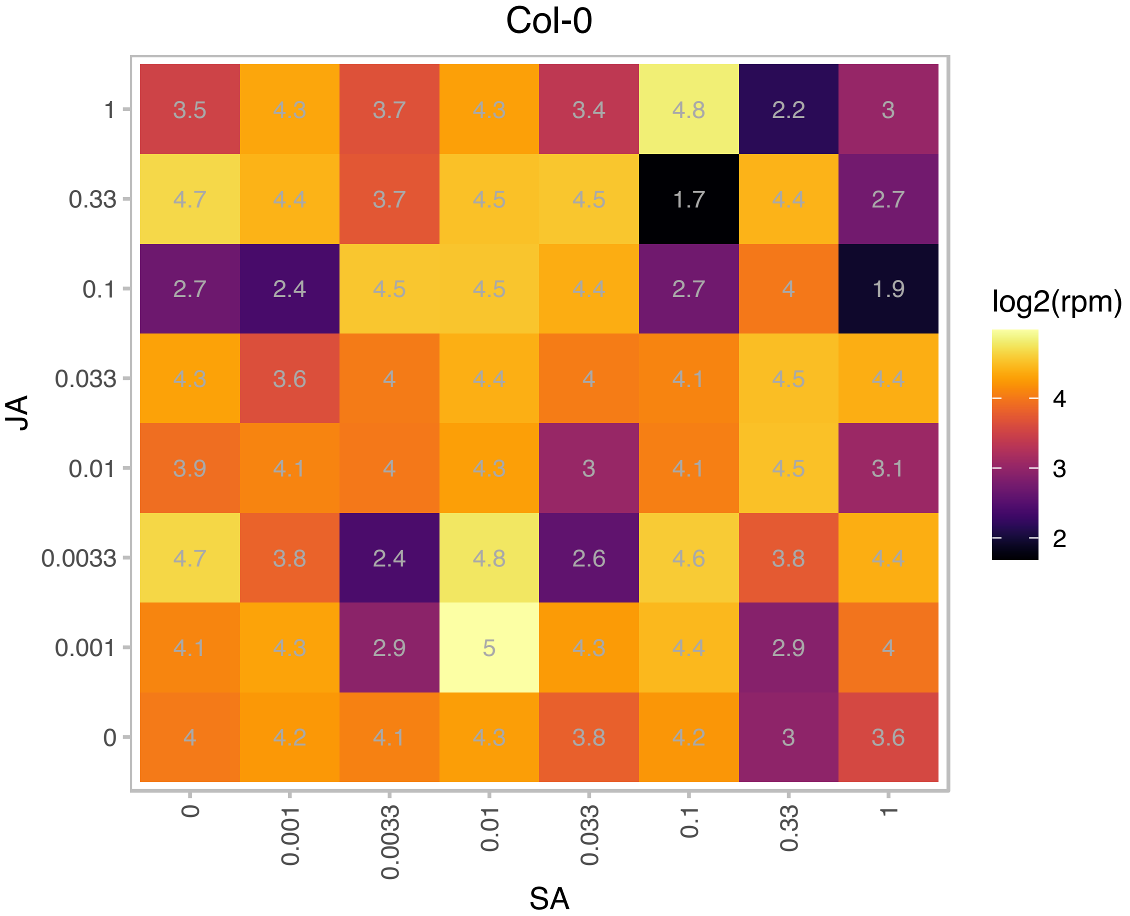


| | |
|---|---|
| Gene | AT4G19150 |
| is.expressed_genes | TRUE |
| is.responsive_genes | TRUE |
| is.apcluster | F-03 |

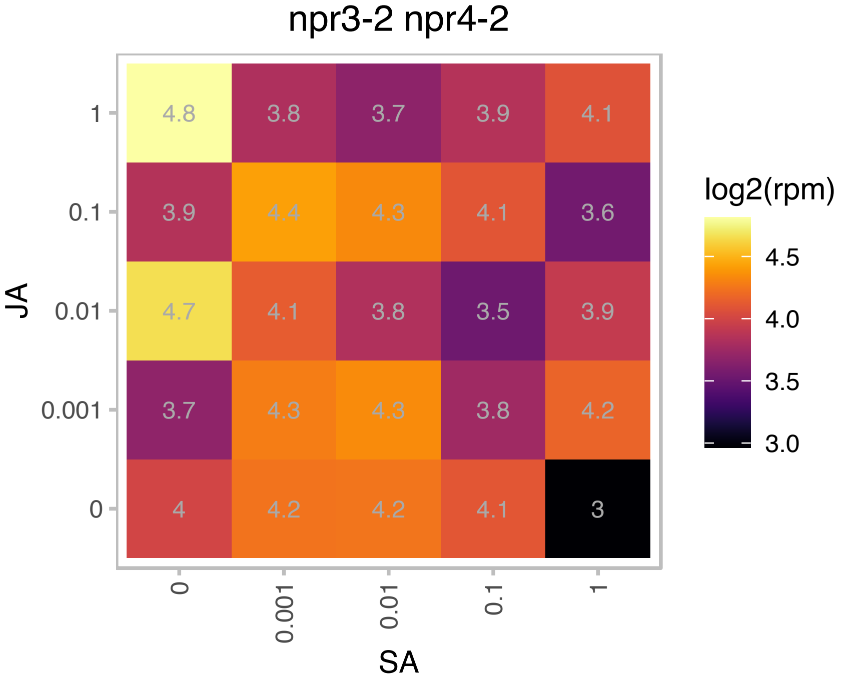


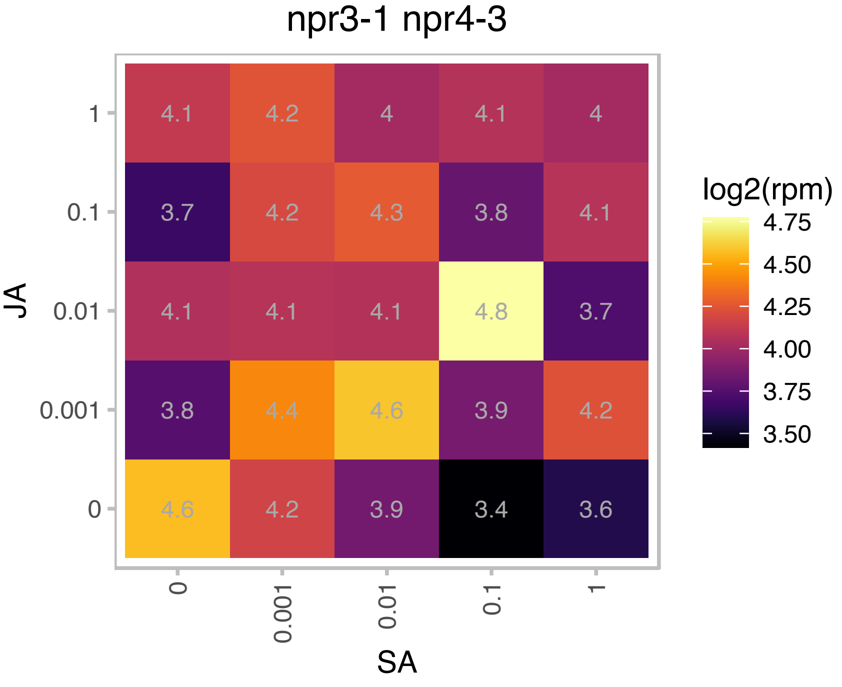

### AT4G20960.1 : PyrD

Cytidine/deoxycytidylate deaminase family protein

encodes diaminohydroxyphosphoribosylaminopyrimidine deaminase catalyzing the second step in the riboflavin biosynthesis

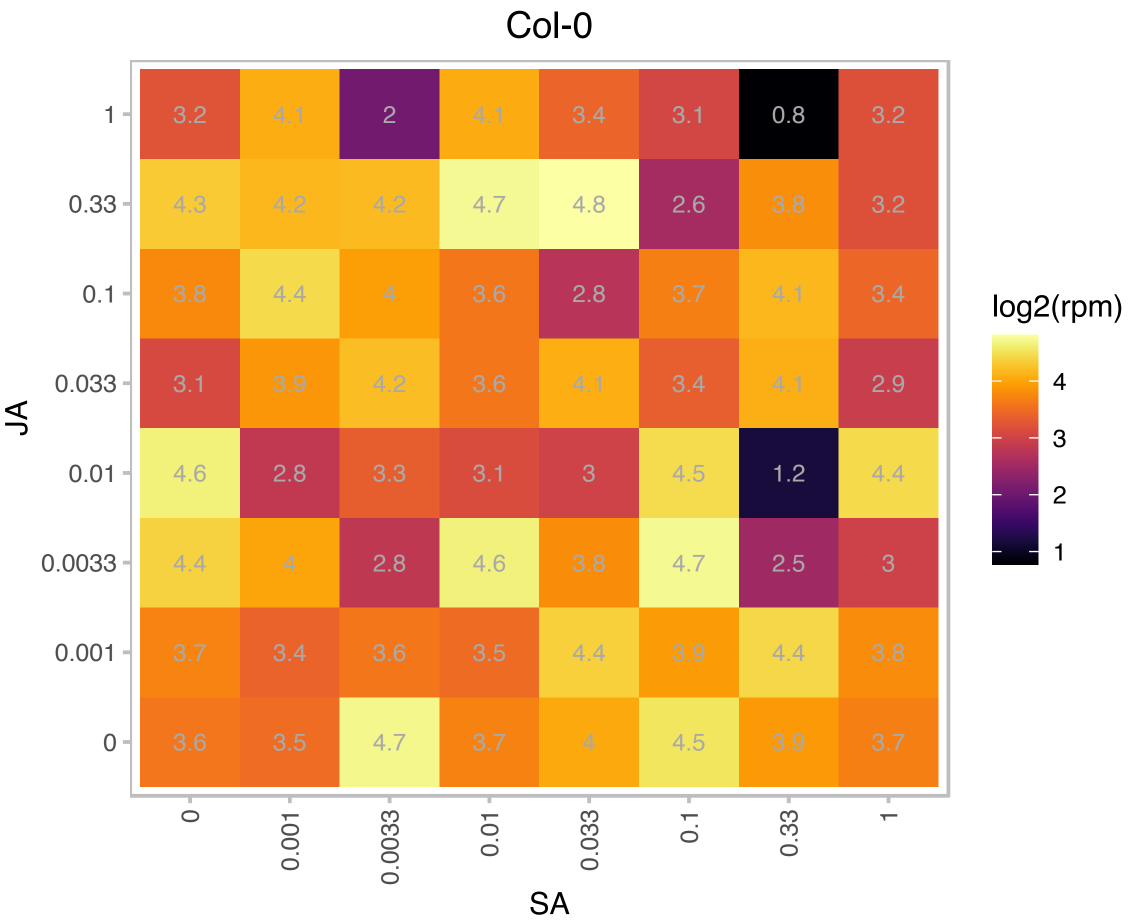


| | |
|---|---|
| Gene | AT4G20960 |
| is.expressed_genes | TRUE |
| is.responsive_genes | TRUE |
| is.apcluster | F-03 |

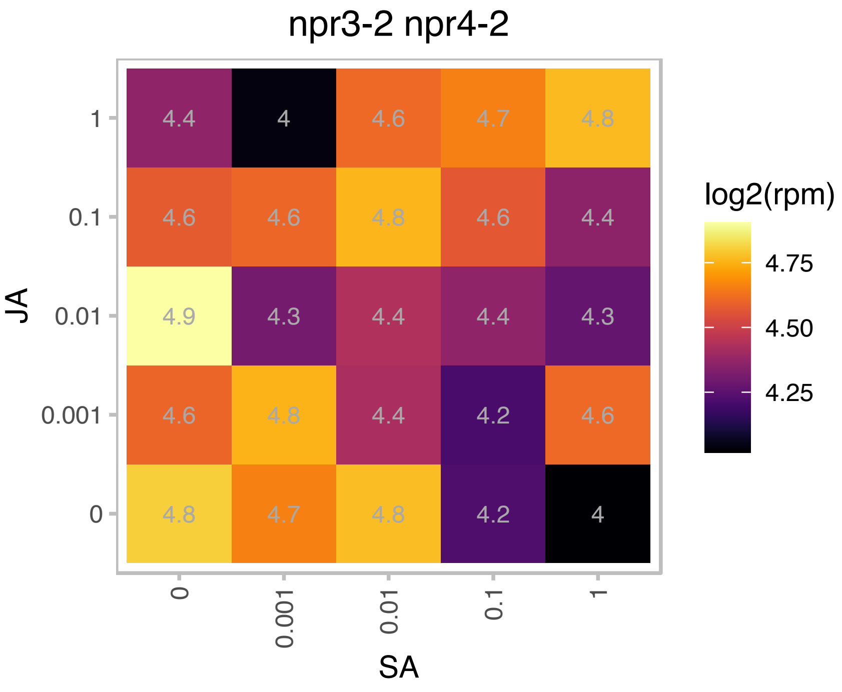


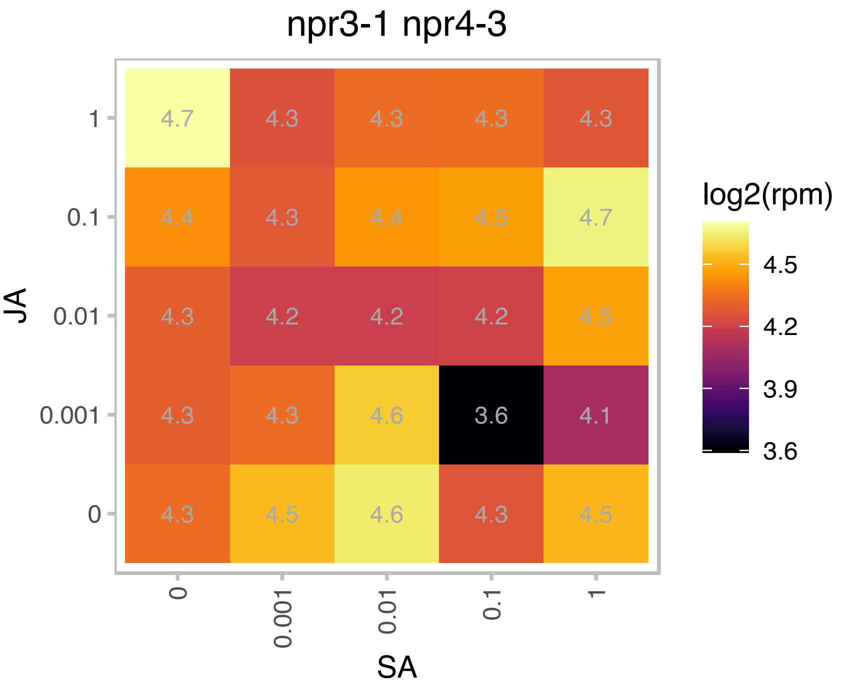

### AT4G27520.1 : AtENODL2, ENODL2

early nodulin-like protein 2

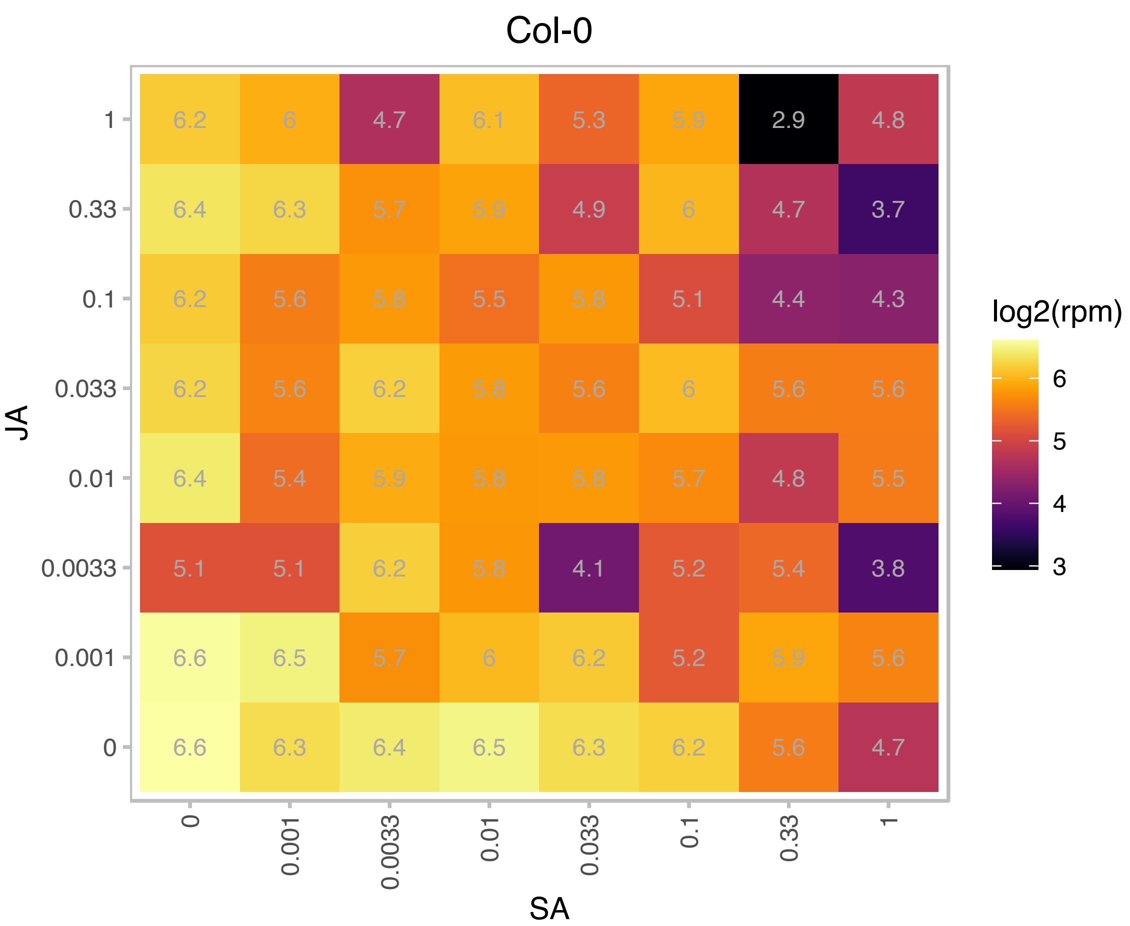


| | |
|---|---|
| Gene | AT4G27520 |
| is.expressed_genes | TRUE |
| is.responsive_genes | TRUE |
| is.apcluster | F-03 |

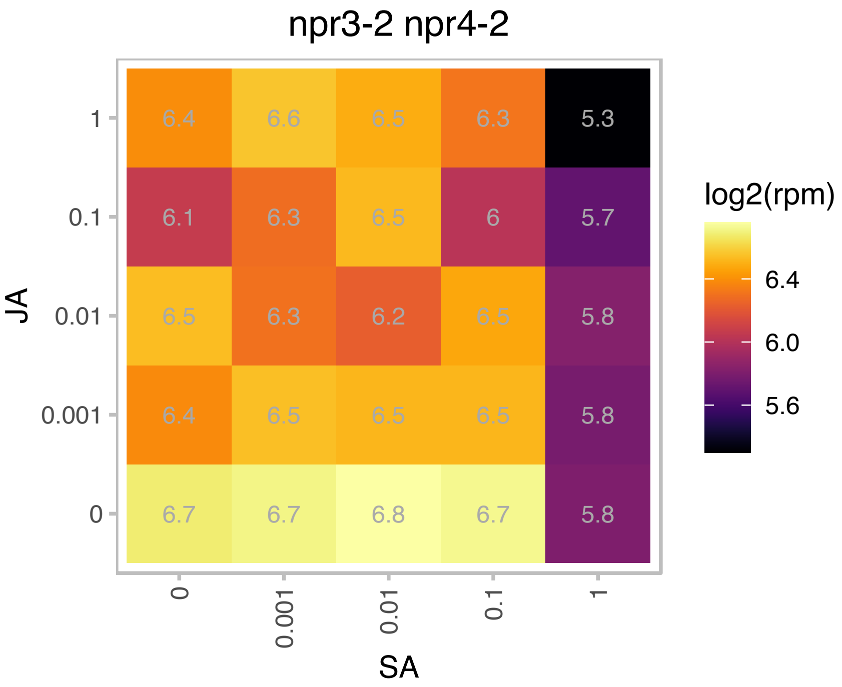


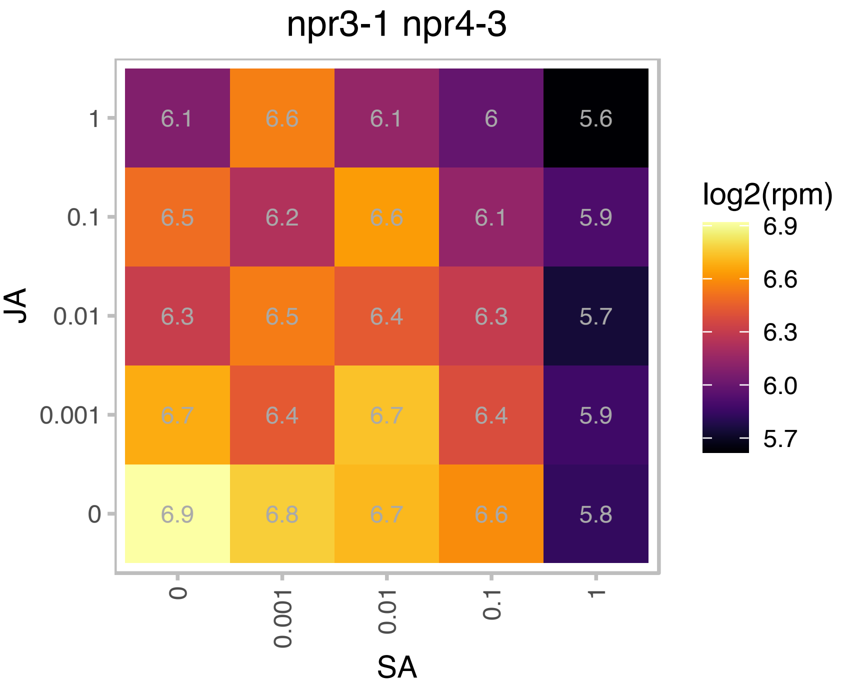

# AT4G28830.1 : NA

S-adenosyl-L-methionine-dependent methyltransferases superfamily protein

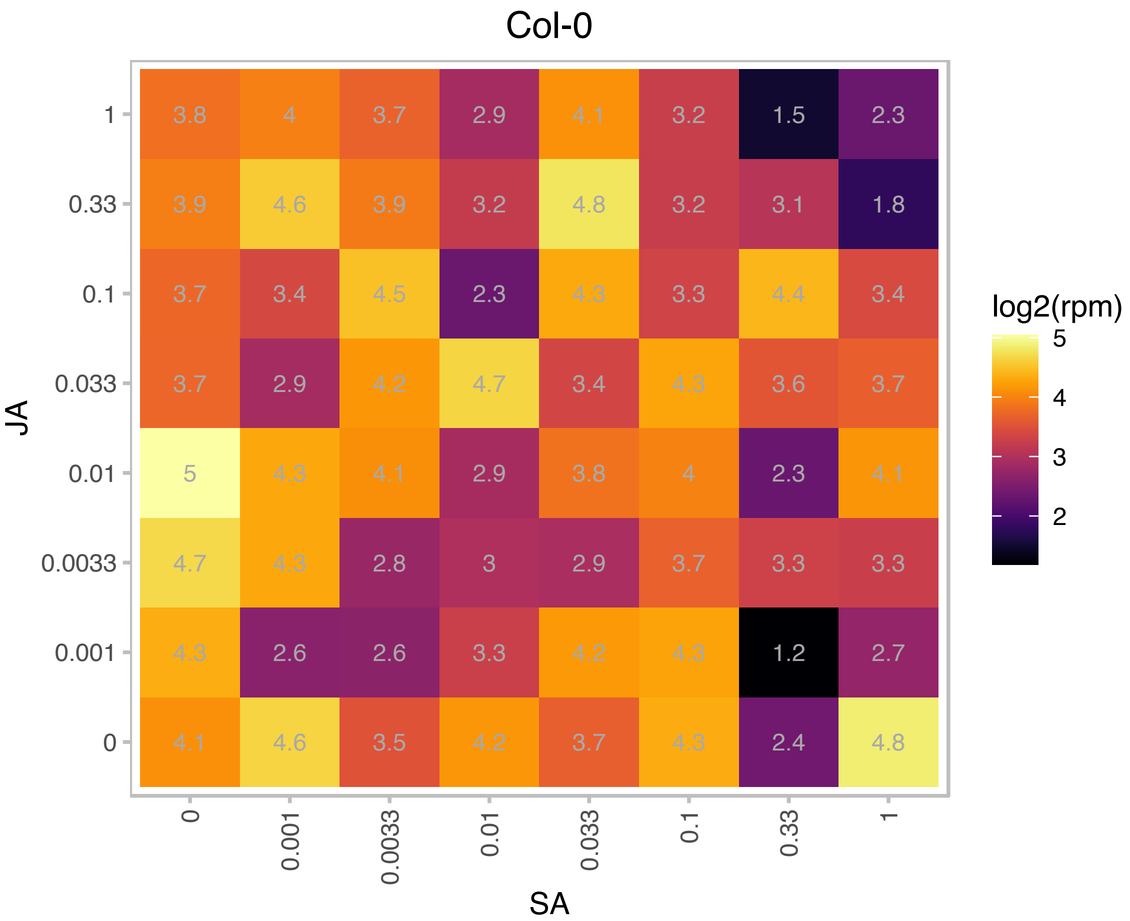


| | |
|---|---|
| Gene | AT4G28830 |
| is.expressed_genes | TRUE |
| is.responsive_genes | TRUE |
| is.apcluster | F-03 |

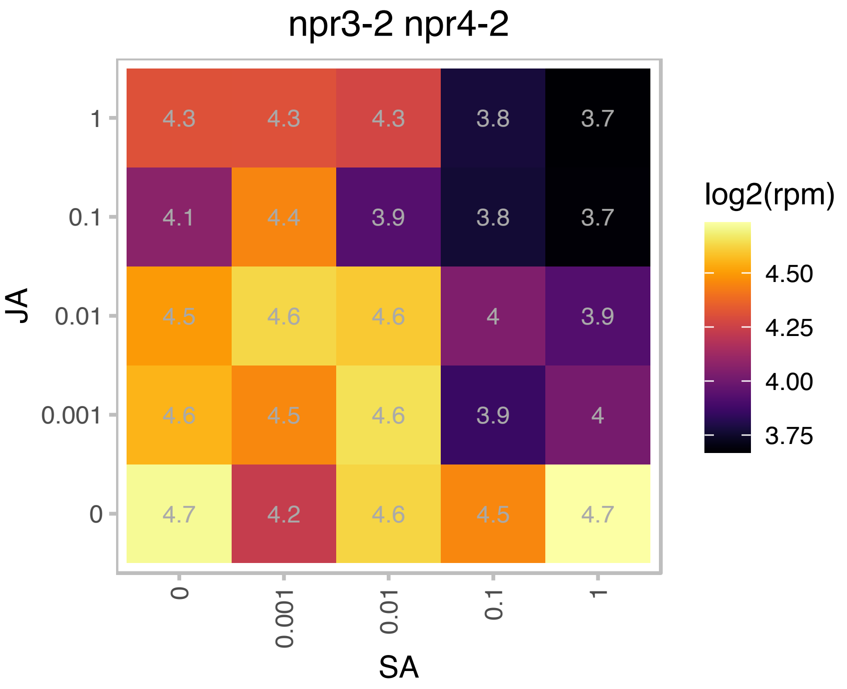


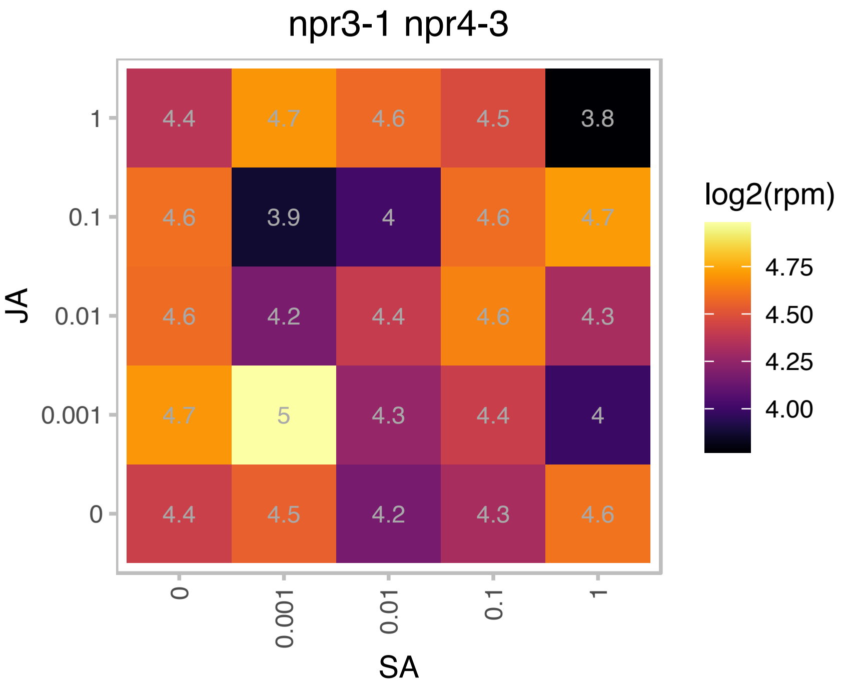

# AT4G29870.1 : NA

Oligosaccharyltransferase complex/magnesium transporter family protein

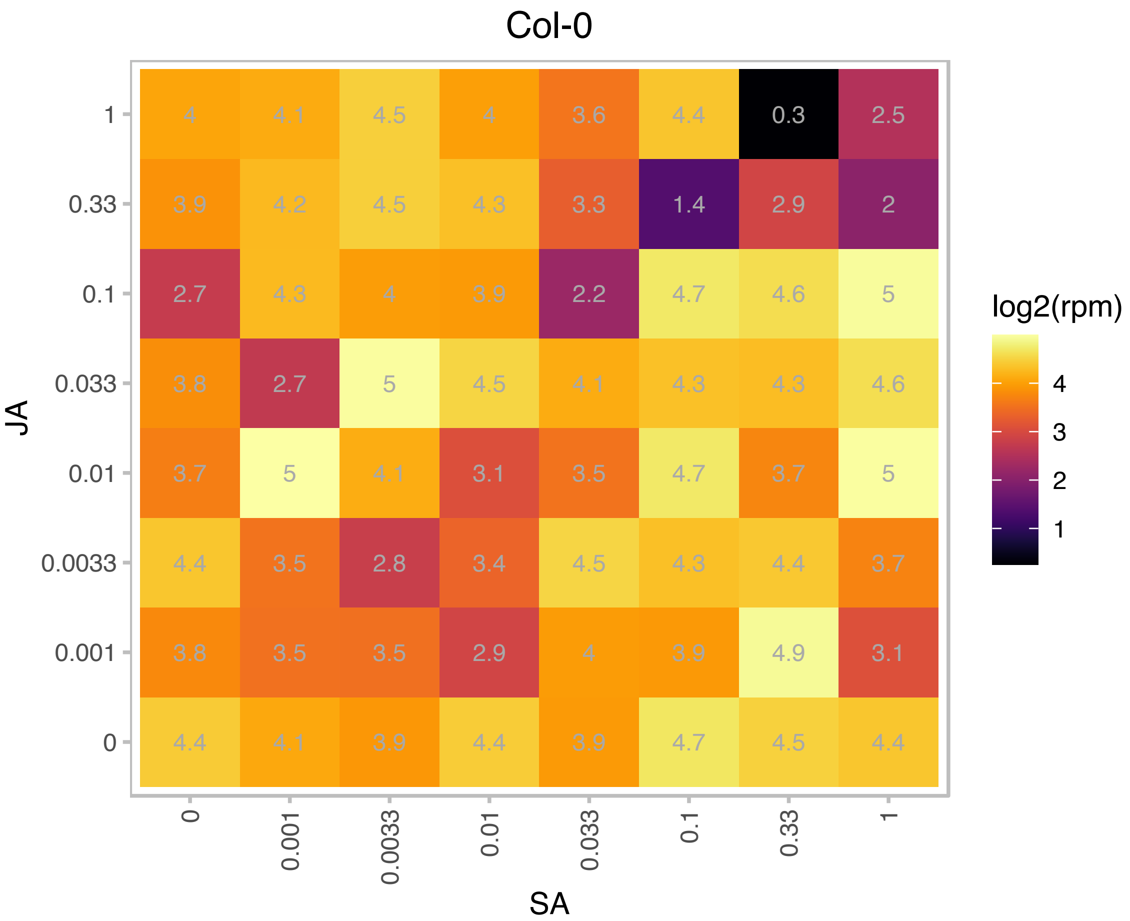


| | |
|---|---|
| Gene | AT4G29870 |
| is.expressed_genes | TRUE |
| is.responsive_genes | TRUE |
| is.apcluster | F-03 |

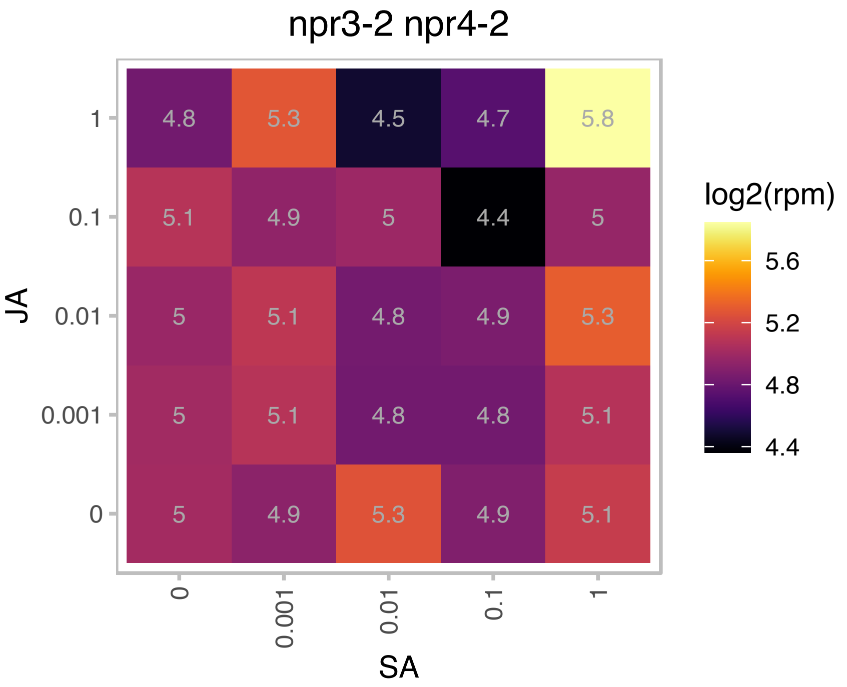


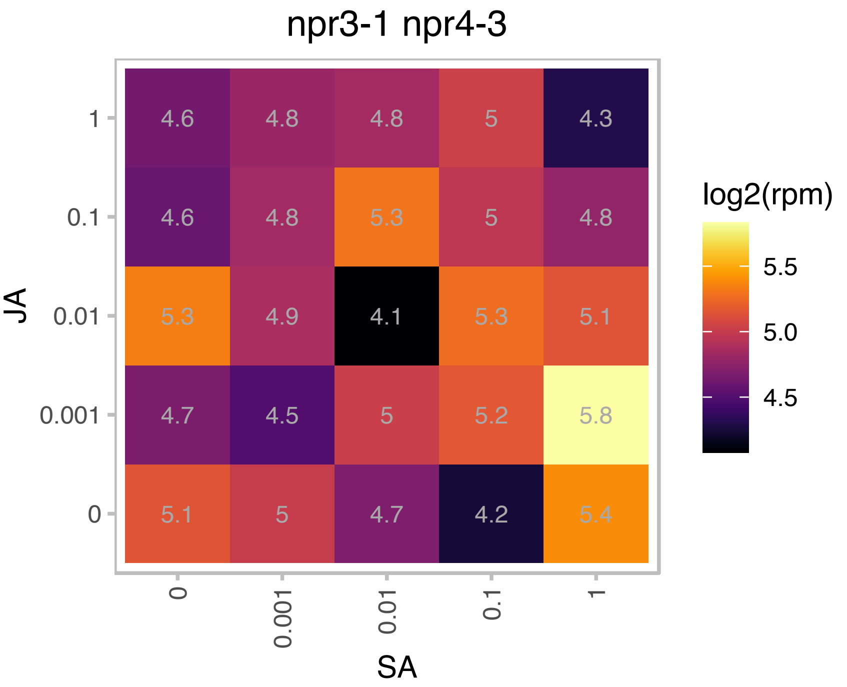

# AT4G33080.1 : NA

AGC (cAMP-dependent, cGMP-dependent and protein kinase C) kinase family protein

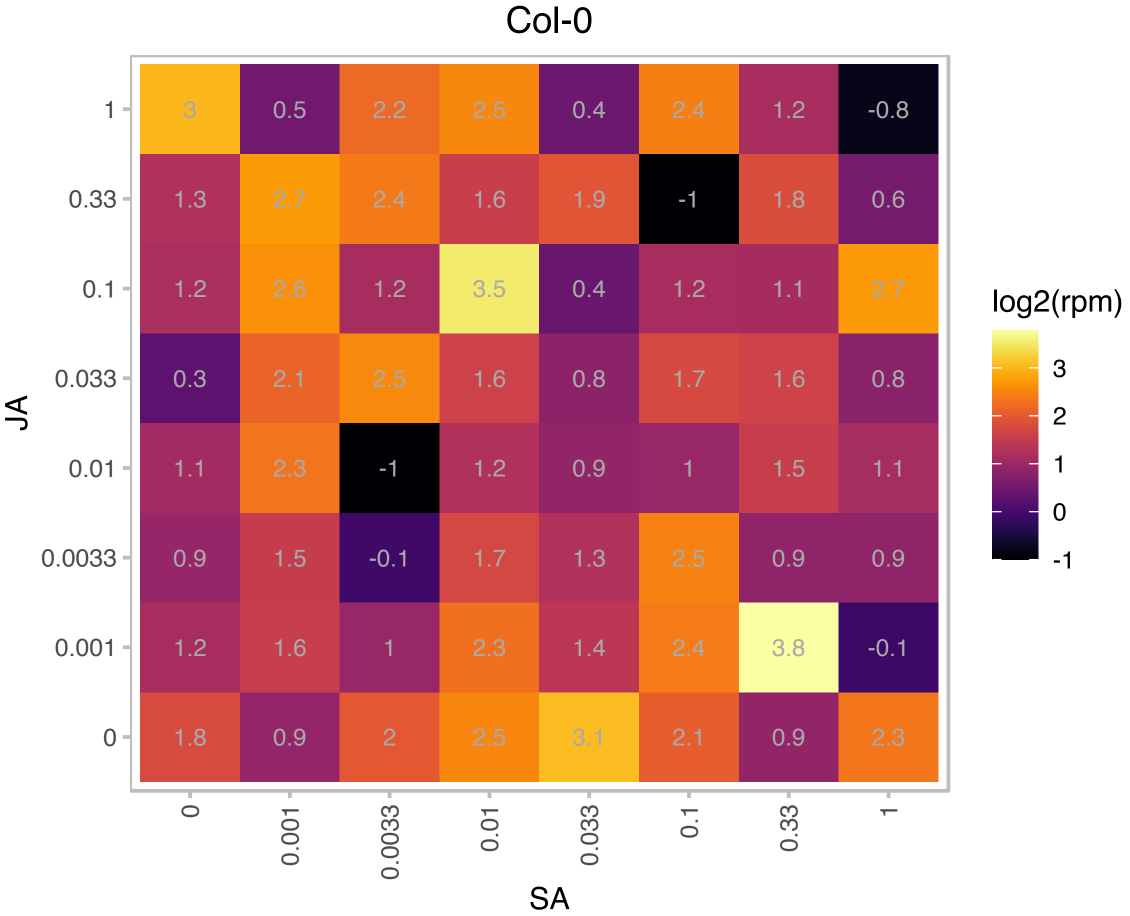


| | |
|---|---|
| Gene | AT4G33080 |
| is.expressed_genes | TRUE |
| is.responsive_genes | TRUE |
| is.apcluster | F-03 |

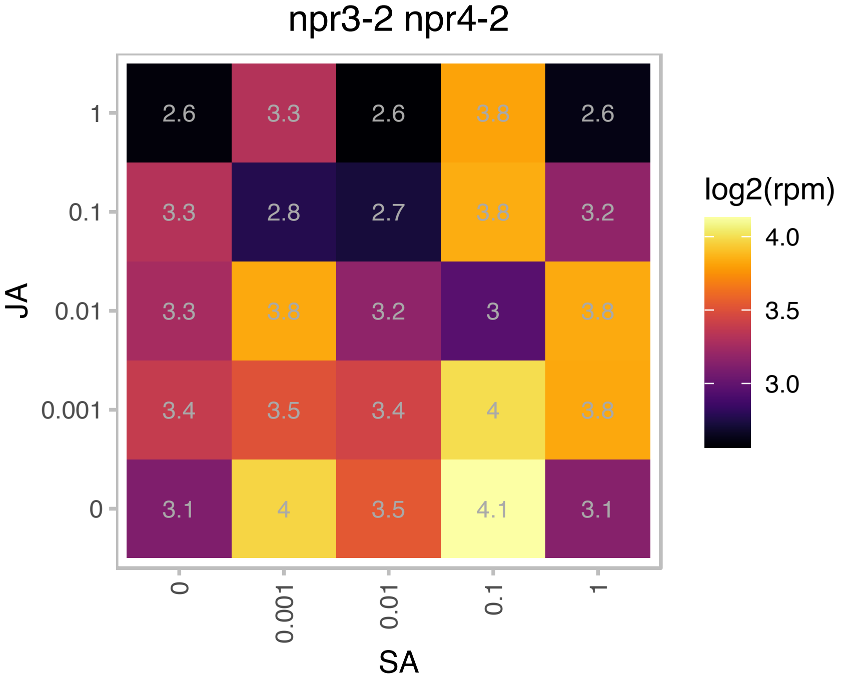


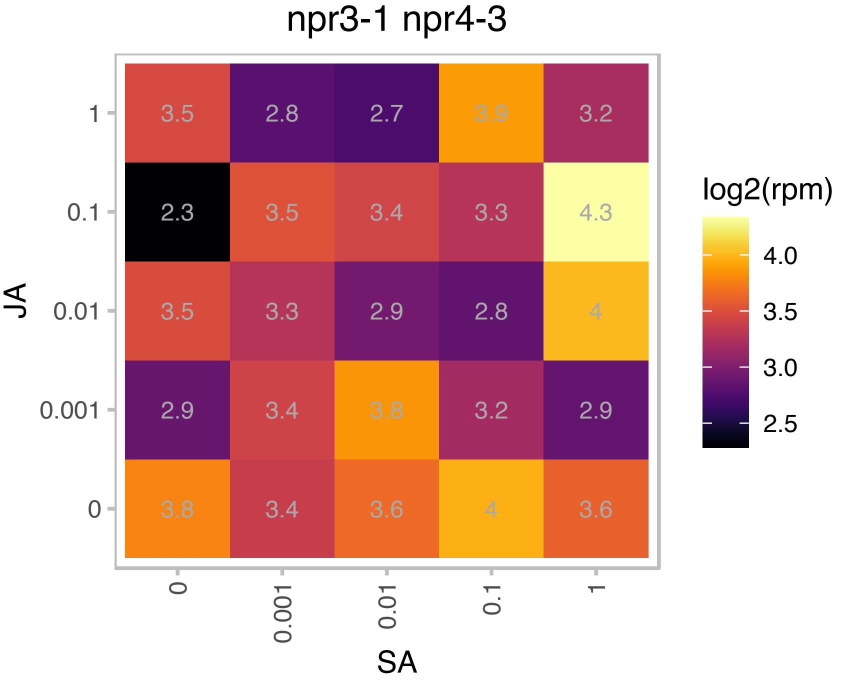

### AT5G03300.1 : ADK2

adenosine kinase 2

Encodes adenosine kinase 2 (ADK2), a typical, constitutively expressed housekeeping enzyme. Shows a high sequence identity with ADK1. Involved in salvage synthesis of adenylates and methyl recycling. Enzyme activity is substantially inhibited in roots, siliques and dry seeds by an unknown compound. May contribute to cytokinin interconversion. The mRNA is cell-to-cell mobile.

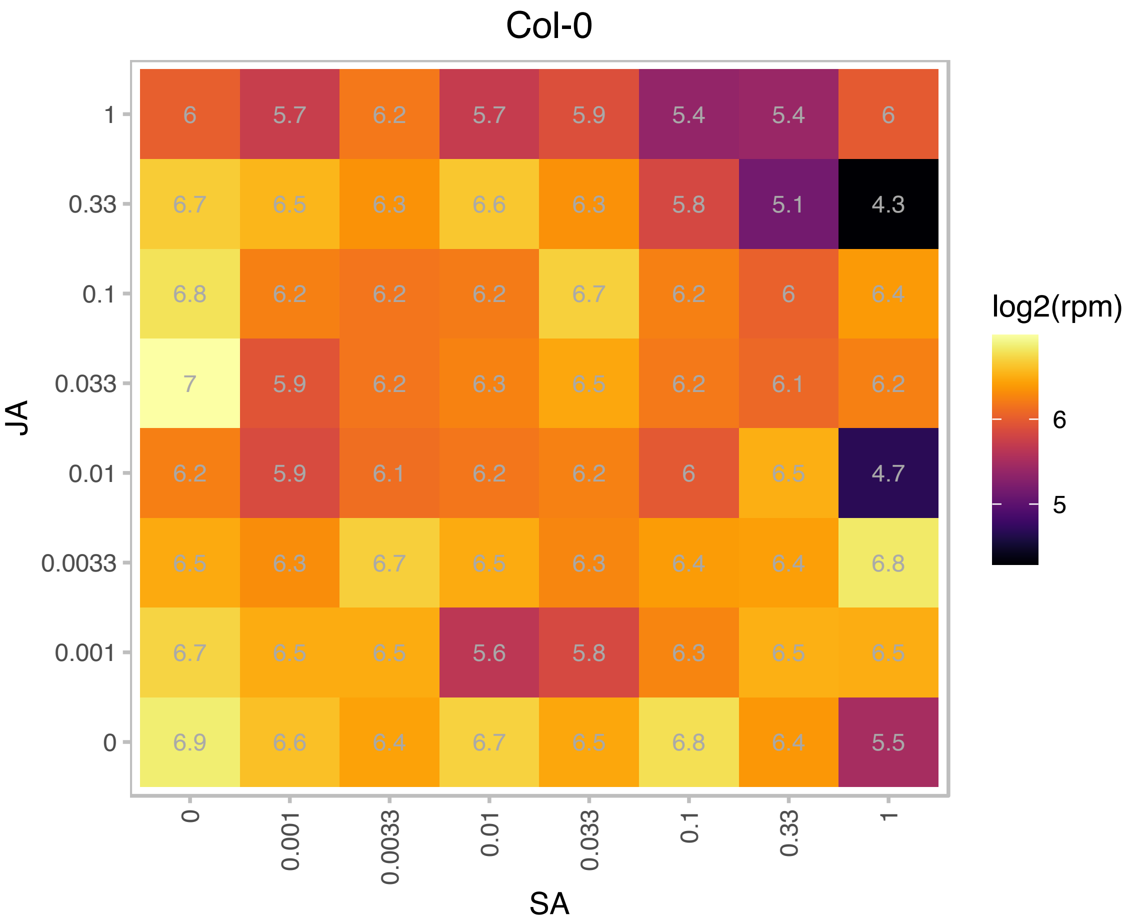


| | |
|---|---|
| Gene | AT5G03300 |
| is.expressed_genes | TRUE |
| is.responsive_genes | TRUE |
| is.apcluster | F-03 |

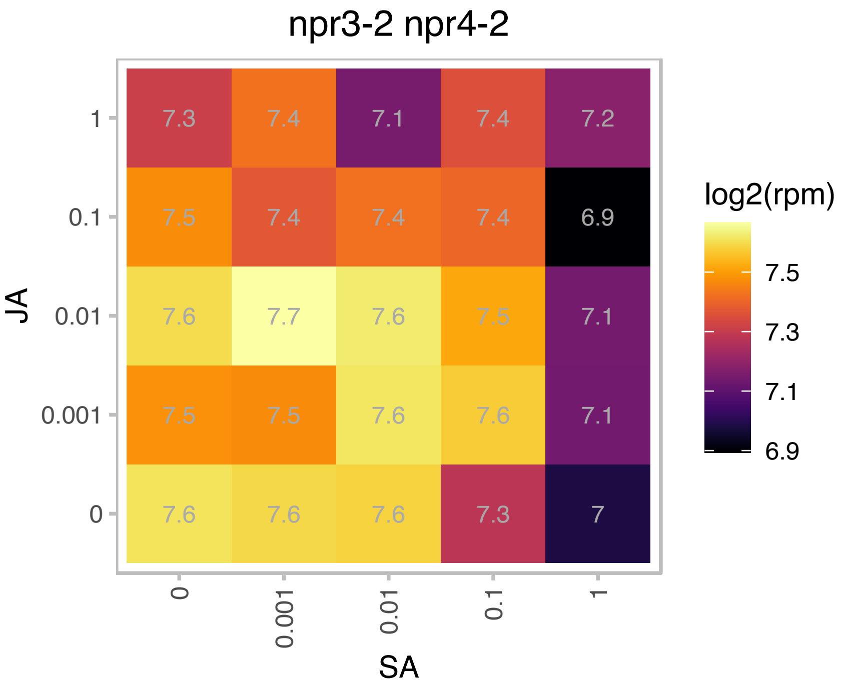


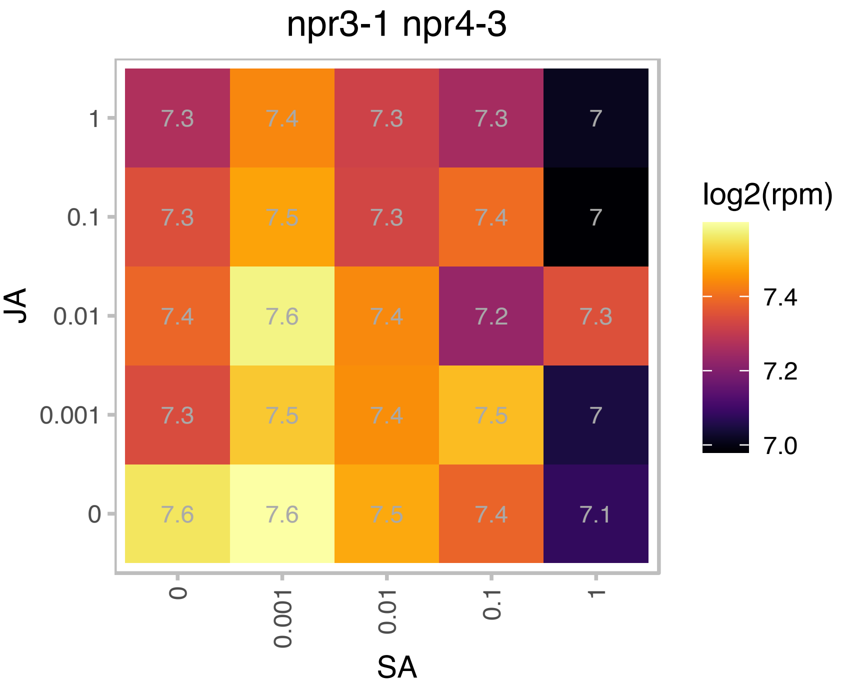

### AT5G13930.1 : ATCHS, CHS, TT4

Chalcone and stilbene synthase family protein
Encodes chalcone synthase (CHS), a key enzyme involved in the biosynthesis of flavonoids. Required for the accumulation of purple anthocyanins in leaves and stems. Also involved in the regulation of auxin transport and the modulation of root gravitropism. The mRNA is cell-to-cell mobile.

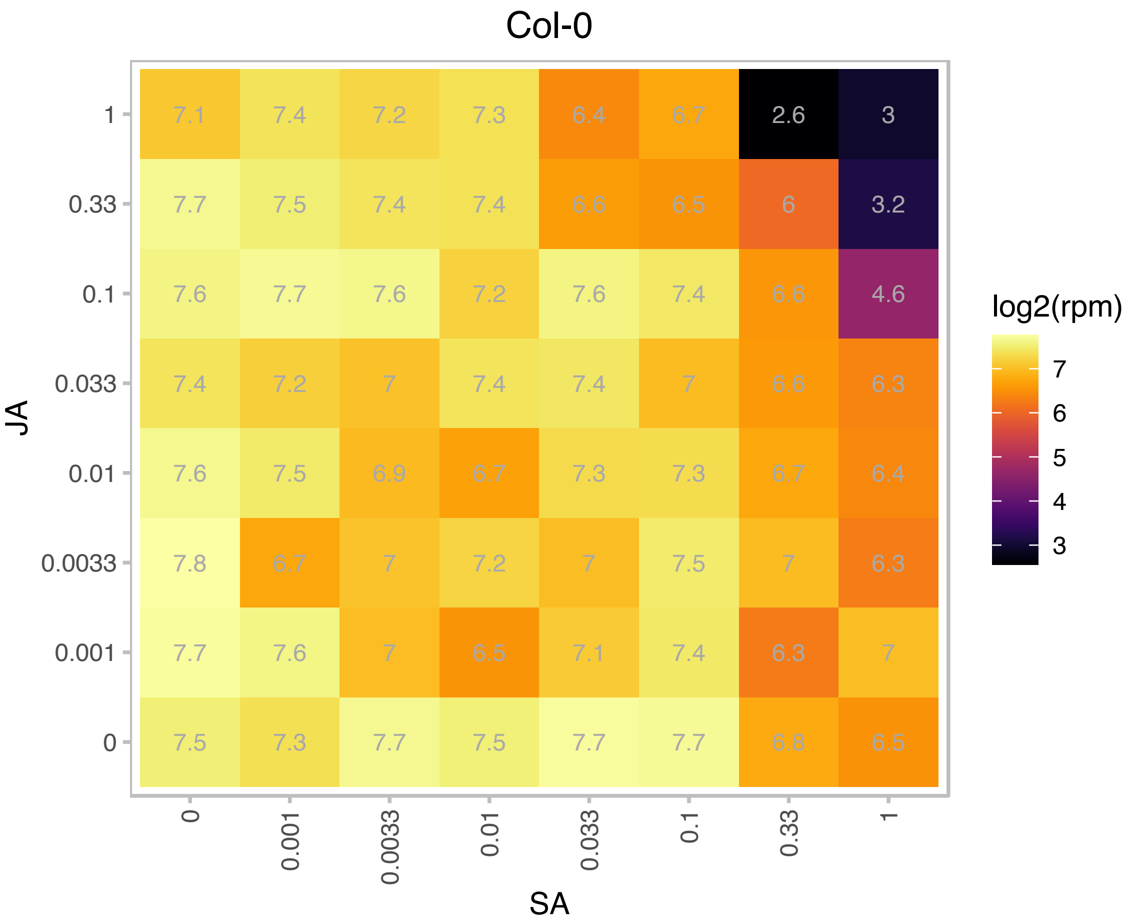


| | |
|---|---|
| Gene | AT5G13930 |
| is.expressed_genes | TRUE |
| is.responsive_genes | TRUE |
| is.apcluster | F-03 |

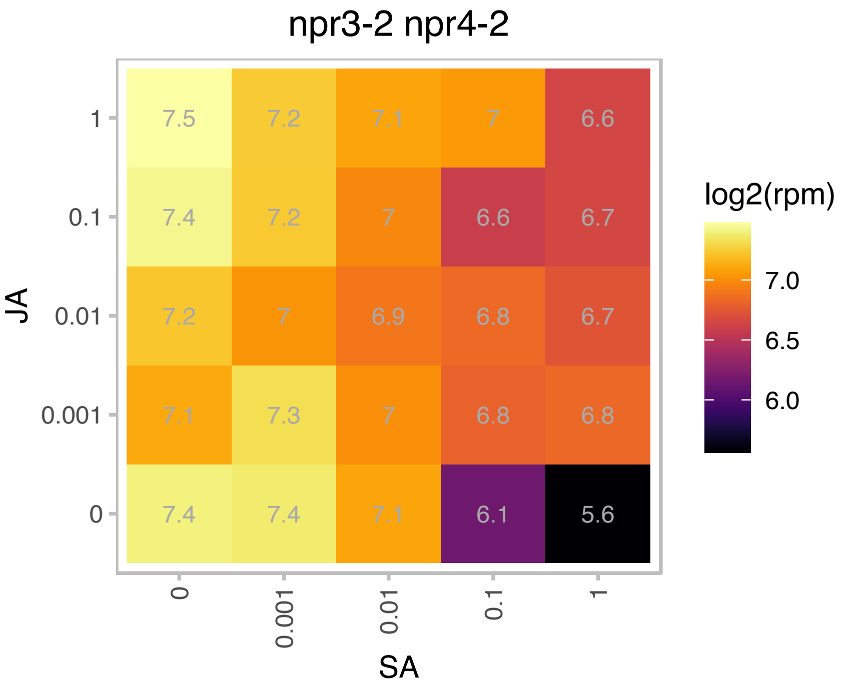


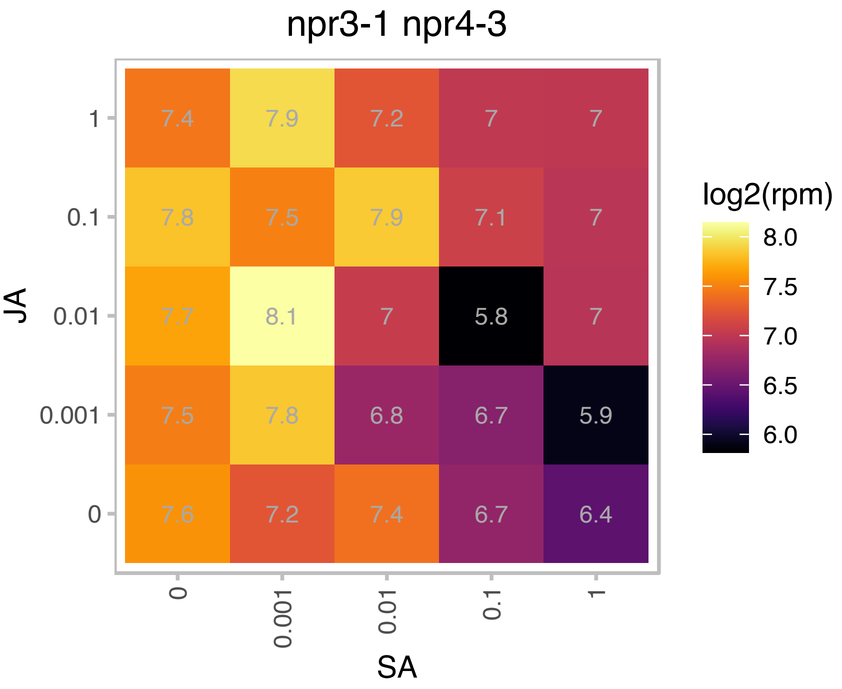

### AT5G15230.1 : GASA4

GAST1 protein homolog 4

Encodes gibberellin-regulated protein GASA4. Promotes GA responses and exhibits redox activity.

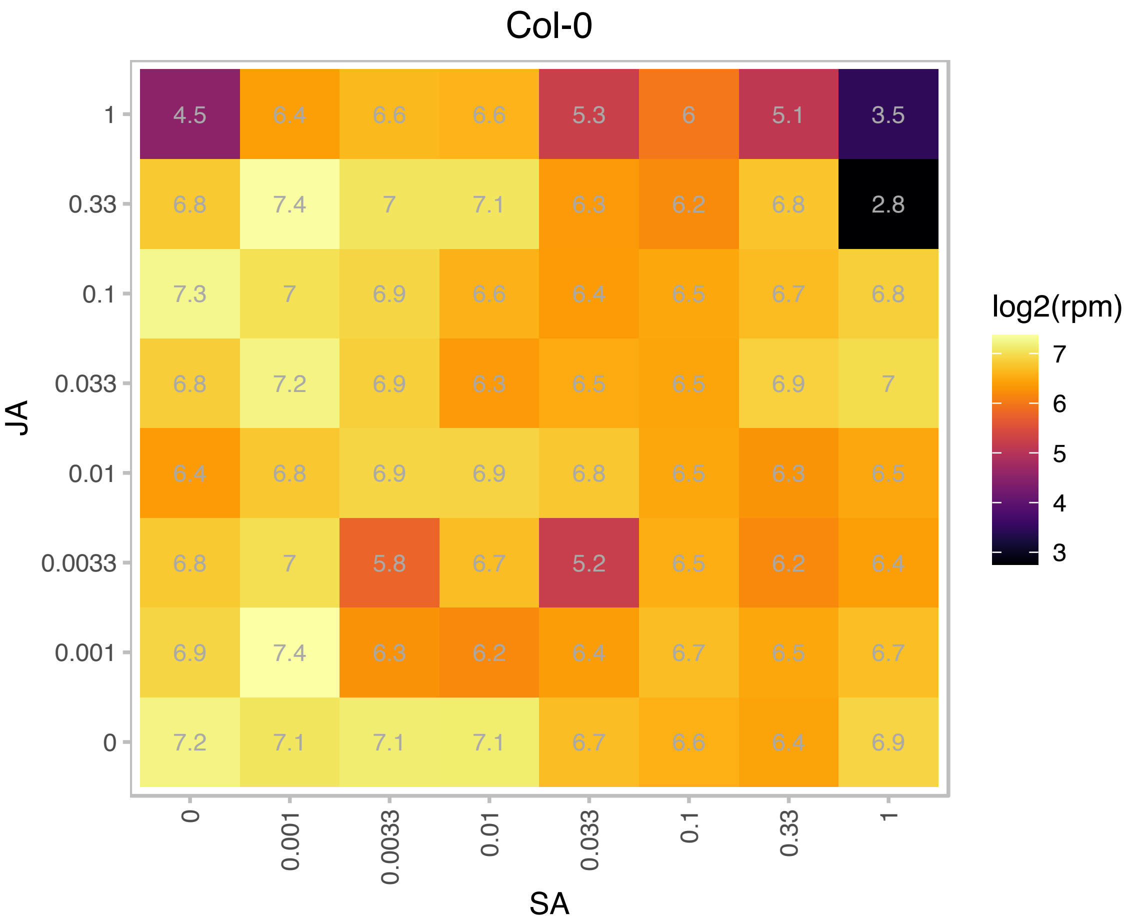


| | |
|---|---|
| Gene | AT5G15230 |
| is.expressed_genes | TRUE |
| is.responsive_genes | TRUE |
| is.apcluster | F-03 |

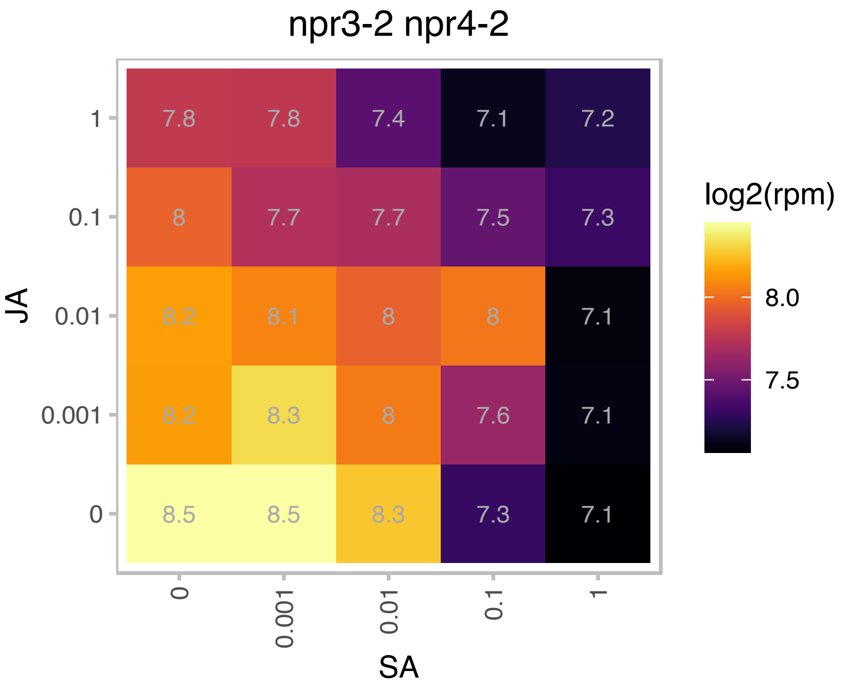


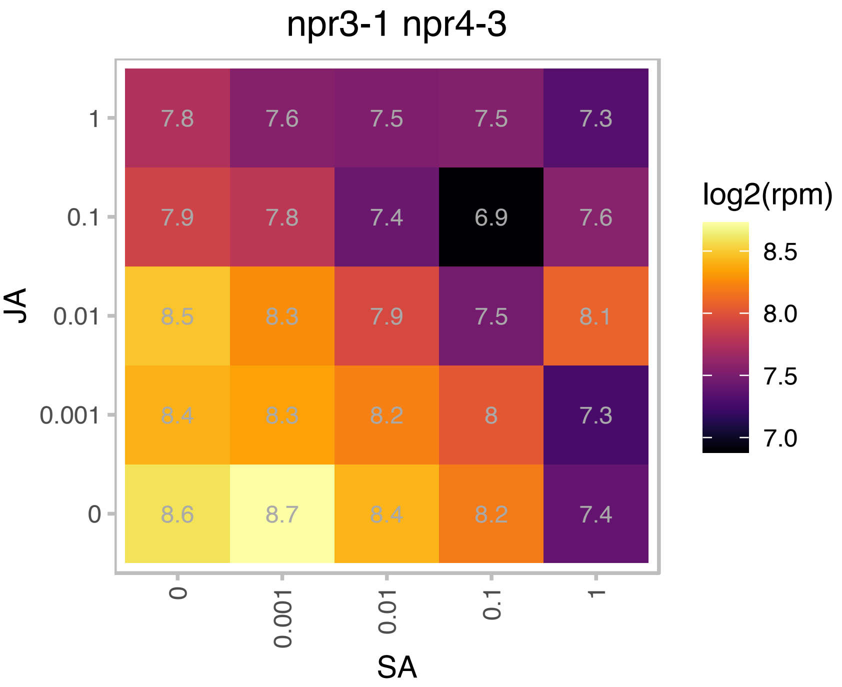

### AT5G20070.1 : ATNUDT19, ATNUDX19, NUDX19

nudix hydrolase homolog 19

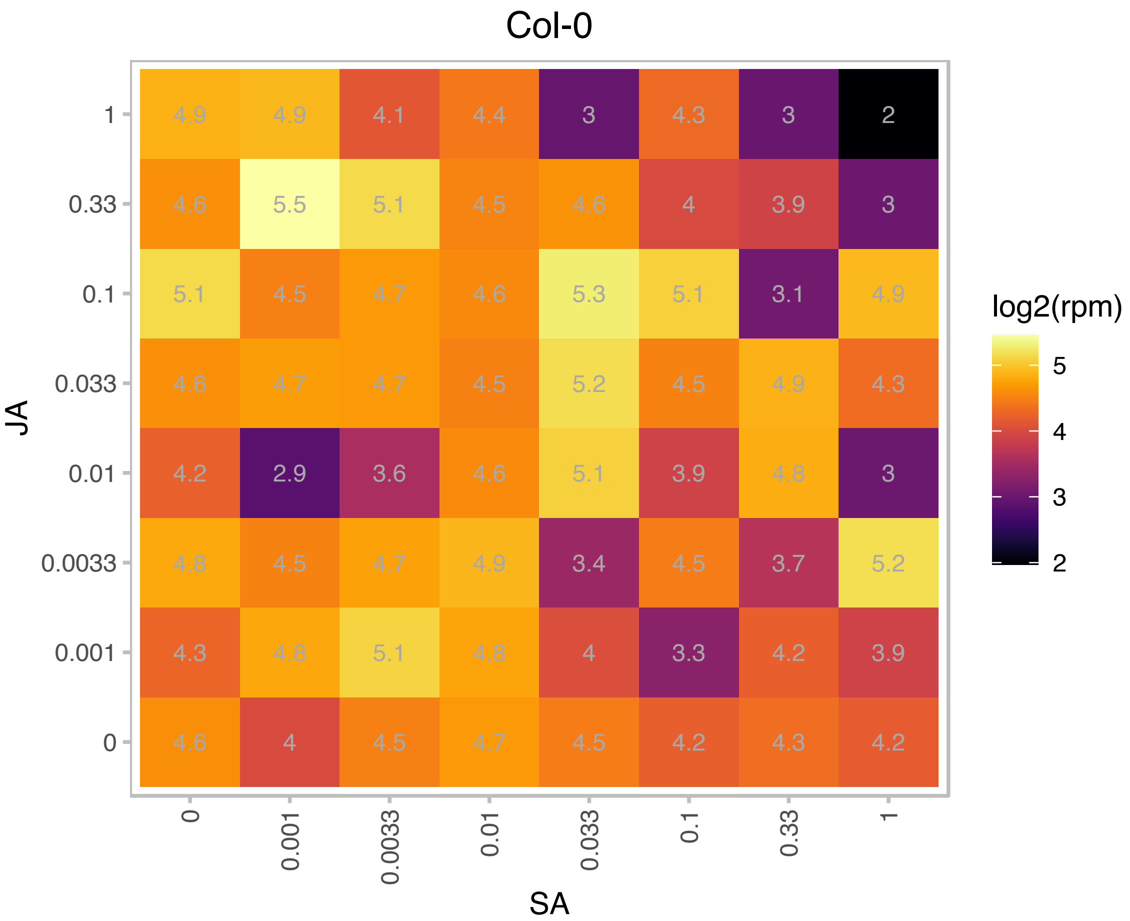


| | |
|---|---|
| Gene | AT5G20070 |
| is.expressed_genes | TRUE |
| is.responsive_genes | TRUE |
| is.apcluster | F-03 |

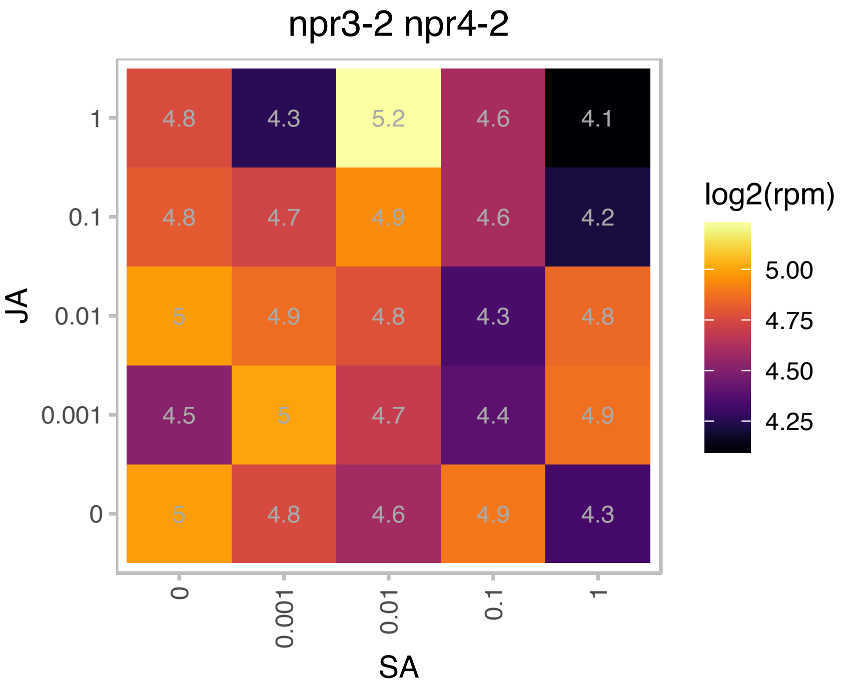


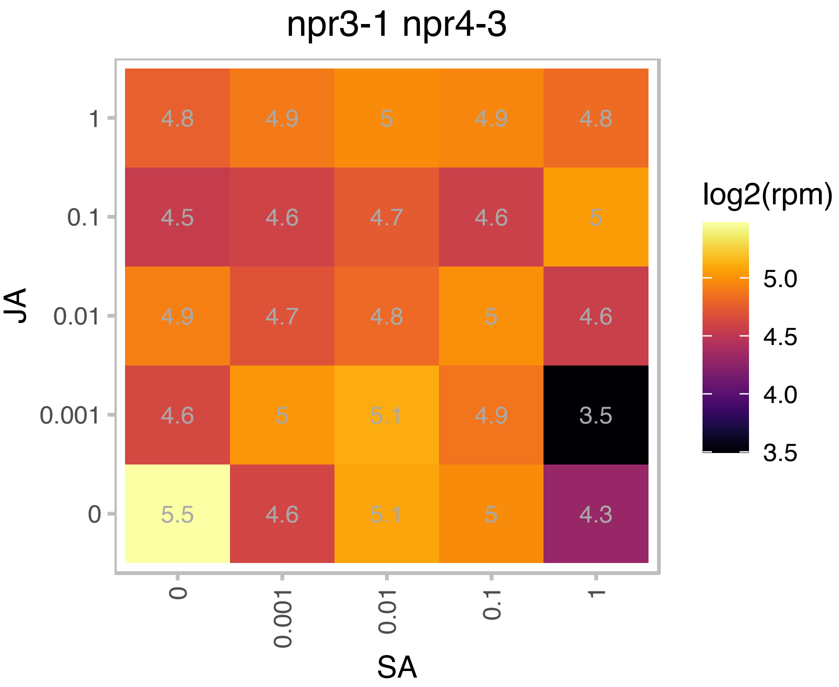

# AT5G22320.1 : NA

Leucine-rich repeat (LRR) family protein

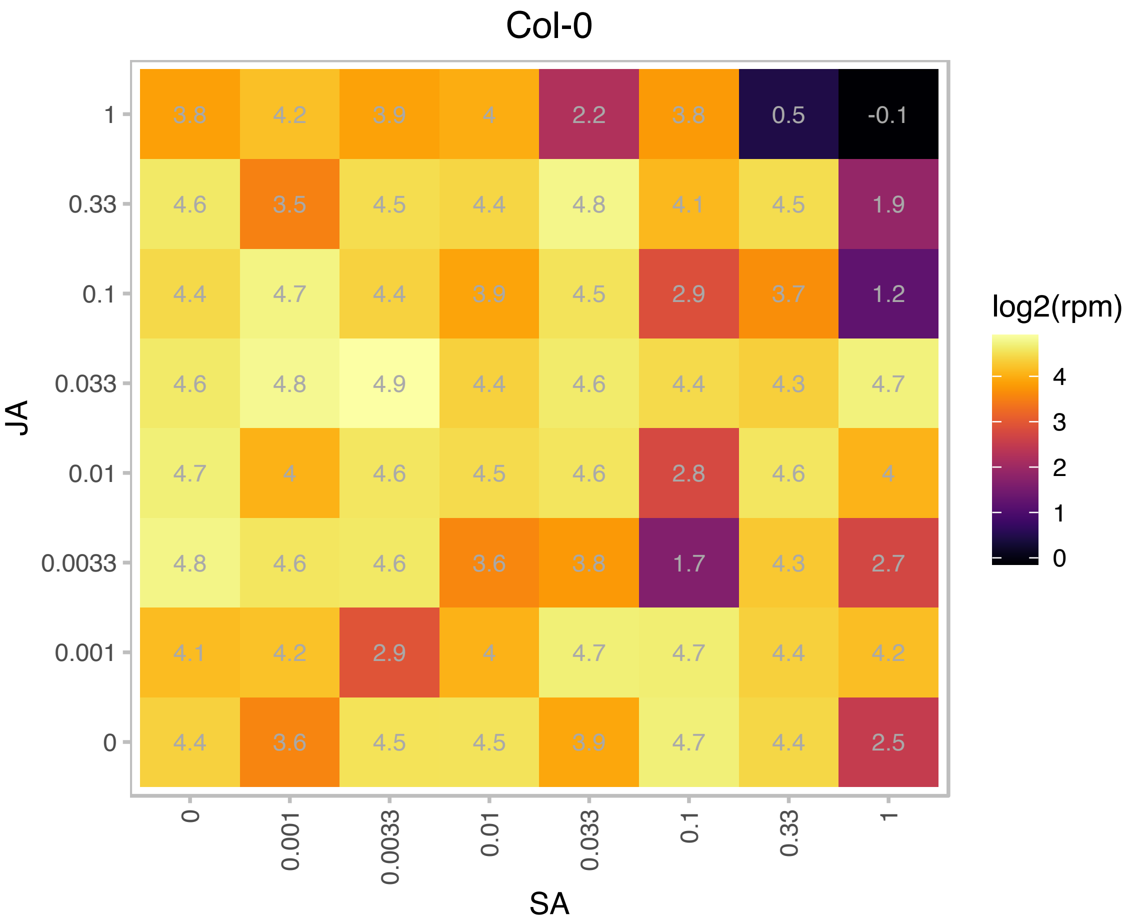


| | |
|---|---|
| Gene | AT5G22320 |
| is.expressed_genes | TRUE |
| is.responsive_genes | TRUE |
| is.apcluster | F-03 |

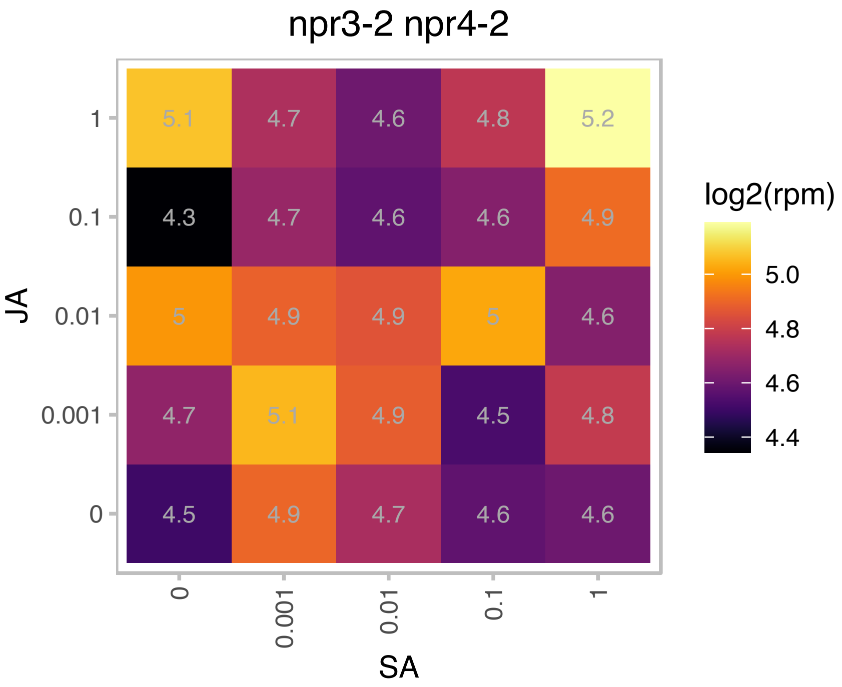


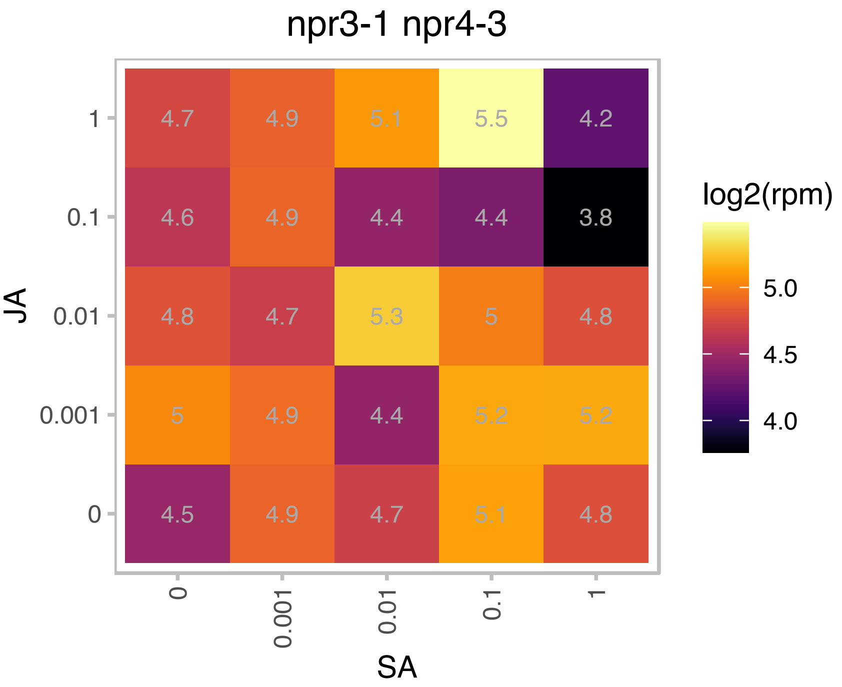

# AT5G44730.1 : NA

Haloacid dehalogenase-like hydrolase (HAD) superfamily protein

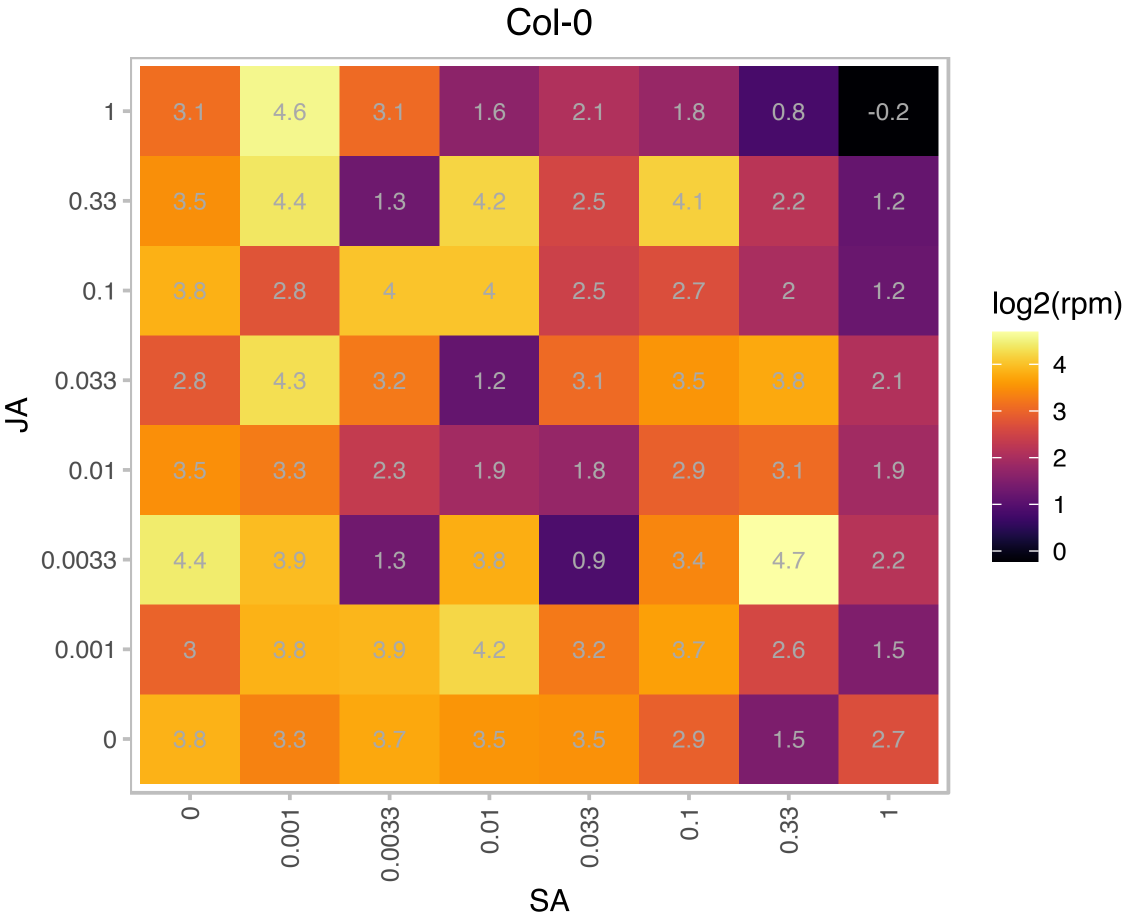


| | |
|---|---|
| Gene | AT5G44730 |
| is.expressed_genes | TRUE |
| is.responsive_genes | TRUE |
| is.apcluster | F-03 |

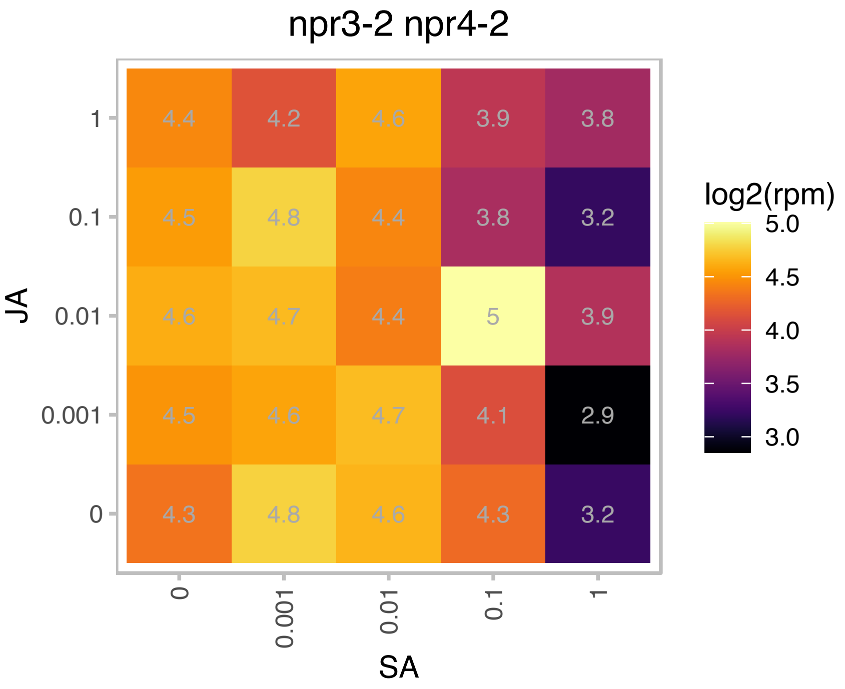


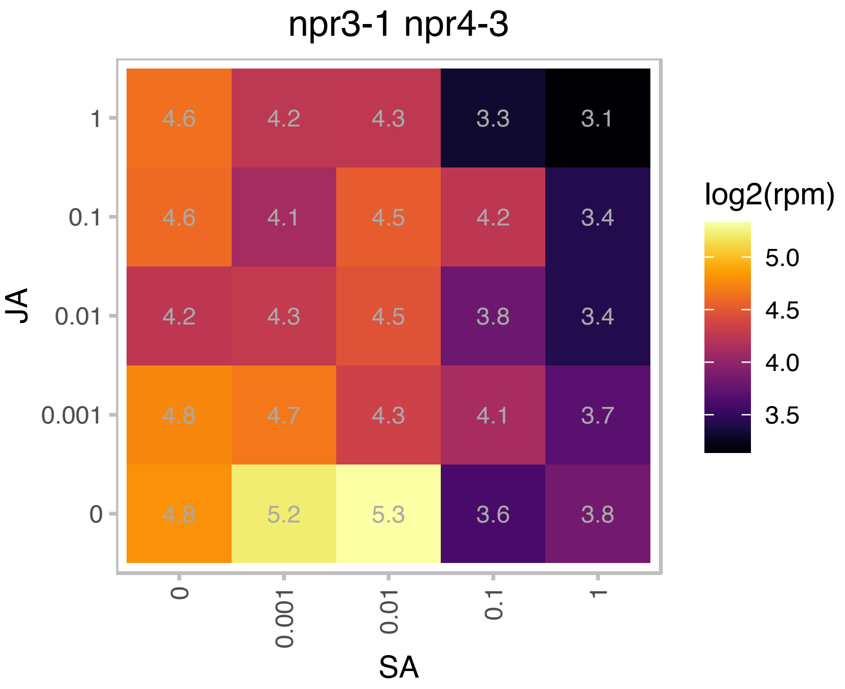

### AT5G47100.1 : ATCBL9, CBL9

calcineurin B-like protein 9

member of AtCBLs (Calcineurin B-like Calcium Sensor Proteins. CBL9 interacts with and targets CIPK23 to the plasma membrane in vivo.

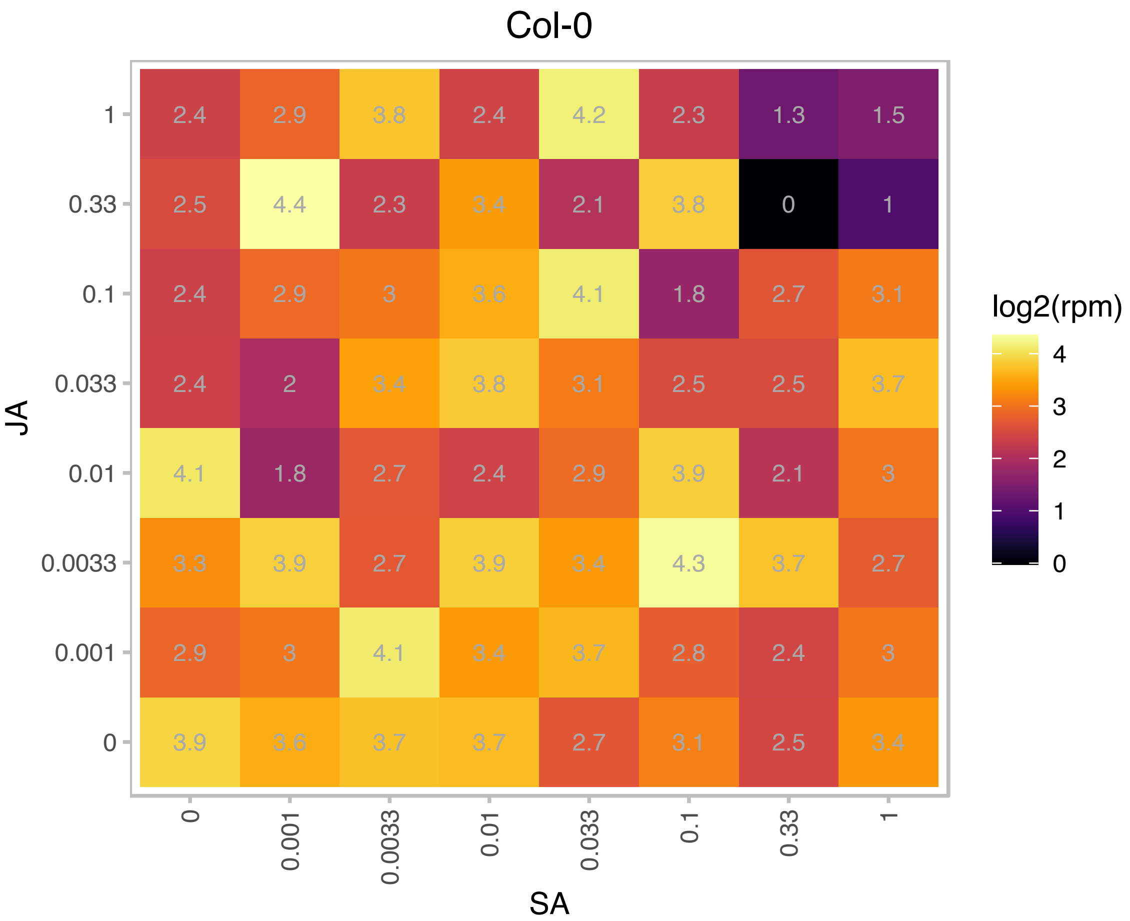


| | |
|---|---|
| Gene | AT5G47100 |
| is.expressed_genes | TRUE |
| is.responsive_genes | TRUE |
| is.apcluster | F-03 |

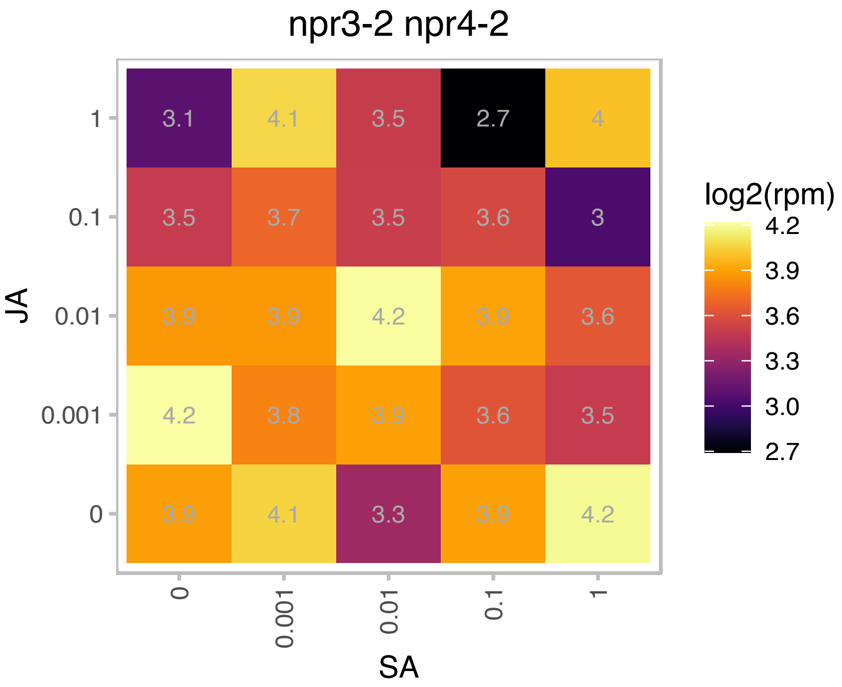


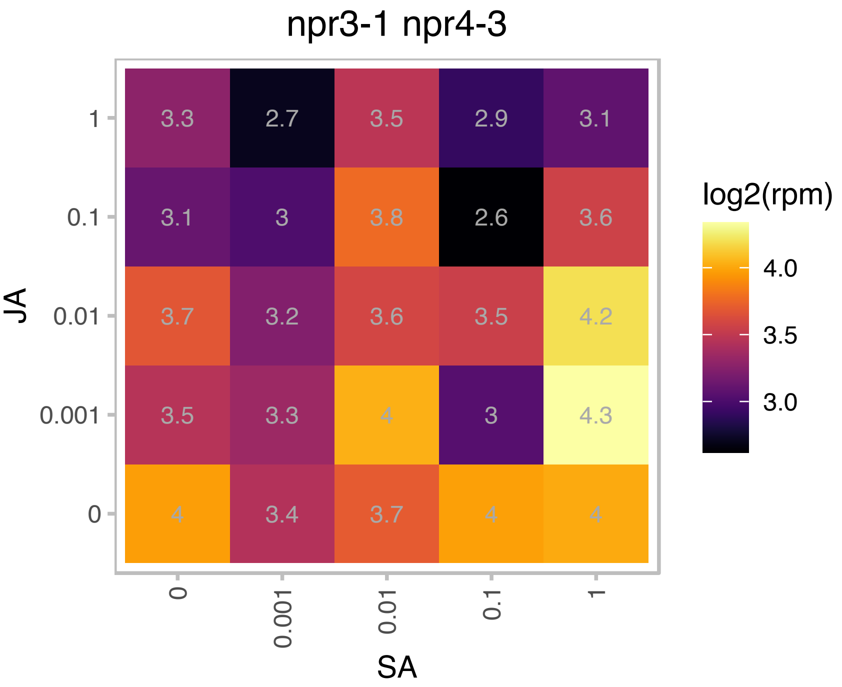

### AT5G48485.1 : DIR1

Encodes a putative apoplastic lipid transfer protein that is involved in systemic acquired resistance. Mutants in this gene exhibit wild-type local resistance to avirulent and virulent Pseudomonas syringae, but pathogenesis-related gene expression is abolished in uninoculated distant leaves and fail to develop SAR to virulent Pseudomonas or Peronospora parasitica. DIR1 protein is cell-to-cell mobile and is transported via phloem sap.

Bifunctional inhibitor/lipid-transfer protein/seed storage 2S albumin superfamily protein

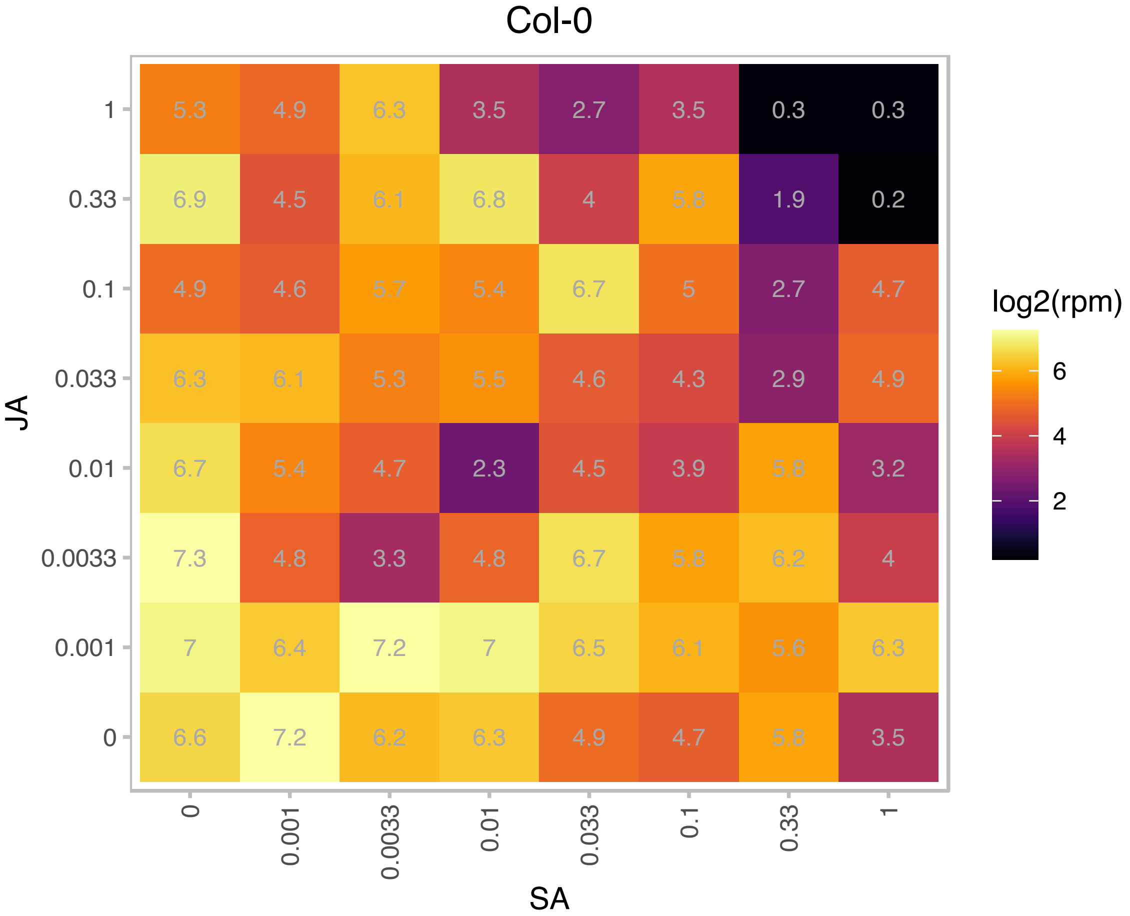


| | |
|---|---|
| Gene | AT5G48485 |
| is.expressed_genes | TRUE |
| is.responsive_genes | TRUE |
| is.apcluster | F-03 |

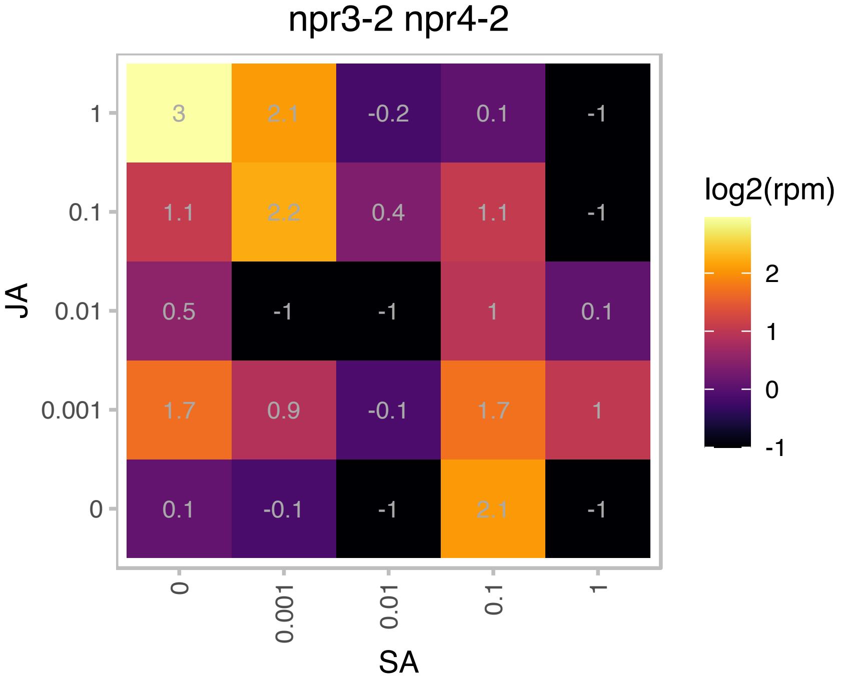


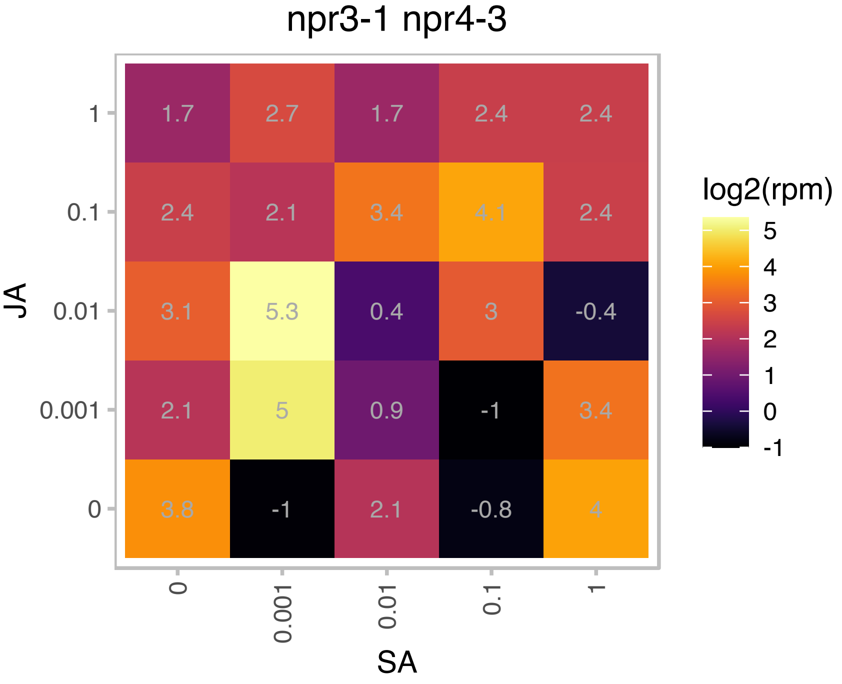

# AT5G48630.1 : NA

Cyclin family protein

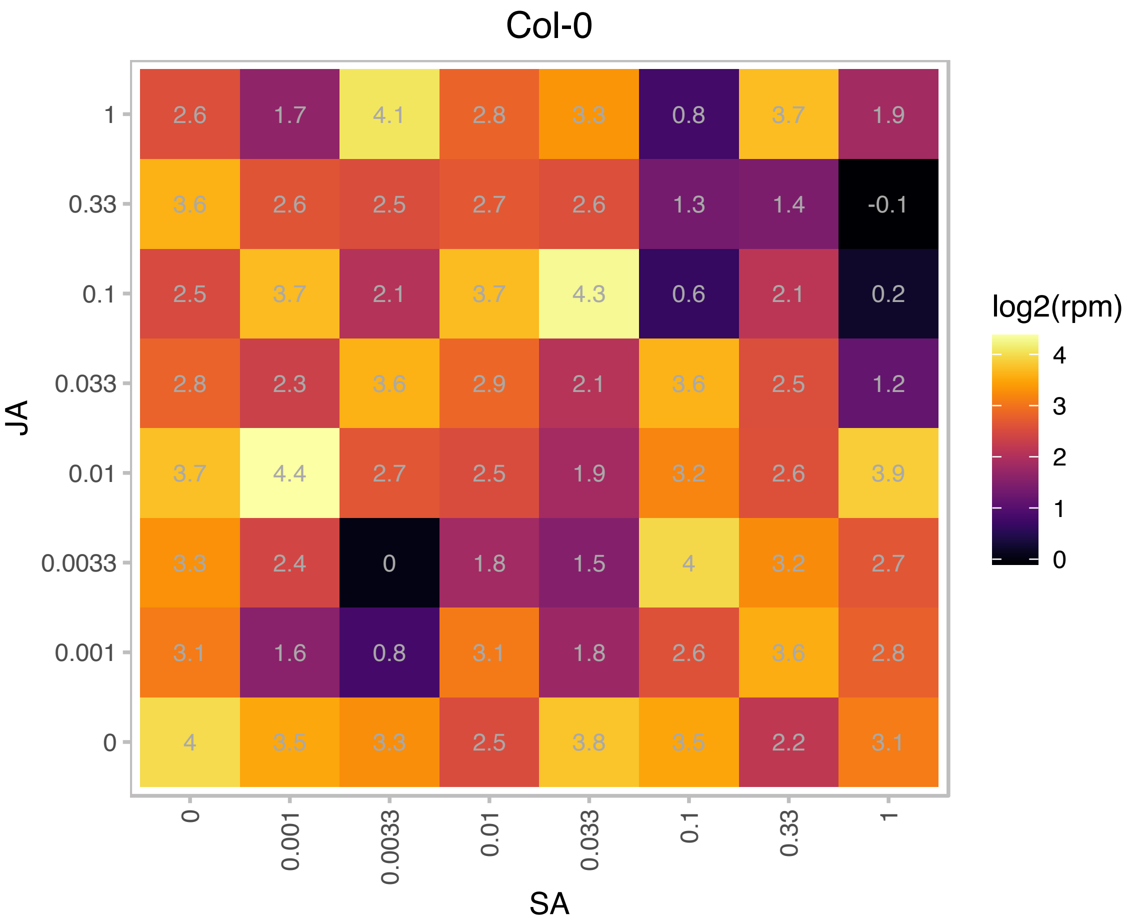


| | |
|---|---|
| Gene | AT5G48630 |
| is.expressed_genes | TRUE |
| is.responsive_genes | TRUE |
| is.apcluster | F-03 |

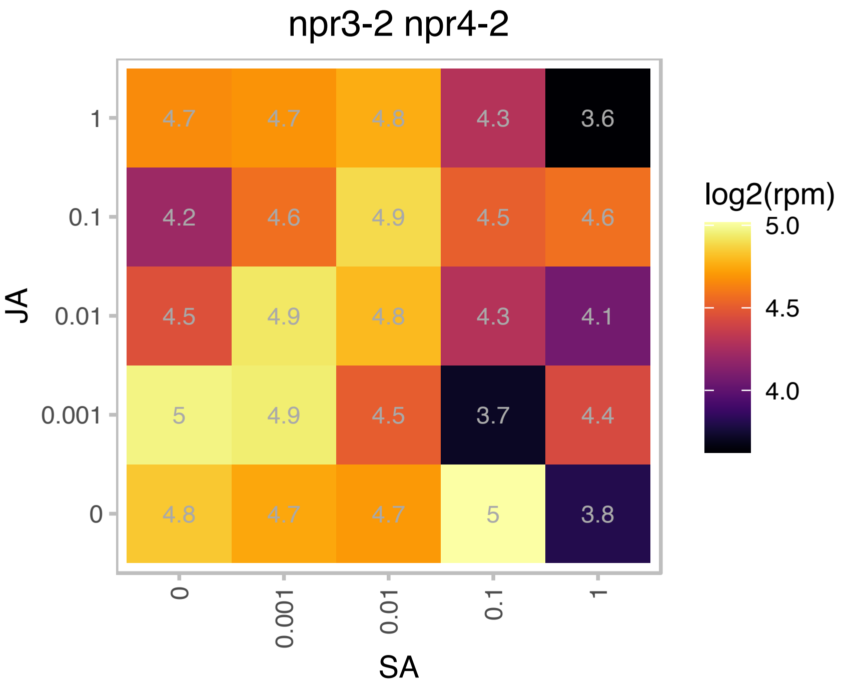


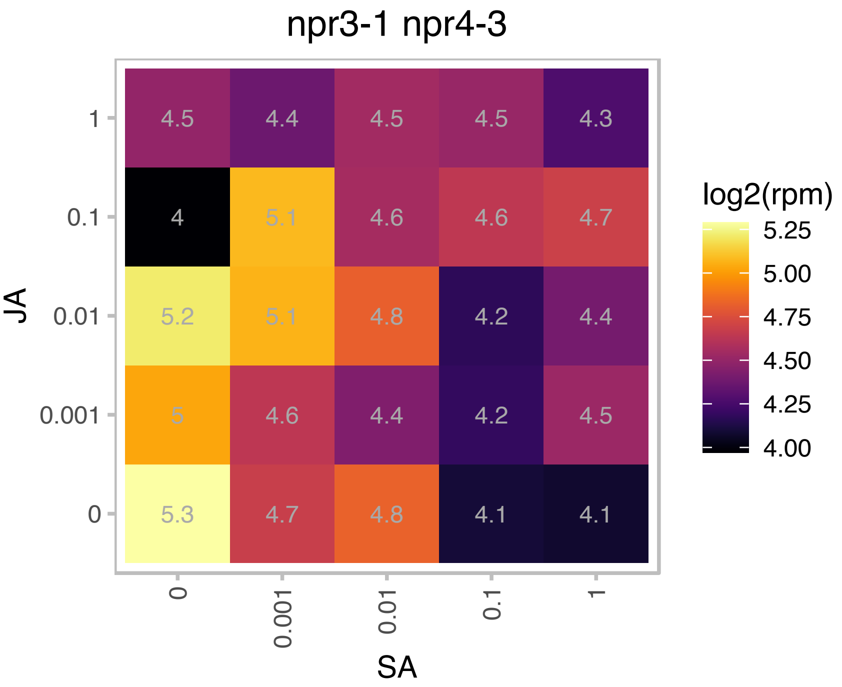

### AT5G49510.1 : PFD3

prefoldin 3

| | |
|---|---|
| Gene | AT5G49510 |
| is.expressed_genes | TRUE |
| is.responsive_genes | TRUE |
| is.apcluster | F-03 |

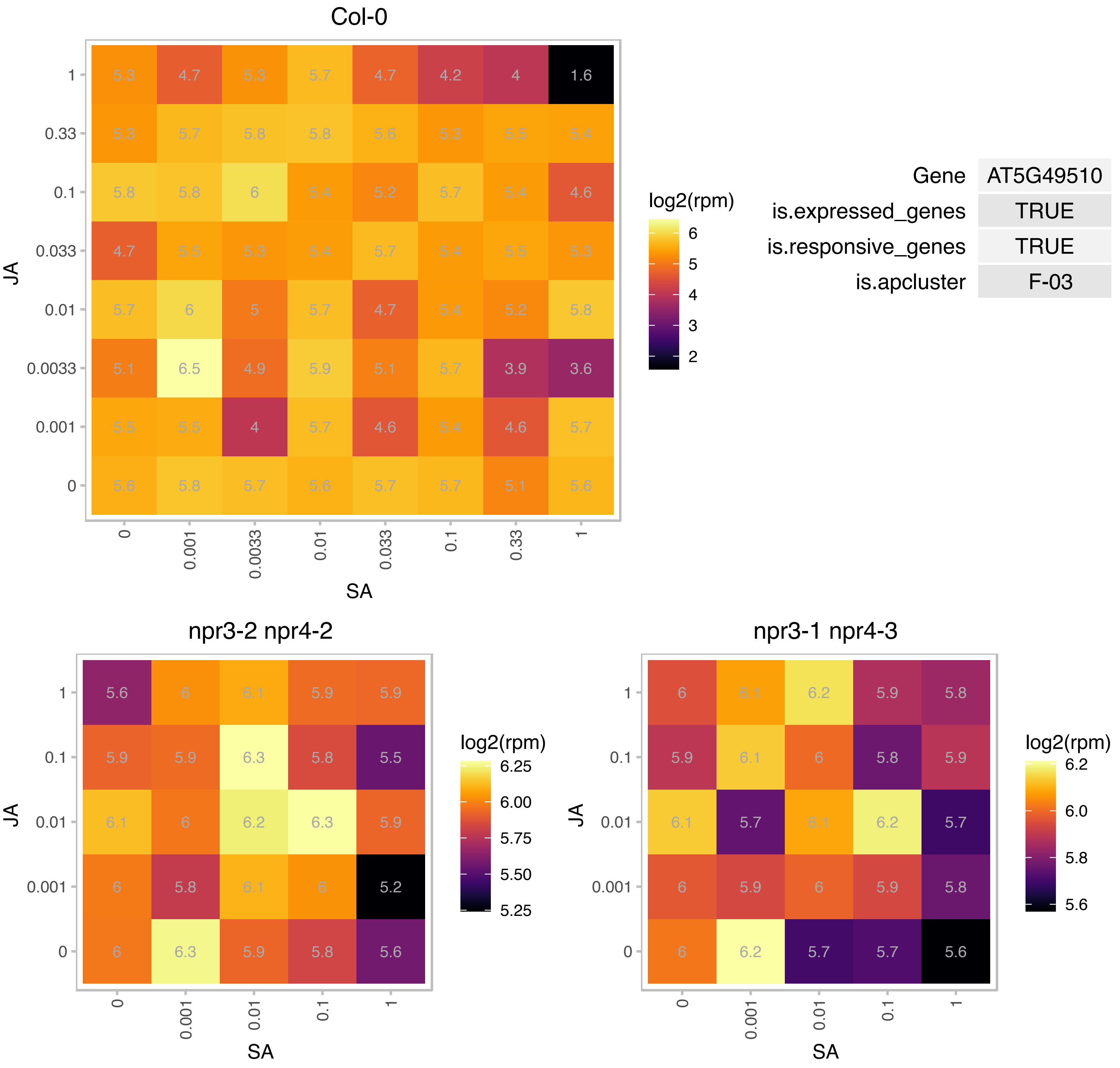

# AT5G54855.1 : NA

Pollen Ole e 1 allergen and extensin family protein

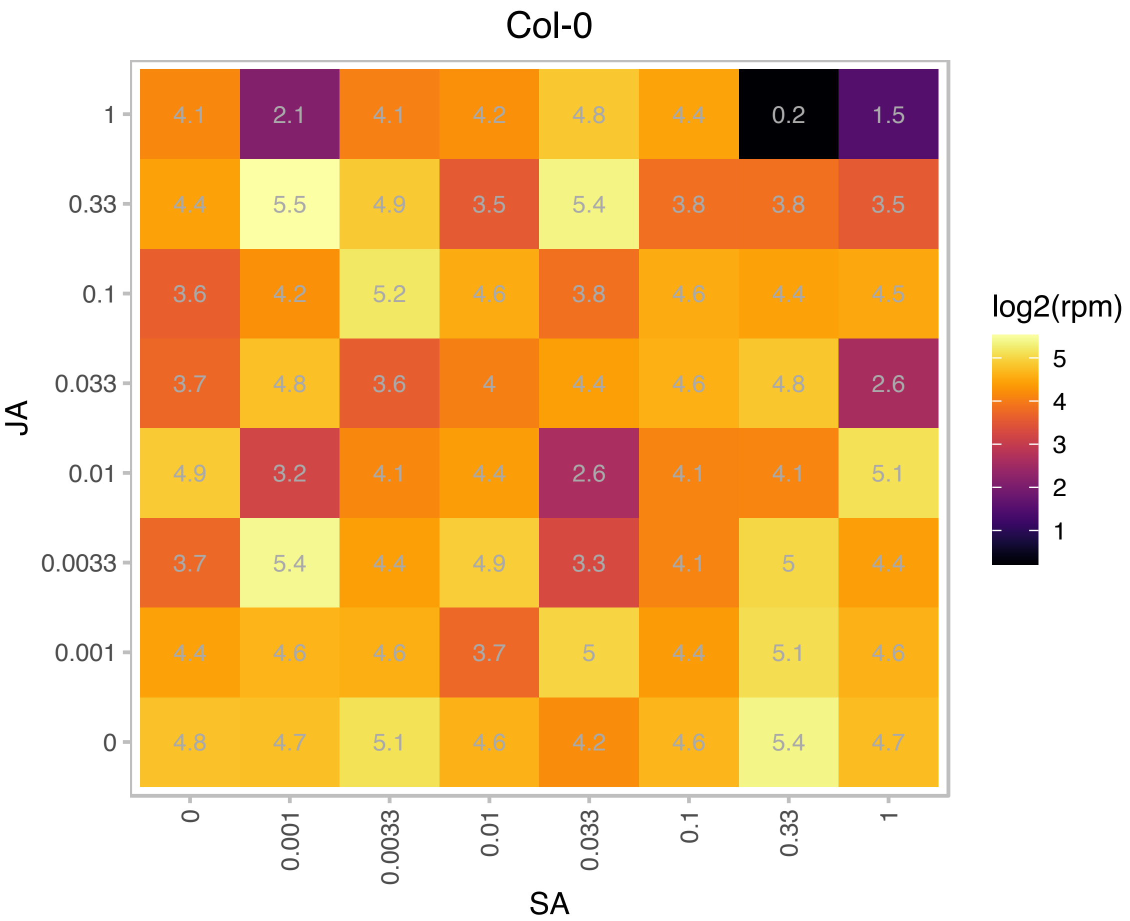


| Gene | AT5G54855 |
|---|---|
| is.expressed_genes | TRUE |
| is.responsive_genes | TRUE |
| is.apcluster | F-03 |

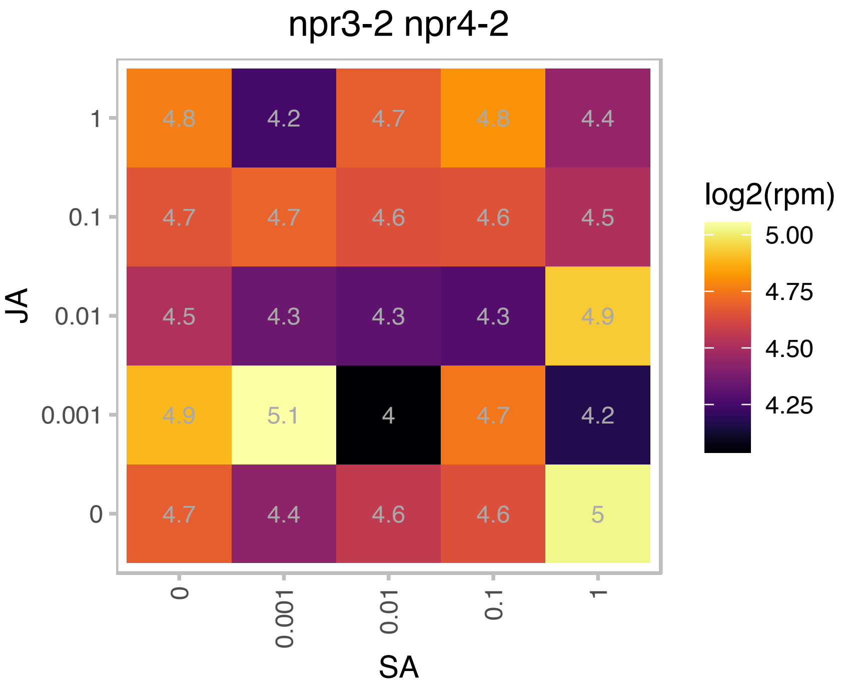


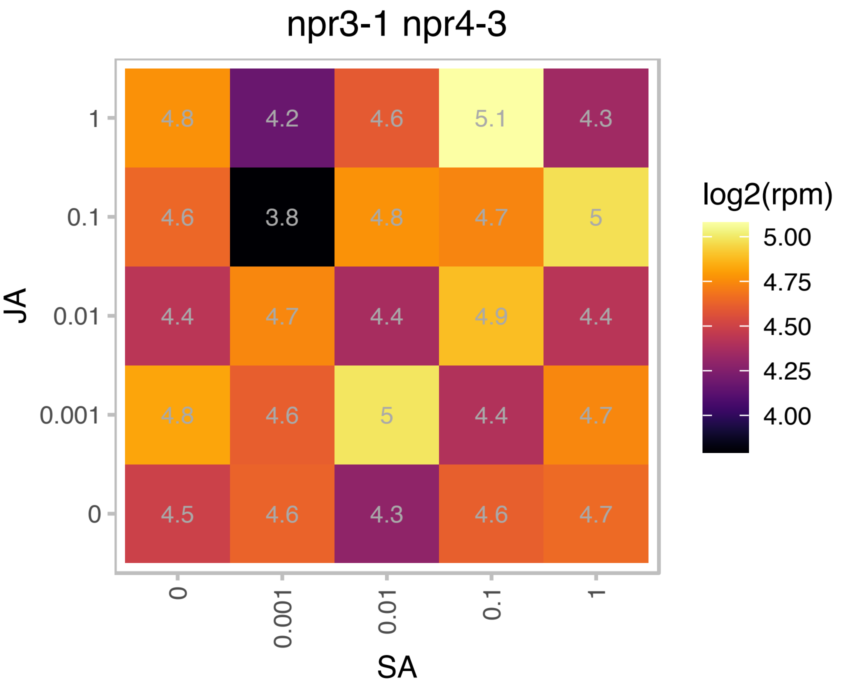

# AT5G61830.1 : NA

NAD(P)-binding Rossmann-fold superfamily protein

| | |
|---|---|
| Gene | AT5G61830 |
| is.expressed_genes | TRUE |
| is.responsive_genes | TRUE |
| is.apcluster | F-03 |

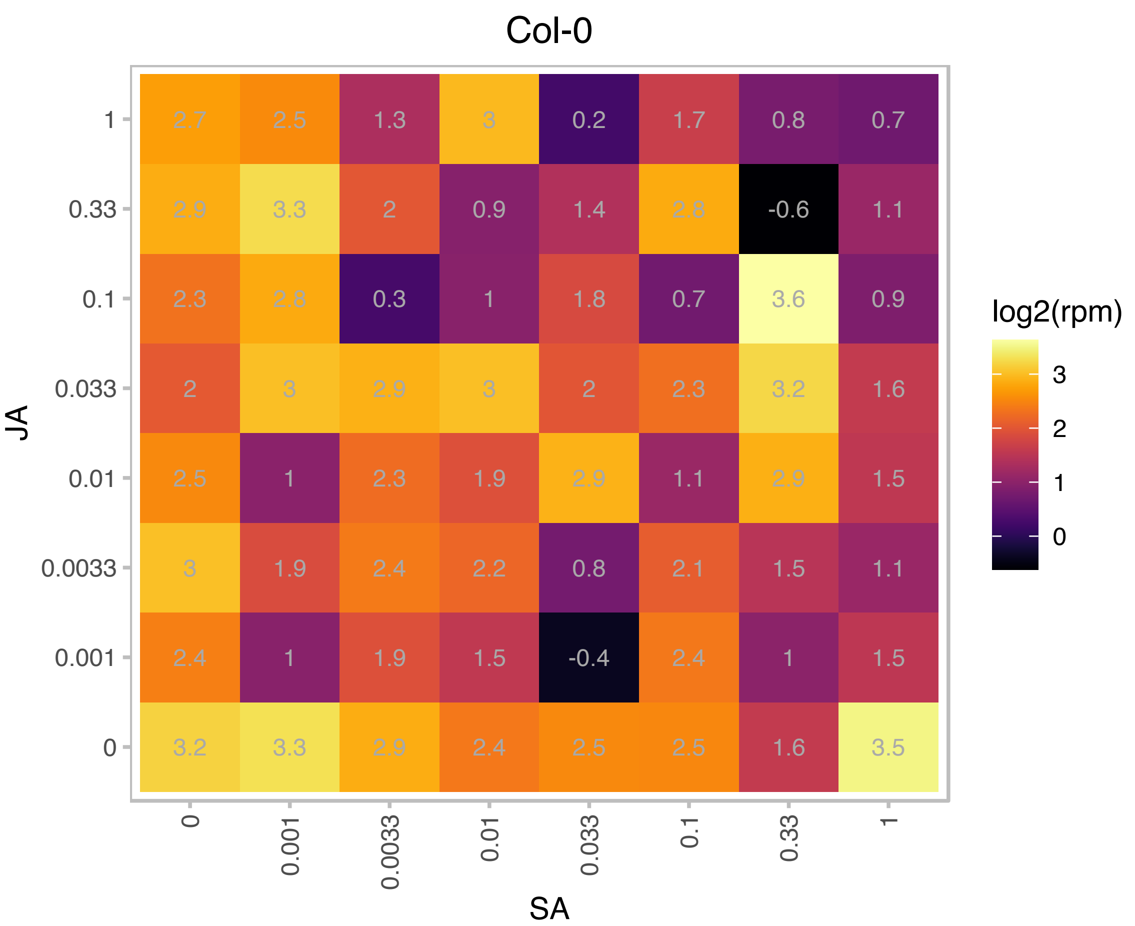


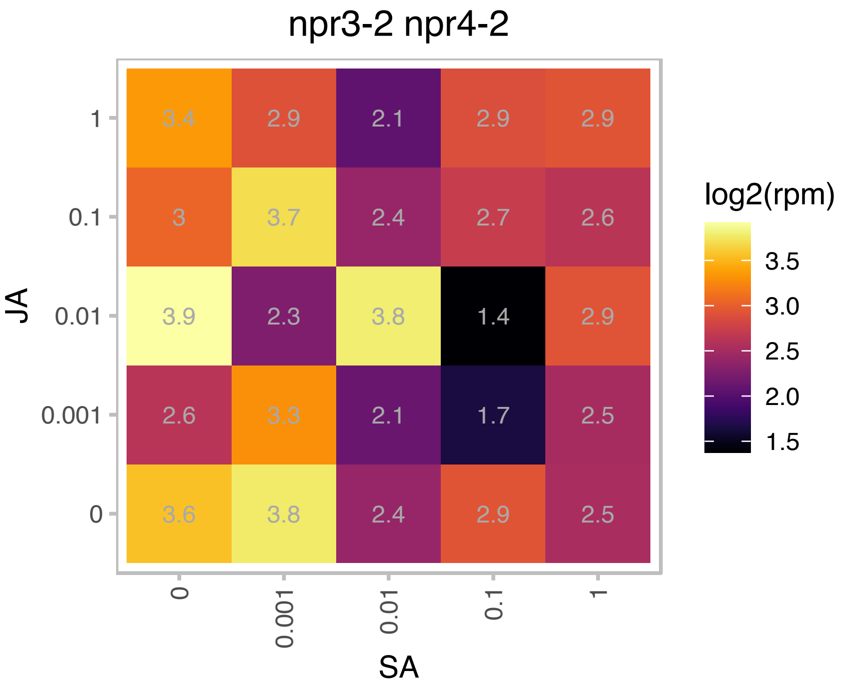


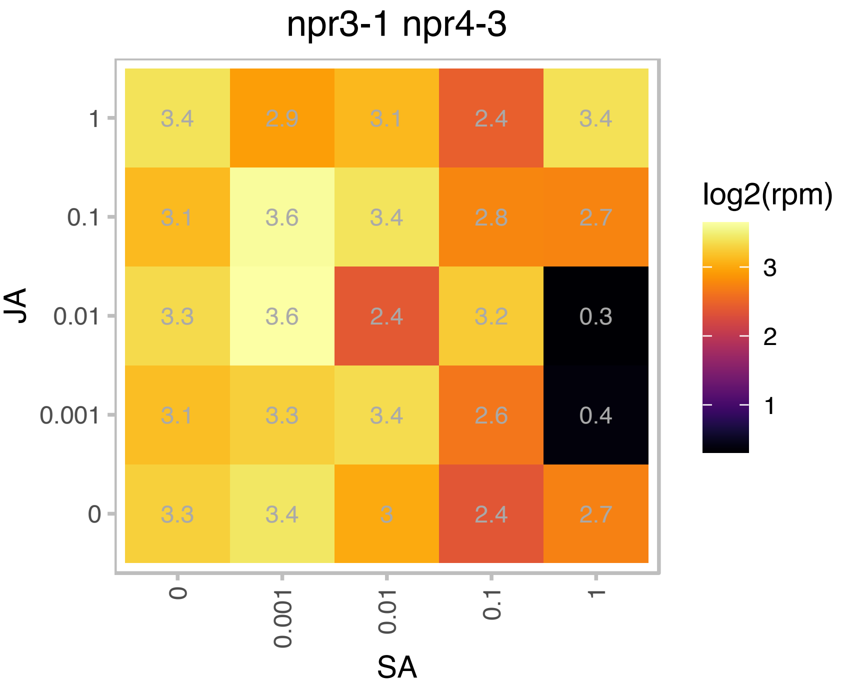

### ATCG00220.1 : PSBM

photosystem II reaction center protein M

PSII low MW protein

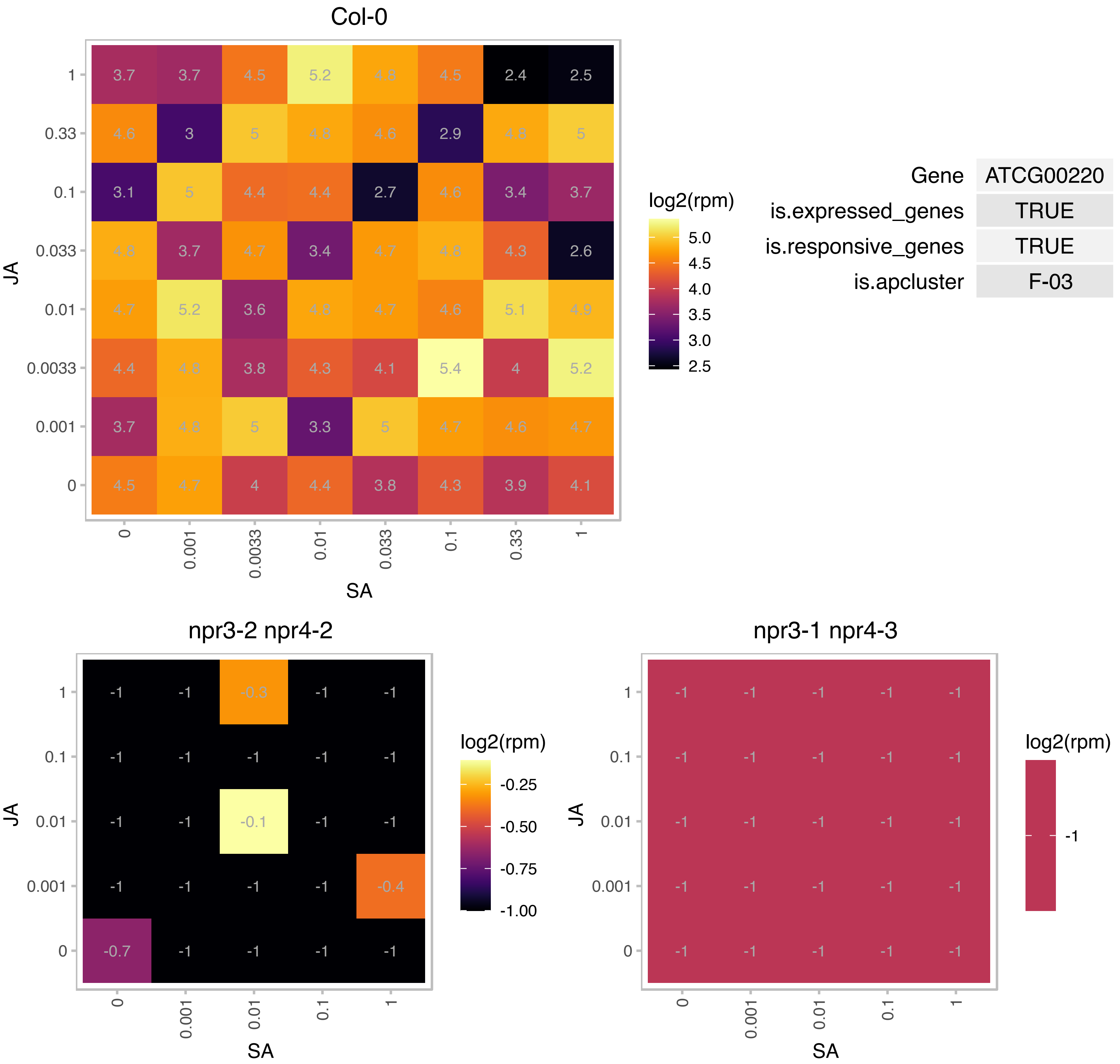


| | |
|---|---|
| Gene | ATCG00220 |
| is.expressed_genes | TRUE |
| is.responsive_genes | TRUE |
| is.apcluster | F-03 |

# AT1G01725.1 : NA

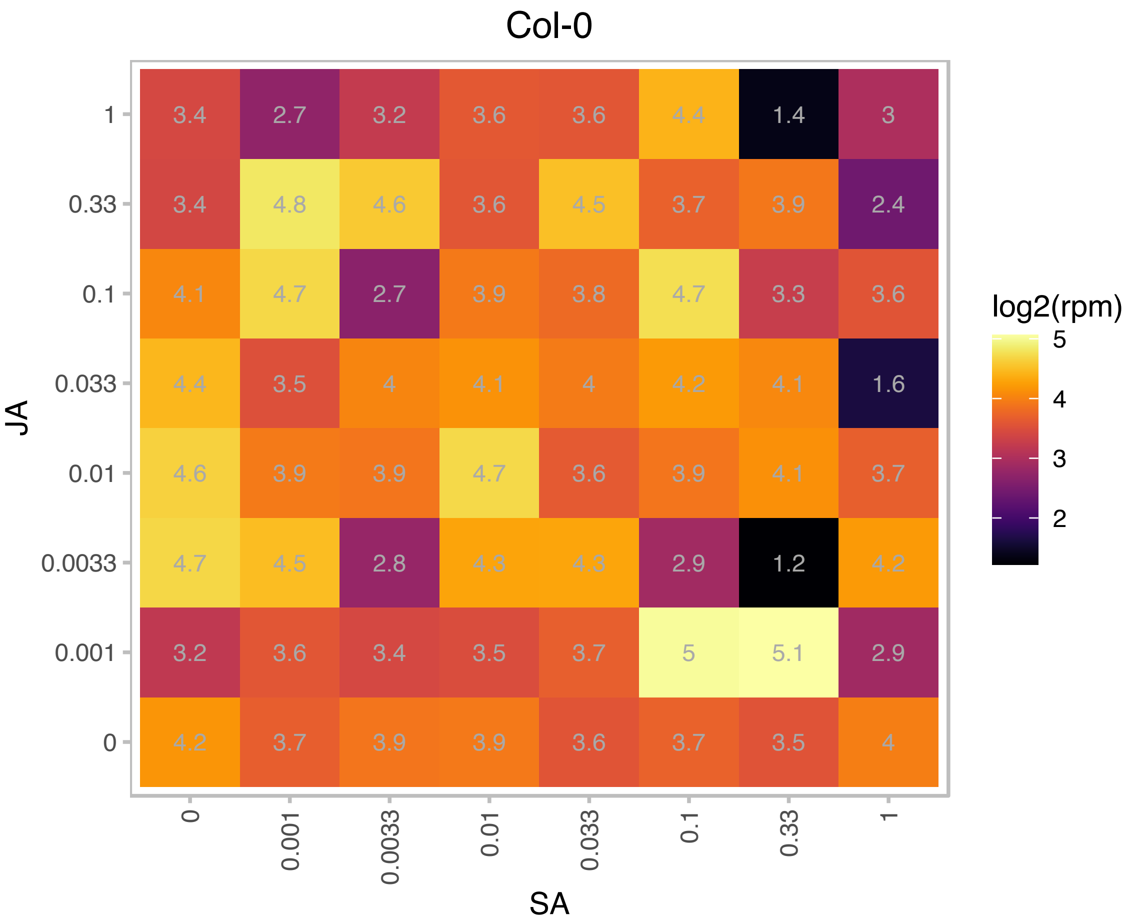

| | |
|---|---|
| Gene | AT1G01725 |
| is.expressed_genes | TRUE |
| is.responsive_genes | TRUE |
| is.apcluster | F-04 |

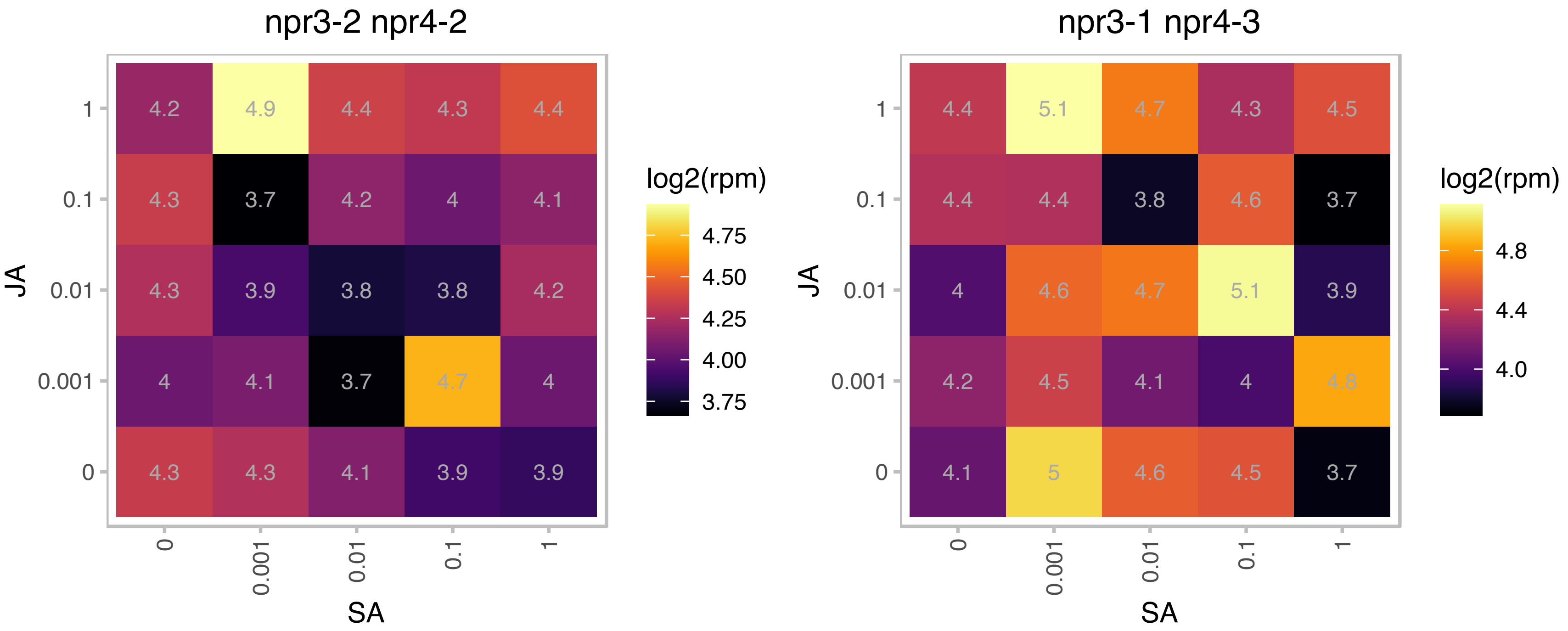

# AT1G05205.1 : NA

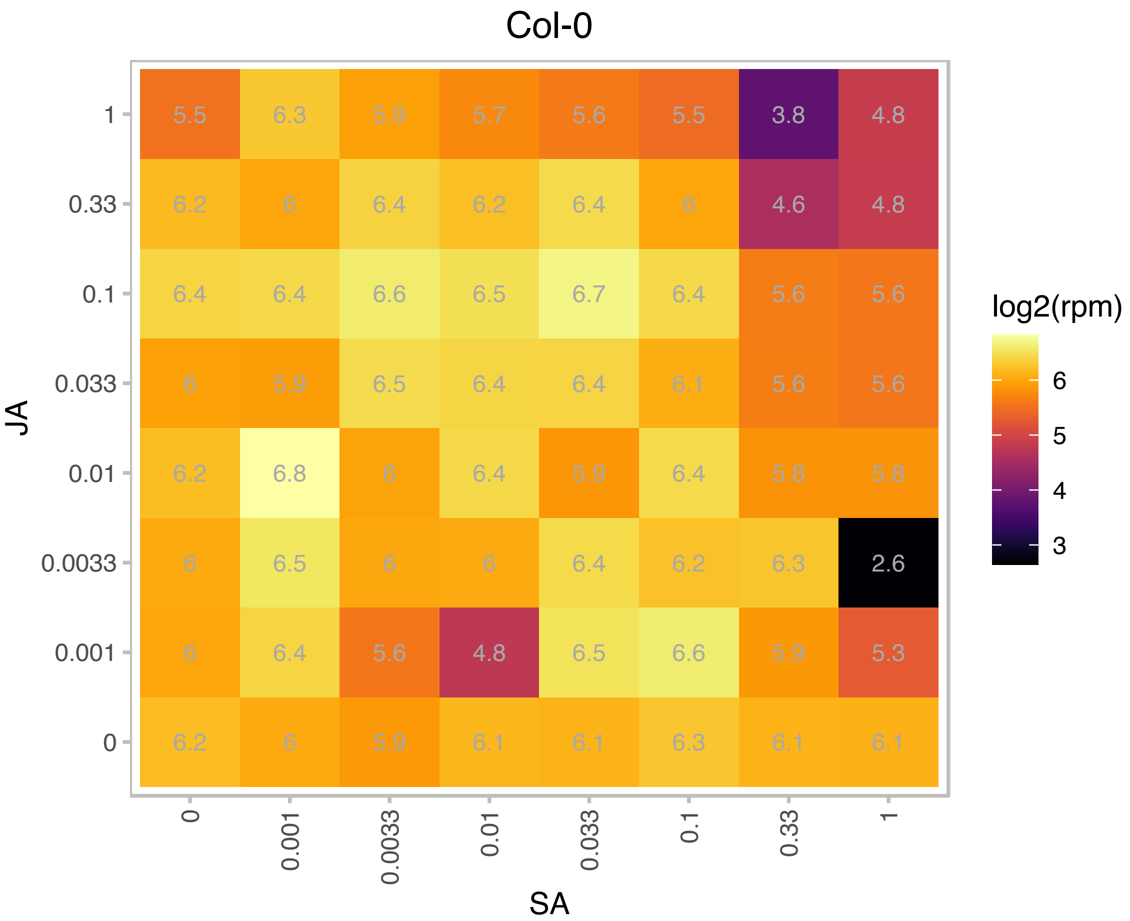


| | |
|---|---|
| Gene | AT1G05205 |
| is.expressed_genes | TRUE |
| is.responsive_genes | TRUE |
| is.apcluster | F-04 |

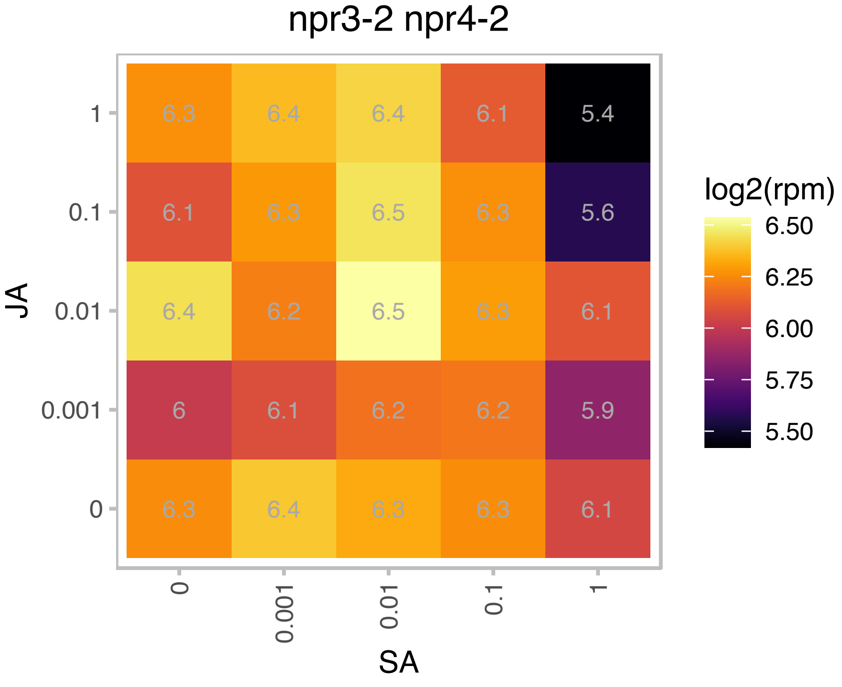


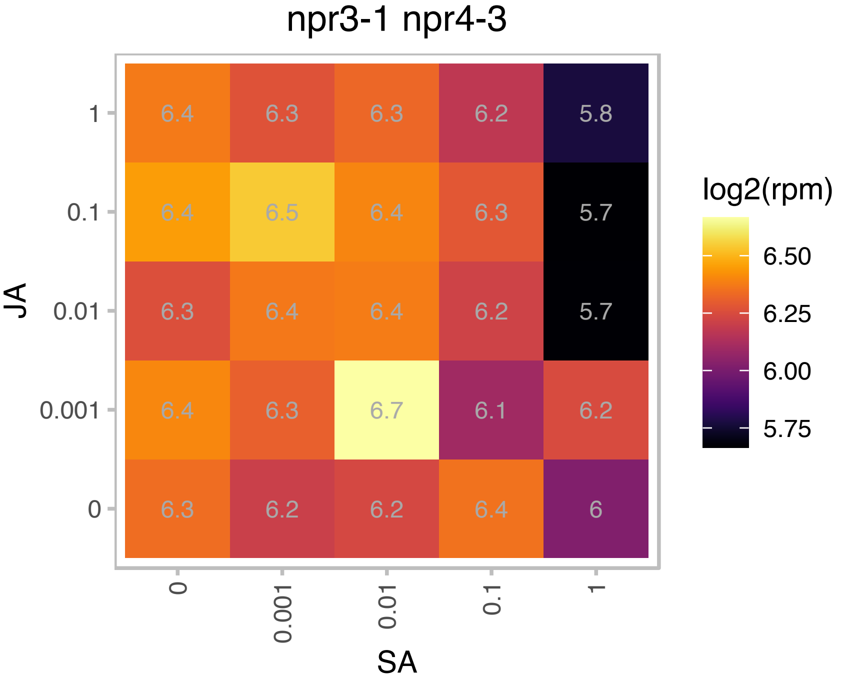

# AT1G06550.1 : NA

ATP-dependent caseinolytic (Clp) protease/crotonase family protein

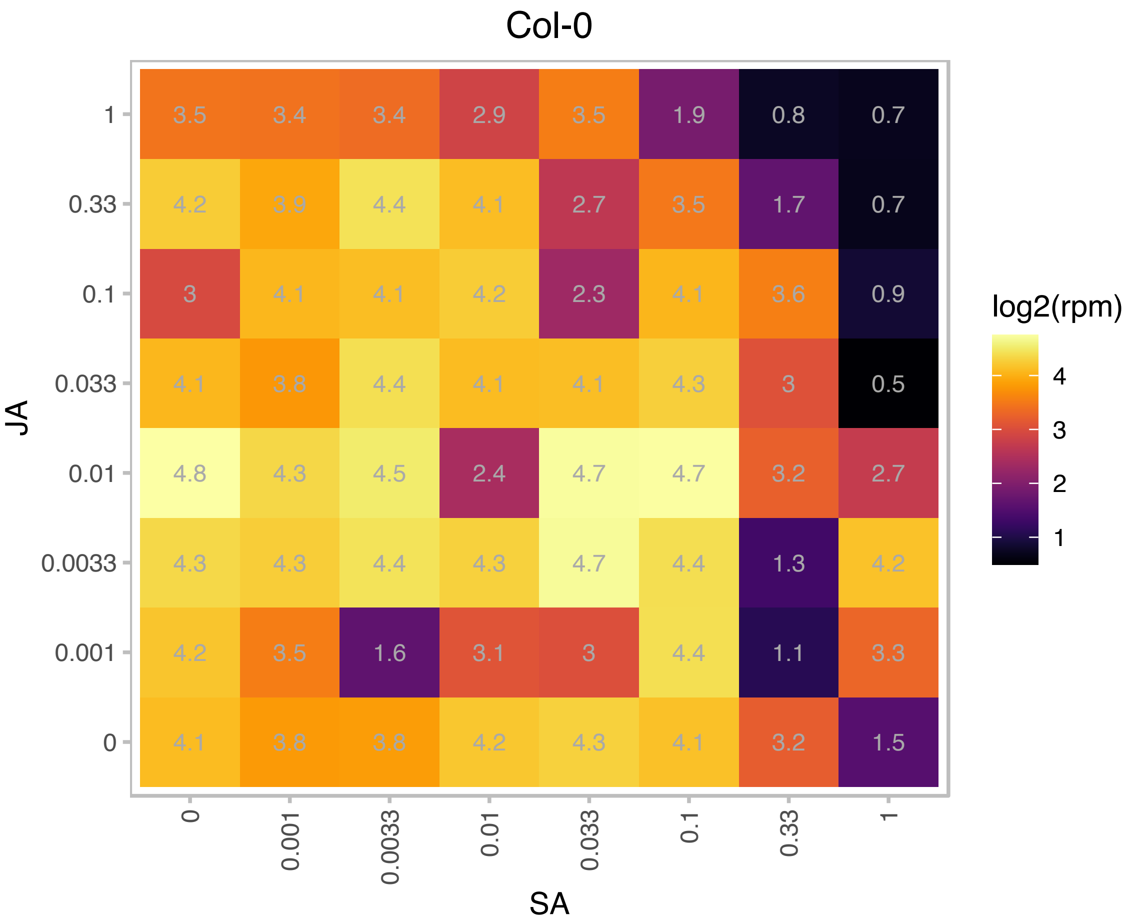


| | |
|---|---|
| Gene | AT1G06550 |
| is.expressed_genes | TRUE |
| is.responsive_genes | TRUE |
| is.apcluster | F-04 |

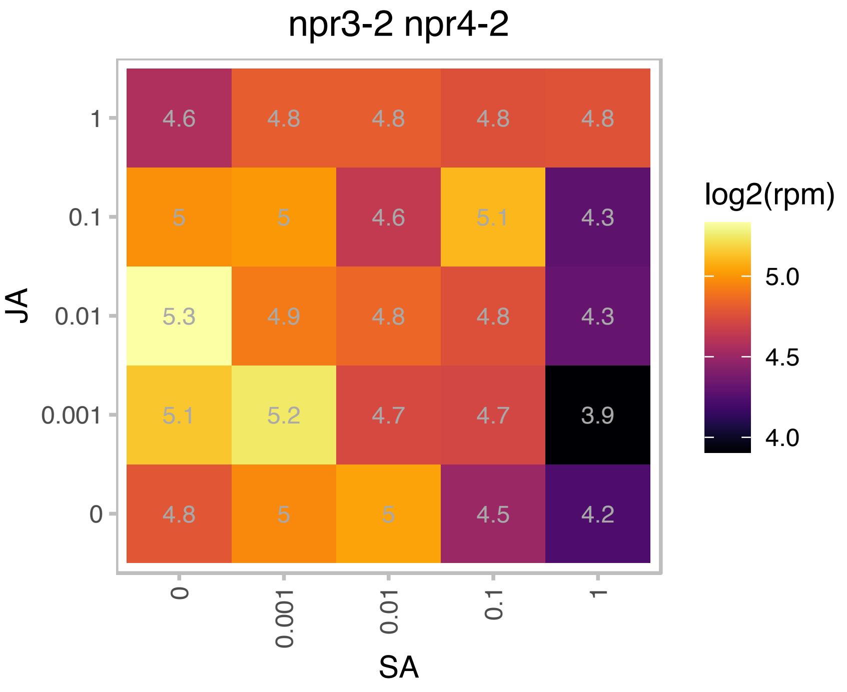


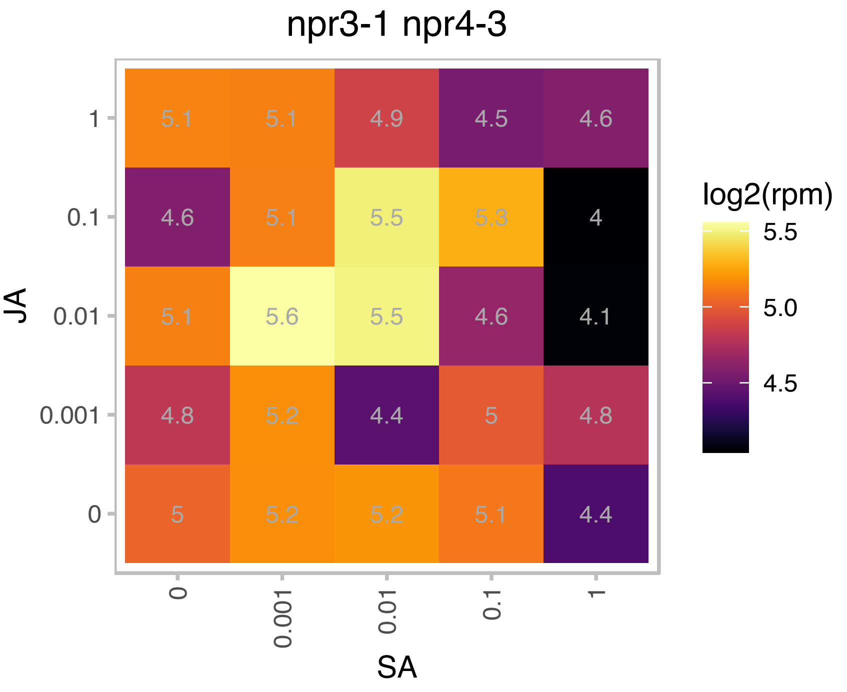

### AT1G09780.1 : iPGAM1

Phosphoglycerate mutase, 2,3-bisphosphoglycerate-independent

Encodes a 2,3-biphosphoglycerate-independent phosphoglycerate mutase that is involved in pollen development and stomatal movement.

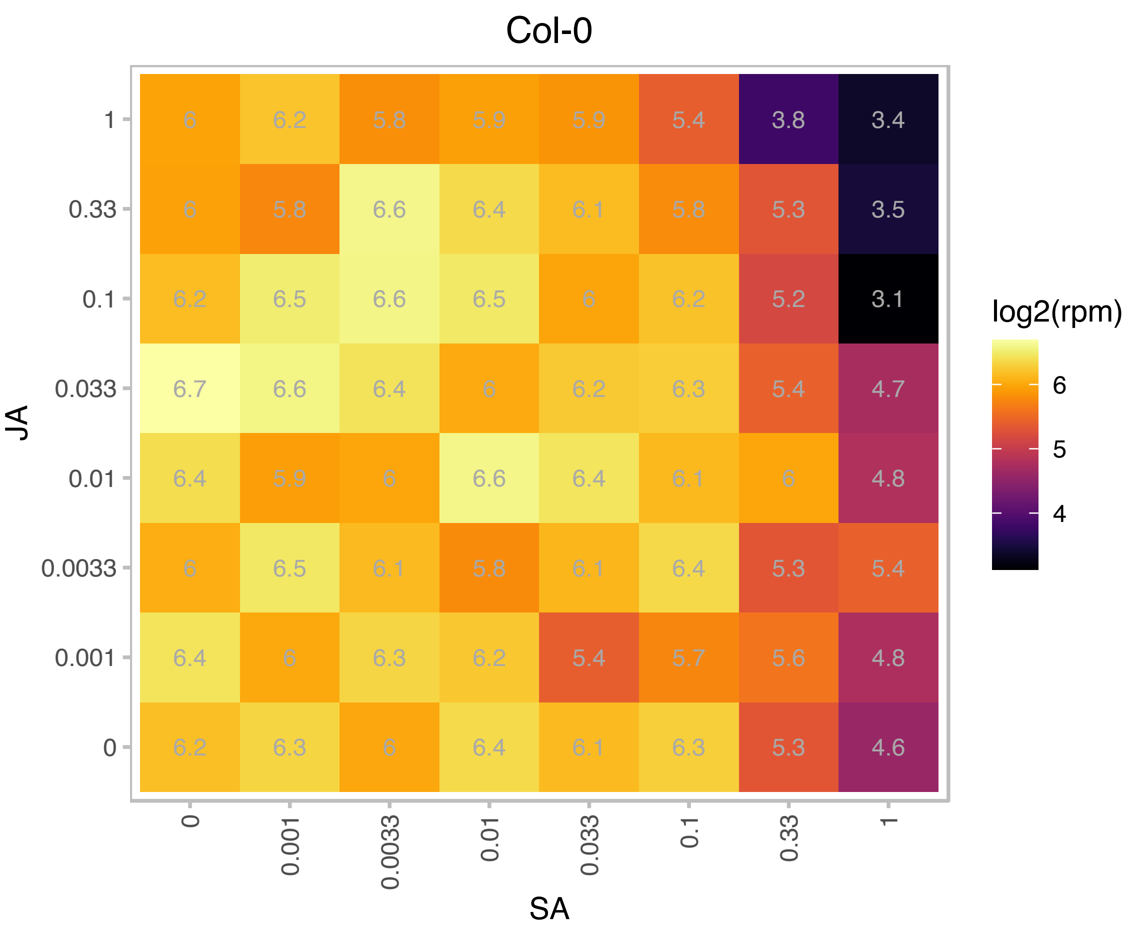


| | |
|---|---|
| Gene | AT1G09780 |
| is.expressed_genes | TRUE |
| is.responsive_genes | TRUE |
| is.apcluster | F-04 |

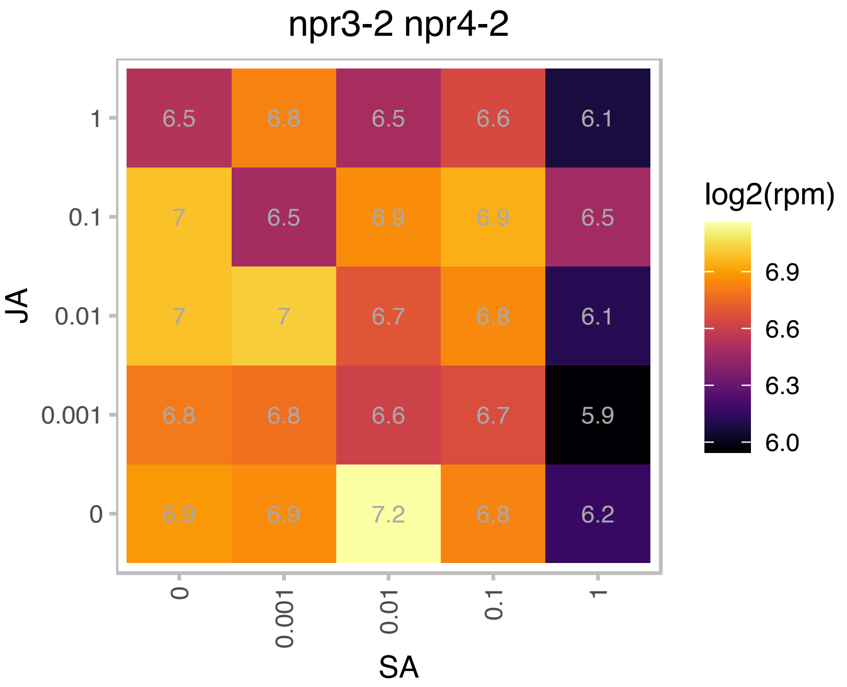


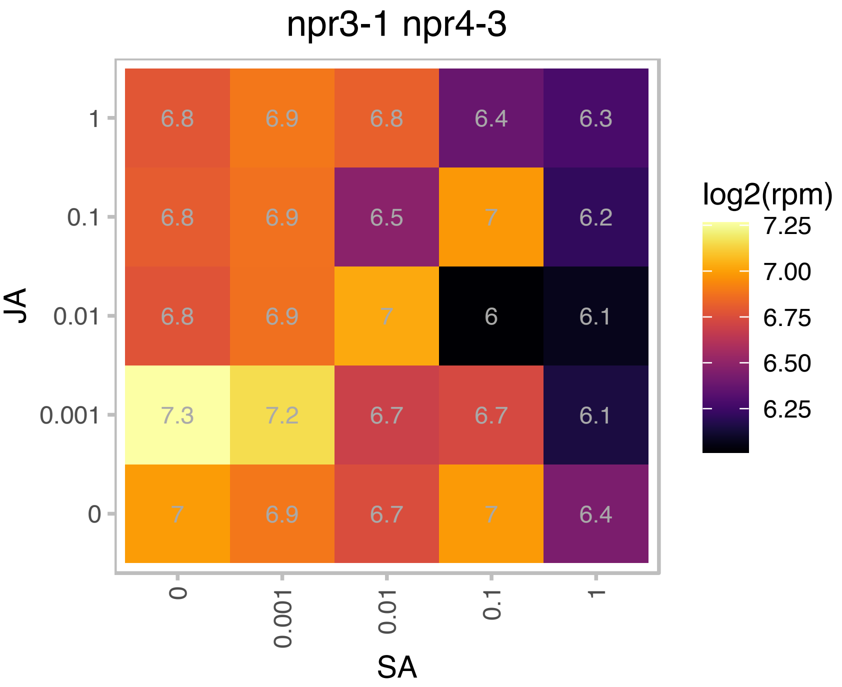

### AT1G10200.1 : AtWLIM1, WLIM1

GATA type zinc finger transcription factor family protein
Encodes a member of the Arabidopsis LIM proteins: a family of actin bundlers with distinct expression patterns. WLIM1, WLIM2a, and WLIM2b are widely expressed, whereas PLIM2a, PLIM2b, and PLIM2c are predominantly expressed in pollen. Regulates actin cytoskeleton organization.

Col-0

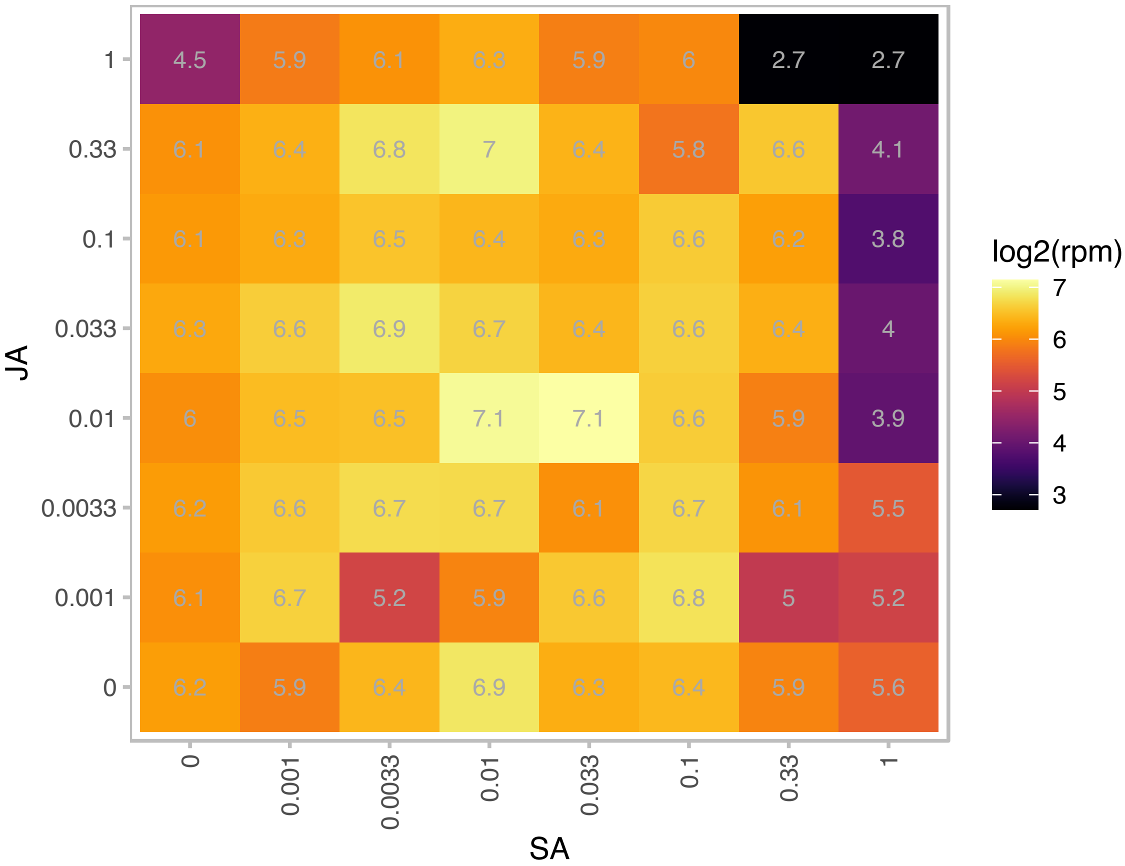


| | |
|---|---|
| Gene | AT1G10200 |
| is.expressed_genes | TRUE |
| is.responsive_genes | TRUE |
| is.apcluster | F-04 |

npr3-2 npr4-2

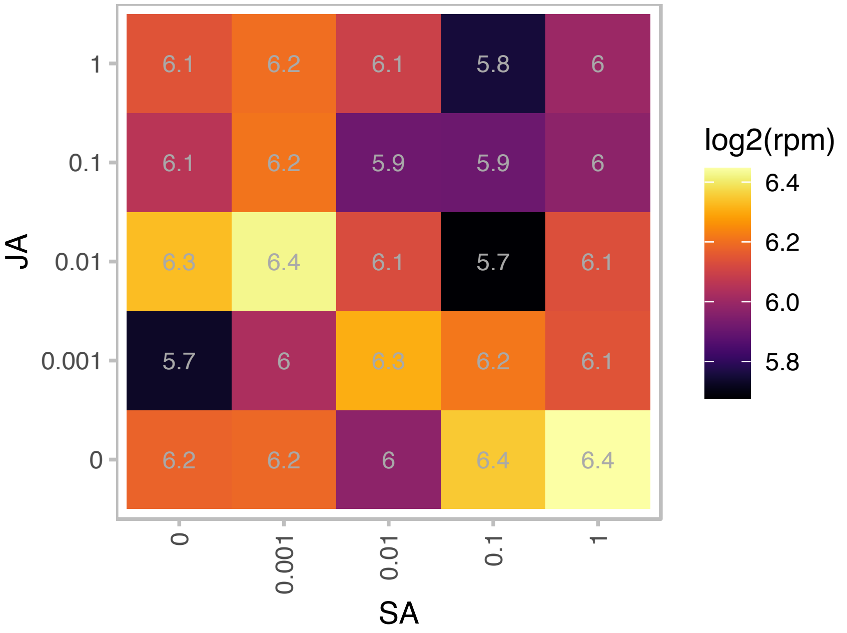


npr3-1 npr4-3

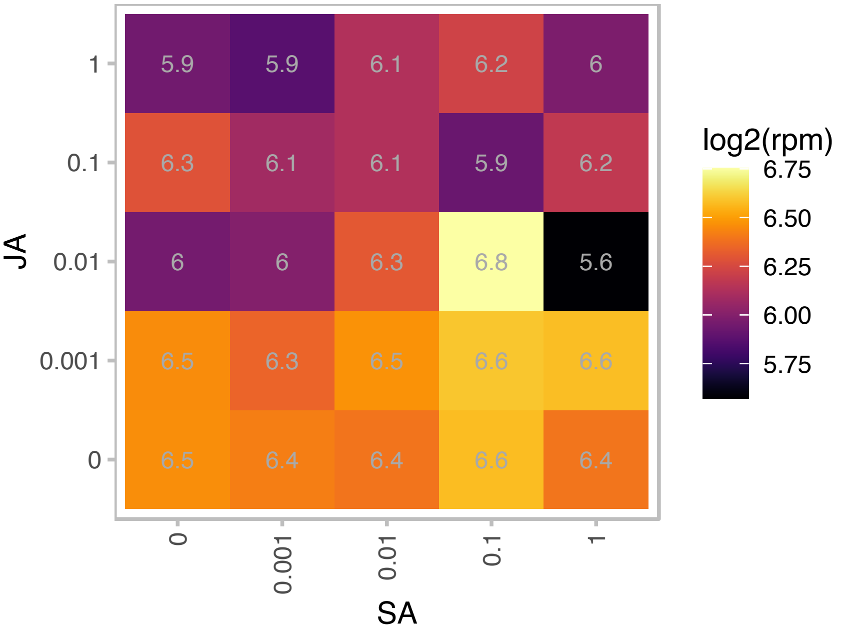

### AT1G12530.1 : MOS9

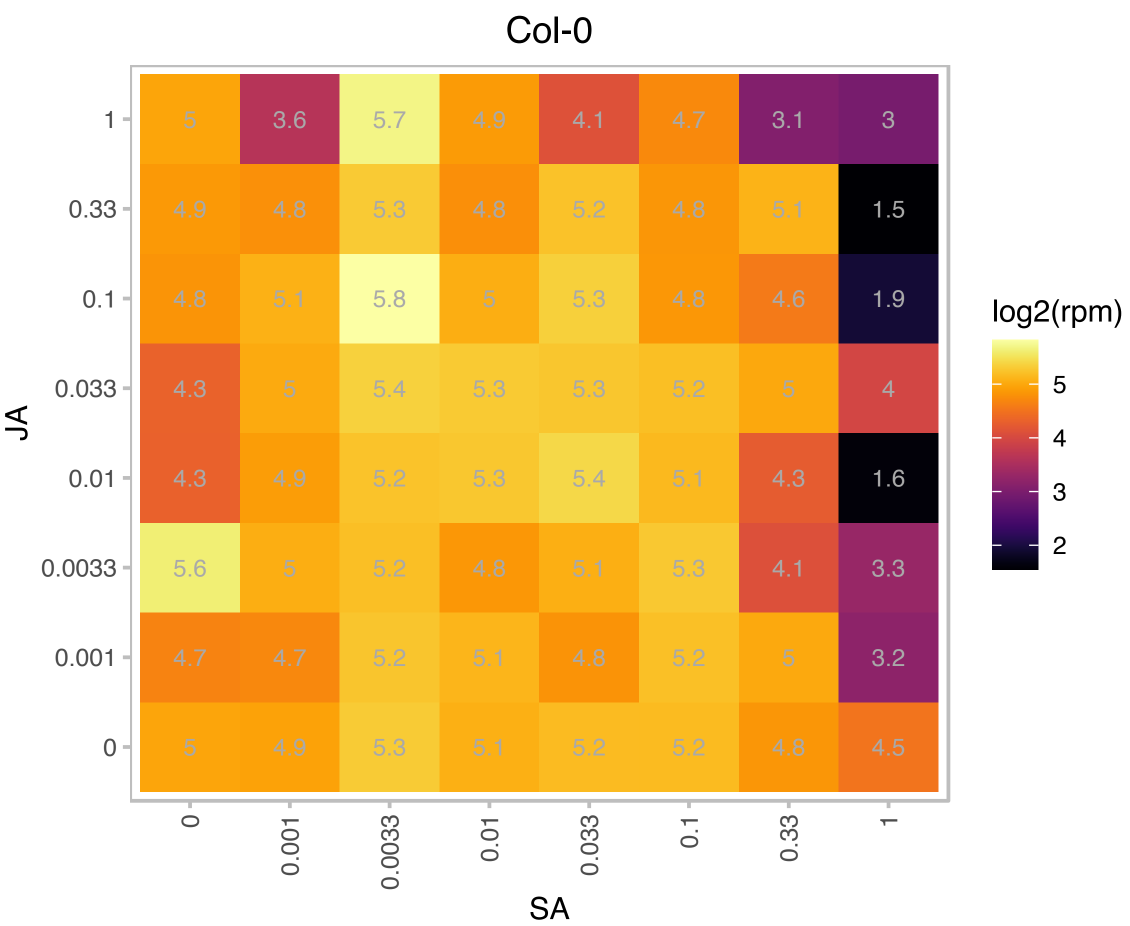


| | |
|---|---|
| Gene | AT1G12530 |
| is.expressed_genes | TRUE |
| is.responsive_genes | TRUE |
| is.apcluster | F-04 |

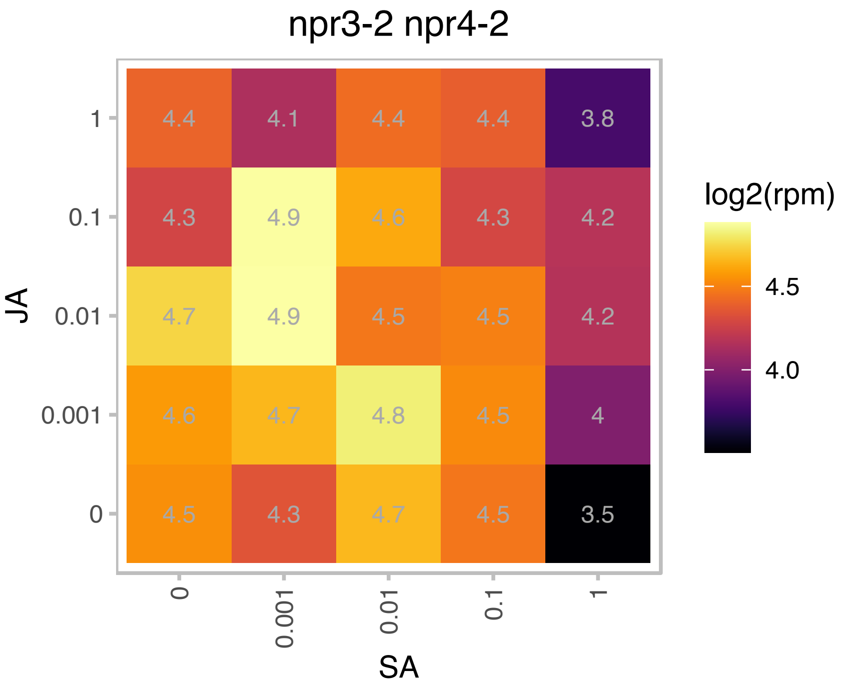


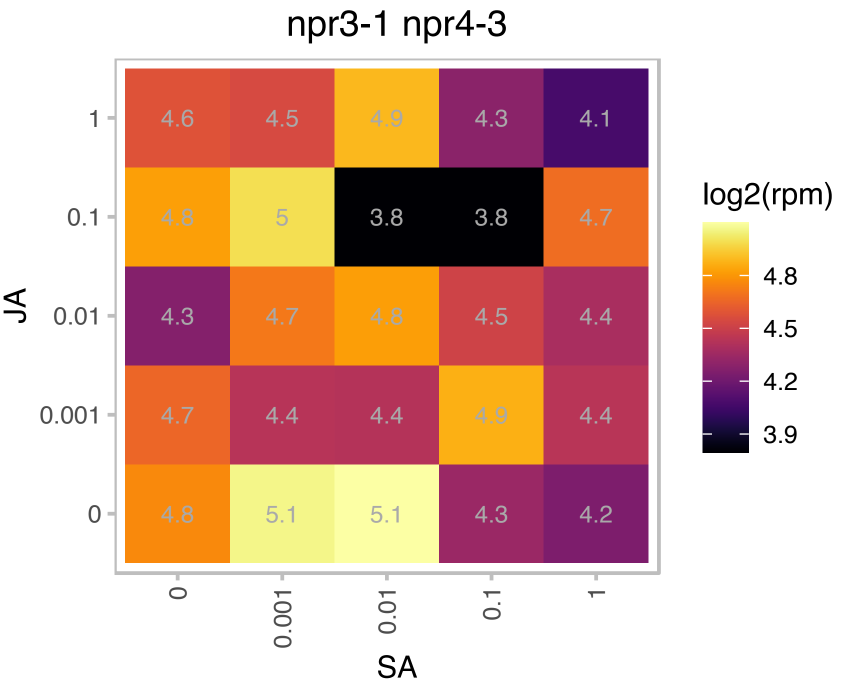

# AT1G18950.1 : NA

DDT domain superfamily

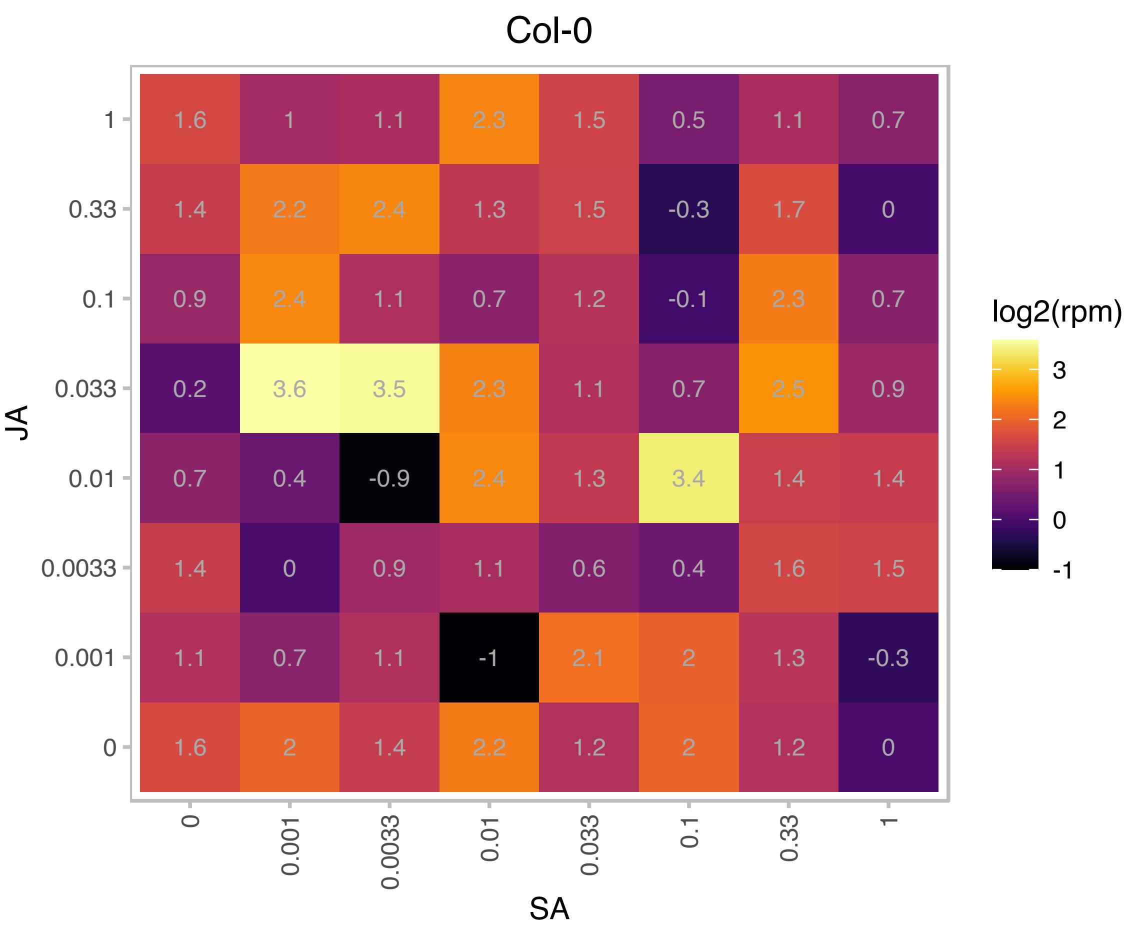


| | |
|---|---|
| Gene | AT1G18950 |
| is.expressed_genes | TRUE |
| is.responsive_genes | TRUE |
| is.apcluster | F-04 |

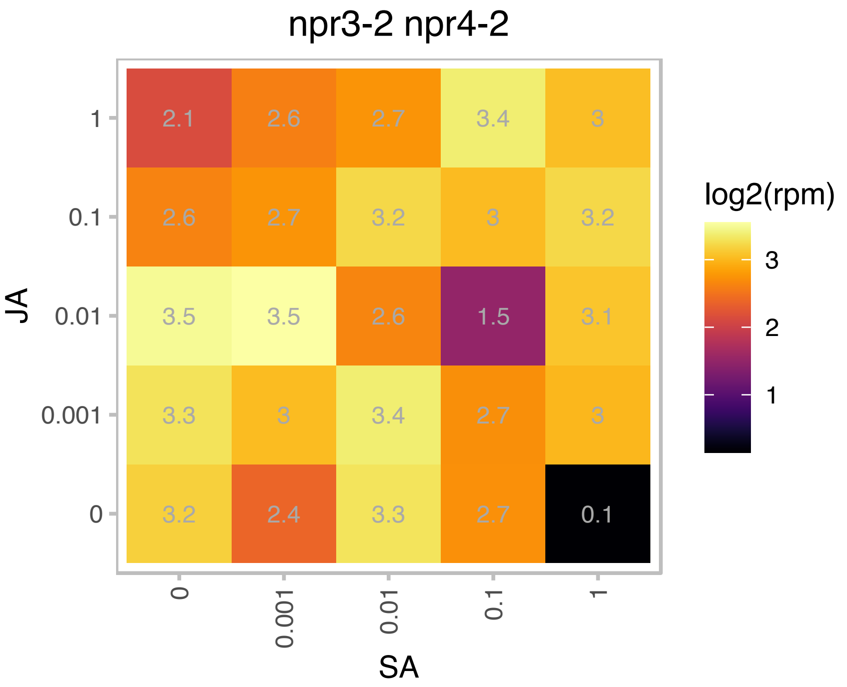


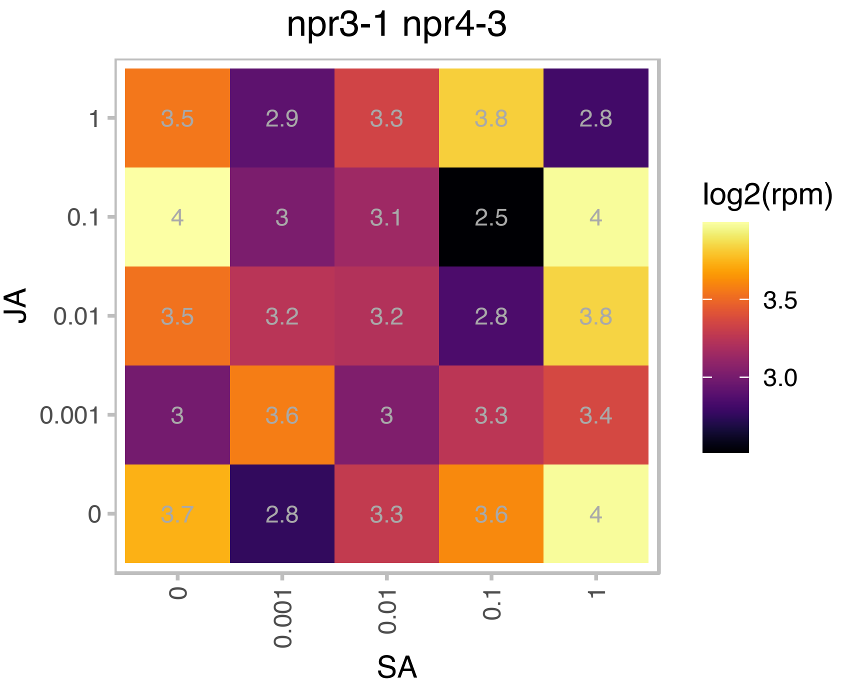

# AT1G19000.1 : NA

Homeodomain-like superfamily protein

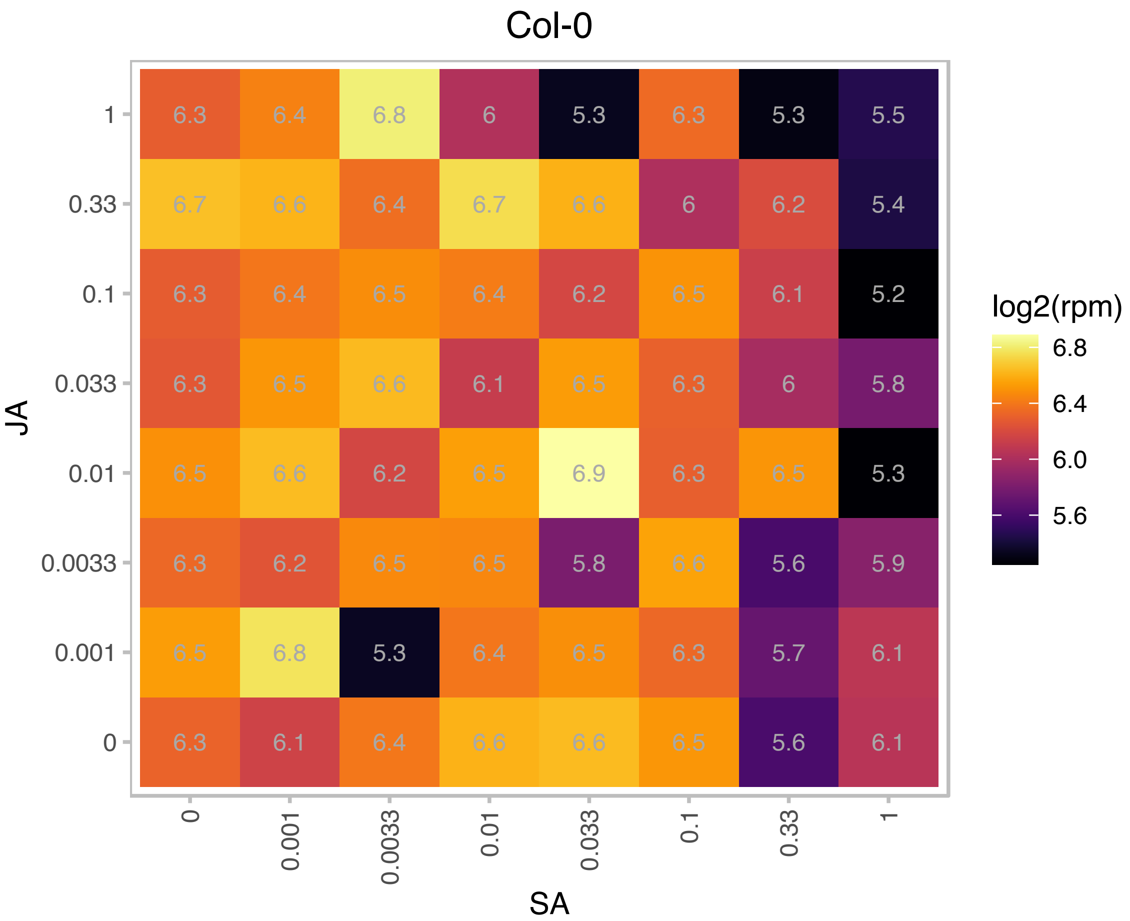


| | |
|---|---|
| Gene | AT1G19000 |
| is.expressed_genes | TRUE |
| is.responsive_genes | TRUE |
| is.apcluster | F-04 |

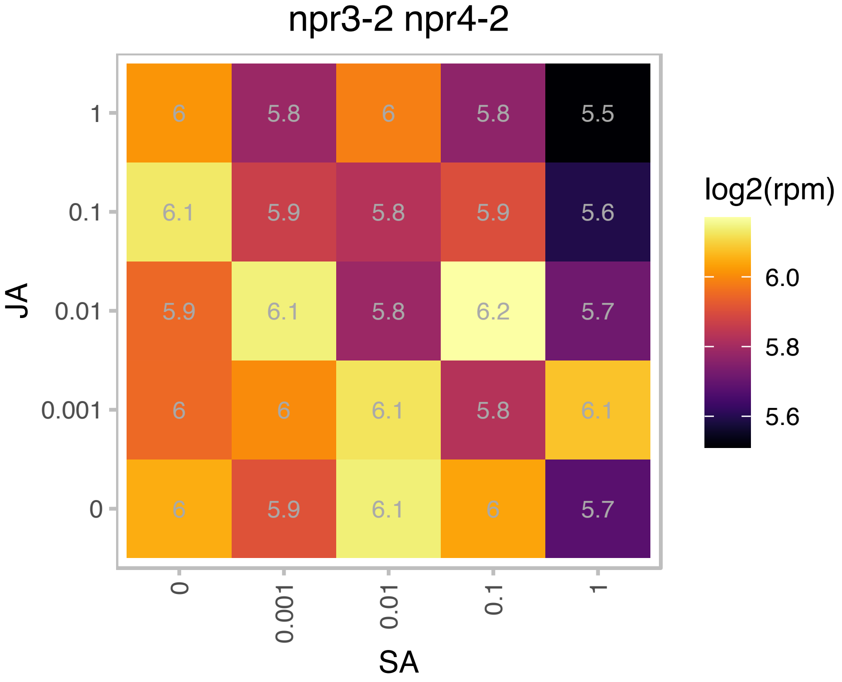


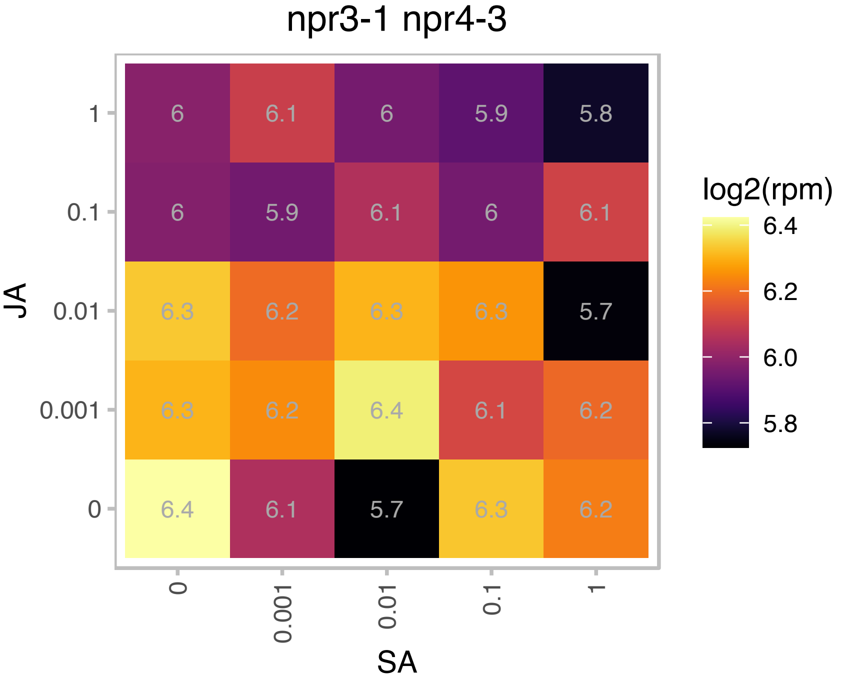

### AT1G19920.1 : APS2, ASA1, ATPS2

Pseudouridine synthase/archaeosine transglycosylase-like family protein

encodes a chloroplast form of ATP sulfurylase.

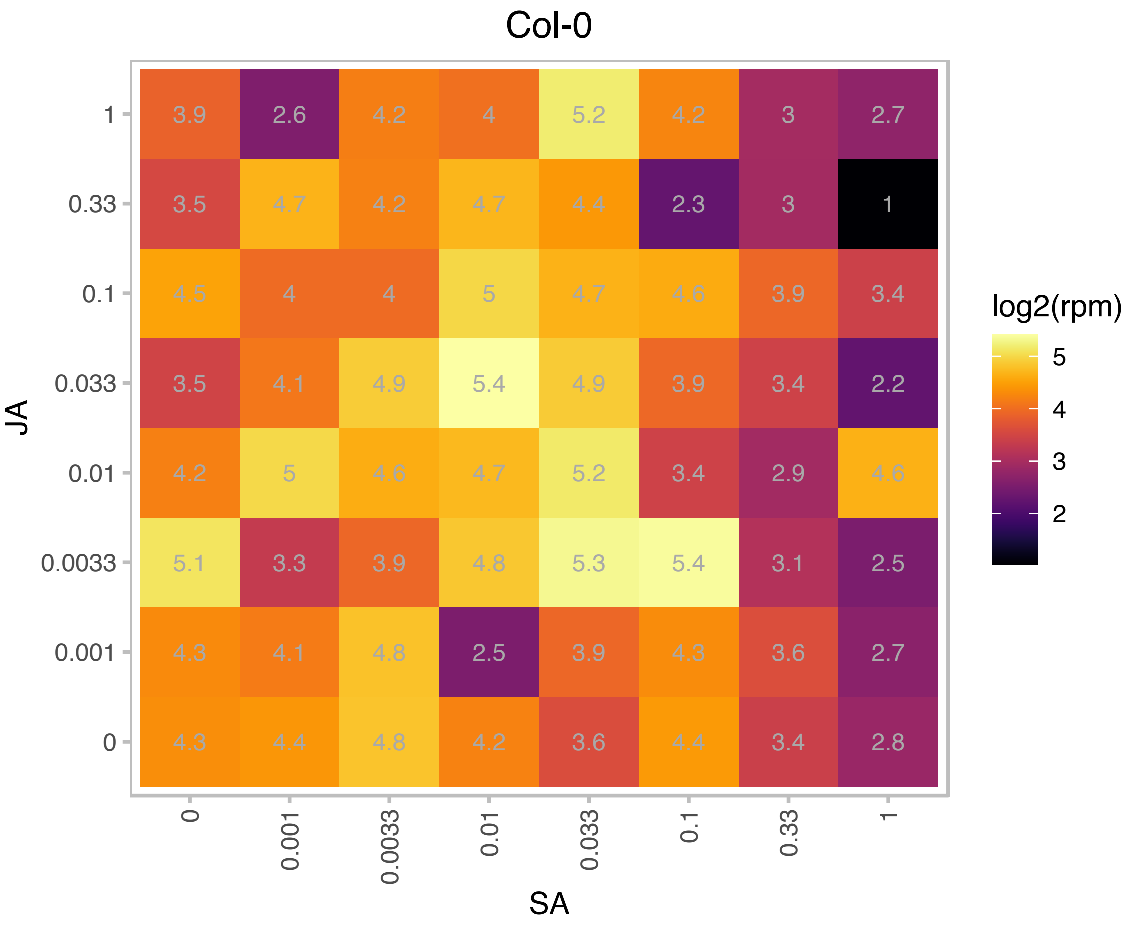


| | |
|---|---|
| Gene | AT1G19920 |
| is.expressed_genes | TRUE |
| is.responsive_genes | TRUE |
| is.apcluster | F-04 |

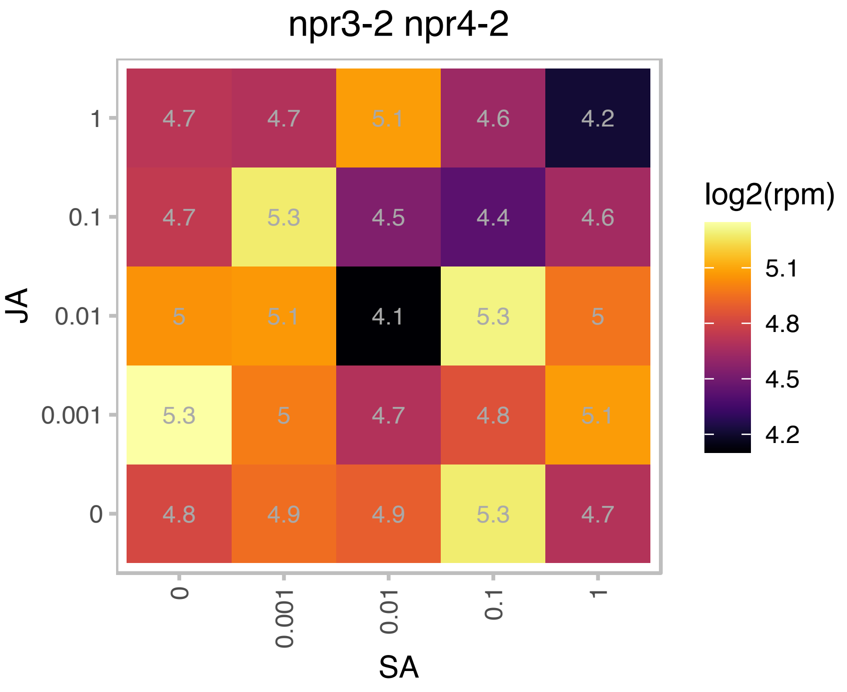


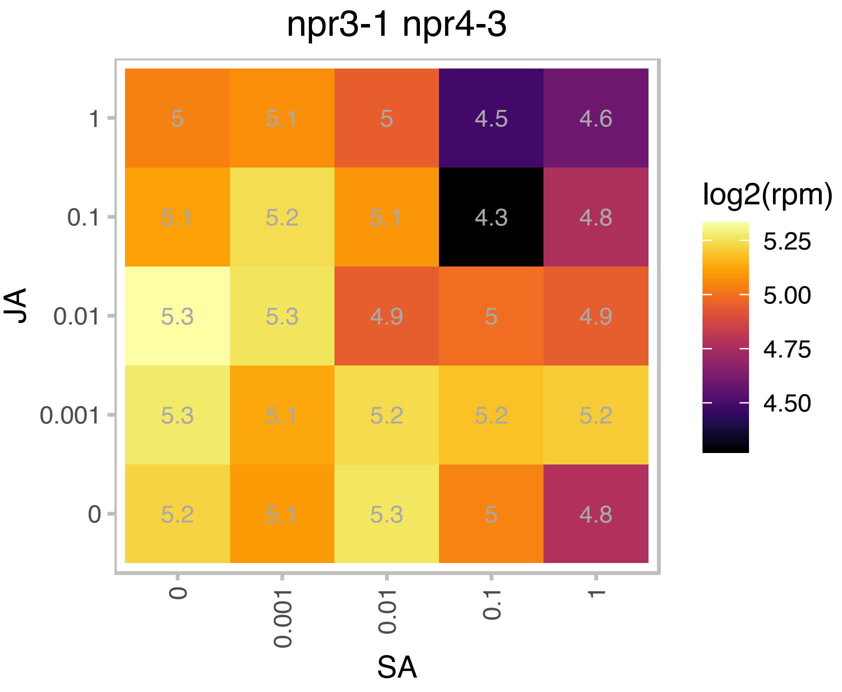

### AT1G22450.1 : ATCOX6B2, COX6B

cytochrome C oxidase 6B

subunit 6b of cytochrome c oxidase

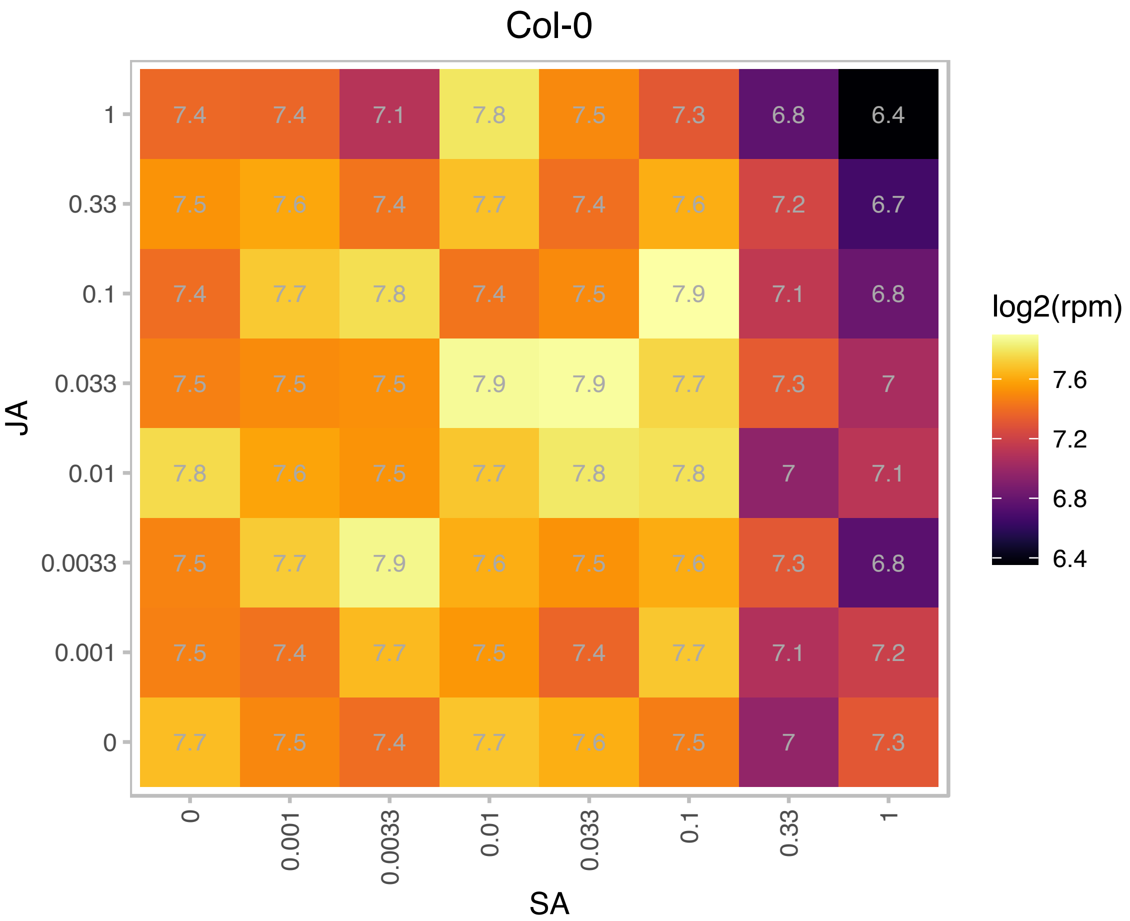


| | |
|---|---|
| Gene | AT1G22450 |
| is.expressed_genes | TRUE |
| is.responsive_genes | TRUE |
| is.apcluster | F-04 |

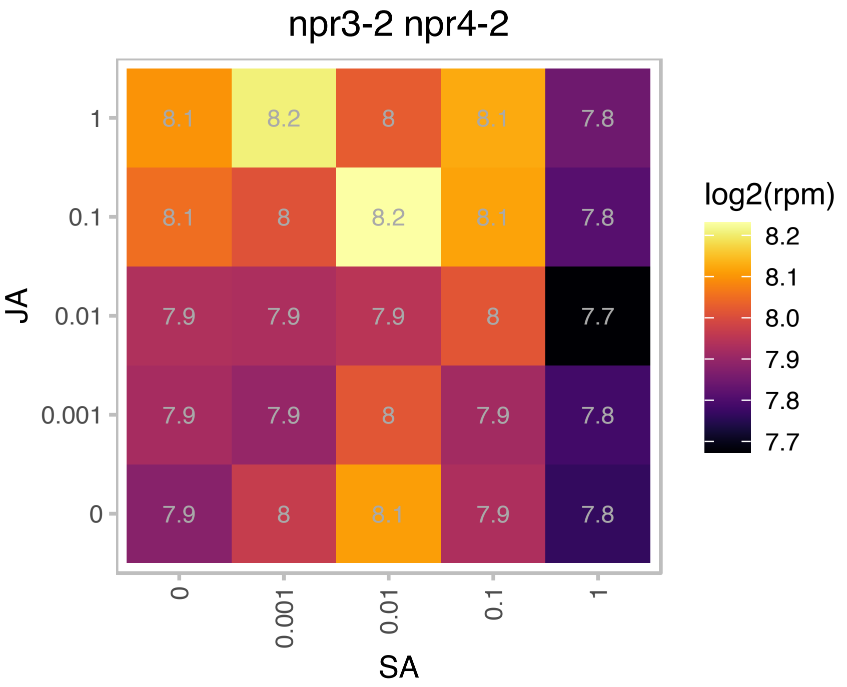


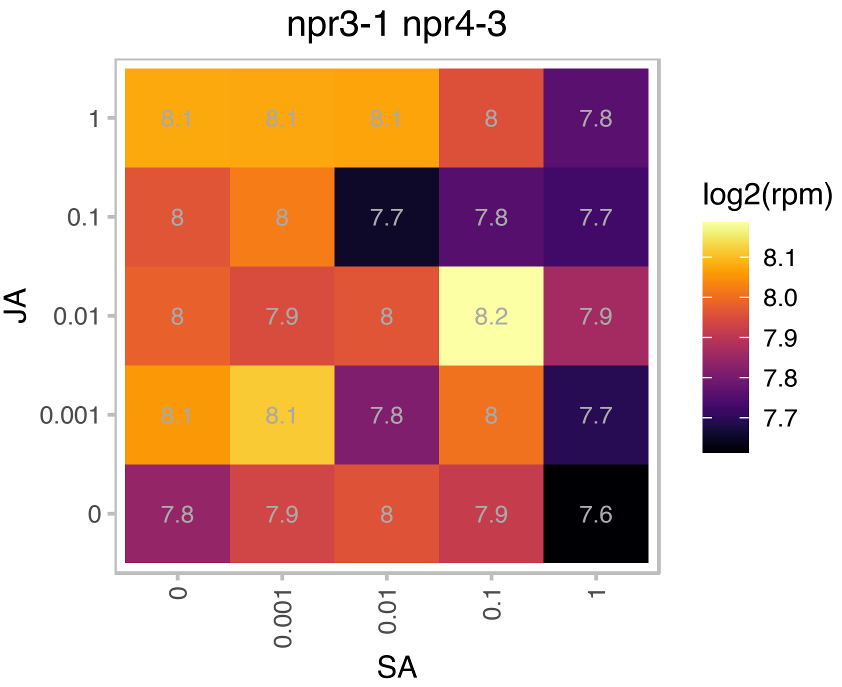

### AT1G25440.1 : BBX15

B-box type zinc finger protein with CCT domain

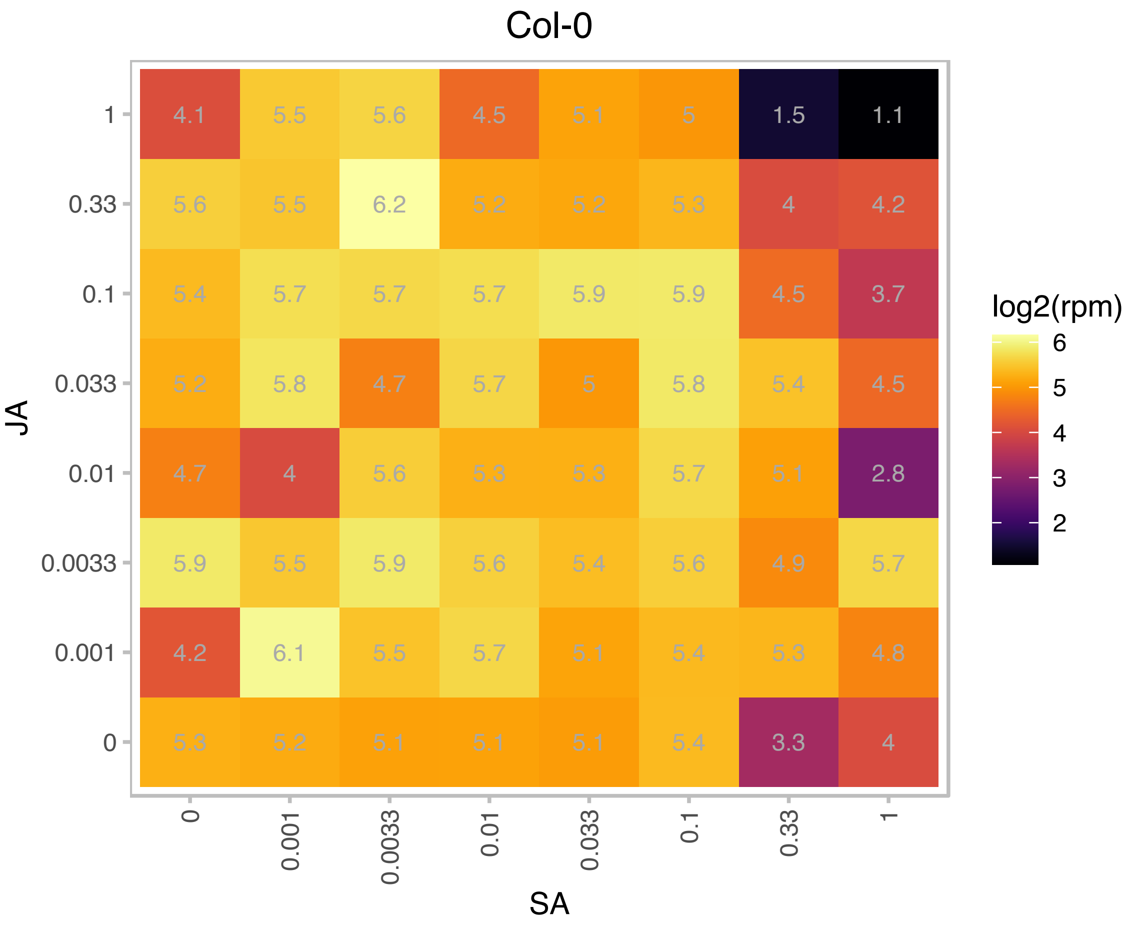


| | |
|---|---|
| Gene | AT1G25440 |
| is.expressed_genes | TRUE |
| is.responsive_genes | TRUE |
| is.apcluster | F-04 |

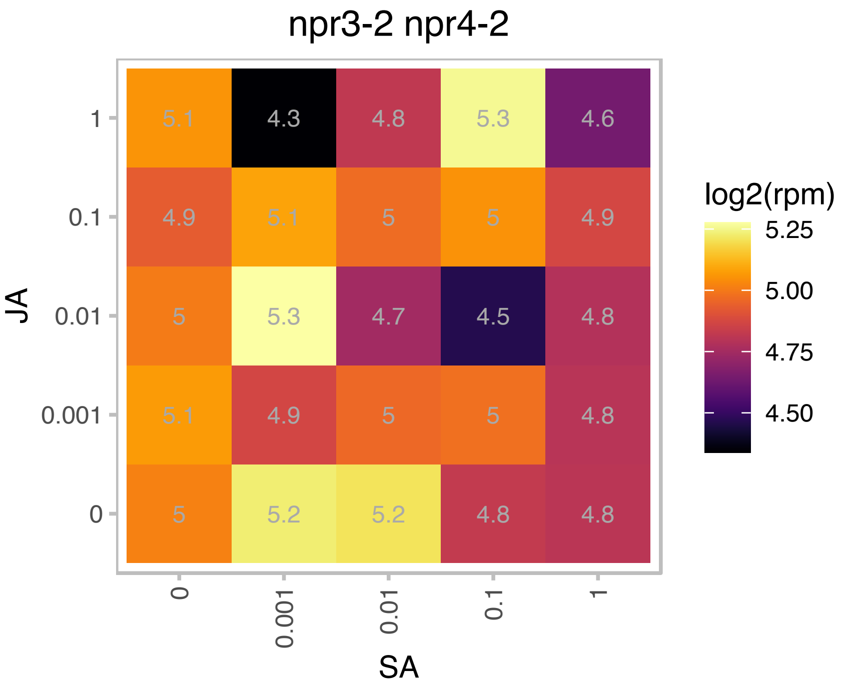


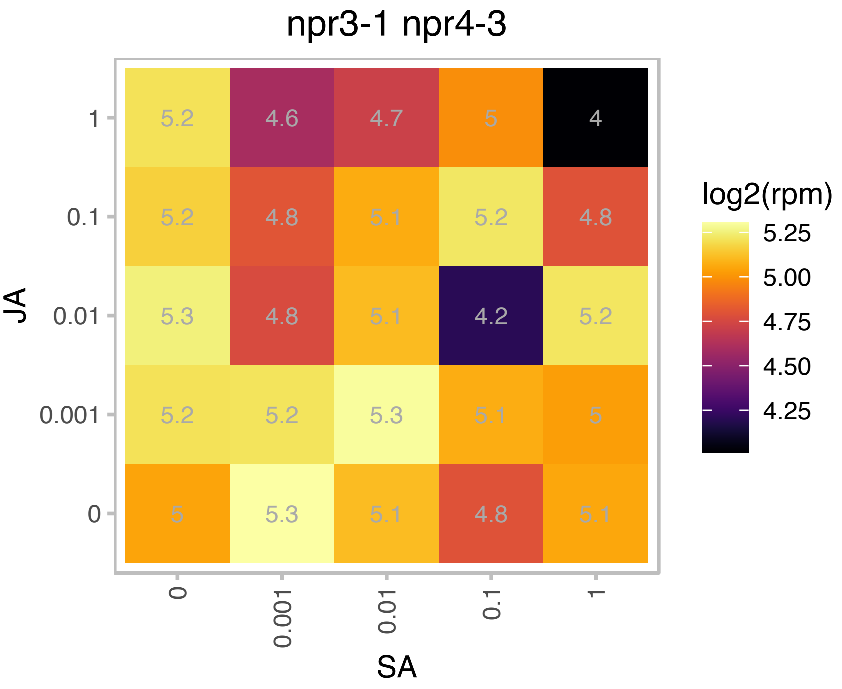

### AT1G27390.1 : TOM20-2

translocase outer membrane 20-2

Form of TOM20, which is a component of the TOM complex, involved in transport of nuclear-encoded mitochondrial proteins

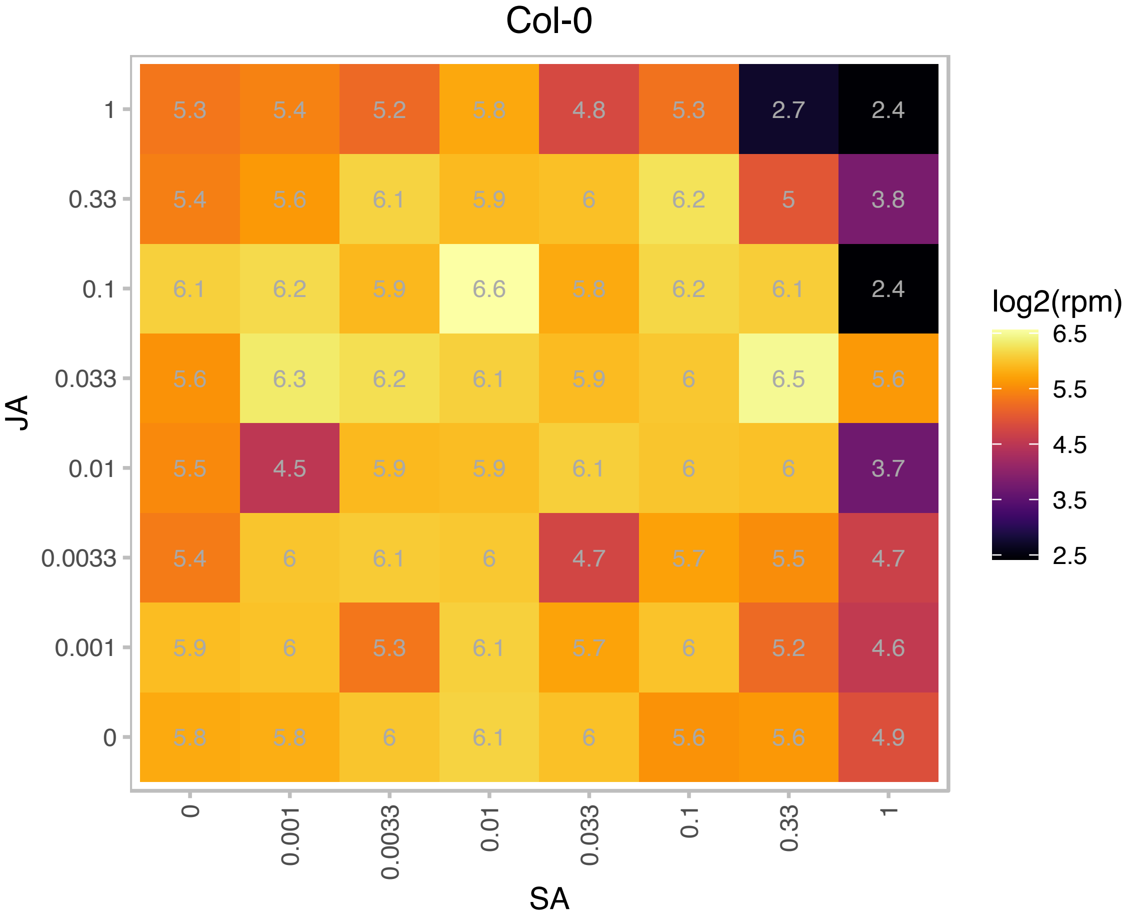


| | |
|---|---|
| Gene | AT1G27390 |
| is.expressed_genes | TRUE |
| is.responsive_genes | TRUE |
| is.apcluster | F-04 |

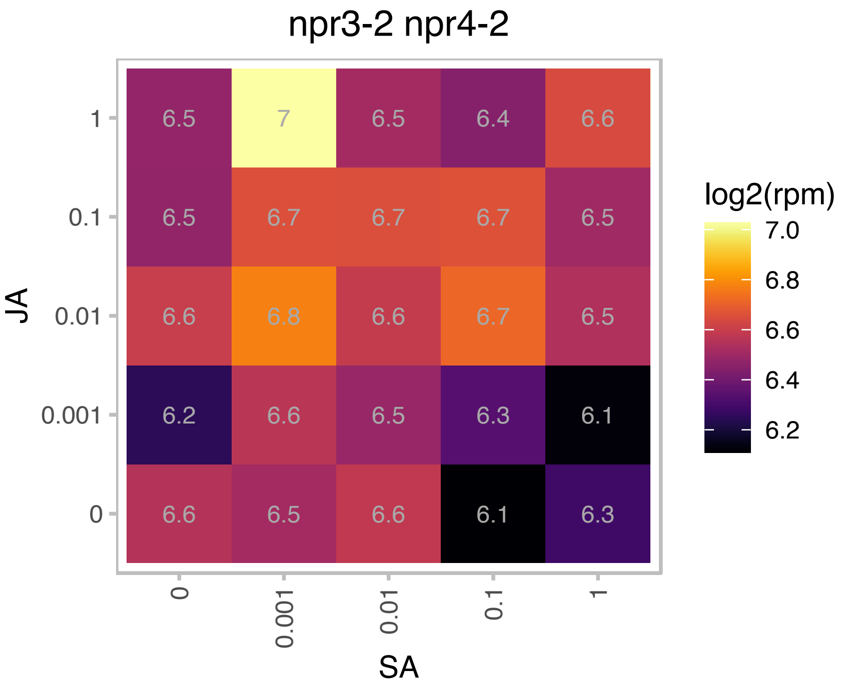


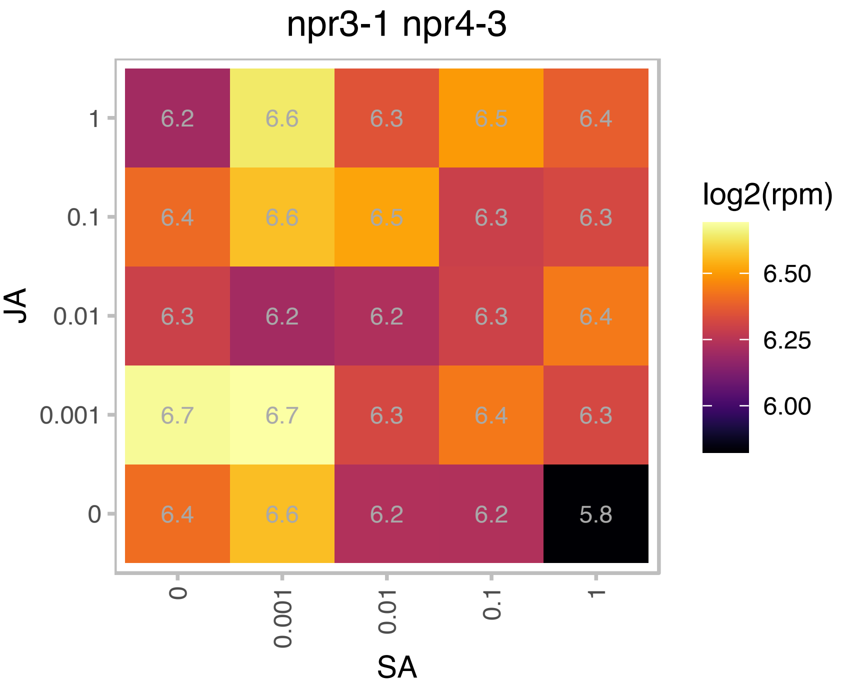

# AT1G30590.1 : NA

RNA polymerase I specific transcription initiation factor RRN3 protein

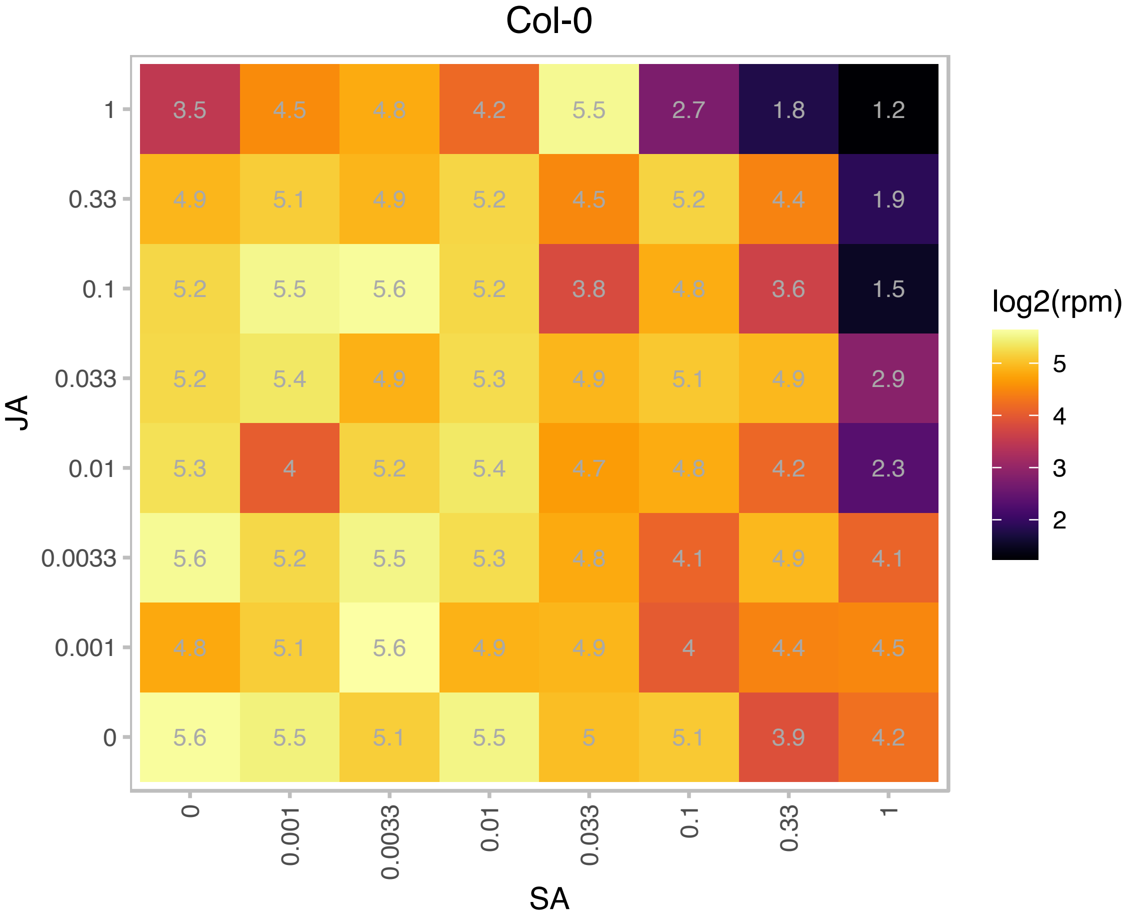


| | |
|---|---|
| Gene | AT1G30590 |
| is.expressed_genes | TRUE |
| is.responsive_genes | TRUE |
| is.apcluster | F-04 |

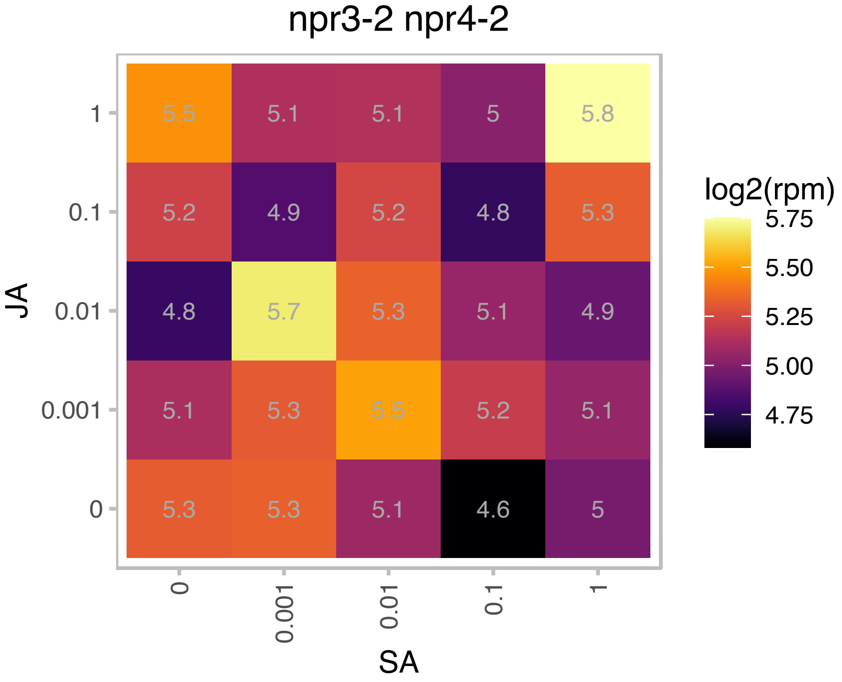


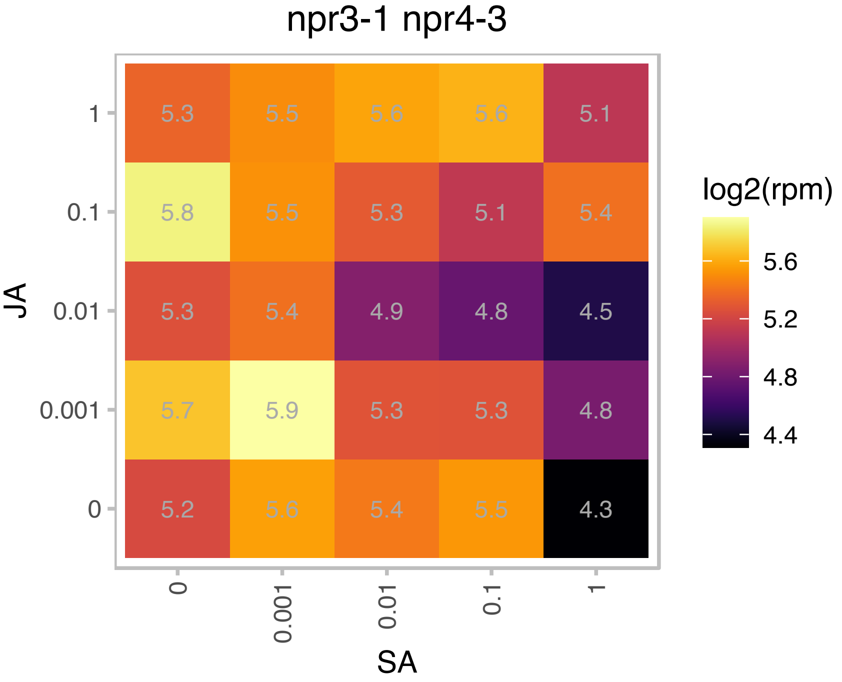

### AT1G56110.1 : NOP56

homolog of nucleolar protein NOP56

NOP56-like protein

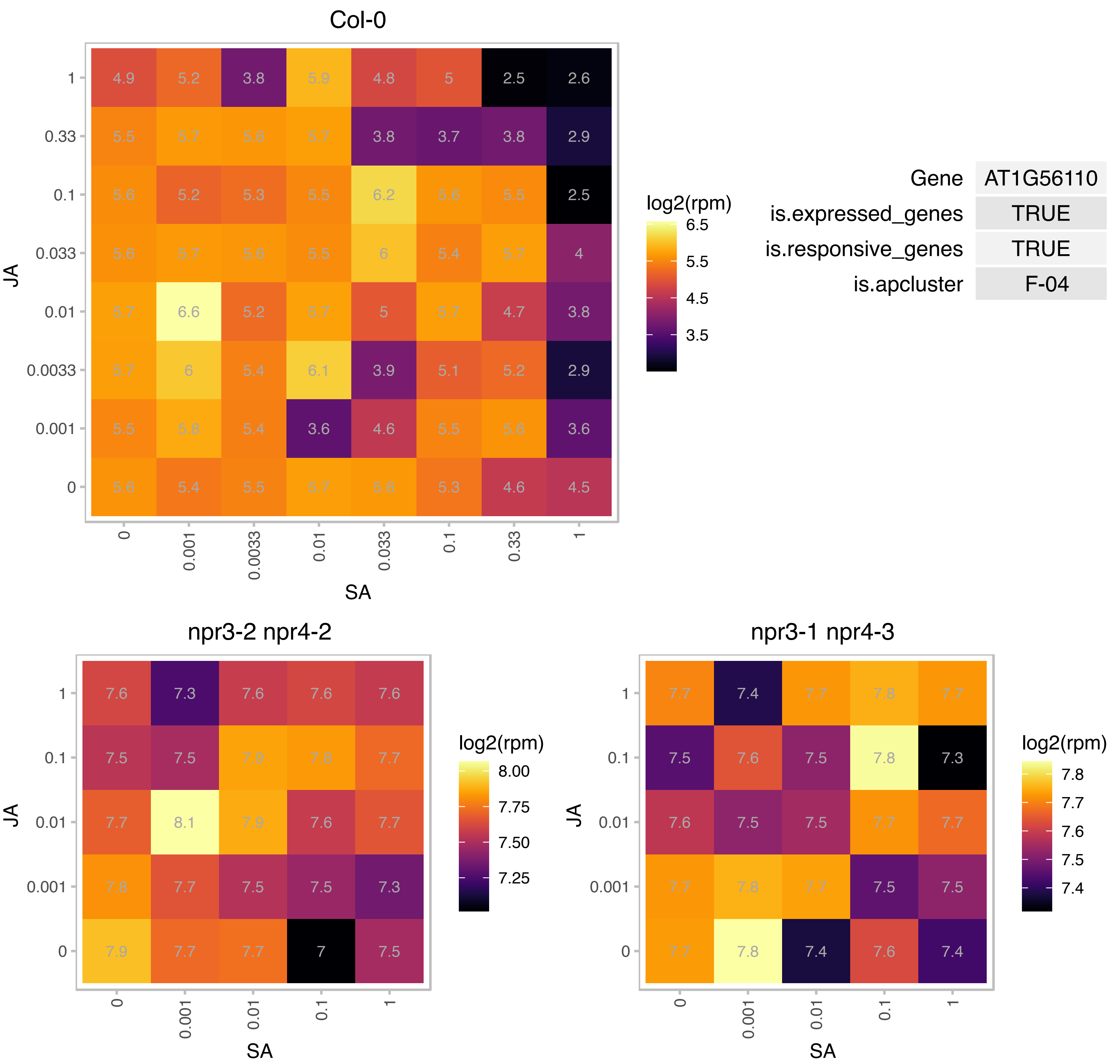


| | |
|---|---|
| Gene | AT1G56110 |
| is.expressed_genes | TRUE |
| is.responsive_genes | TRUE |
| is.apcluster | F-04 |

### AT1G61580.1 : ARP2, RPL3B

R-protein L3 B

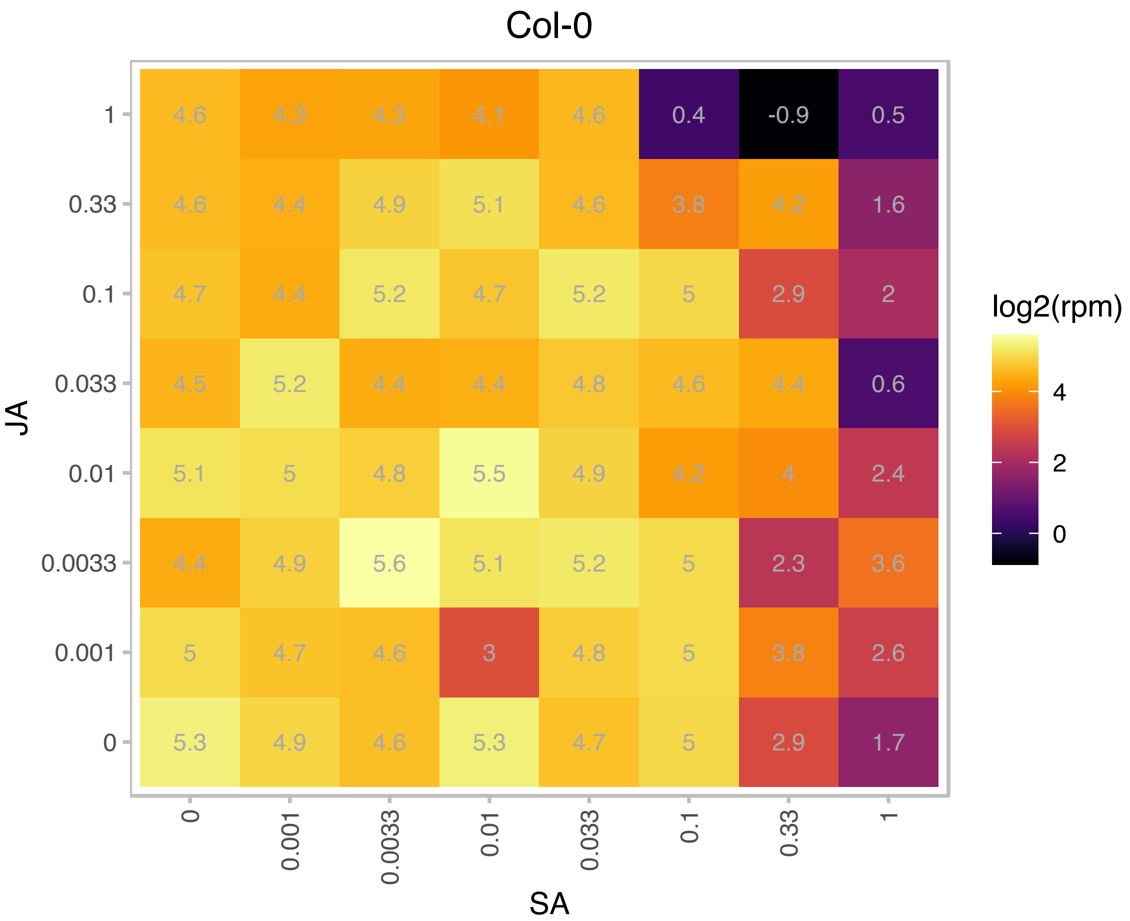


| | |
|---|---|
| Gene | AT1G61580 |
| is.expressed_genes | TRUE |
| is.responsive_genes | TRUE |
| is.apcluster | F-04 |

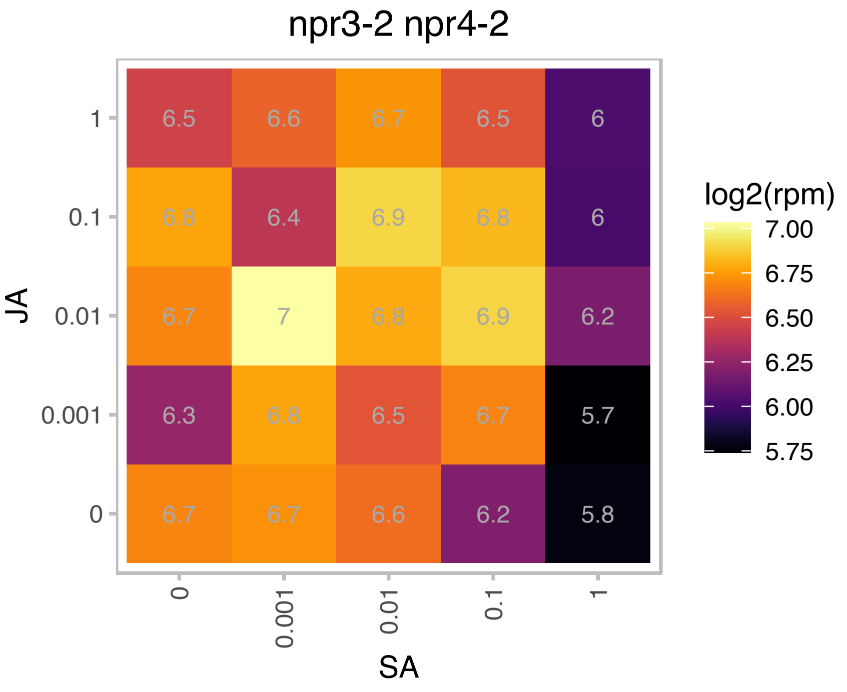


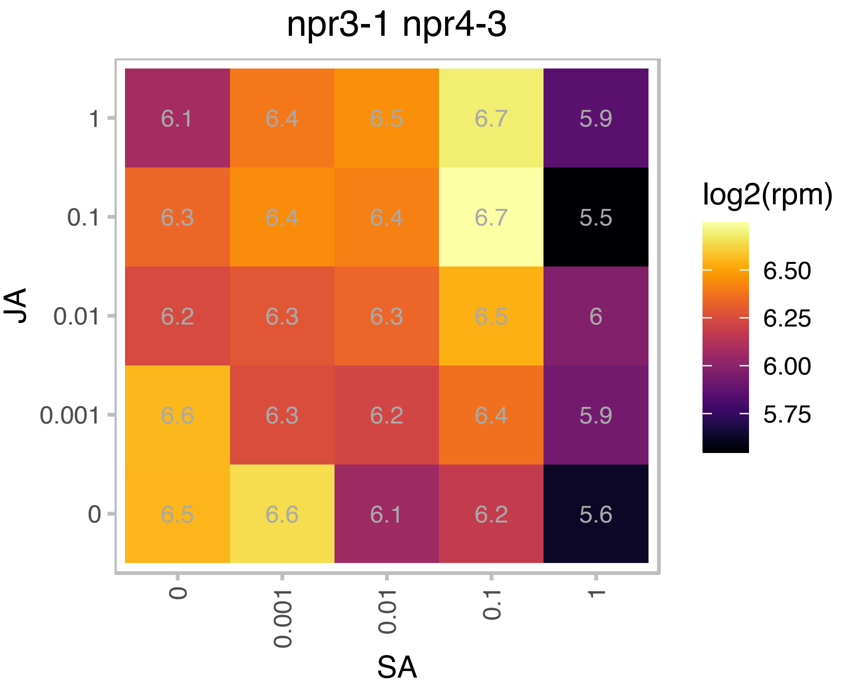

### AT1G64690.1 : BLT

branchless trichome
Encodes BRANCHLESS TRICHOME (BLT) involved in trichome development.  A large portion of the internal amino acid sequence of BLT is predicted to form a coiled-coil domain.  BLT mutants form branchless trichomes with blunt tips.

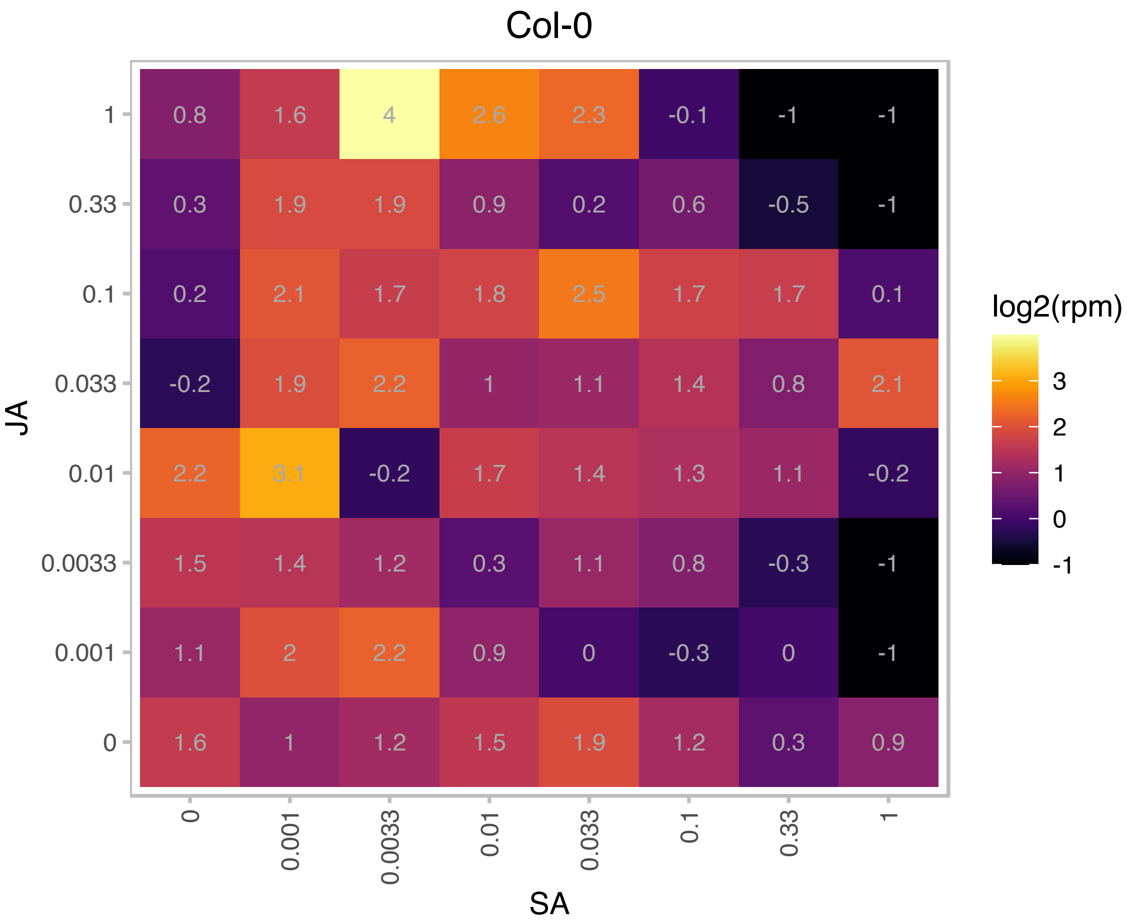


| | |
|---|---|
| Gene | AT1G64690 |
| is.expressed_genes | TRUE |
| is.responsive_genes | TRUE |
| is.apcluster | F-04 |

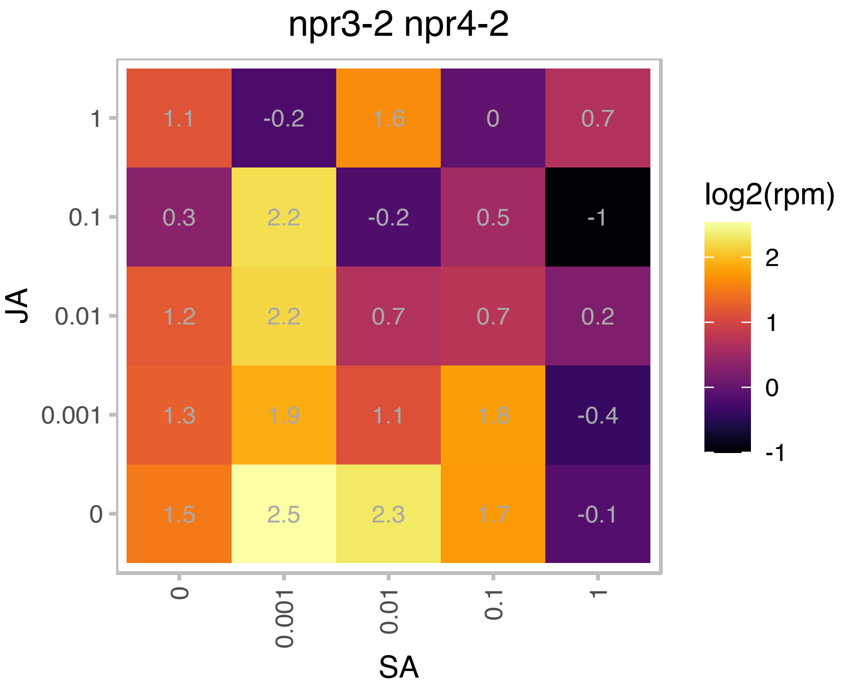


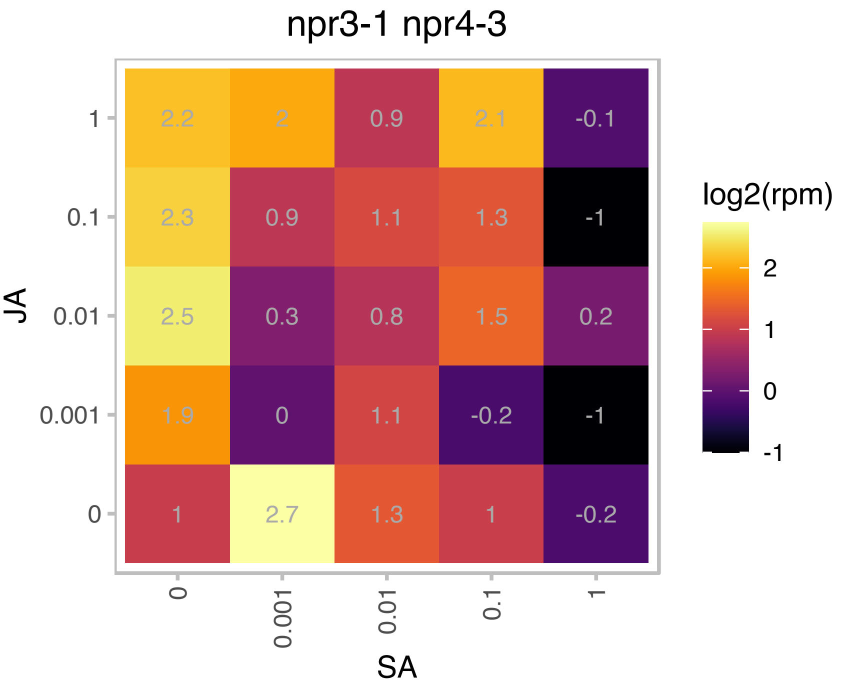

### AT1G68790.1 : CRWN3, LINC3

little nuclei3

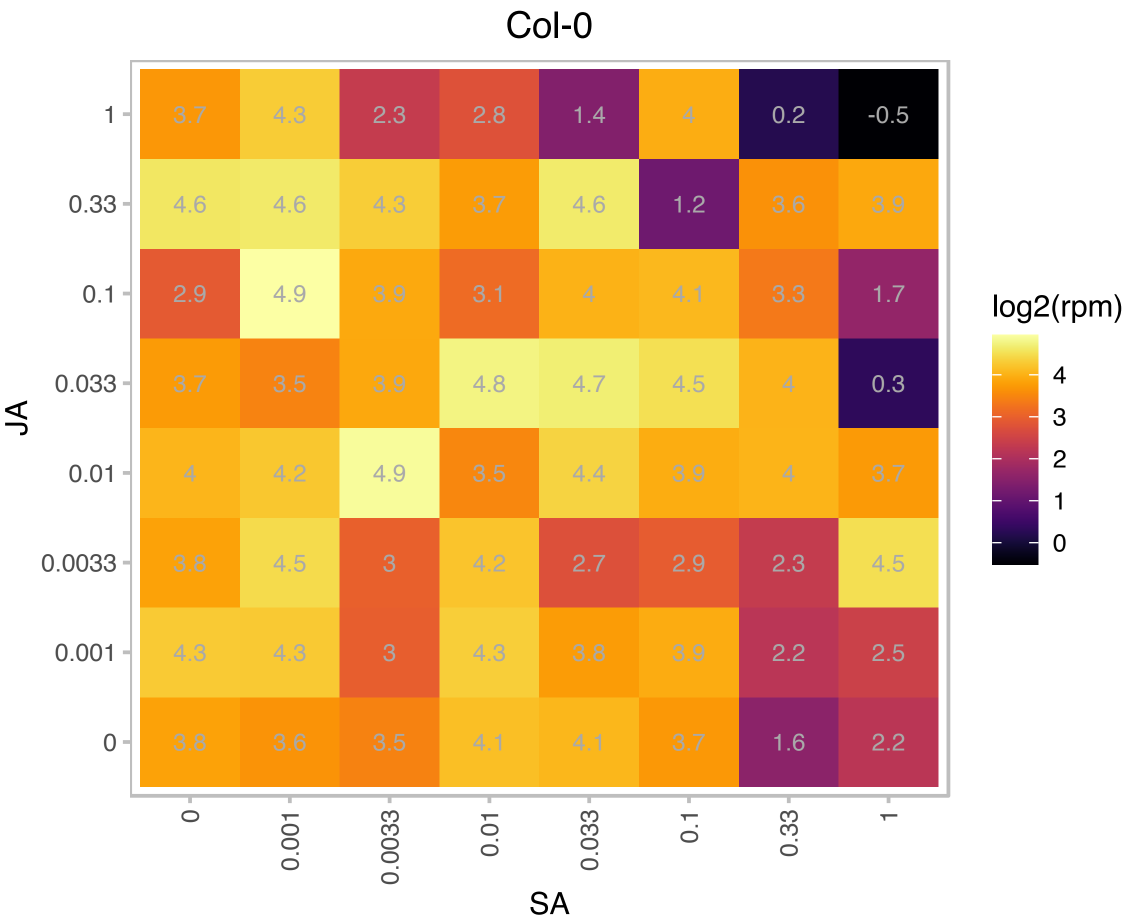


| | |
|---|---|
| Gene | AT1G68790 |
| is.expressed_genes | TRUE |
| is.responsive_genes | TRUE |
| is.apcluster | F-04 |

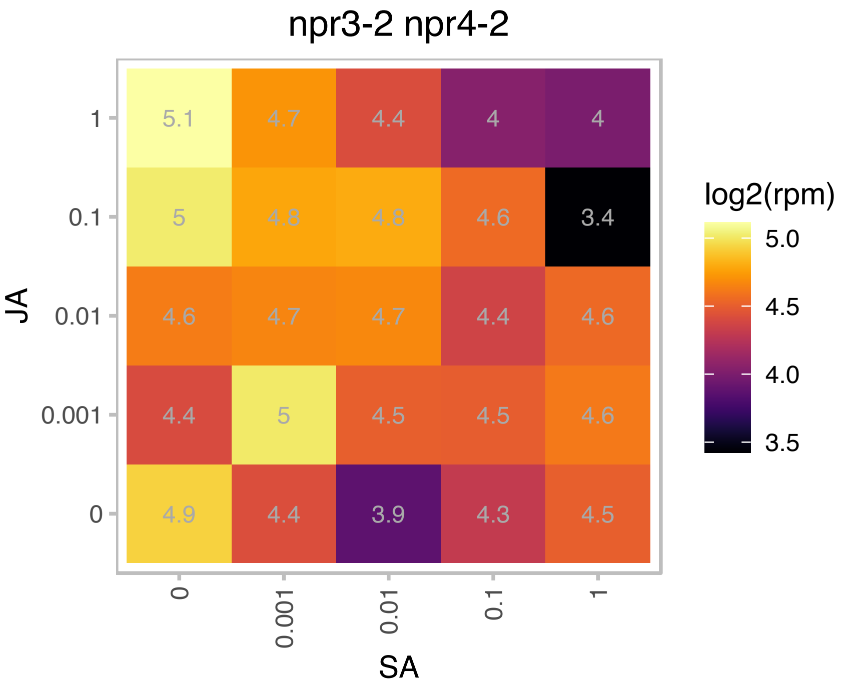


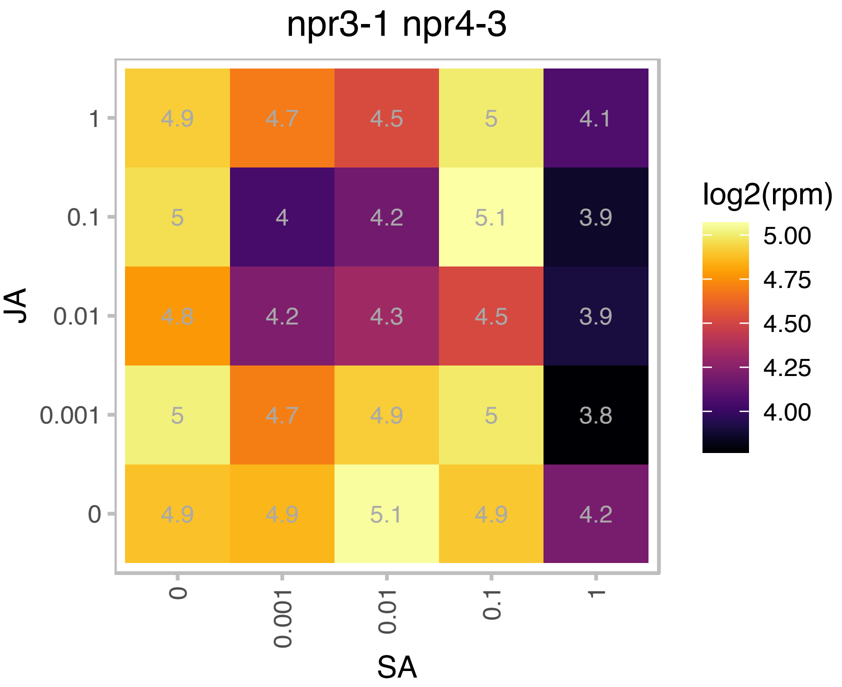

# AT1G73490.1 : NA

RNA-binding (RRM/RBD/RNP motifs) family protein

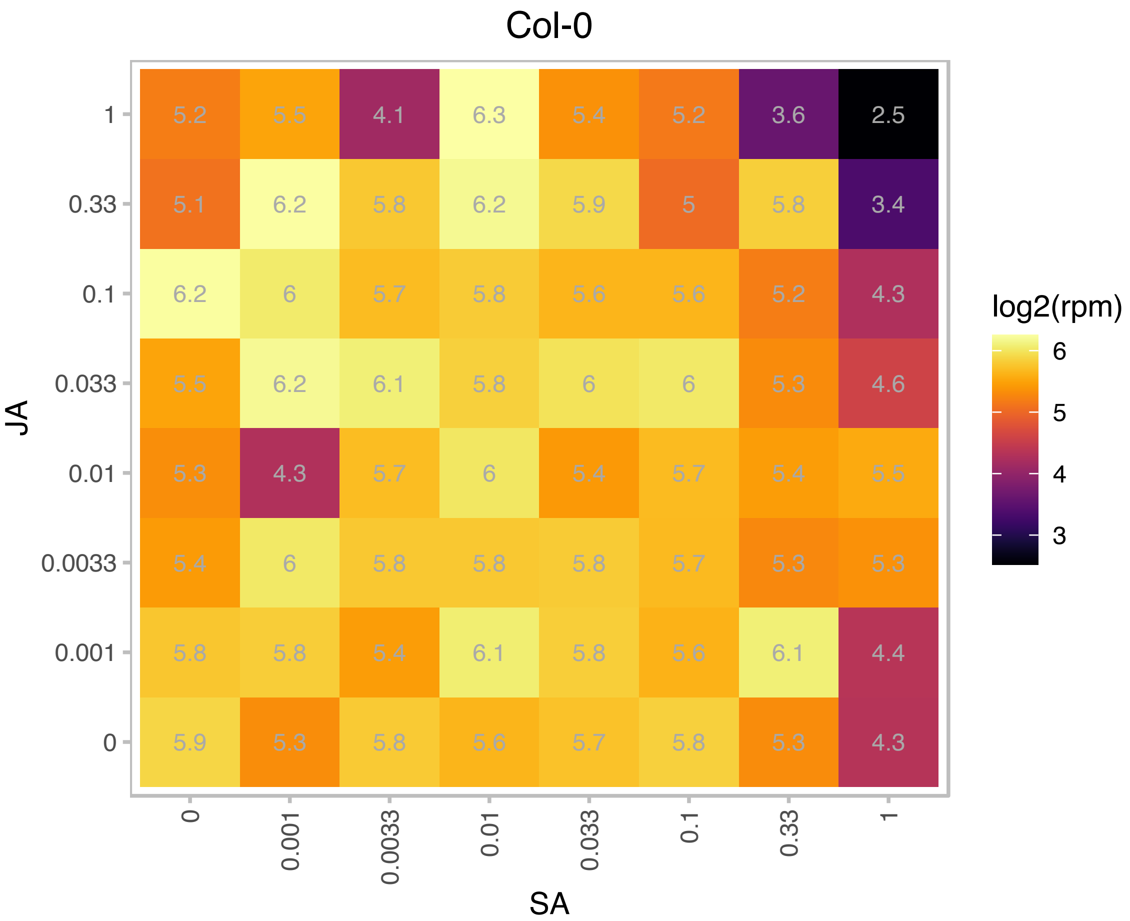


| | |
|---|---|
| Gene | AT1G73490 |
| is.expressed_genes | TRUE |
| is.responsive_genes | TRUE |
| is.apcluster | F-04 |

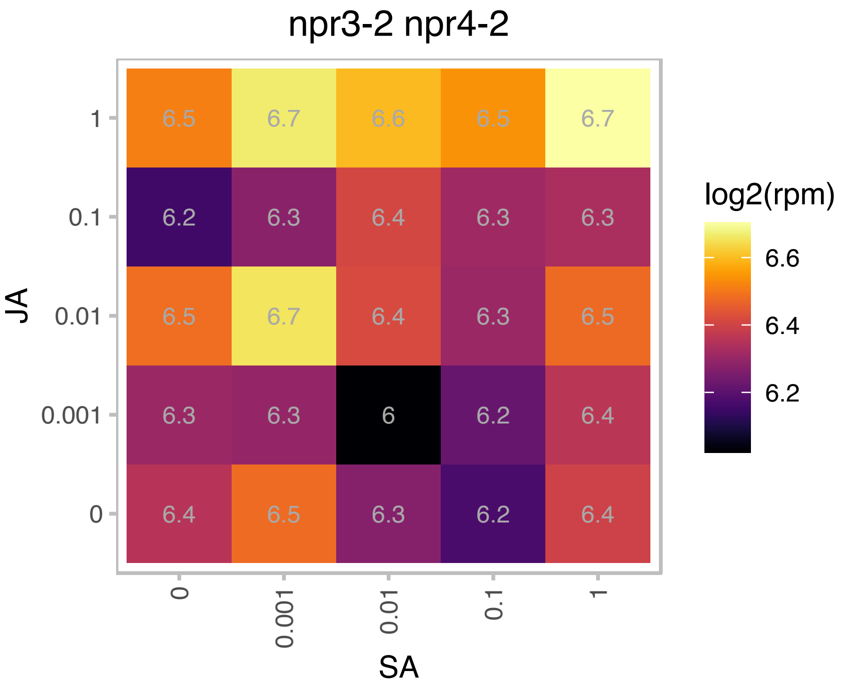


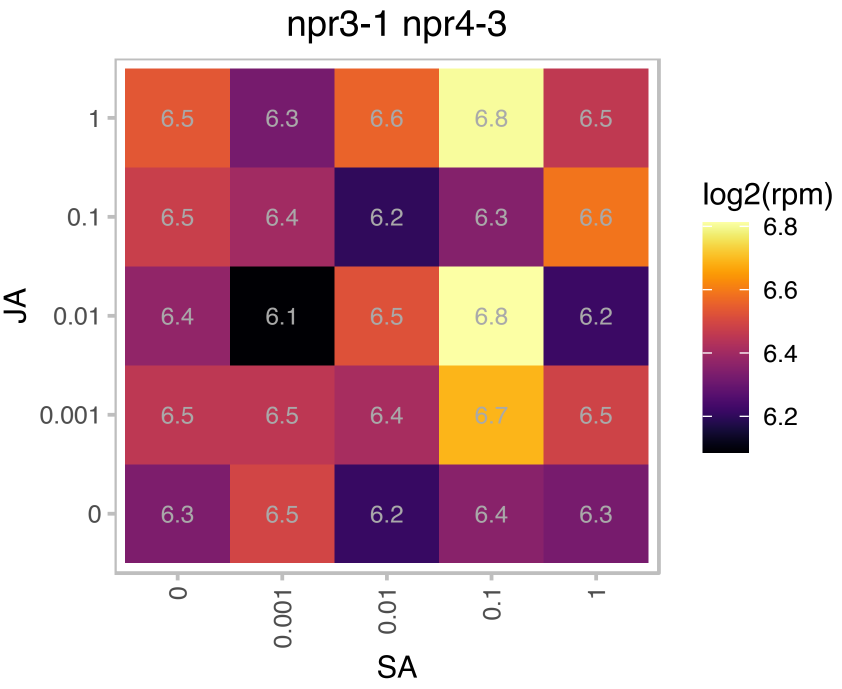

# AT1G74060.1 : NA

Ribosomal protein L6 family protein

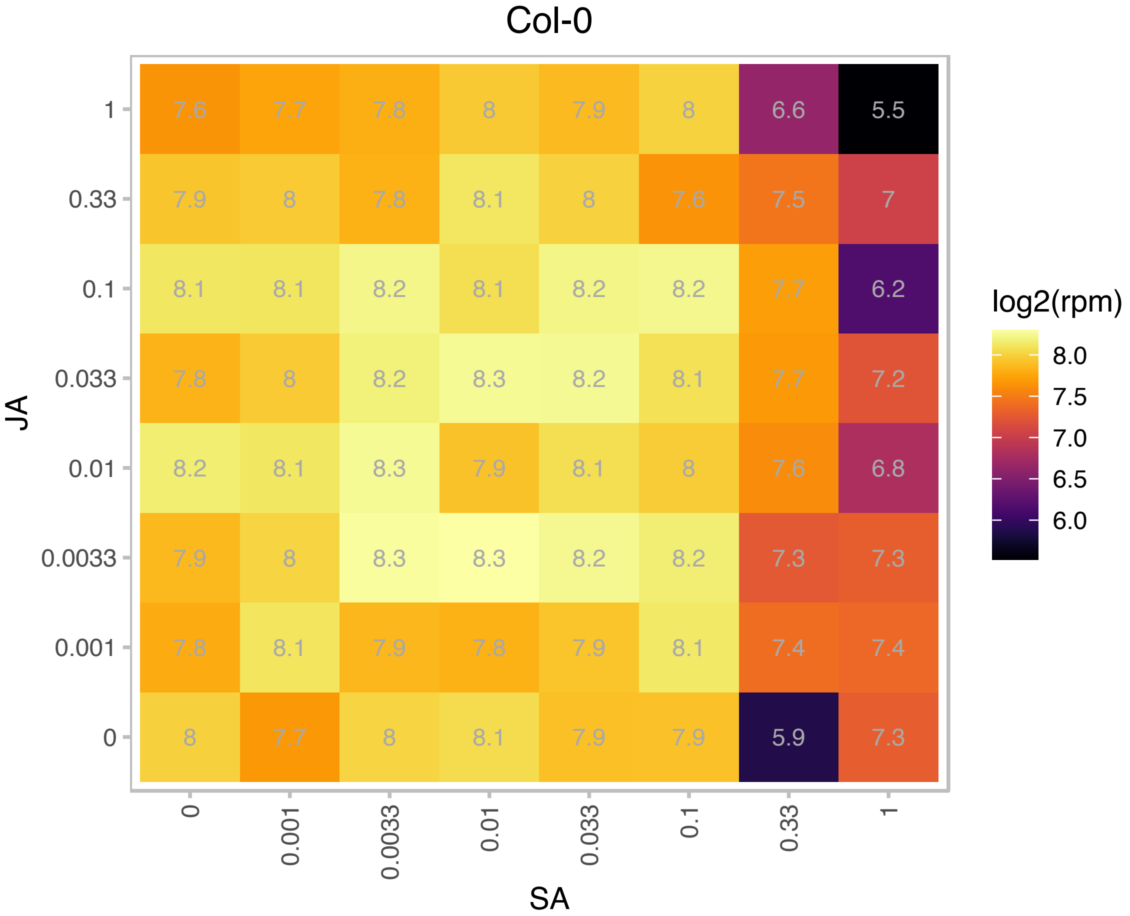


| | |
|---|---|
| Gene | AT1G74060 |
| is.expressed_genes | TRUE |
| is.responsive_genes | TRUE |
| is.apcluster | F-04 |

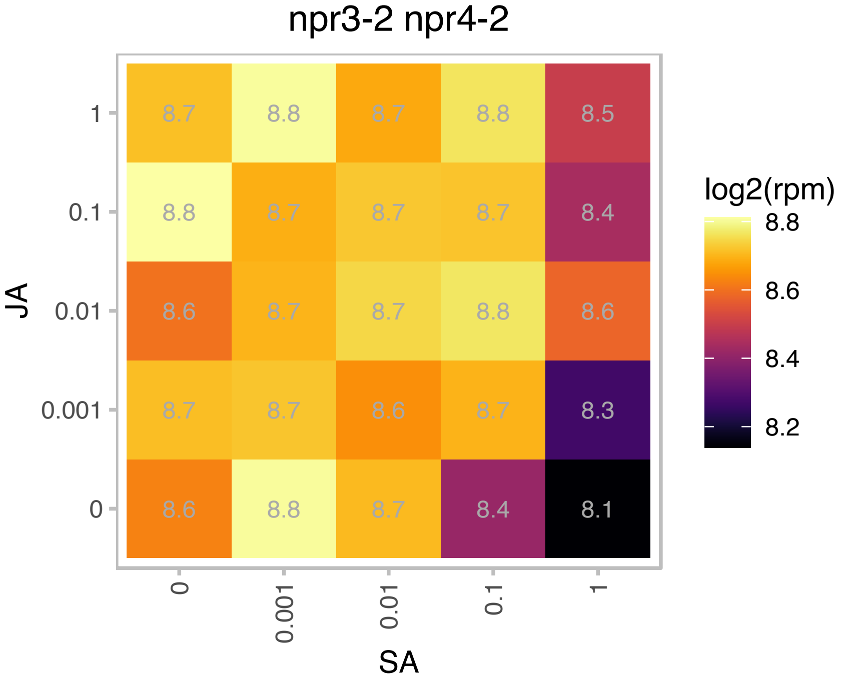


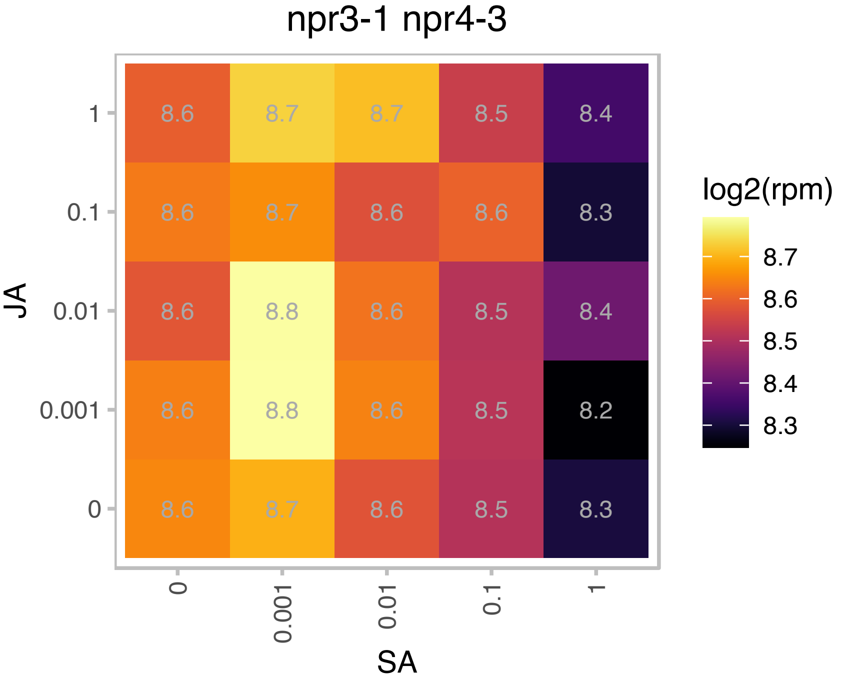

# AT2G16930.1 : NA

Ribosomal protein L27 family protein

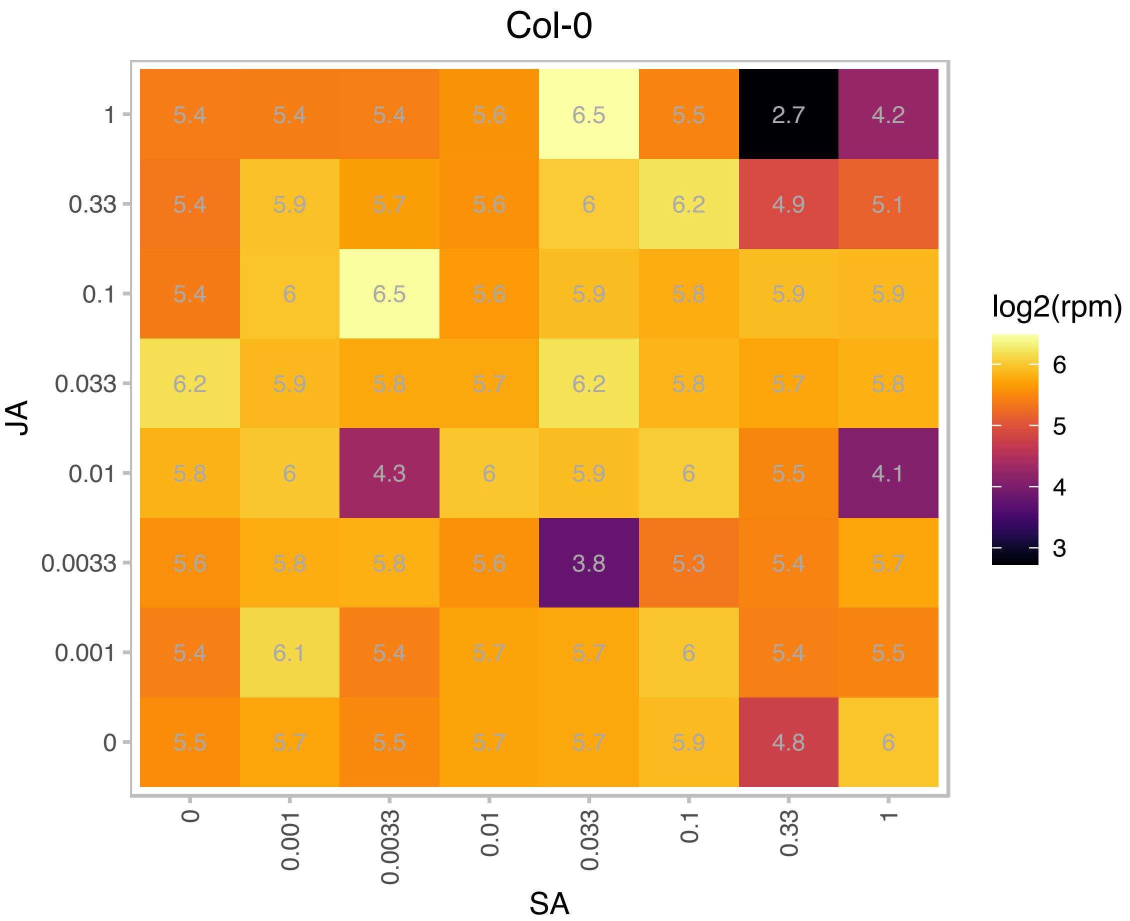


| | |
|---|---|
| Gene | AT2G16930 |
| is.expressed_genes | TRUE |
| is.responsive_genes | TRUE |
| is.apcluster | F-04 |

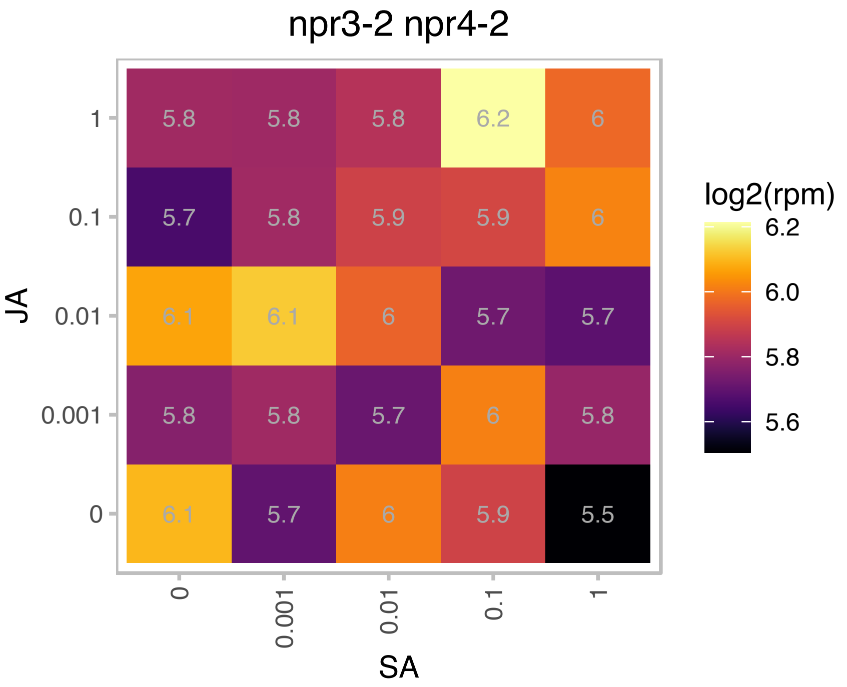


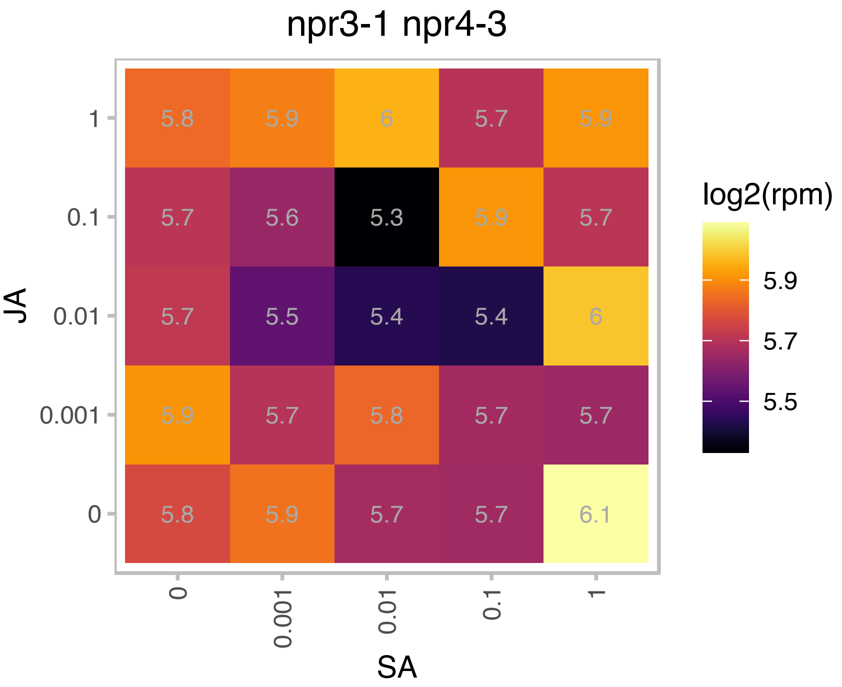

### AT2G17265.1 : DMR1, HSK

homoserine kinase
Encodes a homoserine kinase (HSK) which produces O-phospho-L-homoserine (HserP), a compound at the branch point of methionine and threonine biosynthesis. HSK is found in the stromal fraction of chloroplasts. Mutation of this gene results in higher level of the amino acid homoserine and resistance to downy mildew pathogen Hyaloperonospora arabidopsidis.

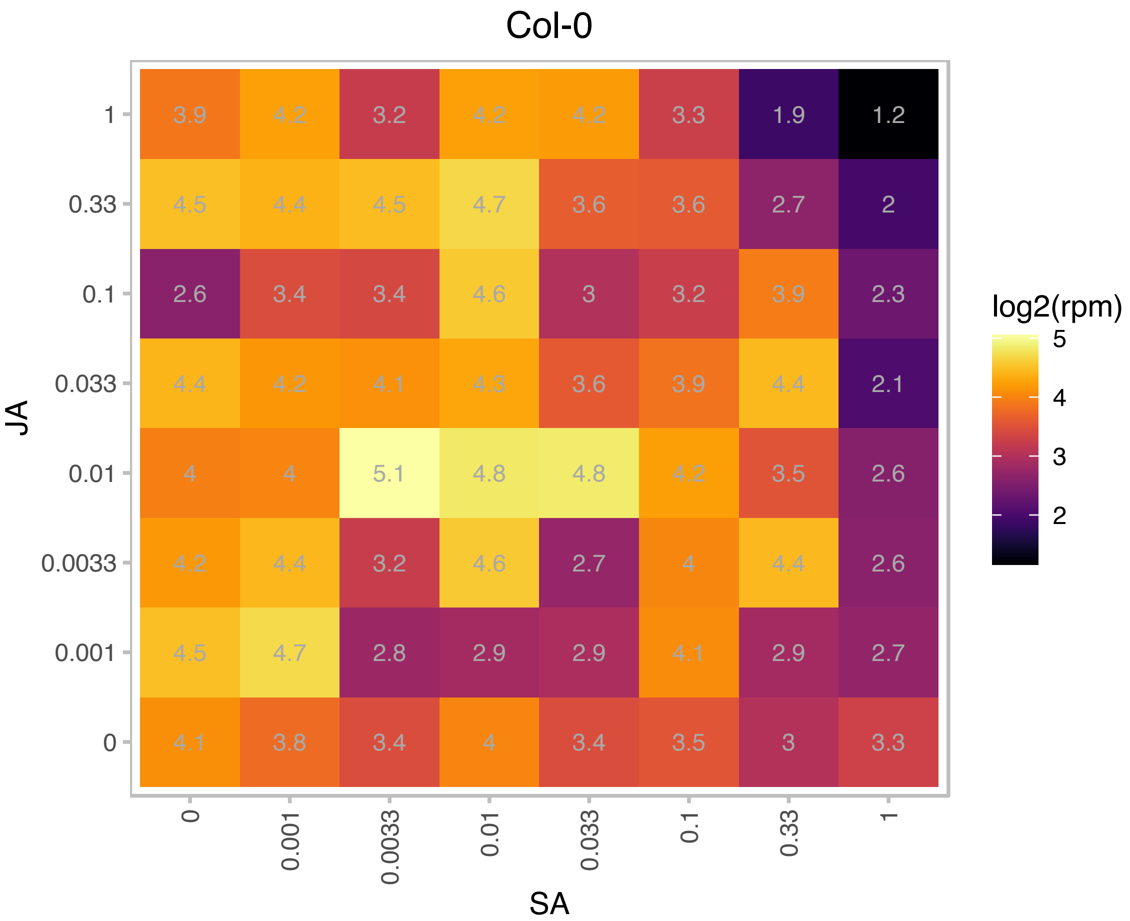


| | |
|---|---|
| Gene | AT2G17265 |
| is.expressed_genes | TRUE |
| is.responsive_genes | TRUE |
| is.apcluster | F-04 |

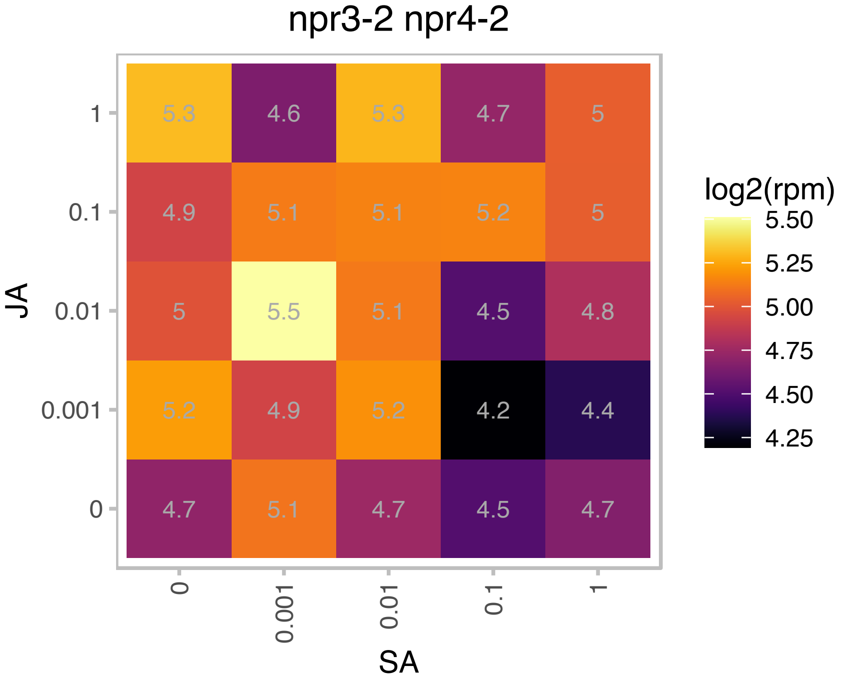


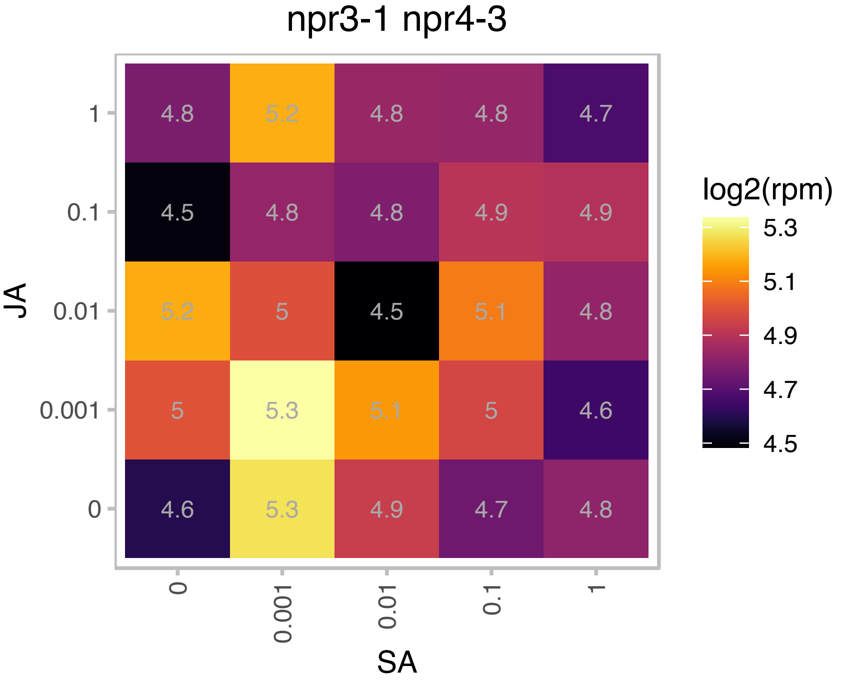

### AT2G19520.1 : ACG1, ATMSI4, FVE, MSI4, NFC04, NFC4

Transducin family protein / WD-40 repeat family protein

Controls flowering.

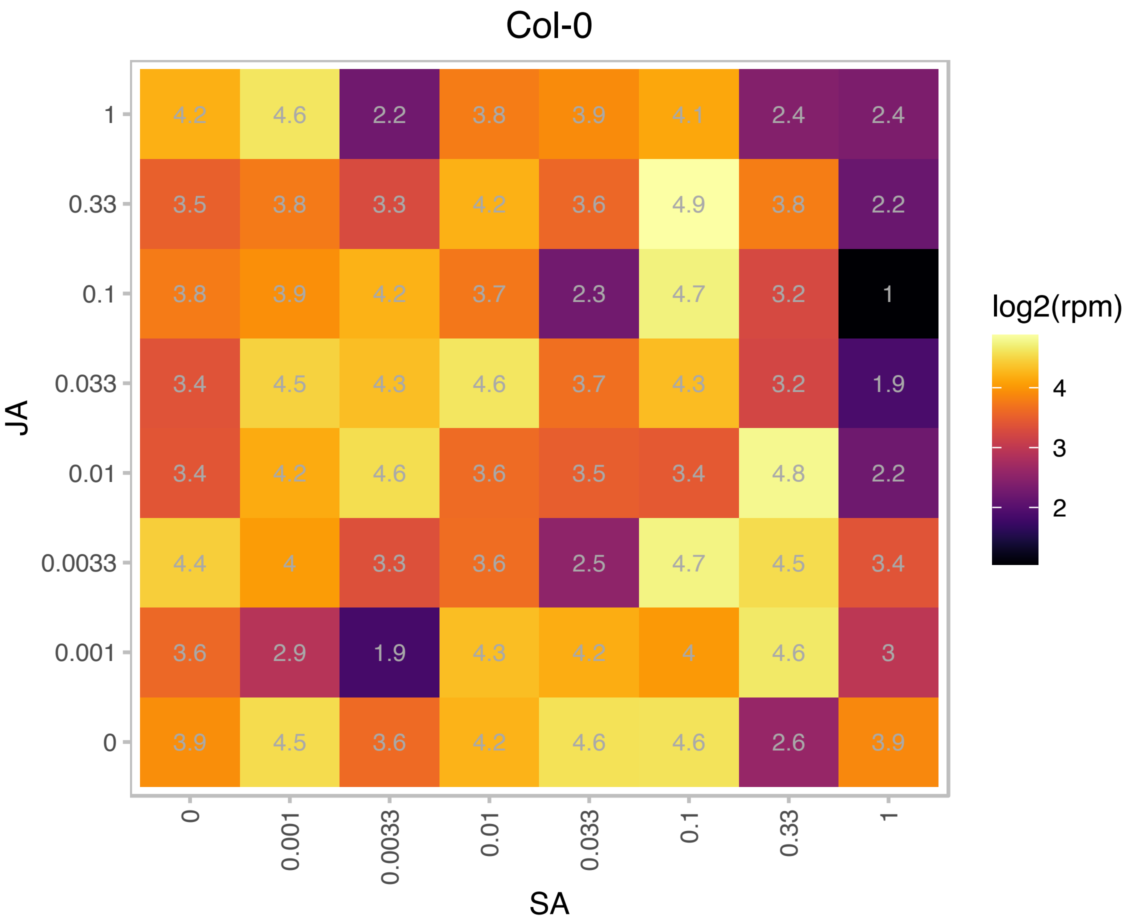


| | |
|---|---|
| Gene | AT2G19520 |
| is.expressed_genes | TRUE |
| is.responsive_genes | TRUE |
| is.apcluster | F-04 |

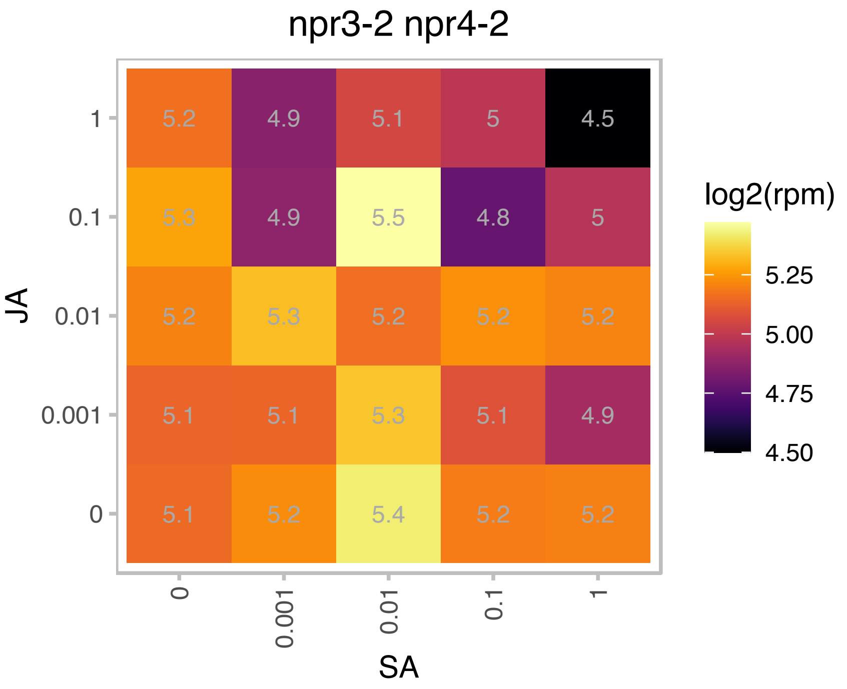


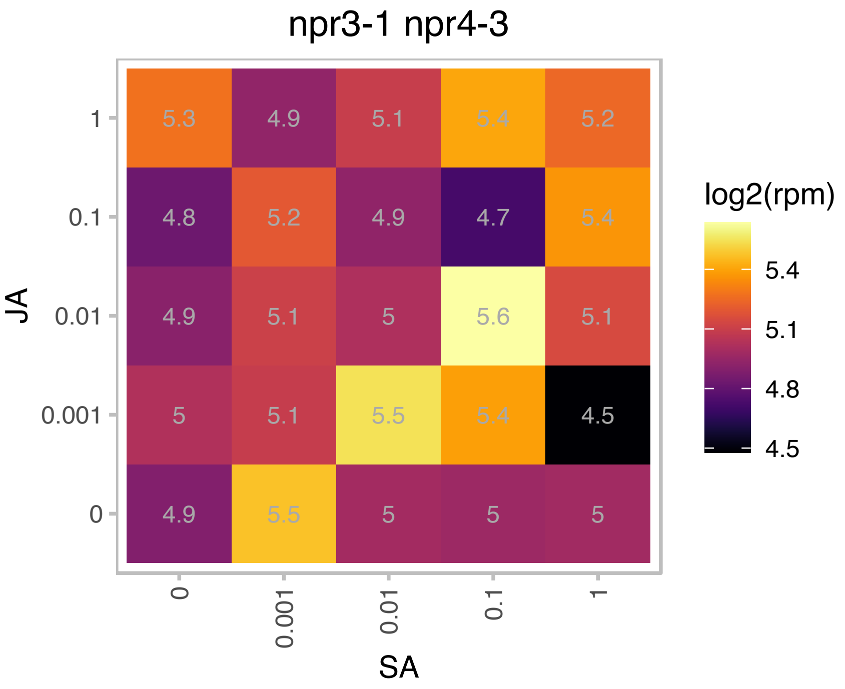

### AT2G20630.1 : PIA1

PP2C induced by AVRRPM1

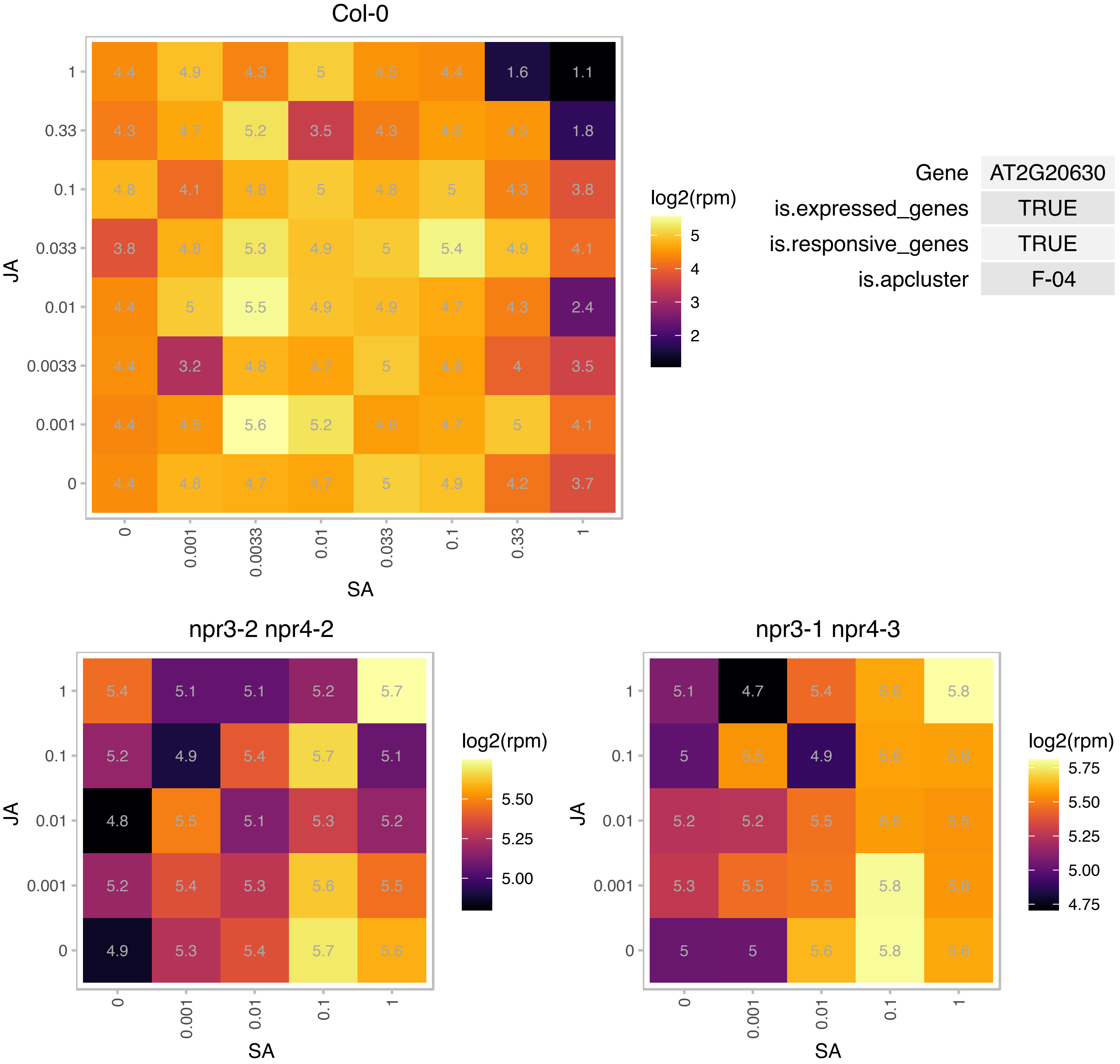


| | |
|---|---|
| Gene | AT2G20630 |
| is.expressed_genes | TRUE |
| is.responsive_genes | TRUE |
| is.apcluster | F-04 |

### AT2G23350.1 : PAB4, PABP4

poly(A) binding protein 4

polyadenylate-binding protein, putative / PABP, putative.Member of the Class II family of PABP proteins. Highly and ubiquitously expressed. The mRNA is cell-to-cell mobile.

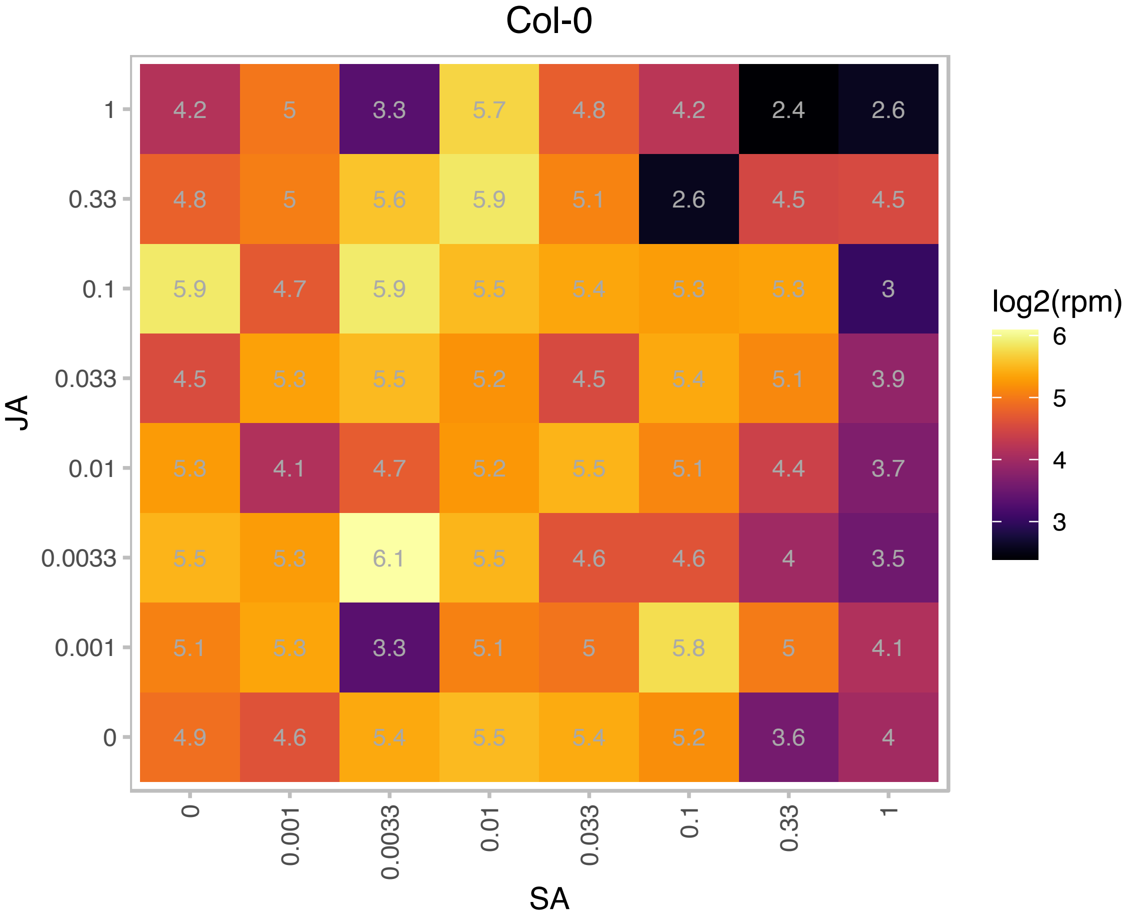


| Gene | AT2G23350 |
|---|---|
| is.expressed_genes | TRUE |
| is.responsive_genes | TRUE |
| is.apcluster | F-04 |

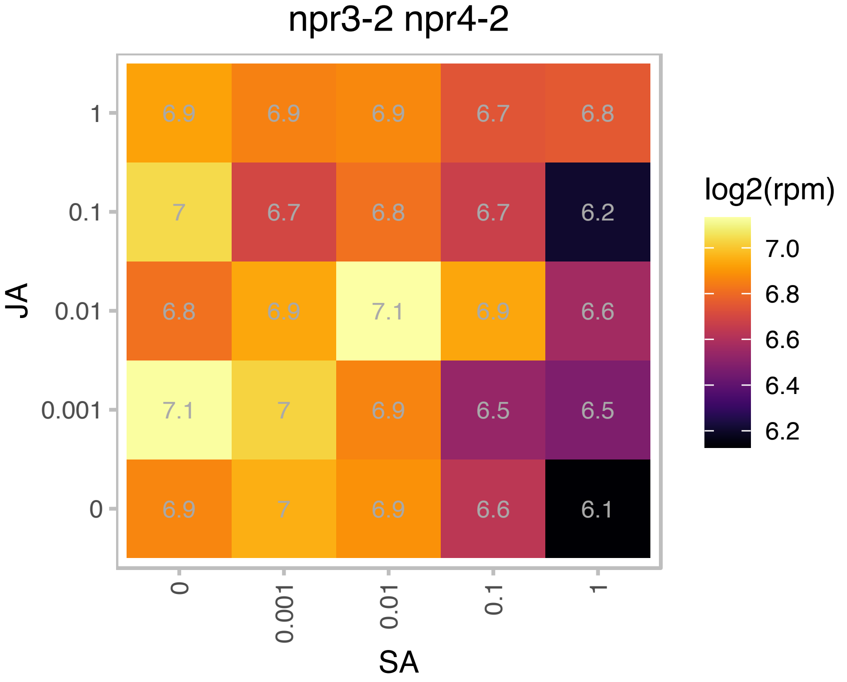


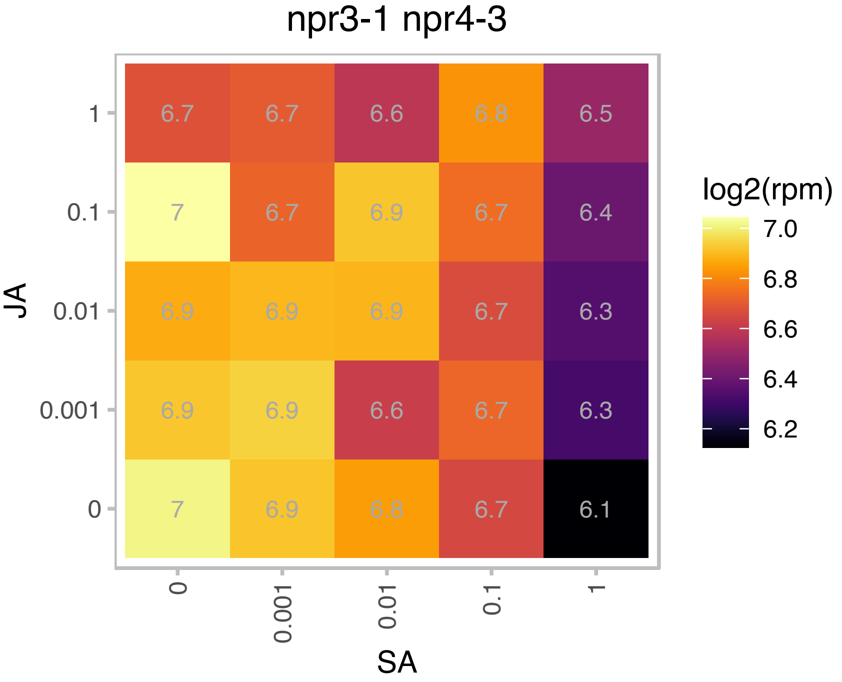

### AT2G23610.1 : ATMES3, MES3

methyl esterase 3

Encodes a protein shown to have carboxylesterase activity, methyl IAA esterase activity, and methyl jasmonate esterase activity in vitro. This protein does not act on methyl salicylate, MeGA4, or MEGA9 in vitro.

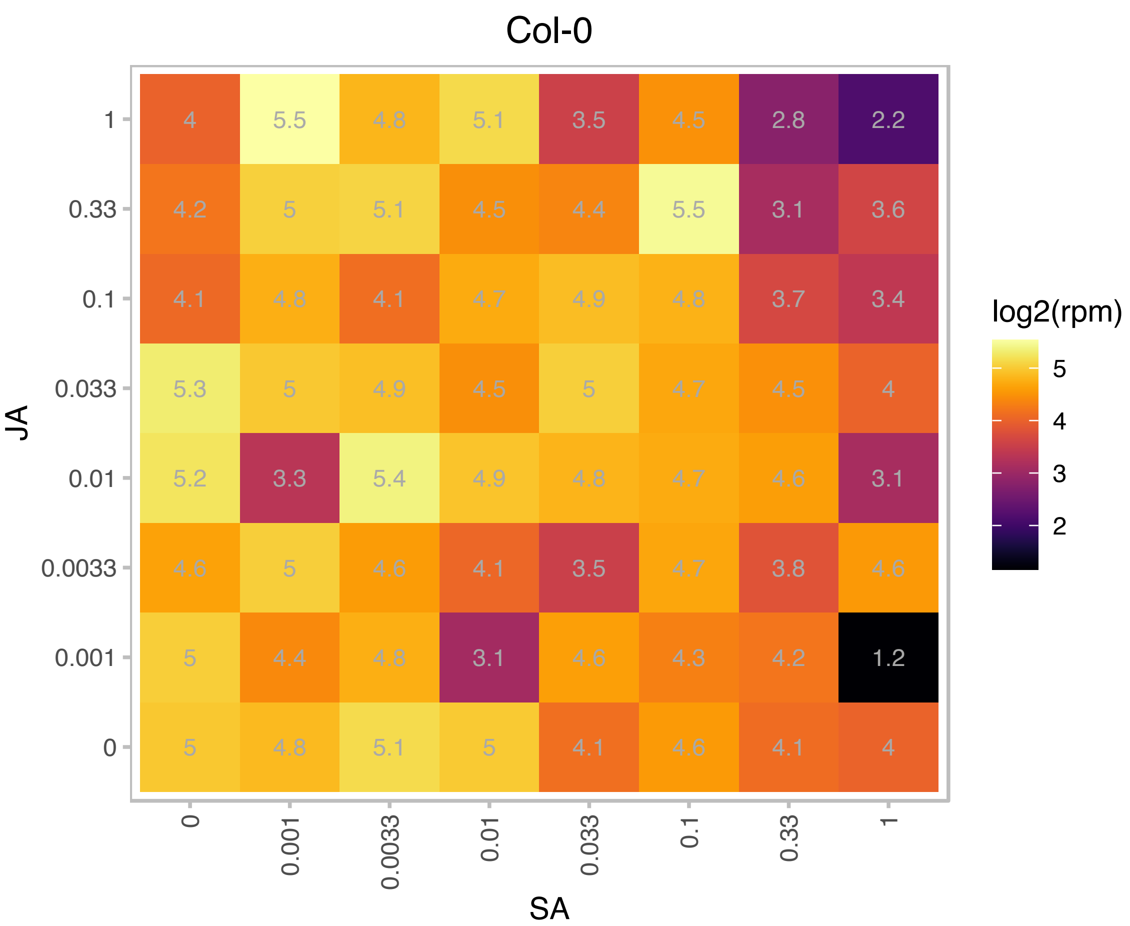


| | |
|---|---|
| Gene | AT2G23610 |
| is.expressed_genes | TRUE |
| is.responsive_genes | TRUE |
| is.apcluster | F-04 |

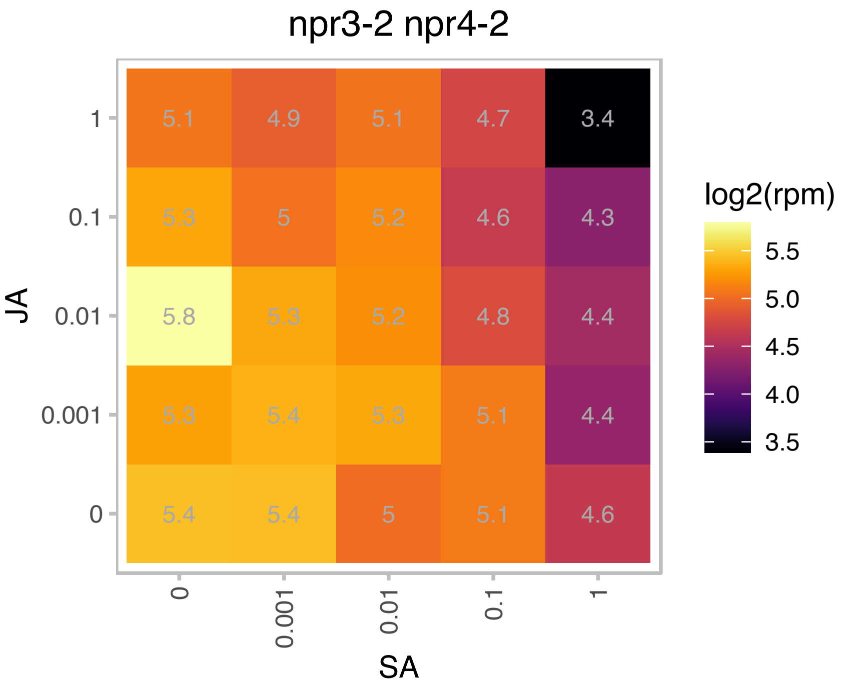


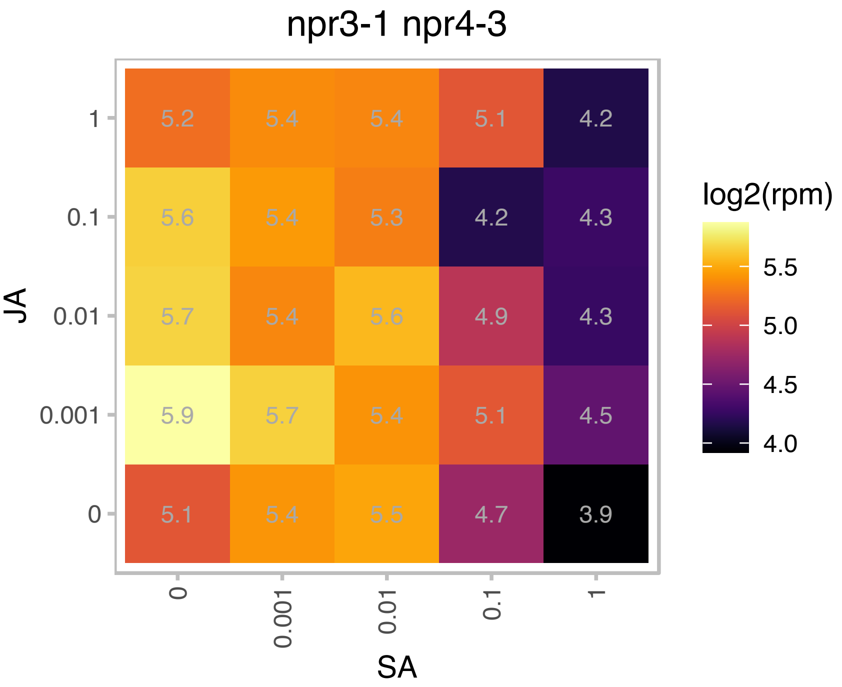

### AT2G24050.1 : eIFiso4G2

MIF4G domain-containing protein / MA3 domain-containing protein

Encodes a putative eukaryotic translation initiation factor.

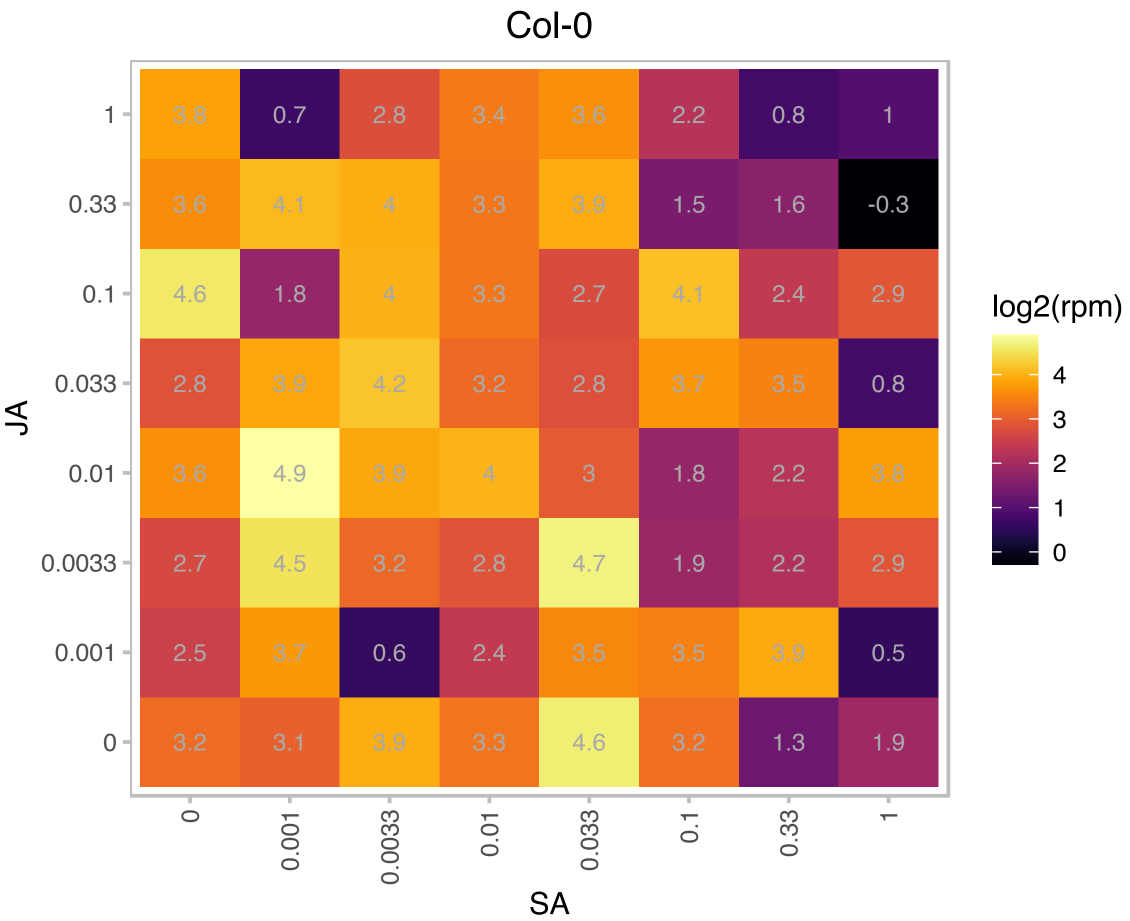


| | |
|---|---|
| Gene | AT2G24050 |
| is.expressed_genes | TRUE |
| is.responsive_genes | TRUE |
| is.apcluster | F-04 |

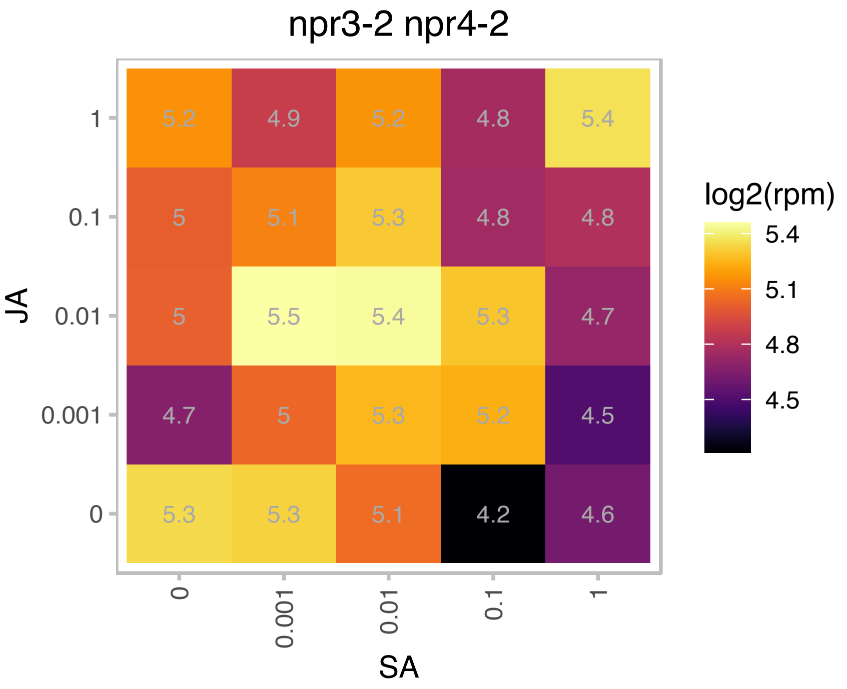


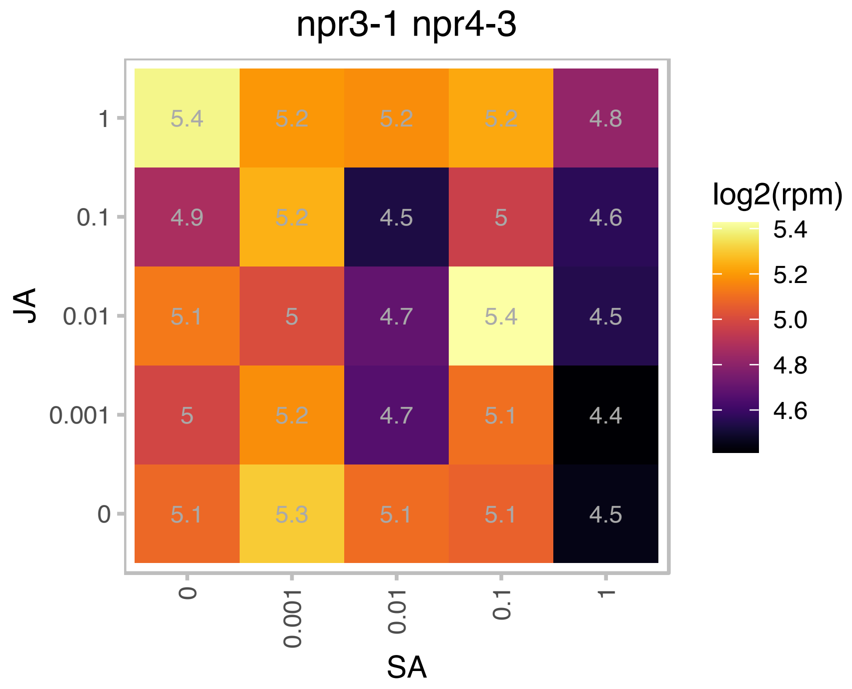

# AT2G27470.1 : NF-YB11

nuclear factor Y, subunit B11

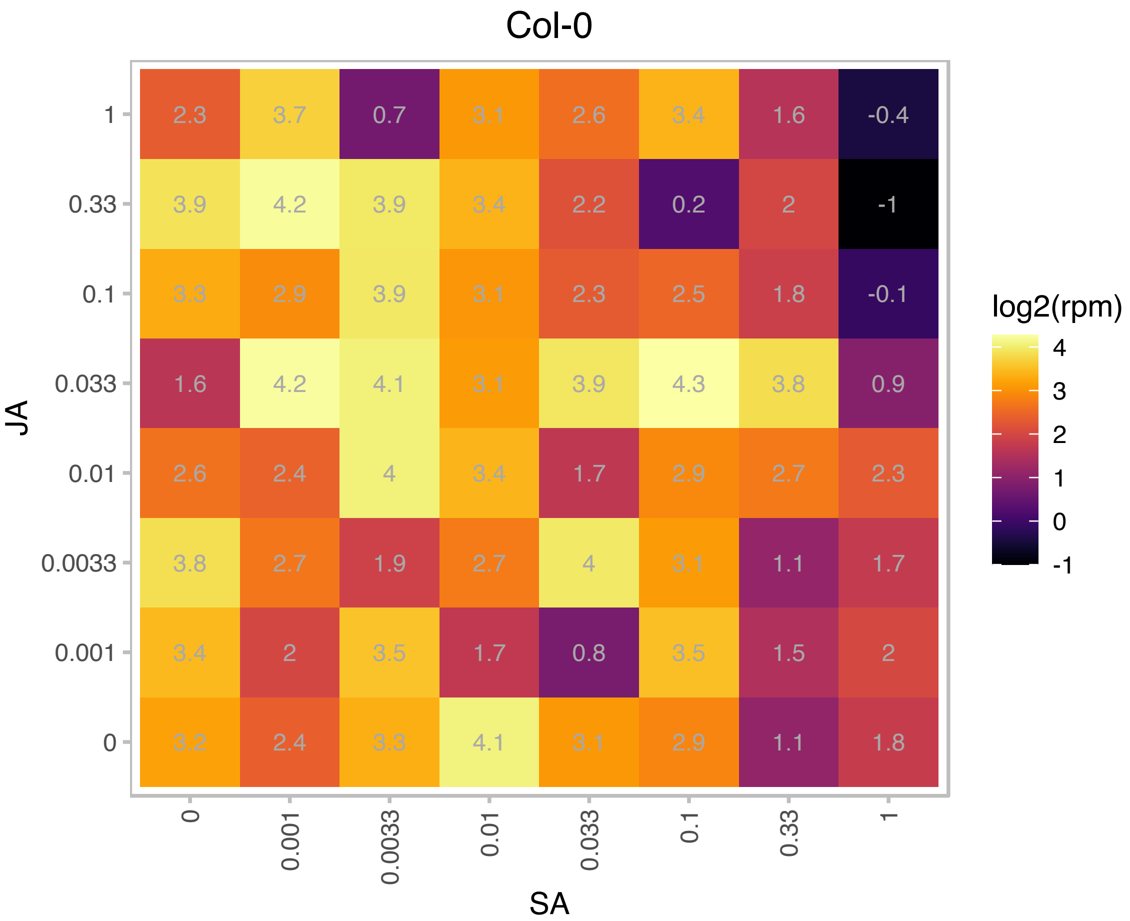


| | |
|---|---|
| Gene | AT2G27470 |
| is.expressed_genes | TRUE |
| is.responsive_genes | TRUE |
| is.apcluster | F-04 |

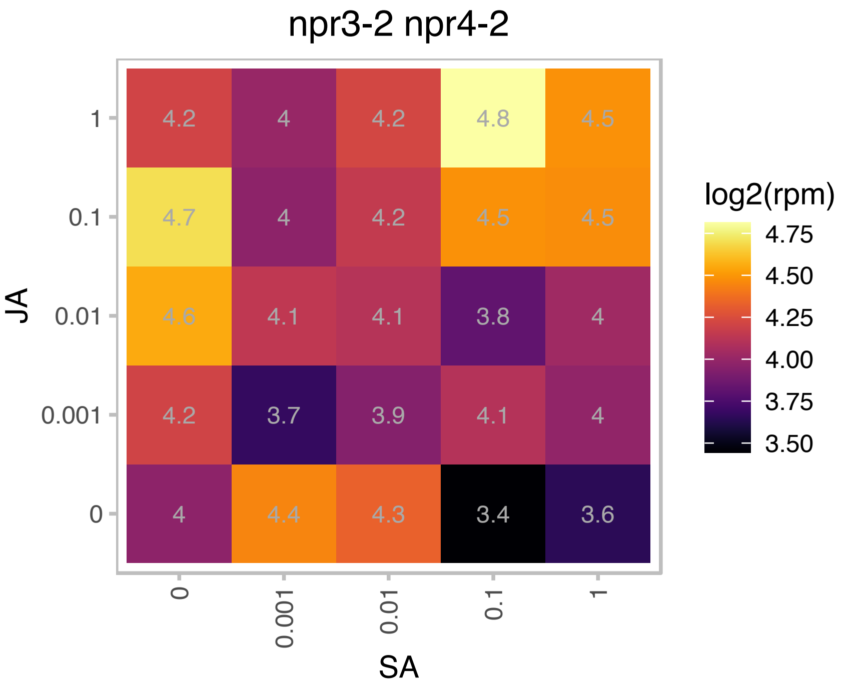


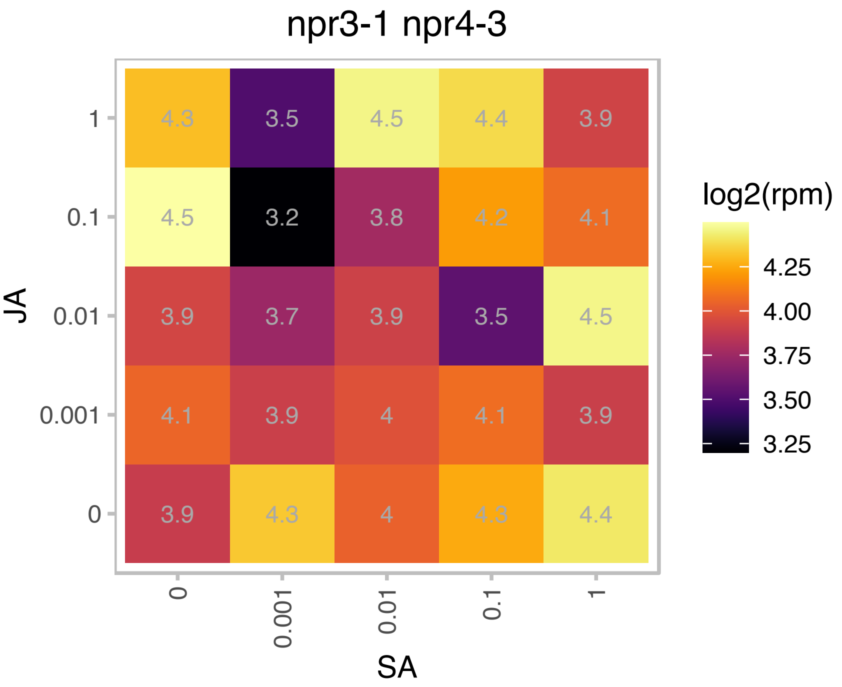

# AT2G35040.1 : NA

AICARFT/IMPCHase bienzyme family protein

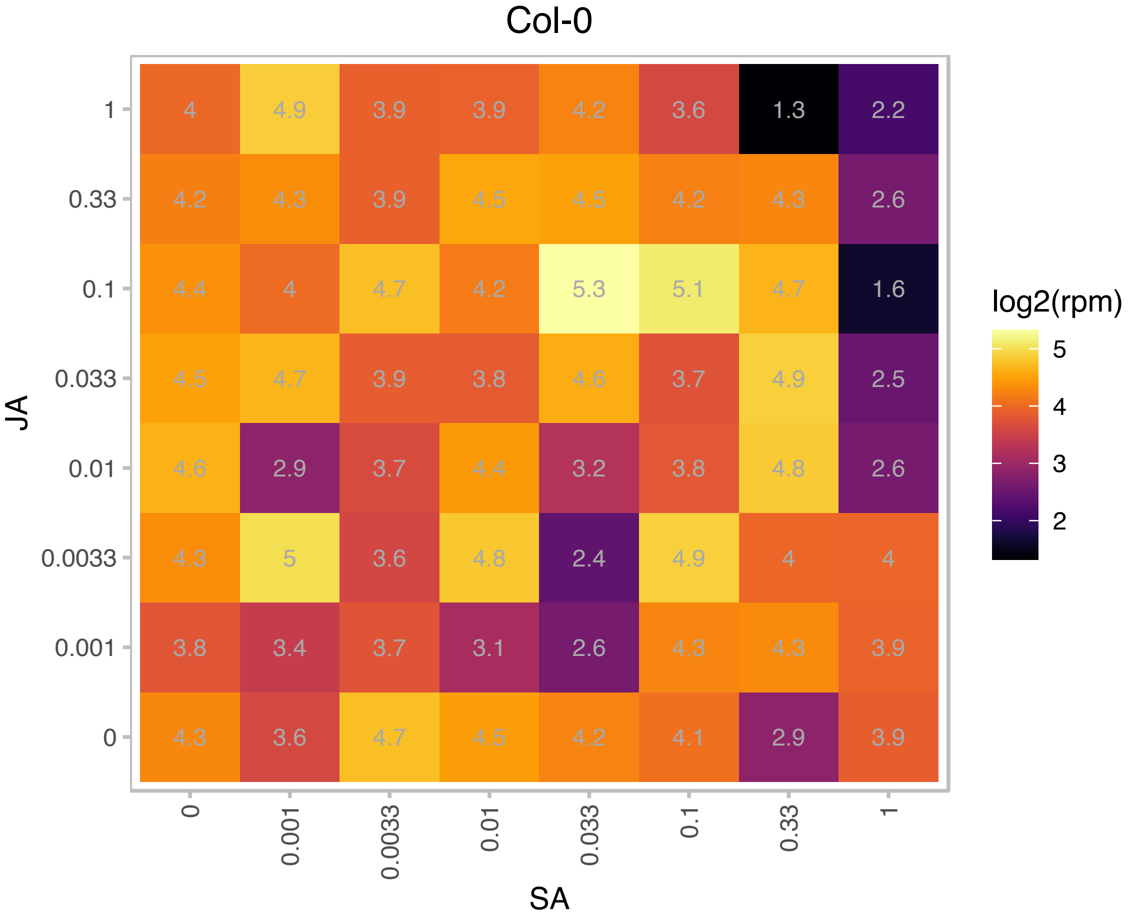


| | |
|---|---|
| Gene | AT2G35040 |
| is.expressed_genes | TRUE |
| is.responsive_genes | TRUE |
| is.apcluster | F-04 |

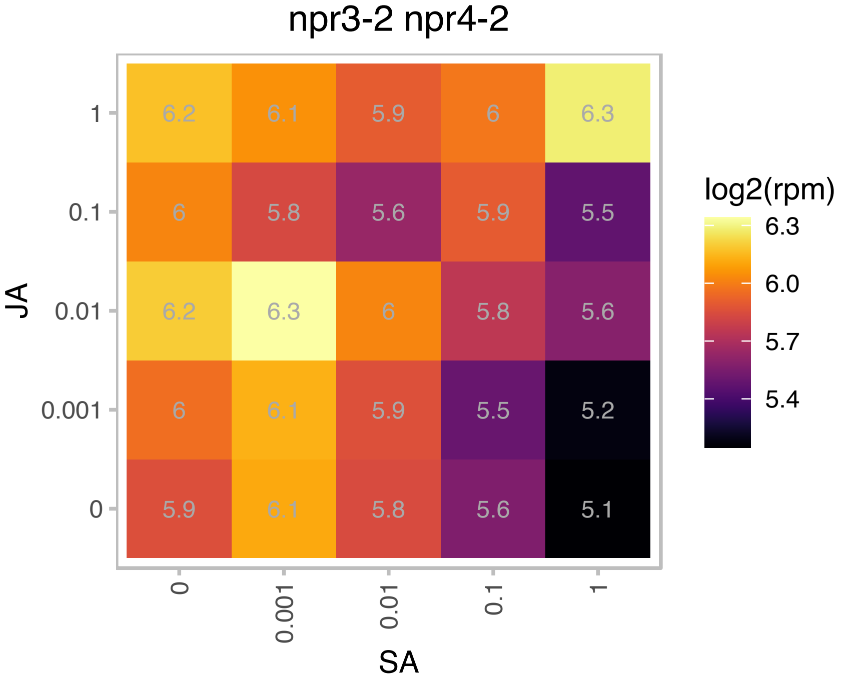


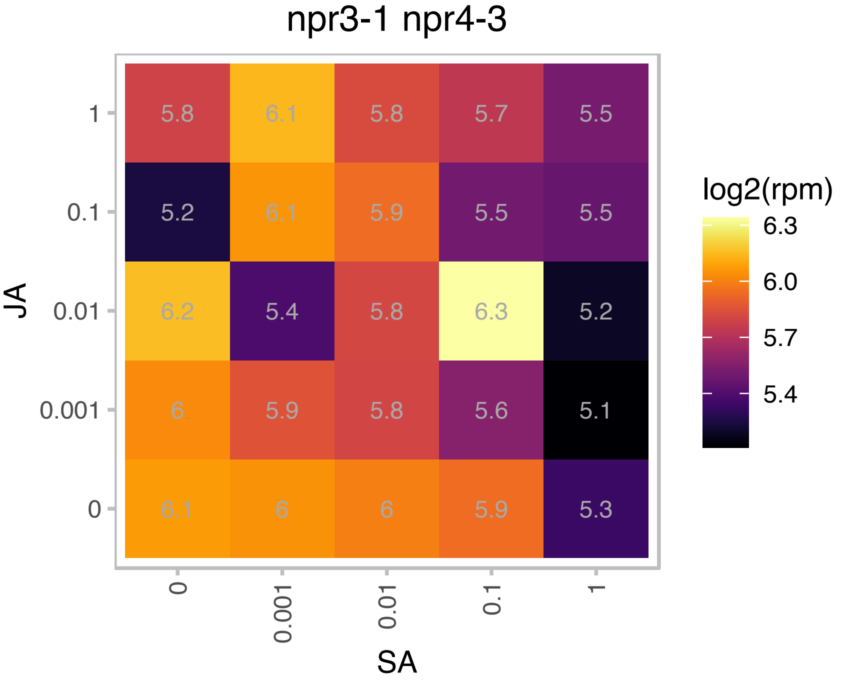

# AT2G35120.1 : NA

Single hybrid motif superfamily protein

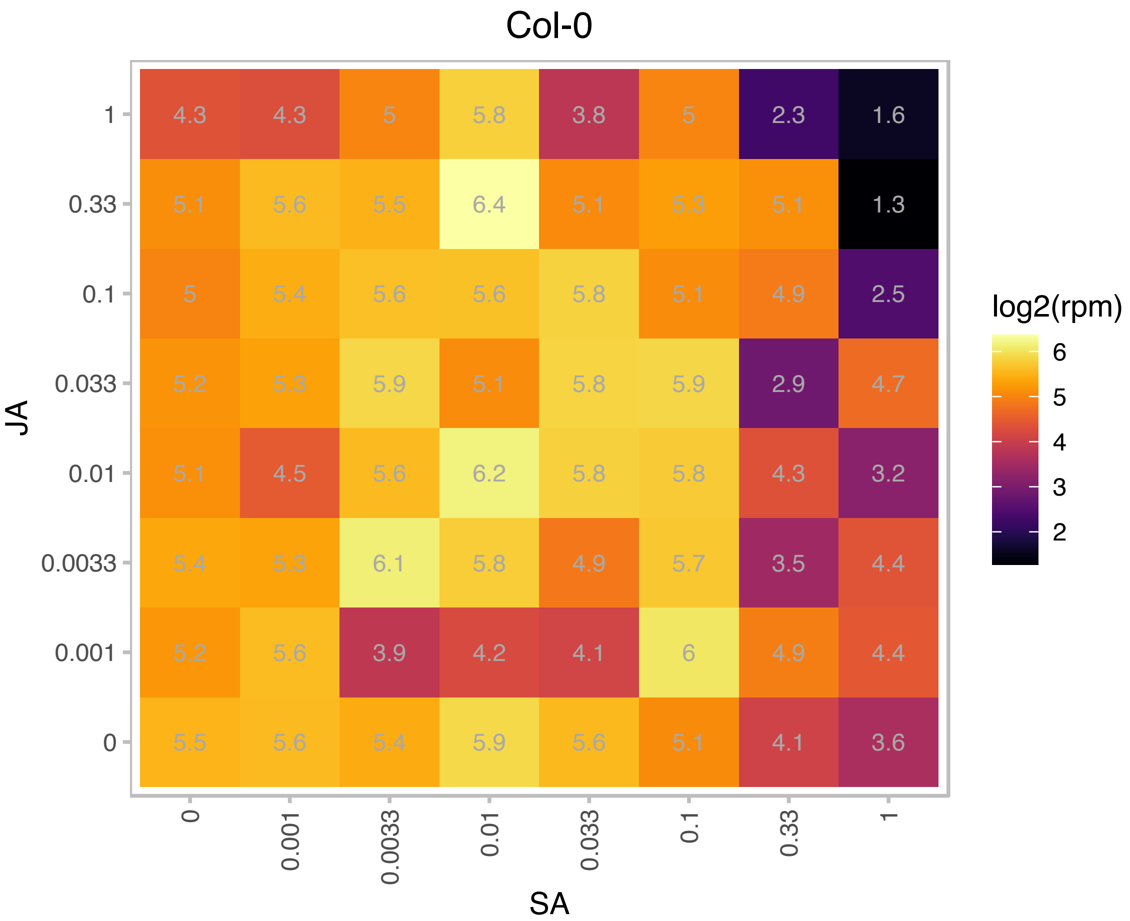


| | |
|---|---|
| Gene | AT2G35120 |
| is.expressed_genes | TRUE |
| is.responsive_genes | TRUE |
| is.apcluster | F-04 |

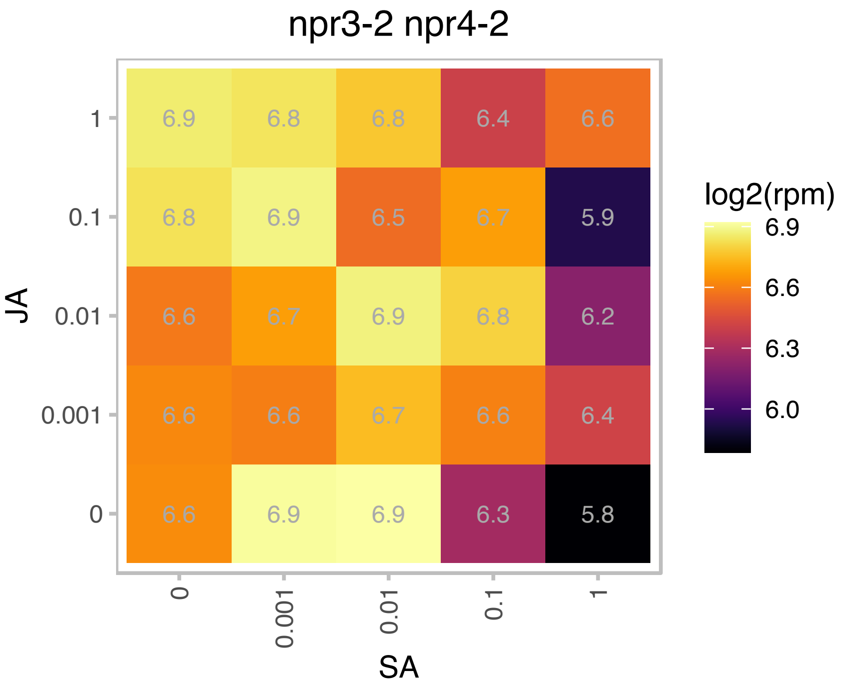


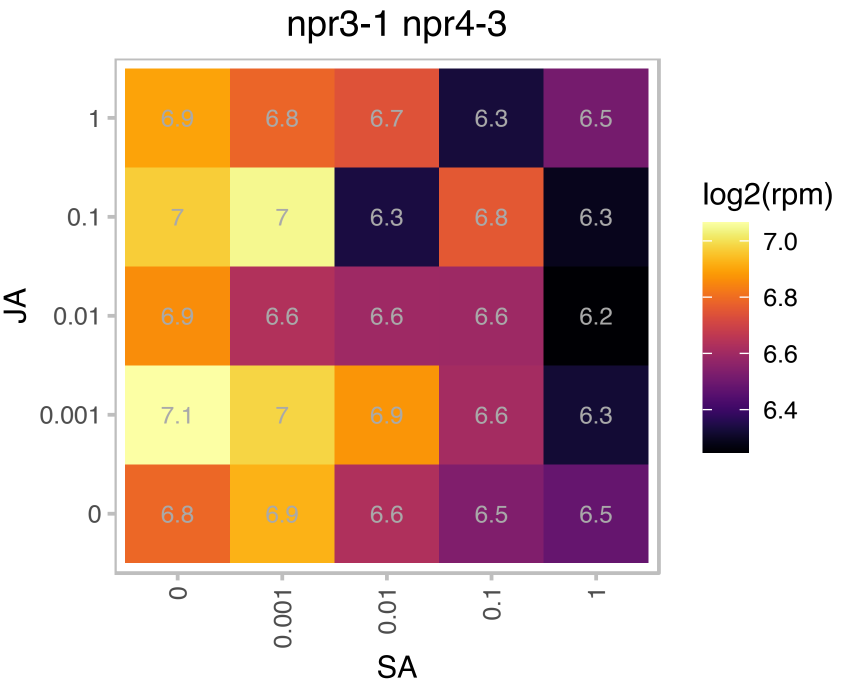

### AT2G36880.1 : MAT3

methionine adenosyltransferase 3

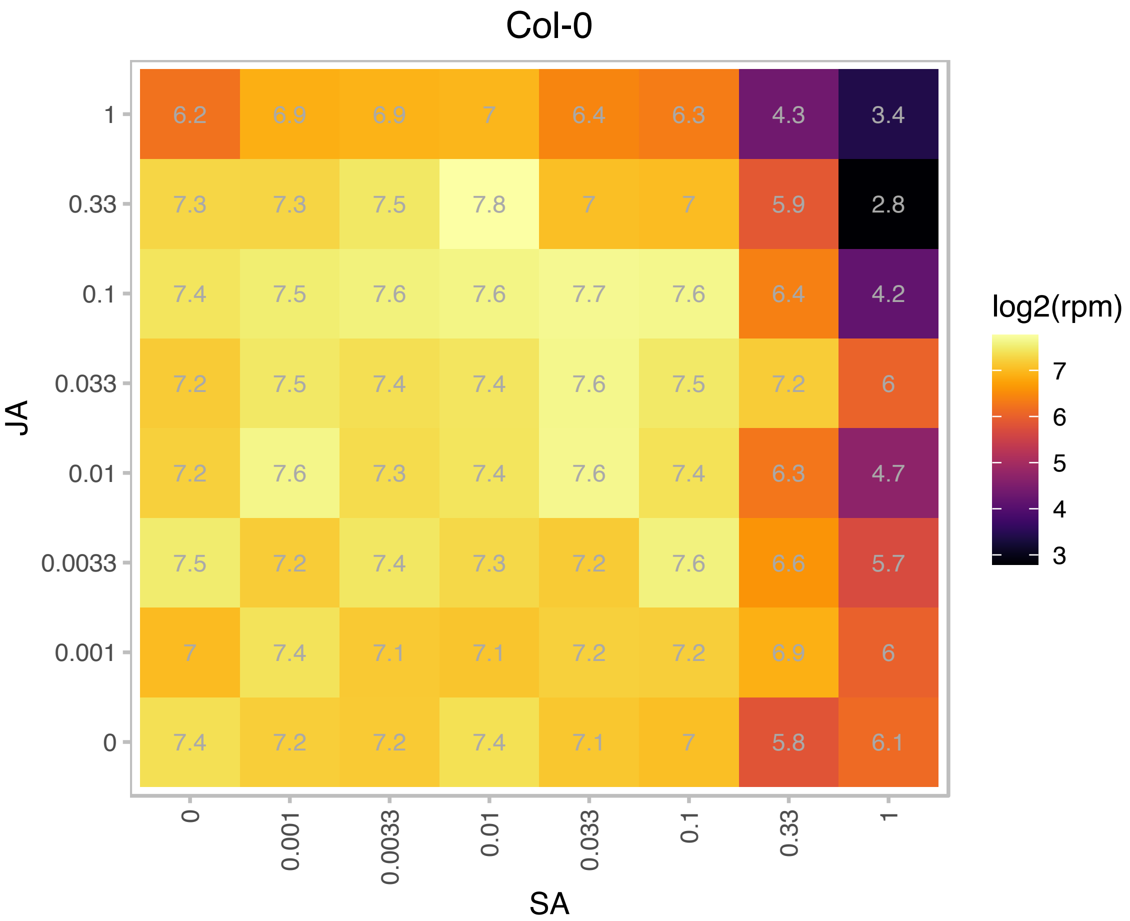


| | |
|---|---|
| Gene | AT2G36880 |
| is.expressed_genes | TRUE |
| is.responsive_genes | TRUE |
| is.apcluster | F-04 |

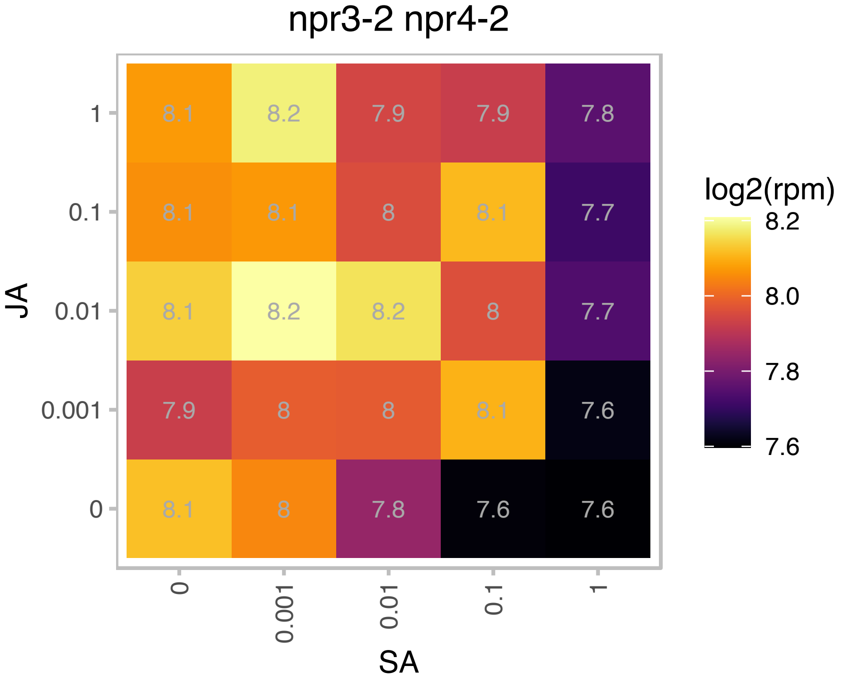


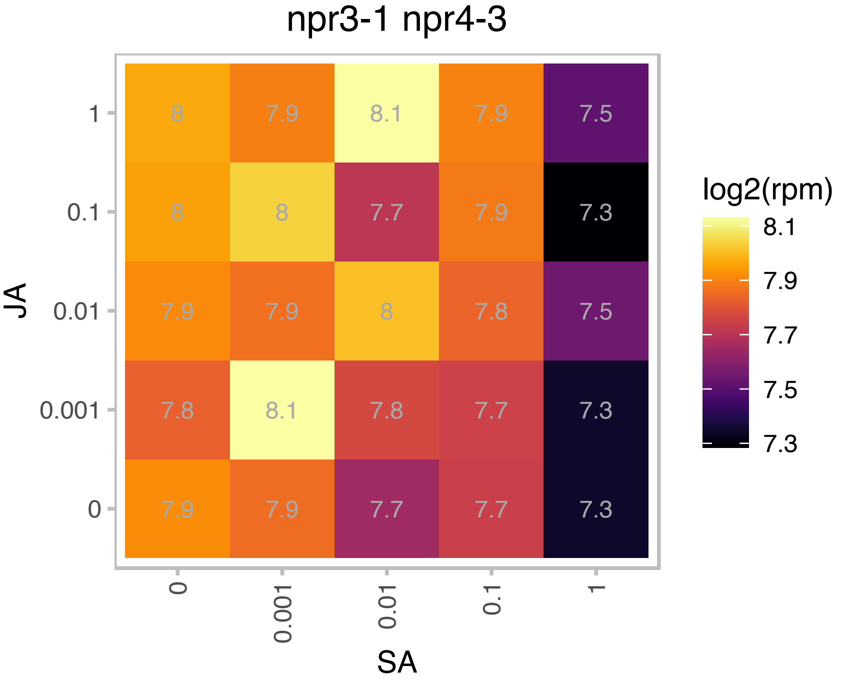

# AT2G43630.1 : NA

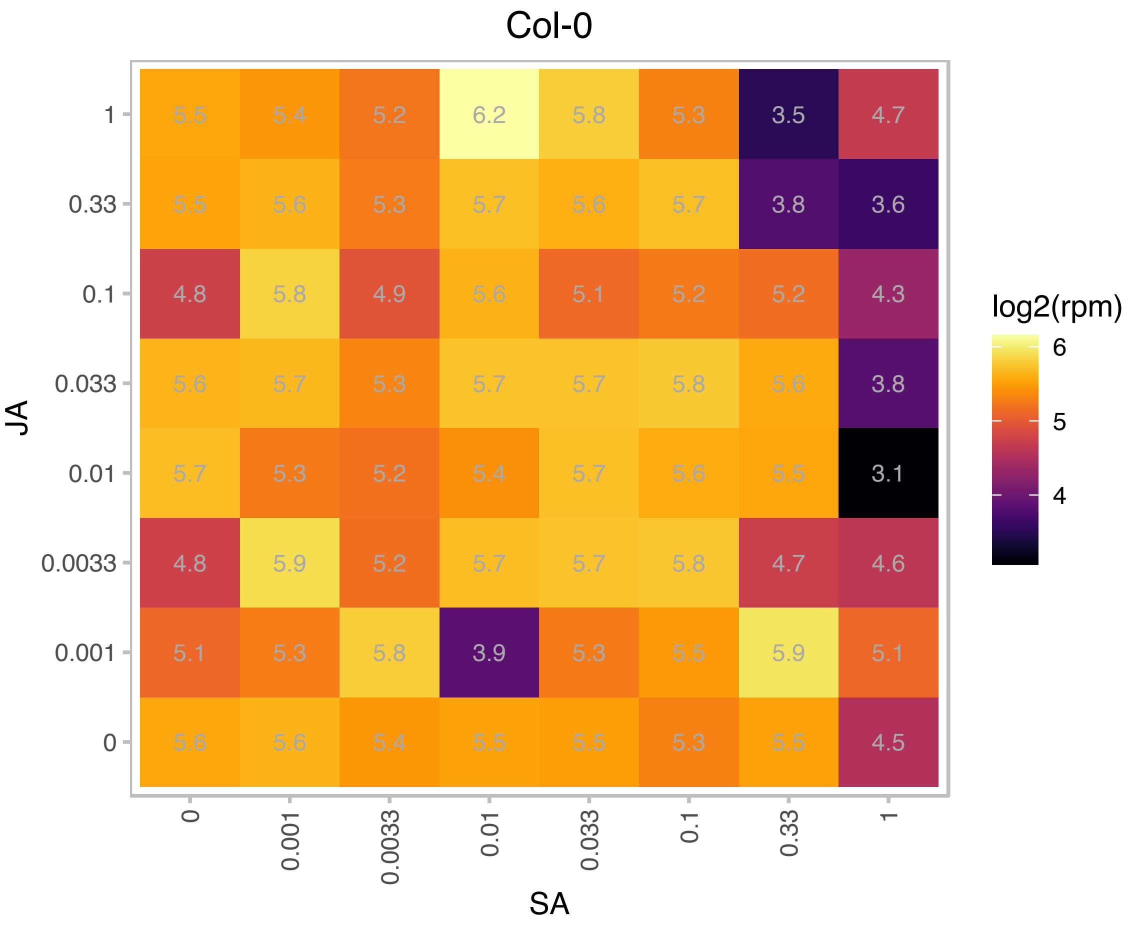

| | |
|---|---|
| Gene | AT2G43630 |
| is.expressed_genes | TRUE |
| is.responsive_genes | TRUE |
| is.apcluster | F-04 |

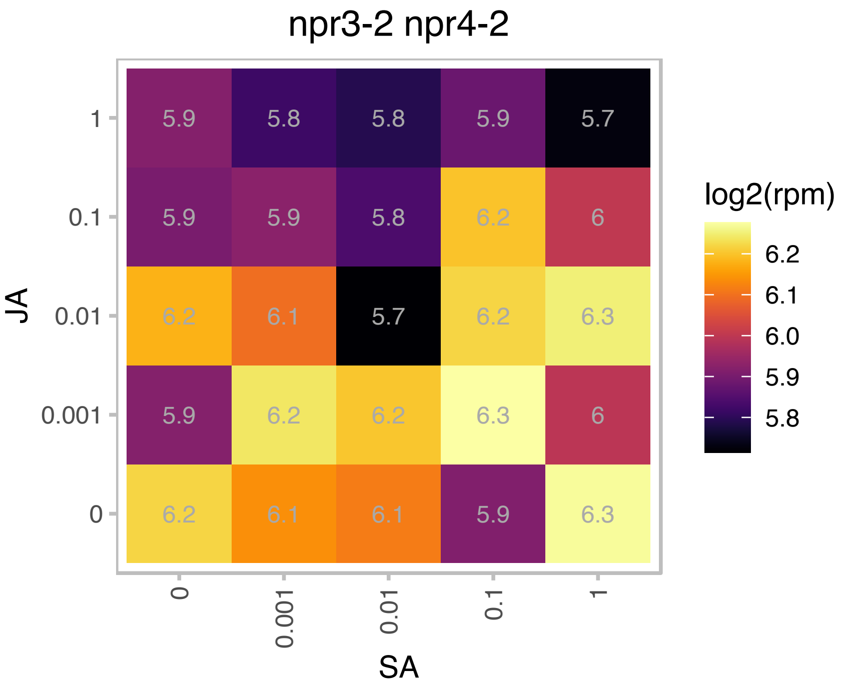

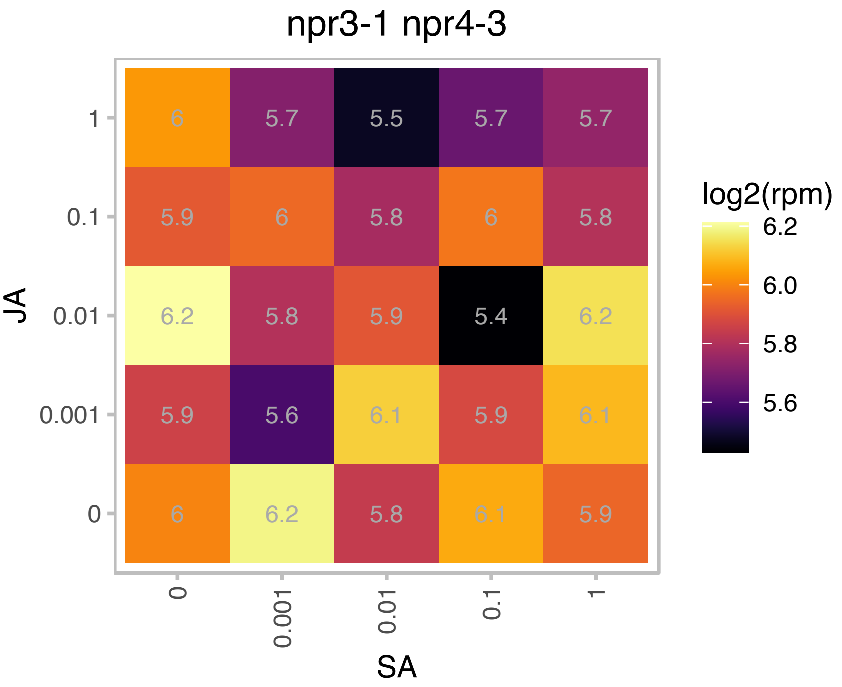

# AT2G45520.1 : NA

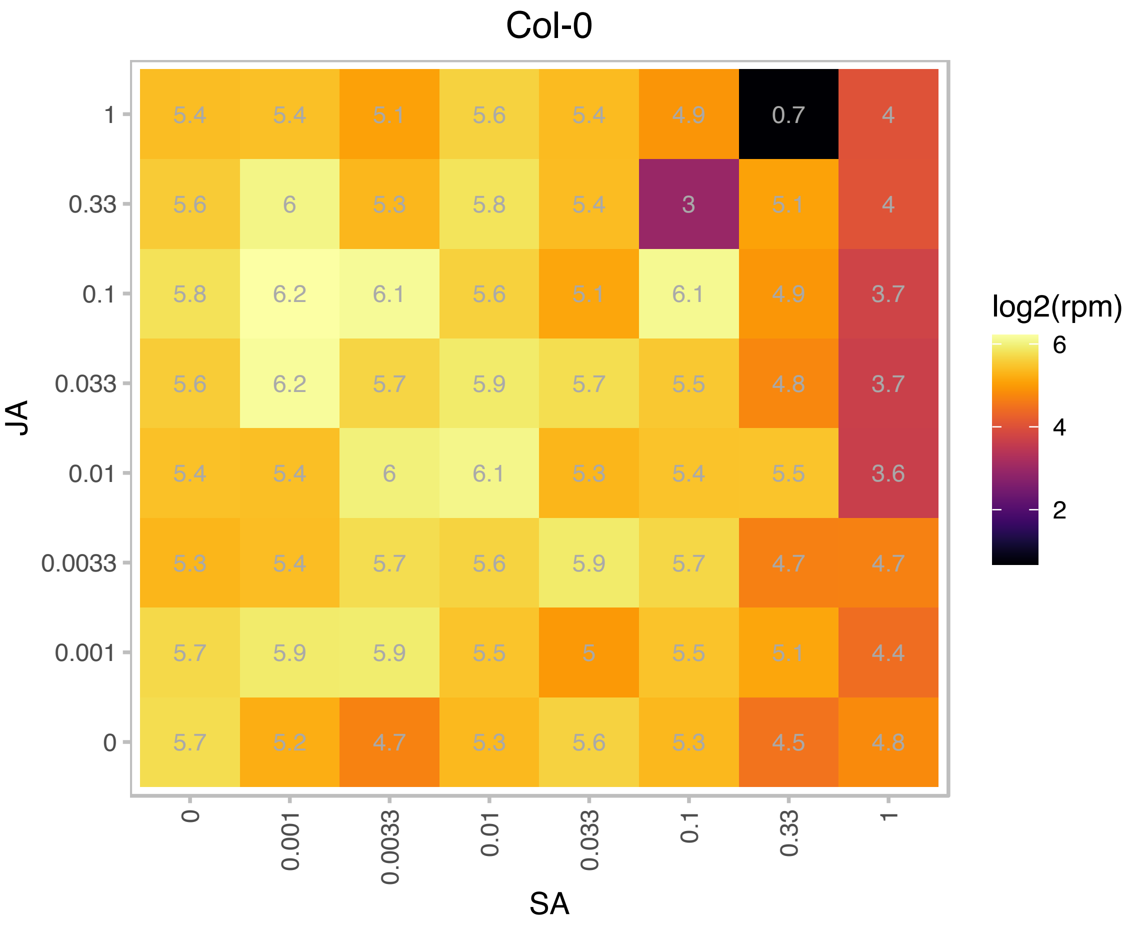


| | |
|---|---|
| Gene | AT2G45520 |
| is.expressed_genes | TRUE |
| is.responsive_genes | TRUE |
| is.apcluster | F-04 |

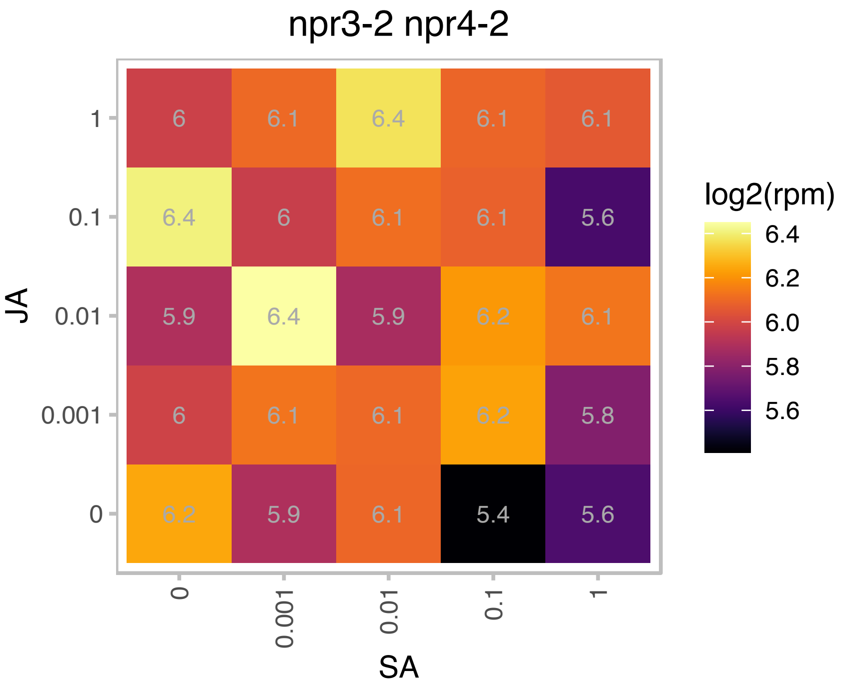


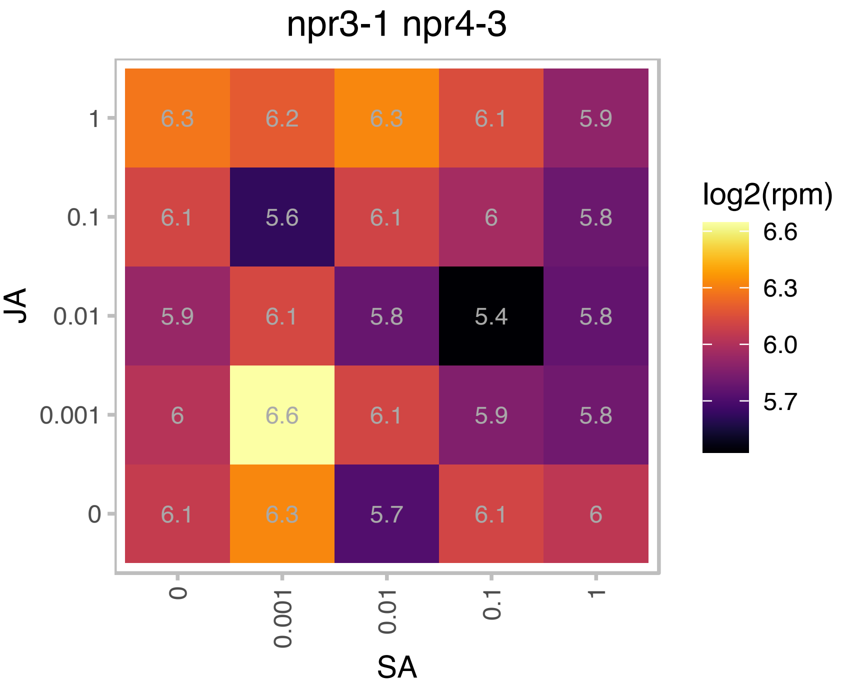

### AT2G47650.1 : UXS4

UDP-xylose synthase 4
encodes a protein similar to UDP-glucuronic acid decarboxylase. UDP-glucuronic acid decarboxylase produces UDP-xylose, which is a substrate for many cell wall carbohydrates including hemicellulose and pectin. UDP-xylose is also known to feedback regulate several cell wall biosynthetic enzymes.

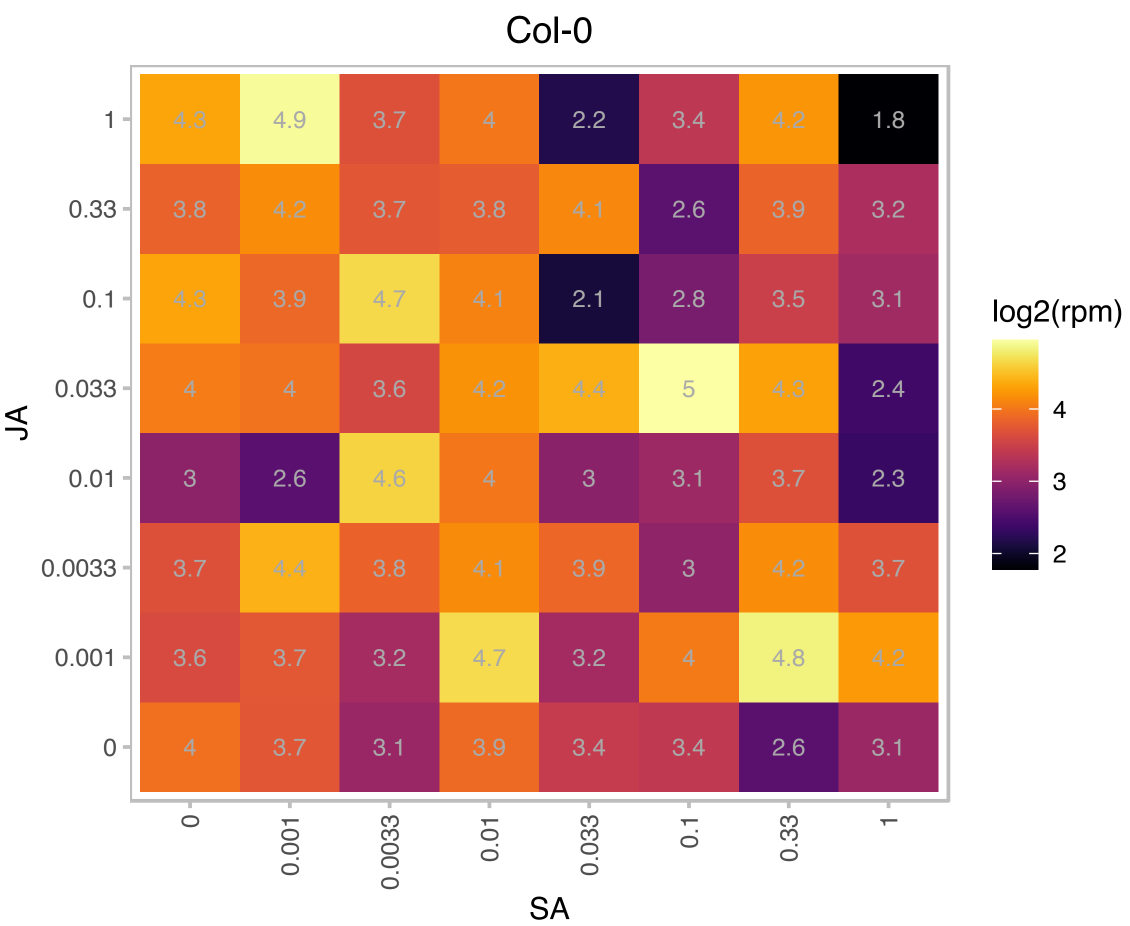


| | |
|---|---|
| Gene | AT2G47650 |
| is.expressed_genes | TRUE |
| is.responsive_genes | TRUE |
| is.apcluster | F-04 |

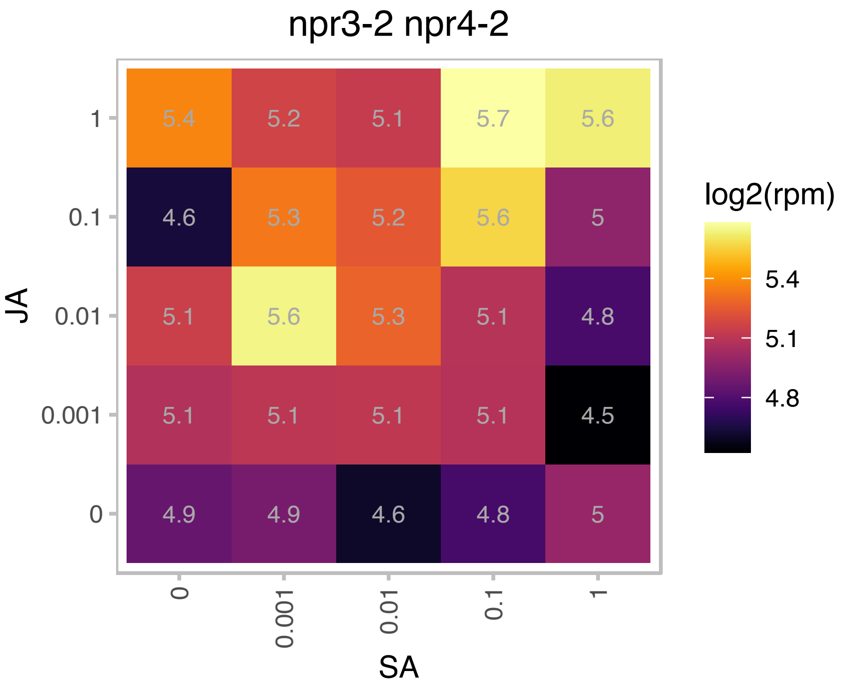


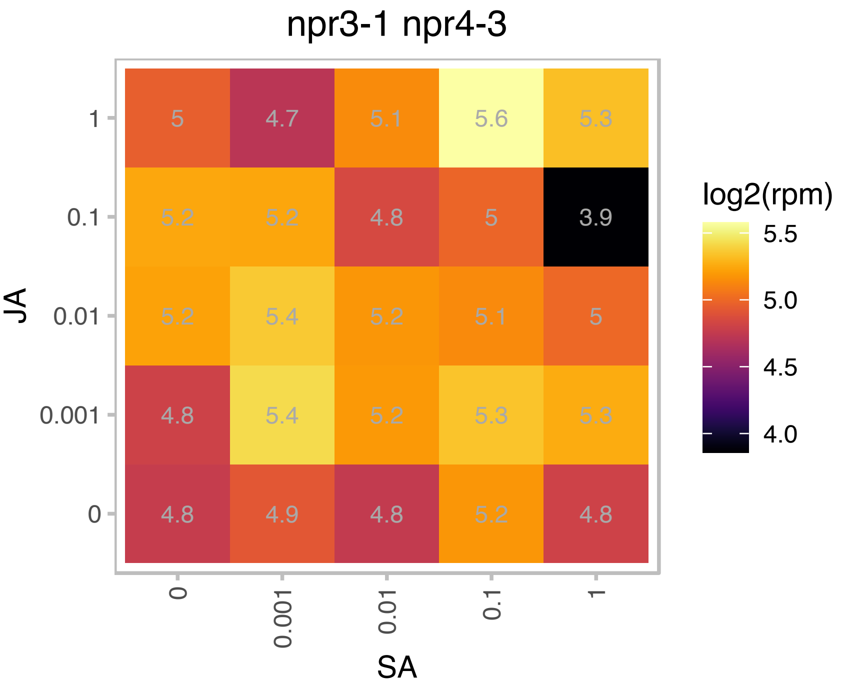

# AT2G48090.1 : NA

| | |
|---|---|
| Gene | AT2G48090 |
| is.expressed_genes | TRUE |
| is.responsive_genes | TRUE |
| is.apcluster | F-04 |

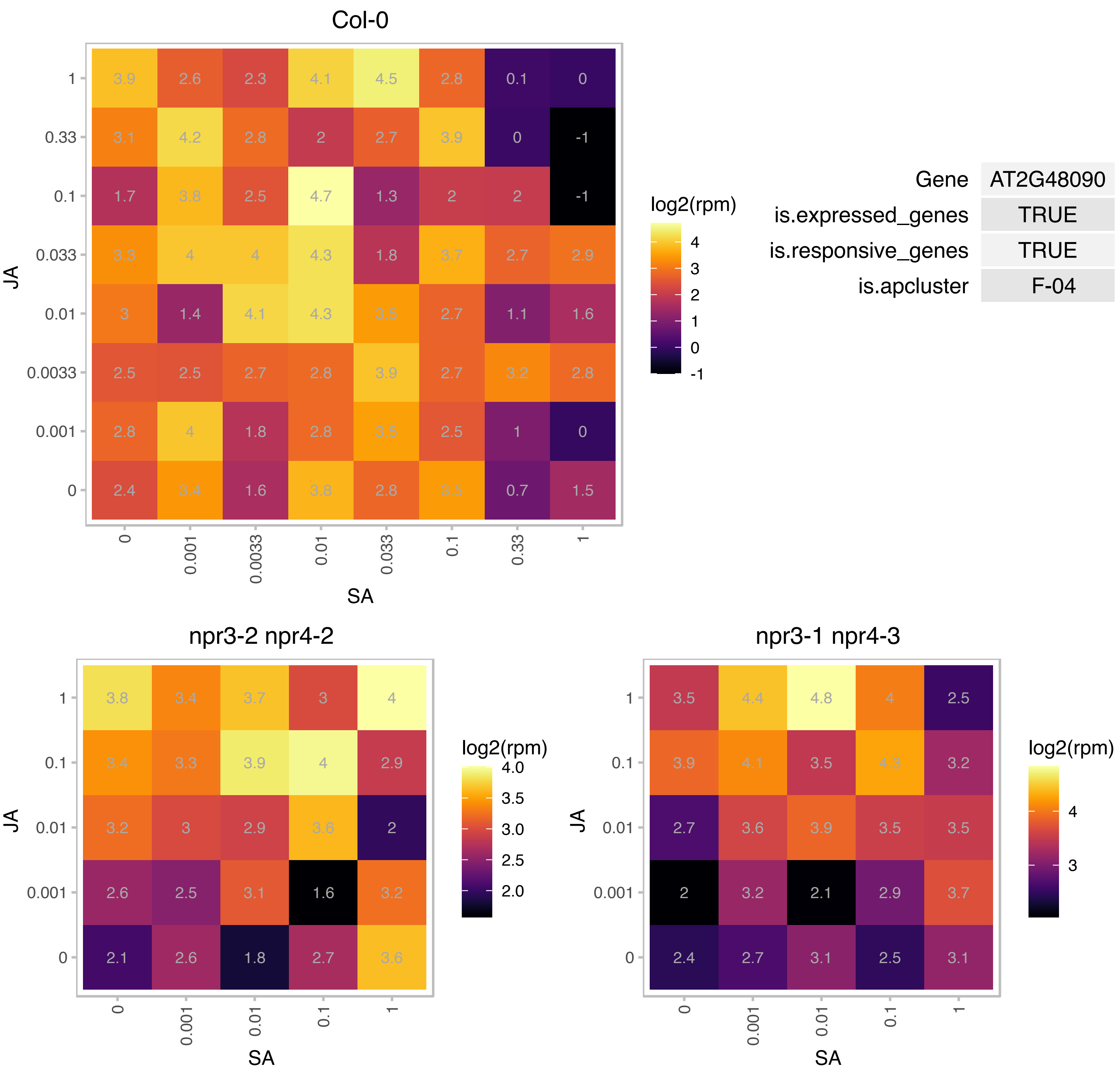

# AT3G01160.1 : NA

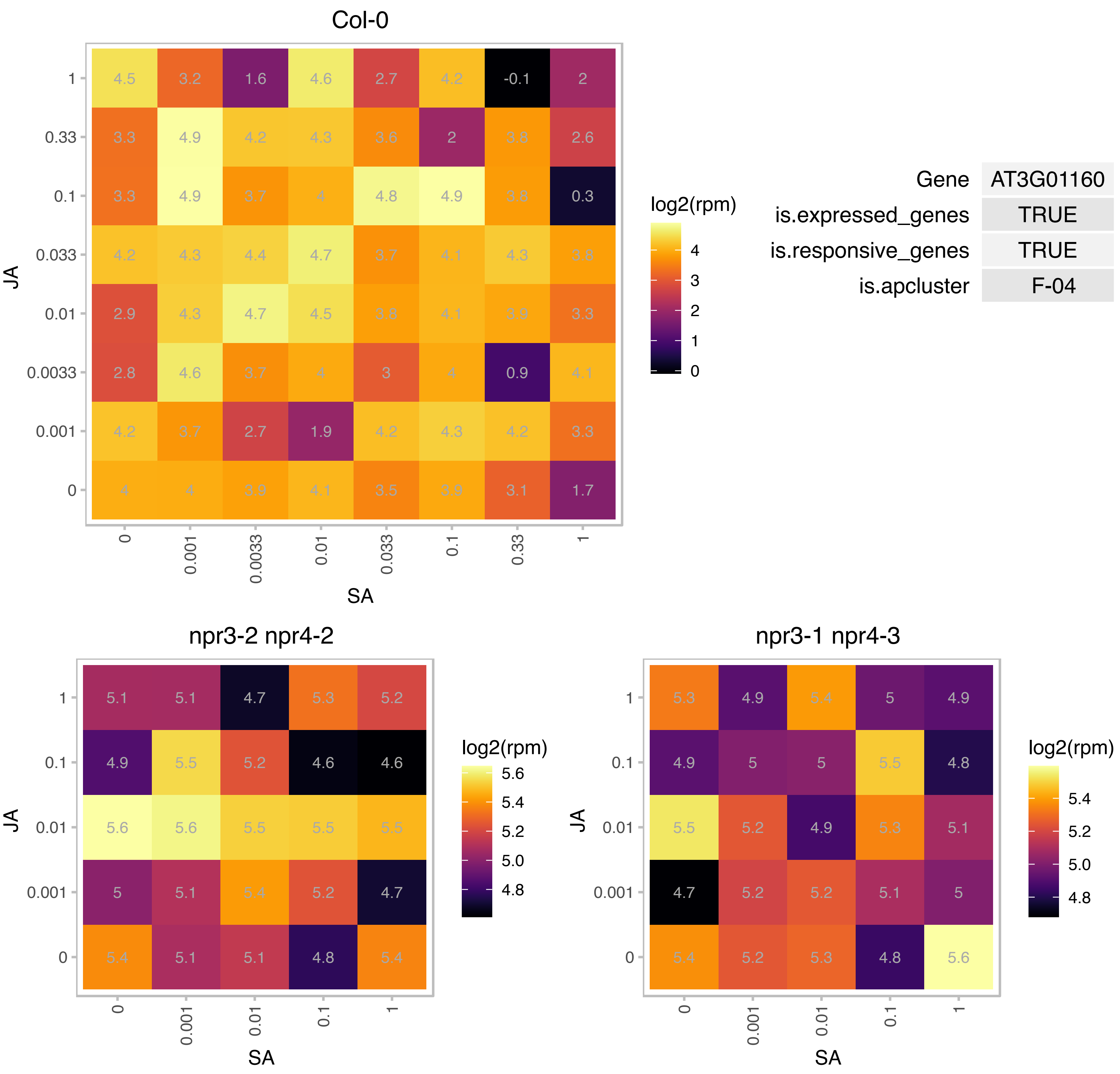

# AT3G06355.1 : NA

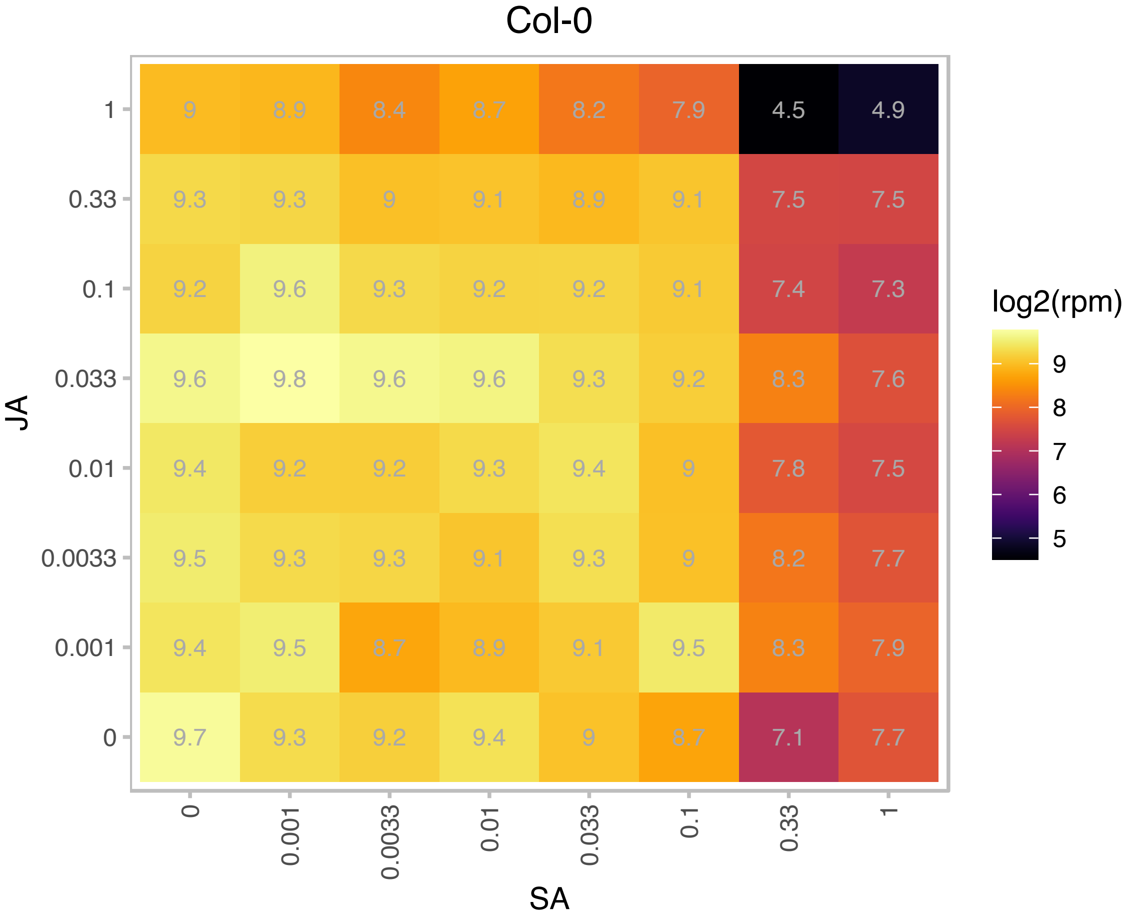


| Gene | AT3G06355 |
|---|---|
| is.expressed_genes | TRUE |
| is.responsive_genes | TRUE |
| is.apcluster | F-04 |

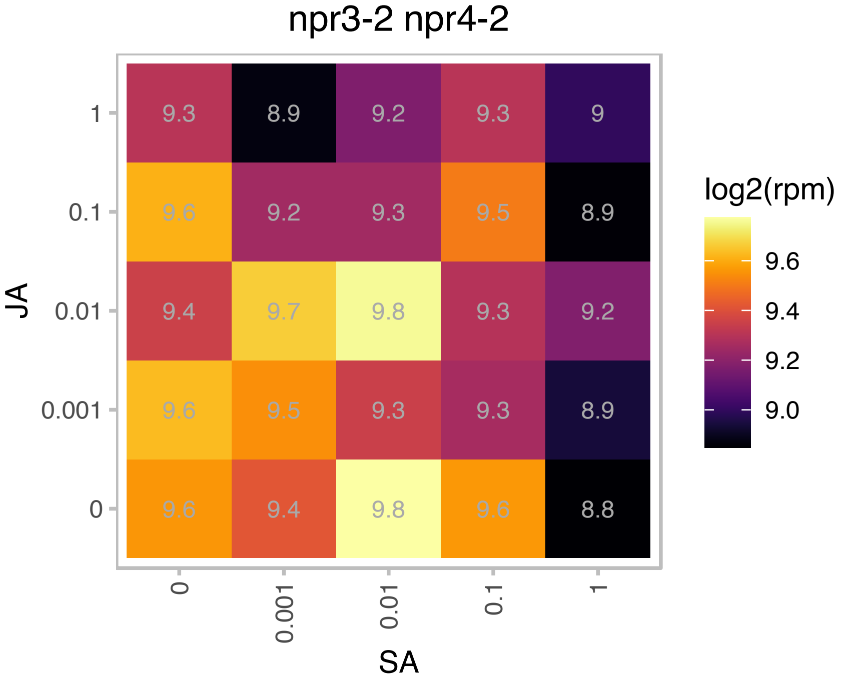


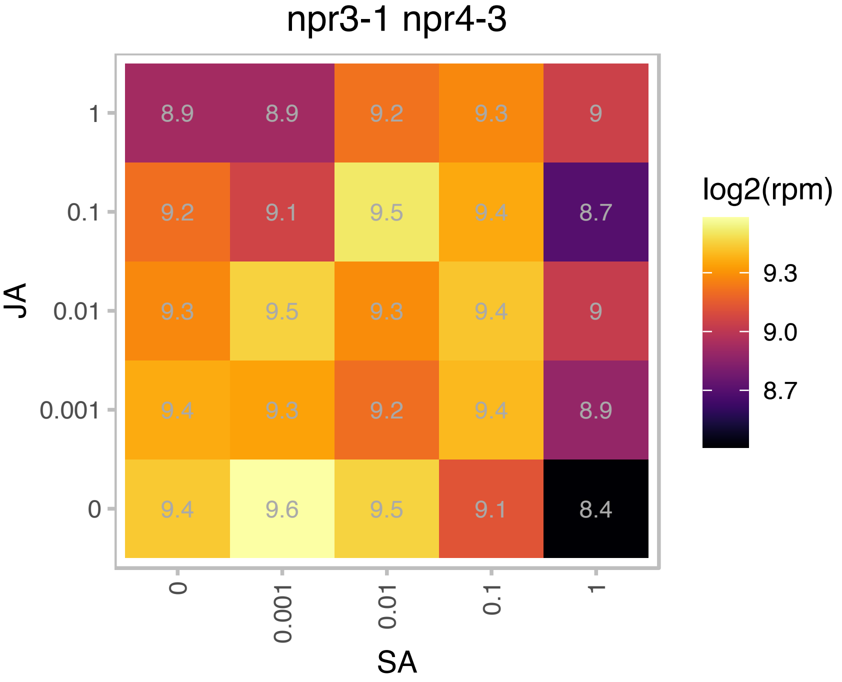

# AT3G08020.1 : NA

PHD finger family protein

Col-0

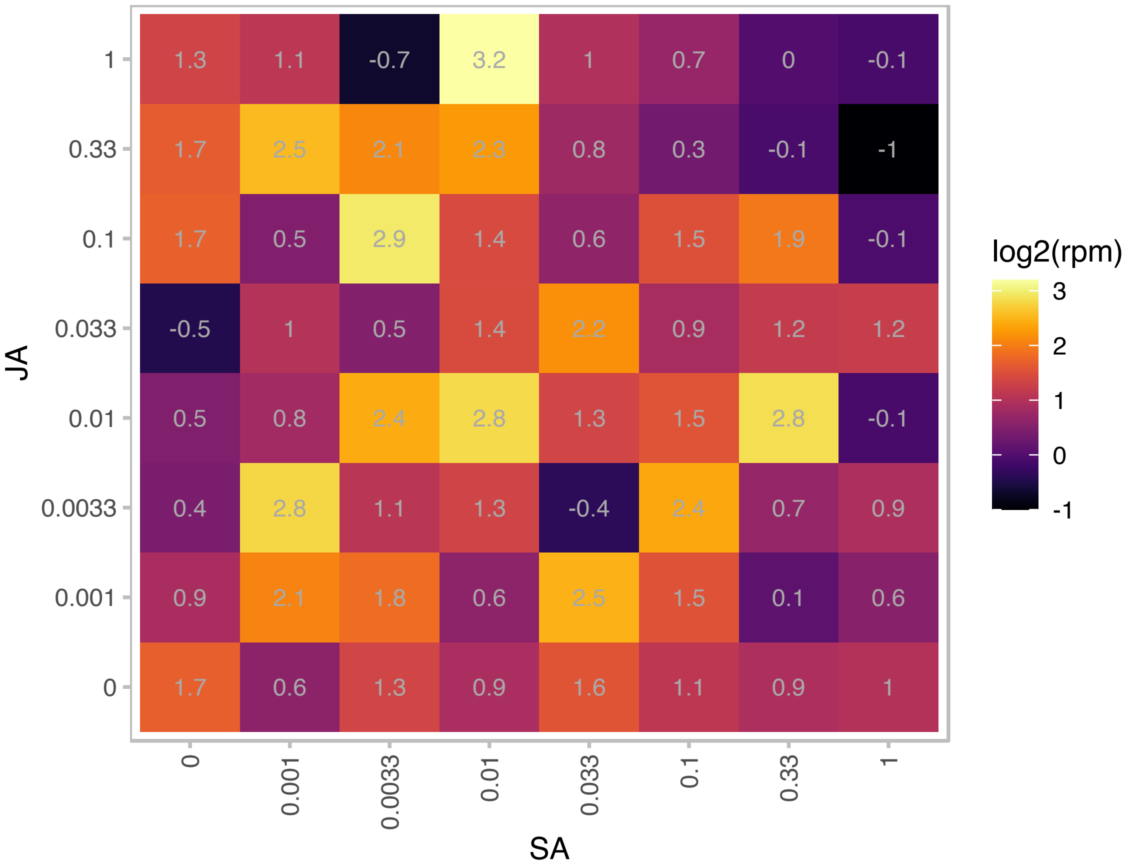


| | |
|---|---|
| Gene | AT3G08020 |
| is.expressed_genes | TRUE |
| is.responsive_genes | TRUE |
| is.apcluster | F-04 |

npr3-2 npr4-2

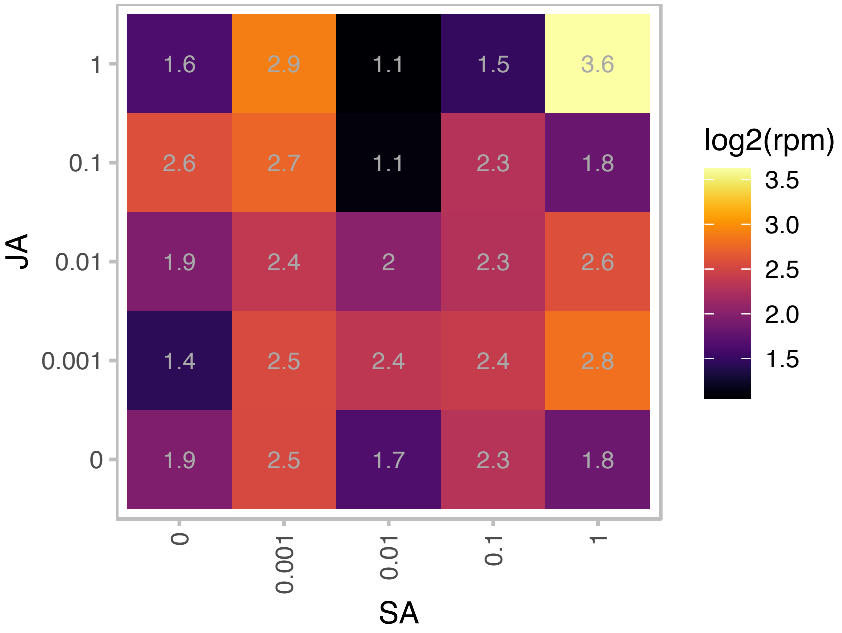


npr3-1 npr4-3

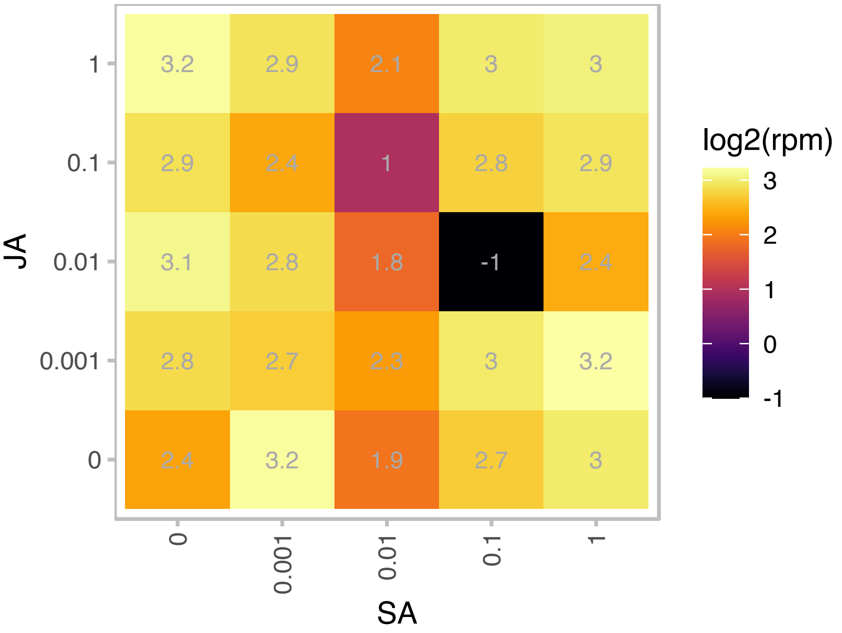

### AT3G08580.1 : AAC1

ADP/ATP carrier 1

mitochondrial ADP/ATP carrier

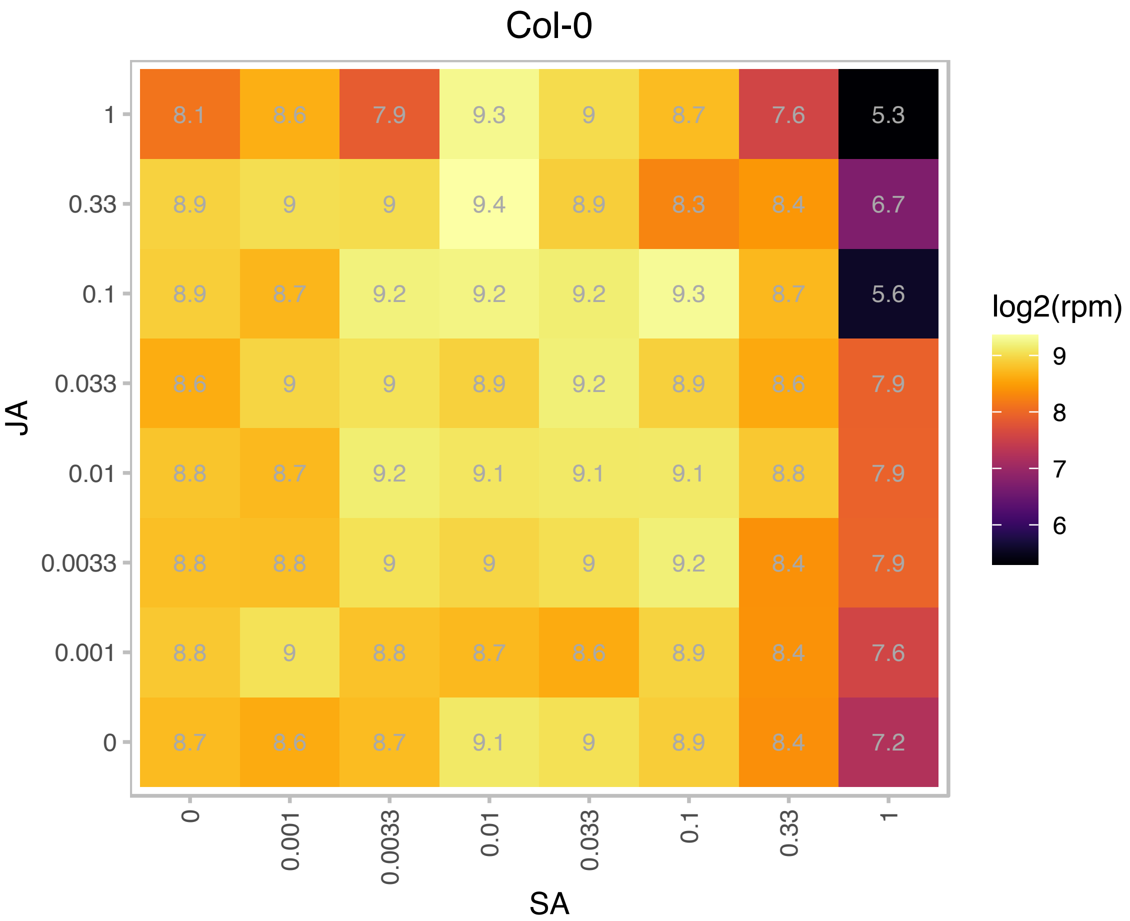


| | |
|---|---|
| Gene | AT3G08580 |
| is.expressed_genes | TRUE |
| is.responsive_genes | TRUE |
| is.apcluster | F-04 |

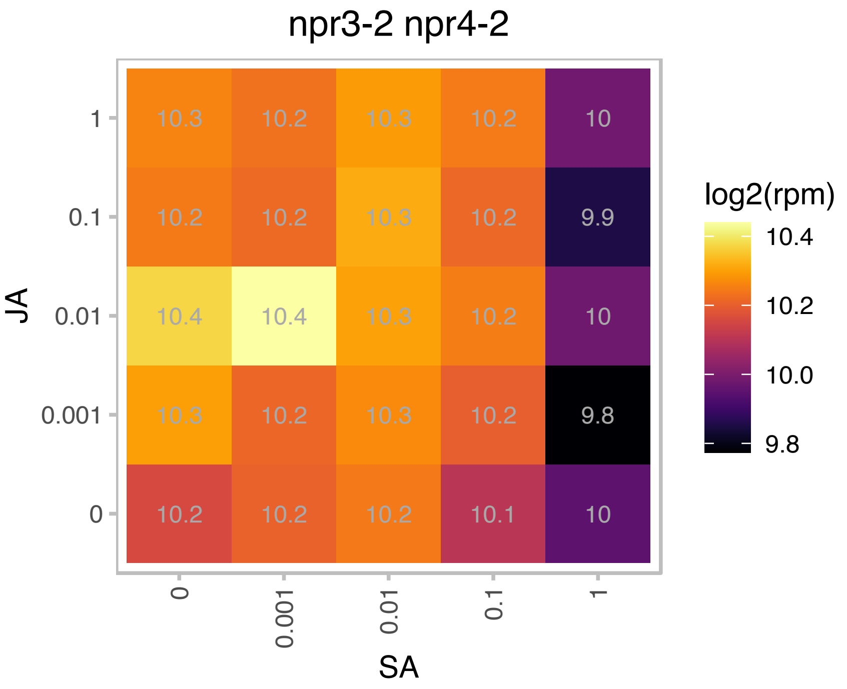


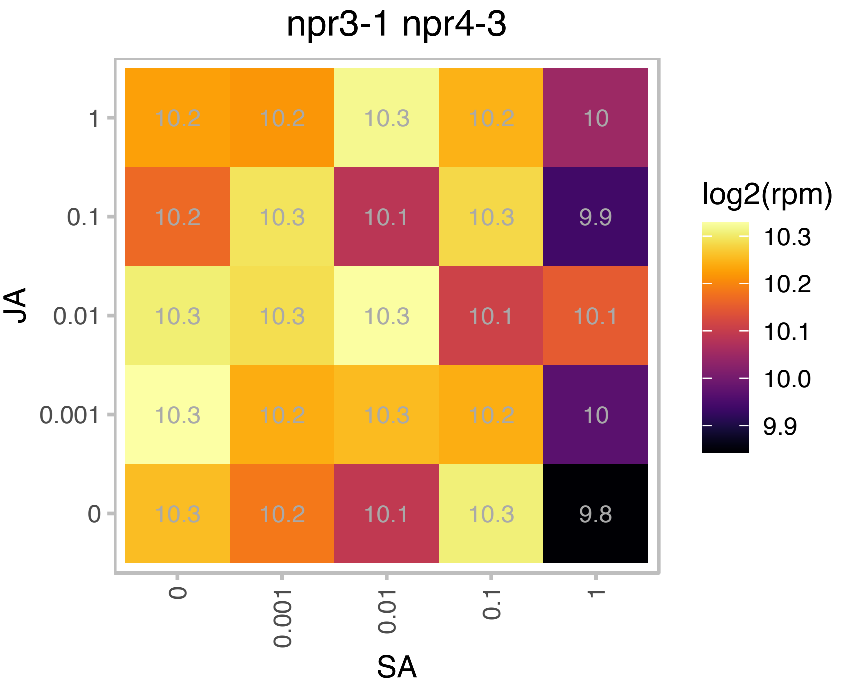

# AT3G09455.1 : NA

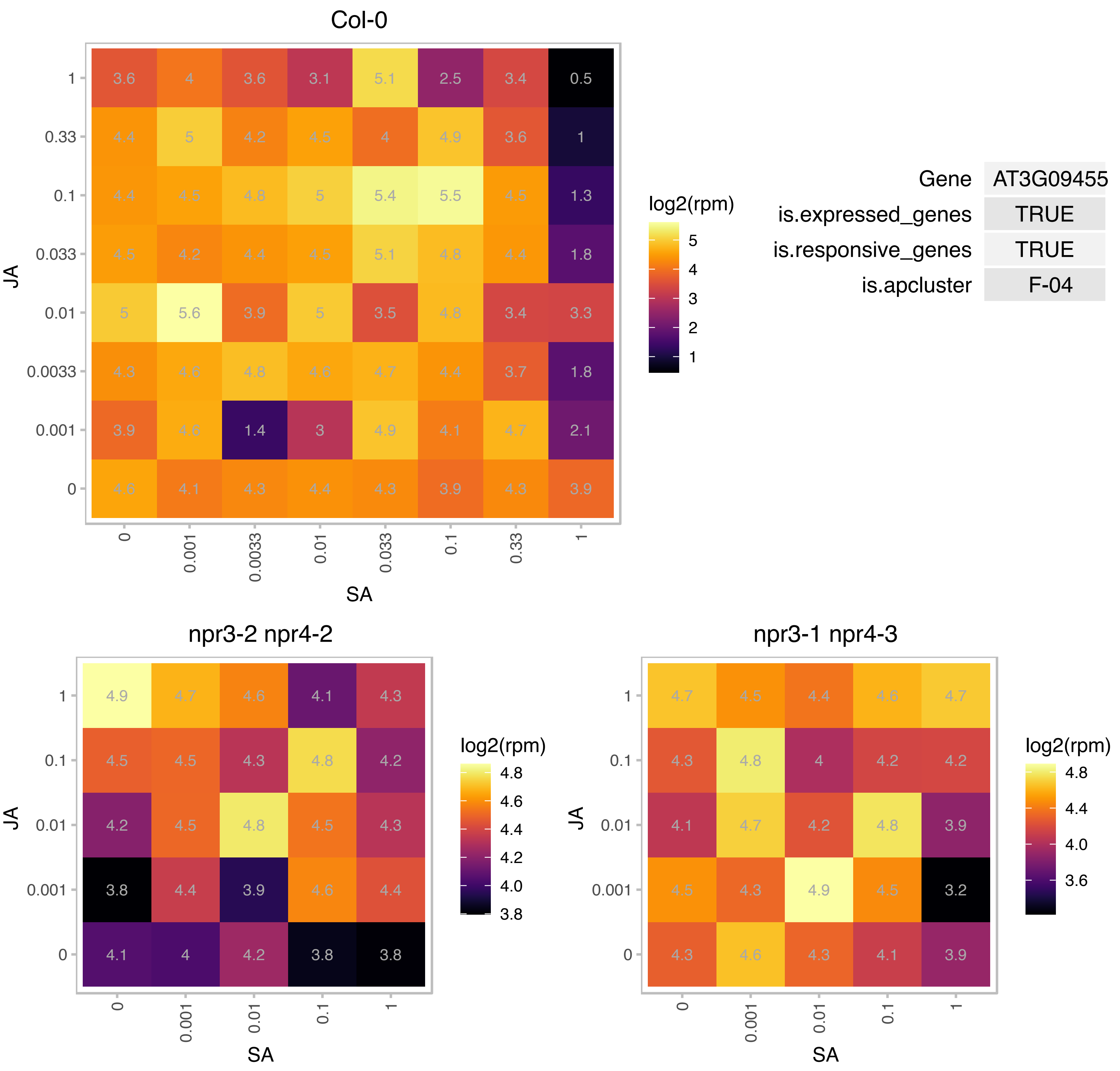


| | |
|---|---|
| Gene | AT3G09455 |
| is.expressed_genes | TRUE |
| is.responsive_genes | TRUE |
| is.apcluster | F-04 |

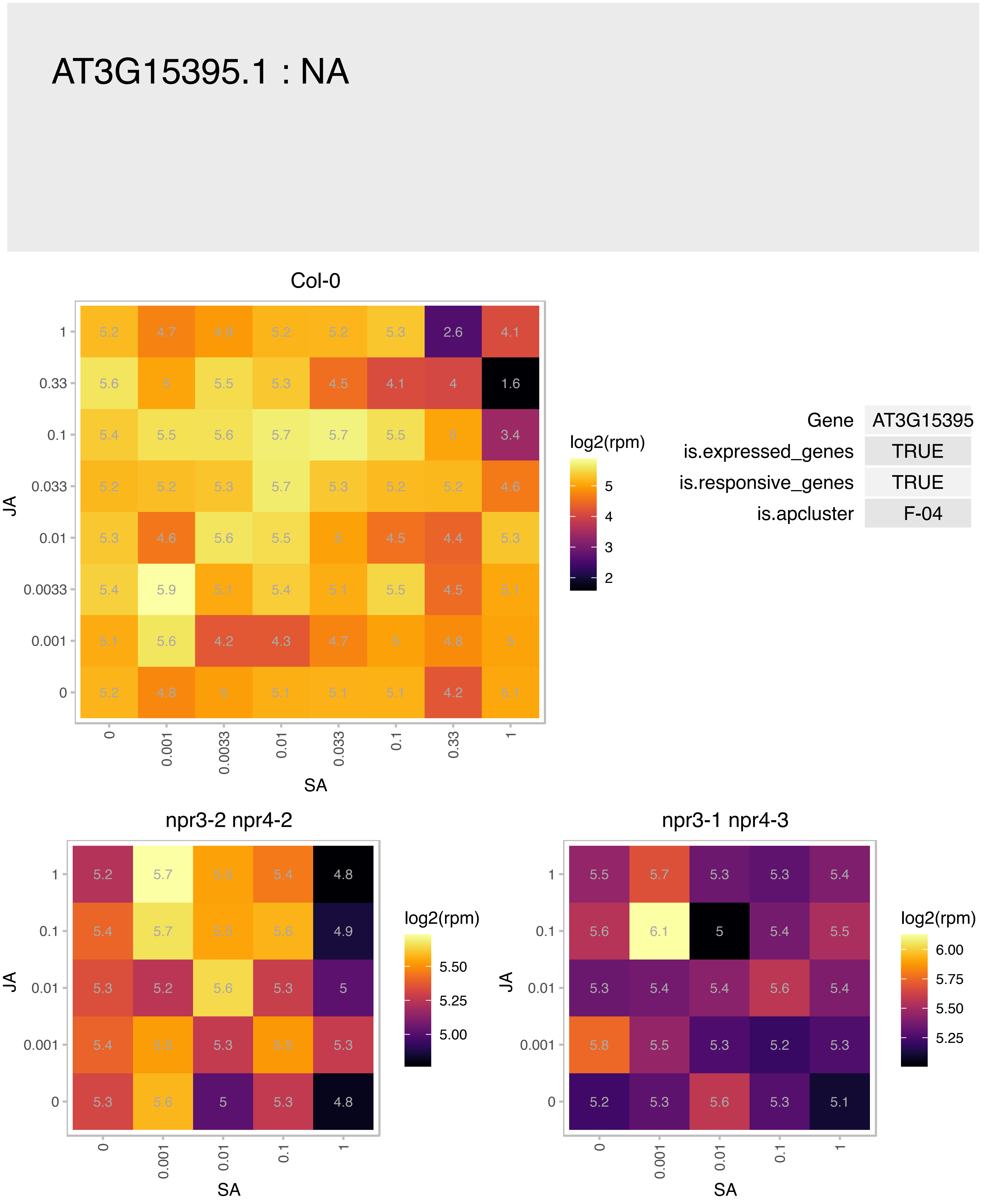
AT3G15395.1 : NA
Col-0
log2(rpm)
JA
SA
Gene AT3G15395
is.expressed_genes TRUE
is.responsive_genes TRUE
is.apcluster F-04
npr3-2 npr4-2
npr3-1 npr4-3

### AT3G16290.1 : EMB2083

AAA-type ATPase family protein

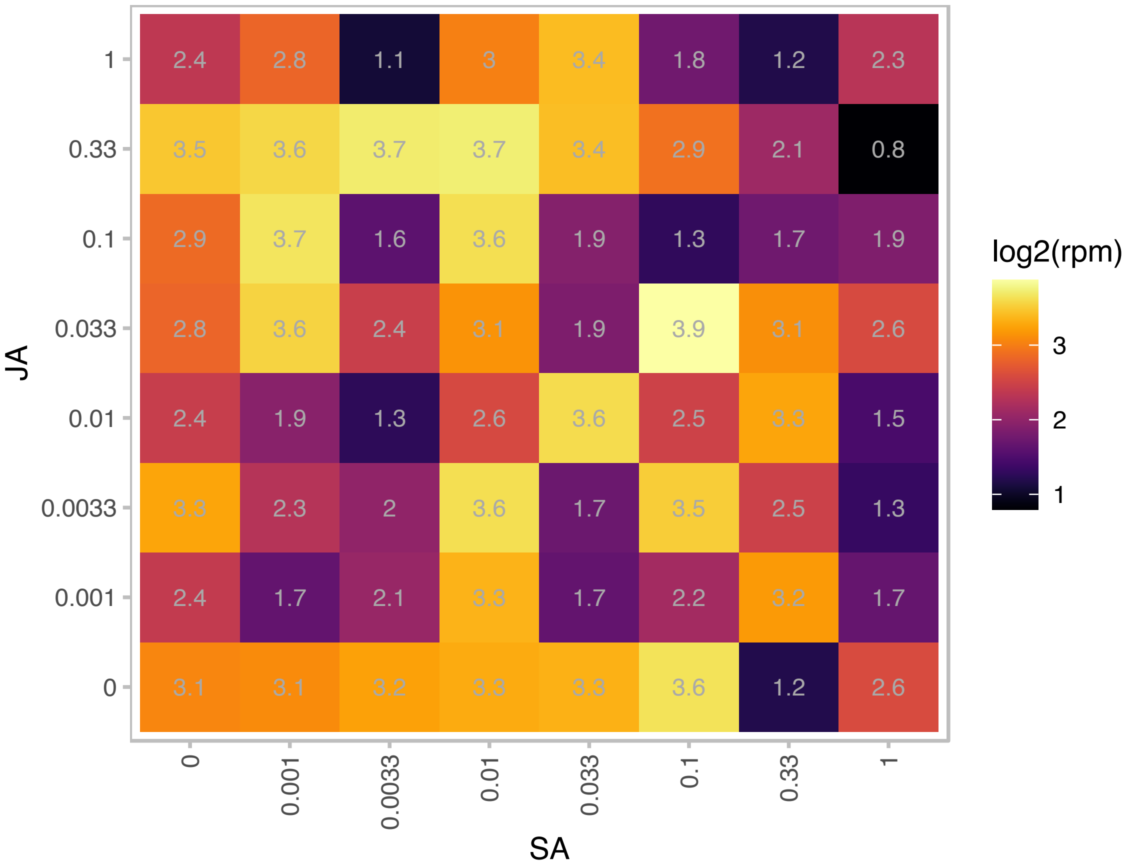


| | |
|---|---|
| Gene | AT3G16290 |
| is.expressed_genes | TRUE |
| is.responsive_genes | TRUE |
| is.apcluster | F-04 |

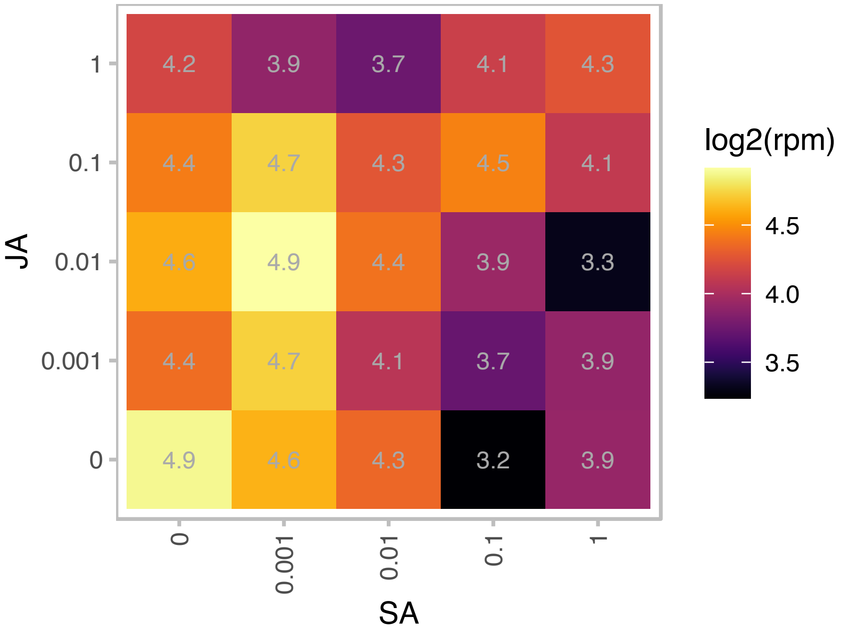


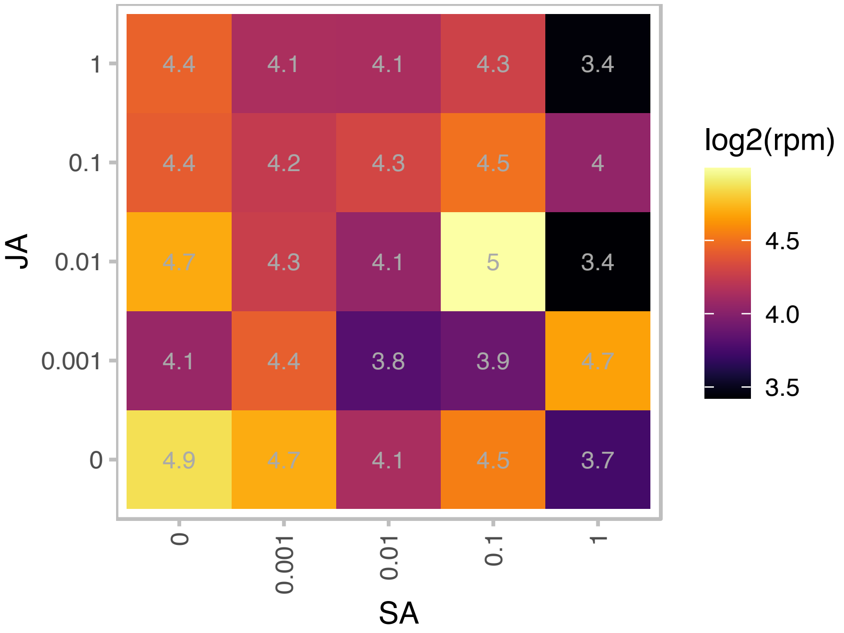

### AT3G16780.1 : RPL19B

Ribosomal protein L19e family protein

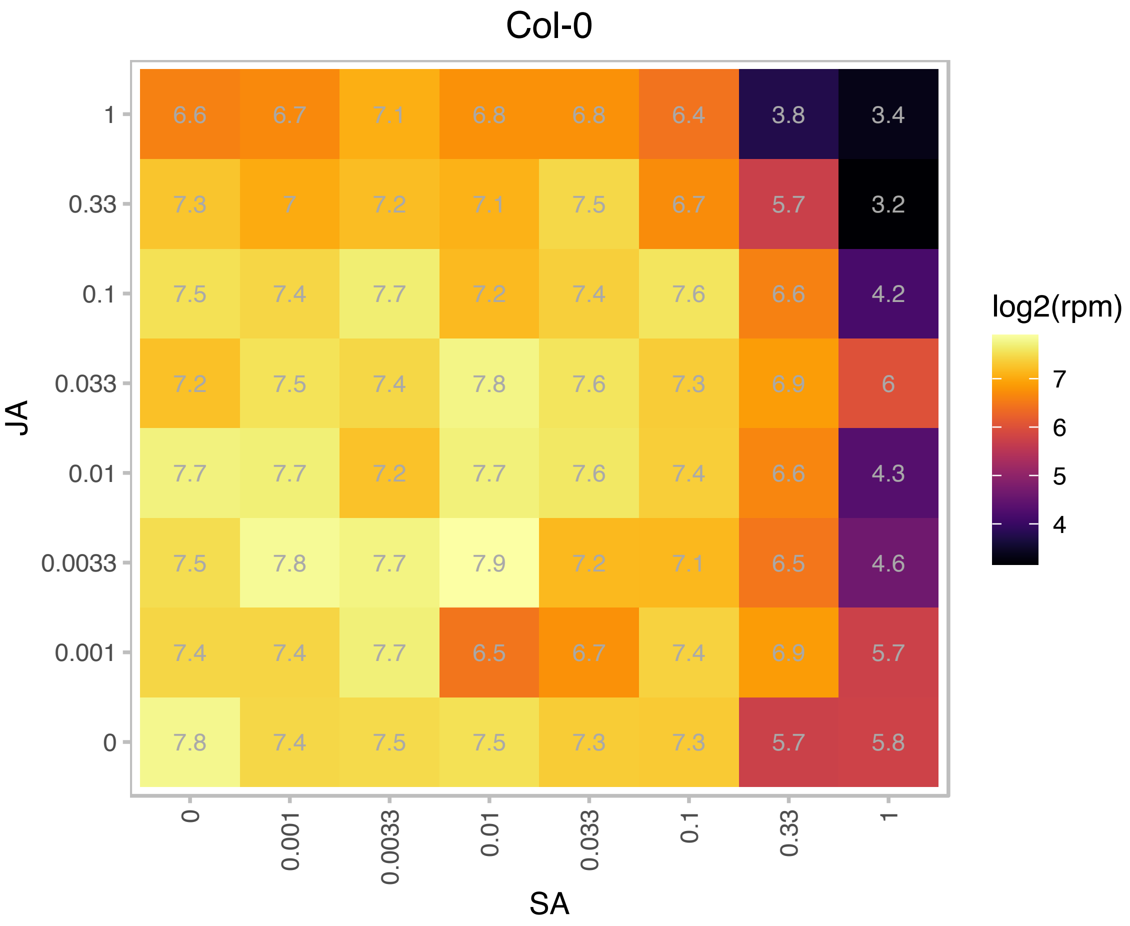


| | |
|---|---|
| Gene | AT3G16780 |
| is.expressed_genes | TRUE |
| is.responsive_genes | TRUE |
| is.apcluster | F-04 |

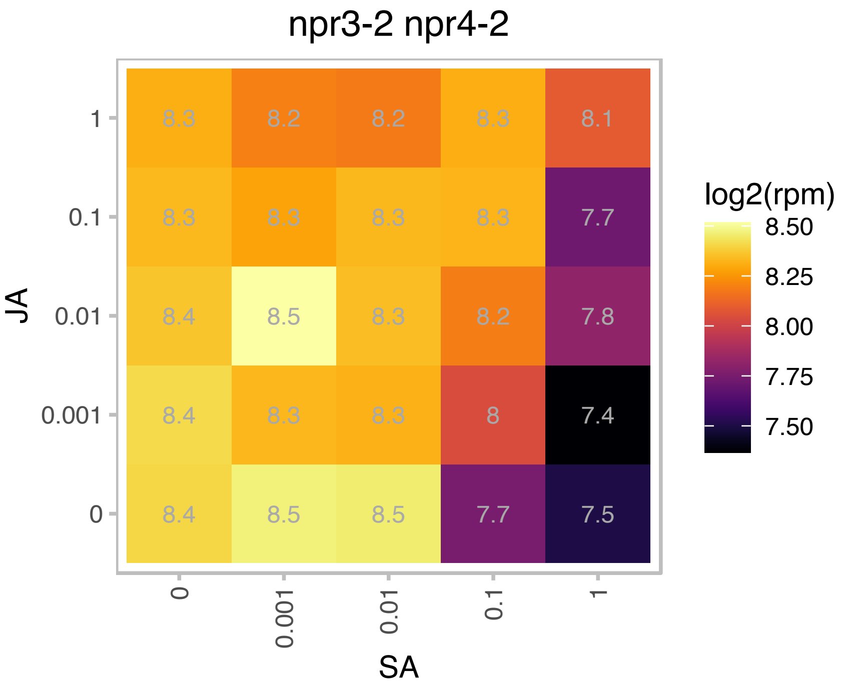


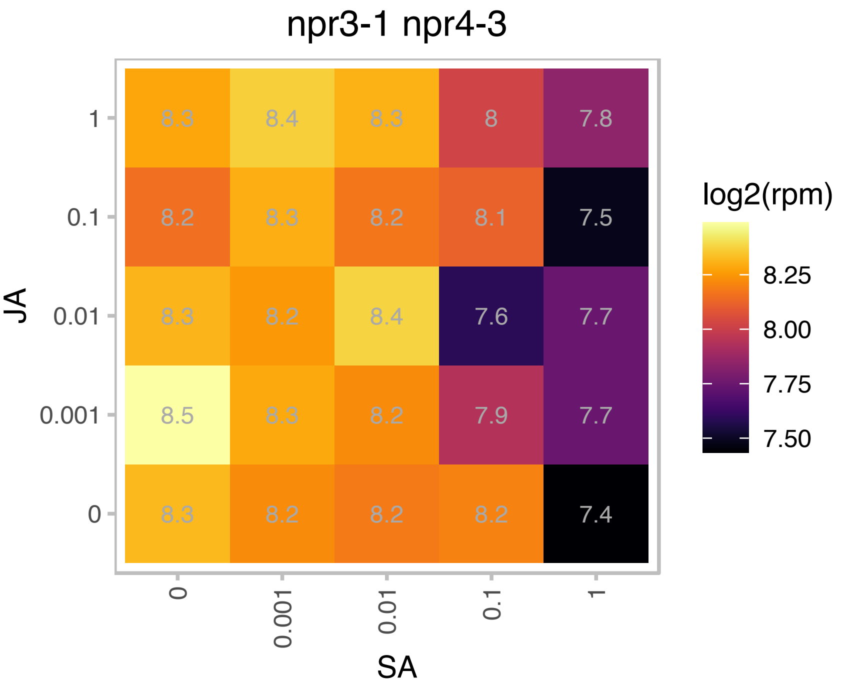

# AT3G18190.1 : NA

TCP-1/cpn60 chaperonin family protein

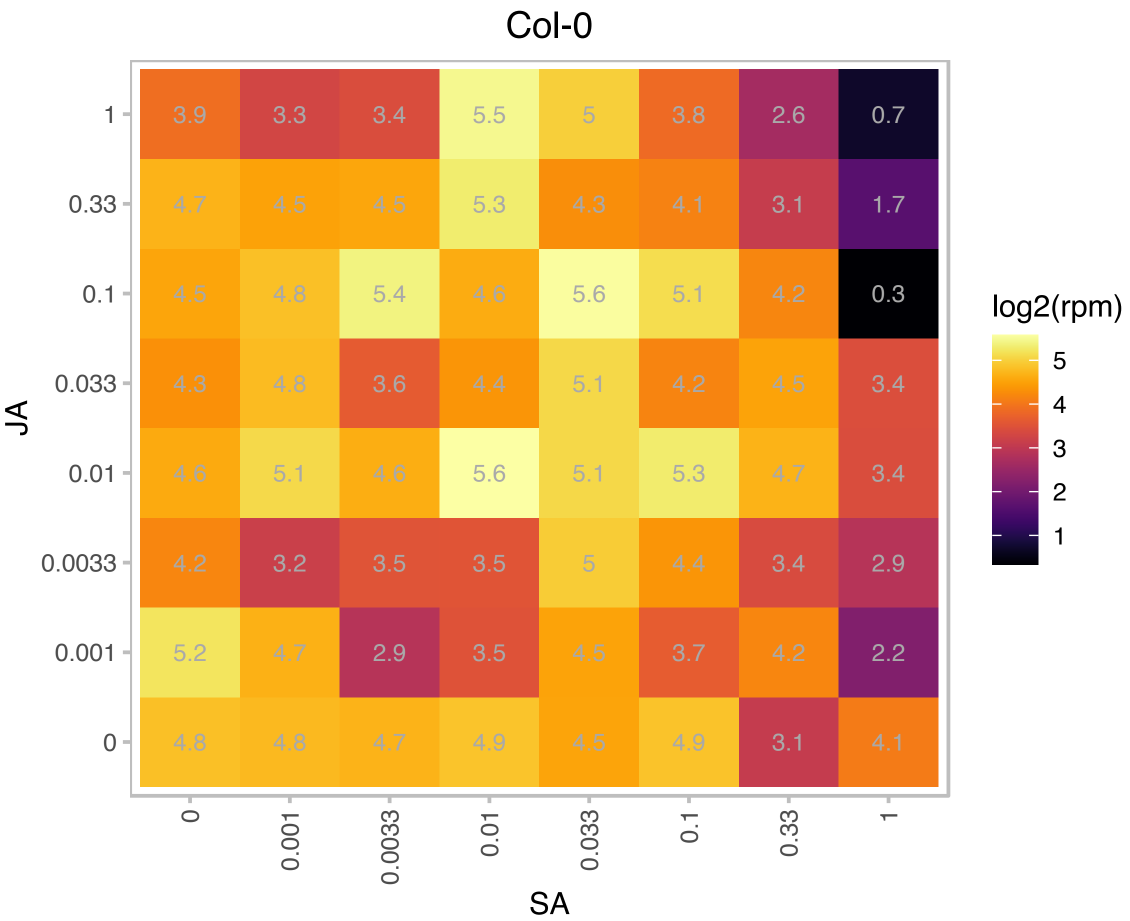


| | |
|---|---|
| Gene | AT3G18190 |
| is.expressed_genes | TRUE |
| is.responsive_genes | TRUE |
| is.apcluster | F-04 |

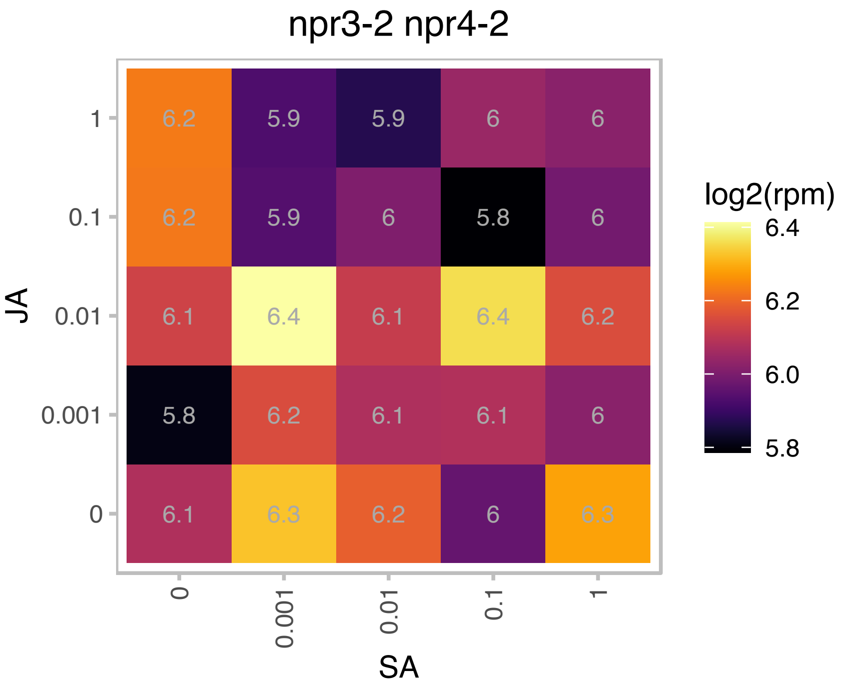


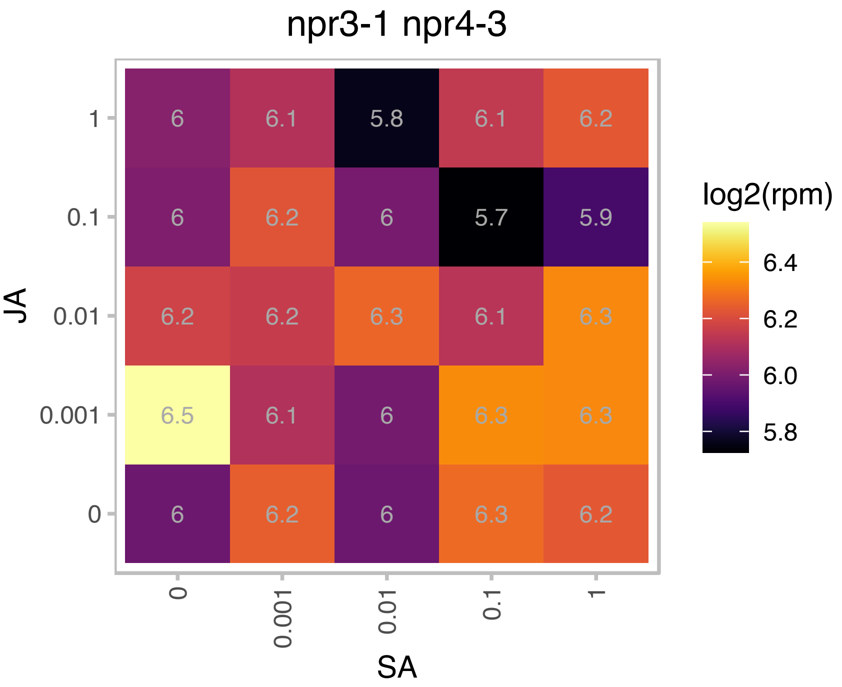

### AT3G20970.1 : ATNFU2, NFU4

NFU domain protein 4

Encodes a protein containing the NFU domain that may be involved in iron-sulfur cluster assembly. Part of a five member gene family, more closely related to NFU5 than to NFU1,2, and 3. Targeted to the mitochondrion.

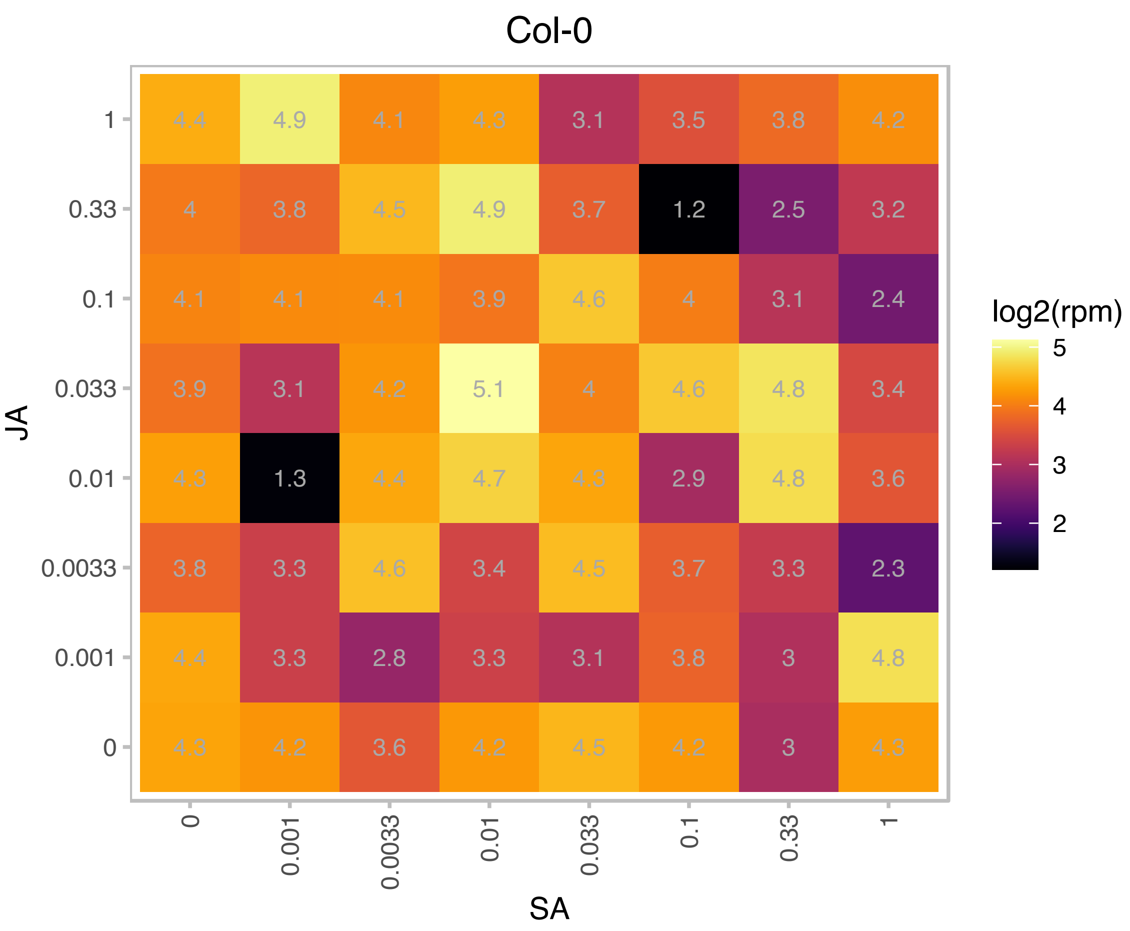


| | |
|---|---|
| Gene | AT3G20970 |
| is.expressed_genes | TRUE |
| is.responsive_genes | TRUE |
| is.apcluster | F-04 |

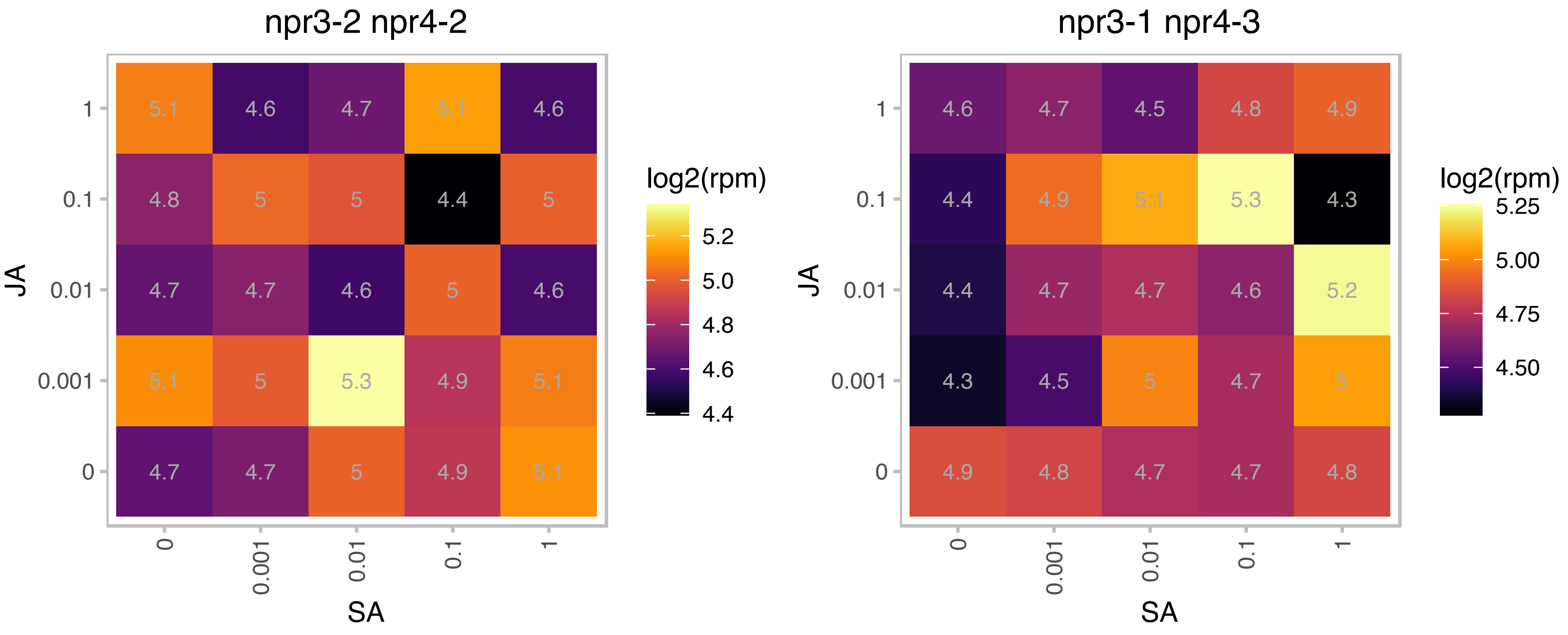

### AT3G21760.1 : HYR1

UDP-Glycosyltransferase superfamily protein

Encodes HYR1, a UDP glycosyltransferase (UGT).  HYR1 glucosylates hypostatin, an inhibitor of cell expansion in vivo to form a bioactive glucoside.

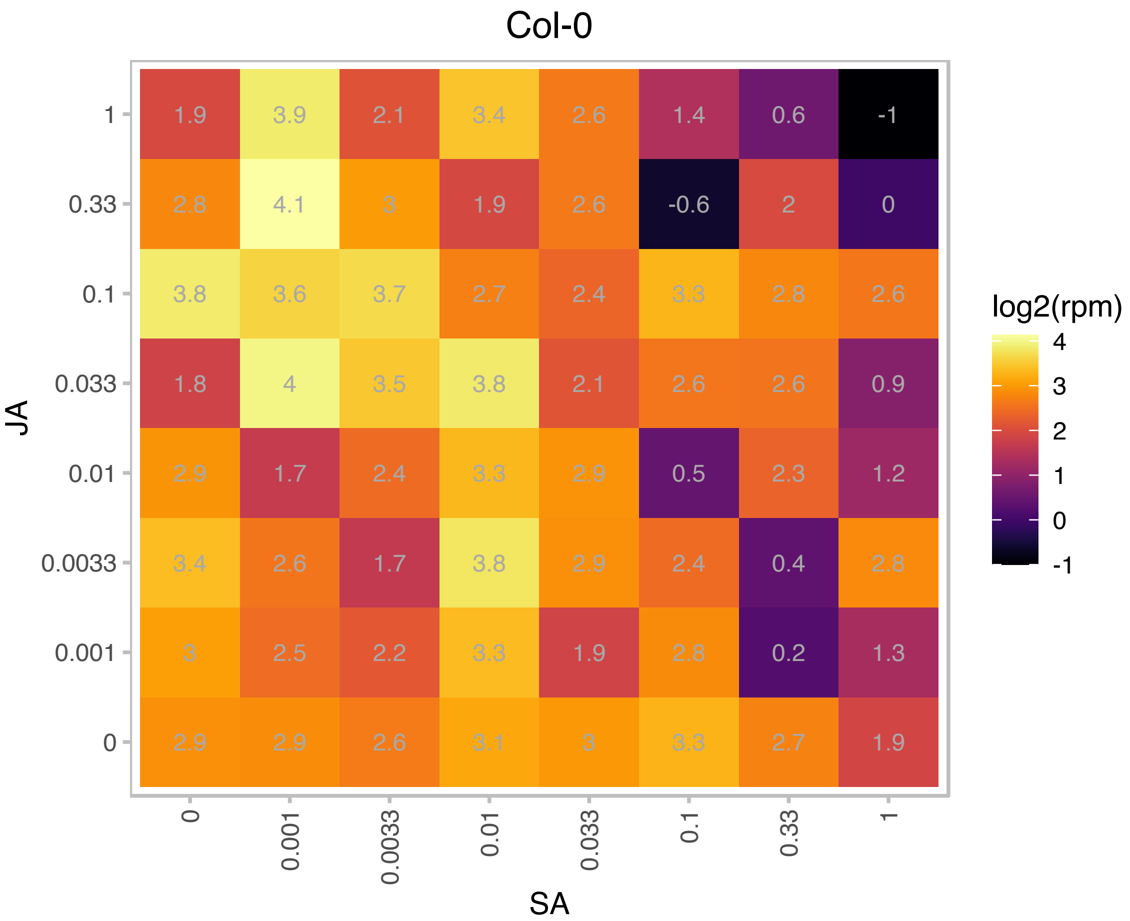


| | |
|---|---|
| Gene | AT3G21760 |
| is.expressed_genes | TRUE |
| is.responsive_genes | TRUE |
| is.apcluster | F-04 |

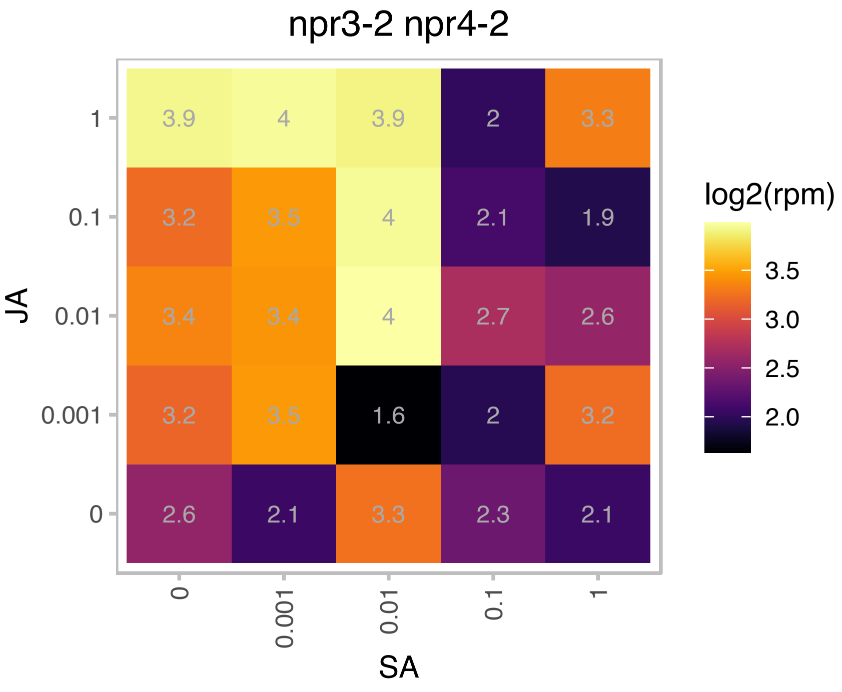


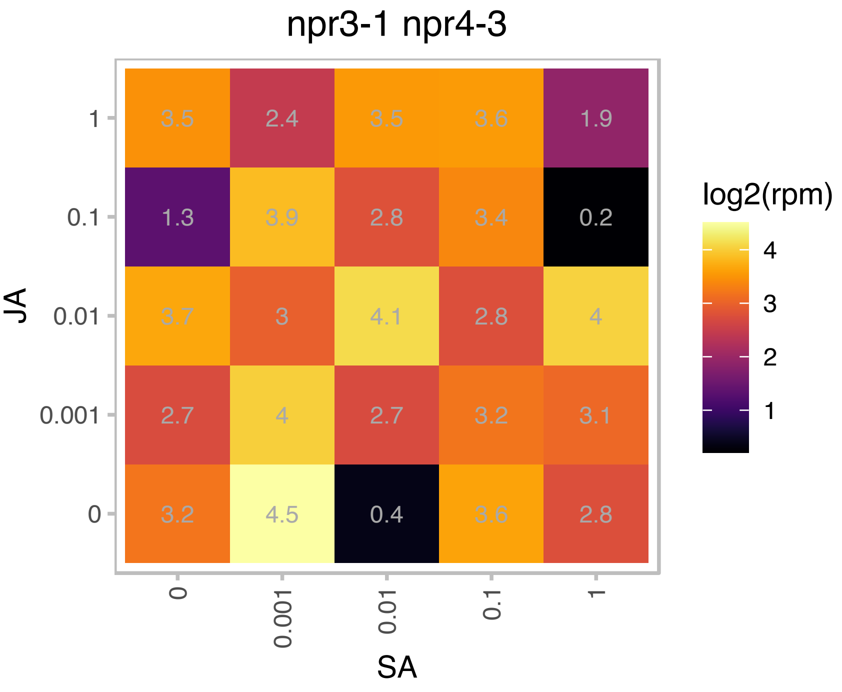

### AT3G28920.1 : AtHB34, HB34, ZHD9

homeobox protein 34

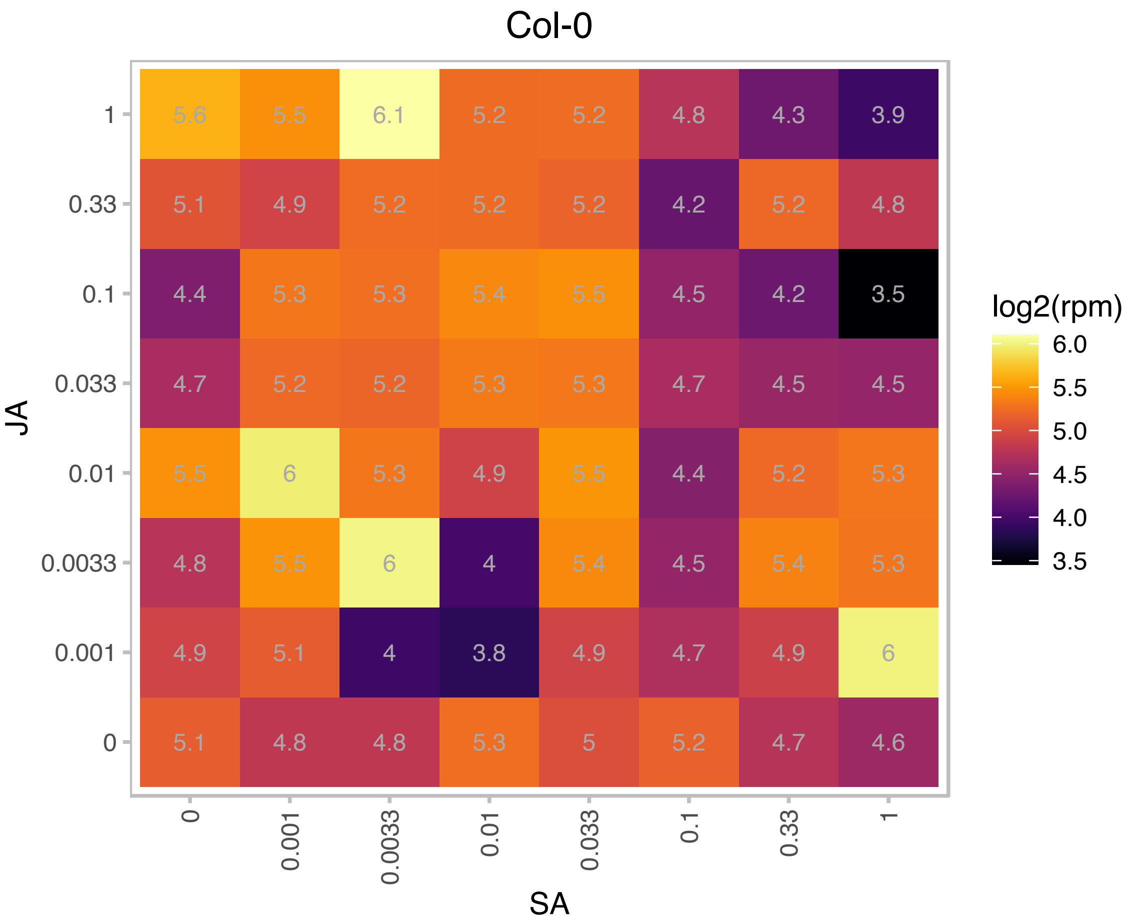


| | |
|---|---|
| Gene | AT3G28920 |
| is.expressed_genes | TRUE |
| is.responsive_genes | TRUE |
| is.apcluster | F-04 |

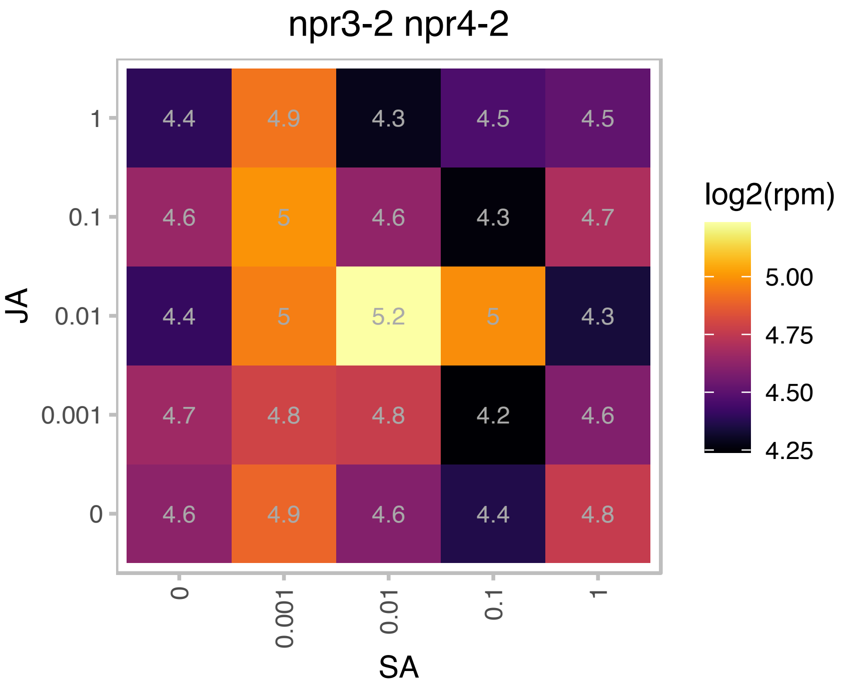


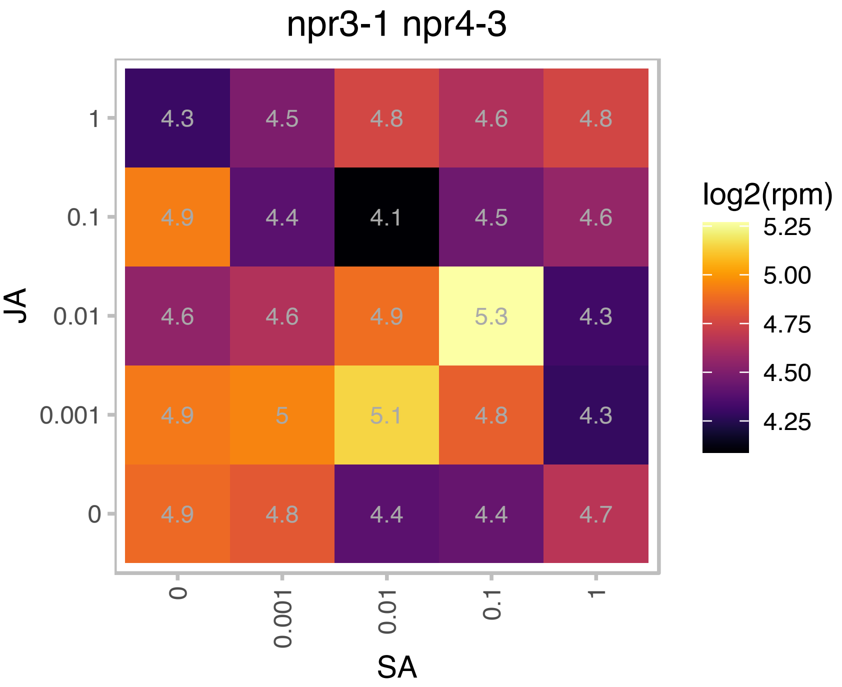

# AT3G51950.1 : NA

Zinc finger (CCCH-type) family protein / RNA recognition motif (RRM)-containing protein

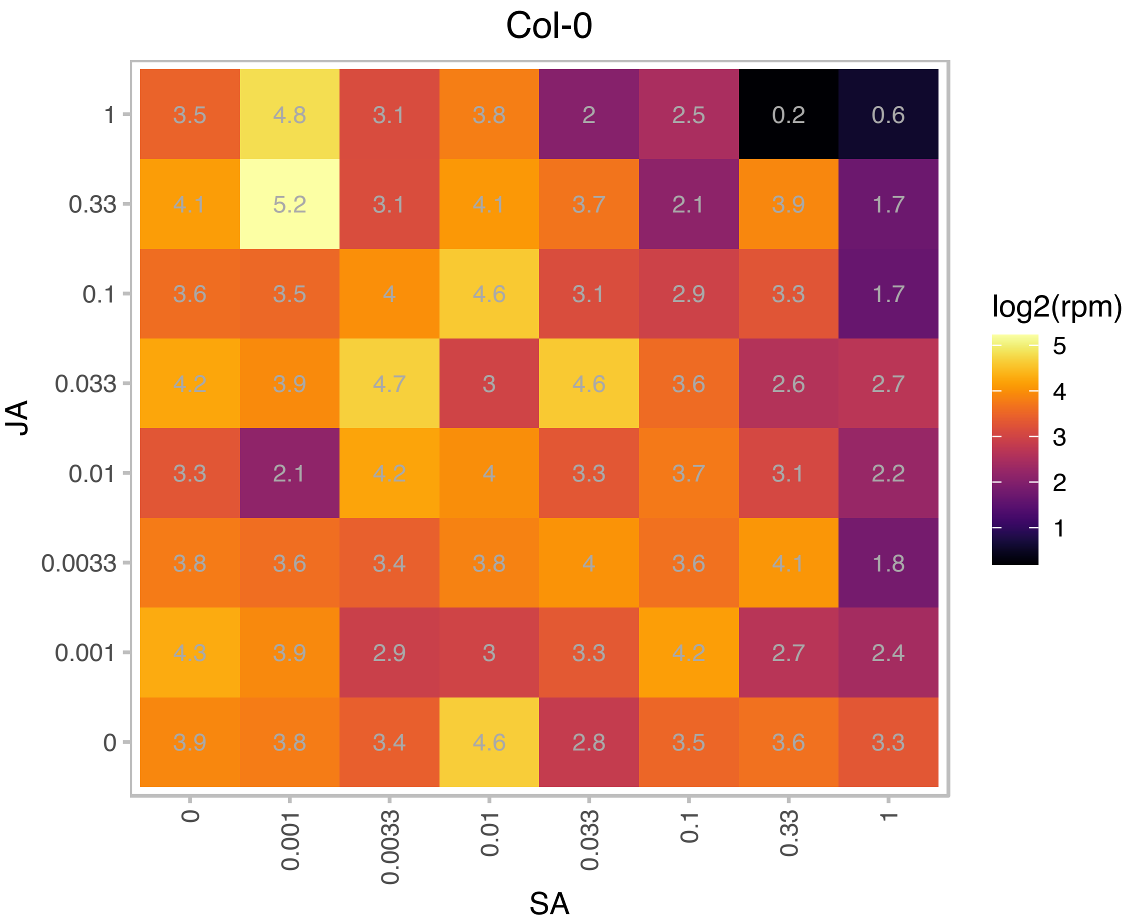


| | |
|---|---|
| Gene | AT3G51950 |
| is.expressed_genes | TRUE |
| is.responsive_genes | TRUE |
| is.apcluster | F-04 |

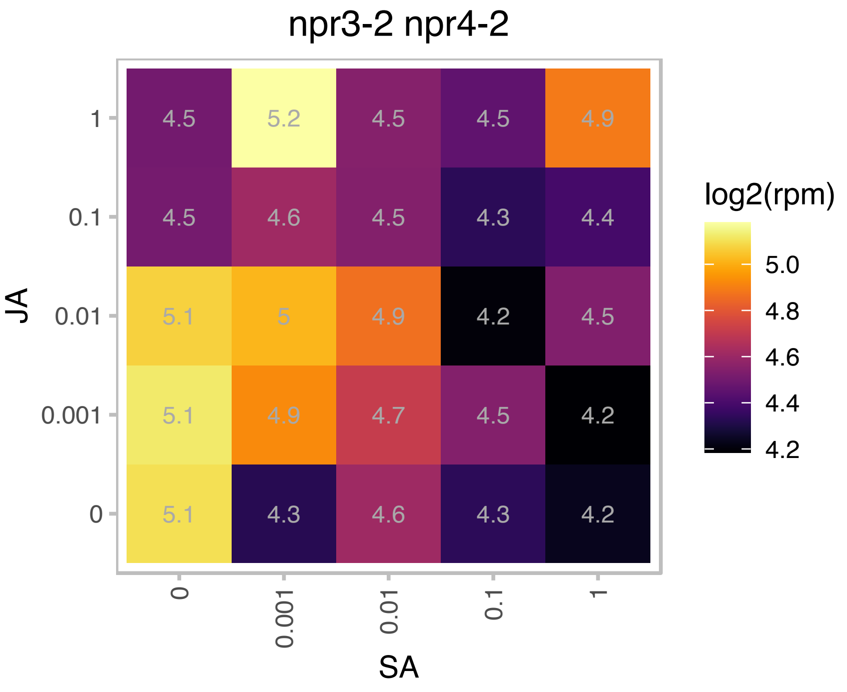


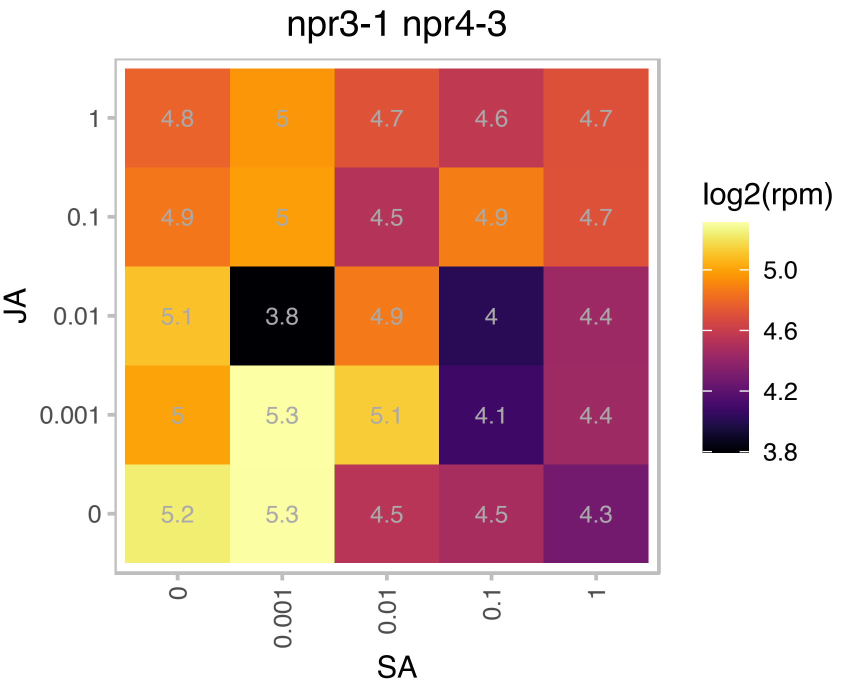

### AT3G54090.1 : FLN1

fructokinase-like 1

Encodes a fructokinase-like protein (AT3G54090/FLN1, AT1G69200/FLN2), a member of the pfkB-carbohydrate kinase family. FLN1 and FLN2 are potential plastidial thioredoxin z (TRX z) targets. Mutants display mutant chloroplast development, general plant growth and development defects and defects in PEP-dependent transcription.

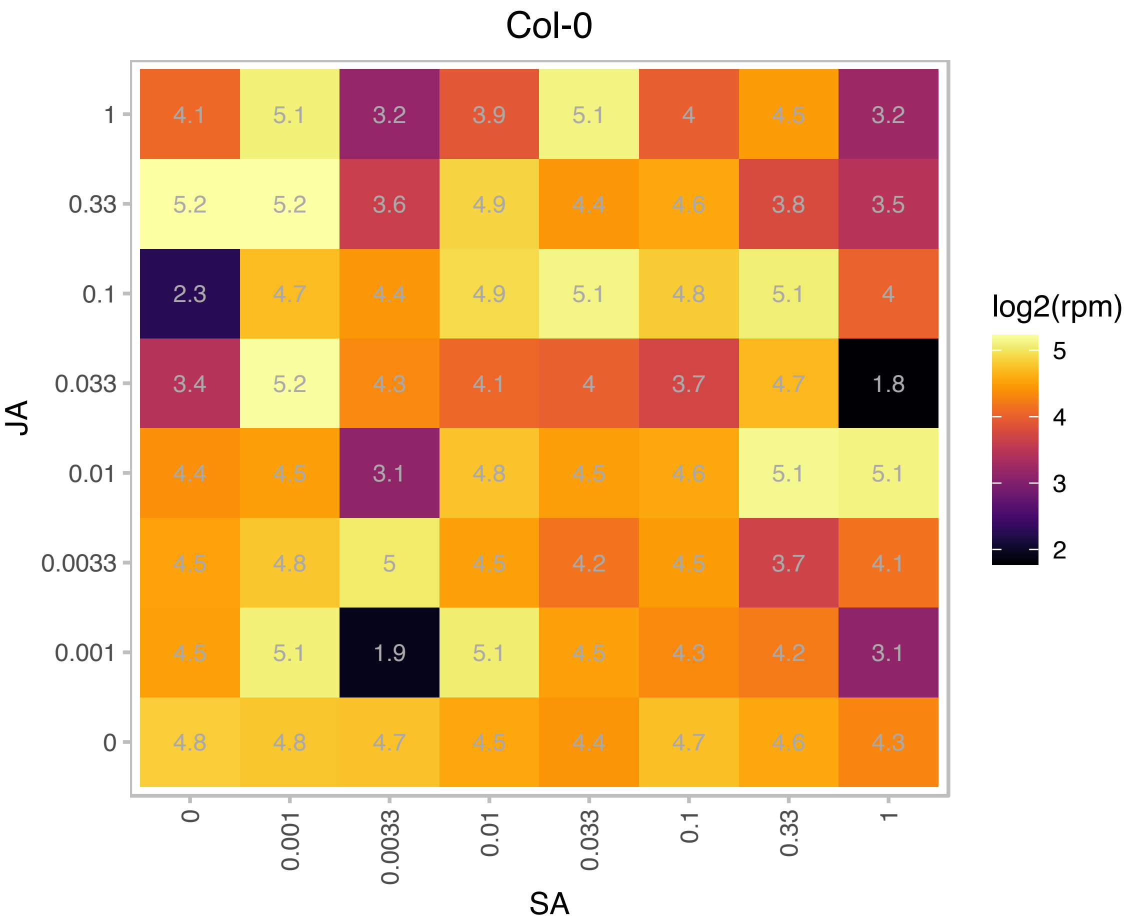


| | |
|---|---|
| Gene | AT3G54090 |
| is.expressed_genes | TRUE |
| is.responsive_genes | TRUE |
| is.apcluster | F-04 |

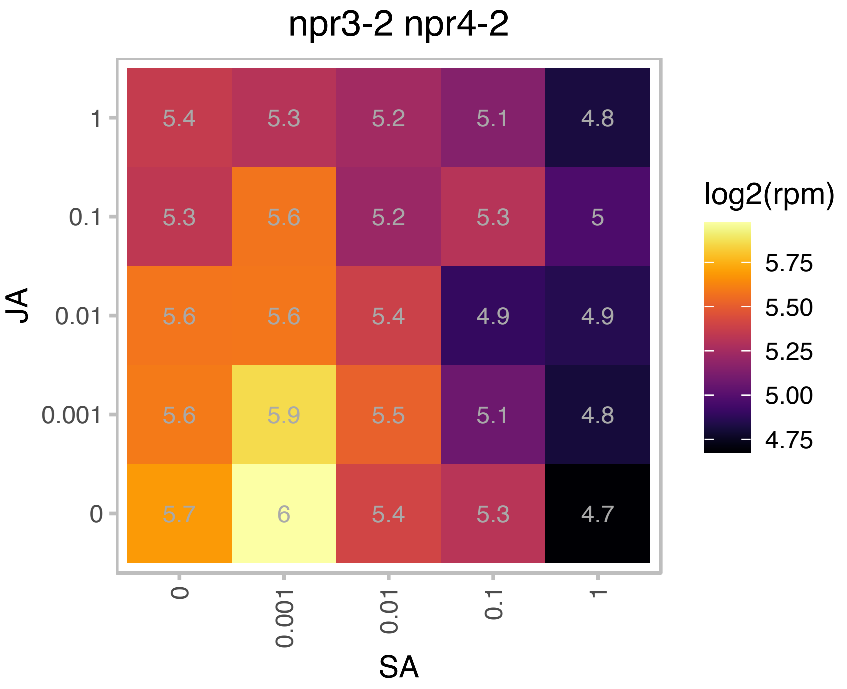


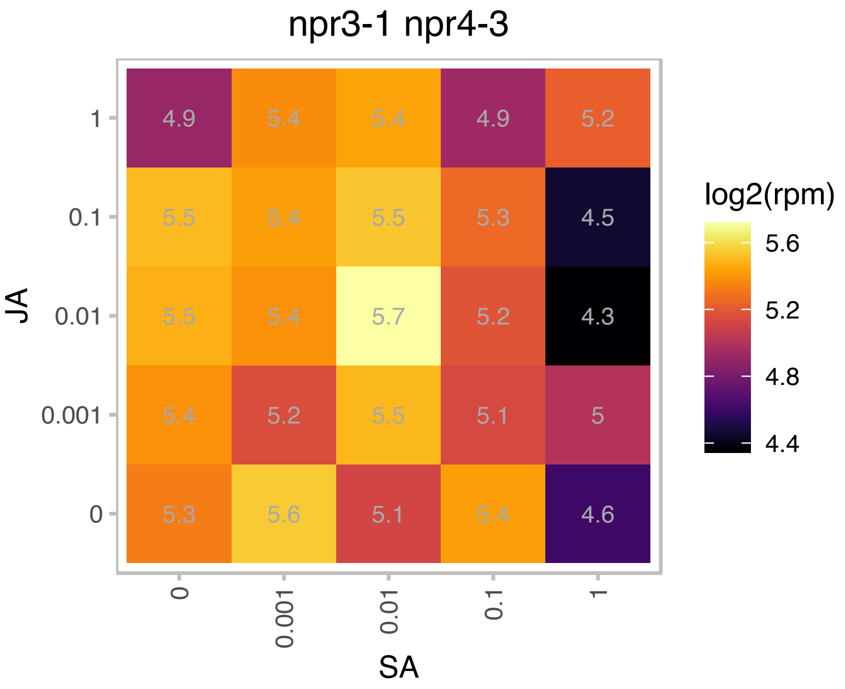

### AT3G55340.1 : PHIP1

phragmoplastin interacting protein 1

Plant-specific protein. Interacts with phragmoplastin, Rop1 and Rop2. Involved in cell plate formation.

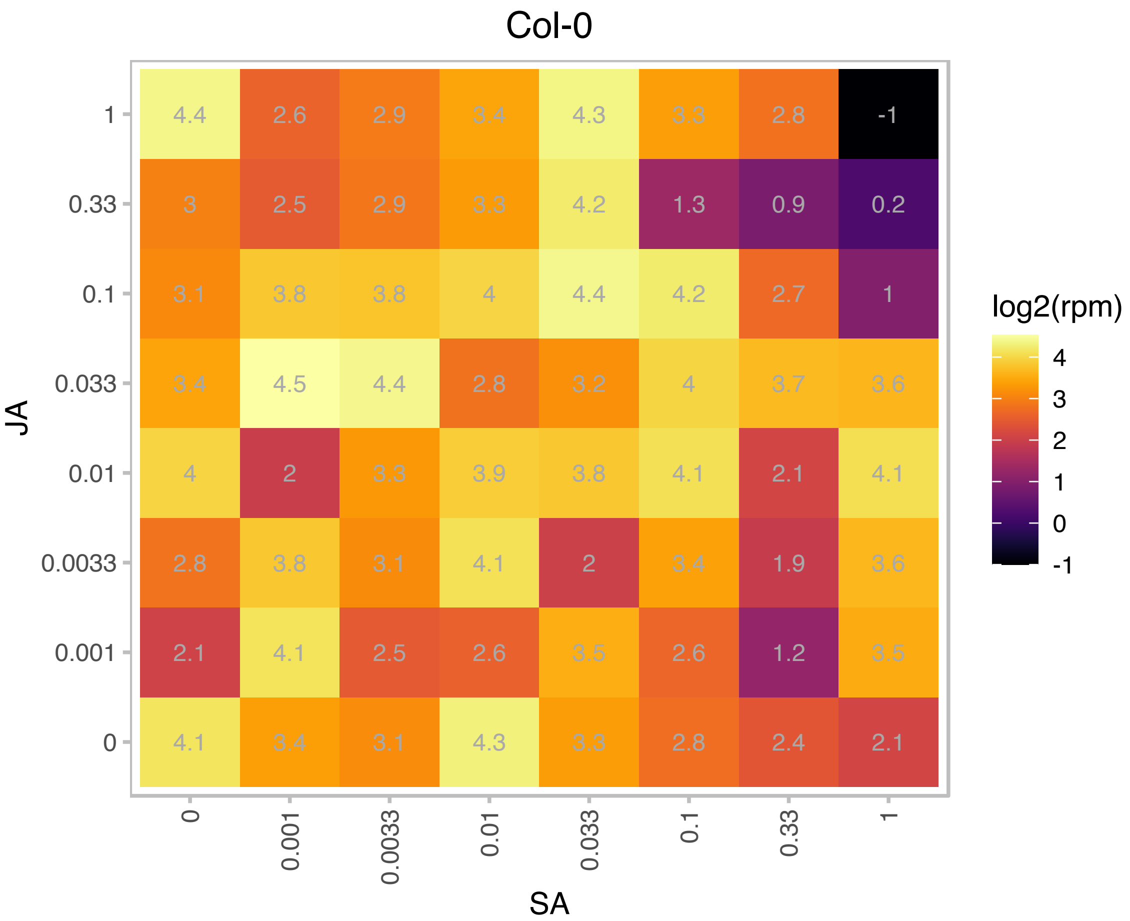


| | |
|---|---|
| Gene | AT3G55340 |
| is.expressed_genes | TRUE |
| is.responsive_genes | TRUE |
| is.apcluster | F-04 |

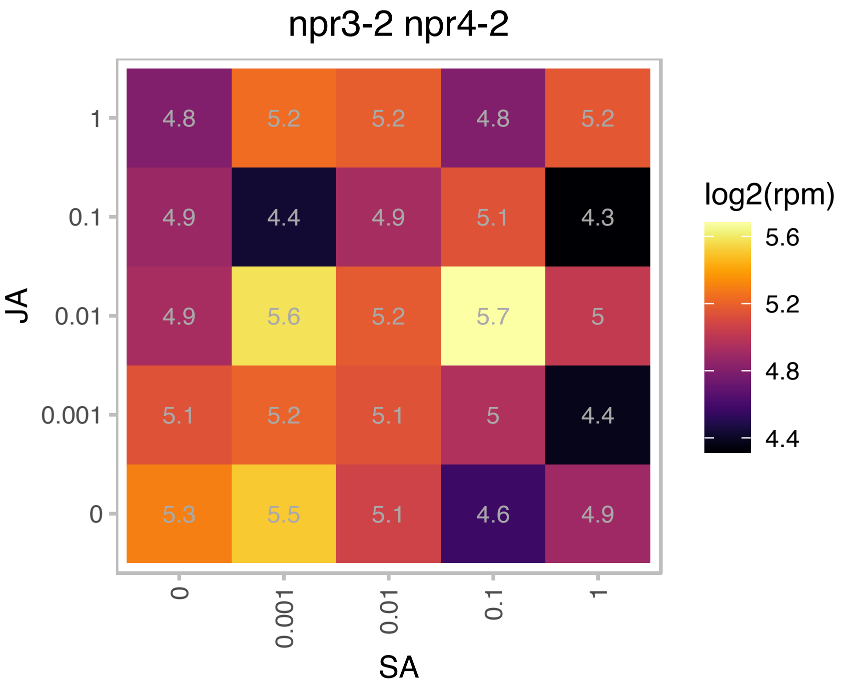


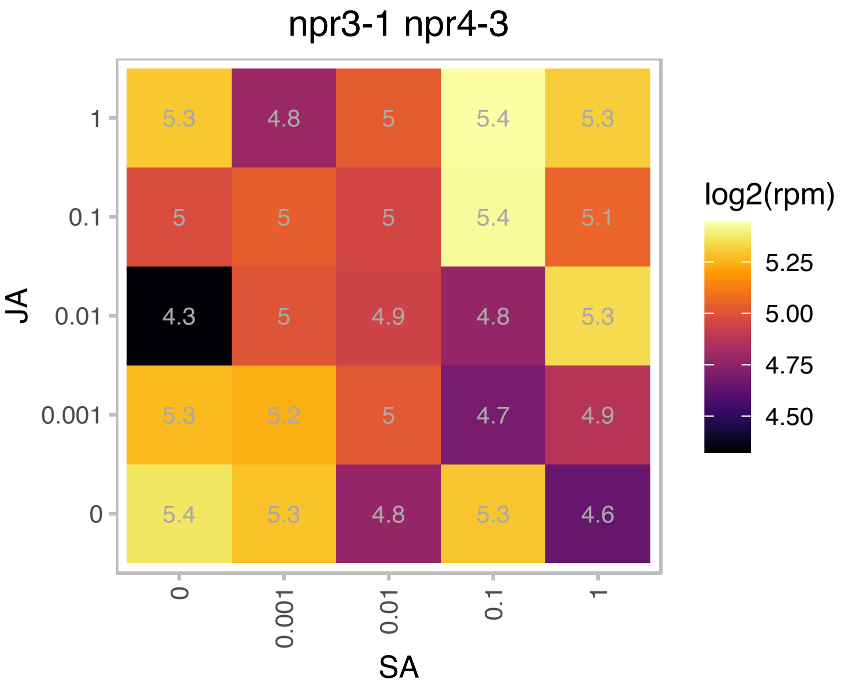

### AT3G55460.1 : At-SCL30, SCL30

SC35-like splicing factor 30
encodes an SC35-like splicing factor that is localized to nuclear specks. Barta et al (2010) have proposed a nomenclature for Serine/Arginine-Rich Protein Splicing Factors (SR proteins): Plant Cell. 2010, 22:2926. The mRNA is cell-to-cell mobile.

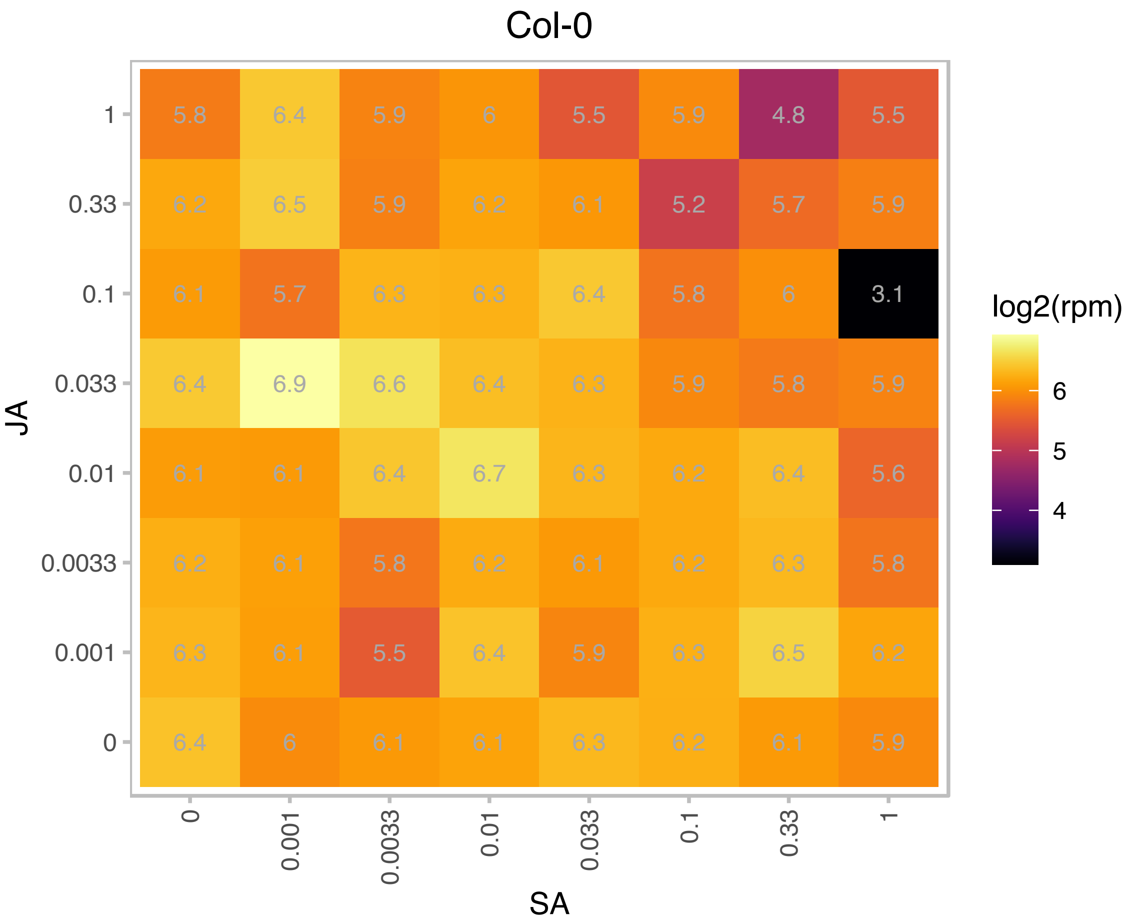


| | |
|---|---|
| Gene | AT3G55460 |
| is.expressed_genes | TRUE |
| is.responsive_genes | TRUE |
| is.apcluster | F-04 |

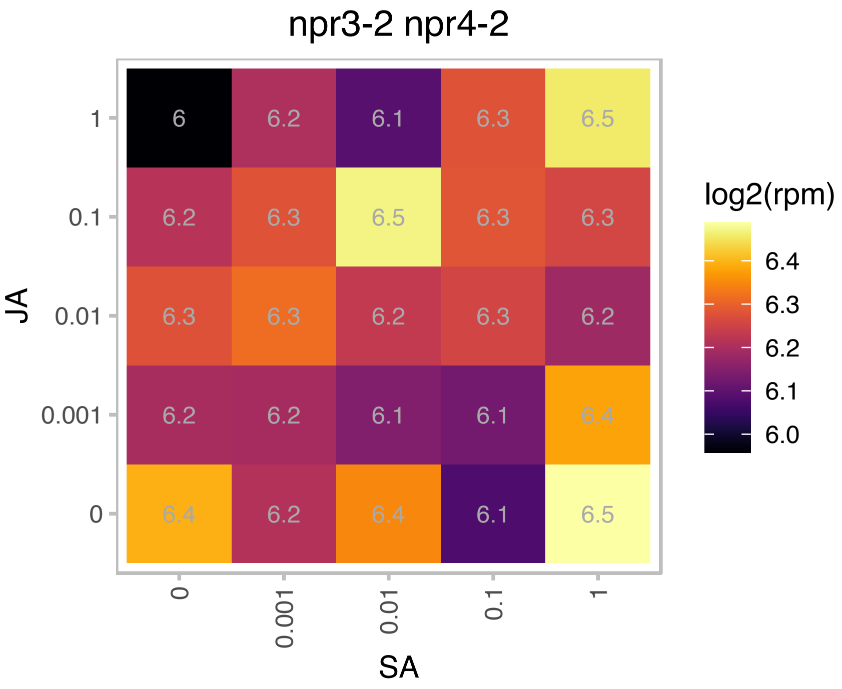


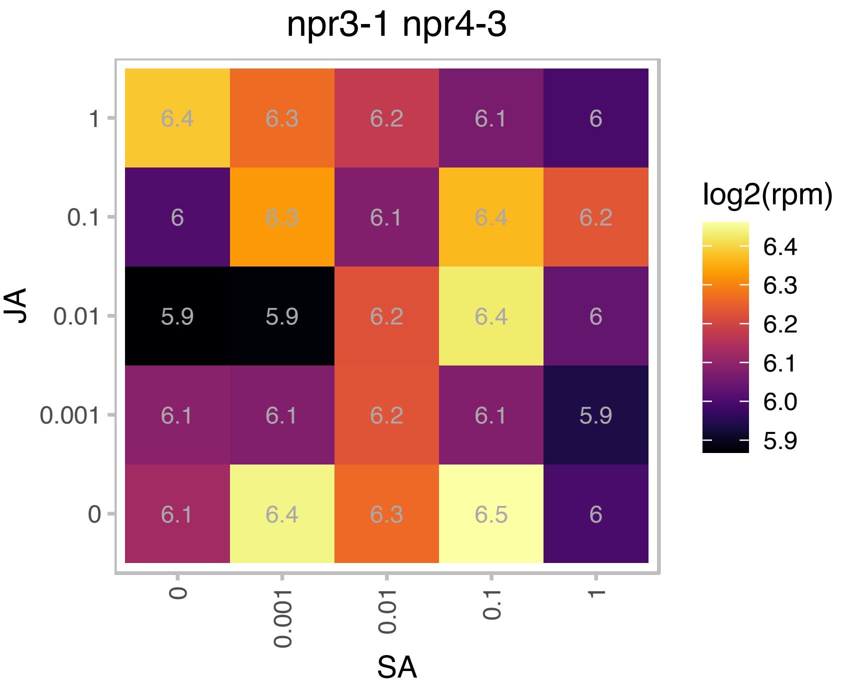

### AT3G58840.1 : PMD1

Tropomyosin-related

Encodes PEROXISOMAL AND MITOCHONDRIAL DIVISION FACTOR1. Involved in the morphogenesis and prolifer
peroxisomes and mitochondria.

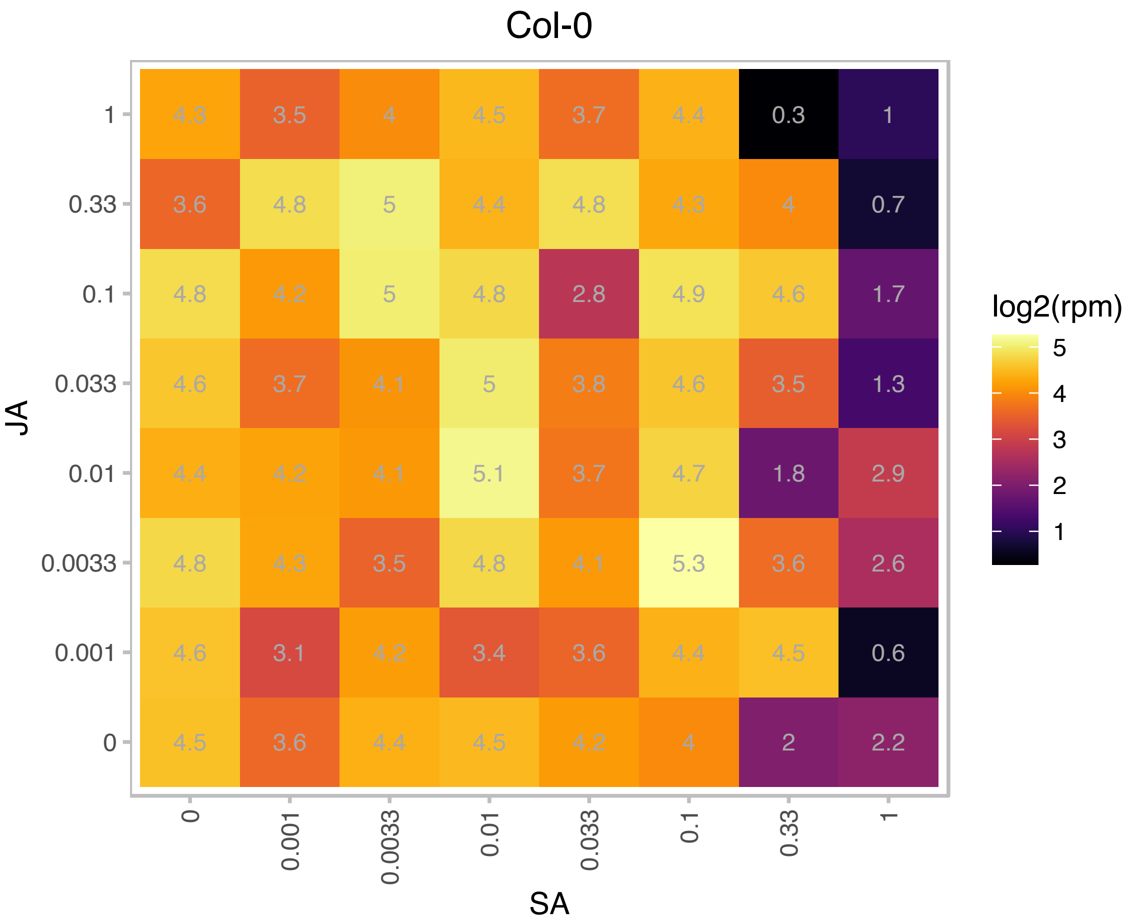


| | |
|---|---|
| Gene | AT3G58840 |
| is.expressed_genes | TRUE |
| is.responsive_genes | TRUE |
| is.apcluster | F-04 |

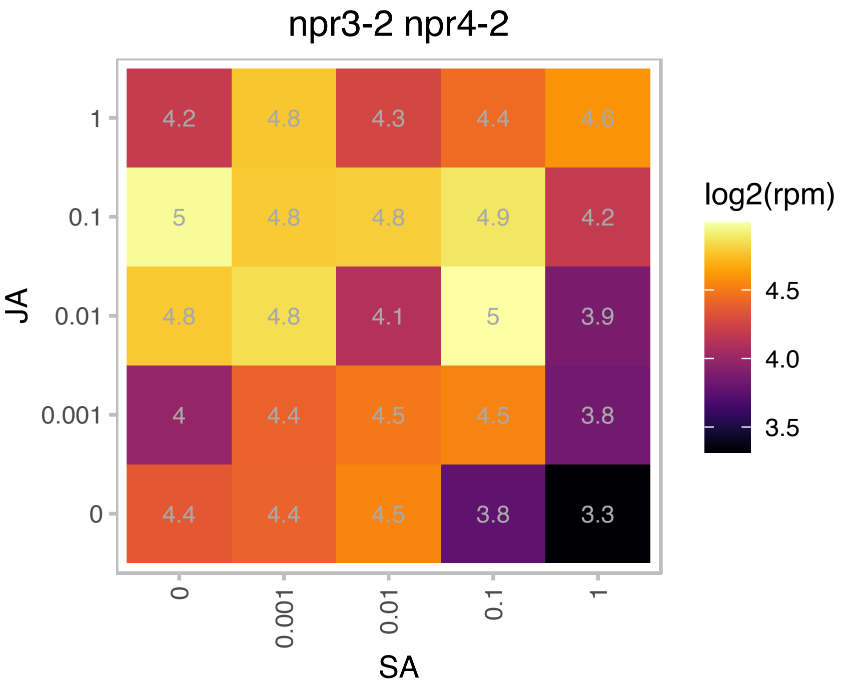


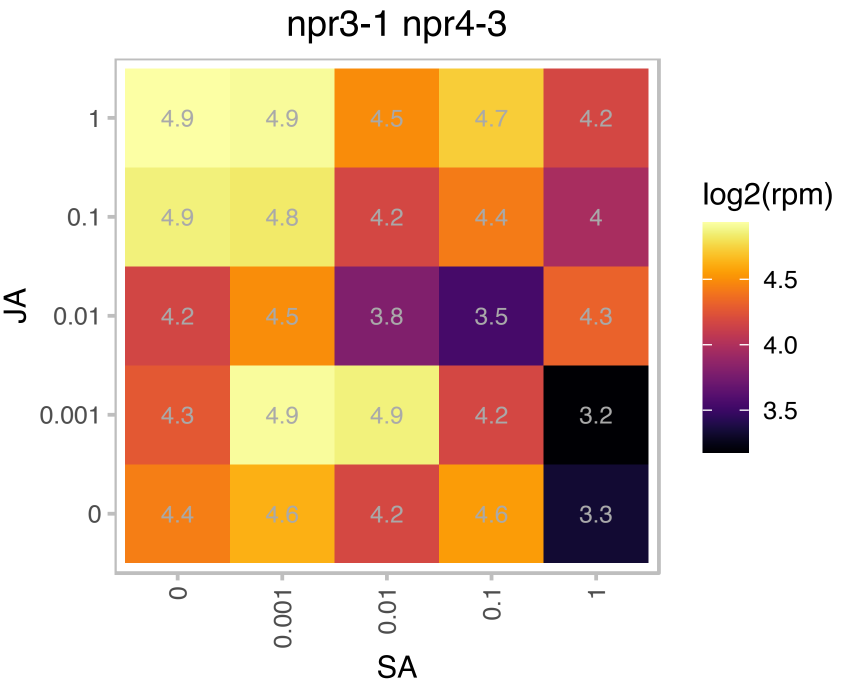

### AT3G59970.1 : MTHFR1

methylenetetrahydrofolate reductase 1

methylenetetrahydrofolate reductase MTHFR1 mRNA, complete

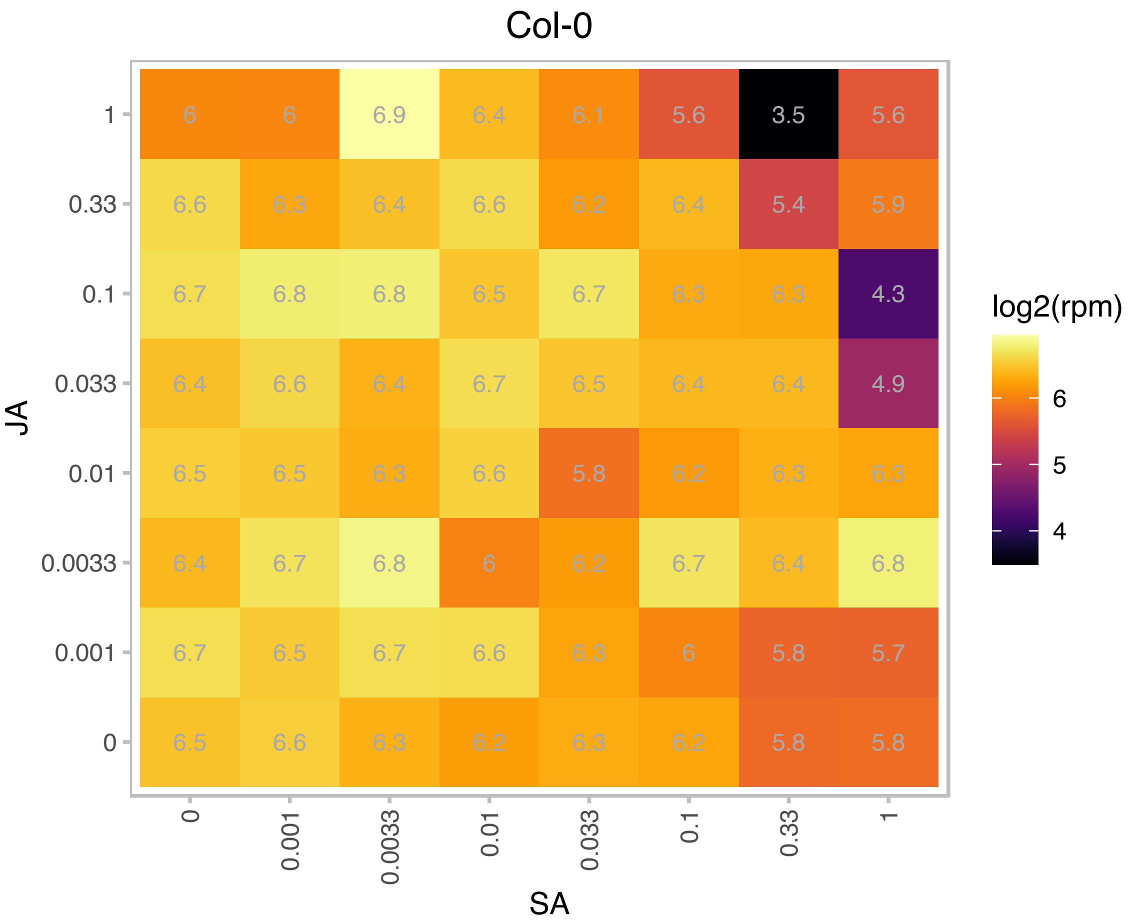


| | |
|---|---|
| Gene | AT3G59970 |
| is.expressed_genes | TRUE |
| is.responsive_genes | TRUE |
| is.apcluster | F-04 |

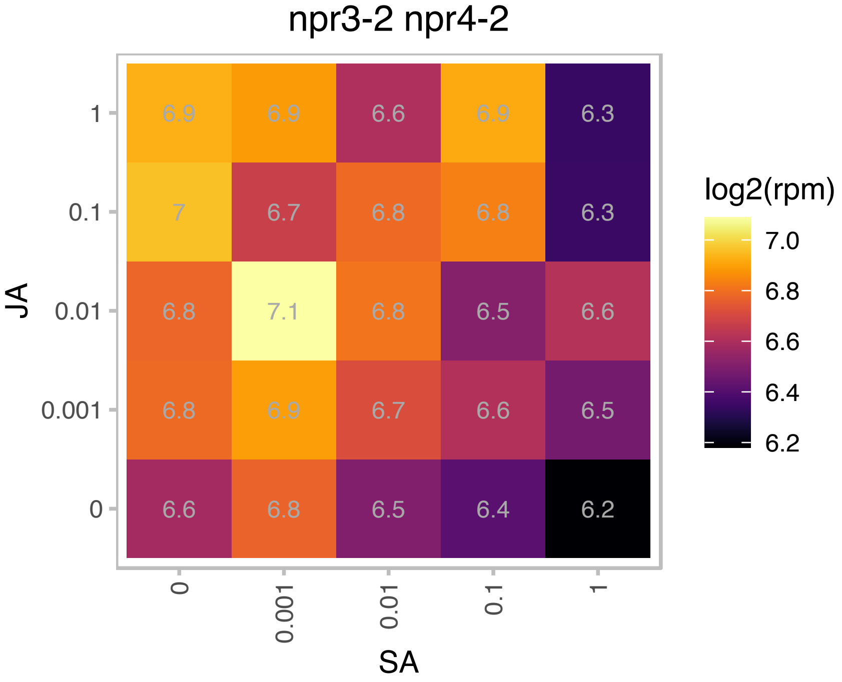


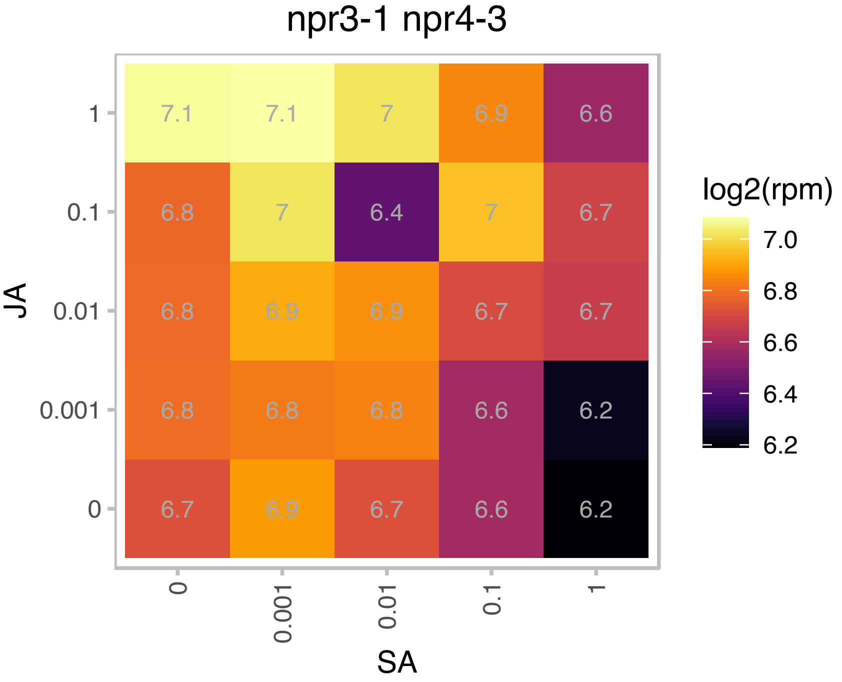

# AT3G62650.1 : NA

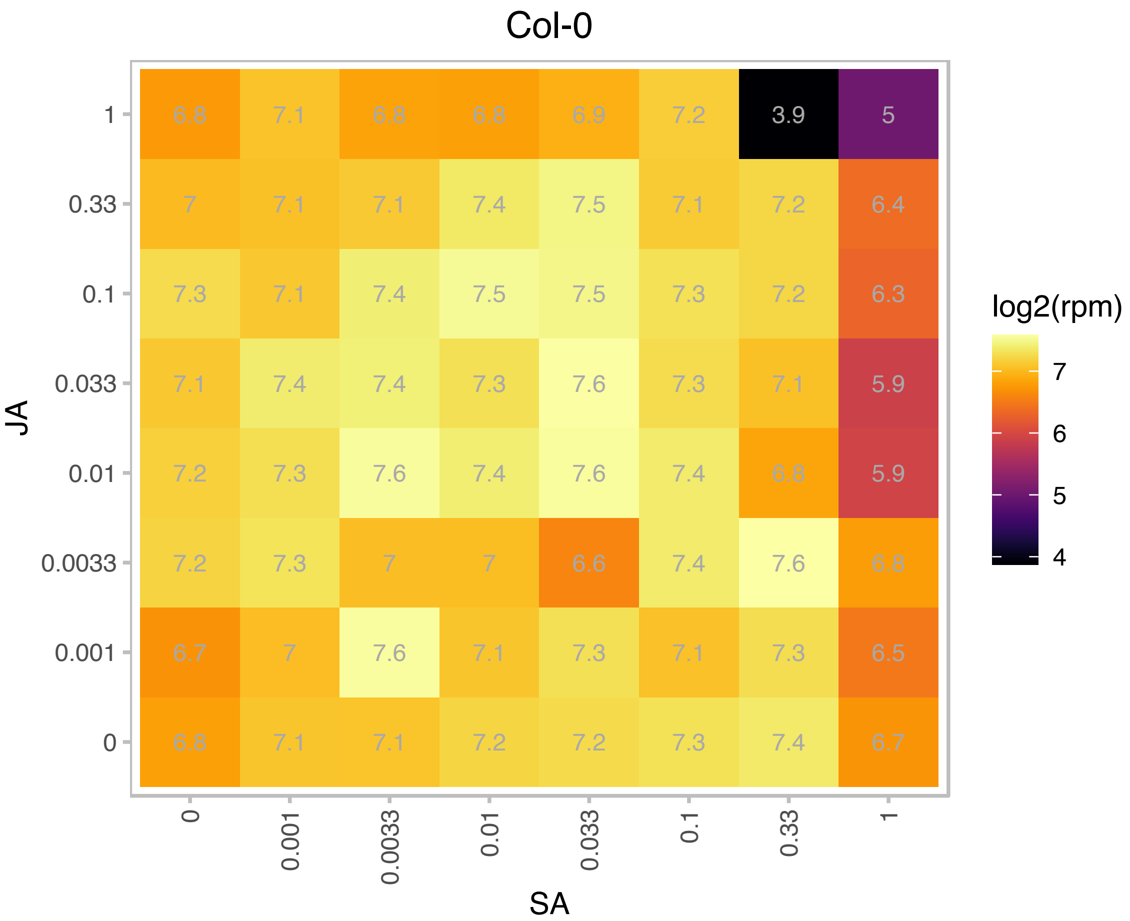


| | |
|---|---|
| Gene | AT3G62650 |
| is.expressed_genes | TRUE |
| is.responsive_genes | TRUE |
| is.apcluster | F-04 |

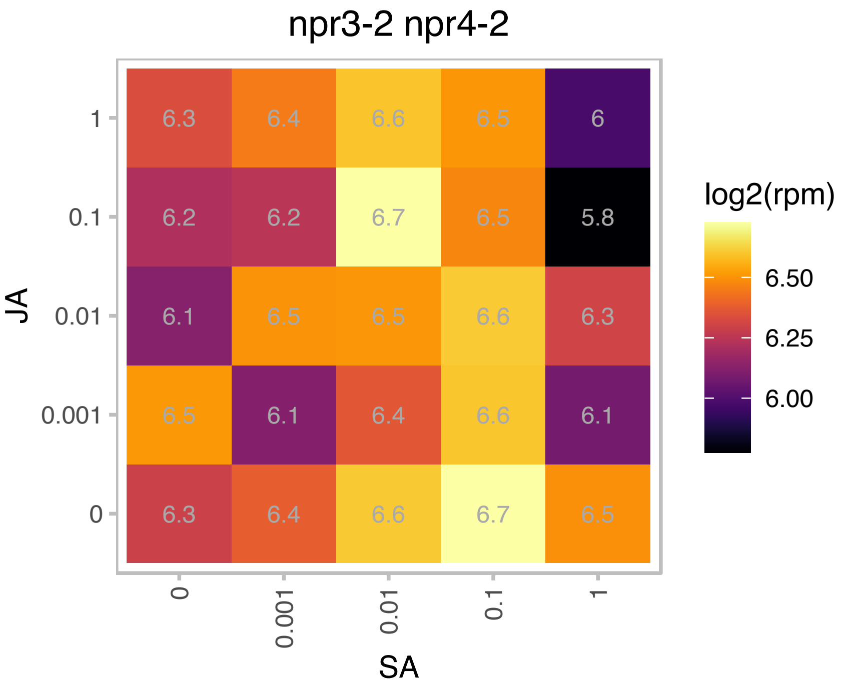


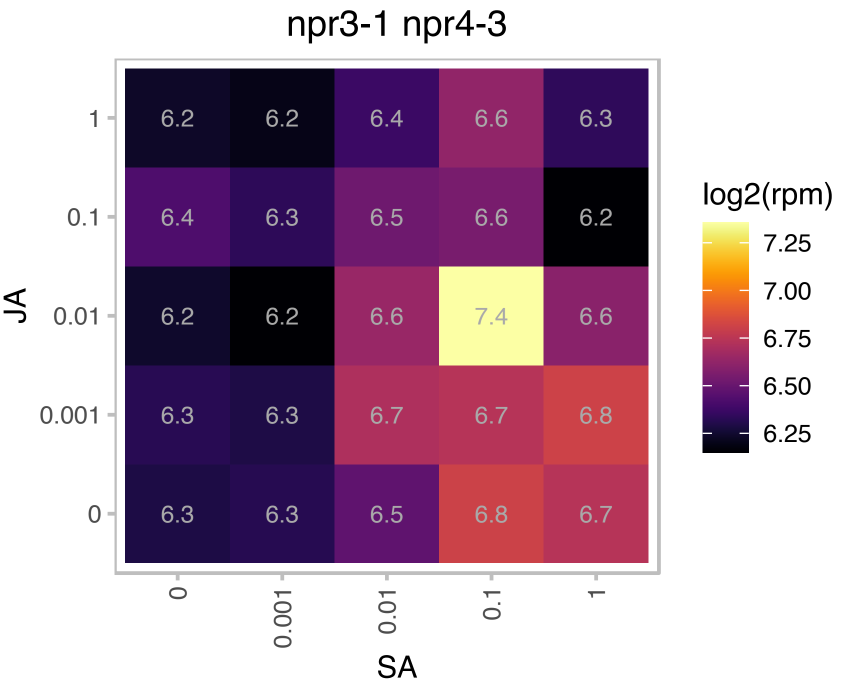

# AT3G62810.1 : NA

complex 1 family protein / LVR family protein

| | |
|---|---|
| Gene | AT3G62810 |
| is.expressed_genes | TRUE |
| is.responsive_genes | TRUE |
| is.apcluster | F-04 |

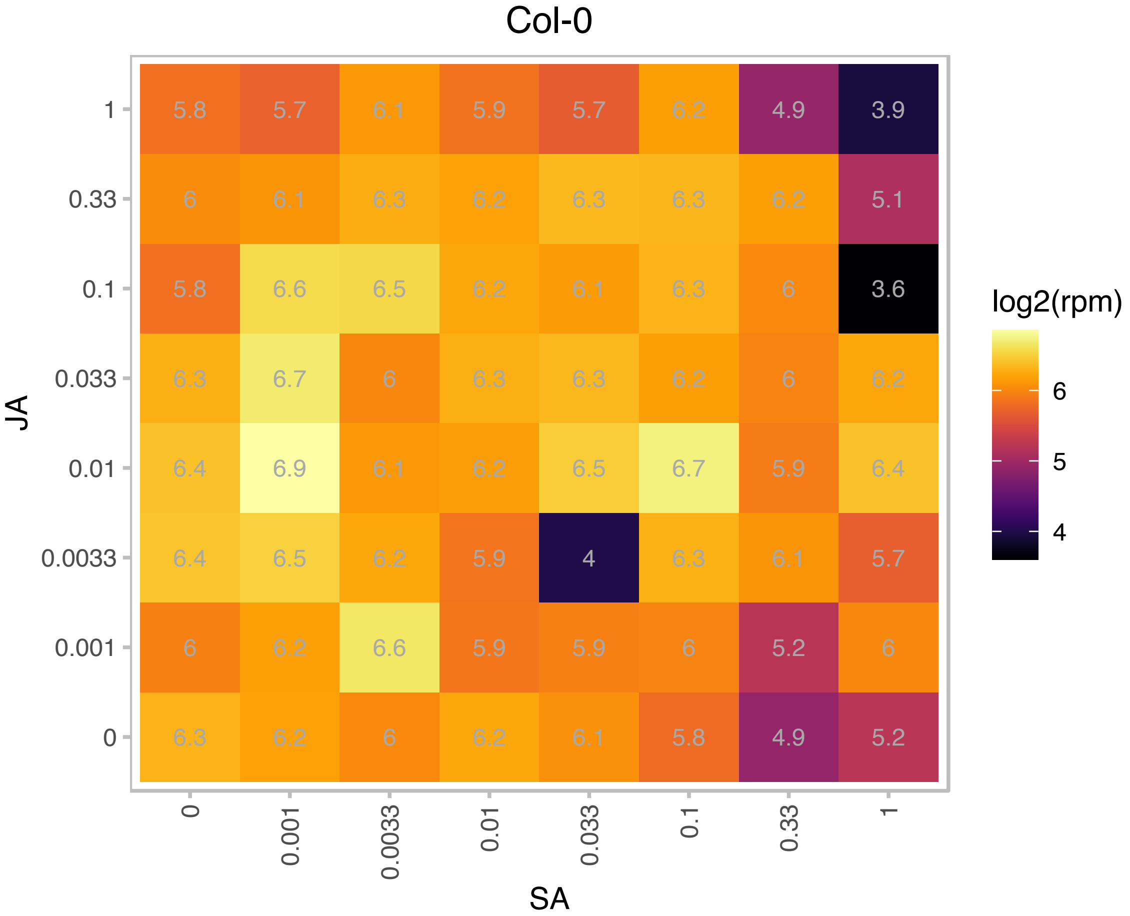


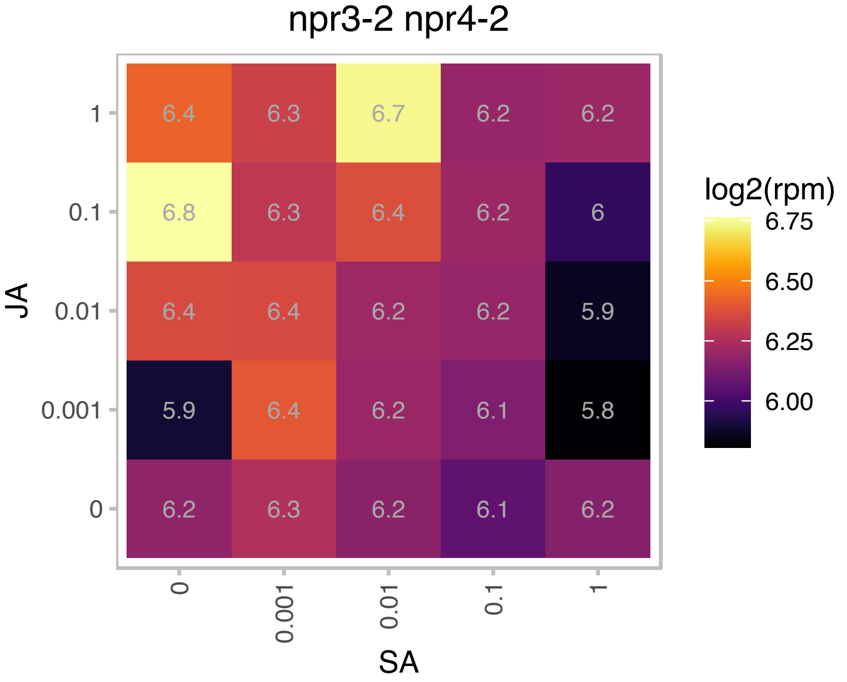


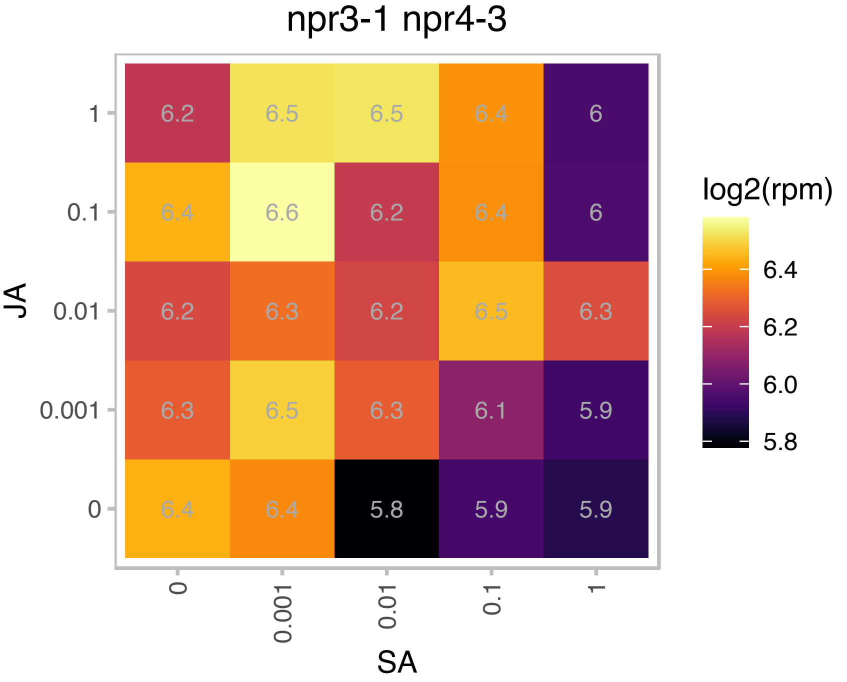

# AT3G62970.1 : NA

zinc finger (C3HC4-type RING finger) family protein

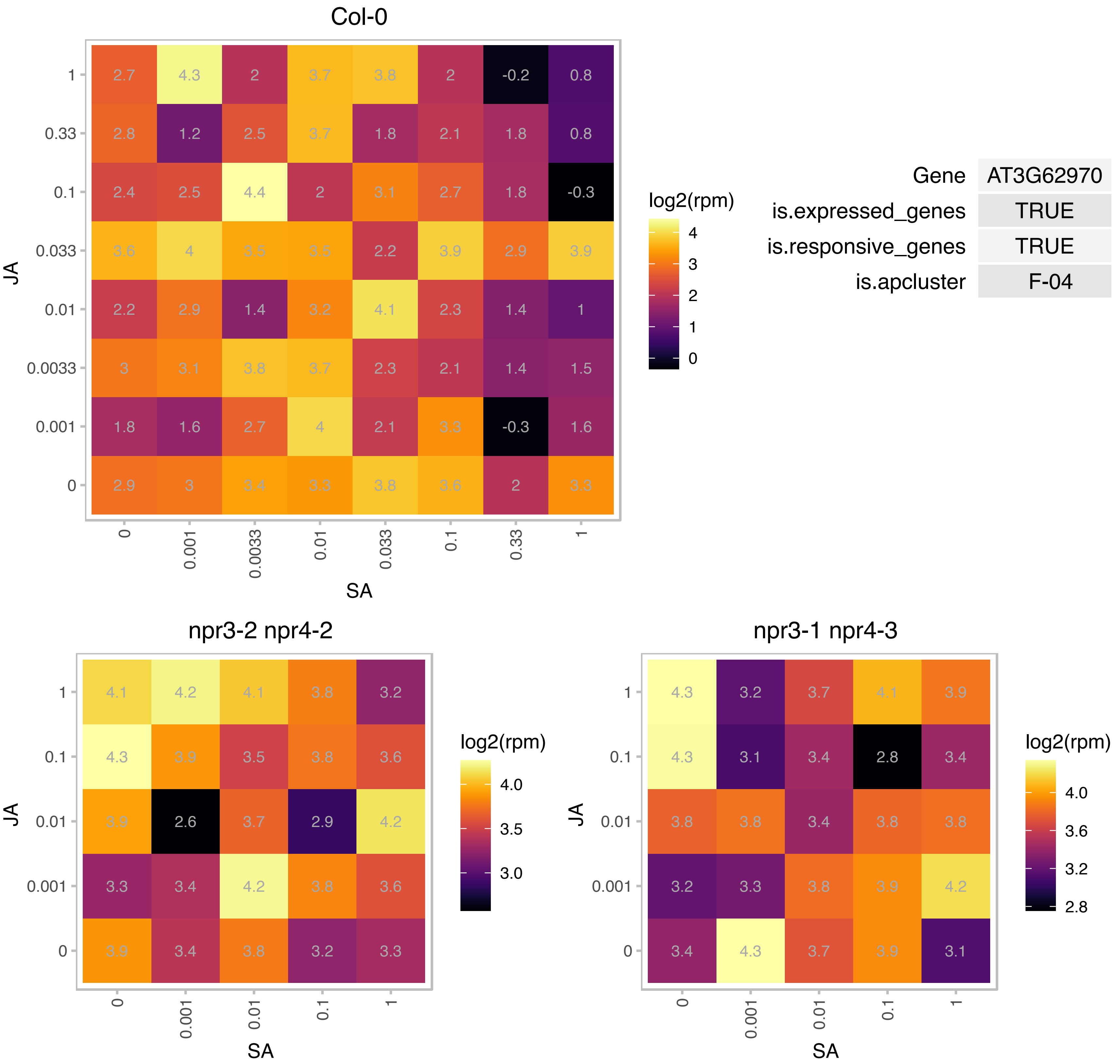


| | |
|---|---|
| Gene | AT3G62970 |
| is.expressed_genes | TRUE |
| is.responsive_genes | TRUE |
| is.apcluster | F-04 |

# AT4G00530.1 : NA

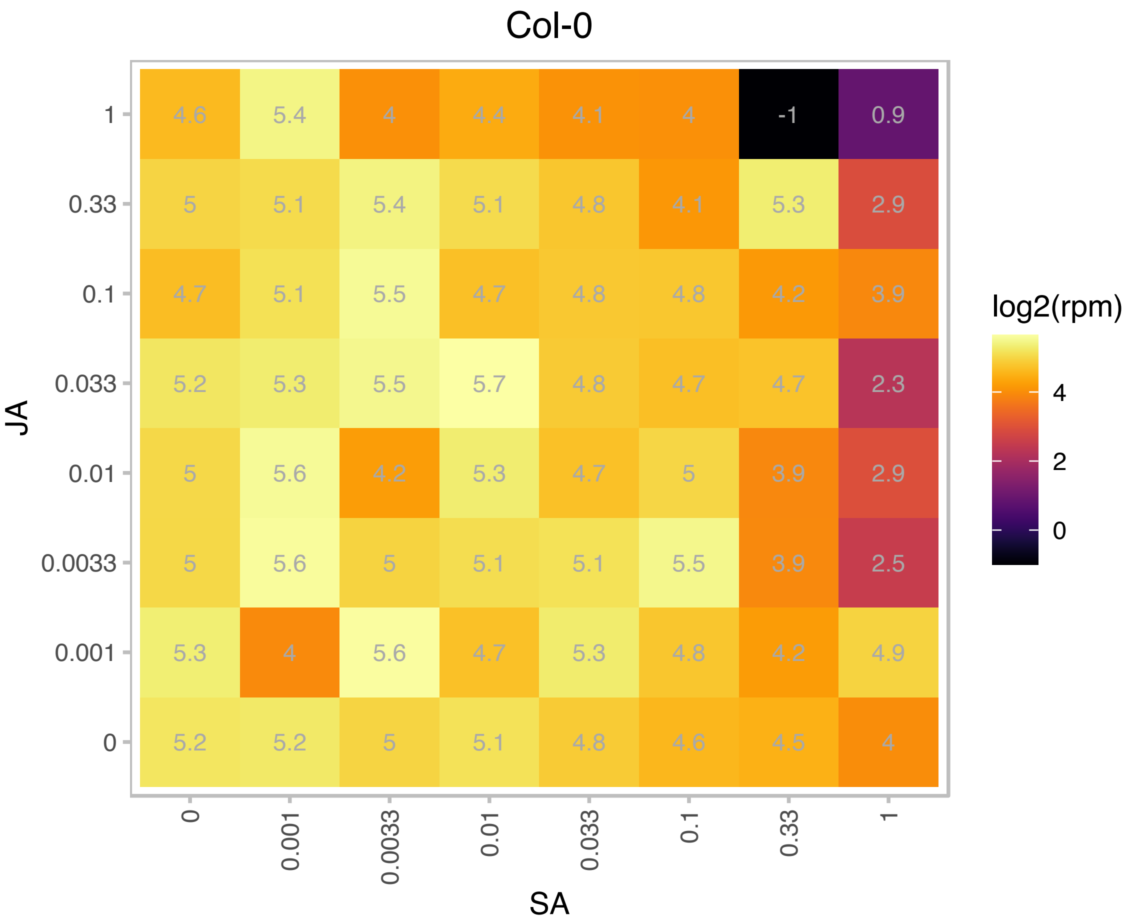


| | |
|---|---|
| Gene | AT4G00530 |
| is.expressed_genes | TRUE |
| is.responsive_genes | TRUE |
| is.apcluster | F-04 |

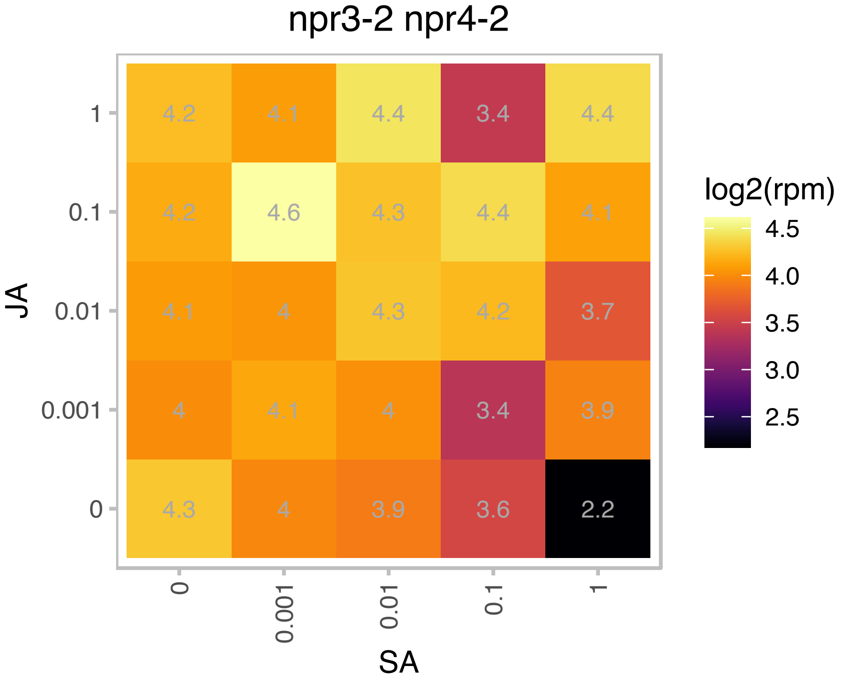


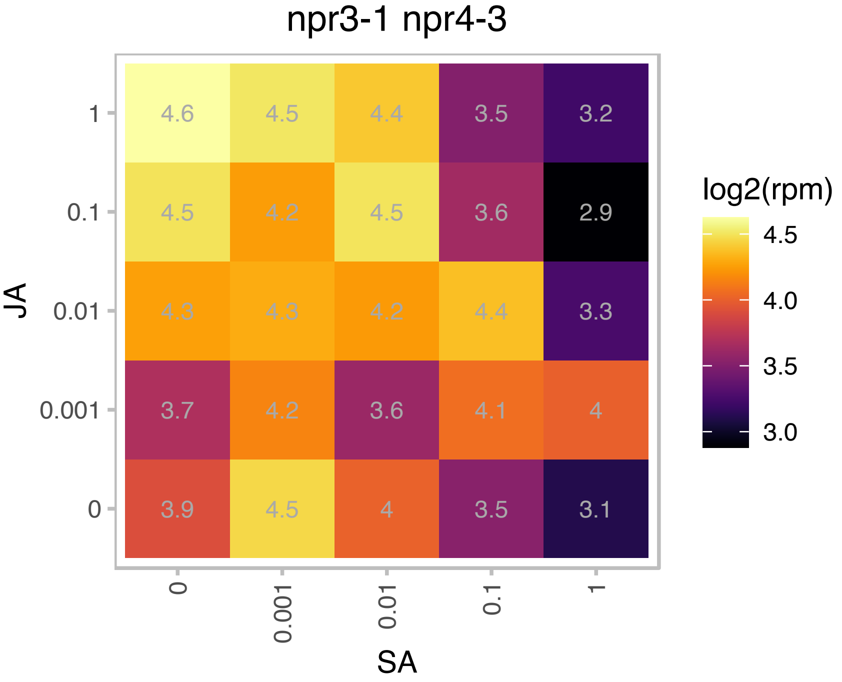

### AT4G00830.1 : LIF2

RNA-binding (RRM/RBD/RNP motifs) family protein

Encodes a heterogeneous nuclear ribonucleoprotein (hnRNP-Q) that is involved in the plant innate immune response and may function as a suppressor of cell-autonomous immunity.

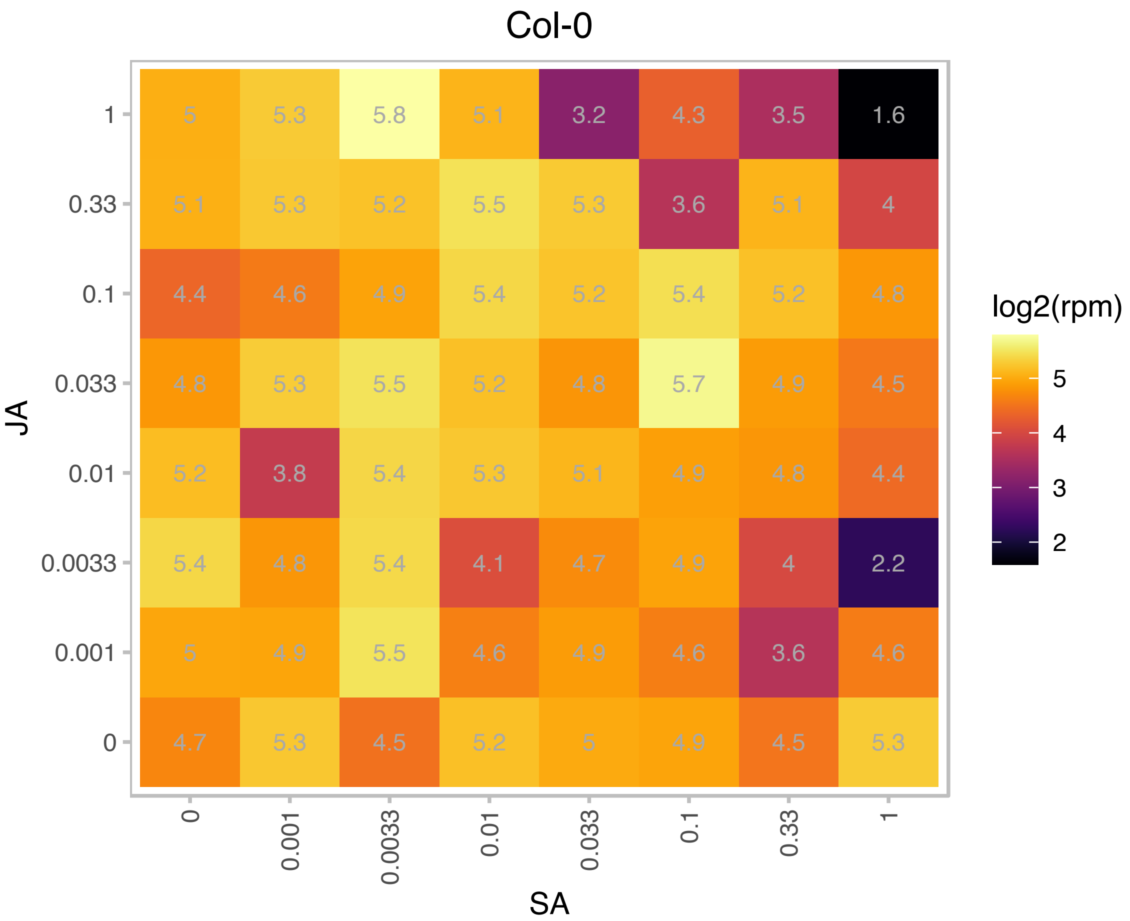


| | |
|---|---|
| Gene | AT4G00830 |
| is.expressed_genes | TRUE |
| is.responsive_genes | TRUE |
| is.apcluster | F-04 |

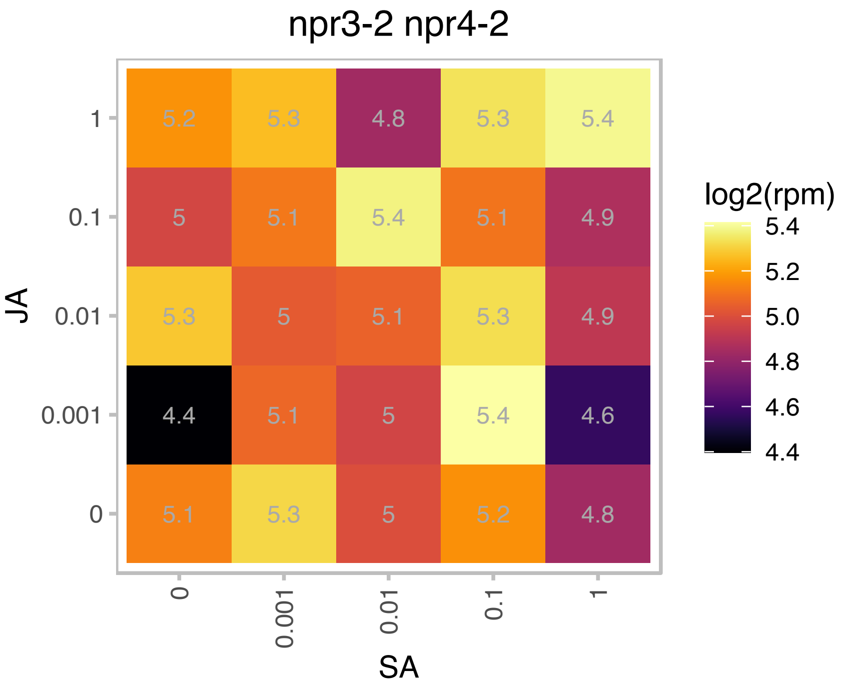


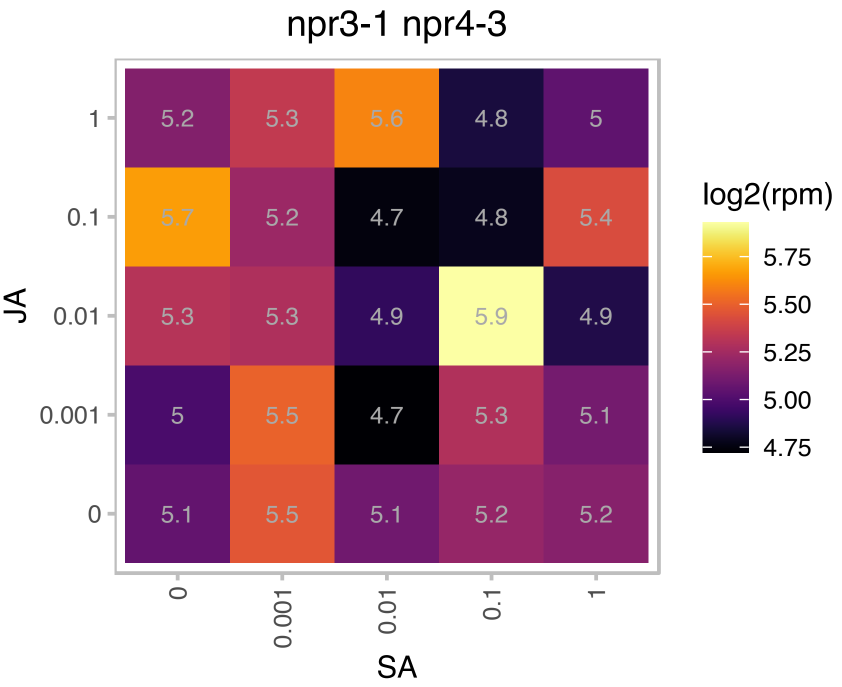

### AT4G04830.1 : ATMSRB5, MSRB5

methionine sulfoxide reductase B5

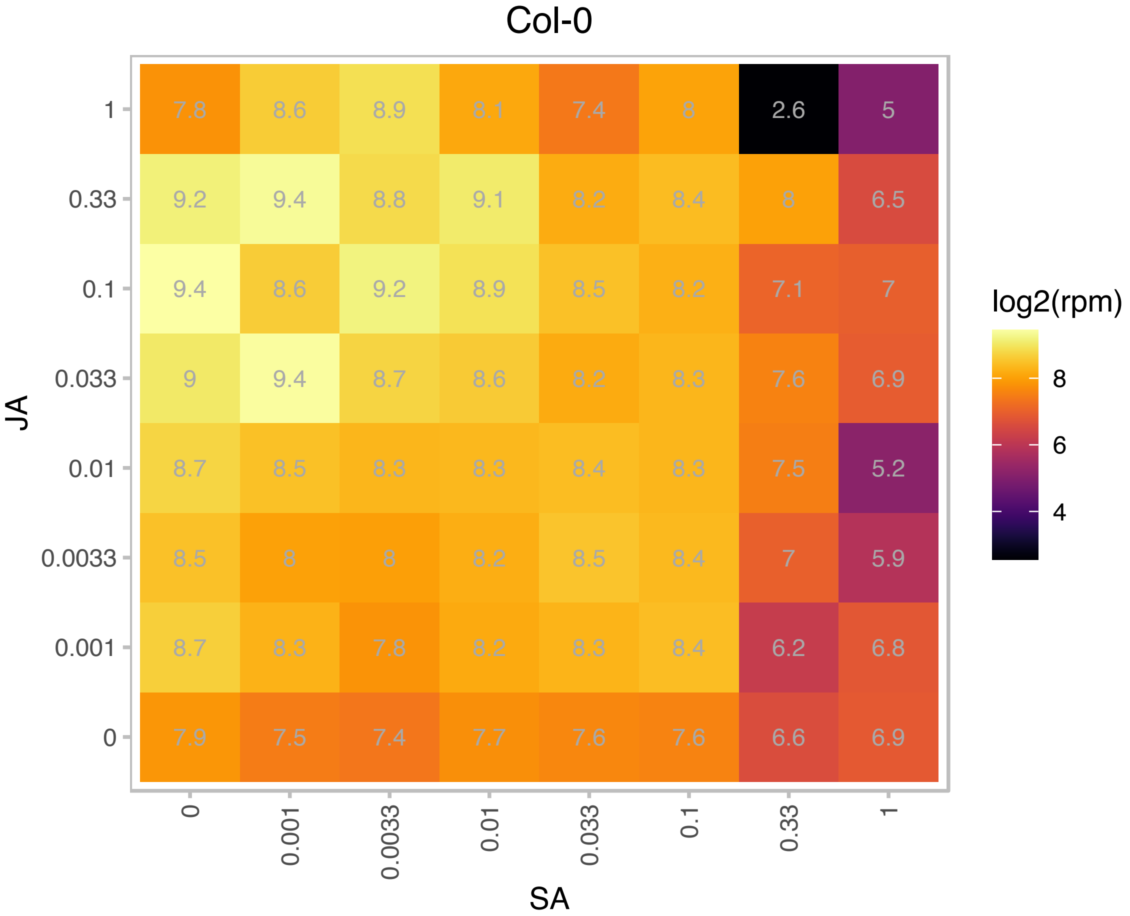


| | |
|---|---|
| Gene | AT4G04830 |
| is.expressed_genes | TRUE |
| is.responsive_genes | TRUE |
| is.apcluster | F-04 |

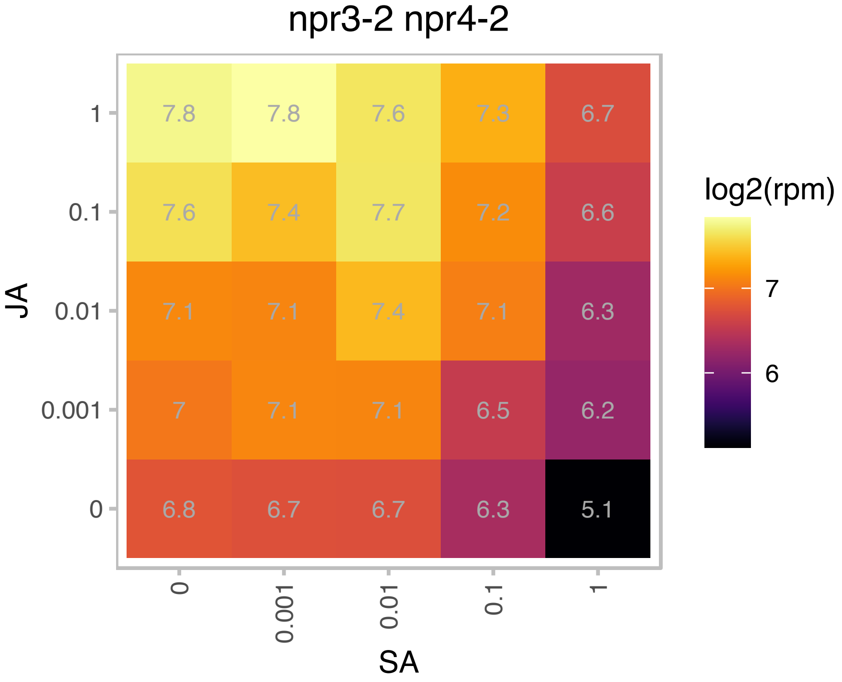


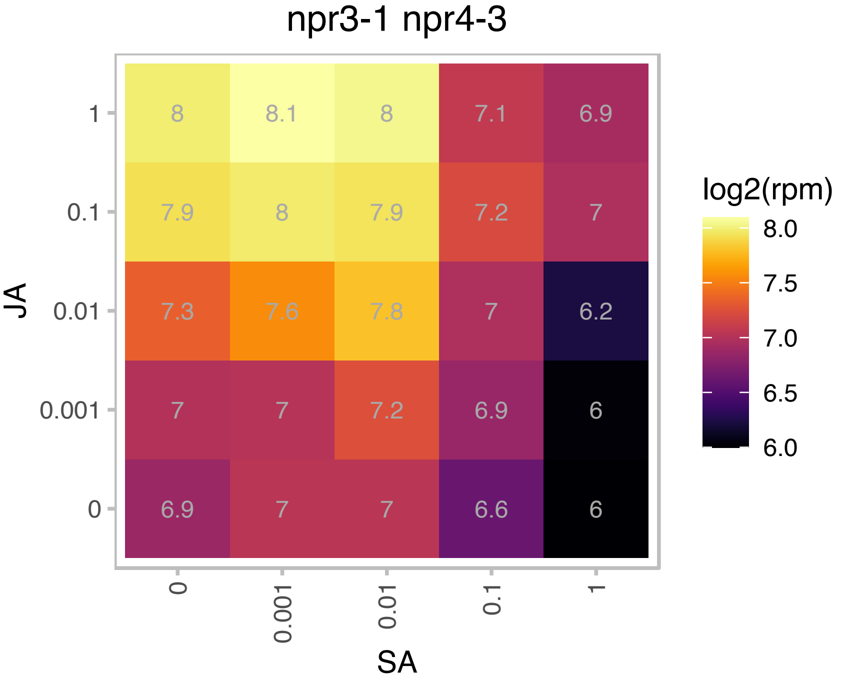

# AT4G08310.1 : NA

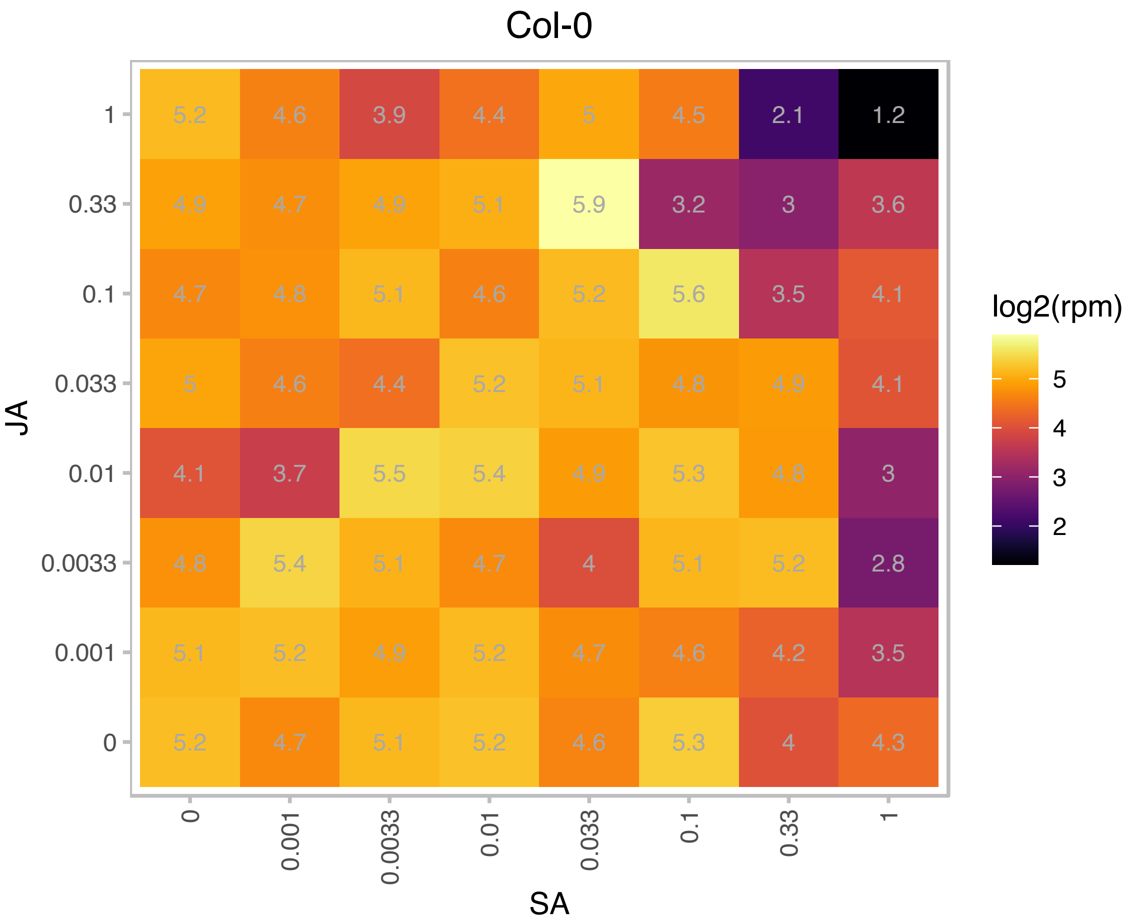


| | |
|---|---|
| Gene | AT4G08310 |
| is.expressed_genes | TRUE |
| is.responsive_genes | TRUE |
| is.apcluster | F-04 |

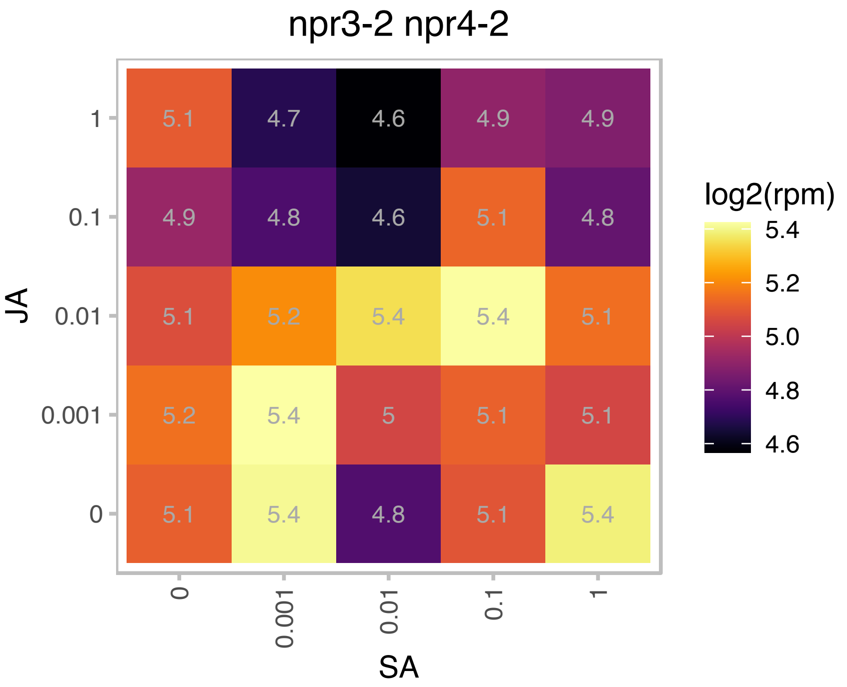


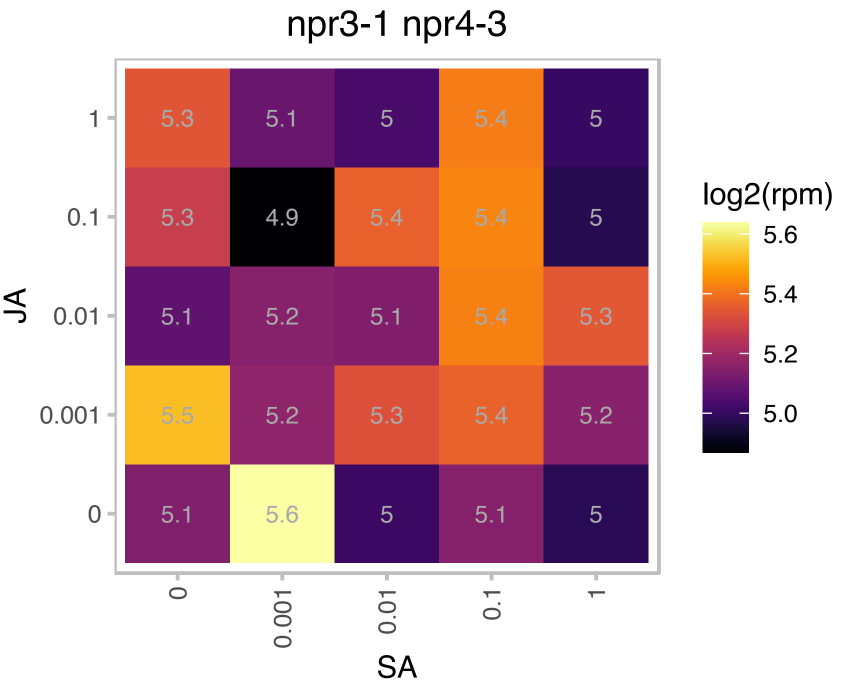

# AT4G08580.1 : NA

microfibrillar-associated protein-related

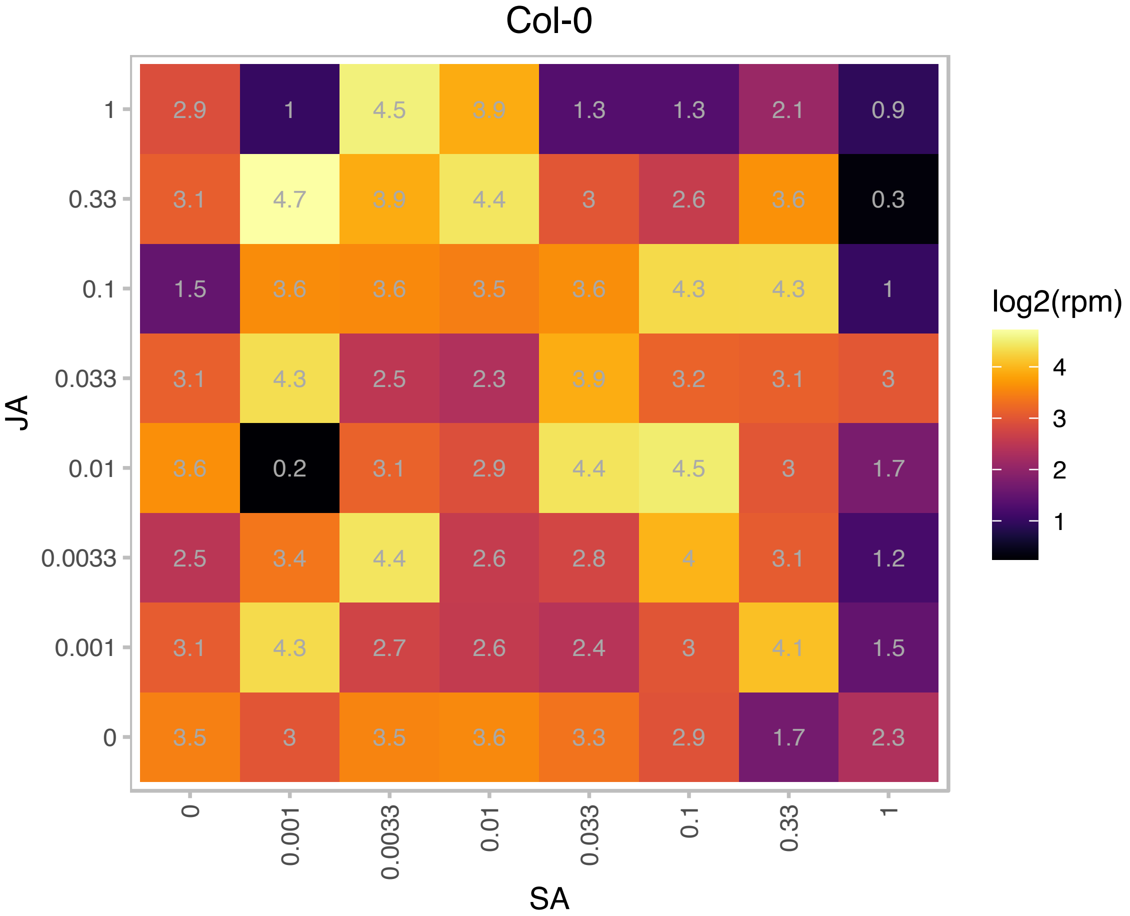


| | |
|---|---|
| Gene | AT4G08580 |
| is.expressed_genes | TRUE |
| is.responsive_genes | TRUE |
| is.apcluster | F-04 |

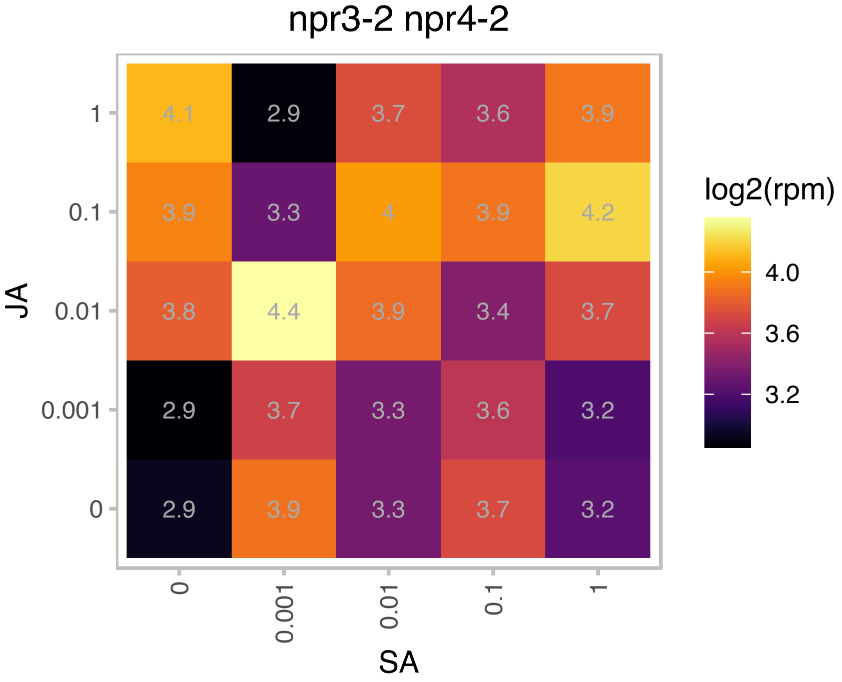


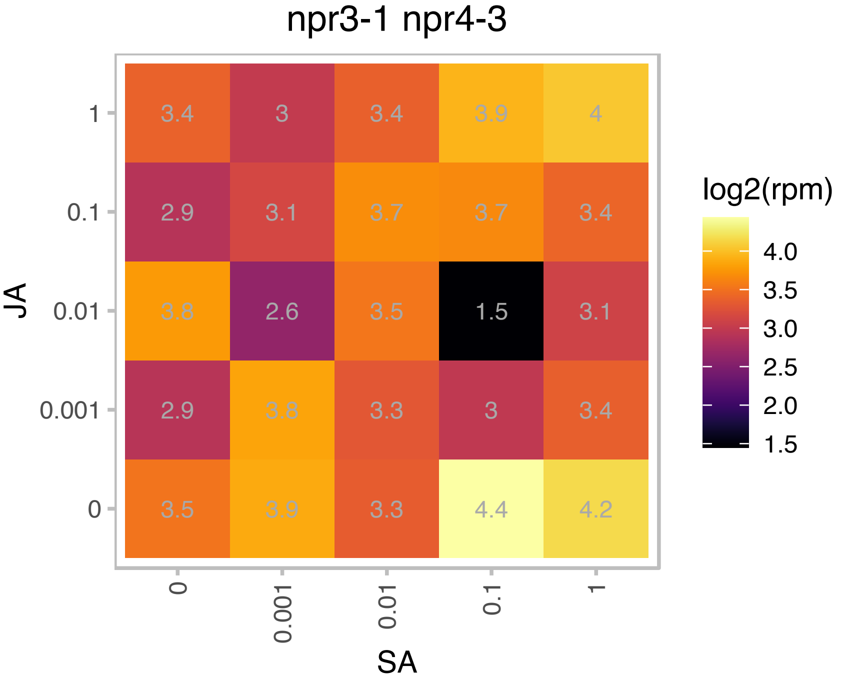

### AT4G13930.1 : SHM4

serine hydroxymethyltransferase 4

Encodes a serine hydroxymethyltransferase maximally expressed in root

#### Col-0

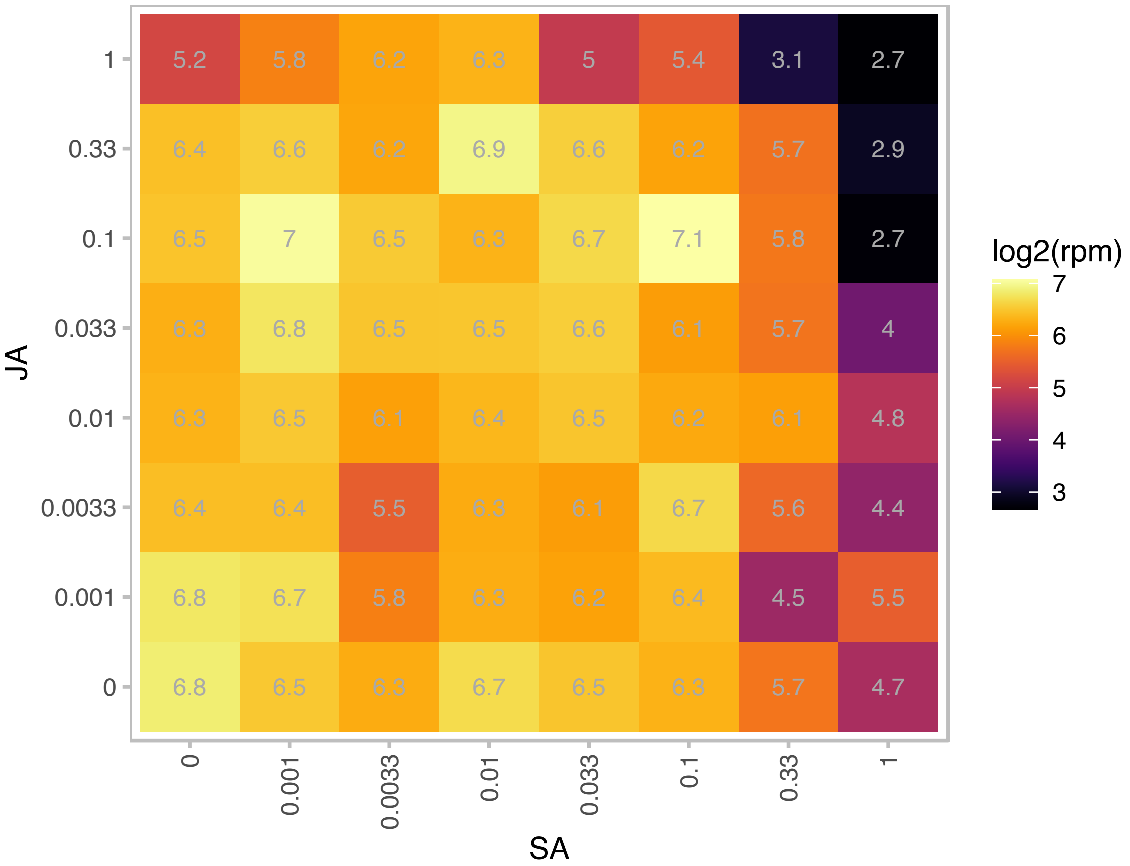


| | |
|---|---|
| Gene | AT4G13930 |
| is.expressed_genes | TRUE |
| is.responsive_genes | TRUE |
| is.apcluster | F-04 |

#### npr3-2 npr4-2

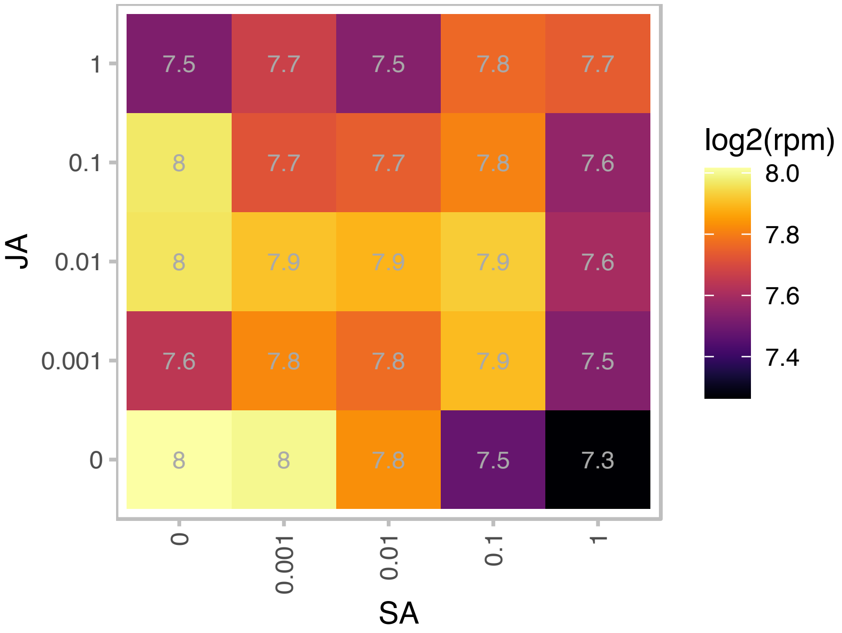


#### npr3-1 npr4-3

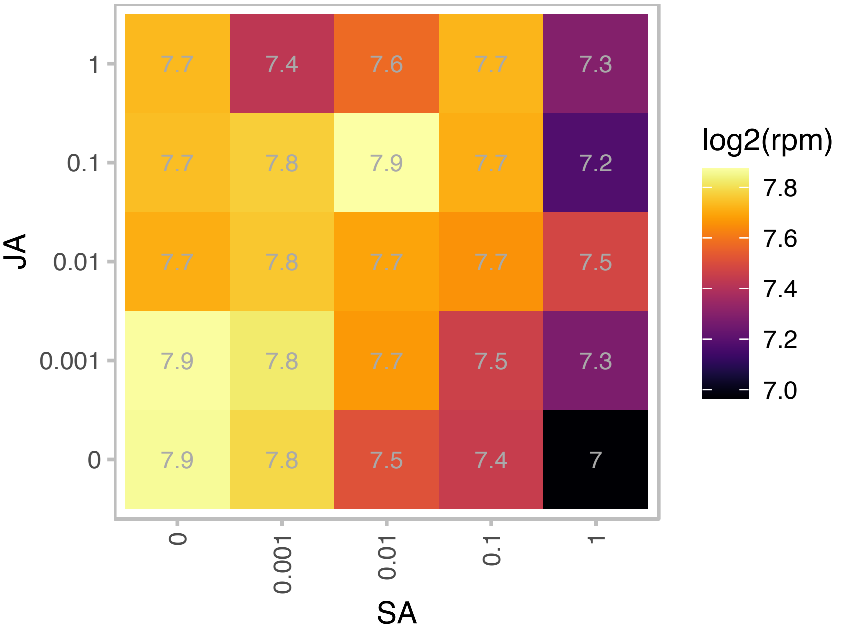

# AT4G20980.1 : NA

Eukaryotic translation initiation factor 3 subunit 7 (eIF-3)

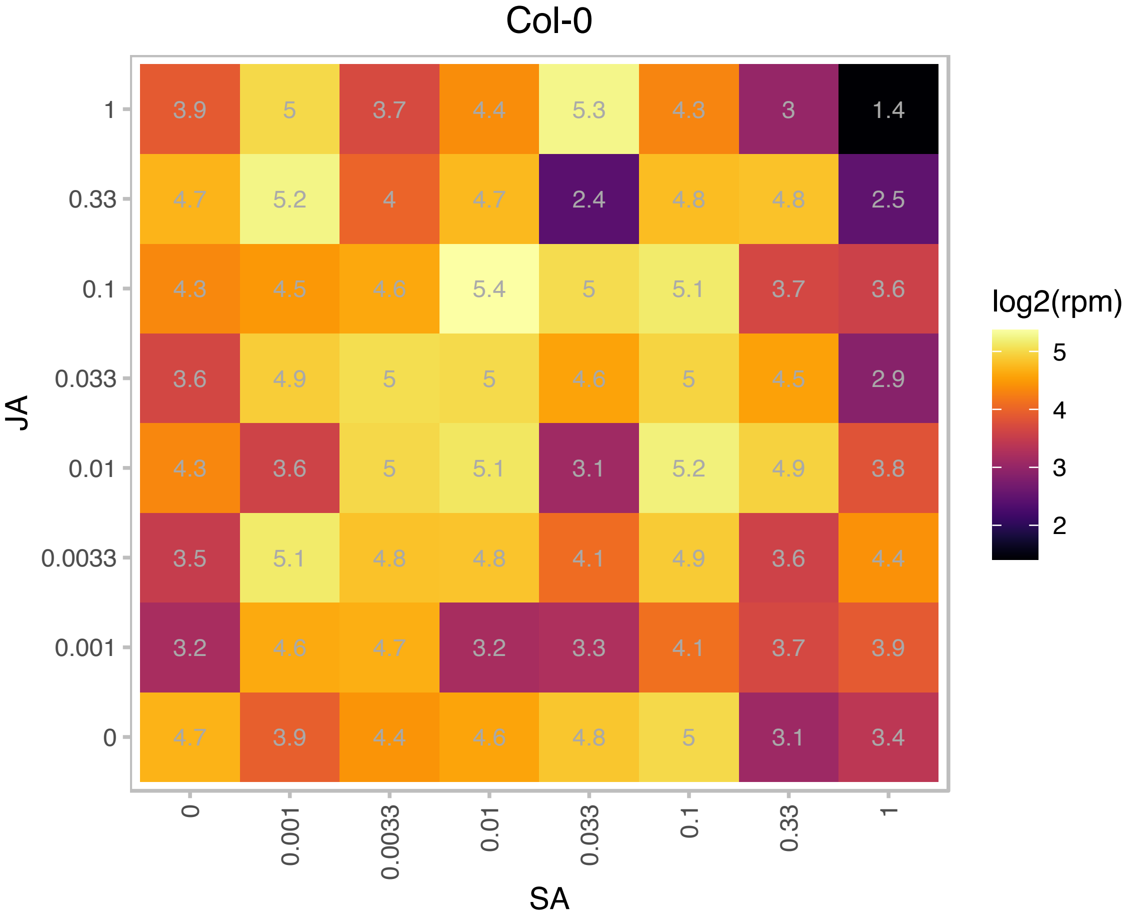


| | |
|---|---|
| Gene | AT4G20980 |
| is.expressed_genes | TRUE |
| is.responsive_genes | TRUE |
| is.apcluster | F-04 |

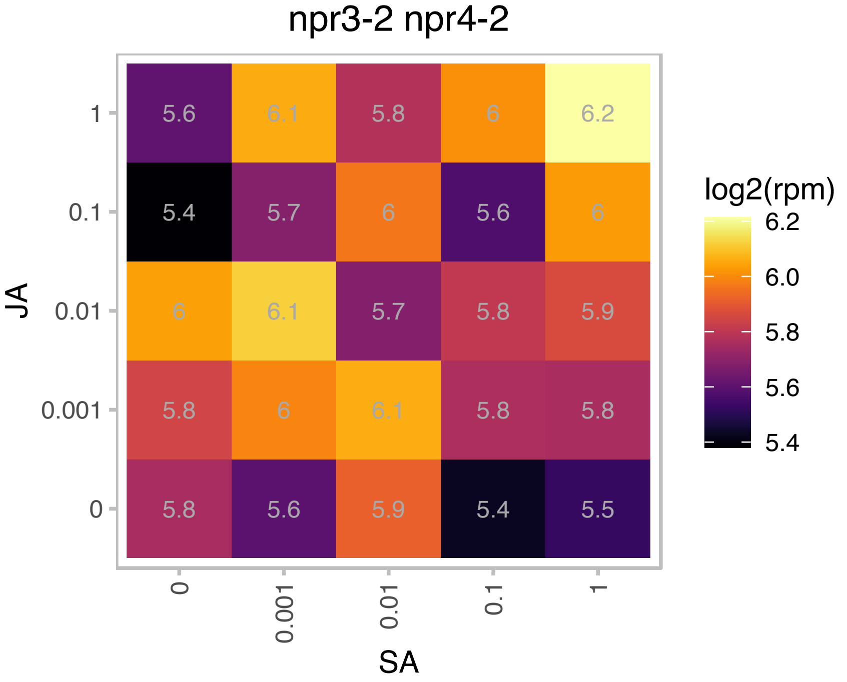


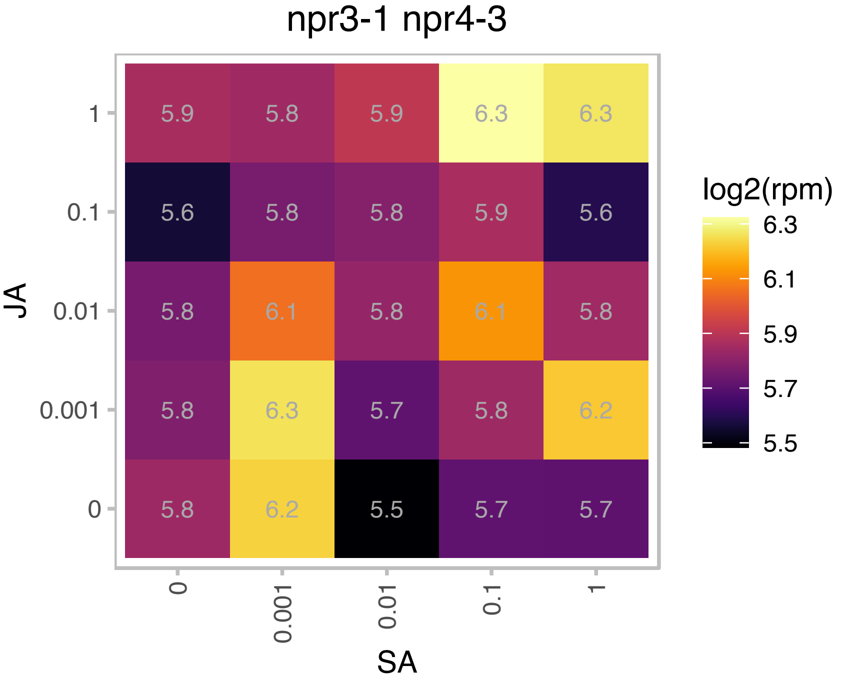

# AT4G32420.1 : NA

Cyclophilin-like peptidyl-prolyl cis-trans isomerase family protein

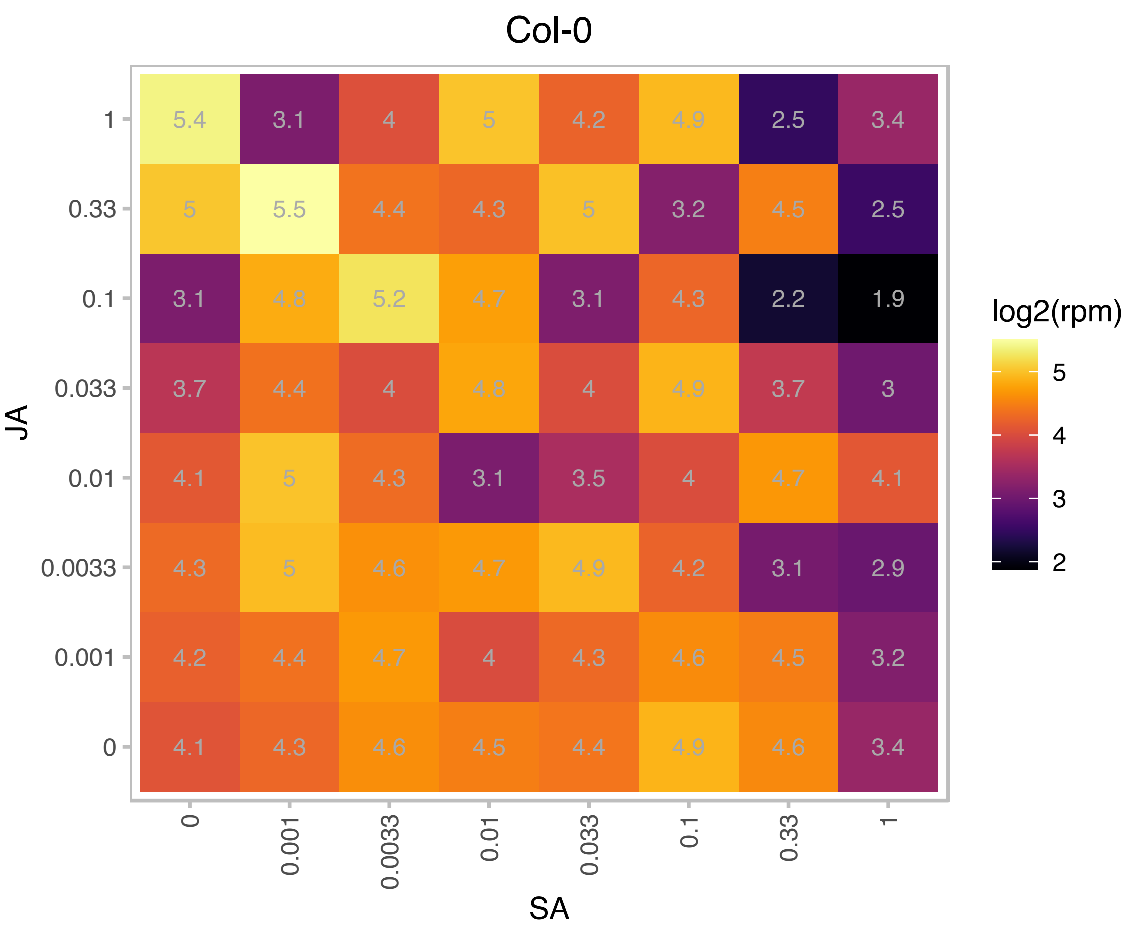


| | |
|---|---|
| Gene | AT4G32420 |
| is.expressed_genes | TRUE |
| is.responsive_genes | TRUE |
| is.apcluster | F-04 |

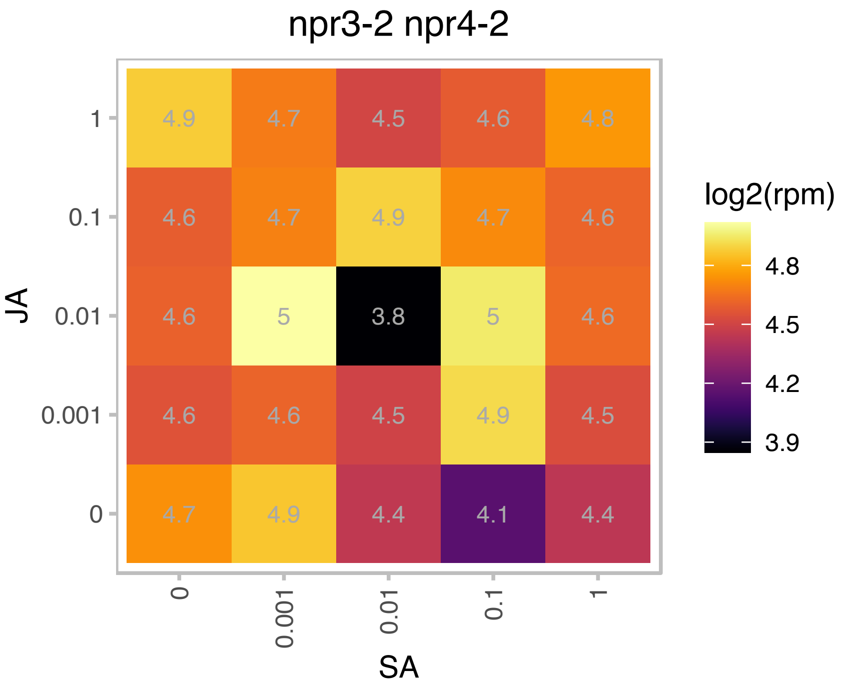


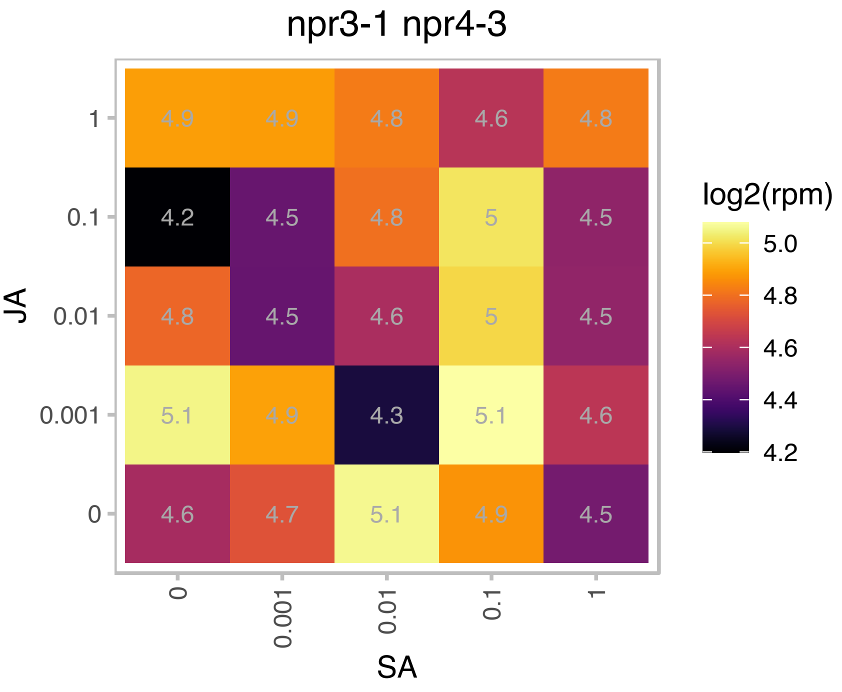

# AT4G35980.1 : NA

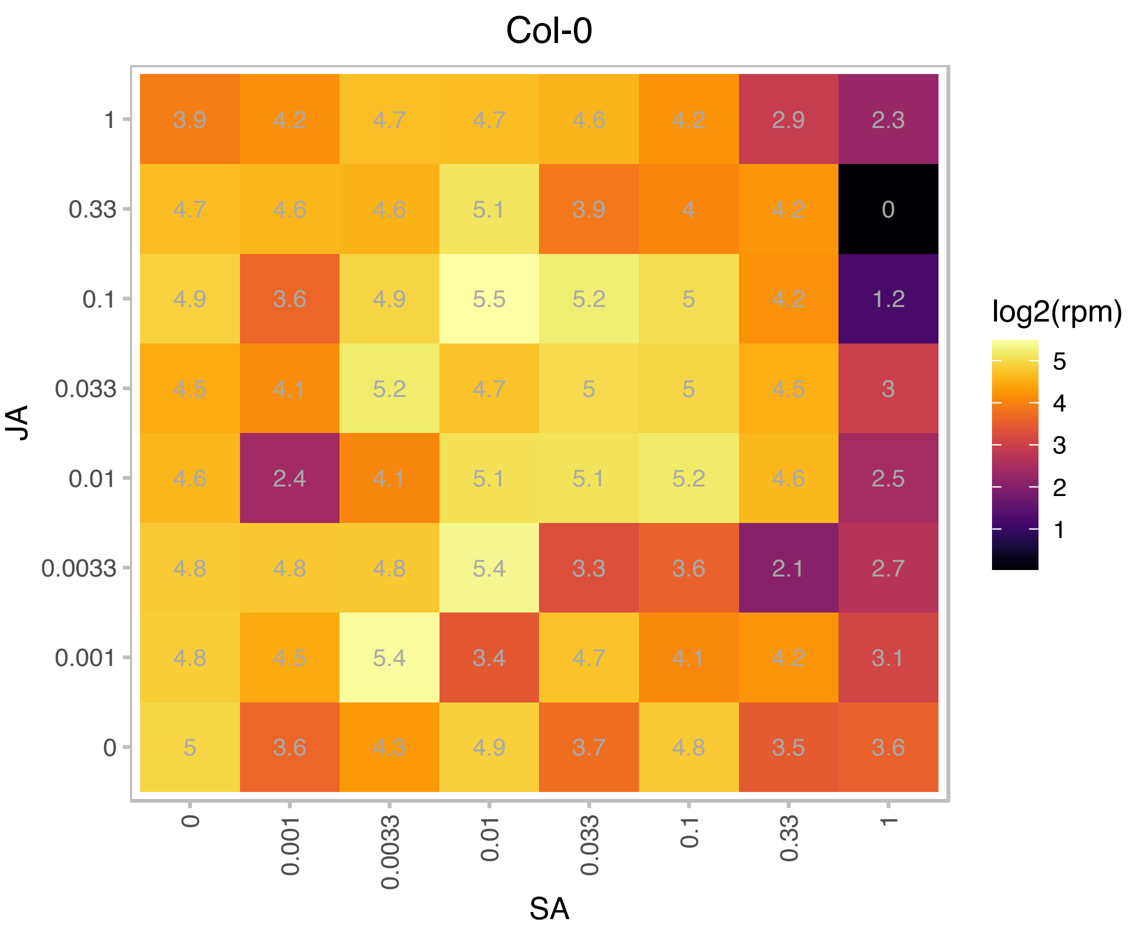


| | |
|---|---|
| Gene | AT4G35980 |
| is.expressed_genes | TRUE |
| is.responsive_genes | TRUE |
| is.apcluster | F-04 |

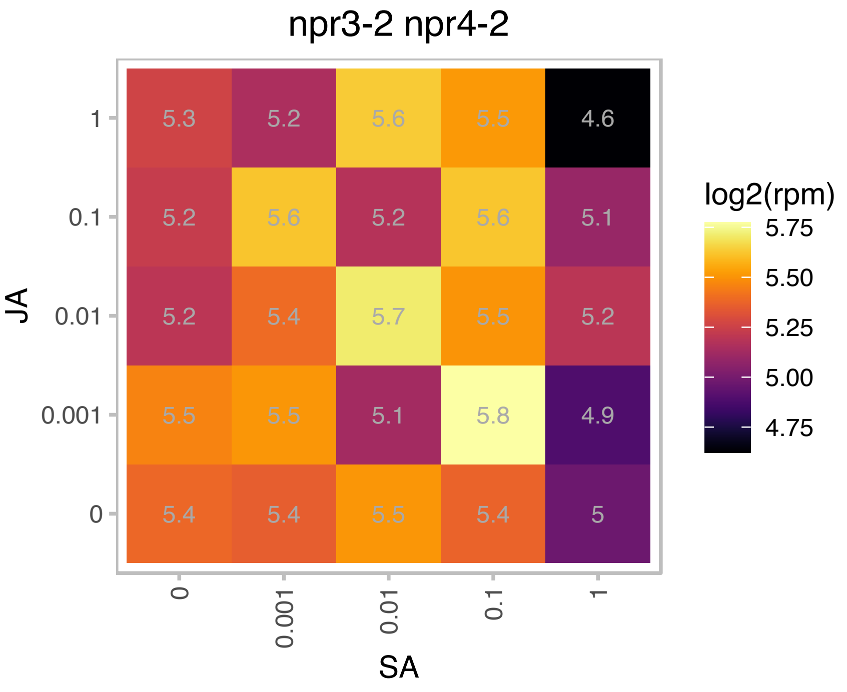


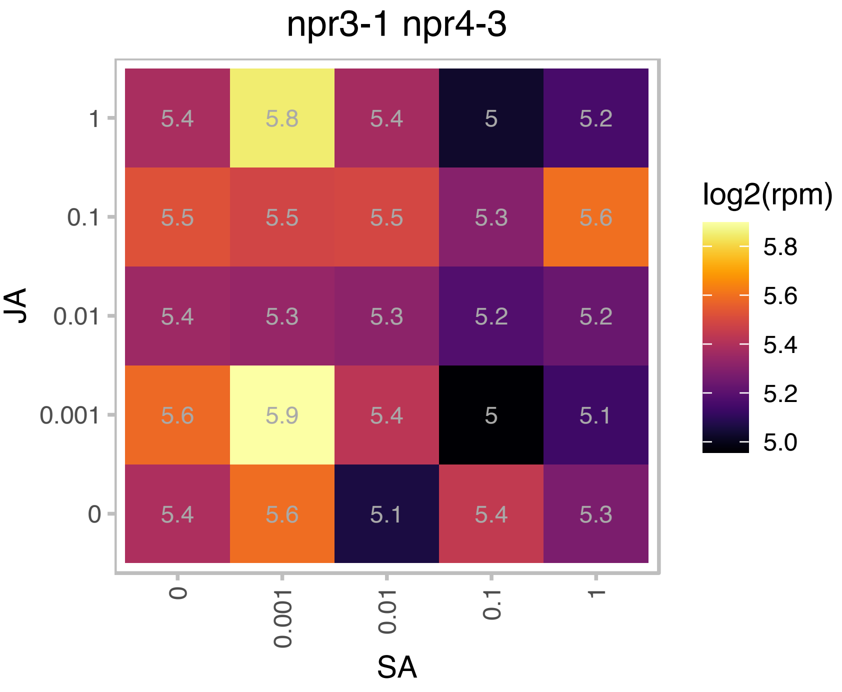

### AT4G36020.1 : AtCSP1, CSDP1, CSP1

cold shock domain protein 1

Encodes a cold shock domain protein. Involved in cold acclimation by blocking the secondary structure of mRNA which in turn facilitates translation at cold temperature.

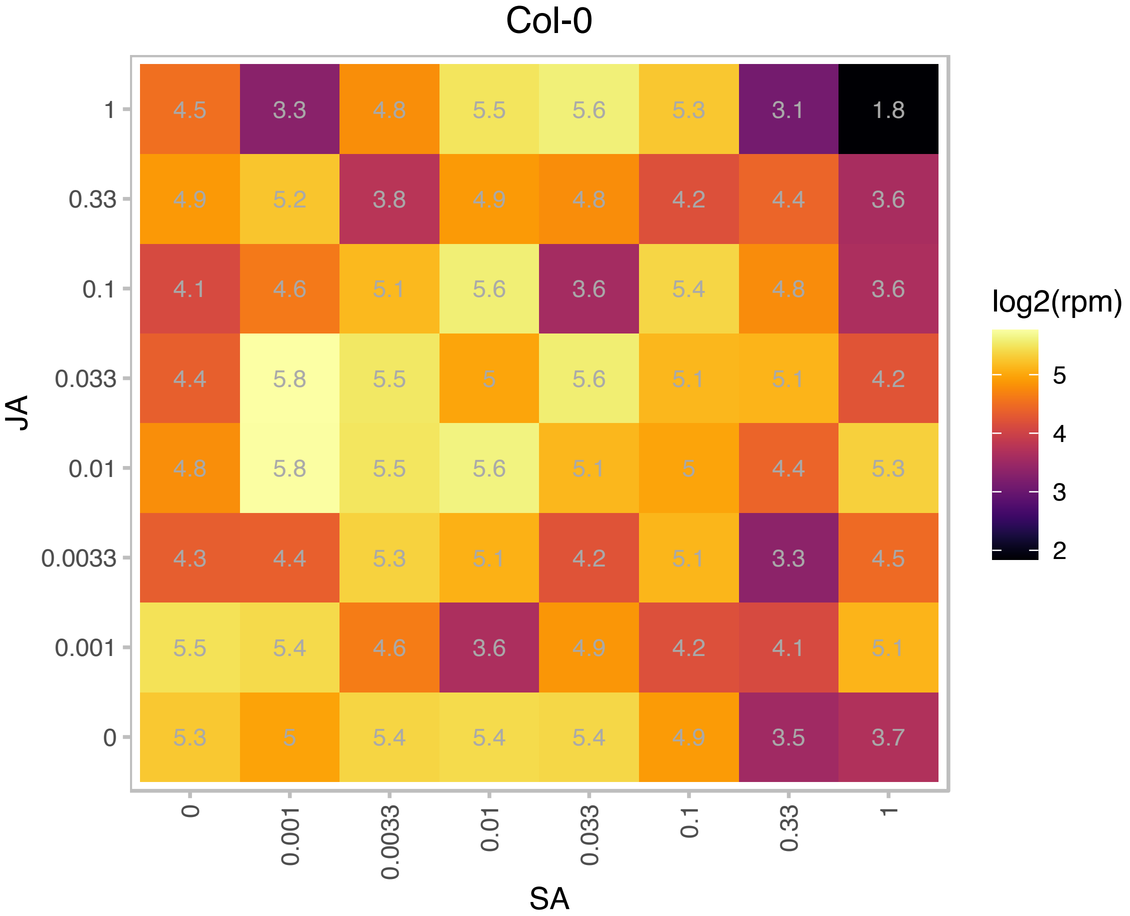


| | |
|---|---|
| Gene | AT4G36020 |
| is.expressed_genes | TRUE |
| is.responsive_genes | TRUE |
| is.apcluster | F-04 |

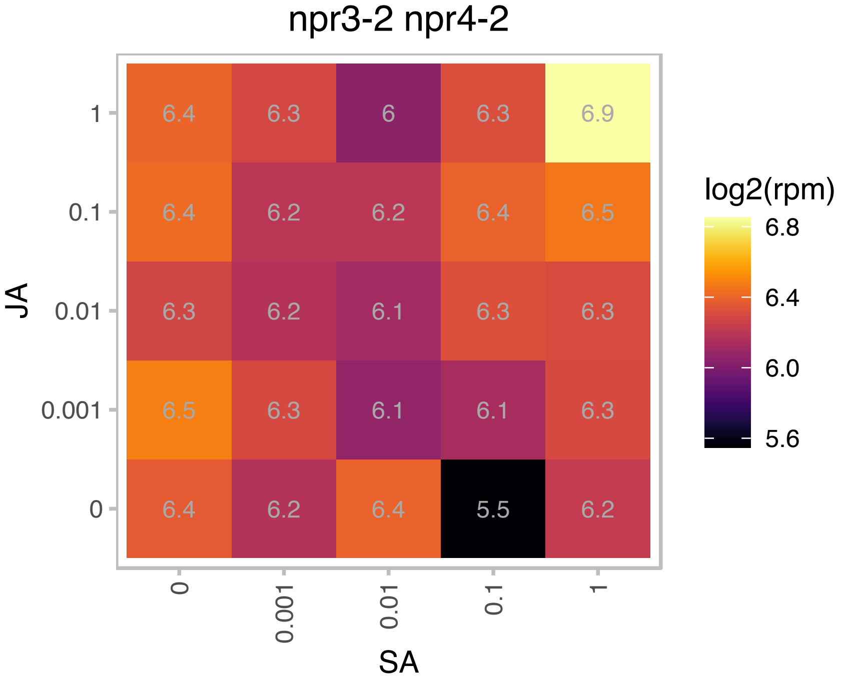


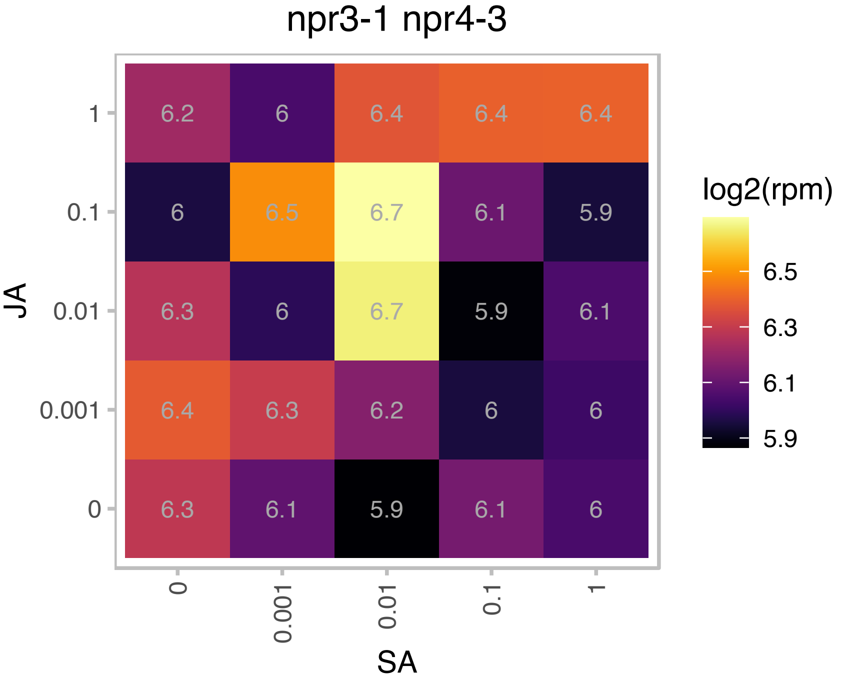

### AT5G04430.1 : BTR1, BTR1L, BTR1S

binding to TOMV RNA 1L (long form)

Gene model AT5G04430.1 produces active protein. (BTS1S). Binds to ToMV genomic RNA and prevents viral multiplication.

Col-0

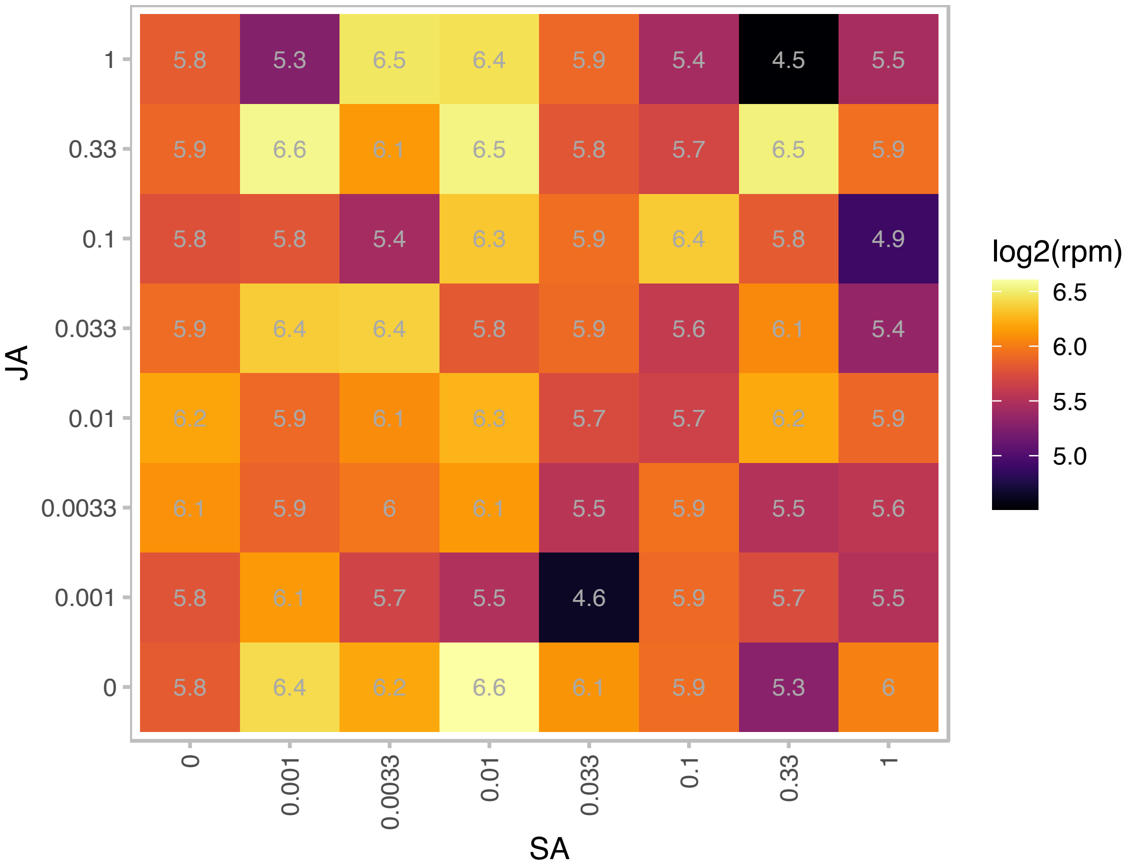


| | |
|---|---|
| Gene | AT5G04430 |
| is.expressed_genes | TRUE |
| is.responsive_genes | TRUE |
| is.apcluster | F-04 |

npr3-2 npr4-2

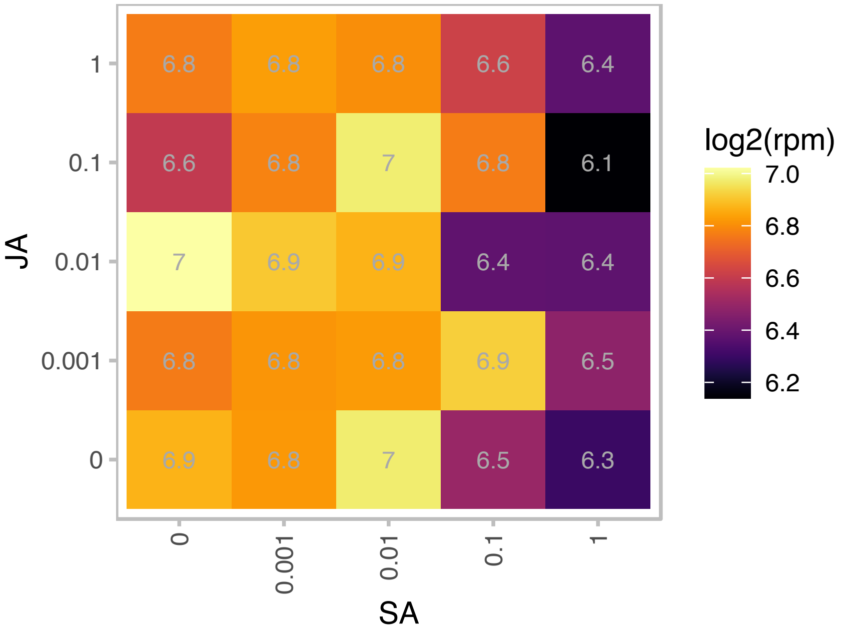


npr3-1 npr4-3

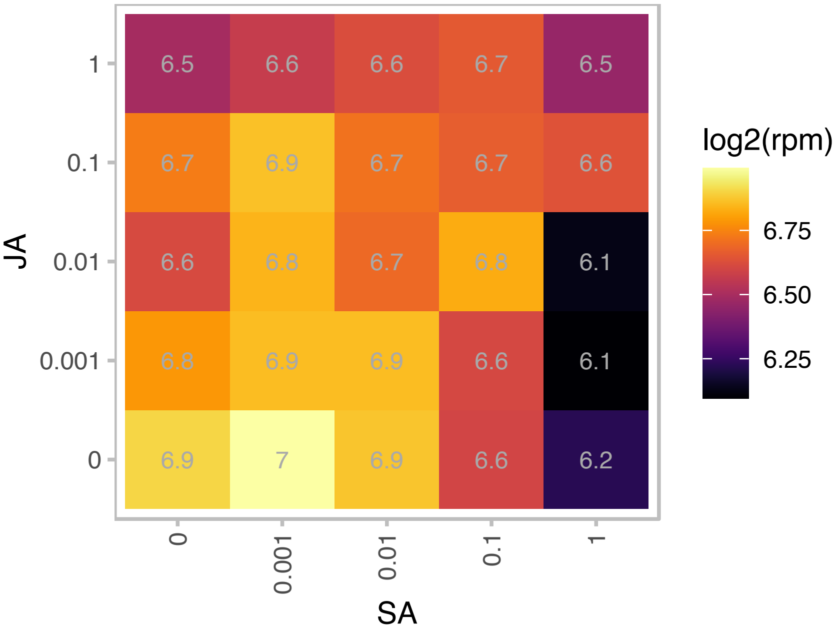

# AT5G07940.1 : NA

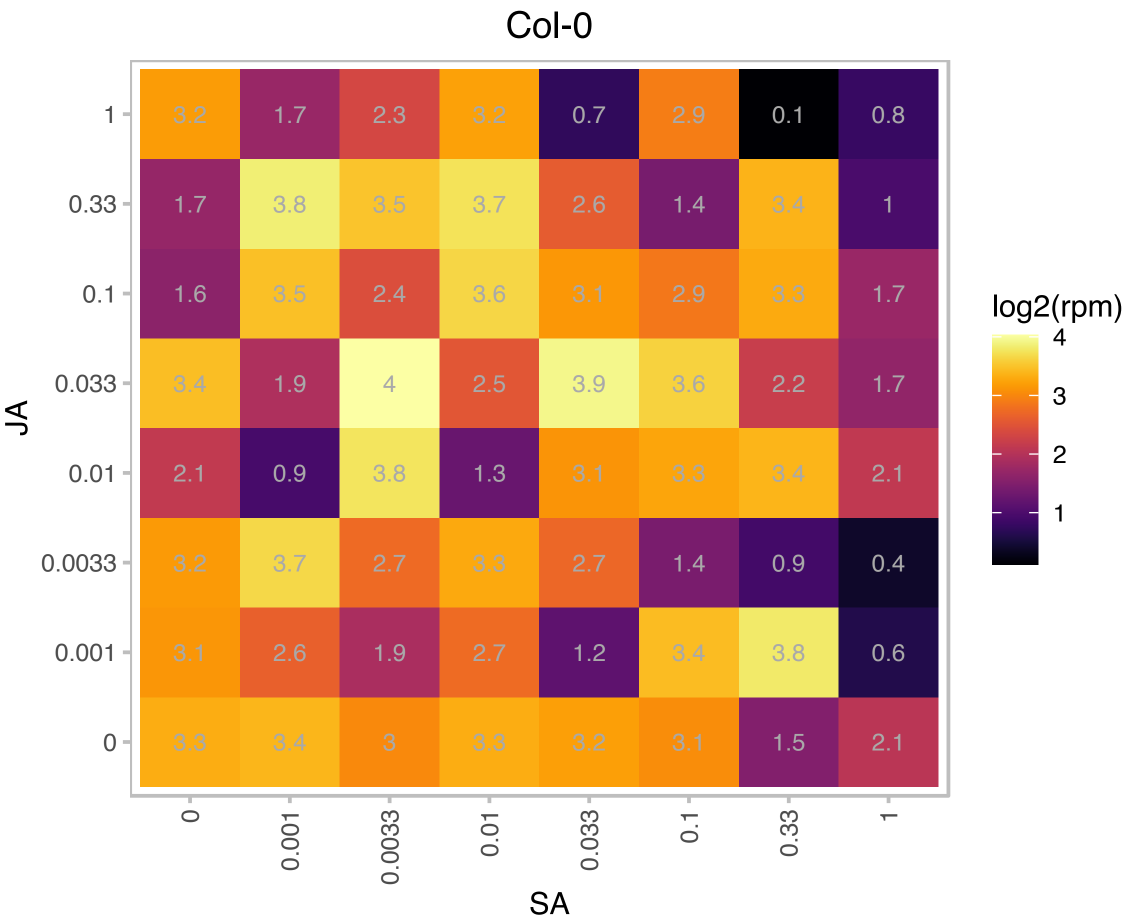


| | |
|---|---|
| Gene | AT5G07940 |
| is.expressed_genes | TRUE |
| is.responsive_genes | TRUE |
| is.apcluster | F-04 |

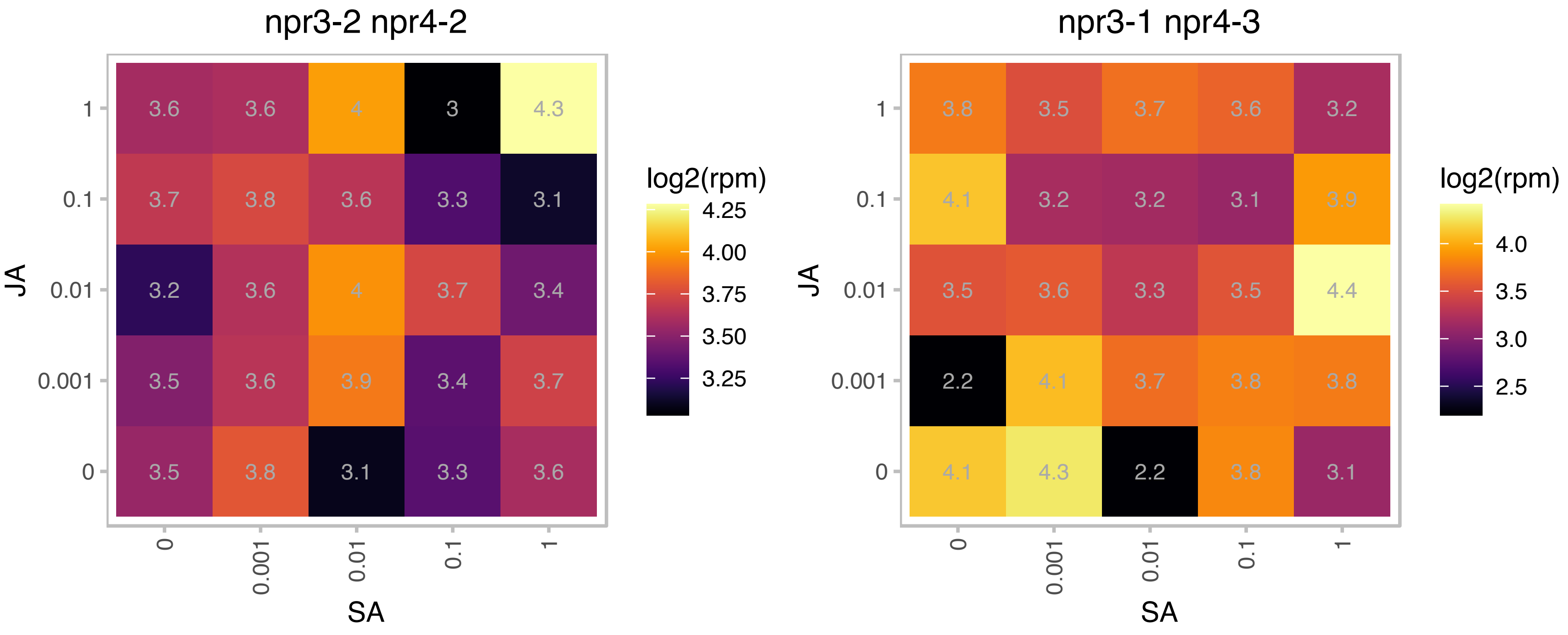

# AT5G08420.1 : NA

RNA-binding KH domain-containing protein

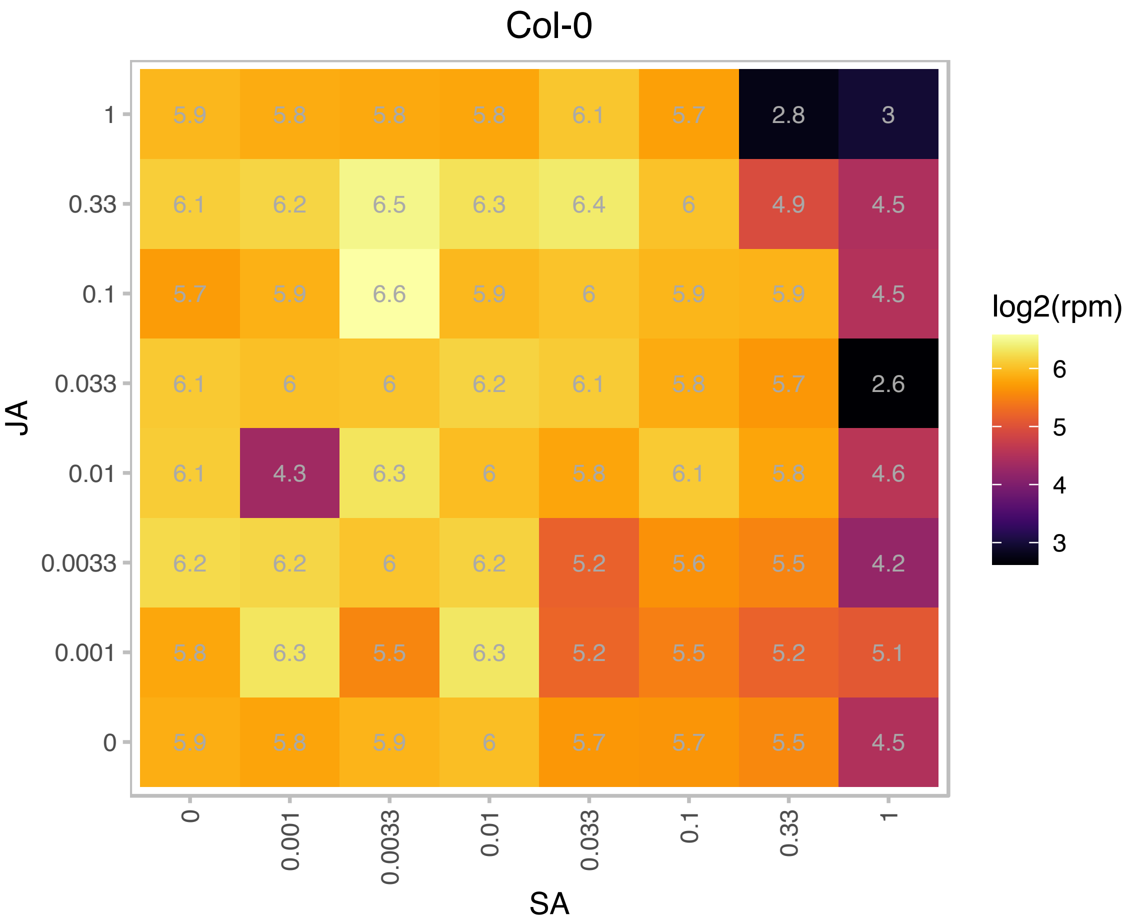


| | |
|---|---|
| Gene | AT5G08420 |
| is.expressed_genes | TRUE |
| is.responsive_genes | TRUE |
| is.apcluster | F-04 |

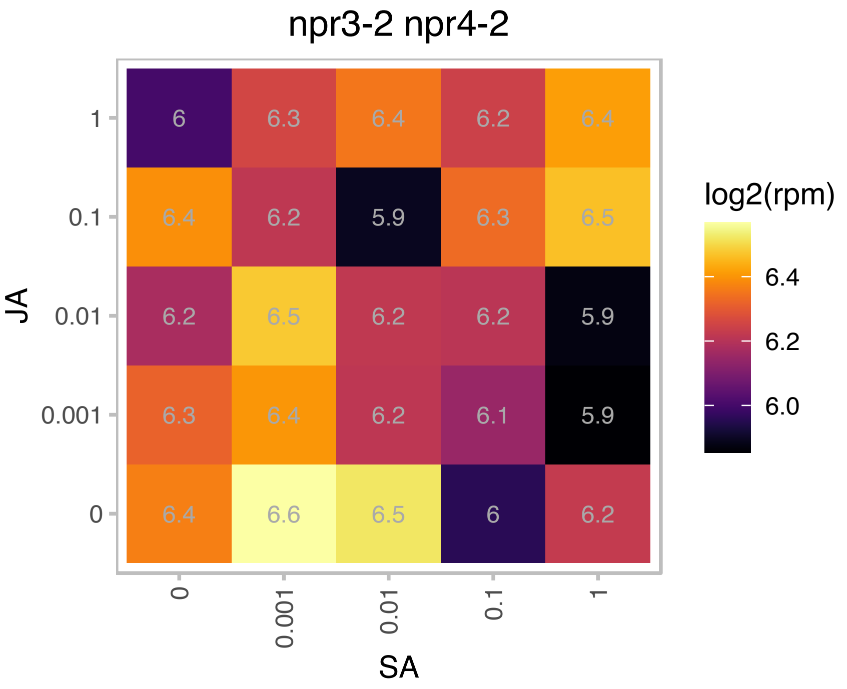


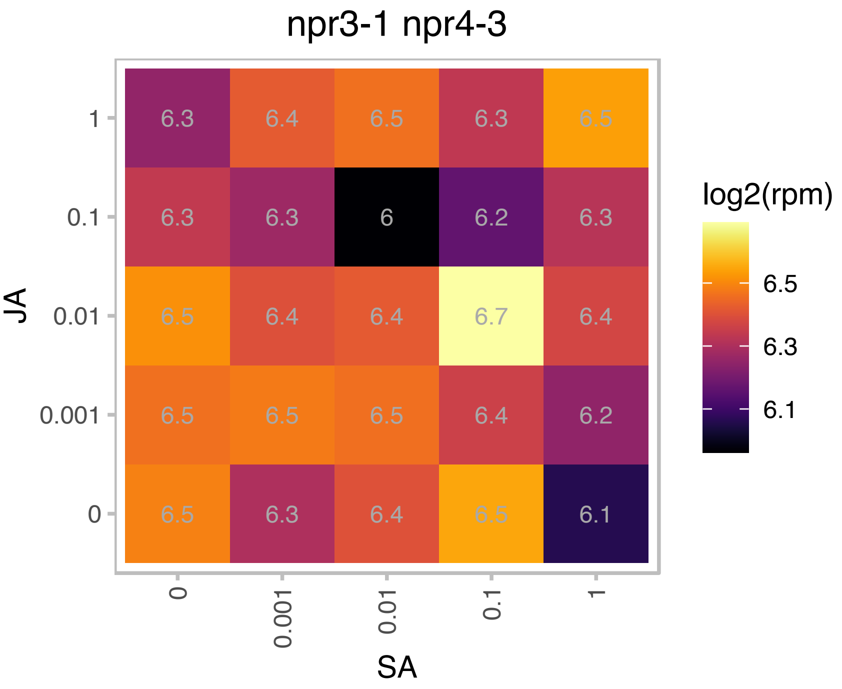

### AT5G10350.1 : RRM

RNA-binding (RRM/RBD/RNP motifs) family protein
Encodes a nuclear-localized RNA recognition motif-containing protein that forms homodimers.  It interacts with telomere and non-telomere associated proteins. Null mutation does not result in changes in telomerase activity or telomere length.

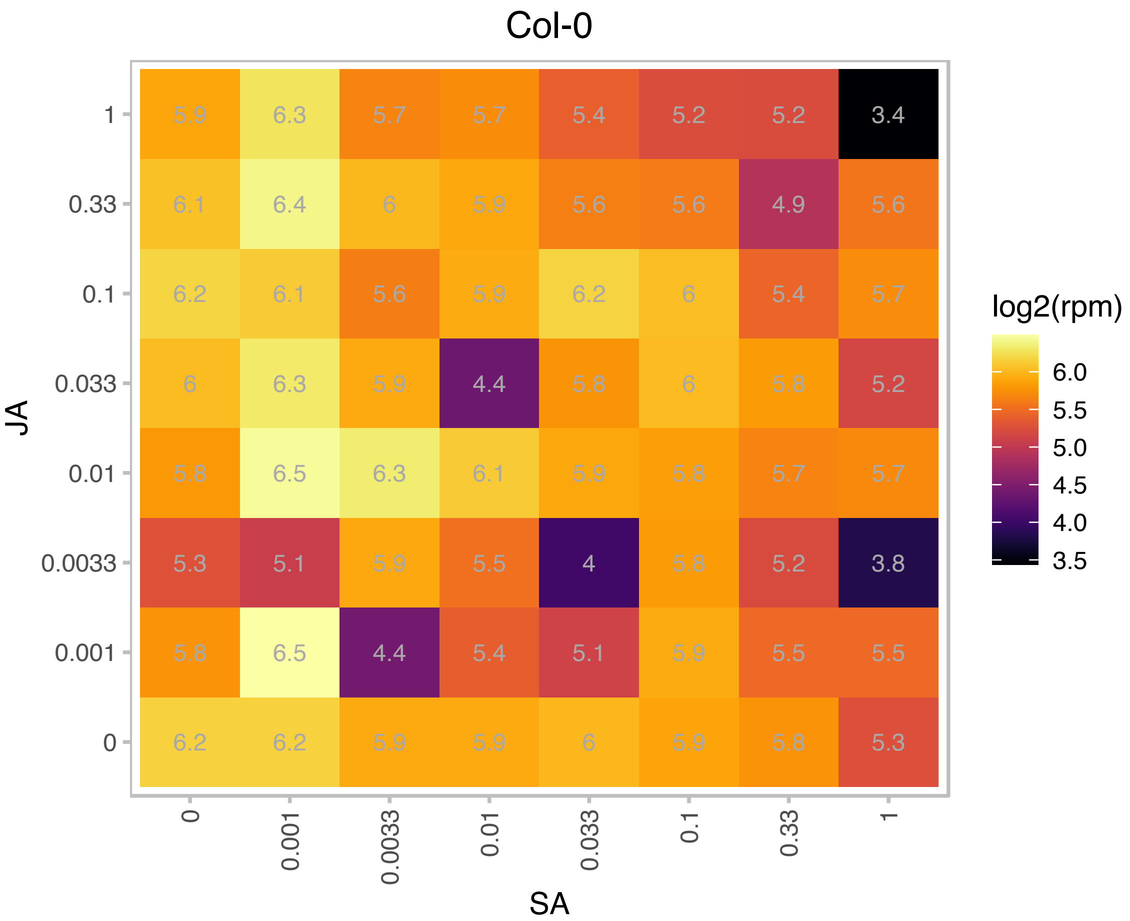


| | |
|---|---|
| Gene | AT5G10350 |
| is.expressed_genes | TRUE |
| is.responsive_genes | TRUE |
| is.apcluster | F-04 |

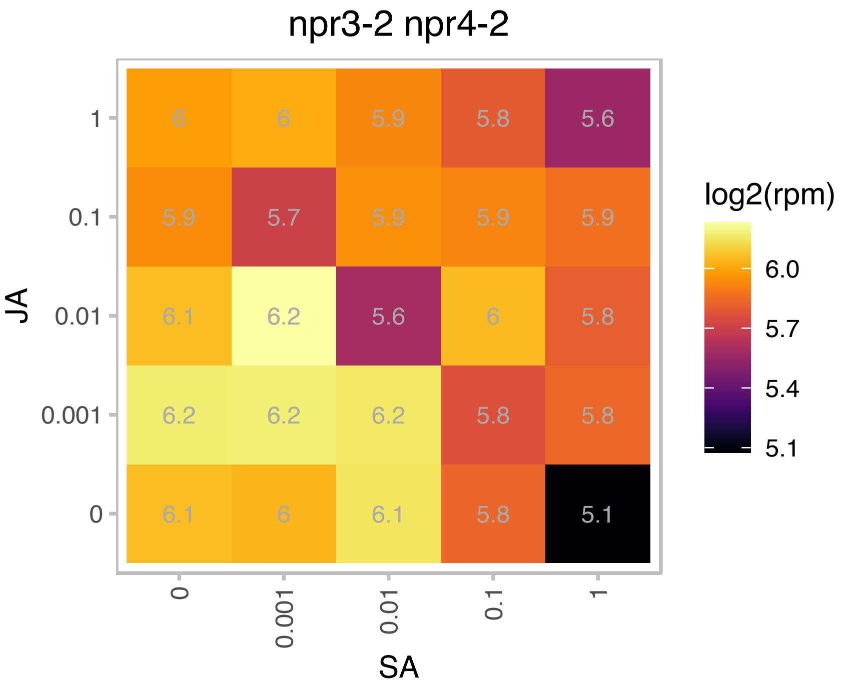


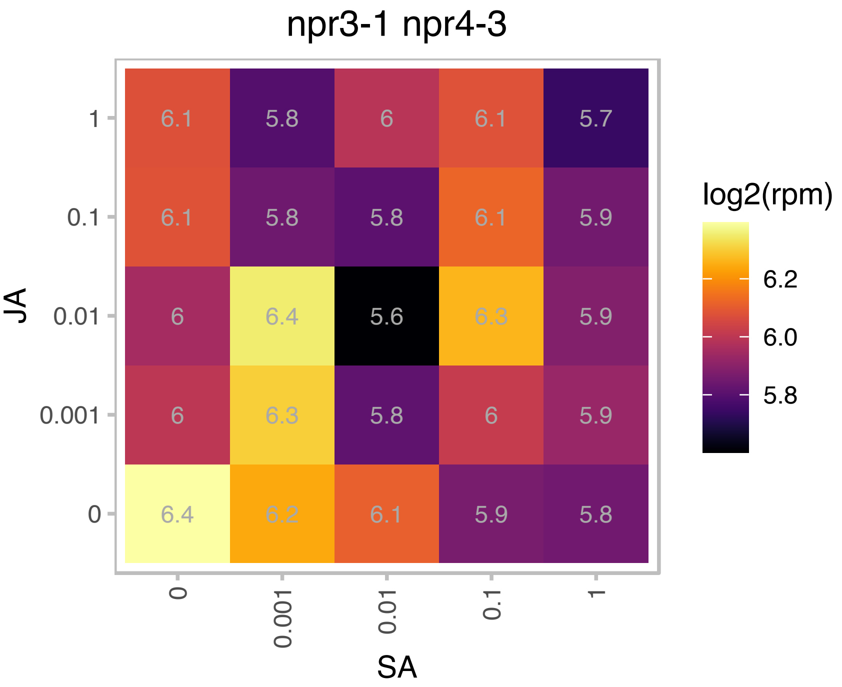

# AT5G12410.1 : NA

THUMP domain-containing protein

| | |
|---|---|
| Gene | AT5G12410 |
| is.expressed_genes | TRUE |
| is.responsive_genes | TRUE |
| is.apcluster | F-04 |

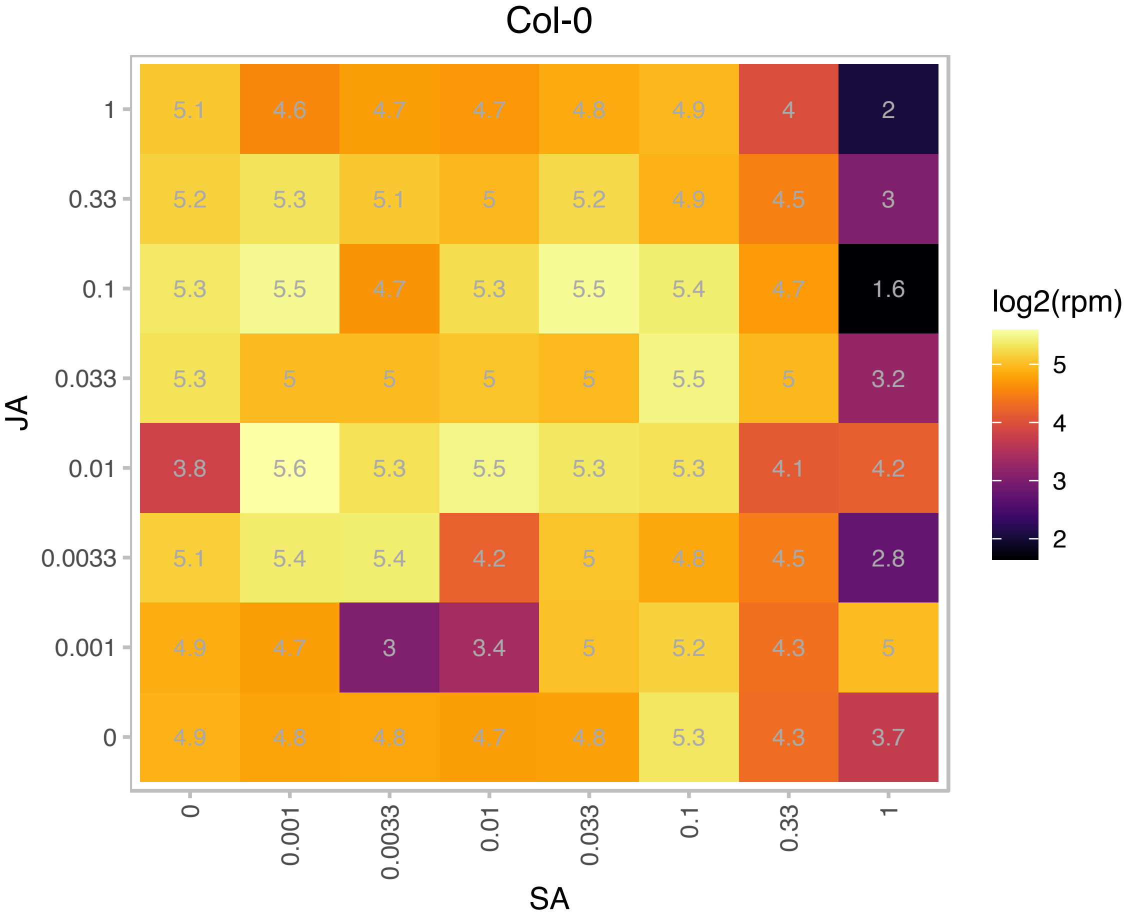


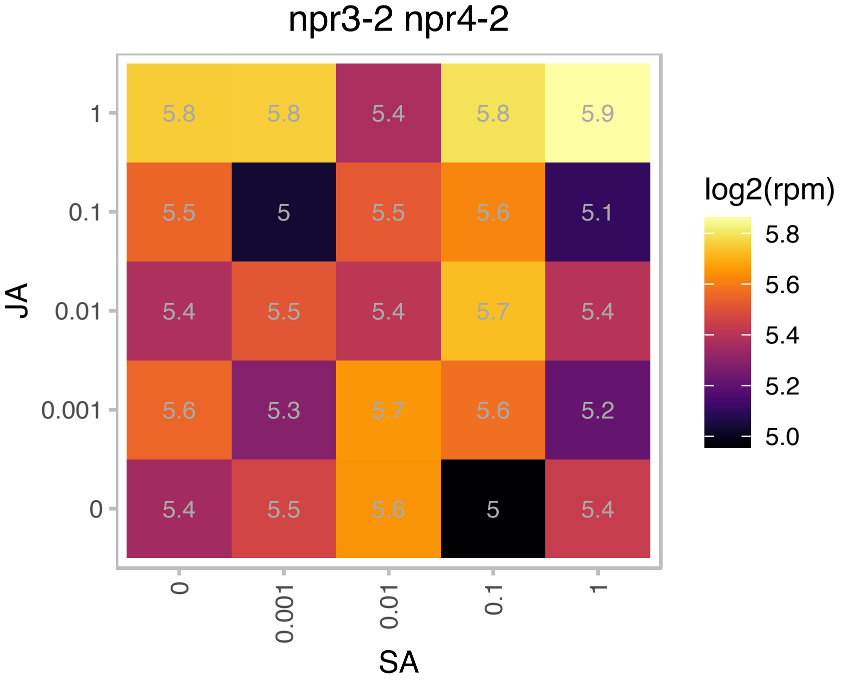


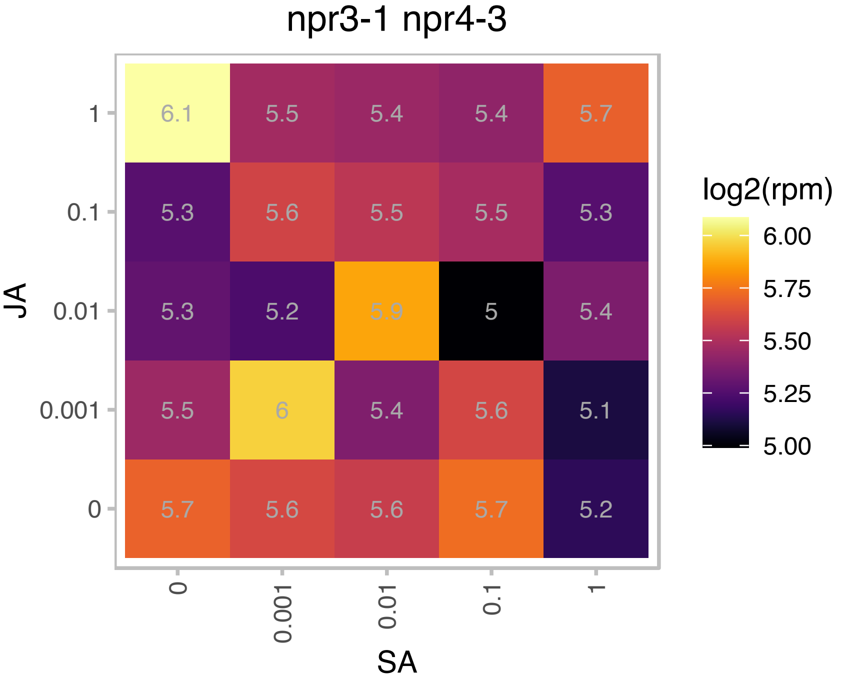

# AT5G13980.1 : NA

Glycosyl hydrolase family 38 protein

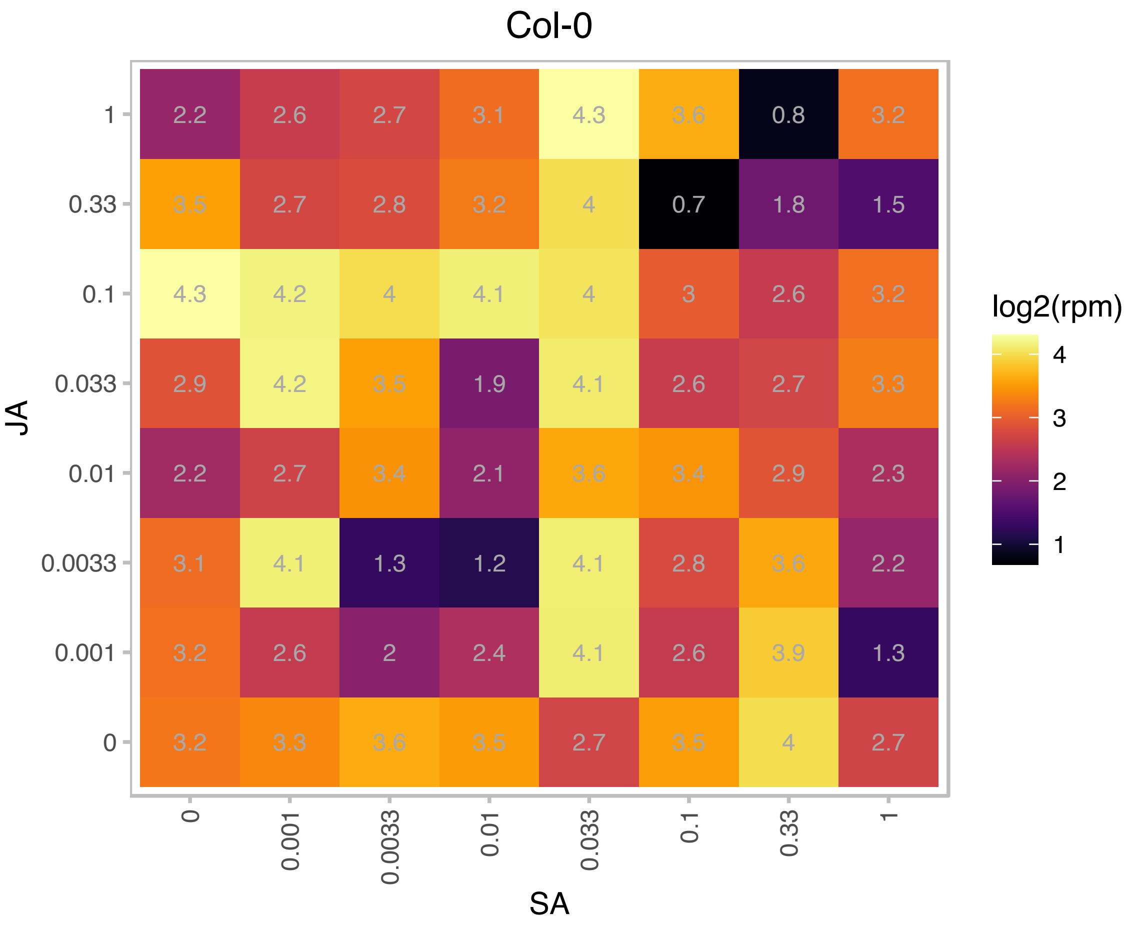


| | |
|---|---|
| Gene | AT5G13980 |
| is.expressed_genes | TRUE |
| is.responsive_genes | TRUE |
| is.apcluster | F-04 |

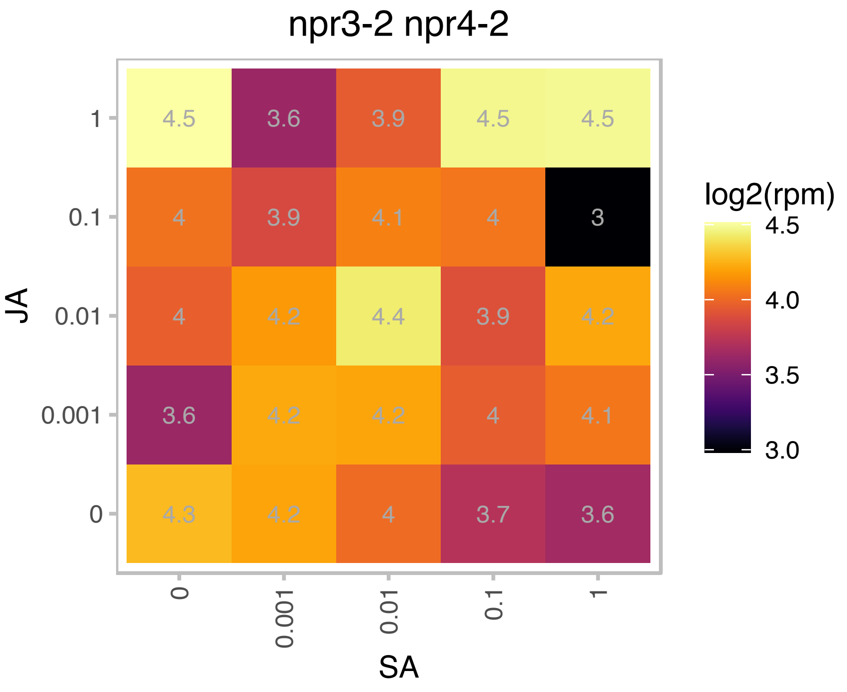


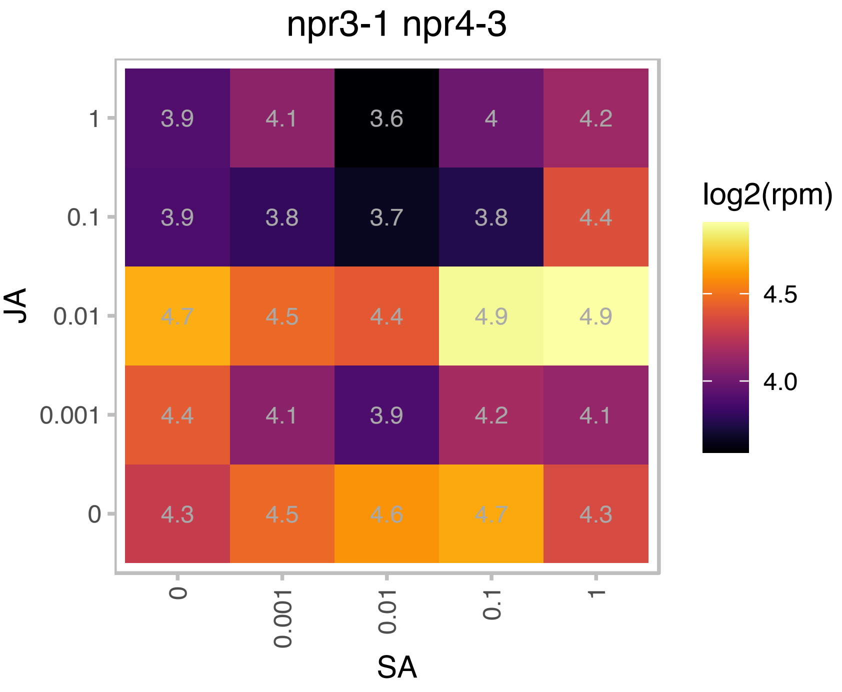

### AT5G17050.1 : UGT78D2

UDP-glucosyl transferase 78D2
The At5g17050 encodes a anthocyanidin 3-O-glucosyltransferase which specifically glucosylates the 3-position of the flavonoid C-ring. Anthocyanidins such as cyanidin and pelargonidin as well as flavonols such as kaempferol and quercetin are accepted substrates.

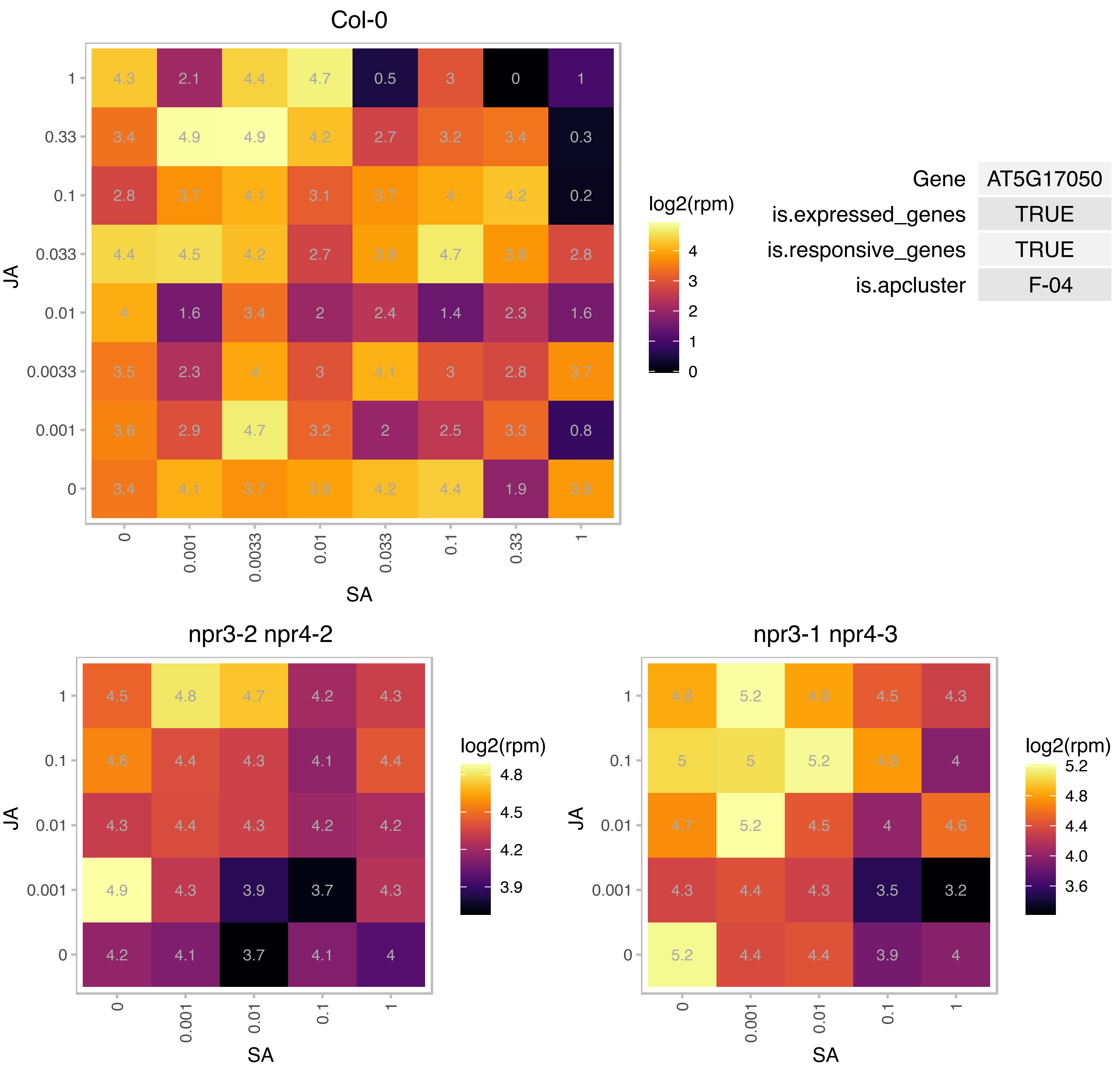


| | |
|---|---|
| Gene | AT5G17050 |
| is.expressed_genes | TRUE |
| is.responsive_genes | TRUE |
| is.apcluster | F-04 |

### AT5G17920.1 : ATCIMS, ATMETS, ATMS1

Cobalamin-independent synthase family protein
Encodes a cytosolic cobalamin-independent methionine synthase, involved in methionine regeneration via the activated methyl cycle (SAM cycle). The protein undergoes thiolation following treatment with the oxidant tert-butylhydroperoxide. The mRNA is cell-to-cell mobile.

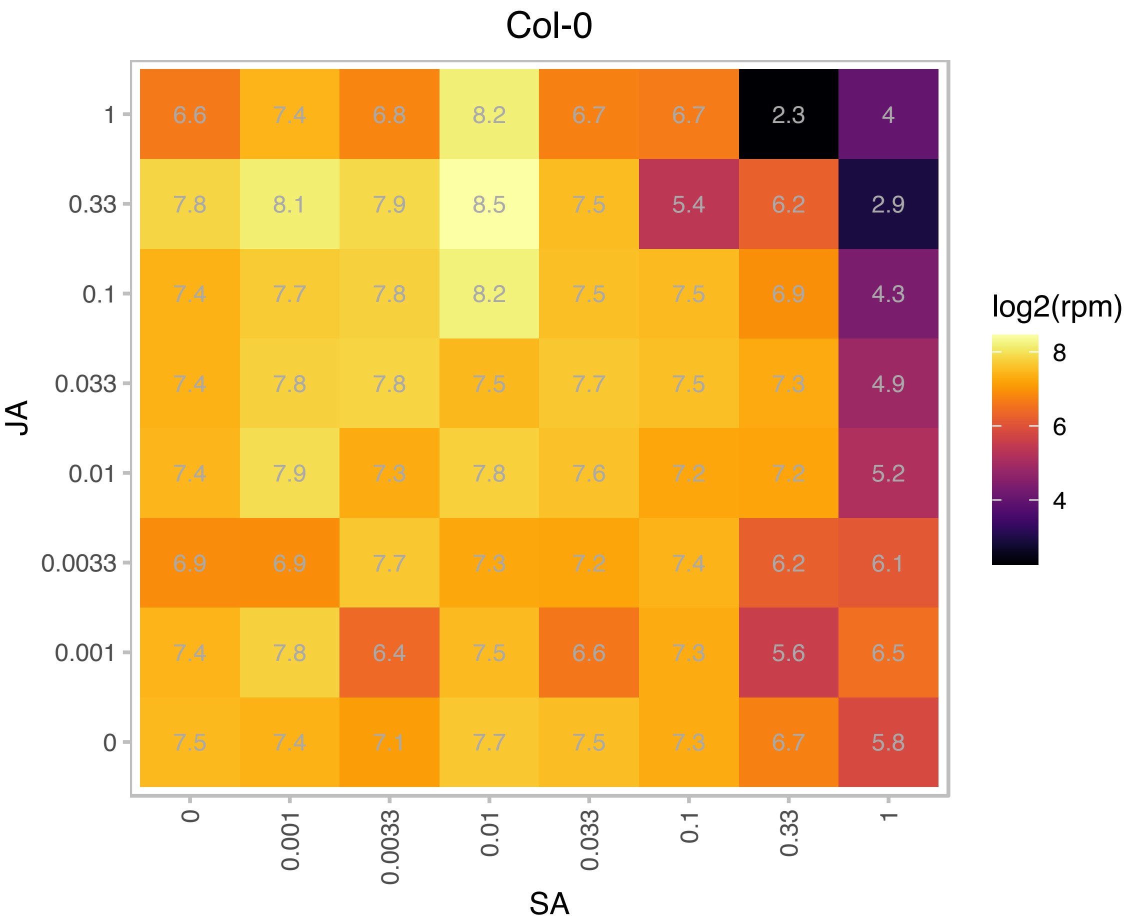


| | |
|---|---|
| Gene | AT5G17920 |
| is.expressed_genes | TRUE |
| is.responsive_genes | TRUE |
| is.apcluster | F-04 |

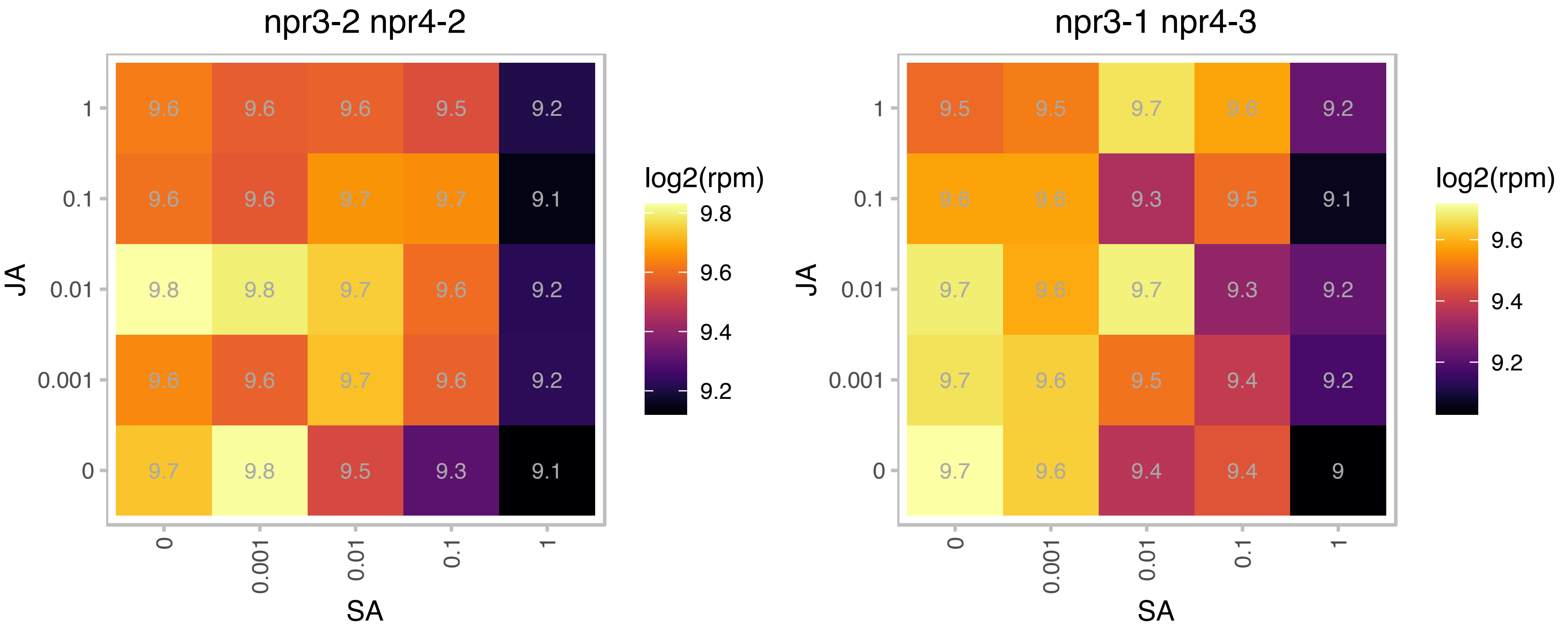

### AT5G19820.1 : EMB2734, KETCH1

ARM repeat superfamily protein

Encodes an importin that transports HYL1, a component of the microprocessor, from the cytoplasm to the nucleus to constitute functional microprocessor, thereby affecting miRNA processing. Knockdown amiR mutants significantly reduced nuclear portion of HYL1 protein and correspondingly compromised the pri-miRNA processing in the nucleus.

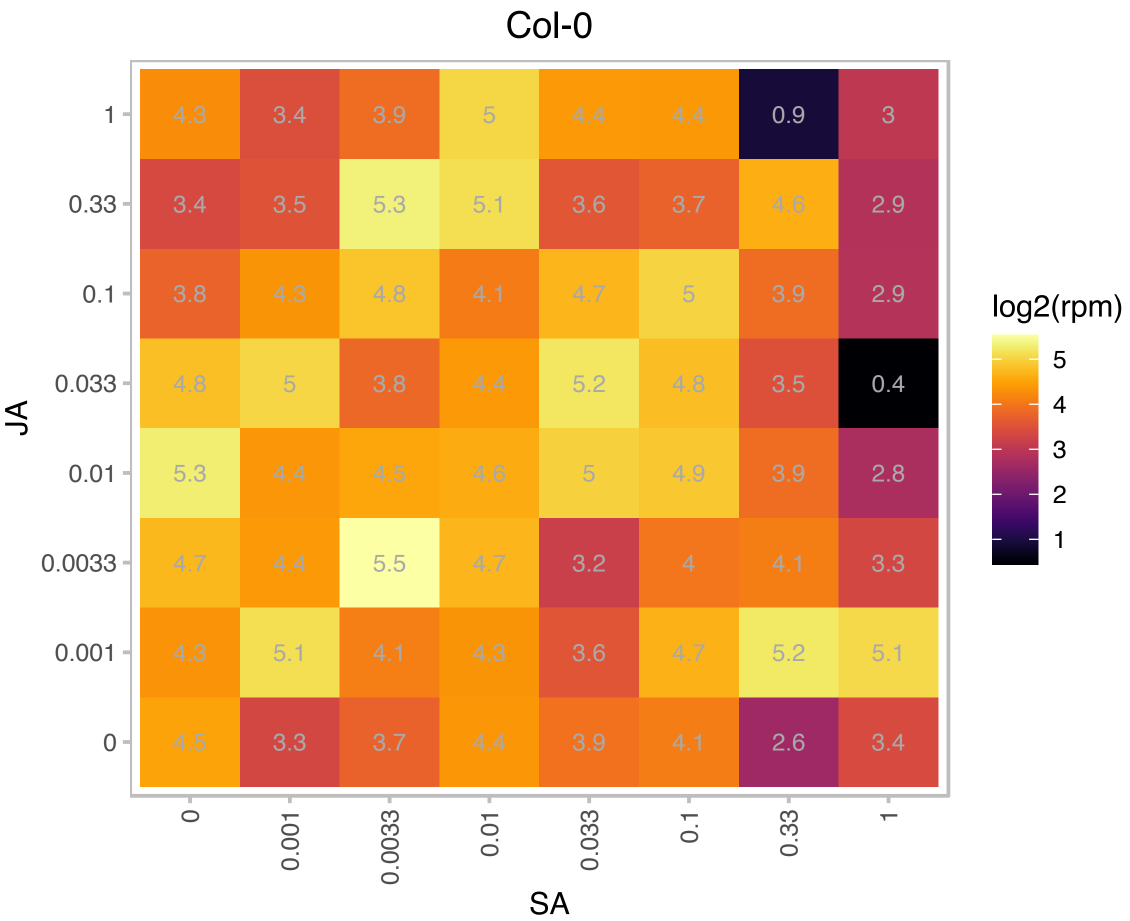


| Gene | AT5G19820 |
|---|---|
| is.expressed_genes | TRUE |
| is.responsive_genes | TRUE |
| is.apcluster | F-04 |

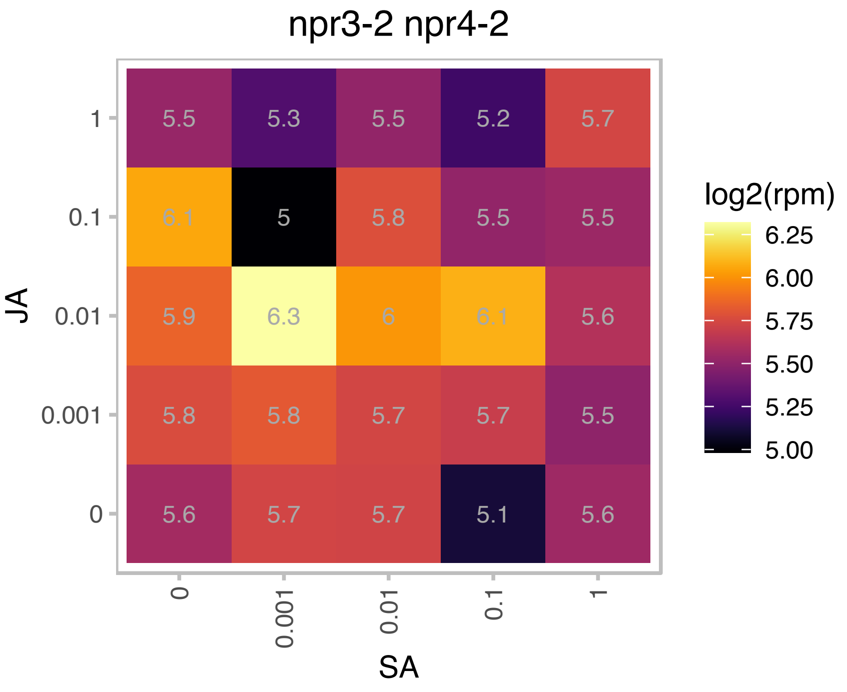


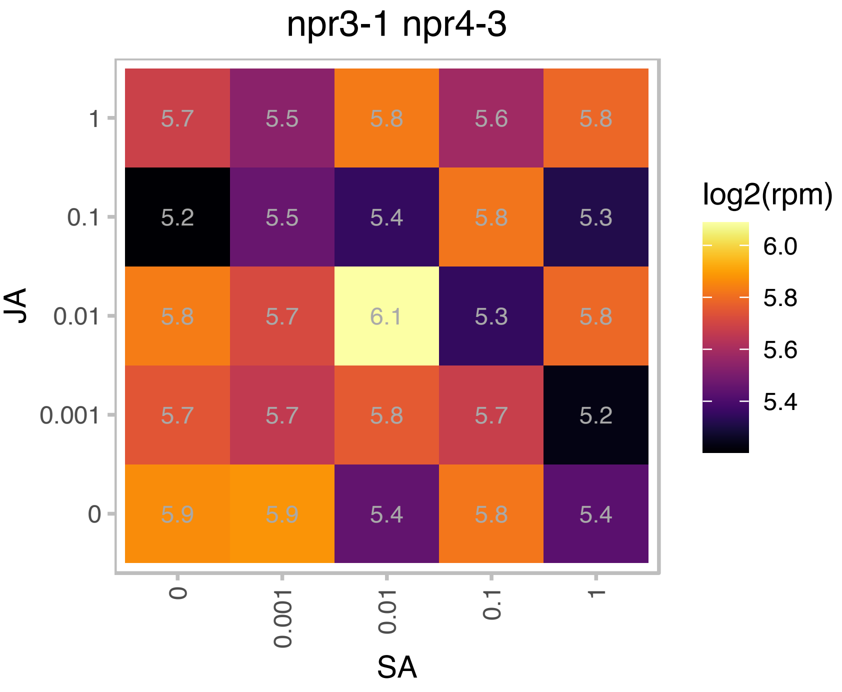

### AT5G22650.1 : ATHD2, ATHD2B, HD2, HD2B, HDA4, HDT

histone deacetylase 2B

Encodes a member of a plant-specific class of histone deacetylases. Controls the development of adaxial/abaxial leaf polarity. Its mRNA is widely expressed in stems, leaves, flowers and young siliques. Plant lines expressing RNAi constructs directed against this gene showed a marked reduction in agrobacterium-mediated root transformation.

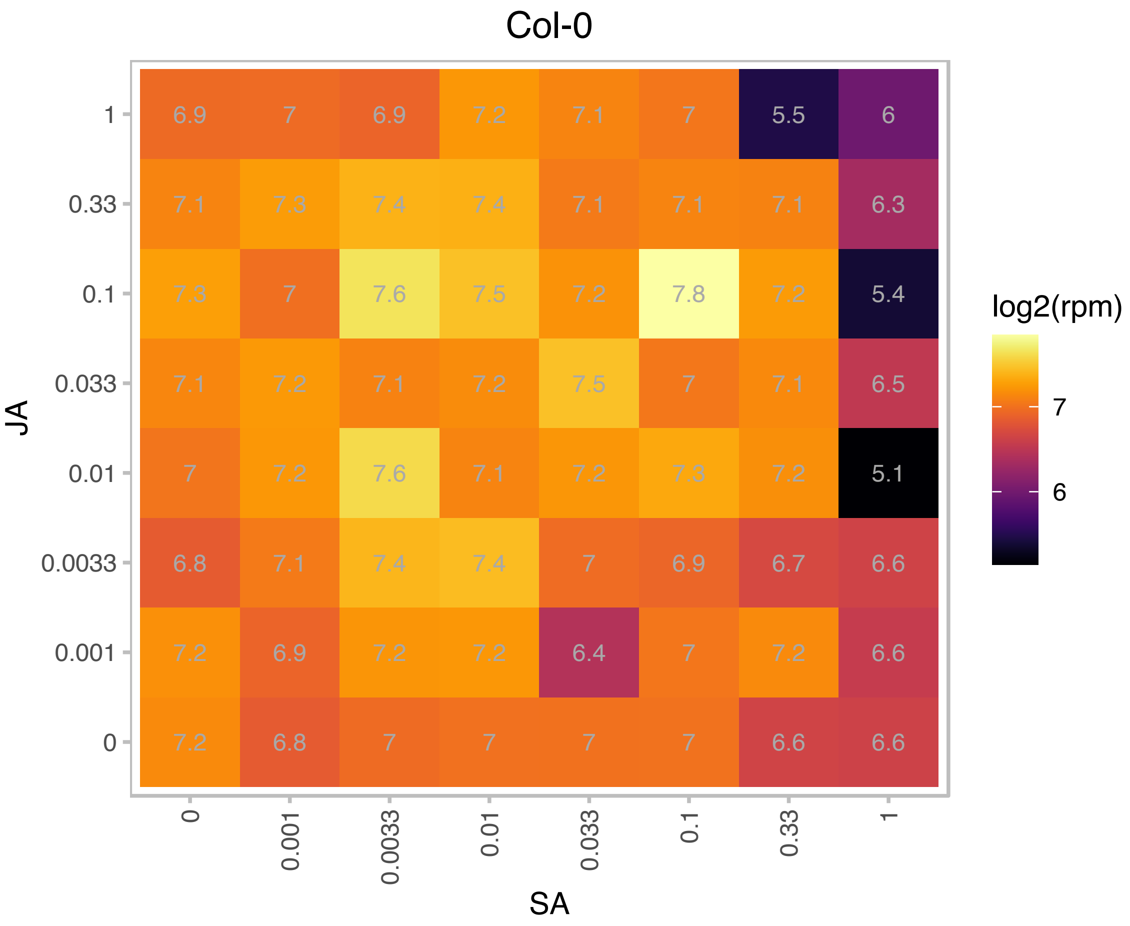


| | |
|---|---|
| Gene | AT5G22650 |
| is.expressed_genes | TRUE |
| is.responsive_genes | TRUE |
| is.apcluster | F-04 |

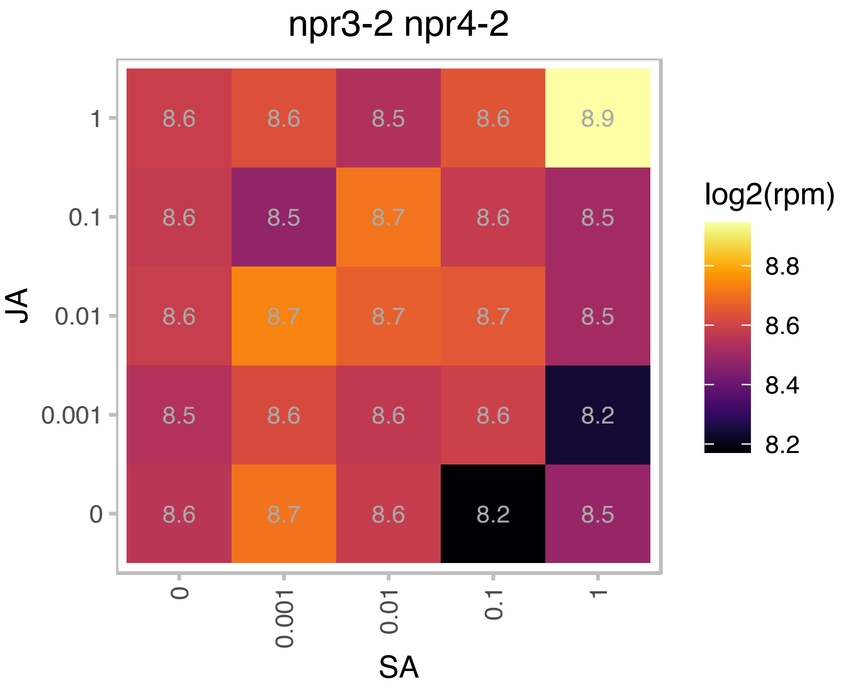


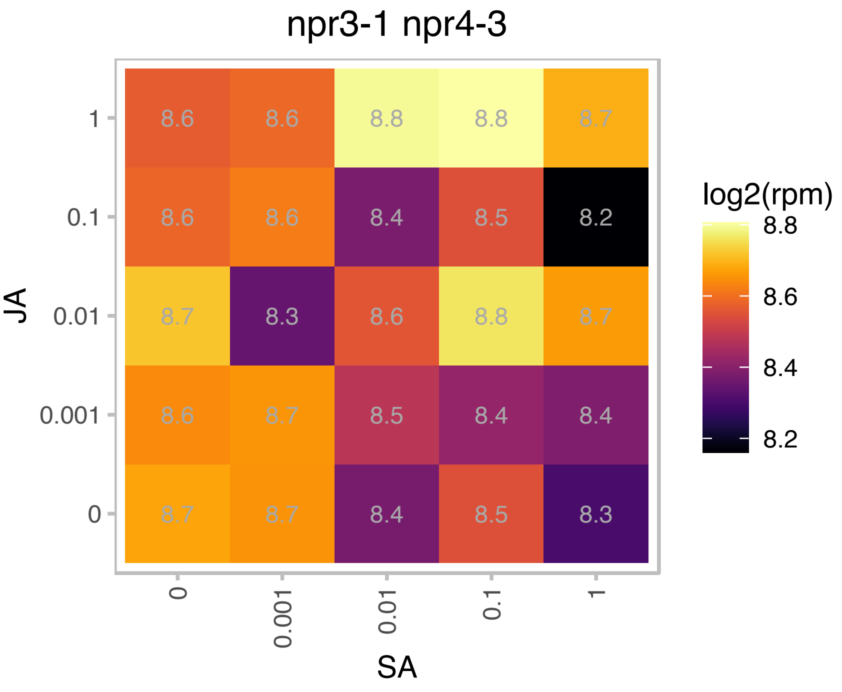

### AT5G25890.1 : IAA28, IAR2

indole-3-acetic acid inducible 28
encodes a protein that may be a negative regulator of lateral root formation in response to auxin. It is a member of IAA/ARF gene family and is plant-specific. Gain of function mutations in this gene suppresses lateral root formation and is resistant to inhibition of root elongation by auxin, cytokinin, and ethylene.

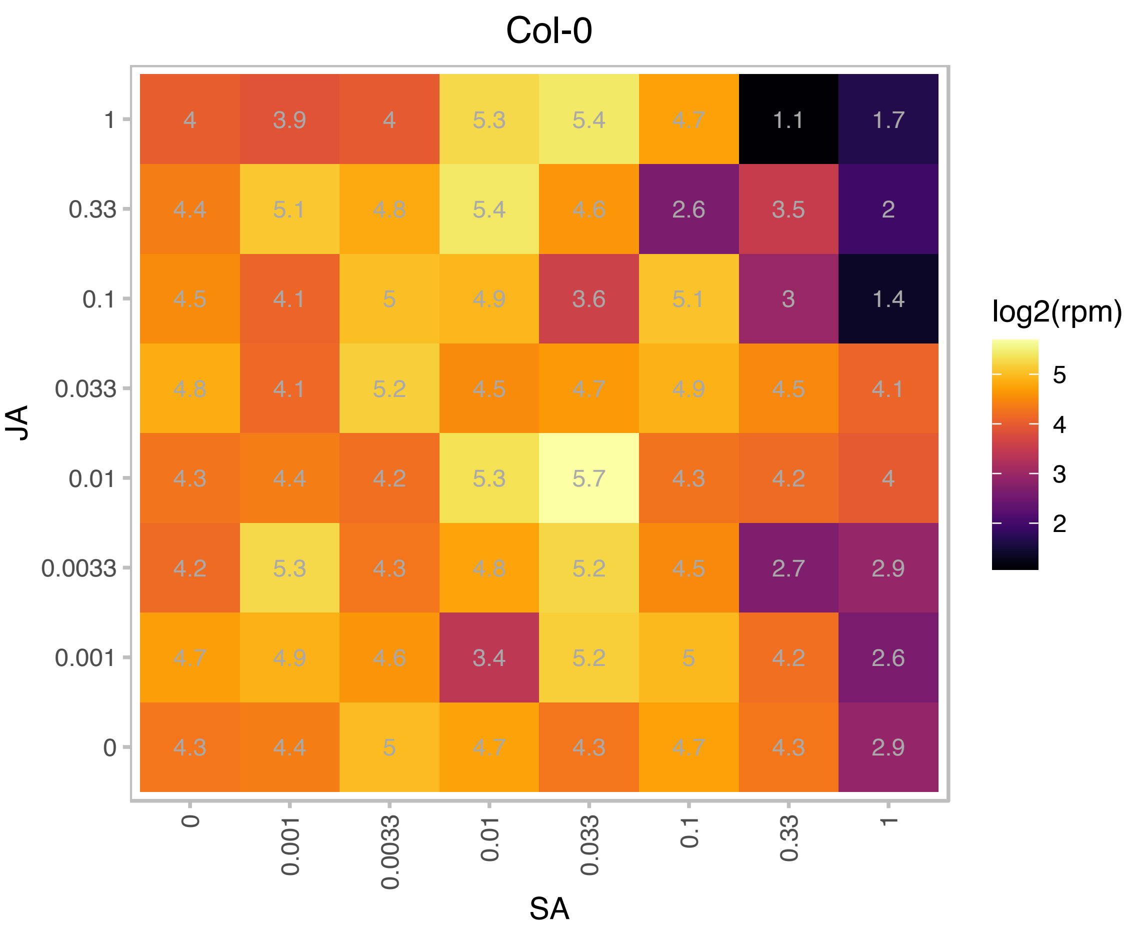


| | |
|---|---|
| Gene | AT5G25890 |
| is.expressed_genes | TRUE |
| is.responsive_genes | TRUE |
| is.apcluster | F-04 |

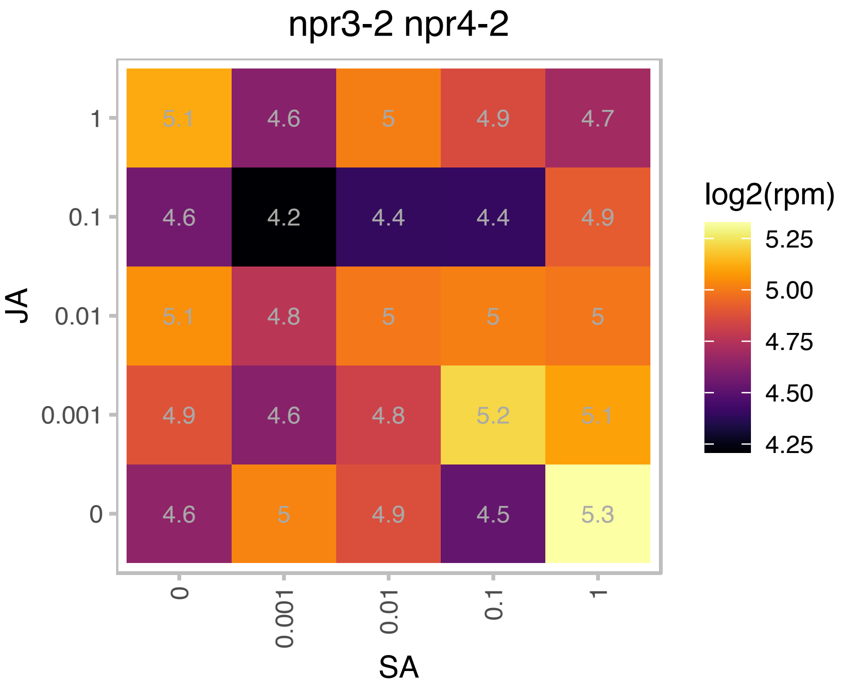


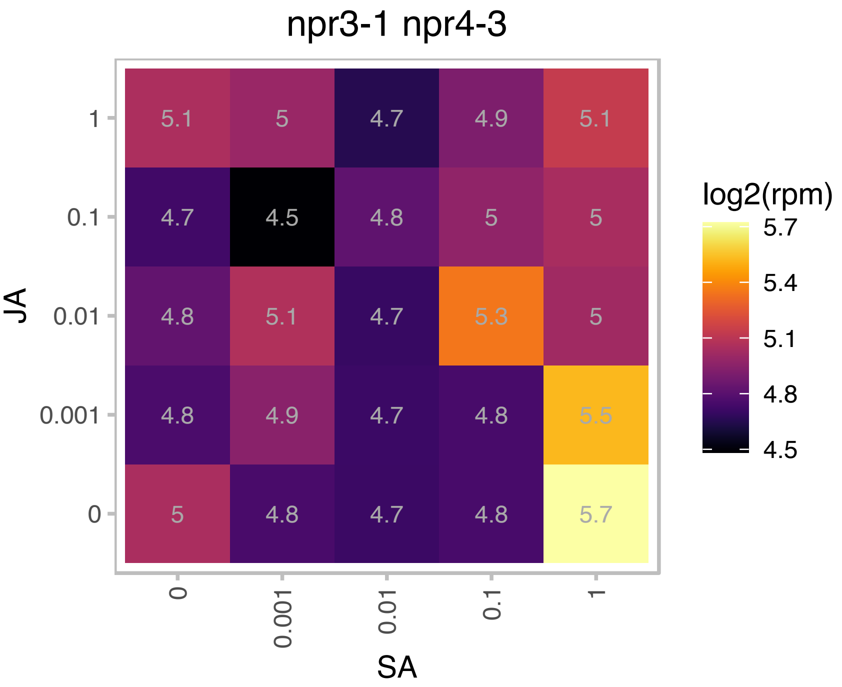

# AT5G28850.1 : NA

Calcium-binding EF-hand family protein

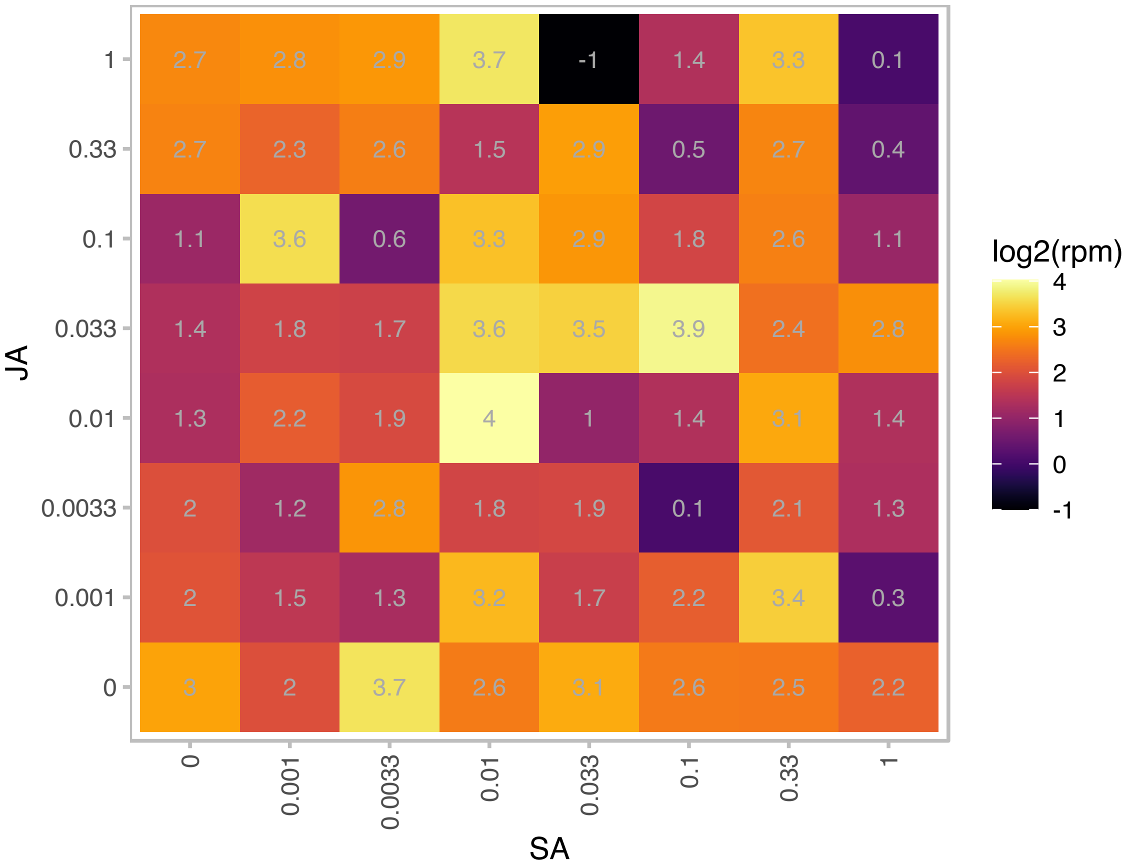


| | |
|---|---|
| Gene | AT5G28850 |
| is.expressed_genes | TRUE |
| is.responsive_genes | TRUE |
| is.apcluster | F-04 |

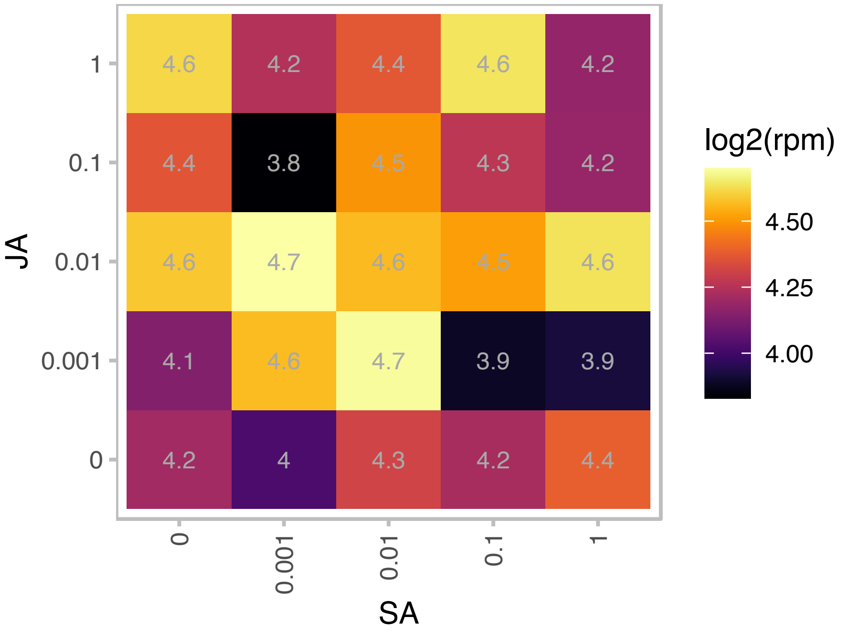


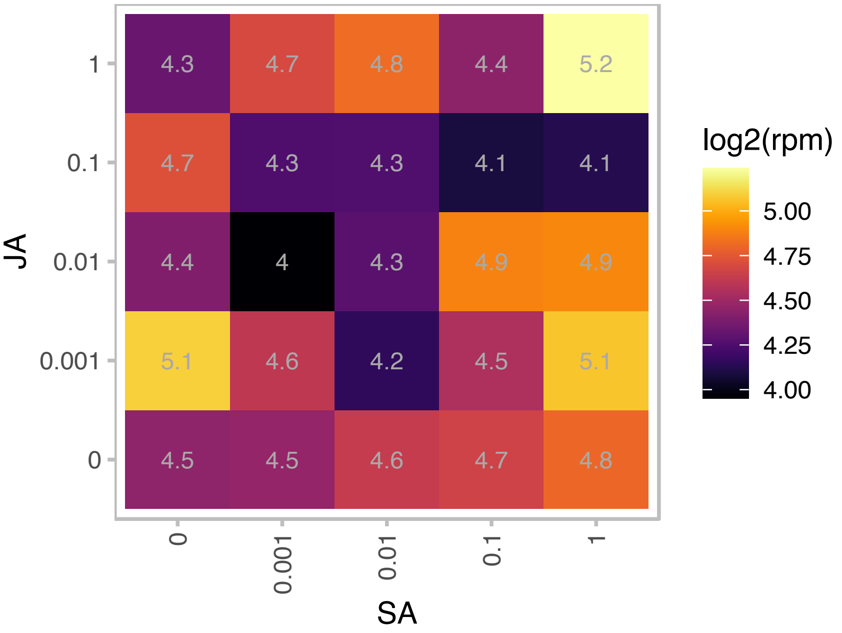

# AT5G42060.1 : NA

DEK, chromatin associated protein

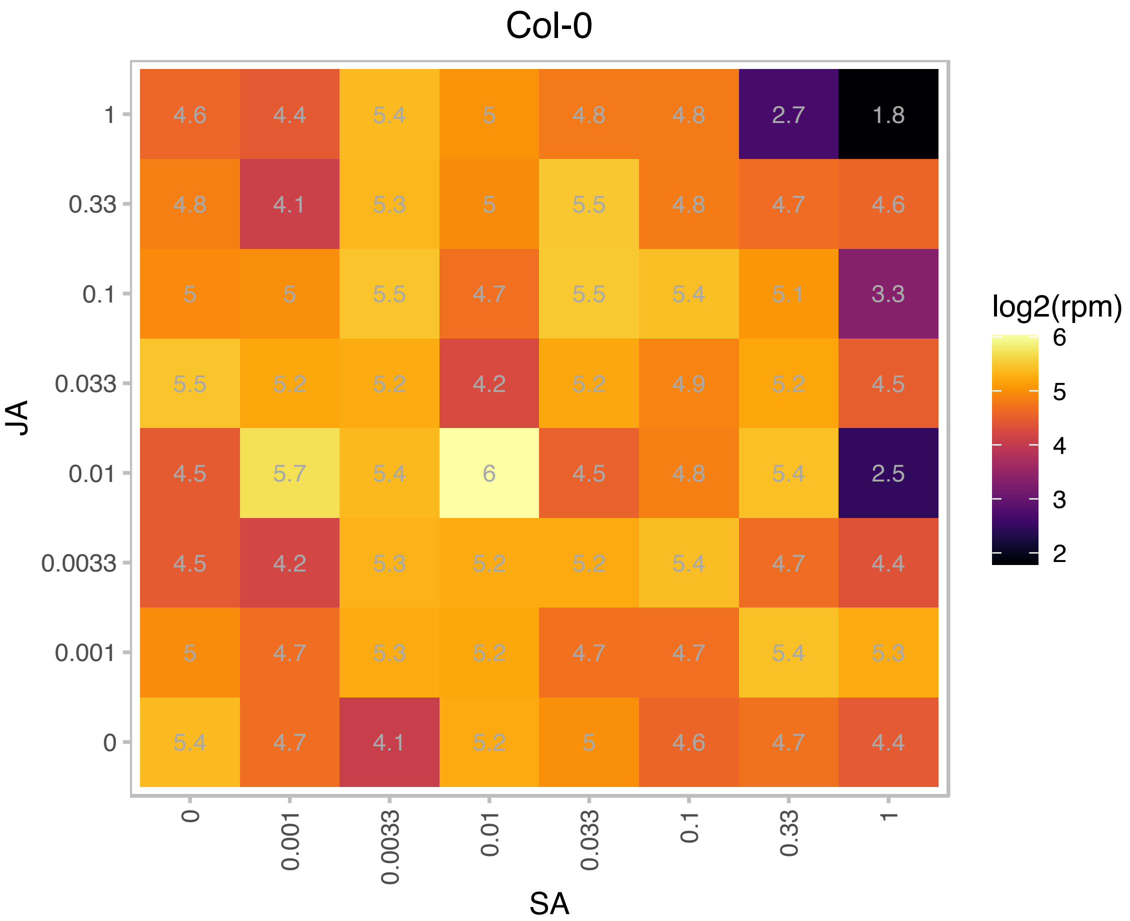


| | |
|---|---|
| Gene | AT5G42060 |
| is.expressed_genes | TRUE |
| is.responsive_genes | TRUE |
| is.apcluster | F-04 |

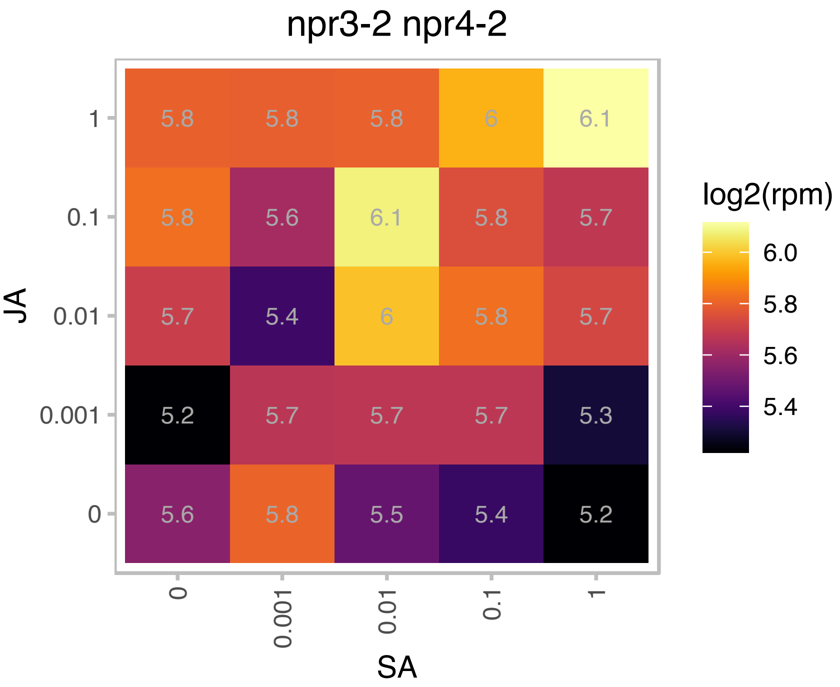


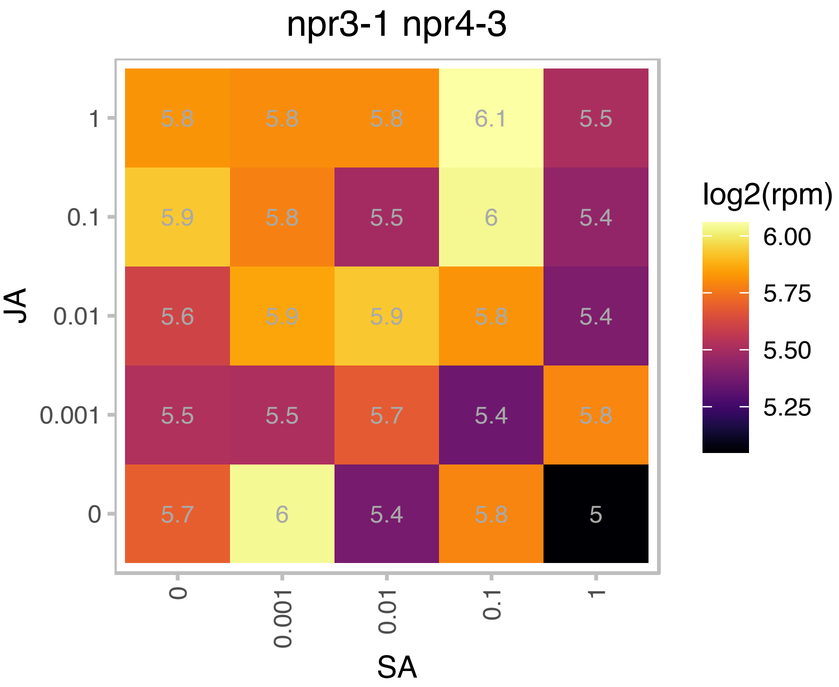

### AT5G44110.1 : ABCI21, ATNAP2, ATPOP1, POP1

P-loop containing nucleoside triphosphate hydrolases superfamily protein

Encodes a member of the NAP subfamily of ABC transporters whose expression pattern is regulated by light and sucrose.

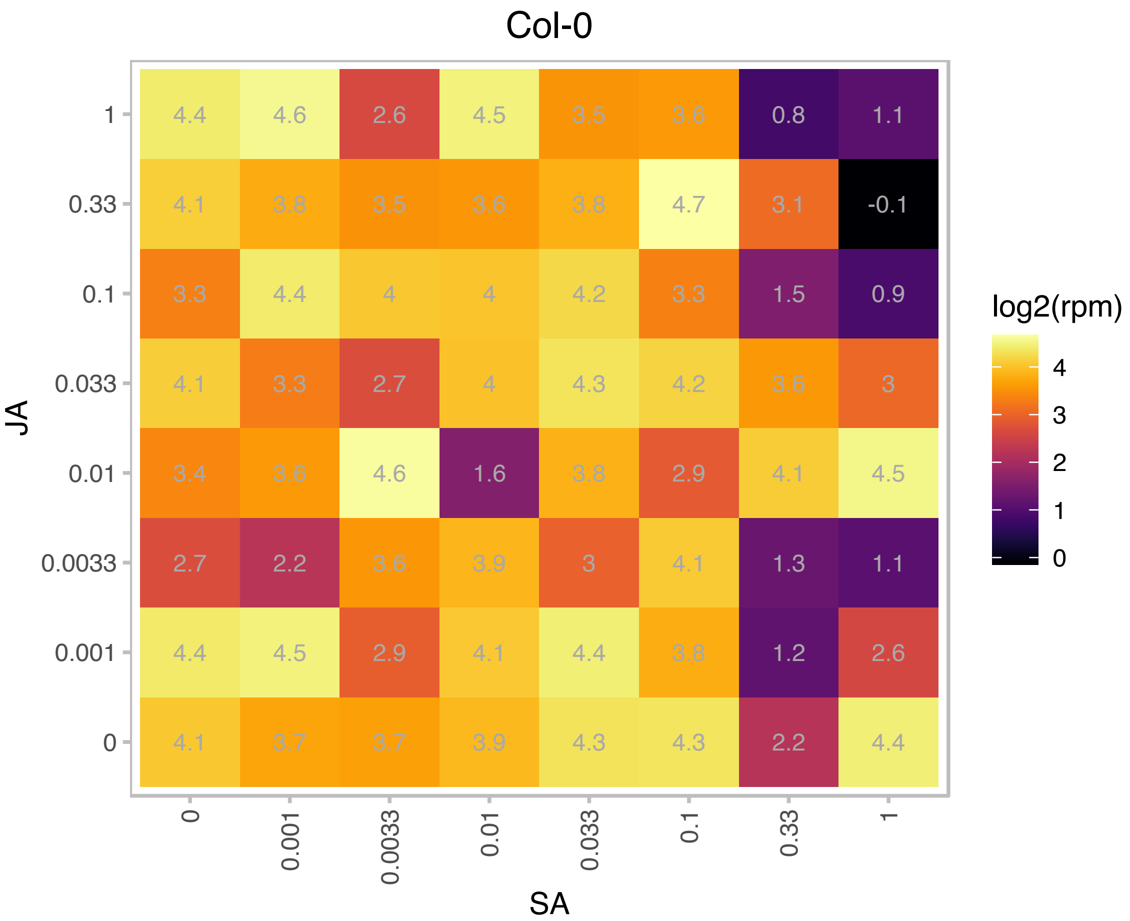


| Gene | AT5G44110 |
|---|---|
| is.expressed_genes | TRUE |
| is.responsive_genes | TRUE |
| is.apcluster | F-04 |

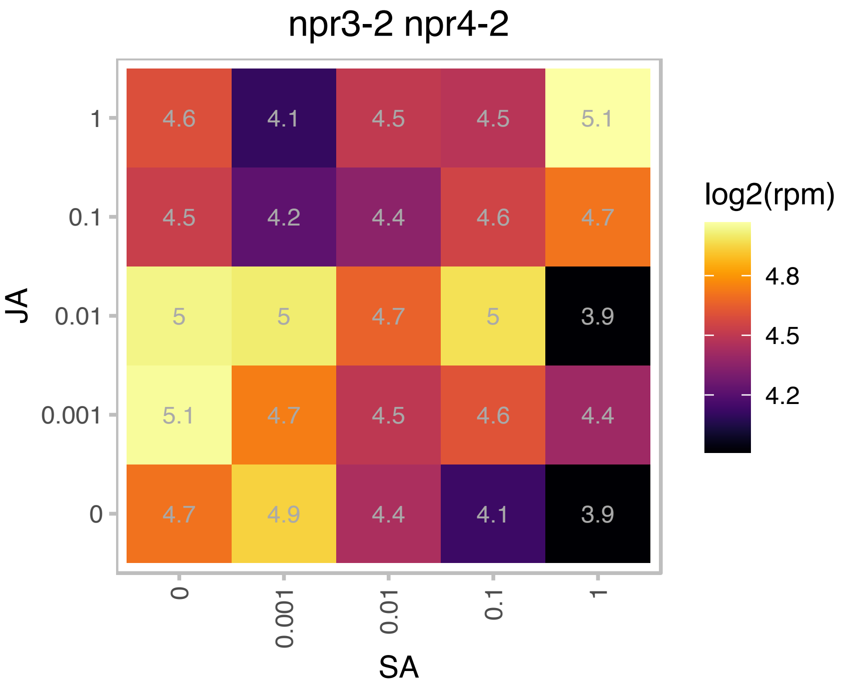


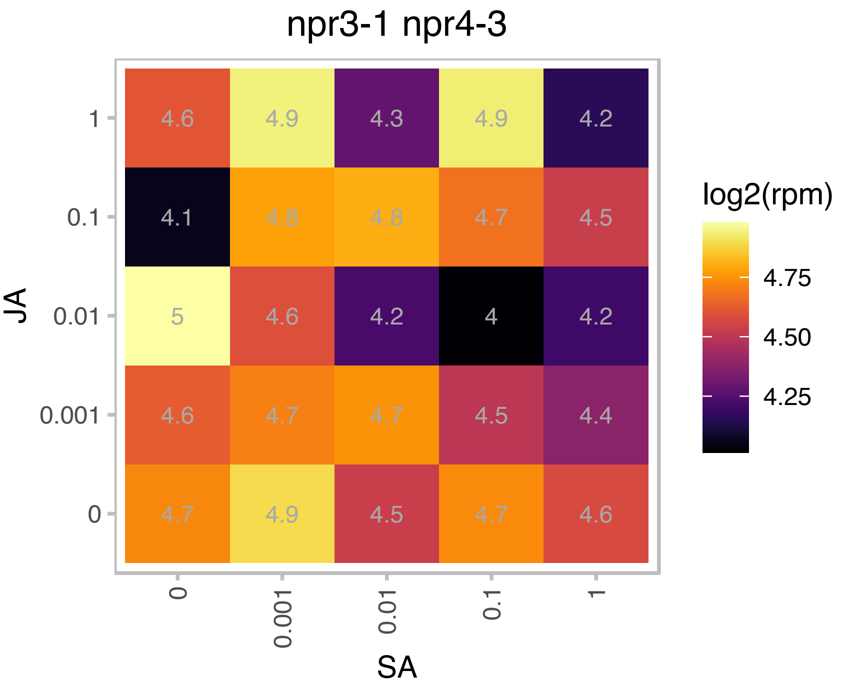

# AT5G44450.1 : NA

methyltransferases

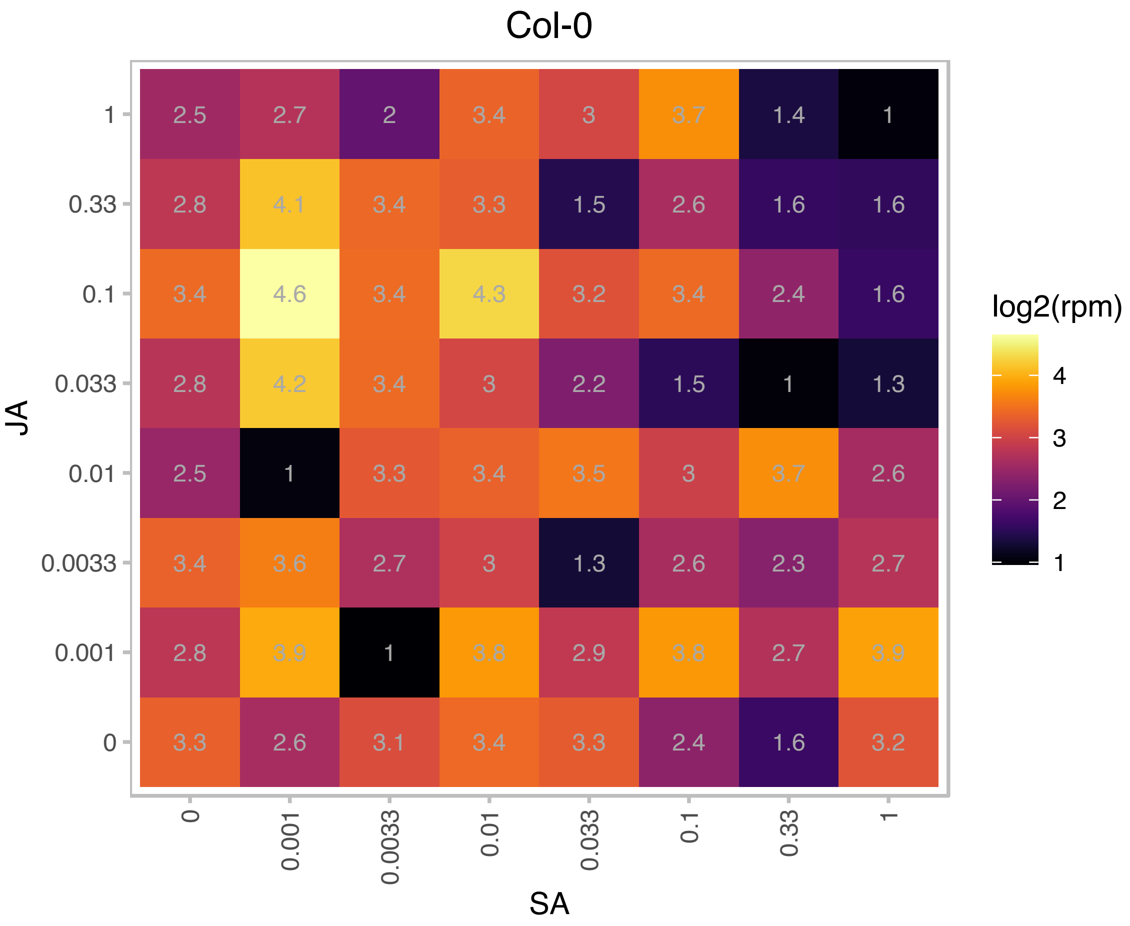


| | |
|---|---|
| Gene | AT5G44450 |
| is.expressed_genes | TRUE |
| is.responsive_genes | TRUE |
| is.apcluster | F-04 |

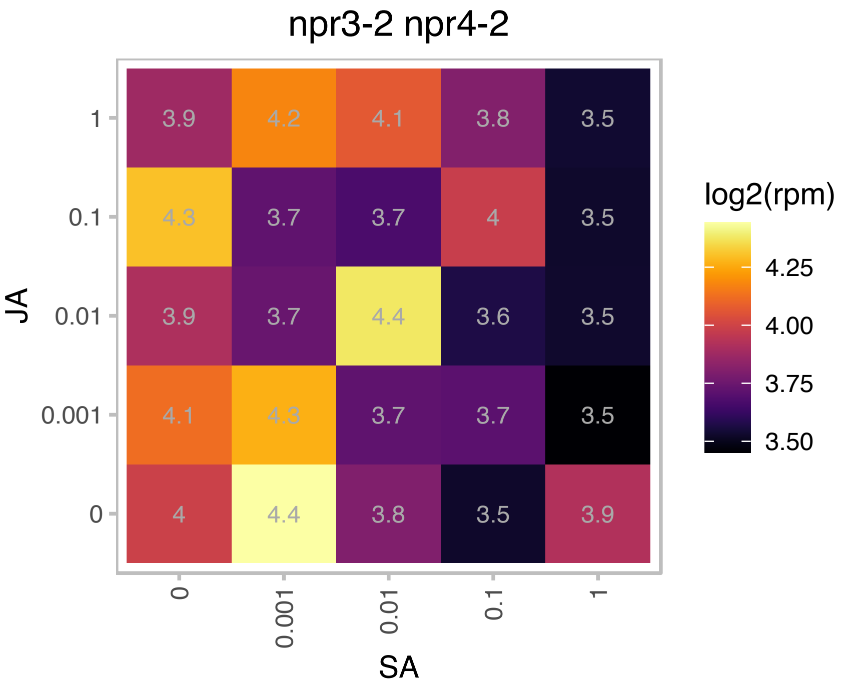


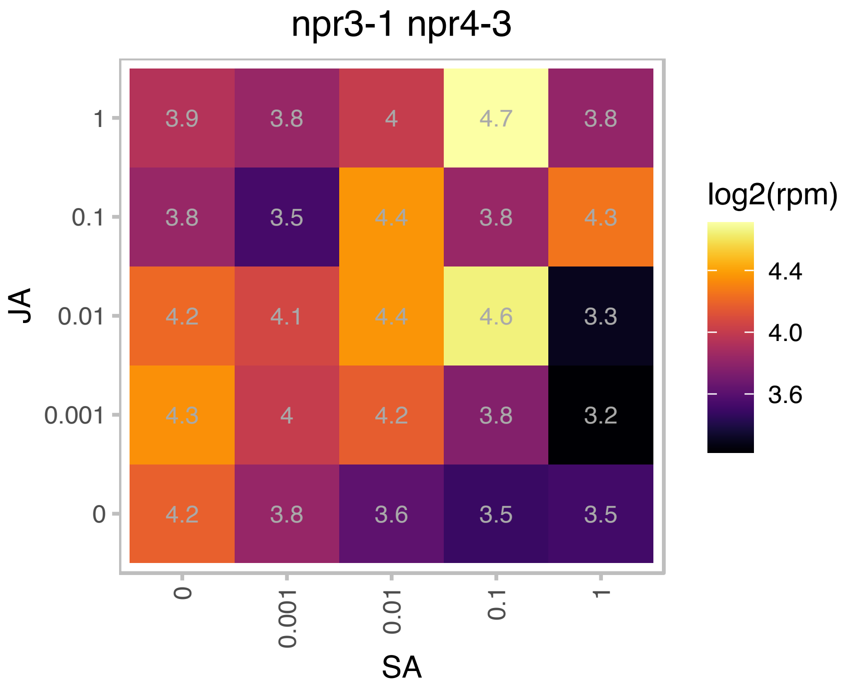

# AT5G46430.1 : NA

Ribosomal protein L32e

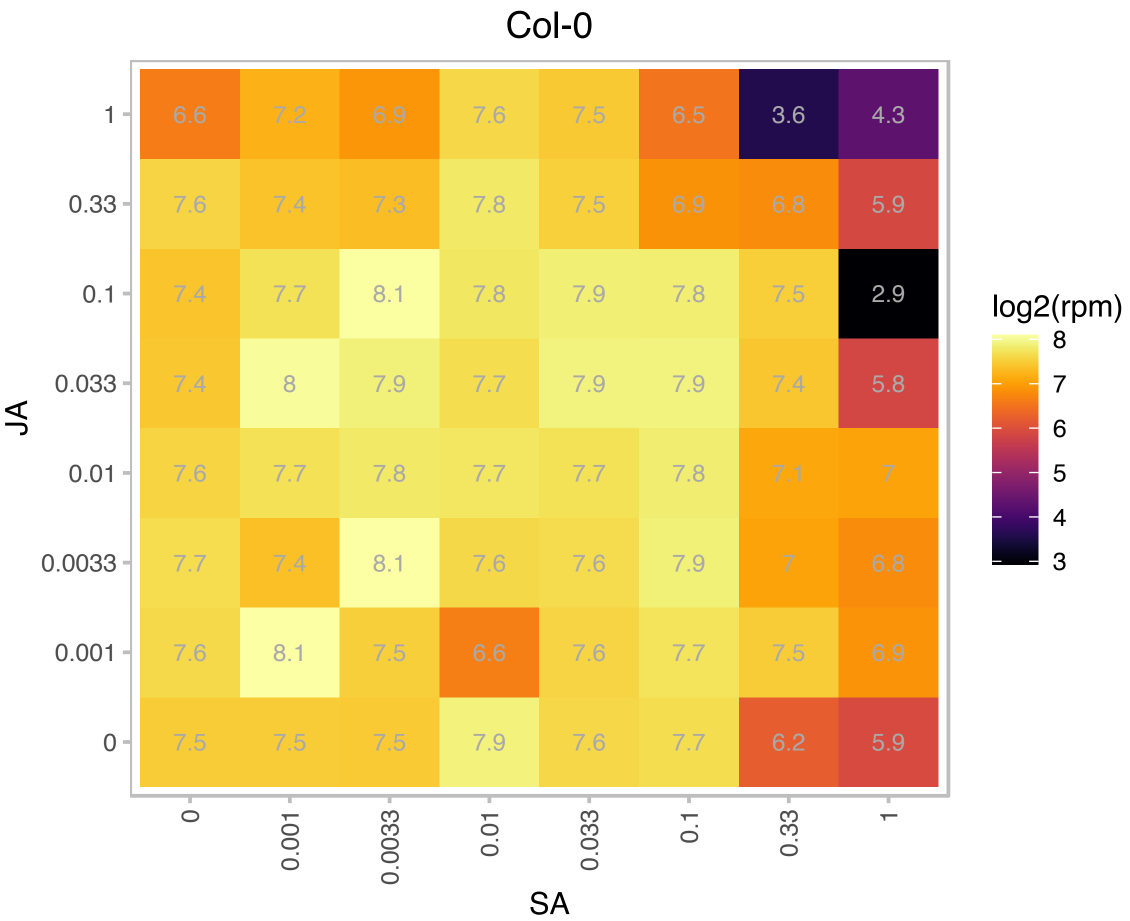


| | |
|---|---|
| Gene | AT5G46430 |
| is.expressed_genes | TRUE |
| is.responsive_genes | TRUE |
| is.apcluster | F-04 |

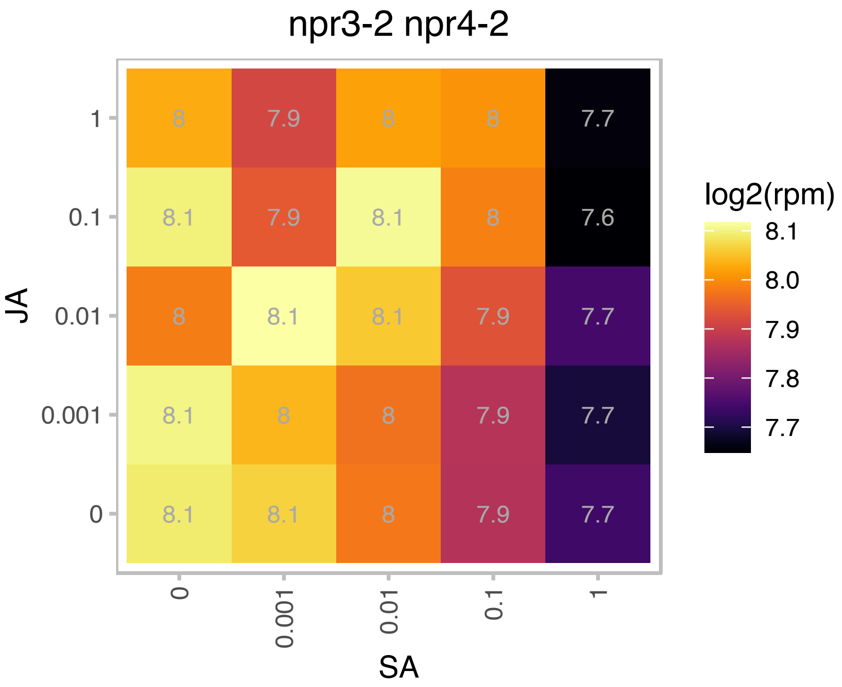


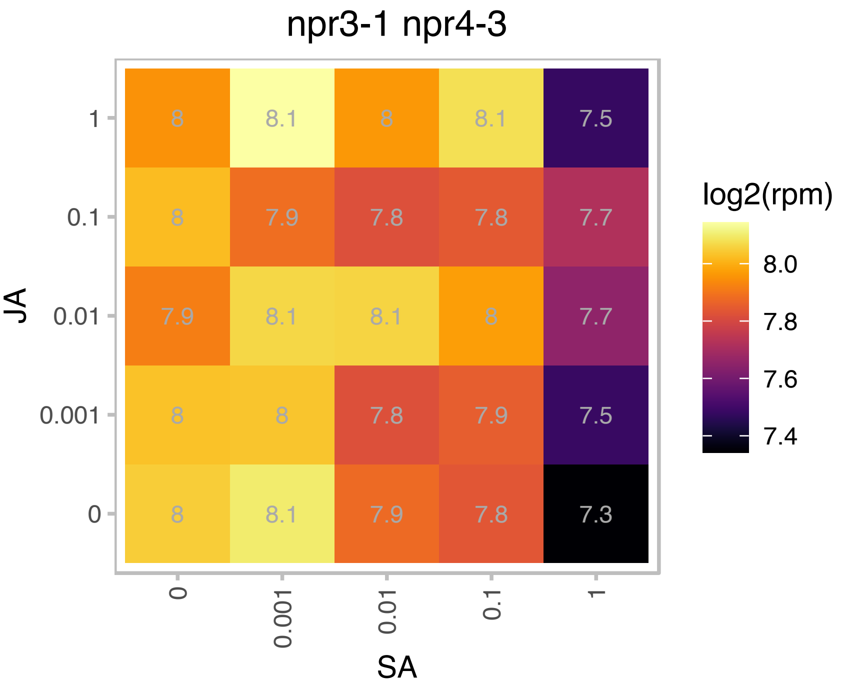

### AT5G47680.1 : AtTRM10, TRM10

Encodes a protein involved in modification of nucleosides in tRNA. Mutants have 50% less 1-methylguanosine than wt counterparts.

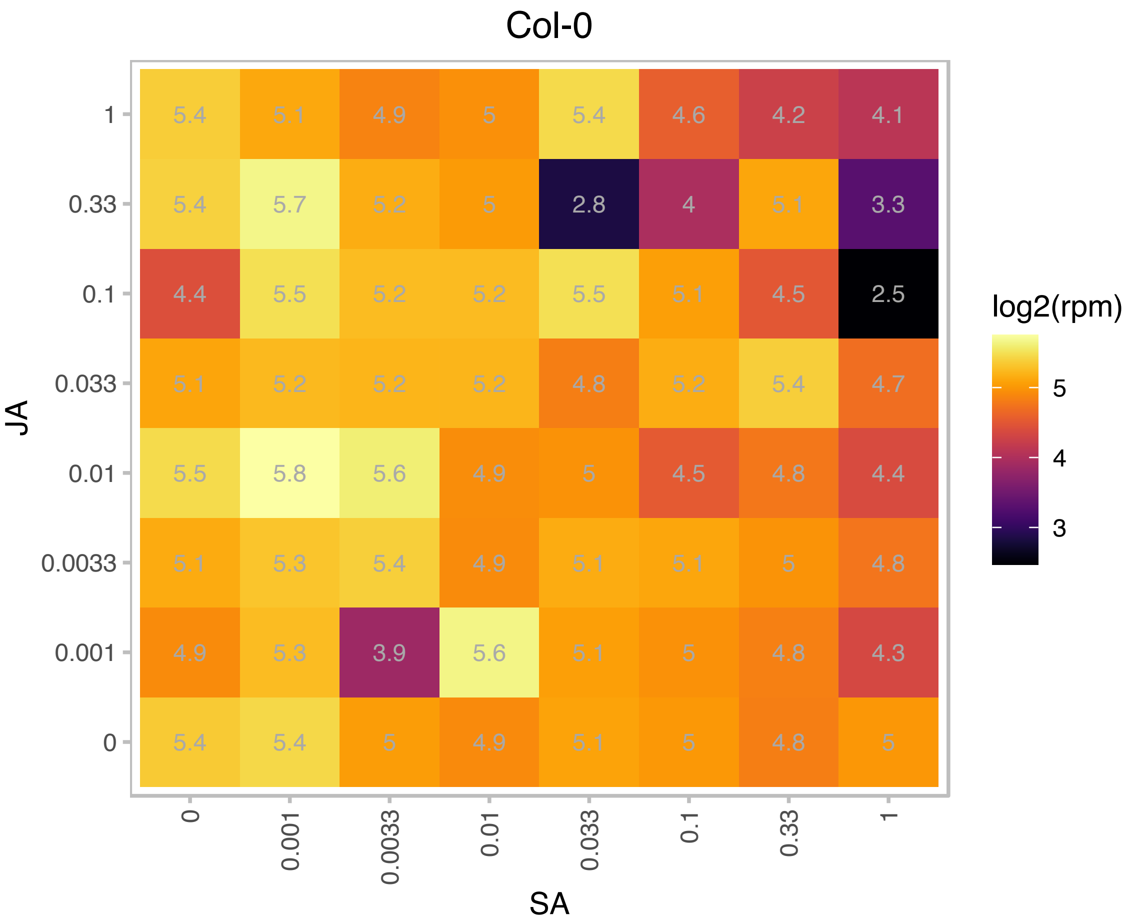


| | |
|---|---|
| Gene | AT5G47680 |
| is.expressed_genes | TRUE |
| is.responsive_genes | TRUE |
| is.apcluster | F-04 |

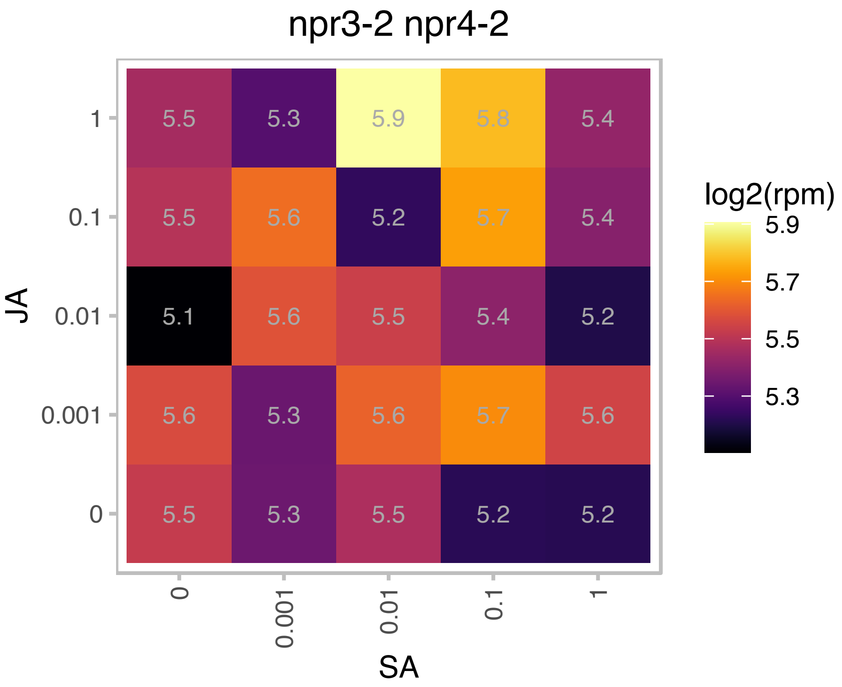


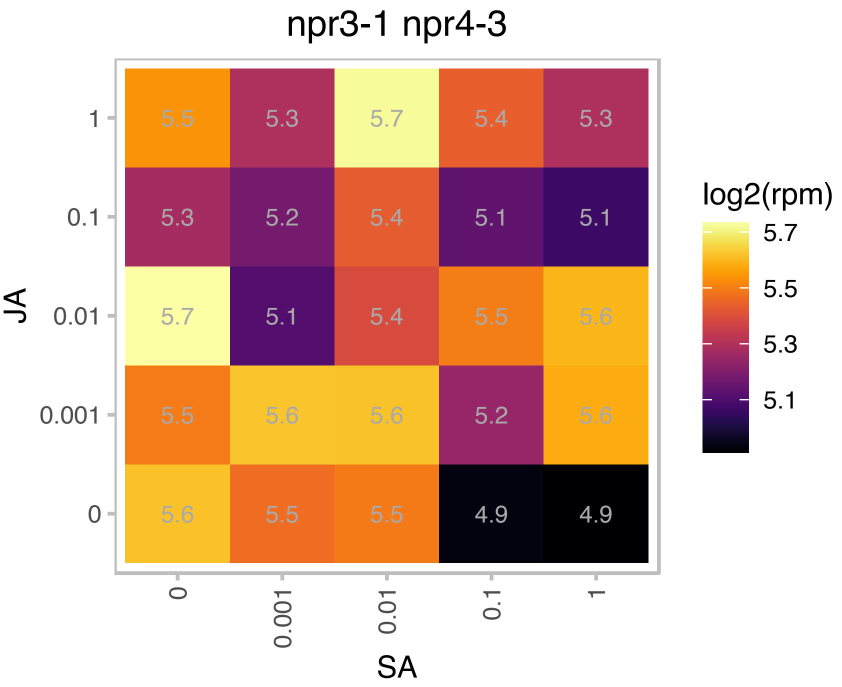

### AT5G51550.1 : EXL3

EXORDIUM like 3

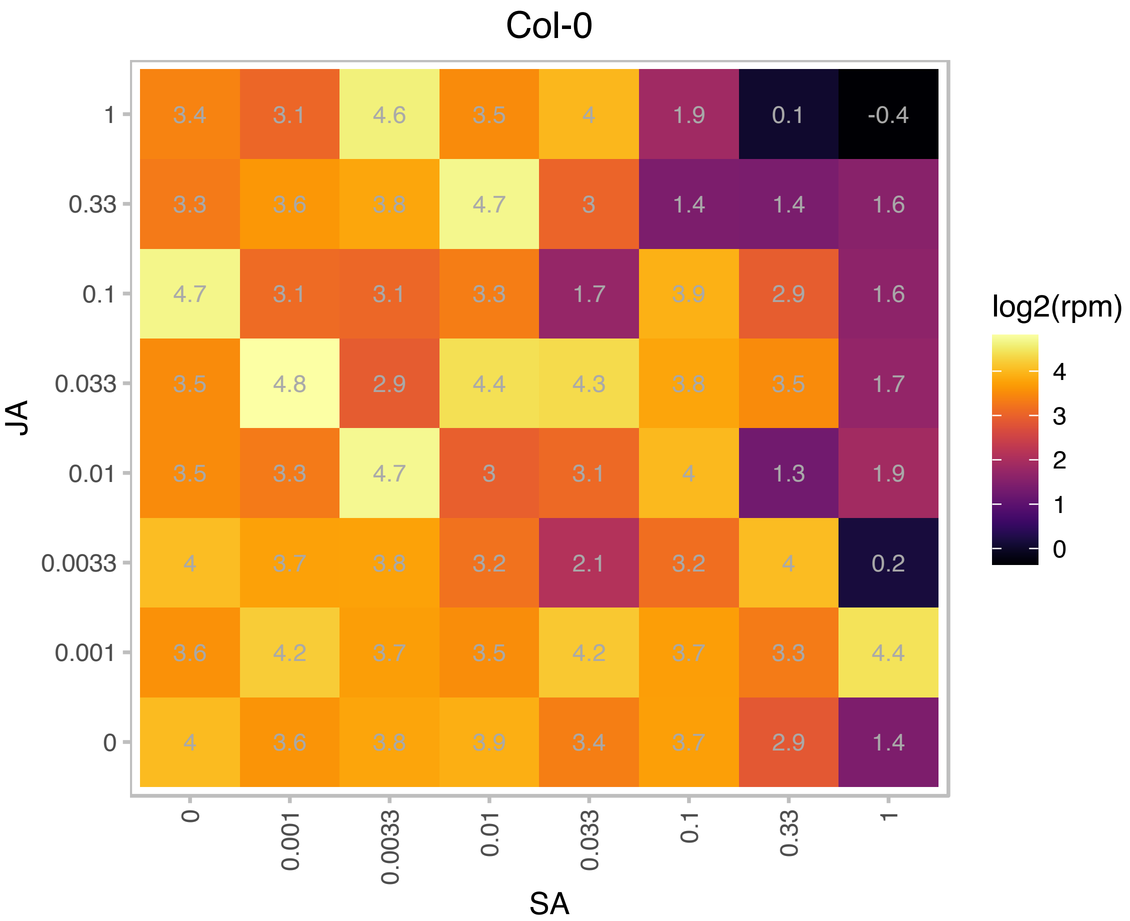


| | |
|---|---|
| Gene | AT5G51550 |
| is.expressed_genes | TRUE |
| is.responsive_genes | TRUE |
| is.apcluster | F-04 |

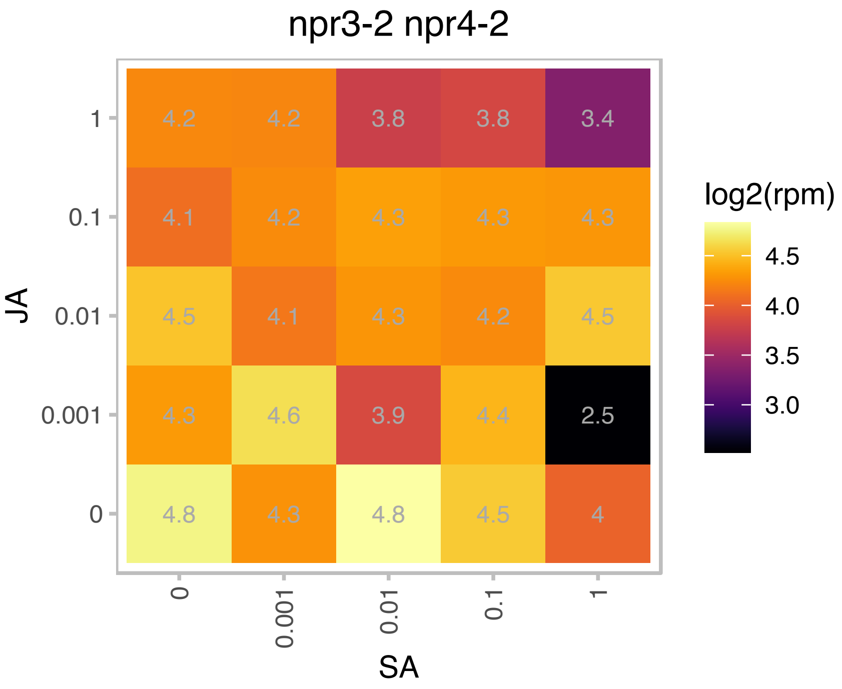


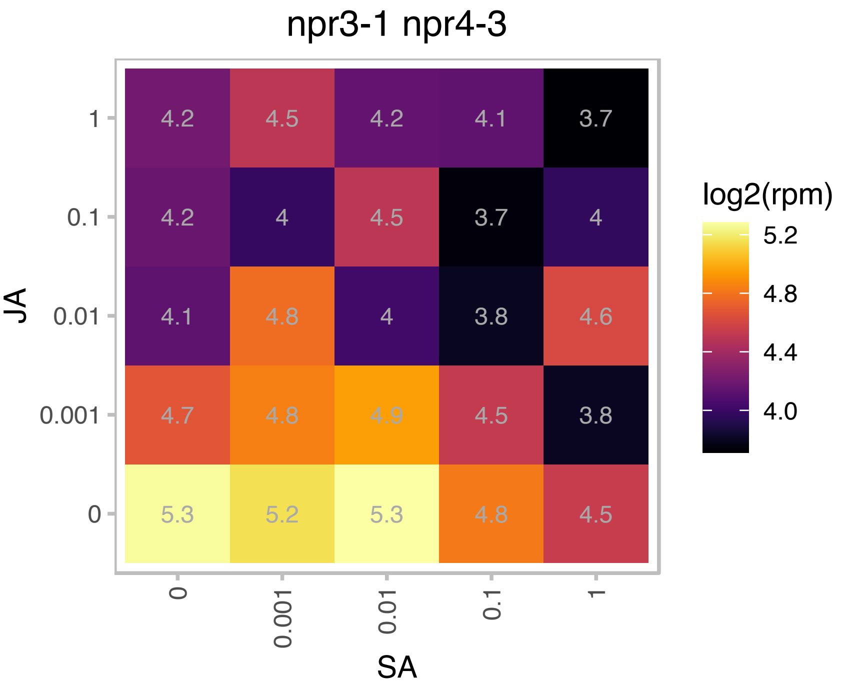

# AT5G53140.1 : NA

Protein phosphatase 2C family protein

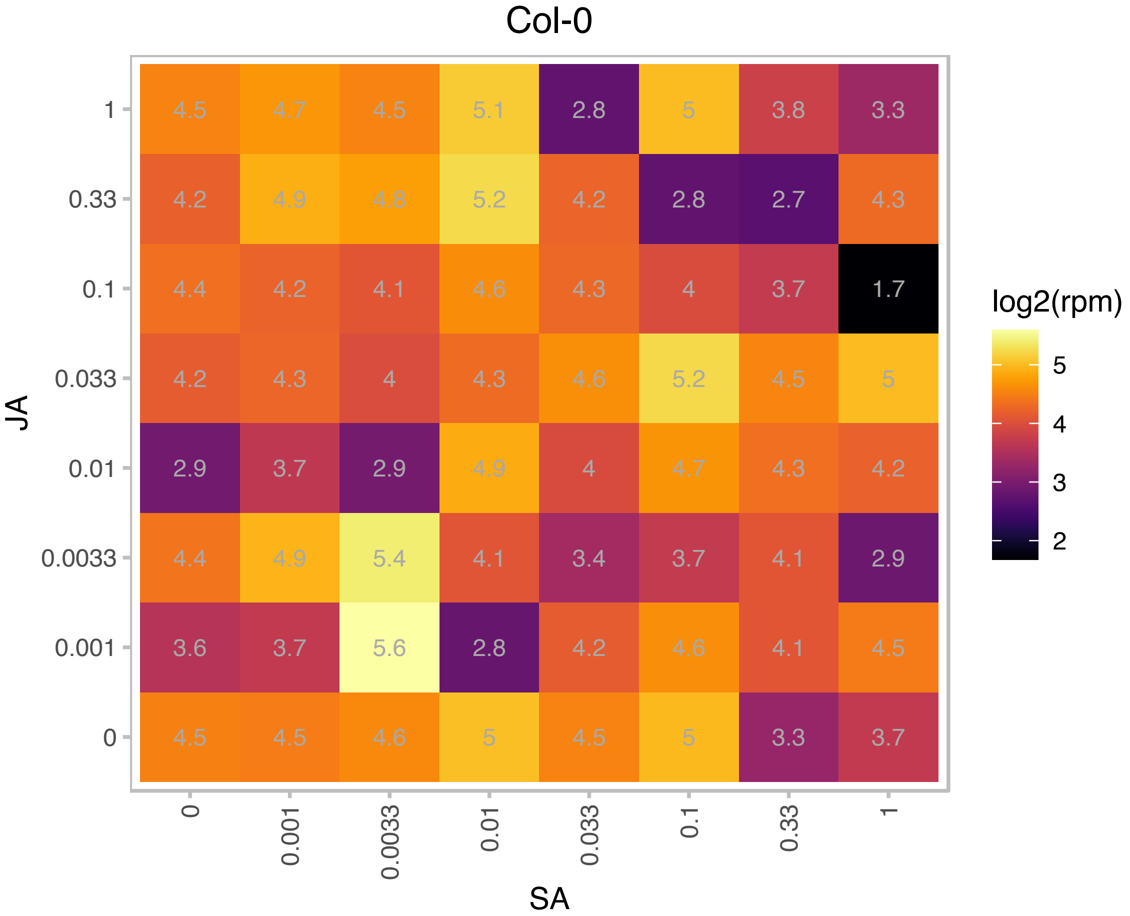


| | |
|---|---|
| Gene | AT5G53140 |
| is.expressed_genes | TRUE |
| is.responsive_genes | TRUE |
| is.apcluster | F-04 |

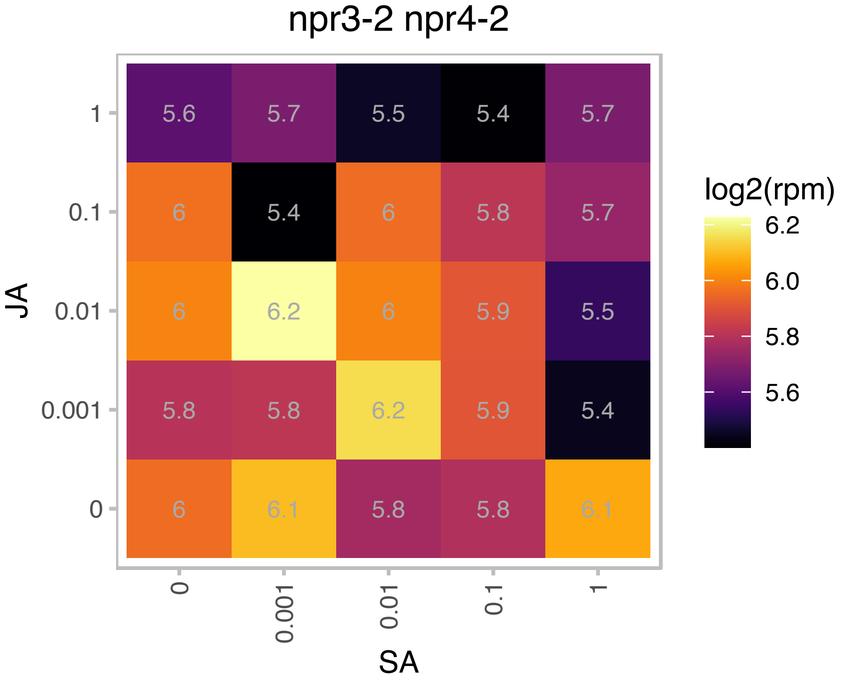


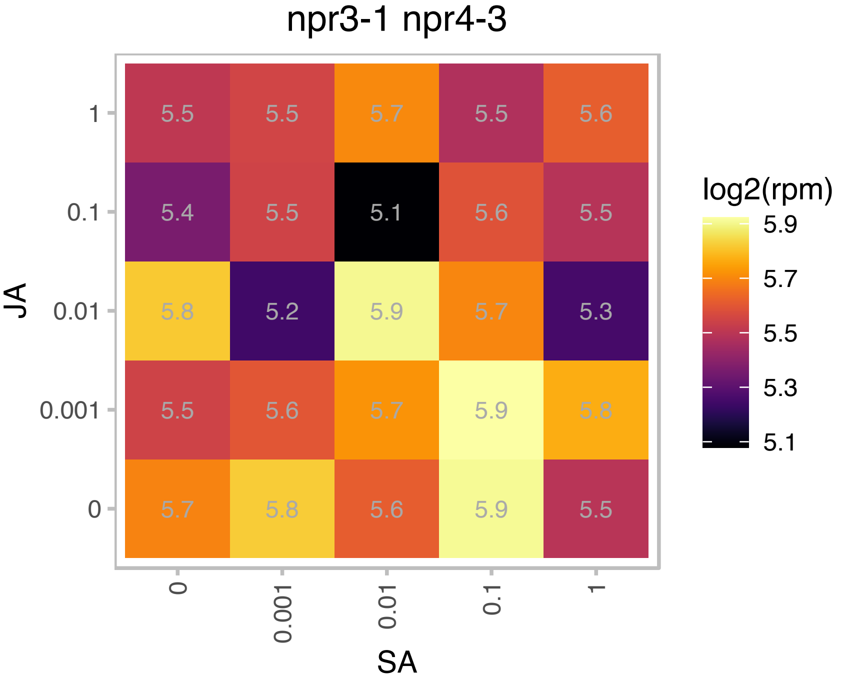

### AT5G56130.1 : AtTEX1, TEX1, THO3

Transducin/WD40 repeat-like superfamily protein

Encodes a component of the putative Arabidopsis THO/TREX complex: THO1 or HPR1 (At5g09860), THO2 (At1g24 THO3 or TEX1 (At5g56130), THO5 (At5g42920, At1g45233), THO6 (At2g19430), and THO7 (At5g16790, At3g0295 THO/TREX complexes in animals have been implicated in the transport of mRNA precursors. Mutants of THO3/TEX1, THO1, THO6 accumulate reduced amount of small interfering (si)RNA, suggesting a role of the putative Arabidopsis THO/TREX in siRNA biosynthesis.

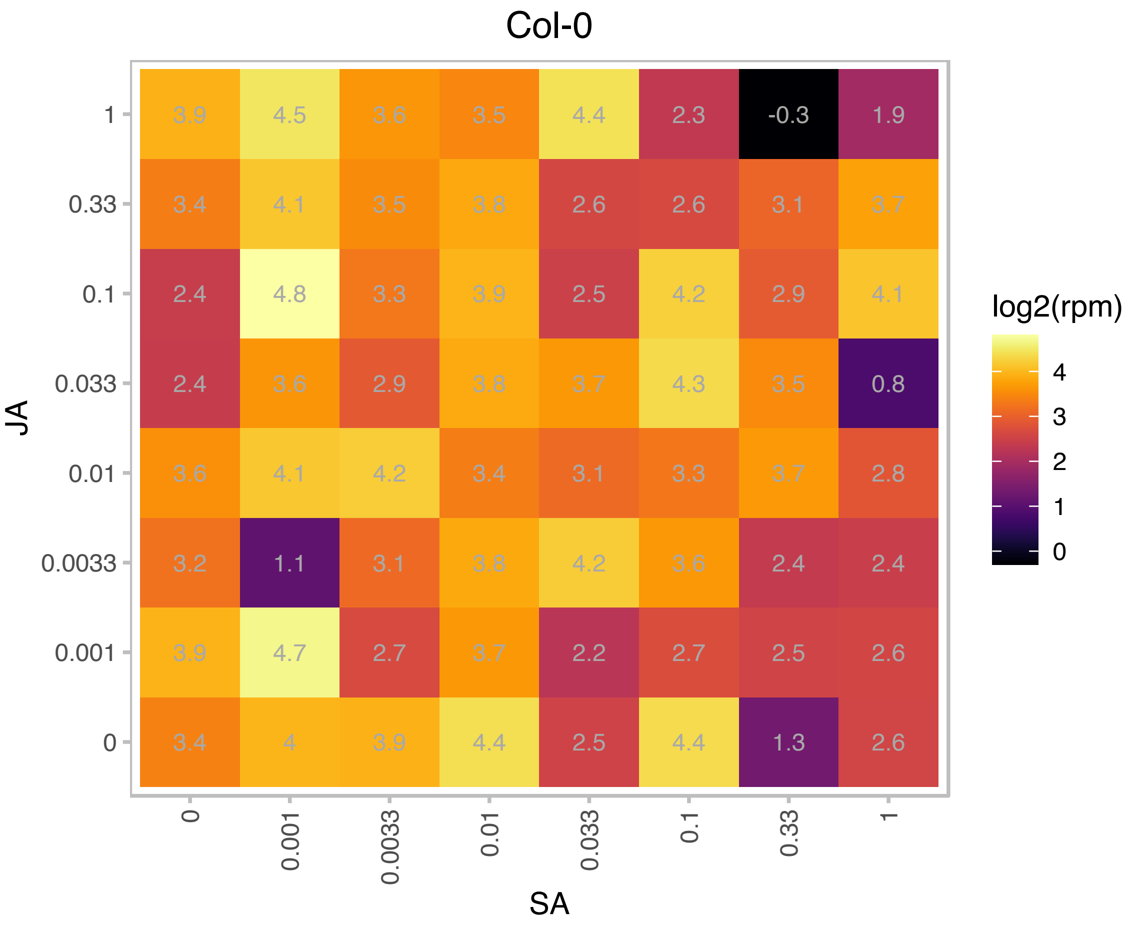


| | |
|---|---|
| Gene | AT5G56130 |
| is.expressed_genes | TRUE |
| is.responsive_genes | TRUE |
| is.apcluster | F-04 |

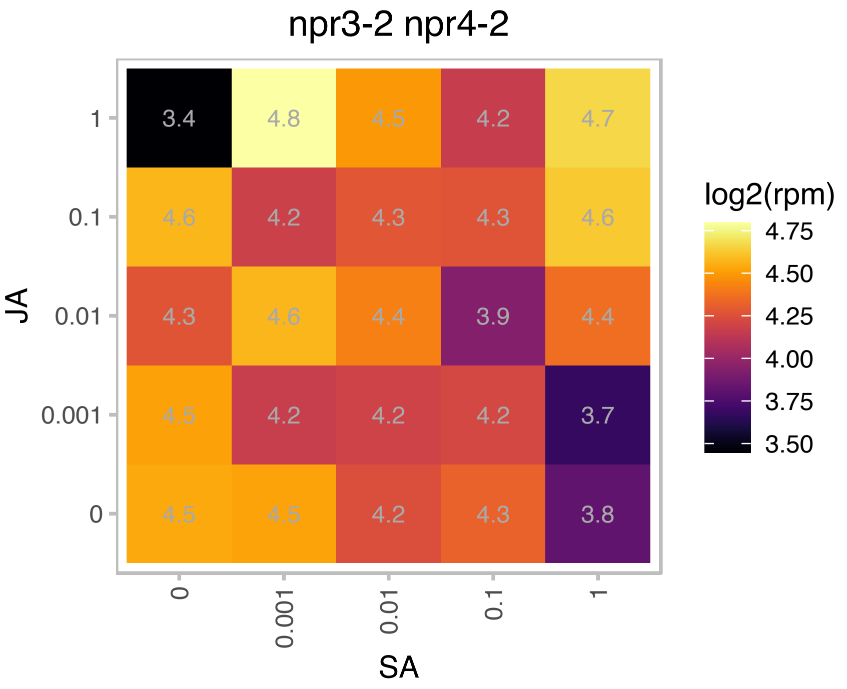


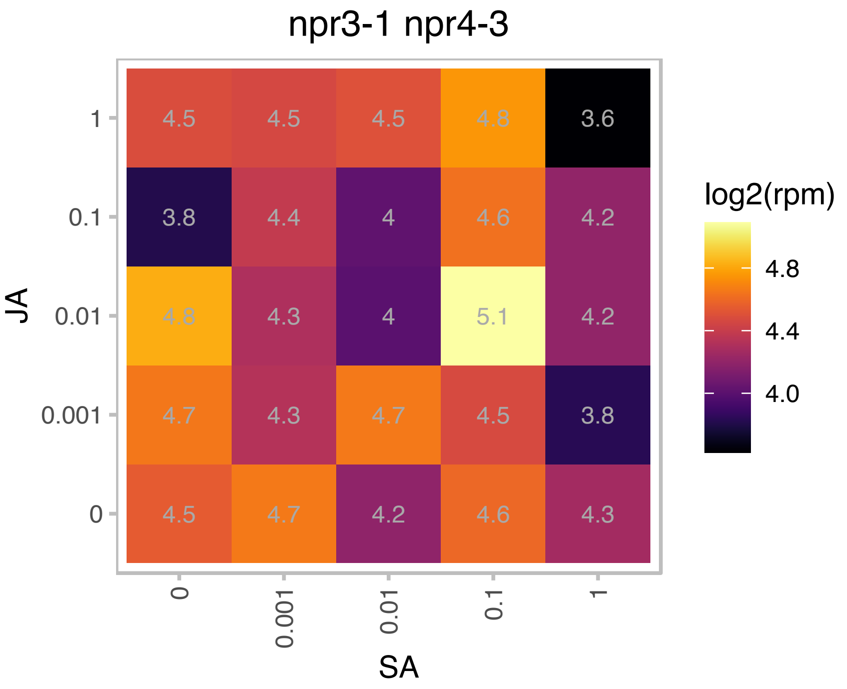

### AT5G56650.1 : ILL1

IAA-leucine resistant (ILR)-like 1

encodes a protein similar to IAA conjugate hydrolases. Enzyme assays with purified GST-tagged protein shows some activity in vitro but too low to be physiologically significant.

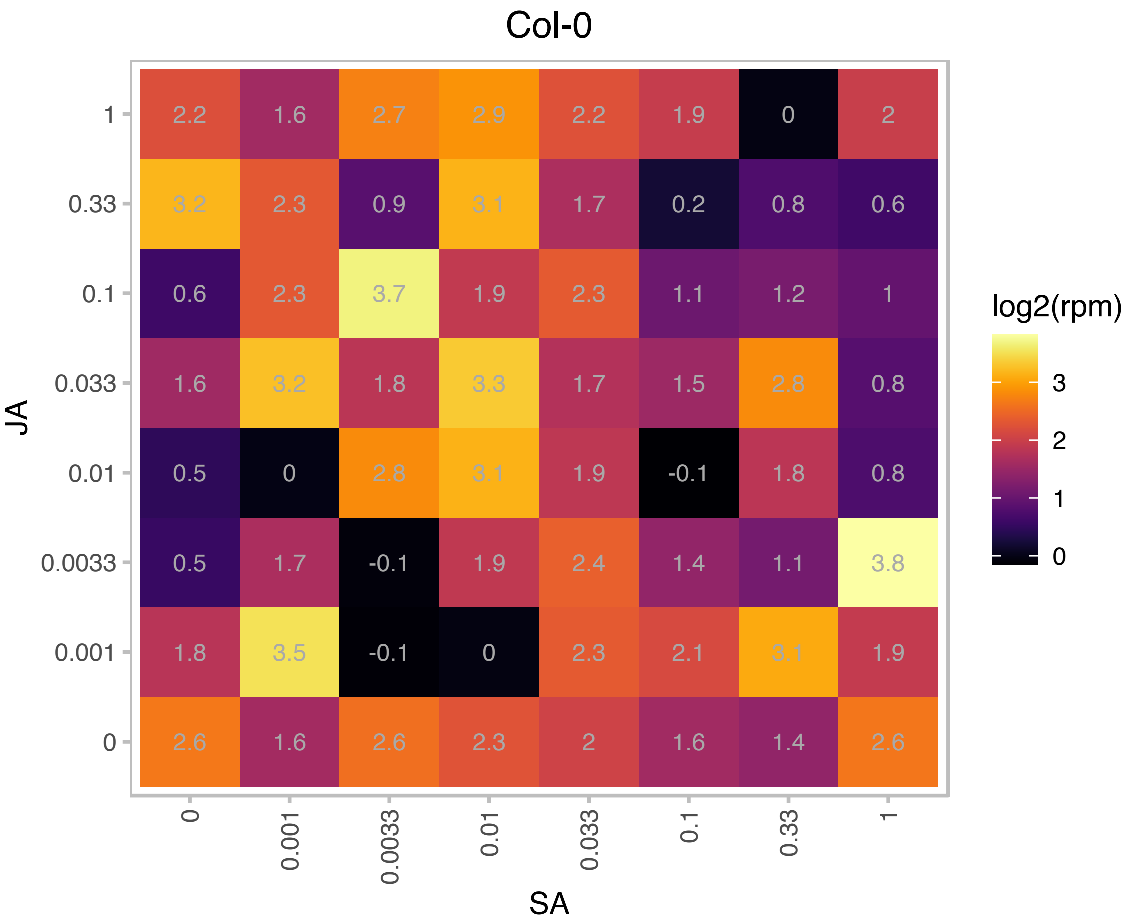


| | |
|---|---|
| Gene | AT5G56650 |
| is.expressed_genes | TRUE |
| is.responsive_genes | TRUE |
| is.apcluster | F-04 |

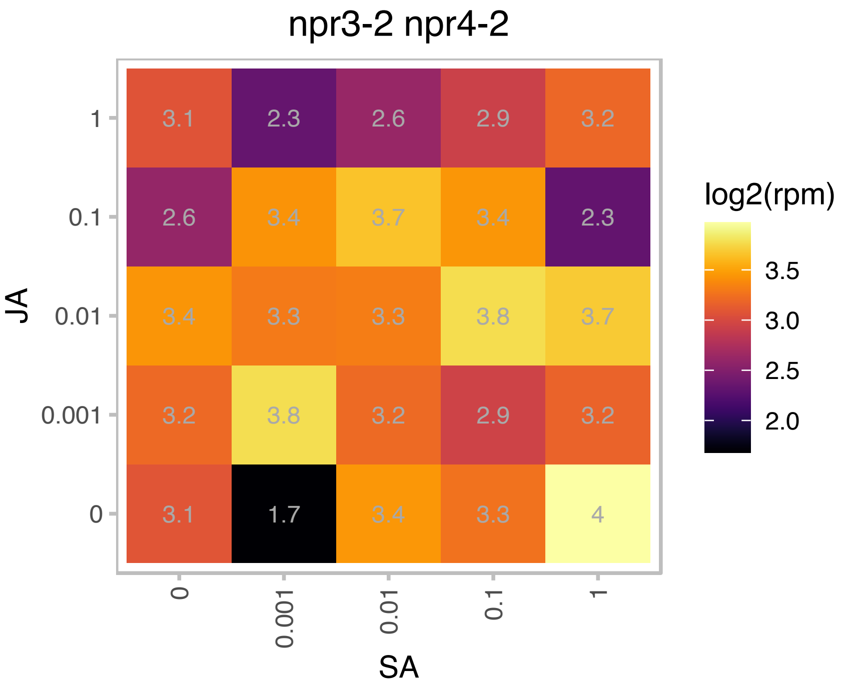


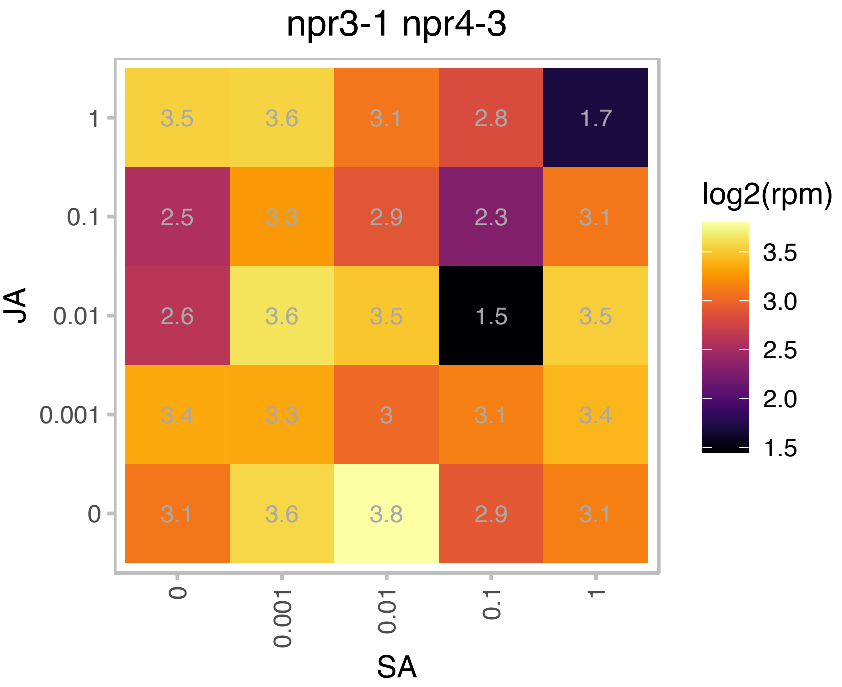

### AT5G61770.1 : PPAN, SNAIL1

PETER PAN-like protein

A single-copy gene encoding a 346 aa protein with a single Brix domain. Similar to yeast ribosome biogenesis proteins Ssf1/2.

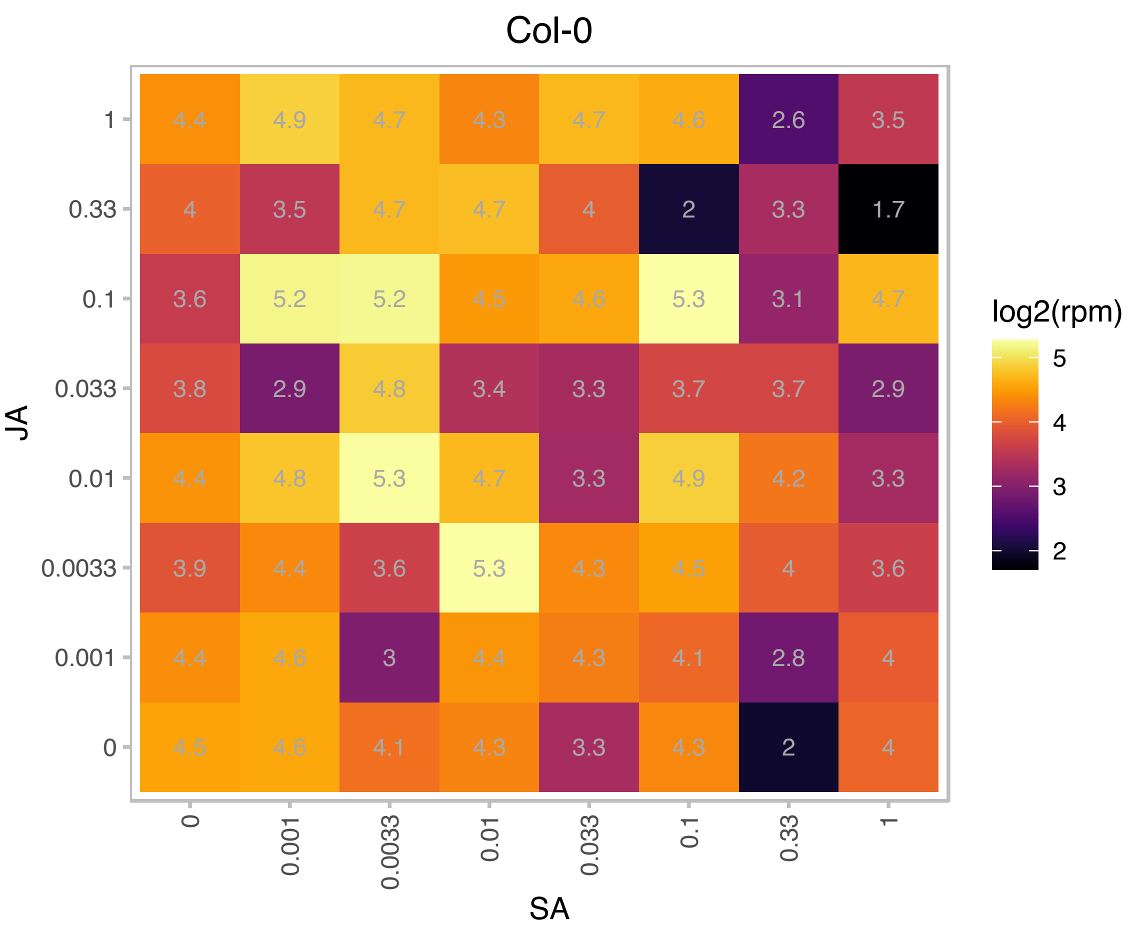


| | |
|---|---|
| Gene | AT5G61770 |
| is.expressed_genes | TRUE |
| is.responsive_genes | TRUE |
| is.apcluster | F-04 |

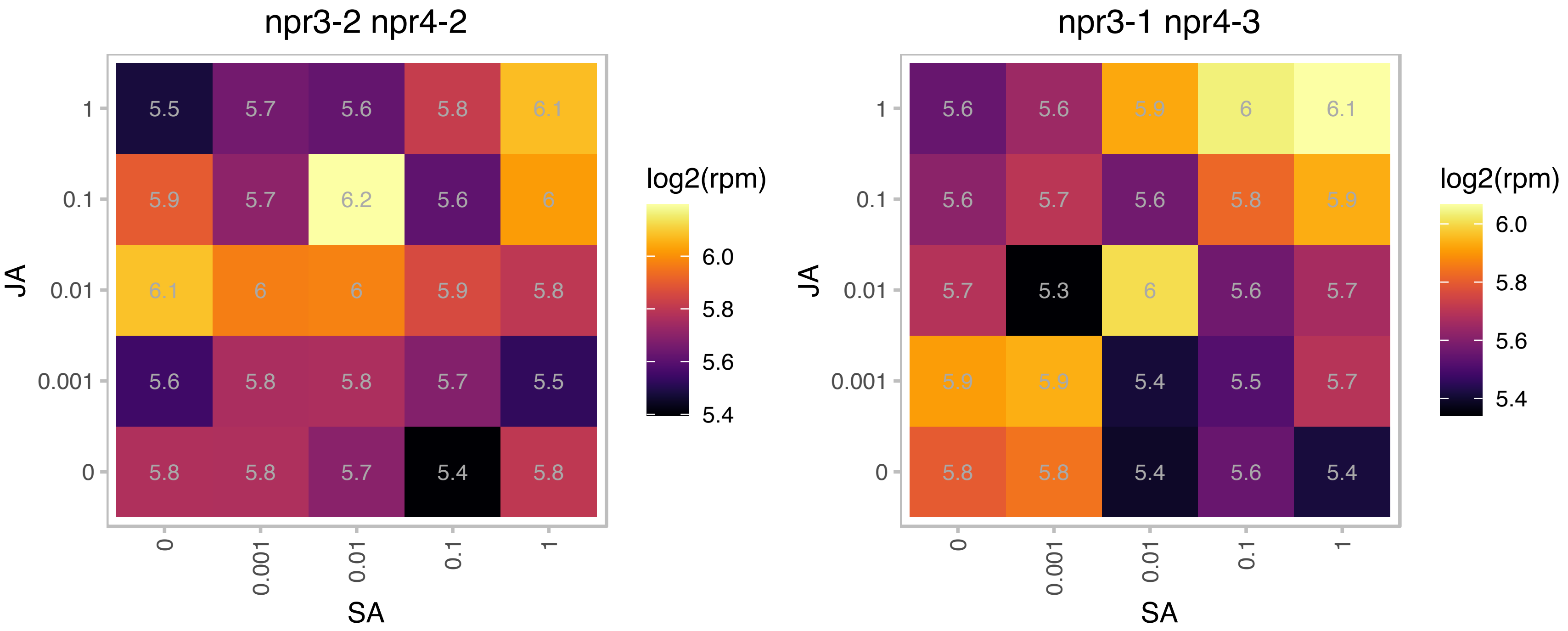

### AT5G62190.1 : AtRH7, PRH75

DEAD box RNA helicase (PRH75)

DEAD/DEAH box RNA helicase PRH75

| | |
|---|---|
| Gene | AT5G62190 |
| is.expressed_genes | TRUE |
| is.responsive_genes | TRUE |
| is.apcluster | F-04 |

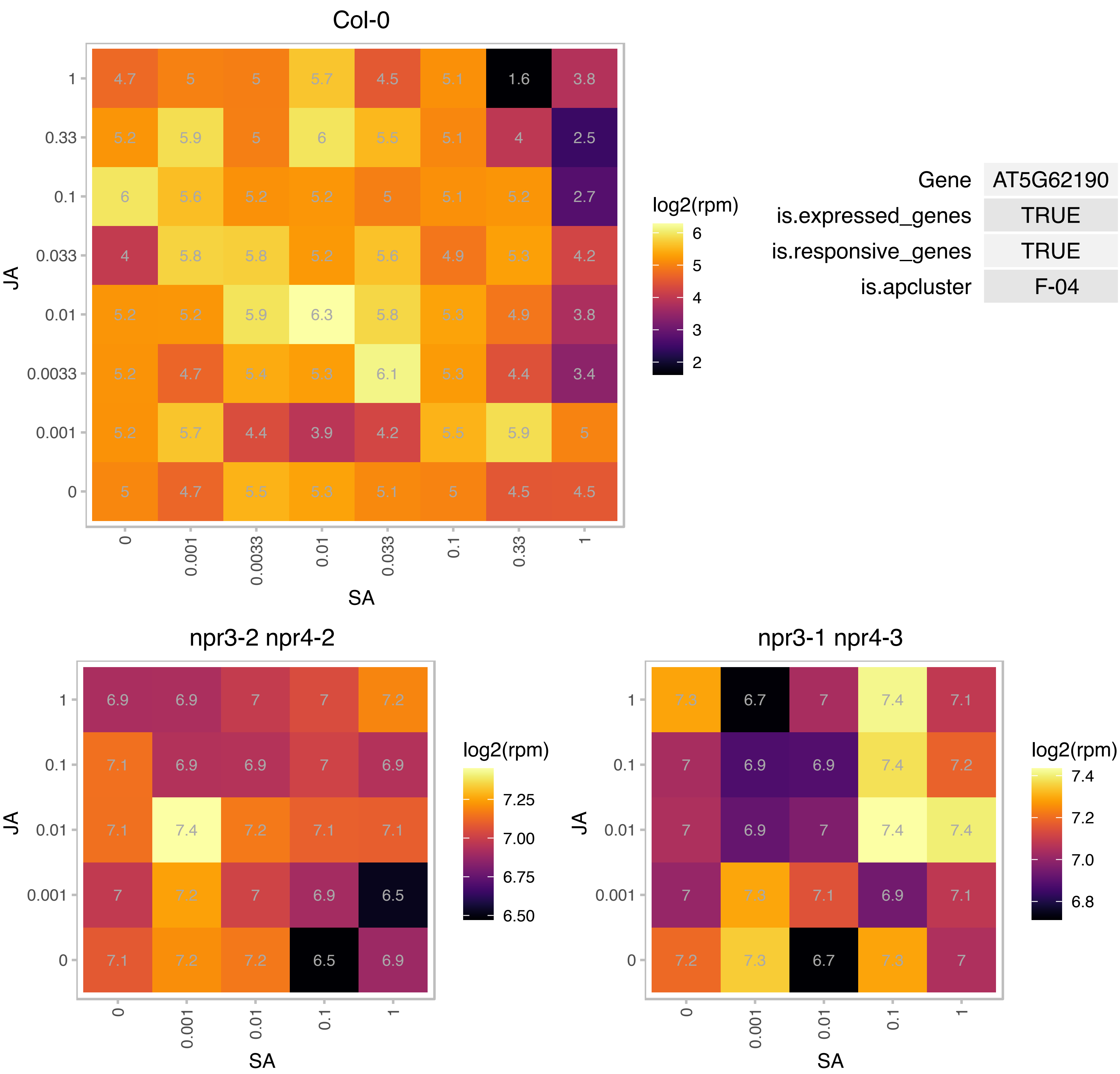

### AT5G62670.1 : AHA11, HA11

H(+)-ATPase 11

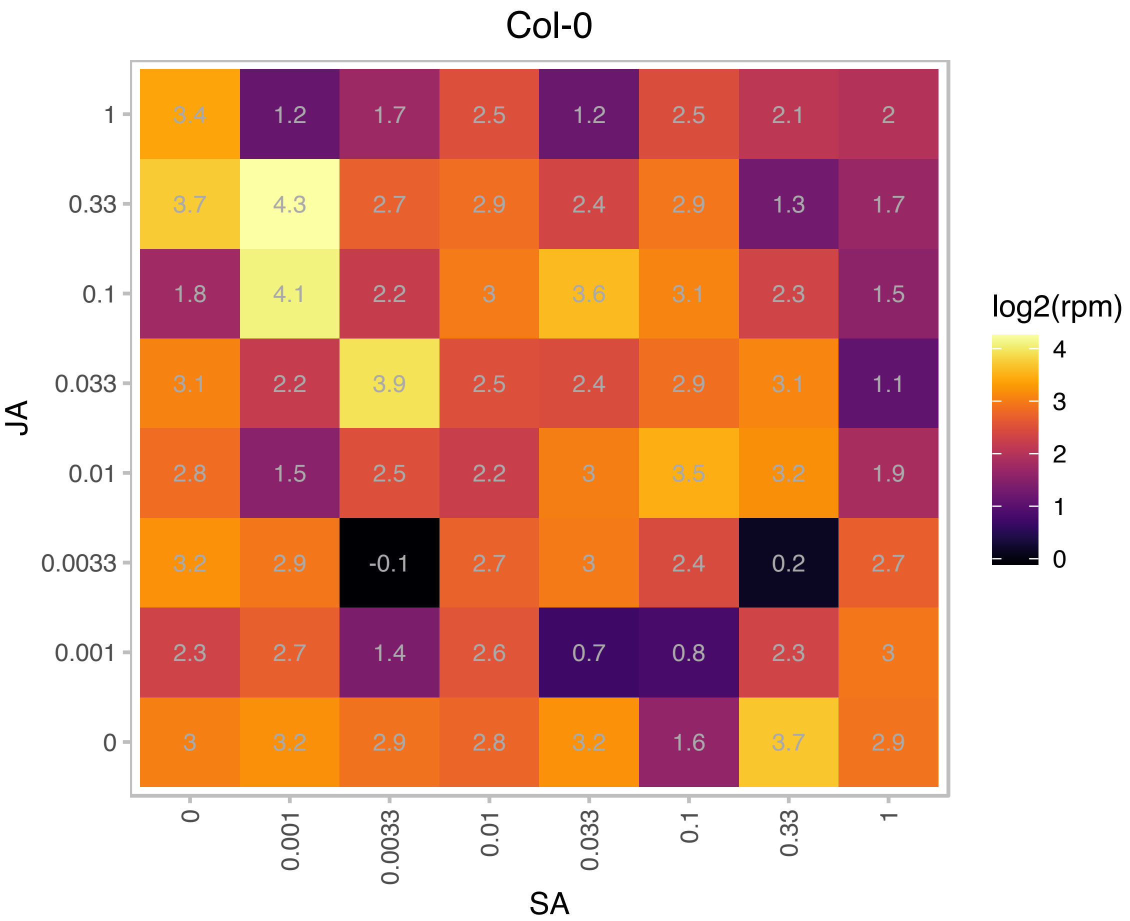


| | |
|---|---|
| Gene | AT5G62670 |
| is.expressed_genes | TRUE |
| is.responsive_genes | TRUE |
| is.apcluster | F-04 |

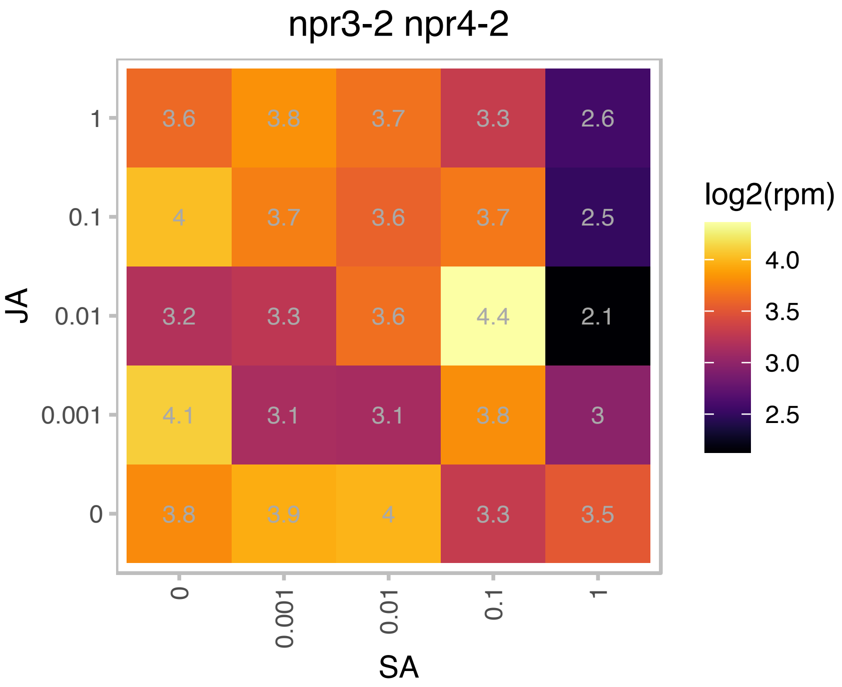


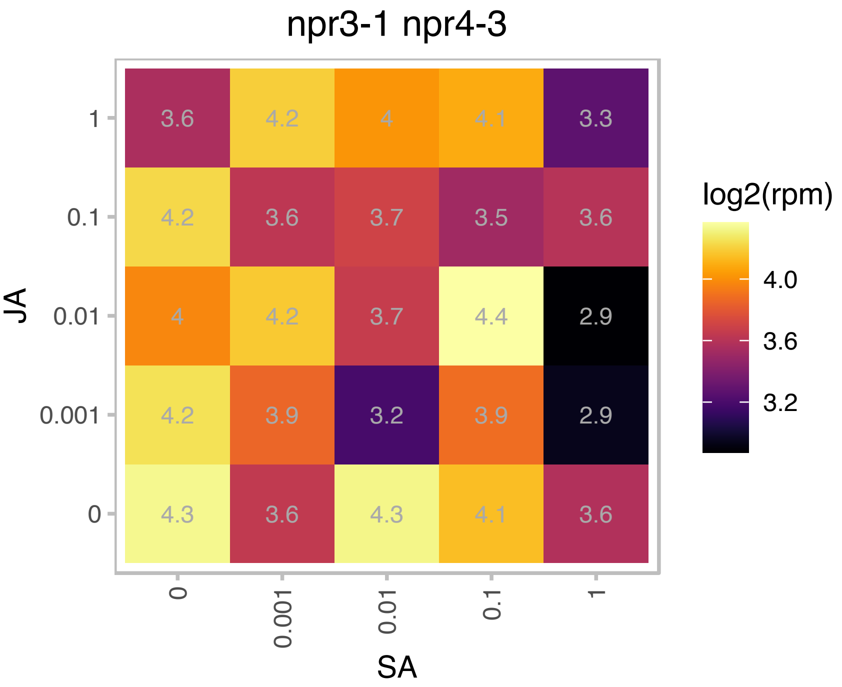

# AT5G63550.1 : NA

DEK domain-containing chromatin associated protein

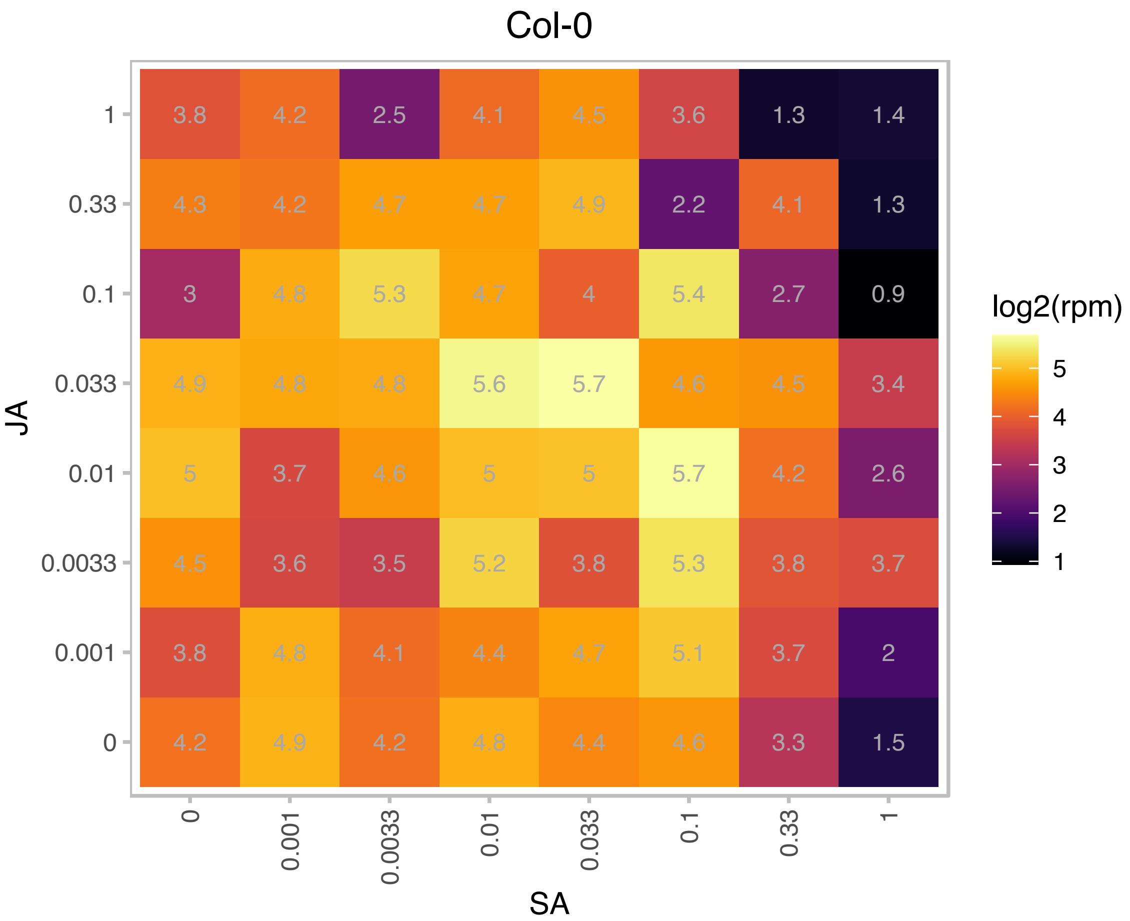


| Gene | AT5G63550 |
|---|---|
| is.expressed_genes | TRUE |
| is.responsive_genes | TRUE |
| is.apcluster | F-04 |

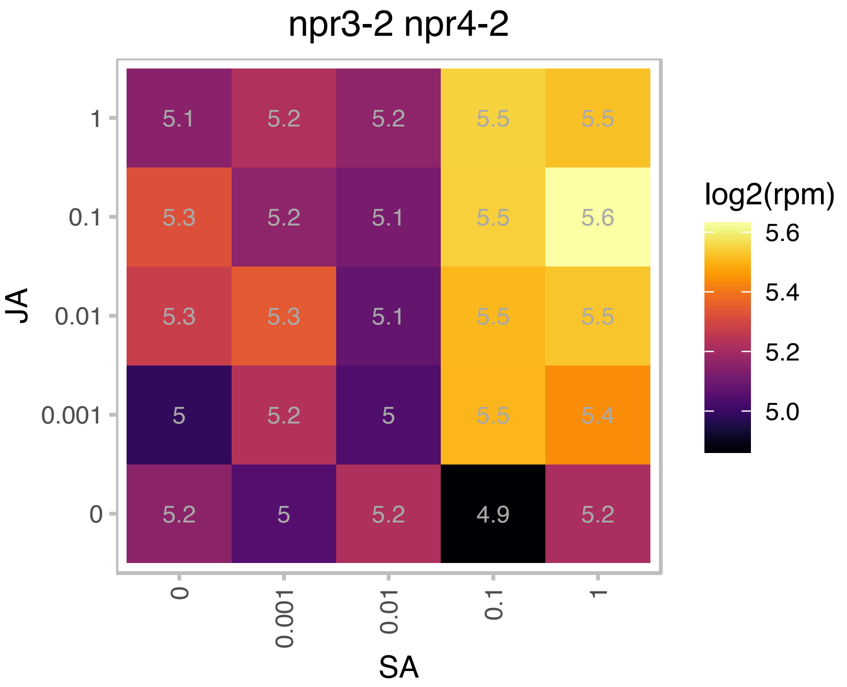


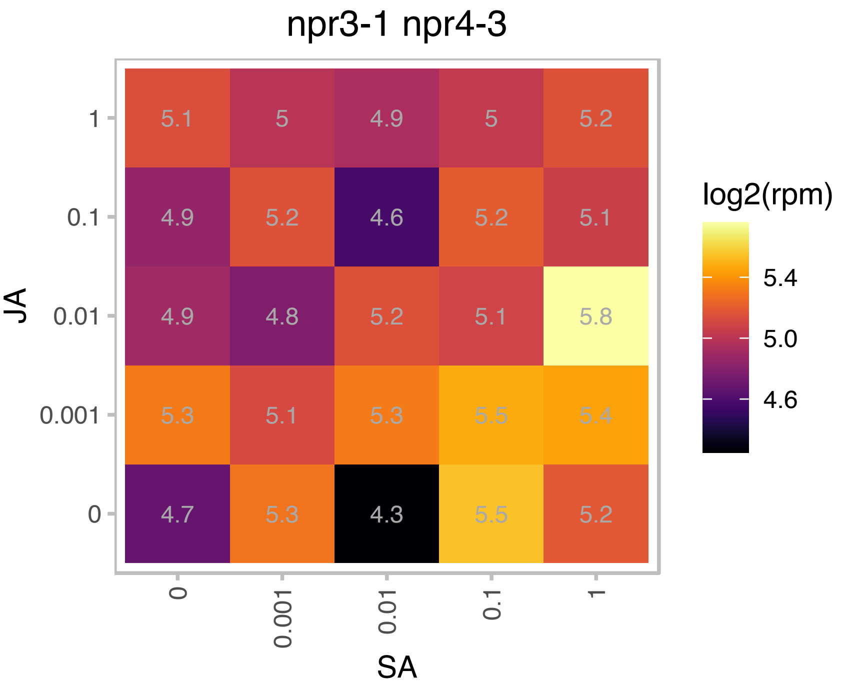

### ATCG00790.1 : RPL16

ribosomal protein L16

chloroplast gene encoding a ribosomal protein L16, which is a constituent of 50S large ribosomal subunit

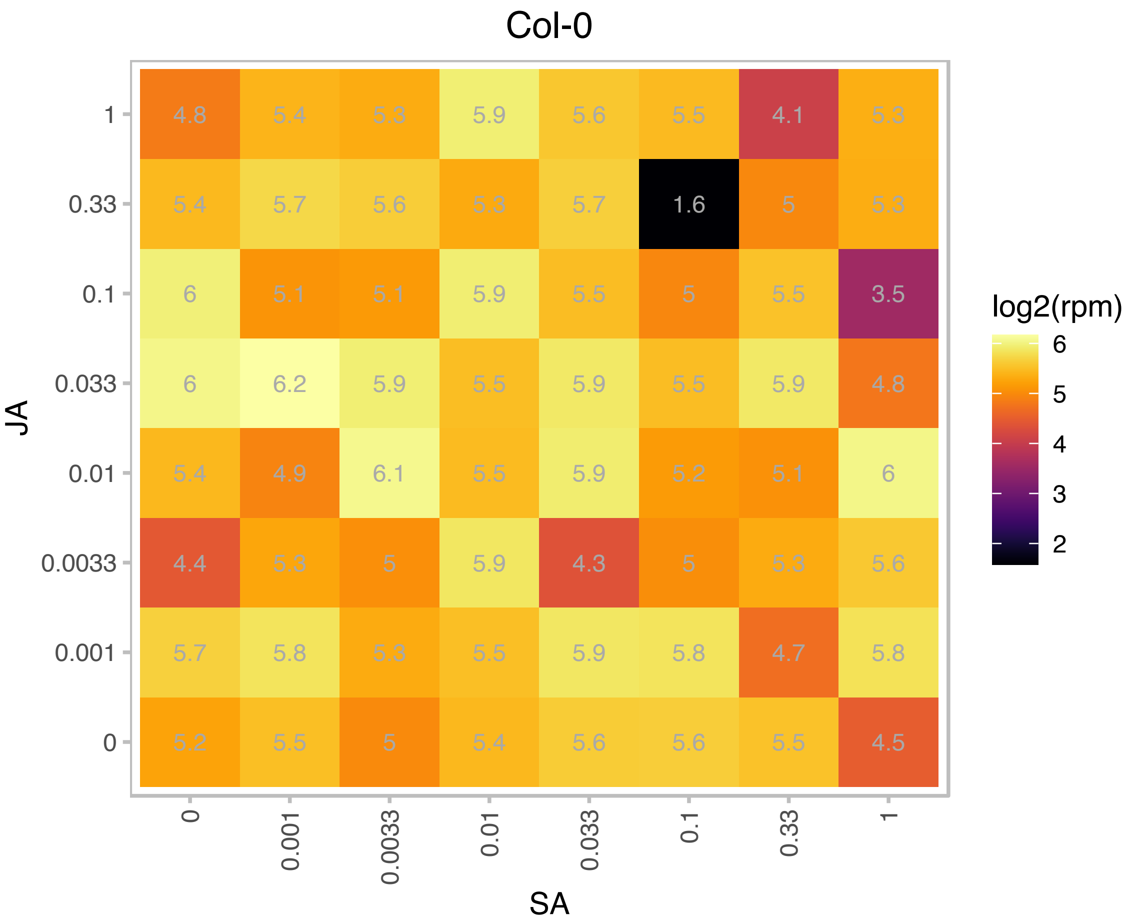


| | |
|---|---|
| Gene | ATCG00790 |
| is.expressed_genes | TRUE |
| is.responsive_genes | TRUE |
| is.apcluster | F-04 |

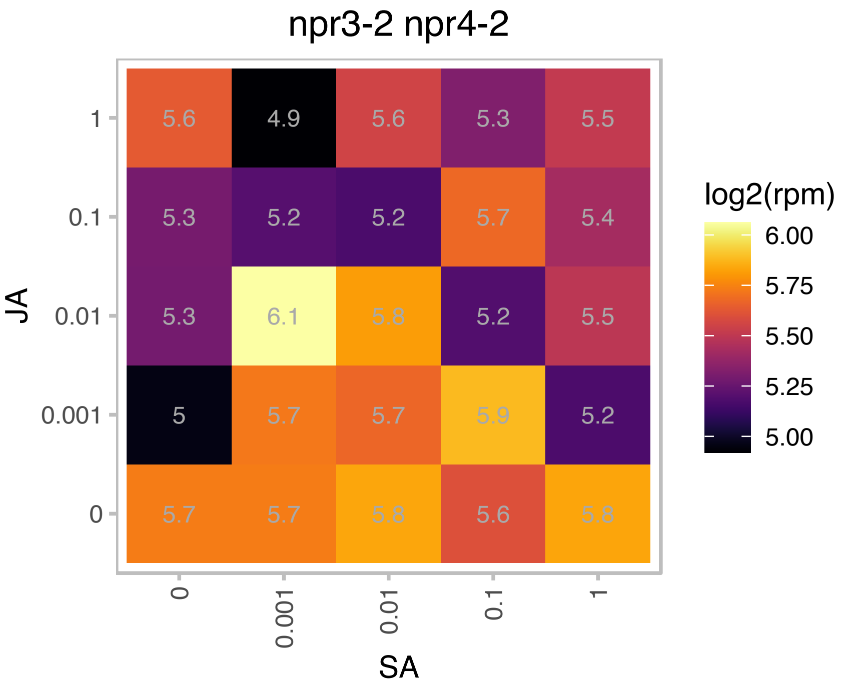


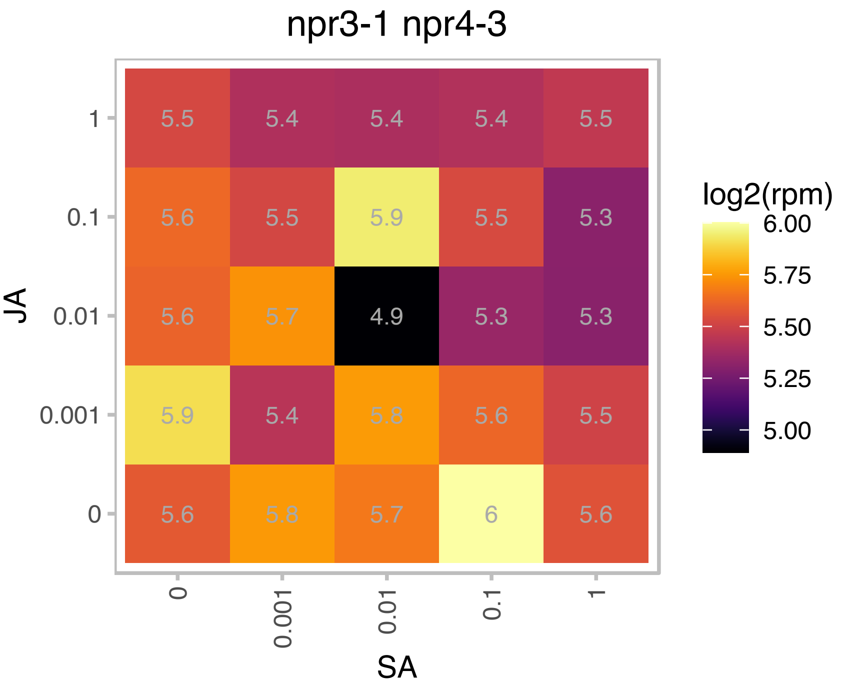

### AT1G01050.1 : AtPPa1, PPa1

pyrophosphorylase 1

Encodes a soluble protein with inorganic pyrophosphatase activity that is highly specific for Mg-inorganic pyrophosphate.

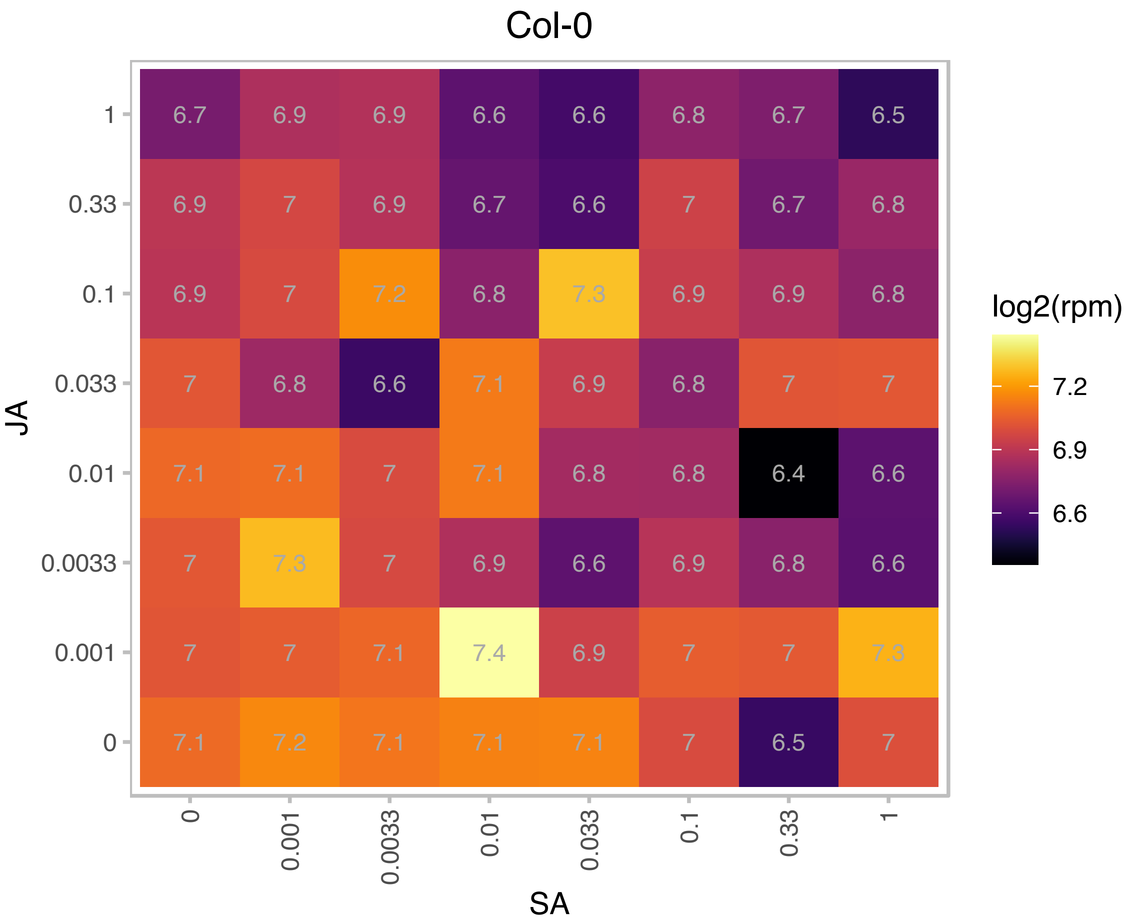


| | |
|---|---|
| Gene | AT1G01050 |
| is.expressed_genes | TRUE |
| is.responsive_genes | TRUE |
| is.apcluster | F-05 |

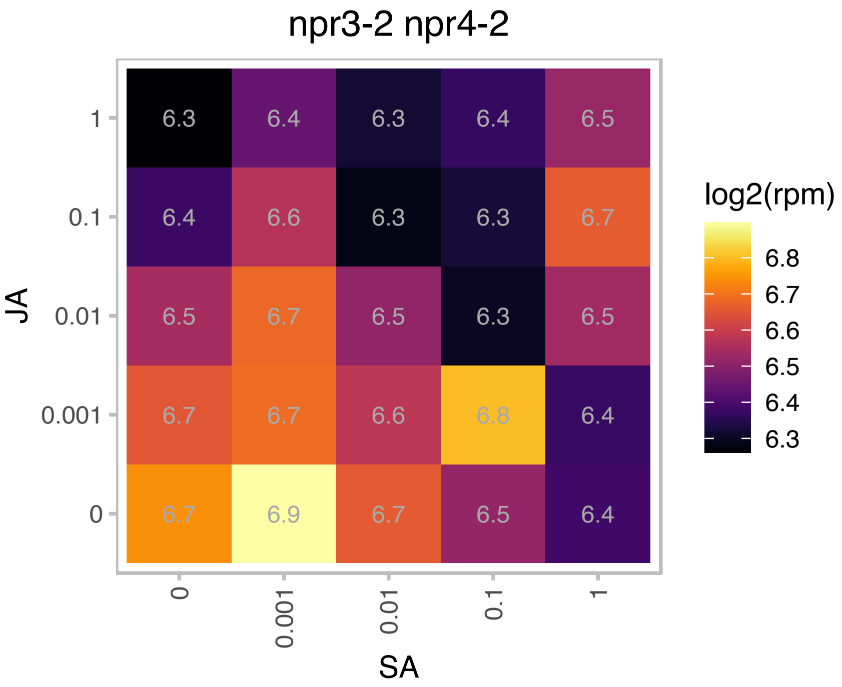


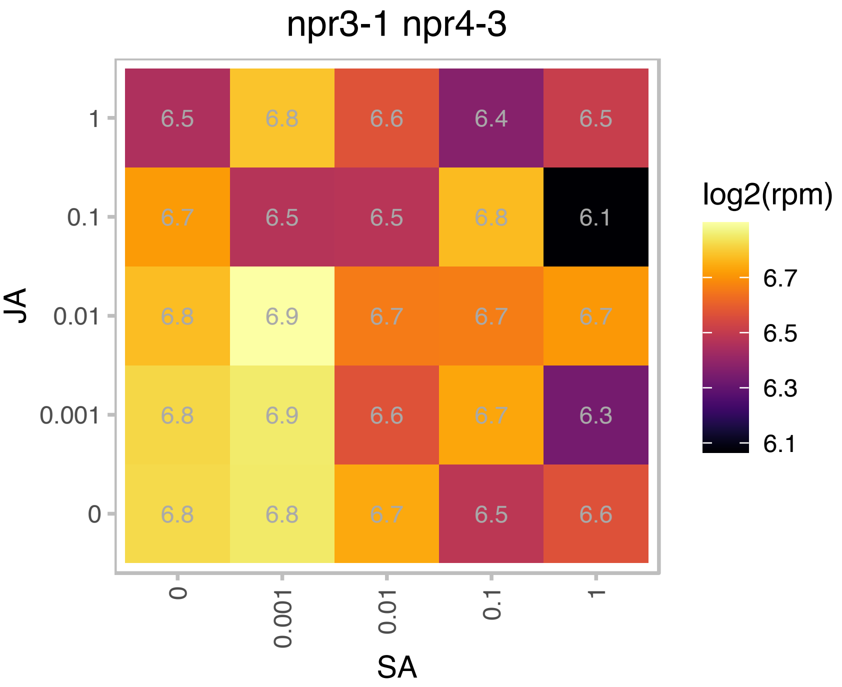

# AT1G01170.1 : NA

Protein of unknown function (DUF1138)

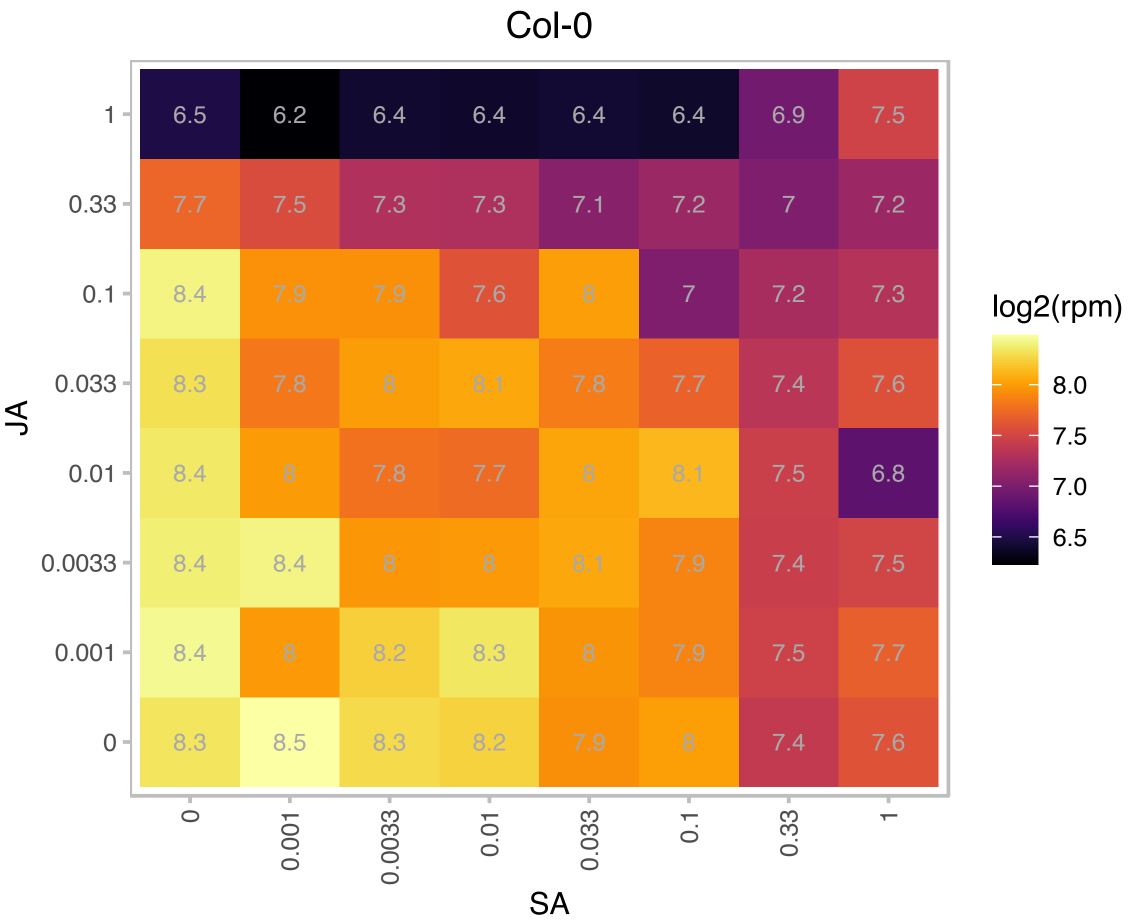


| | |
|---|---|
| Gene | AT1G01170 |
| is.expressed_genes | TRUE |
| is.responsive_genes | TRUE |
| is.apcluster | F-05 |

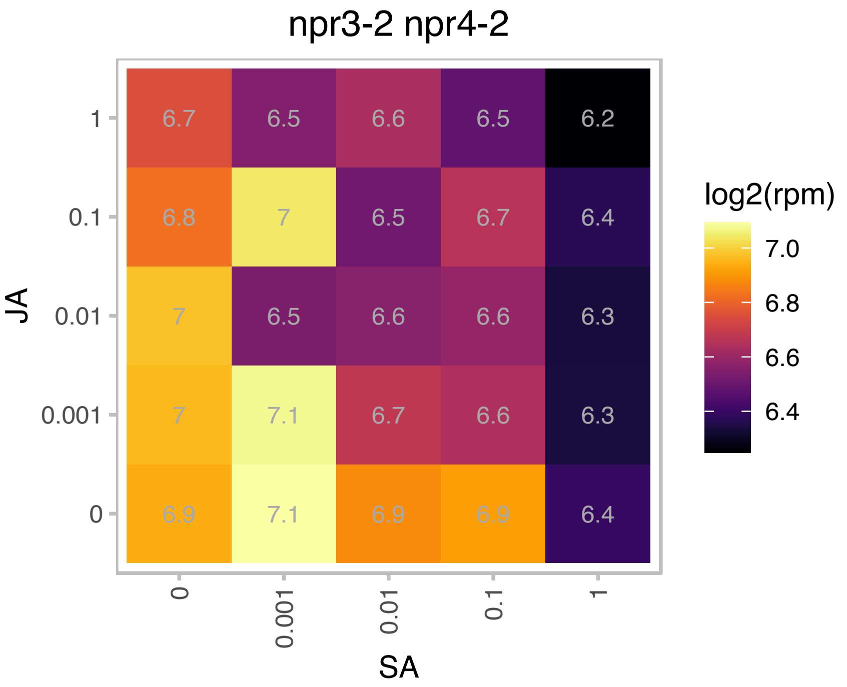


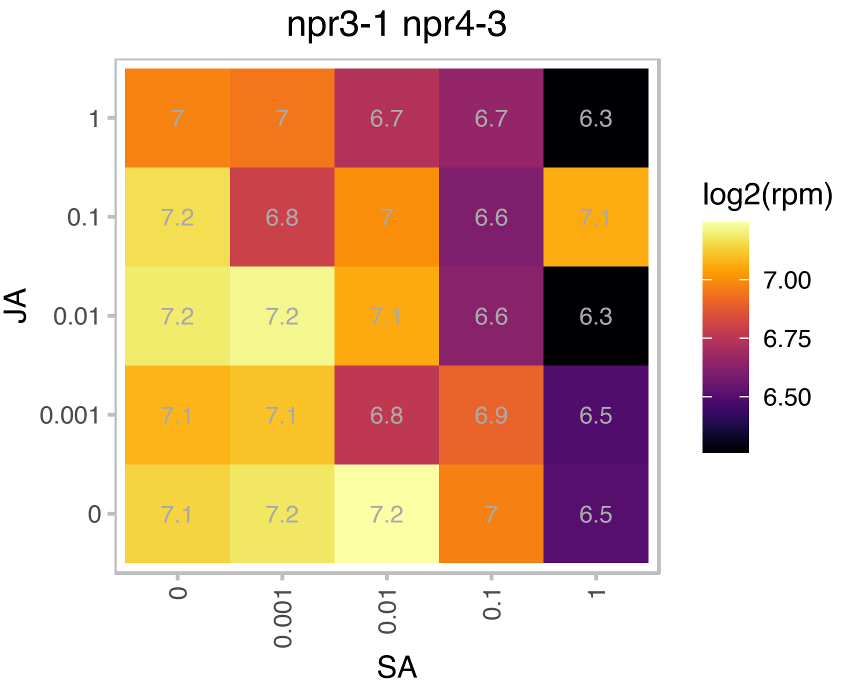

### AT1G01780.1 : PLIM2b

GATA type zinc finger transcription factor family protein

Encodes a member of the Arabidopsis LIM proteins: a family of actin bundlers with distinct expression patterns. WLIM1, WLIM2a, and WLIM2b are widely expressed, whereas PLIM2a, PLIM2b, and PLIM2c are predominantly expressed in pollen. Regulates actin cytoskeleton organization. The mRNA is cell-to-cell mobile.

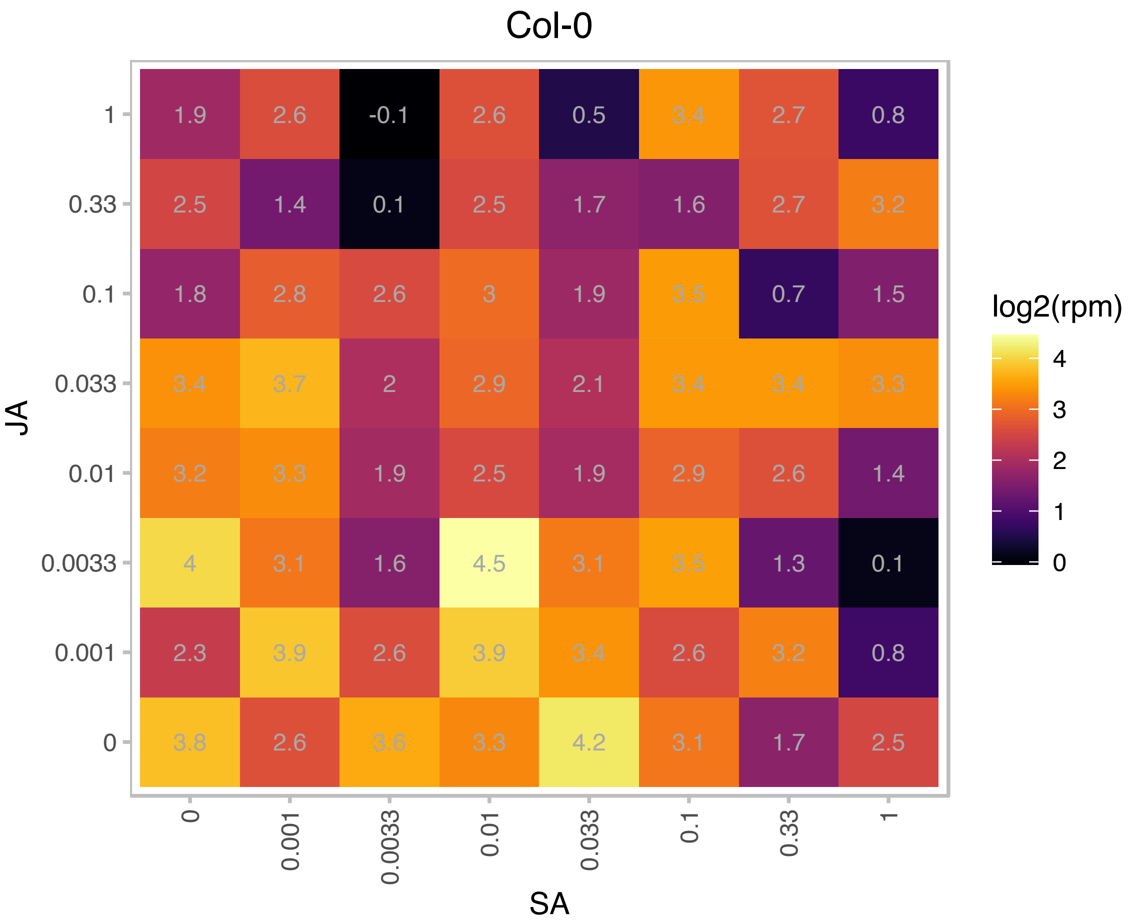


| | |
|---|---|
| Gene | AT1G01780 |
| is.expressed_genes | TRUE |
| is.responsive_genes | TRUE |
| is.apcluster | F-05 |

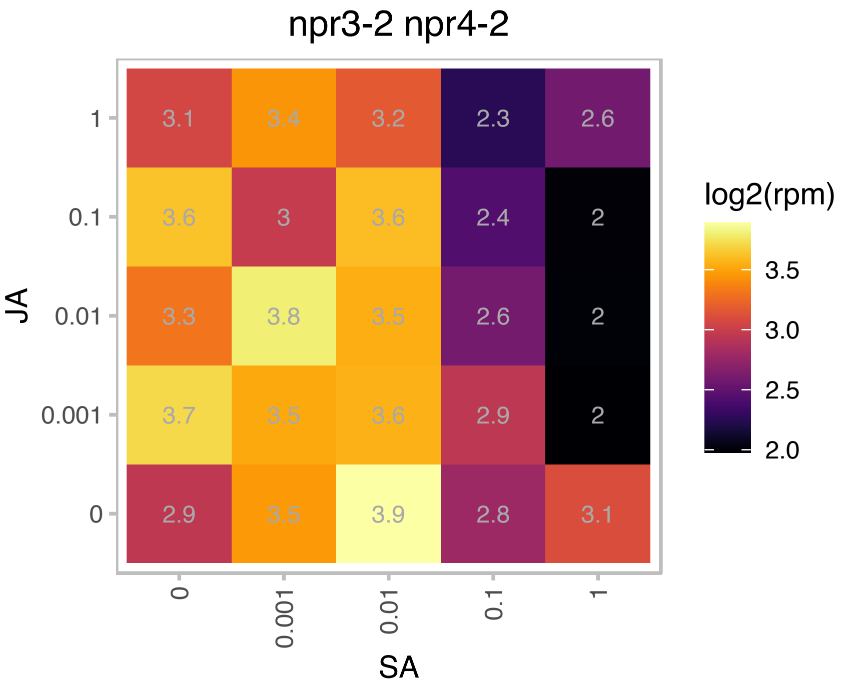


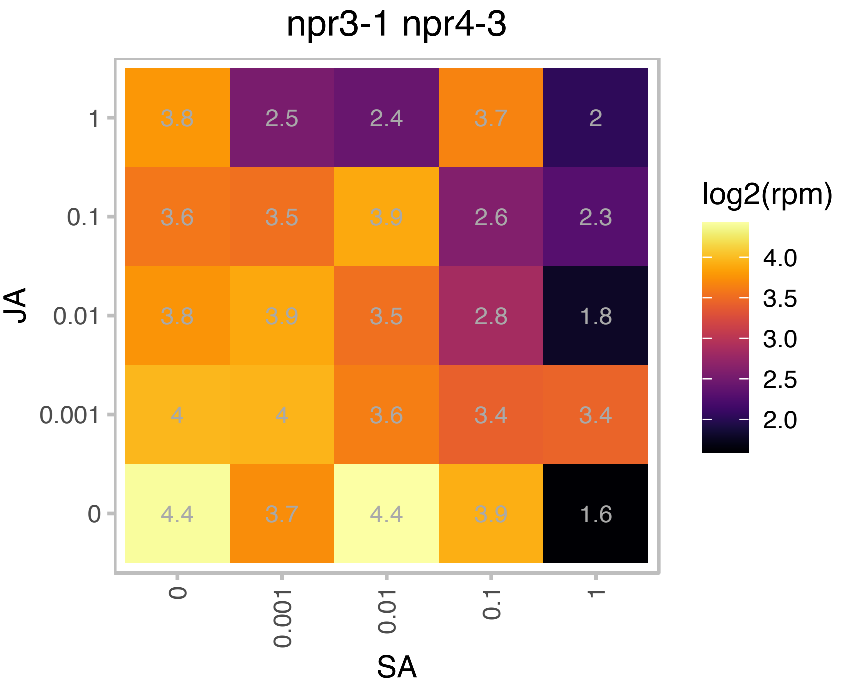

### AT1G03630.1 : POR C, PORC

protochlorophyllide oxidoreductase C

Encodes for a protein with protochlorophyllide oxidoreductase activity. The enzyme is NADPH- and light-dependent.

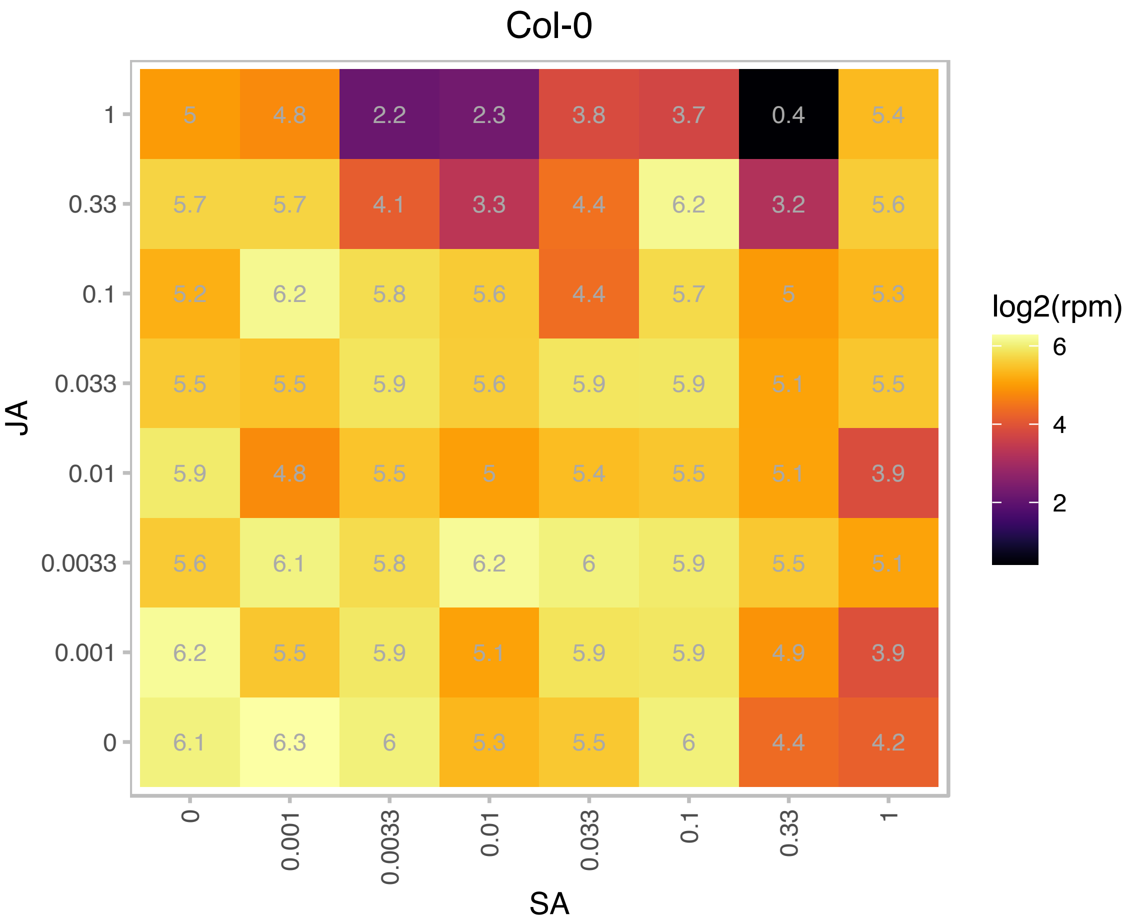


| | |
|---|---|
| Gene | AT1G03630 |
| is.expressed_genes | TRUE |
| is.responsive_genes | TRUE |
| is.apcluster | F-05 |

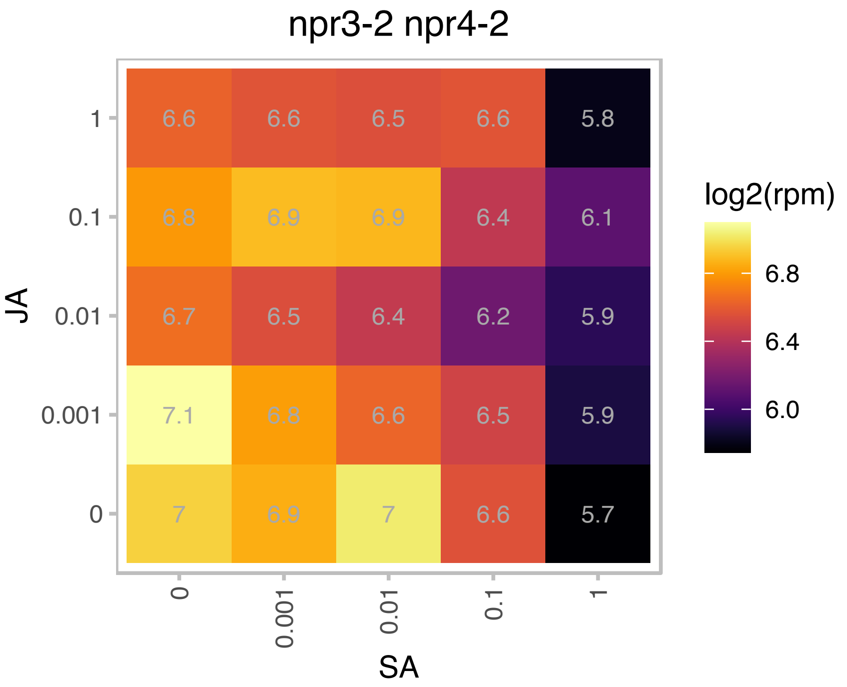


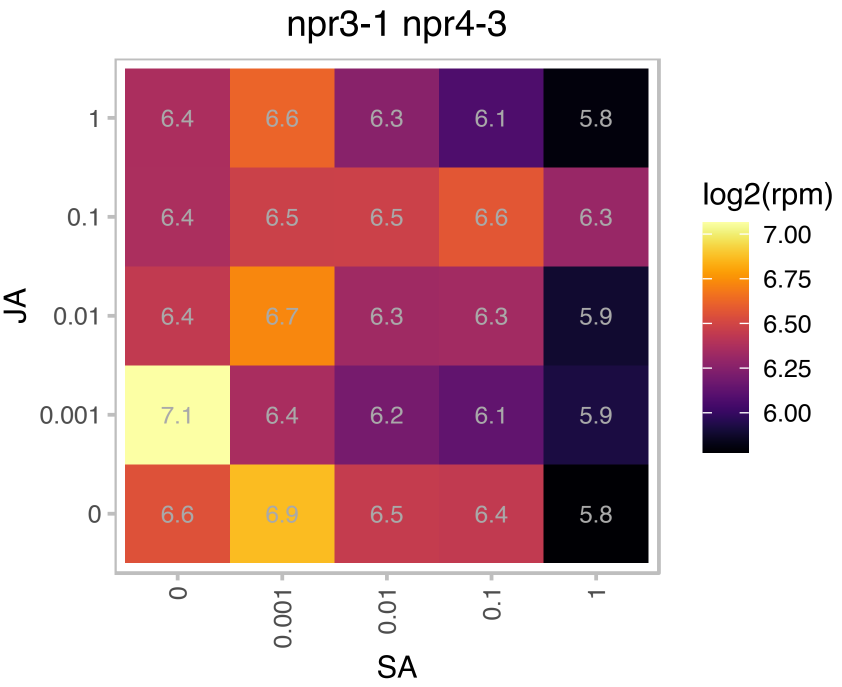

### AT1G04250.1 : AtIAA17, AXR3, IAA17

Transcription regulator acting as repressor of auxin-inducible gene expression. Auxin-inducible AUX/IAA gene. Short-lived nuclear protein with four conserved domains. Domain III has homology to beta alpha alpha dimerization and DNA binding domains. Involved in auxin signaling and is a positive modulator of natural leaf senescence. Auxin induces the degradation of the protein in a dosage-dependent manner in a process mediated by AtRac1. Auxin induced the relocalization of the protein within the nucleus from a diffused nucleoplasmic pattern to a discrete particulated pattern named nuclear protein bodies or NPB in a process also mediated by [illegible]

AUX/IAA transcriptional regulator family protein

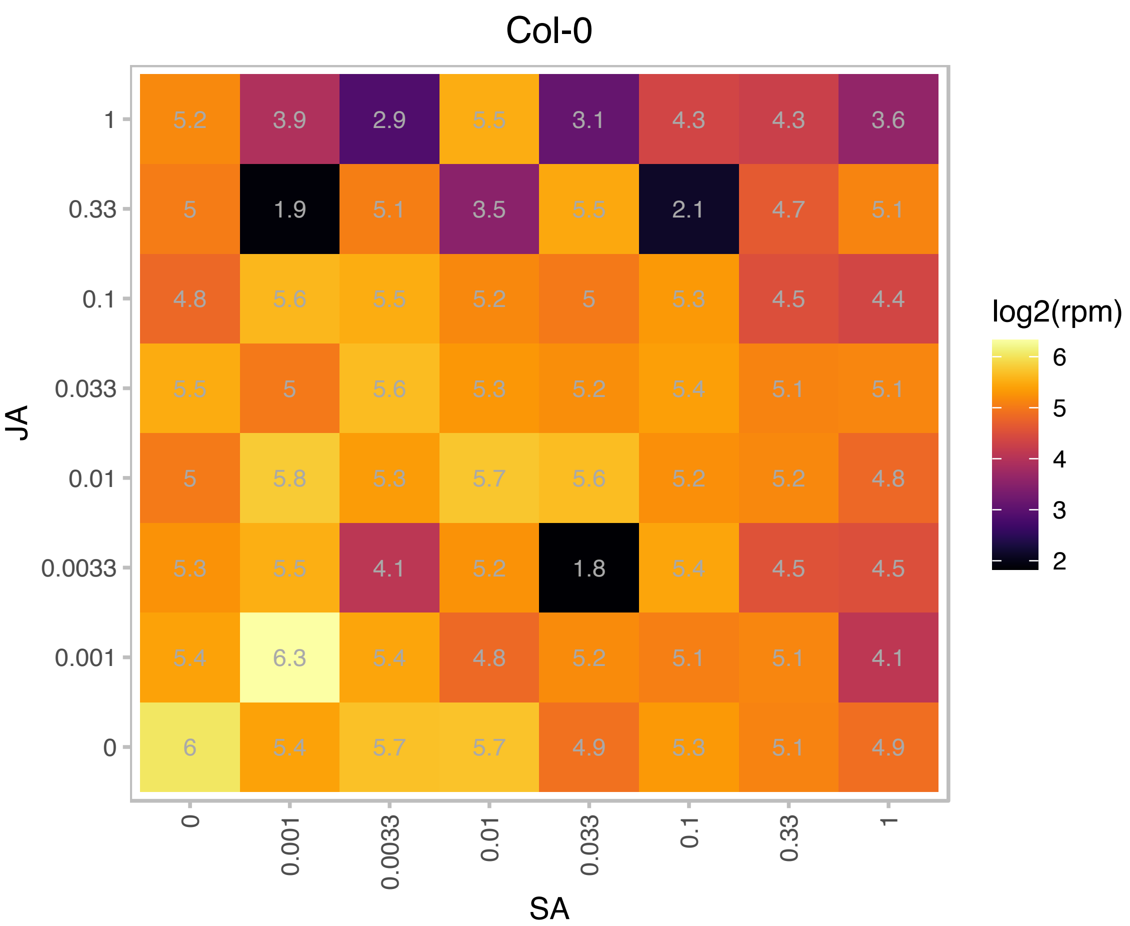


| | |
|---|---|
| Gene | AT1G04250 |
| is.expressed_genes | TRUE |
| is.responsive_genes | TRUE |
| is.apcluster | F-05 |

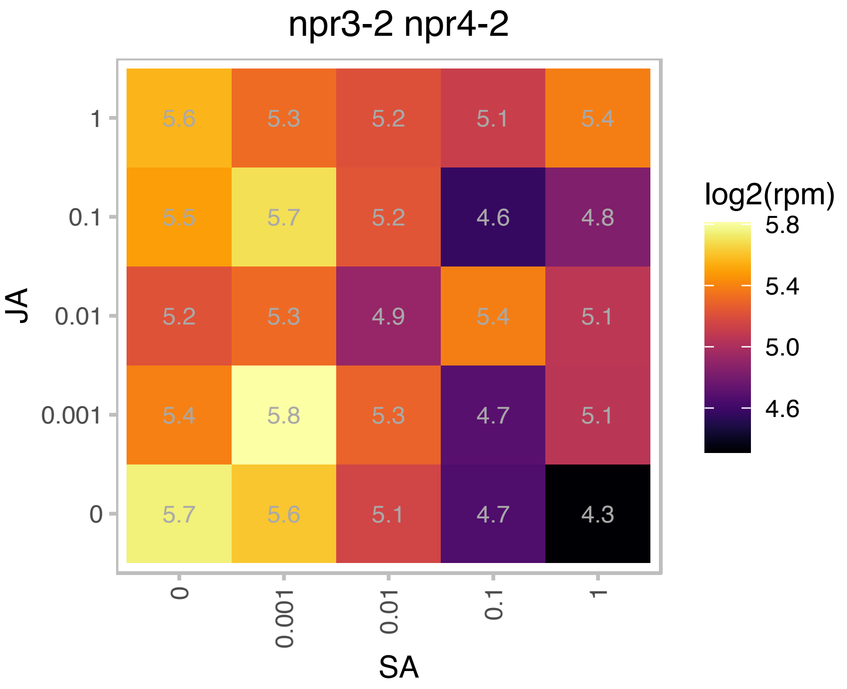


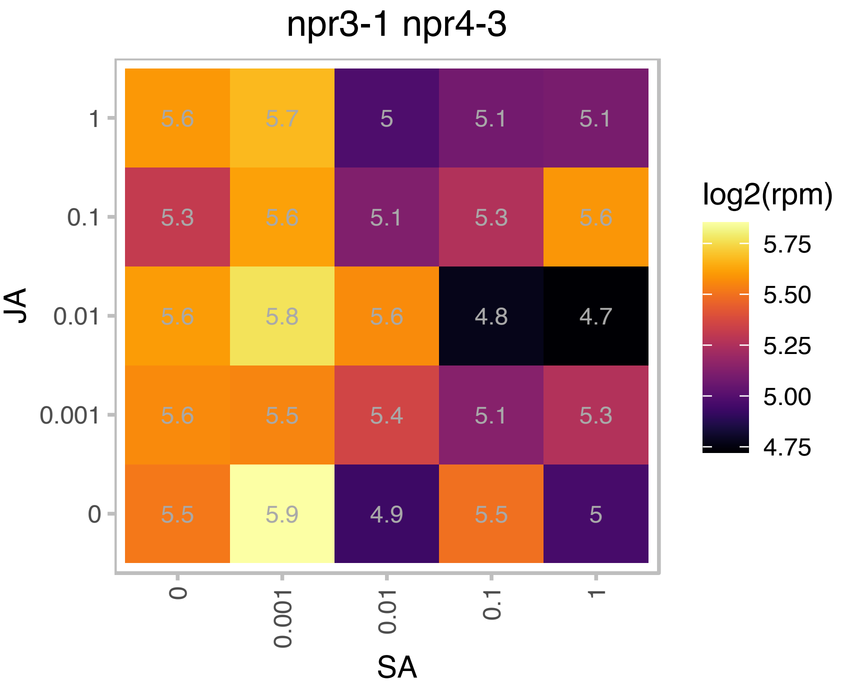

### AT1G05385.1 : LPA19, Psb27-H1

photosystem II 11 kDa protein-related

Encodes a Psb27 homolog involved in photosystem II biogenesis.

| | |
|---|---|
| Gene | AT1G05385 |
| is.expressed_genes | TRUE |
| is.responsive_genes | TRUE |
| is.apcluster | F-05 |

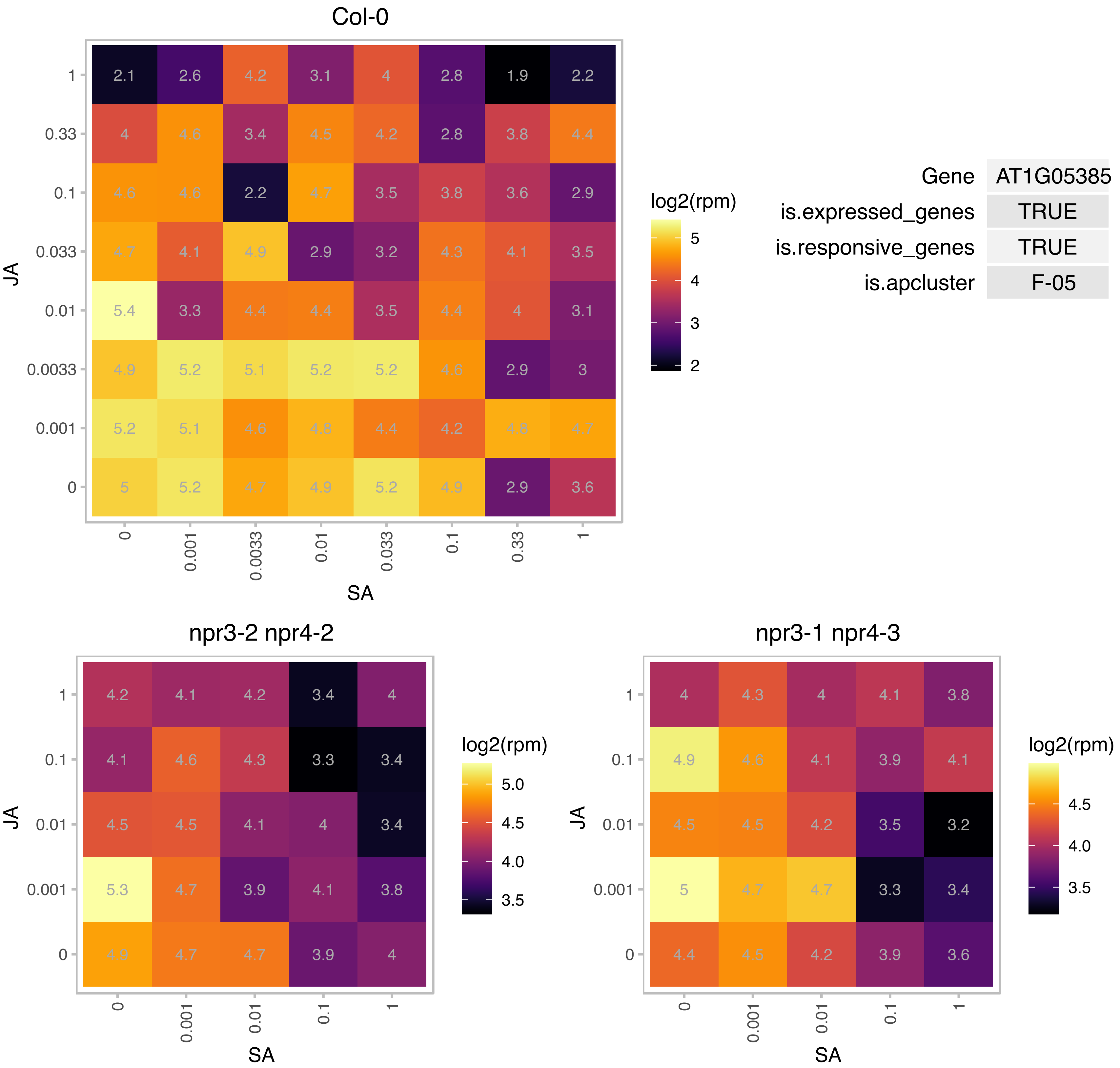

### AT1G10522.1 : PRIN2

Encodes PRIN2 (plastid redox insensitive 2). PRIN2 mutants are impaired in PEP (plastid-encoded RNA polymerase) activity and high light-dependent plastid redox signalling to the nucleus.

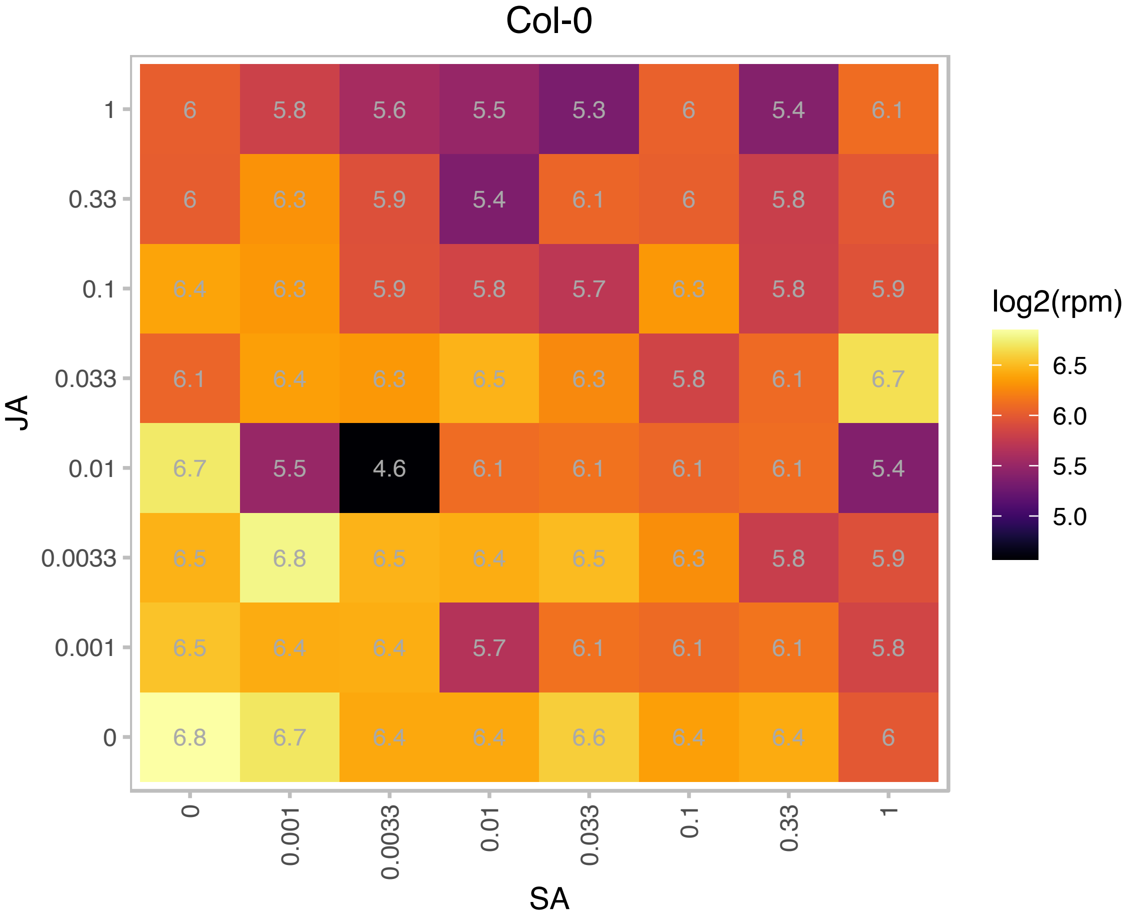


| Gene | AT1G10522 |
|---|---|
| is.expressed_genes | TRUE |
| is.responsive_genes | TRUE |
| is.apcluster | F-05 |

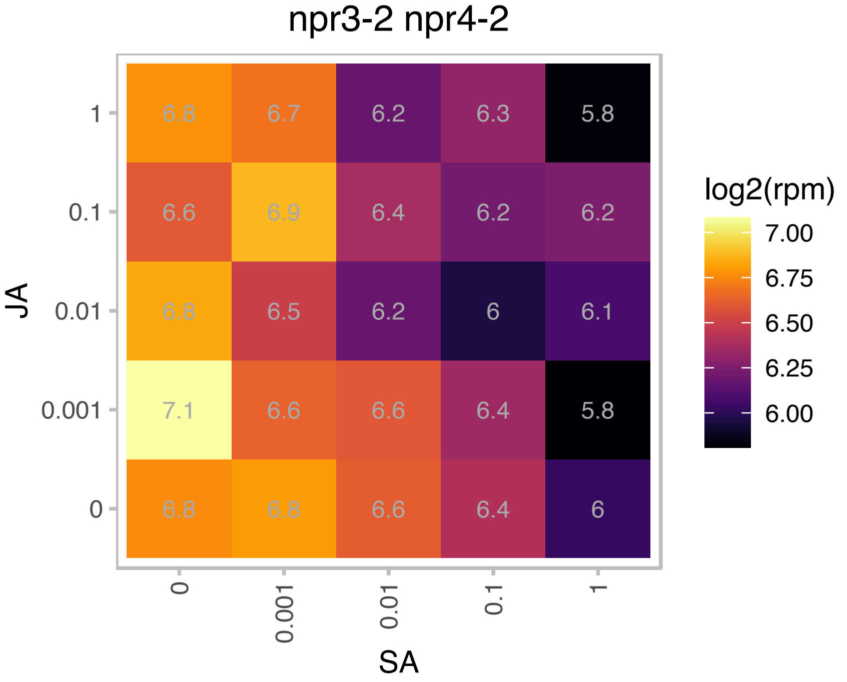


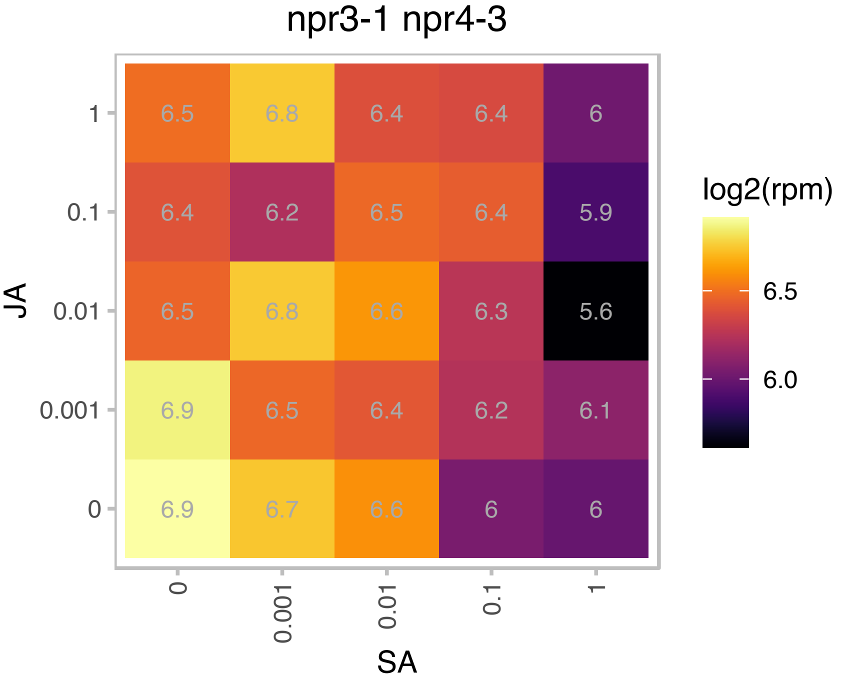

### AT1G12860.1 : ICE2, SCRM2

basic helix-loop-helix (bHLH) DNA-binding superfamily protein
Encodes ICE2 (Inducer of CBF Expression 2), a transcription factor of the bHLH family that participates in the response to deep freezing through the cold acclimation-dependent pathway. Overexpression of ICE2 results in increased tolerance to deep freezing stress after cold acclimation.

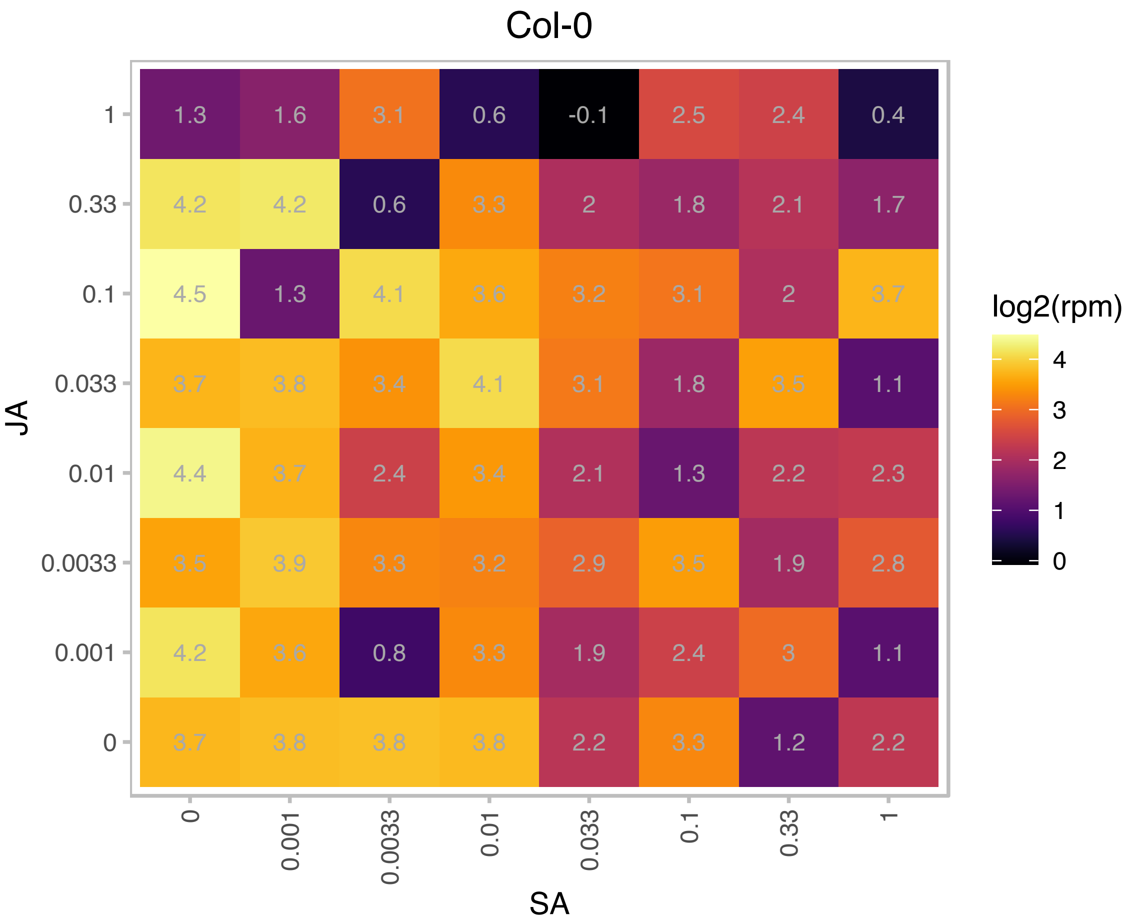


| | |
|---|---|
| Gene | AT1G12860 |
| is.expressed_genes | TRUE |
| is.responsive_genes | TRUE |
| is.apcluster | F-05 |

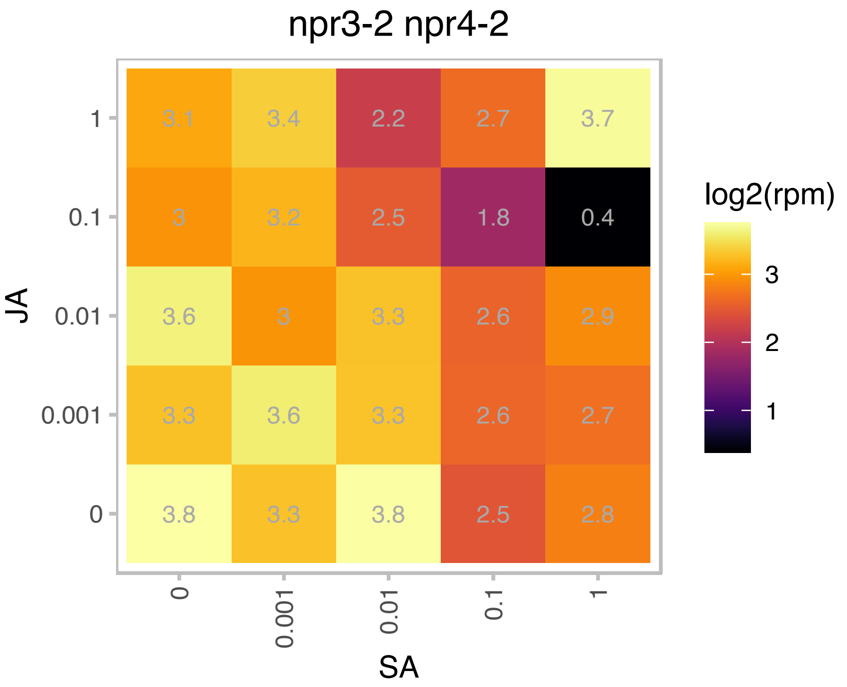


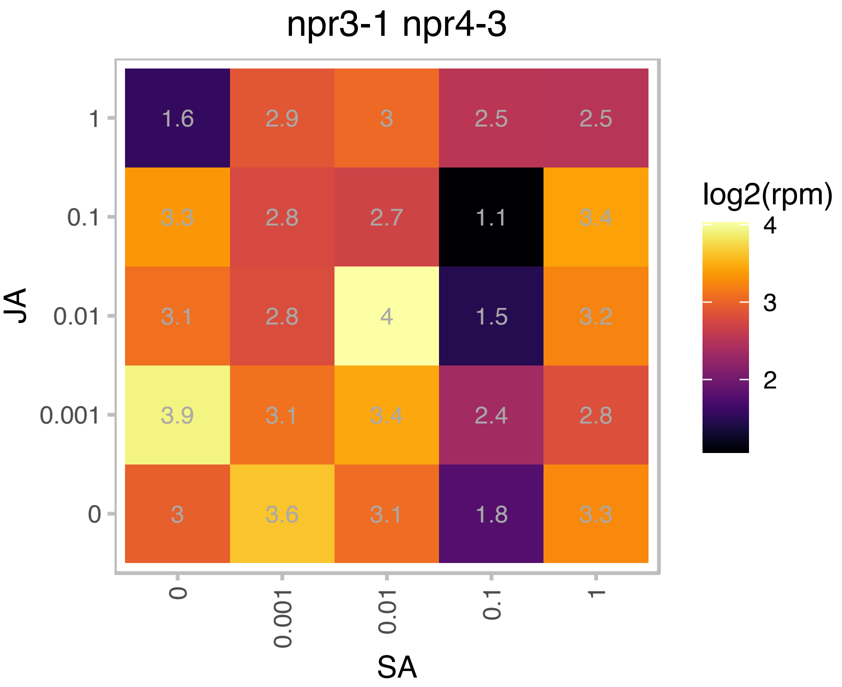

### AT1G12900.1 : GAPA-2

glyceraldehyde 3-phosphate dehydrogenase A subunit 2

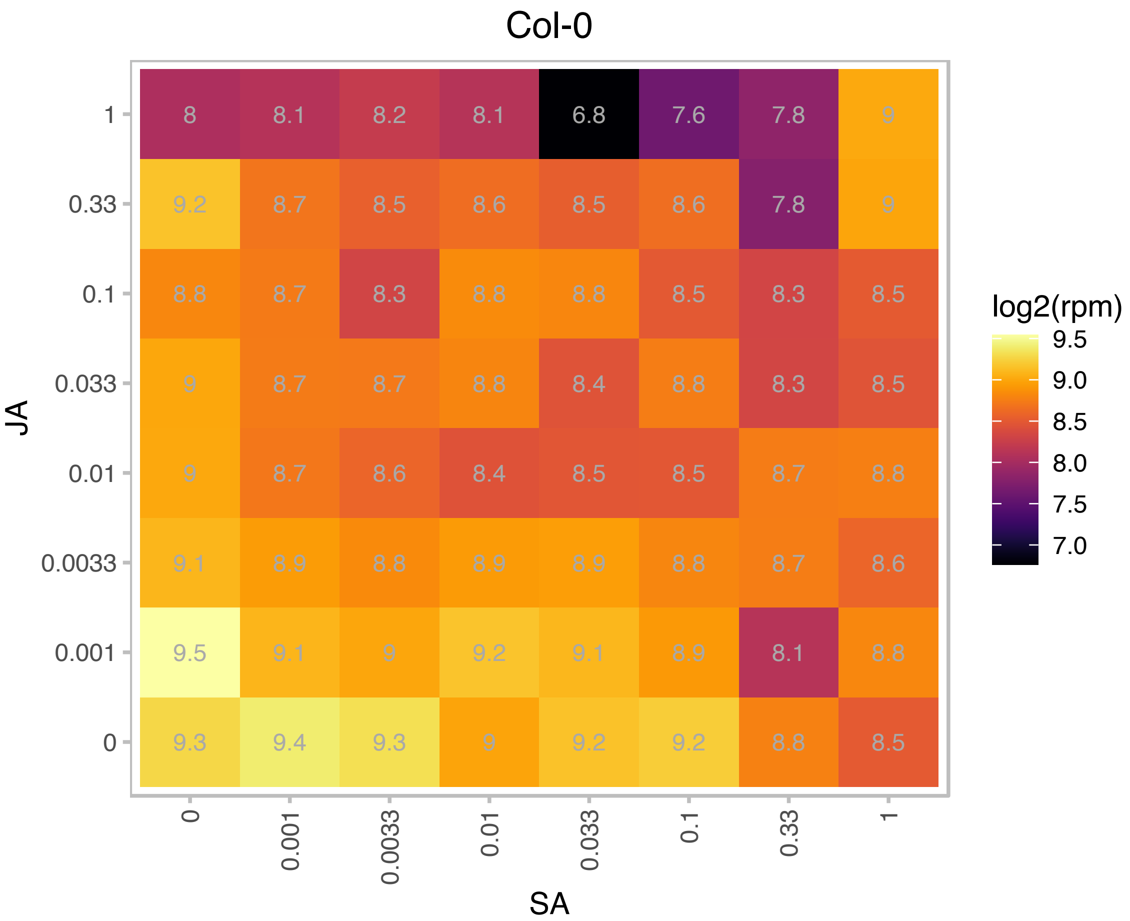


| | |
|---|---|
| Gene | AT1G12900 |
| is.expressed_genes | TRUE |
| is.responsive_genes | TRUE |
| is.apcluster | F-05 |

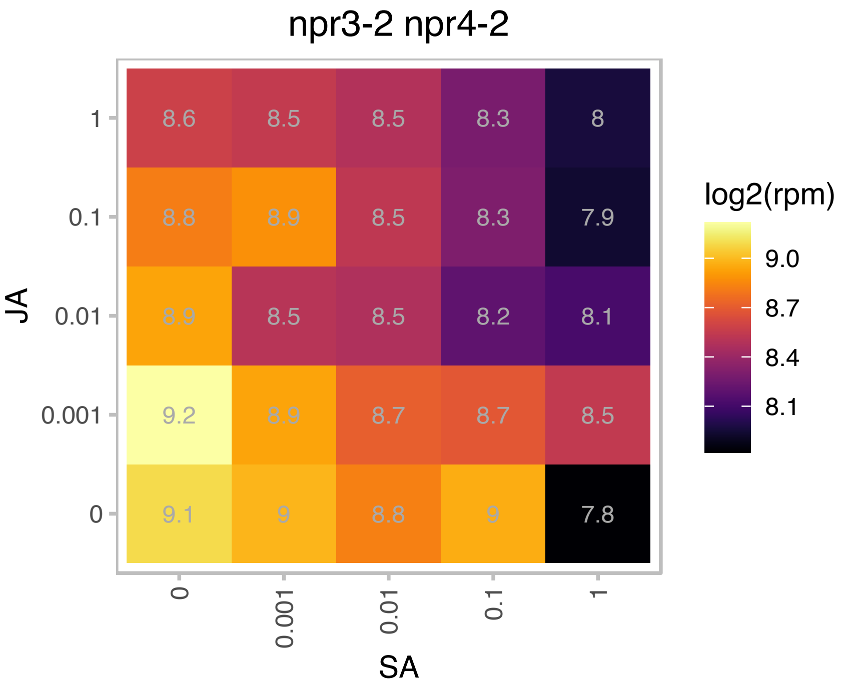


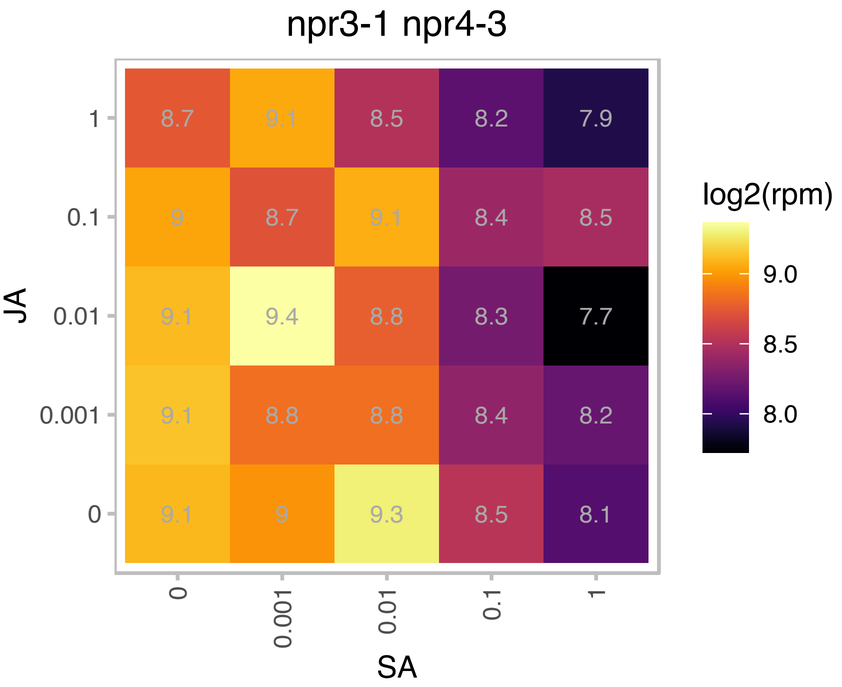

# AT1G13380.1 : NA

Protein of unknown function (DUF1218)

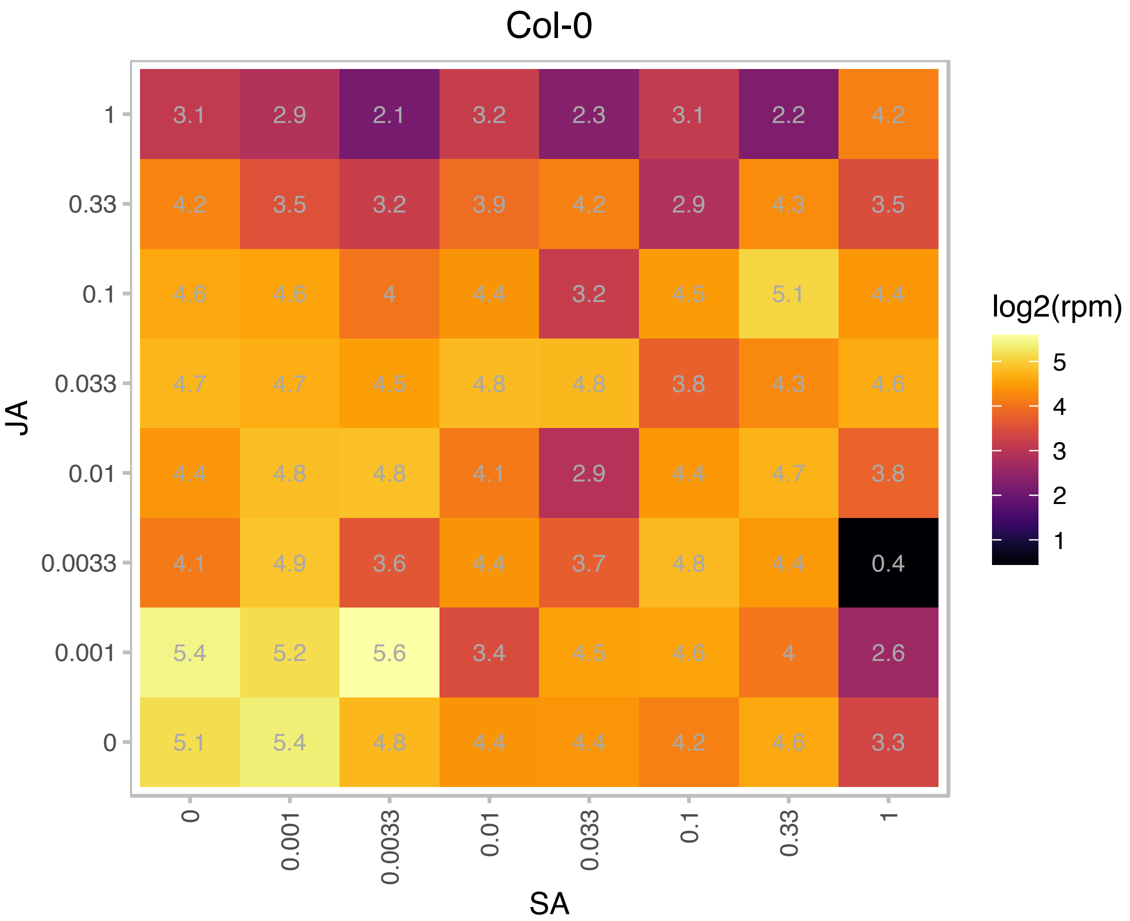


| | |
|---|---|
| Gene | AT1G13380 |
| is.expressed_genes | TRUE |
| is.responsive_genes | TRUE |
| is.apcluster | F-05 |

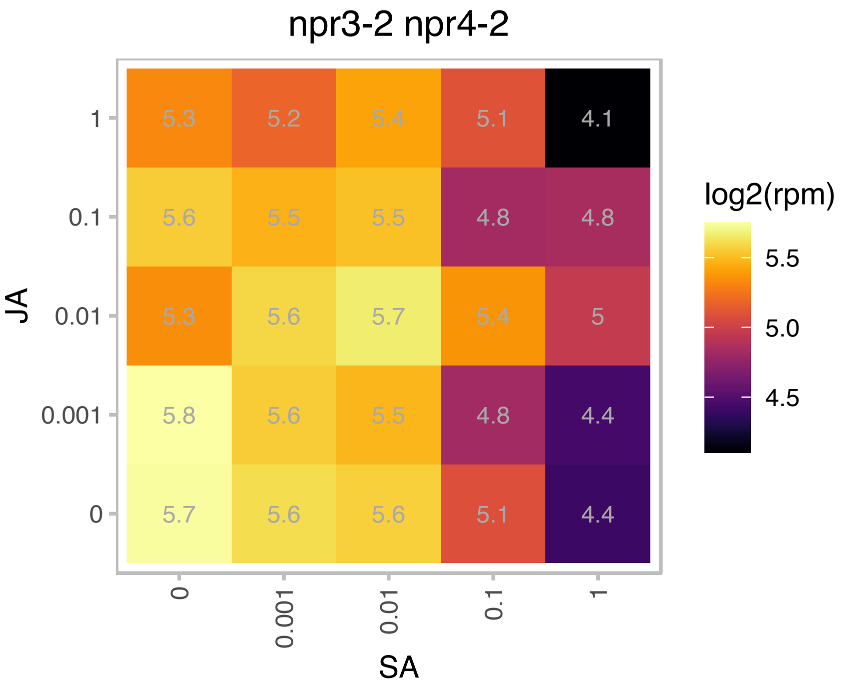


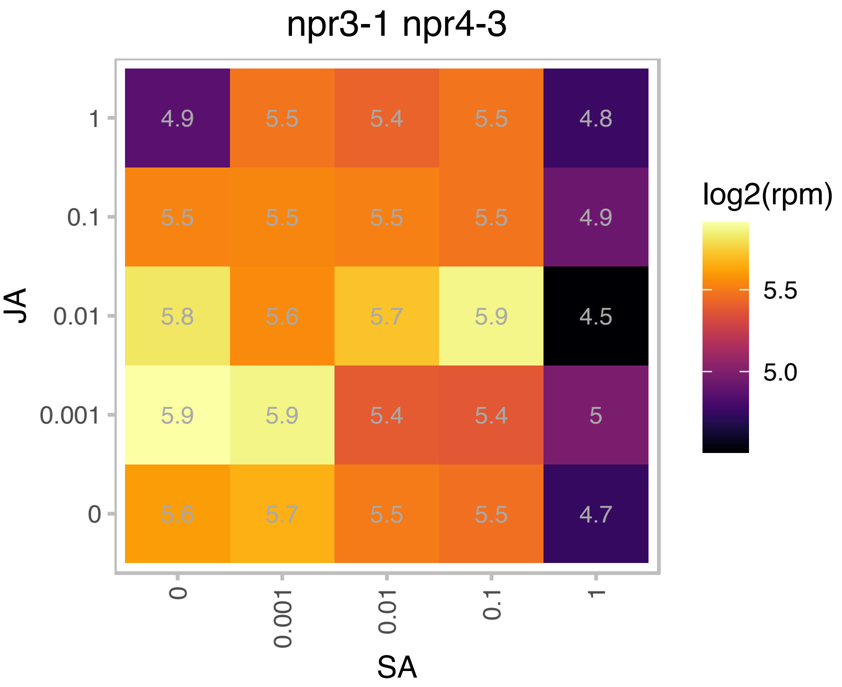

### AT1G20010.1 : TUB5

tubulin beta-5 chain

beta tubulin

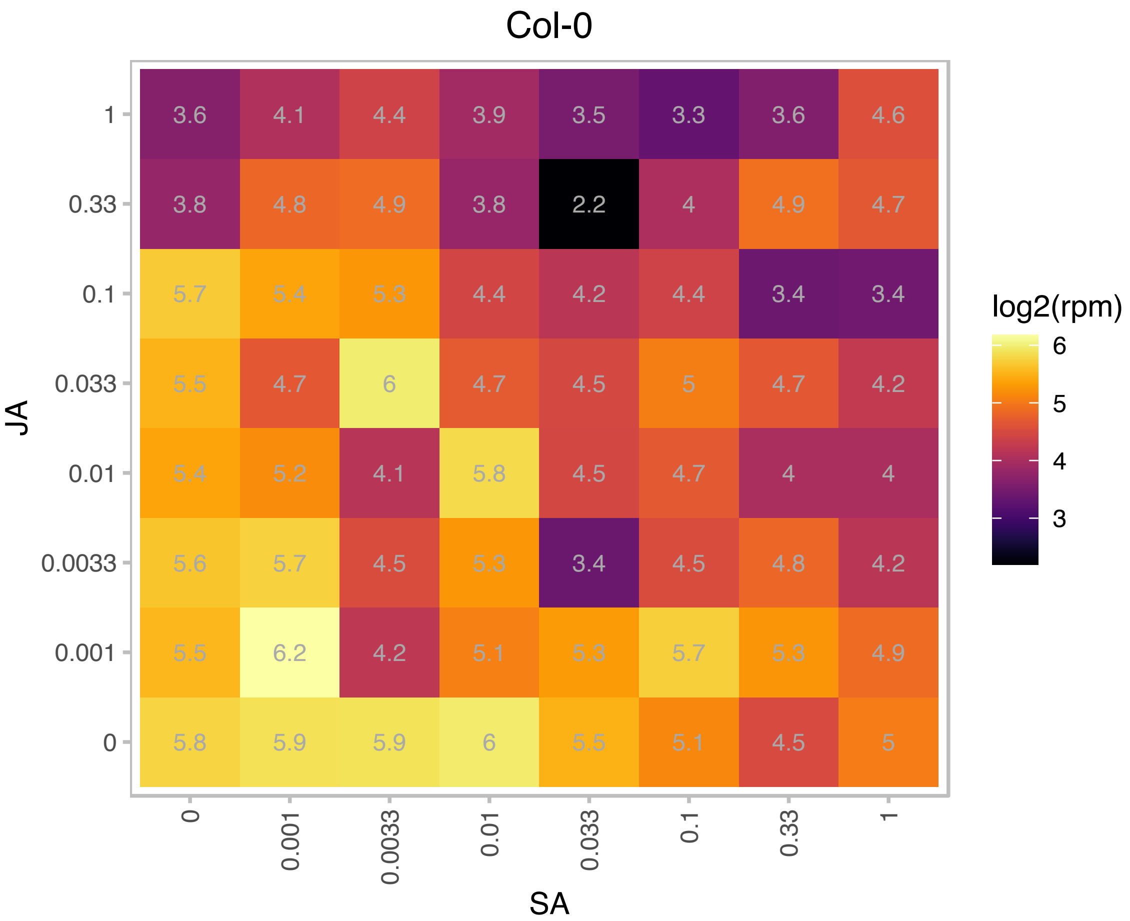


| | |
|---|---|
| Gene | AT1G20010 |
| is.expressed_genes | TRUE |
| is.responsive_genes | TRUE |
| is.apcluster | F-05 |

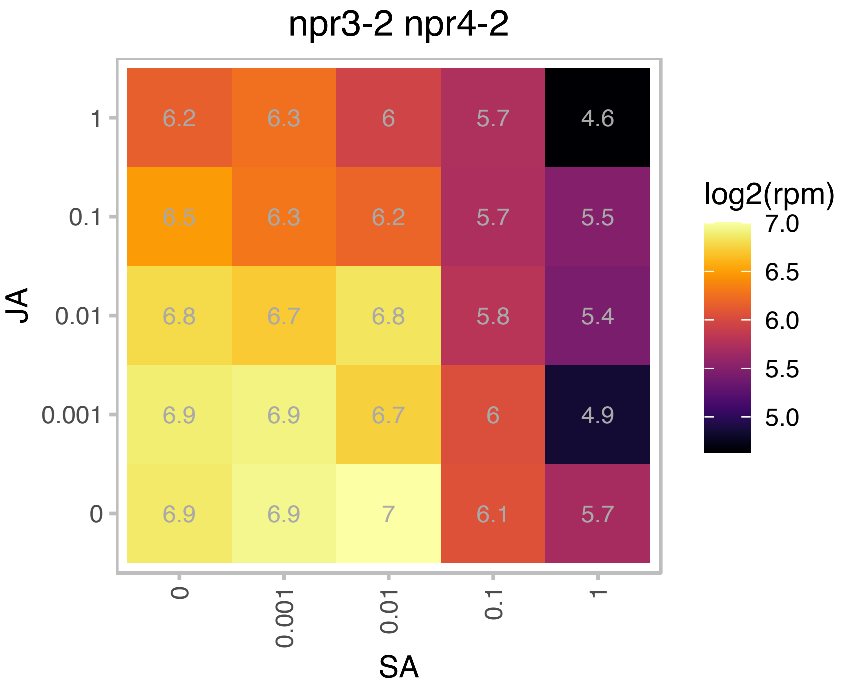


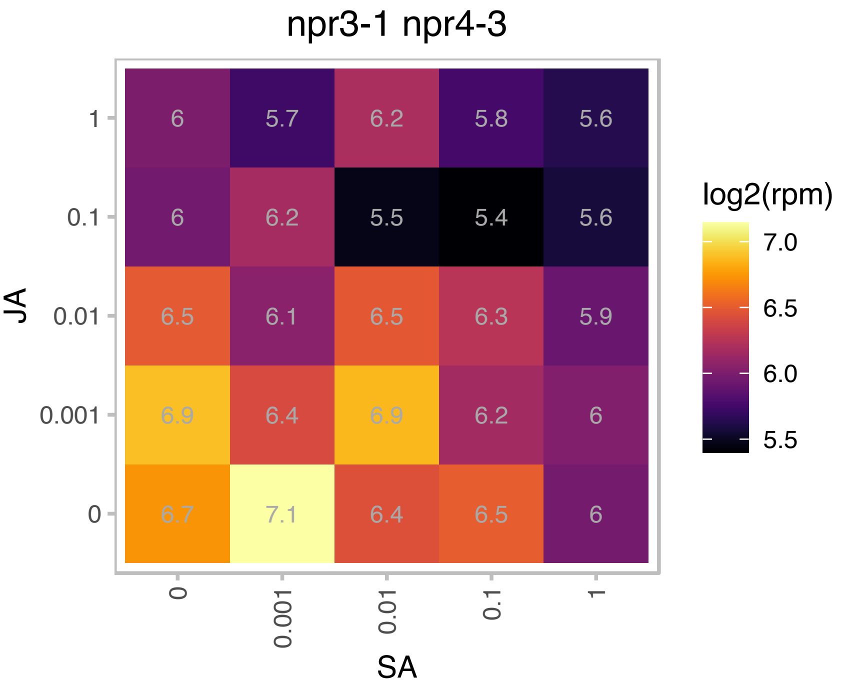

# AT1G20810.1 : NA

FKBP-like peptidyl-prolyl cis-trans isomerase family protein

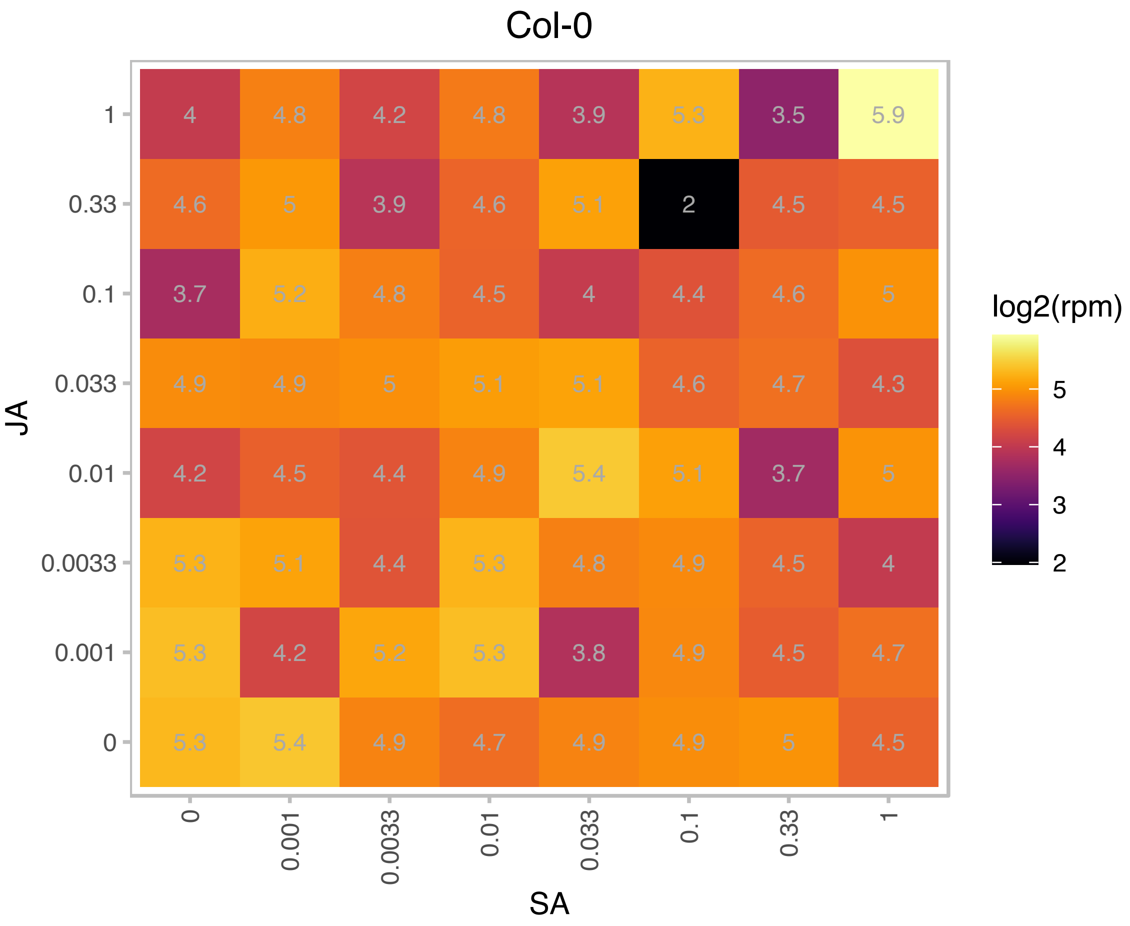


| | |
|---|---|
| Gene | AT1G20810 |
| is.expressed_genes | TRUE |
| is.responsive_genes | TRUE |
| is.apcluster | F-05 |

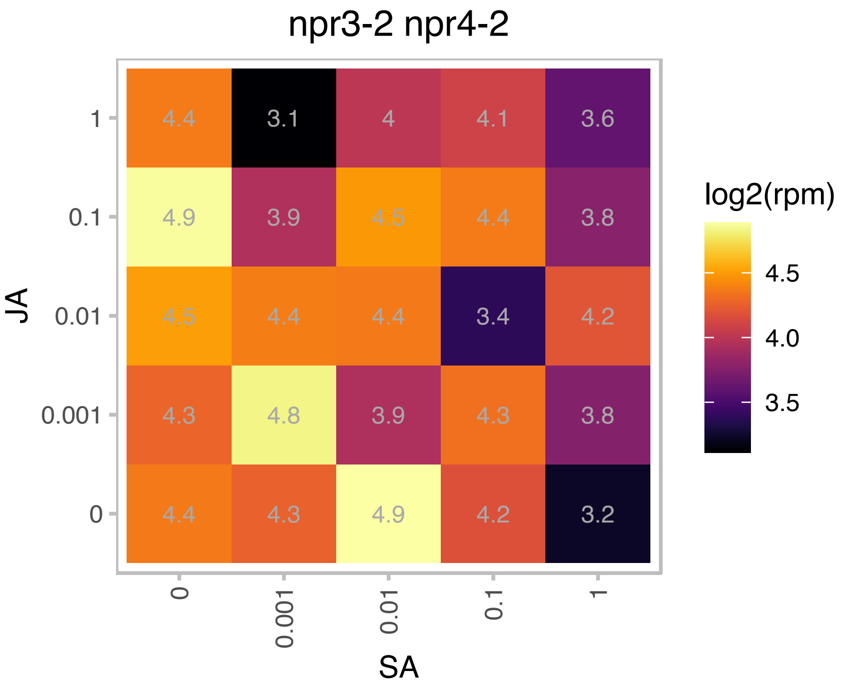


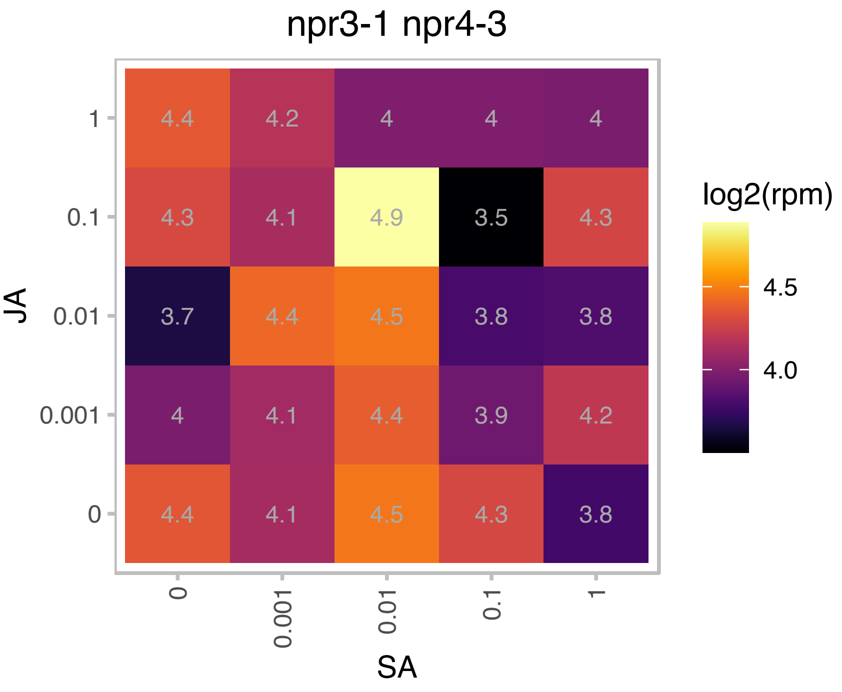

# AT1G29670.1 : NA

GDSL-like Lipase/Acylhydrolase superfamily protein

GDSL-motif esterase/acyltransferase/lipase. Enzyme group with broad substrate specificity that may catalyze acyltransfer or hydrolase reactions with lipid and non-lipid substrates. The mRNA is cell-to-cell mobile.

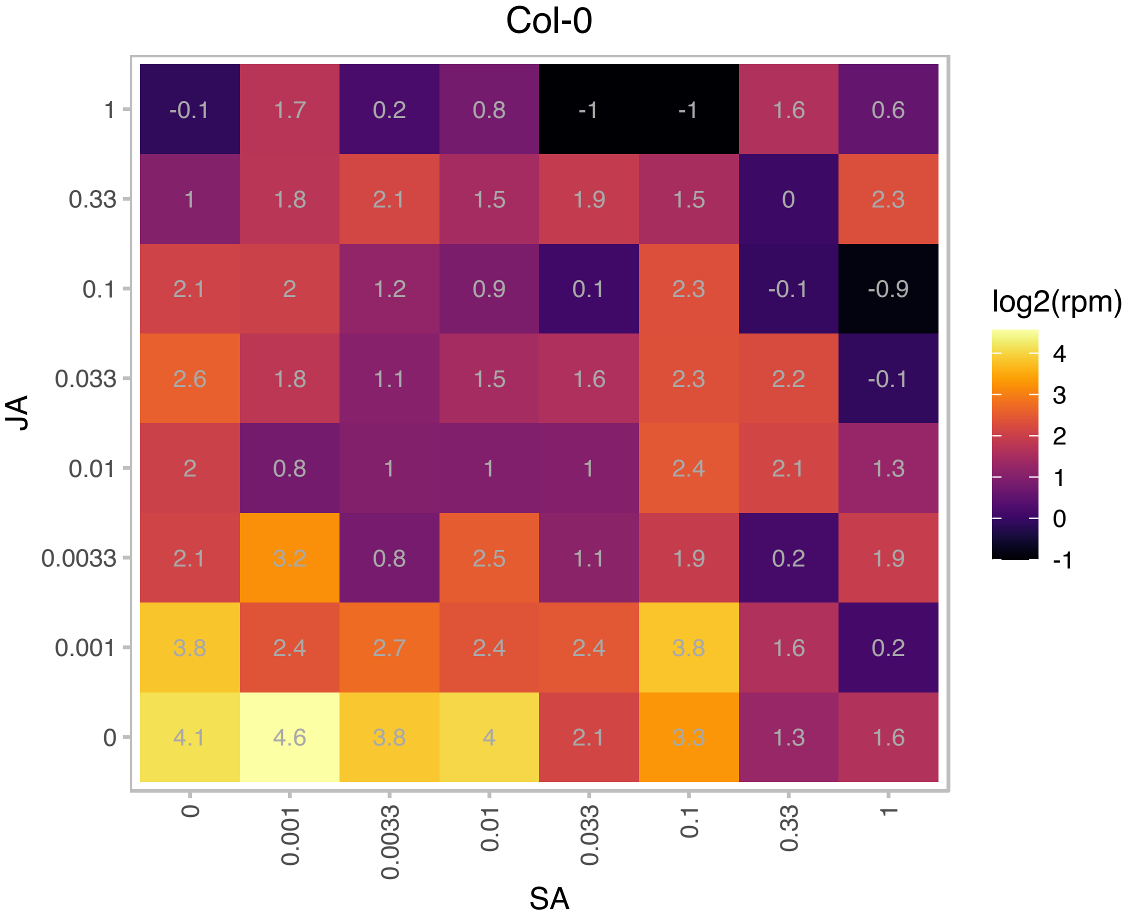


| | |
|---|---|
| Gene | AT1G29670 |
| is.expressed_genes | TRUE |
| is.responsive_genes | TRUE |
| is.apcluster | F-05 |

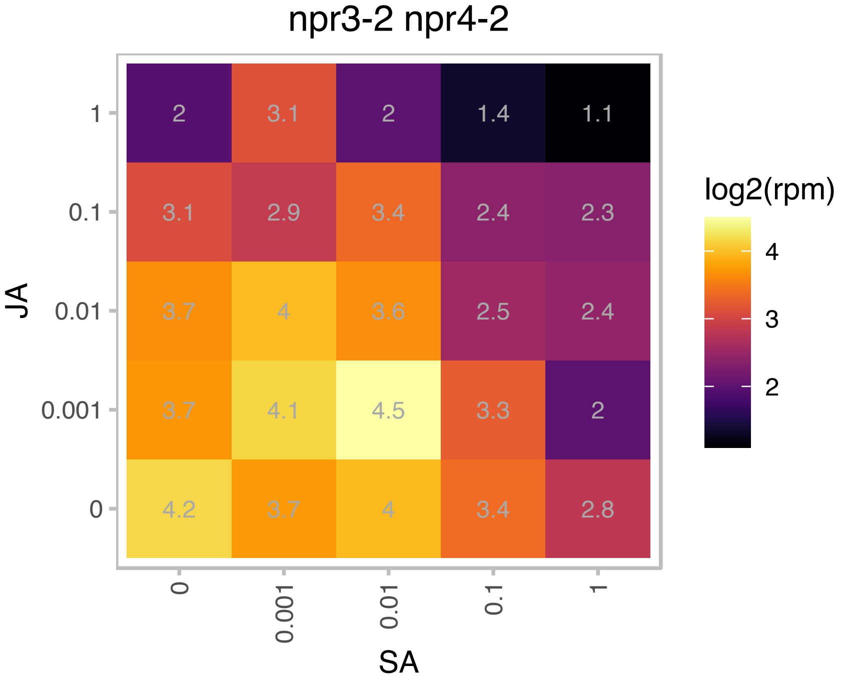


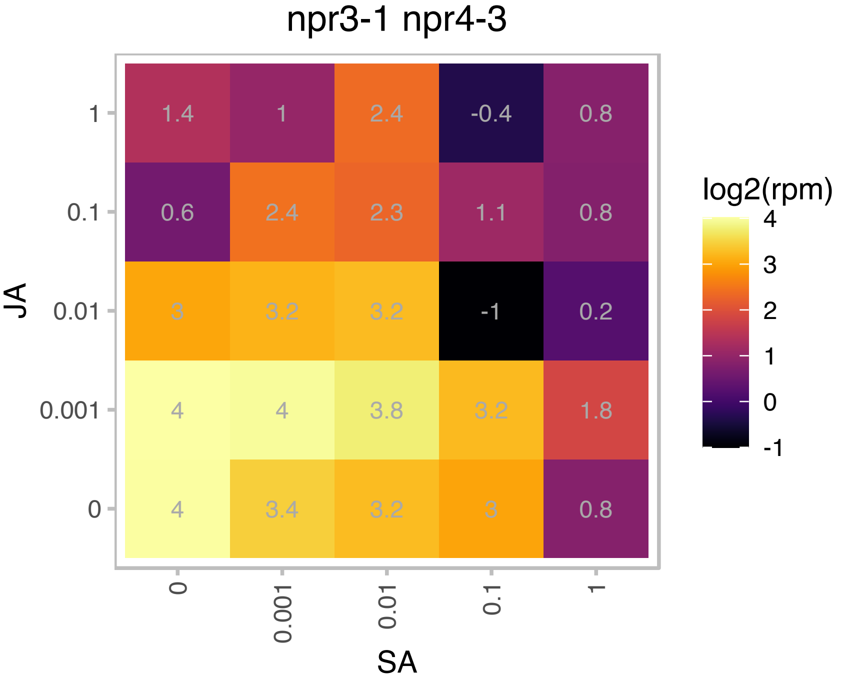

# AT1G32470.1 : NA

Single hybrid motif superfamily protein

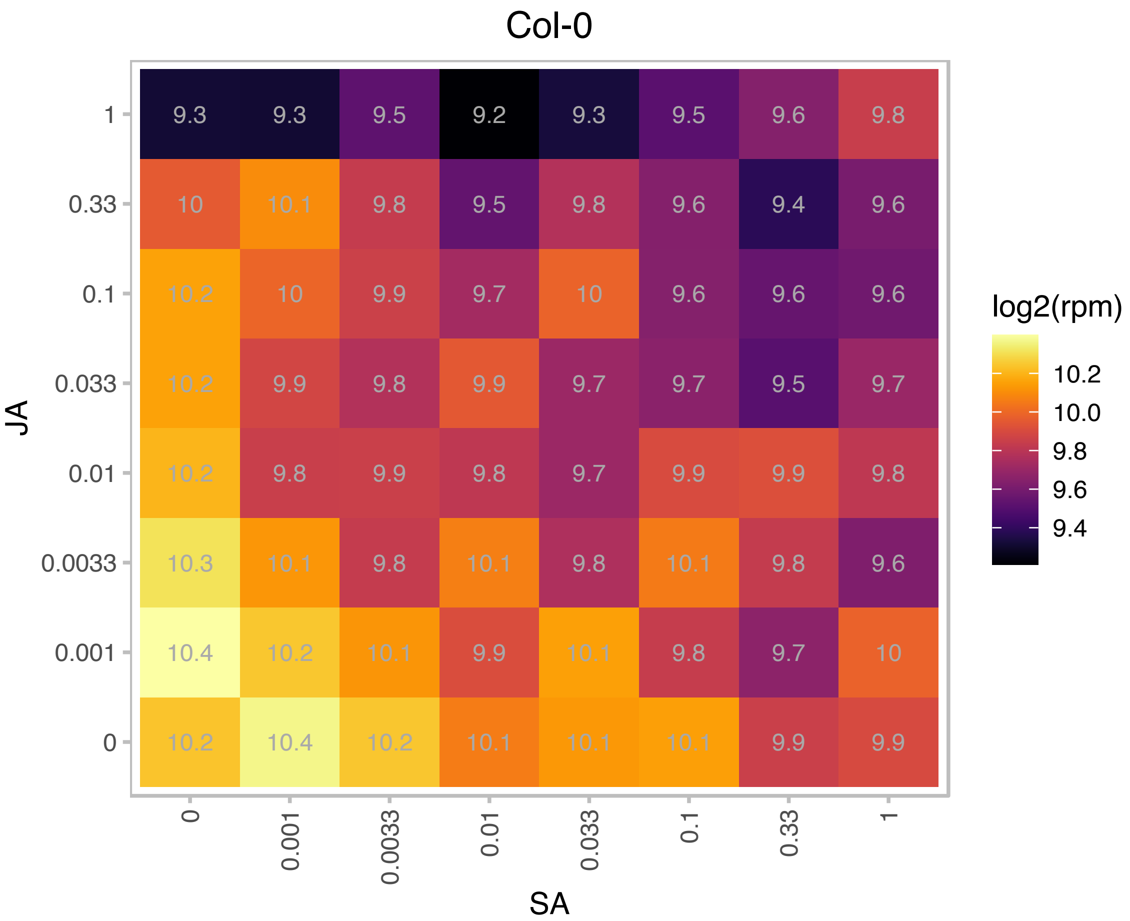


| | |
|---|---|
| Gene | AT1G32470 |
| is.expressed_genes | TRUE |
| is.responsive_genes | TRUE |
| is.apcluster | F-05 |

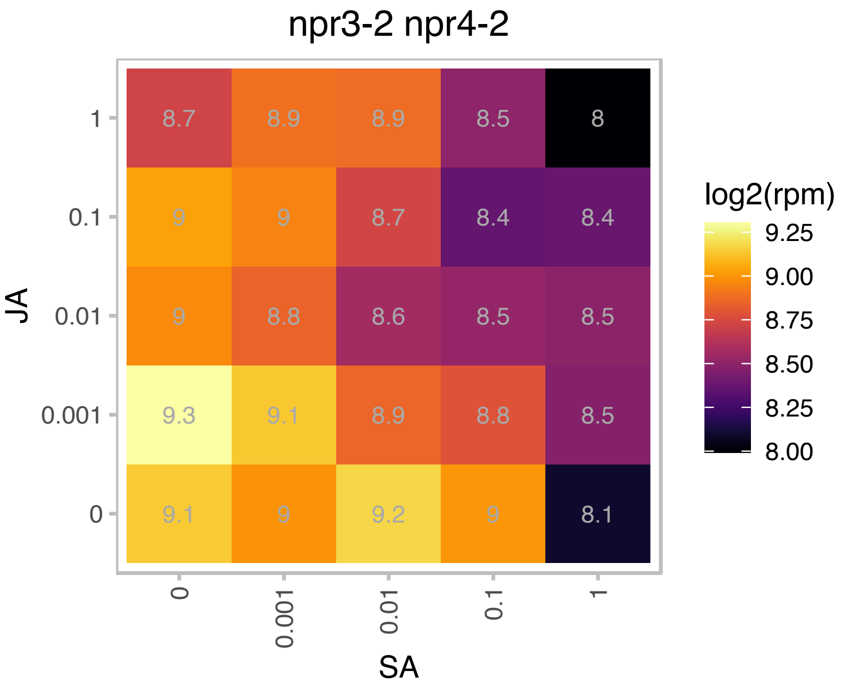


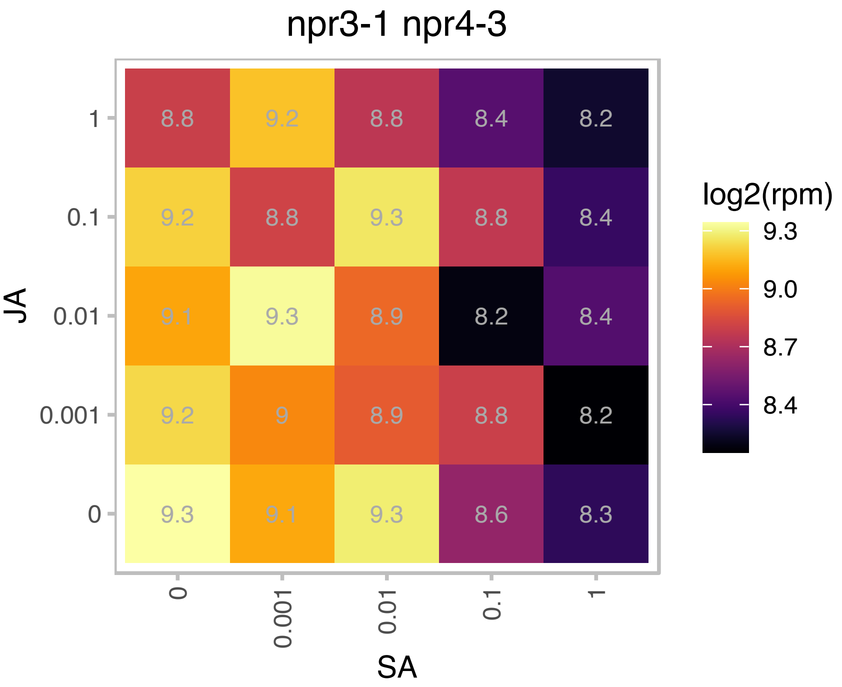

### AT1G33040.1 : NACA5

nascent polypeptide-associated complex subunit alpha-like protein 5

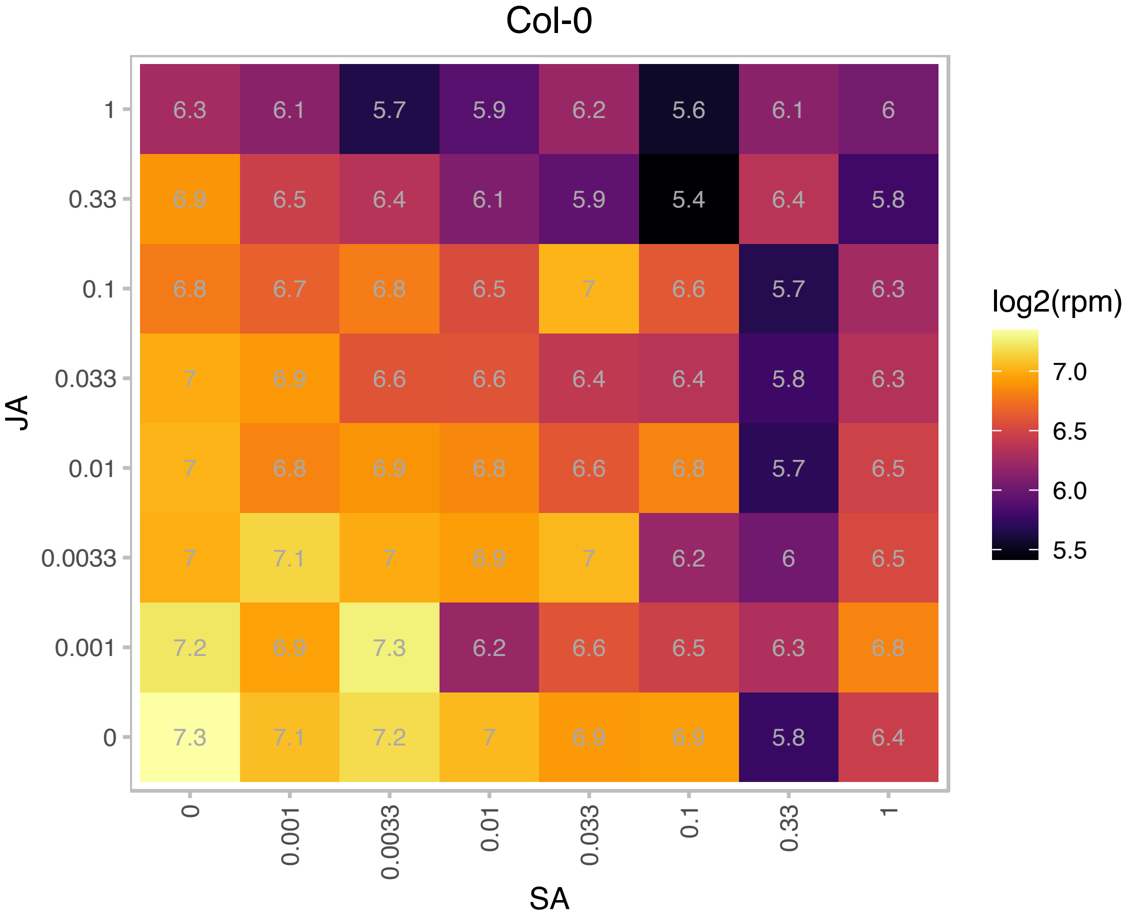


| | |
|---|---|
| Gene | AT1G33040 |
| is.expressed_genes | TRUE |
| is.responsive_genes | TRUE |
| is.apcluster | F-05 |

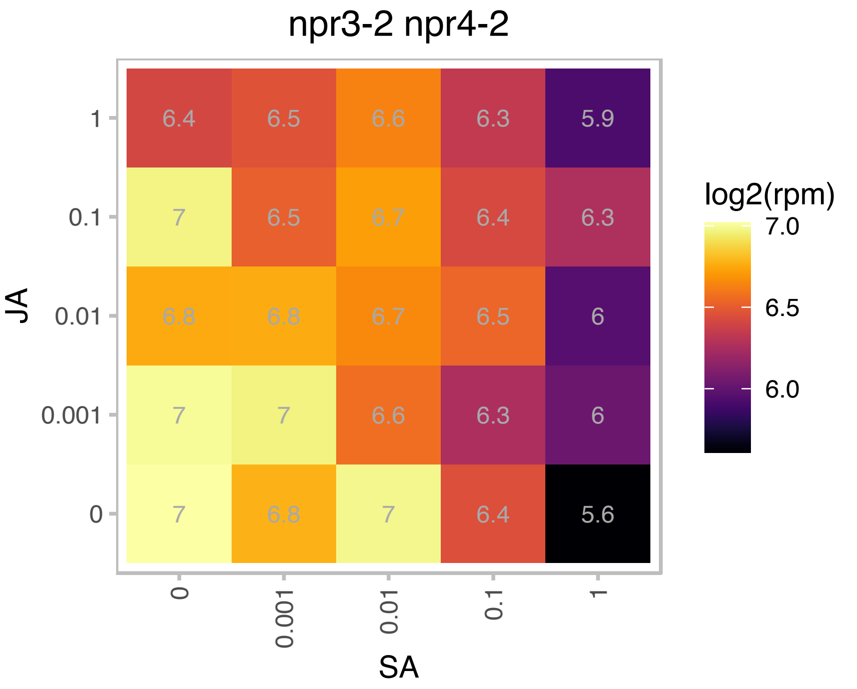


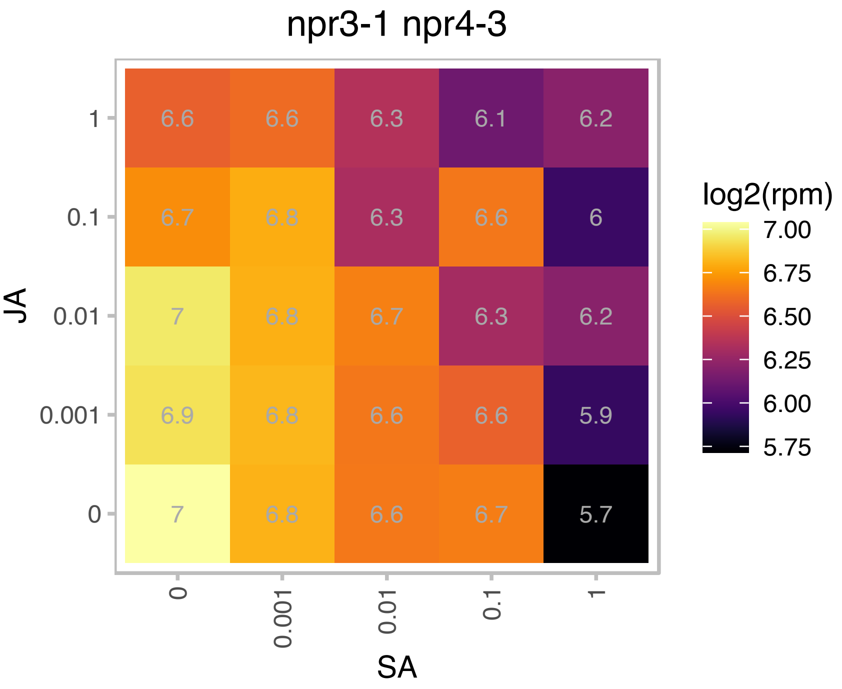

# AT1G33810.1 : NA

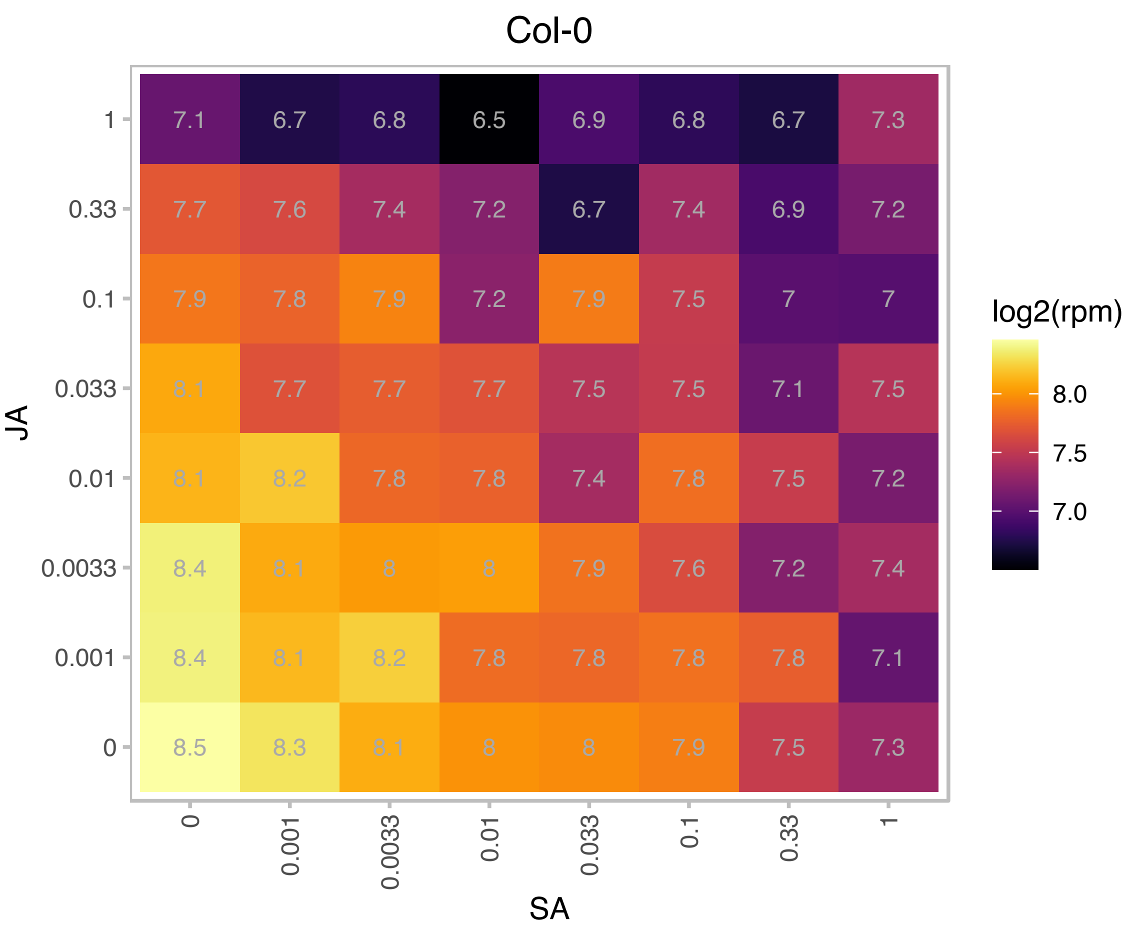


| | |
|---|---|
| Gene | AT1G33810 |
| is.expressed_genes | TRUE |
| is.responsive_genes | TRUE |
| is.apcluster | F-05 |

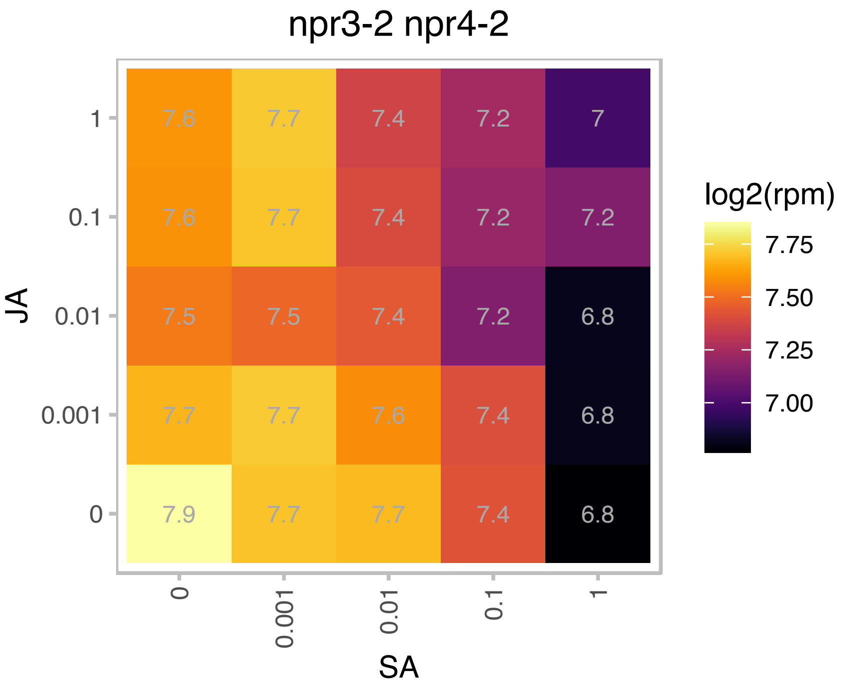


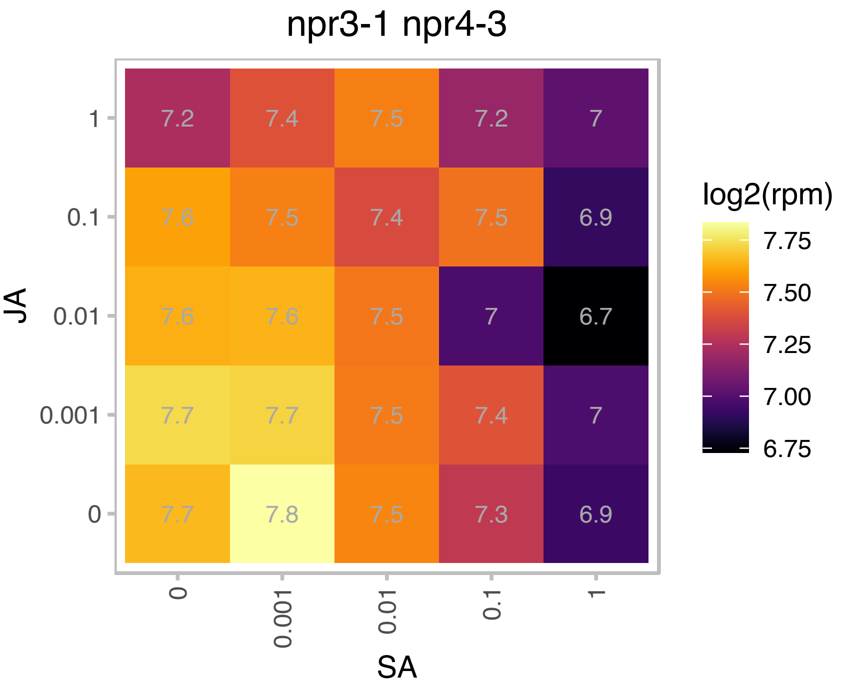

# AT1G35420.1 : NA

alpha/beta-Hydrolases superfamily protein

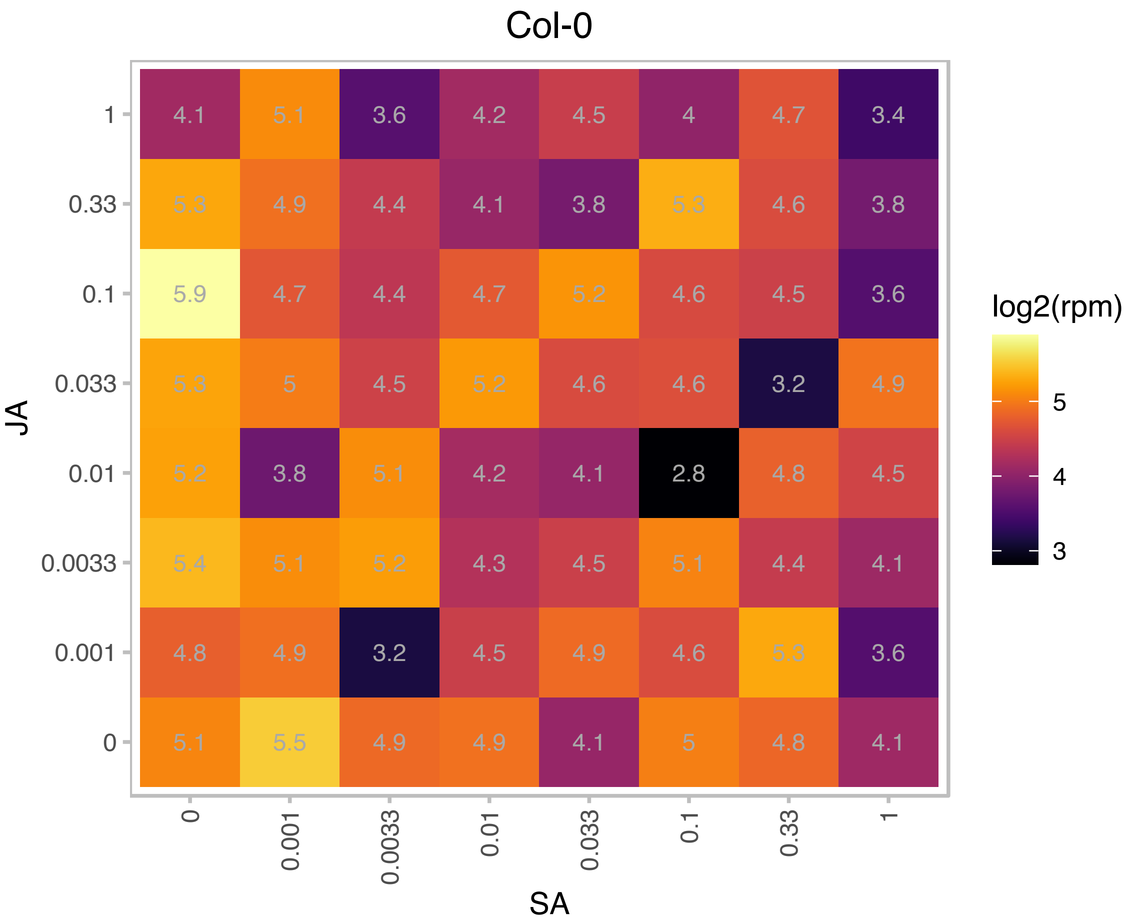


| | |
|---|---|
| Gene | AT1G35420 |
| is.expressed_genes | TRUE |
| is.responsive_genes | TRUE |
| is.apcluster | F-05 |

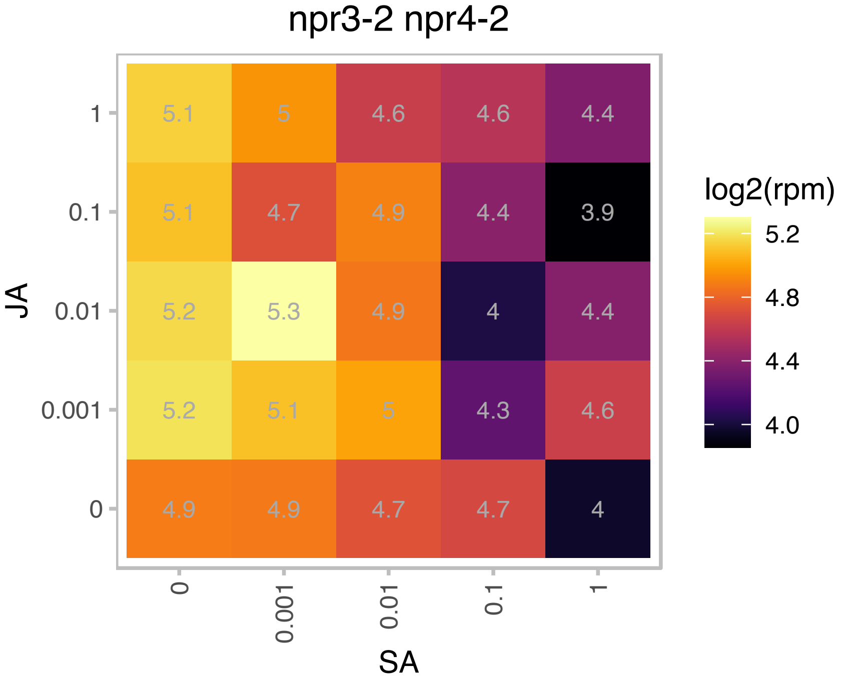


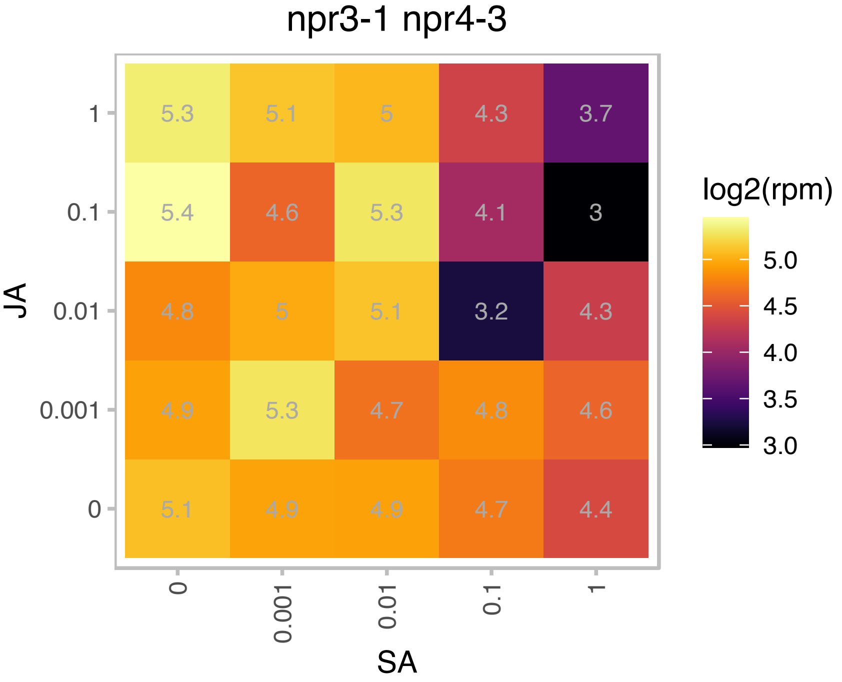

# AT1G50290.1 : NA

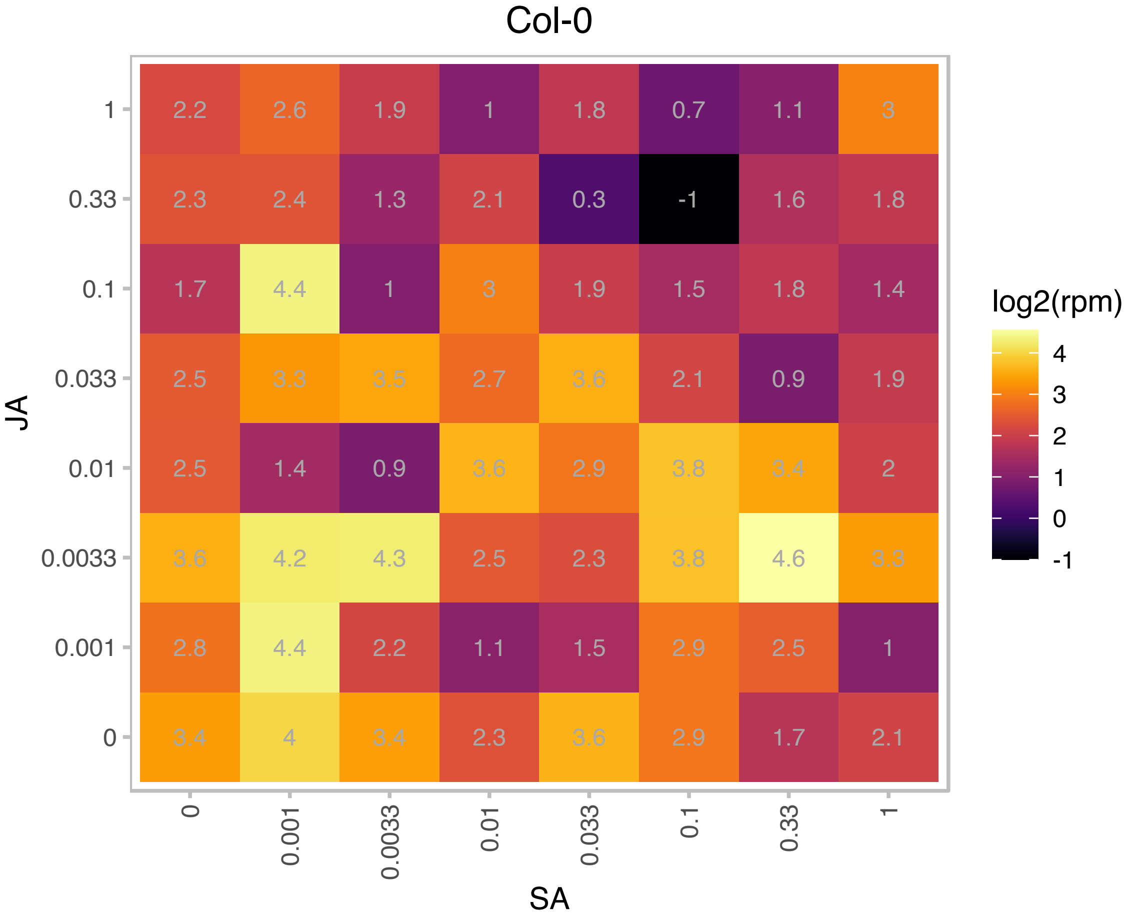


| | |
|---|---|
| Gene | AT1G50290 |
| is.expressed_genes | TRUE |
| is.responsive_genes | TRUE |
| is.apcluster | F-05 |

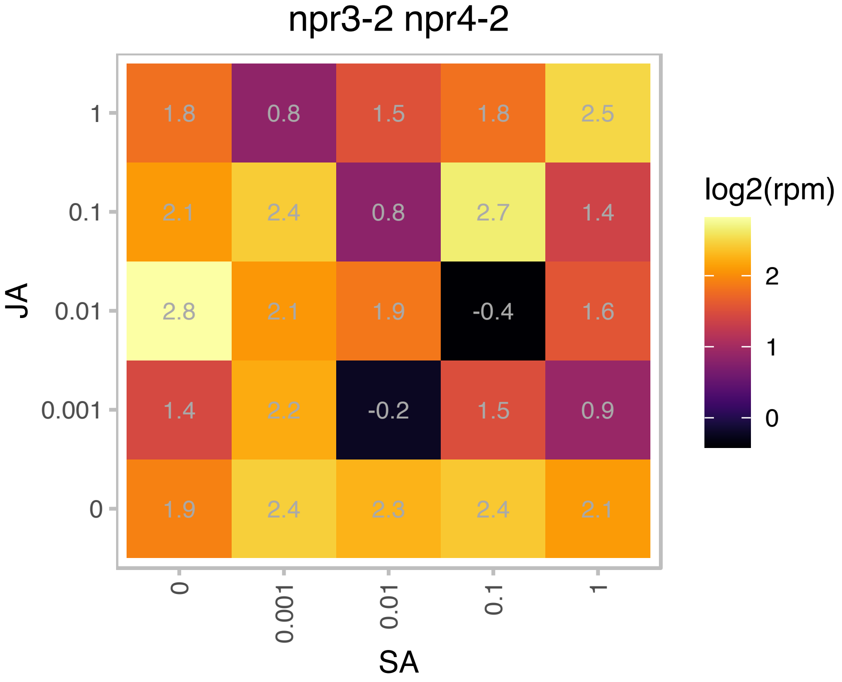


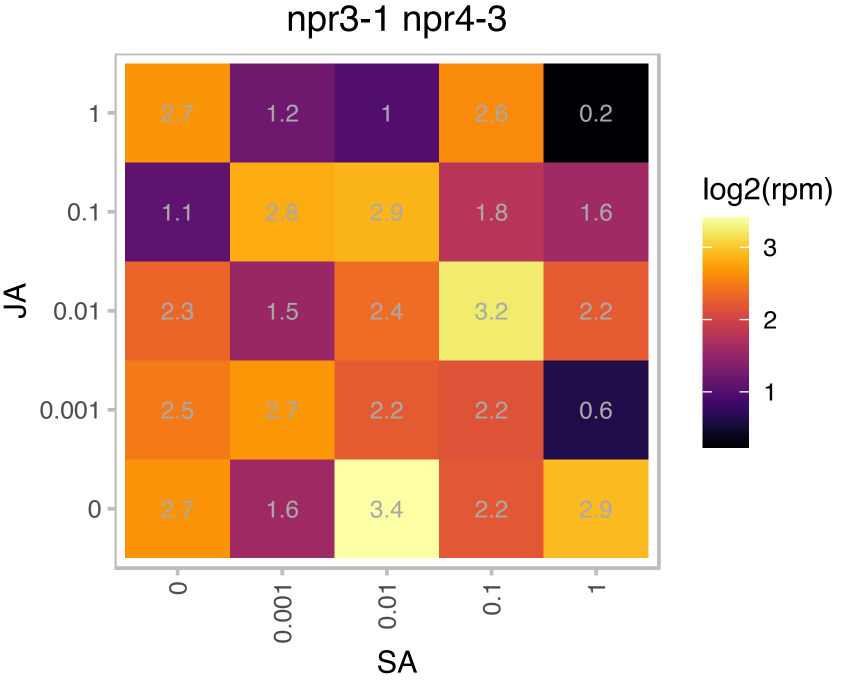

### AT1G74690.1 : IQD31

IQ-domain 31

Encodes a microtubule-associated protein.

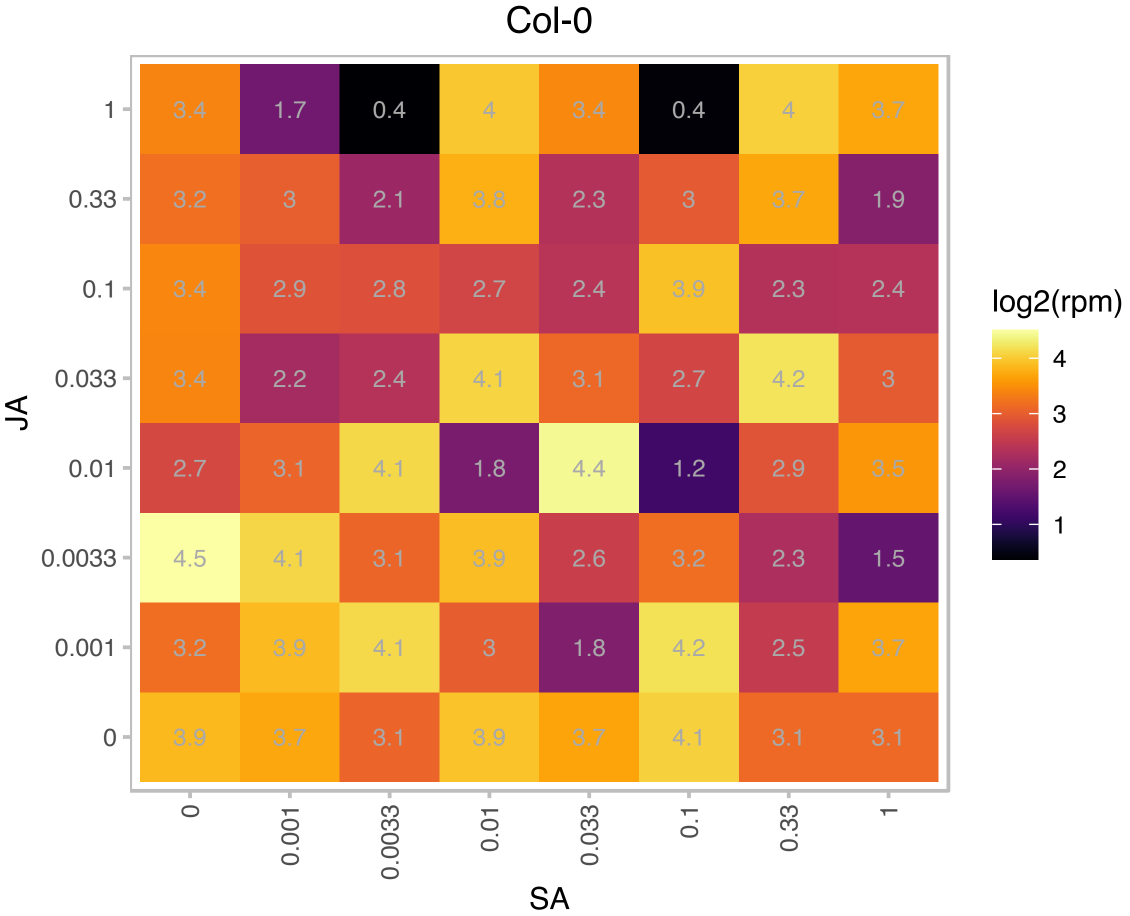


| | |
|---|---|
| Gene | AT1G74690 |
| is.expressed_genes | TRUE |
| is.responsive_genes | TRUE |
| is.apcluster | F-05 |

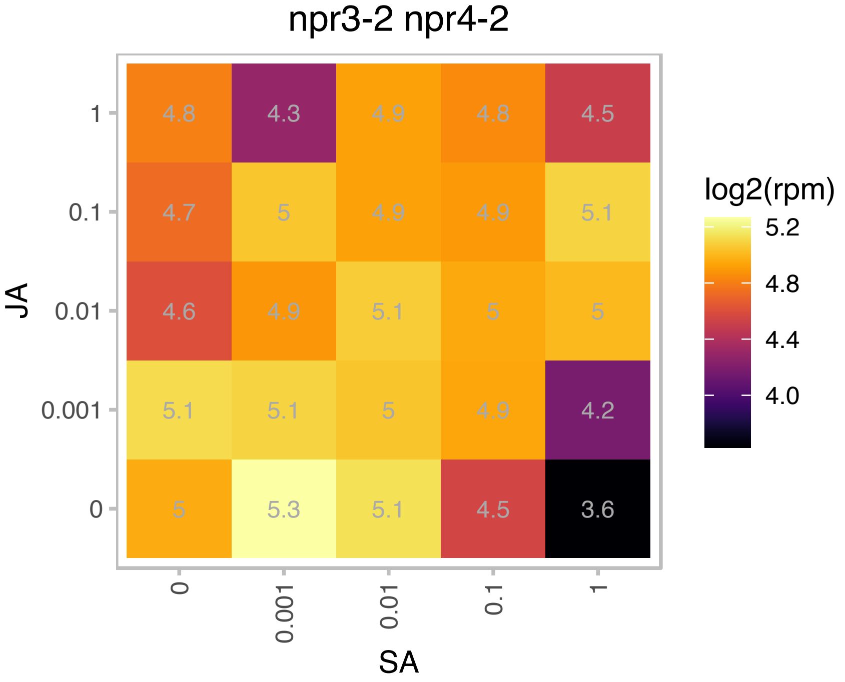


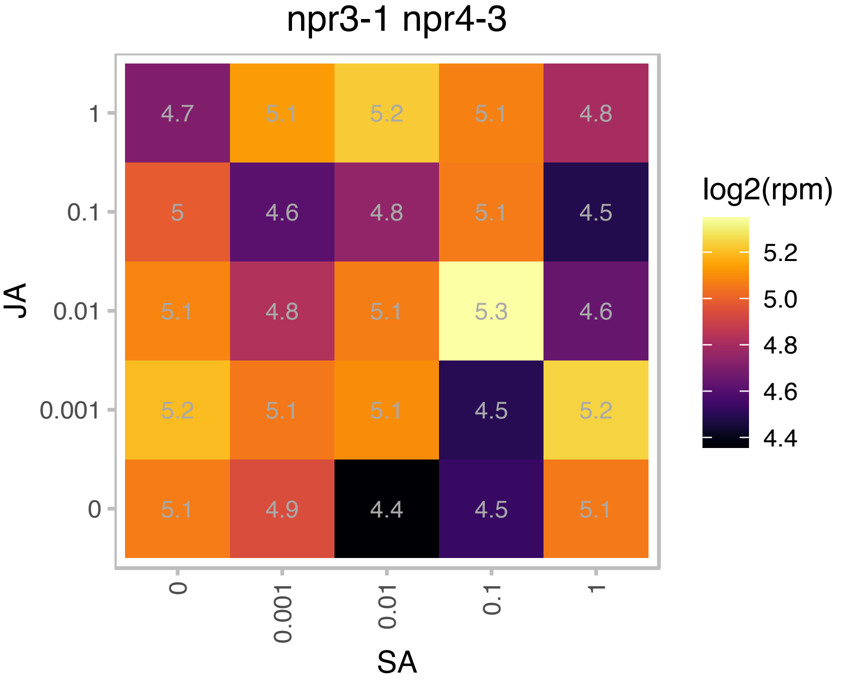

### AT1G75690.1 : LQY1

DnaJ/Hsp40 cysteine-rich domain superfamily protein

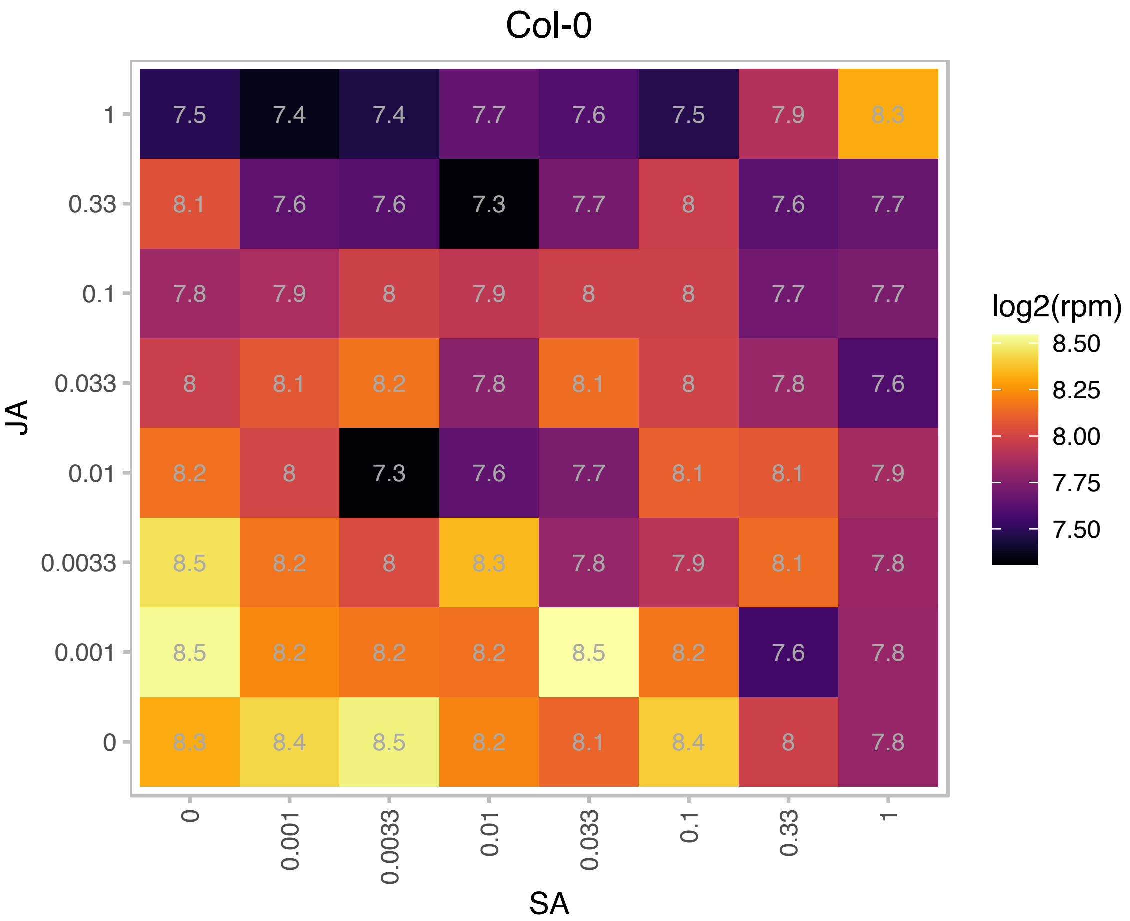


| | |
|---|---|
| Gene | AT1G75690 |
| is.expressed_genes | TRUE |
| is.responsive_genes | TRUE |
| is.apcluster | F-05 |

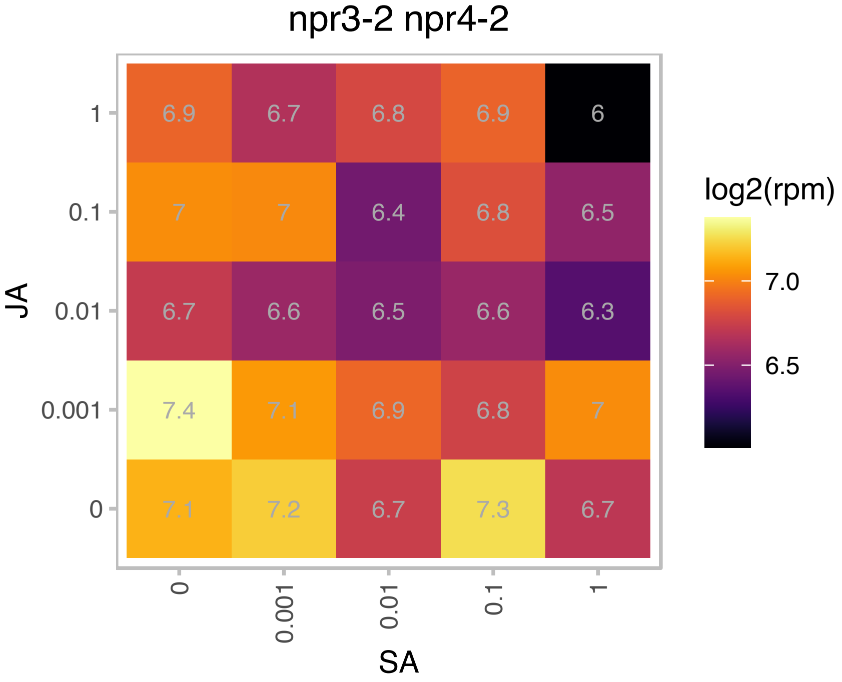


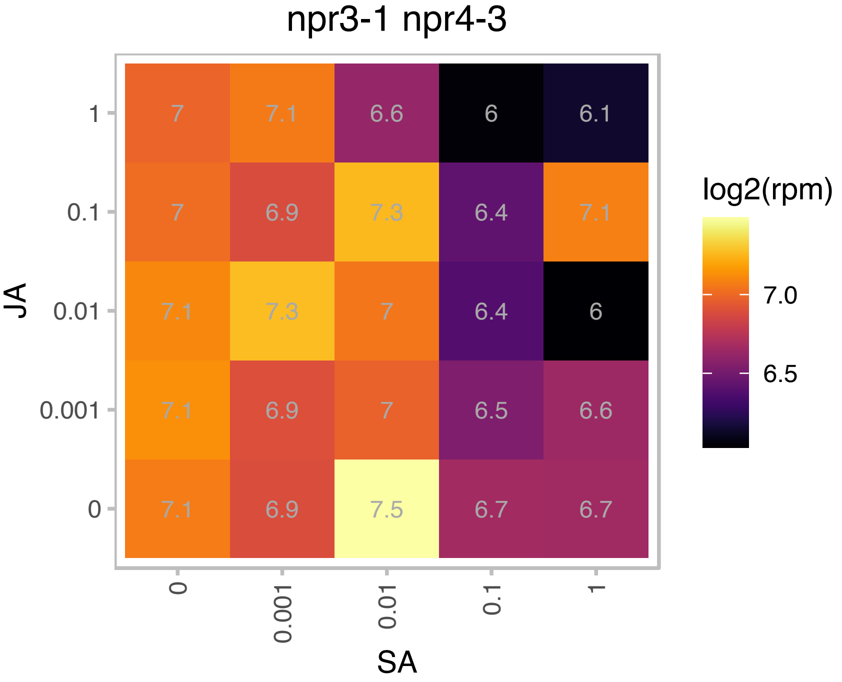

### AT2G04780.1 : FLA7

FASCICLIN-like arabinoogalactan 7

fasciclin-like arabinogalactan-protein 7 (Fla7)

| | |
|---|---|
| Gene | AT2G04780 |
| is.expressed_genes | TRUE |
| is.responsive_genes | TRUE |
| is.apcluster | F-05 |

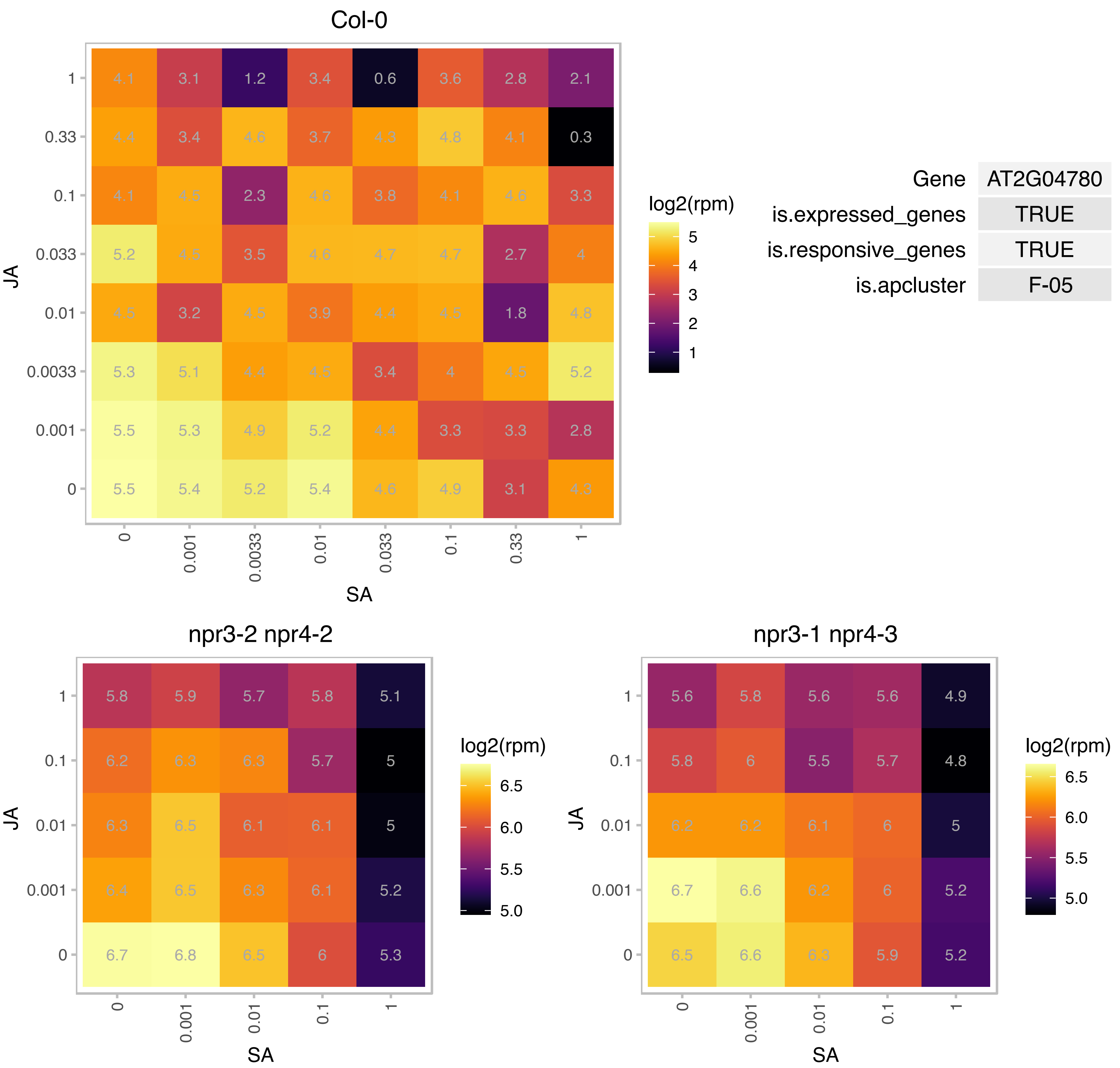

### AT2G18300.1 : HBI1

basic helix-loop-helix (bHLH) DNA-binding superfamily protein

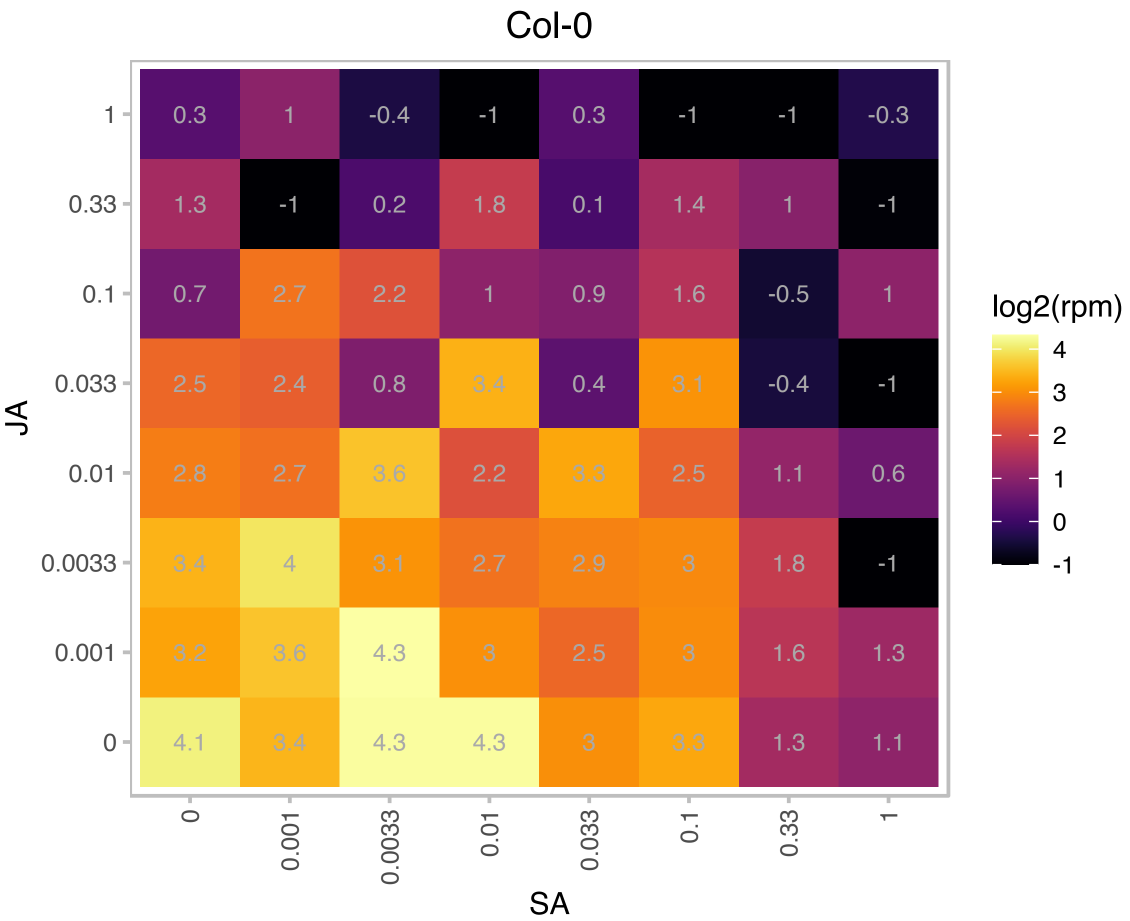


| | |
|---|---|
| Gene | AT2G18300 |
| is.expressed_genes | TRUE |
| is.responsive_genes | TRUE |
| is.apcluster | F-05 |

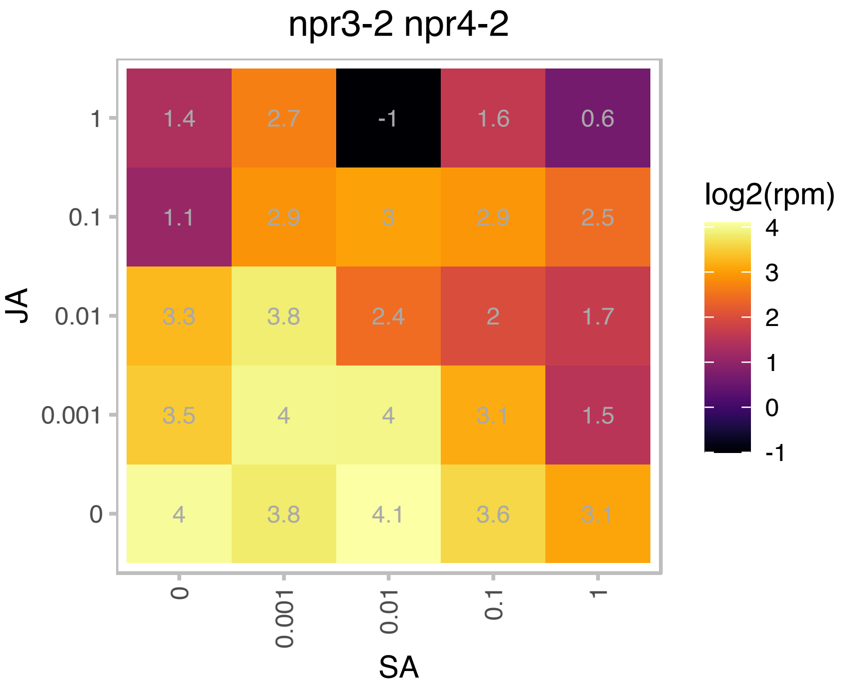


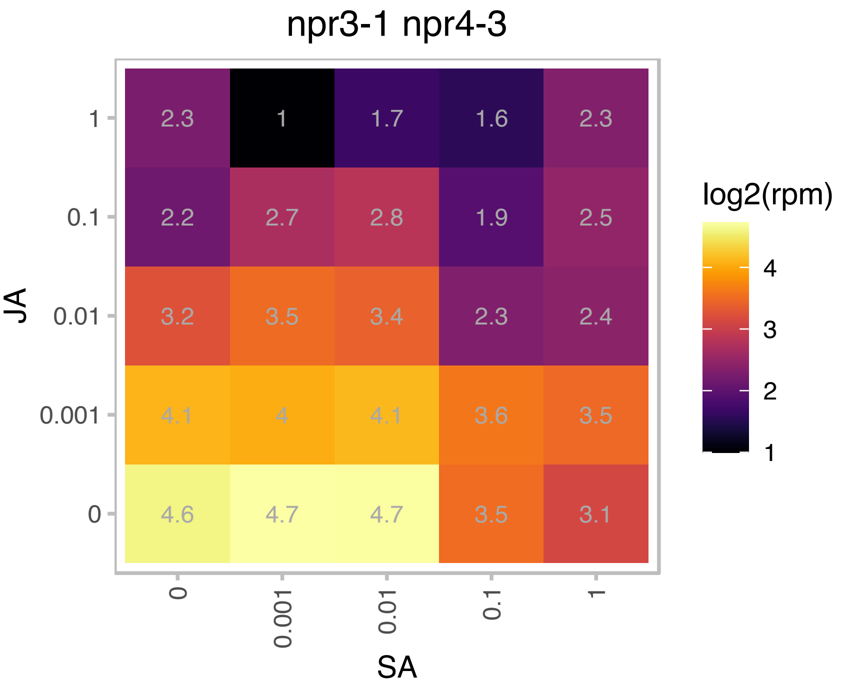

# AT2G21195.1 : NA

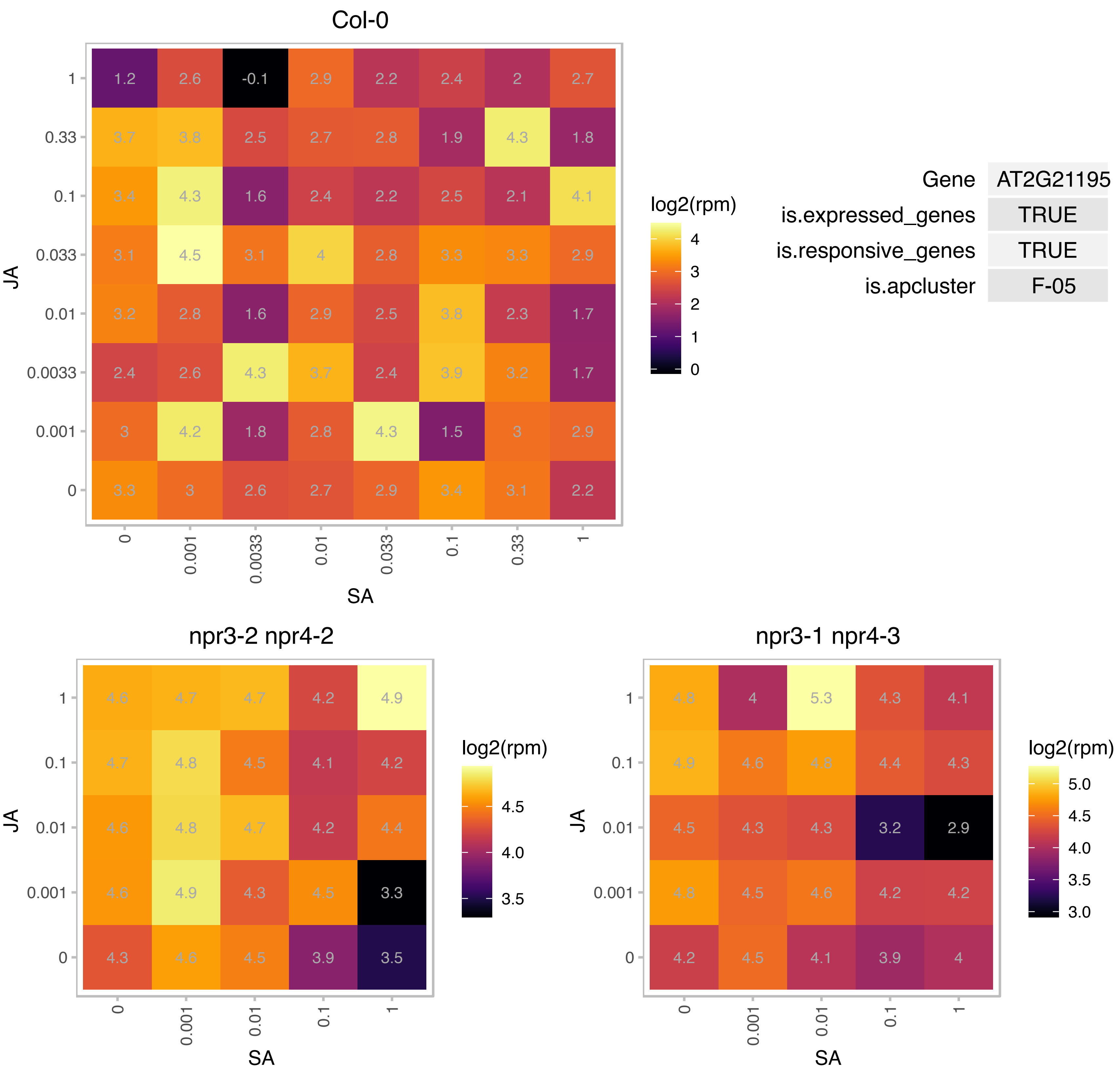


| | |
|---|---|
| Gene | AT2G21195 |
| is.expressed_genes | TRUE |
| is.responsive_genes | TRUE |
| is.apcluster | F-05 |

### AT2G22170.1 : PLAT2

Lipase/lipooxygenase, PLAT/LH2 family protein

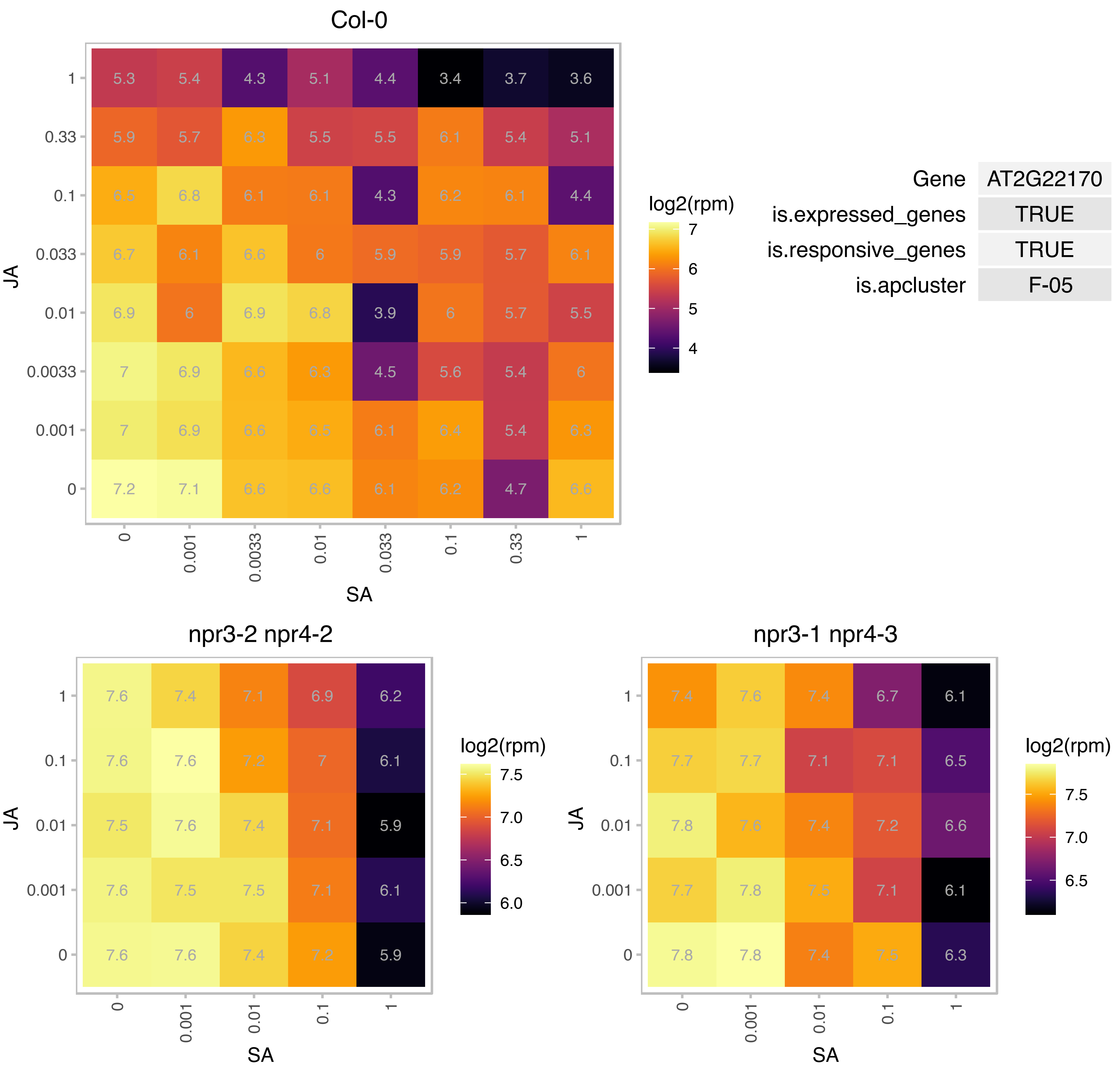


| | |
|---|---|
| Gene | AT2G22170 |
| is.expressed_genes | TRUE |
| is.responsive_genes | TRUE |
| is.apcluster | F-05 |

### AT2G25900.1 : ATCTH, ATTZF1

Zinc finger C-x8-C-x5-C-x3-H type family protein

Encodes a protein with two tandem-arrayed CCCH-type zinc fingers that binds RNA and is involved in RNA turnover. The mRNA is cell-to-cell mobile.

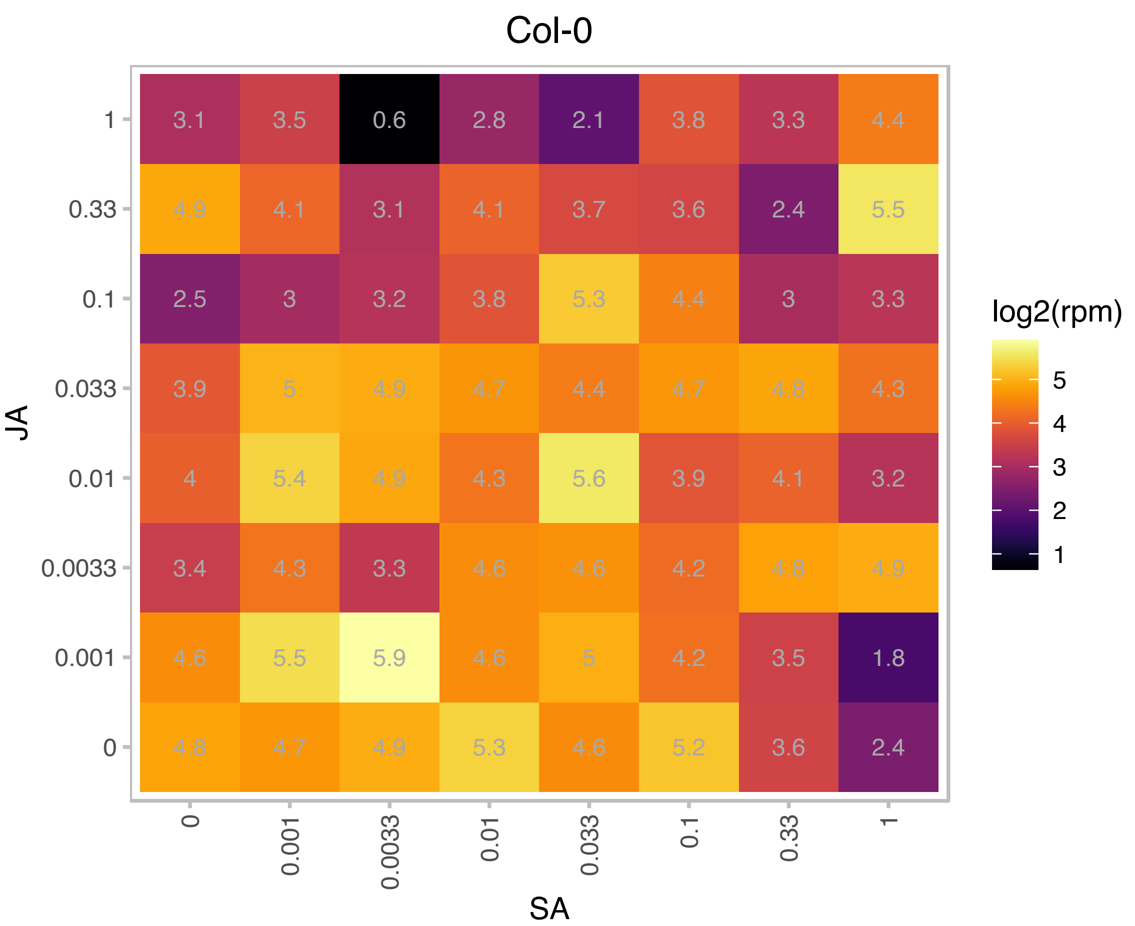


| | |
|---|---|
| Gene | AT2G25900 |
| is.expressed_genes | TRUE |
| is.responsive_genes | TRUE |
| is.apcluster | F-05 |

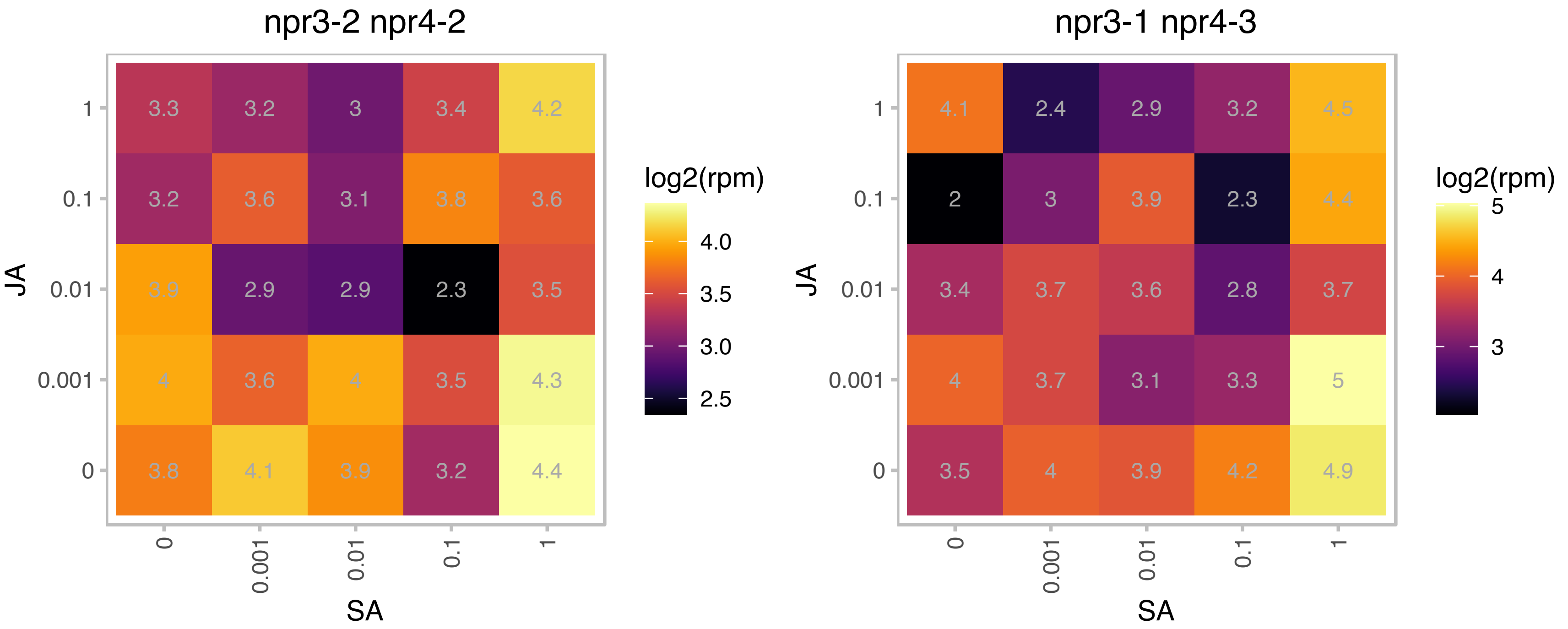

# AT2G27402.1 : NA

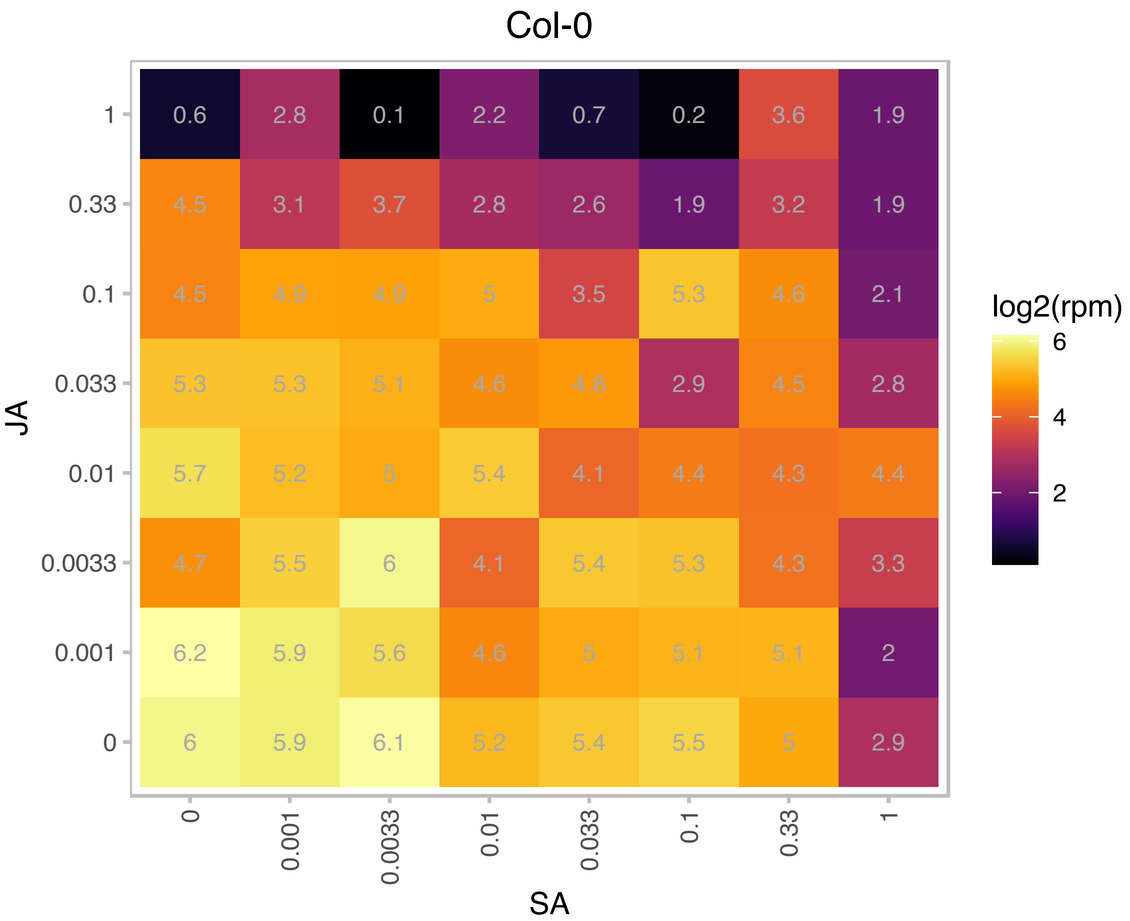


| | |
|---|---|
| Gene | AT2G27402 |
| is.expressed_genes | TRUE |
| is.responsive_genes | TRUE |
| is.apcluster | F-05 |

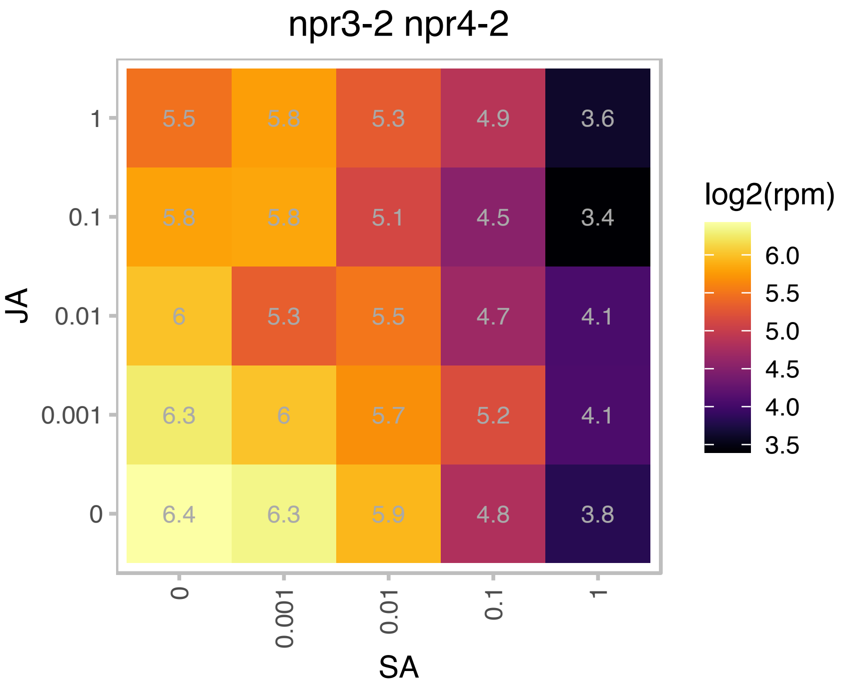


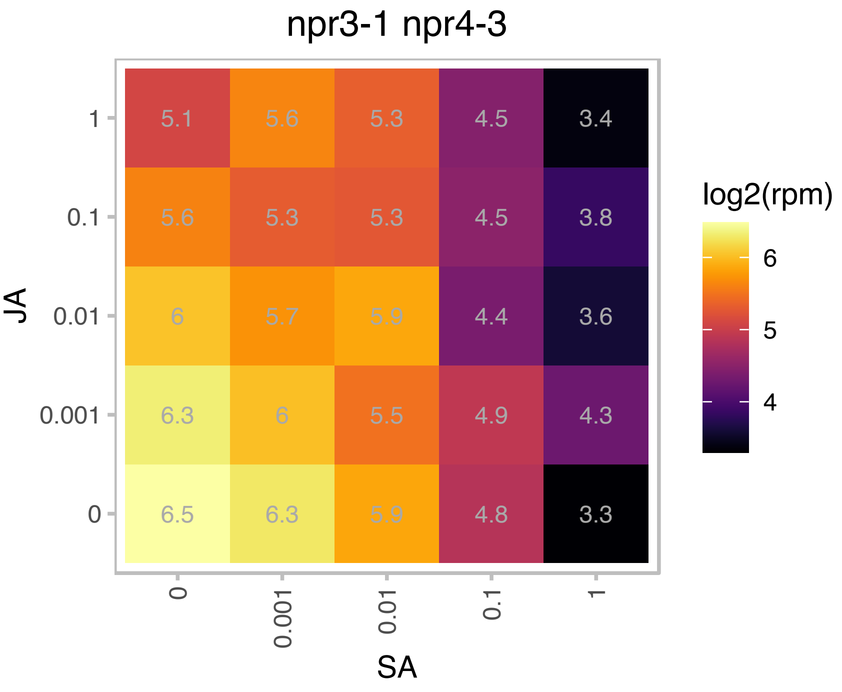

# AT2G29180.1 : NA

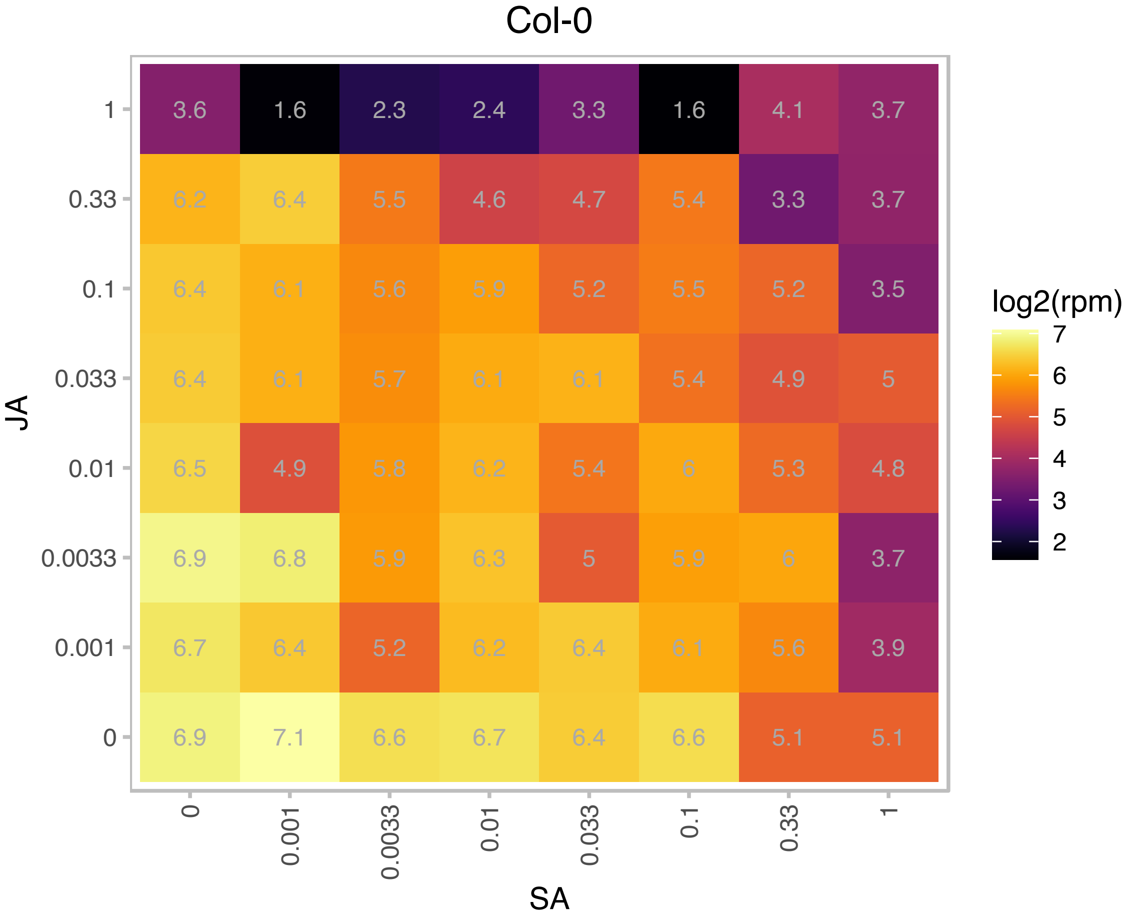


| | |
|---|---|
| Gene | AT2G29180 |
| is.expressed_genes | TRUE |
| is.responsive_genes | TRUE |
| is.apcluster | F-05 |

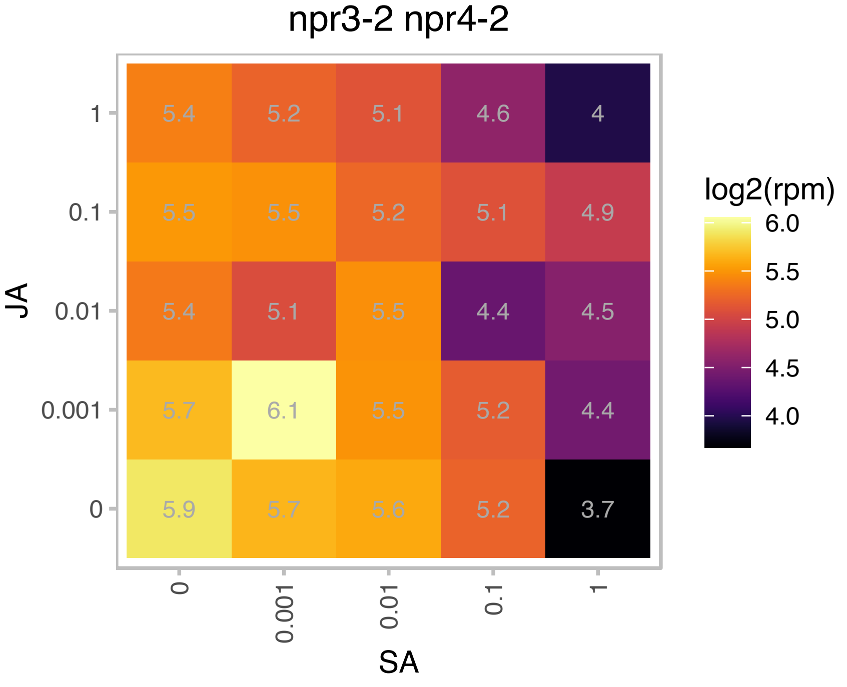


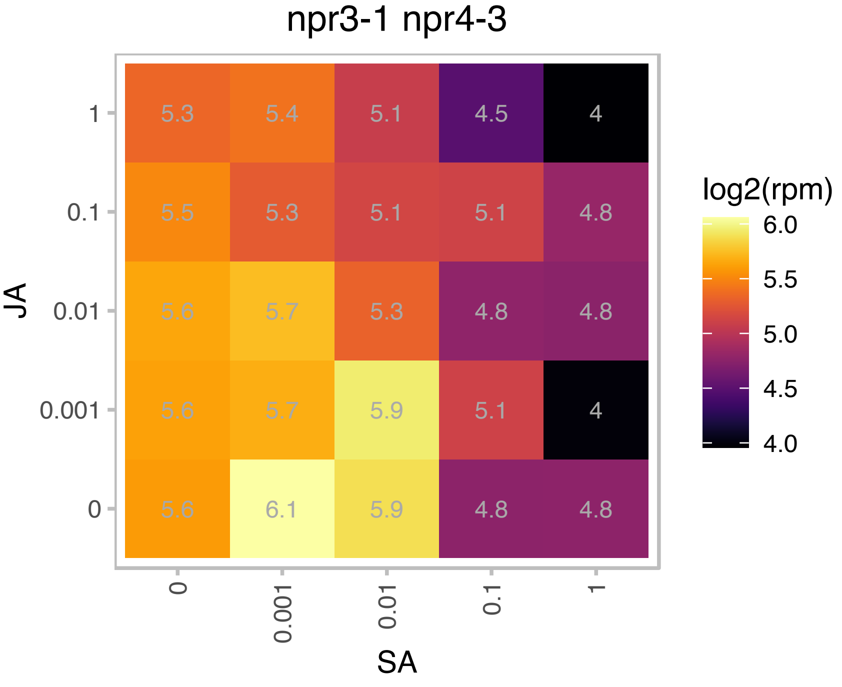

# AT2G29290.1 : NA

NAD(P)-binding Rossmann-fold superfamily protein

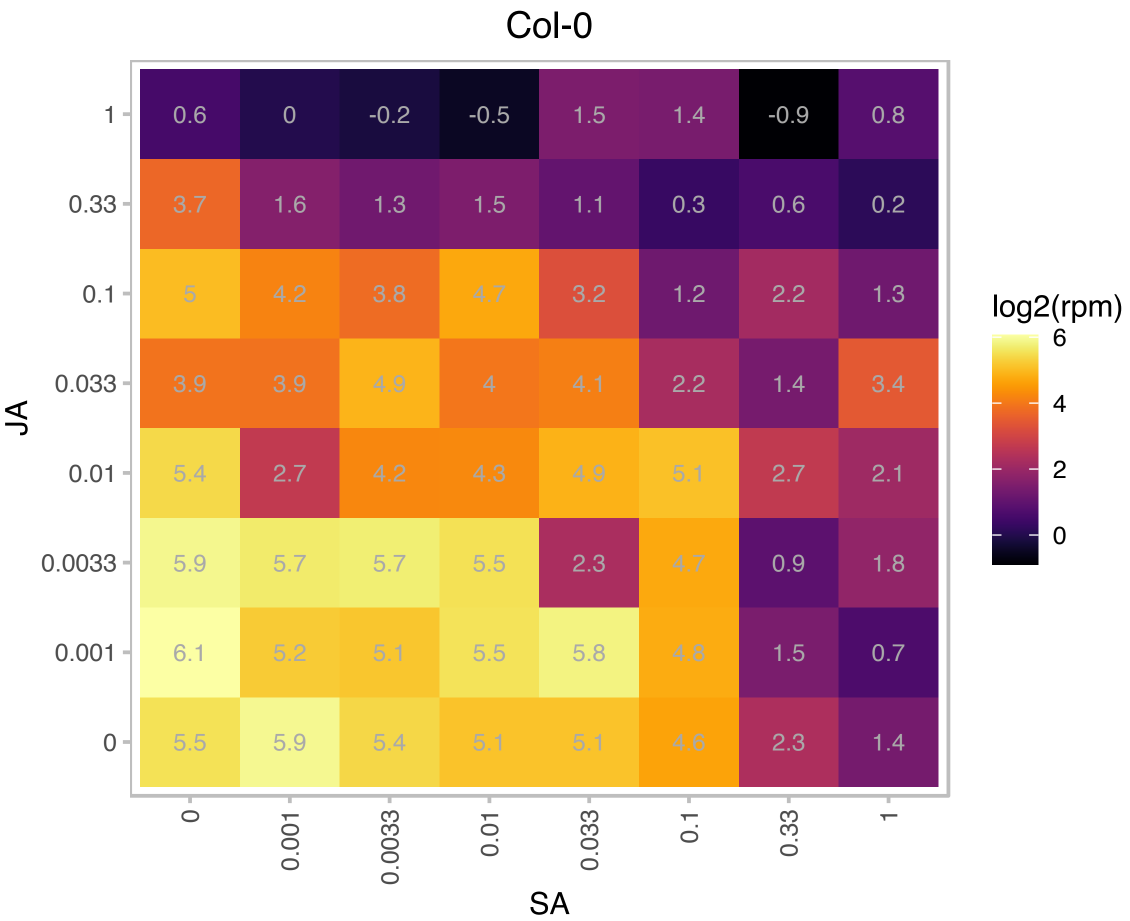


| | |
|---|---|
| Gene | AT2G29290 |
| is.expressed_genes | TRUE |
| is.responsive_genes | TRUE |
| is.apcluster | F-05 |

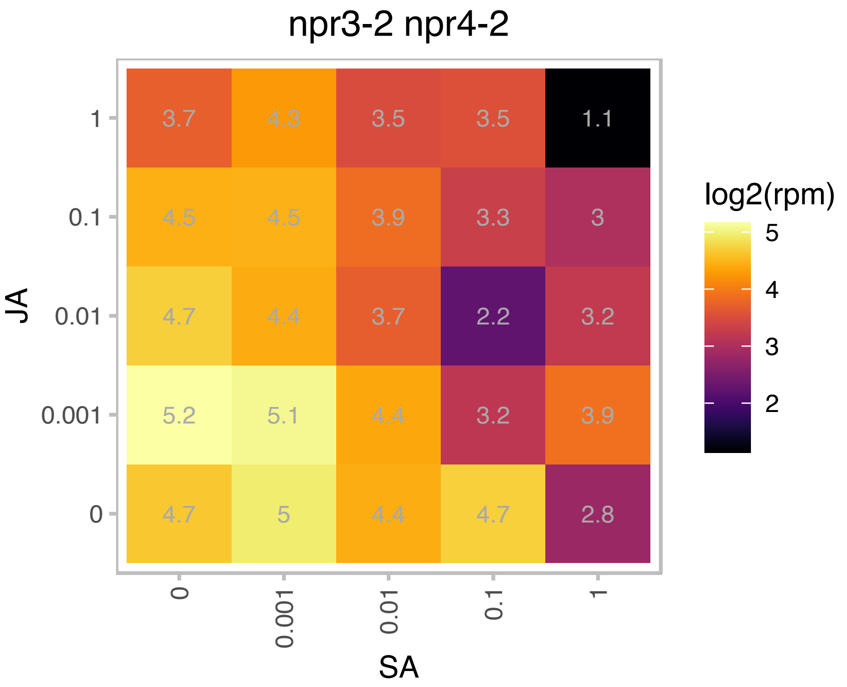


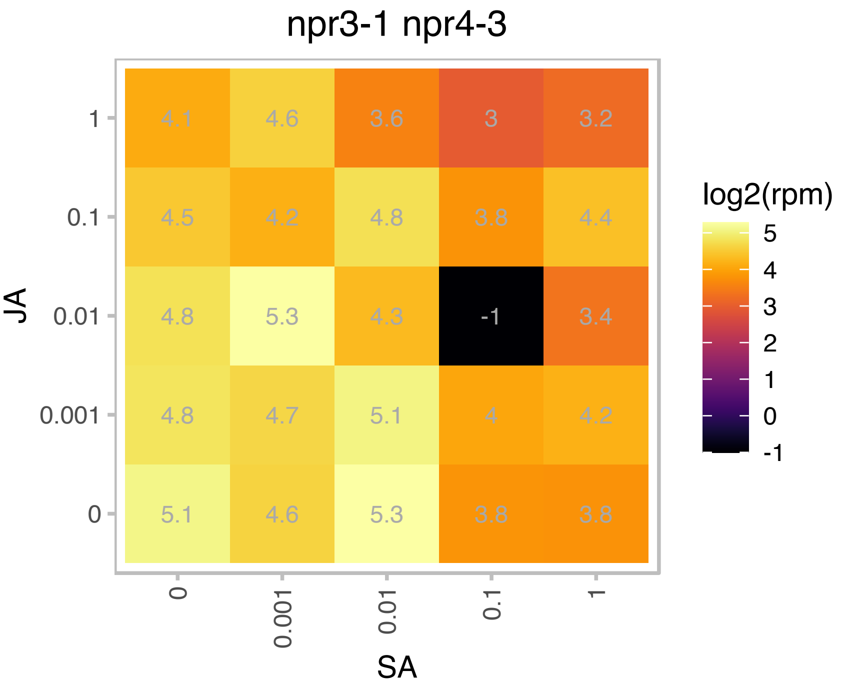

### AT2G32230.1 : PRORP1

proteinaceous RNase P 1
Encodes a protein-only RNase P that is involved in the 5? cleavage of the precursor tRNAs and is able to cleave tRNA-like structures involved in the maturation of plant mitochondrial mRNAs. Mutants show a drastic reduction in the levels of mature plastid tRNA-Phe(GAA) and tRNA-Arg(ACG), limiting plastid gene expression.

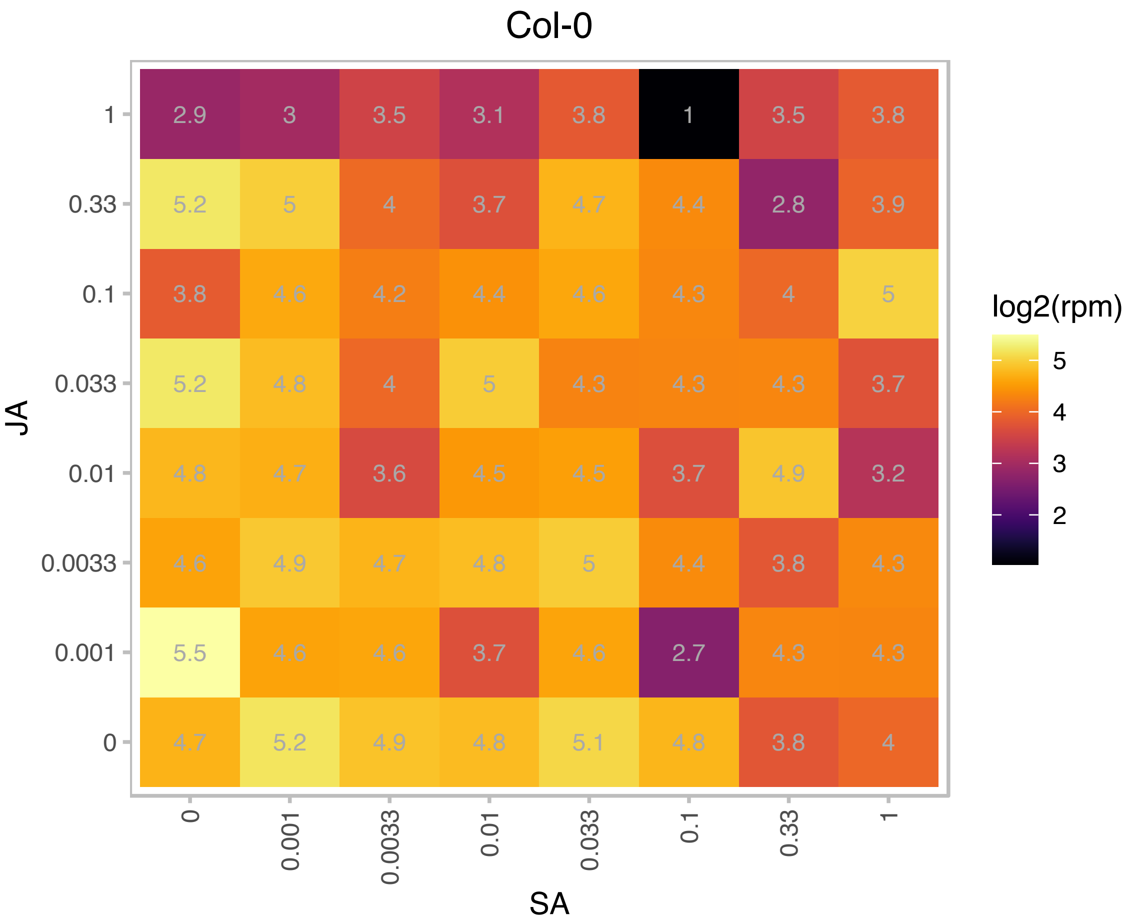


| | |
|---|---|
| Gene | AT2G32230 |
| is.expressed_genes | TRUE |
| is.responsive_genes | TRUE |
| is.apcluster | F-05 |

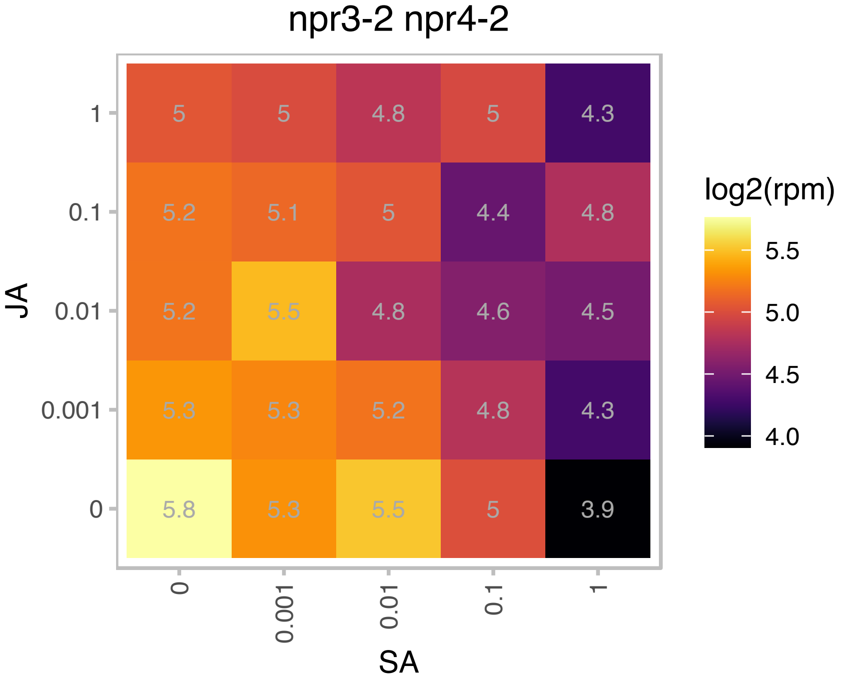


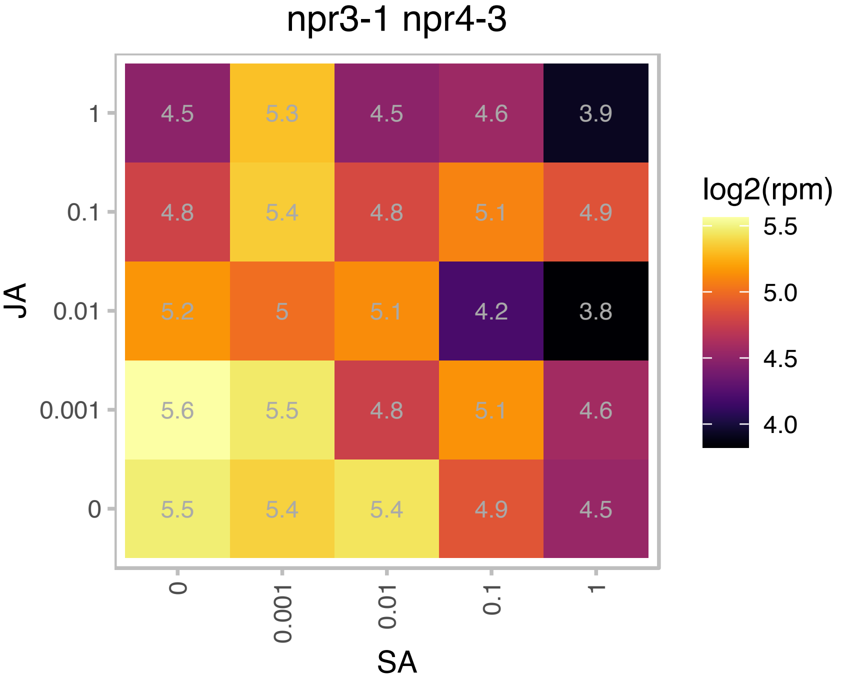

# AT3G02690.1 : NA

nodulin MtN21 /EamA-like transporter family protein

Nucleotide/sugar transporter family protein

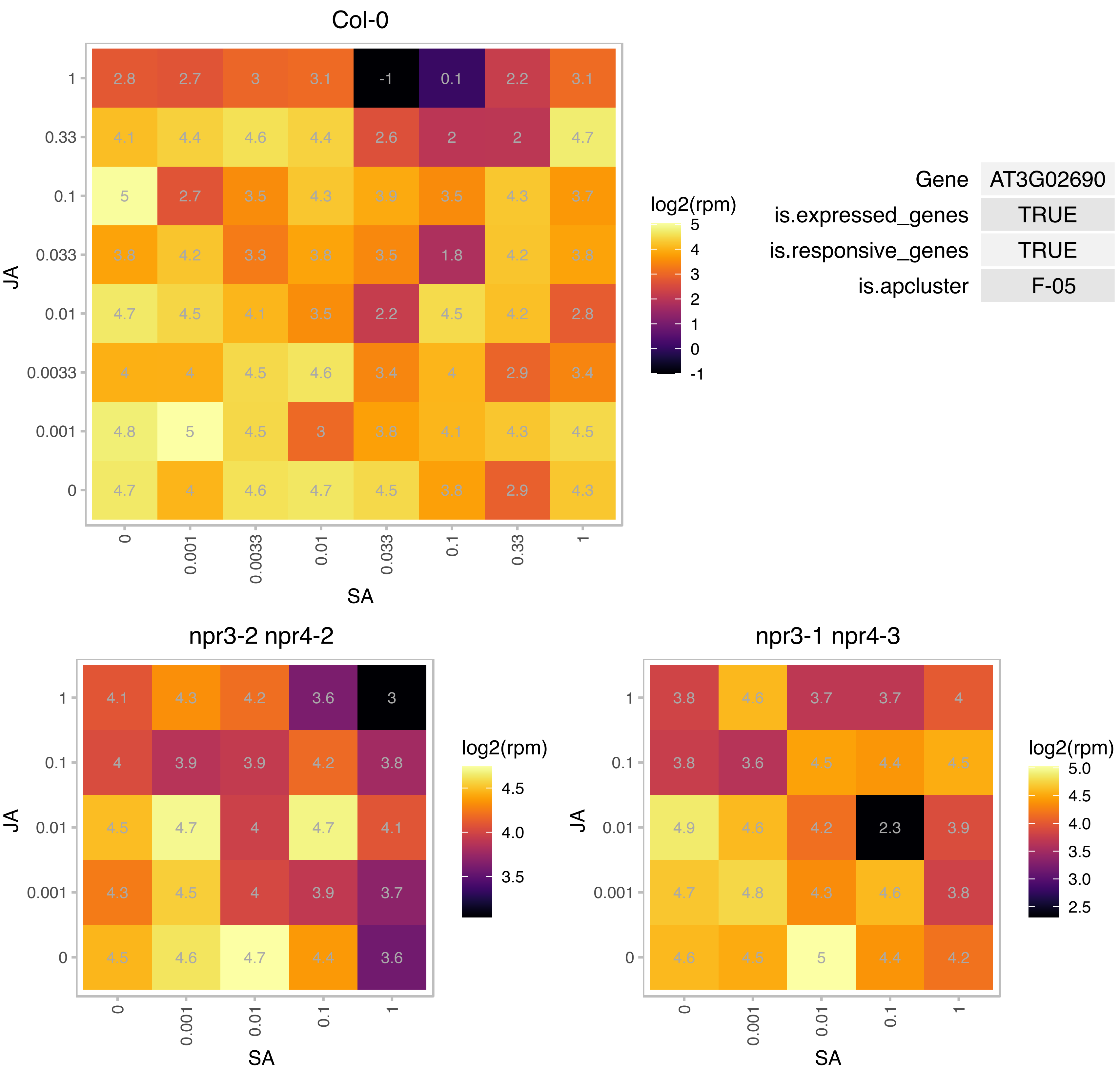


| | |
|---|---|
| Gene | AT3G02690 |
| is.expressed_genes | TRUE |
| is.responsive_genes | TRUE |
| is.apcluster | F-05 |

# AT3G05410.1 : NA

Photosystem II reaction center PsbP family protein

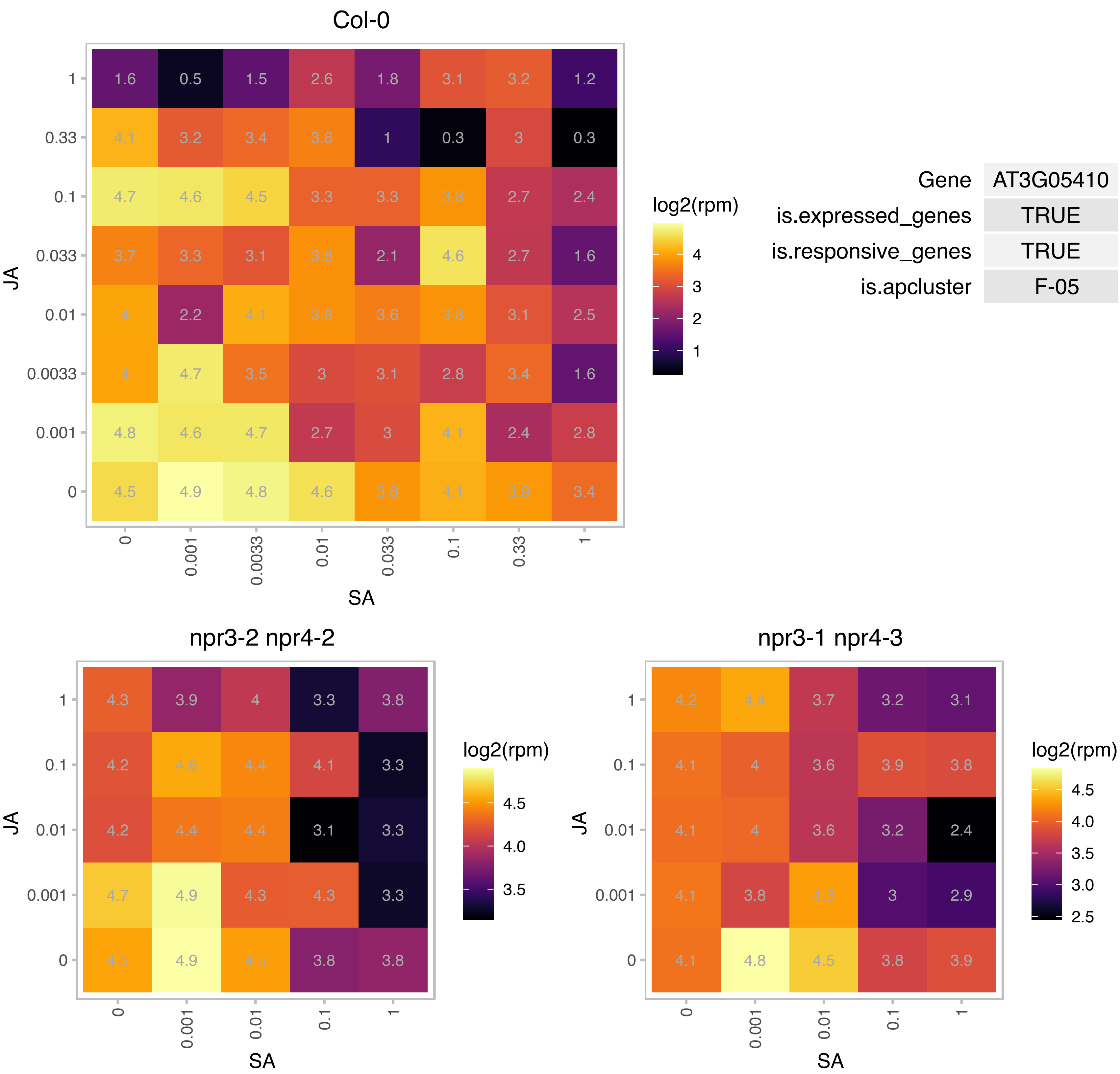


| | |
|---|---|
| Gene | AT3G05410 |
| is.expressed_genes | TRUE |
| is.responsive_genes | TRUE |
| is.apcluster | F-05 |

### AT3G06430.1 : AtPPR2, EMB2750, PPR2

Tetratricopeptide repeat (TPR)-like superfamily protein

Encodes PPR2, a pentatricopeptide repeat protein. Binds to plastid 23S rRNA and plays an important role in the first mitotic division during gametogenesis and in cell proliferation during embryogenesis.

| | |
|---|---|
| Gene | AT3G06430 |
| is.expressed_genes | TRUE |
| is.responsive_genes | TRUE |
| is.apcluster | F-05 |

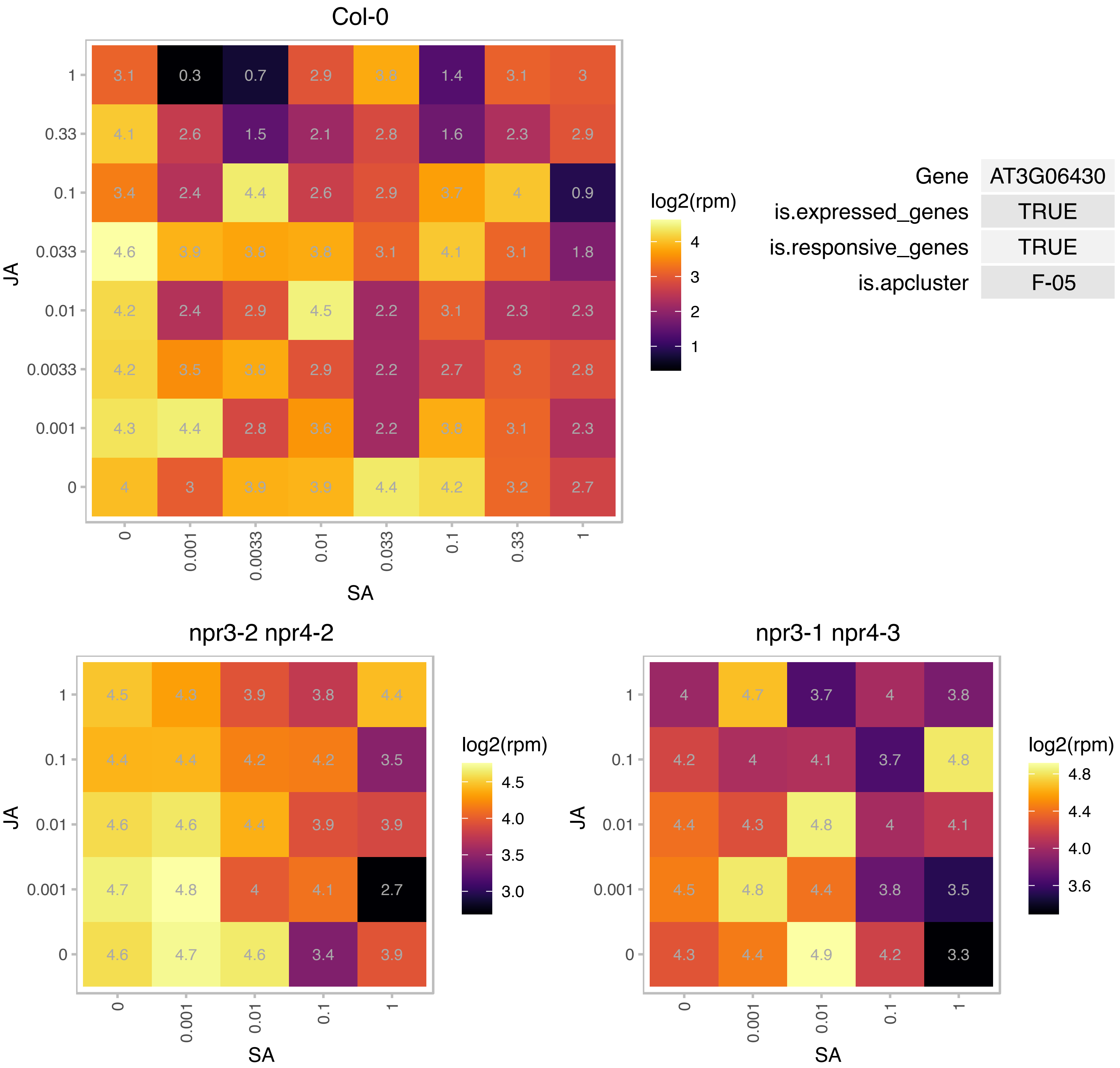

# AT3G07200.1 : NA

RING/U-box superfamily protein

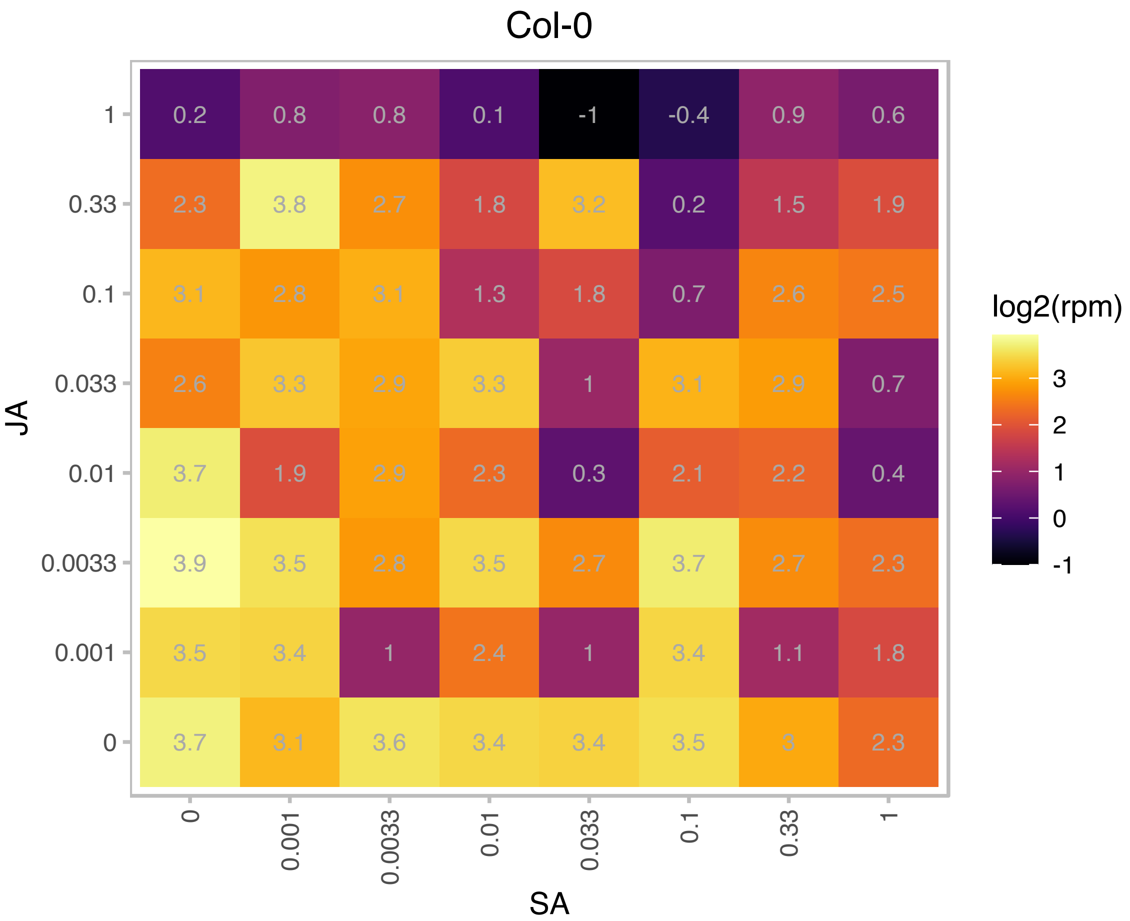


| | |
|---|---|
| Gene | AT3G07200 |
| is.expressed_genes | TRUE |
| is.responsive_genes | TRUE |
| is.apcluster | F-05 |

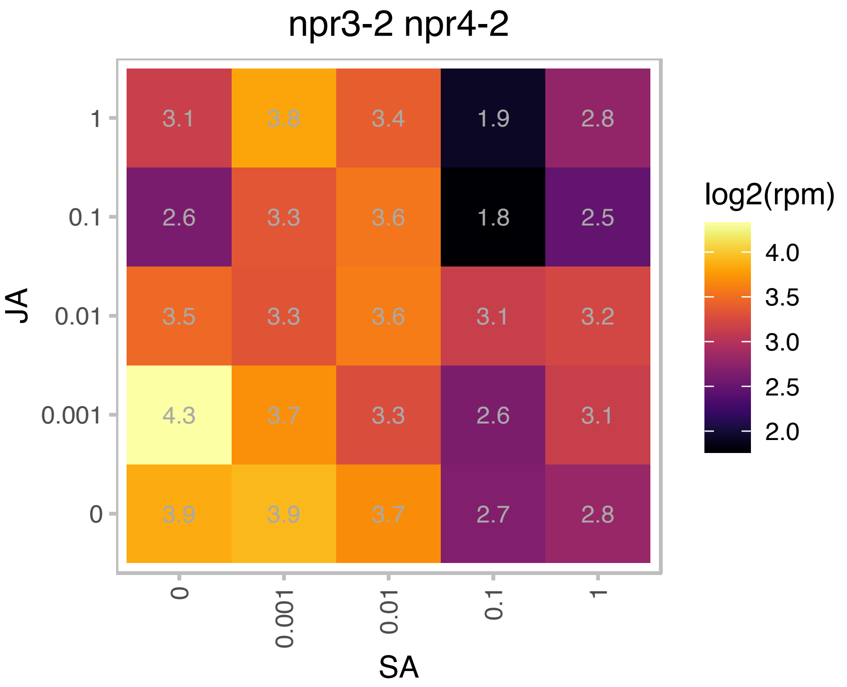


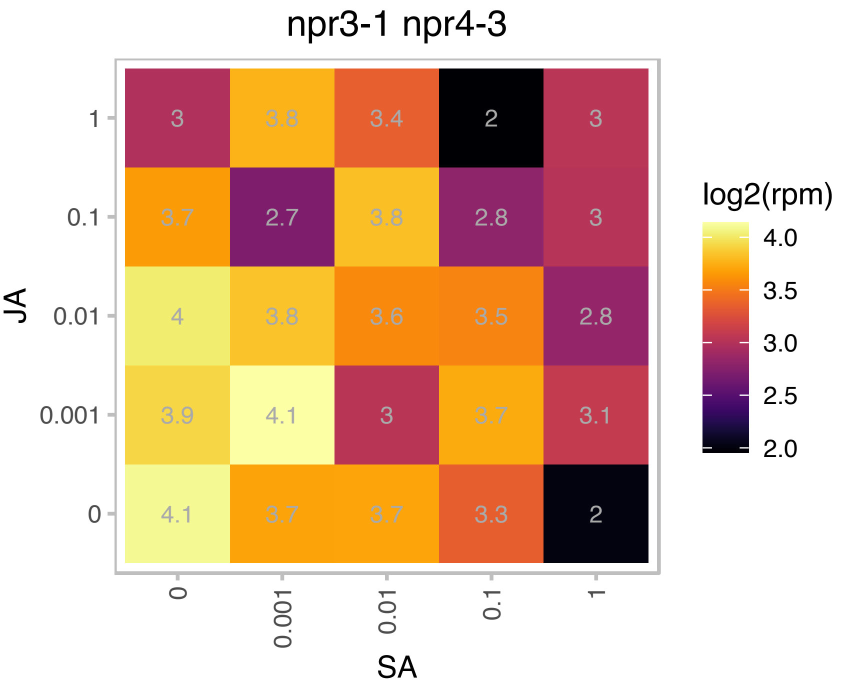

### AT3G09210.1 : PTAC13

plastid transcriptionally active 13

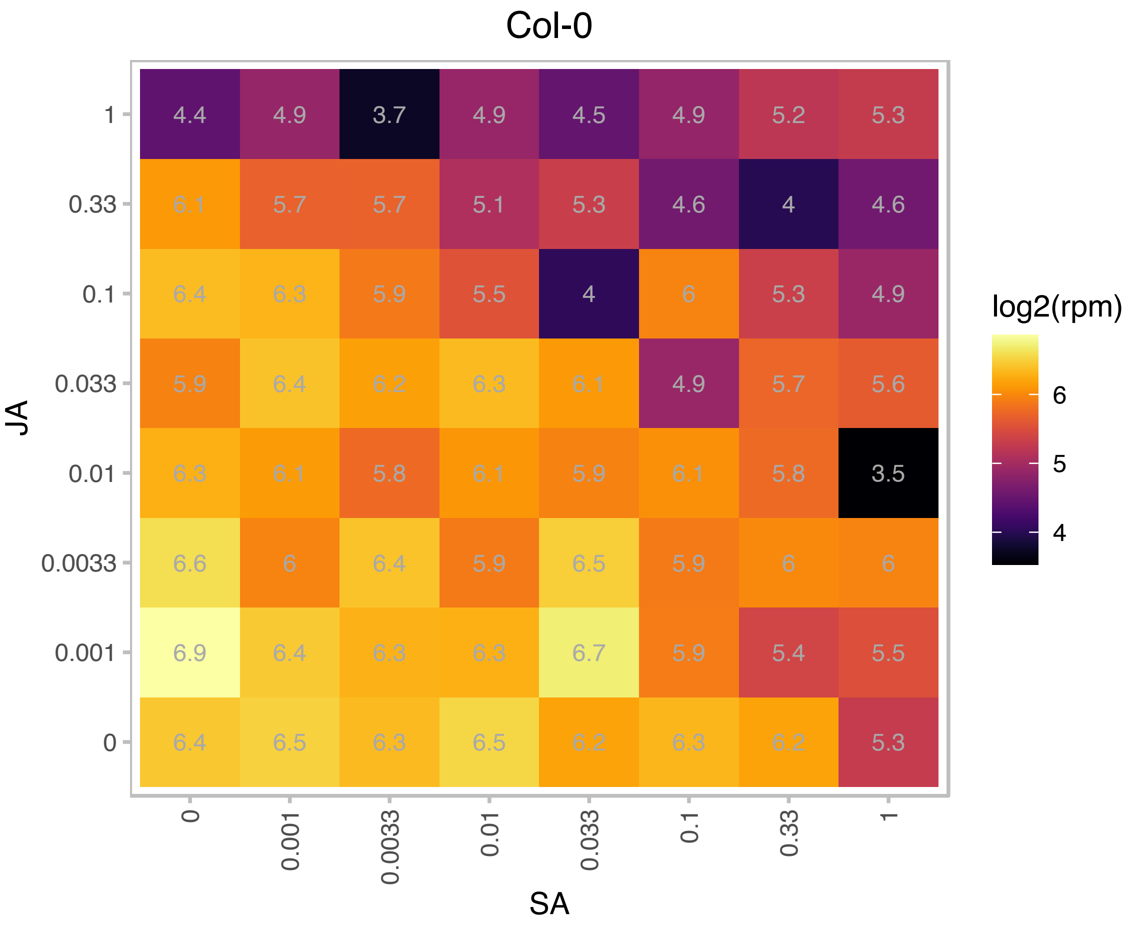


| | |
|---|---|
| Gene | AT3G09210 |
| is.expressed_genes | TRUE |
| is.responsive_genes | TRUE |
| is.apcluster | F-05 |

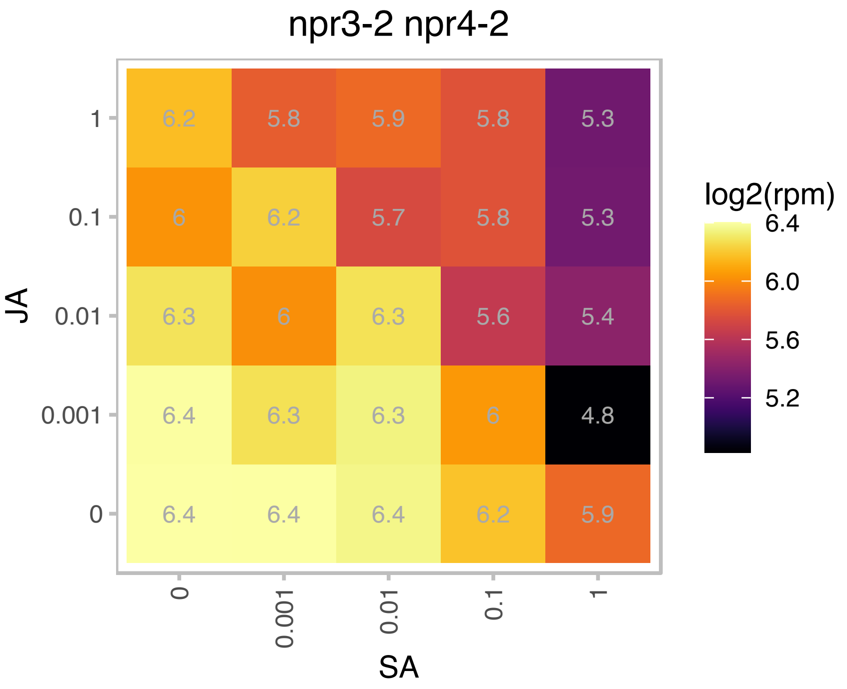


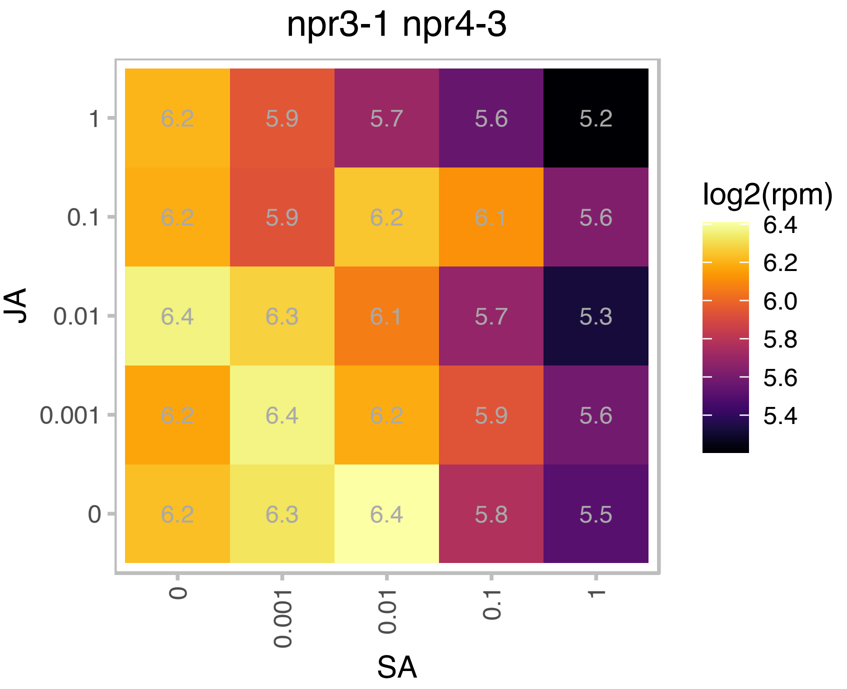

# AT3G09310.1 : NA

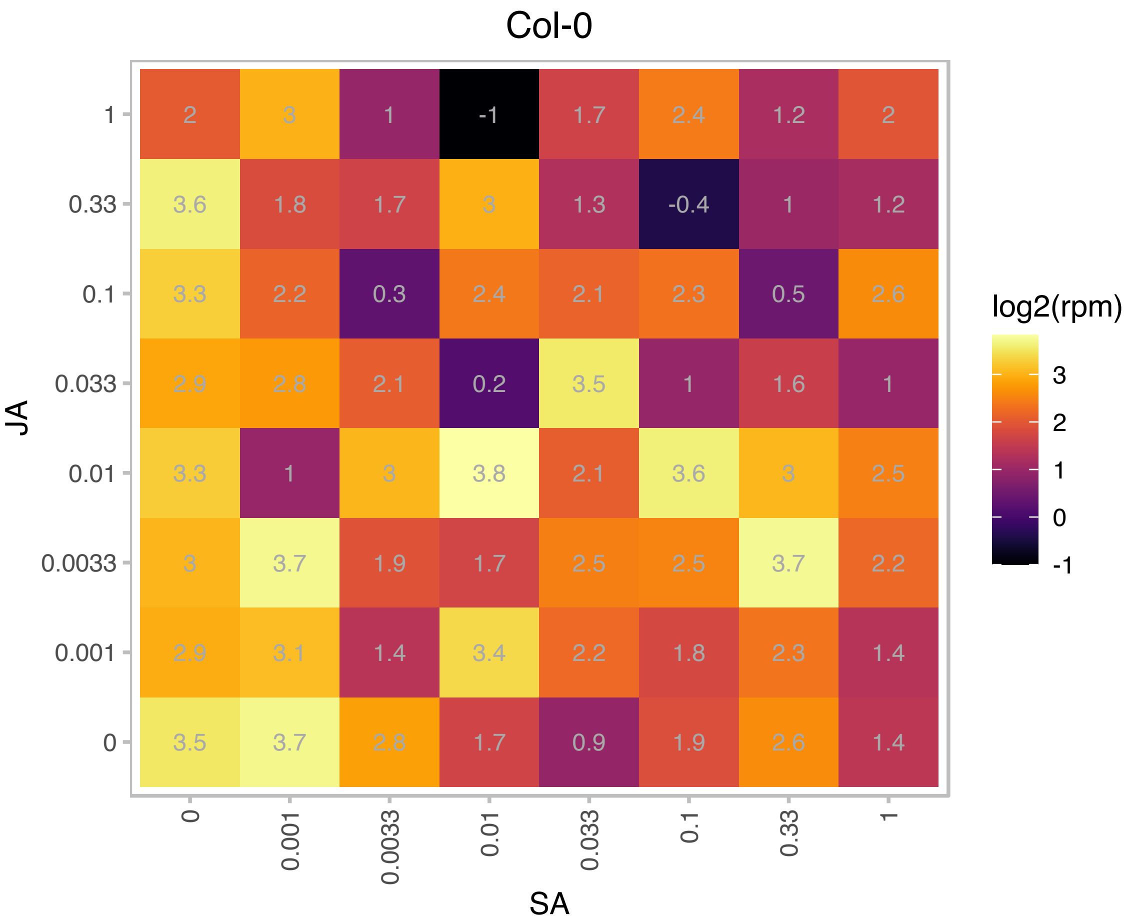


| | |
|---|---|
| Gene | AT3G09310 |
| is.expressed_genes | TRUE |
| is.responsive_genes | TRUE |
| is.apcluster | F-05 |

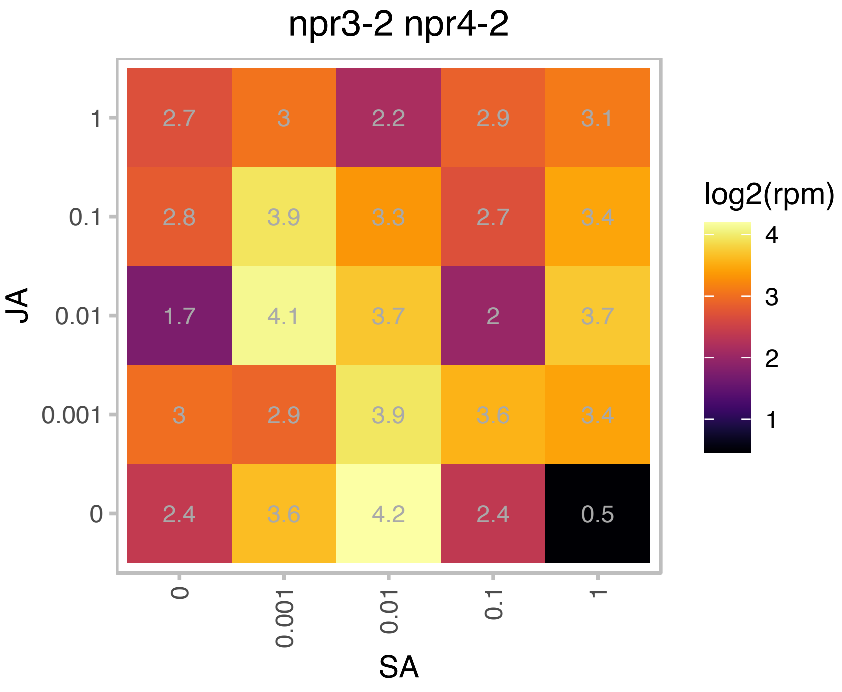


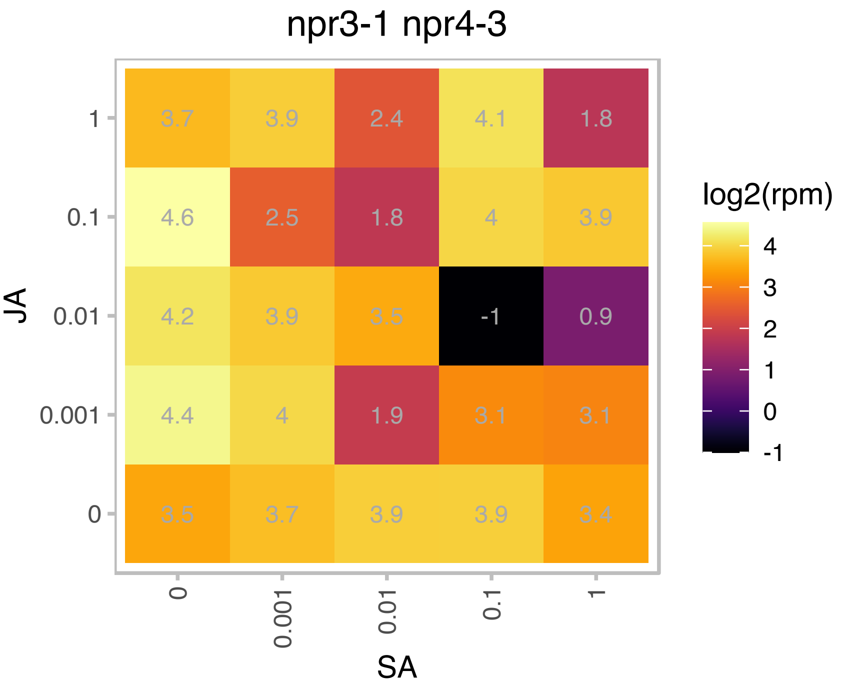

# AT3G12870.1 : NA

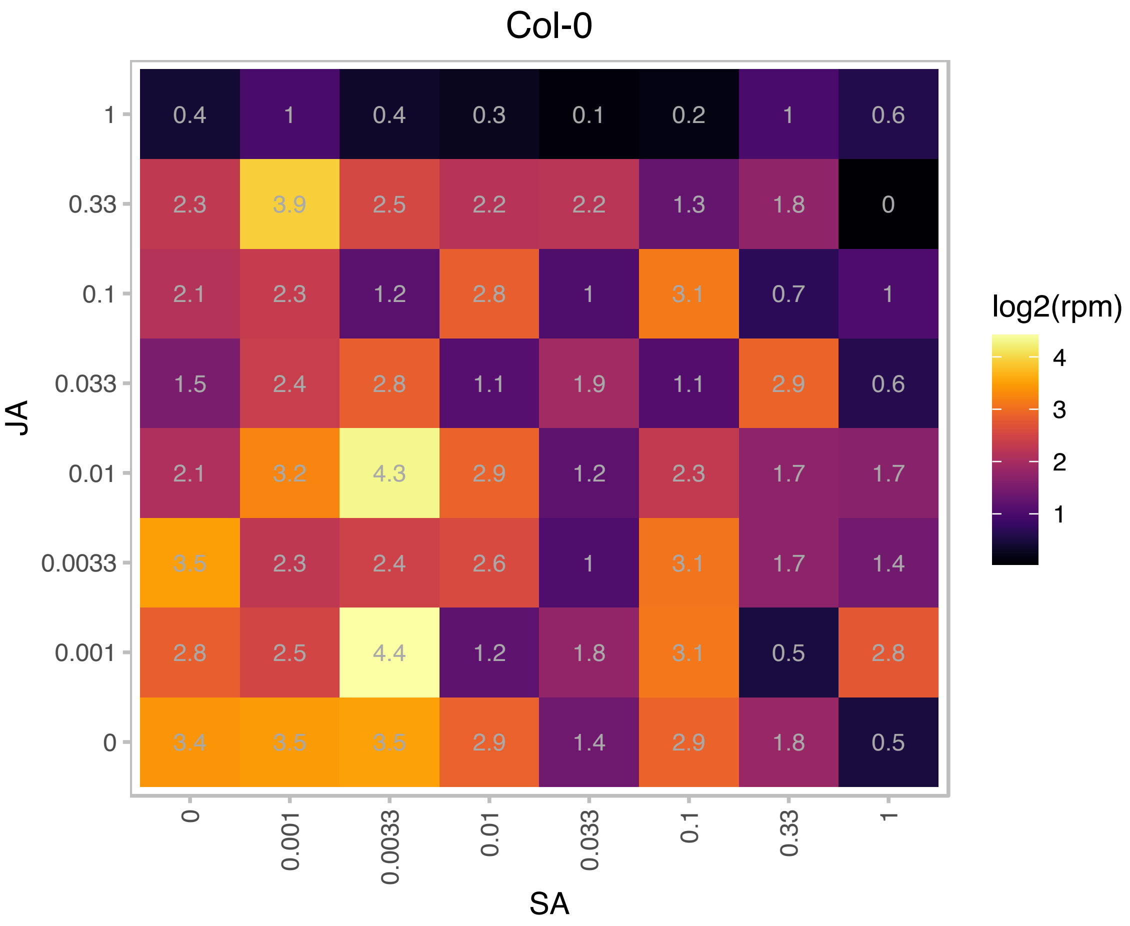


| | |
|---|---|
| Gene | AT3G12870 |
| is.expressed_genes | TRUE |
| is.responsive_genes | TRUE |
| is.apcluster | F-05 |

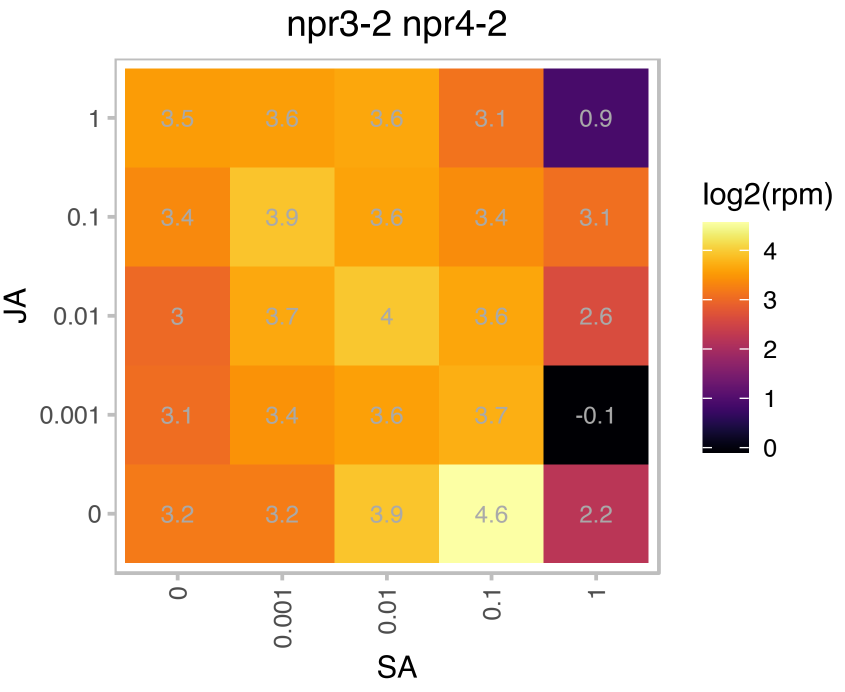


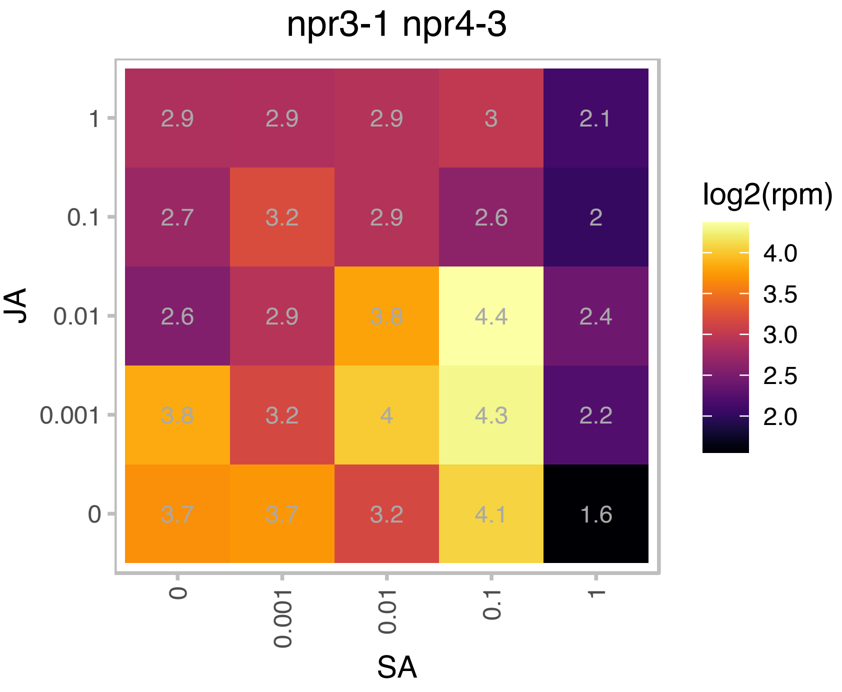

### AT3G15850.1 : ADS3, AtADS3, FAD5, FADB, JB67

fatty acid desaturase 5

Chloroplastic enzyme responsible for the synthesis of 16:1 fatty acids from galactolipids and sulpholipids. Uses ferredoxin as electron donor. The mRNA is cell-to-cell mobile.

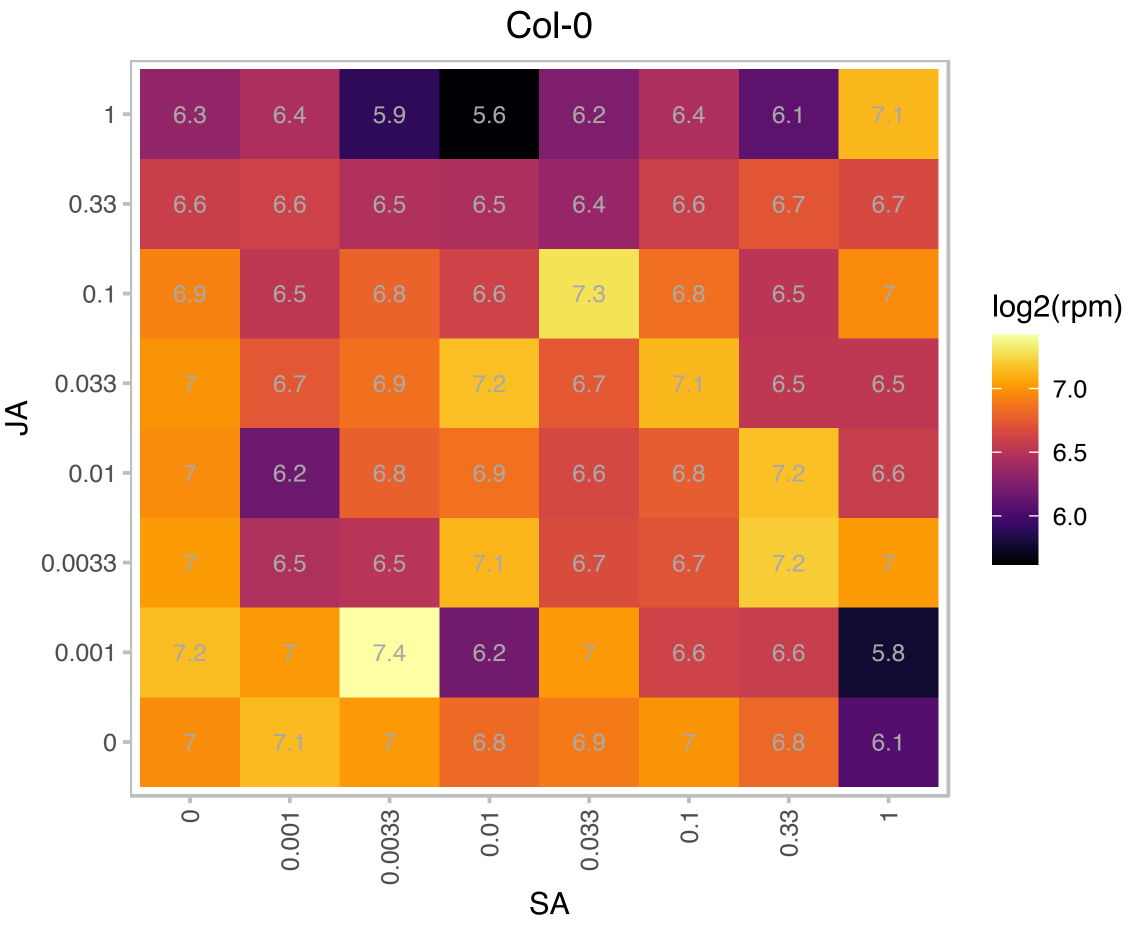


| | |
|---|---|
| Gene | AT3G15850 |
| is.expressed_genes | TRUE |
| is.responsive_genes | TRUE |
| is.apcluster | F-05 |

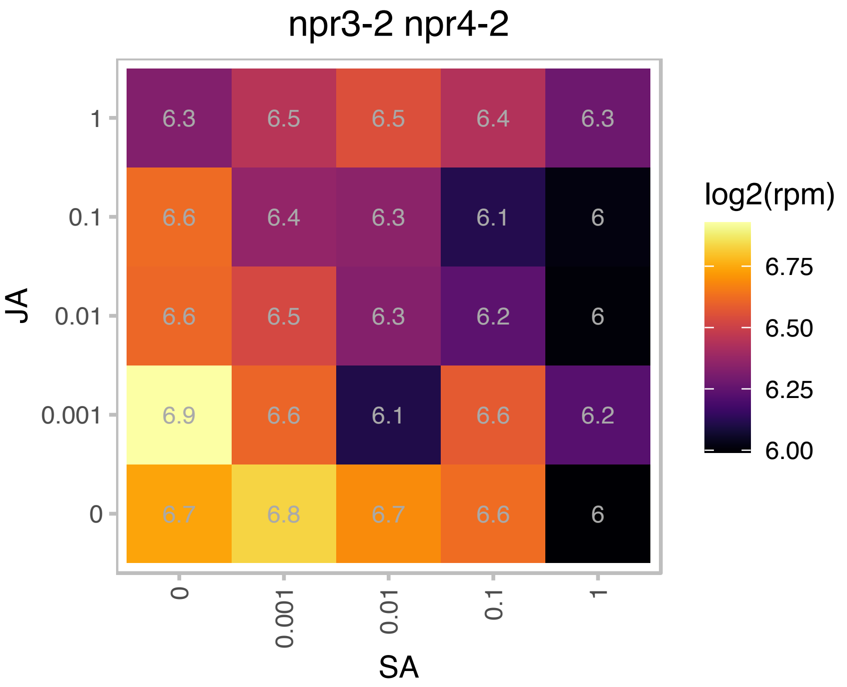


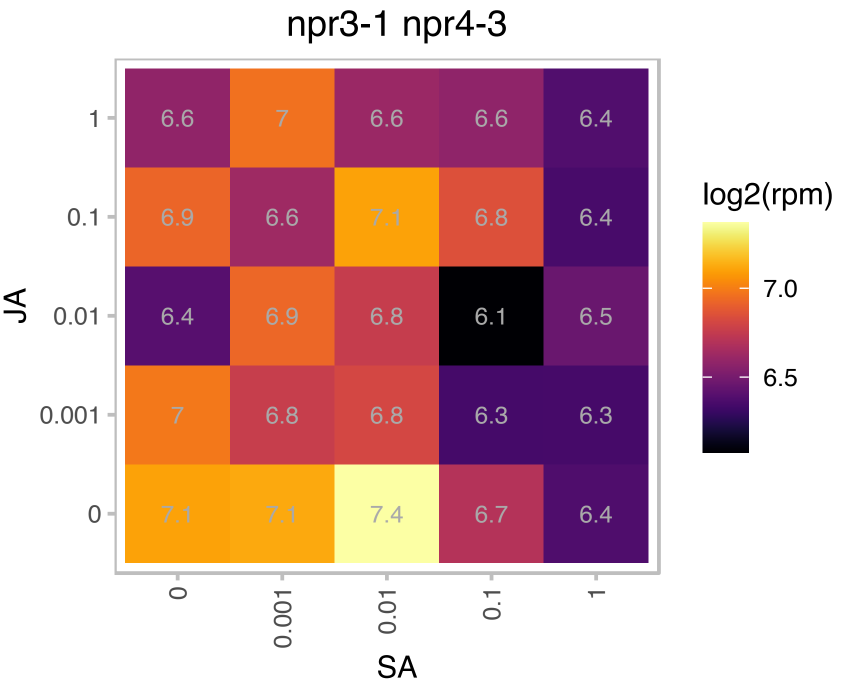

### AT3G17185.1 : ATTAS3, sORF24, TAS3, TASIR-ARF

TAS3/TASIR-ARF (TRANS-ACTING SIRNA3); other RNA
Encodes a trans-acting siRNA (tasi-RNA) that regulates the expression of auxin response factor genes (ARF2, ARF4, ETT). One of 3 genomic loci that encode the TAS3 siRNA. Has been identified as a translated small open reading frame by ribosome profiling.

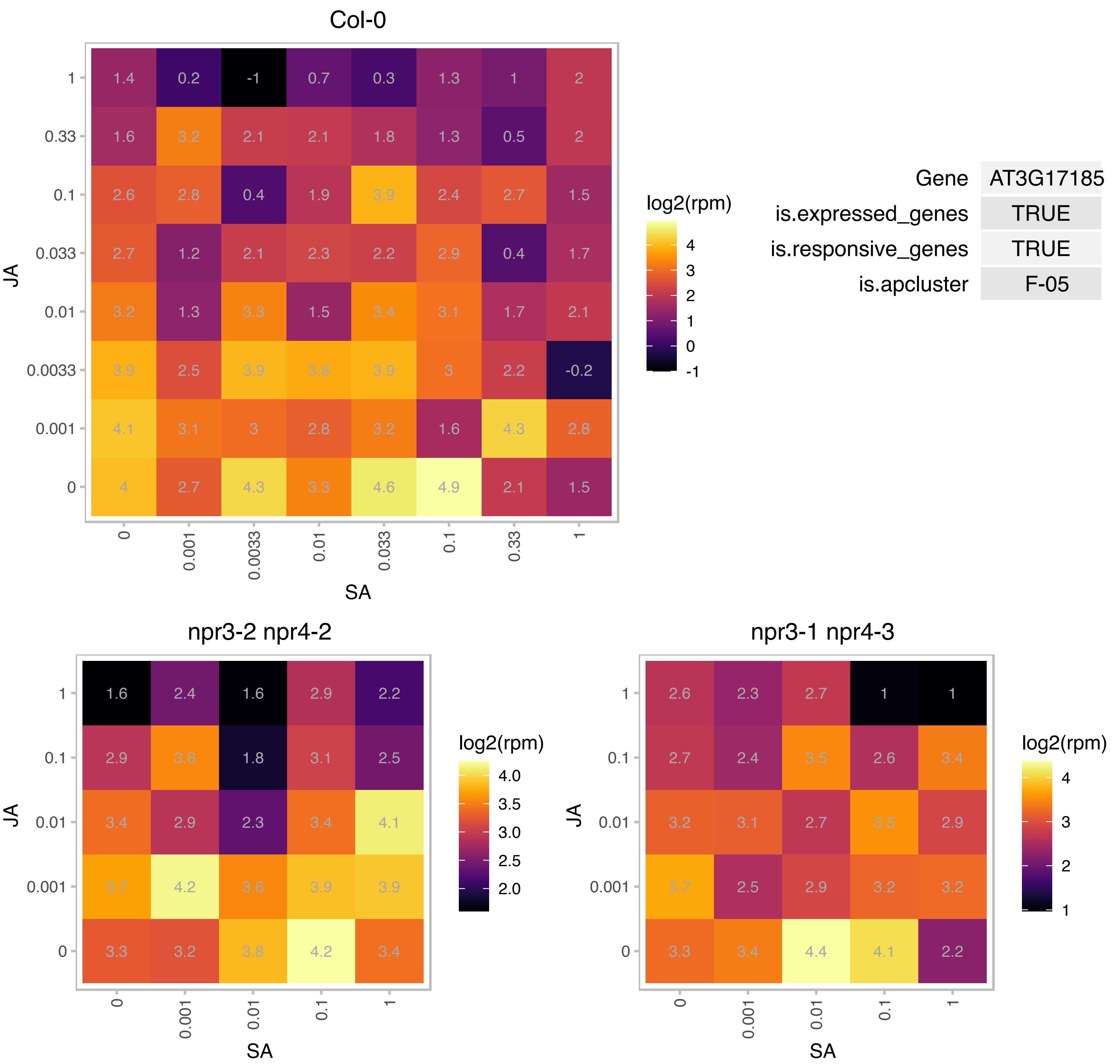


| | |
|---|---|
| Gene | AT3G17185 |
| is.expressed_genes | TRUE |
| is.responsive_genes | TRUE |
| is.apcluster | F-05 |

# AT3G20230.1 : NA

Ribosomal L18p/L5e family protein

| | |
|---|---|
| Gene | AT3G20230 |
| is.expressed_genes | TRUE |
| is.responsive_genes | TRUE |
| is.apcluster | F-05 |

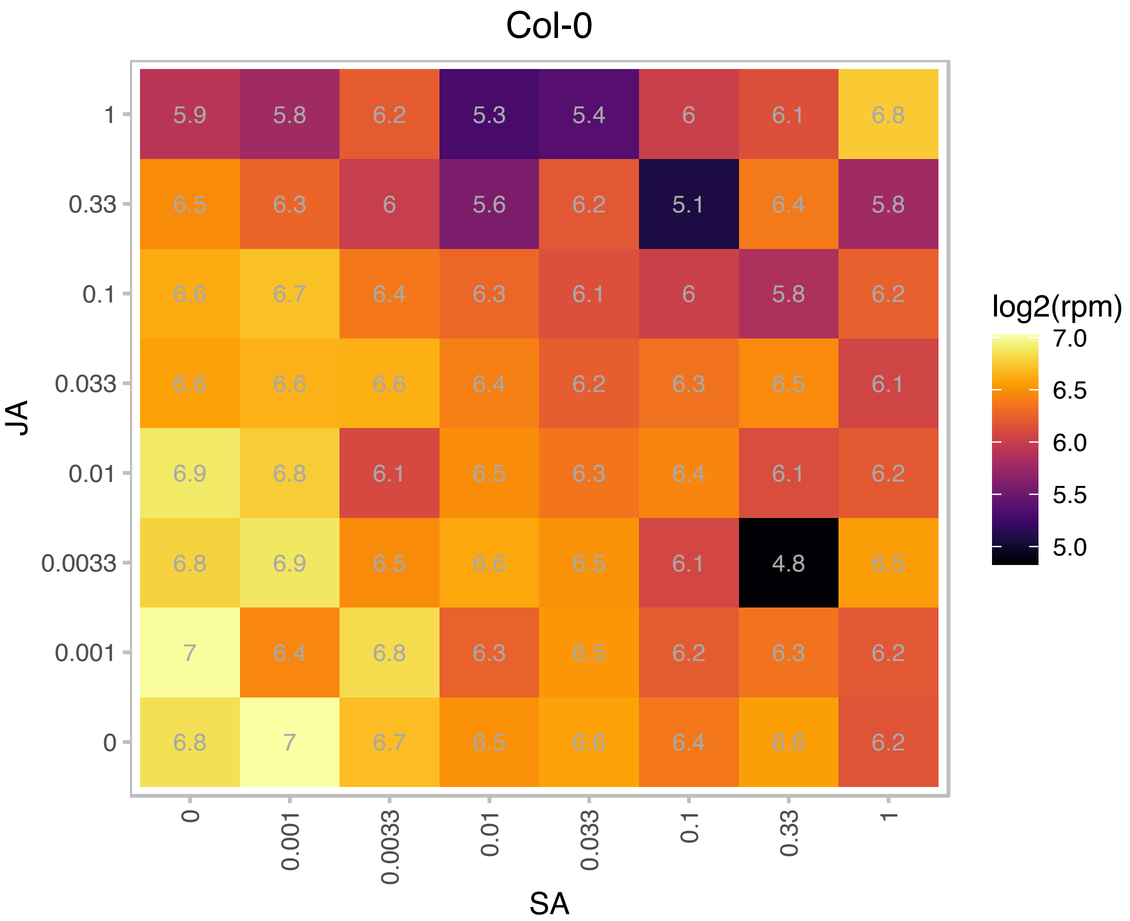


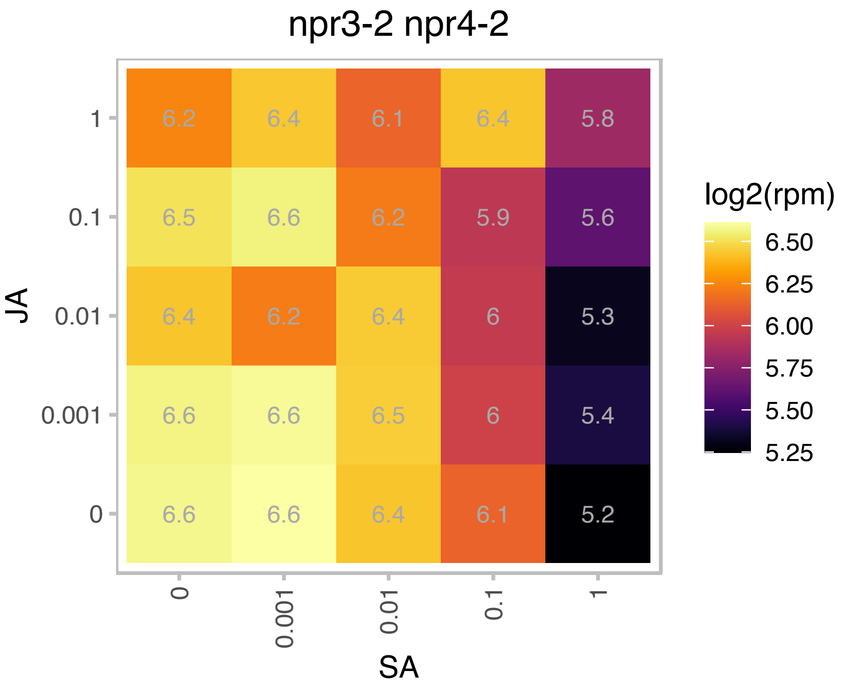


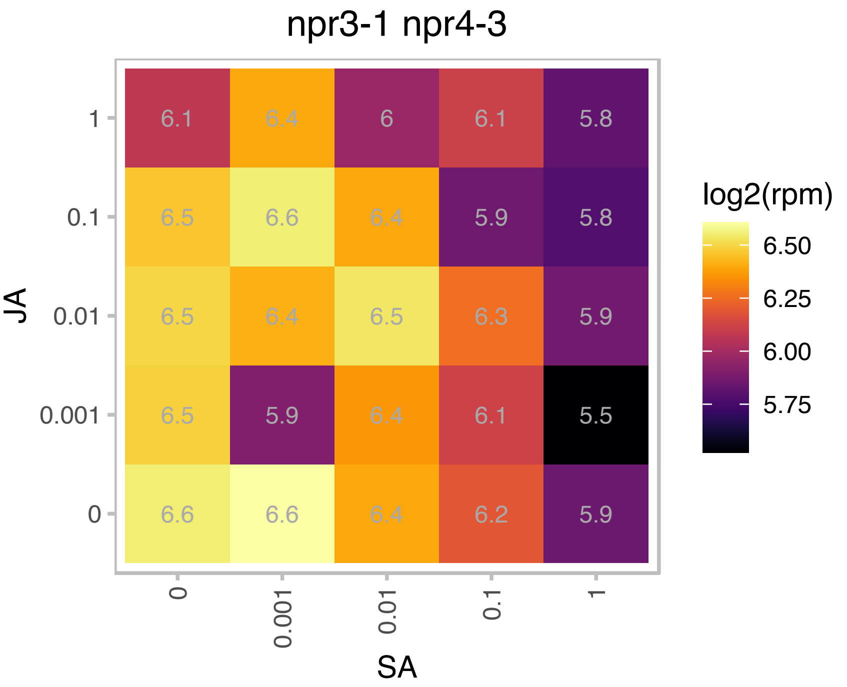

# AT3G21351.1 : NA

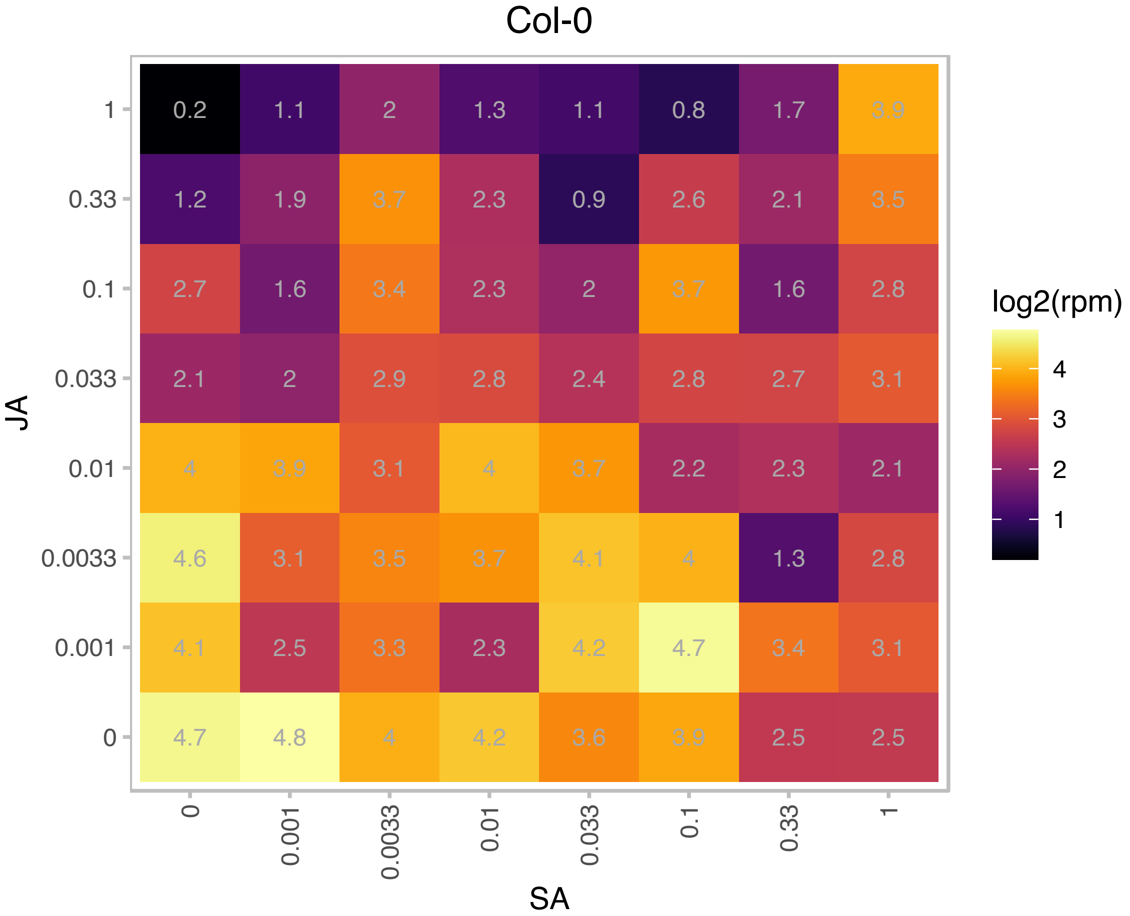


| | |
|---|---|
| Gene | AT3G21351 |
| is.expressed_genes | TRUE |
| is.responsive_genes | TRUE |
| is.apcluster | F-05 |

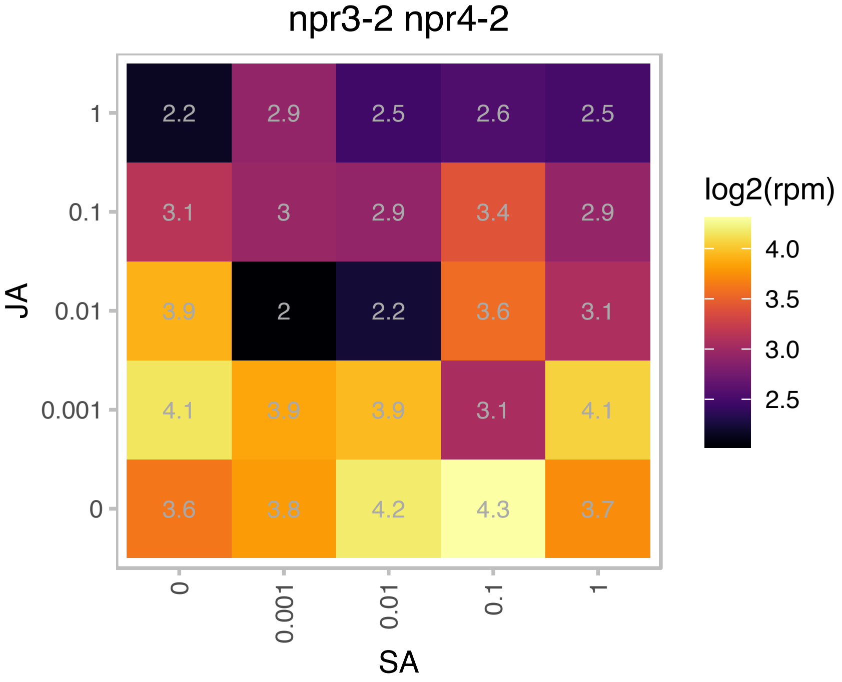


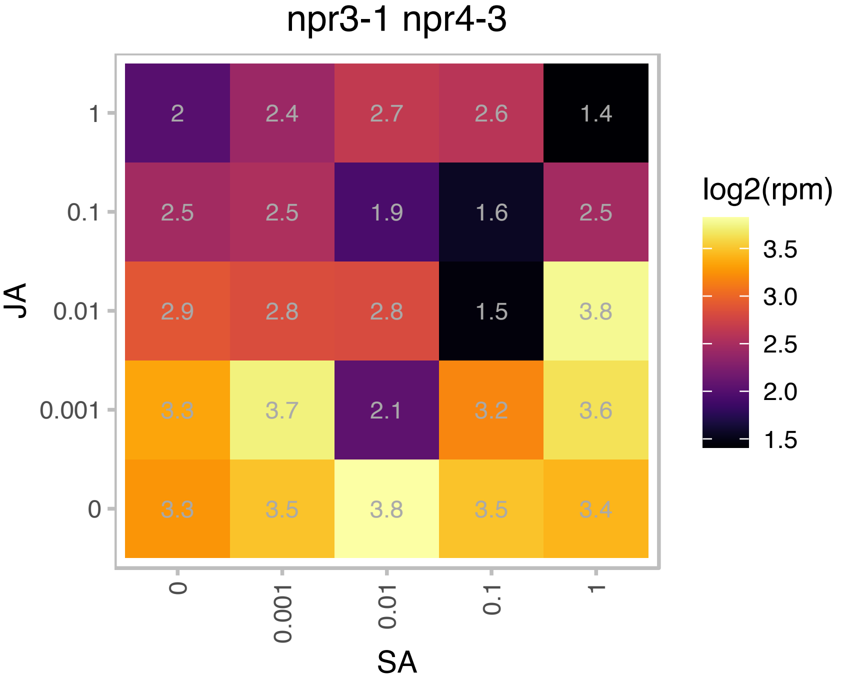

### AT3G21870.1 : CYCP2;1

cyclin p2;1

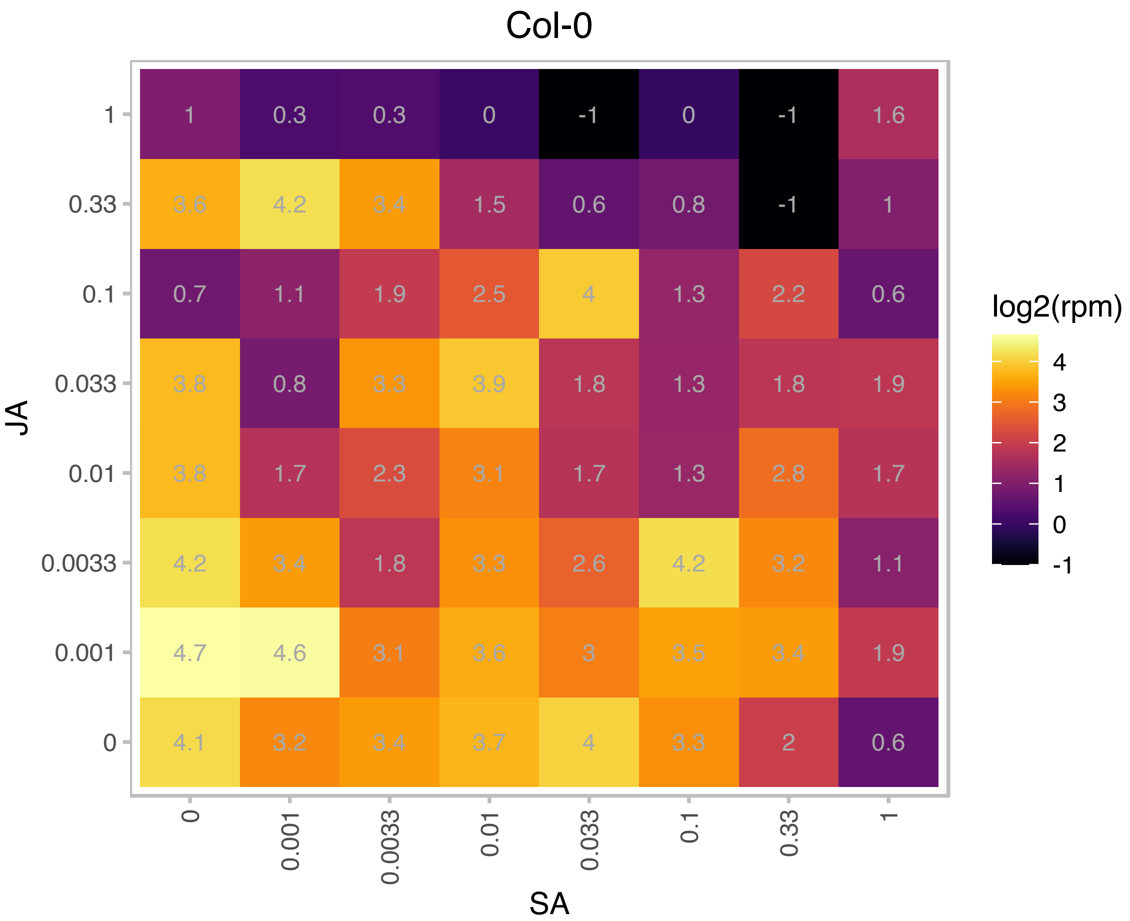


| | |
|---|---|
| Gene | AT3G21870 |
| is.expressed_genes | TRUE |
| is.responsive_genes | TRUE |
| is.apcluster | F-05 |

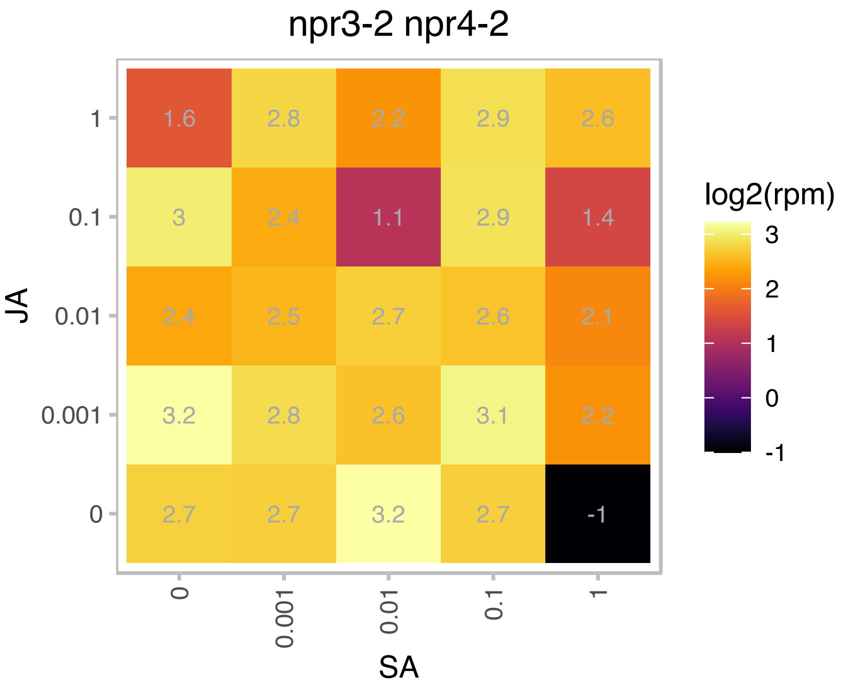


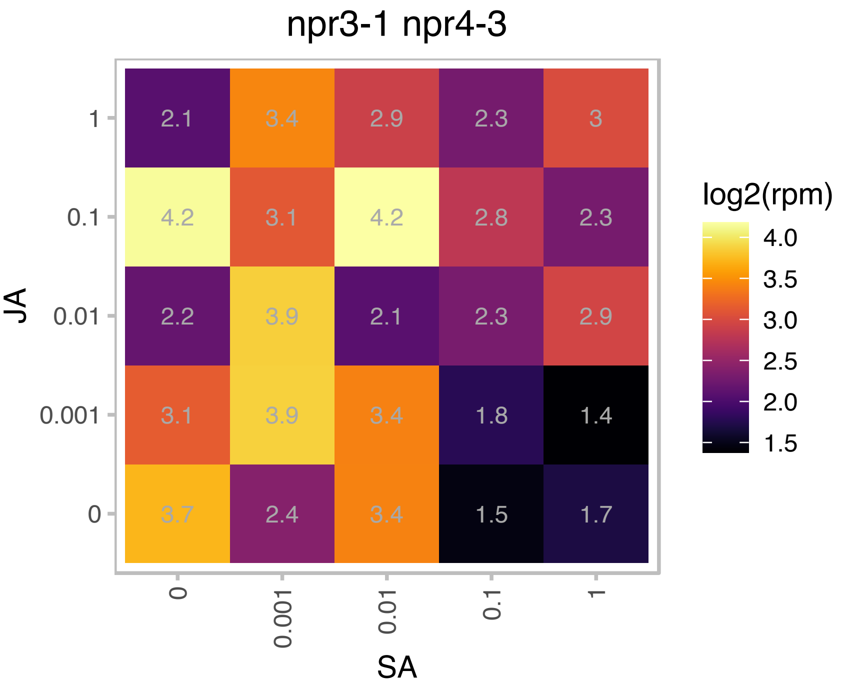

### AT3G23030.1 : IAA2

indole-3-acetic acid inducible 2

auxin inducible gene expressed in the nucleus

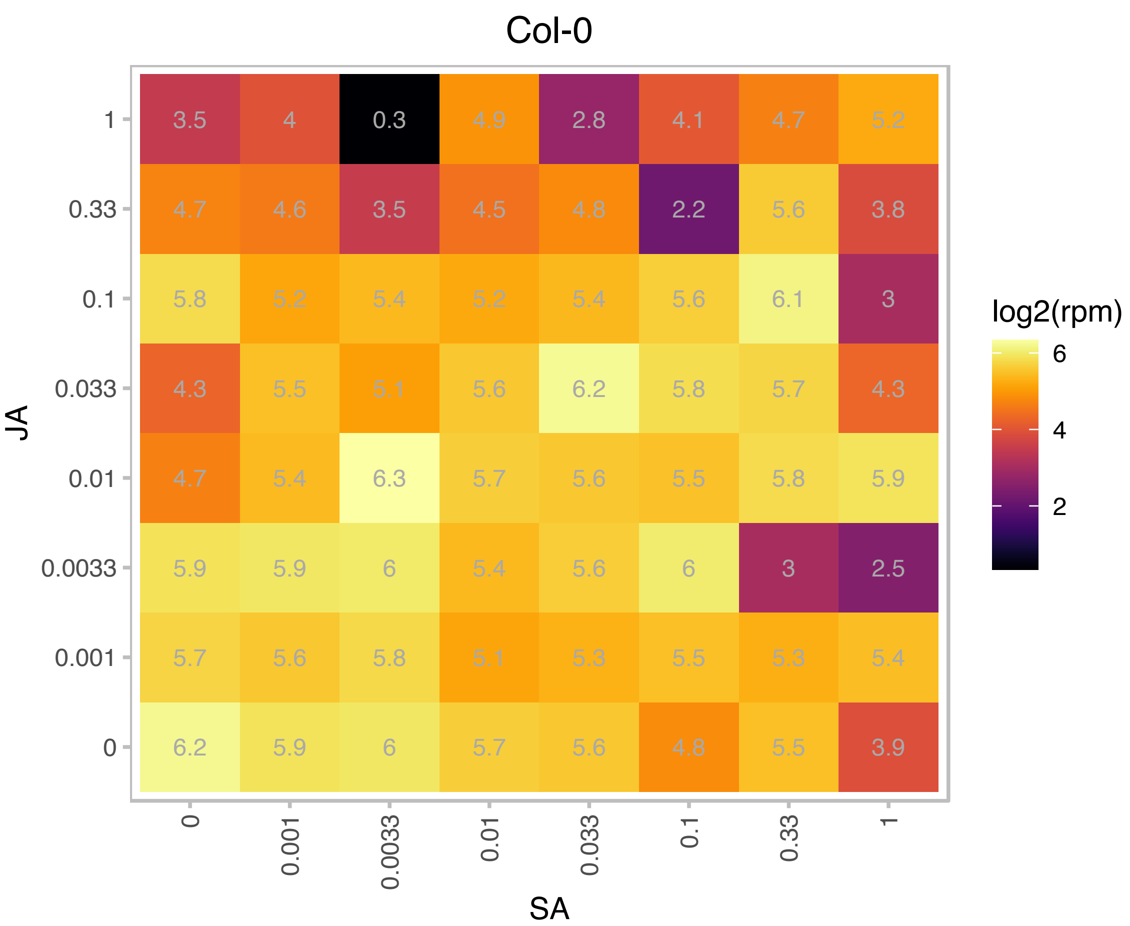


| | |
|---|---|
| Gene | AT3G23030 |
| is.expressed_genes | TRUE |
| is.responsive_genes | TRUE |
| is.apcluster | F-05 |

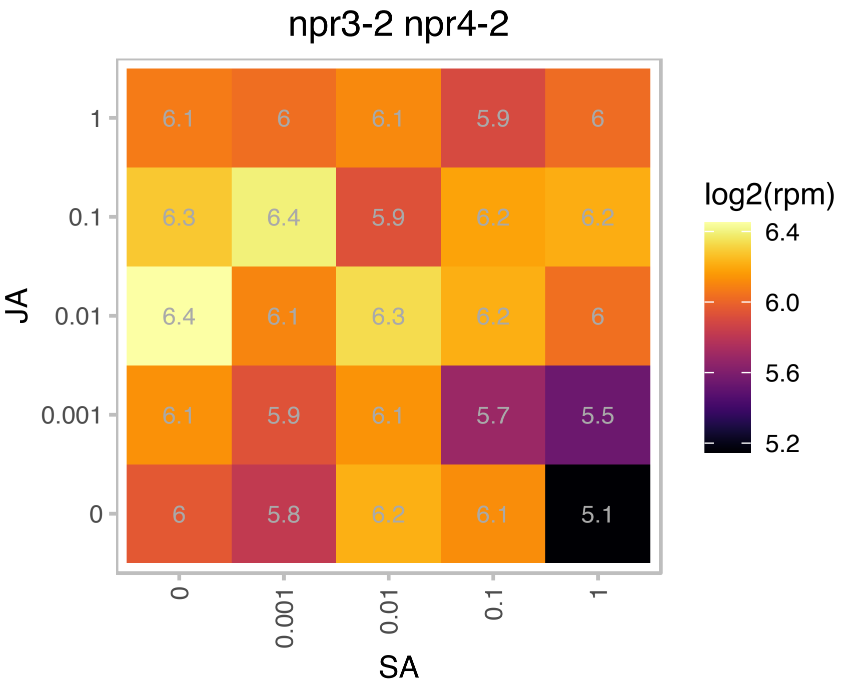


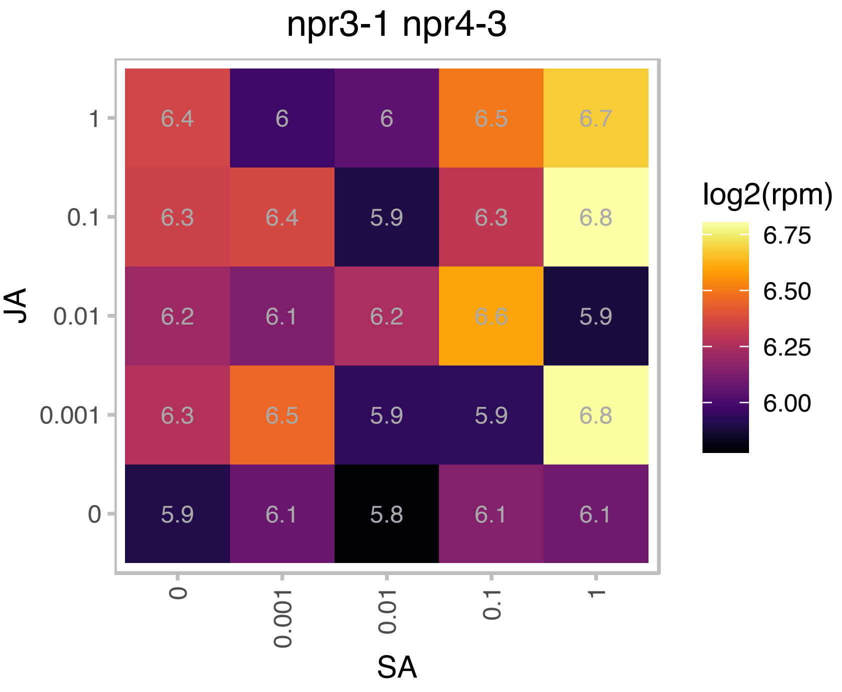

### AT3G24590.1 : PLSP1

plastidic type i signal peptidase 1

Encodes a signal peptidase Plsp1 (plastidic type I signal peptidase 1). Required for thylakoid development. Functions in the maturation of the 75-kD component of the translocon at the outer envelope membrane of chloroplasts and oxygen evolving complex subunit 33 (OE33). Involved in the maturation of PsbO, plastocyanin and FtsH2/8. The mRNA is cell-to-cell mobile.

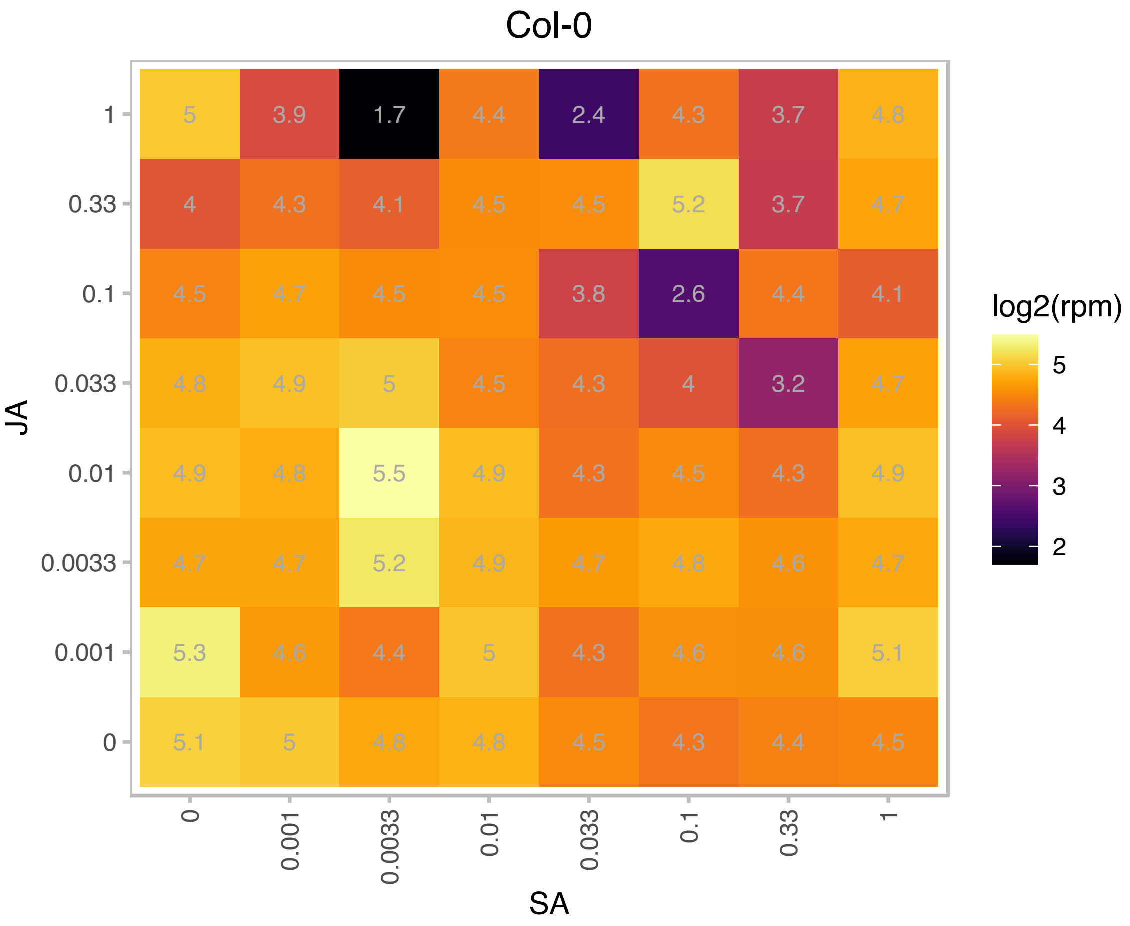


| | |
|---|---|
| Gene | AT3G24590 |
| is.expressed_genes | TRUE |
| is.responsive_genes | TRUE |
| is.apcluster | F-05 |

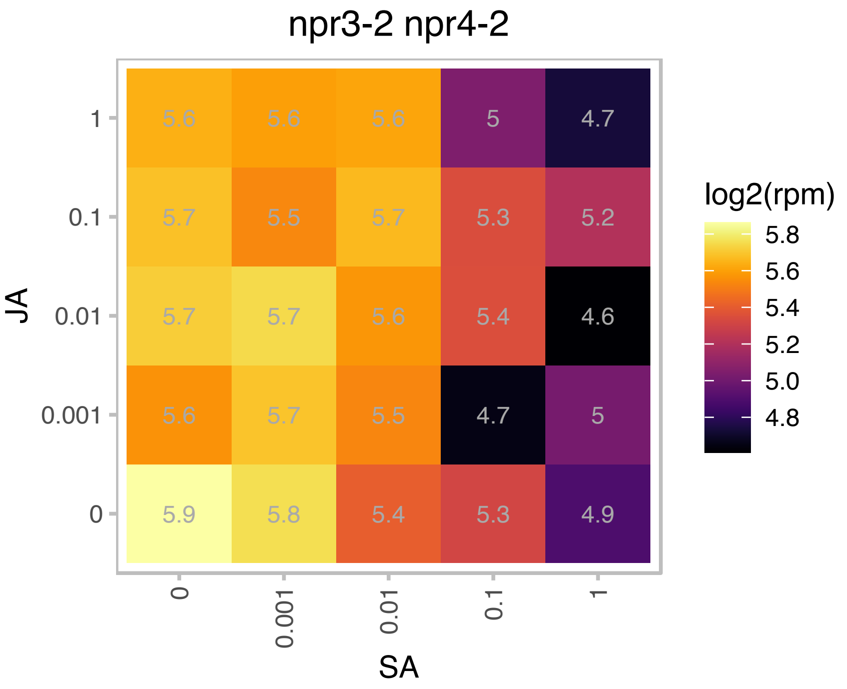


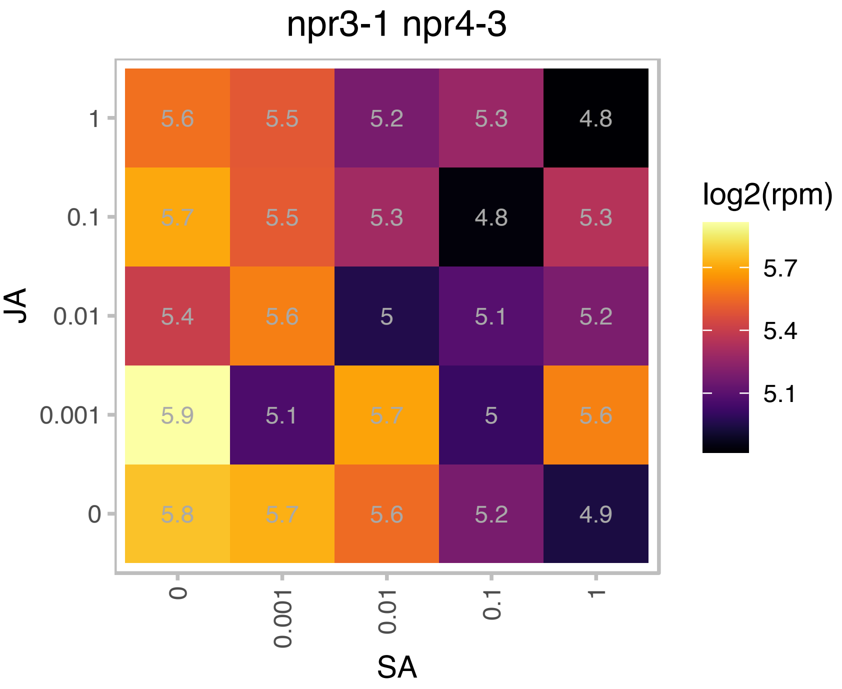

### AT3G27850.1 : RPL12-C

ribosomal protein L12-C

50S ribosomal protein L12-C The mRNA is cell-to-cell mobile.

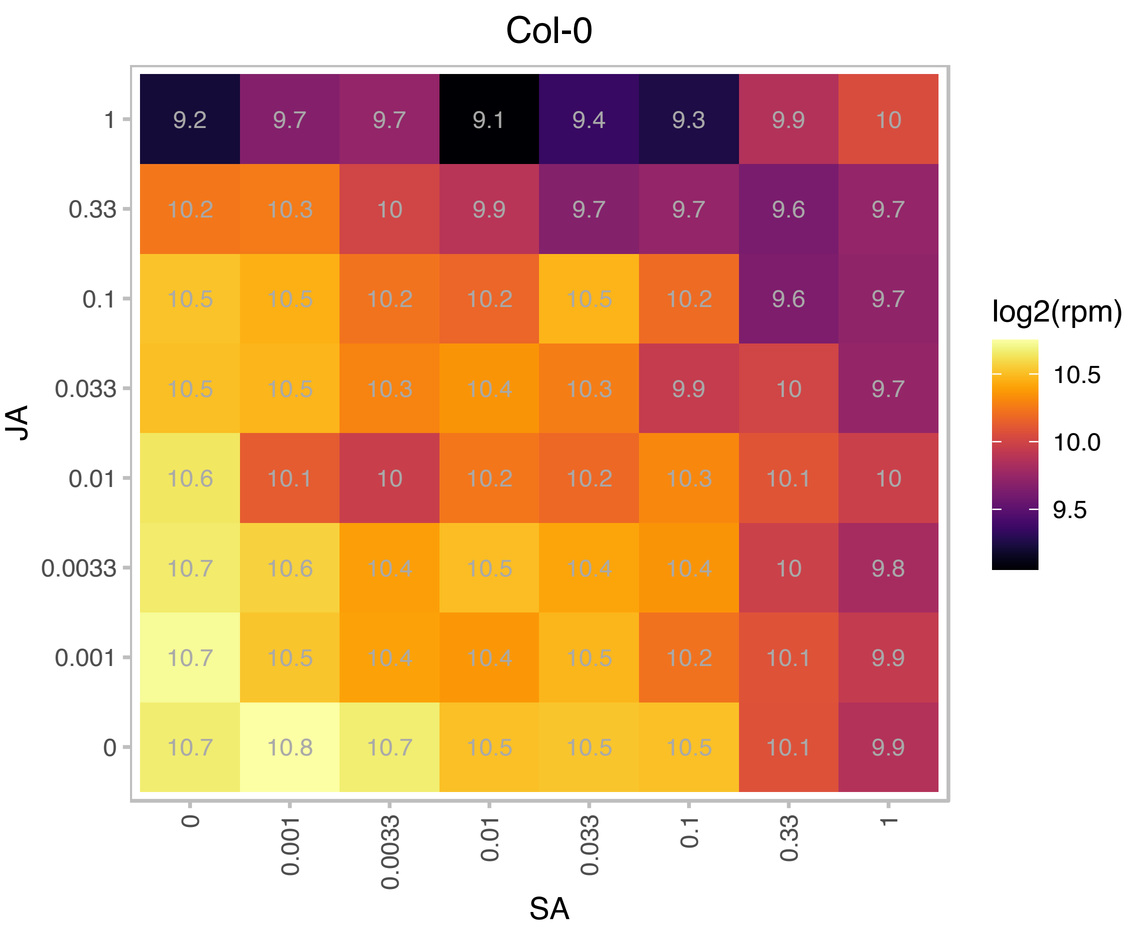


| | |
|---|---|
| Gene | AT3G27850 |
| is.expressed_genes | TRUE |
| is.responsive_genes | TRUE |
| is.apcluster | F-05 |

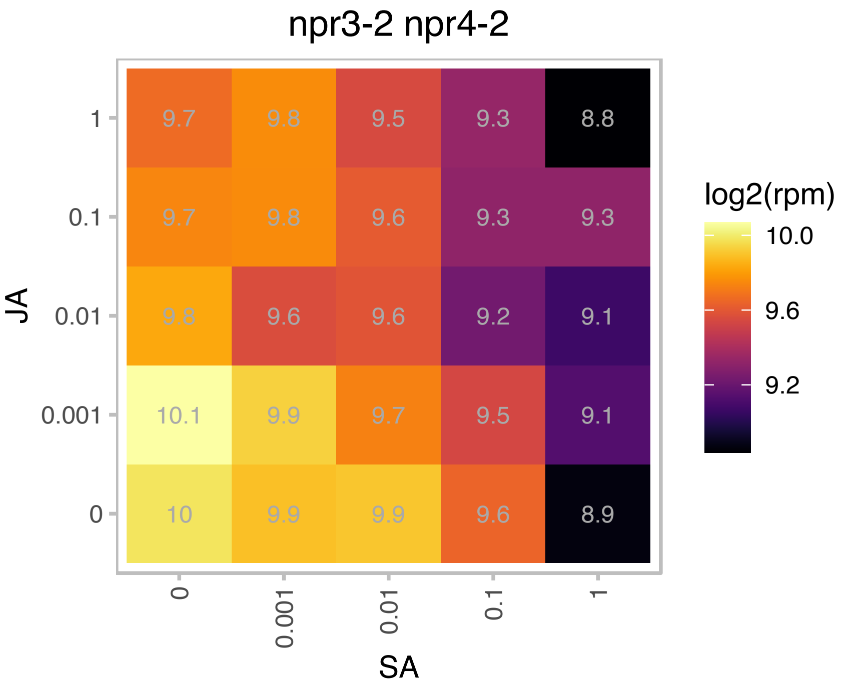


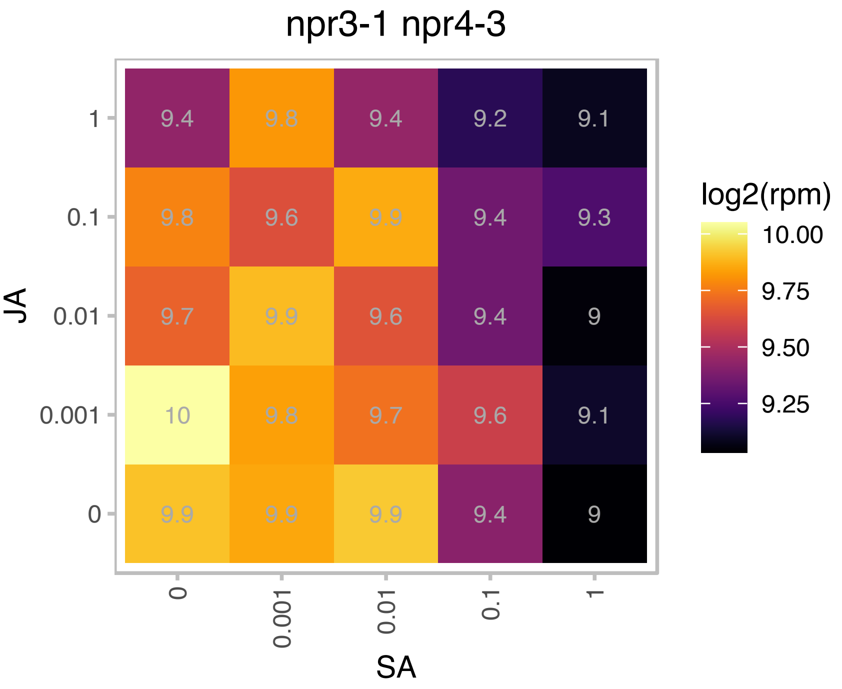

# AT3G32930.1 : NA

Protein of unknown function. Locus is correlated with bacterial hypersensitive response, expression is reduced after injection with avrRpm1.

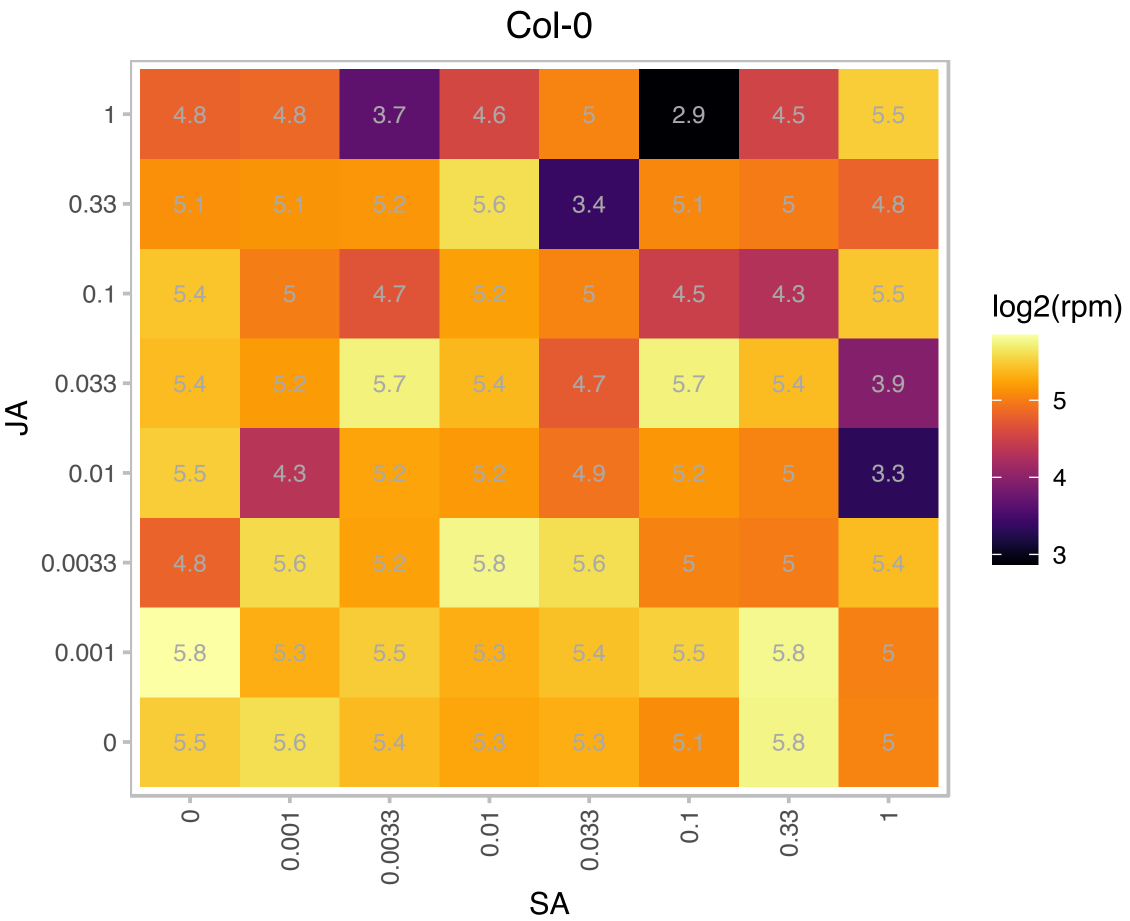


| | |
|---|---|
| Gene | AT3G32930 |
| is.expressed_genes | TRUE |
| is.responsive_genes | TRUE |
| is.apcluster | F-05 |

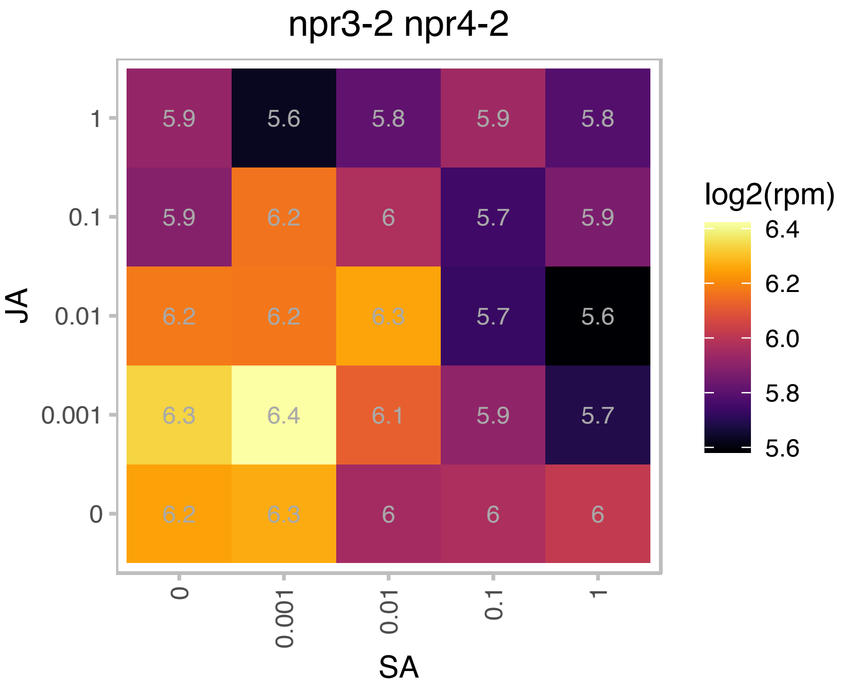


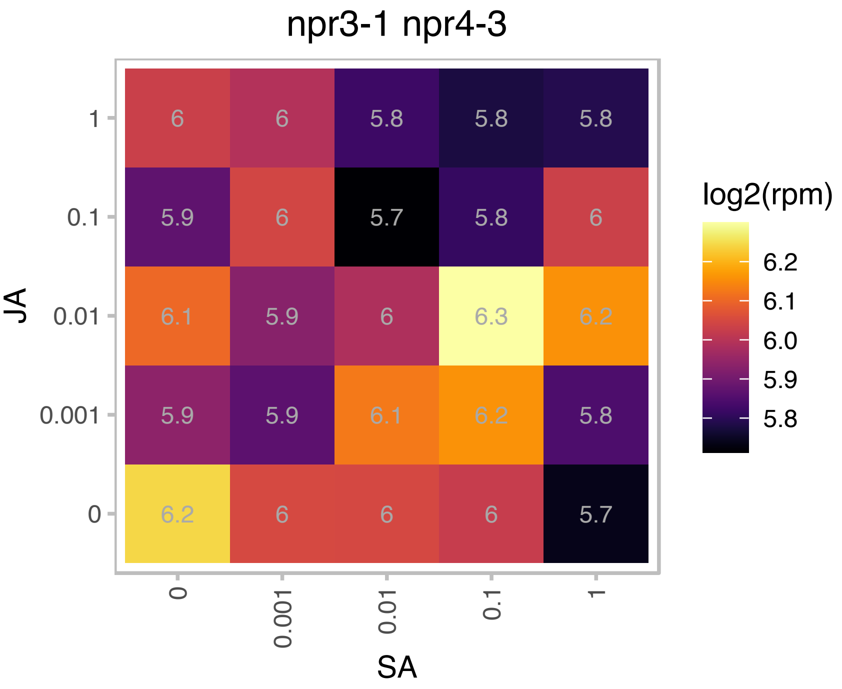

# AT3G47070.1 : NA

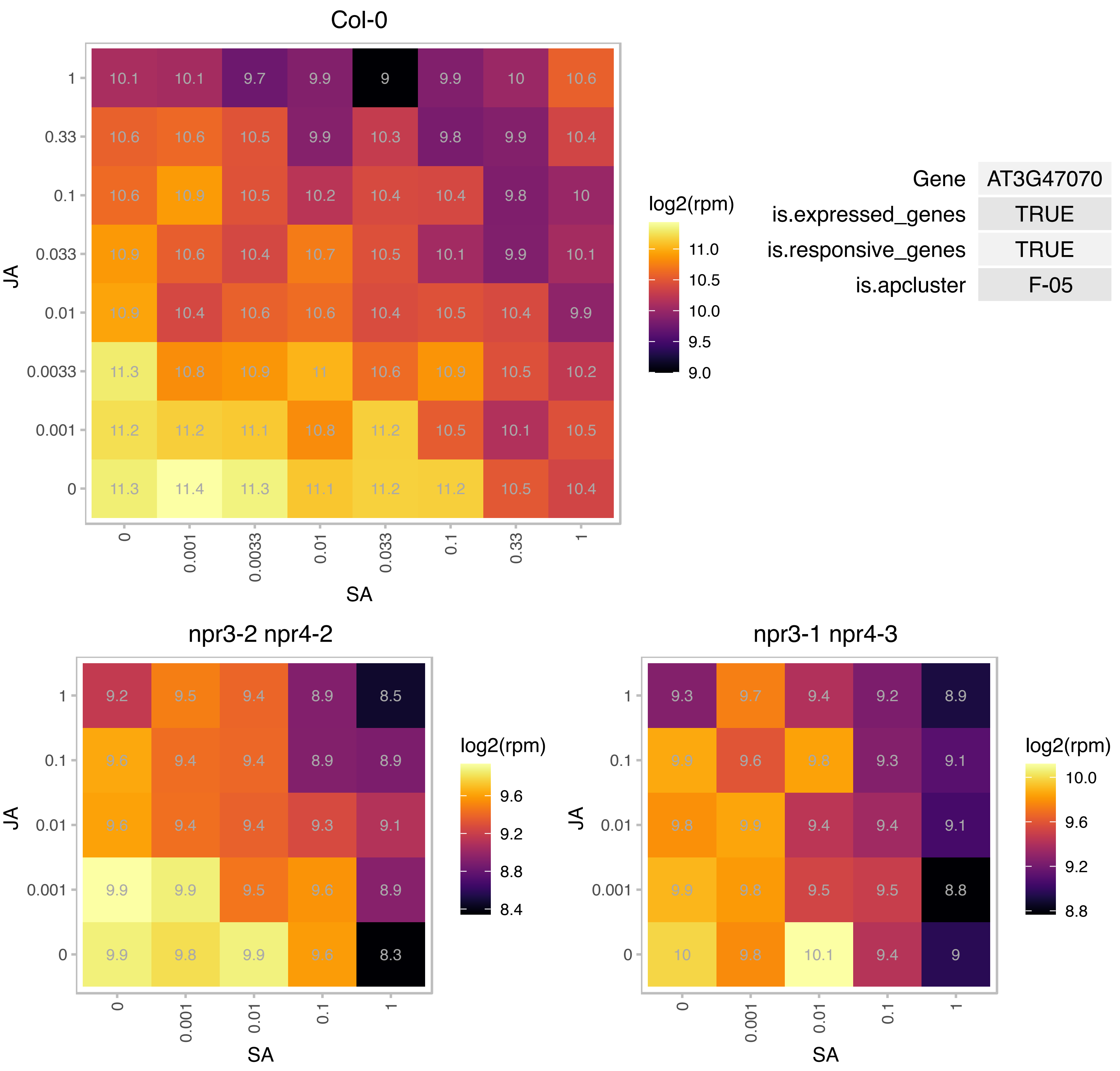

### AT3G47650.1 : BDS2

DnaJ/Hsp40 cysteine-rich domain superfamily protein

Col-0

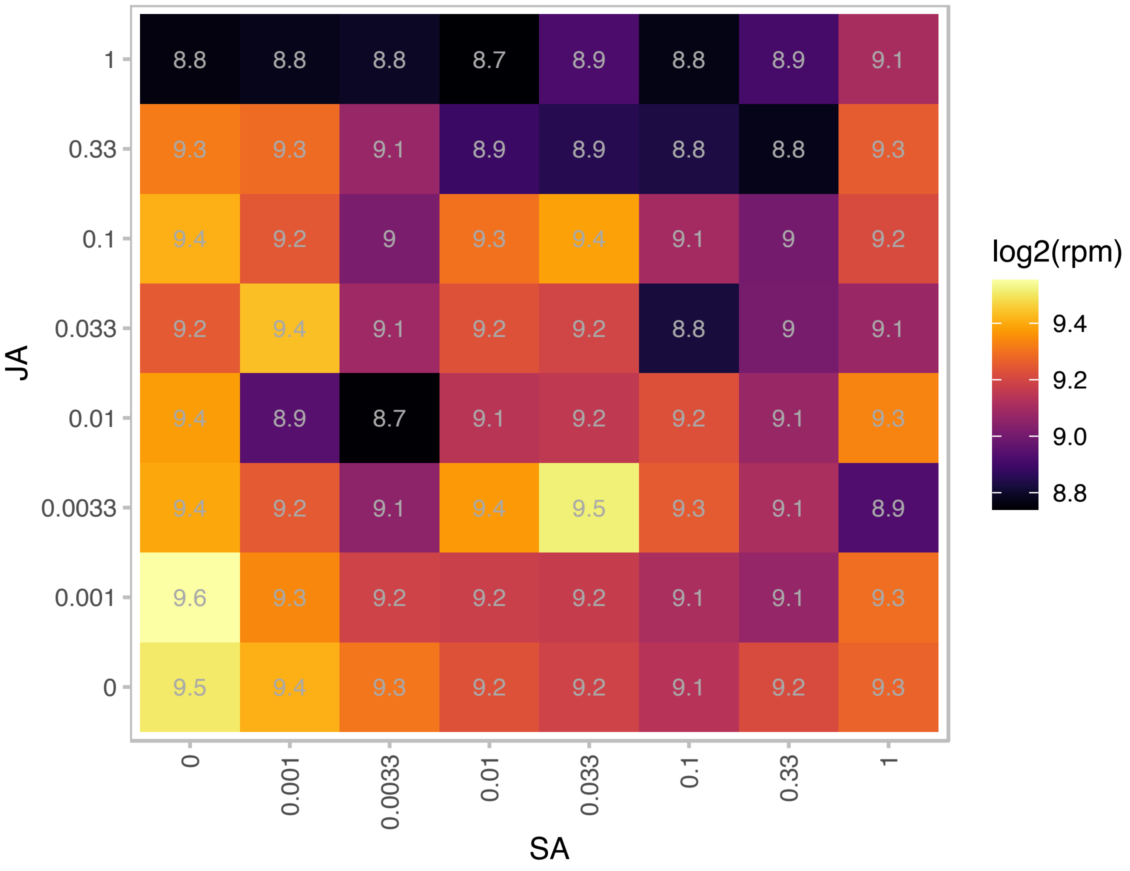


| | |
|---|---|
| Gene | AT3G47650 |
| is.expressed_genes | TRUE |
| is.responsive_genes | TRUE |
| is.apcluster | F-05 |

npr3-2 npr4-2

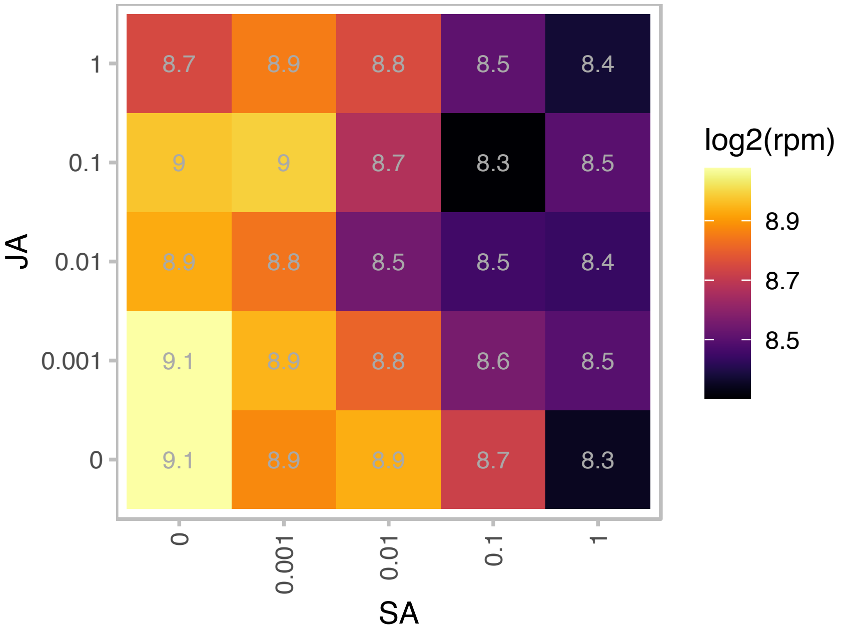


npr3-1 npr4-3

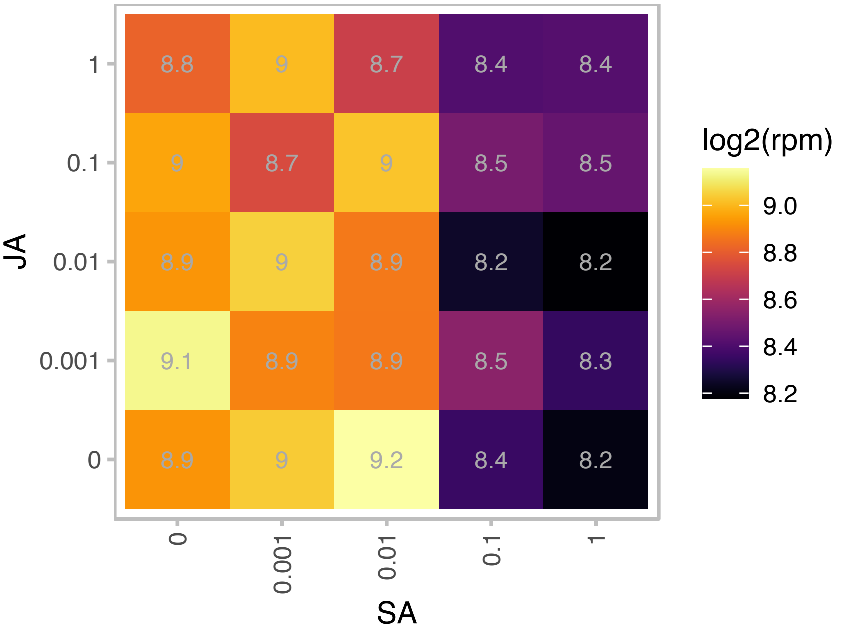

### AT3G48110.1 : EDD, EDD1

glycine-tRNA ligases

glycine-tRNA ligase

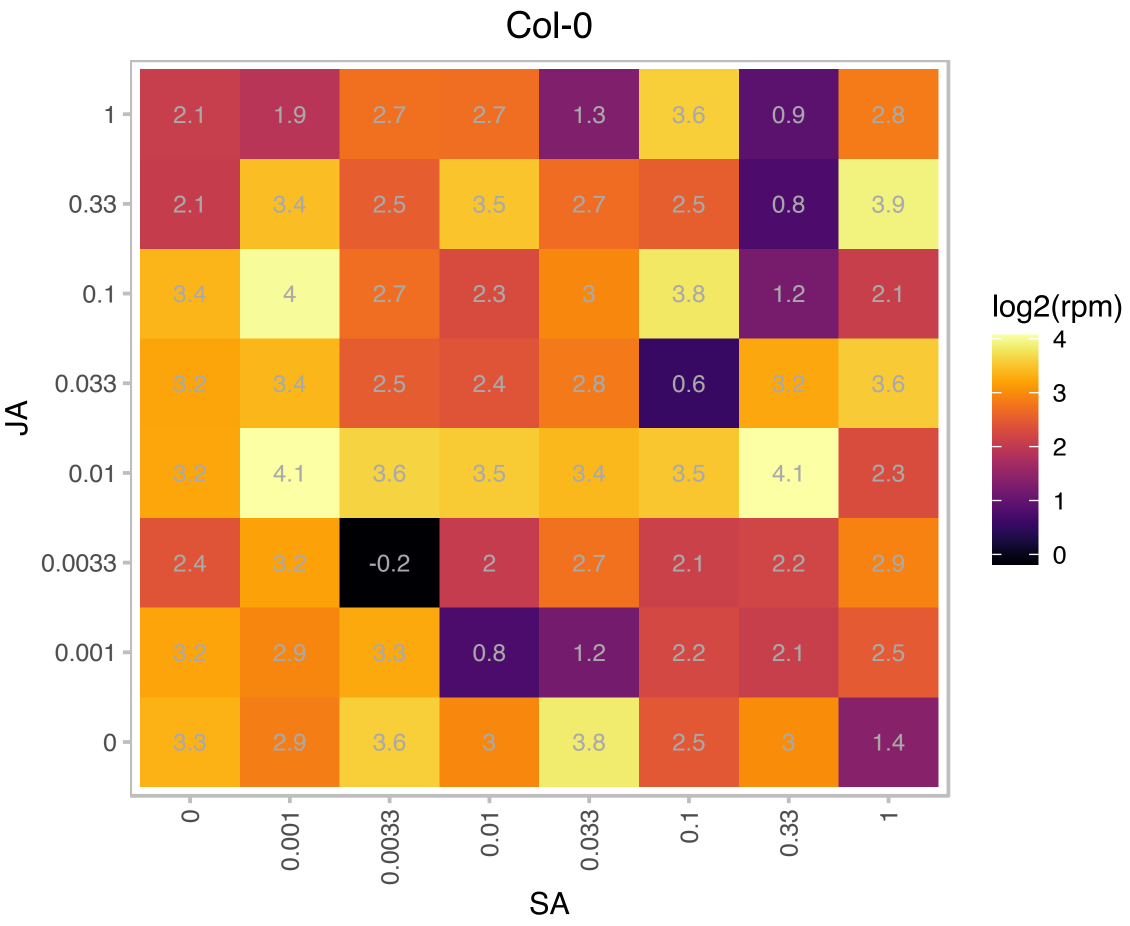


| | |
|---|---|
| Gene | AT3G48110 |
| is.expressed_genes | TRUE |
| is.responsive_genes | TRUE |
| is.apcluster | F-05 |

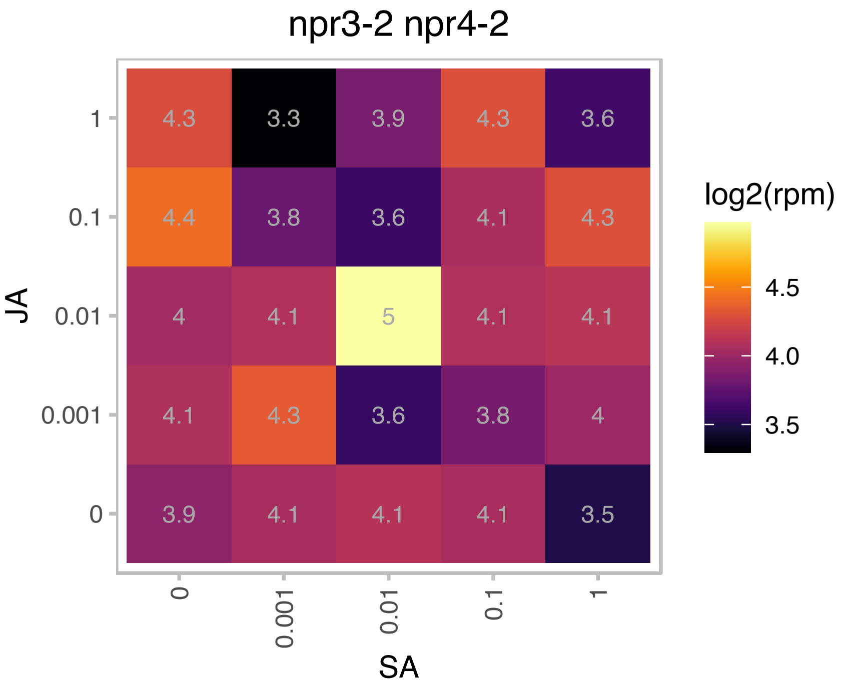


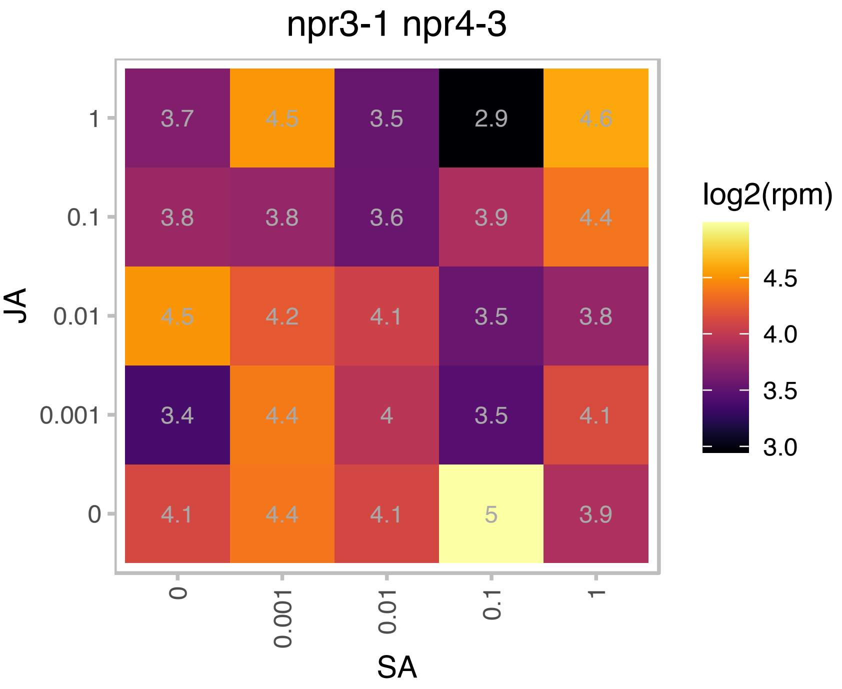

# AT3G49140.1 : NA

Pentatricopeptide repeat (PPR) superfamily protein

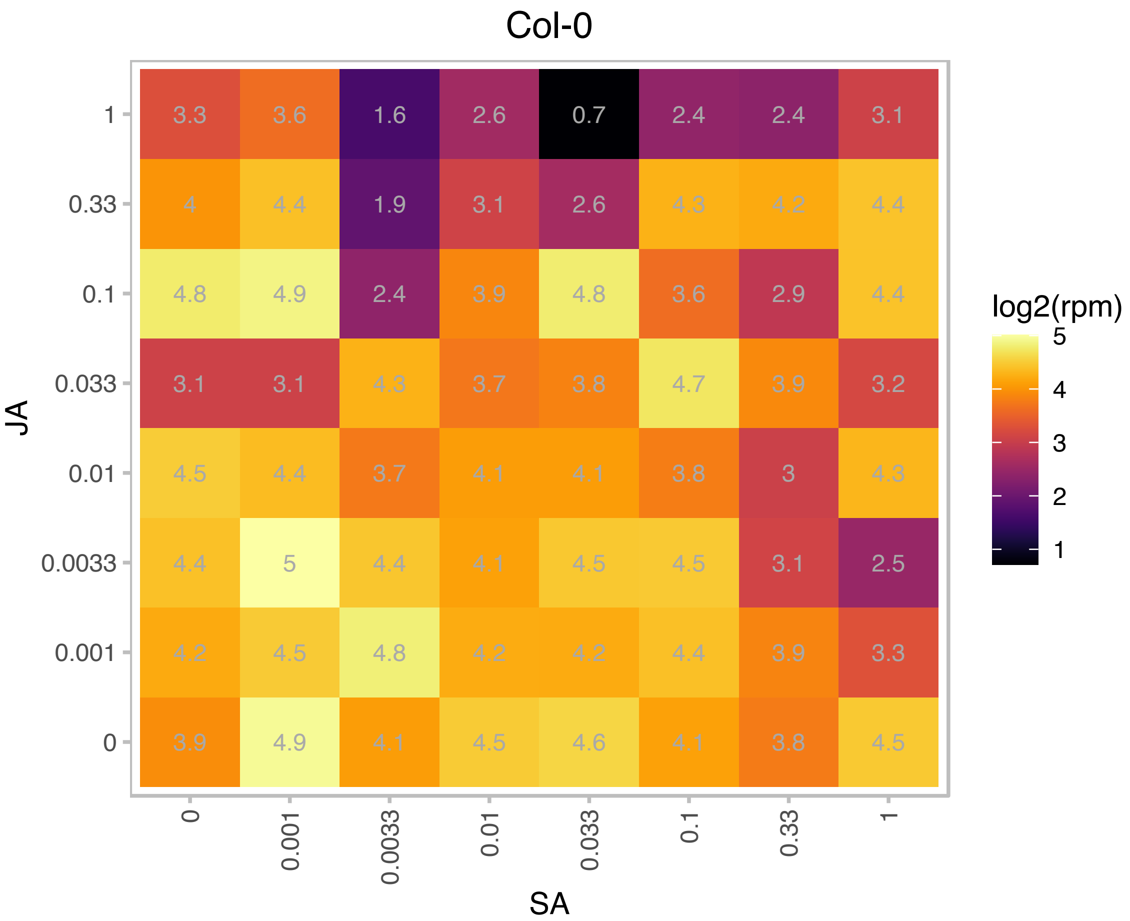


| | |
|---|---|
| Gene | AT3G49140 |
| is.expressed_genes | TRUE |
| is.responsive_genes | TRUE |
| is.apcluster | F-05 |

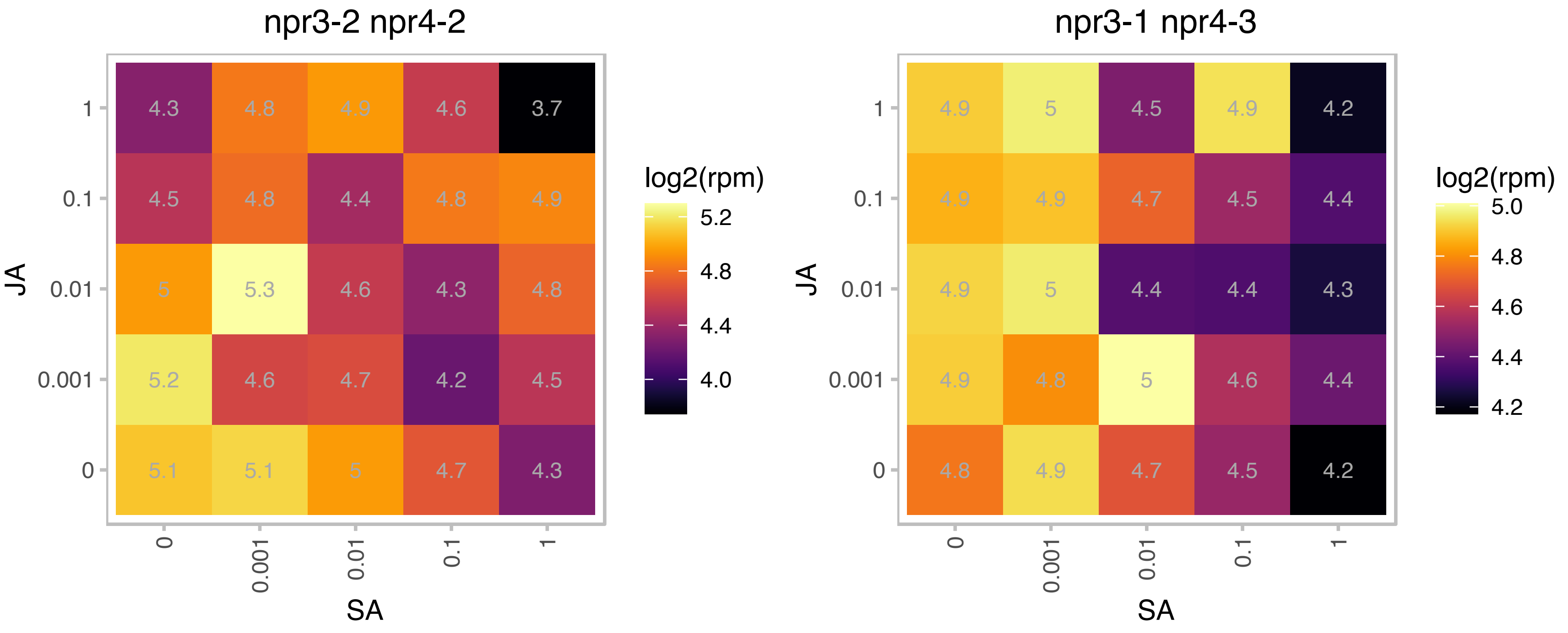

# AT3G50685.1 : NA

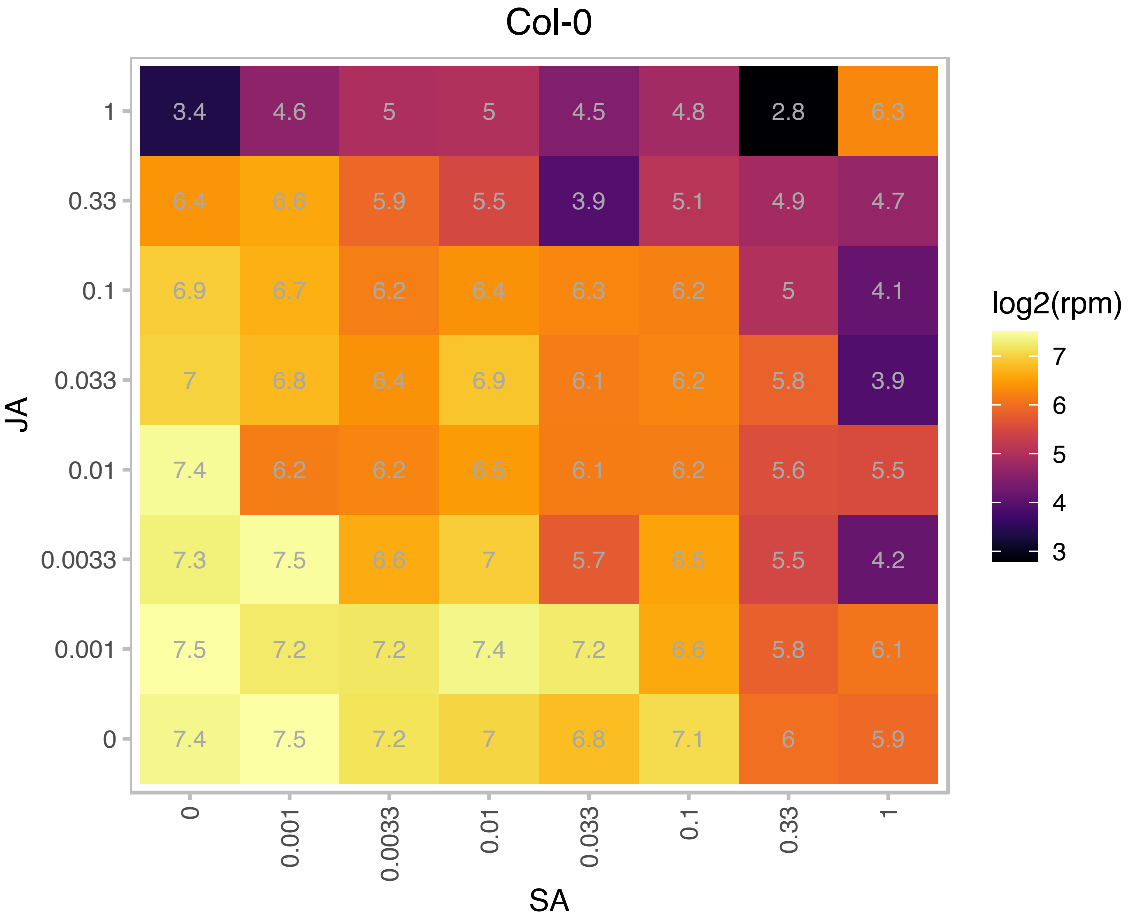


| | |
|---|---|
| Gene | AT3G50685 |
| is.expressed_genes | TRUE |
| is.responsive_genes | TRUE |
| is.apcluster | F-05 |

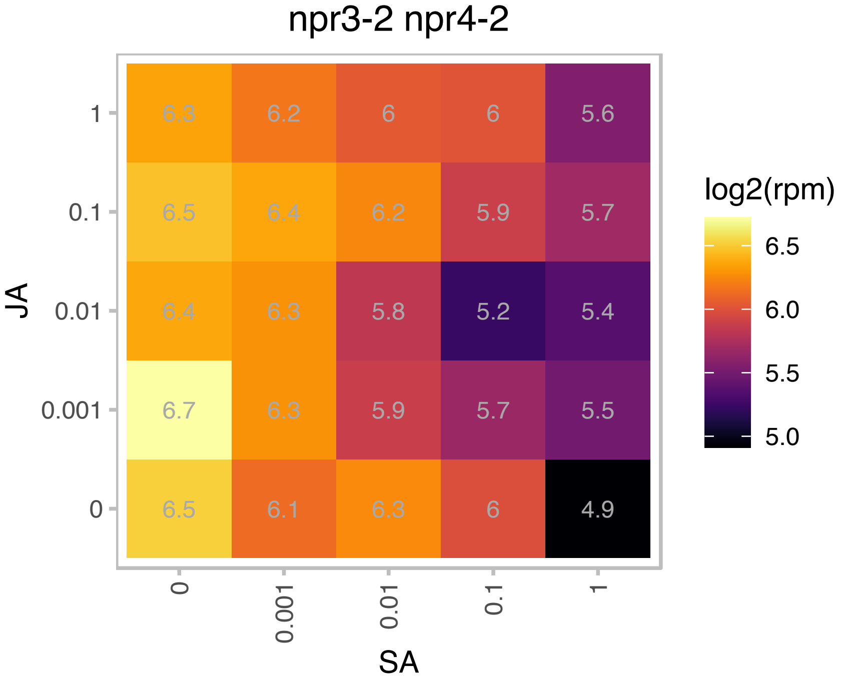


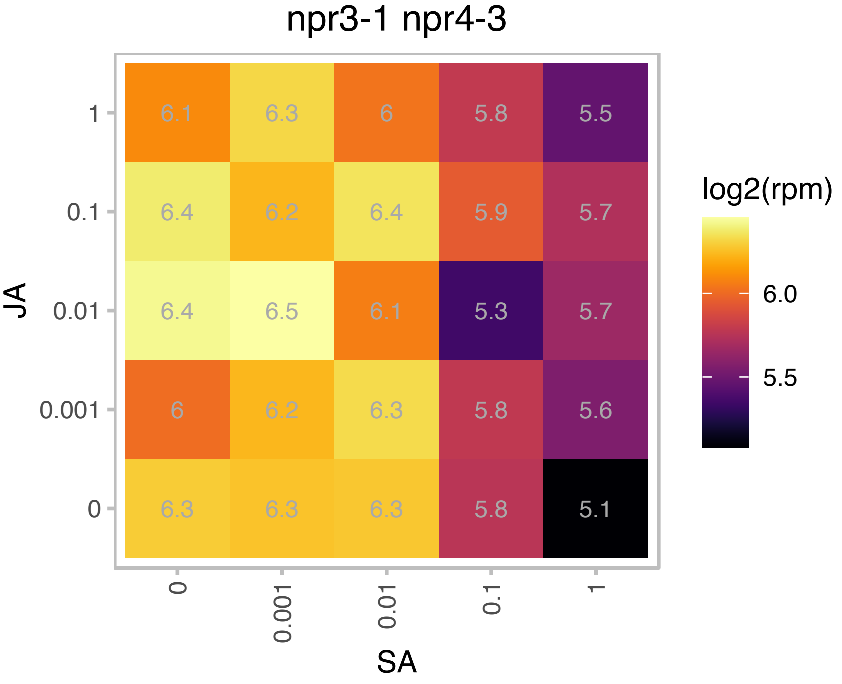

# AT3G52155.1 : NA

Phosphoglycerate mutase family protein

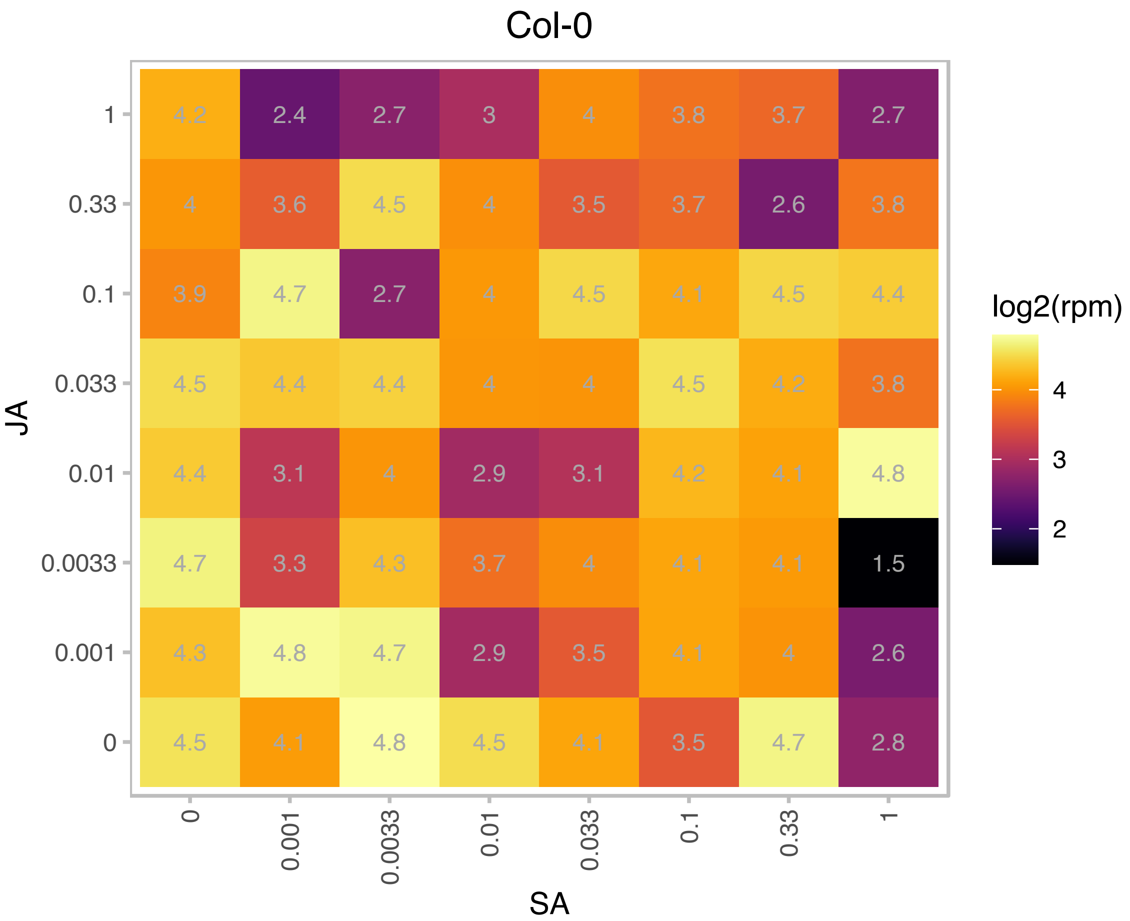


| | |
|---|---|
| Gene | AT3G52155 |
| is.expressed_genes | TRUE |
| is.responsive_genes | TRUE |
| is.apcluster | F-05 |

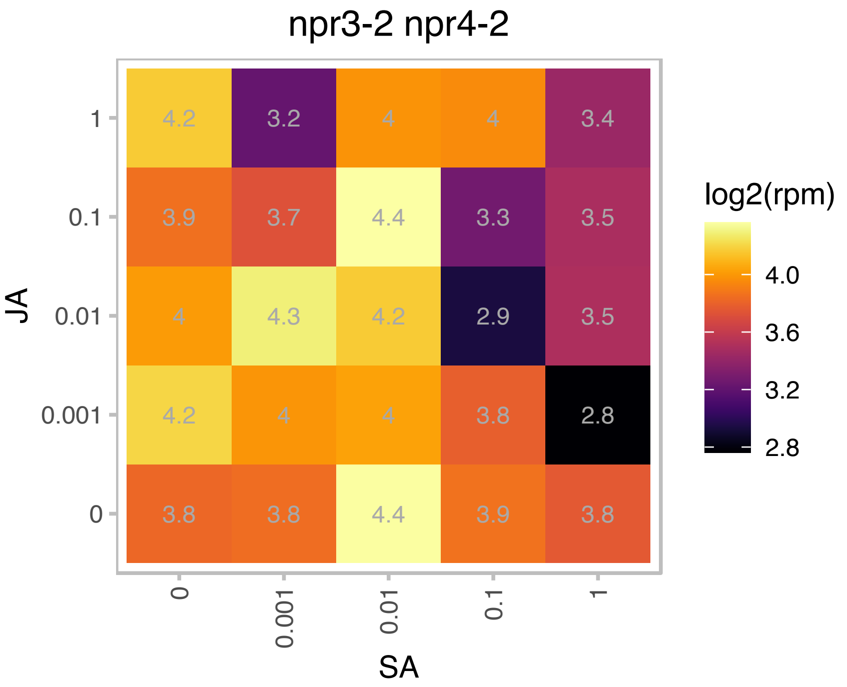


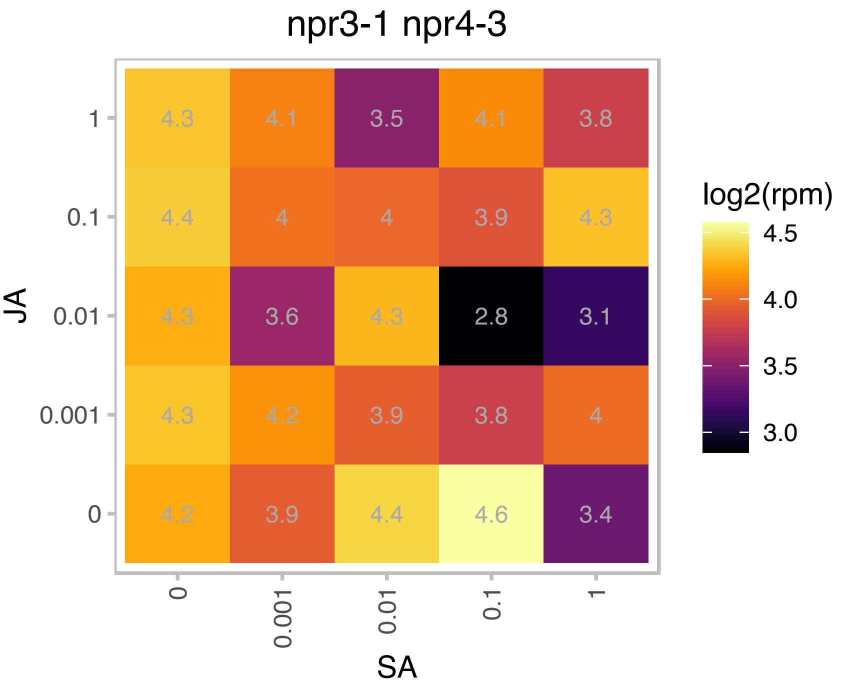

### AT3G53130.1 : CYP97C1, LUT1

Cytochrome P450 superfamily protein

Lutein-deficient 1 (LUT1) required for lutein biosynthesis, member of the xanthophyll class of carotenoids. Involved in epsilon ring hydroxylation. Maps at 67.3 cM on chromosome 3.

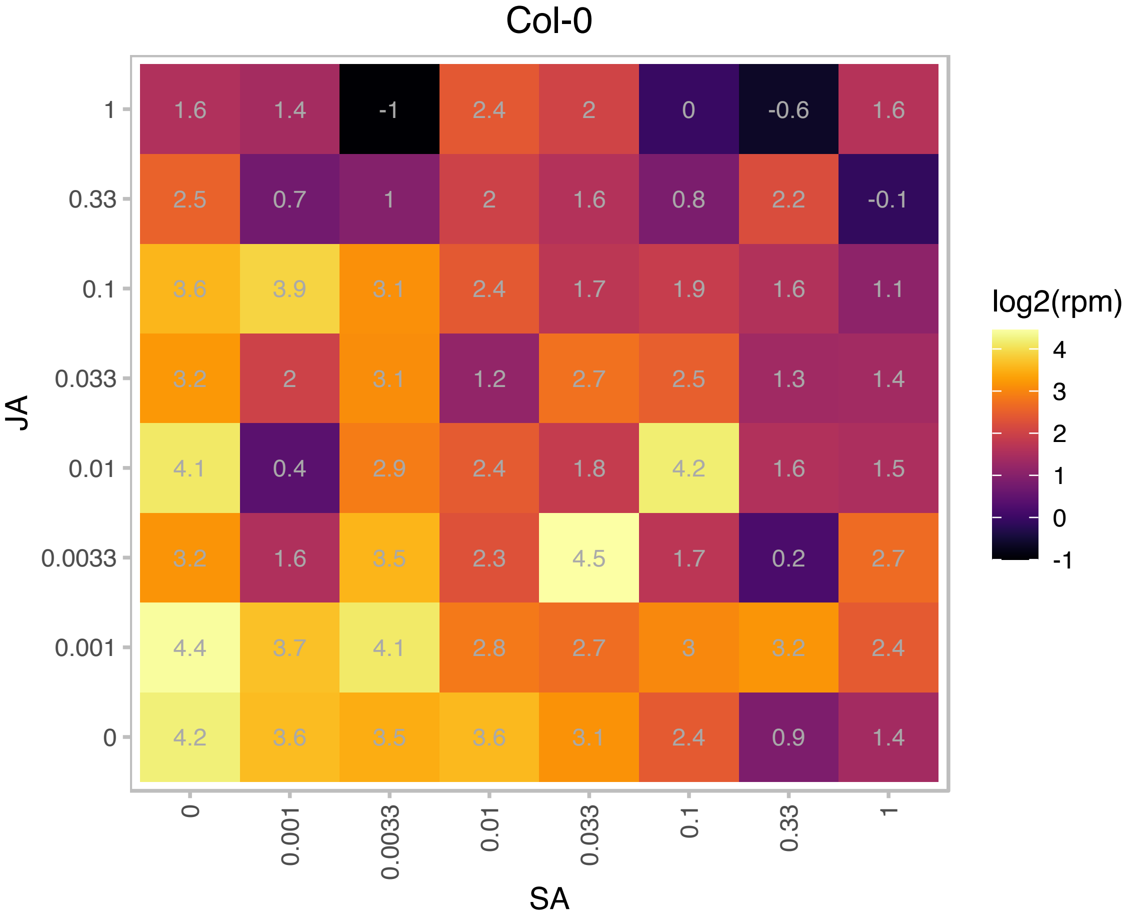


| | |
|---|---|
| Gene | AT3G53130 |
| is.expressed_genes | TRUE |
| is.responsive_genes | TRUE |
| is.apcluster | F-05 |

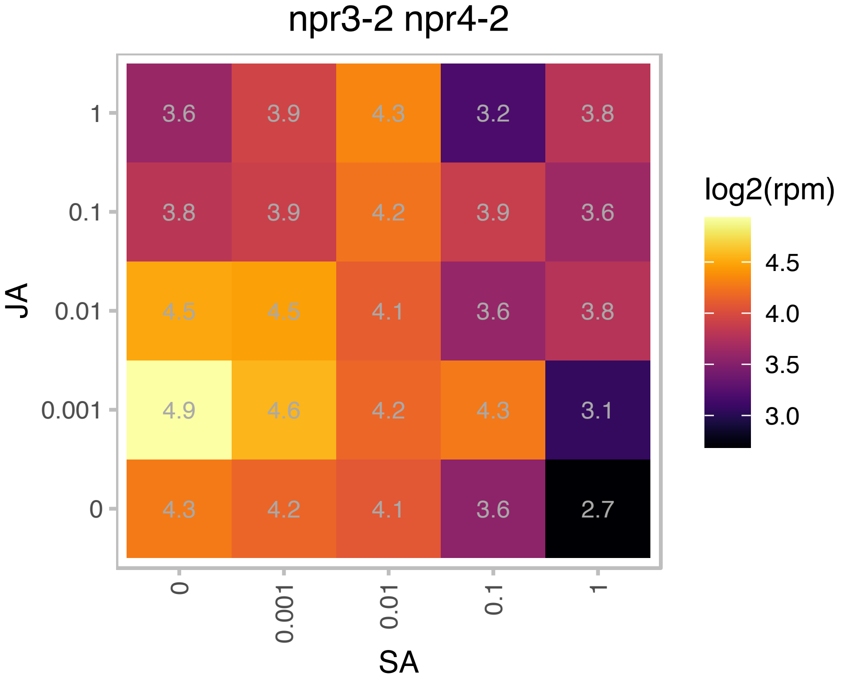


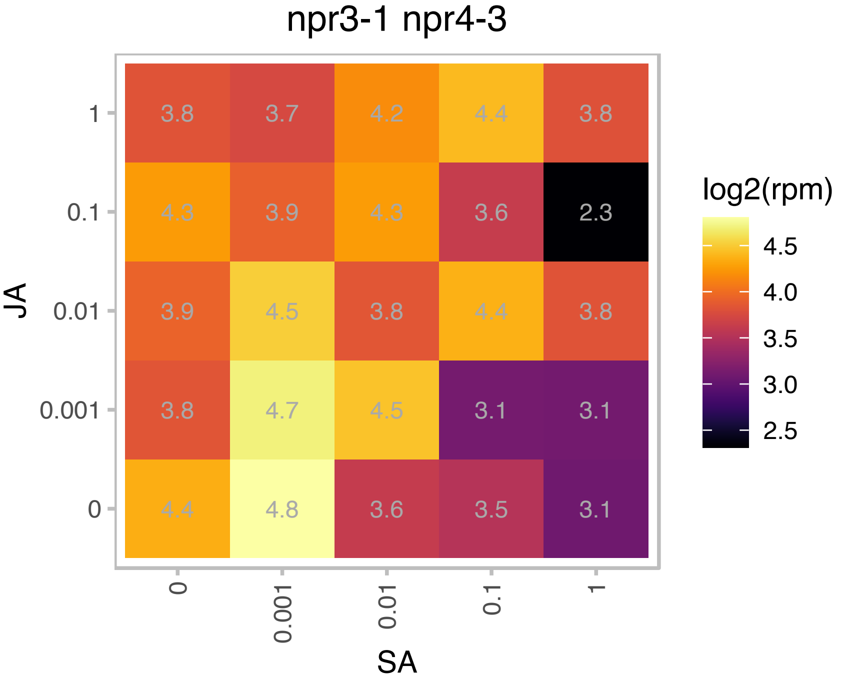

### AT3G53530.1 : NAKR3

Chloroplast-targeted copper chaperone protein

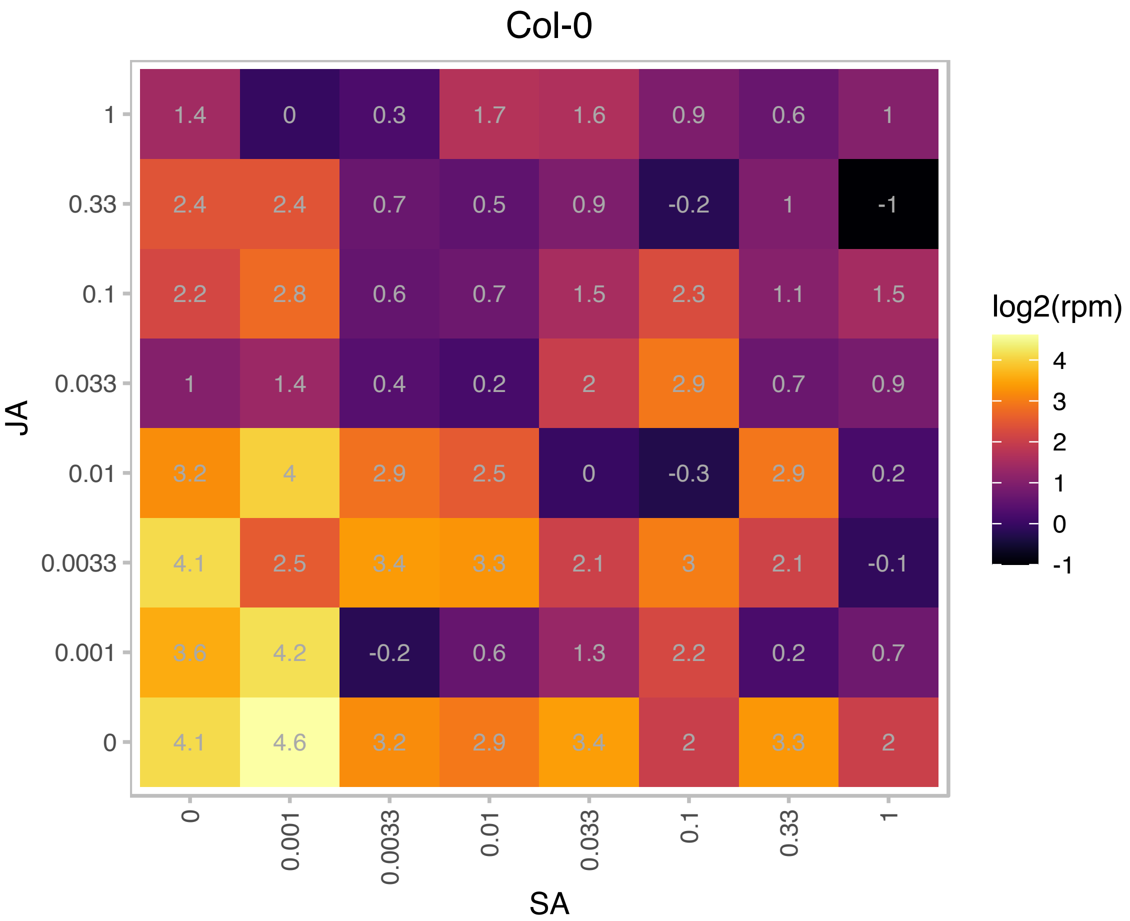


| | |
|---|---|
| Gene | AT3G53530 |
| is.expressed_genes | TRUE |
| is.responsive_genes | TRUE |
| is.apcluster | F-05 |

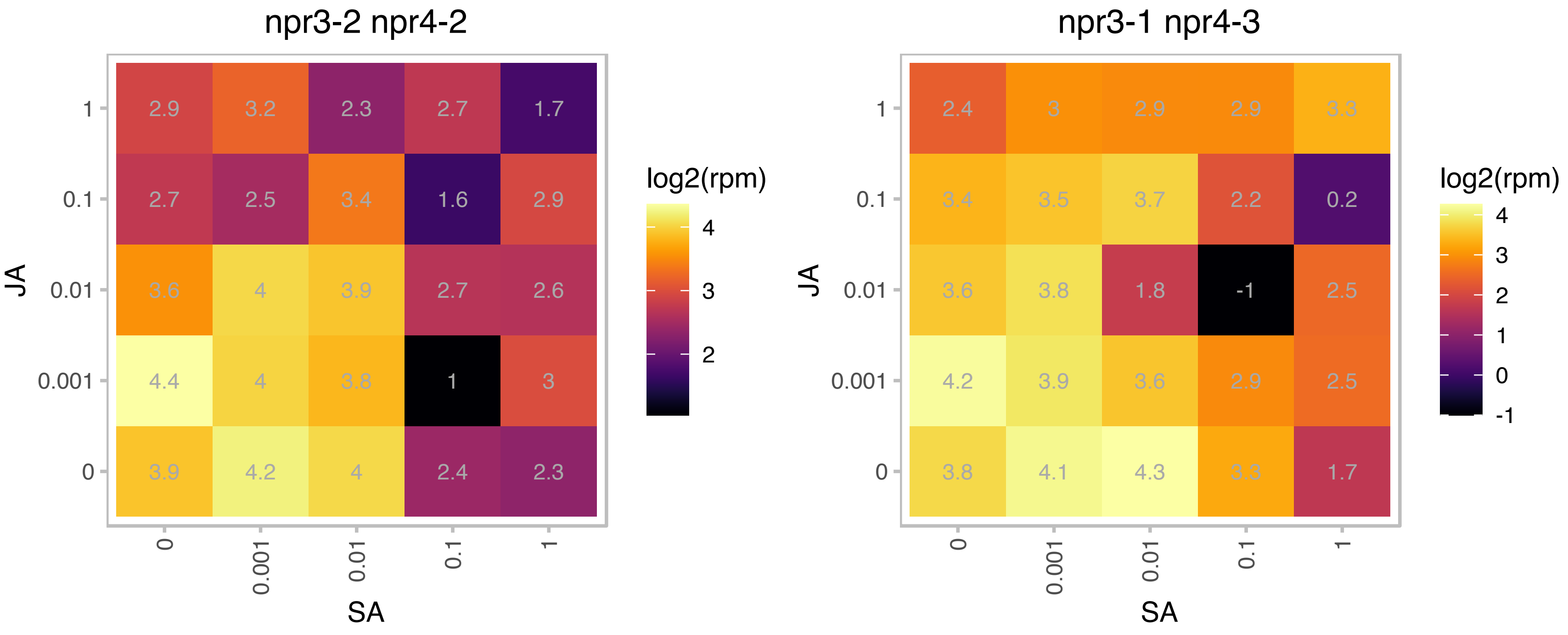

# AT3G56010.1 : NA

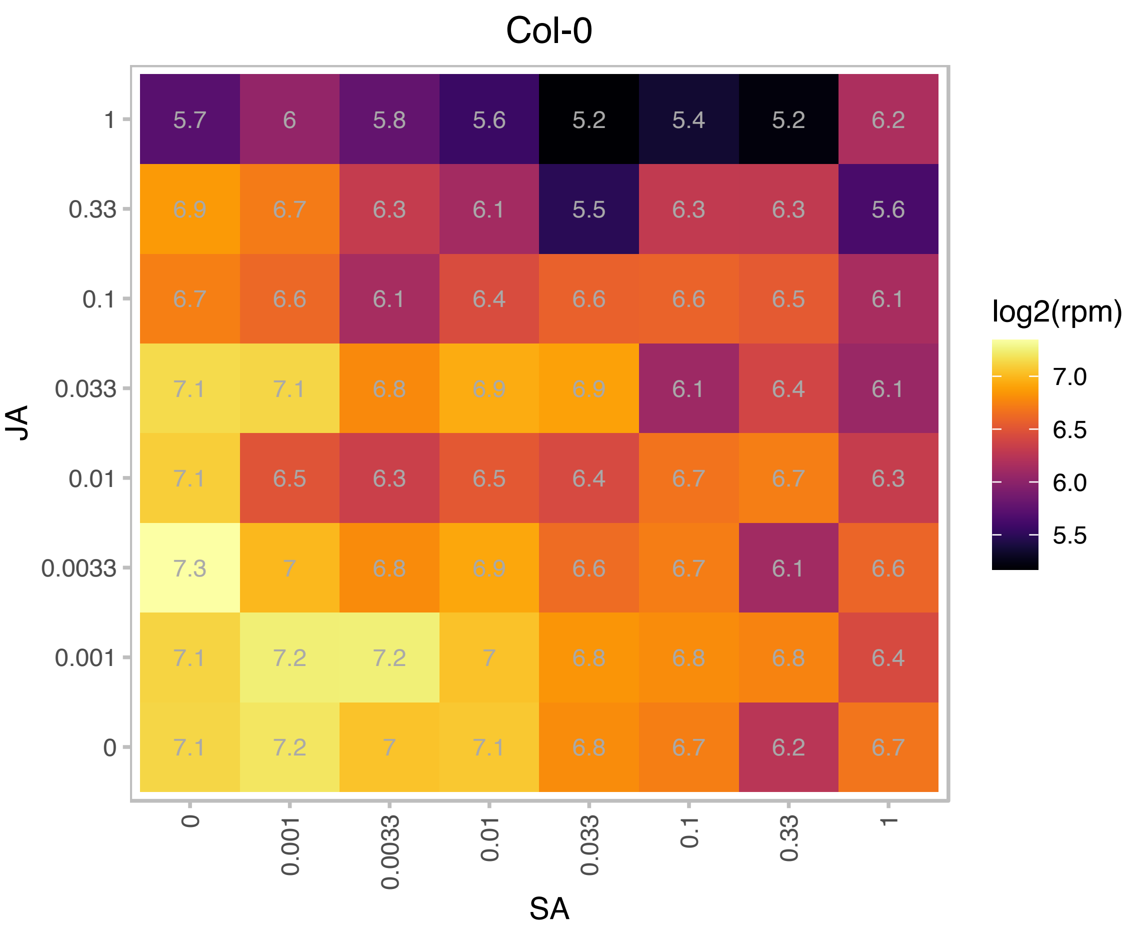


| | |
|---|---|
| Gene | AT3G56010 |
| is.expressed_genes | TRUE |
| is.responsive_genes | TRUE |
| is.apcluster | F-05 |

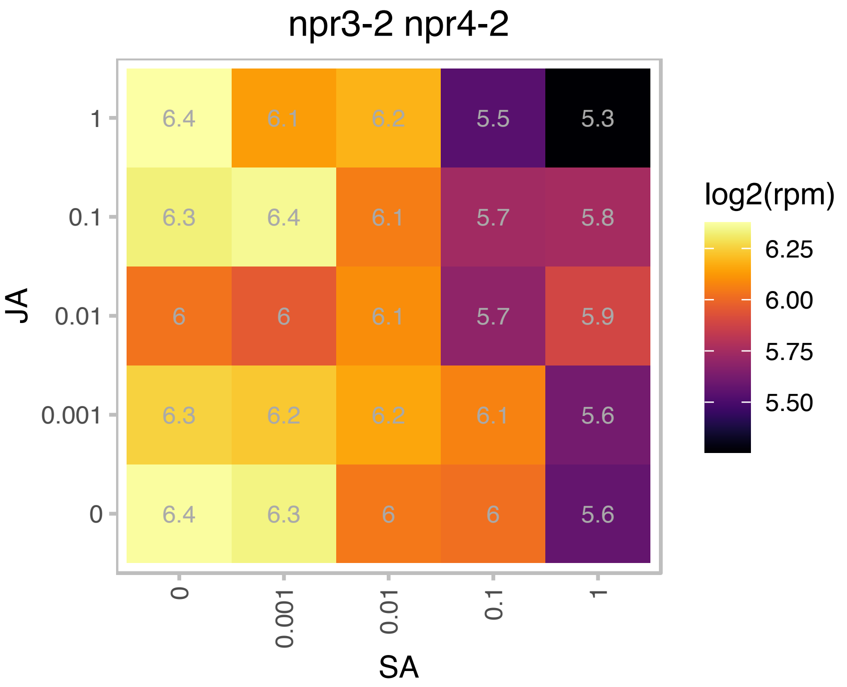


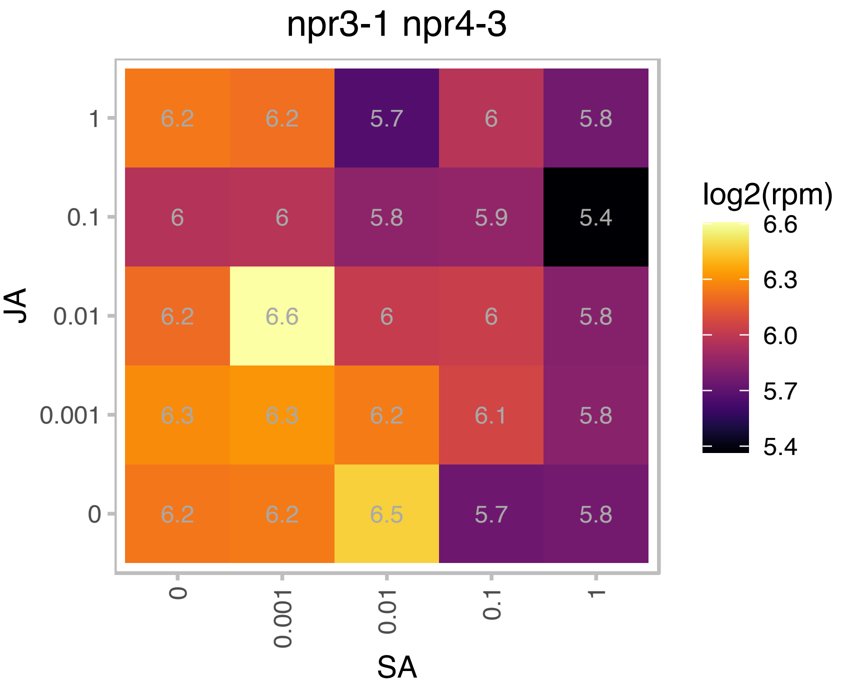

### AT3G56910.1 : PSRP5

plastid-specific 50S ribosomal protein 5

Encodes PSRP5 (PLASTID-SPECIFIC 50S RIBOSOMAL PROTEIN 5). Functions in plastid translation.

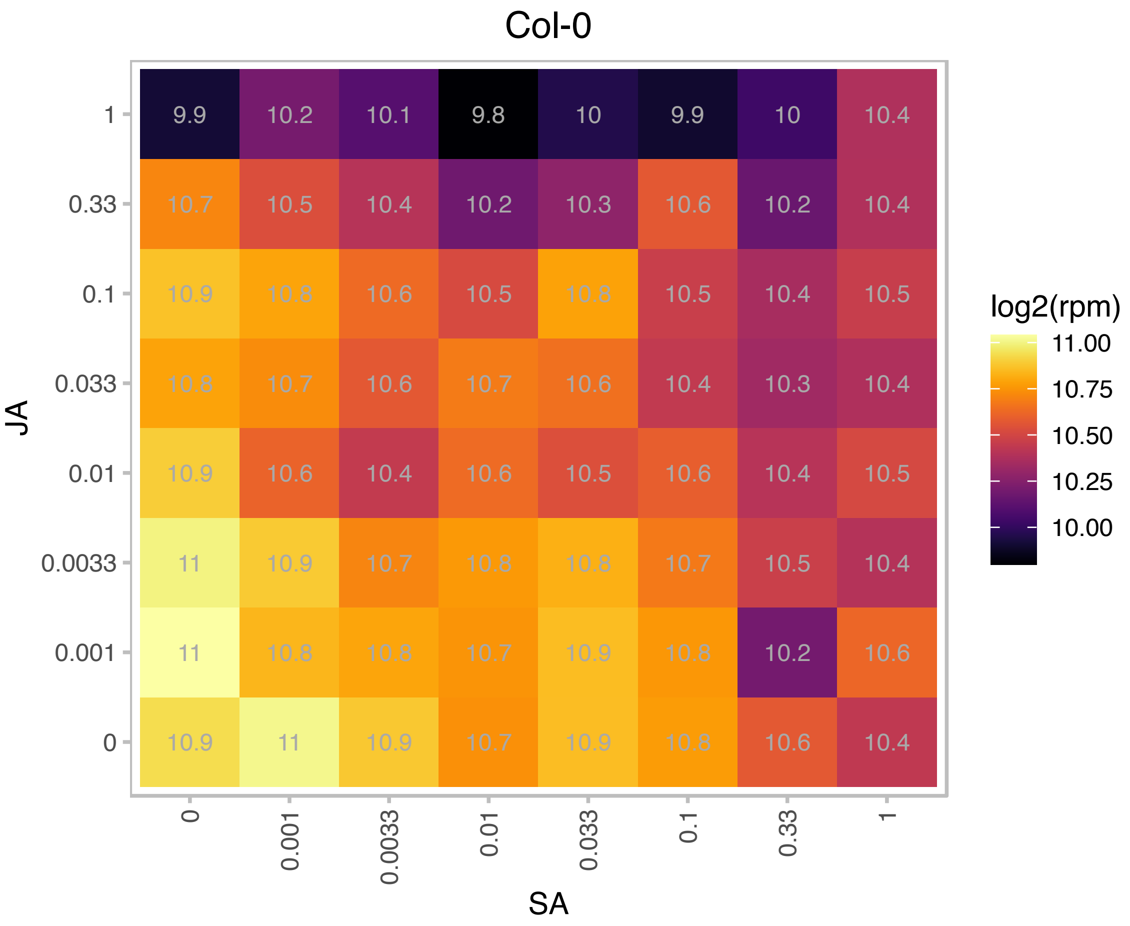


| | |
|---|---|
| Gene | AT3G56910 |
| is.expressed_genes | TRUE |
| is.responsive_genes | TRUE |
| is.apcluster | F-05 |

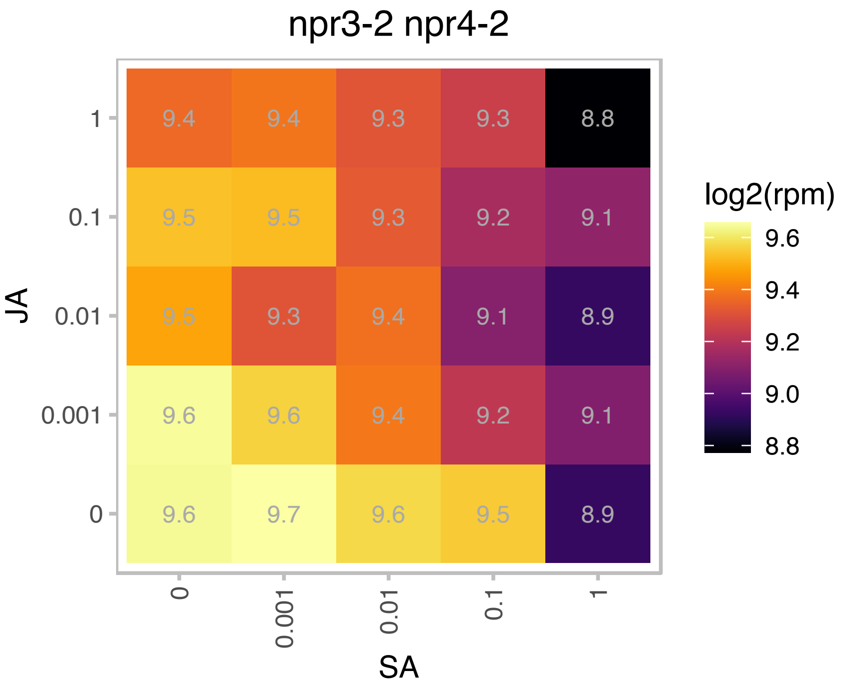


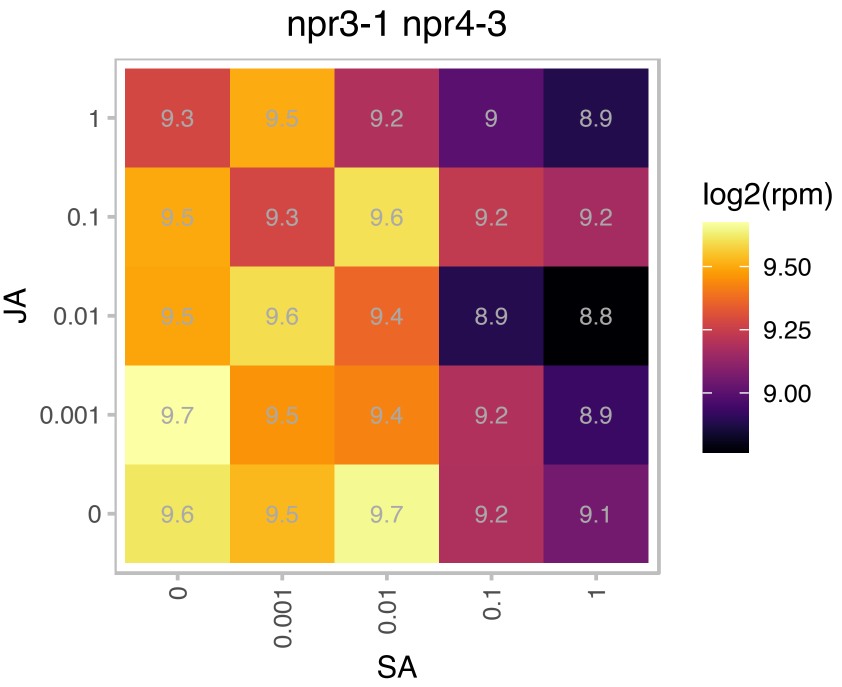

### AT4G00050.1 : UNE10

basic helix-loop-helix (bHLH) DNA-binding superfamily protein

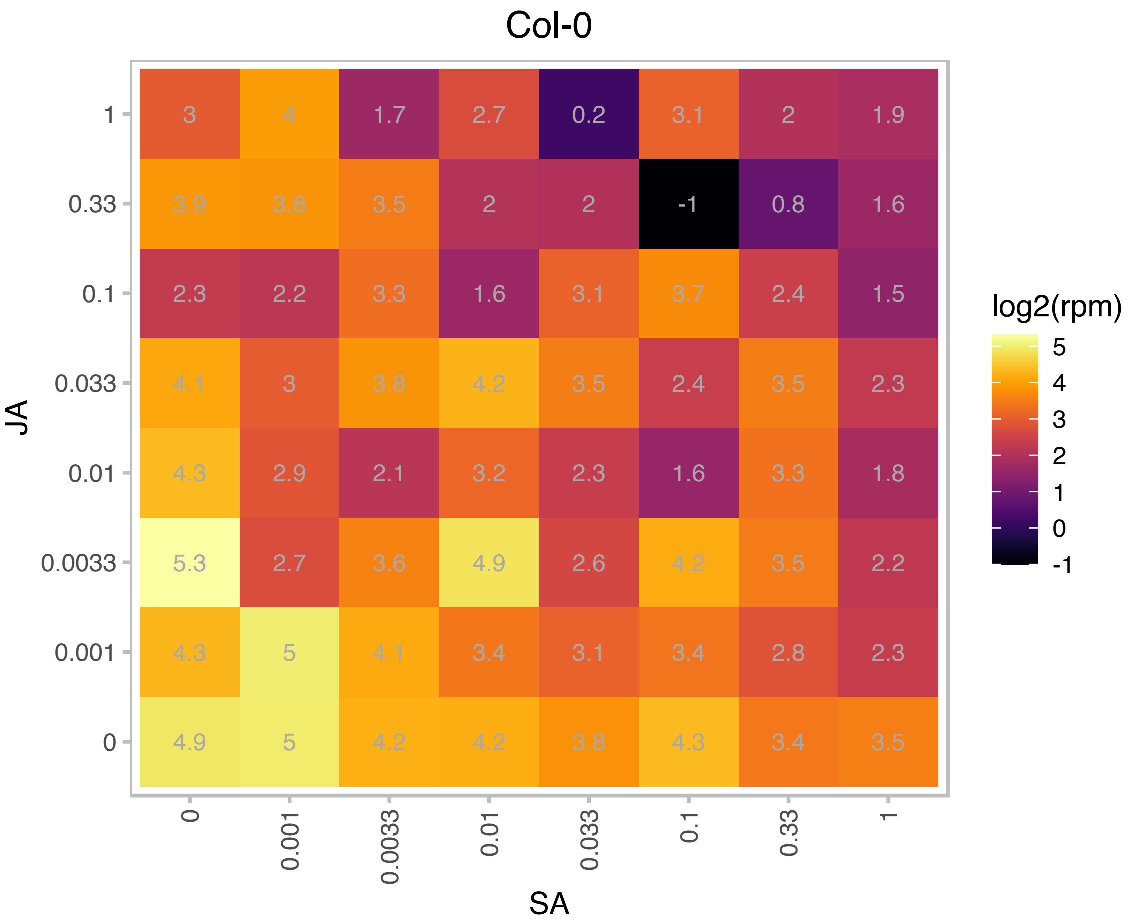


| | |
|---|---|
| Gene | AT4G00050 |
| is.expressed_genes | TRUE |
| is.responsive_genes | TRUE |
| is.apcluster | F-05 |

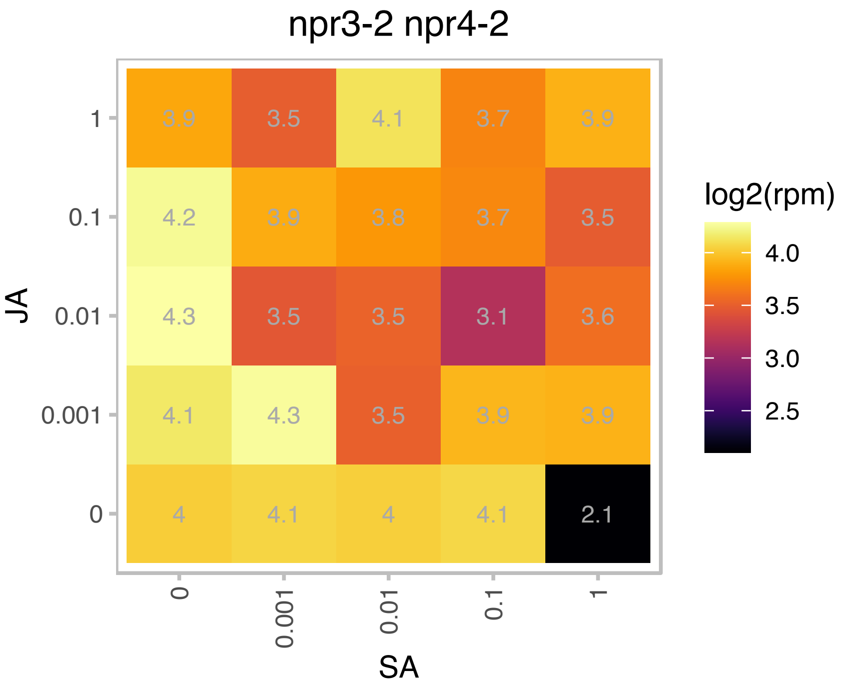


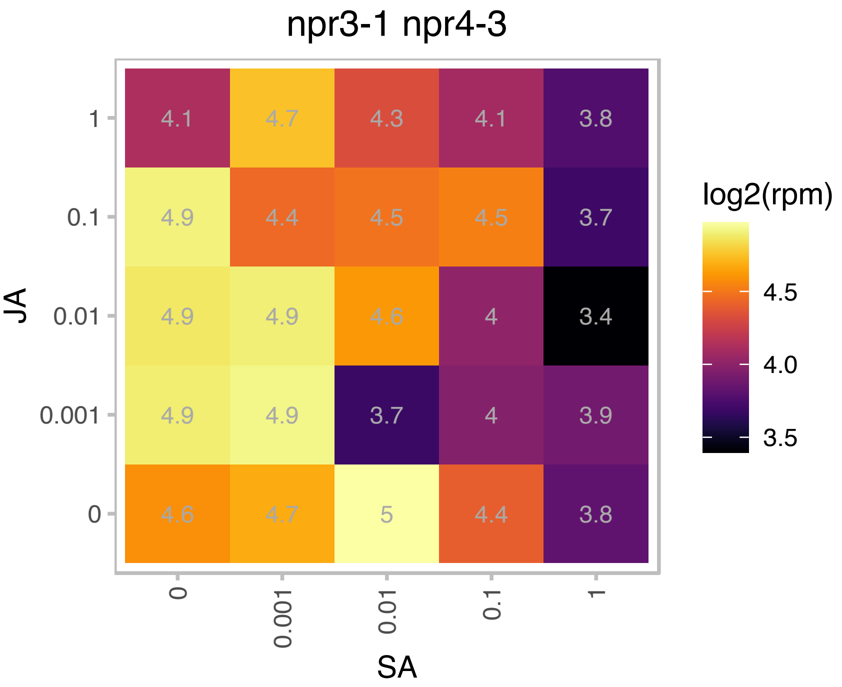

### AT4G01037.1 : AtWTF1, WTF1

Ubiquitin carboxyl-terminal hydrolase family protein

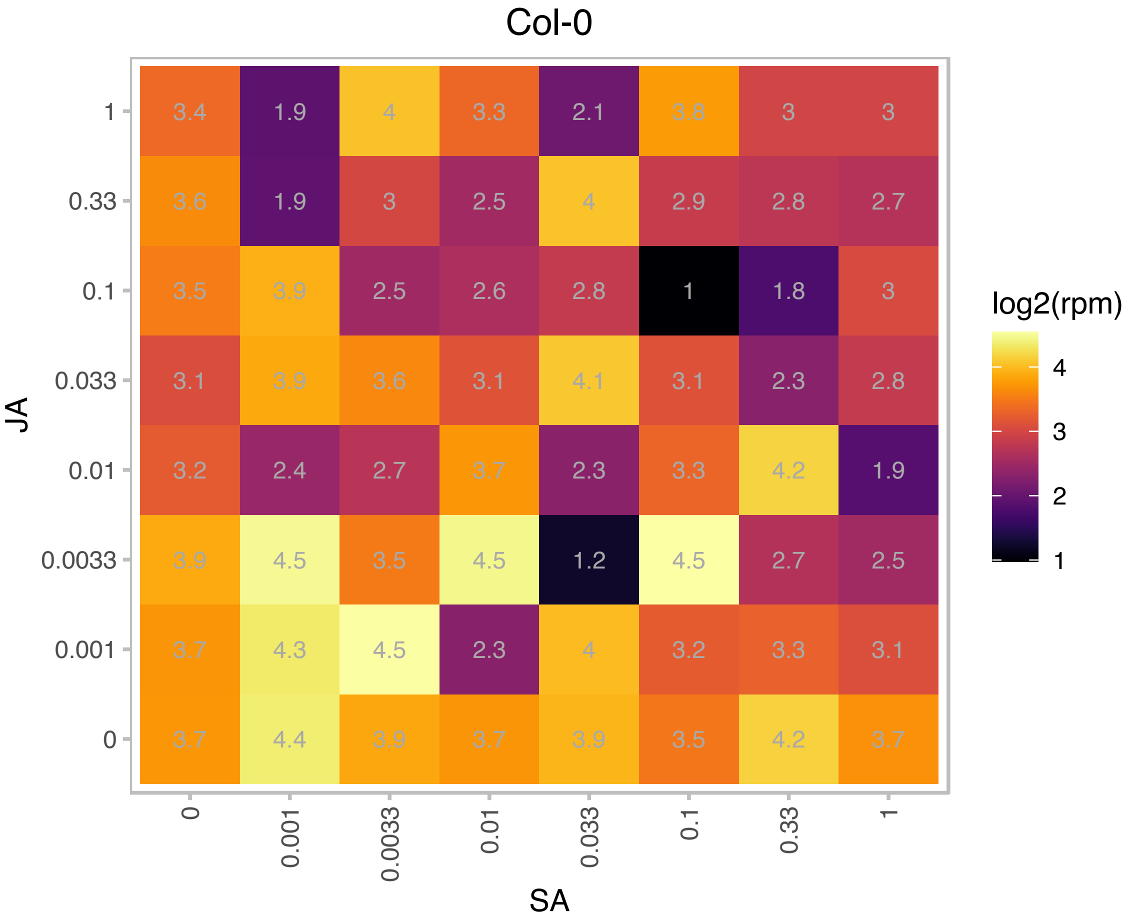


| | |
|---|---|
| Gene | AT4G01037 |
| is.expressed_genes | TRUE |
| is.responsive_genes | TRUE |
| is.apcluster | F-05 |

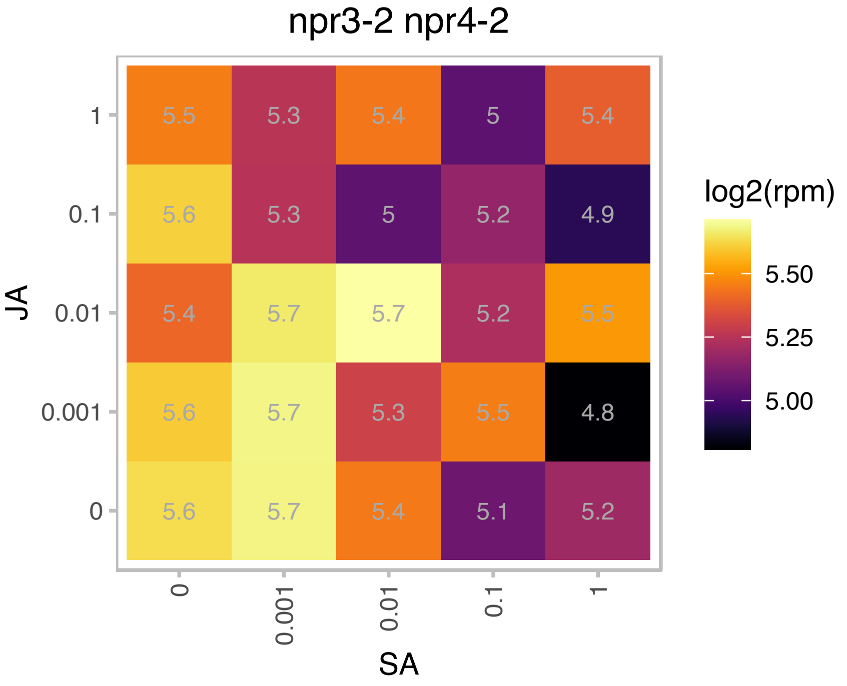


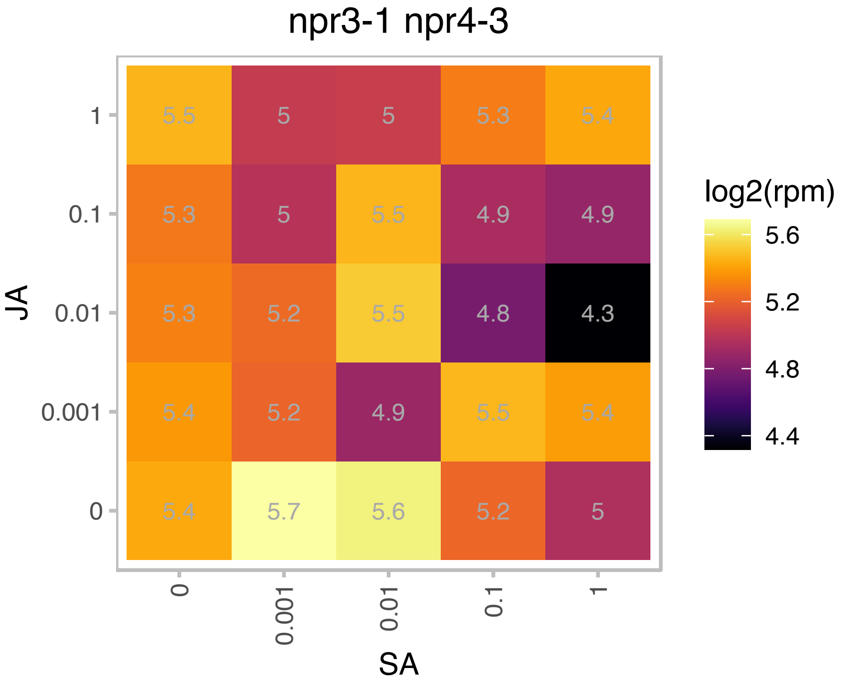

# AT4G01935.1 : NA

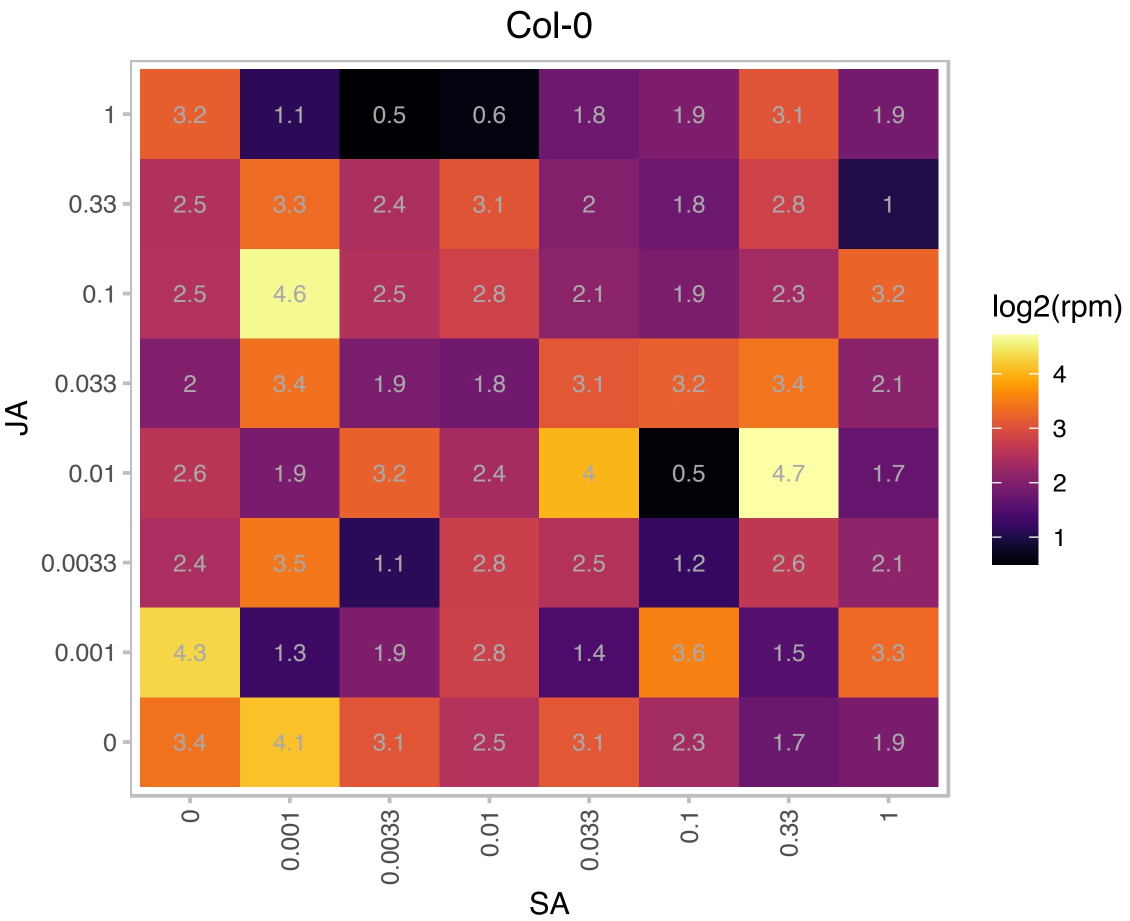


| | |
|---|---|
| Gene | AT4G01935 |
| is.expressed_genes | TRUE |
| is.responsive_genes | TRUE |
| is.apcluster | F-05 |

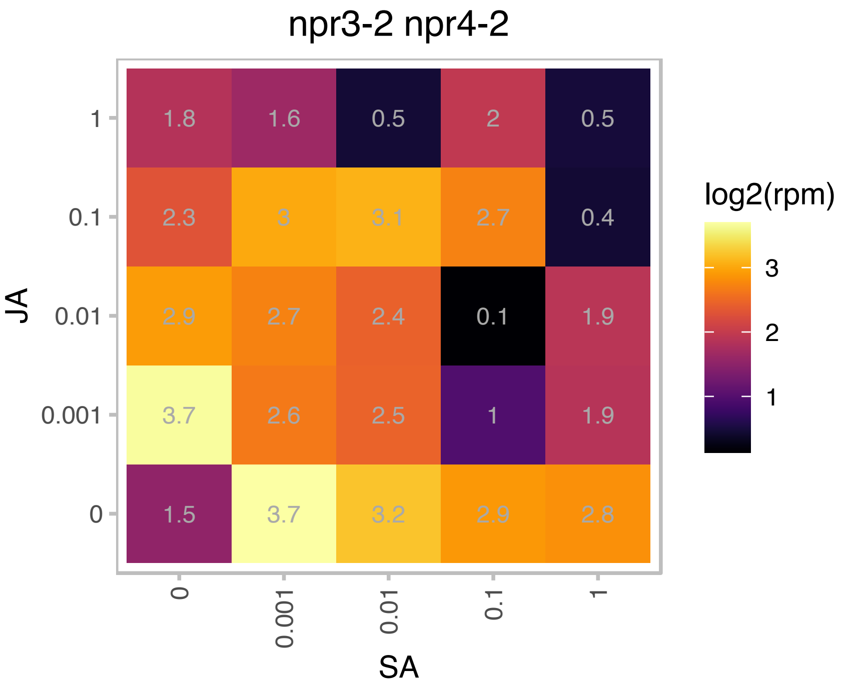


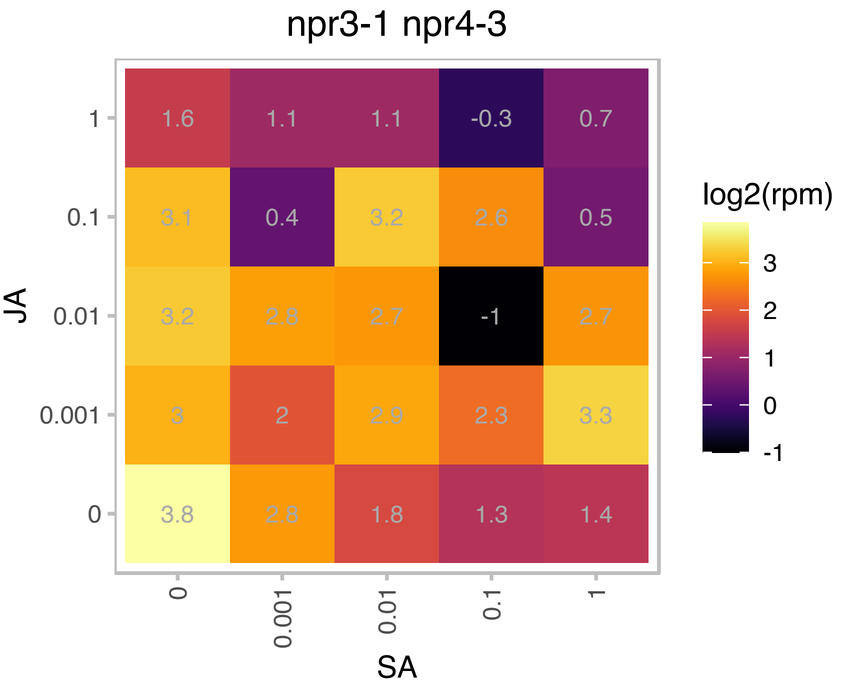

### AT4G04780.1 : MED21

mediator 21
Encodes the med21 subunit of the mediator complex which is involved in transcriptional regulation. MED21 interacts physically with the E3 ligase HUB1 and this interaction may be important in mediation defense responses to fungal pathogens.

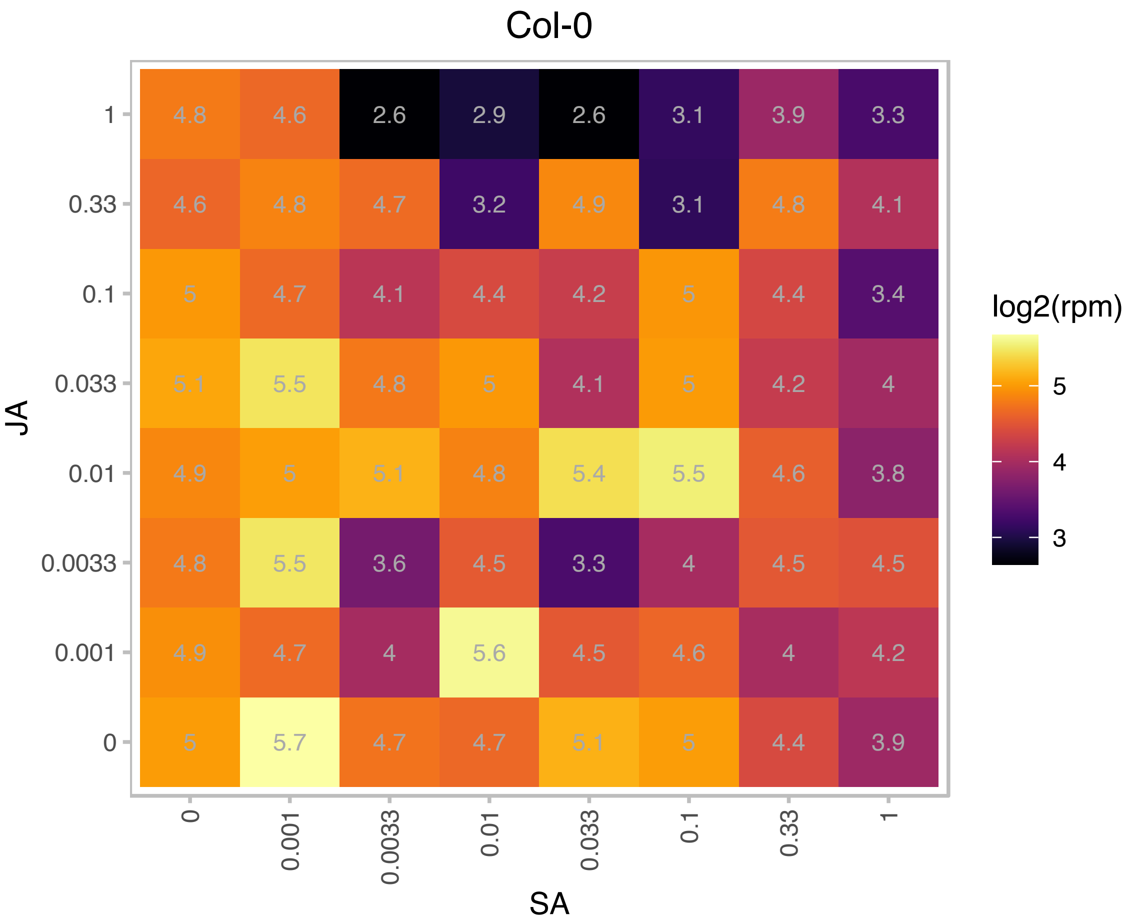


| | |
|---|---|
| Gene | AT4G04780 |
| is.expressed_genes | TRUE |
| is.responsive_genes | TRUE |
| is.apcluster | F-05 |

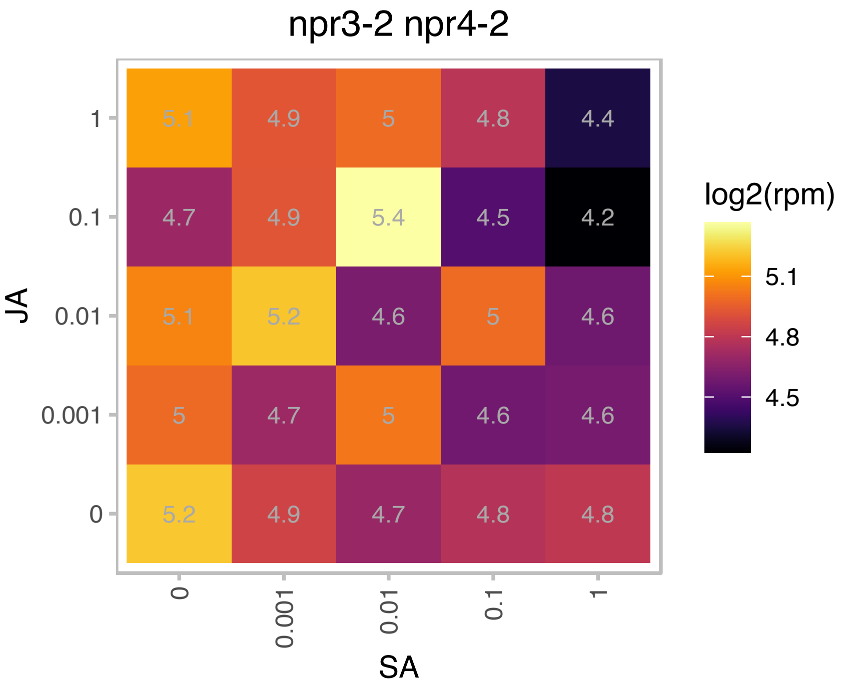


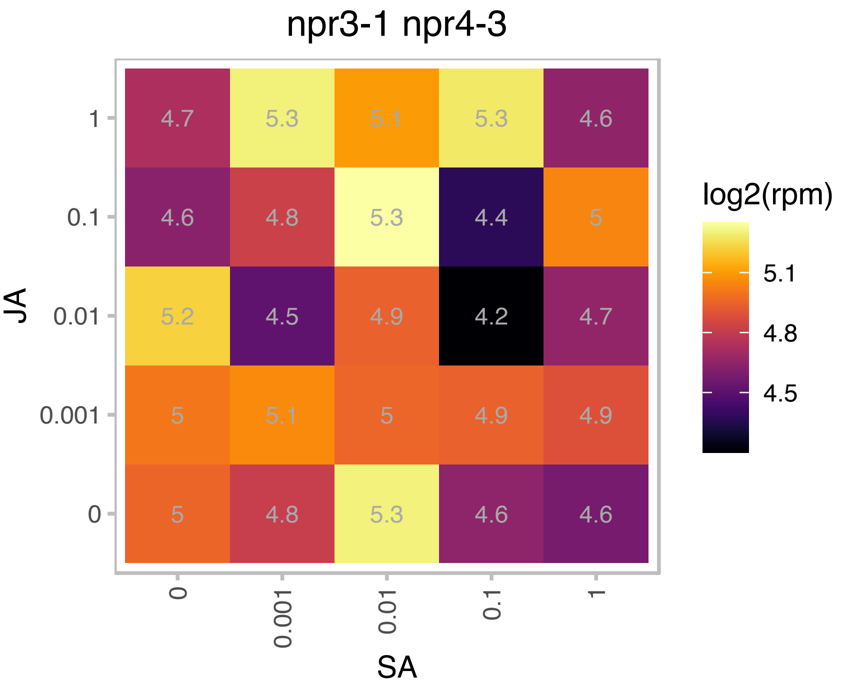

# AT4G13200.1 : NA

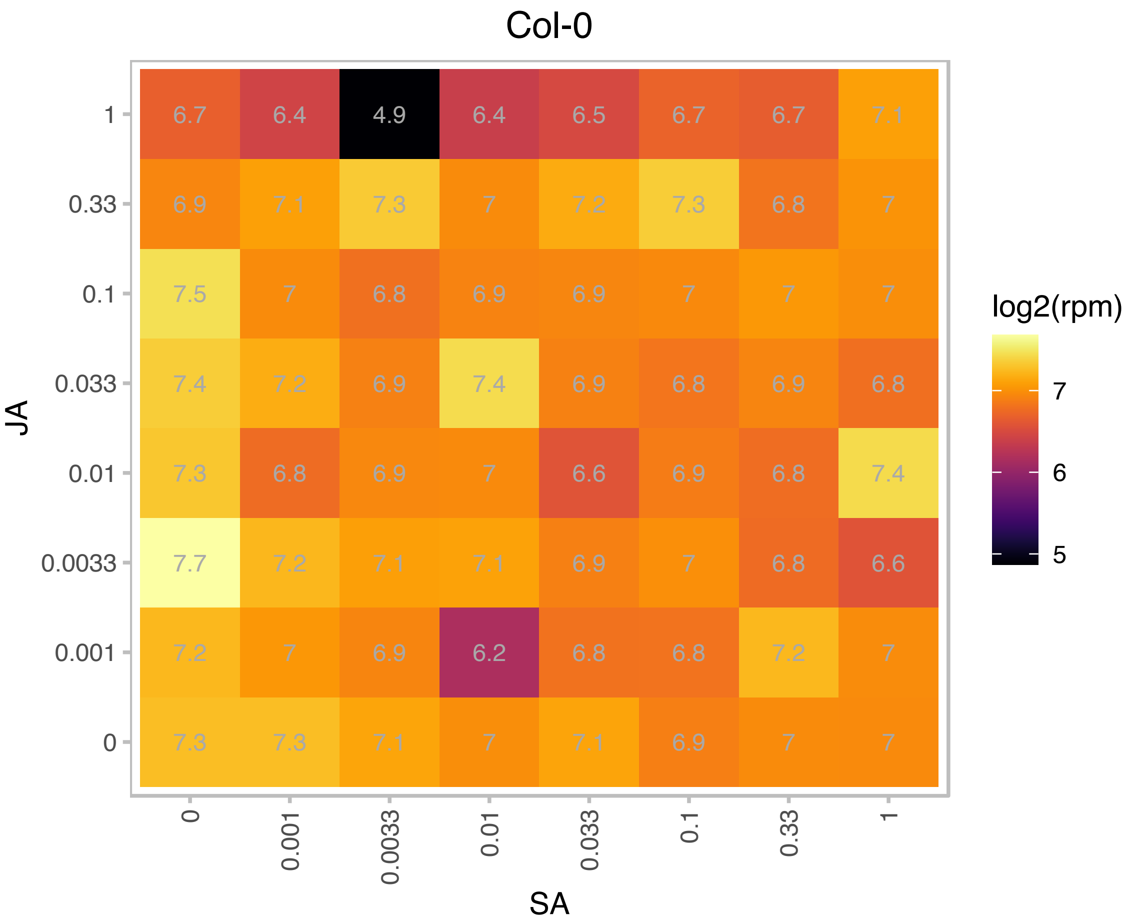


| | |
|---|---|
| Gene | AT4G13200 |
| is.expressed_genes | TRUE |
| is.responsive_genes | TRUE |
| is.apcluster | F-05 |

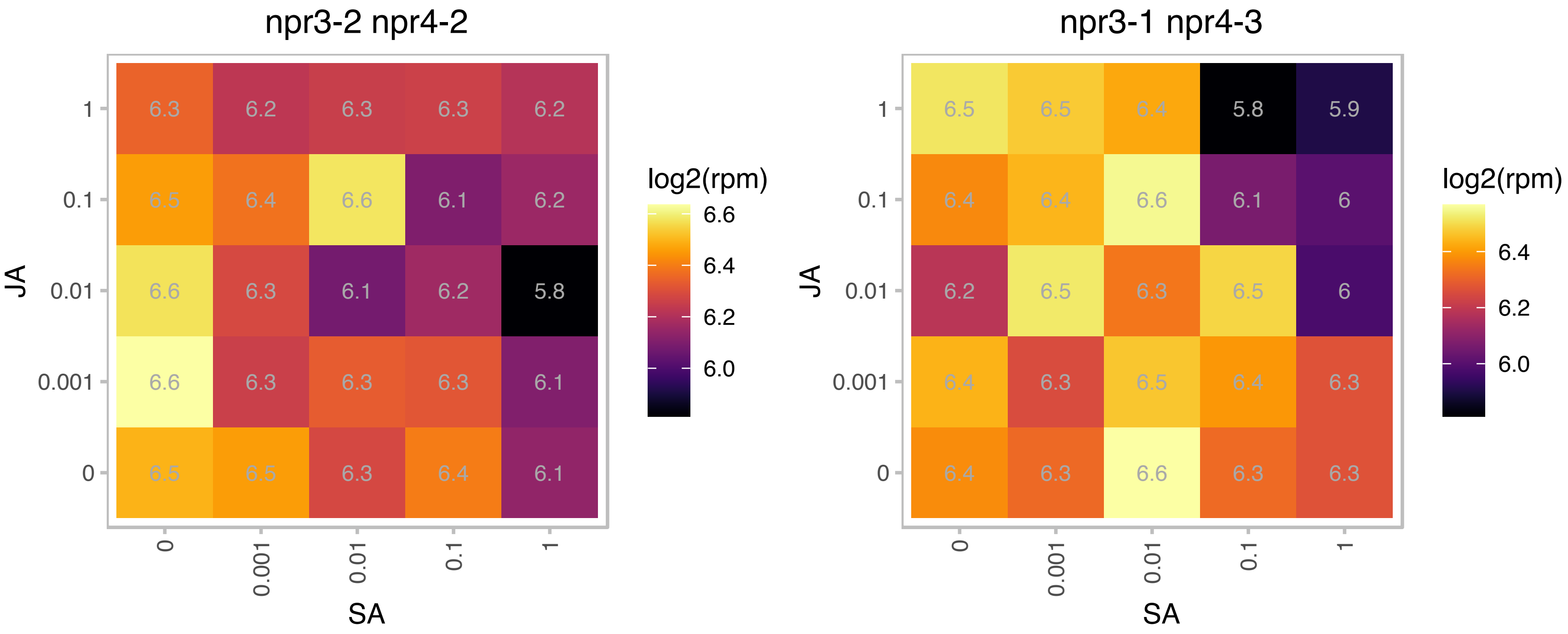

# AT4G20030.1 : NA

RNA-binding (RRM/RBD/RNP motifs) family protein

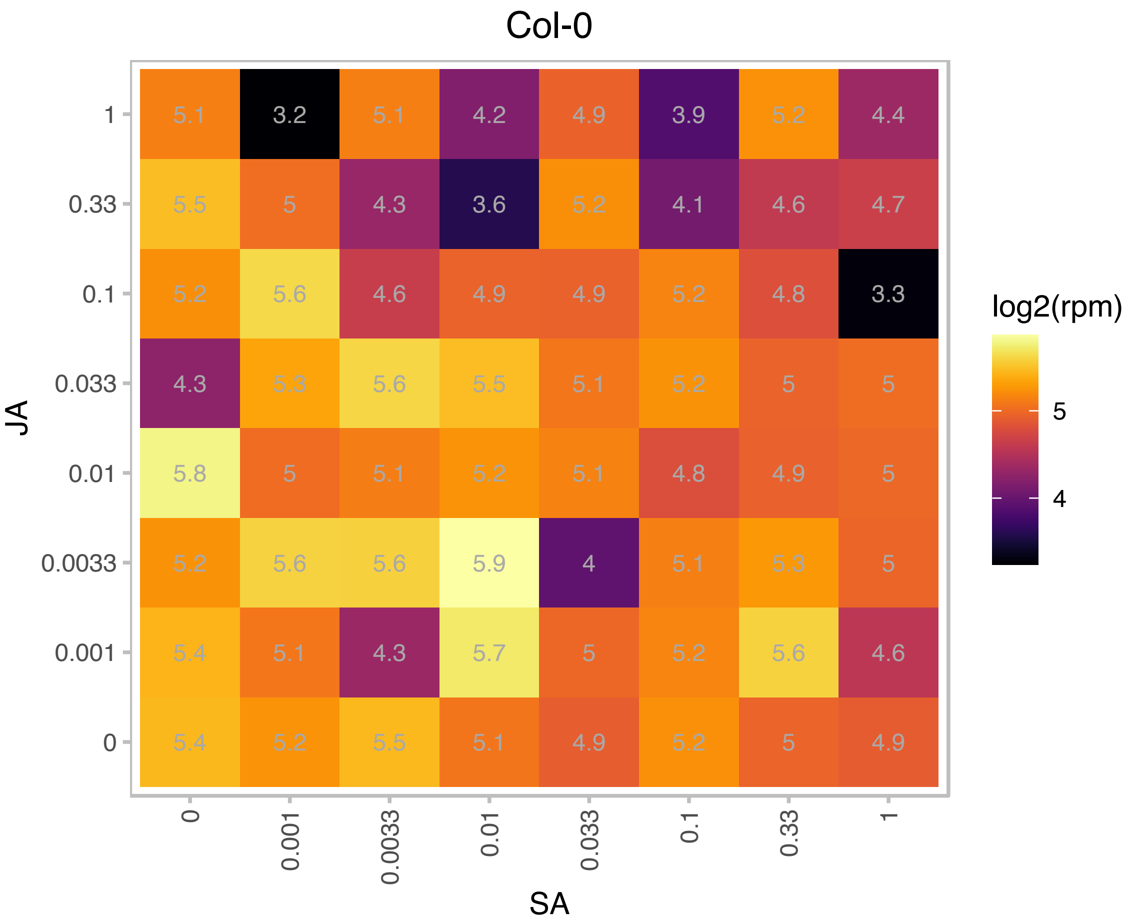


| | |
|---|---|
| Gene | AT4G20030 |
| is.expressed_genes | TRUE |
| is.responsive_genes | TRUE |
| is.apcluster | F-05 |

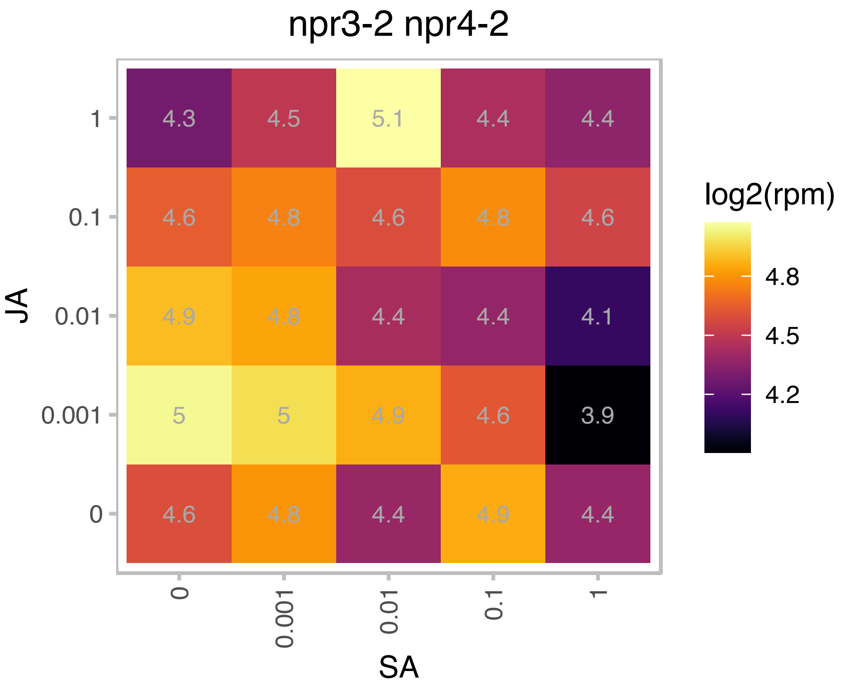


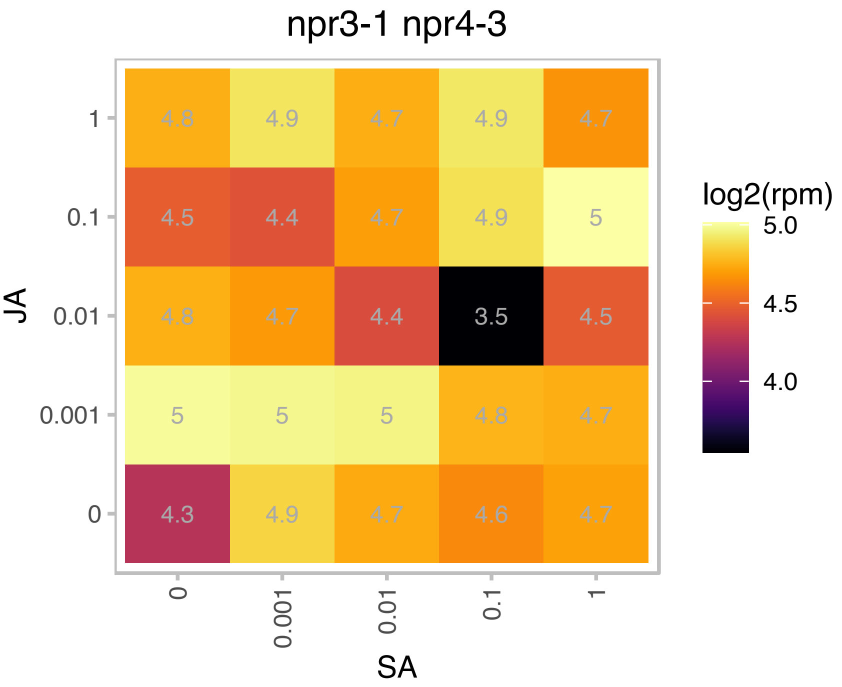

### AT4G21280.1 : PSBQ, PSBQ-1, PSBQA

photosystem II subunit QA

Encodes the PsbQ subunit of the oxygen evolving complex of photosystem II.

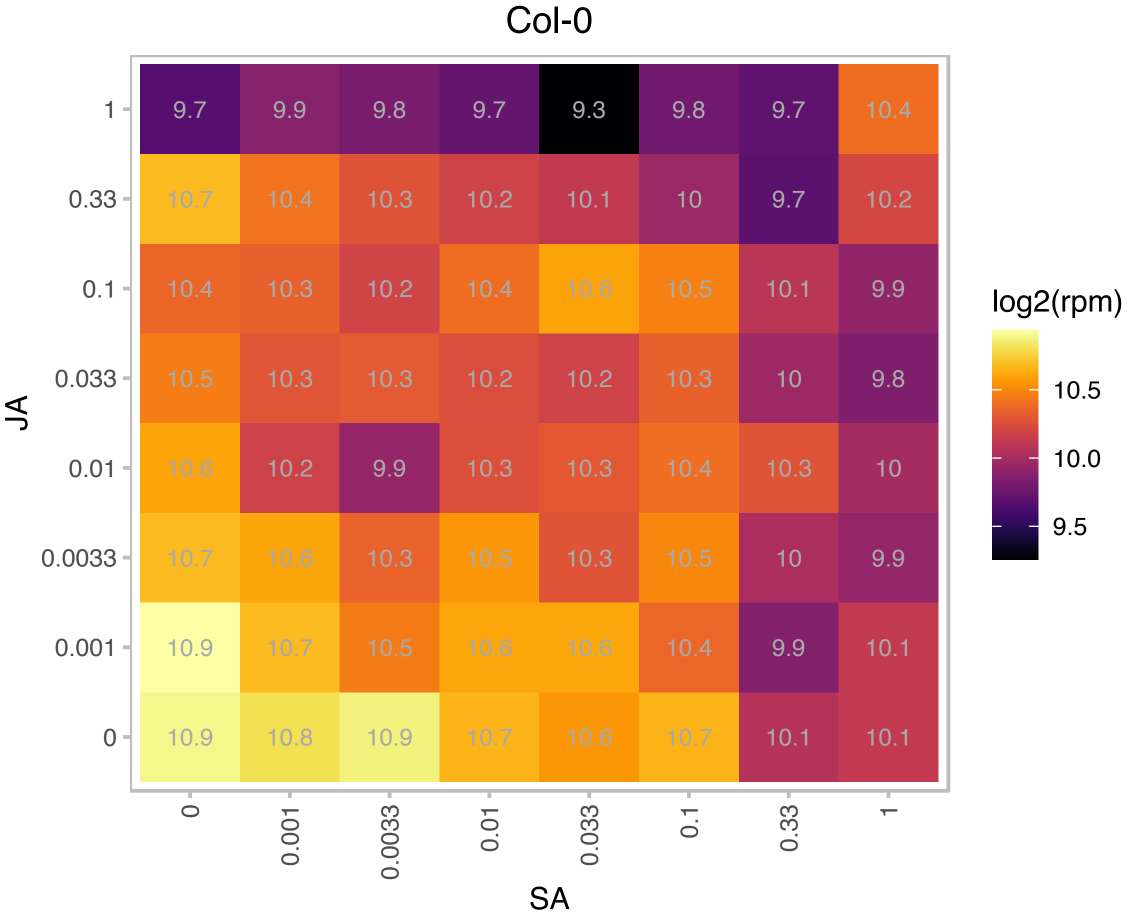


| | |
|---|---|
| Gene | AT4G21280 |
| is.expressed_genes | TRUE |
| is.responsive_genes | TRUE |
| is.apcluster | F-05 |

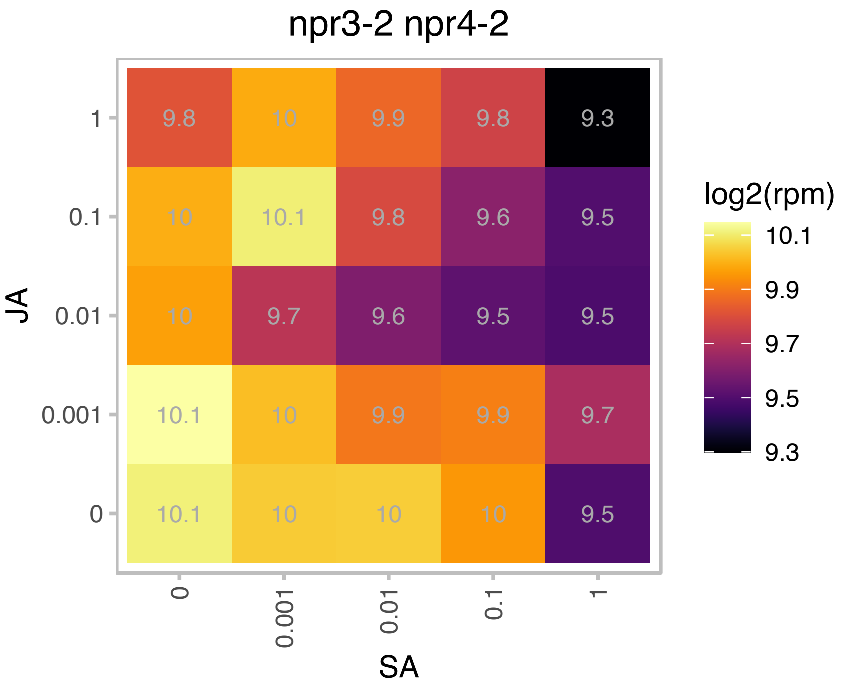


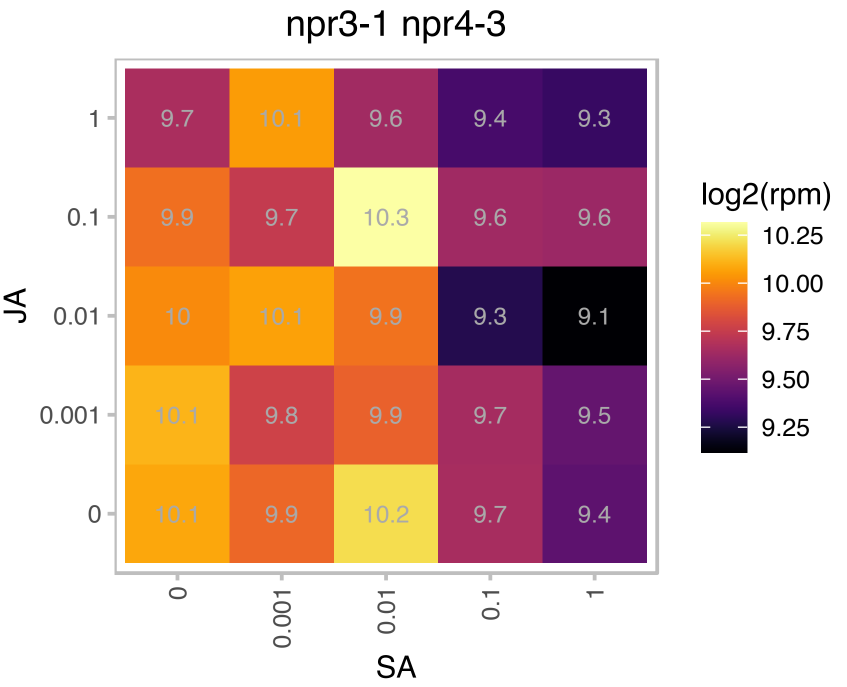

# AT4G26370.1 : NA

antitermination NusB domain-containing protein

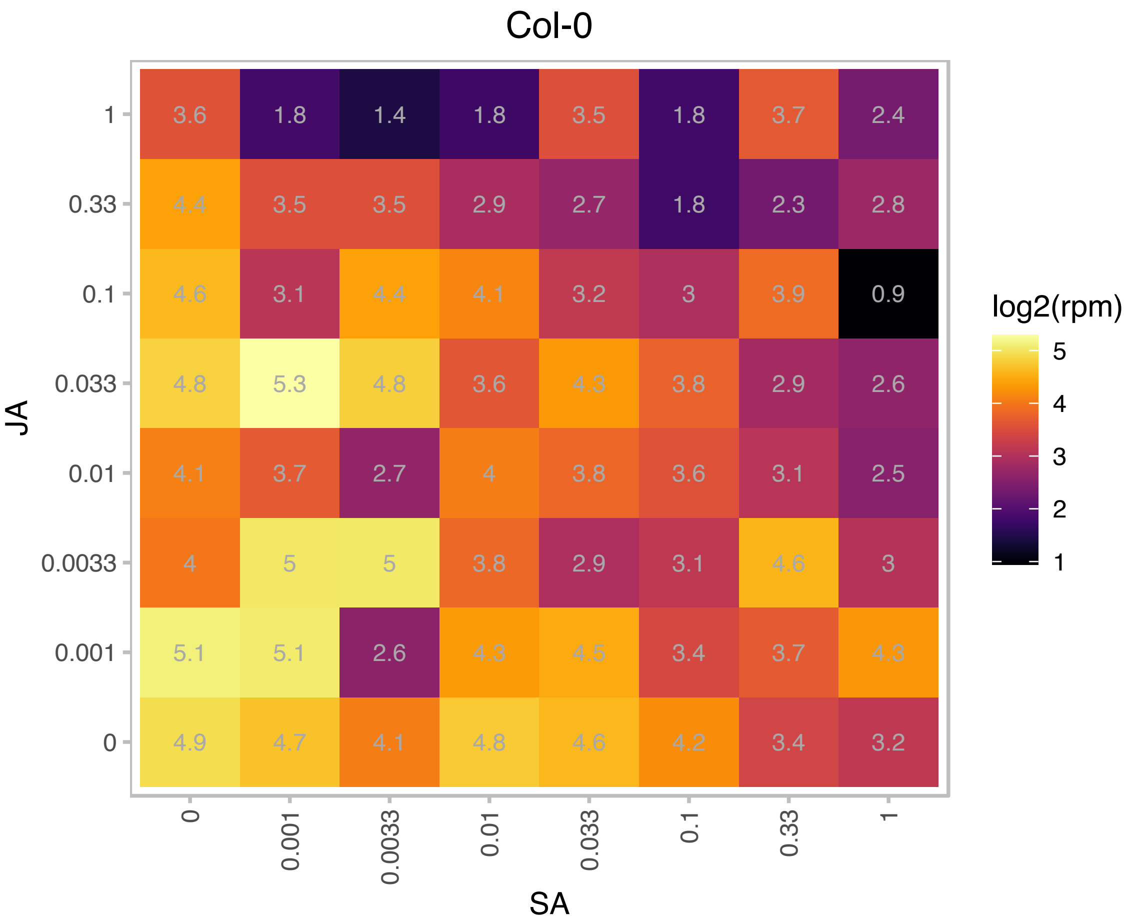


| | |
|---|---|
| Gene | AT4G26370 |
| is.expressed_genes | TRUE |
| is.responsive_genes | TRUE |
| is.apcluster | F-05 |

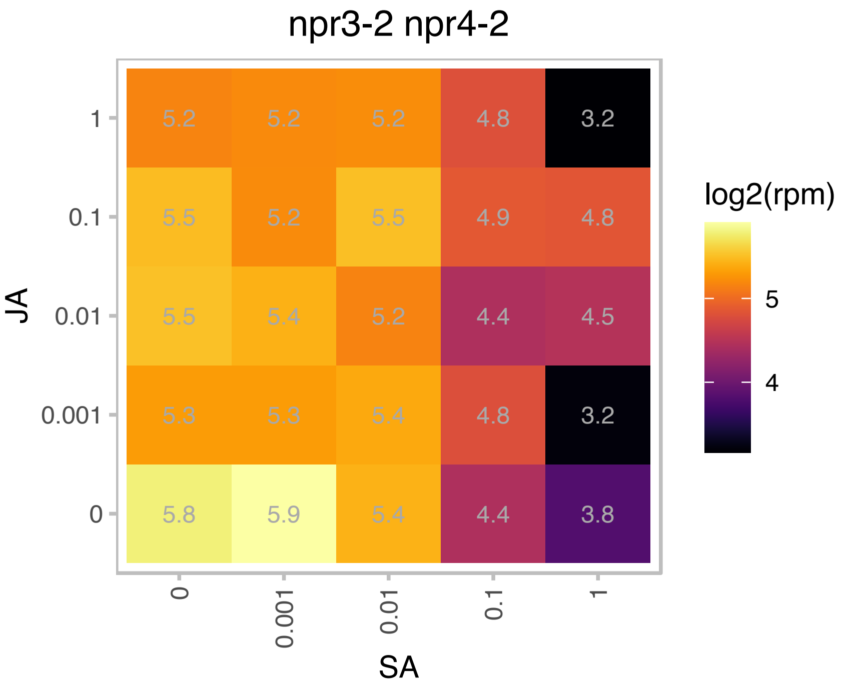


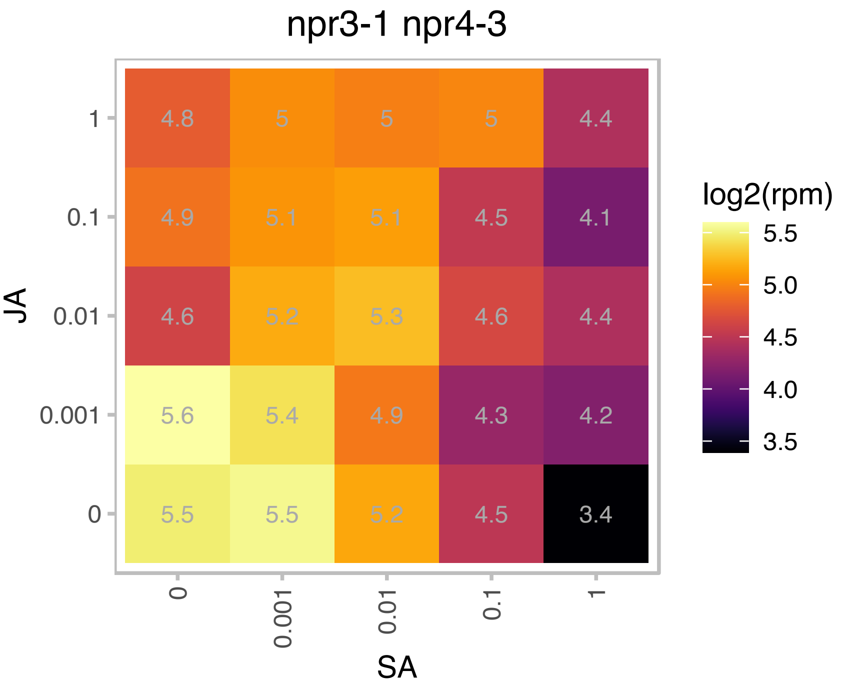

### AT4G26530.1 : AtFBA5, FBA5

Aldolase superfamily protein

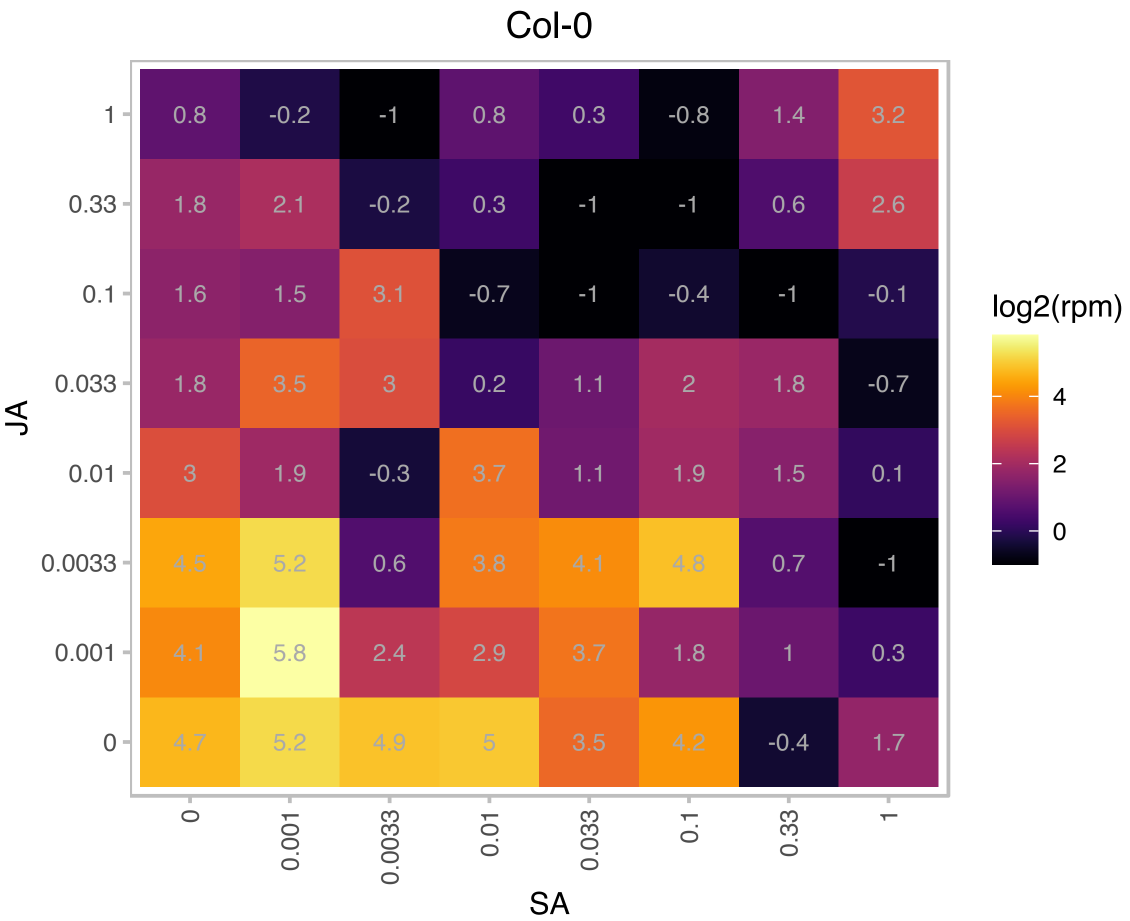


| | |
|---|---|
| Gene | AT4G26530 |
| is.expressed_genes | TRUE |
| is.responsive_genes | TRUE |
| is.apcluster | F-05 |

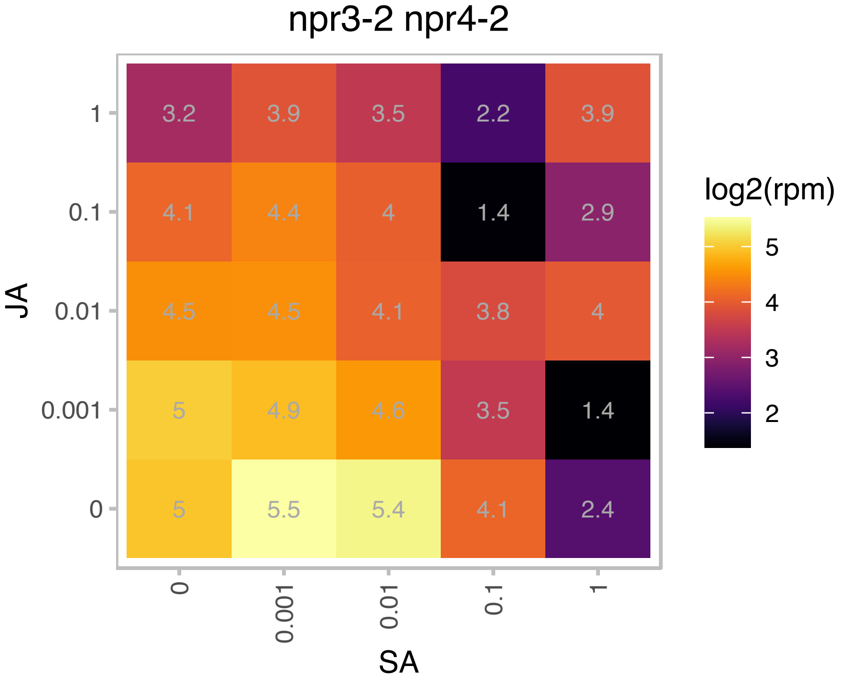


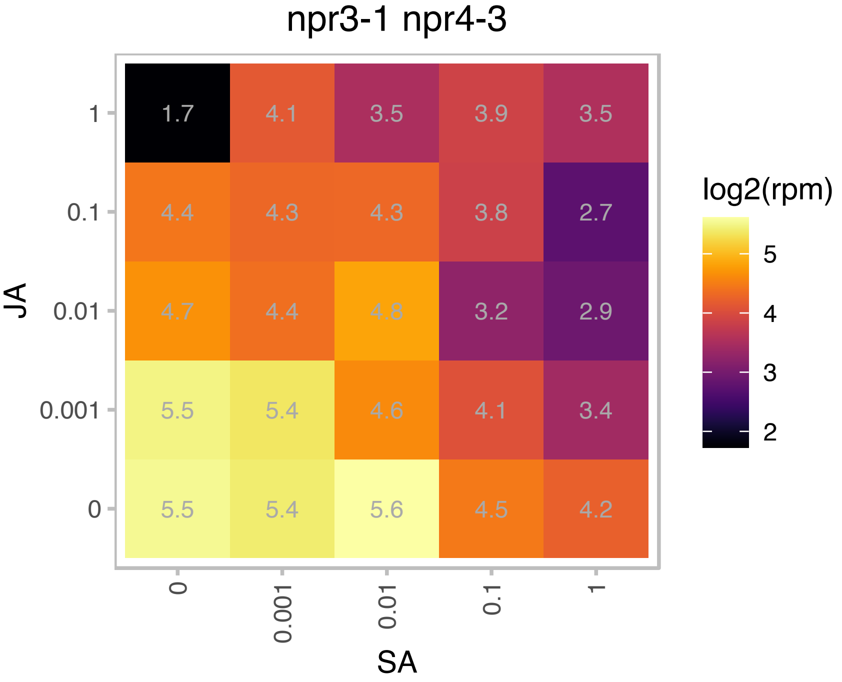

### AT4G28210.1 : EMB1923

embryo defective 1923

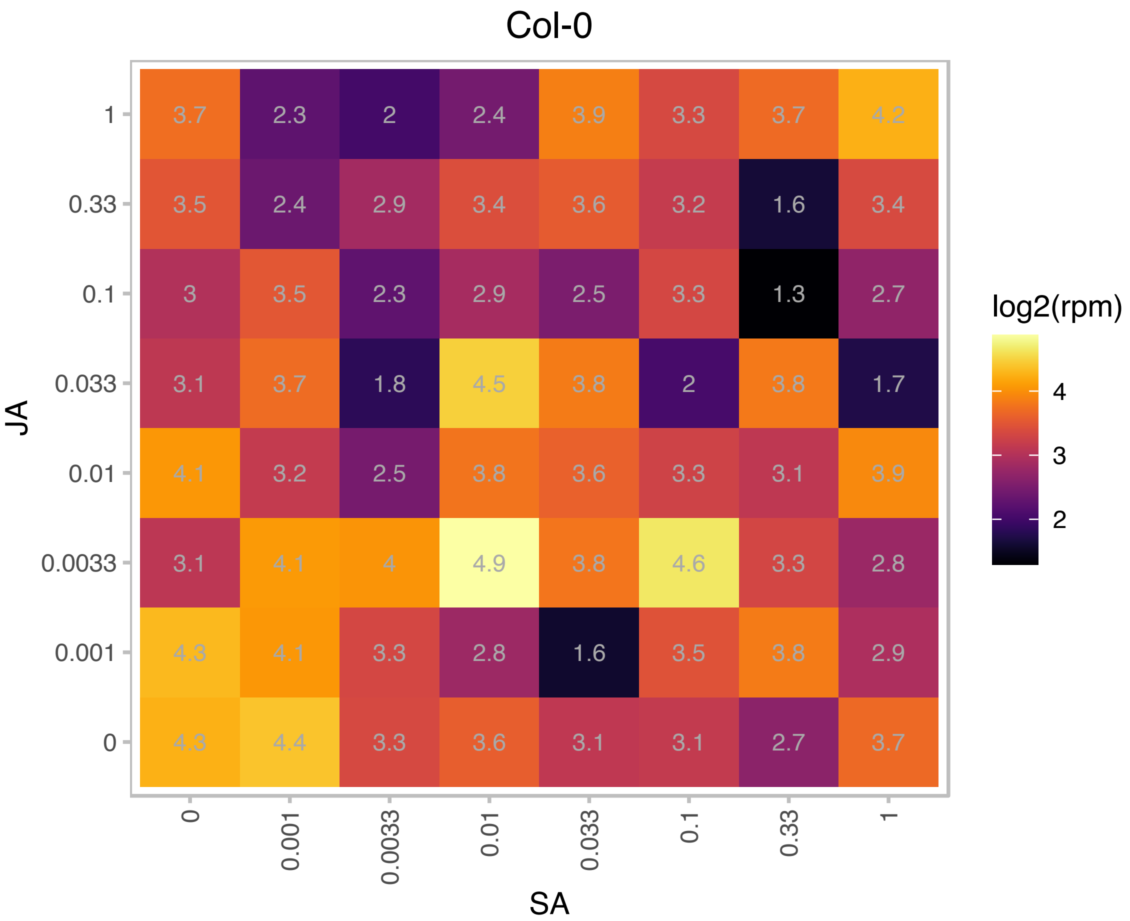


| | |
|---|---|
| Gene | AT4G28210 |
| is.expressed_genes | TRUE |
| is.responsive_genes | TRUE |
| is.apcluster | F-05 |

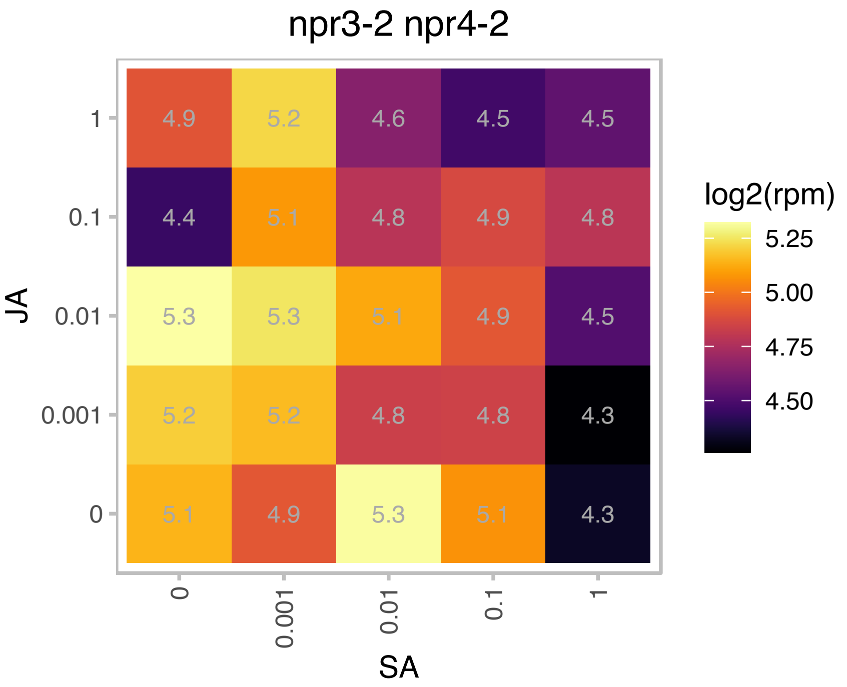


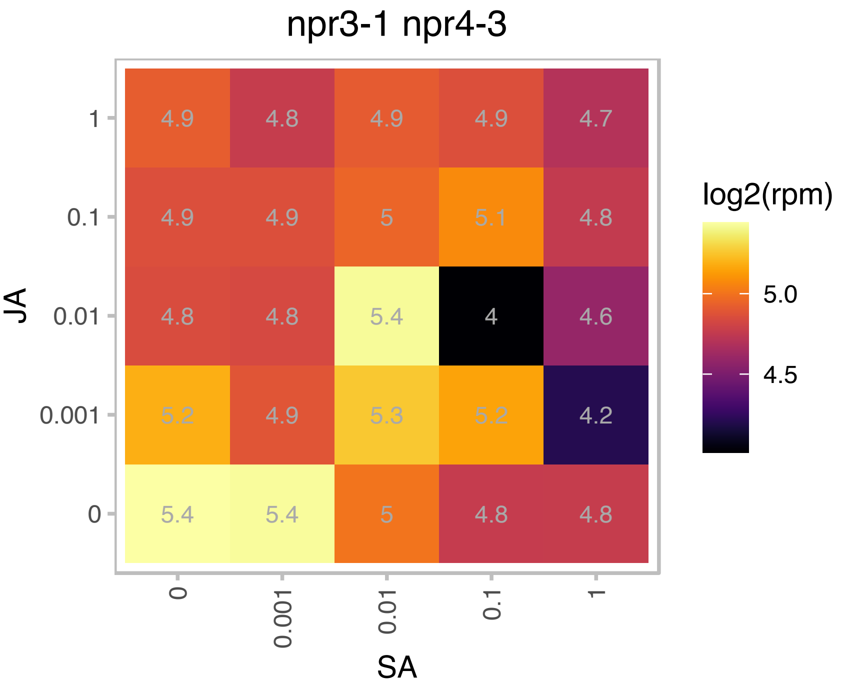

### AT4G28660.1 : PSB28

photosystem II reaction center PSB28 protein

Similar to PsbW subunit of photosystem II.

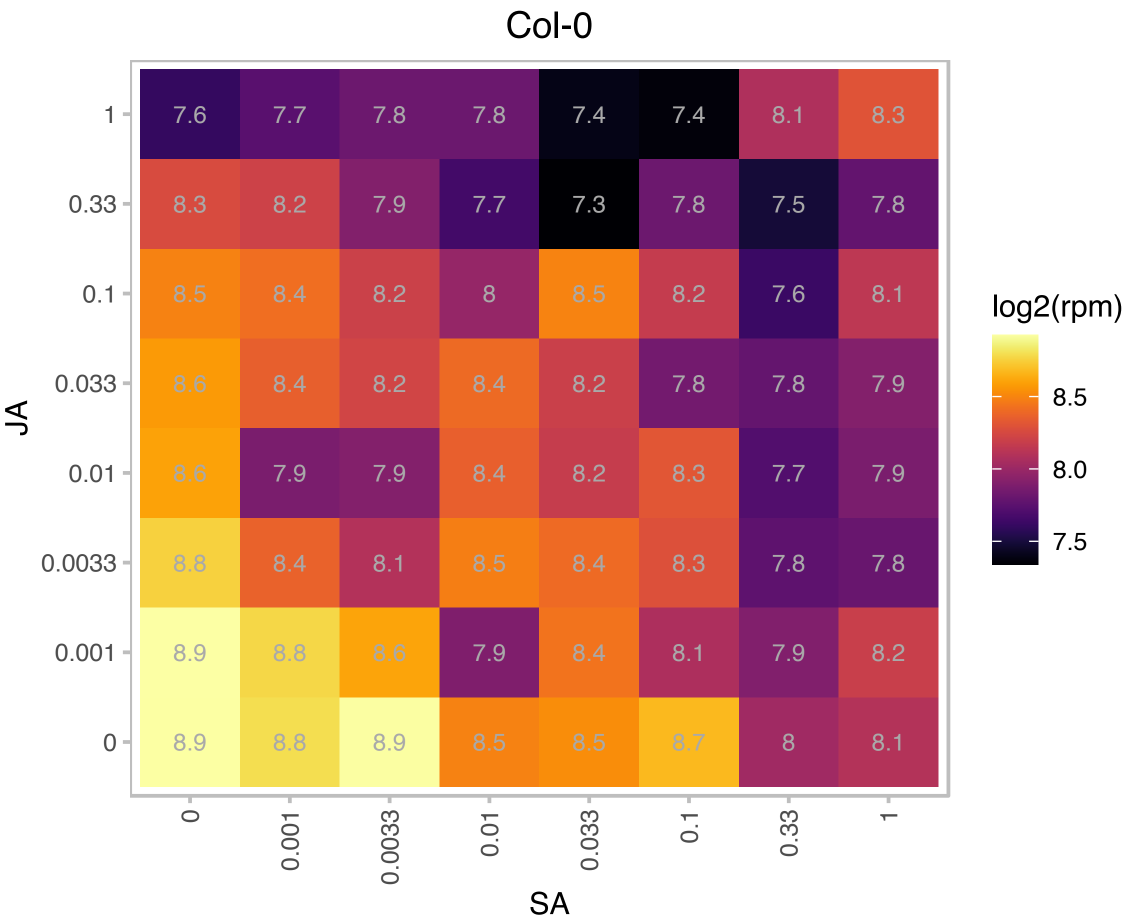


| | |
|---|---|
| Gene | AT4G28660 |
| is.expressed_genes | TRUE |
| is.responsive_genes | TRUE |
| is.apcluster | F-05 |

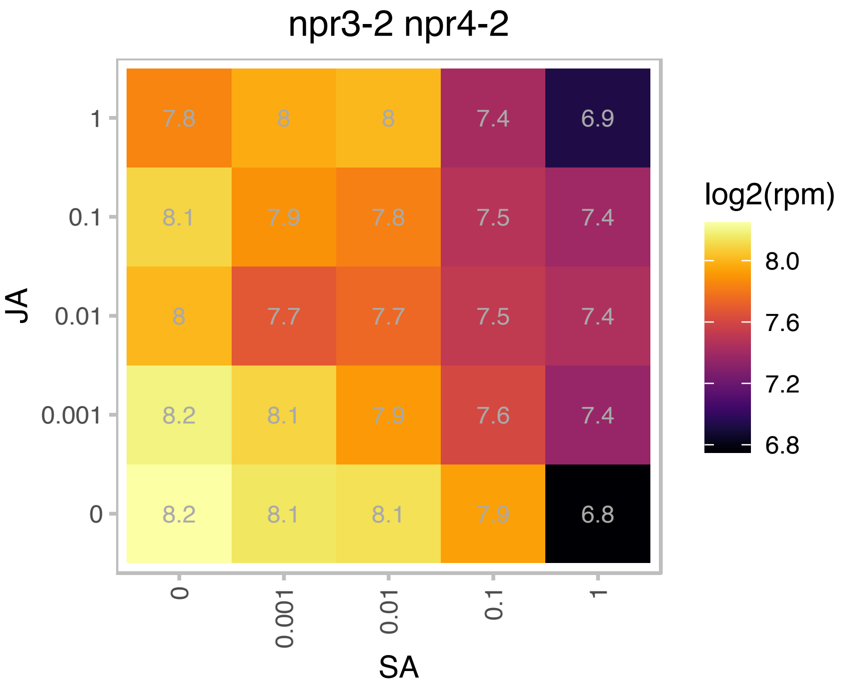


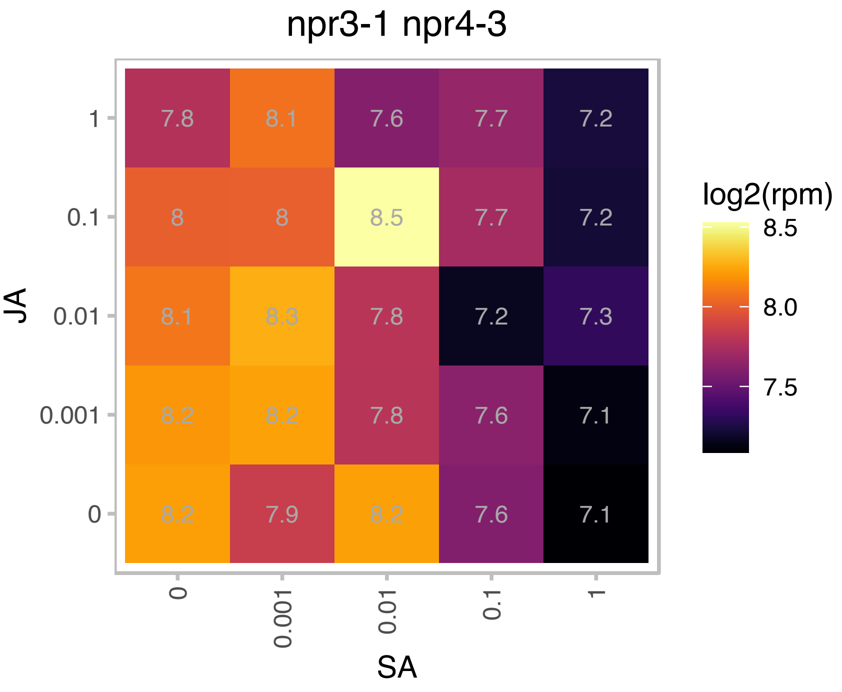

### AT4G30720.1 : PDE327

FAD/NAD(P)-binding oxidoreductase family protein
Encodes a putative oxidoreductase/electron carrier detected in the chloroplast stroma that is essential to ensure a correct electron flow through the photosynthetic chain and, hence, photosynthesis efficiency and normal growth. Mutations in the Col-0 allele result in pale green pigmentation and defective growth.

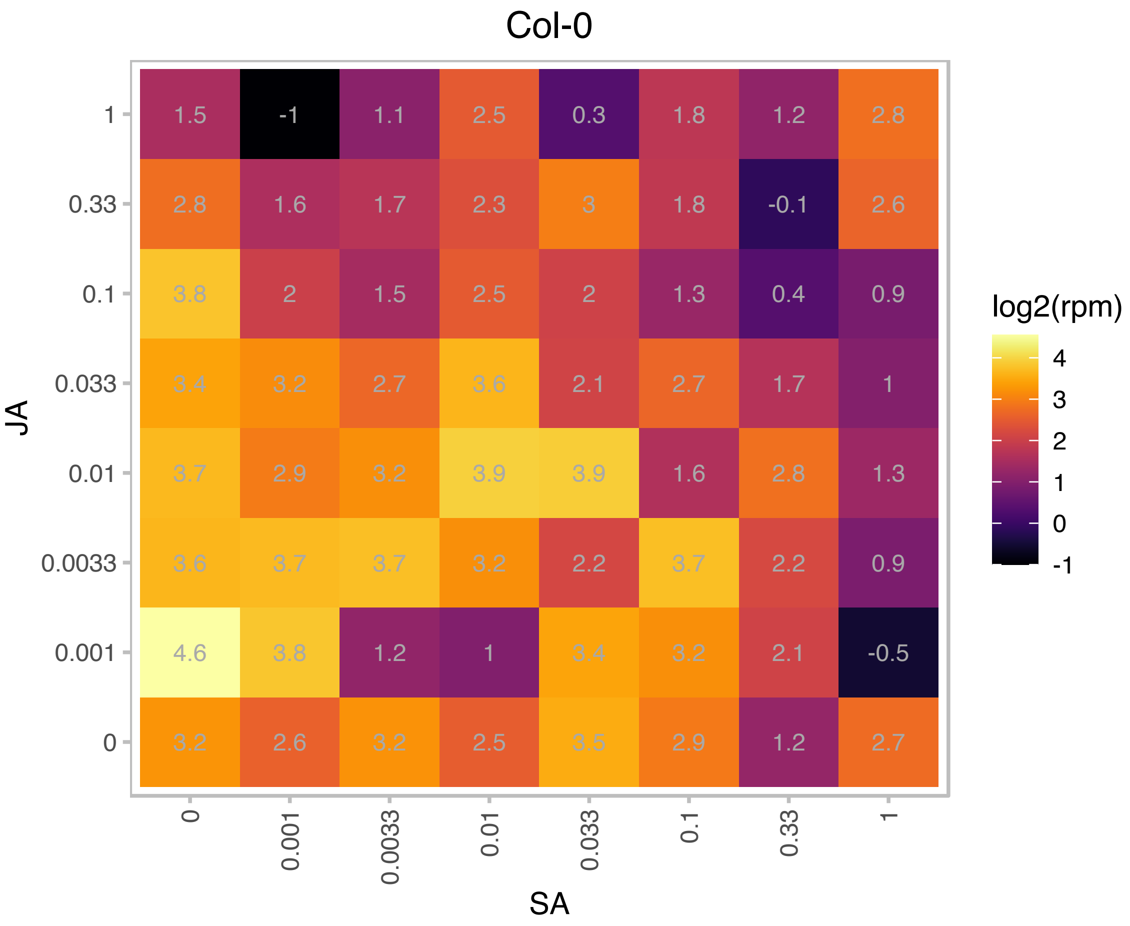


| | |
|---|---|
| Gene | AT4G30720 |
| is.expressed_genes | TRUE |
| is.responsive_genes | TRUE |
| is.apcluster | F-05 |

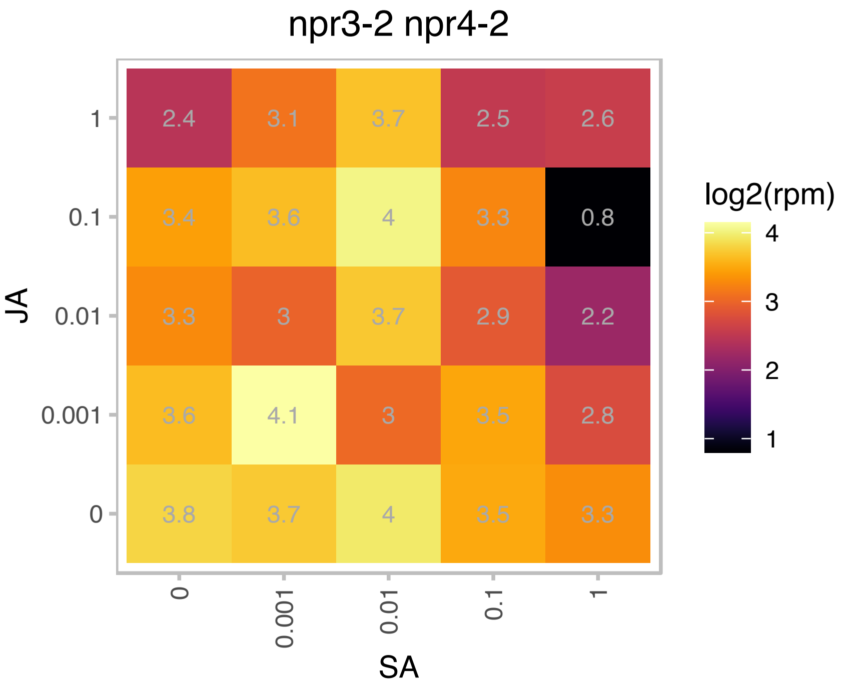


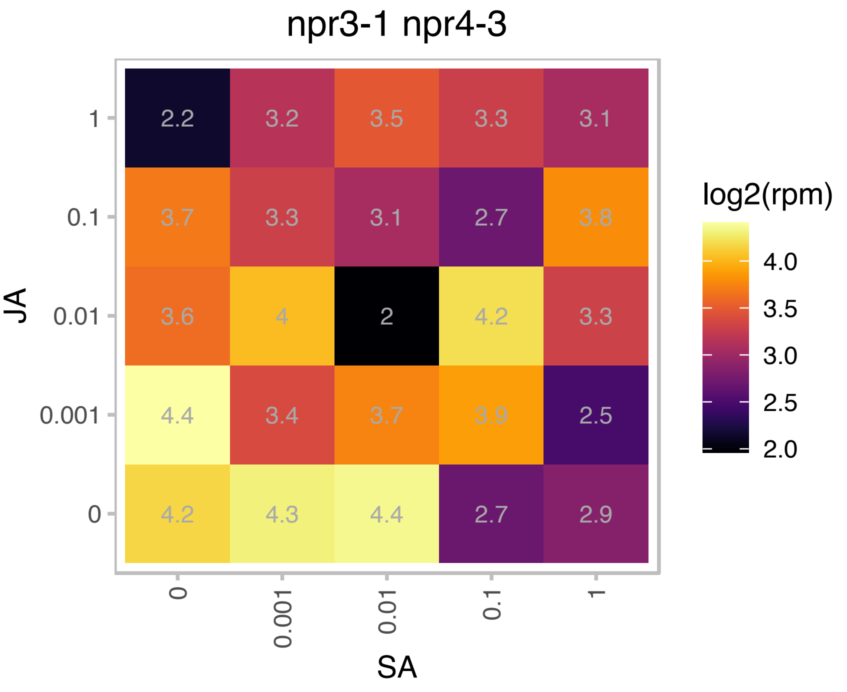

# AT4G31115.1 : NA

Protein of unknown function (DUF1997)

#### Col-0

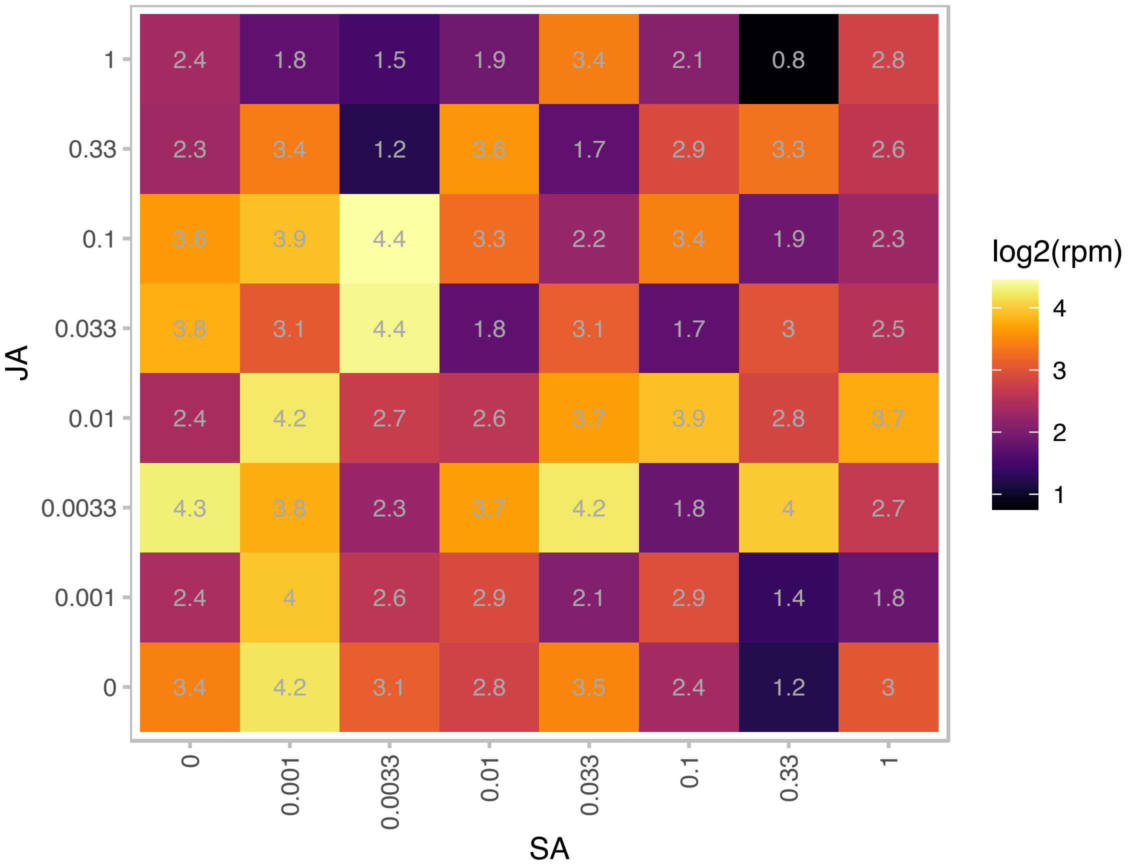


| | |
|---|---|
| Gene | AT4G31115 |
| is.expressed_genes | TRUE |
| is.responsive_genes | TRUE |
| is.apcluster | F-05 |

#### npr3-2 npr4-2

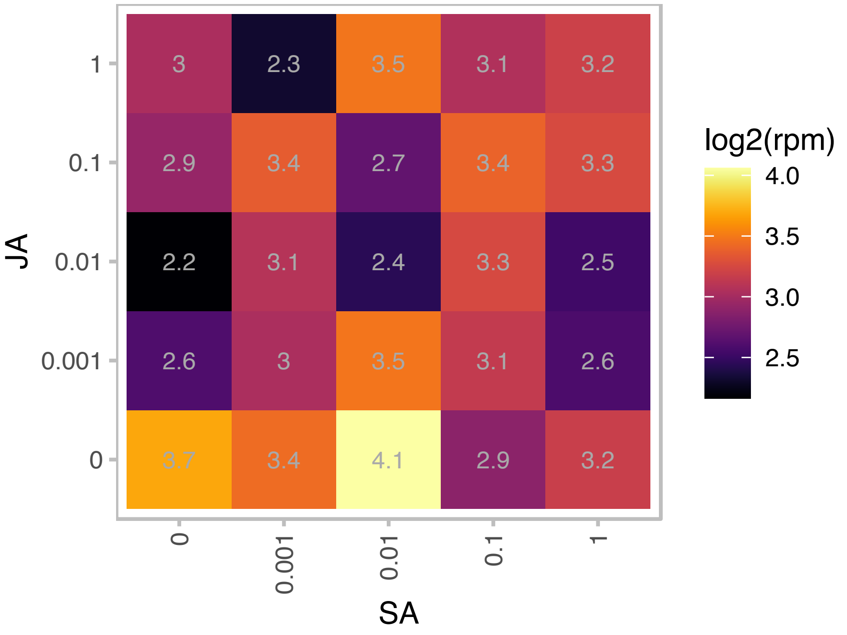


#### npr3-1 npr4-3

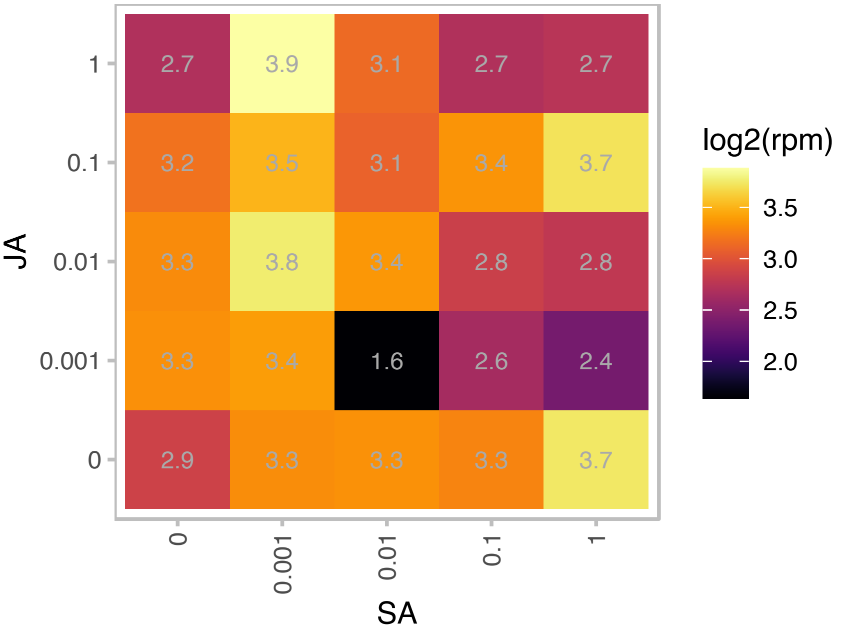

# AT5G01015.1 : NA

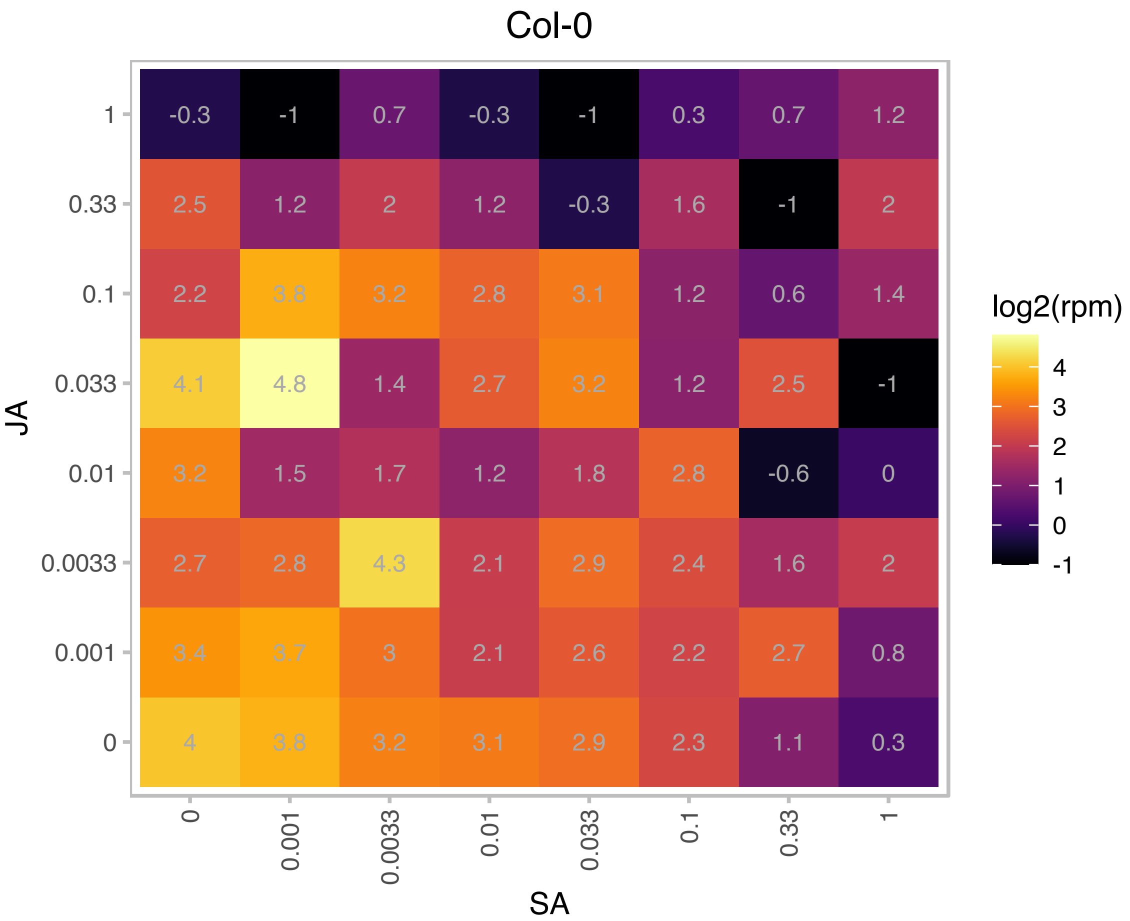


| | |
|---|---|
| Gene | AT5G01015 |
| is.expressed_genes | TRUE |
| is.responsive_genes | TRUE |
| is.apcluster | F-05 |

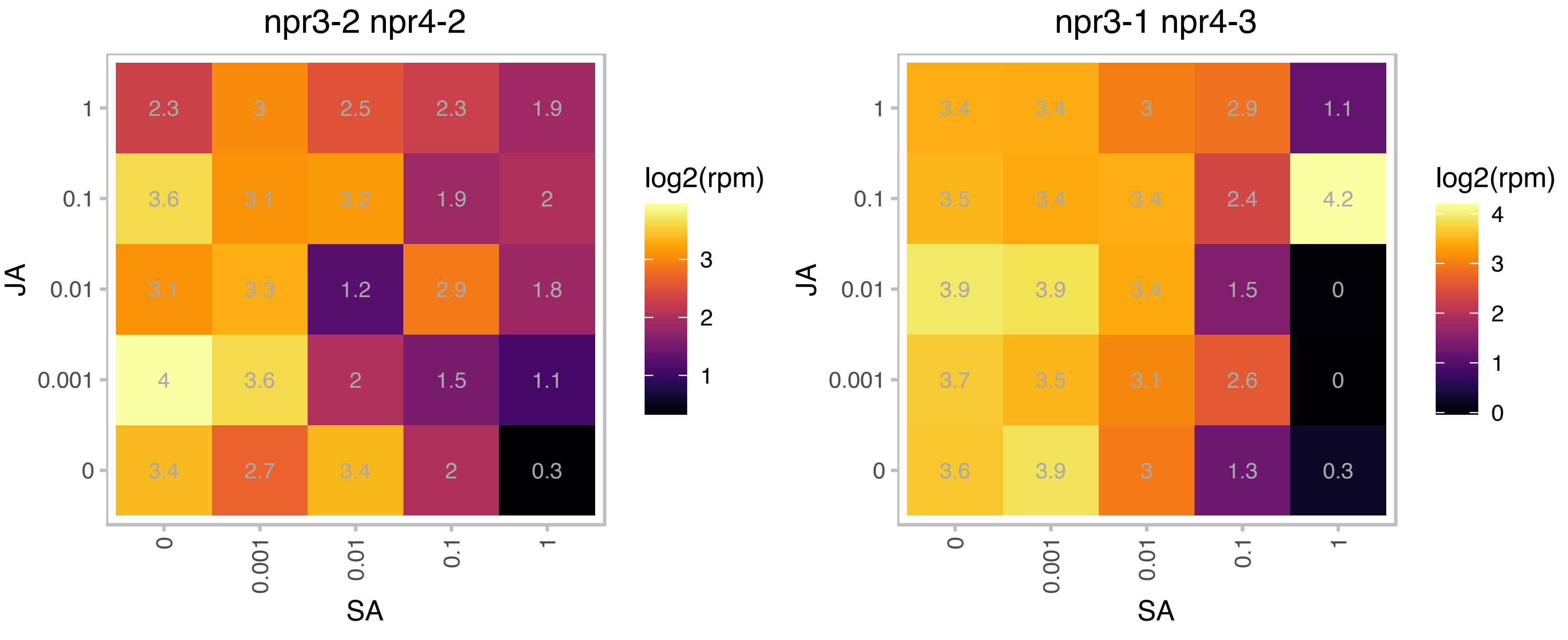

# AT5G01881.1 : NA

| | |
|---|---|
| Gene | AT5G01881 |
| is.expressed_genes | TRUE |
| is.responsive_genes | TRUE |
| is.apcluster | F-05 |

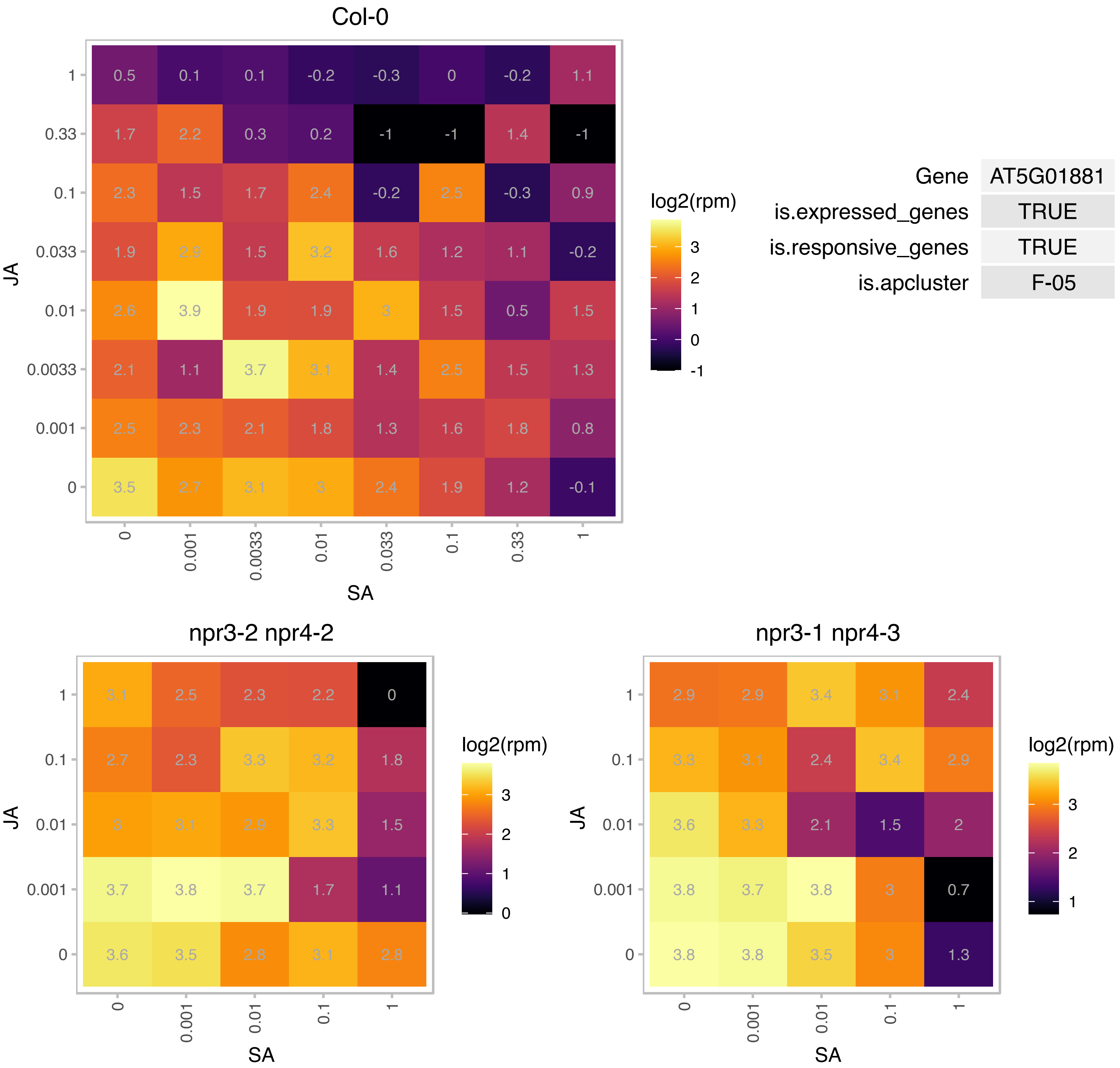

### AT5G13510.1 : EMB3136

Ribosomal protein L10 family protein

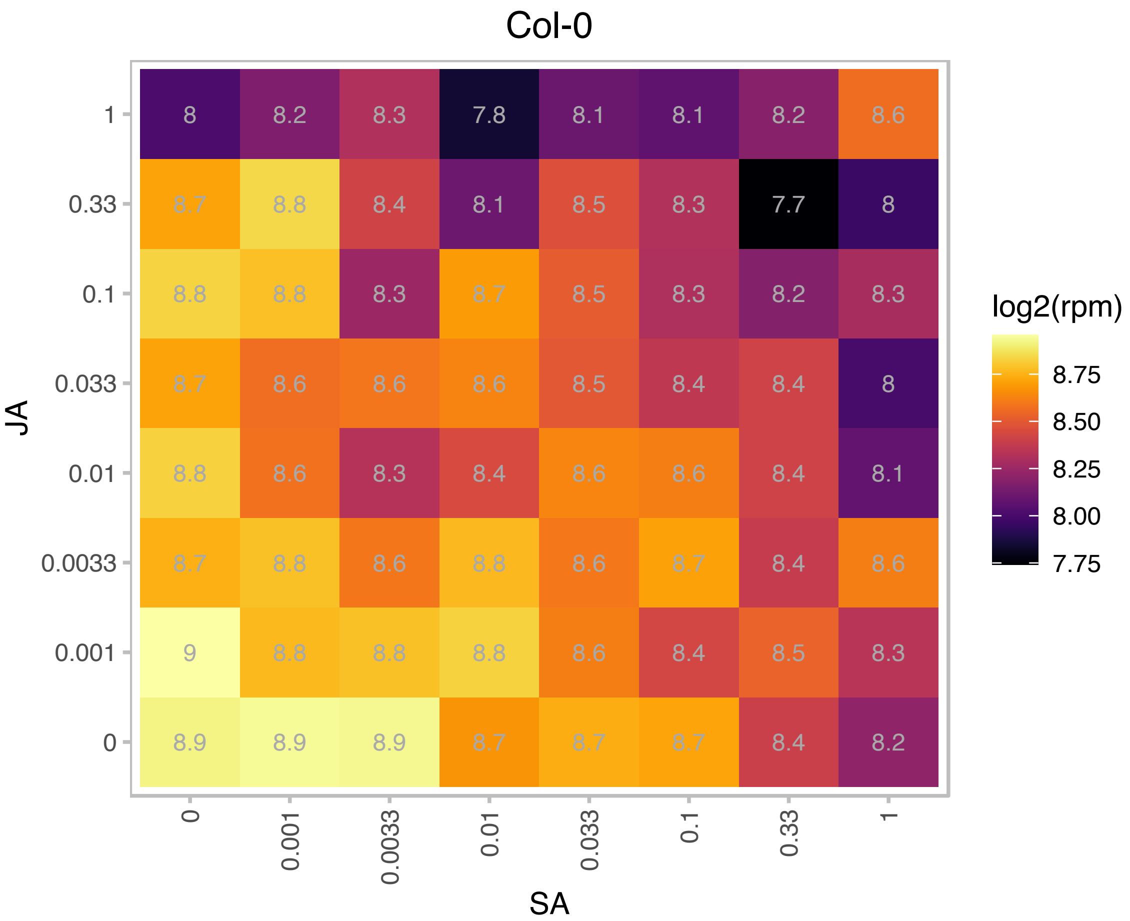


| | |
|---|---|
| Gene | AT5G13510 |
| is.expressed_genes | TRUE |
| is.responsive_genes | TRUE |
| is.apcluster | F-05 |

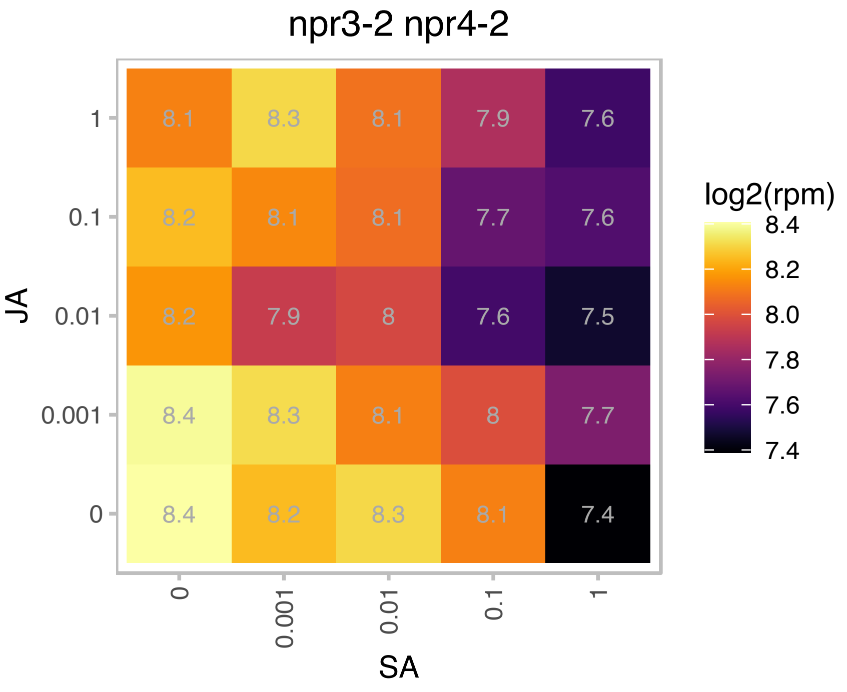


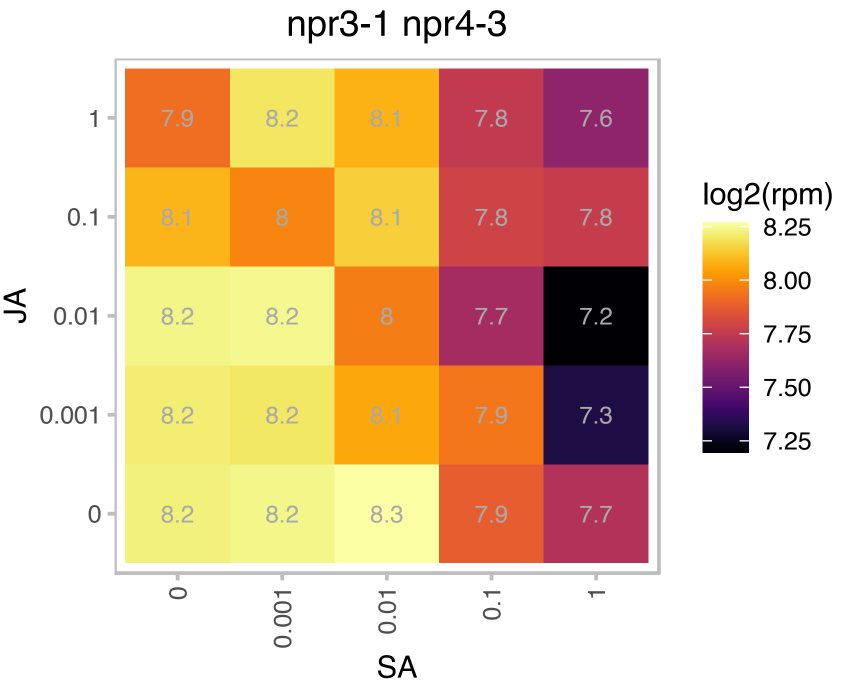

# AT5G27730.1 : NA

Protein of unknown function (DUF1624)

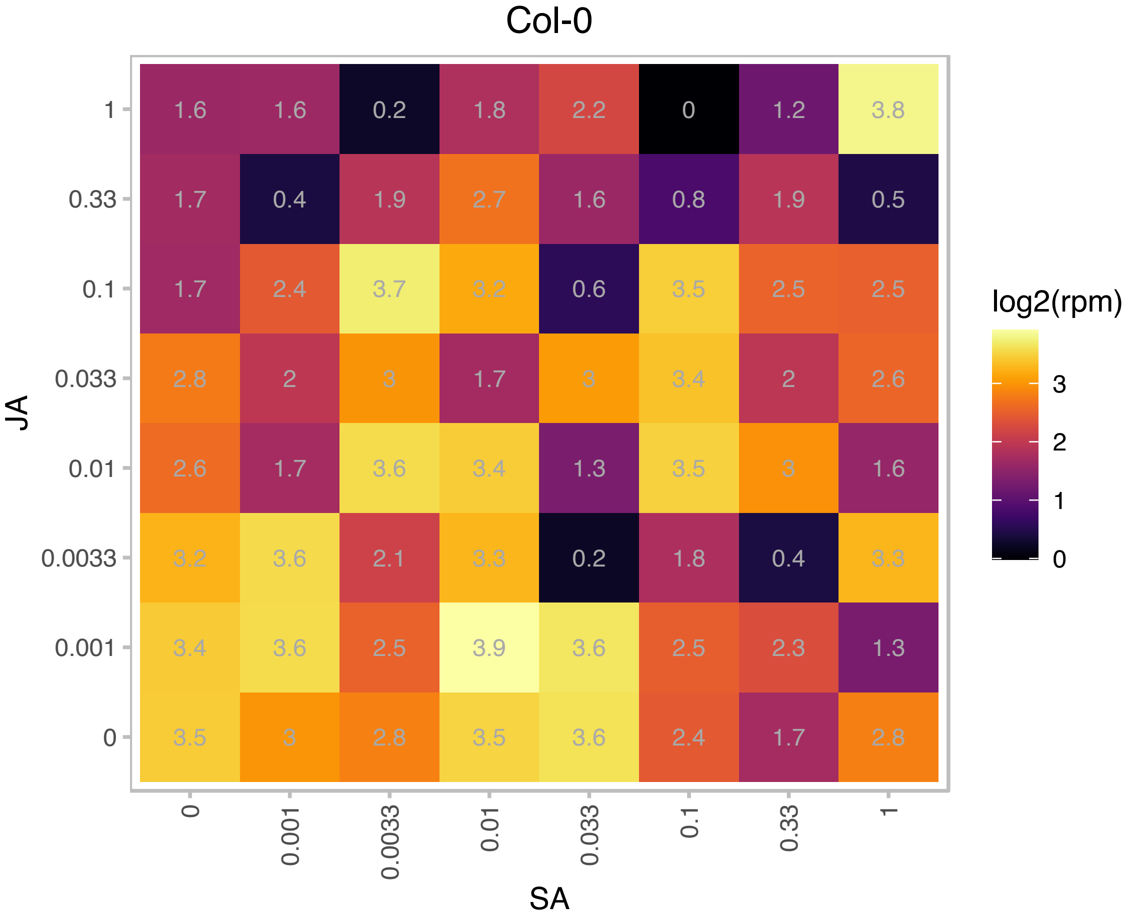


| | |
|---|---|
| Gene | AT5G27730 |
| is.expressed_genes | TRUE |
| is.responsive_genes | TRUE |
| is.apcluster | F-05 |

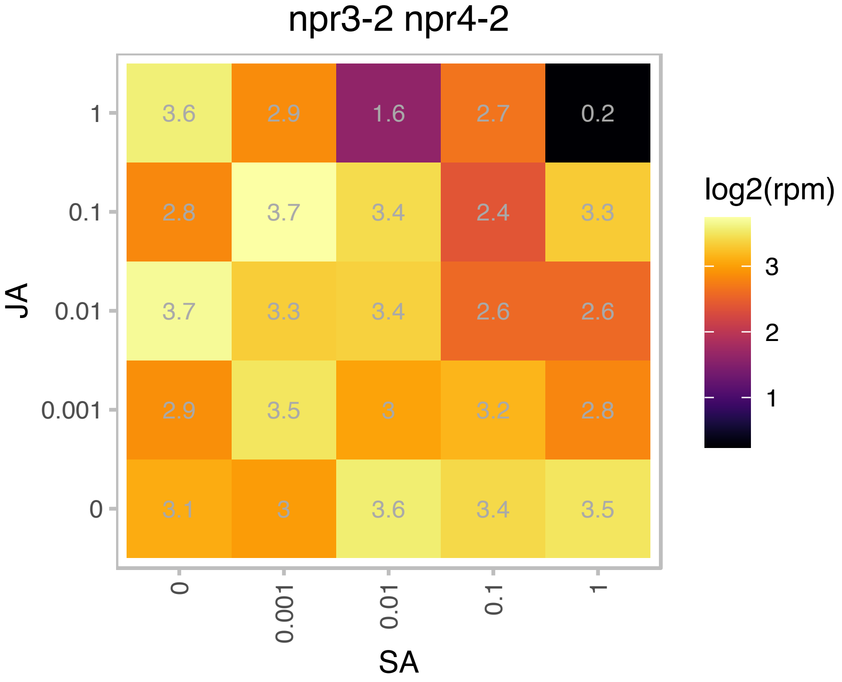


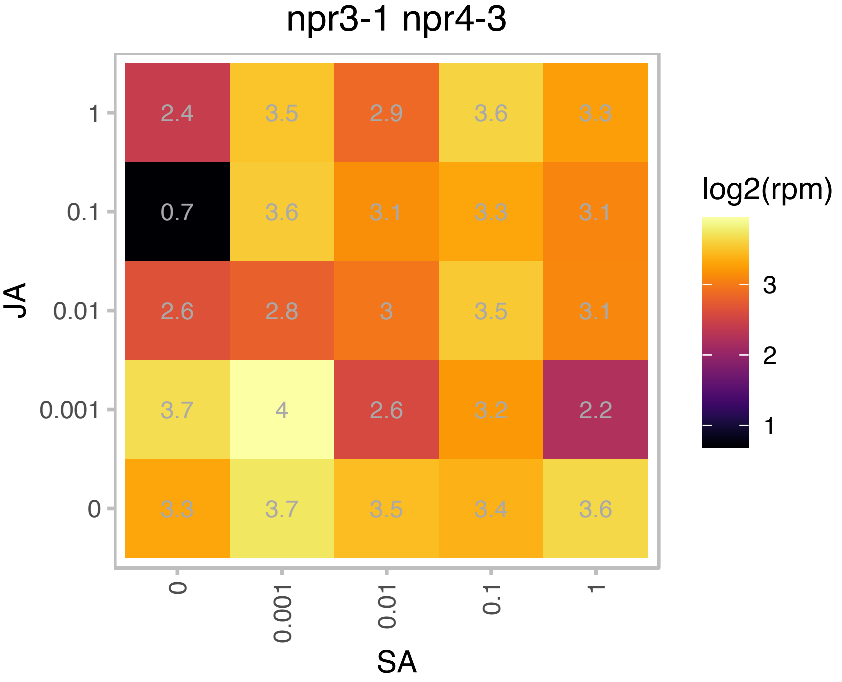

### AT5G37260.1 : CIR1, RVE2

Homeodomain-like superfamily protein

Encodes a MYB family transcription factor Circadian 1 (CIR1). Involved in circadian regulation in Arabidopsis.

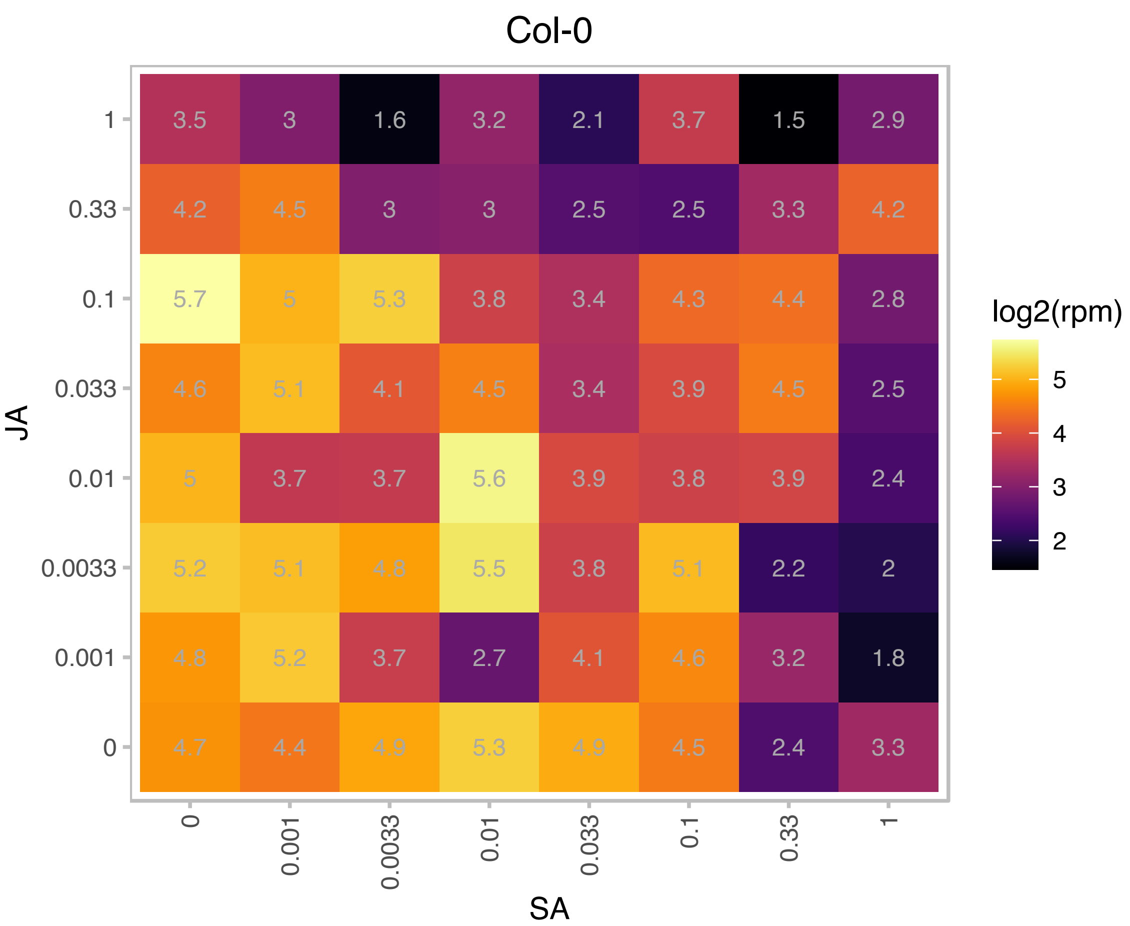


| | |
|---|---|
| Gene | AT5G37260 |
| is.expressed_genes | TRUE |
| is.responsive_genes | TRUE |
| is.apcluster | F-05 |

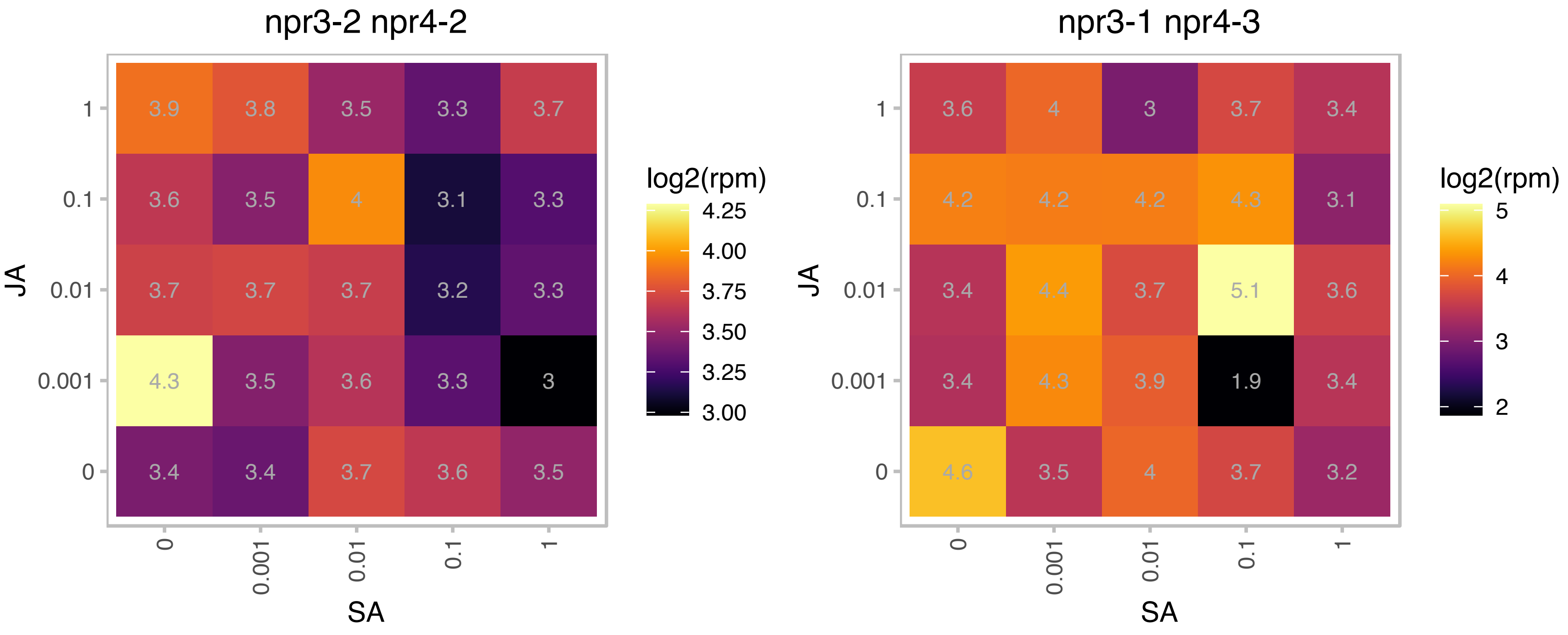

### AT5G38420.1 : RBCS2B

Ribulose bisphosphate carboxylase (small chain) family protein

Encodes a member of the Rubisco small subunit (RBCS) multigene family: RBCS1A (At1g67090), RBCS1B (At5g38430), RBCS2B (At5g38420), and RBCS3B (At5g38410).

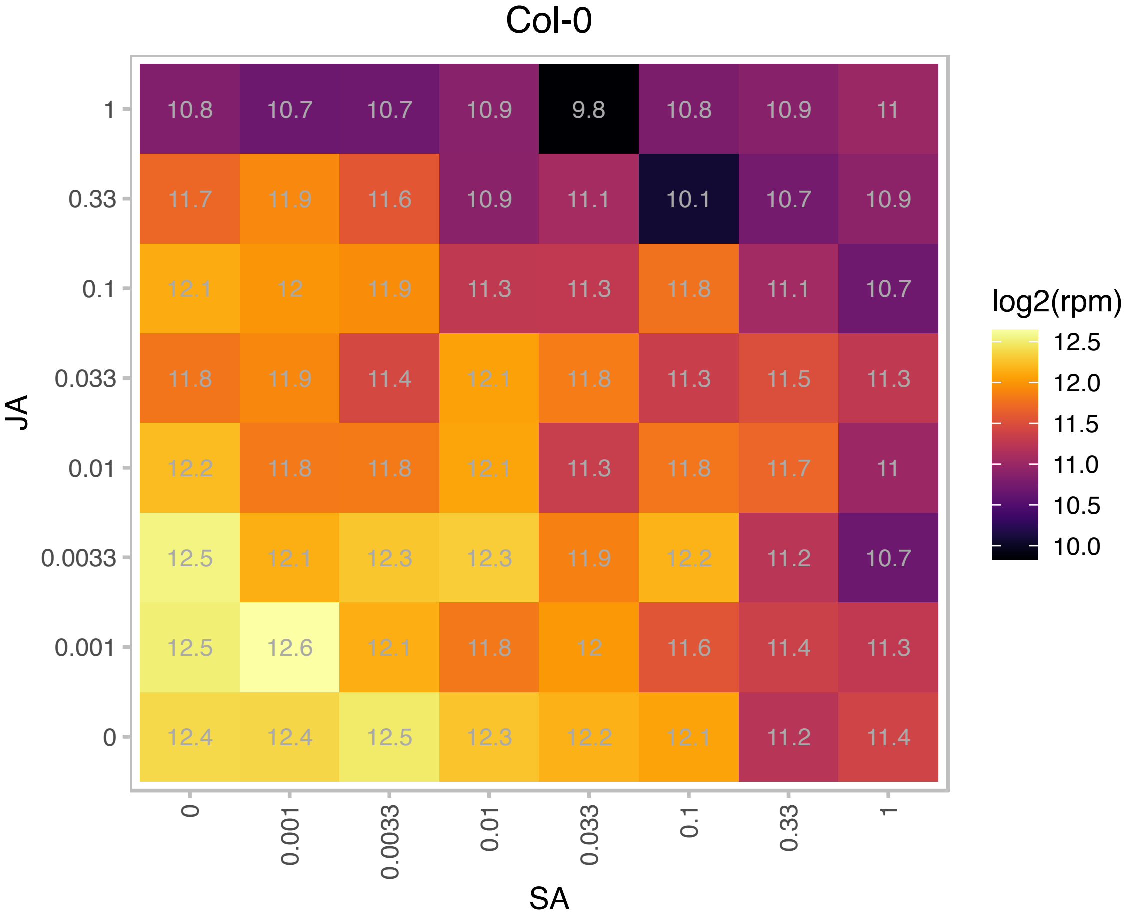


| | |
|---|---|
| Gene | AT5G38420 |
| is.expressed_genes | TRUE |
| is.responsive_genes | TRUE |
| is.apcluster | F-05 |

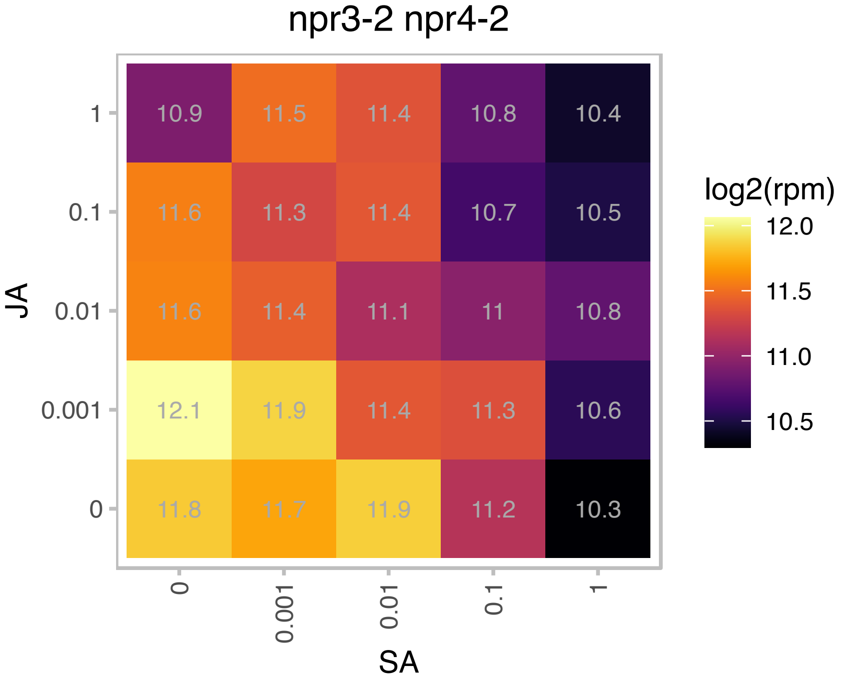


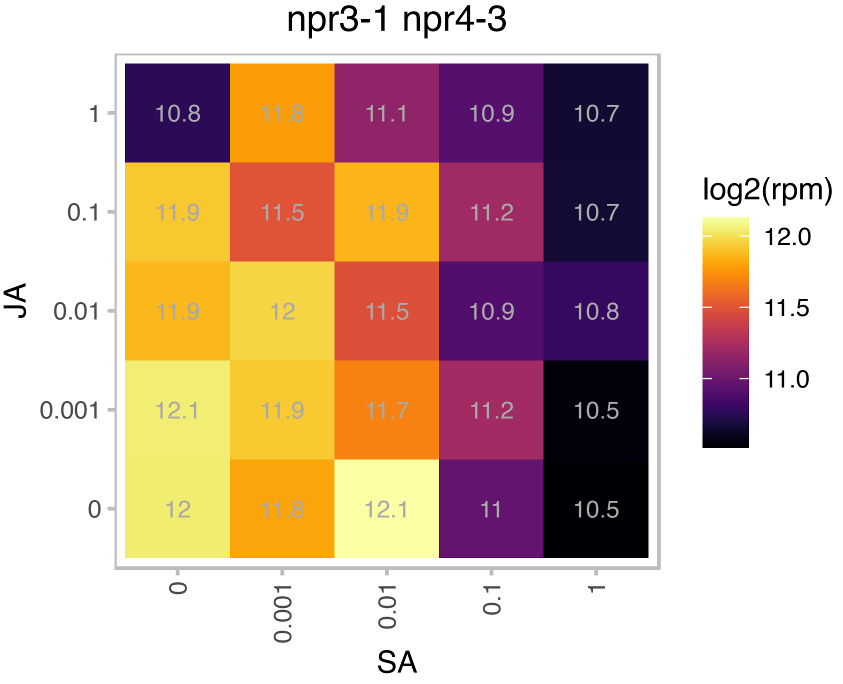

### AT5G40730.1 : AGP24, ATAGP24

arabinogalactan protein 24

Encodes an arabinogalactan-protein (AGP24).

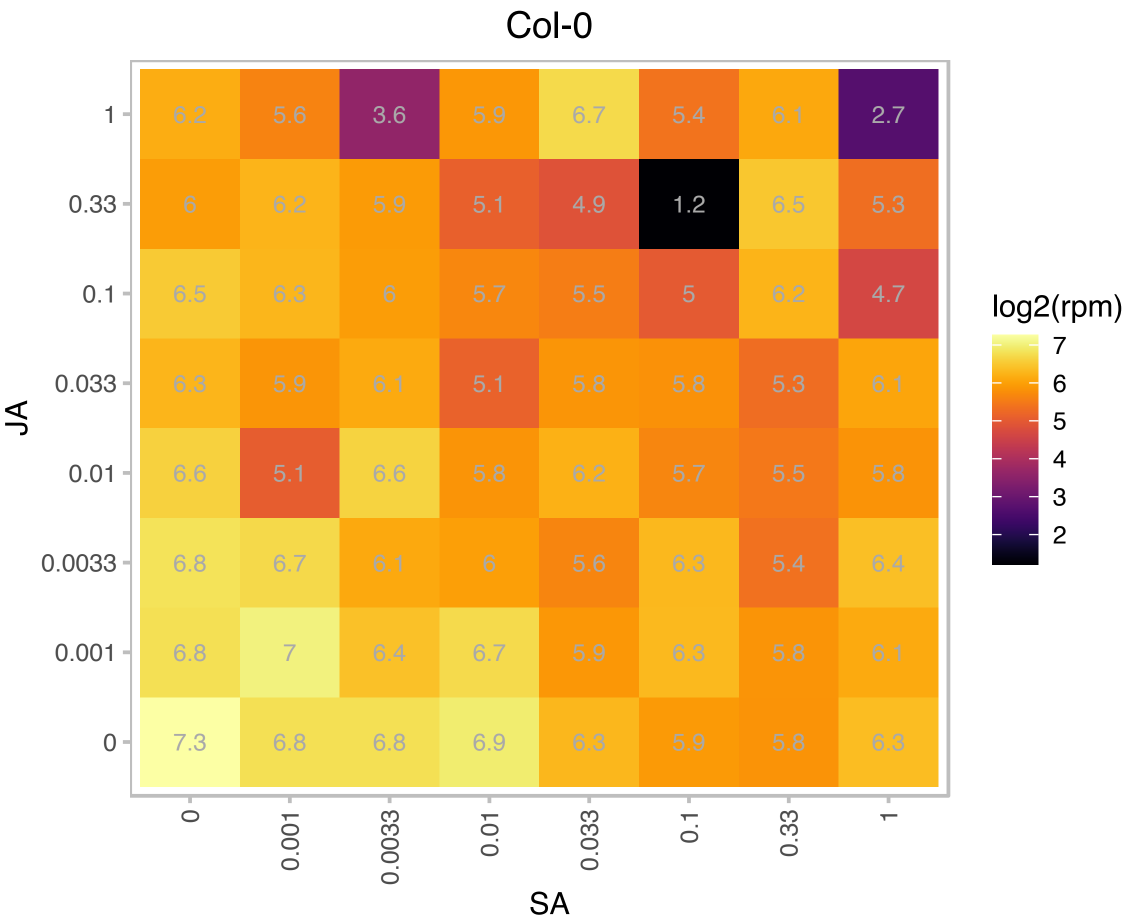


| | |
|---|---|
| Gene | AT5G40730 |
| is.expressed_genes | TRUE |
| is.responsive_genes | TRUE |
| is.apcluster | F-05 |

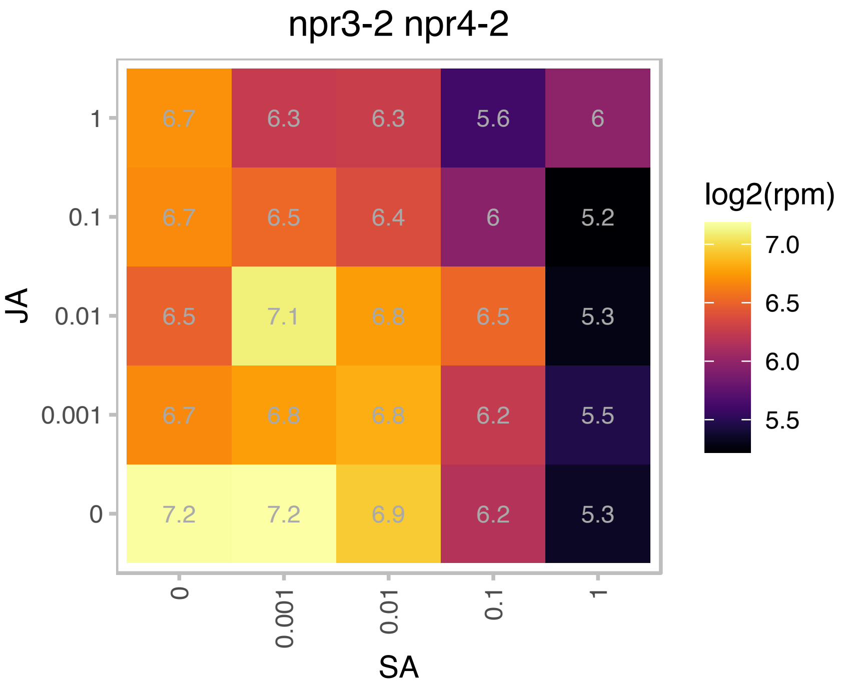


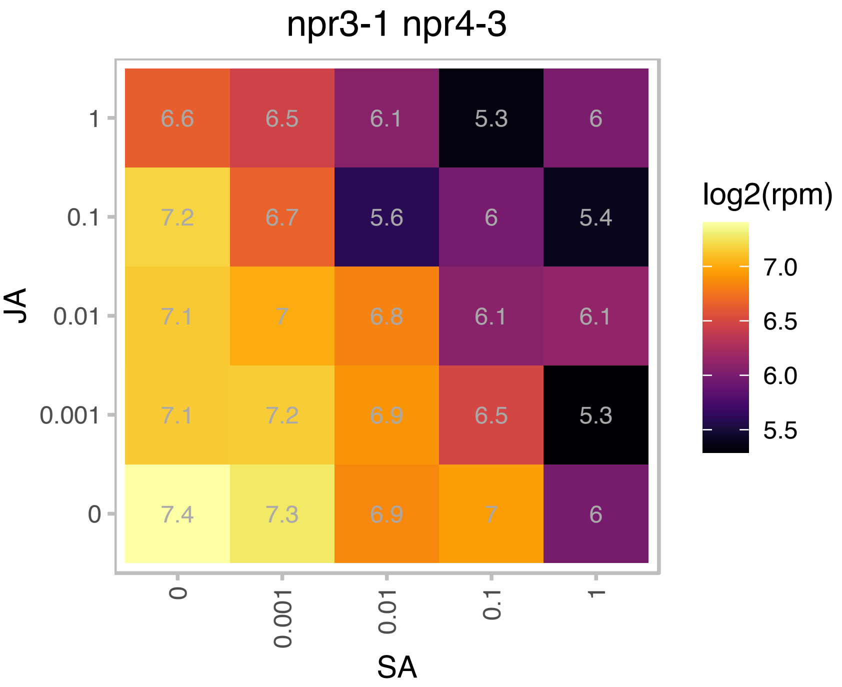

# AT5G41761.1 : NA

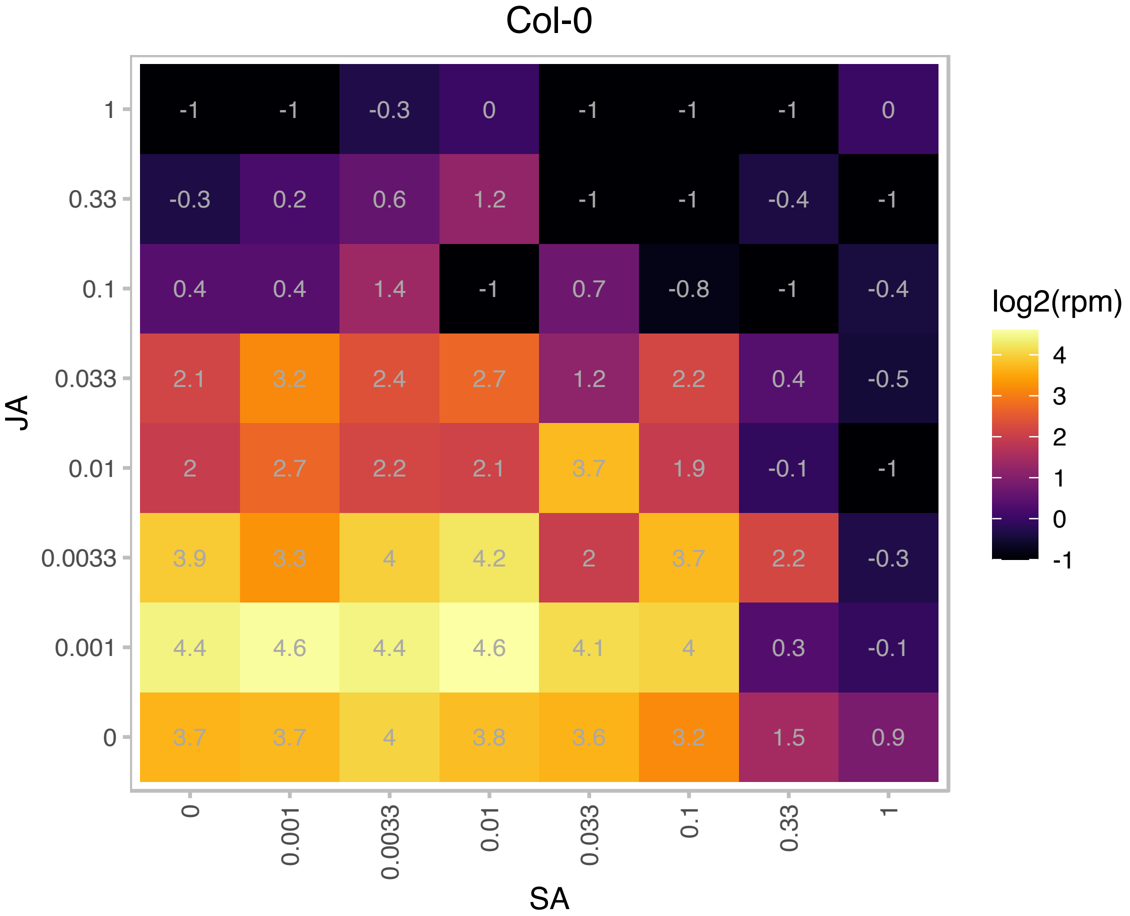


| | |
|---|---|
| Gene | AT5G41761 |
| is.expressed_genes | TRUE |
| is.responsive_genes | TRUE |
| is.apcluster | F-05 |

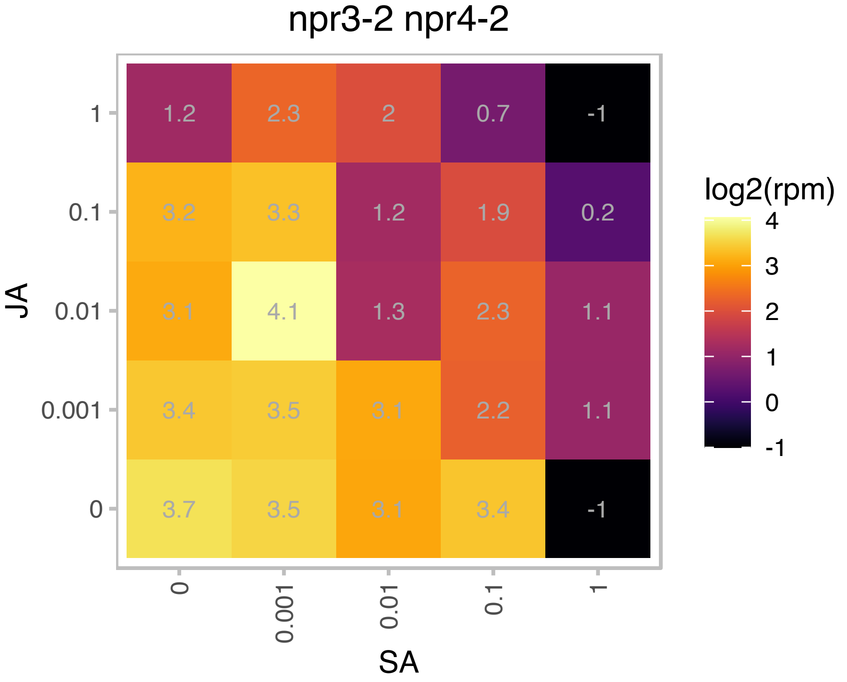


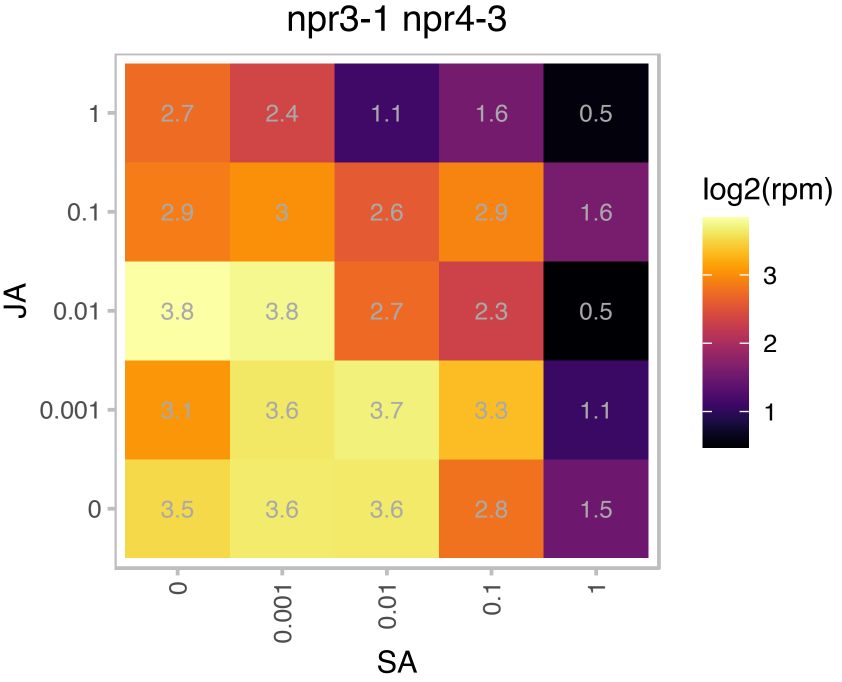

# AT5G44020.1 : NA

HAD superfamily, subfamily IIIB acid phosphatase

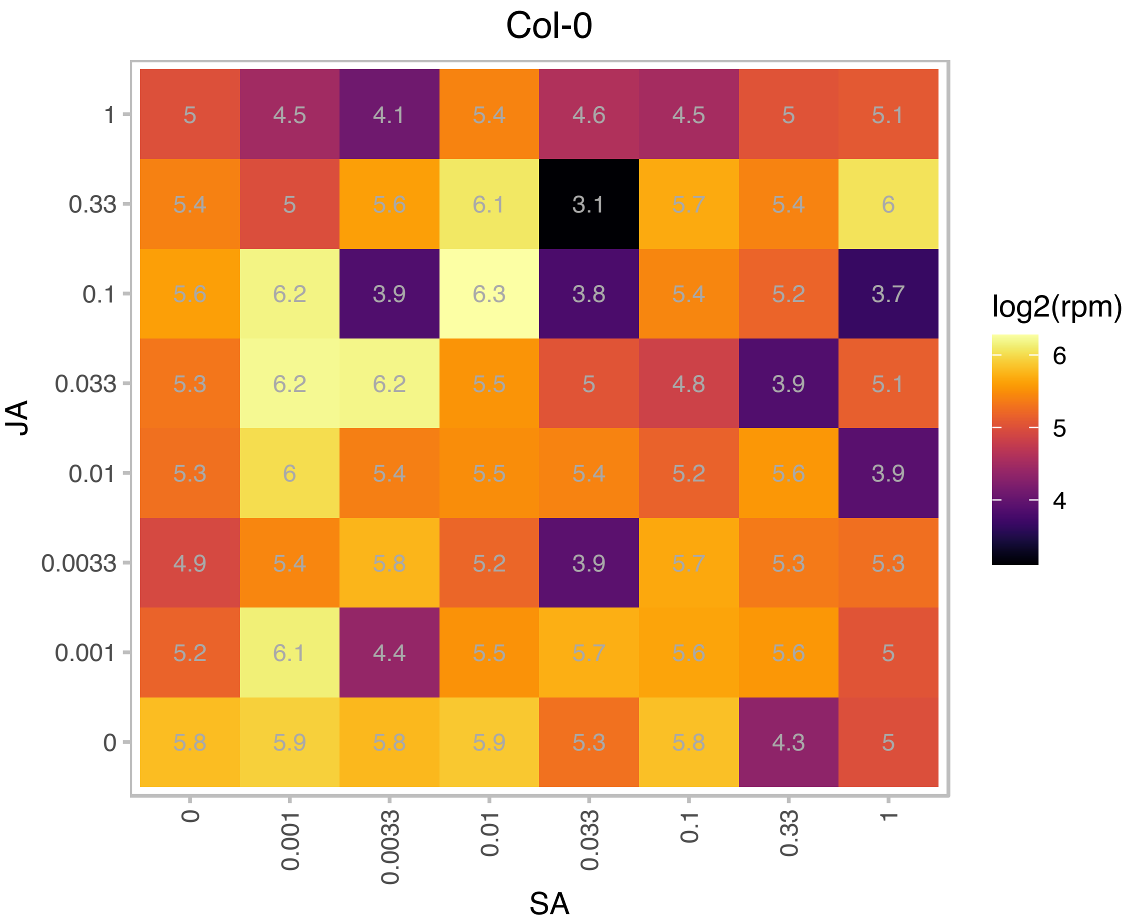


| | |
|---|---|
| Gene | AT5G44020 |
| is.expressed_genes | TRUE |
| is.responsive_genes | TRUE |
| is.apcluster | F-05 |

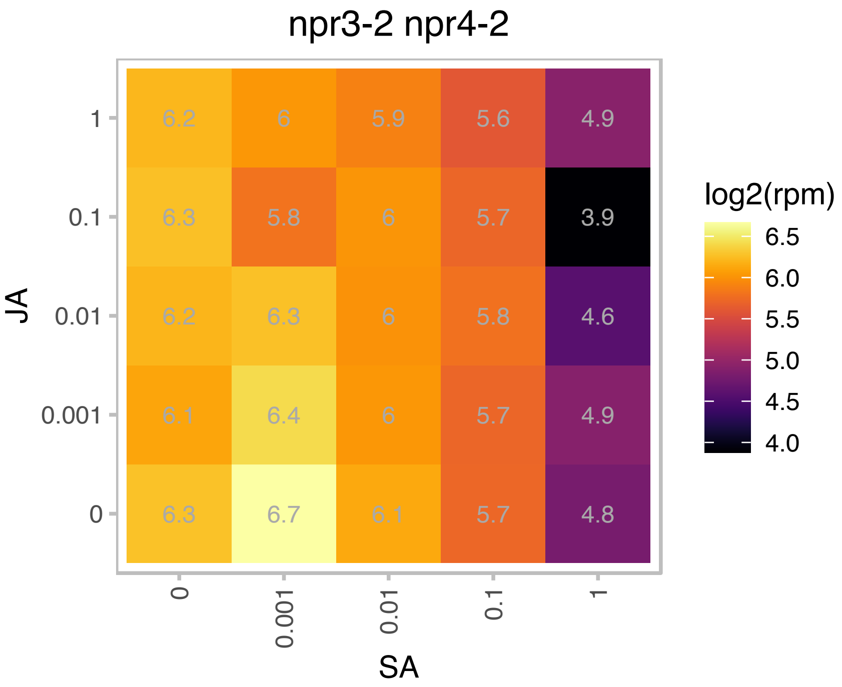


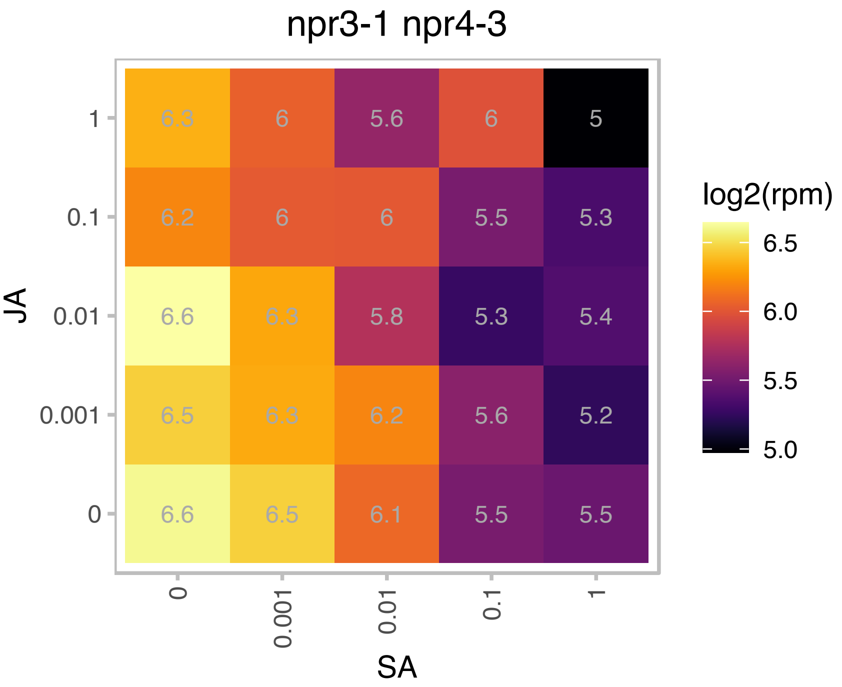

### AT5G44785.1 : OSB3

organellar single-stranded DNA binding protein 3

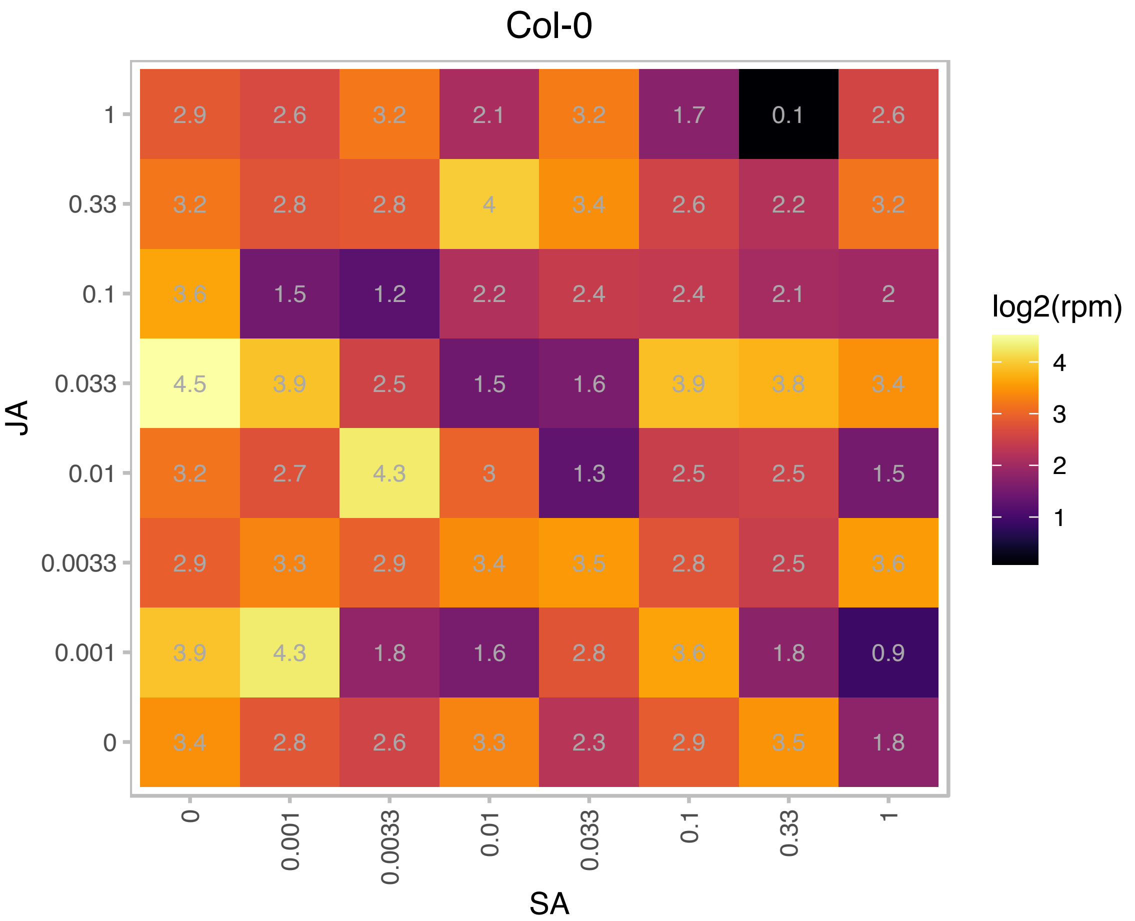


| | |
|---|---|
| Gene | AT5G44785 |
| is.expressed_genes | TRUE |
| is.responsive_genes | TRUE |
| is.apcluster | F-05 |

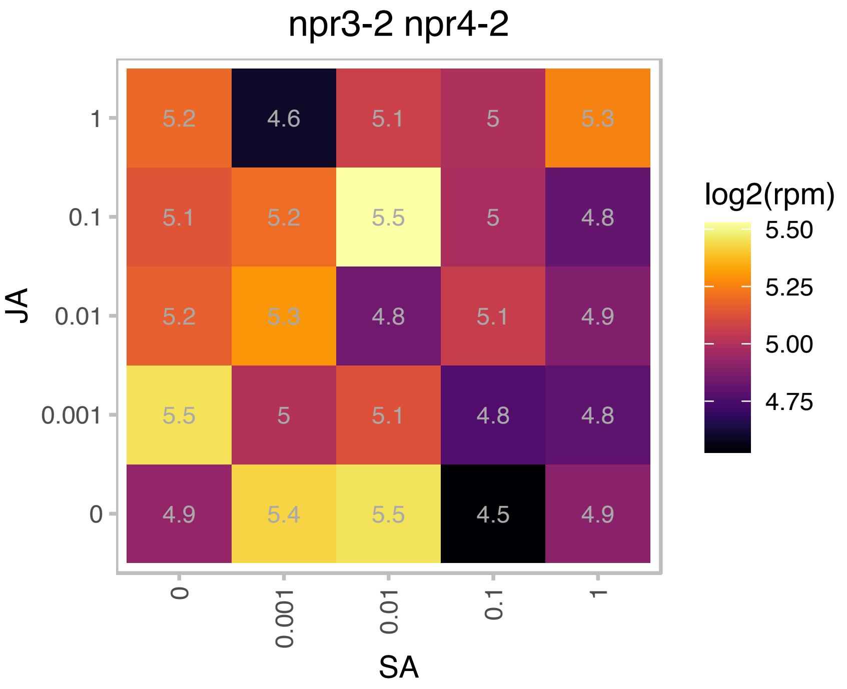


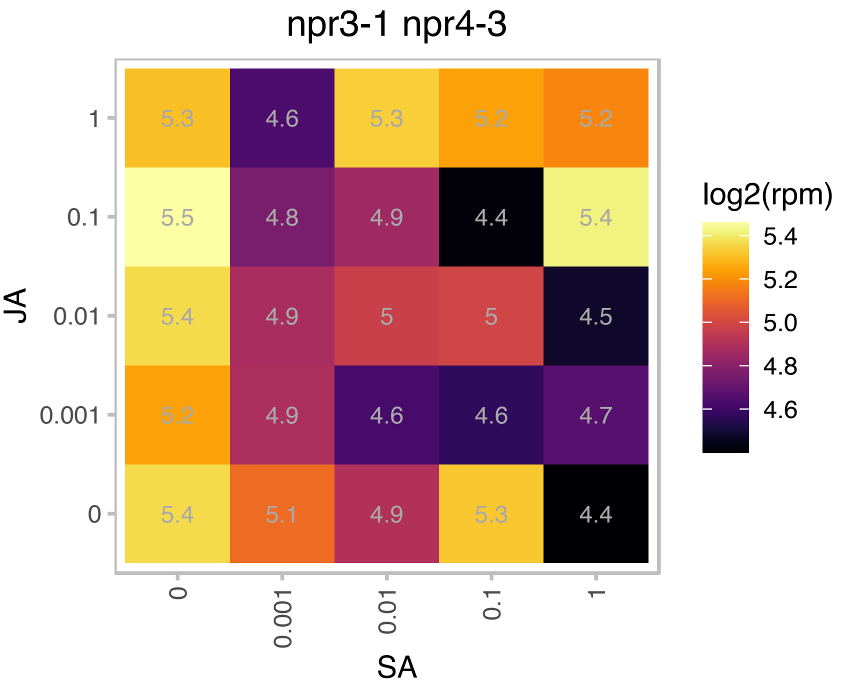

### AT5G45040.1 : CYTC6A

Cytochrome c
Encodes a Class I cytochrome c family member possessing a high structural homology with photosynthetic cytochrome c(6) from cyanobacteria, but structurally and functionally distinct through the presence of a disulfide bond.

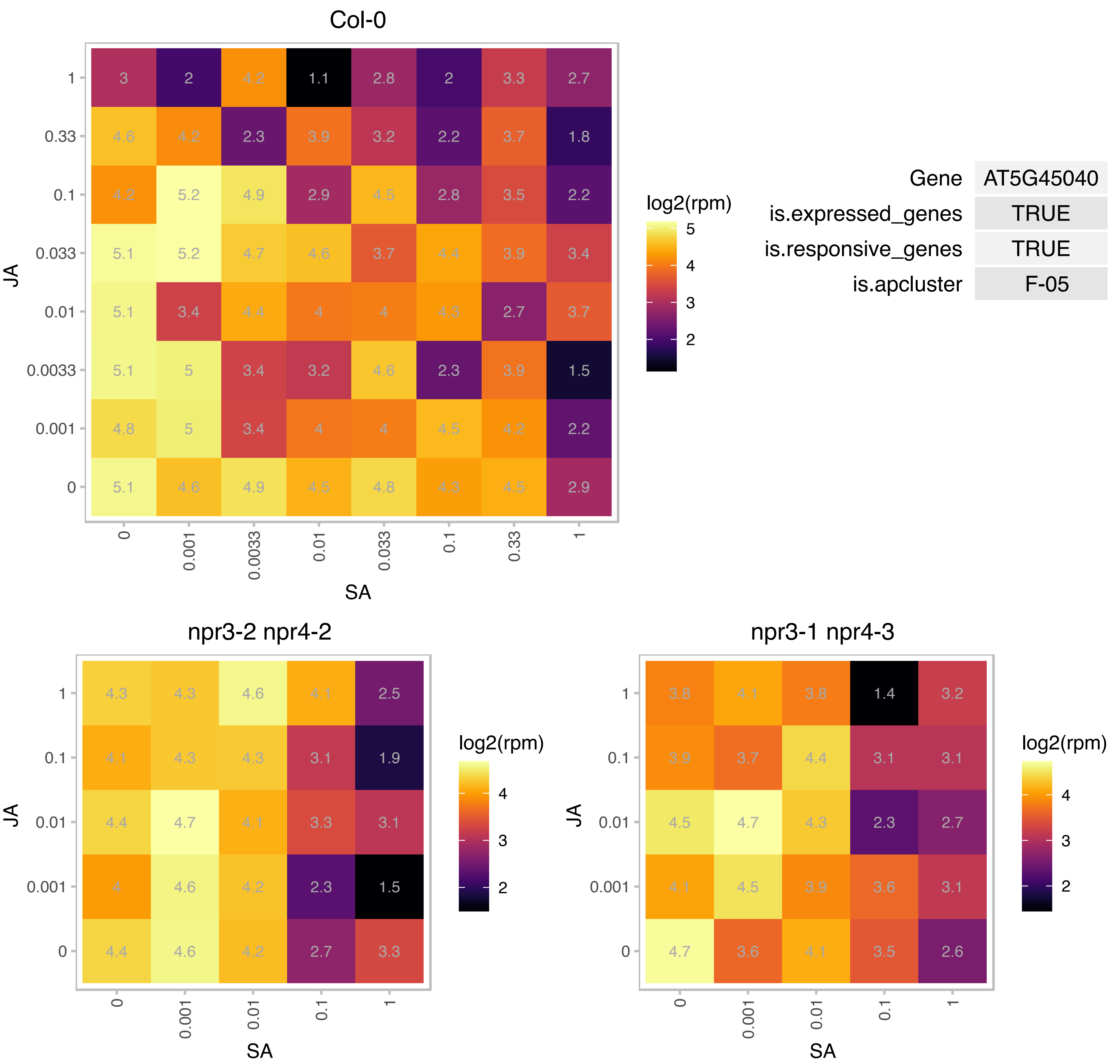


| | |
|---|---|
| Gene | AT5G45040 |
| is.expressed_genes | TRUE |
| is.responsive_genes | TRUE |
| is.apcluster | F-05 |

# AT5G46890.1 : NA

Bifunctional inhibitor/lipid-transfer protein/seed storage 2S albumin superfamily protein

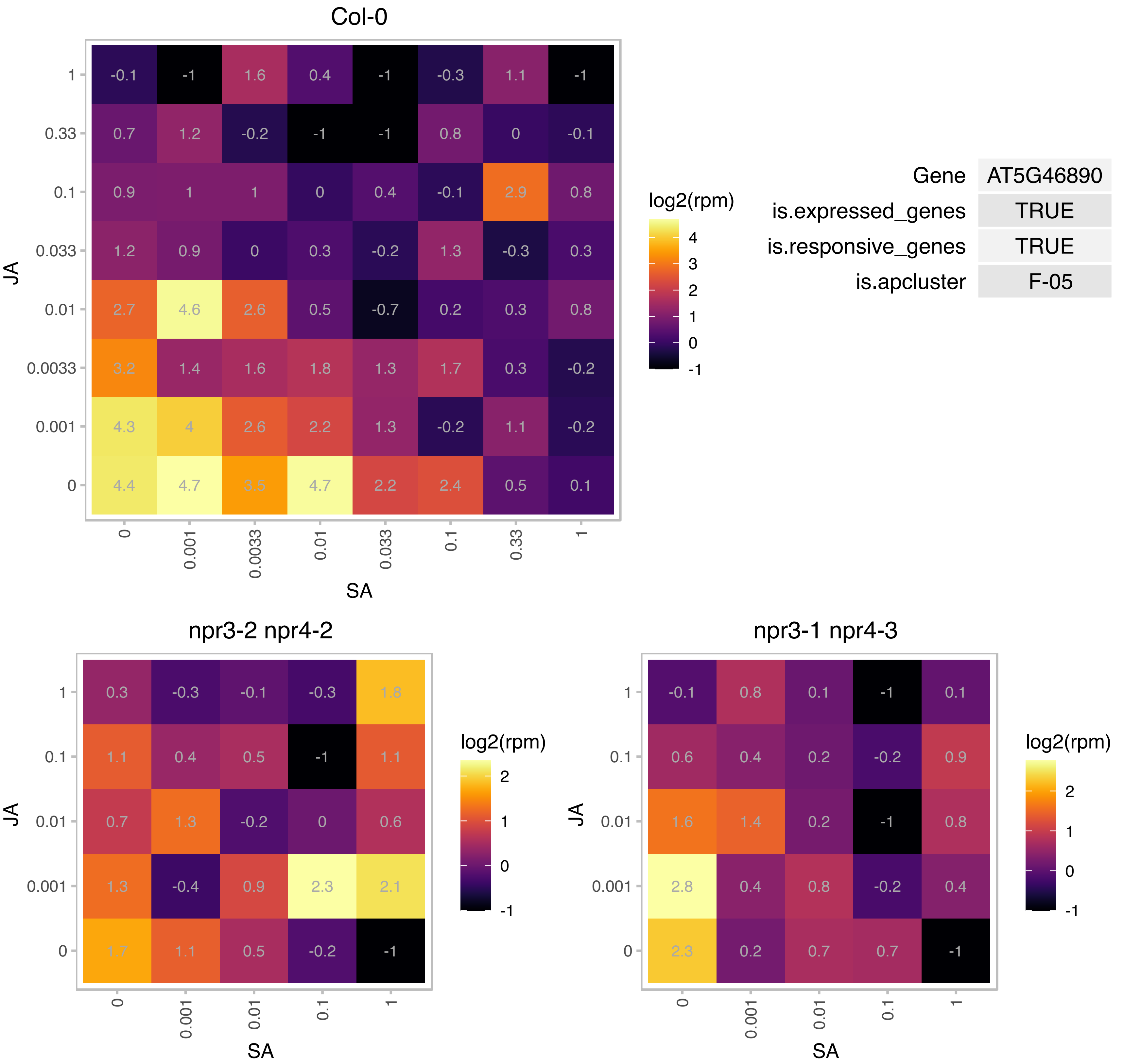


| | |
|---|---|
| Gene | AT5G46890 |
| is.expressed_genes | TRUE |
| is.responsive_genes | TRUE |
| is.apcluster | F-05 |

# AT5G49560.1 : NA

Putative methyltransferase family protein

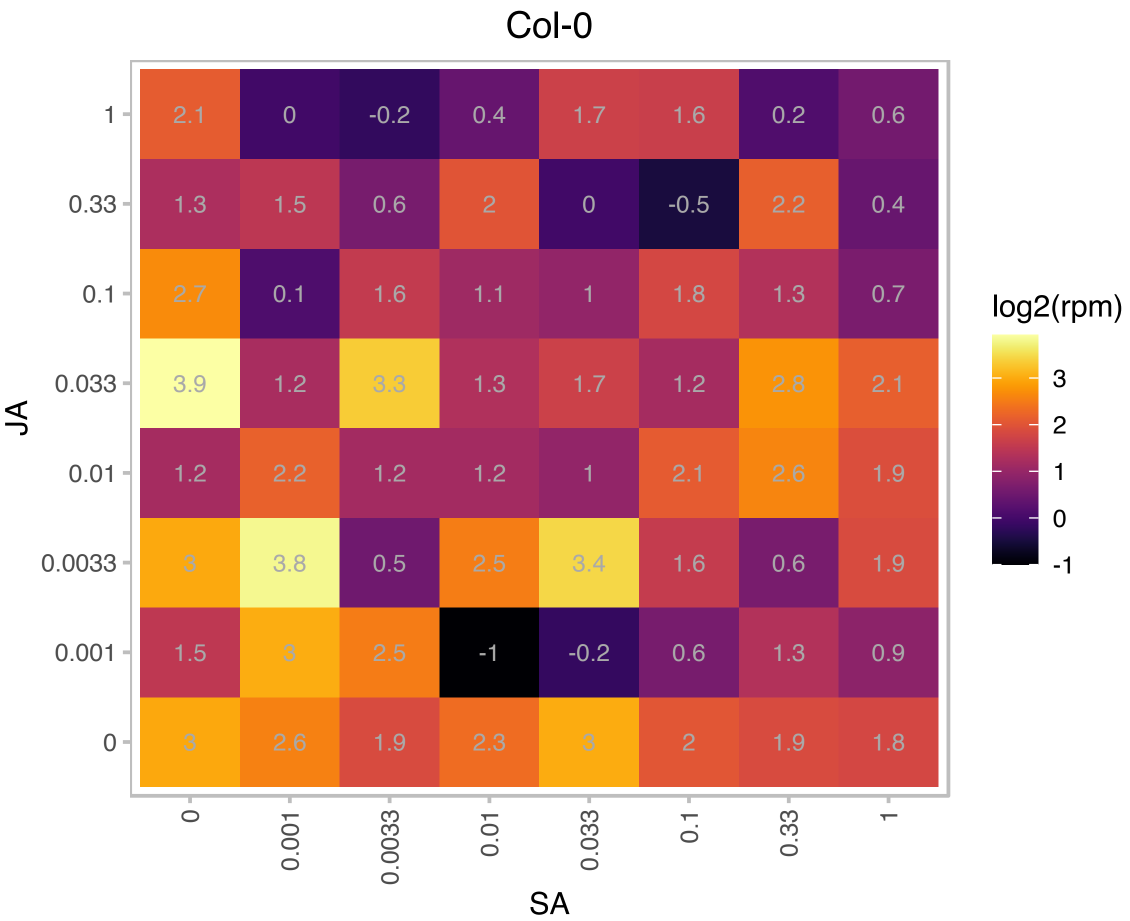


| | |
|---|---|
| Gene | AT5G49560 |
| is.expressed_genes | TRUE |
| is.responsive_genes | TRUE |
| is.apcluster | F-05 |

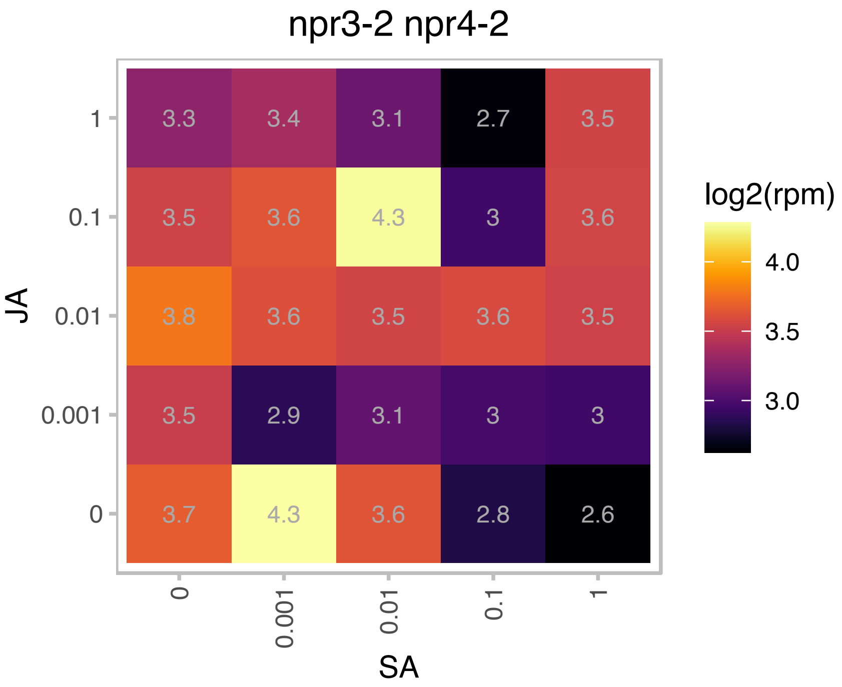


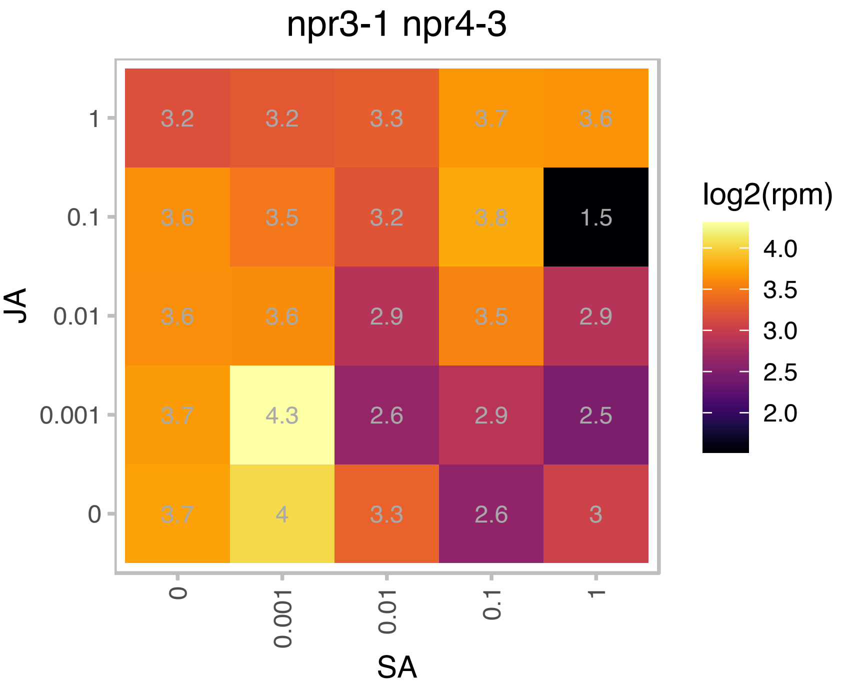

# AT5G50560.1 : NA

Protein of unknown function (DUF 3339)

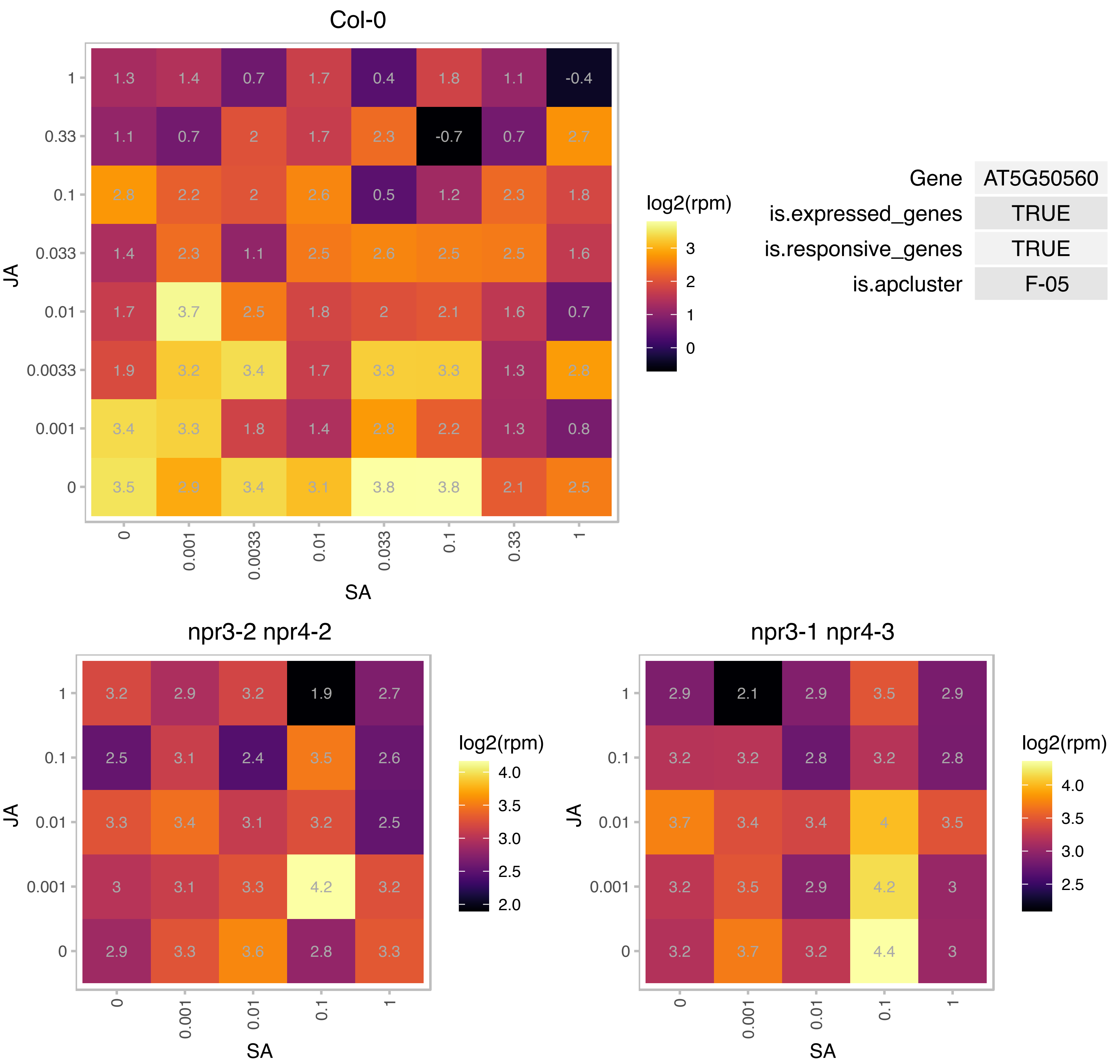


| | |
|---|---|
| Gene | AT5G50560 |
| is.expressed_genes | TRUE |
| is.responsive_genes | TRUE |
| is.apcluster | F-05 |

# AT5G50660.1 : NA

Protein of unknown function (DUF 3339)

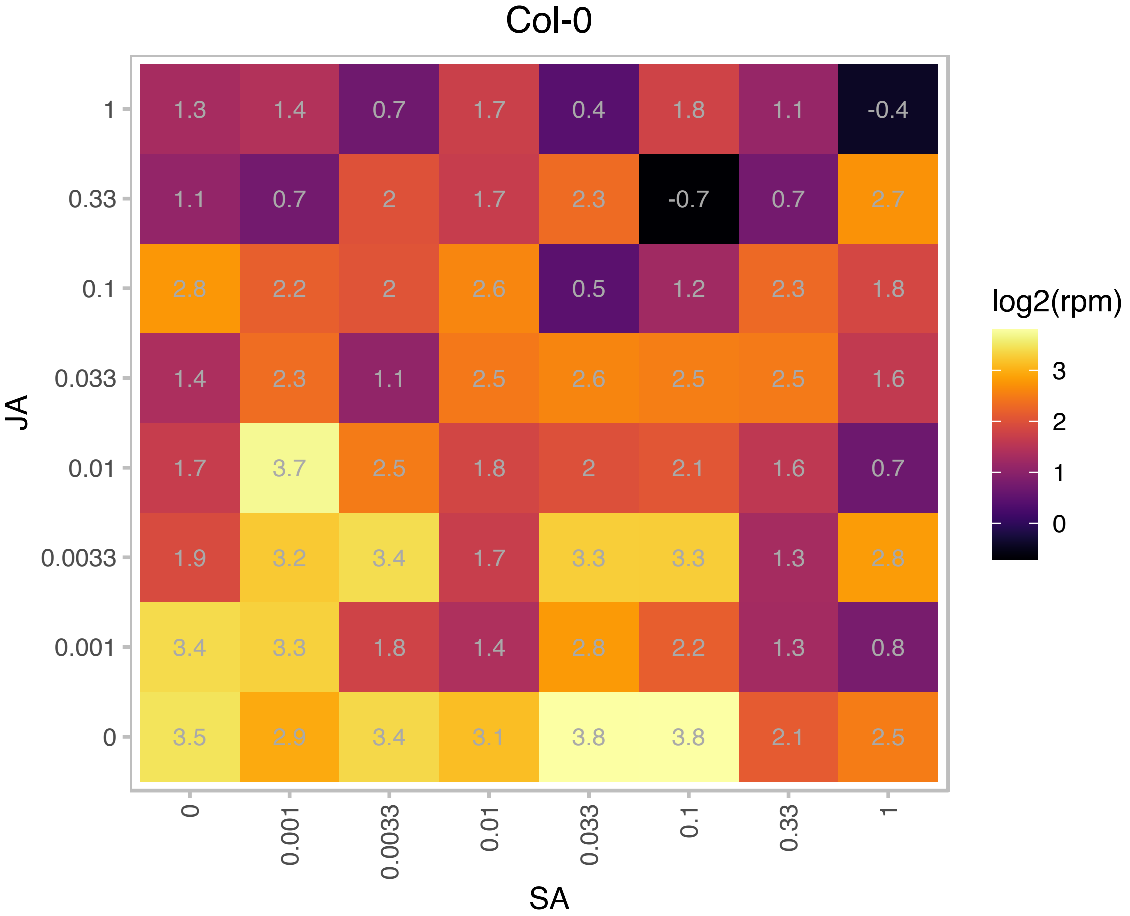


| | |
|---|---|
| Gene | AT5G50660 |
| is.expressed_genes | TRUE |
| is.responsive_genes | TRUE |
| is.apcluster | F-05 |

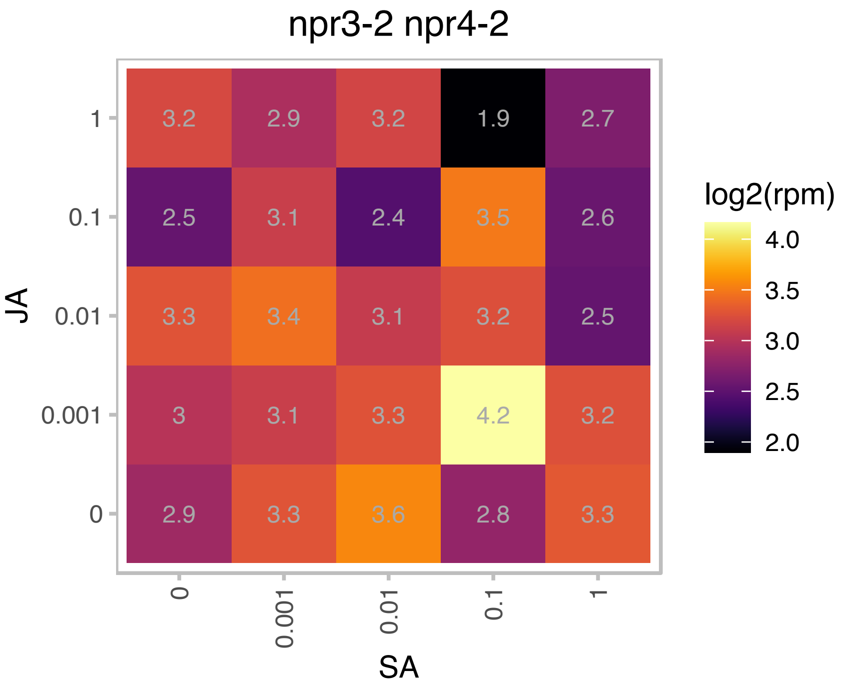


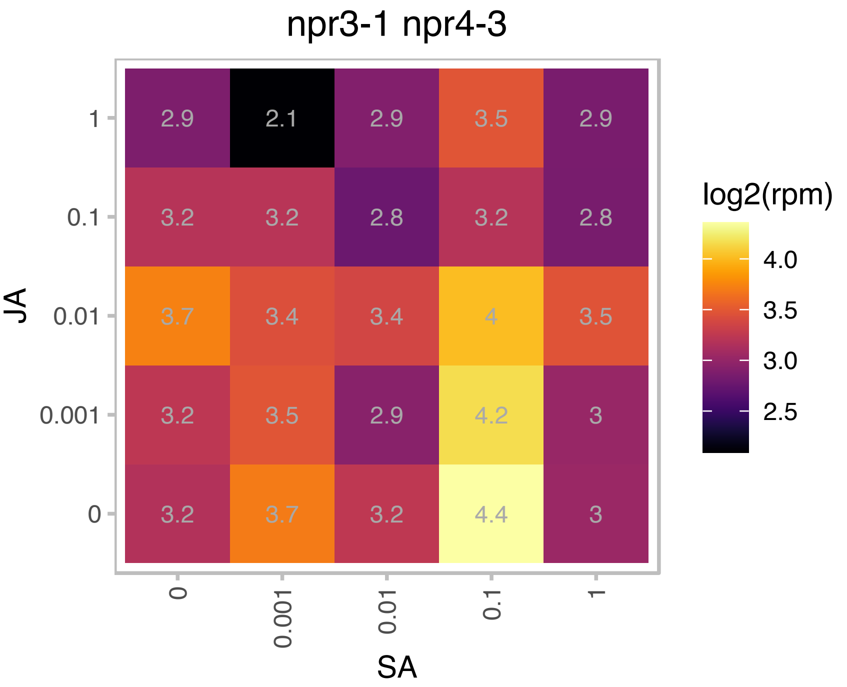

### AT5G51545.1 : LPA2

low psii accumulation2

Encodes LPA2 (low psii accumulation2), an intrinsic thylakoid membrane protein required for efficient assembly of Photosystem II.

| | |
|---|---|
| Gene | AT5G51545 |
| is.expressed_genes | TRUE |
| is.responsive_genes | TRUE |
| is.apcluster | F-05 |

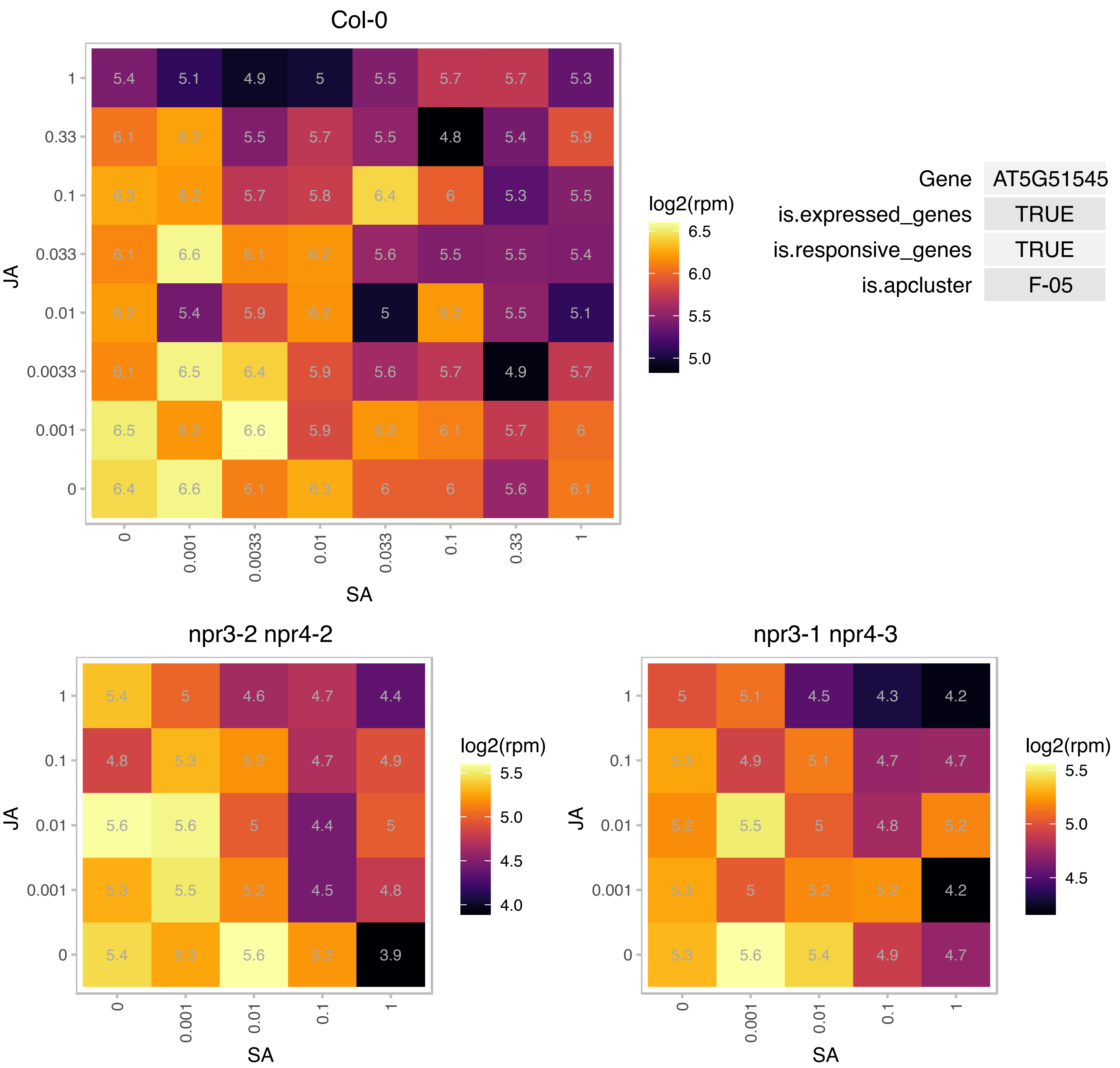

# AT5G56100.1 : NA

glycine-rich protein / oleosin

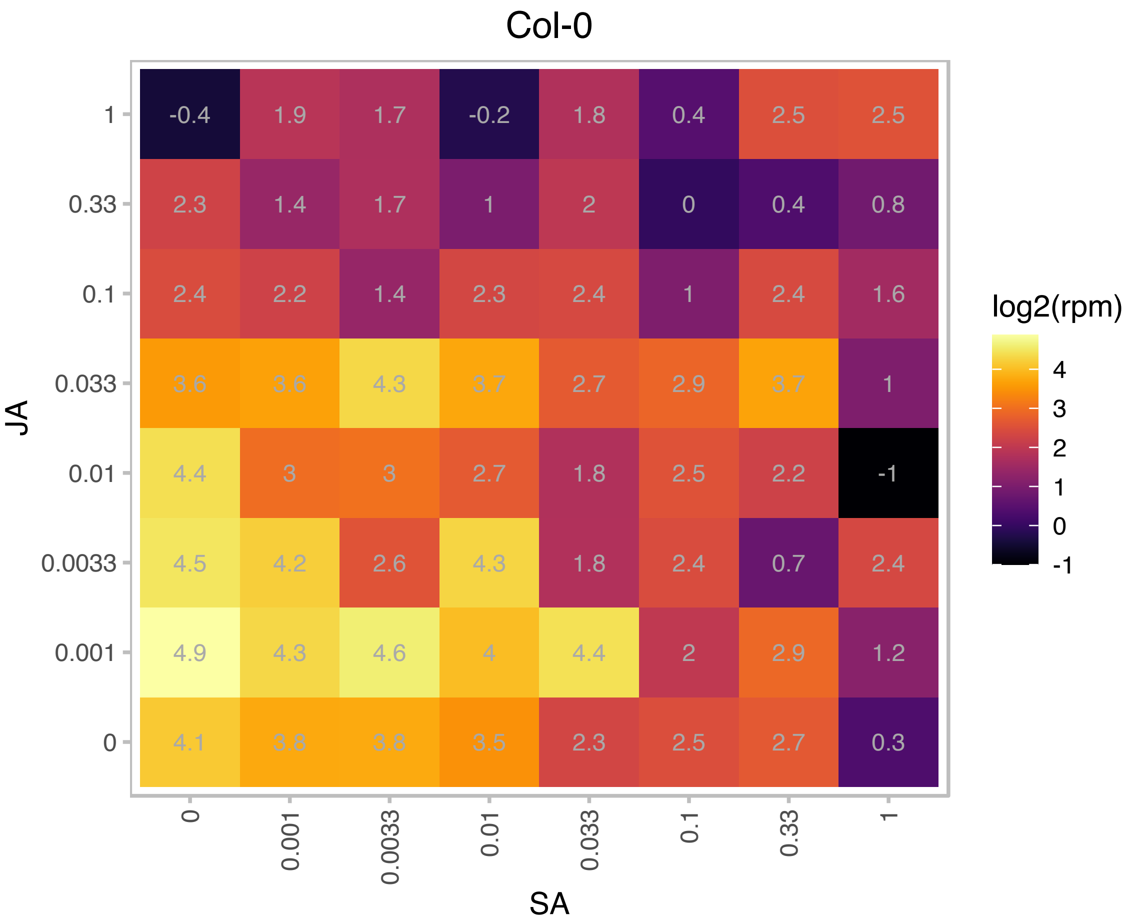


| | |
|---|---|
| Gene | AT5G56100 |
| is.expressed_genes | TRUE |
| is.responsive_genes | TRUE |
| is.apcluster | F-05 |

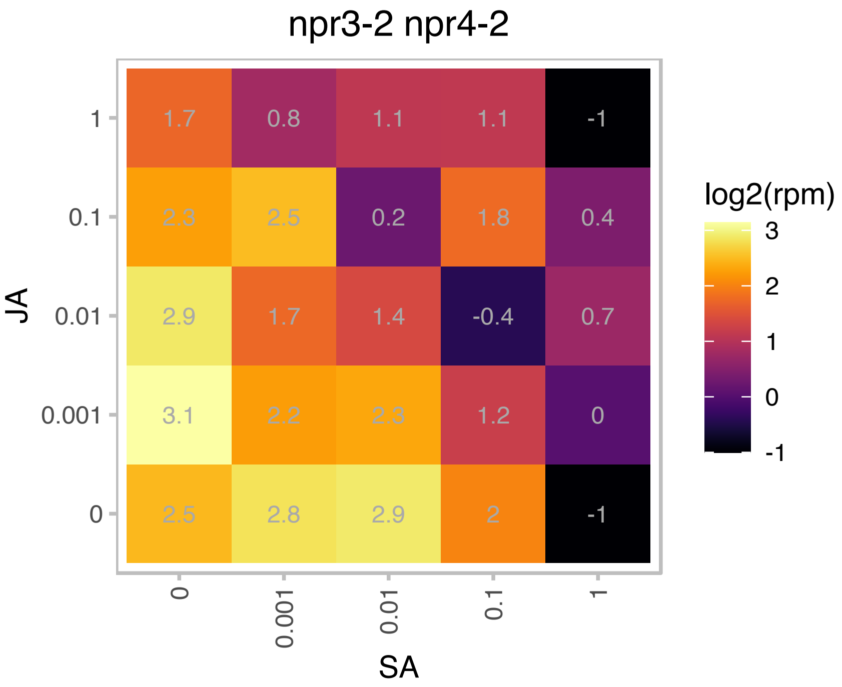


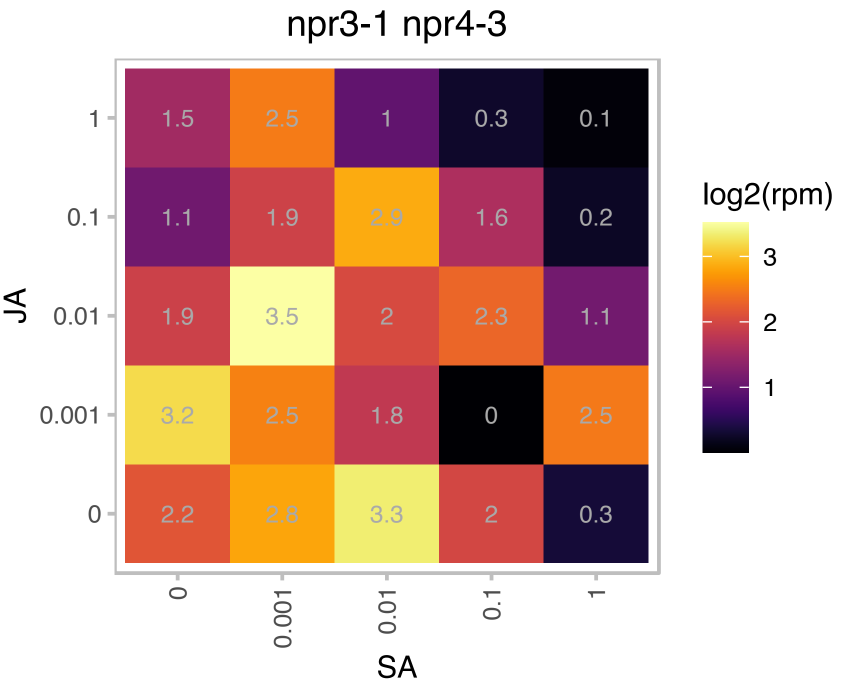

# AT5G57760.1 : NA

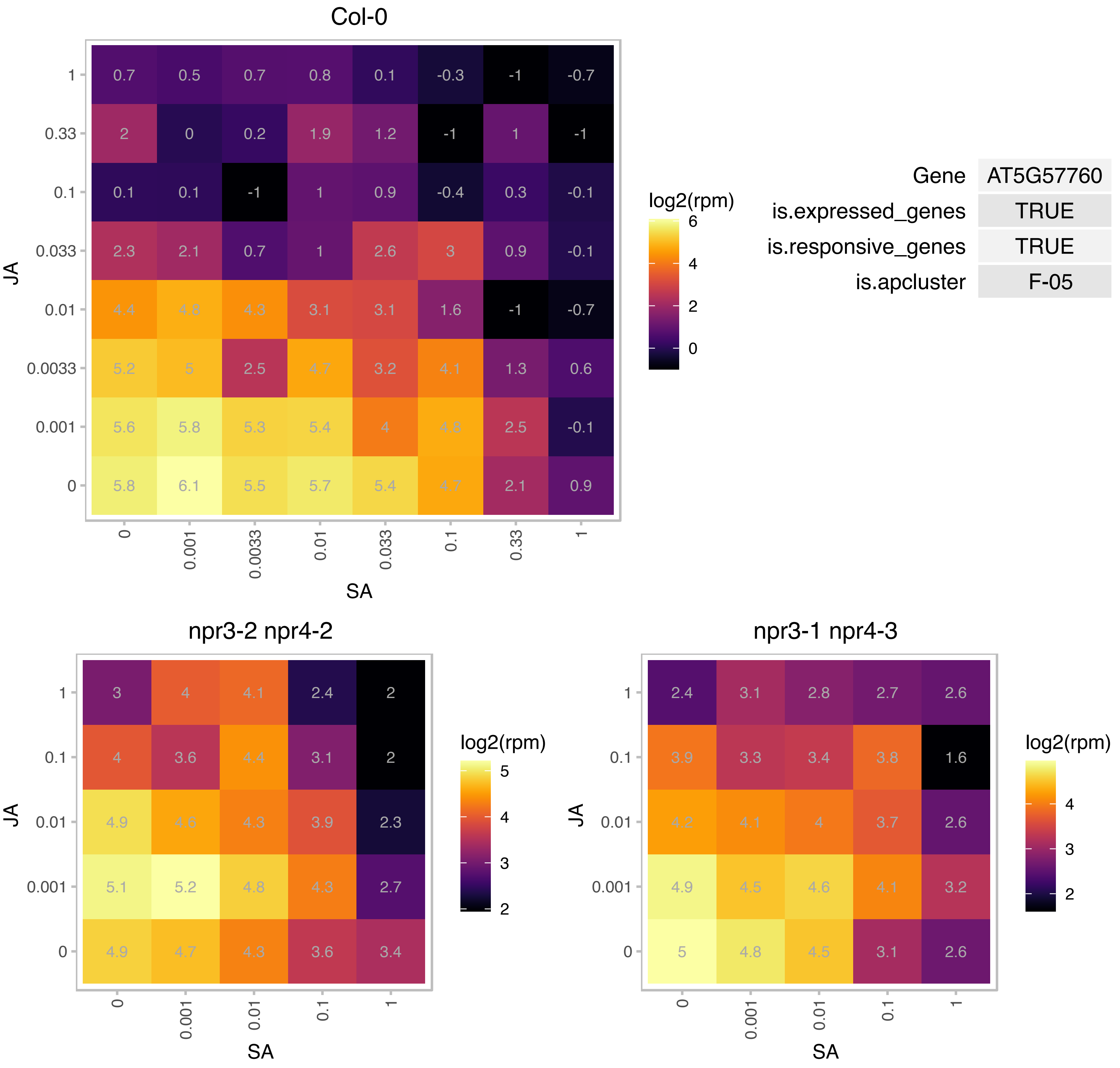

### AT5G58250.1 : EMB3143, LCAA/YCF54

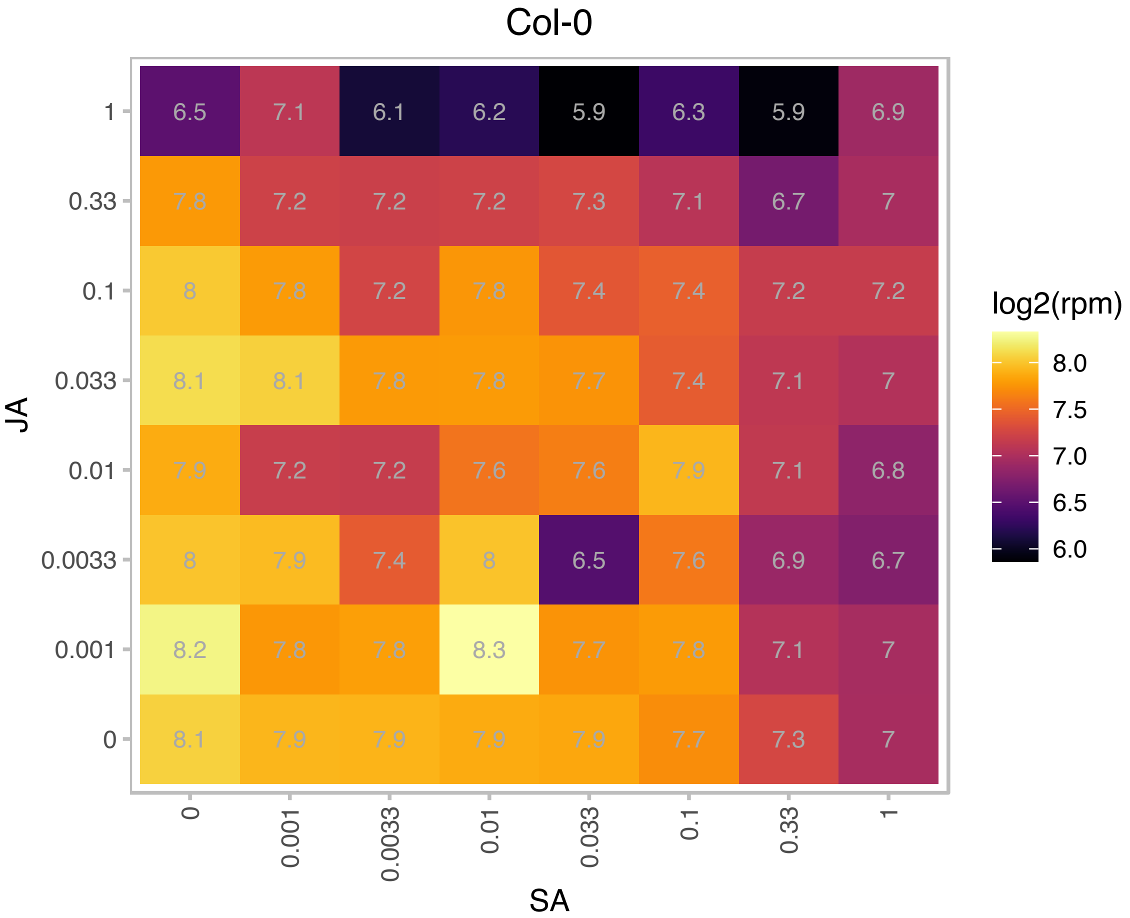


| | |
|---|---|
| Gene | AT5G58250 |
| is.expressed_genes | TRUE |
| is.responsive_genes | TRUE |
| is.apcluster | F-05 |

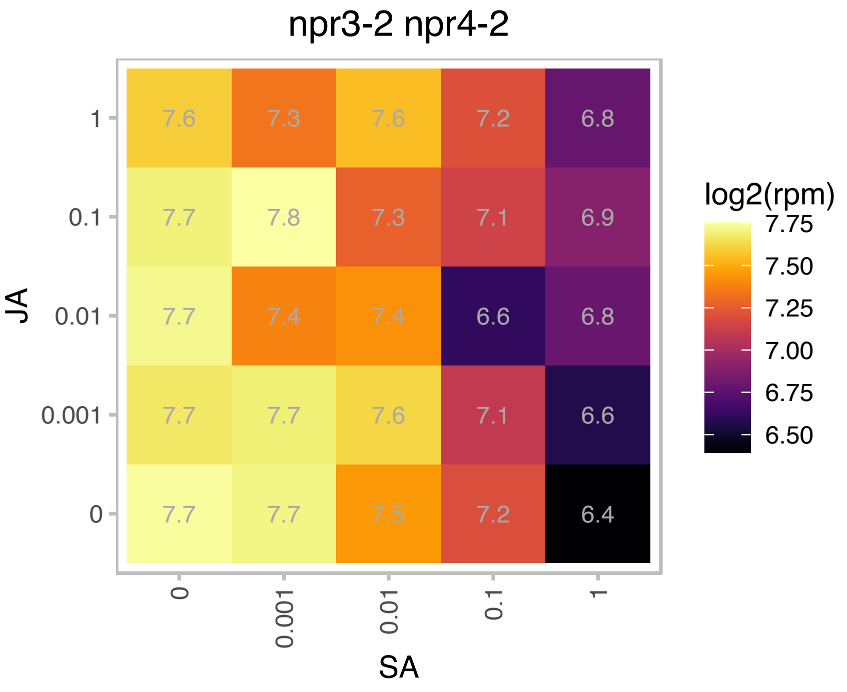


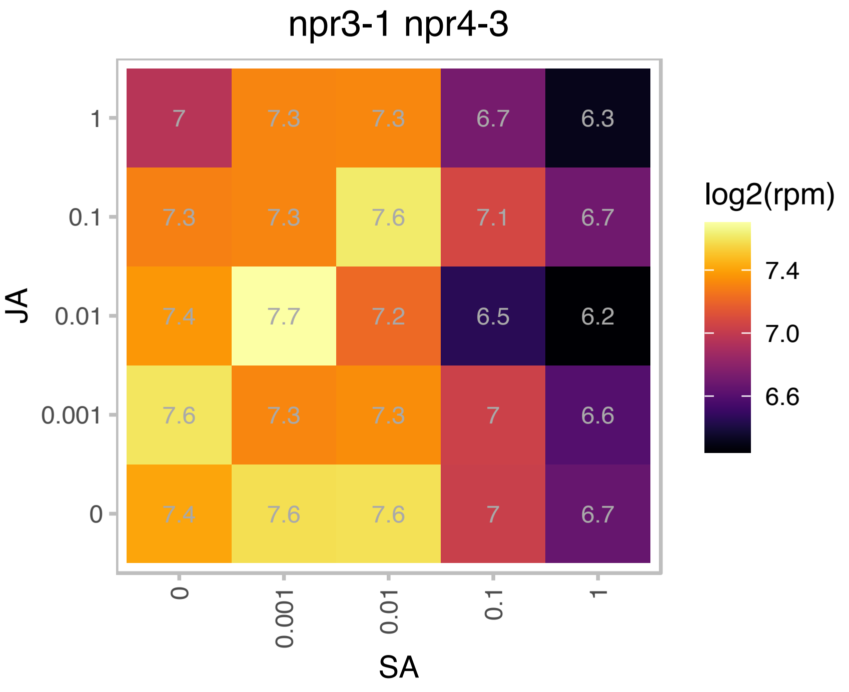

### AT5G58260.1 : NdhN

oxidoreductases, acting on NADH or NADPH, quinone or similar compound as acceptor

Encodes subunit NDH-N of NAD(P)H:plastoquinone dehydrogenase complex (Ndh complex) present in the thylakoid membrane of chloroplasts. This subunit is thought to be required for Ndh complex assembly.

| | |
|---|---|
| Gene | AT5G58260 |
| is.expressed_genes | TRUE |
| is.responsive_genes | TRUE |
| is.apcluster | F-05 |

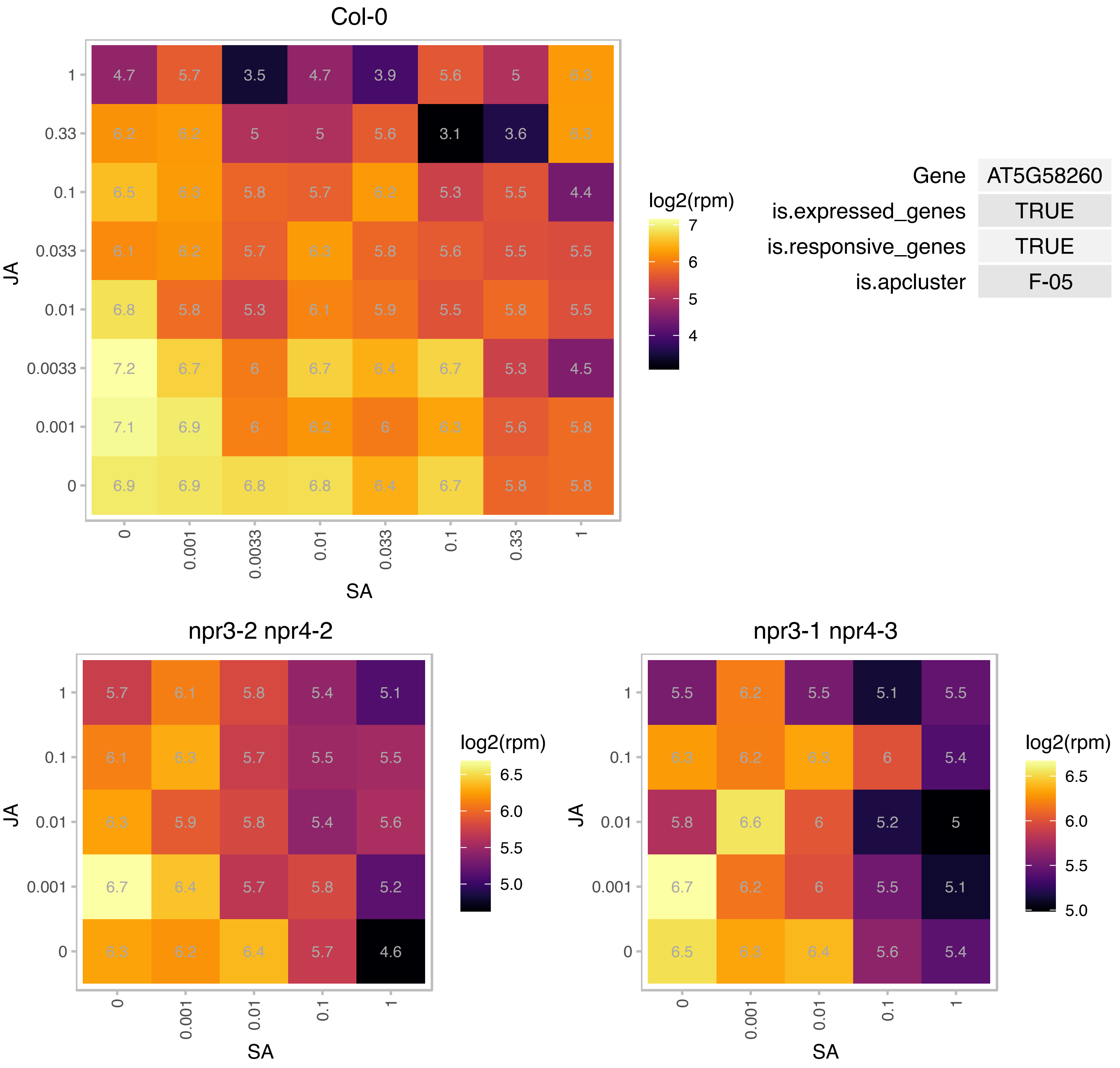

# AT5G58800.1 : NA

Quinone reductase family protein

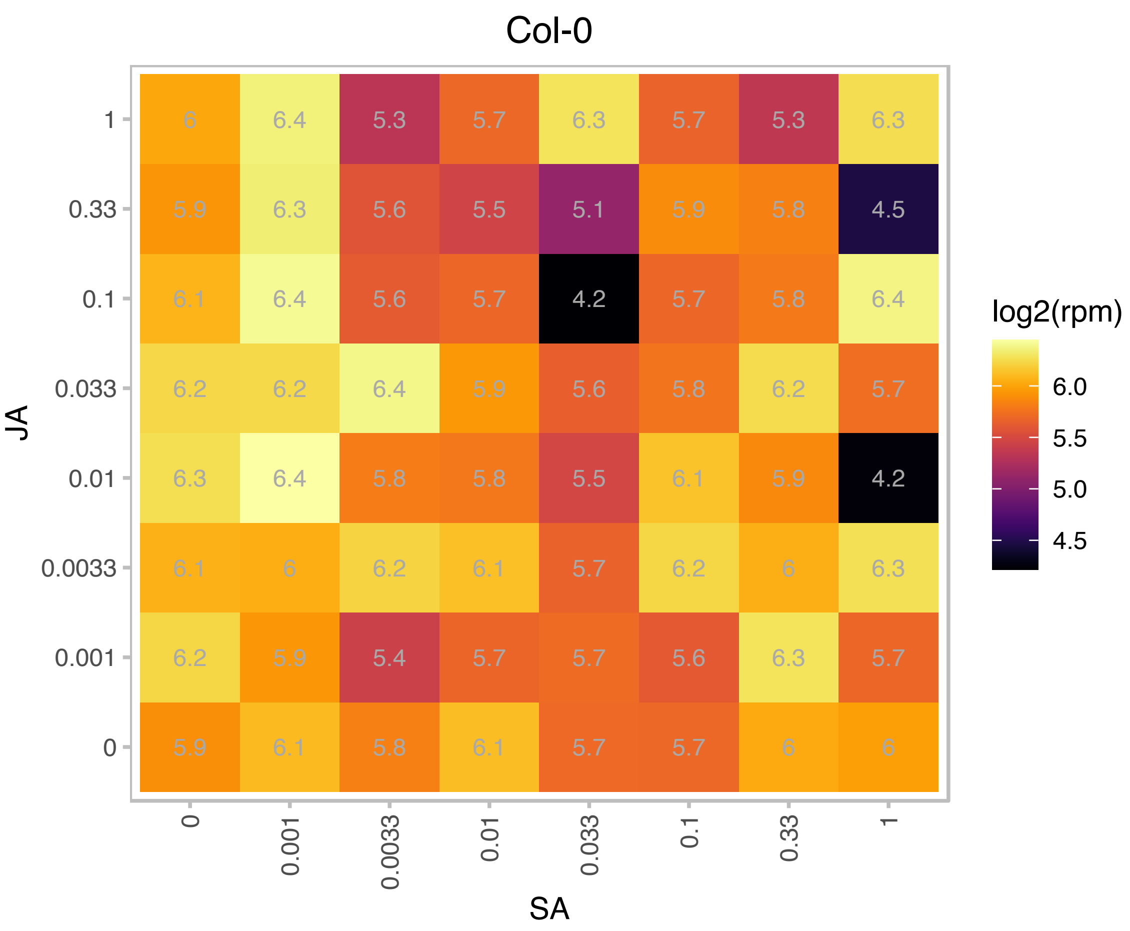


| | |
|---|---|
| Gene | AT5G58800 |
| is.expressed_genes | TRUE |
| is.responsive_genes | TRUE |
| is.apcluster | F-05 |

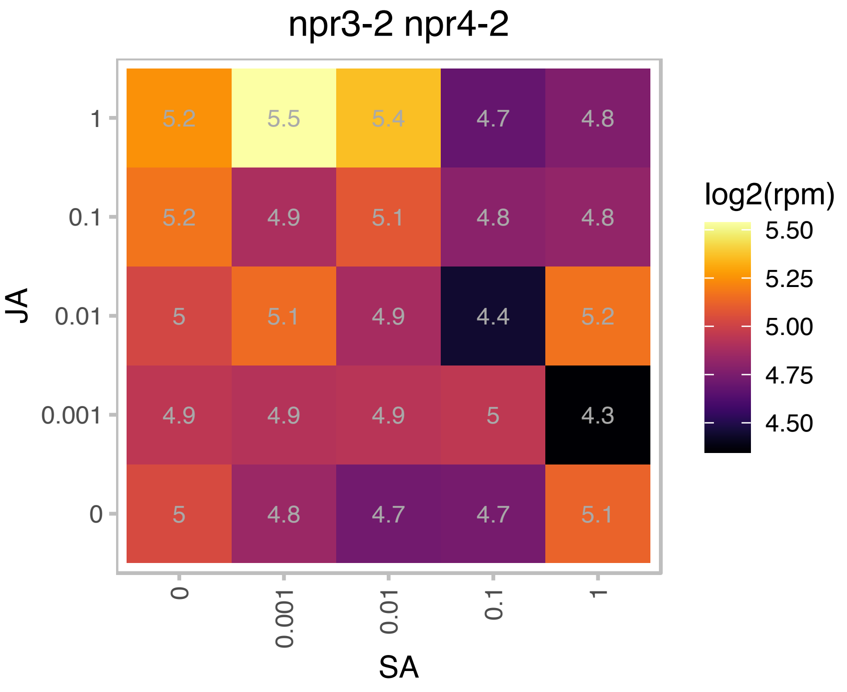


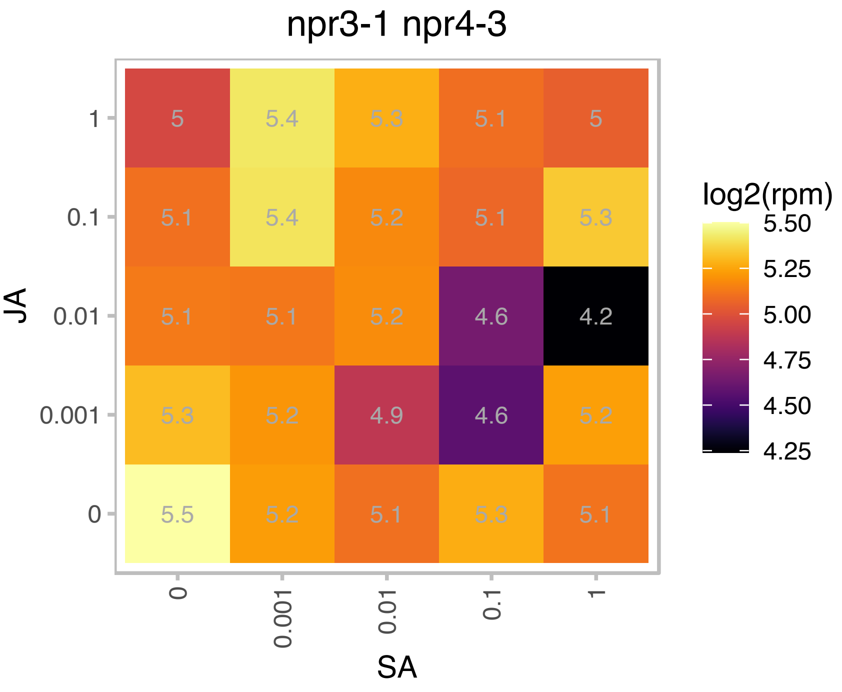

### AT5G64040.1 : PSAN

photosystem I reaction center subunit PSI-N, chloroplast, putative / PSI-N, putative (PSAN)
Encodes the only subunit of photosystem I located entirely in the thylakoid lumen. May be involved in the interaction between plastocyanin and the photosystem I complex. Phosphorylation of this protein is dependent on calcium.

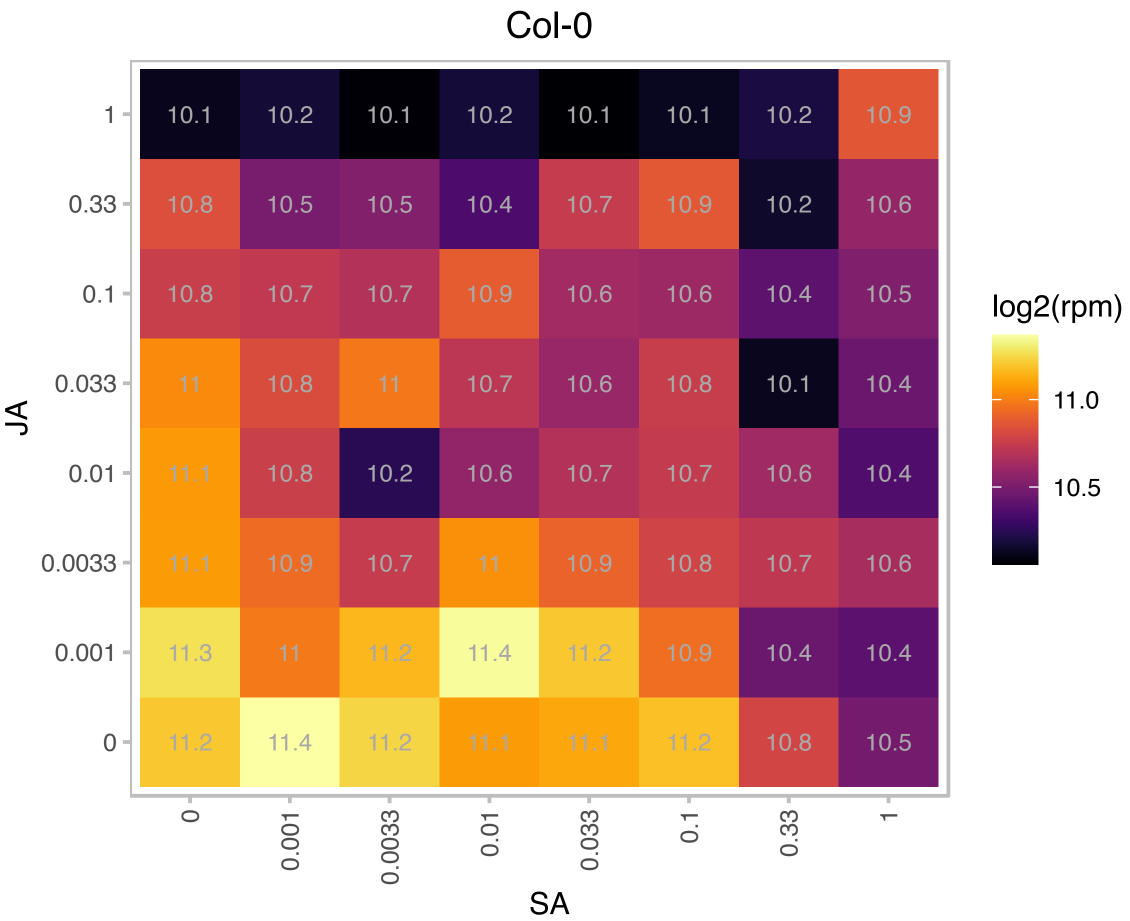


| | |
|---|---|
| Gene | AT5G64040 |
| is.expressed_genes | TRUE |
| is.responsive_genes | TRUE |
| is.apcluster | F-05 |

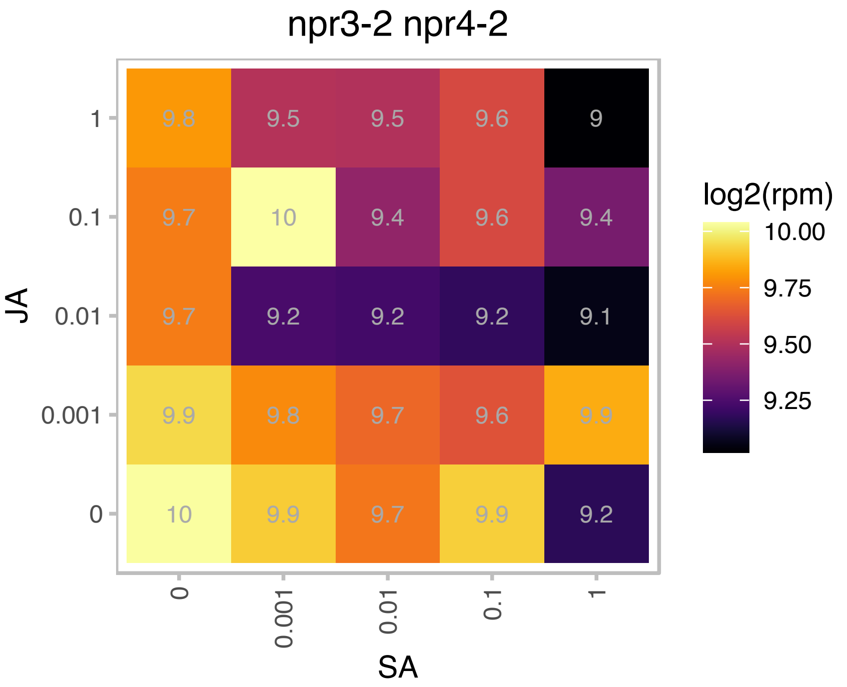


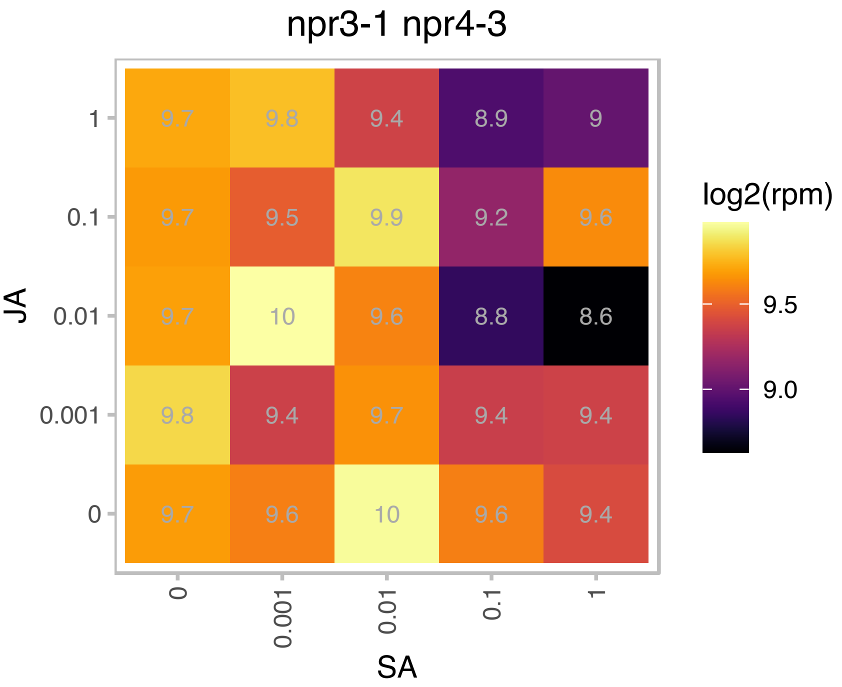

### AT5G67270.1 : ATEB1C, ATEB1H1, EB1C

end binding protein 1C

encodes a homolog of animal microtubule-end-binding protein. There are two other members of this family. EB1 forms foci at regions where the minus ends of microtubules are gathered during mitosis and early cytokinesis.

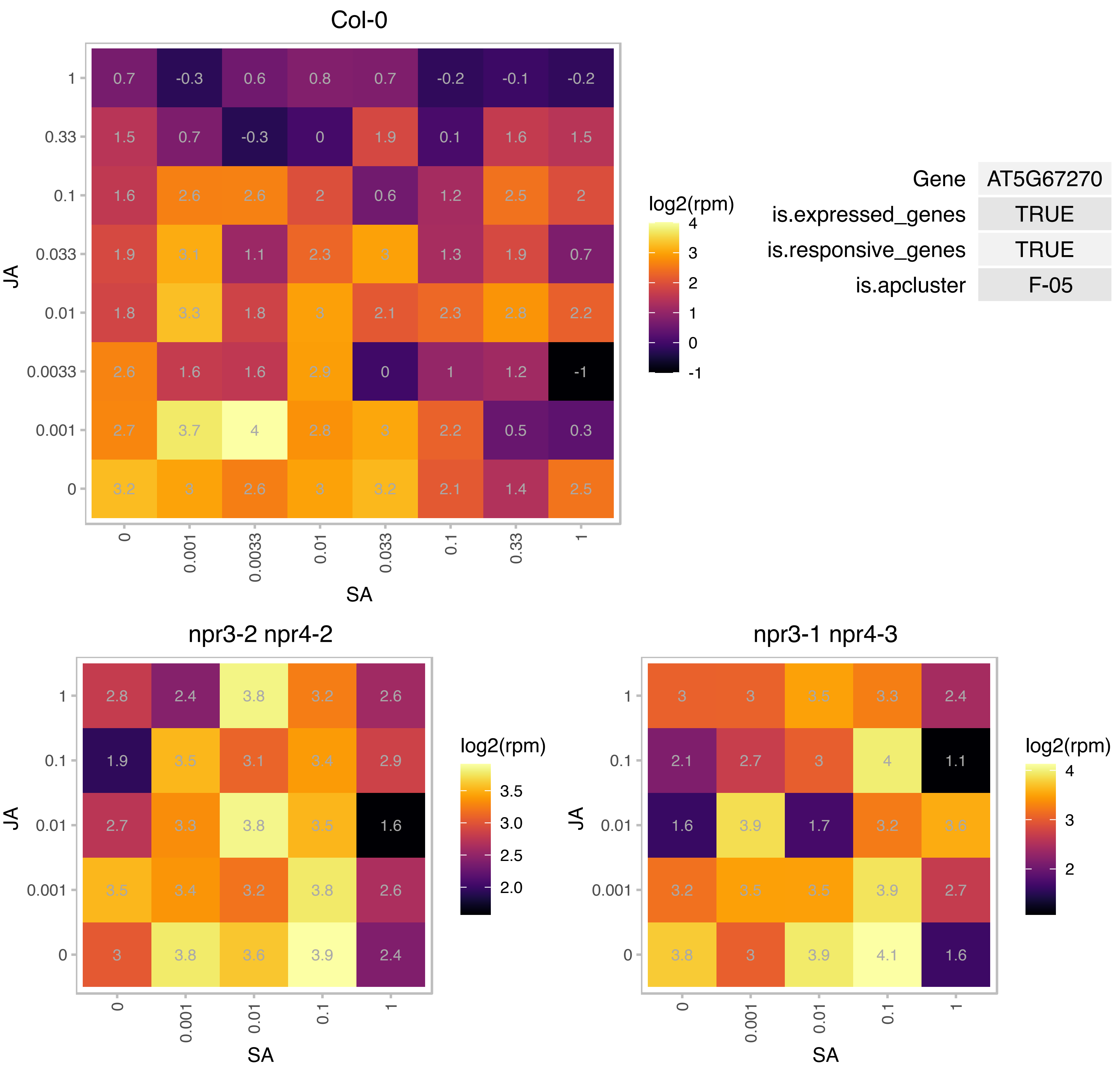


| | |
|---|---|
| Gene | AT5G67270 |
| is.expressed_genes | TRUE |
| is.responsive_genes | TRUE |
| is.apcluster | F-05 |

### AT1G02280.1 : ATTOC33, PPI1, TOC33

translocon at the outer envelope membrane of chloroplasts 33

Encodes a GTP-binding GTP-ase. Component of the chloroplast protein import machinery. Required for import of POR B into plastids. Toc33 phosphorylation may not play an important role in vivo.

Col-0

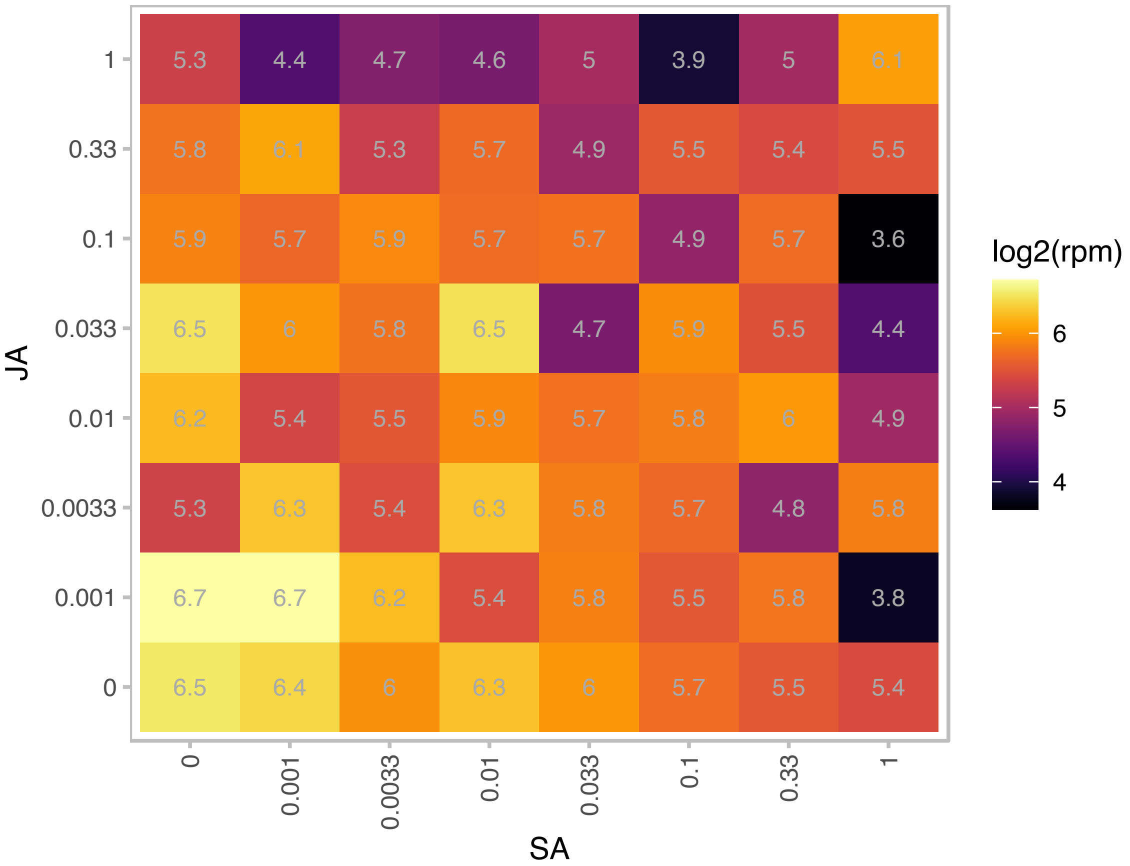


| | |
|---|---|
| Gene | AT1G02280 |
| is.expressed_genes | TRUE |
| is.responsive_genes | TRUE |
| is.apcluster | F-06 |

npr3-2 npr4-2

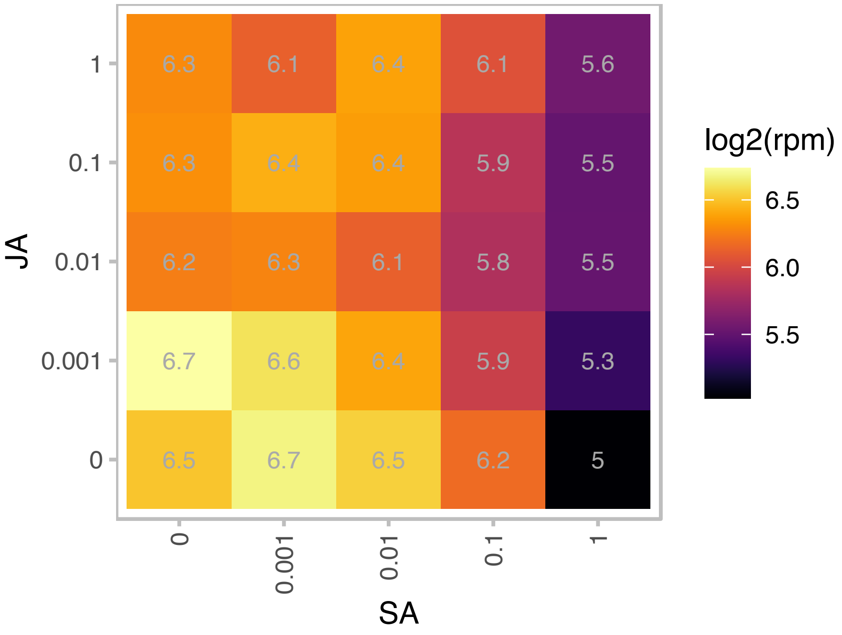


npr3-1 npr4-3

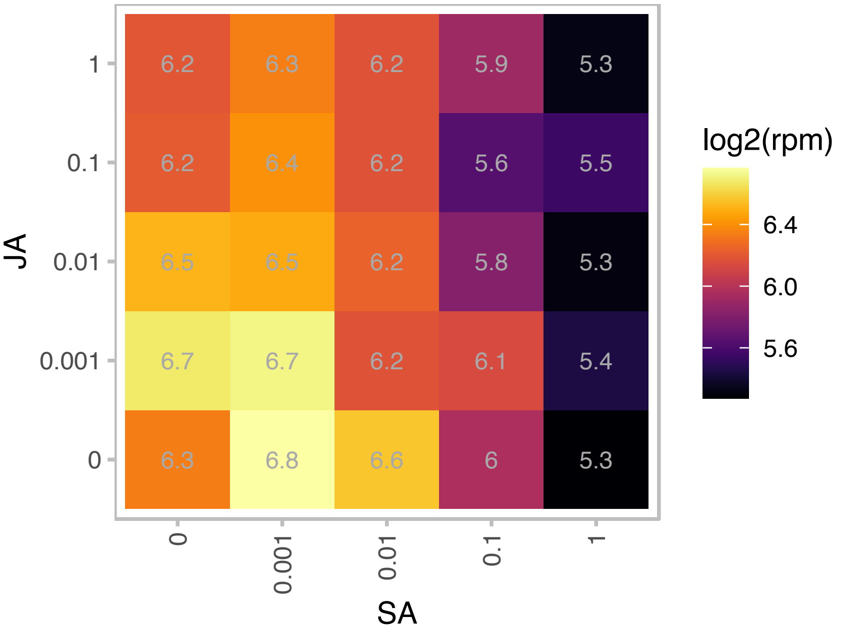

### AT1G03600.1 : PSB27

photosystem II family protein

PSB27 is a chloroplast lumen localized protein that is involved in adaptation to changes in light intensity.

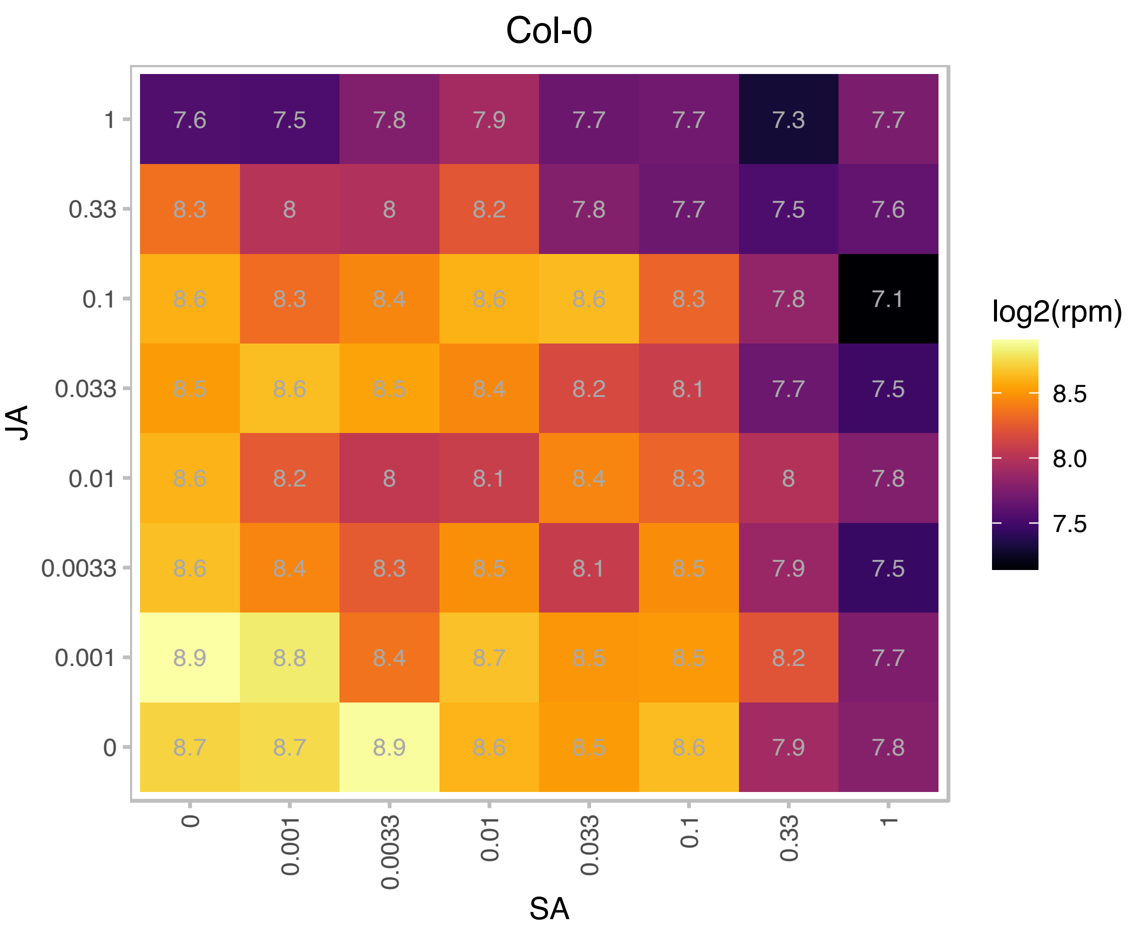


| Gene | AT1G03600 |
|---|---|
| is.expressed_genes | TRUE |
| is.responsive_genes | TRUE |
| is.apcluster | F-06 |

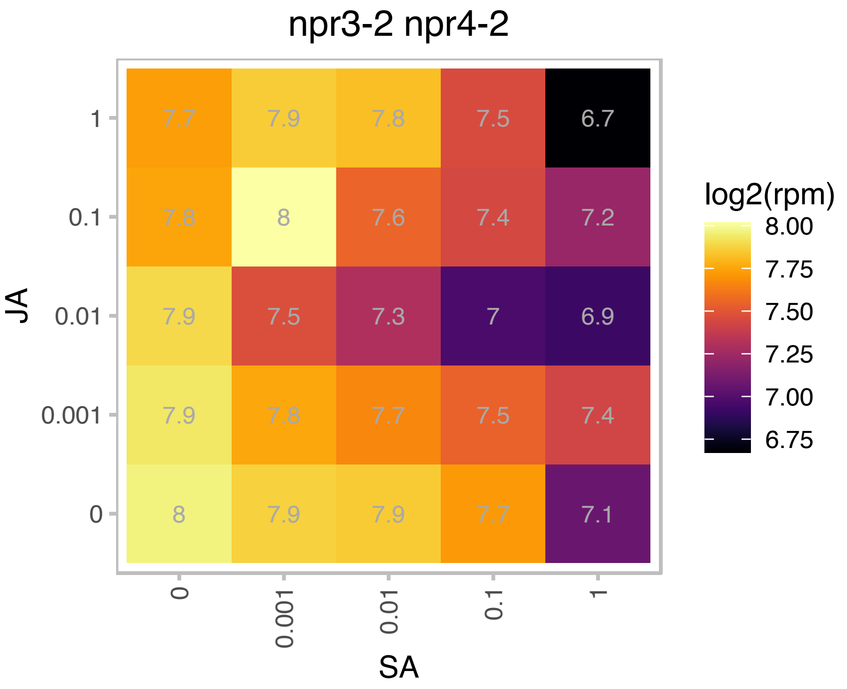


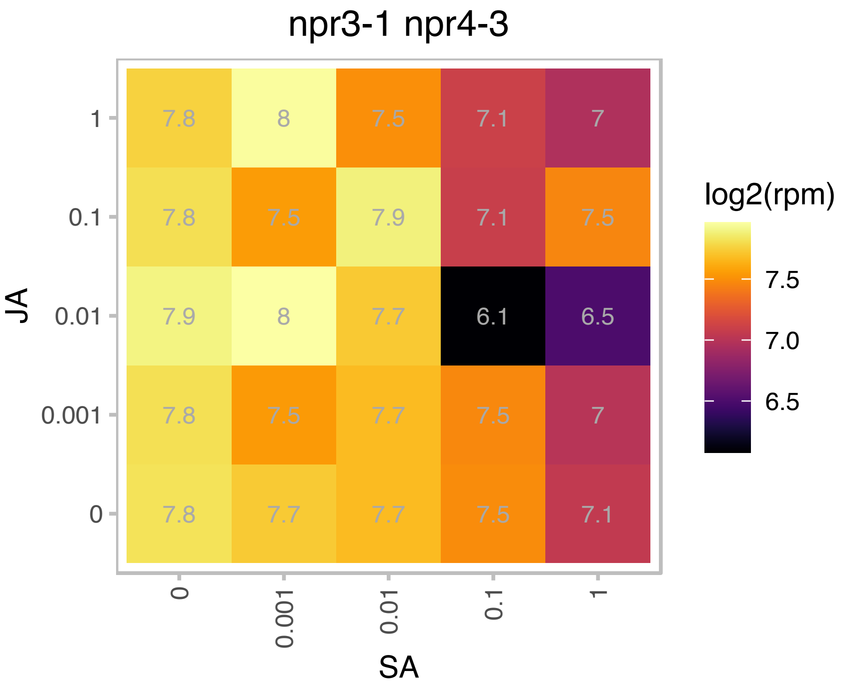

# AT1G04040.1 : NA

HAD superfamily, subfamily IIIB acid phosphatase

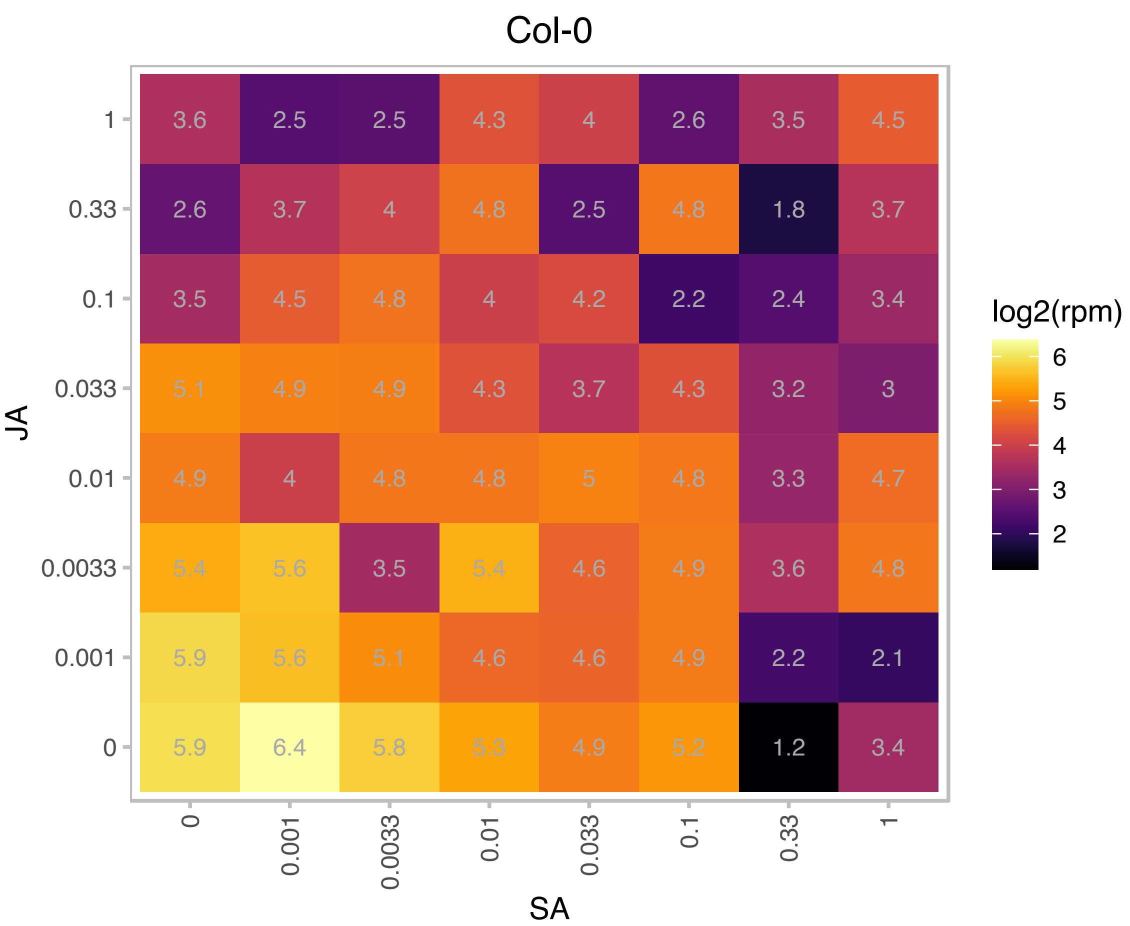


| | |
|---|---|
| Gene | AT1G04040 |
| is.expressed_genes | TRUE |
| is.responsive_genes | TRUE |
| is.apcluster | F-06 |

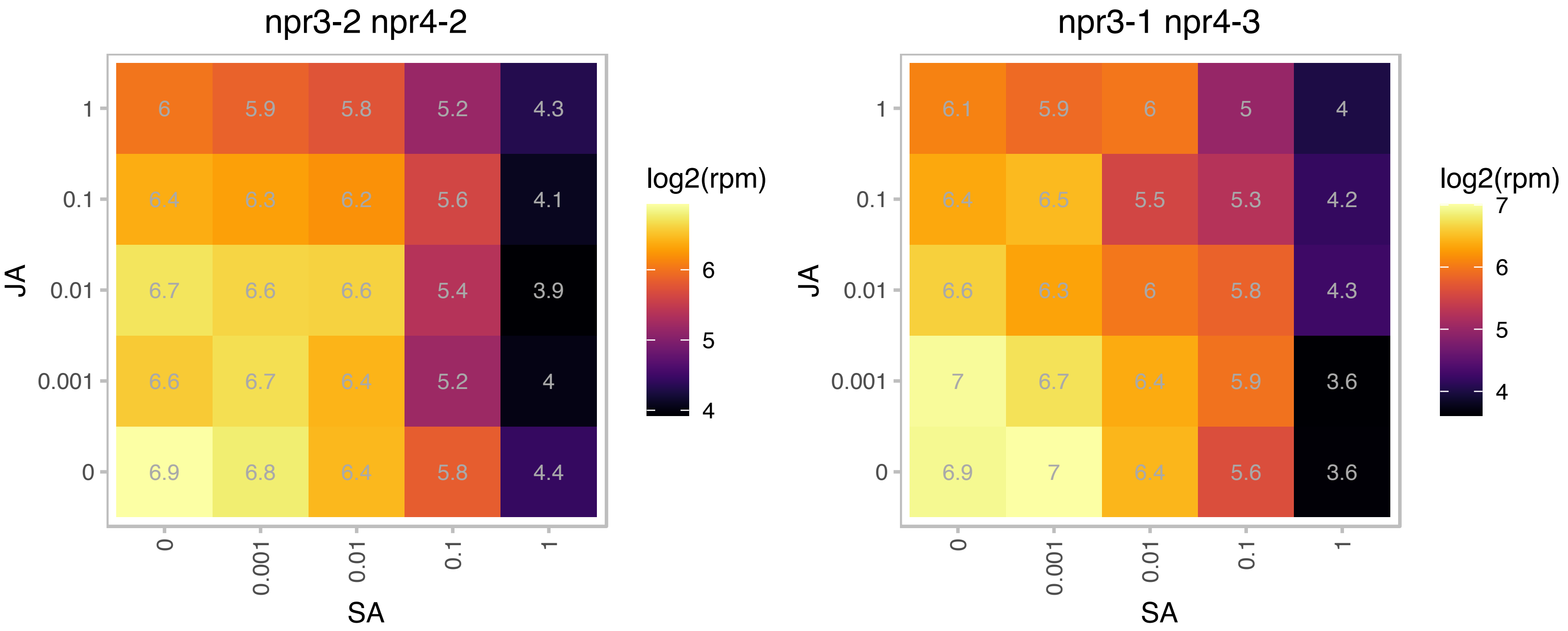

### AT1G06190.1 : RHON1

Rho termination factor

Encodes a novel ribonucleic acid-binding protein that interacts with the endonuclease RNase E and supports its function in processing plastid ribonucleic acids.

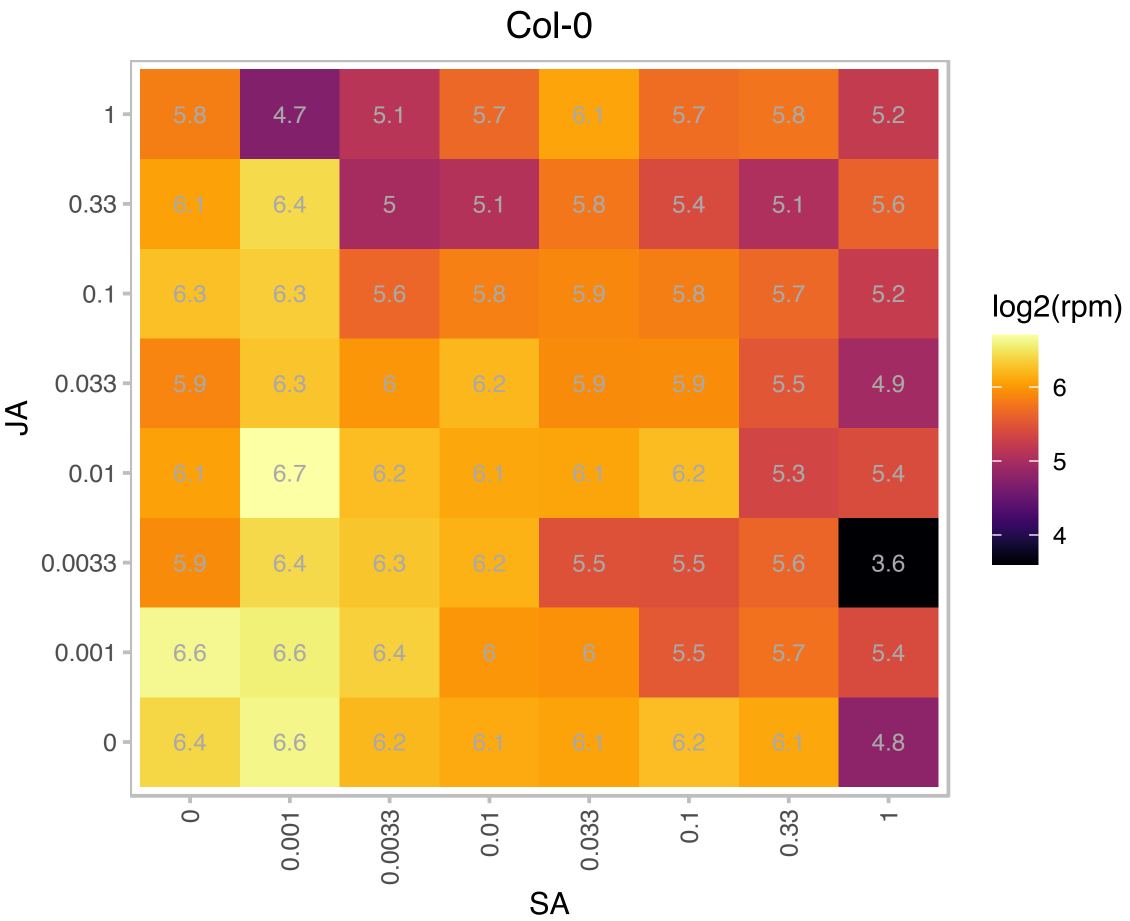


| | |
|---|---|
| Gene | AT1G06190 |
| is.expressed_genes | TRUE |
| is.responsive_genes | TRUE |
| is.apcluster | F-06 |

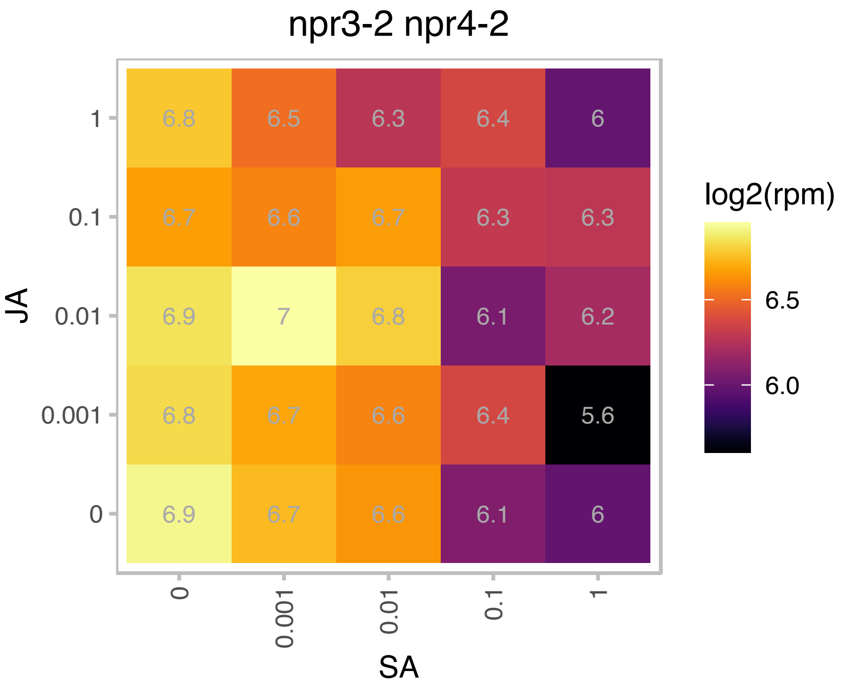


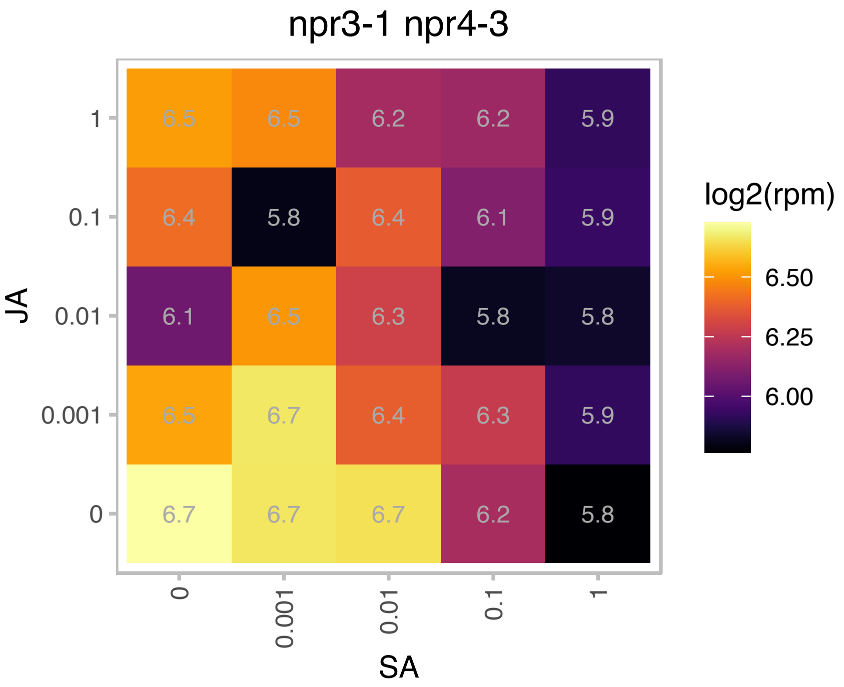

### AT1G07320.1 : EMB2784, PRPL4, RPL4

ribosomal protein L4

encodes a plastid ribosomal protein L4

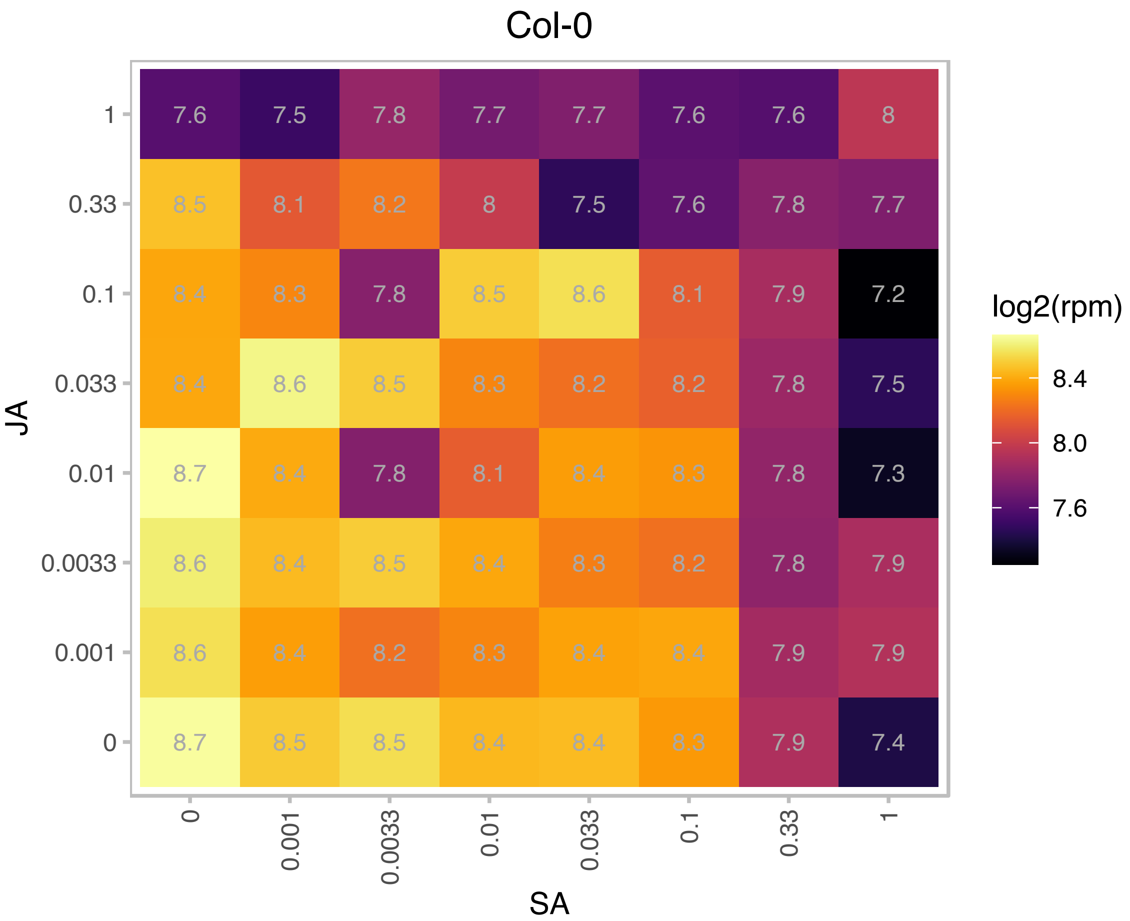


| | |
|---|---|
| Gene | AT1G07320 |
| is.expressed_genes | TRUE |
| is.responsive_genes | TRUE |
| is.apcluster | F-06 |

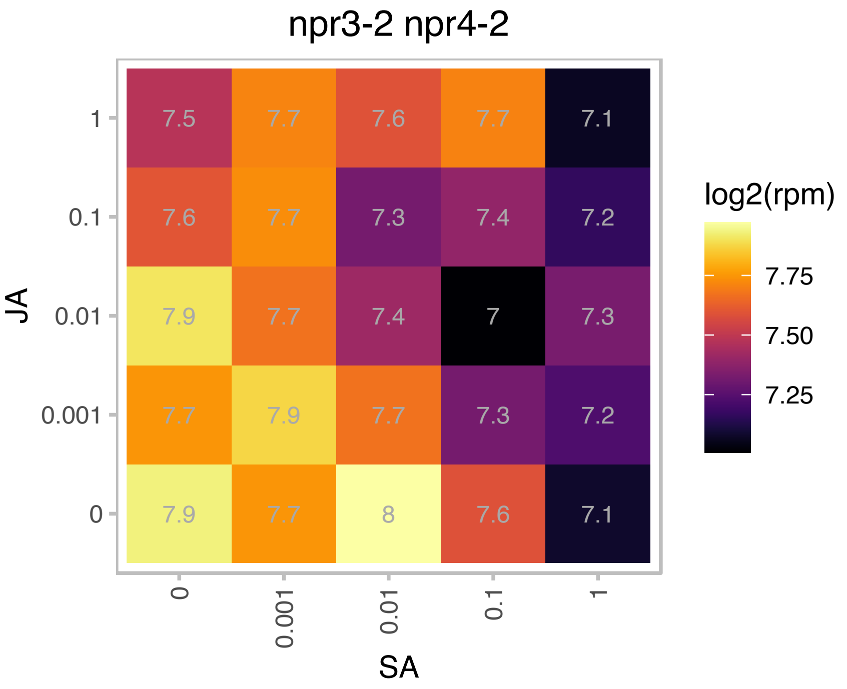


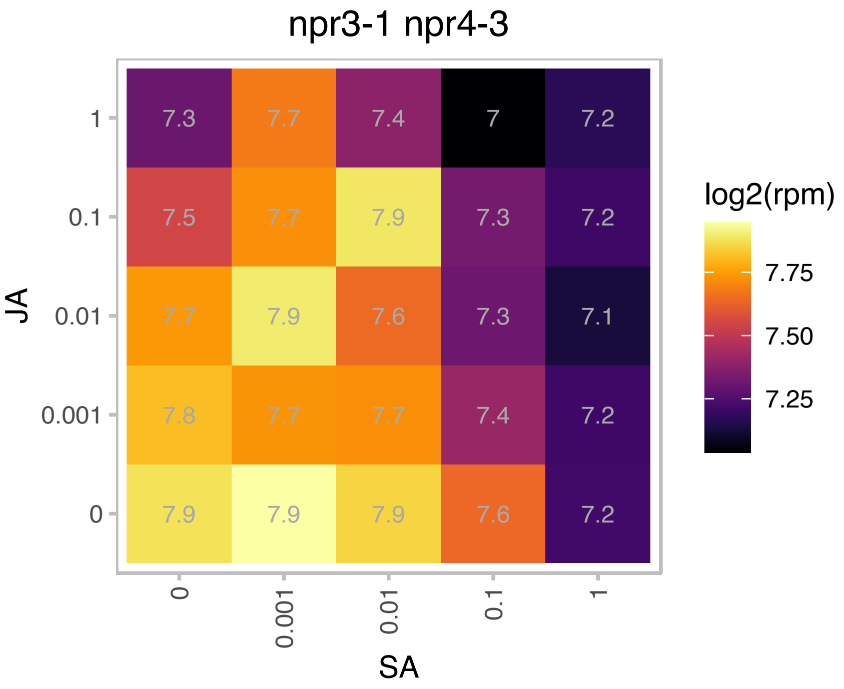

# AT1G07440.1 : NA

NAD(P)-binding Rossmann-fold superfamily protein

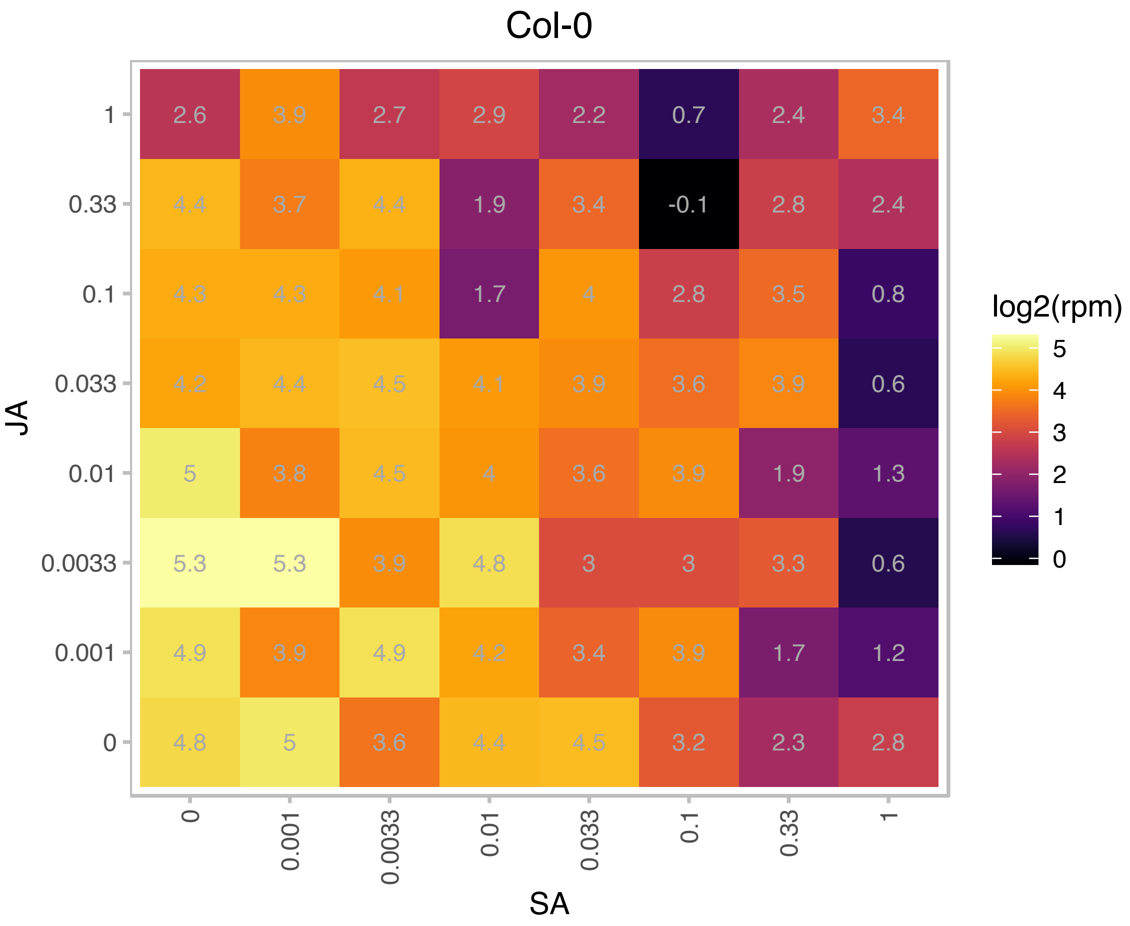


| | |
|---|---|
| Gene | AT1G07440 |
| is.expressed_genes | TRUE |
| is.responsive_genes | TRUE |
| is.apcluster | F-06 |

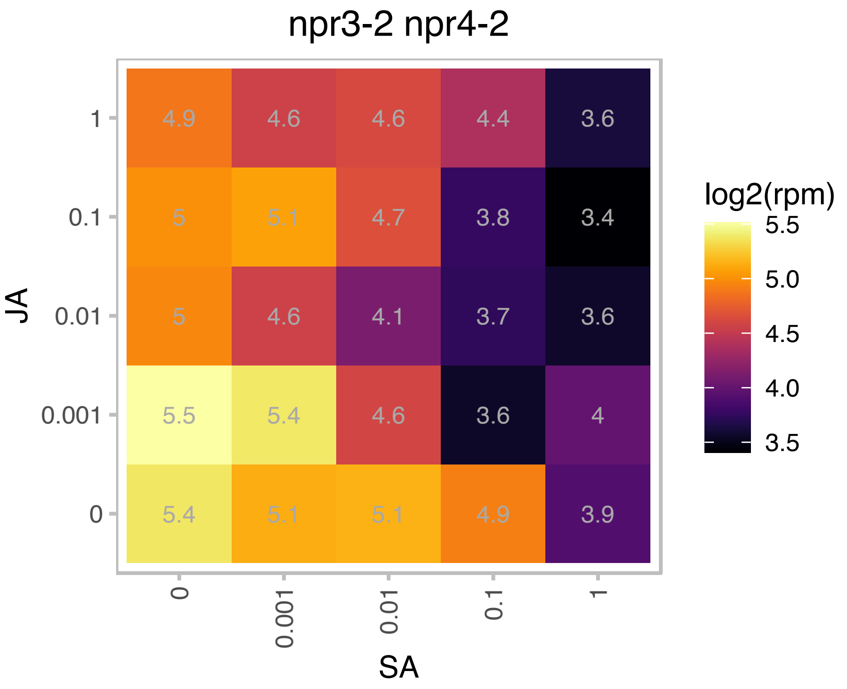


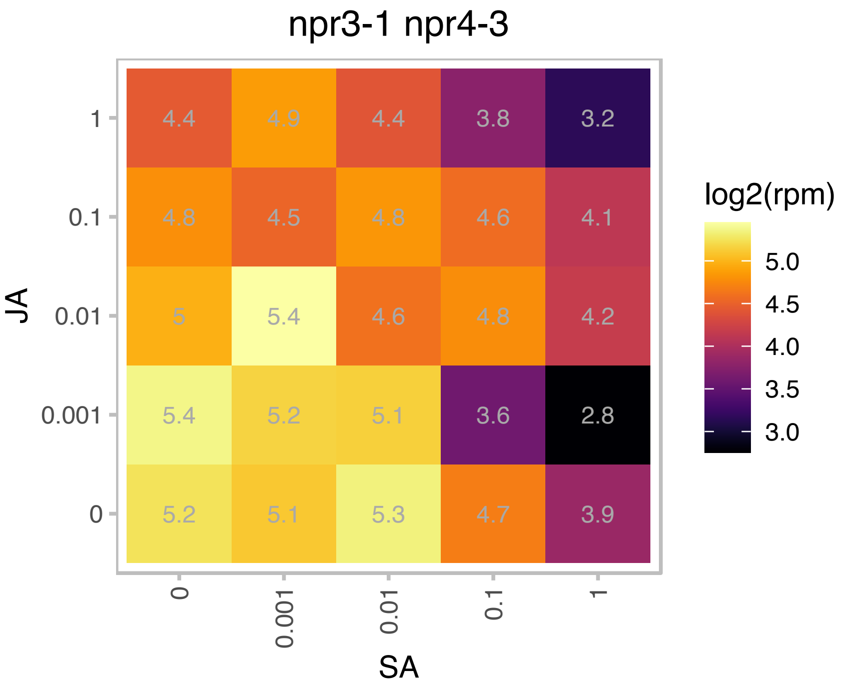

### AT1G08970.1 : HAP5C, NF-YC9

nuclear factor Y, subunit C9

Encodes a NUCLEAR FACTOR-Y C (NF-YC) homologue NF-YC9. NF-YC3., NF-YC4 and NF-YC9 redundantly mod
and ABA-mediated seed germination.

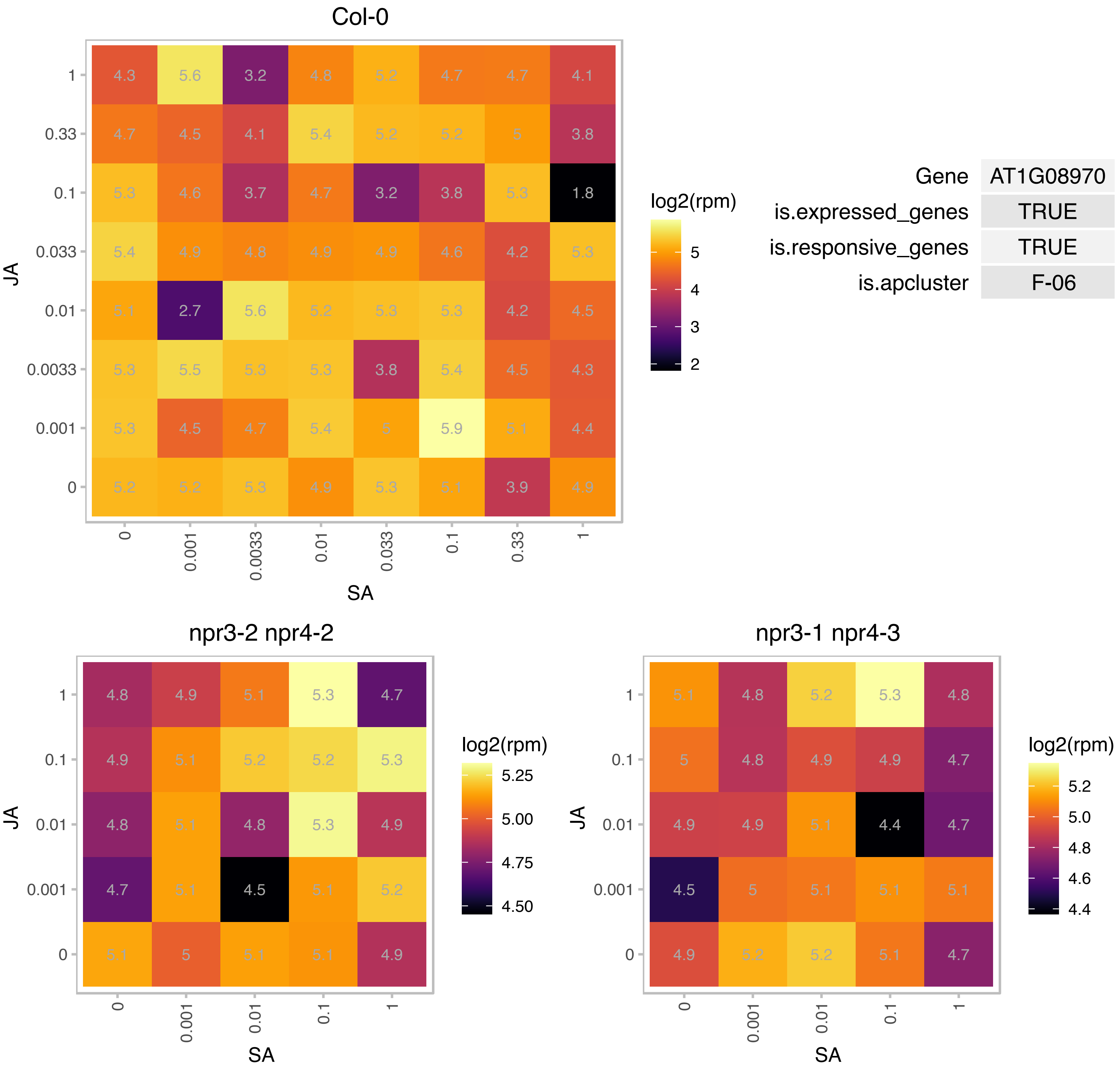


| | |
|---|---|
| Gene | AT1G08970 |
| is.expressed_genes | TRUE |
| is.responsive_genes | TRUE |
| is.apcluster | F-06 |

# AT1G12250.1 : TL20.3

Pentapeptide repeat-containing protein

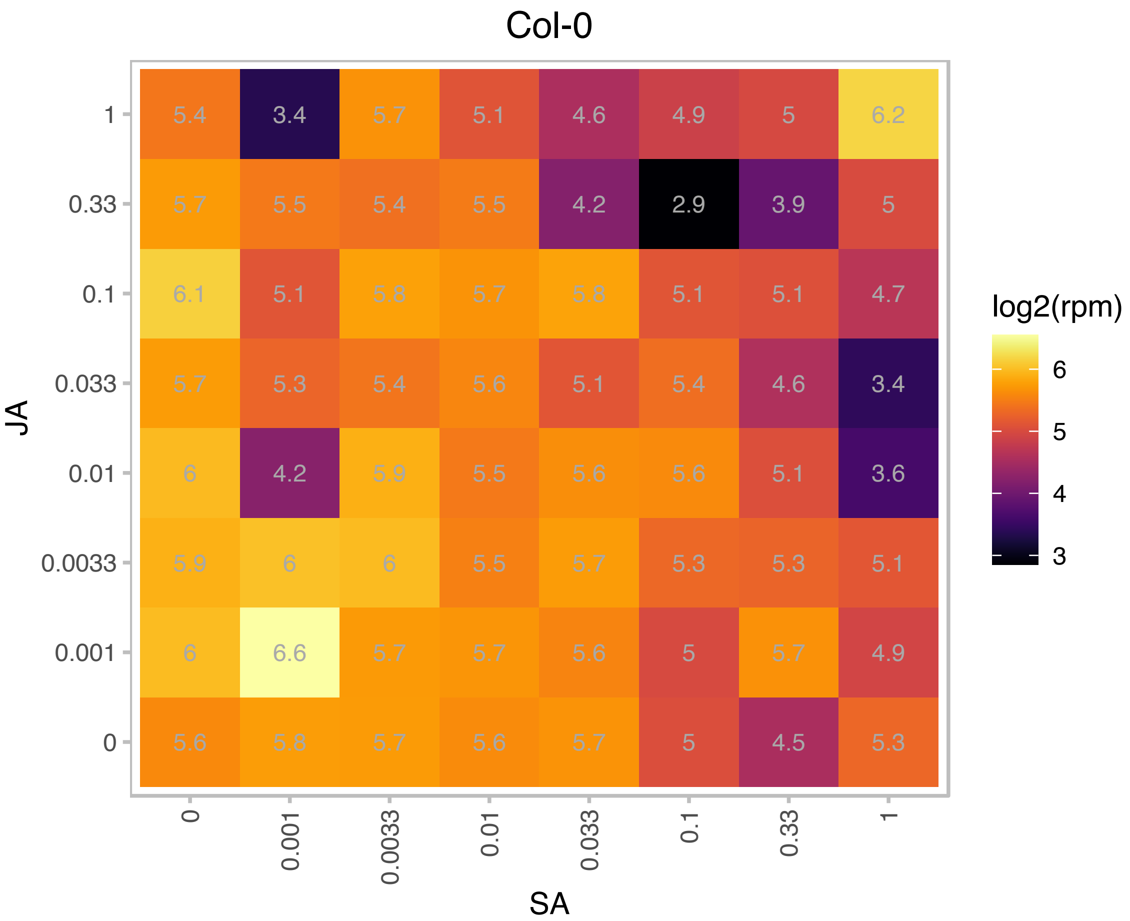


| | |
|---|---|
| Gene | AT1G12250 |
| is.expressed_genes | TRUE |
| is.responsive_genes | TRUE |
| is.apcluster | F-06 |

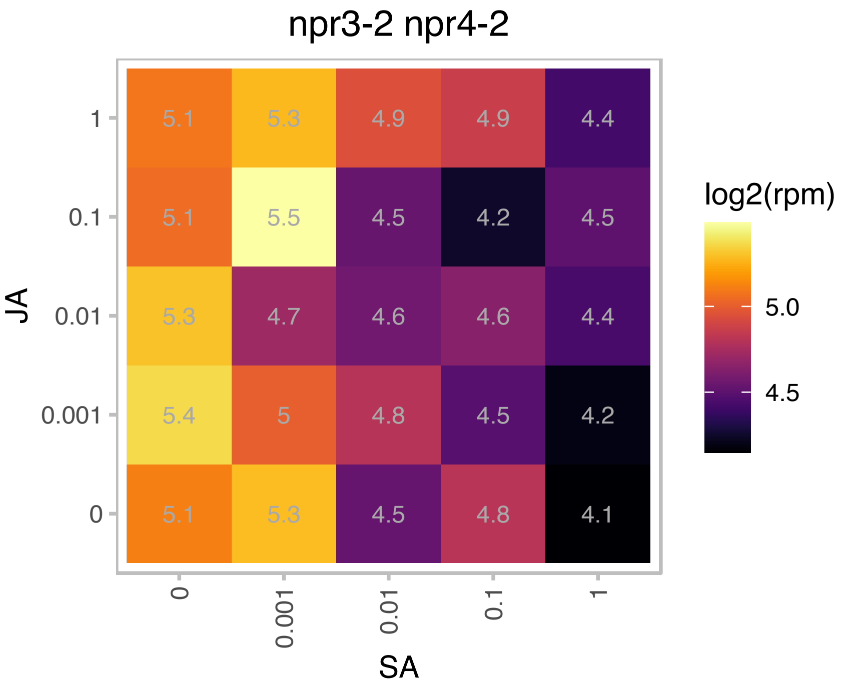


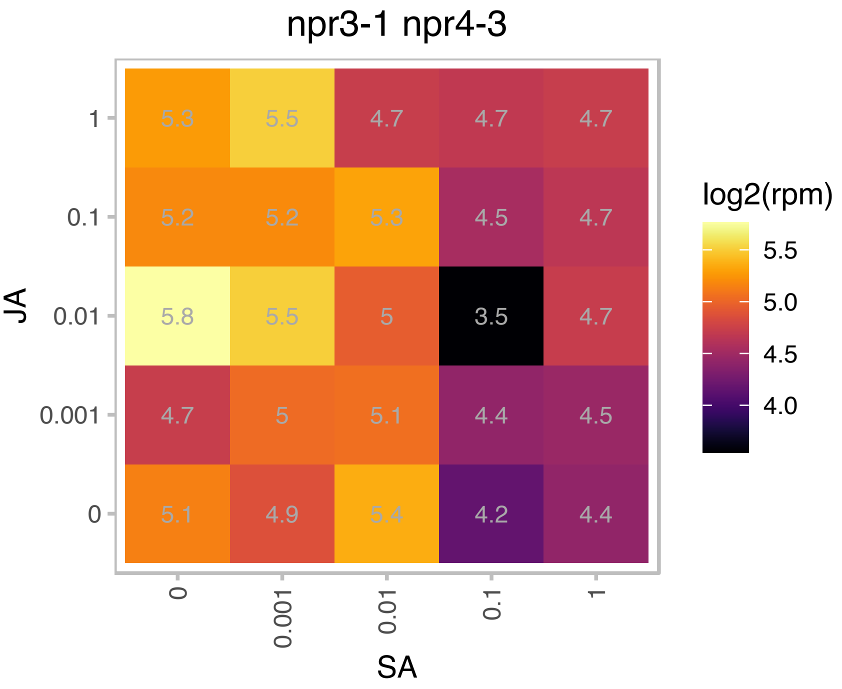

# AT1G13640.1 : NA

Phosphatidylinositol 3- and 4-kinase family protein

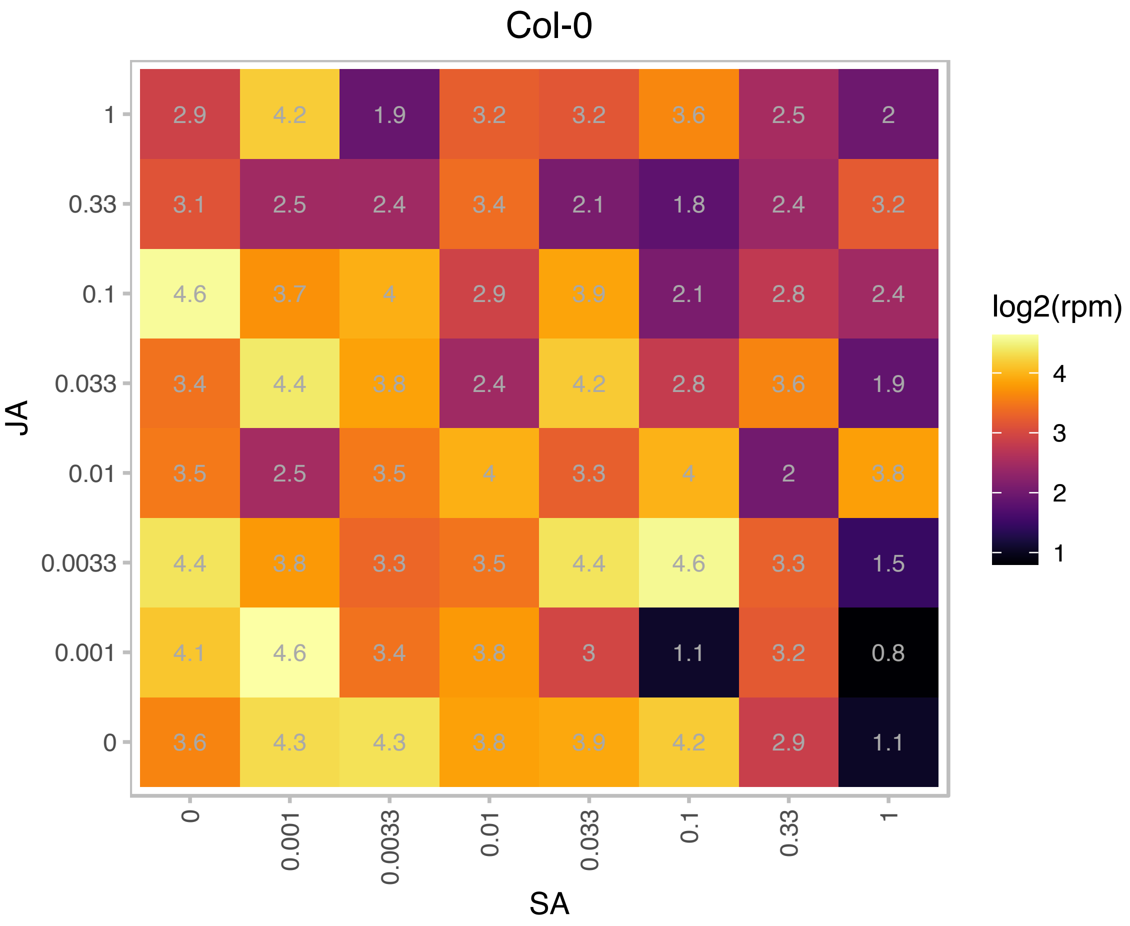


| | |
|---|---|
| Gene | AT1G13640 |
| is.expressed_genes | TRUE |
| is.responsive_genes | TRUE |
| is.apcluster | F-06 |

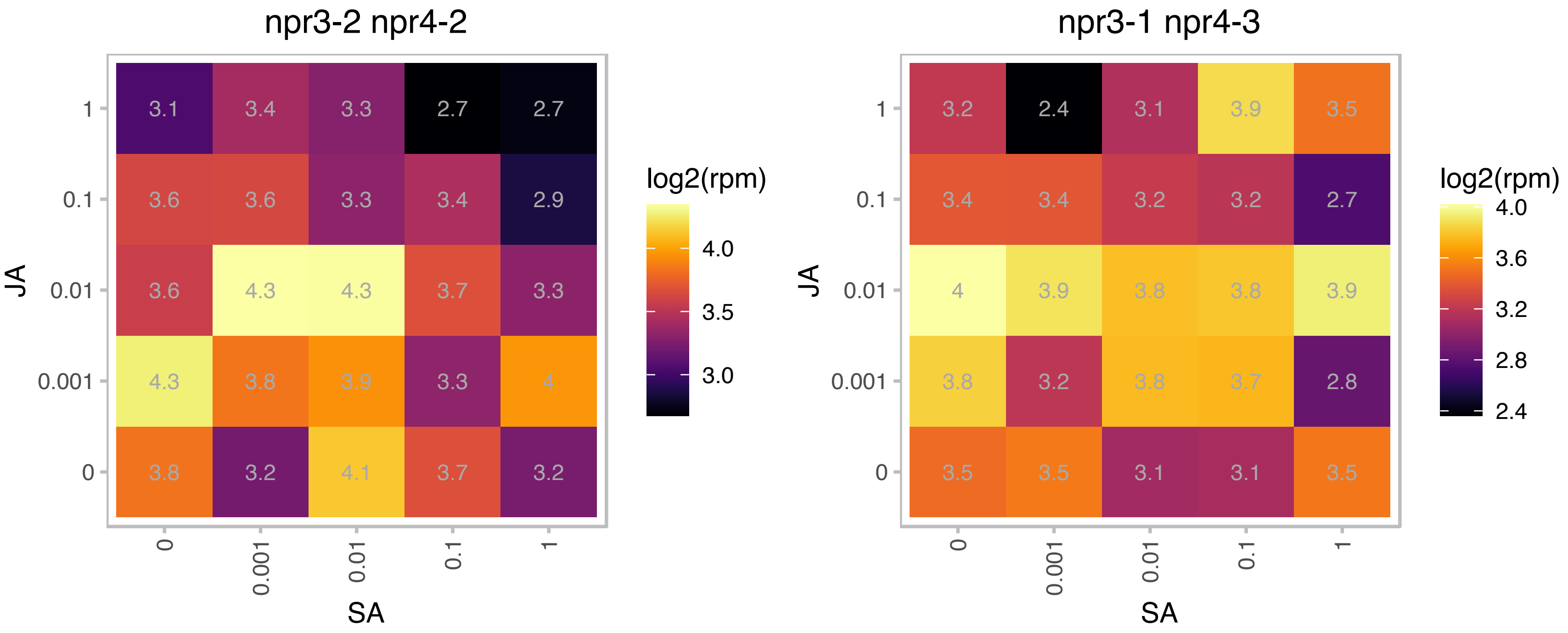

### AT1G14150.1 : PnsL2, PQL1, PQL2

PsbQ-like 2

Encodes a subunit of the NAD(P)H dehydrogenase complex located in the chloroplast thylakoid lumen.

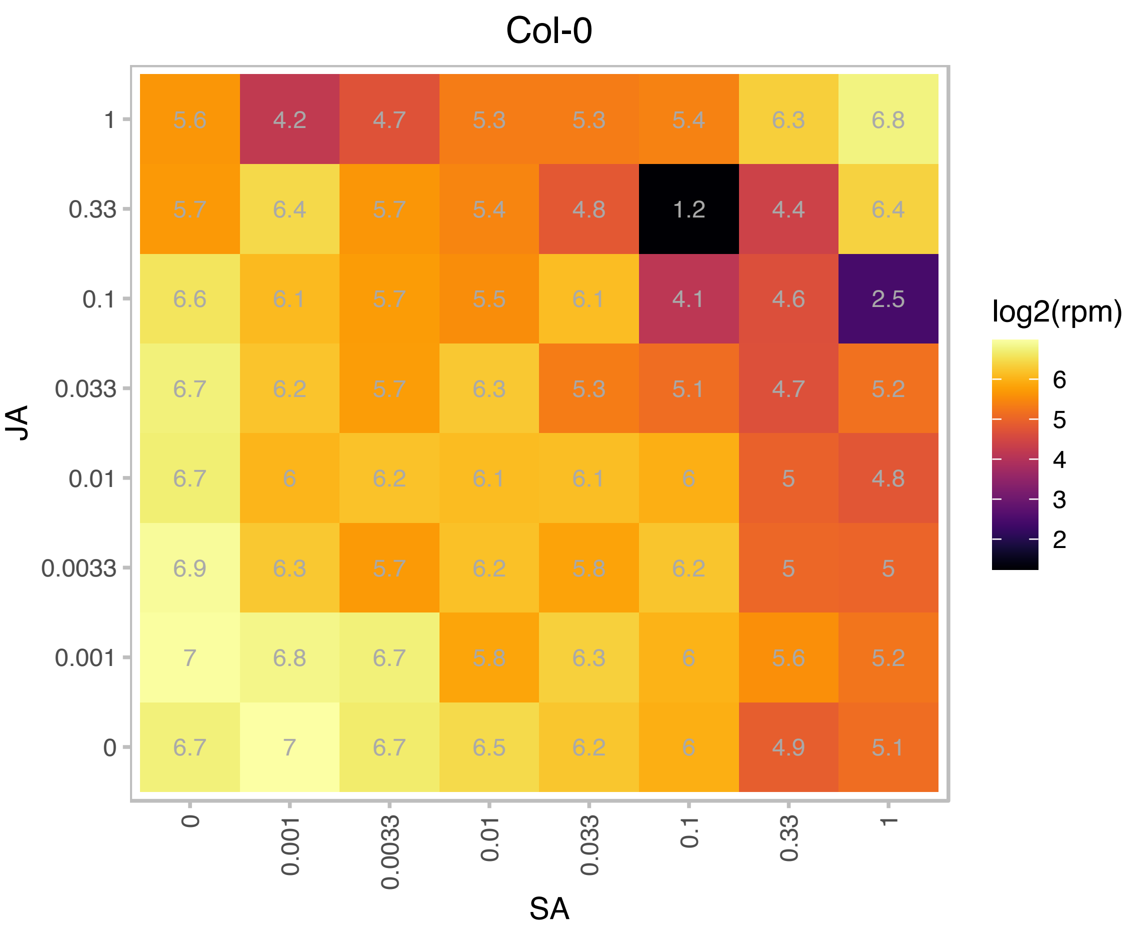


| | |
|---|---|
| Gene | AT1G14150 |
| is.expressed_genes | TRUE |
| is.responsive_genes | TRUE |
| is.apcluster | F-06 |

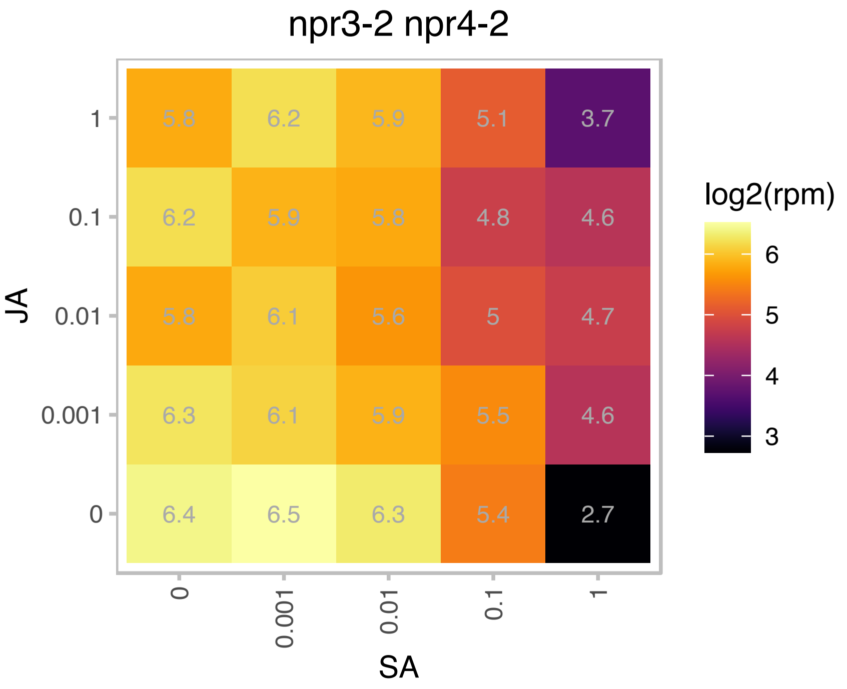


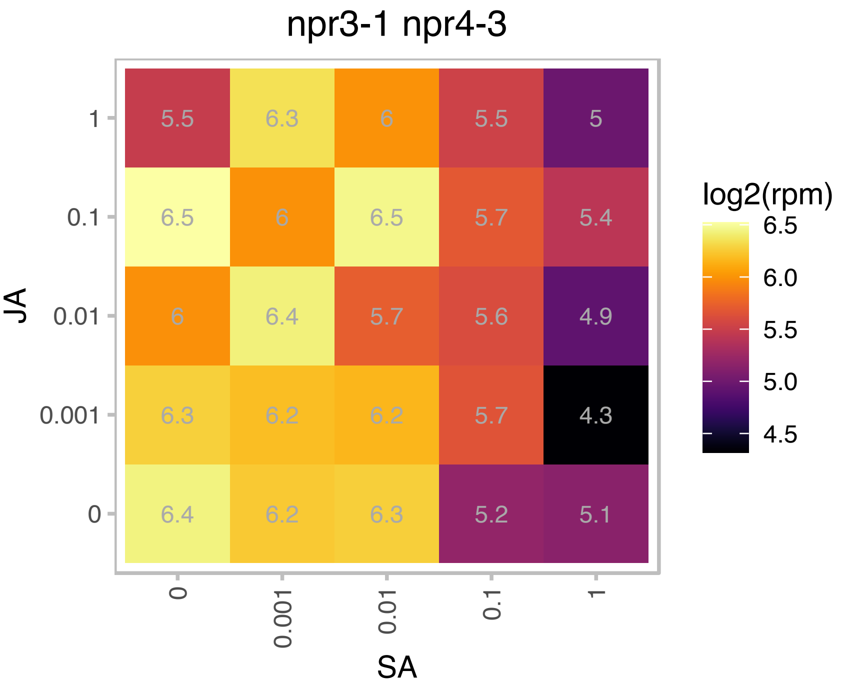

# AT1G16170.1 : NA

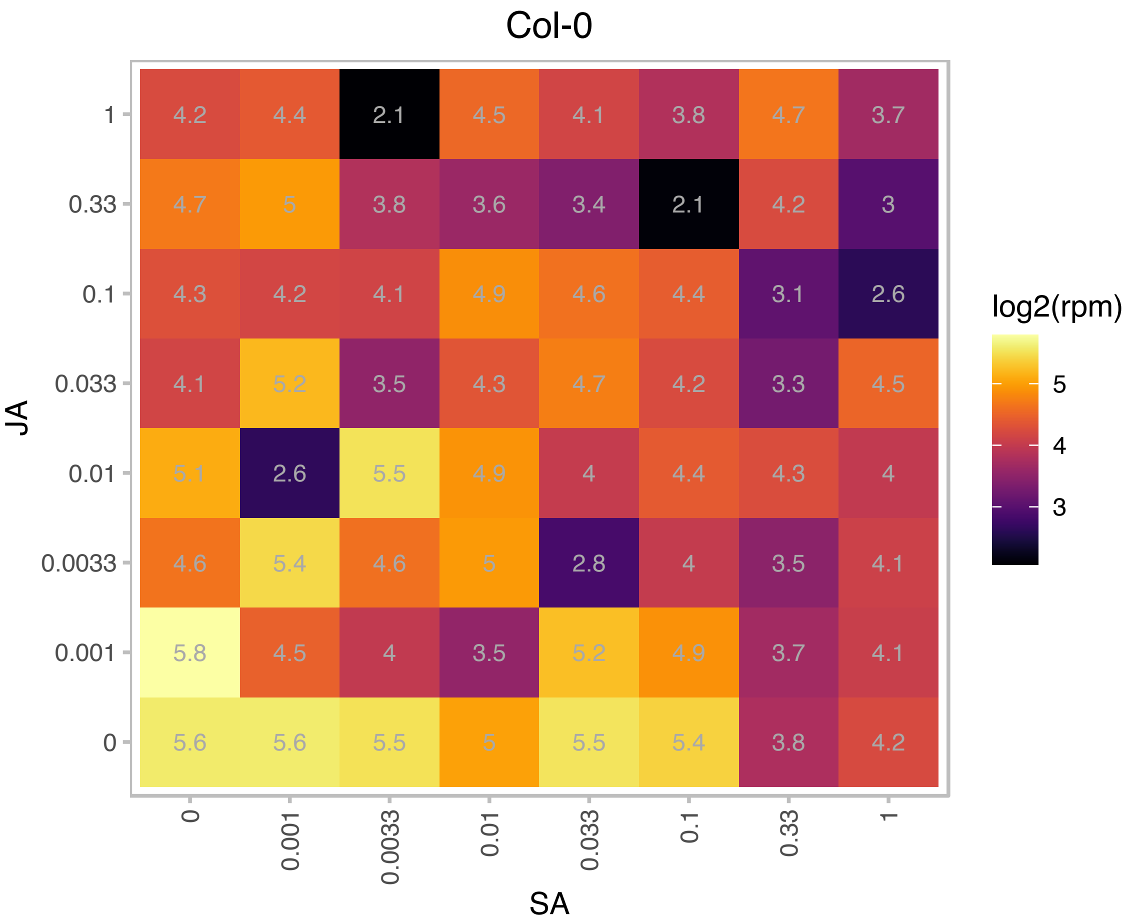


| | |
|---|---|
| Gene | AT1G16170 |
| is.expressed_genes | TRUE |
| is.responsive_genes | TRUE |
| is.apcluster | F-06 |

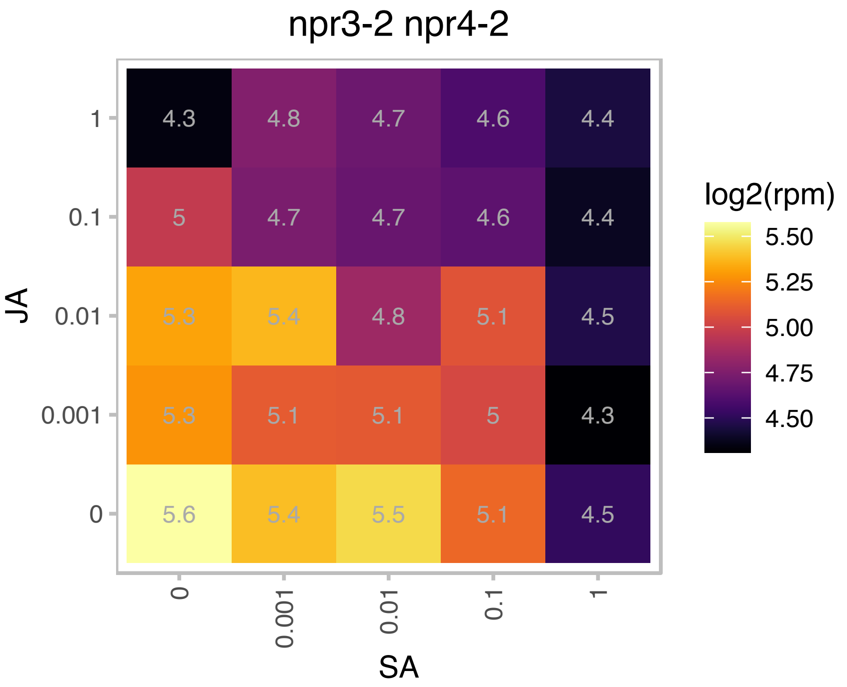


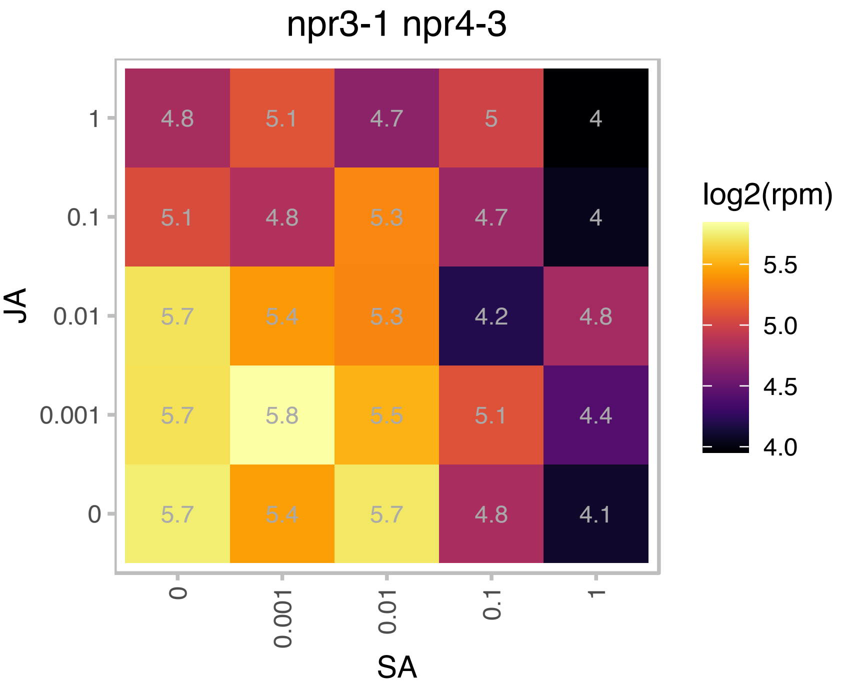

### AT1G19150.1 : LHCA2*1, Lhca6

photosystem I light harvesting complex gene 6

PSI type II chlorophyll a/b-binding protein (Lhca2*1) mRNA, The mRNA is cell-to-cell mobile.

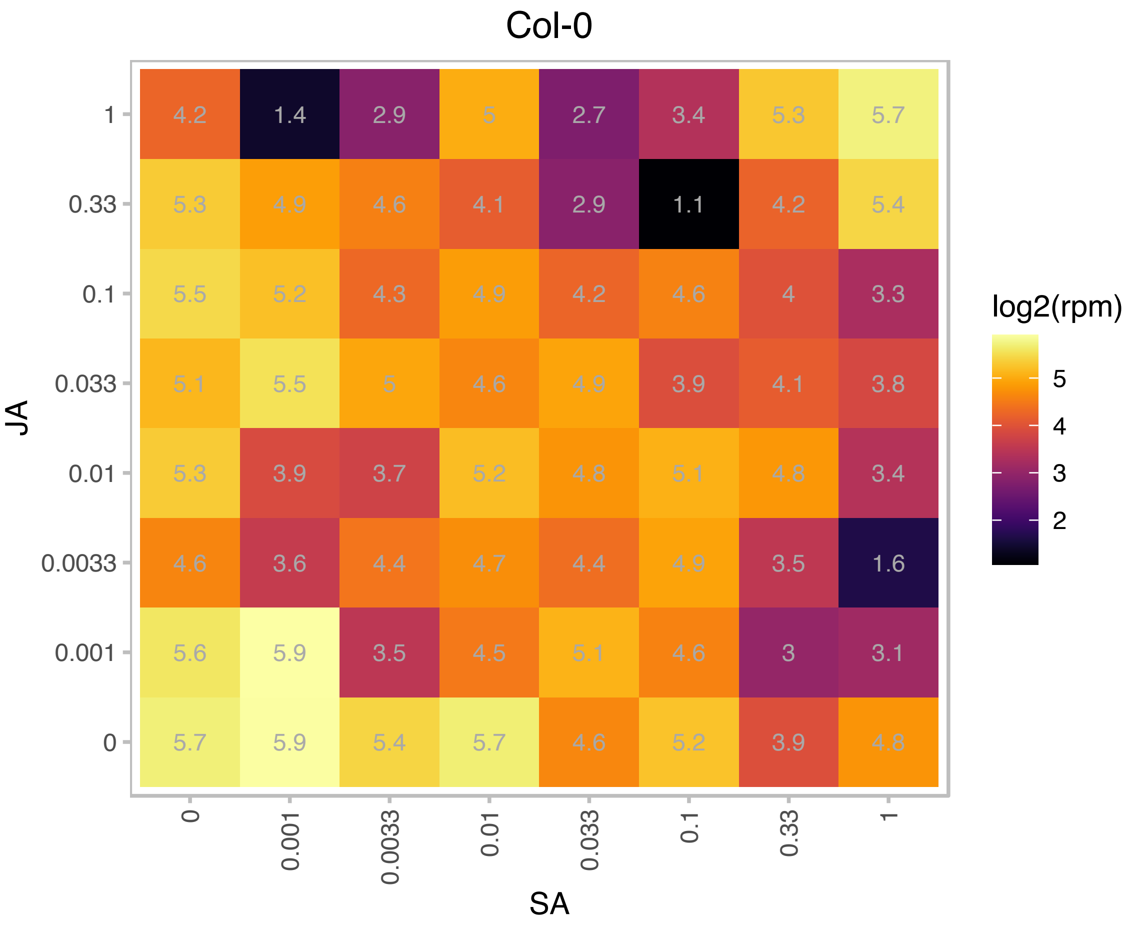


| | |
|---|---|
| Gene | AT1G19150 |
| is.expressed_genes | TRUE |
| is.responsive_genes | TRUE |
| is.apcluster | F-06 |

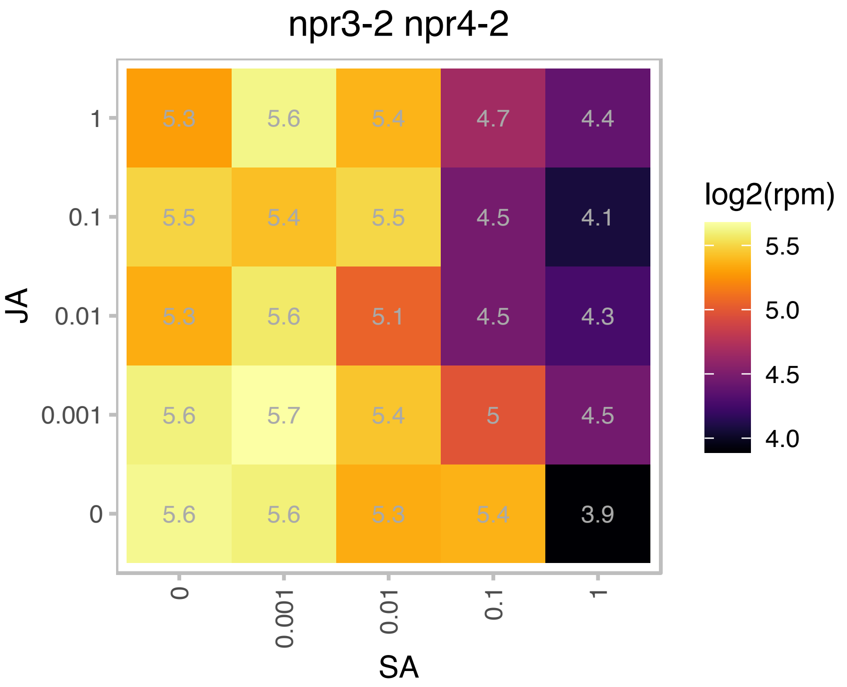


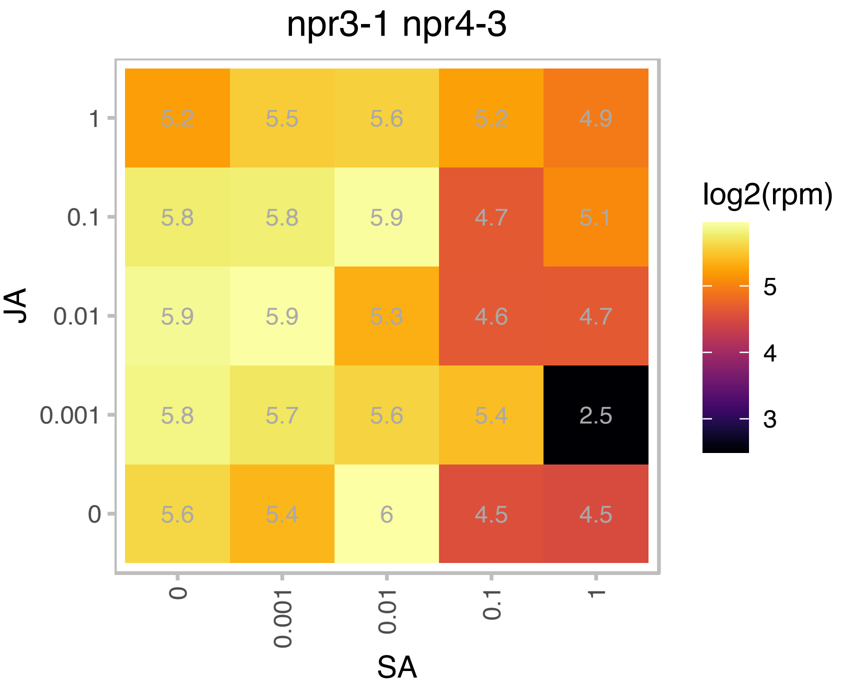

### AT1G21130.1 : IGMT4

O-methyltransferase family protein

| | |
|---|---|
| Gene | AT1G21130 |
| is.expressed_genes | TRUE |
| is.responsive_genes | TRUE |
| is.apcluster | F-06 |

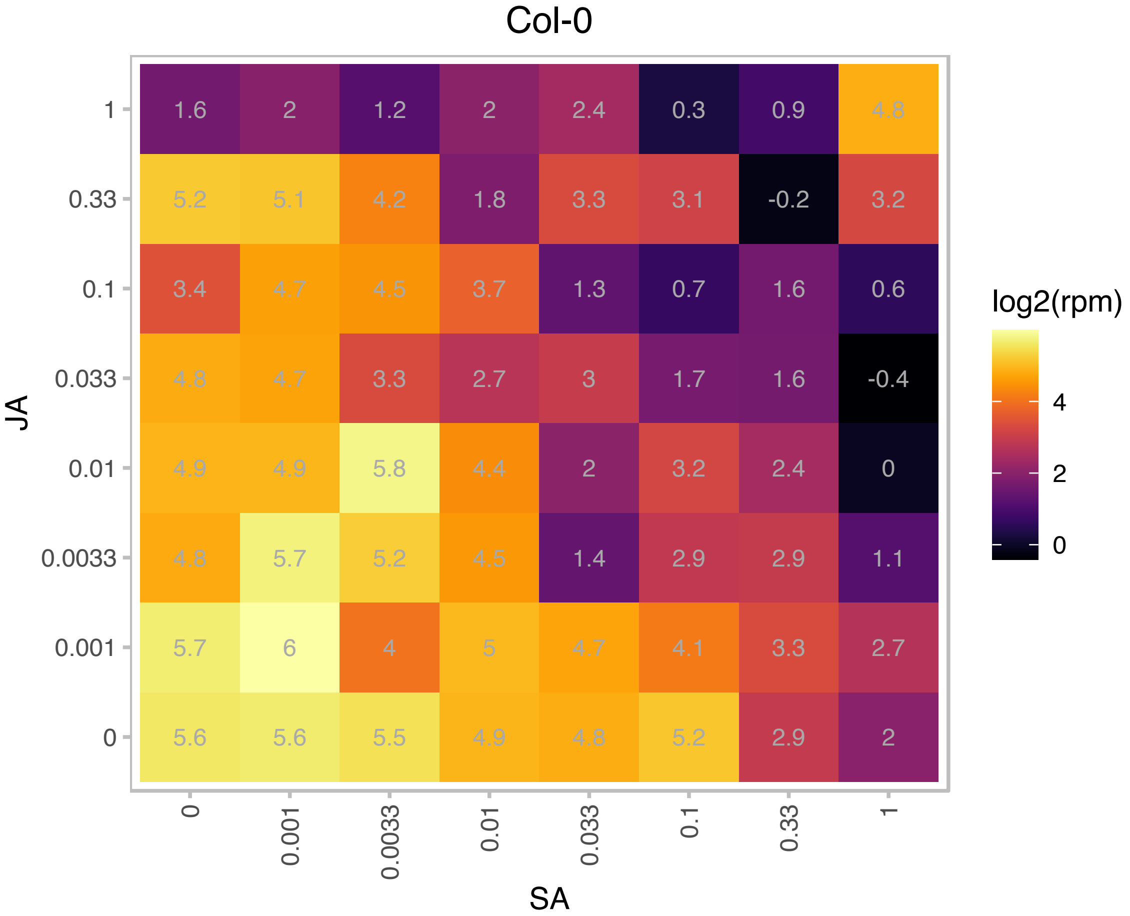


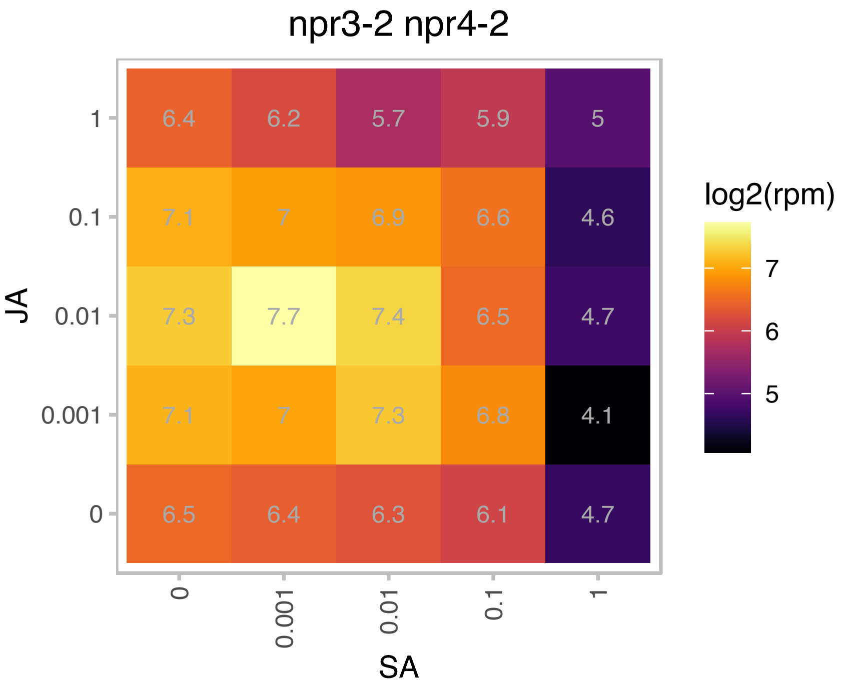


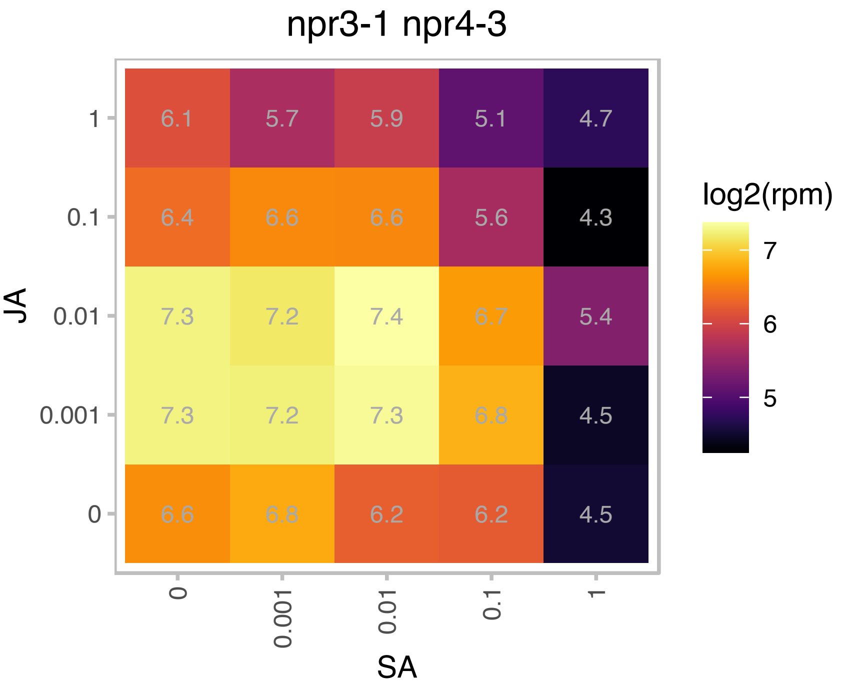

### AT1G21910.1 : DREB26

Integrase-type DNA-binding superfamily protein

encodes a member of the DREB subfamily A-5 of ERF/AP2 transcription factor family. The protein contains one AP2 domain. There are 15 members in this subfamily including RAP2.1, RAP2.9 and RAP2.10.

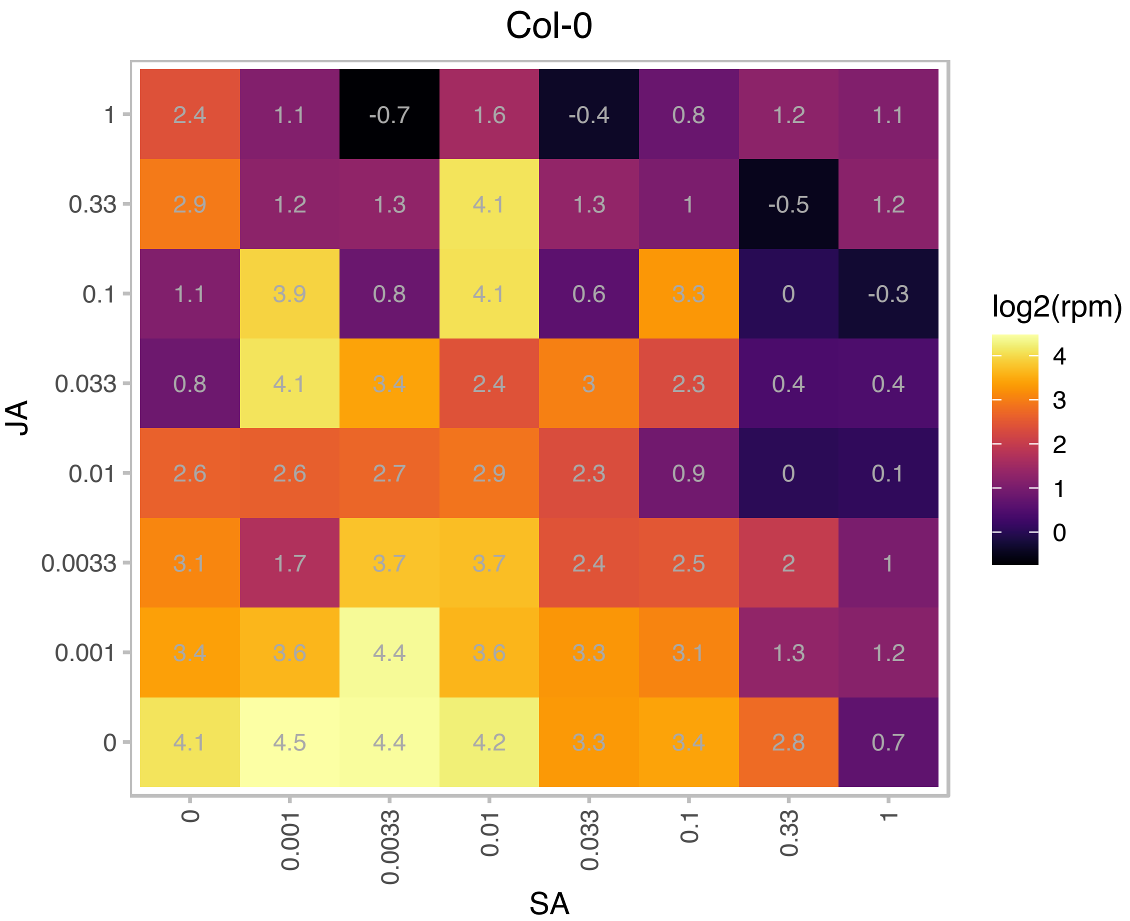


| | |
|---|---|
| Gene | AT1G21910 |
| is.expressed_genes | TRUE |
| is.responsive_genes | TRUE |
| is.apcluster | F-06 |

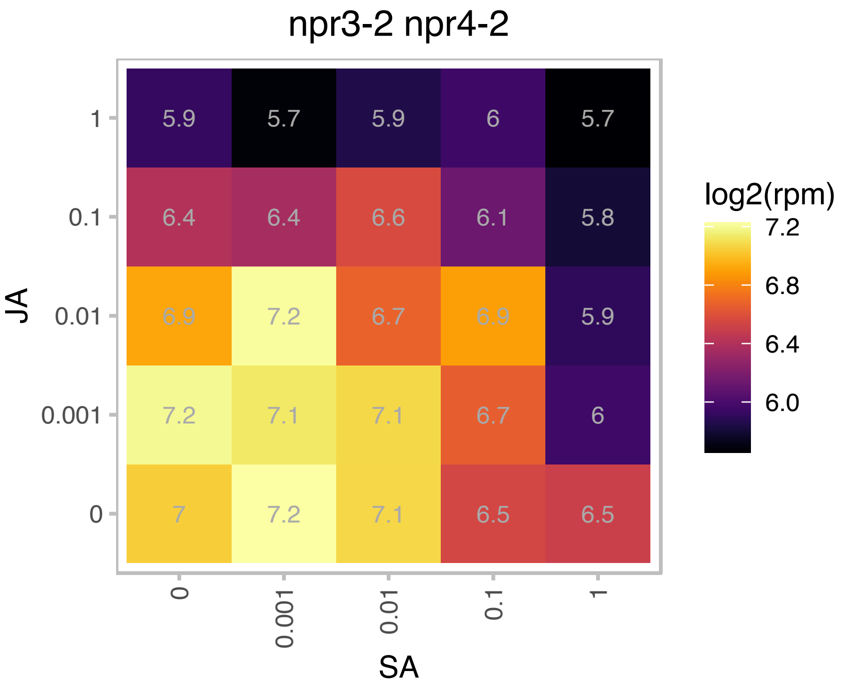


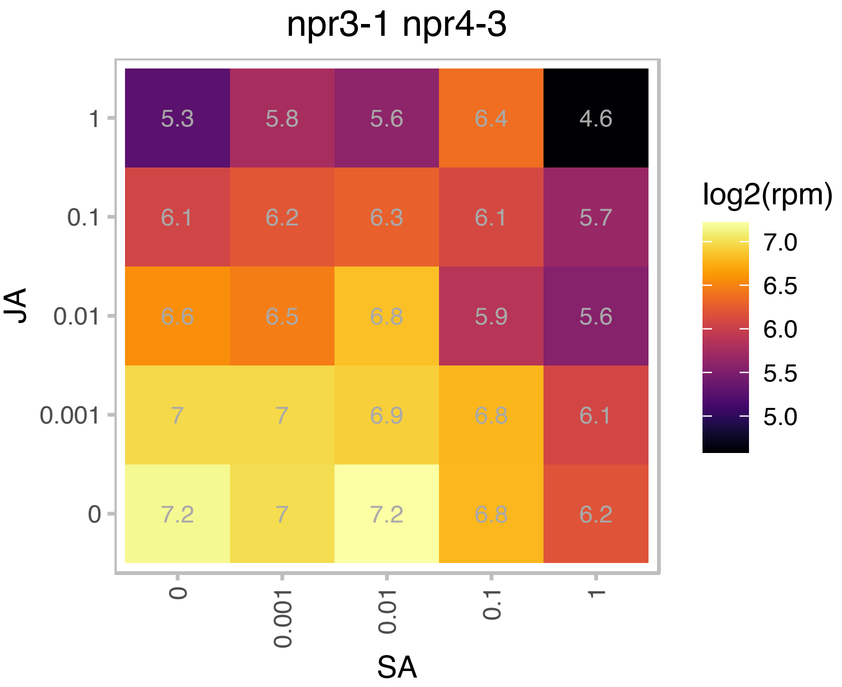

### AT1G23740.1 : AOR

Oxidoreductase, zinc-binding dehydrogenase family protein

AOR is an alkenal/one oxidoreductase that acts on compounds with unsaturated alpha,beta-carbonyls. The activity of this enzyme with a number of substrates, including acrolein and 3-buten-2-one, was demonstrated in vitro using a truncated form of the protein that lacked approximately 80 of the first amino acids. This protein appears to localize to the chloroplast where it likely helps to maintain the photosynthetic process by detoxifying reactive carbonyls formed during lipid peroxidation.

Col-0

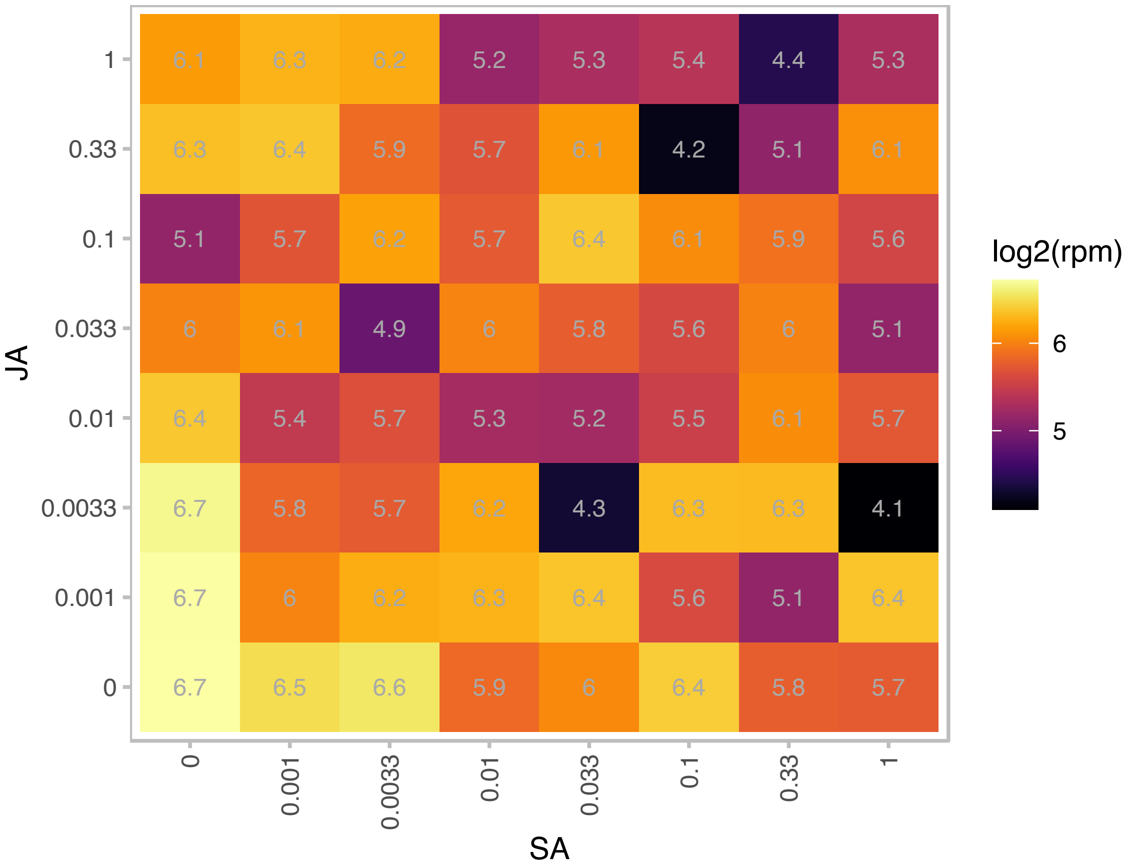


| | |
|---|---|
| Gene | AT1G23740 |
| is.expressed_genes | TRUE |
| is.responsive_genes | TRUE |
| is.apcluster | F-06 |

npr3-2 npr4-2

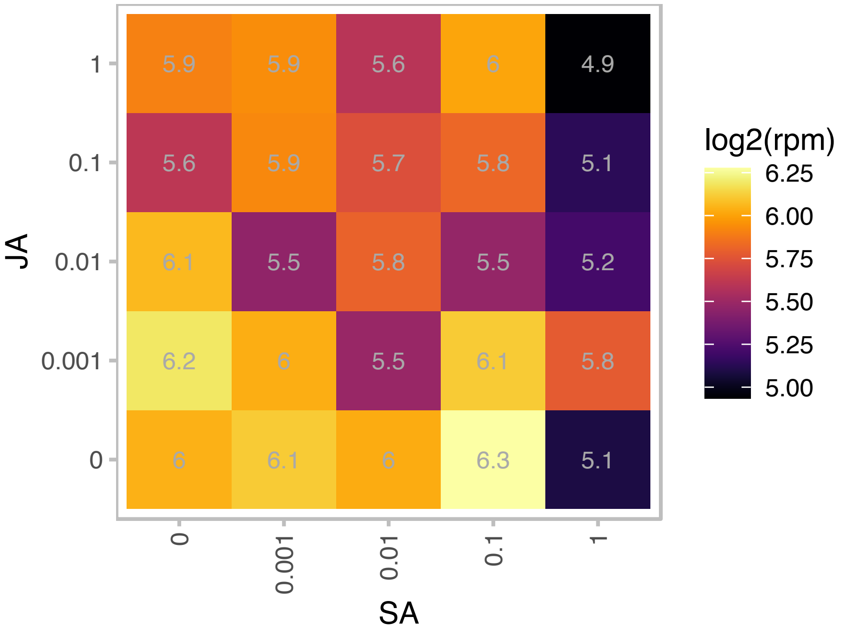


npr3-1 npr4-3

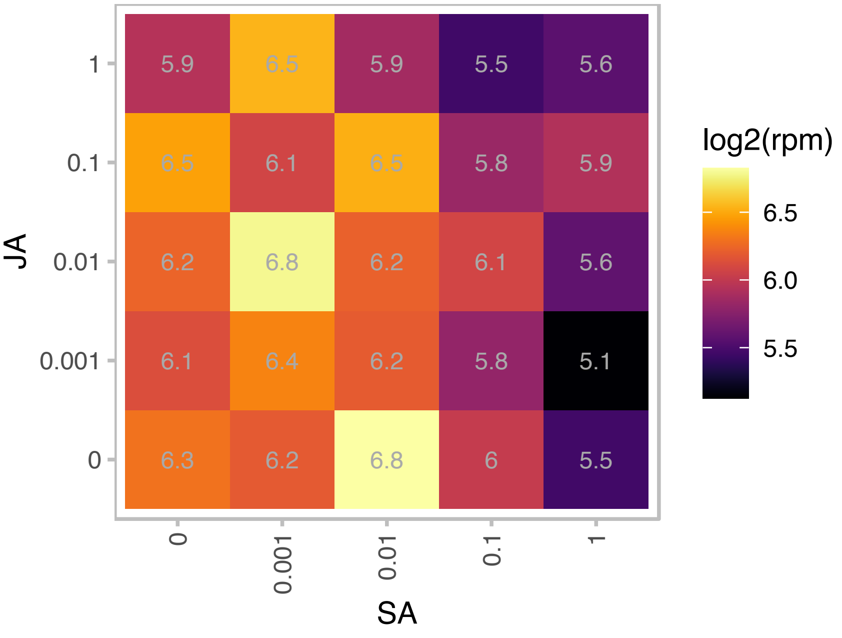

# AT1G27480.1 : NA

alpha/beta-Hydrolases superfamily protein

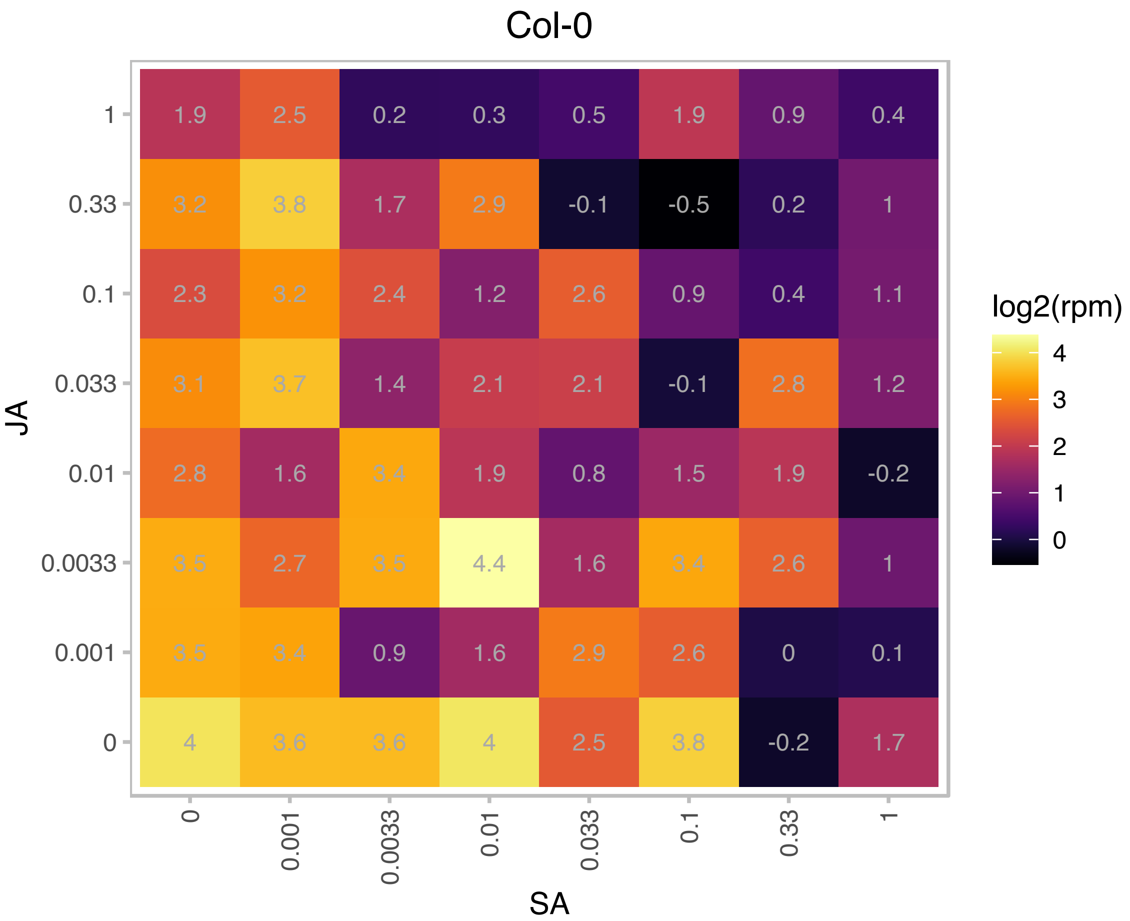


| | |
|---|---|
| Gene | AT1G27480 |
| is.expressed_genes | TRUE |
| is.responsive_genes | TRUE |
| is.apcluster | F-06 |

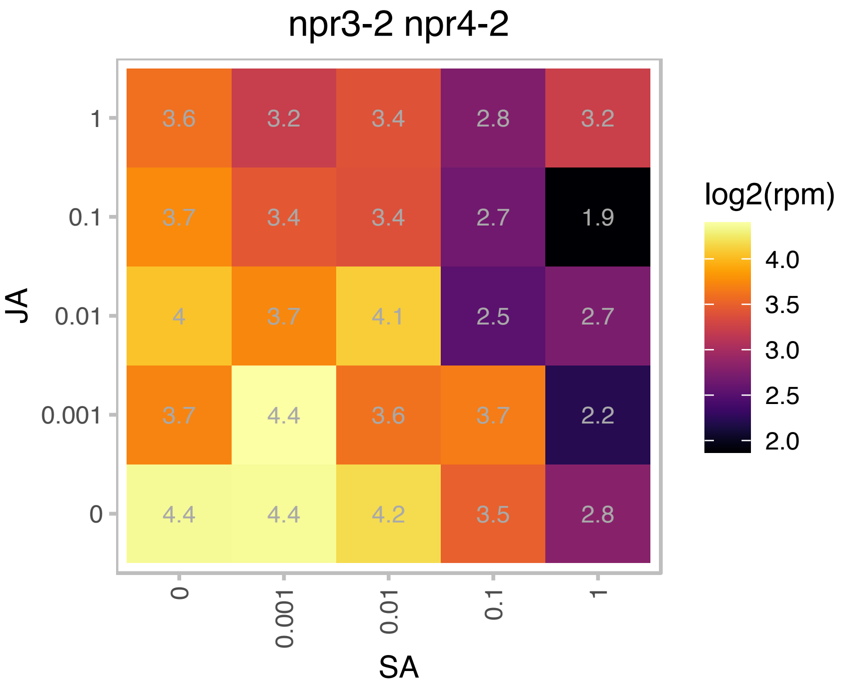


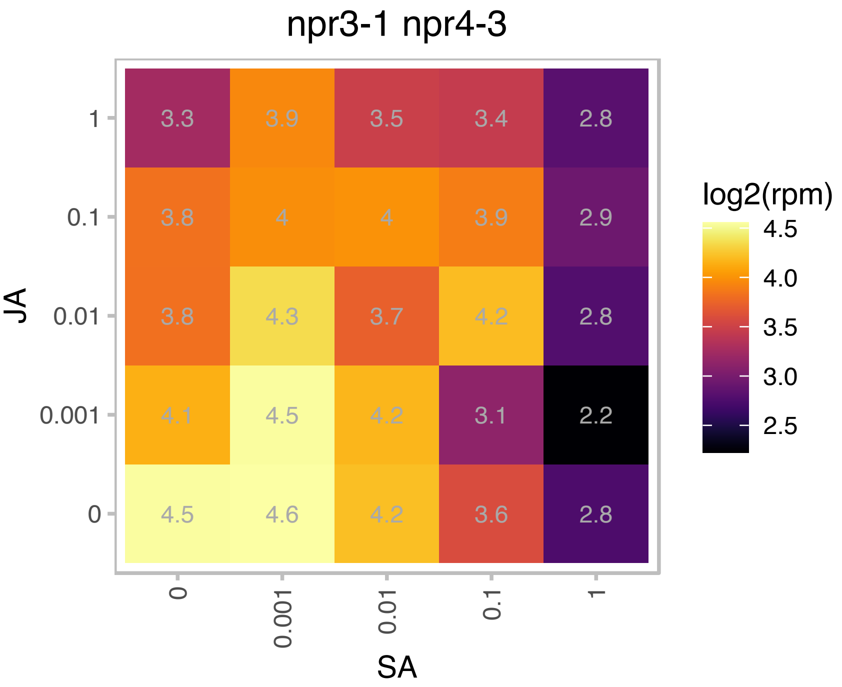

# AT1G45170.1 : NA

| | |
|---|---|
| Gene | AT1G45170 |
| is.expressed_genes | TRUE |
| is.responsive_genes | TRUE |
| is.apcluster | F-06 |

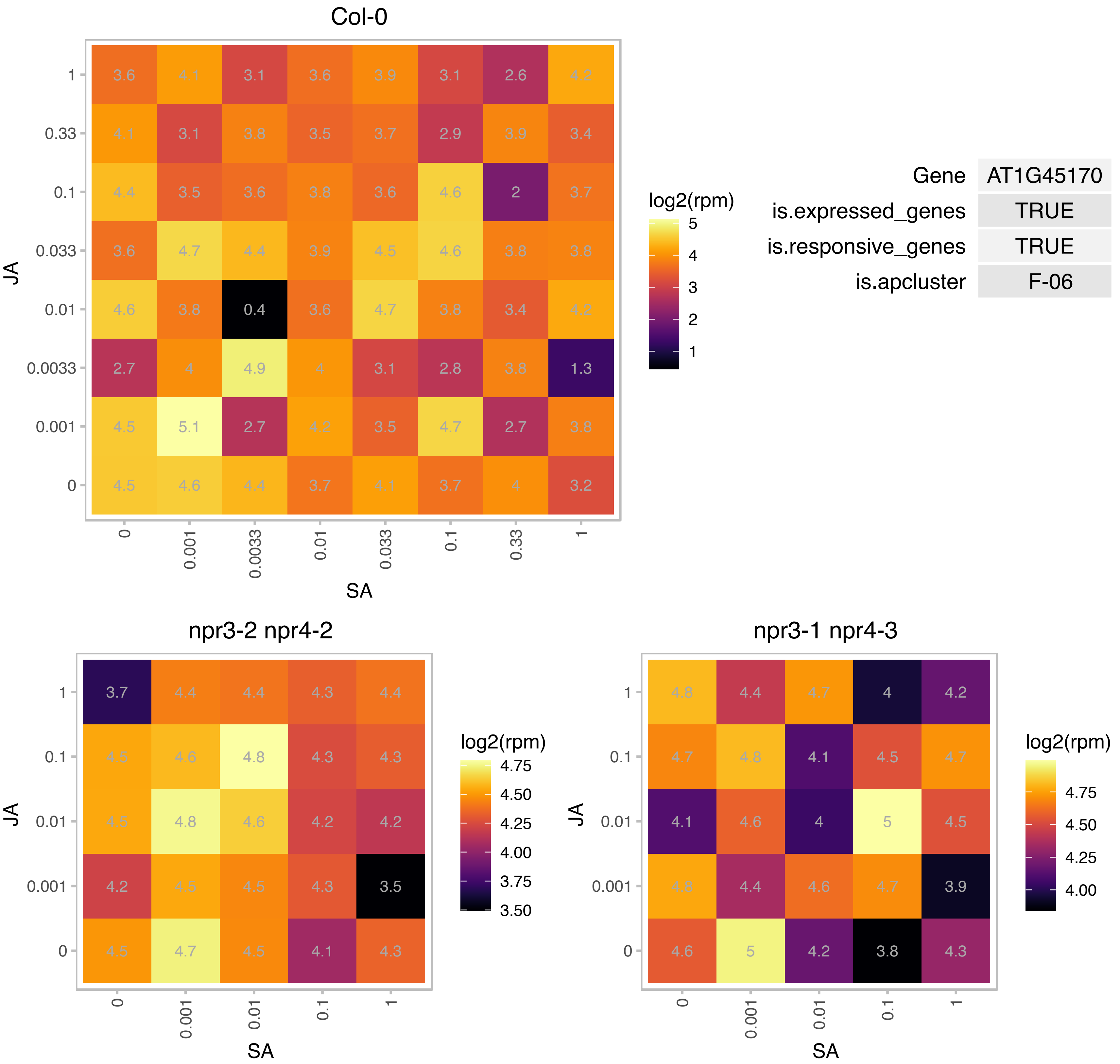

# AT1G48750.1 : NA

Bifunctional inhibitor/lipid-transfer protein/seed storage 2S albumin superfamily protein

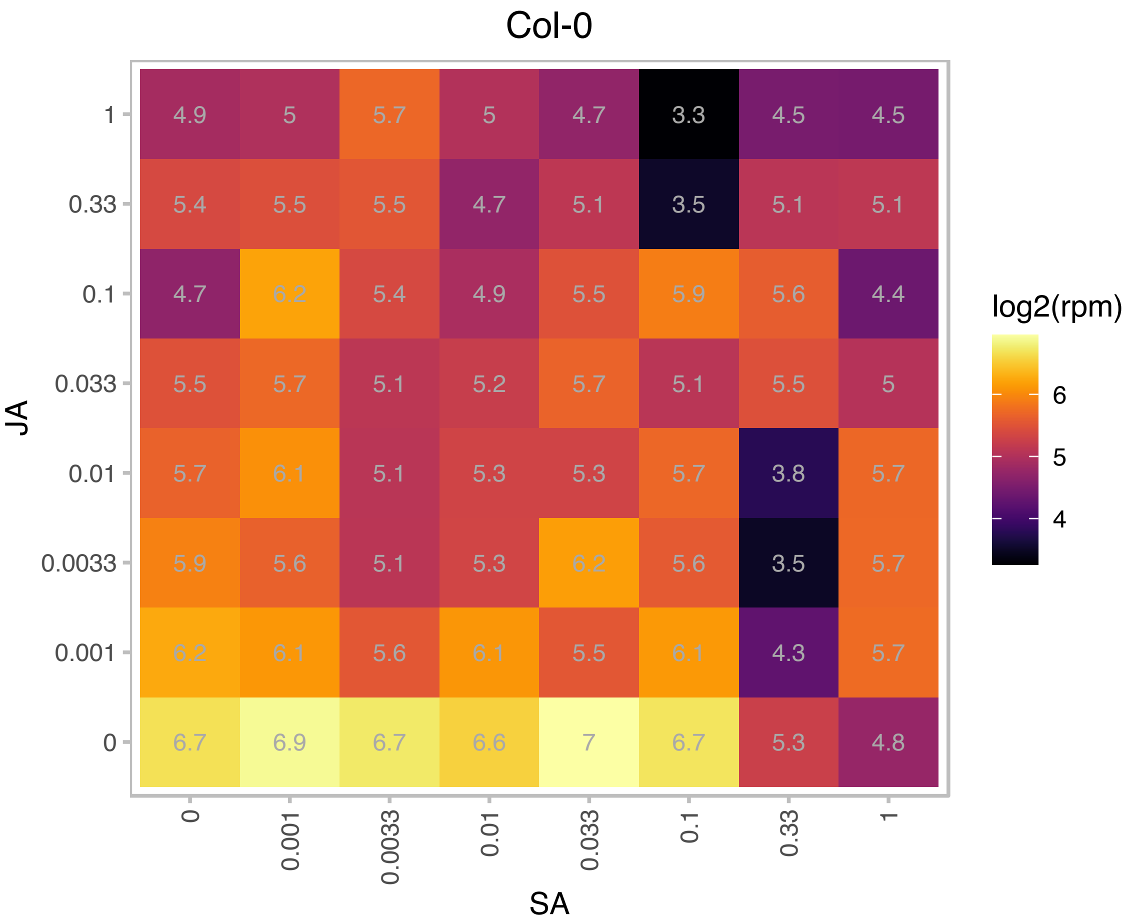


| | |
|---|---|
| Gene | AT1G48750 |
| is.expressed_genes | TRUE |
| is.responsive_genes | TRUE |
| is.apcluster | F-06 |

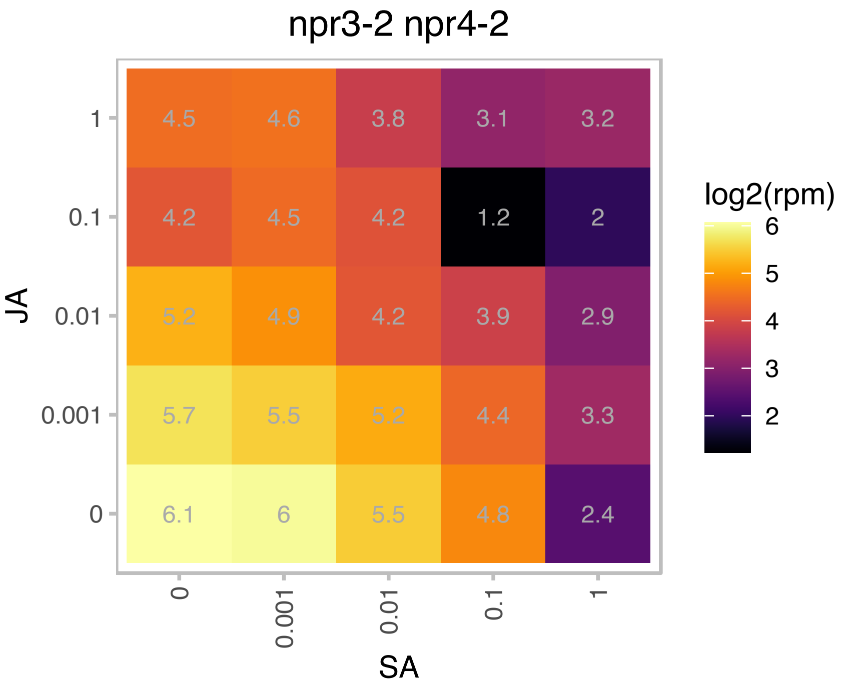


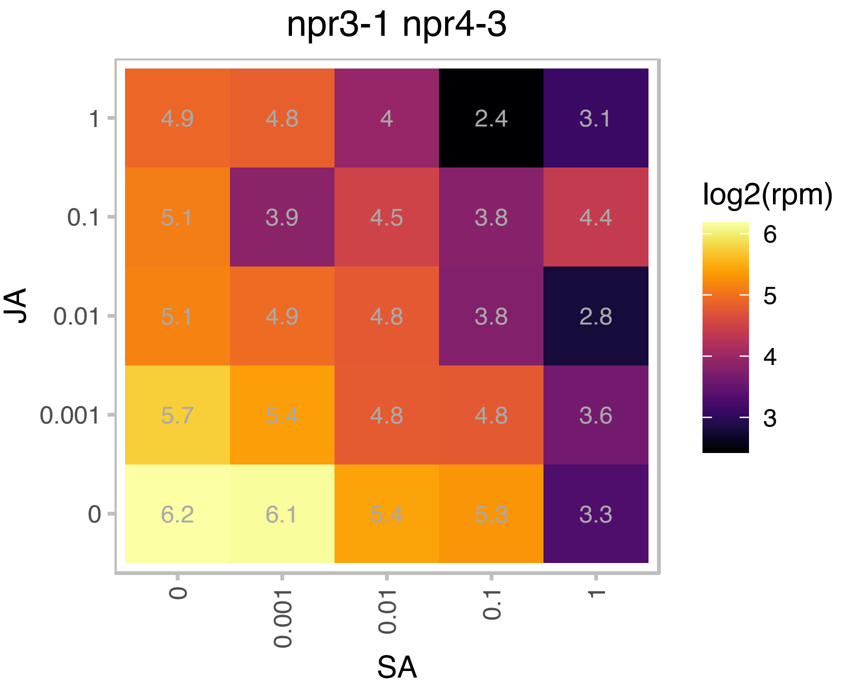

# AT1G50732.1 : NA

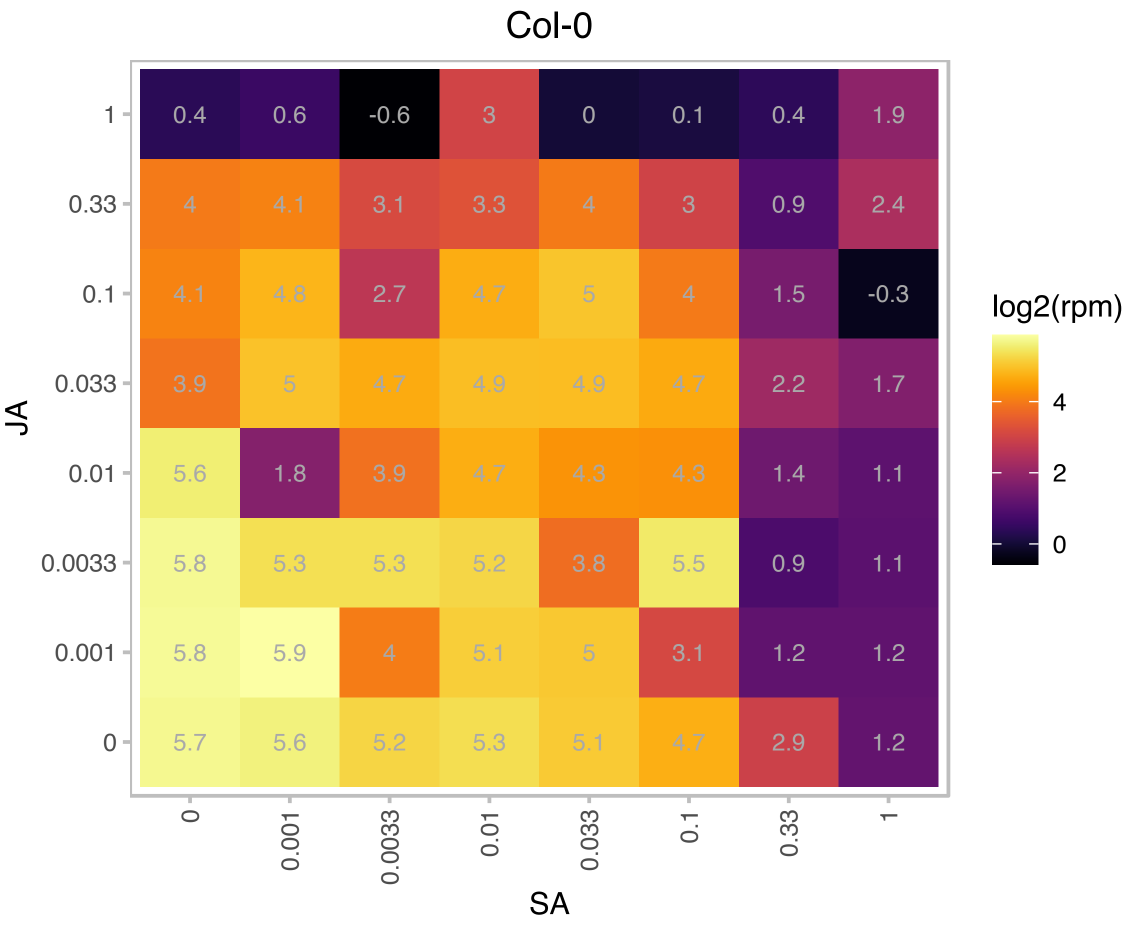


| | |
|---|---|
| Gene | AT1G50732 |
| is.expressed_genes | TRUE |
| is.responsive_genes | TRUE |
| is.apcluster | F-06 |

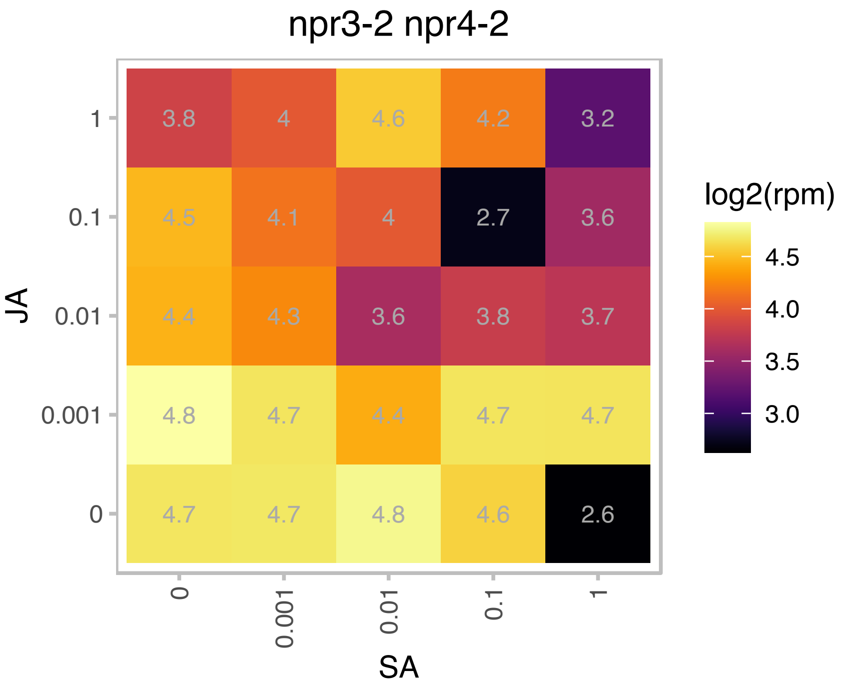


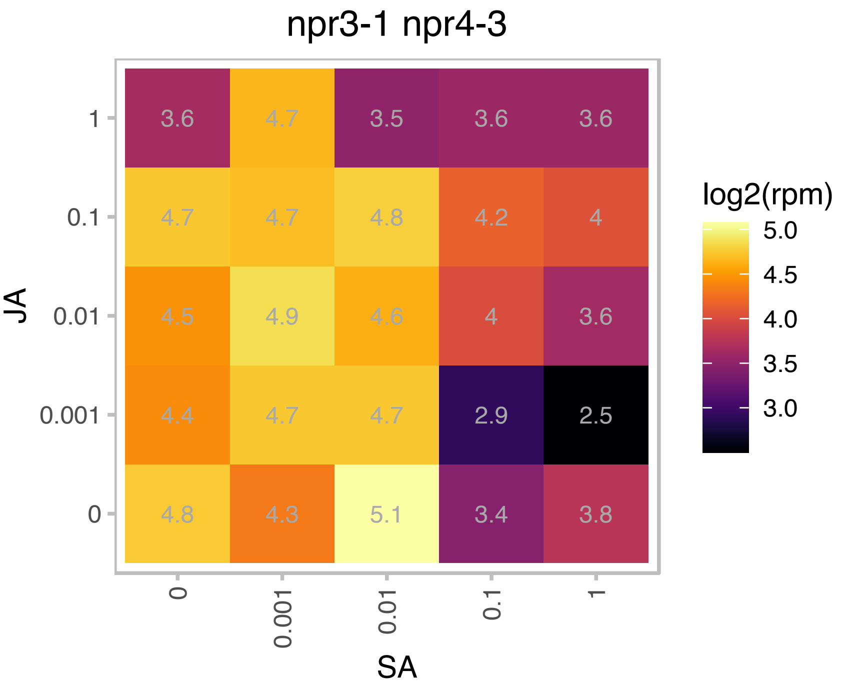

### AT1G51100.1 : CRR41

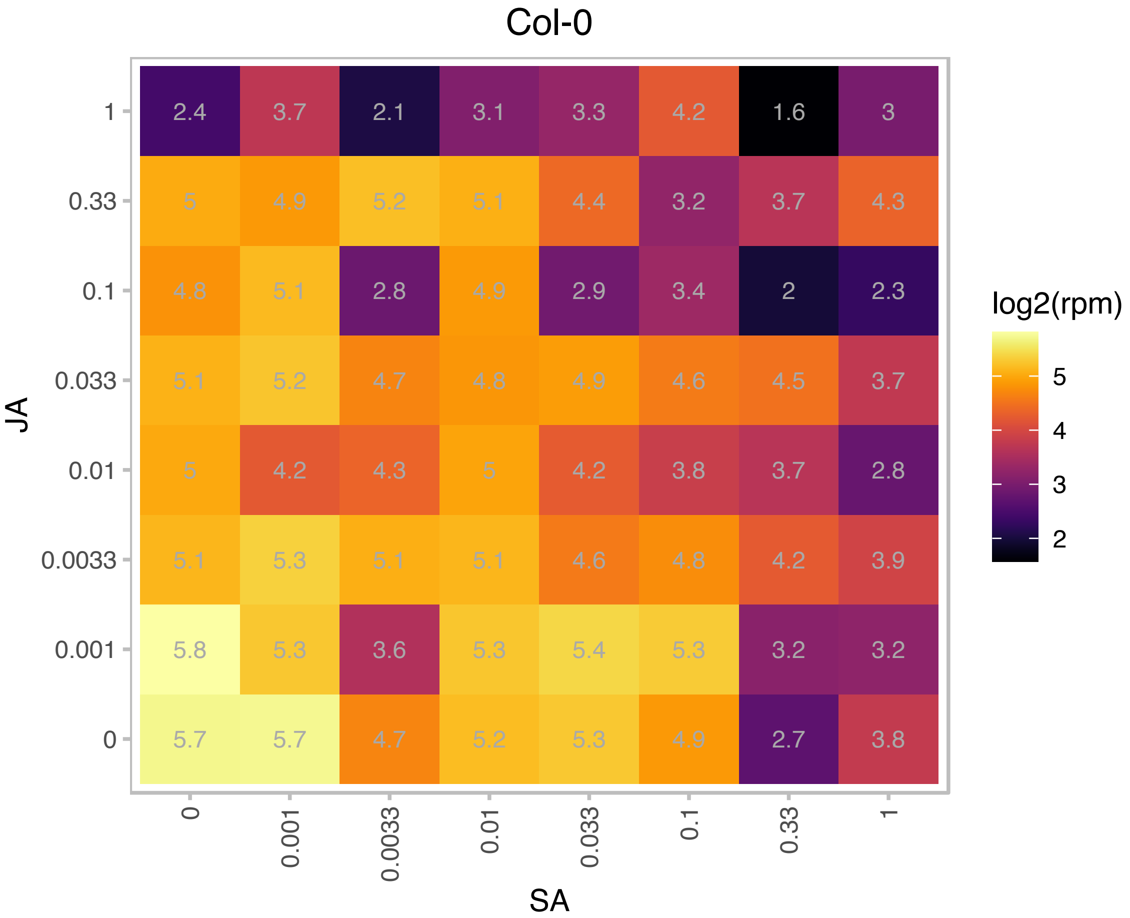


| | |
|---|---|
| Gene | AT1G51100 |
| is.expressed_genes | TRUE |
| is.responsive_genes | TRUE |
| is.apcluster | F-06 |

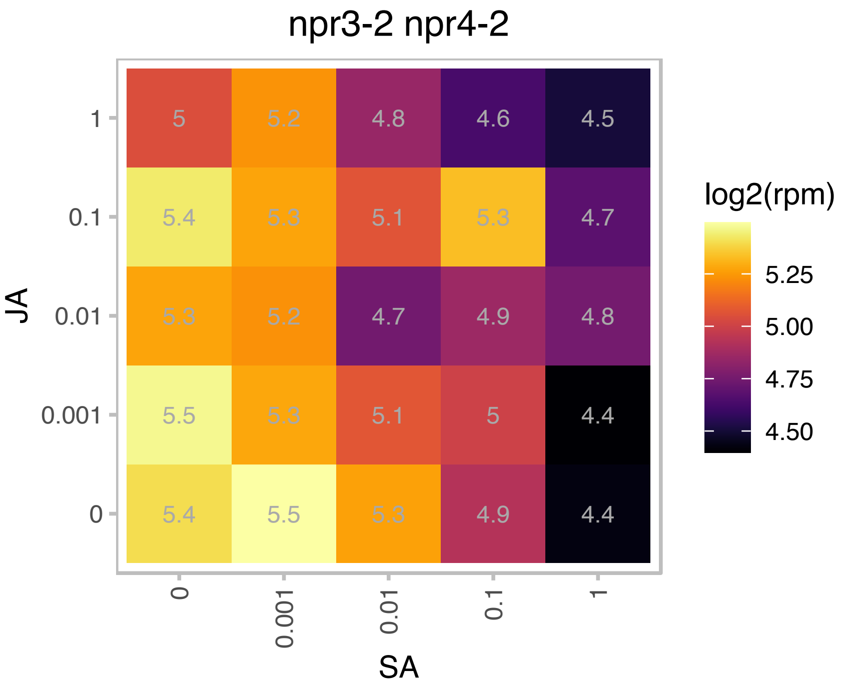


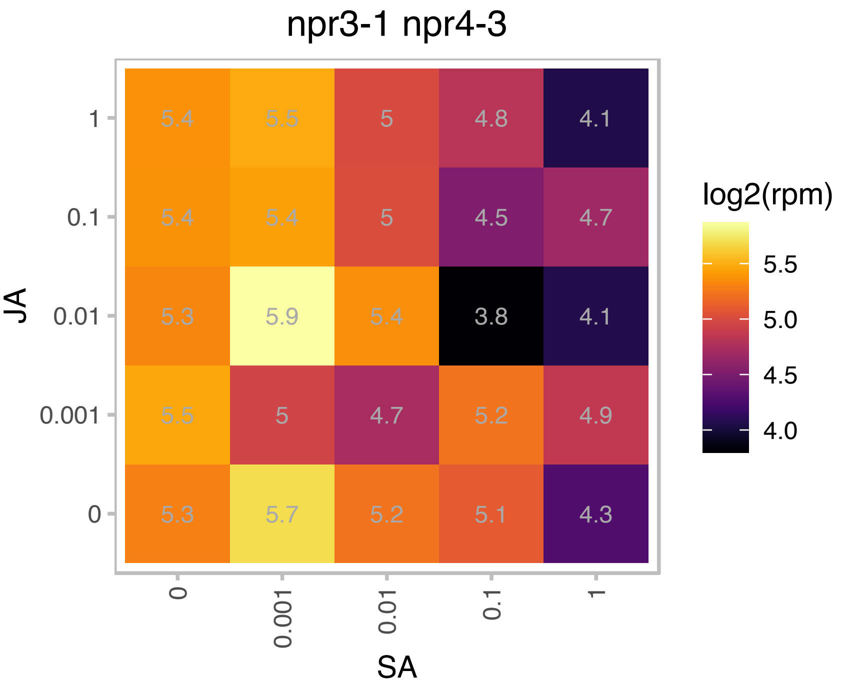

### AT1G60950.1 : ATFD2, FD2, FED A

2Fe-2S ferredoxin-like superfamily protein

encodes a major leaf ferredoxin

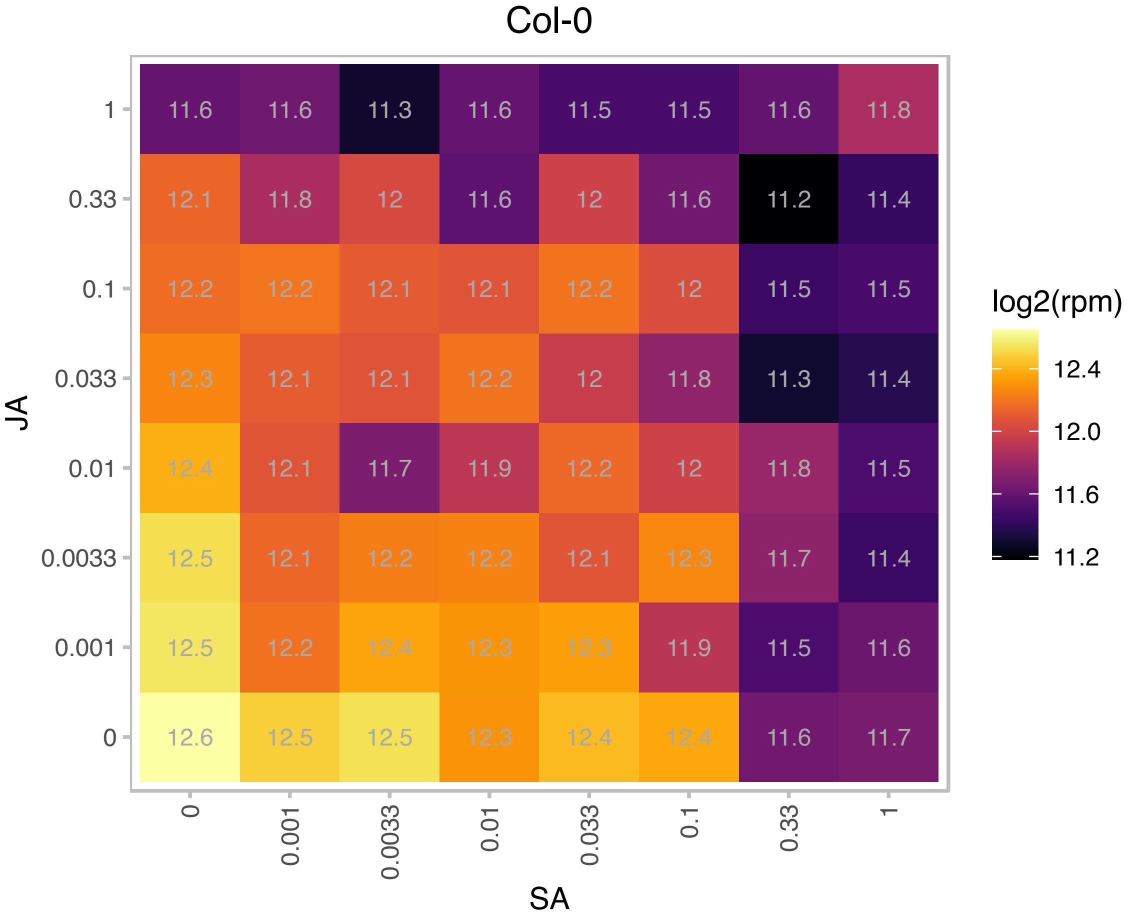


| | |
|---|---|
| Gene | AT1G60950 |
| is.expressed_genes | TRUE |
| is.responsive_genes | TRUE |
| is.apcluster | F-06 |

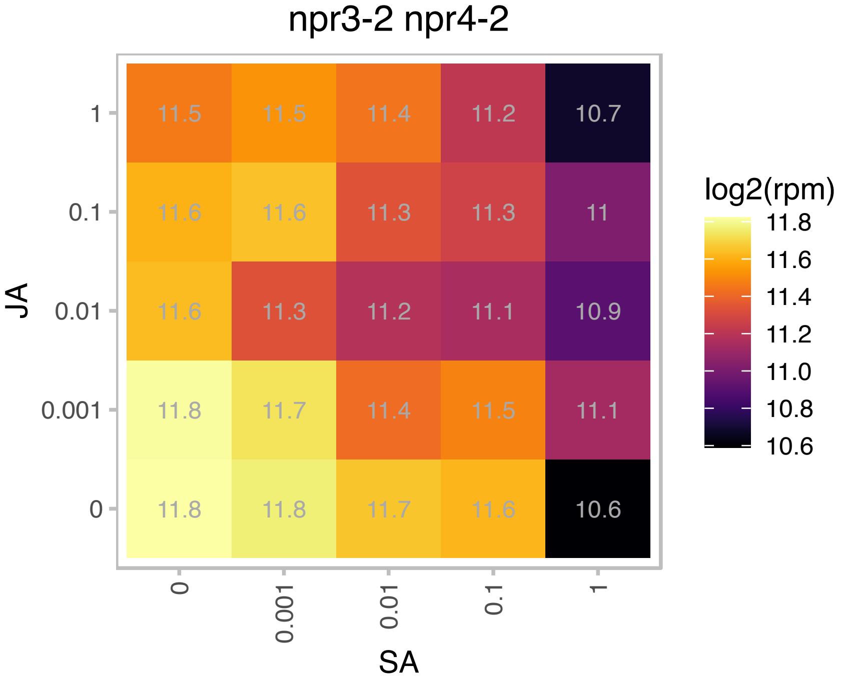


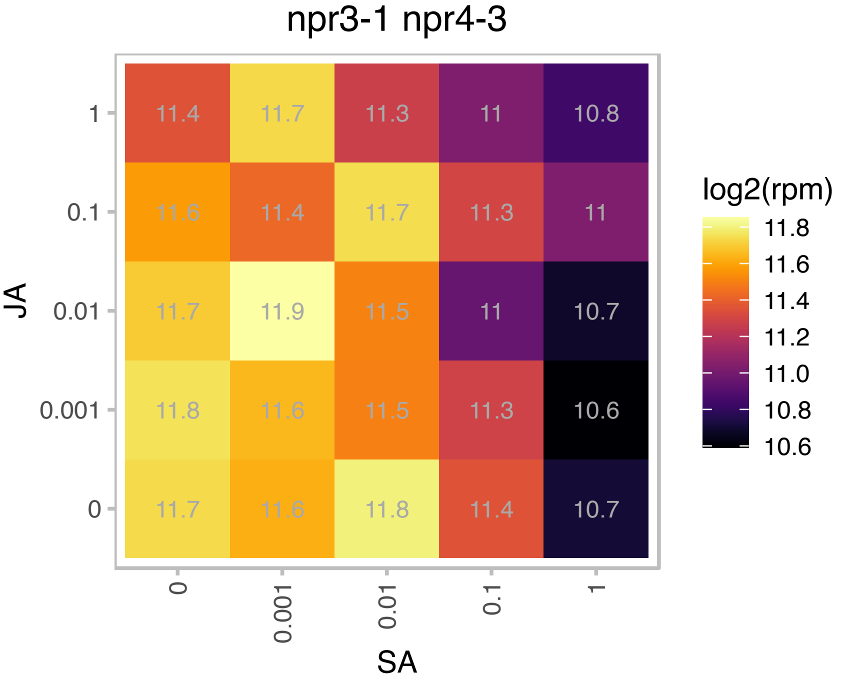

### AT1G62750.1 : ATSCO1, ATSCO1/CPEF-G, SCO1

Translation elongation factor EFG/EF2 protein

Nuclear-encoded protein consists of the five domains conserved in EF-G proteins, with two GTP-binding sites in the first domain, and an additional transit peptide at the N-terminus. Localized in chloroplasts. Point mutation results in a delay in the onset of germination. At early developmental stage embryos still contain undifferentiated proplastids. The greening of cotyledons is severely impaired in light-grown mutant sco1 seedlings, whereas the following true leaves develop normally as in wild-type plants.

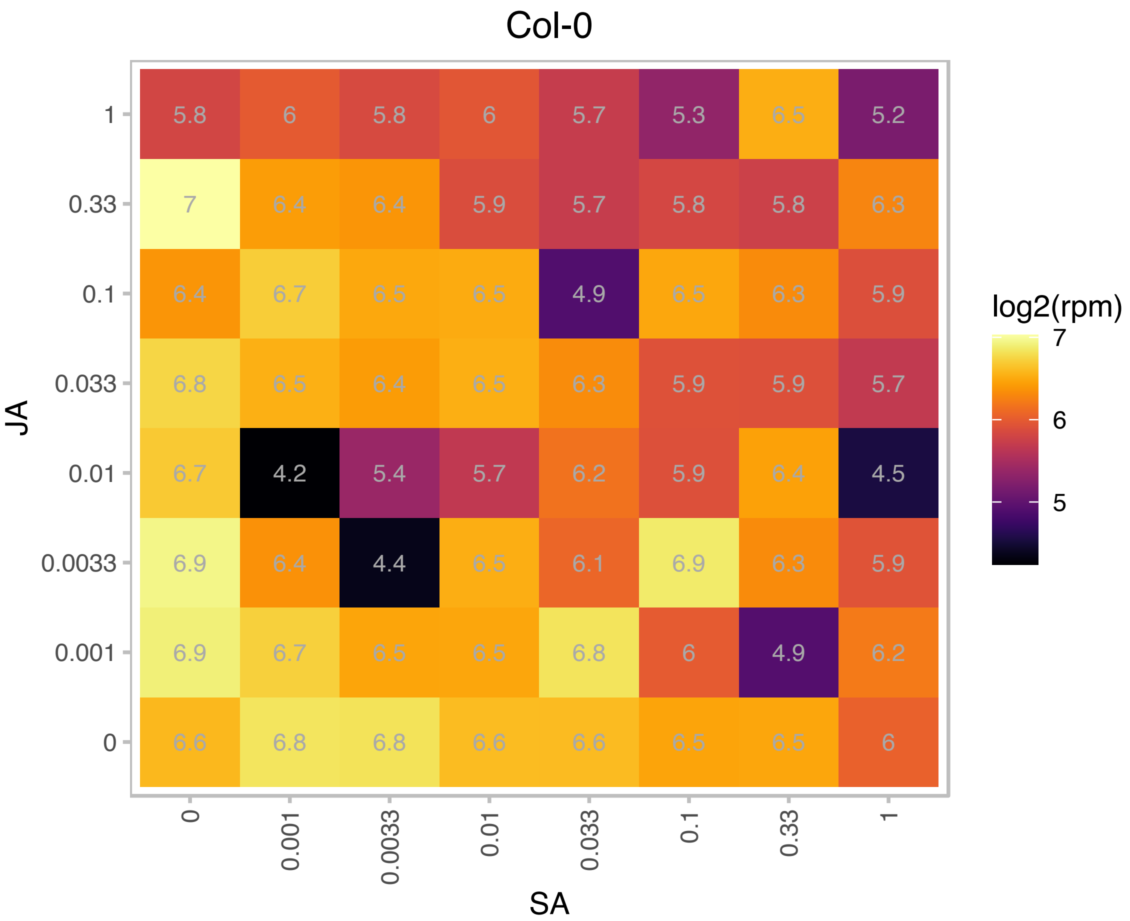


| | |
|---|---|
| Gene | AT1G62750 |
| is.expressed_genes | TRUE |
| is.responsive_genes | TRUE |
| is.apcluster | F-06 |

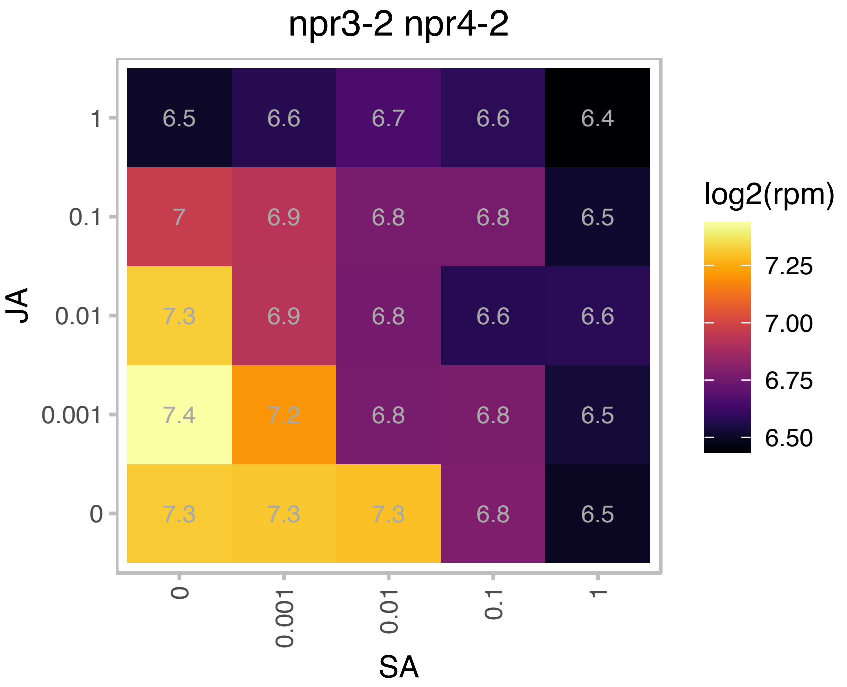


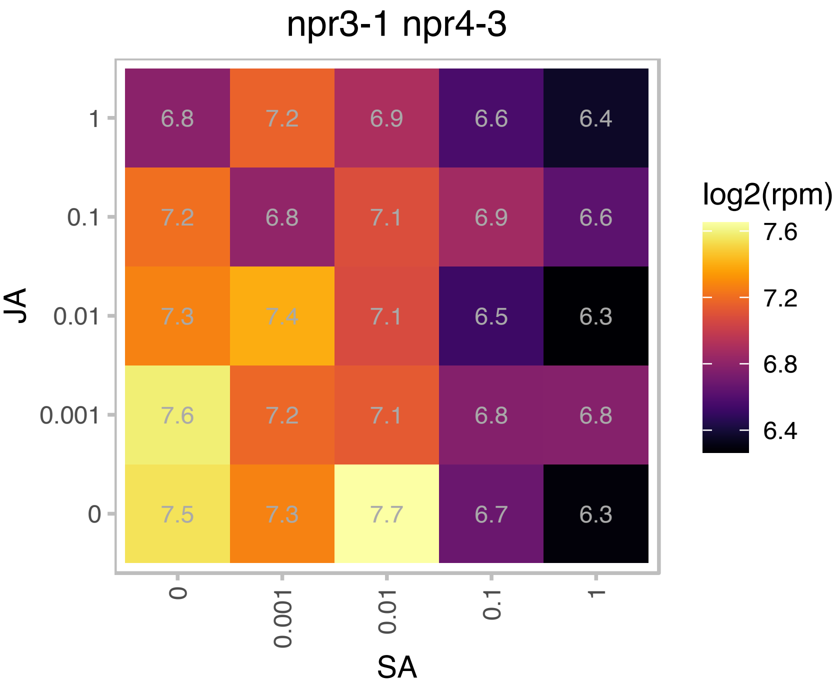

# AT1G65230.1 : NA

Uncharacterized conserved protein (DUF2358)

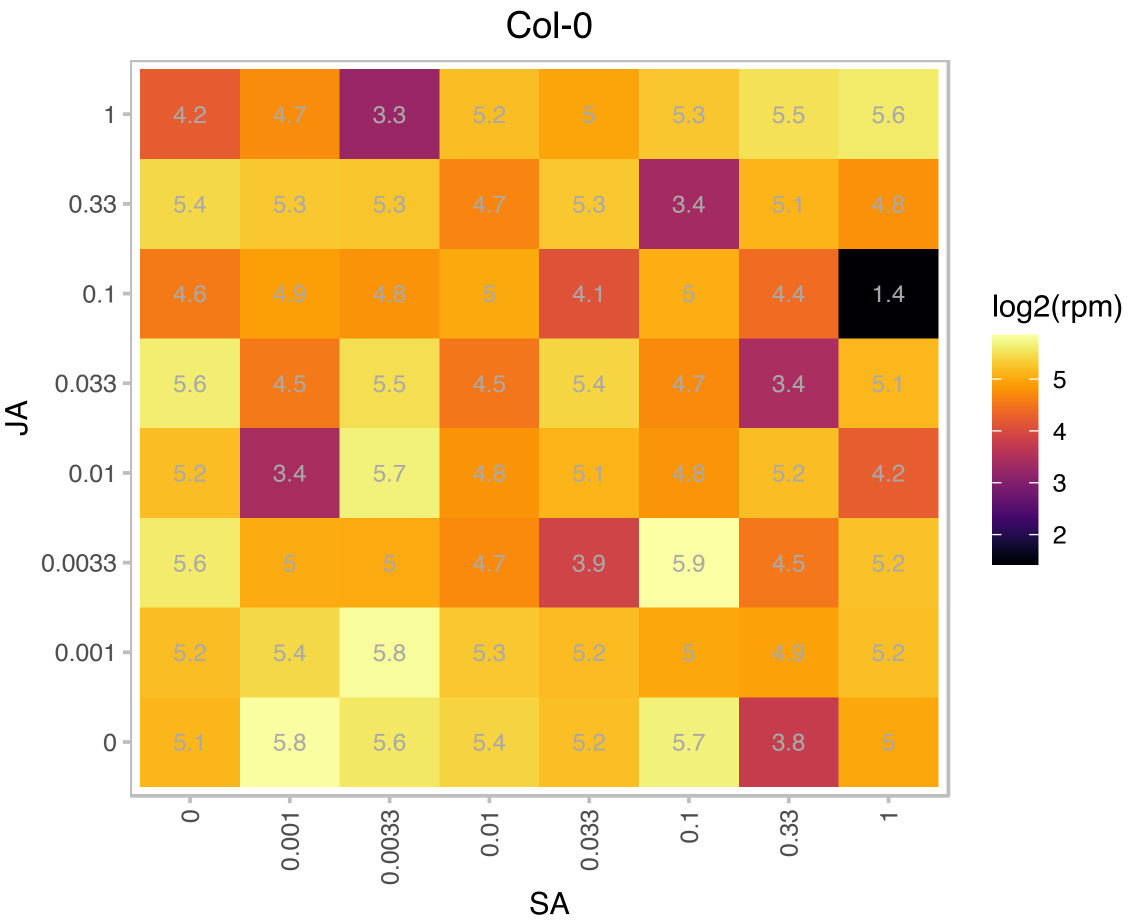


| | |
|---|---|
| Gene | AT1G65230 |
| is.expressed_genes | TRUE |
| is.responsive_genes | TRUE |
| is.apcluster | F-06 |

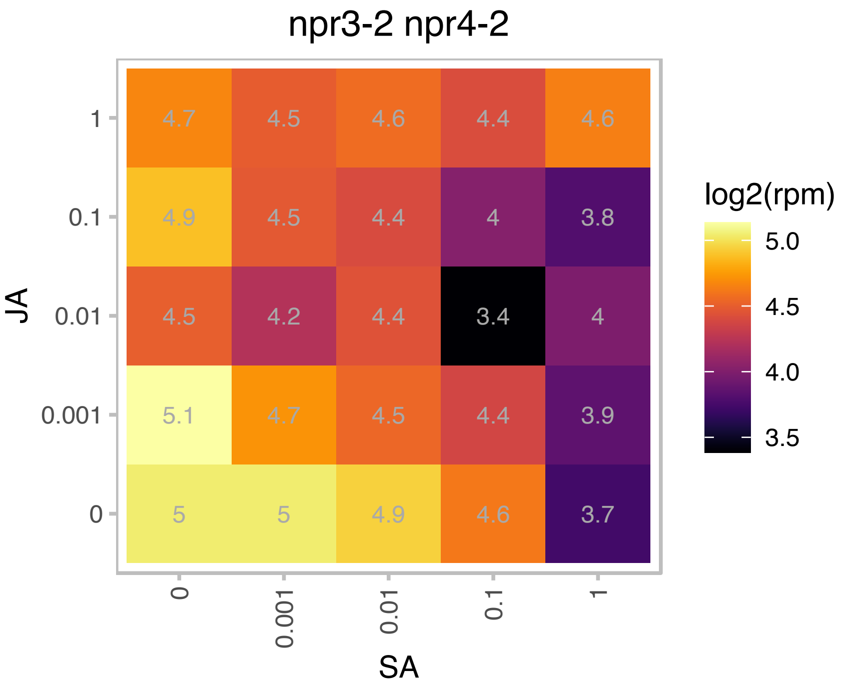


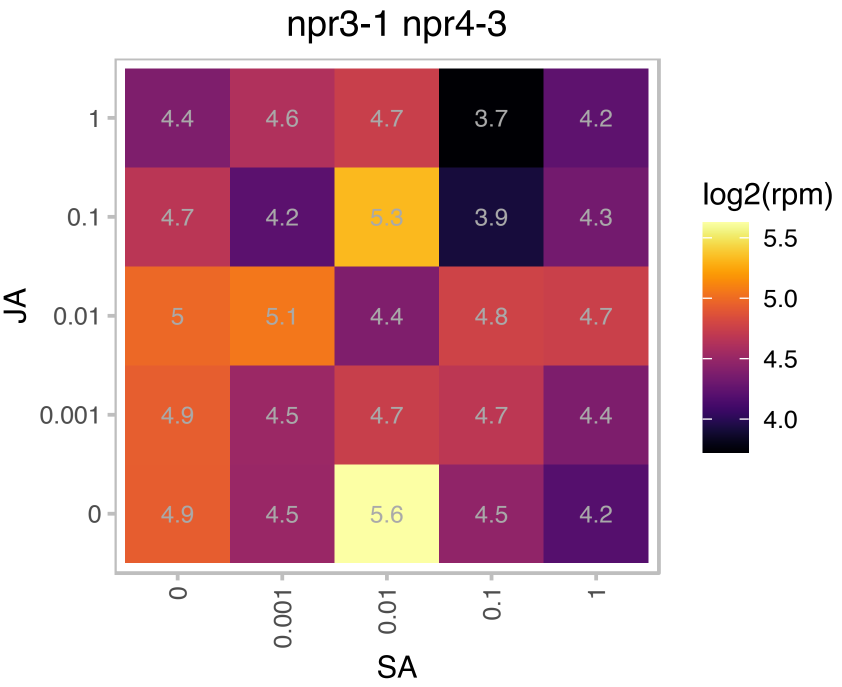

# AT1G66890.1 : NA

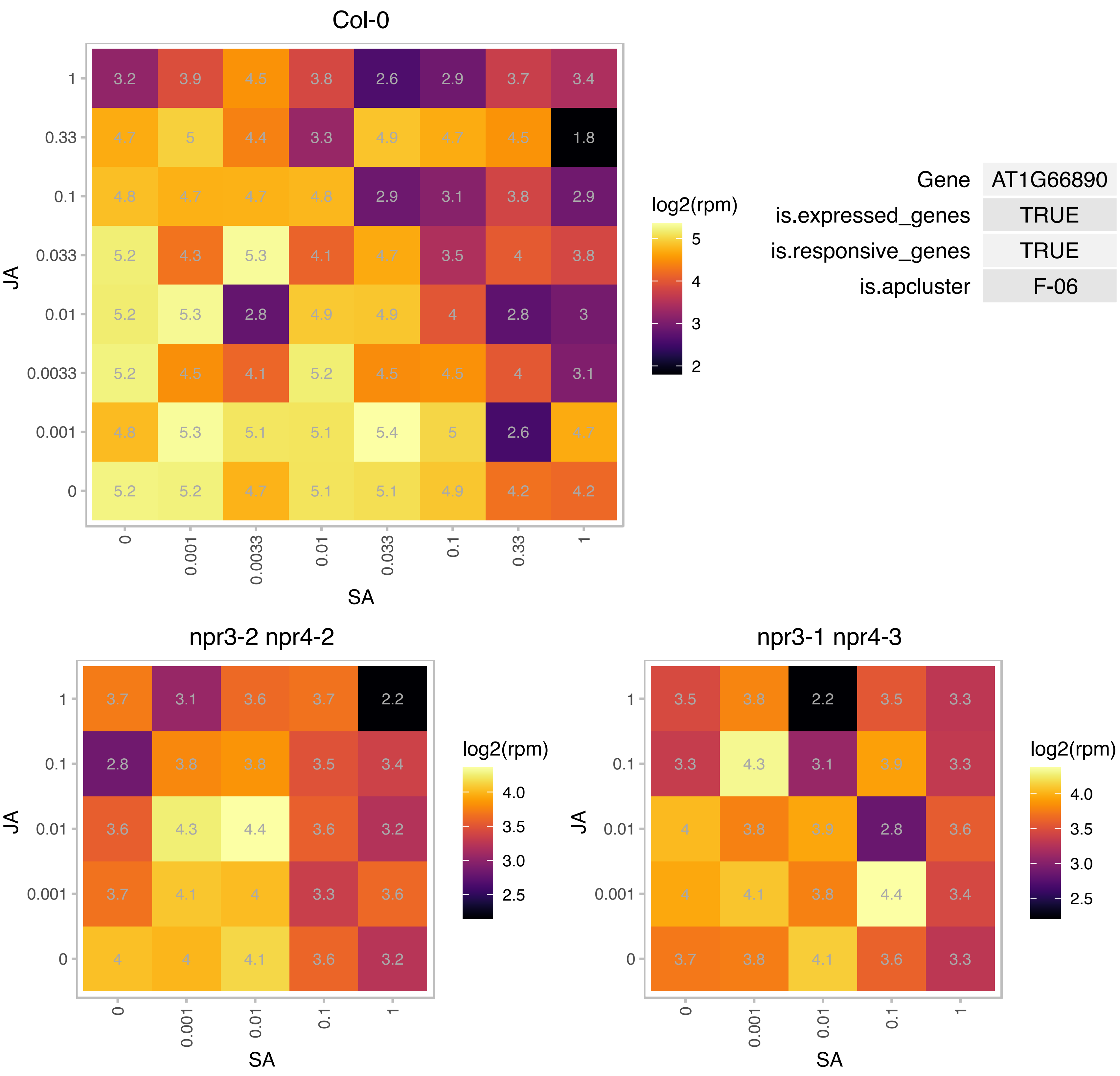

### AT1G68660.1 : AtClpS1, ClpS1

Ribosomal protein L12/ ATP-dependent Clp protease adaptor protein ClpS family protein

ClpS1 is a member of the caseionolytic proteinase S family of N-recognins. It is involved in proteolysis in the chloroplast stroma. An arginine residue (Arg50) controls low-affinity substrate binding.

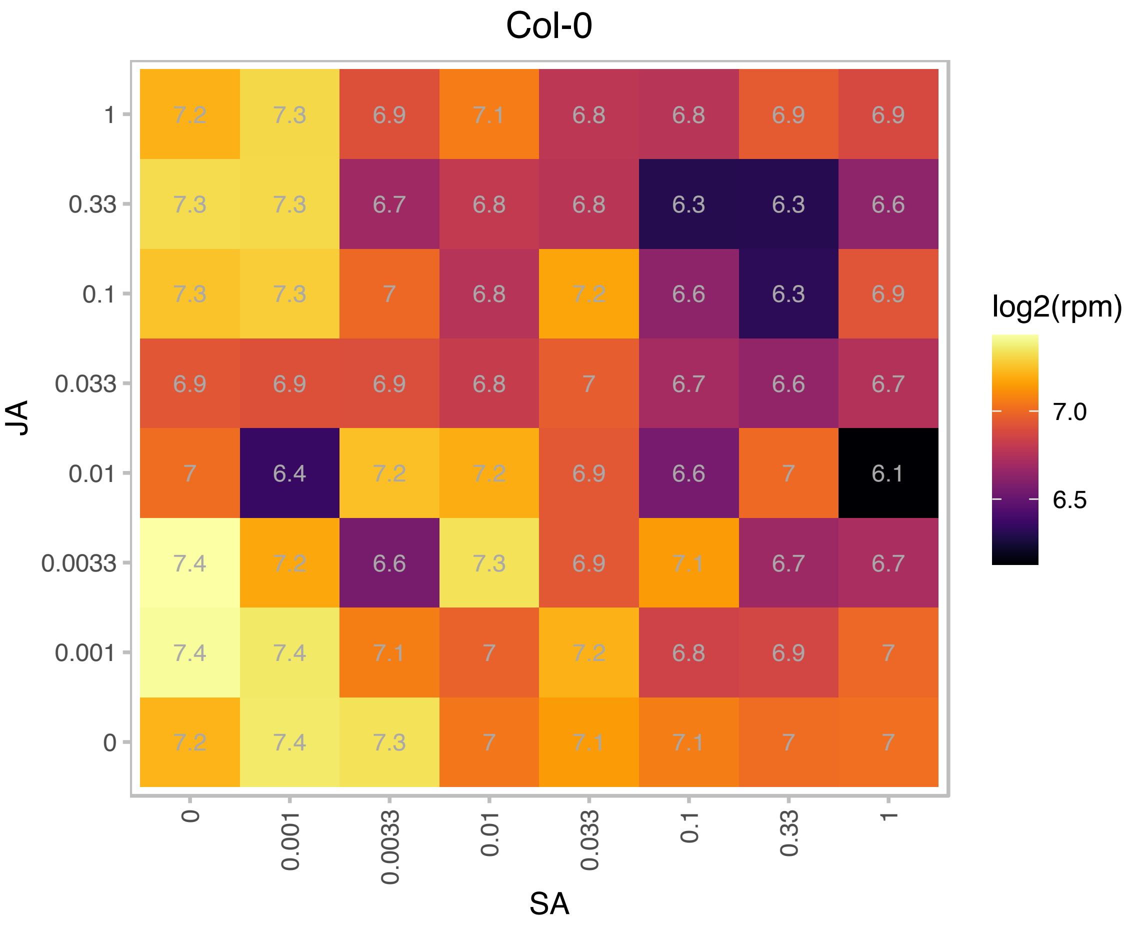


| | |
|---|---|
| Gene | AT1G68660 |
| is.expressed_genes | TRUE |
| is.responsive_genes | TRUE |
| is.apcluster | F-06 |

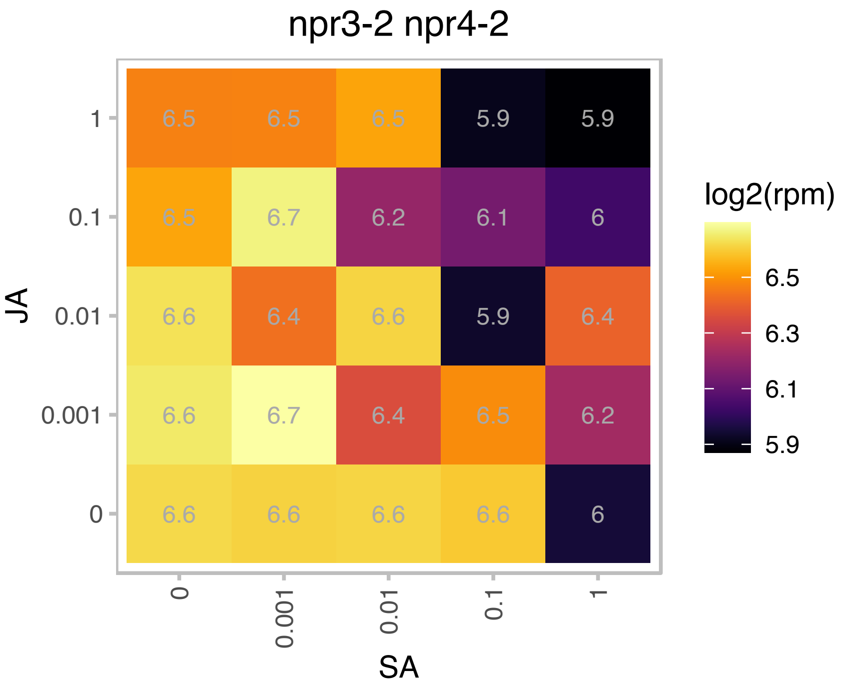


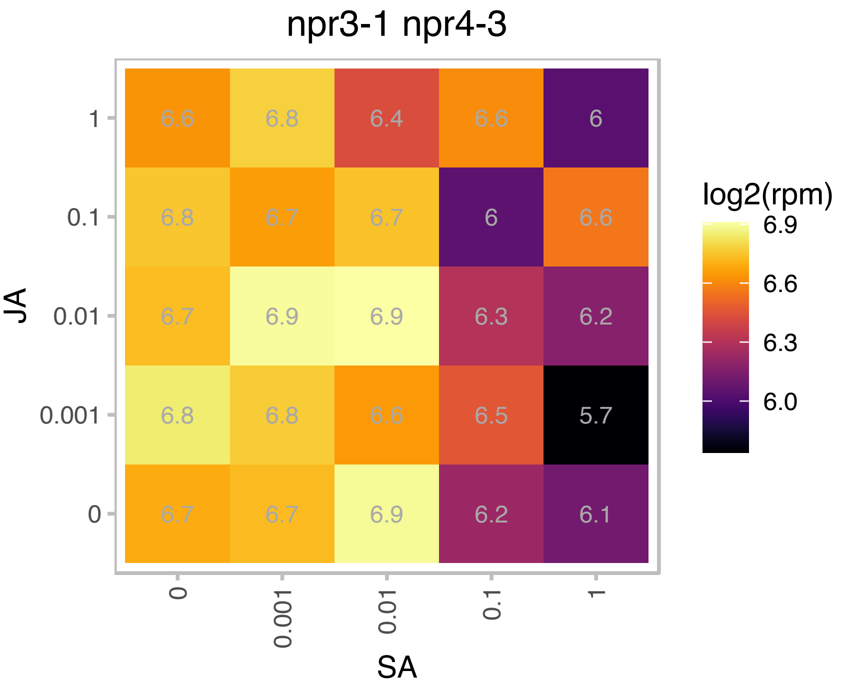

### AT1G73830.1 : BEE3

BR enhanced expression 3

Encodes the brassinosteroid signaling component BEE3 (BR-ENHANCED EXPRESSION 3). Positively modulates t
shade avoidance syndrome in Arabidopsis seedlings.

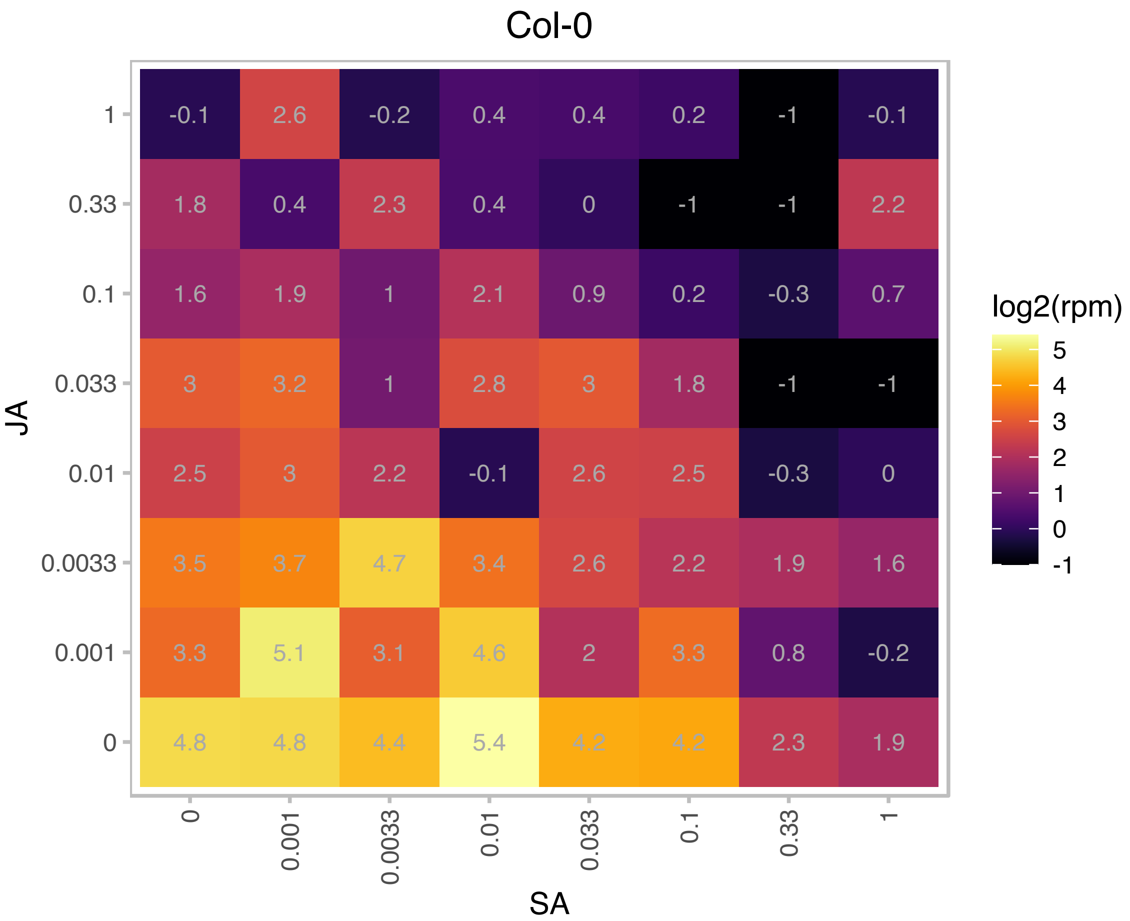


| | |
|---|---|
| Gene | AT1G73830 |
| is.expressed_genes | TRUE |
| is.responsive_genes | TRUE |
| is.apcluster | F-06 |

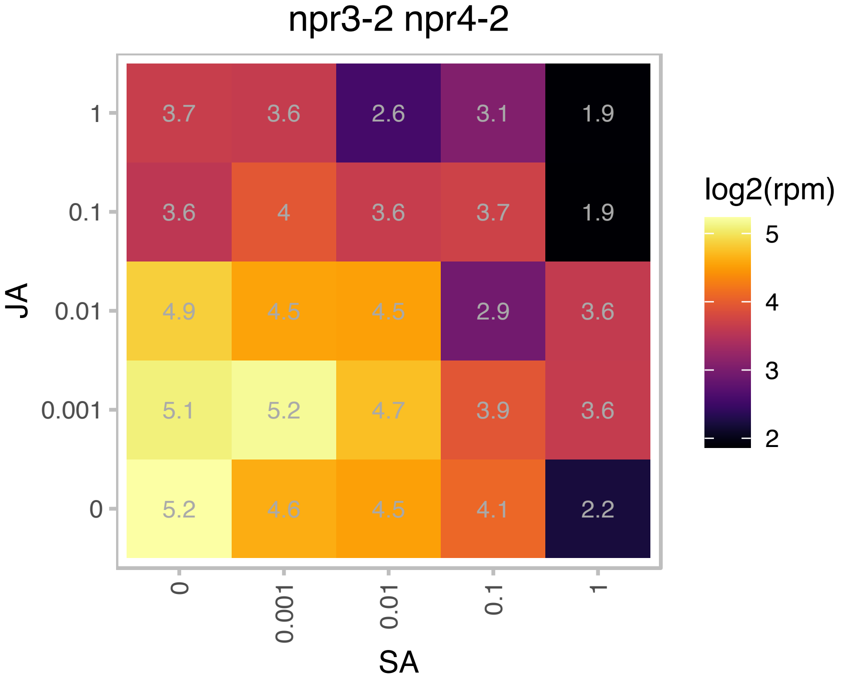


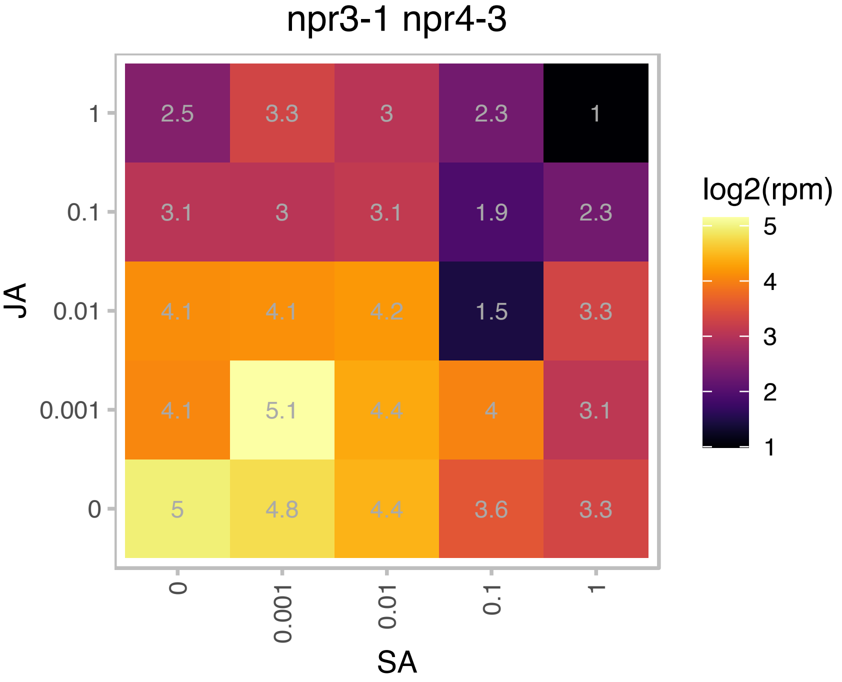

# AT1G73885.1 : NA

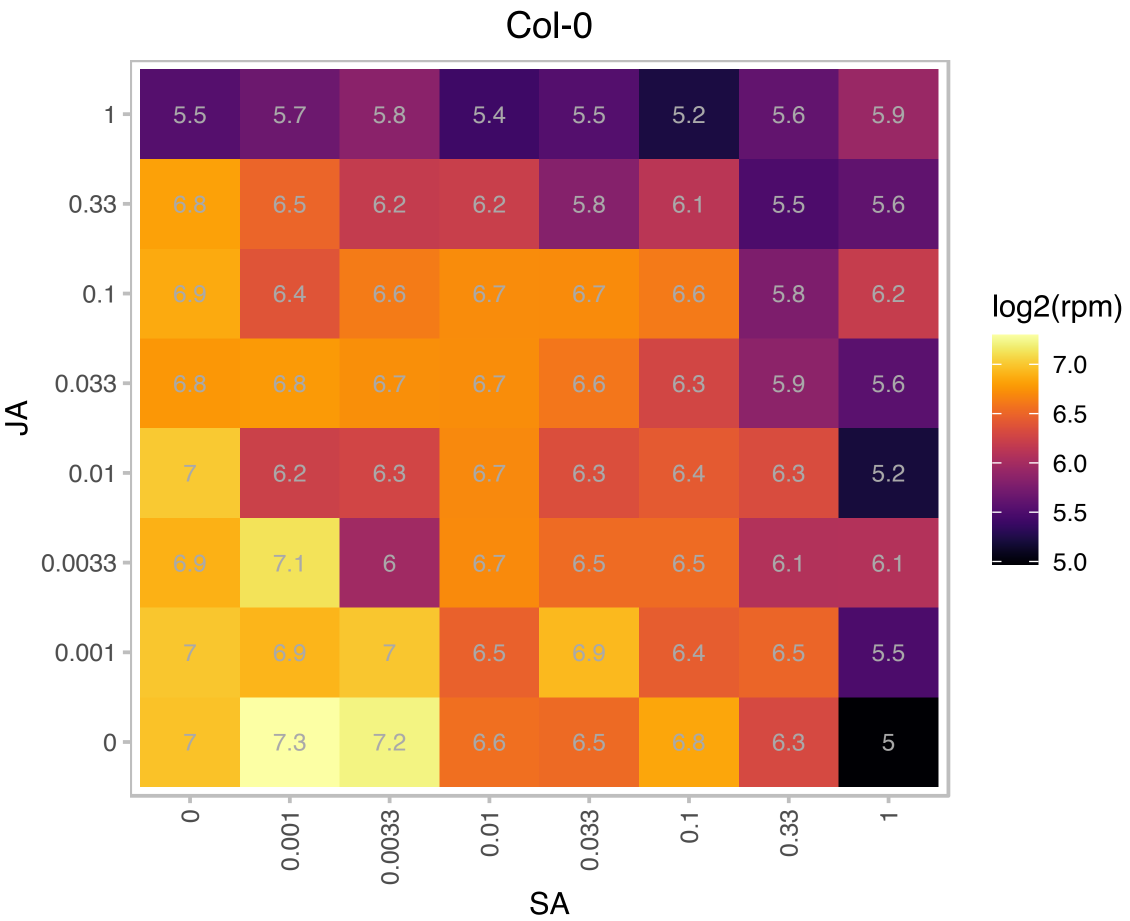


| | |
|---|---|
| Gene | AT1G73885 |
| is.expressed_genes | TRUE |
| is.responsive_genes | TRUE |
| is.apcluster | F-06 |

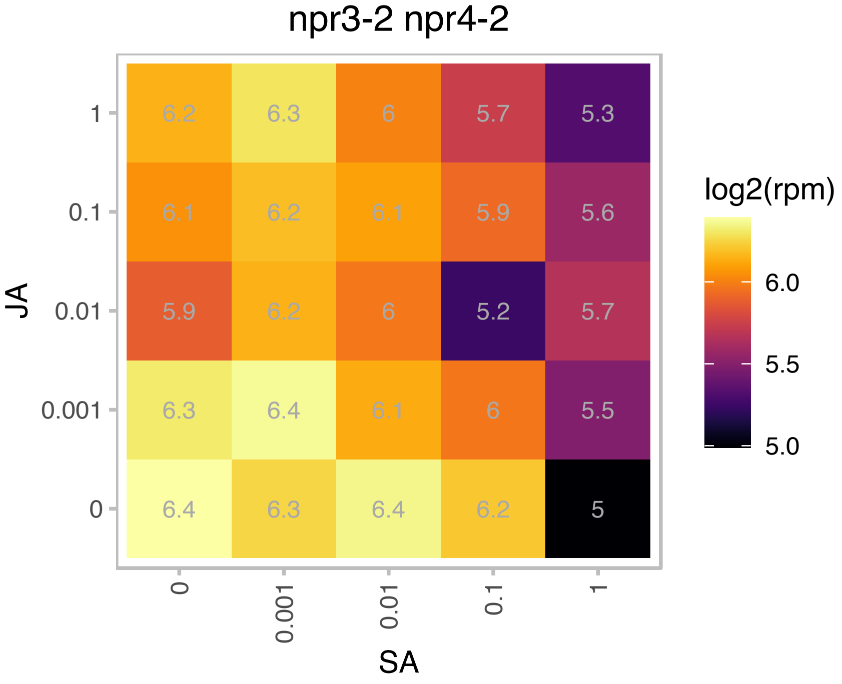


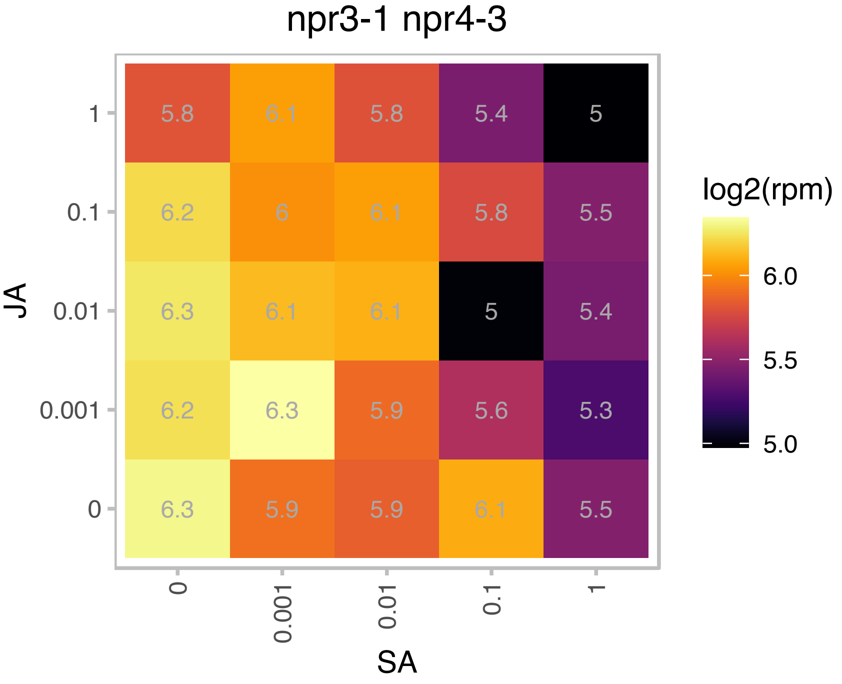

### AT2G01590.1 : CRR3

chlororespiratory reduction 3

Likely a subunit of the chloroplast NAD(P)H dehydrogenase complex, involved in PSI cyclic electron transport. Located in the membrane fraction of chloroplast. Mutant has impaired NAD(P)H dehydrogenase activity.

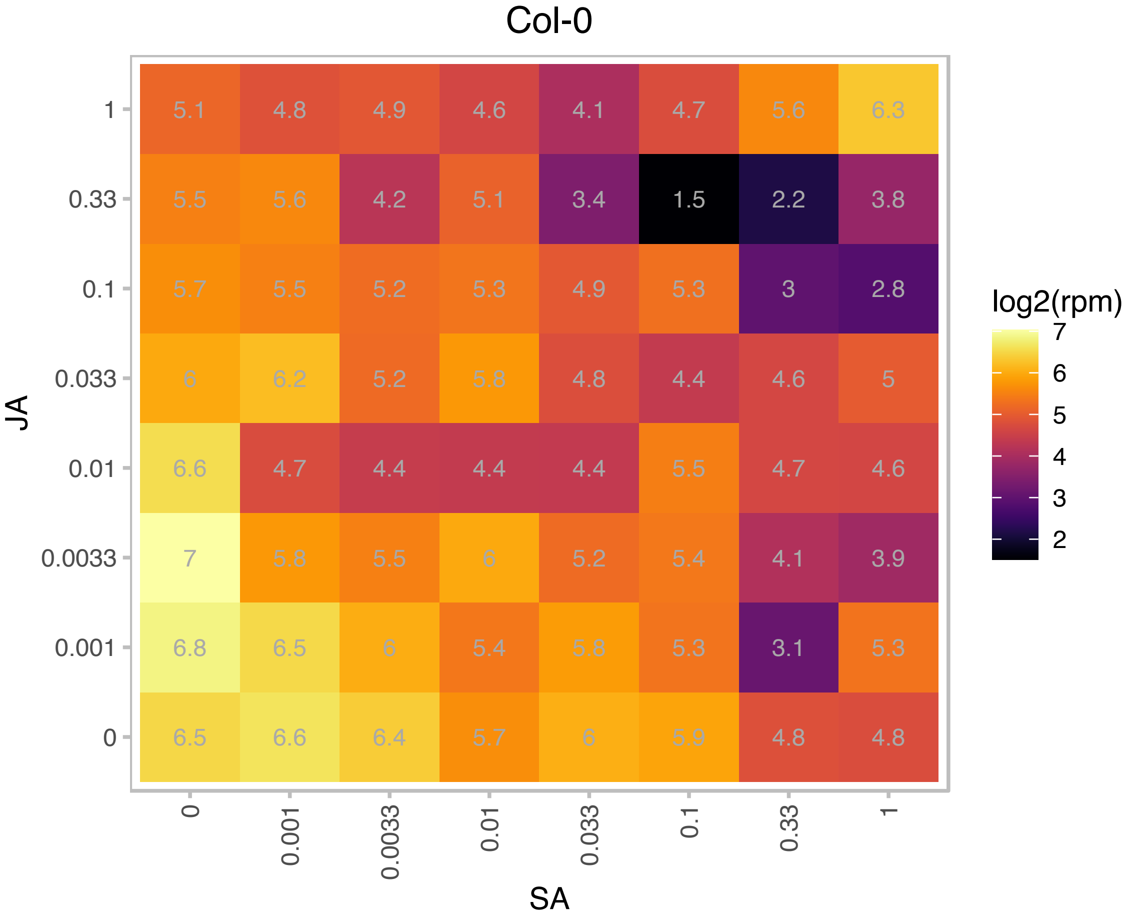


| | |
|---|---|
| Gene | AT2G01590 |
| is.expressed_genes | TRUE |
| is.responsive_genes | TRUE |
| is.apcluster | F-06 |

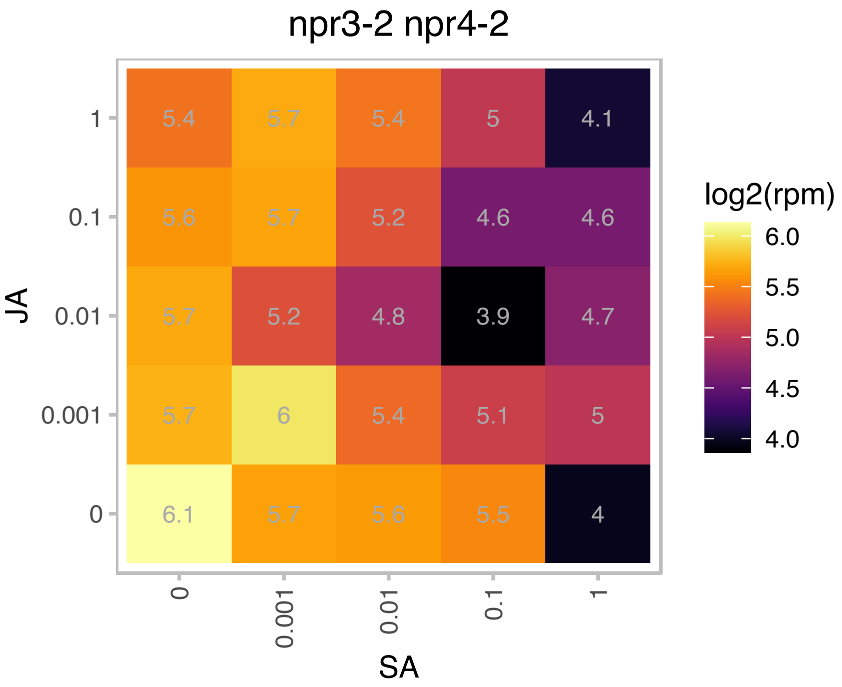


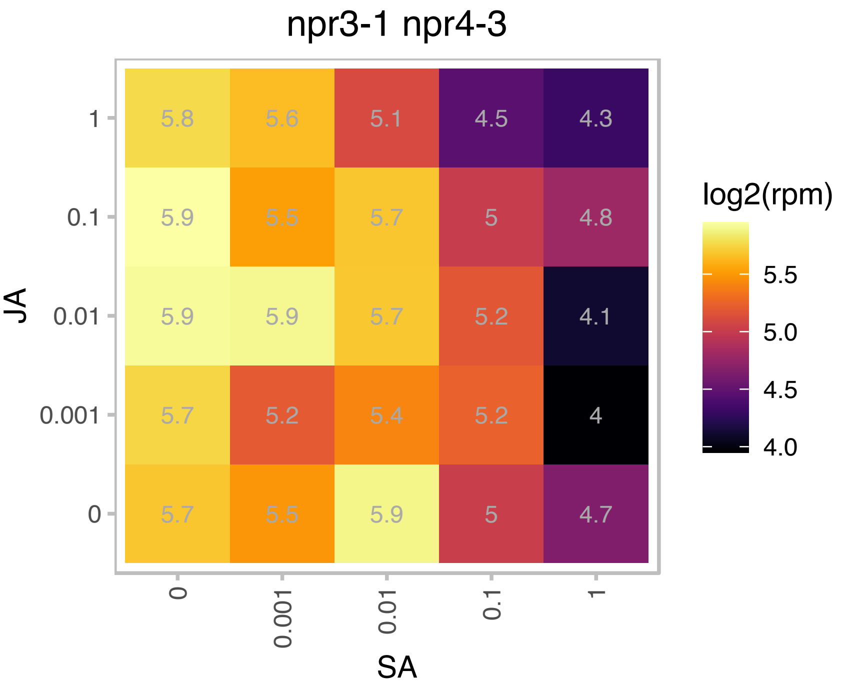

### AT2G04039.1 : NdhV

NdhV is loosely associated with the NDH complex and is required for stabilizing NDH subcomplexes A and E.

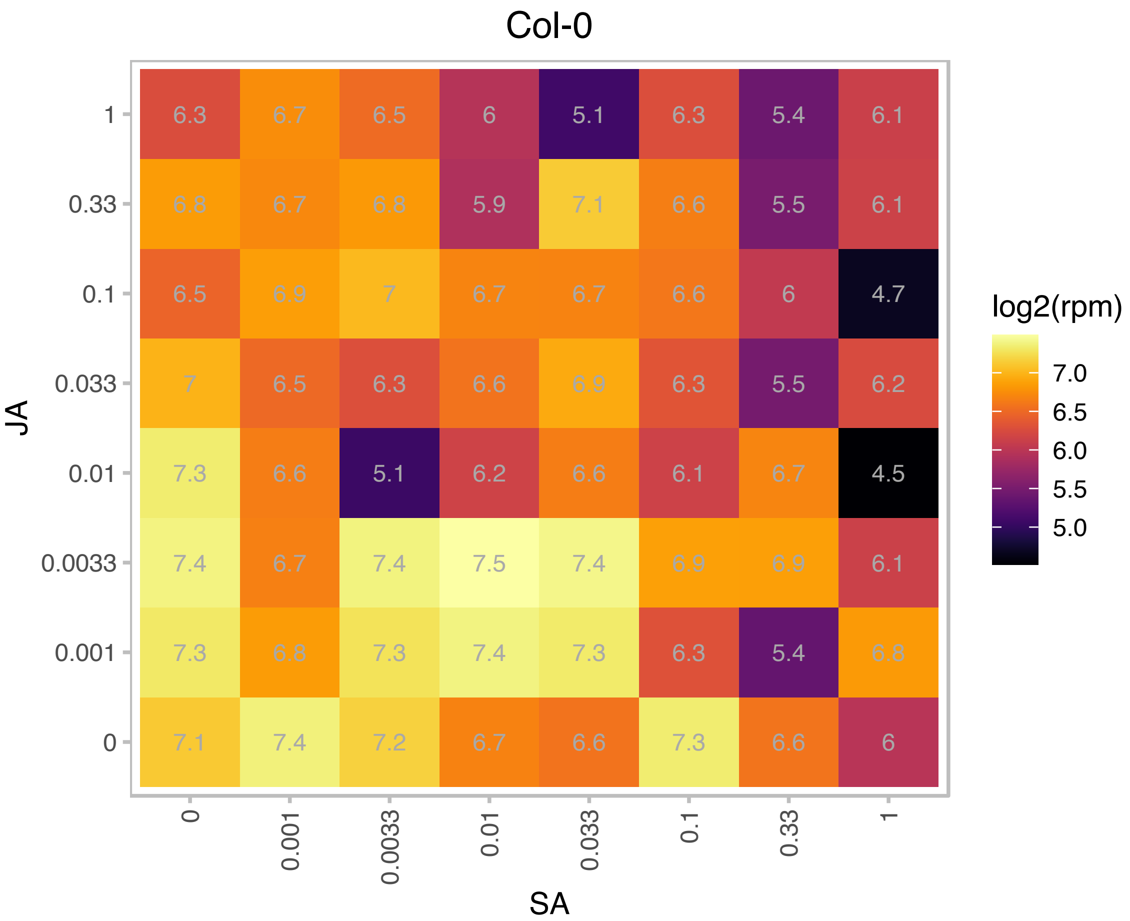


| | |
|---|---|
| Gene | AT2G04039 |
| is.expressed_genes | TRUE |
| is.responsive_genes | TRUE |
| is.apcluster | F-06 |

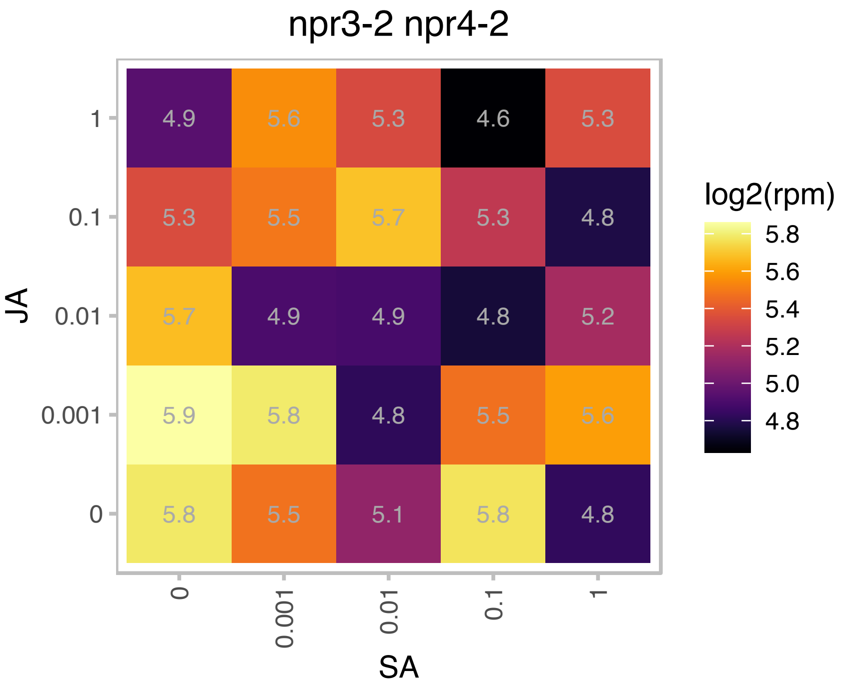


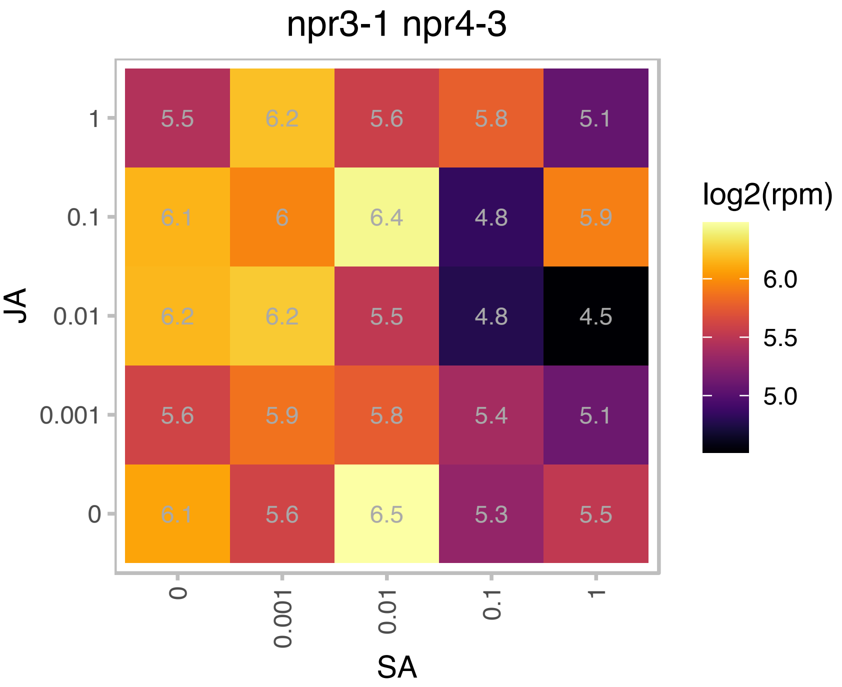

# AT2G04790.1 : NA

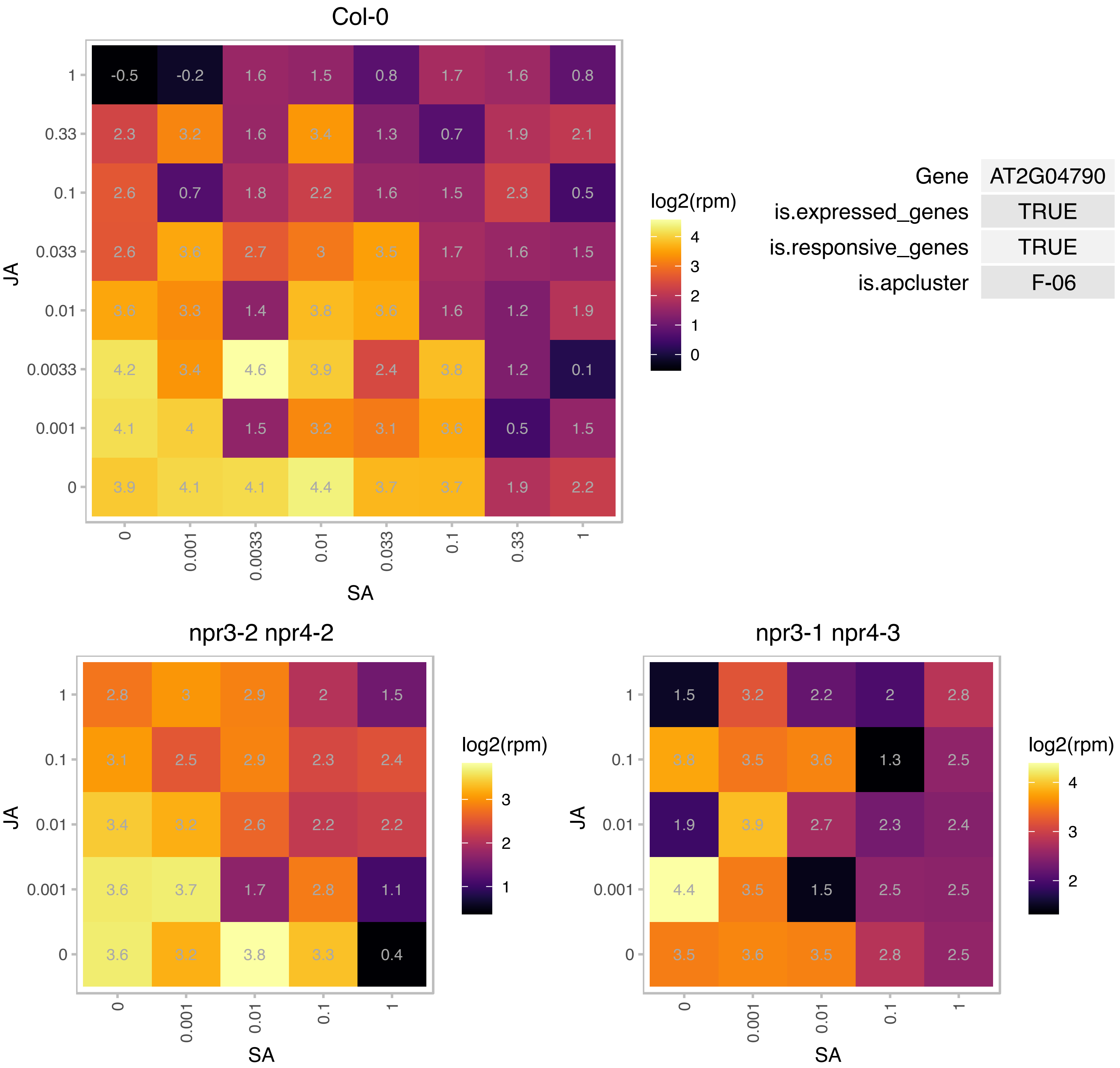


| | |
|---|---|
| Gene | AT2G04790 |
| is.expressed_genes | TRUE |
| is.responsive_genes | TRUE |
| is.apcluster | F-06 |

### AT2G19590.1 : ACO1, ATACO1

ACC oxidase 1

encodes a protein whose sequence is similar to 1-aminocyclopropane-1-carboxylate oxidase

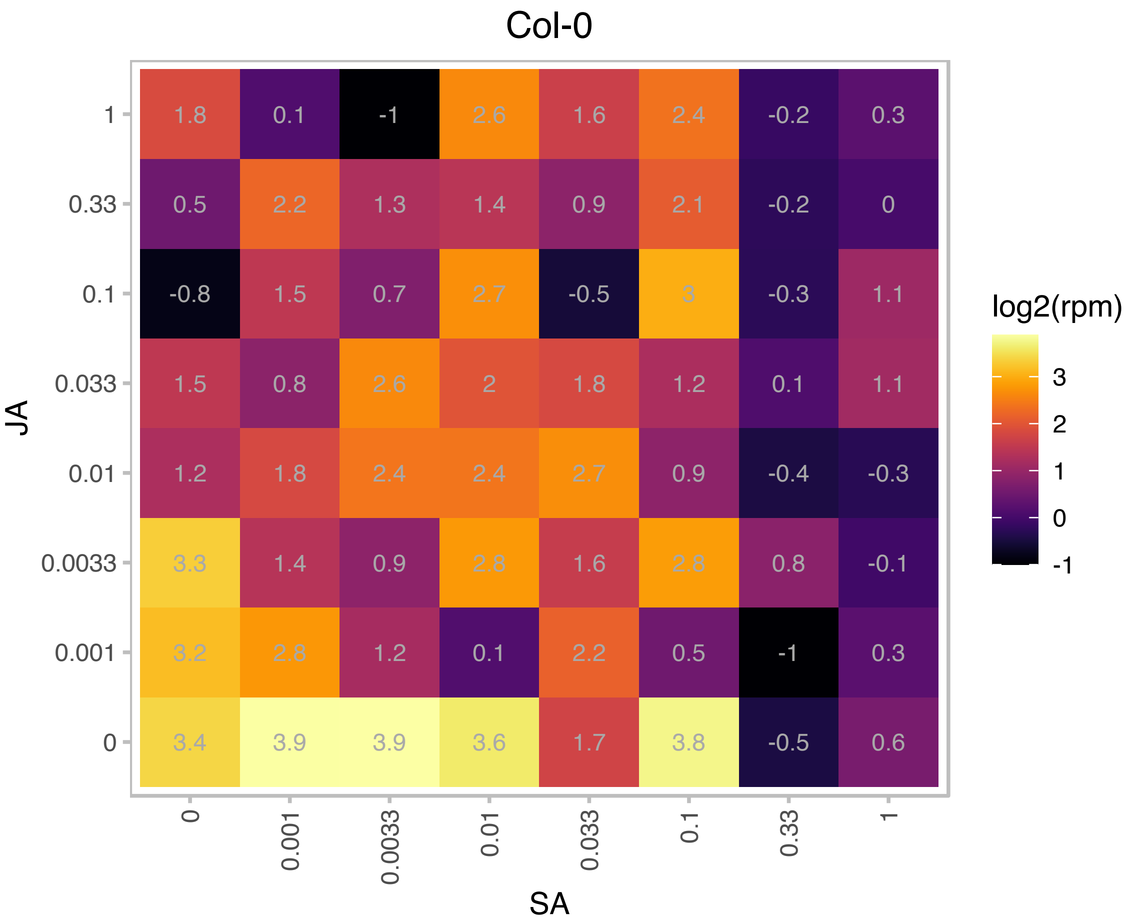


| | |
|---|---|
| Gene | AT2G19590 |
| is.expressed_genes | TRUE |
| is.responsive_genes | TRUE |
| is.apcluster | F-06 |

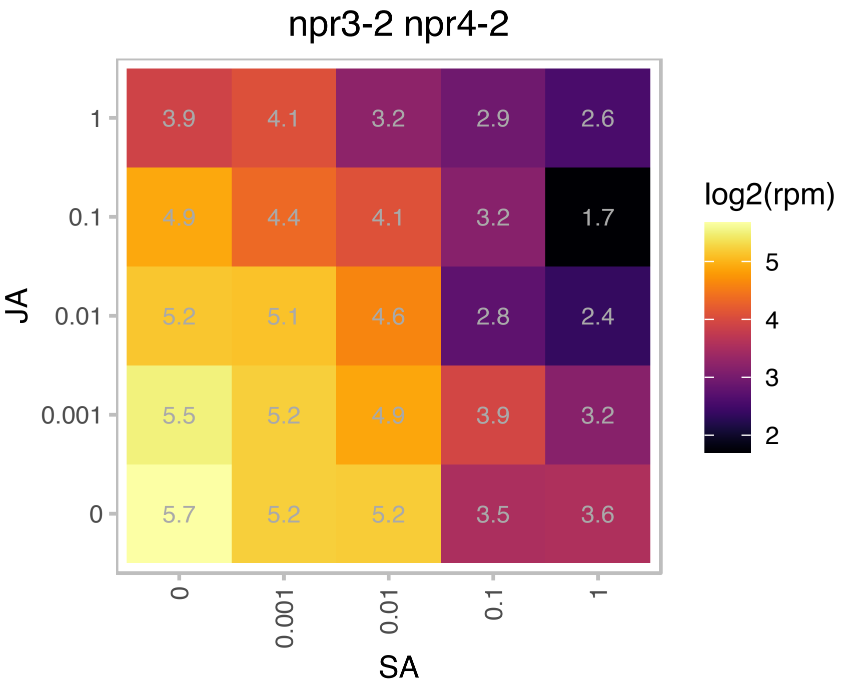


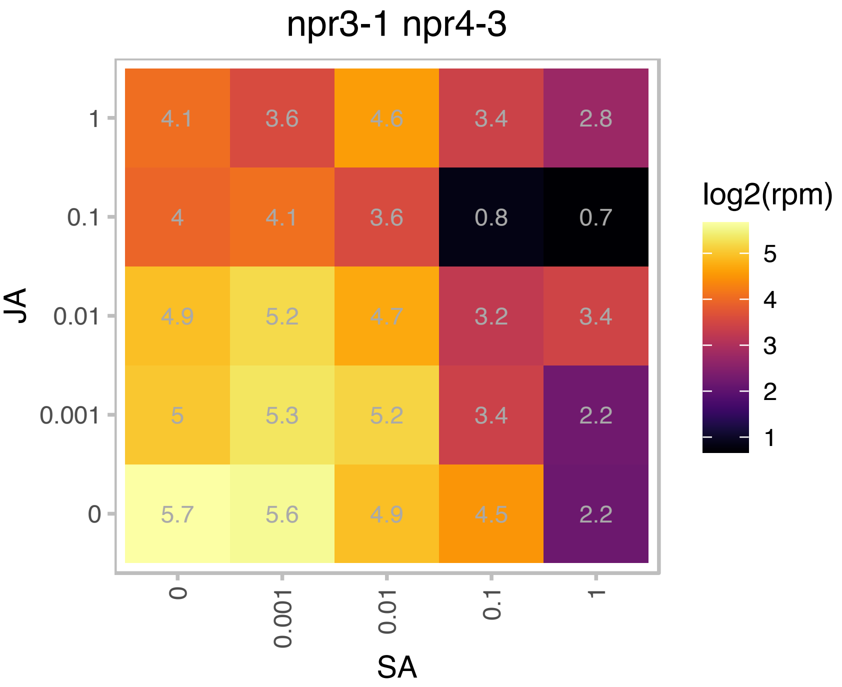

# AT2G21185.1 : NA

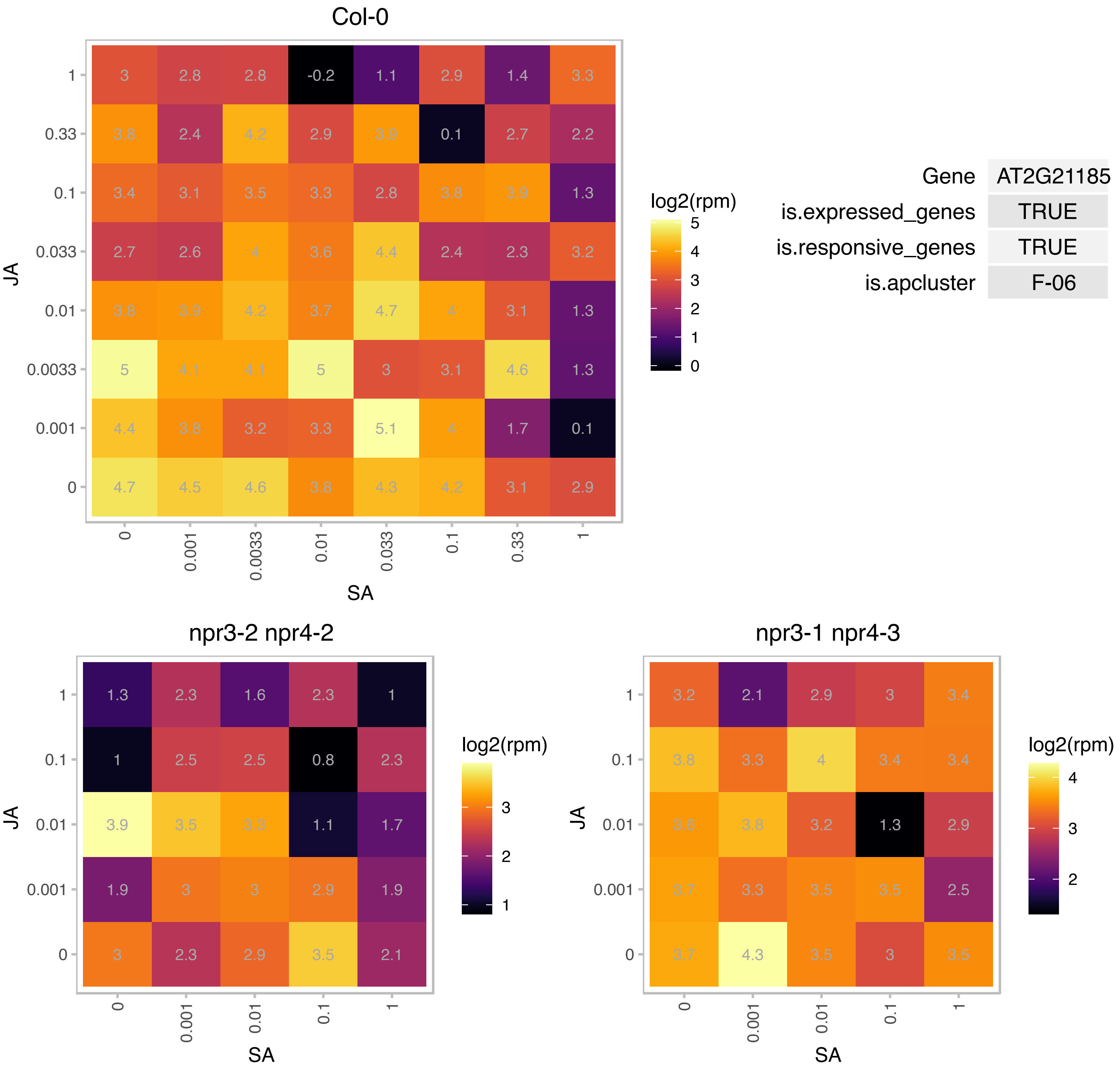


| | |
|---|---|
| Gene | AT2G21185 |
| is.expressed_genes | TRUE |
| is.responsive_genes | TRUE |
| is.apcluster | F-06 |

### AT2G21330.1 : AtFBA1, FBA1

fructose-bisphosphate aldolase 1

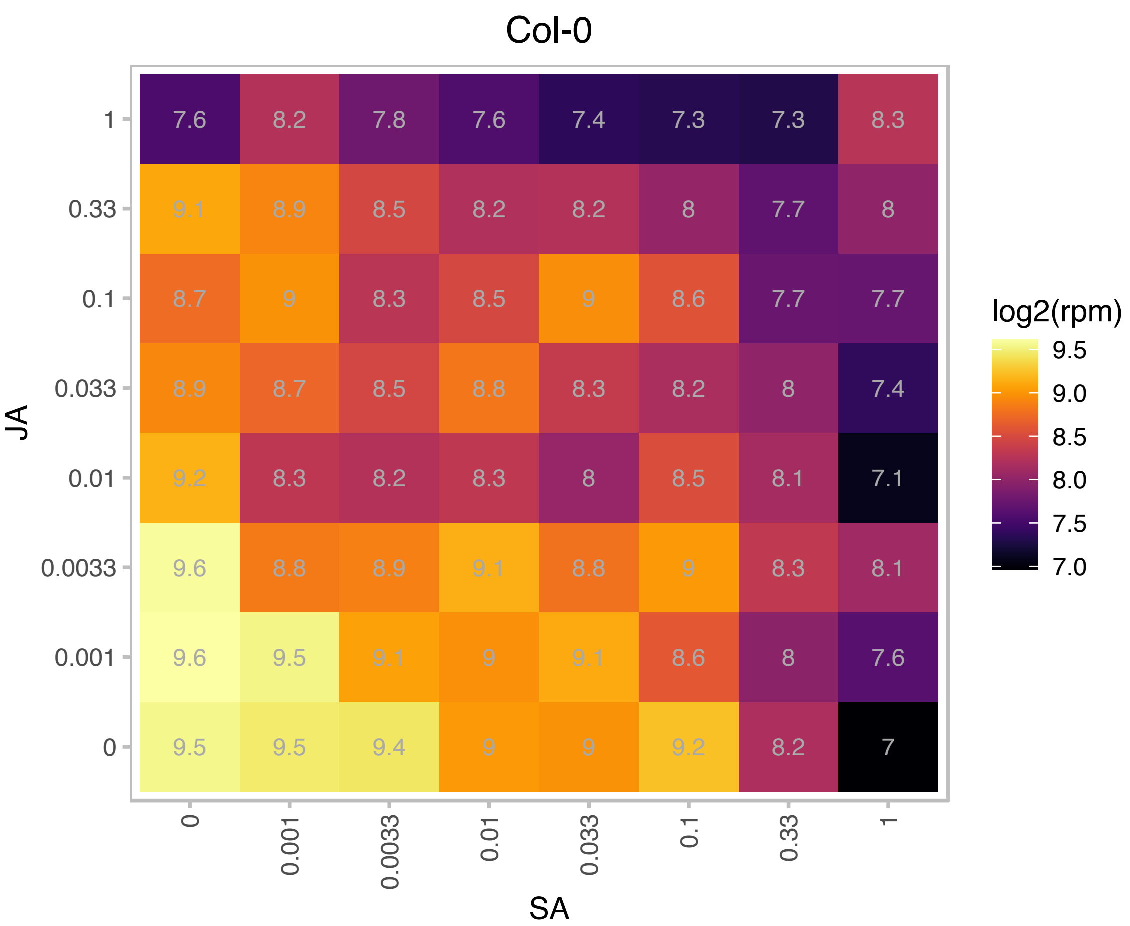


| | |
|---|---|
| Gene | AT2G21330 |
| is.expressed_genes | TRUE |
| is.responsive_genes | TRUE |
| is.apcluster | F-06 |

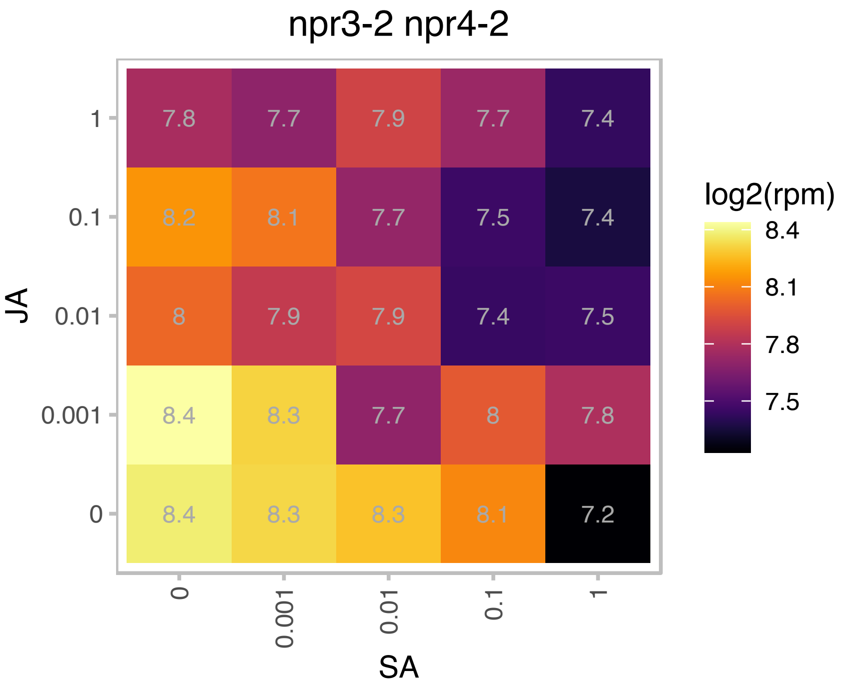


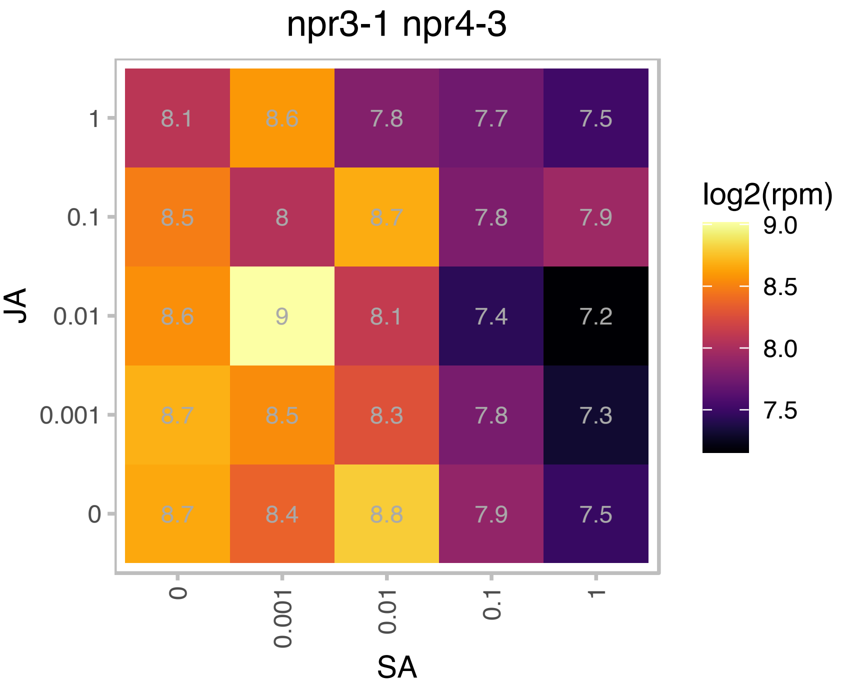

# AT2G21530.1 : NA

SMAD/FHA domain-containing protein

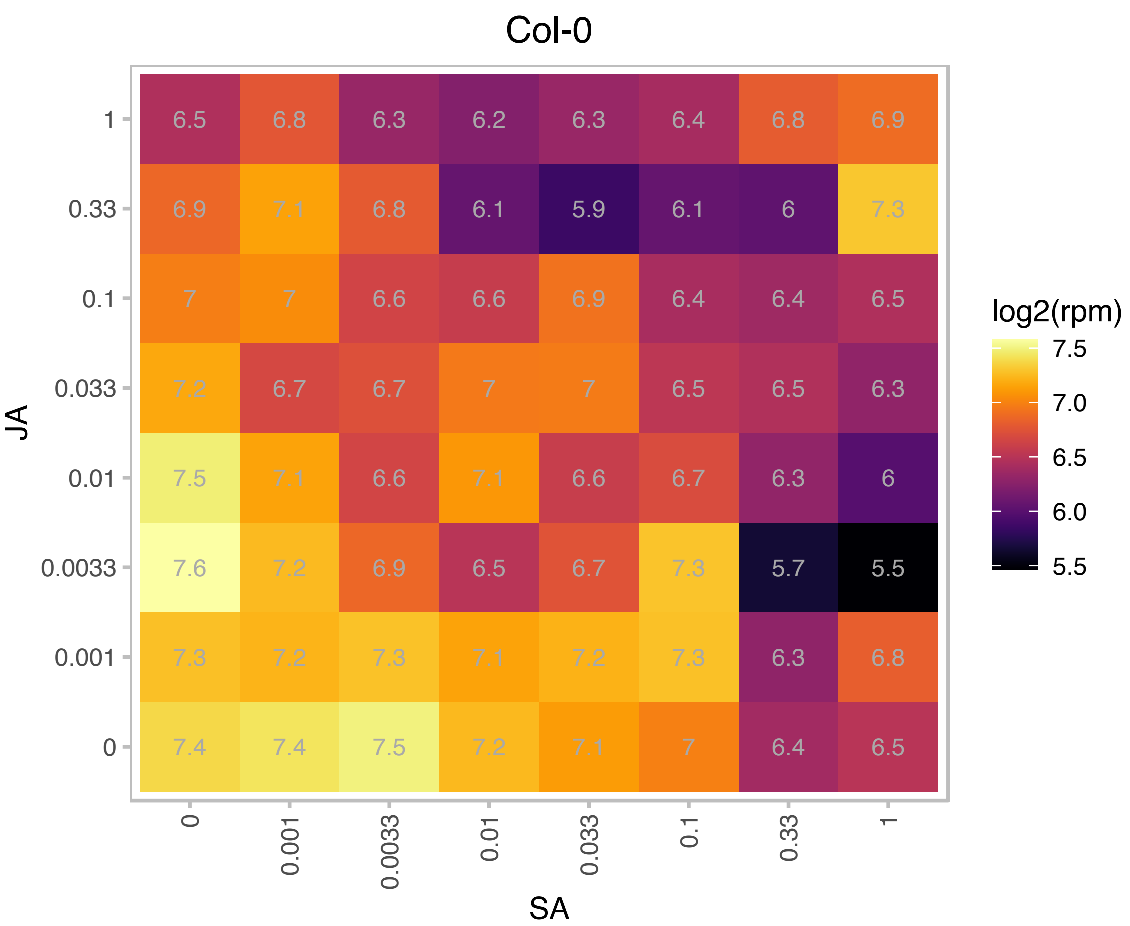


| | |
|---|---|
| Gene | AT2G21530 |
| is.expressed_genes | TRUE |
| is.responsive_genes | TRUE |
| is.apcluster | F-06 |

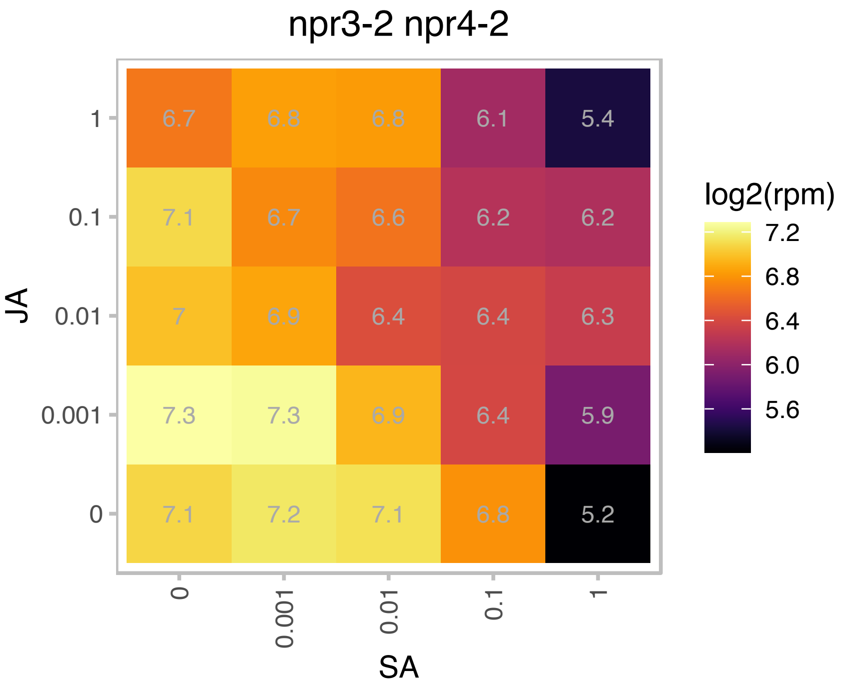


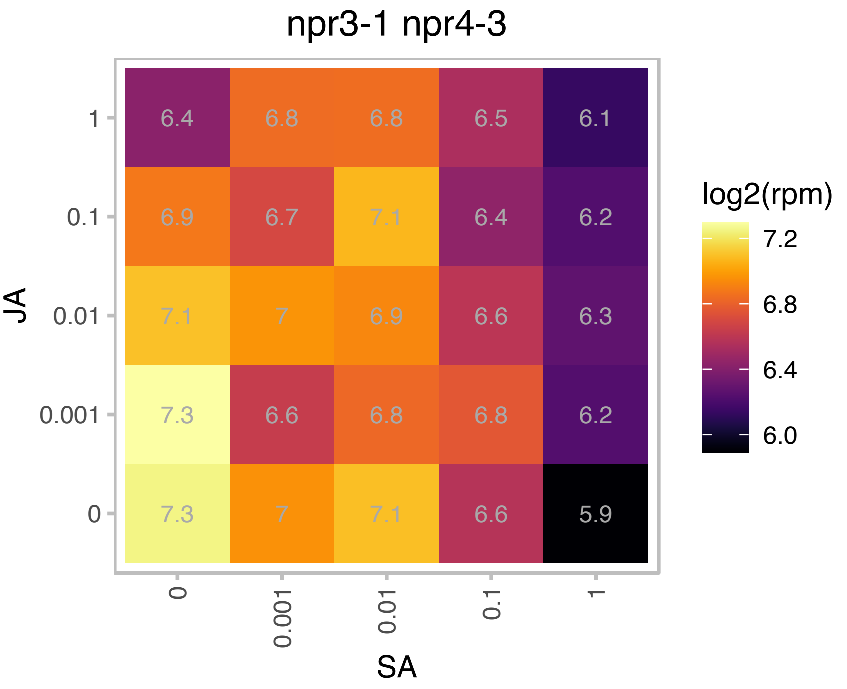

### AT2G24060.1 : AtIF3-2, AtINFC-2, SVR9

Translation initiation factor 3 protein

SUPPRESSOR OF VARIEGATION9 (SVR9),encodes a chloroplast-localized prokaryotic type translation initiation factor 3. Mutant plants shows both chloroplast development defect, and a series of leaf developmental abnormalities including more serrated leaf margin, disorganized mesophyll cells, and altered cotyledon venation patterns.

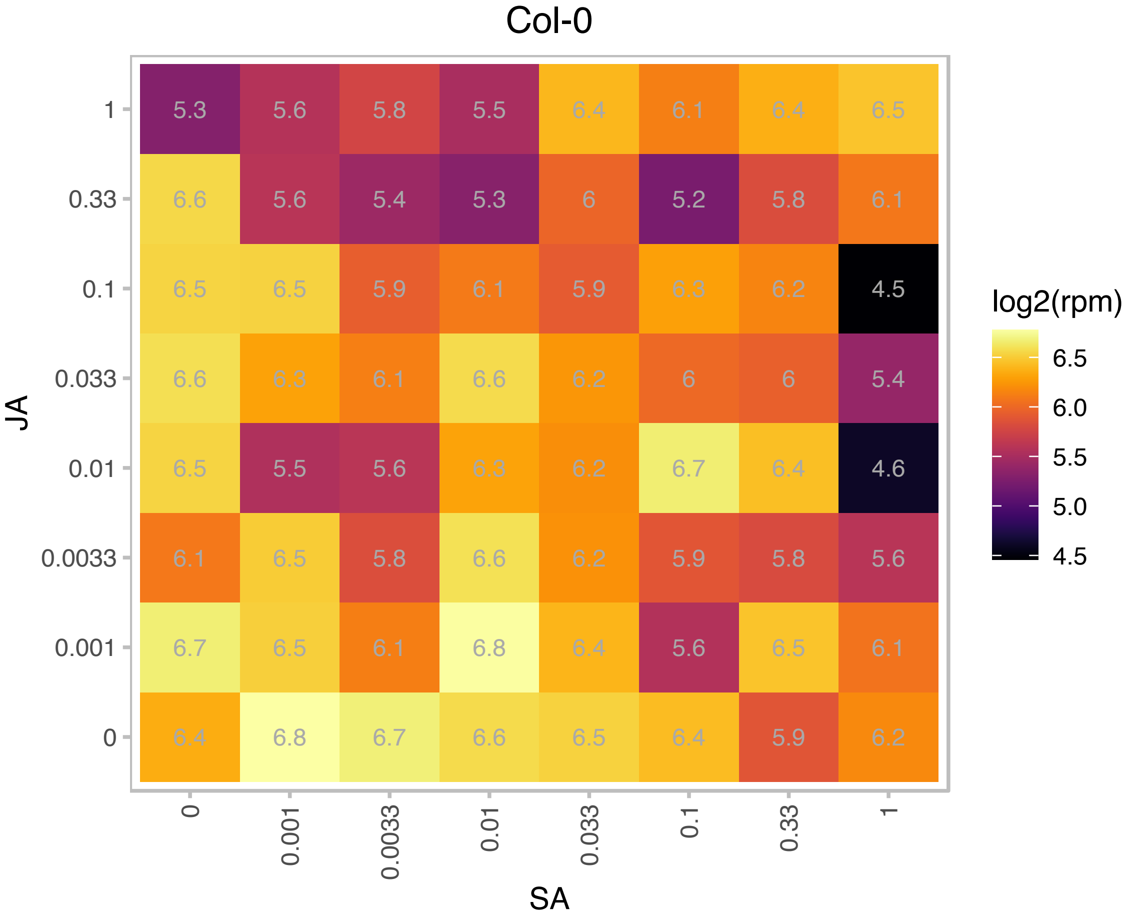


| | |
|---|---|
| Gene | AT2G24060 |
| is.expressed_genes | TRUE |
| is.responsive_genes | TRUE |
| is.apcluster | F-06 |

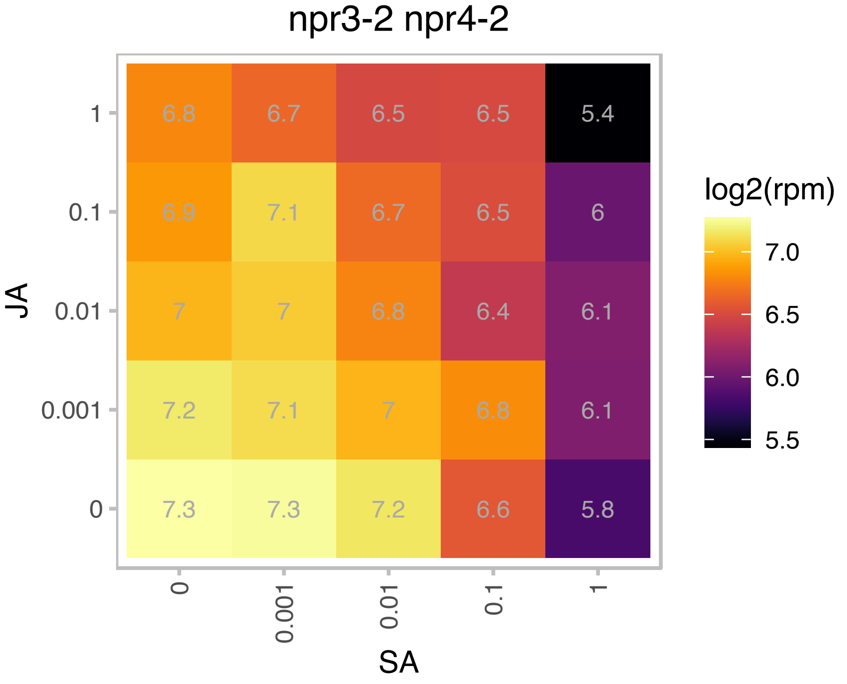


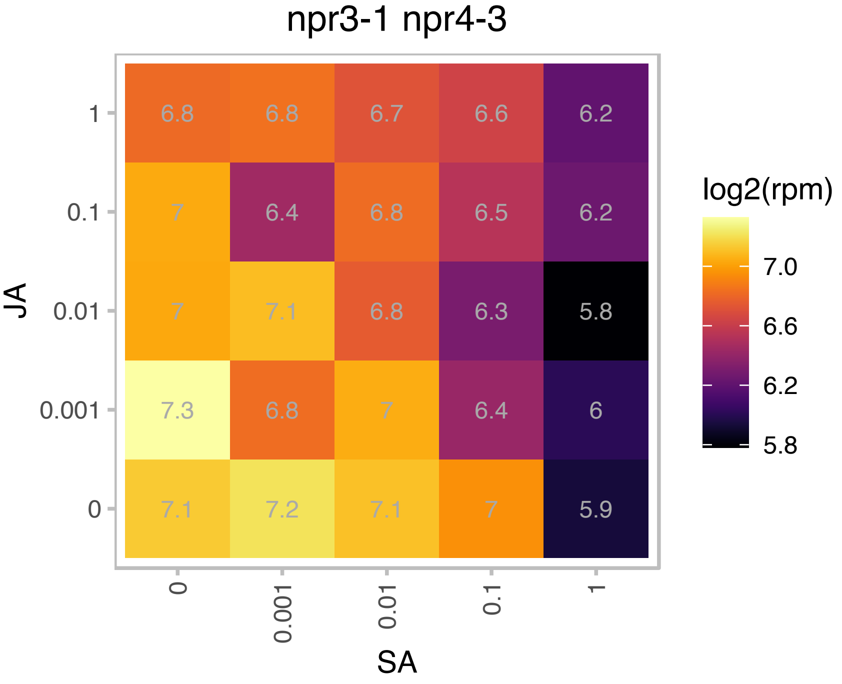

# AT2G25830.1 : NA

YebC-related

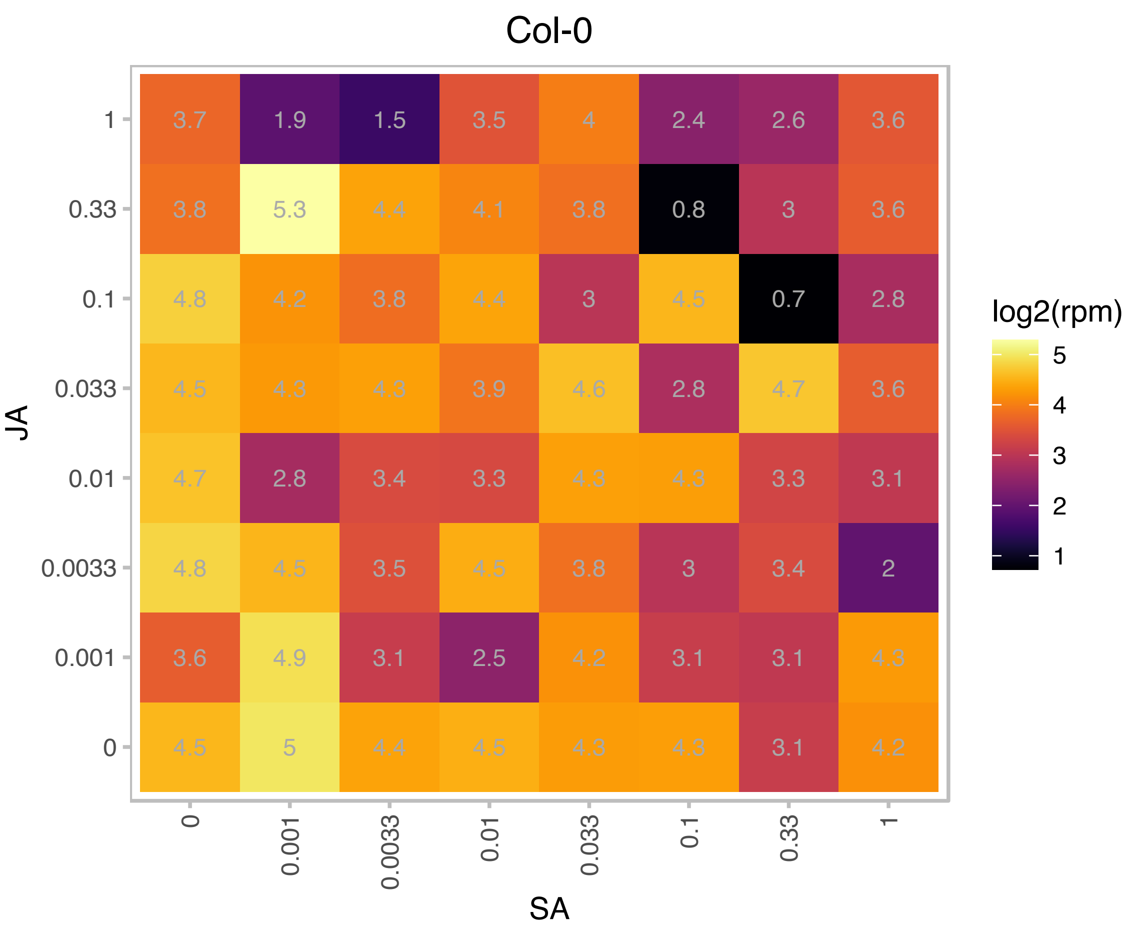


| | |
|---|---|
| Gene | AT2G25830 |
| is.expressed_genes | TRUE |
| is.responsive_genes | TRUE |
| is.apcluster | F-06 |

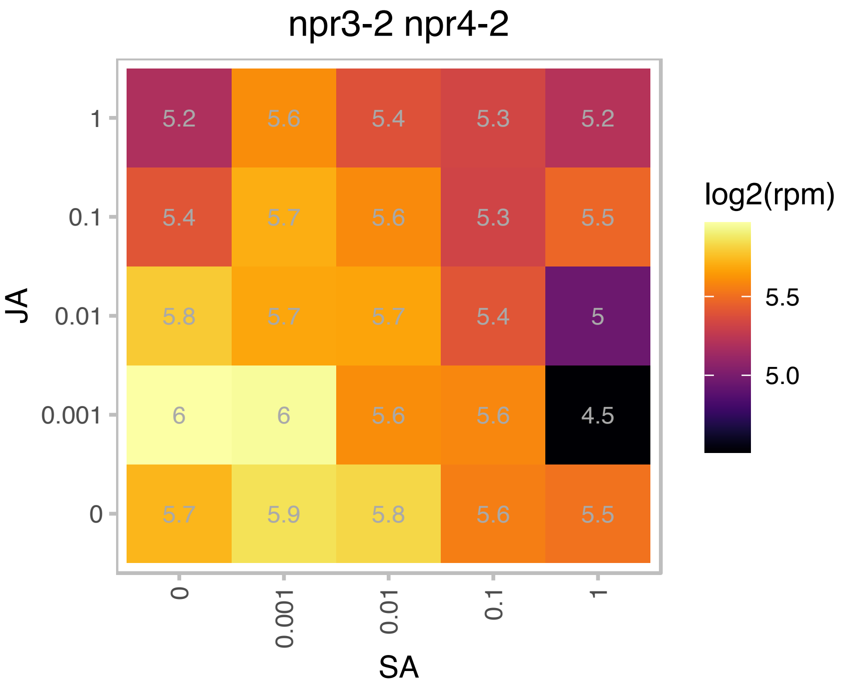


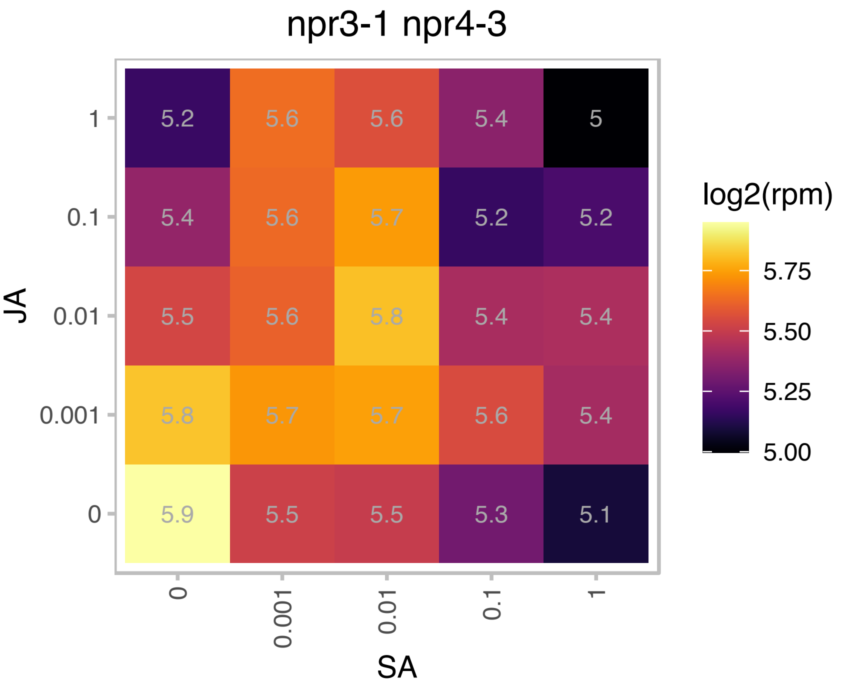

# AT2G29340.1 : NA

NAD-dependent epimerase/dehydratase family protein

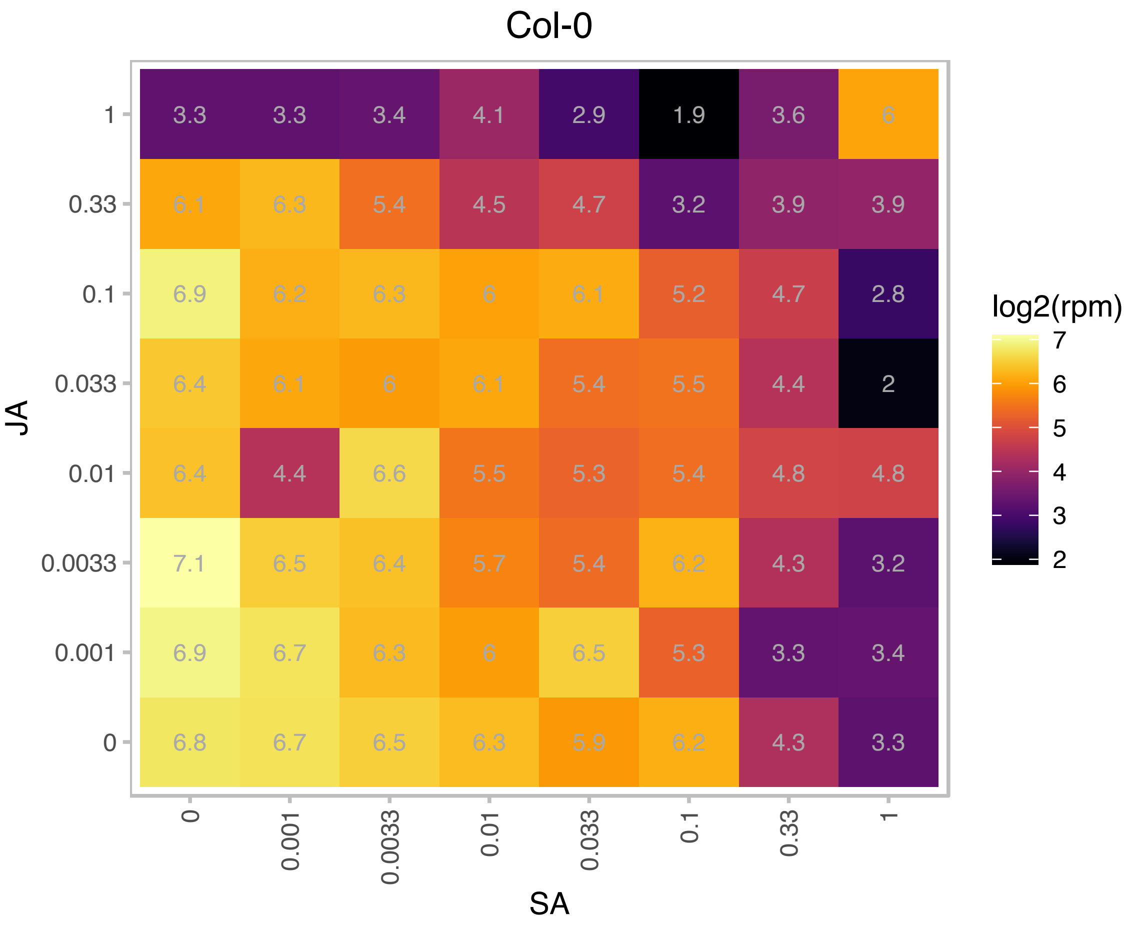


| | |
|---|---|
| Gene | AT2G29340 |
| is.expressed_genes | TRUE |
| is.responsive_genes | TRUE |
| is.apcluster | F-06 |

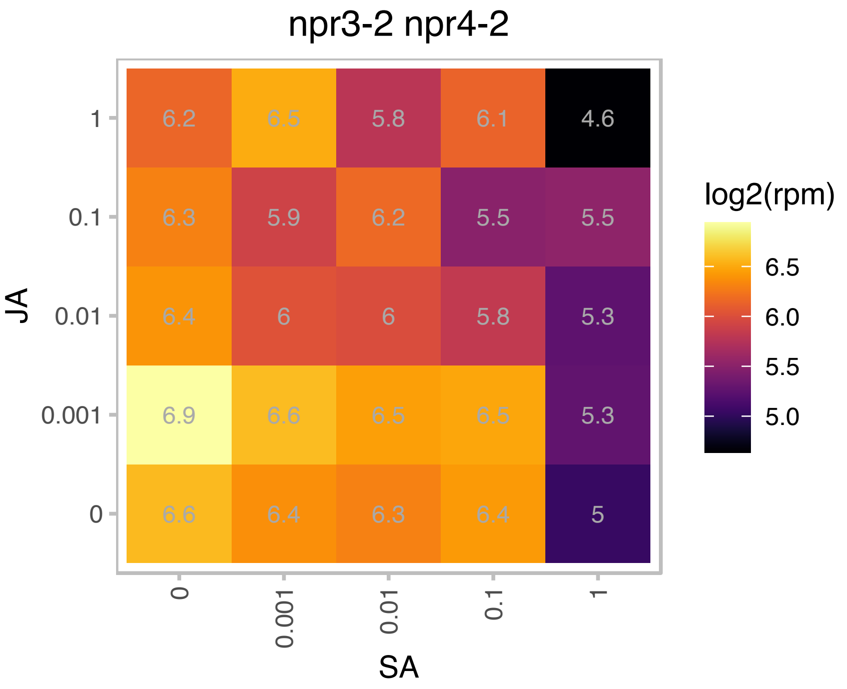


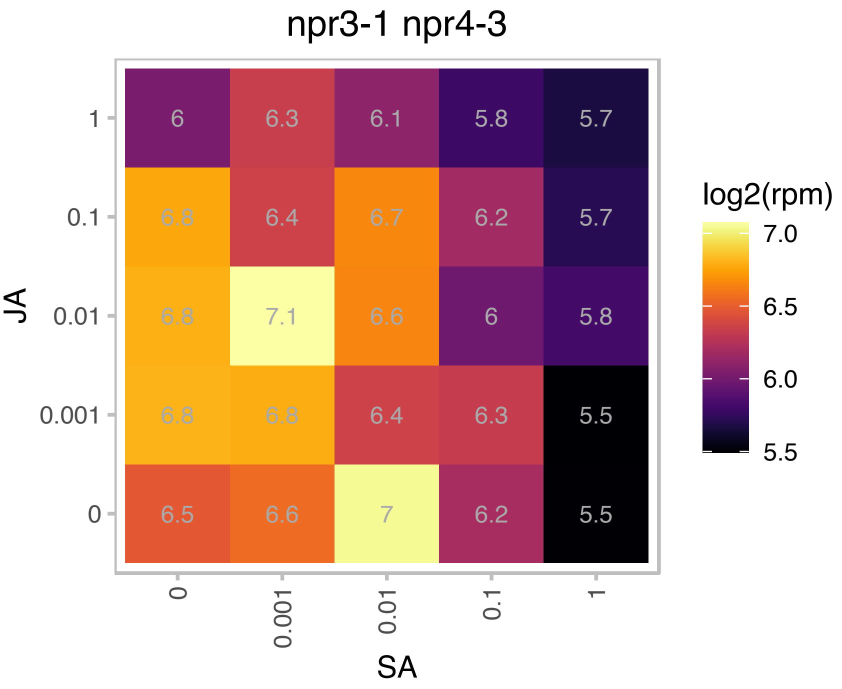

# AT2G40010.1 : NA

Ribosomal protein L10 family protein

| | |
|---|---|
| Gene | AT2G40010 |
| is.expressed_genes | TRUE |
| is.responsive_genes | TRUE |
| is.apcluster | F-06 |

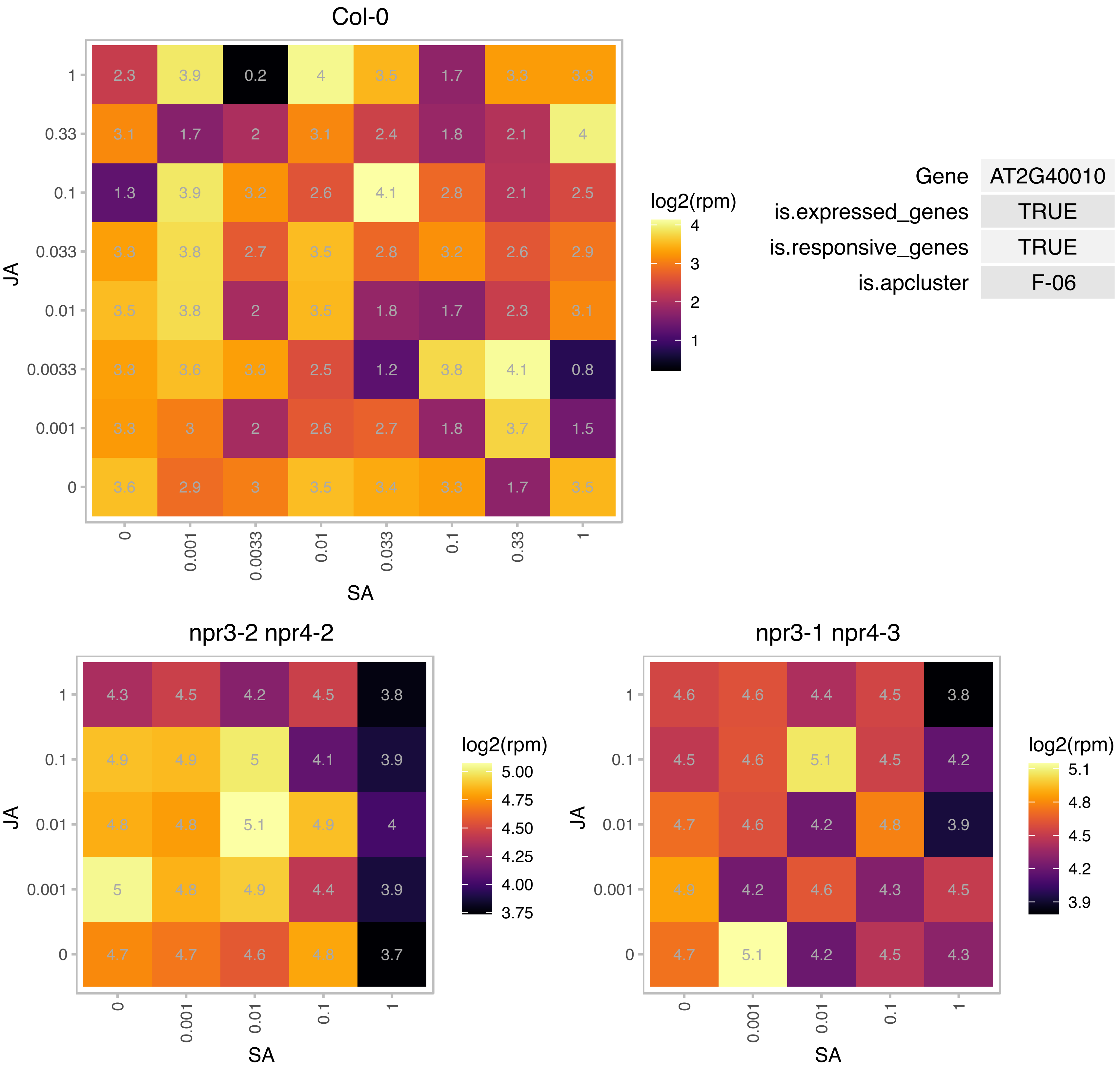

### AT2G43280.1 : FRF4

Far-red impaired responsive (FAR1) family protein

Encodes one of four FRS (FAR1-RELATED SEQUENCE) factor-like genes in Arabidopsis. FRS factors are characterized by having an N-terminal C2H2-type chelating motif of the WRKY- Glial Cell Missing1 family, a central core transposase domain of Mutator-like element transposases, and a C-terminal SWIM domain. The four FRF-like genes in Arabidopsis share only the N-terminal motif with FRS proteins.

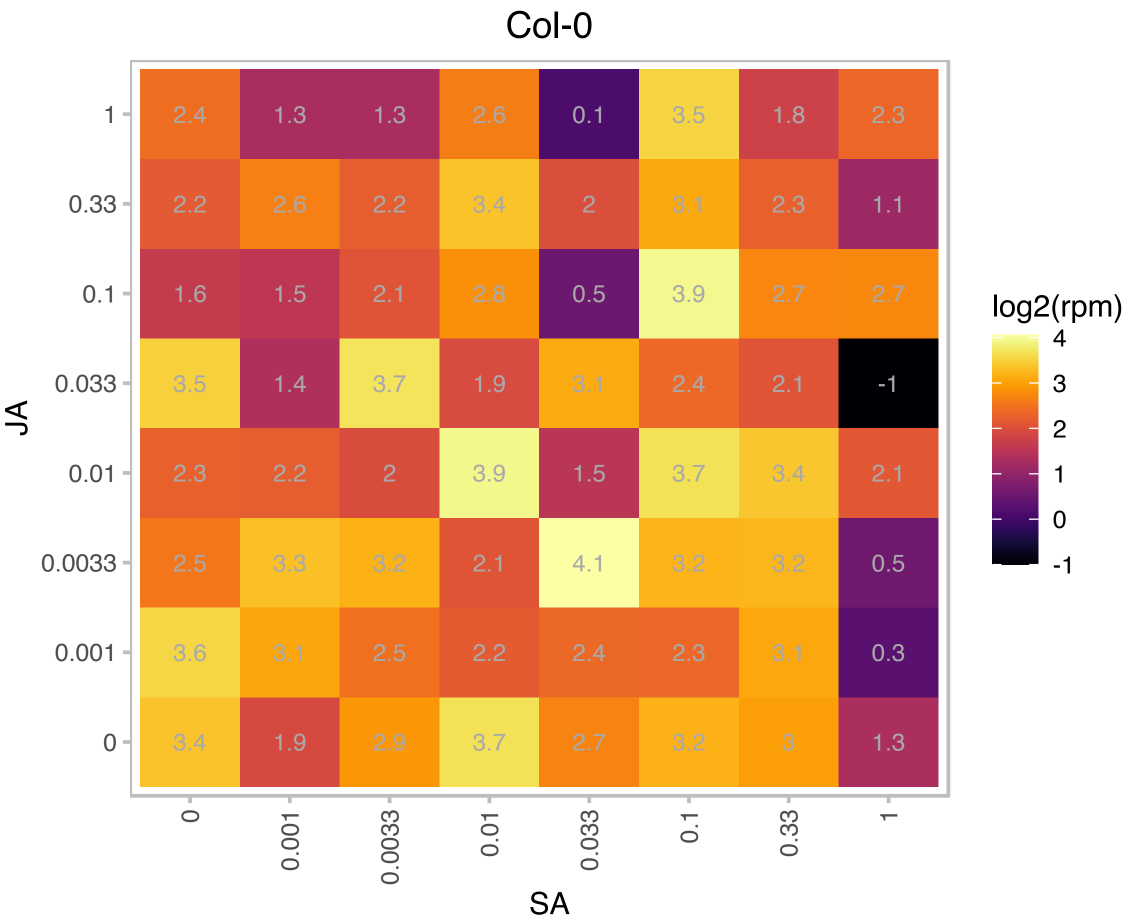


| | |
|---|---|
| Gene | AT2G43280 |
| is.expressed_genes | TRUE |
| is.responsive_genes | TRUE |
| is.apcluster | F-06 |

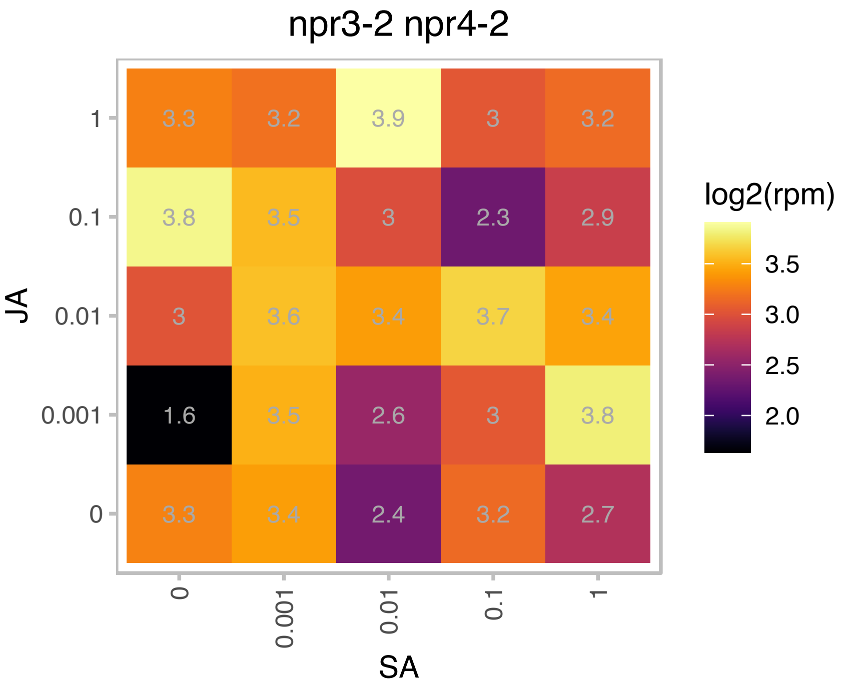


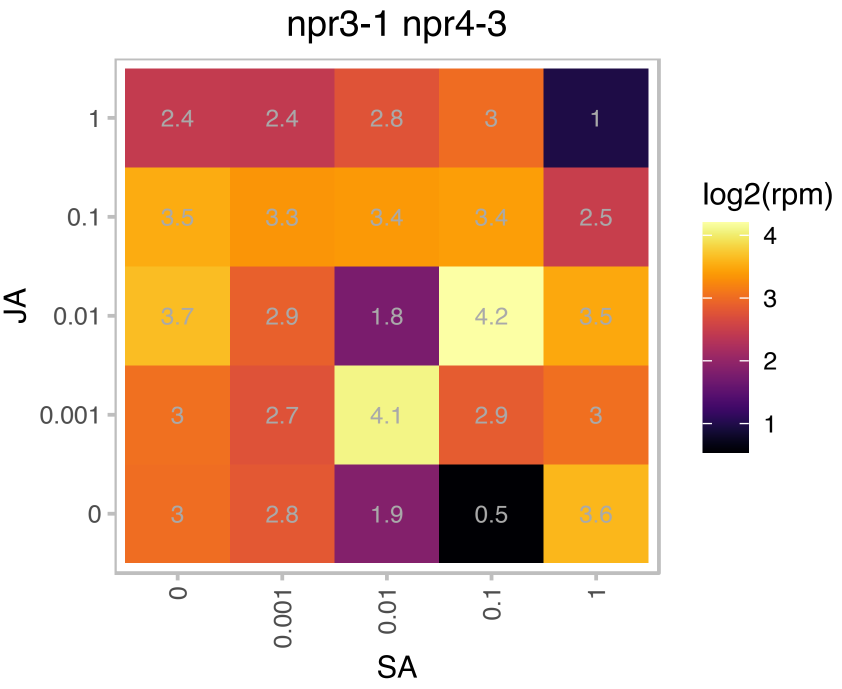

### AT2G45470.1 : AGP8, FLA8

FASCICLIN-like arabinogalactan protein 8

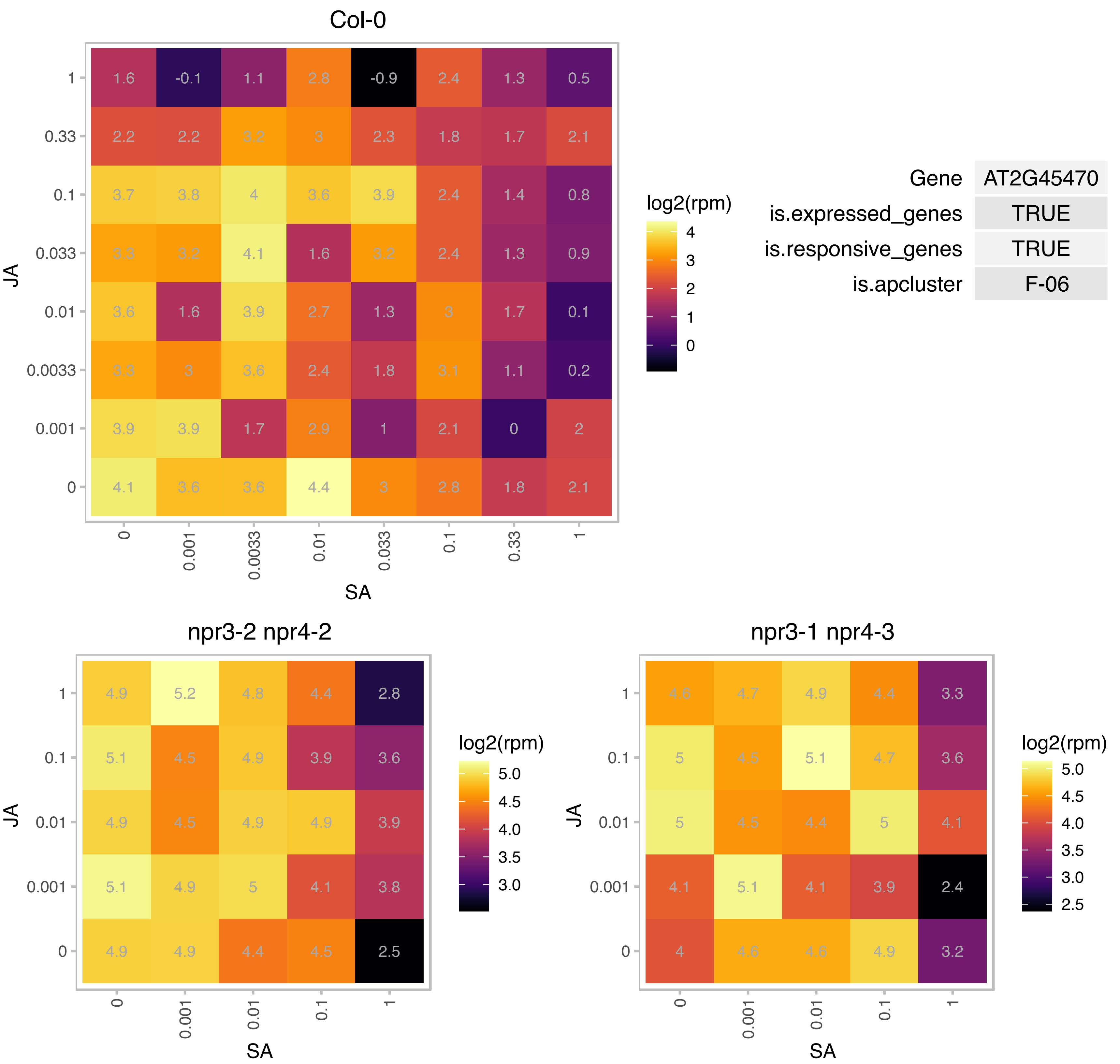


| | |
|---|---|
| Gene | AT2G45470 |
| is.expressed_genes | TRUE |
| is.responsive_genes | TRUE |
| is.apcluster | F-06 |

### AT2G47930.1 : AGP26, ATAGP26

arabinogalactan protein 26

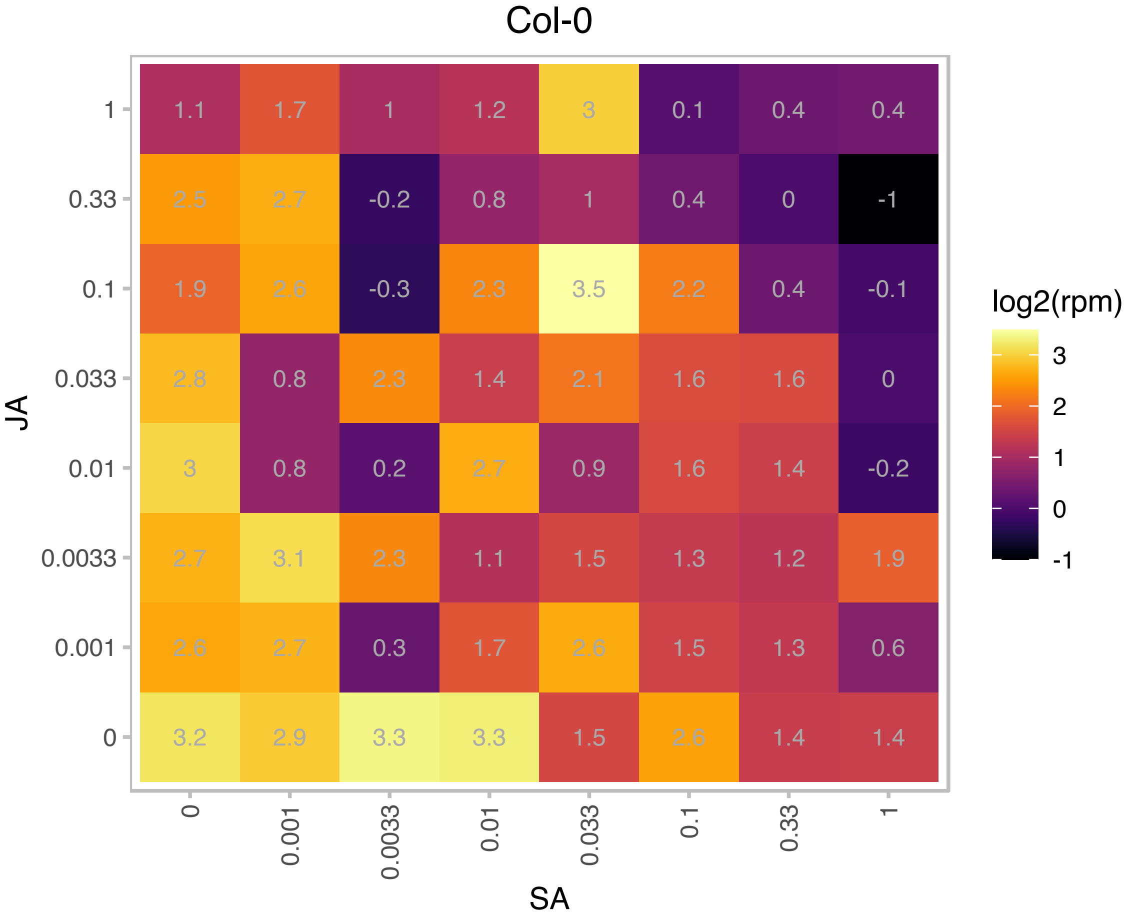


| | |
|---|---|
| Gene | AT2G47930 |
| is.expressed_genes | TRUE |
| is.responsive_genes | TRUE |
| is.apcluster | F-06 |

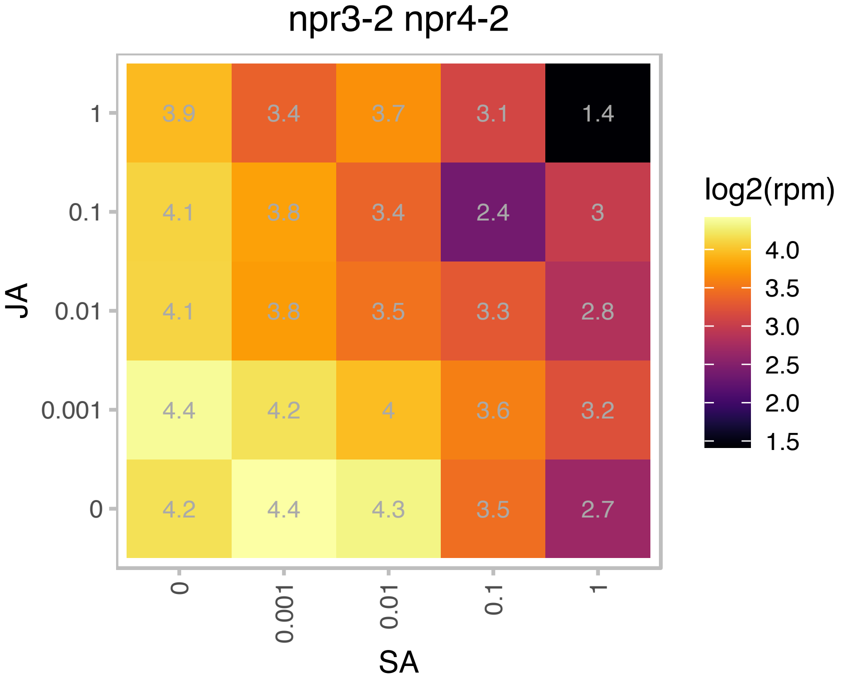


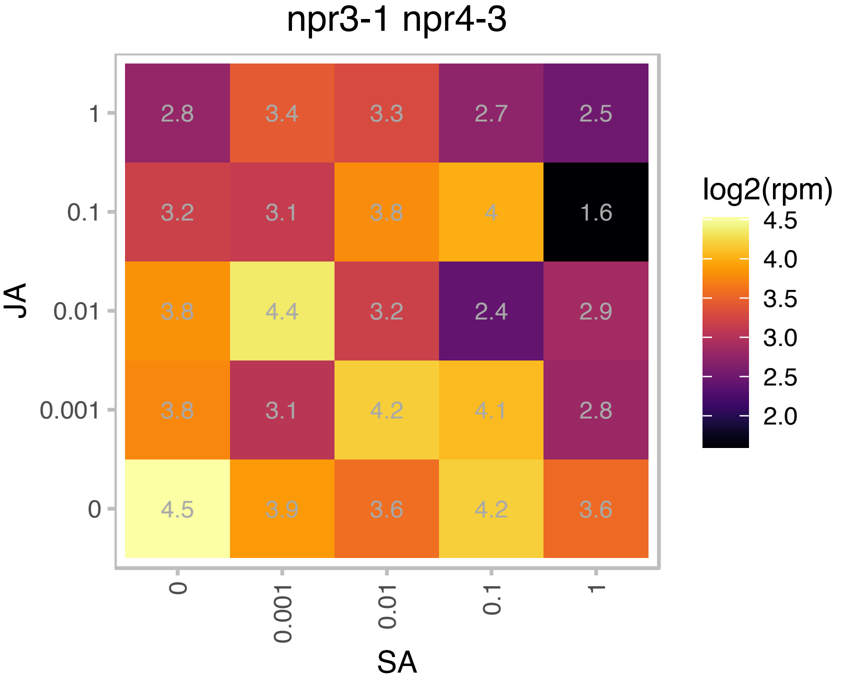

### AT2G48070.1 : RPH1

resistance to phytophthora 1

Encodes a chloroplast protein RPH1 (resistance to Phytophthora 1). Involved in immune response to Phytophthora brassicae. rph1 mutant plants (in Ws background) are susceptible to P. brassicae.

Col-0

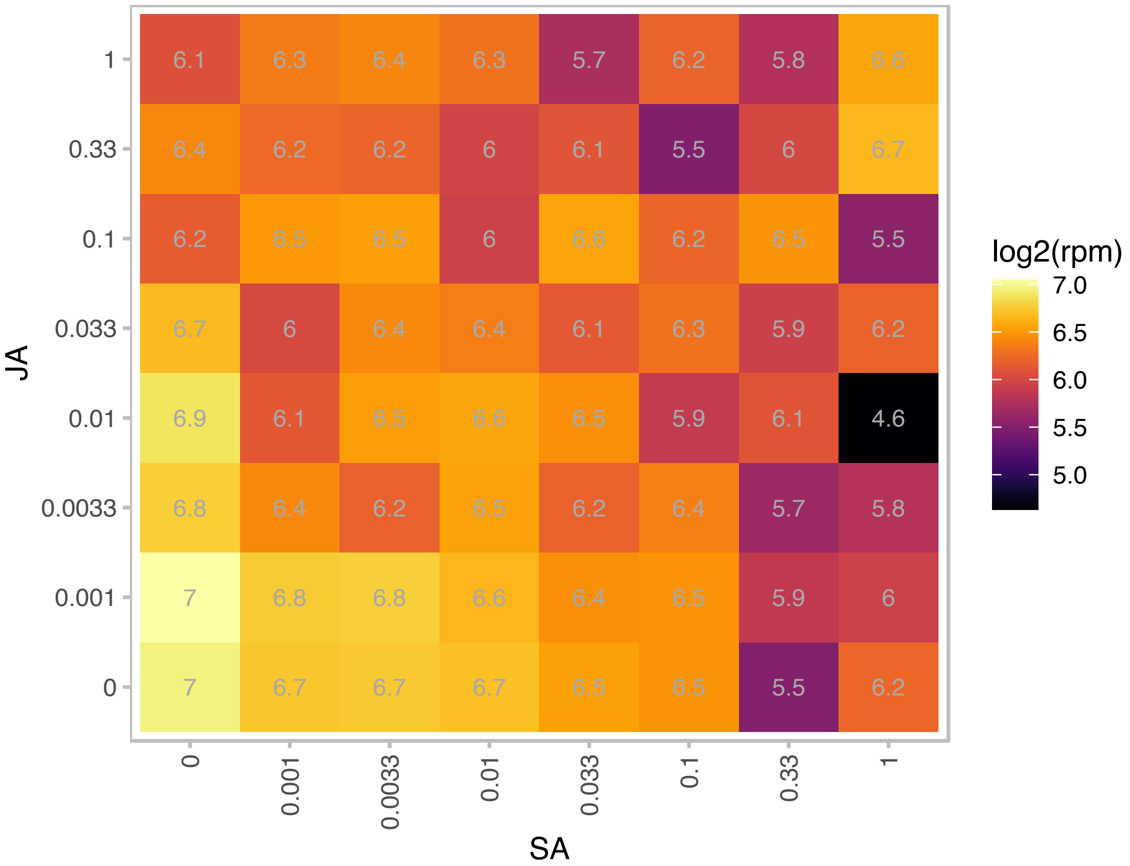


| | |
|---|---|
| Gene | AT2G48070 |
| is.expressed_genes | TRUE |
| is.responsive_genes | TRUE |
| is.apcluster | F-06 |

npr3-2 npr4-2

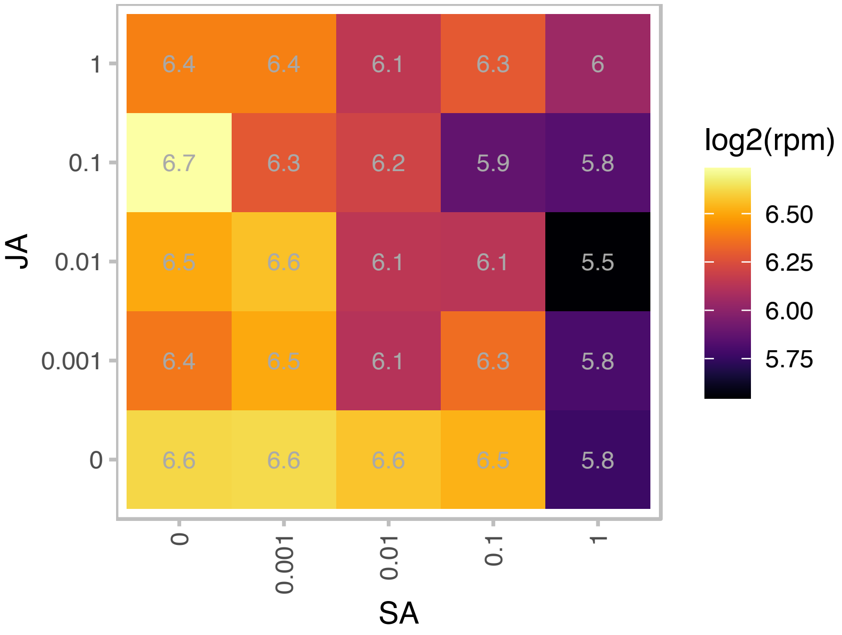


npr3-1 npr4-3

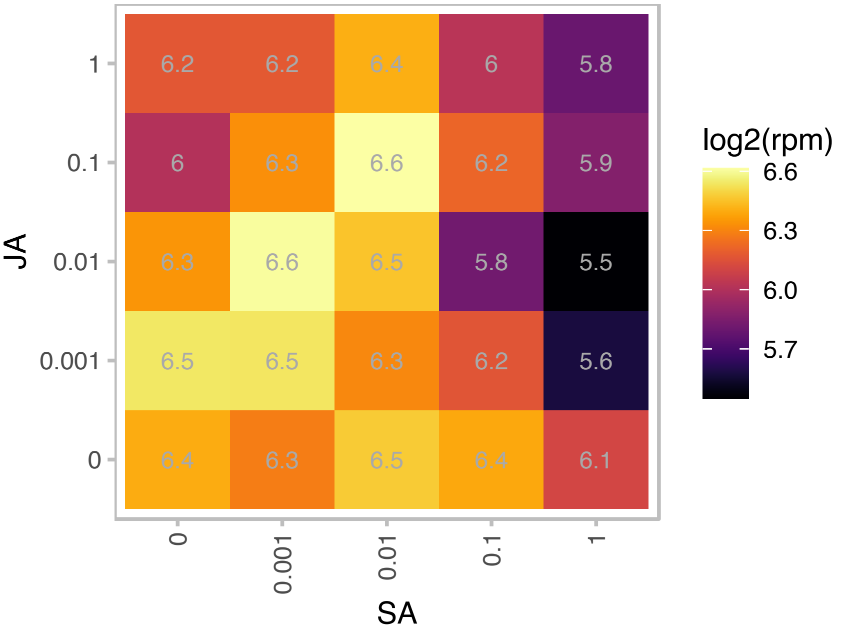

# AT3G02450.1 : NA

cell division protein ftsH, putative

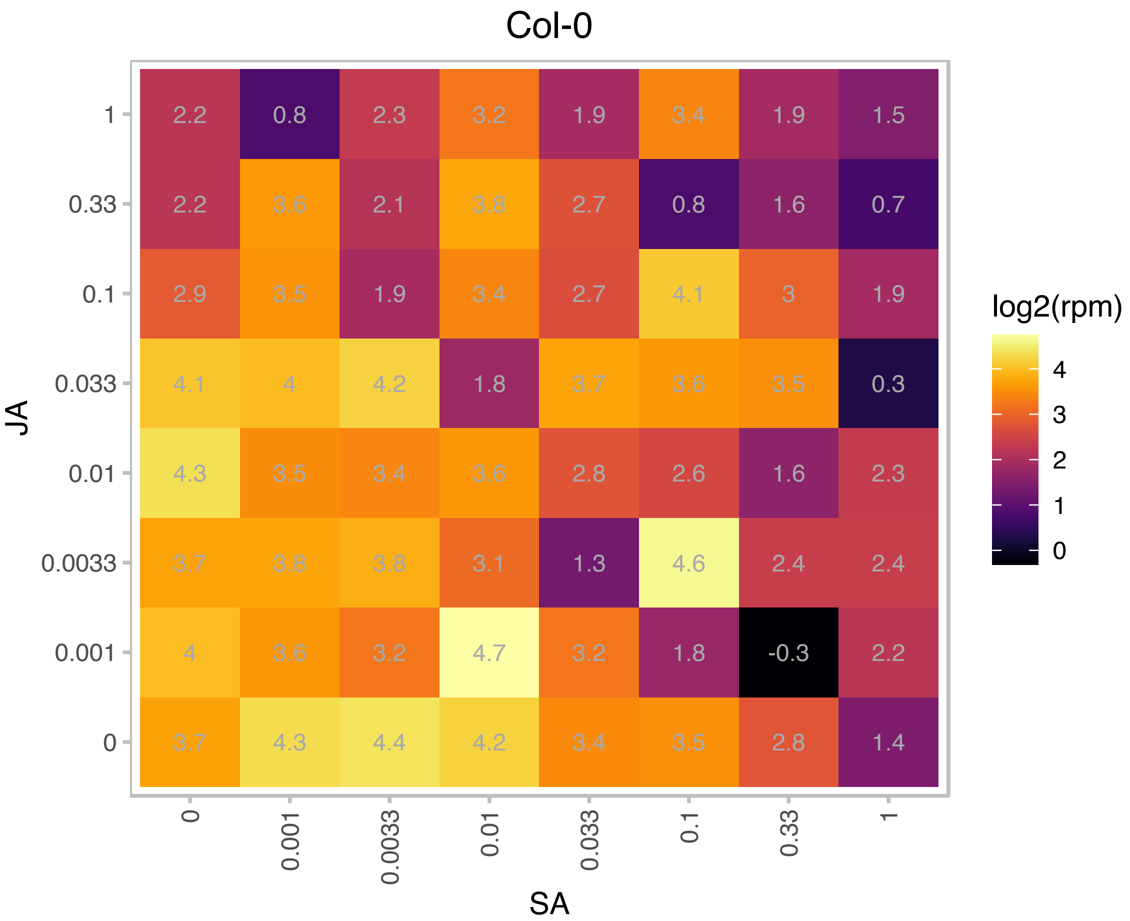


| | |
|---|---|
| Gene | AT3G02450 |
| is.expressed_genes | TRUE |
| is.responsive_genes | TRUE |
| is.apcluster | F-06 |

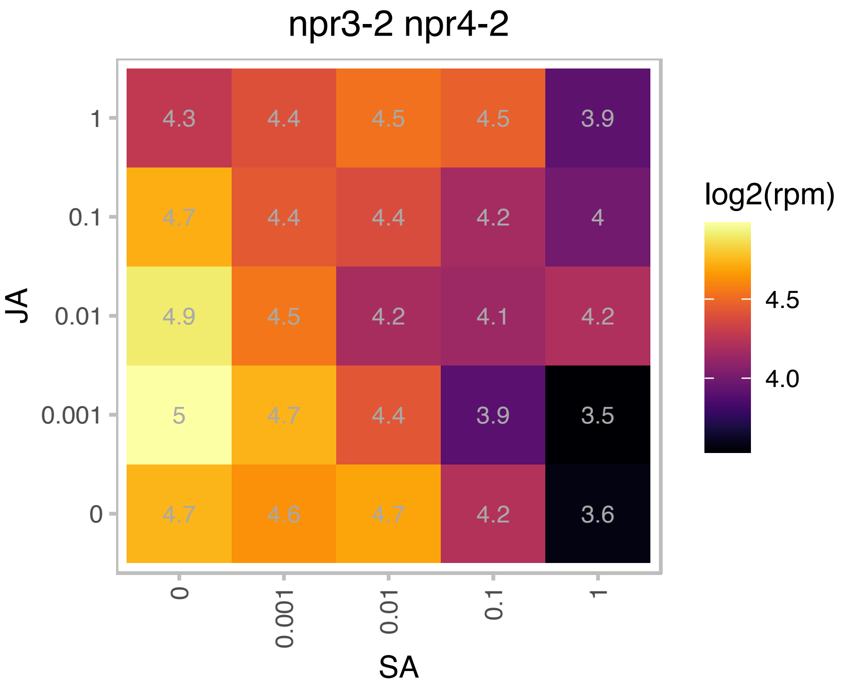


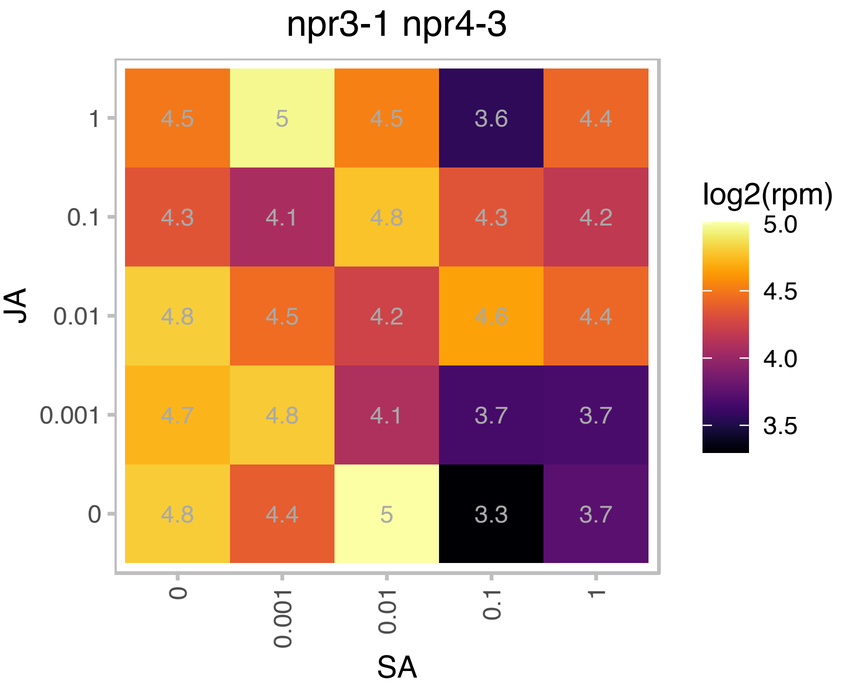

# AT3G06145.1 : NA

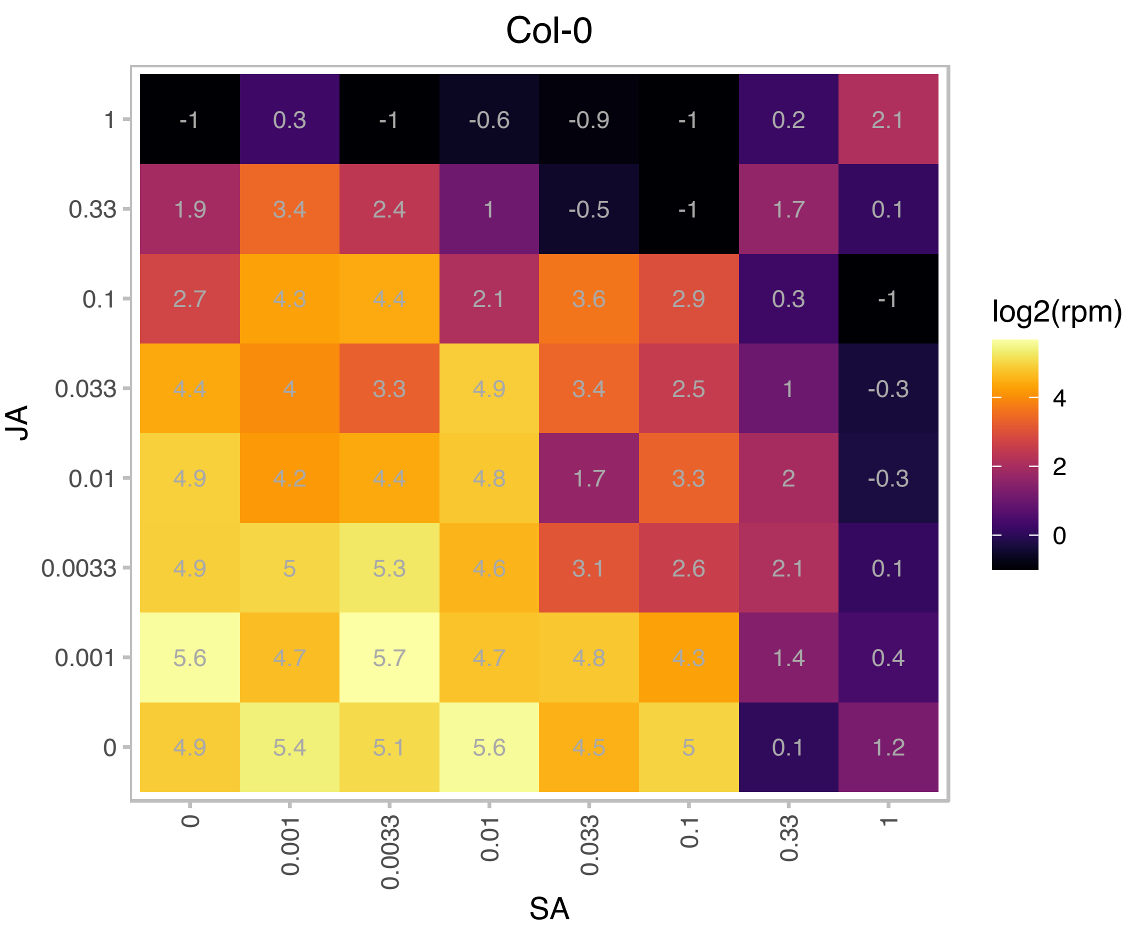


| | |
|---|---|
| Gene | AT3G06145 |
| is.expressed_genes | TRUE |
| is.responsive_genes | TRUE |
| is.apcluster | F-06 |

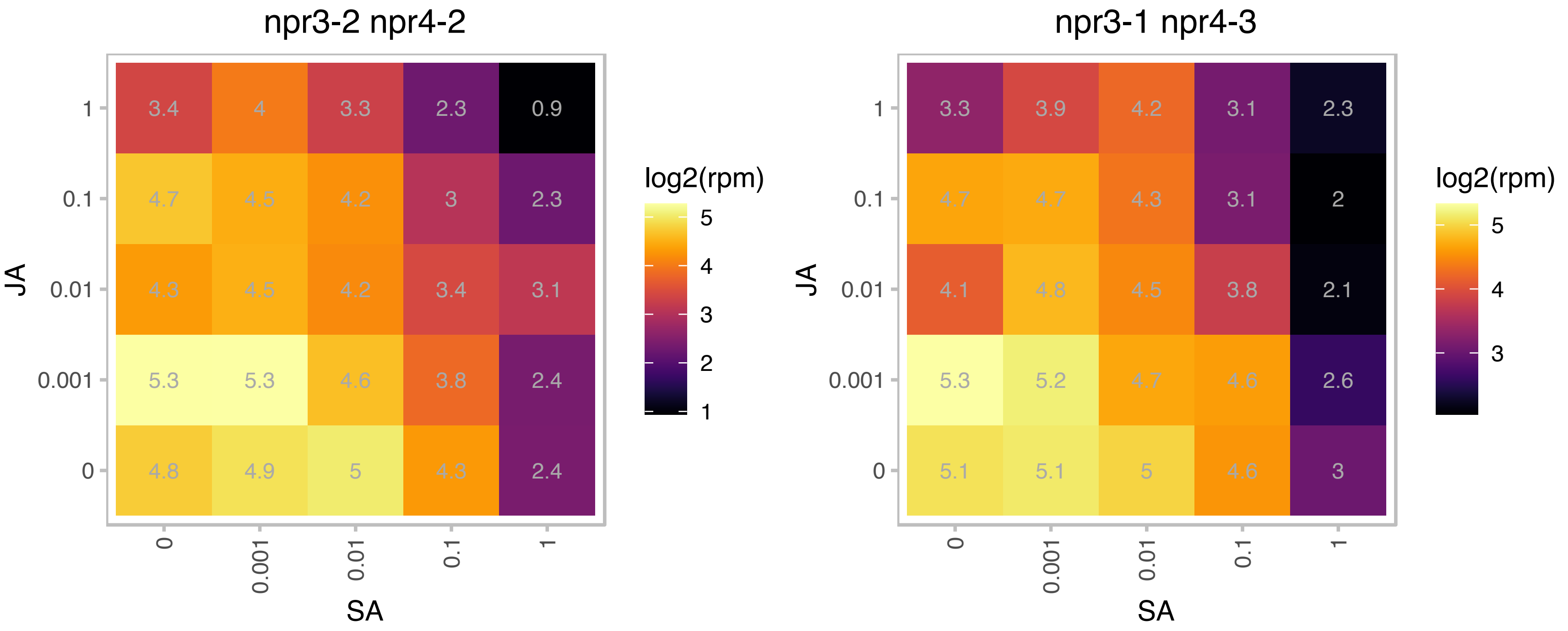

# AT3G08920.1 : NA

Rhodanese/Cell cycle control phosphatase superfamily protein

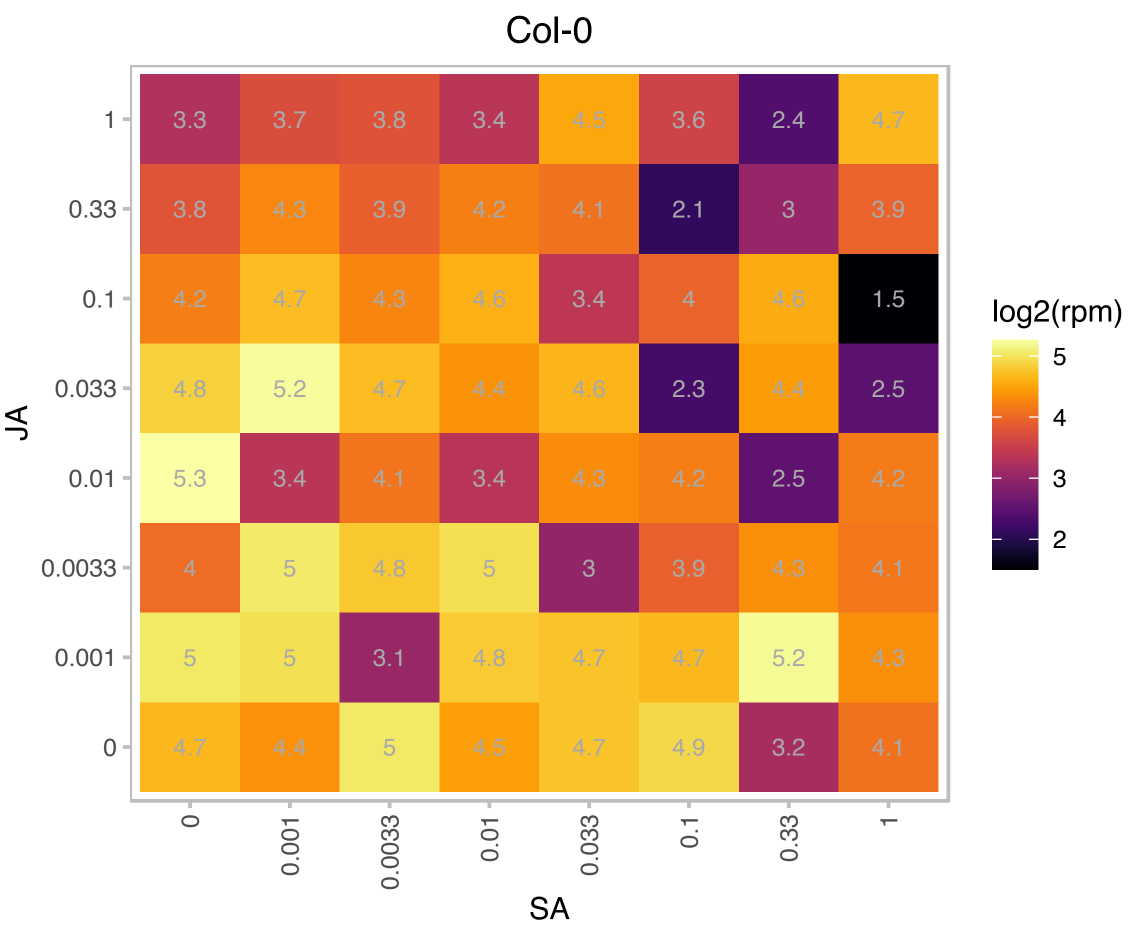


| | |
|---|---|
| Gene | AT3G08920 |
| is.expressed_genes | TRUE |
| is.responsive_genes | TRUE |
| is.apcluster | F-06 |

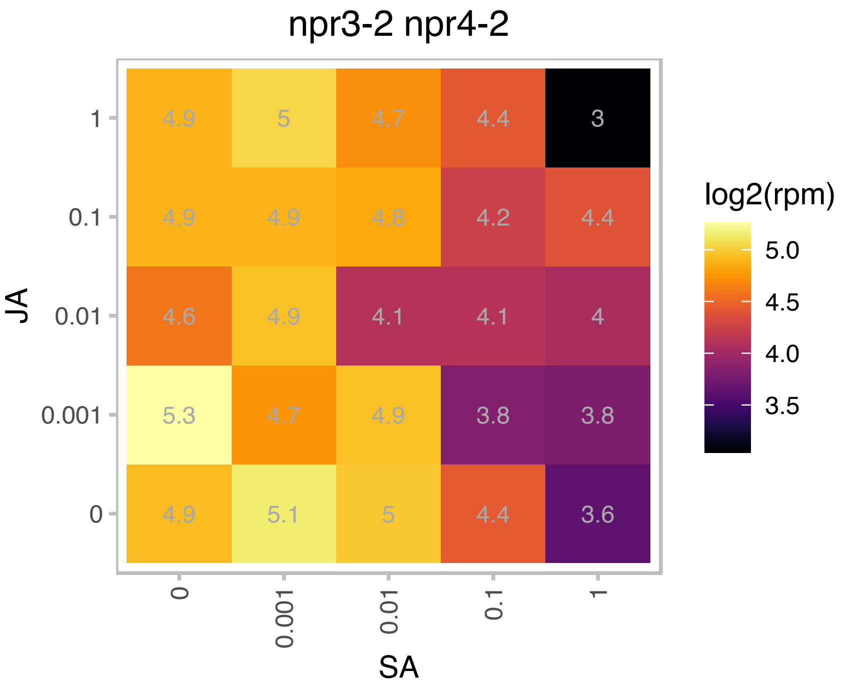


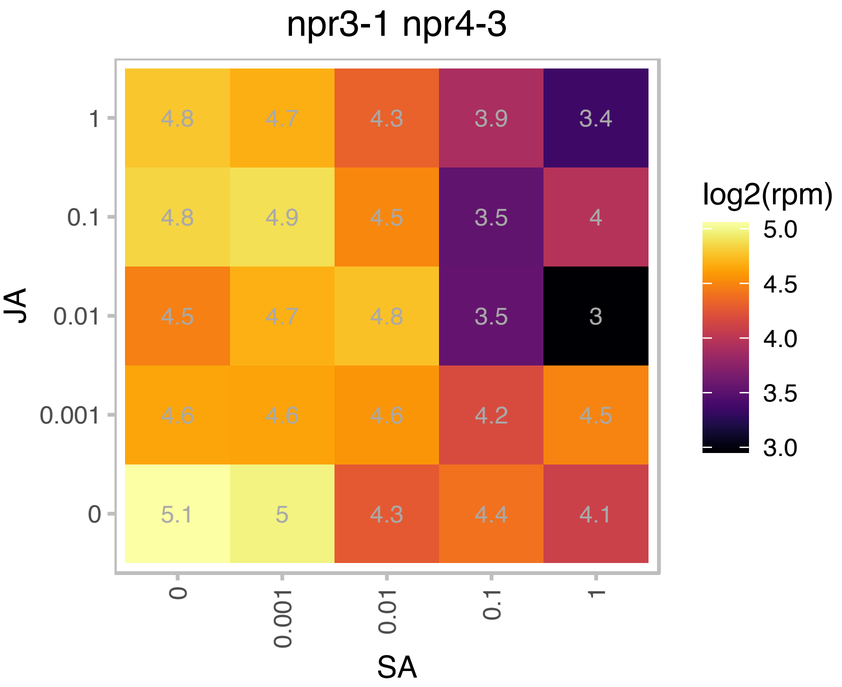

### AT3G13470.1 : Cpn60beta2, CPNB2

TCP-1/cpn60 chaperonin family protein

Encodes a subunit of chloroplasts chaperonins that are involved in mediating the folding of newly synthesized, translocated, or stress-denatured proteins. Cpn60 subunits are: Cpn60alpha1 (At2g28000), AtCpn60alpha2 (At5g18820), AtCpn60beta1 (At1g55490), AtCpn60beta2 (At3g13470), AtCpn60beta3 (At5g56500), AtCpn60beta4 (At1g26230).

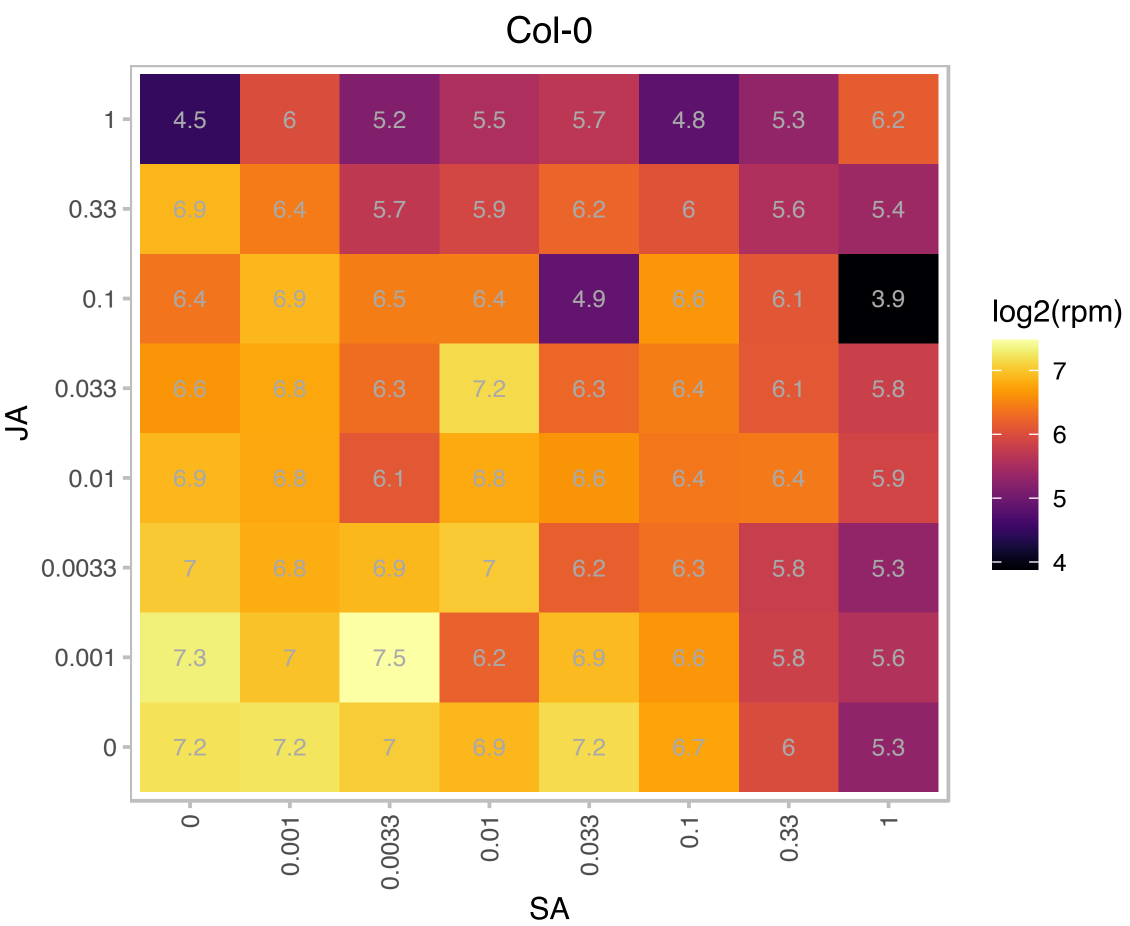


| | |
|---|---|
| Gene | AT3G13470 |
| is.expressed_genes | TRUE |
| is.responsive_genes | TRUE |
| is.apcluster | F-06 |

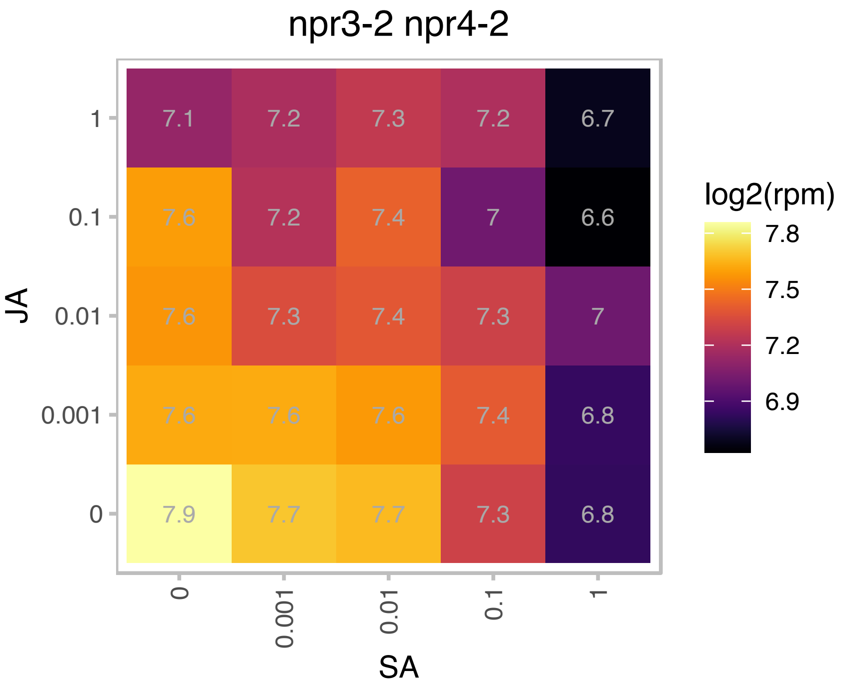


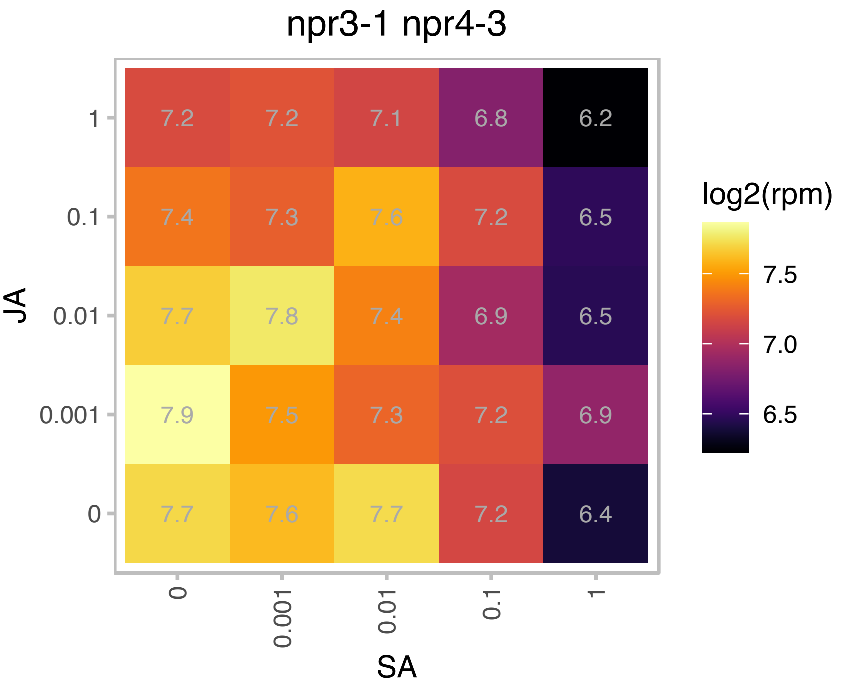

### AT3G13740.1 : RNC4

Ribonuclease III family protein

Encodes one of two chloroplast Mini-RNase III-like enzymes in Arabidopsis.  Double mutants display imprecise maturation of 23S rRNA and other rRNAs.

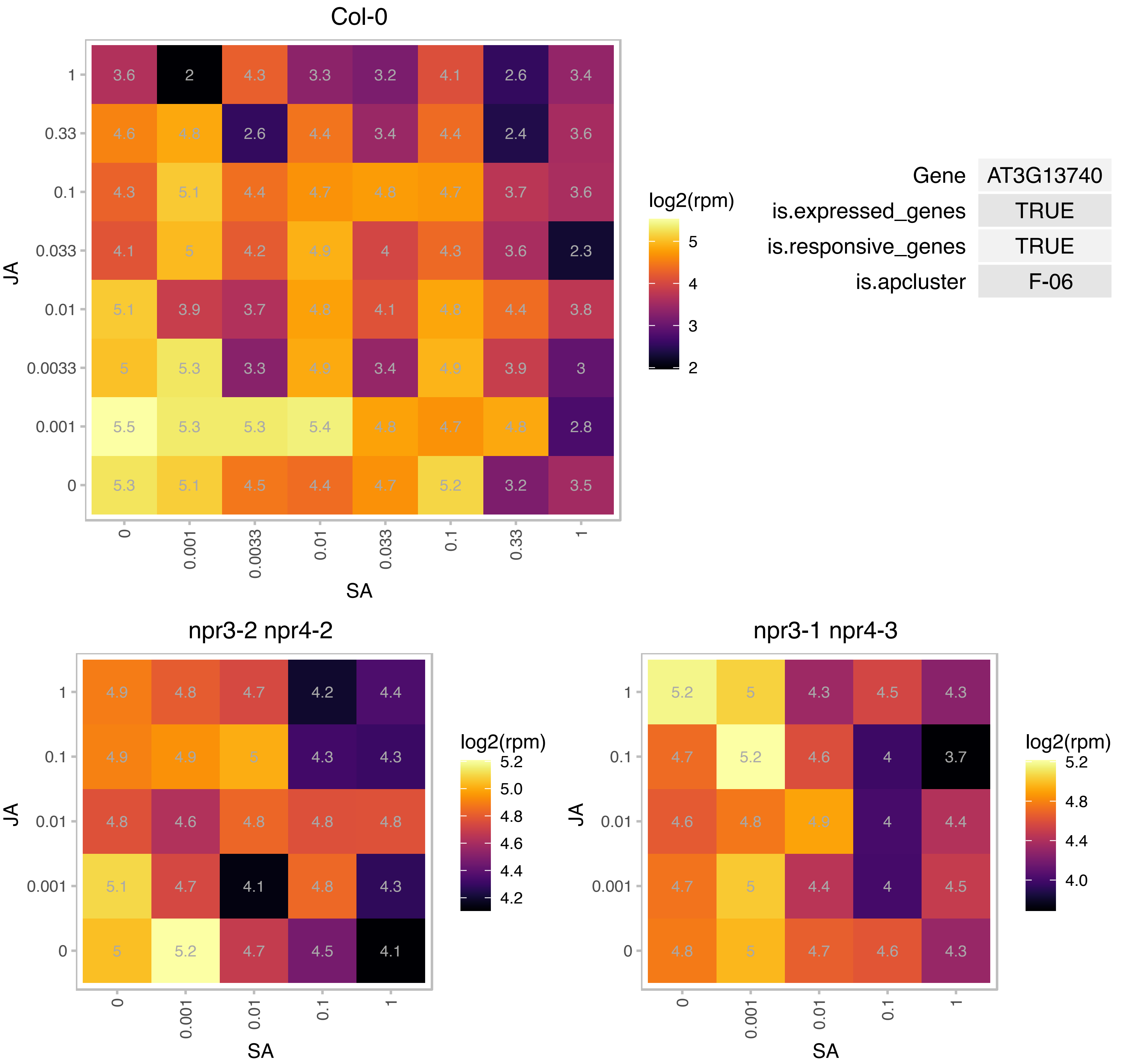


| | |
|---|---|
| Gene | AT3G13740 |
| is.expressed_genes | TRUE |
| is.responsive_genes | TRUE |
| is.apcluster | F-06 |

### AT3G14350.1 : SRF7

STRUBBELIG-receptor family 7

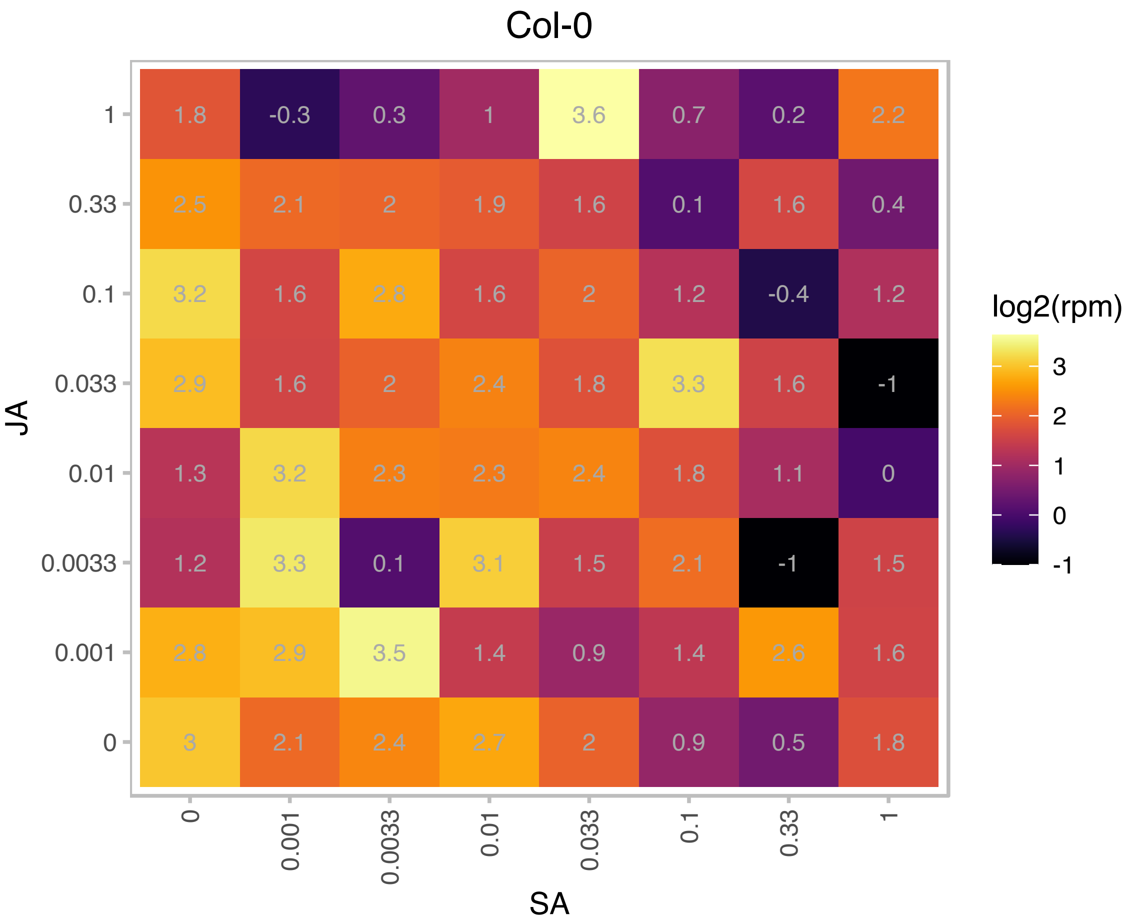


| | |
|---|---|
| Gene | AT3G14350 |
| is.expressed_genes | TRUE |
| is.responsive_genes | TRUE |
| is.apcluster | F-06 |

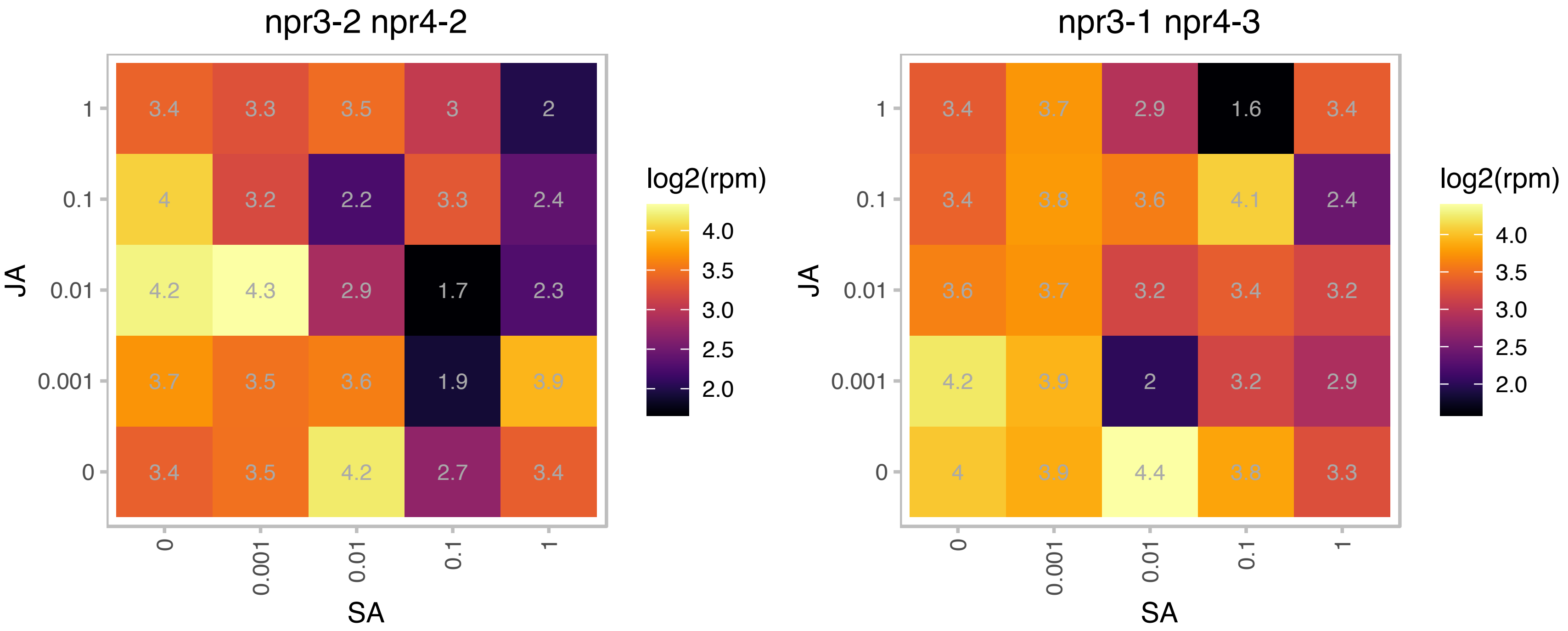

### AT3G16250.1 : NDF4, PnsB3

NDH-dependent cyclic electron flow 1

encodes a novel subunit of the chloroplast NAD(P)H dehydrogenase complex, involved in cyclic electron flow around photosystem I to produce ATP. Contains a 4Fe-4S cluster.

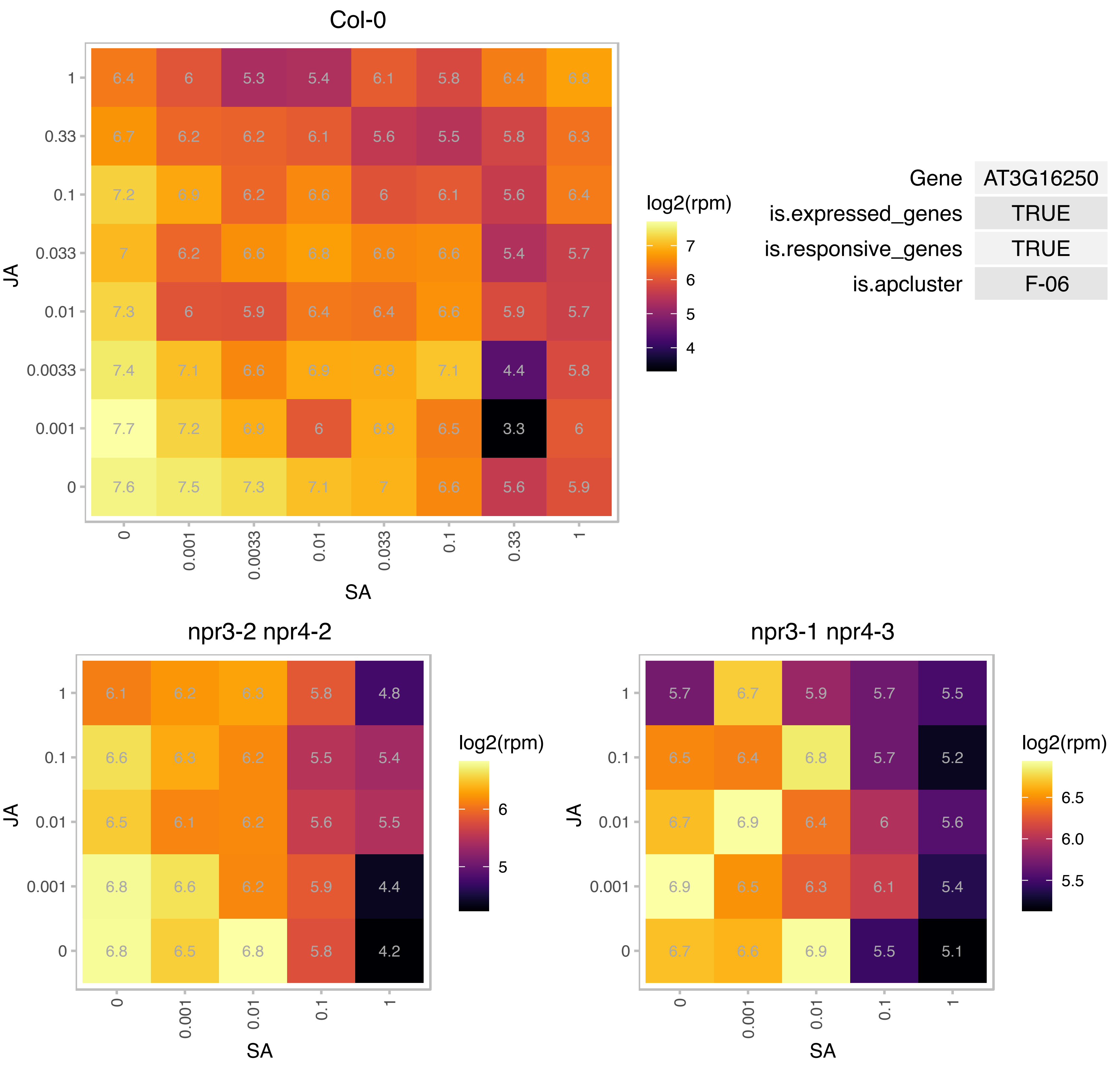


| | |
|---|---|
| Gene | AT3G16250 |
| is.expressed_genes | TRUE |
| is.responsive_genes | TRUE |
| is.apcluster | F-06 |

### AT3G18390.1 : EMB1865

CRS1 / YhbY (CRM) domain-containing protein

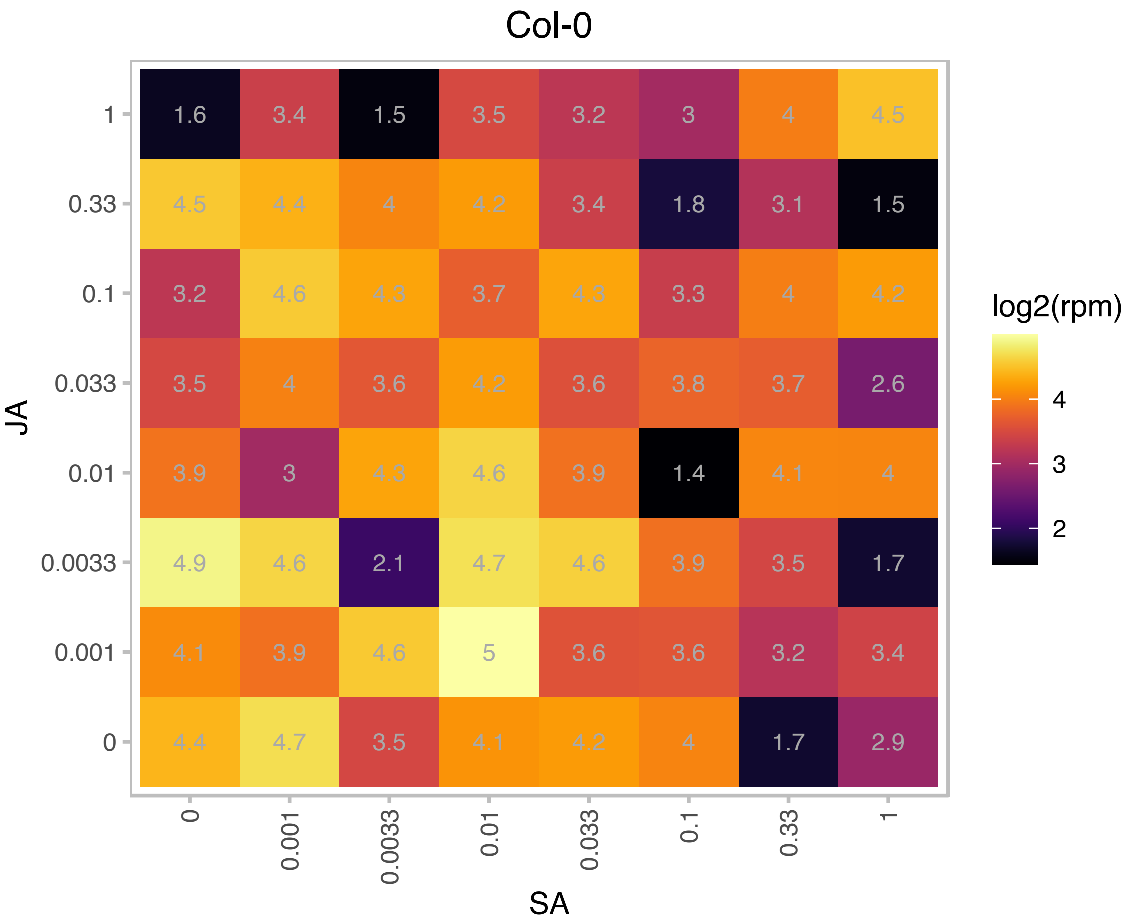


| | |
|---|---|
| Gene | AT3G18390 |
| is.expressed_genes | TRUE |
| is.responsive_genes | TRUE |
| is.apcluster | F-06 |

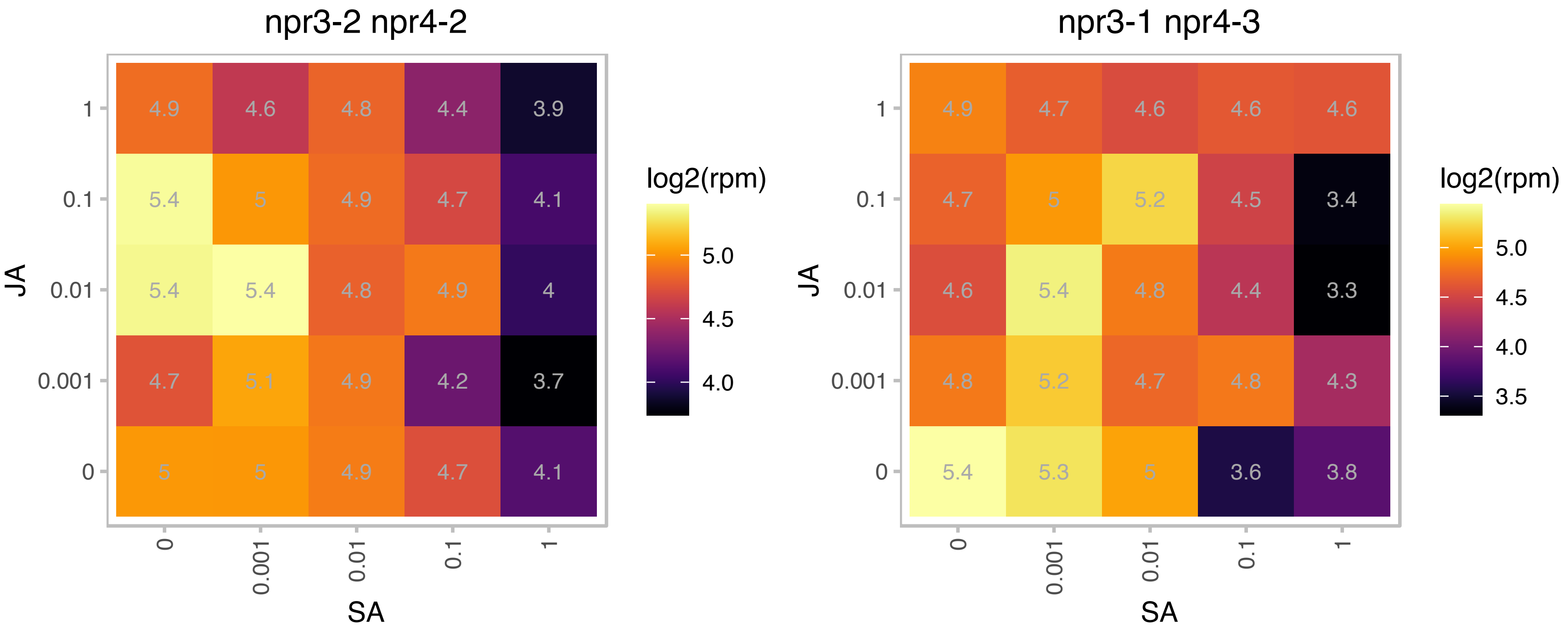

# AT3G25660.1 : NA

Amidase family protein

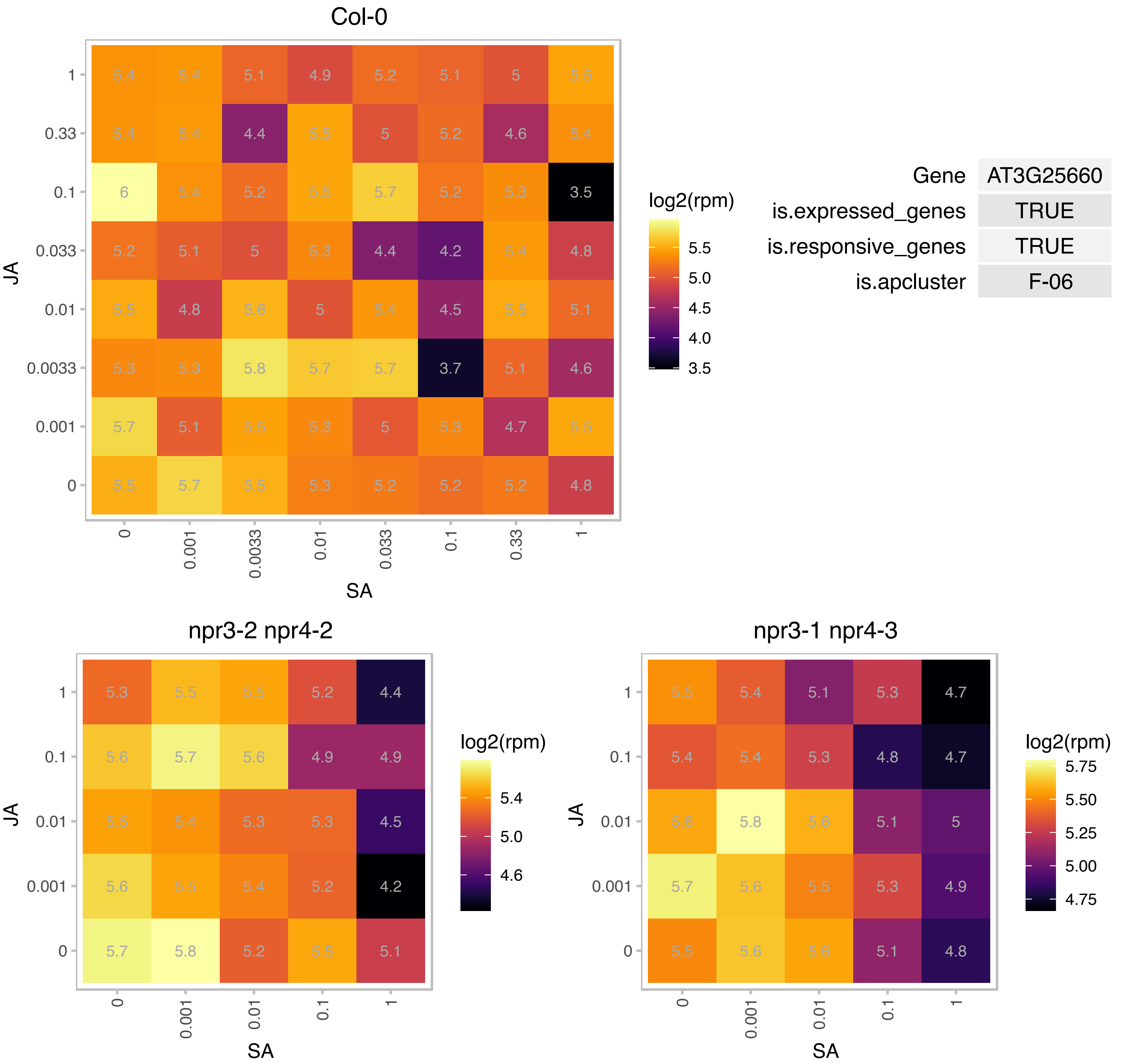


| | |
|---|---|
| Gene | AT3G25660 |
| is.expressed_genes | TRUE |
| is.responsive_genes | TRUE |
| is.apcluster | F-06 |

### AT3G26300.1 : CYP71B34

cytochrome P450, family 71, subfamily B, polypeptide 34

putative cytochrome P450

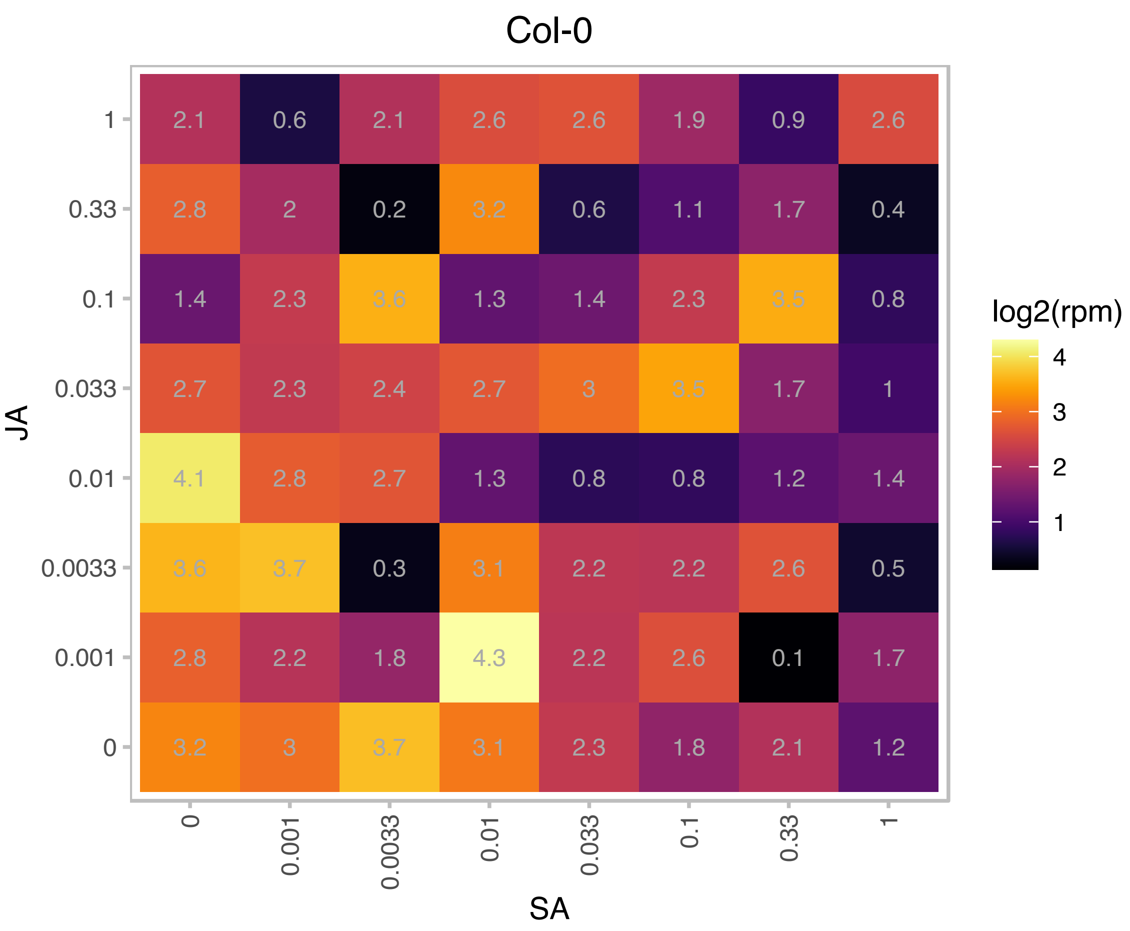


| | |
|---|---|
| Gene | AT3G26300 |
| is.expressed_genes | TRUE |
| is.responsive_genes | TRUE |
| is.apcluster | F-06 |

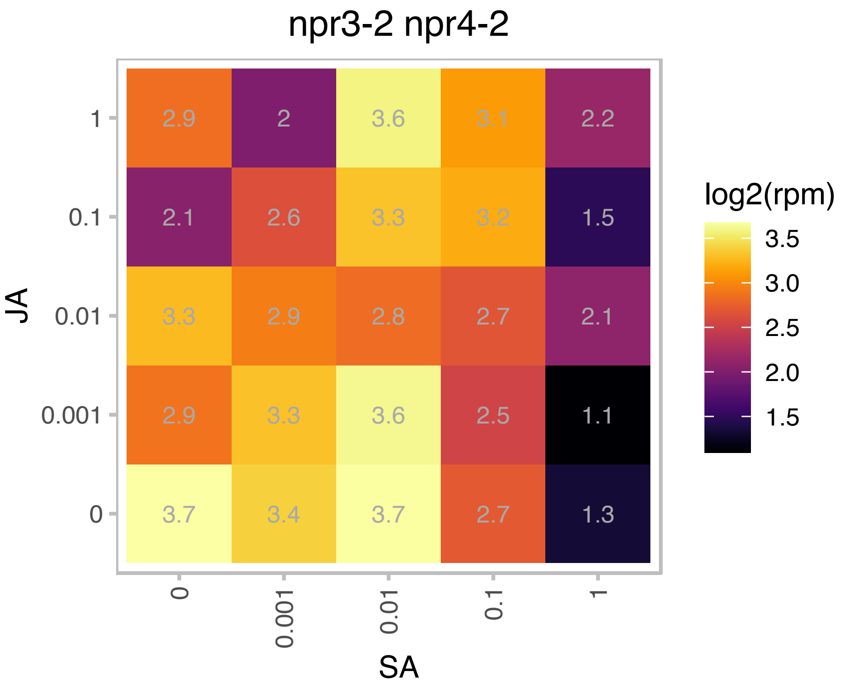


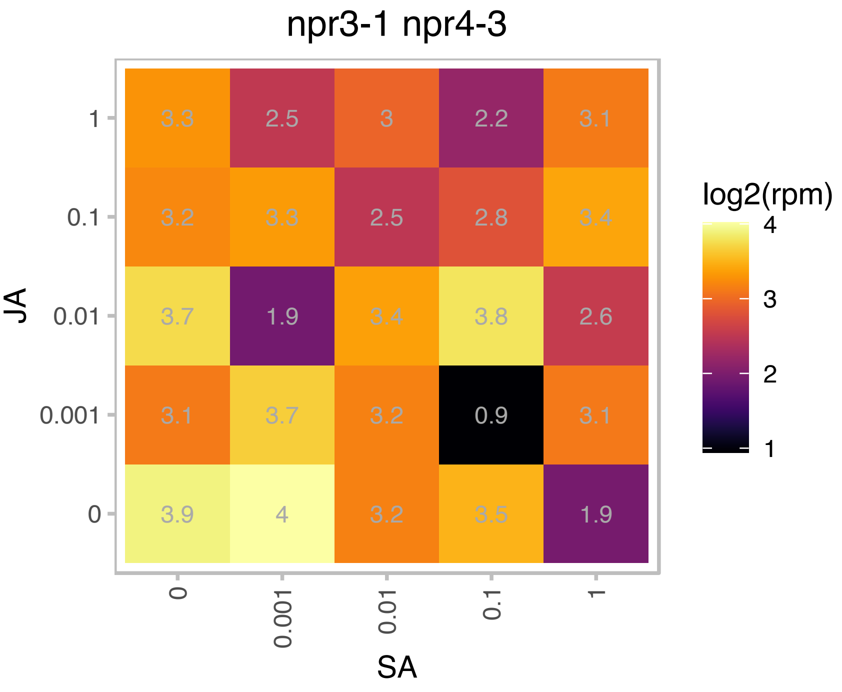

### AT3G27690.1 : LHCB2, LHCB2.3, LHCB2.4

photosystem II light harvesting complex gene 2.3
Encodes Lhcb2.4. Belongs to the Lhc super-gene family encodes the light-harvesting chlorophyll a/b-binding (LHC) proteins that constitute the antenna system of the photosynthetic apparatus. The mRNA is cell-to-cell mobile.

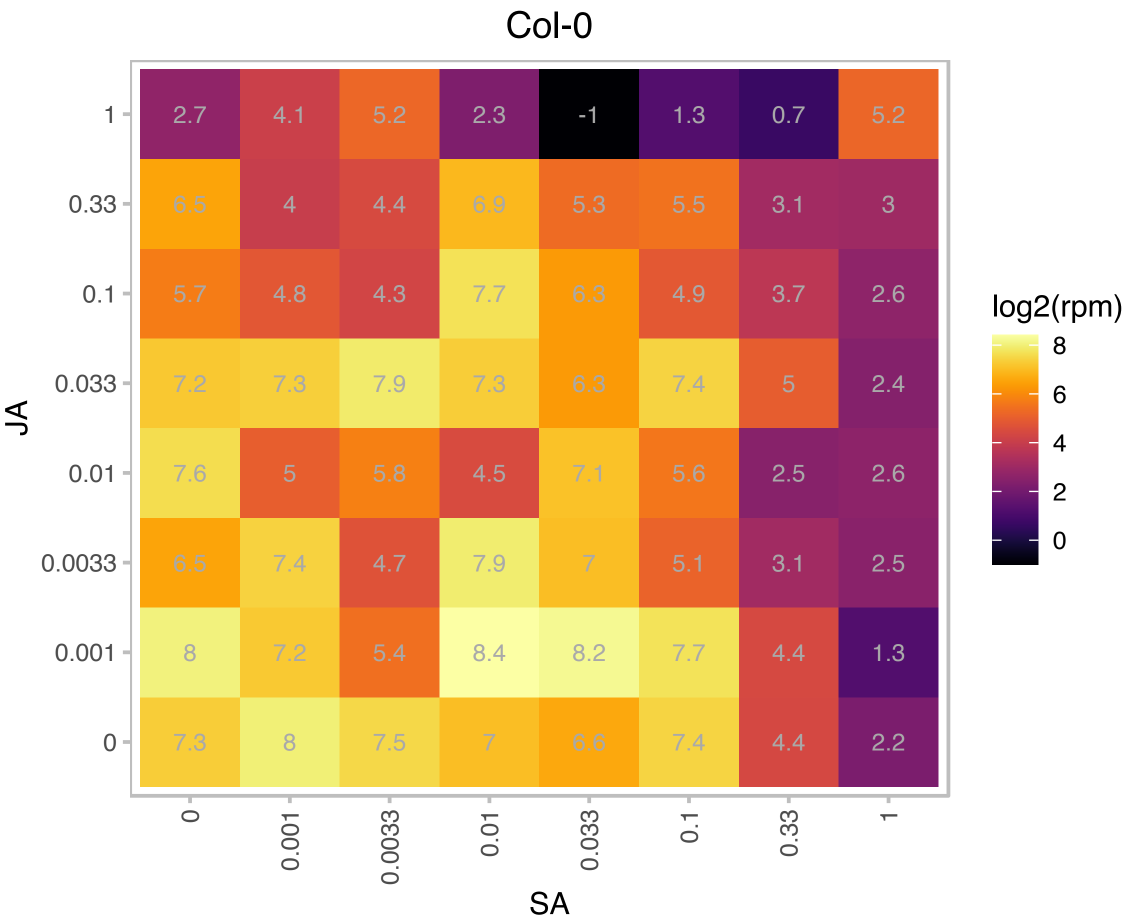


| | |
|---|---|
| Gene | AT3G27690 |
| is.expressed_genes | TRUE |
| is.responsive_genes | TRUE |
| is.apcluster | F-06 |

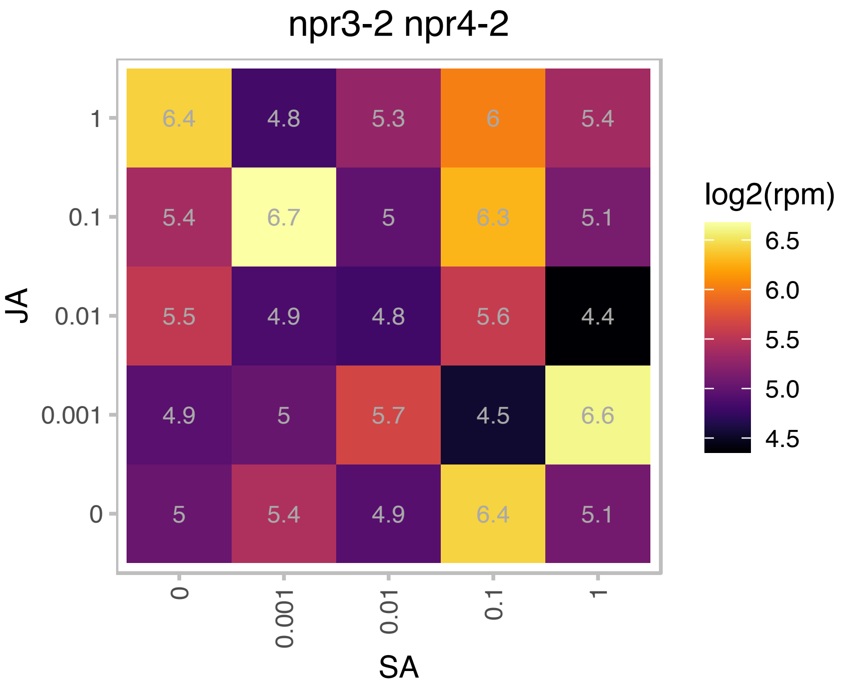


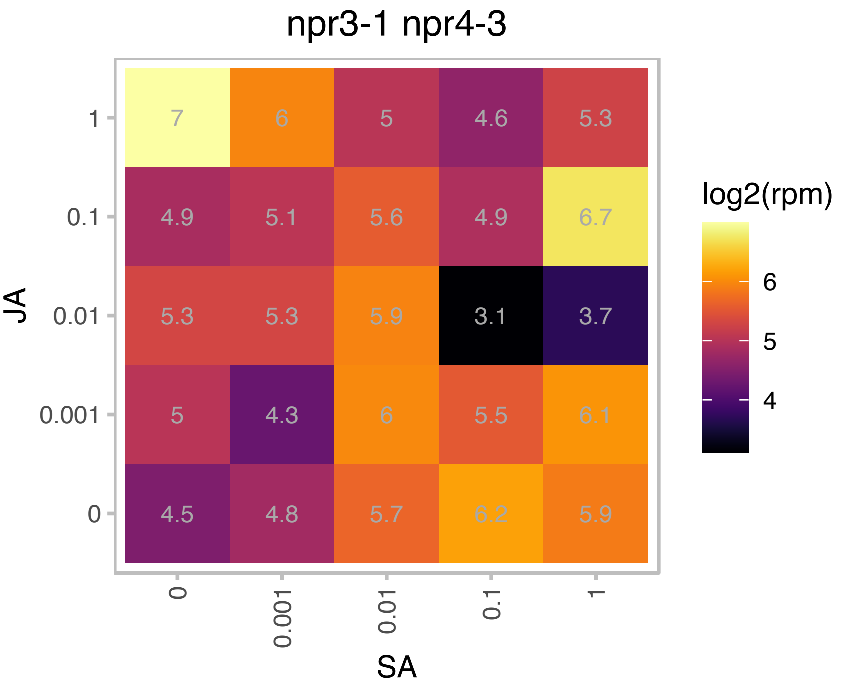

### AT3G44990.1 : AtXTH31, ATXTR8, XTH31, XTR8

xyloglucan endo-transglycosylase-related 8

Encodes a xyloglucan endotransglycosylase/hydrolase. Protein sequence and phylogenetic analysis indicates that this enzyme resides in Group III-A of the XTH family, with high similarity to Tropaeolum majus (nasturtium) xyloglucanase 1 (TmNXG1).Enzyme kinetic analysis indicates predominant xyloglucan endo-hydrolase activity (EC 3.2.1.151) with only limited potential to act as a xyloglucan endo-transglycosylase (EC 2.4.1.207).

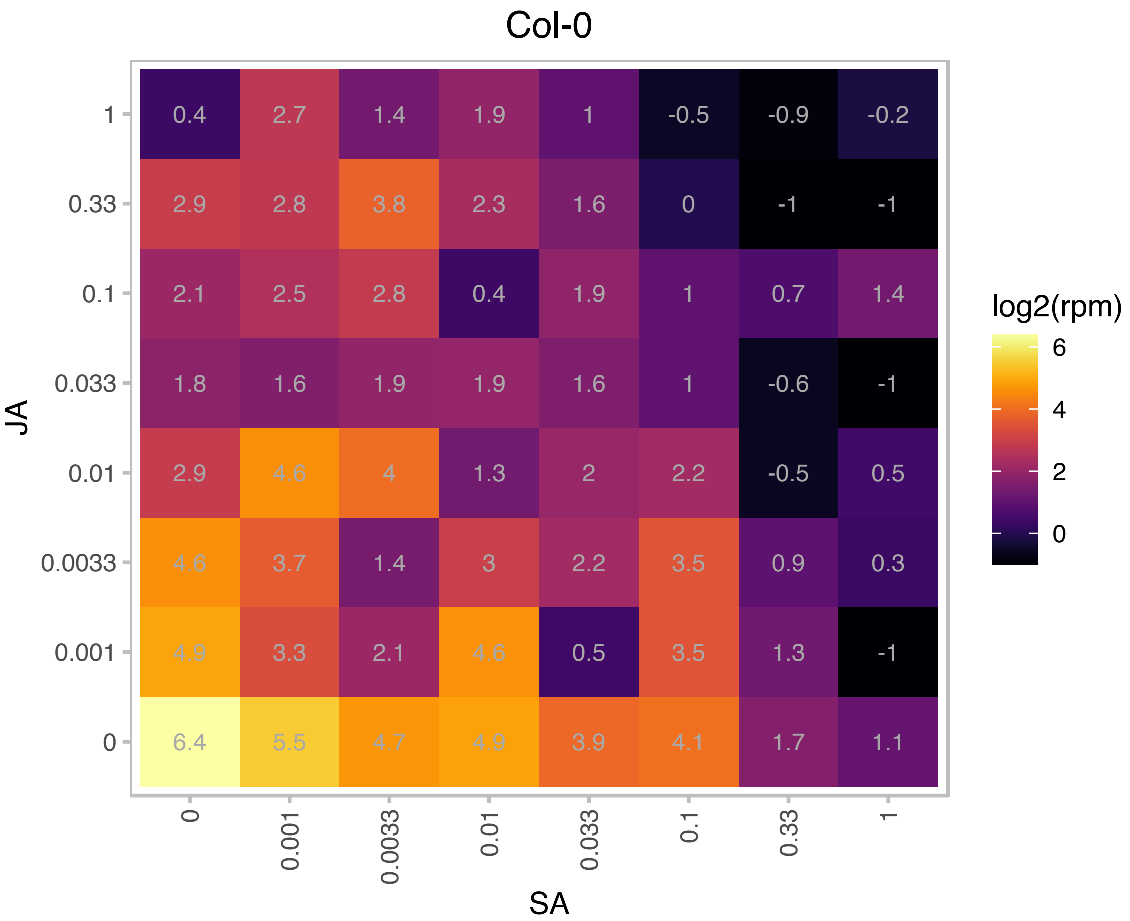


| | |
|---|---|
| Gene | AT3G44990 |
| is.expressed_genes | TRUE |
| is.responsive_genes | TRUE |
| is.apcluster | F-06 |

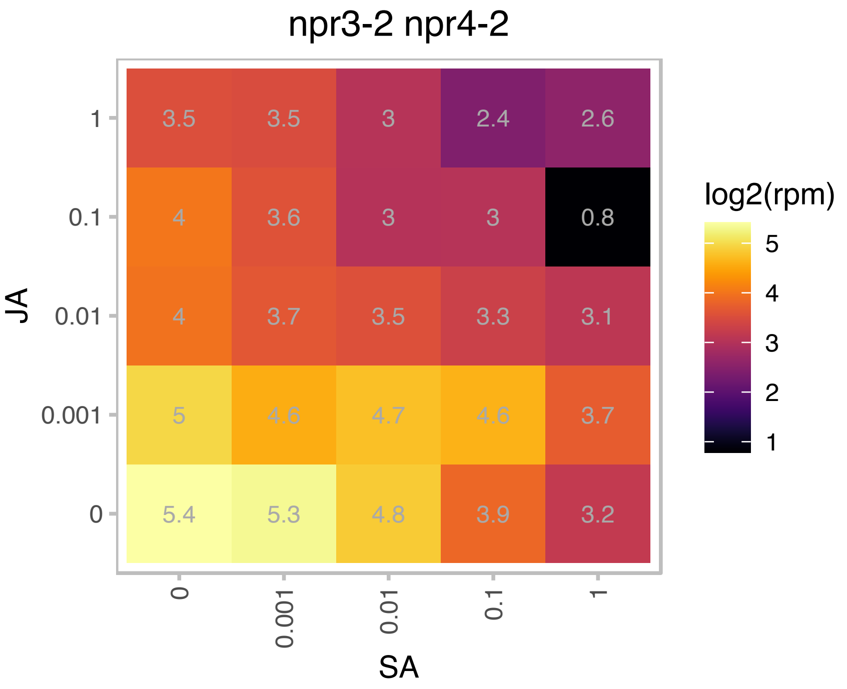


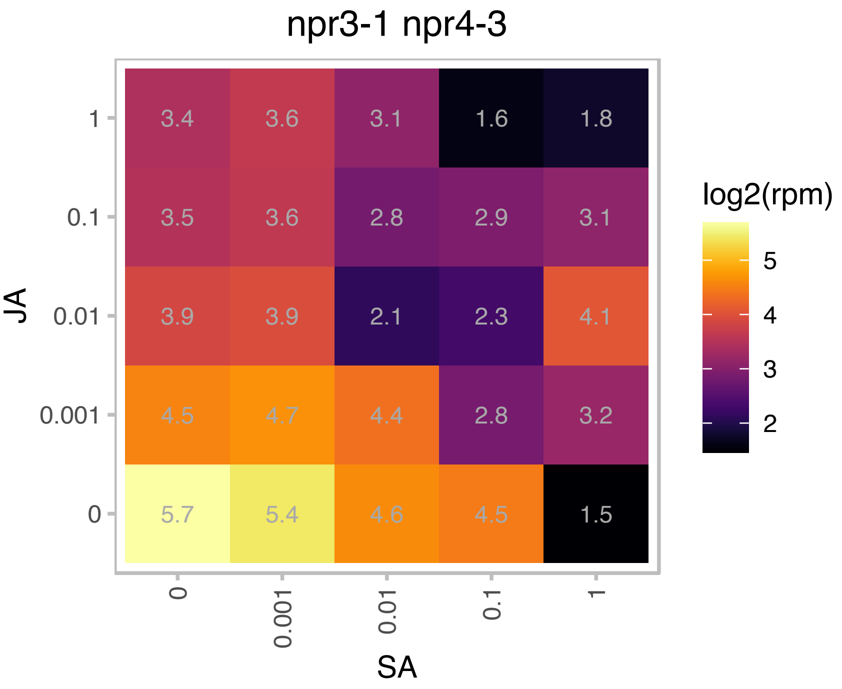

### AT3G50060.1 : MYB77

myb domain protein 77
Encodes a member of the R2R3 transcription factor gene family. Expressed in response to potassium deprivation and auxin. Involved in lateral root development. Interacts with ARF7 and regulates the expression of some auxin responsive genes.

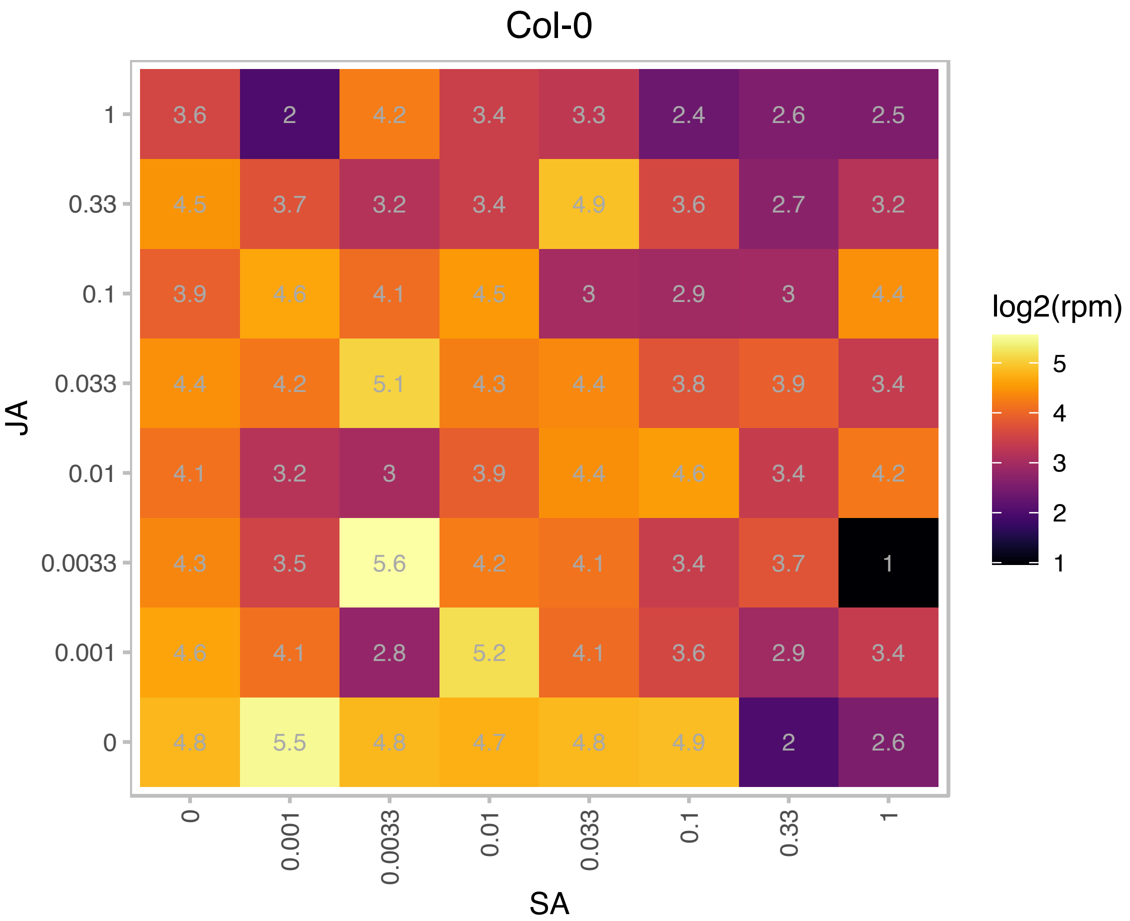


| | |
|---|---|
| Gene | AT3G50060 |
| is.expressed_genes | TRUE |
| is.responsive_genes | TRUE |
| is.apcluster | F-06 |

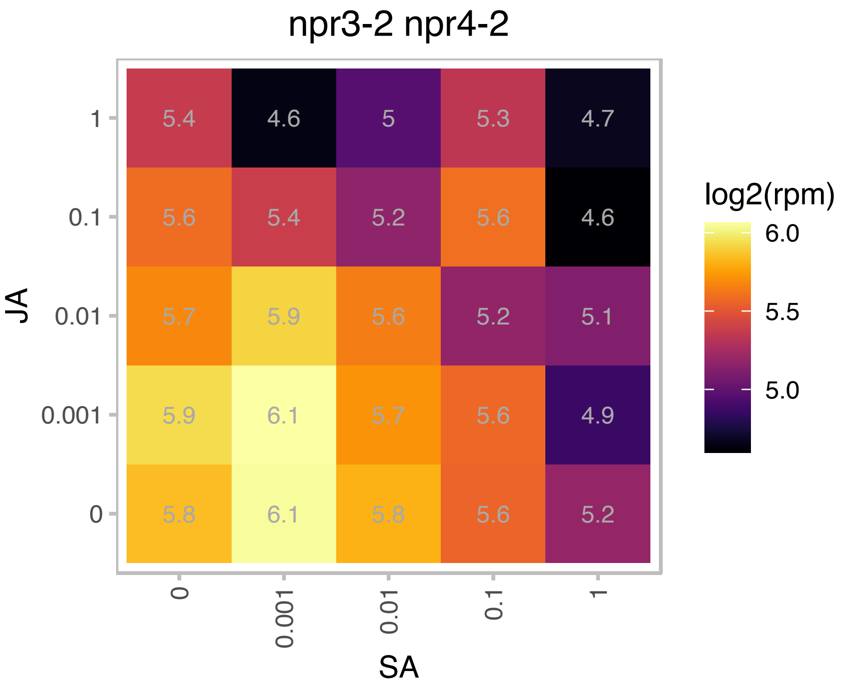


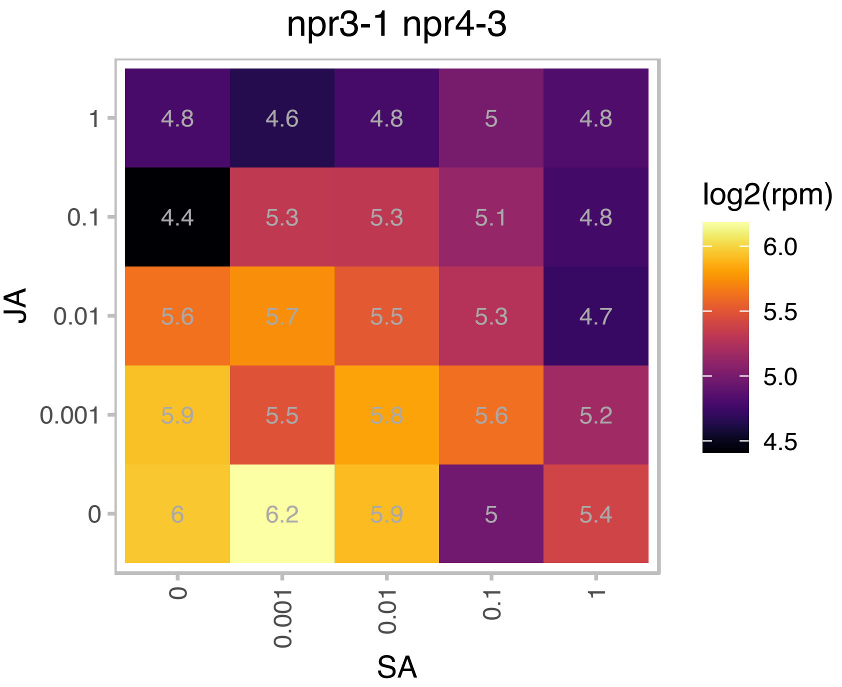

### AT3G52150.1 : PSRP2

RNA-binding (RRM/RBD/RNP motifs) family protein

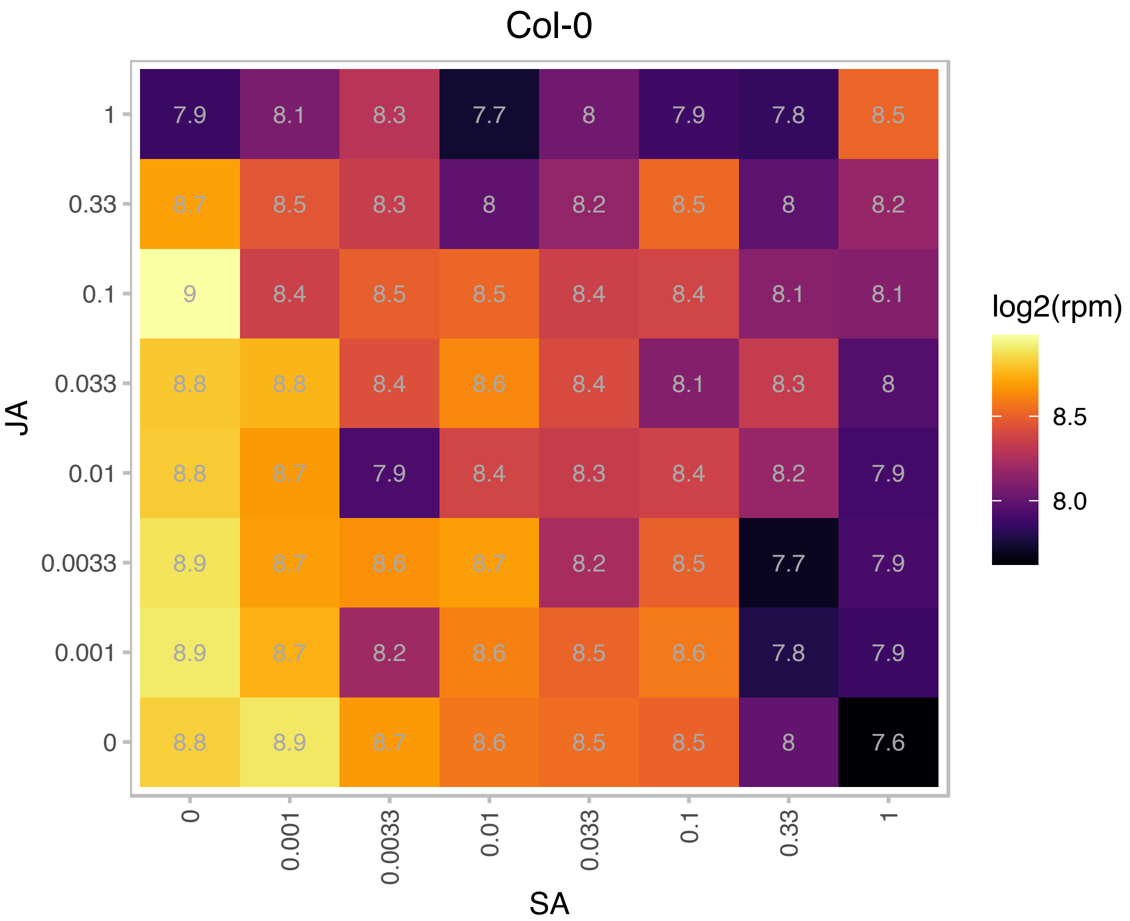


| | |
|---|---|
| Gene | AT3G52150 |
| is.expressed_genes | TRUE |
| is.responsive_genes | TRUE |
| is.apcluster | F-06 |

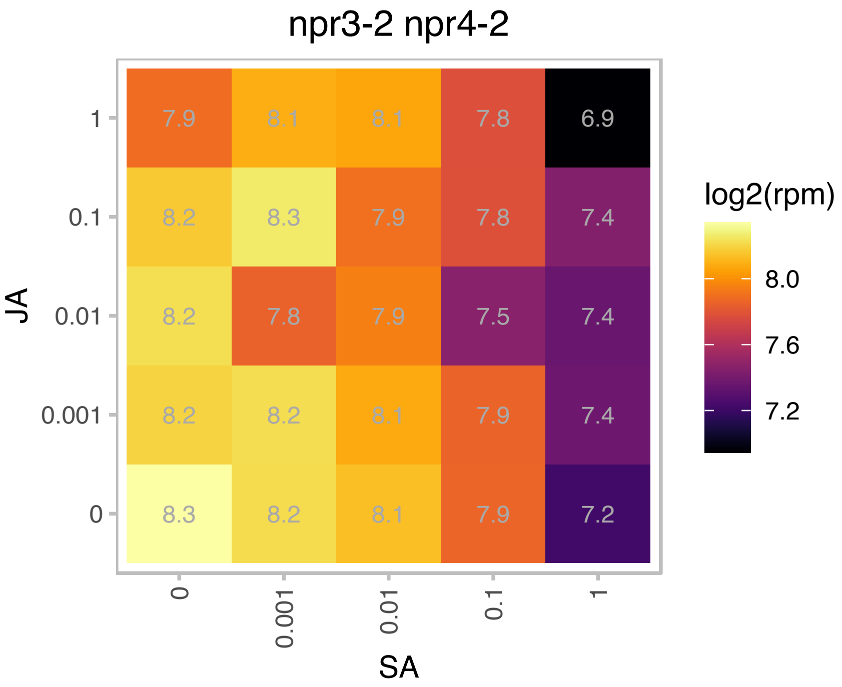


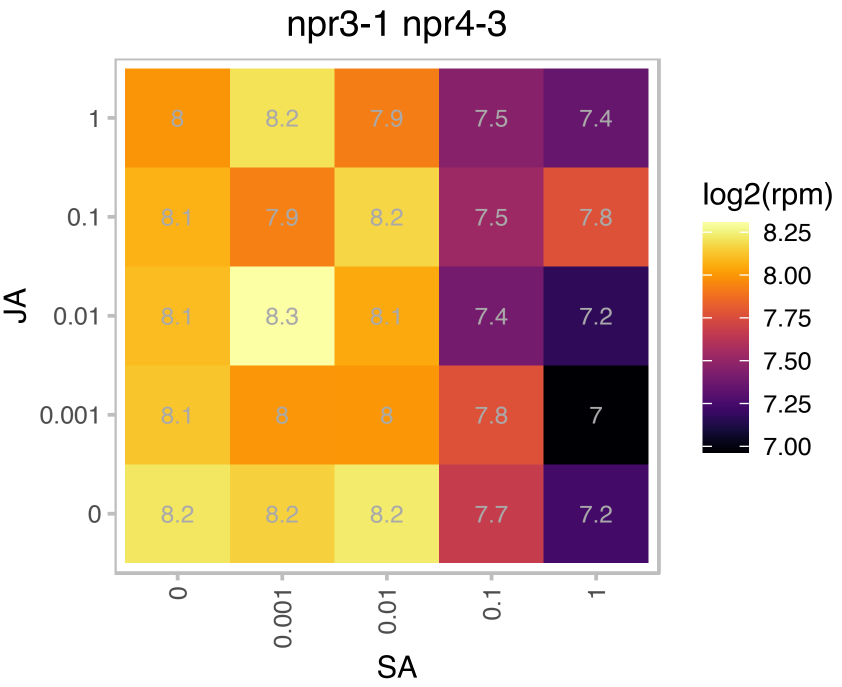

### AT3G52380.1 : CP33, PDE322

chloroplast RNA-binding protein 33
Encodes a chloroplast RNA-binding protein that stabilizes chloroplast RNAs as evidenced by analyses of transcript accumulation in null mutants. Essential for seedling development (albino, strongly retarded growth even on sucrose-containing medium).

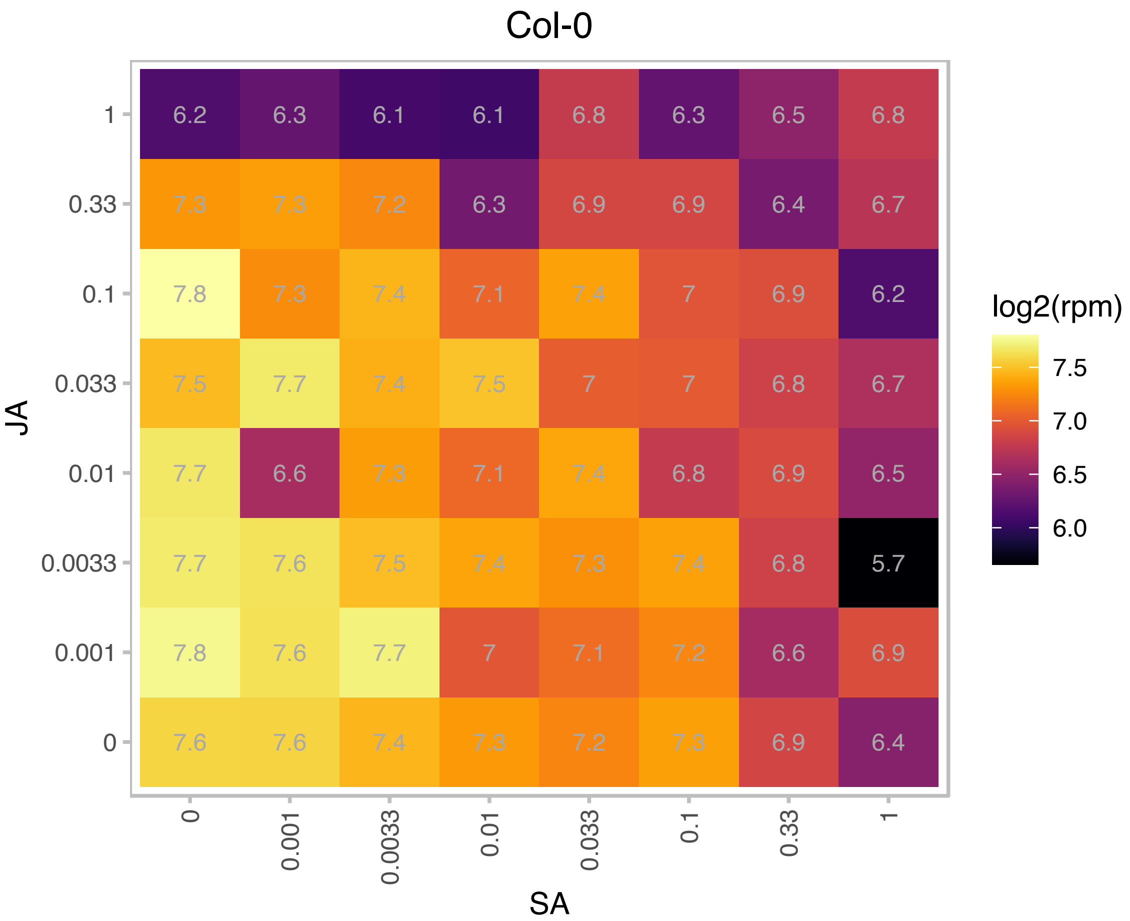


| | |
|---|---|
| Gene | AT3G52380 |
| is.expressed_genes | TRUE |
| is.responsive_genes | TRUE |
| is.apcluster | F-06 |

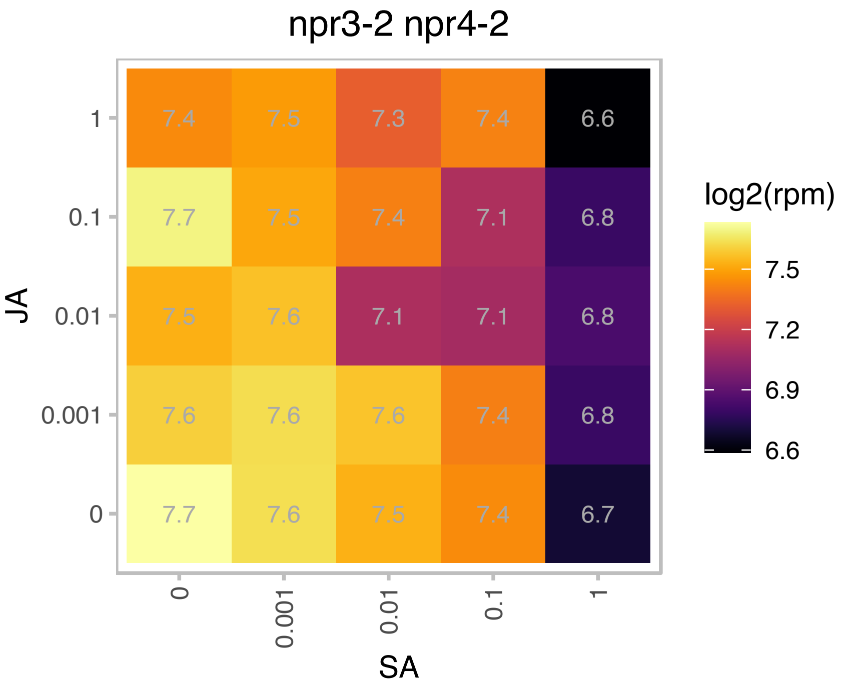


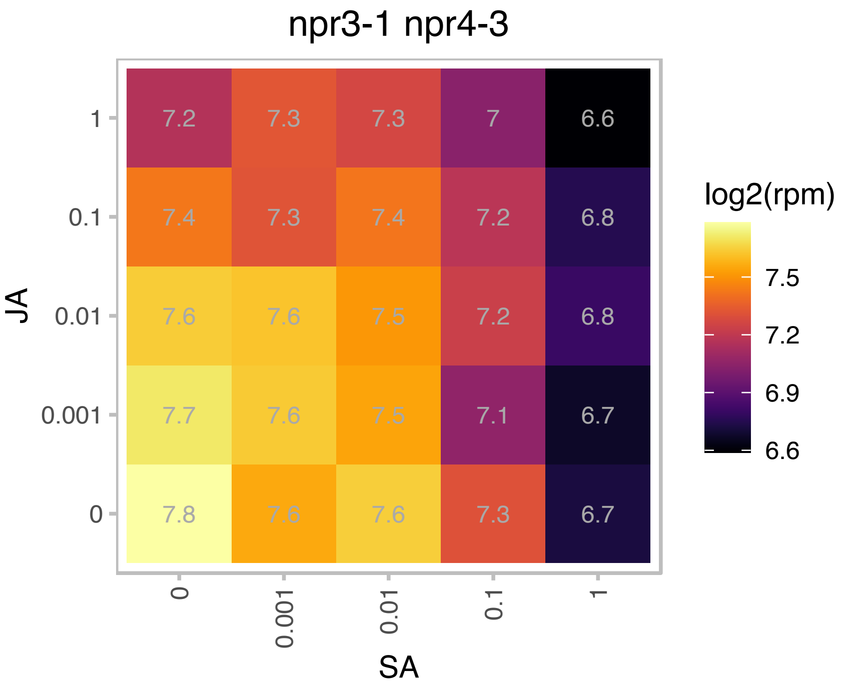

### AT3G52960.1 : PrxIIE

Thioredoxin superfamily protein

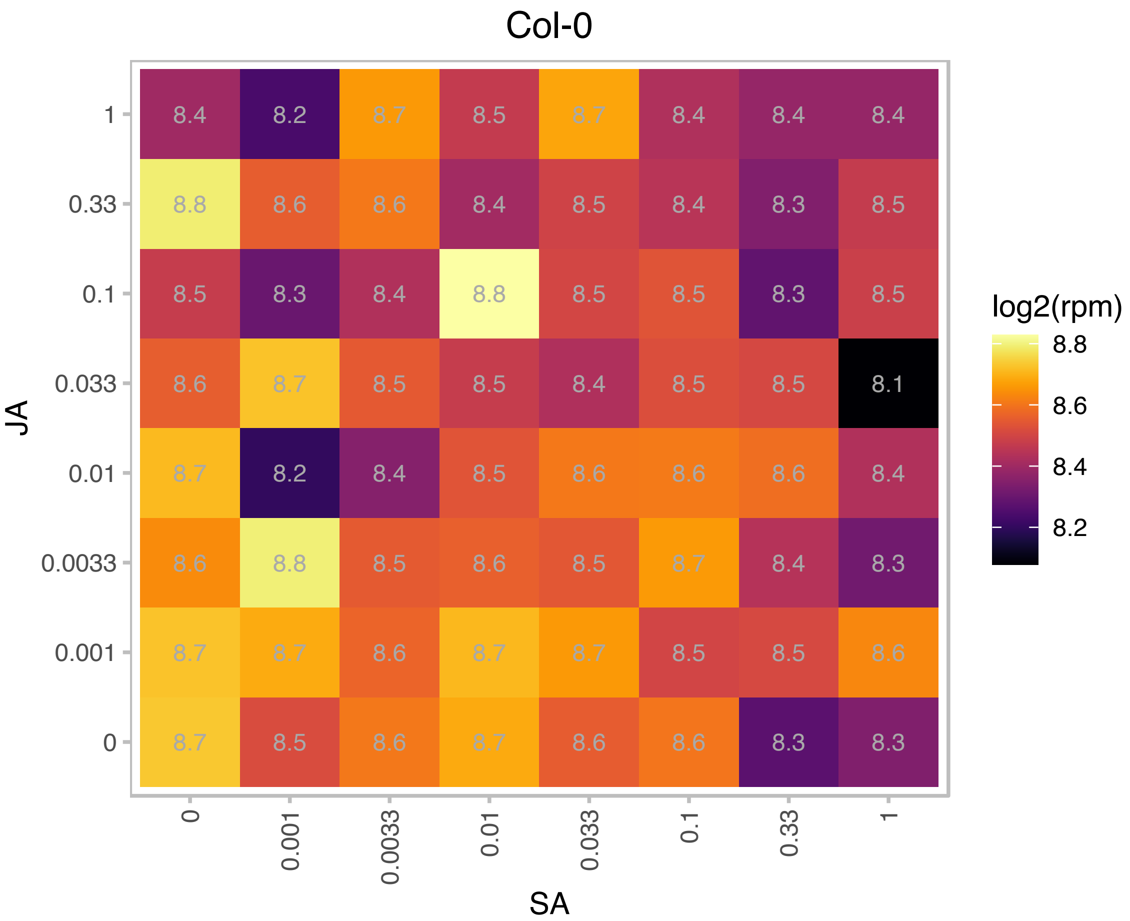


| | |
|---|---|
| Gene | AT3G52960 |
| is.expressed_genes | TRUE |
| is.responsive_genes | TRUE |
| is.apcluster | F-06 |

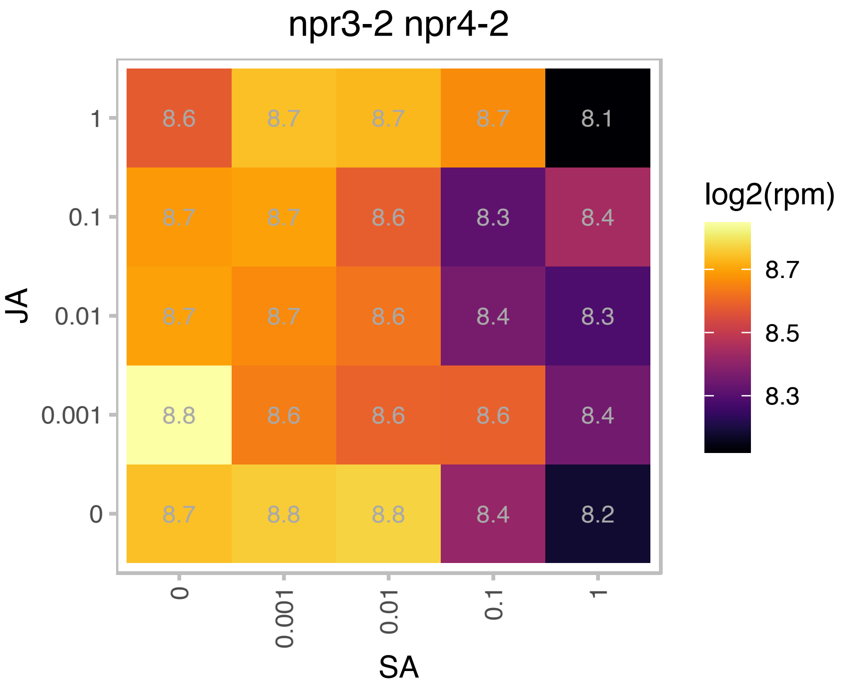


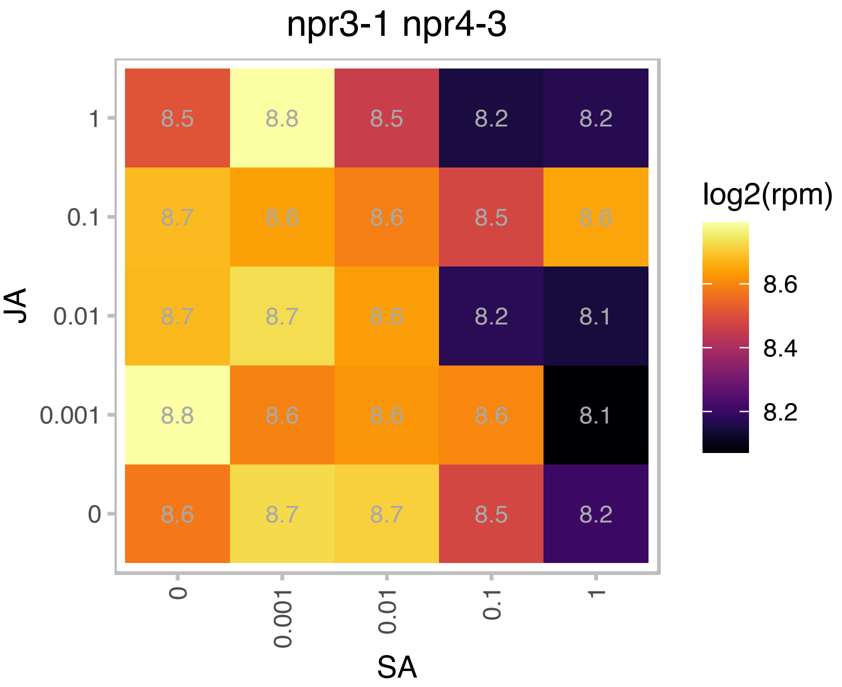

# AT3G58610.1 : NA

ketol-acid reductoisomerase

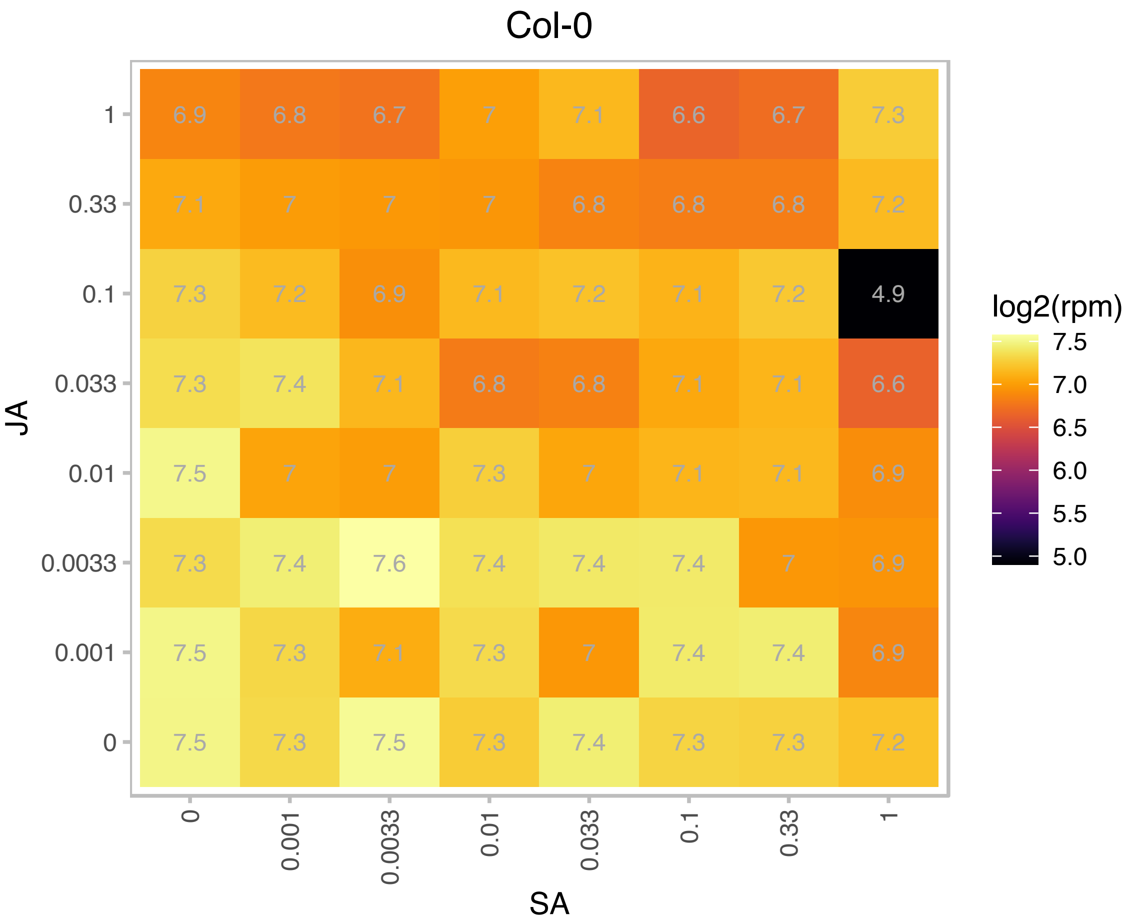


| | |
|---|---|
| Gene | AT3G58610 |
| is.expressed_genes | TRUE |
| is.responsive_genes | TRUE |
| is.apcluster | F-06 |

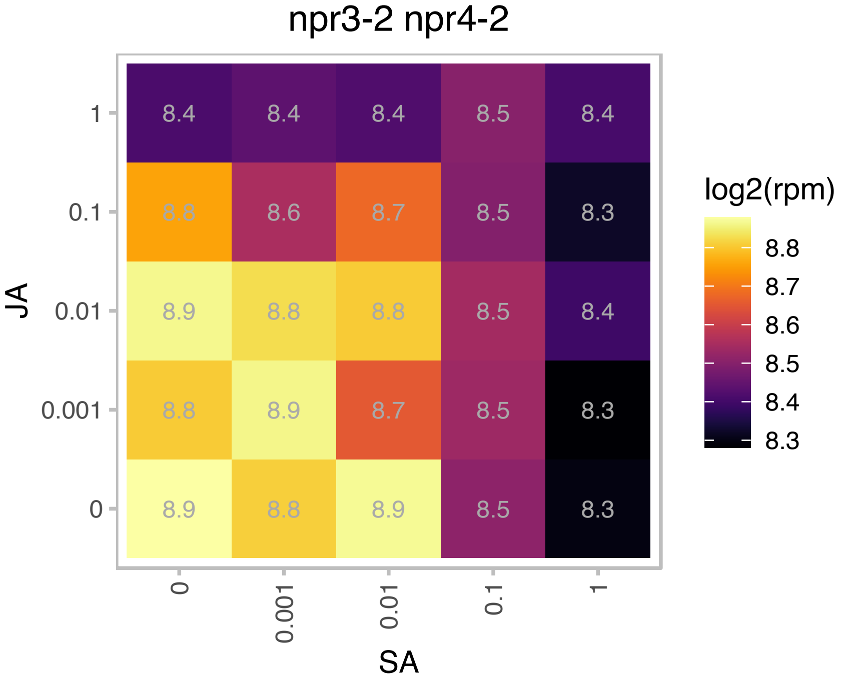


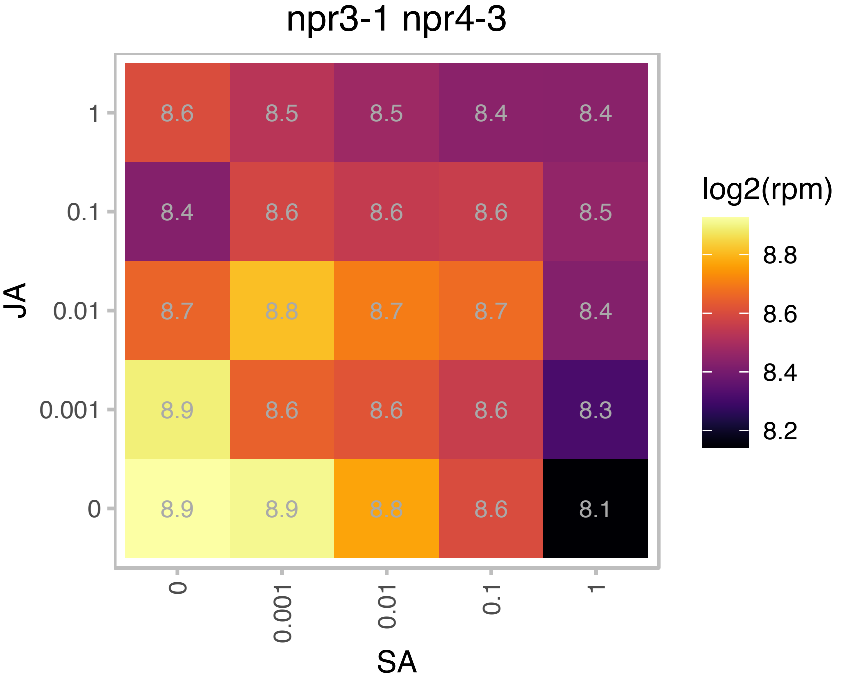

### AT4G02770.1 : PSAD-1

photosystem I subunit D-1

Encodes a protein predicted by sequence similarity with spinach PsaD to be photosystem I reaction center subunit II (PsaD1)

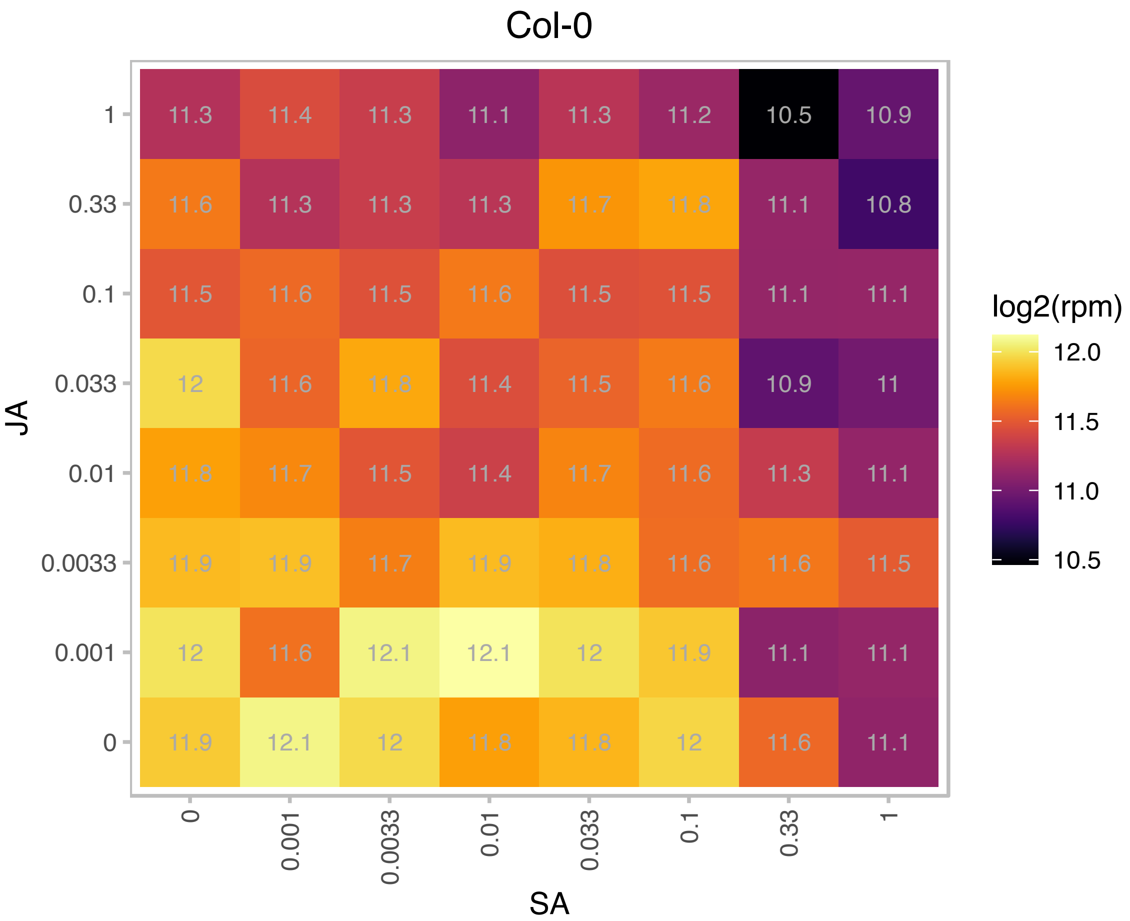


| | |
|---|---|
| Gene | AT4G02770 |
| is.expressed_genes | TRUE |
| is.responsive_genes | TRUE |
| is.apcluster | F-06 |

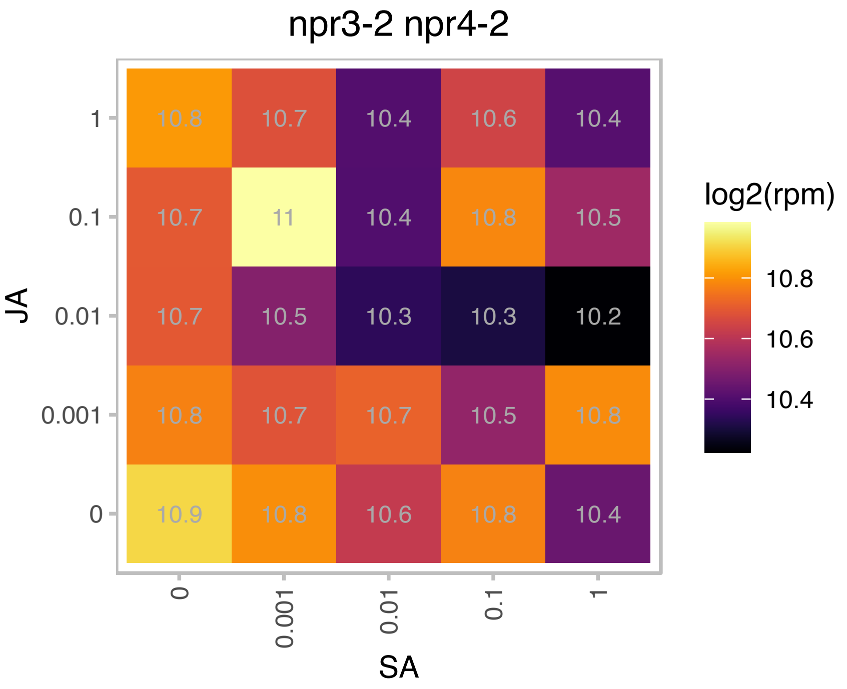


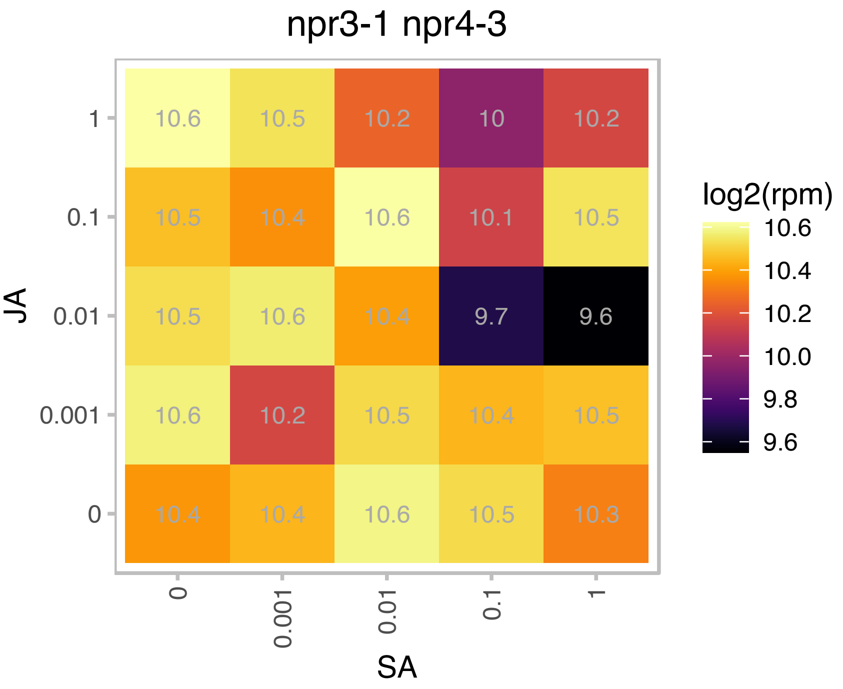

### AT4G02790.1 : AtRBGA, EMB3129, RBGA

GTP-binding family protein

Encodes a GTPase that is targeted to chloroplasts and co-fractionated with chloroplast ribosomes. Mutants are embryo lethal due to this essential function being lost.

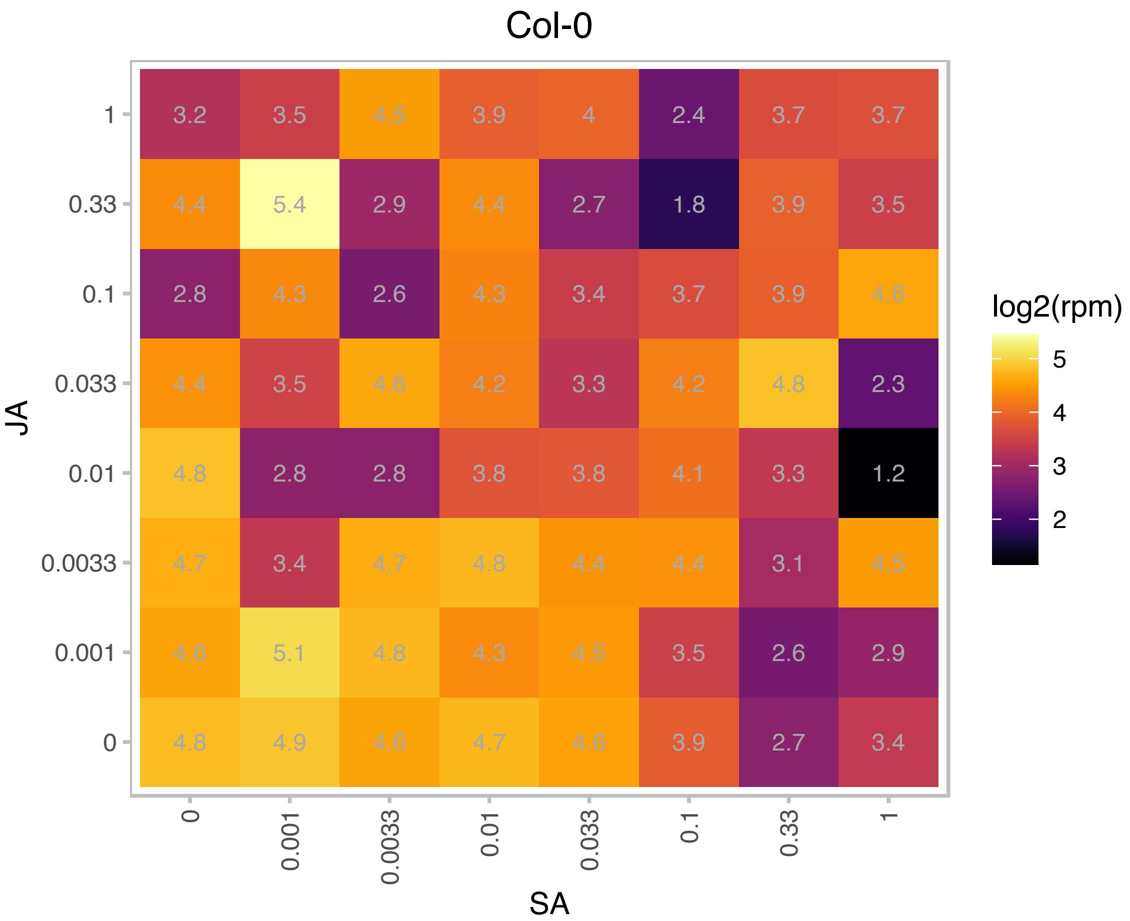


| | |
|---|---|
| Gene | AT4G02790 |
| is.expressed_genes | TRUE |
| is.responsive_genes | TRUE |
| is.apcluster | F-06 |

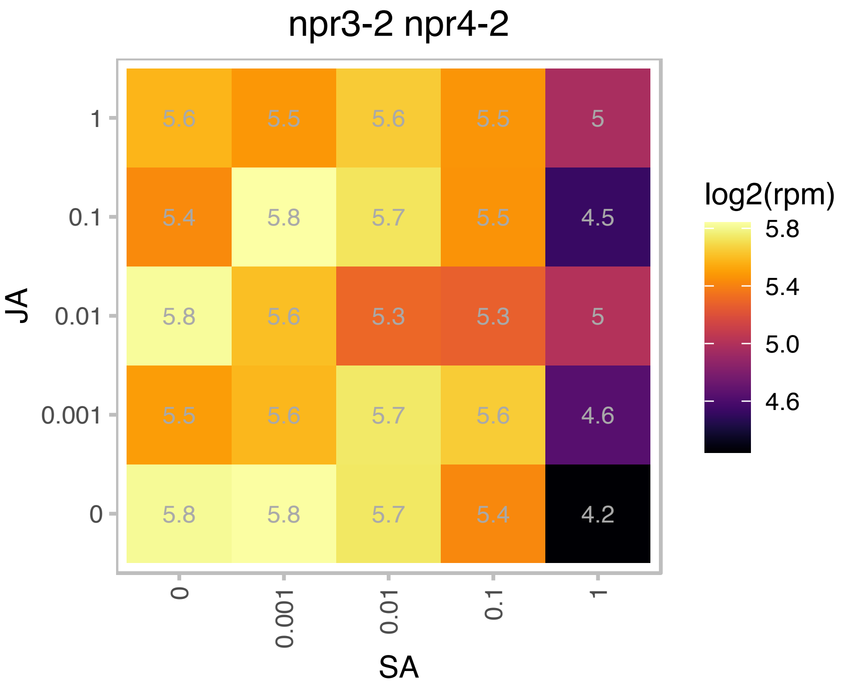


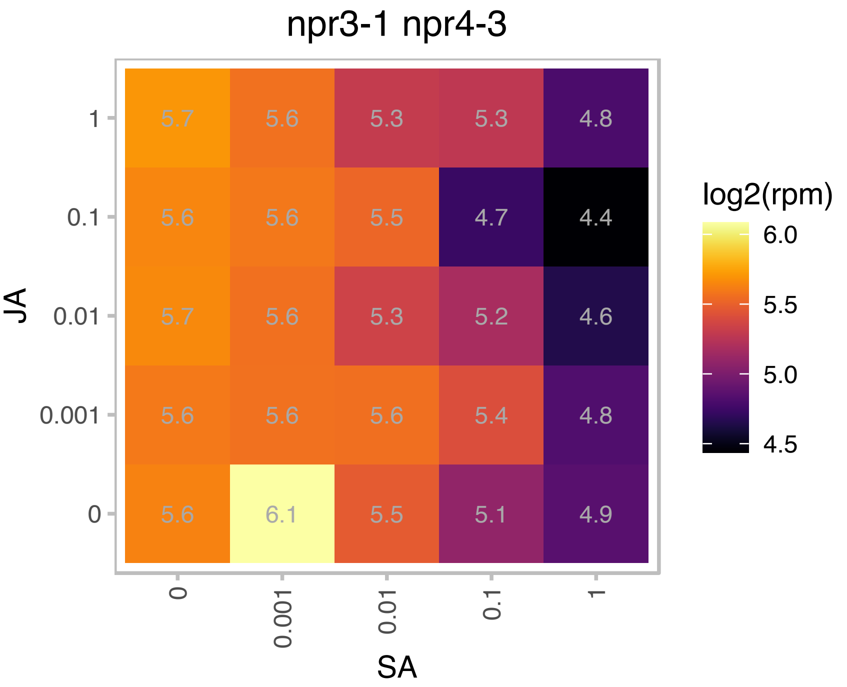

# AT4G02970.1 : AT7SL-1

7SL RNA1

Signal recognition particle. Type 4 of RNA polymerase III dependent genes.

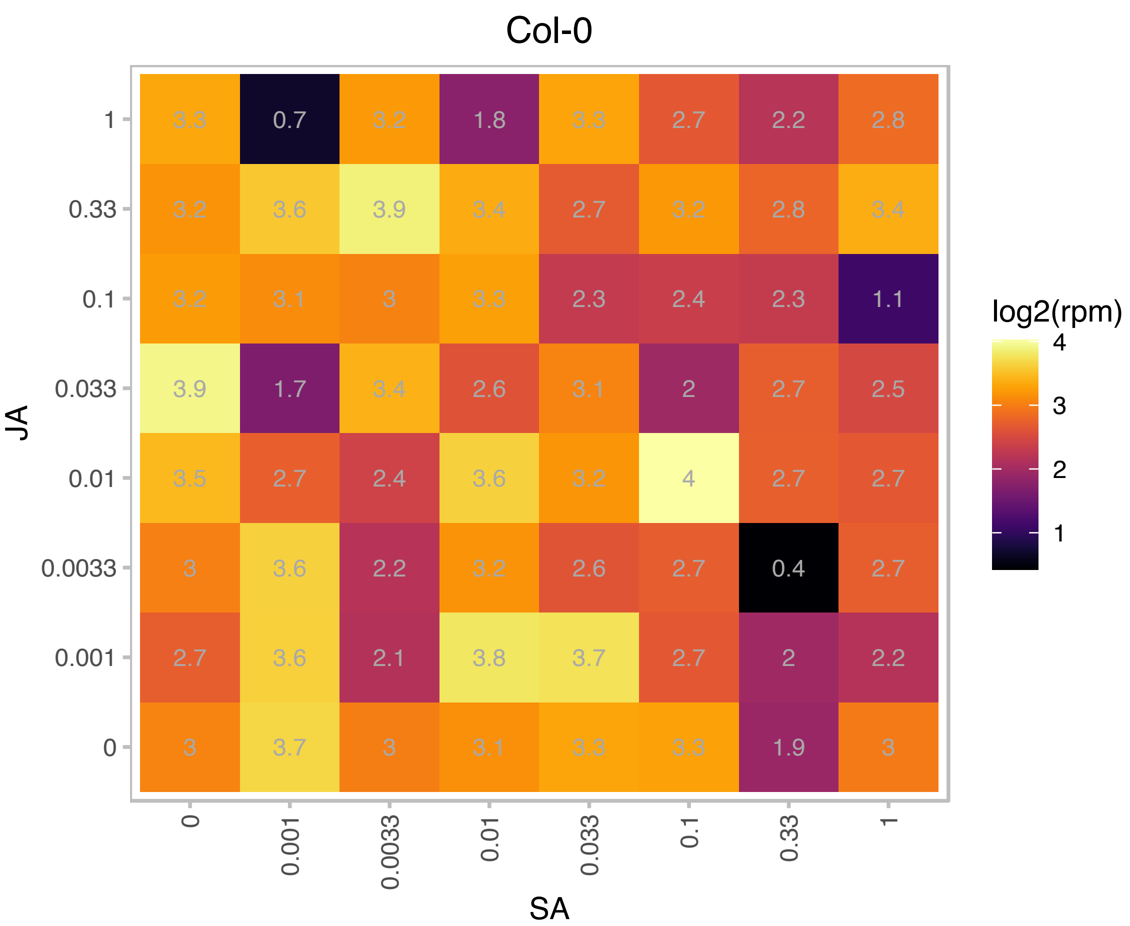


| | |
|---|---|
| Gene | AT4G02970 |
| is.expressed_genes | TRUE |
| is.responsive_genes | TRUE |
| is.apcluster | F-06 |

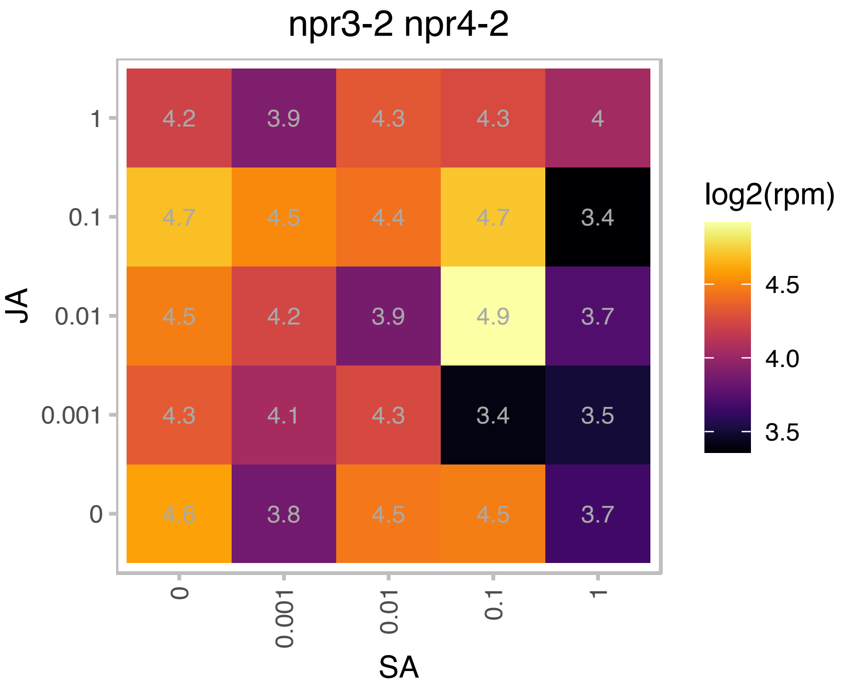


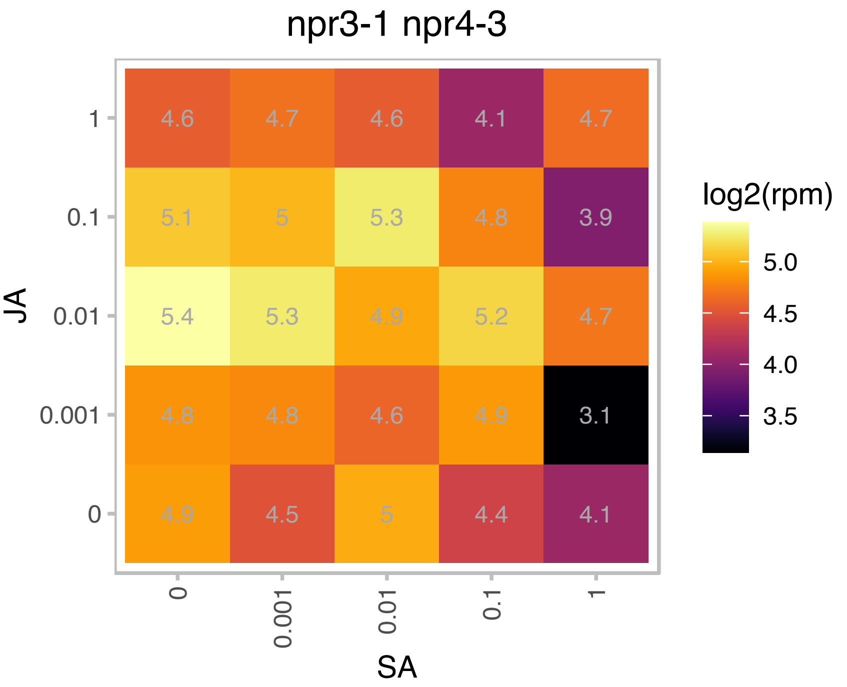

### AT4G08685.1 : SAH7

Pollen Ole e 1 allergen and extensin family protein

Encodes a protein, expressed in leaves, with similarity to pollen allergens. The mRNA is cell-to-cell mobile.

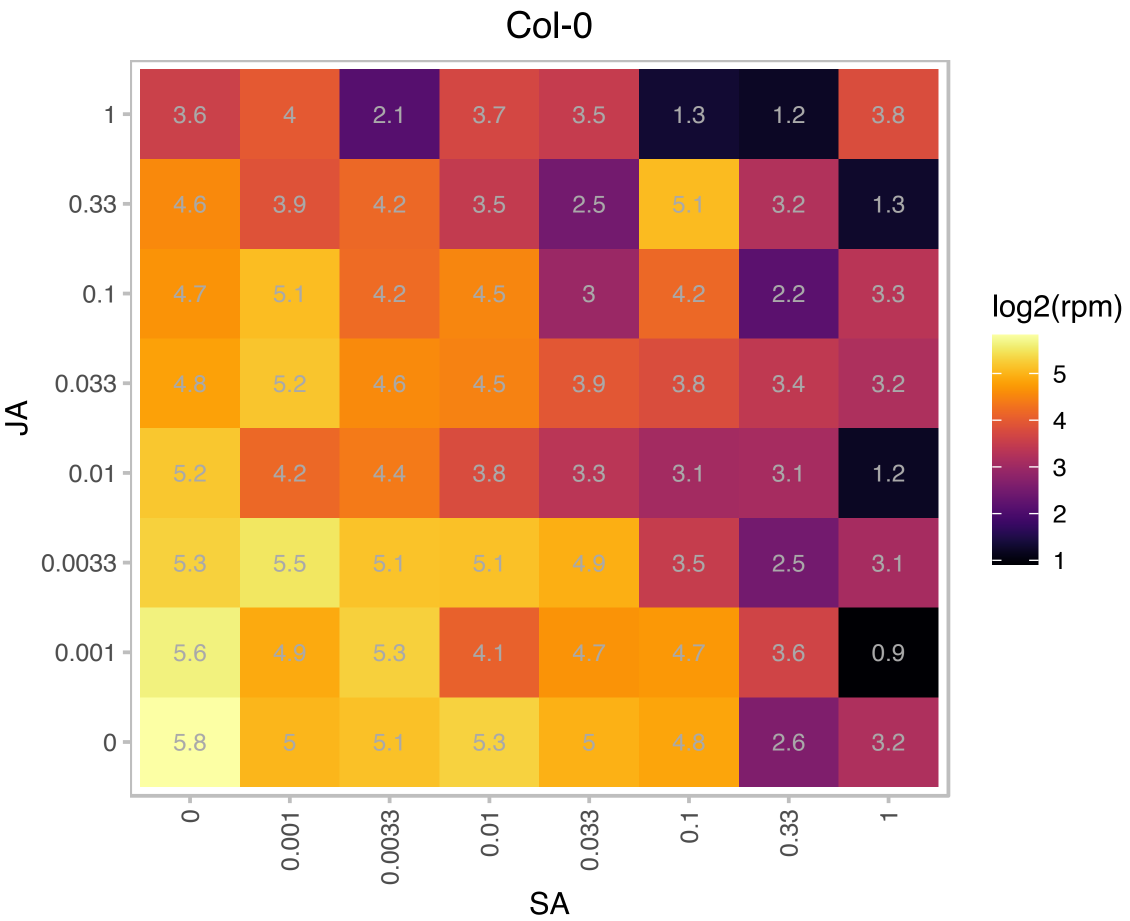


| | |
|---|---|
| Gene | AT4G08685 |
| is.expressed_genes | TRUE |
| is.responsive_genes | TRUE |
| is.apcluster | F-06 |

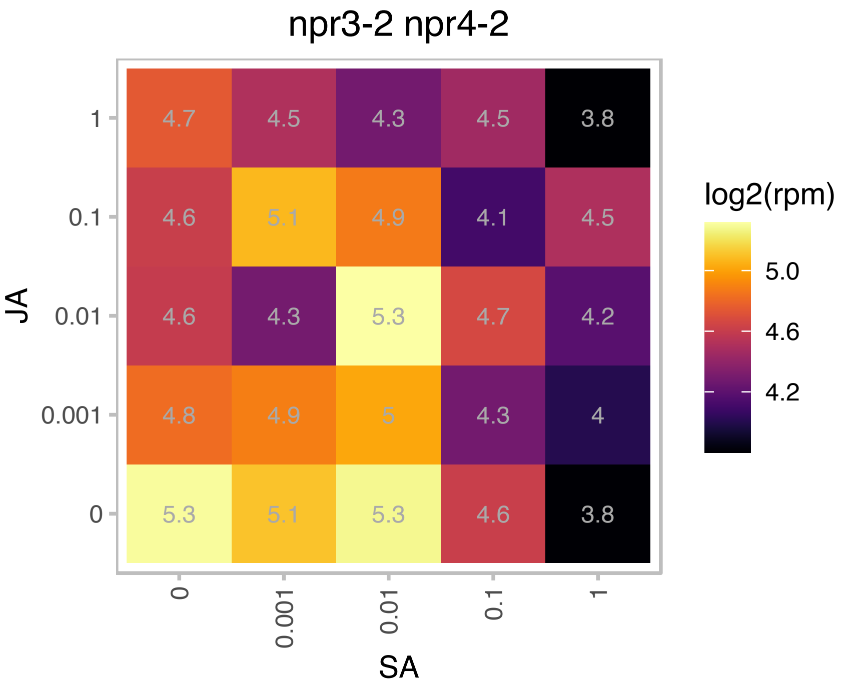


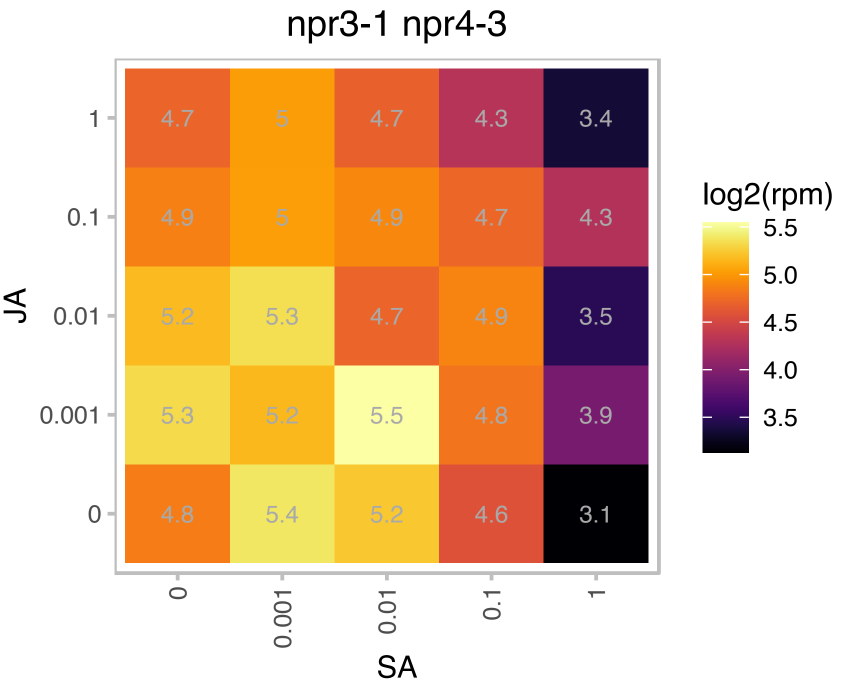

### AT4G12390.1 : PME1

pectin methylesterase inhibitor 1

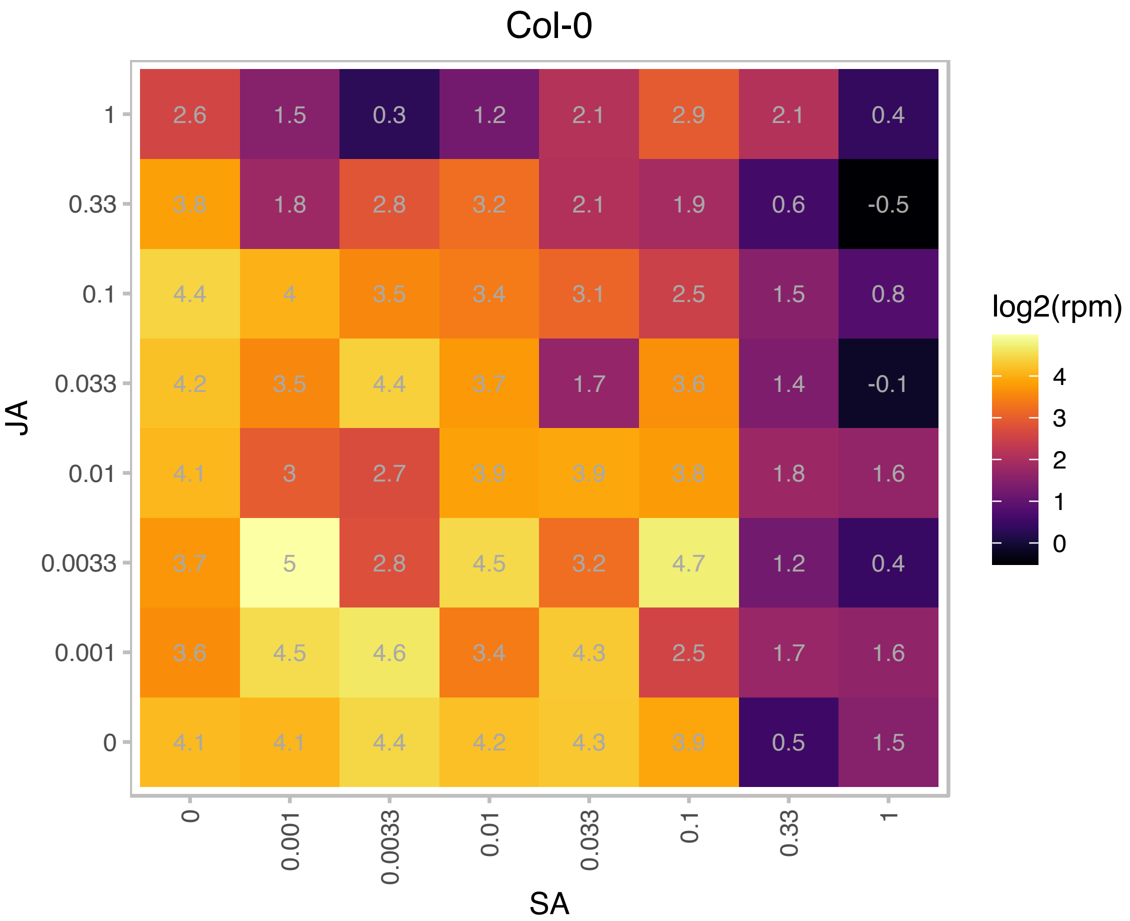


| | |
|---|---|
| Gene | AT4G12390 |
| is.expressed_genes | TRUE |
| is.responsive_genes | TRUE |
| is.apcluster | F-06 |

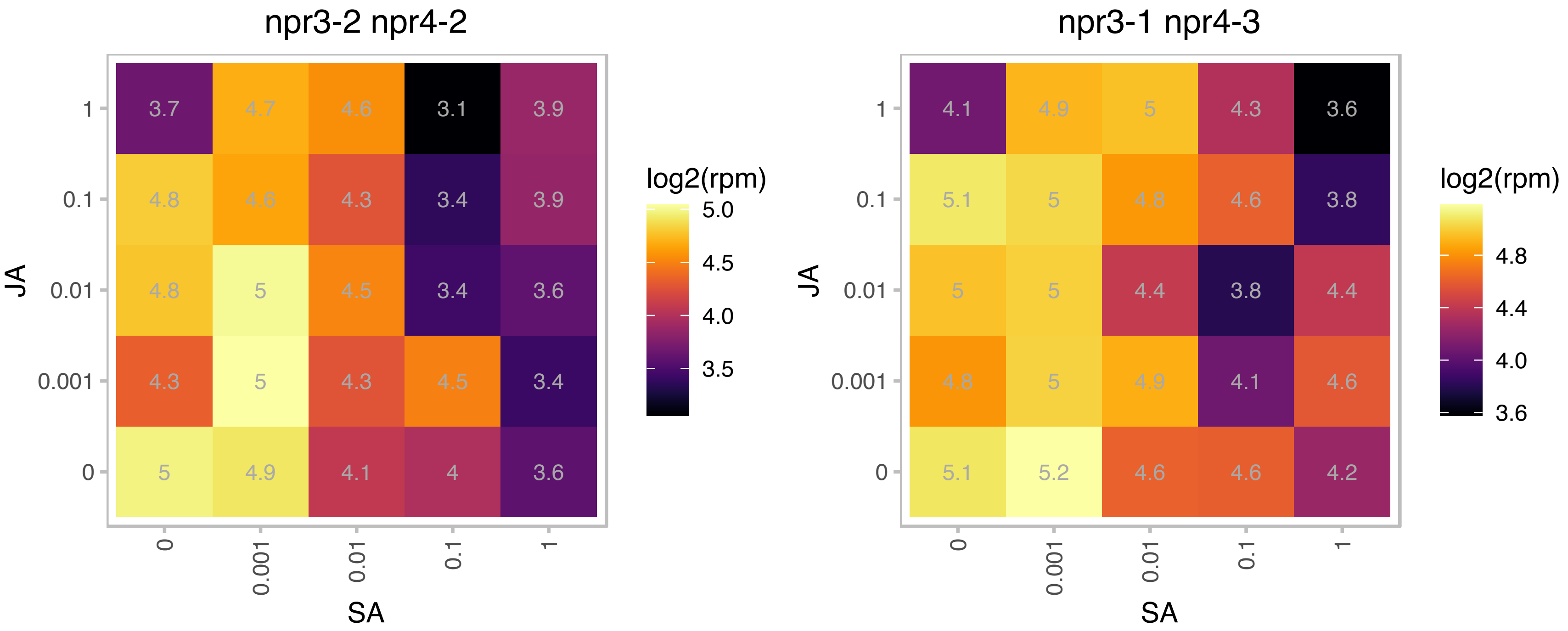

### AT4G14870.1 : SECE1

secE/sec61-gamma protein transport protein

Encodes a component of the thylakoid-localized Sec system involved in the translocation of cytoplasmic proteins into plastid. The mRNA is cell-to-cell mobile.

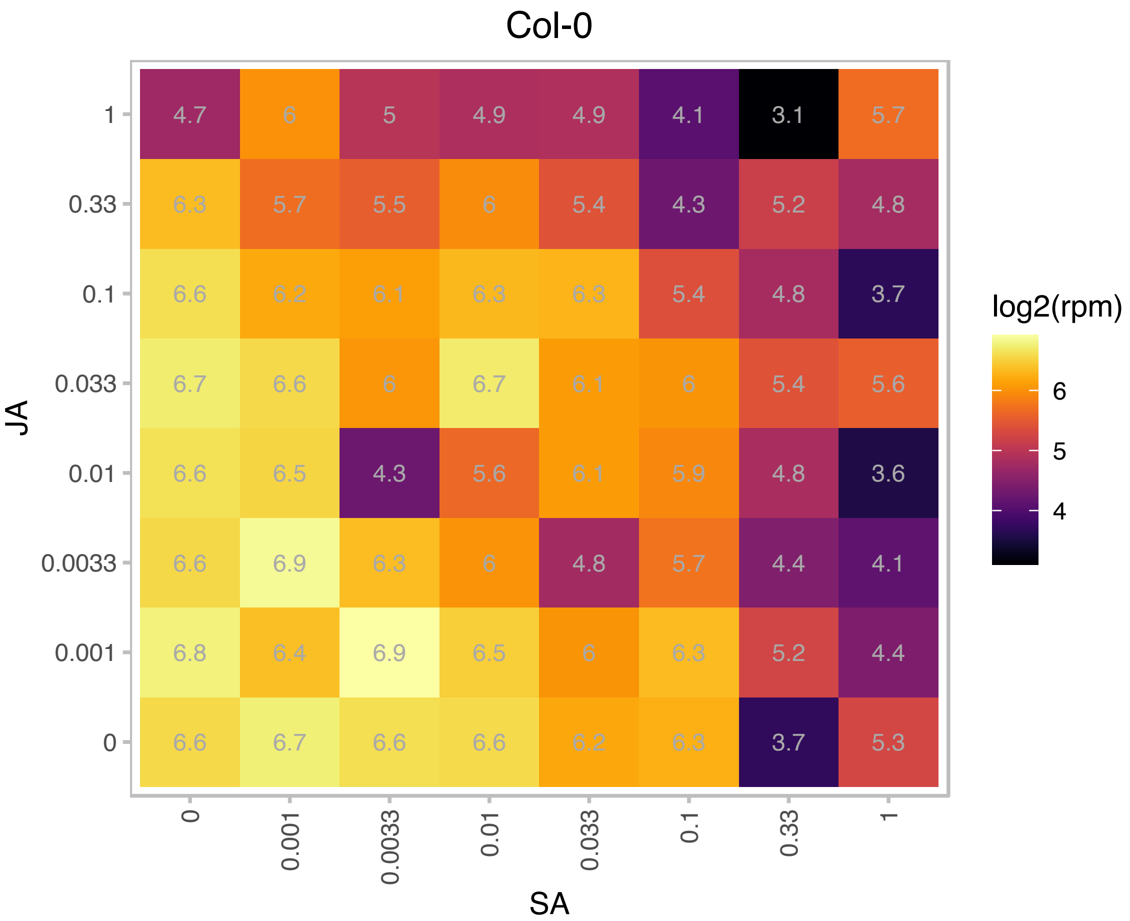


| | |
|---|---|
| Gene | AT4G14870 |
| is.expressed_genes | TRUE |
| is.responsive_genes | TRUE |
| is.apcluster | F-06 |

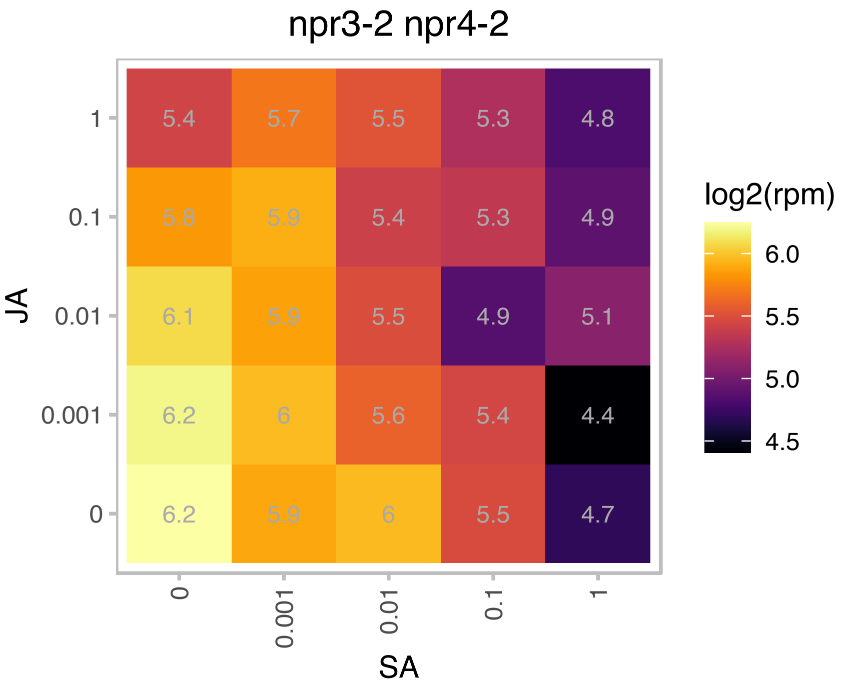


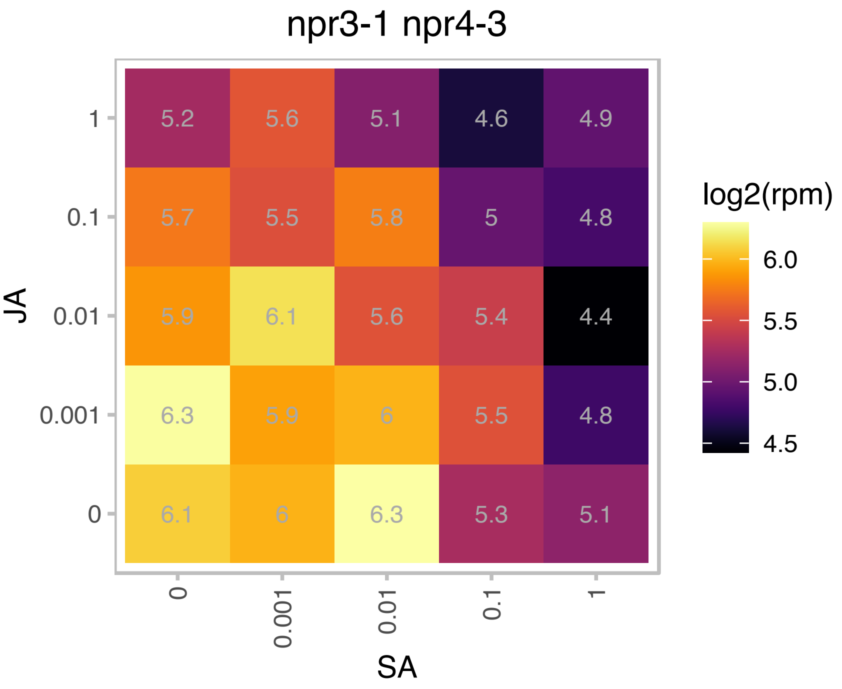

### AT4G15510.1 : PPD1

Photosystem II reaction center PsbP family protein

A nuclear-encoded thylakoid lumenal protein that is associated with photosystem I (PSI) but is not an integral subunit of PSI. It is an PSI assembly factor that assists the proper folding and integration of PsaB and PsaA into the thylakoid membrane. It is involved in the assembly of PSI reaction center heterodimer core.

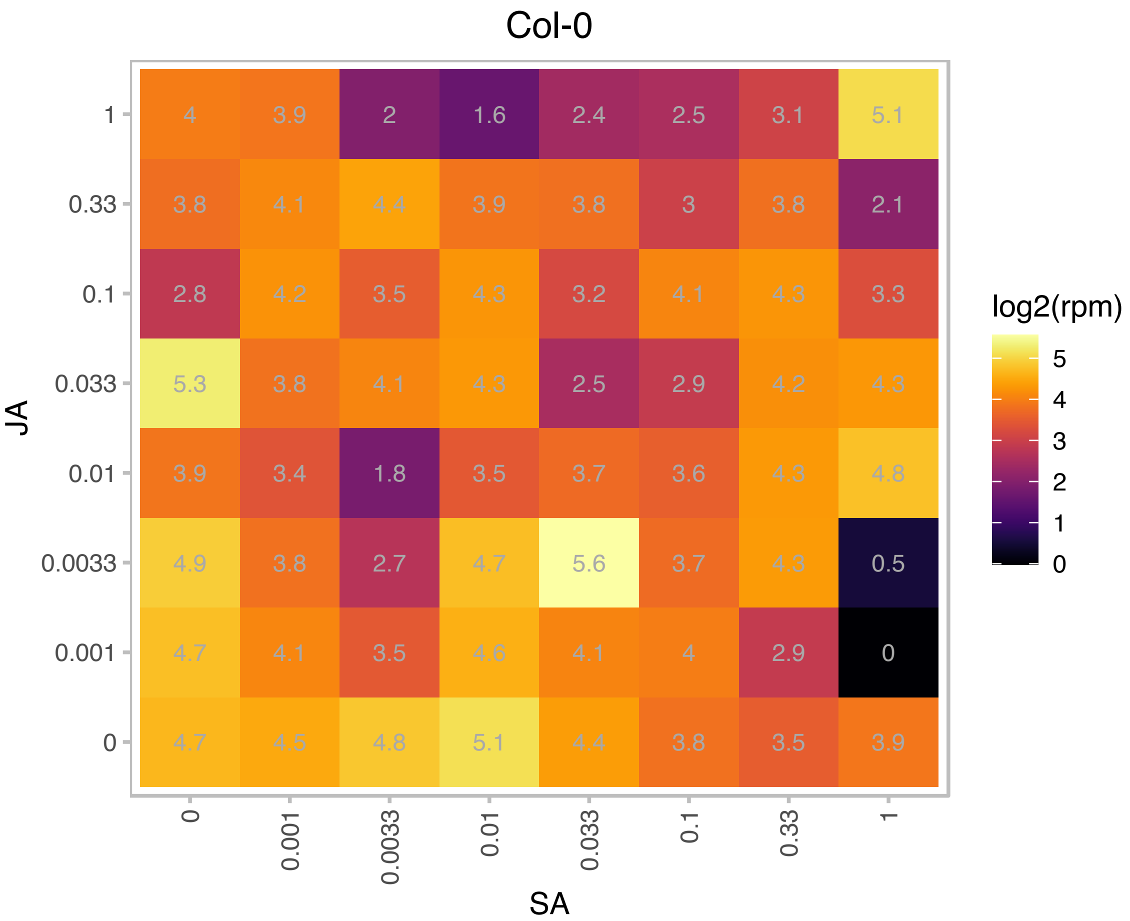


| | |
|---|---|
| Gene | AT4G15510 |
| is.expressed_genes | TRUE |
| is.responsive_genes | TRUE |
| is.apcluster | F-06 |

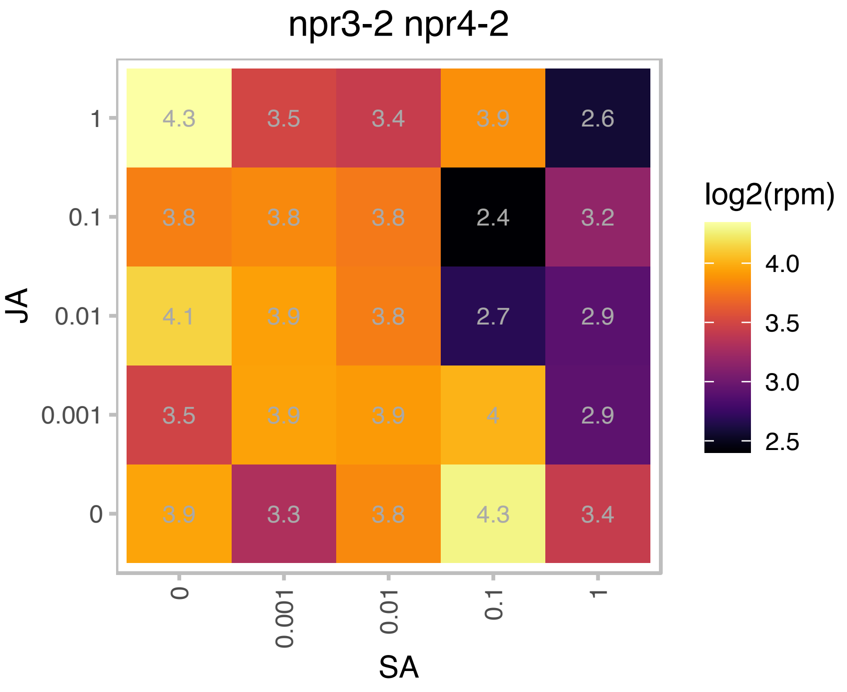


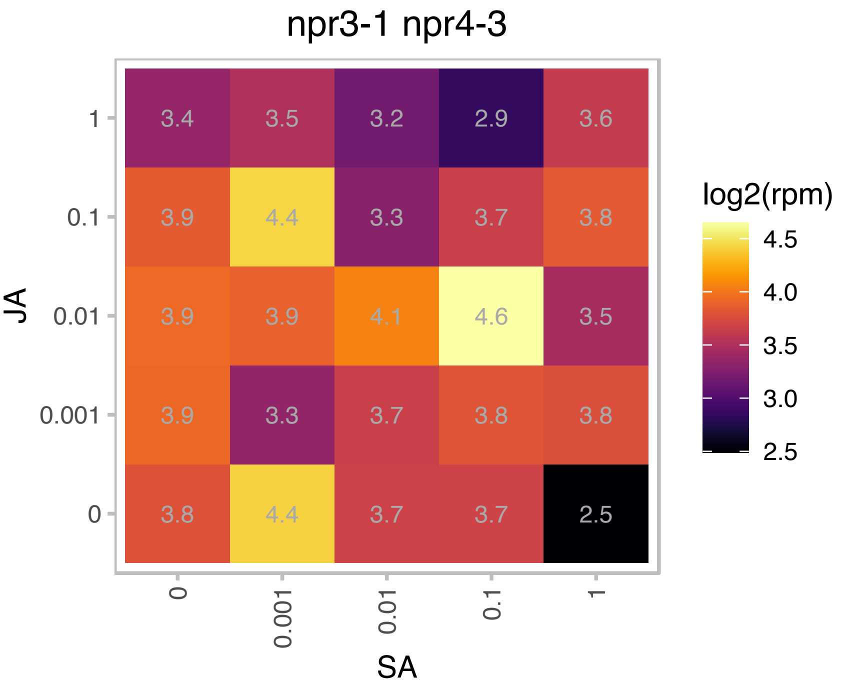

# AT4G17540.1 : NA

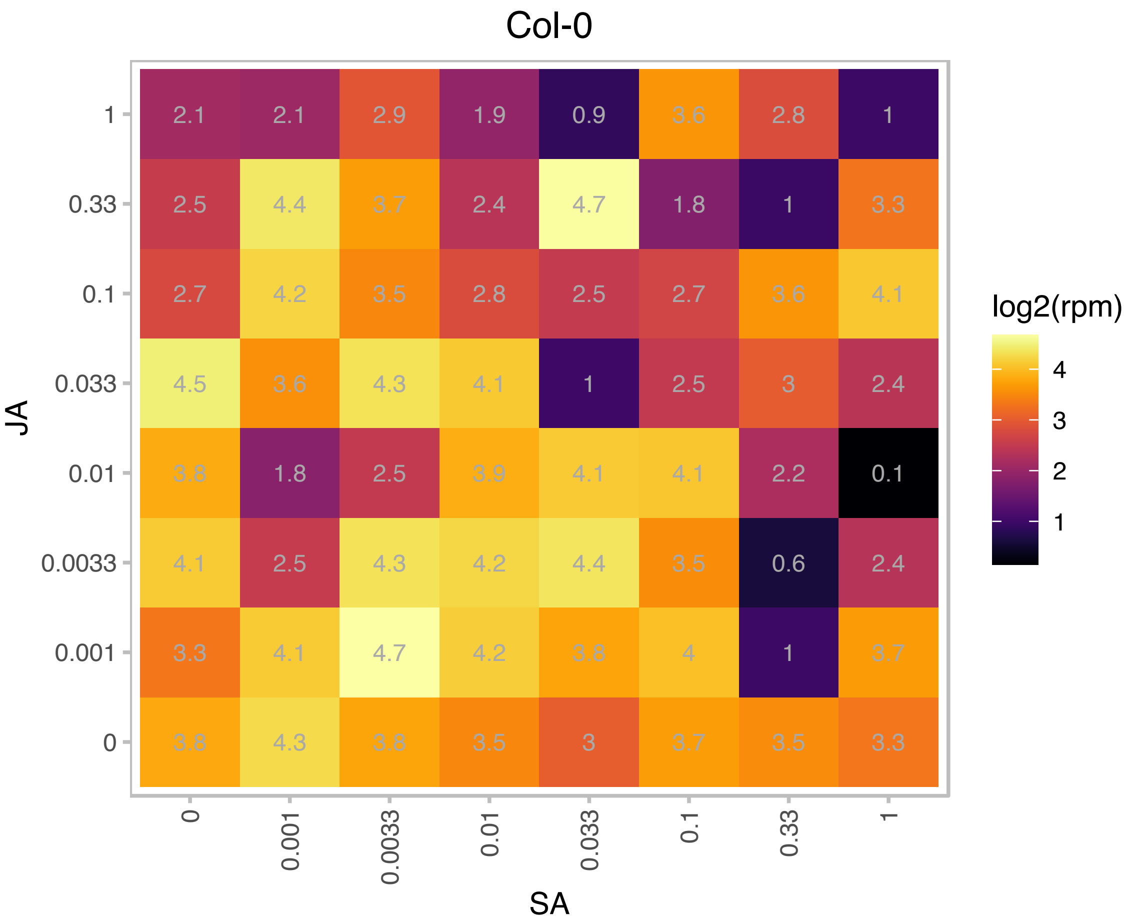


| | |
|---|---|
| Gene | AT4G17540 |
| is.expressed_genes | TRUE |
| is.responsive_genes | TRUE |
| is.apcluster | F-06 |

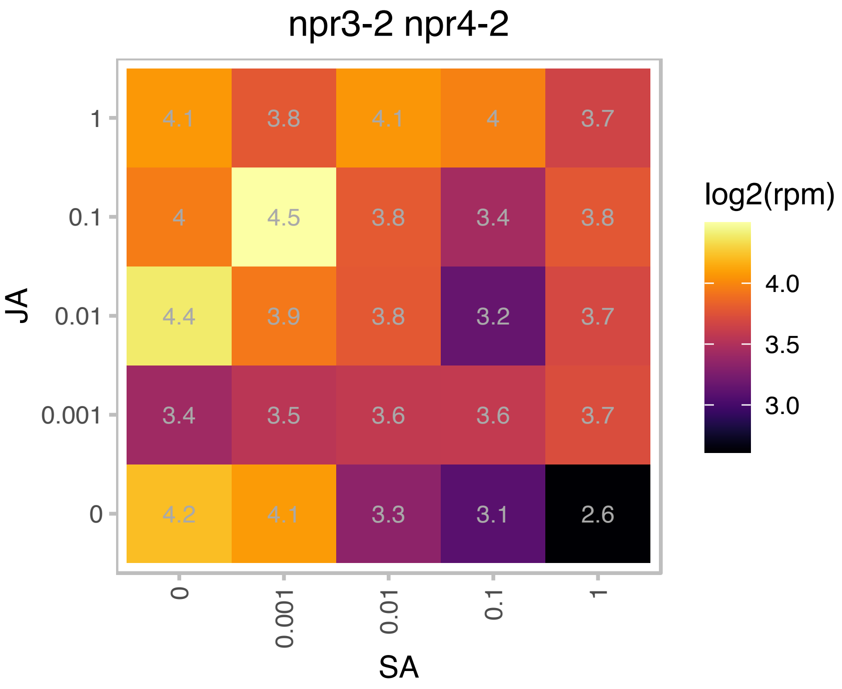


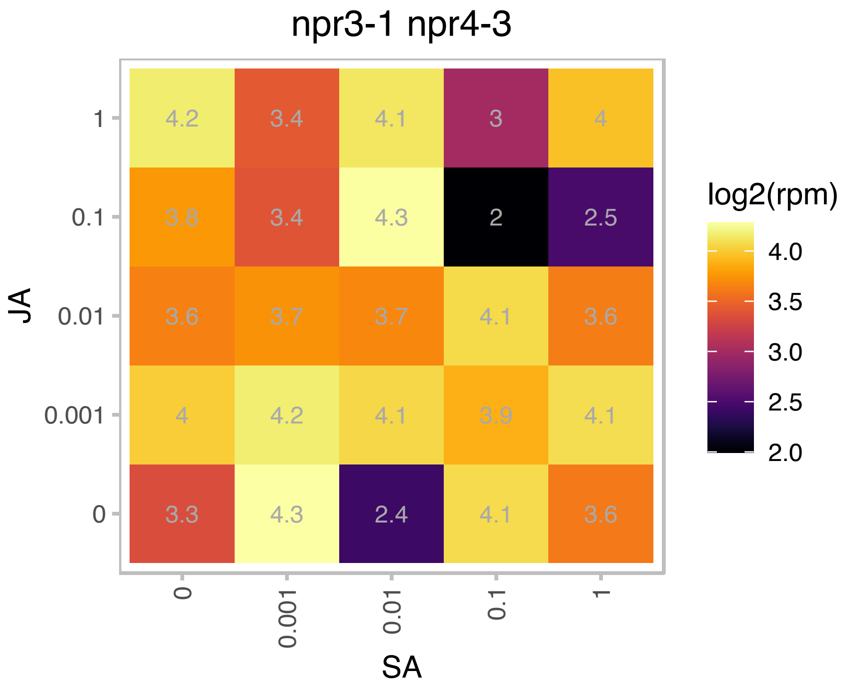

### AT4G25050.1 : ACP4, AtACP4

acyl carrier protein 4

encodes an acyl carrier protein predominantly expressed in leaves. Gene expression is upregulated by light.

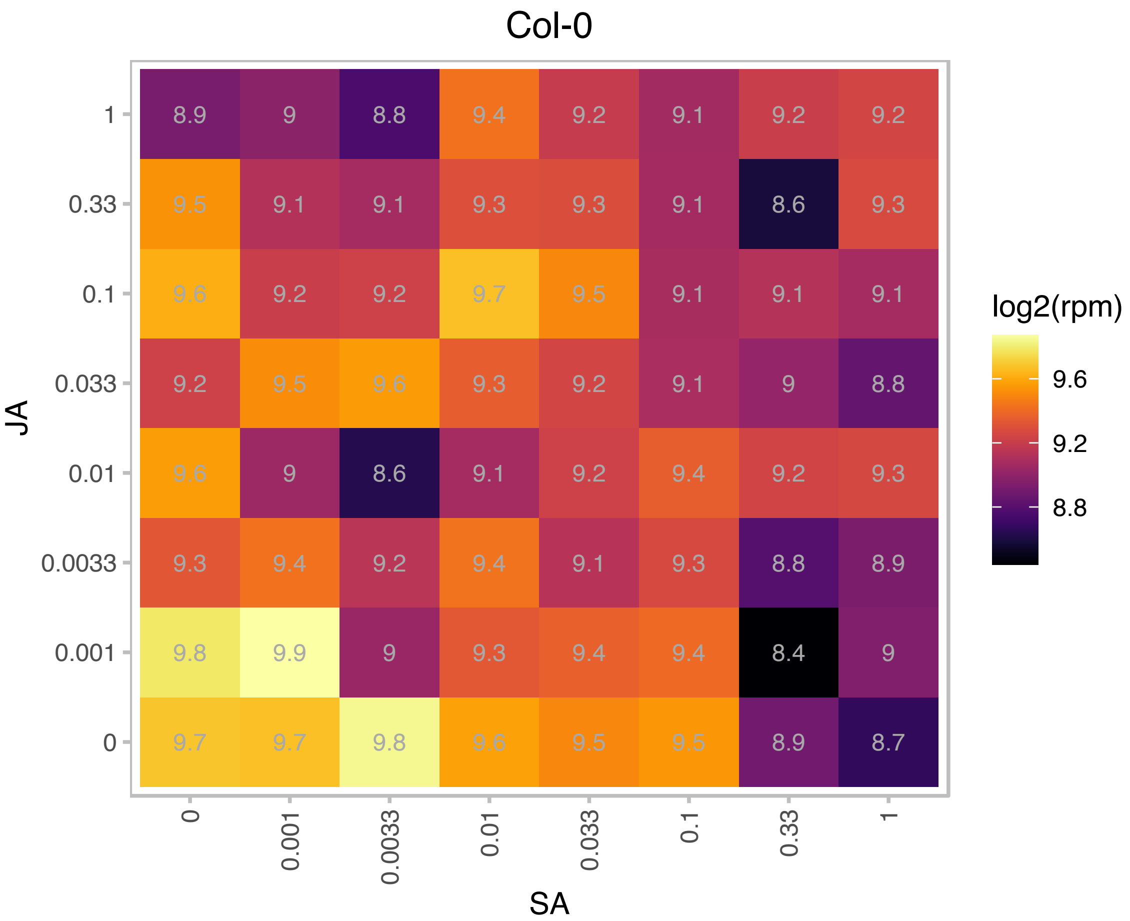


| | |
|---|---|
| Gene | AT4G25050 |
| is.expressed_genes | TRUE |
| is.responsive_genes | TRUE |
| is.apcluster | F-06 |

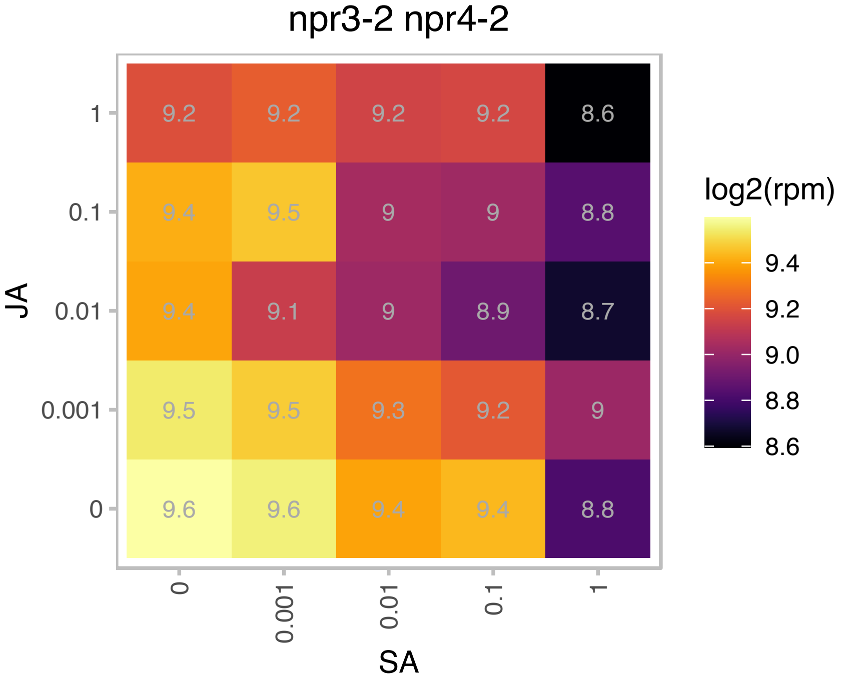


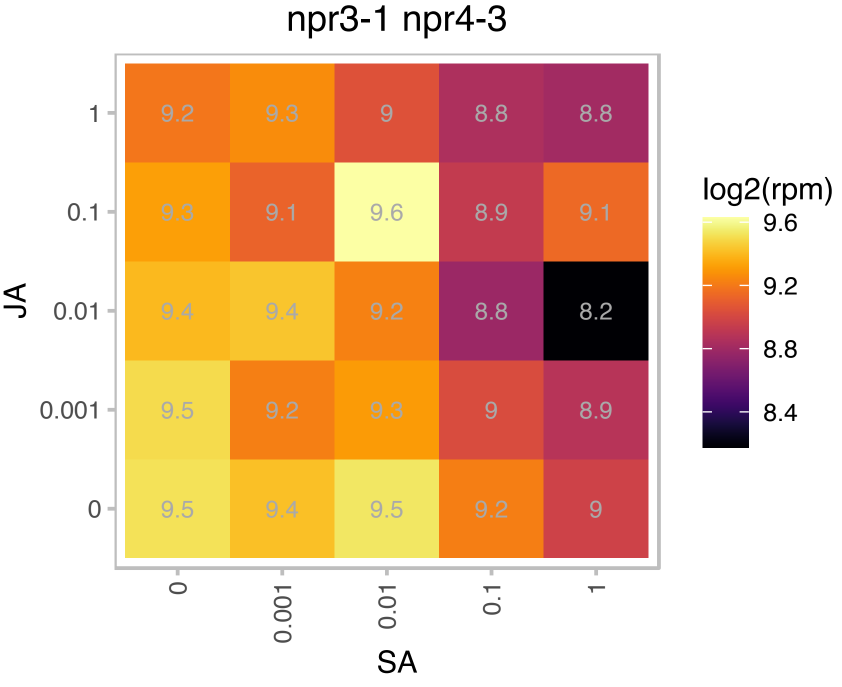

### AT4G25080.1 : CHLM

magnesium-protoporphyrin IX methyltransferase

Encodes a protein with methyltransferase activity responsible for the methylation of magnesium protoporphyrin IX. Mutants defective in this gene are affected in chlorophyll biosynthesis and show a reduction in the accumulation of a number of major thylakoid-associated proteins including components of PSI (LHCI), PSII (LHCII, D1, CP43) and the cytochrome b6f complex (Cytf). By contrast, no significant changes were detected for the proteins of the stroma and the chloroplast envelope.

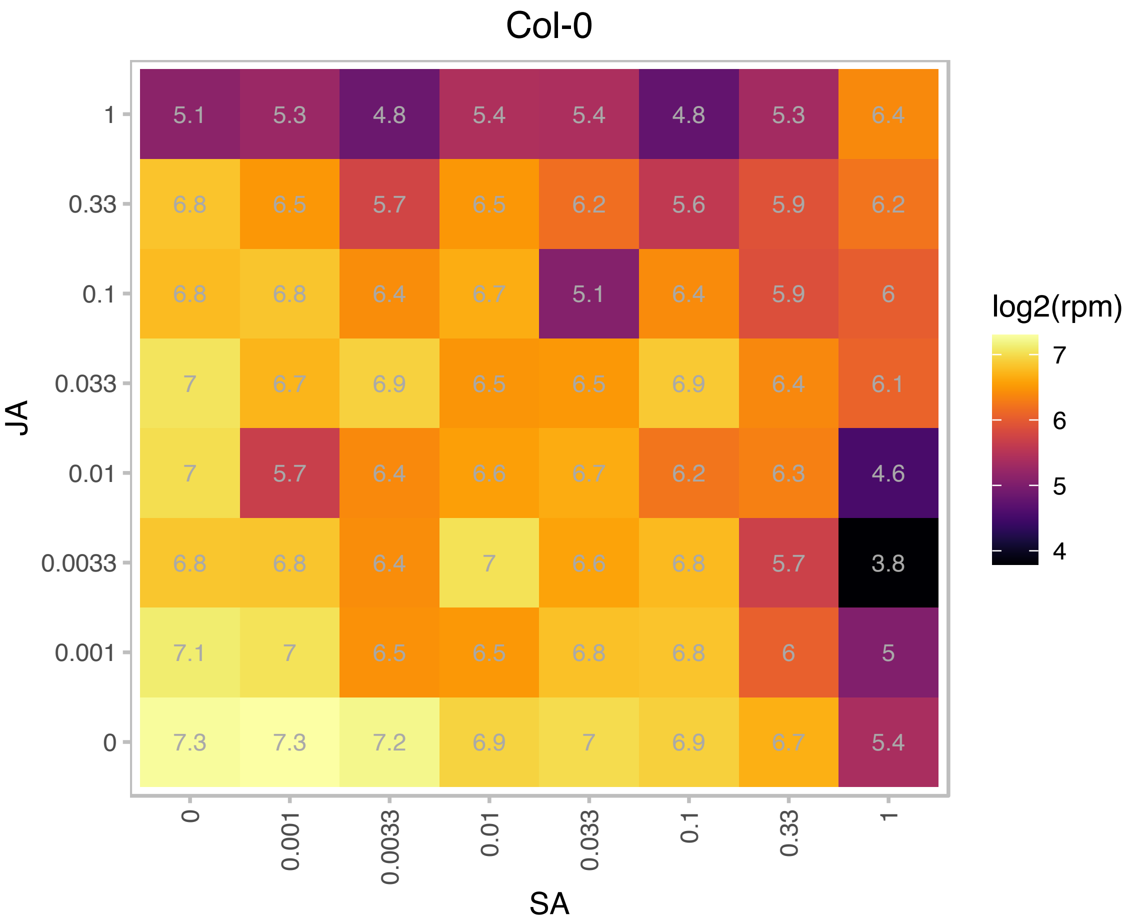


| | |
|---|---|
| Gene | AT4G25080 |
| is.expressed_genes | TRUE |
| is.responsive_genes | TRUE |
| is.apcluster | F-06 |

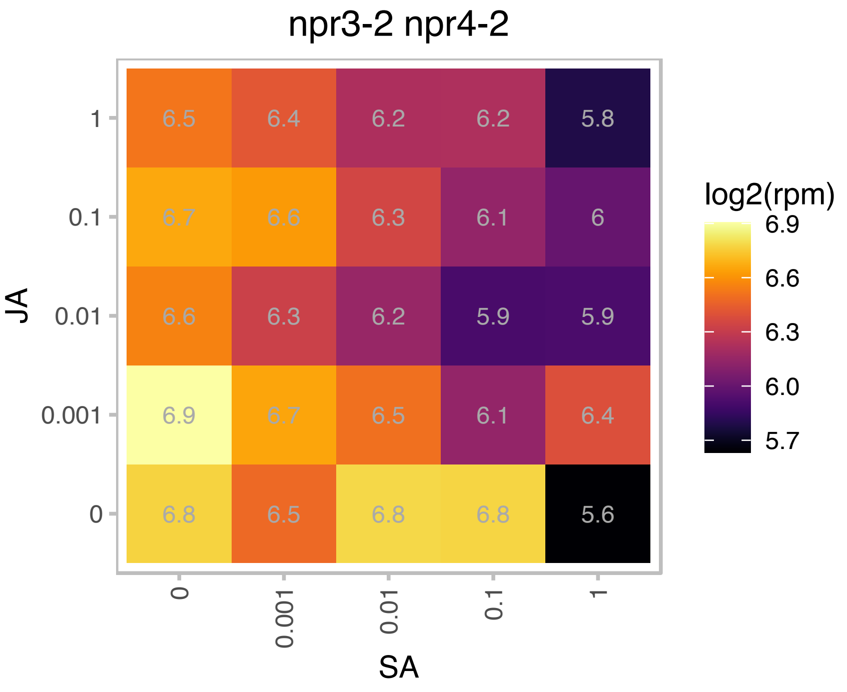


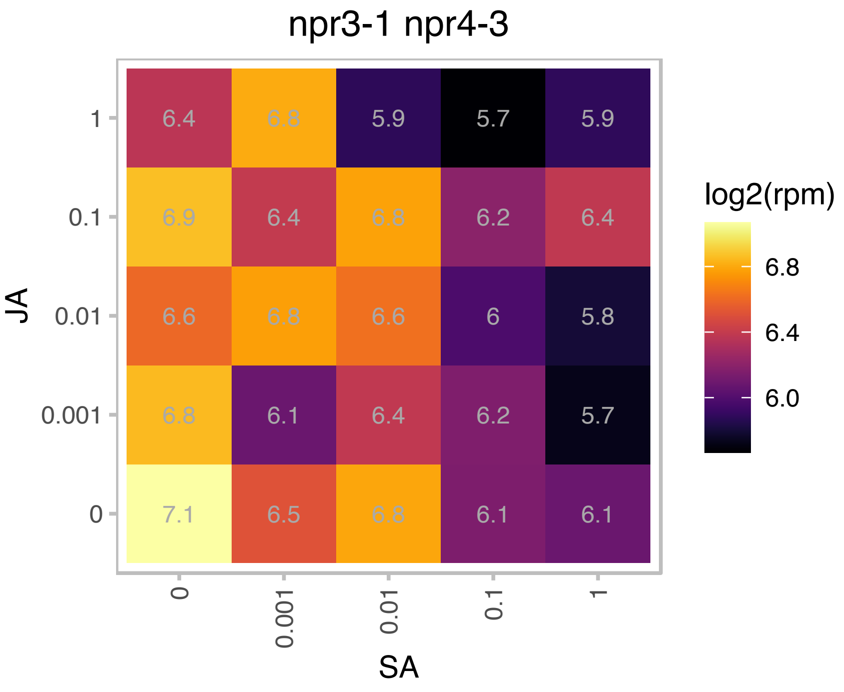

# AT4G33560.1 : NA

Wound-responsive family protein

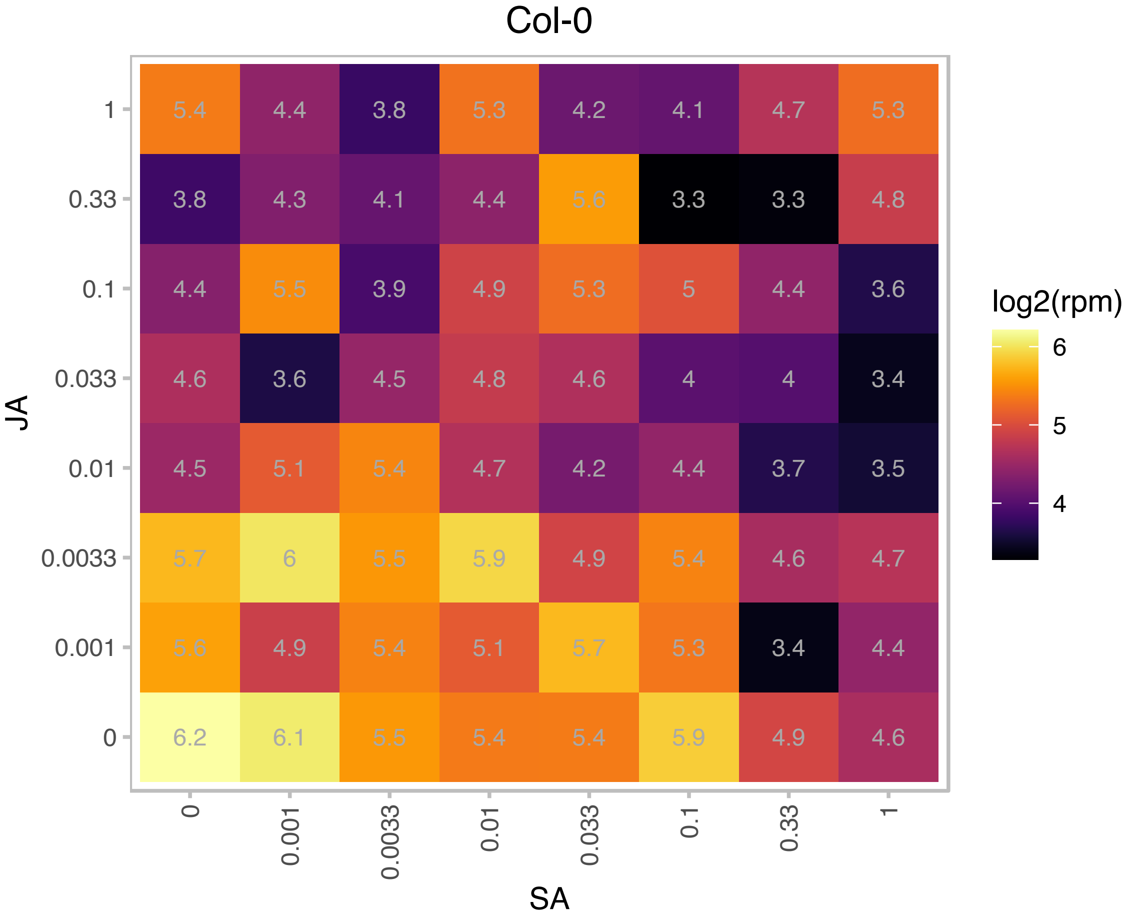


| | |
|---|---|
| Gene | AT4G33560 |
| is.expressed_genes | TRUE |
| is.responsive_genes | TRUE |
| is.apcluster | F-06 |

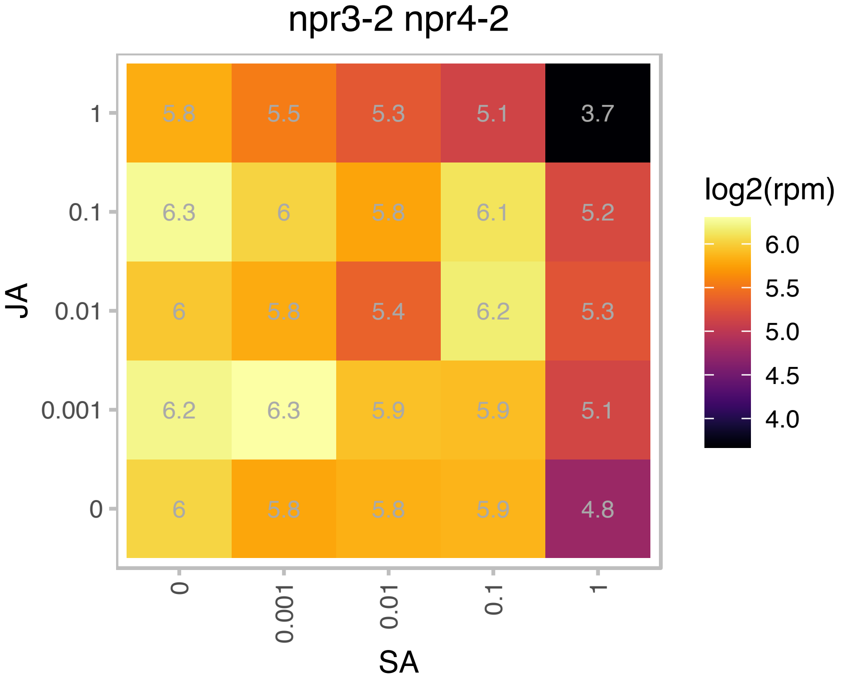


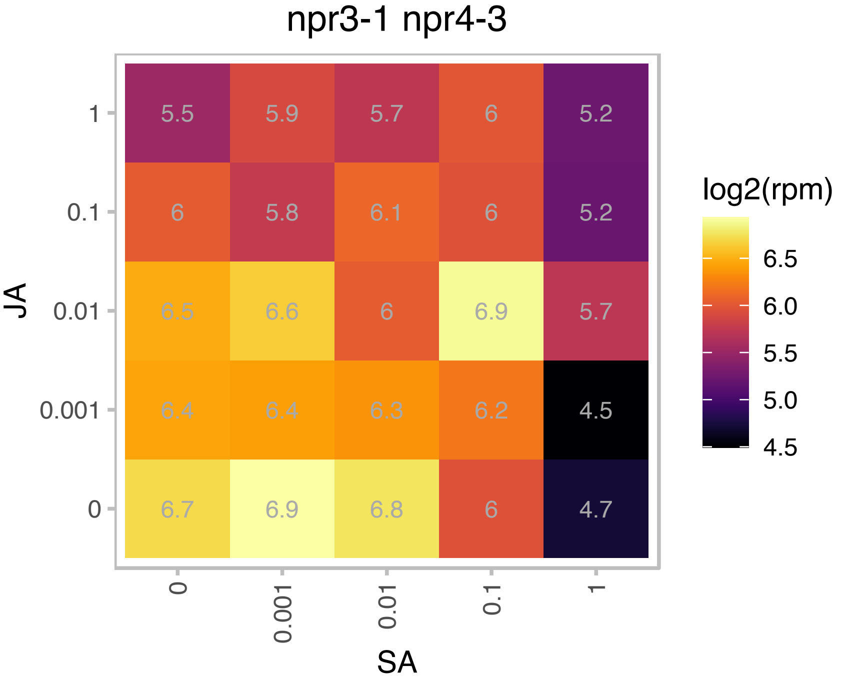

# AT4G33670.1 : NA

NAD(P)-linked oxidoreductase superfamily protein

Encodes a L-galactose dehydrogenase, involved in ascorbate biosynthesis

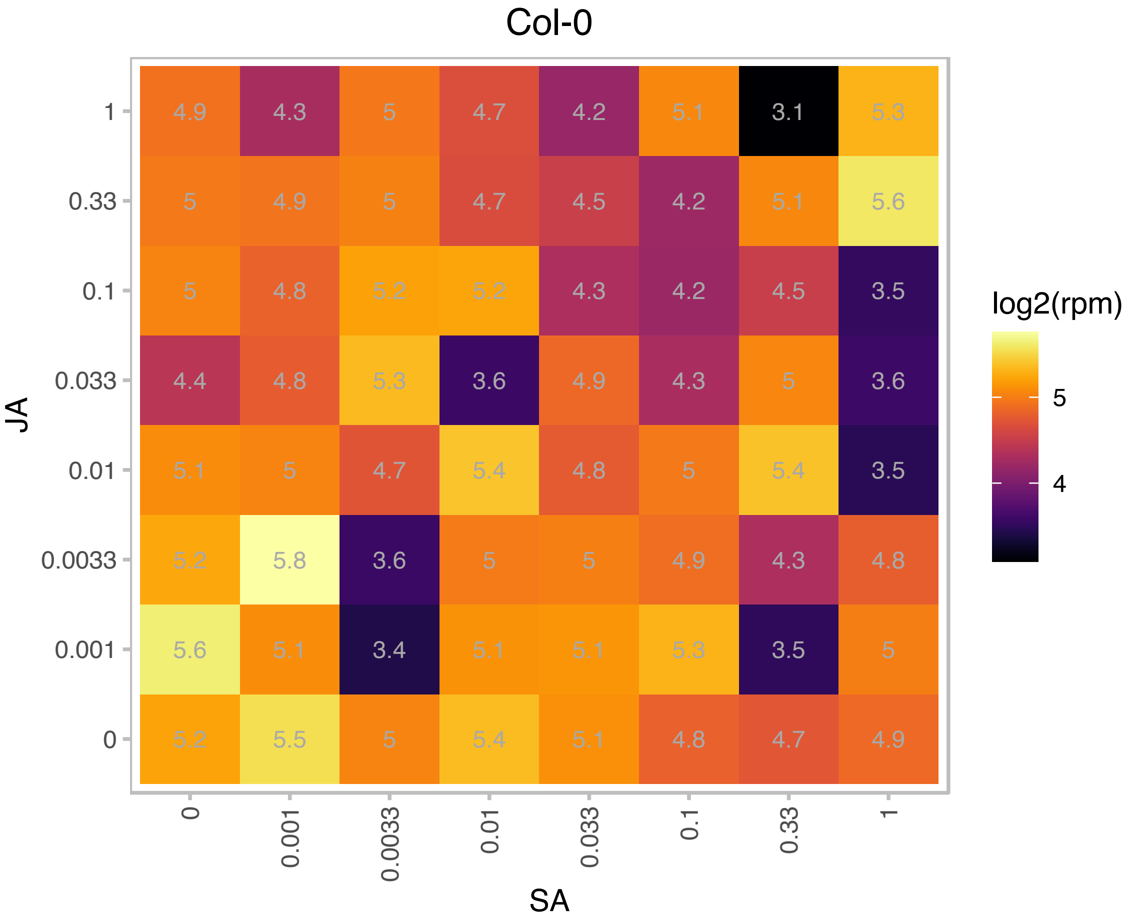


| | |
|---|---|
| Gene | AT4G33670 |
| is.expressed_genes | TRUE |
| is.responsive_genes | TRUE |
| is.apcluster | F-06 |

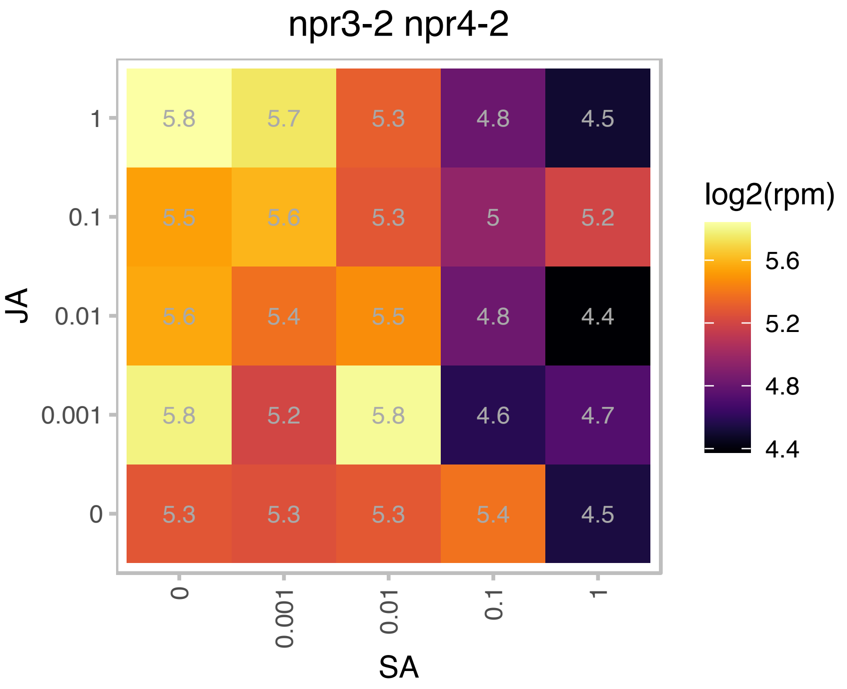


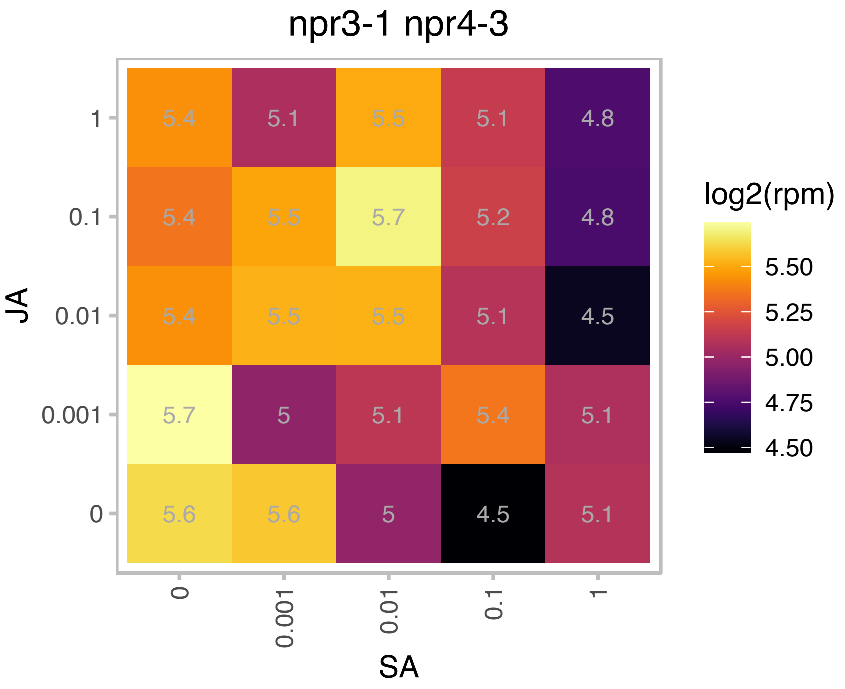

### AT4G34090.1 : PAB

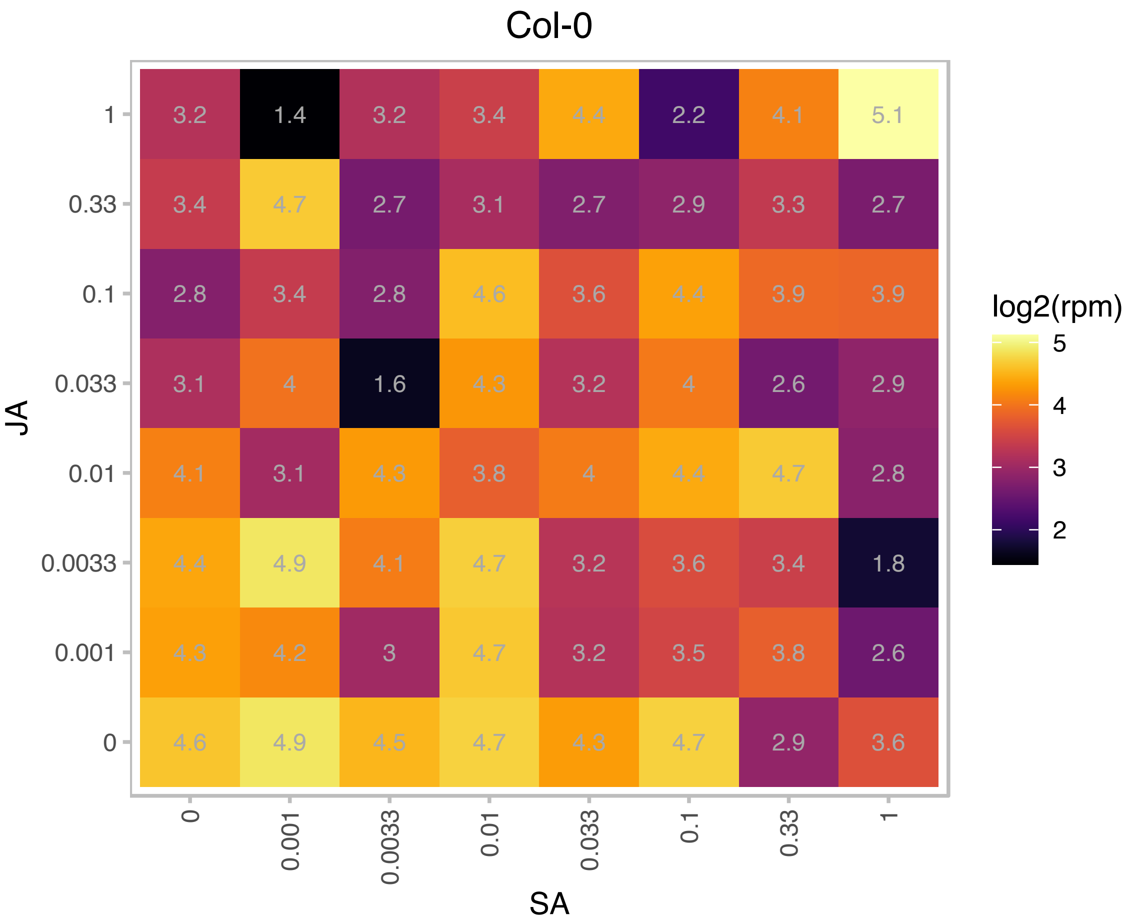


| | |
|---|---|
| Gene | AT4G34090 |
| is.expressed_genes | TRUE |
| is.responsive_genes | TRUE |
| is.apcluster | F-06 |

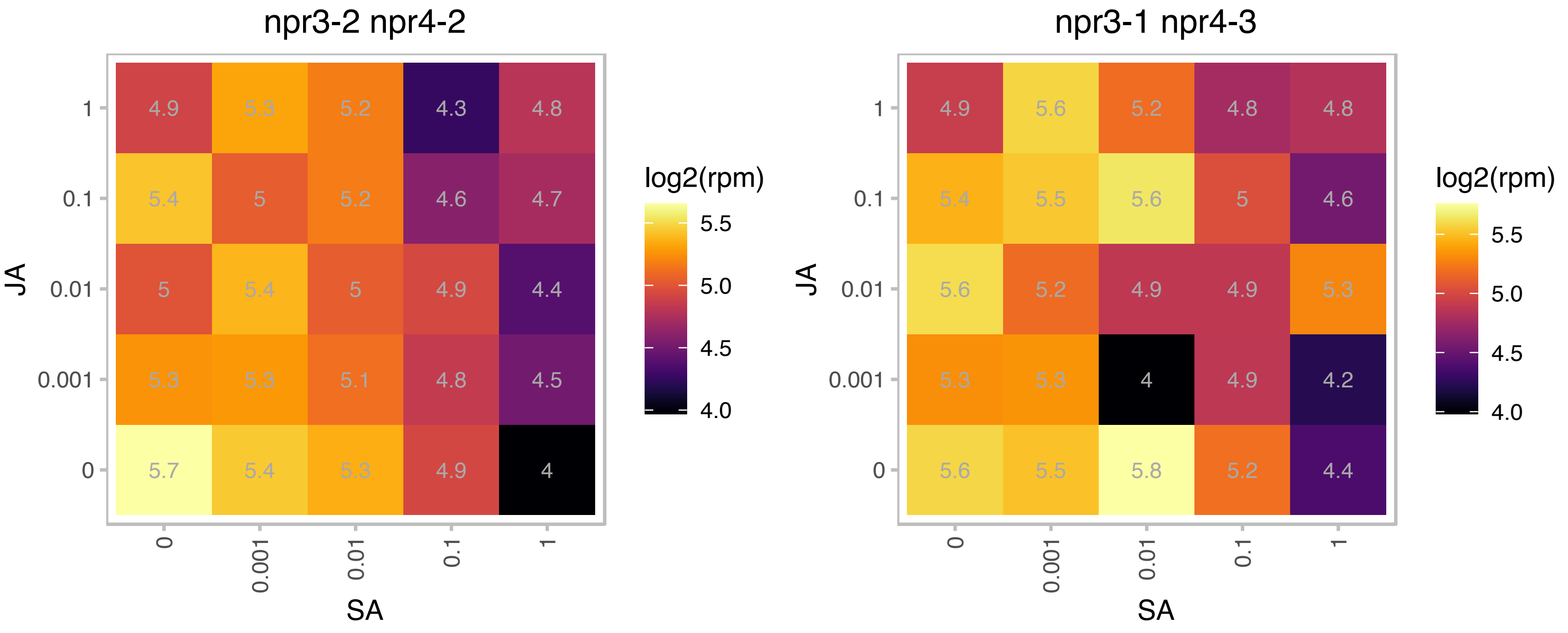

### AT4G34770.1 : SAUR1

SAUR-like auxin-responsive protein family

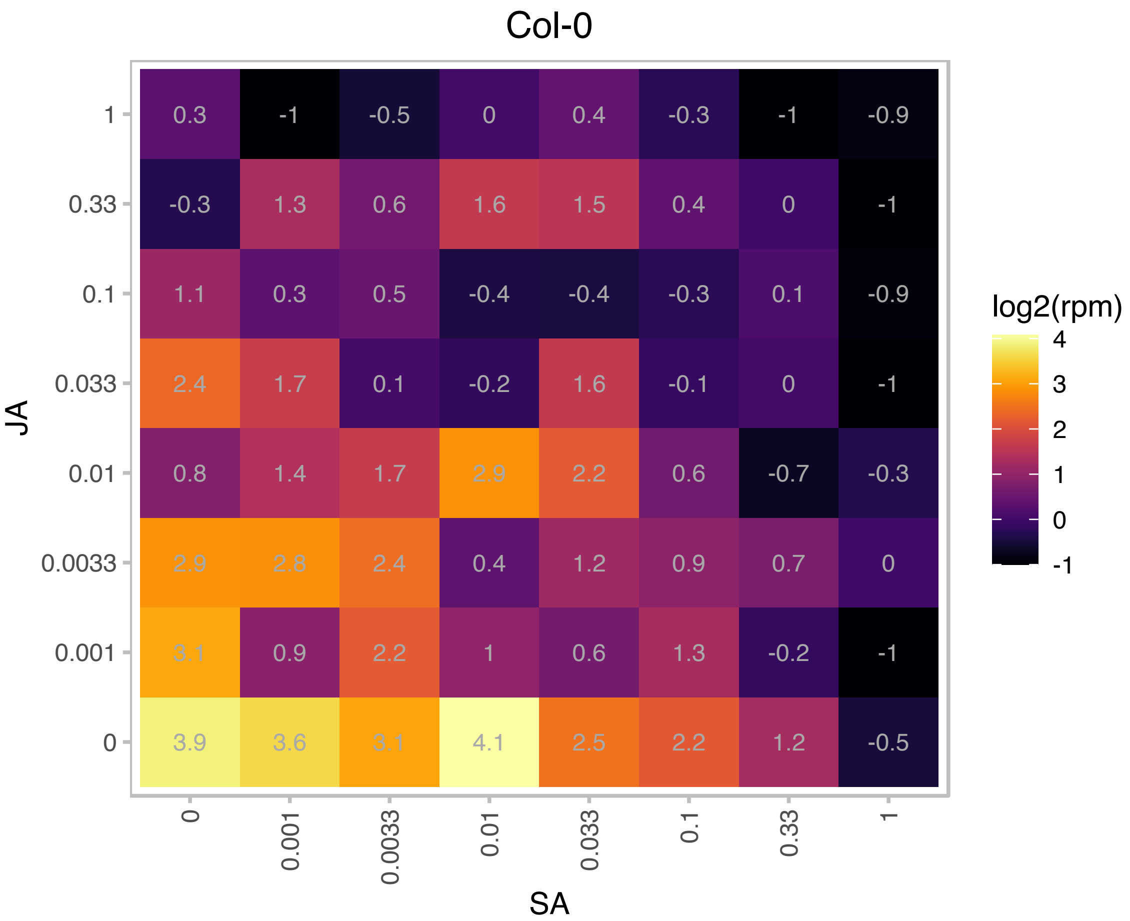


| | |
|---|---|
| Gene | AT4G34770 |
| is.expressed_genes | TRUE |
| is.responsive_genes | TRUE |
| is.apcluster | F-06 |

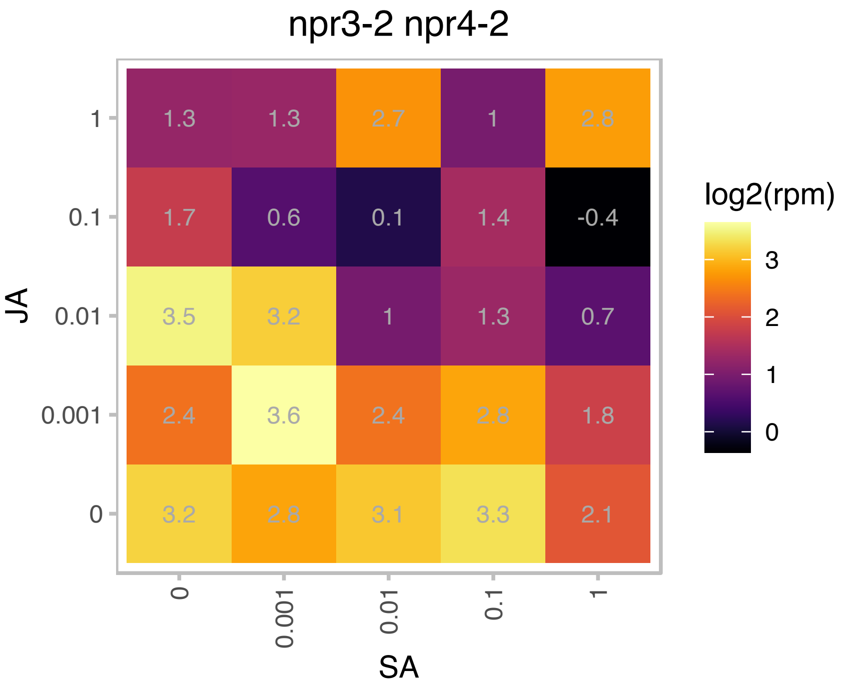


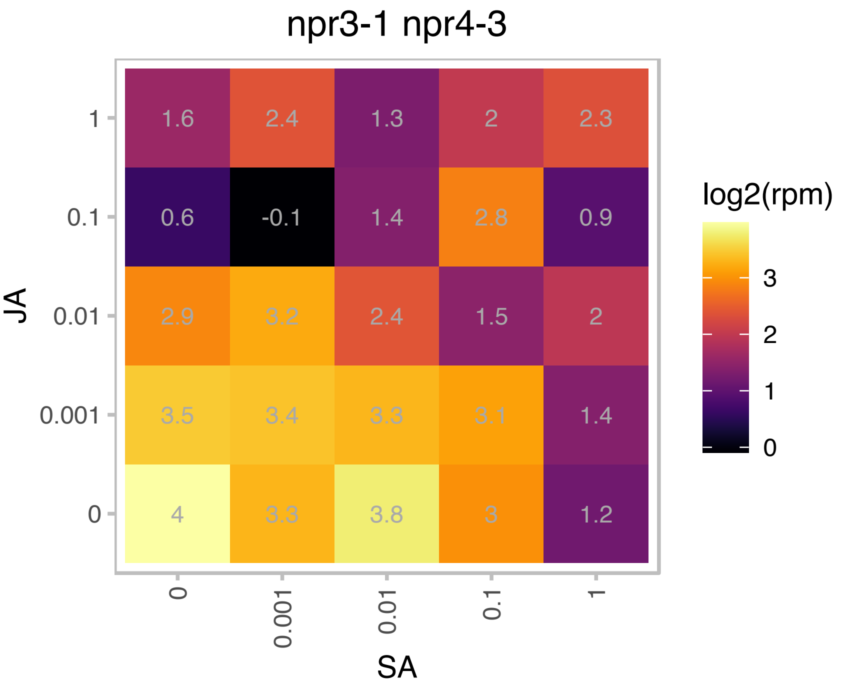

### AT4G37800.1 : XTH7

xyloglucan endotransglucosylase/hydrolase 7

Col-0

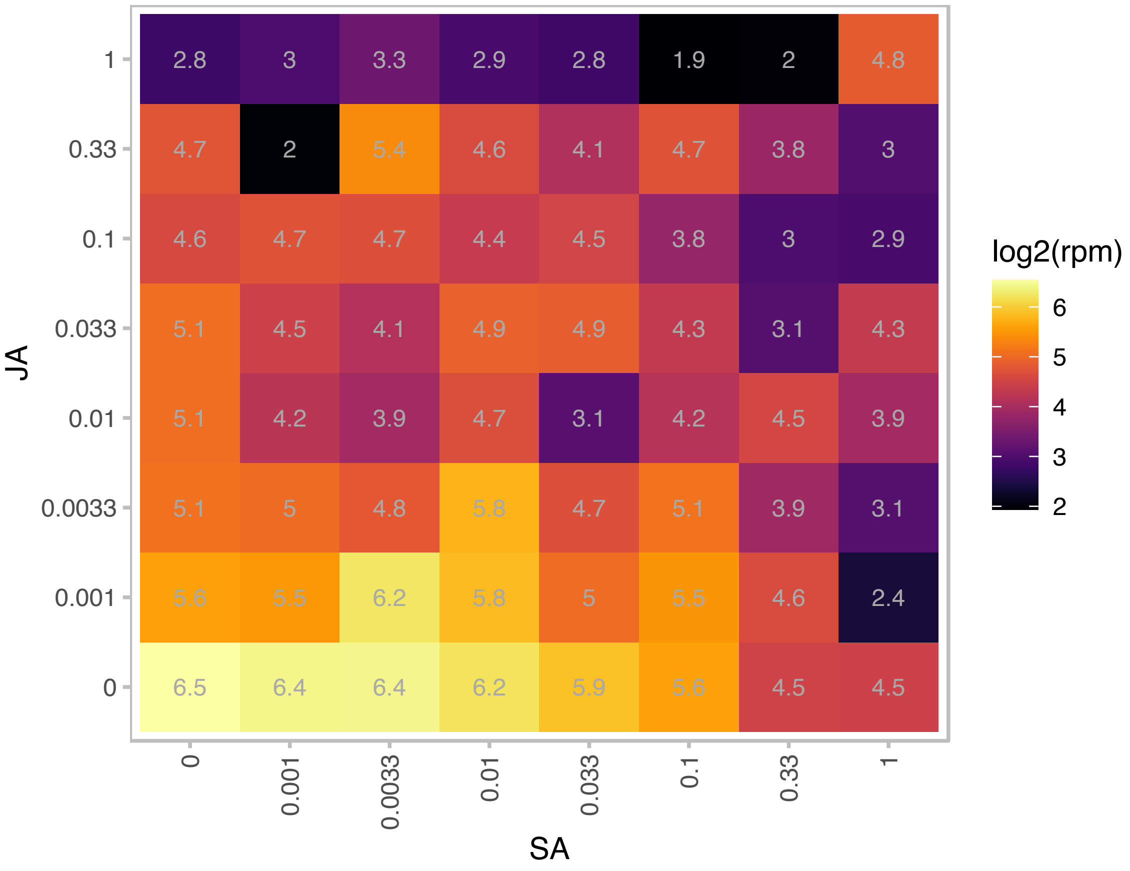


| | |
|---|---|
| Gene | AT4G37800 |
| is.expressed_genes | TRUE |
| is.responsive_genes | TRUE |
| is.apcluster | F-06 |

npr3-2 npr4-2

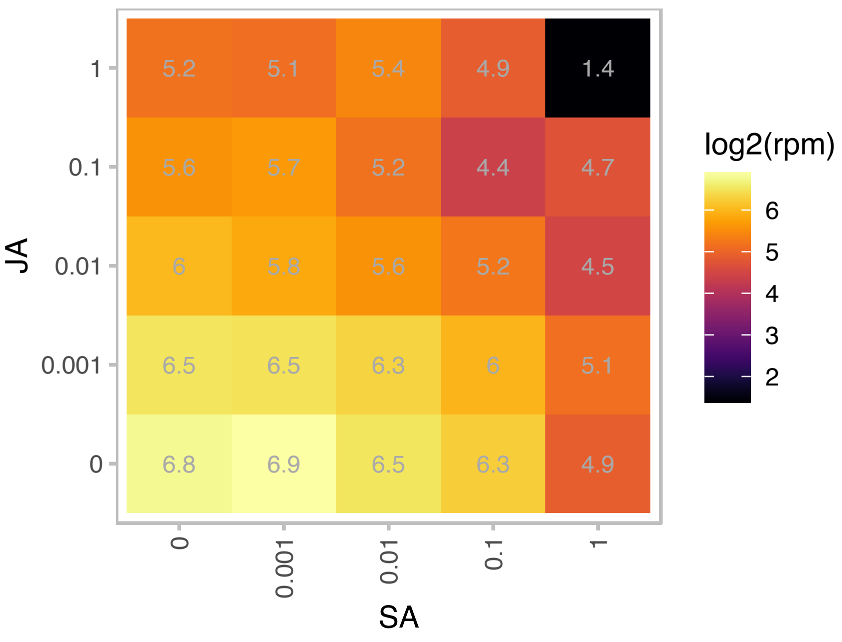


npr3-1 npr4-3

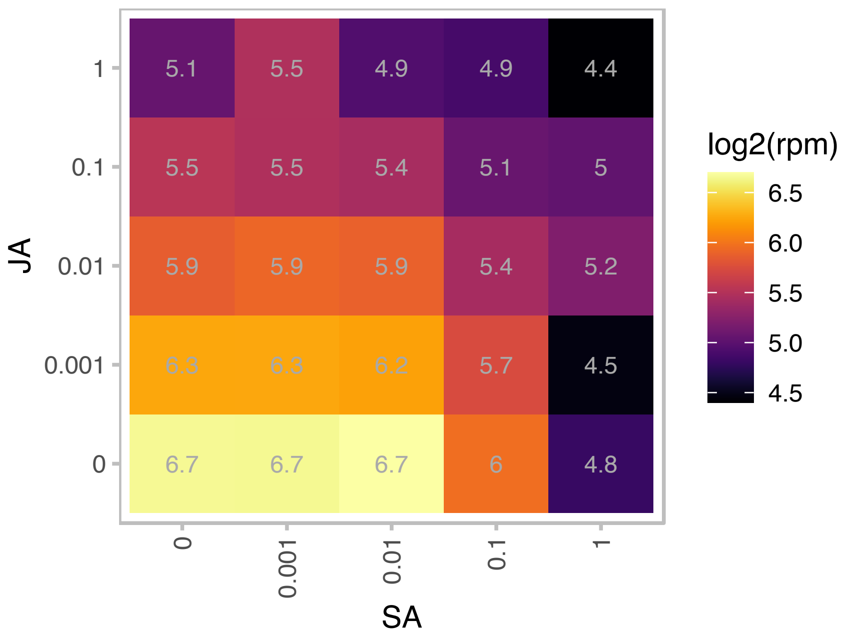

# AT4G37920.1 : NA

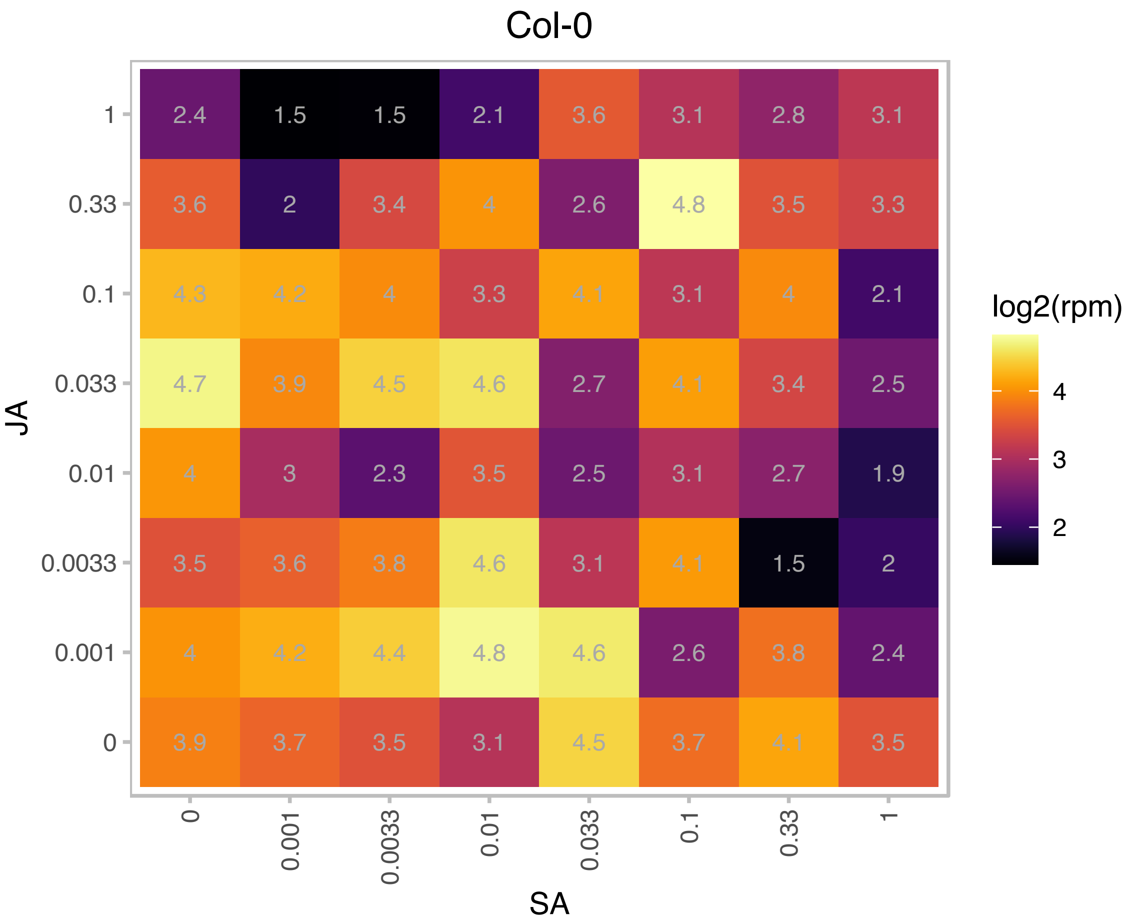


| | |
|---|---|
| Gene | AT4G37920 |
| is.expressed_genes | TRUE |
| is.responsive_genes | TRUE |
| is.apcluster | F-06 |

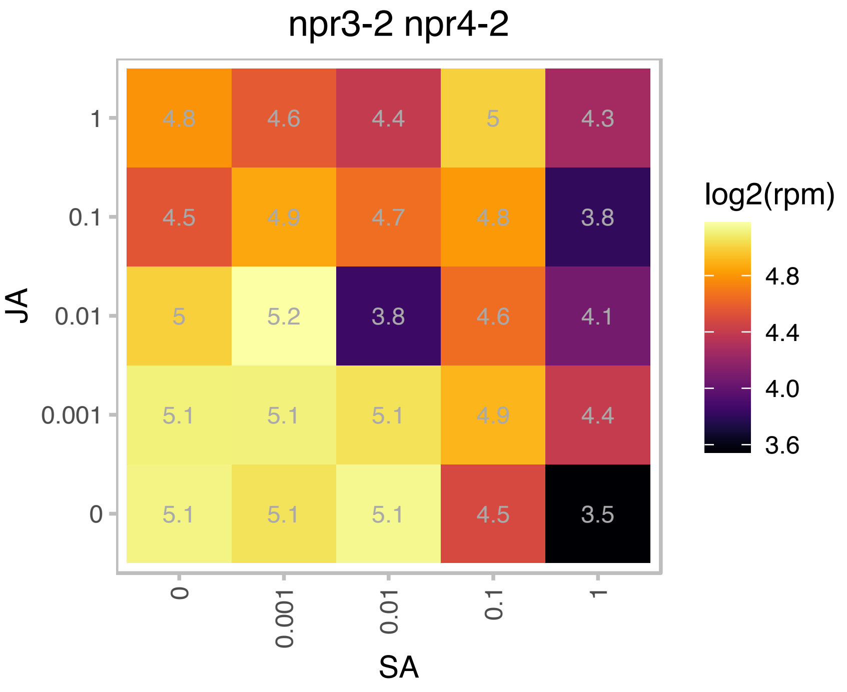


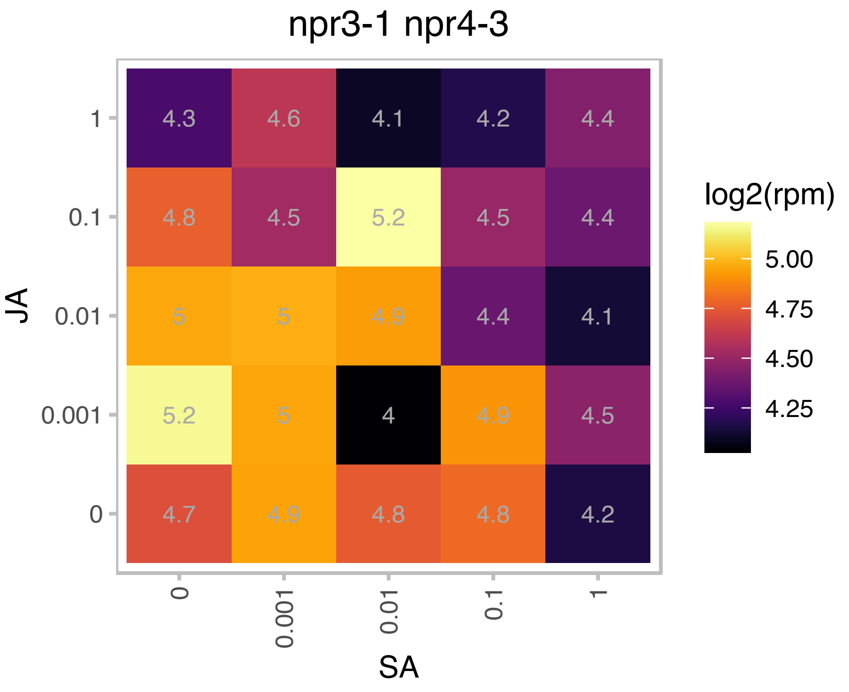

### AT5G04950.1 : ATNAS1, NAS1

nicotianamine synthase 1

Encodes a nicotianamide synthase.

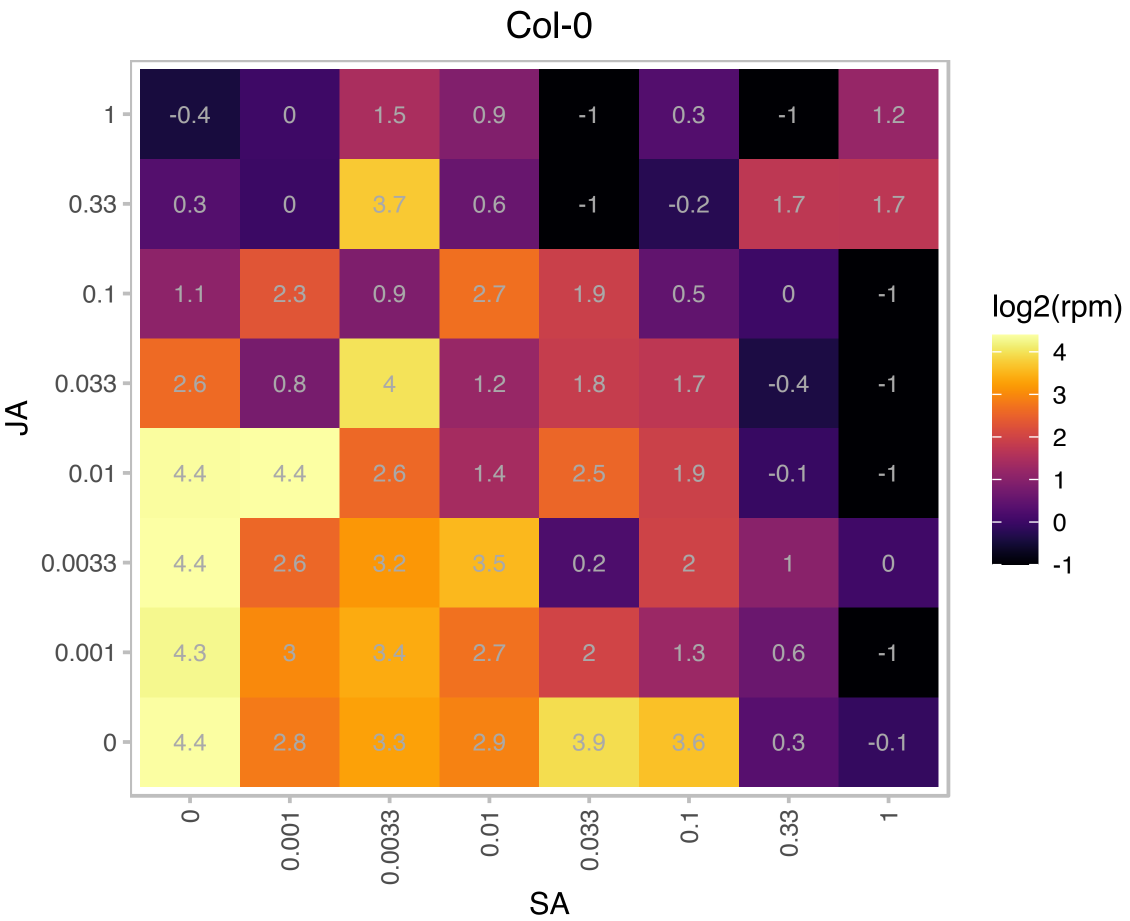


| | |
|---|---|
| Gene | AT5G04950 |
| is.expressed_genes | TRUE |
| is.responsive_genes | TRUE |
| is.apcluster | F-06 |

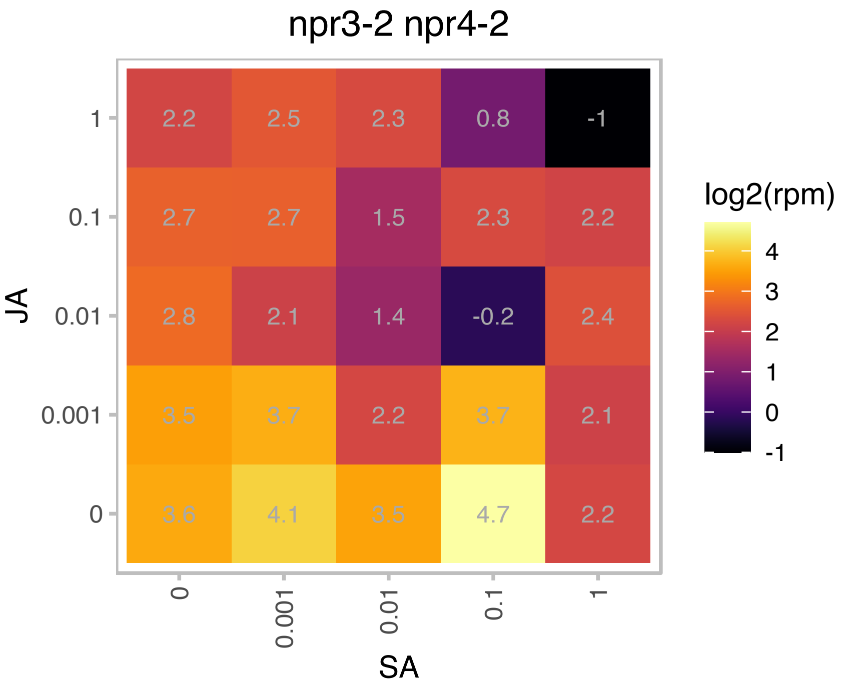


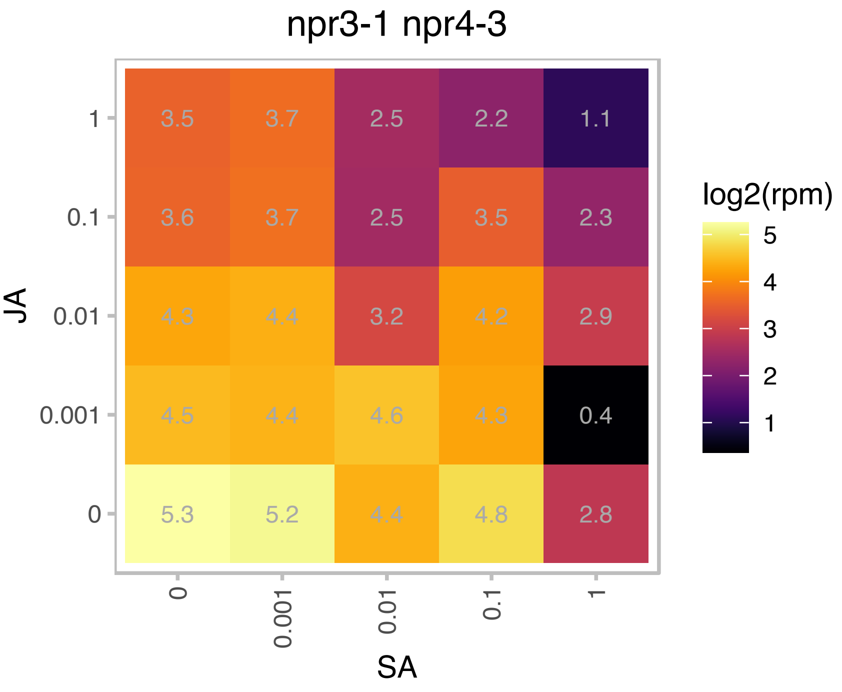

# AT5G05960.1 : NA

Bifunctional inhibitor/lipid-transfer protein/seed storage 2S albumin superfamily protein

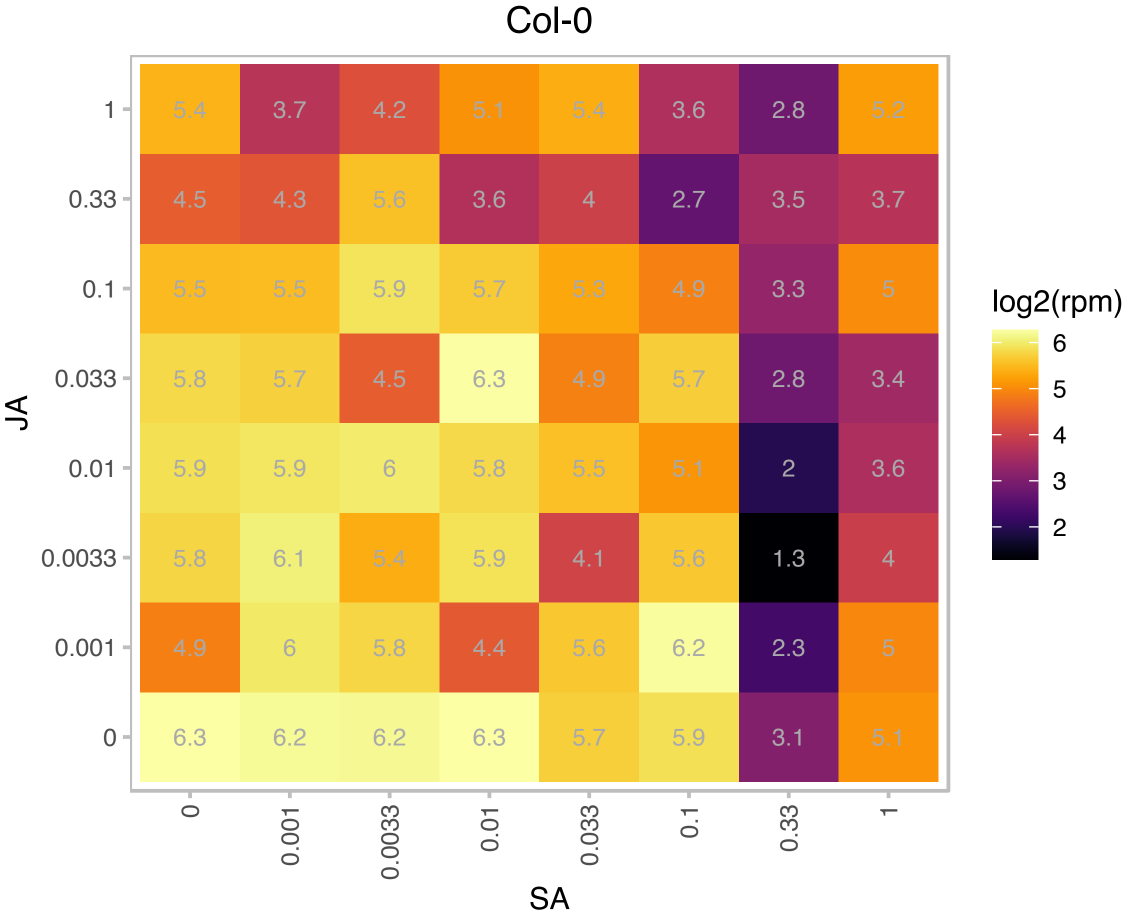


| | |
|---|---|
| Gene | AT5G05960 |
| is.expressed_genes | TRUE |
| is.responsive_genes | TRUE |
| is.apcluster | F-06 |

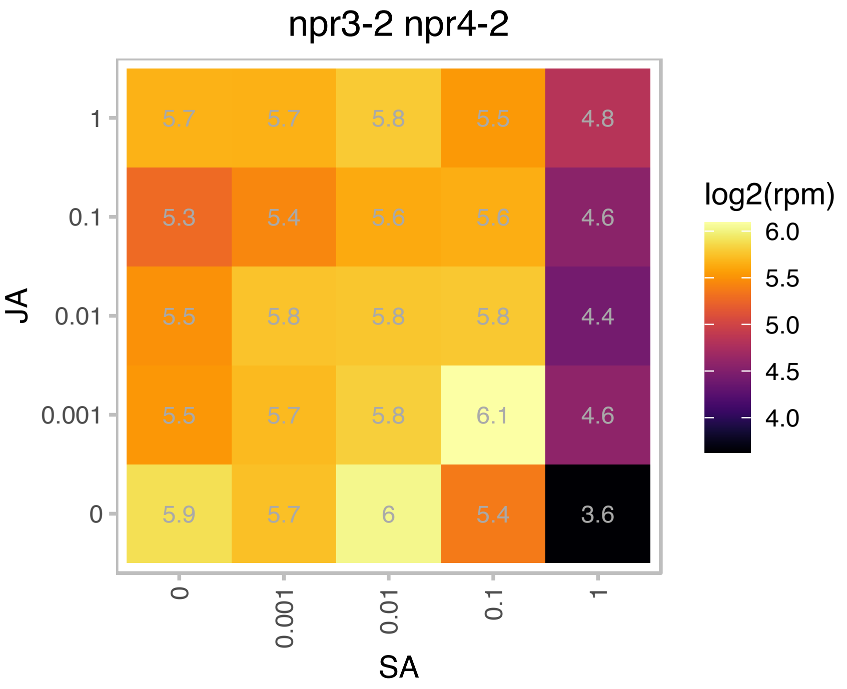


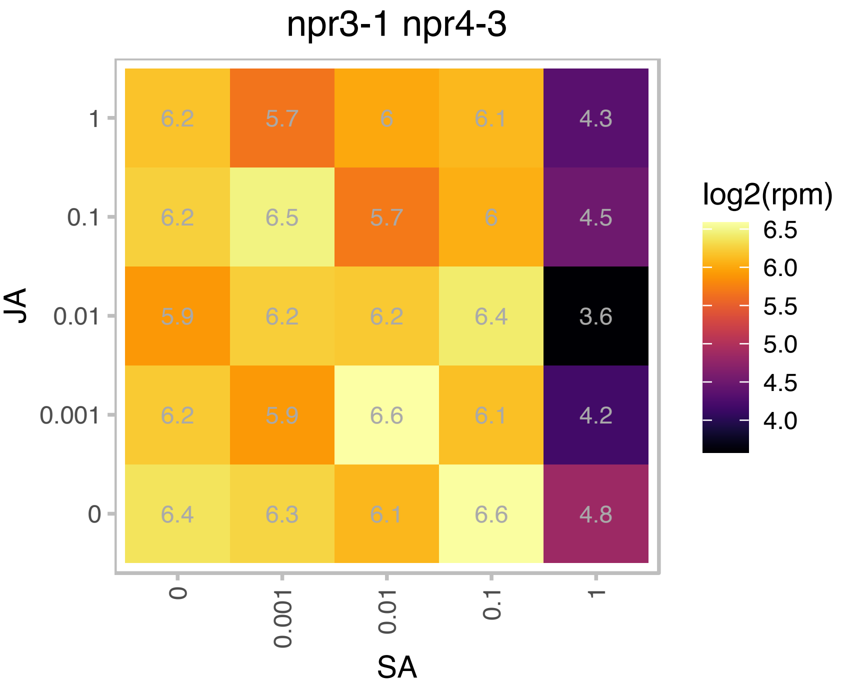

### AT5G17170.1 : ENH1

rubredoxin family protein

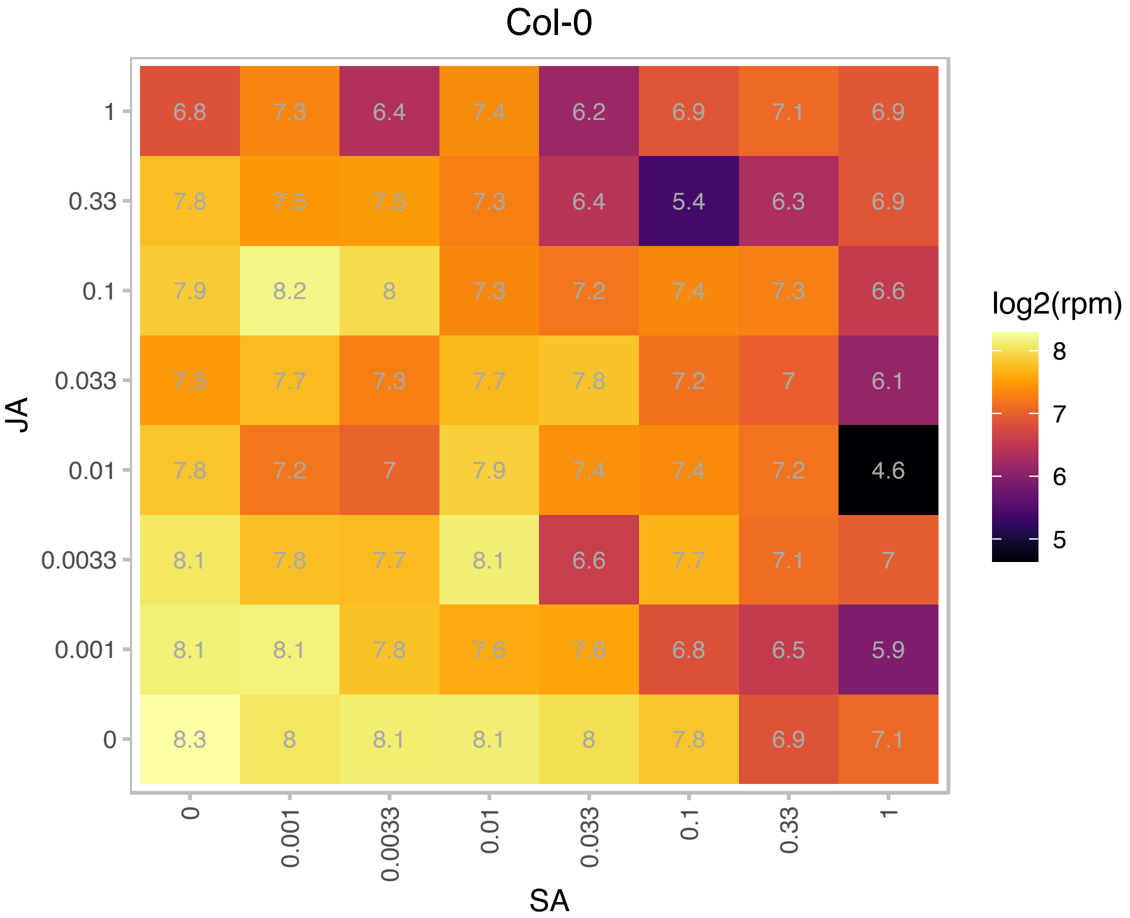


| | |
|---|---|
| Gene | AT5G17170 |
| is.expressed_genes | TRUE |
| is.responsive_genes | TRUE |
| is.apcluster | F-06 |

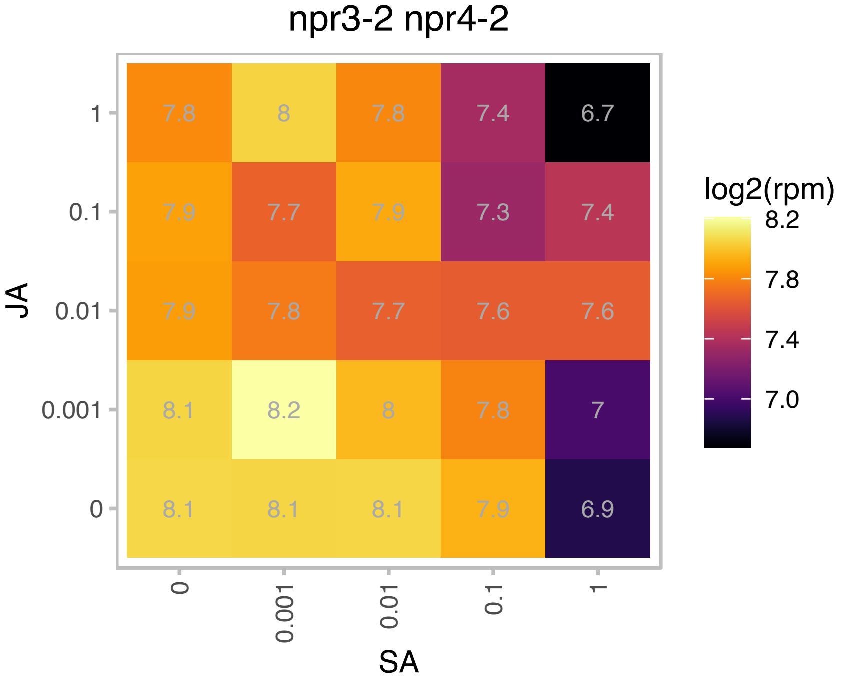


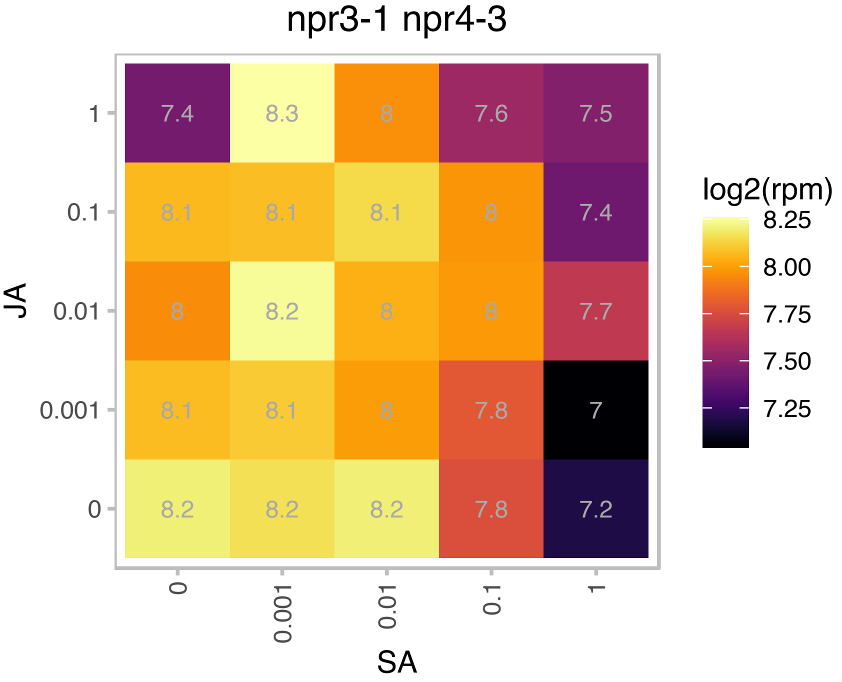

# AT5G17460.1 : NA

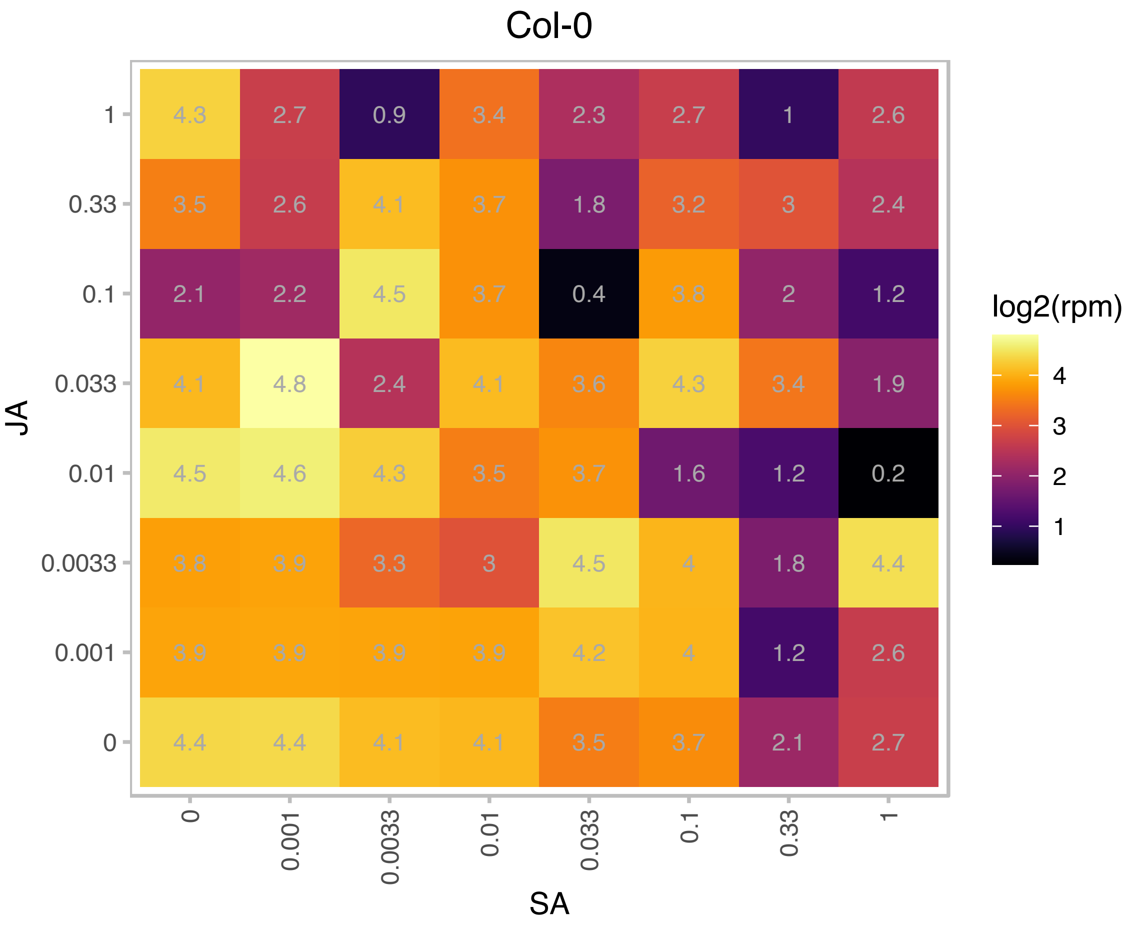


| | |
|---|---|
| Gene | AT5G17460 |
| is.expressed_genes | TRUE |
| is.responsive_genes | TRUE |
| is.apcluster | F-06 |

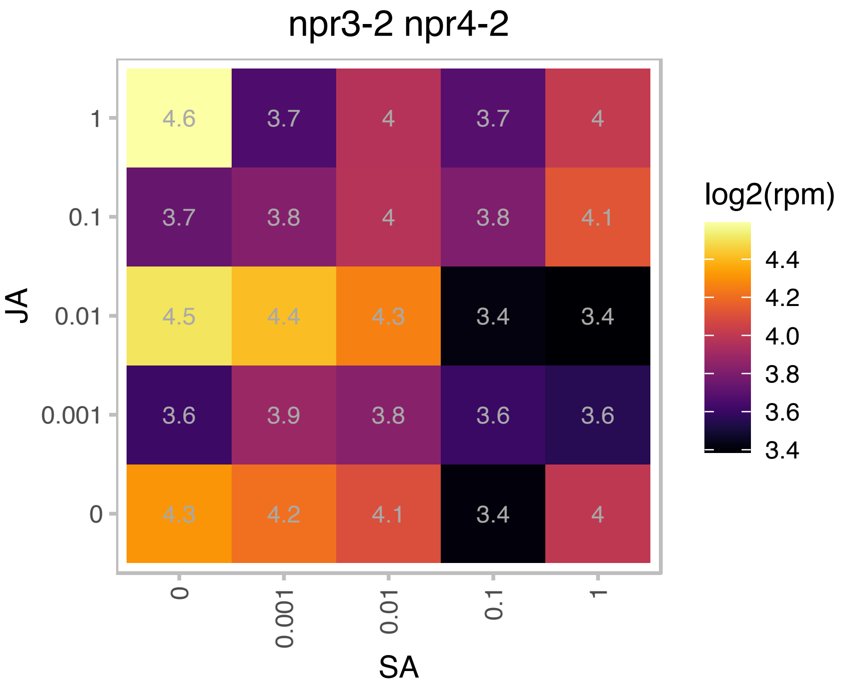


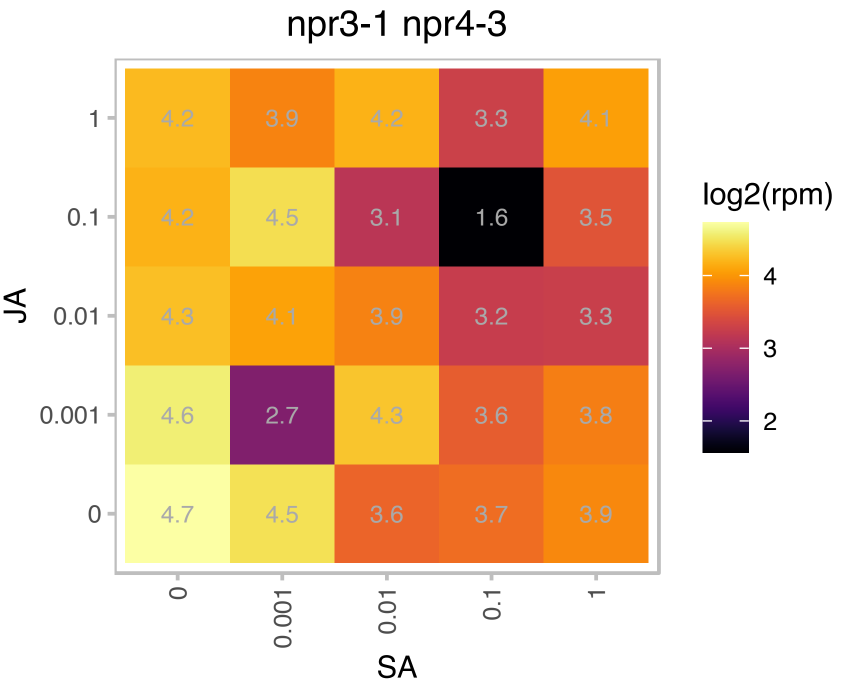

### AT5G19530.1 : ACL5

Encodes a spermine synthase. Required for internode elongation and vascular development, specifically in the mechanism that defines the boundaries between veins and nonvein regions. This mechanism may be mediated by polar auxin transport. Though ACL5 has been shown to function as a spermine synthase in E. coli, an ACL5 knockout has no effect on the endogenous levels of free and conjugated polyamines in Arabidopsis, suggesting that ACL5 may have a very specific or altogether different in vivo function.

S-adenosyl-L-methionine-dependent methyltransferases superfamily protein

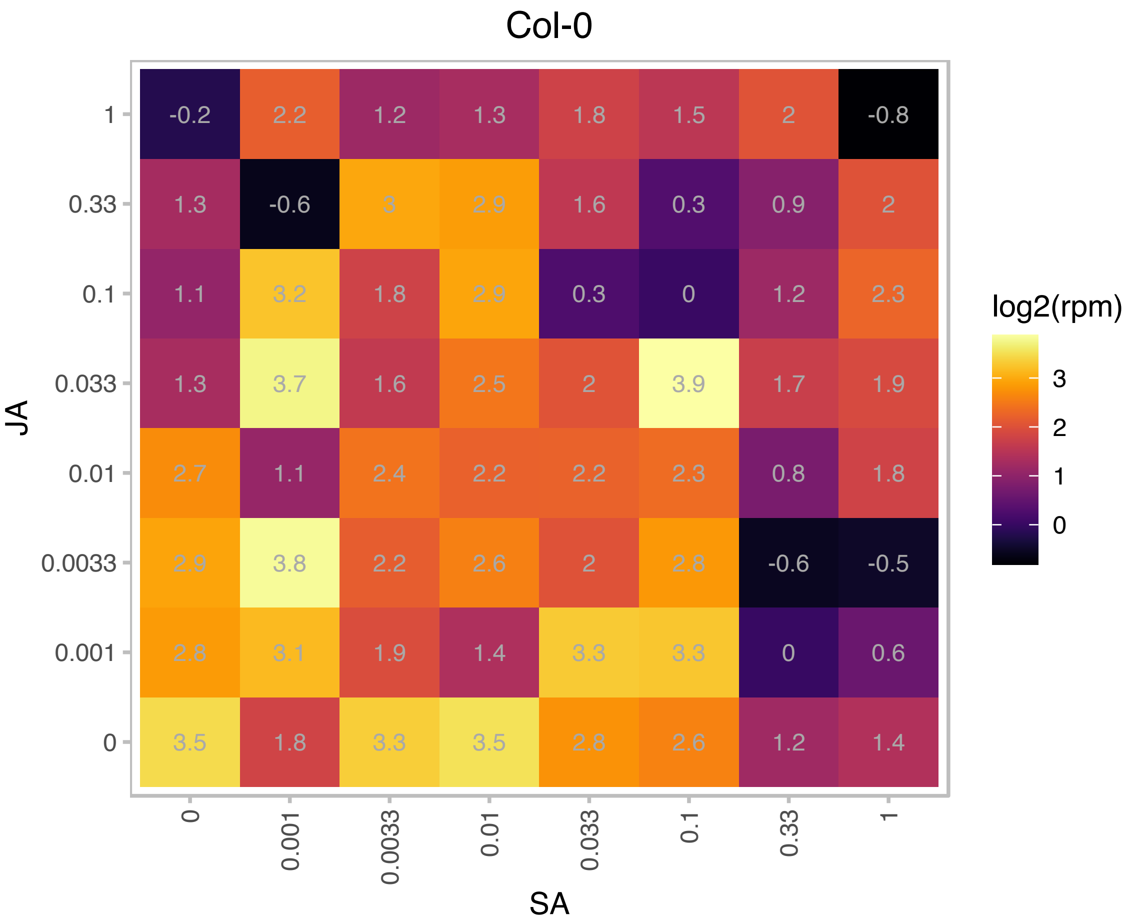


| | |
|---|---|
| Gene | AT5G19530 |
| is.expressed_genes | TRUE |
| is.responsive_genes | TRUE |
| is.apcluster | F-06 |

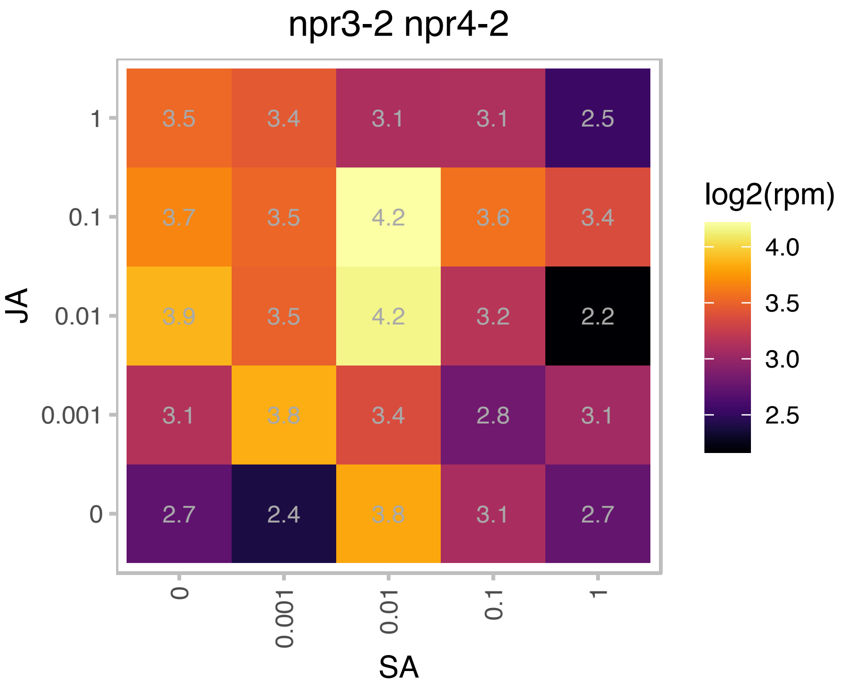


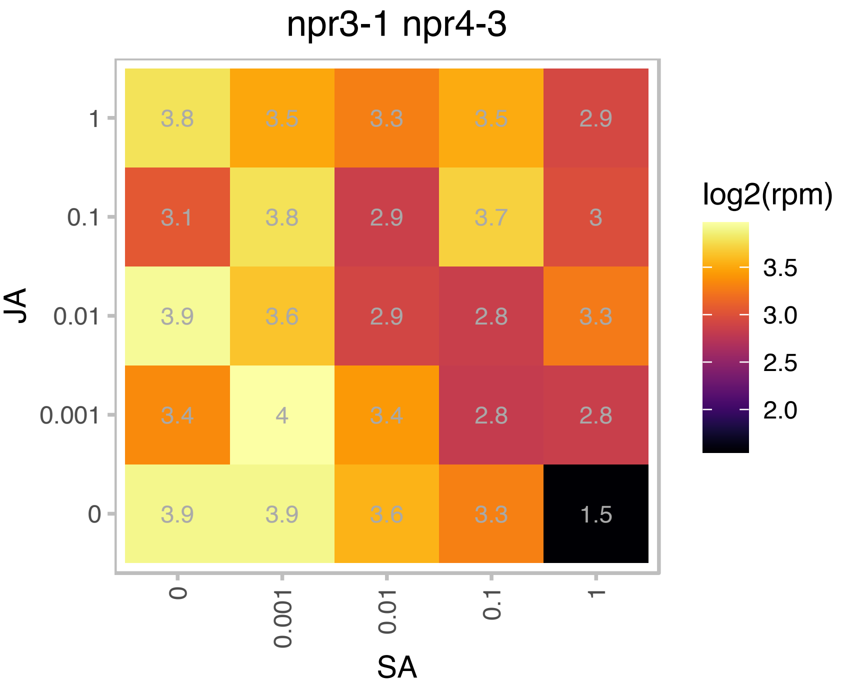

### AT5G23070.1 : AtTK1b, TK1b, TK2

Thymidine kinase

Encodes a thymidine kinase that salvages DNA precursors.

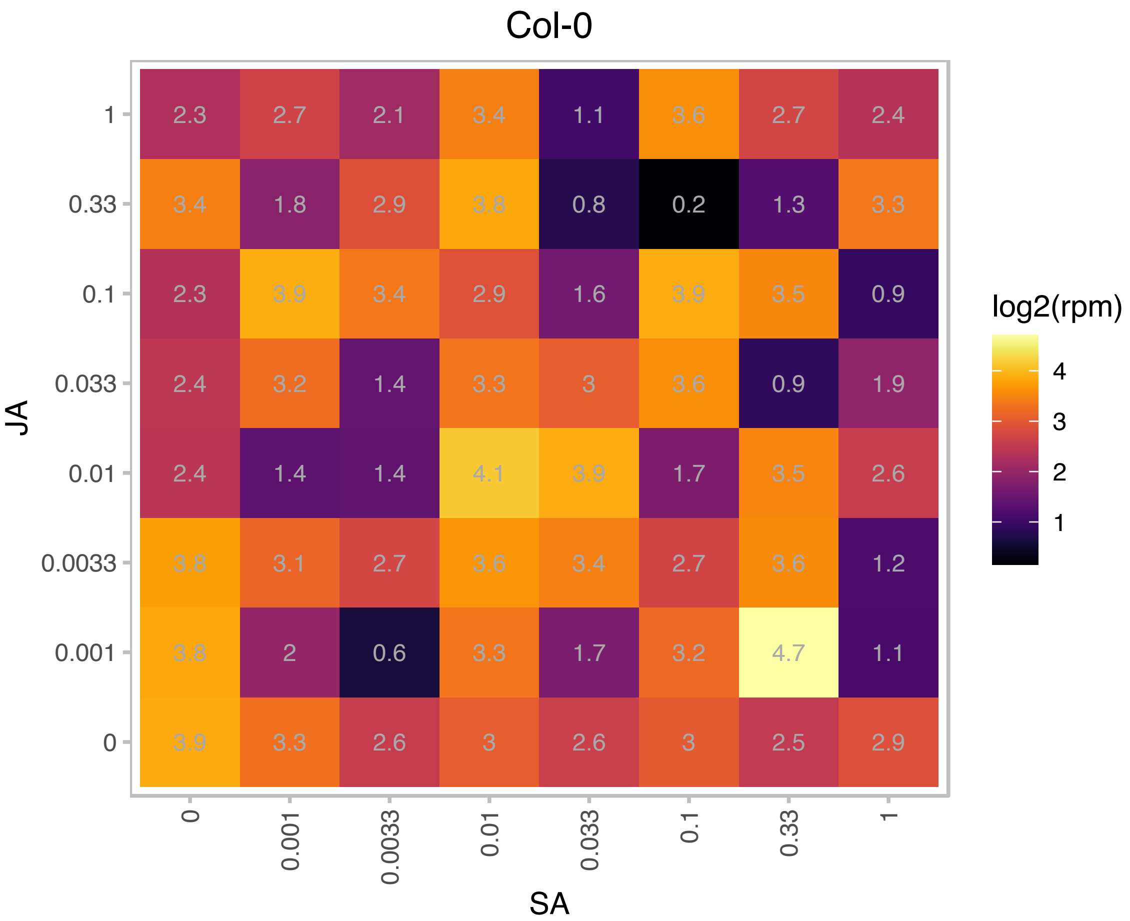


| | |
|---|---|
| Gene | AT5G23070 |
| is.expressed_genes | TRUE |
| is.responsive_genes | TRUE |
| is.apcluster | F-06 |

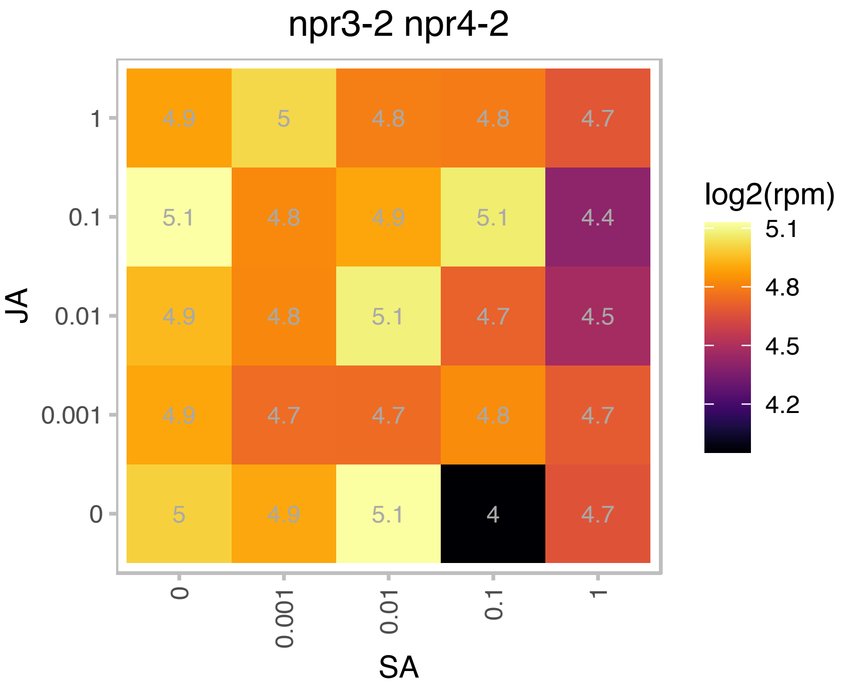


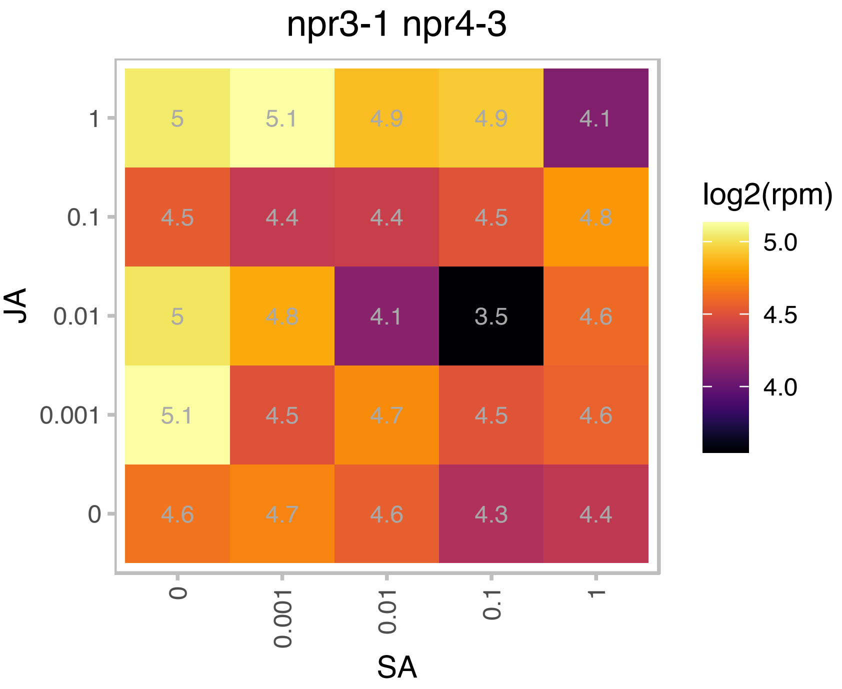

# AT5G35740.1 : NA

Carbohydrate-binding X8 domain superfamily protein

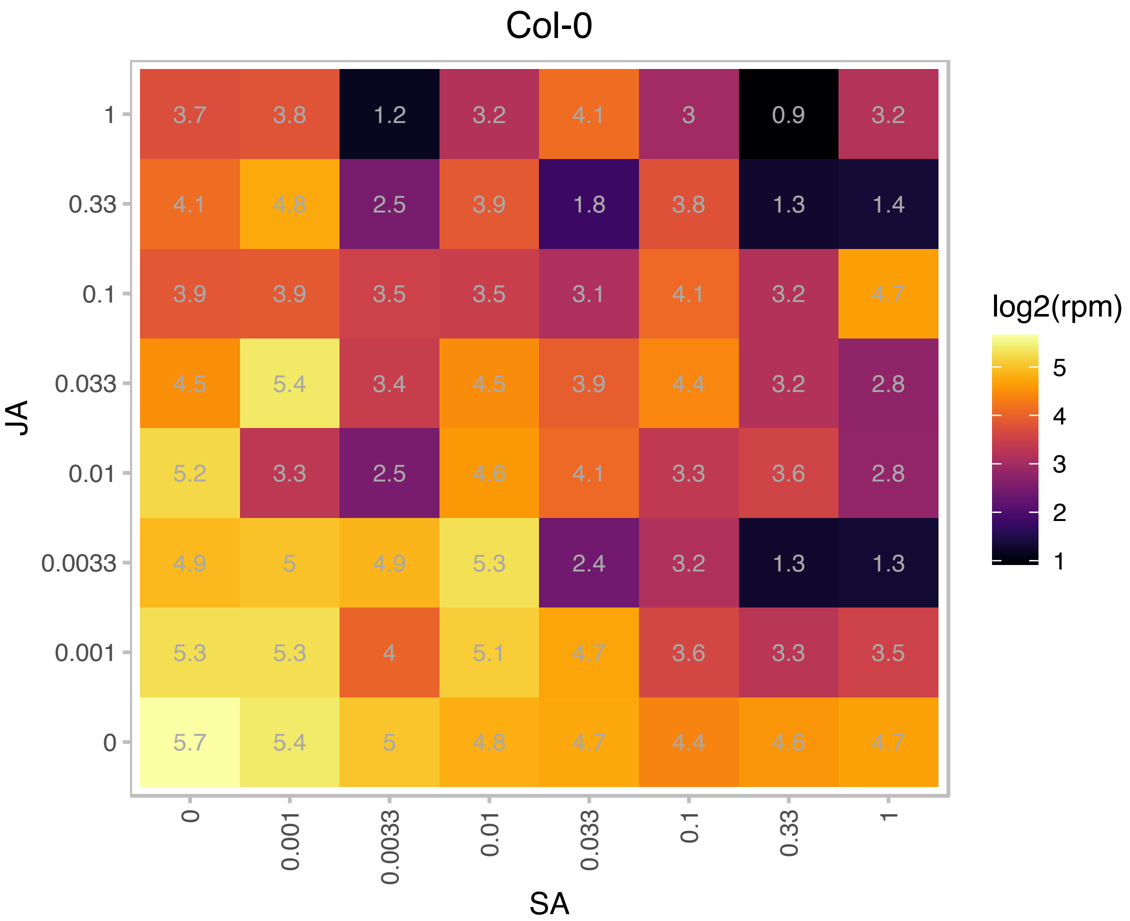


| | |
|---|---|
| Gene | AT5G35740 |
| is.expressed_genes | TRUE |
| is.responsive_genes | TRUE |
| is.apcluster | F-06 |

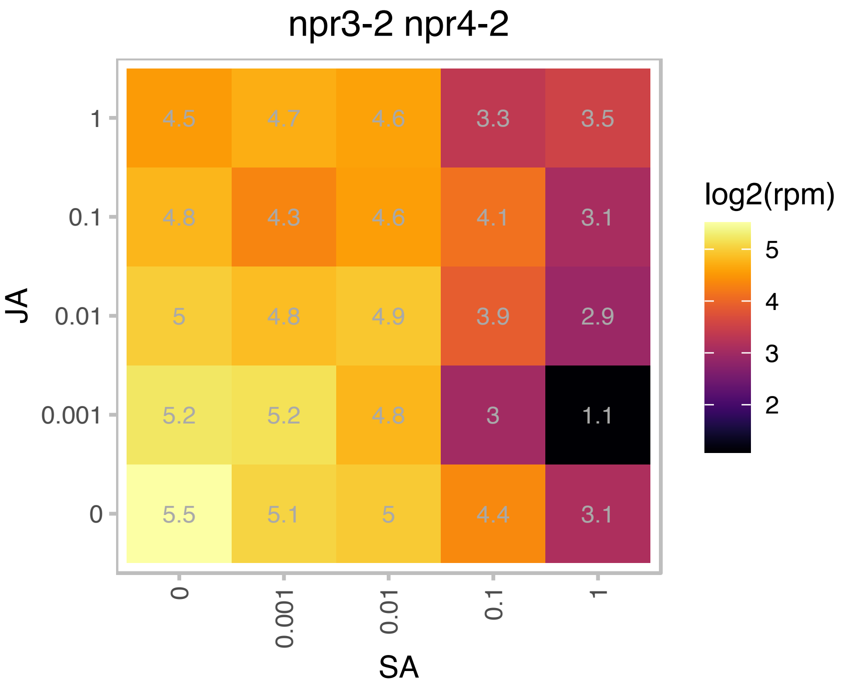


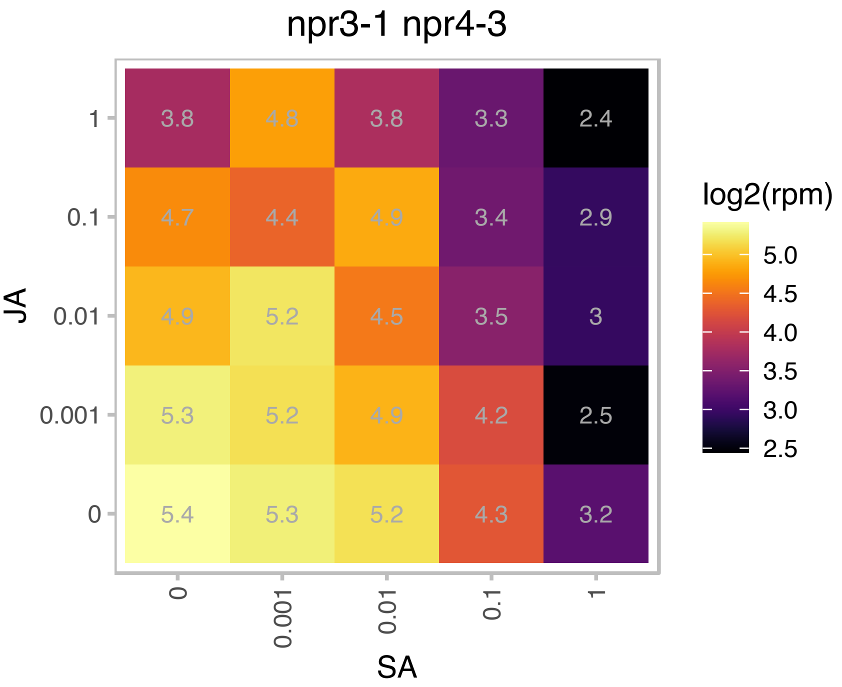

# AT5G42110.1 : NA

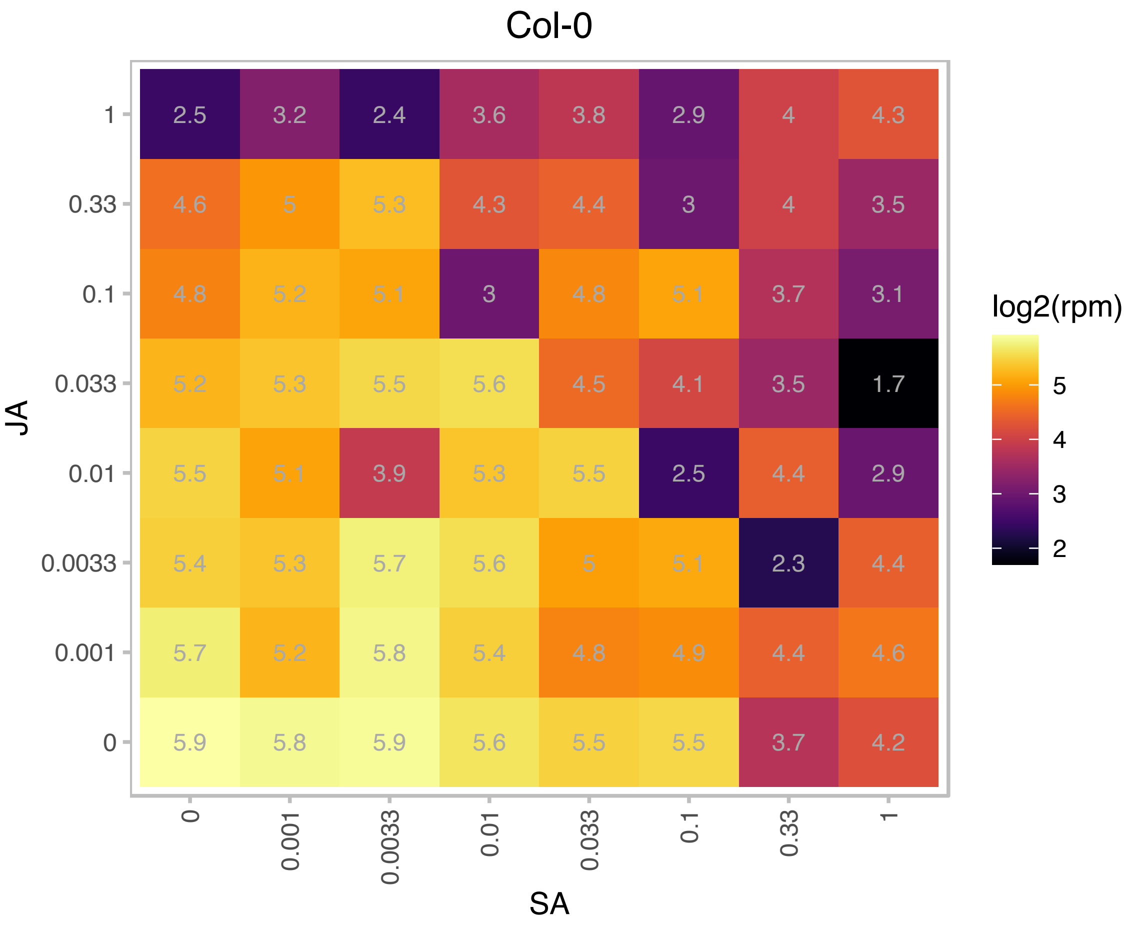


| | |
|---|---|
| Gene | AT5G42110 |
| is.expressed_genes | TRUE |
| is.responsive_genes | TRUE |
| is.apcluster | F-06 |

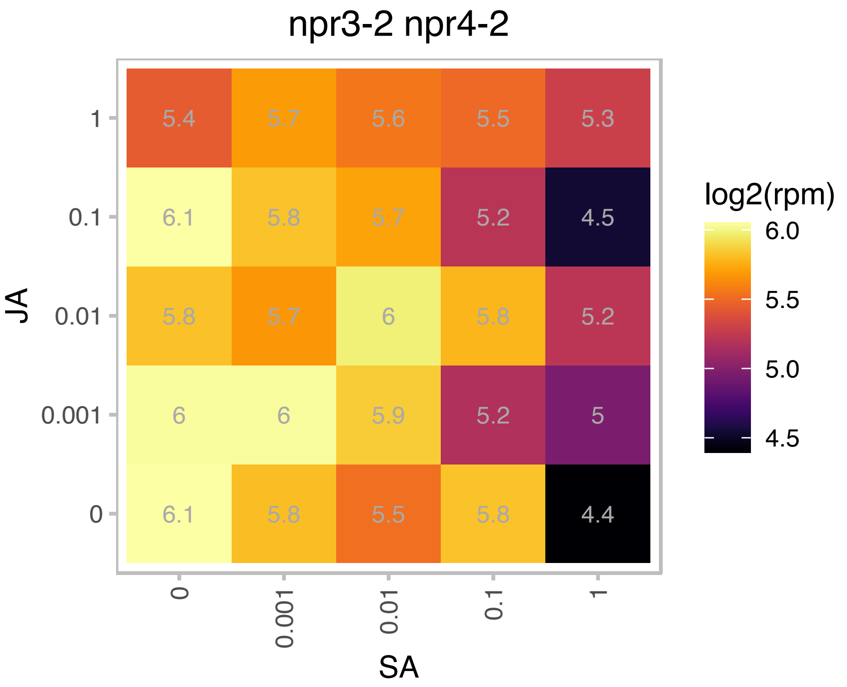


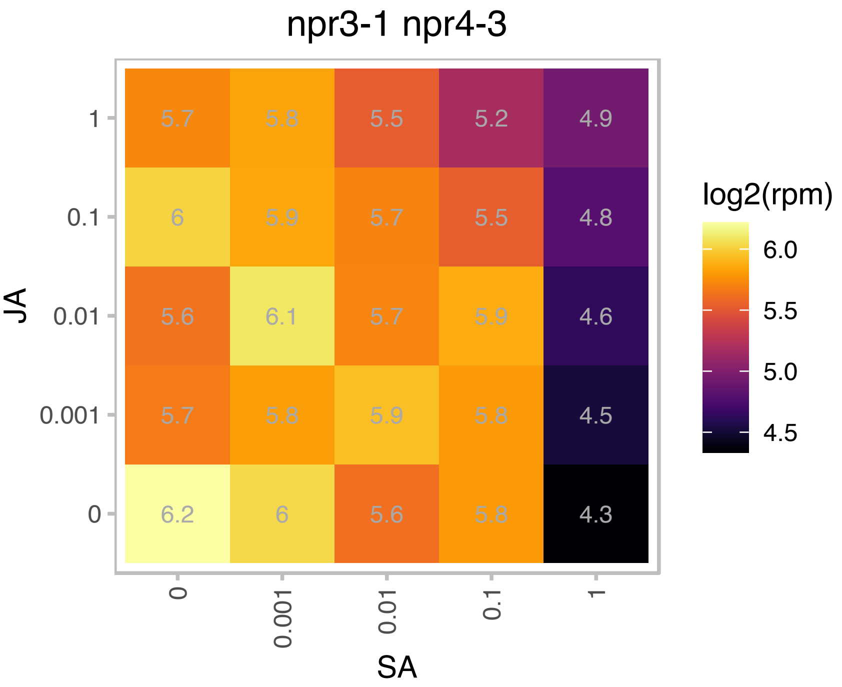

### AT5G44650.1 : AtCEST, CEST, Y3IP1

Encodes a nucleus-encoded thylakoid protein, cooperates with the plastid-encoded Ycf3 protein in photosystem i assembly. Also induces tolerance to multiple environmental stresses and reduces photooxidative damage.

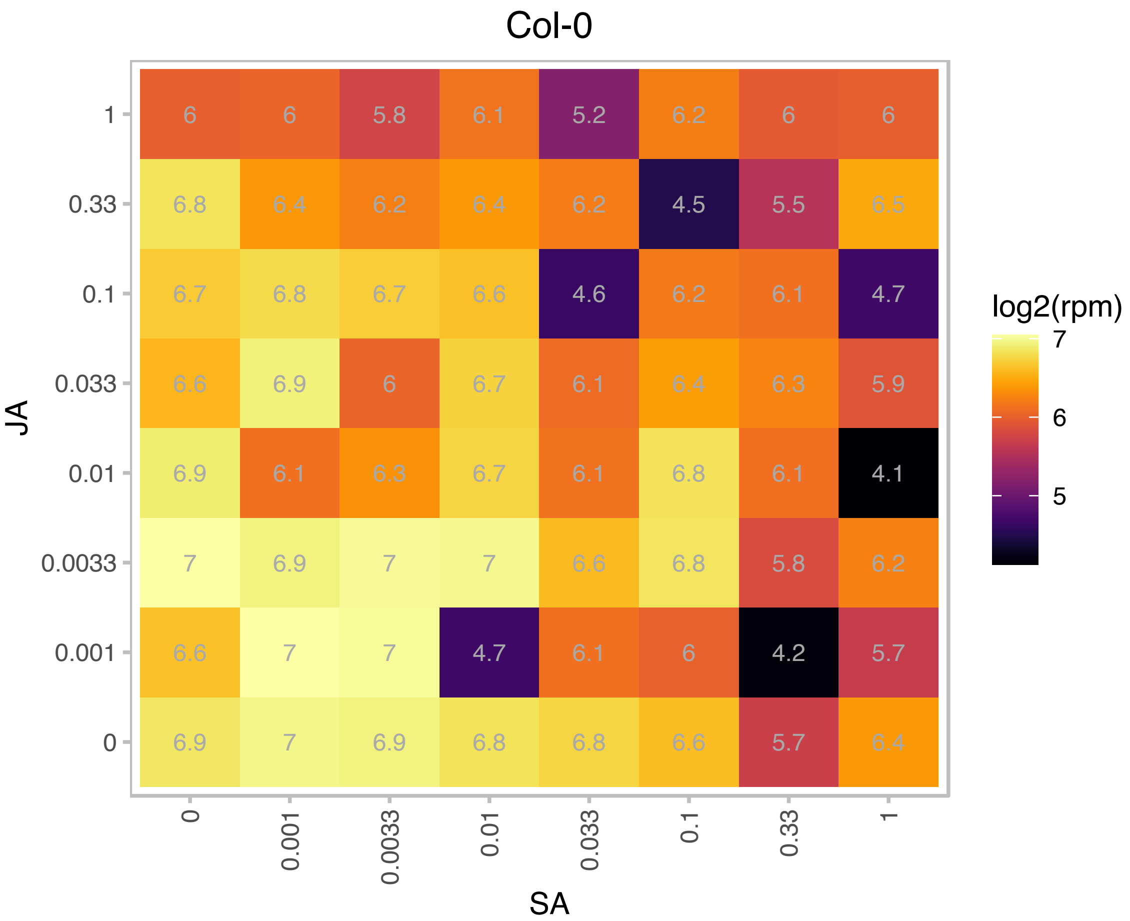


| | |
|---|---|
| Gene | AT5G44650 |
| is.expressed_genes | TRUE |
| is.responsive_genes | TRUE |
| is.apcluster | F-06 |

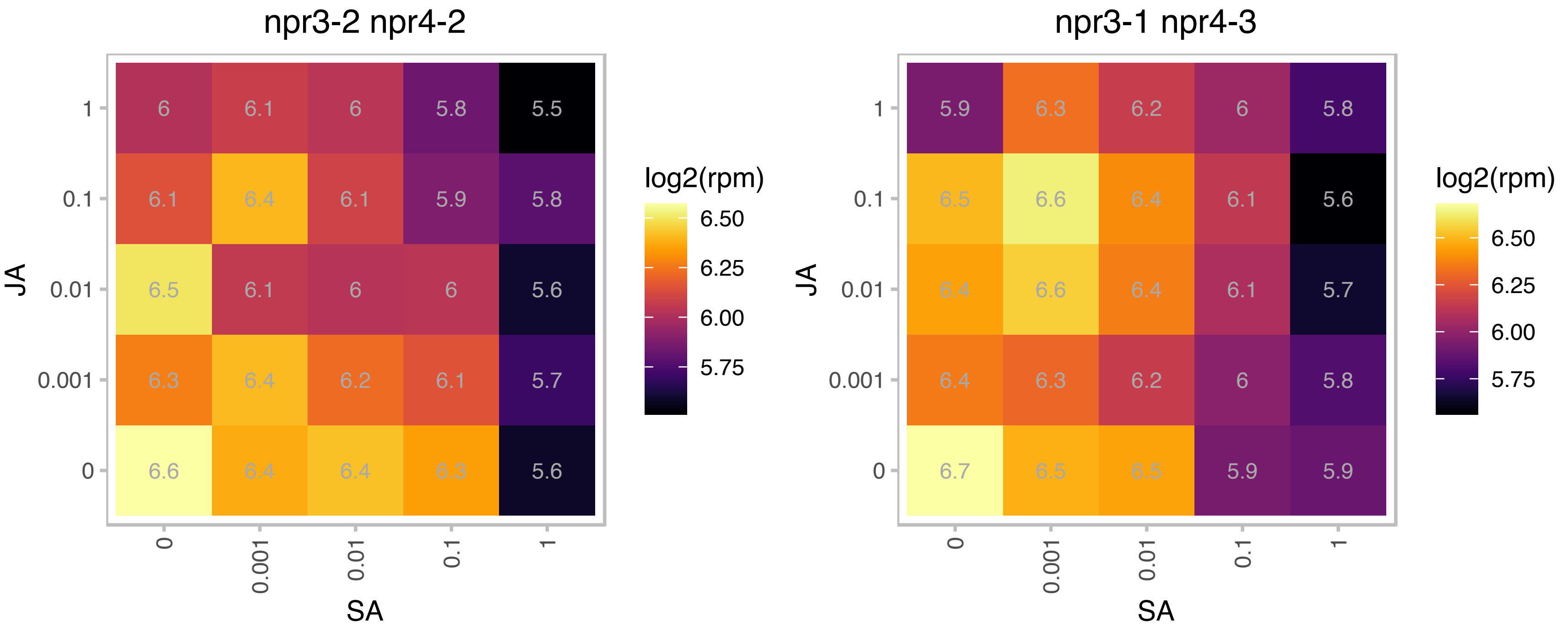

# AT5G51010.1 : NA

Rubredoxin-like superfamily protein

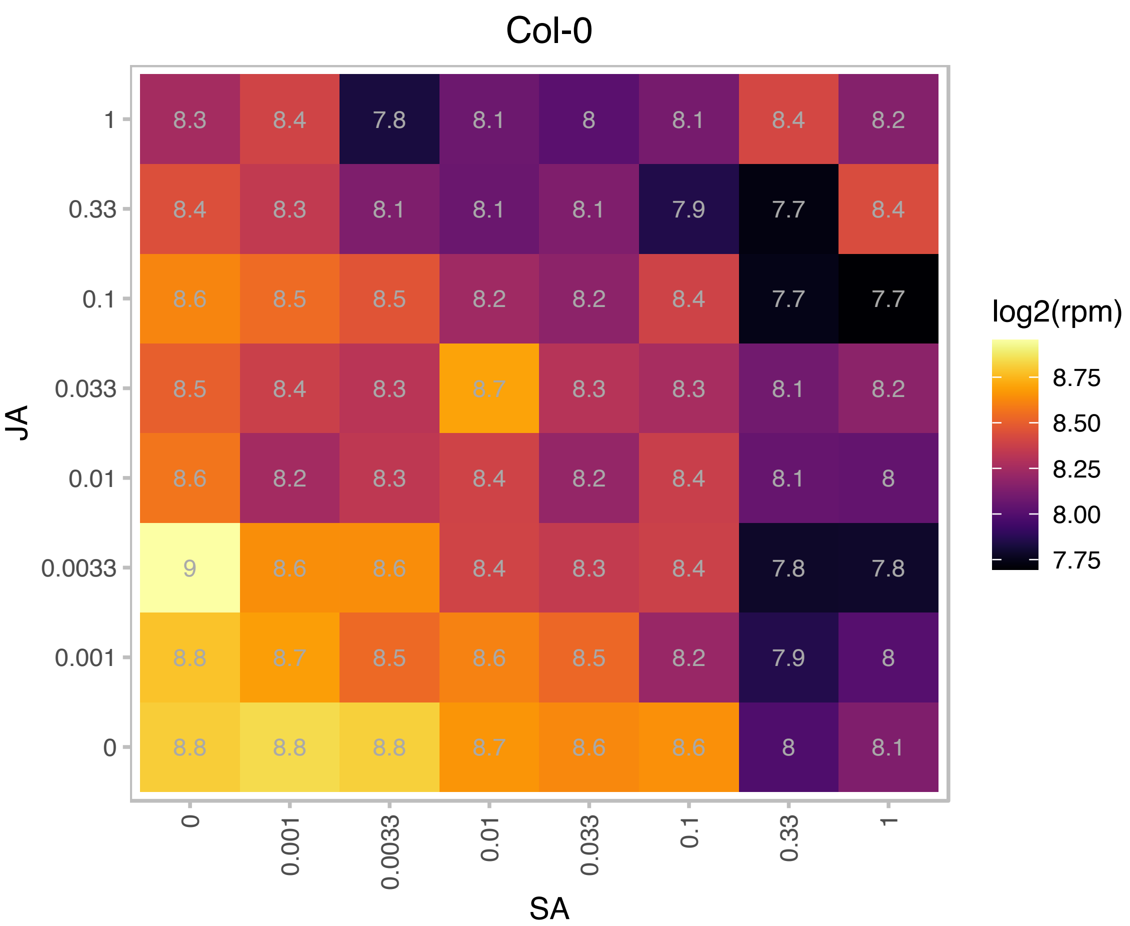


| | |
|---|---|
| Gene | AT5G51010 |
| is.expressed_genes | TRUE |
| is.responsive_genes | TRUE |
| is.apcluster | F-06 |

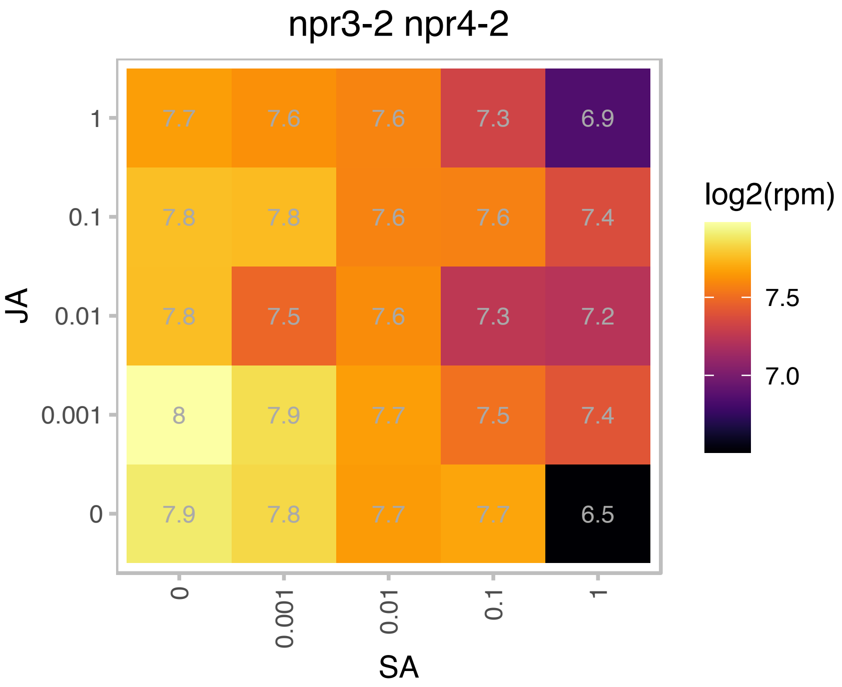


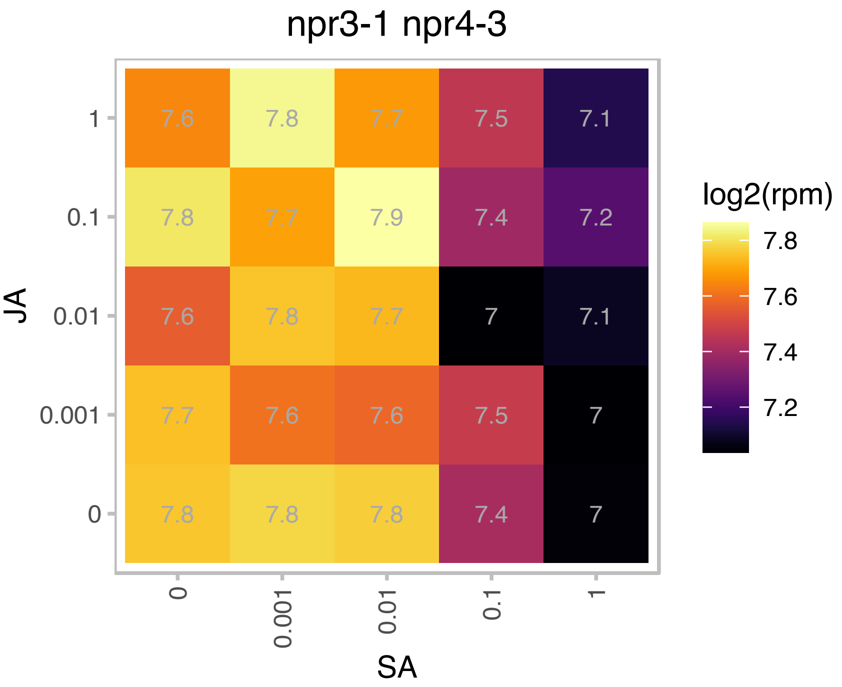

### AT5G51720.1 : At-NEET, NEET

2 iron, 2 sulfur cluster binding

Encodes a protein with biochemical, structural, and biophysical characteristics of a NEET protein. It plays a key role in plant development, senescence, reactive oxygen homeostasis, and Fe metabolism.

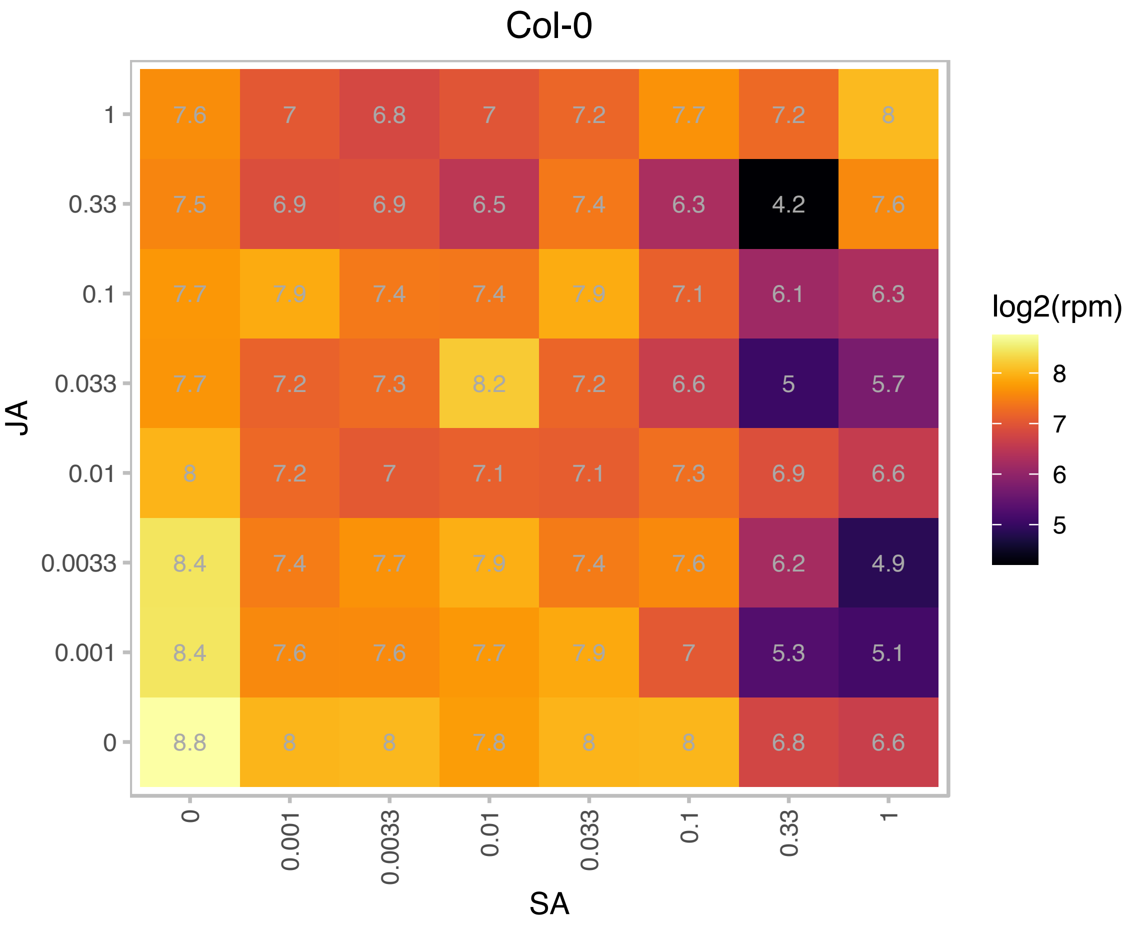


| | |
|---|---|
| Gene | AT5G51720 |
| is.expressed_genes | TRUE |
| is.responsive_genes | TRUE |
| is.apcluster | F-06 |

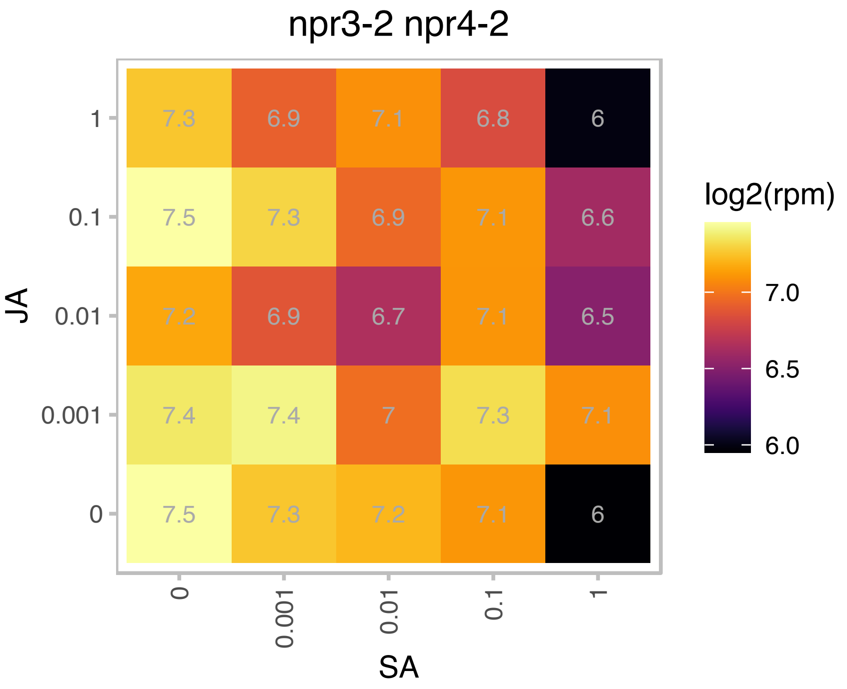


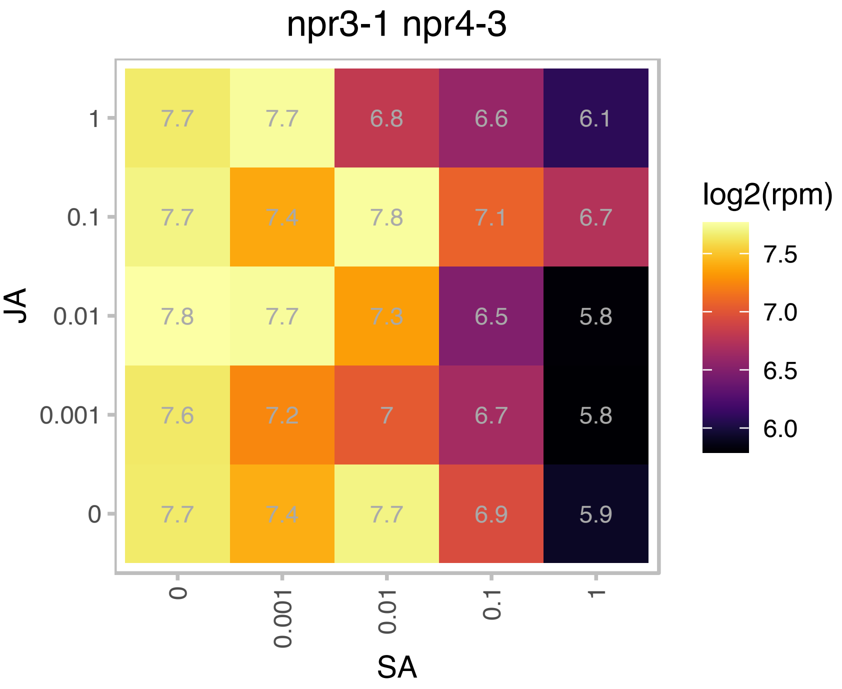

# AT5G52960.1 : NA

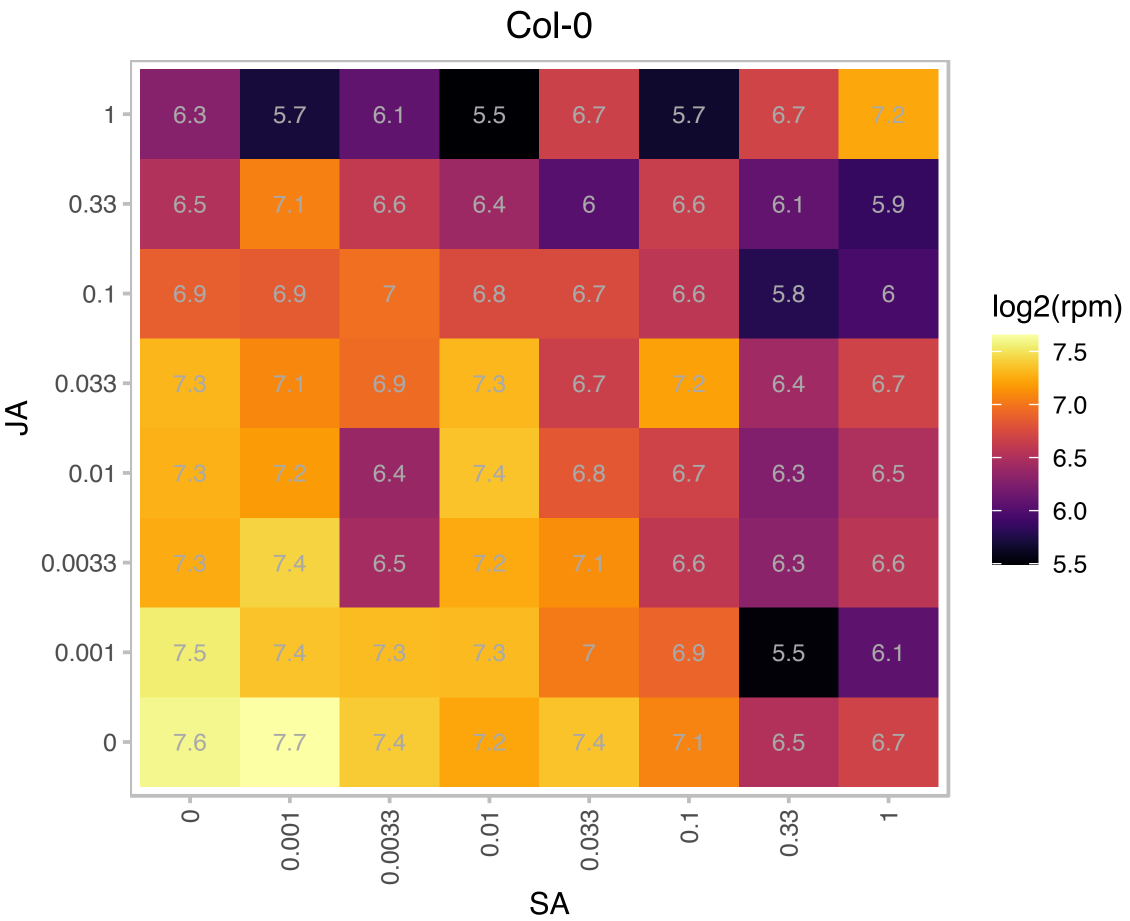


| | |
|---|---|
| Gene | AT5G52960 |
| is.expressed_genes | TRUE |
| is.responsive_genes | TRUE |
| is.apcluster | F-06 |

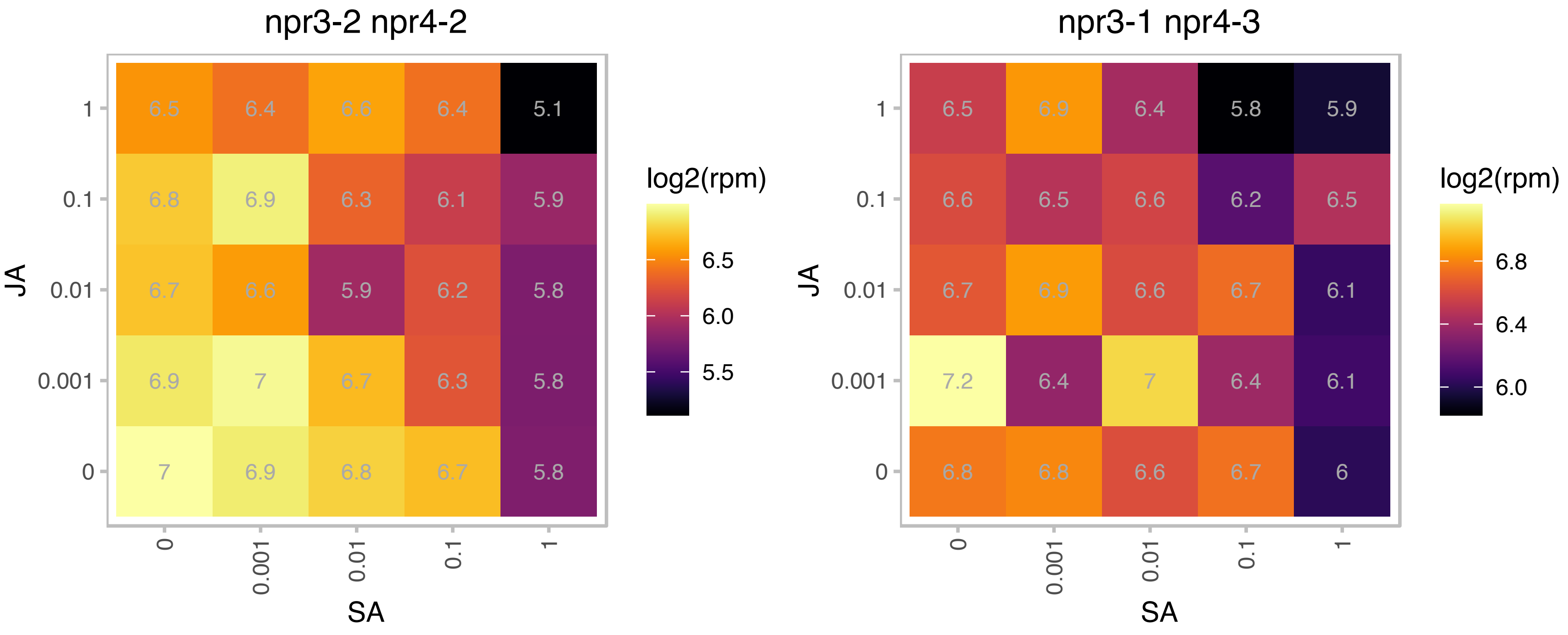

### AT5G53580.1 : AtPLR1, PLR1

NAD(P)-linked oxidoreductase superfamily protein

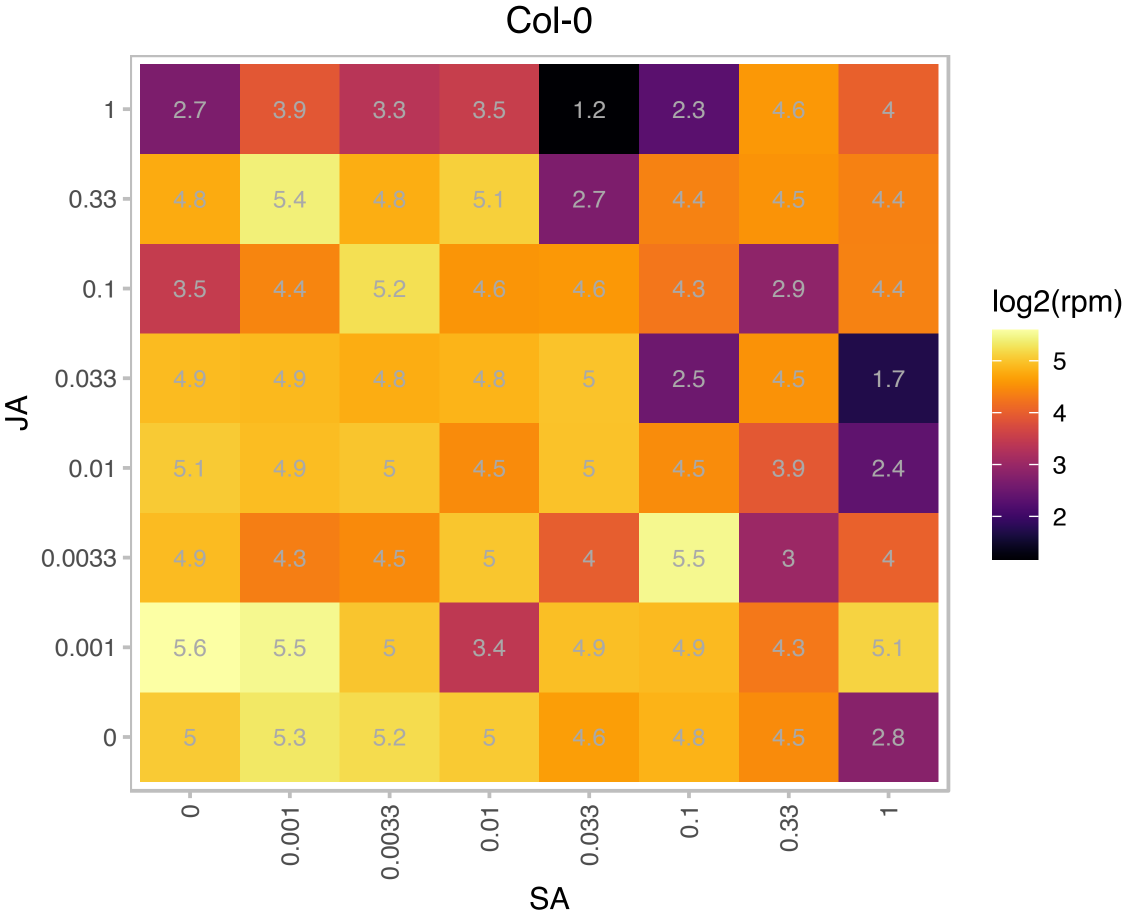


| | |
|---|---|
| Gene | AT5G53580 |
| is.expressed_genes | TRUE |
| is.responsive_genes | TRUE |
| is.apcluster | F-06 |

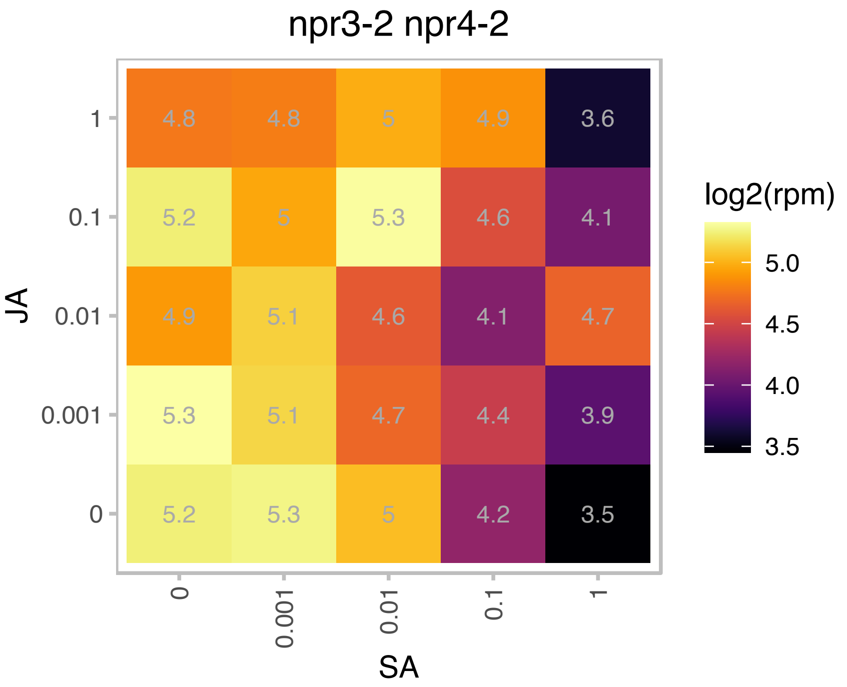


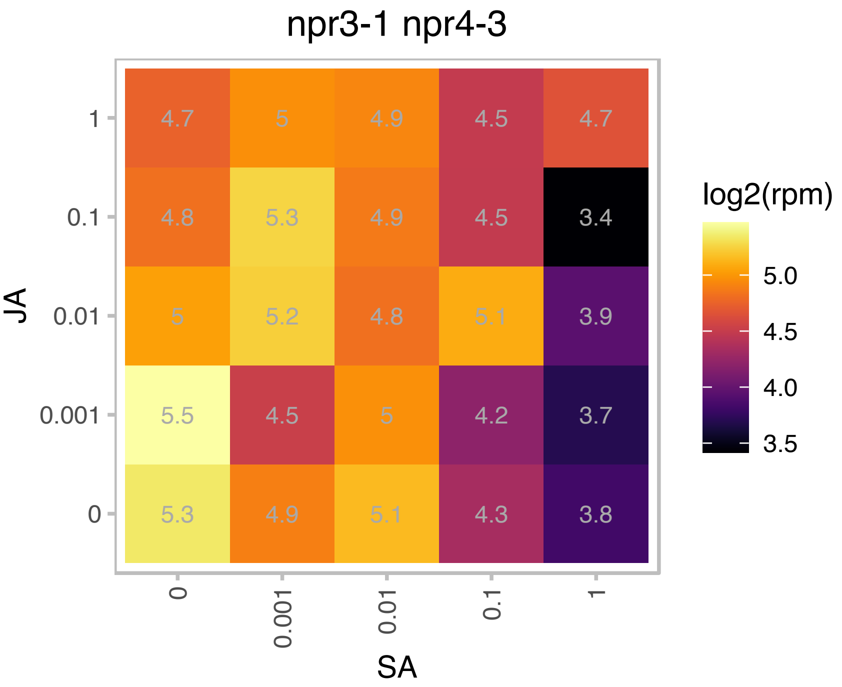

# AT5G62720.1 : NA

Integral membrane HPP family protein

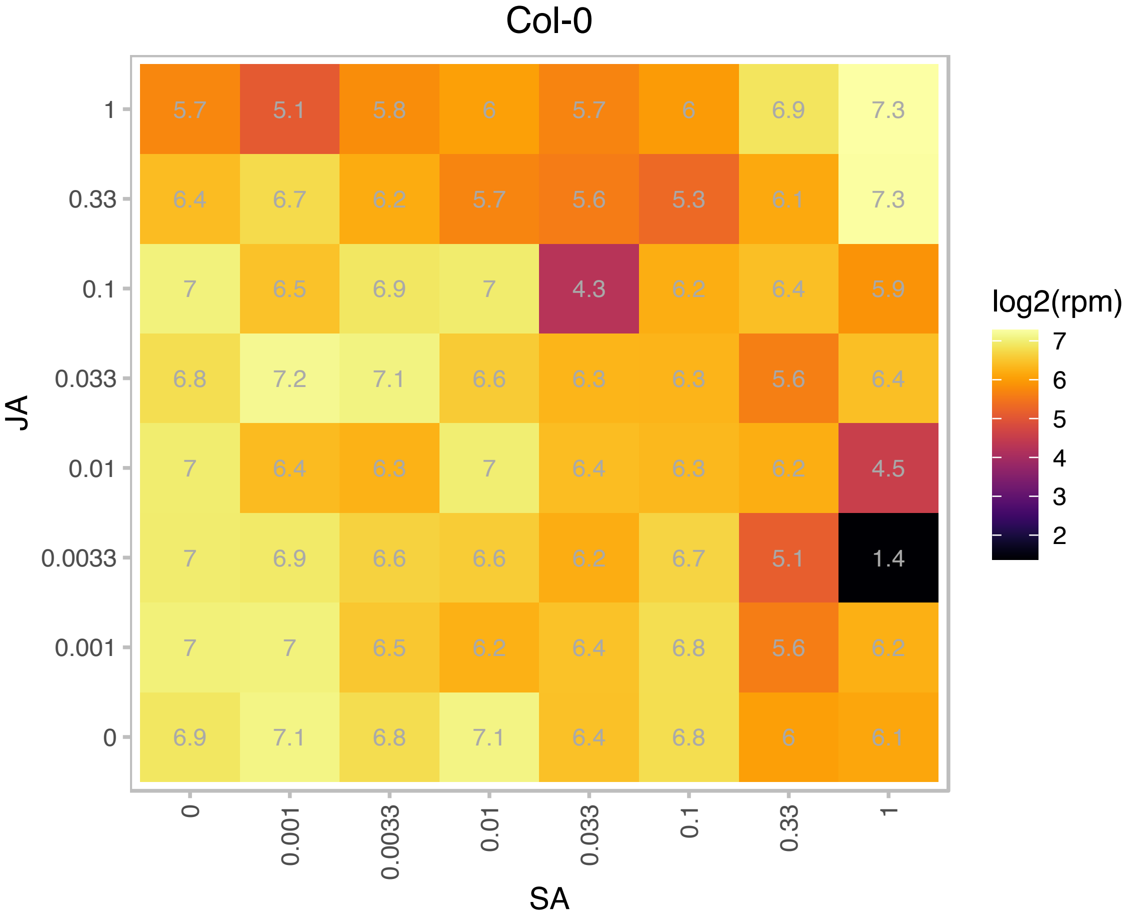


| | |
|---|---|
| Gene | AT5G62720 |
| is.expressed_genes | TRUE |
| is.responsive_genes | TRUE |
| is.apcluster | F-06 |

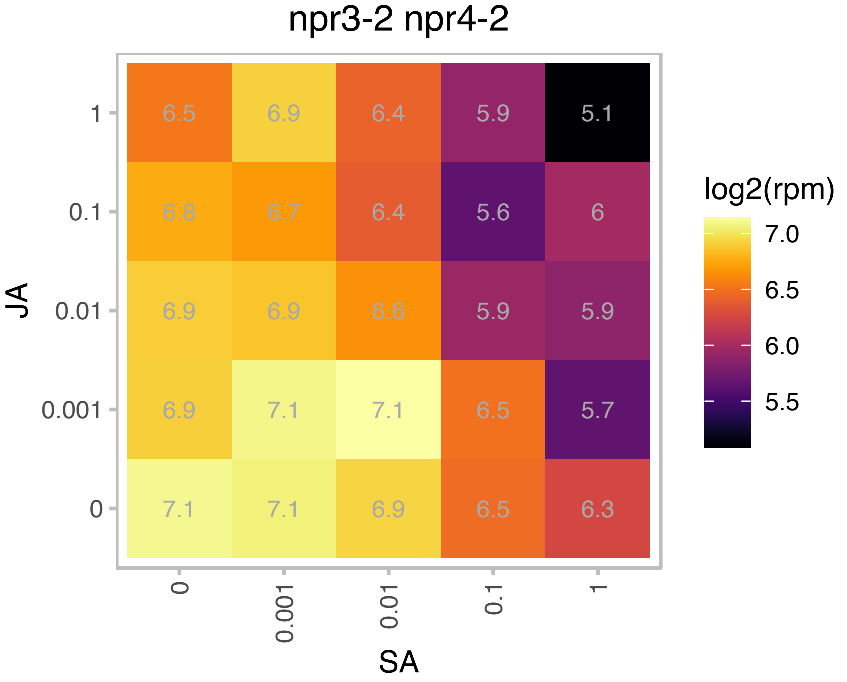


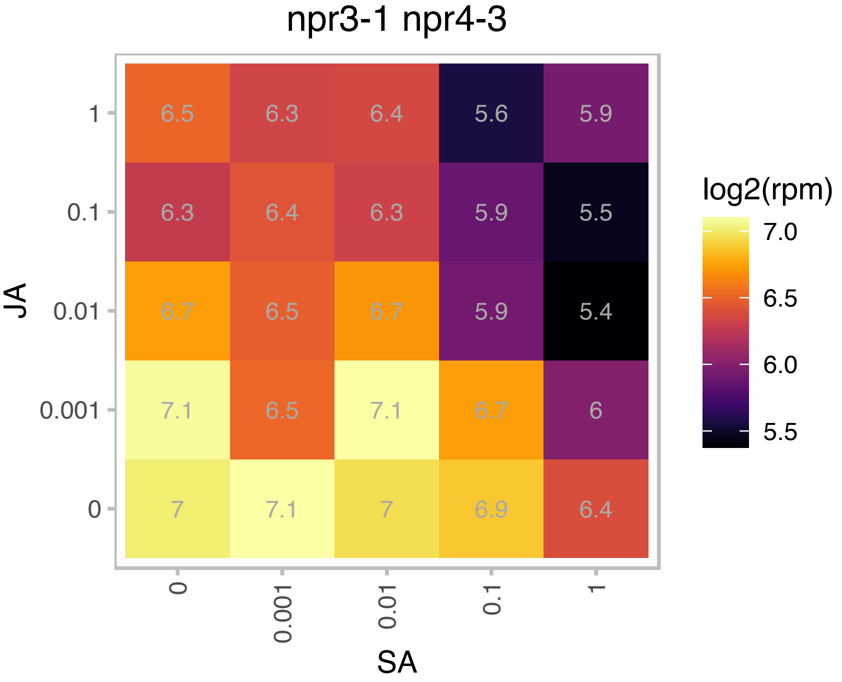

### AT5G65010.1 : ASN2

asparagine synthetase 2

Encodes asparagine synthetase (ASN2).

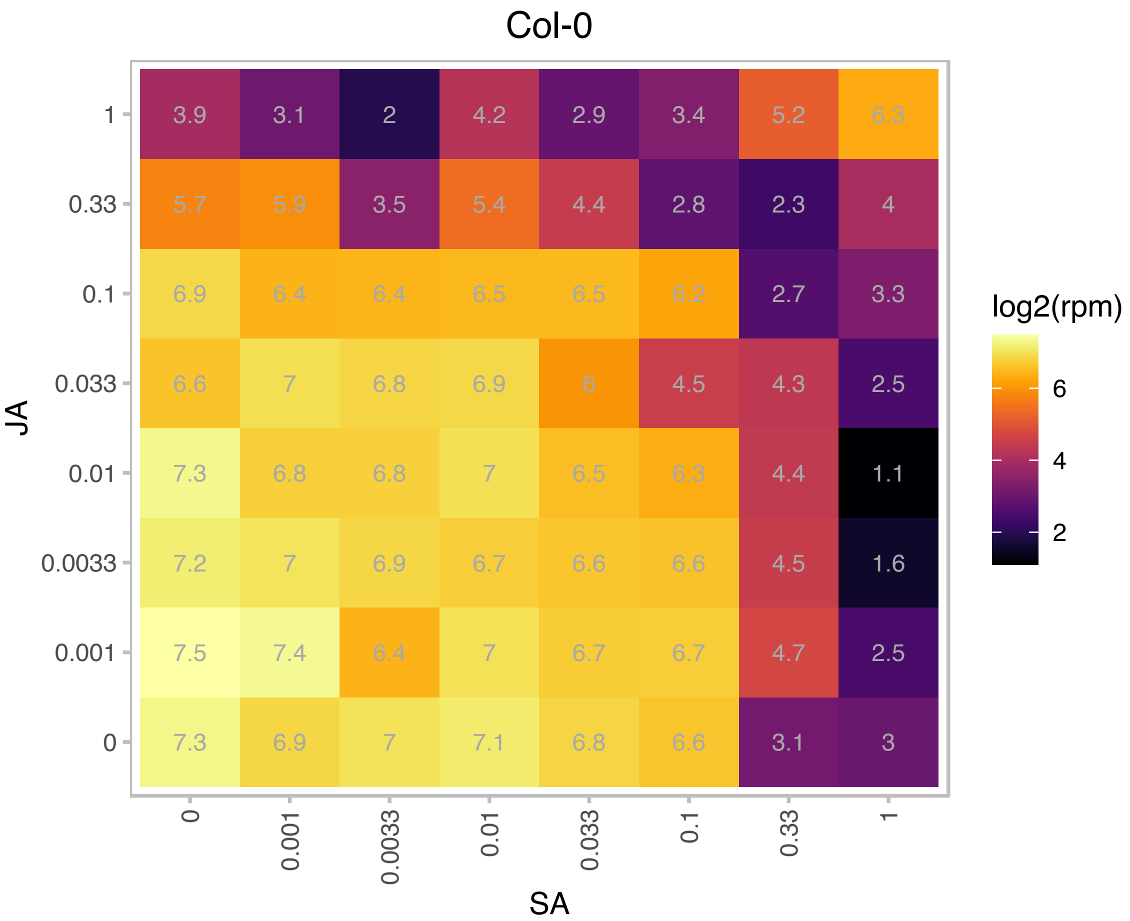


| | |
|---|---|
| Gene | AT5G65010 |
| is.expressed_genes | TRUE |
| is.responsive_genes | TRUE |
| is.apcluster | F-06 |

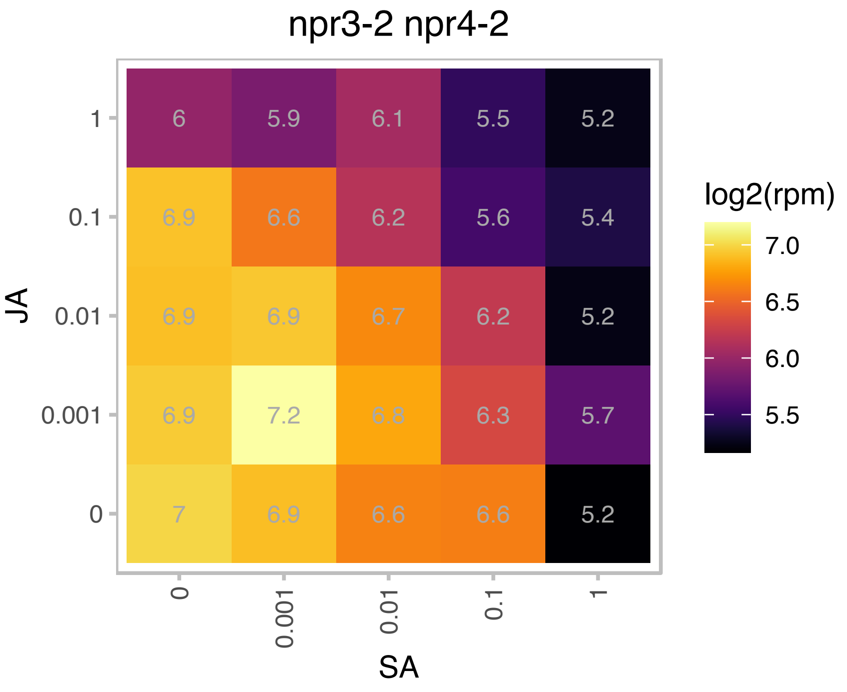


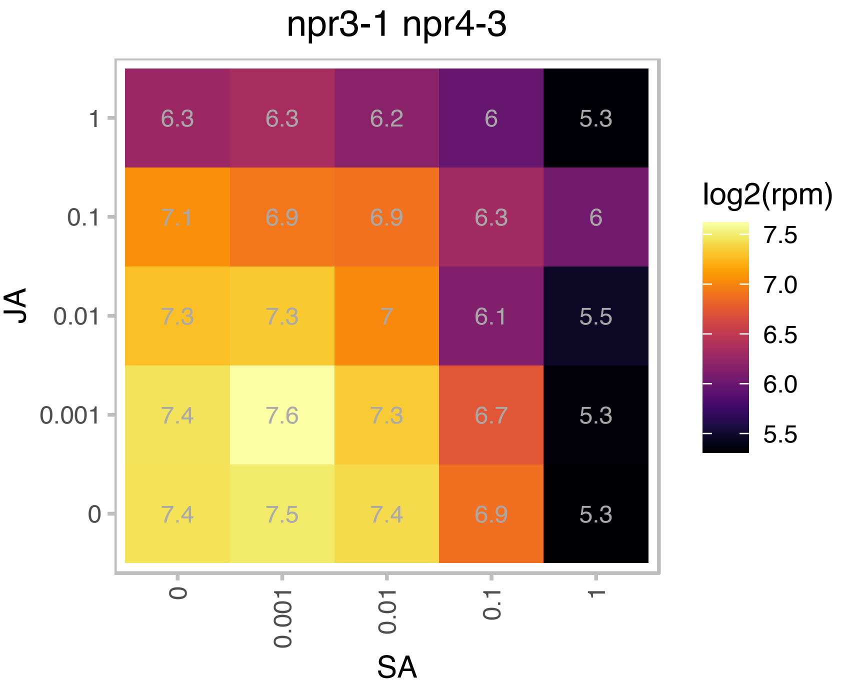

### ATCG00840.1 : RPL23, RPL23.1

ribosomal protein L23.1

One of two chloroplast genes that encode chloroplast ribosomal protein L23, a constituent of the large subunit of the ribosomal complex

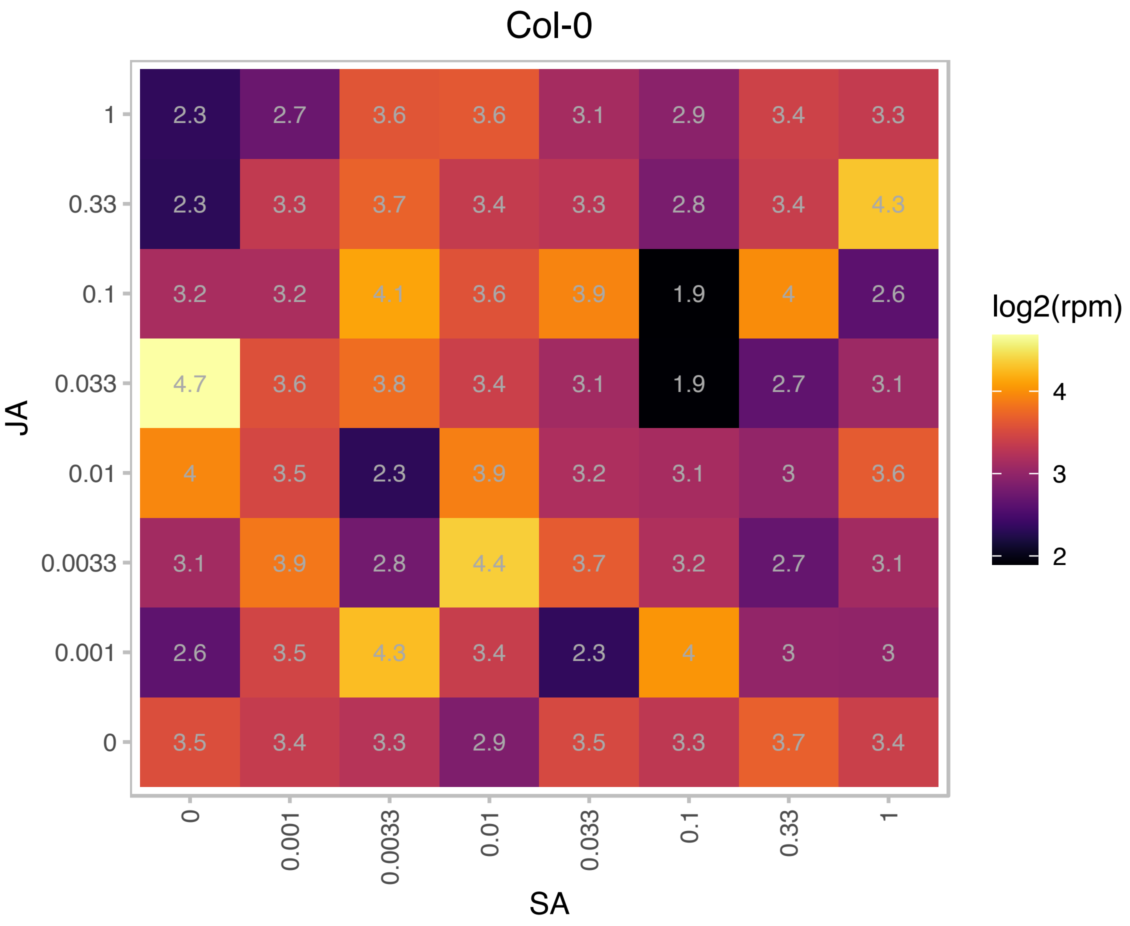


| | |
|---|---|
| Gene | ATCG00840 |
| is.expressed_genes | TRUE |
| is.responsive_genes | TRUE |
| is.apcluster | F-06 |

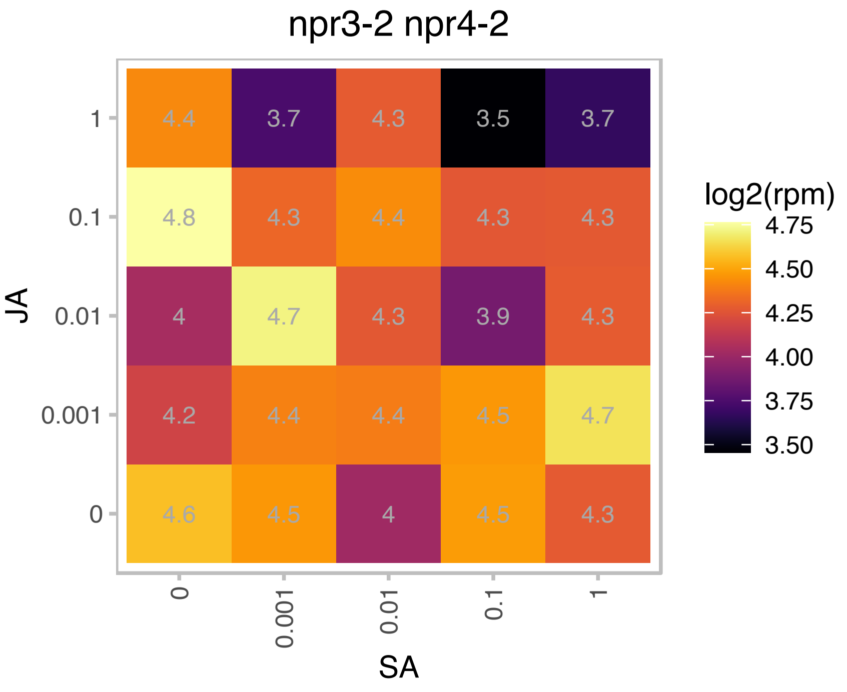


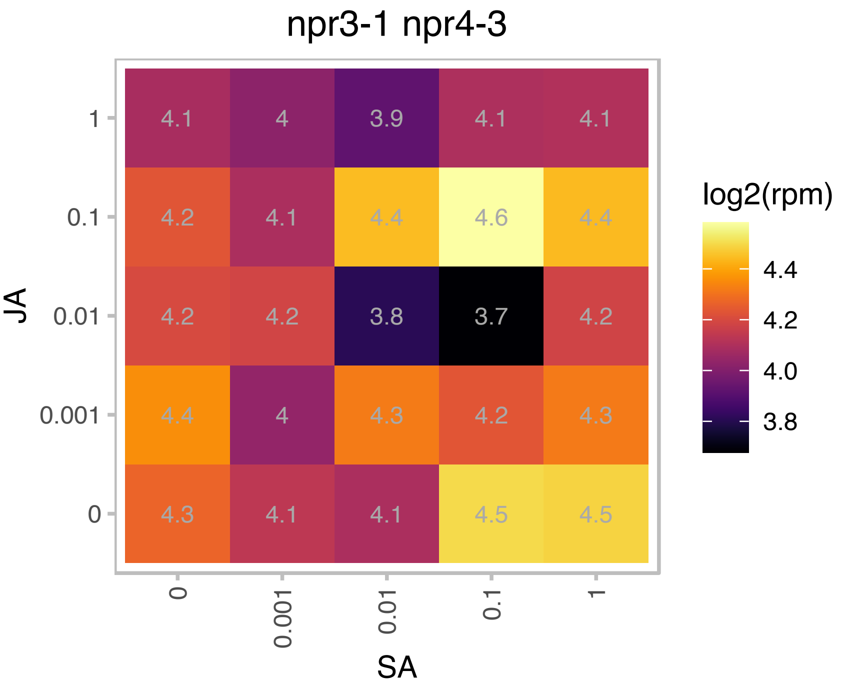

### ATCG01300.1 : RPL23.2

ribosomal protein L23

One of two chloroplast genes that encode chloroplast ribosomal protein L23, a constituent of the large subunit of the ribosomal complex

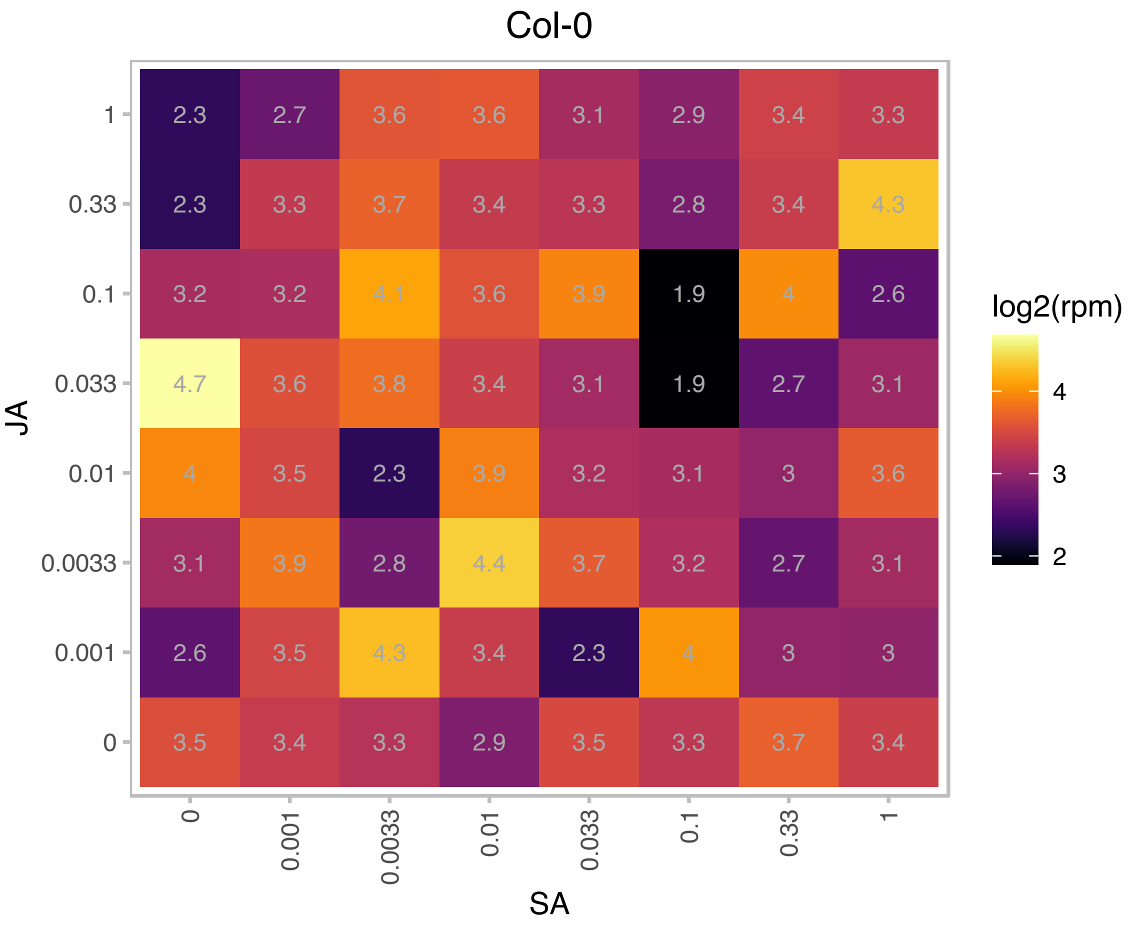


| | |
|---|---|
| Gene | ATCG01300 |
| is.expressed_genes | TRUE |
| is.responsive_genes | TRUE |
| is.apcluster | F-06 |

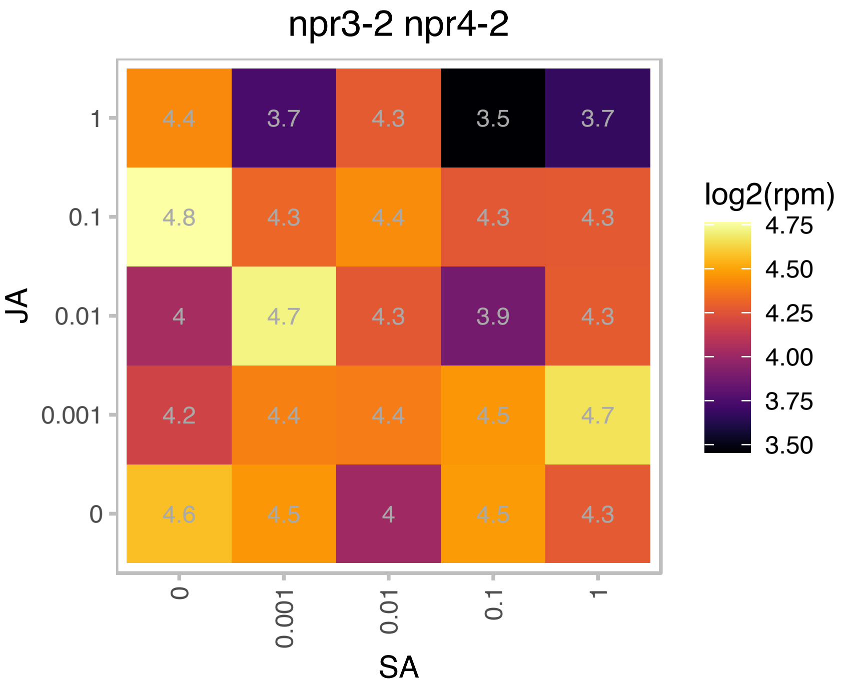


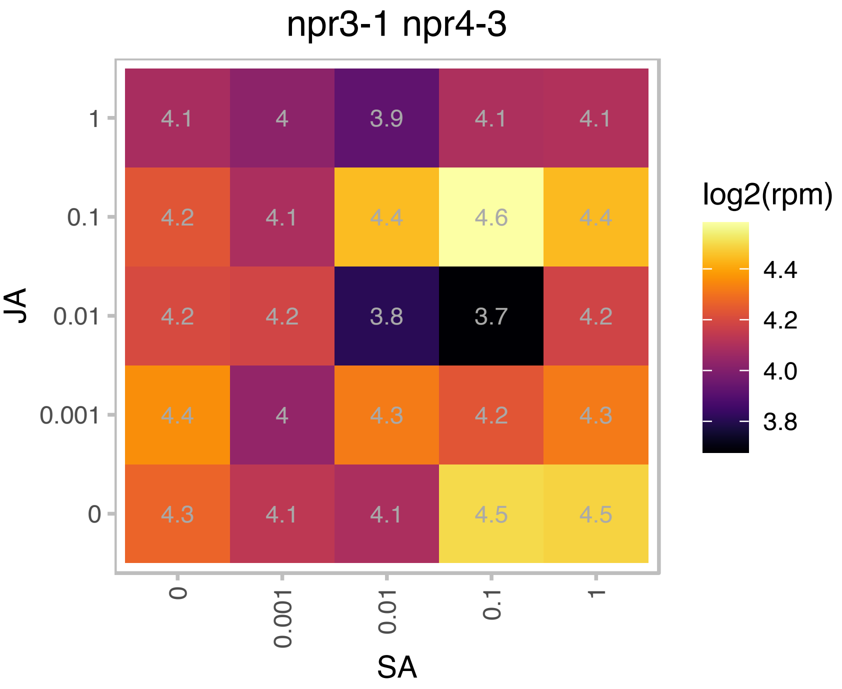

# AT1G01100.1 : NA

60S acidic ribosomal protein family

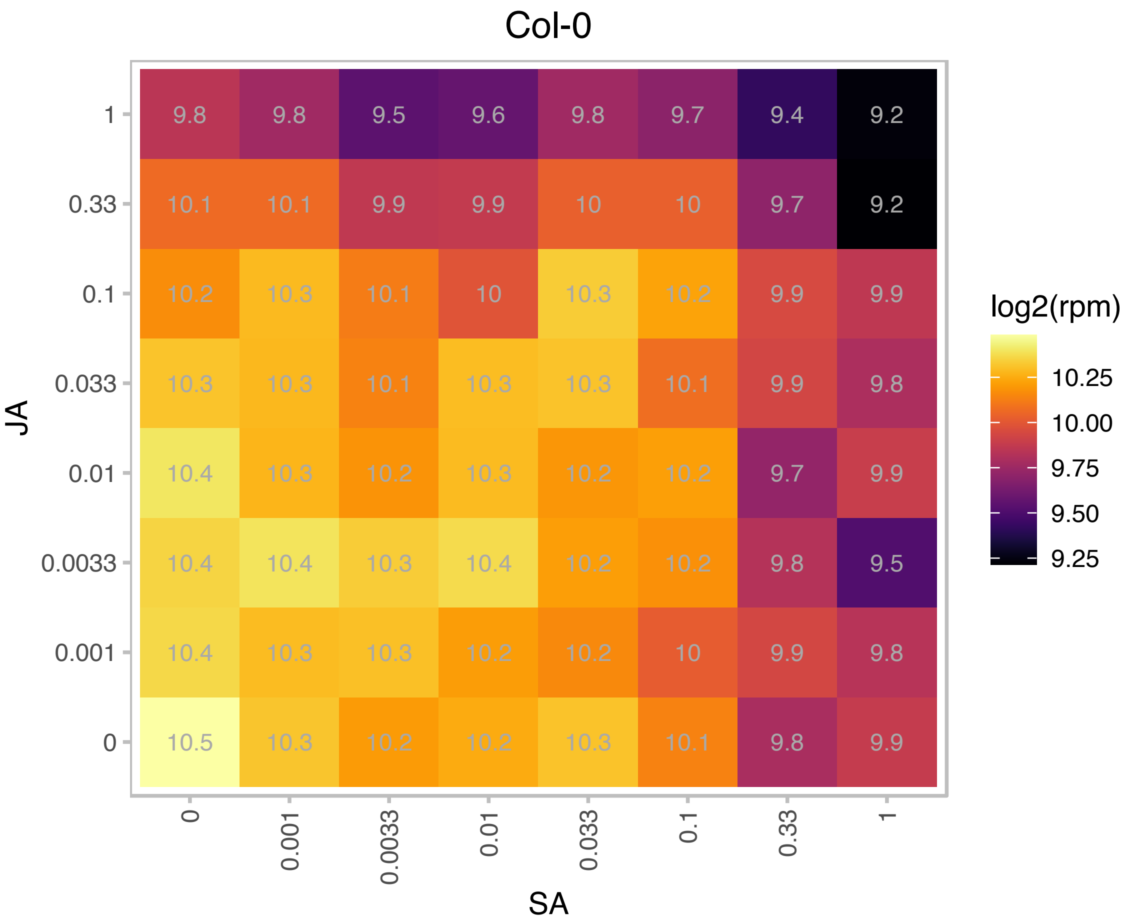


| | |
|---|---|
| Gene | AT1G01100 |
| is.expressed_genes | TRUE |
| is.responsive_genes | TRUE |
| is.apcluster | F-07 |

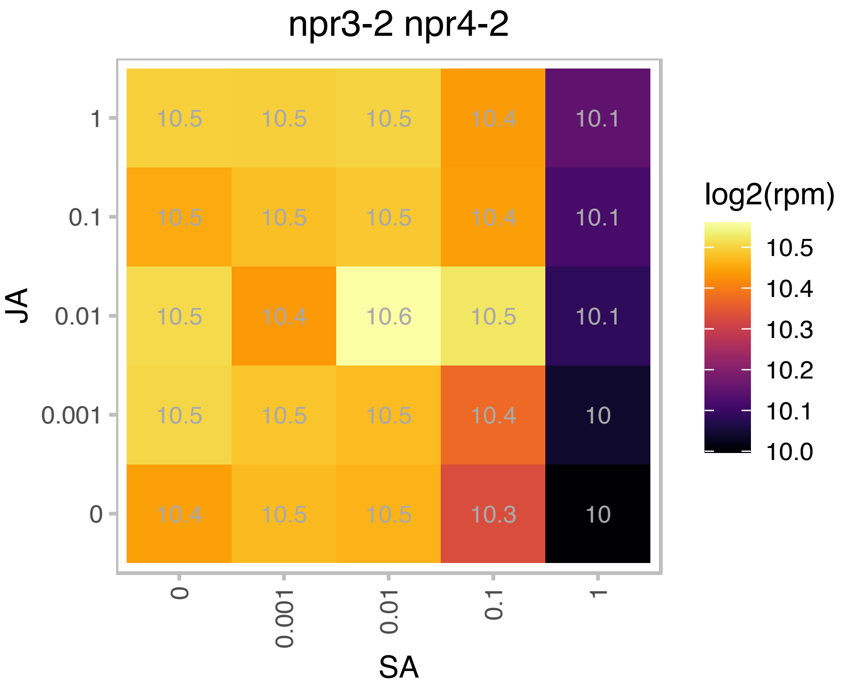


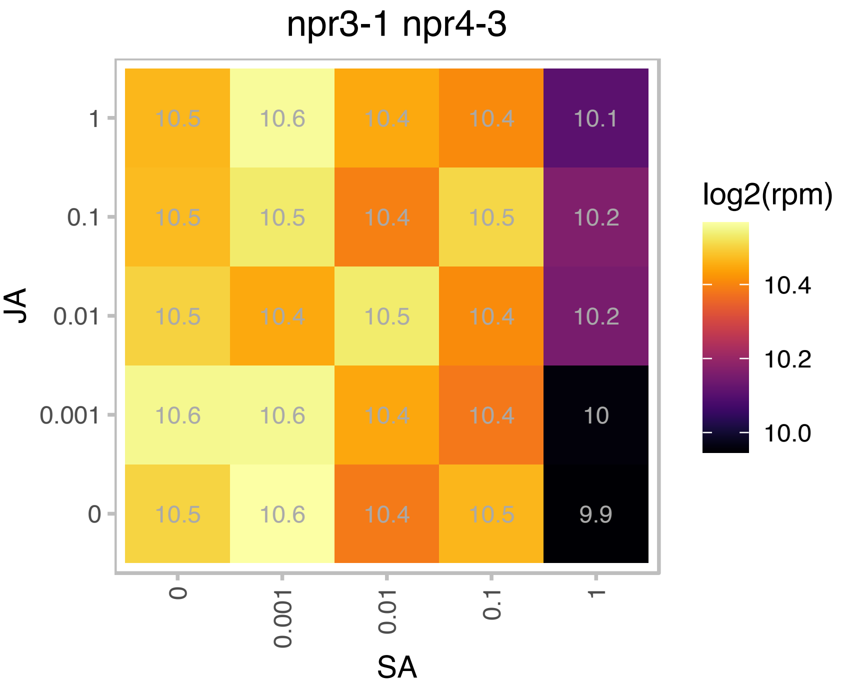

### AT1G01320.1 : REC1

Tetratricopeptide repeat (TPR)-like superfamily protein

Encodes REDUCED CHLOROPLAST COVERAGE 1 (REC1). Located in the nucleus and cytosol. Contributes to establishing the size of the chloroplast compartment.

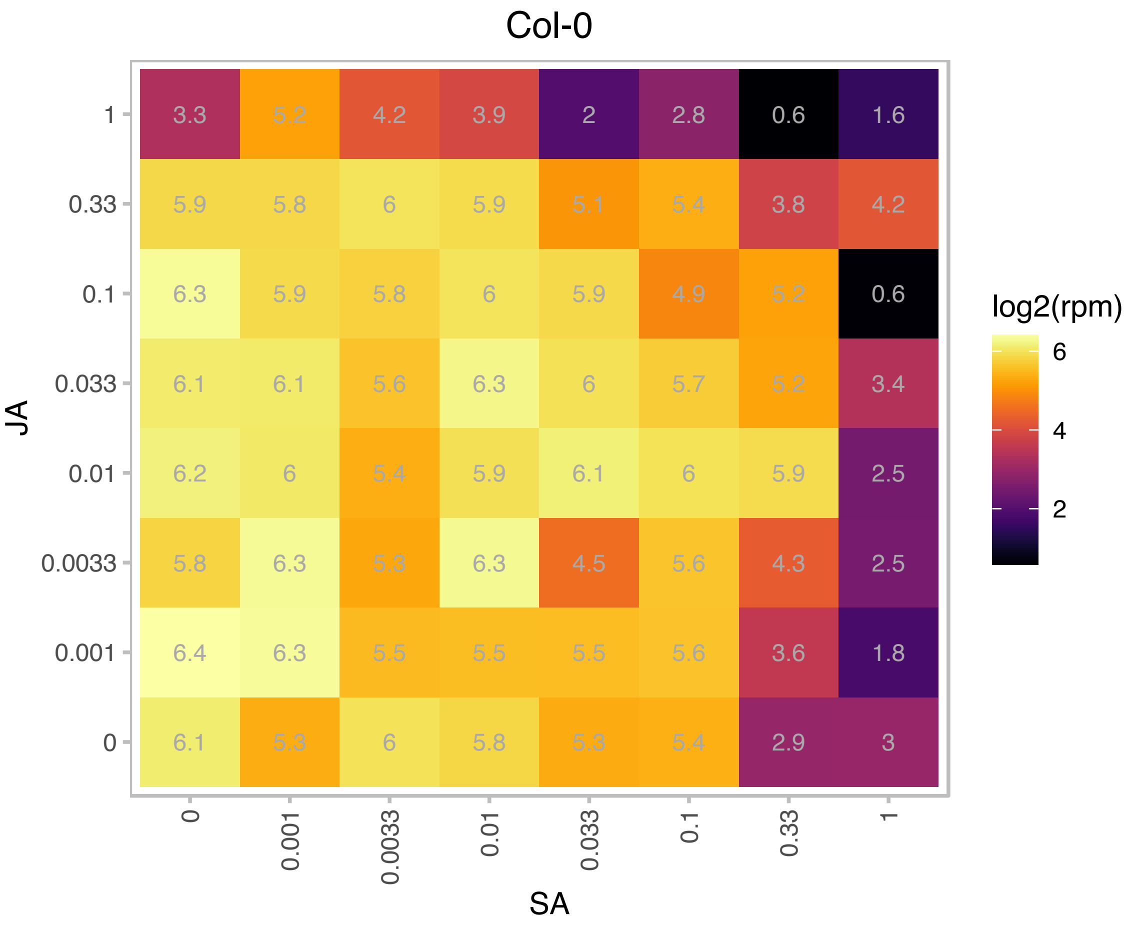


| | |
|---|---|
| Gene | AT1G01320 |
| is.expressed_genes | TRUE |
| is.responsive_genes | TRUE |
| is.apcluster | F-07 |

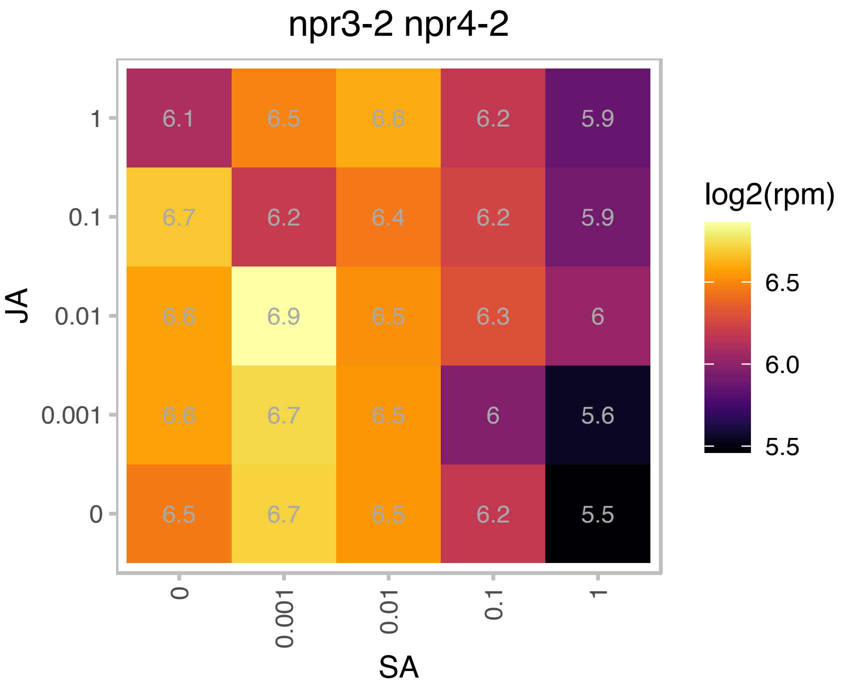


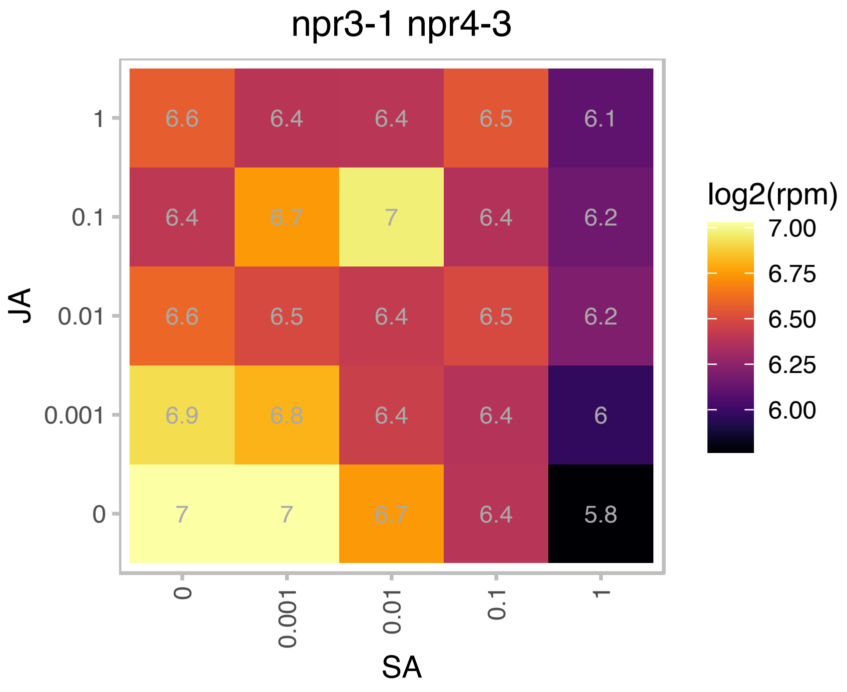

# AT1G02150.1 : NA

Tetratricopeptide repeat (TPR)-like superfamily protein

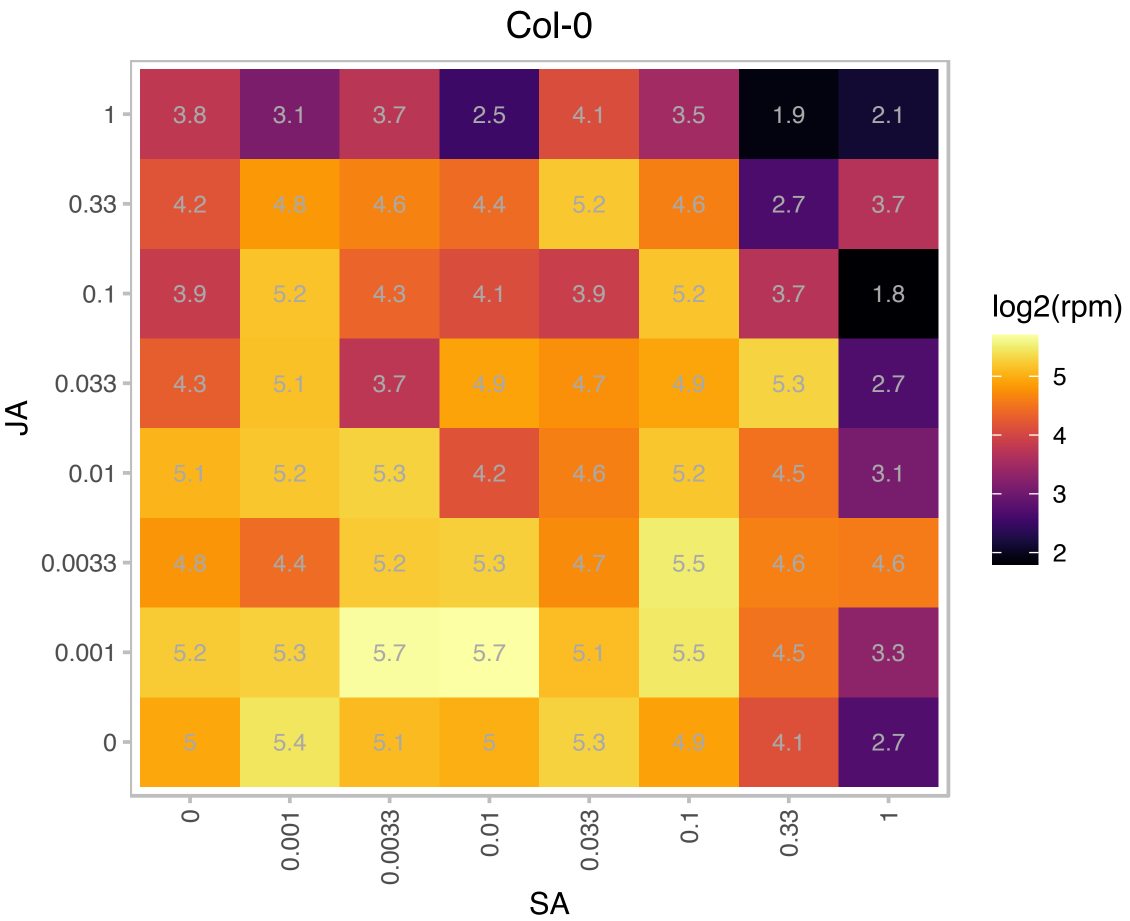


| | |
|---|---|
| Gene | AT1G02150 |
| is.expressed_genes | TRUE |
| is.responsive_genes | TRUE |
| is.apcluster | F-07 |

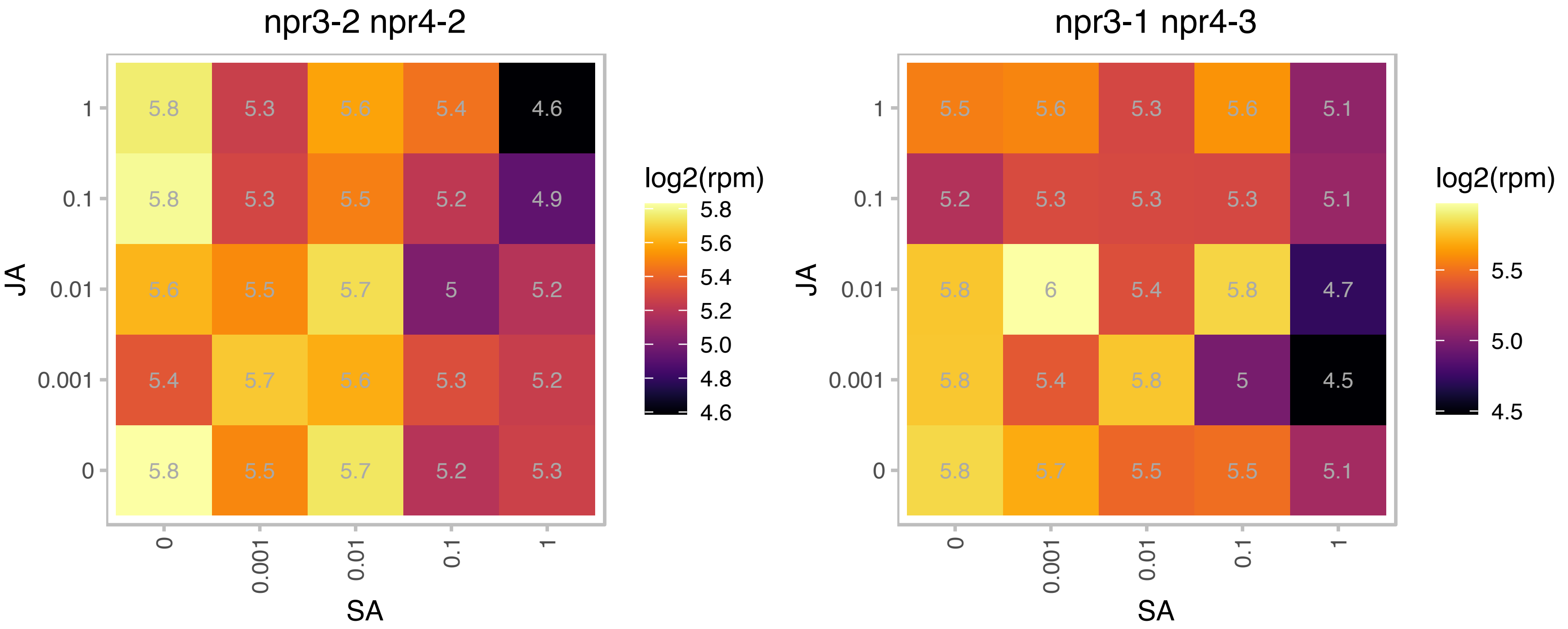

### AT1G02780.1 : emb2386

Ribosomal protein L19e family protein

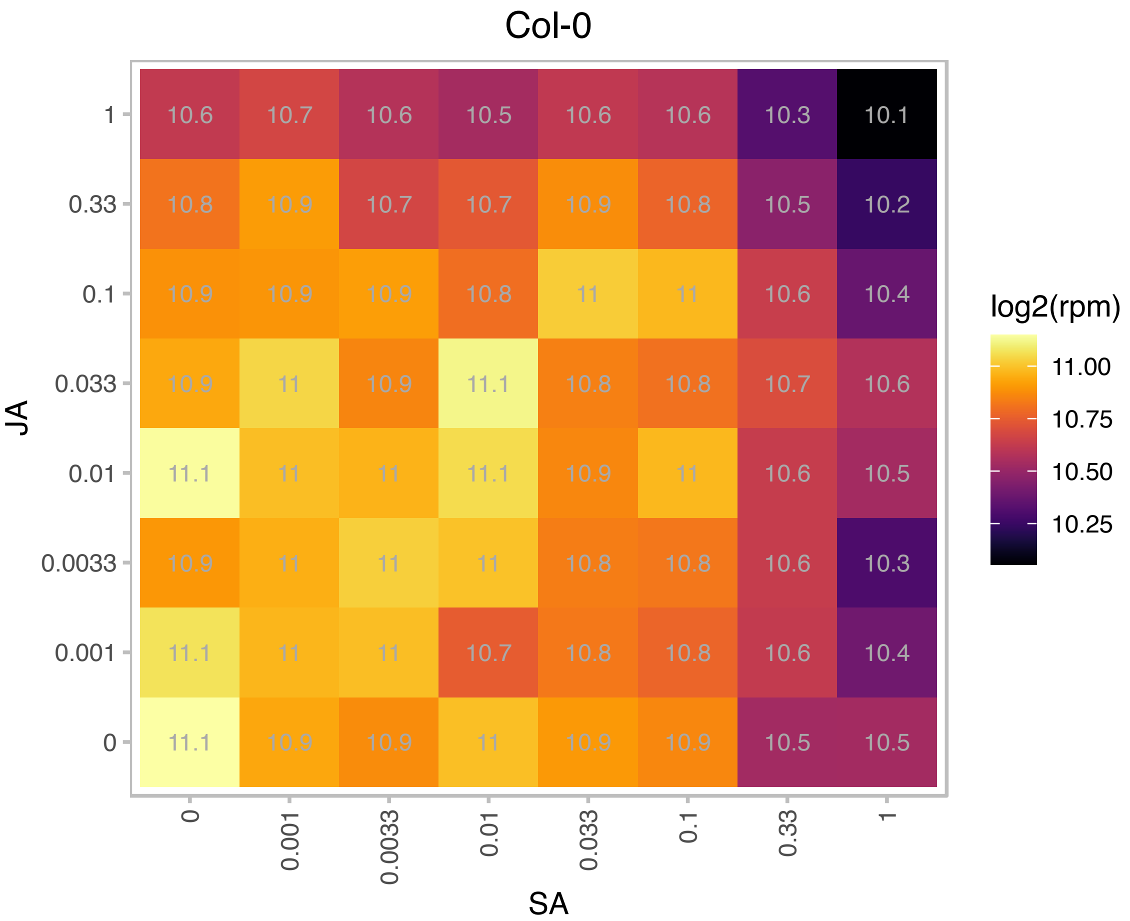


| | |
|---|---|
| Gene | AT1G02780 |
| is.expressed_genes | TRUE |
| is.responsive_genes | TRUE |
| is.apcluster | F-07 |

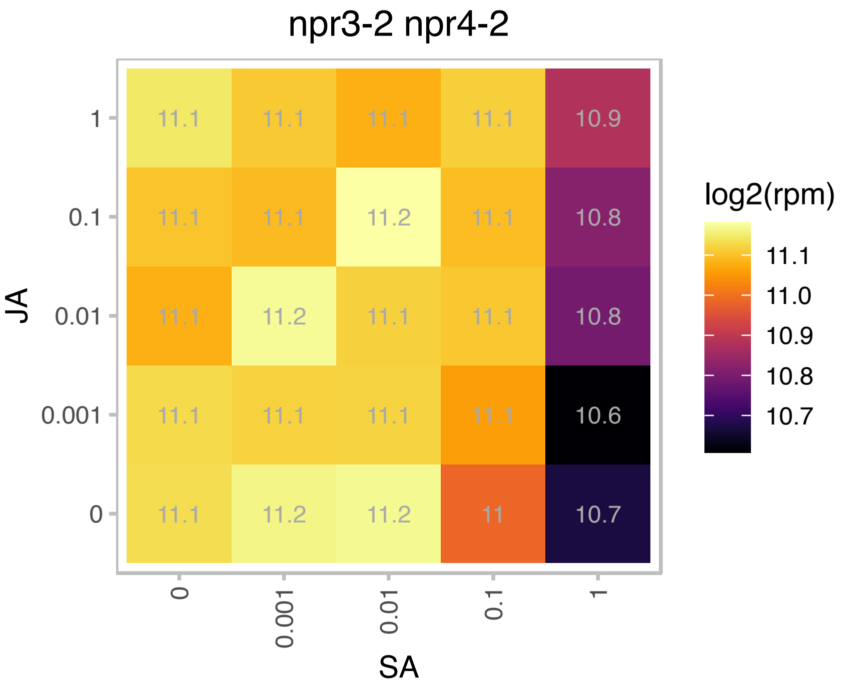


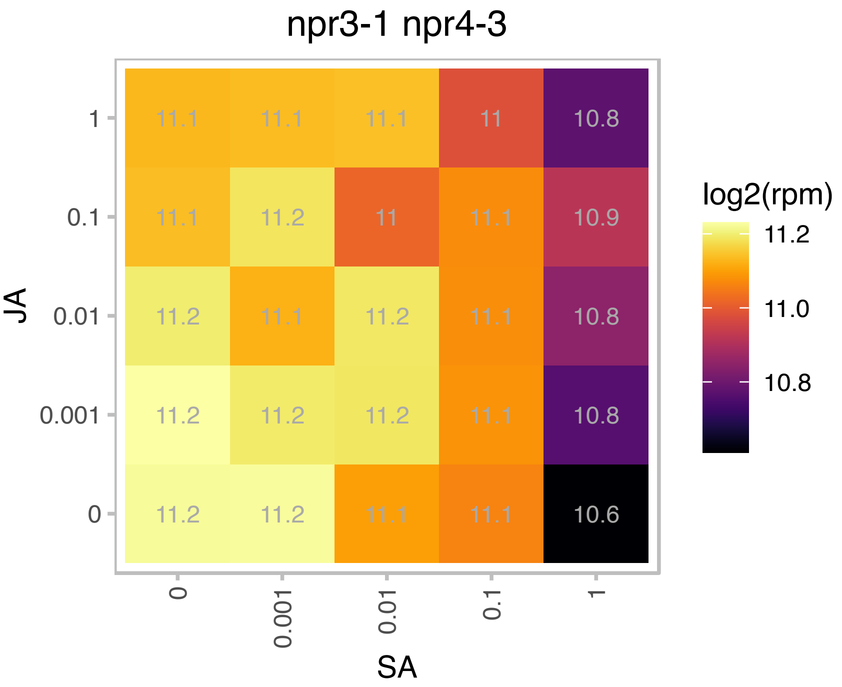

# AT1G05320.1 : NA

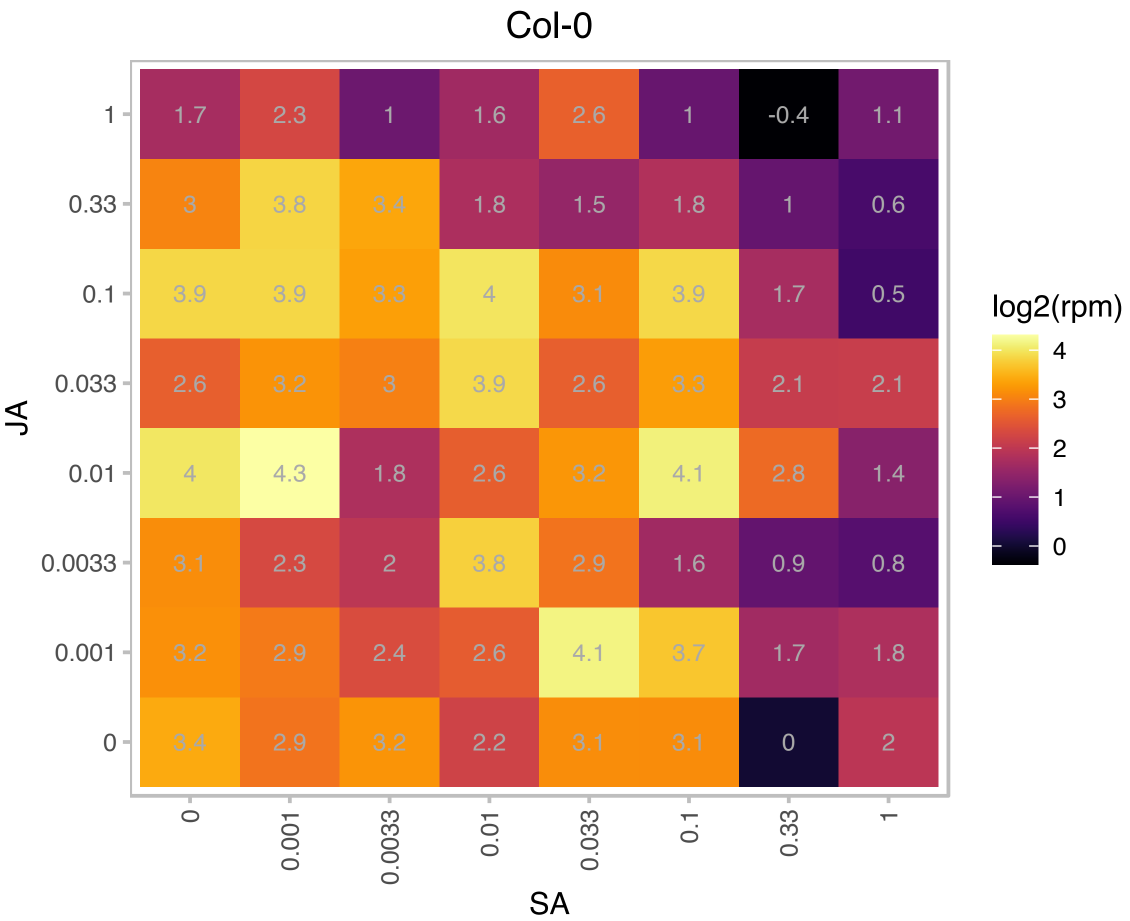


| | |
|---|---|
| Gene | AT1G05320 |
| is.expressed_genes | TRUE |
| is.responsive_genes | TRUE |
| is.apcluster | F-07 |

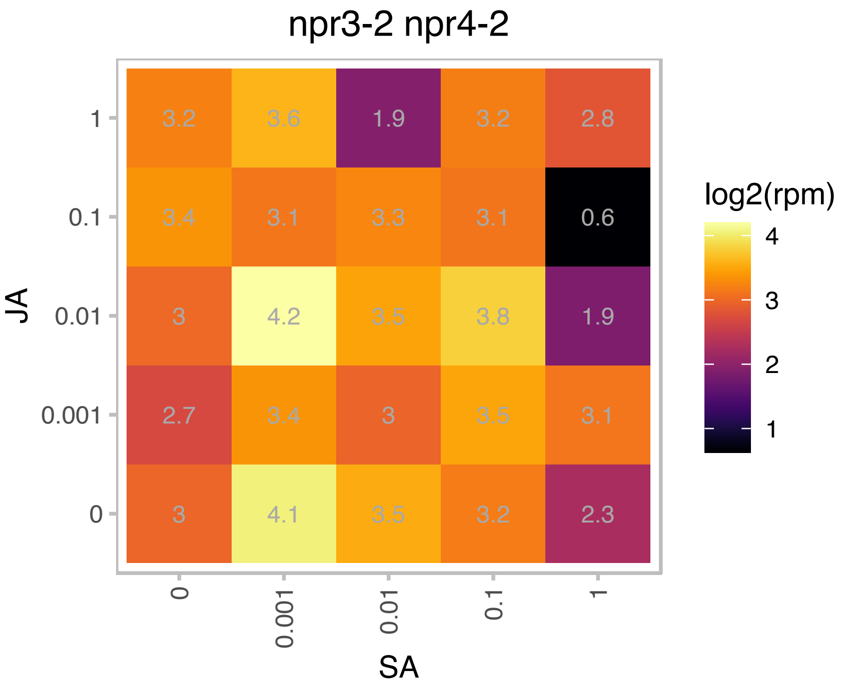


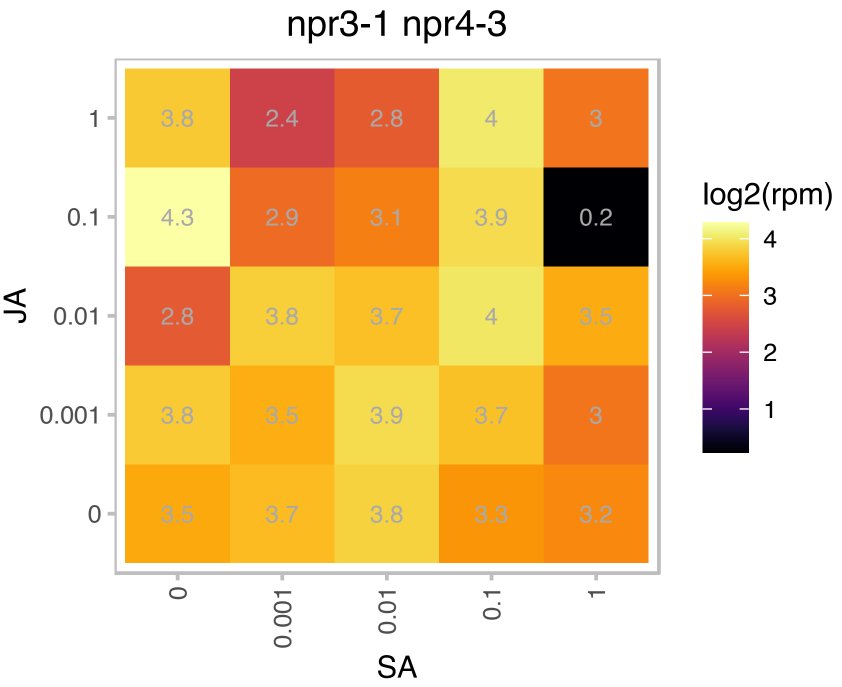

# AT1G07070.1 : NA

Ribosomal protein L35Ae family protein

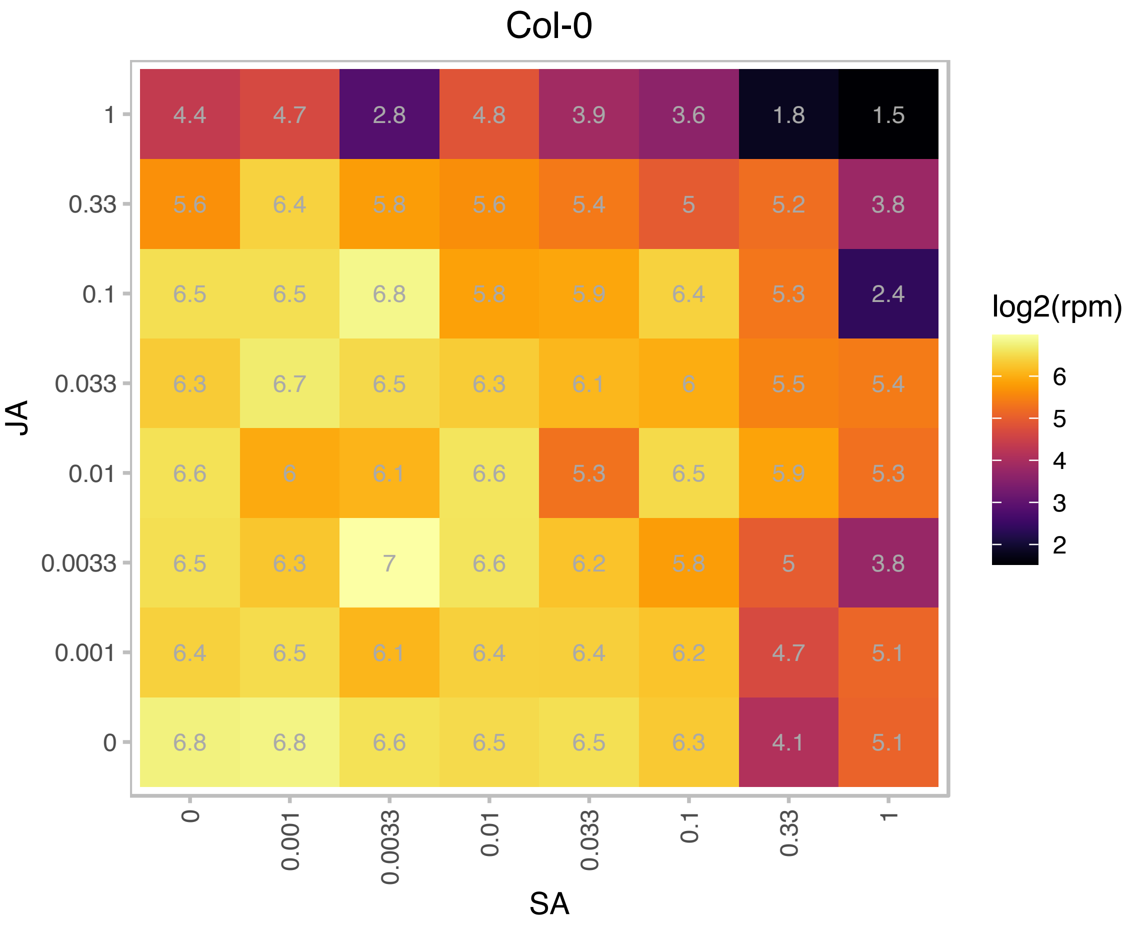


| | |
|---|---|
| Gene | AT1G07070 |
| is.expressed_genes | TRUE |
| is.responsive_genes | TRUE |
| is.apcluster | F-07 |

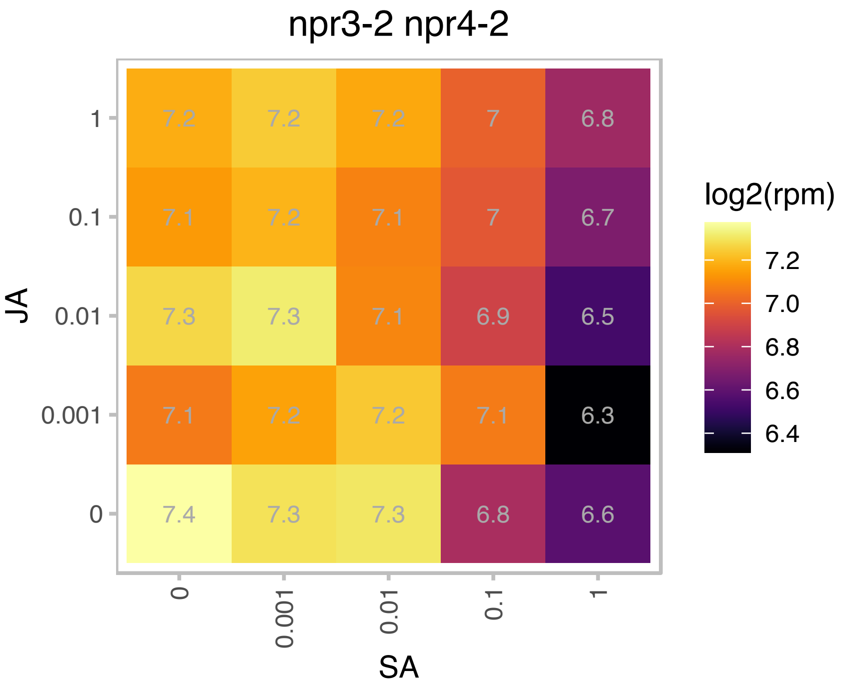


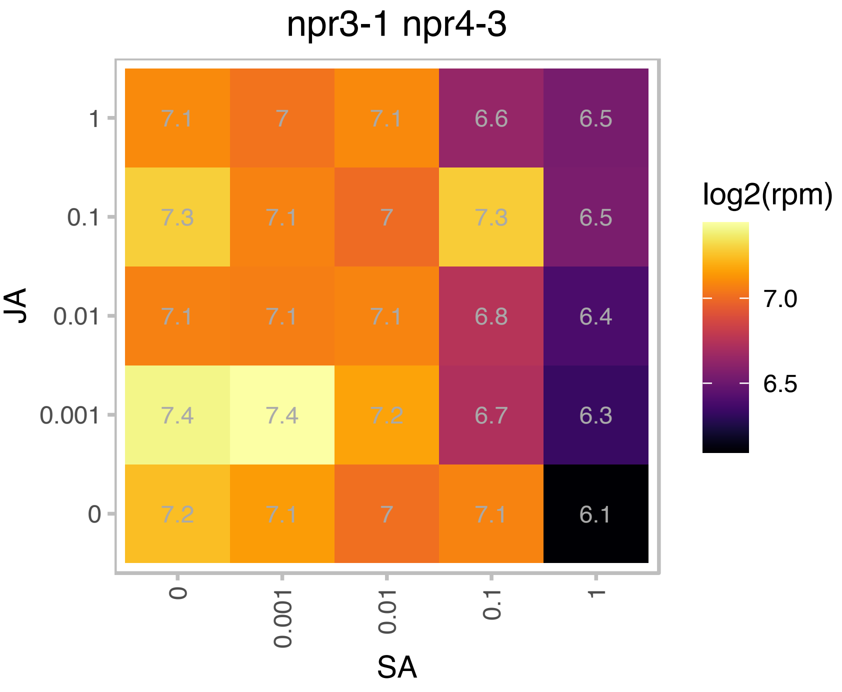

### AT1G07370.1 : ATPCNA1, PCNA1

proliferating cellular nuclear antigen 1

Encodes putative proliferating cell nuclear antigen involved in cell cycle regulation. May be sumoylated.

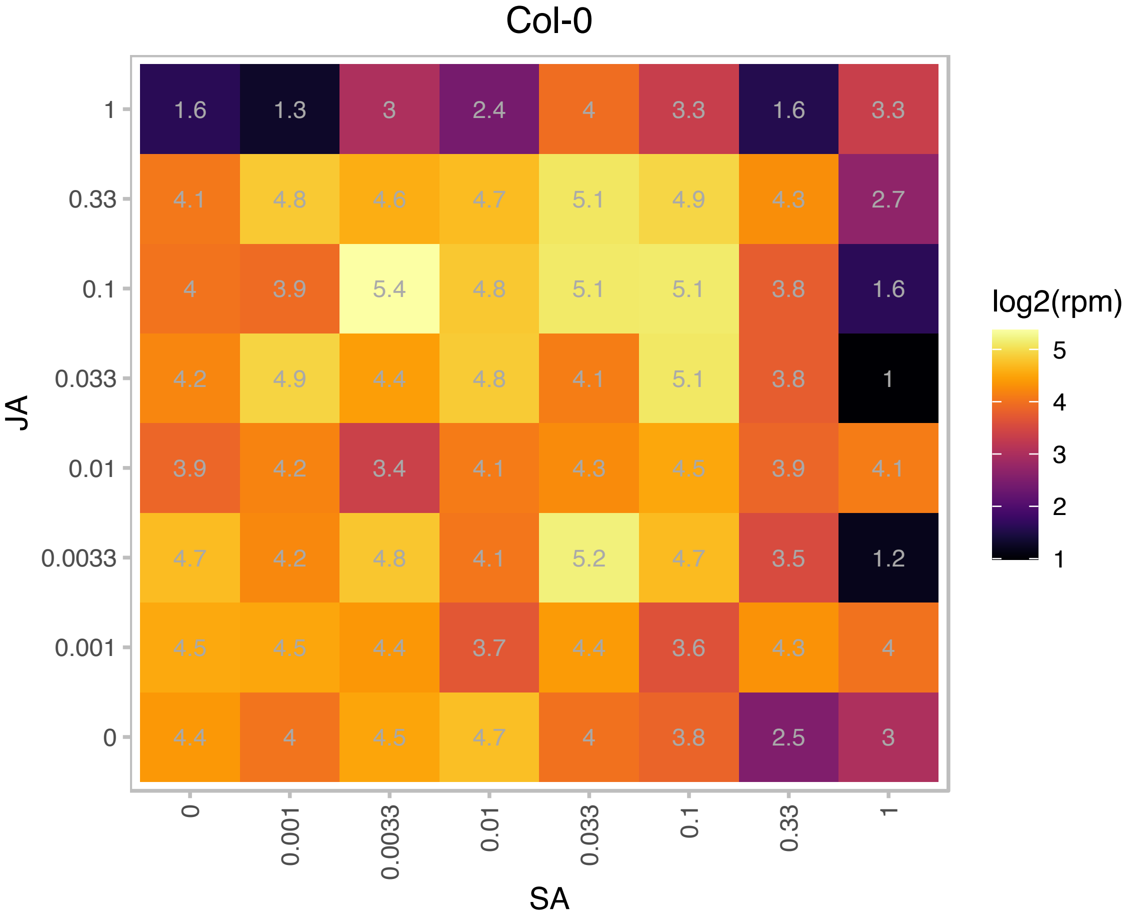


| | |
|---|---|
| Gene | AT1G07370 |
| is.expressed_genes | TRUE |
| is.responsive_genes | TRUE |
| is.apcluster | F-07 |

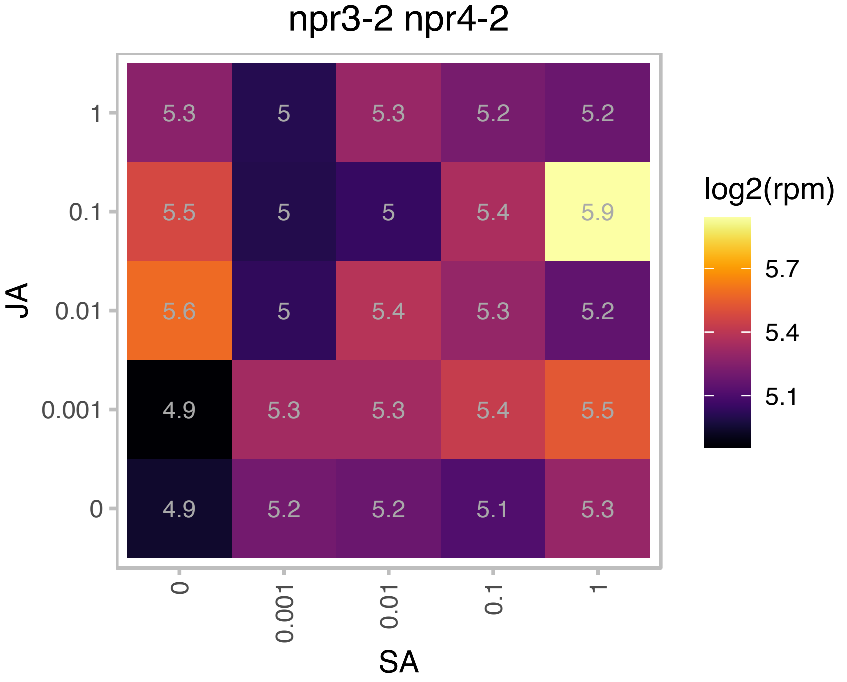


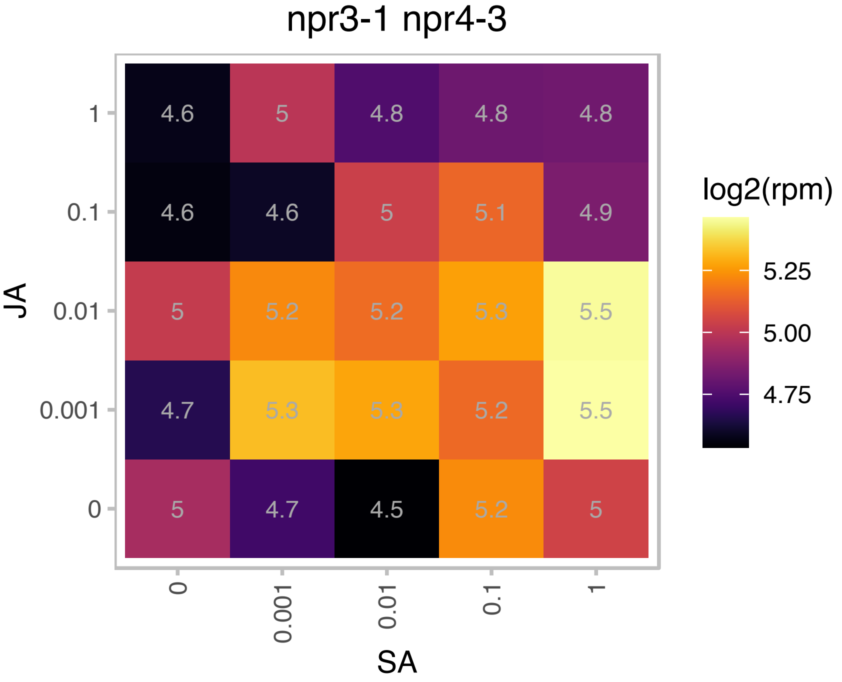

# AT1G08360.1 : NA

Ribosomal protein L1p/L10e family

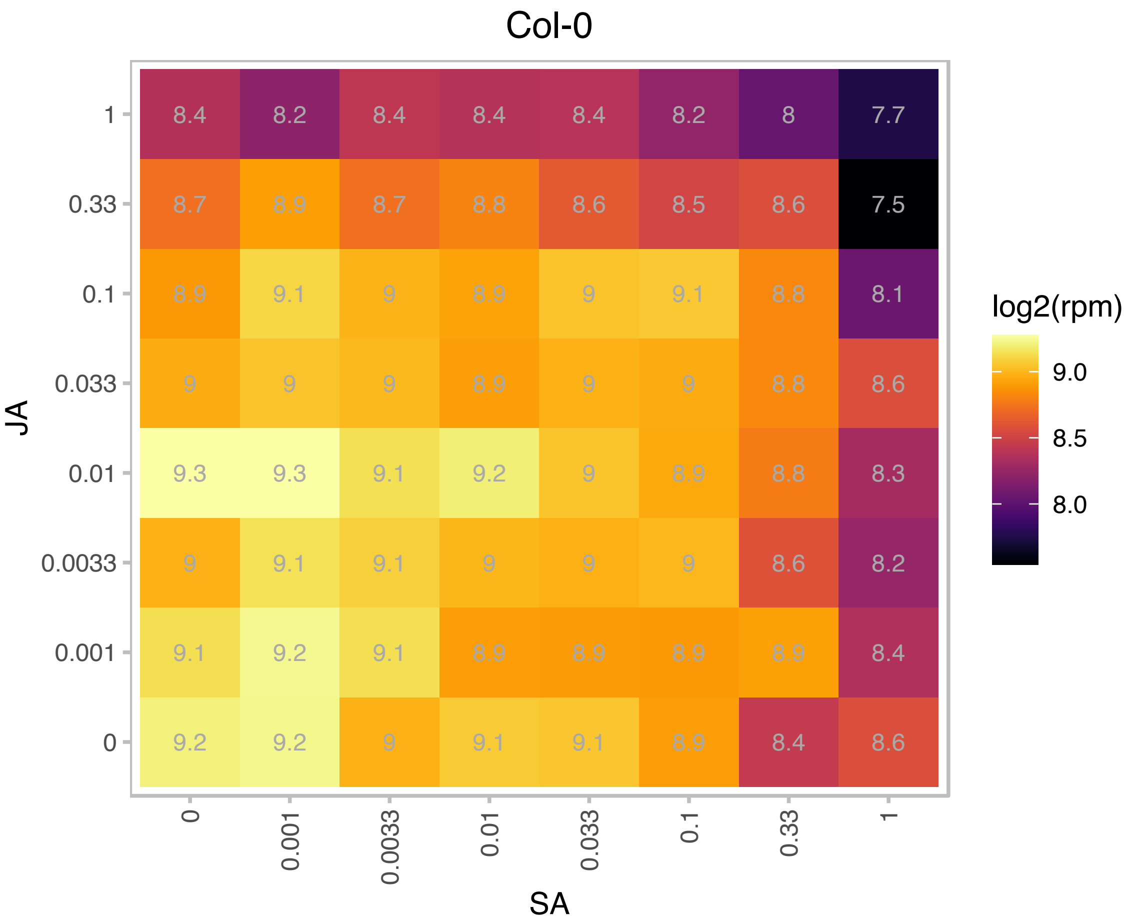


| | |
|---|---|
| Gene | AT1G08360 |
| is.expressed_genes | TRUE |
| is.responsive_genes | TRUE |
| is.apcluster | F-07 |

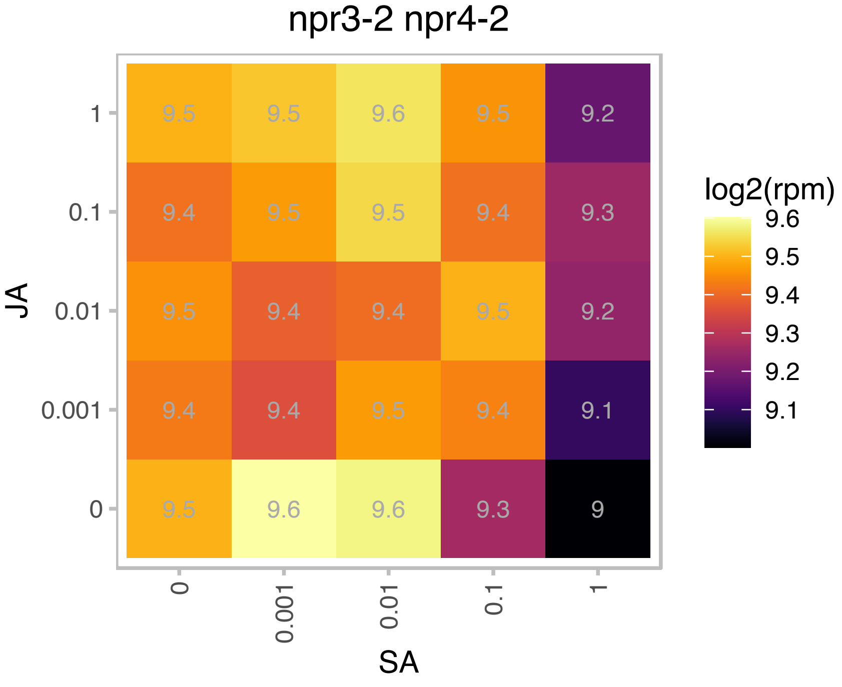


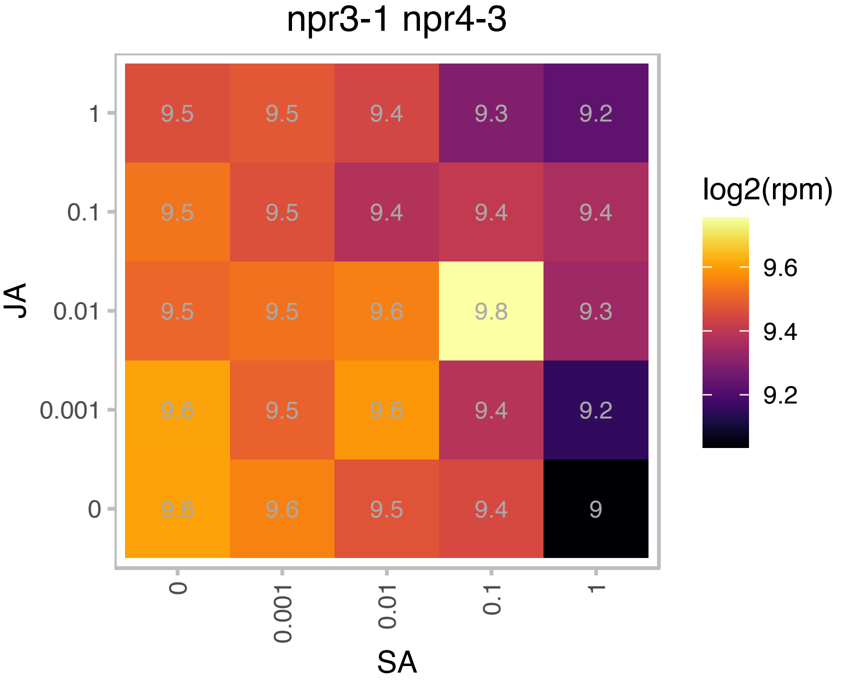

# AT1G08580.1 : NA

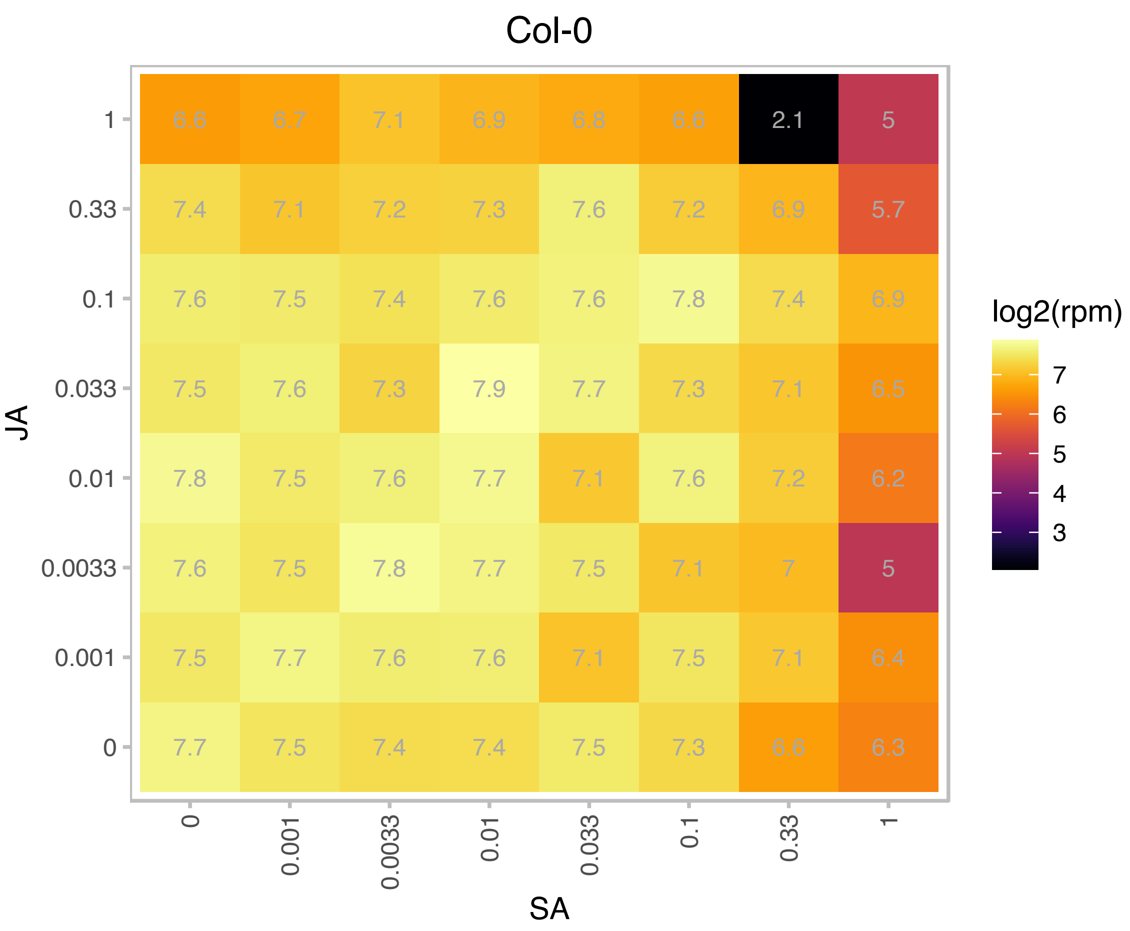


| | |
|---|---|
| Gene | AT1G08580 |
| is.expressed_genes | TRUE |
| is.responsive_genes | TRUE |
| is.apcluster | F-07 |

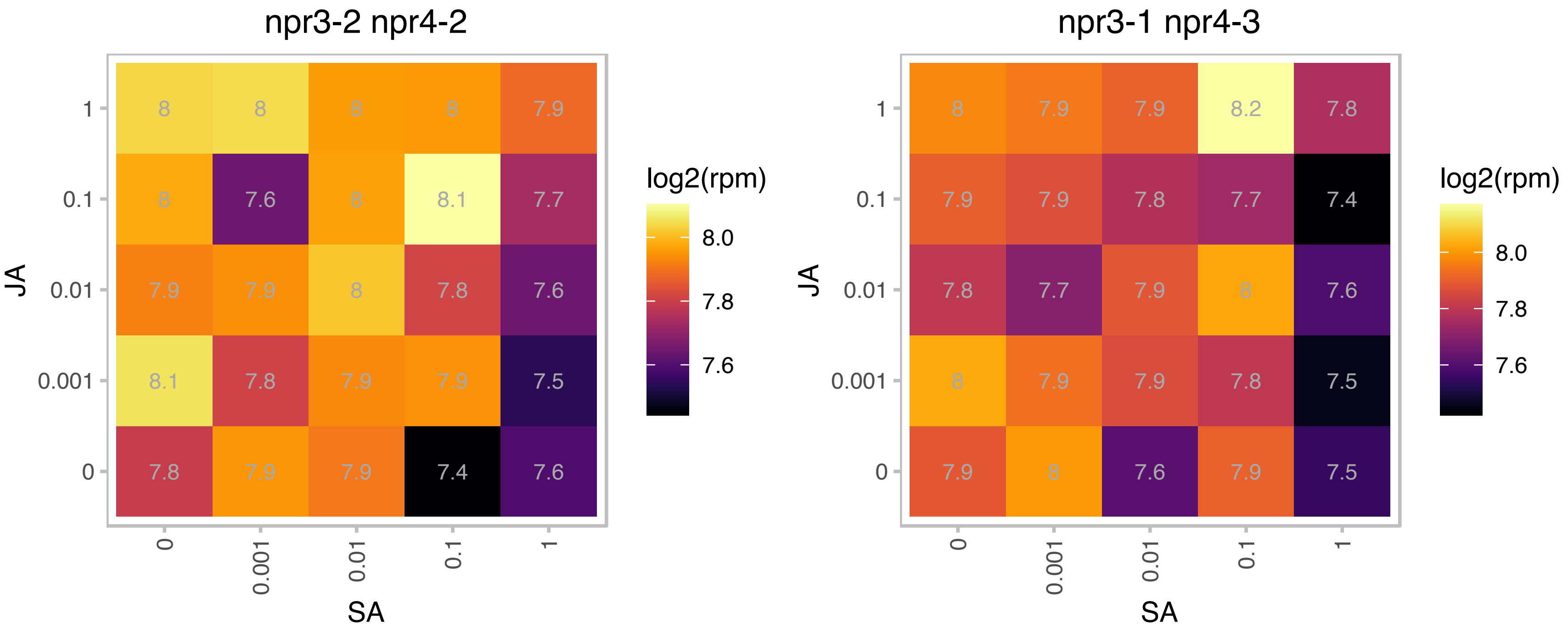

### AT1G09415.1 : NIMIN-3

NIM1-interacting 3

encodes a kinase that physically interacts with NPR1/NIM1

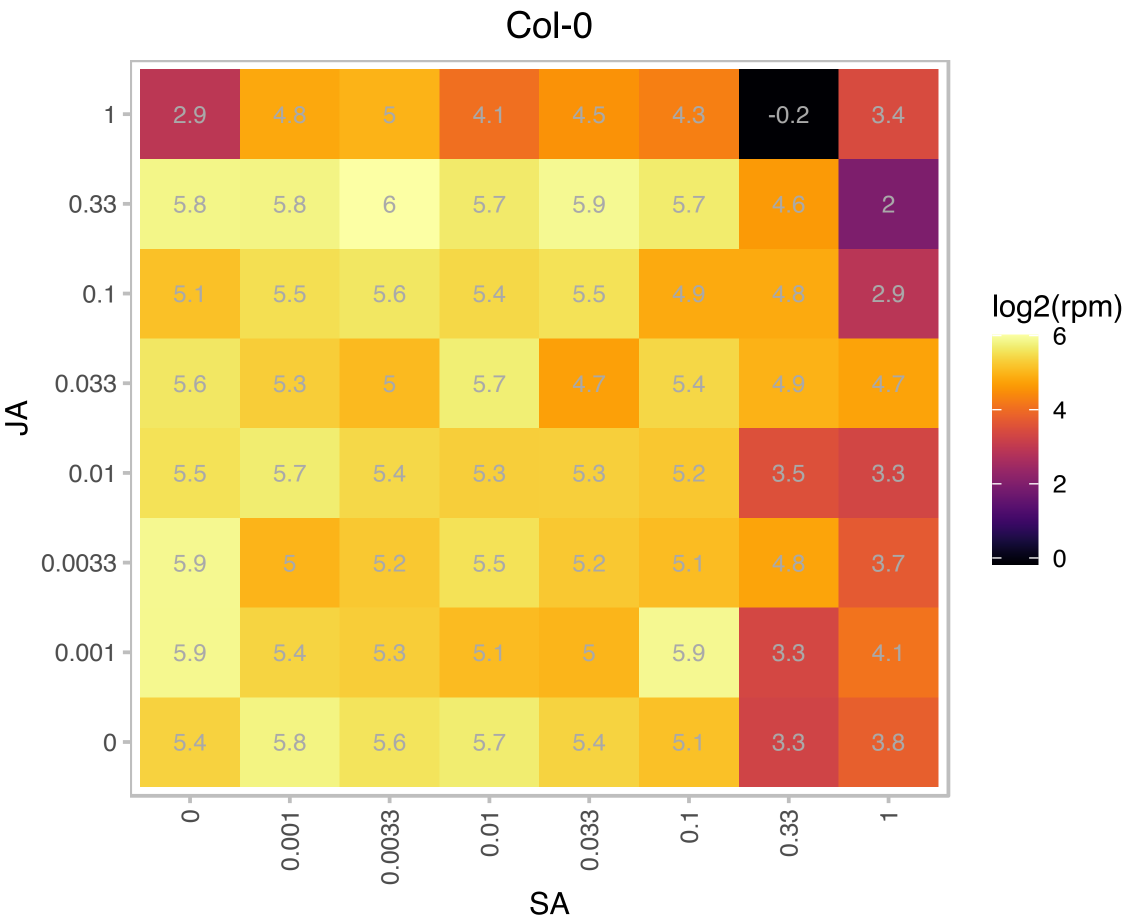


| | |
|---|---|
| Gene | AT1G09415 |
| is.expressed_genes | TRUE |
| is.responsive_genes | TRUE |
| is.apcluster | F-07 |

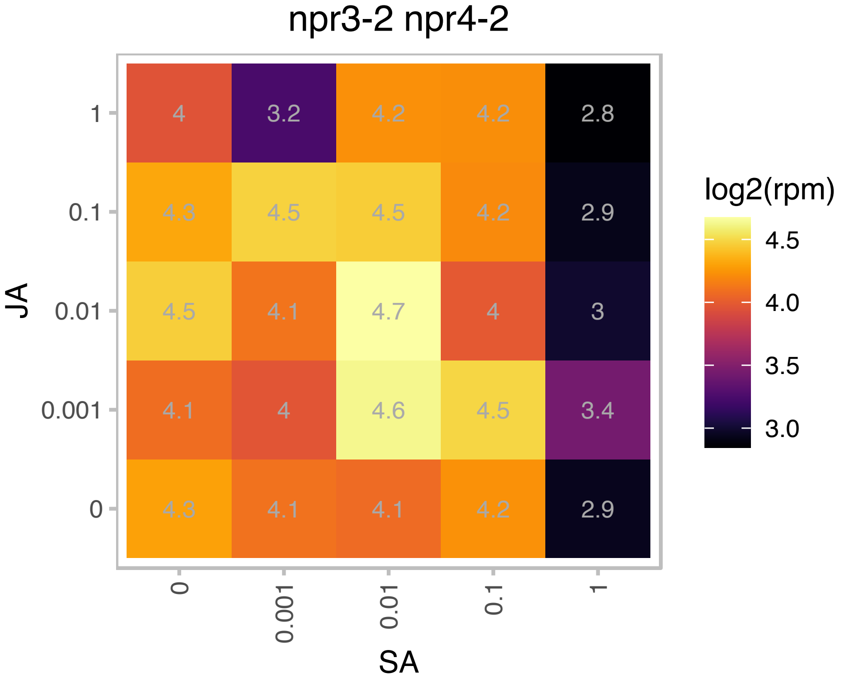


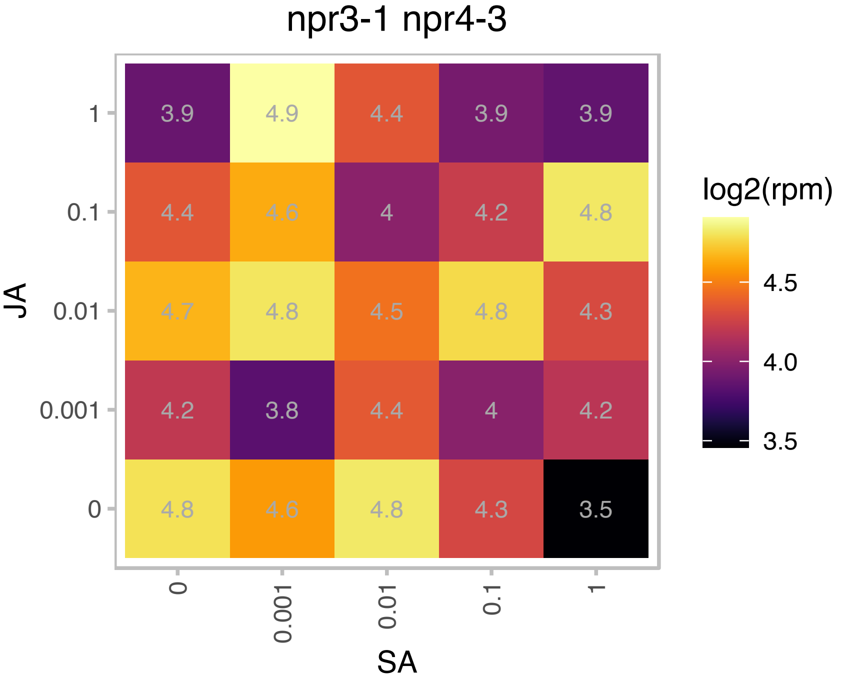

# AT1G09590.1 : NA

Translation protein SH3-like family protein

| | |
|---|---|
| Gene | AT1G09590 |
| is.expressed_genes | TRUE |
| is.responsive_genes | TRUE |
| is.apcluster | F-07 |

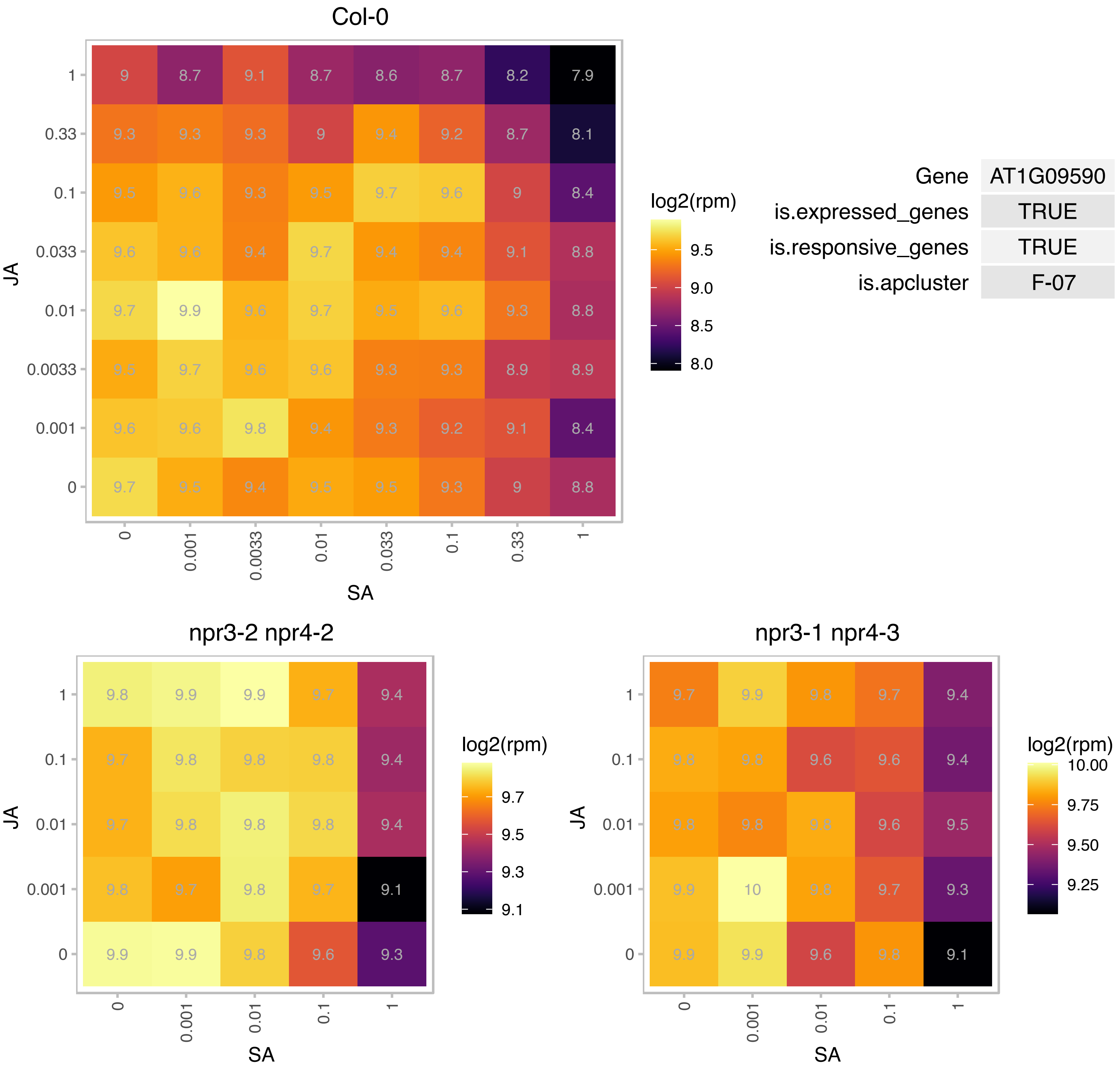

# AT1G09660.1 : NA

RNA-binding KH domain-containing protein

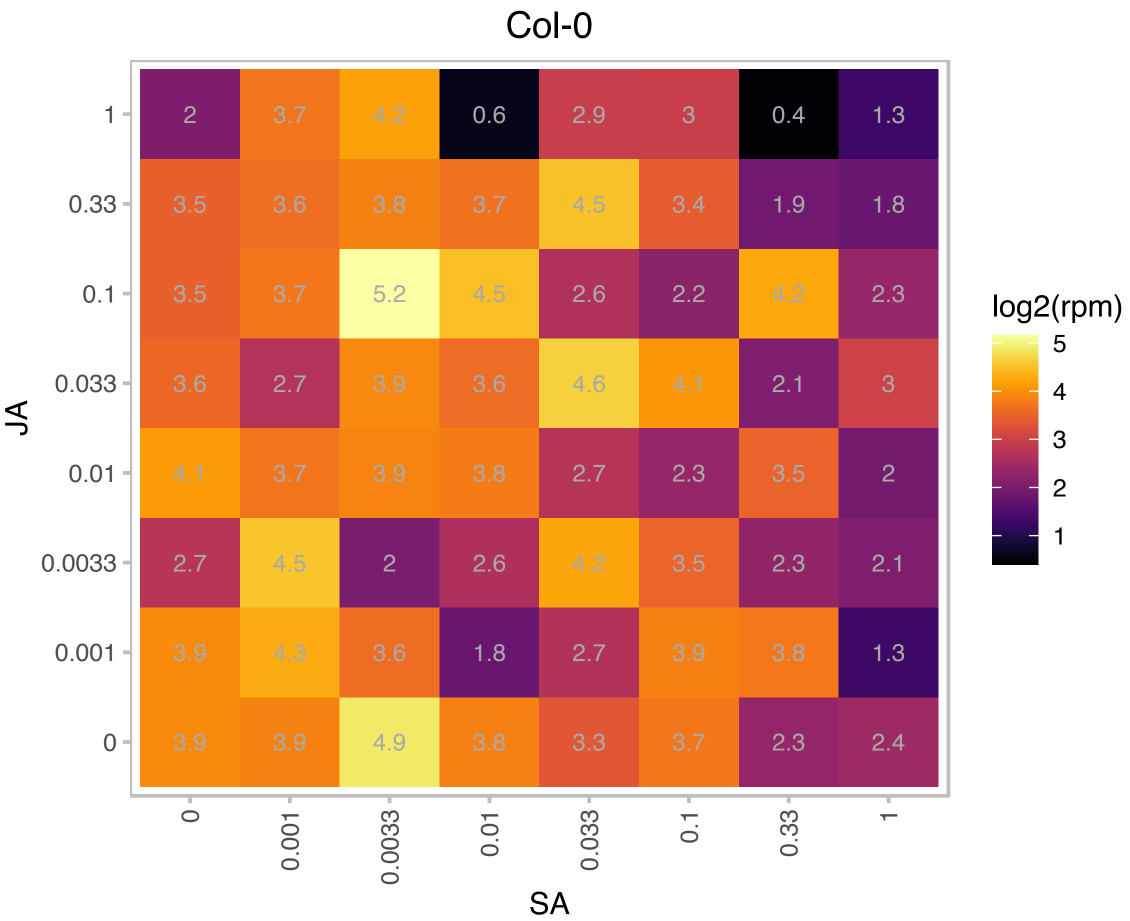


| | |
|---|---|
| Gene | AT1G09660 |
| is.expressed_genes | TRUE |
| is.responsive_genes | TRUE |
| is.apcluster | F-07 |

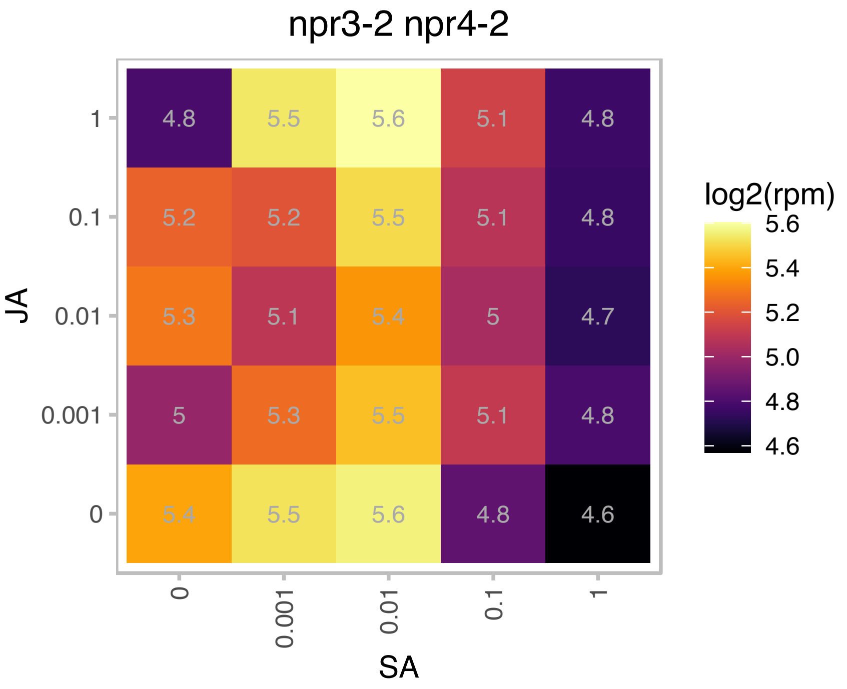


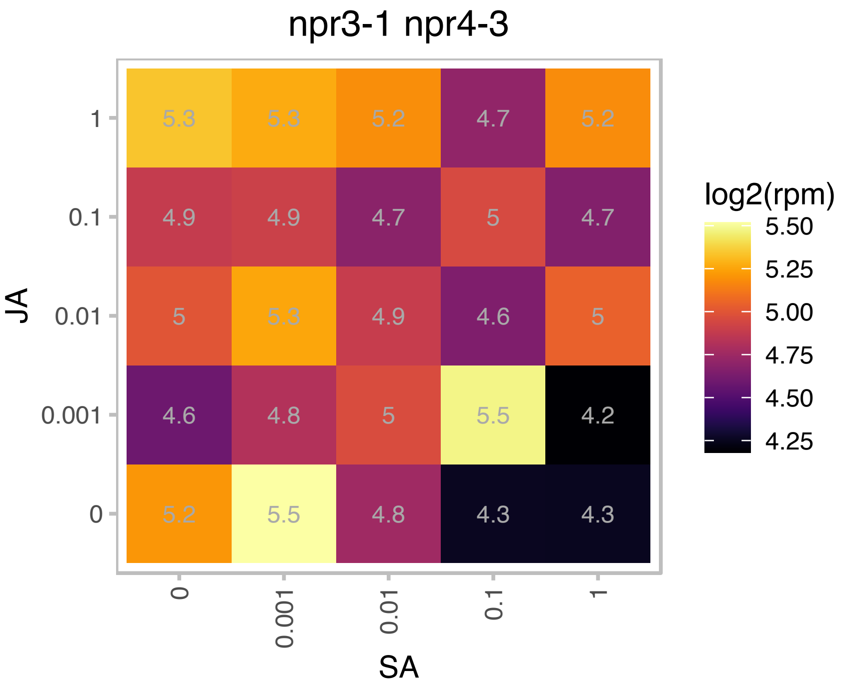

# AT1G11240.1 : NA

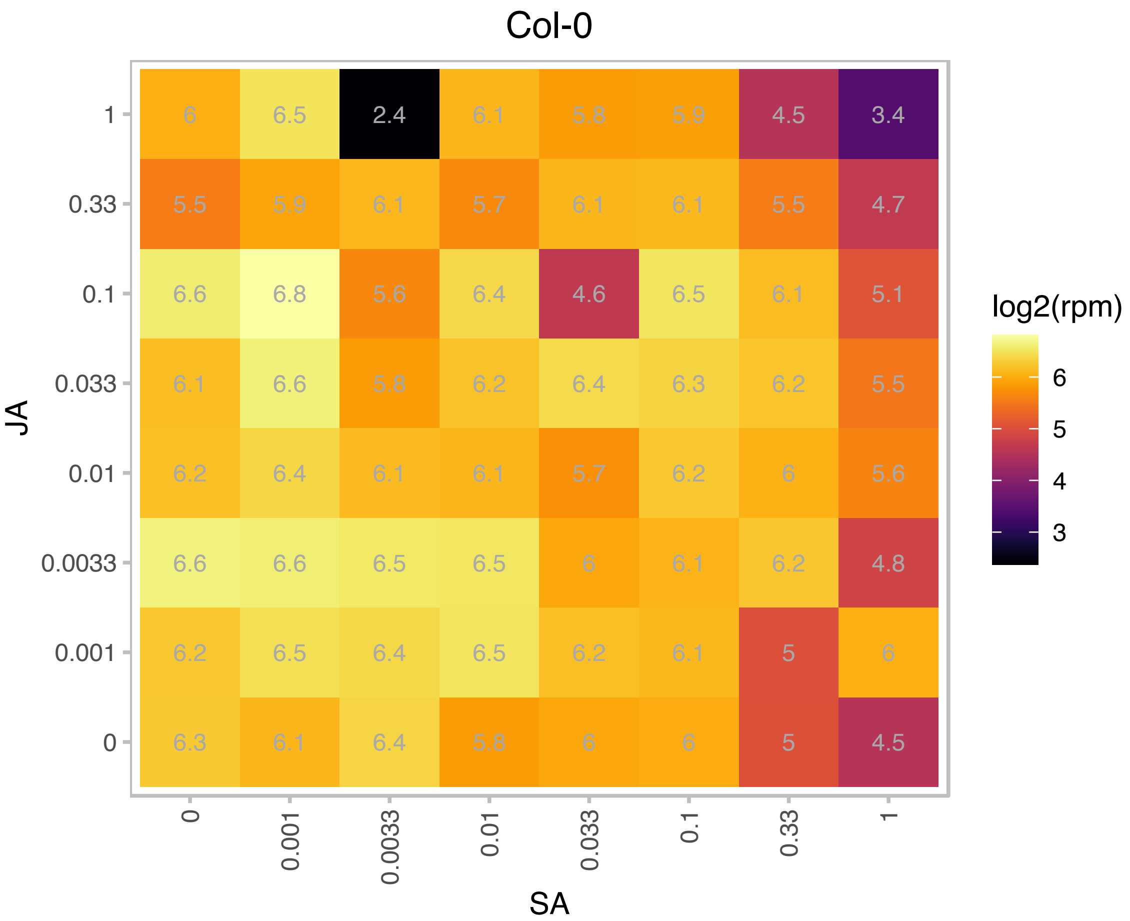


| | |
|---|---|
| Gene | AT1G11240 |
| is.expressed_genes | TRUE |
| is.responsive_genes | TRUE |
| is.apcluster | F-07 |

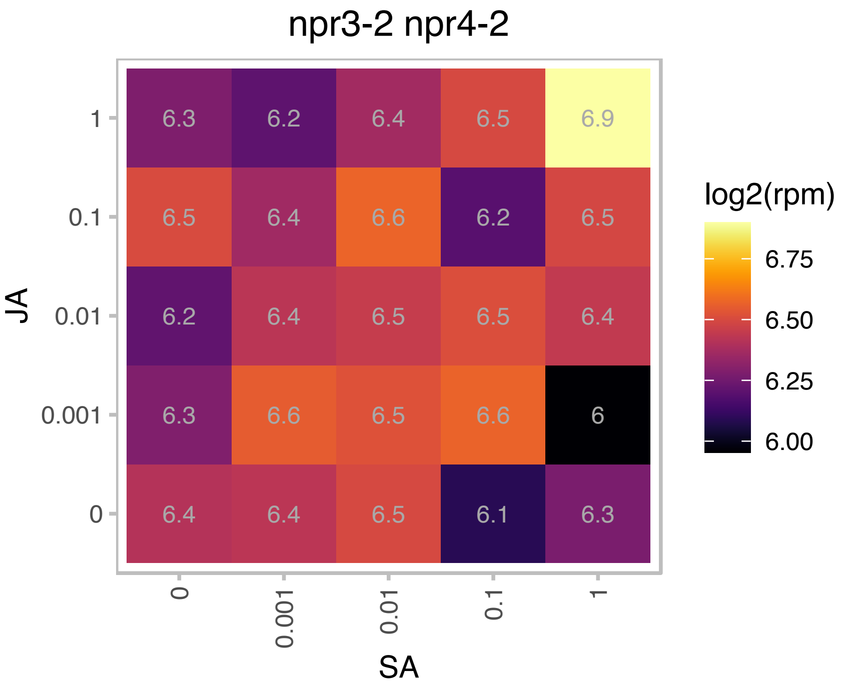


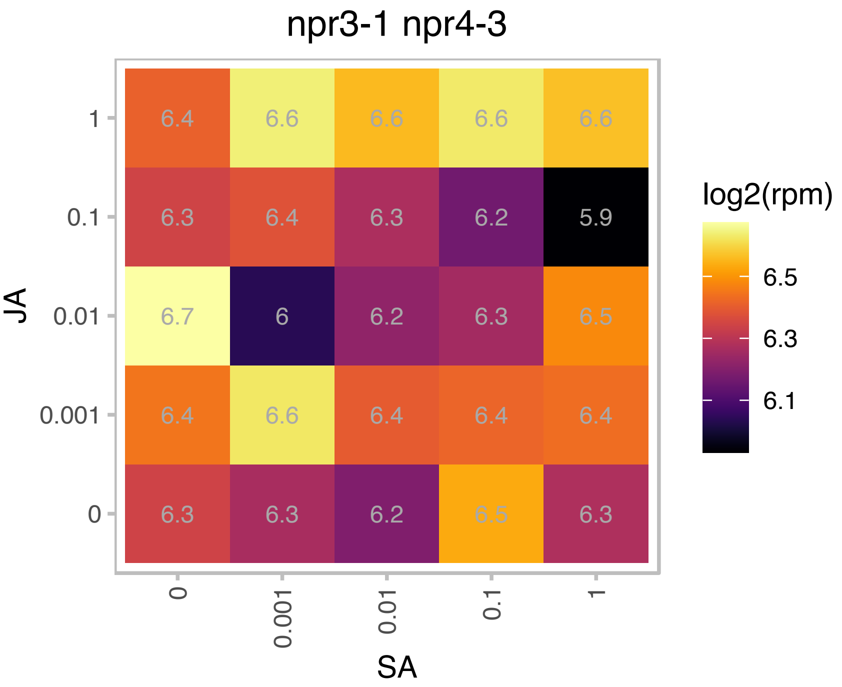

# AT1G12020.1 : NA

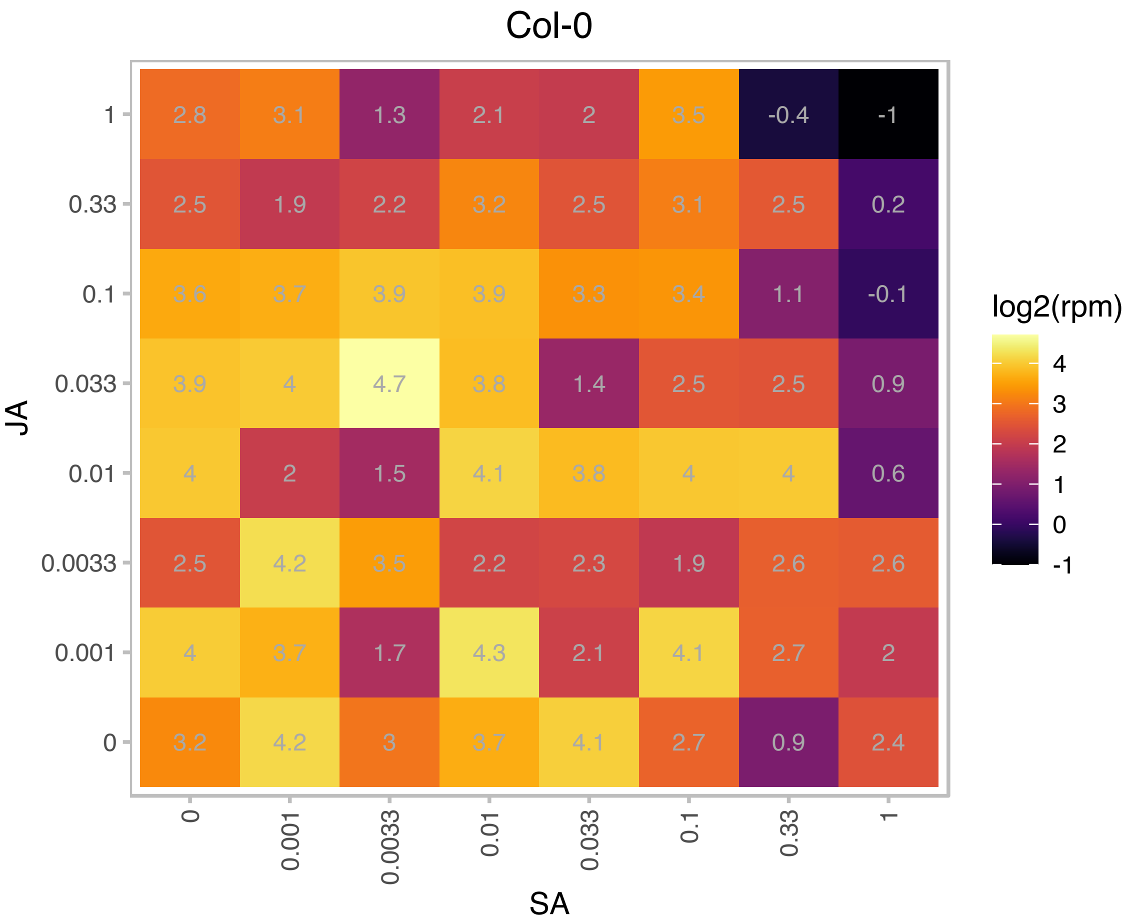


| | |
|---|---|
| Gene | AT1G12020 |
| is.expressed_genes | TRUE |
| is.responsive_genes | TRUE |
| is.apcluster | F-07 |

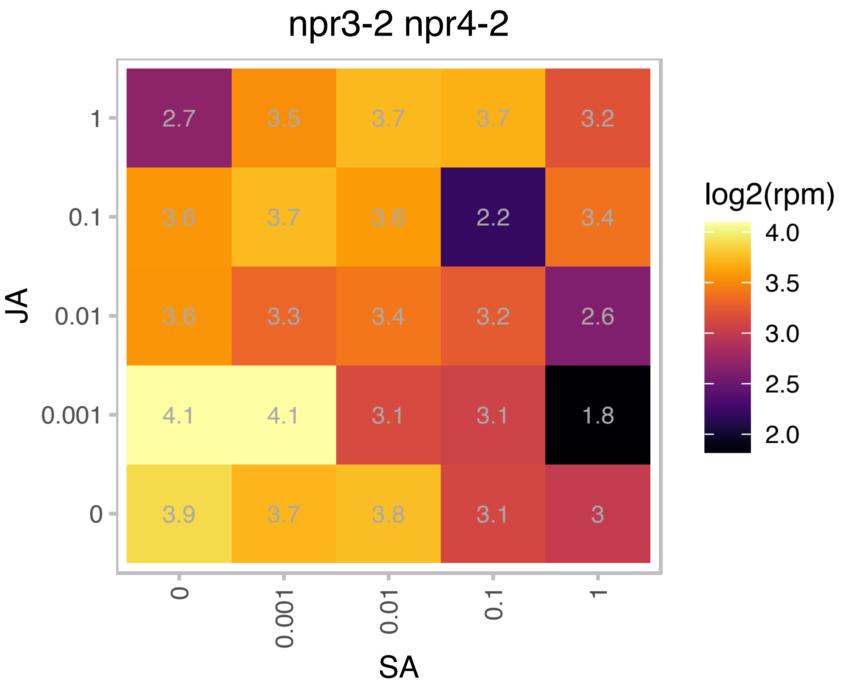


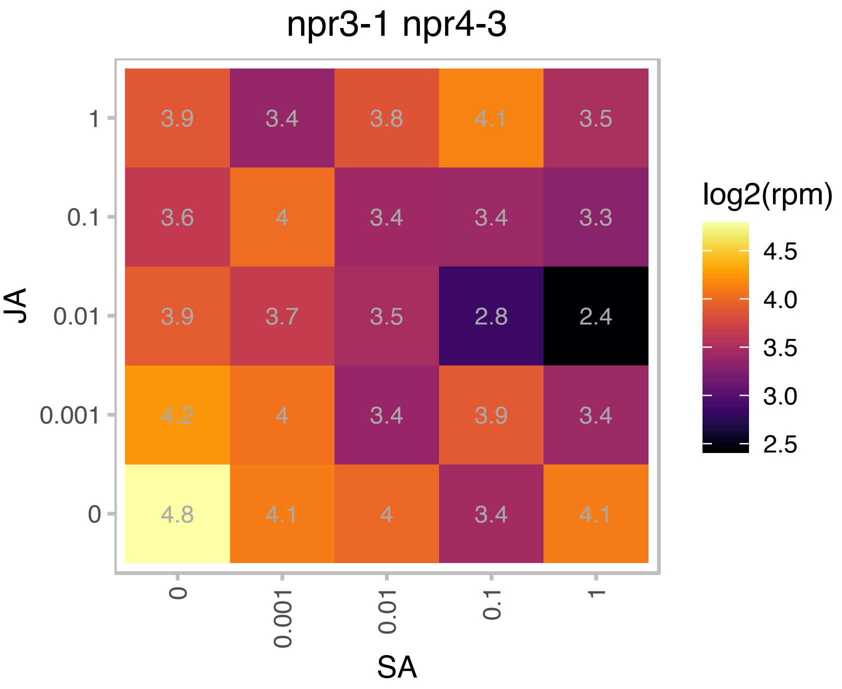

### AT1G13300.1 : AtNIGT1, HRS1, NIGT1.4

myb-like transcription factor family protein
Encodes a nuclear localized member of the GARP family of transcription factors. Involved in nitrate/phosphate signaling in roots. It is transcriptionally regulated by nitrate and post transcriptionally by phospate and functions to integrate these two nutrient signaling pathways in the root.

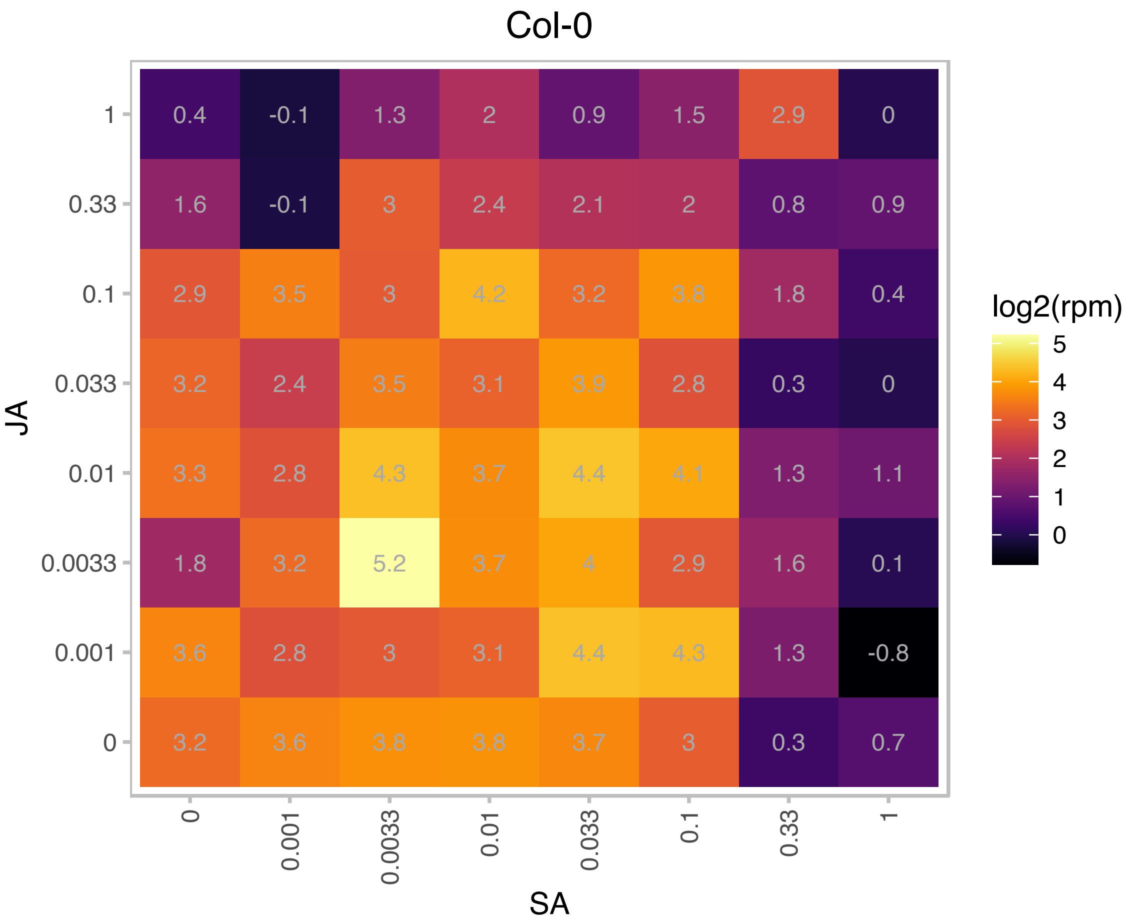


| JA \ SA | 0 | 0.001 | 0.0033 | 0.01 | 0.033 | 0.1 | 0.33 | 1 |
|---|---|---|---|---|---|---|---|---|
| 1 | 0.4 | -0.1 | 1.3 | 2 | 0.9 | 1.5 | 2.9 | 0 |
| 0.33 | 1.6 | -0.1 | 3 | 2.4 | 2.1 | 2 | 0.8 | 0.9 |
| 0.1 | 2.9 | 3.5 | 3 | 4.2 | 3.2 | 3.8 | 1.8 | 0.4 |
| 0.033 | 3.2 | 2.4 | 3.5 | 3.1 | 3.9 | 2.8 | 0.3 | 0 |
| 0.01 | 3.3 | 2.8 | 4.3 | 3.7 | 4.4 | 4.1 | 1.3 | 1.1 |
| 0.0033 | 1.8 | 3.2 | 5.2 | 3.7 | 4 | 2.9 | 1.6 | 0.1 |
| 0.001 | 3.6 | 2.8 | 3 | 3.1 | 4.4 | 4.3 | 1.3 | -0.8 |
| 0 | 3.2 | 3.6 | 3.8 | 3.8 | 3.7 | 3 | 0.3 | 0.7 |

| | |
|---|---|
| Gene | AT1G13300 |
| is.expressed_genes | TRUE |
| is.responsive_genes | TRUE |
| is.apcluster | F-07 |

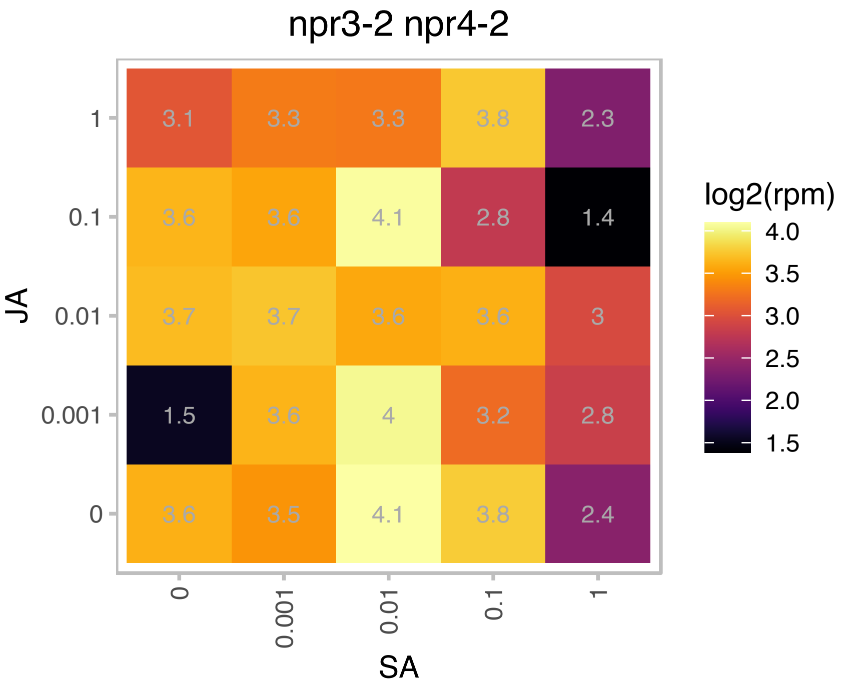


| JA \ SA | 0 | 0.001 | 0.01 | 0.1 | 1 |
|---|---|---|---|---|---|
| 1 | 3.1 | 3.3 | 3.3 | 3.8 | 2.3 |
| 0.1 | 3.6 | 3.6 | 4.1 | 2.8 | 1.4 |
| 0.01 | 3.7 | 3.7 | 3.6 | 3.6 | 3 |
| 0.001 | 1.5 | 3.6 | 4 | 3.2 | 2.8 |
| 0 | 3.6 | 3.5 | 4.1 | 3.8 | 2.4 |

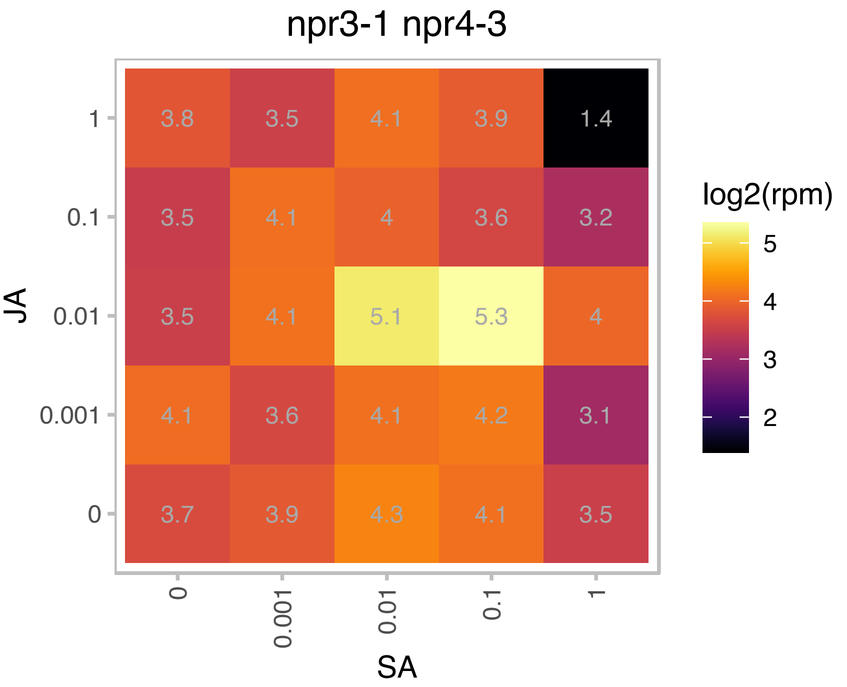


| JA \ SA | 0 | 0.001 | 0.01 | 0.1 | 1 |
|---|---|---|---|---|---|
| 1 | 3.8 | 3.5 | 4.1 | 3.9 | 1.4 |
| 0.1 | 3.5 | 4.1 | 4 | 3.6 | 3.2 |
| 0.01 | 3.5 | 4.1 | 5.1 | 5.3 | 4 |
| 0.001 | 4.1 | 3.6 | 4.1 | 4.2 | 3.1 |
| 0 | 3.7 | 3.9 | 4.3 | 4.1 | 3.5 |

### AT1G14280.1 : PKS2

phytochrome kinase substrate 2

Encodes phytochrome kinase substrate 2.  PKS proteins are critical for hypocotyl phototropism. Forms a complex with Phot1, Phot2 and NPH3.

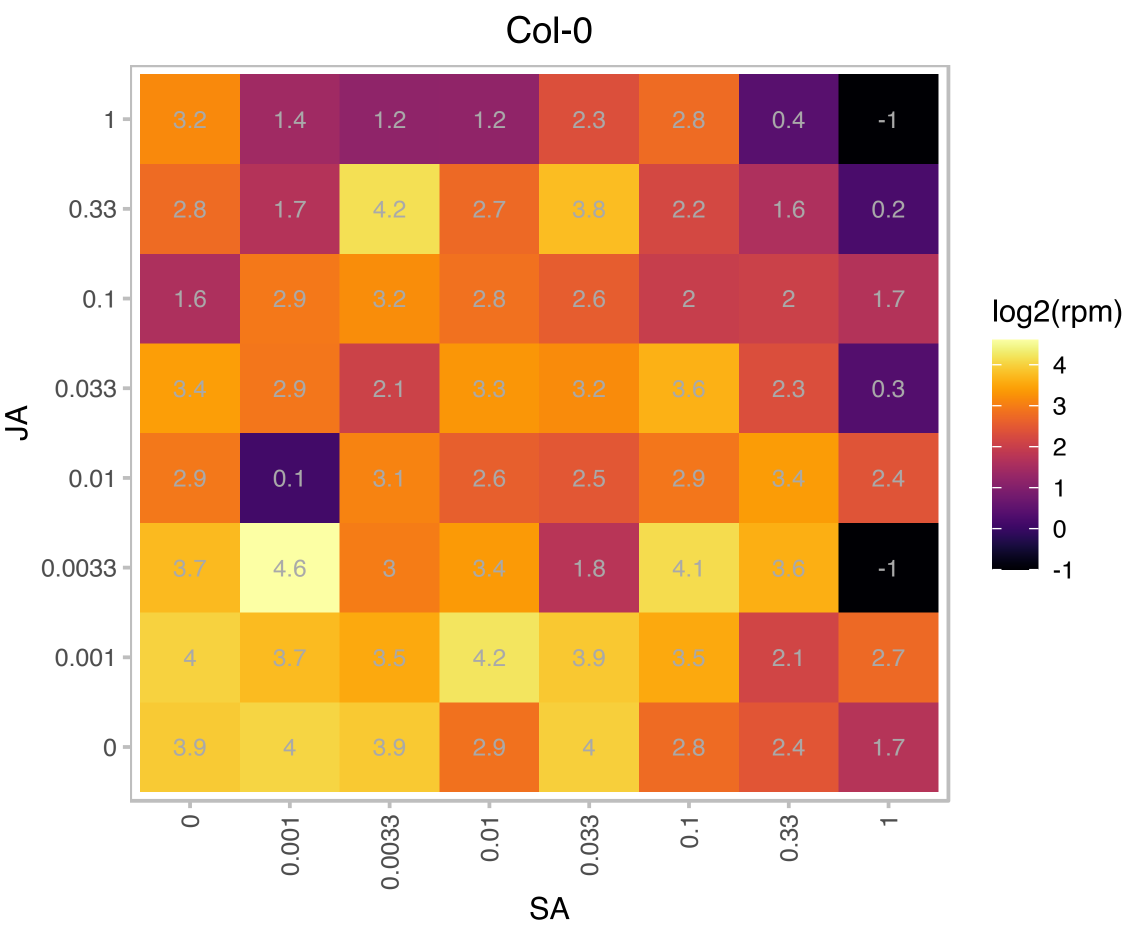


| | |
|---|---|
| Gene | AT1G14280 |
| is.expressed_genes | TRUE |
| is.responsive_genes | TRUE |
| is.apcluster | F-07 |

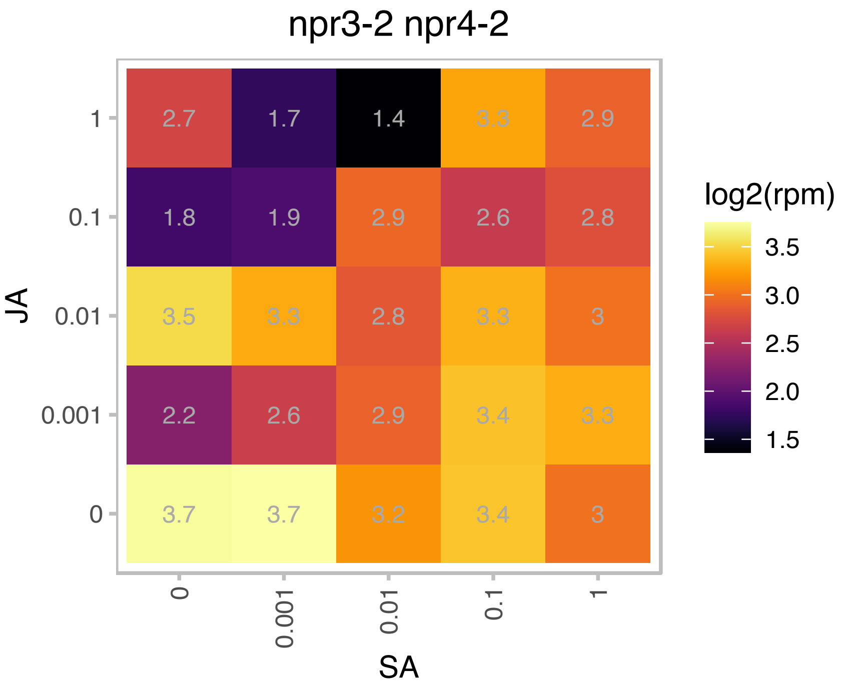


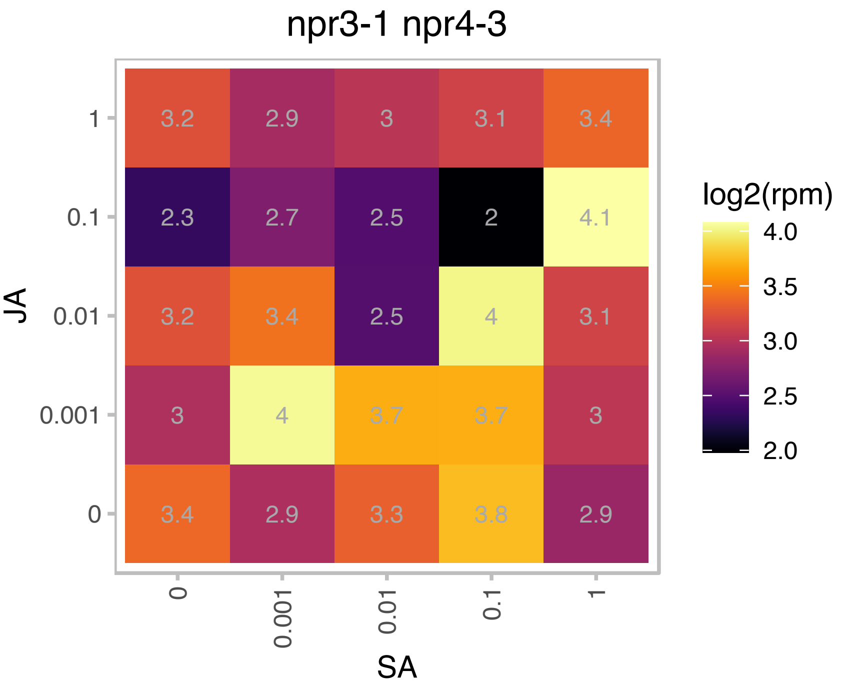

### AT1G14320.1 : RPL10, RPL10A, SAC52

Ribosomal protein L16p/L10e family protein
Encodes a ribosomal protein L10 and may be involved in translation regulation. Semi-dominant mutations in SAC552 can suppress defects in acaulis5, which encodes a thermospermine synthase, by enhancing translation of acl5 and itself.

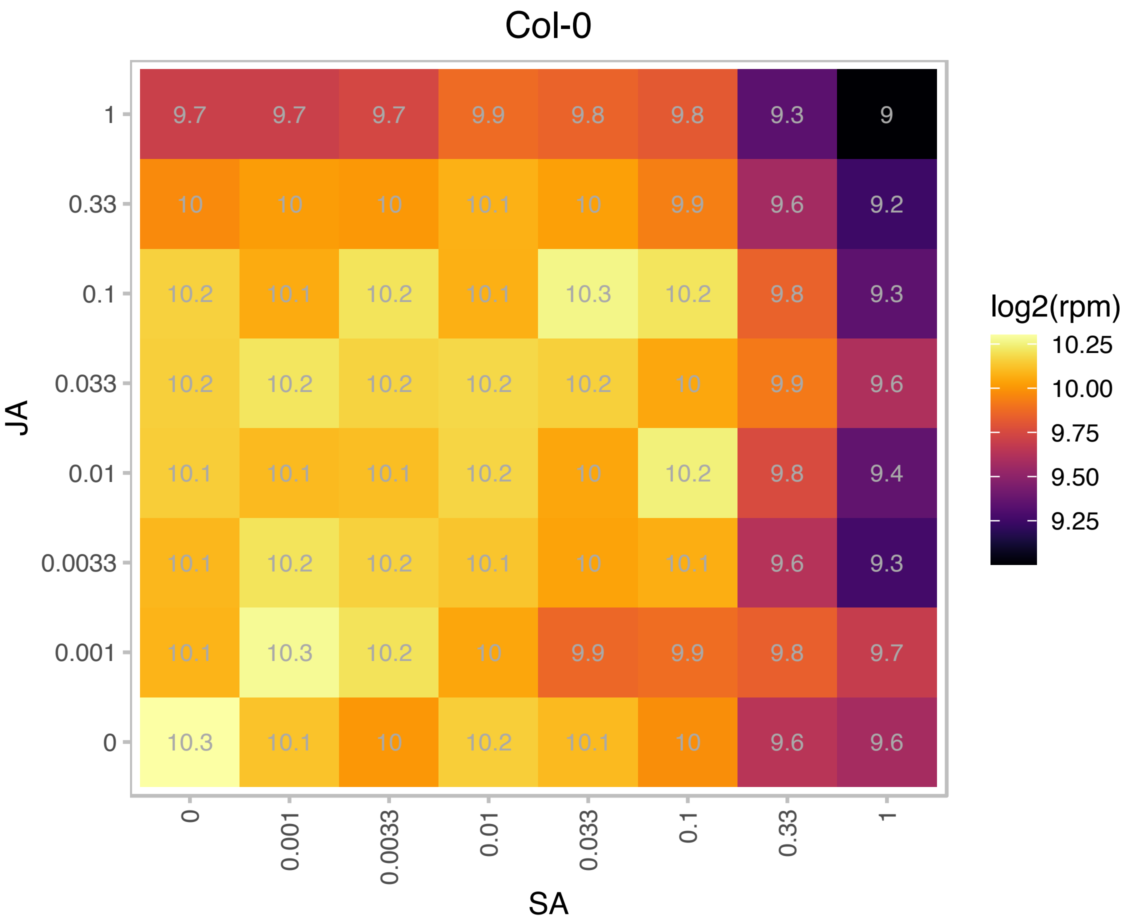


| | |
|---|---|
| Gene | AT1G14320 |
| is.expressed_genes | TRUE |
| is.responsive_genes | TRUE |
| is.apcluster | F-07 |

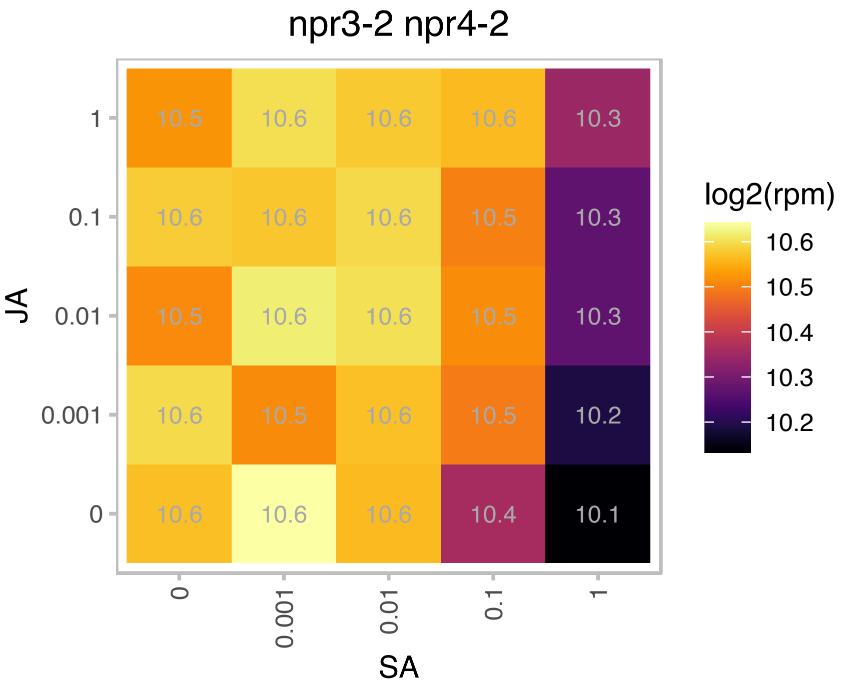


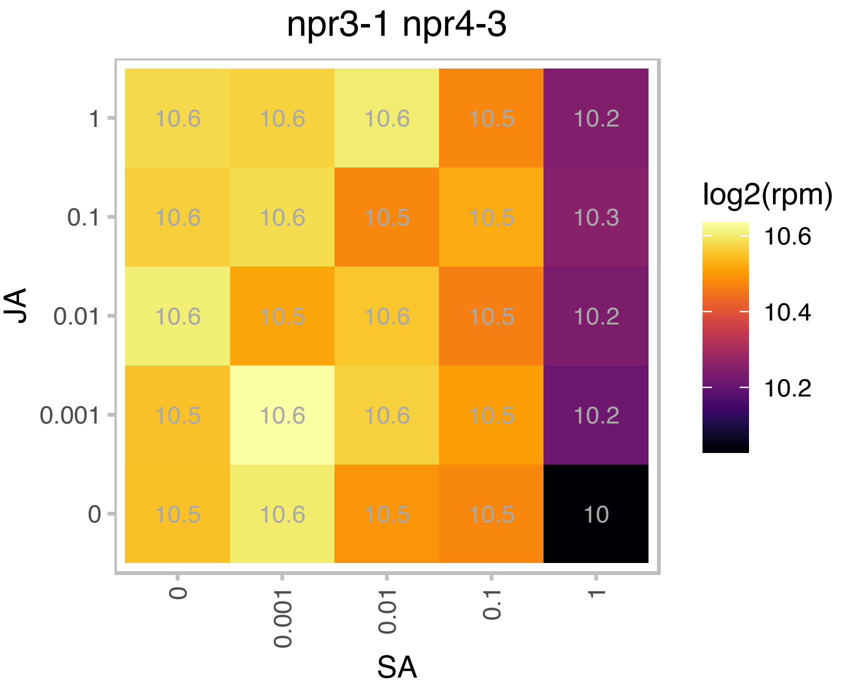

### AT1G14410.1 : ATWHY1, PTAC1, WHY1

ssDNA-binding transcriptional regulator

Encodes a homolog of the potato p24 protein. Binds single strand telomeric repeats. Negatively regulates telomerase activity and telomere length.

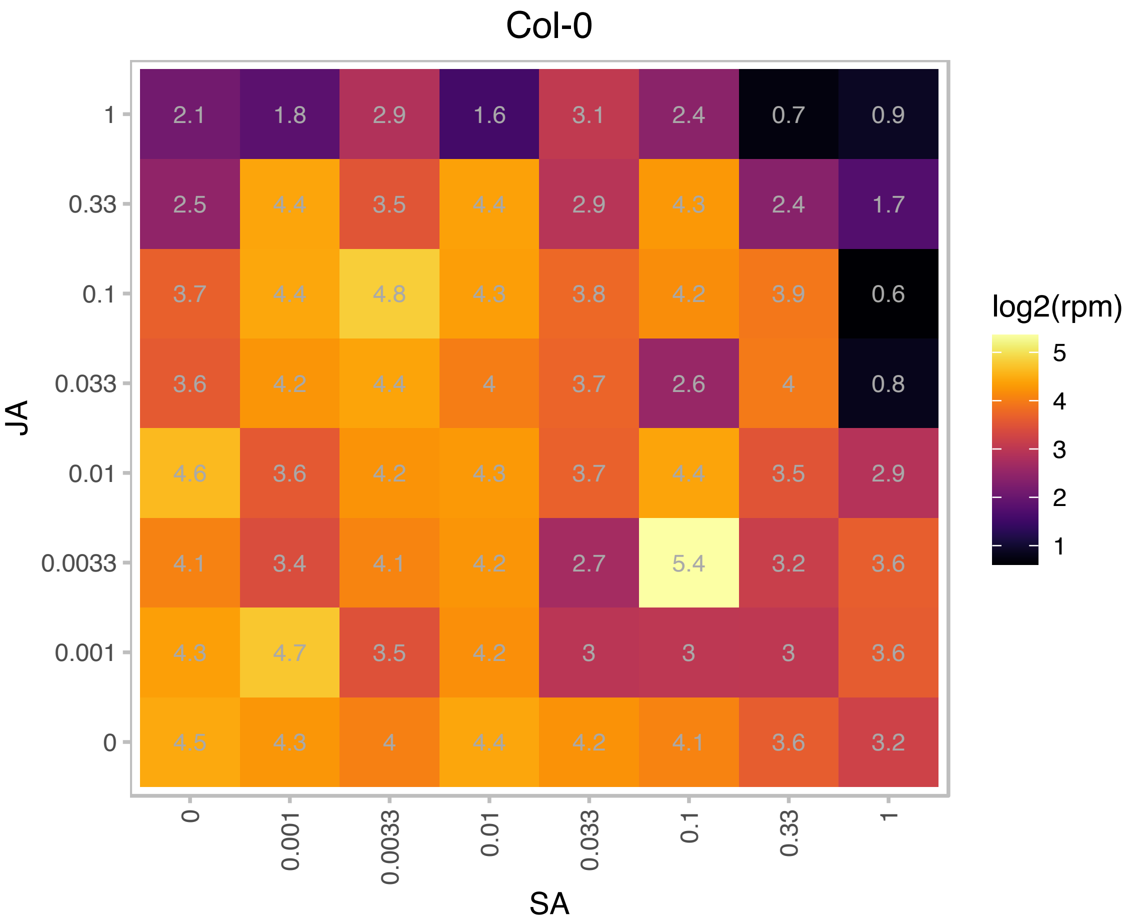


| | |
|---|---|
| Gene | AT1G14410 |
| is.expressed_genes | TRUE |
| is.responsive_genes | TRUE |
| is.apcluster | F-07 |

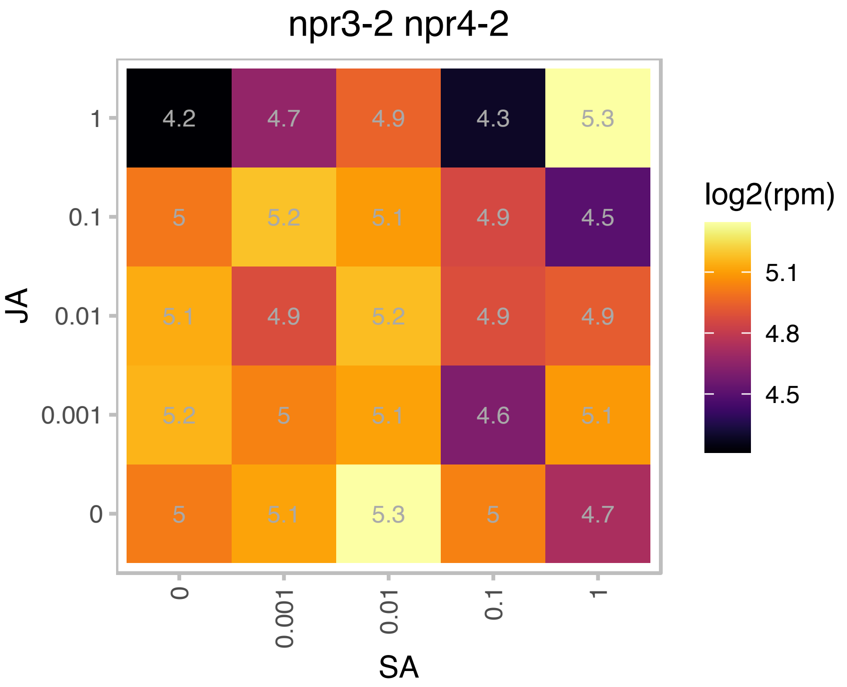


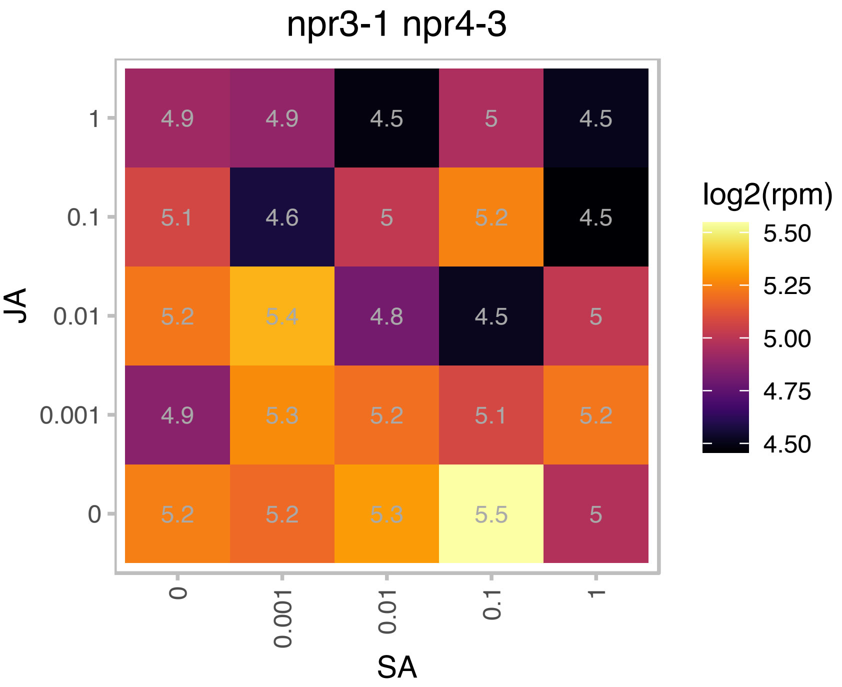

### AT1G14900.1 : HMGA

high mobility group A

Encodes a protein belonging to the subgroup of HMGA (high mobility group A) proteins that interact with A/T-rich stretches of DNA.

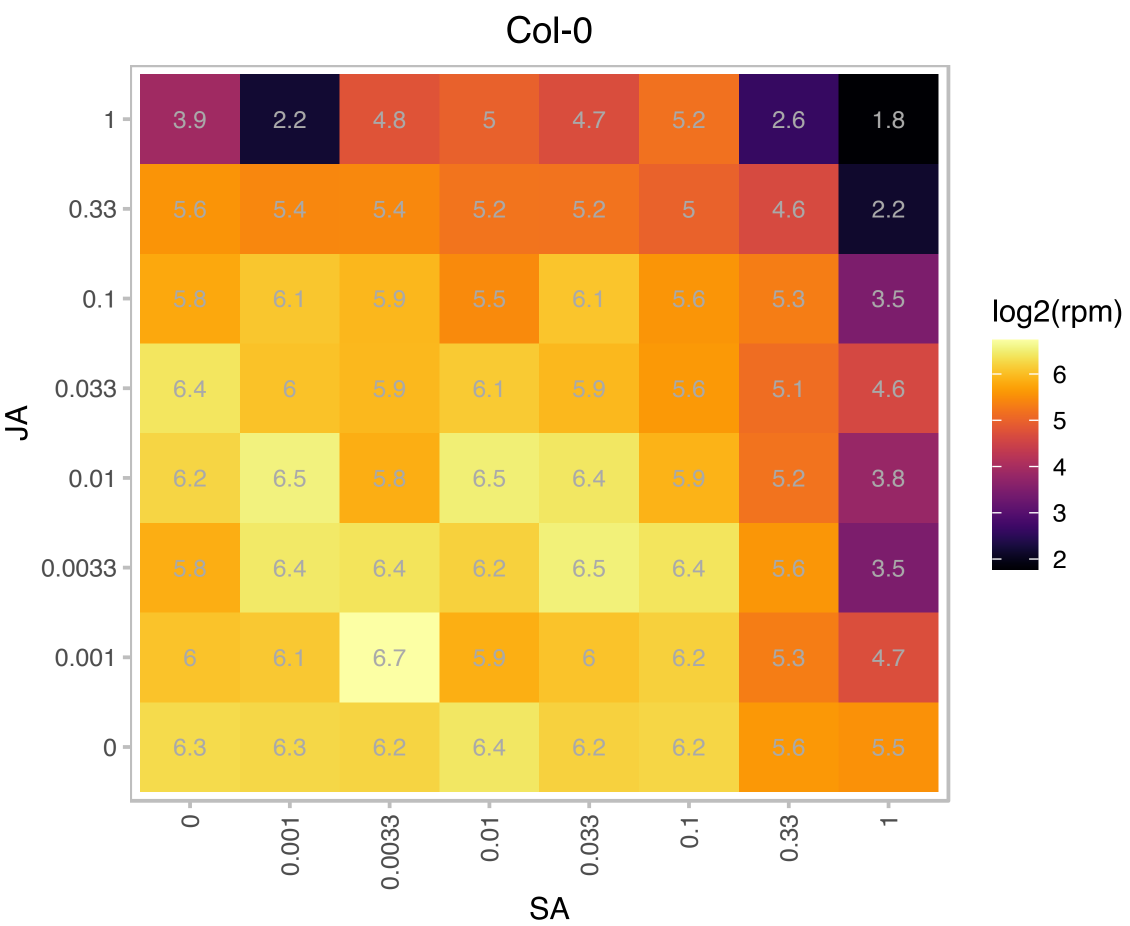


| | |
|---|---|
| Gene | AT1G14900 |
| is.expressed_genes | TRUE |
| is.responsive_genes | TRUE |
| is.apcluster | F-07 |

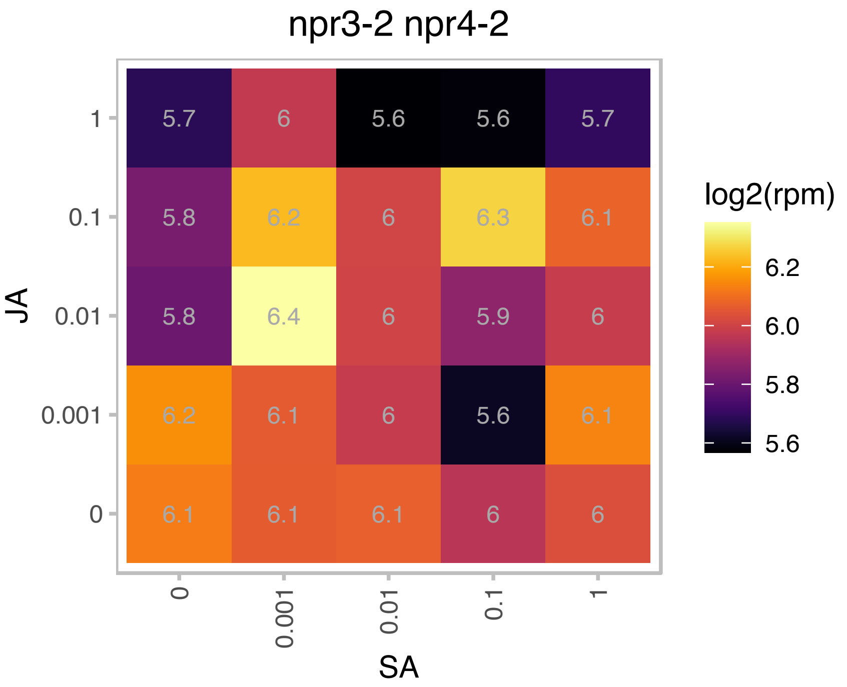


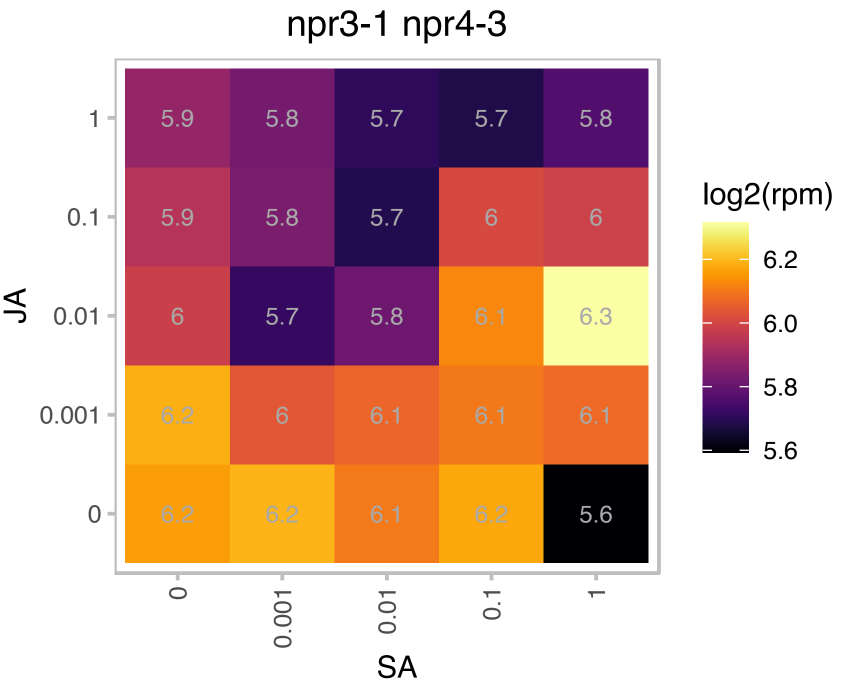

# AT1G15030.1 : NA

Protein of unknown function (DUF789)

Encodes a Cysteine-rich peptide (CRP) family protein

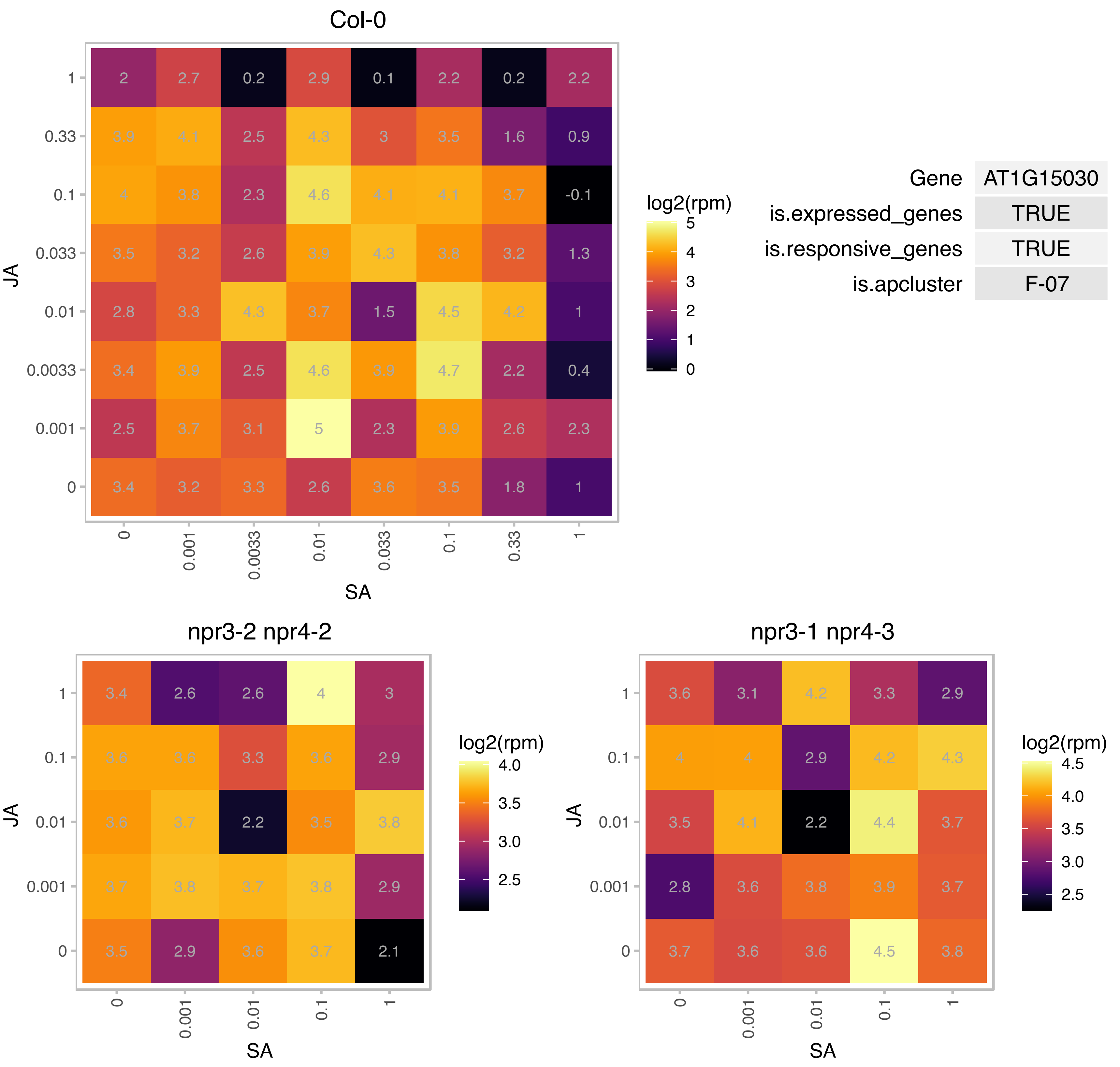


| | |
|---|---|
| Gene | AT1G15030 |
| is.expressed_genes | TRUE |
| is.responsive_genes | TRUE |
| is.apcluster | F-07 |

### AT1G15500.1 : ATNTT2

TLC ATP/ADP transporter

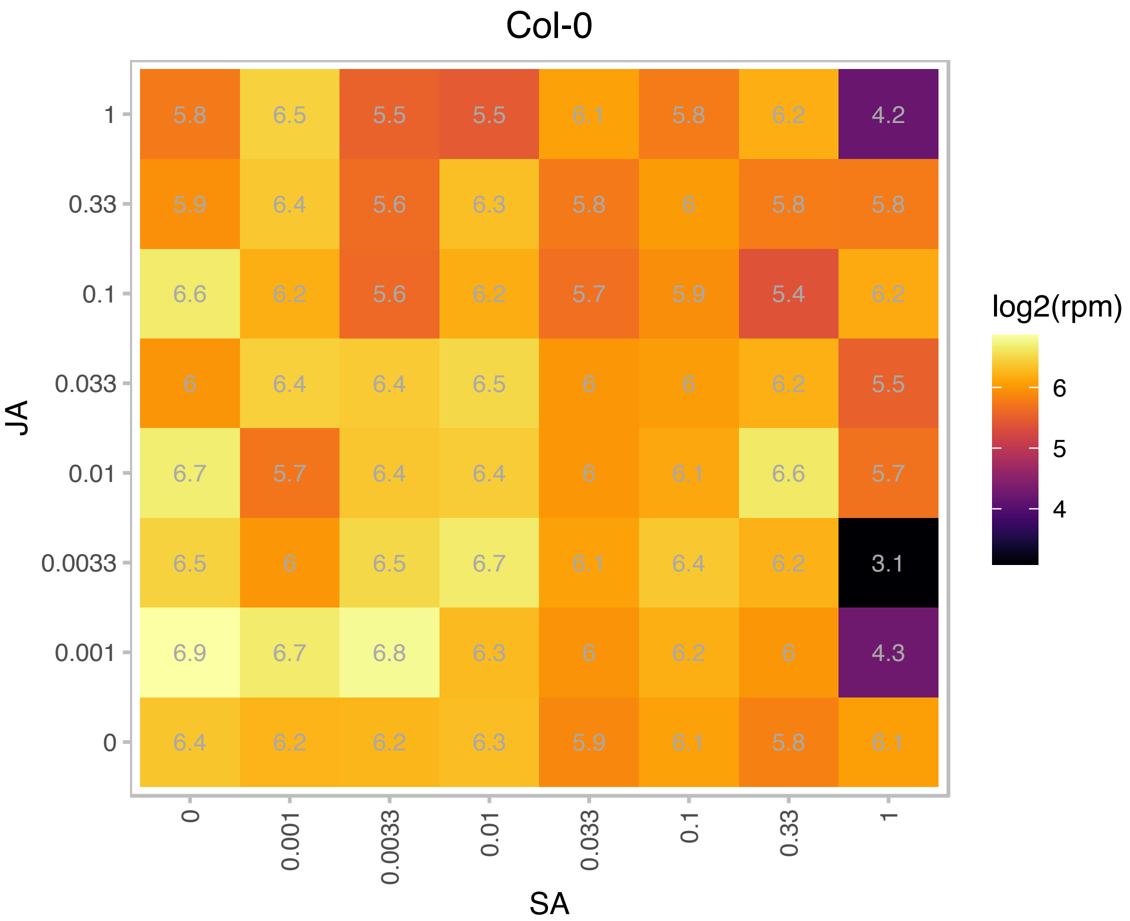


| | |
|---|---|
| Gene | AT1G15500 |
| is.expressed_genes | TRUE |
| is.responsive_genes | TRUE |
| is.apcluster | F-07 |

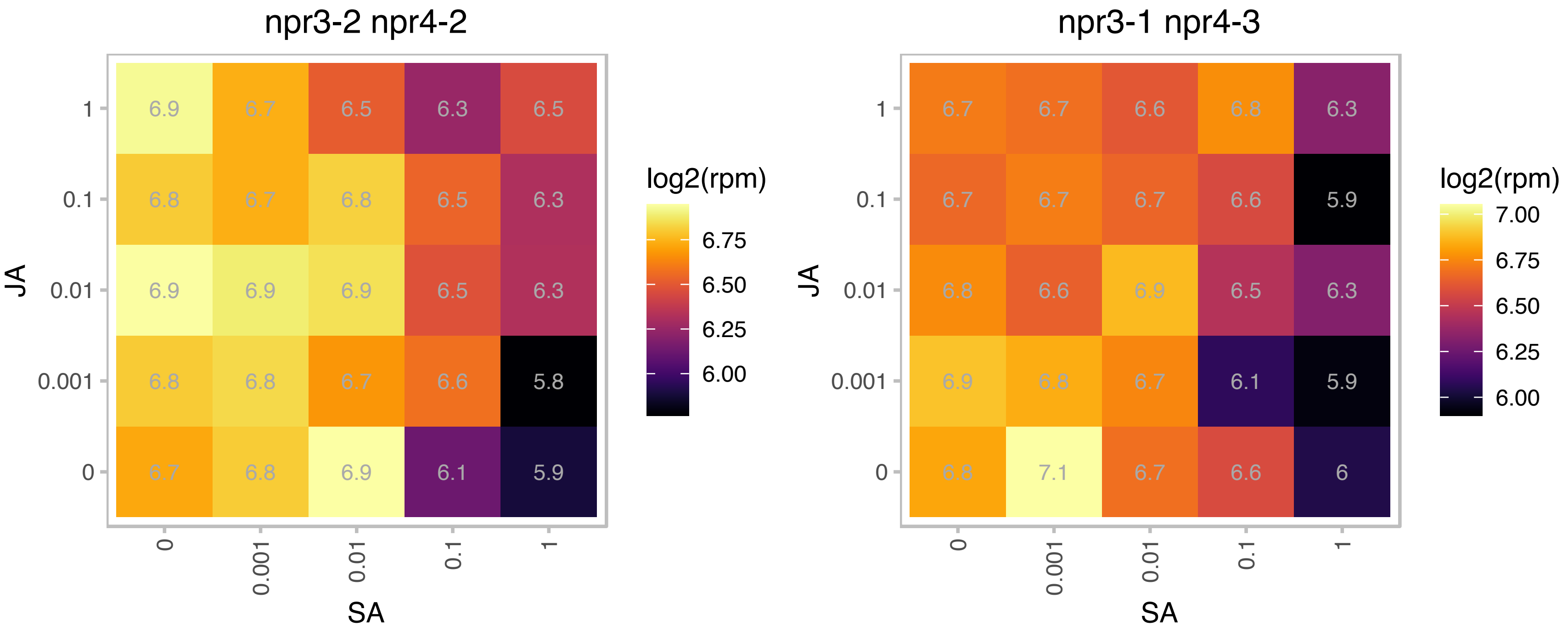

# AT1G15930.1 : NA

Ribosomal protein L7Ae/L30e/S12e/Gadd45 family protein

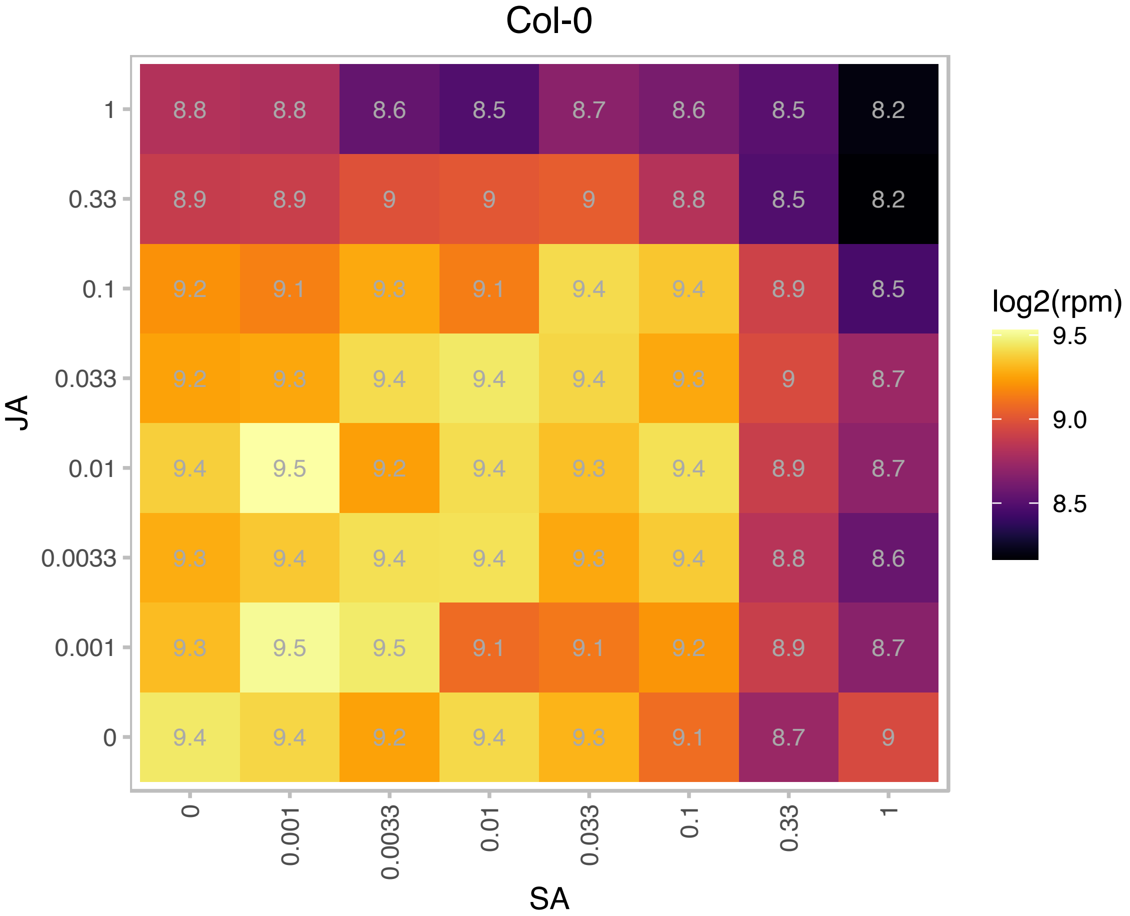


| | |
|---|---|
| Gene | AT1G15930 |
| is.expressed_genes | TRUE |
| is.responsive_genes | TRUE |
| is.apcluster | F-07 |

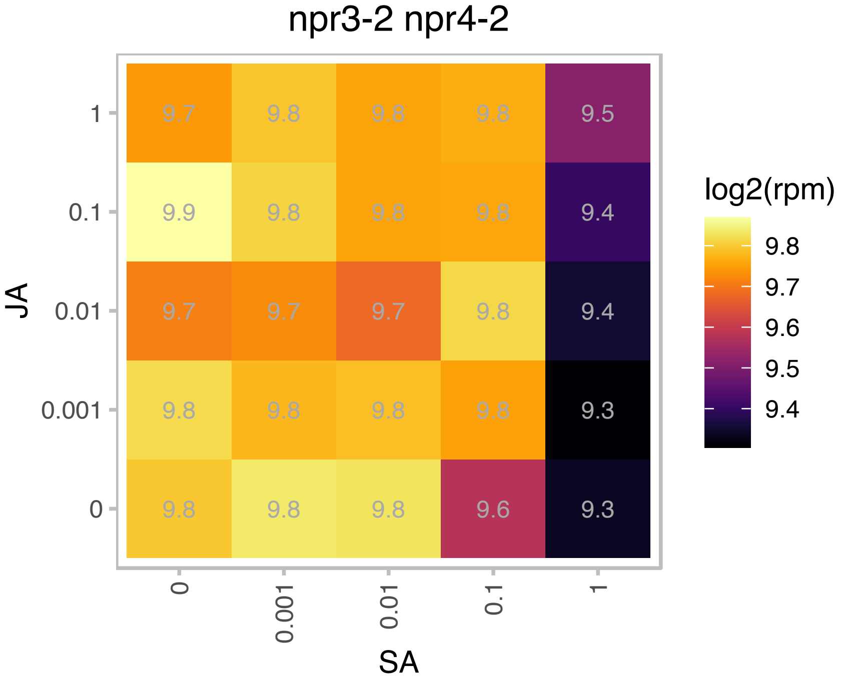


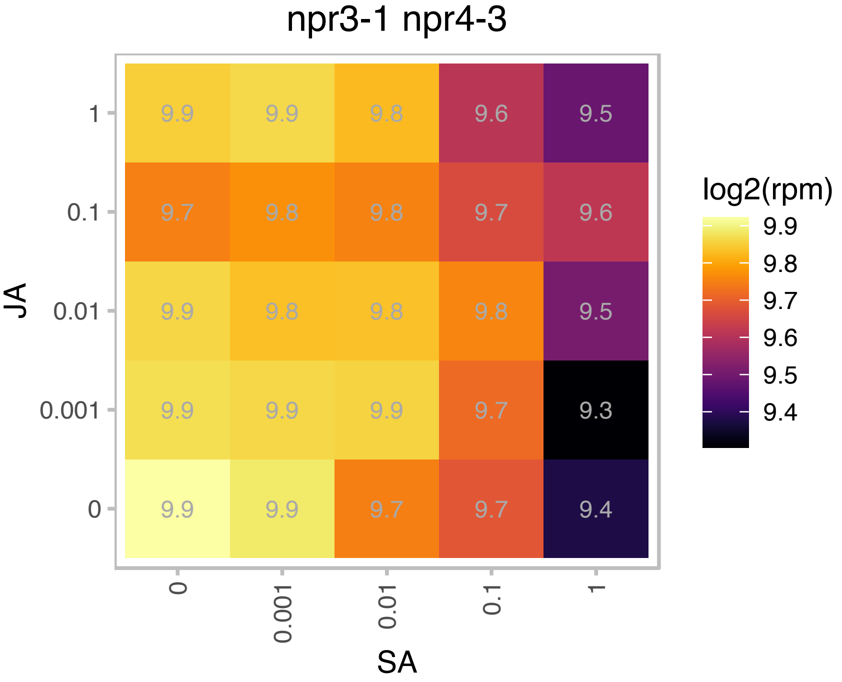

### AT1G16410.1 : BUS1, CYP79F1, SPS1

cytochrome p450 79f1

member of CYP79F The mRNA is cell-to-cell mobile.

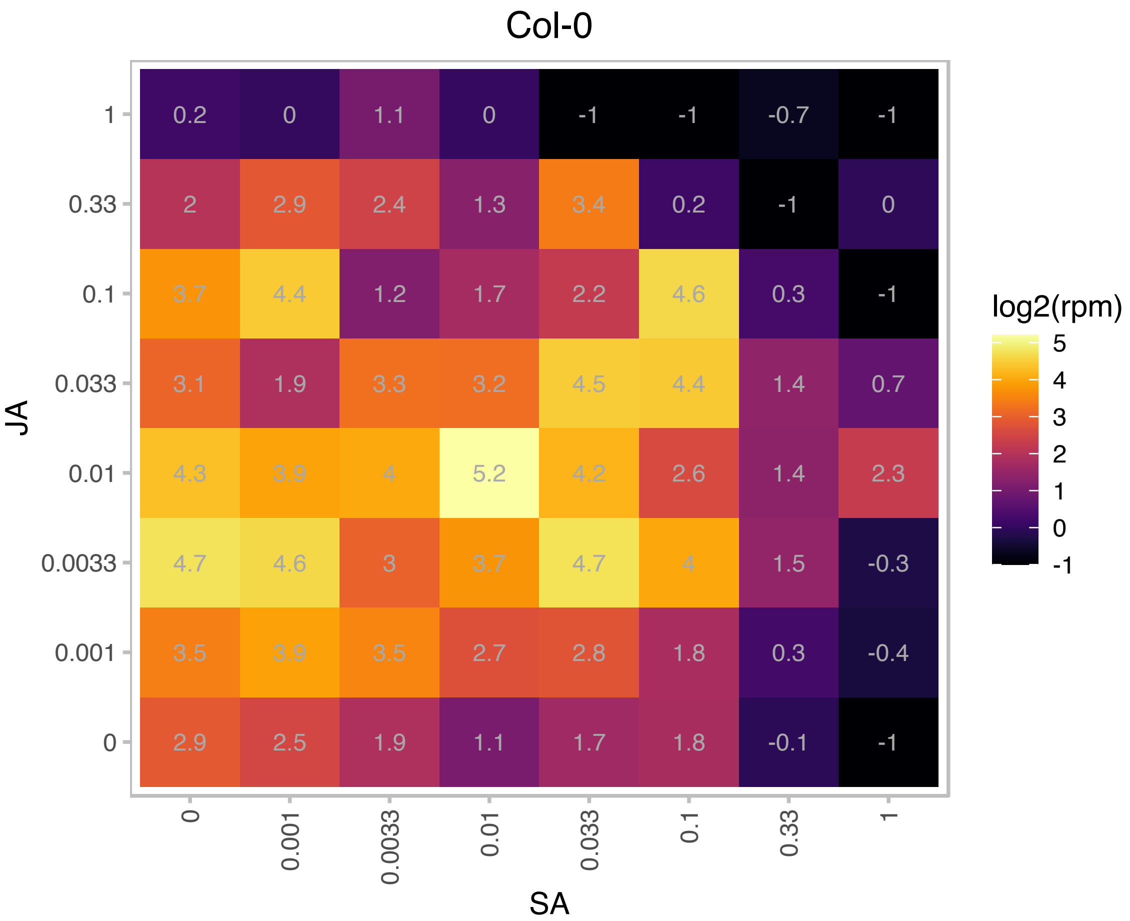


| | |
|---|---|
| Gene | AT1G16410 |
| is.expressed_genes | TRUE |
| is.responsive_genes | TRUE |
| is.apcluster | F-07 |

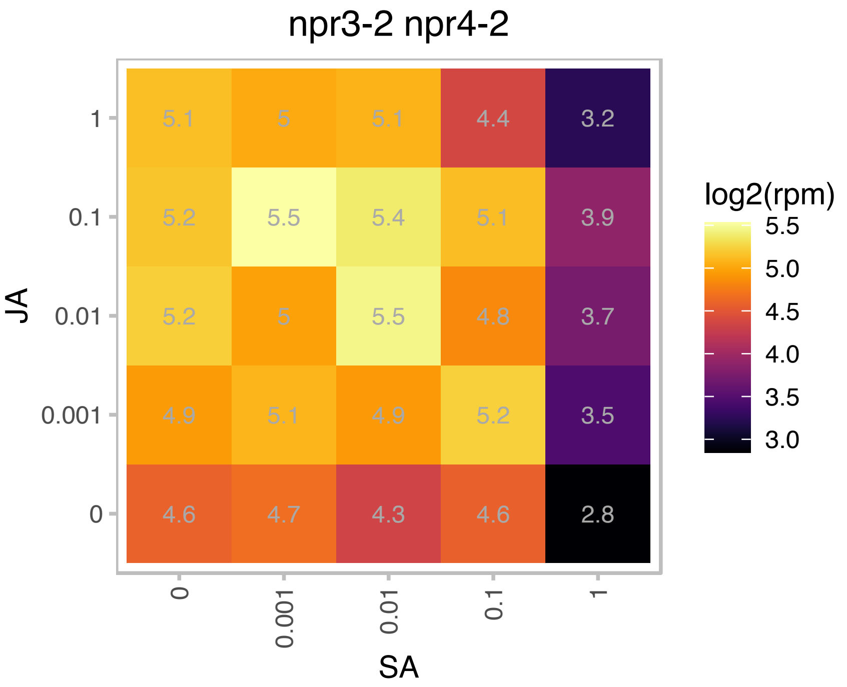


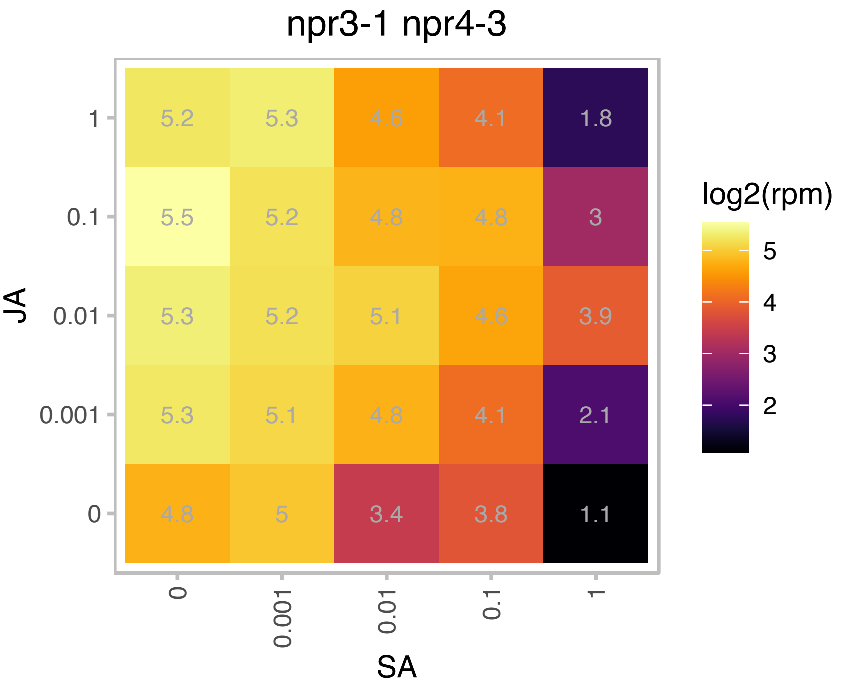

# AT1G16790.1 : NA

ribosomal protein-related

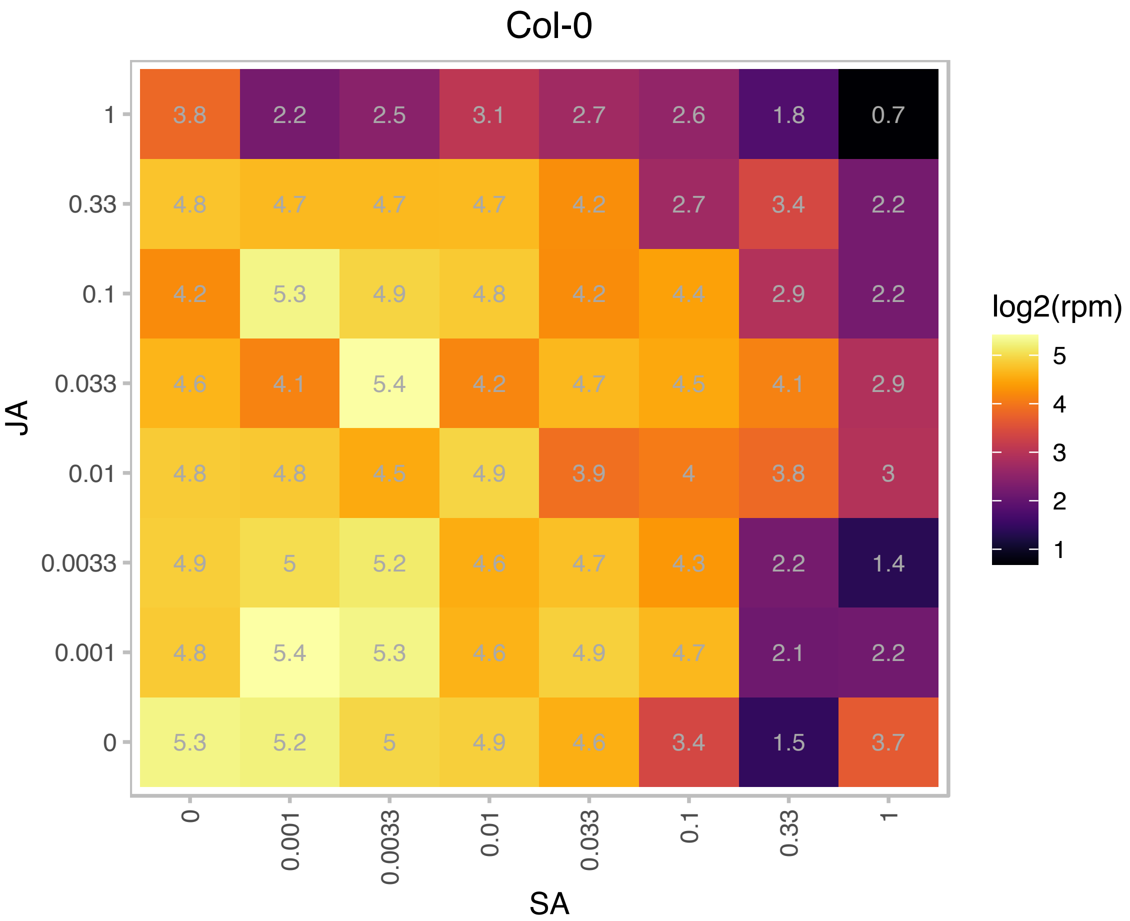


| | |
|---|---|
| Gene | AT1G16790 |
| is.expressed_genes | TRUE |
| is.responsive_genes | TRUE |
| is.apcluster | F-07 |

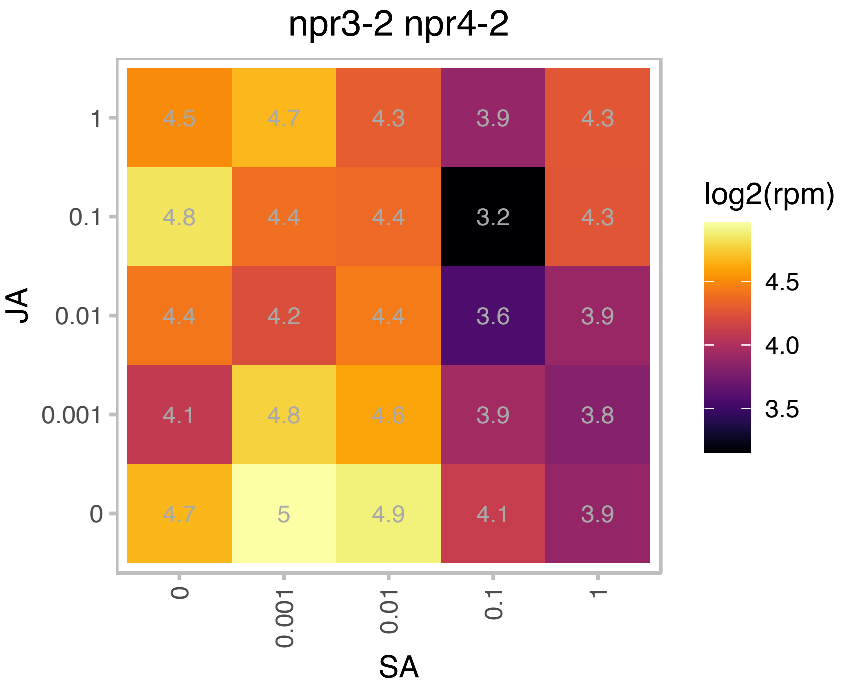


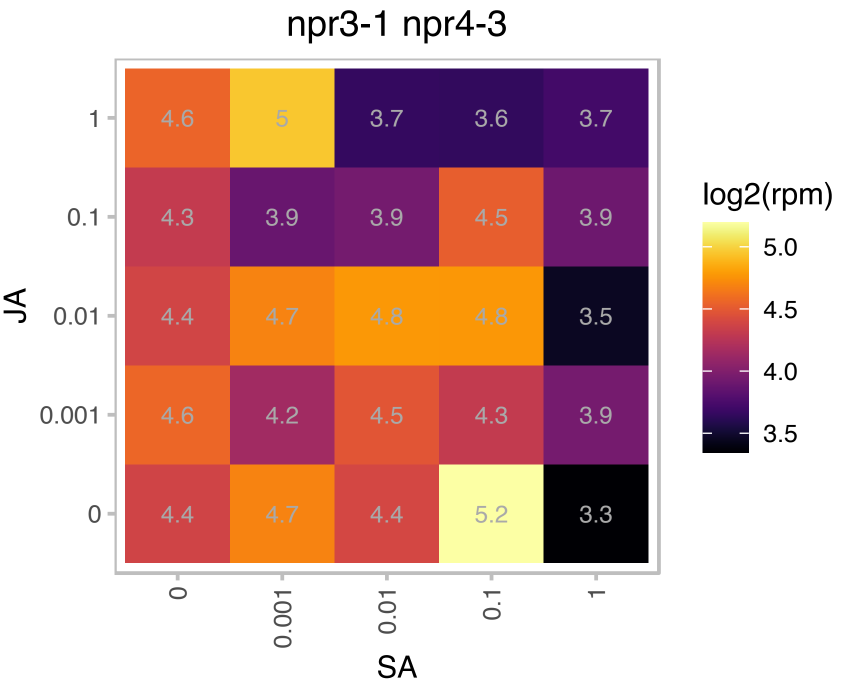

### AT1G17140.1 : ICR1, RIP1

interactor of constitutive active rops 1

Encodes a ROP/RAC effector, designated interactor of constitutive active ROPs 1 (ICR1), that interacts with GTP-bound ROPs. ICR1 is a scaffold mediating formation of protein complexes that are required for cell polarity.  ICR1 is comprised of coiled-coil domains and forms complexes with itself and the exocyst vesicle-tethering complex subunit SEC3.

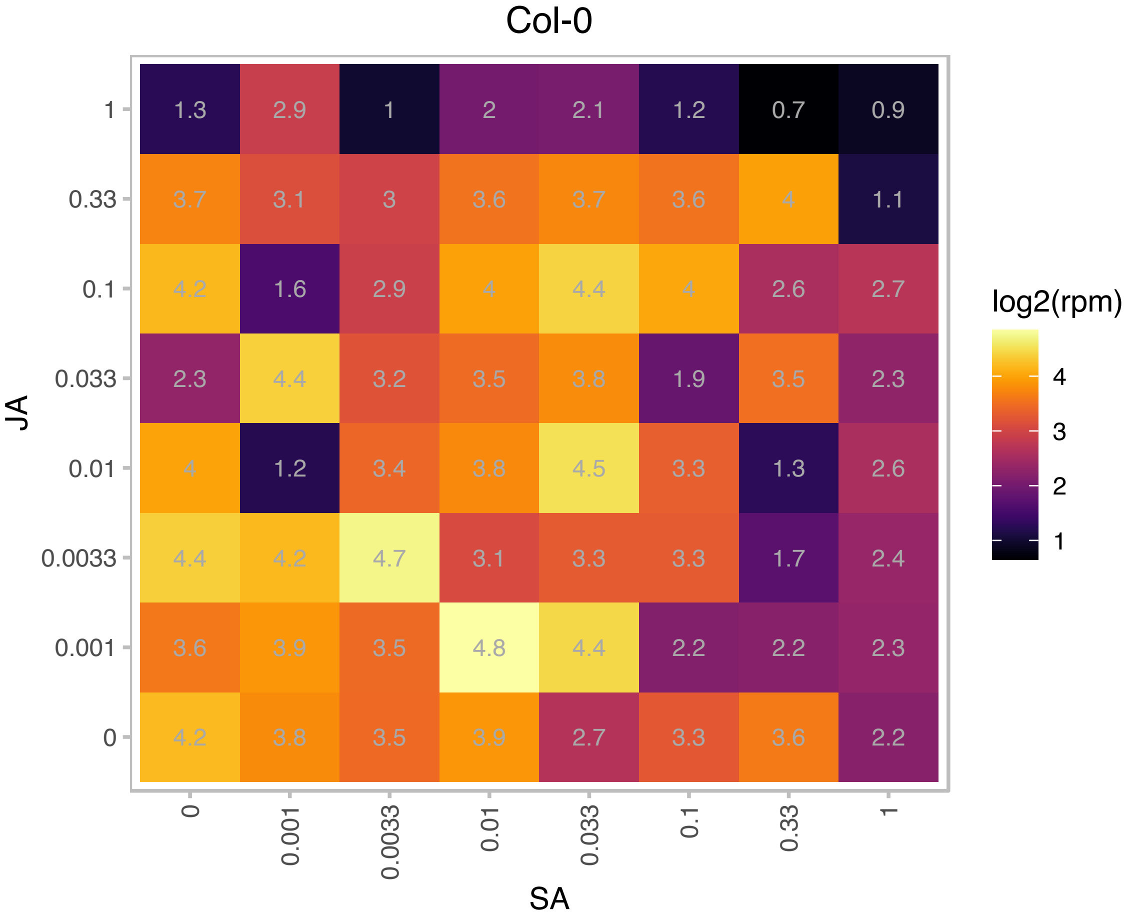


| | |
|---|---|
| Gene | AT1G17140 |
| is.expressed_genes | TRUE |
| is.responsive_genes | TRUE |
| is.apcluster | F-07 |

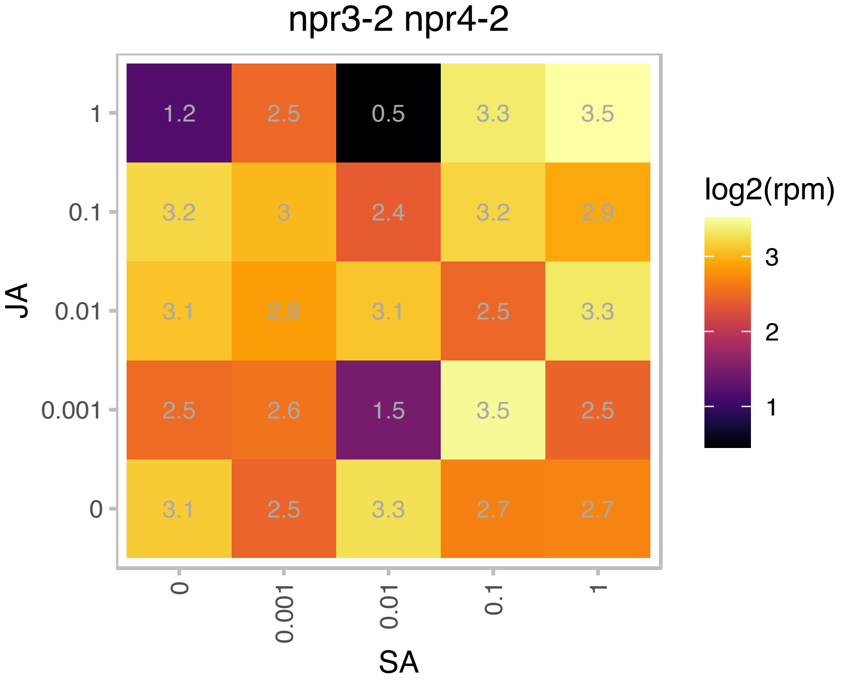


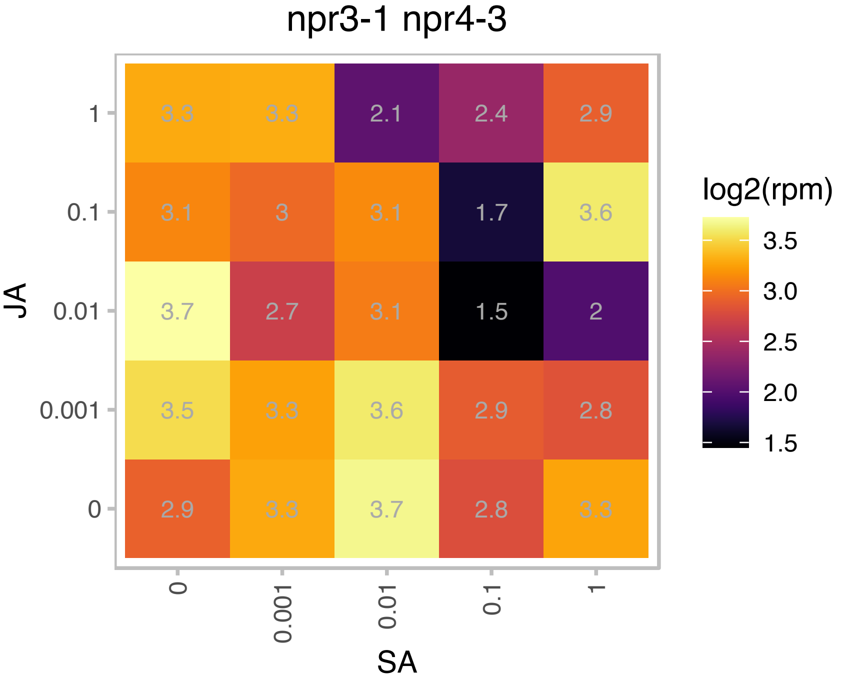

# AT1G18540.1 : NA

Ribosomal protein L6 family protein

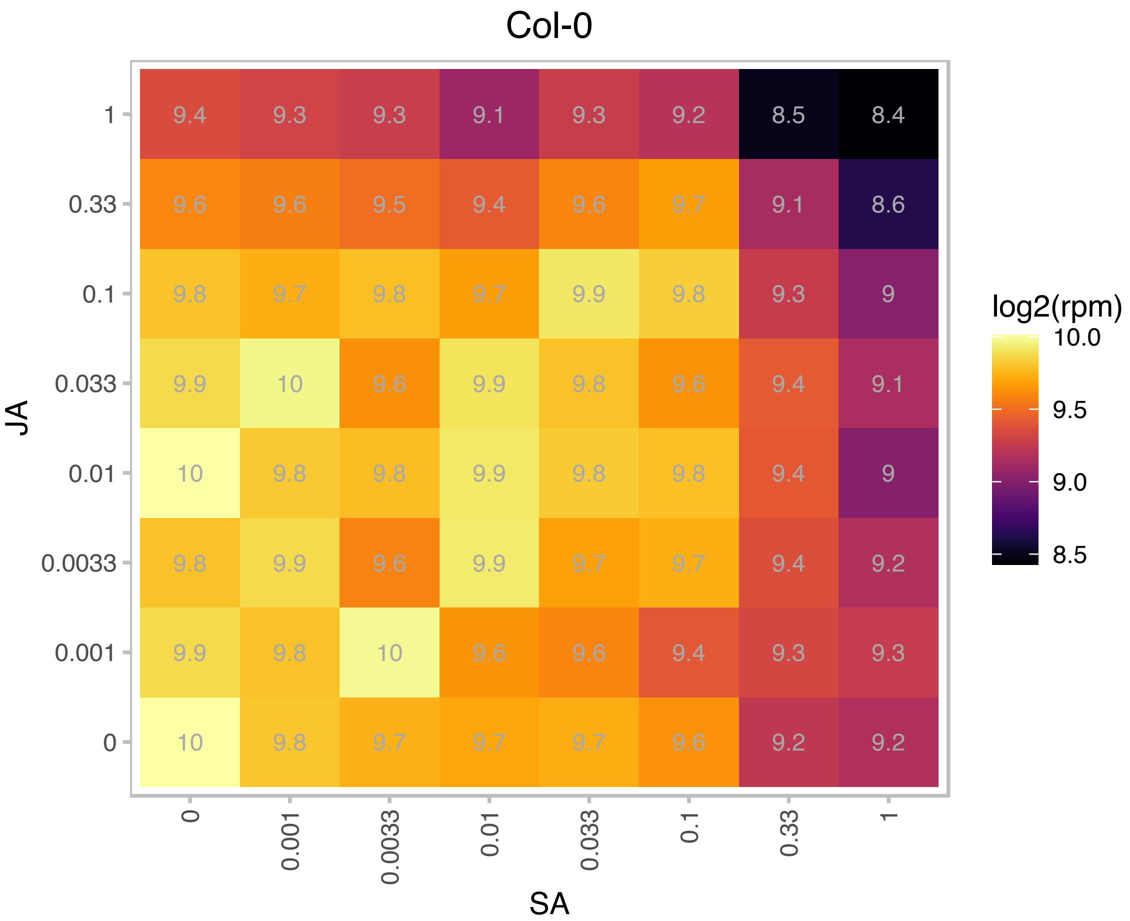


| | |
|---|---|
| Gene | AT1G18540 |
| is.expressed_genes | TRUE |
| is.responsive_genes | TRUE |
| is.apcluster | F-07 |

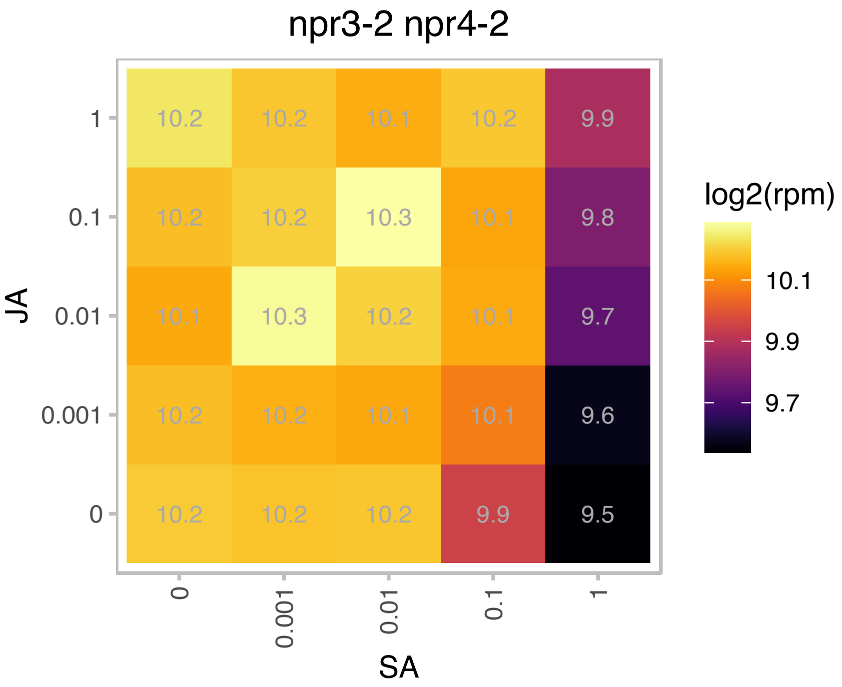


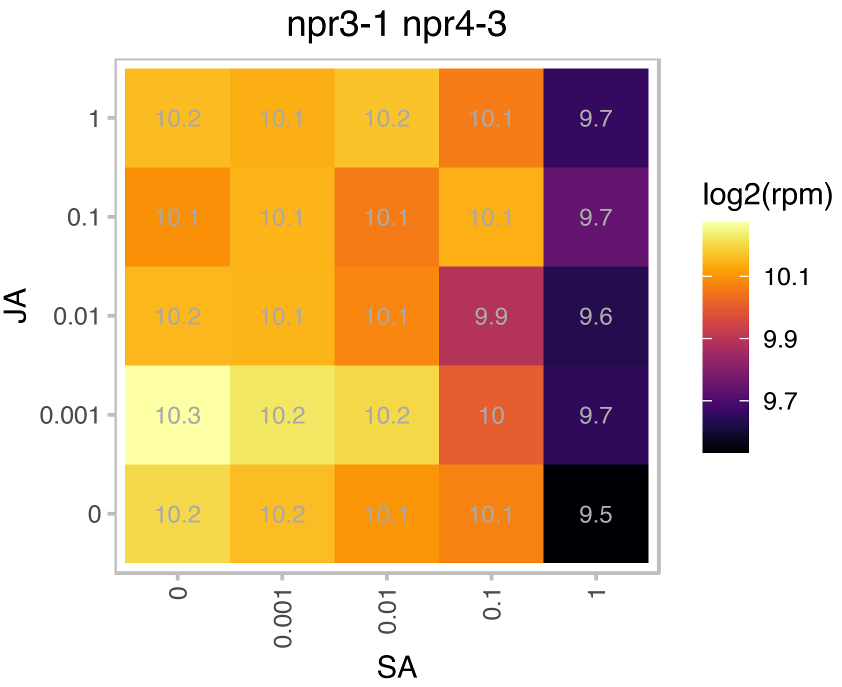

### AT1G19800.1 : ABCI14, TGD1

Encodes a permease-like protein involved in lipid transfer from the ER to the chloroplast, more specifically,
trigalactosyldiacylglycerol 1
transfer of phosphatidate across the chloroplast inner membrane. Mutant leaves accumulate trigalactosyldiacylglycerol, triacylglycerol and phosphatidate. Chloroplast lipids are altered in their fatty acid composition and as a consequence the development of chloroplasts in the mutants are impacted. The mutant seeds has a higher abortion rate. Mutations in this gene suppress the low temperature-induced phenotype of Arabidopsis tocopherol-deficient mutant vte2.

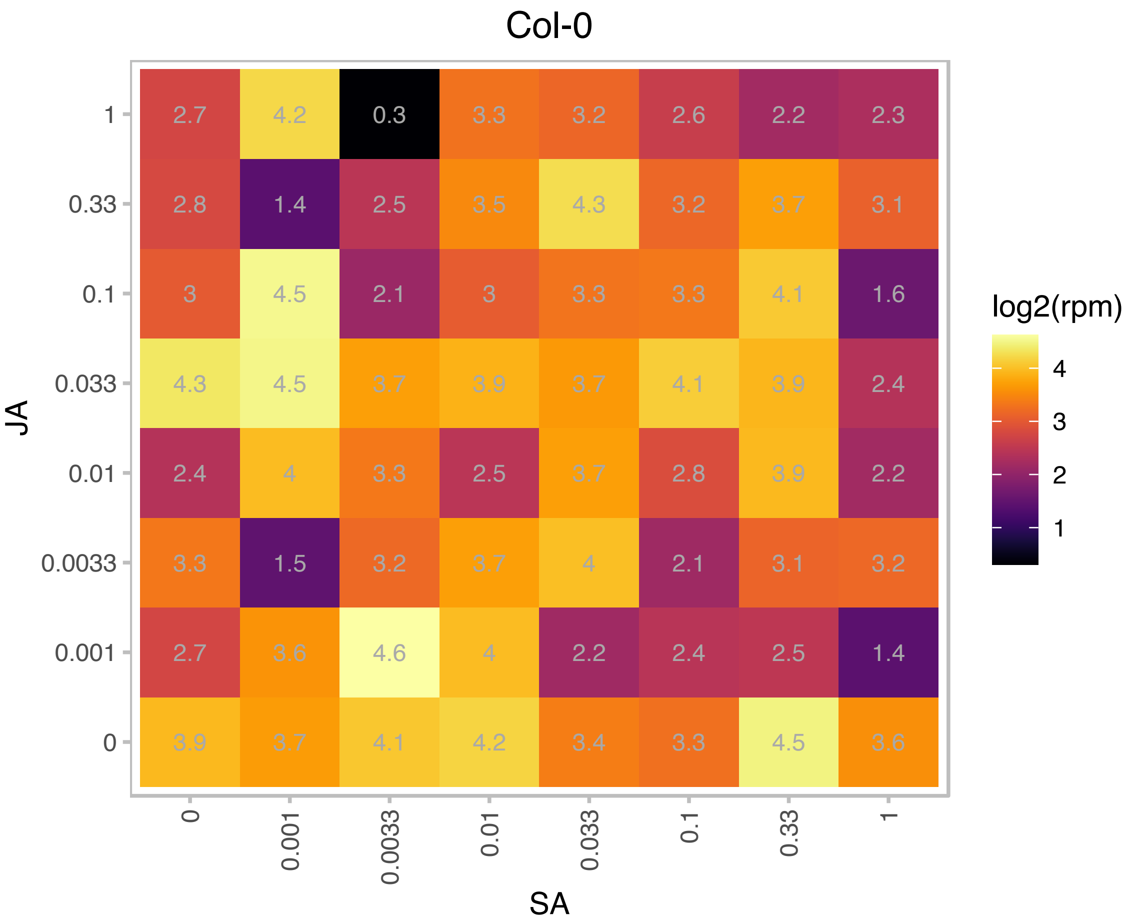


| | |
|---|---|
| Gene | AT1G19800 |
| is.expressed_genes | TRUE |
| is.responsive_genes | TRUE |
| is.apcluster | F-07 |

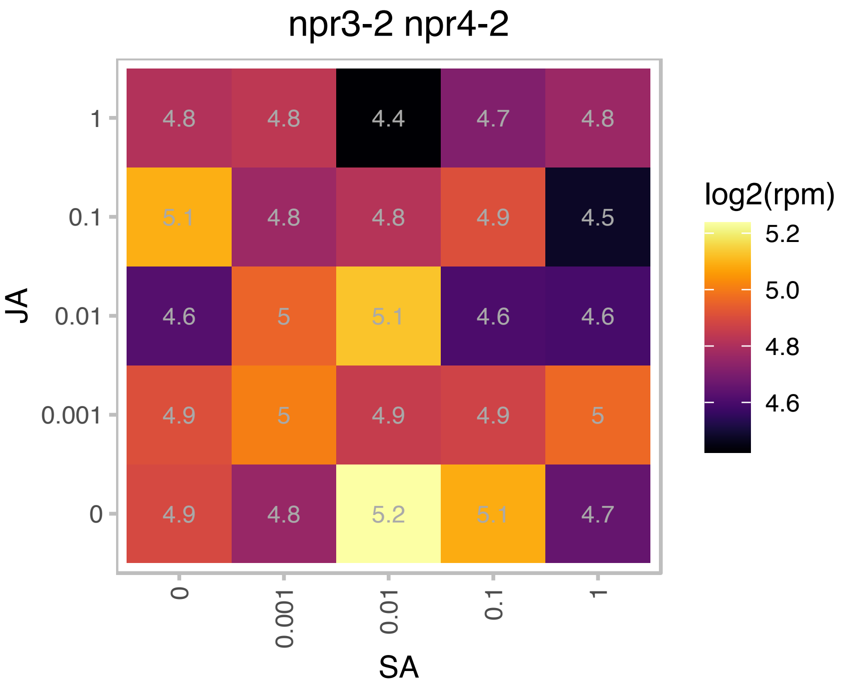


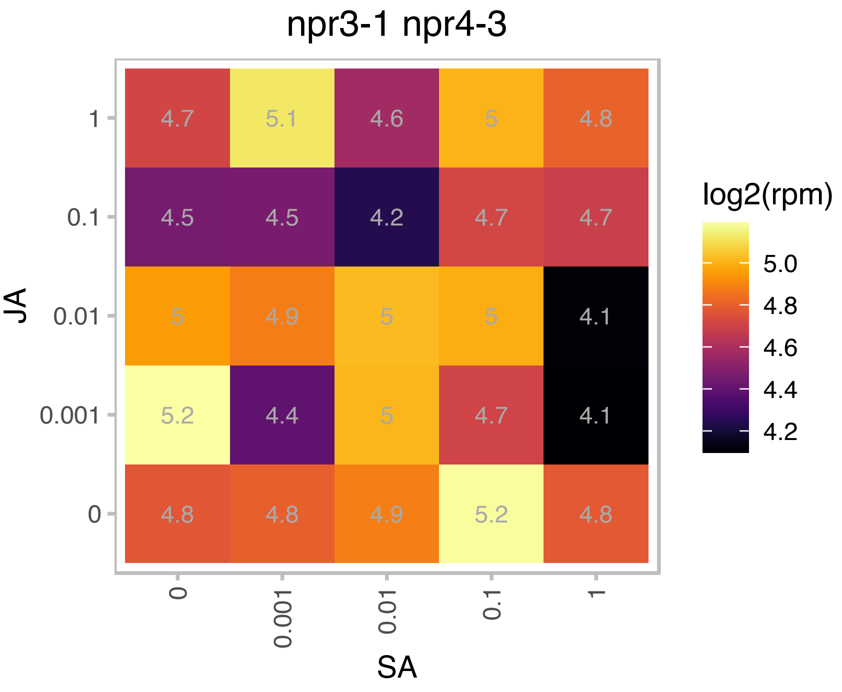

# AT1G19990.1 : NA

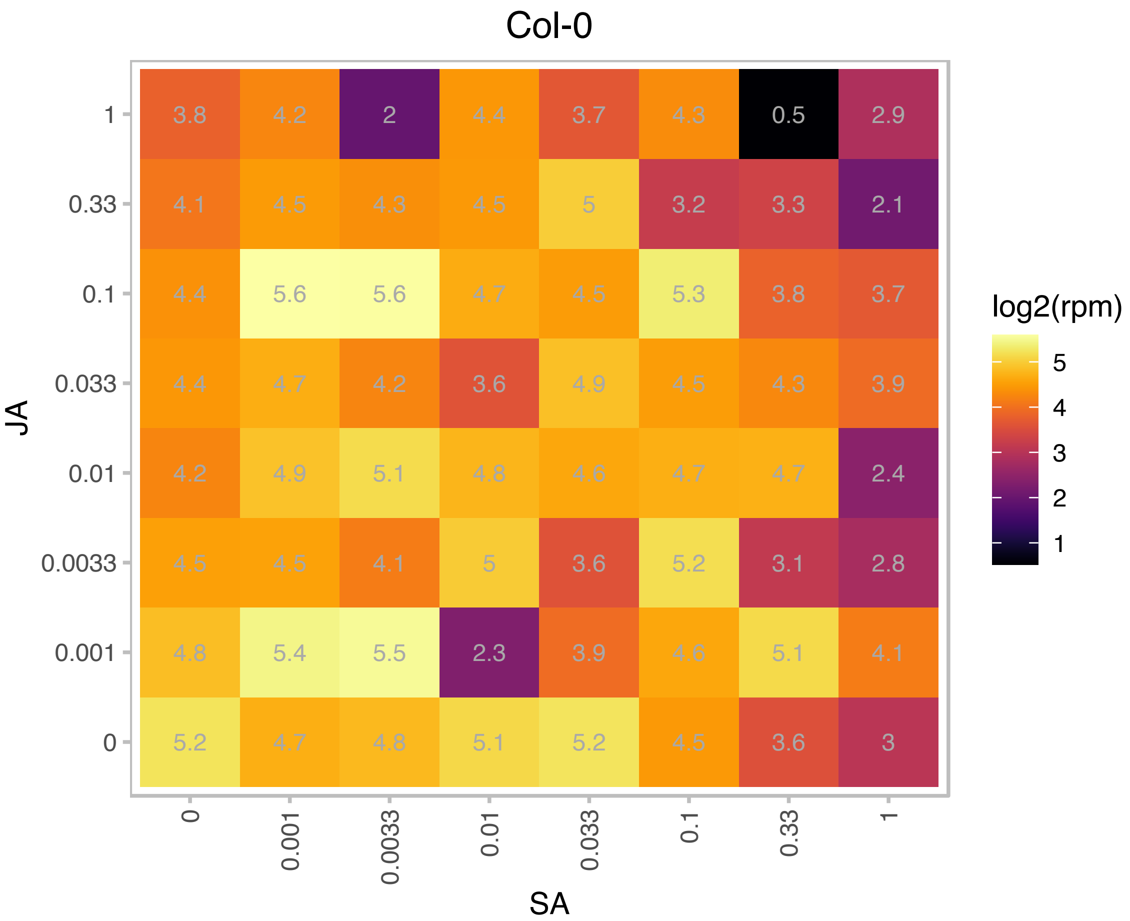


| | |
|---|---|
| Gene | AT1G19990 |
| is.expressed_genes | TRUE |
| is.responsive_genes | TRUE |
| is.apcluster | F-07 |

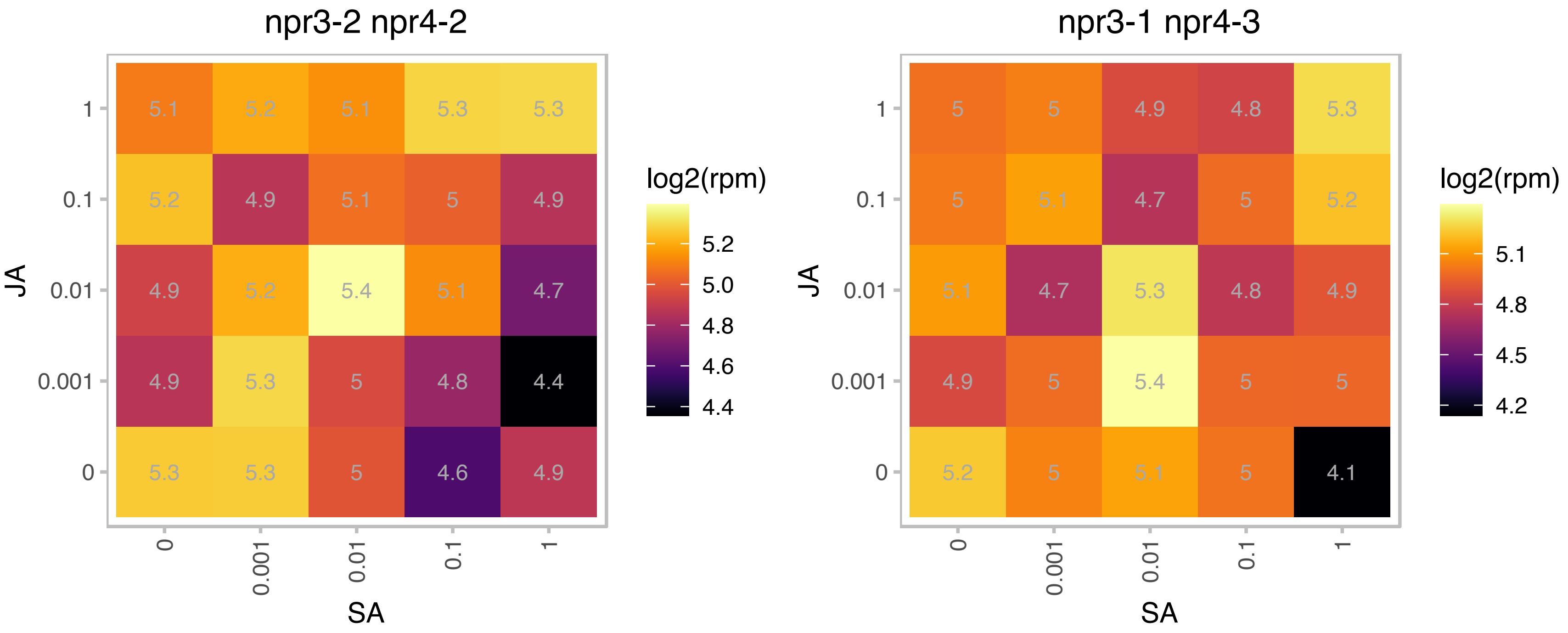

### AT1G21710.1 : ATOGG1, OGG1

8-oxoguanine-DNA glycosylase 1

Encodes 8-oxoguanine-DNA glycosylase. DNA repair enzyme.

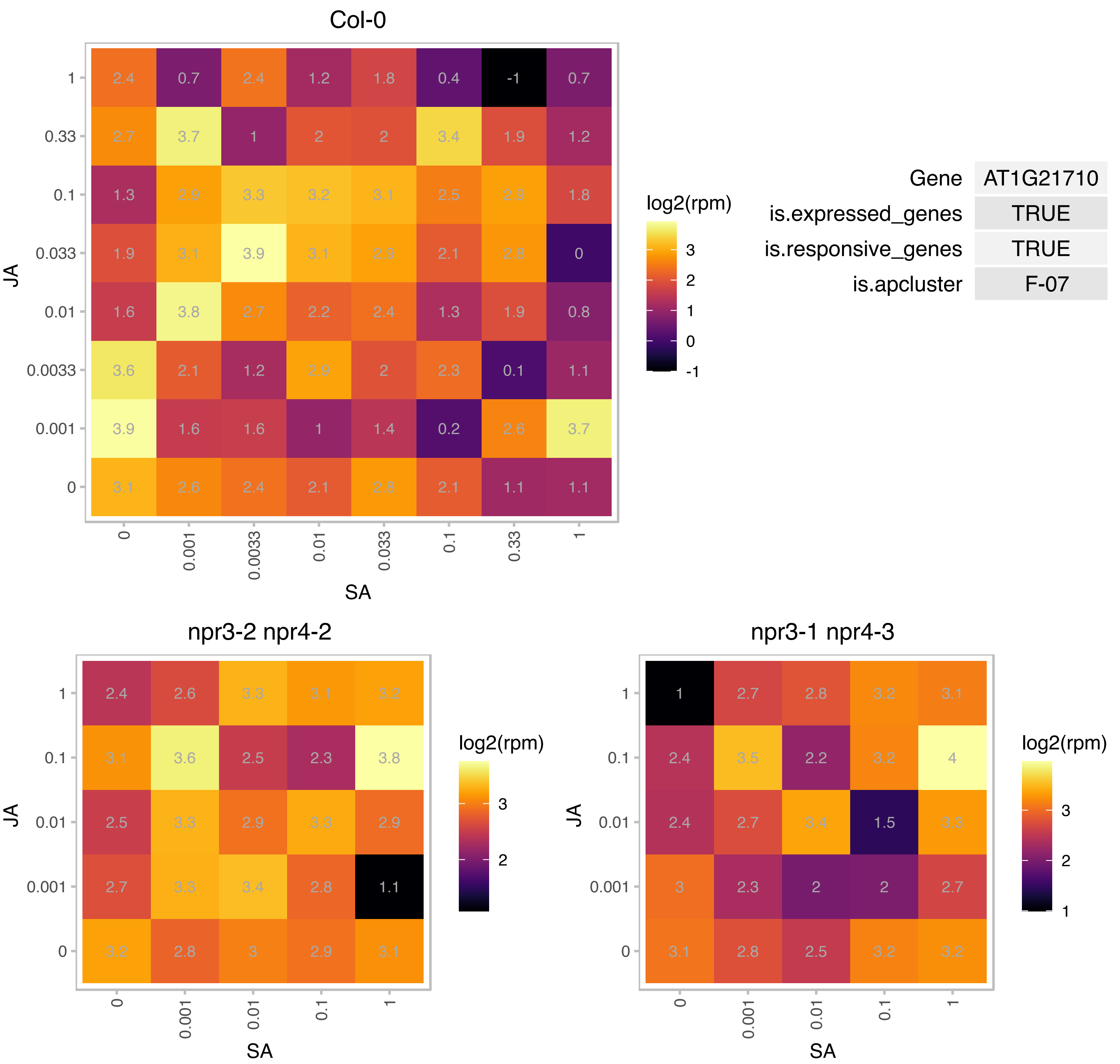


| | |
|---|---|
| Gene | AT1G21710 |
| is.expressed_genes | TRUE |
| is.responsive_genes | TRUE |
| is.apcluster | F-07 |

### AT1G23400.1 : ATCAF2, CAF2

RNA-binding CRS1 / YhbY (CRM) domain-containing protein

Promotes the splicing of chloroplast group II introns.

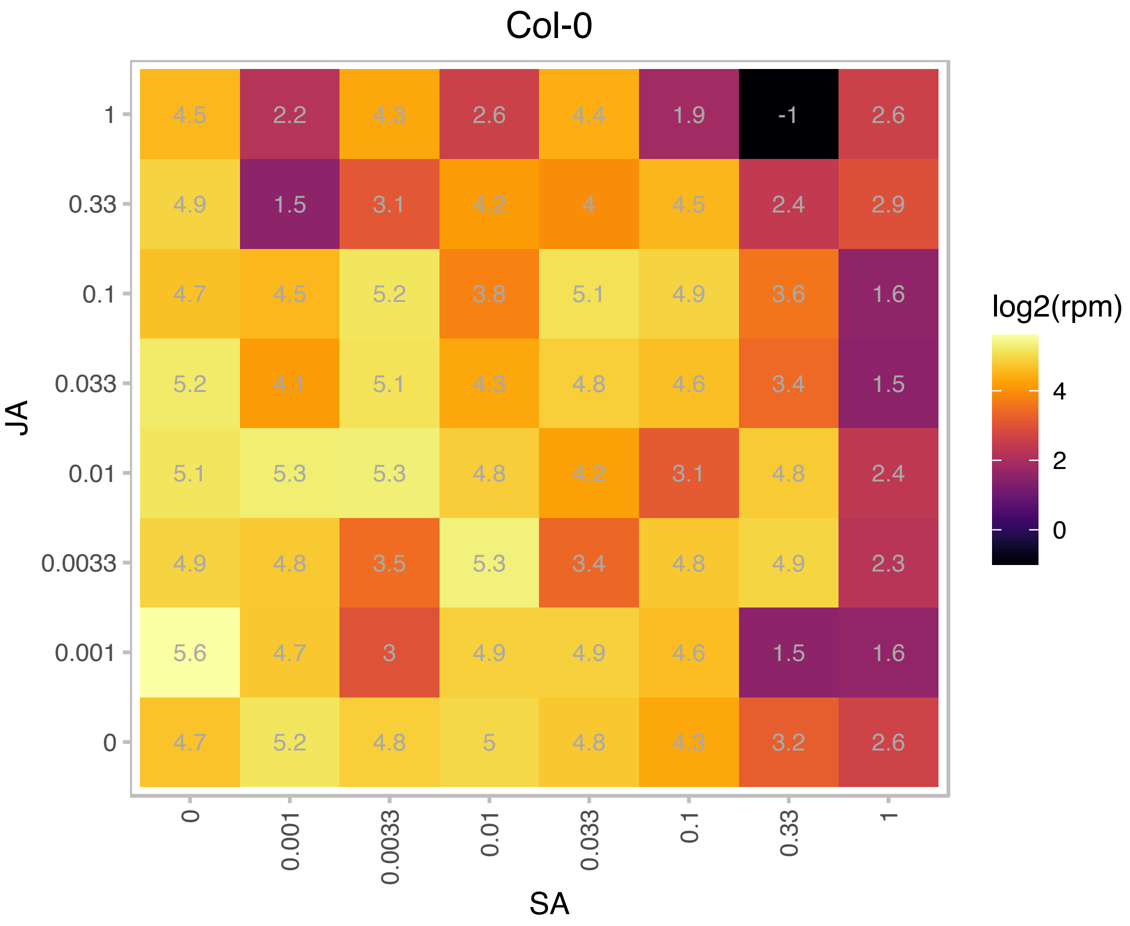


| | |
|---|---|
| Gene | AT1G23400 |
| is.expressed_genes | TRUE |
| is.responsive_genes | TRUE |
| is.apcluster | F-07 |

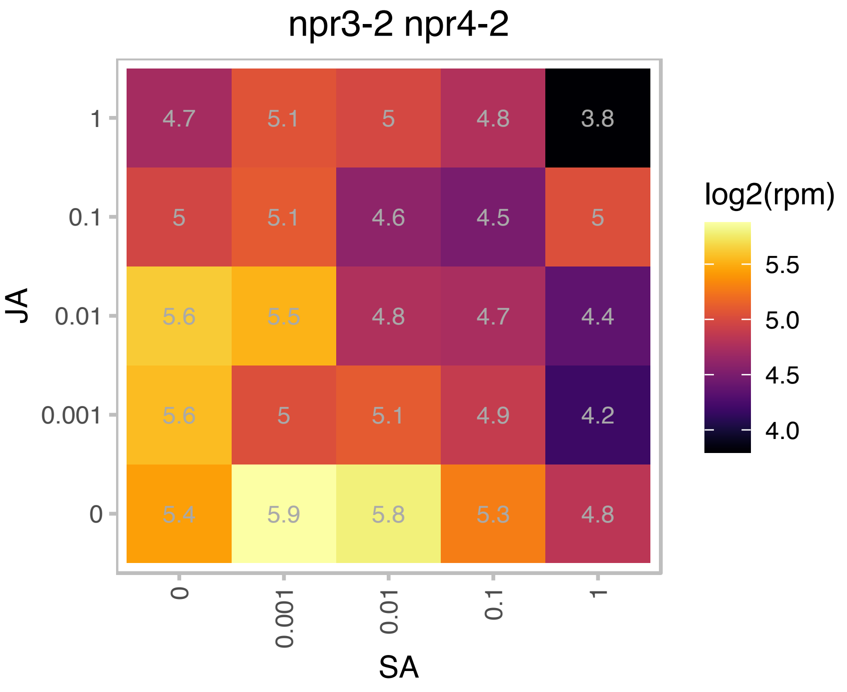


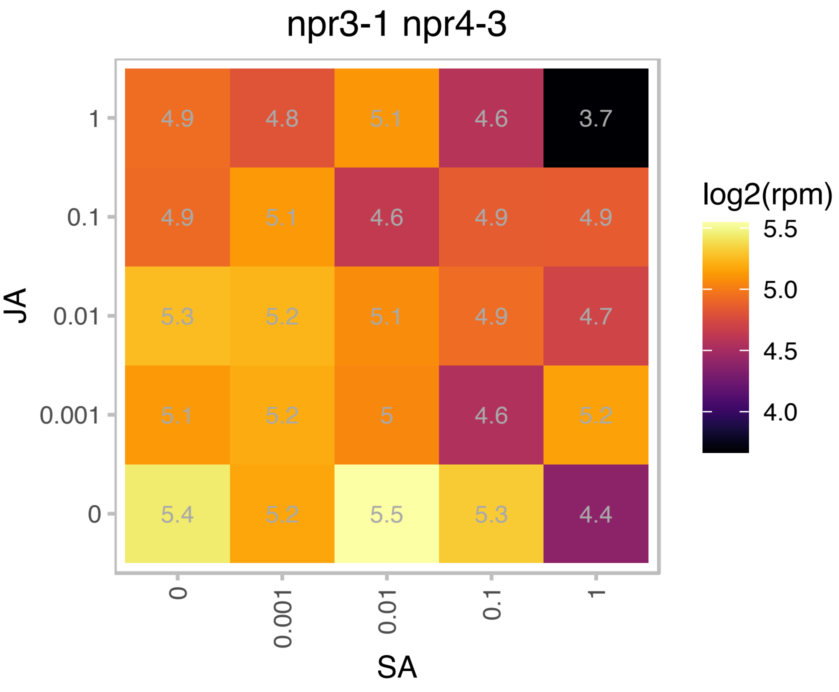

### AT1G24020.1 : MLP423

MLP-like protein 423

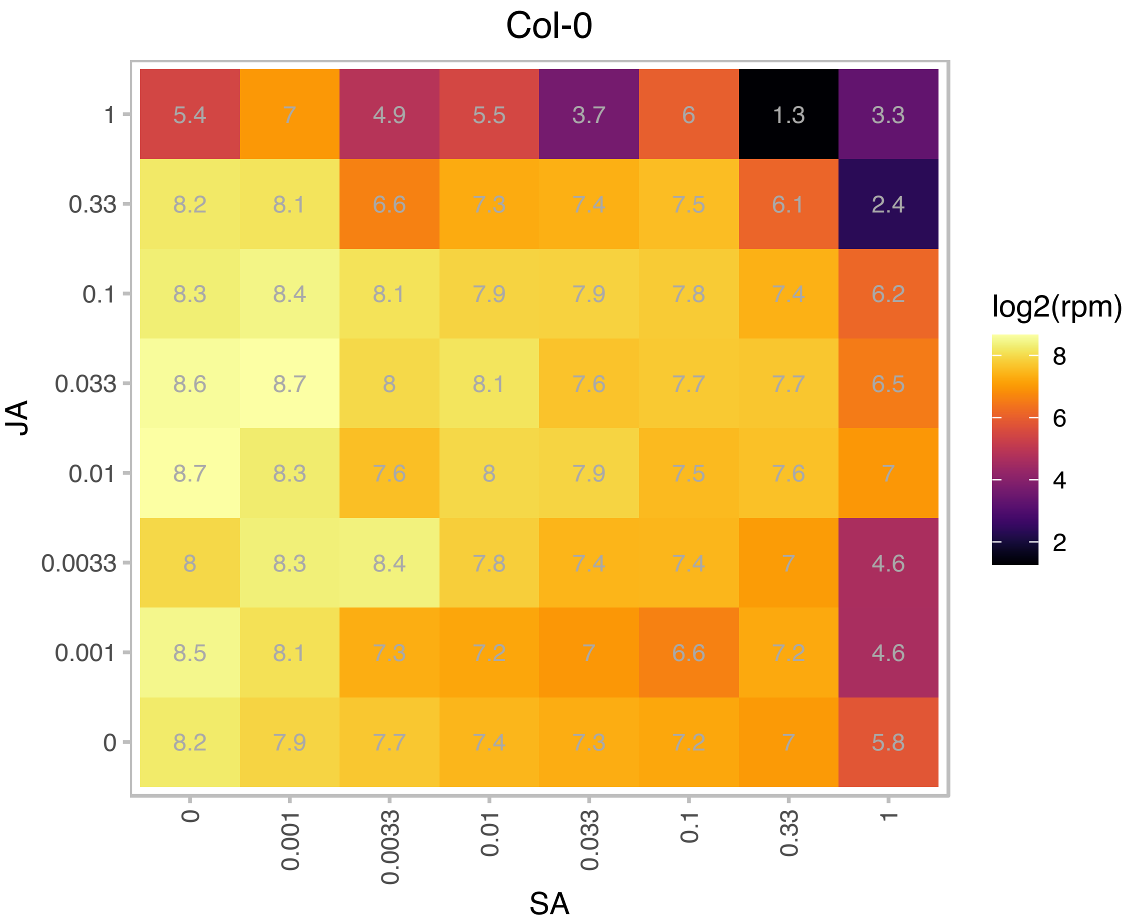


| | |
|---|---|
| Gene | AT1G24020 |
| is.expressed_genes | TRUE |
| is.responsive_genes | TRUE |
| is.apcluster | F-07 |

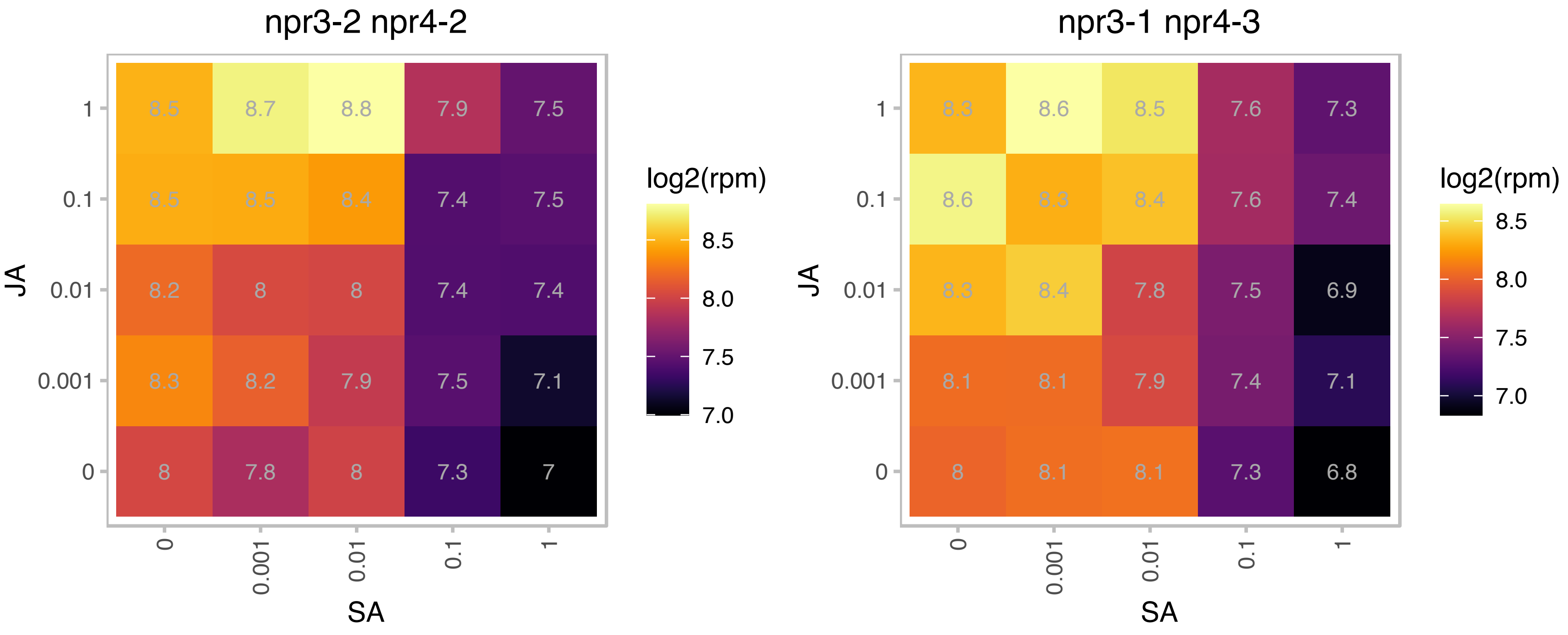

# AT1G26880.1 : NA

Ribosomal protein L34e superfamily protein

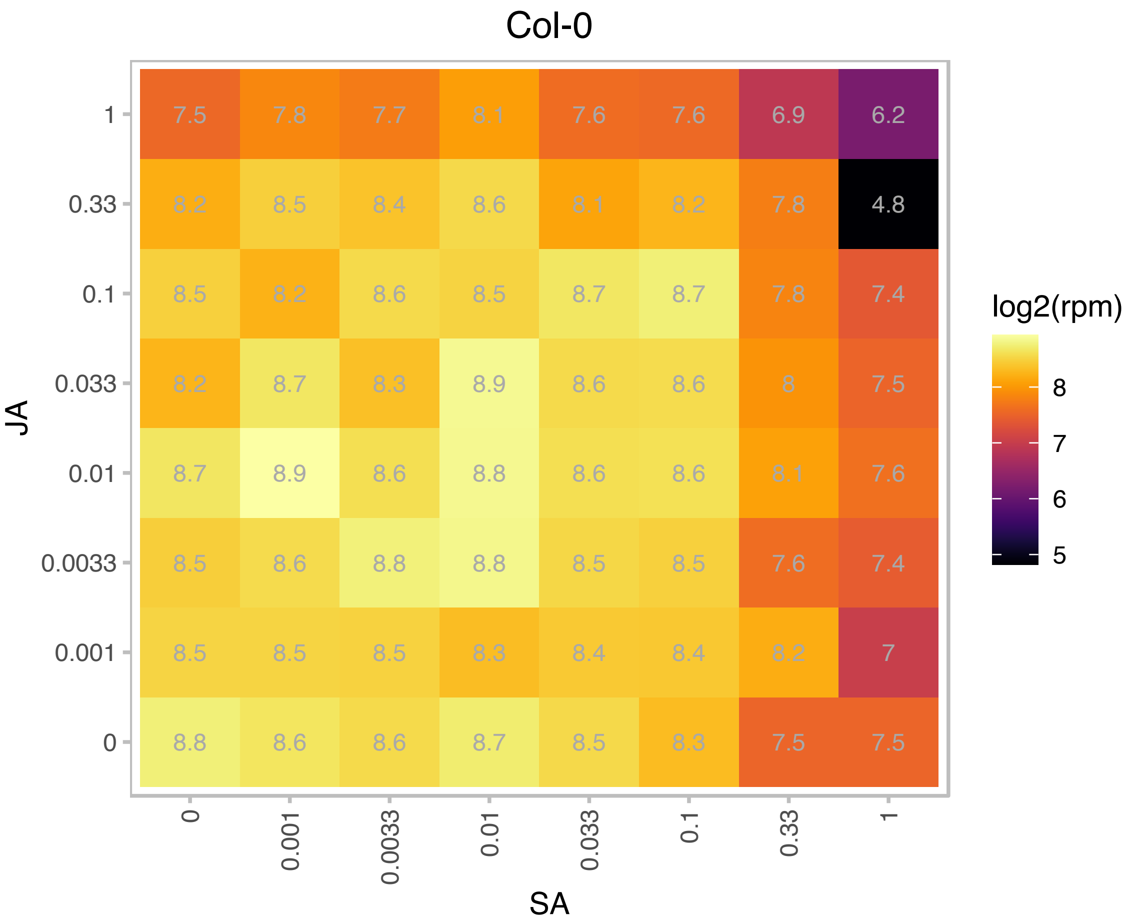


| | |
|---|---|
| Gene | AT1G26880 |
| is.expressed_genes | TRUE |
| is.responsive_genes | TRUE |
| is.apcluster | F-07 |

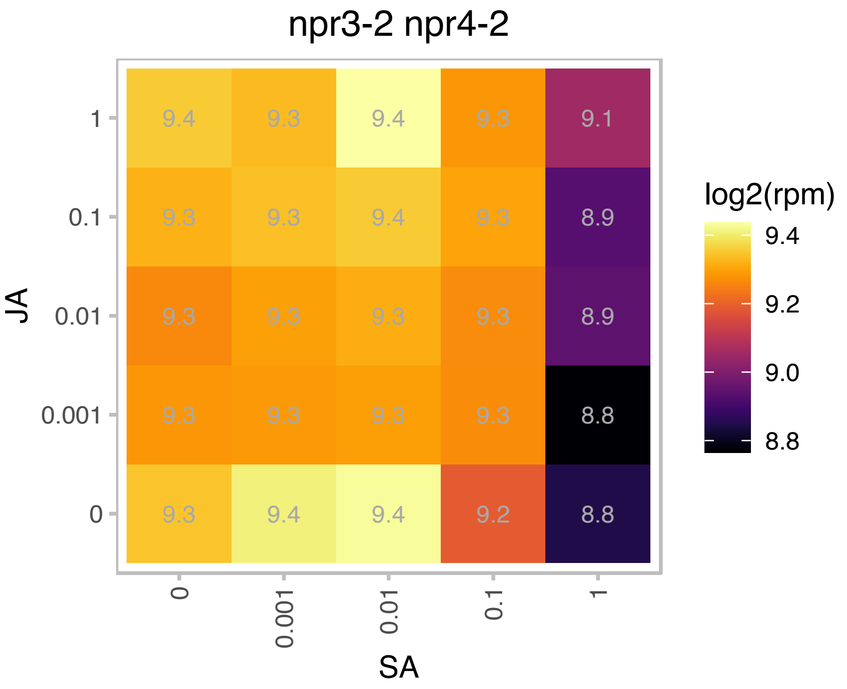


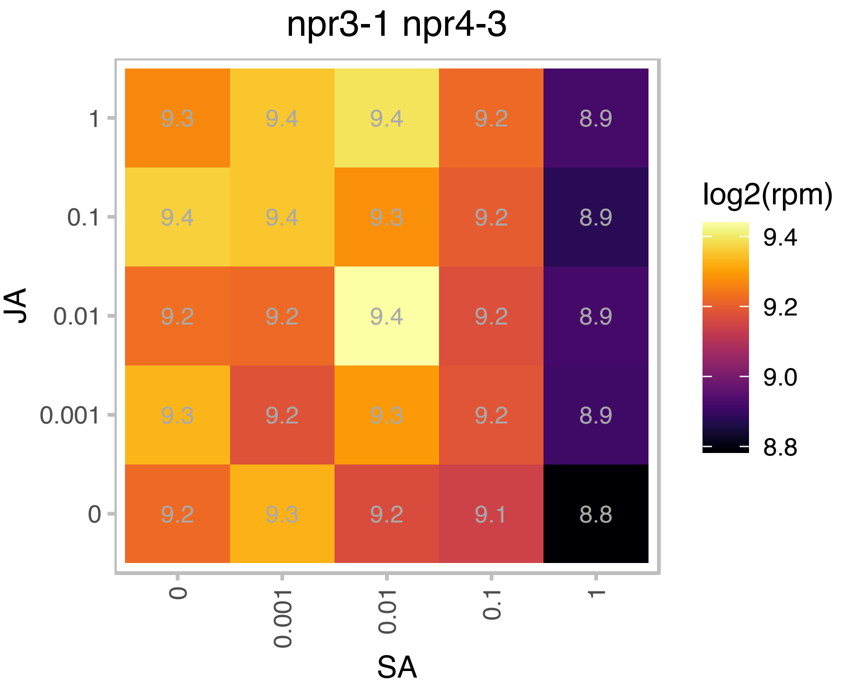

# AT1G27400.1 : NA

Ribosomal protein L22p/L17e family protein

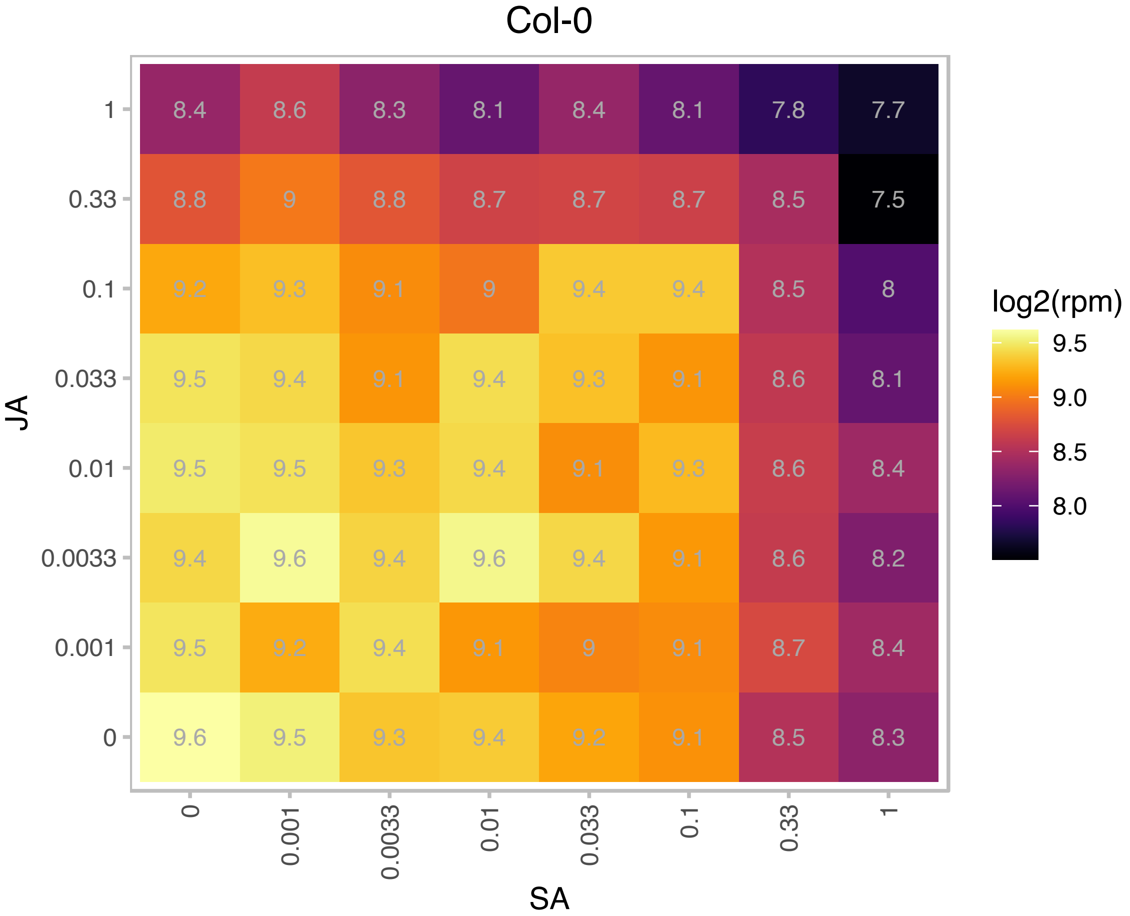


| | |
|---|---|
| Gene | AT1G27400 |
| is.expressed_genes | TRUE |
| is.responsive_genes | TRUE |
| is.apcluster | F-07 |

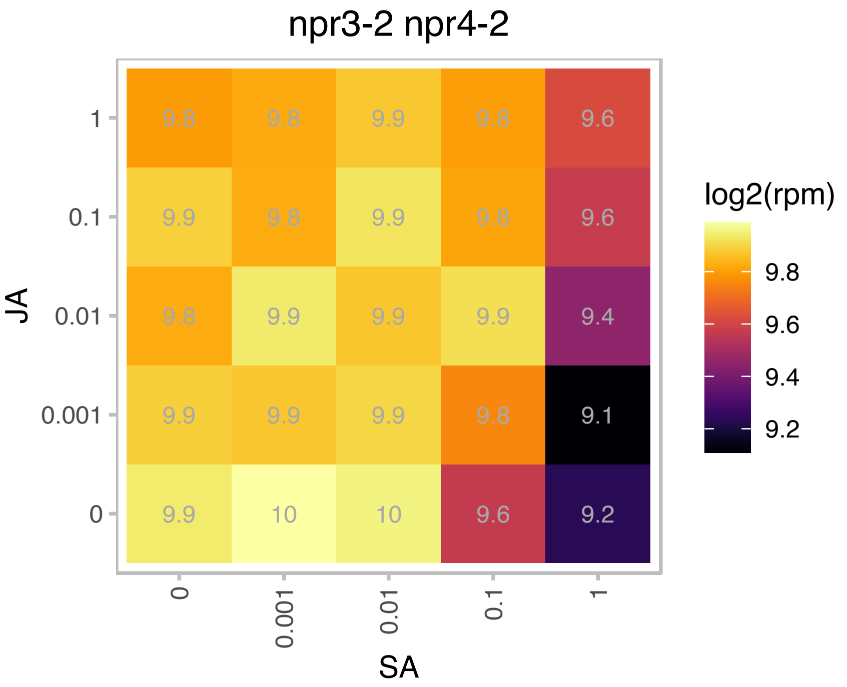


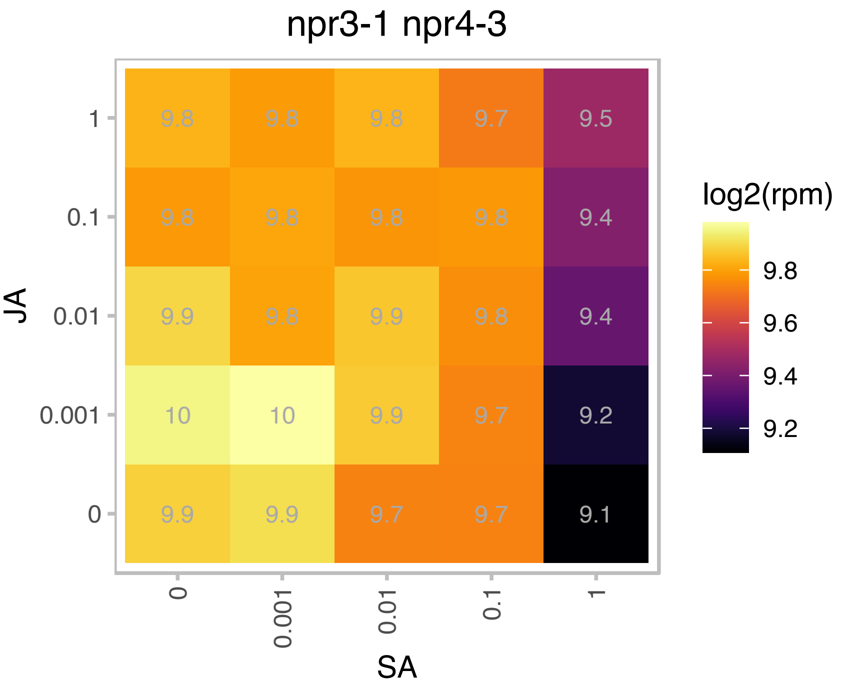

### AT1G28290.1 : AGP31

arabinogalactan protein 31
Encodes an atypical arabinogalactan protein that is localized to the plasma membrange. AGP31 is highly expressed in flowers and vascular tissue and is repressed by jasmonic acid. AGP31 may play a role in vascular tissue function during defense and development.

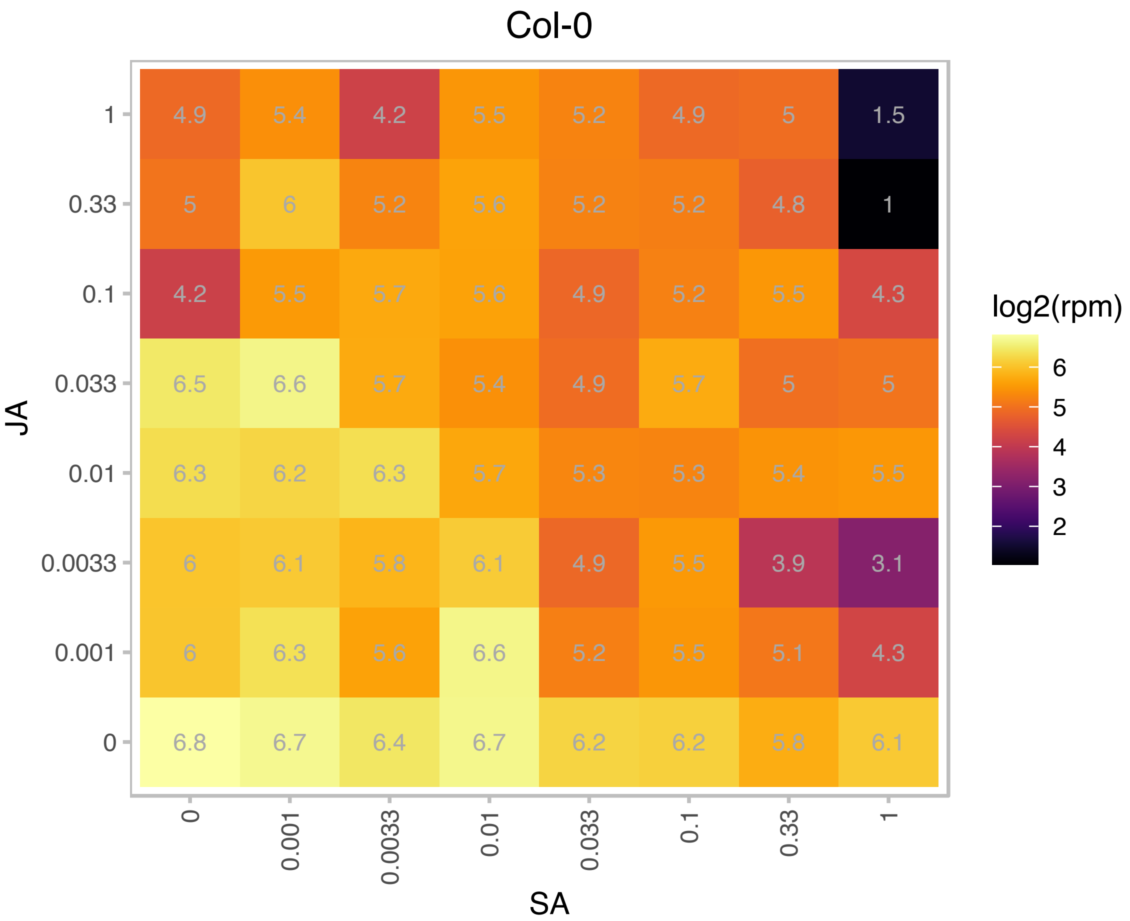


| | |
|---|---|
| Gene | AT1G28290 |
| is.expressed_genes | TRUE |
| is.responsive_genes | TRUE |
| is.apcluster | F-07 |

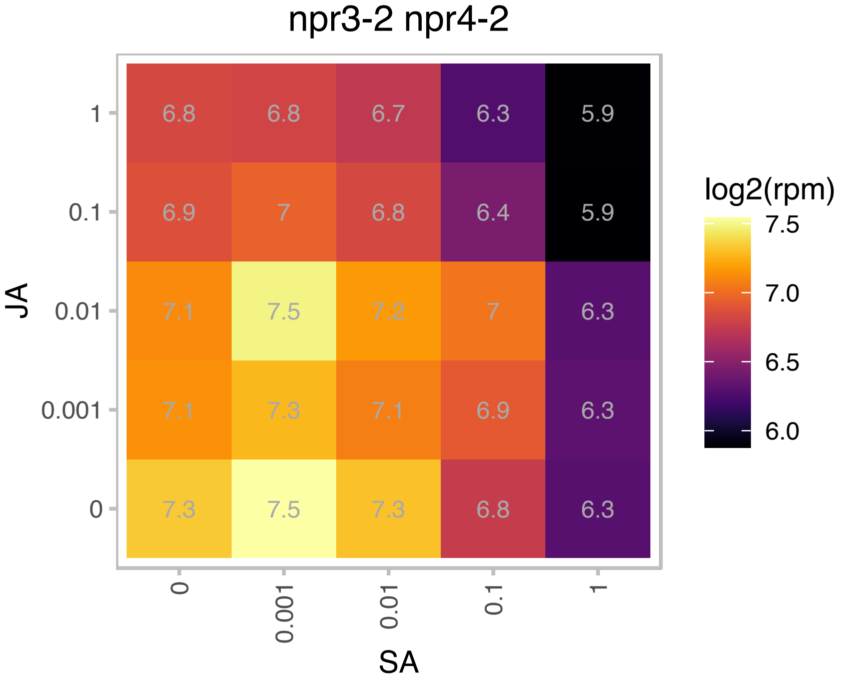


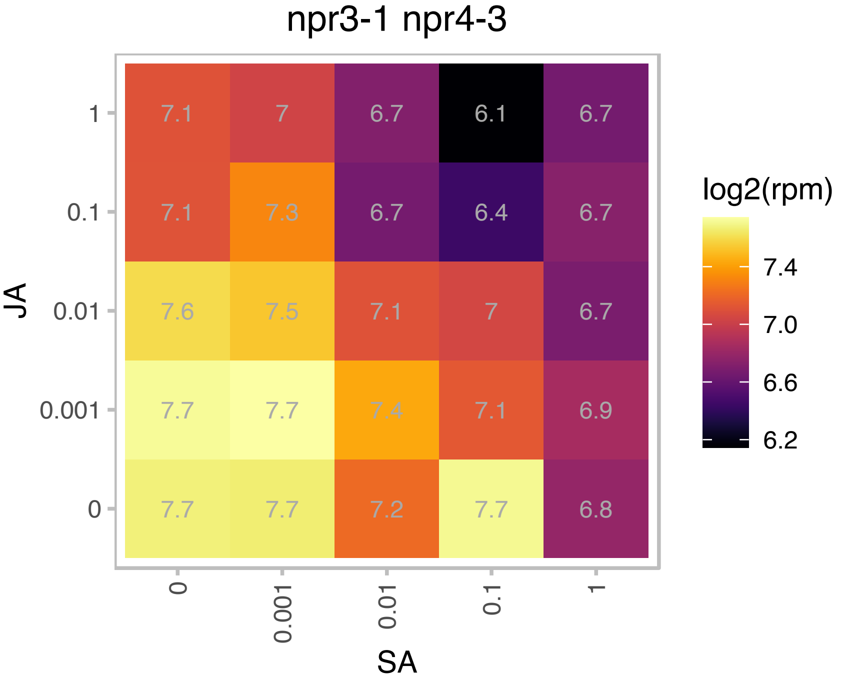

# AT1G29250.1 : NA

Alba DNA/RNA-binding protein

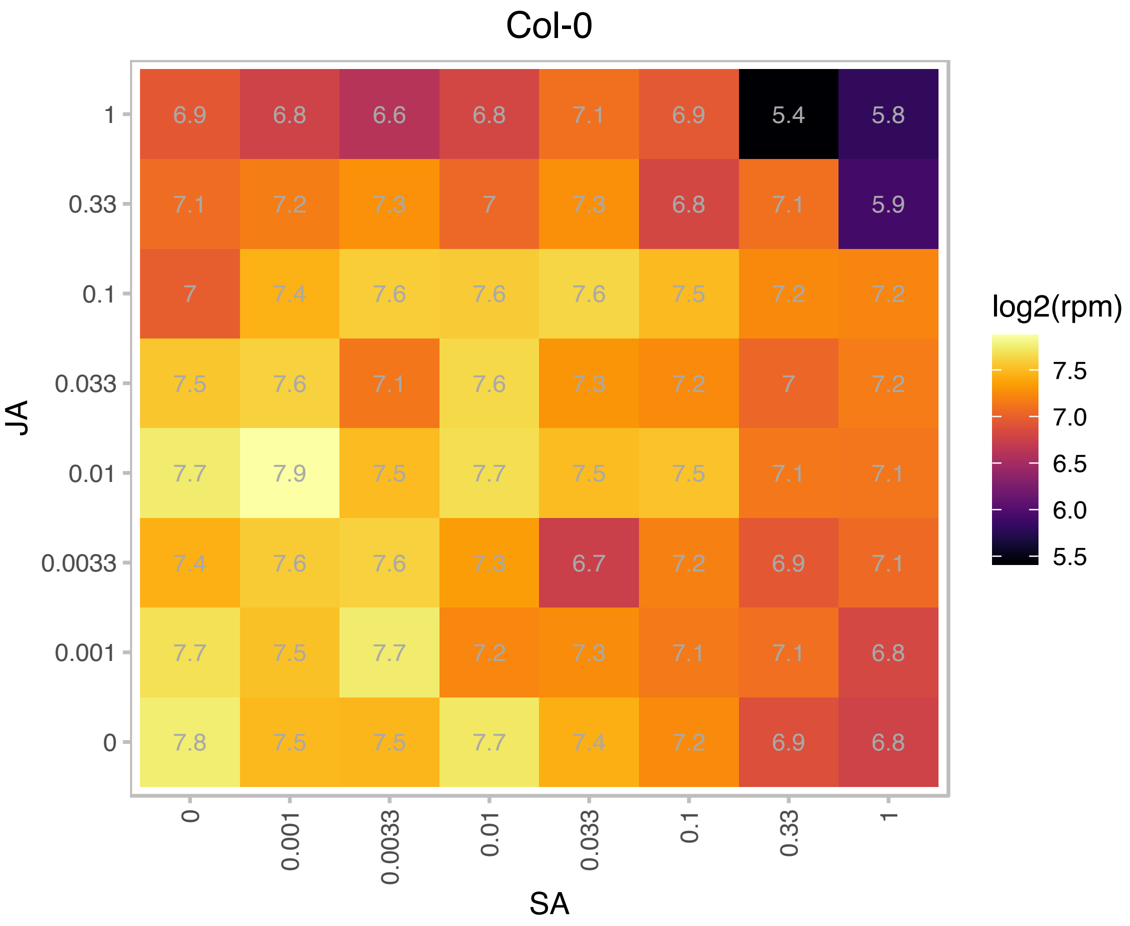


| | |
|---|---|
| Gene | AT1G29250 |
| is.expressed_genes | TRUE |
| is.responsive_genes | TRUE |
| is.apcluster | F-07 |

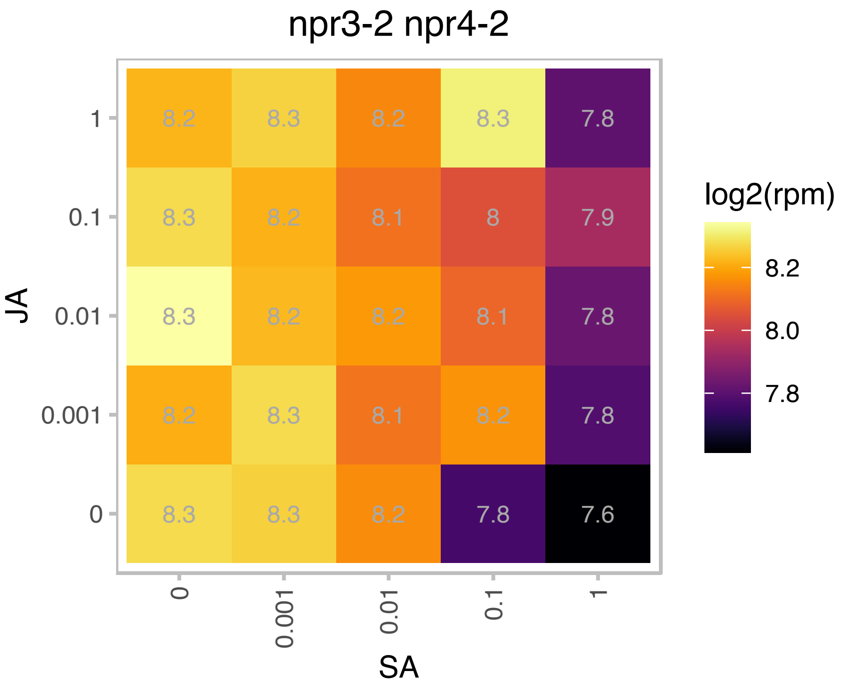


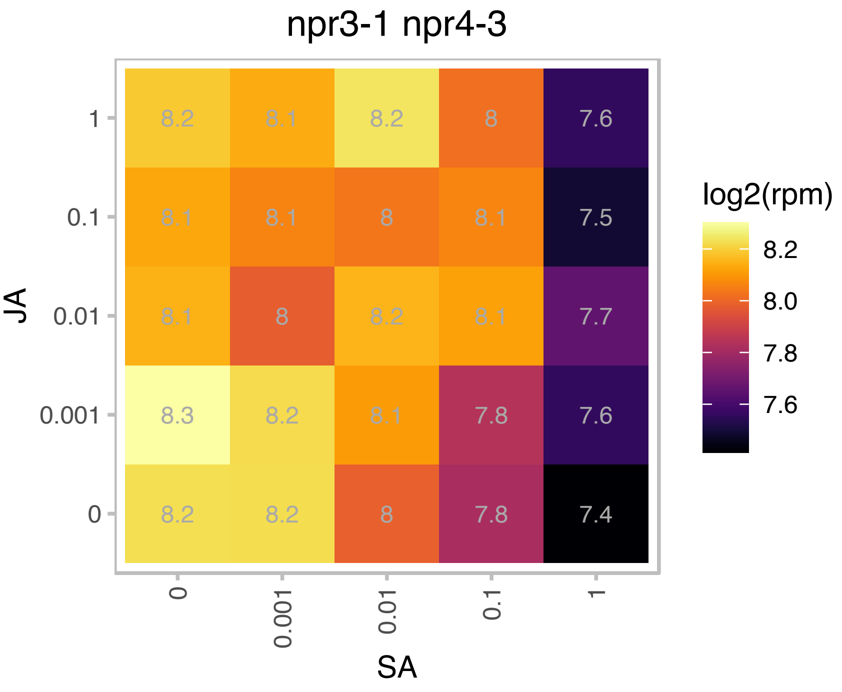

# AT1G29418.1 : NA

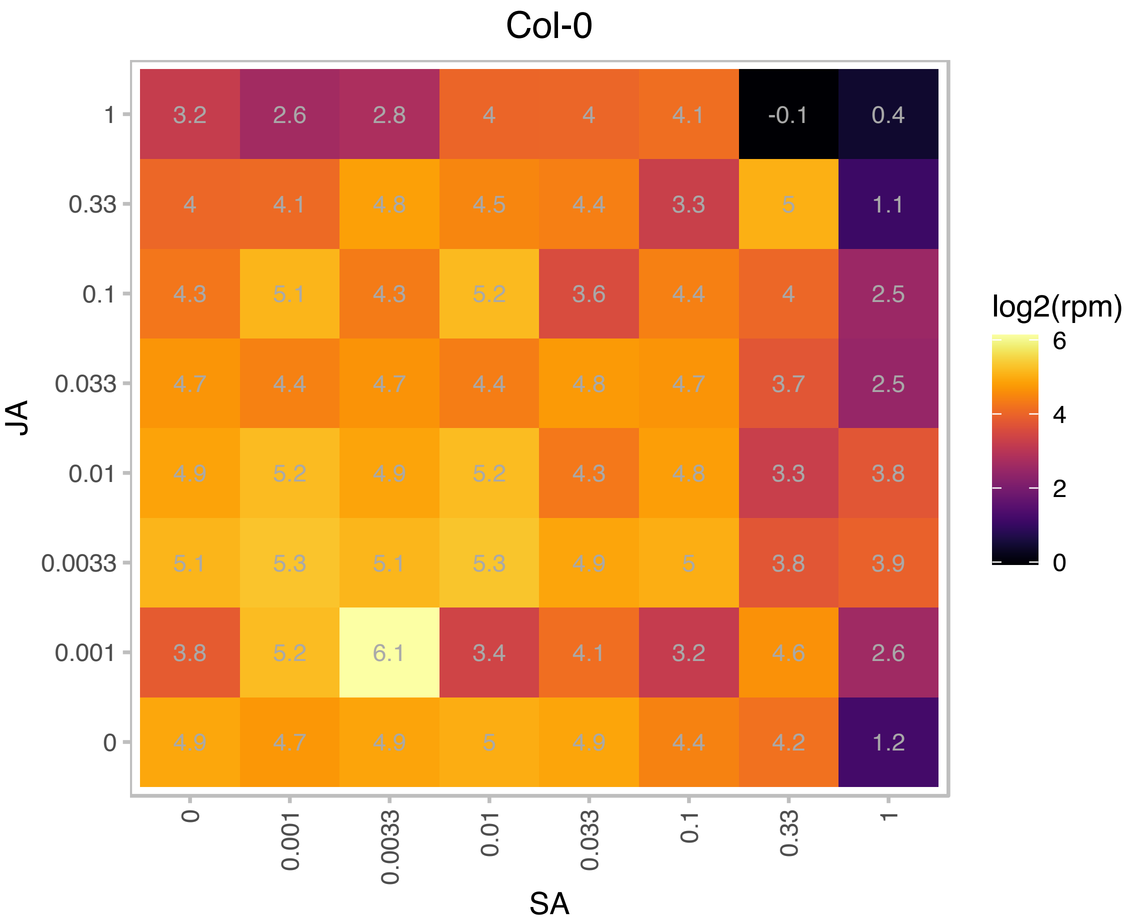


| | |
|---|---|
| Gene | AT1G29418 |
| is.expressed_genes | TRUE |
| is.responsive_genes | TRUE |
| is.apcluster | F-07 |

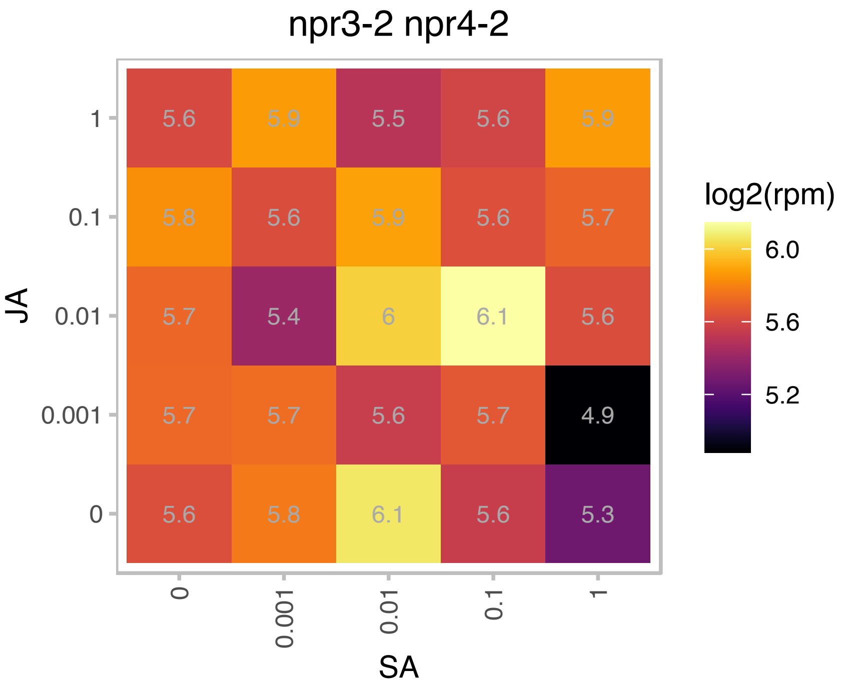


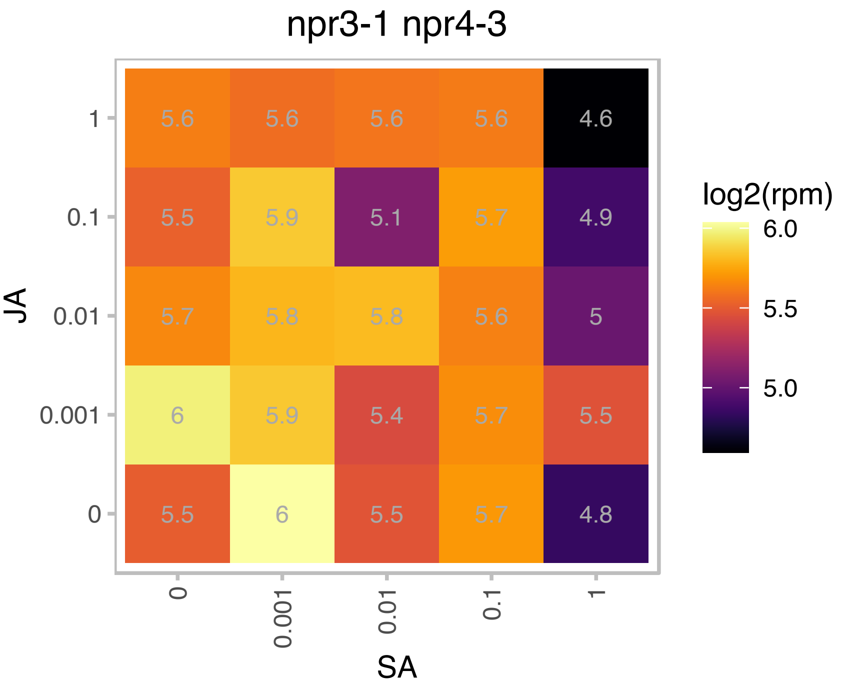

# AT1G29530.1 : NA

| | |
|---|---|
| Gene | AT1G29530 |
| is.expressed_genes | TRUE |
| is.responsive_genes | TRUE |
| is.apcluster | F-07 |

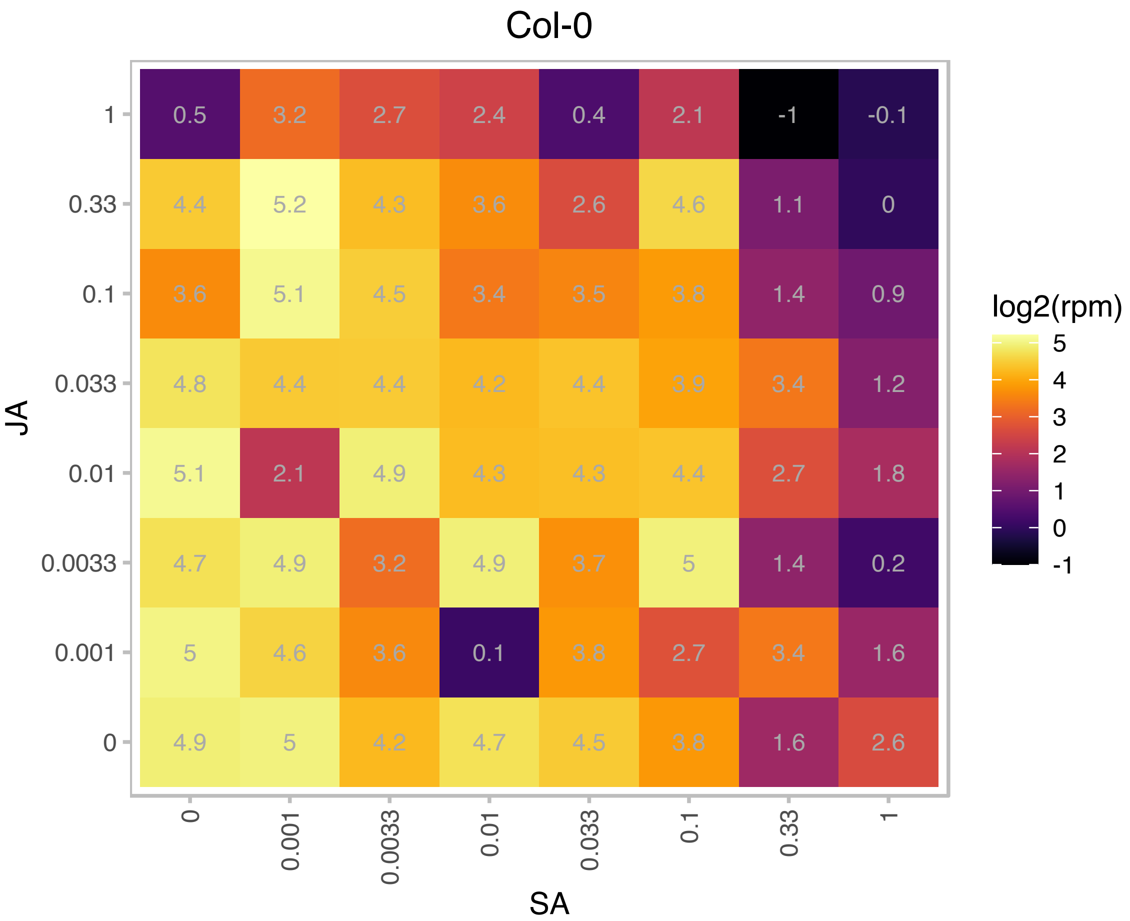


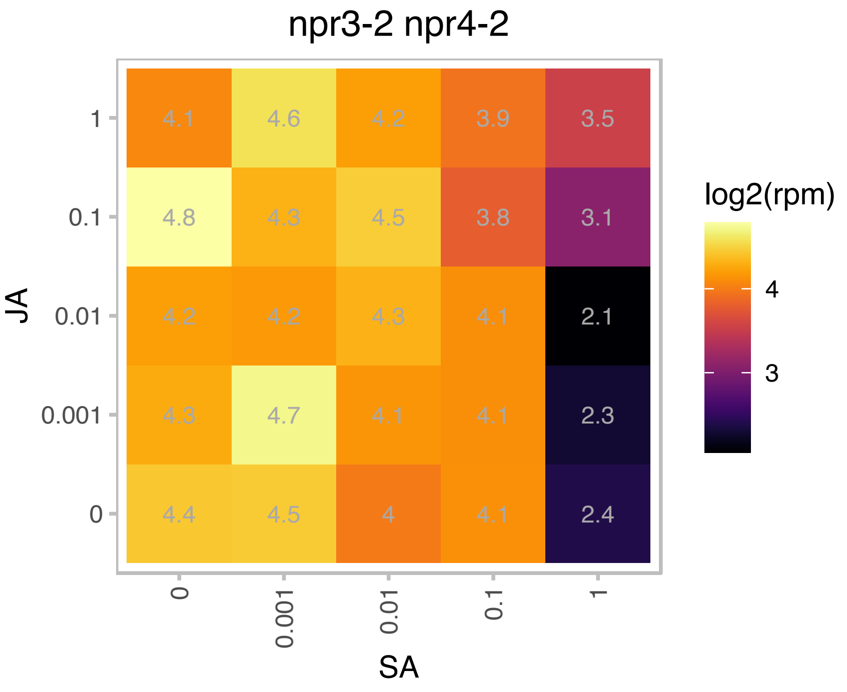


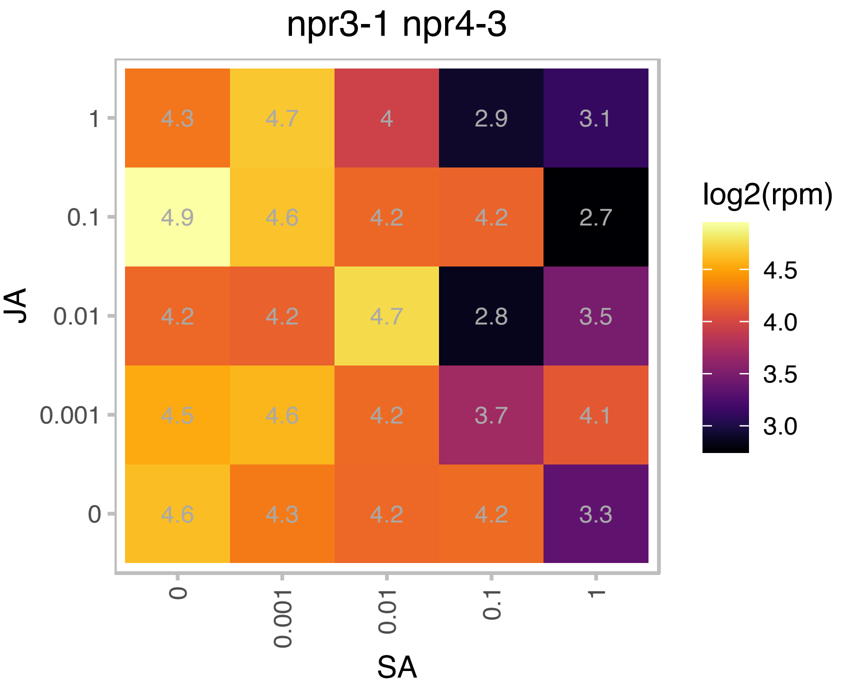

### AT1G32580.1 : MORF5

plastid developmental protein DAG, putative

Member of MORF family consisting of of nine full-length proteins encoded in the nuclear genome. MORF proteins are required for all RNA editing events in plastids and for many, possibly also all, sites in mitochondria. Potential link between the RNA binding PPR protein and the protein contributing the enzymatic activity in RNA editing.

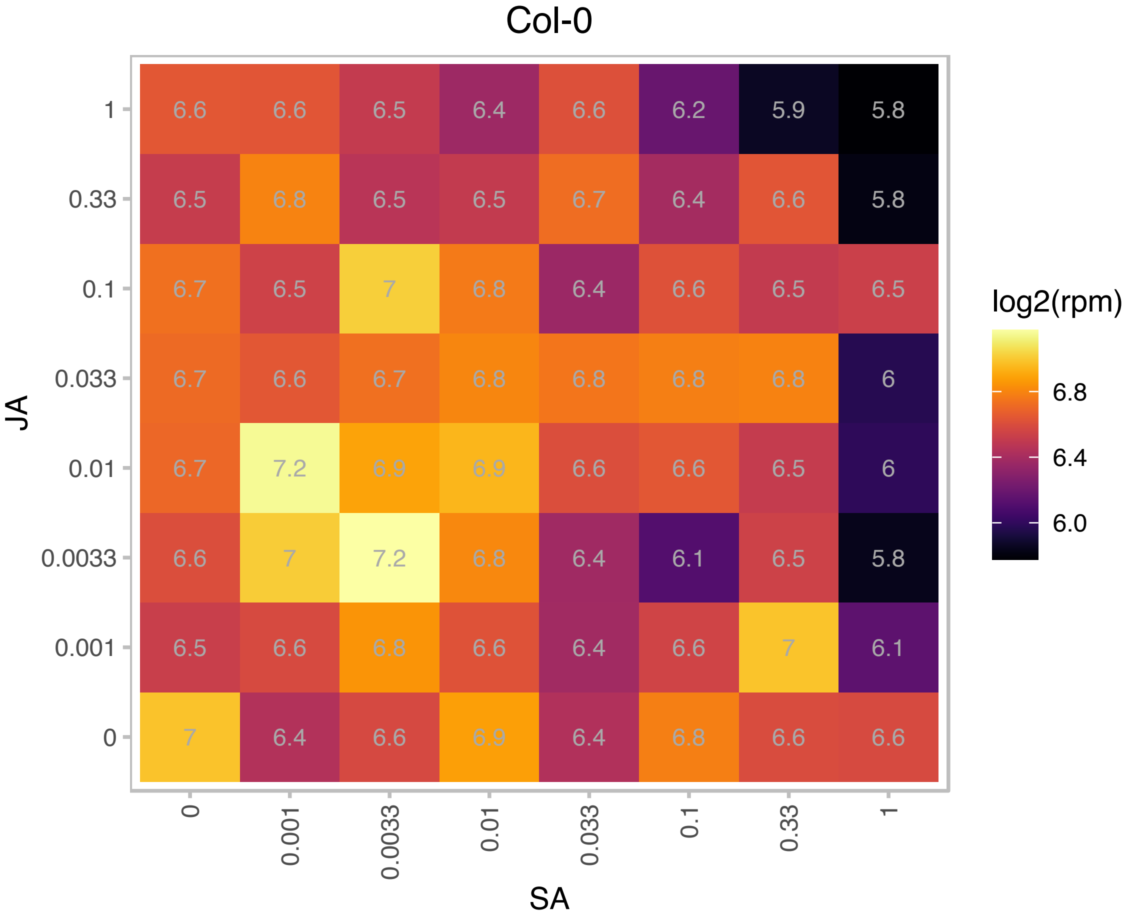


| Gene | AT1G32580 |
|---|---|
| is.expressed_genes | TRUE |
| is.responsive_genes | TRUE |
| is.apcluster | F-07 |

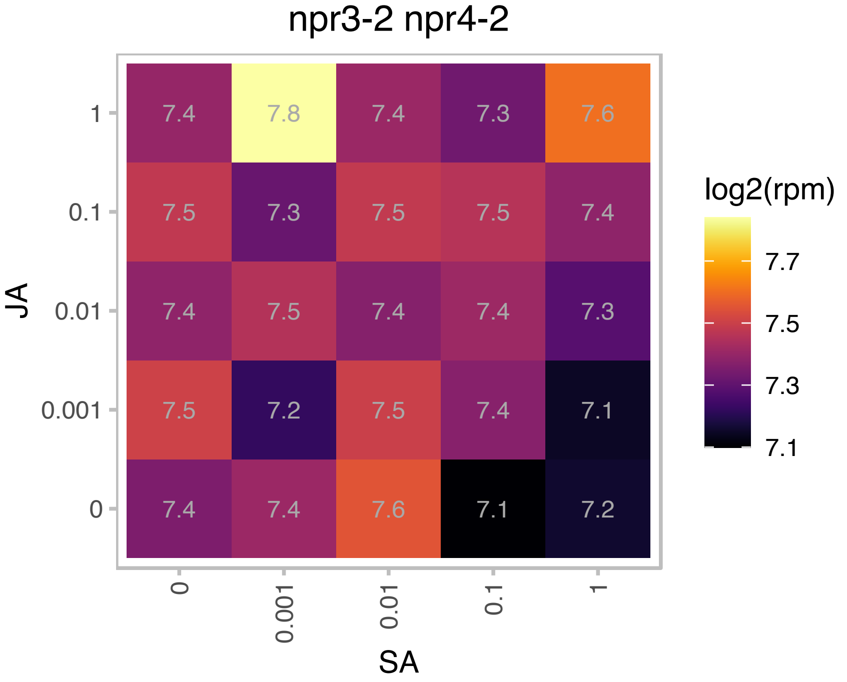


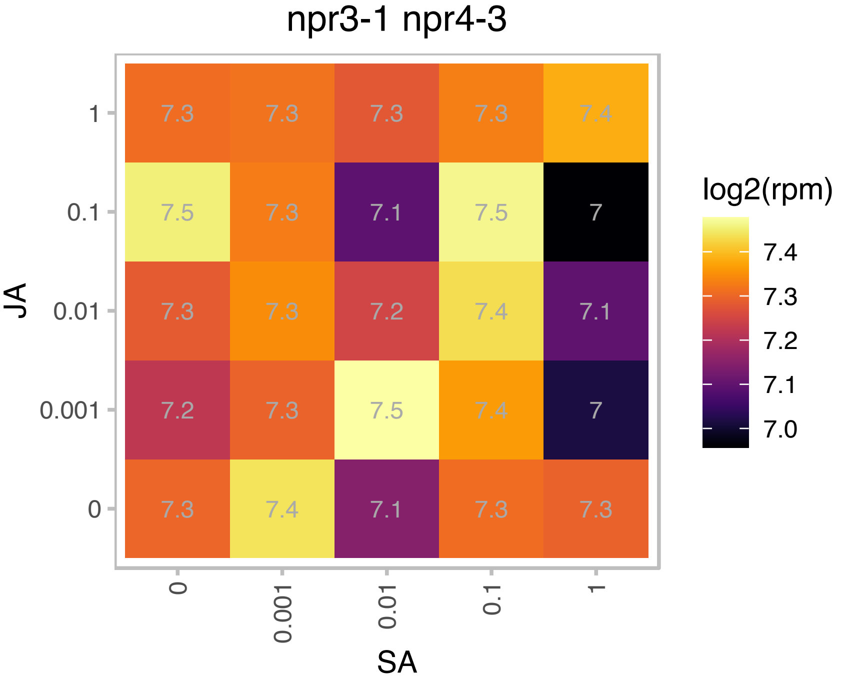

# AT1G33120.1 : NA

Ribosomal protein L6 family

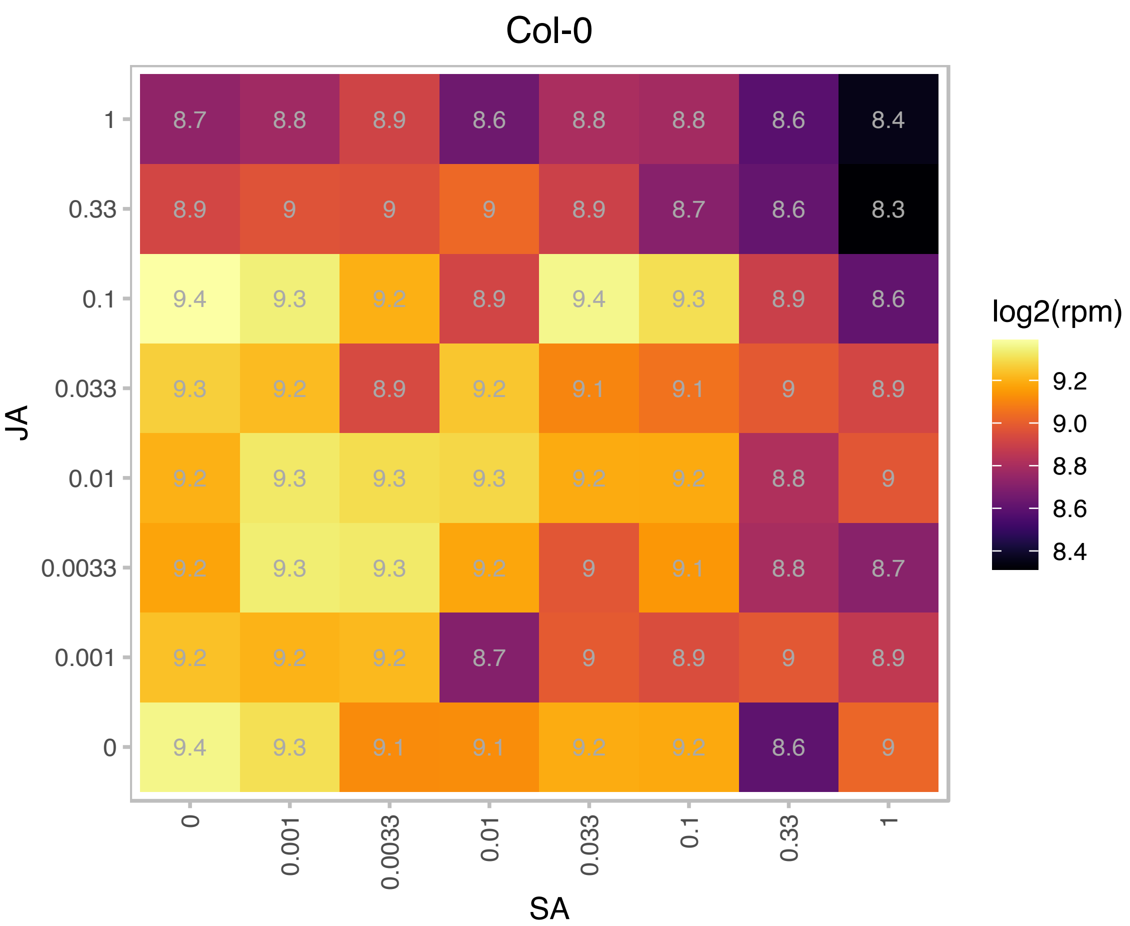


| | |
|---|---|
| Gene | AT1G33120 |
| is.expressed_genes | TRUE |
| is.responsive_genes | TRUE |
| is.apcluster | F-07 |

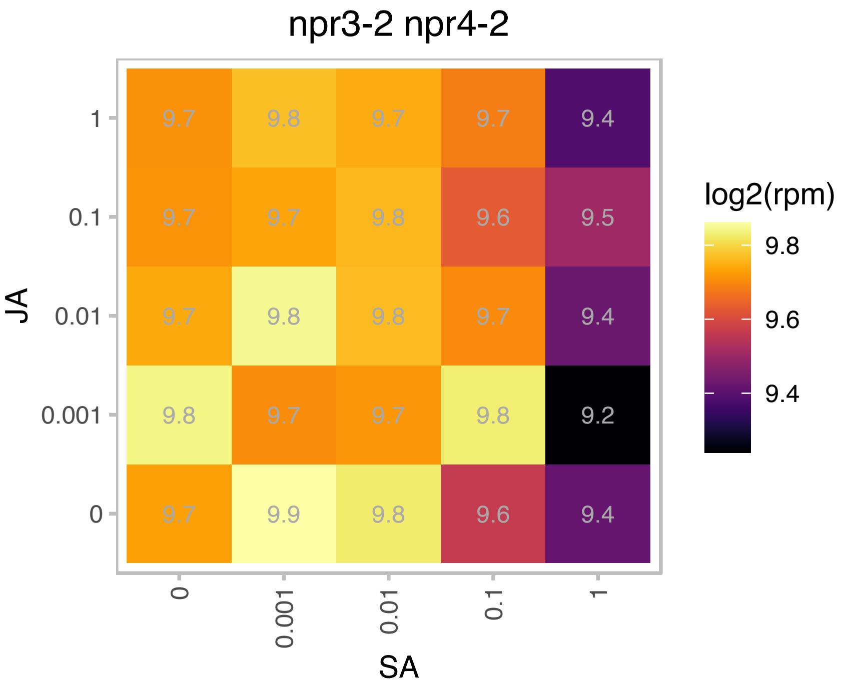


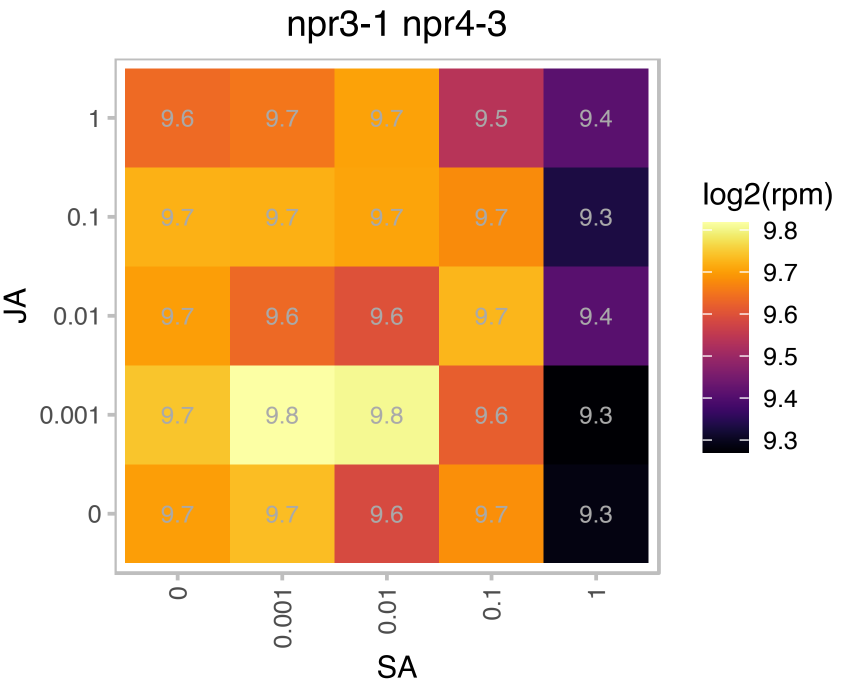

# AT1G33265.1 : NA

Transmembrane proteins 14C

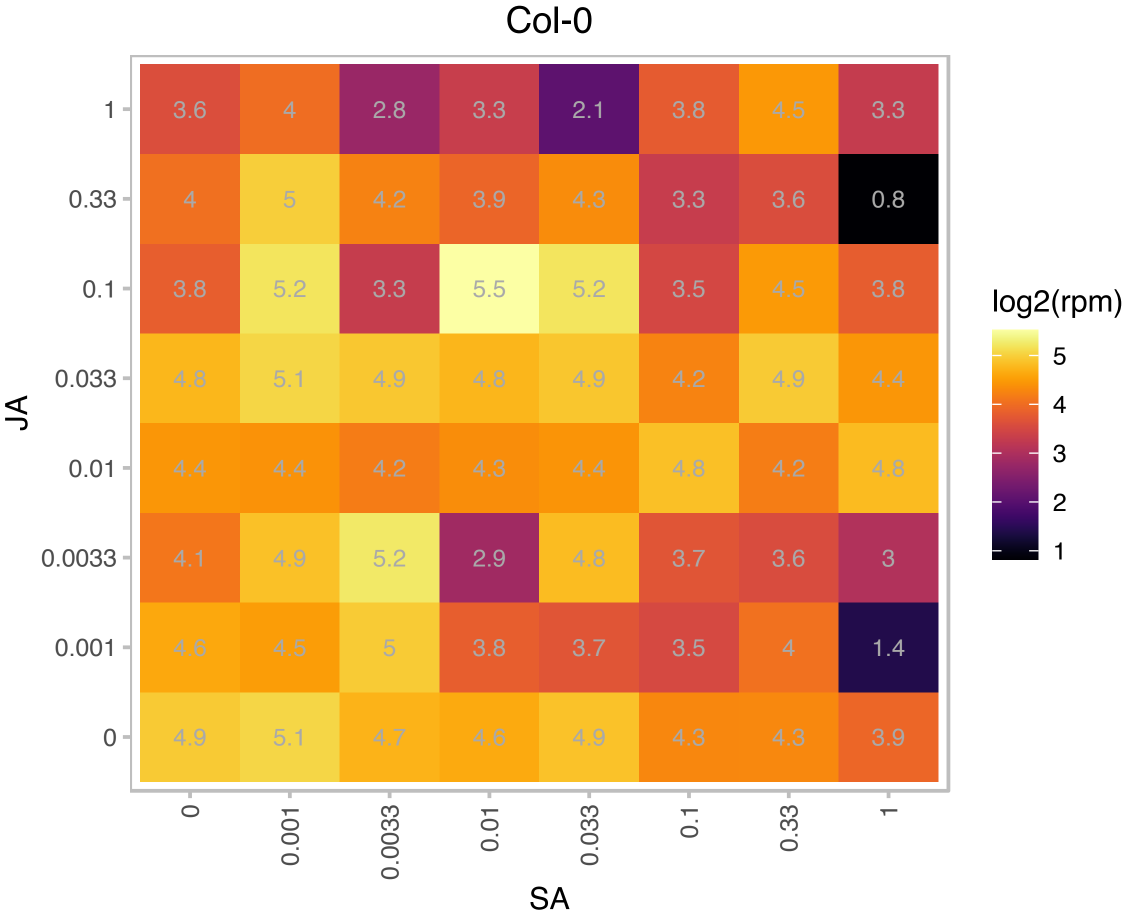


| | |
|---|---|
| Gene | AT1G33265 |
| is.expressed_genes | TRUE |
| is.responsive_genes | TRUE |
| is.apcluster | F-07 |

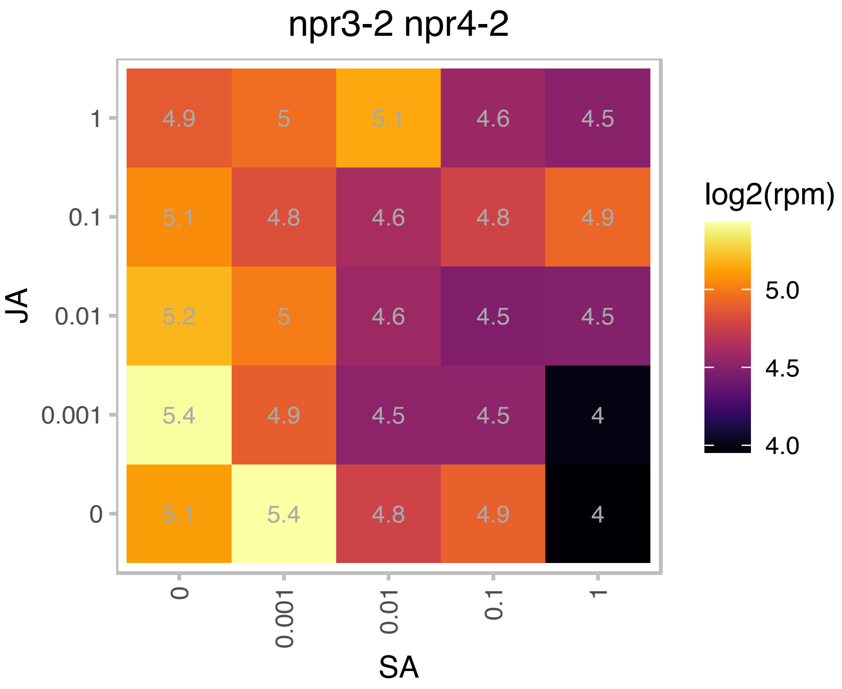


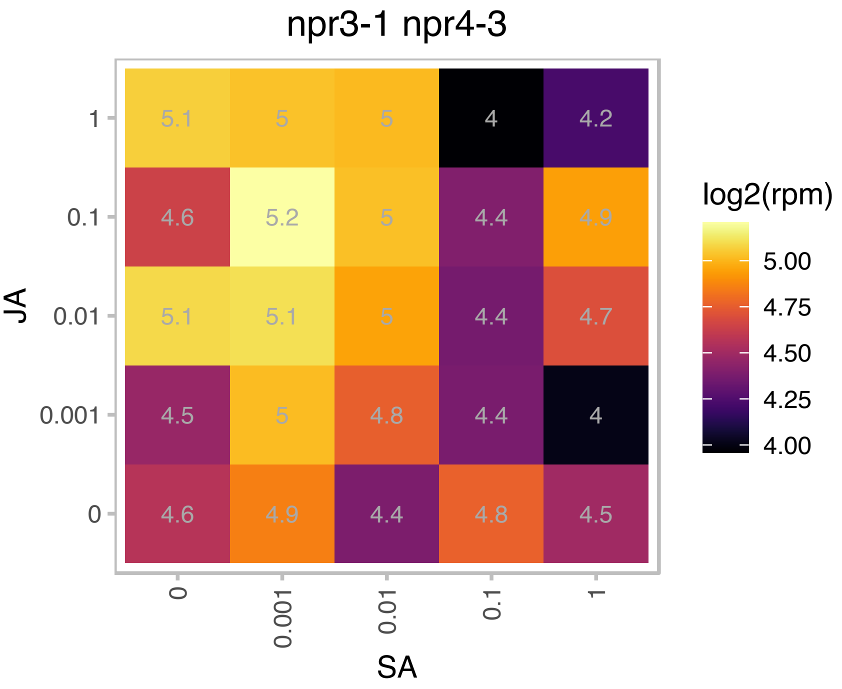

# AT1G34030.1 : NA

Ribosomal protein S13/S18 family

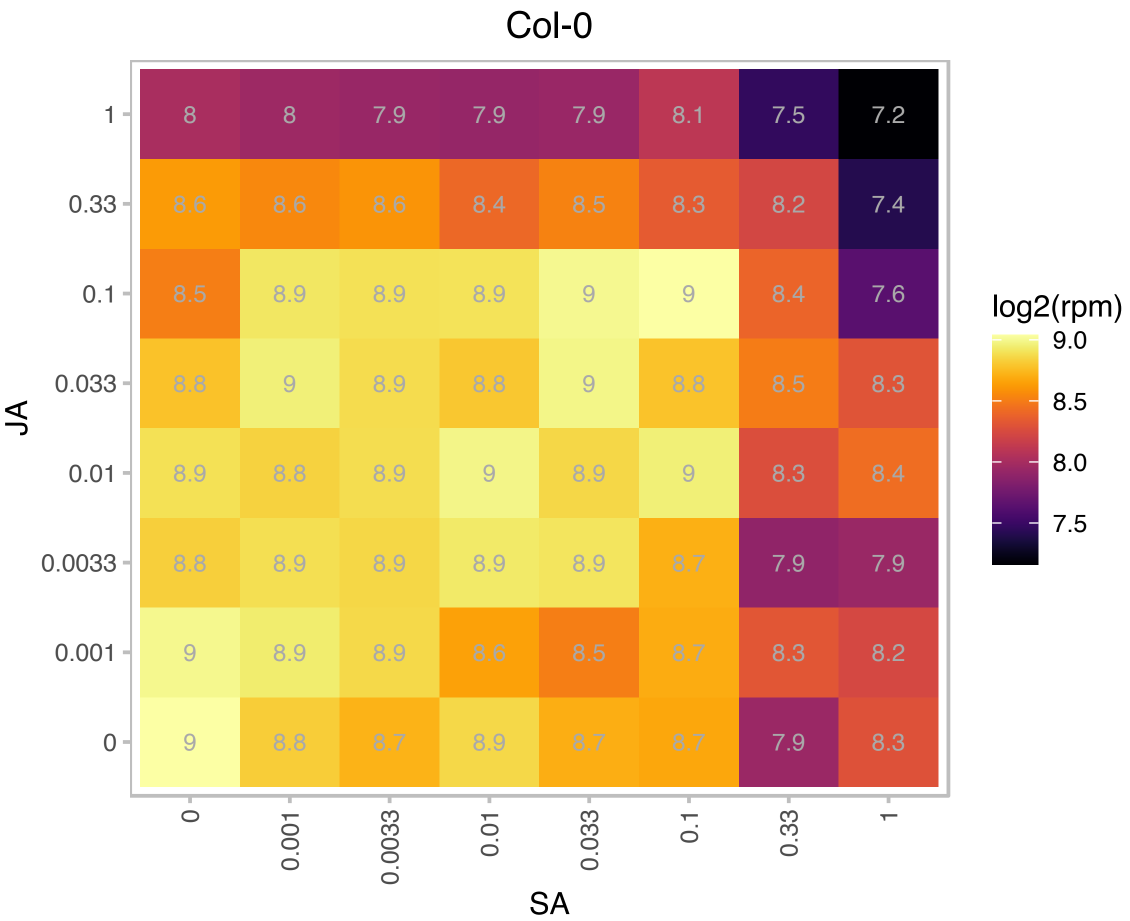


| | |
|---|---|
| Gene | AT1G34030 |
| is.expressed_genes | TRUE |
| is.responsive_genes | TRUE |
| is.apcluster | F-07 |

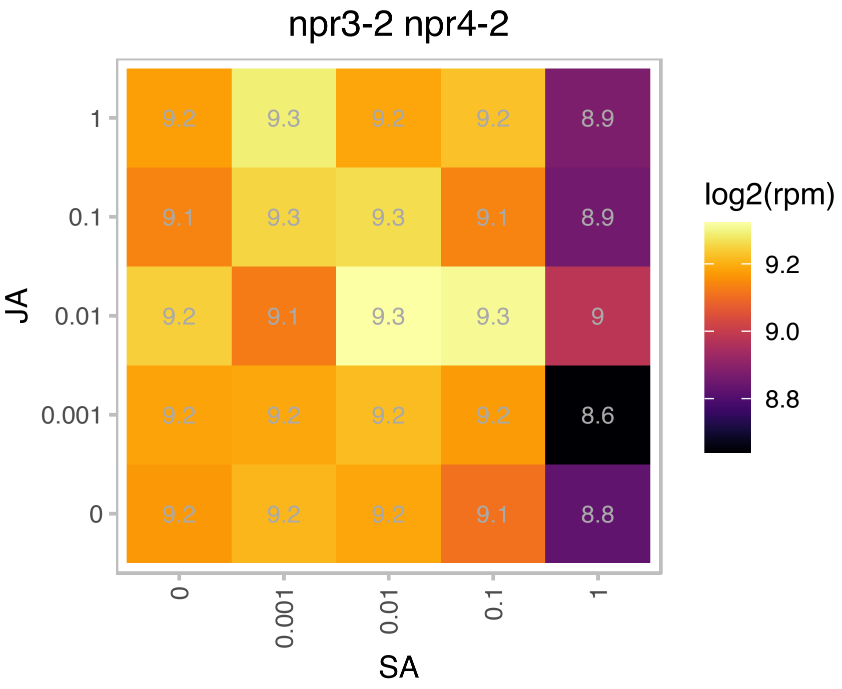


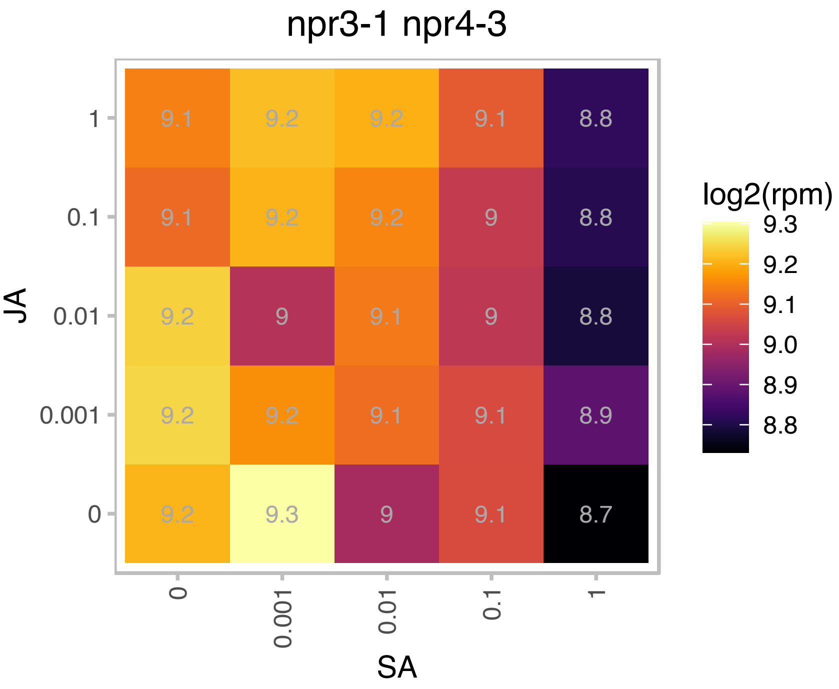

### AT1G44000.1 : SGRL

Col-0

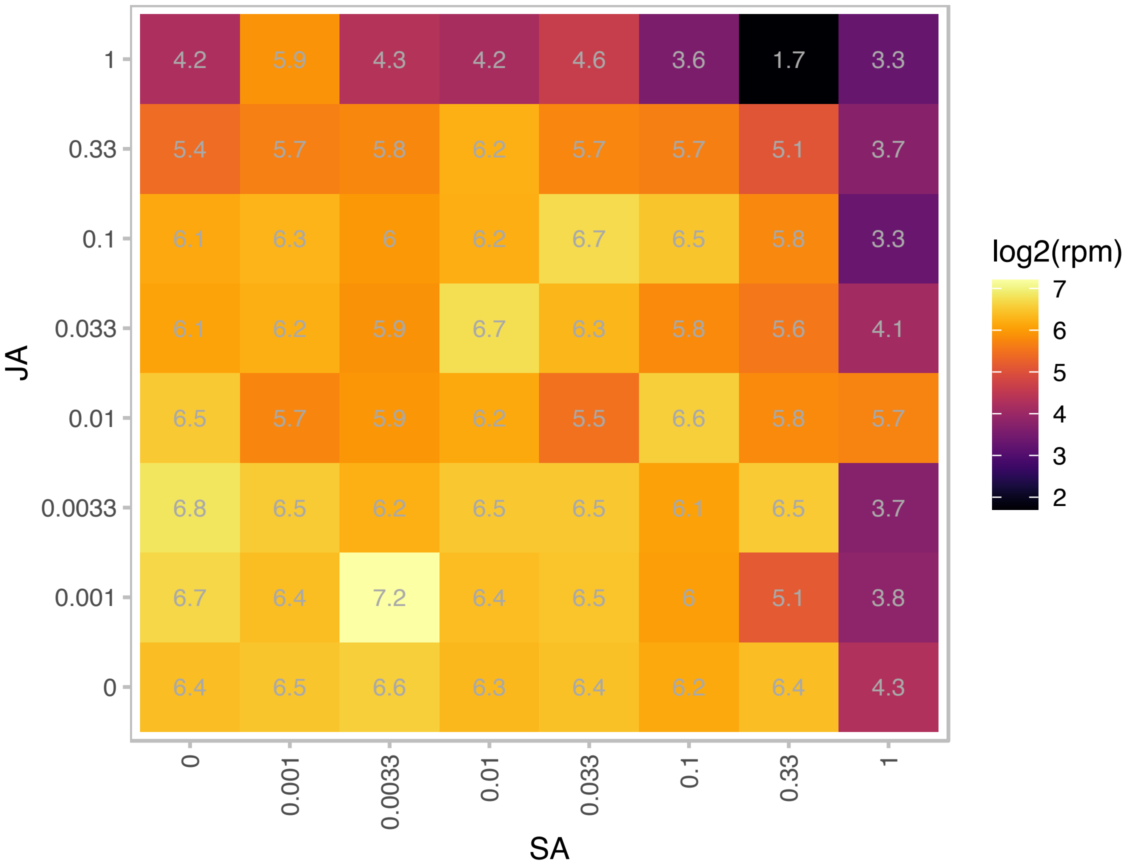


| | |
|---|---|
| Gene | AT1G44000 |
| is.expressed_genes | TRUE |
| is.responsive_genes | TRUE |
| is.apcluster | F-07 |

npr3-2 npr4-2

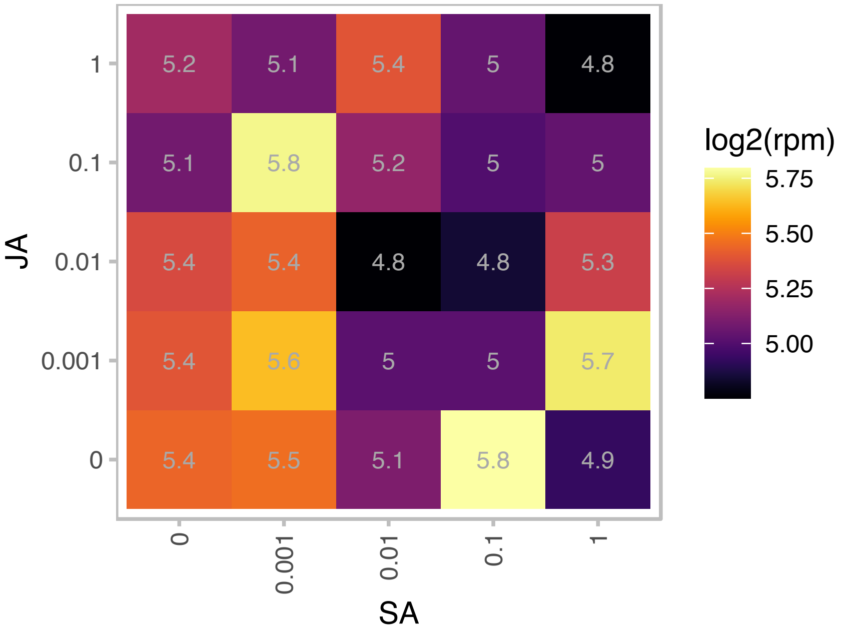


npr3-1 npr4-3

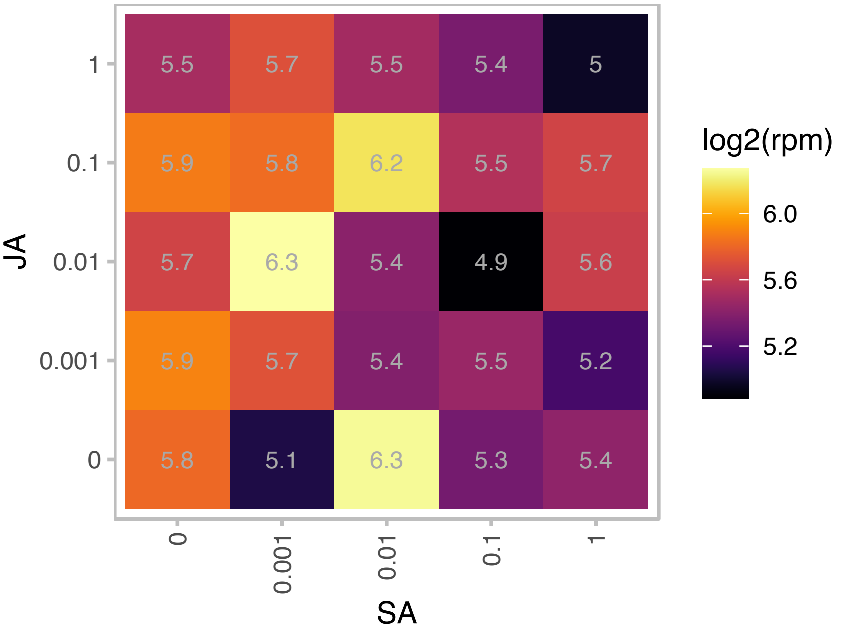

# AT1G48830.1 : NA

Ribosomal protein S7e family protein

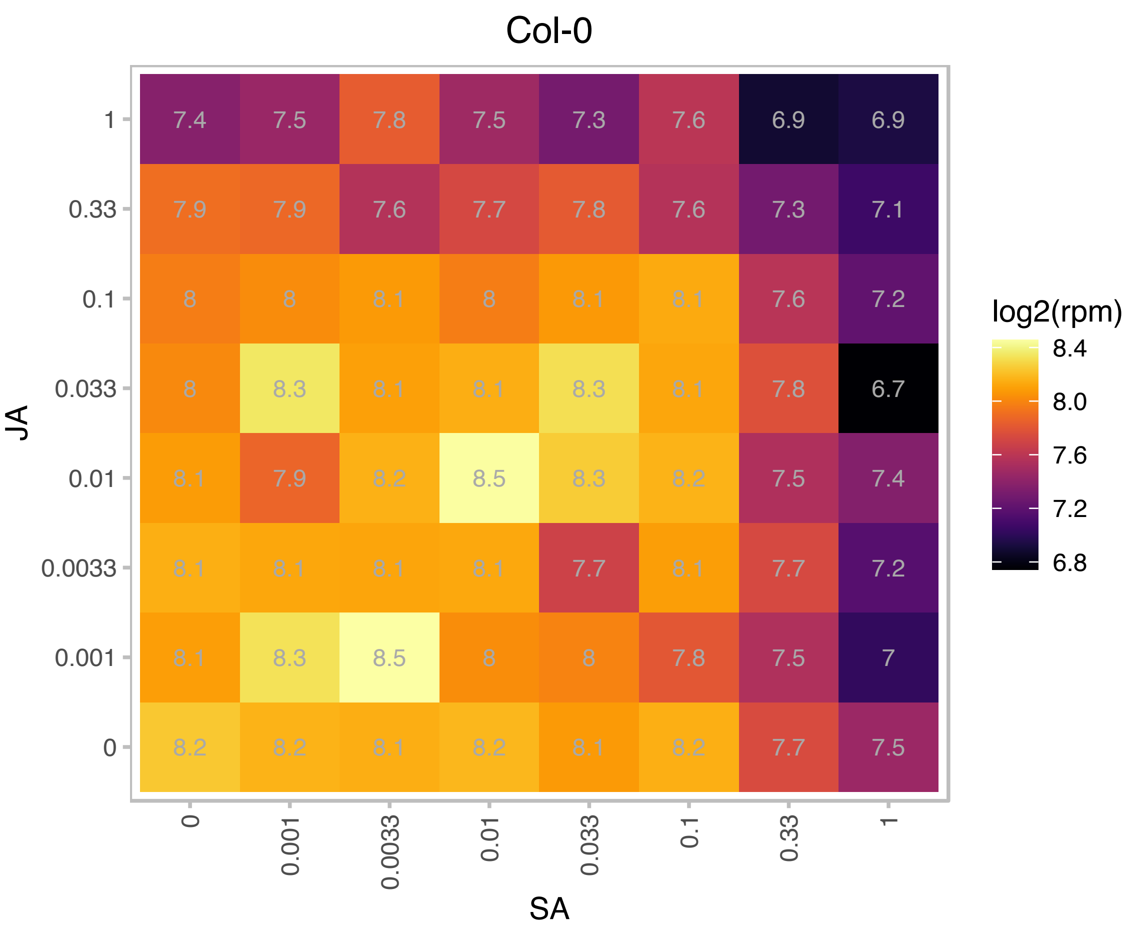


| | |
|---|---|
| Gene | AT1G48830 |
| is.expressed_genes | TRUE |
| is.responsive_genes | TRUE |
| is.apcluster | F-07 |

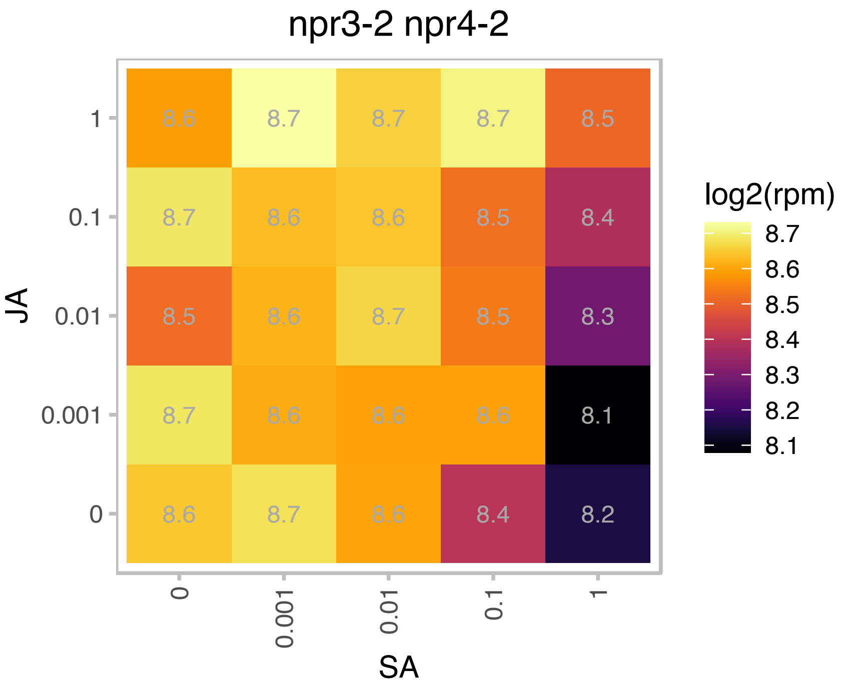


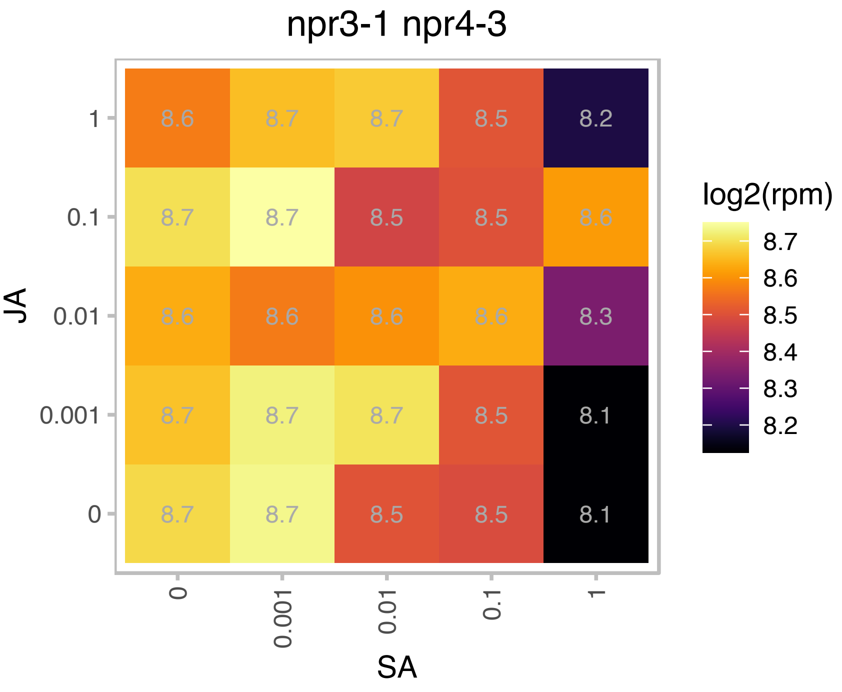

### AT1G49130.1 : BBX17

B-box type zinc finger protein with CCT domain

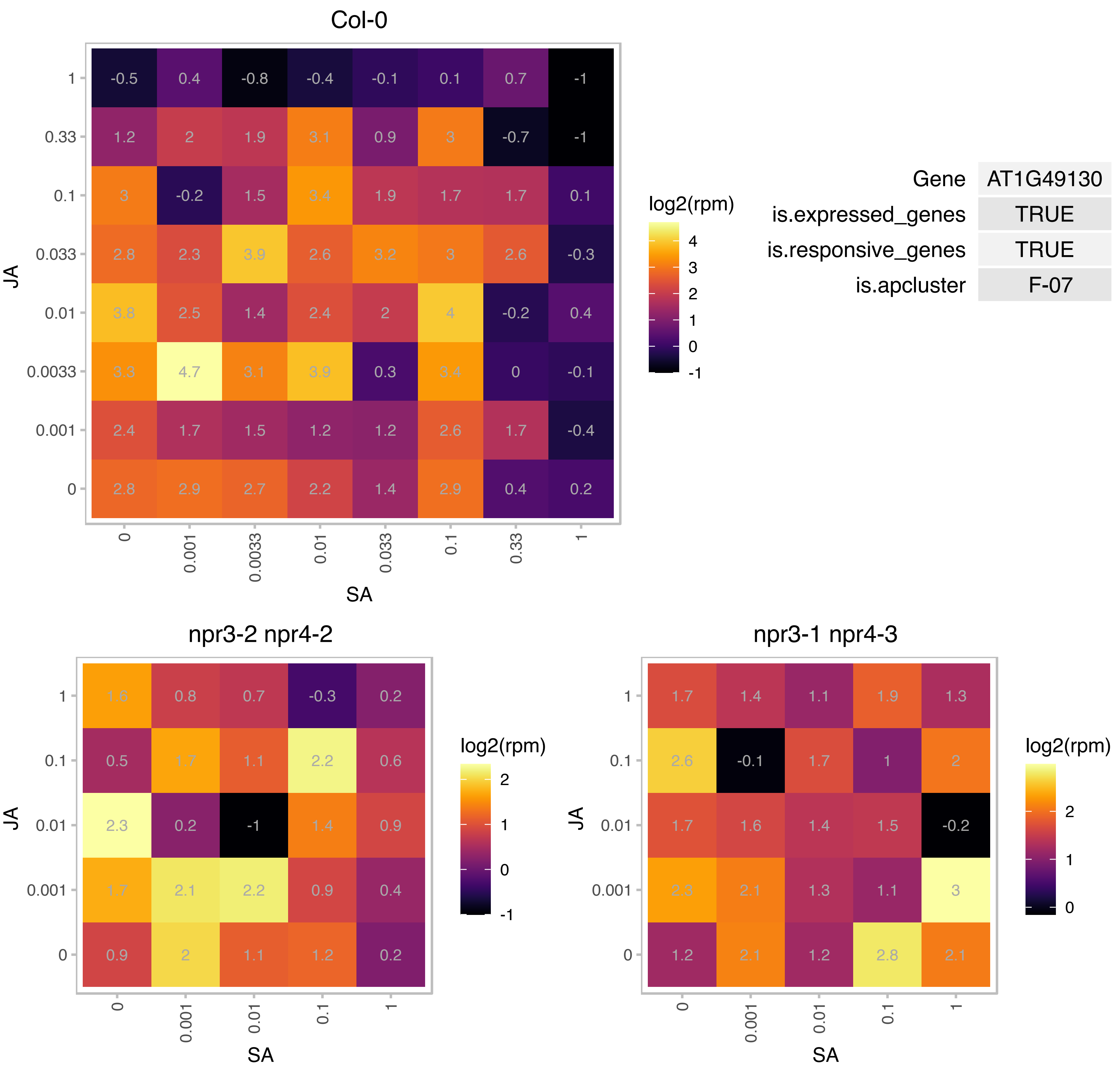


| | |
|---|---|
| Gene | AT1G49130 |
| is.expressed_genes | TRUE |
| is.responsive_genes | TRUE |
| is.apcluster | F-07 |

### AT1G49400.1 : emb1129

Nucleic acid-binding, OB-fold-like protein

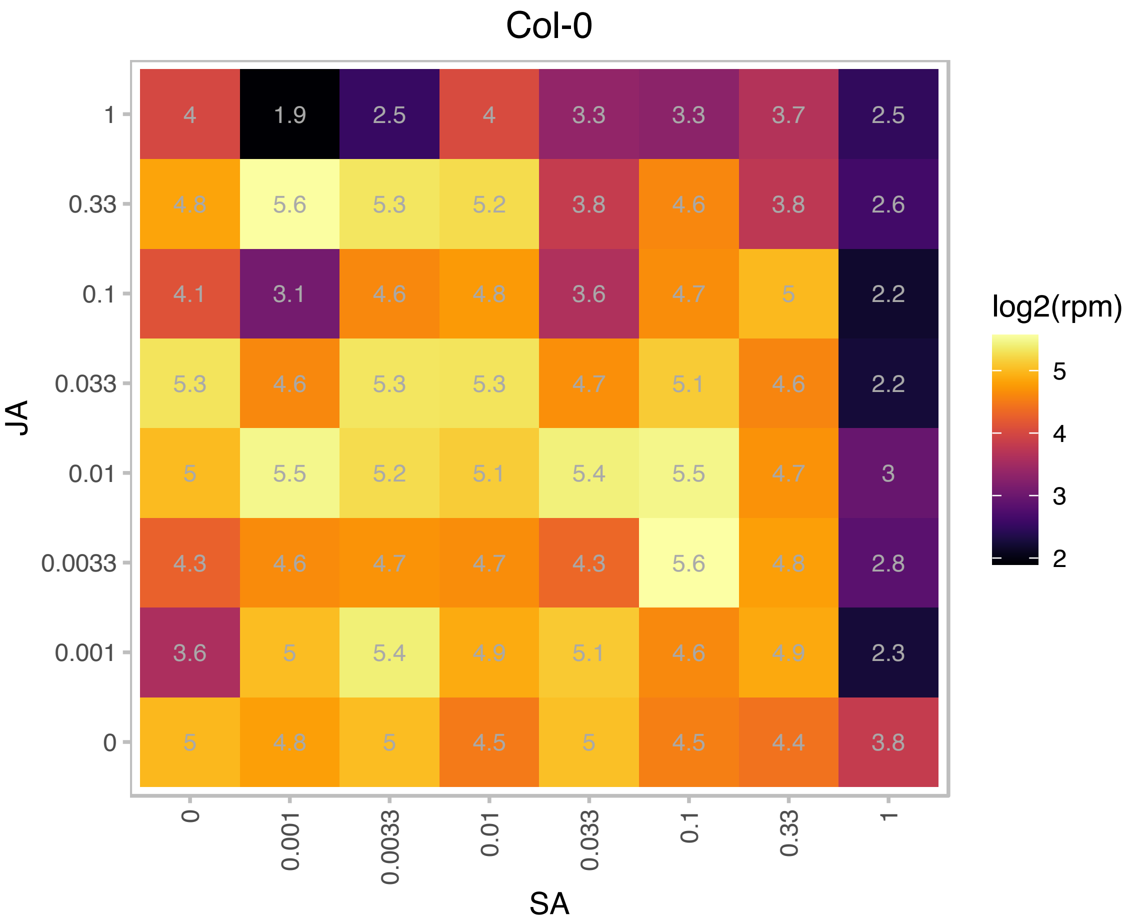


| | |
|---|---|
| Gene | AT1G49400 |
| is.expressed_genes | TRUE |
| is.responsive_genes | TRUE |
| is.apcluster | F-07 |

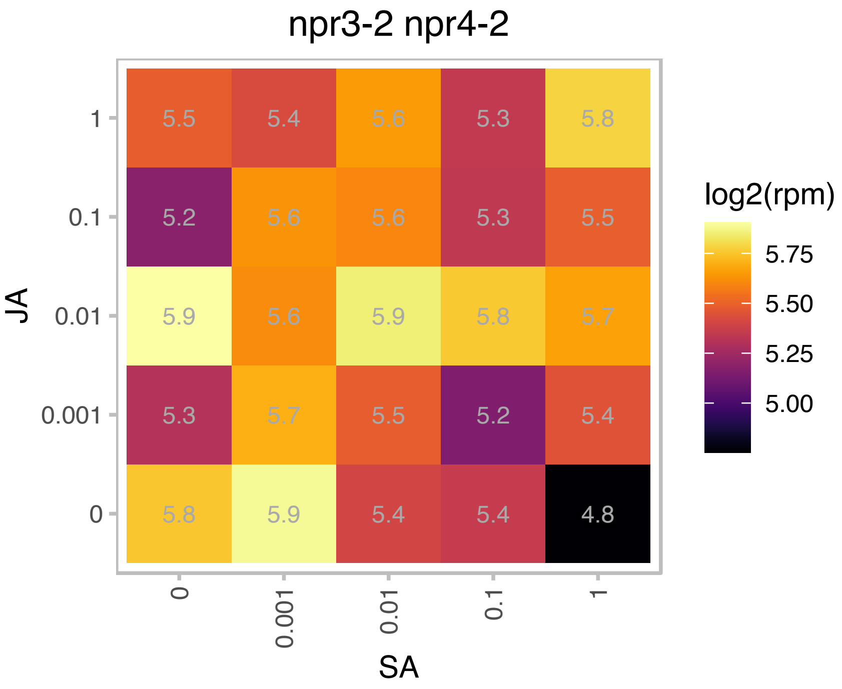


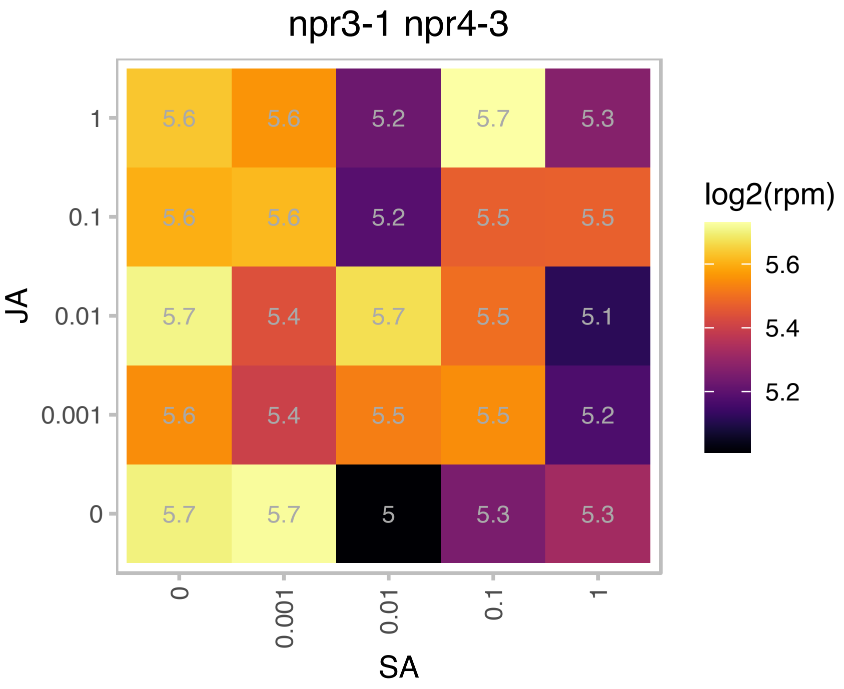

### AT1G49510.1 : DPG1, emb1273

embryo defective 1273

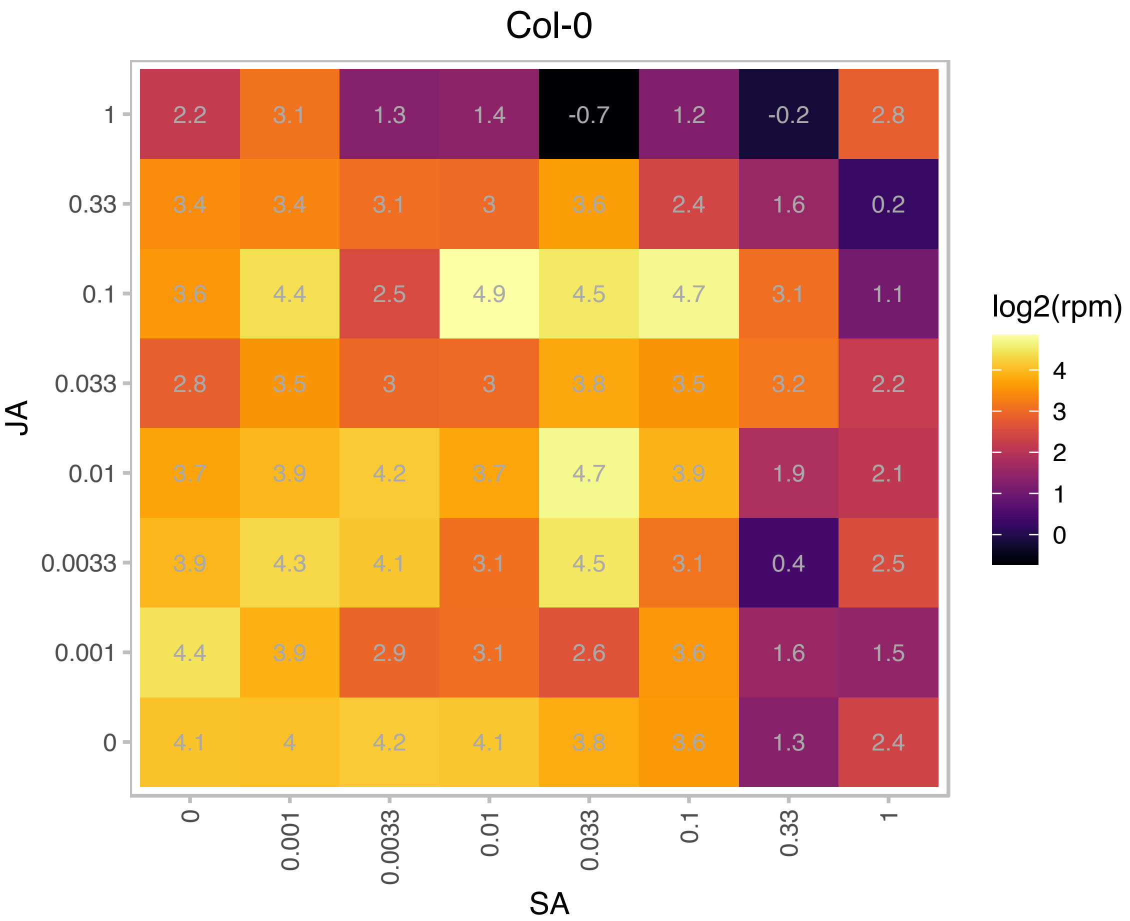


| | |
|---|---|
| Gene | AT1G49510 |
| is.expressed_genes | TRUE |
| is.responsive_genes | TRUE |
| is.apcluster | F-07 |

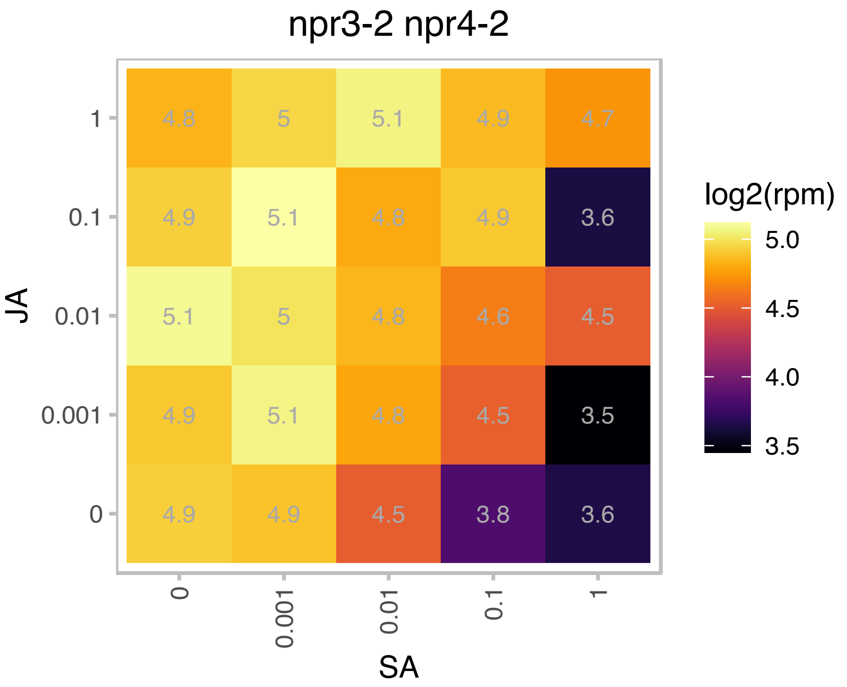


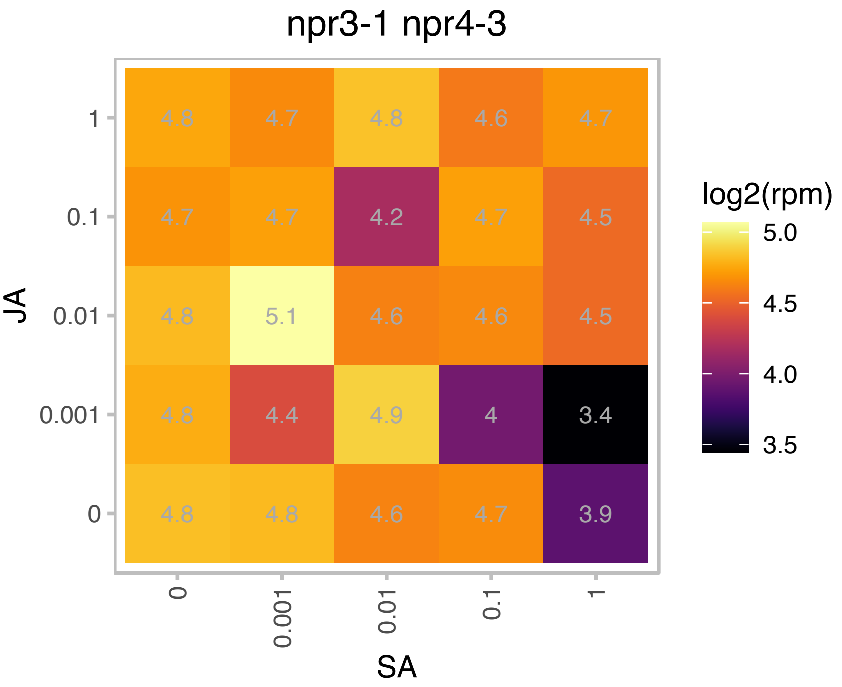

### AT1G50010.1 : TUA2

tubulin alpha-2 chain

Encodes alpha-2,4 tubulin.  TUA2 and TUA4 encode identical proteins. The mRNA is cell-to-cell mobile.

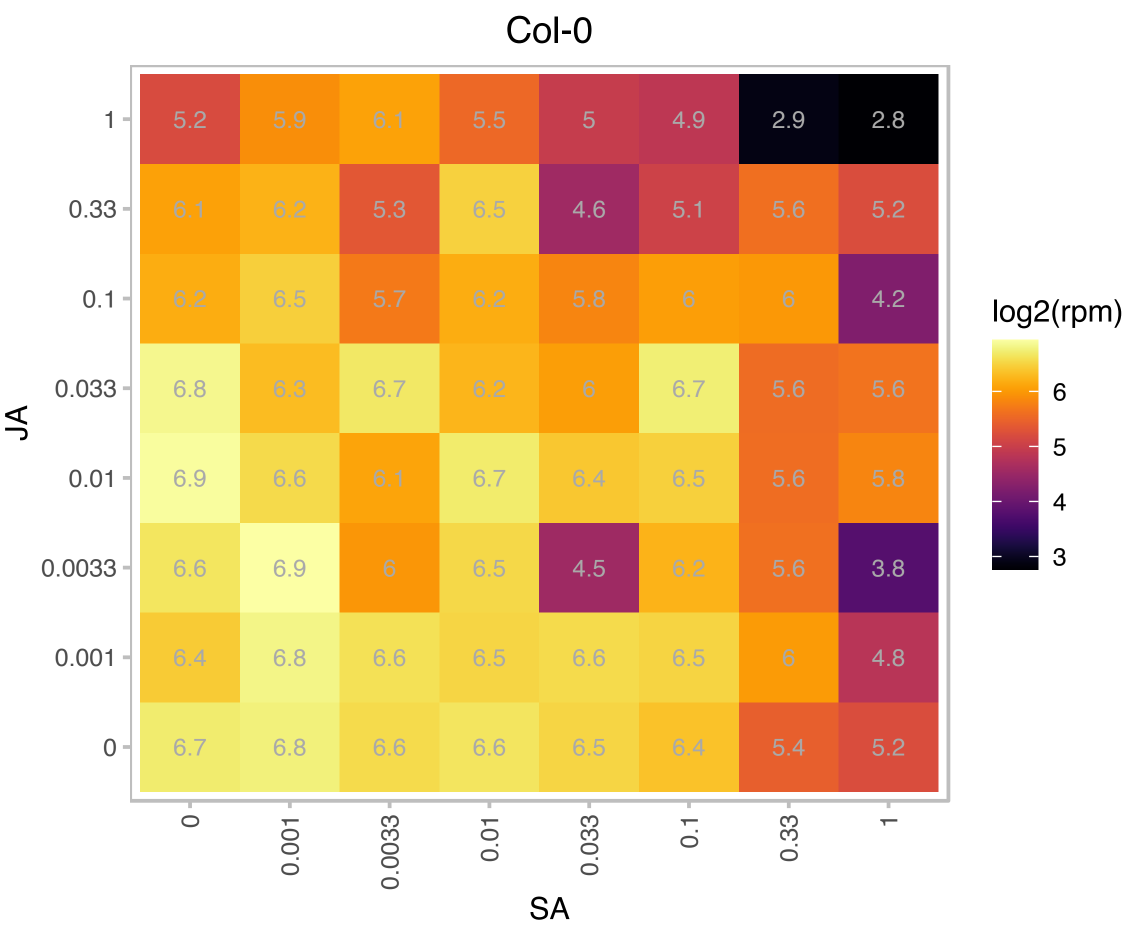


| | |
|---|---|
| Gene | AT1G50010 |
| is.expressed_genes | TRUE |
| is.responsive_genes | TRUE |
| is.apcluster | F-07 |

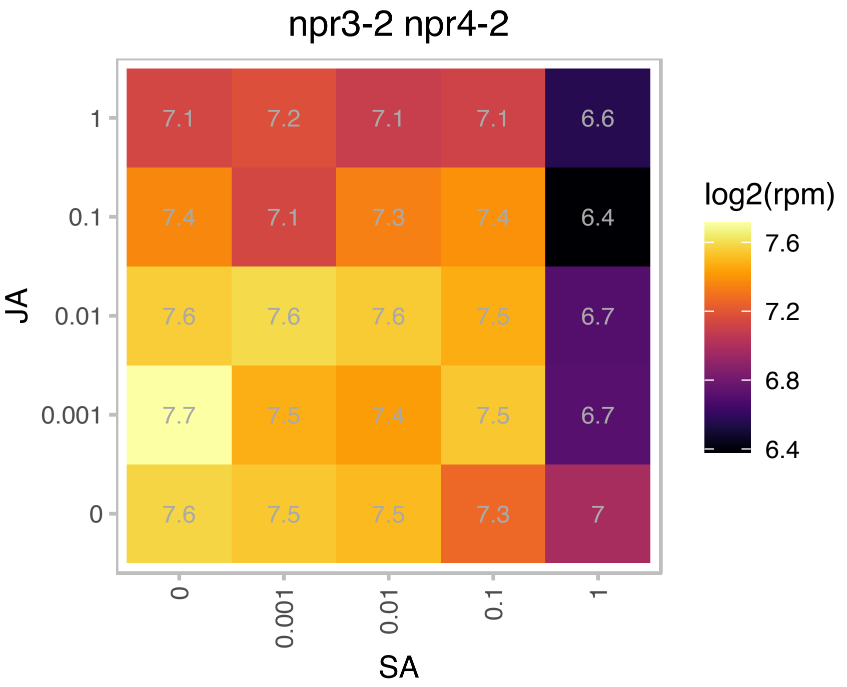


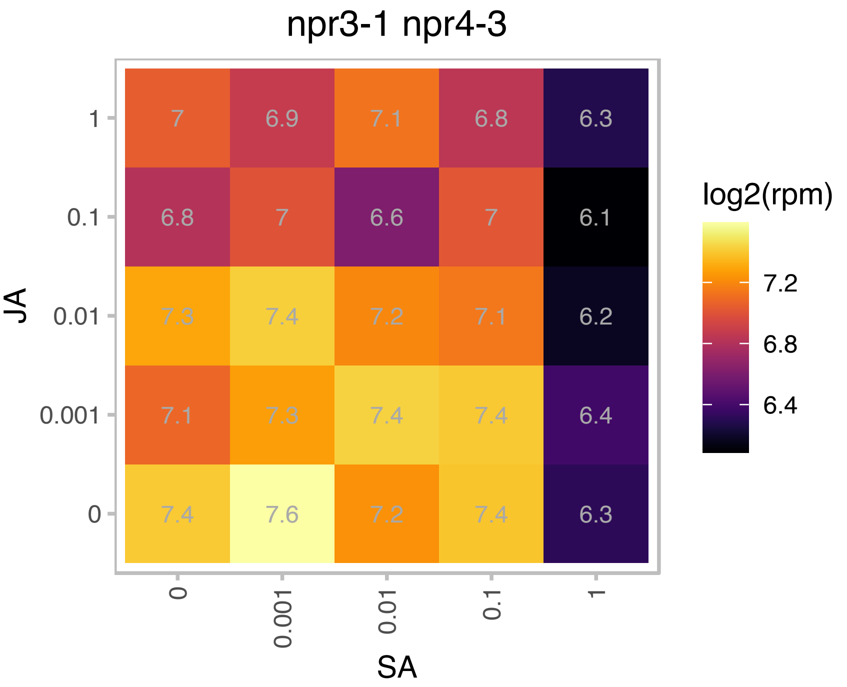

### AT1G51060.1 : HTA10

histone H2A 10

Encodes HTA10, a histone H2A protein. The mRNA is cell-to-cell mobile.

Col-0

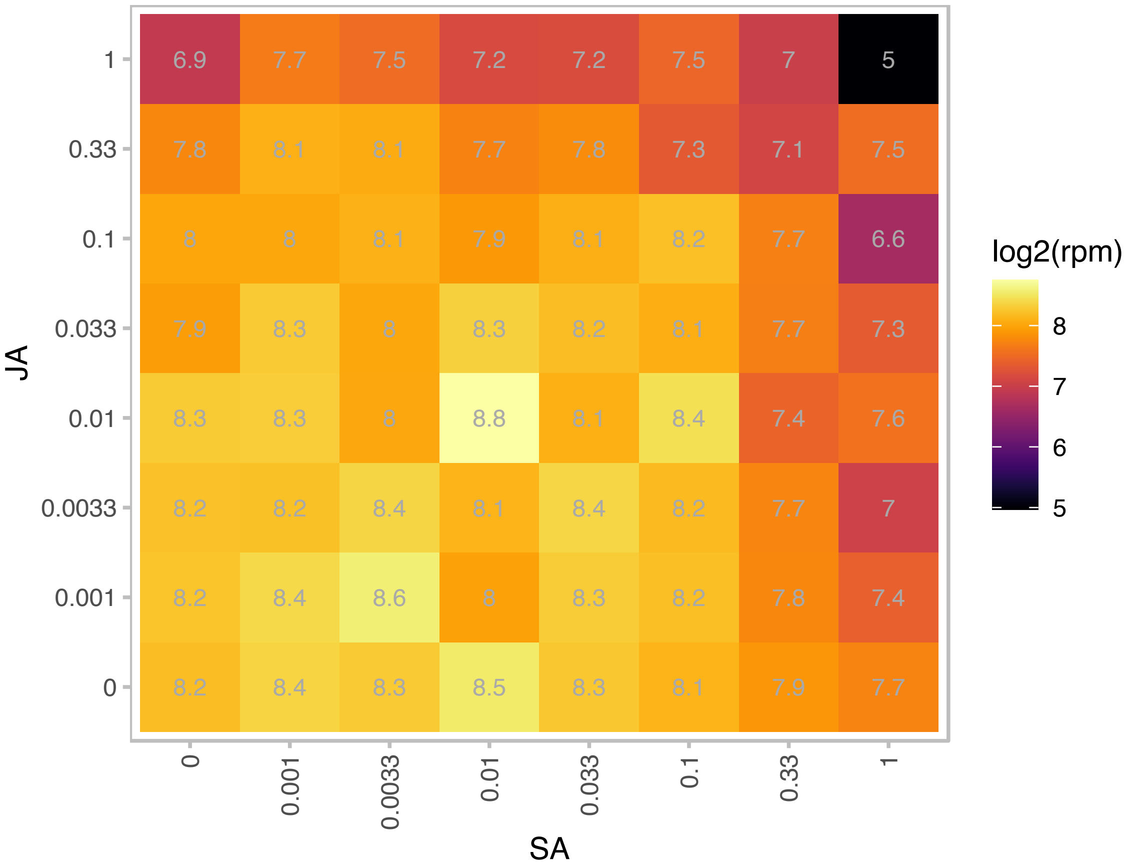


| | |
|---|---|
| Gene | AT1G51060 |
| is.expressed_genes | TRUE |
| is.responsive_genes | TRUE |
| is.apcluster | F-07 |

npr3-2 npr4-2

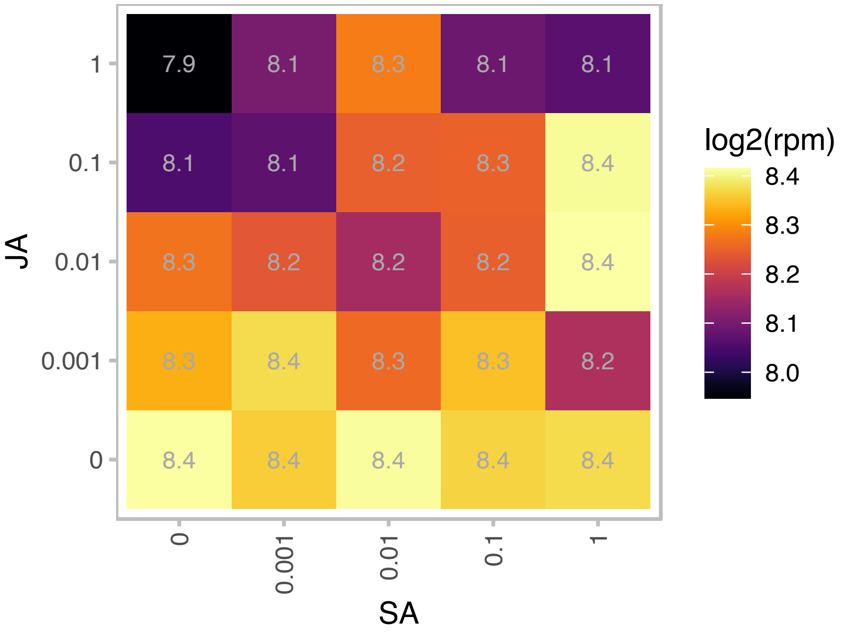


npr3-1 npr4-3

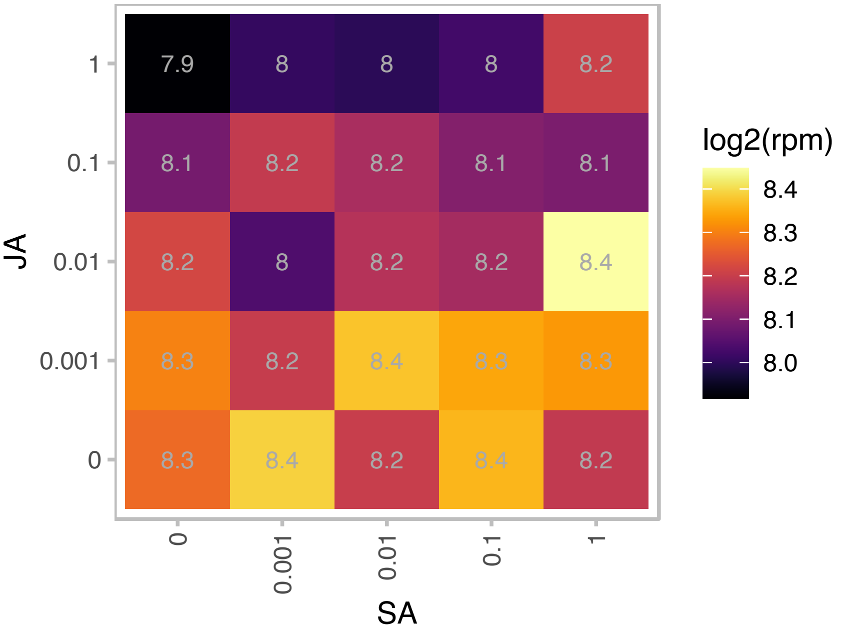

# AT1G51680.1 : 4CL.1, 4CL1, AT4CL1

4-coumarate:CoA ligase 1

encodes an isoform of 4-coumarate:CoA ligase (4CL), which is involved in the last step of the general phenylpropanoid pathway. In addition to 4-coumarate, it also converts ferulate. The catalytic efficiency was in the following (descending) order: p-coumaric acid, ferulic acid, caffeic acid and 5-OH-ferulic acid. At4CL1 was unable to use sinapic acid as substrate.

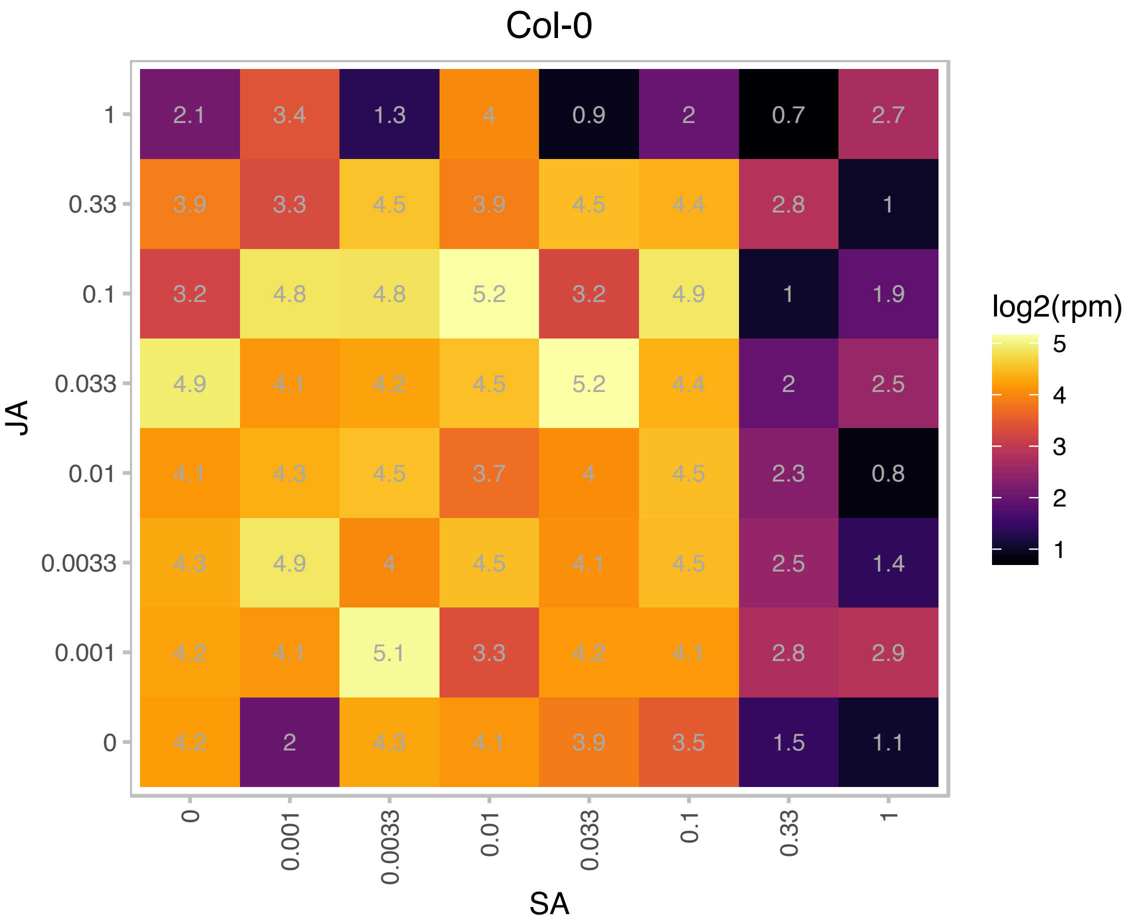


| | |
|---|---|
| Gene | AT1G51680 |
| is.expressed_genes | TRUE |
| is.responsive_genes | TRUE |
| is.apcluster | F-07 |

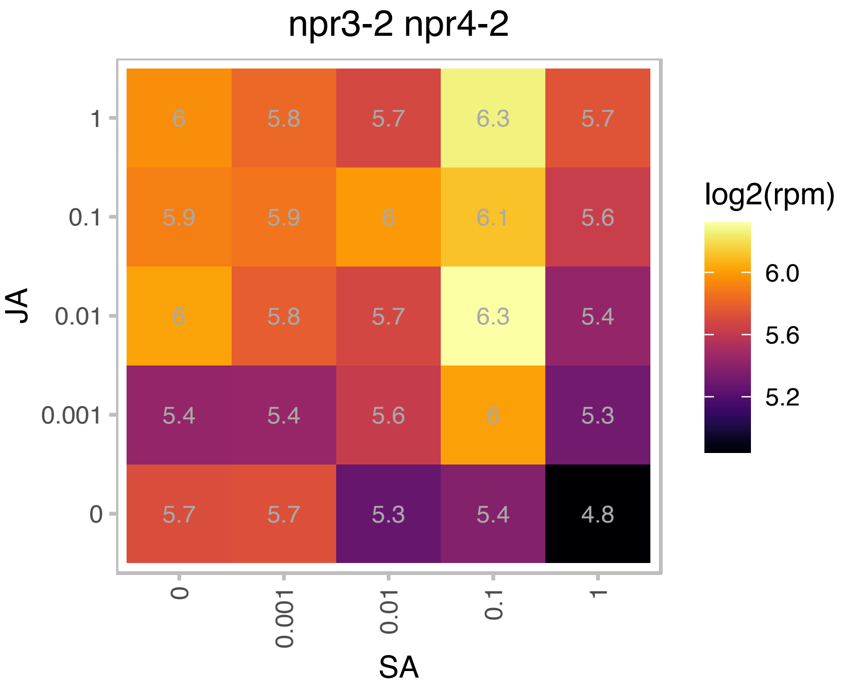


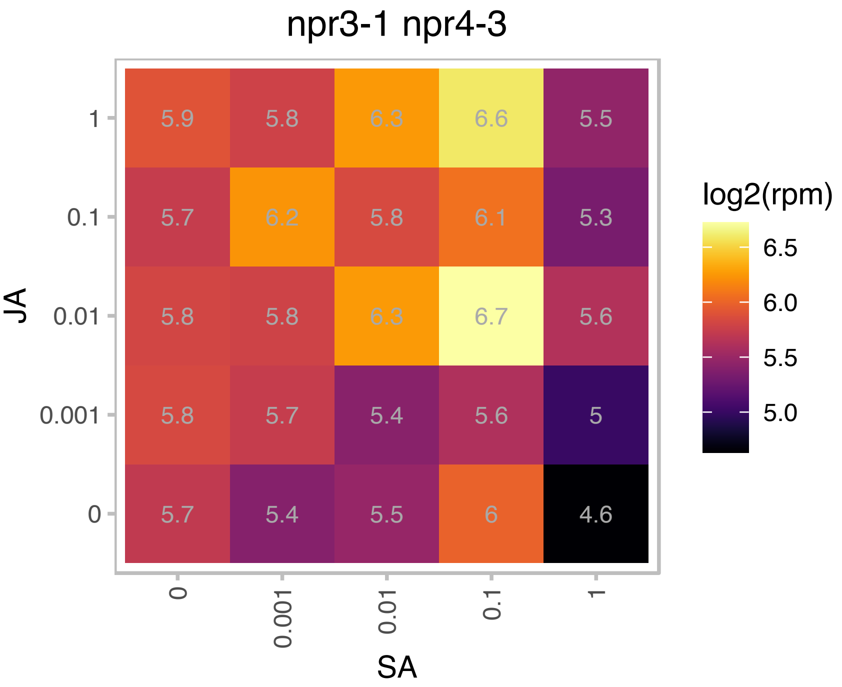

### AT1G51700.1 : ADOF1, DOF1

DOF zinc finger protein 1

Encodes dof zinc finger protein (adof1). The mRNA is cell-to-cell mobile.

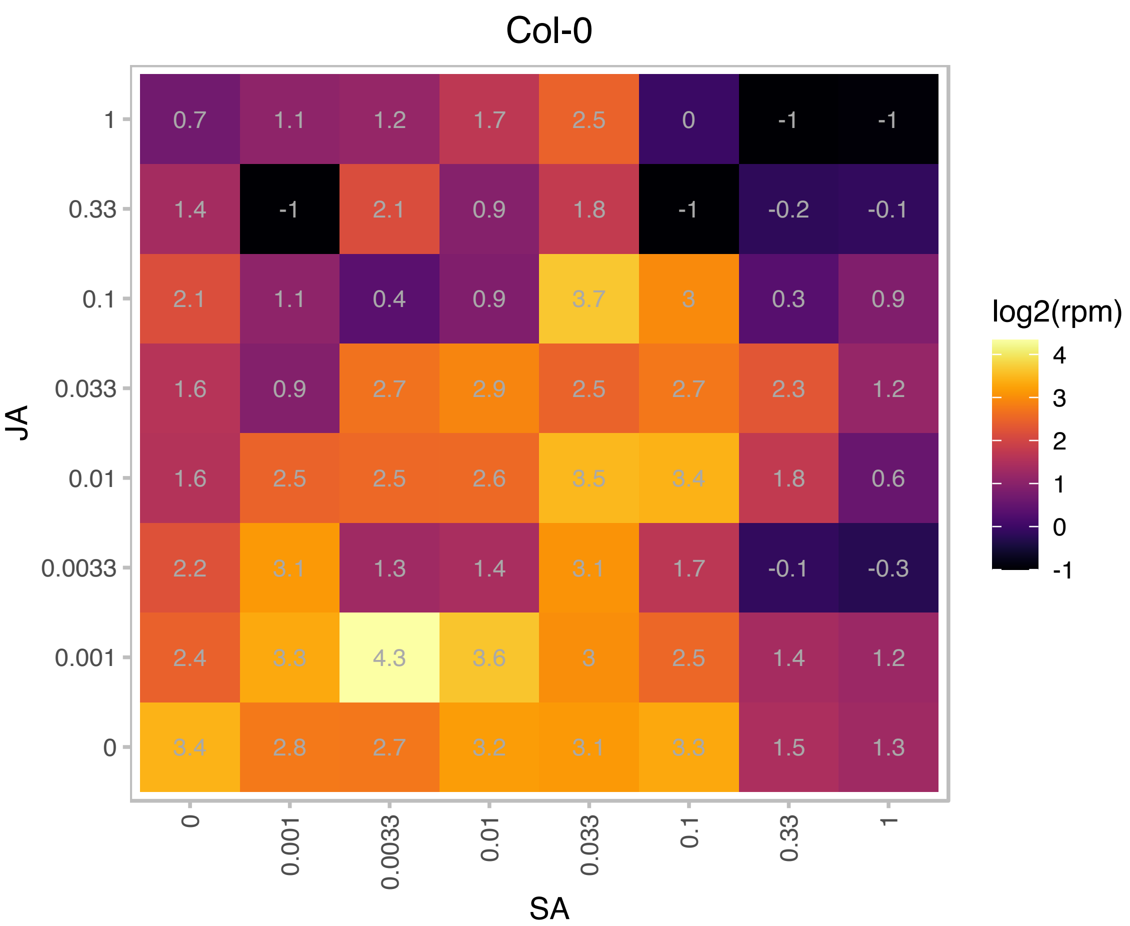


| | |
|---|---|
| Gene | AT1G51700 |
| is.expressed_genes | TRUE |
| is.responsive_genes | TRUE |
| is.apcluster | F-07 |

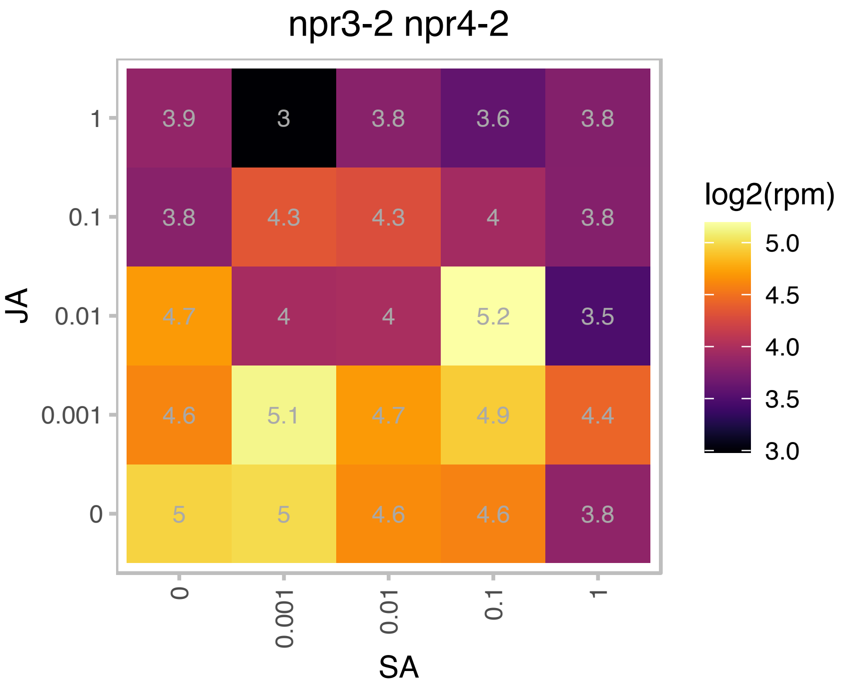


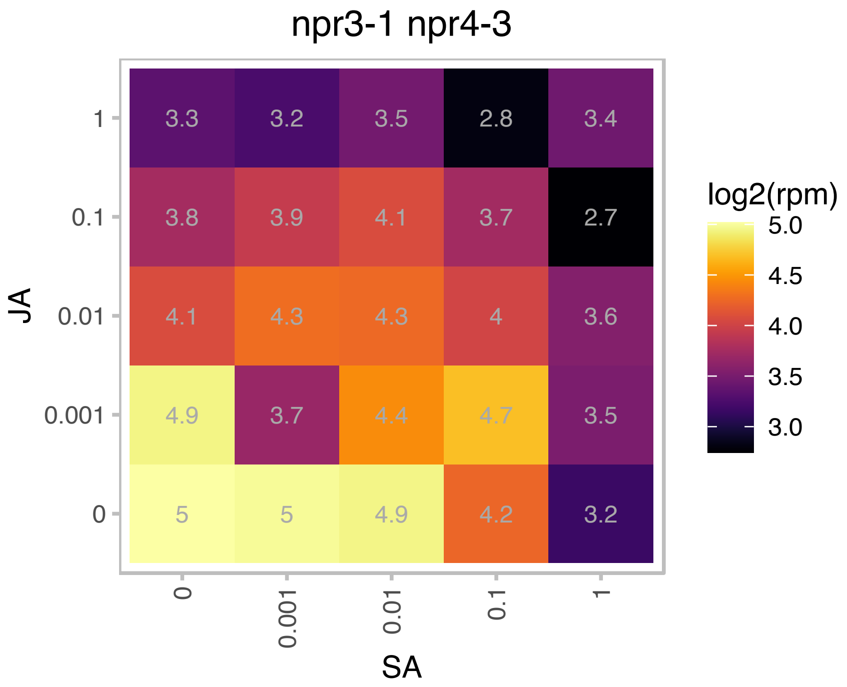

# AT1G52300.1 : NA

Zinc-binding ribosomal protein family protein

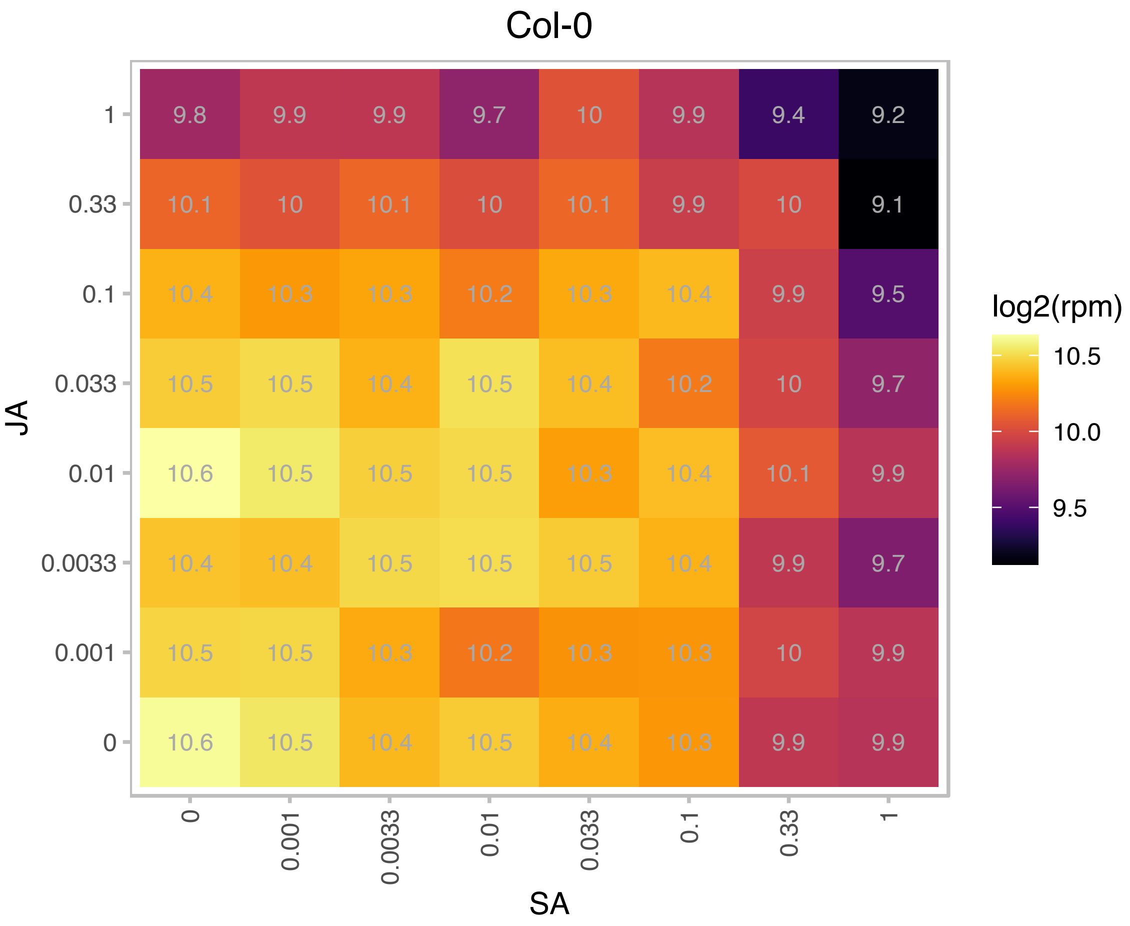


| | |
|---|---|
| Gene | AT1G52300 |
| is.expressed_genes | TRUE |
| is.responsive_genes | TRUE |
| is.apcluster | F-07 |

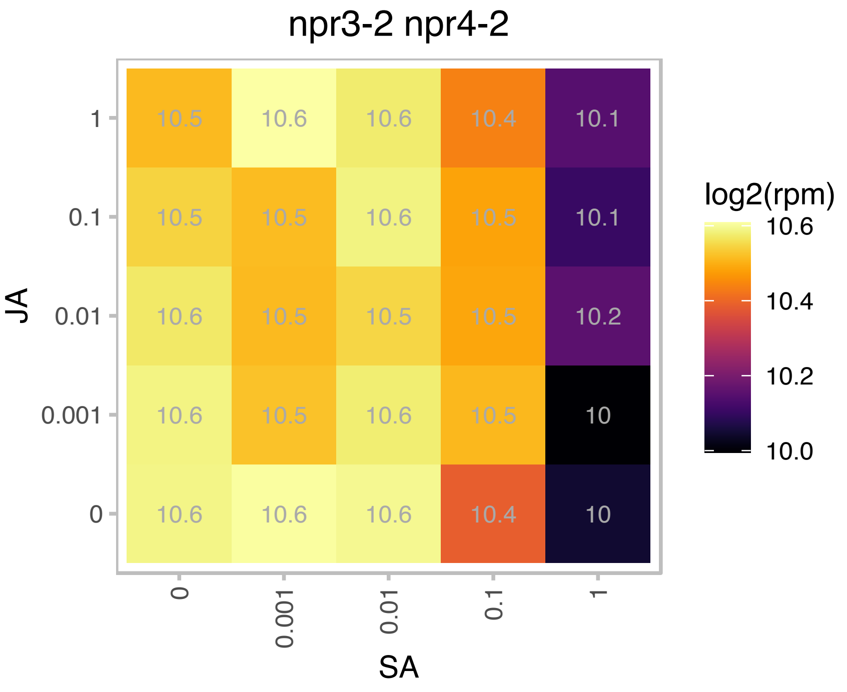


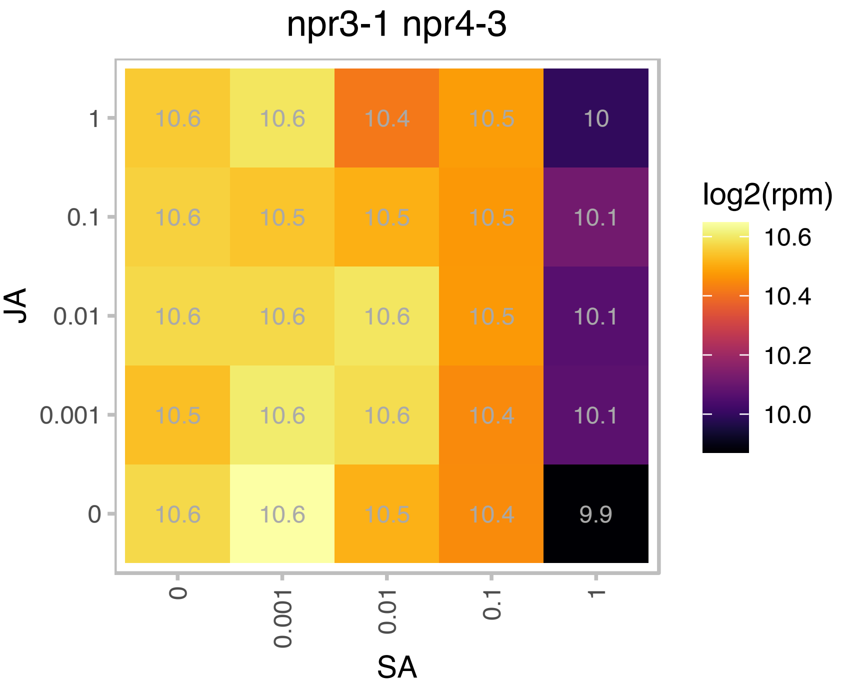

### AT1G54690.1 ; G-H2AX, GAMMA-H2AX, H2AXB, HTA3

gamma histone variant H2AX

Encodes HTA3, a histone H2A protein. H2AX is a meiosis-specific isoform of histone H2A. Upon DSB formation, rapid accumulation of phosphorylated H2AX (γ-H2AX) occurs around the break site. □ H2AX foci accumulate early G2. Immunolocalization studies in spread preparations of wild-type meiocytes at G2/early leptotene revealed the accumulation of numerous rather diffuse γ-H2AX foci throughout the chromatin. However, their accumulation is not contemporaneous with that of AtSPO11-1. At 3 h post-S, no γ-H2AX foci are detected. During the 3- to 5-h window when AtSPO11-1 foci rapidly disappear, there is an equally swift accumulation of γ-H2AX to a maximum of >50 diffuse foci. The level of γH2AX then remains constant [illegible]

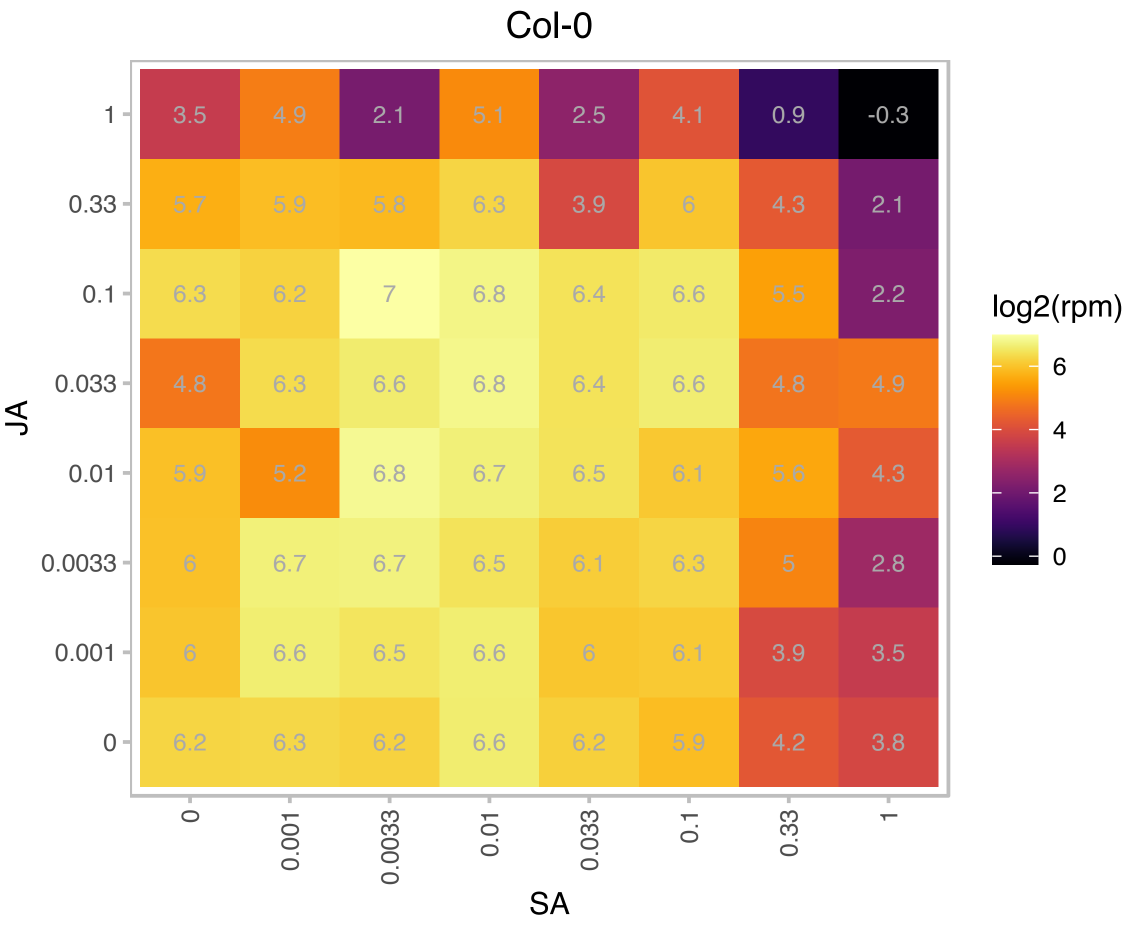


| | |
|---|---|
| Gene | AT1G54690 |
| is.expressed_genes | TRUE |
| is.responsive_genes | TRUE |
| is.apcluster | F-07 |

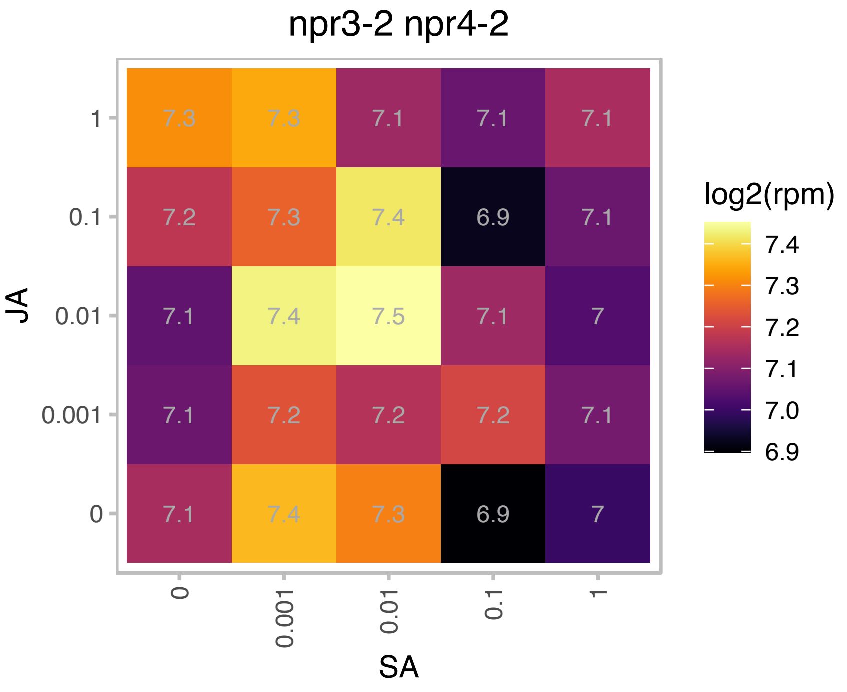


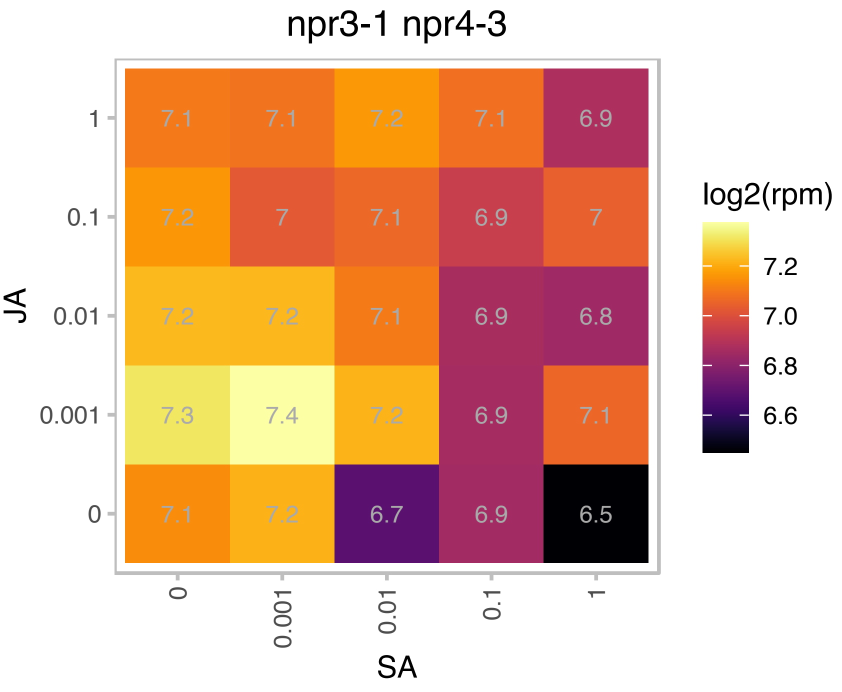

# AT1G57660.1 : NA

Translation protein SH3-like family protein

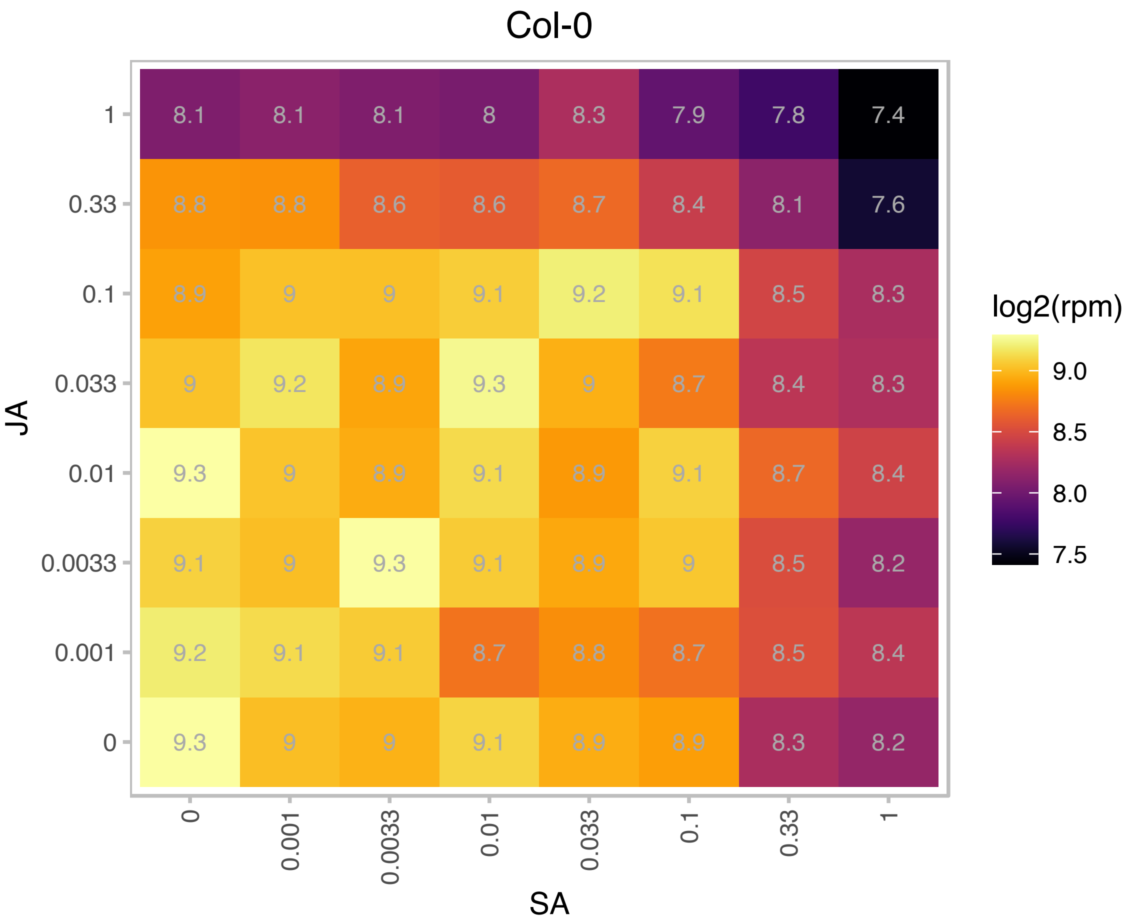


| | |
|---|---|
| Gene | AT1G57660 |
| is.expressed_genes | TRUE |
| is.responsive_genes | TRUE |
| is.apcluster | F-07 |

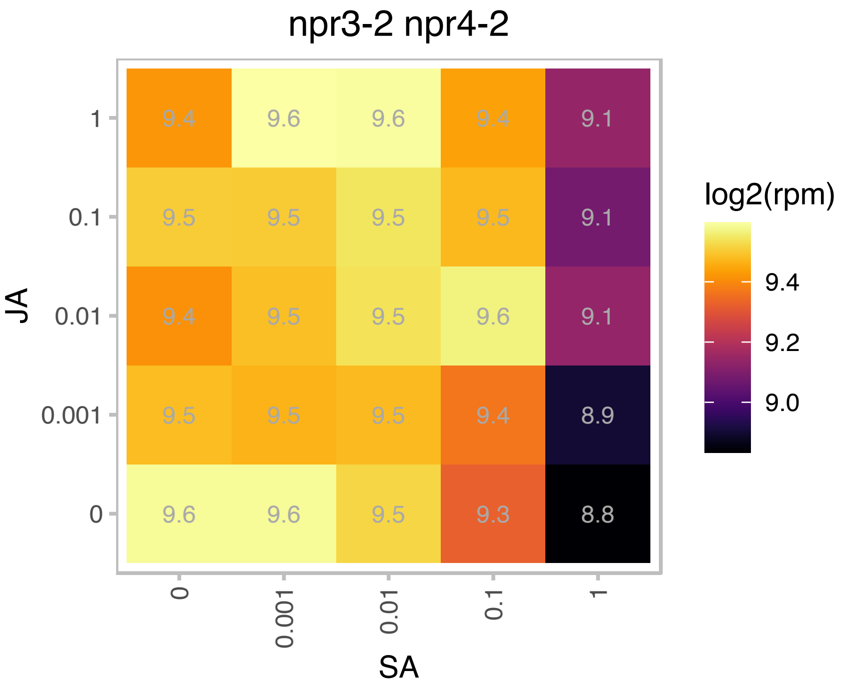


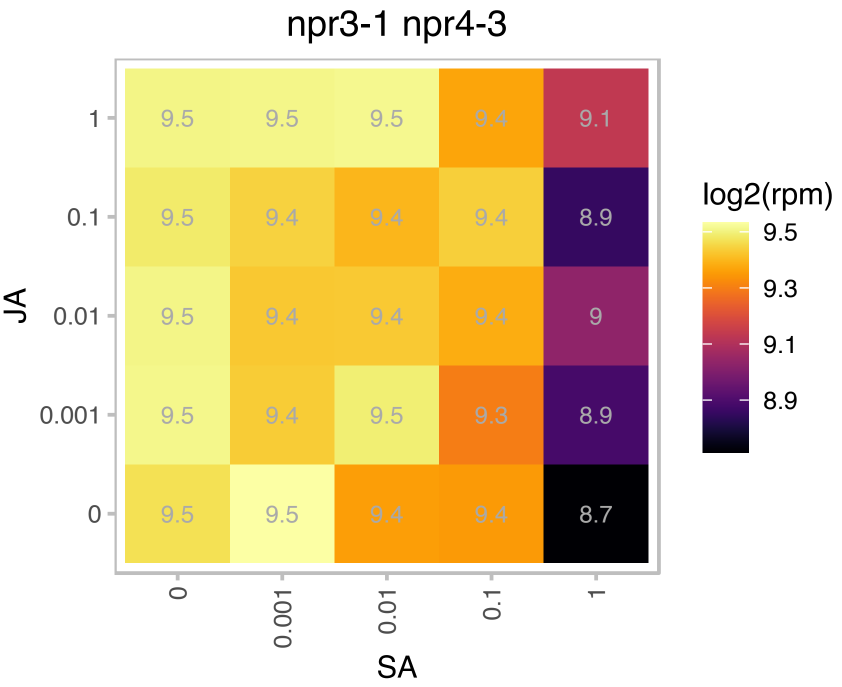

# AT1G57860.1 : NA

Translation protein SH3-like family protein

| | |
|---|---|
| Gene | AT1G57860 |
| is.expressed_genes | TRUE |
| is.responsive_genes | TRUE |
| is.apcluster | F-07 |

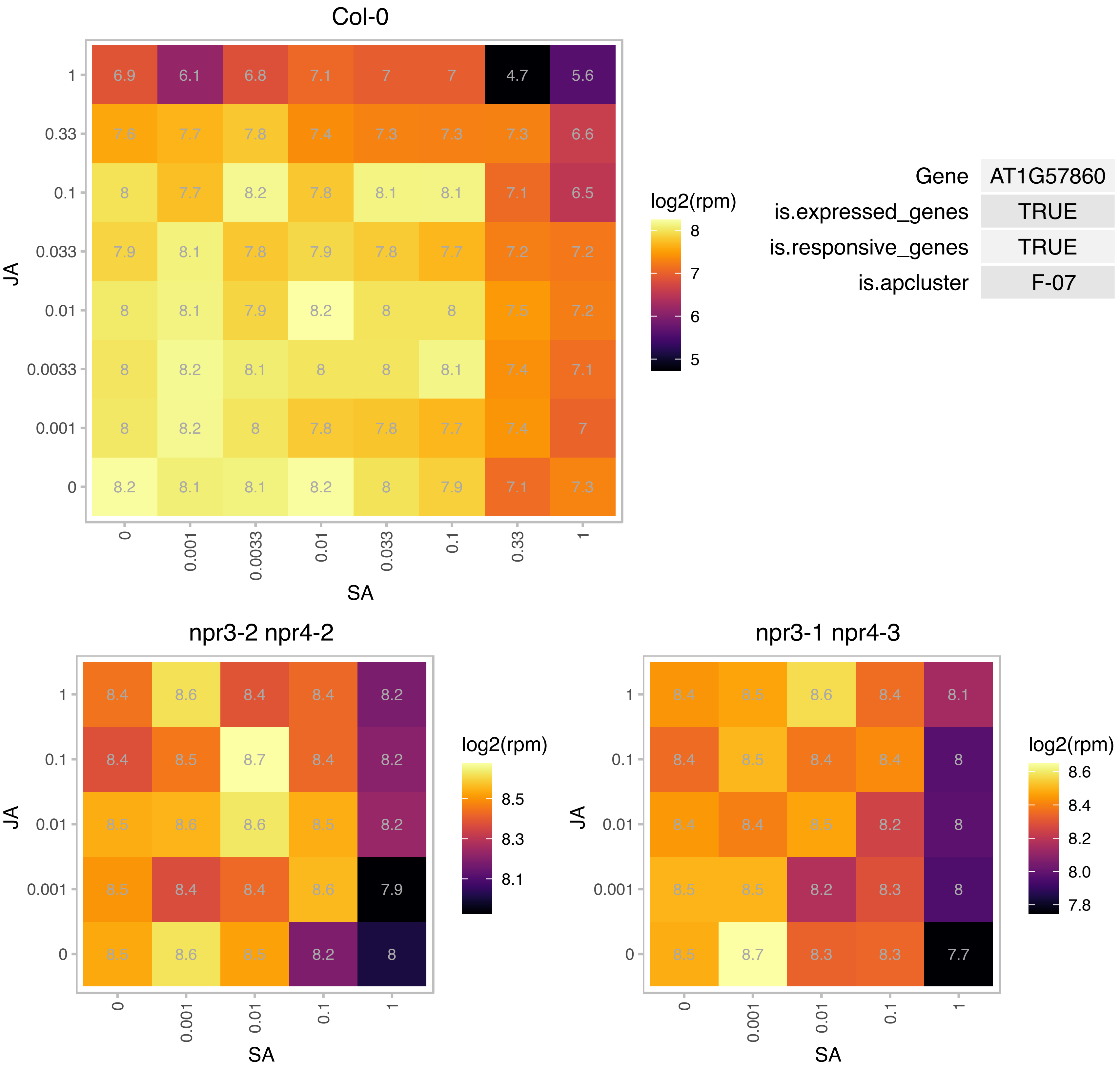

# AT1G58380.1 : XW6

Ribosomal protein S5 family protein

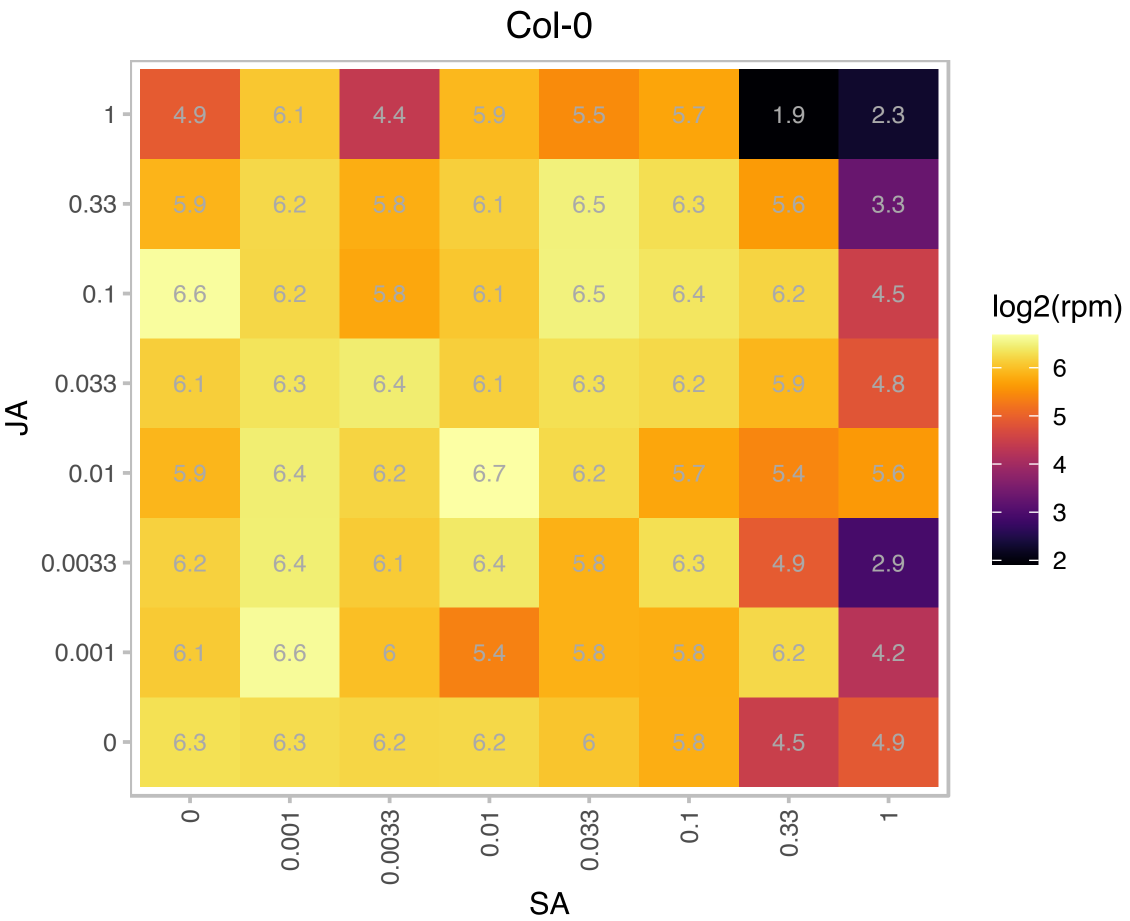


| | |
|---|---|
| Gene | AT1G58380 |
| is.expressed_genes | TRUE |
| is.responsive_genes | TRUE |
| is.apcluster | F-07 |

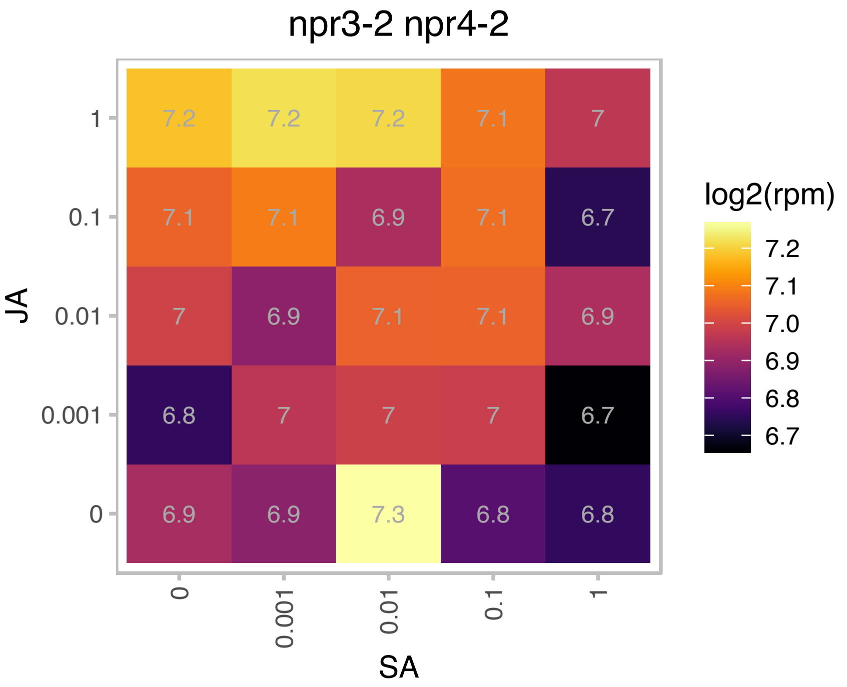


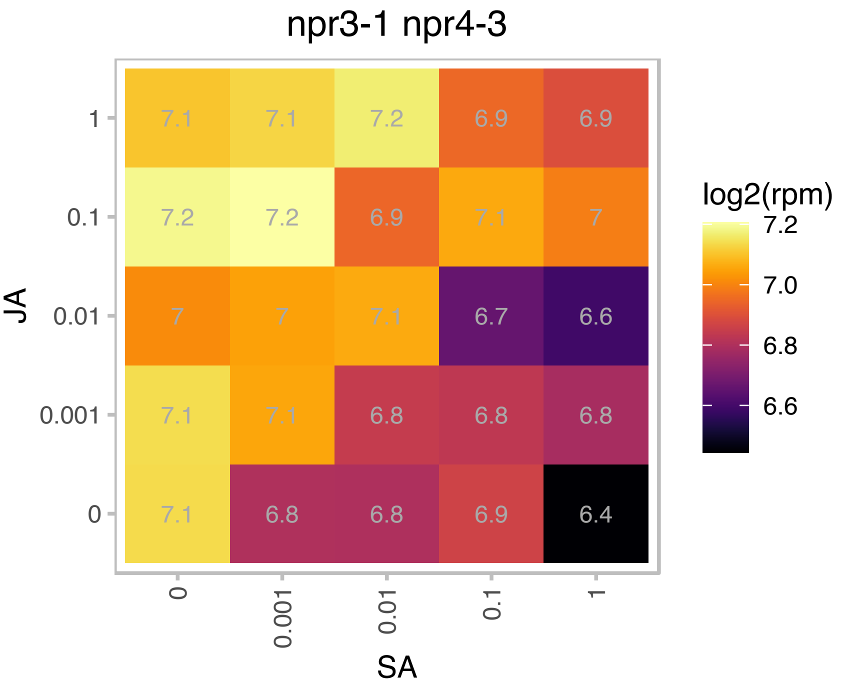

# AT1G58983.1 : NA

Ribosomal protein S5 family protein

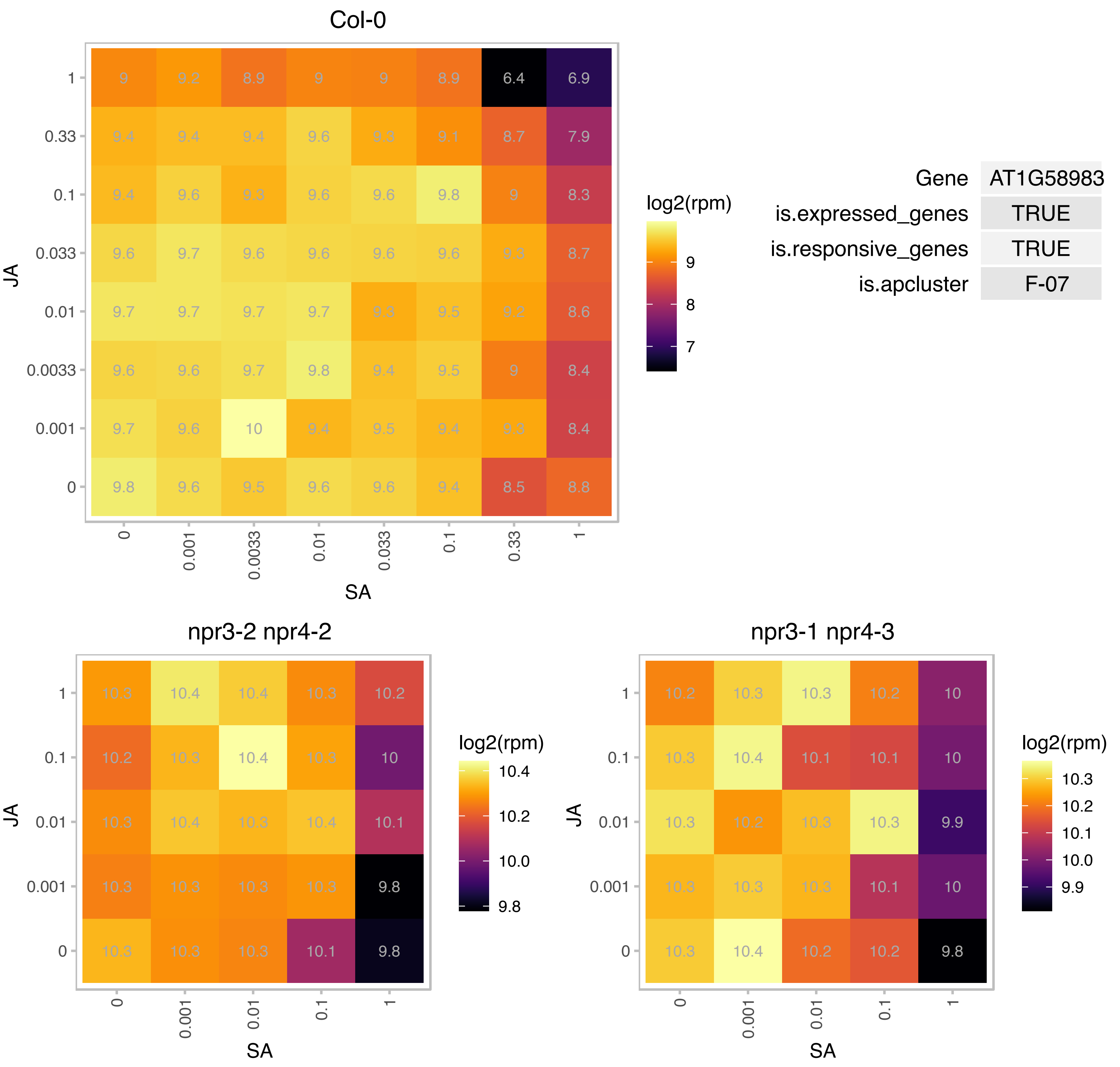


| | |
|---|---|
| Gene | AT1G58983 |
| is.expressed_genes | TRUE |
| is.responsive_genes | TRUE |
| is.apcluster | F-07 |

# AT1G60000.1 : NA

RNA-binding (RRM/RBD/RNP motifs) family protein

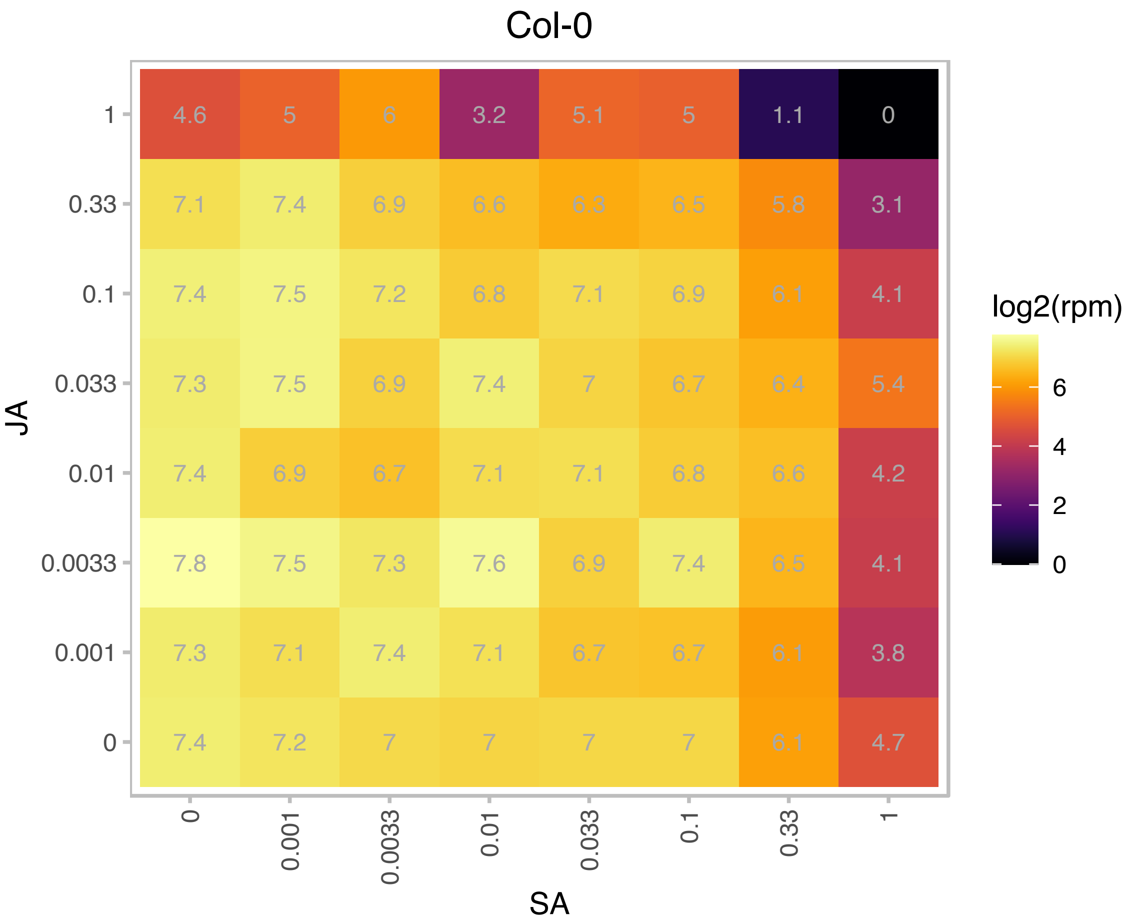


| | |
|---|---|
| Gene | AT1G60000 |
| is.expressed_genes | TRUE |
| is.responsive_genes | TRUE |
| is.apcluster | F-07 |

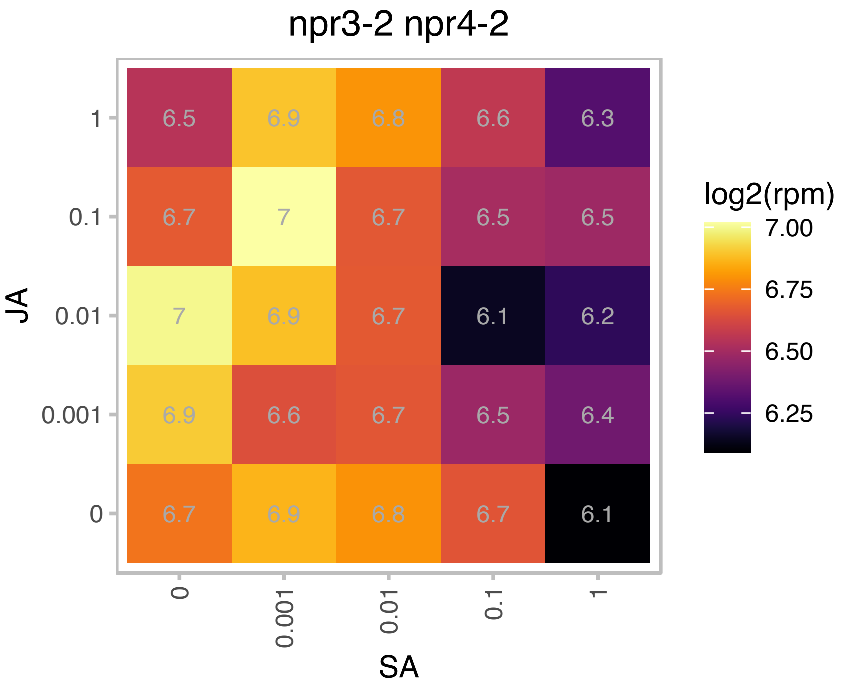


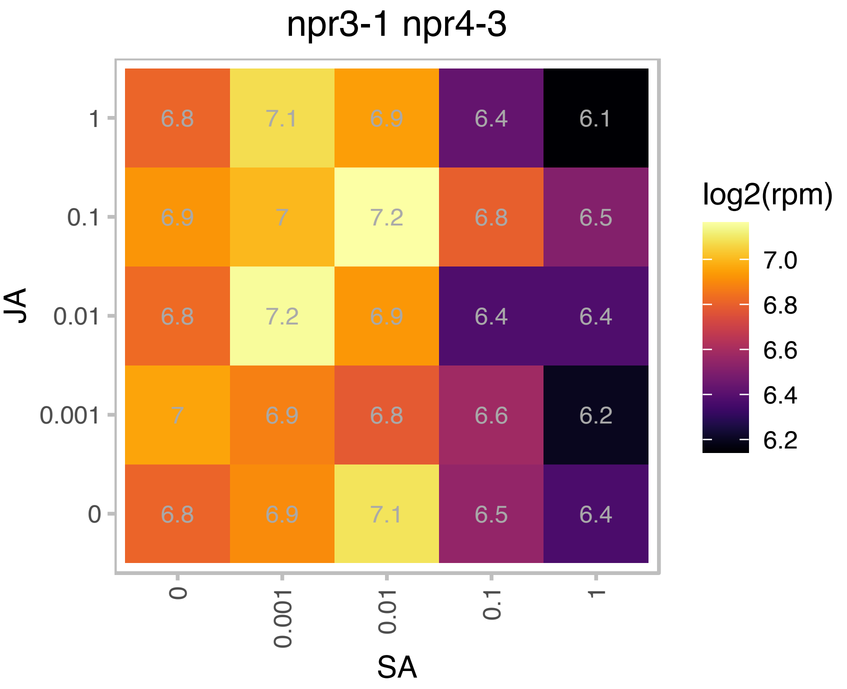

### AT1G60550.1 : DHNS, ECHID

enoyl-CoA hydratase/isomerase D

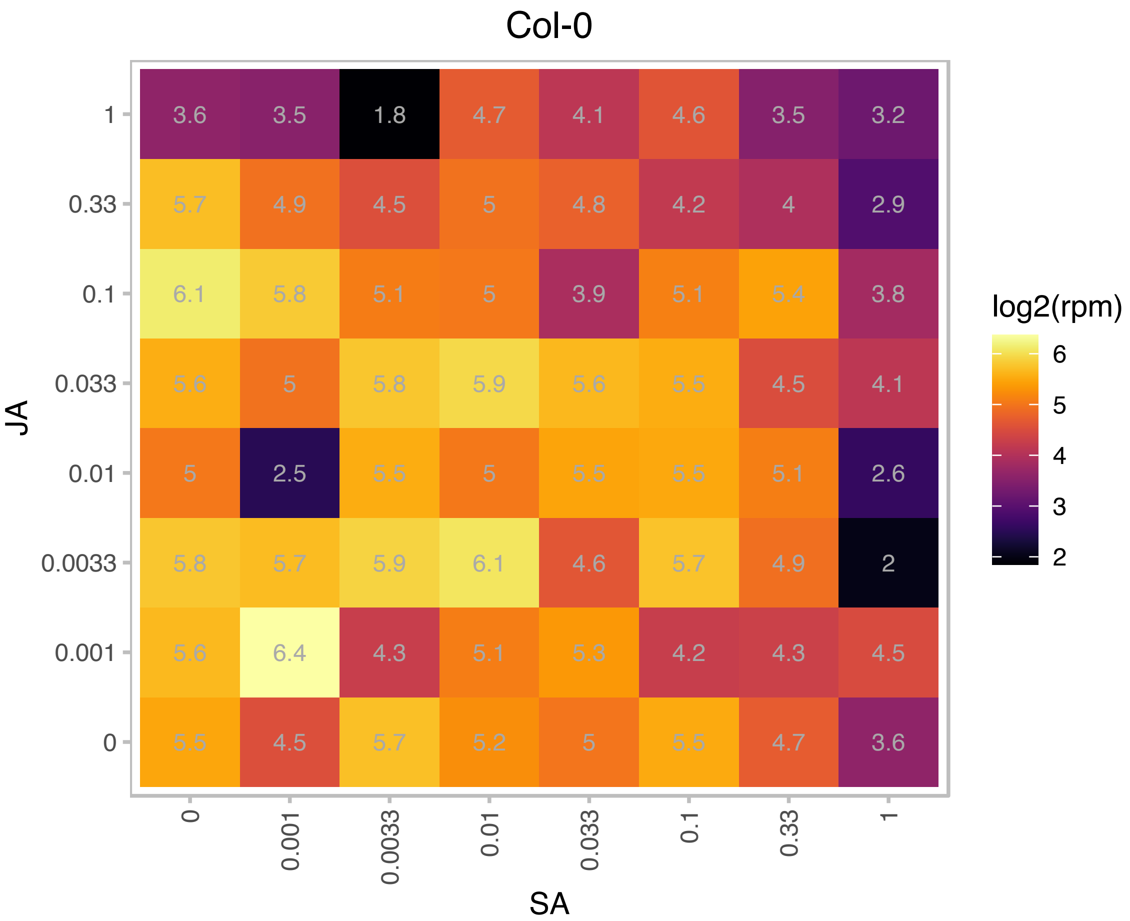


| | |
|---|---|
| Gene | AT1G60550 |
| is.expressed_genes | TRUE |
| is.responsive_genes | TRUE |
| is.apcluster | F-07 |

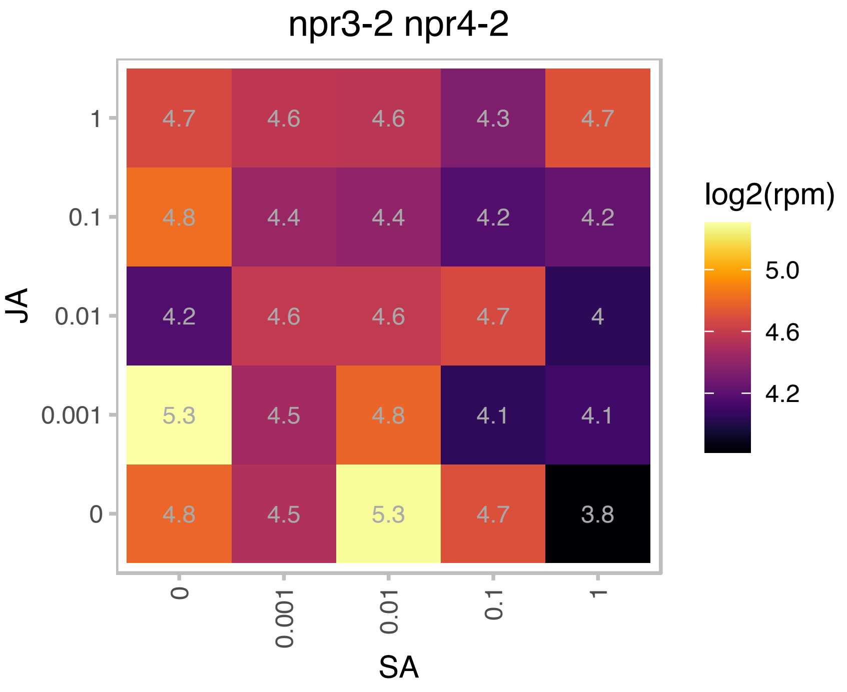


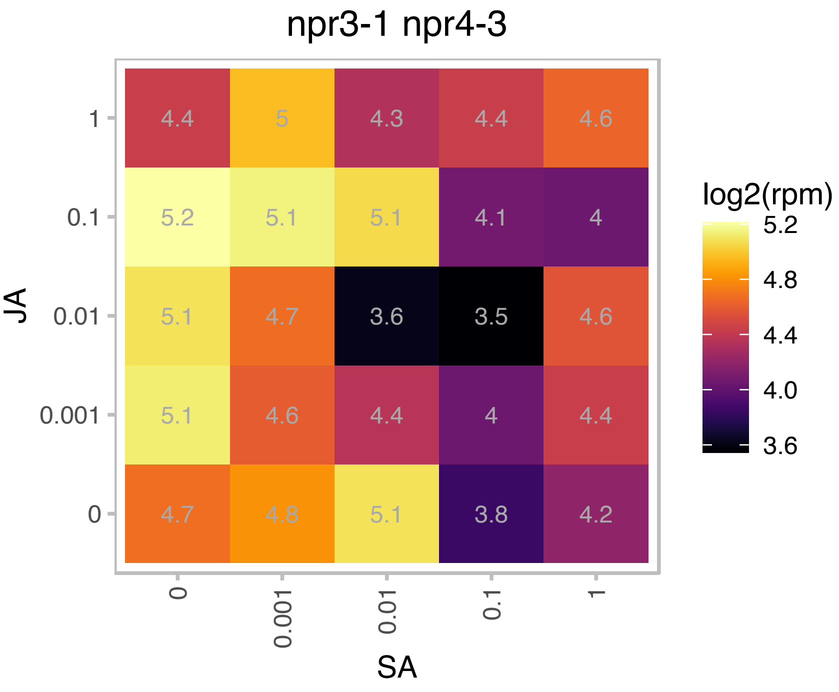

# AT1G67430.1 : NA

Ribosomal protein L22p/L17e family protein

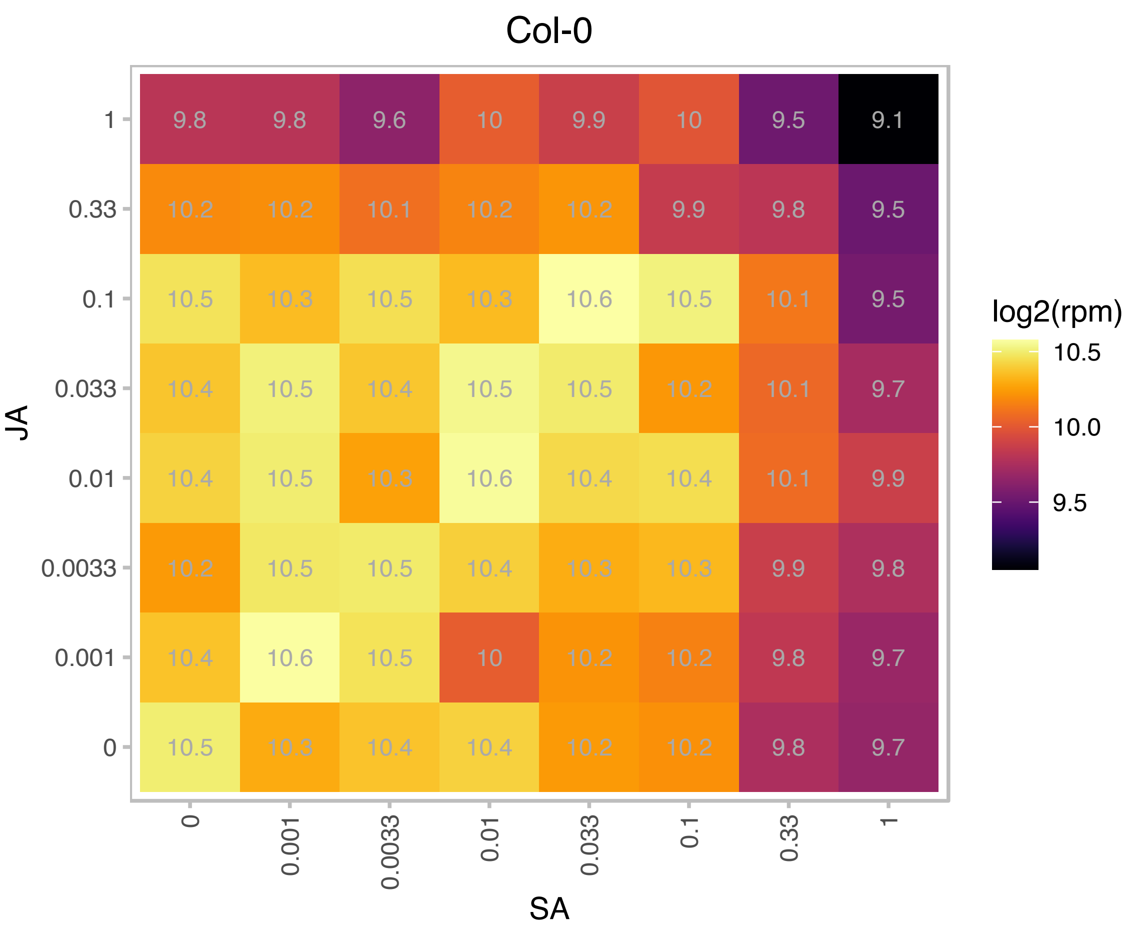


| | |
|---|---|
| Gene | AT1G67430 |
| is.expressed_genes | TRUE |
| is.responsive_genes | TRUE |
| is.apcluster | F-07 |

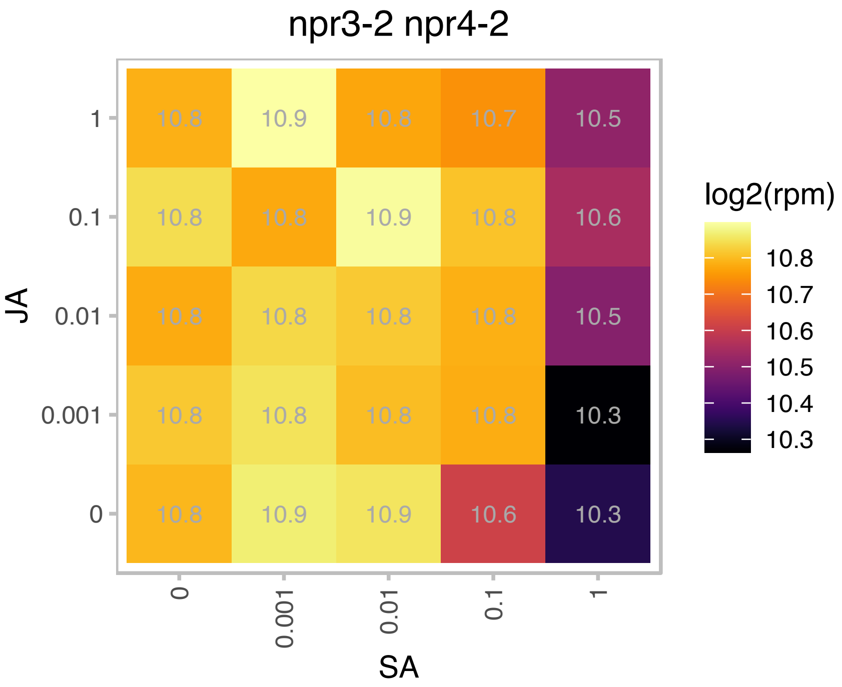


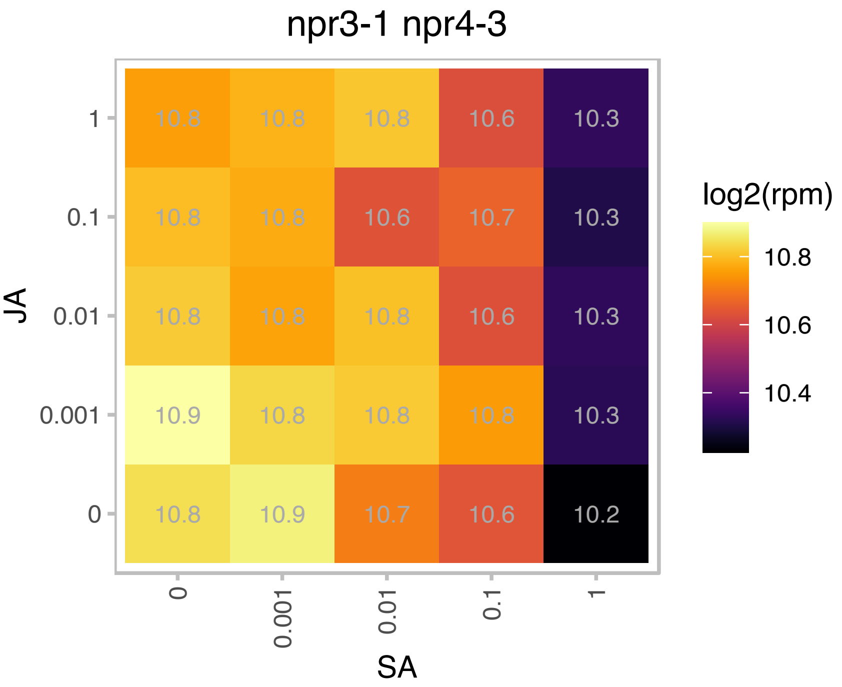

### AT1G68520.1 : BBX14

B-box type zinc finger protein with CCT domain

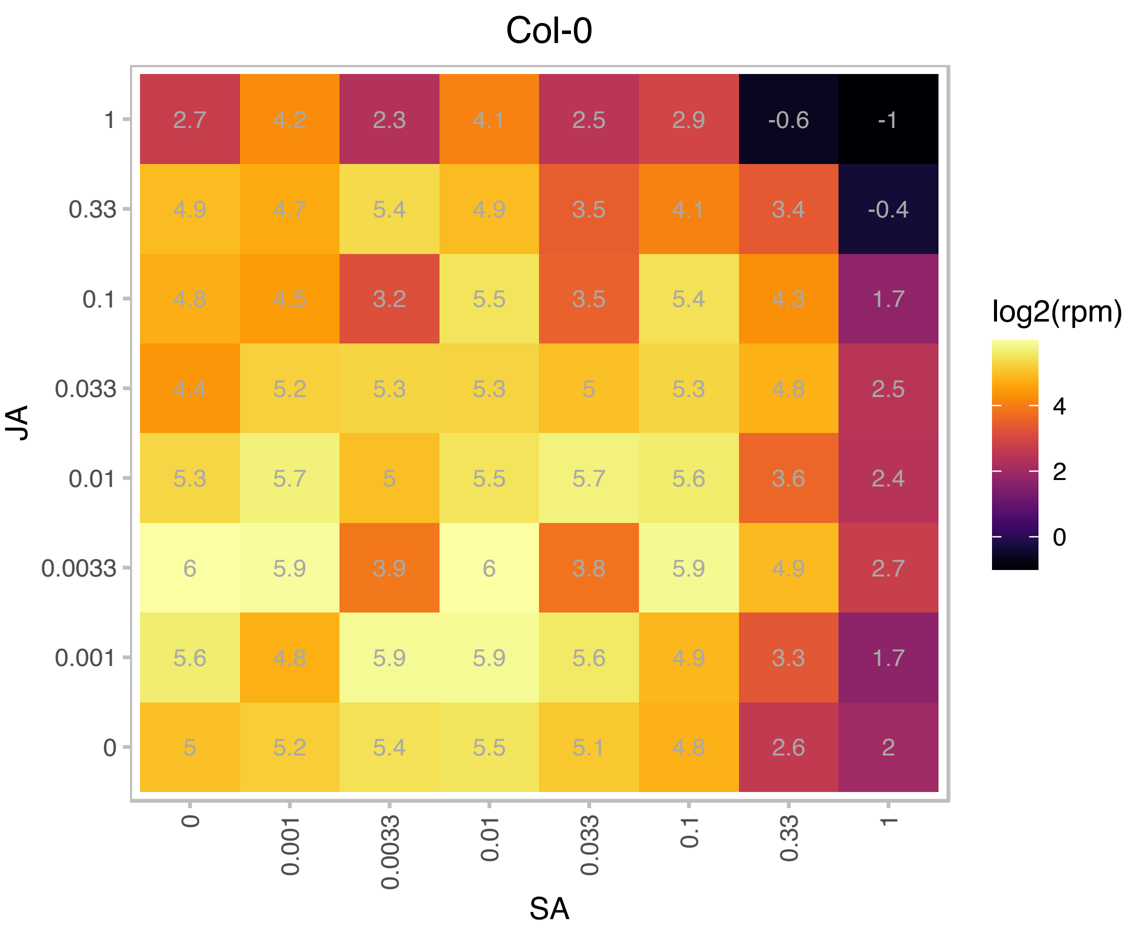


| | |
|---|---|
| Gene | AT1G68520 |
| is.expressed_genes | TRUE |
| is.responsive_genes | TRUE |
| is.apcluster | F-07 |

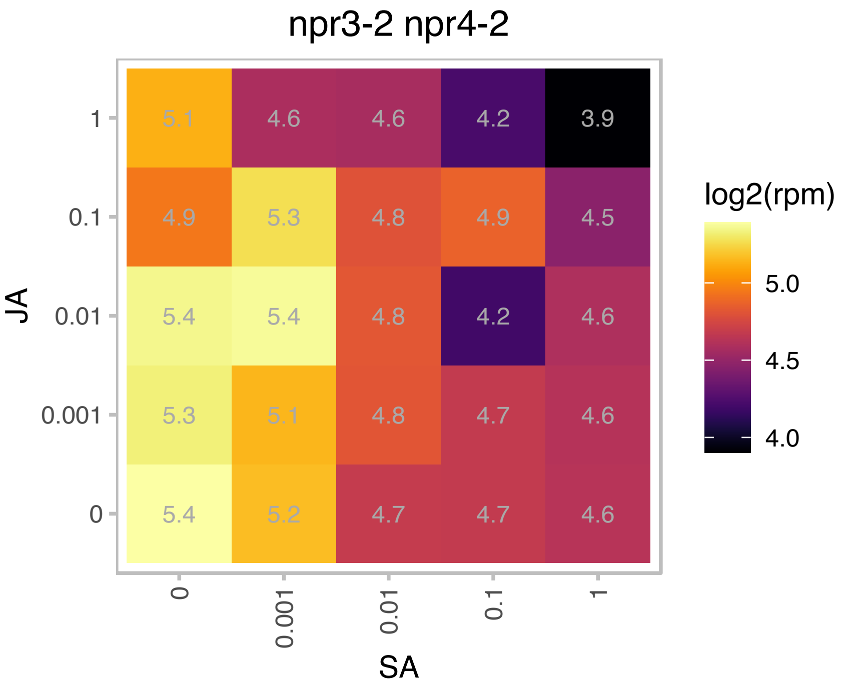


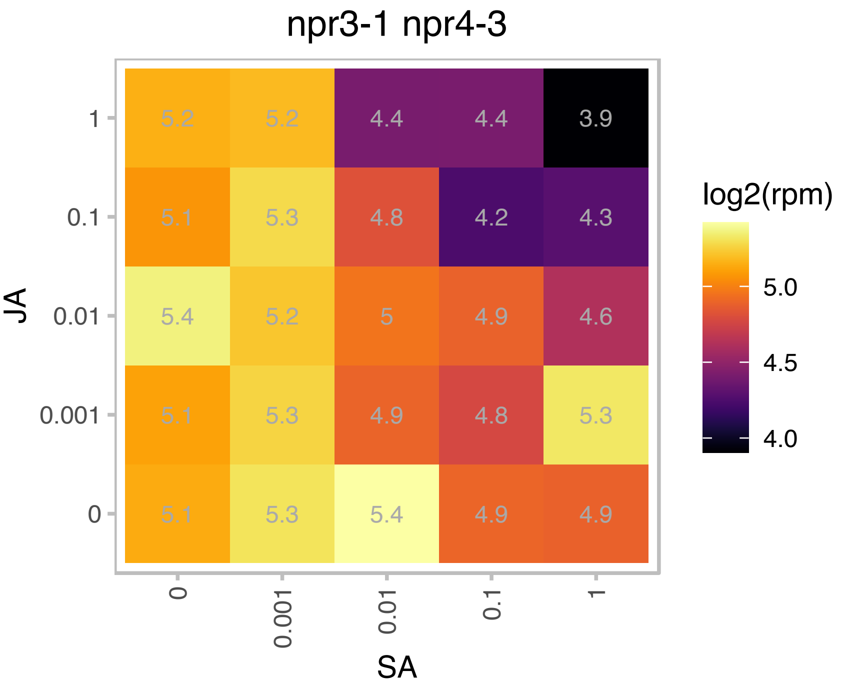

# AT1G68730.1 : NA

Zim17-type zinc finger protein

Col-0

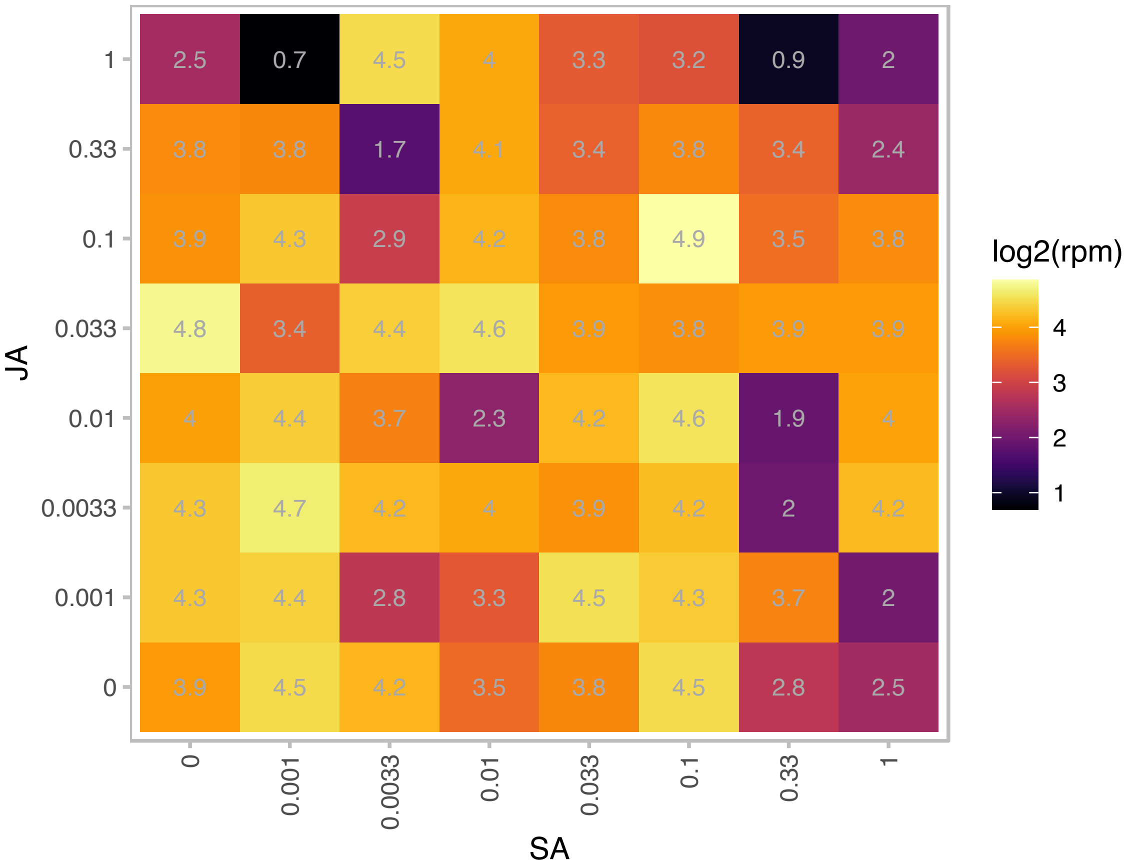


| Gene | AT1G68730 |
|---|---|
| is.expressed_genes | TRUE |
| is.responsive_genes | TRUE |
| is.apcluster | F-07 |

npr3-2 npr4-2

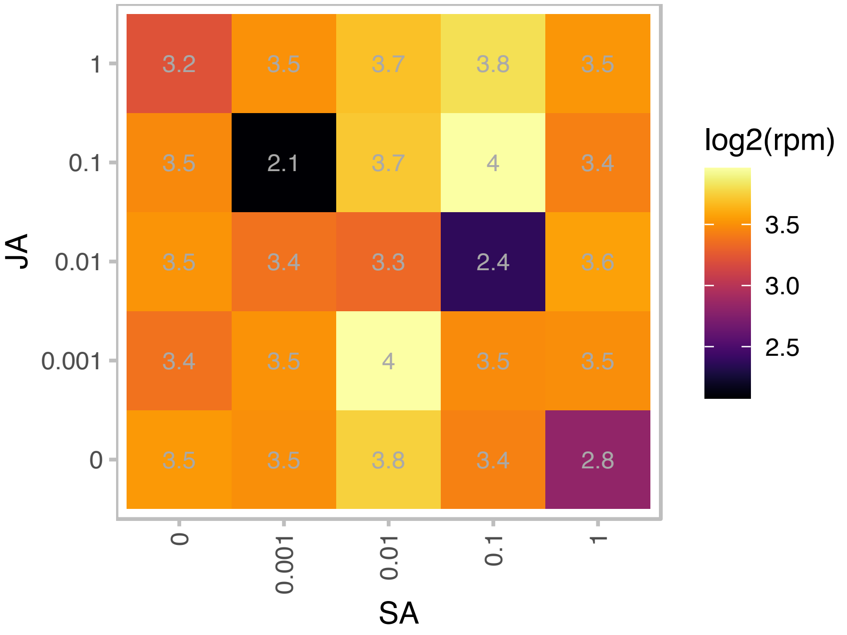


npr3-1 npr4-3

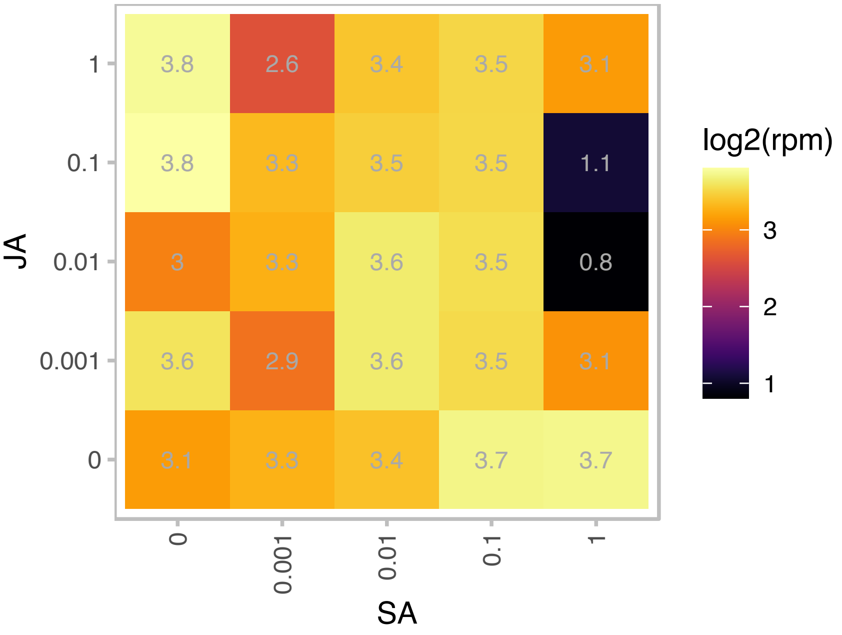

# AT1G68945.1 : NA

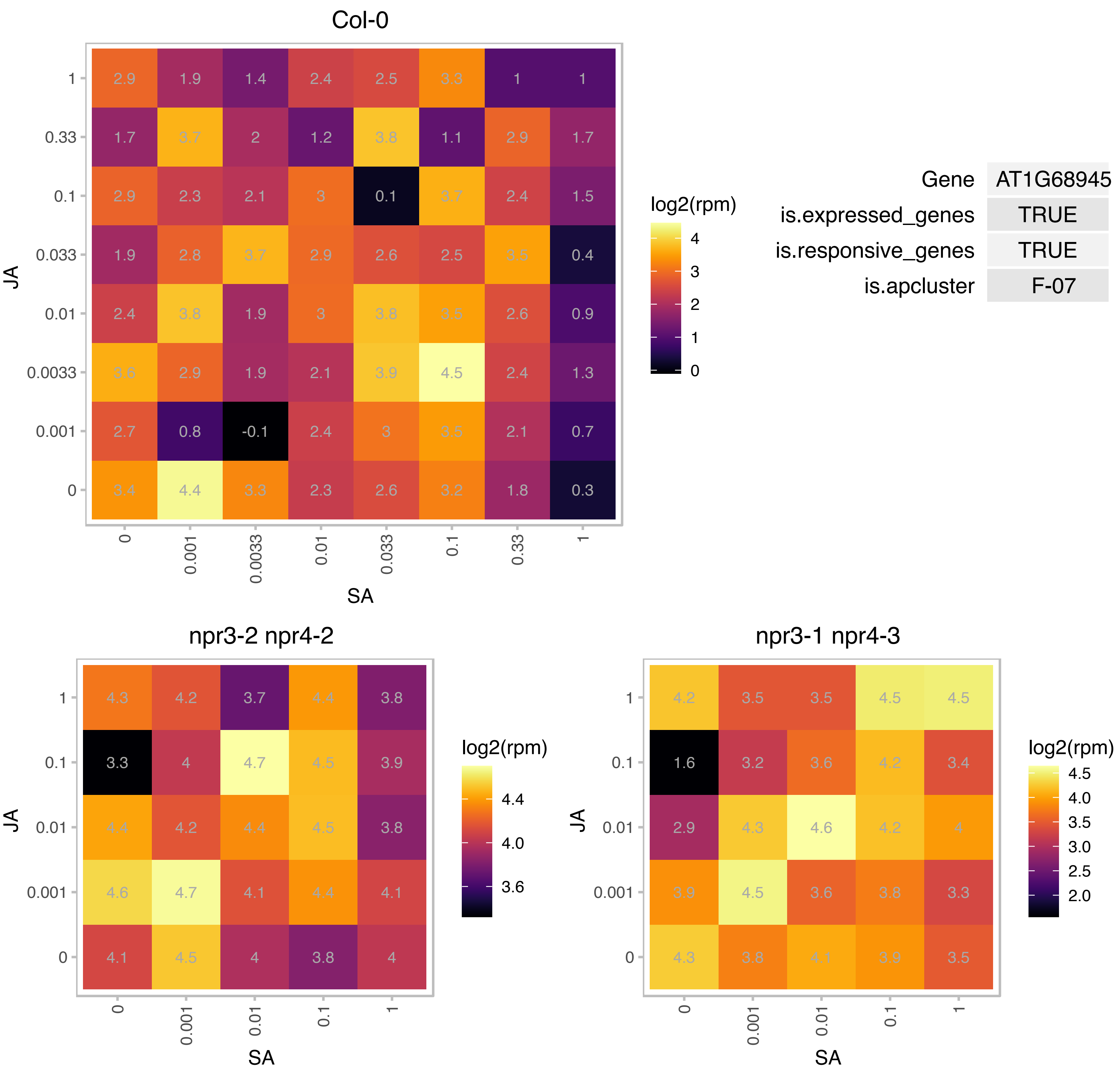

### AT1G69620.1 : RPL34

ribosomal protein L34

putative 60S ribosomal protein L34 The mRNA is cell-to-cell mobile.

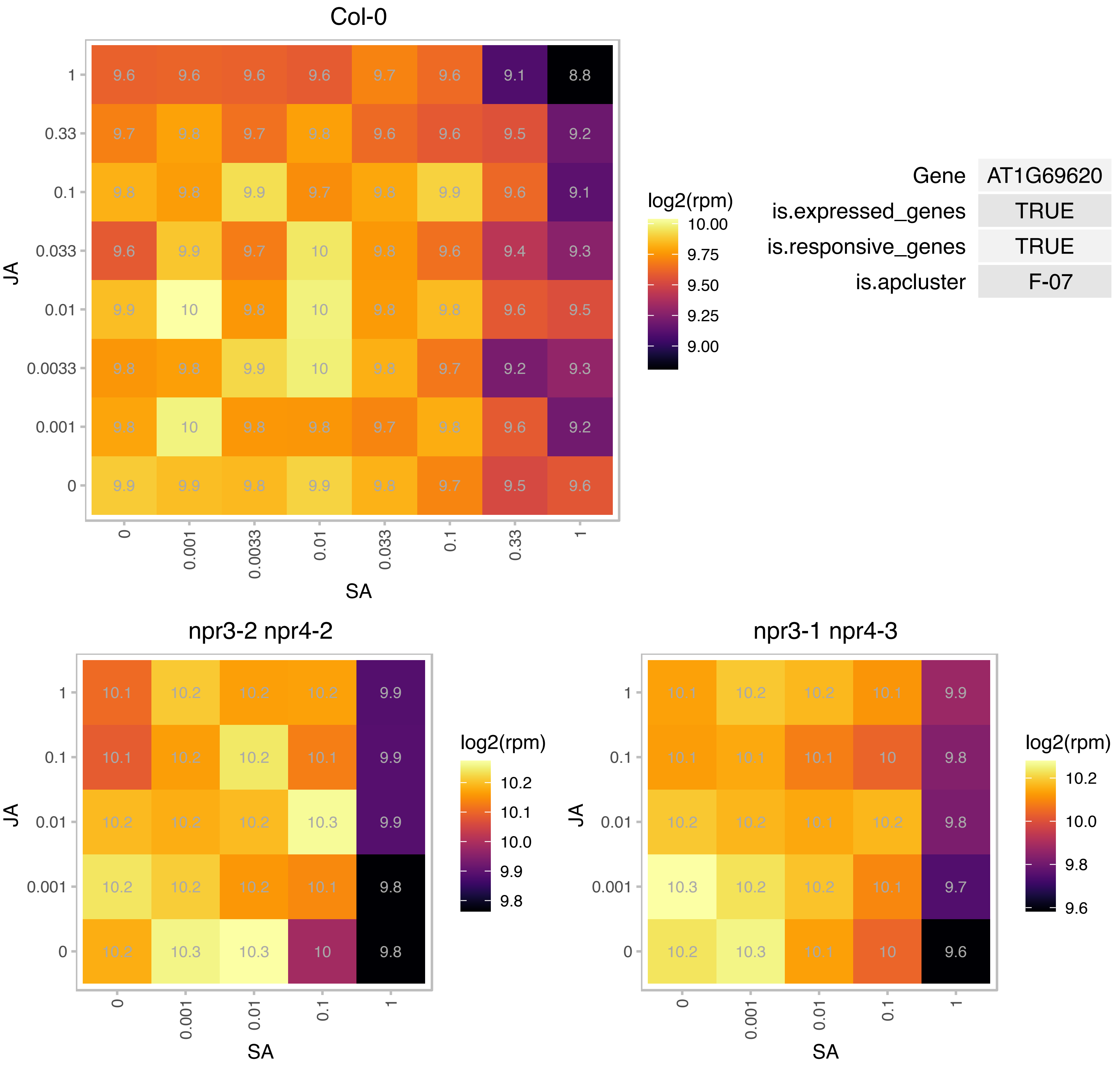

### AT1G72040.1 : AtdNK, dNK

P-loop containing nucleoside triphosphate hydrolases superfamily protein

Encodes a multisubstrate deoxyribonucleoside kinase that salvages DNA precursors.

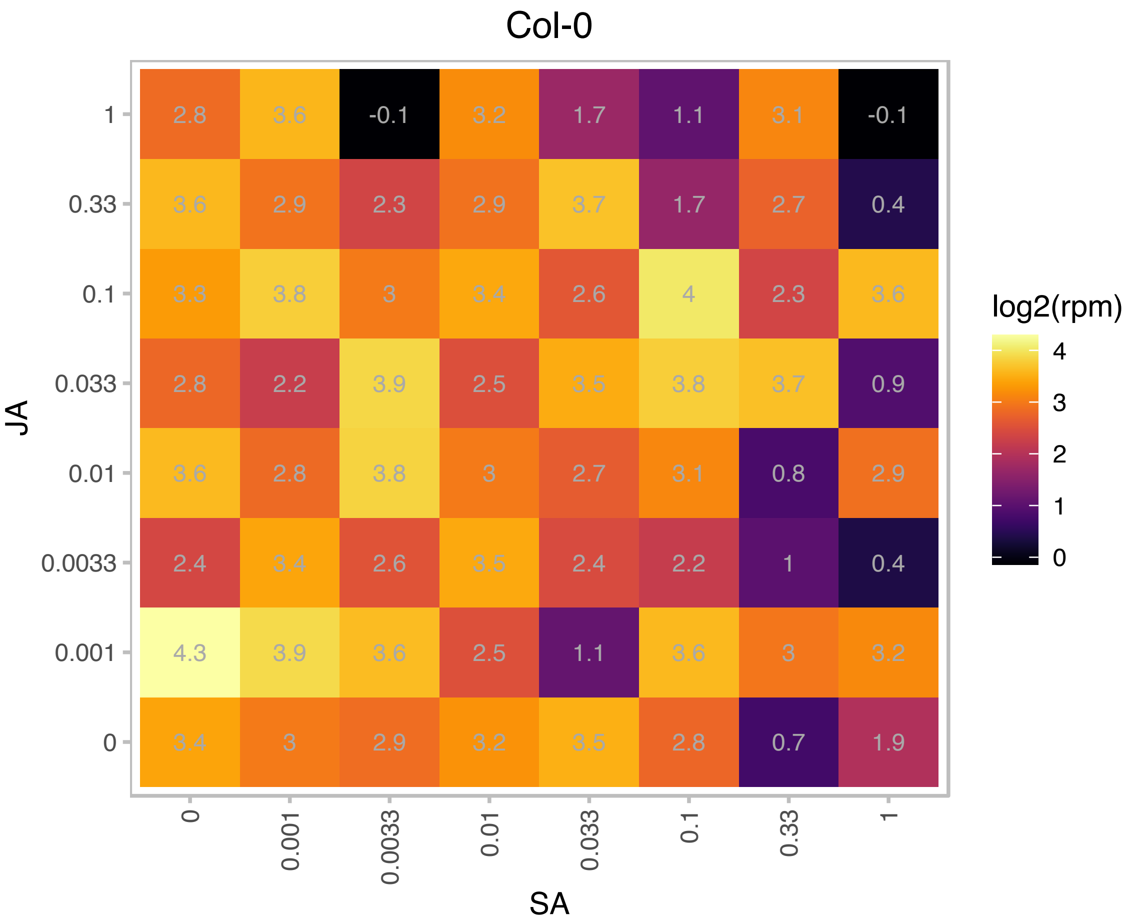


| | |
|---|---|
| Gene | AT1G72040 |
| is.expressed_genes | TRUE |
| is.responsive_genes | TRUE |
| is.apcluster | F-07 |

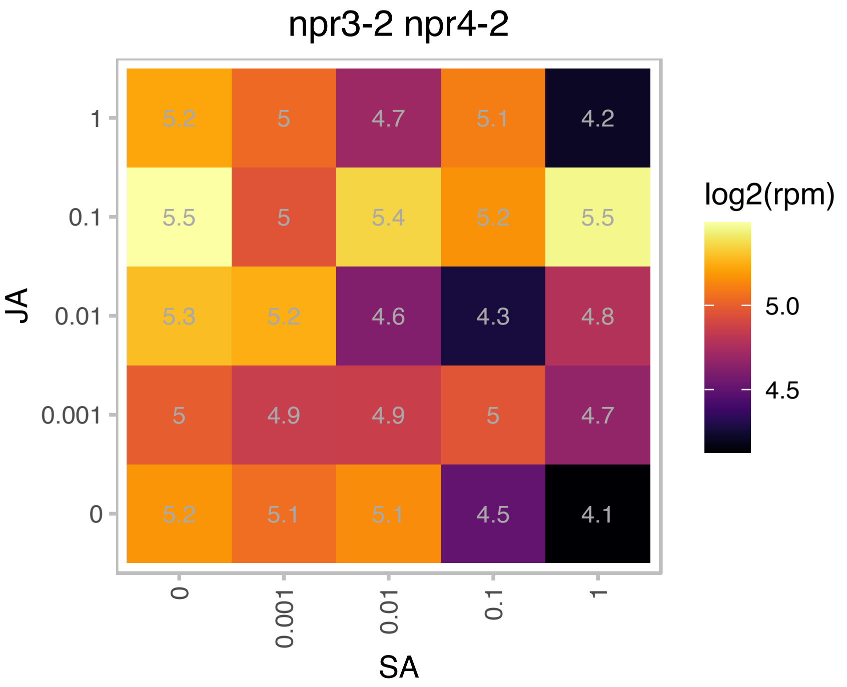


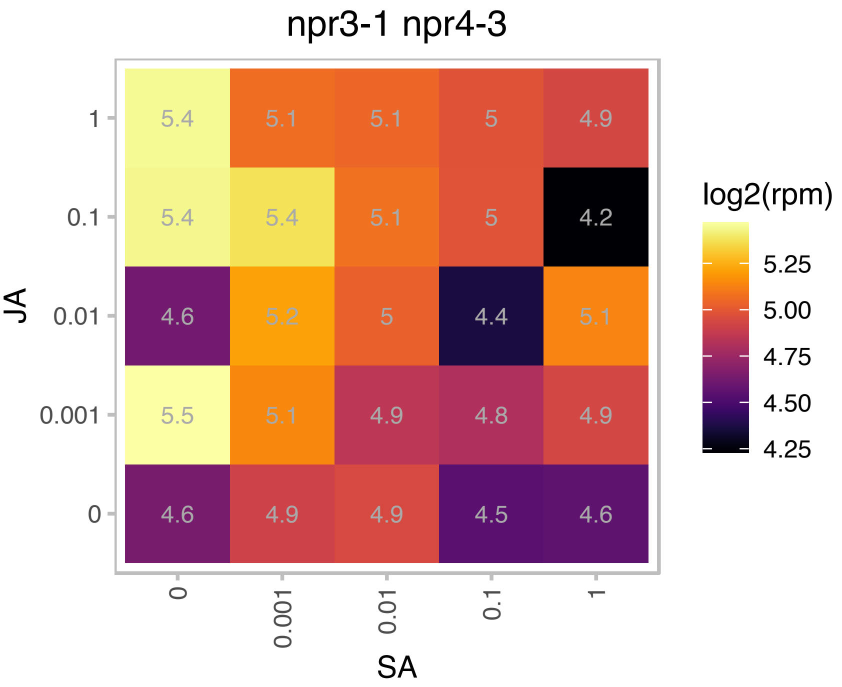

# AT1G72170.1 : NA

Domain of unknown function (DUF543)

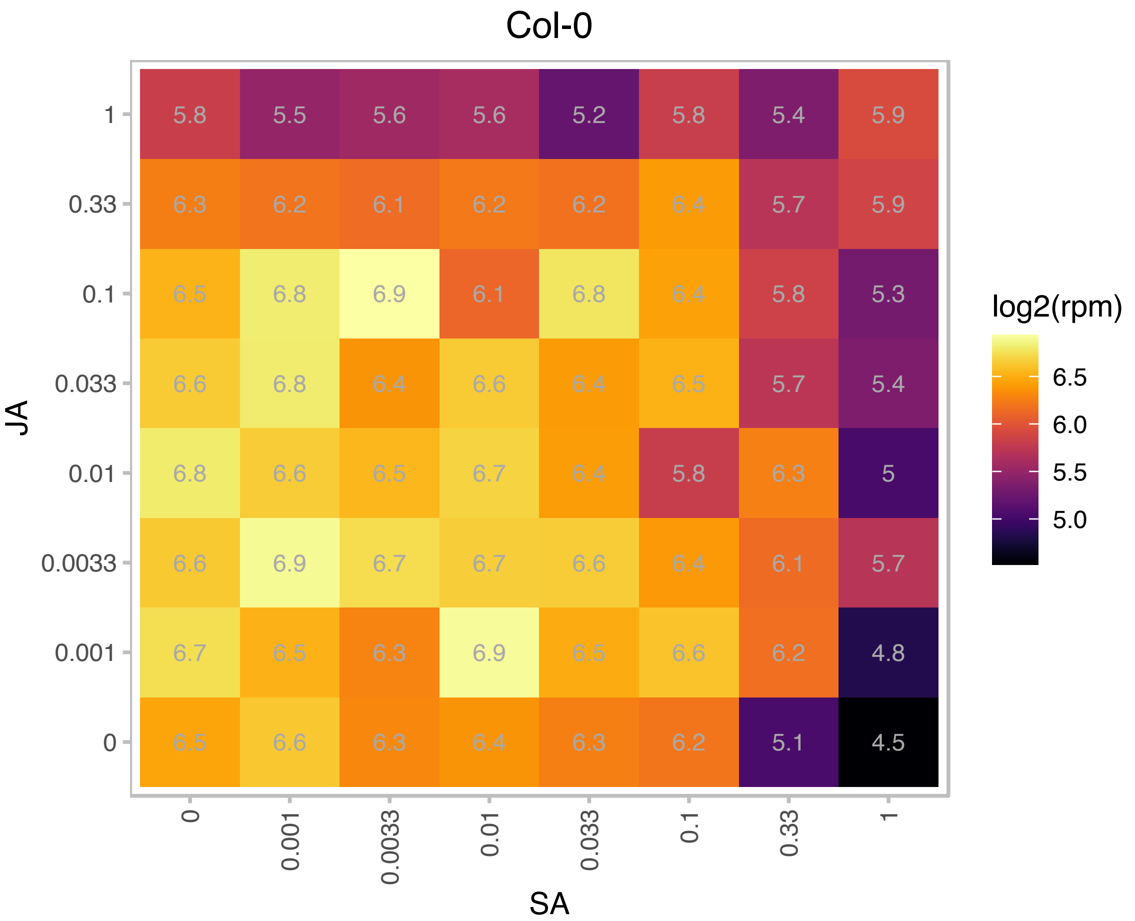


| | |
|---|---|
| Gene | AT1G72170 |
| is.expressed_genes | TRUE |
| is.responsive_genes | TRUE |
| is.apcluster | F-07 |

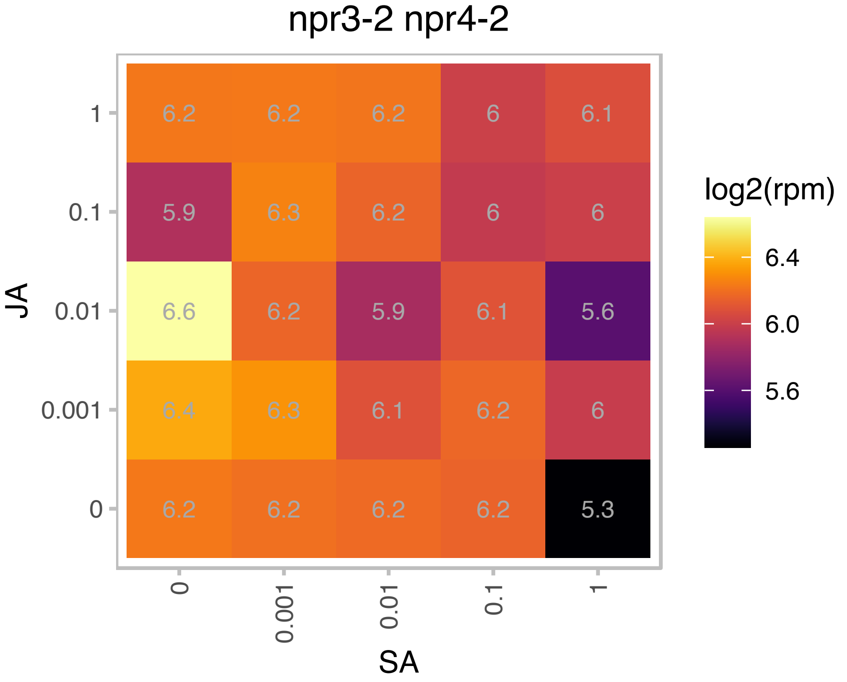


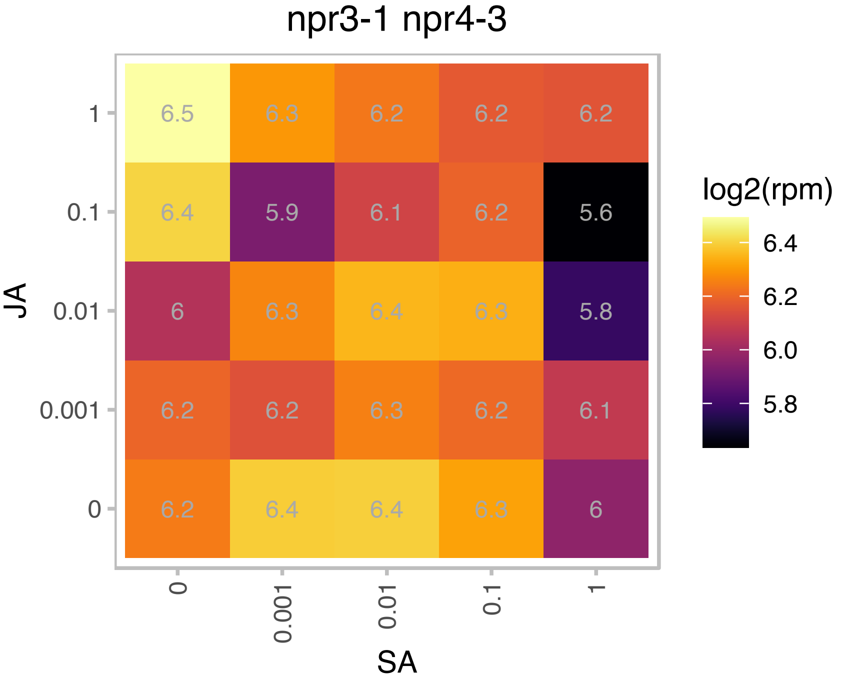

### AT1G72370.1 : AP40, P40, RP40, RPSAA

40s ribosomal protein SA

acidic protein associated to 40S ribosomal subunit of ribosomes. Involved in polysome formation during active protein synthesis. Expressed in actively growing tissue.

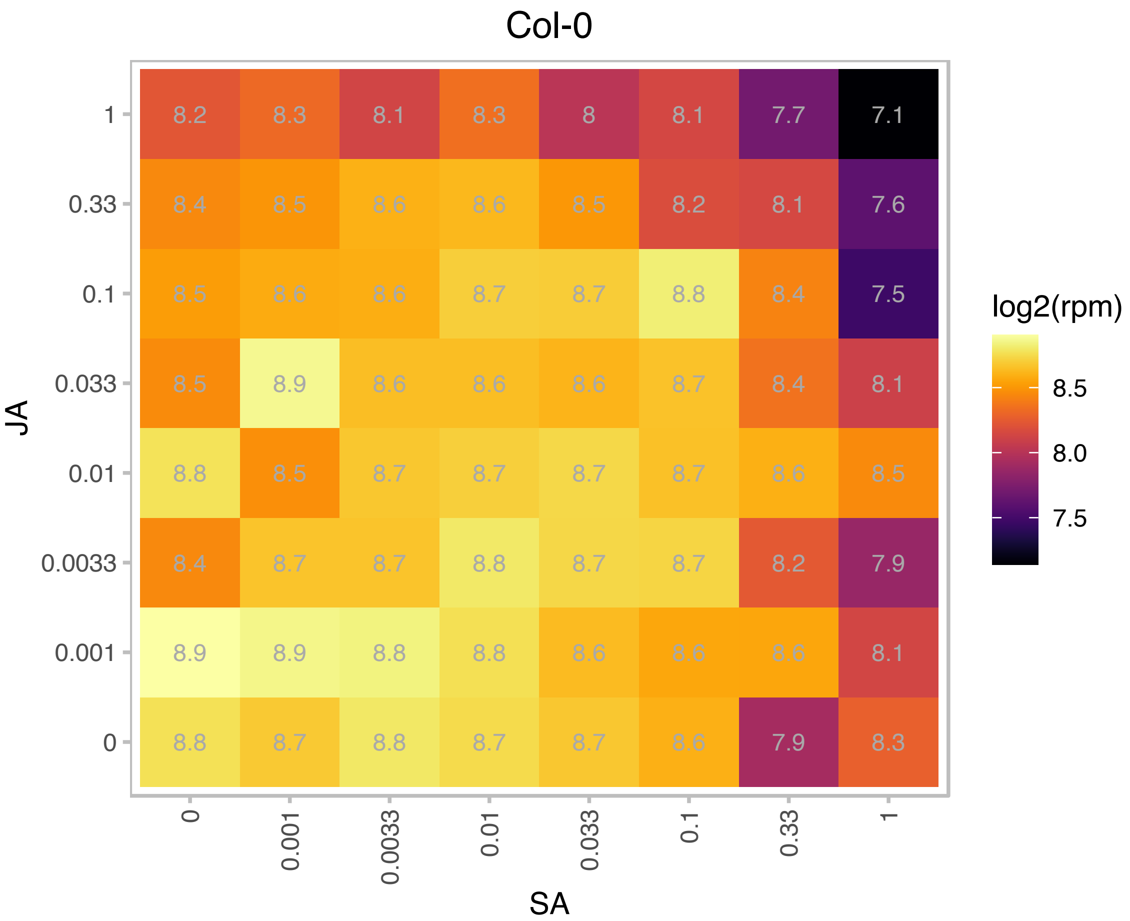


| | |
|---|---|
| Gene | AT1G72370 |
| is.expressed_genes | TRUE |
| is.responsive_genes | TRUE |
| is.apcluster | F-07 |

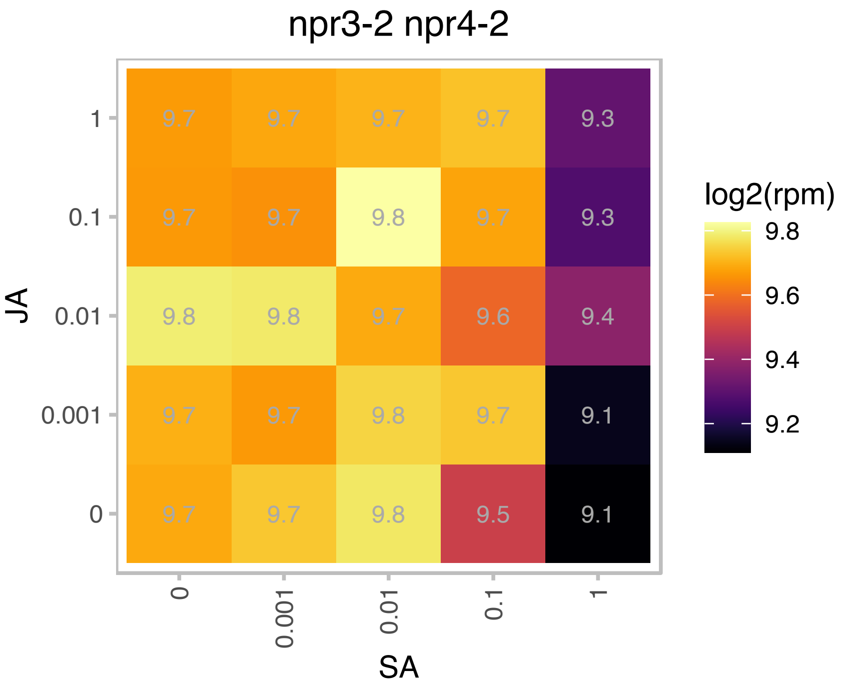


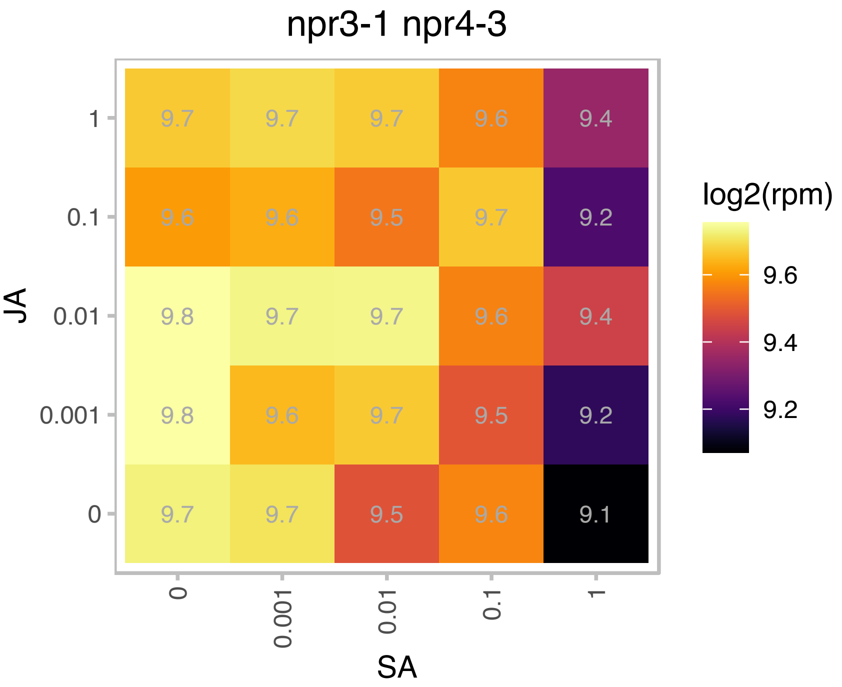

# AT1G73230.1 : NA

Nascent polypeptide-associated complex NAC

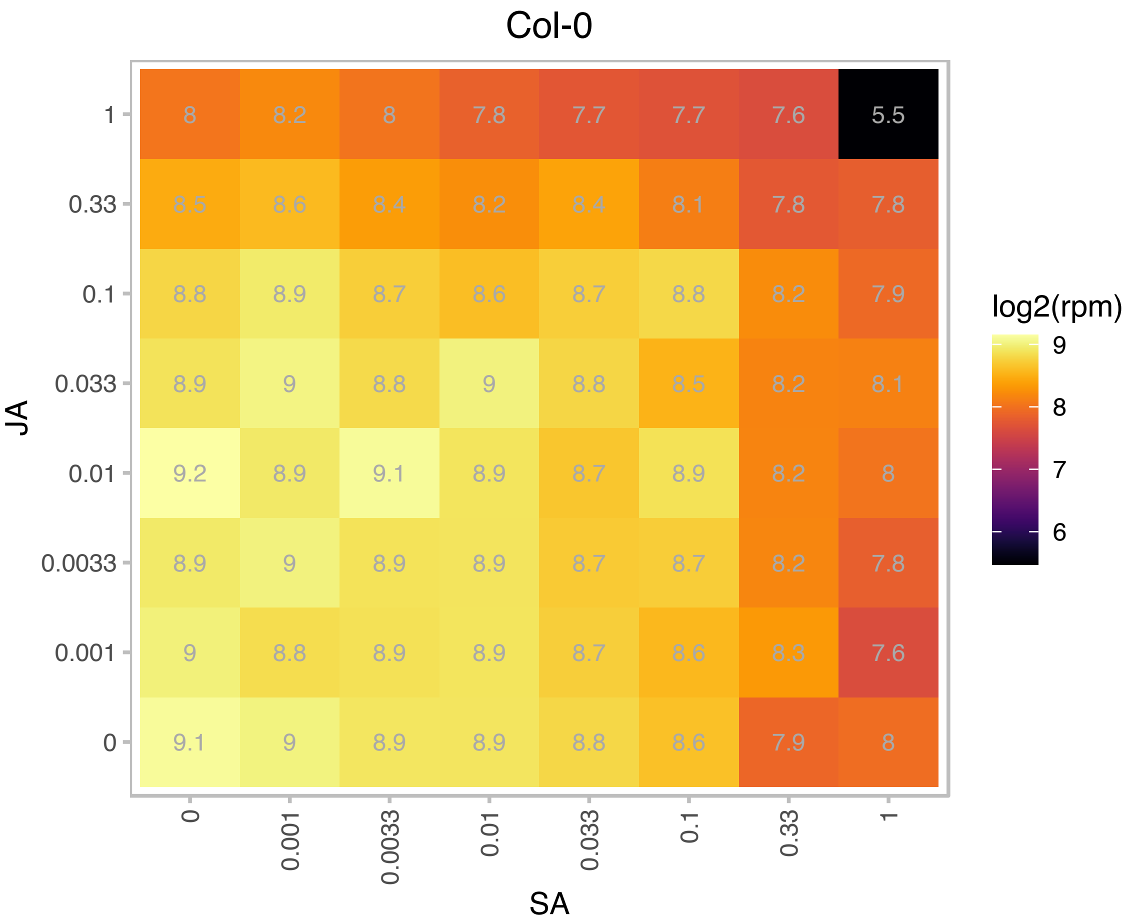


| | |
|---|---|
| Gene | AT1G73230 |
| is.expressed_genes | TRUE |
| is.responsive_genes | TRUE |
| is.apcluster | F-07 |

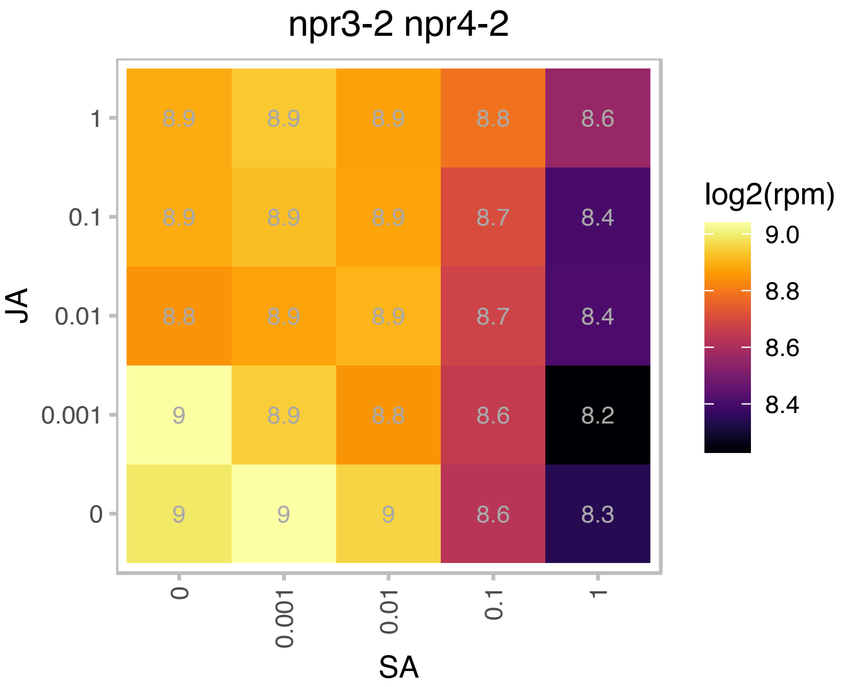


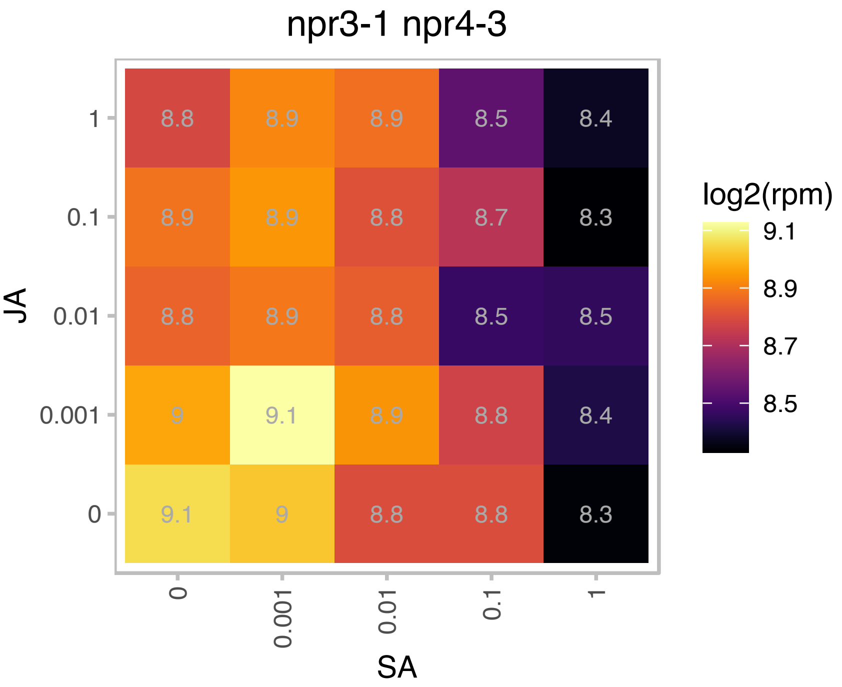

# AT1G74050.1 : NA

Ribosomal protein L6 family protein

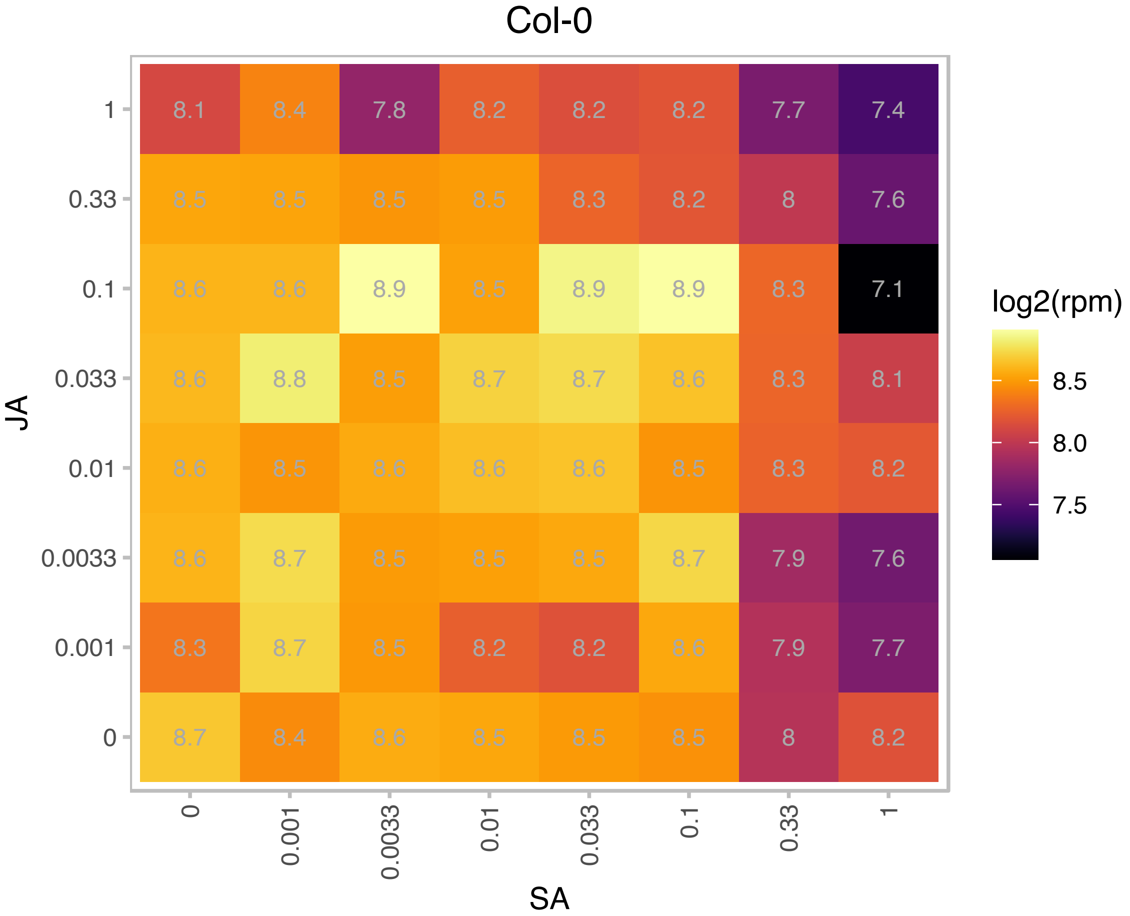


| | |
|---|---|
| Gene | AT1G74050 |
| is.expressed_genes | TRUE |
| is.responsive_genes | TRUE |
| is.apcluster | F-07 |

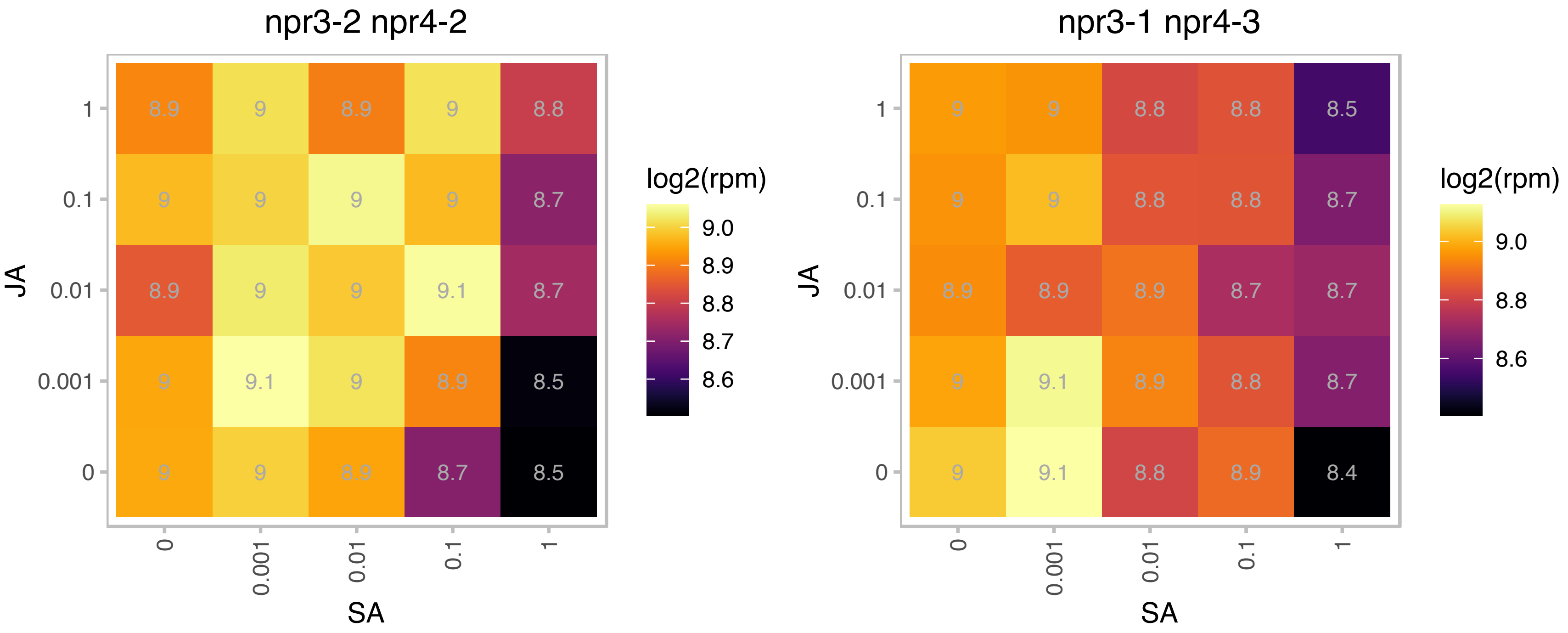

### AT1G74560.1 : NRP1

NAP1-related protein 1

Double nrp1-1 nrp2-1 mutants show arrest of cell cycle progression at G2/M and disordered cellular organization occurred in root tips. Localize in the nucleus and can form homomeric and heteromeric protein complexes with NRP2. Bind histones Histone2A and Histone2B and associate with chromatin in vivo. Plant mutated in both NRP1 and NRP2 genes show hypersensitivity to genotoxic stresses including UV and DSB-inducing agent Bleomycin. NRP genes act synergistically with NAP1 genes in promoting somatic homologous recombination.

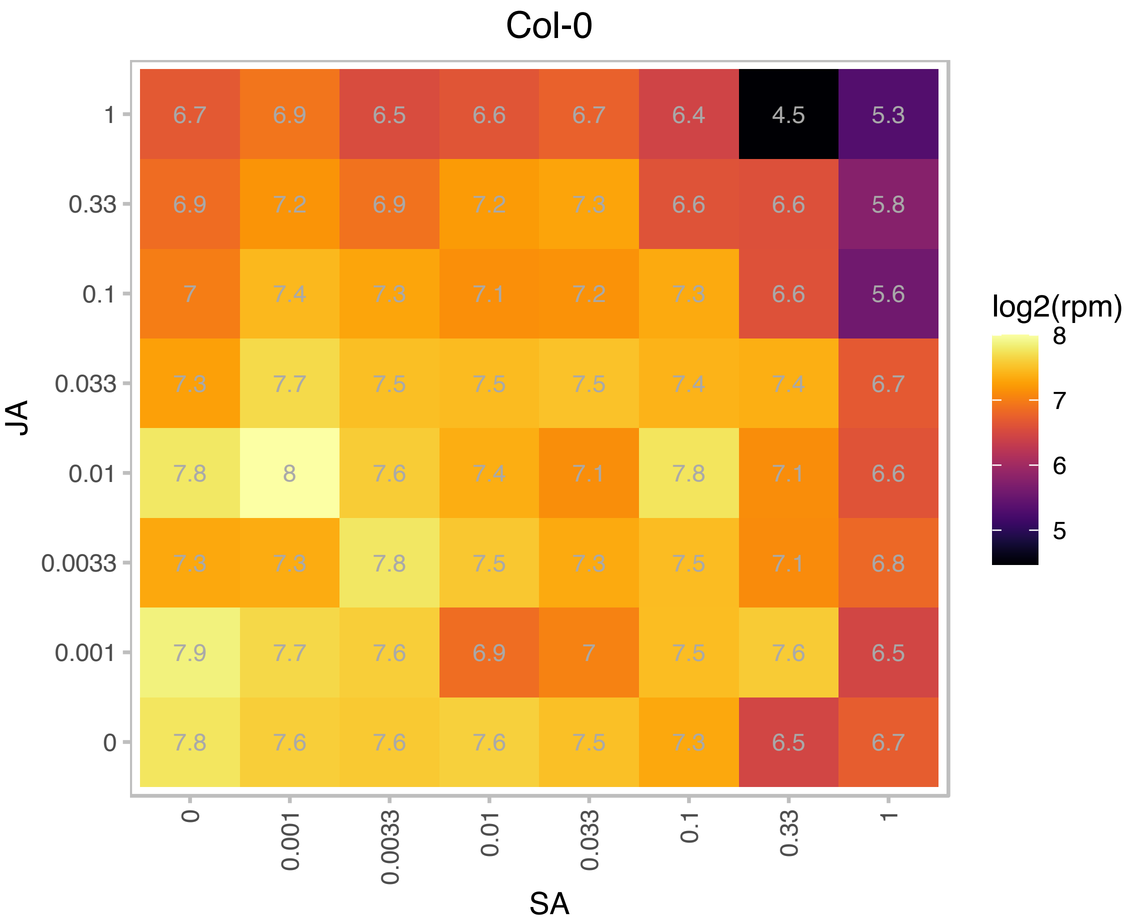


| | |
|---|---|
| Gene | AT1G74560 |
| is.expressed_genes | TRUE |
| is.responsive_genes | TRUE |
| is.apcluster | F-07 |

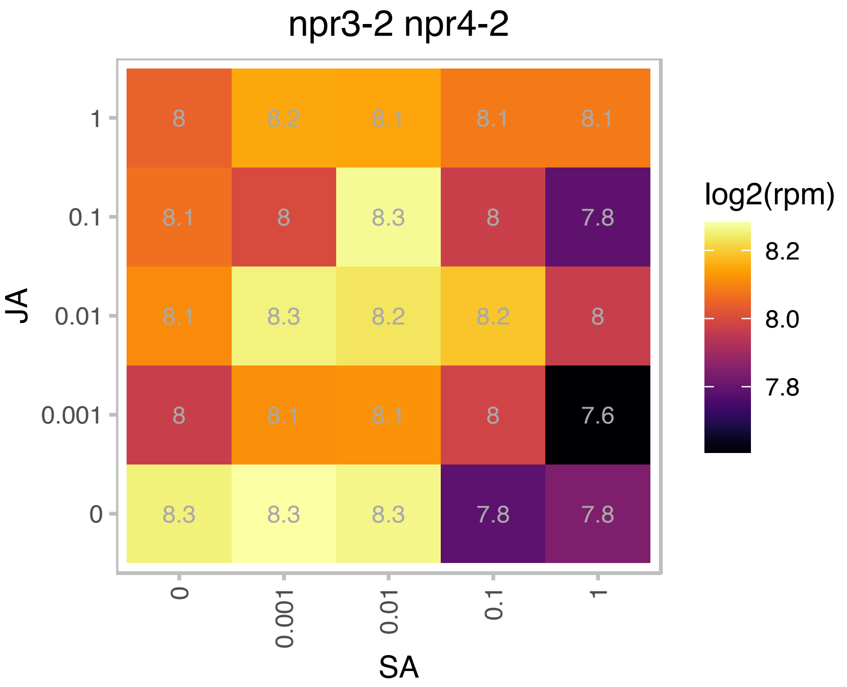


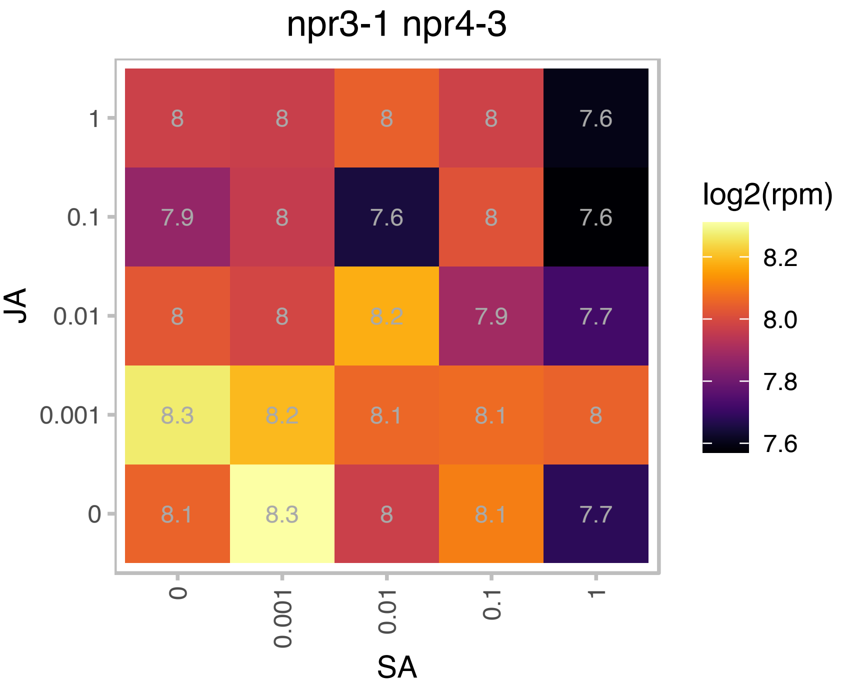

# AT1G75335.1 : NA

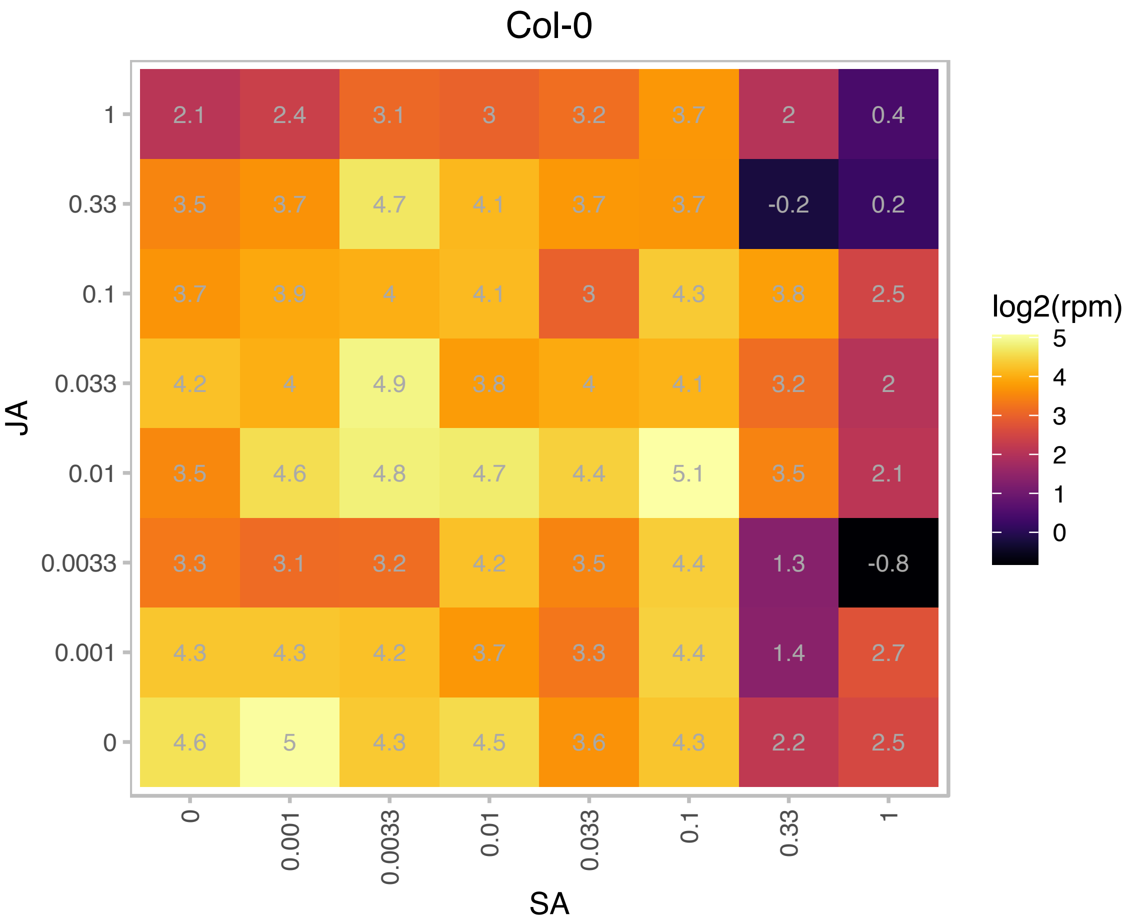


| | |
|---|---|
| Gene | AT1G75335 |
| is.expressed_genes | TRUE |
| is.responsive_genes | TRUE |
| is.apcluster | F-07 |

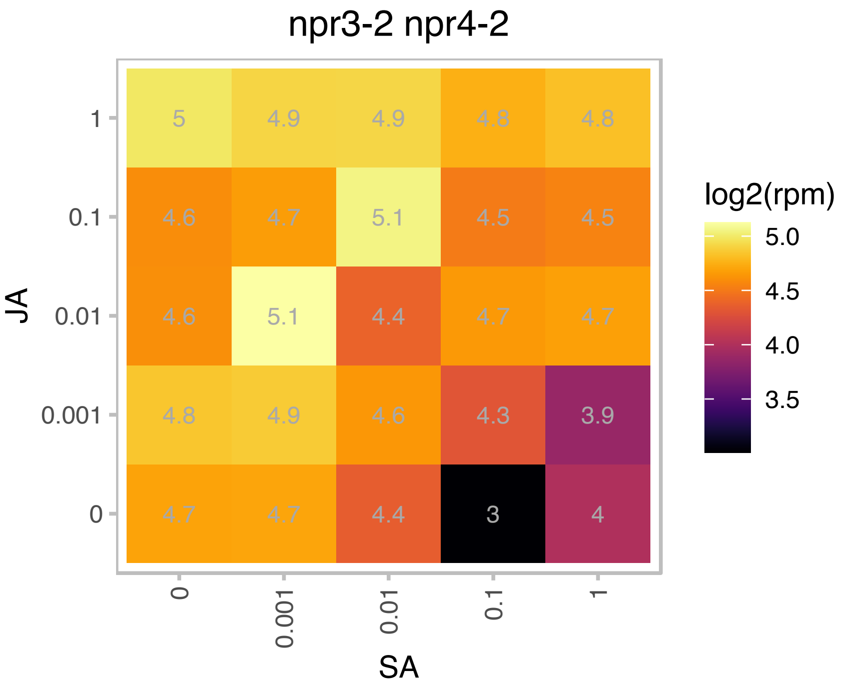


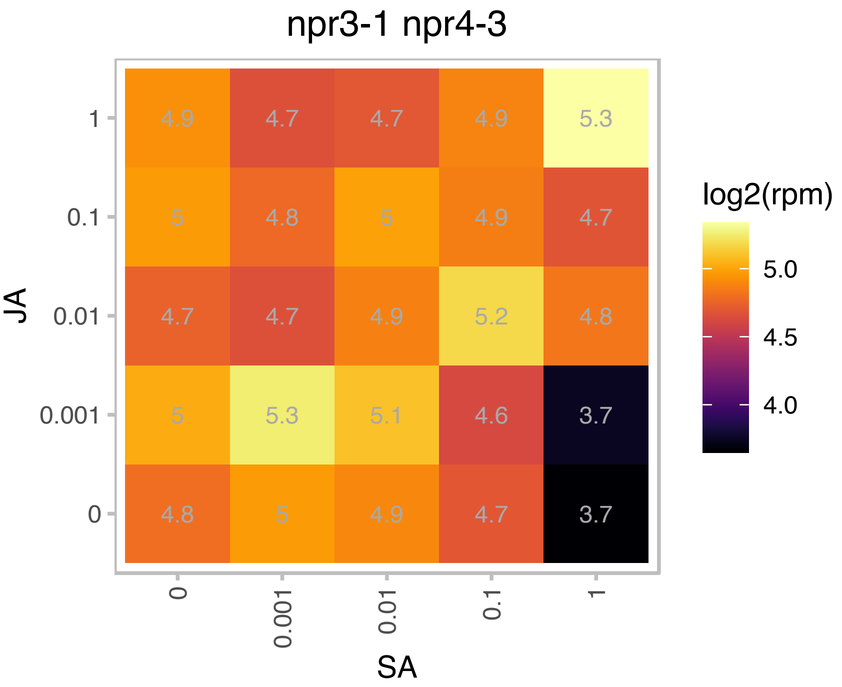

### AT1G76100.1 : PETE1

plastocyanin 1
One of two Arabidopsis plastocyanin genes. Expressed at 1/10th level of PETE2. Does not respond to increased copper levels and is thought to be the isoform that participates in electron transport under copper-limiting conditions. Mutation of this gene does not have obvious effect on photosynthesis.

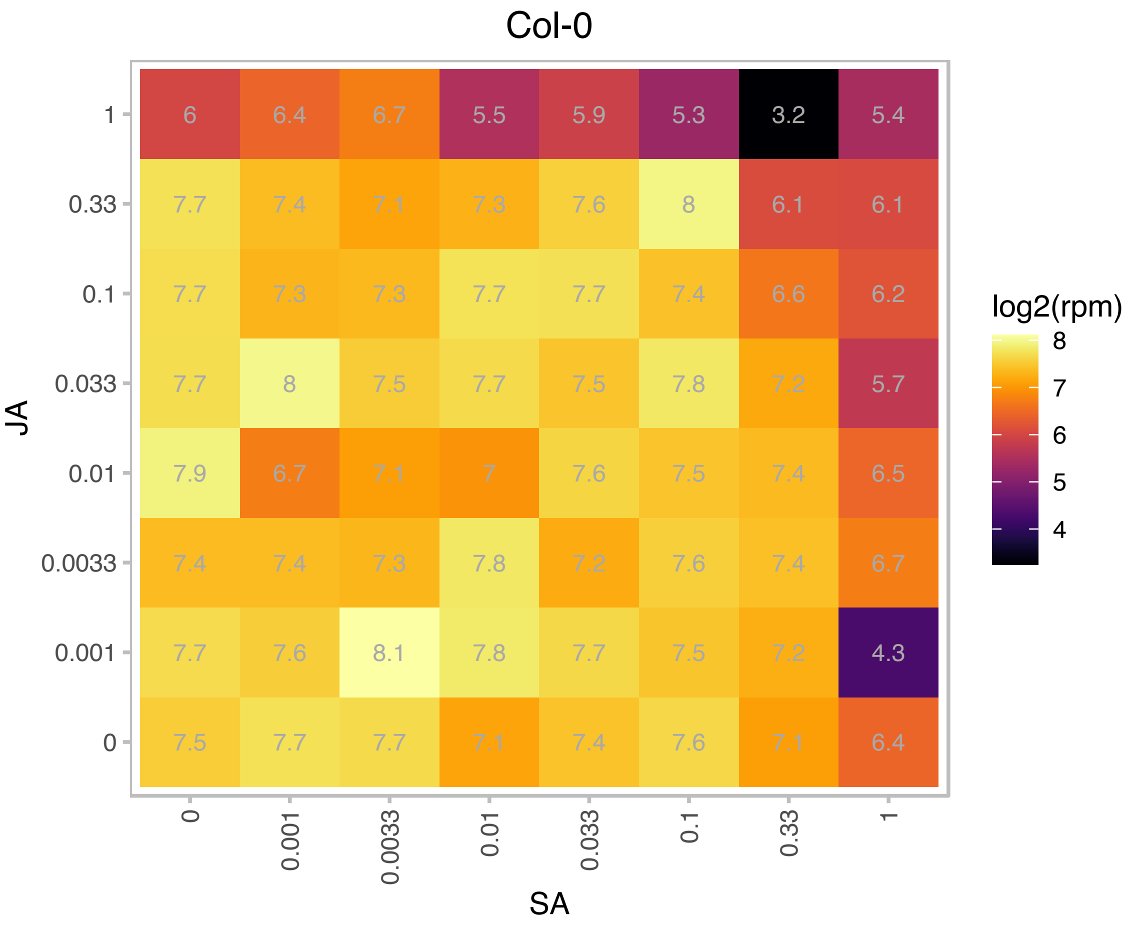


| | |
|---|---|
| Gene | AT1G76100 |
| is.expressed_genes | TRUE |
| is.responsive_genes | TRUE |
| is.apcluster | F-07 |

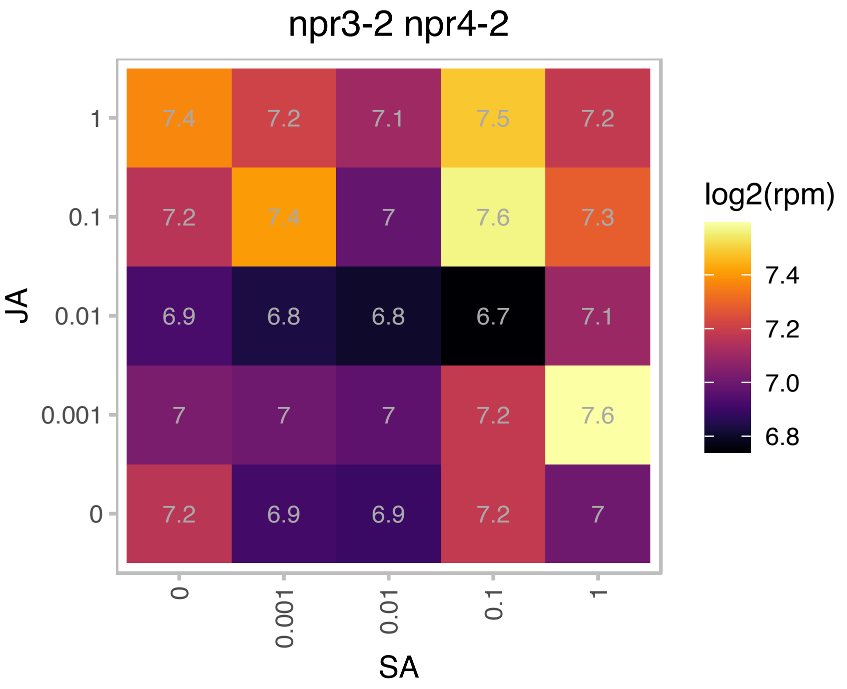


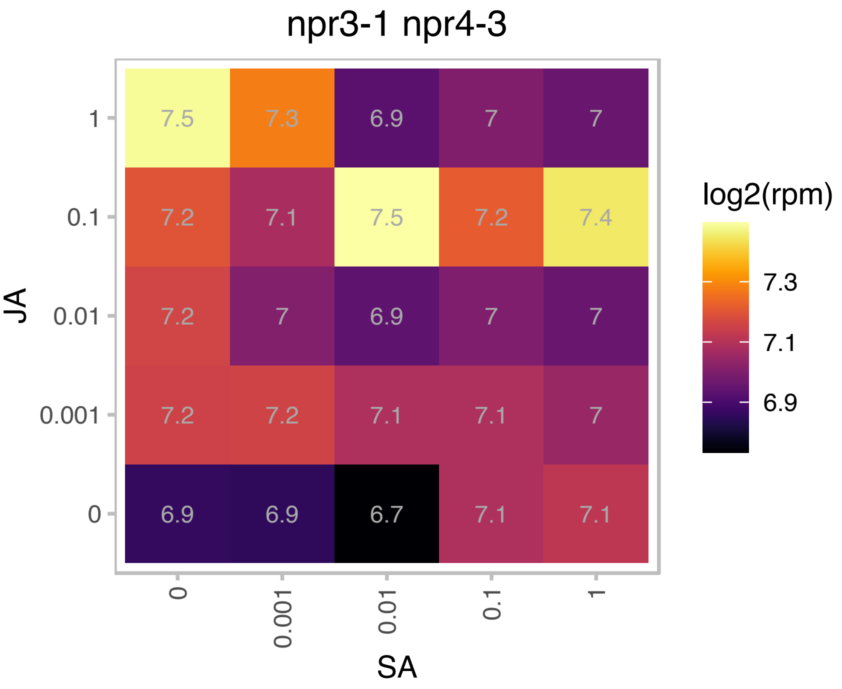

# AT1G77940.1 : NA

Ribosomal protein L7Ae/L30e/S12e/Gadd45 family protein

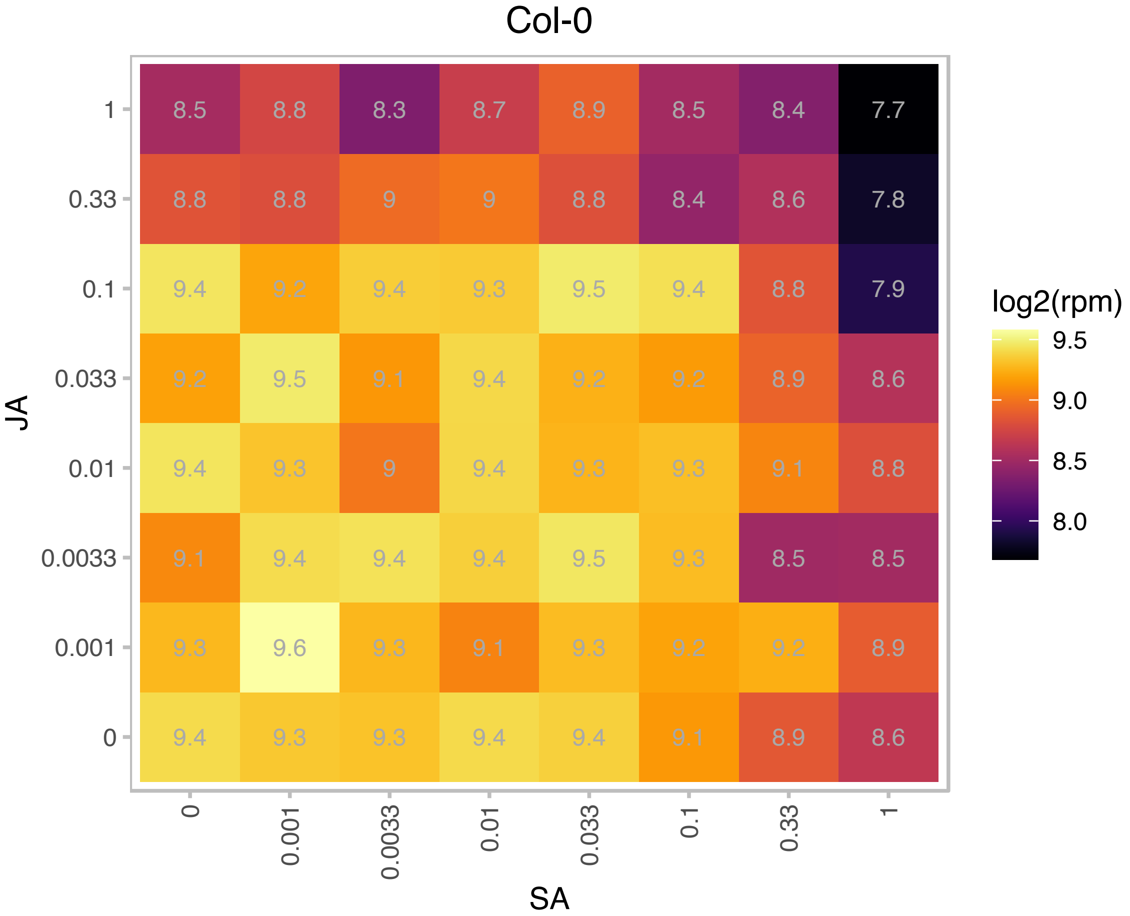


| | |
|---|---|
| Gene | AT1G77940 |
| is.expressed_genes | TRUE |
| is.responsive_genes | TRUE |
| is.apcluster | F-07 |

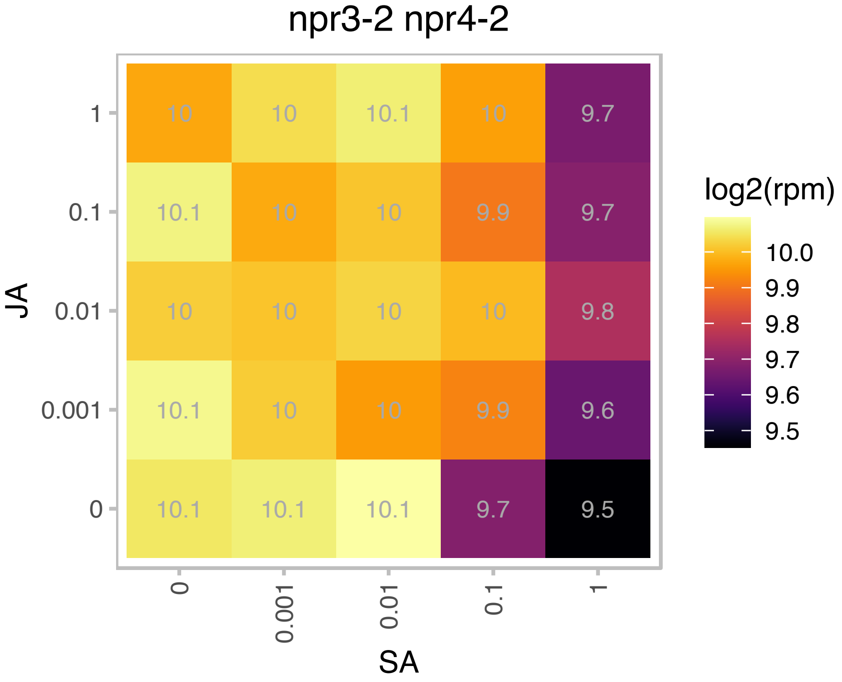


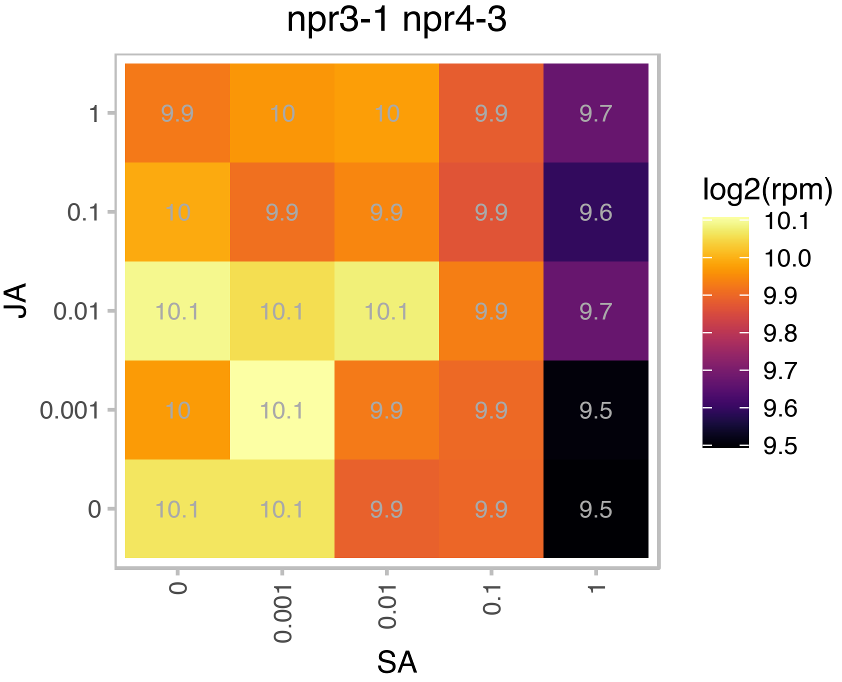

# AT1G78460.1 : NA

SOUL heme-binding family protein

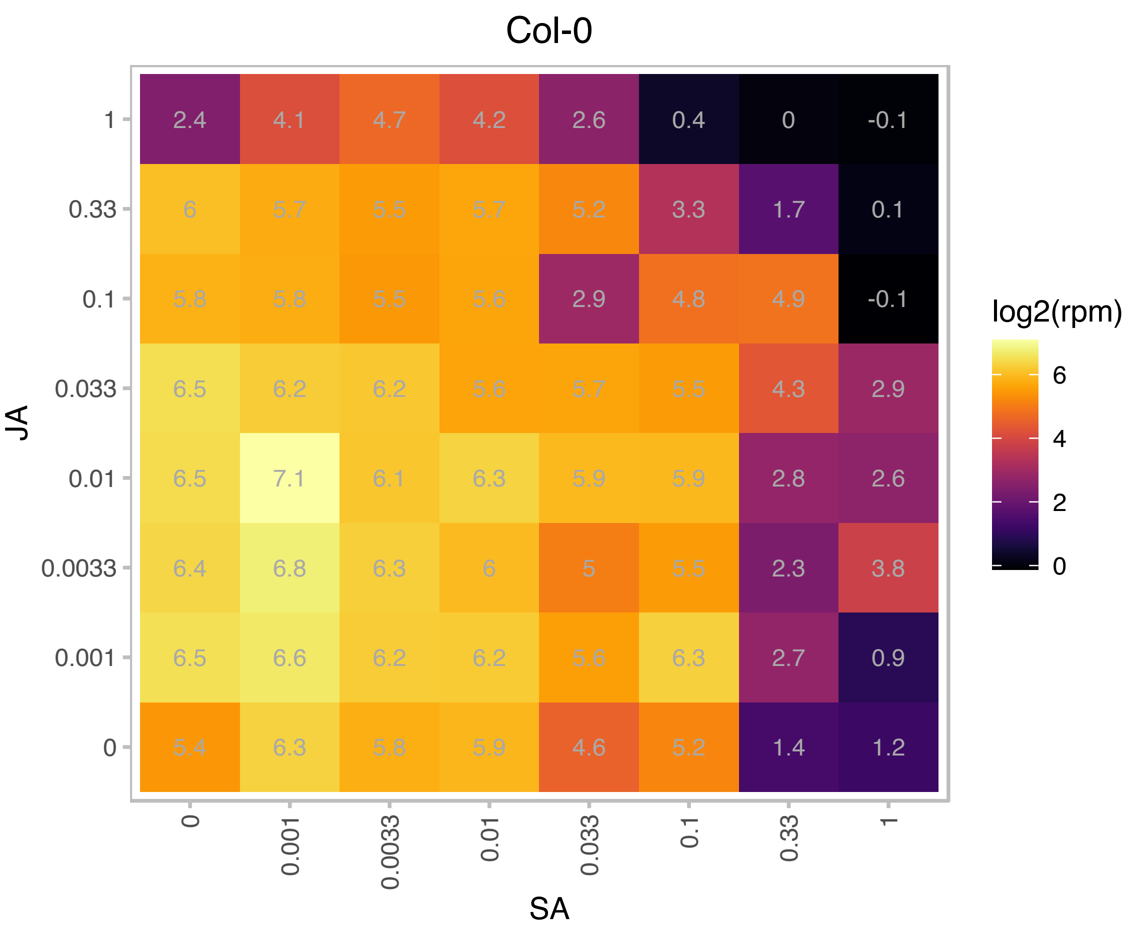


| | |
|---|---|
| Gene | AT1G78460 |
| is.expressed_genes | TRUE |
| is.responsive_genes | TRUE |
| is.apcluster | F-07 |

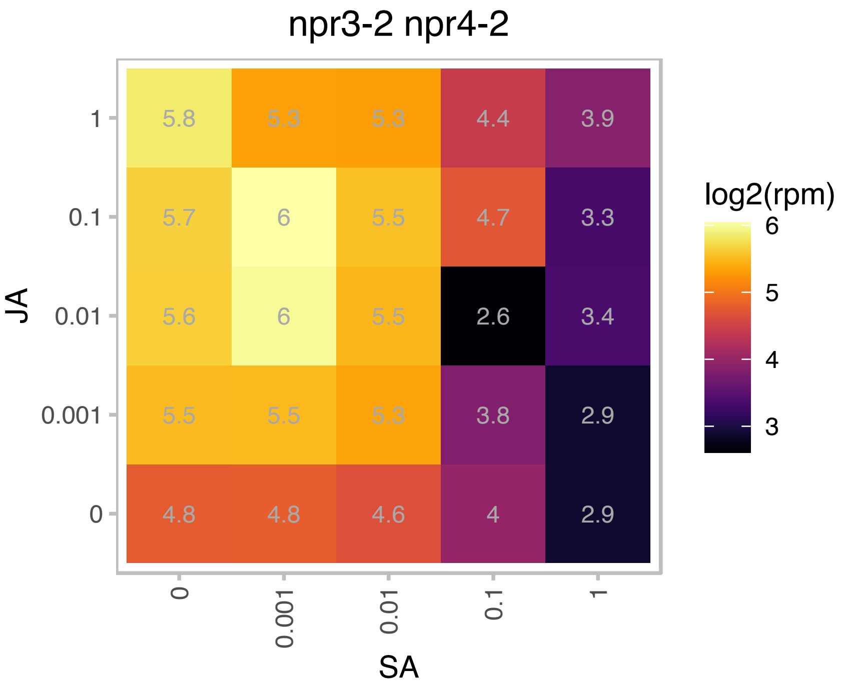


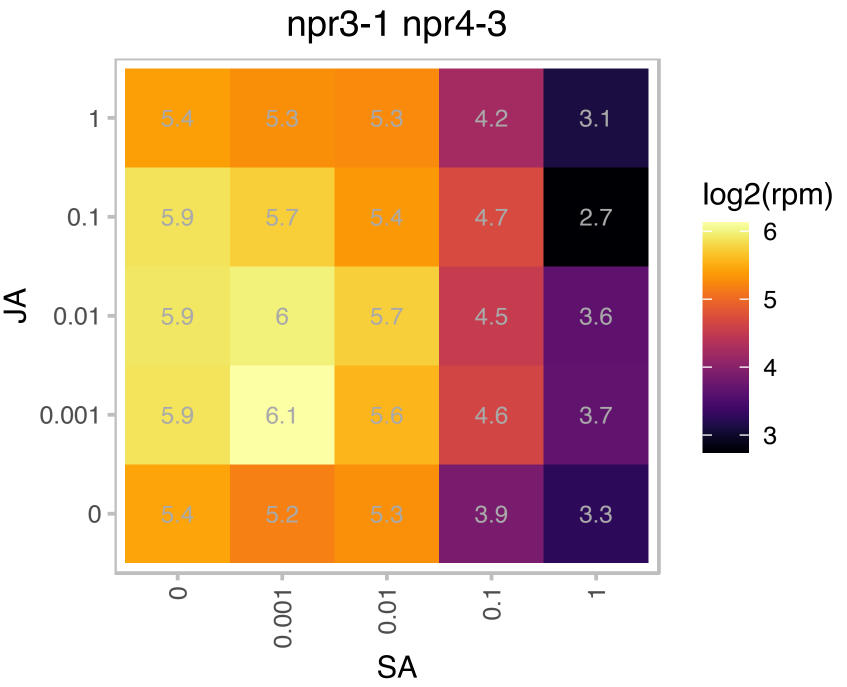

### AT1G78850.1 : MBL1

D-mannose binding lectin protein with Apple-like carbohydrate-binding domain
curculin-like (mannose-binding) lectin family protein, low similarity to ser/thr protein kinase from Zea mays (GI:2598067); contains Pfam lectin (probable mannose binding) domain PF01453 but not the protein kinase domain of the Z. mays protein

Col-0

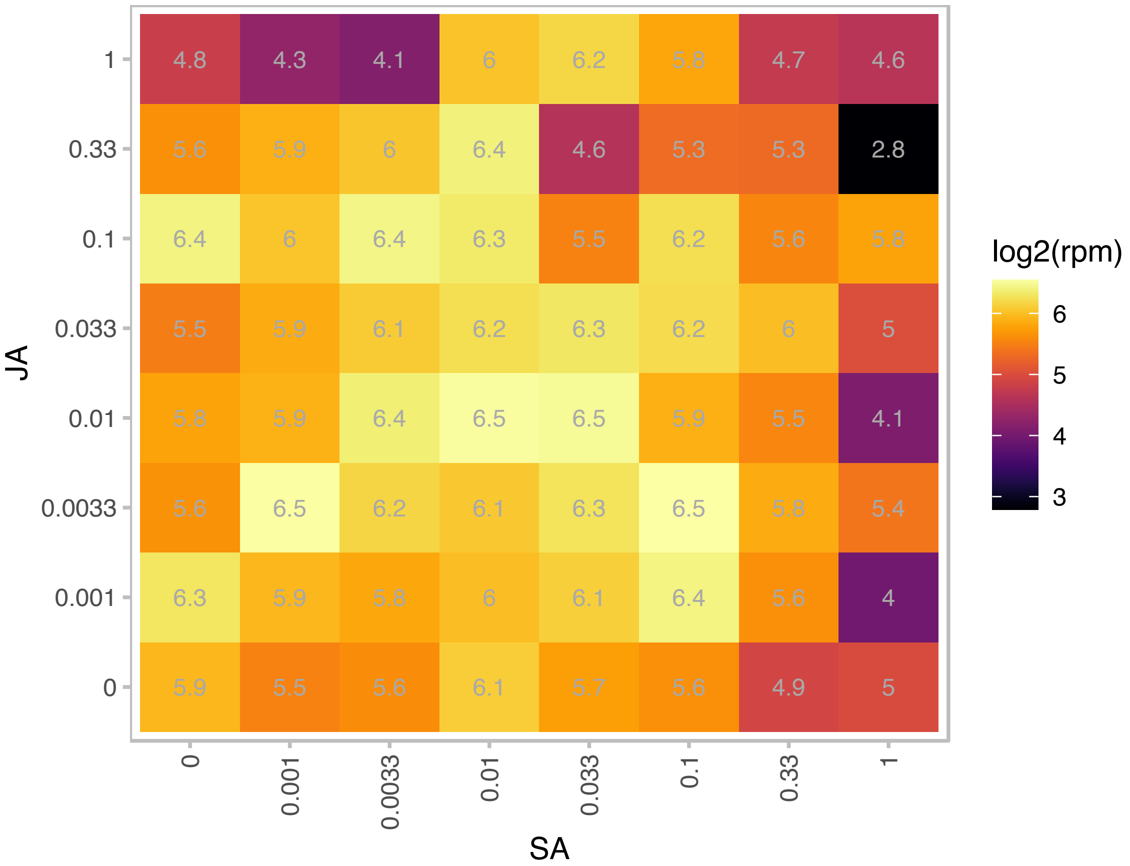


| | |
|---|---|
| Gene | AT1G78850 |
| is.expressed_genes | TRUE |
| is.responsive_genes | TRUE |
| is.apcluster | F-07 |

npr3-2 npr4-2

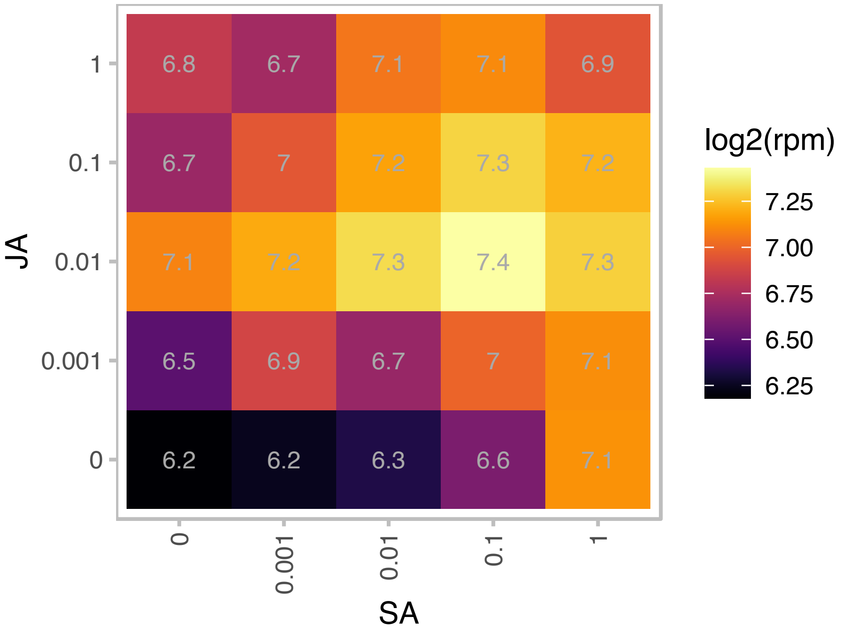


npr3-1 npr4-3

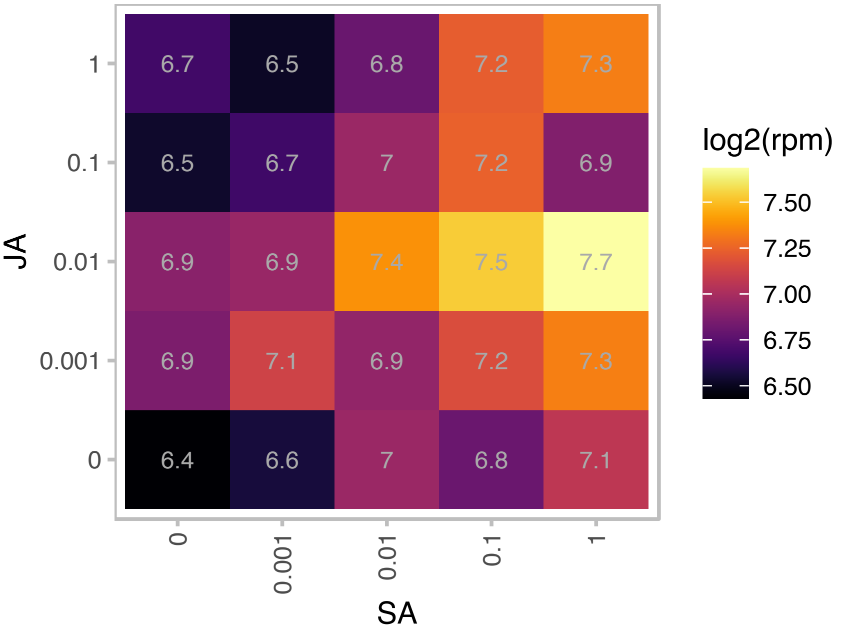

# AT1G79390.1 : NA

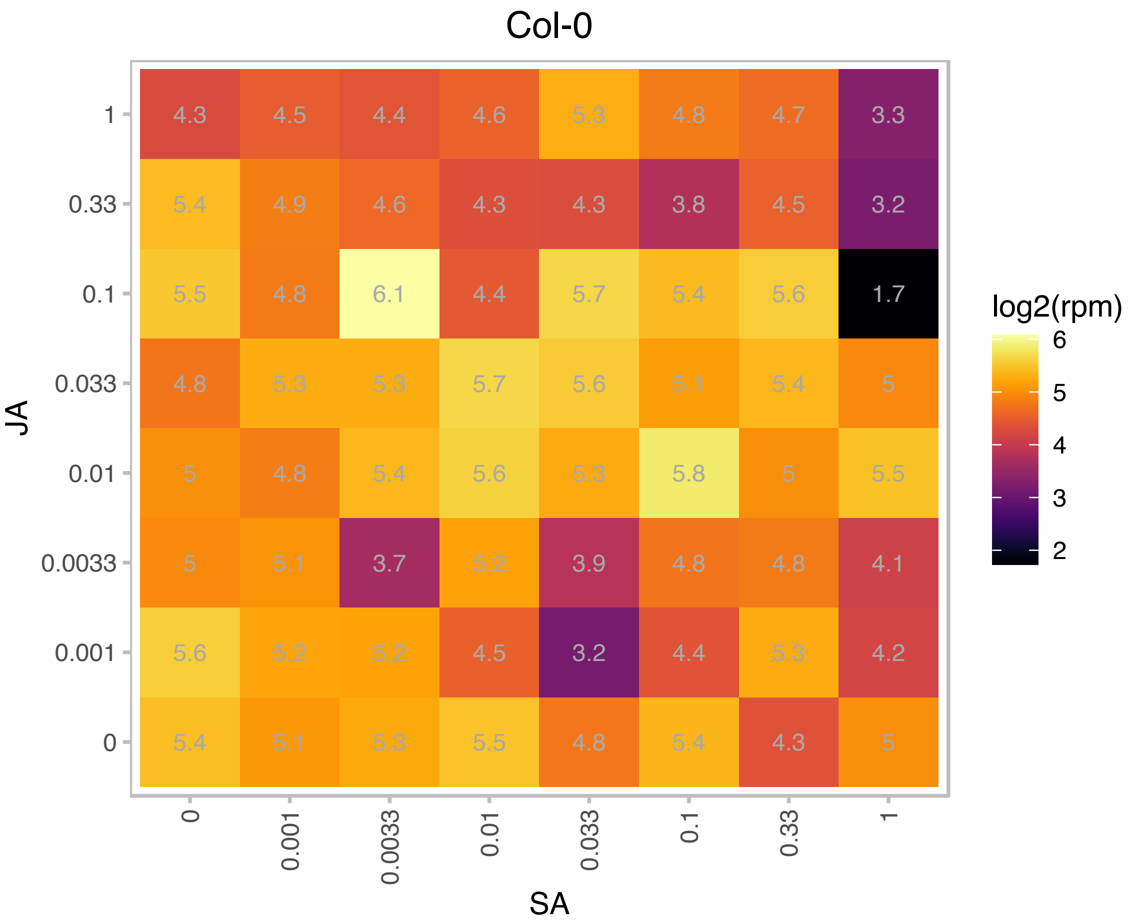


| | |
|---|---|
| Gene | AT1G79390 |
| is.expressed_genes | TRUE |
| is.responsive_genes | TRUE |
| is.apcluster | F-07 |

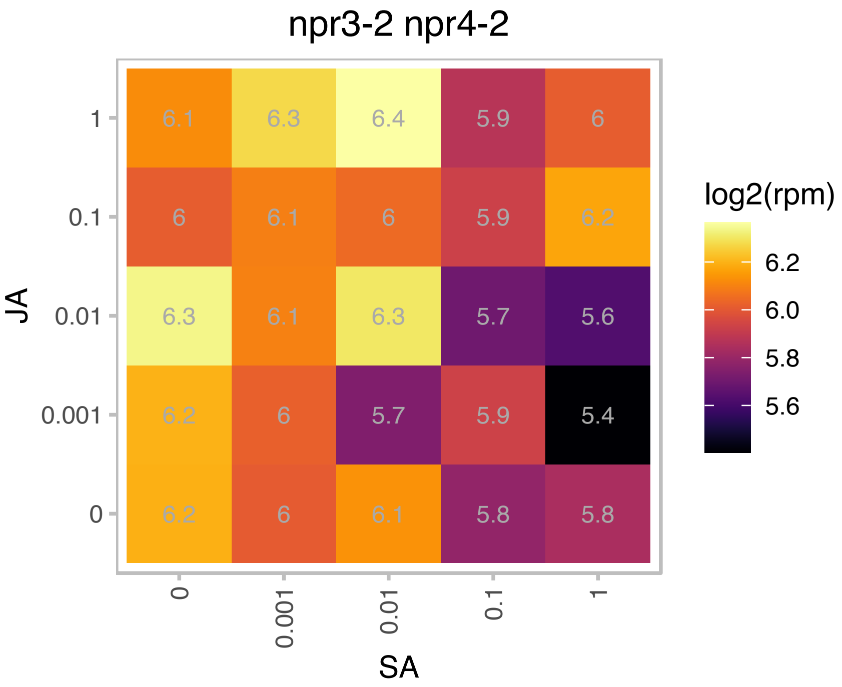


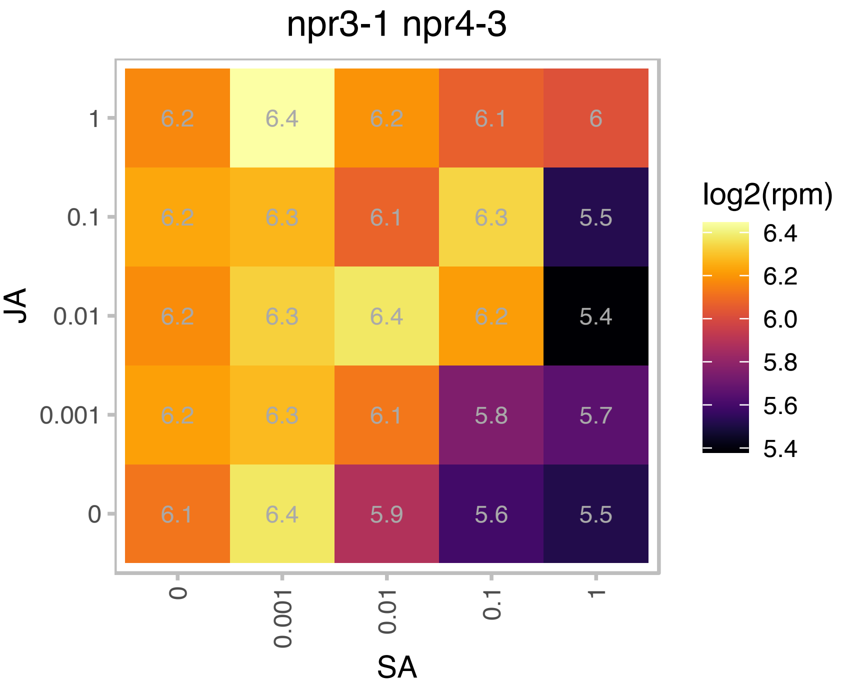

# AT1G80180.1 : NA

Encodes a substrate of the MAPK kinases. Phenotypic analyses of Arabidopsis expressing phosphorylation site mutant forms of At1g80180.1 showed clustered stomata and higher stomatal index in cotyledons expressing the phosphomimetic form of At1g80180.1.

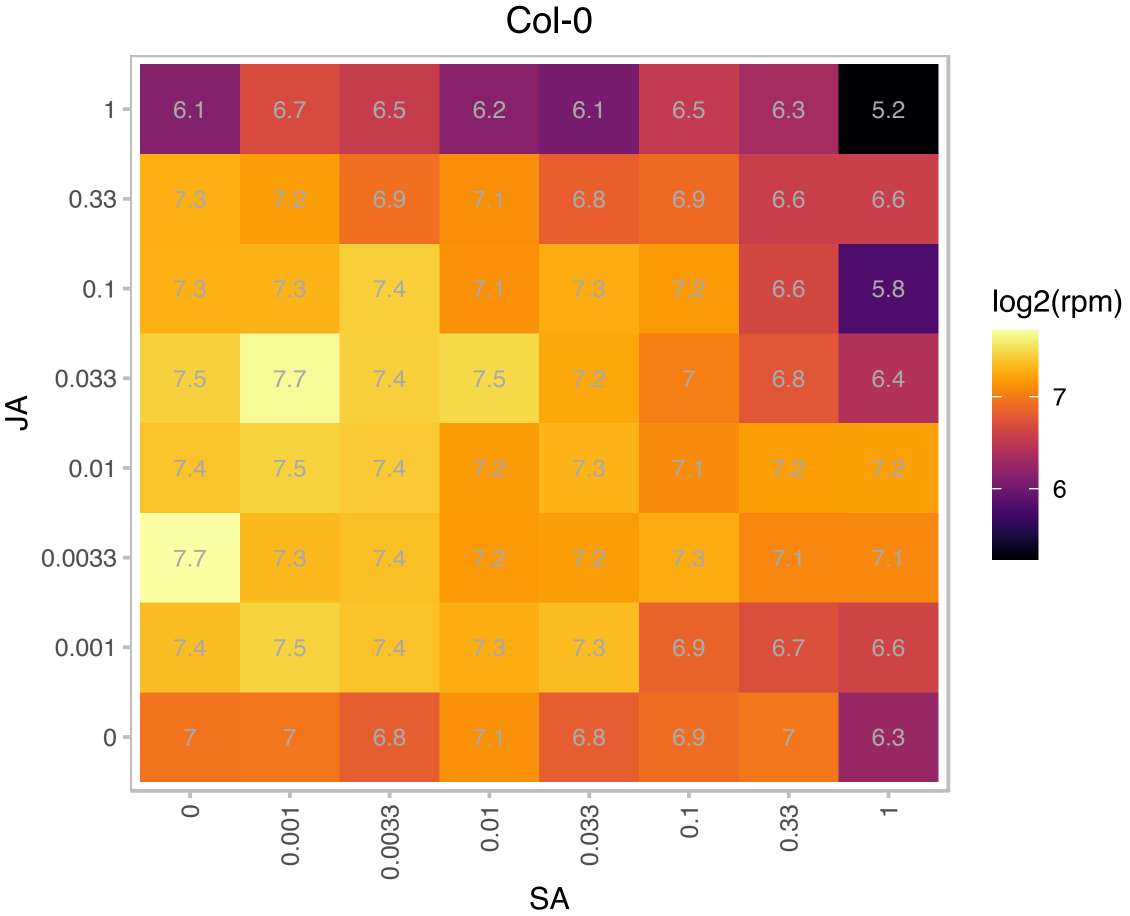


| | |
|---|---|
| Gene | AT1G80180 |
| is.expressed_genes | TRUE |
| is.responsive_genes | TRUE |
| is.apcluster | F-07 |

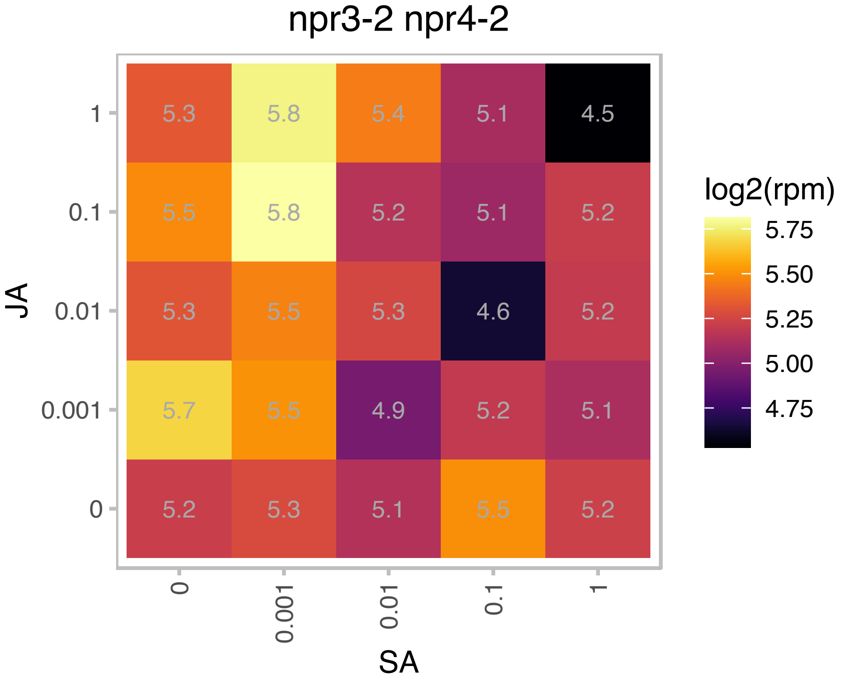


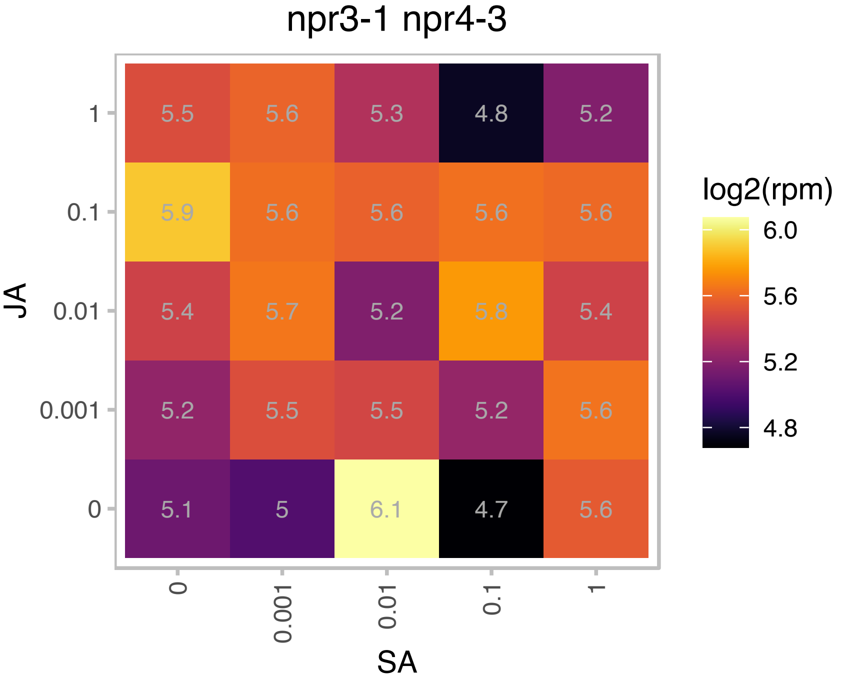

### AT1G80190.1 : PSF1

partner of SLD five 1

Similar to the PSF1 component of GINS complex, which in other organism was shown to be involved in the initiation of DNA replication.

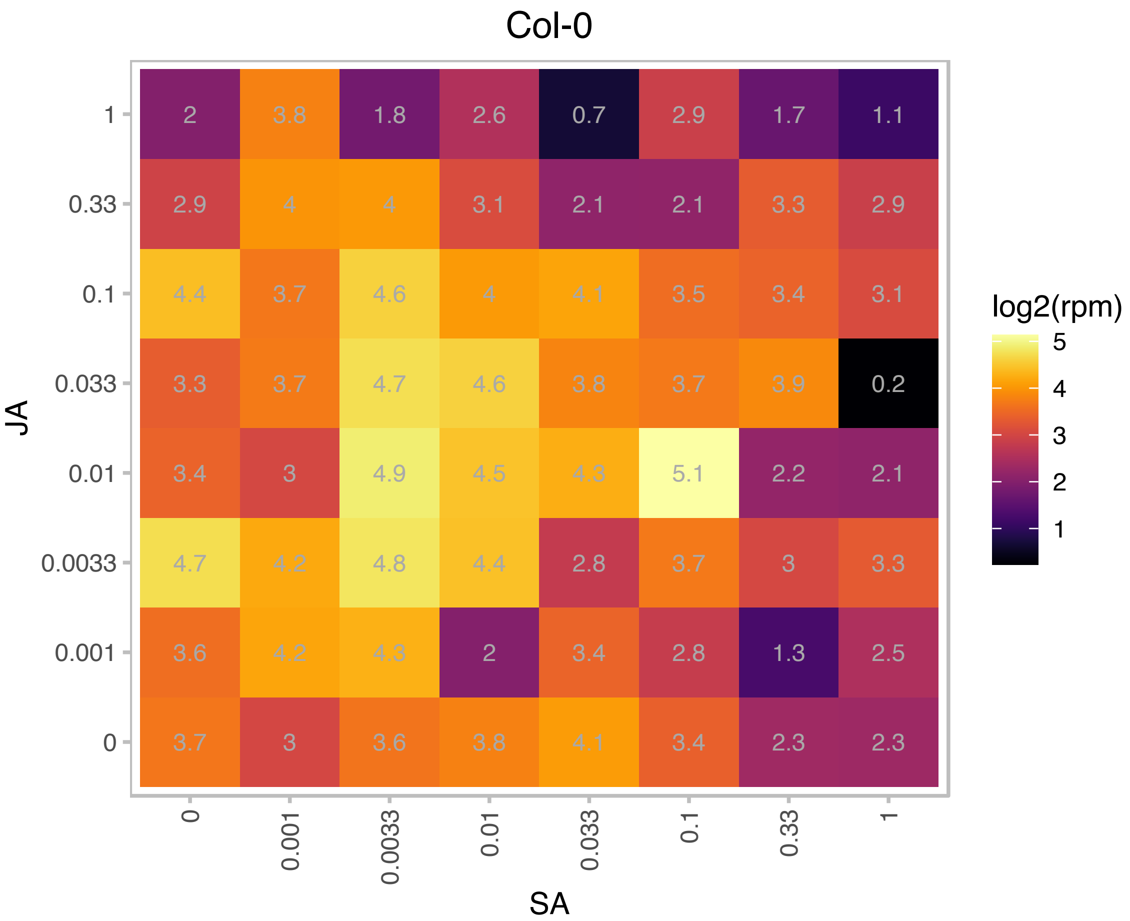


| | |
|---|---|
| Gene | AT1G80190 |
| is.expressed_genes | TRUE |
| is.responsive_genes | TRUE |
| is.apcluster | F-07 |

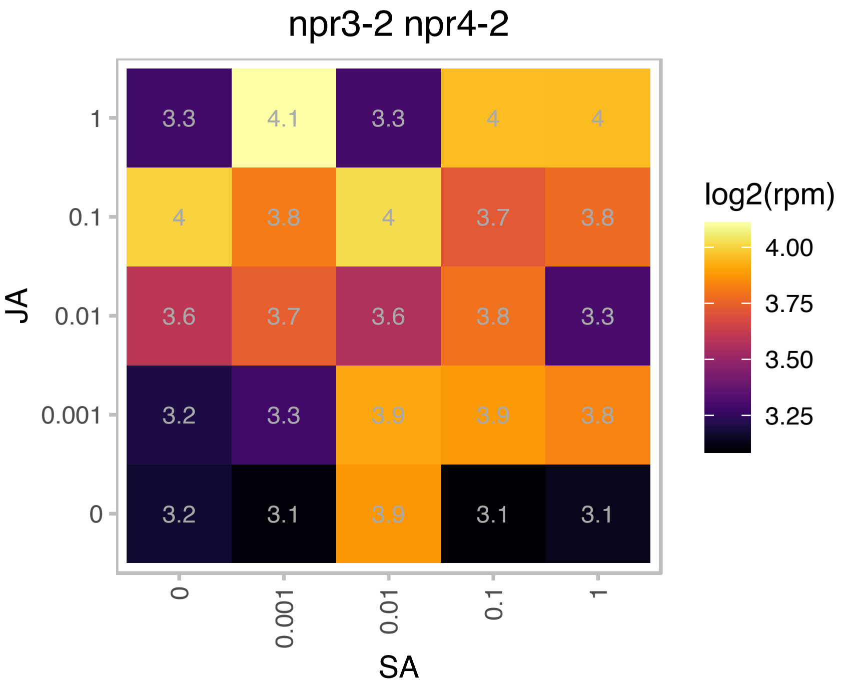


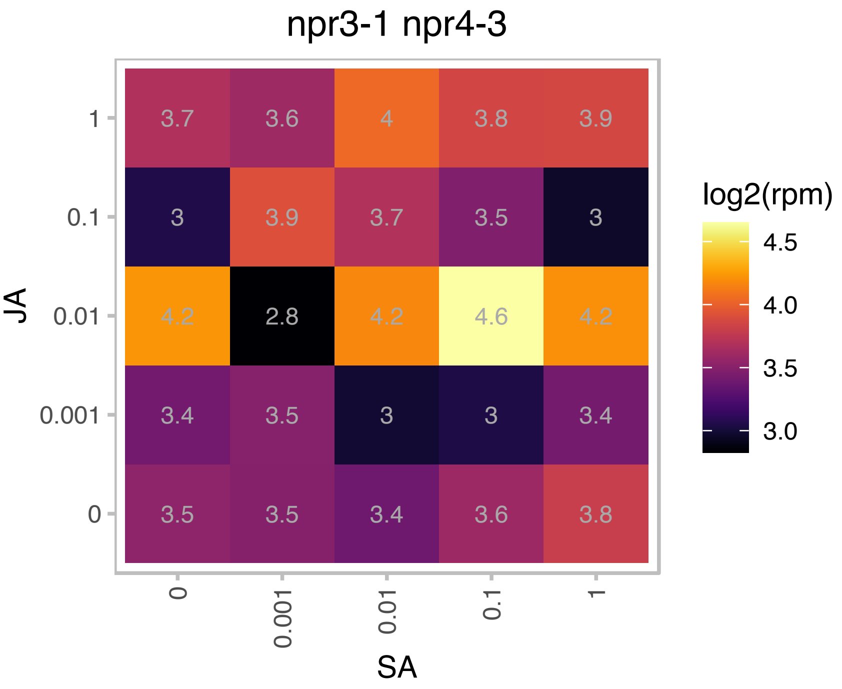

### AT1G80770.1 : PDE318

P-loop containing nucleoside triphosphate hydrolases superfamily protein

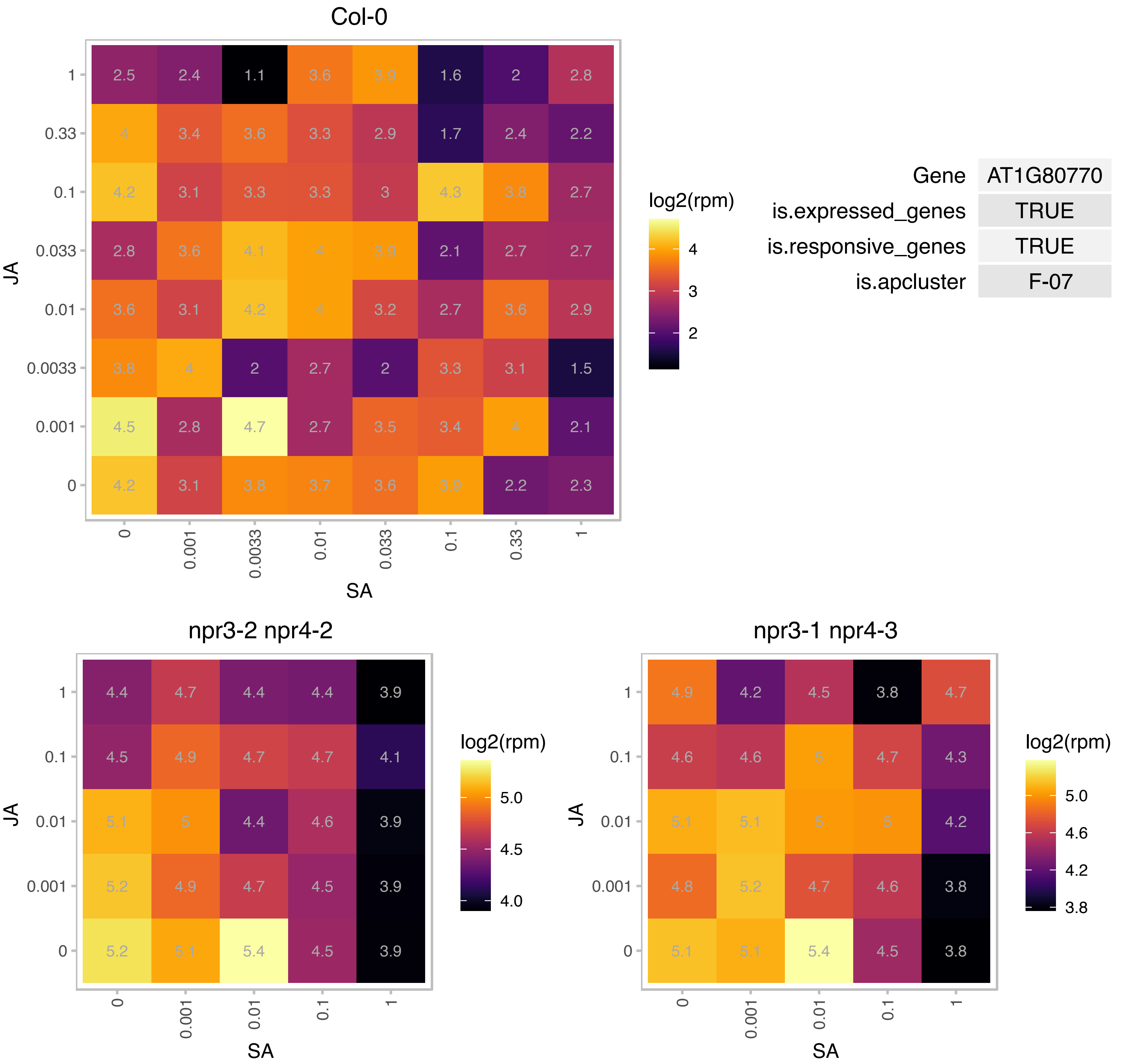


| | |
|---|---|
| Gene | AT1G80770 |
| is.expressed_genes | TRUE |
| is.responsive_genes | TRUE |
| is.apcluster | F-07 |

### AT1G80860.1 : ATPLMT, PLMT

phospholipid N-methyltransferase
Encodes a single-copy phospholipid N-methyltransferase, involved in phosphatidylcholine biosynthesis. Has specific activity towards phosphatidylmonomethylethanolamine and phosphatidyldimethylethanolamine, but not phosphatidylethanolamine.

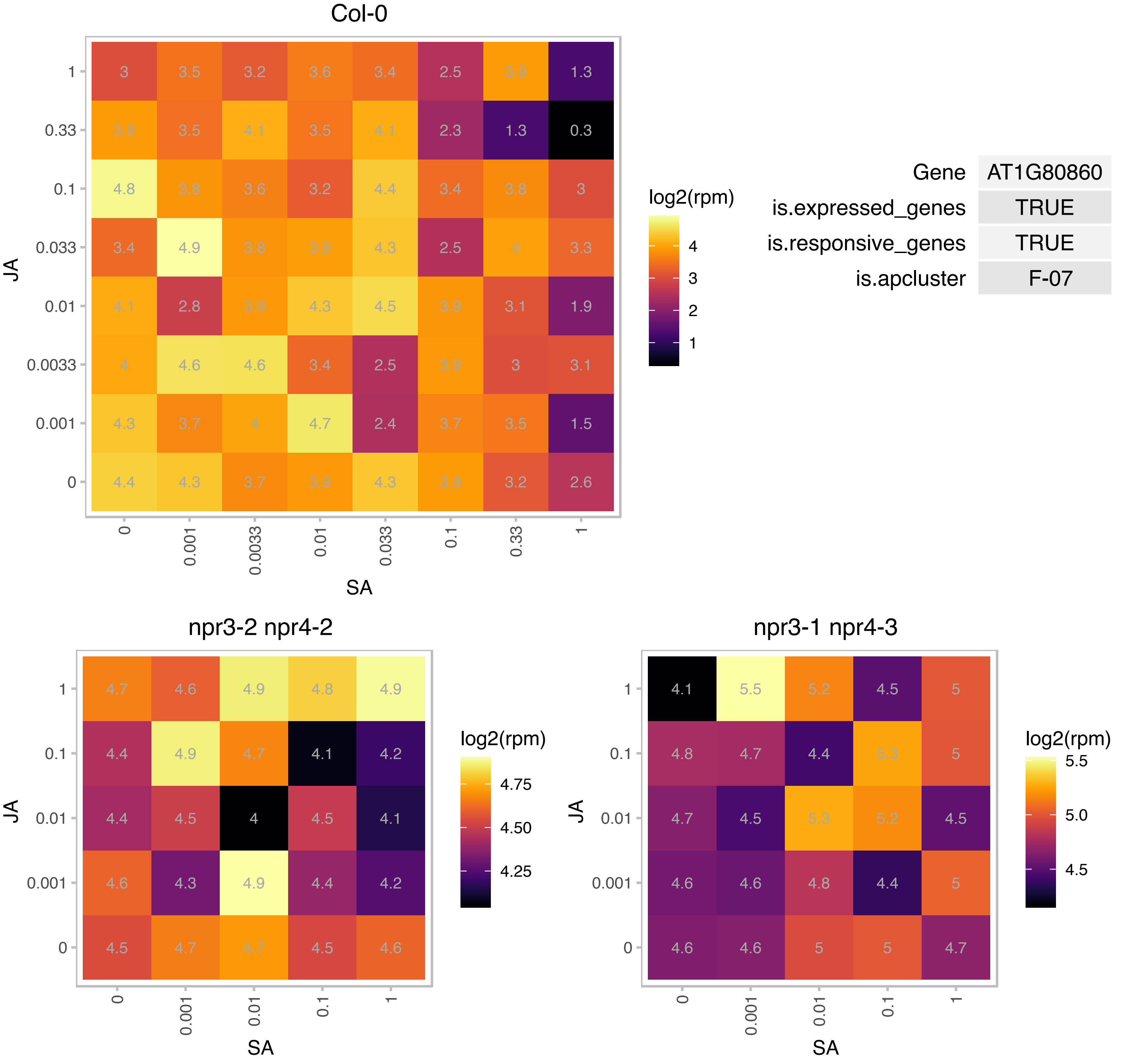


| | |
|---|---|
| Gene | AT1G80860 |
| is.expressed_genes | TRUE |
| is.responsive_genes | TRUE |
| is.apcluster | F-07 |

### AT2G01250.1 : RPL7B

Ribosomal protein L30/L7 family protein

Cytosolic ribosomal 60S subunit protein.

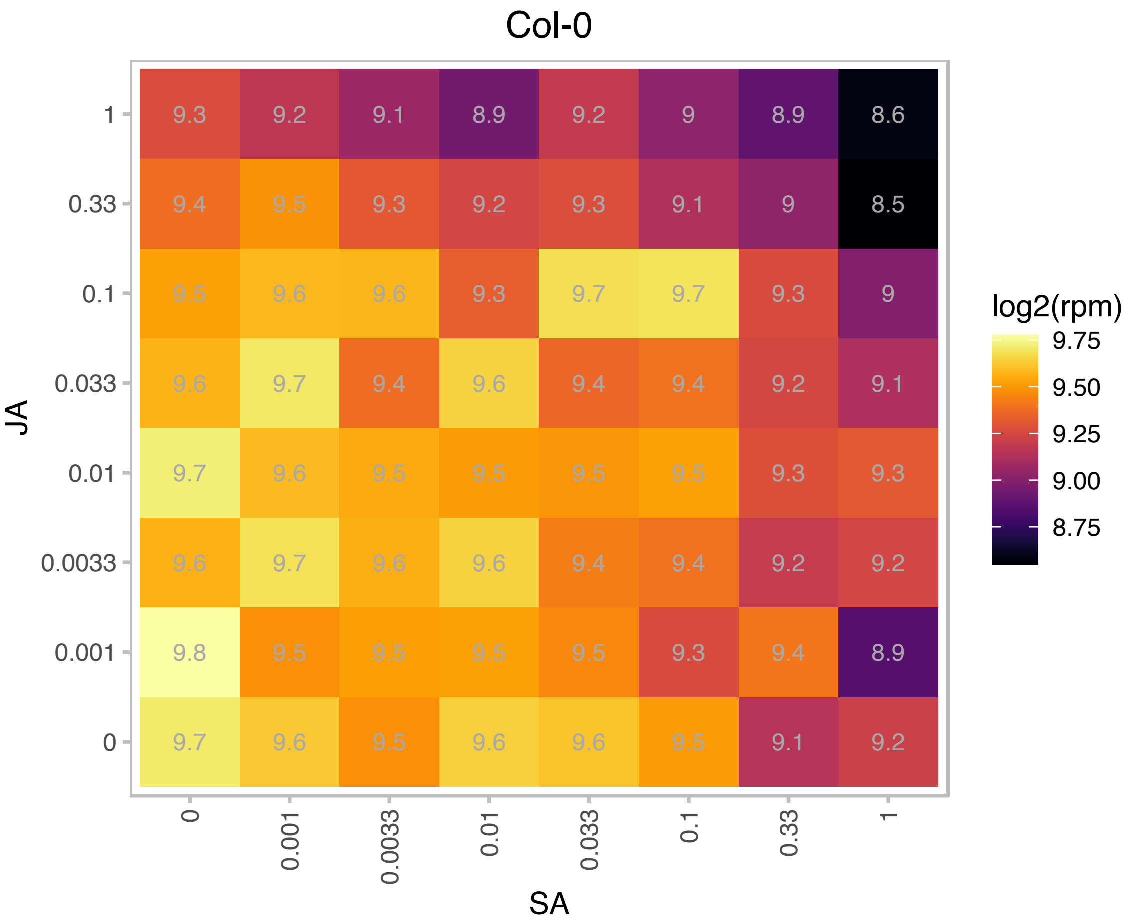


| | |
|---|---|
| Gene | AT2G01250 |
| is.expressed_genes | TRUE |
| is.responsive_genes | TRUE |
| is.apcluster | F-07 |

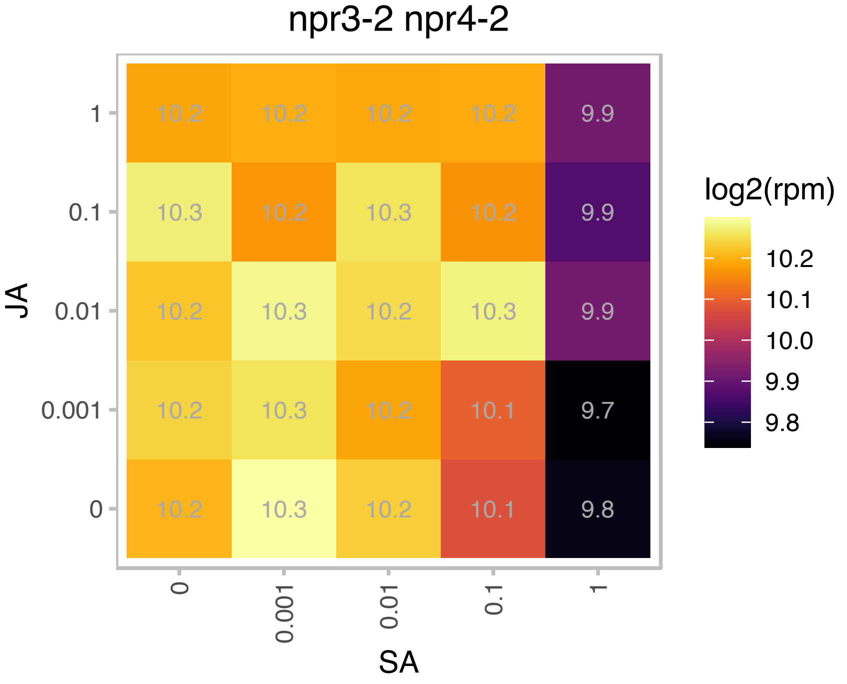


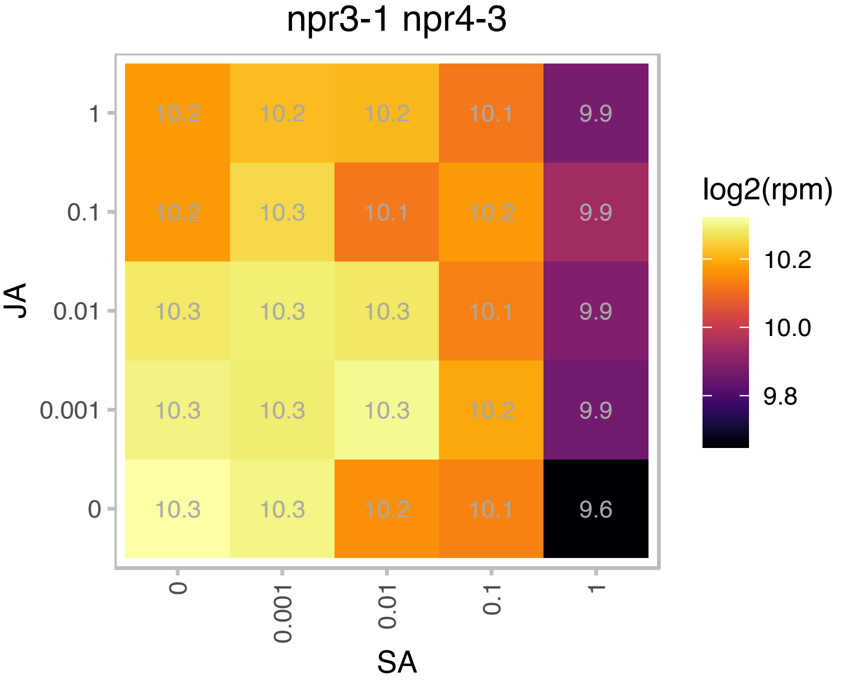

### AT2G02100.1 : LCR69, PDF2.2

Predicted to encode a PR (pathogenesis-related) protein. Belongs to the plant defensin (PDF) family with the following members: At1g75830/PDF1.1, At5g44420/PDF1.2a, At2g26020/PDF1.2b, At5g44430/PDF1.2c, At2g26010/PDF1.3, At1g19610/PDF1.4, At1g55010/PDF1.5, At2g02120/PDF2.1, At2g02100/PDF2.2, At2g02130/PD At1g61070/PDF2.4, At5g63660/PDF2.5, At2g02140/PDF2.6, At5g38330/PDF3.1 and At4g30070/PDF3.2. The mRN cell-to-cell mobile.

low-molecular-weight cysteine-rich 69

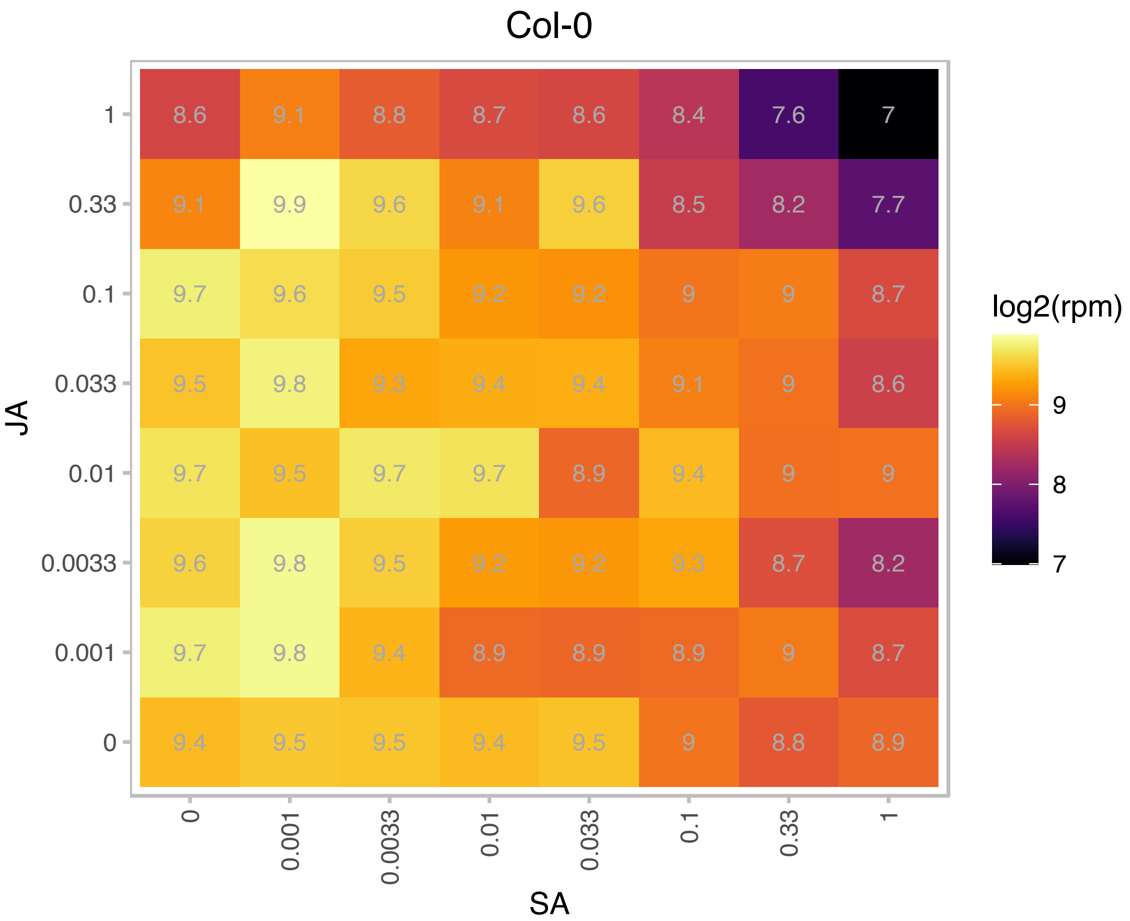


| | |
|---|---|
| Gene | AT2G02100 |
| is.expressed_genes | TRUE |
| is.responsive_genes | TRUE |
| is.apcluster | F-07 |

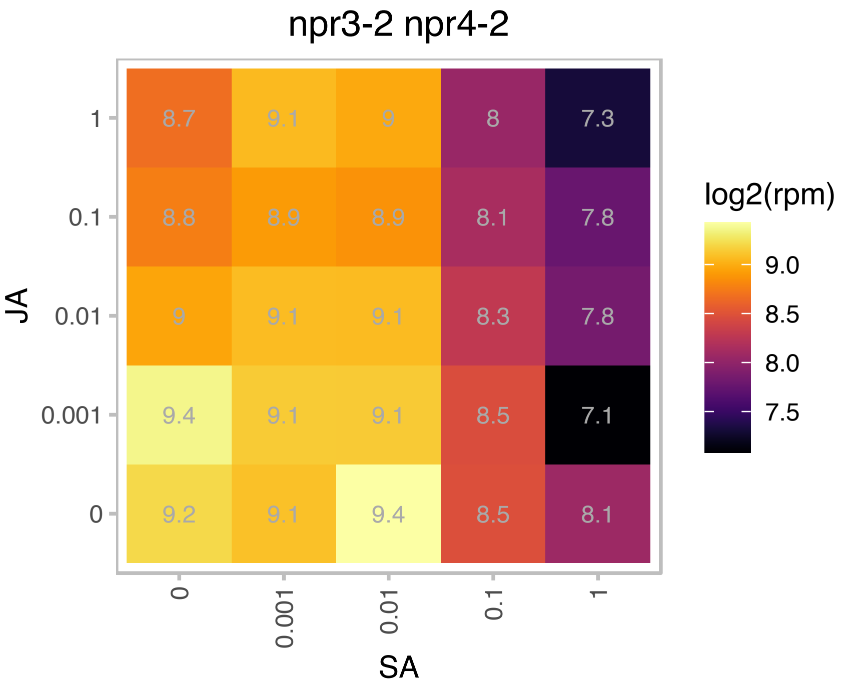


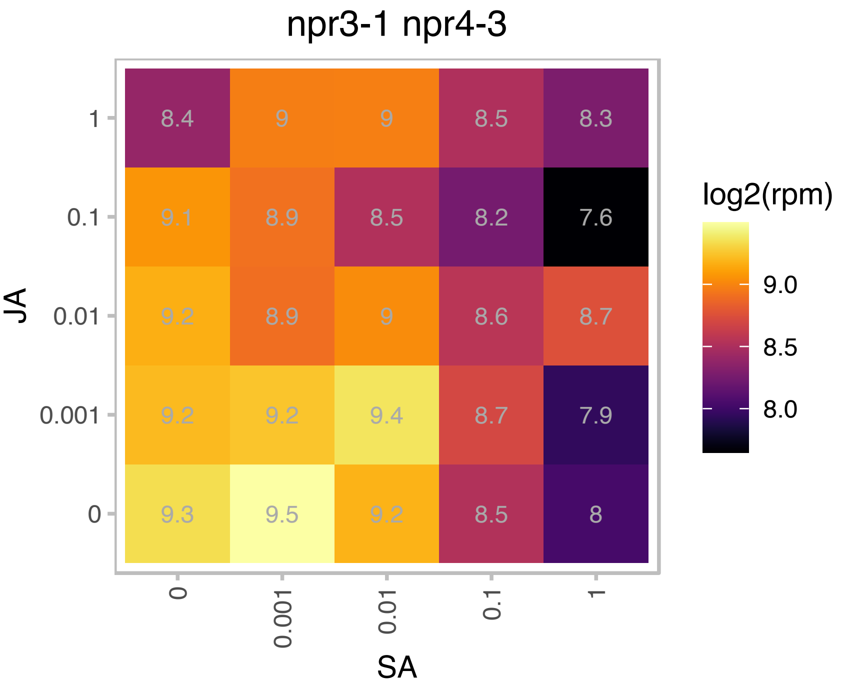

# AT2G03310.1 : NA

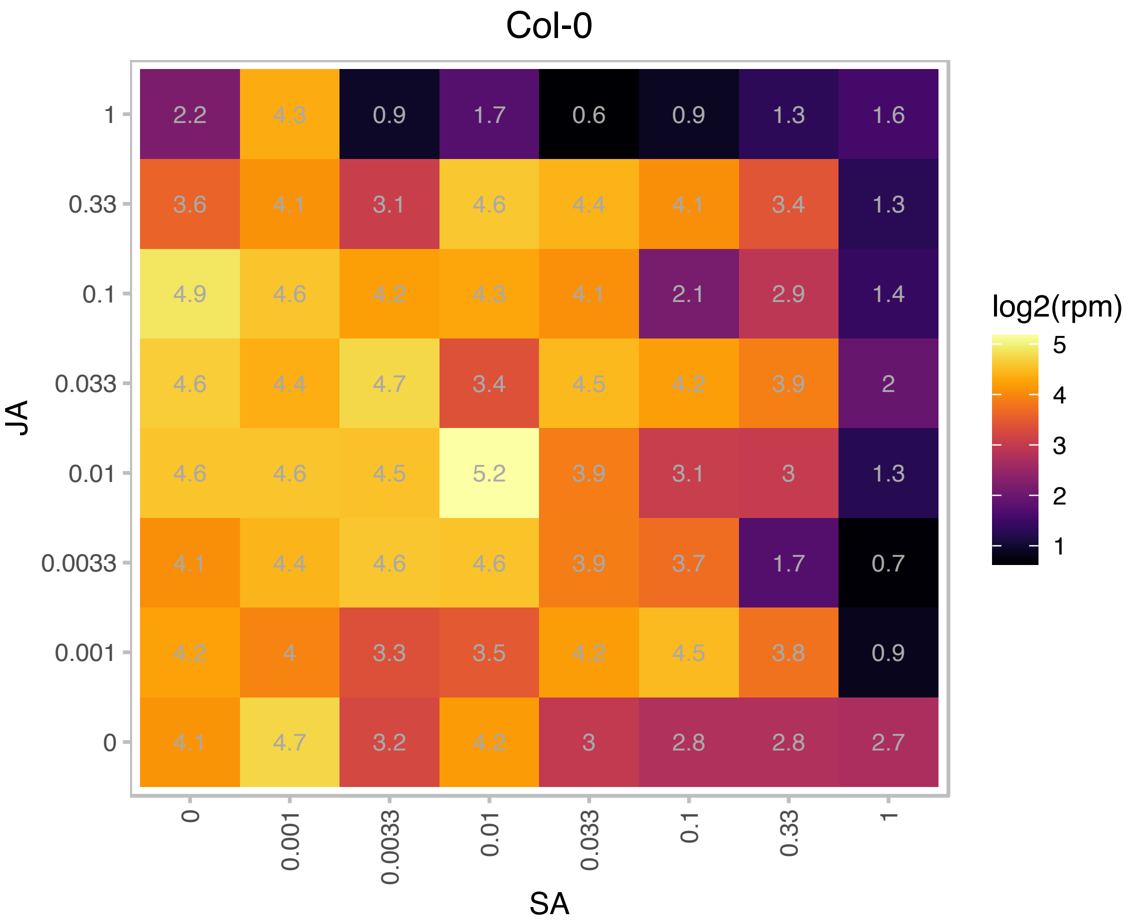


| | |
|---|---|
| Gene | AT2G03310 |
| is.expressed_genes | TRUE |
| is.responsive_genes | TRUE |
| is.apcluster | F-07 |

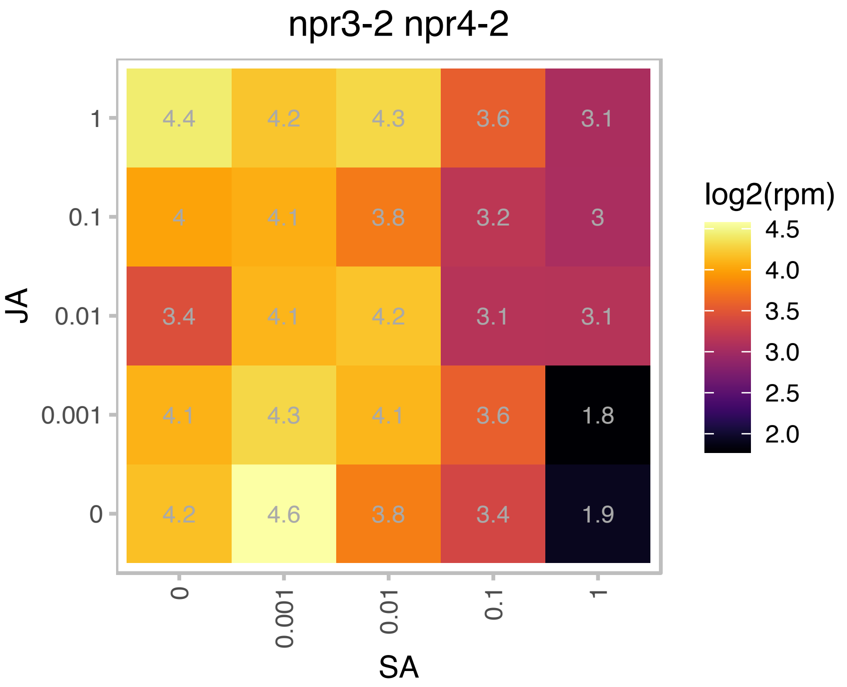


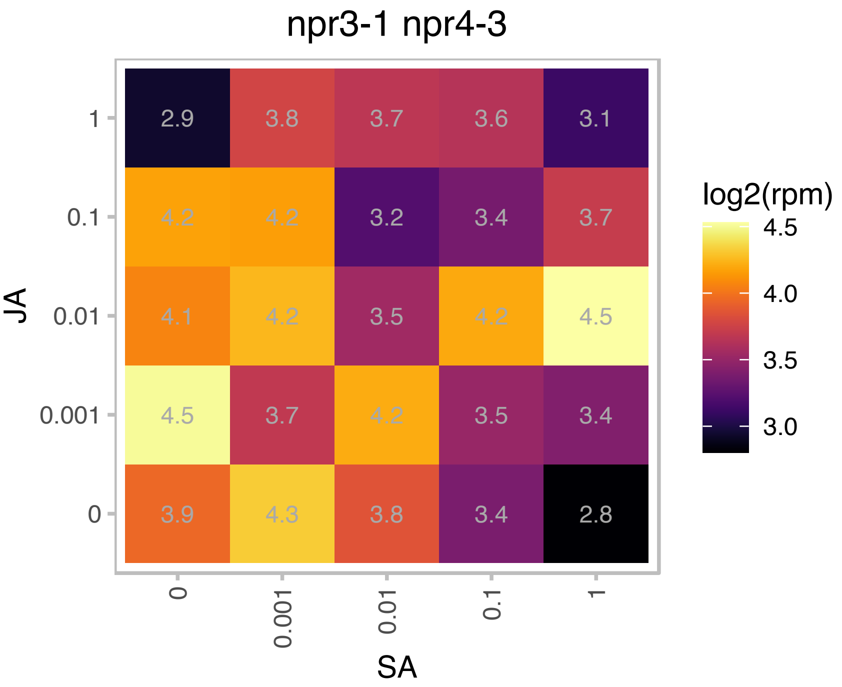

# AT2G05220.1 : NA

Ribosomal S17 family protein

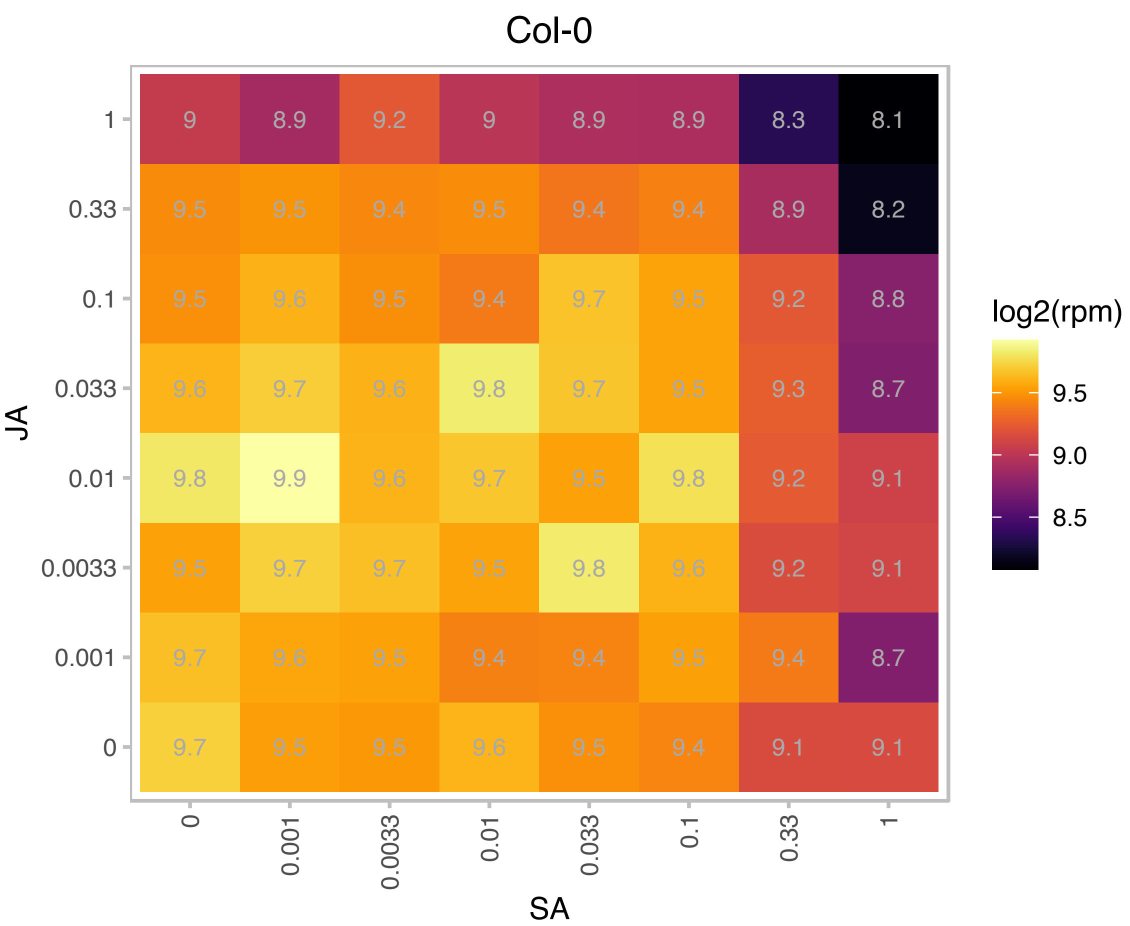


| | |
|---|---|
| Gene | AT2G05220 |
| is.expressed_genes | TRUE |
| is.responsive_genes | TRUE |
| is.apcluster | F-07 |

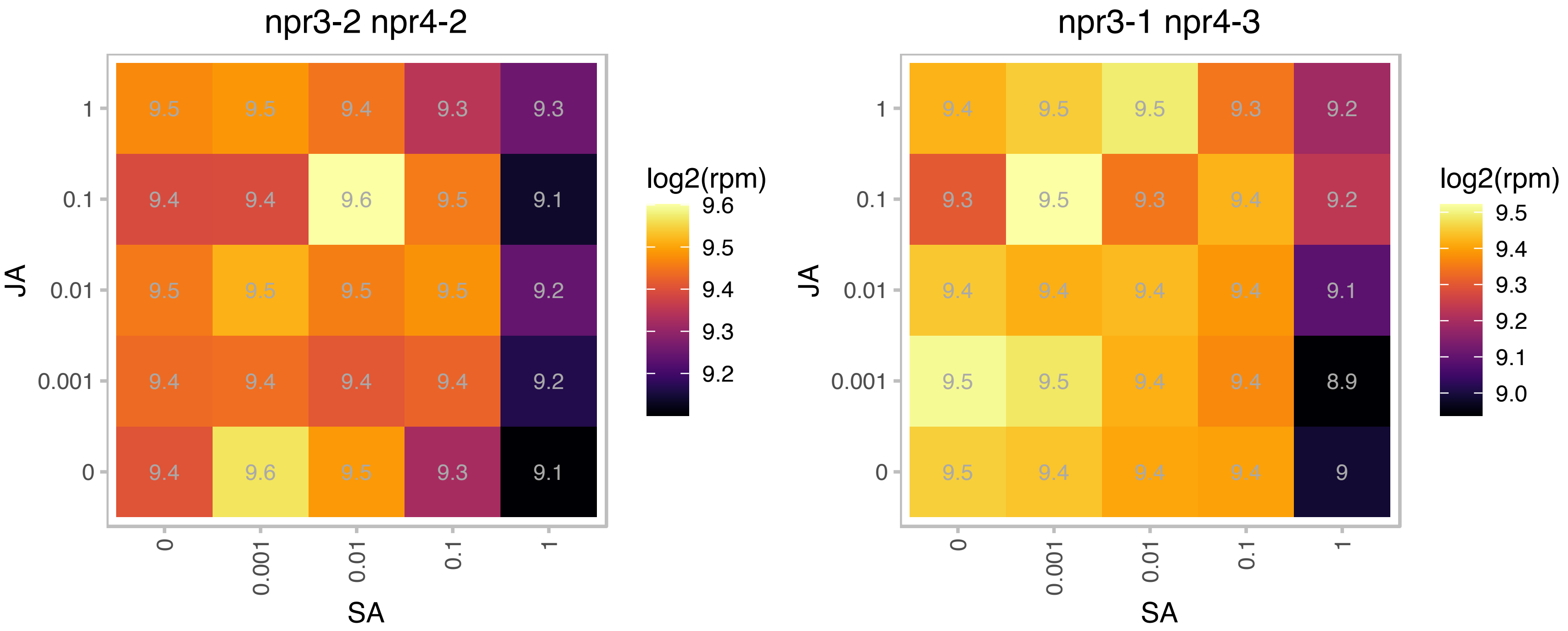

### AT2G07340.1 : PFD1

PREFOLDIN 1

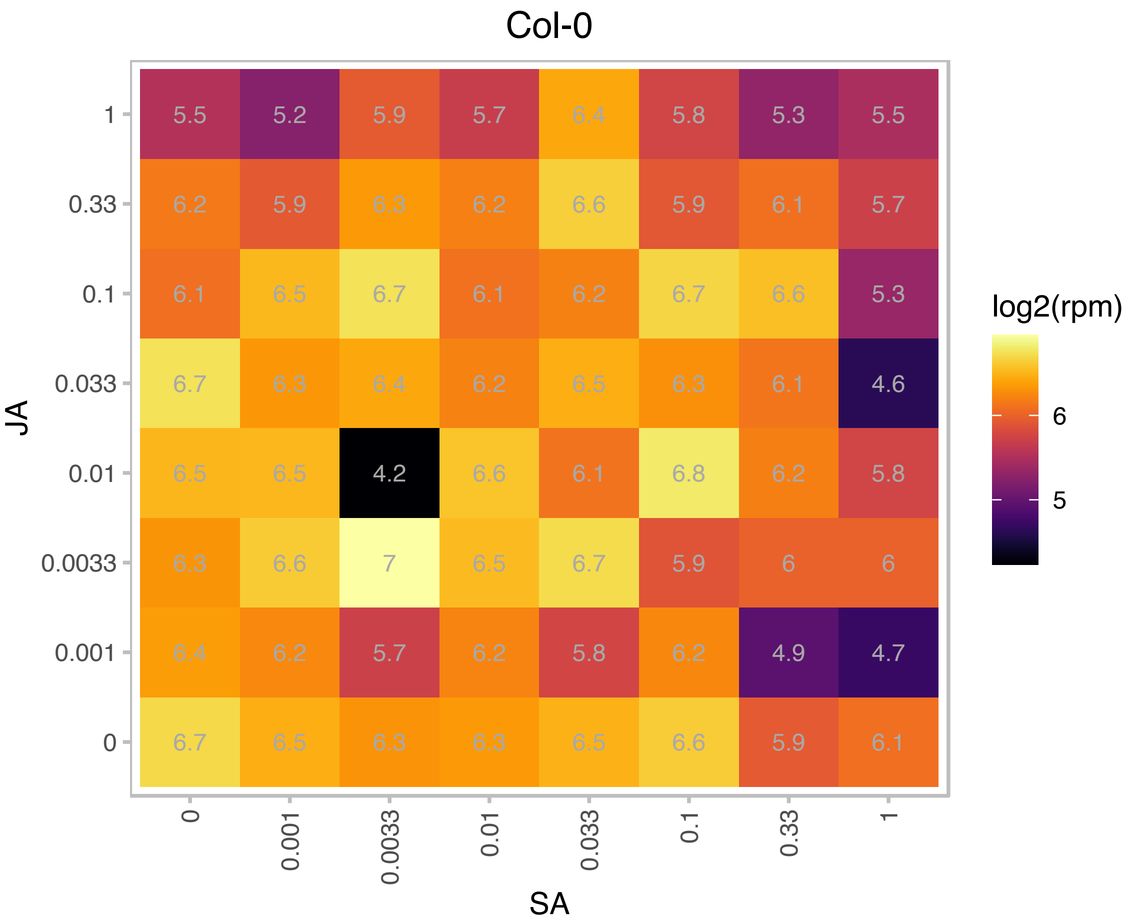


| | |
|---|---|
| Gene | AT2G07340 |
| is.expressed_genes | TRUE |
| is.responsive_genes | TRUE |
| is.apcluster | F-07 |

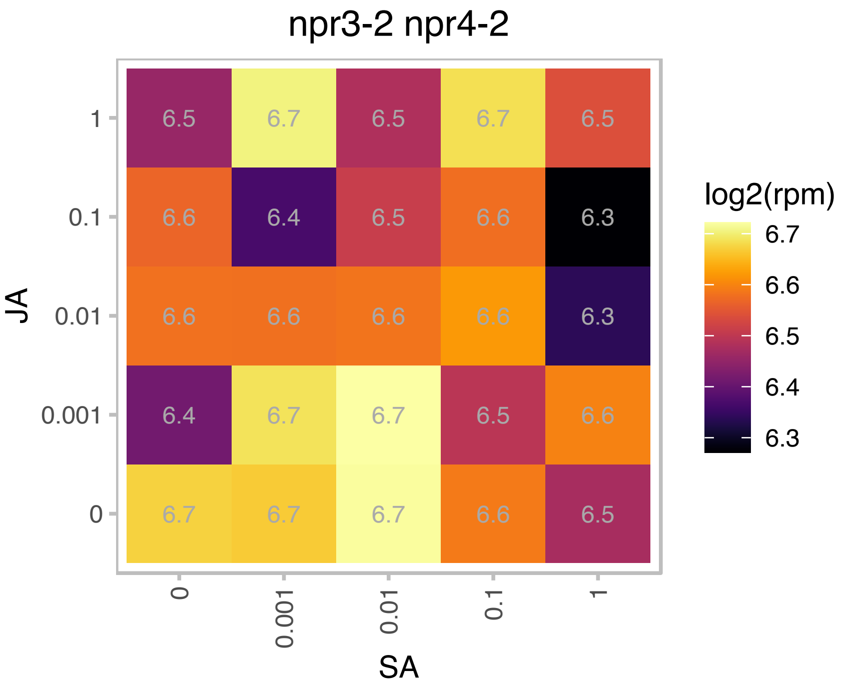


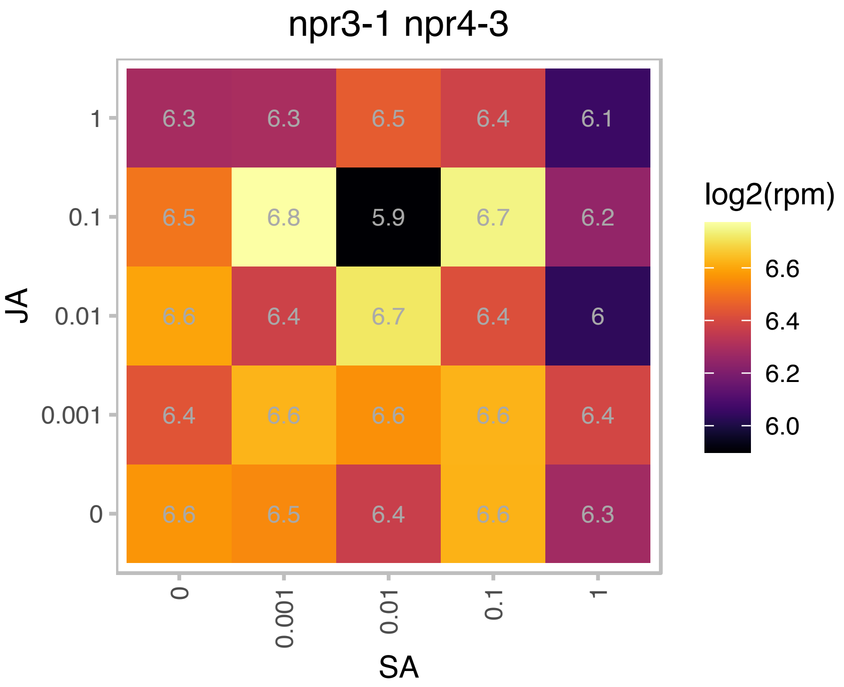

# AT2G17240.1 : NA

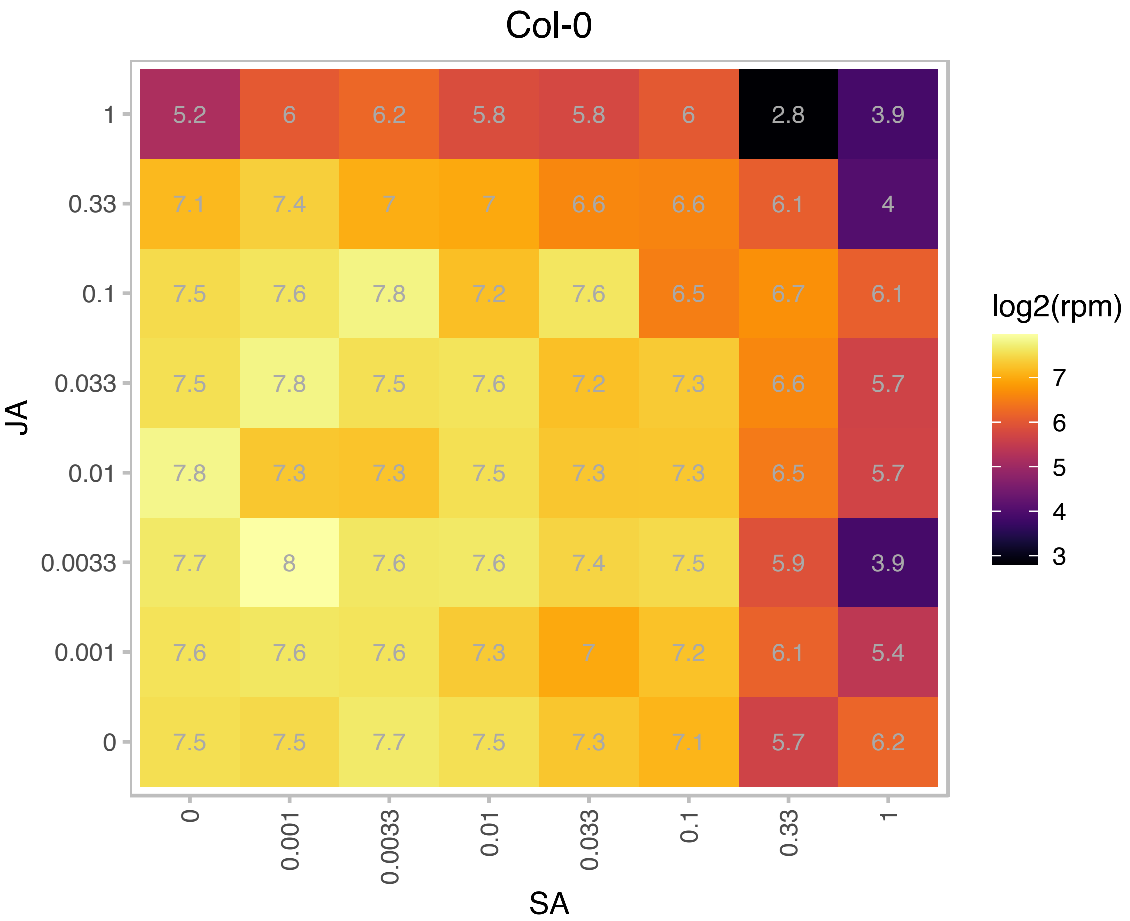


| | |
|---|---|
| Gene | AT2G17240 |
| is.expressed_genes | TRUE |
| is.responsive_genes | TRUE |
| is.apcluster | F-07 |

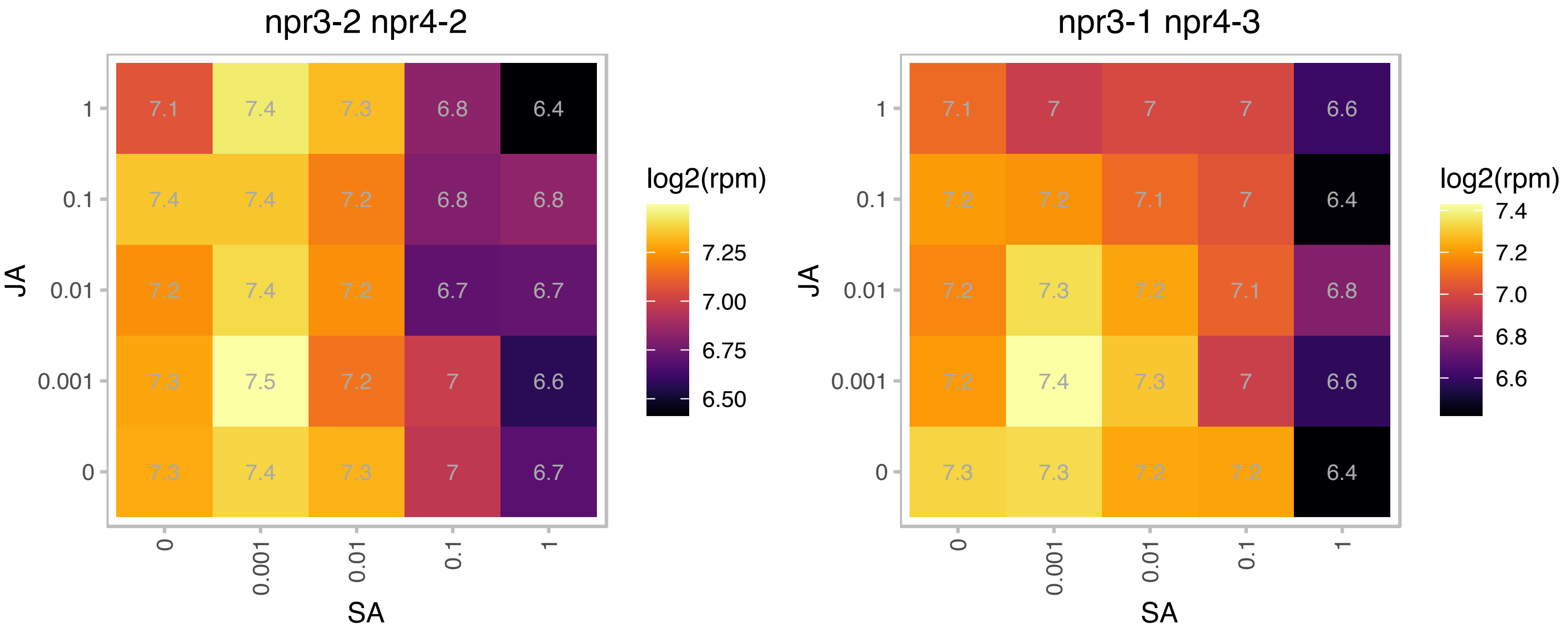

# AT2G17972.1 : NA

| | |
|---|---|
| Gene | AT2G17972 |
| is.expressed_genes | TRUE |
| is.responsive_genes | TRUE |
| is.apcluster | F-07 |

Col-0

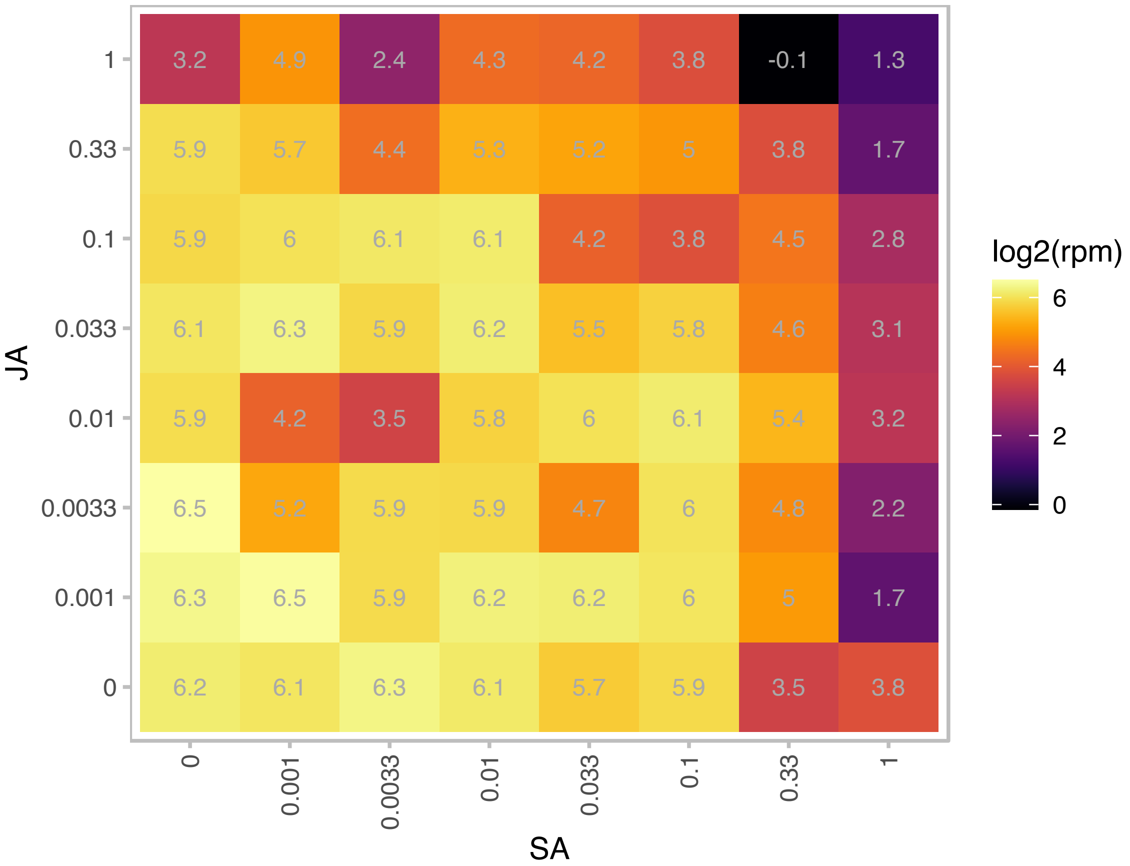


npr3-2 npr4-2

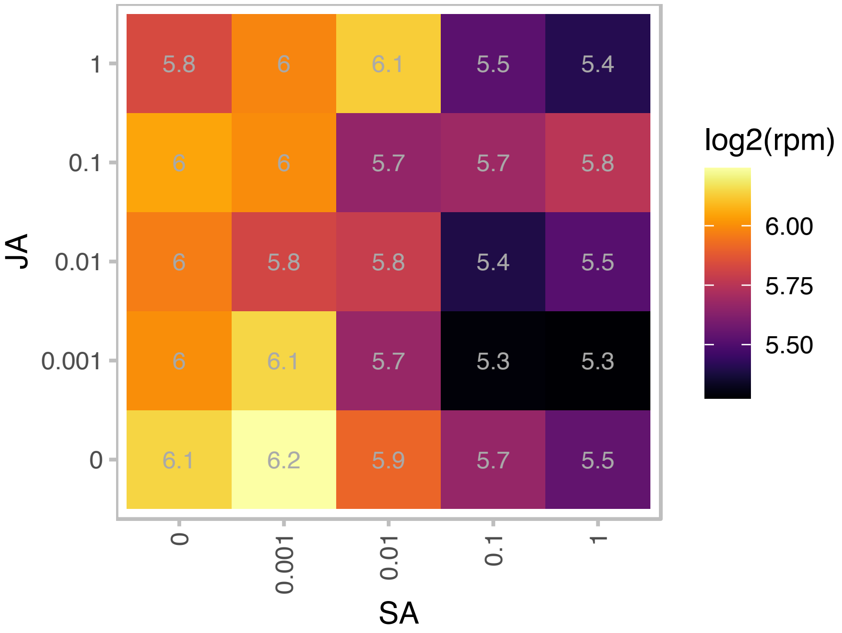


npr3-1 npr4-3

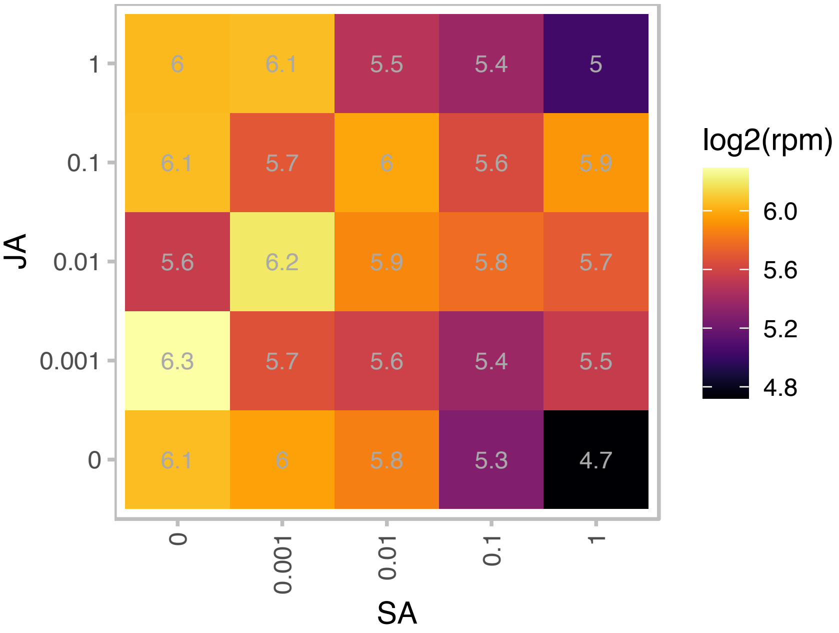

### AT2G18020.1 : EMB2296

Ribosomal protein L2 family

Col-0

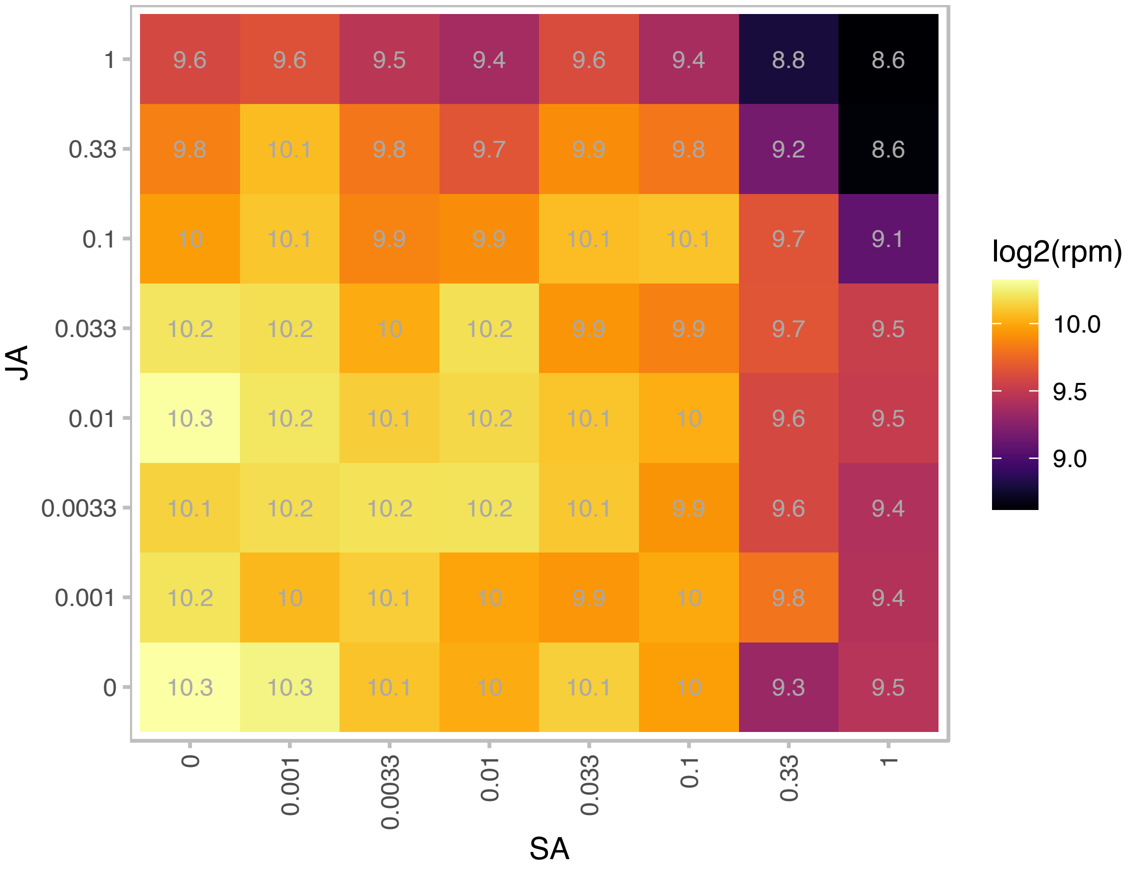


| | |
|---|---|
| Gene | AT2G18020 |
| is.expressed_genes | TRUE |
| is.responsive_genes | TRUE |
| is.apcluster | F-07 |

npr3-2 npr4-2

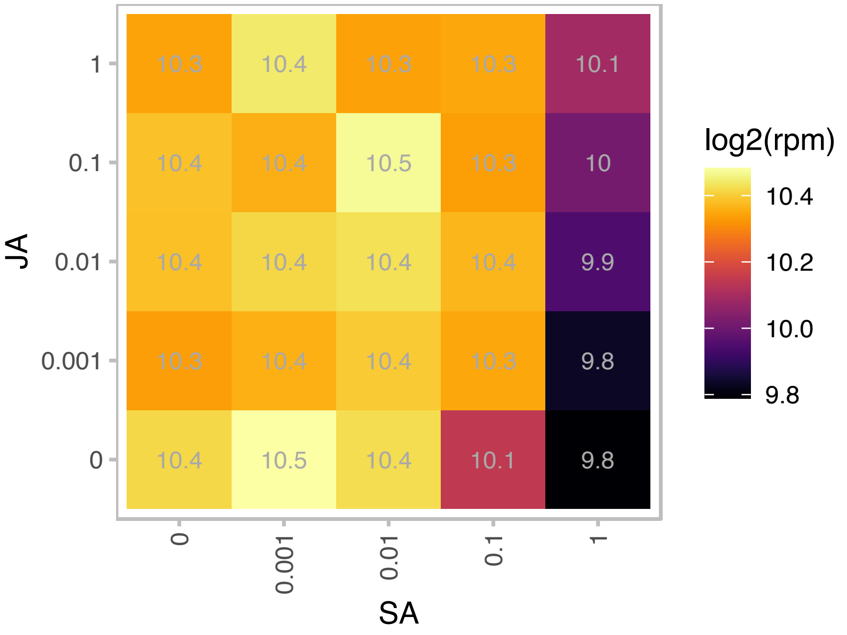


npr3-1 npr4-3

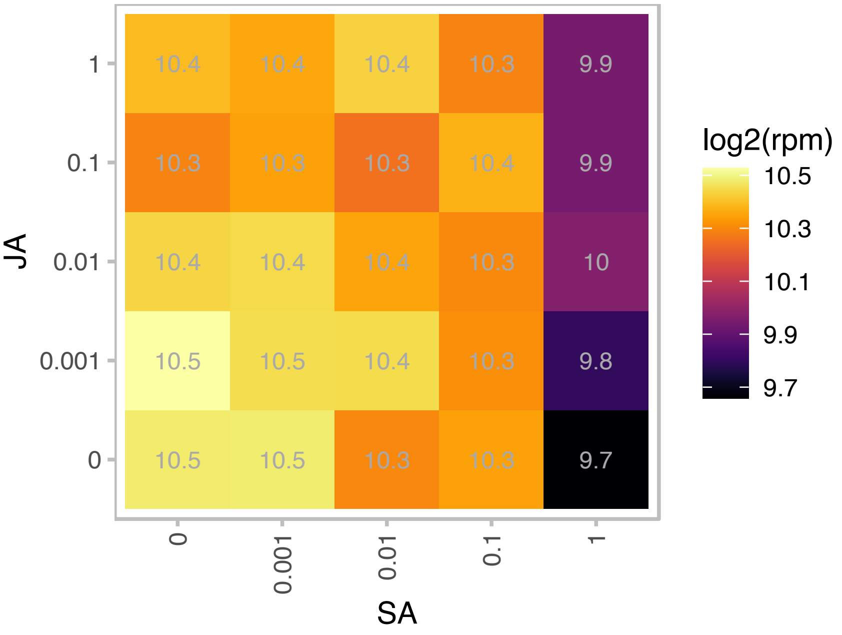

# AT2G19730.1 : NA

Ribosomal L28e protein family

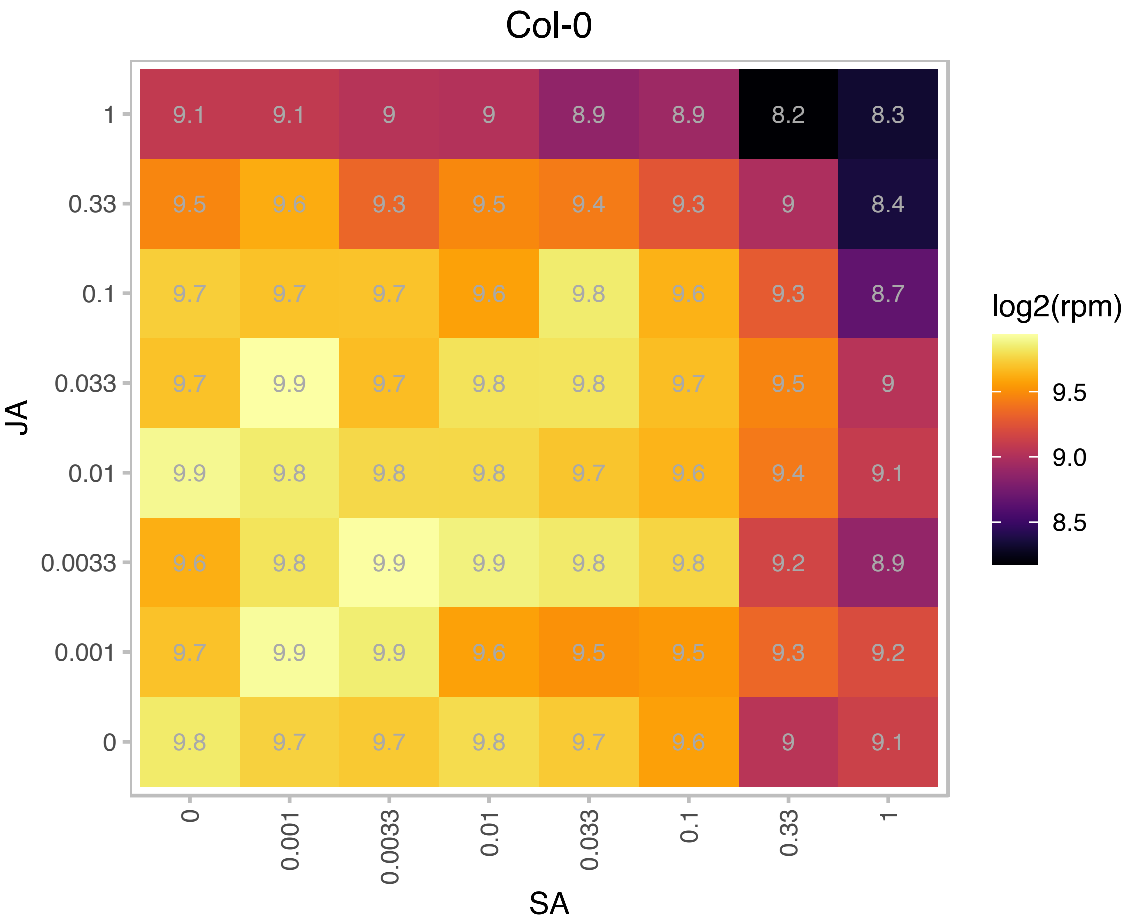


| | |
|---|---|
| Gene | AT2G19730 |
| is.expressed_genes | TRUE |
| is.responsive_genes | TRUE |
| is.apcluster | F-07 |

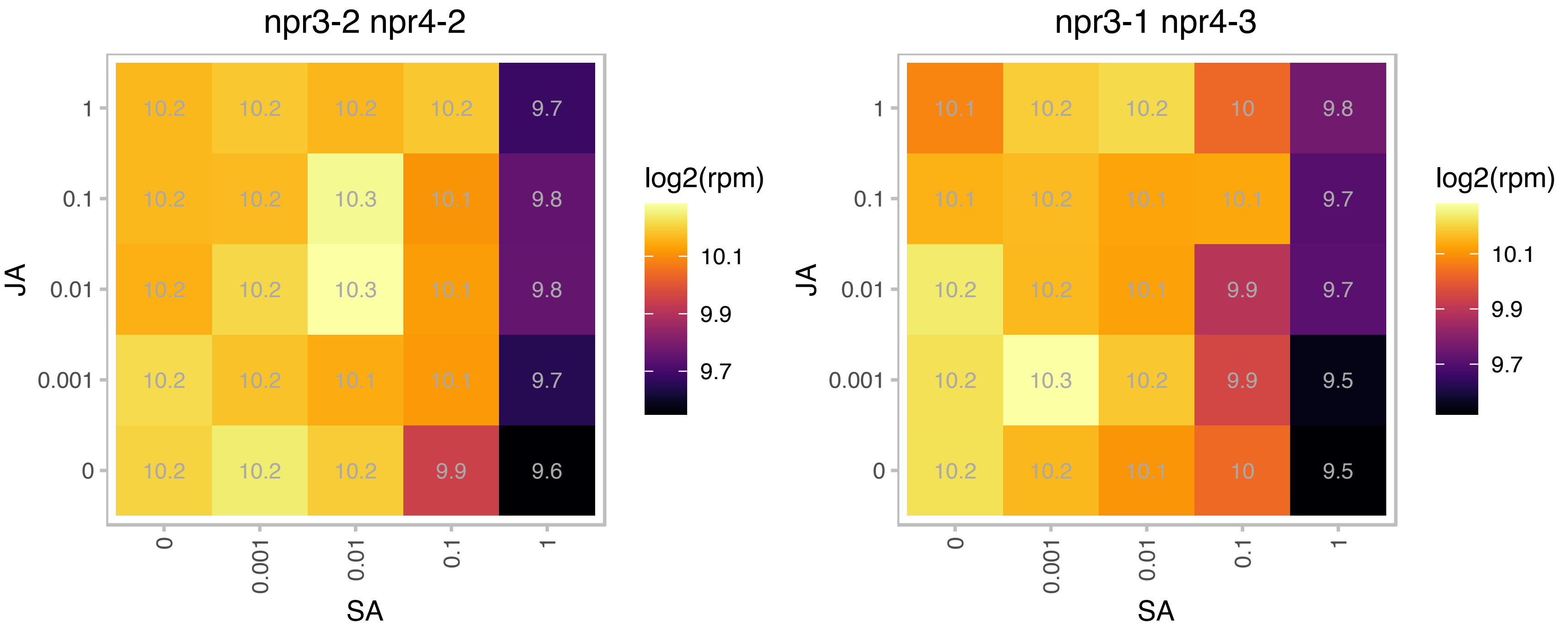

# AT2G19750.1 : NA

Ribosomal protein S30 family protein

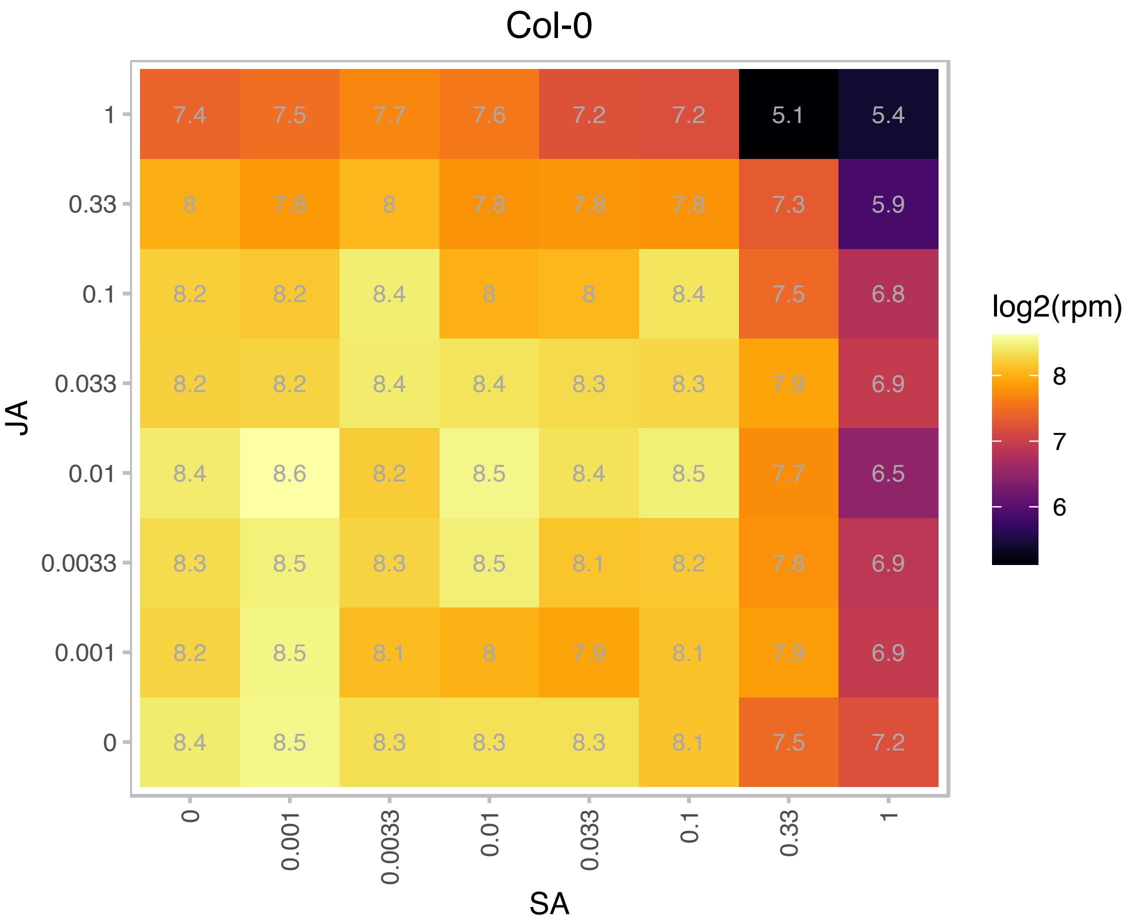


| | |
|---|---|
| Gene | AT2G19750 |
| is.expressed_genes | TRUE |
| is.responsive_genes | TRUE |
| is.apcluster | F-07 |

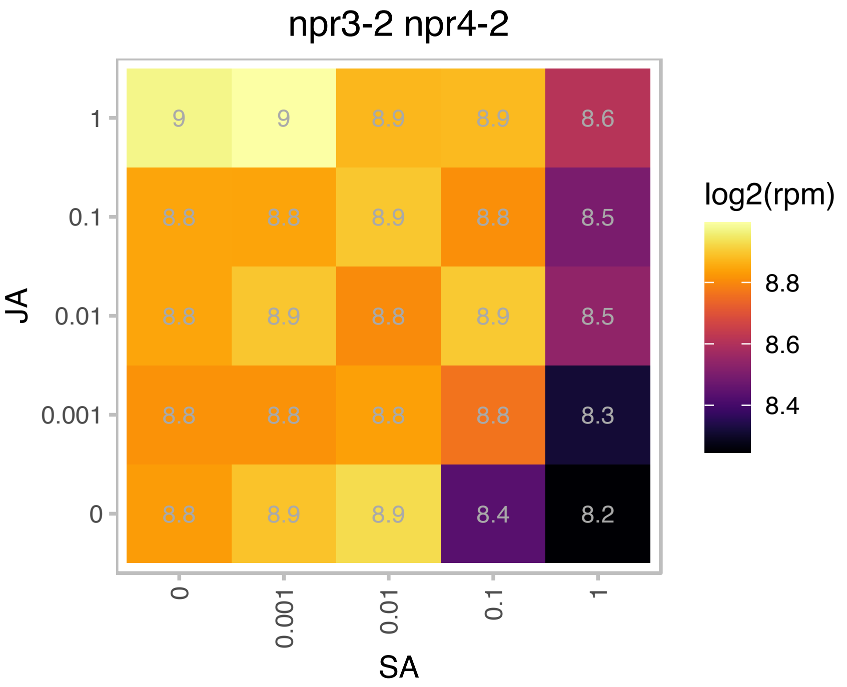


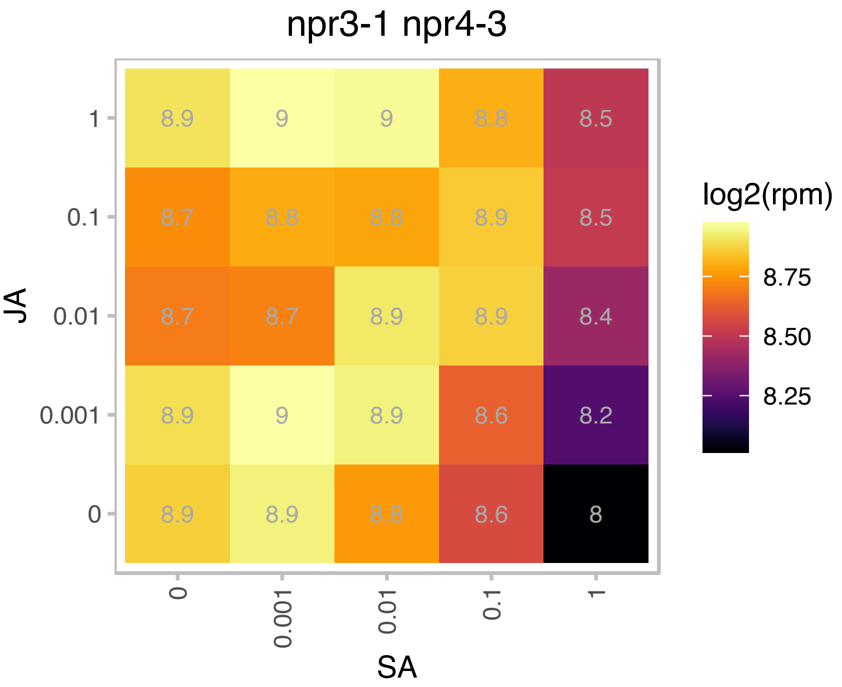

# AT2G20450.1 : NA

Ribosomal protein L14

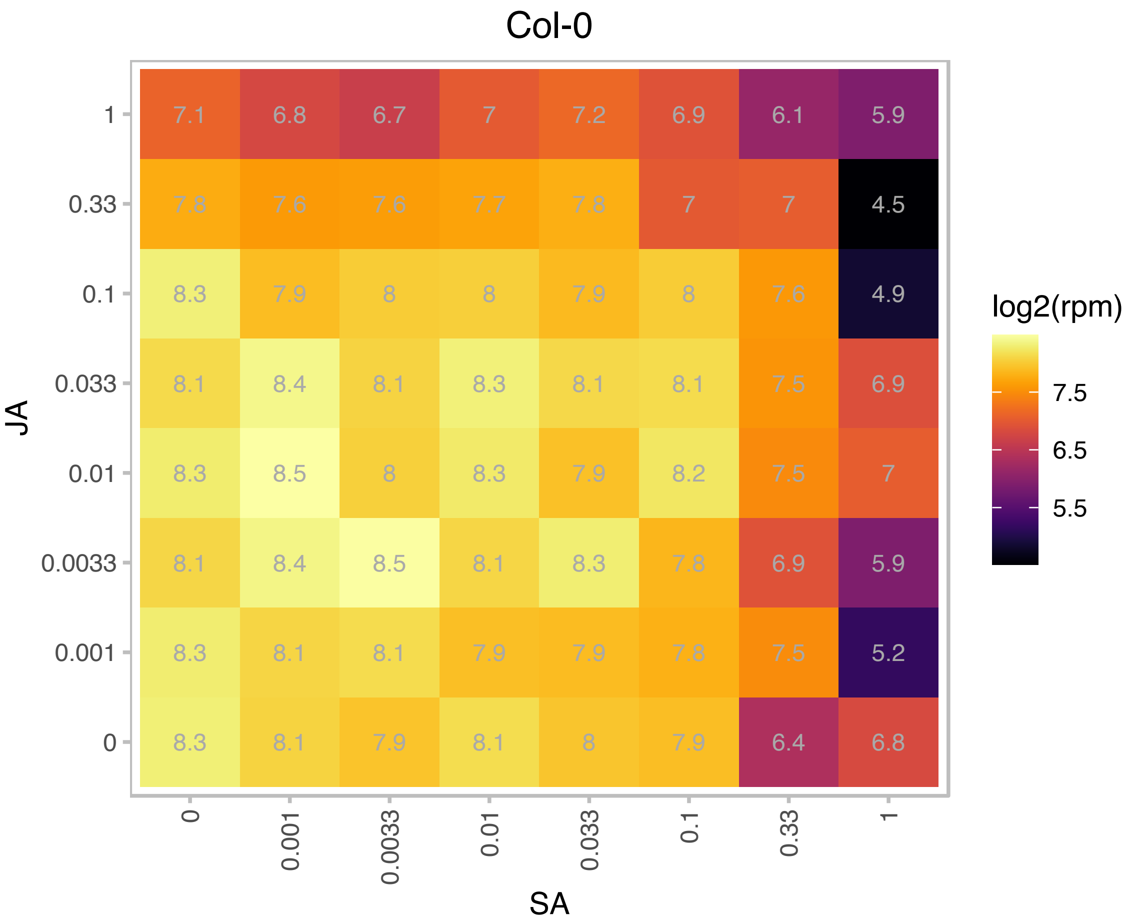


| | |
|---|---|
| Gene | AT2G20450 |
| is.expressed_genes | TRUE |
| is.responsive_genes | TRUE |
| is.apcluster | F-07 |

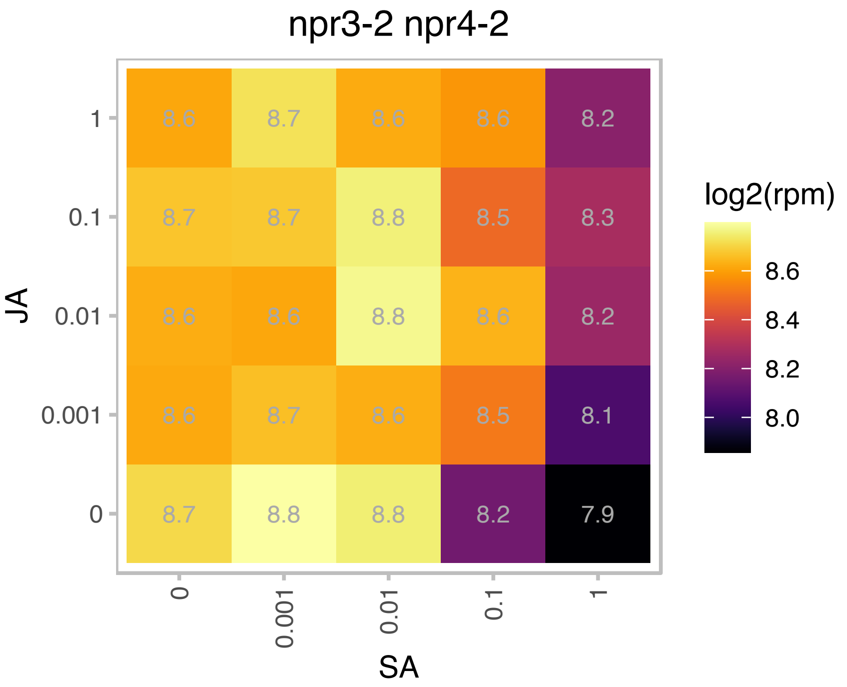


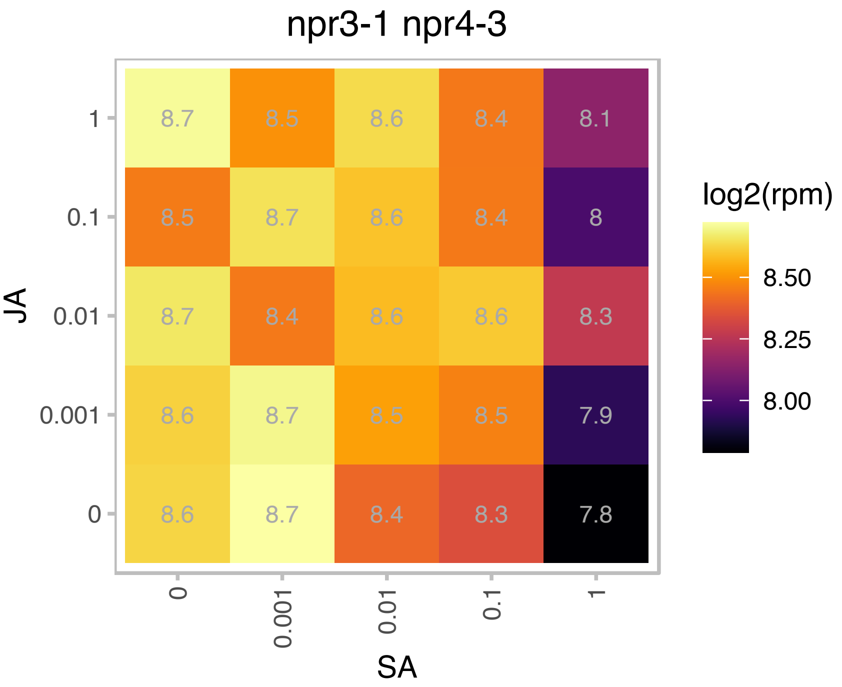

# AT2G20690.1 : NA

lumazine-binding family protein
A synthetic gene encoding the catalytic domain of the Arabidopsis thaliana gene At2g20690 was recombinant expressed in E. coli demonstrating the molecular function of riboflavin synthase. The mRNA is cell-to-cell mobile.

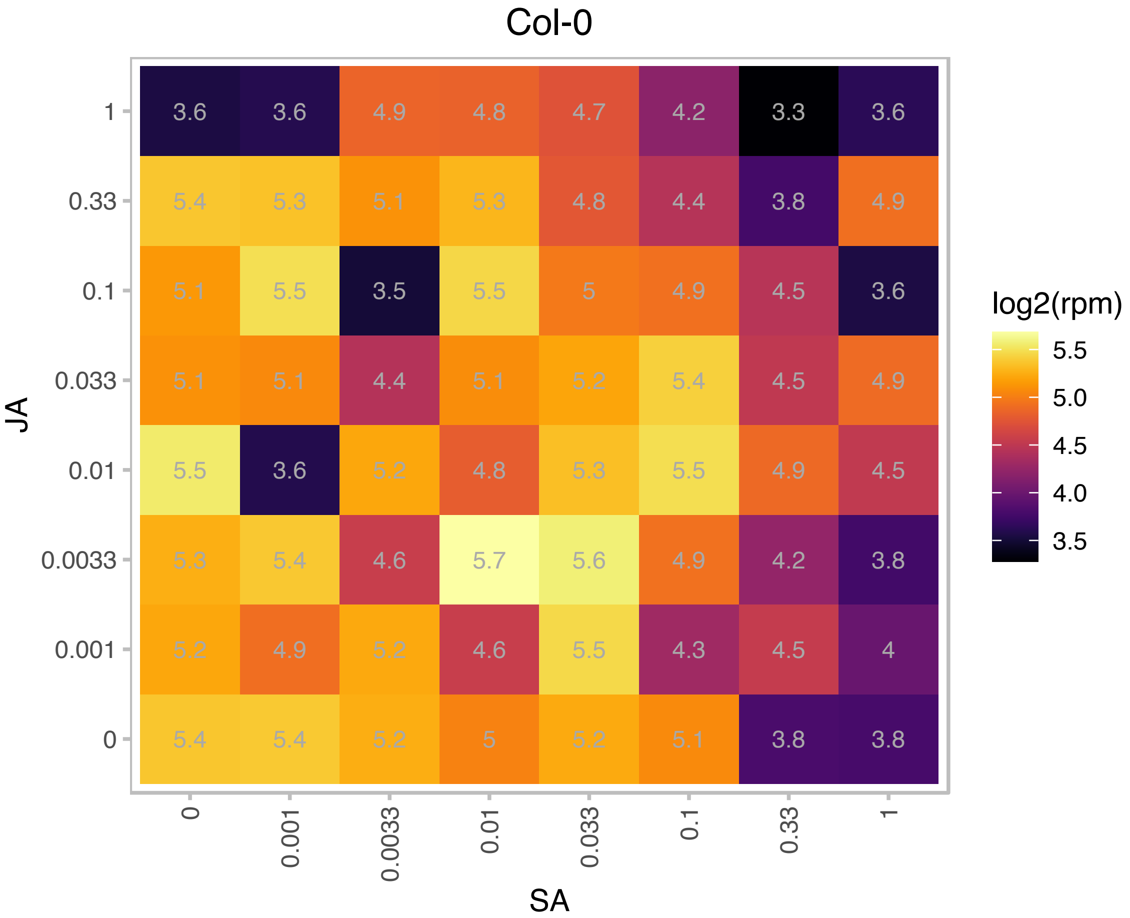


| | |
|---|---|
| Gene | AT2G20690 |
| is.expressed_genes | TRUE |
| is.responsive_genes | TRUE |
| is.apcluster | F-07 |

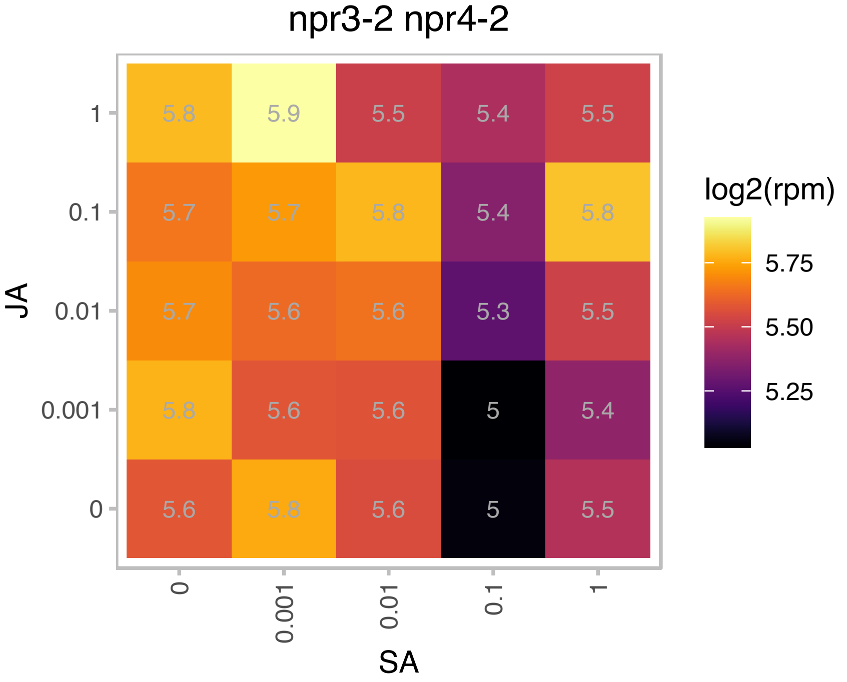


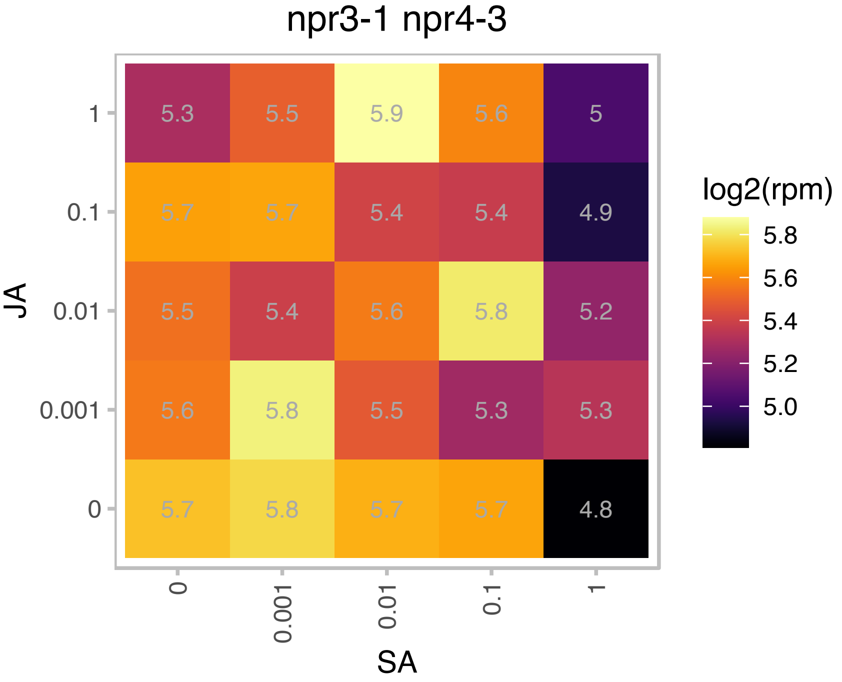

# AT2G21580.1 : NA

Ribosomal protein S25 family protein

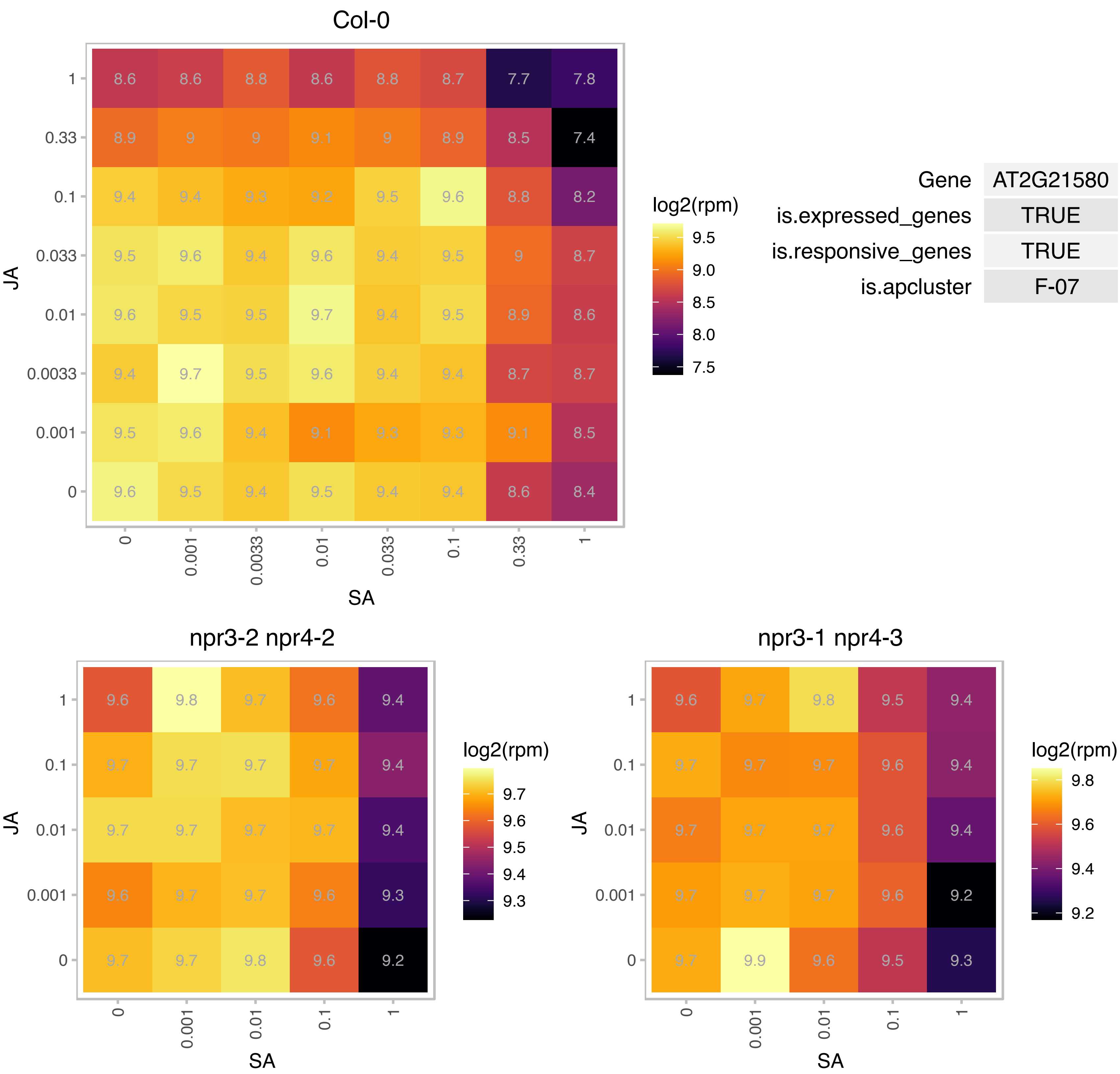


| | |
|---|---|
| Gene | AT2G21580 |
| is.expressed_genes | TRUE |
| is.responsive_genes | TRUE |
| is.apcluster | F-07 |

### AT2G21660.1 : ATGRP7, CCR2, GR-RBP7, GRP7, RBGA3

cold, circadian rhythm, and rna binding 2

Encodes a small glycine-rich RNA binding protein that is part of a negative-feedback loop through which AtGRP7 regulates the circadian oscillations of its own transcript. Gene expression is induced by cold. GRP7 appears to promote stomatal opening and reduce tolerance under salt and dehydration stress conditions, but, promotes stomatal closing and thereby increases stress tolerance under conditions of cold tolerance. Loss of function mutations have increased susceptibility to pathogens suggesting a role in mediating innate immune response. Mutants are also late flowering in a non-photoperiodic manner and are responsive to vernalization suggesting an interaction with the autonomous flowering pathway. There is a reduction of mRNA export from [illegible]

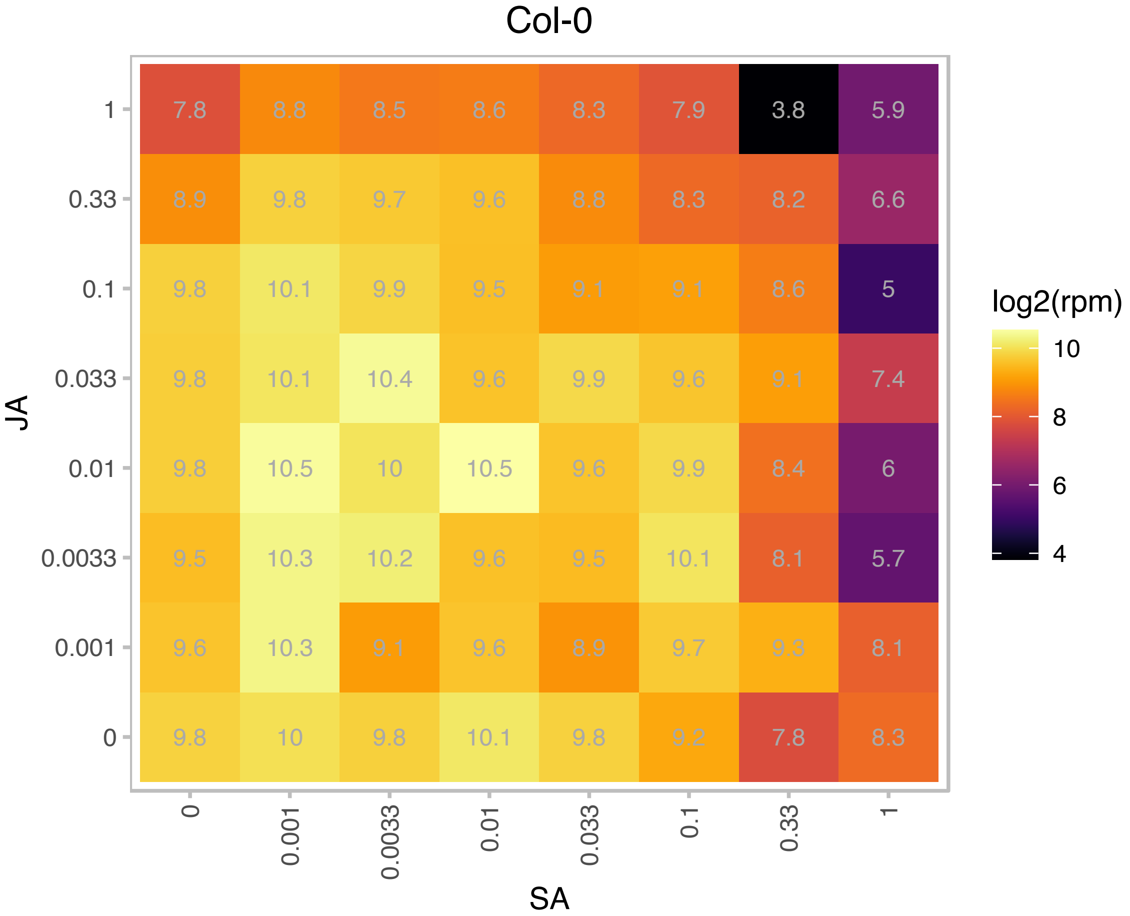


| | |
|---|---|
| Gene | AT2G21660 |
| is.expressed_genes | TRUE |
| is.responsive_genes | TRUE |
| is.apcluster | F-07 |

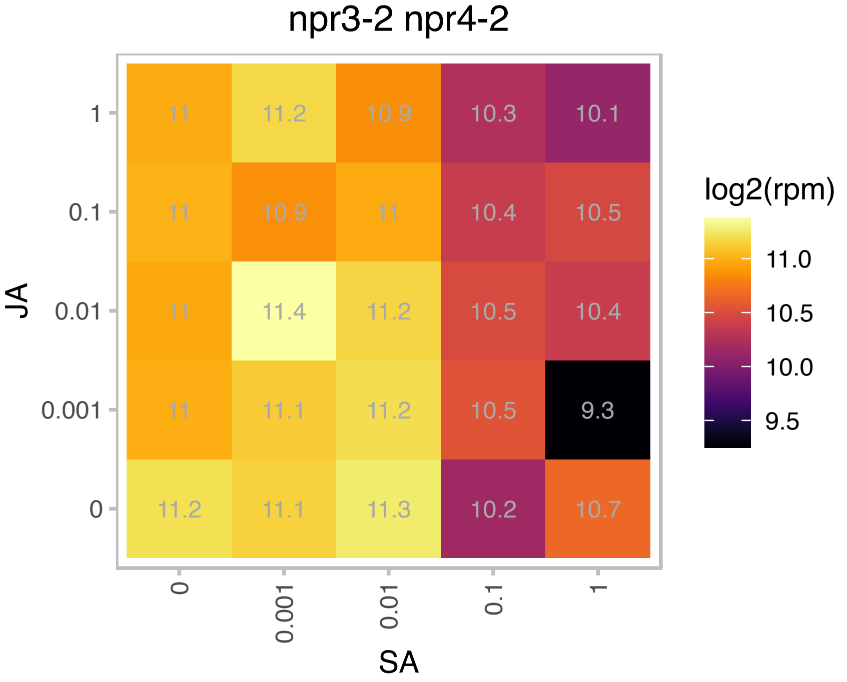


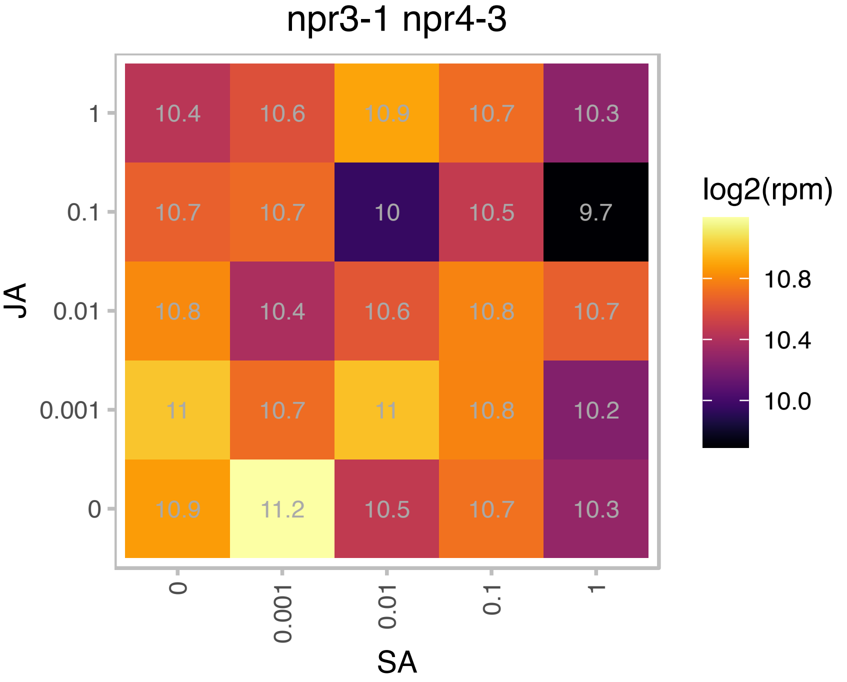

# AT2G23672.1 : NA

other RNA

Potential natural antisense gene, locus overlaps with AT2G23670

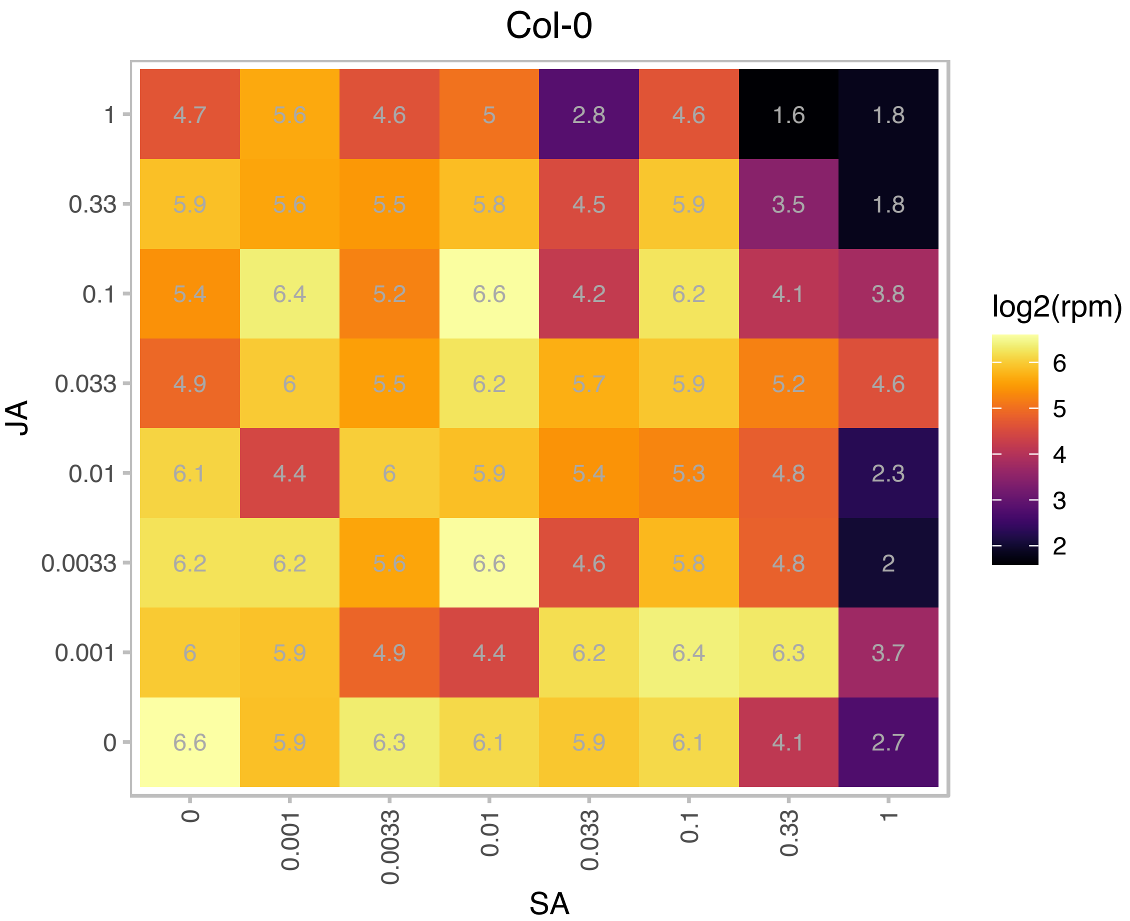


| | |
|---|---|
| Gene | AT2G23672 |
| is.expressed_genes | TRUE |
| is.responsive_genes | TRUE |
| is.apcluster | F-07 |

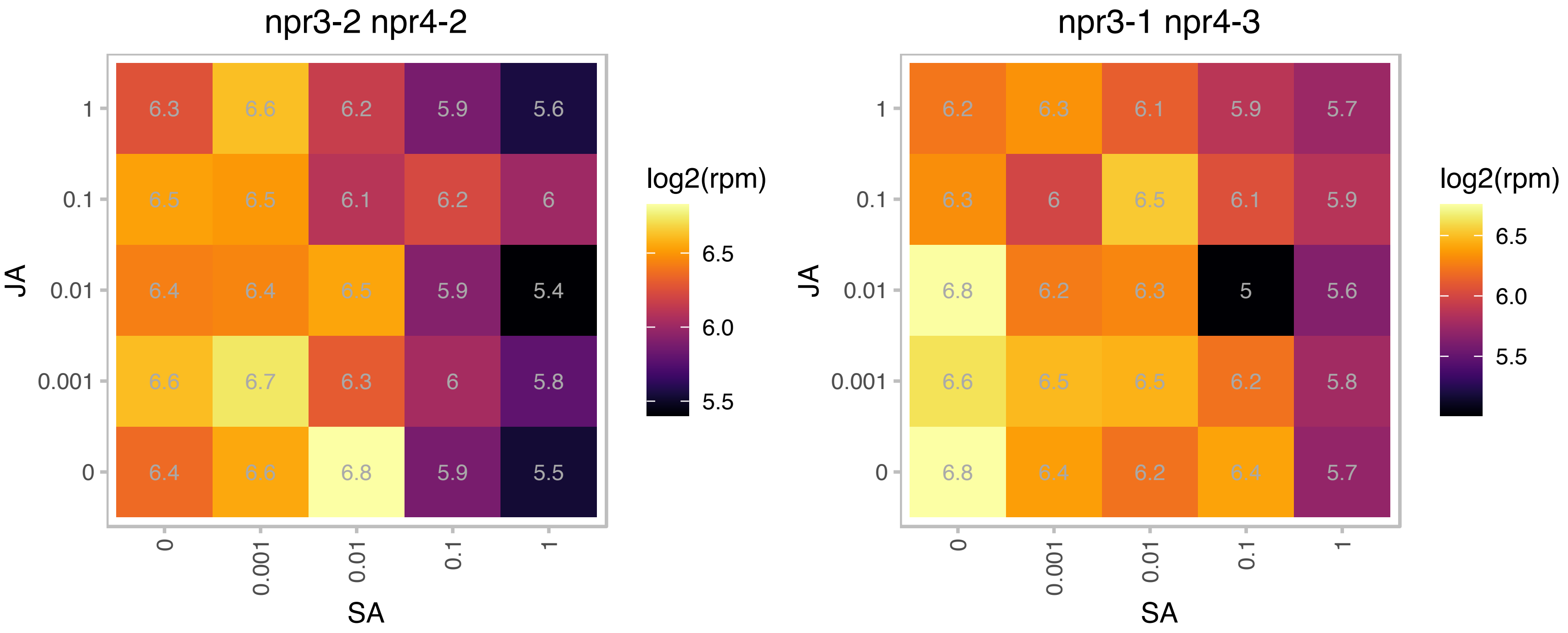

# AT2G25480.1 : NA

TPX2 (targeting protein for Xklp2) protein family

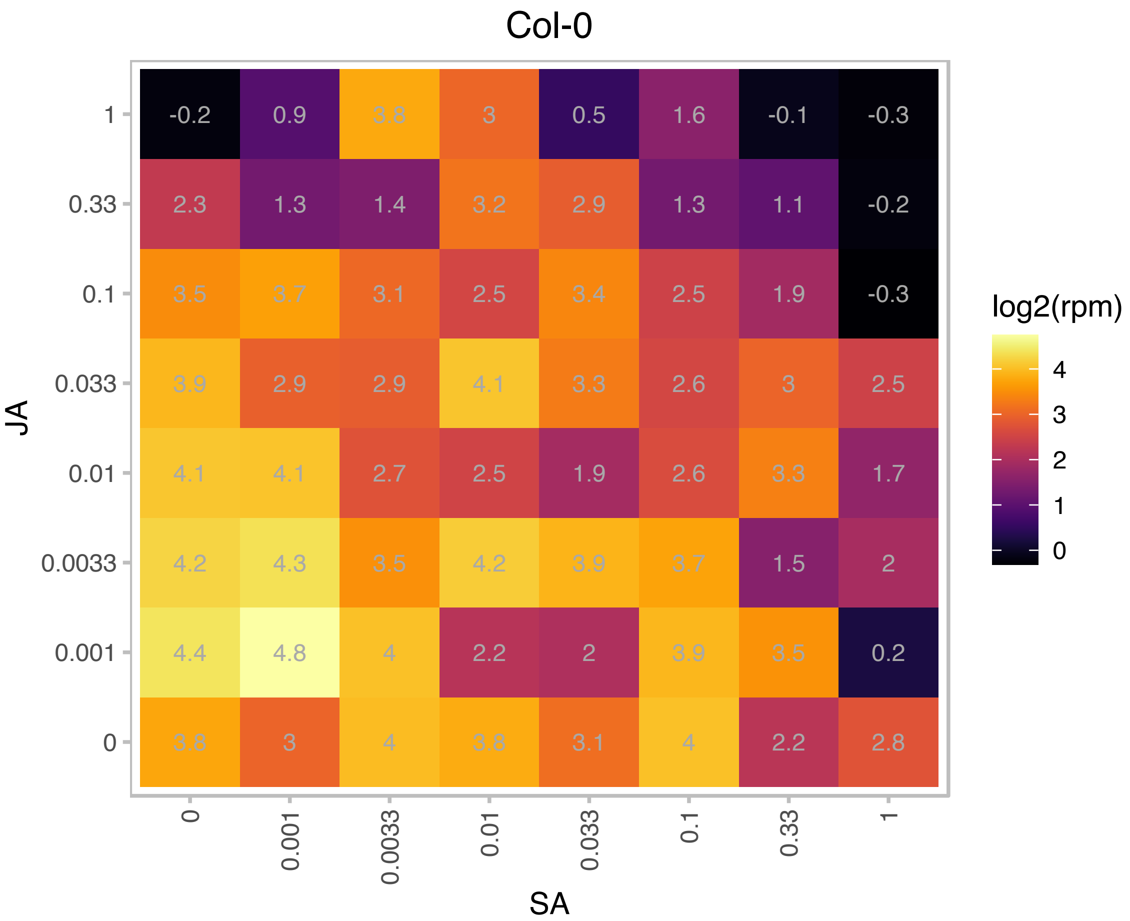


| | |
|---|---|
| Gene | AT2G25480 |
| is.expressed_genes | TRUE |
| is.responsive_genes | TRUE |
| is.apcluster | F-07 |

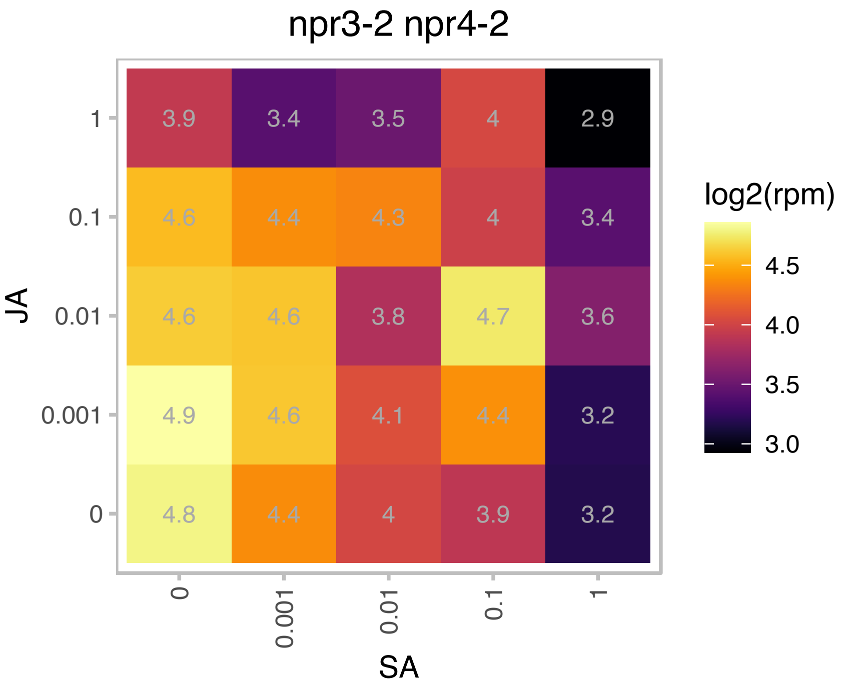


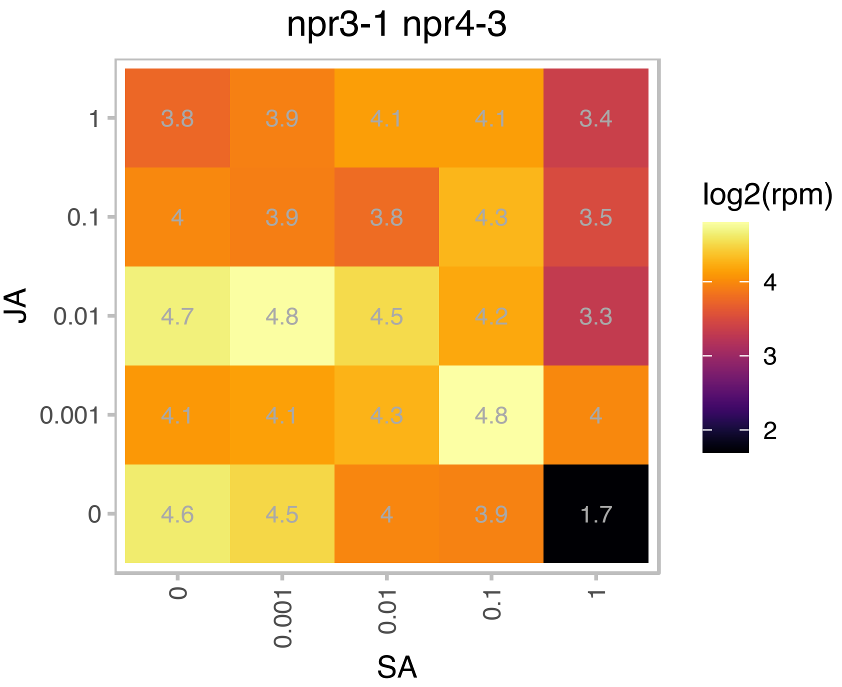

### AT2G26580.1 : YAB5

plant-specific transcription factor YABBY family protein

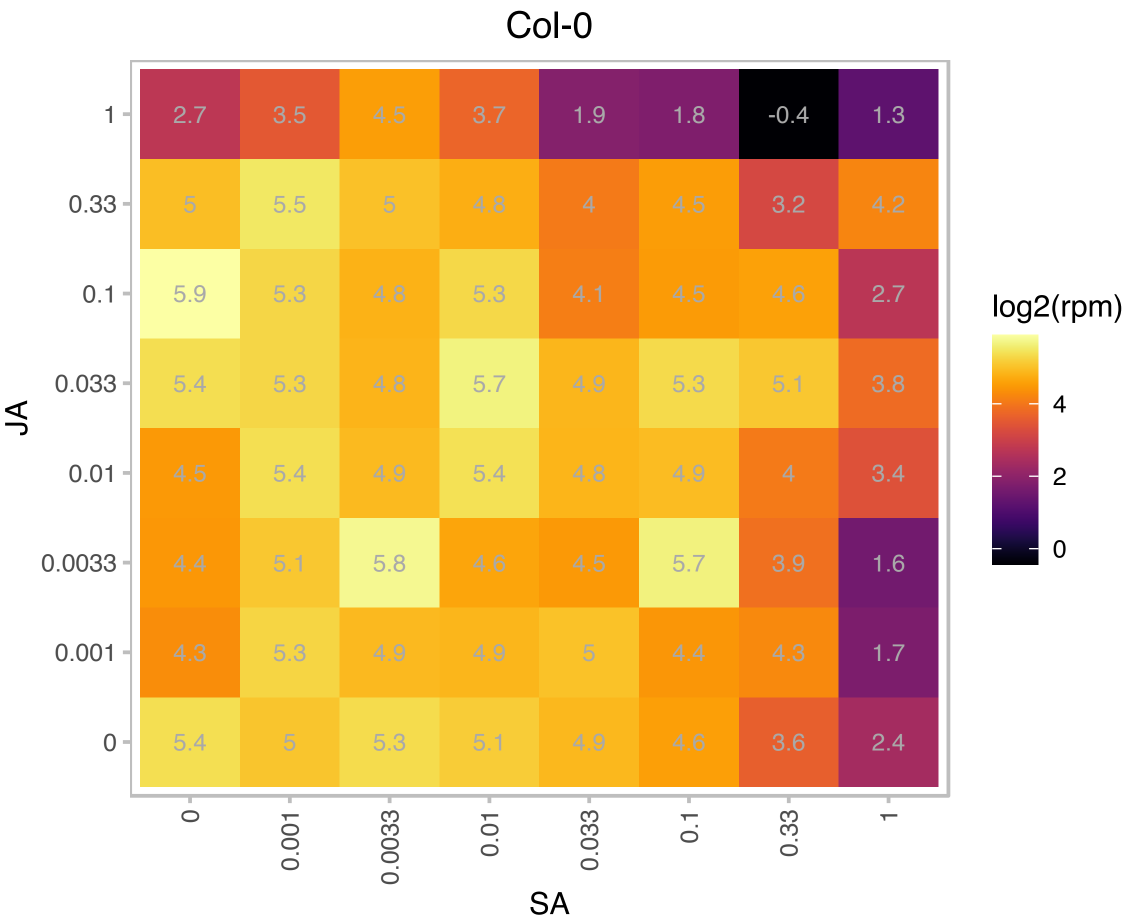


| | |
|---|---|
| Gene | AT2G26580 |
| is.expressed_genes | TRUE |
| is.responsive_genes | TRUE |
| is.apcluster | F-07 |

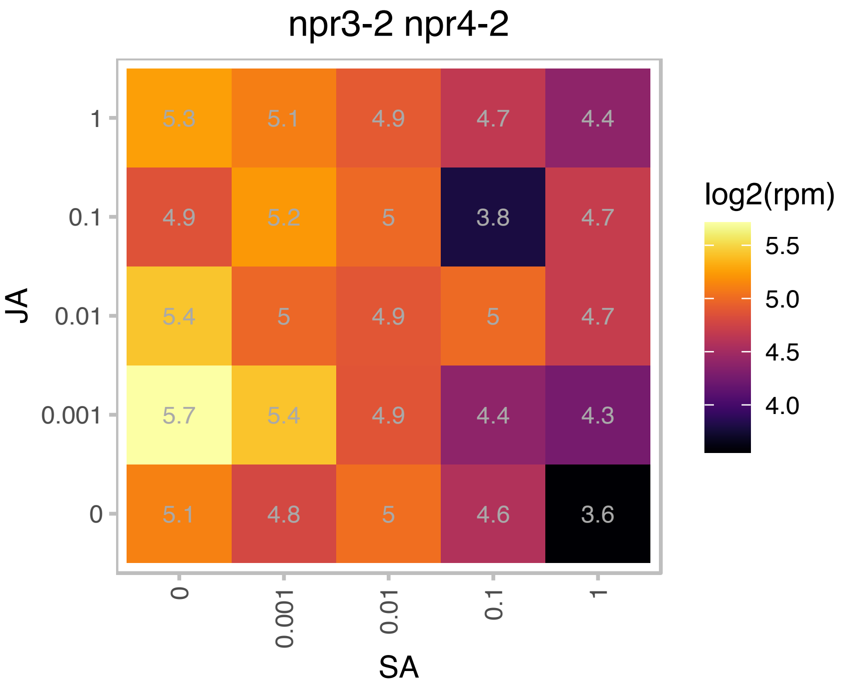


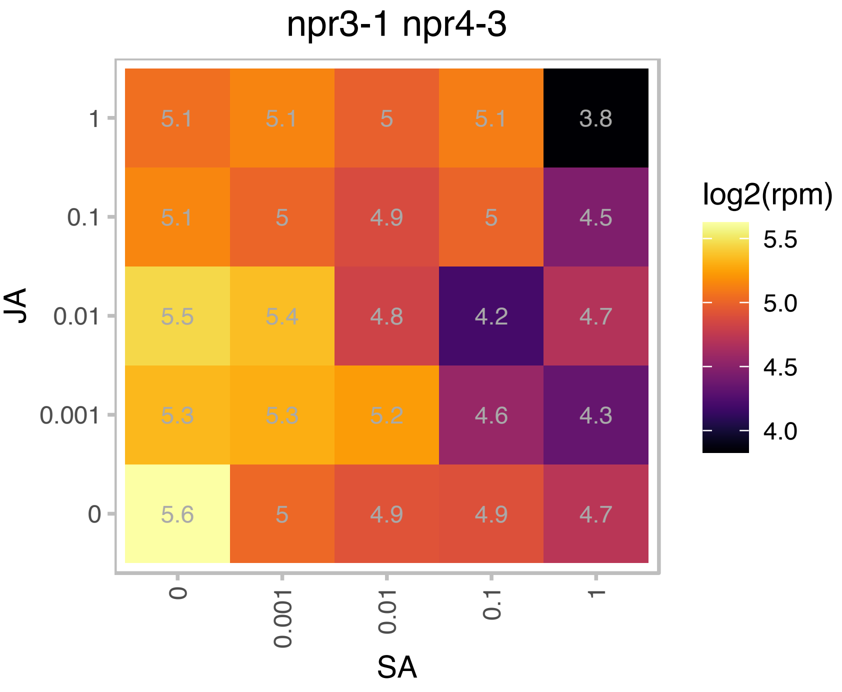

# AT2G26695.1 : NA

Ran BP2/NZF zinc finger-like superfamily protein

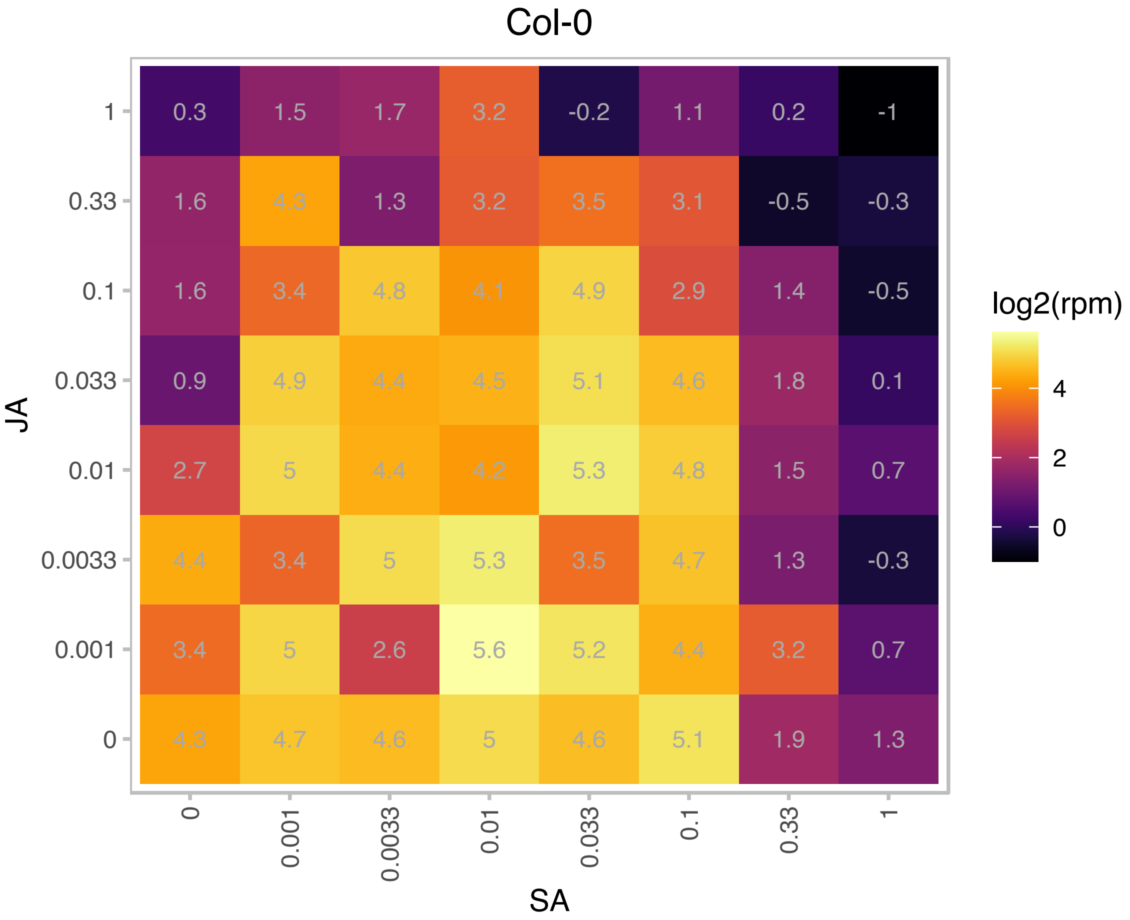


| | |
|---|---|
| Gene | AT2G26695 |
| is.expressed_genes | TRUE |
| is.responsive_genes | TRUE |
| is.apcluster | F-07 |

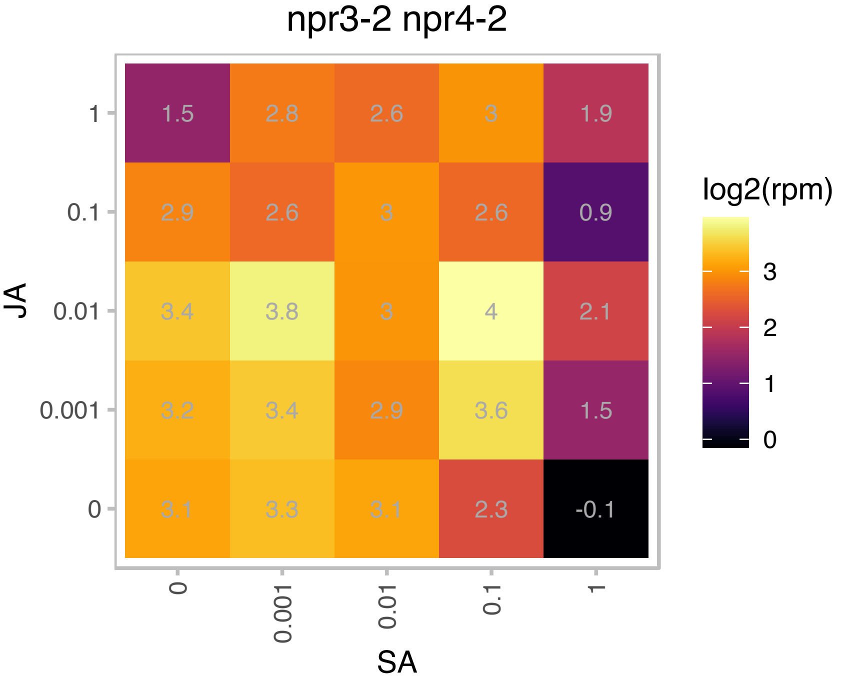


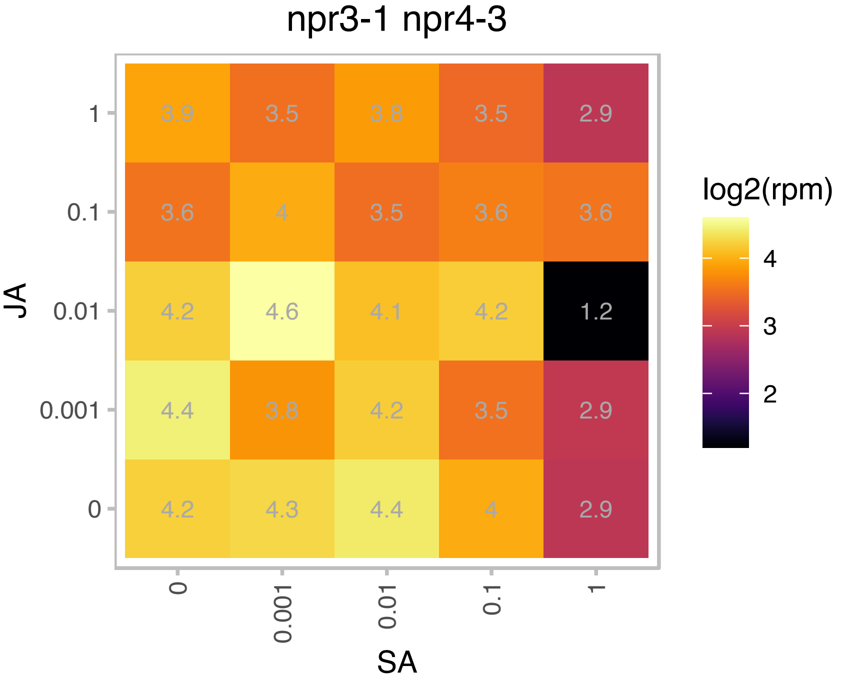

### AT2G26980.1 : CIPK3, SnRK3.17

CBL-interacting protein kinase 3
encodes a serine-threonine protein kinase whose expression increases in response to abscisic acid, cold, drought, high salt, and wounding conditions. The gene is expressed in developing seeds and seedlings. Lines carrying a T-DNA insertions have reduced germination efficiency and expression of cold, high-salt, and abscisic acid marker genes are altered, but not drought-response markers.

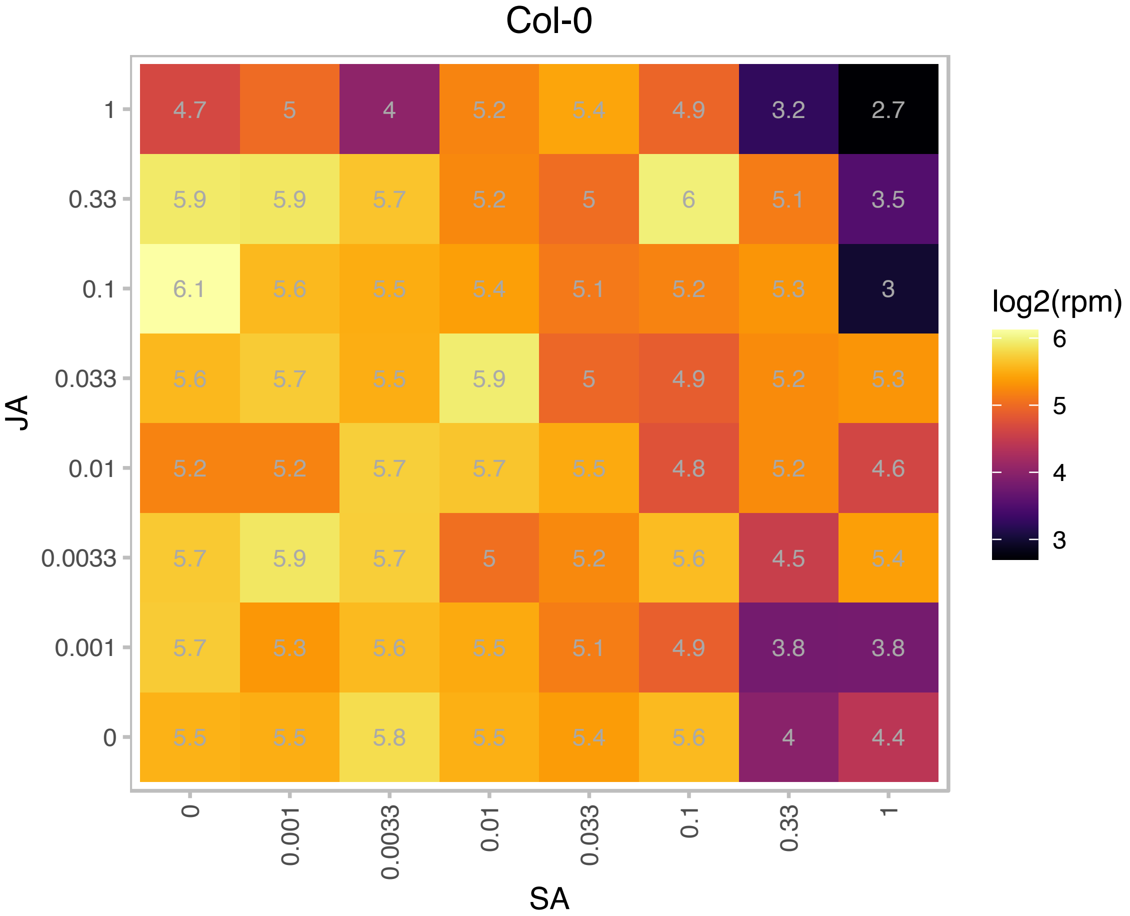


| Gene | AT2G26980 |
|---|---|
| is.expressed_genes | TRUE |
| is.responsive_genes | TRUE |
| is.apcluster | F-07 |

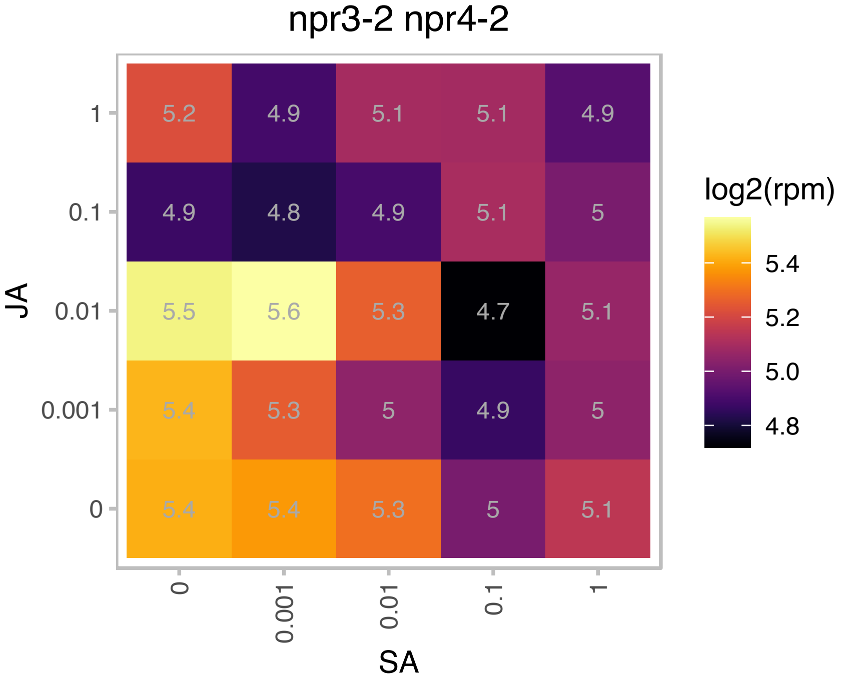


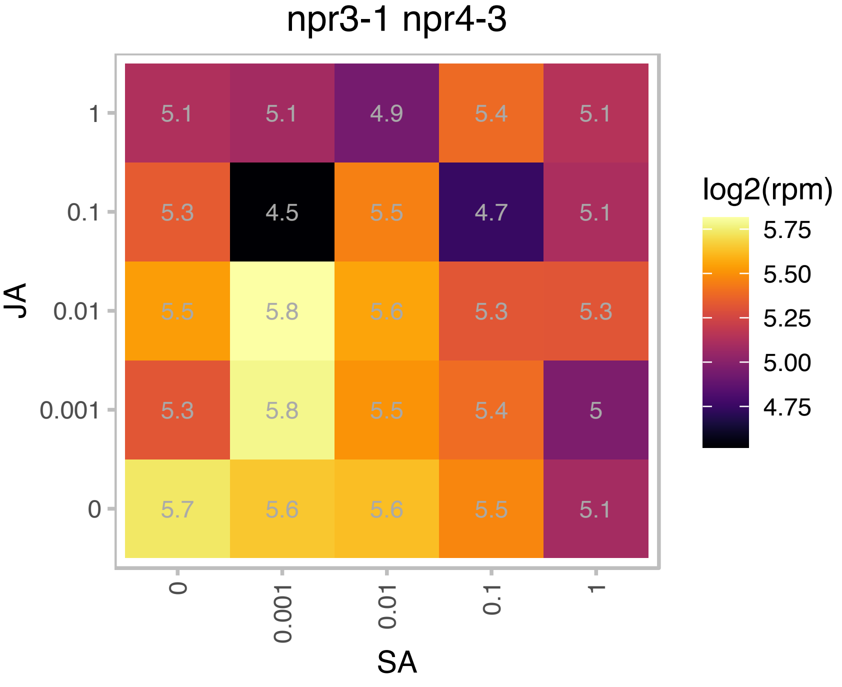

### AT2G27530.1 : PGY1

Ribosomal protein L1p/L10e family
Encodes ribosomal protein L10aP. Identified in a screen for enhancers of as1. as1/pgy double mutants show defects in leaf vascular patterning and adaxial cell fate. Double mutant analysis indicates pgy genes function in the same pathway as REV, KAN1 and KAN2.

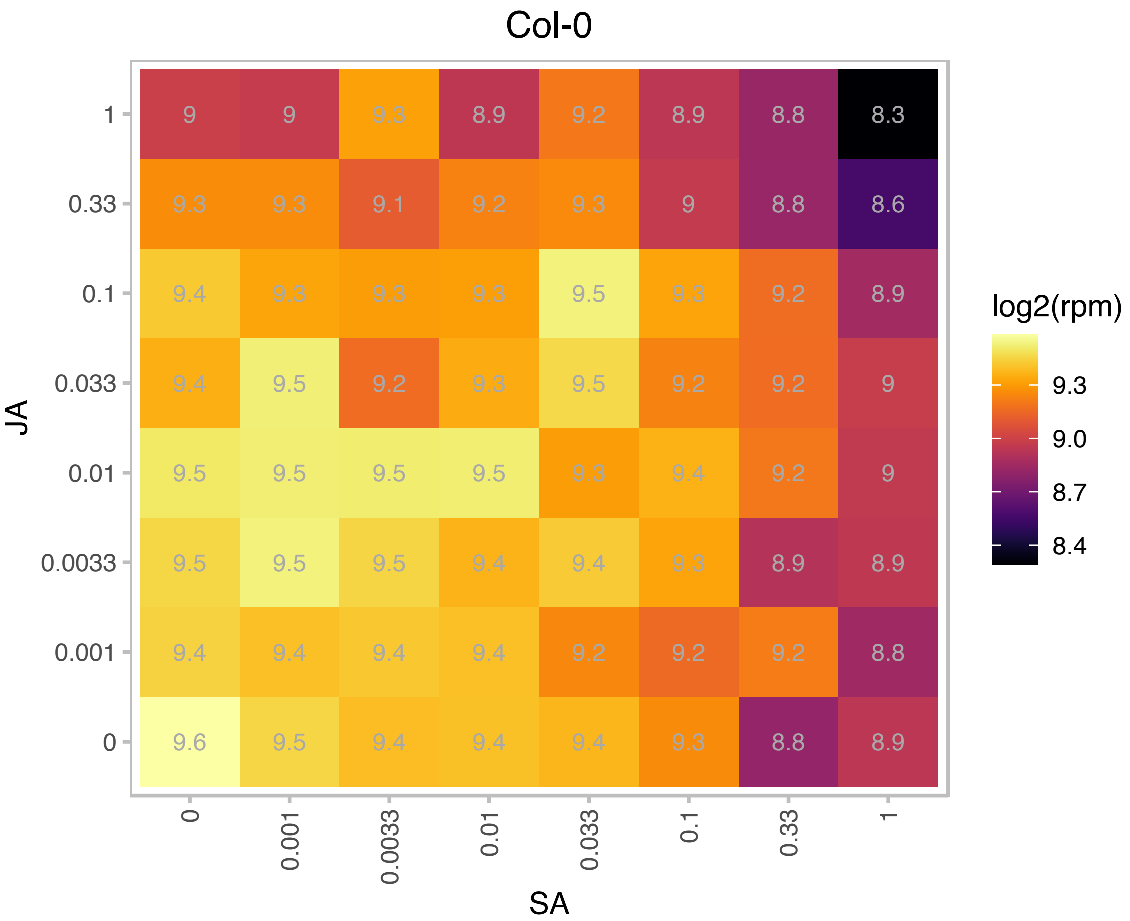


| | |
|---|---|
| Gene | AT2G27530 |
| is.expressed_genes | TRUE |
| is.responsive_genes | TRUE |
| is.apcluster | F-07 |

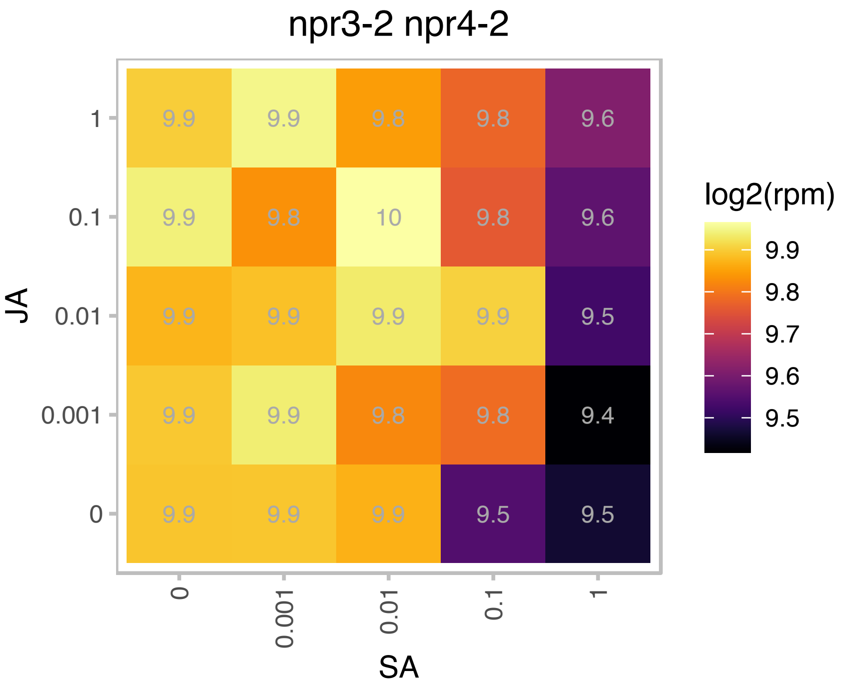


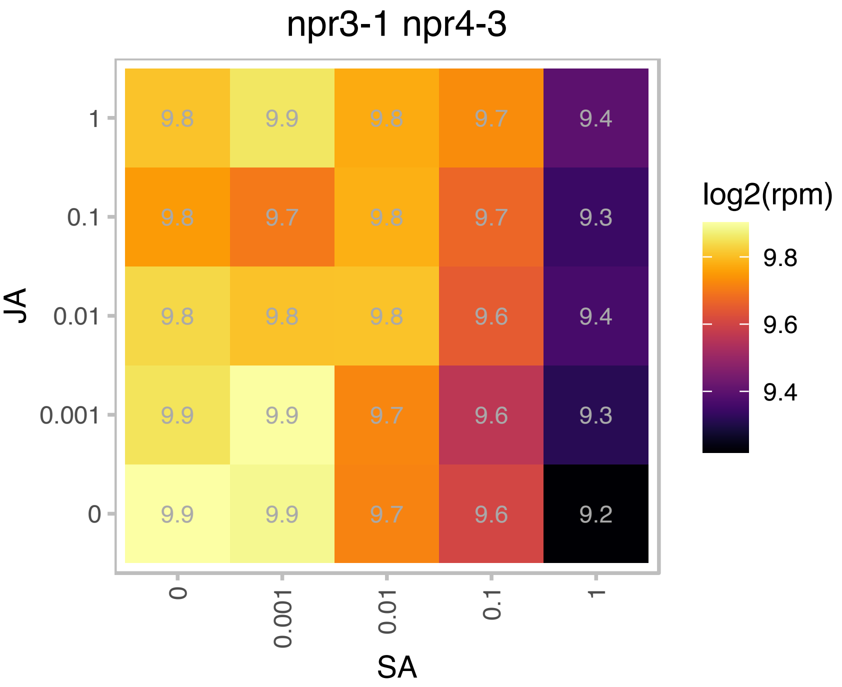

# AT2G27710.1 : NA

60S acidic ribosomal protein family

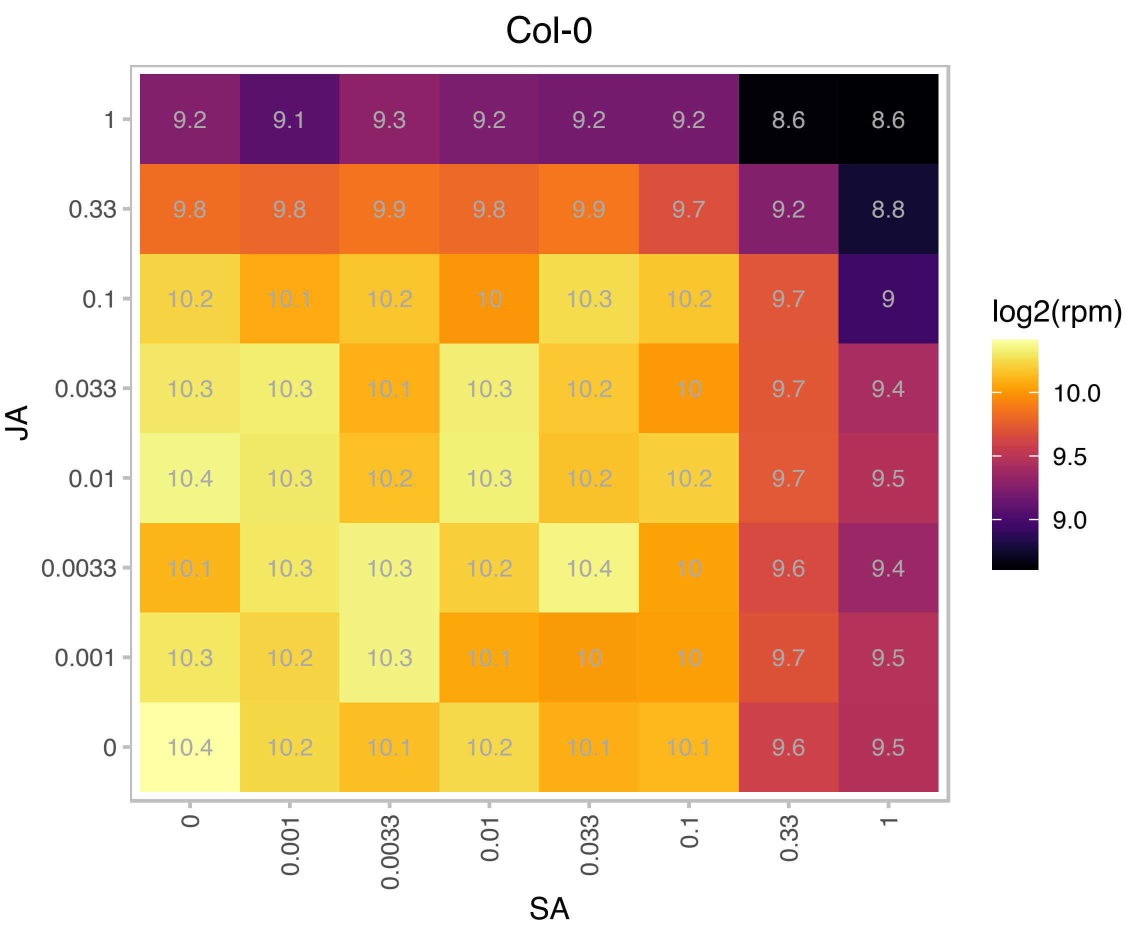


| | |
|---|---|
| Gene | AT2G27710 |
| is.expressed_genes | TRUE |
| is.responsive_genes | TRUE |
| is.apcluster | F-07 |

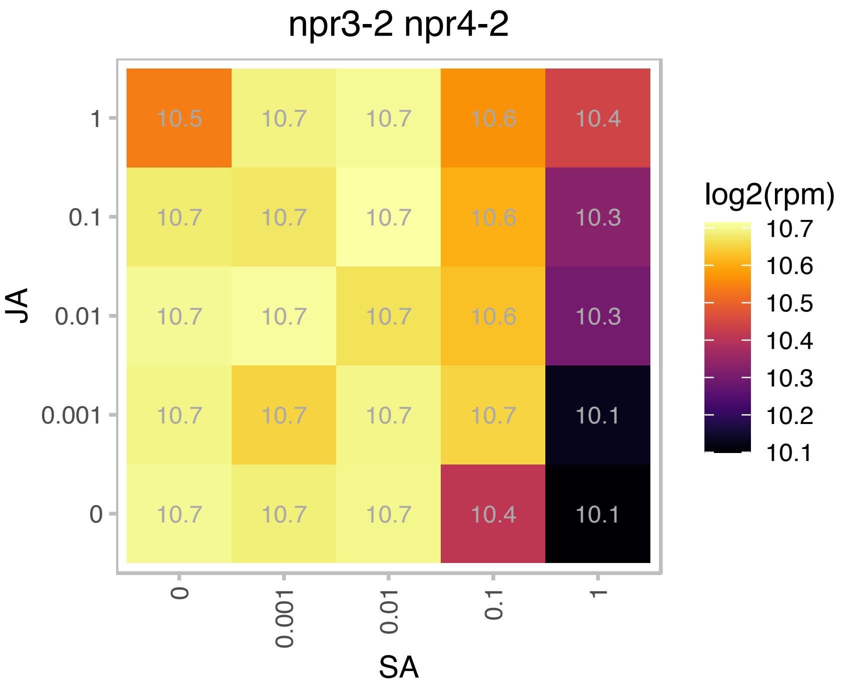


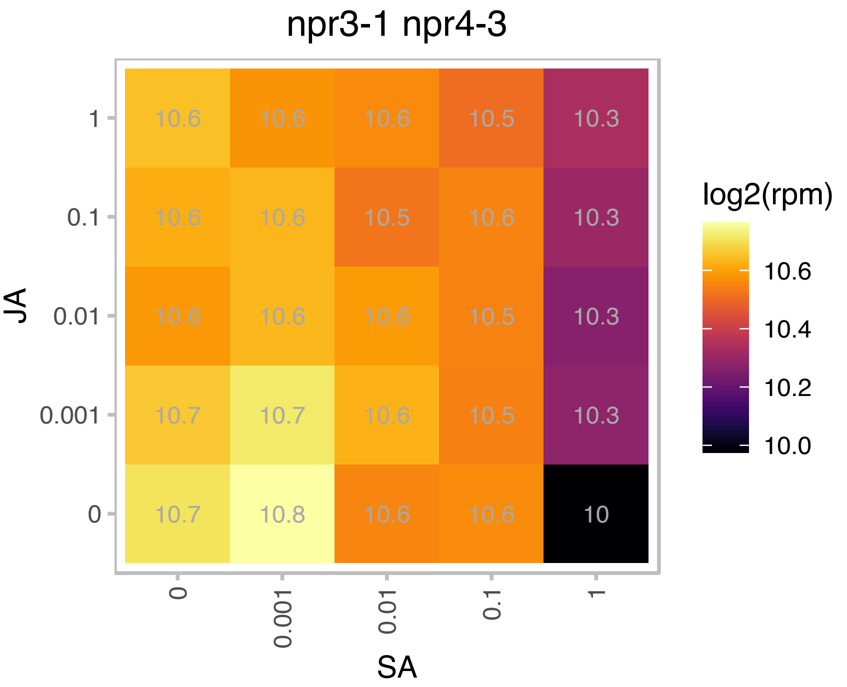

### AT2G27840.1 : HD2D, HDA13, HDT04, HDT4

histone deacetylase-related / HD-related

Belongs to the plant specific HD2 type proteins; similar to nucleolar Zea mays histone deacetylase; HD2-p39

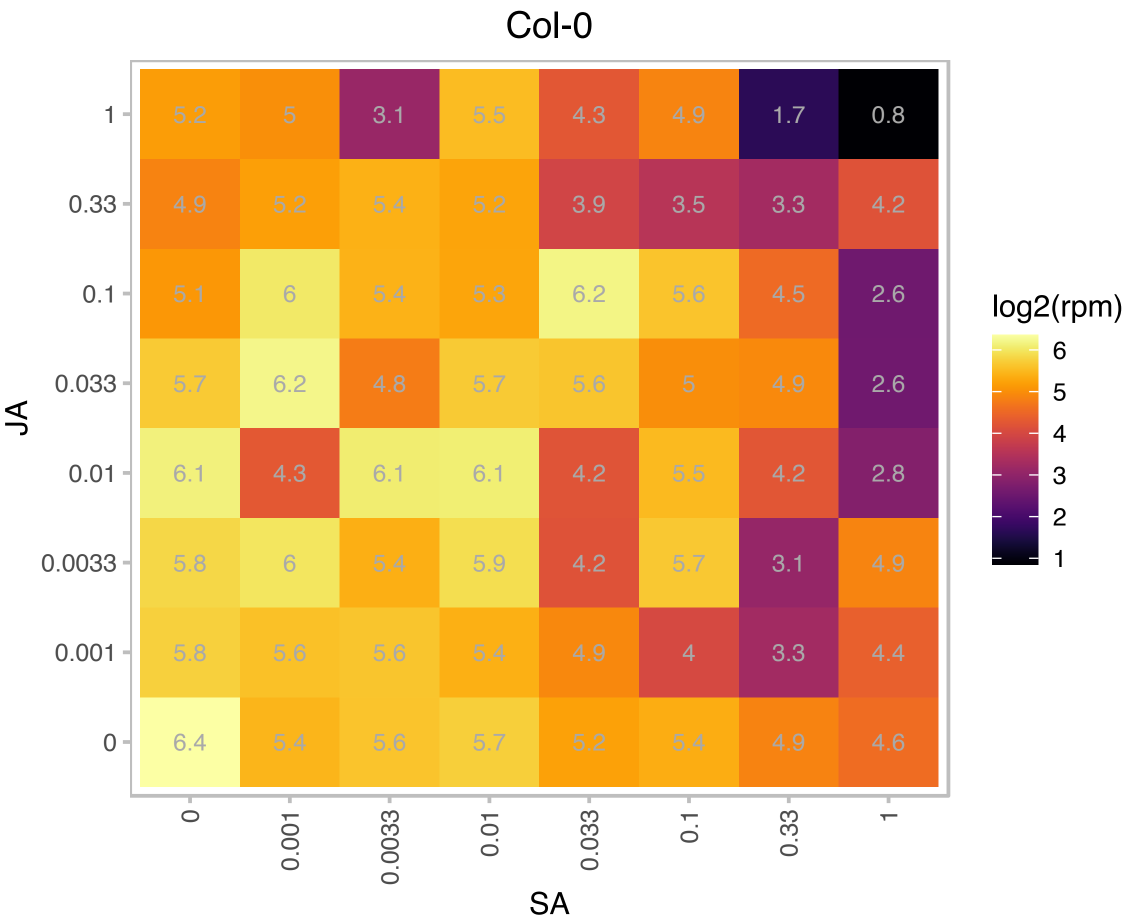


| | |
|---|---|
| Gene | AT2G27840 |
| is.expressed_genes | TRUE |
| is.responsive_genes | TRUE |
| is.apcluster | F-07 |

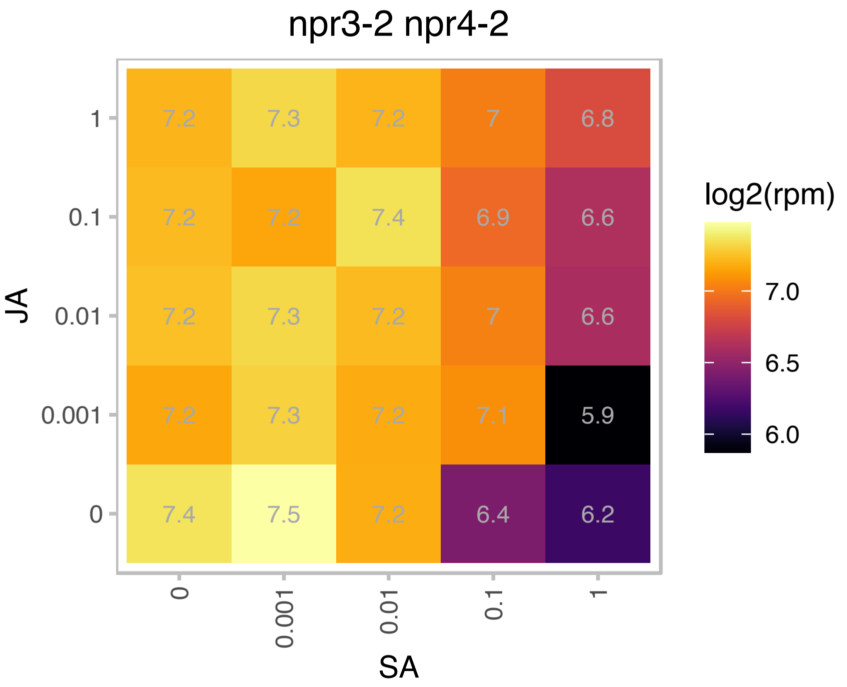


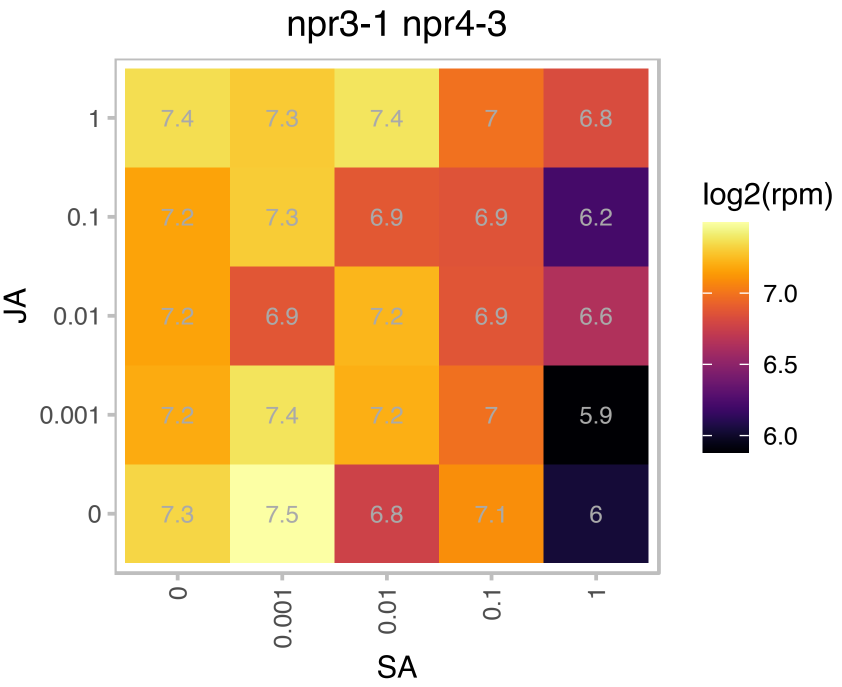

### AT2G28190.1 : AtSOD2, CSD2, CZSOD2, SOD2

copper/zinc superoxide dismutase 2

Encodes a chloroplastic copper/zinc superoxide dismutase CSD2 that can detoxify superoxide radicals. Its expression is affected by miR398-directed mRNA cleavage. Activation depends totally on CCS. Overexpression of a miR398-resistant form of CSD2 leads to more dramatic improvements in stress (hight light, Cu2+ and methyl viologen) tolerance than overexpression of wild-type CSD2. The mRNA is cell-to-cell mobile.

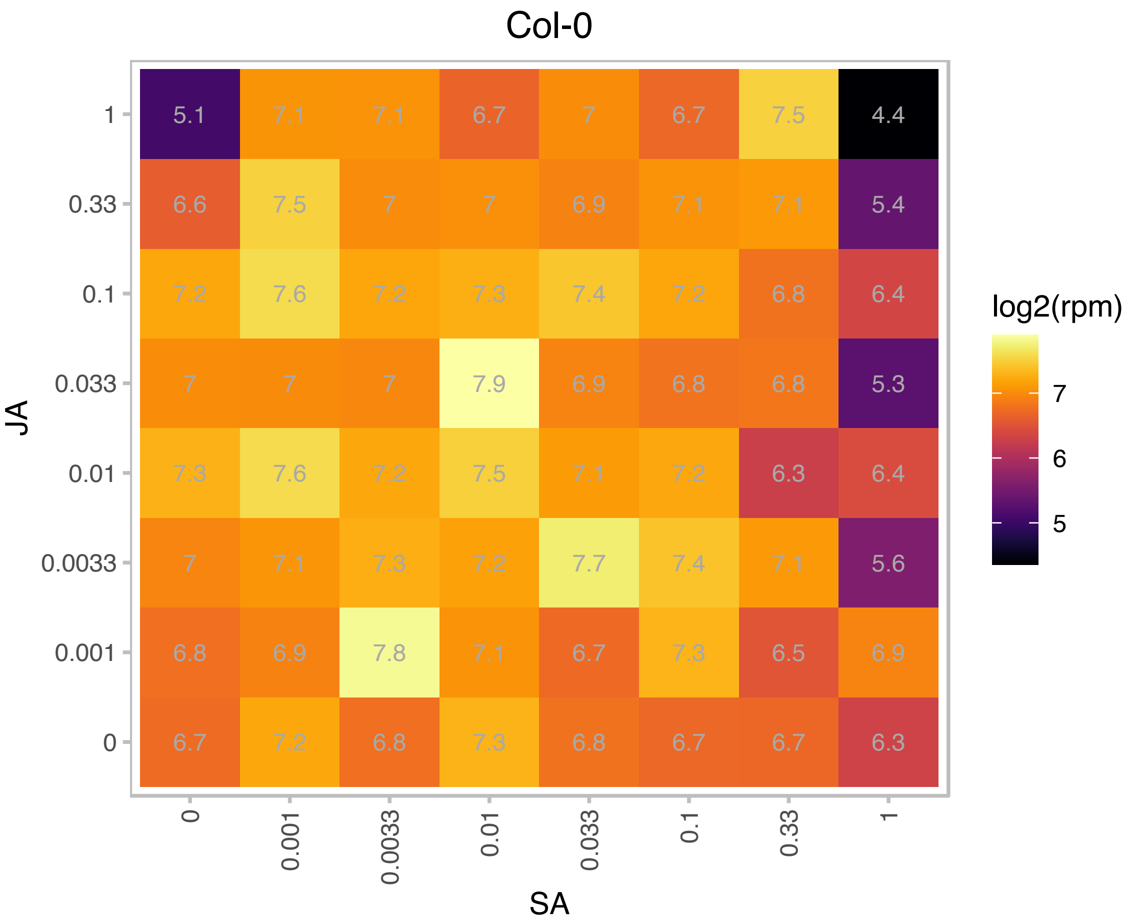


| | |
|---|---|
| Gene | AT2G28190 |
| is.expressed_genes | TRUE |
| is.responsive_genes | TRUE |
| is.apcluster | F-07 |

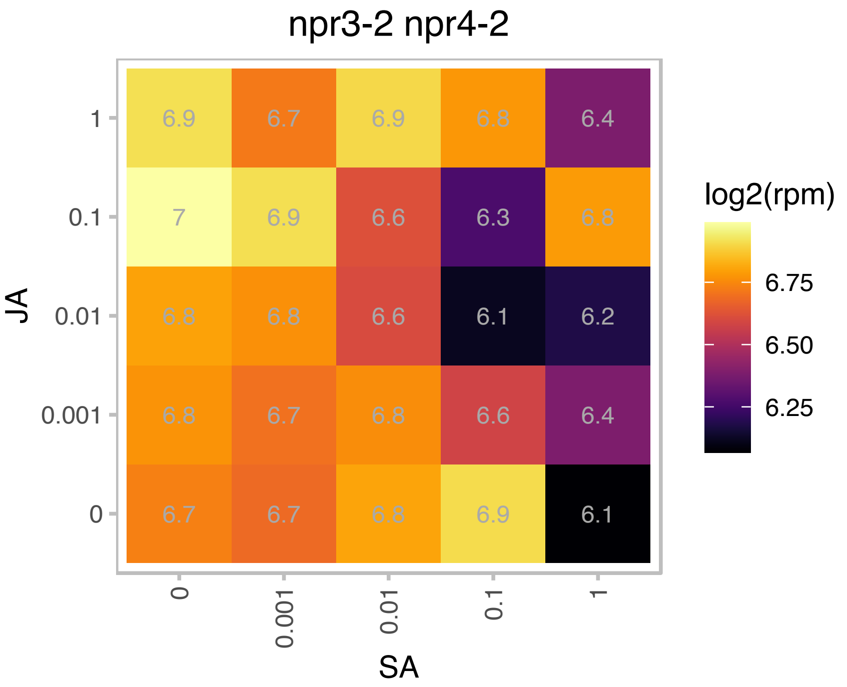


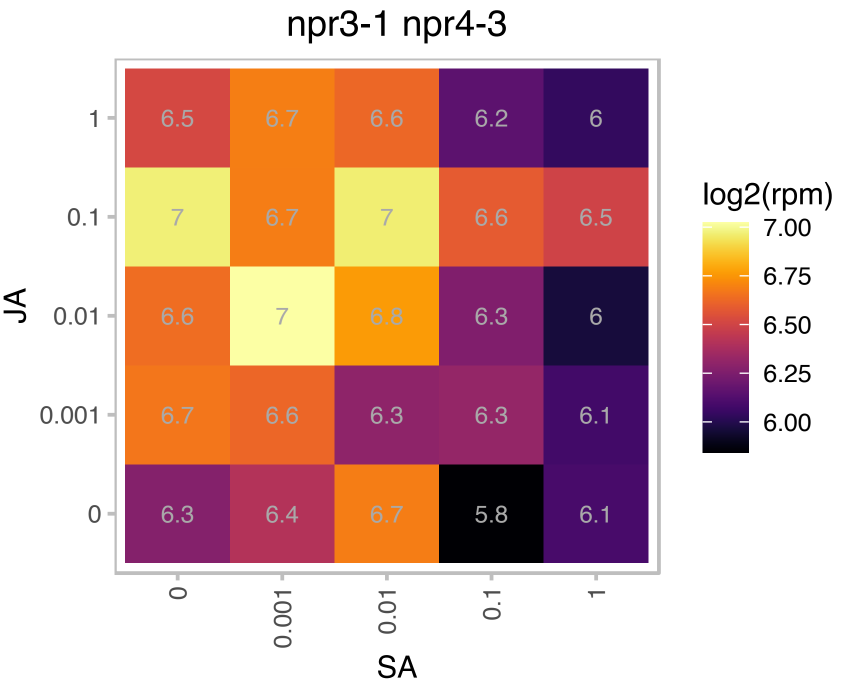

# AT2G28200.1 : NA

C2H2-type zinc finger family protein

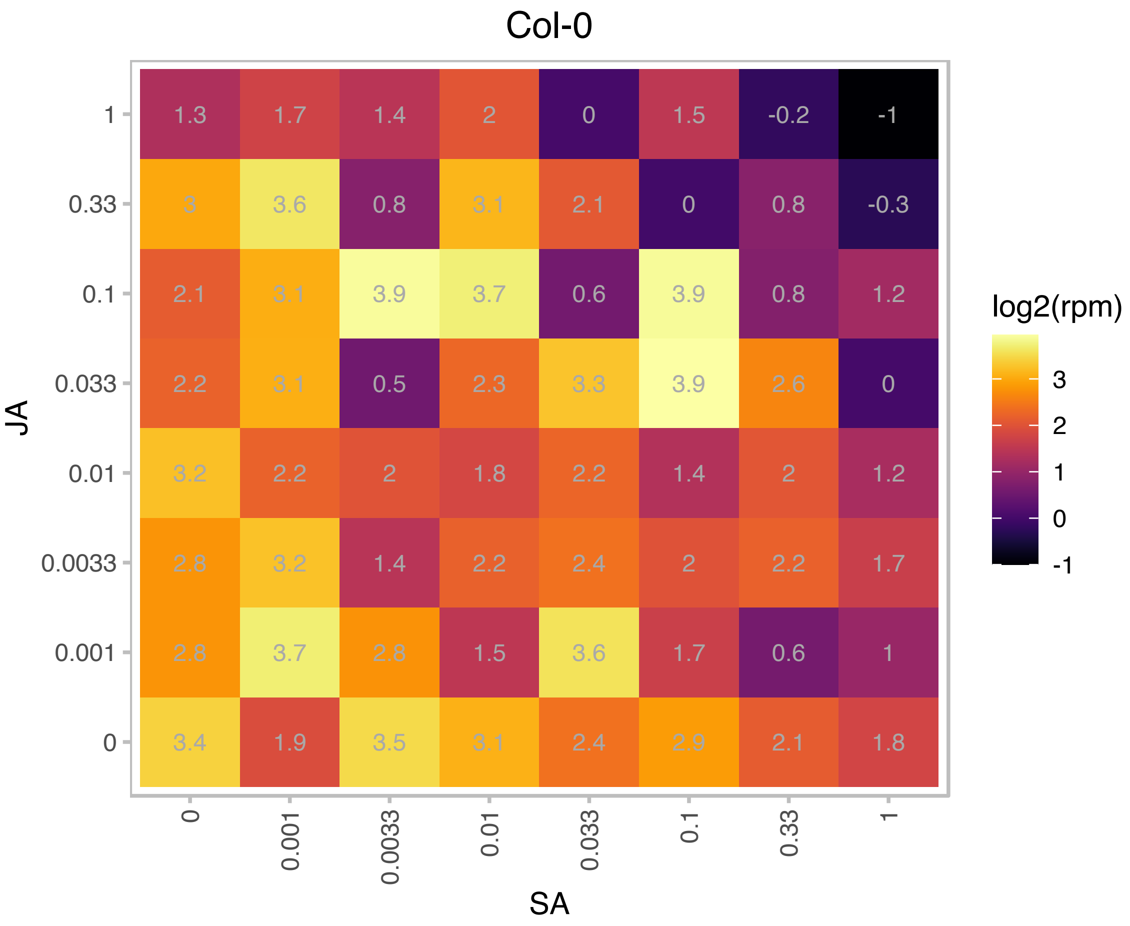


| | |
|---|---|
| Gene | AT2G28200 |
| is.expressed_genes | TRUE |
| is.responsive_genes | TRUE |
| is.apcluster | F-07 |

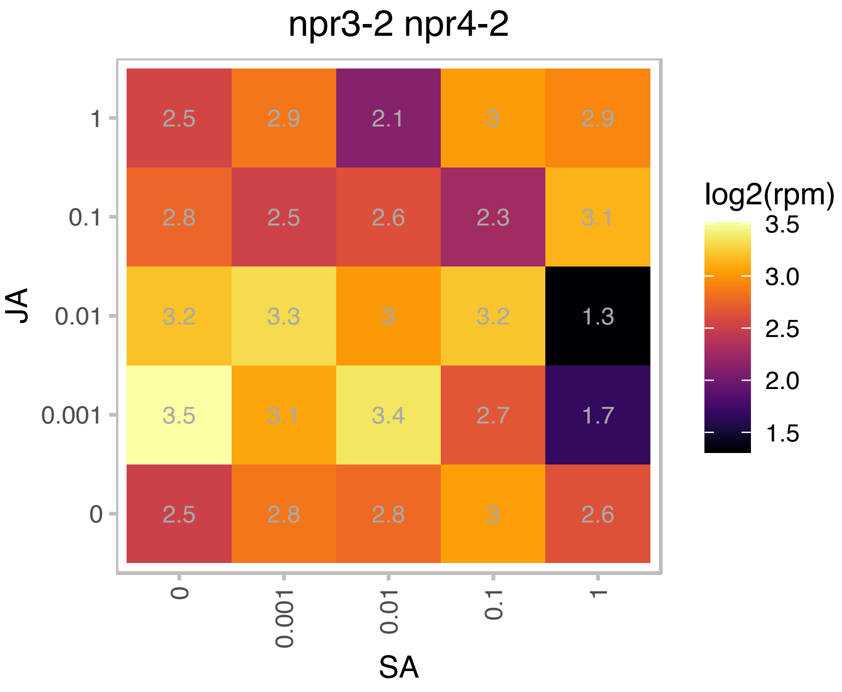


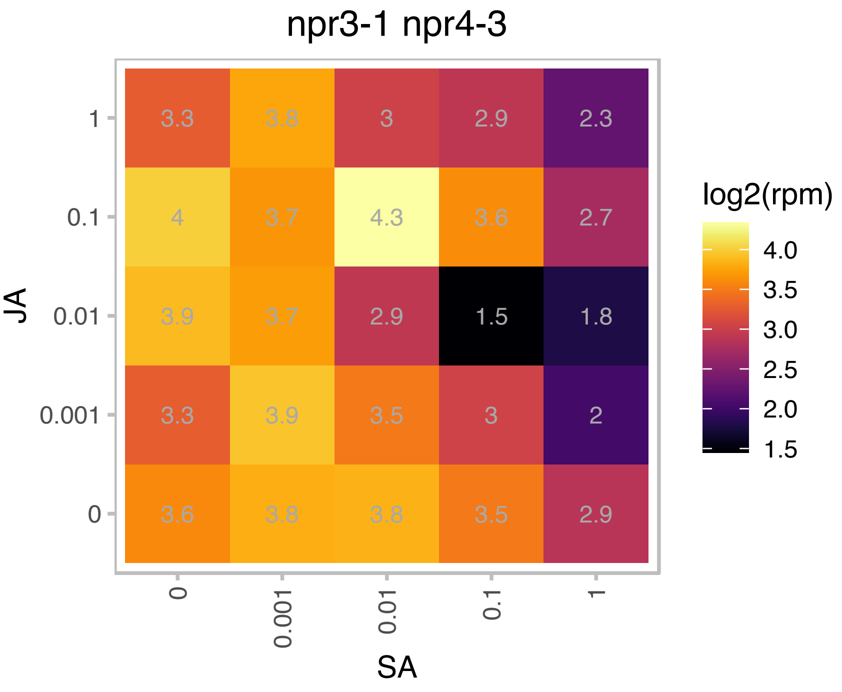

### AT2G28305.1 : ATLOG1, LOG1

Putative lysine decarboxylase family protein

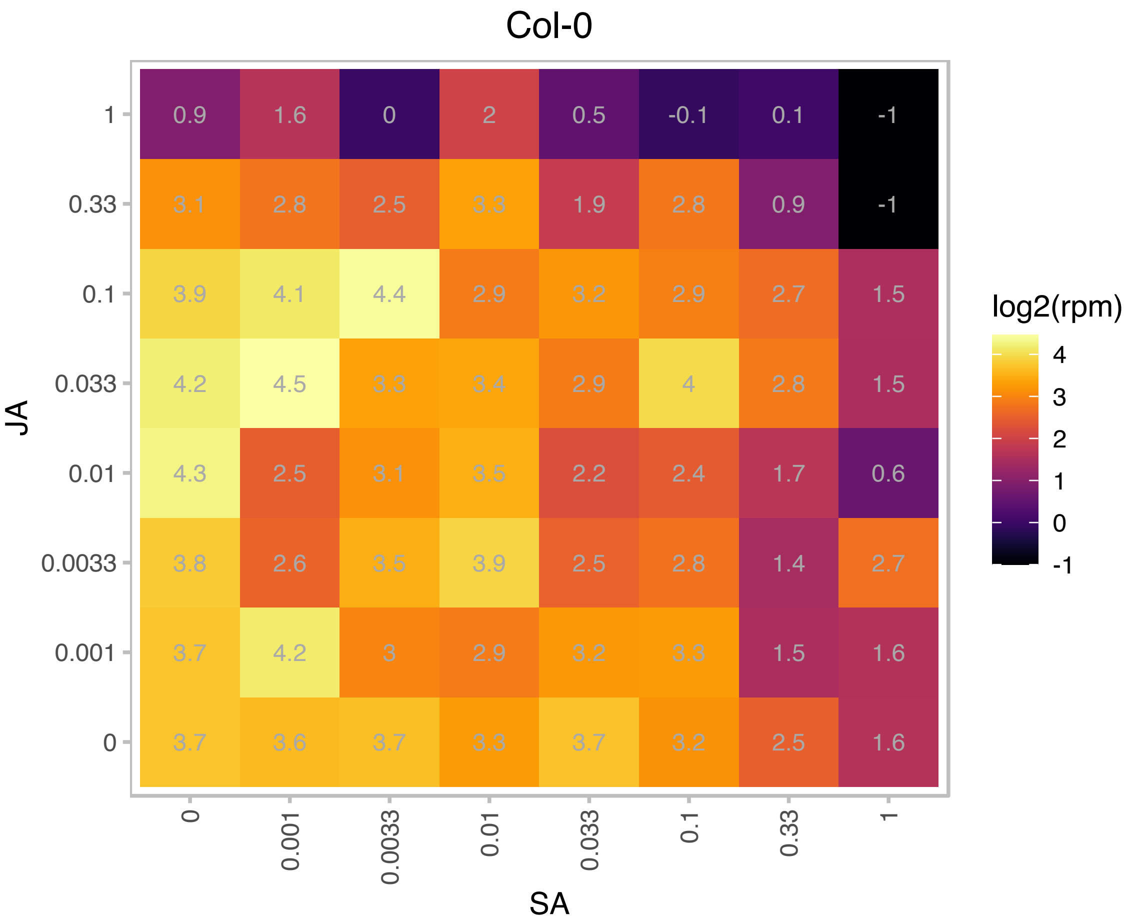


| | |
|---|---|
| Gene | AT2G28305 |
| is.expressed_genes | TRUE |
| is.responsive_genes | TRUE |
| is.apcluster | F-07 |

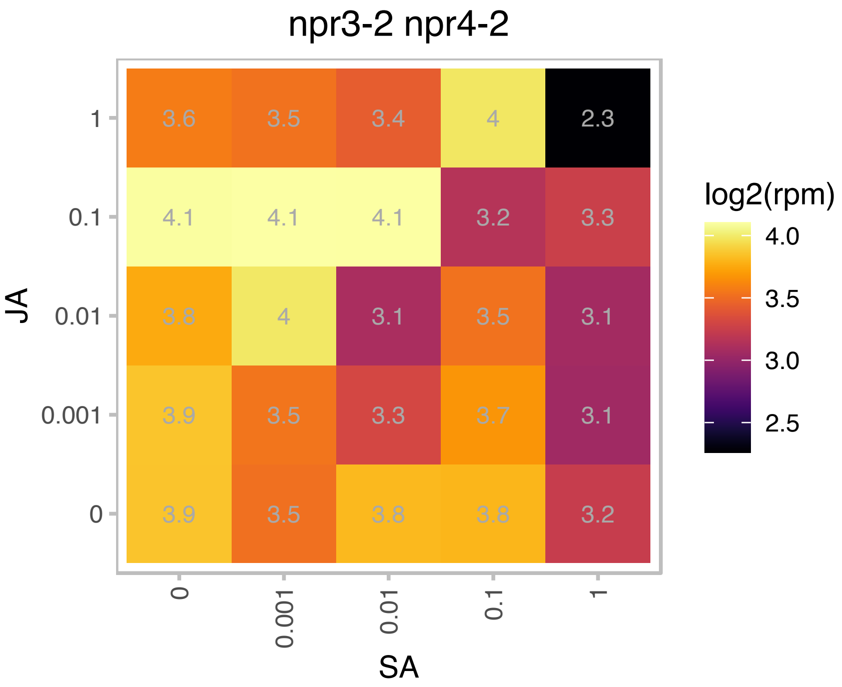


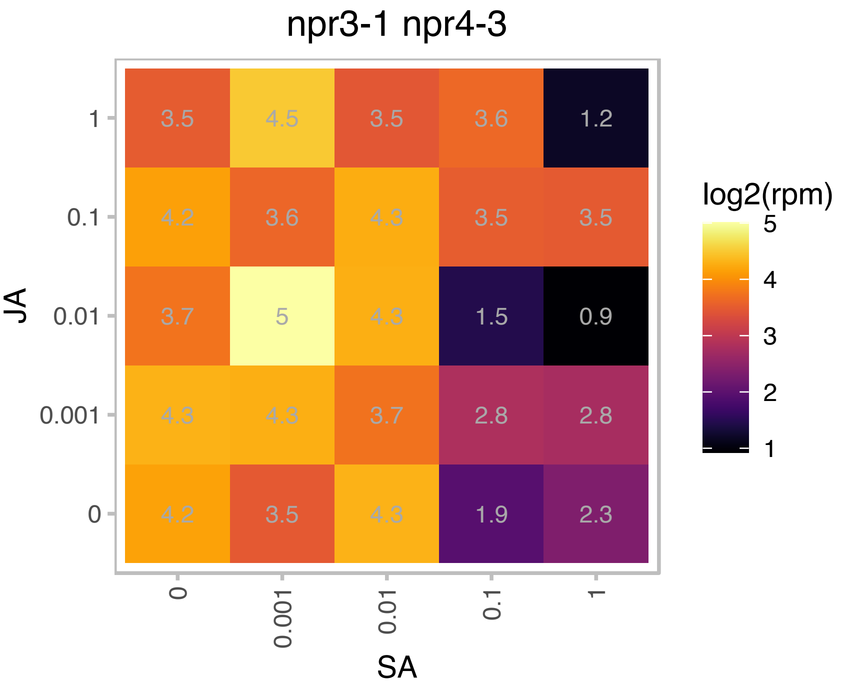

# AT2G28660.1 : NA

Chloroplast-targeted copper chaperone protein

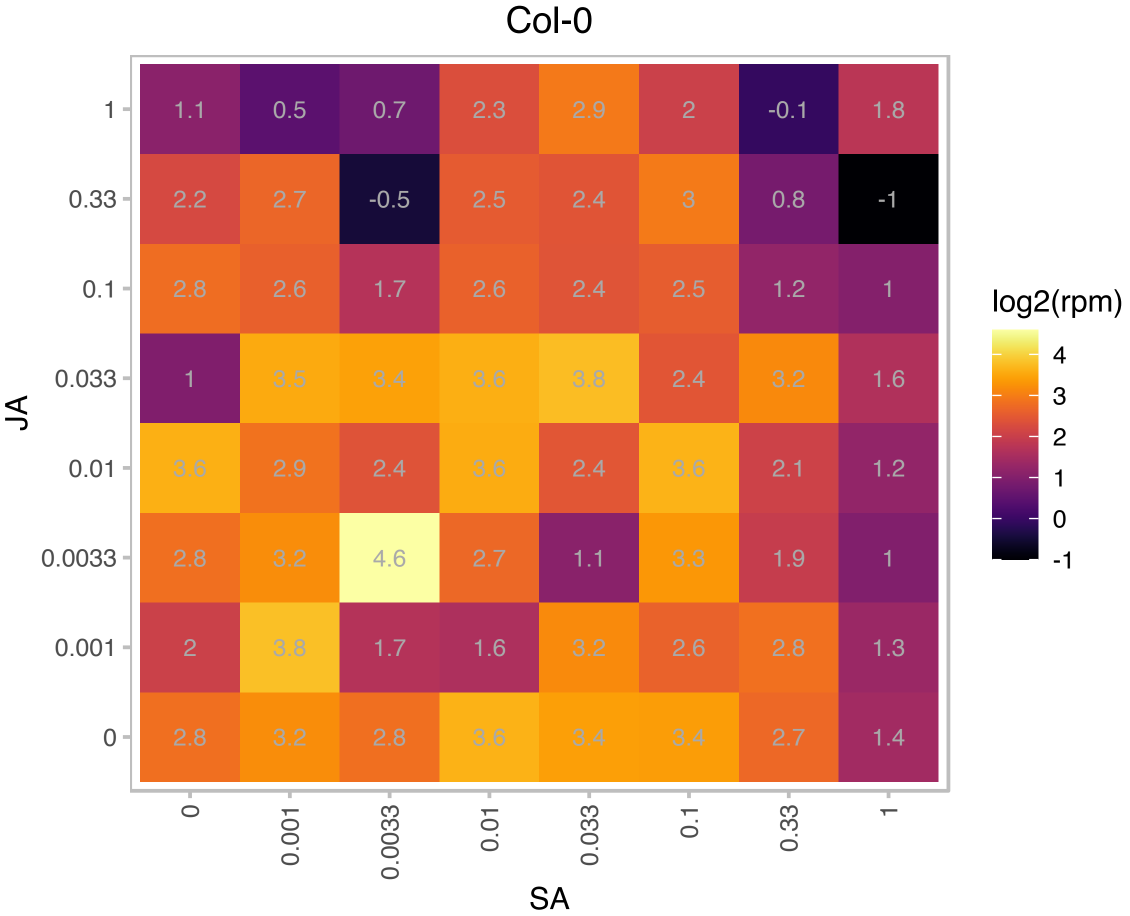


| | |
|---|---|
| Gene | AT2G28660 |
| is.expressed_genes | TRUE |
| is.responsive_genes | TRUE |
| is.apcluster | F-07 |

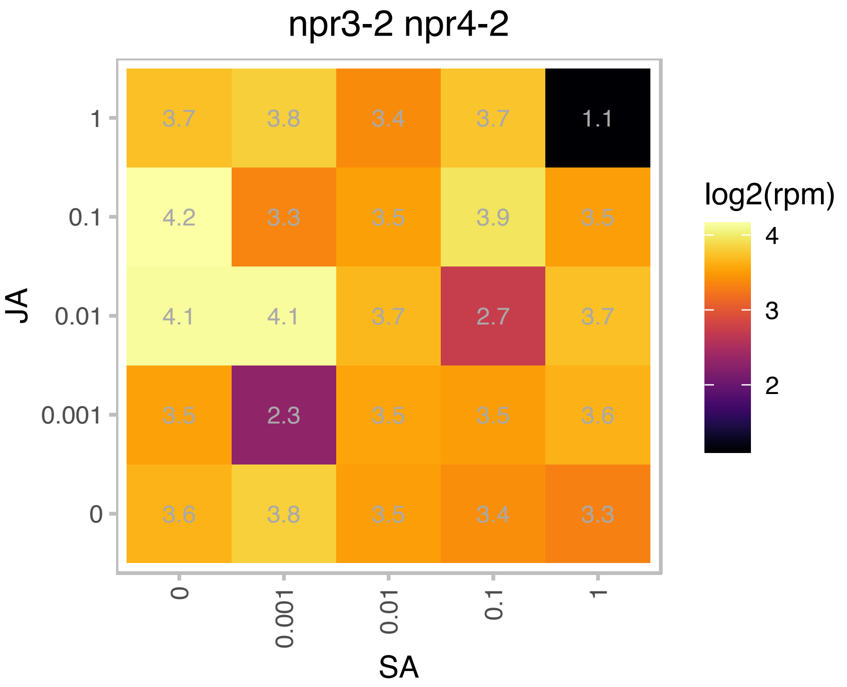


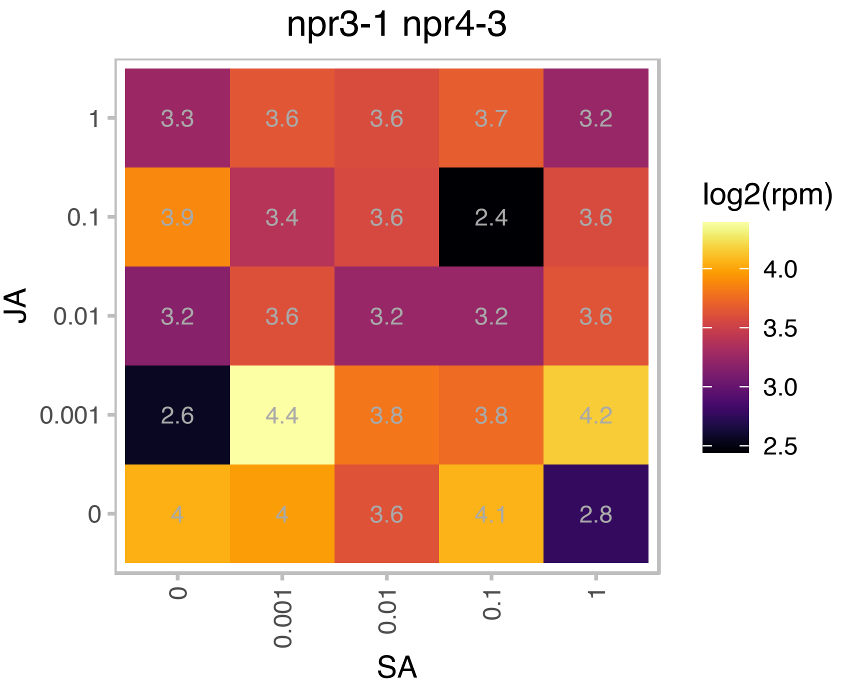

# AT2G31790.1 : NA

UDP-Glycosyltransferase superfamily protein

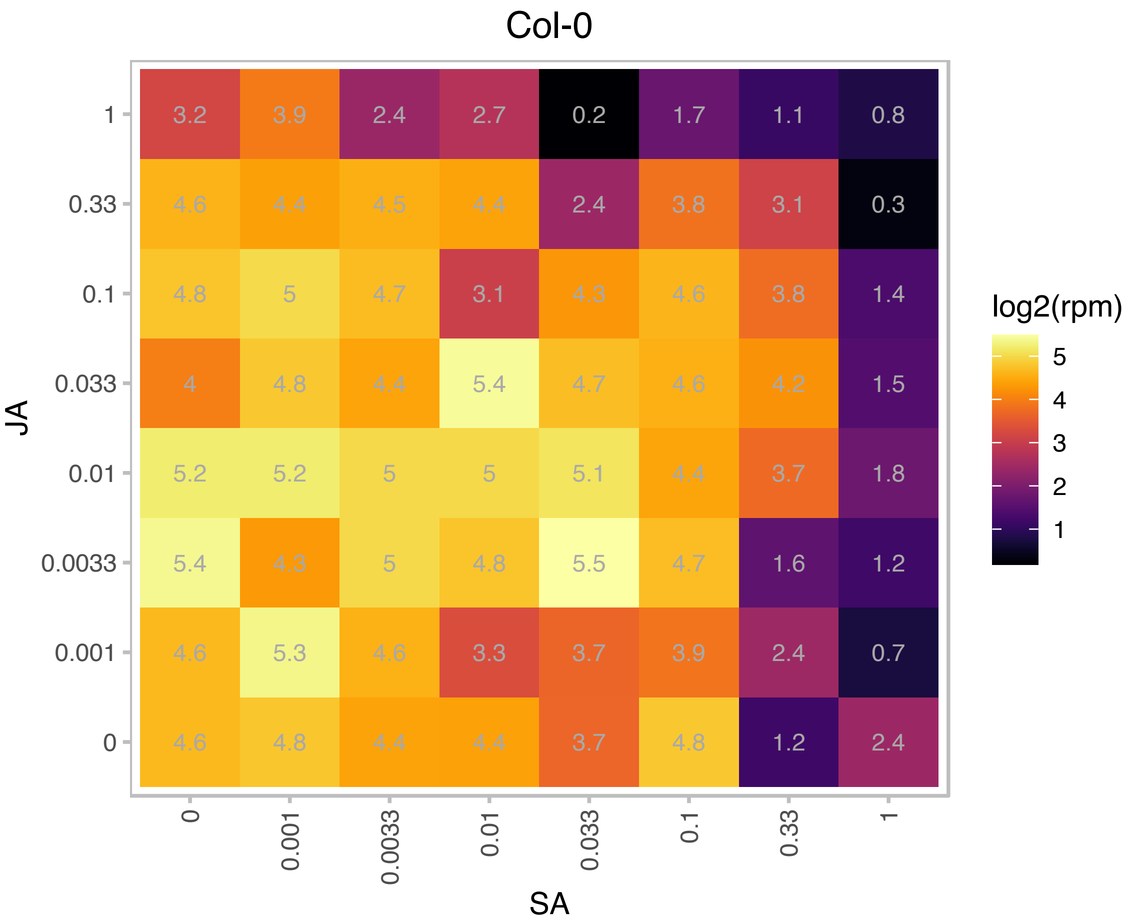


| | |
|---|---|
| Gene | AT2G31790 |
| is.expressed_genes | TRUE |
| is.responsive_genes | TRUE |
| is.apcluster | F-07 |

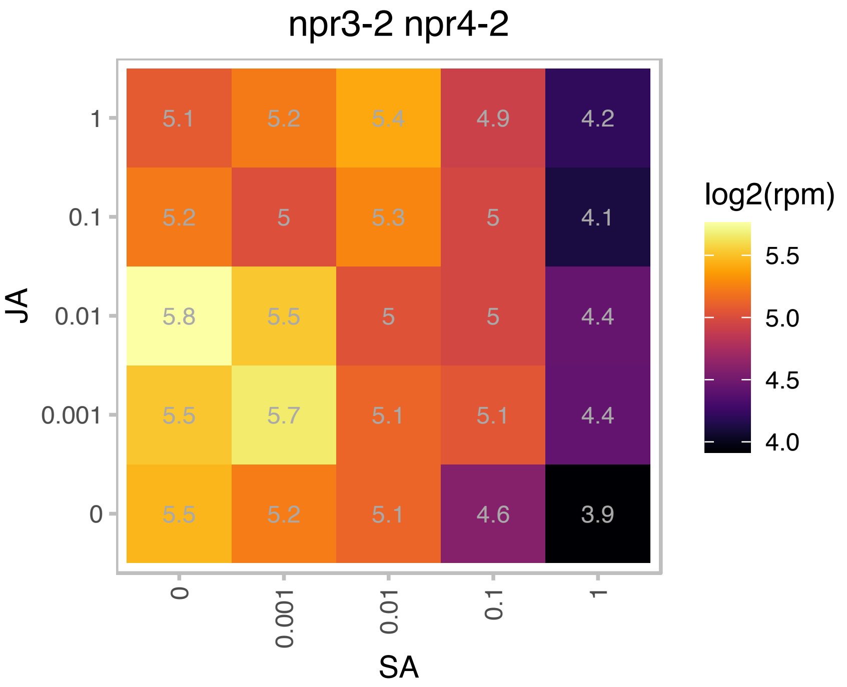


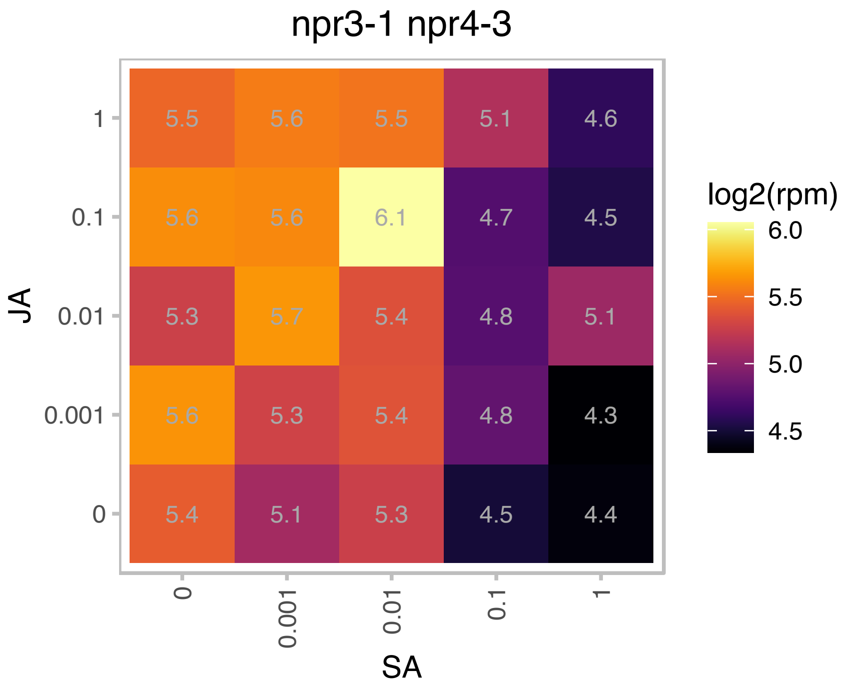

# AT2G32060.1 : NA

Ribosomal protein L7Ae/L30e/S12e/Gadd45 family protein

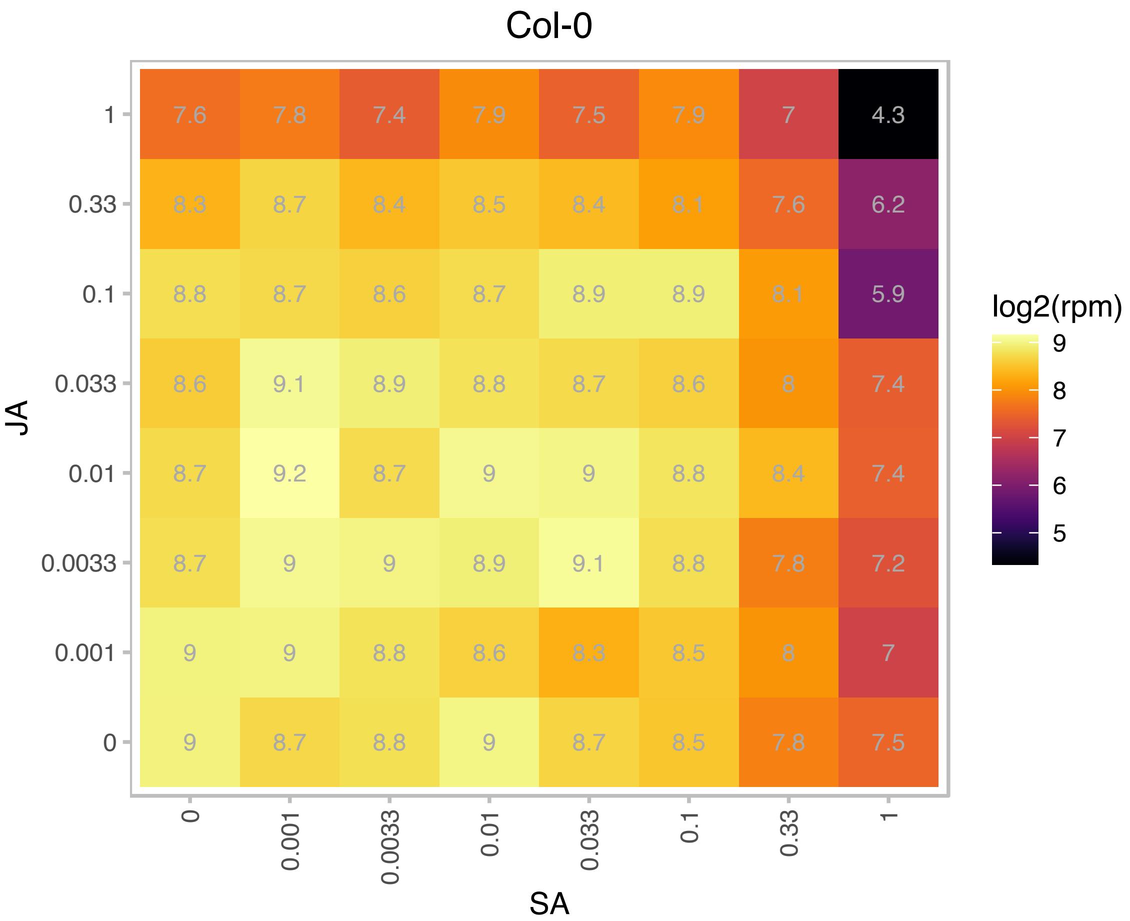


| | |
|---|---|
| Gene | AT2G32060 |
| is.expressed_genes | TRUE |
| is.responsive_genes | TRUE |
| is.apcluster | F-07 |

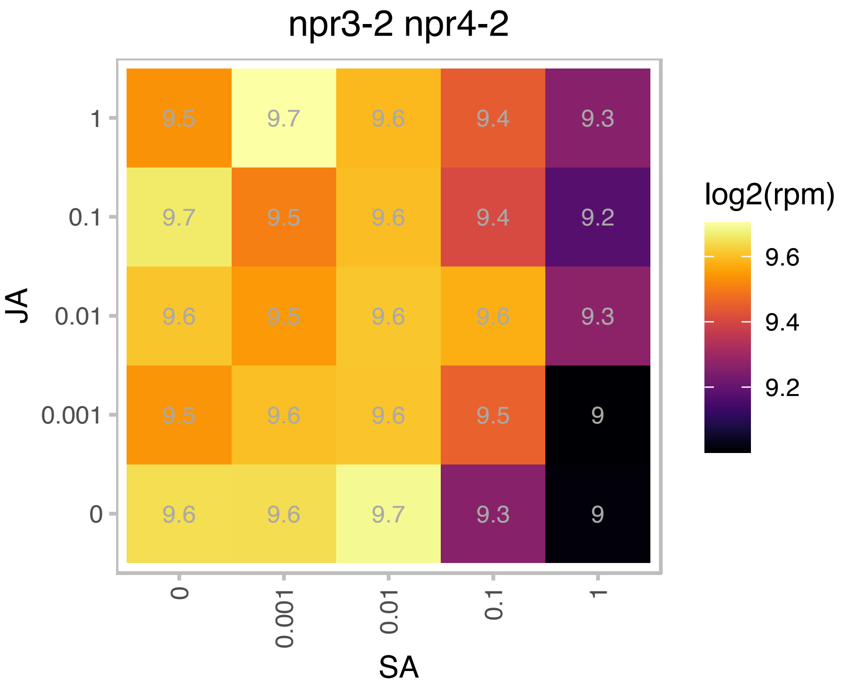


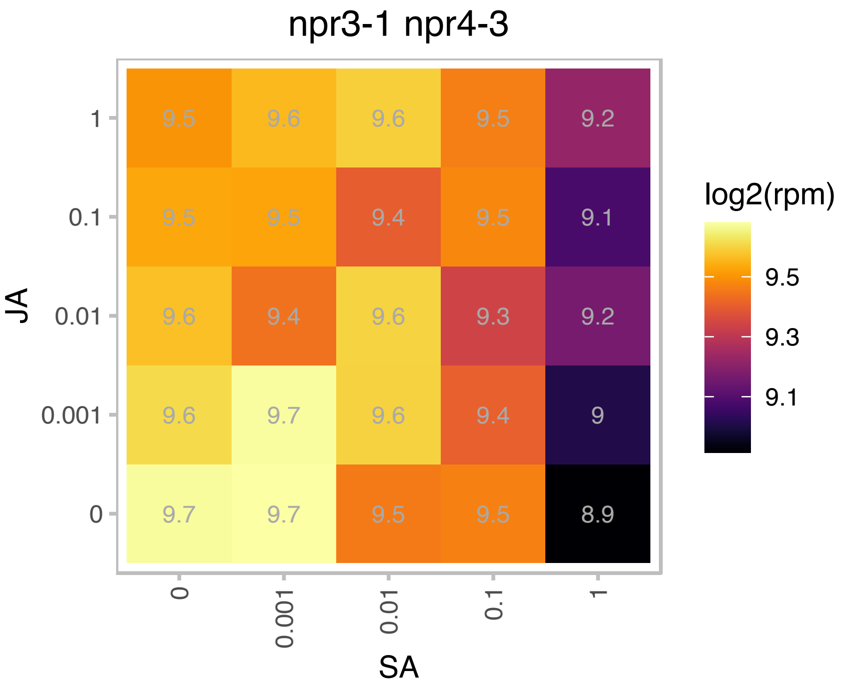

# AT2G33255.1 : NA

Haloacid dehalogenase-like hydrolase (HAD) superfamily protein

| | |
|---|---|
| Gene | AT2G33255 |
| is.expressed_genes | TRUE |
| is.responsive_genes | TRUE |
| is.apcluster | F-07 |

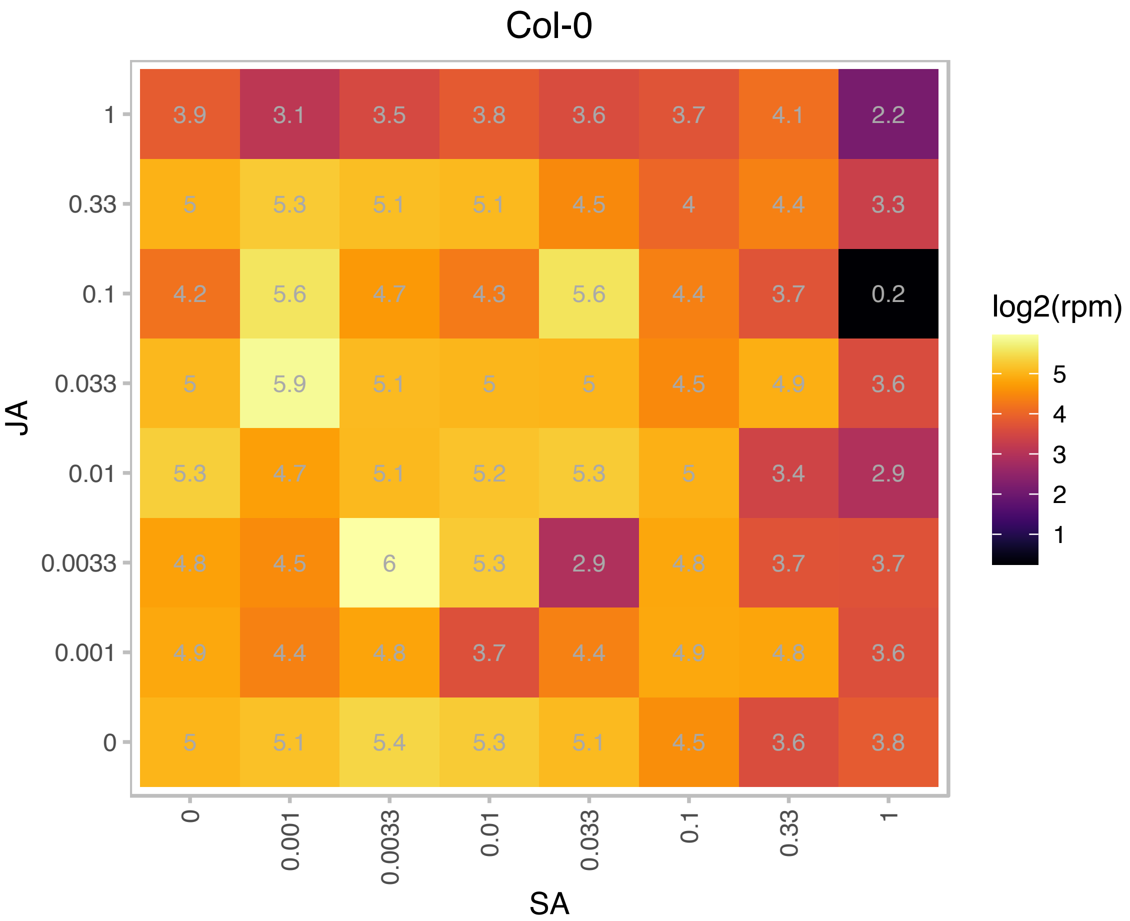


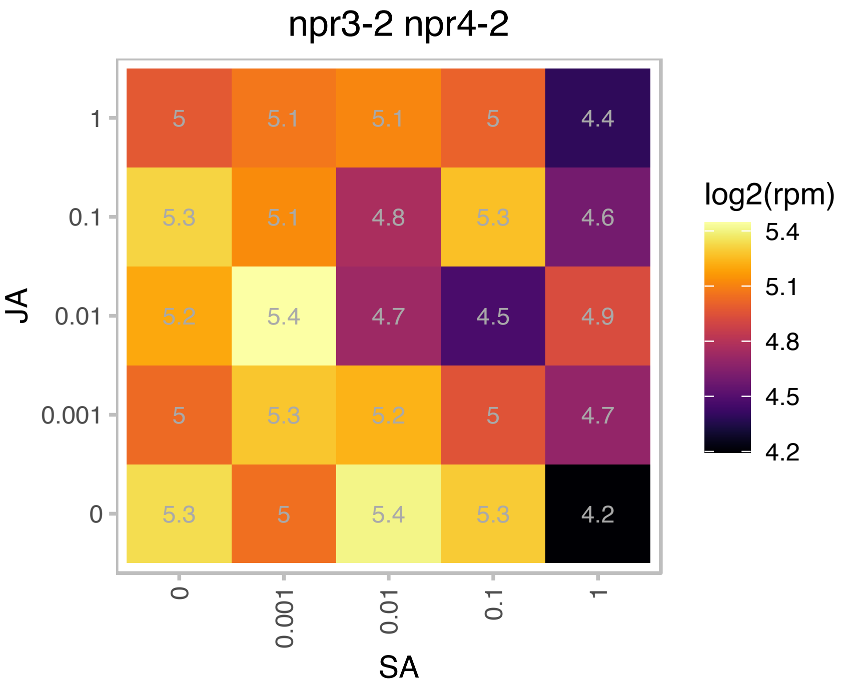


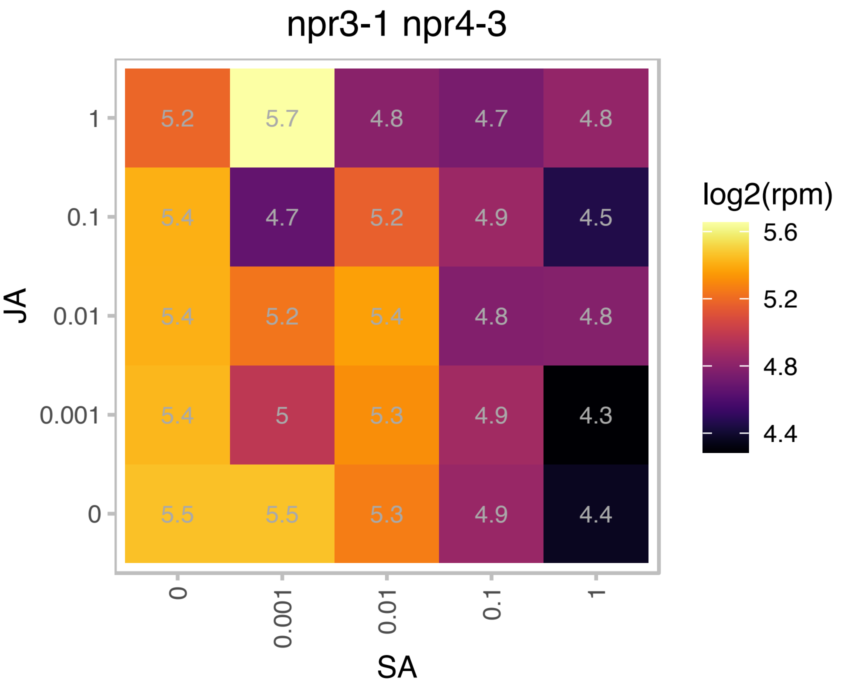

# AT2G34480.1 : L18aB

Ribosomal protein L18ae/LX family protein

Encodes a nuclear localized member of the ribosomal L18ae/LX protein family.  Loss of function mutations show reduced transmission through the gametophytes and embryo lethality.

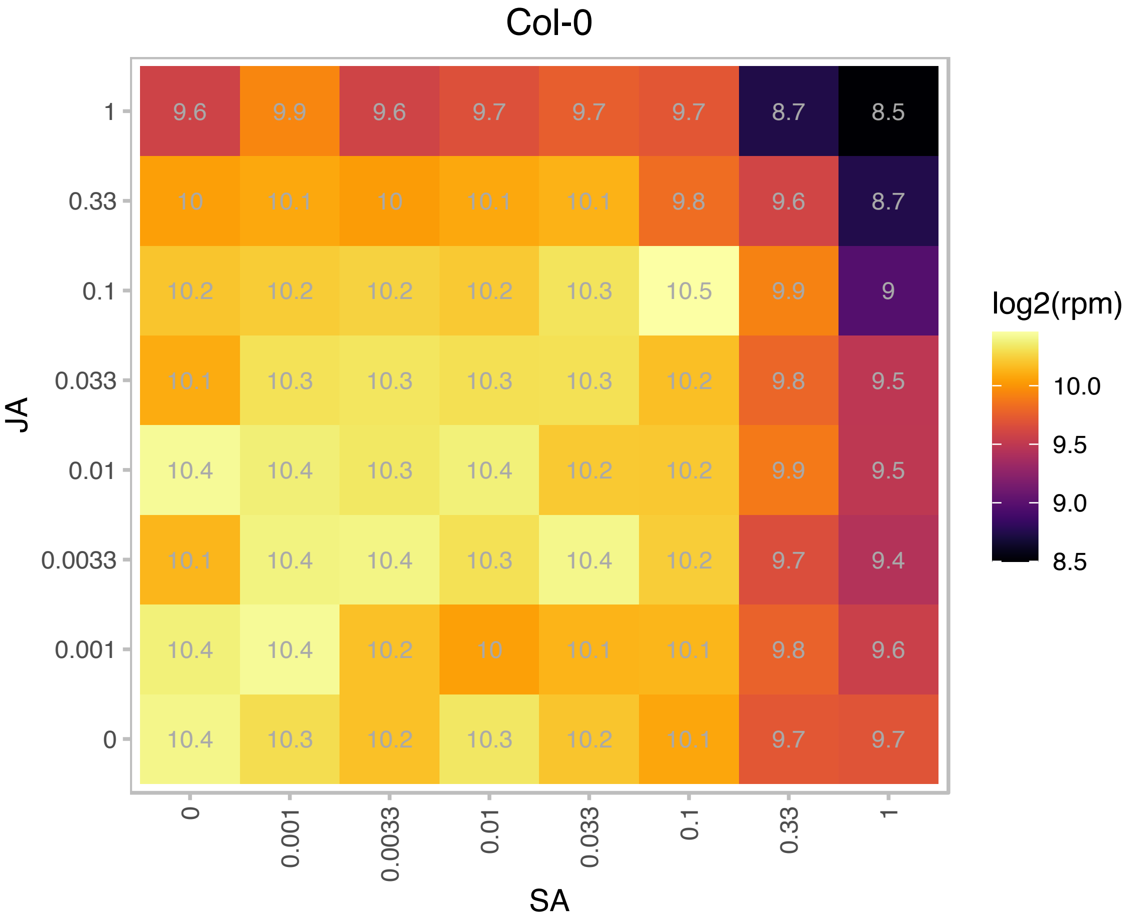


| | |
|---|---|
| Gene | AT2G34480 |
| is.expressed_genes | TRUE |
| is.responsive_genes | TRUE |
| is.apcluster | F-07 |

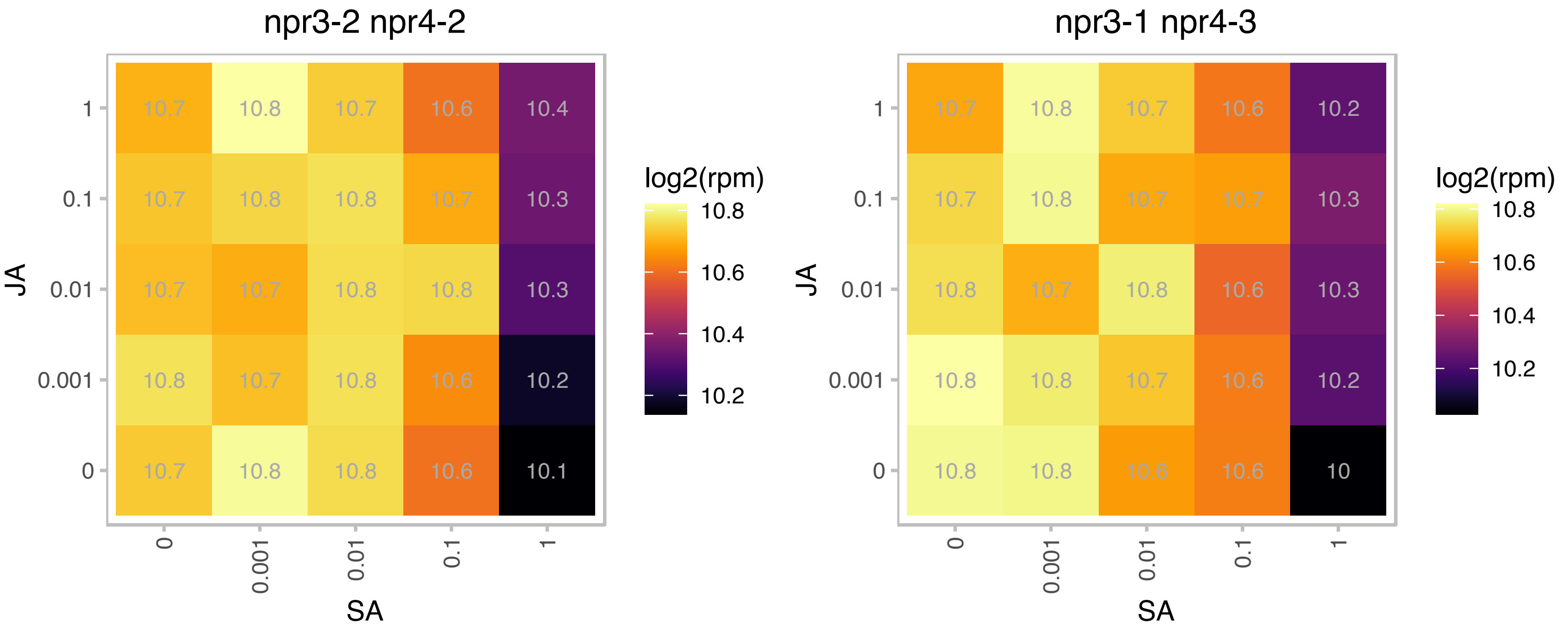

# AT2G35795.1 : NA

Chaperone DnaJ-domain superfamily protein

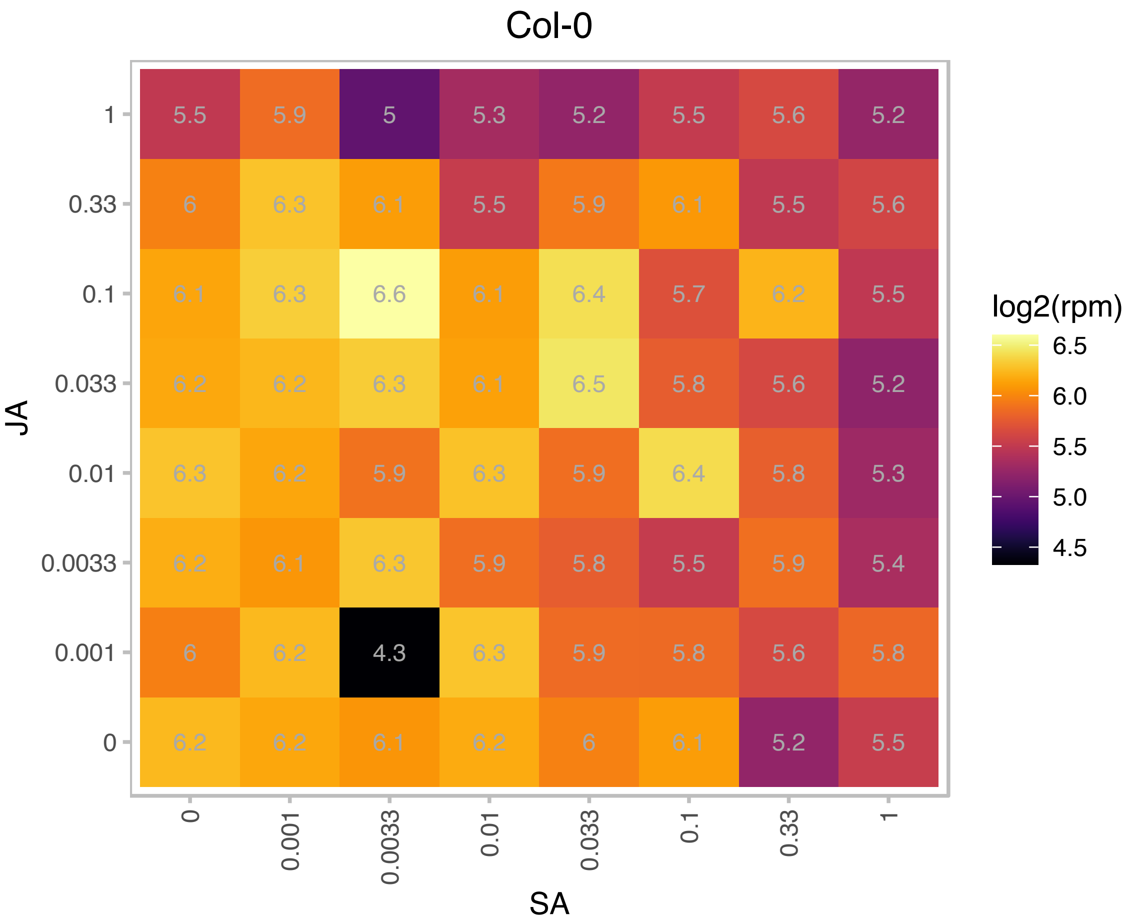


| | |
|---|---|
| Gene | AT2G35795 |
| is.expressed_genes | TRUE |
| is.responsive_genes | TRUE |
| is.apcluster | F-07 |

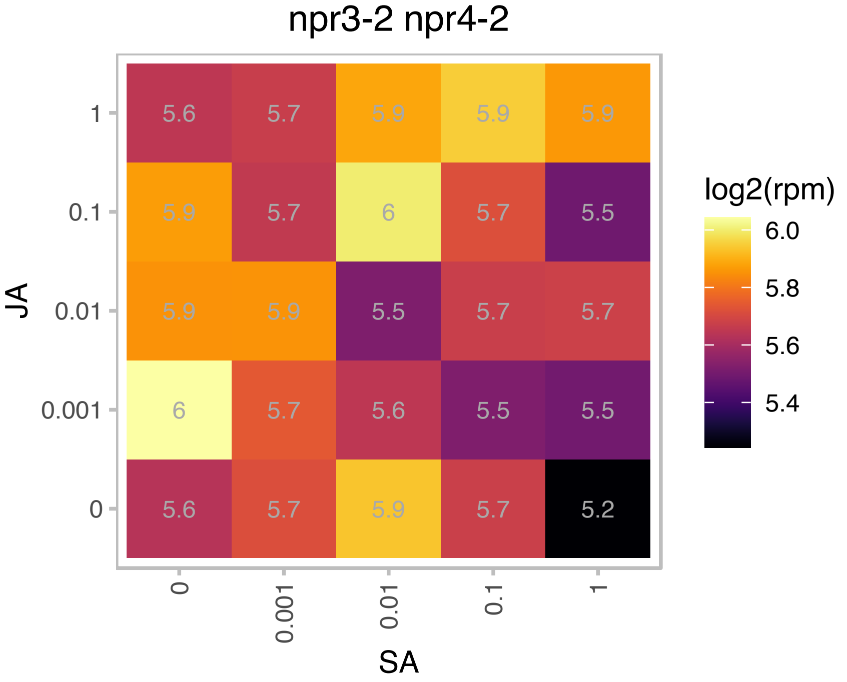


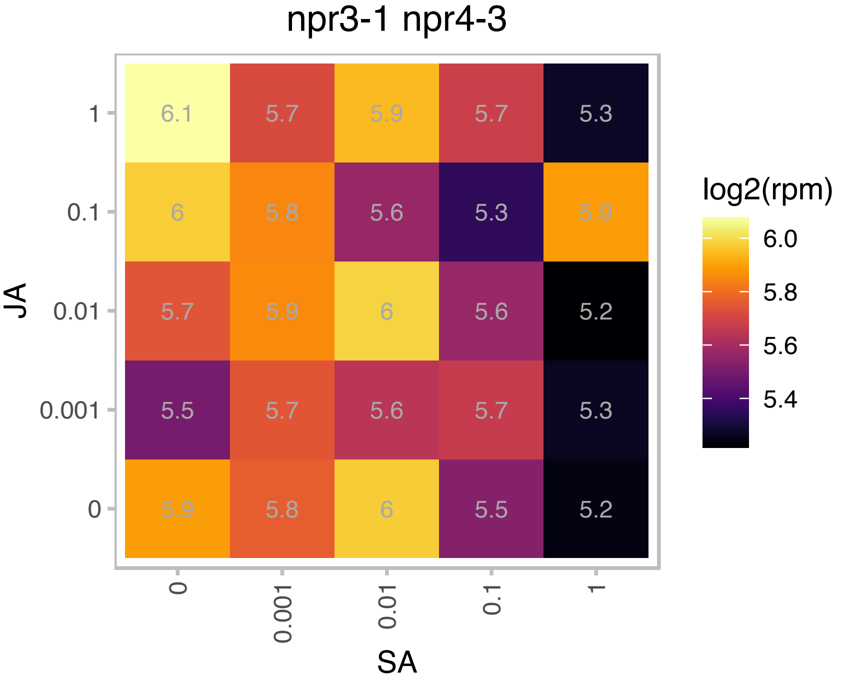

### AT2G36170.1 : RPL40A, UBQ2

Ubiquitin supergroup;Ribosomal protein L40e

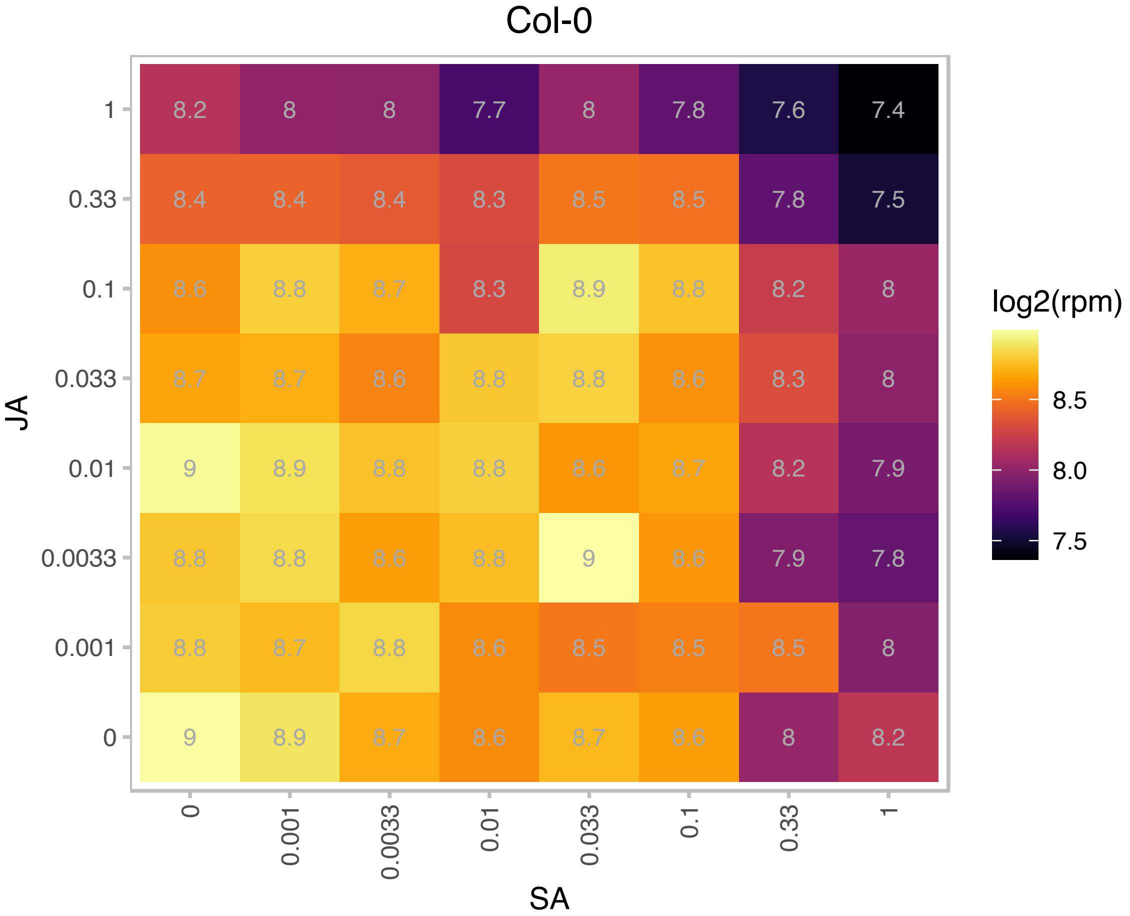


| | |
|---|---|
| Gene | AT2G36170 |
| is.expressed_genes | TRUE |
| is.responsive_genes | TRUE |
| is.apcluster | F-07 |

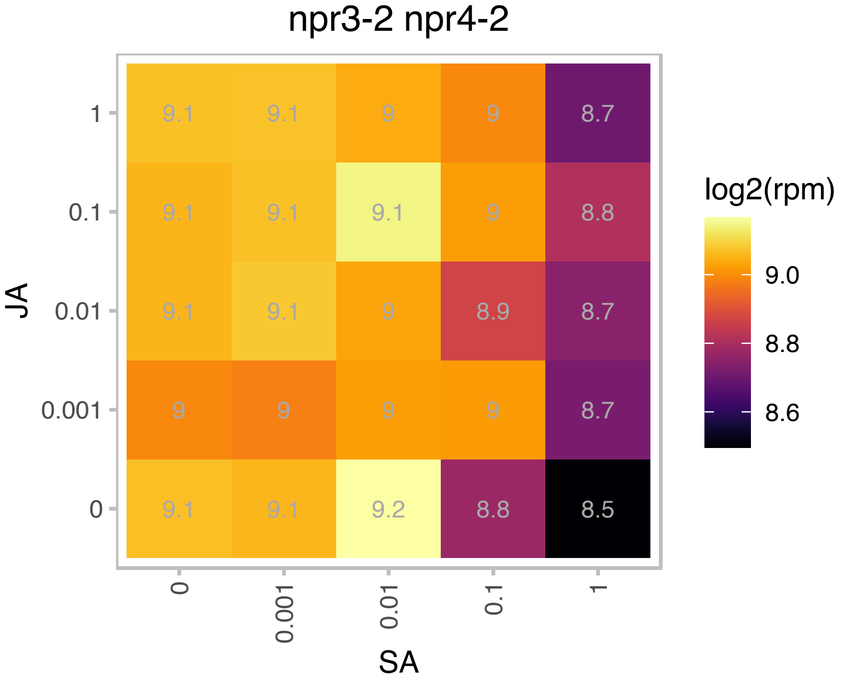


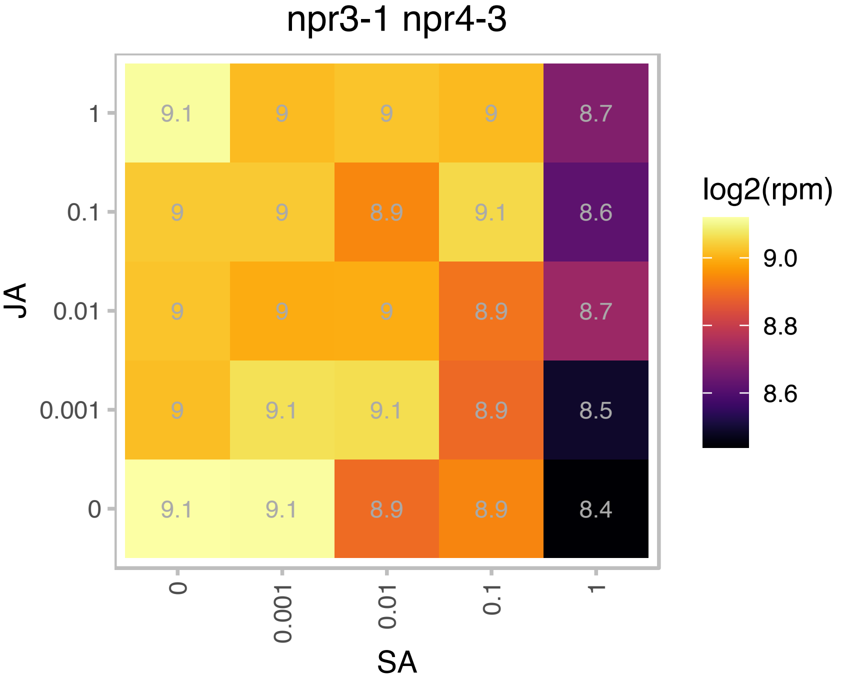

### AT2G36620.1 : RPL24A

ribosomal protein L24

RPL24A encodes ribosomal protein L24, homolog of cytosolic RPL24, found in archaea and higher eukaryotes. Arabidopsis has two RPL24 homologs, RPL24A (AT2G36620) and RPL24B (AT3G53020).

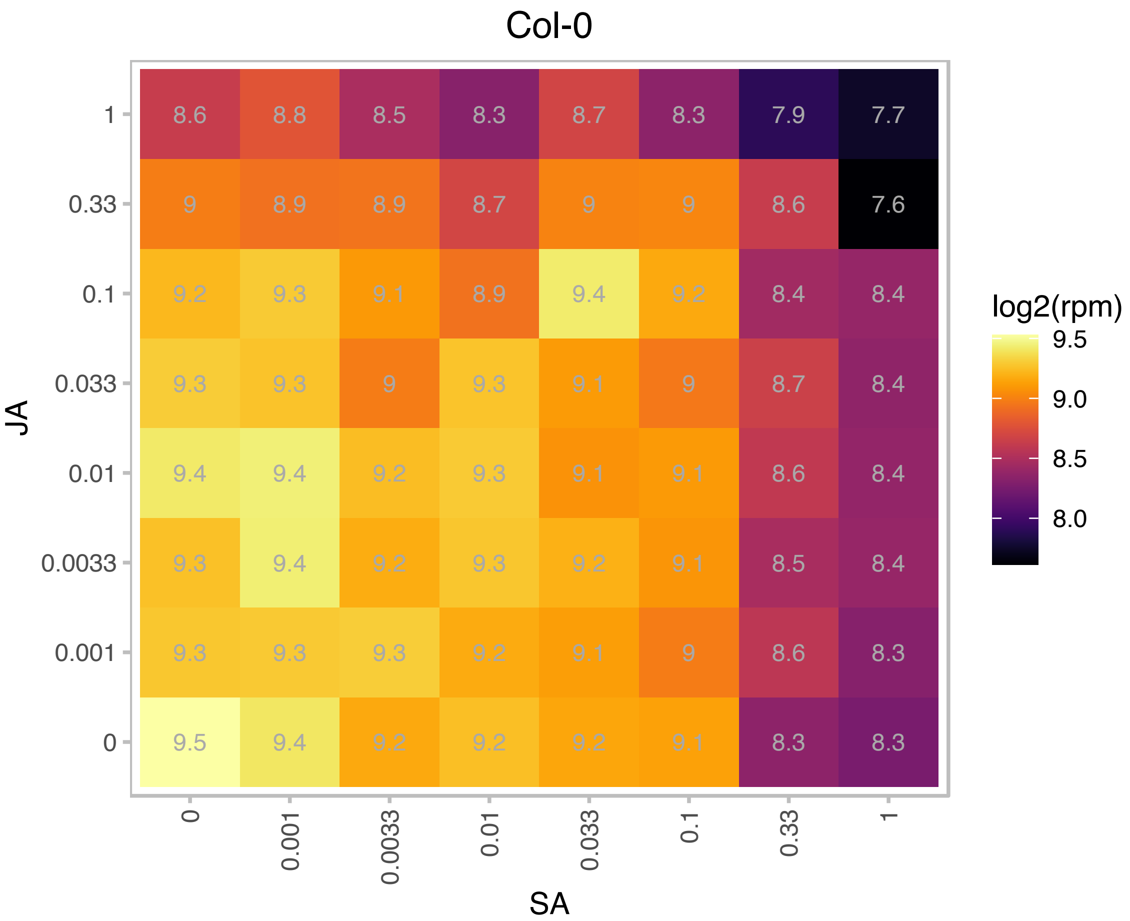


| | |
|---|---|
| Gene | AT2G36620 |
| is.expressed_genes | TRUE |
| is.responsive_genes | TRUE |
| is.apcluster | F-07 |

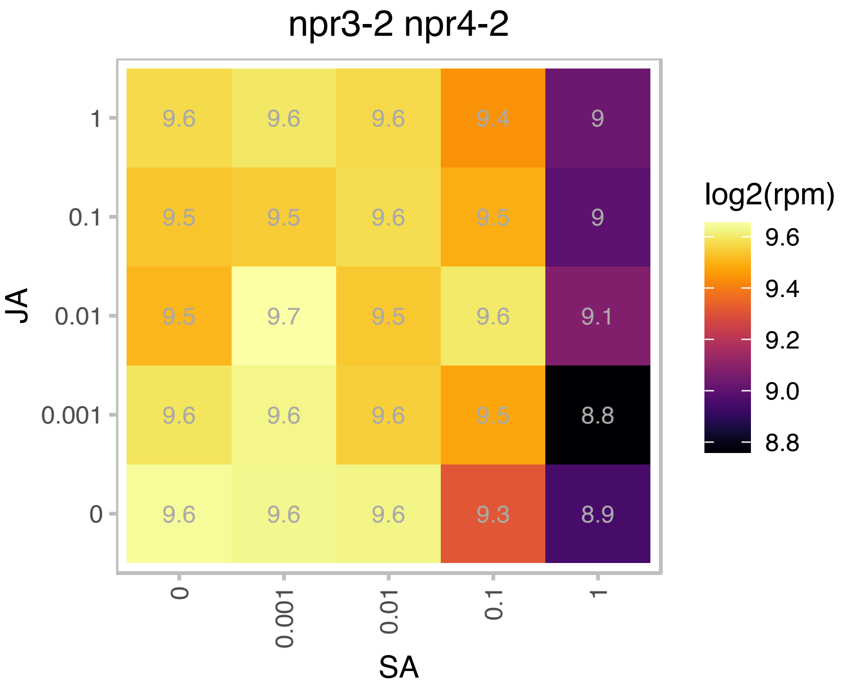


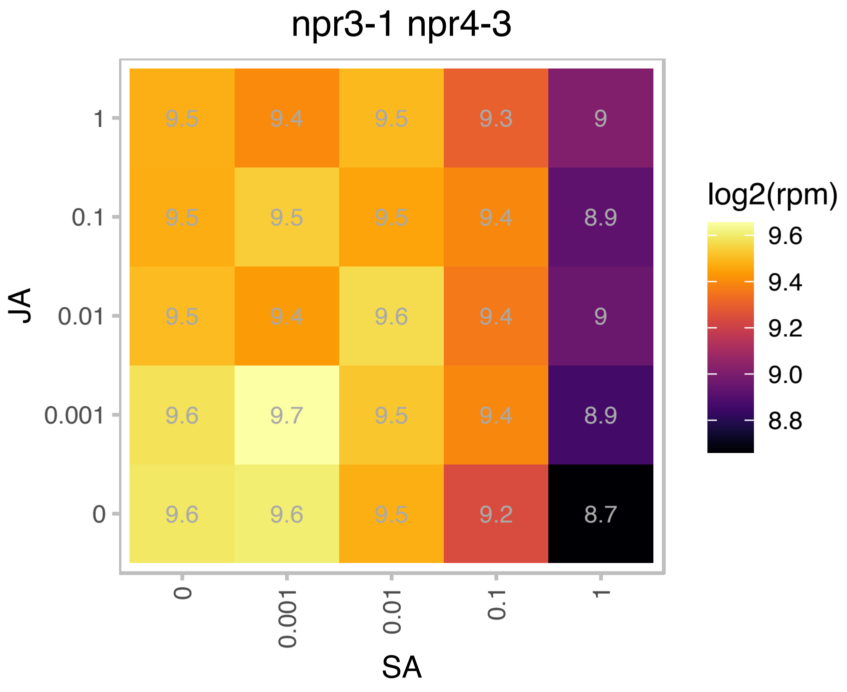

### AT2G37080.1 : RIP2, RIP3

ROP interactive partner 3

Encodes RIP2 (ROP interactive partner 2), a putative Rho protein effector, interacting specifically with the active form of ROPs (Rho proteins of plants).

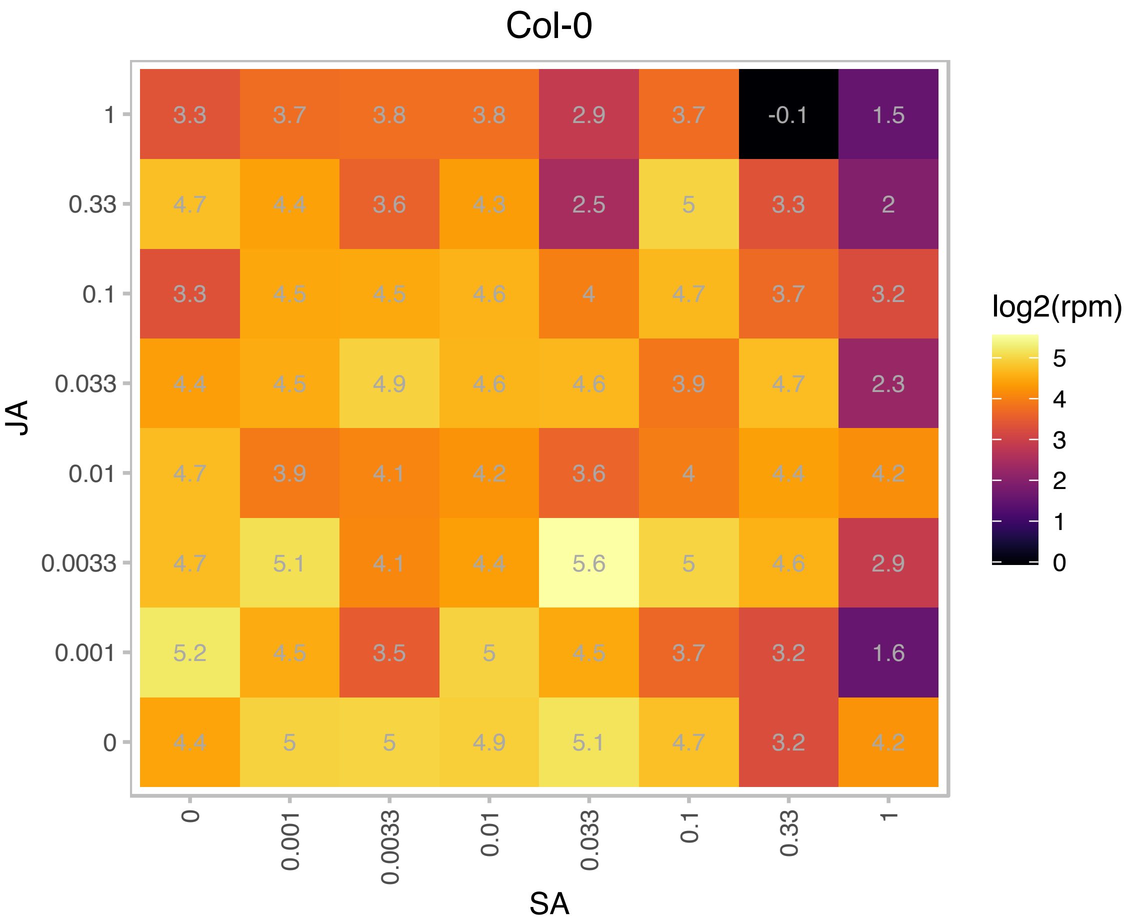


| | |
|---|---|
| Gene | AT2G37080 |
| is.expressed_genes | TRUE |
| is.responsive_genes | TRUE |
| is.apcluster | F-07 |

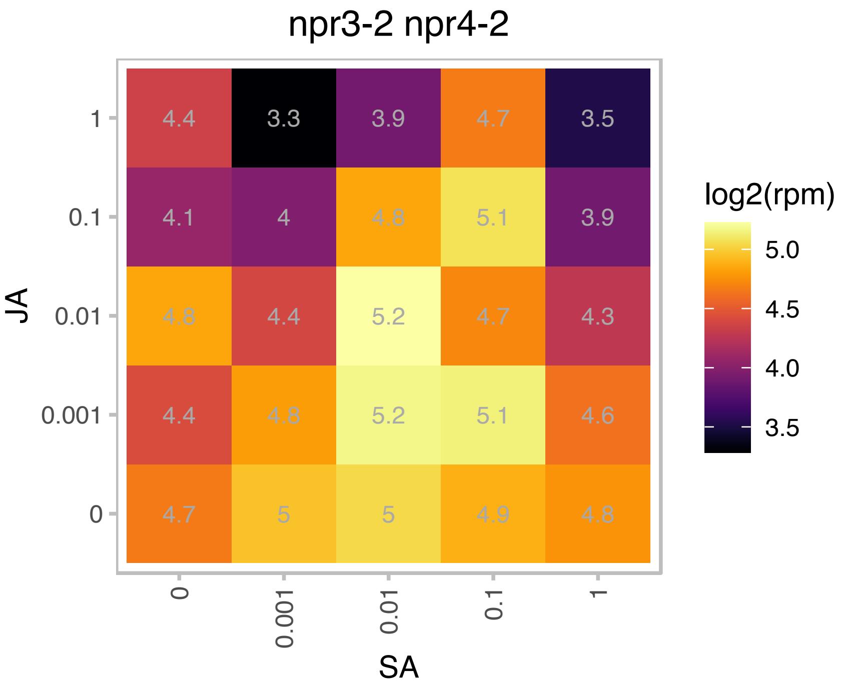


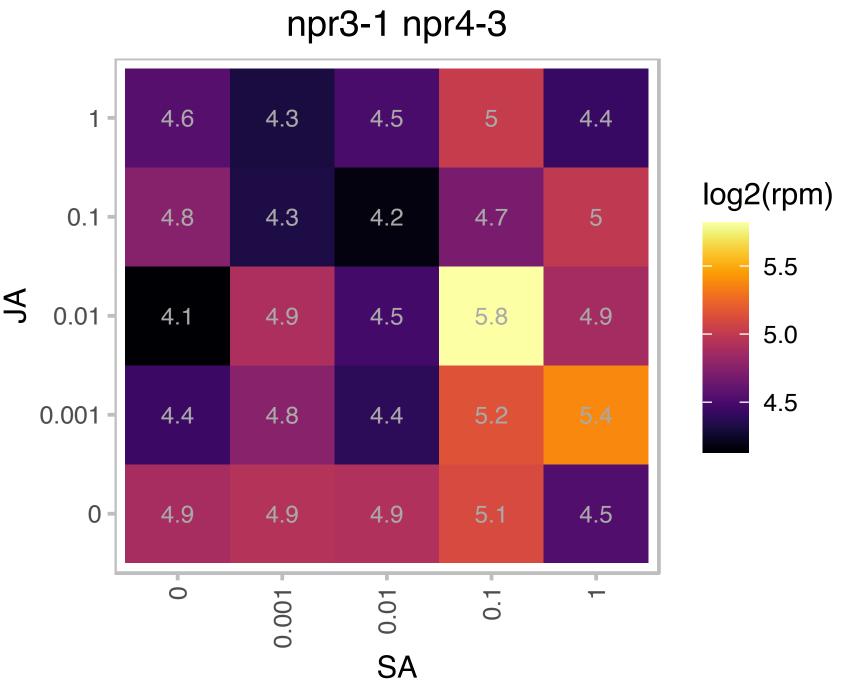

# AT2G37190.1 : NA

Ribosomal protein L11 family protein

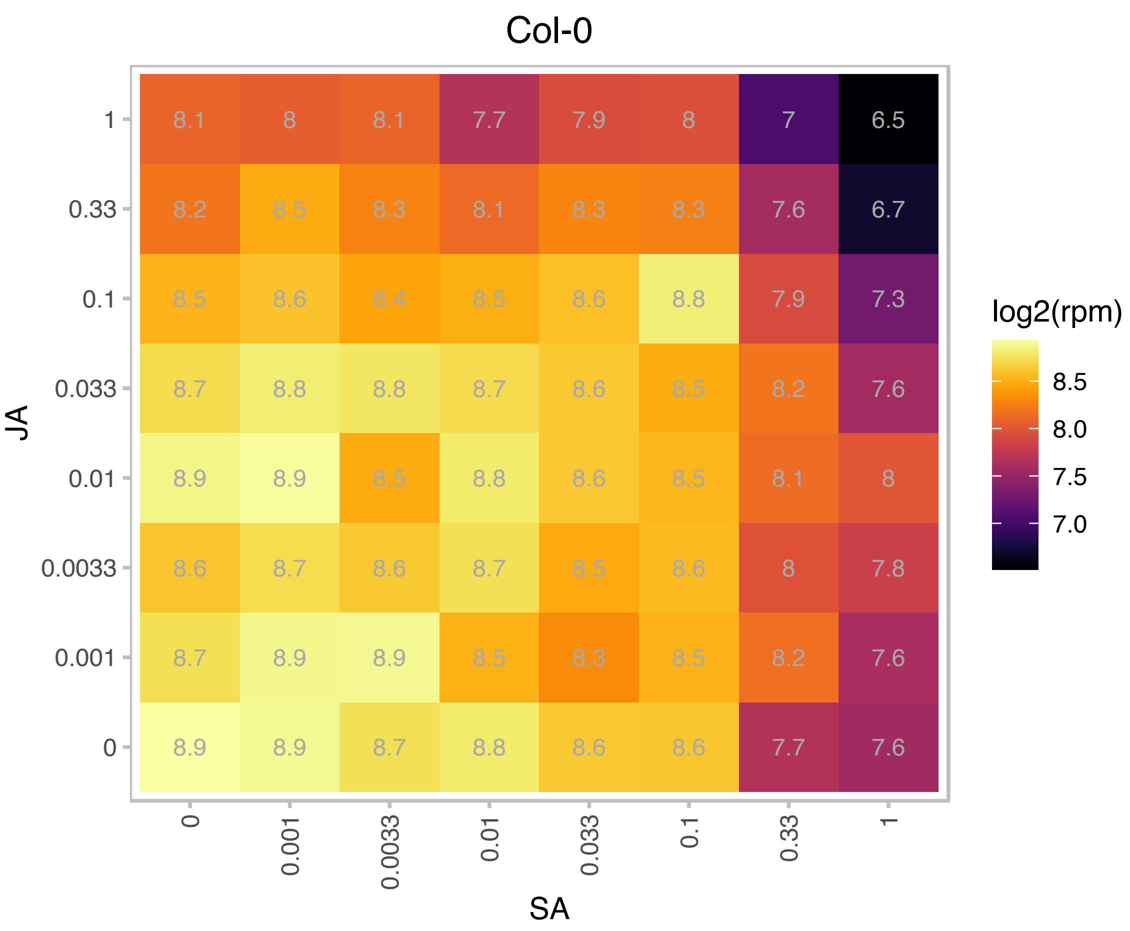


| | |
|---|---|
| Gene | AT2G37190 |
| is.expressed_genes | TRUE |
| is.responsive_genes | TRUE |
| is.apcluster | F-07 |

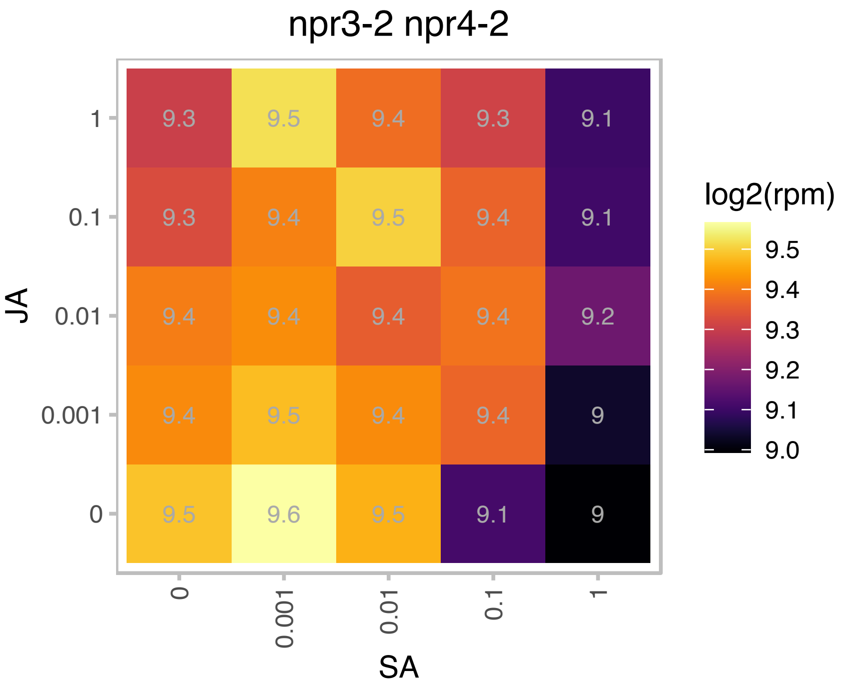


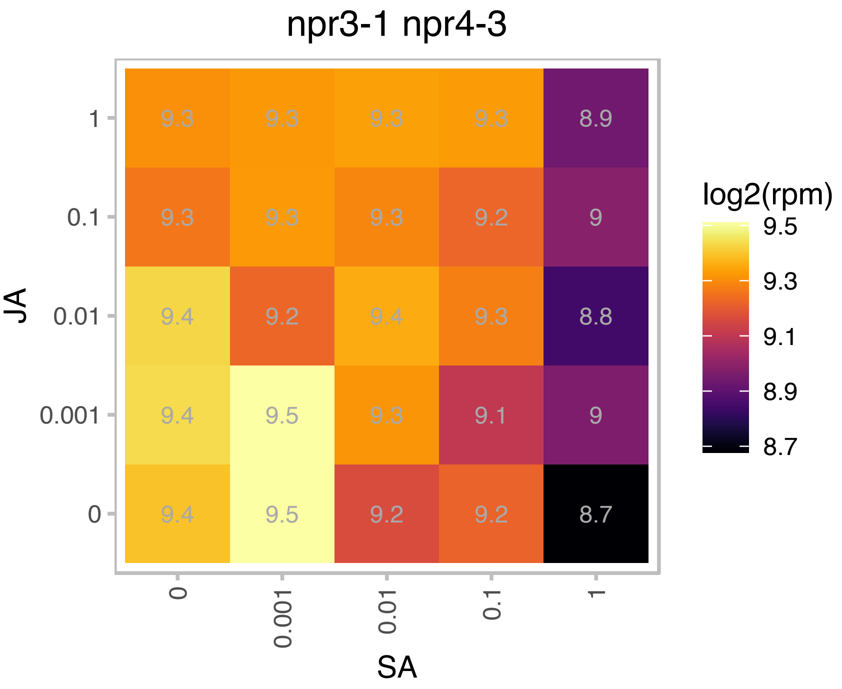

### AT2G37270.1 : ATRPS5B, RPS5B

ribosomal protein 5B

One of two genes encoding the ribosomal protein S5. Expressed at a lower level compared to ATRPS5A.

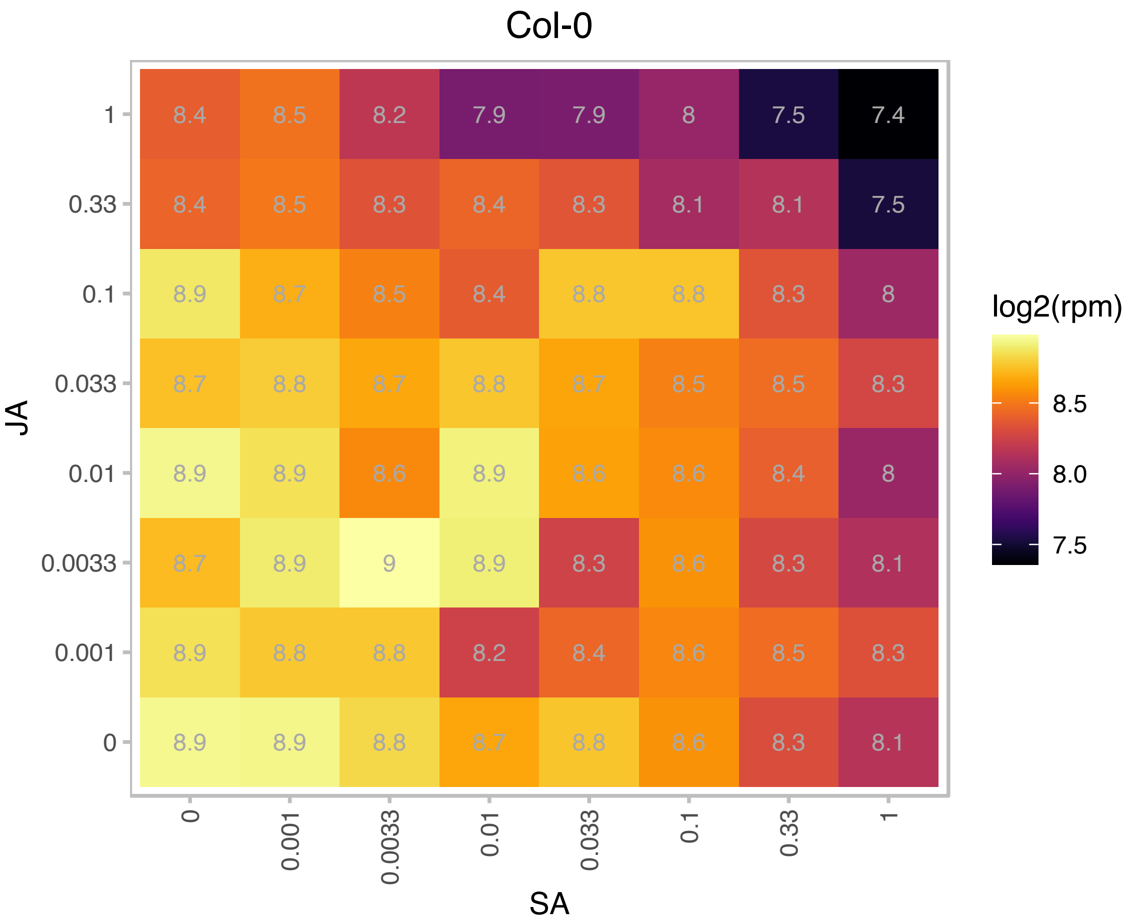


| | |
|---|---|
| Gene | AT2G37270 |
| is.expressed_genes | TRUE |
| is.responsive_genes | TRUE |
| is.apcluster | F-07 |

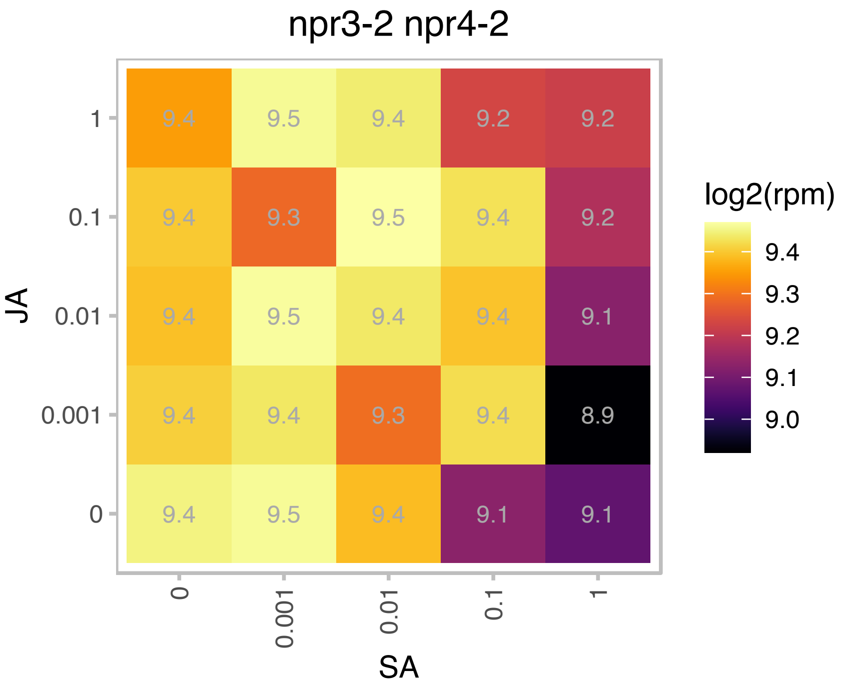


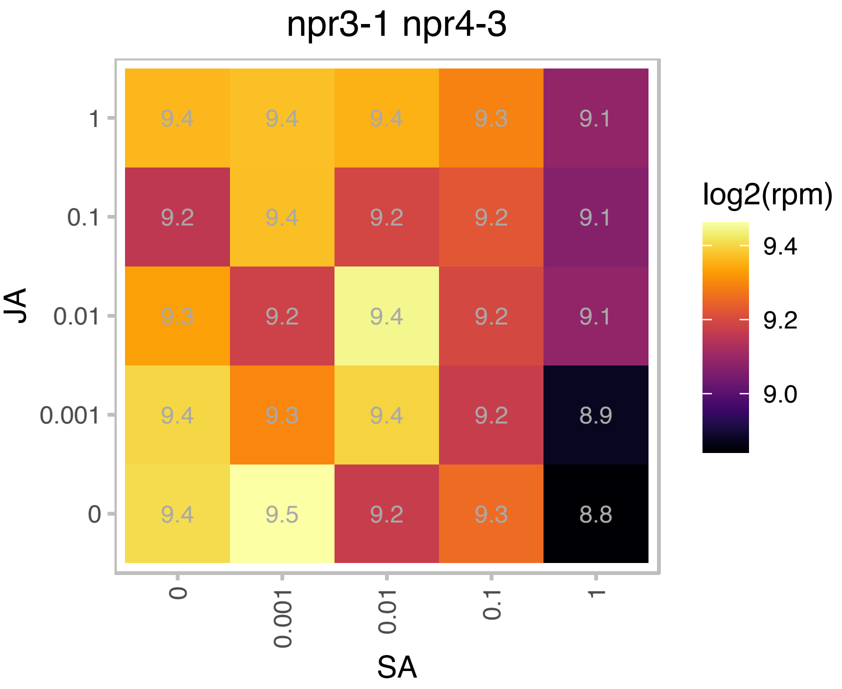

# AT2G37600.1 : NA

Ribosomal protein L36e family protein

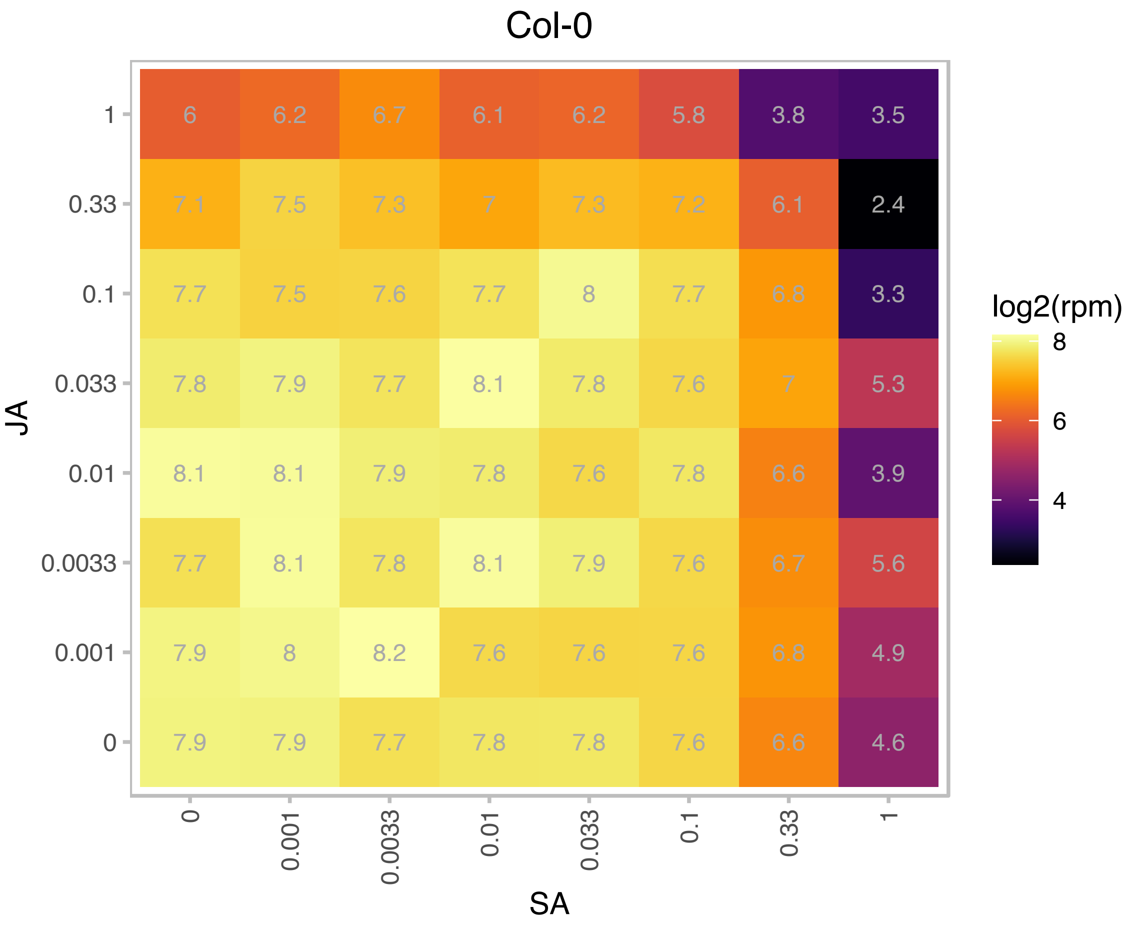


| | |
|---|---|
| Gene | AT2G37600 |
| is.expressed_genes | TRUE |
| is.responsive_genes | TRUE |
| is.apcluster | F-07 |

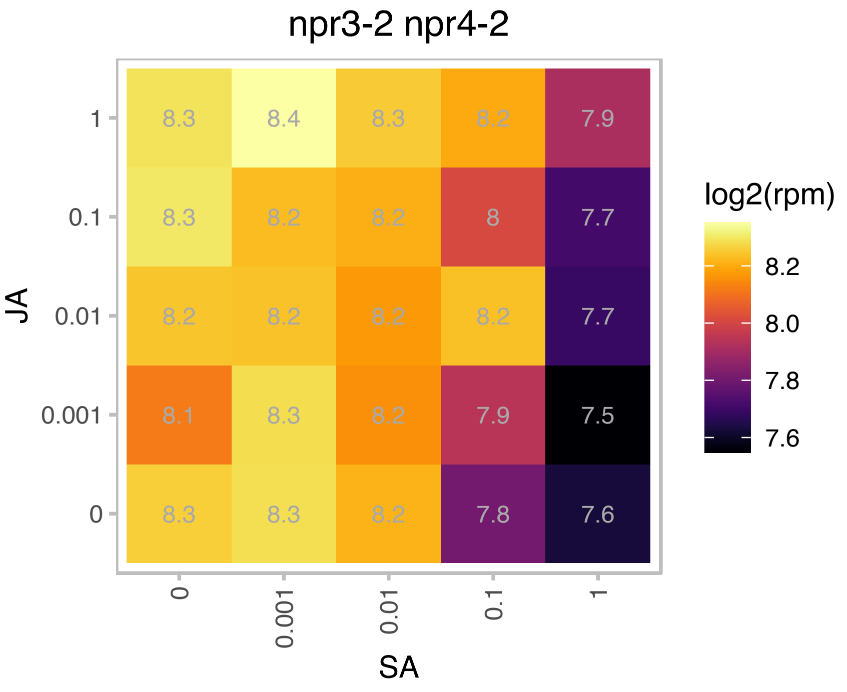


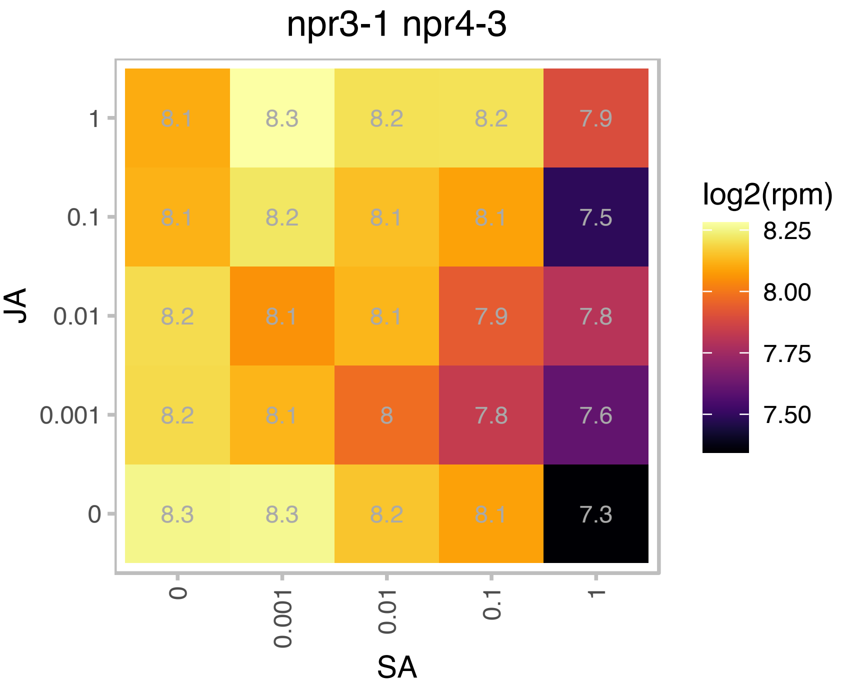

### AT2G38810.1 : H2A.Z, HTA8

histone H2A 8,
Encodes HTA8, a histone H2A protein. Loss of all H2A.Z (triple mutant with HTA9 and HTA11) results in a reduction in DNA methylation of transposons but not that of genes. Loss of H2A.Z causes misregulation of many genes involved in the response to developmental and environmental cues, and that these genes tend to have high levels of gene-body H2A.Z.

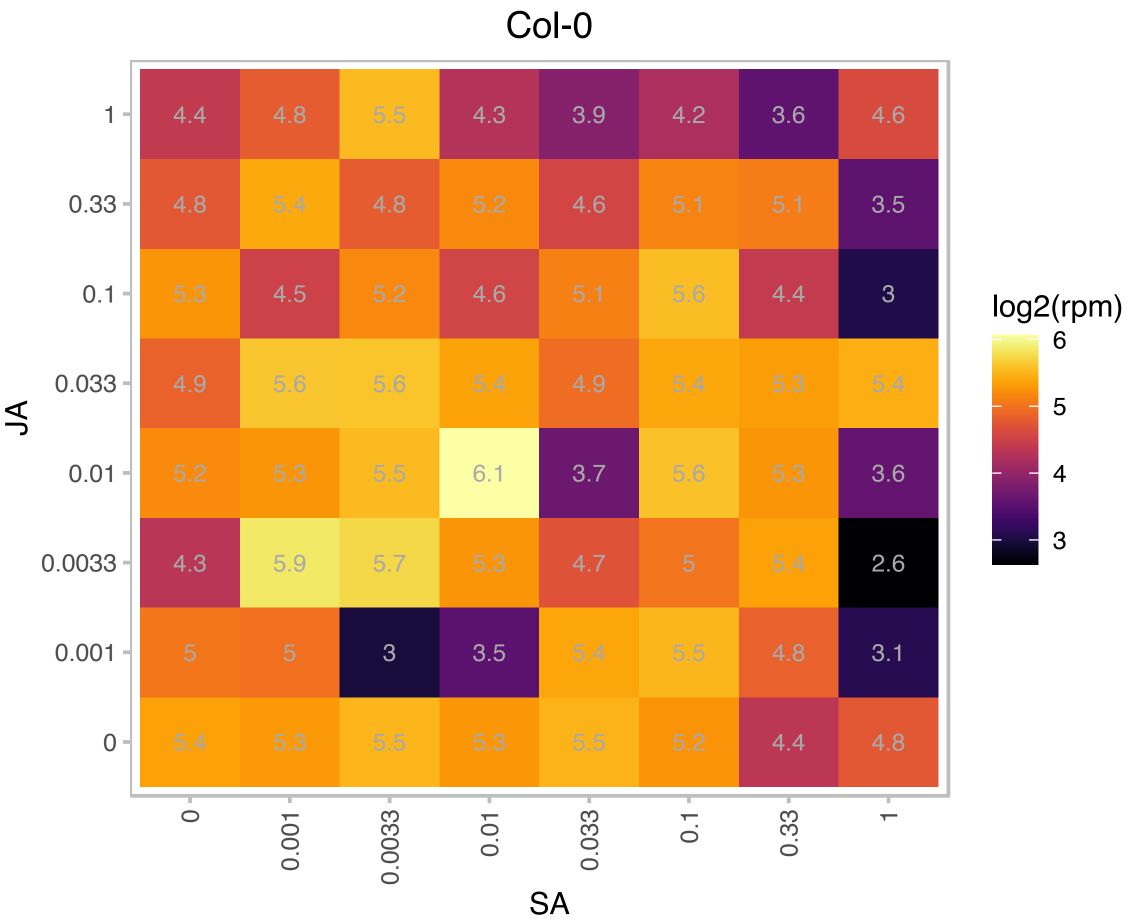


| | |
|---|---|
| Gene | AT2G38810 |
| is.expressed_genes | TRUE |
| is.responsive_genes | TRUE |
| is.apcluster | F-07 |

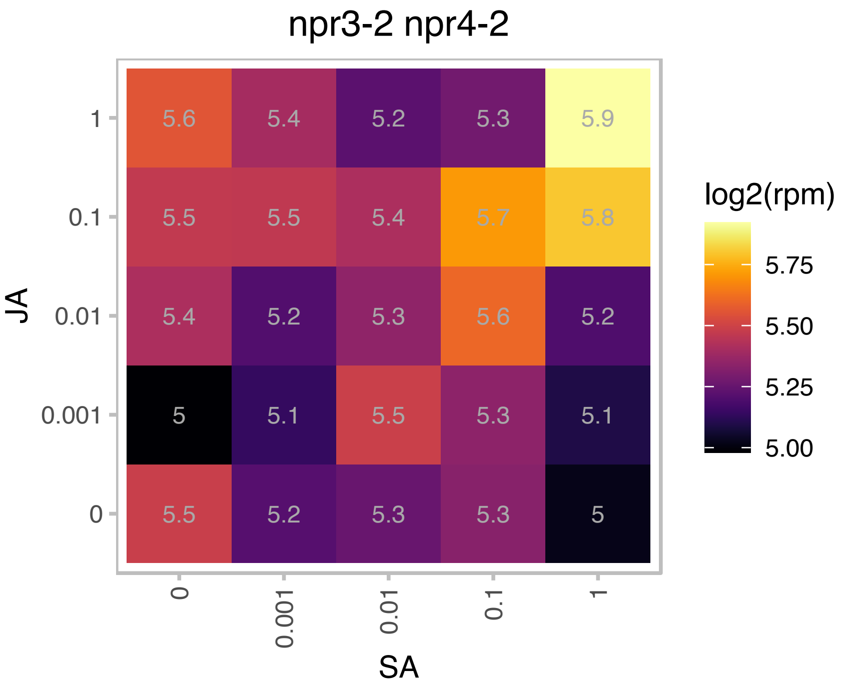


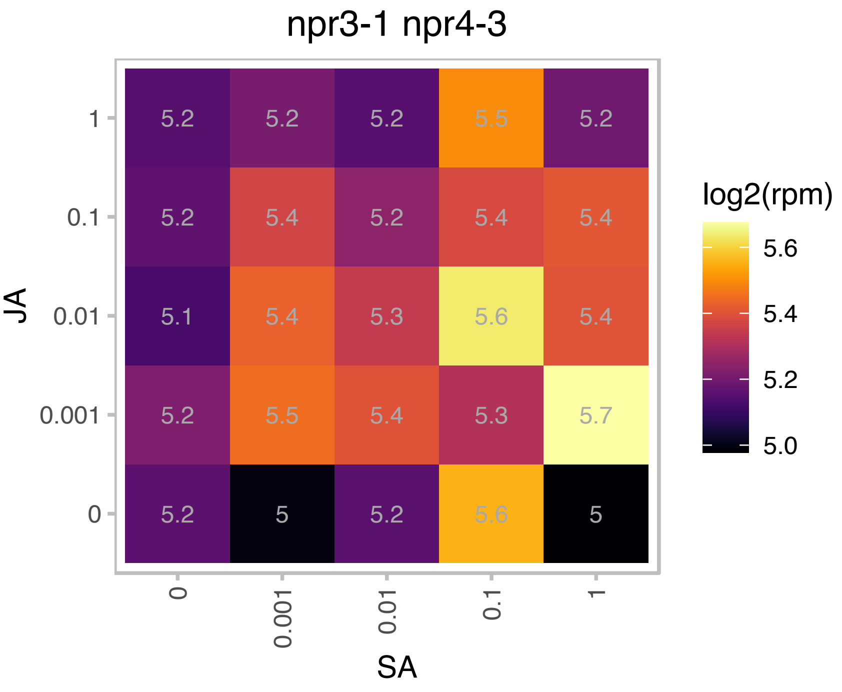

# AT2G39390.1 : NA

Ribosomal L29 family protein

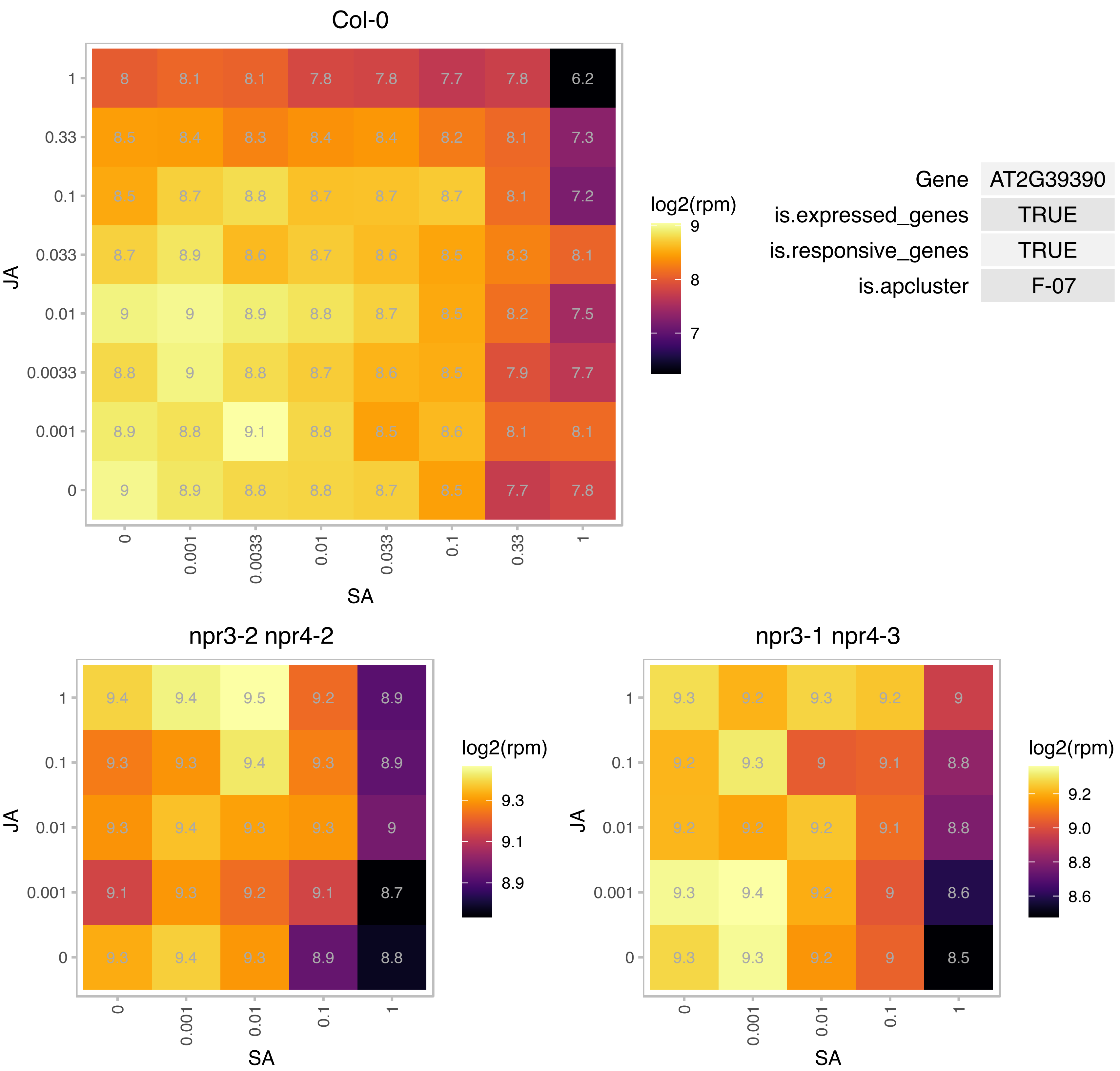

### AT2G39460.1 : ATRPL23A, RPL23A, RPL23A1, RPL23AA

ribosomal protein L23AA

Encodes a 60S ribosomal protein L23aA (AtrpL23aA). Paralog of RLPL23aB.

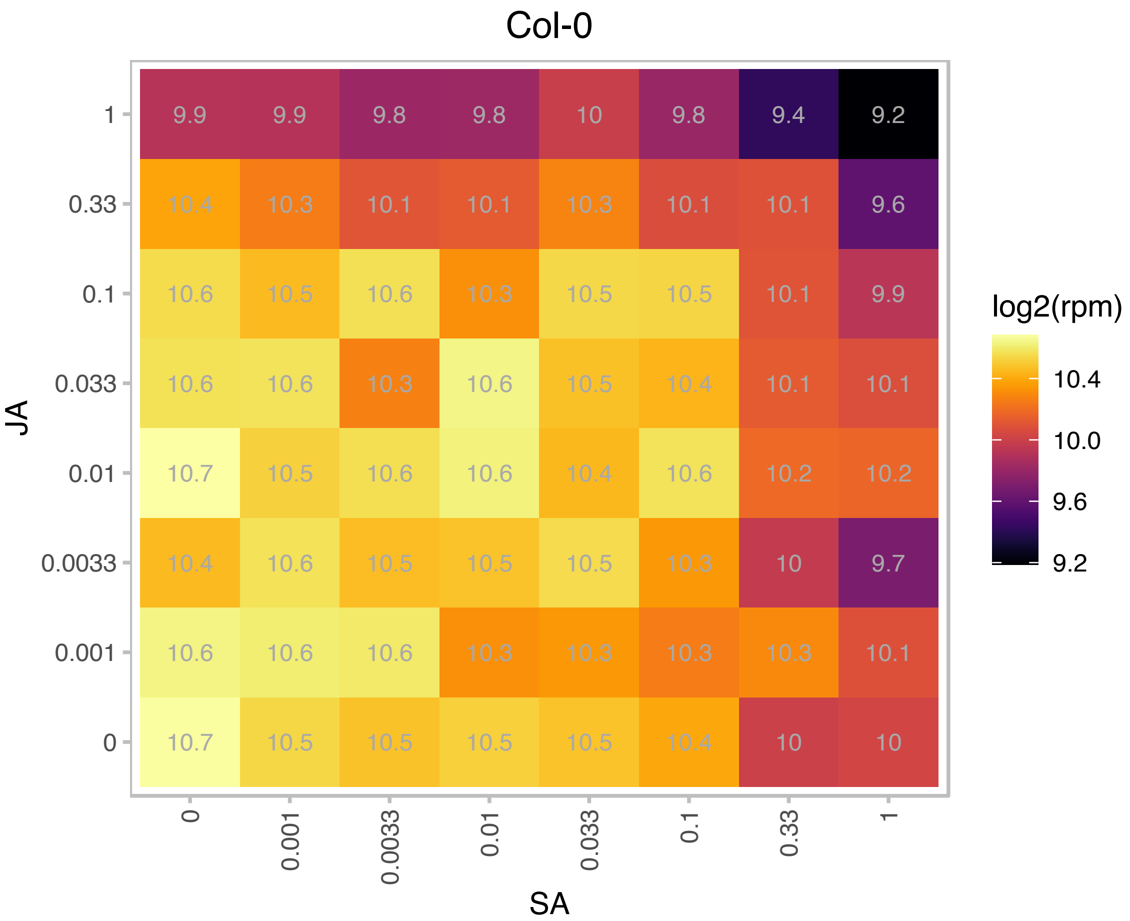


| | |
|---|---|
| Gene | AT2G39460 |
| is.expressed_genes | TRUE |
| is.responsive_genes | TRUE |
| is.apcluster | F-07 |

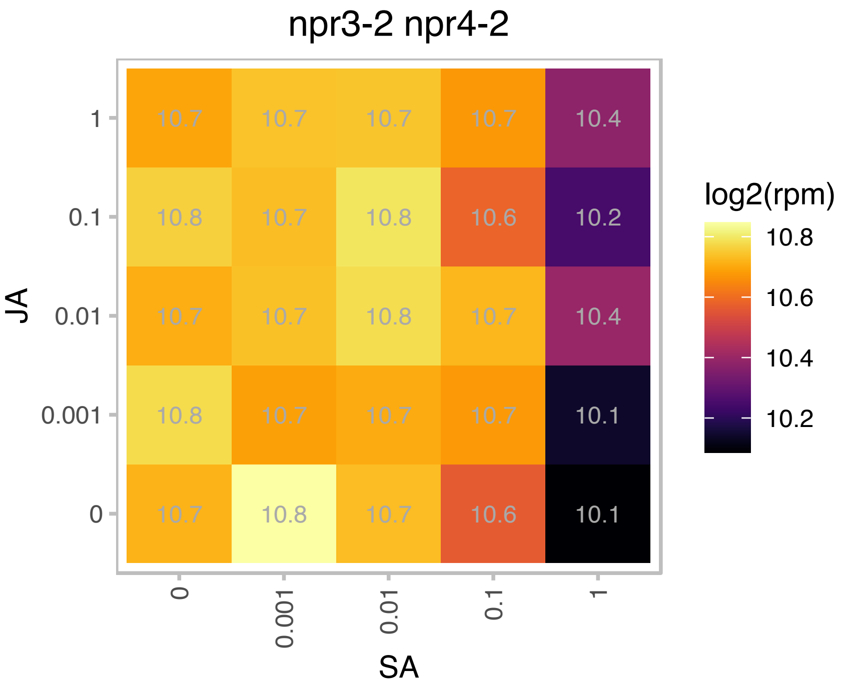


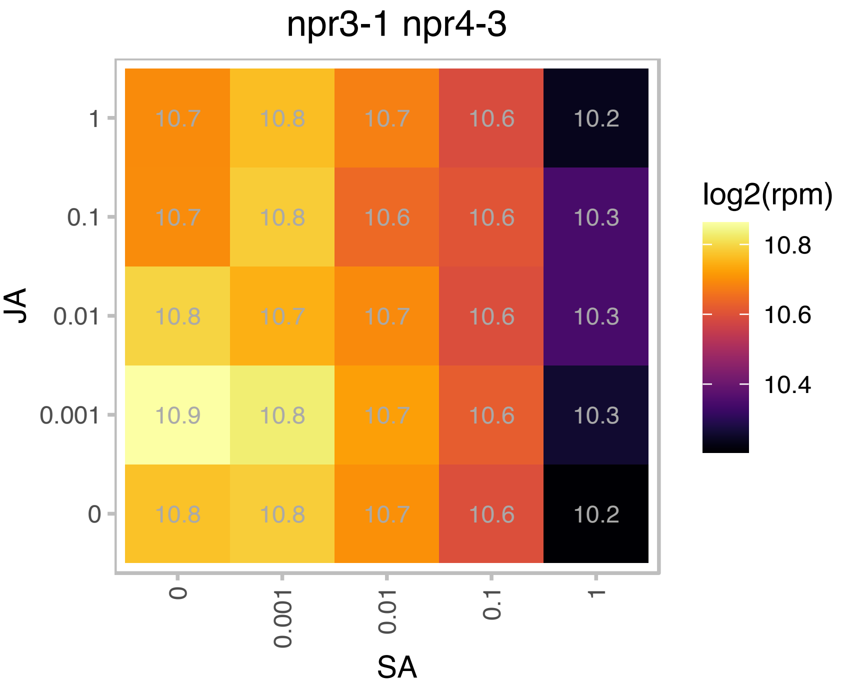

# AT2G40590.1 : NA

Ribosomal protein S26e family protein

| | |
|---|---|
| Gene | AT2G40590 |
| is.expressed_genes | TRUE |
| is.responsive_genes | TRUE |
| is.apcluster | F-07 |

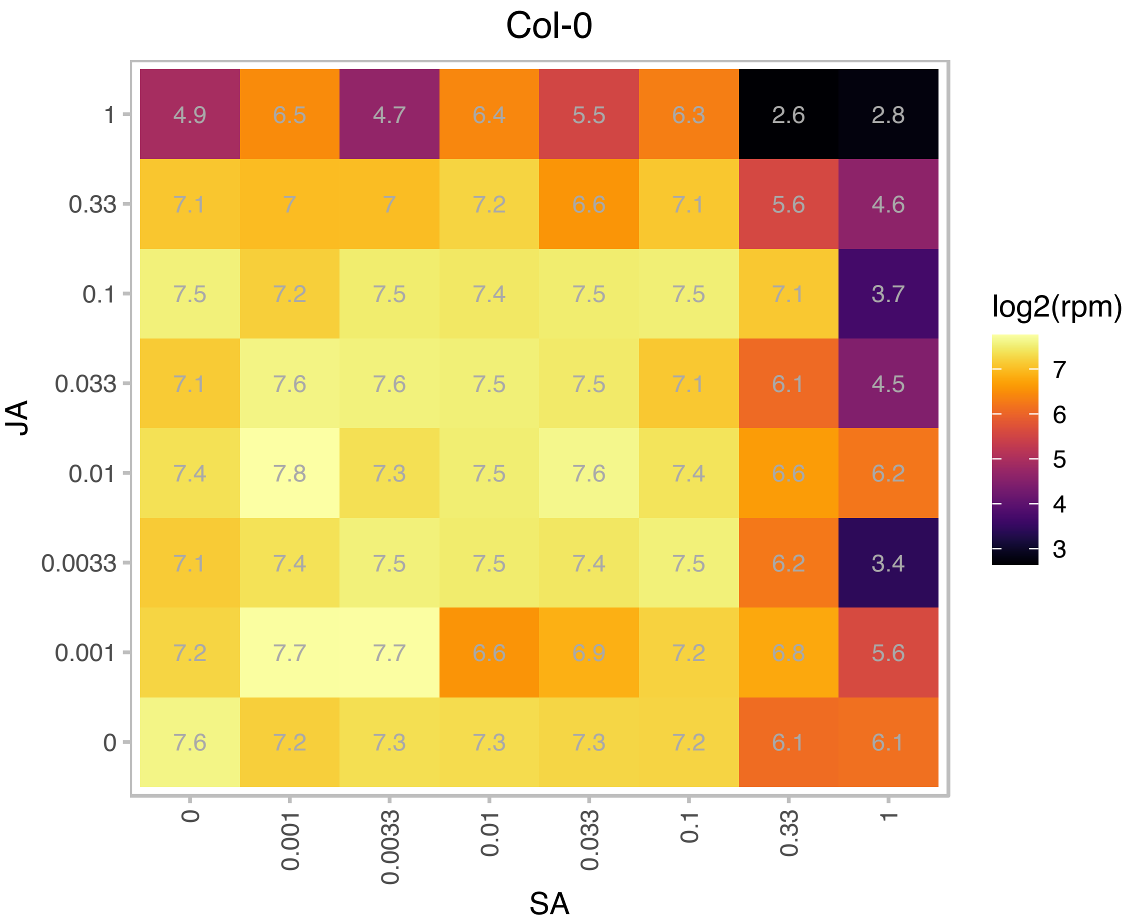


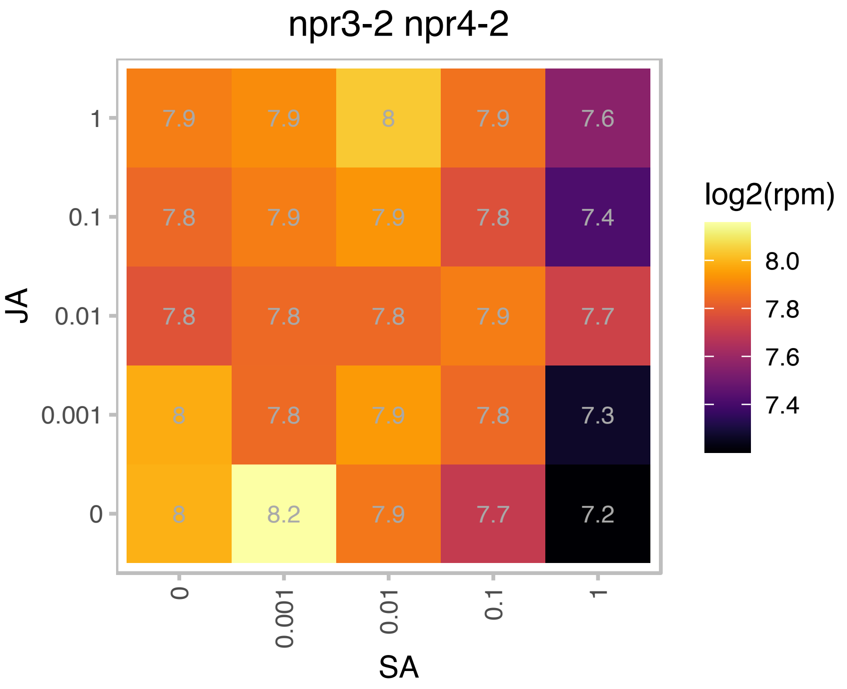


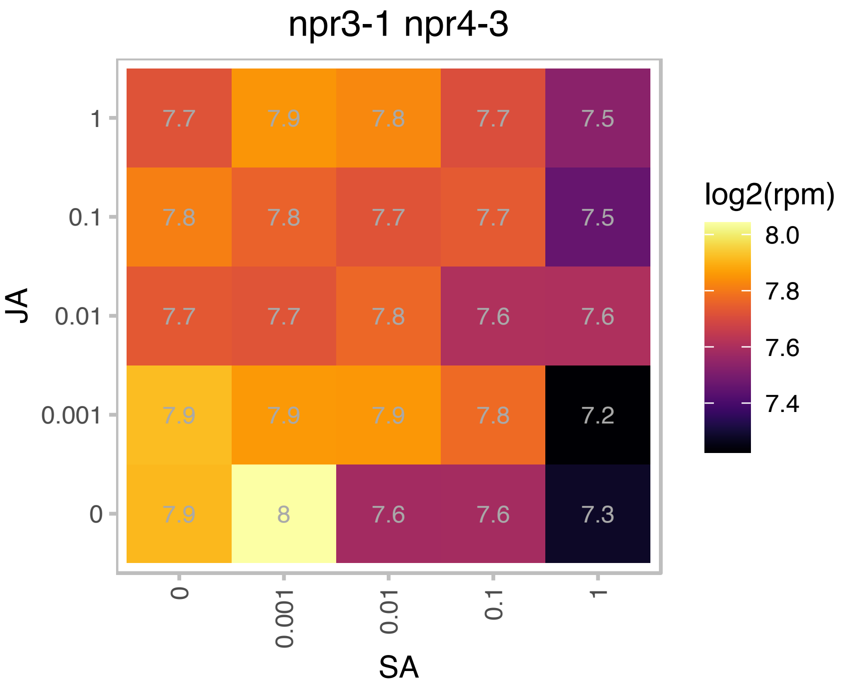

# AT2G41840.1 : NA

Ribosomal protein S5 family protein

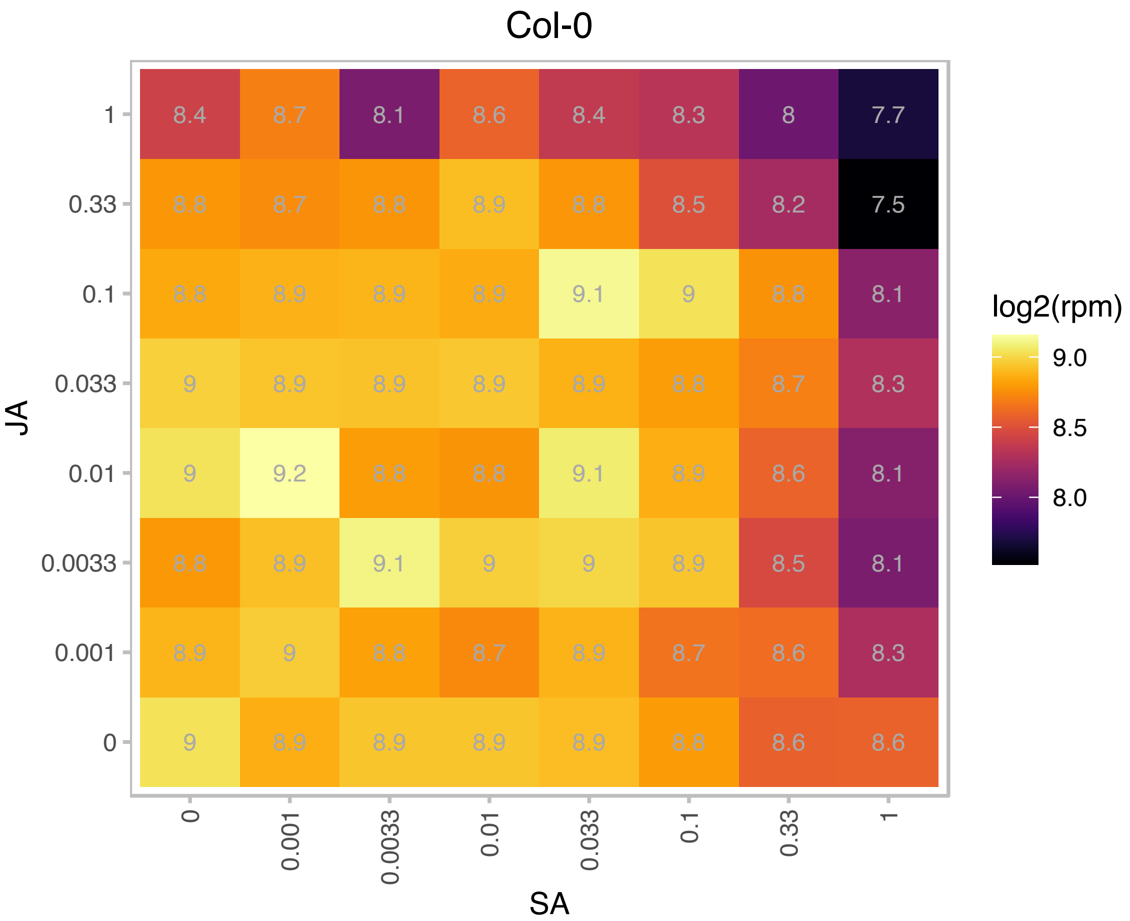


| | |
|---|---|
| Gene | AT2G41840 |
| is.expressed_genes | TRUE |
| is.responsive_genes | TRUE |
| is.apcluster | F-07 |

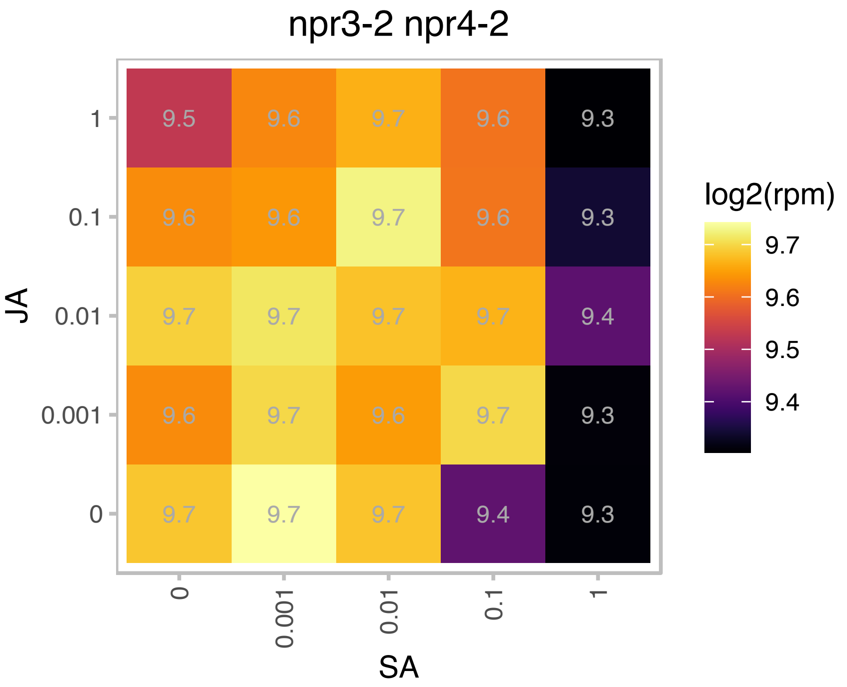


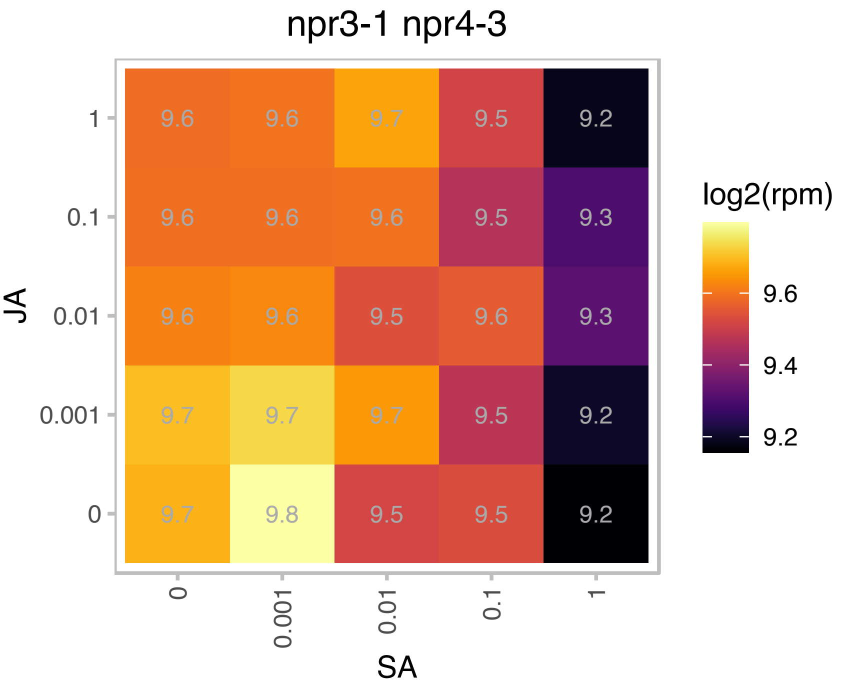

# AT2G43460.1 : NA

Ribosomal L38e protein family

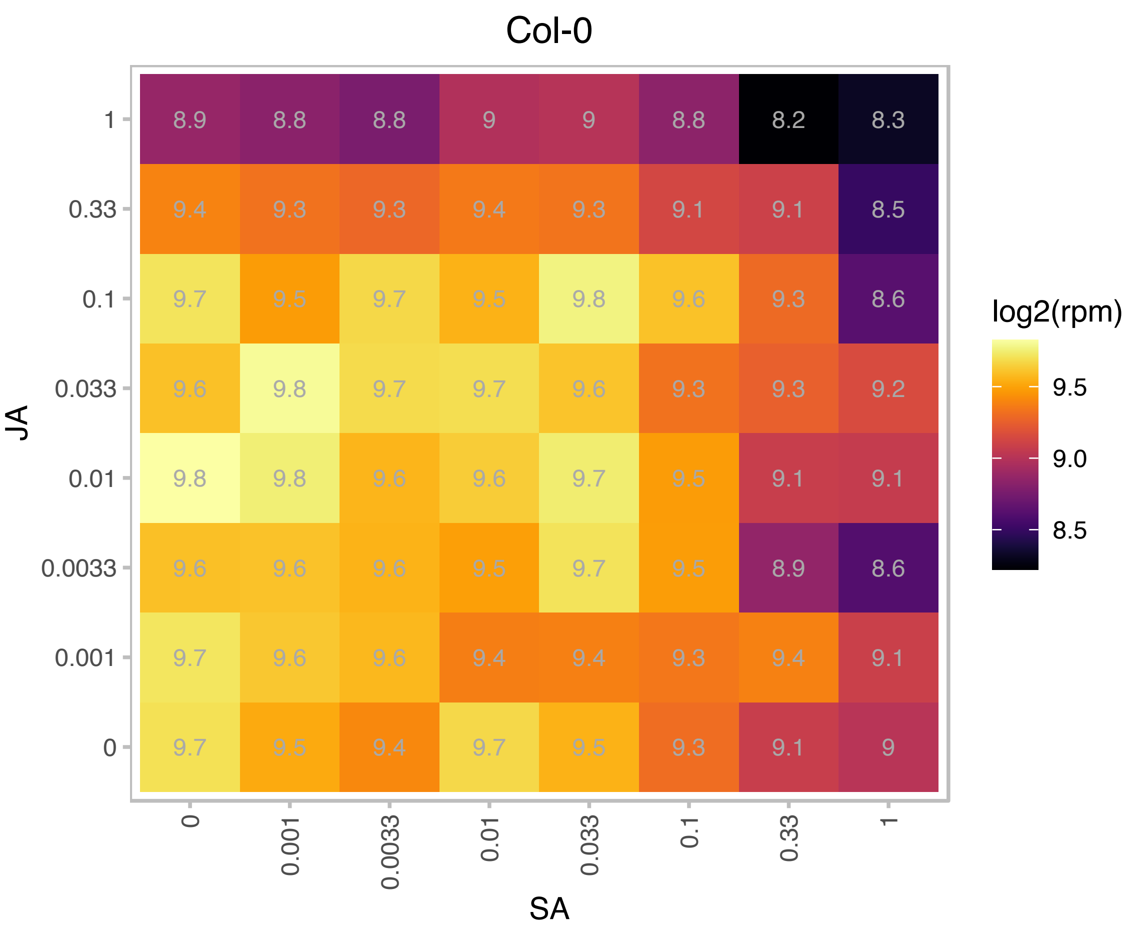


| | |
|---|---|
| Gene | AT2G43460 |
| is.expressed_genes | TRUE |
| is.responsive_genes | TRUE |
| is.apcluster | F-07 |

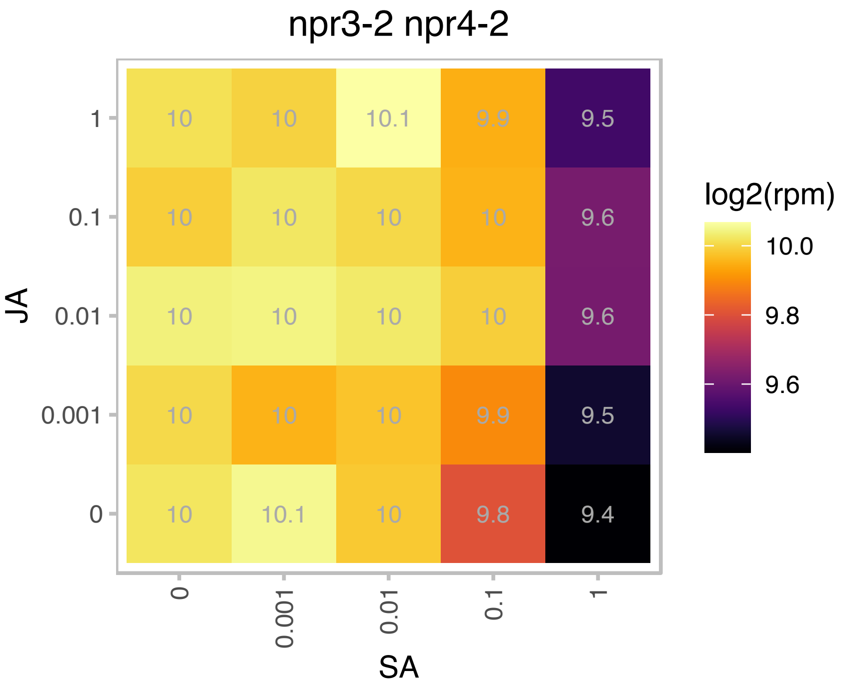


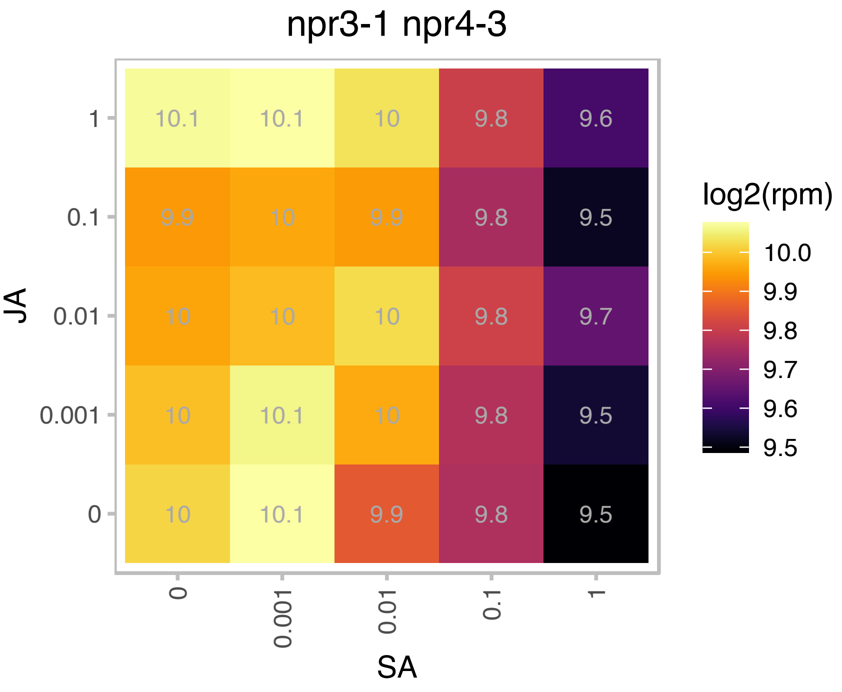

# AT2G44860.1 : NA

Ribosomal protein L24e family protein

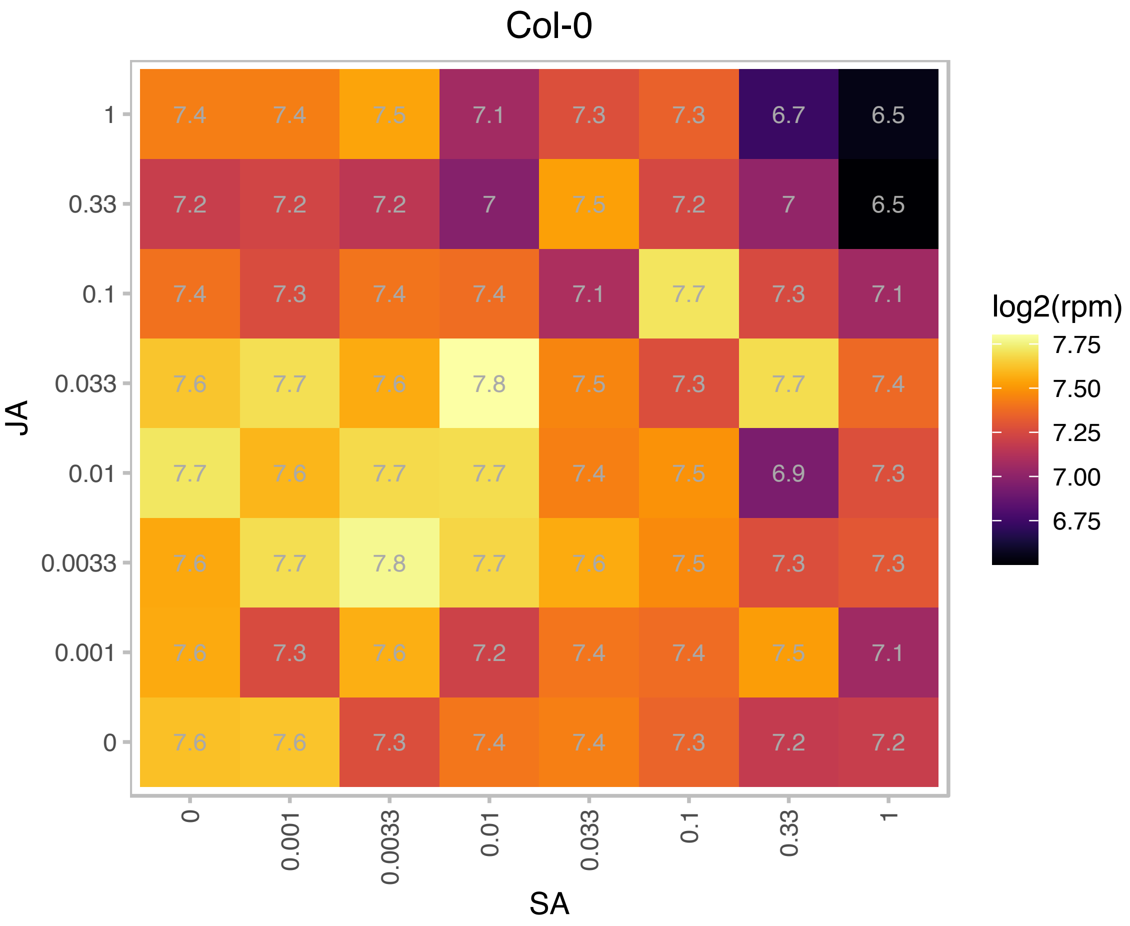


| | |
|---|---|
| Gene | AT2G44860 |
| is.expressed_genes | TRUE |
| is.responsive_genes | TRUE |
| is.apcluster | F-07 |

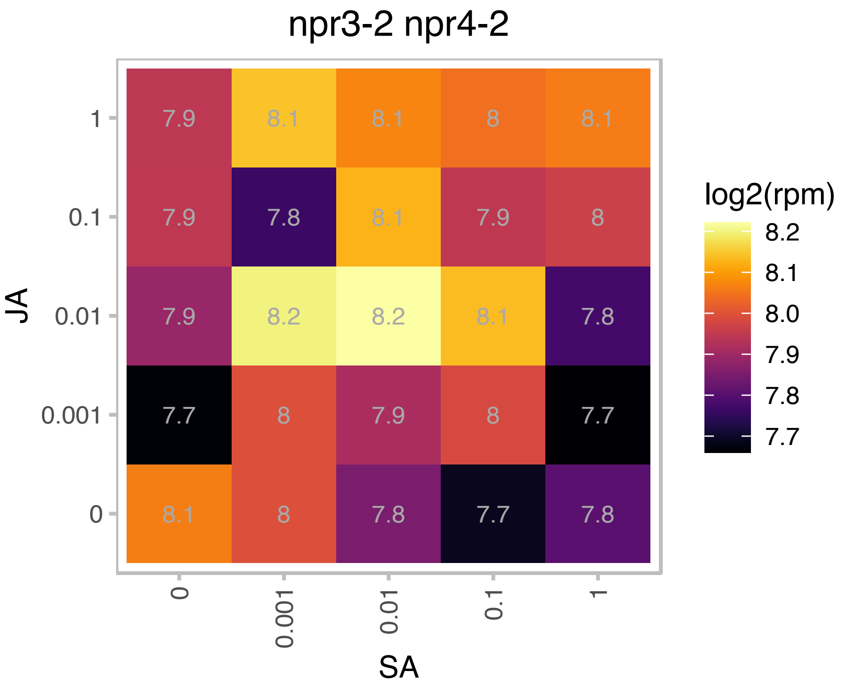


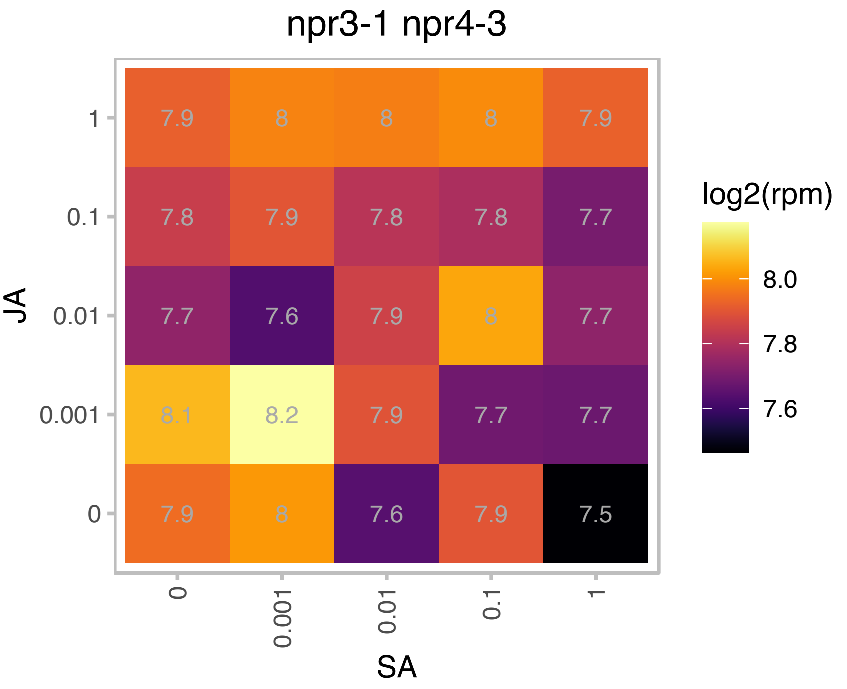

# AT2G45710.1 : NA

Zinc-binding ribosomal protein family protein

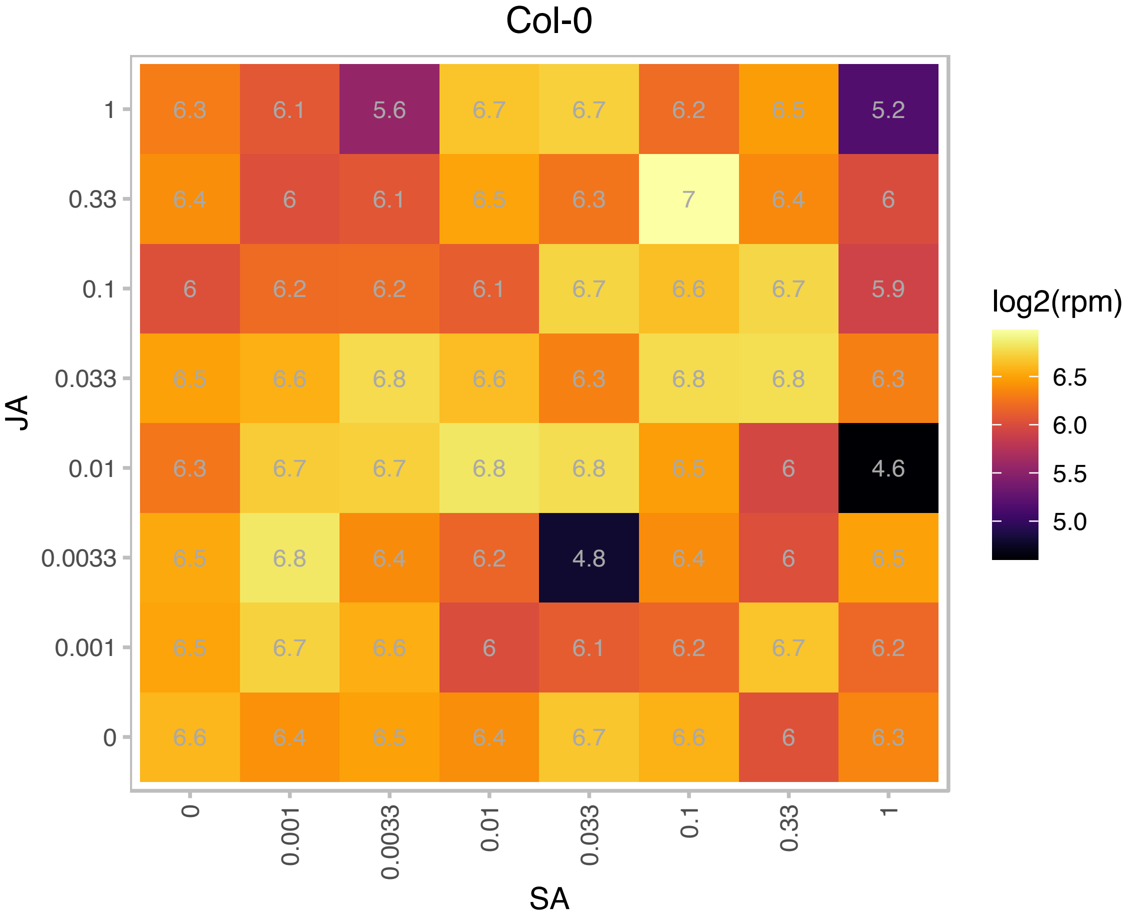


| | |
|---|---|
| Gene | AT2G45710 |
| is.expressed_genes | TRUE |
| is.responsive_genes | TRUE |
| is.apcluster | F-07 |

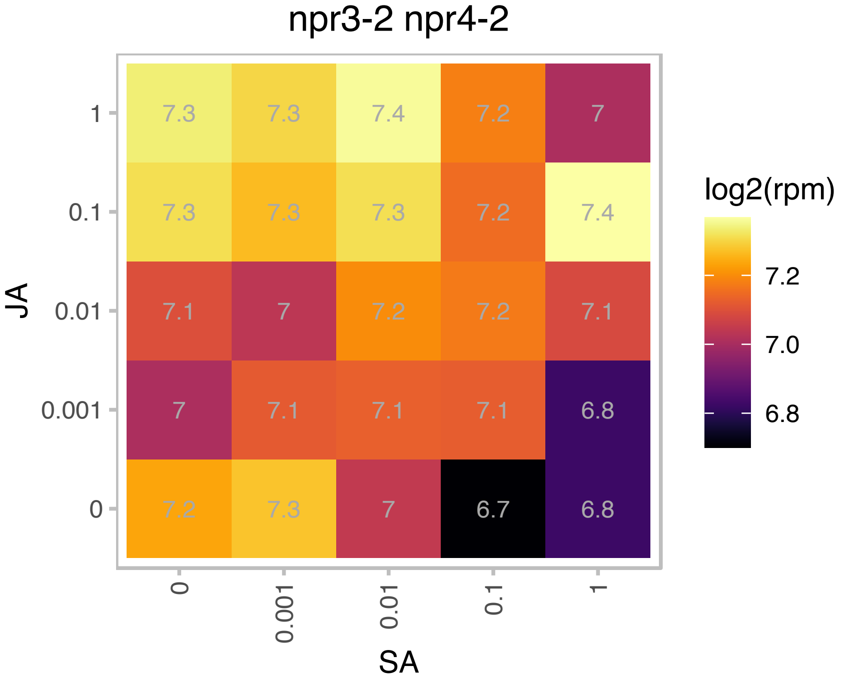


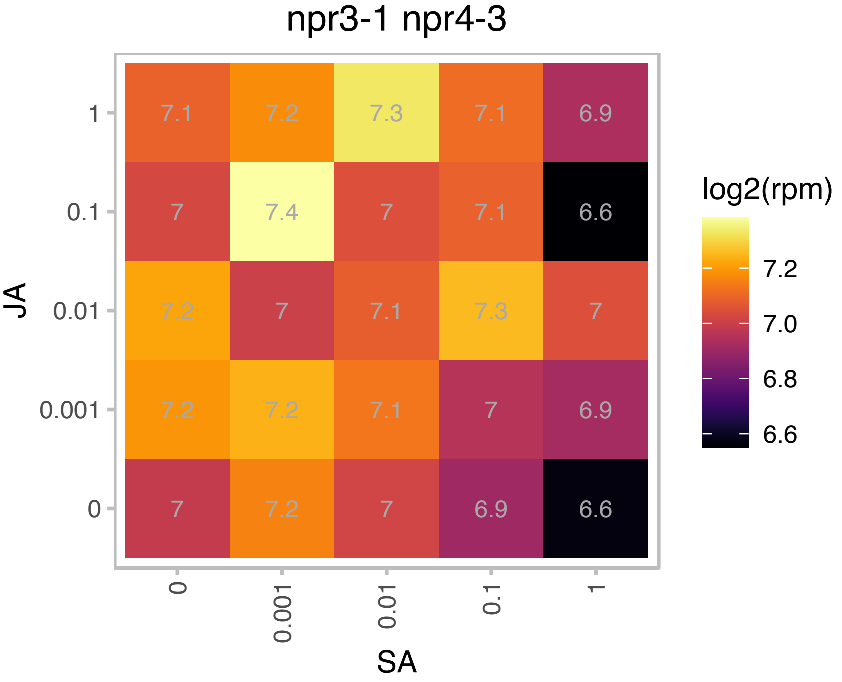

### AT2G47110.1 : UBI6, UBQ6

ubiquitin 6

polyubiquitin gene The mRNA is cell-to-cell mobile.

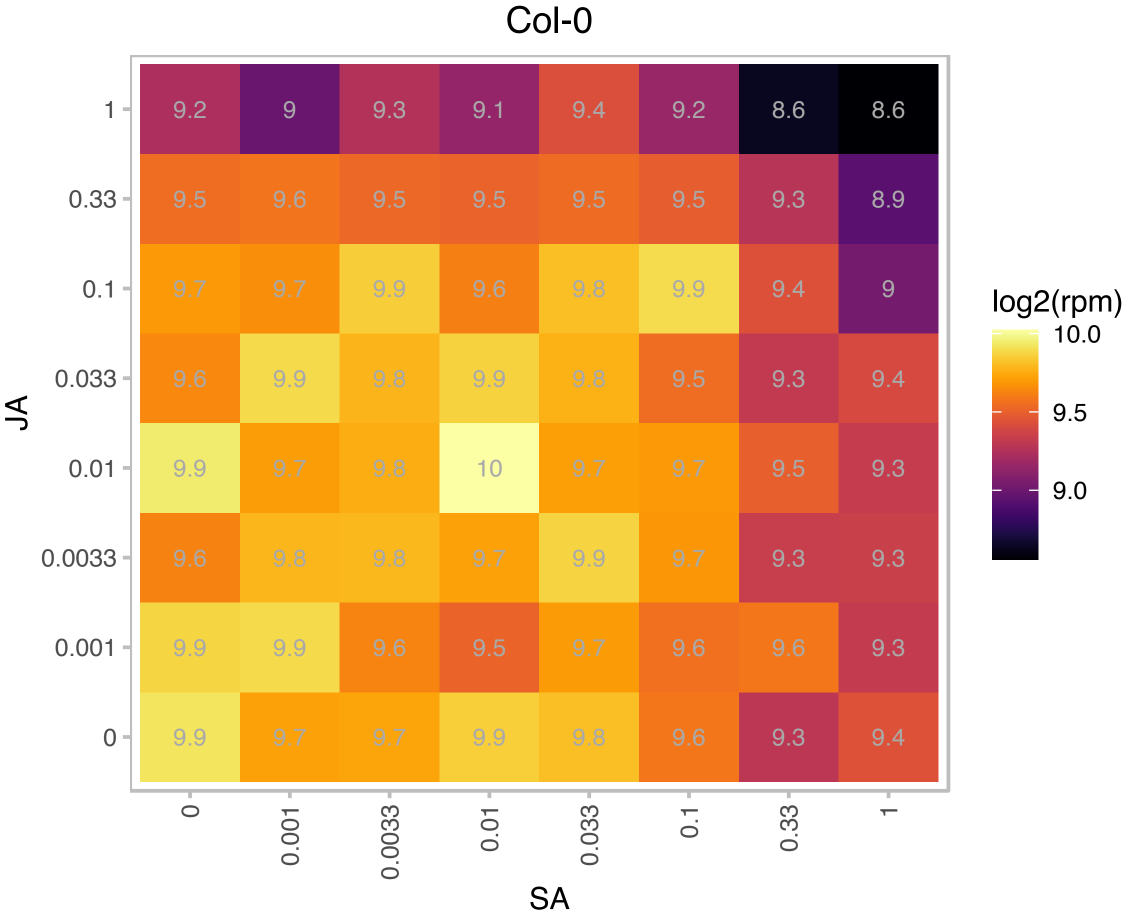


| | |
|---|---|
| Gene | AT2G47110 |
| is.expressed_genes | TRUE |
| is.responsive_genes | TRUE |
| is.apcluster | F-07 |

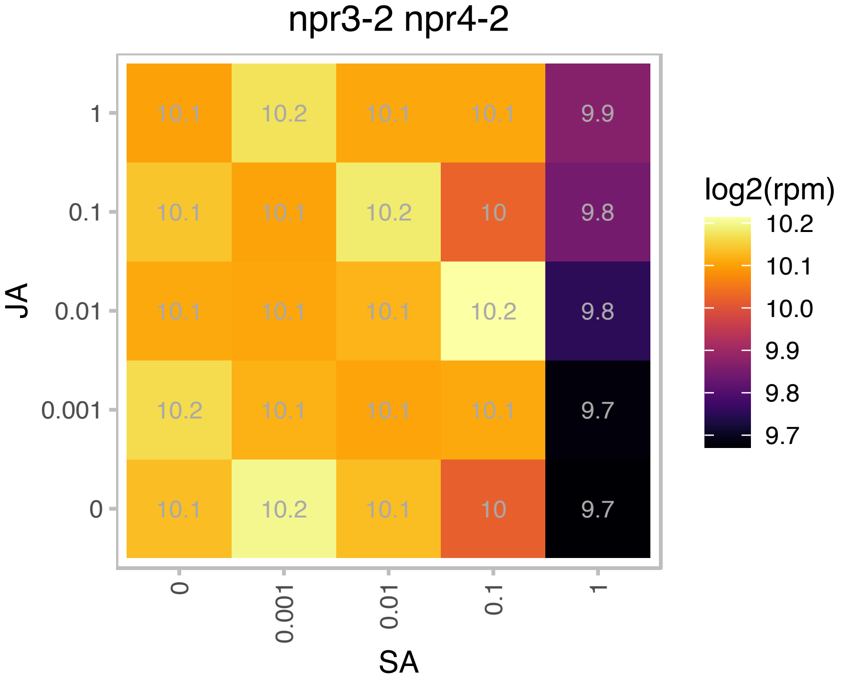


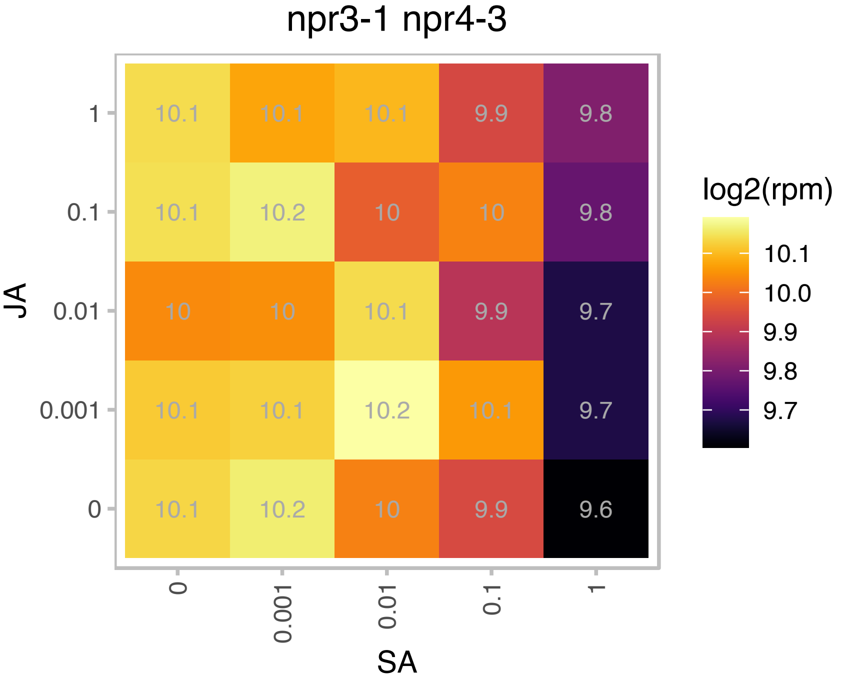

# AT2G47610.1 : NA

Ribosomal protein L7Ae/L30e/S12e/Gadd45 family protein

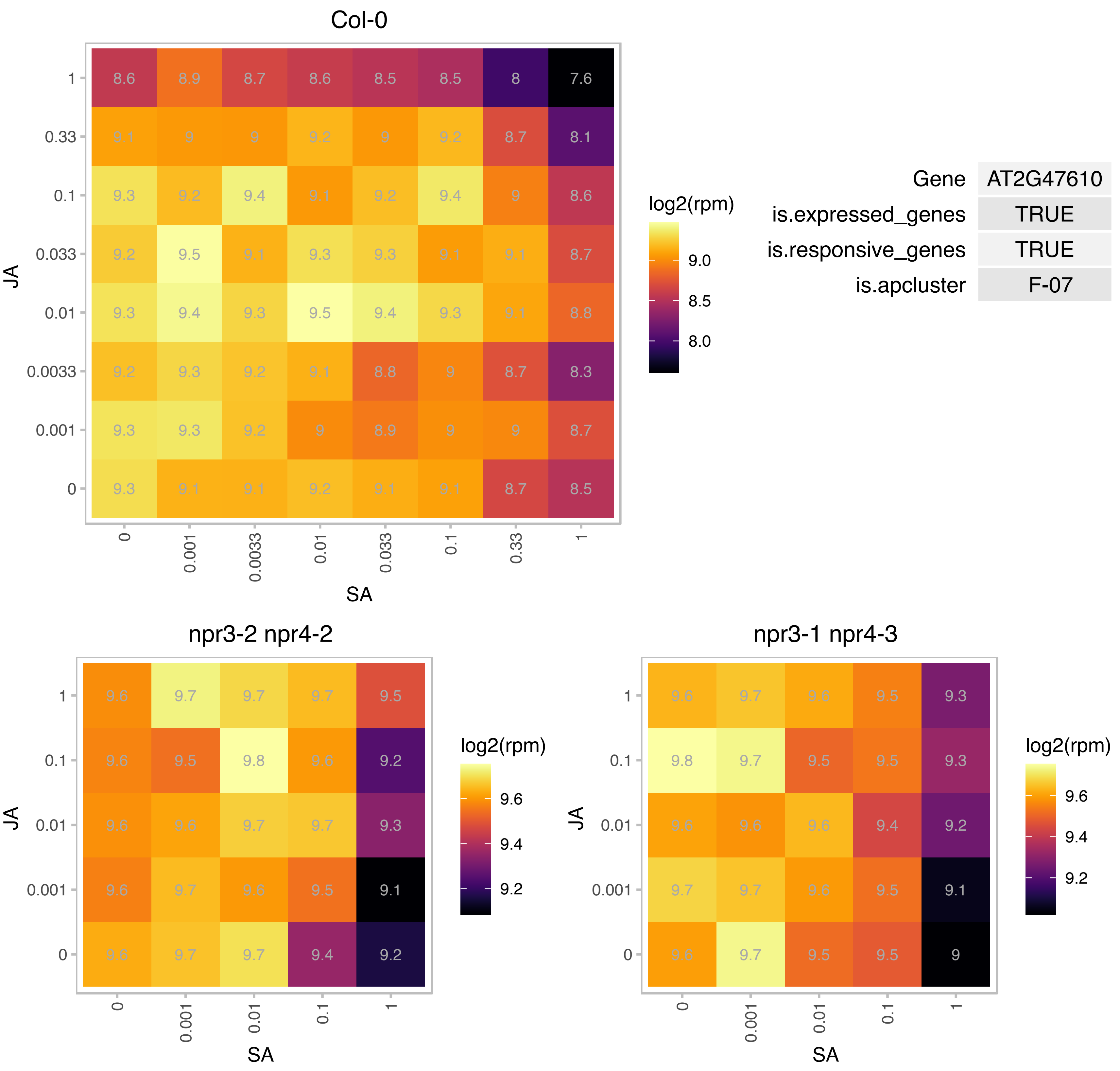


| | |
|---|---|
| Gene | AT2G47610 |
| is.expressed_genes | TRUE |
| is.responsive_genes | TRUE |
| is.apcluster | F-07 |

# AT3G02080.1 : NA

Ribosomal protein S19e family protein

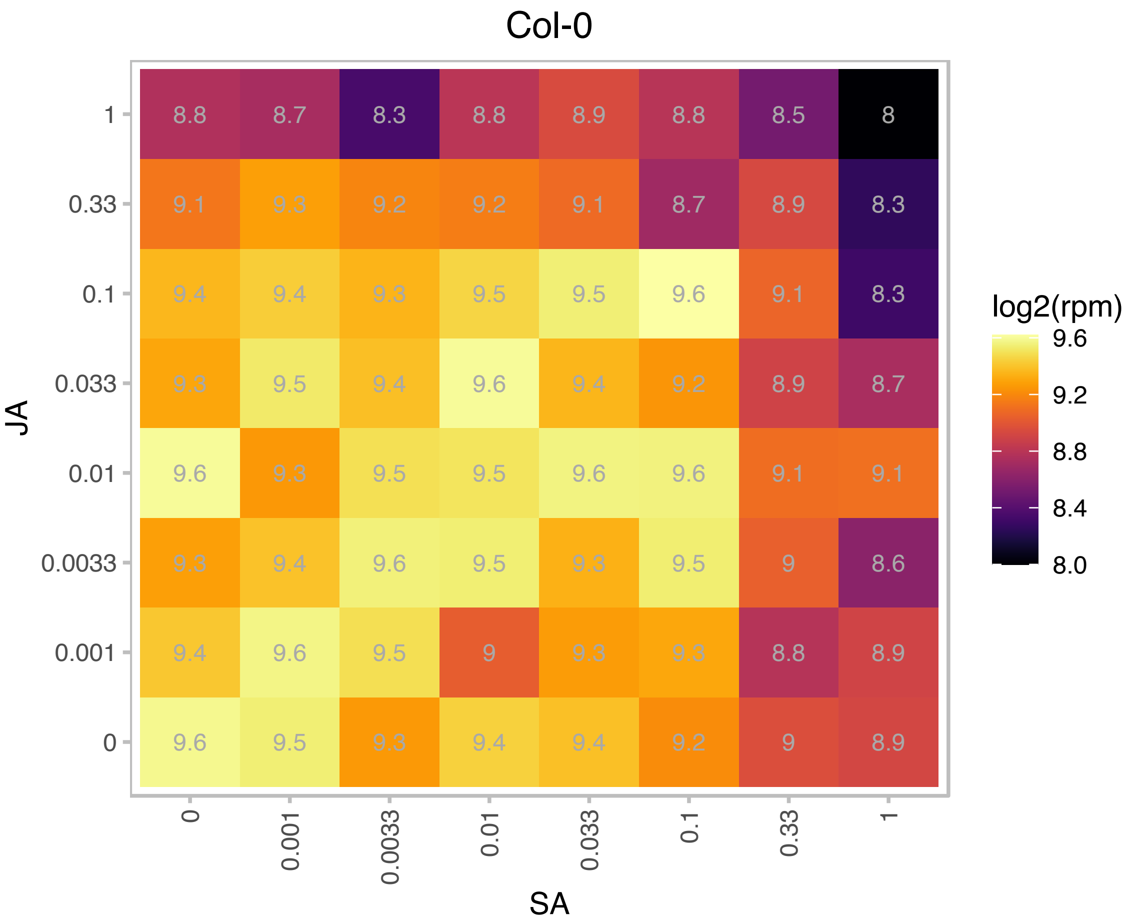


| | |
|---|---|
| Gene | AT3G02080 |
| is.expressed_genes | TRUE |
| is.responsive_genes | TRUE |
| is.apcluster | F-07 |

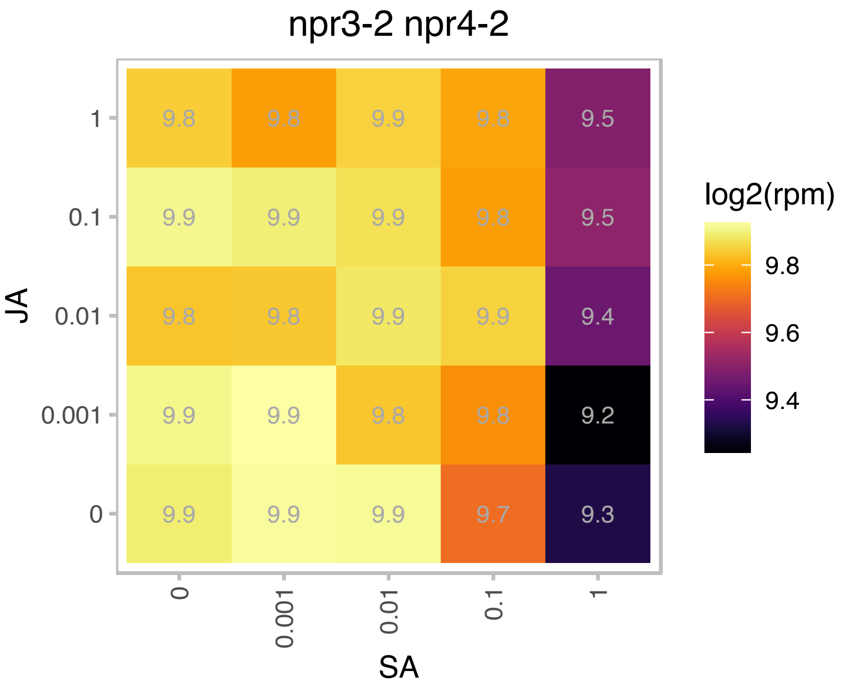


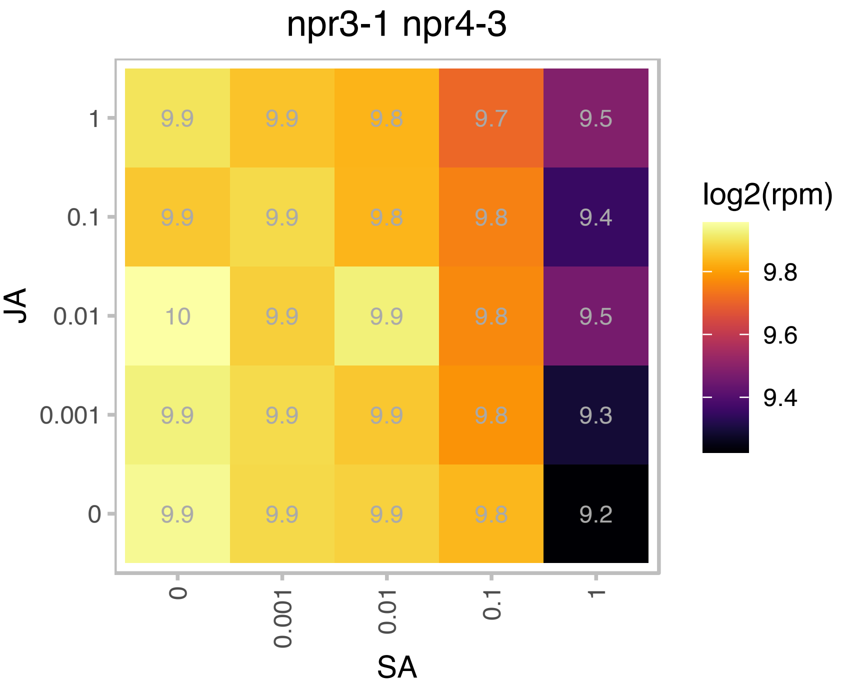

# AT3G02560.1 : NA

Ribosomal protein S7e family protein

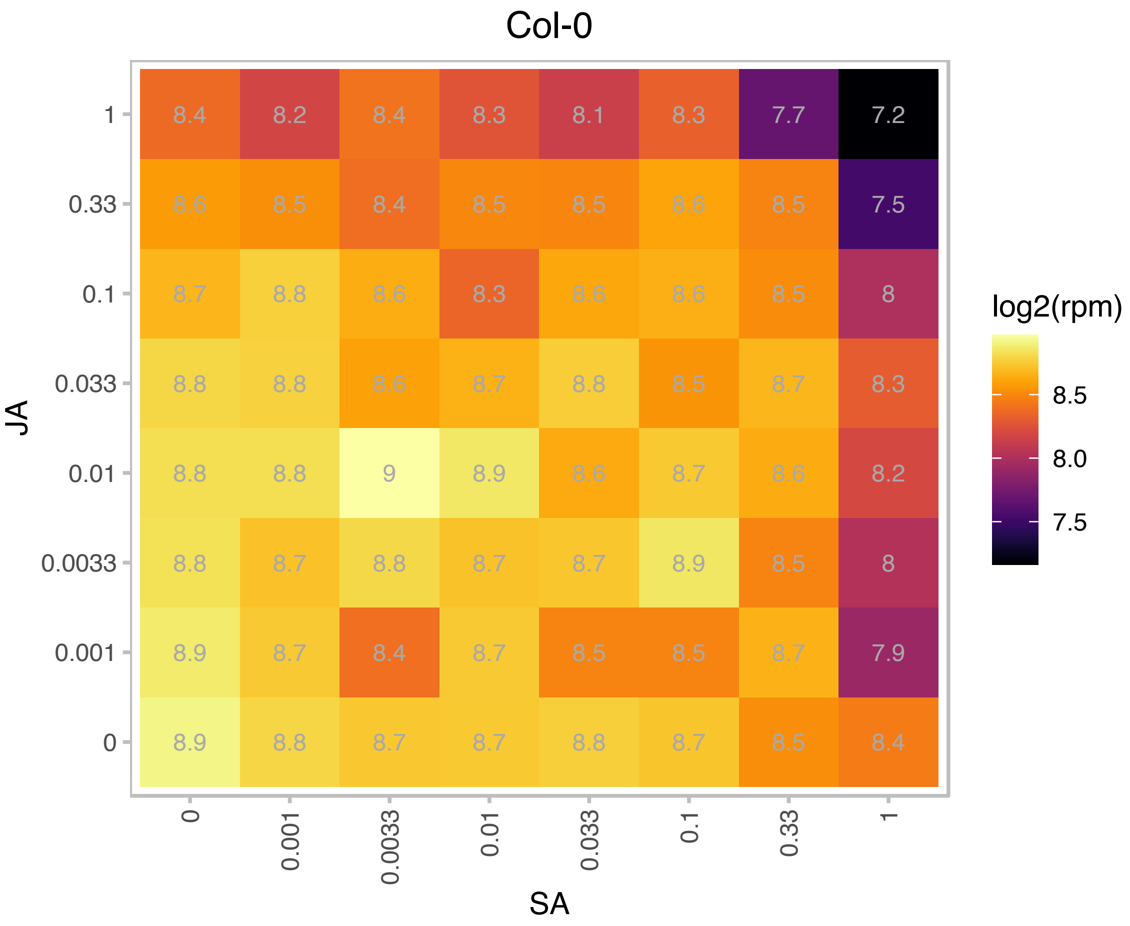


| | |
|---|---|
| Gene | AT3G02560 |
| is.expressed_genes | TRUE |
| is.responsive_genes | TRUE |
| is.apcluster | F-07 |

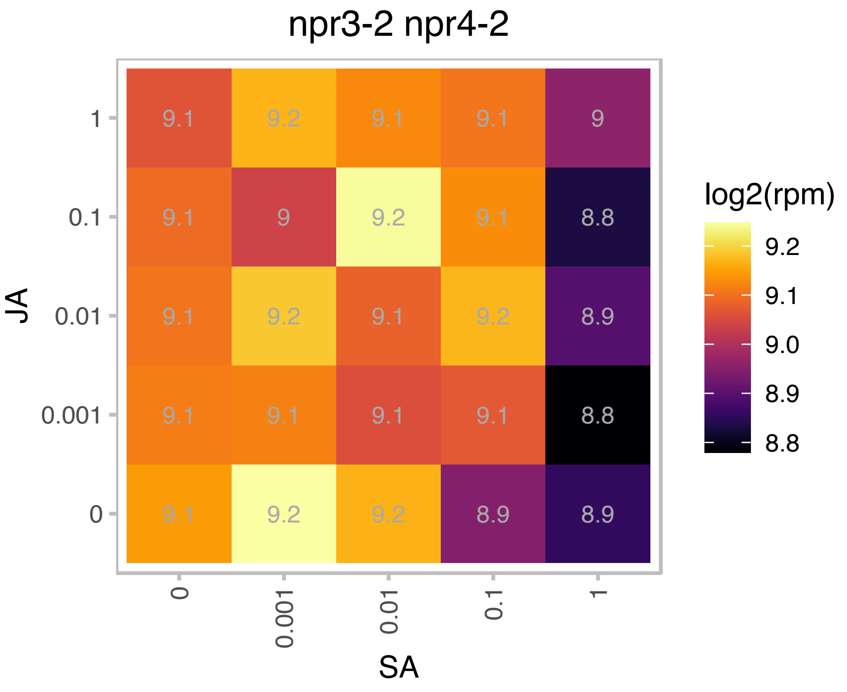


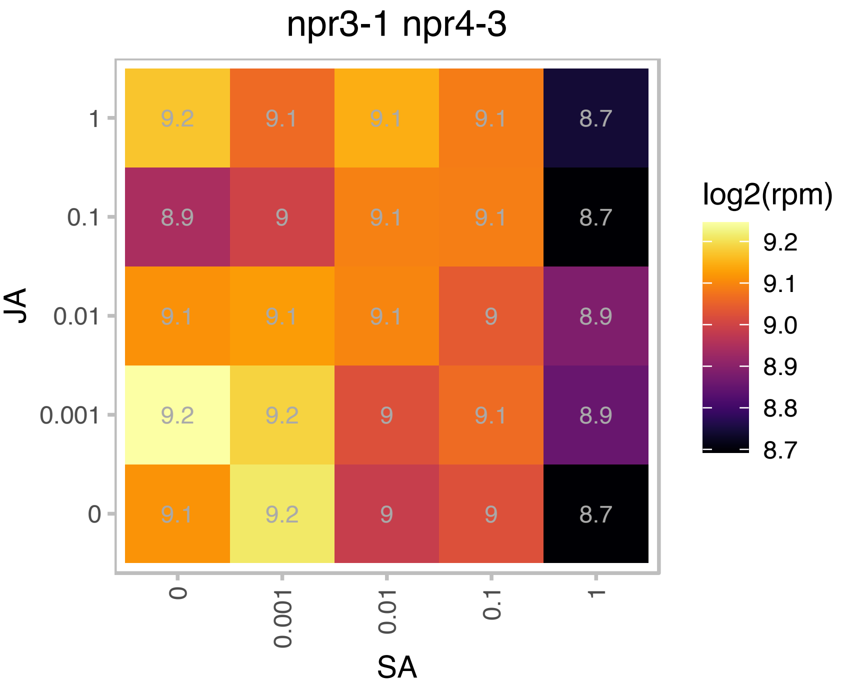

# AT3G02640.1 : NA

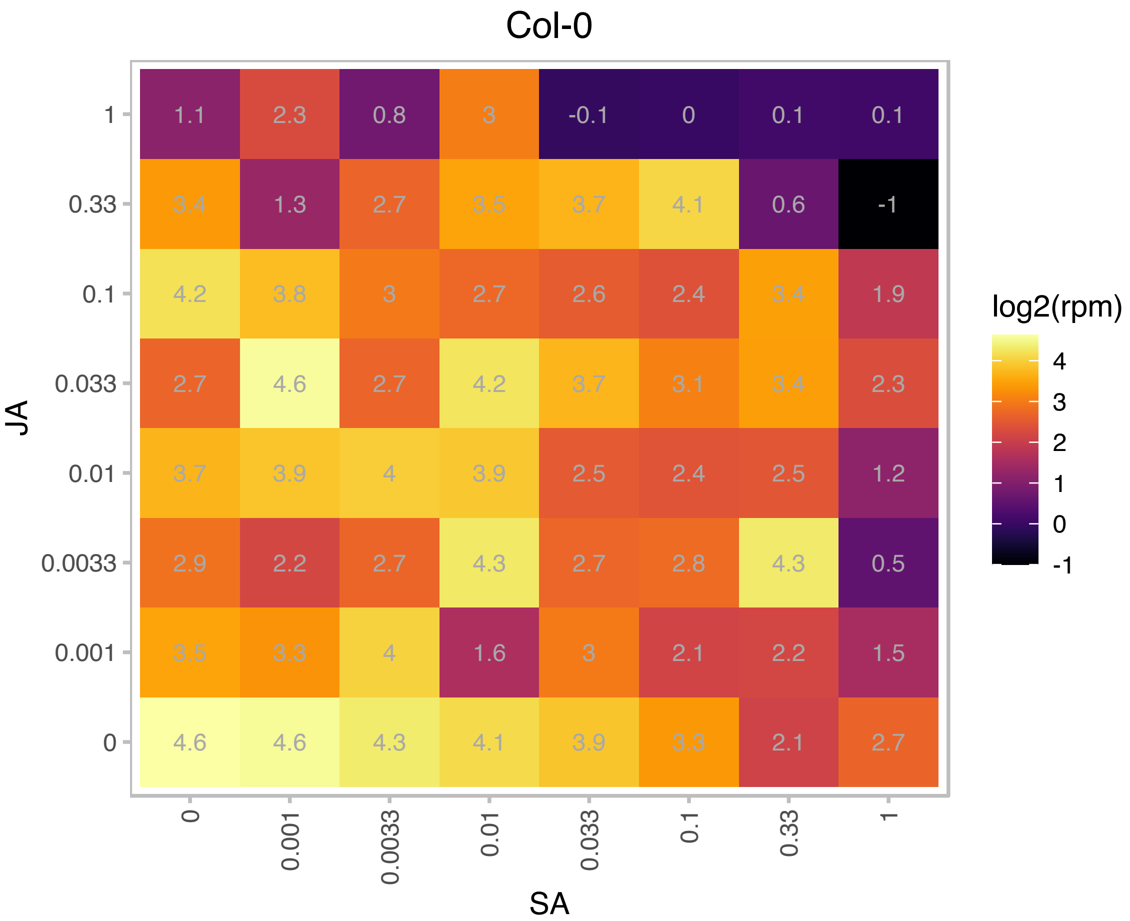


| | |
|---|---|
| Gene | AT3G02640 |
| is.expressed_genes | TRUE |
| is.responsive_genes | TRUE |
| is.apcluster | F-07 |

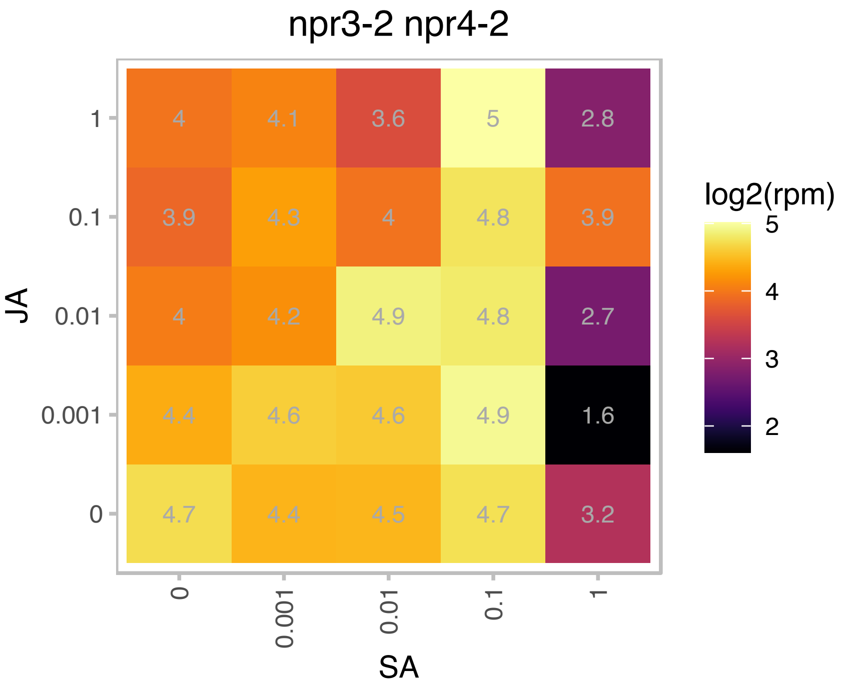


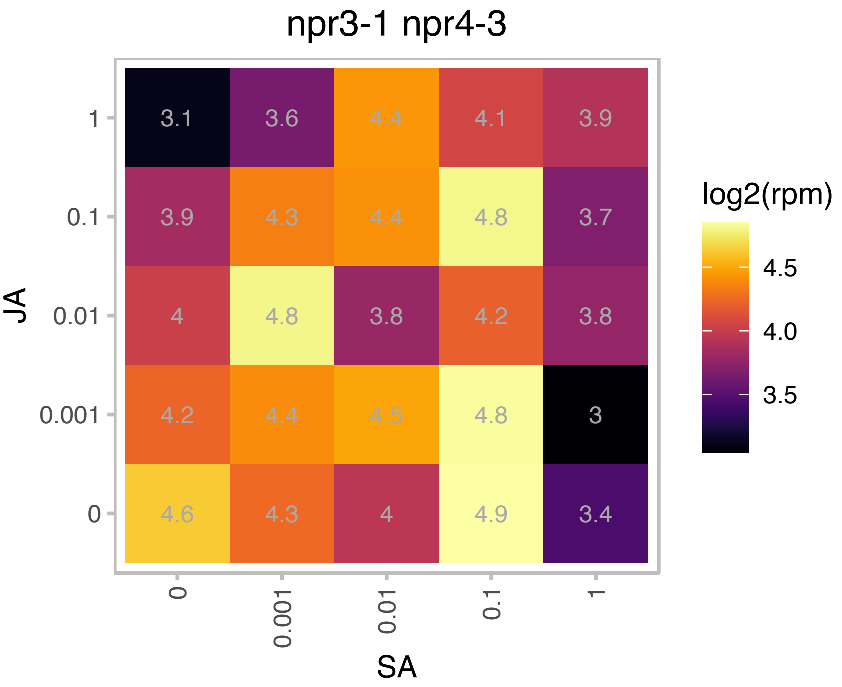

# AT3G02820.1 : NA

zinc knuckle (CCHC-type) family protein

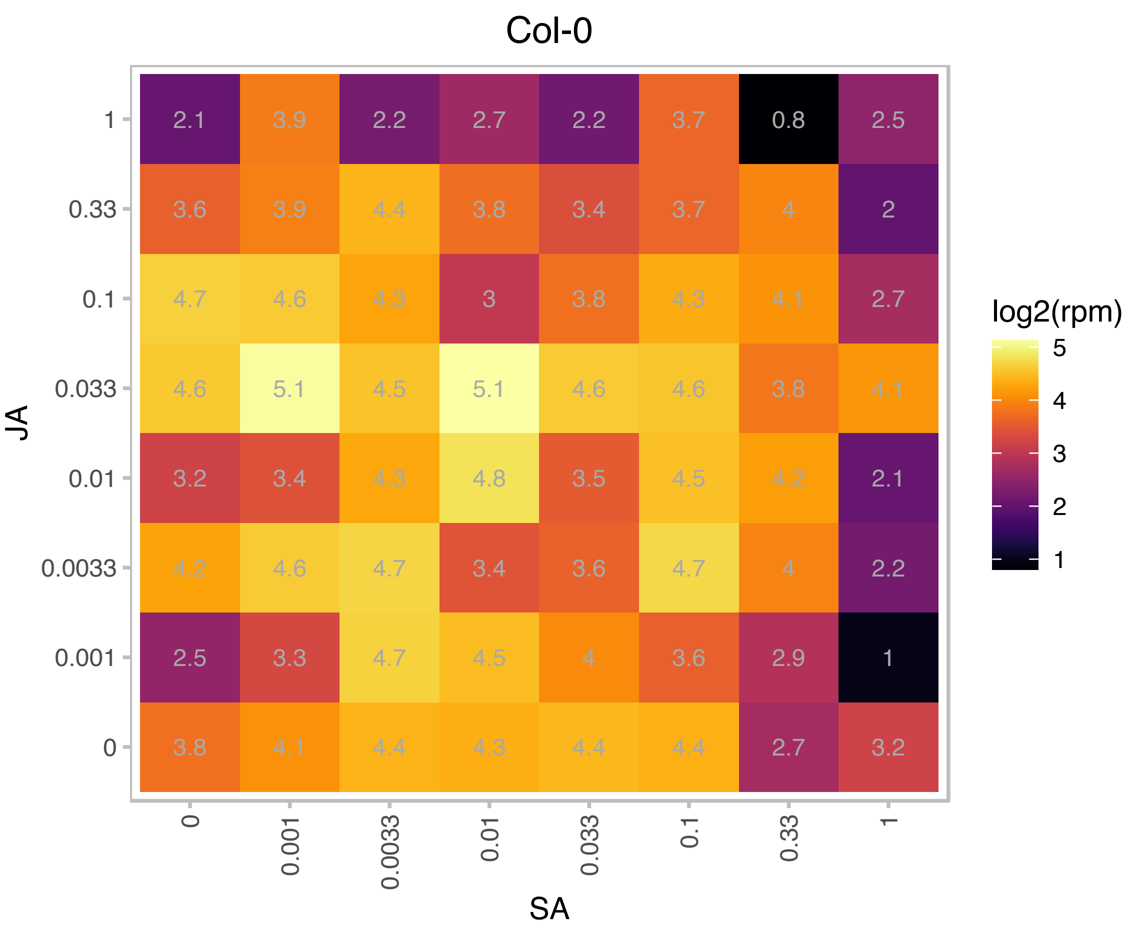


| | |
|---|---|
| Gene | AT3G02820 |
| is.expressed_genes | TRUE |
| is.responsive_genes | TRUE |
| is.apcluster | F-07 |

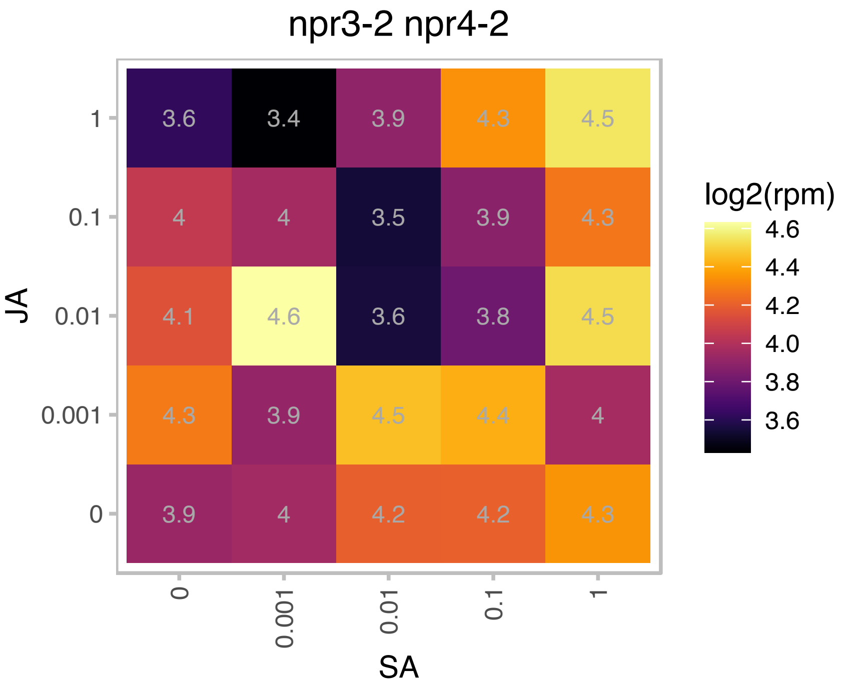


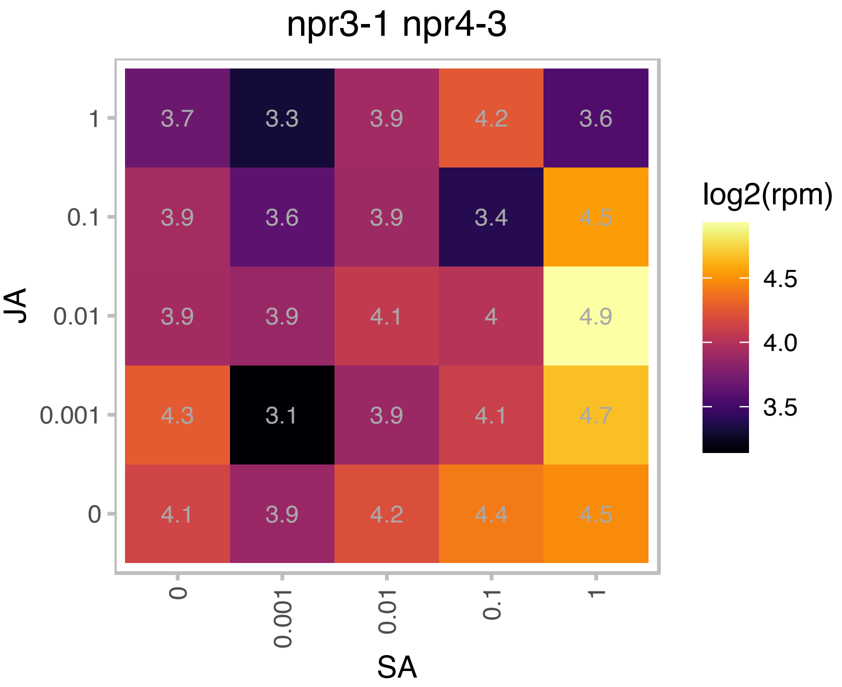

# AT3G03130.1 : NA

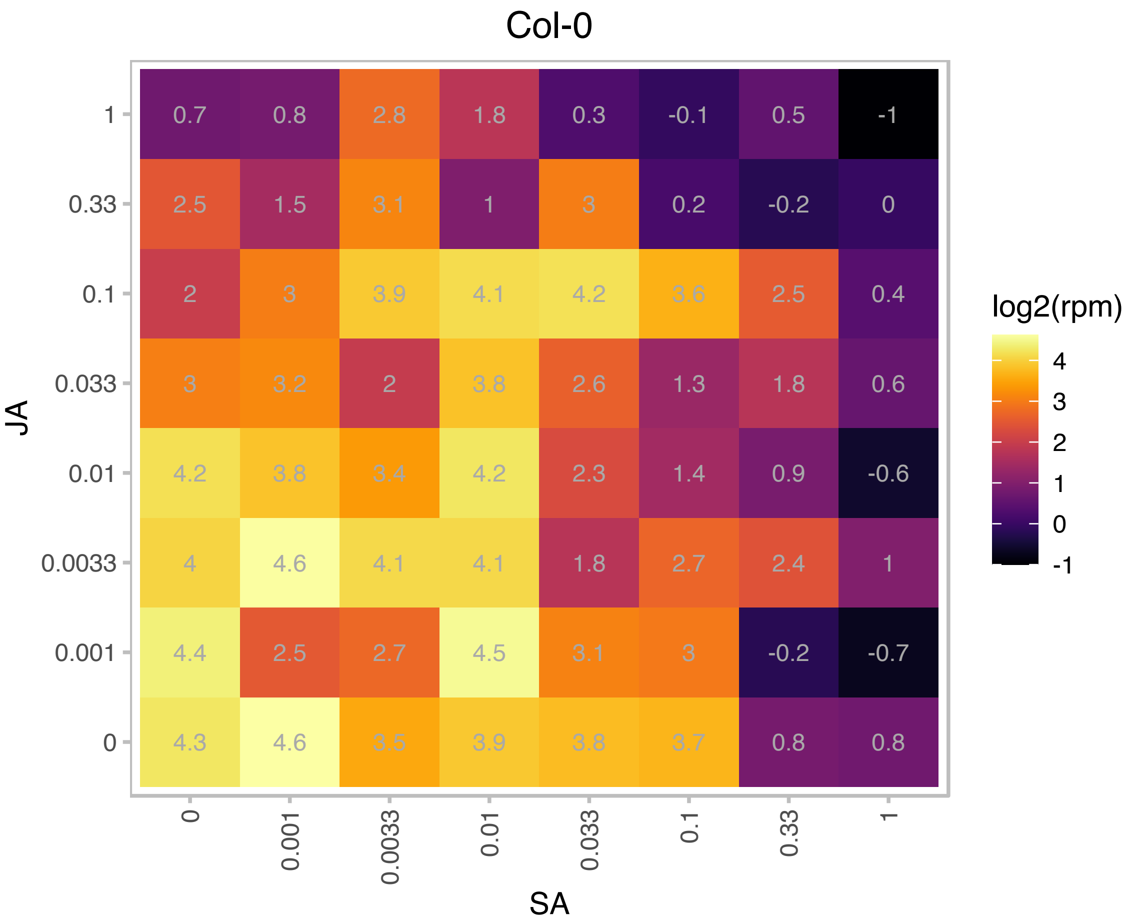


| | |
|---|---|
| Gene | AT3G03130 |
| is.expressed_genes | TRUE |
| is.responsive_genes | TRUE |
| is.apcluster | F-07 |

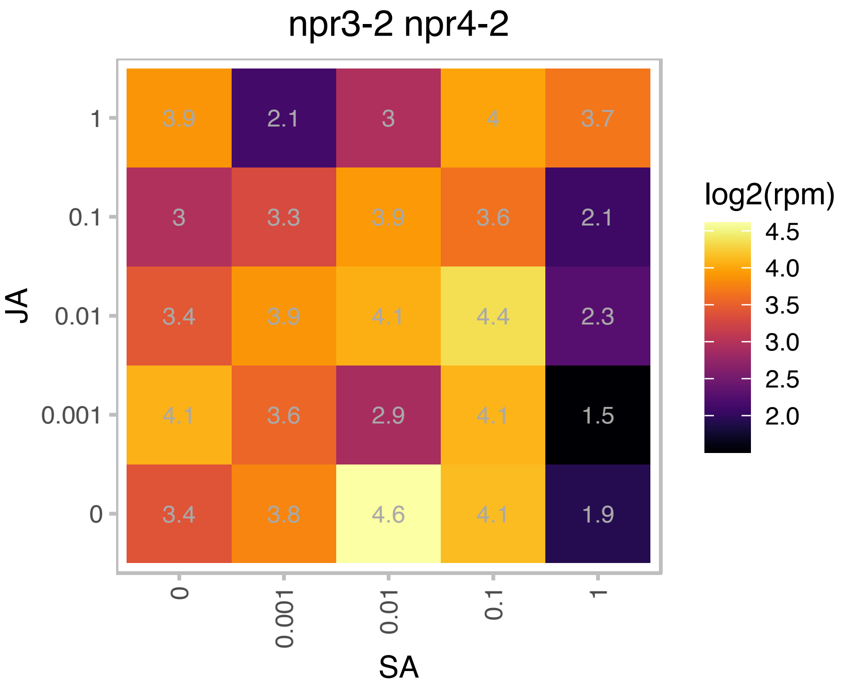


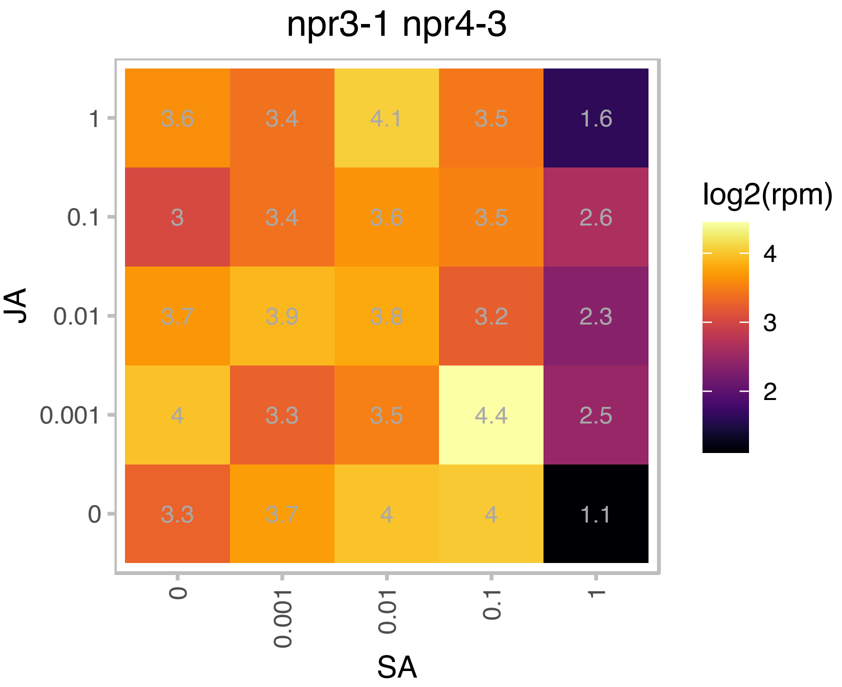

### AT3G03780.1 : ATMS2, MS2

methionine synthase 2

Encodes a cytosolic methionine synthase, involved in methionine regeneration via the activated methyl cycle (or SAM cycle)

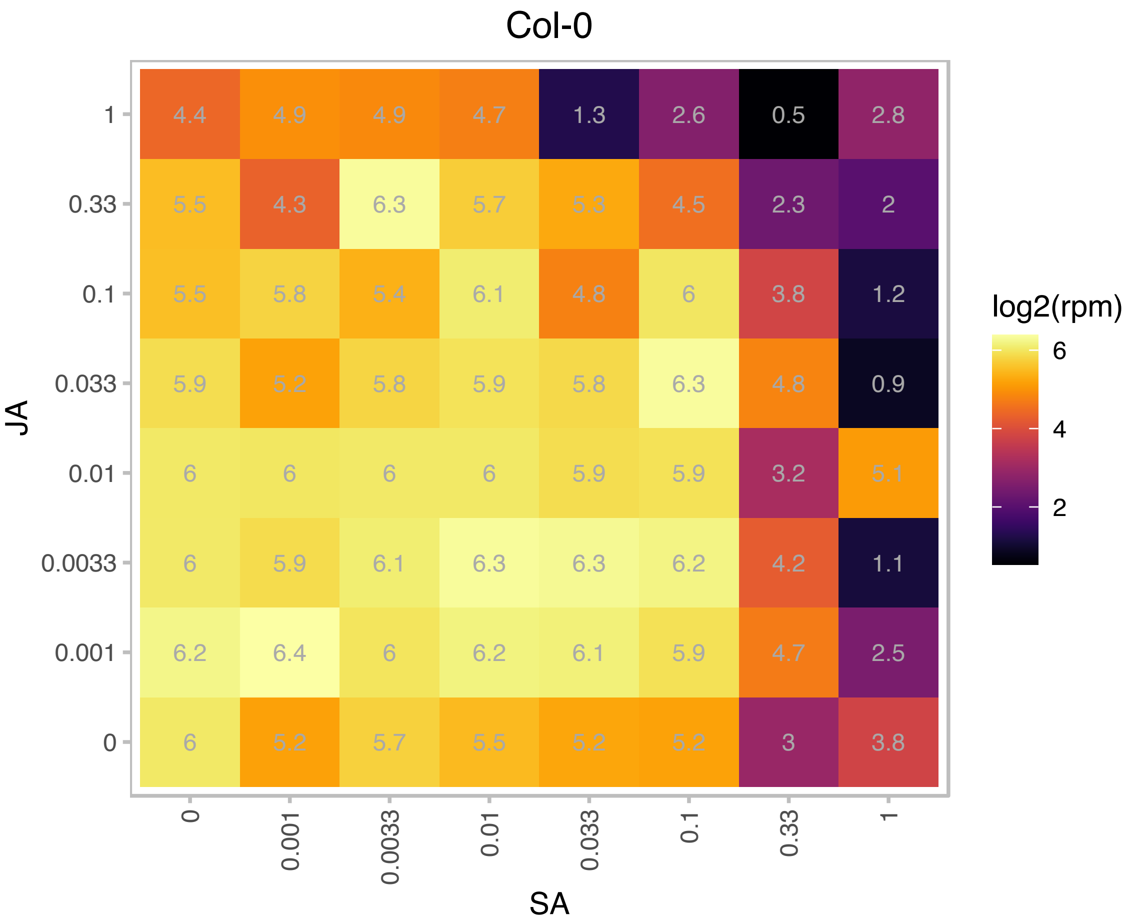


| | |
|---|---|
| Gene | AT3G03780 |
| is.expressed_genes | TRUE |
| is.responsive_genes | TRUE |
| is.apcluster | F-07 |

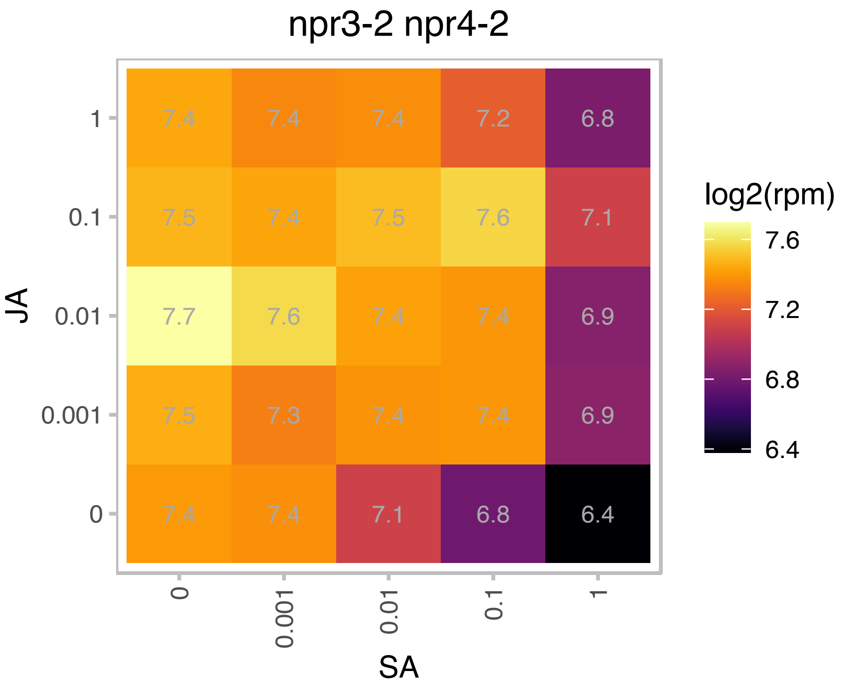


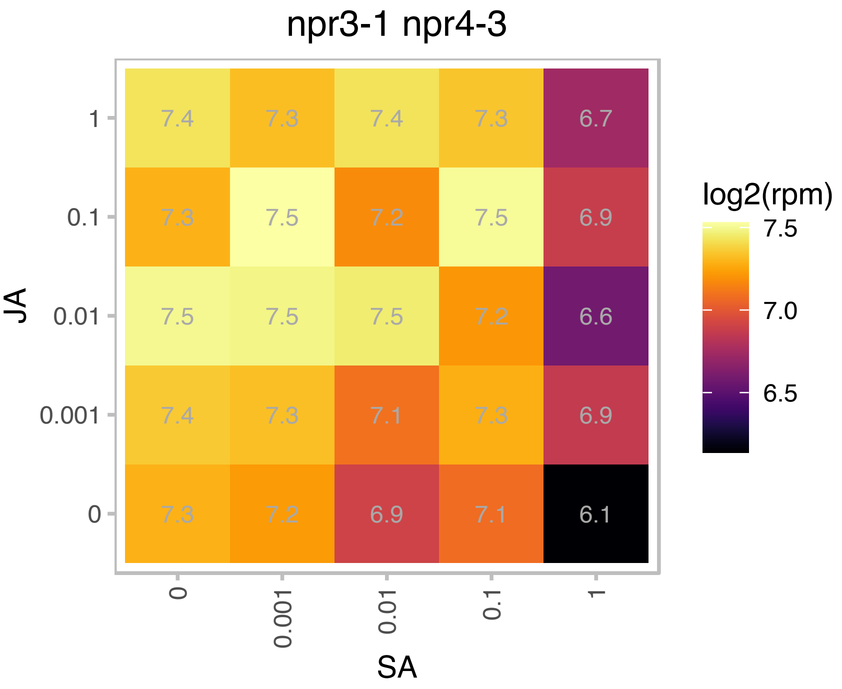

# AT3G04760.1 : NA

Pentatricopeptide repeat (PPR-like) superfamily protein

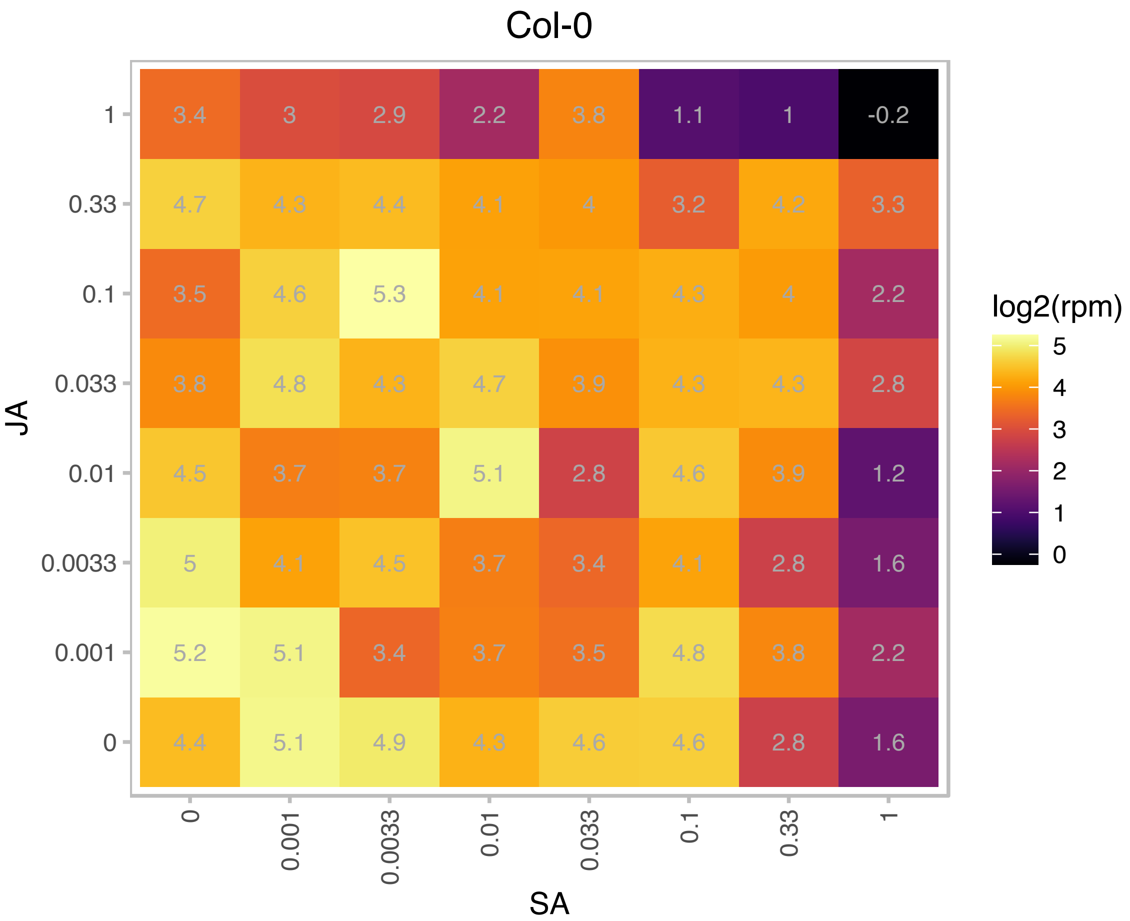


| | |
|---|---|
| Gene | AT3G04760 |
| is.expressed_genes | TRUE |
| is.responsive_genes | TRUE |
| is.apcluster | F-07 |

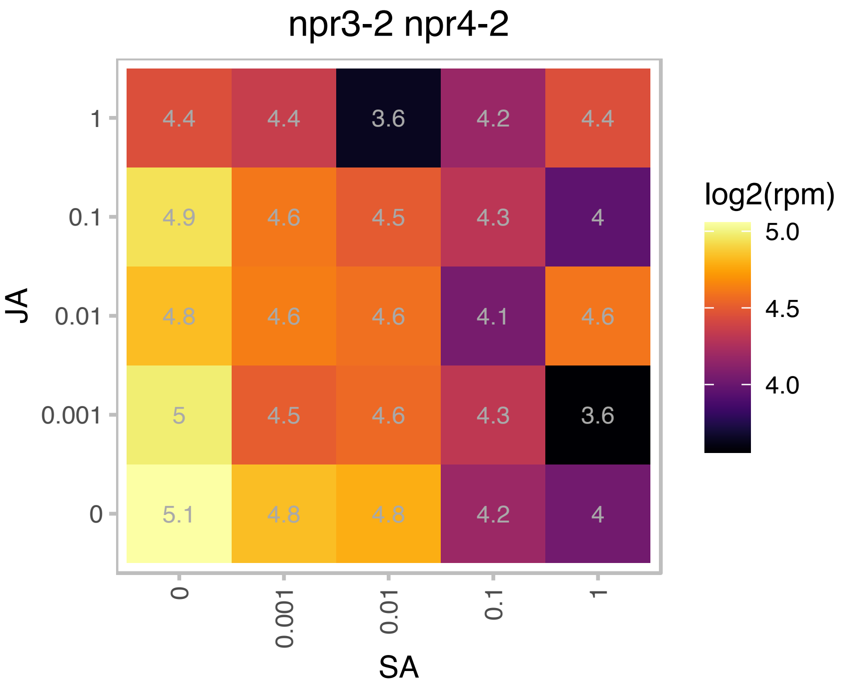


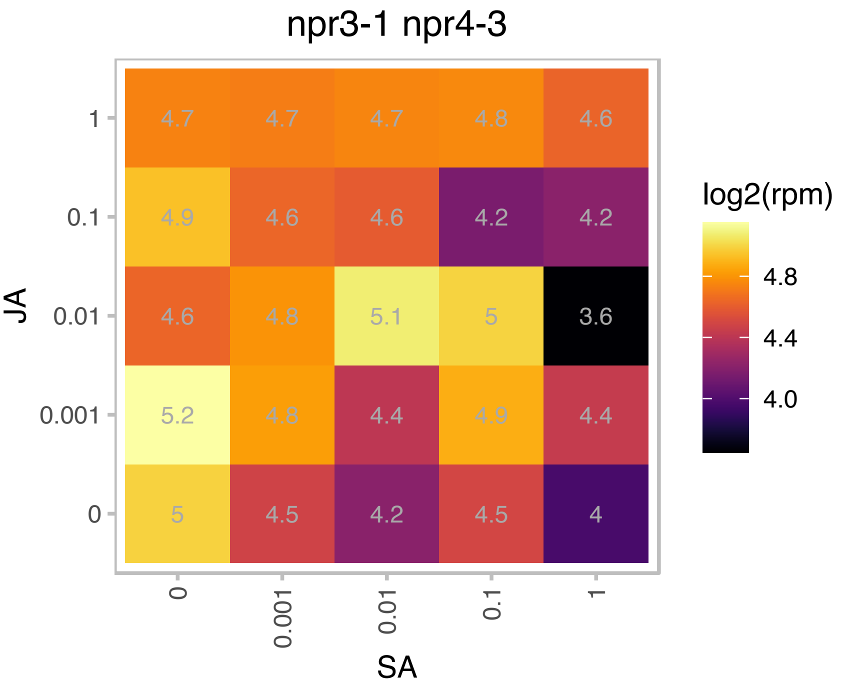

# AT3G04840.1 : NA

Ribosomal protein S3Ae

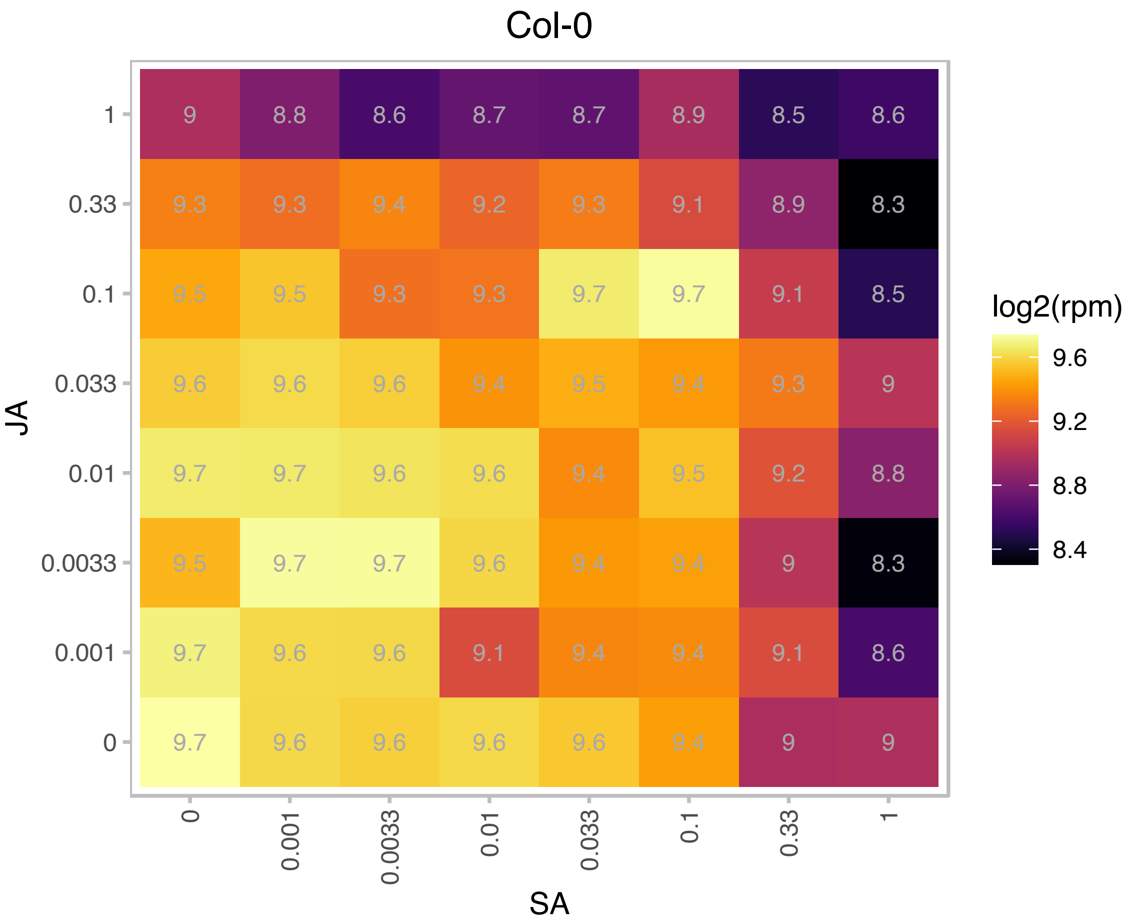


| Gene | AT3G04840 |
|---|---|
| is.expressed_genes | TRUE |
| is.responsive_genes | TRUE |
| is.apcluster | F-07 |

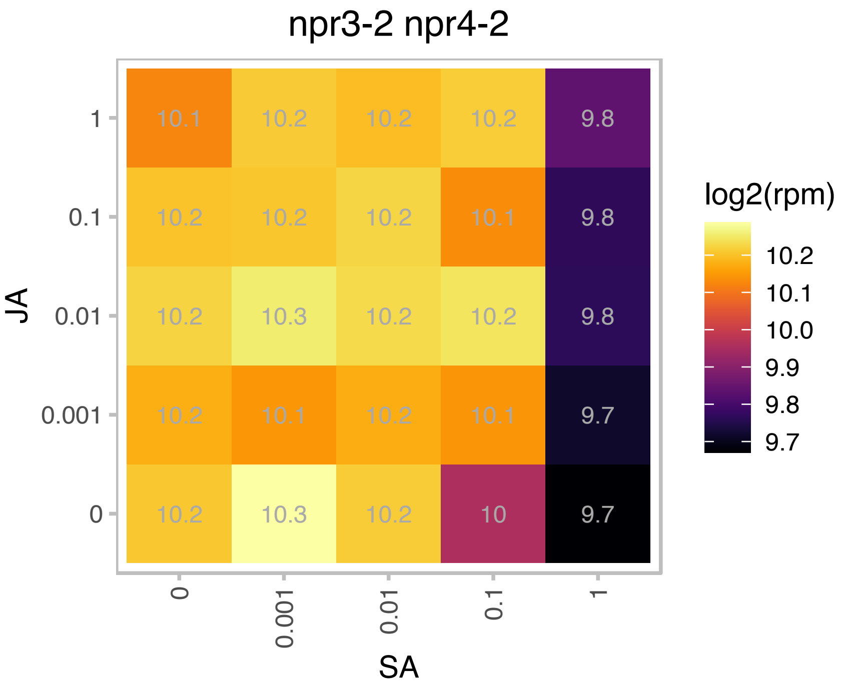


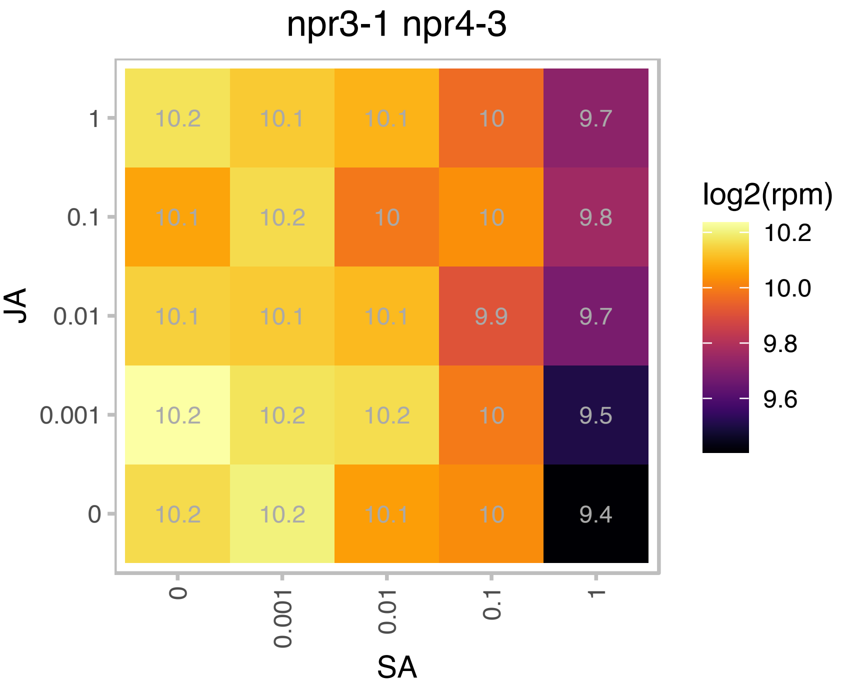

# AT3G04920.1 : NA

Ribosomal protein S24e family protein

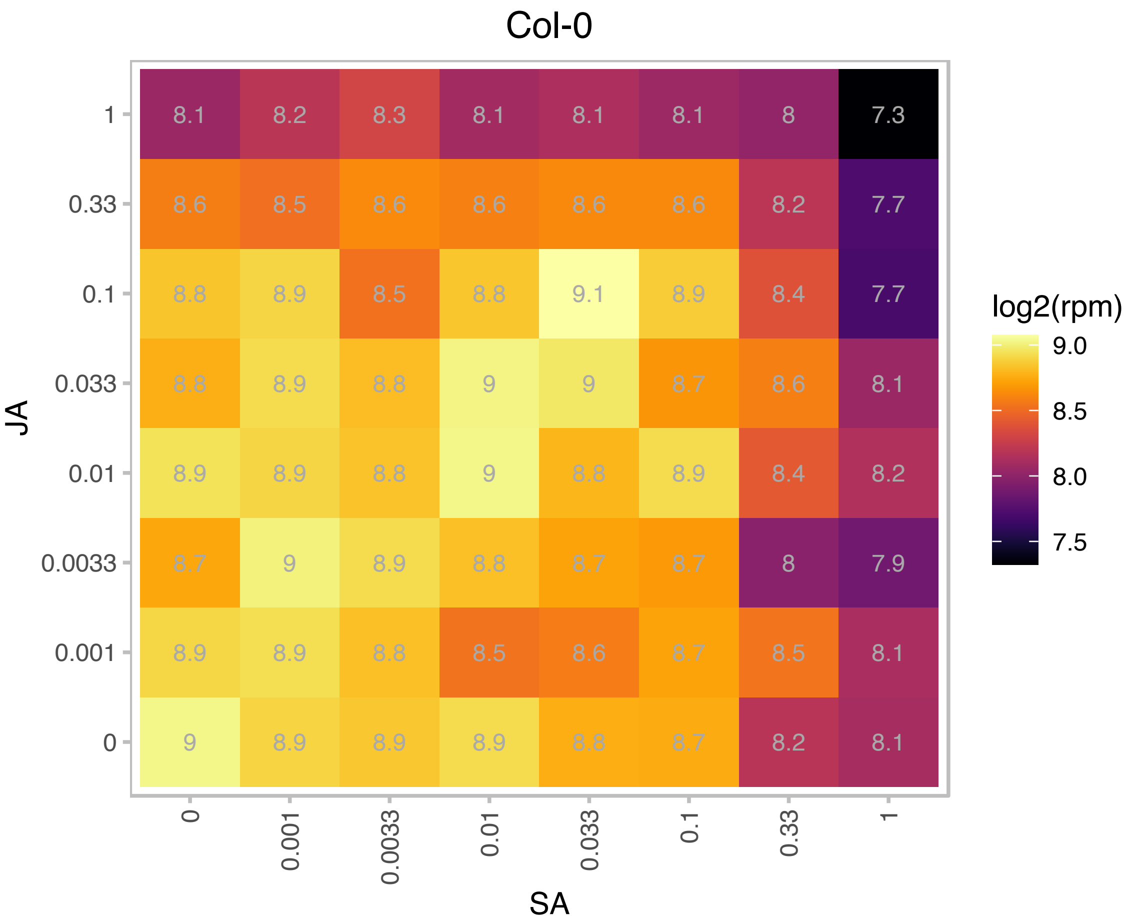


| | |
|---|---|
| Gene | AT3G04920 |
| is.expressed_genes | TRUE |
| is.responsive_genes | TRUE |
| is.apcluster | F-07 |

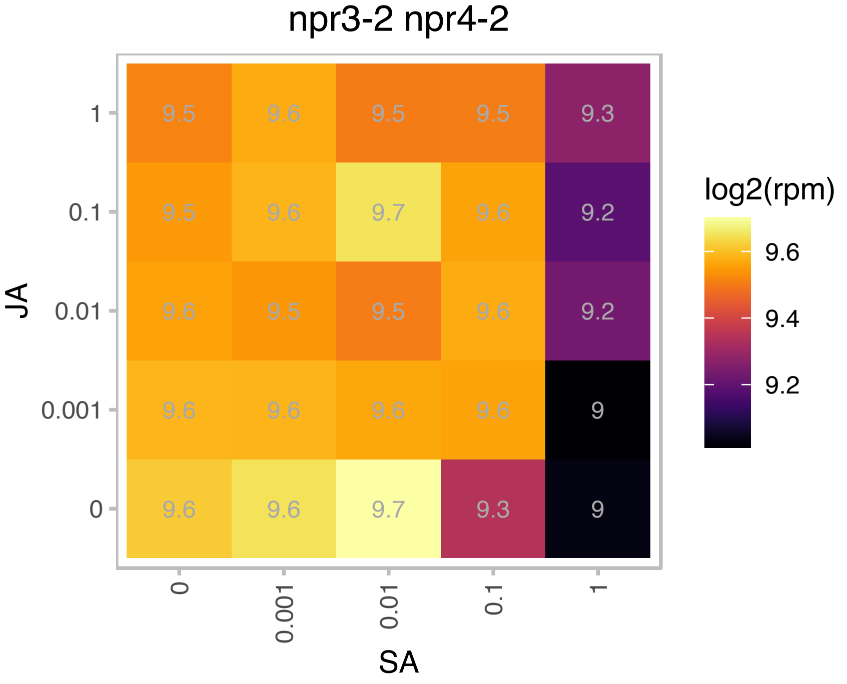


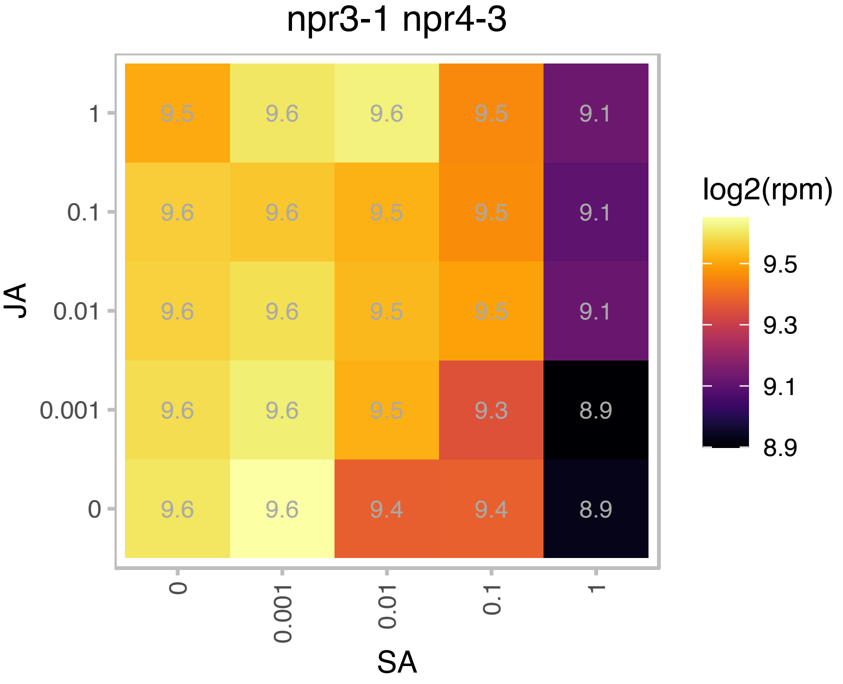

# AT3G05560.1 : NA

Ribosomal L22e protein family

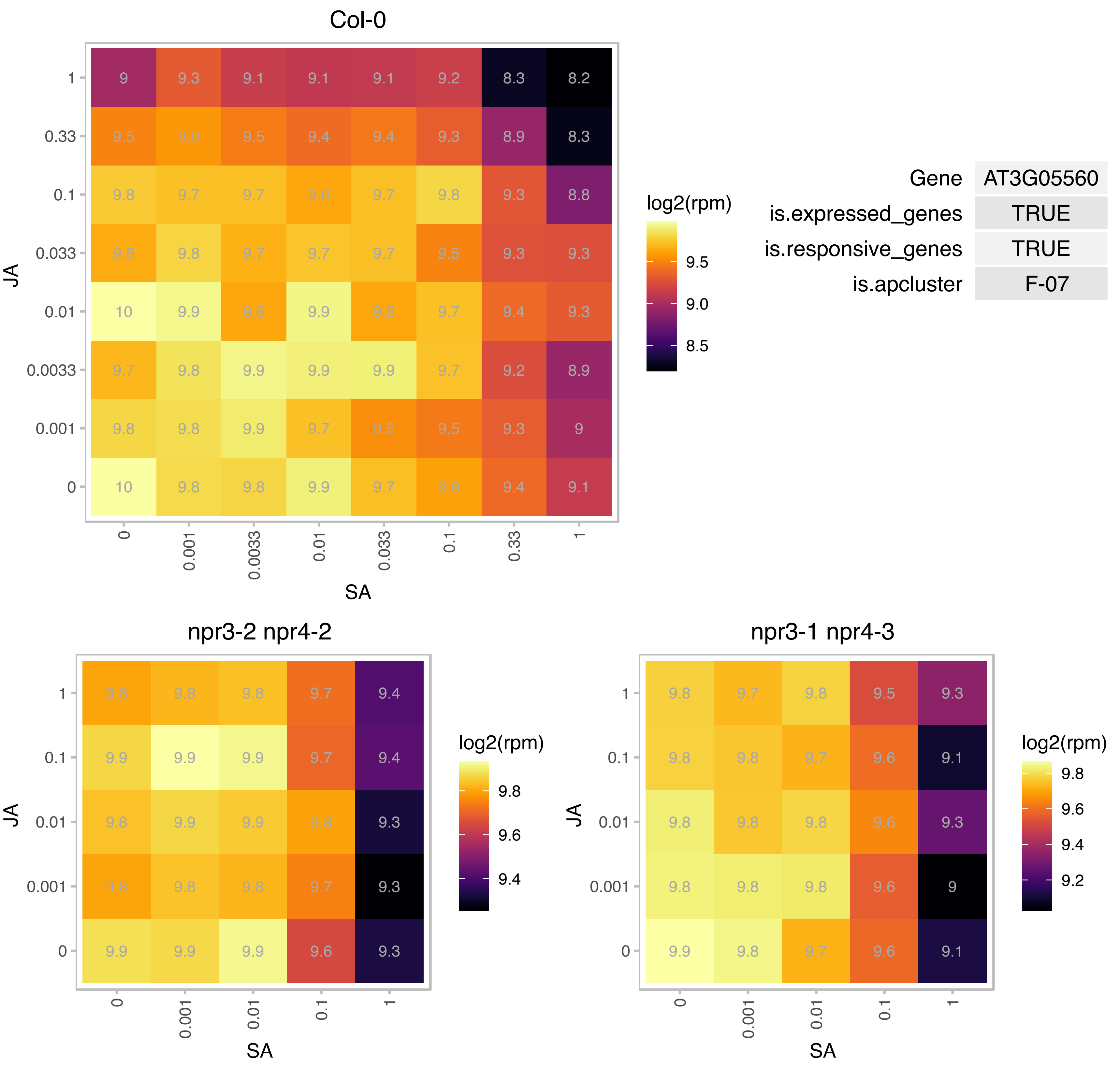


| | |
|---|---|
| Gene | AT3G05560 |
| is.expressed_genes | TRUE |
| is.responsive_genes | TRUE |
| is.apcluster | F-07 |

### AT3G05590.1 : RPL18

ribosomal protein L18

Encodes cytoplasmic ribosomal protein L18.

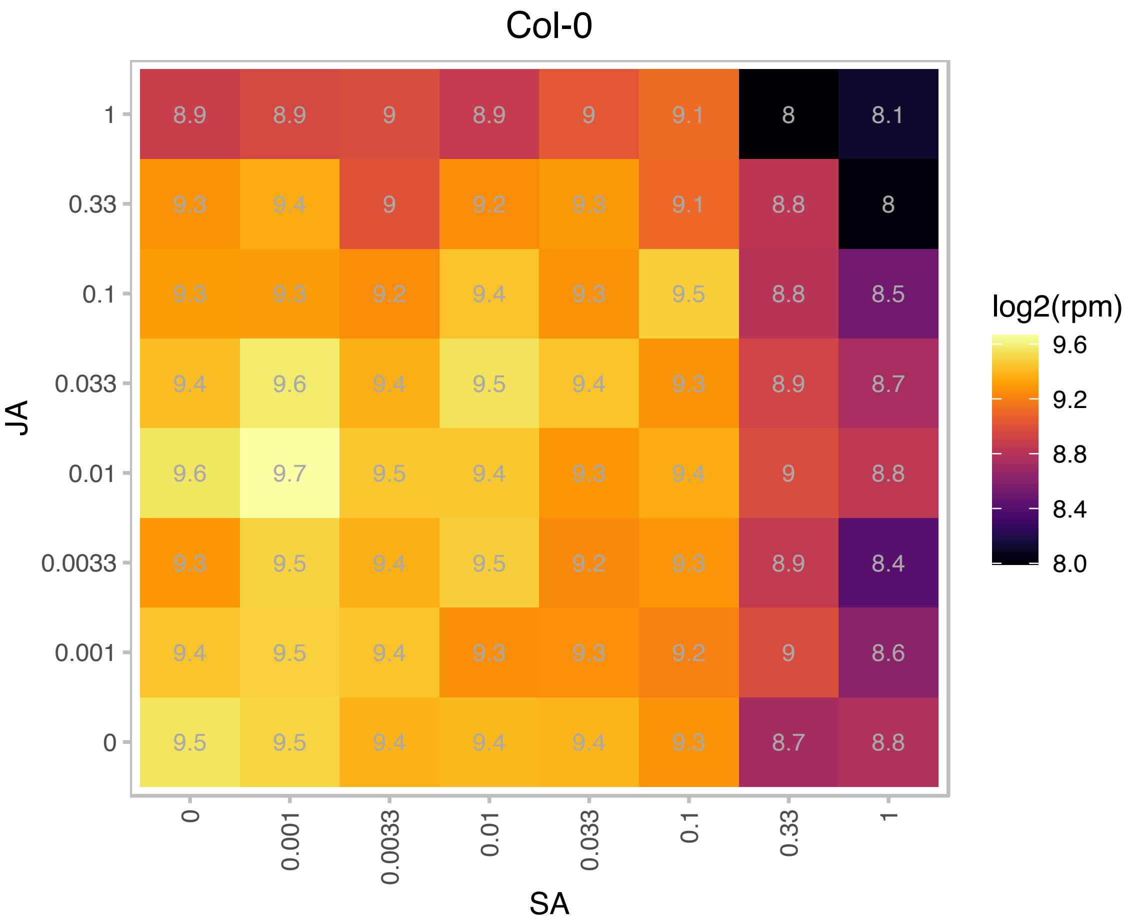


| | |
|---|---|
| Gene | AT3G05590 |
| is.expressed_genes | TRUE |
| is.responsive_genes | TRUE |
| is.apcluster | F-07 |

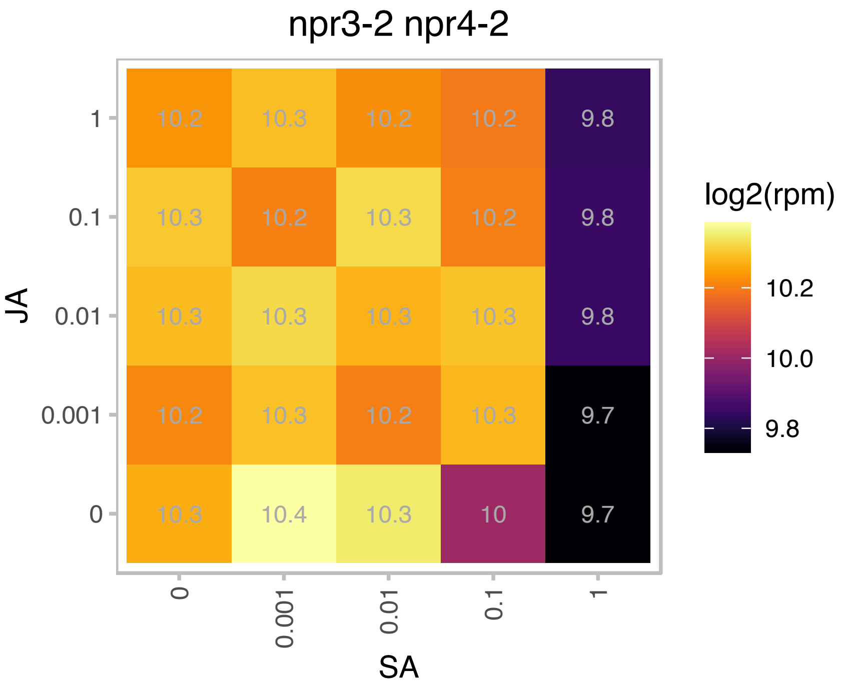


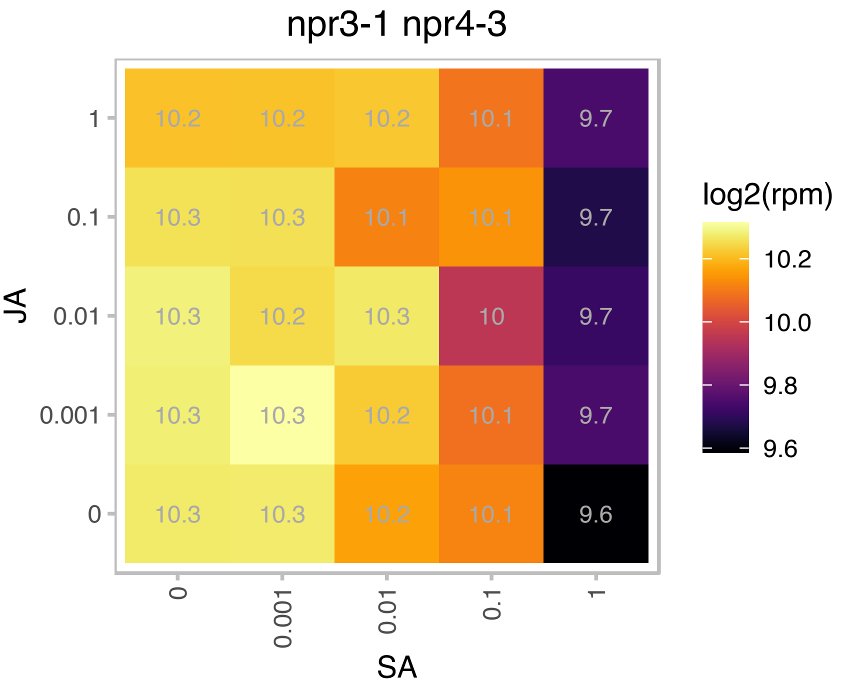

# AT3G06680.1 : NA

Ribosomal L29e protein family

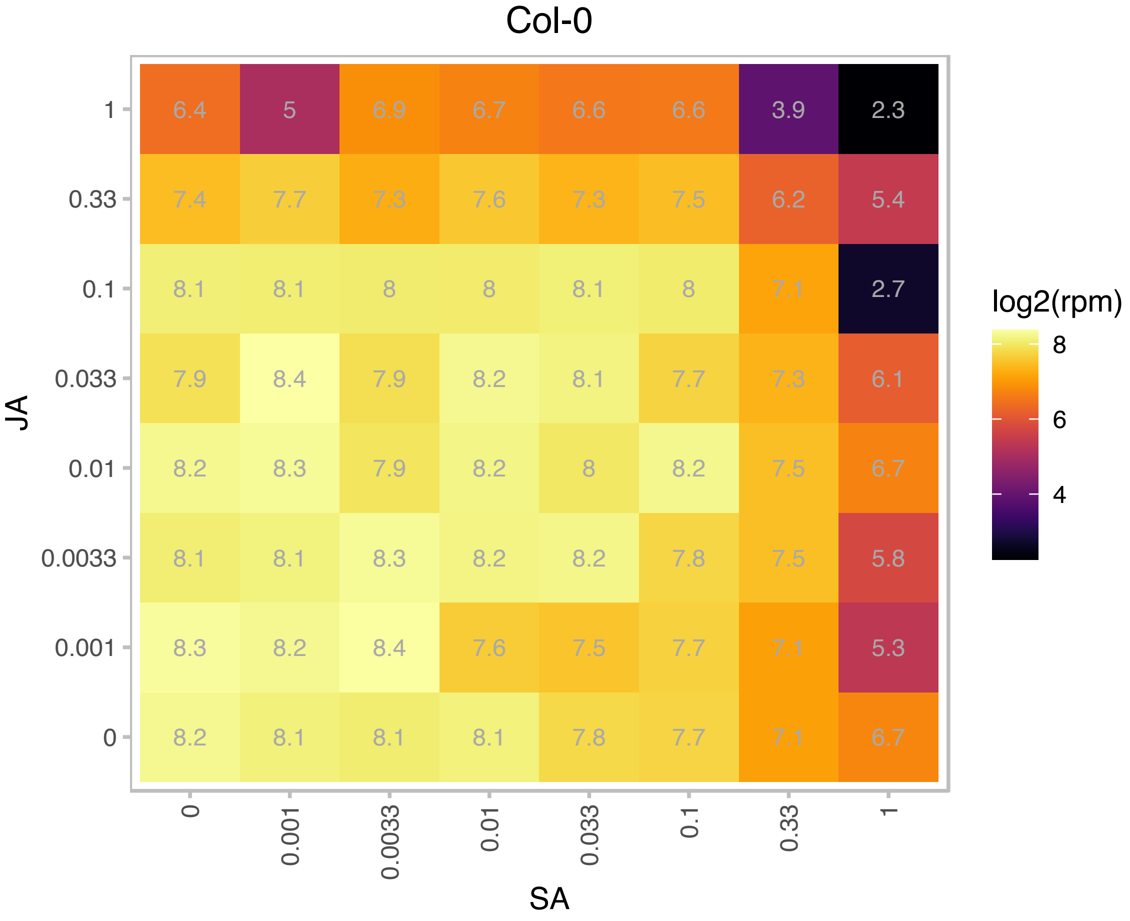


| | |
|---|---|
| Gene | AT3G06680 |
| is.expressed_genes | TRUE |
| is.responsive_genes | TRUE |
| is.apcluster | F-07 |

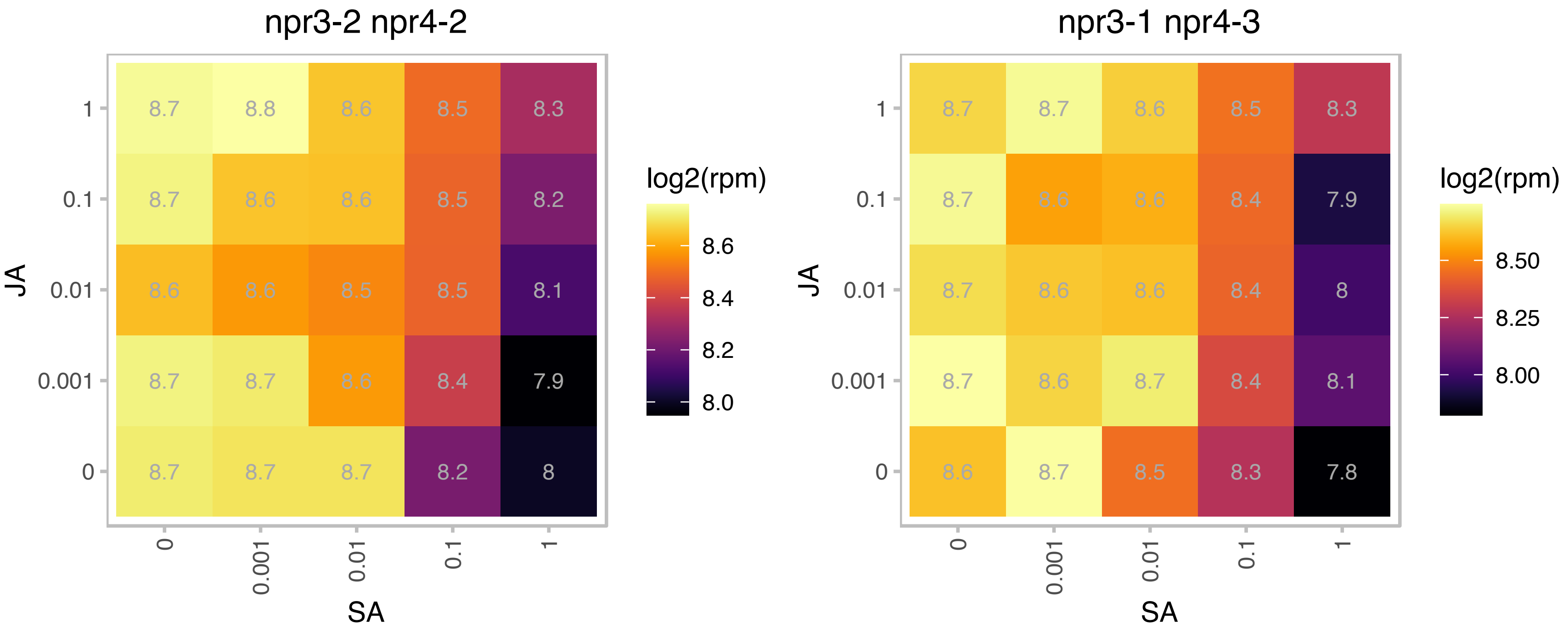

# AT3G06700.1 : NA

Ribosomal L29e protein family

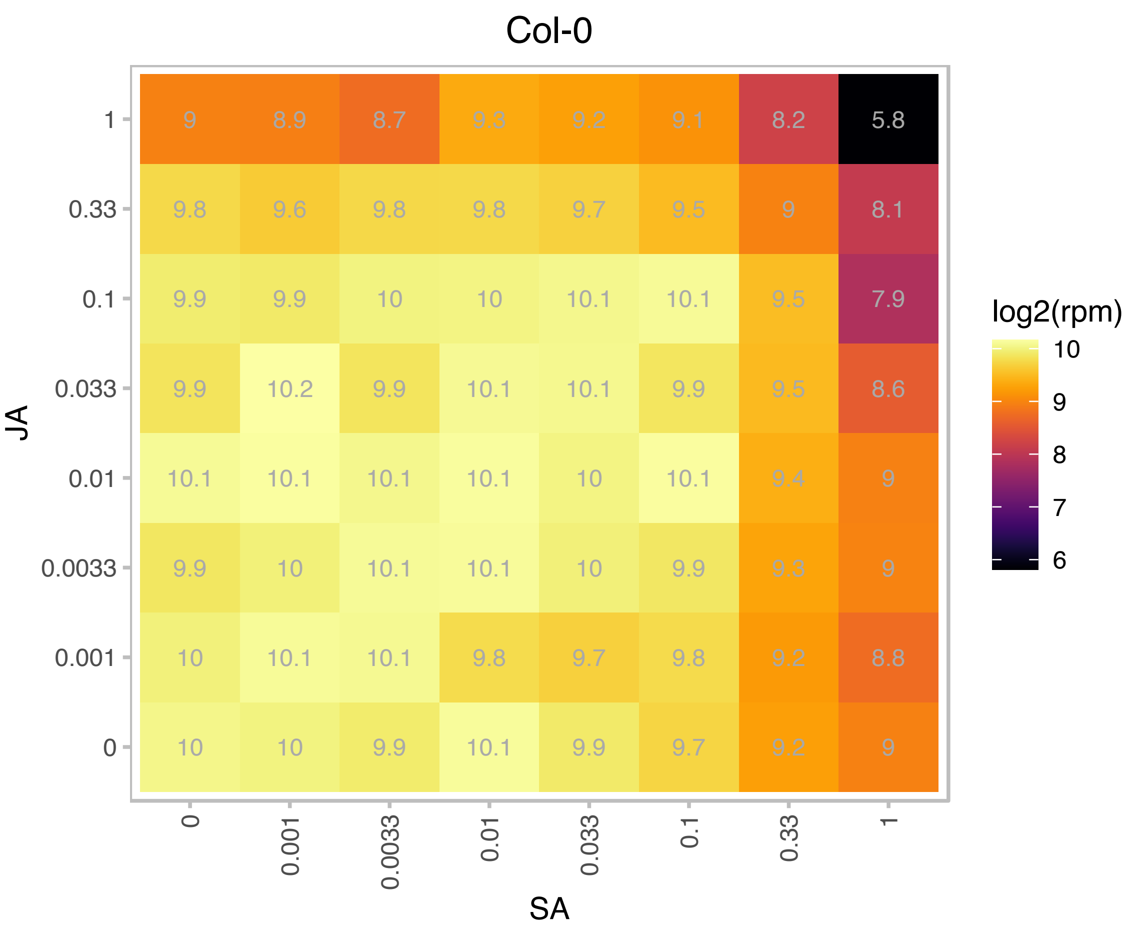


| | |
|---|---|
| Gene | AT3G06700 |
| is.expressed_genes | TRUE |
| is.responsive_genes | TRUE |
| is.apcluster | F-07 |

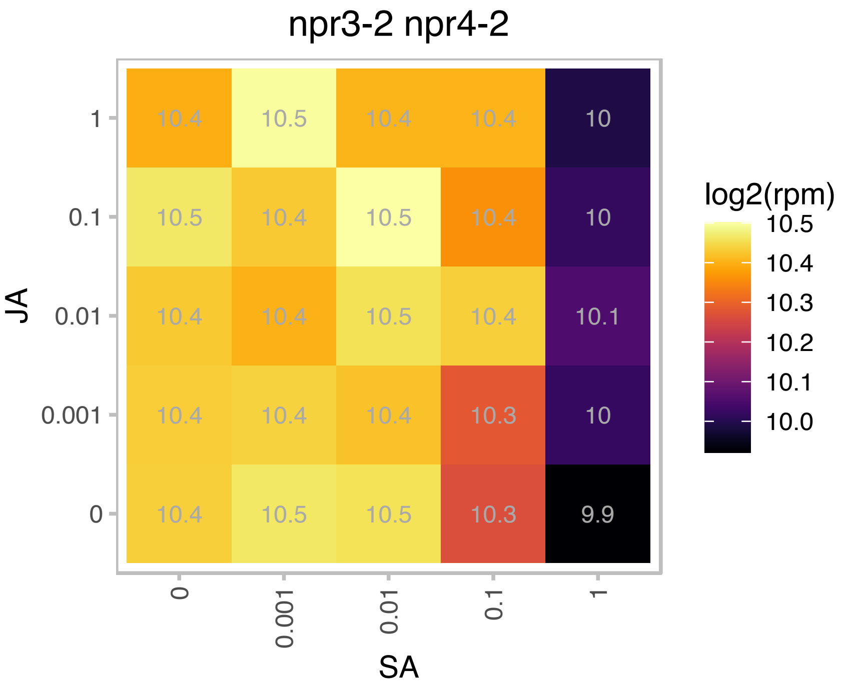


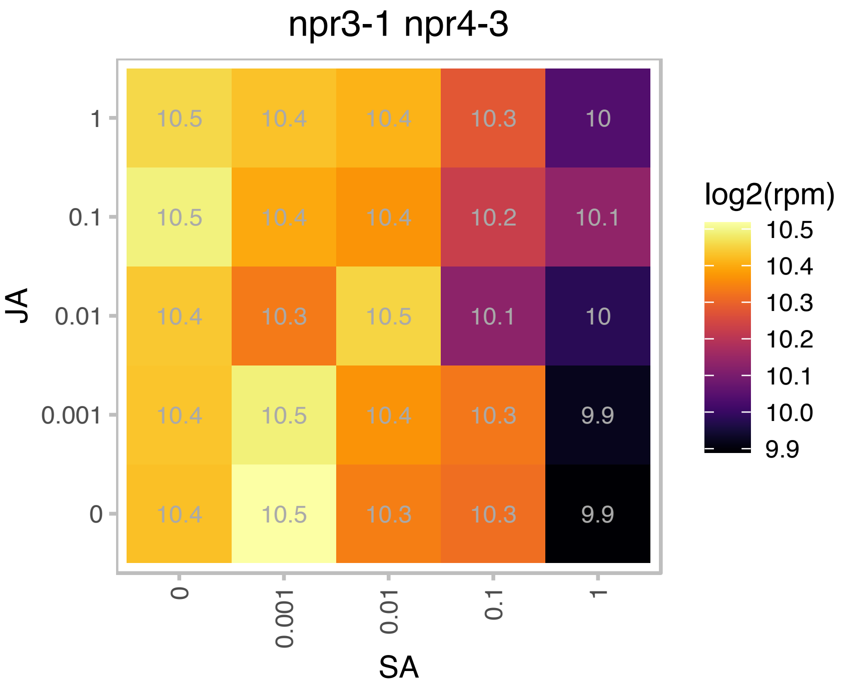

### AT3G06730.1 : TRX P, TRX z

Thioredoxin z

Encodes a plastidial thioredoxin (TRX) isoform that defines a branch of plastidial TRXs lying between x- and y-type TRXs and thus was named TRX z. Possesses disulfide reductase activity in vitro. TRX z was previously named as Trx p (thioredoxin putative and plastidic) before its thioredoxin activity was confirmed (Meng et al., PNAS 2010, 107:3900). Knockout mutant of TRX z has a severe albino phenotype and is inhibited in chloroplast development. Two fructokinase-like proteins (FLN1 and FLN2), members of the pfkB-carbohydrate kinase family, are potential TRX z targets.

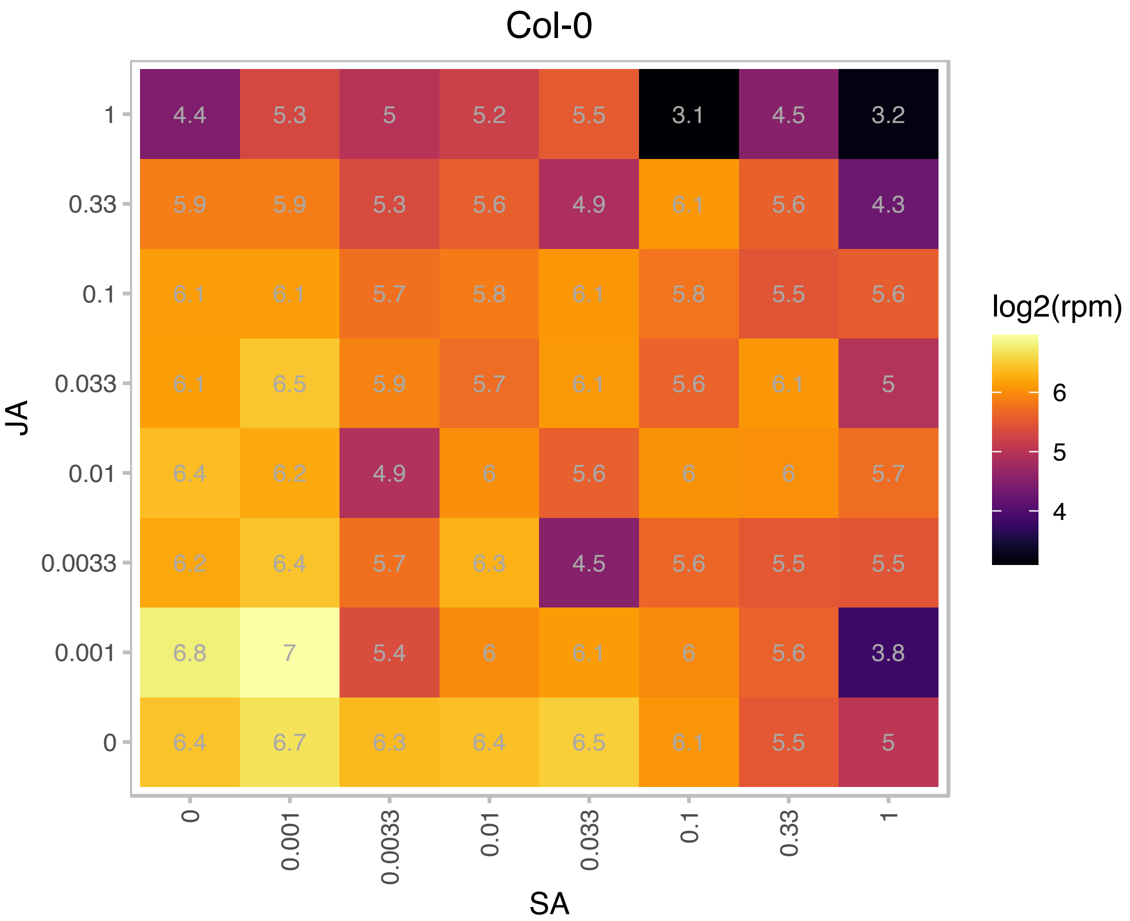


| | |
|---|---|
| Gene | AT3G06730 |
| is.expressed_genes | TRUE |
| is.responsive_genes | TRUE |
| is.apcluster | F-07 |

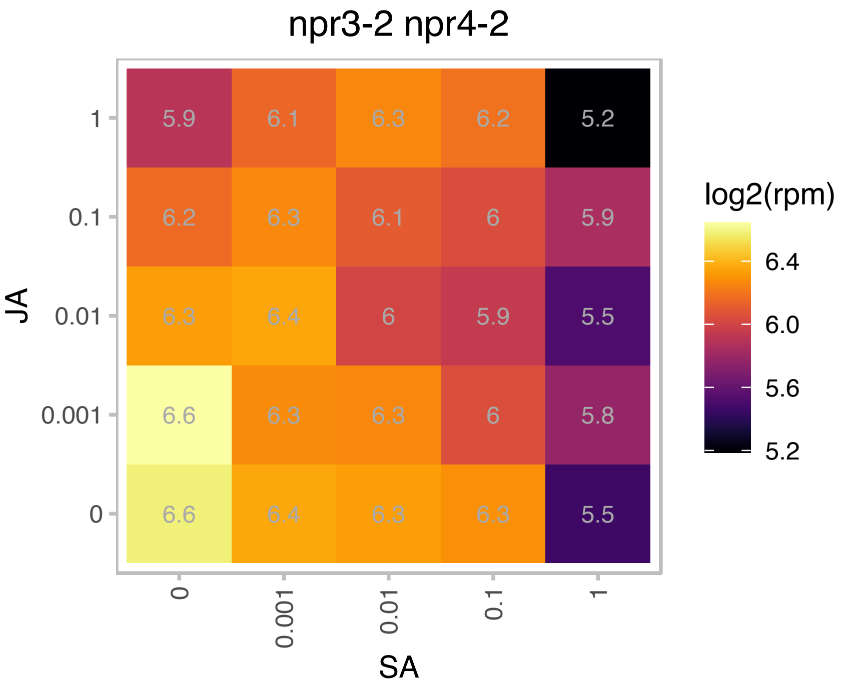


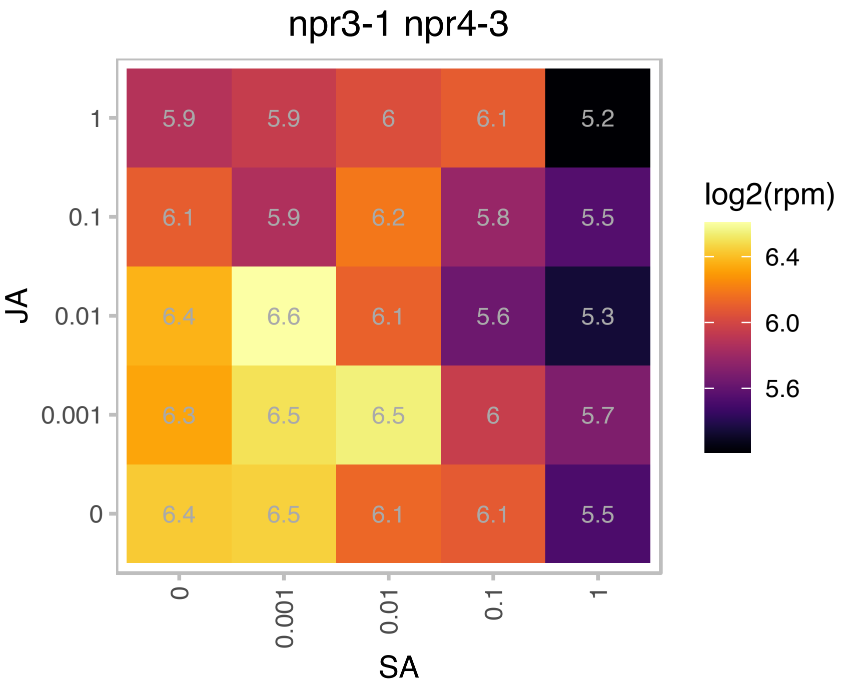

# AT3G07110.1 : NA

Ribosomal protein L13 family protein

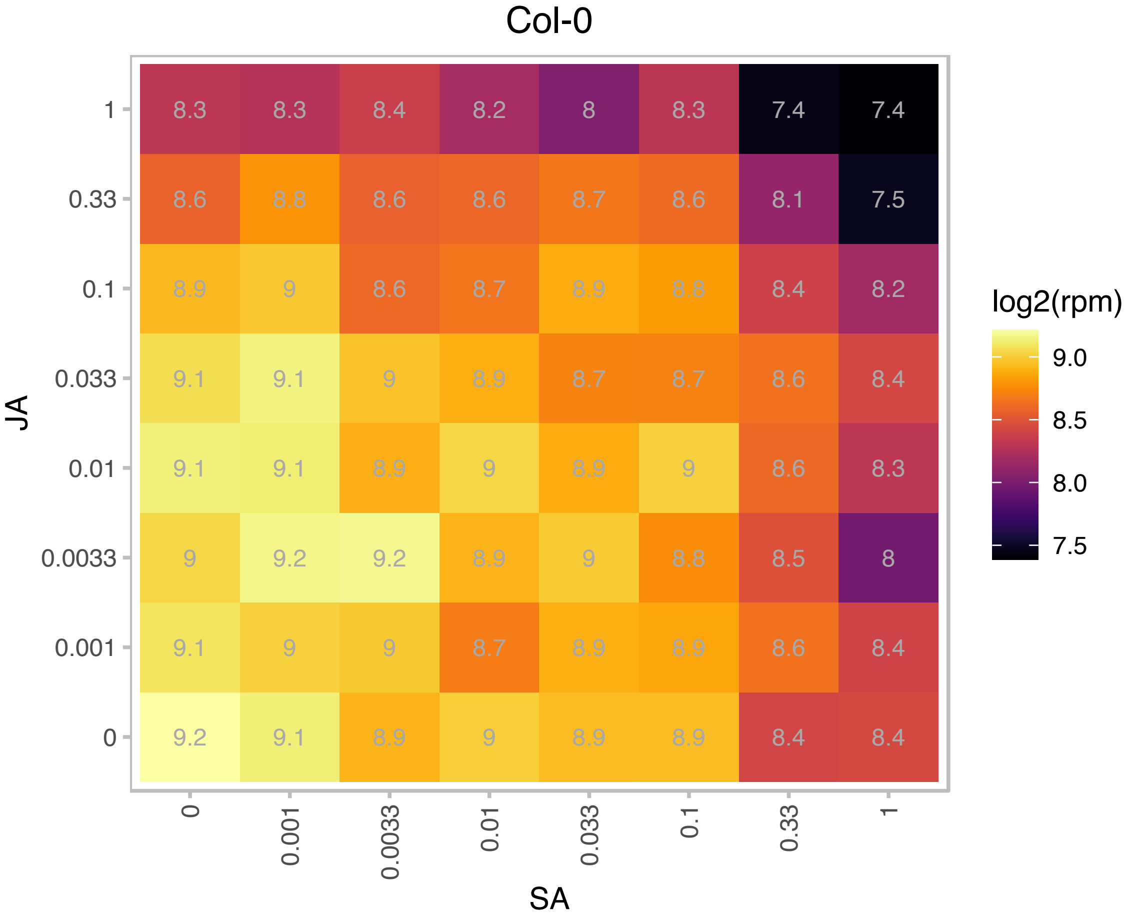


| | |
|---|---|
| Gene | AT3G07110 |
| is.expressed_genes | TRUE |
| is.responsive_genes | TRUE |
| is.apcluster | F-07 |

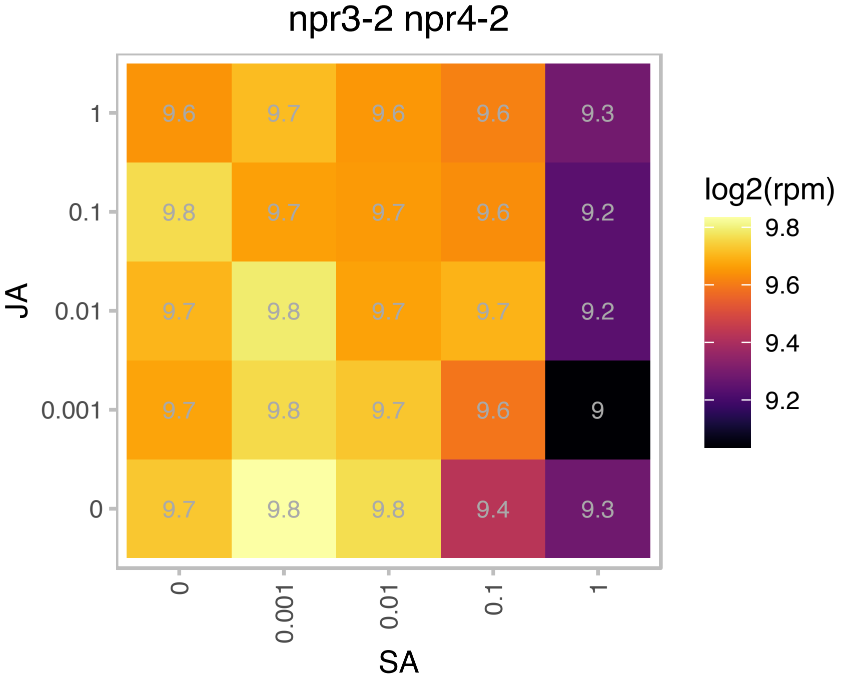


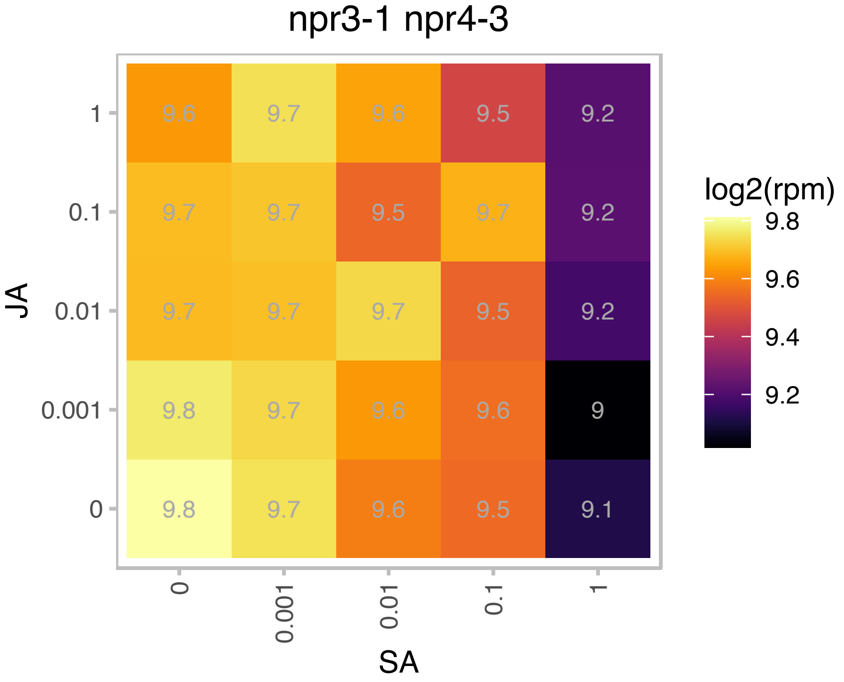

# AT3G08520.1 : NA

Ribosomal protein L41 family

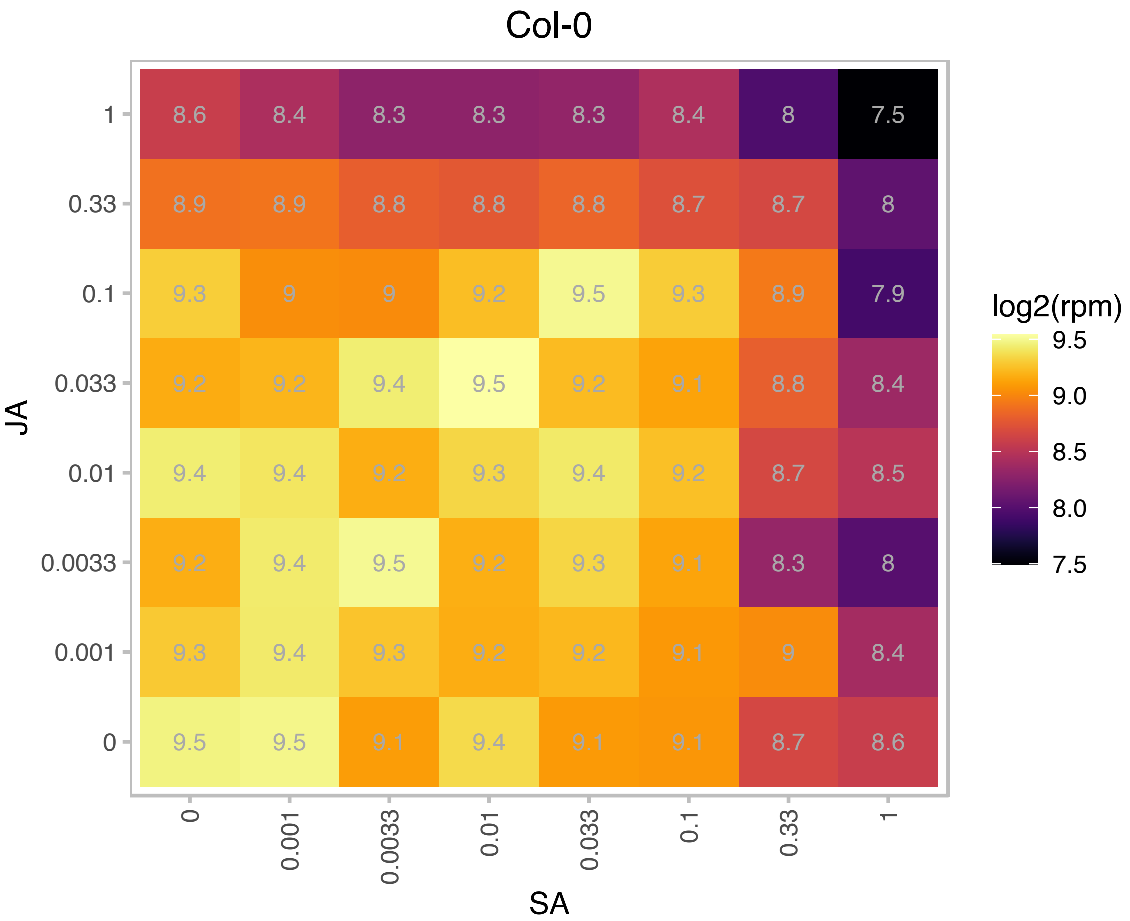


| | |
|---|---|
| Gene | AT3G08520 |
| is.expressed_genes | TRUE |
| is.responsive_genes | TRUE |
| is.apcluster | F-07 |

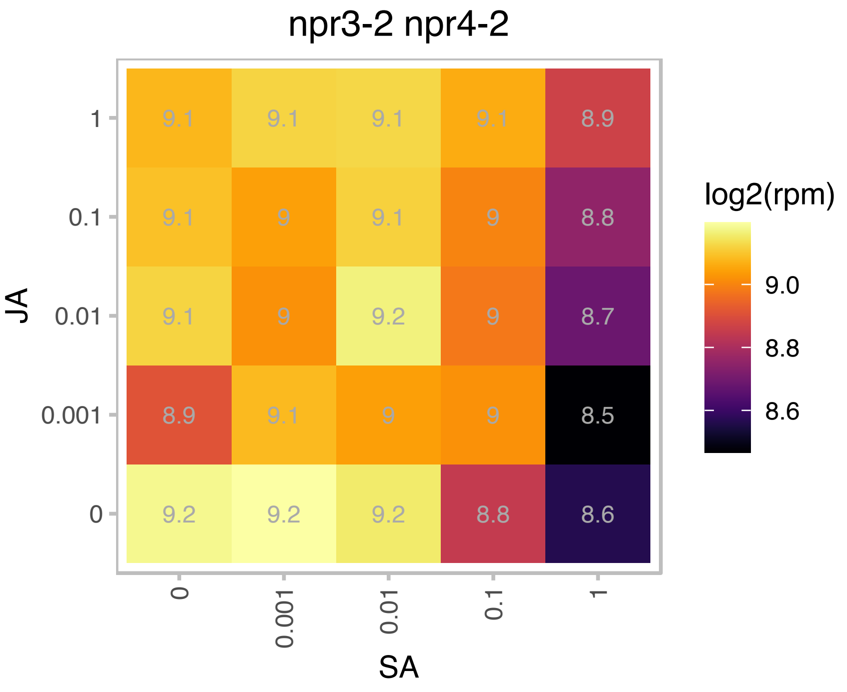


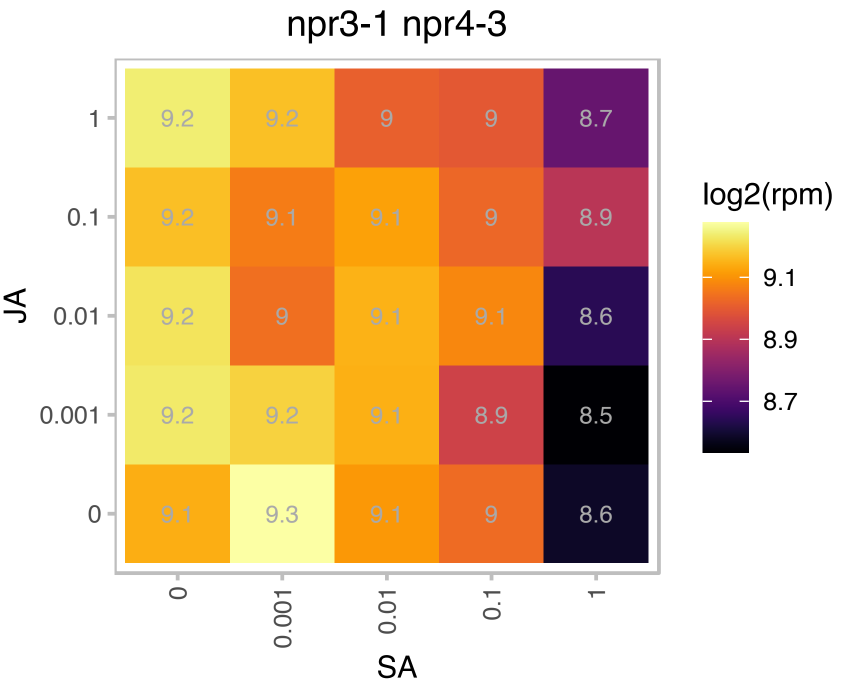

# AT3G09200.1 : NA

Ribosomal protein L10 family protein

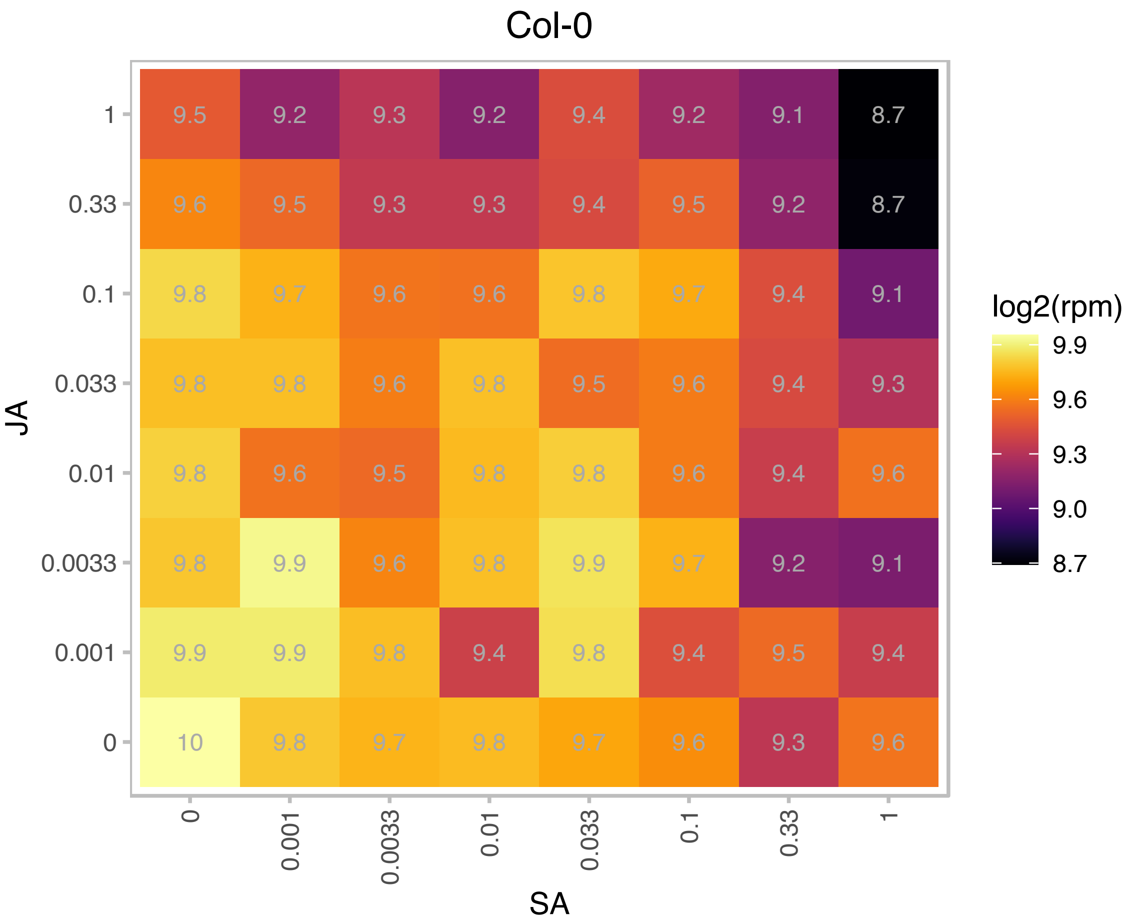


| | |
|---|---|
| Gene | AT3G09200 |
| is.expressed_genes | TRUE |
| is.responsive_genes | TRUE |
| is.apcluster | F-07 |

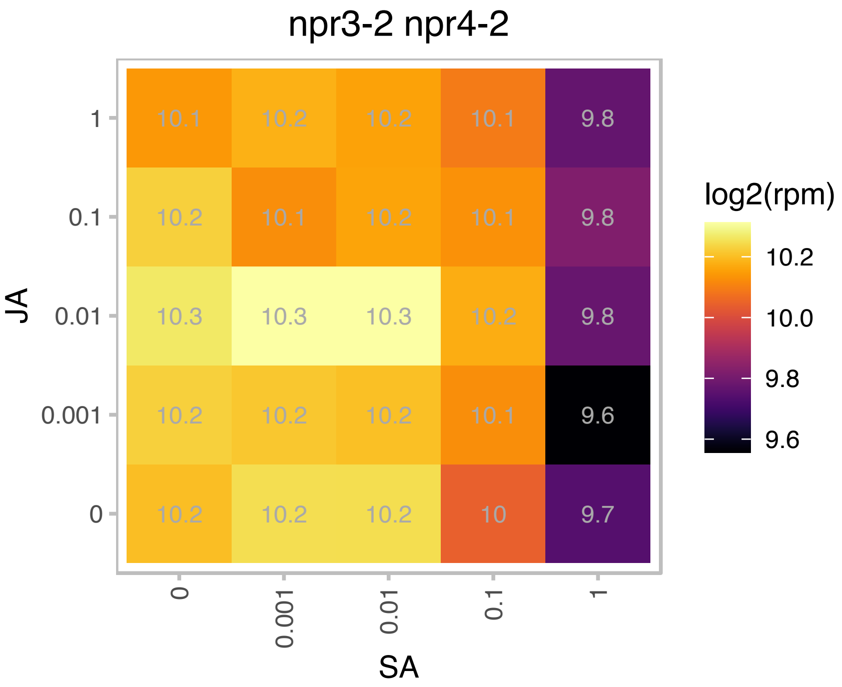


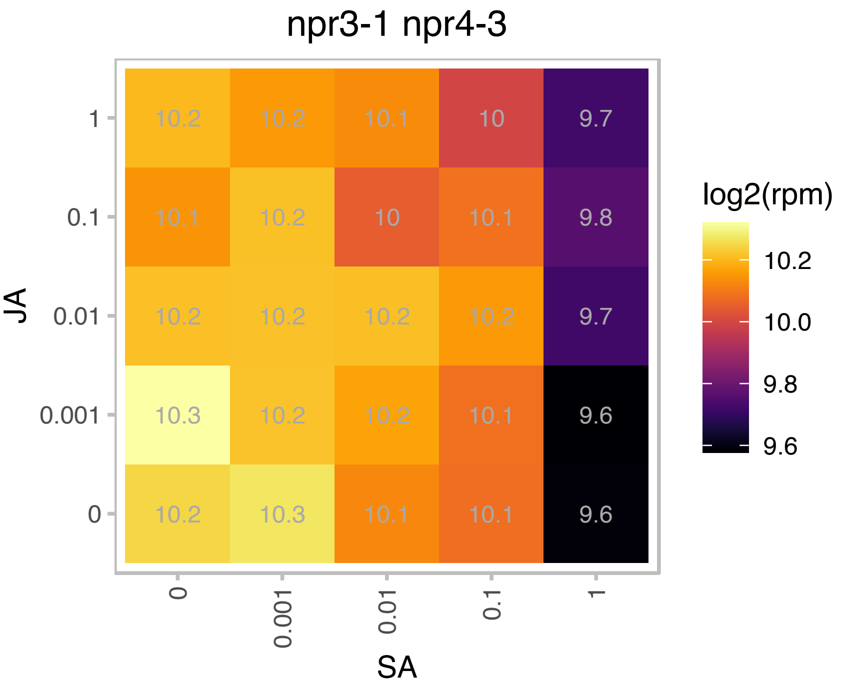

# AT3G10610.1 : NA

Ribosomal S17 family protein

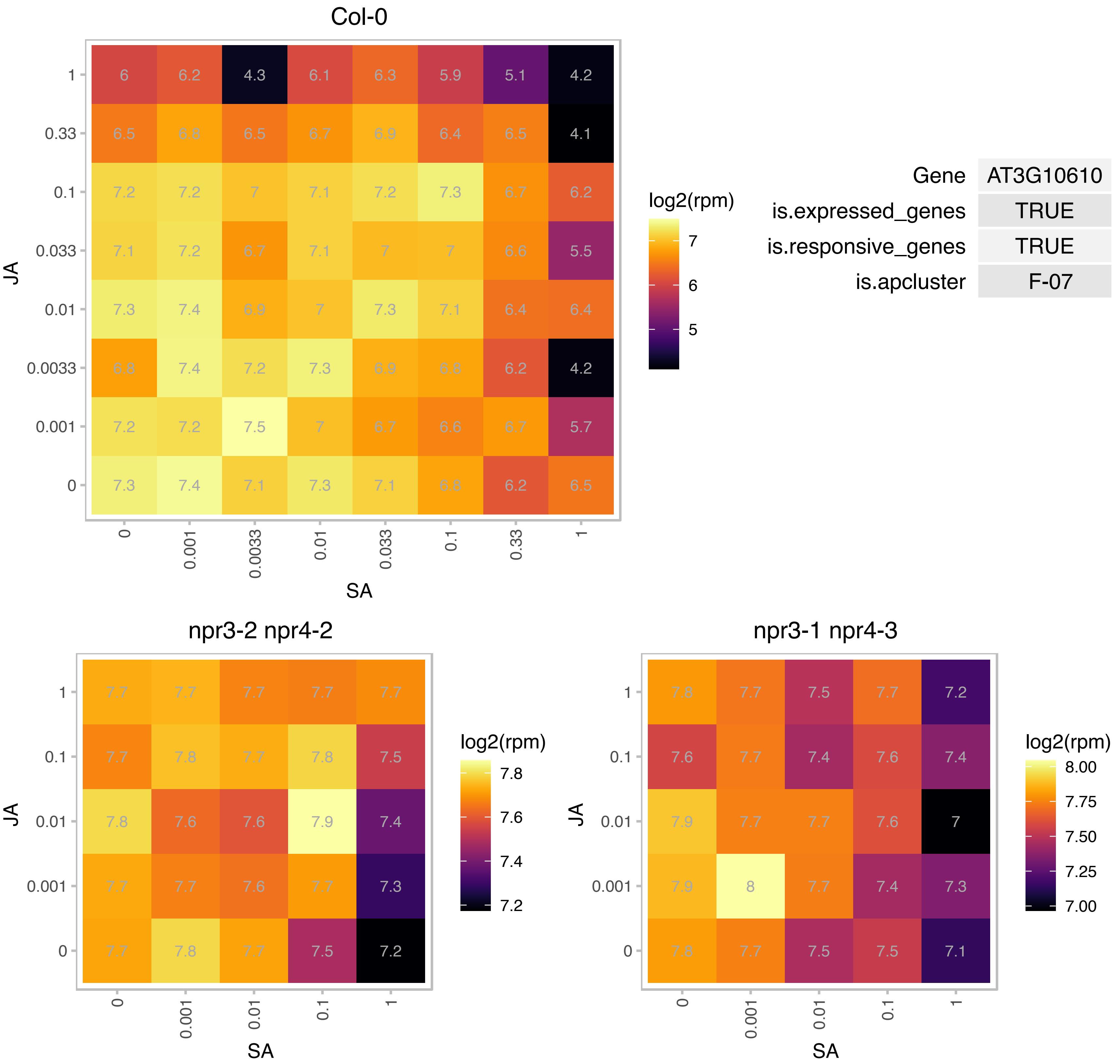


| | |
|---|---|
| Gene | AT3G10610 |
| is.expressed_genes | TRUE |
| is.responsive_genes | TRUE |
| is.apcluster | F-07 |

# AT3G11250.1 : NA

Ribosomal protein L10 family protein

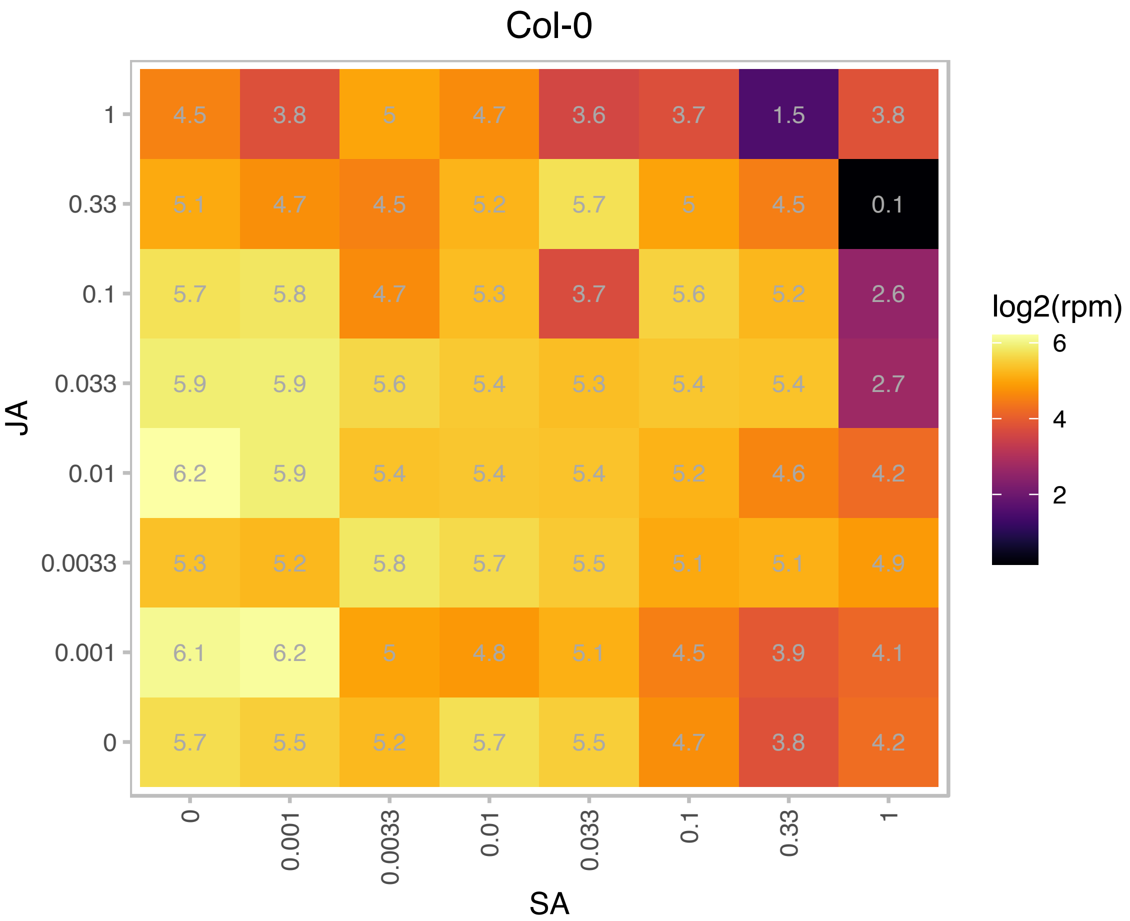


| | |
|---|---|
| Gene | AT3G11250 |
| is.expressed_genes | TRUE |
| is.responsive_genes | TRUE |
| is.apcluster | F-07 |

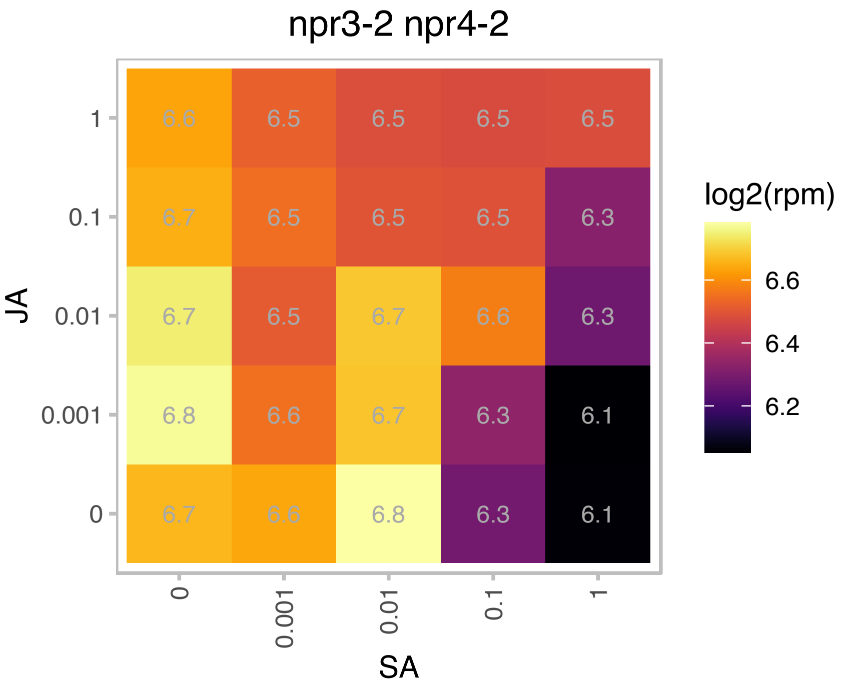


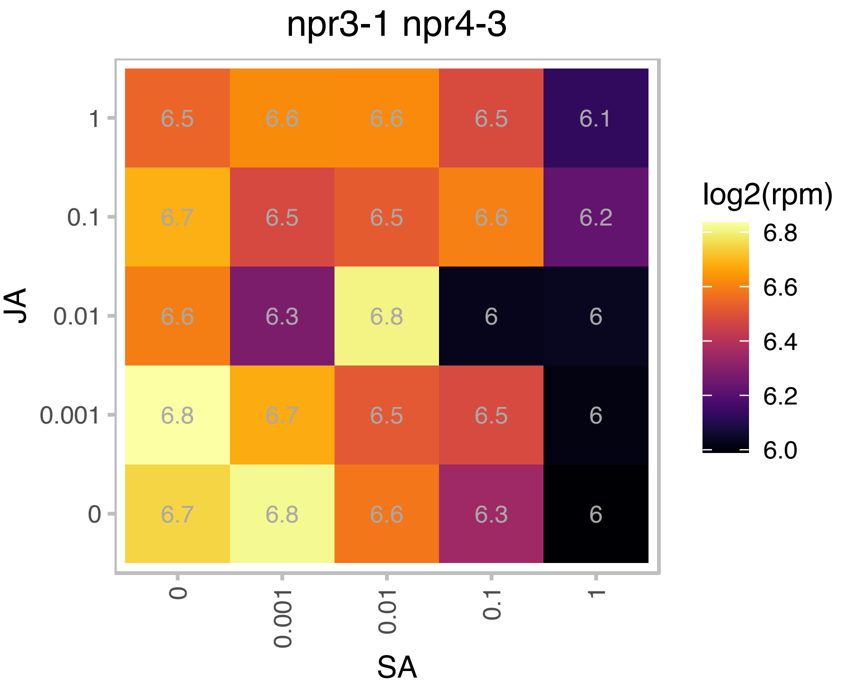

# AT3G11510.1 : NA

Ribosomal protein S11 family protein

| | |
|---|---|
| Gene | AT3G11510 |
| is.expressed_genes | TRUE |
| is.responsive_genes | TRUE |
| is.apcluster | F-07 |

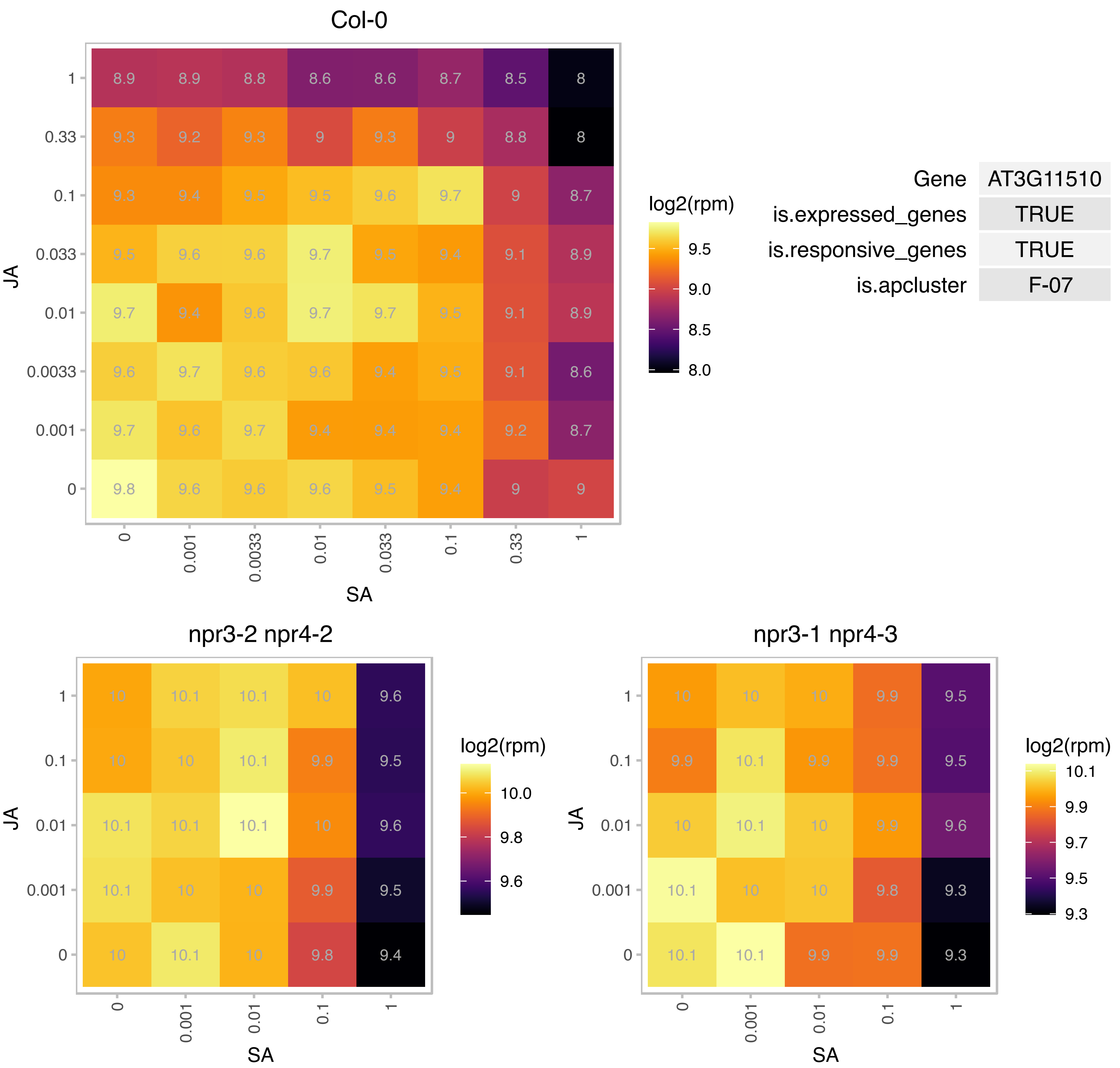

# AT3G12390.1 : NA

Nascent polypeptide-associated complex (NAC), alpha subunit family protein

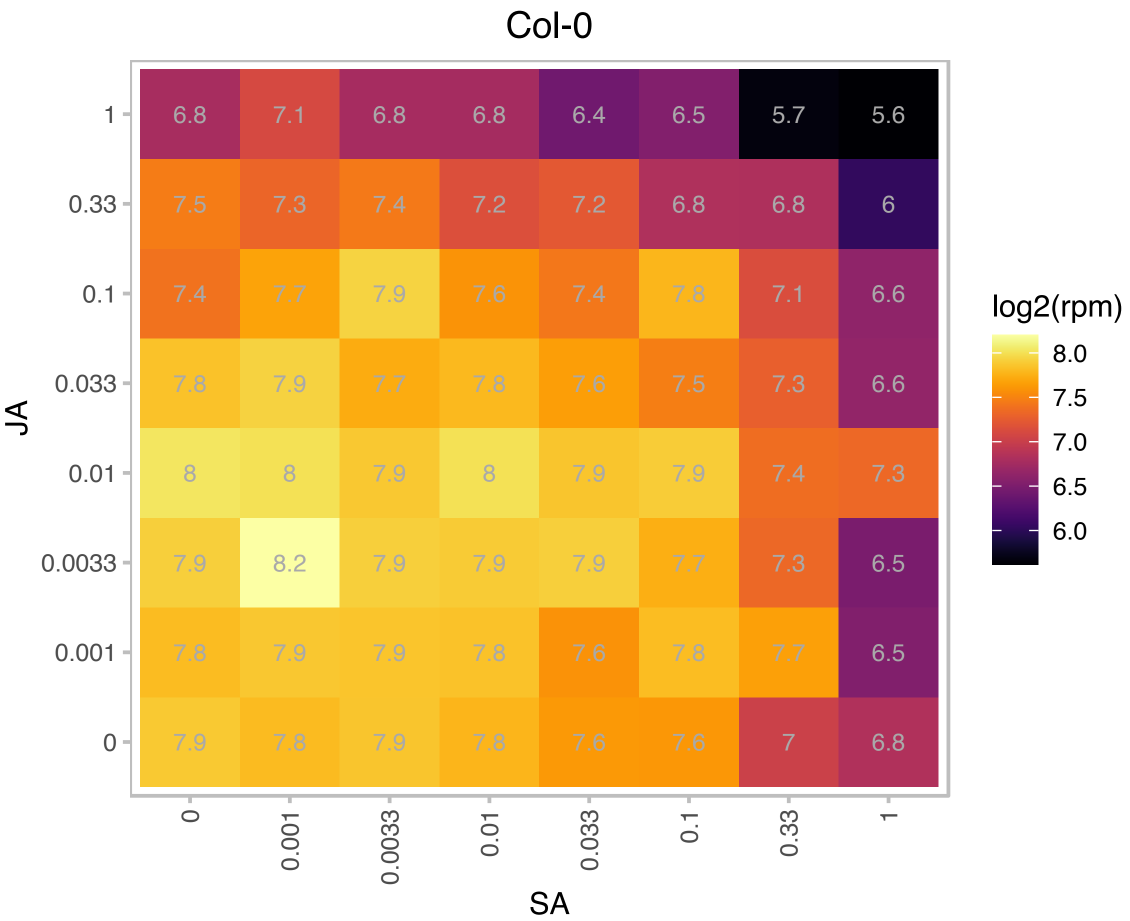


| | |
|---|---|
| Gene | AT3G12390 |
| is.expressed_genes | TRUE |
| is.responsive_genes | TRUE |
| is.apcluster | F-07 |

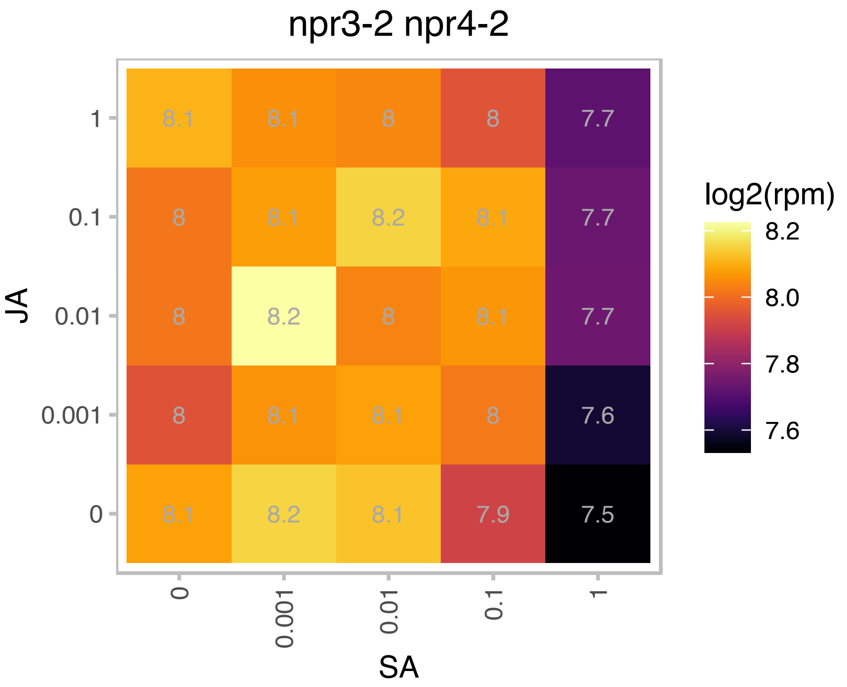


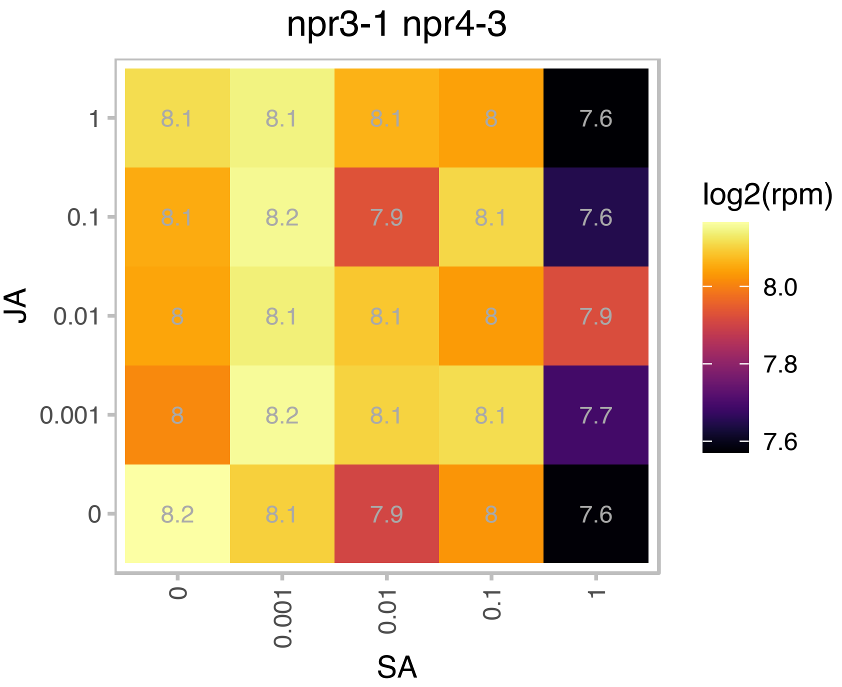

# AT3G12930.1 : DG238

Lojap-related protein

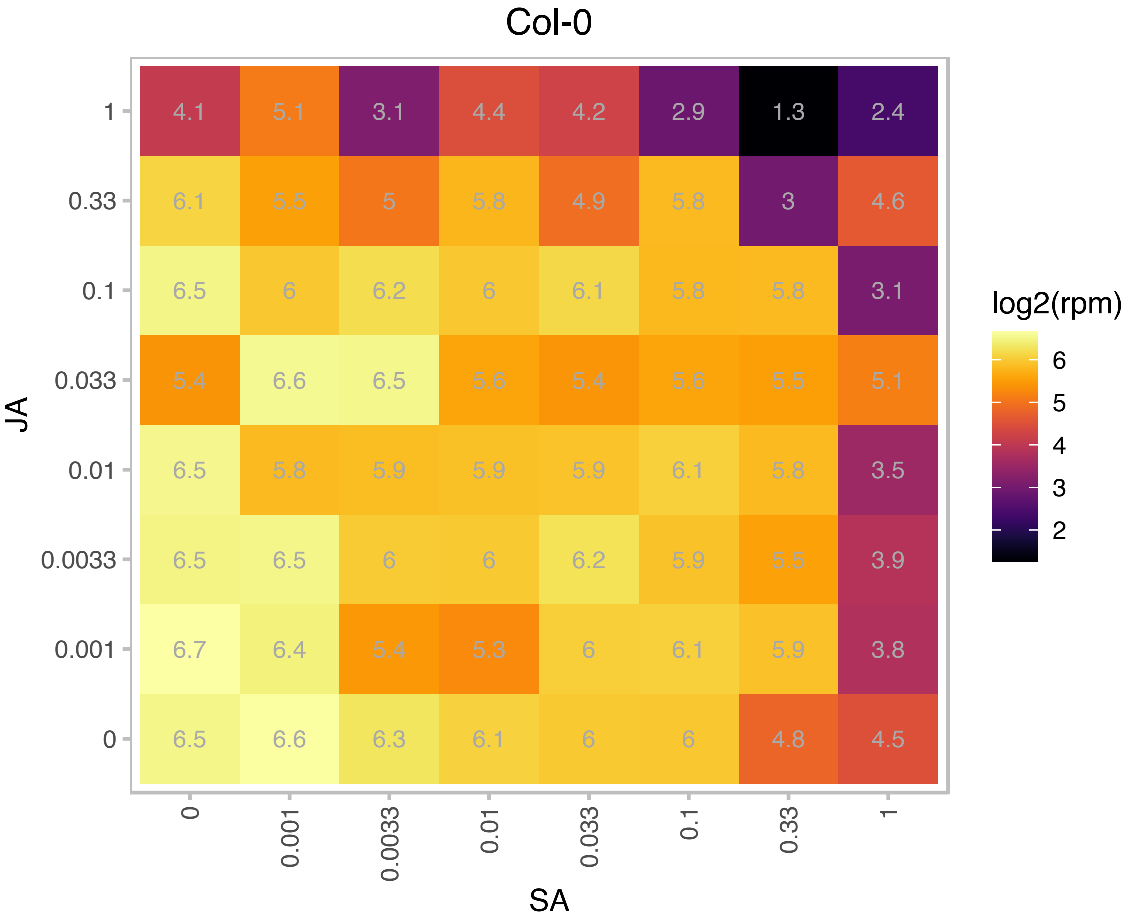


| | |
|---|---|
| Gene | AT3G12930 |
| is.expressed_genes | TRUE |
| is.responsive_genes | TRUE |
| is.apcluster | F-07 |

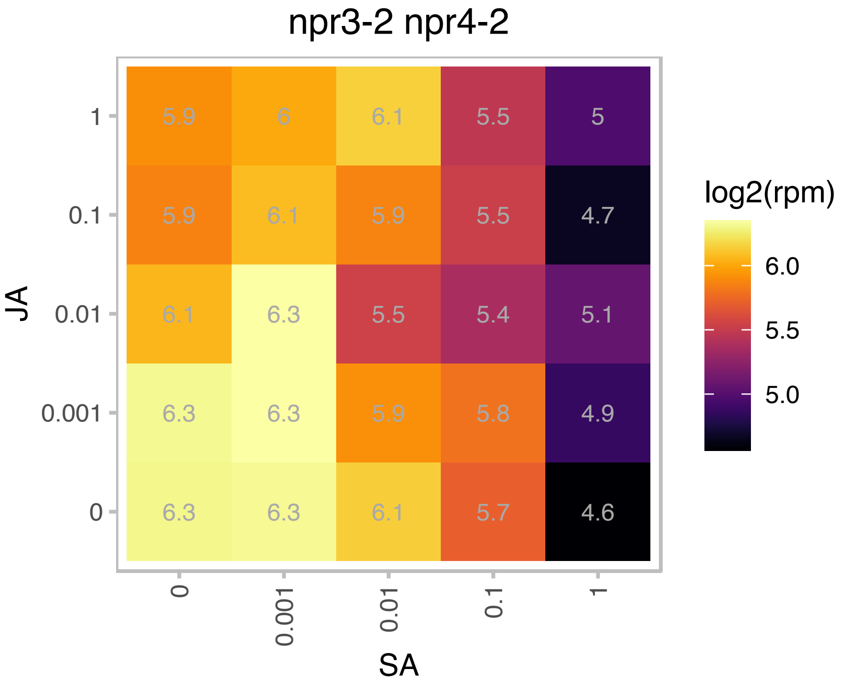


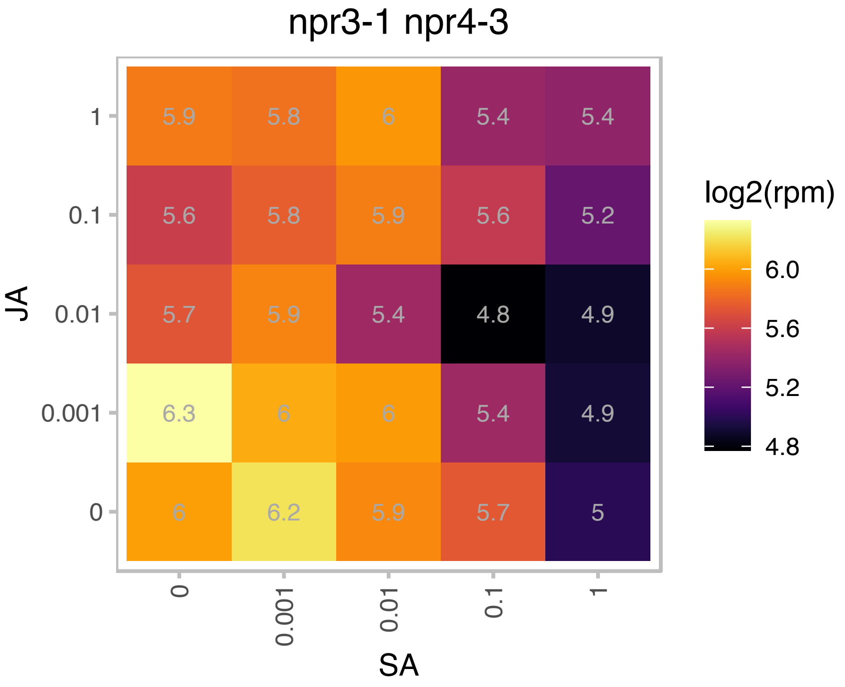

# AT3G13230.1 : NA

RNA-binding KH domain-containing protein

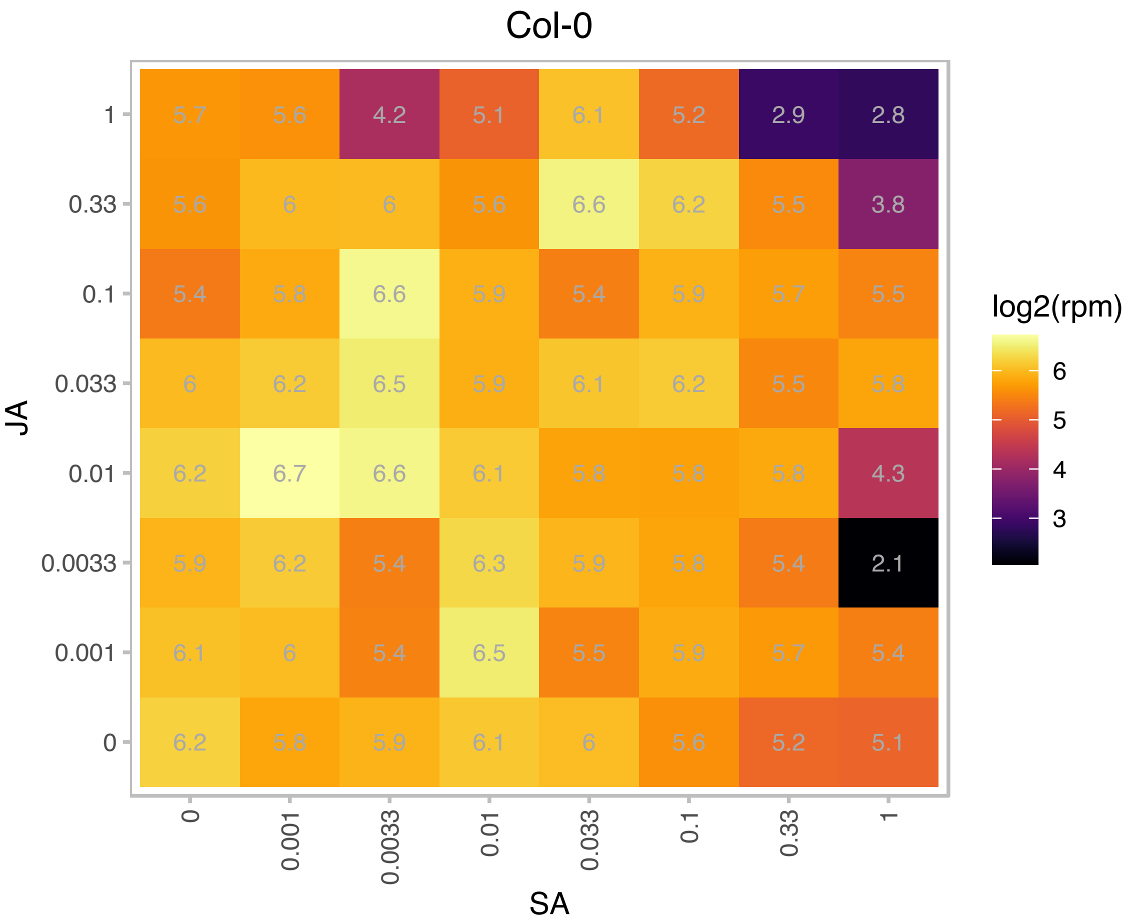


| | |
|---|---|
| Gene | AT3G13230 |
| is.expressed_genes | TRUE |
| is.responsive_genes | TRUE |
| is.apcluster | F-07 |

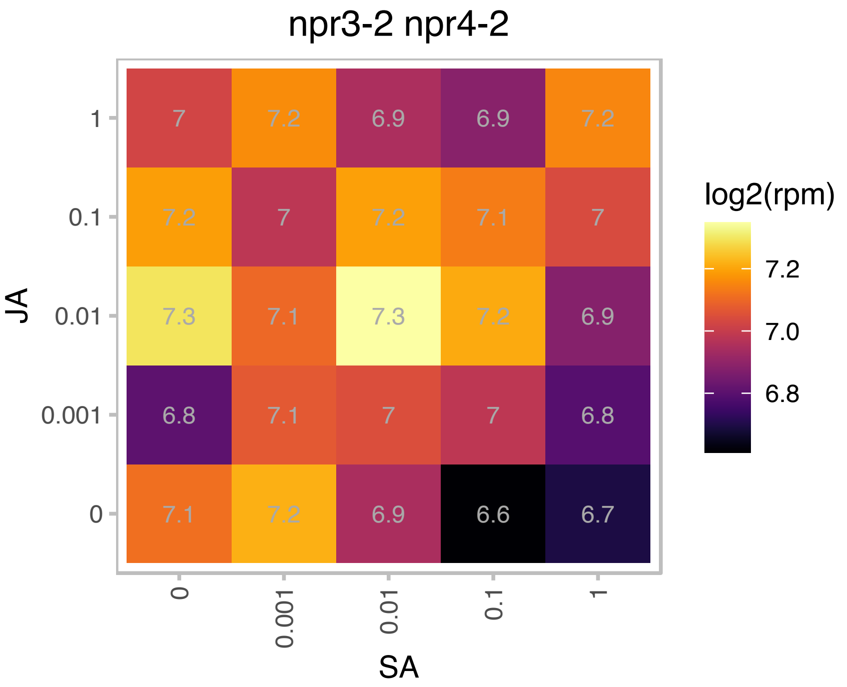


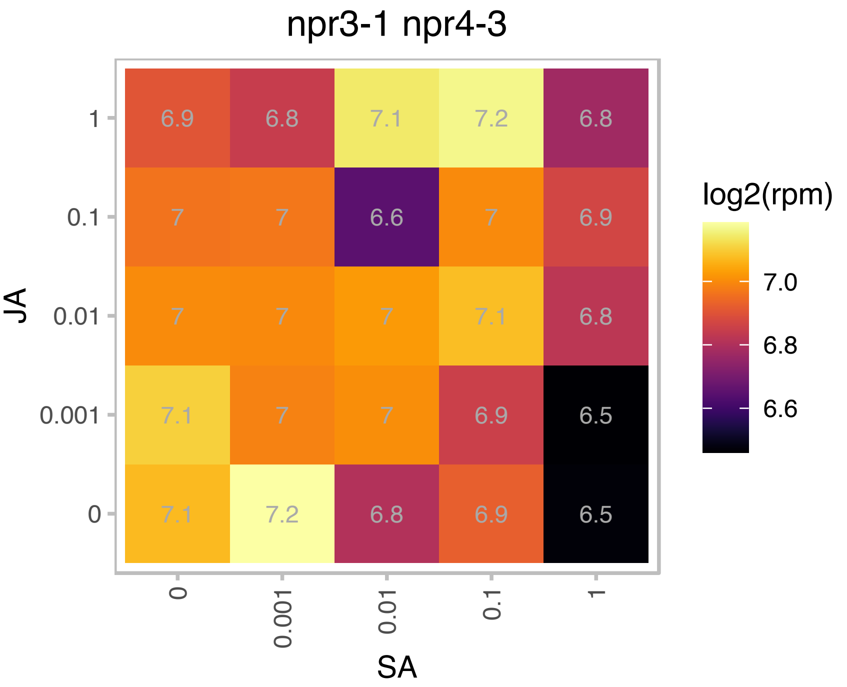

### AT3G13570.1 : At-SCL30A, SCL30A

SC35-like splicing factor 30A
encodes an SC35-like splicing factor of 30 kD that is localized to the nuclear specks. Barta et al (2010) have proposed a nomenclature for Serine/Arginine-Rich Protein Splicing Factors (SR proteins): Plant Cell. 2010, 22:2926.

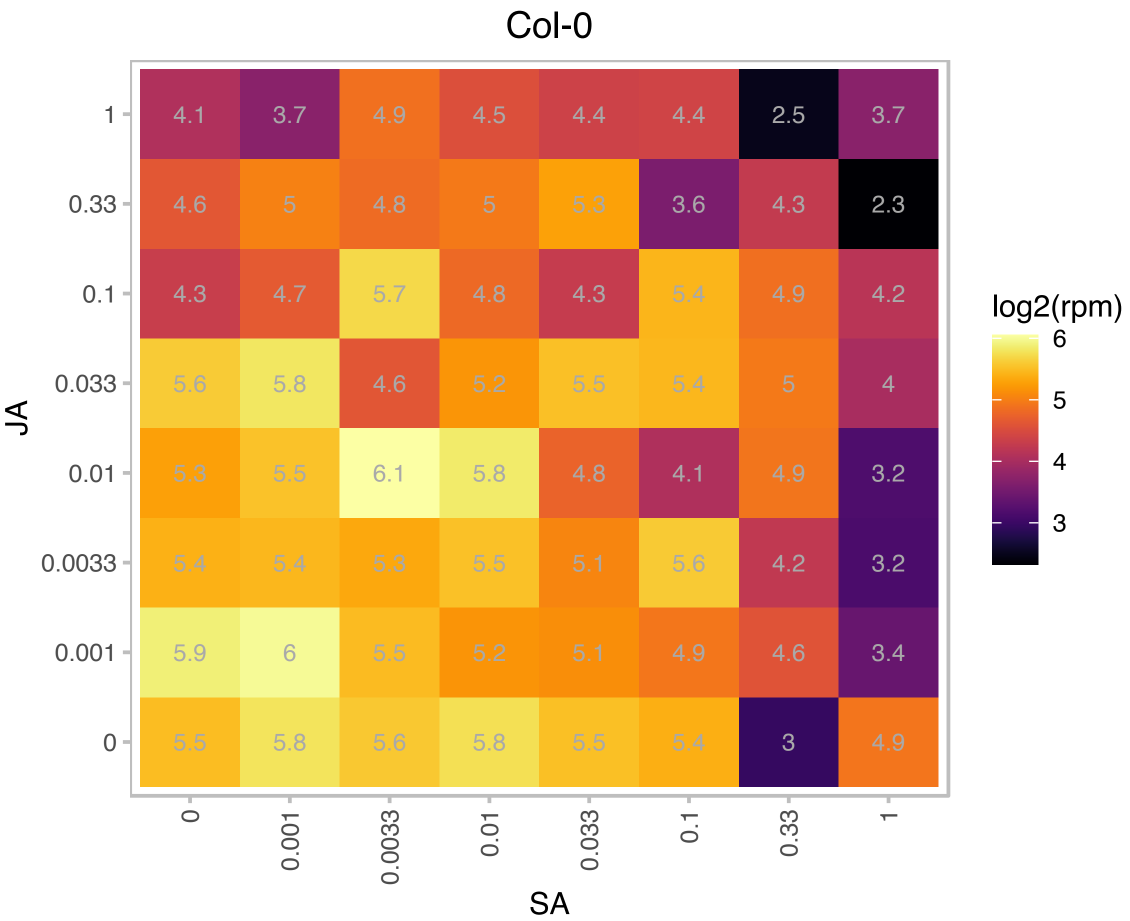


| | |
|---|---|
| Gene | AT3G13570 |
| is.expressed_genes | TRUE |
| is.responsive_genes | TRUE |
| is.apcluster | F-07 |

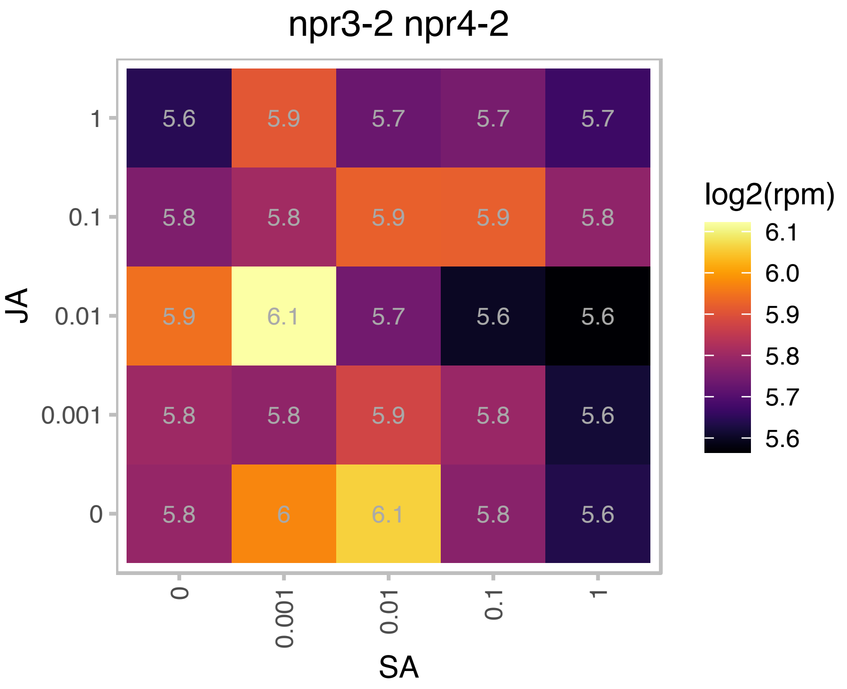


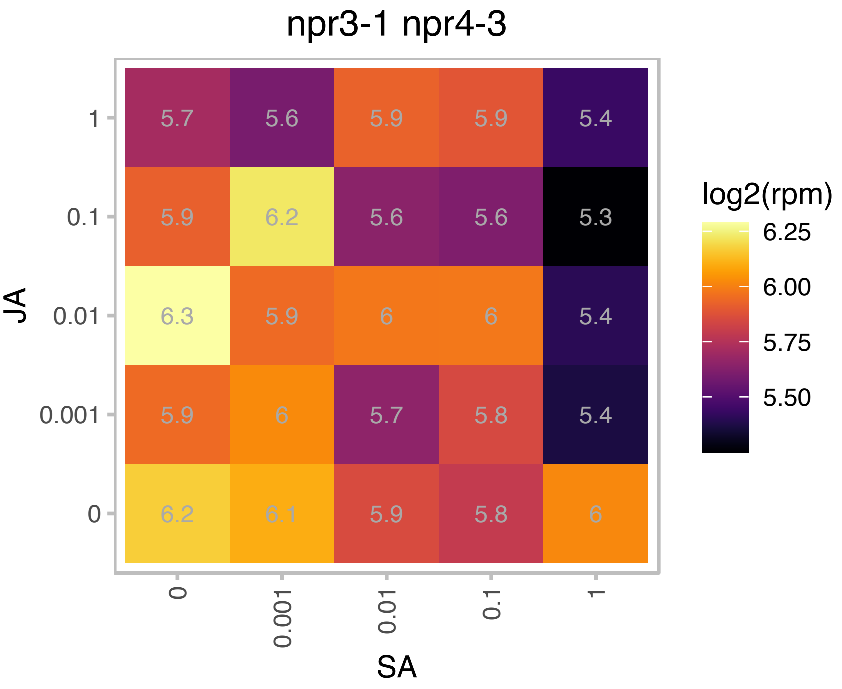

# AT3G15357.1 : NA

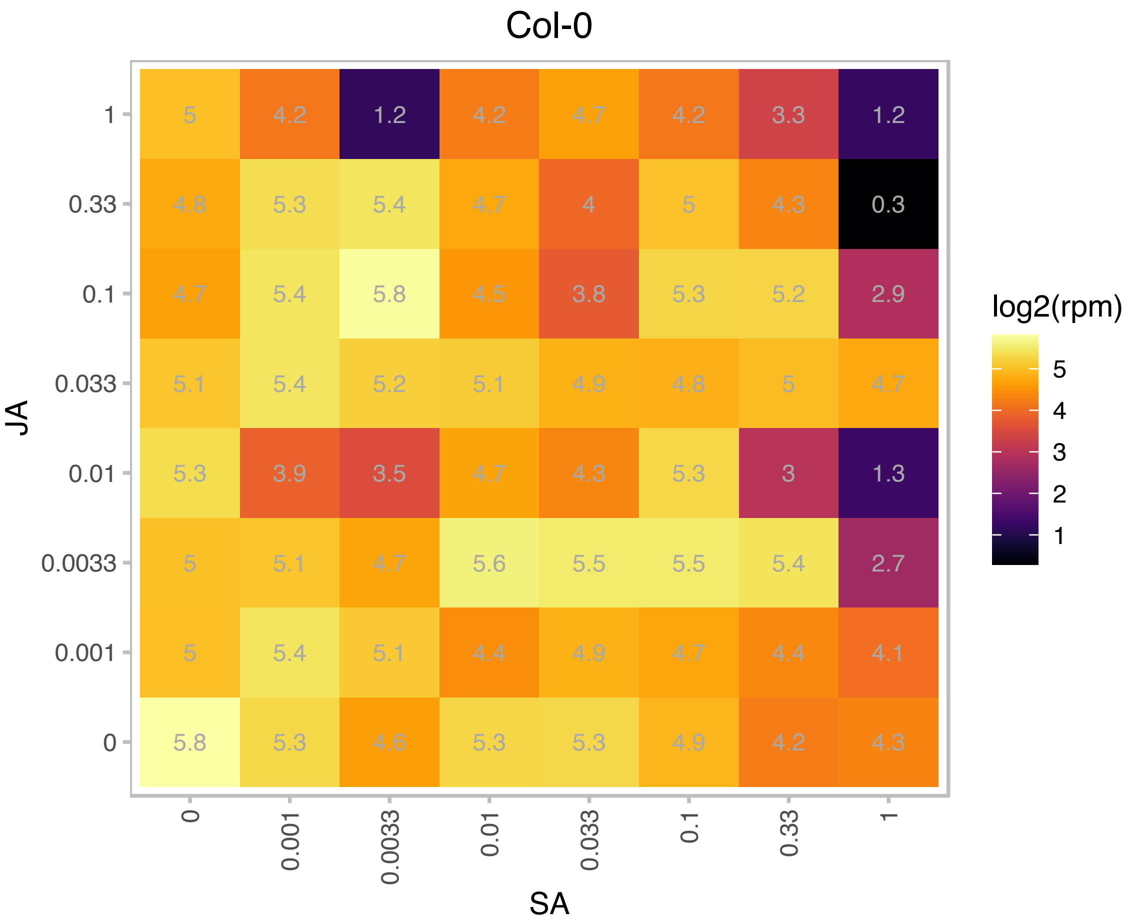


| | |
|---|---|
| Gene | AT3G15357 |
| is.expressed_genes | TRUE |
| is.responsive_genes | TRUE |
| is.apcluster | F-07 |

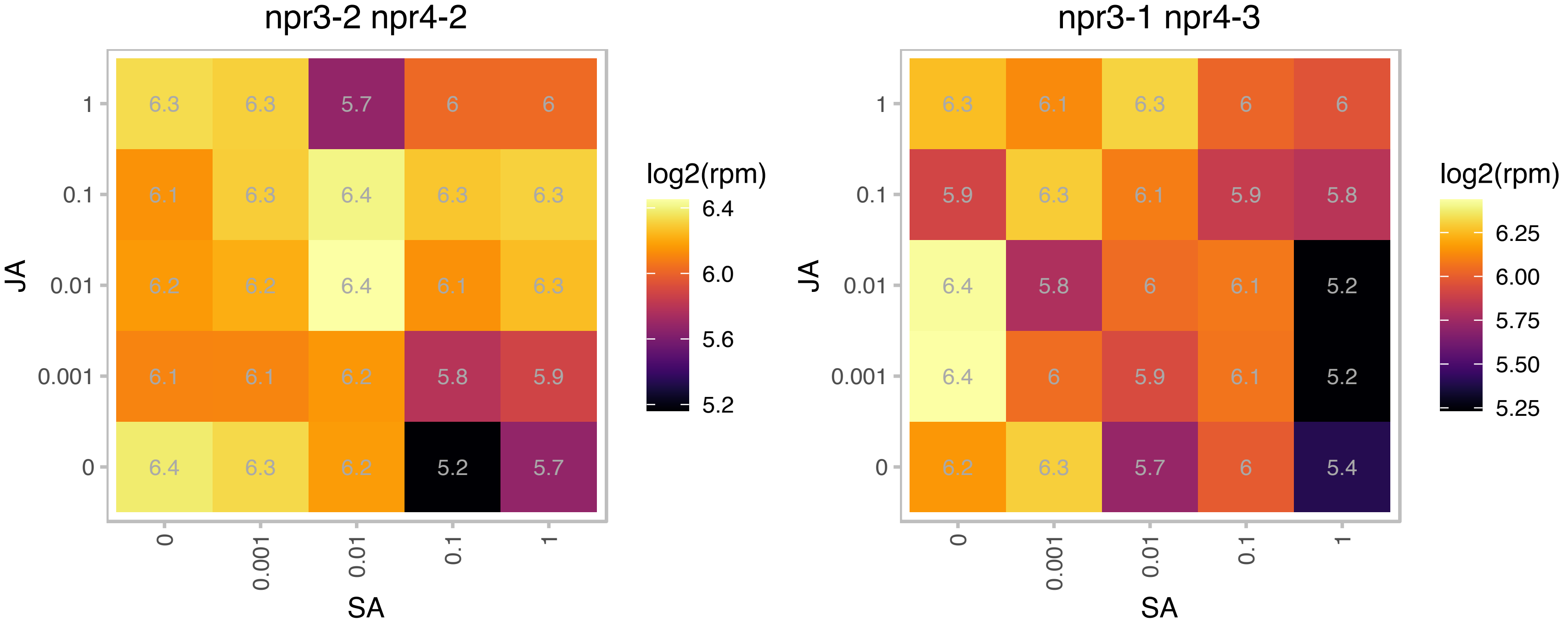

# AT3G15680.1 : NA

Ran BP2/NZF zinc finger-like superfamily protein

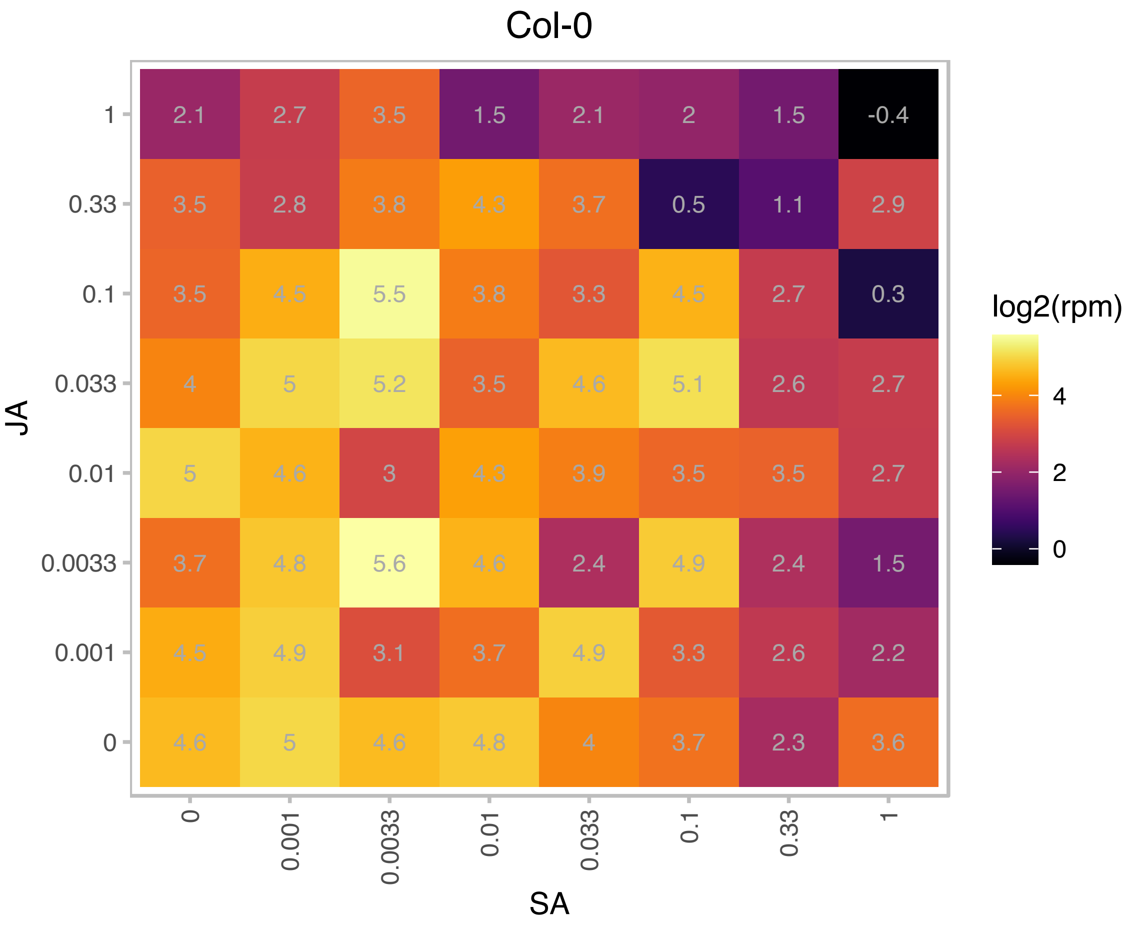


| | |
|---|---|
| Gene | AT3G15680 |
| is.expressed_genes | TRUE |
| is.responsive_genes | TRUE |
| is.apcluster | F-07 |

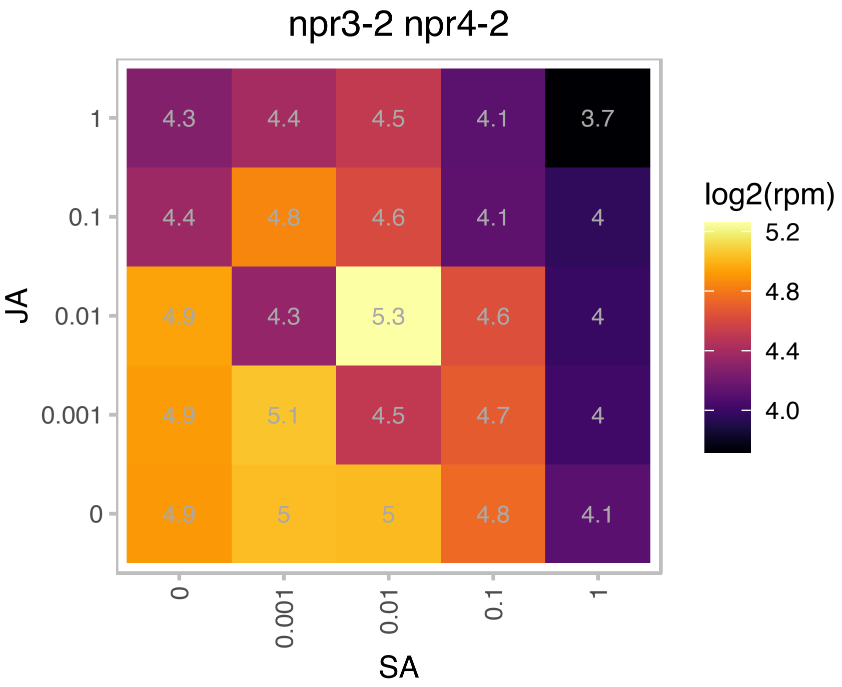


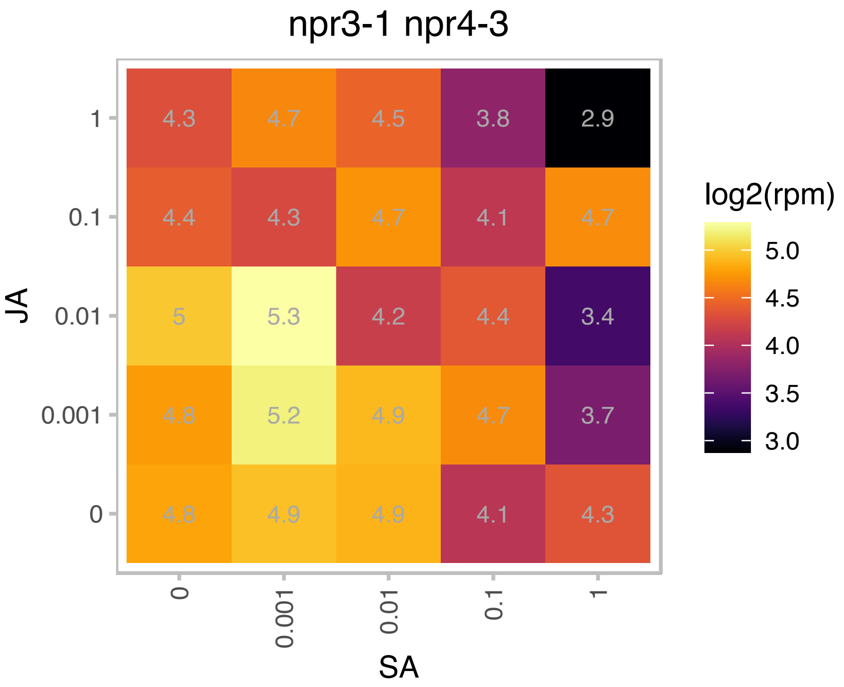

# AT3G16080.1 : NA

Zinc-binding ribosomal protein family protein

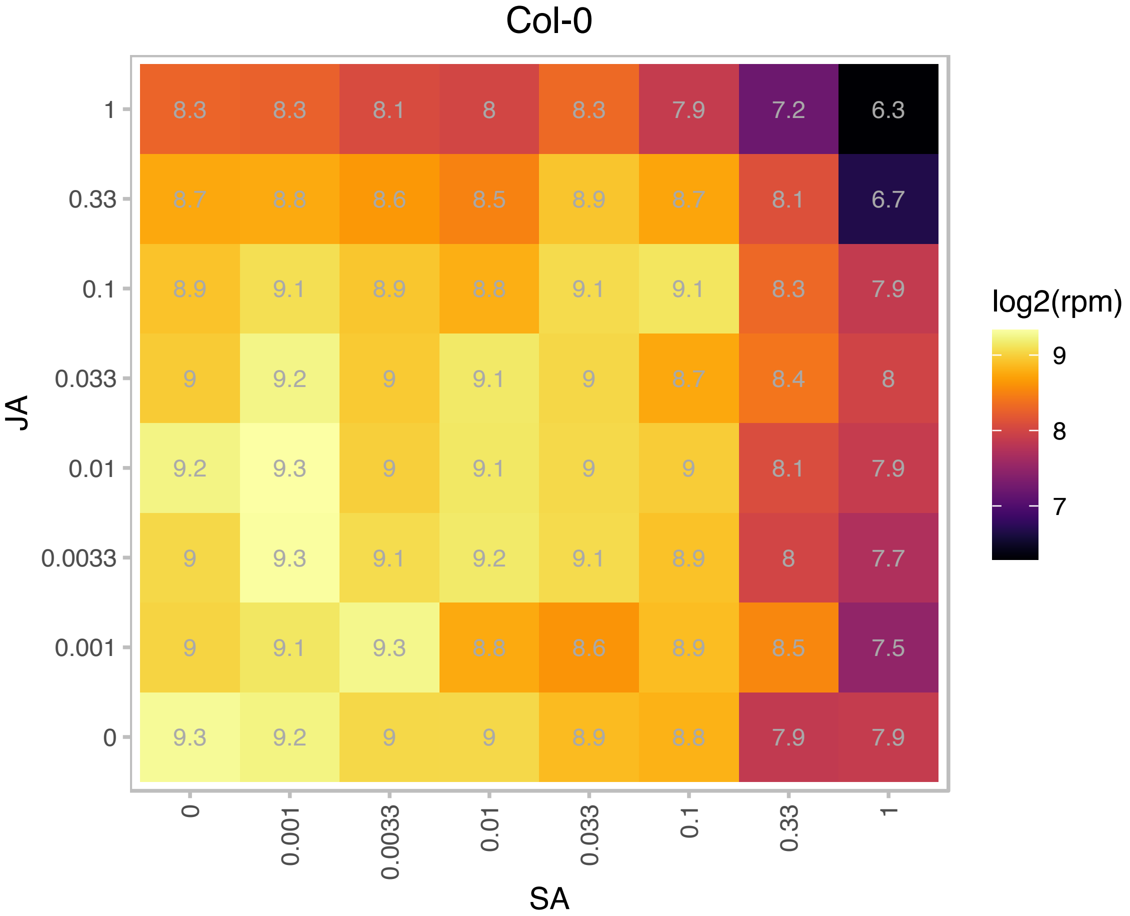


| | |
|---|---|
| Gene | AT3G16080 |
| is.expressed_genes | TRUE |
| is.responsive_genes | TRUE |
| is.apcluster | F-07 |

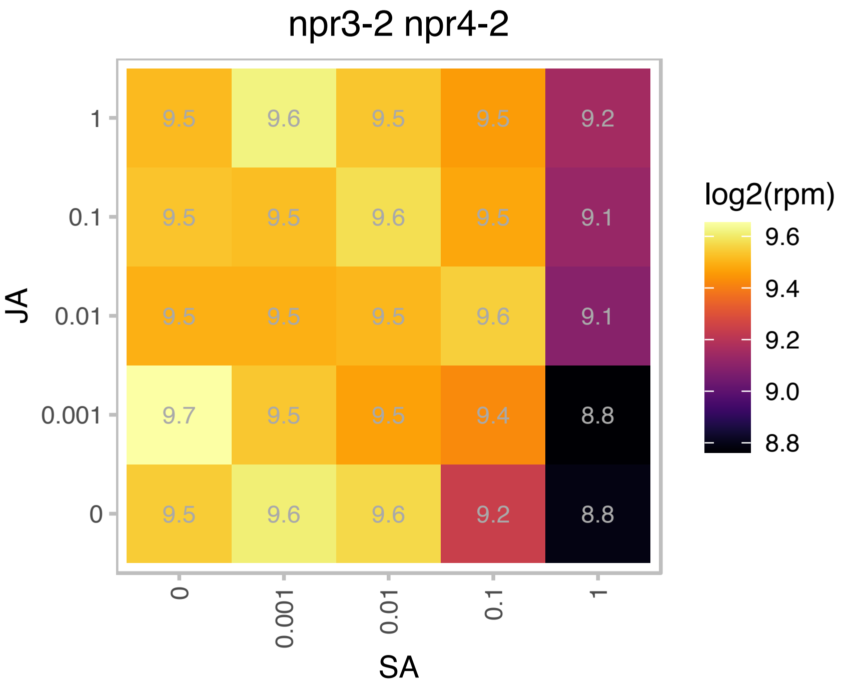


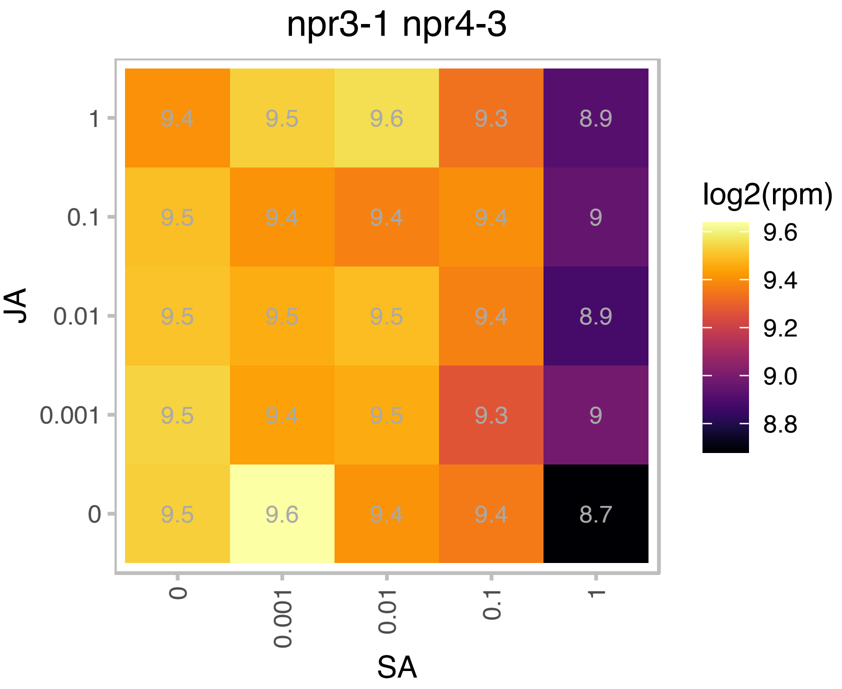

# AT3G17160.1 : NA

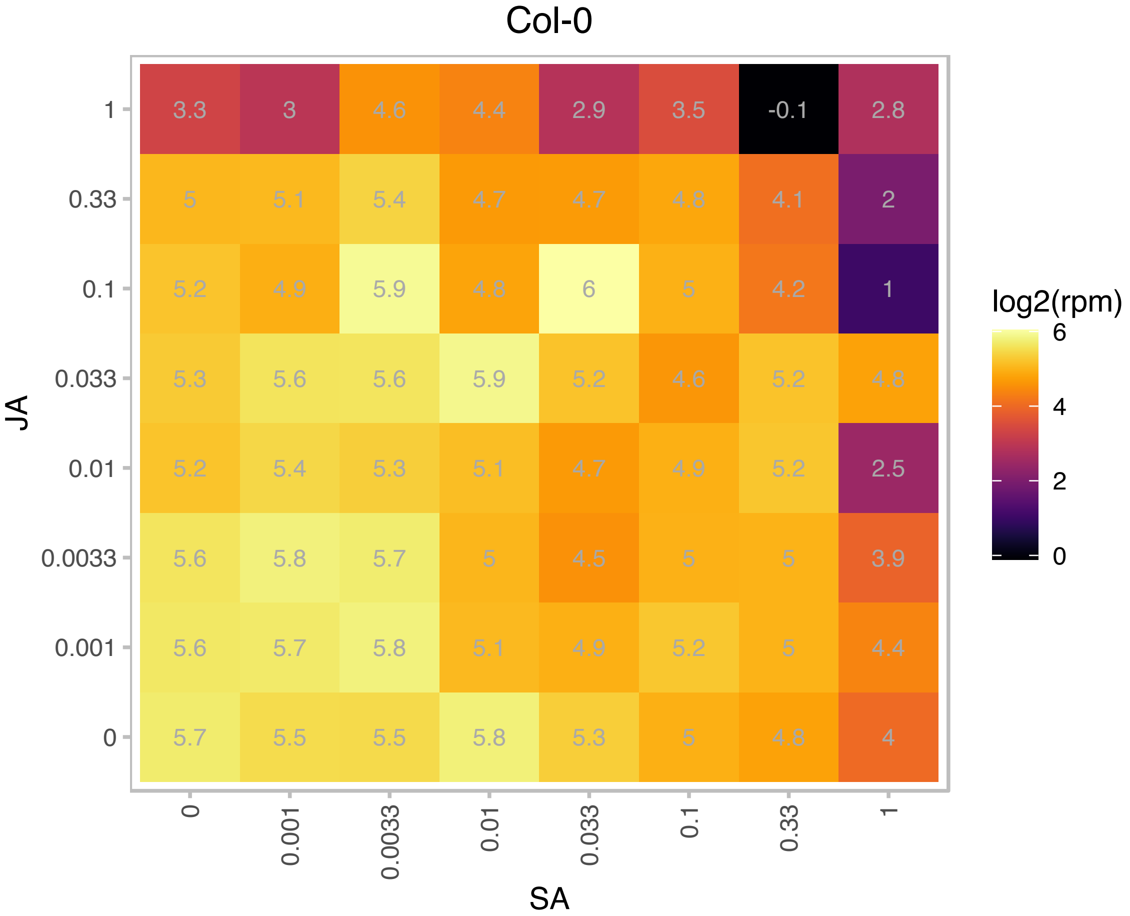


| | |
|---|---|
| Gene | AT3G17160 |
| is.expressed_genes | TRUE |
| is.responsive_genes | TRUE |
| is.apcluster | F-07 |

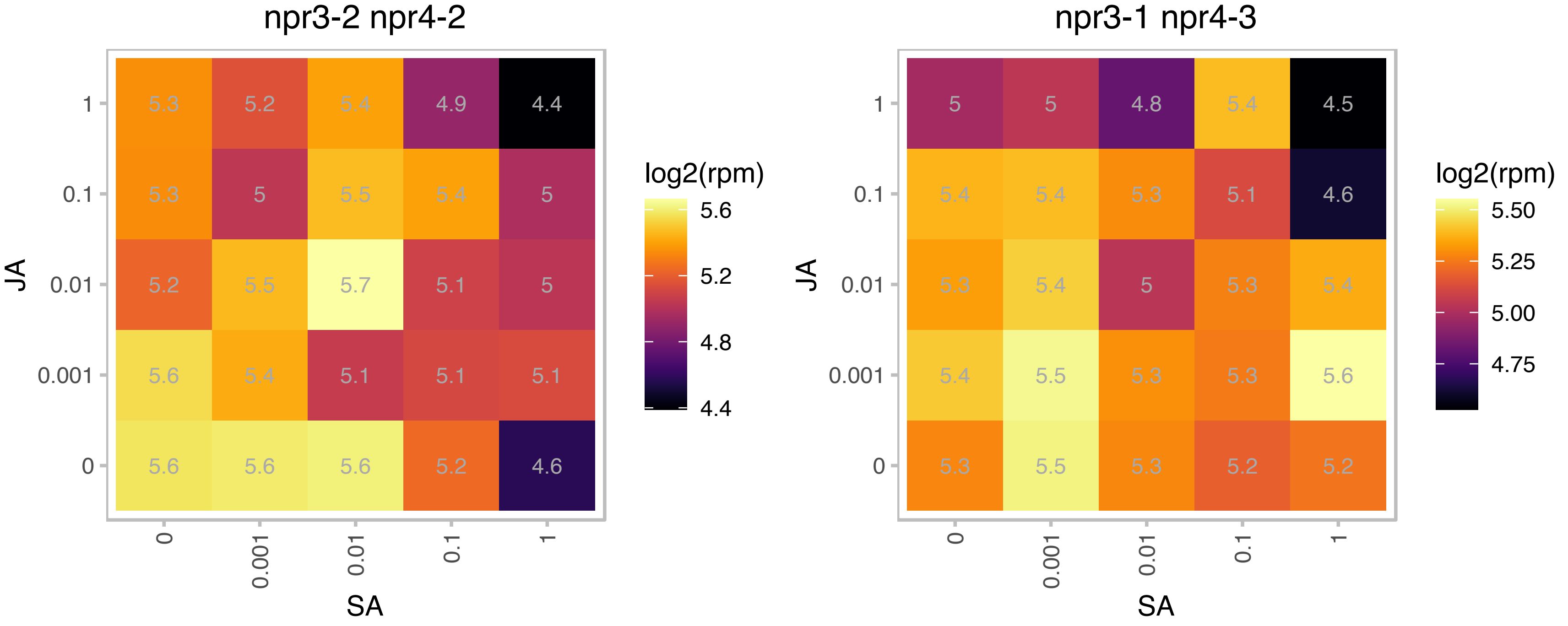

### AT3G18130.1 : RACK1C, RACK1C_AT

receptor for activated C kinase 1C
Encodes a protein with similarity to mammalian RACKs. RACKs function to shuttle activated protein kinase C to different subcellular sites and may also function as a scaffold through physical interactions with other proteins. RACK1C has no phenotype on its own and probably acts redundantly with RACK1A and RACK1B.

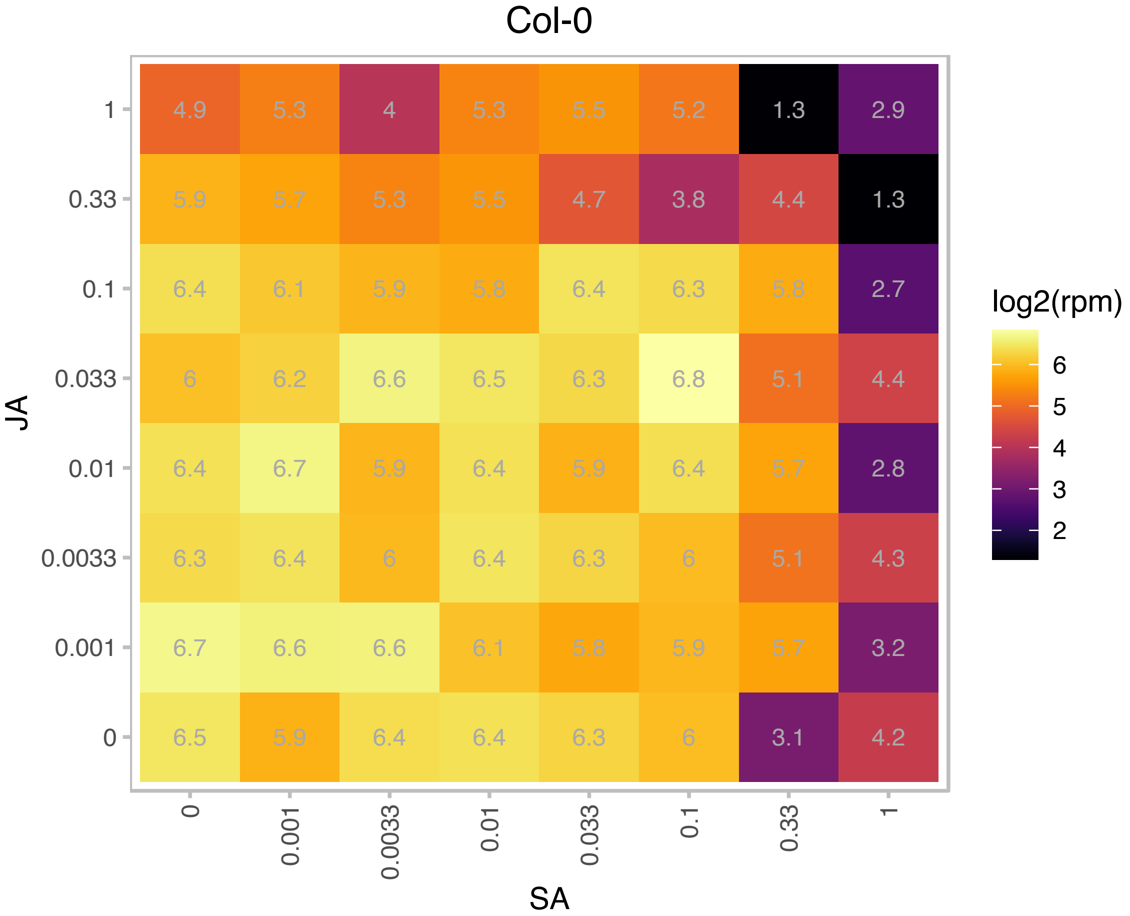


| | |
|---|---|
| Gene | AT3G18130 |
| is.expressed_genes | TRUE |
| is.responsive_genes | TRUE |
| is.apcluster | F-07 |

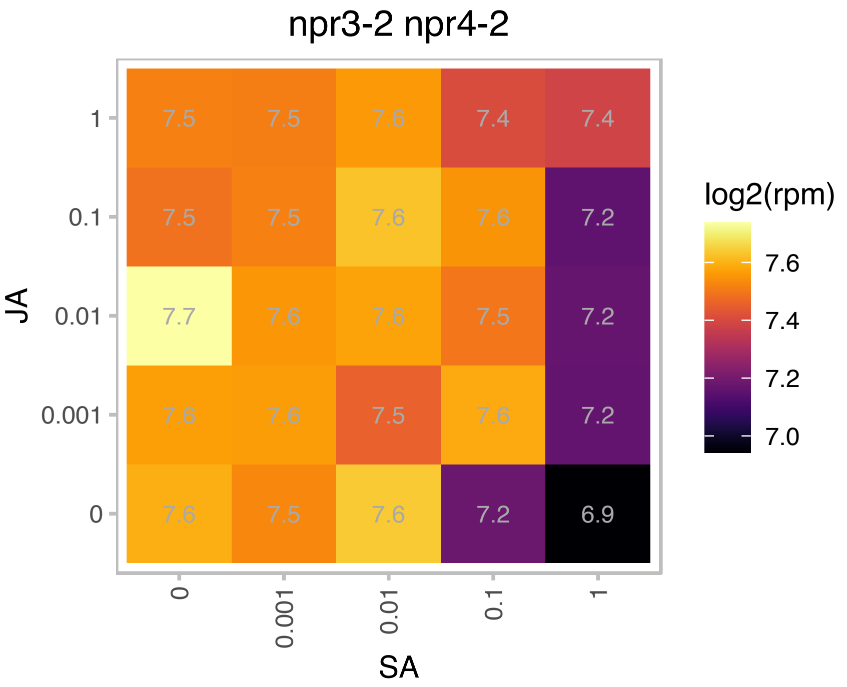


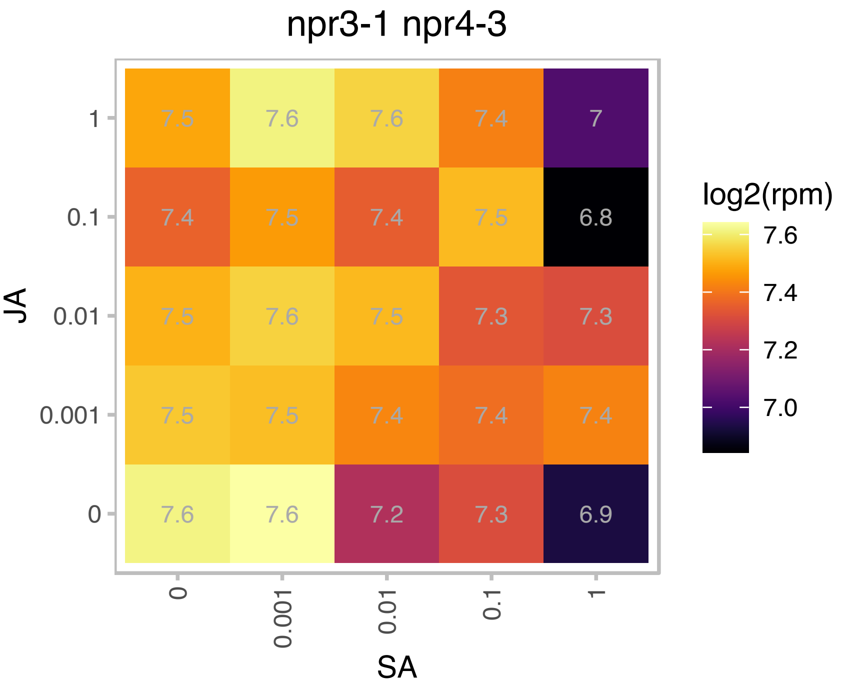

# AT3G18600.1 : NA

P-loop containing nucleoside triphosphate hydrolases superfamily protein

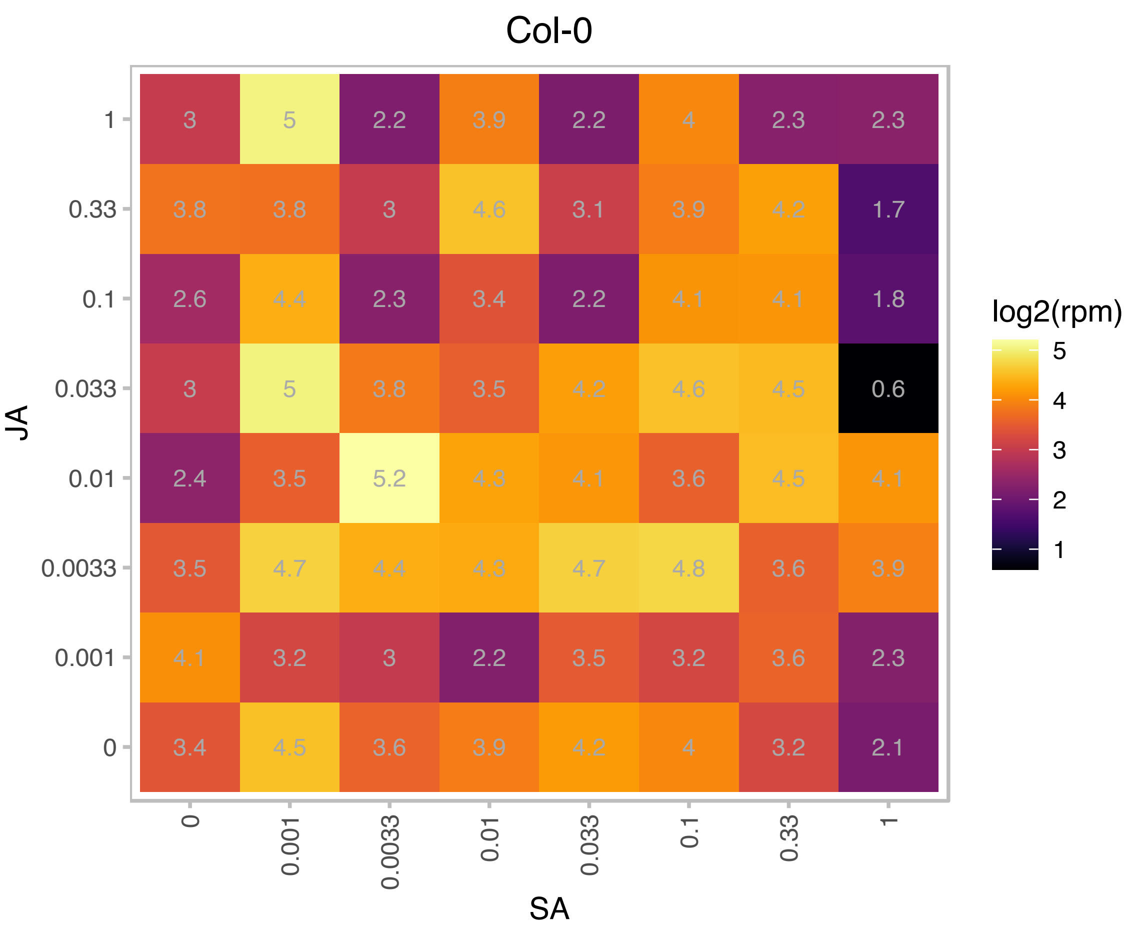


| | |
|---|---|
| Gene | AT3G18600 |
| is.expressed_genes | TRUE |
| is.responsive_genes | TRUE |
| is.apcluster | F-07 |

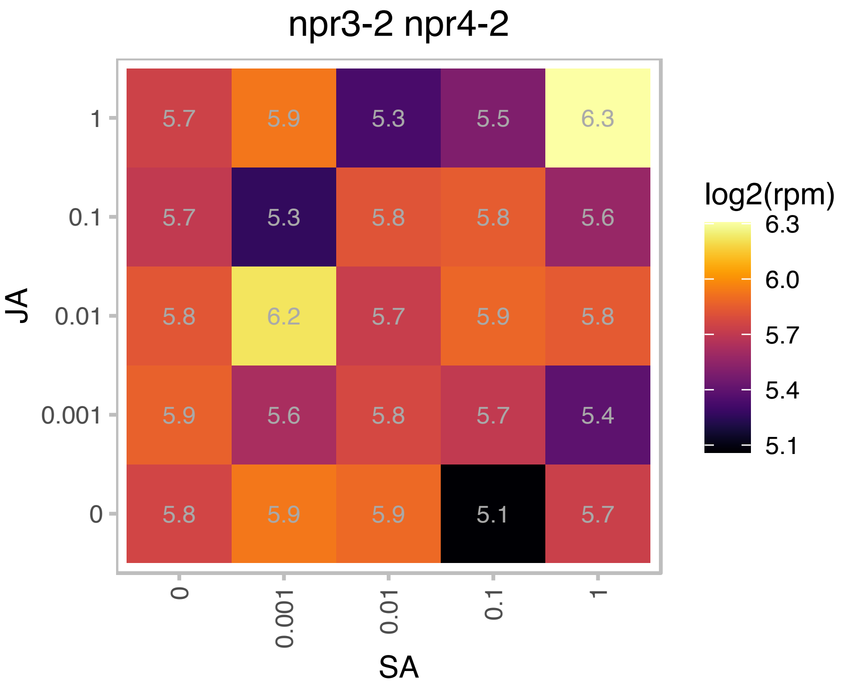


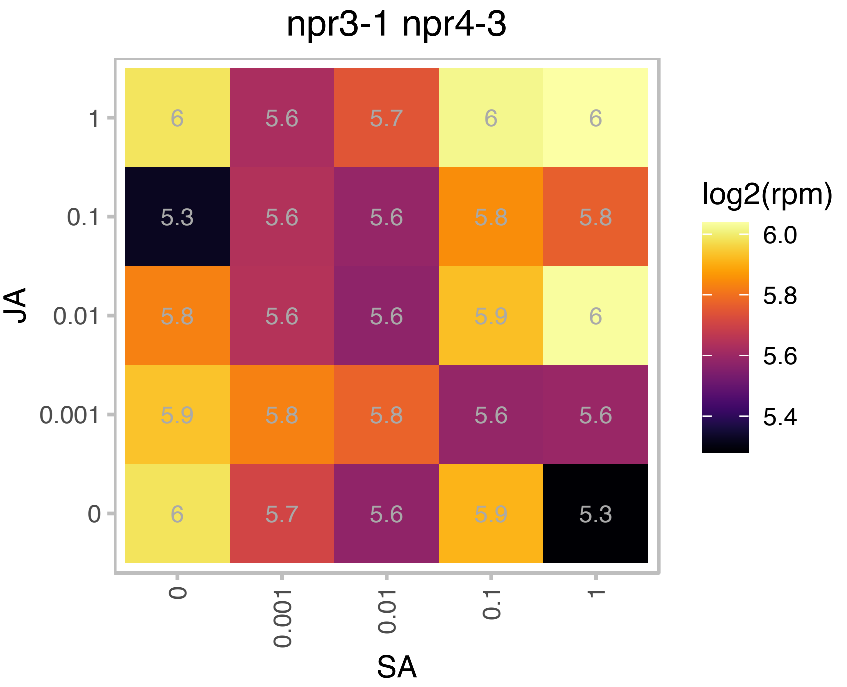

### AT3G18740.1 : RLK902

Ribosomal protein L7Ae/L30e/S12e/Gadd45 family protein

The gene encodes a putative leucine-rich repeat (LRR)Receptor-like kinase (RLK).

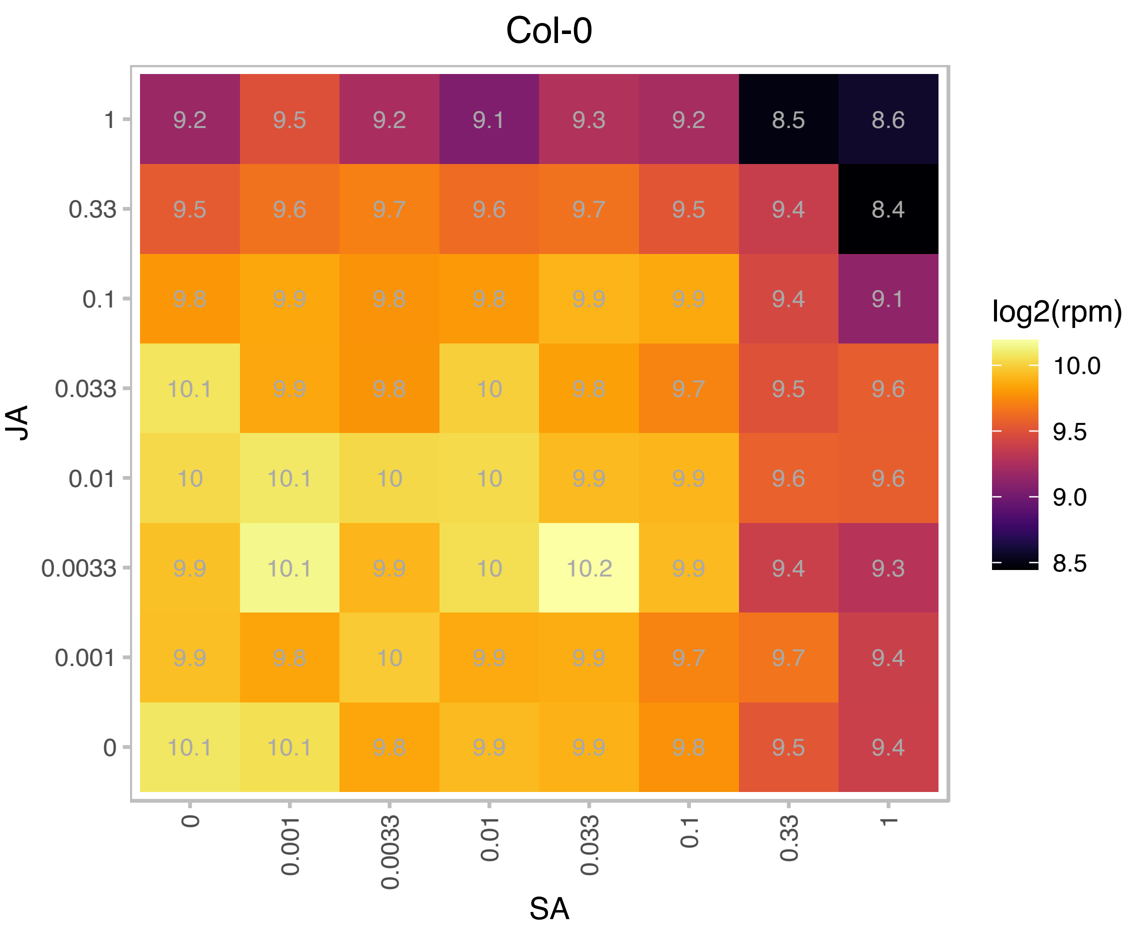


| | |
|---|---|
| Gene | AT3G18740 |
| is.expressed_genes | TRUE |
| is.responsive_genes | TRUE |
| is.apcluster | F-07 |

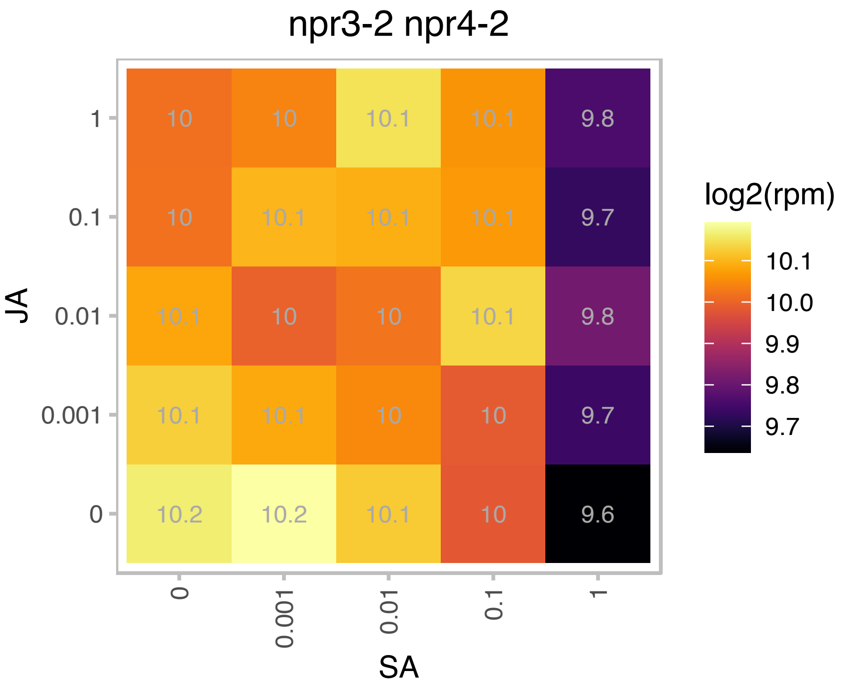


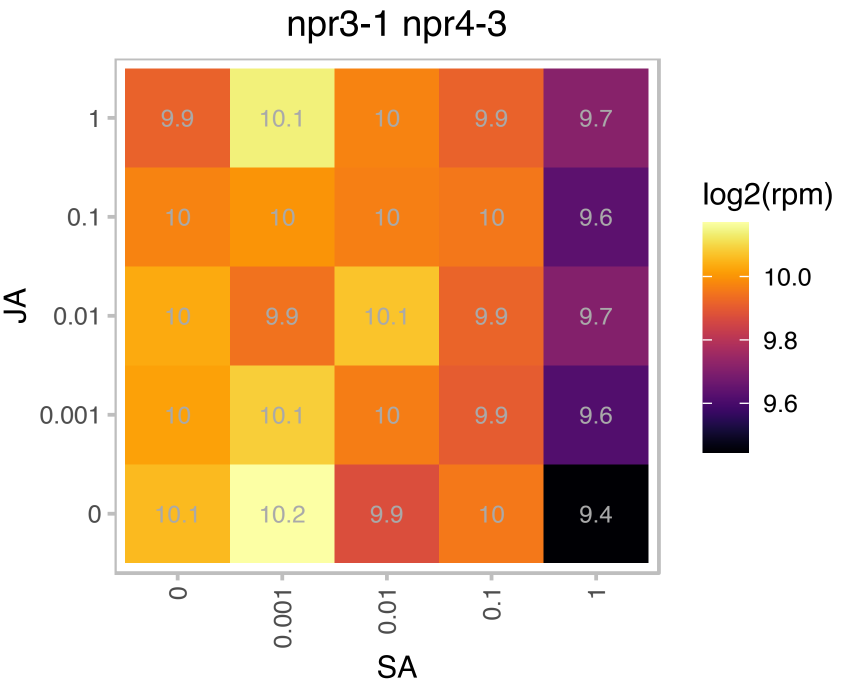

### AT3G21200.1 : GBP, GluTRBP, PGR7

proton gradient regulation 7

Encodes a soluble glutamyl-tRNA reductase (GluTR) binding protein that forms a ternary complex with FLU and GluTR.

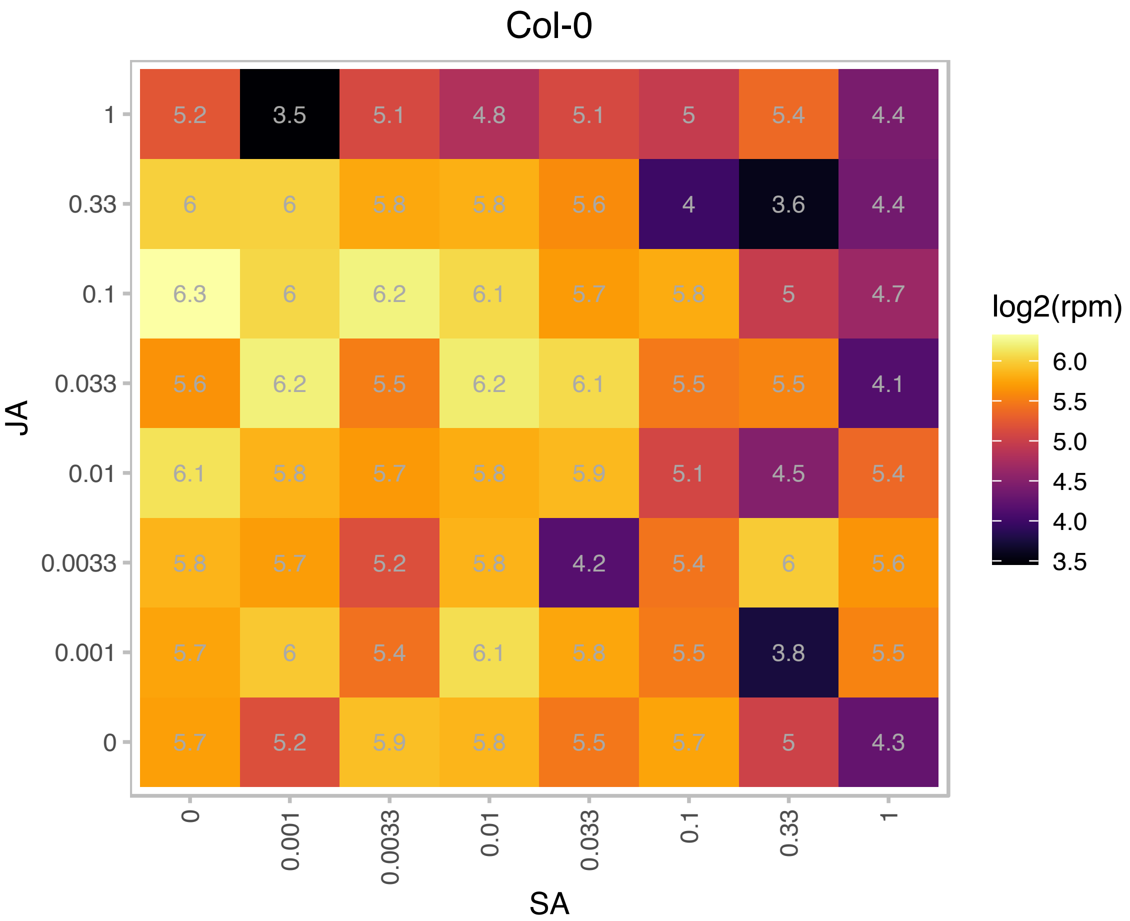


| | |
|---|---|
| Gene | AT3G21200 |
| is.expressed_genes | TRUE |
| is.responsive_genes | TRUE |
| is.apcluster | F-07 |

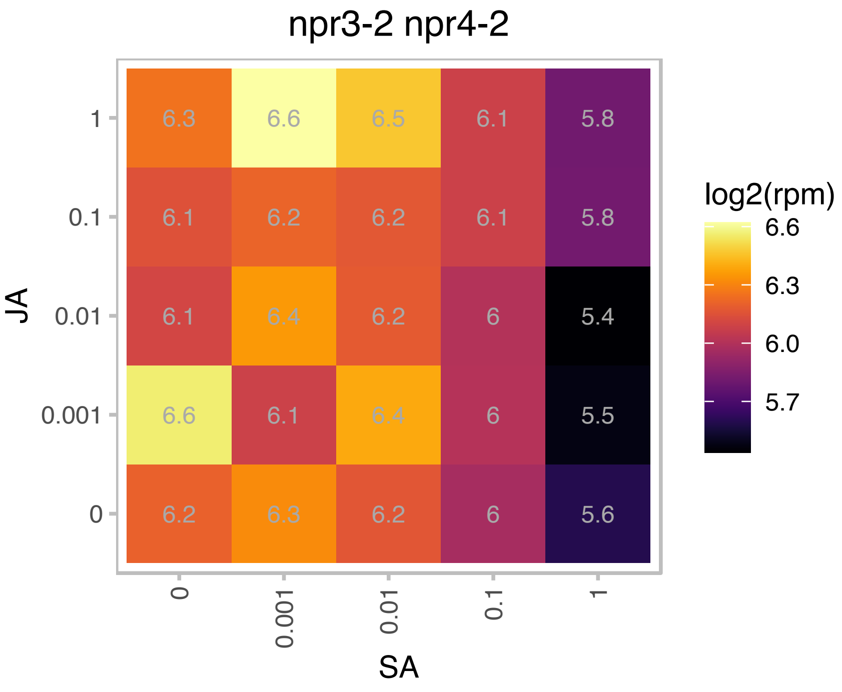


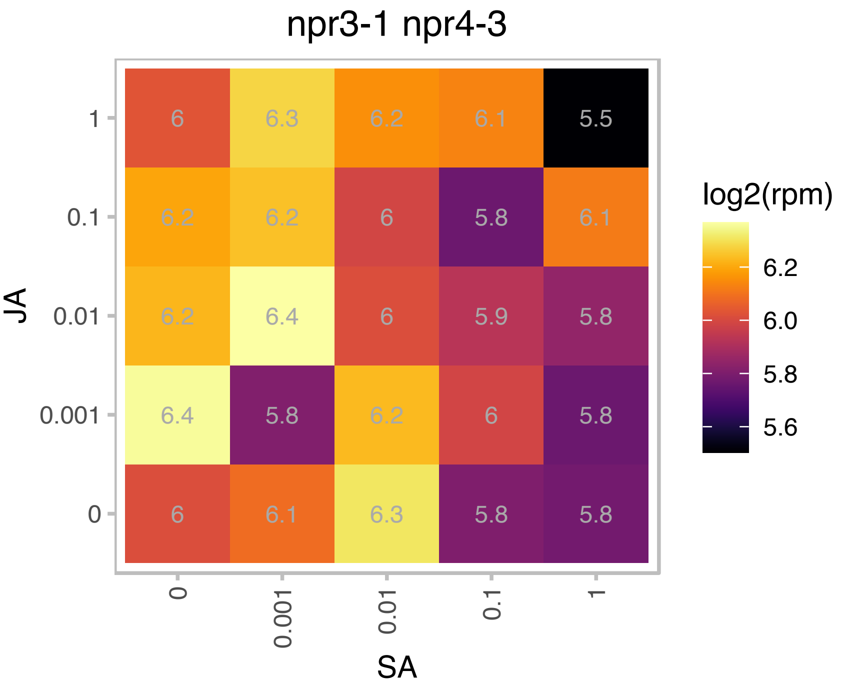

# AT3G21465.1 : NA

| | |
|---|---|
| Gene | AT3G21465 |
| is.expressed_genes | TRUE |
| is.responsive_genes | TRUE |
| is.apcluster | F-07 |

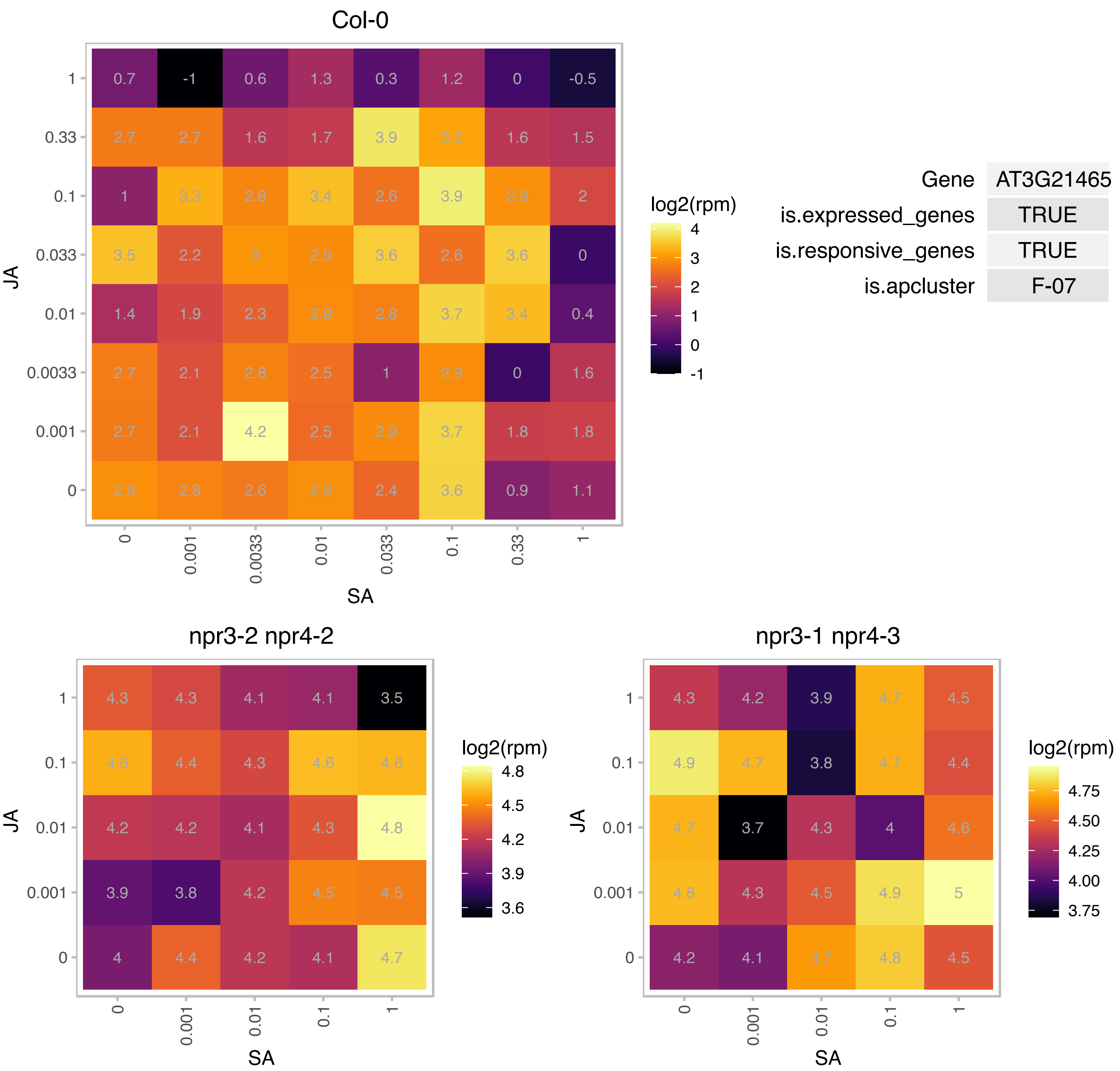

### AT3G22660.1 : EBP2

rRNA processing protein-related

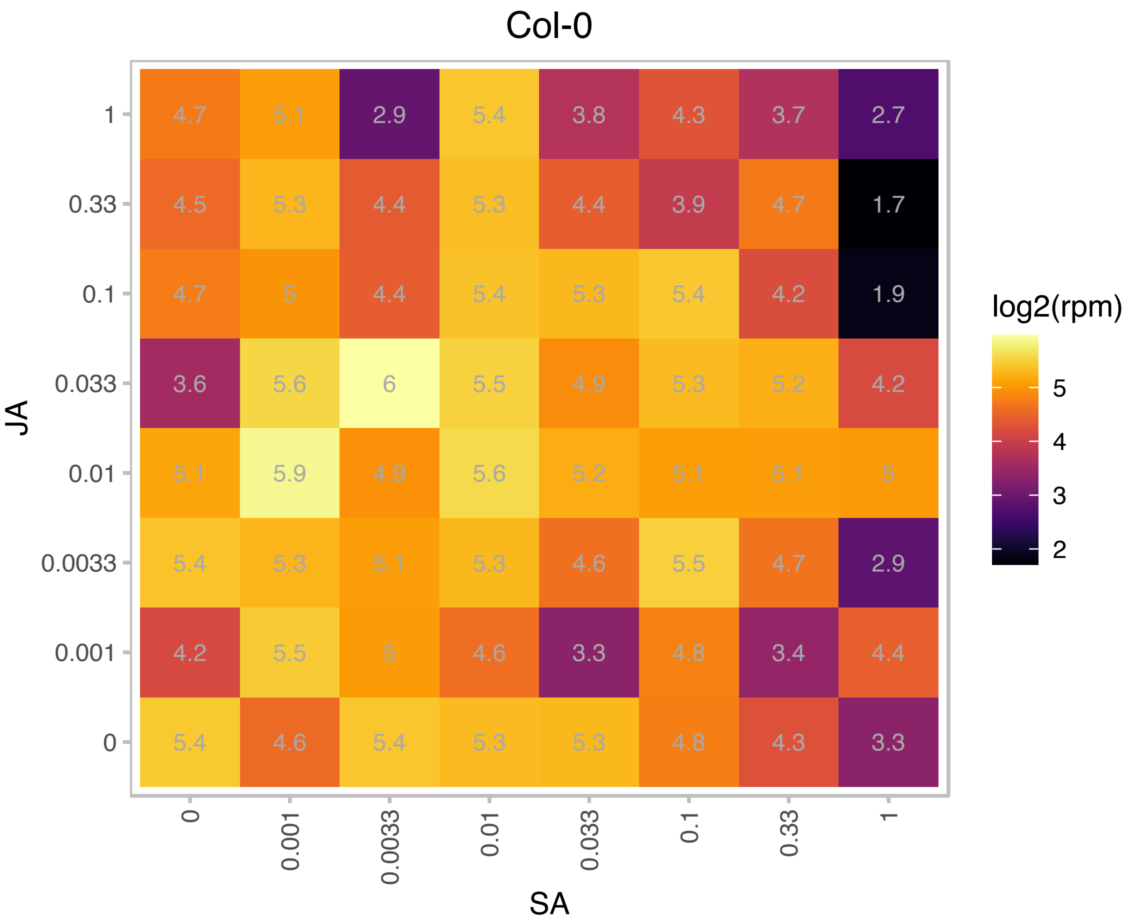


| | |
|---|---|
| Gene | AT3G22660 |
| is.expressed_genes | TRUE |
| is.responsive_genes | TRUE |
| is.apcluster | F-07 |

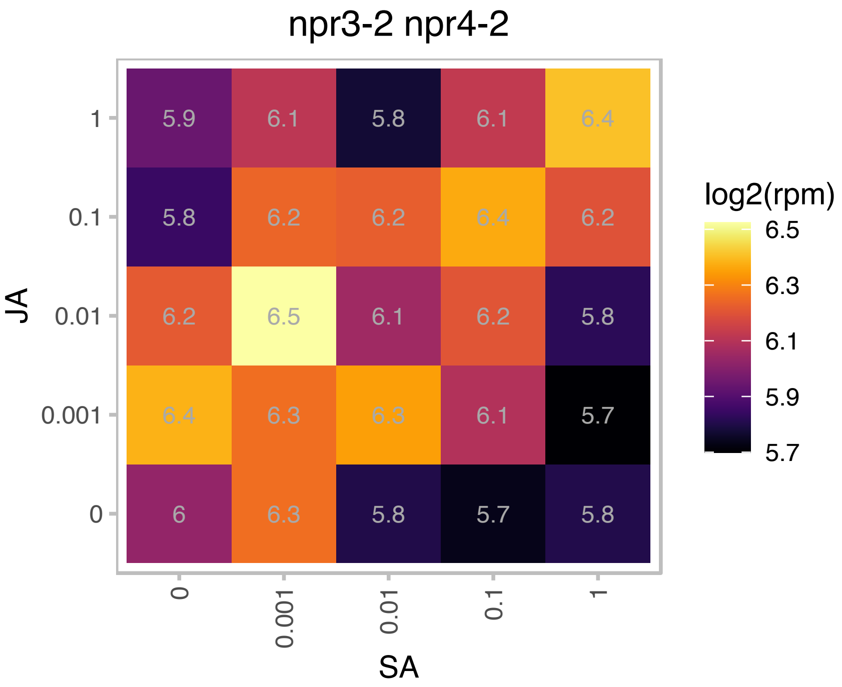


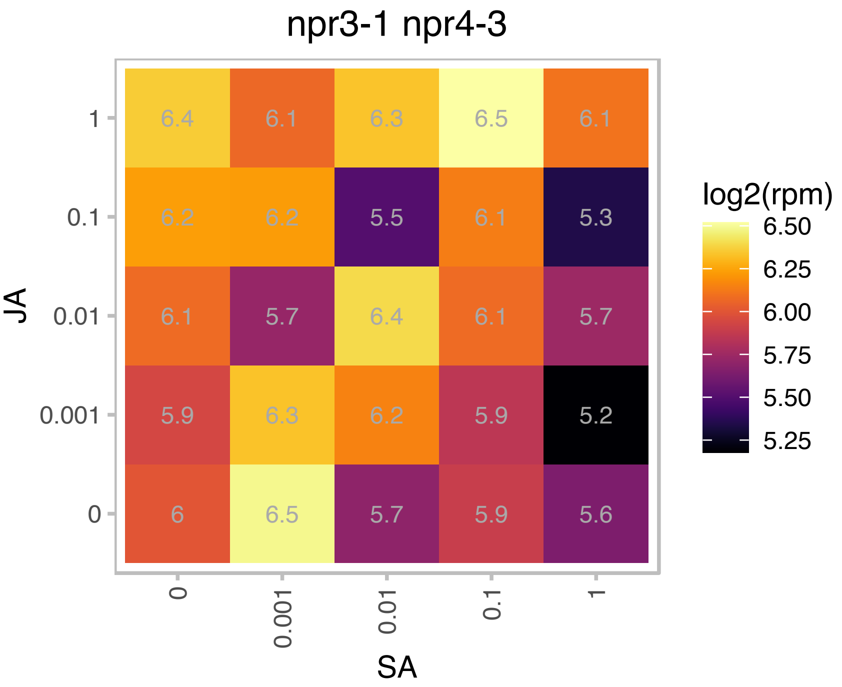

### AT3G23390.1 : RPL36aA

Zinc-binding ribosomal protein family protein

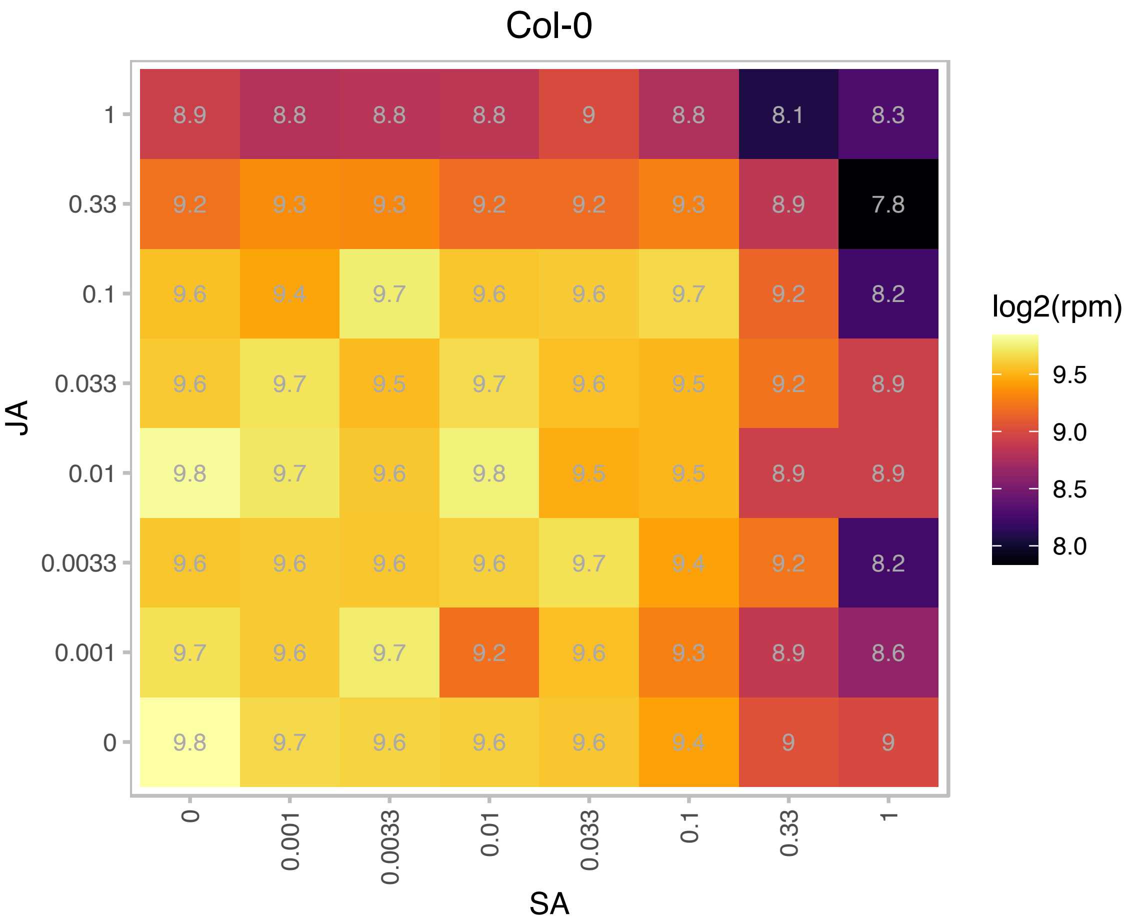


| | |
|---|---|
| Gene | AT3G23390 |
| is.expressed_genes | TRUE |
| is.responsive_genes | TRUE |
| is.apcluster | F-07 |

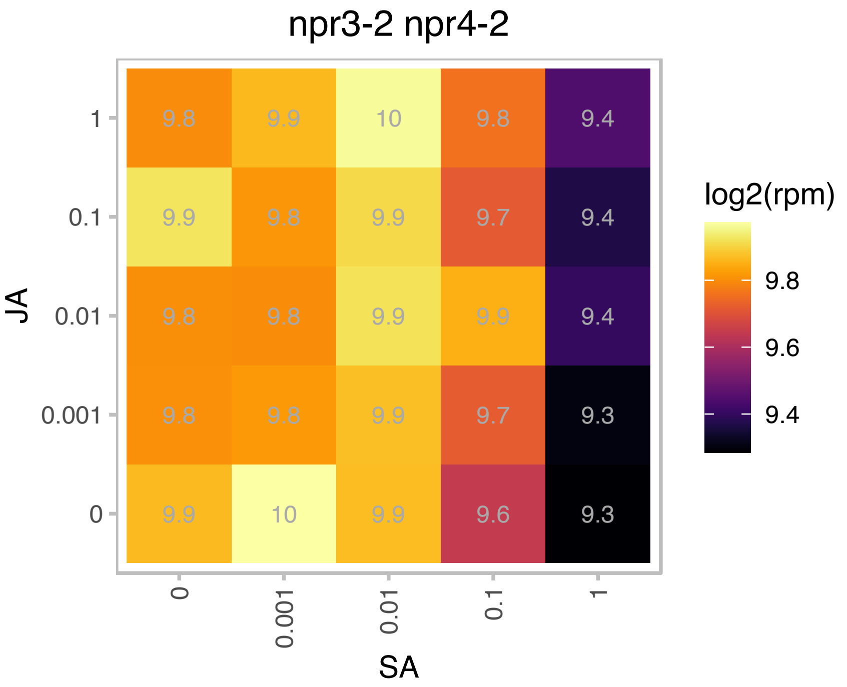


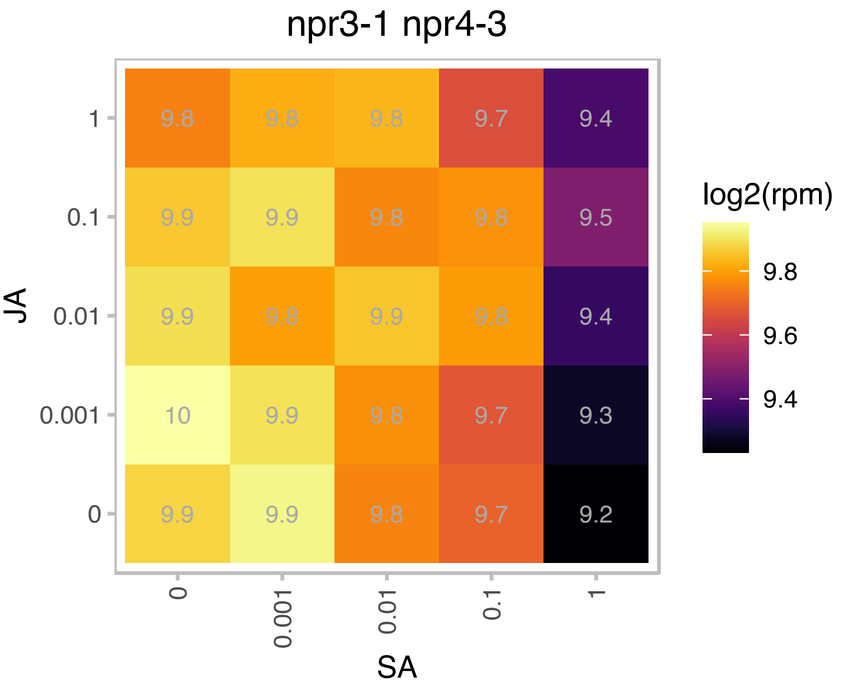

### AT3G23830.1 : AtGRP4, GR-RBP4, GRP4, RBGA4

glycine-rich RNA-binding protein 4
encodes a glycine-rich RNA binding protein. Gene expression is induced by cold and reduced by ionic (salt) and non-ionic (mannitol) osmotic stress. Lines overexpressing the gene are slightly more tolerant to osmotic stress during germination.

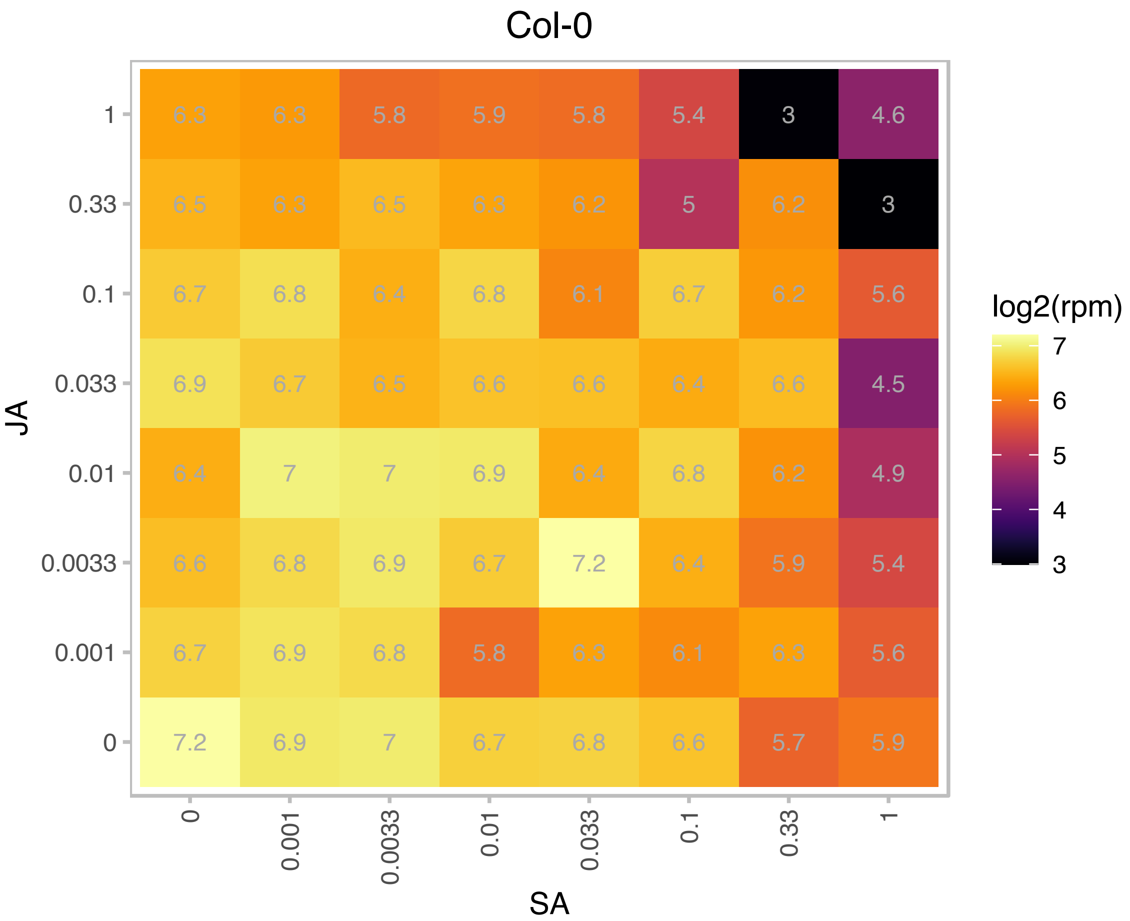


| | |
|---|---|
| Gene | AT3G23830 |
| is.expressed_genes | TRUE |
| is.responsive_genes | TRUE |
| is.apcluster | F-07 |

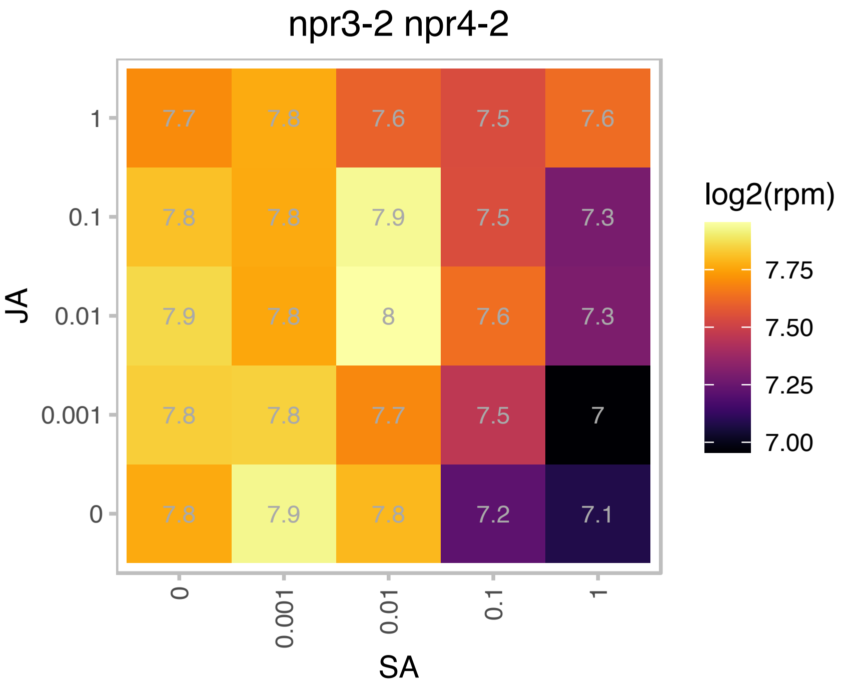


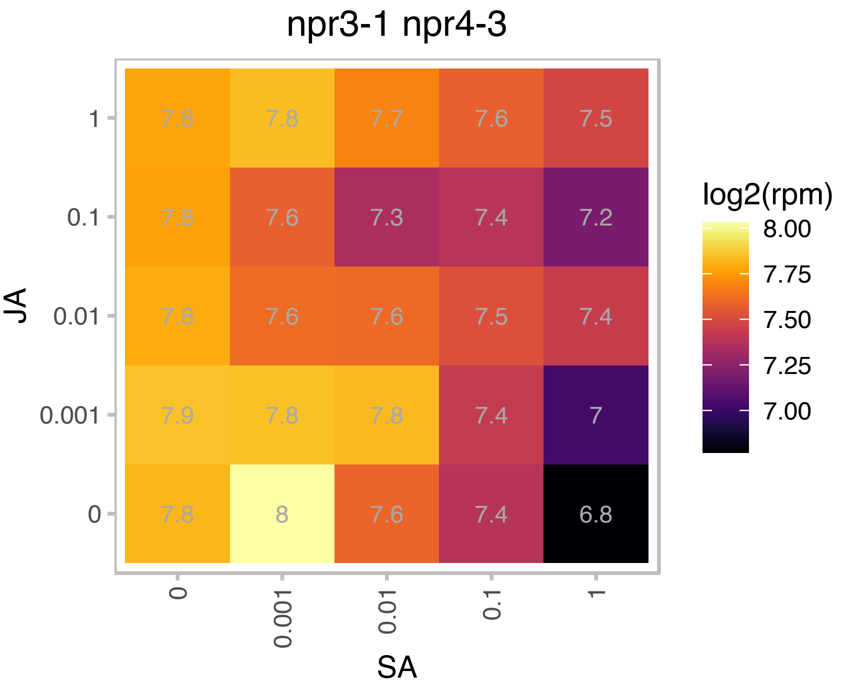

### AT3G25890.1 : CRF11

Integrase-type DNA-binding superfamily protein
encodes a member of the ERF (ethylene response factor) subfamily B-6 of ERF/AP2 transcription factor family. The protein contains one AP2 domain. There are 12 members in this subfamily including RAP2.11. The mRNA is cell-to-cell mobile.

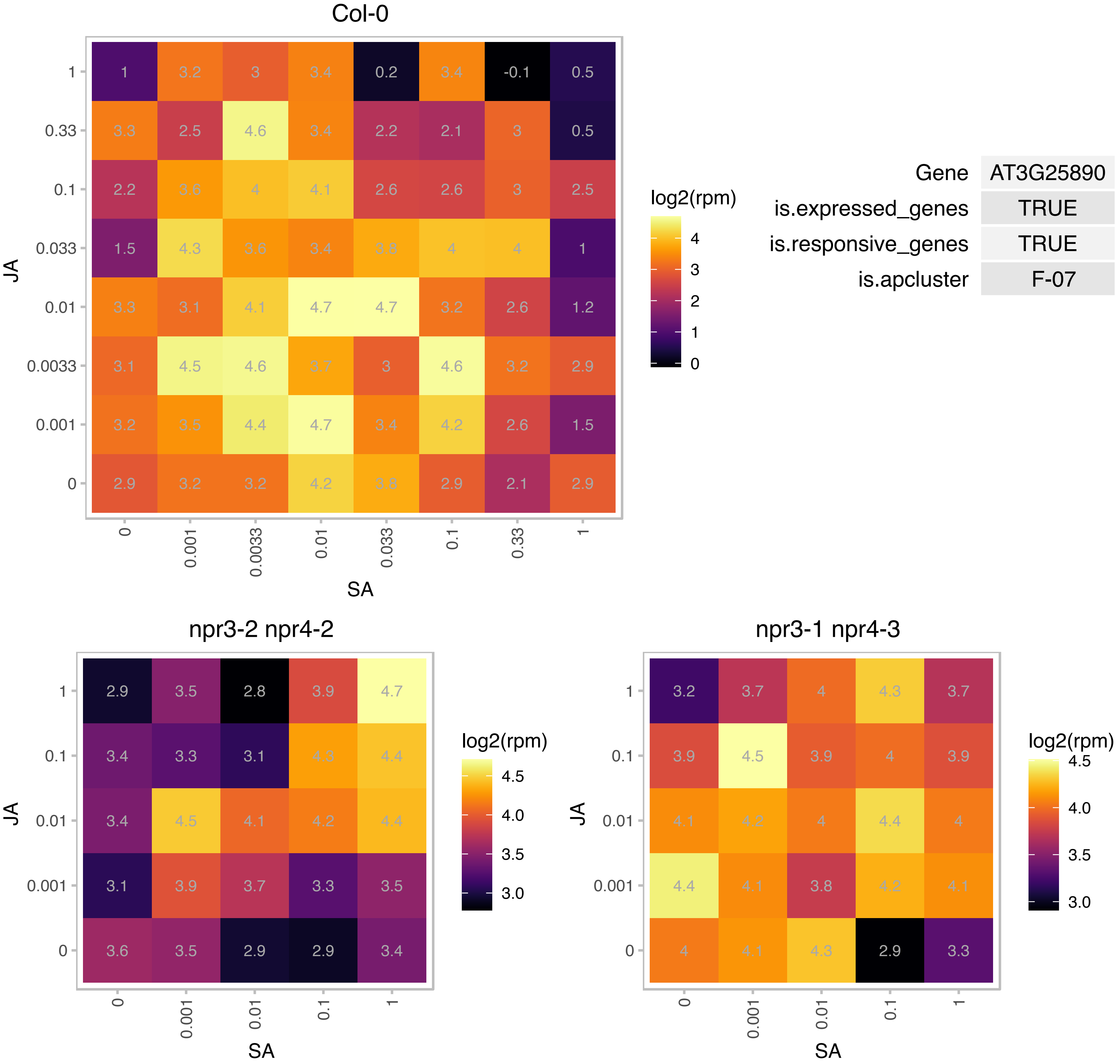


| | |
|---|---|
| Gene | AT3G25890 |
| is.expressed_genes | TRUE |
| is.responsive_genes | TRUE |
| is.apcluster | F-07 |

# AT3G27090.1 : NA

DCD (Development and Cell Death) domain protein

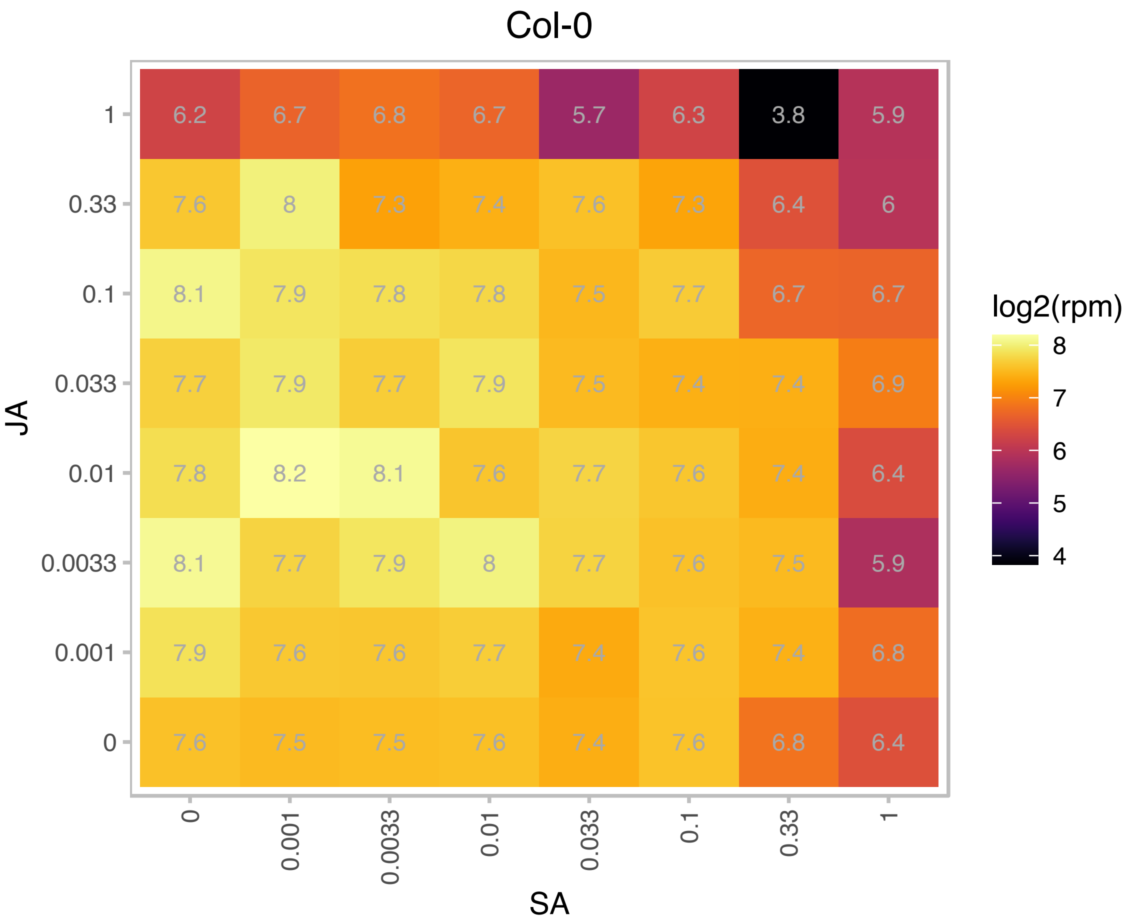


| | |
|---|---|
| Gene | AT3G27090 |
| is.expressed_genes | TRUE |
| is.responsive_genes | TRUE |
| is.apcluster | F-07 |

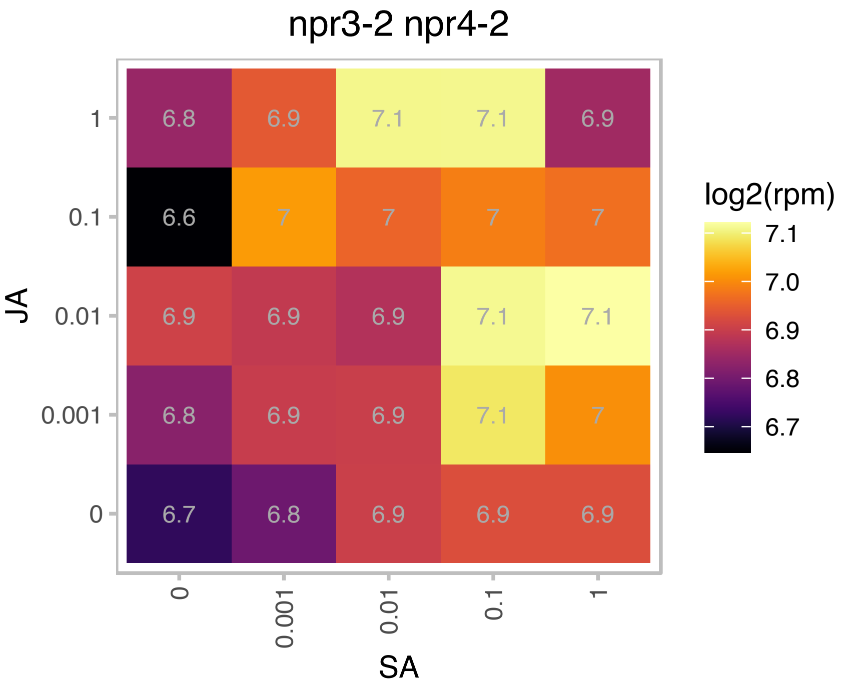


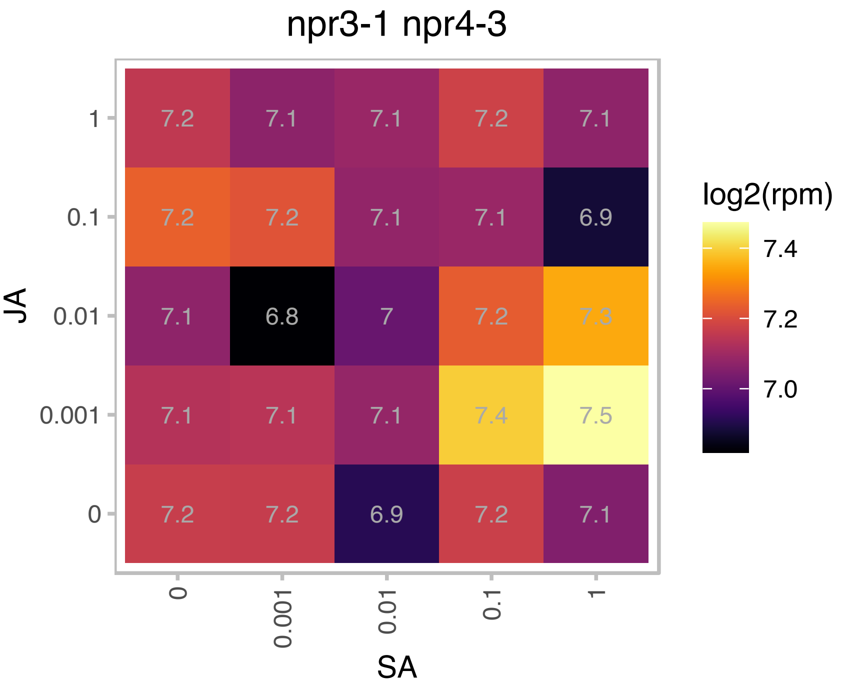

# AT3G28900.1 : NA

Ribosomal protein L34e superfamily protein

| | |
|---|---|
| Gene | AT3G28900 |
| is.expressed_genes | TRUE |
| is.responsive_genes | TRUE |
| is.apcluster | F-07 |

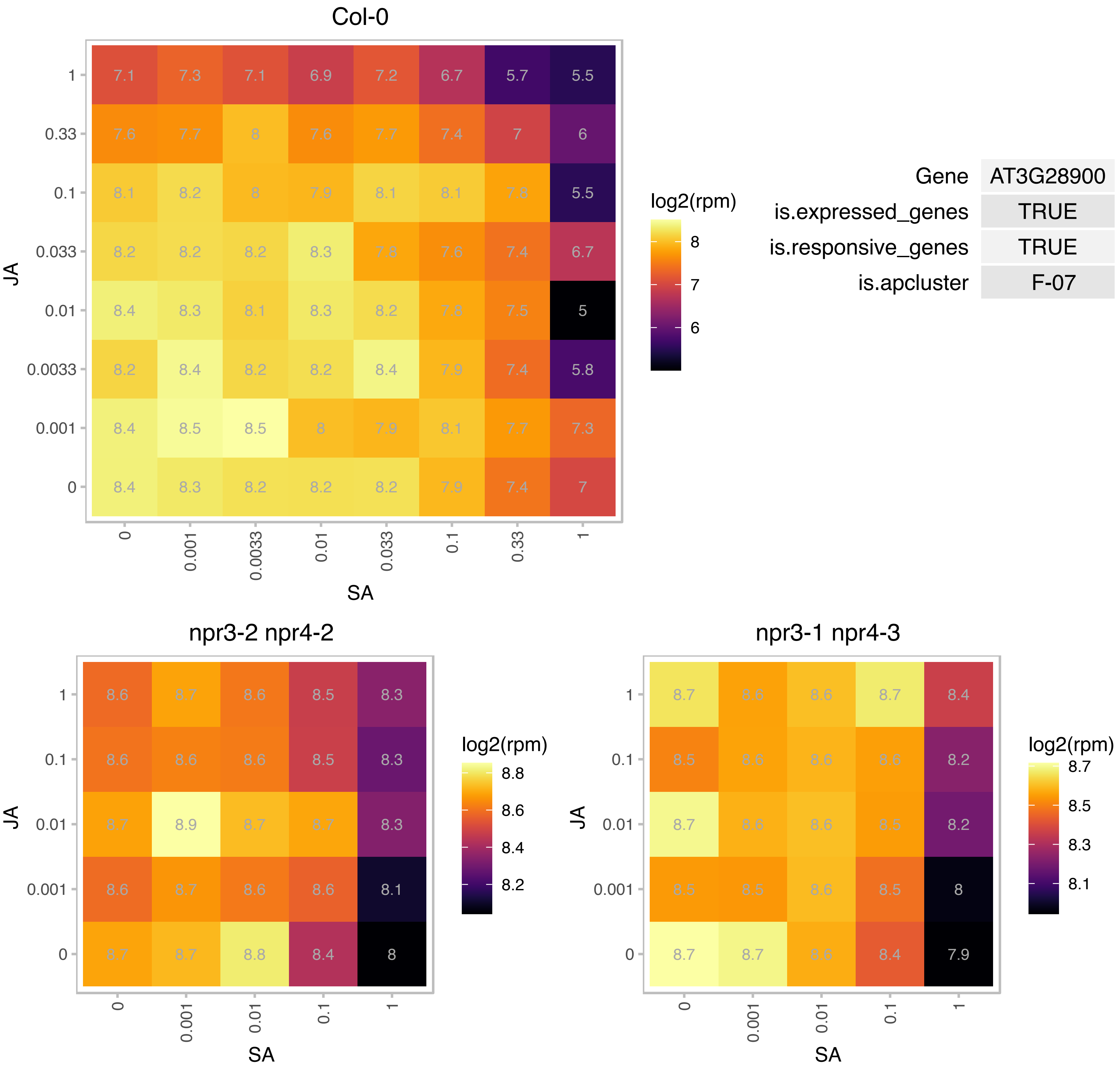

# AT3G28940.1 : NA

AIG2-like (avirulence induced gene) family protein

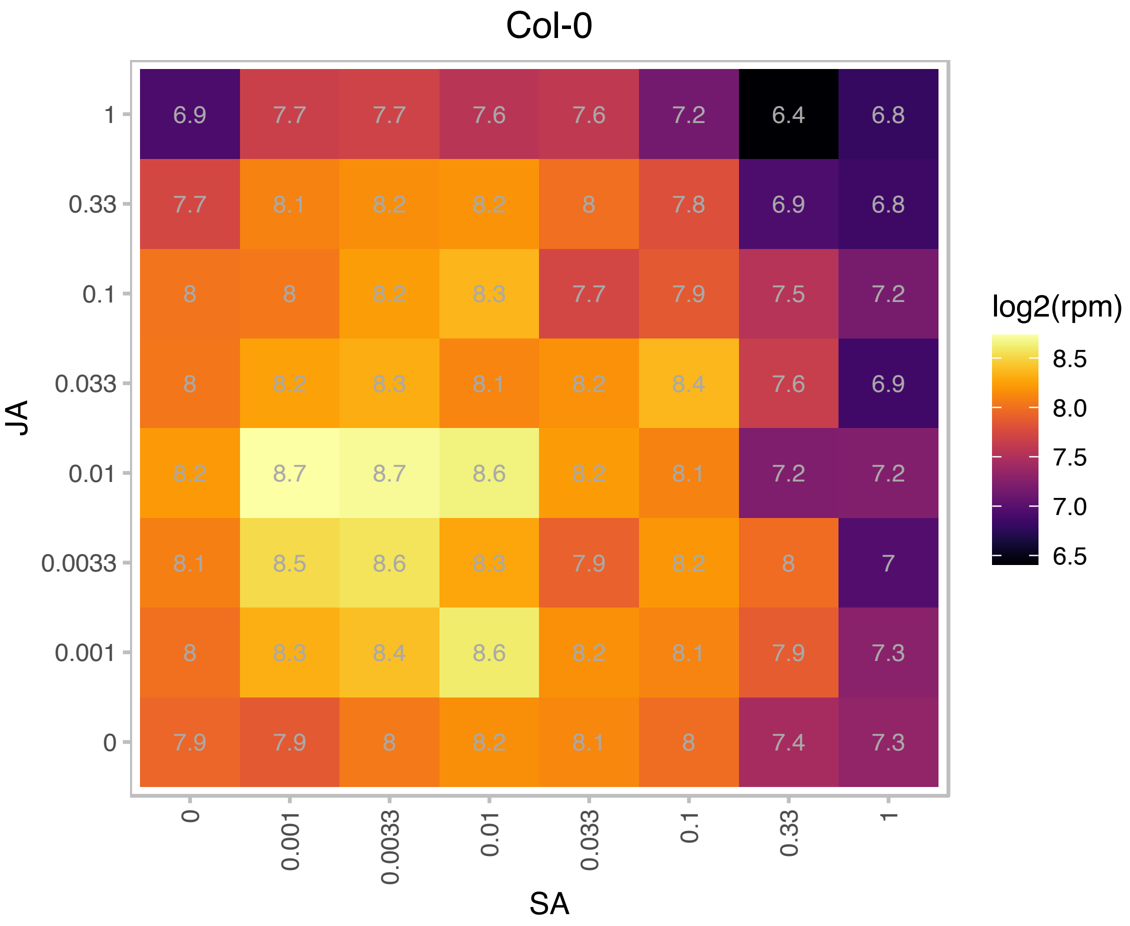


| | |
|---|---|
| Gene | AT3G28940 |
| is.expressed_genes | TRUE |
| is.responsive_genes | TRUE |
| is.apcluster | F-07 |

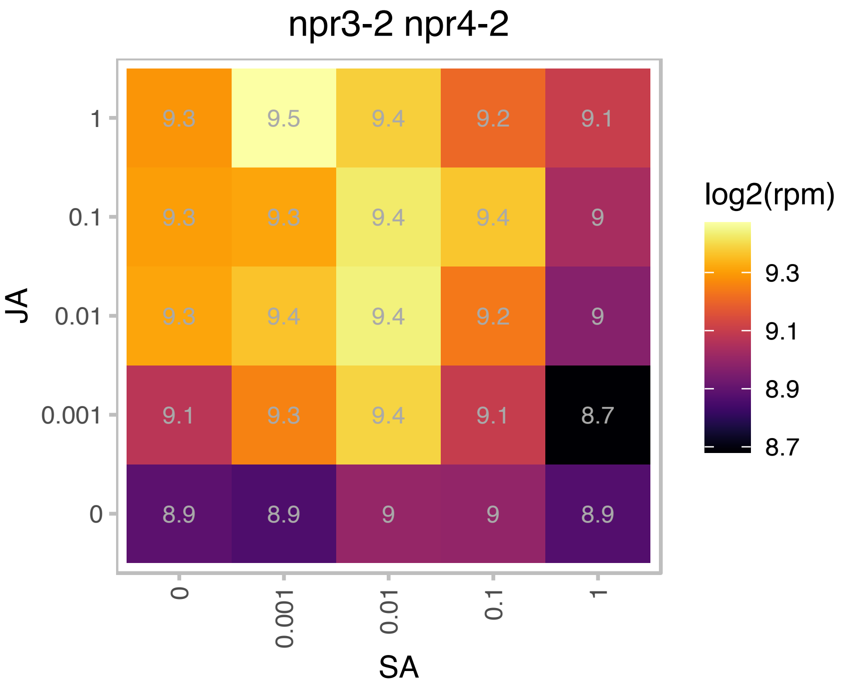


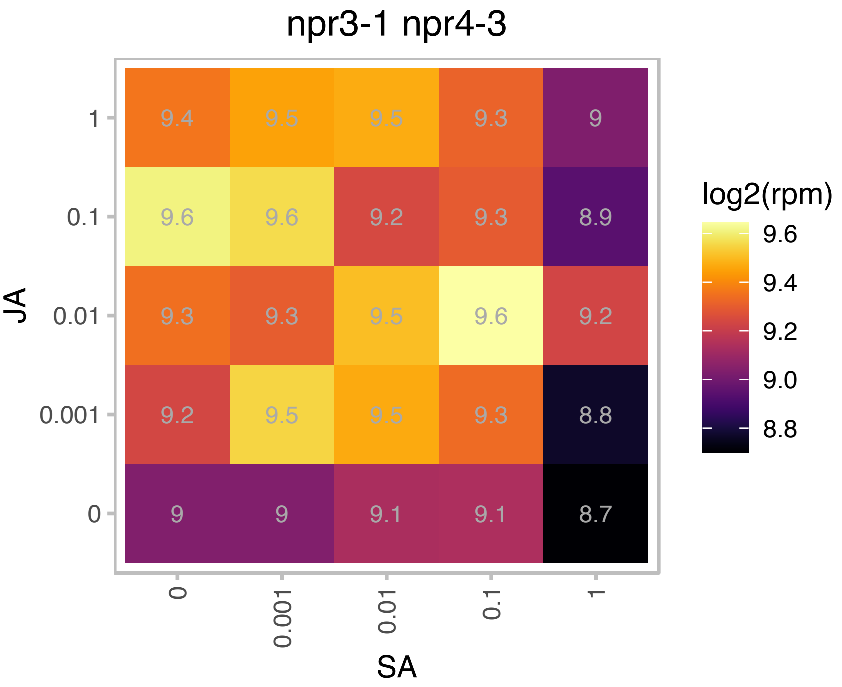

### AT3G44260.1 : AtCAF1a, CAF1a

Polynucleotidyl transferase, ribonuclease H-like superfamily protein

Encodes one of the homologs of the yeast CCR4-associated factor 1: AT3G44260 (CAF1a), AT5G22250 (CAF1b). H
mRNA deadenylation activity.  Also plays a role in plant defense responses.

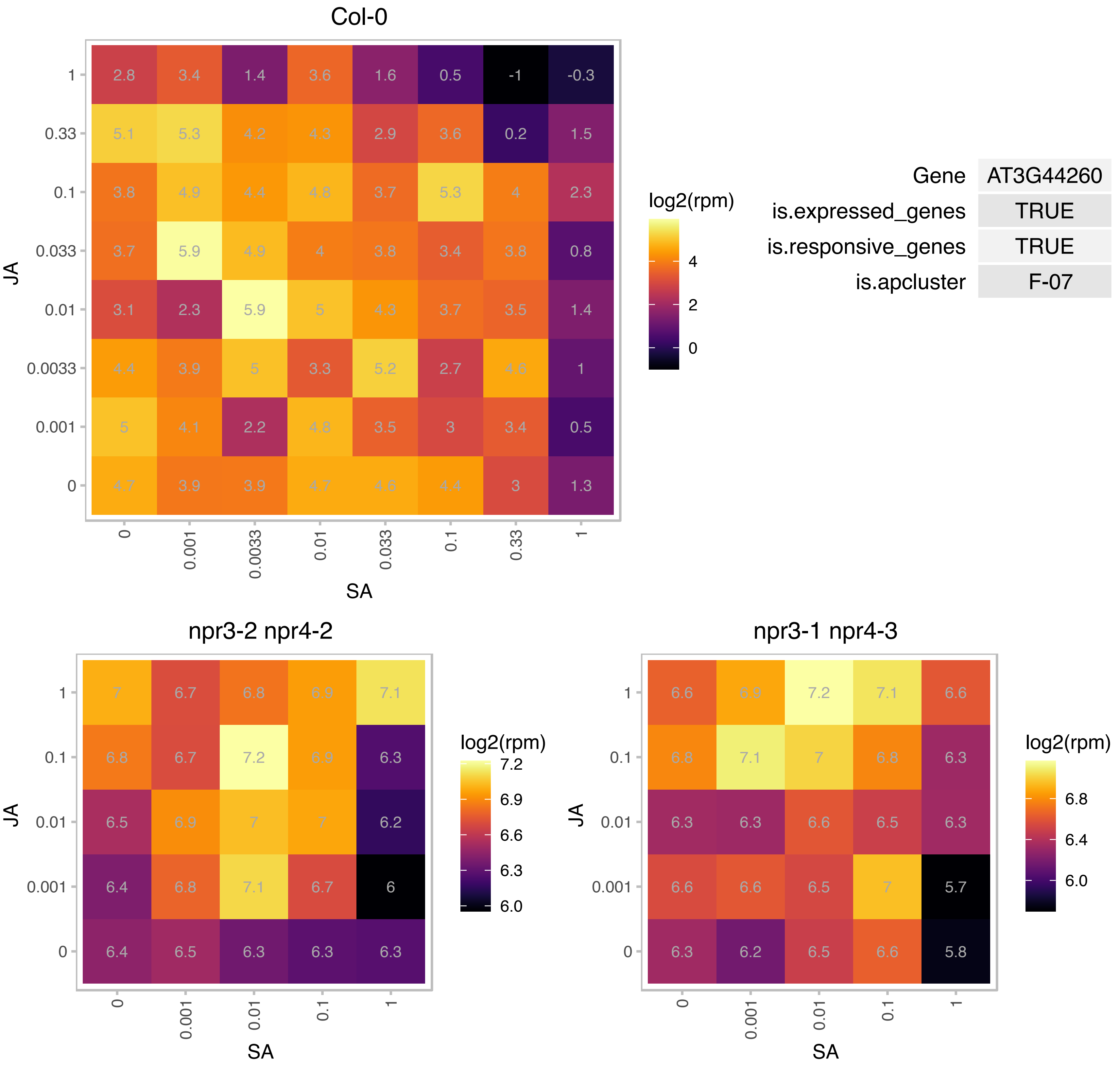


| | |
|---|---|
| Gene | AT3G44260 |
| is.expressed_genes | TRUE |
| is.responsive_genes | TRUE |
| is.apcluster | F-07 |

# AT3G44590.1 : NA

60S acidic ribosomal protein family

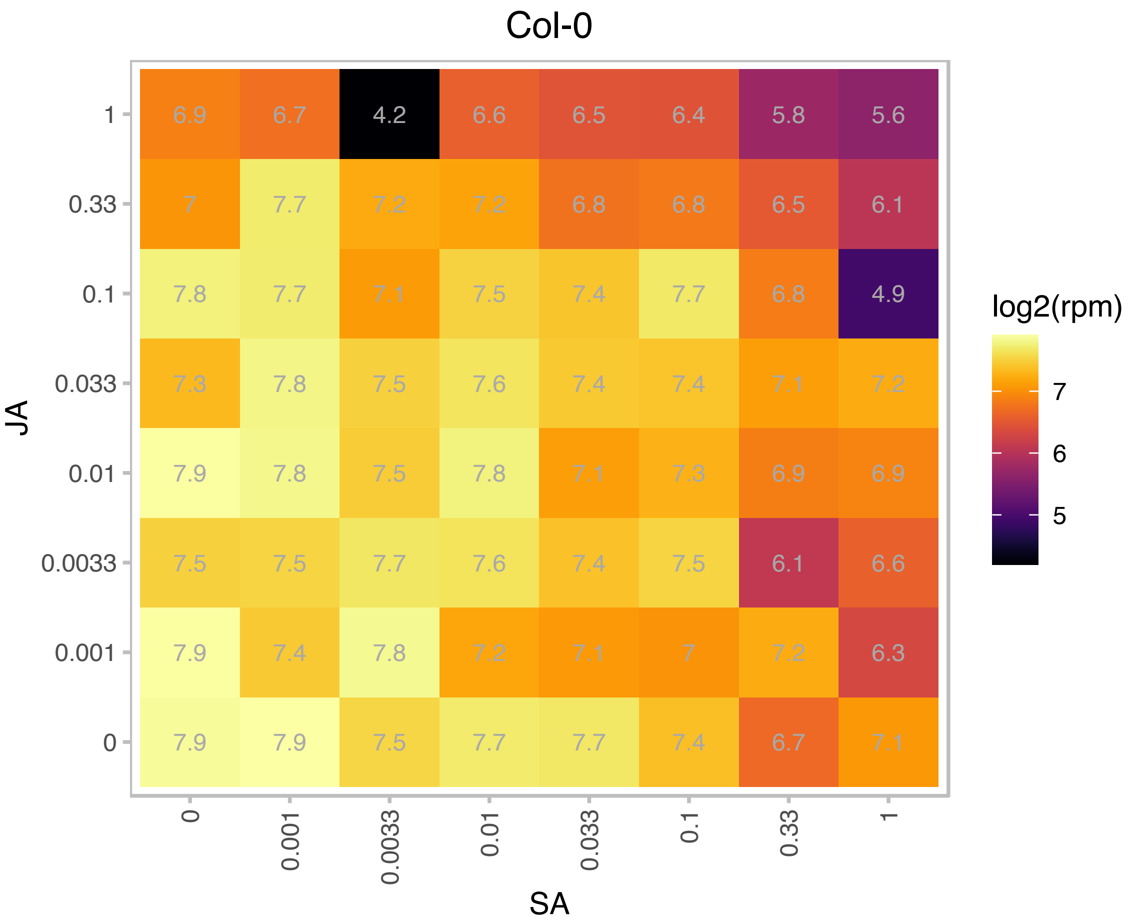


| | |
|---|---|
| Gene | AT3G44590 |
| is.expressed_genes | TRUE |
| is.responsive_genes | TRUE |
| is.apcluster | F-07 |

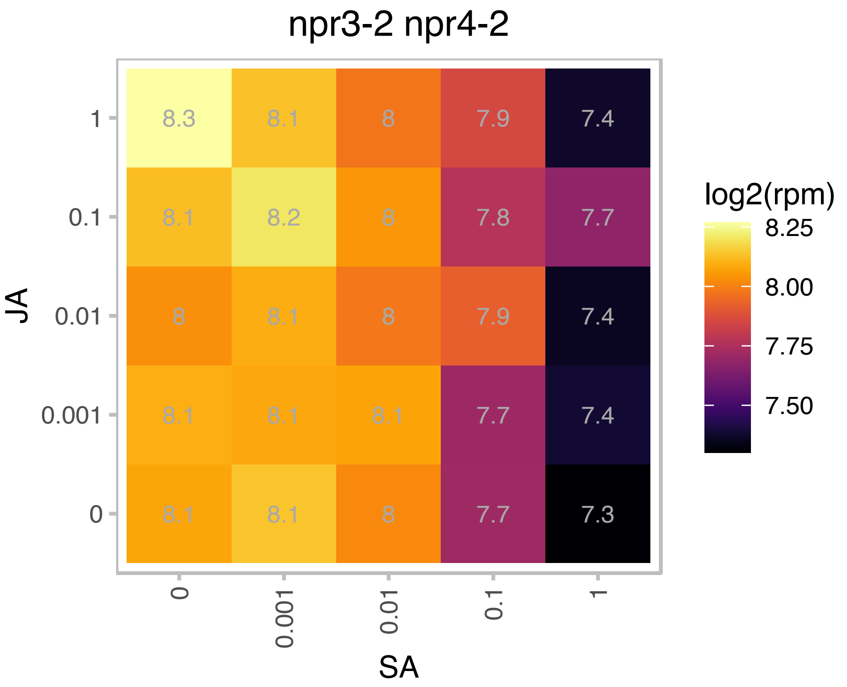


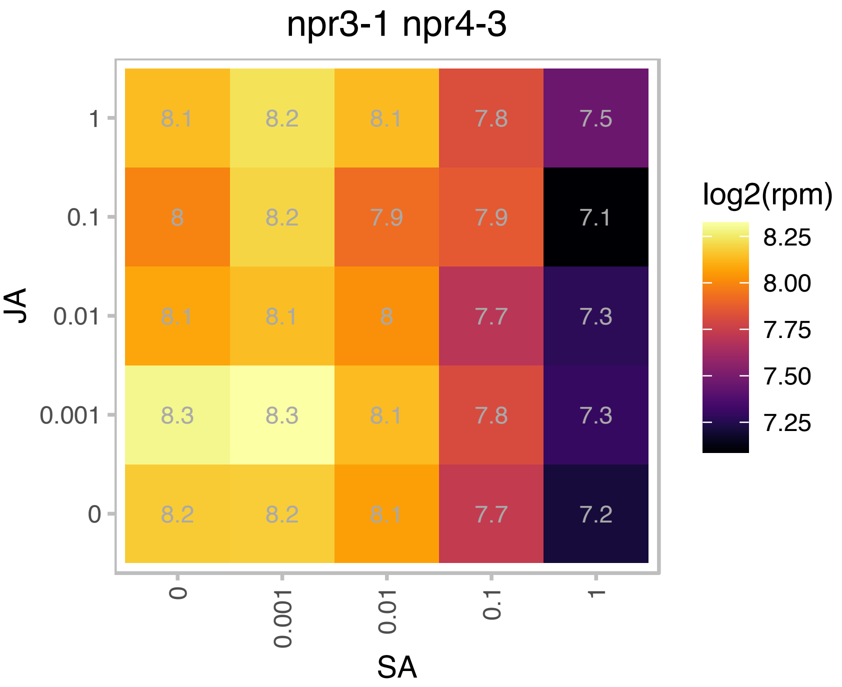

# AT3G45030.1 : NA

Ribosomal protein S10p/S20e family protein

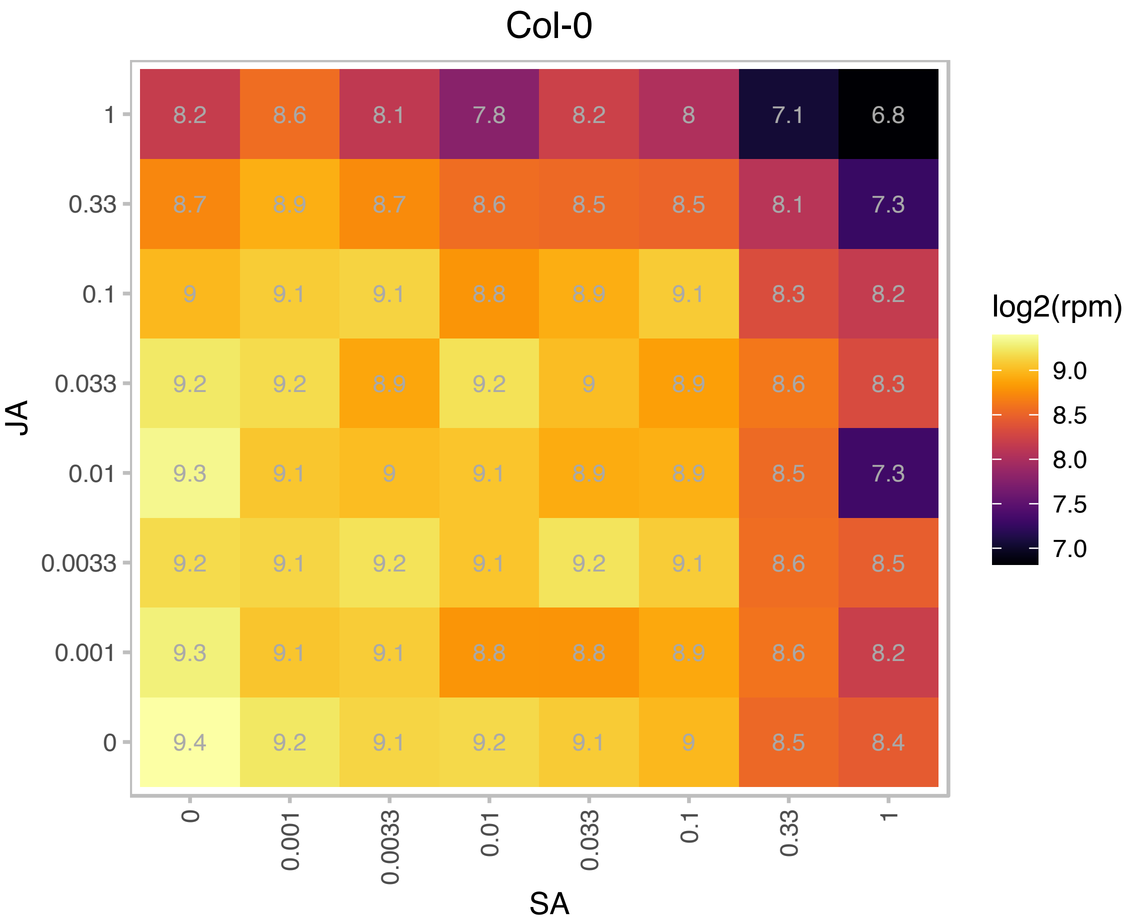


| | |
|---|---|
| Gene | AT3G45030 |
| is.expressed_genes | TRUE |
| is.responsive_genes | TRUE |
| is.apcluster | F-07 |

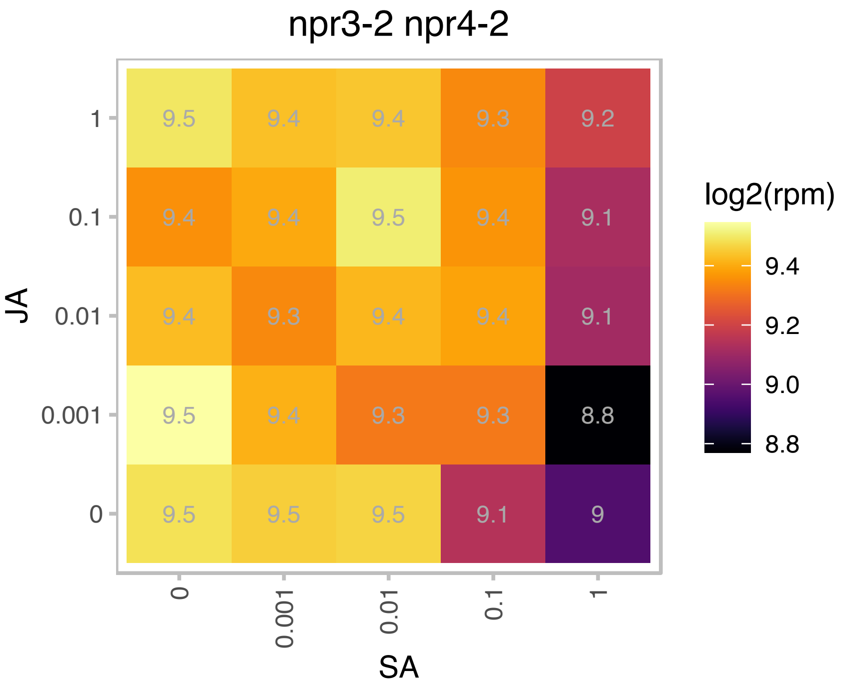


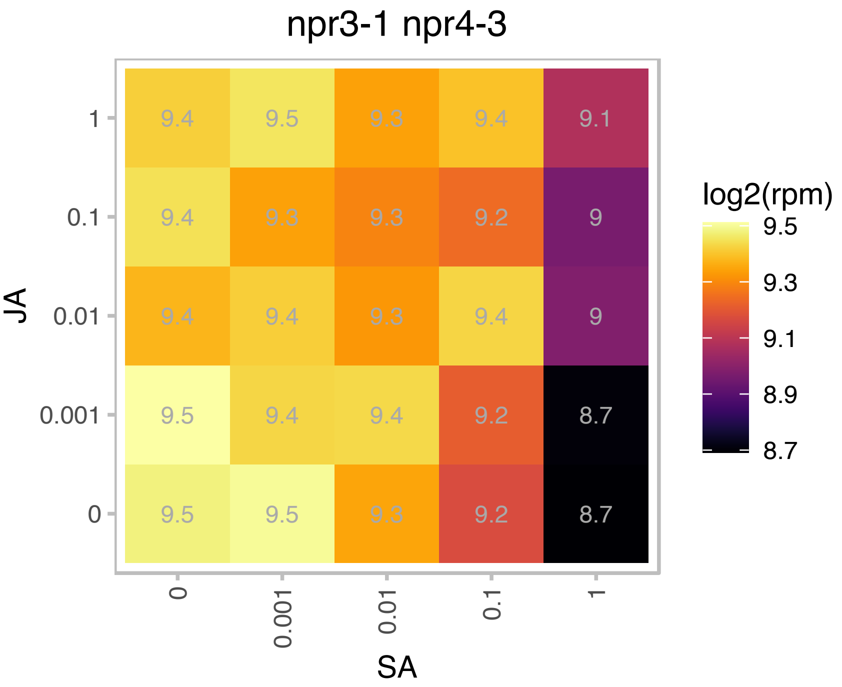

# AT3G46320.1 : NA

Histone superfamily protein

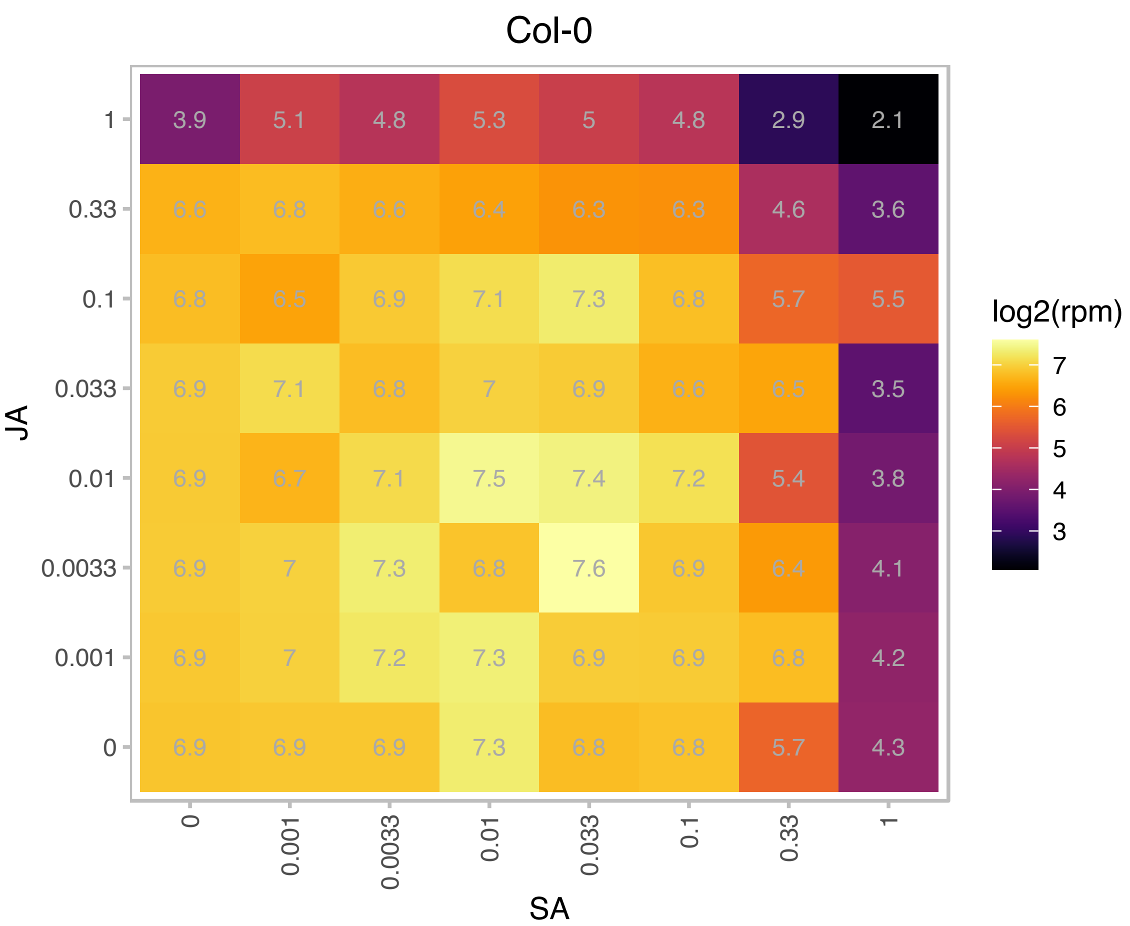


| | |
|---|---|
| Gene | AT3G46320 |
| is.expressed_genes | TRUE |
| is.responsive_genes | TRUE |
| is.apcluster | F-07 |

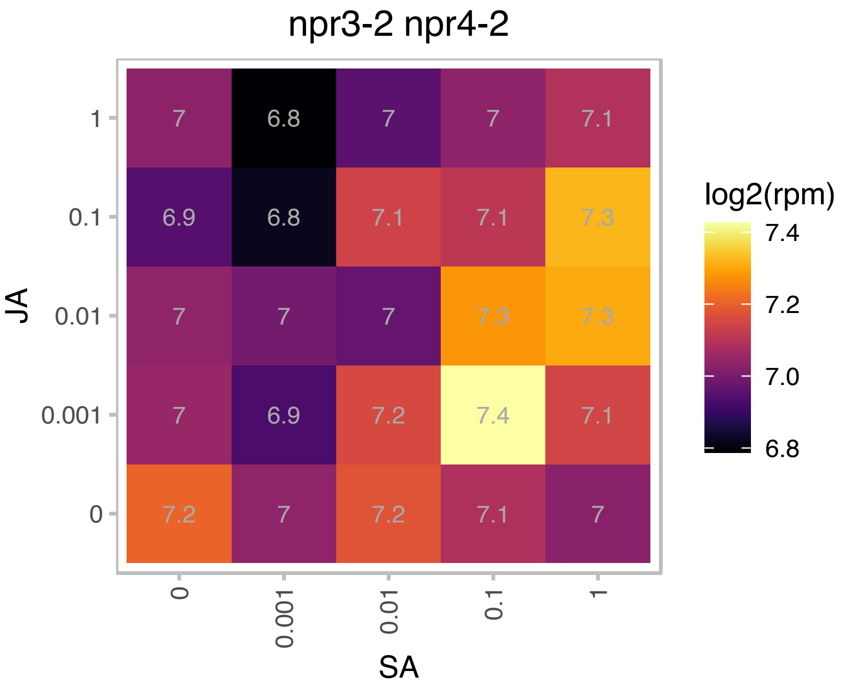


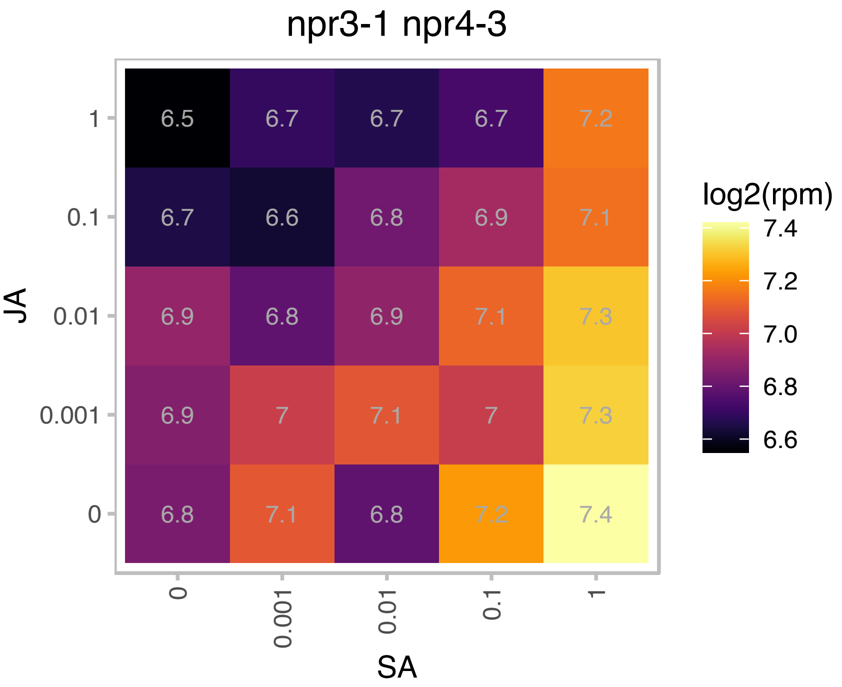

# AT3G47560.1 : NA

alpha/beta-Hydrolases superfamily protein

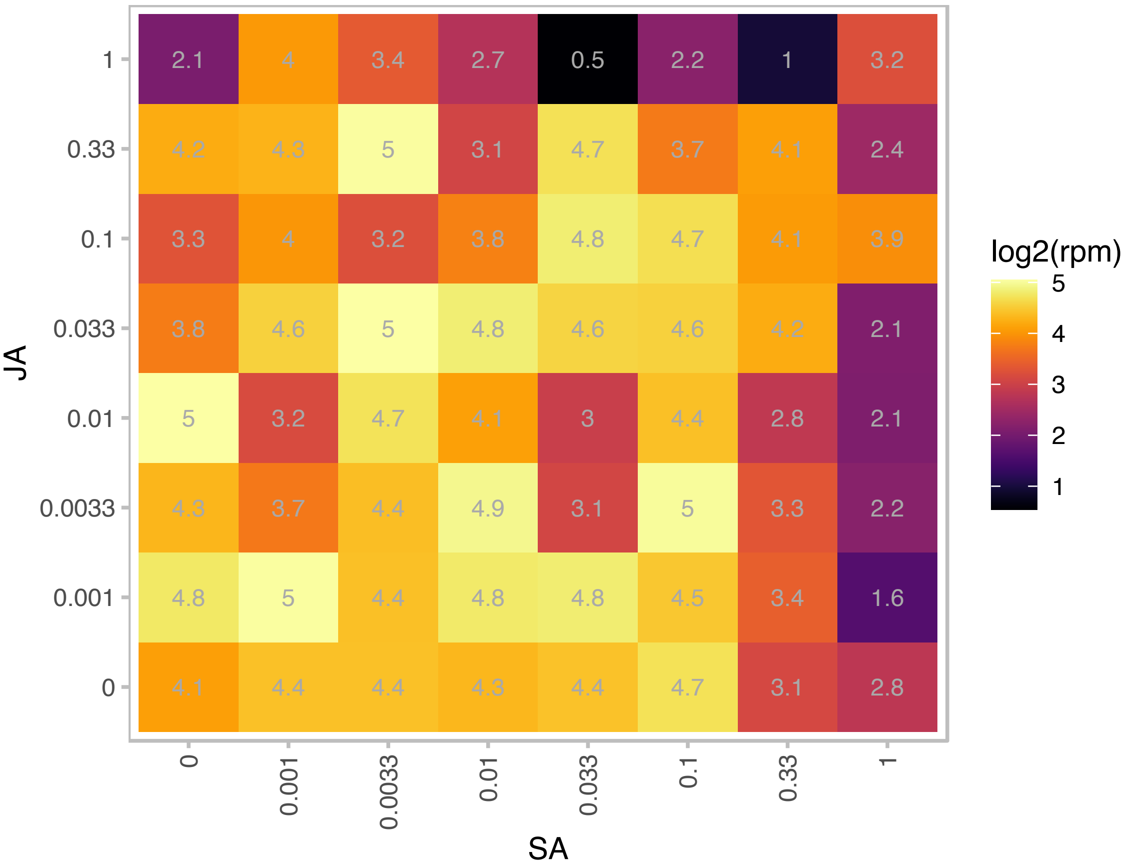


| | |
|---|---|
| Gene | AT3G47560 |
| is.expressed_genes | TRUE |
| is.responsive_genes | TRUE |
| is.apcluster | F-07 |

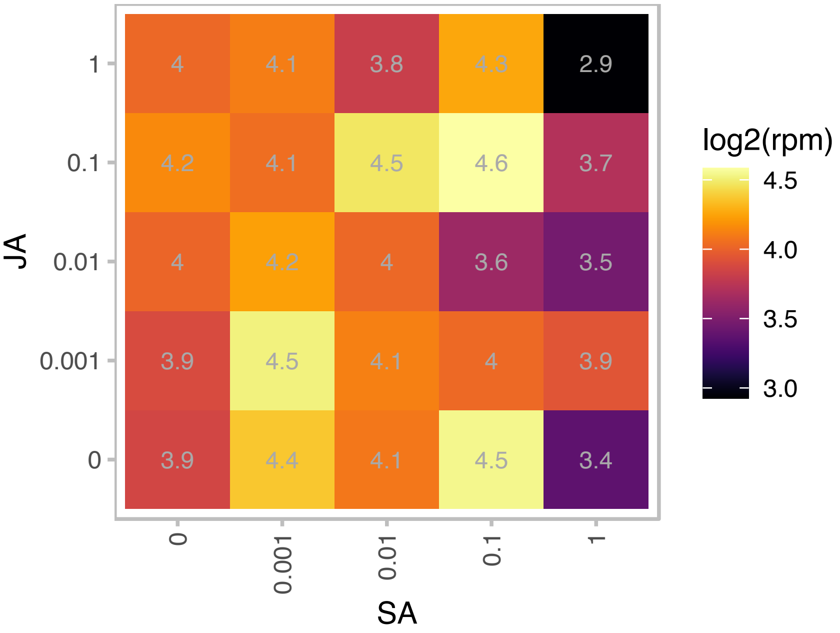


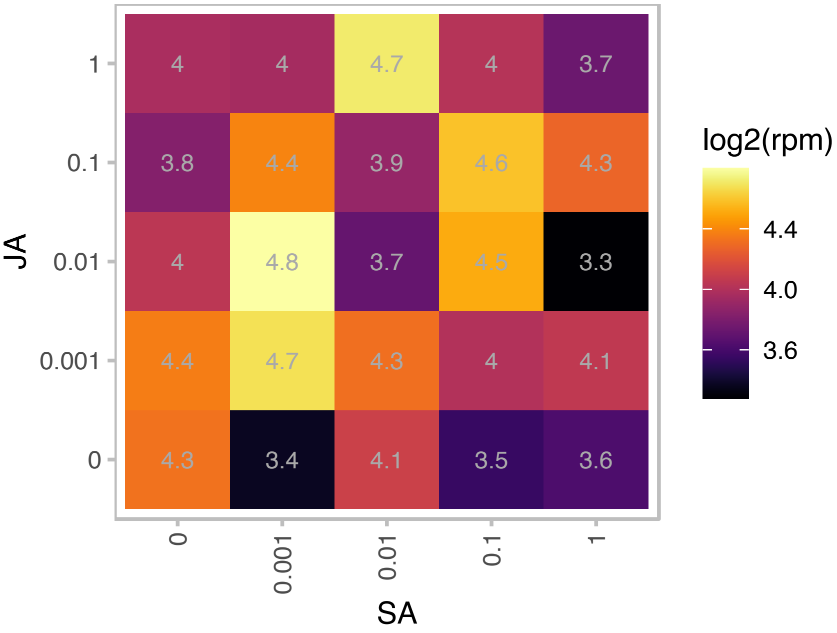

### AT3G48930.1 : EMB1080

Nucleic acid-binding, OB-fold-like protein

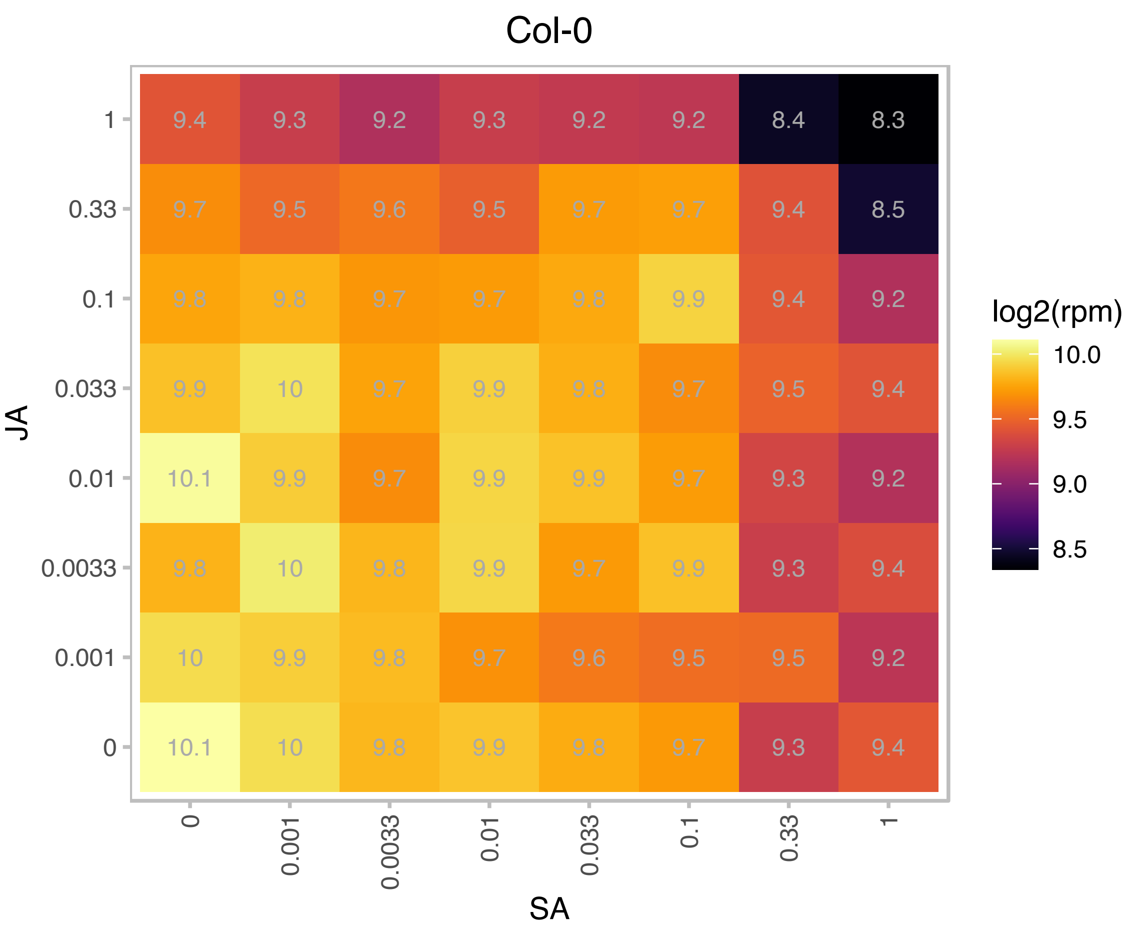


| | |
|---|---|
| Gene | AT3G48930 |
| is.expressed_genes | TRUE |
| is.responsive_genes | TRUE |
| is.apcluster | F-07 |

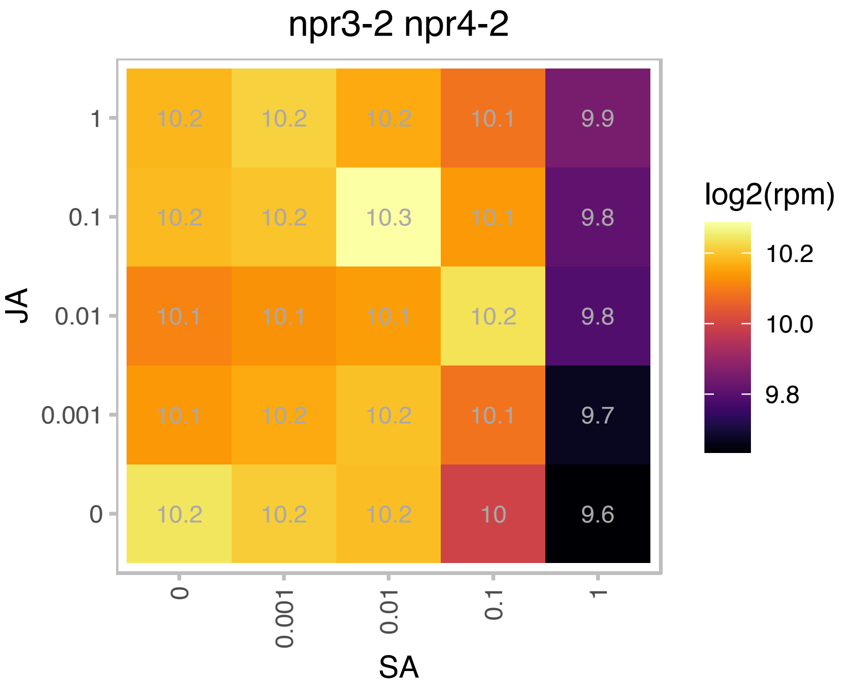


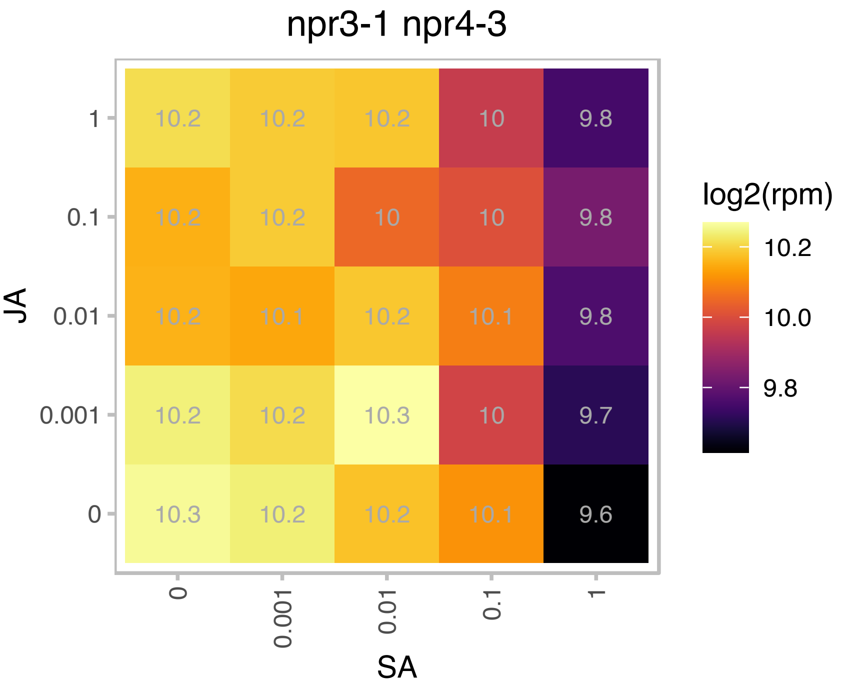

### AT3G49010.1 : ATBBC1, BBC1, RSU2

breast basic conserved 1

Encodes 60S ribosomal protein L13. Homolog of human breast basic conserved 1 (BBC1).

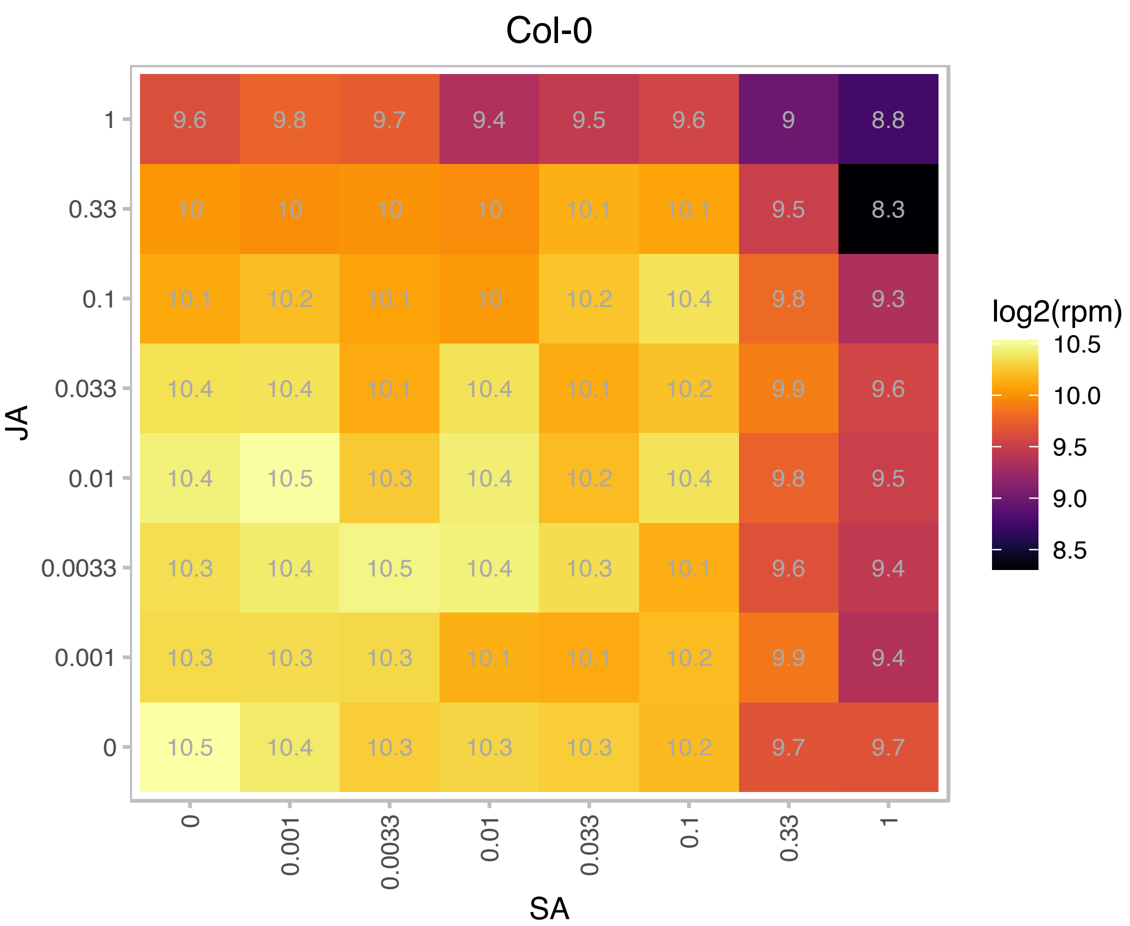


| | |
|---|---|
| Gene | AT3G49010 |
| is.expressed_genes | TRUE |
| is.responsive_genes | TRUE |
| is.apcluster | F-07 |

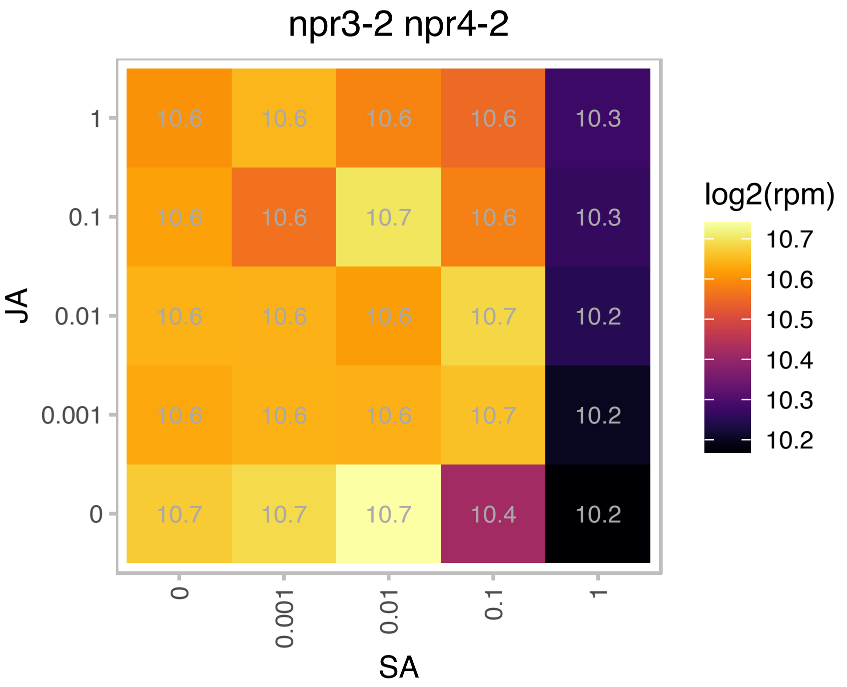


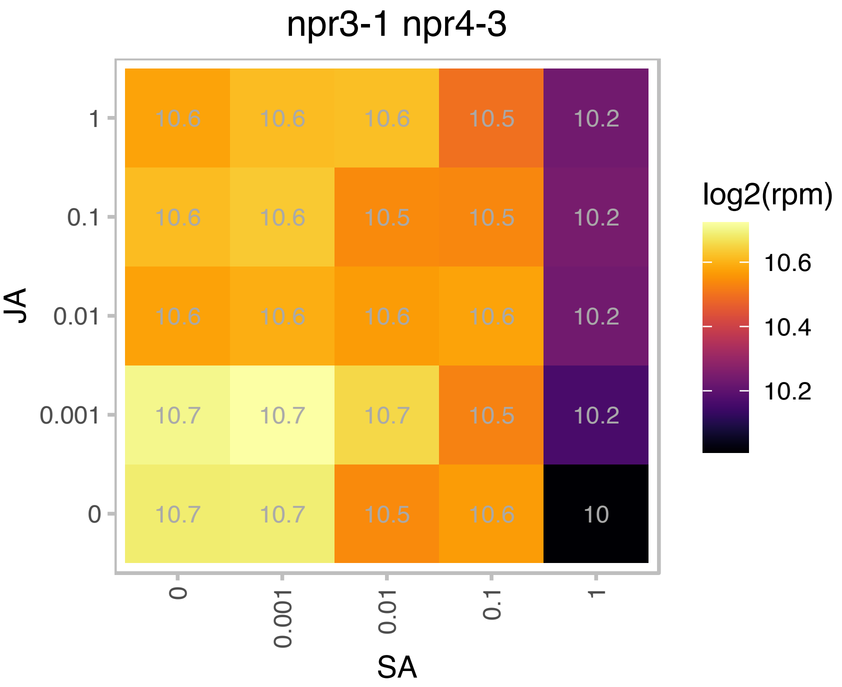

### AT3G49940.1 : LBD38

LOB domain-containing protein 38

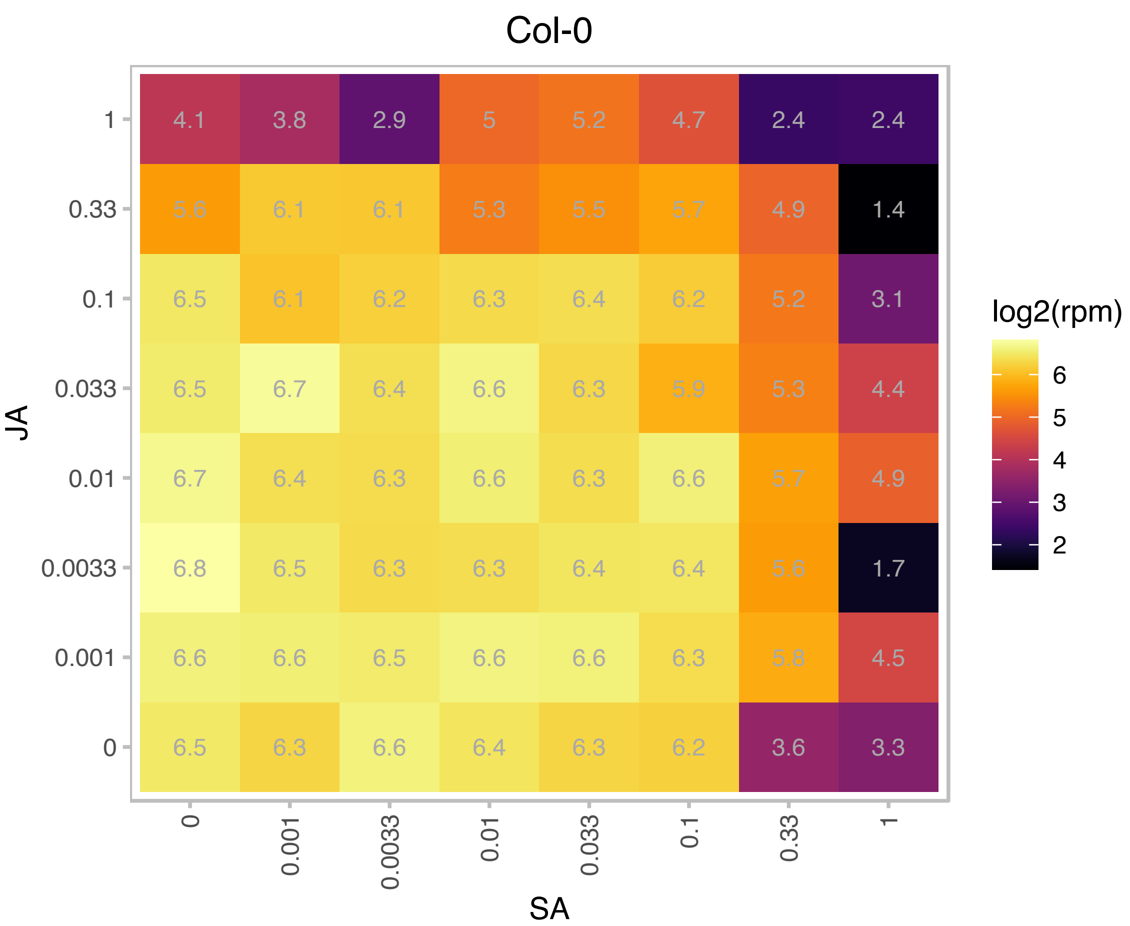


| | |
|---|---|
| Gene | AT3G49940 |
| is.expressed_genes | TRUE |
| is.responsive_genes | TRUE |
| is.apcluster | F-07 |

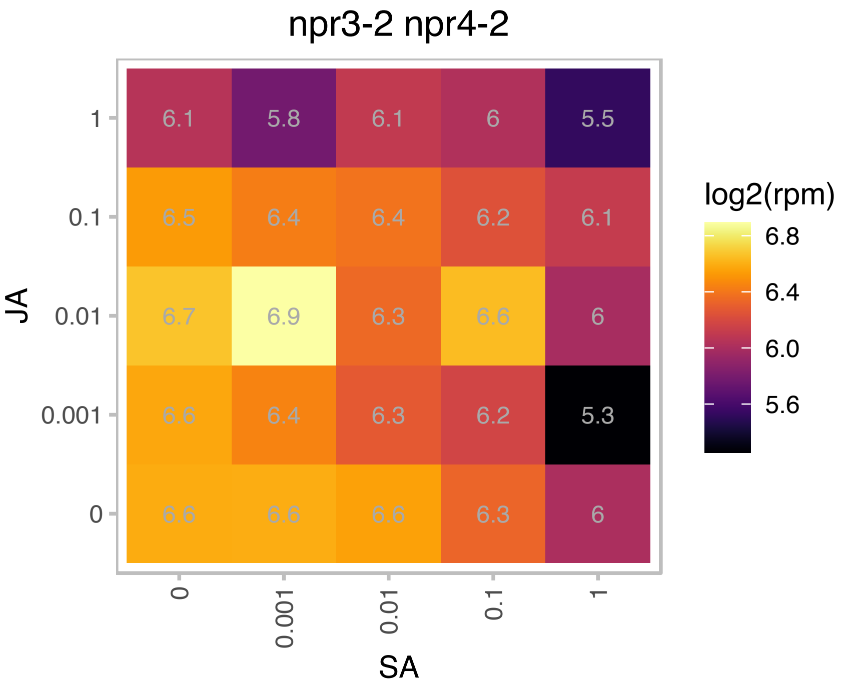


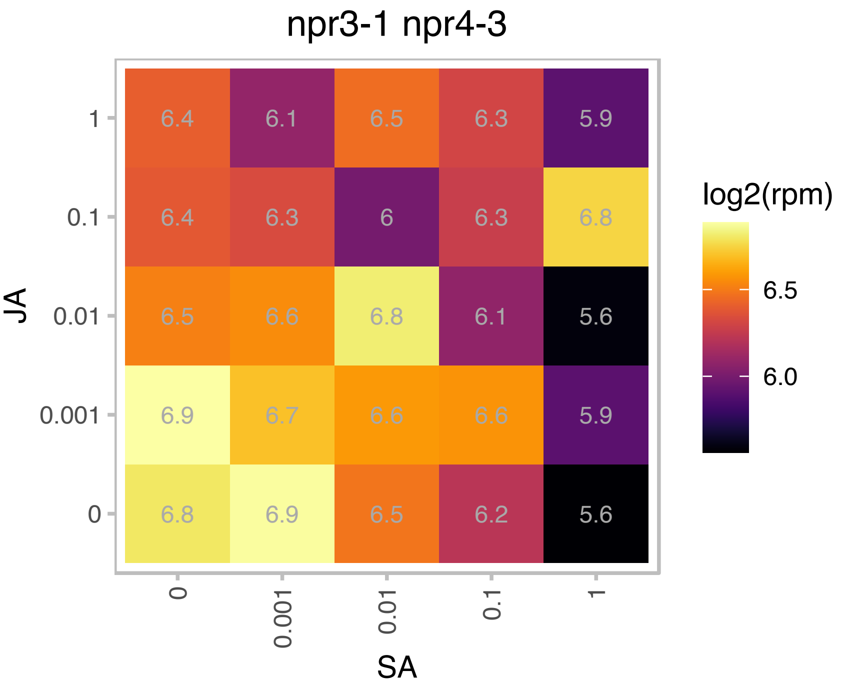

# AT3G52580.1 : NA

Ribosomal protein S11 family protein

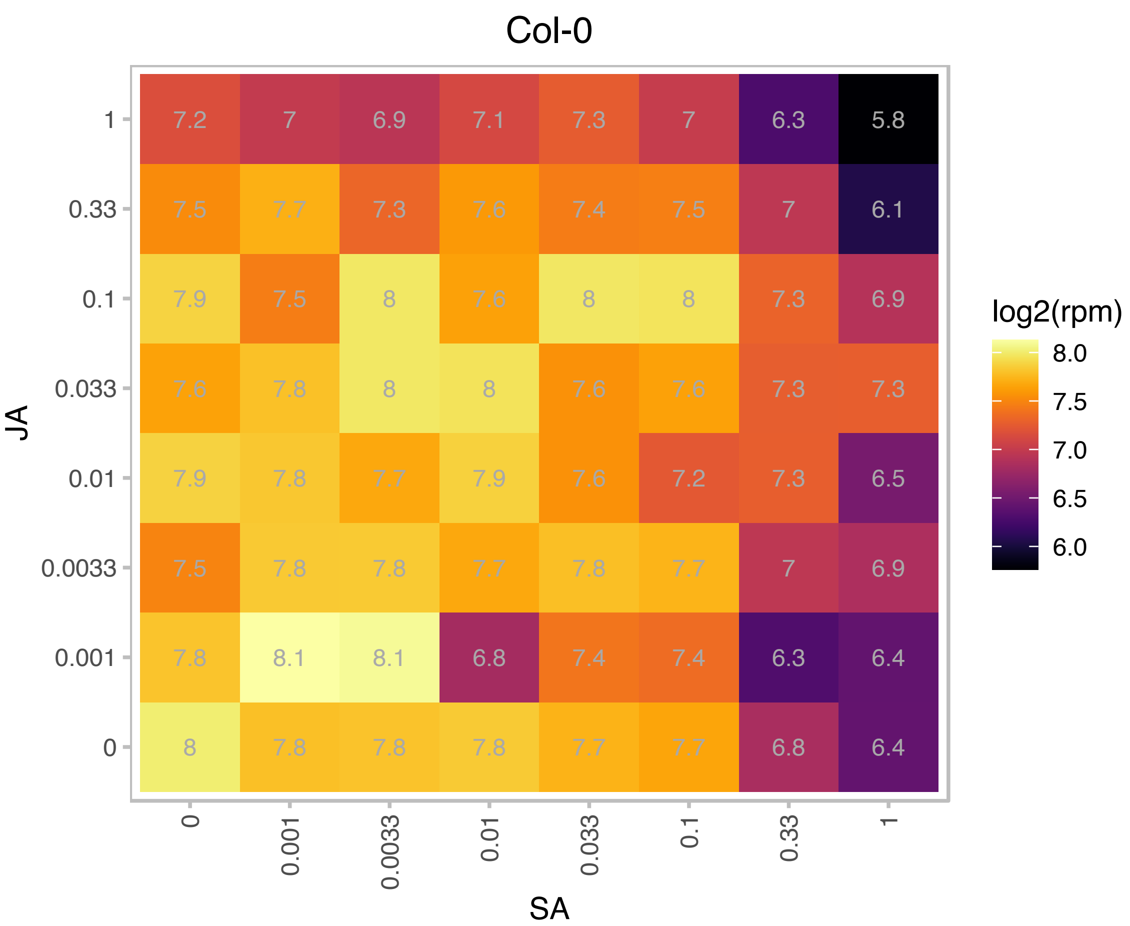


| | |
|---|---|
| Gene | AT3G52580 |
| is.expressed_genes | TRUE |
| is.responsive_genes | TRUE |
| is.apcluster | F-07 |

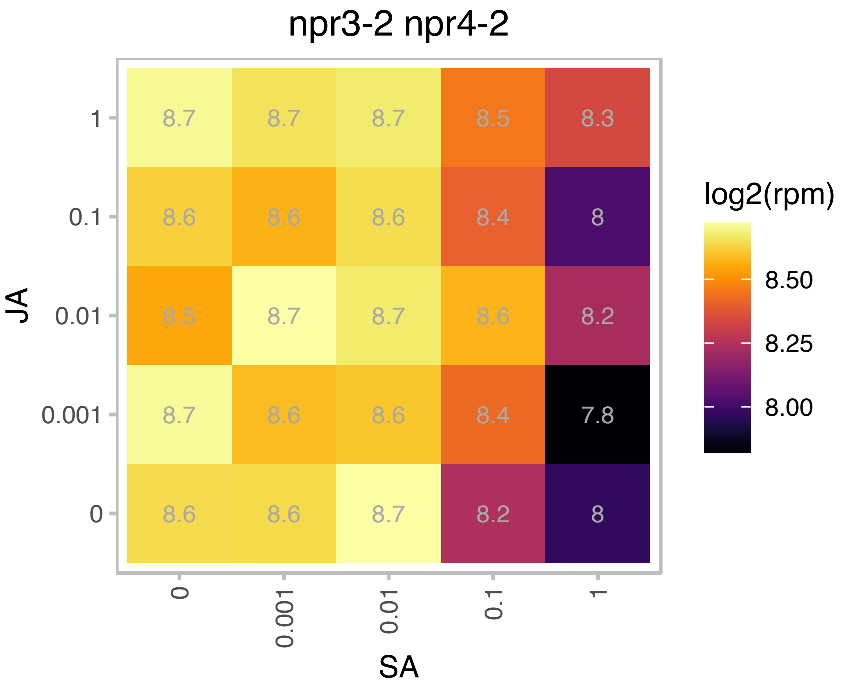


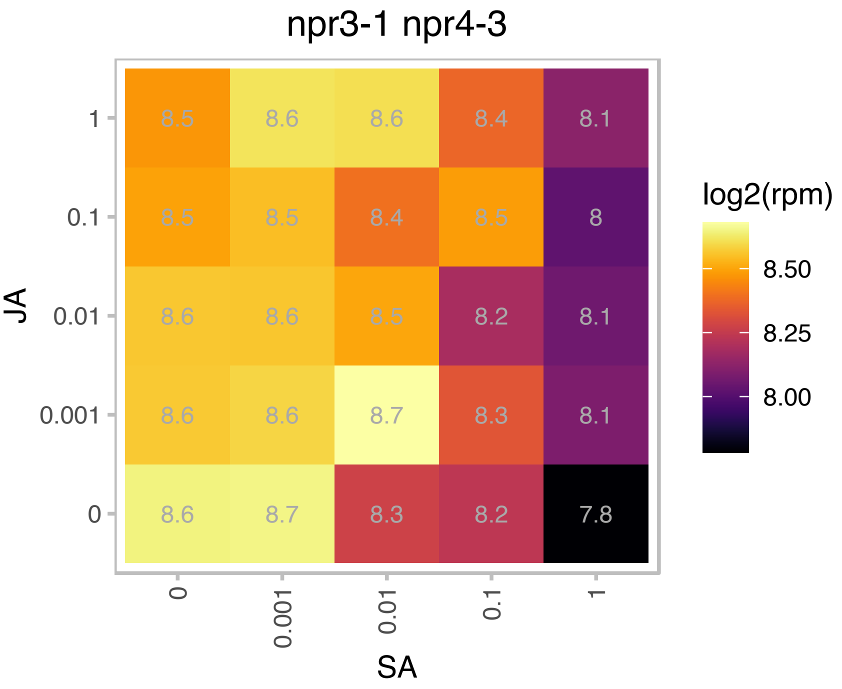

### AT3G53020.1 : RPL24, RPL24B, STV1

RPL24B encodes ribosomal protein L24, homolog of cytosolic RPL24, found in archaea and higher eukaryotes.
Ribosomal protein L24e family protein
Arabidopsis has two RPL24 homologs, RPL24A (AT2G36620) and RPL24B. Mutants showed defects in apical-basa
gynoecium patterning similar to previously described ett and mp mutants. Transformation of stv1-1 mutant with a uORF-eliminated ETT construct partially suppressed the stv1 gynoecium phenotype, implying that STV1 could influence ETT translation through its uORFs. Regulated by TCP20.

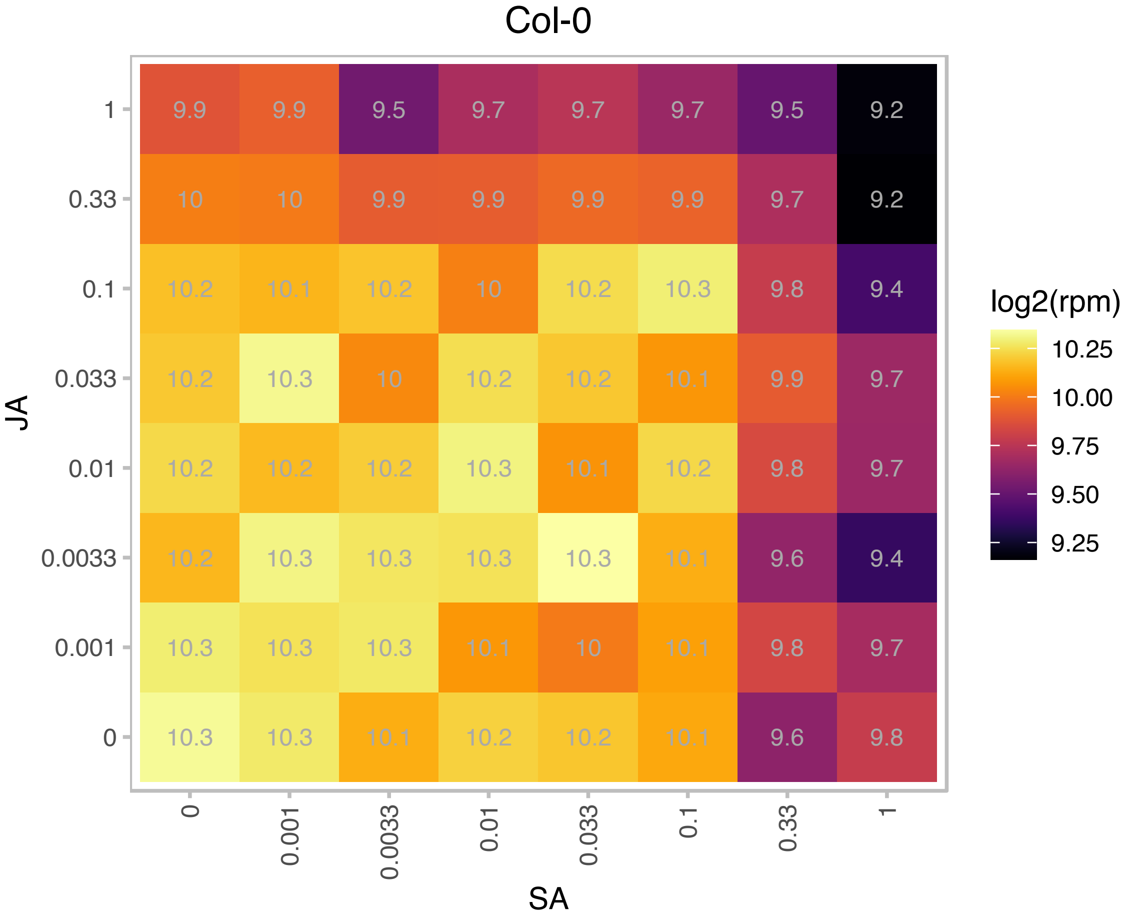


| | |
|---|---|
| Gene | AT3G53020 |
| is.expressed_genes | TRUE |
| is.responsive_genes | TRUE |
| is.apcluster | F-07 |

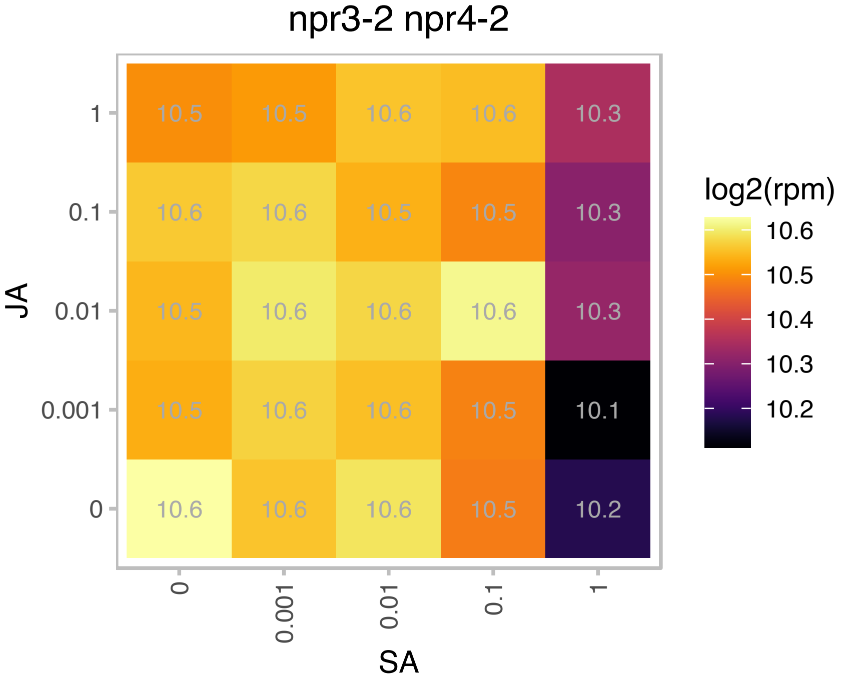


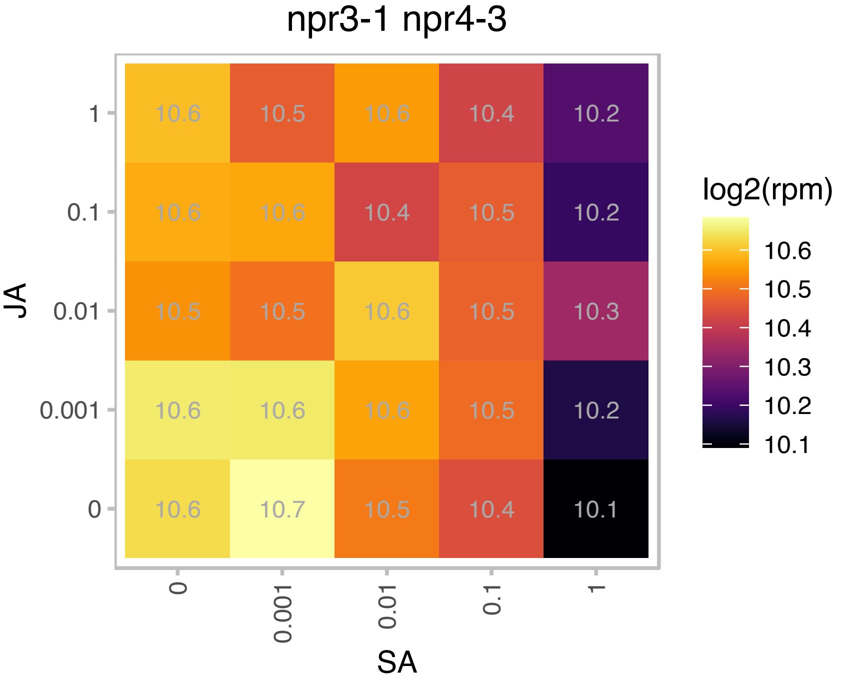

# AT3G53730.1 : NA

Histone superfamily protein

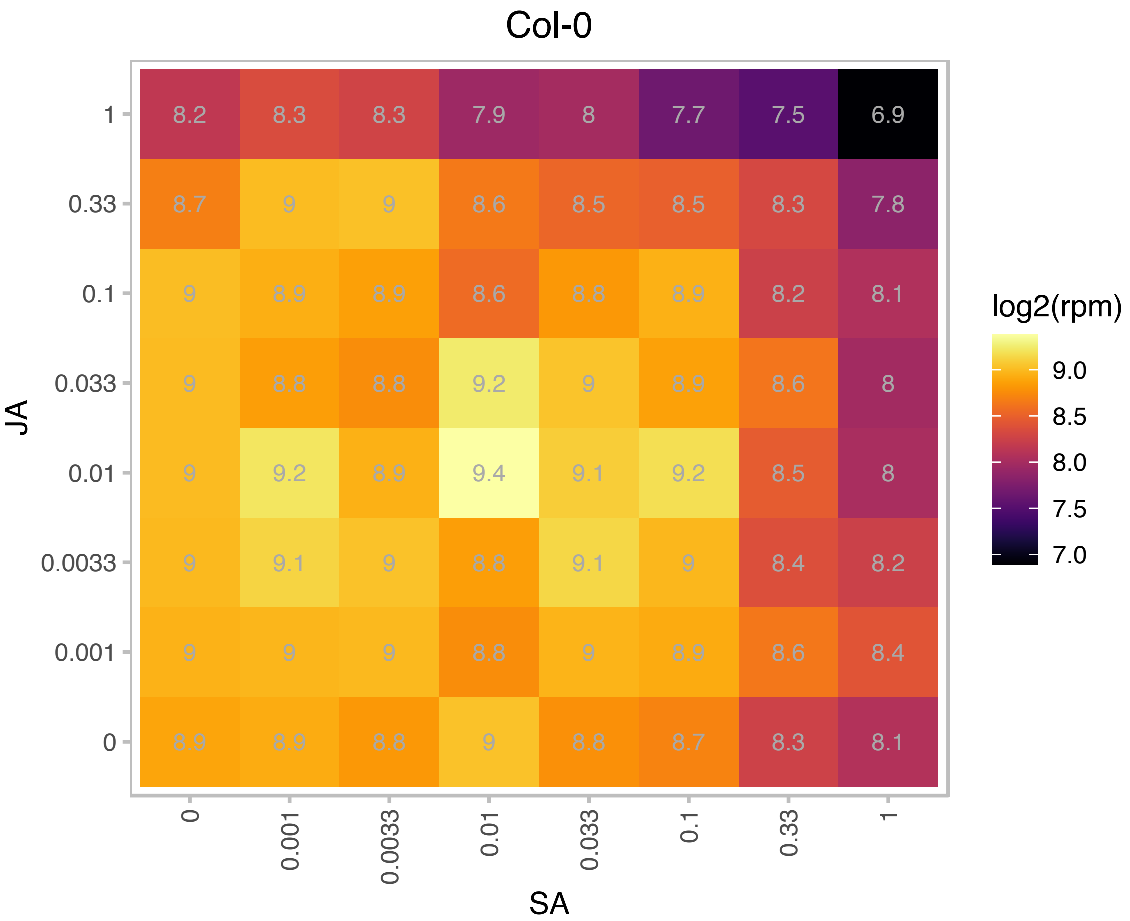


| | |
|---|---|
| Gene | AT3G53730 |
| is.expressed_genes | TRUE |
| is.responsive_genes | TRUE |
| is.apcluster | F-07 |

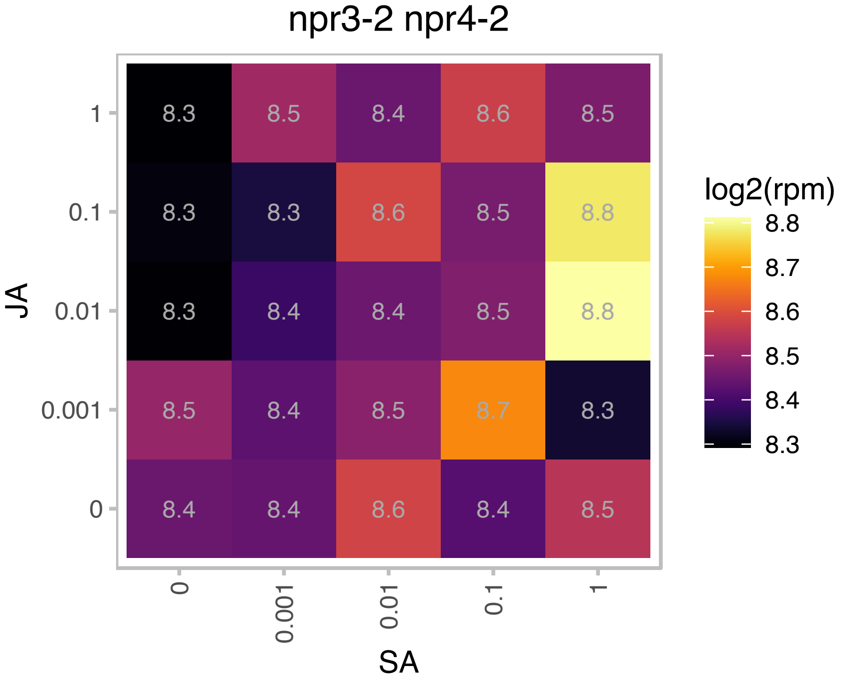


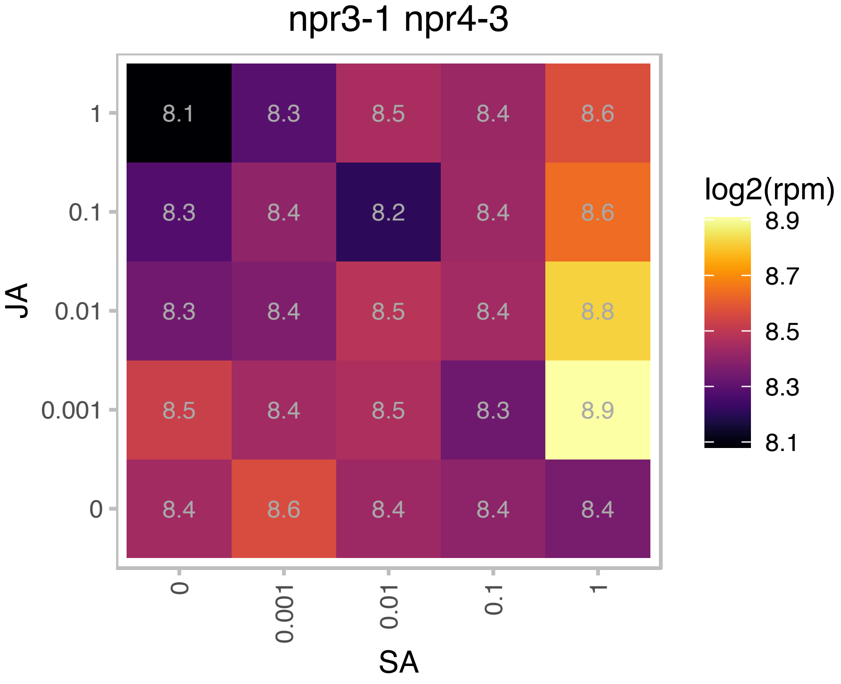

### AT3G53890.1 : EVR1, RPS21B

Ribosomal protein S21e

Cytosolic ribosomal protein. Mutants enhance the varigation effect of var2 mutations suggesting a link between cytosolic translation and chloroplast development.

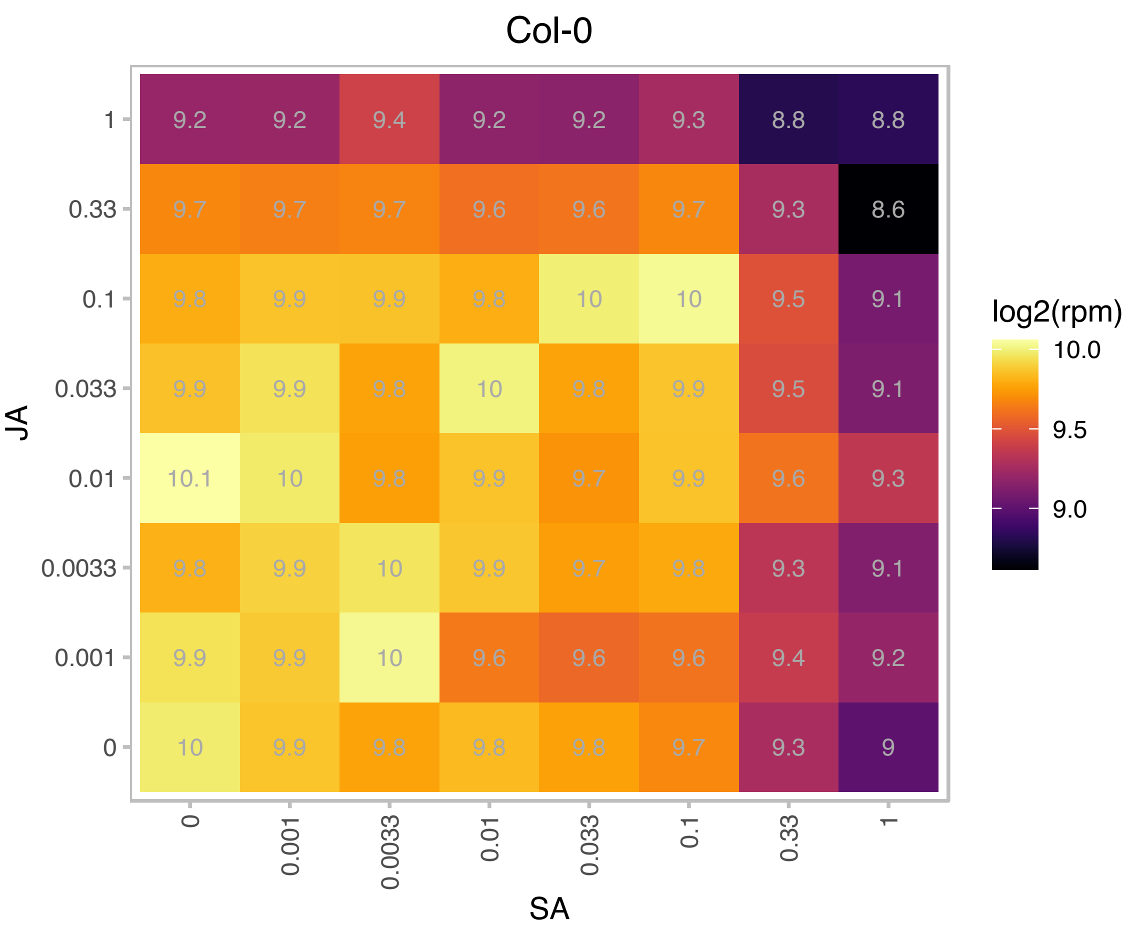


| | |
|---|---|
| Gene | AT3G53890 |
| is.expressed_genes | TRUE |
| is.responsive_genes | TRUE |
| is.apcluster | F-07 |

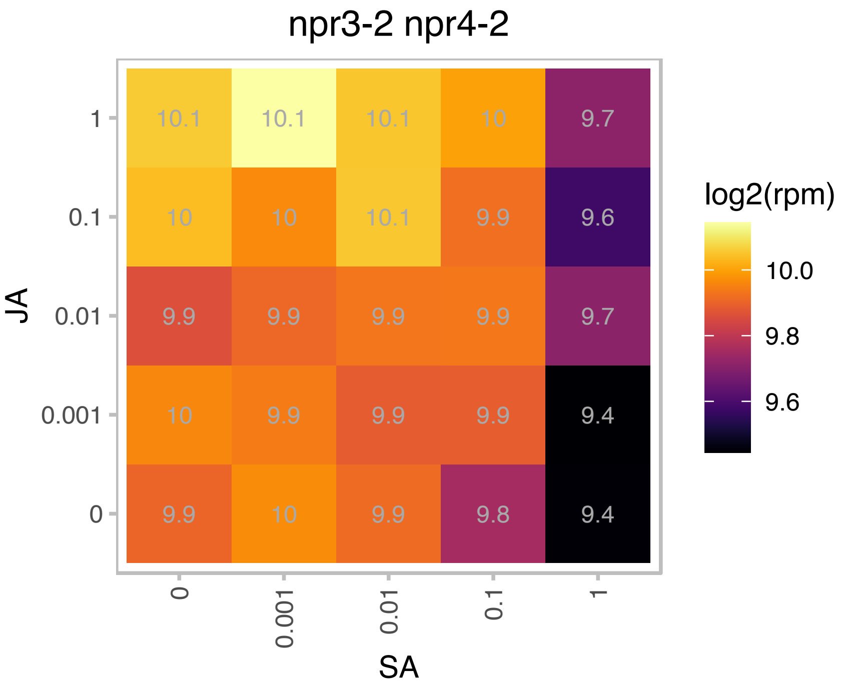


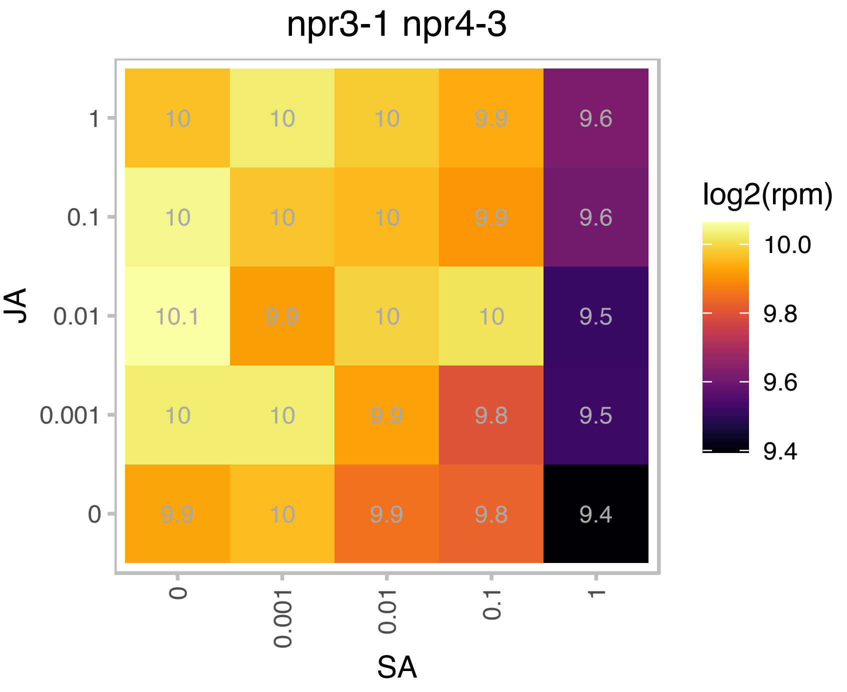

# AT3G54470.1 : NA

uridine 5'-monophosphate synthase / UMP synthase (PYRE-F) (UMPS)

encodes the bi-functional orotate phosphoribosyltransferase/orotidine-5'-phosphate decarboxylase catalyzing the fifth and sixth step in the de novo pyrimidine ribonucleotide biosynthesis

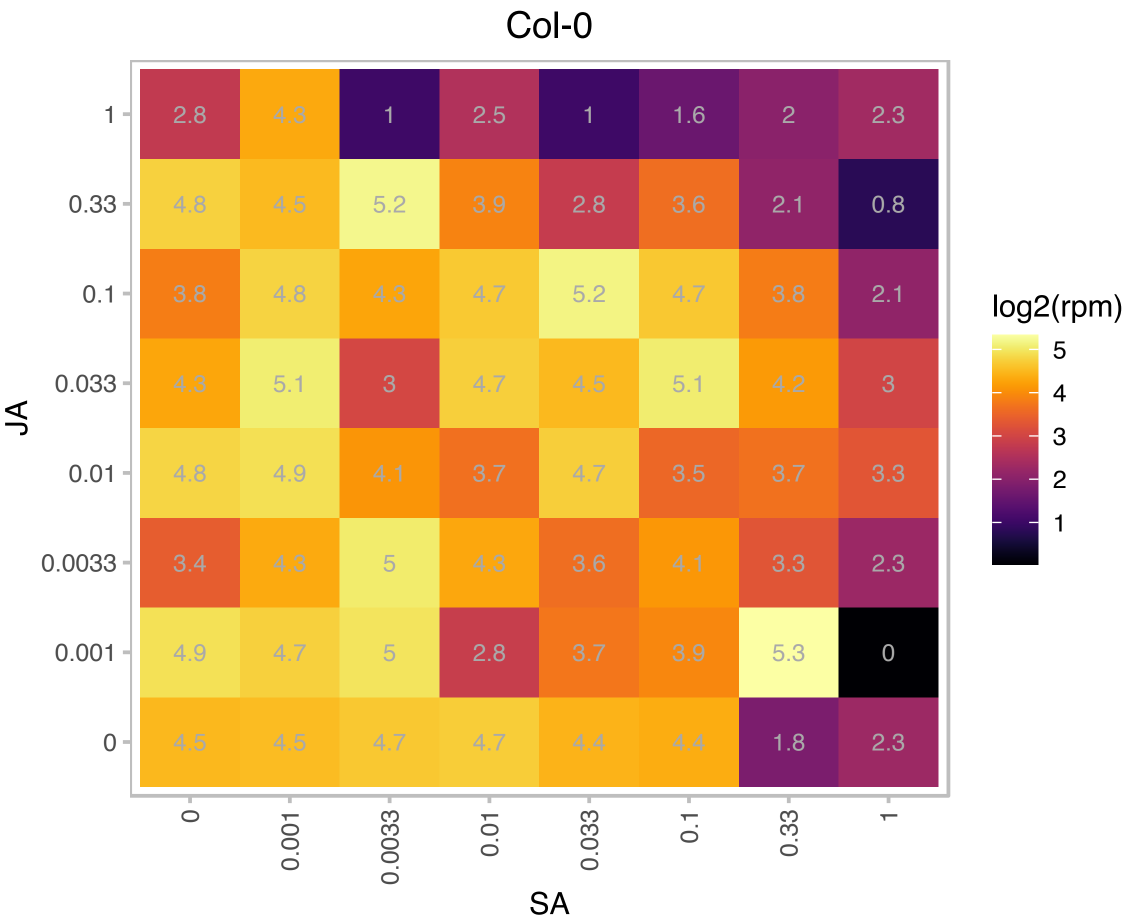


| | |
|---|---|
| Gene | AT3G54470 |
| is.expressed_genes | TRUE |
| is.responsive_genes | TRUE |
| is.apcluster | F-07 |

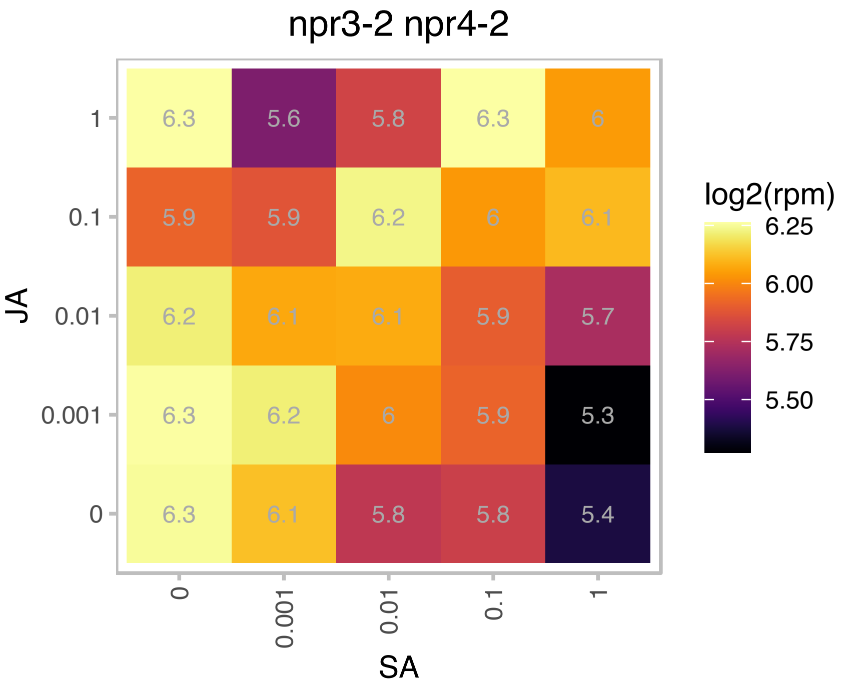


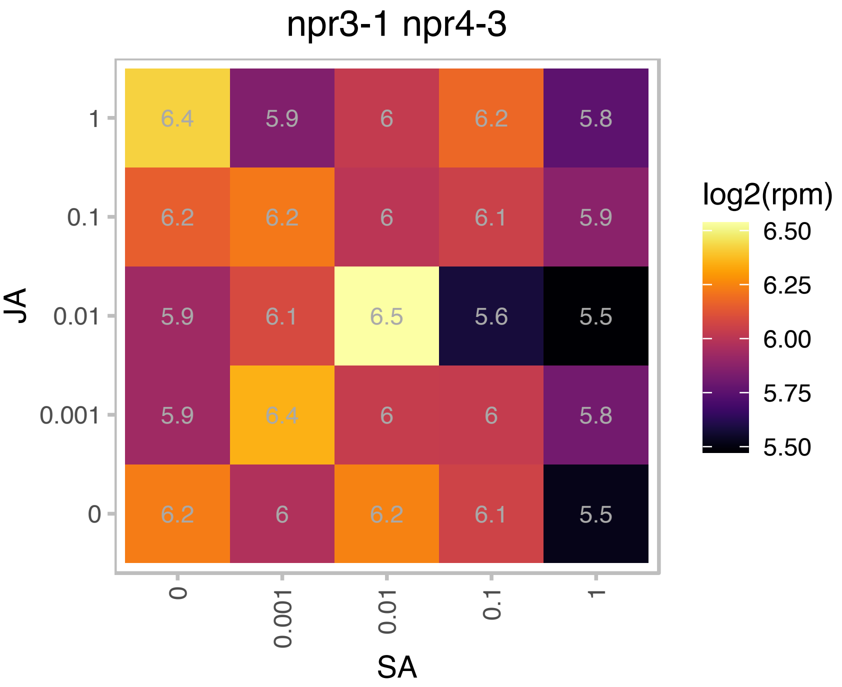

### AT3G55280.1 : RPL23A2, RPL23AB

ribosomal protein L23AB

60S ribosomal protein L23A (RPL23aB). Paralog of RPL23aA

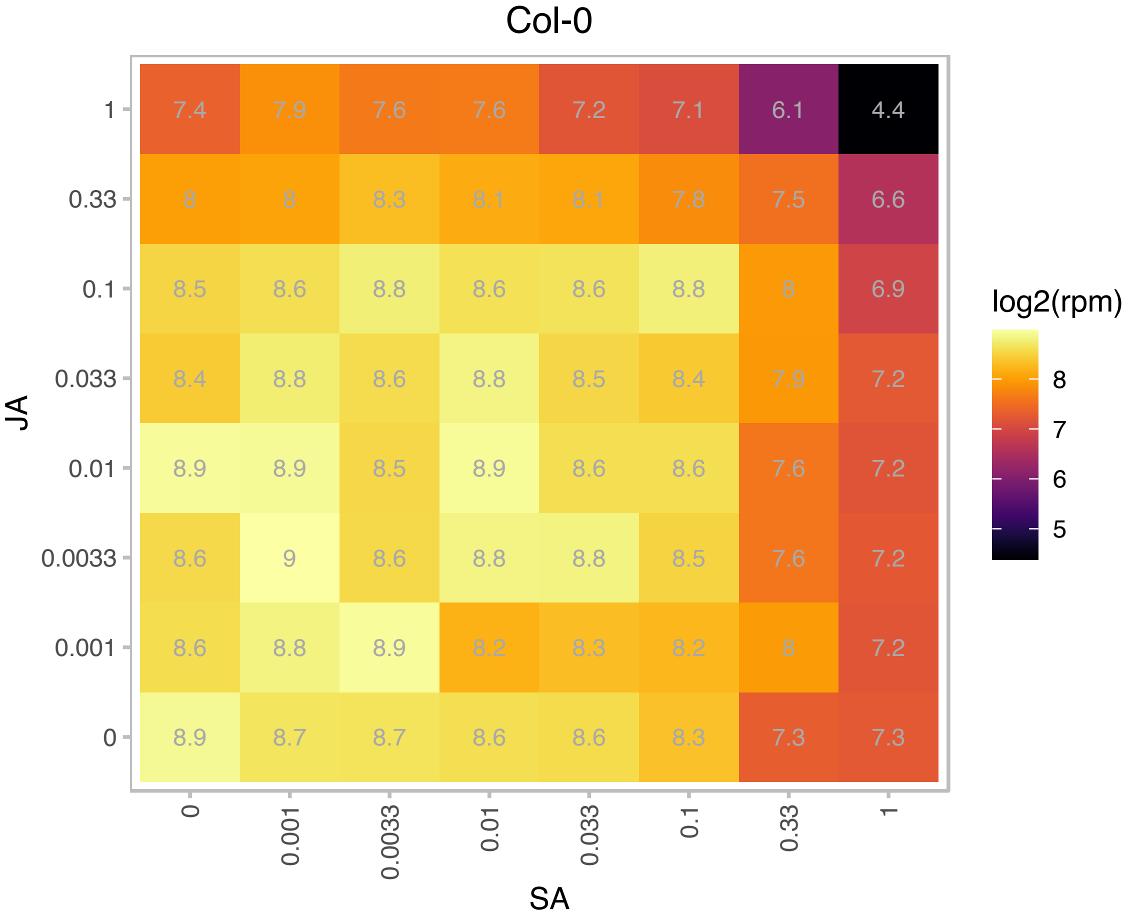


| | |
|---|---|
| Gene | AT3G55280 |
| is.expressed_genes | TRUE |
| is.responsive_genes | TRUE |
| is.apcluster | F-07 |

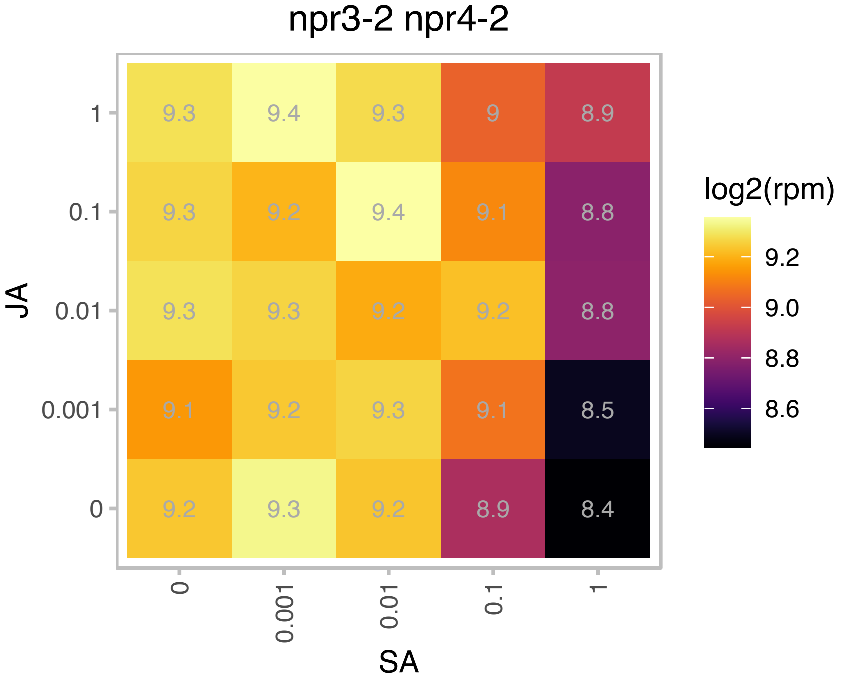


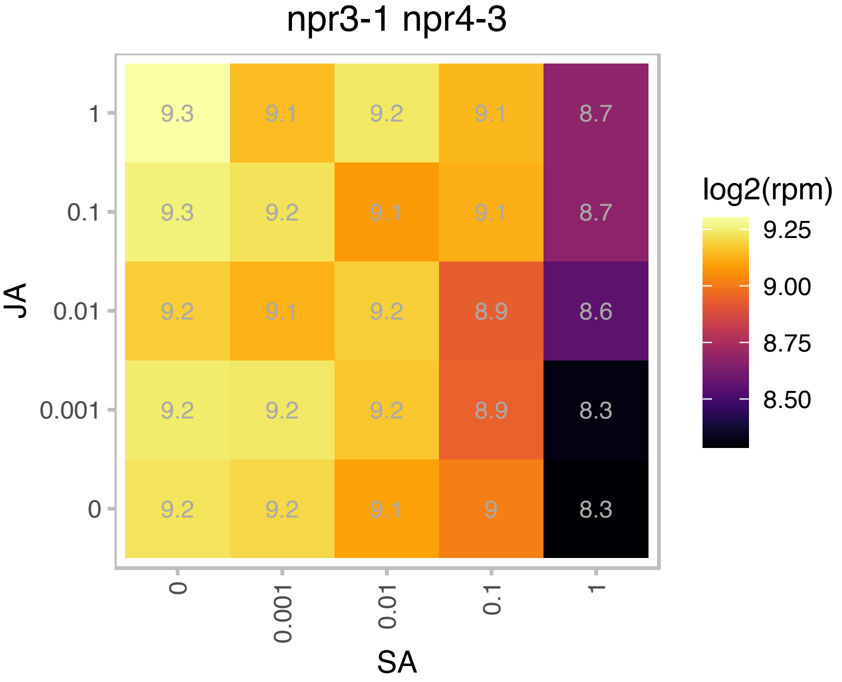

# AT3G55750.1 : NA

Ribosomal protein L35Ae family protein

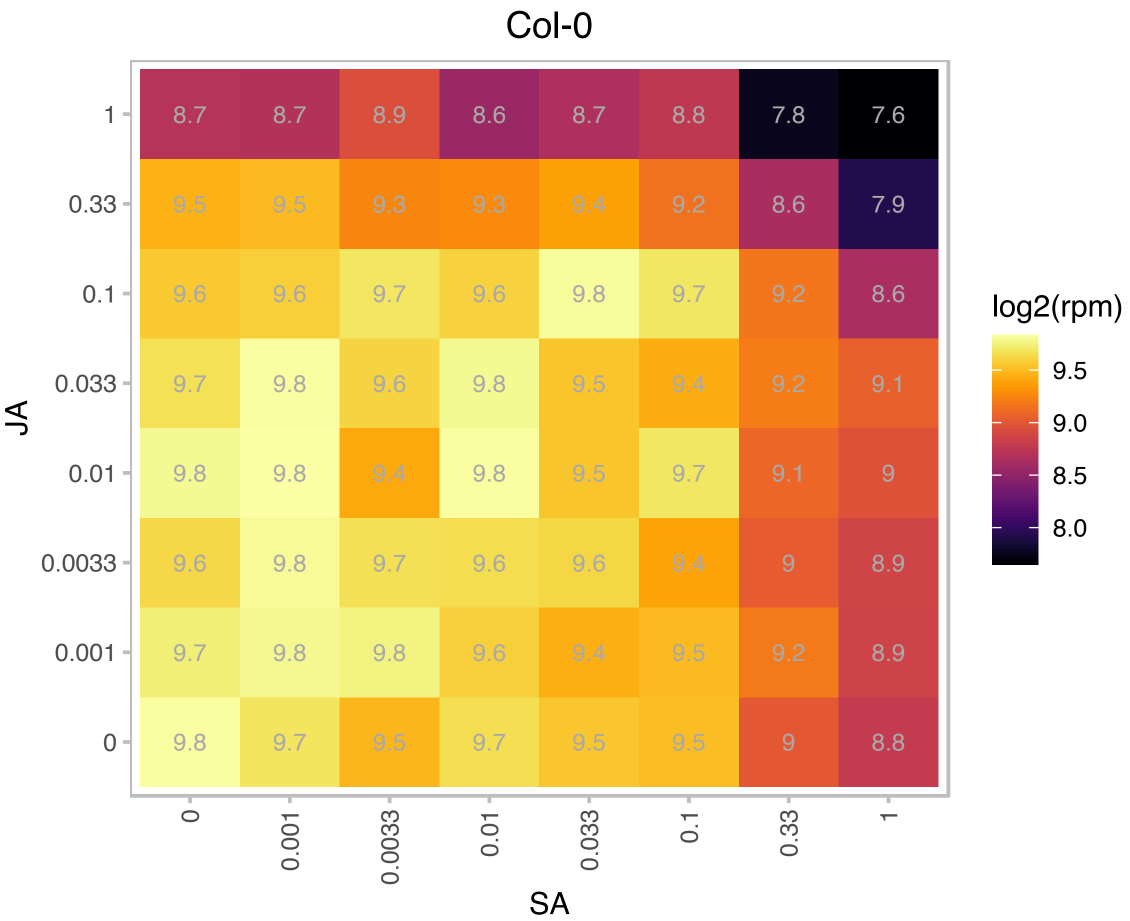


| | |
|---|---|
| Gene | AT3G55750 |
| is.expressed_genes | TRUE |
| is.responsive_genes | TRUE |
| is.apcluster | F-07 |

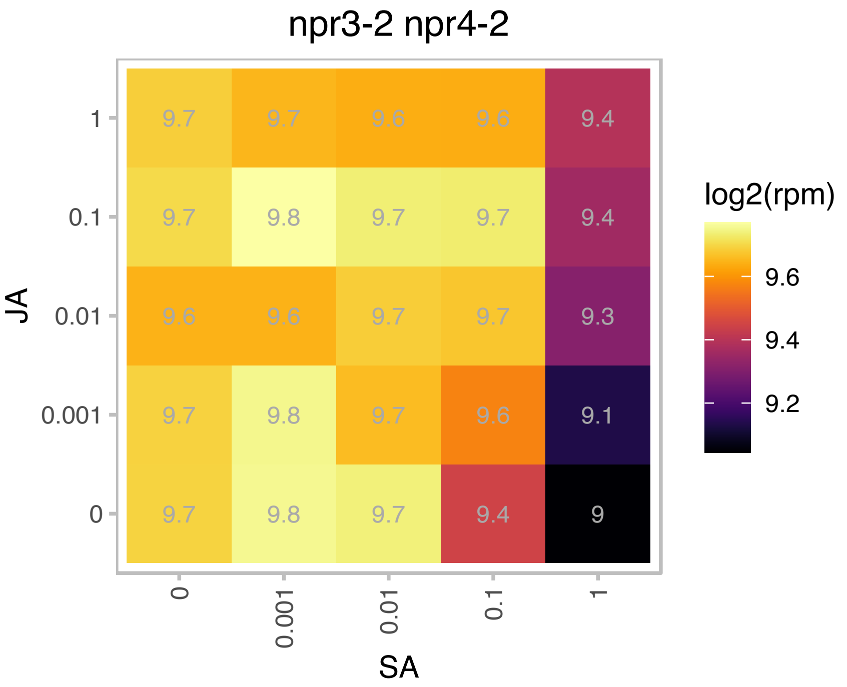


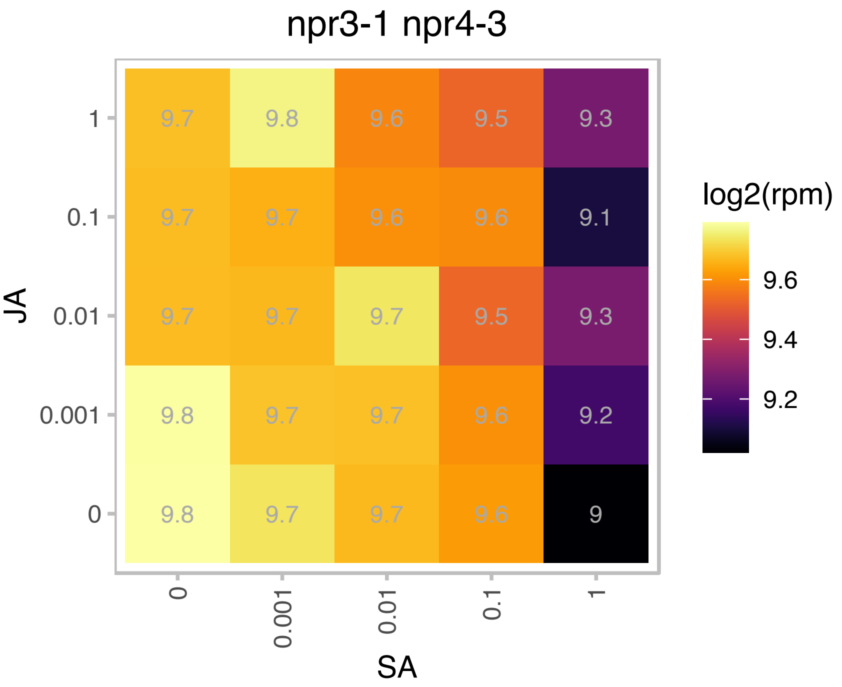

# AT3G56340.1 : NA

Ribosomal protein S26e family protein

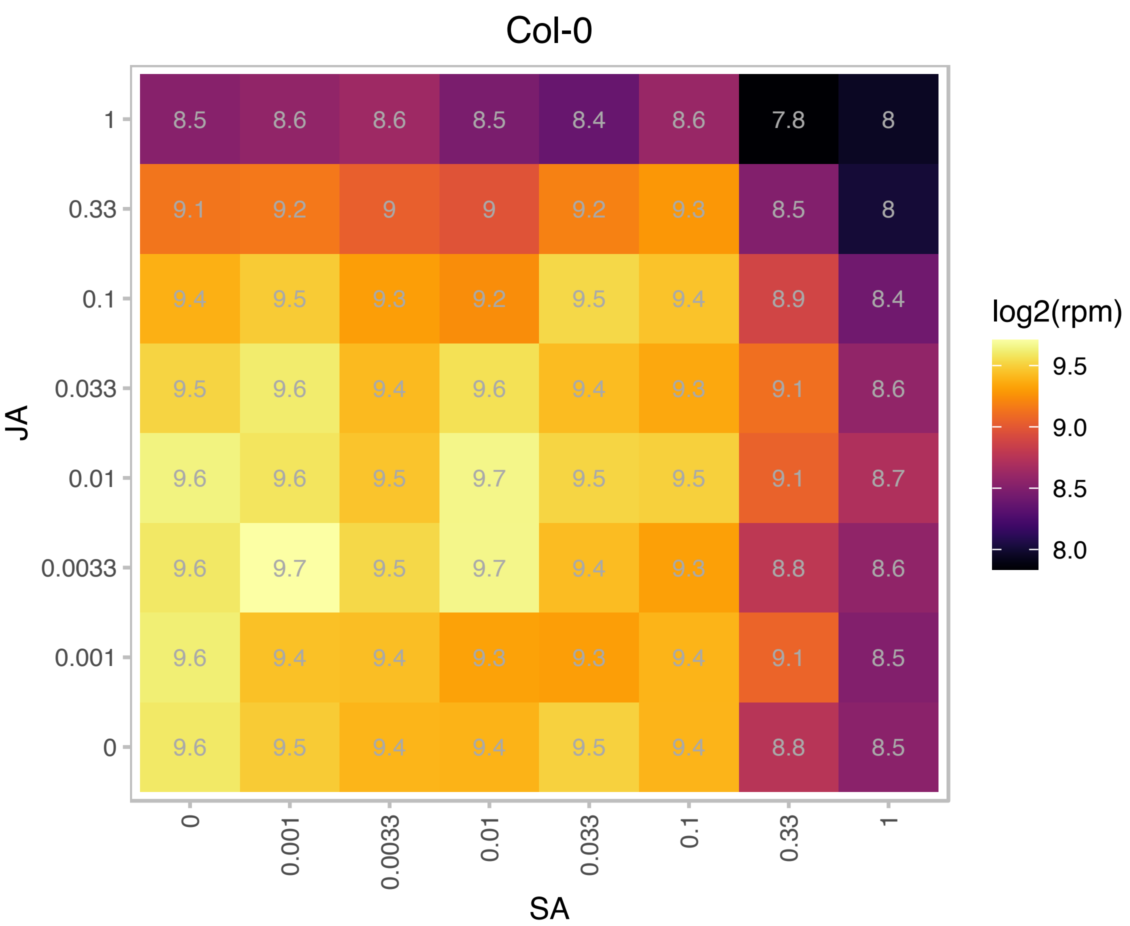


| | |
|---|---|
| Gene | AT3G56340 |
| is.expressed_genes | TRUE |
| is.responsive_genes | TRUE |
| is.apcluster | F-07 |

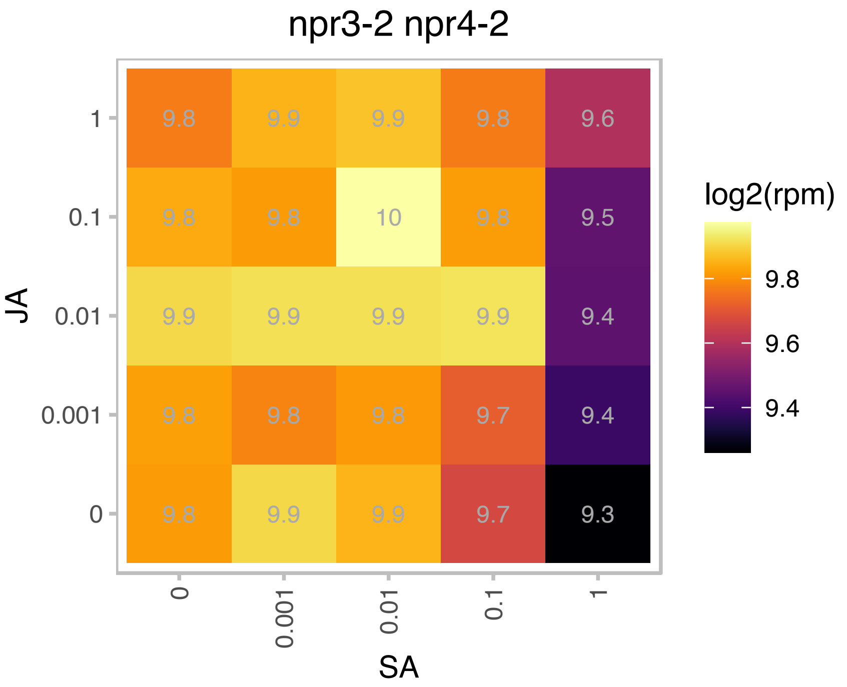


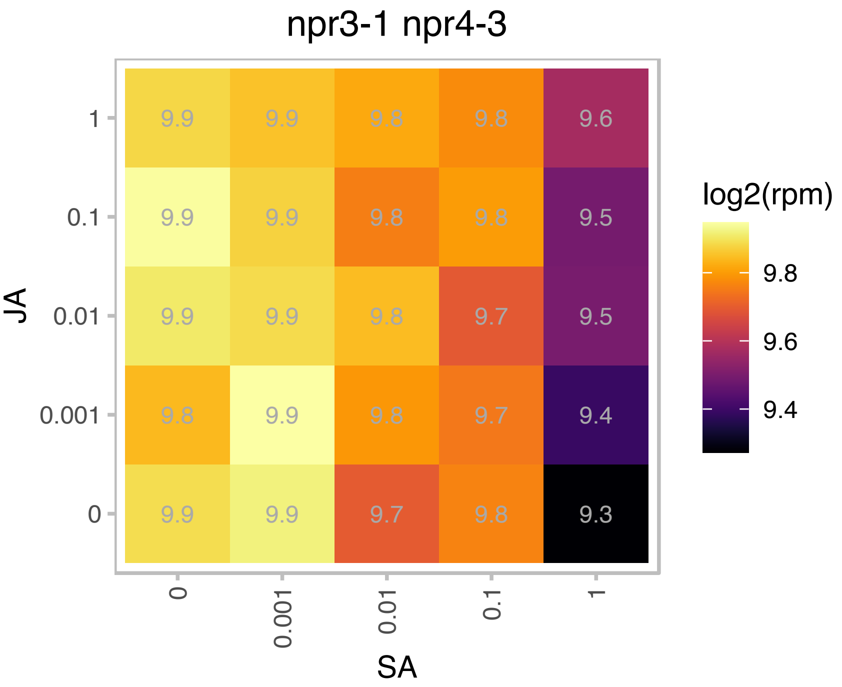

# AT3G57490.1 : NA

Ribosomal protein S5 family protein

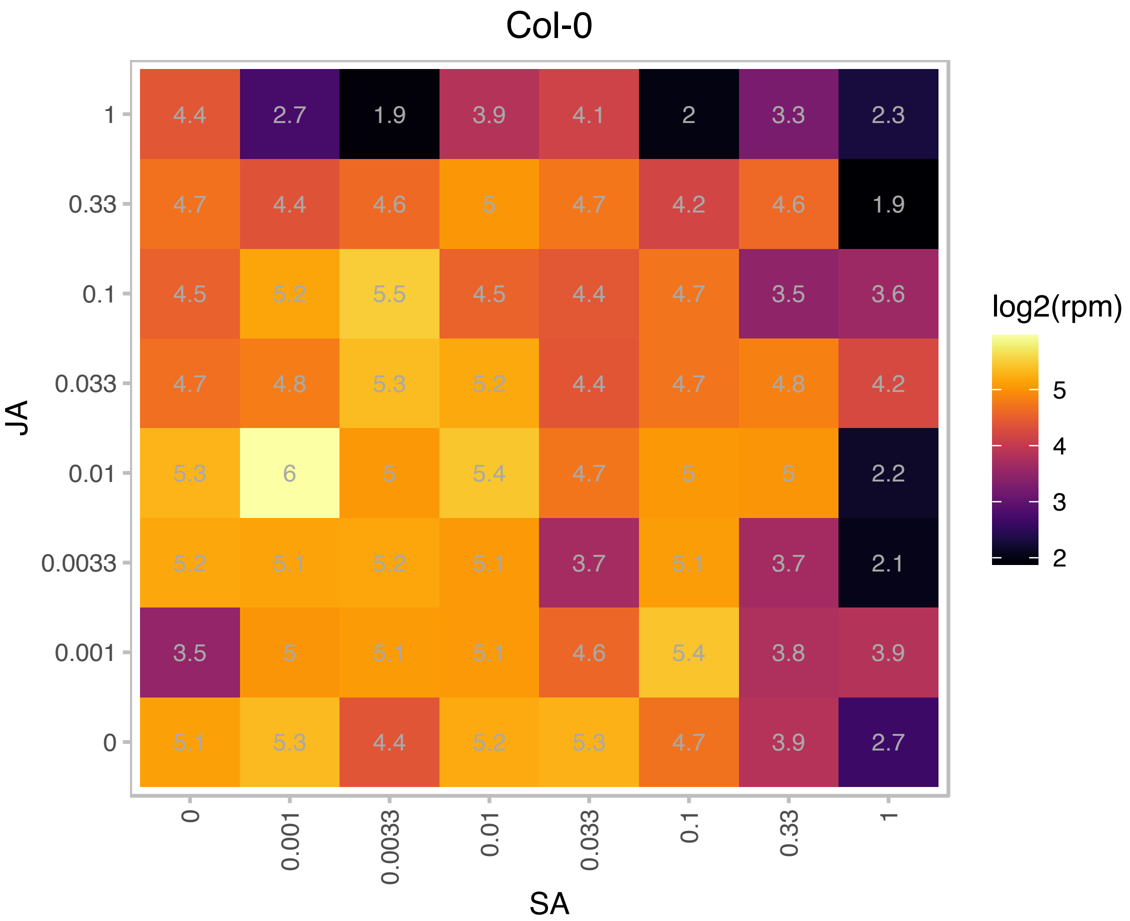


| | |
|---|---|
| Gene | AT3G57490 |
| is.expressed_genes | TRUE |
| is.responsive_genes | TRUE |
| is.apcluster | F-07 |

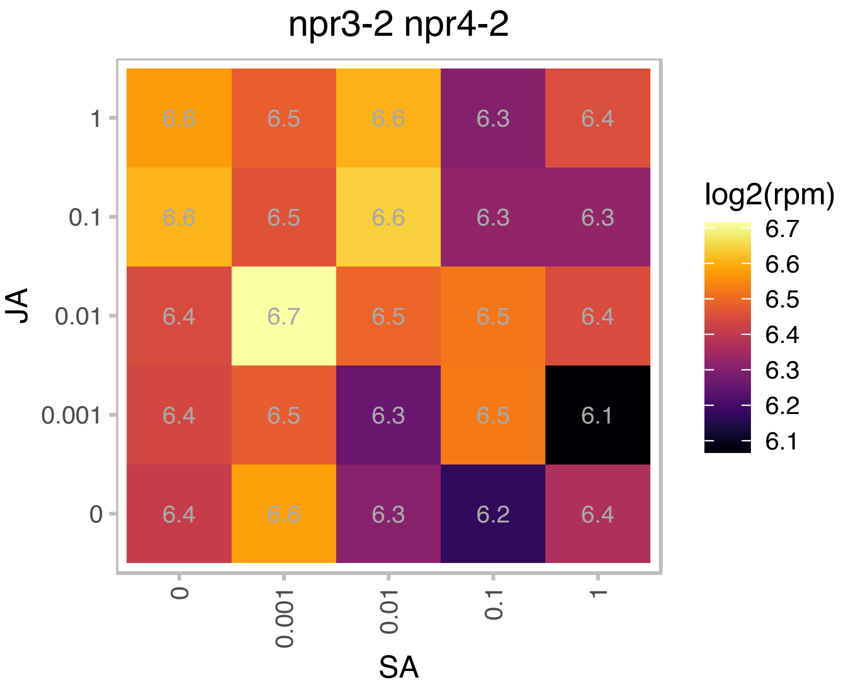


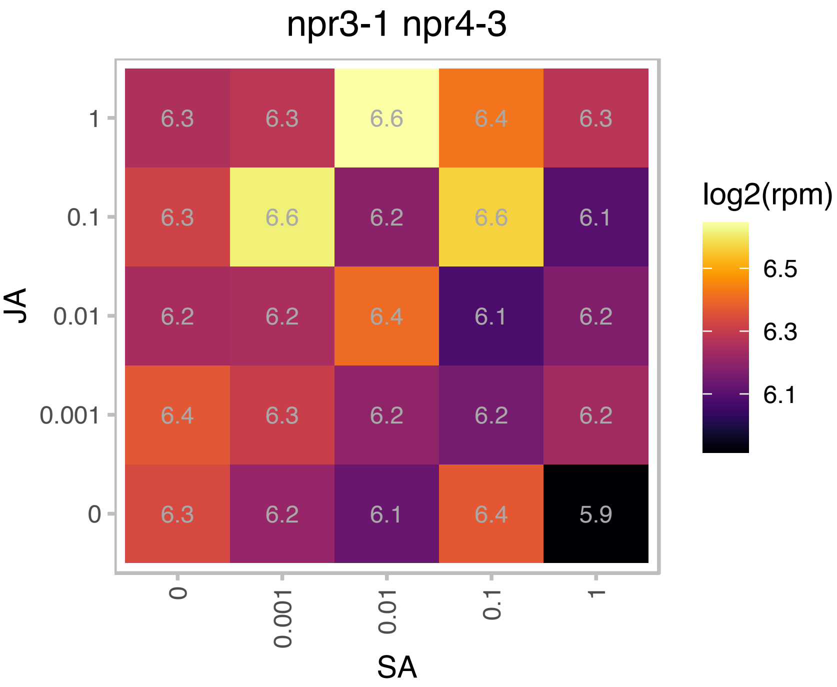

# AT3G58700.1 : NA

Ribosomal L5P family protein

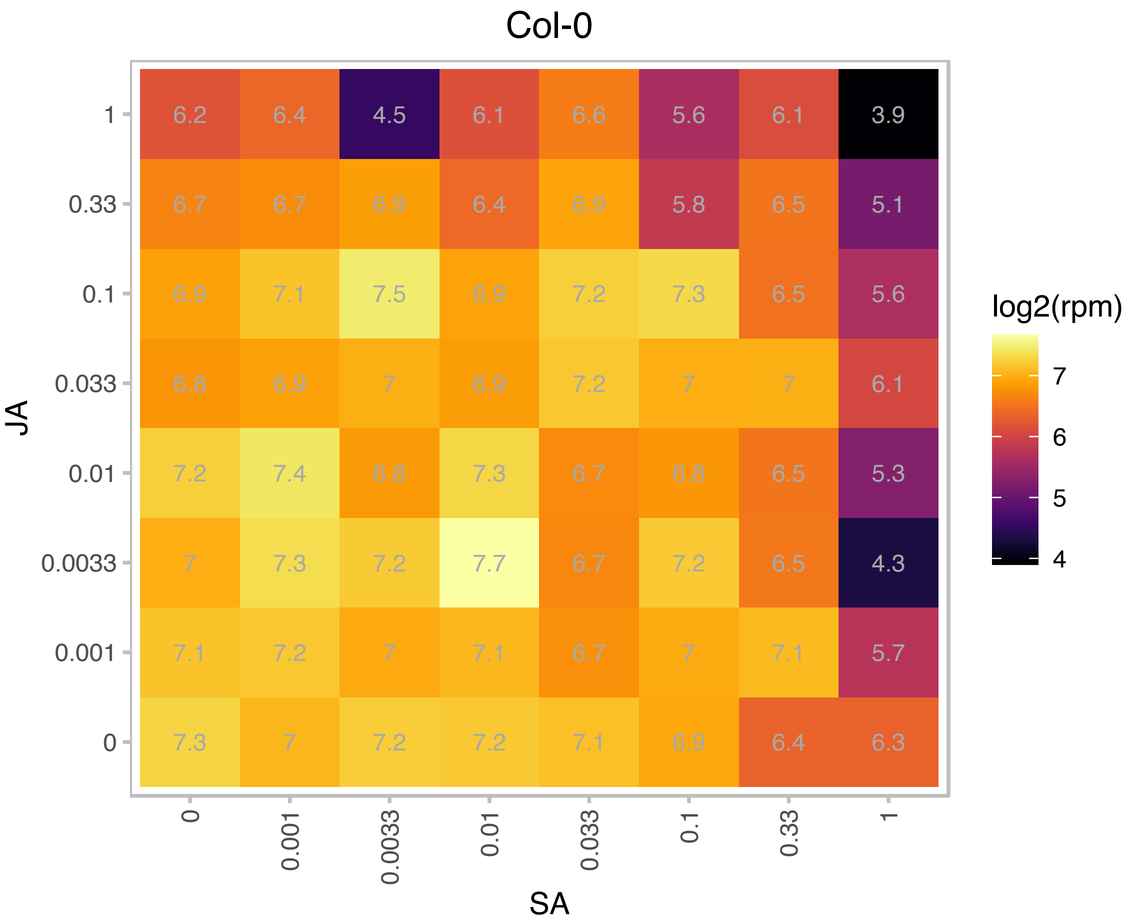


| | |
|---|---|
| Gene | AT3G58700 |
| is.expressed_genes | TRUE |
| is.responsive_genes | TRUE |
| is.apcluster | F-07 |

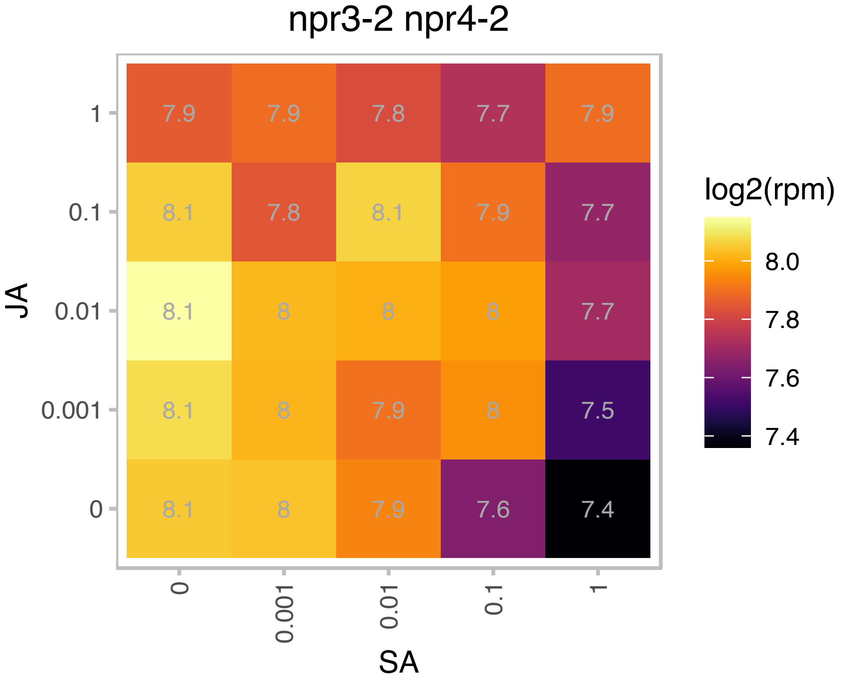


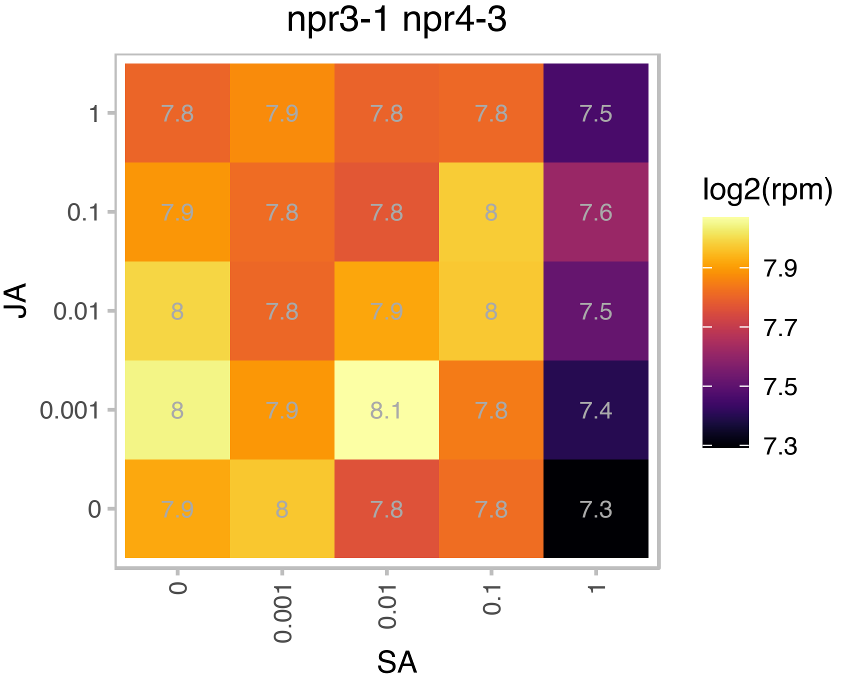

### AT3G58990.1 : IPMI1

isopropylmalate isomerase 1

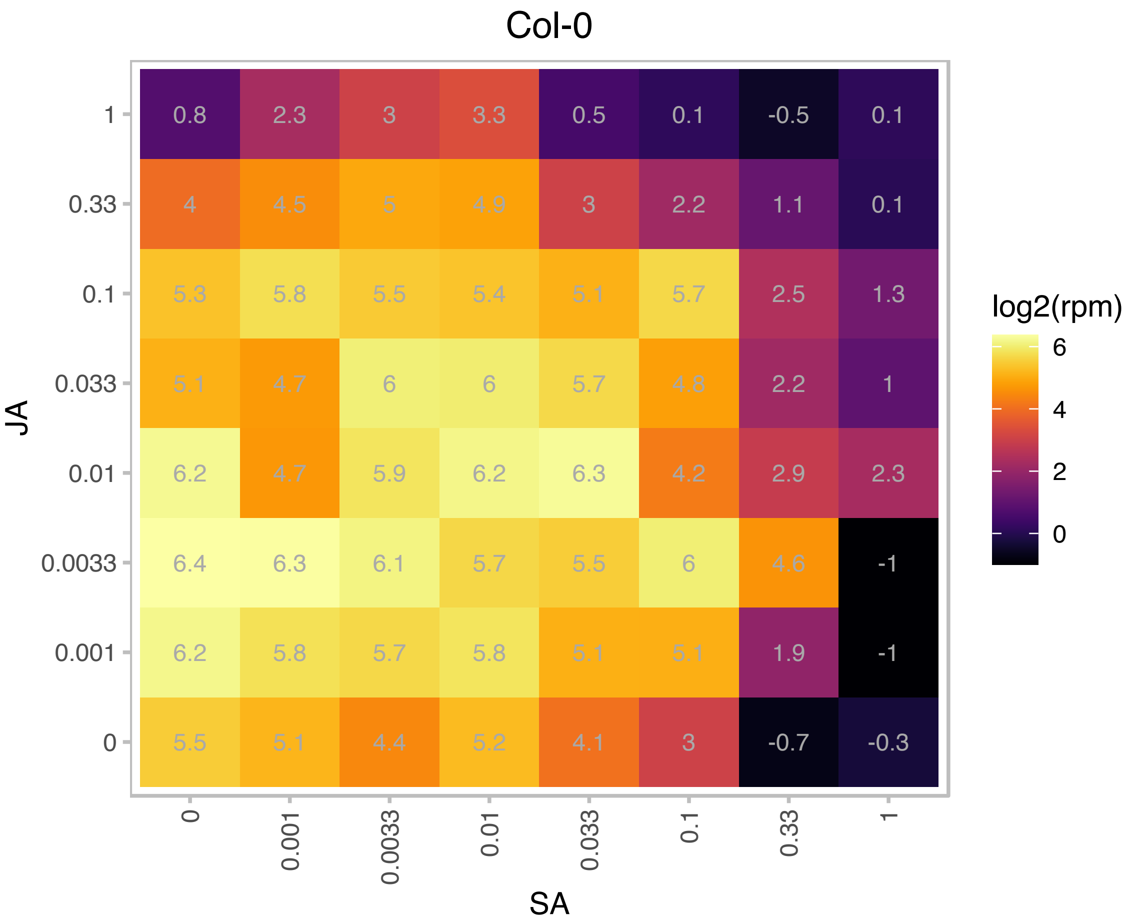


| | |
|---|---|
| Gene | AT3G58990 |
| is.expressed_genes | TRUE |
| is.responsive_genes | TRUE |
| is.apcluster | F-07 |

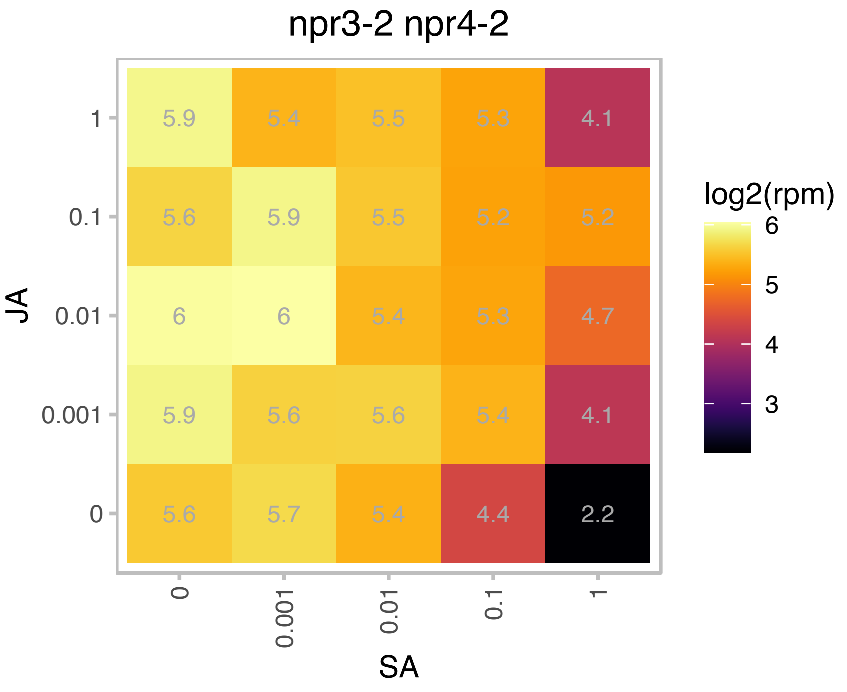


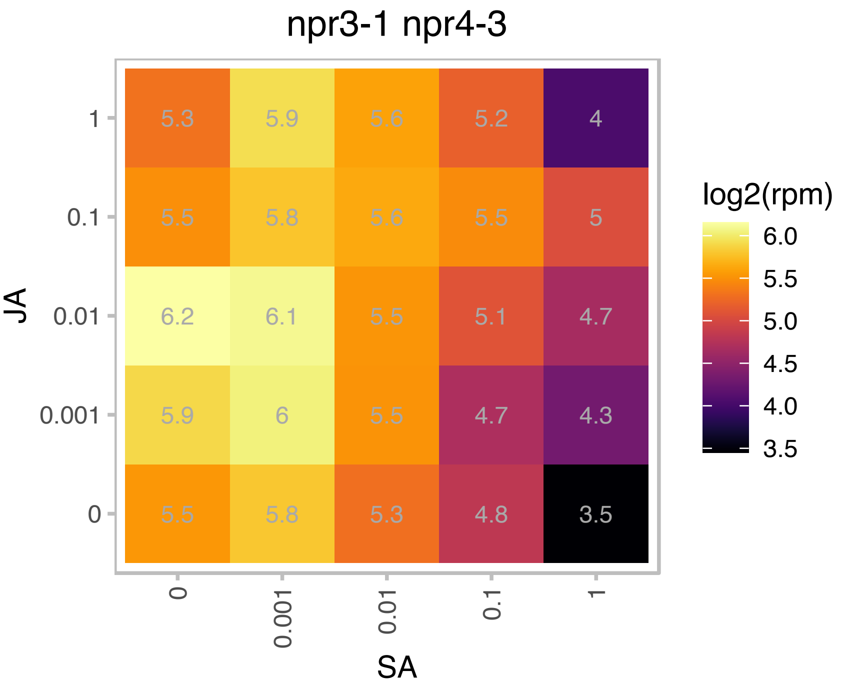

# AT3G59540.1 : NA

Ribosomal L38e protein family

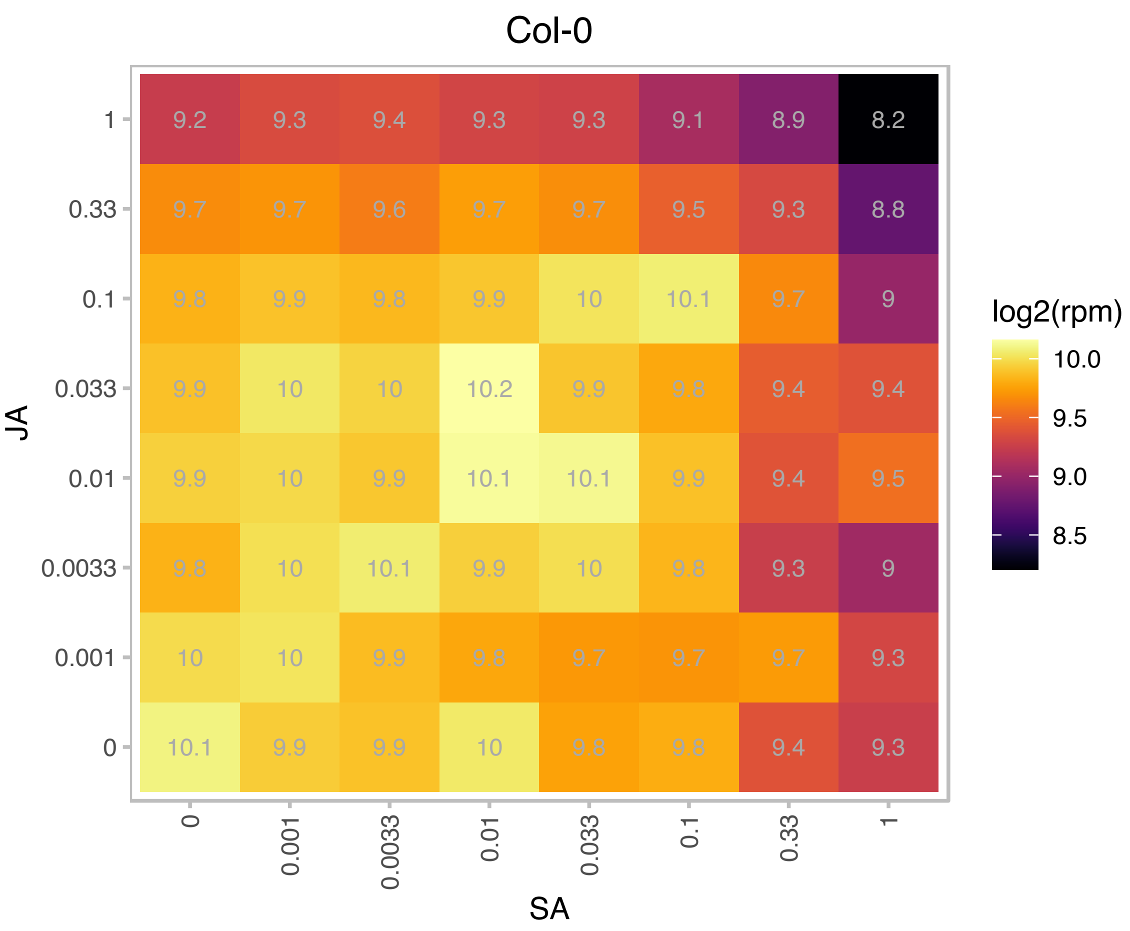


| | |
|---|---|
| Gene | AT3G59540 |
| is.expressed_genes | TRUE |
| is.responsive_genes | TRUE |
| is.apcluster | F-07 |

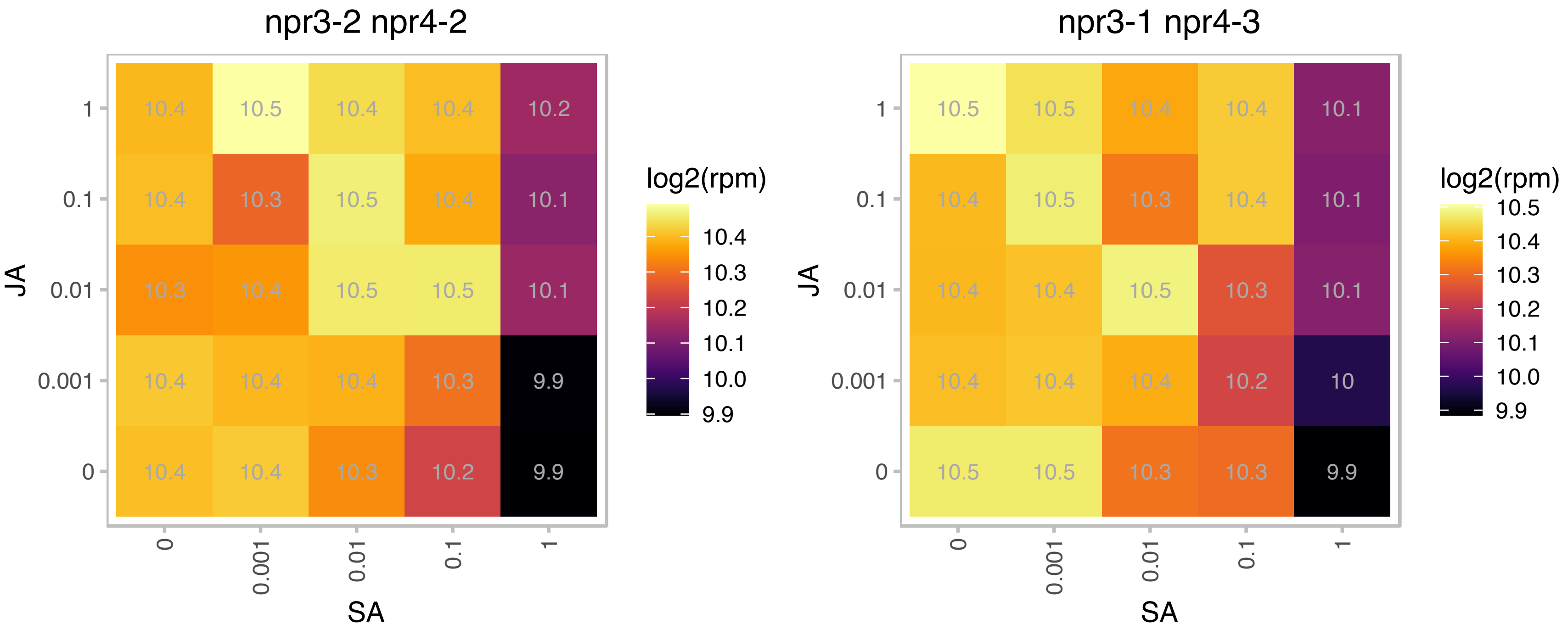

# AT3G59650.1 : NA

mitochondrial ribosomal protein L51/S25/CI-B8 family protein

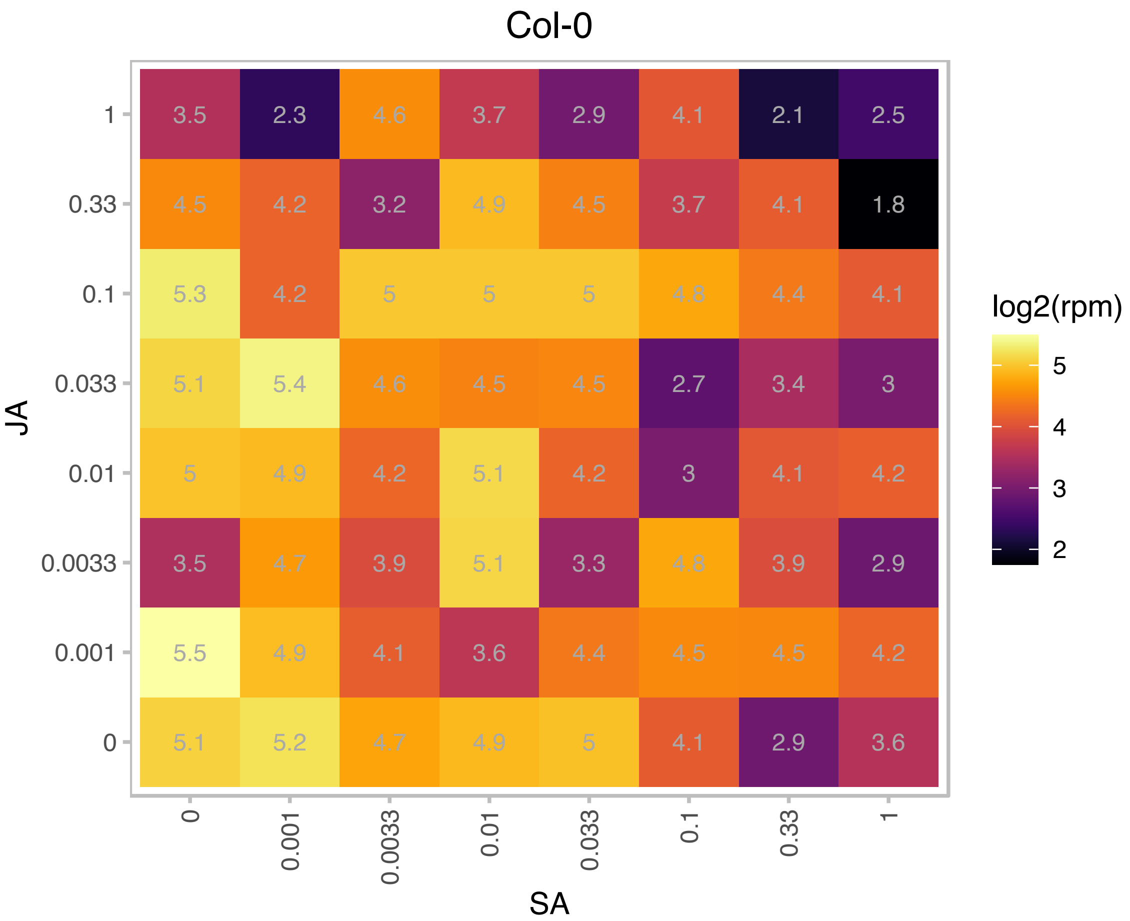


| | |
|---|---|
| Gene | AT3G59650 |
| is.expressed_genes | TRUE |
| is.responsive_genes | TRUE |
| is.apcluster | F-07 |

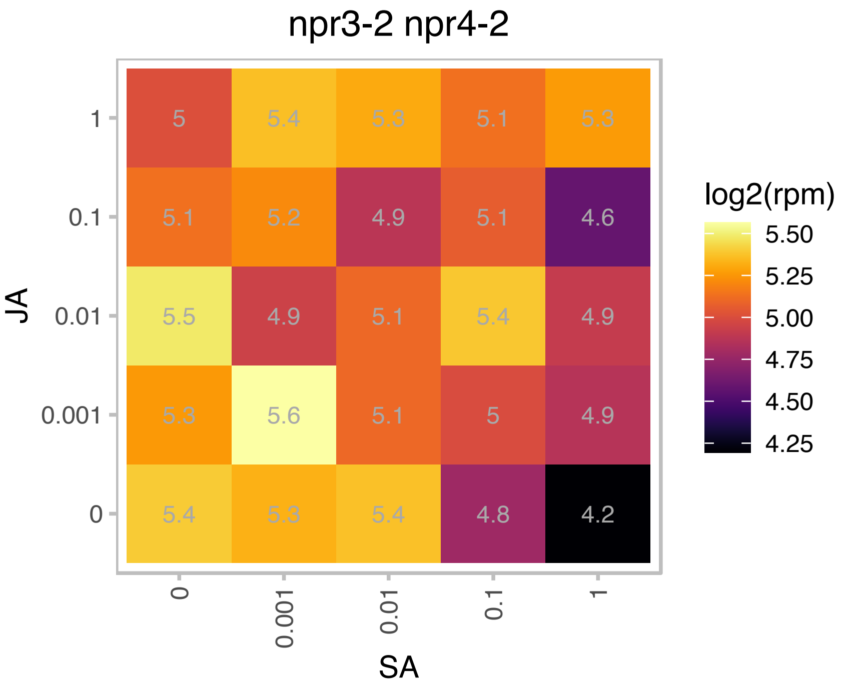


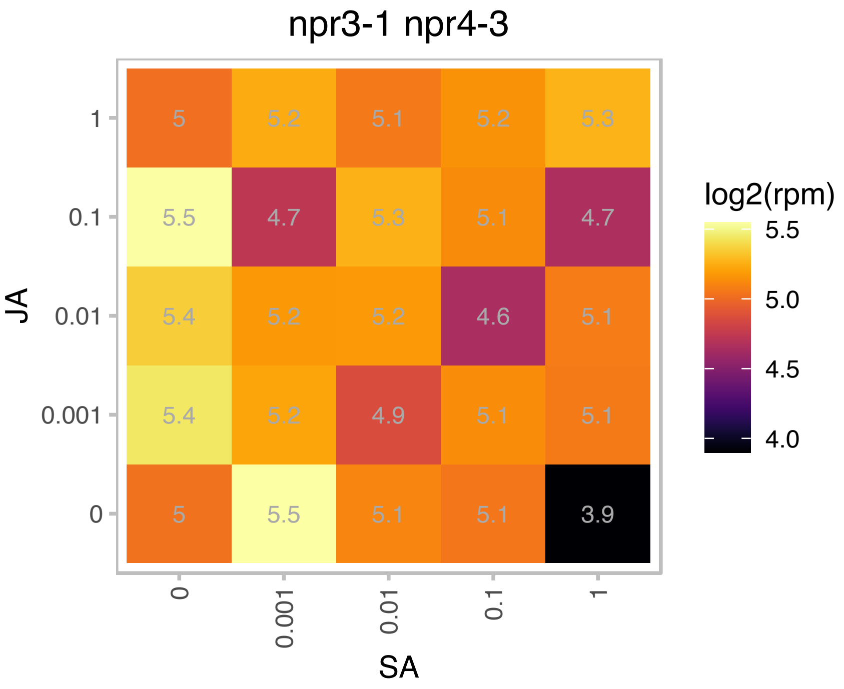

# AT3G60210.1 : NA

GroES-like family protein

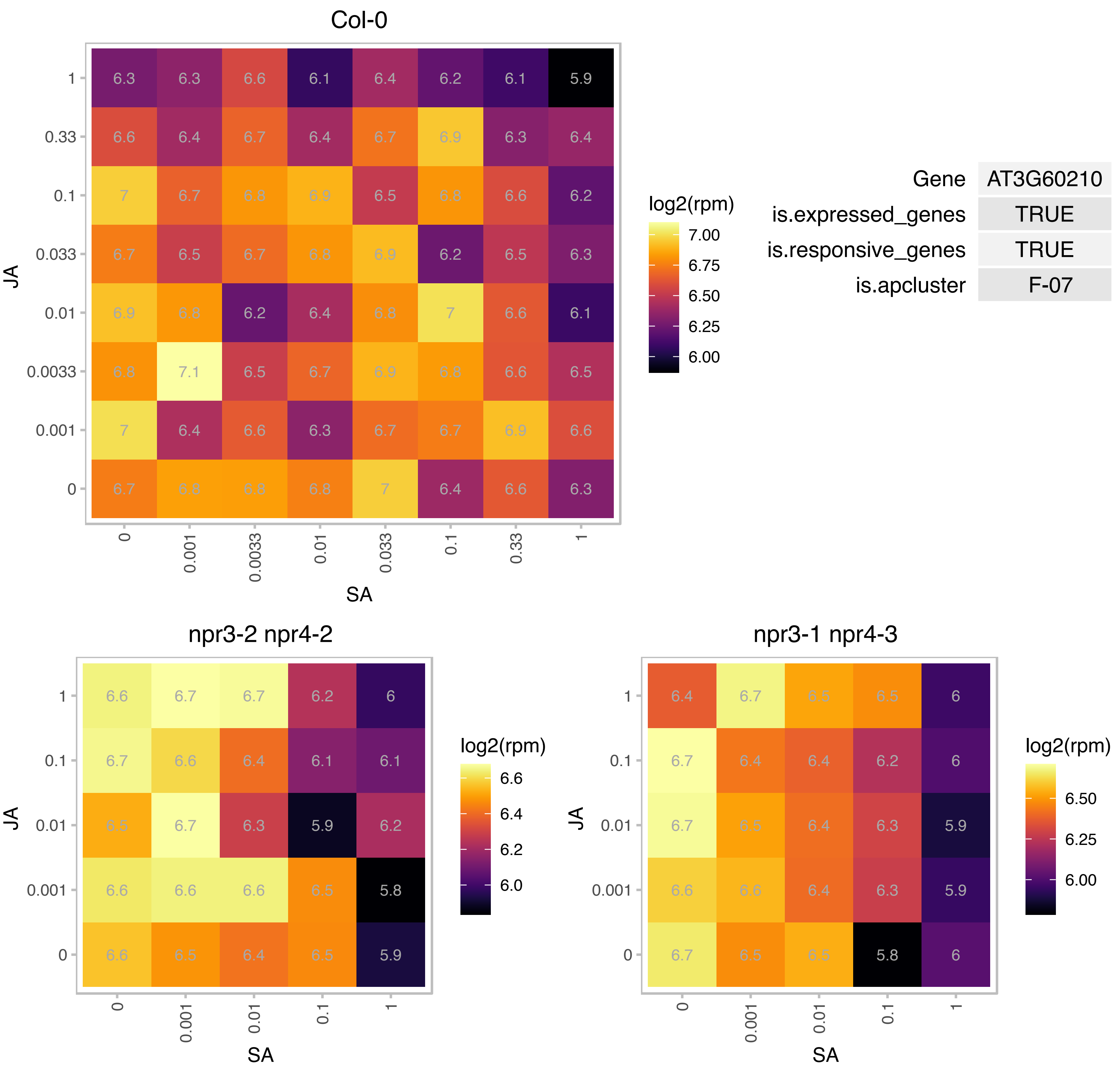


| | |
|---|---|
| Gene | AT3G60210 |
| is.expressed_genes | TRUE |
| is.responsive_genes | TRUE |
| is.apcluster | F-07 |

# AT3G60245.1 : NA

Zinc-binding ribosomal protein family protein

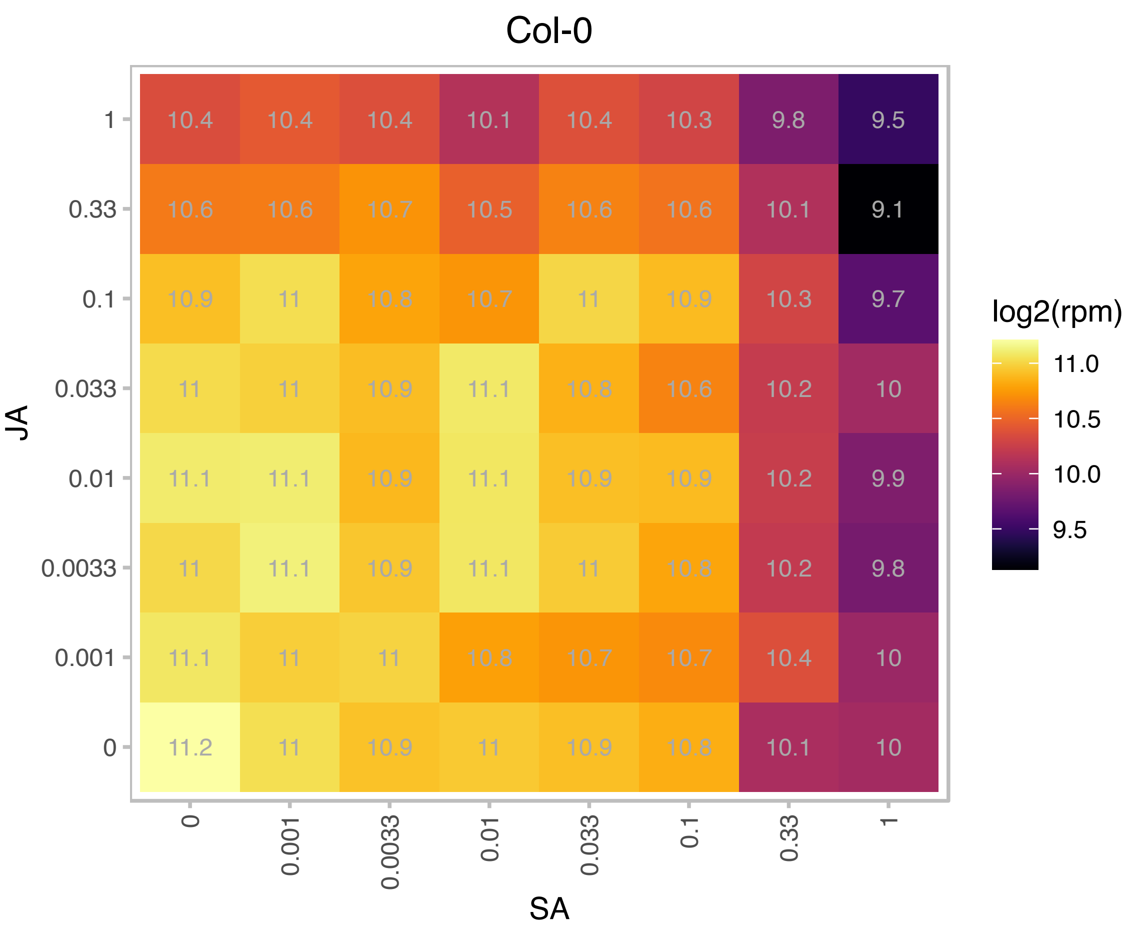


| | |
|---|---|
| Gene | AT3G60245 |
| is.expressed_genes | TRUE |
| is.responsive_genes | TRUE |
| is.apcluster | F-07 |

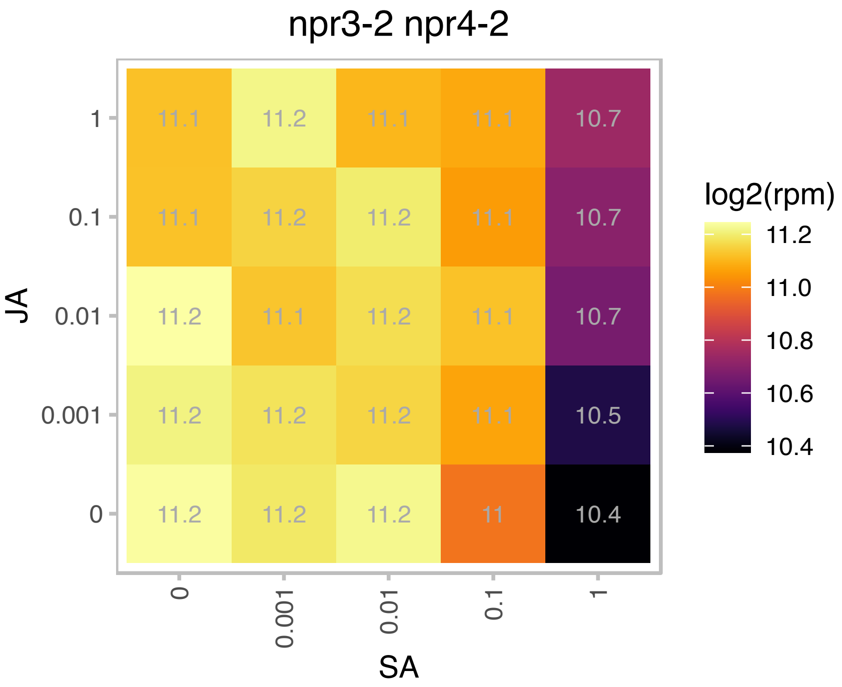


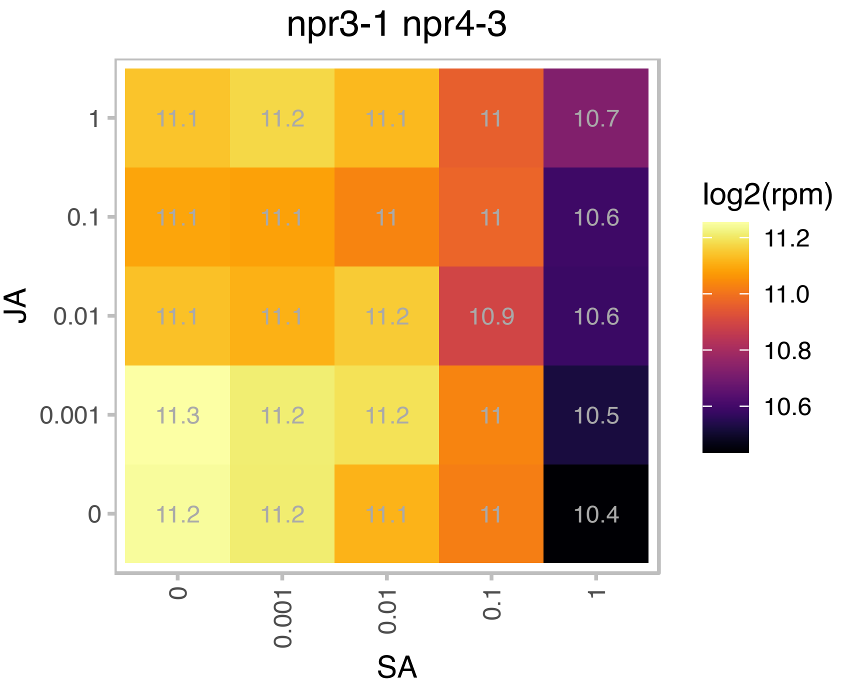

# AT3G60480.1 : NA

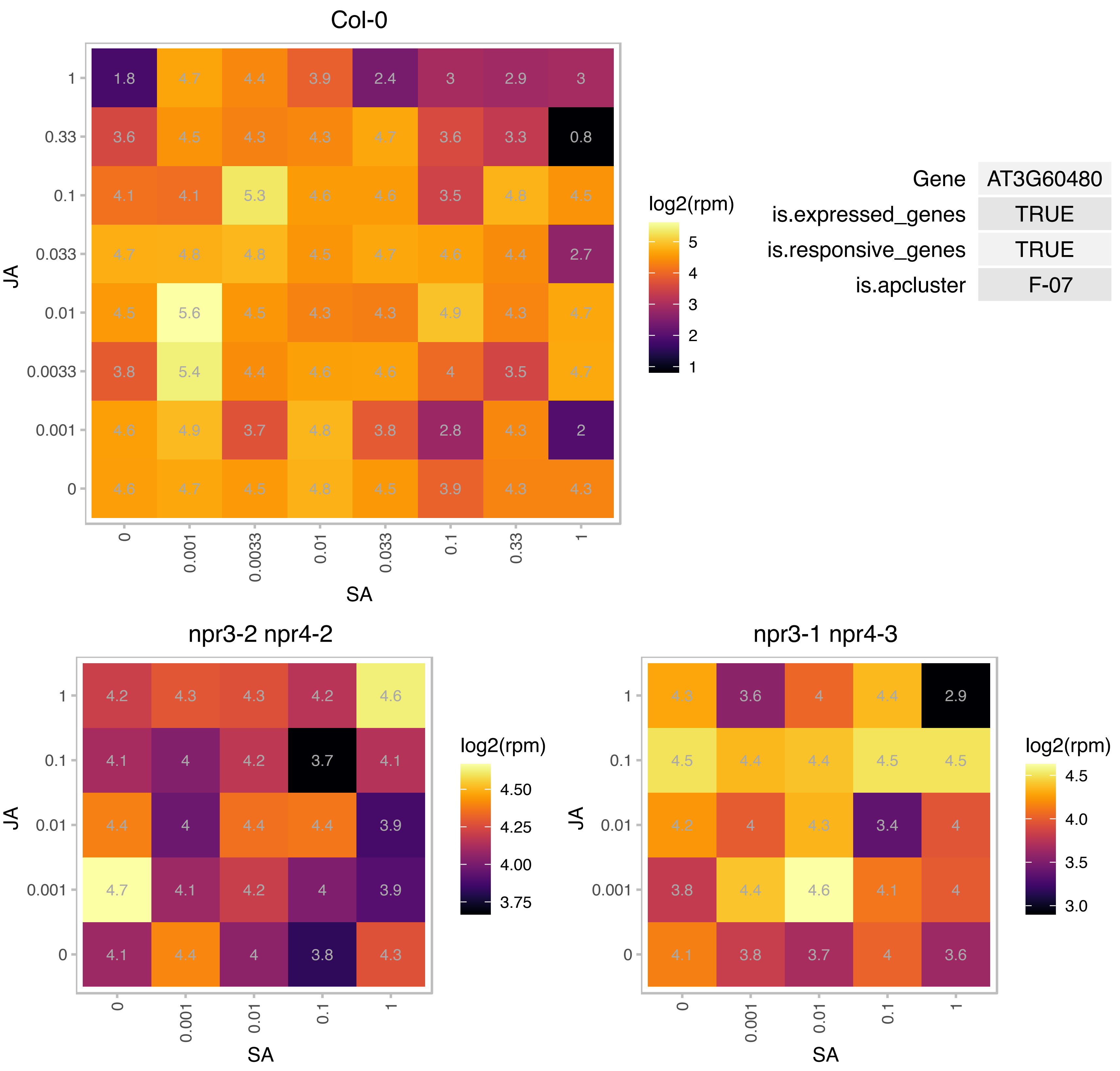


| | |
|---|---|
| Gene | AT3G60480 |
| is.expressed_genes | TRUE |
| is.responsive_genes | TRUE |
| is.apcluster | F-07 |

# AT3G60770.1 : NA

Ribosomal protein S13/S15

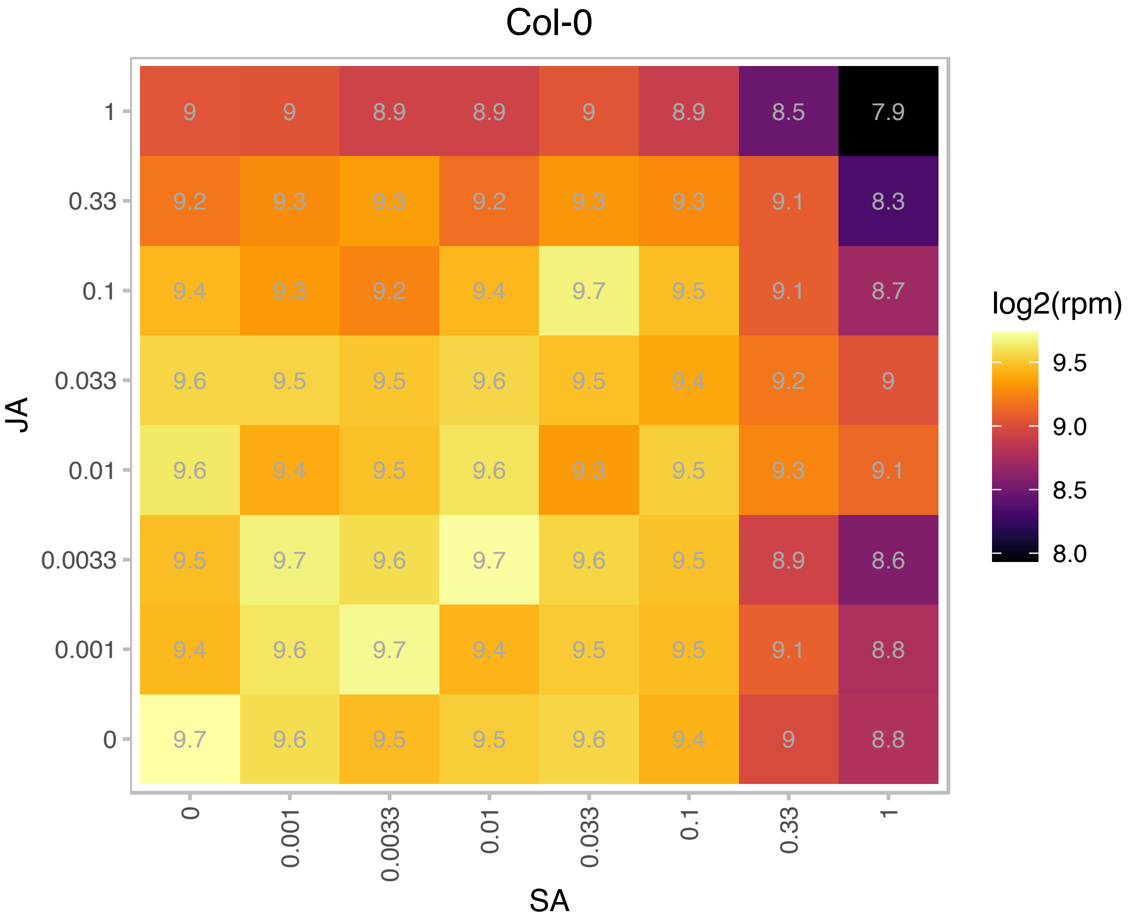


| | |
|---|---|
| Gene | AT3G60770 |
| is.expressed_genes | TRUE |
| is.responsive_genes | TRUE |
| is.apcluster | F-07 |

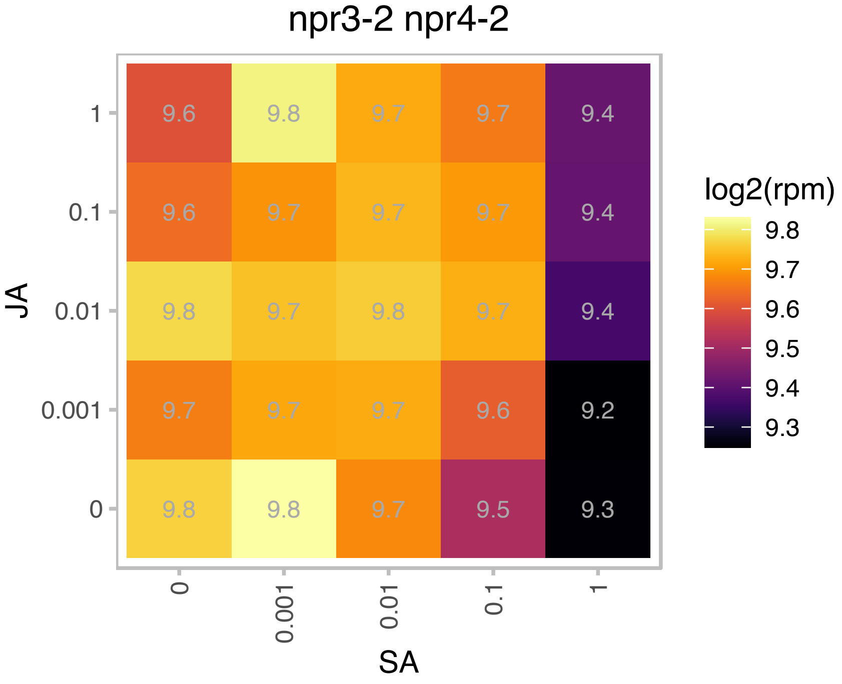


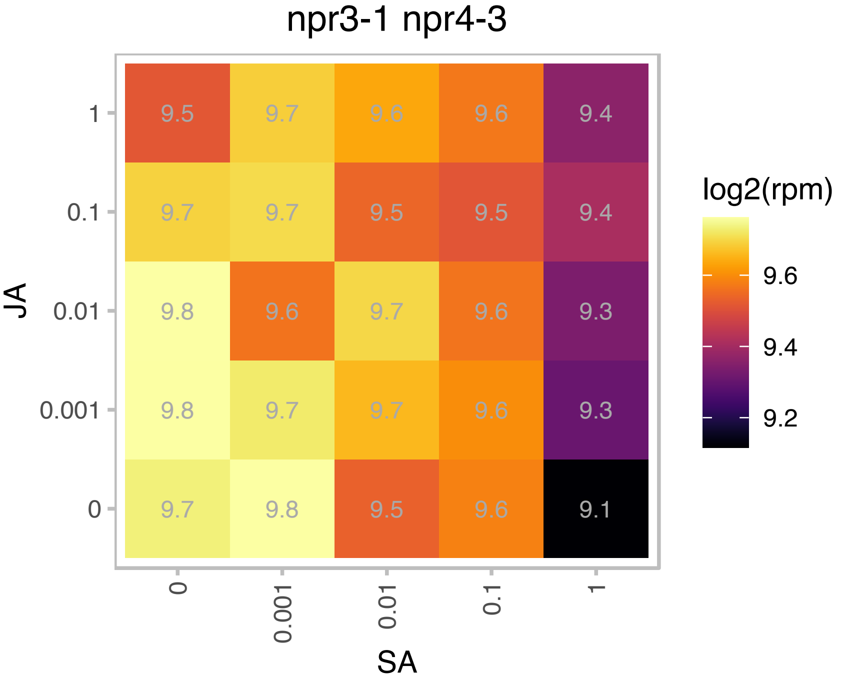

### AT3G61110.1 : ARS27A, RS27A

ribosomal protein S27

Arabidopsis ribosomal protein

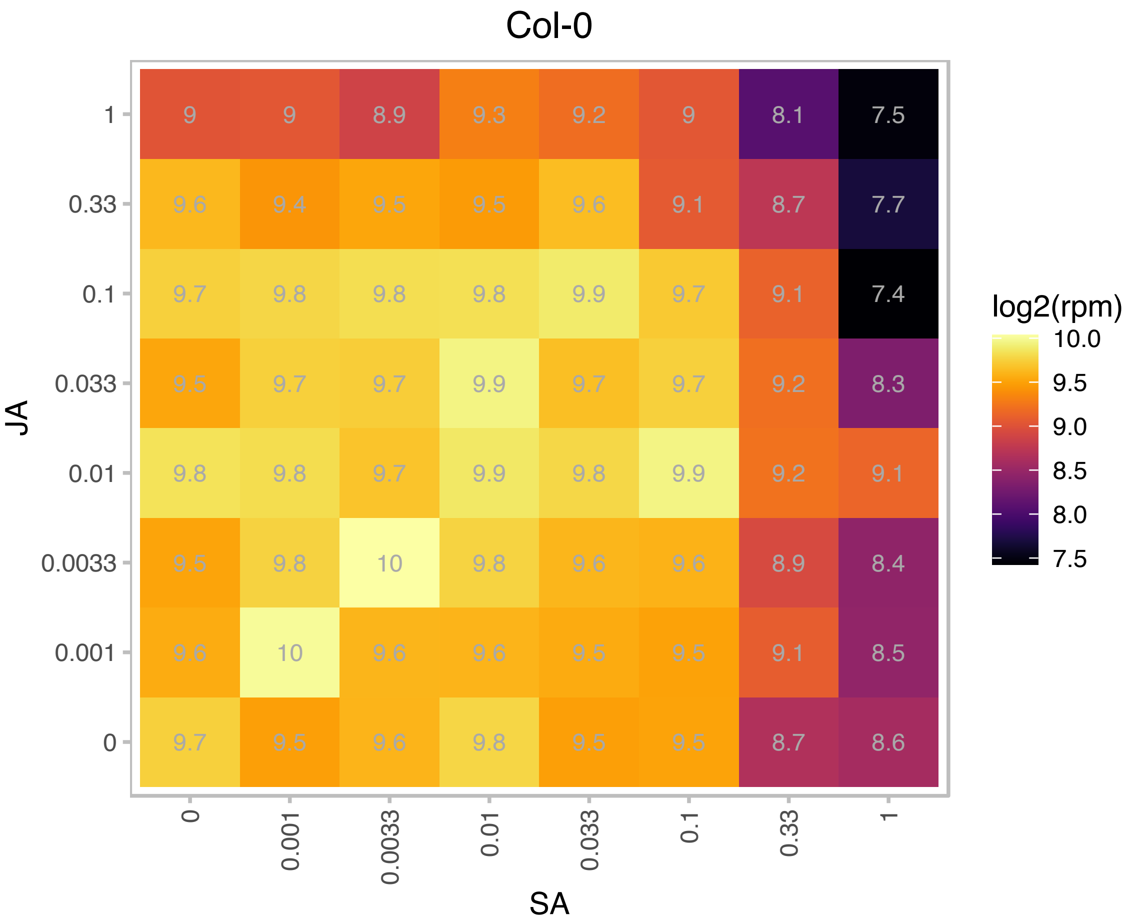


| | |
|---|---|
| Gene | AT3G61110 |
| is.expressed_genes | TRUE |
| is.responsive_genes | TRUE |
| is.apcluster | F-07 |

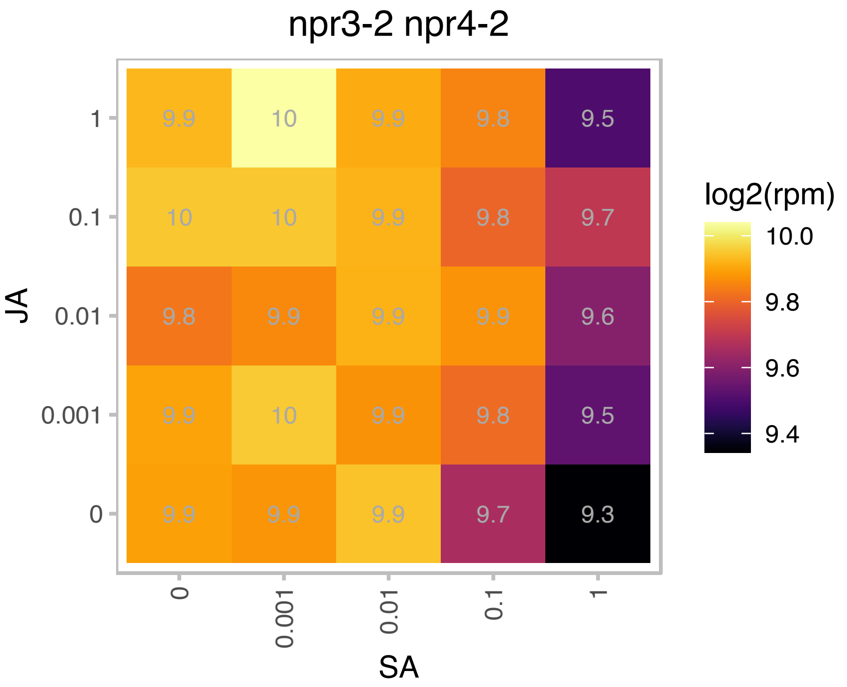


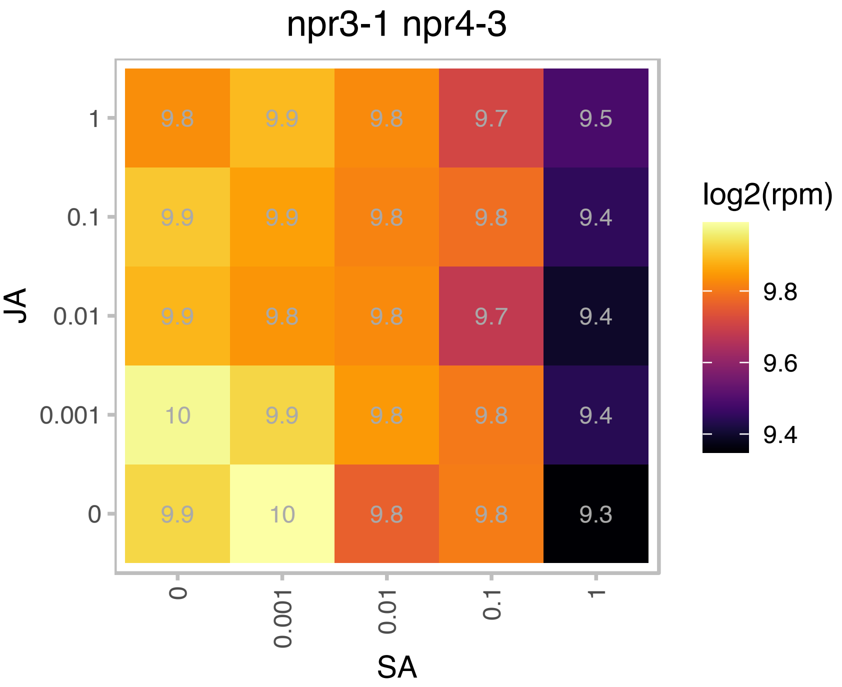

### AT3G61310.1 : AHL11

AT hook motif DNA-binding family protein

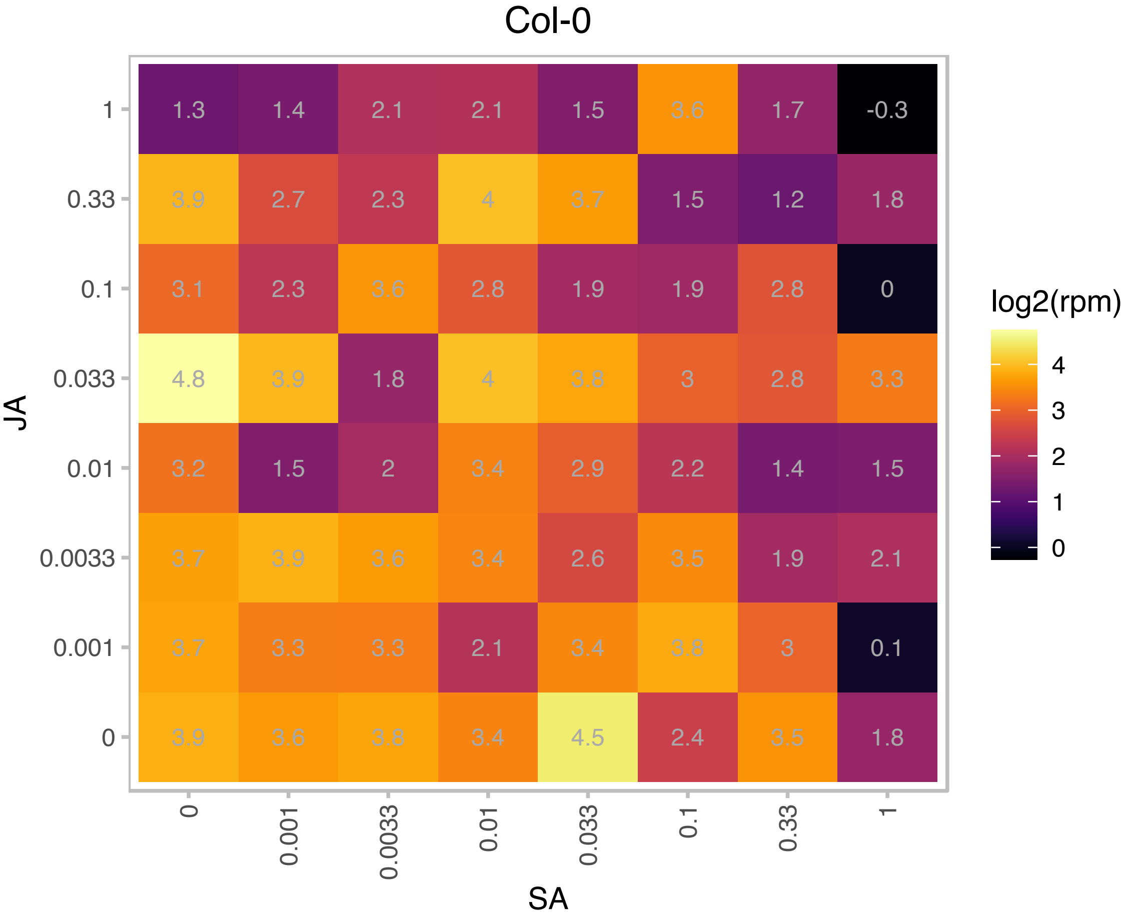


| | |
|---|---|
| Gene | AT3G61310 |
| is.expressed_genes | TRUE |
| is.responsive_genes | TRUE |
| is.apcluster | F-07 |

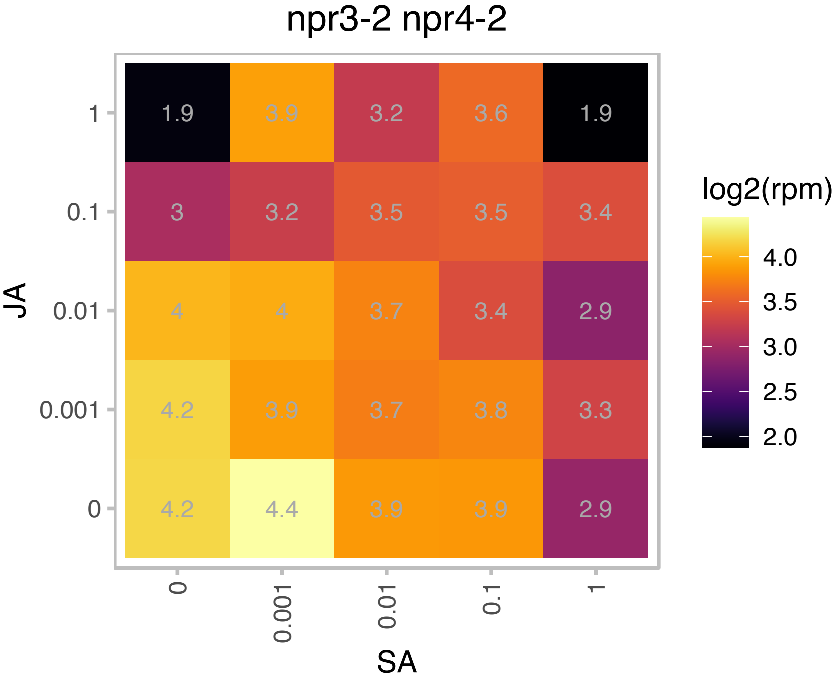


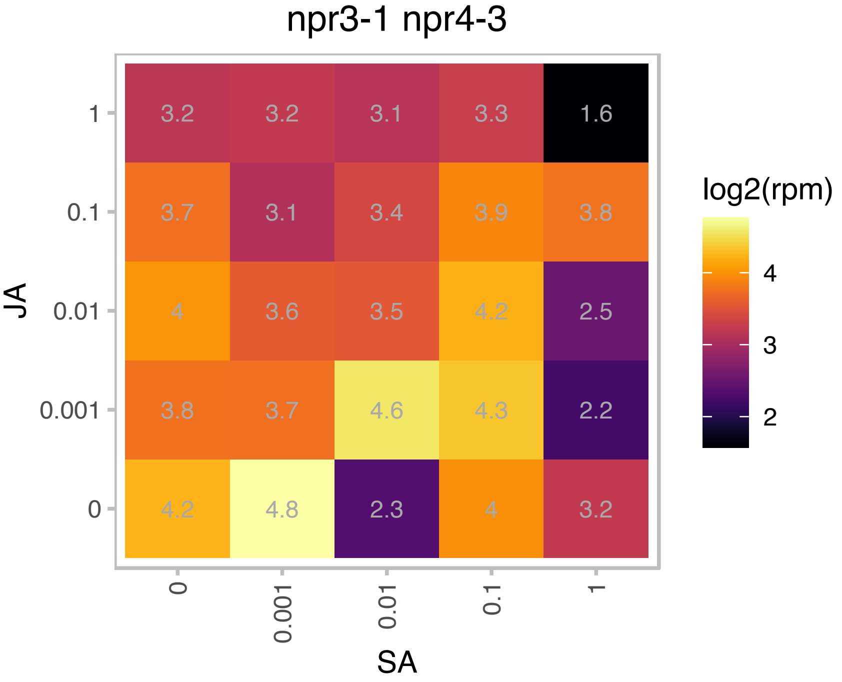

### AT3G62250.1 : UBQ5

ubiquitin 5

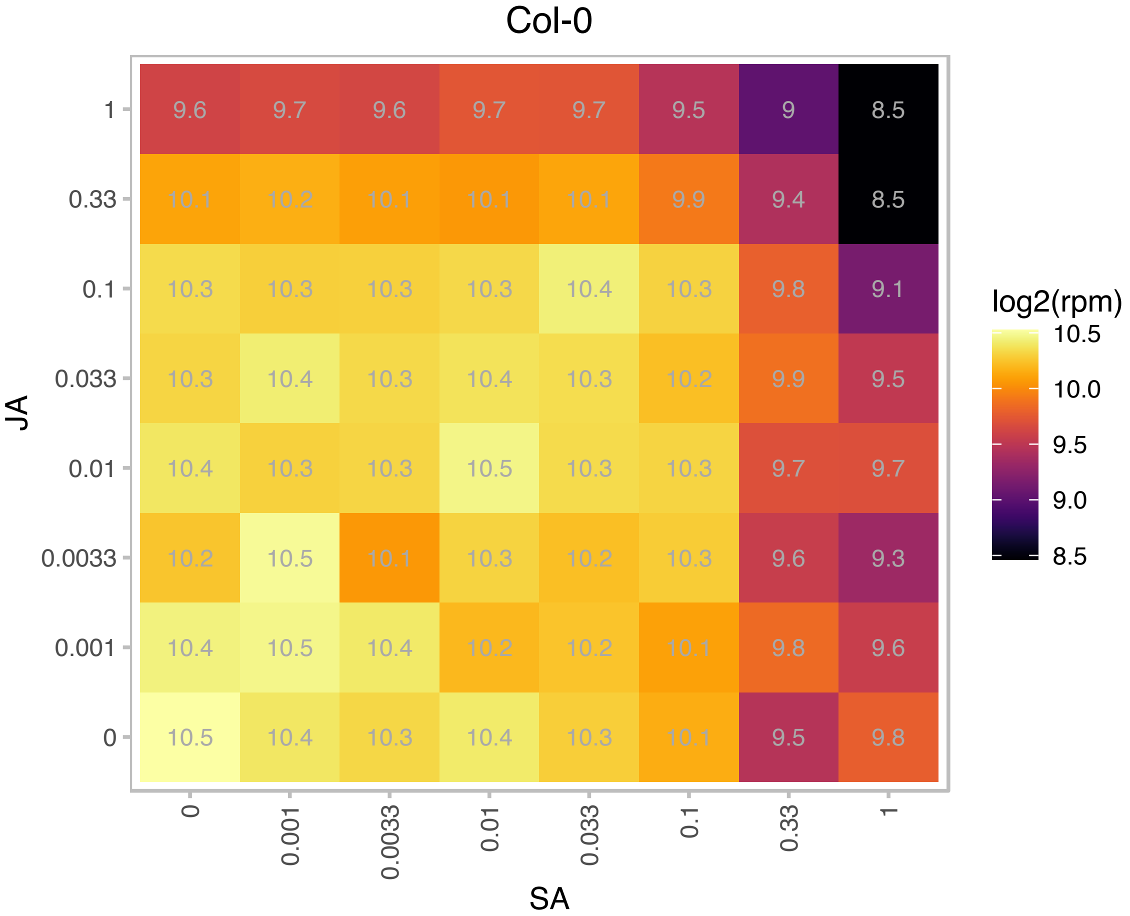


| | |
|---|---|
| Gene | AT3G62250 |
| is.expressed_genes | TRUE |
| is.responsive_genes | TRUE |
| is.apcluster | F-07 |

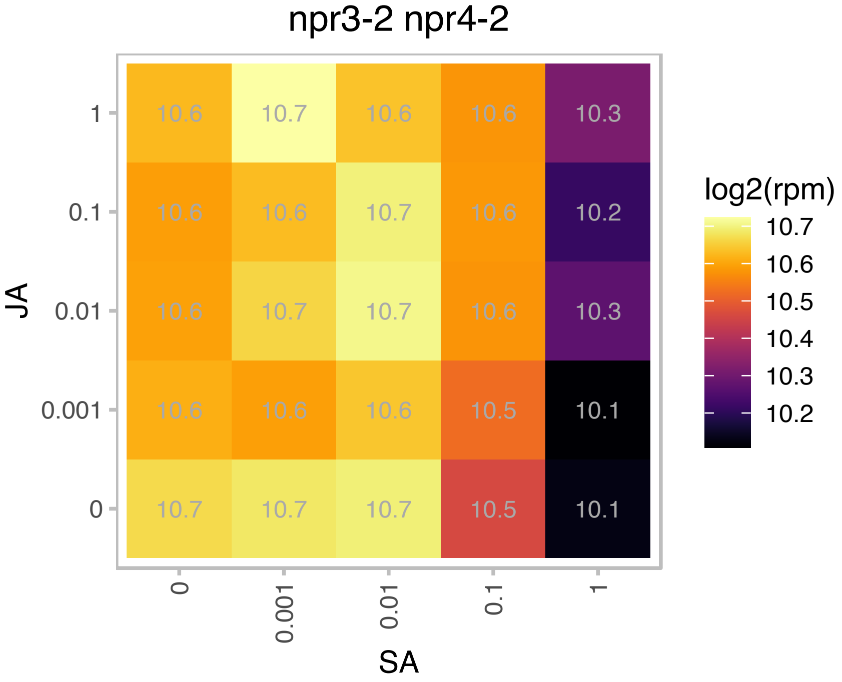


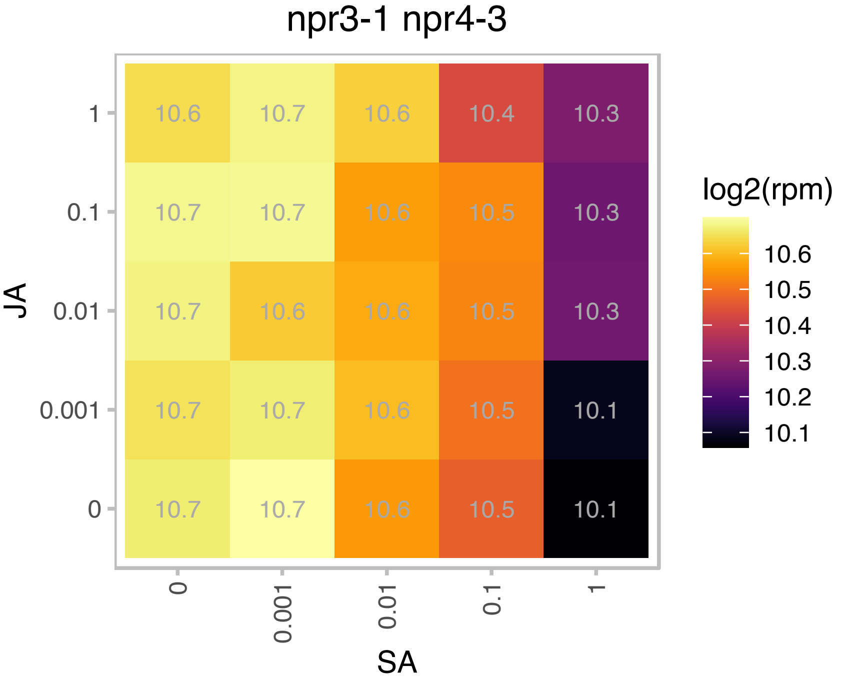

# AT3G62530.1 : NA

ARM repeat superfamily protein

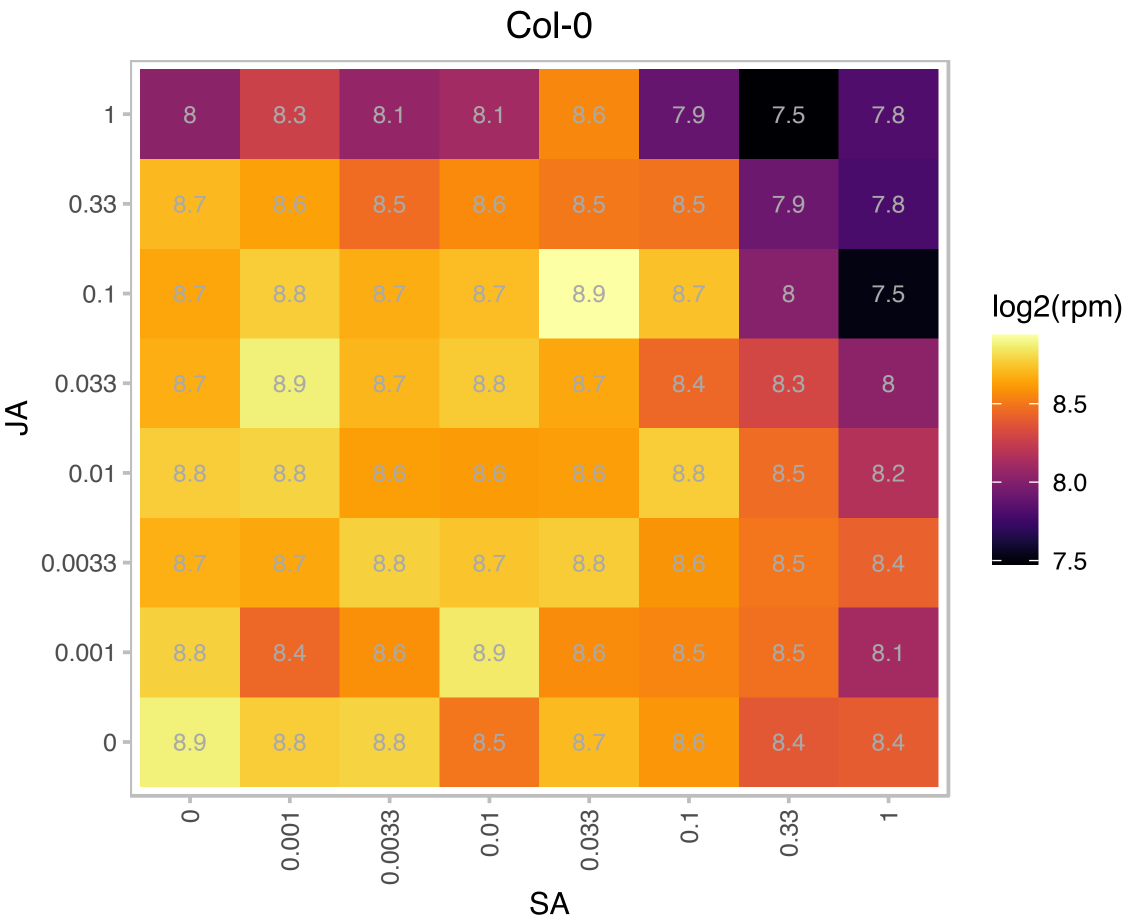


| | |
|---|---|
| Gene | AT3G62530 |
| is.expressed_genes | TRUE |
| is.responsive_genes | TRUE |
| is.apcluster | F-07 |

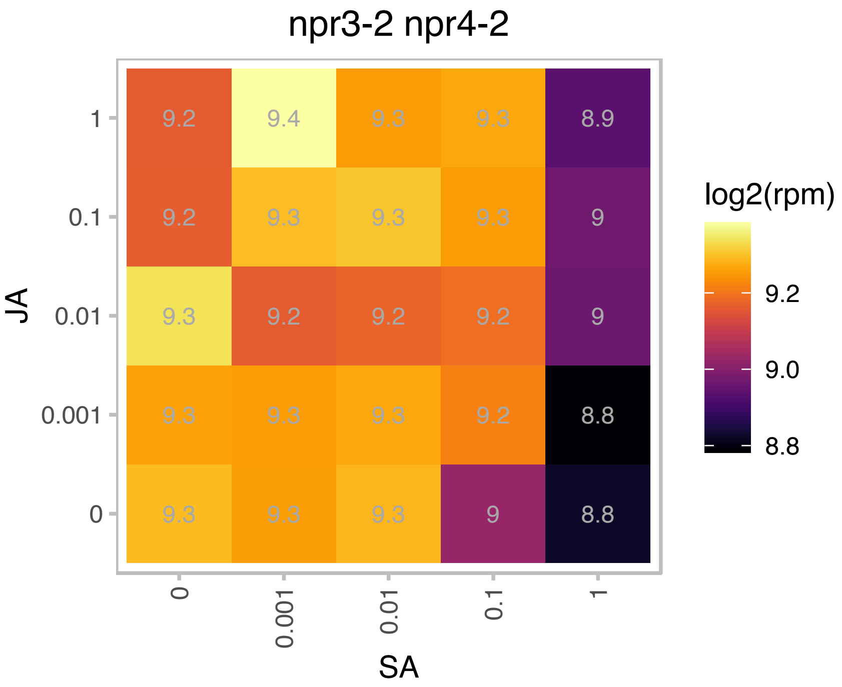


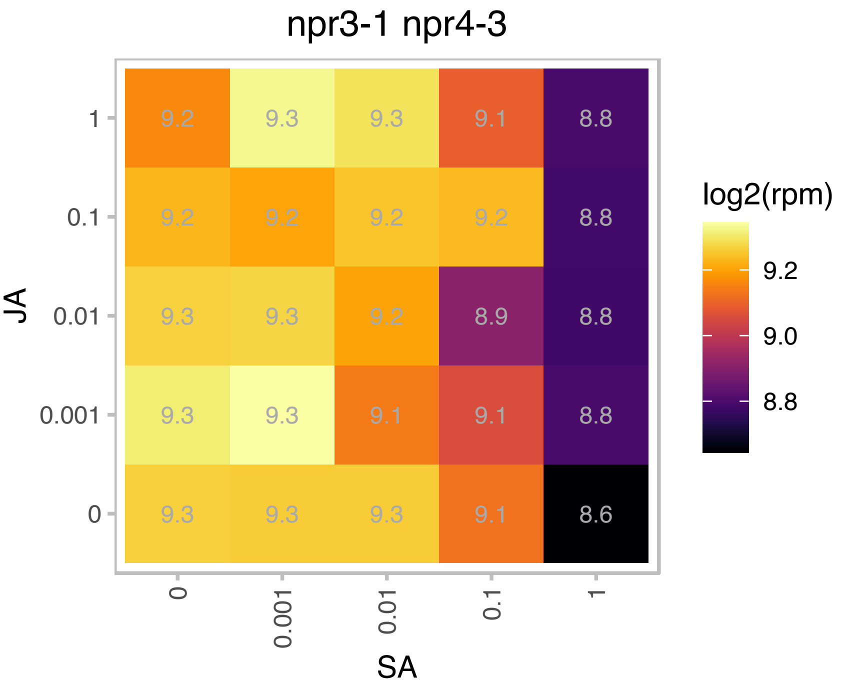

# AT3G62870.1 : NA

Ribosomal protein L7Ae/L30e/S12e/Gadd45 family protein

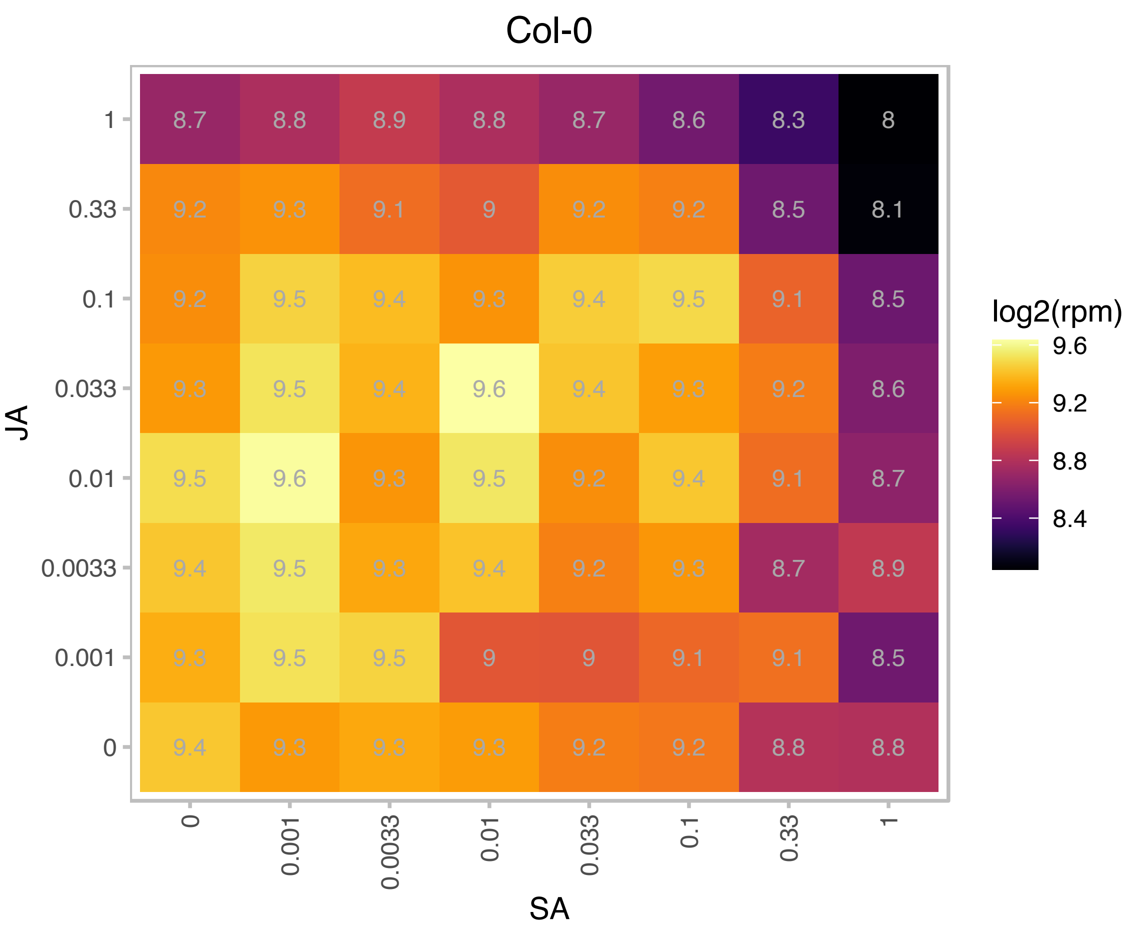


| | |
|---|---|
| Gene | AT3G62870 |
| is.expressed_genes | TRUE |
| is.responsive_genes | TRUE |
| is.apcluster | F-07 |

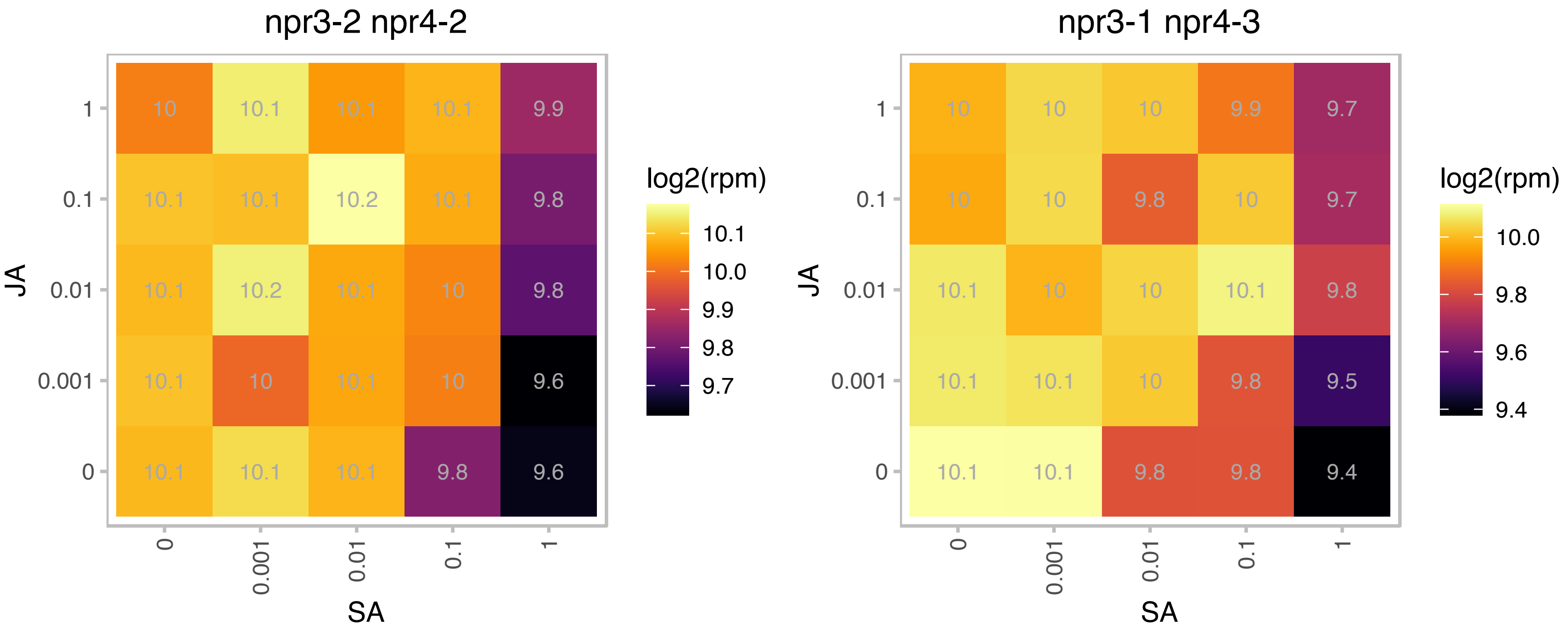

### AT4G00100.1 : ATRPS13A, PFL2, RPS13, RPS13A

ribosomal protein S13A

Encodes a cytoplasmic ribosomal protein S13 homologue involved in early leaf development The mRNA is cell-to-cell mobile.

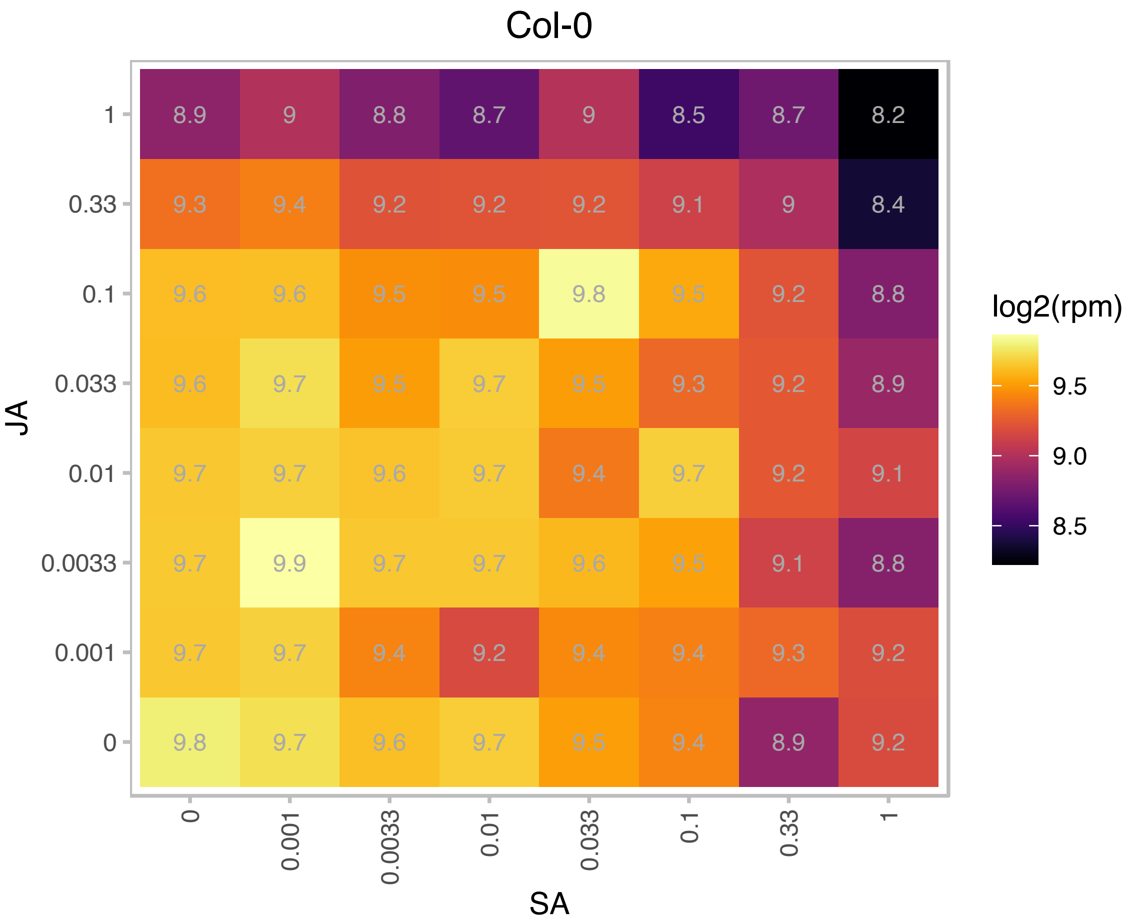


| | |
|---|---|
| Gene | AT4G00100 |
| is.expressed_genes | TRUE |
| is.responsive_genes | TRUE |
| is.apcluster | F-07 |

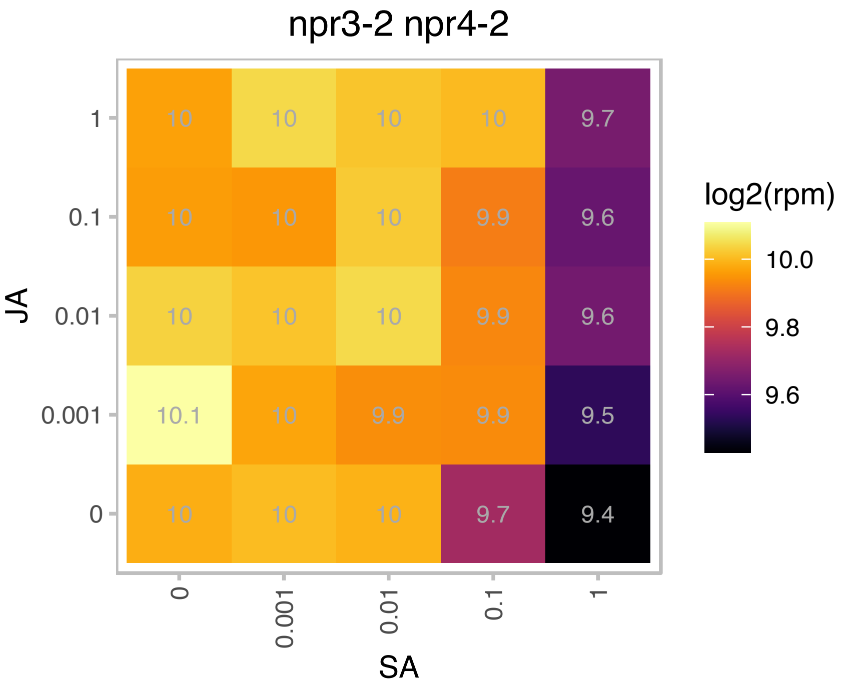


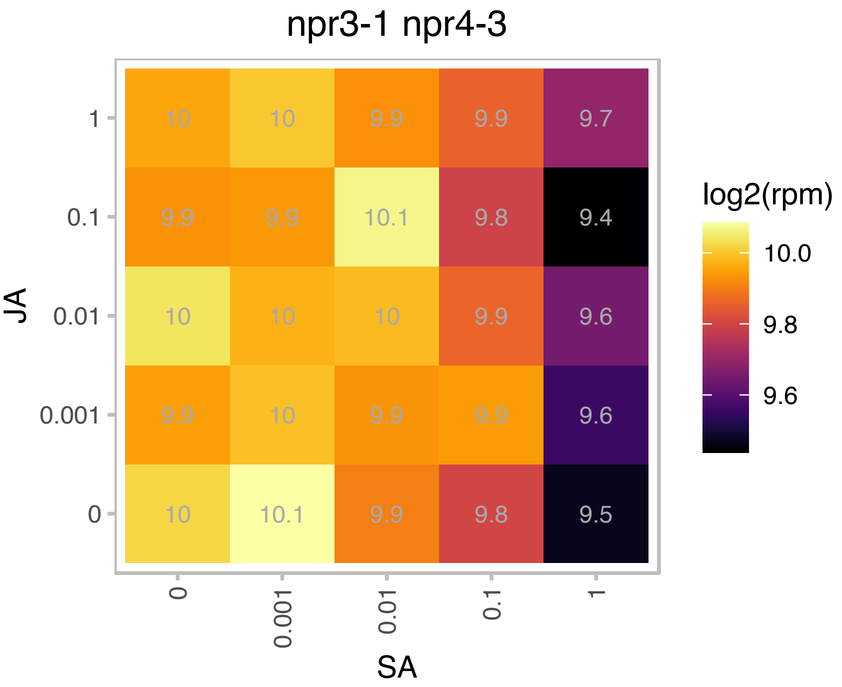

# AT4G05400.1 : NA

copper ion binding

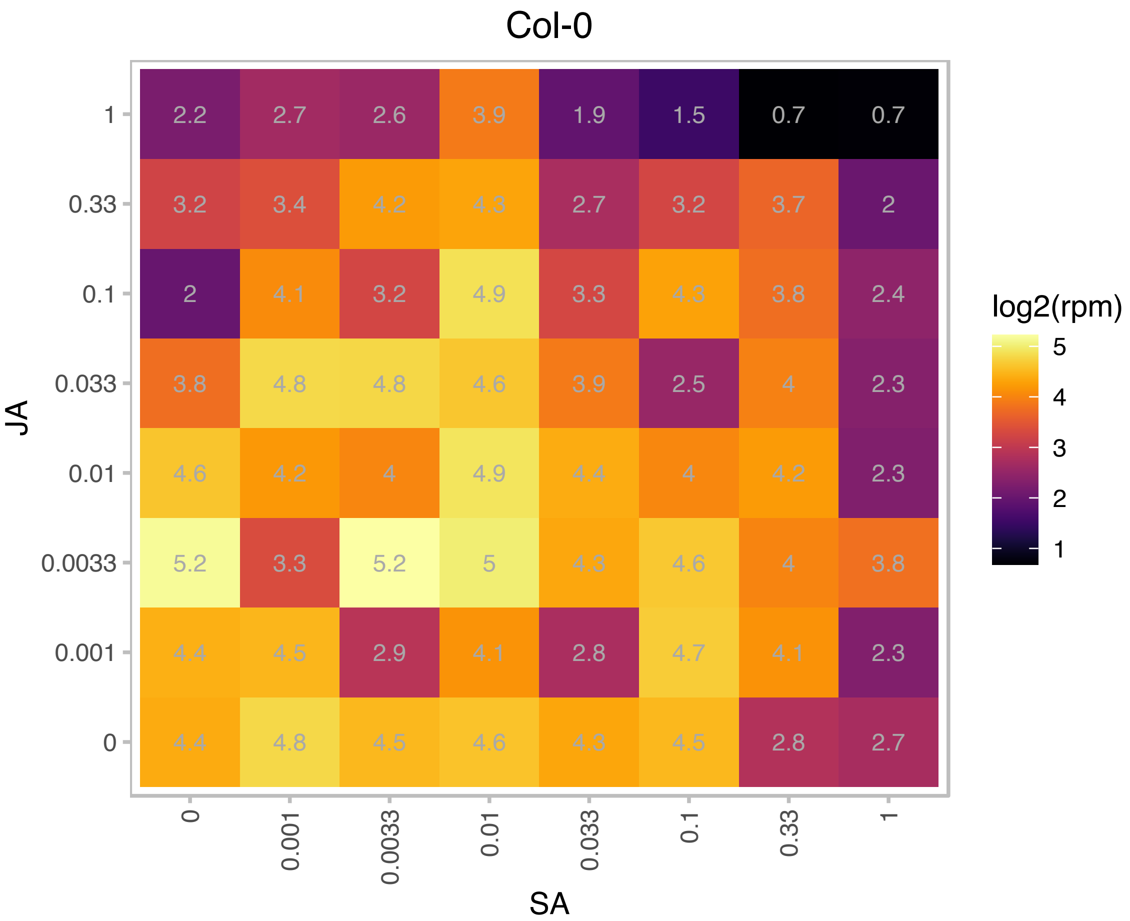


| | |
|---|---|
| Gene | AT4G05400 |
| is.expressed_genes | TRUE |
| is.responsive_genes | TRUE |
| is.apcluster | F-07 |

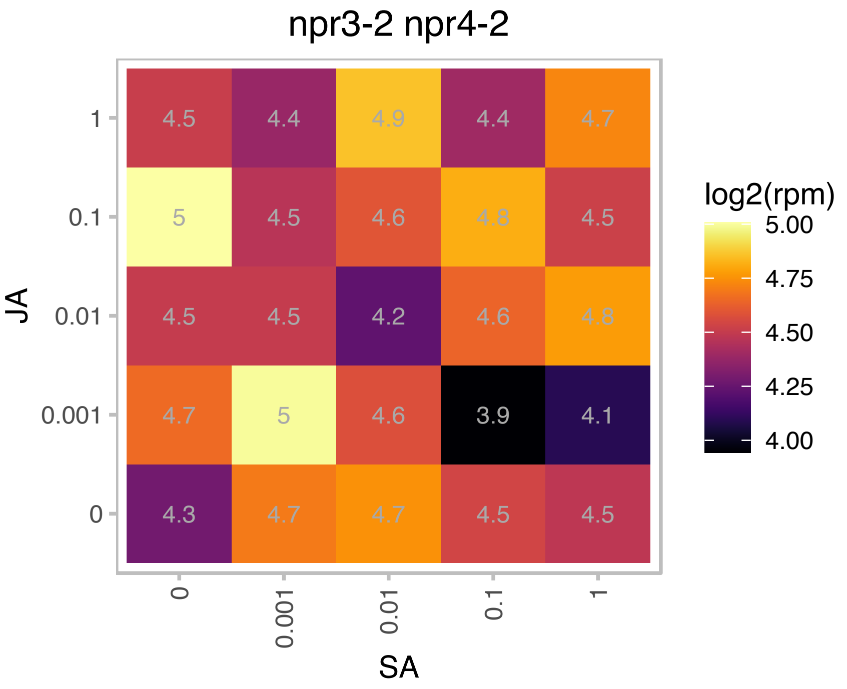


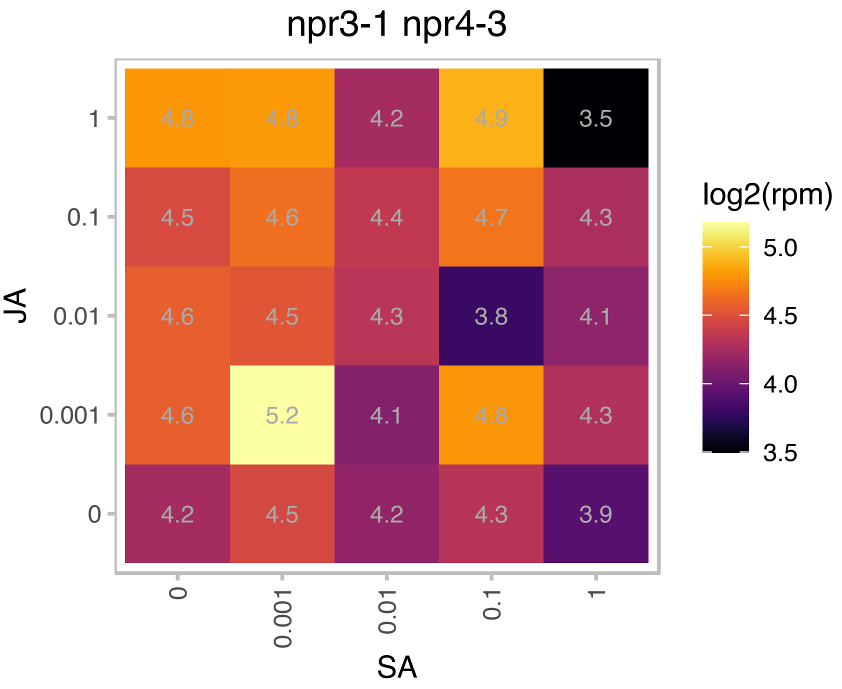

# AT4G09730.1 : RH39

RH39

Encodes RH39, a DEAD-box protein involved in the introduction of the hidden break into the 23S rRNA in the chloroplasts. Recombinant RH39 binds to the 23S rRNA in a segment adjacent to the stem-loop creating the hidden break target loop in a sequence-dependent manner. Has ATP-hydrolyzing activity at a Kcat of 5.3 /min in the presence of rRNA sequence. Mutants have drastically reduced level of level of ribulose 1,5-bisphosphate carboxylase/oxygenase. The mRNA is cell-to-cell mobile.

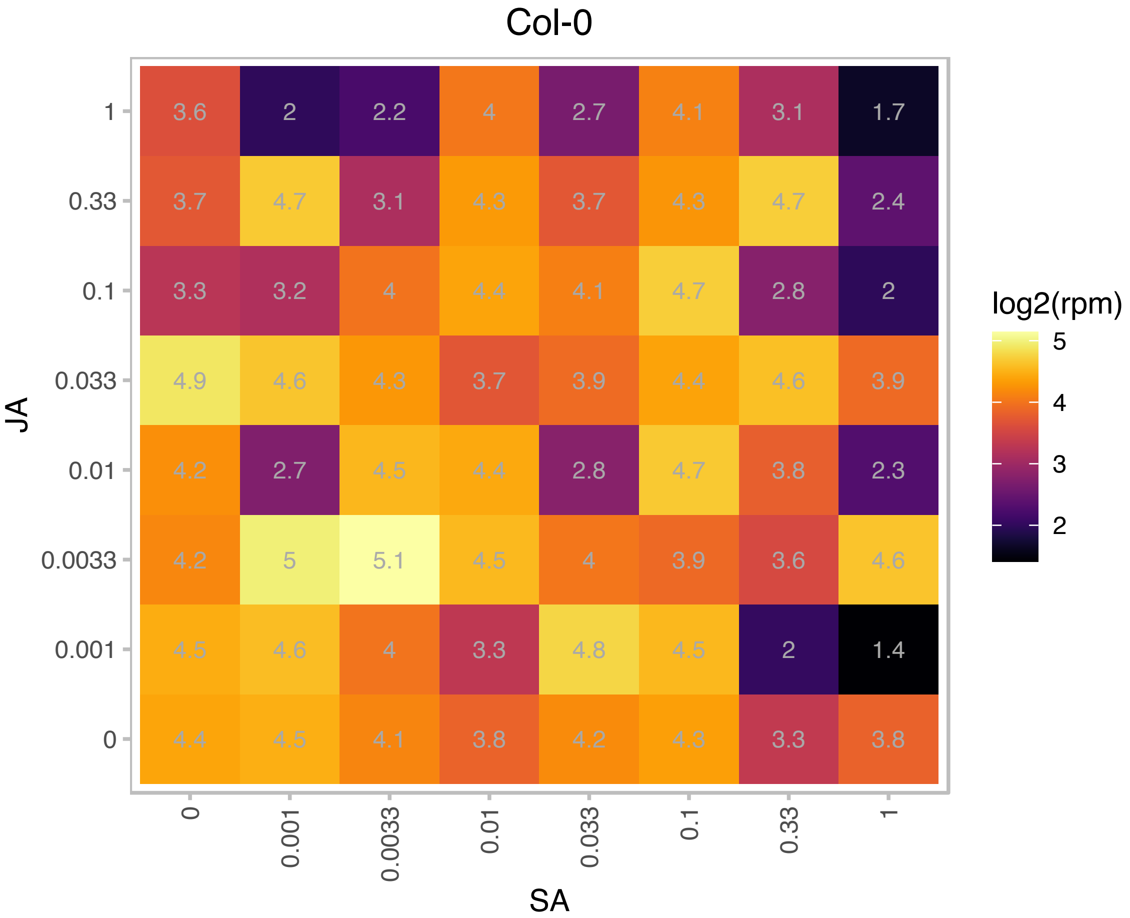


| | |
|---|---|
| Gene | AT4G09730 |
| is.expressed_genes | TRUE |
| is.responsive_genes | TRUE |
| is.apcluster | F-07 |

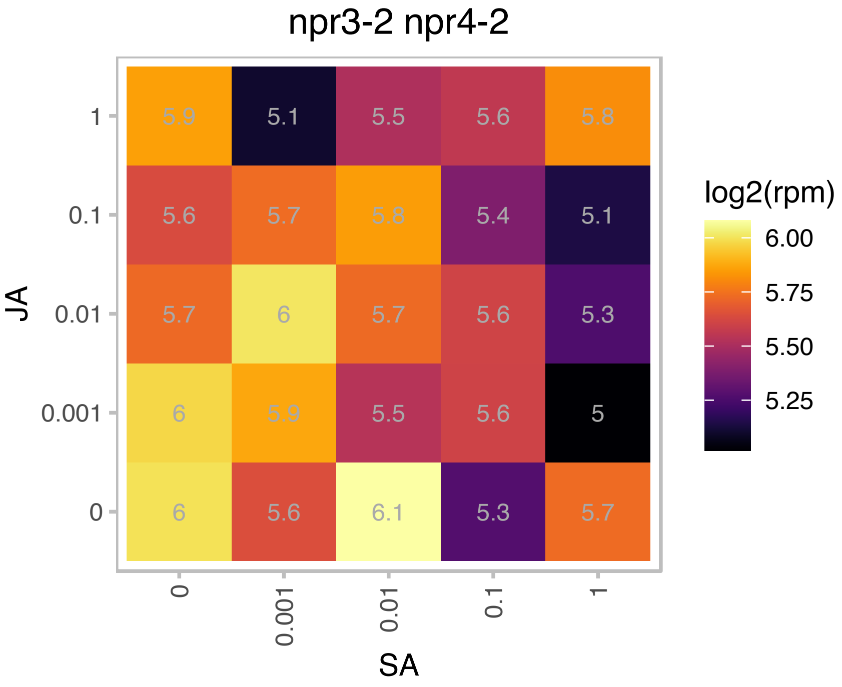


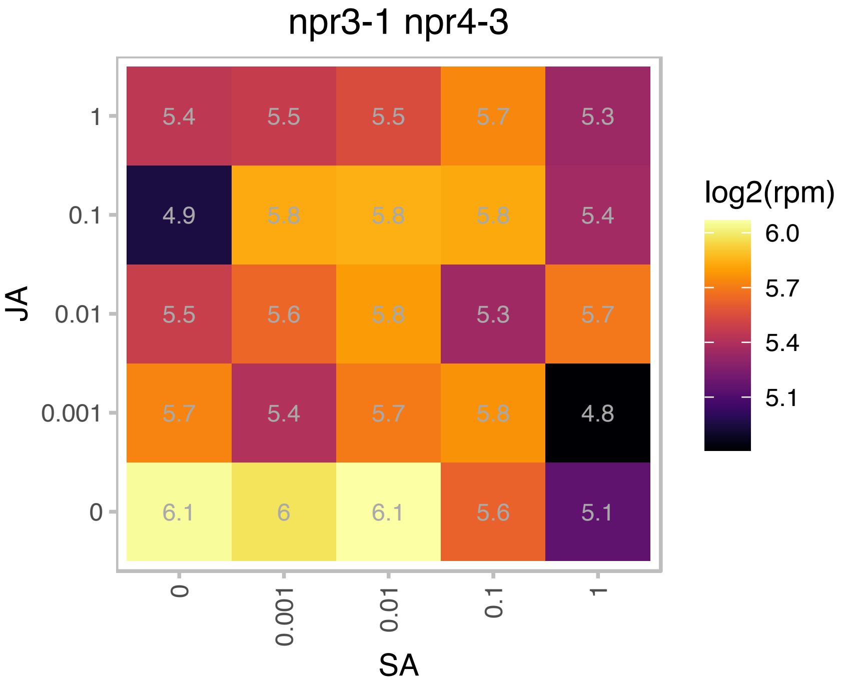

# AT4G09890.1 : NA

Protein of unknown function (DUF3511)

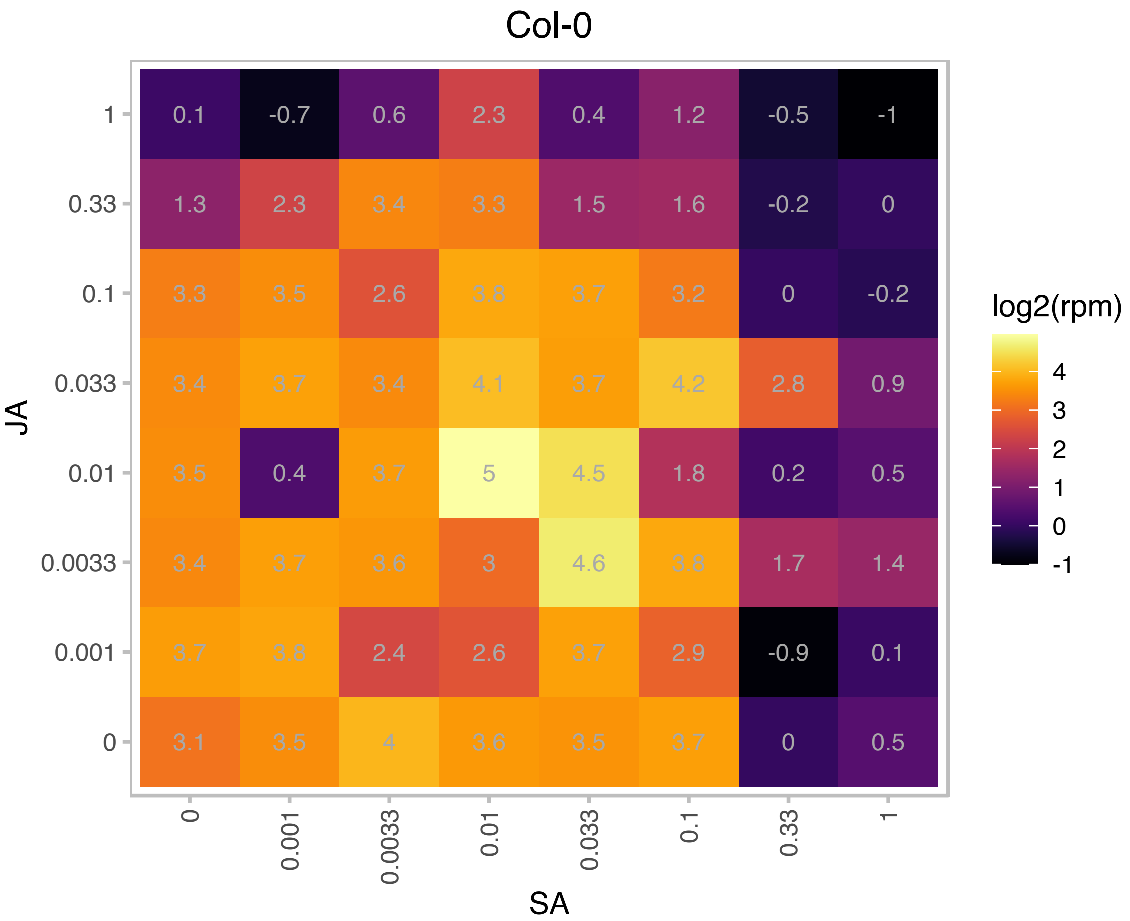


| | |
|---|---|
| Gene | AT4G09890 |
| is.expressed_genes | TRUE |
| is.responsive_genes | TRUE |
| is.apcluster | F-07 |

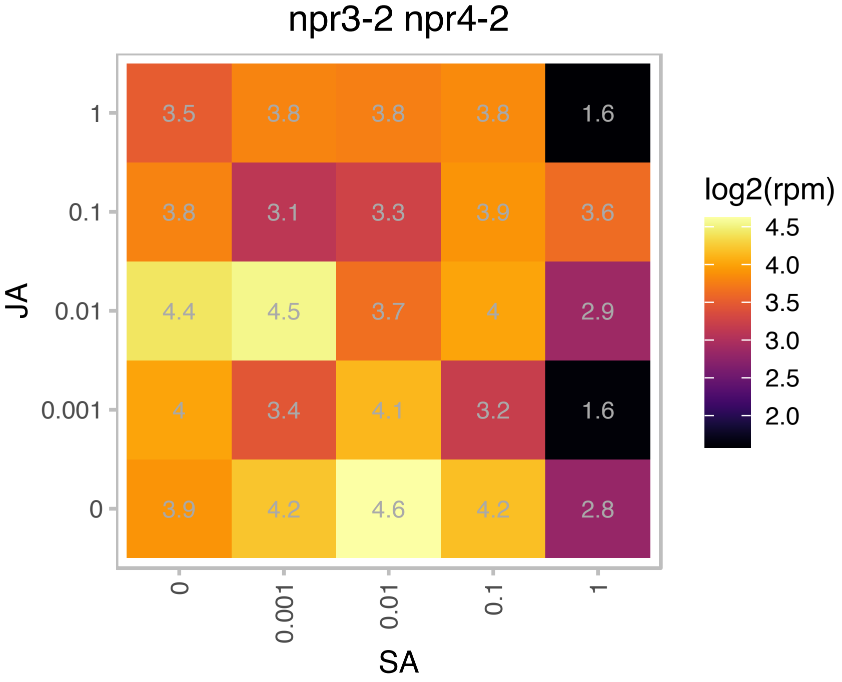


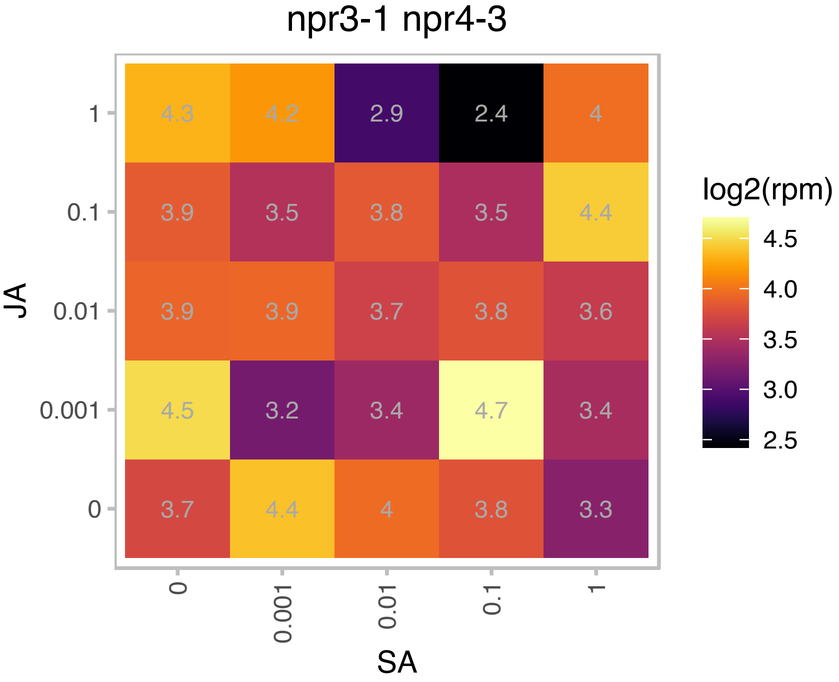

# AT4G10480.1 : NA

Nascent polypeptide-associated complex (NAC), alpha subunit family protein

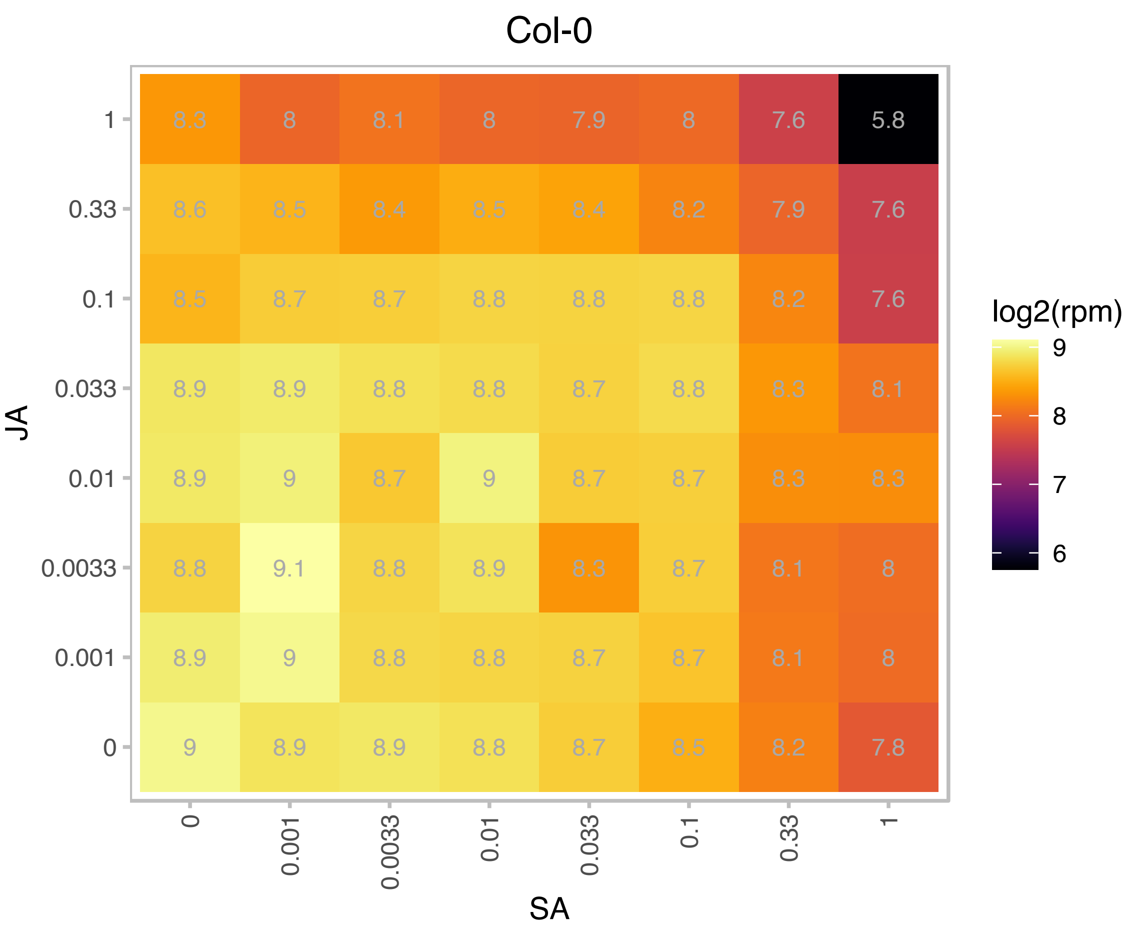


| | |
|---|---|
| Gene | AT4G10480 |
| is.expressed_genes | TRUE |
| is.responsive_genes | TRUE |
| is.apcluster | F-07 |

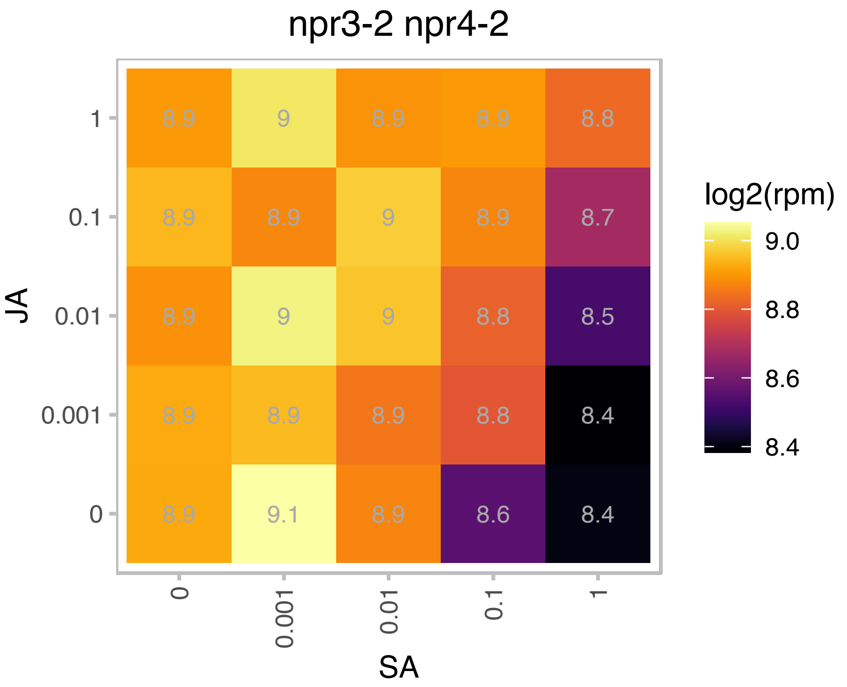


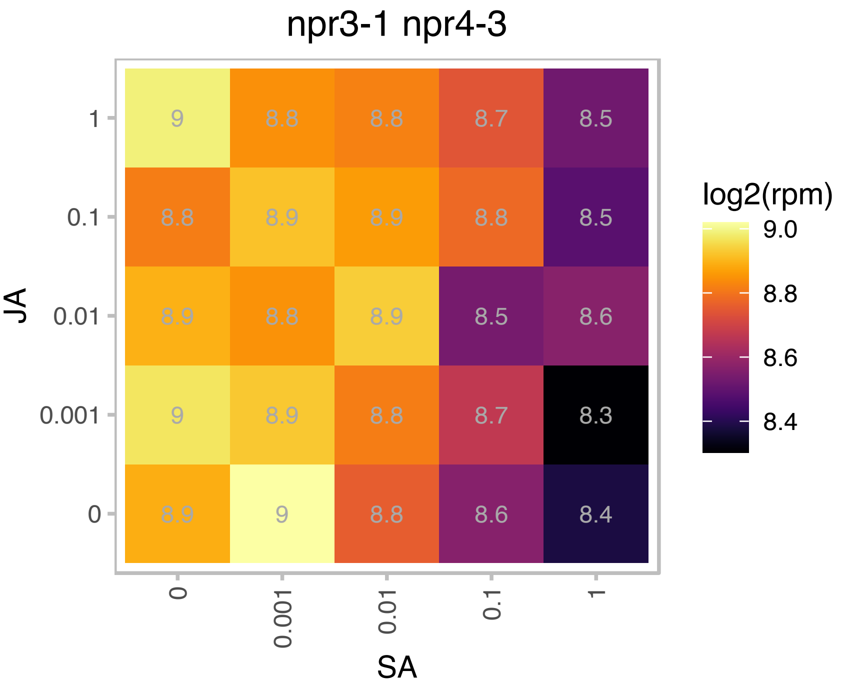

# AT4G11175.1 : NA

Nucleic acid-binding, OB-fold-like protein

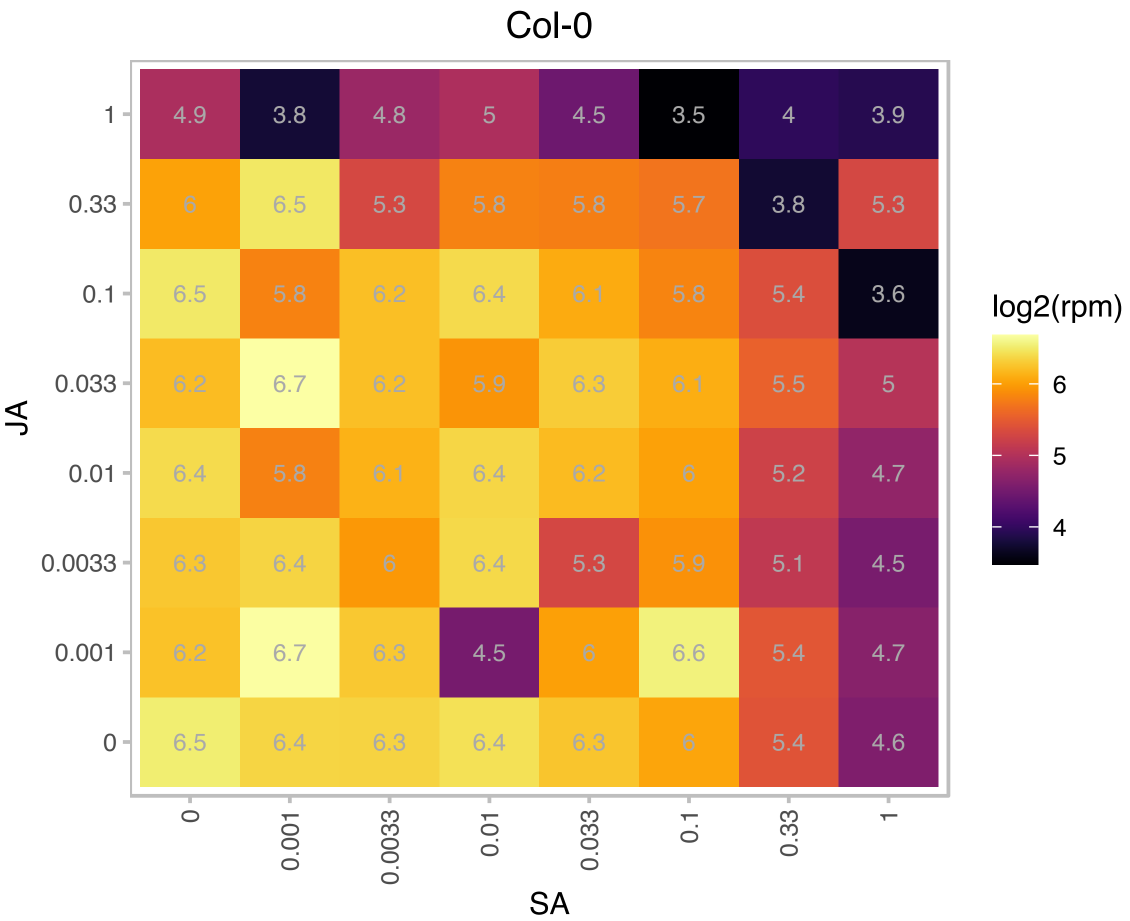


| | |
|---|---|
| Gene | AT4G11175 |
| is.expressed_genes | TRUE |
| is.responsive_genes | TRUE |
| is.apcluster | F-07 |

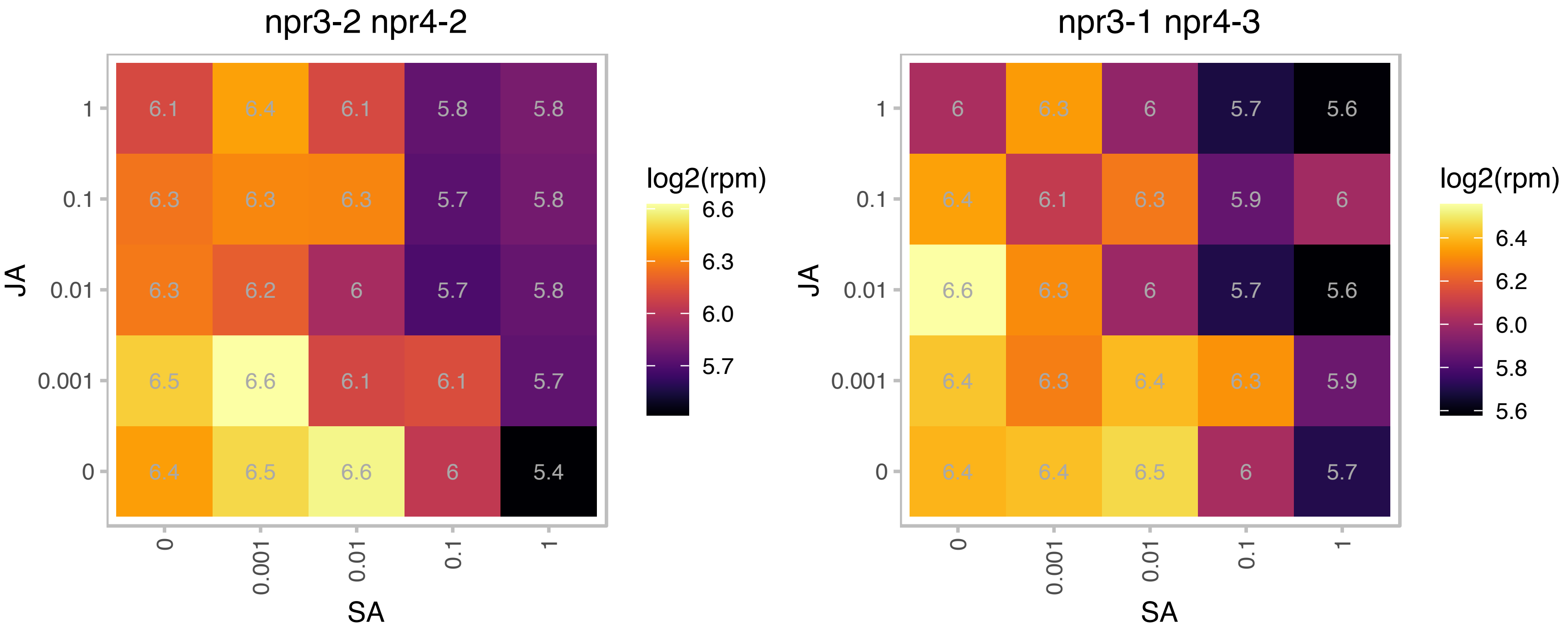

# AT4G13720.1 : NA

Inosine triphosphate pyrophosphatase family protein

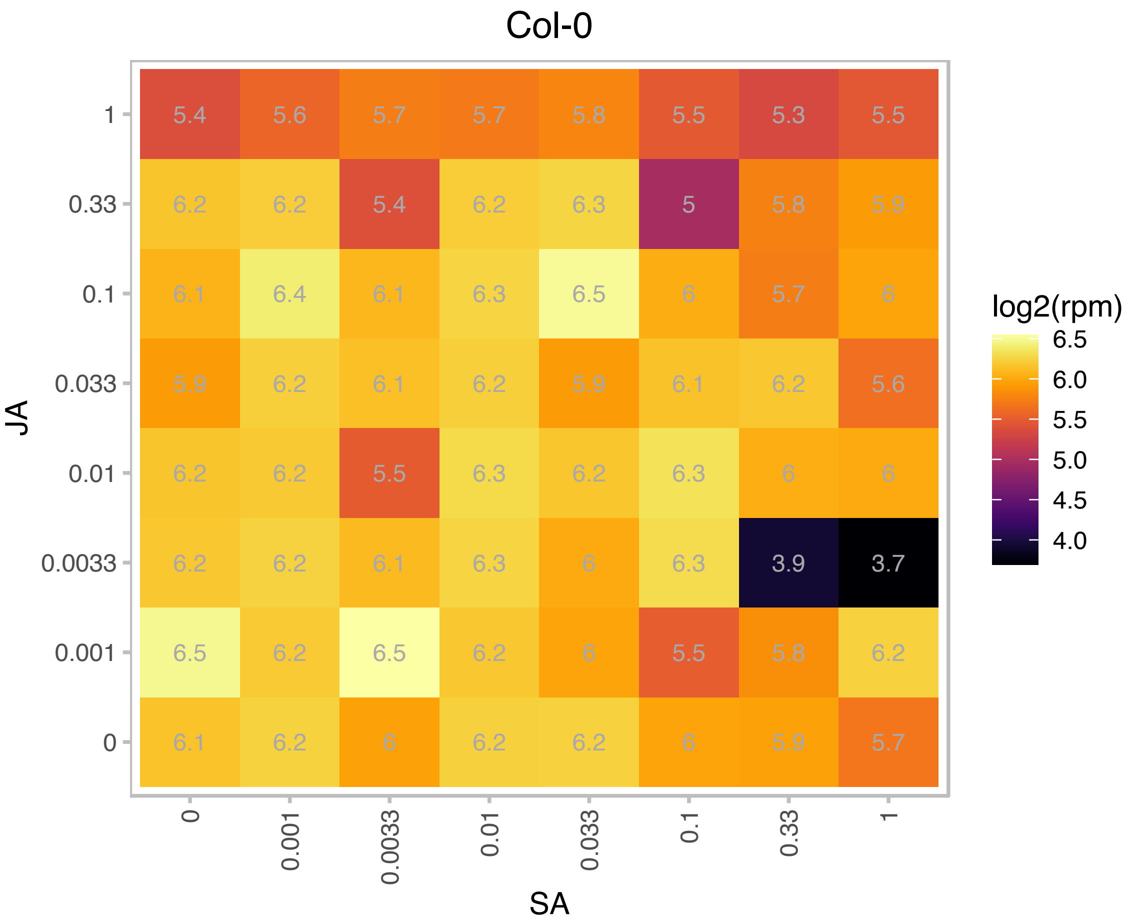


| | |
|---|---|
| Gene | AT4G13720 |
| is.expressed_genes | TRUE |
| is.responsive_genes | TRUE |
| is.apcluster | F-07 |

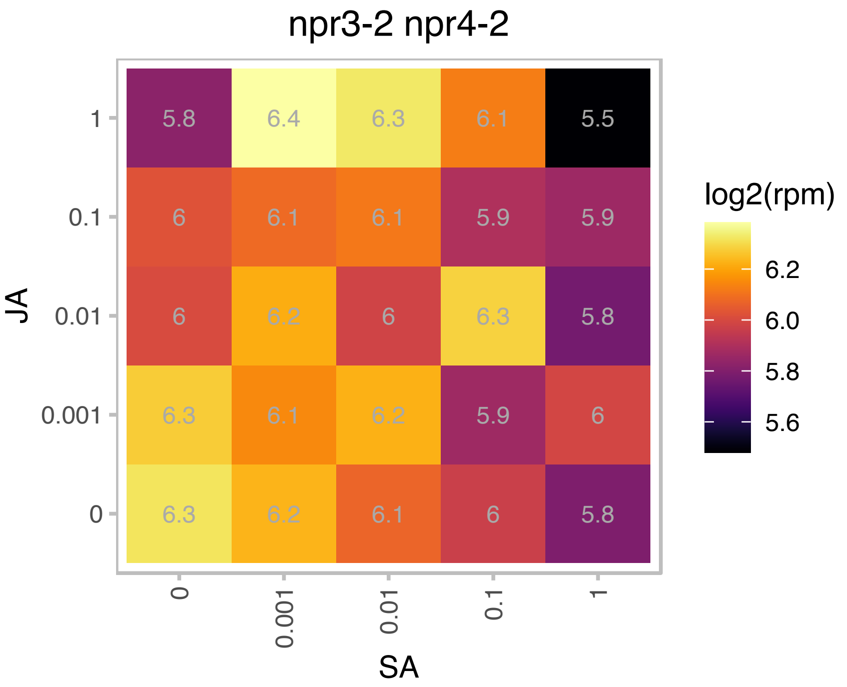


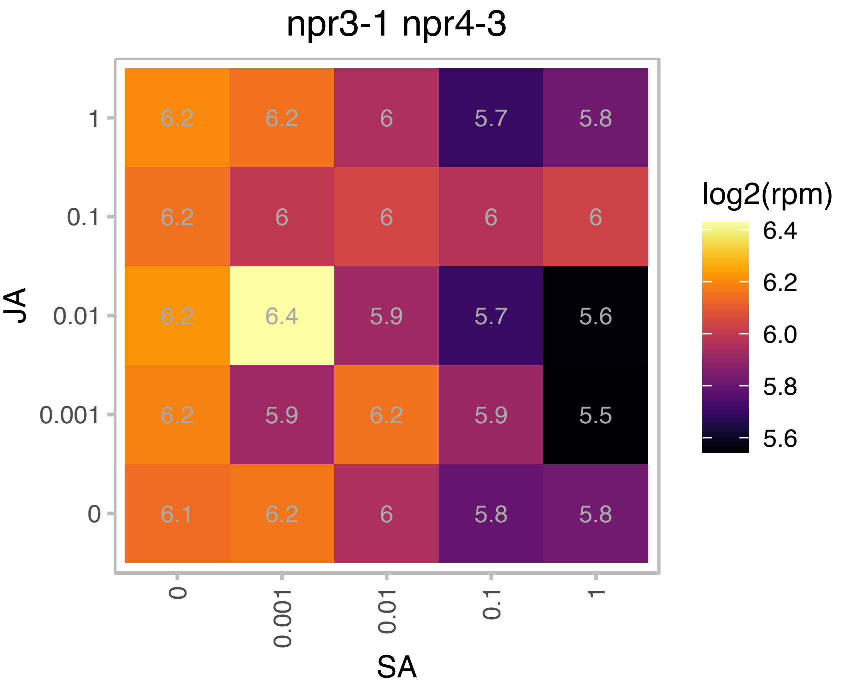

### AT4G13770.1 : CYP83A1, REF2

cytochrome P450, family 83, subfamily A, polypeptide 1

Encodes a cytochrome p450 enzyme that catalyzes the initial conversion of aldoximes to thiohydroximates in the synthesis of glucosinolates not derived from tryptophan.  Also has a role in auxin homeostasis.

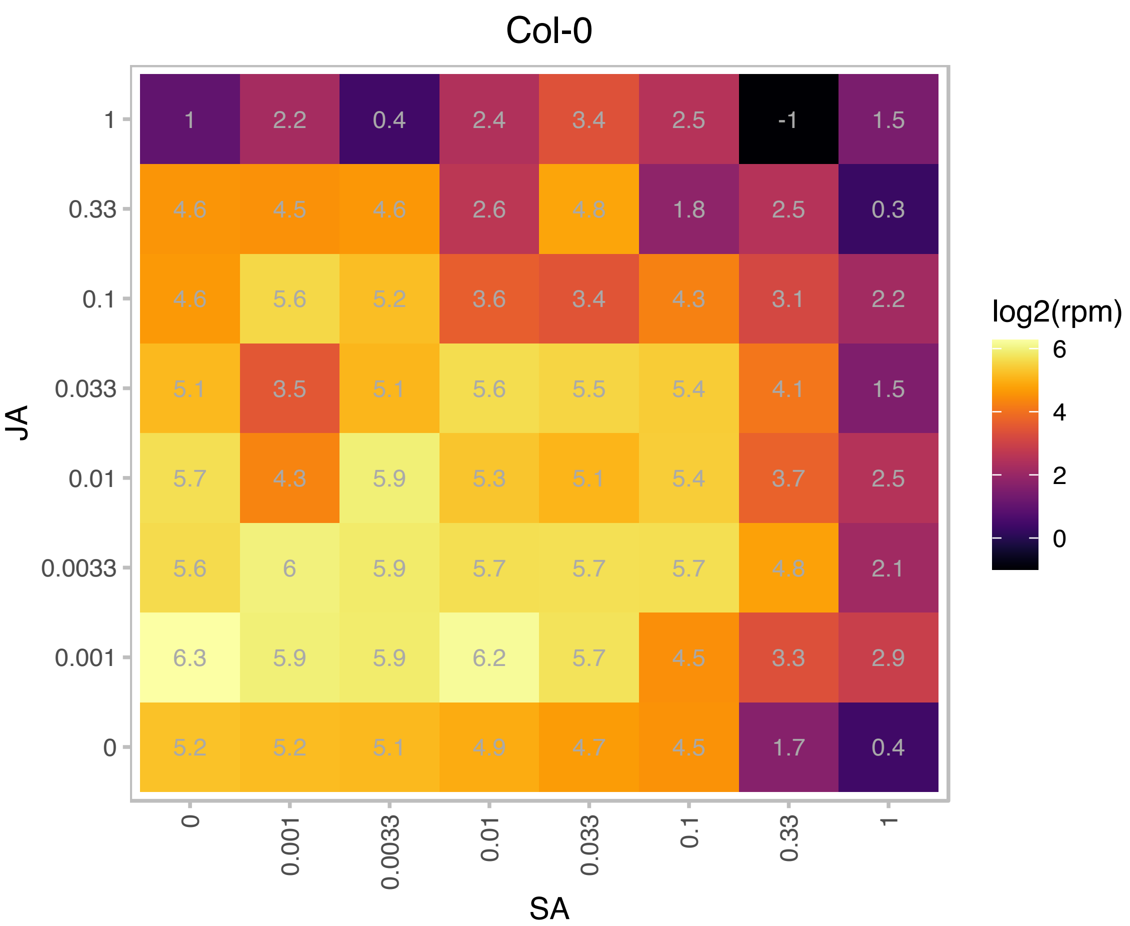


| | |
|---|---|
| Gene | AT4G13770 |
| is.expressed_genes | TRUE |
| is.responsive_genes | TRUE |
| is.apcluster | F-07 |

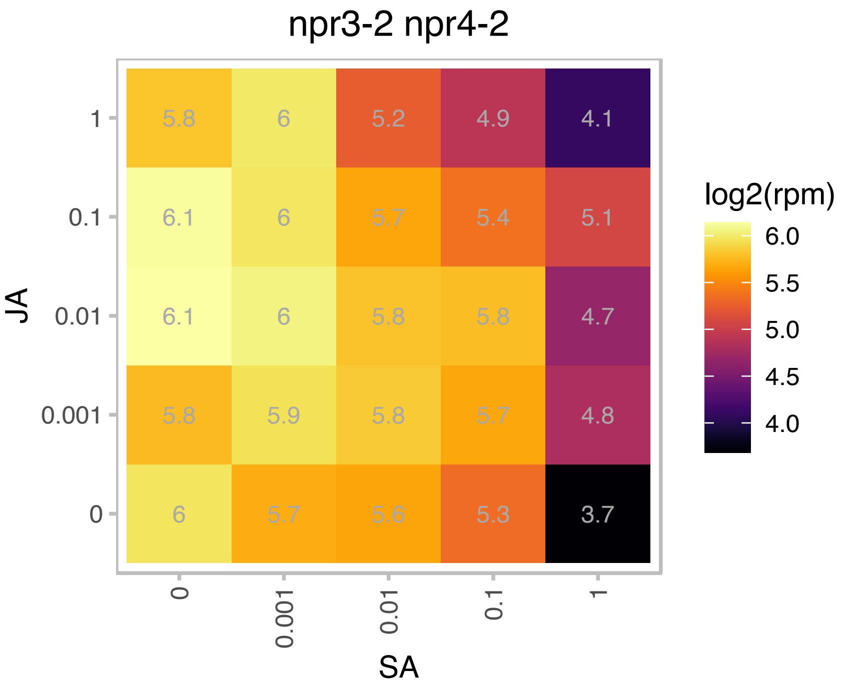


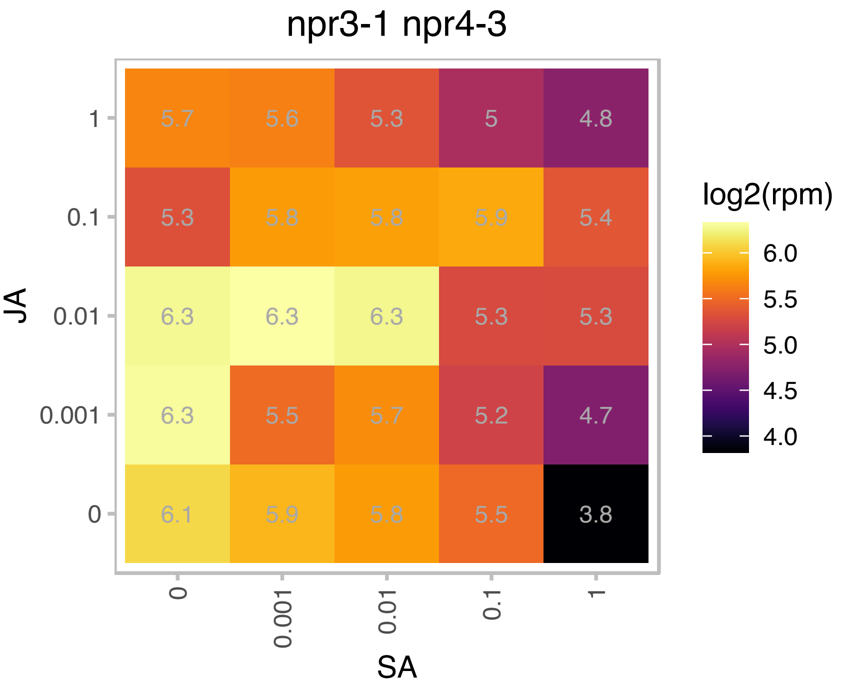

### AT4G14320.1 : API2, RPL36aB

Zinc-binding ribosomal protein family protein

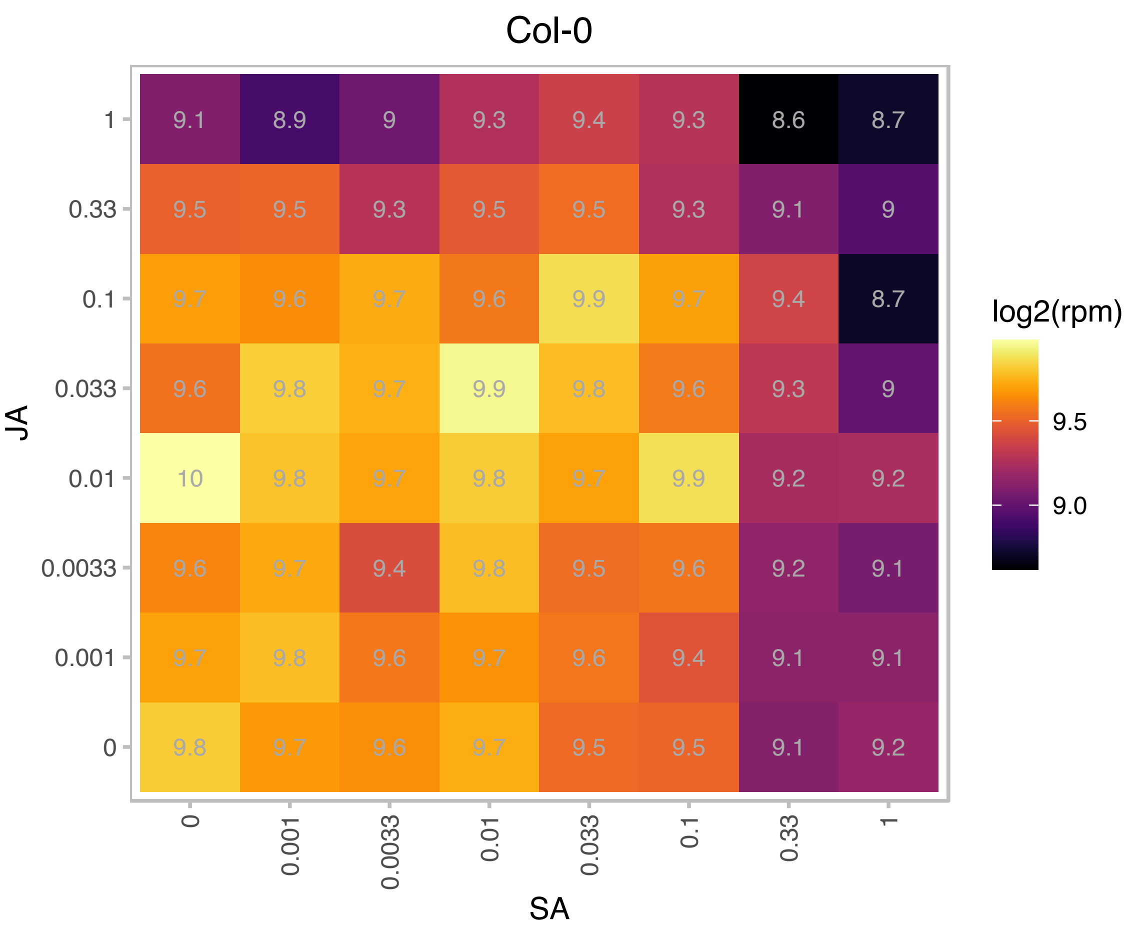


| | |
|---|---|
| Gene | AT4G14320 |
| is.expressed_genes | TRUE |
| is.responsive_genes | TRUE |
| is.apcluster | F-07 |

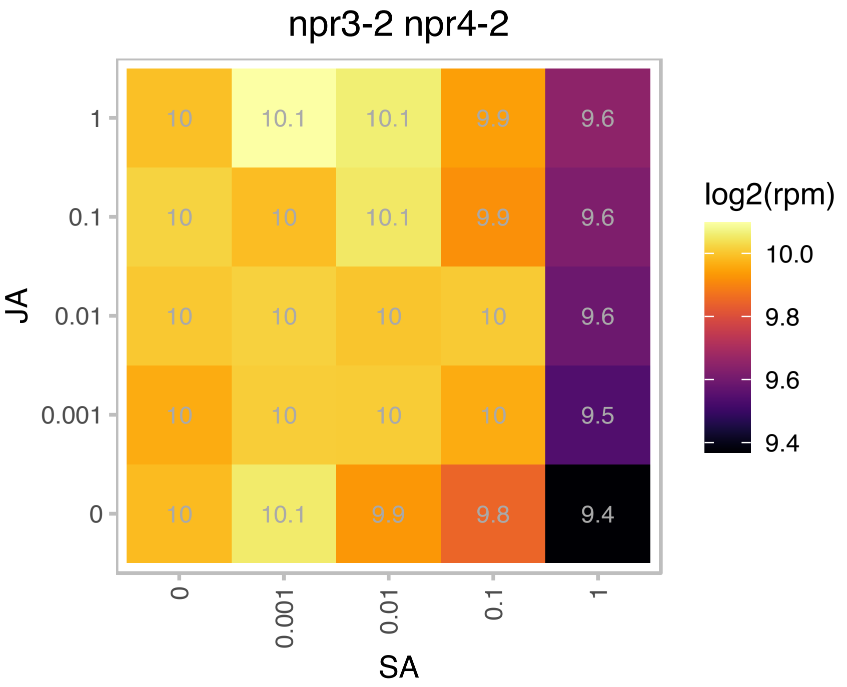


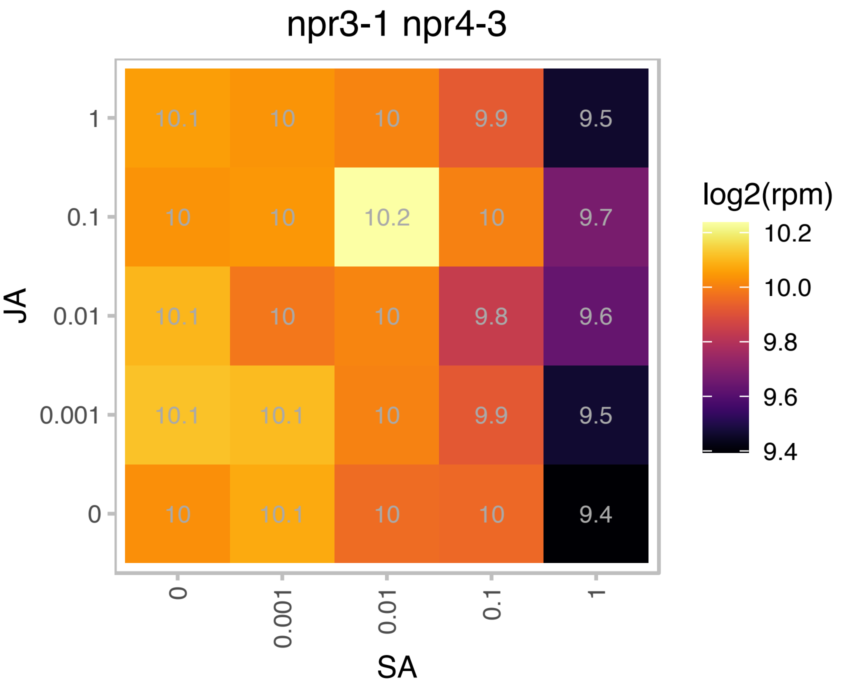

# AT4G15000.1 : NA

Ribosomal L27e protein family

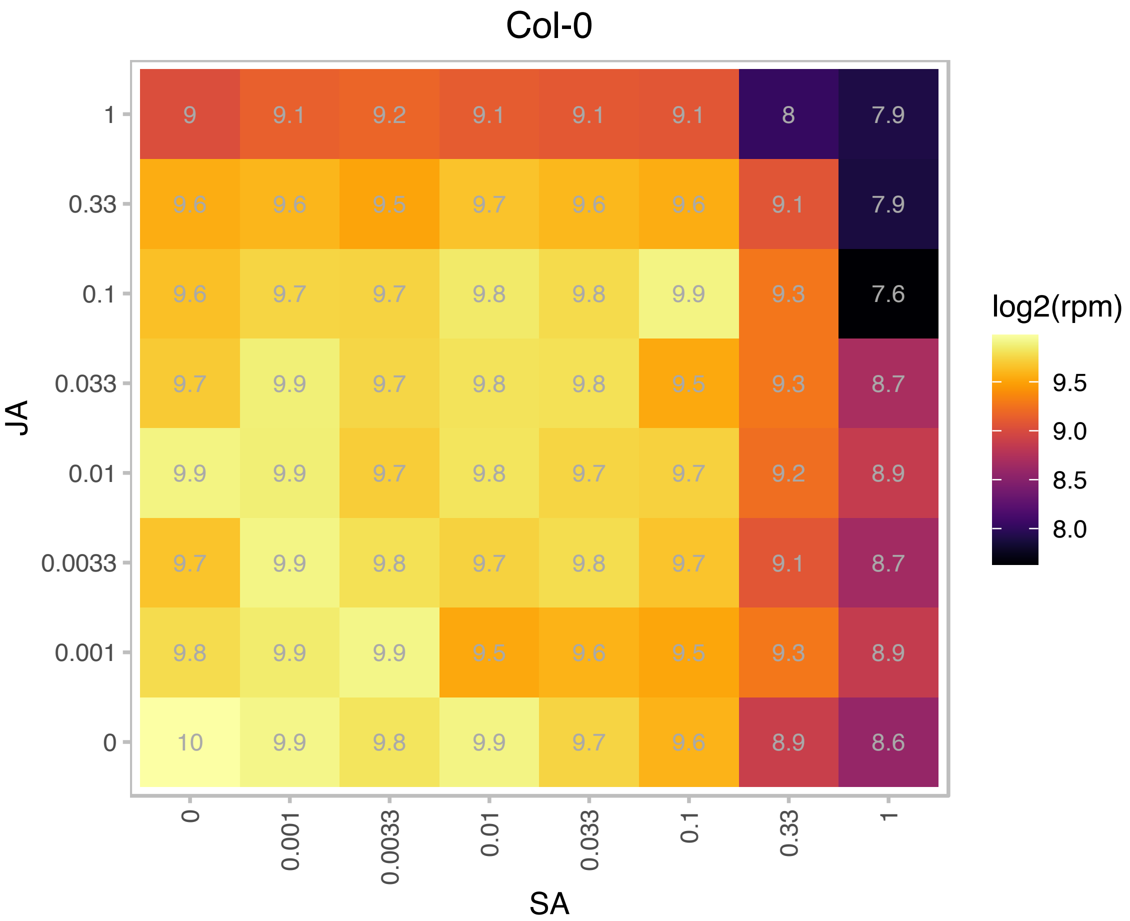


| | |
|---|---|
| Gene | AT4G15000 |
| is.expressed_genes | TRUE |
| is.responsive_genes | TRUE |
| is.apcluster | F-07 |

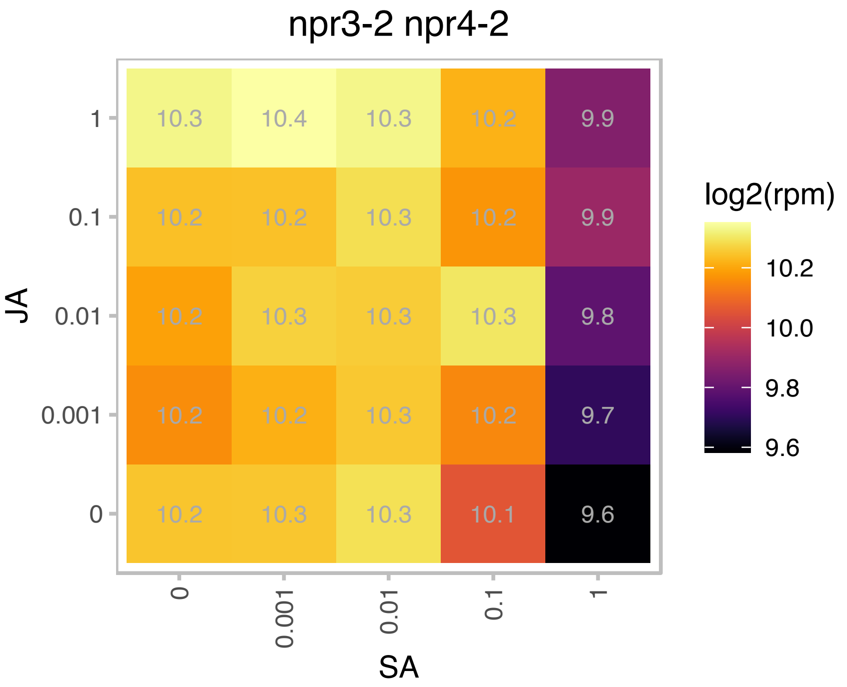


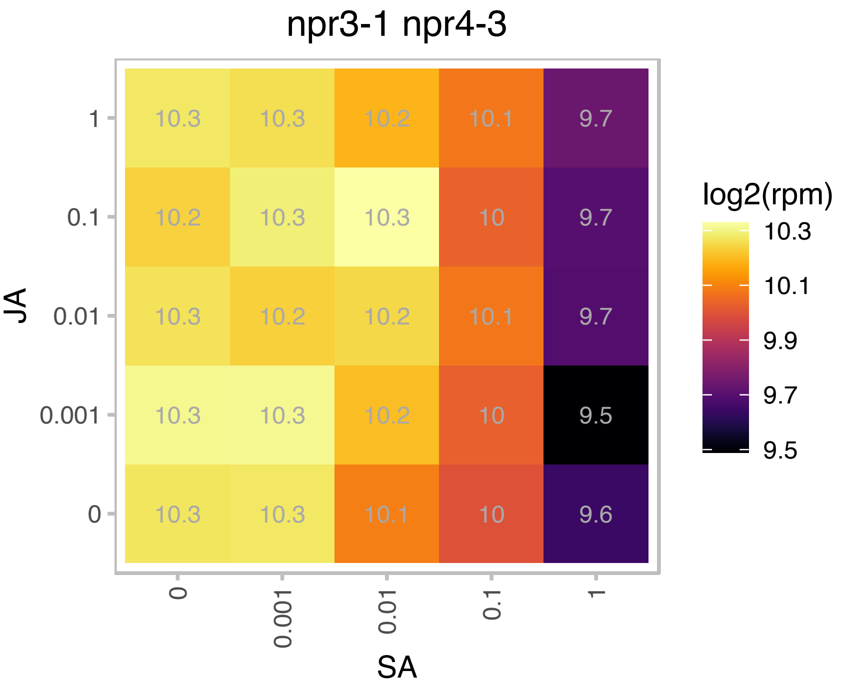

### AT4G16141.1 : AtGATA17L, GATA17L

GATA type zinc finger transcription factor family protein

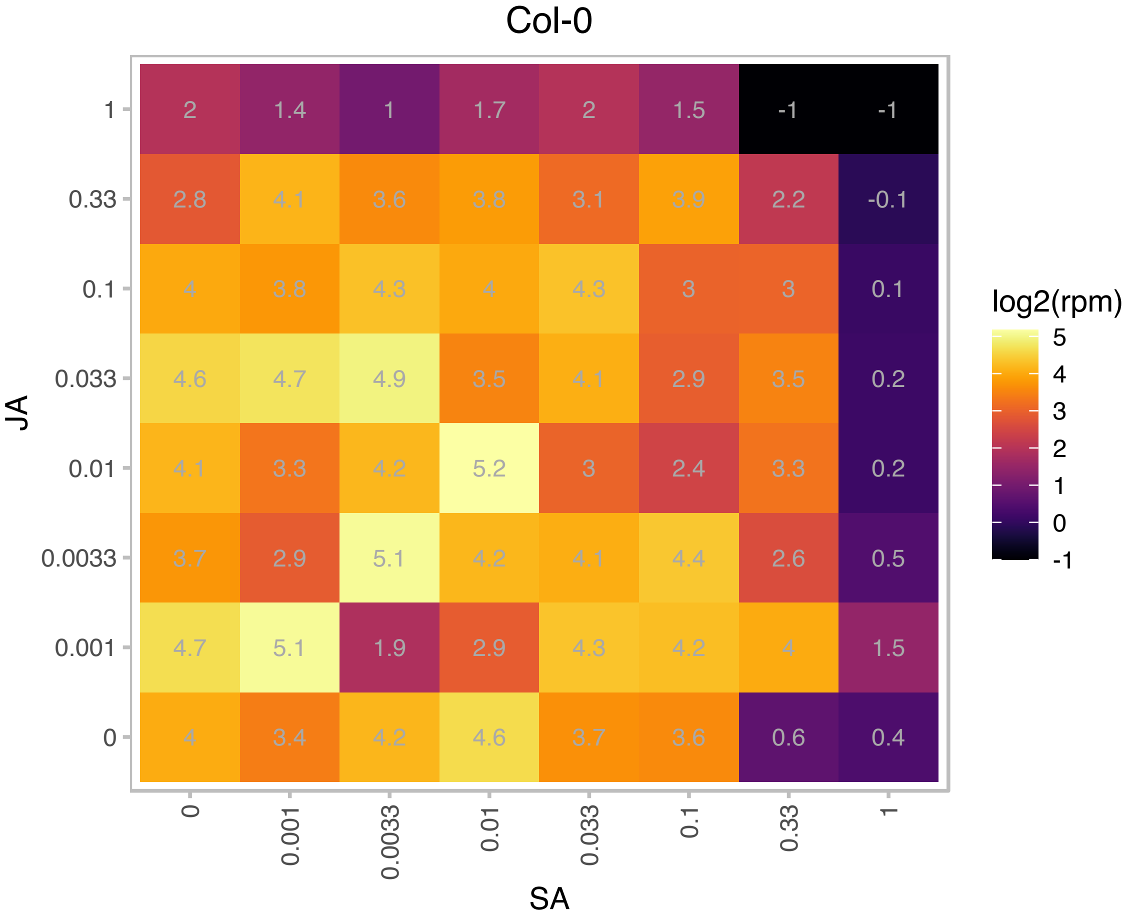


| | |
|---|---|
| Gene | AT4G16141 |
| is.expressed_genes | TRUE |
| is.responsive_genes | TRUE |
| is.apcluster | F-07 |

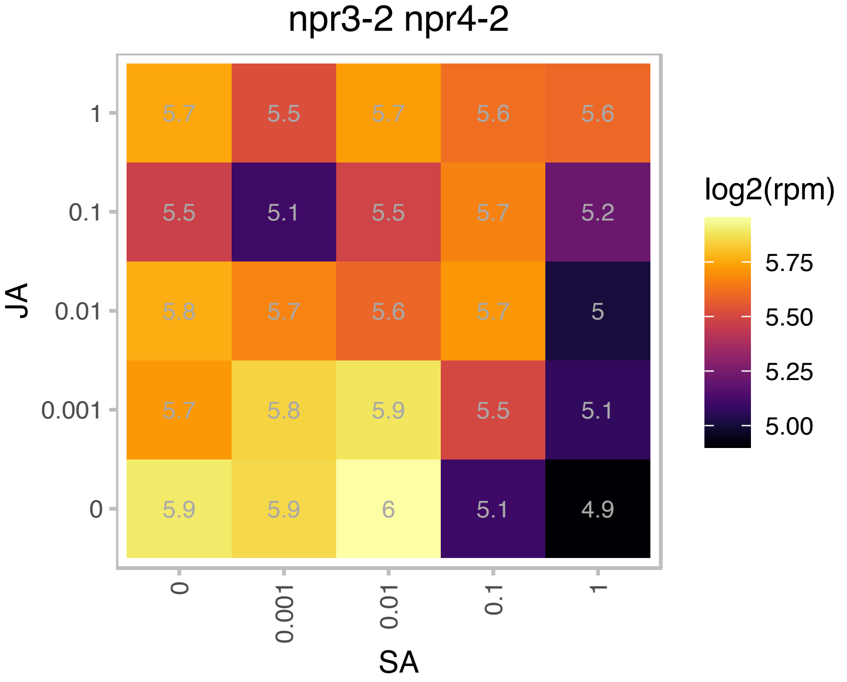


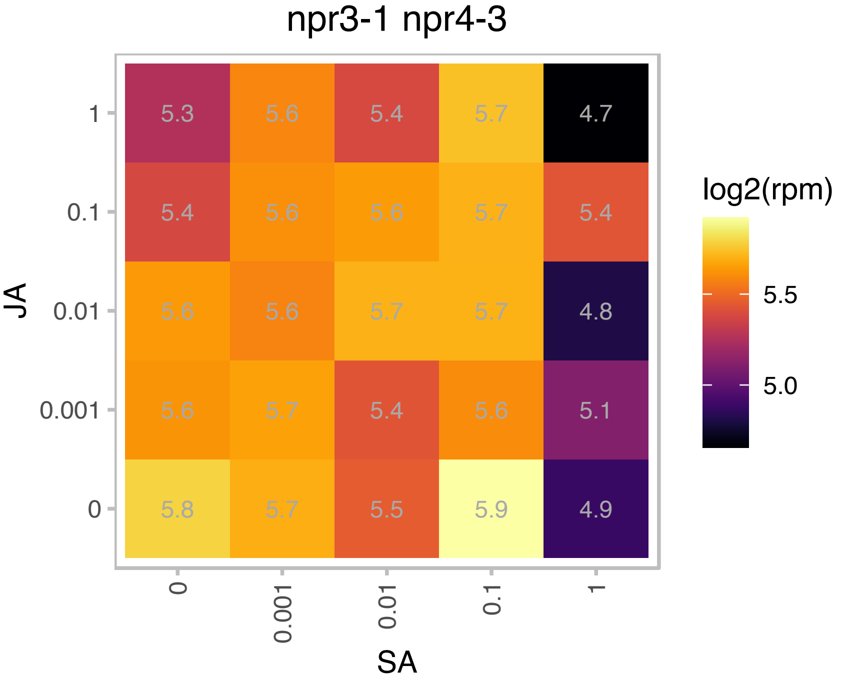

### AT4G16500.1 : AtCYS4, CYS4

Cystatin/monellin superfamily protein

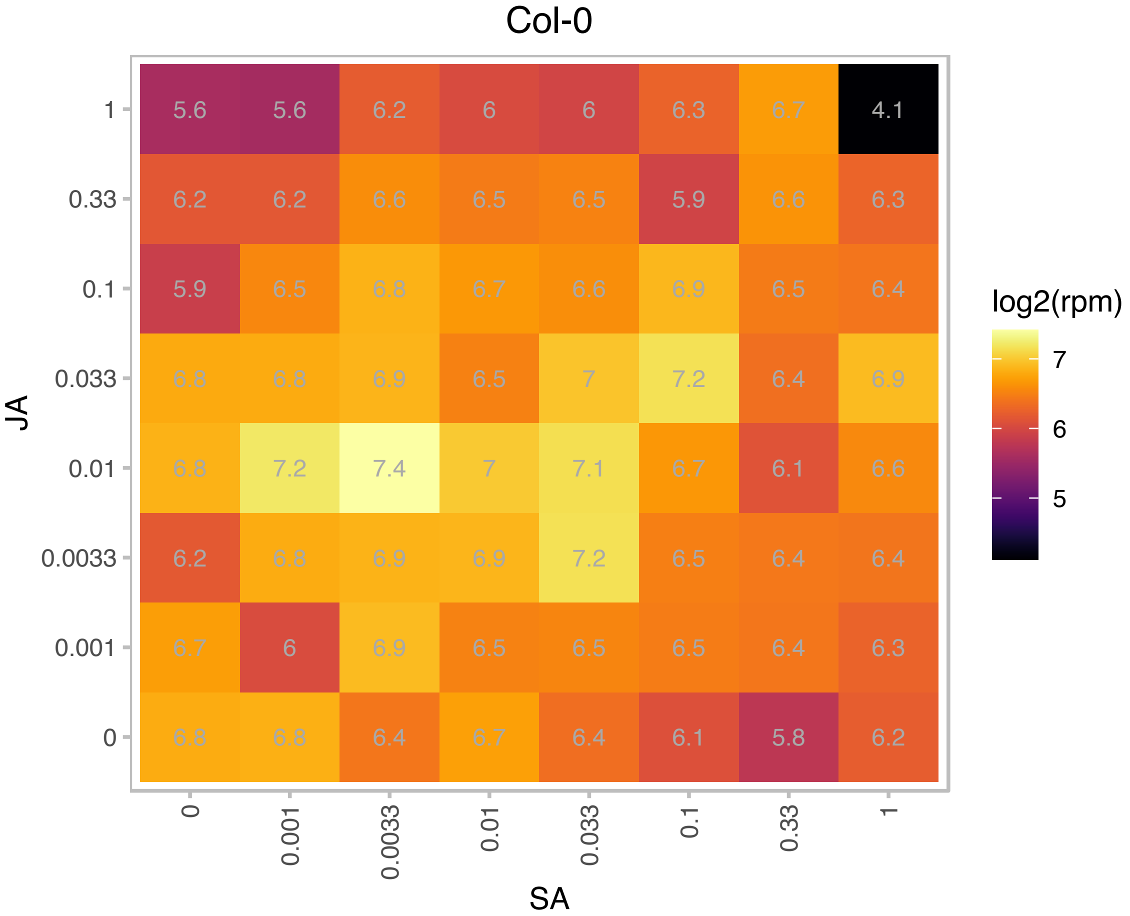


| Gene | AT4G16500 |
|---|---|
| is.expressed_genes | TRUE |
| is.responsive_genes | TRUE |
| is.apcluster | F-07 |

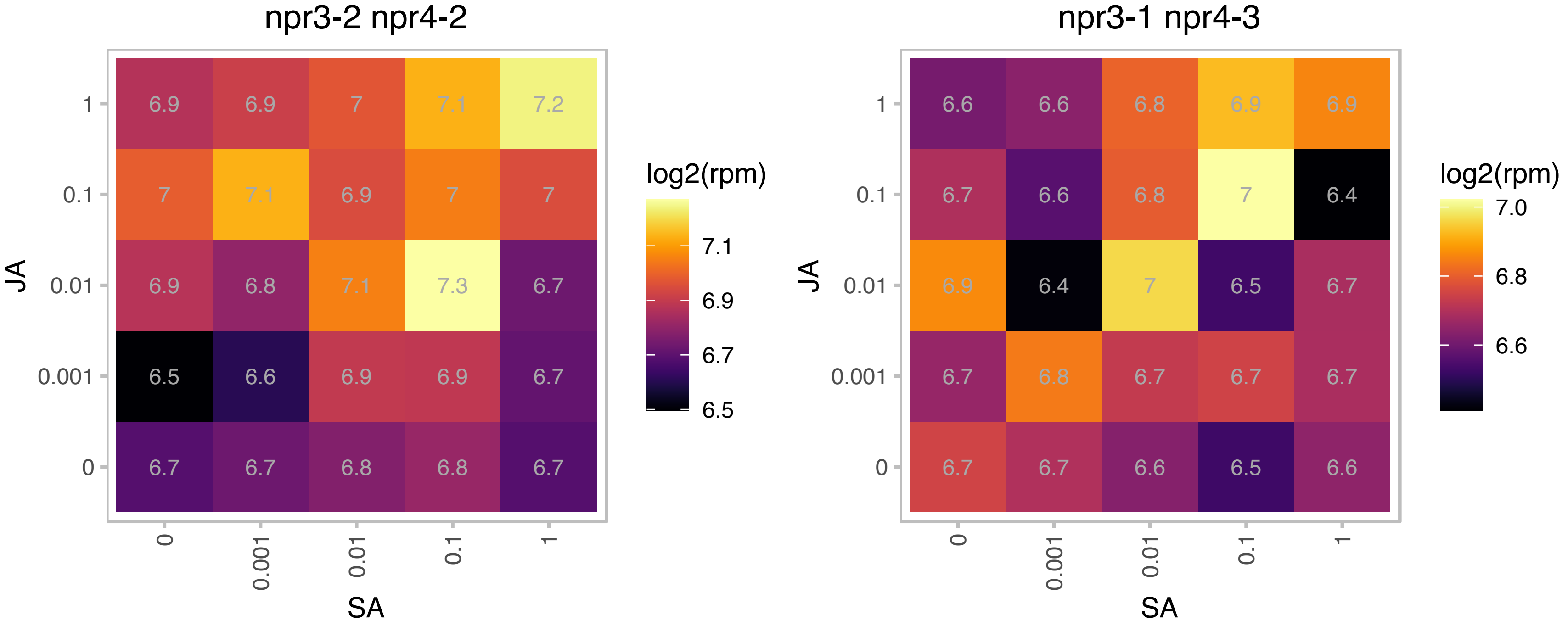

# AT4G16720.1 : NA

Ribosomal protein L23/L15e family protein

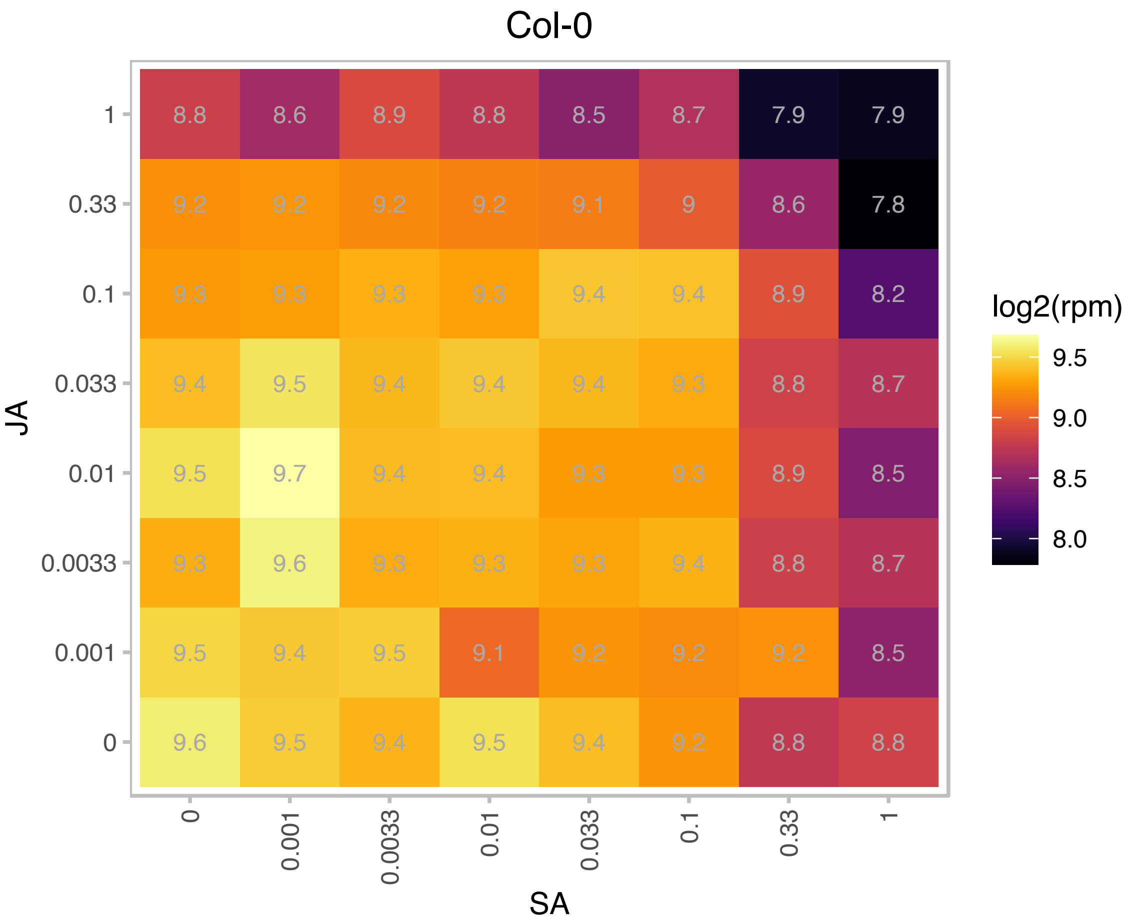


| | |
|---|---|
| Gene | AT4G16720 |
| is.expressed_genes | TRUE |
| is.responsive_genes | TRUE |
| is.apcluster | F-07 |

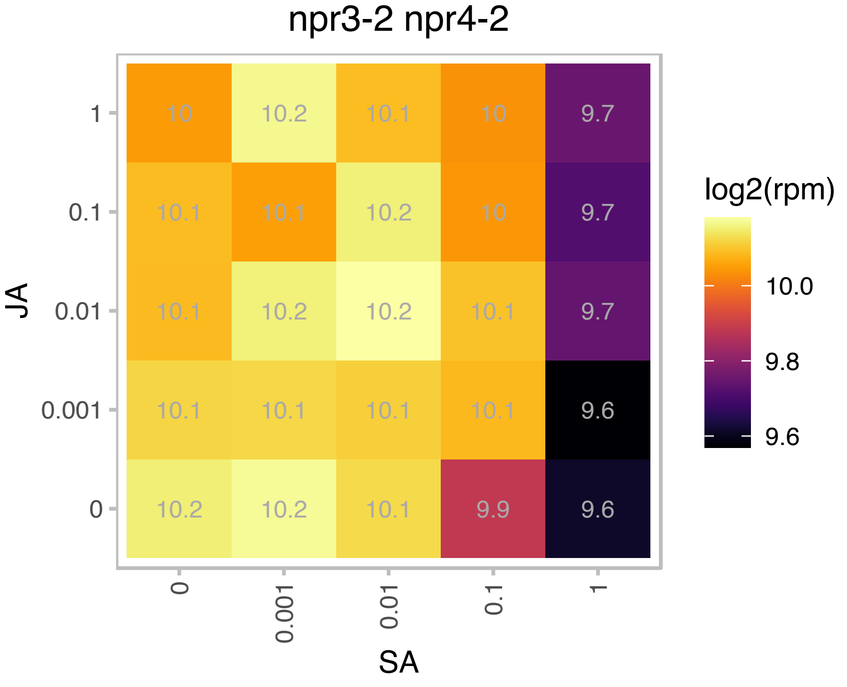


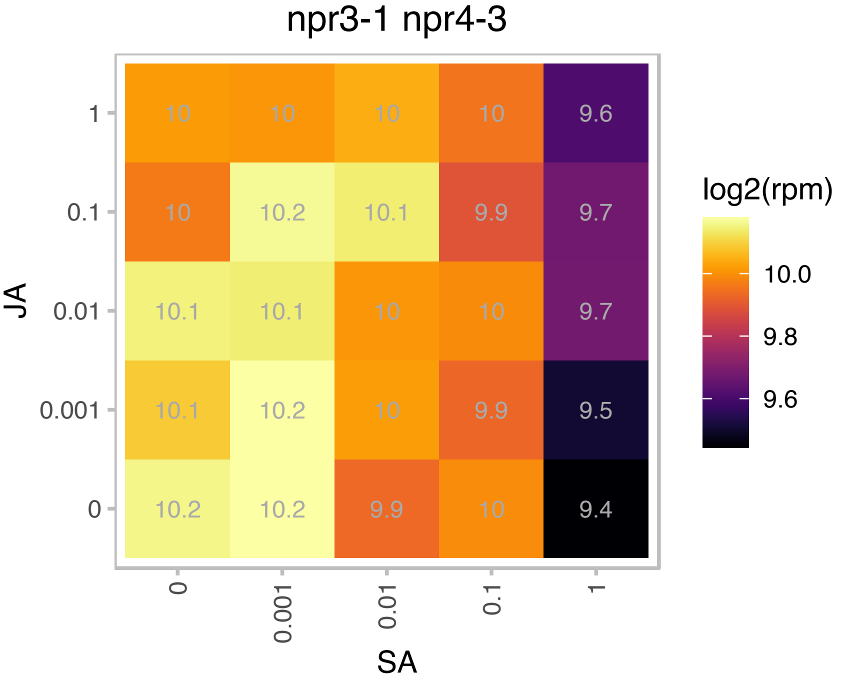

# AT4G17390.1 : NA

Ribosomal protein L23/L15e family protein

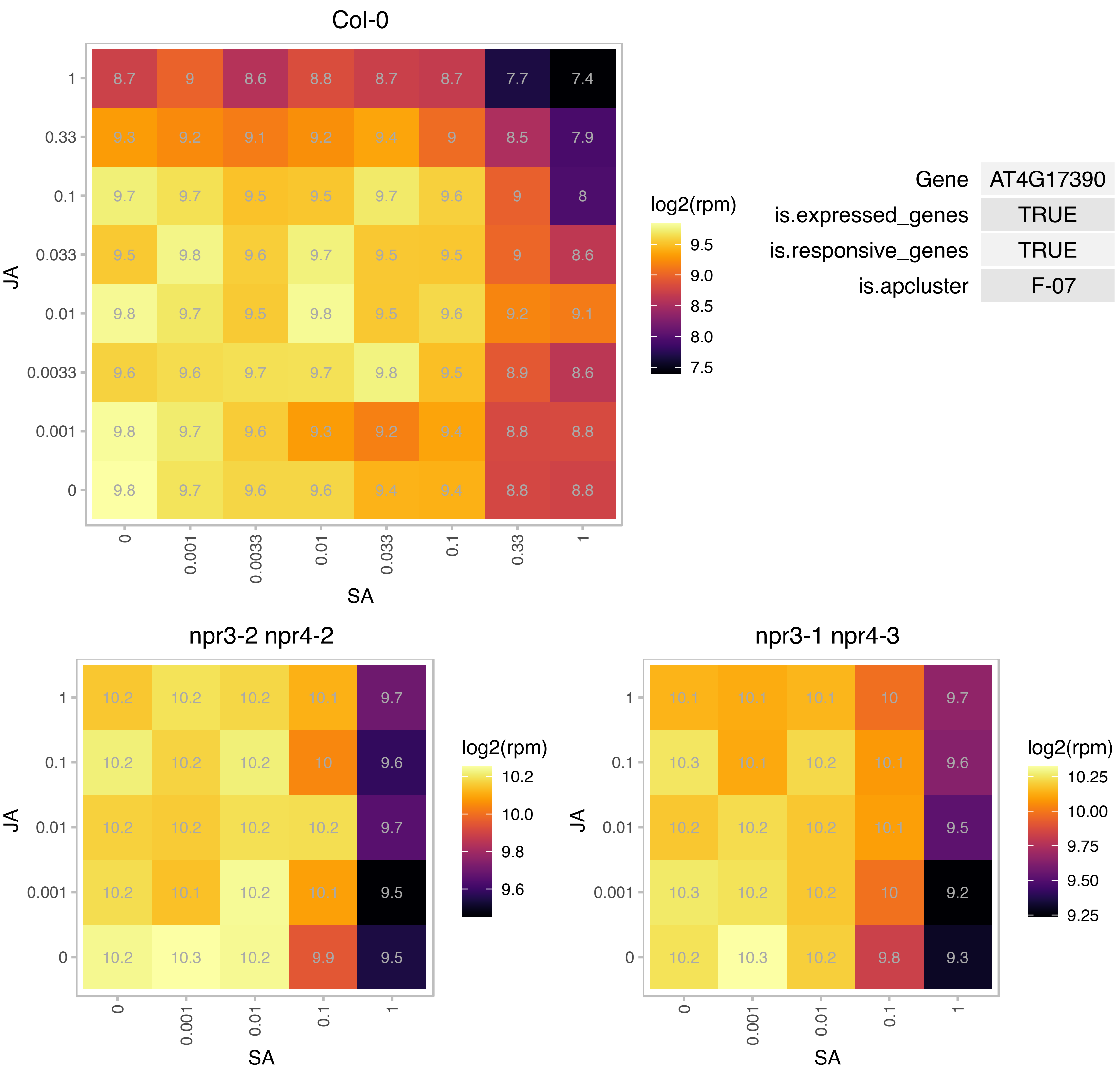


| | |
|---|---|
| Gene | AT4G17390 |
| is.expressed_genes | TRUE |
| is.responsive_genes | TRUE |
| is.apcluster | F-07 |

# AT4G18100.1 : NA

Ribosomal protein L32e

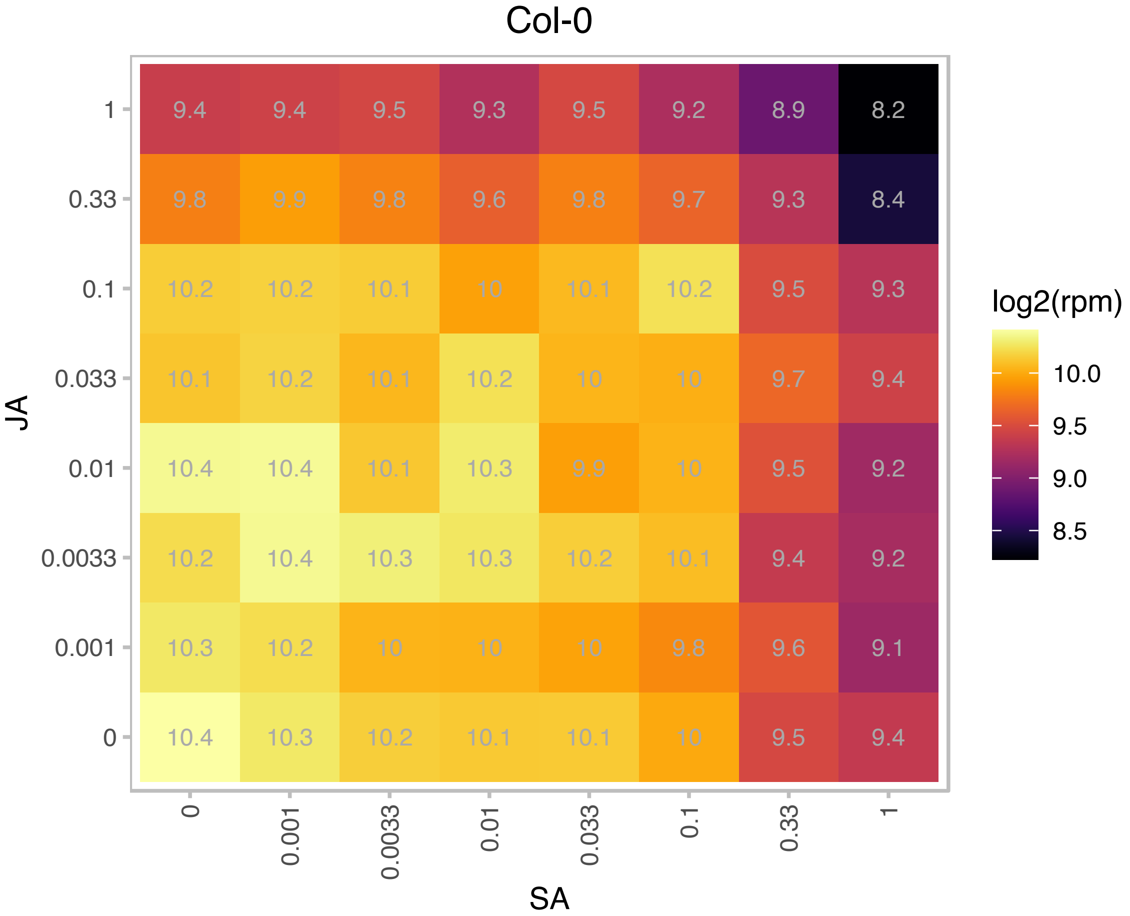


| | |
|---|---|
| Gene | AT4G18100 |
| is.expressed_genes | TRUE |
| is.responsive_genes | TRUE |
| is.apcluster | F-07 |

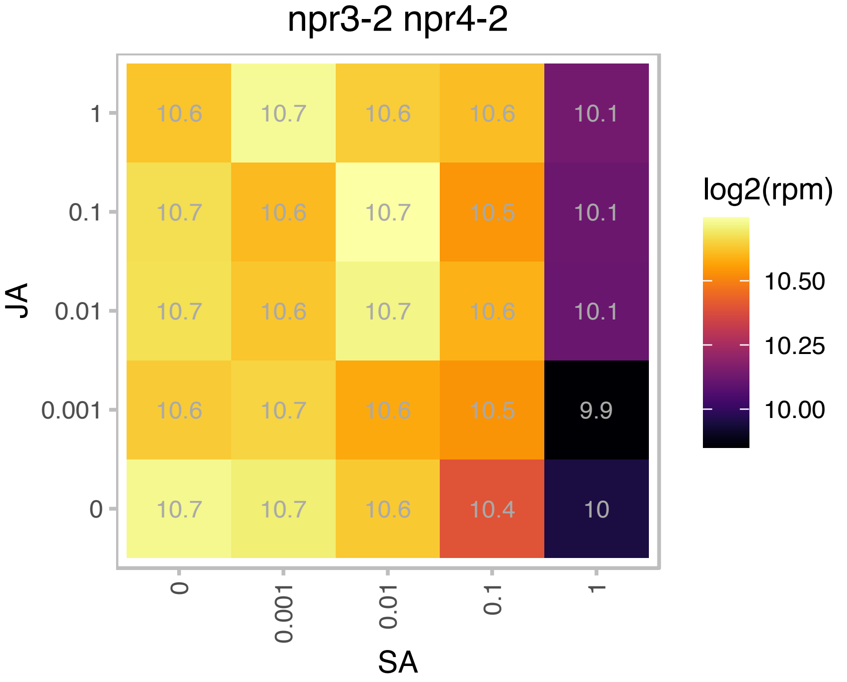


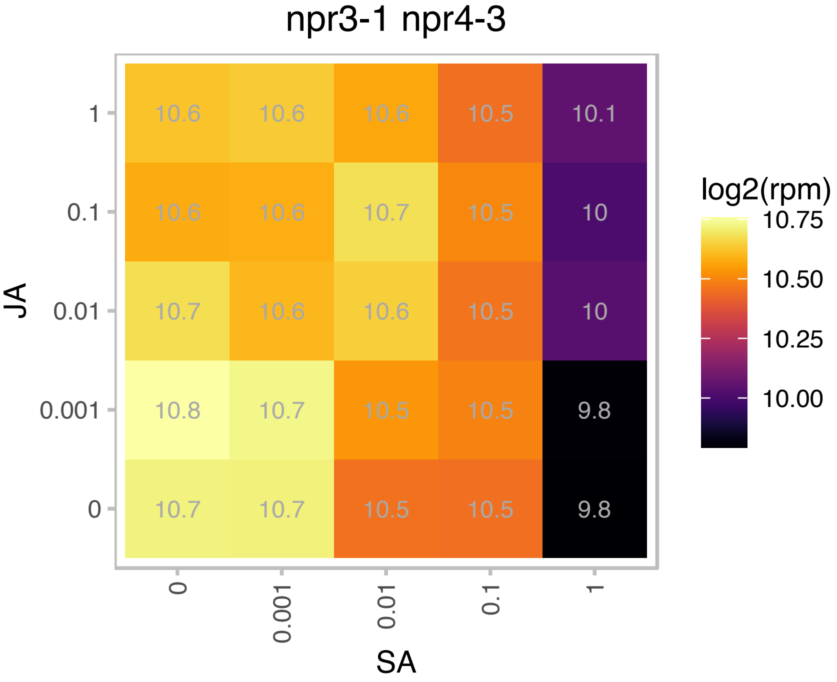

### AT4G18730.1 : RPL16B

ribosomal protein L16B
encodes a cytosolic ribosomal protein L16, which is a constituent of 60S large ribosomal complex. Gene is expressed in root and shoot apical meristems and in lateral root primordia. Expression in lateral root primordia is induced by auxin.

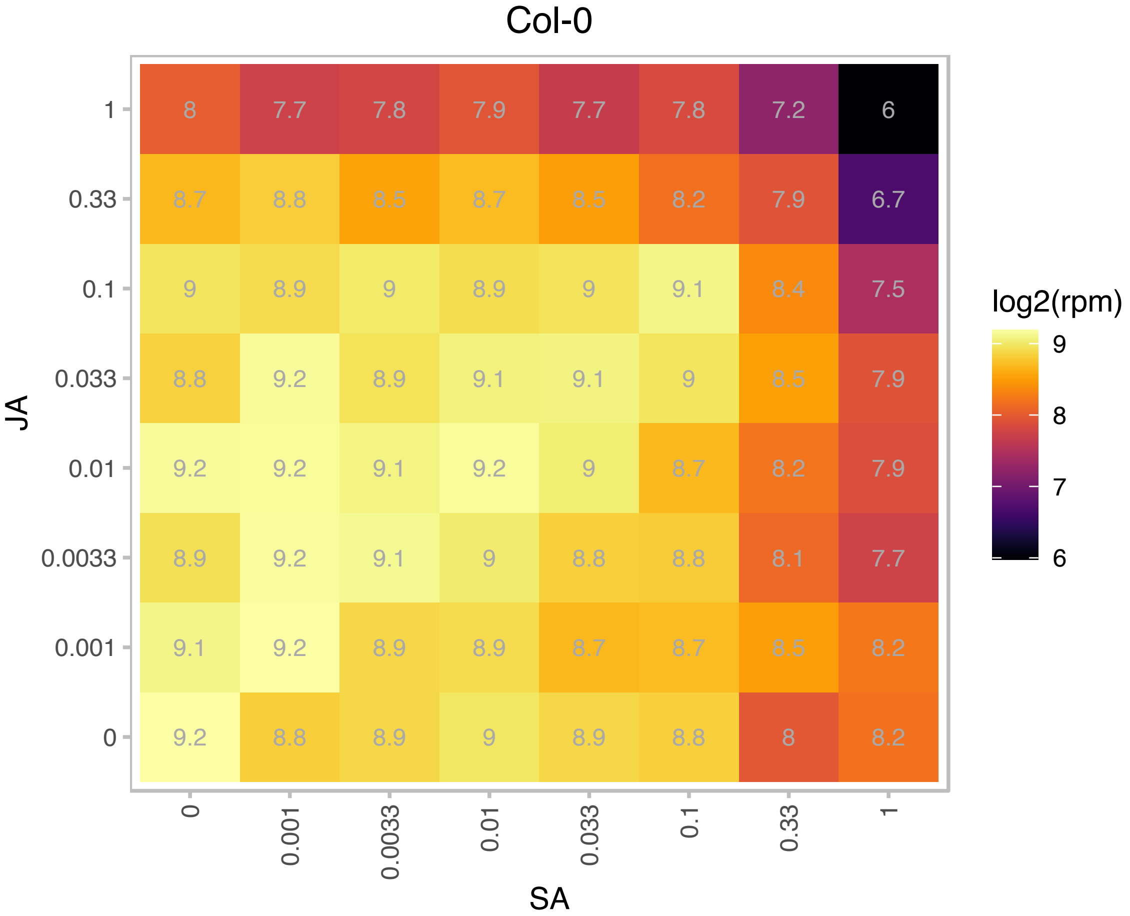


| | |
|---|---|
| Gene | AT4G18730 |
| is.expressed_genes | TRUE |
| is.responsive_genes | TRUE |
| is.apcluster | F-07 |

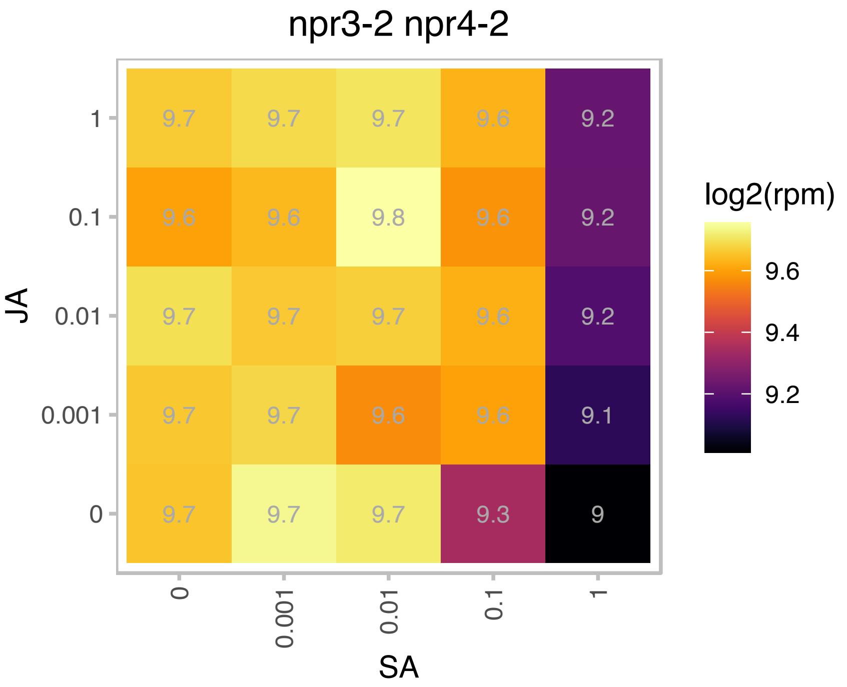


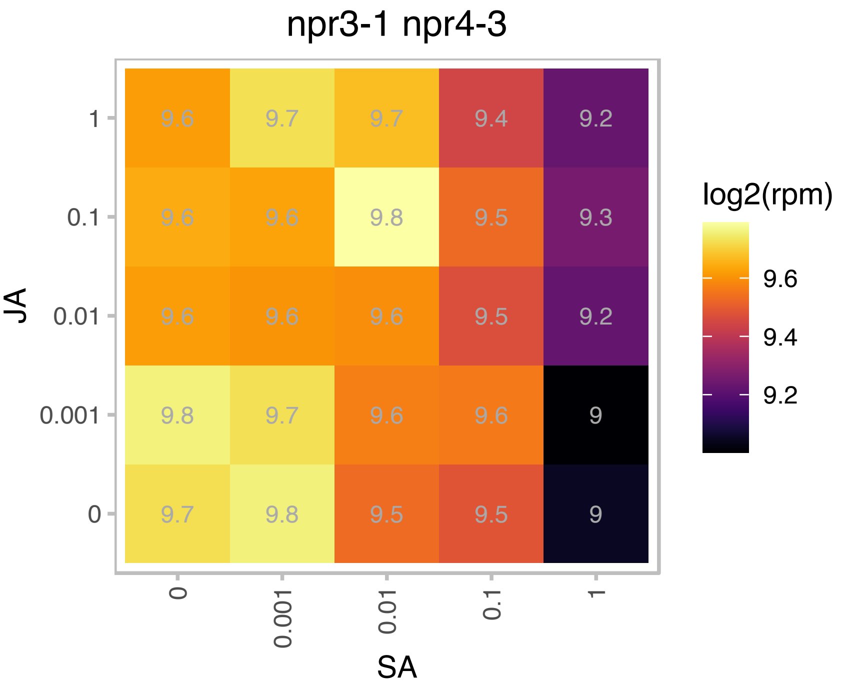

### AT4G20440.1 : smB

small nuclear ribonucleoprotein associated protein B

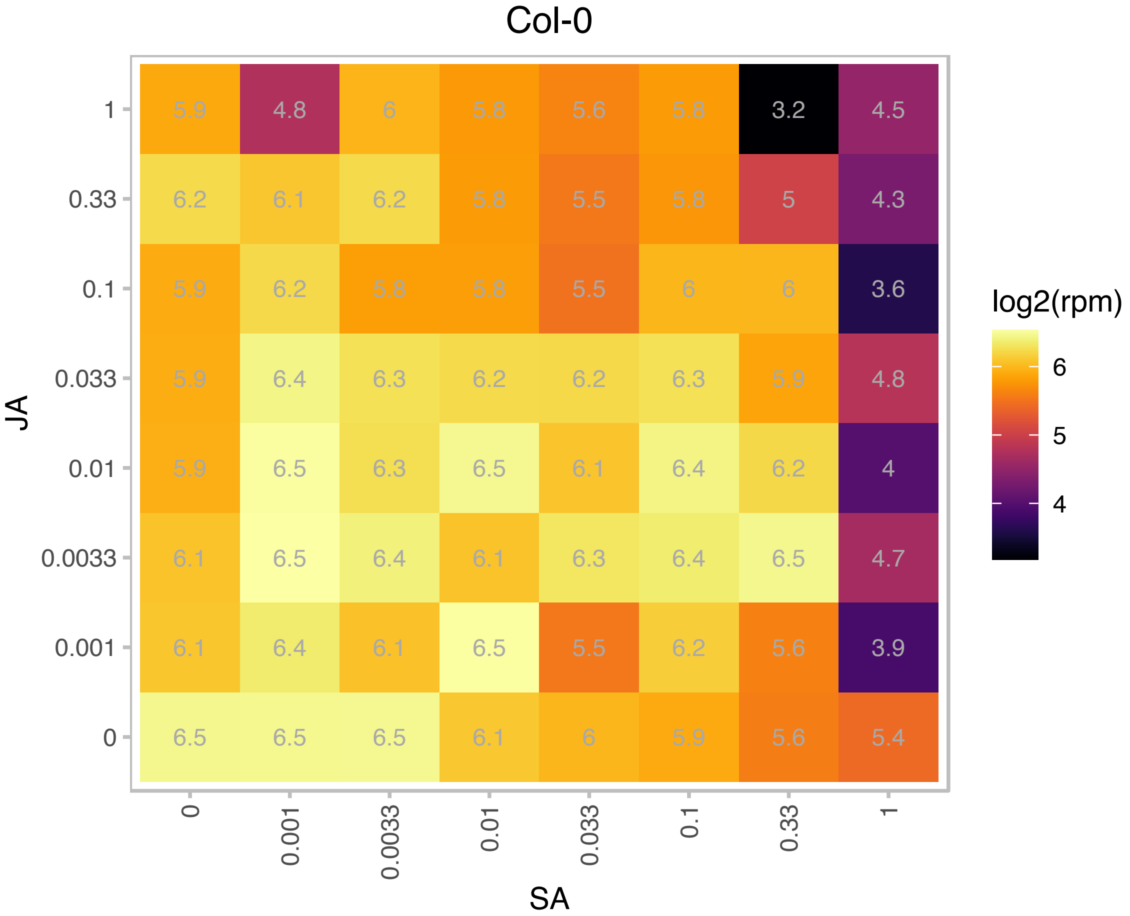


| | |
|---|---|
| Gene | AT4G20440 |
| is.expressed_genes | TRUE |
| is.responsive_genes | TRUE |
| is.apcluster | F-07 |

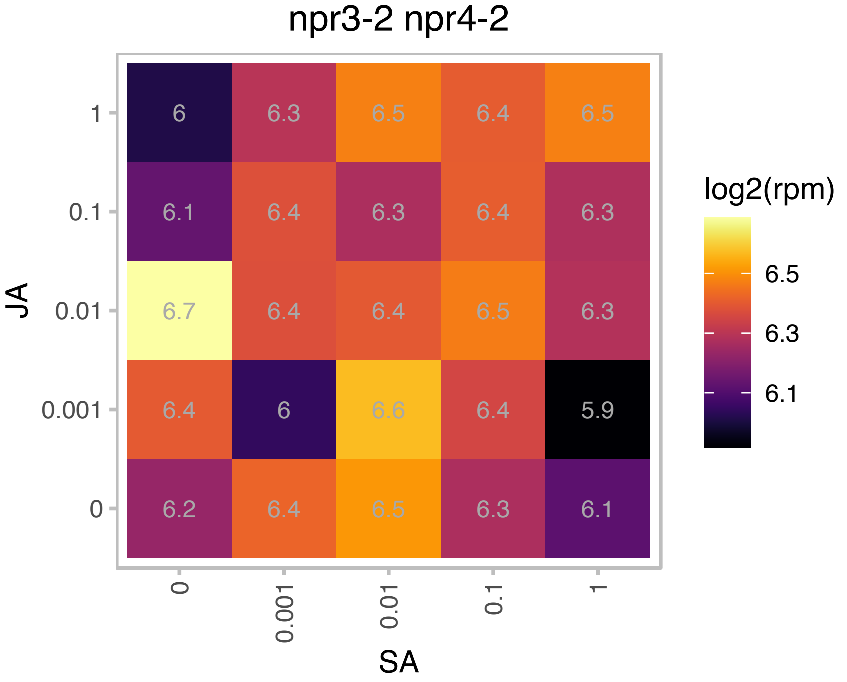


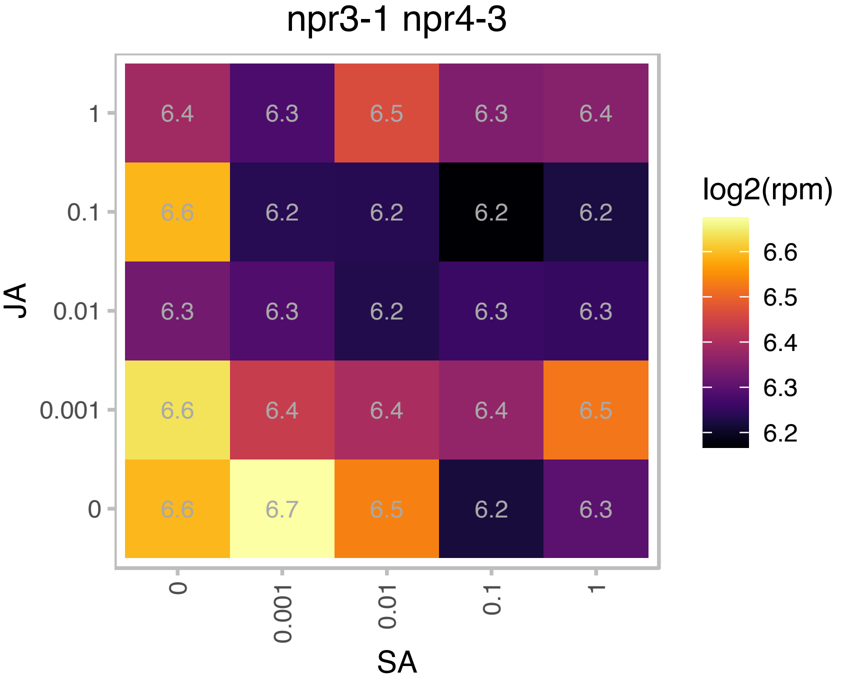

# AT4G21870.1 : NA

HSP20-like chaperones superfamily protein

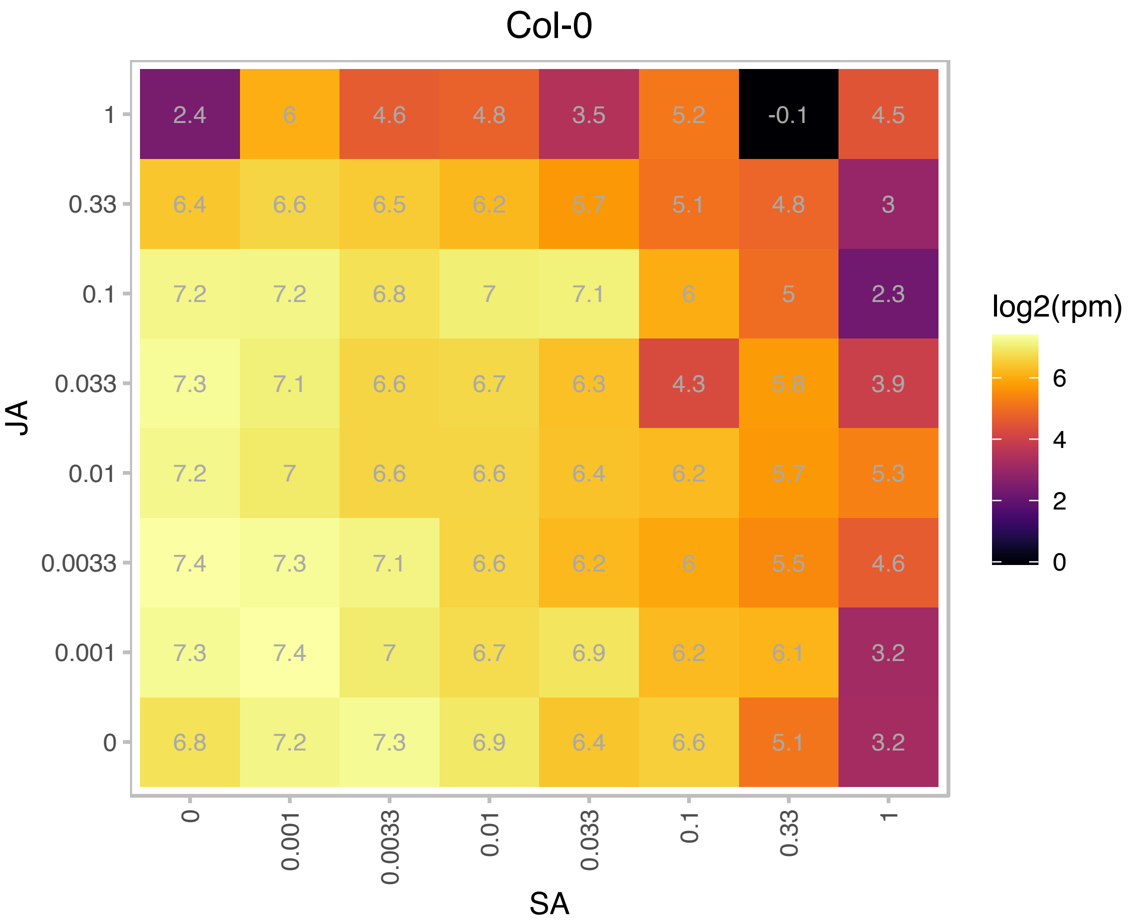


| | |
|---|---|
| Gene | AT4G21870 |
| is.expressed_genes | TRUE |
| is.responsive_genes | TRUE |
| is.apcluster | F-07 |

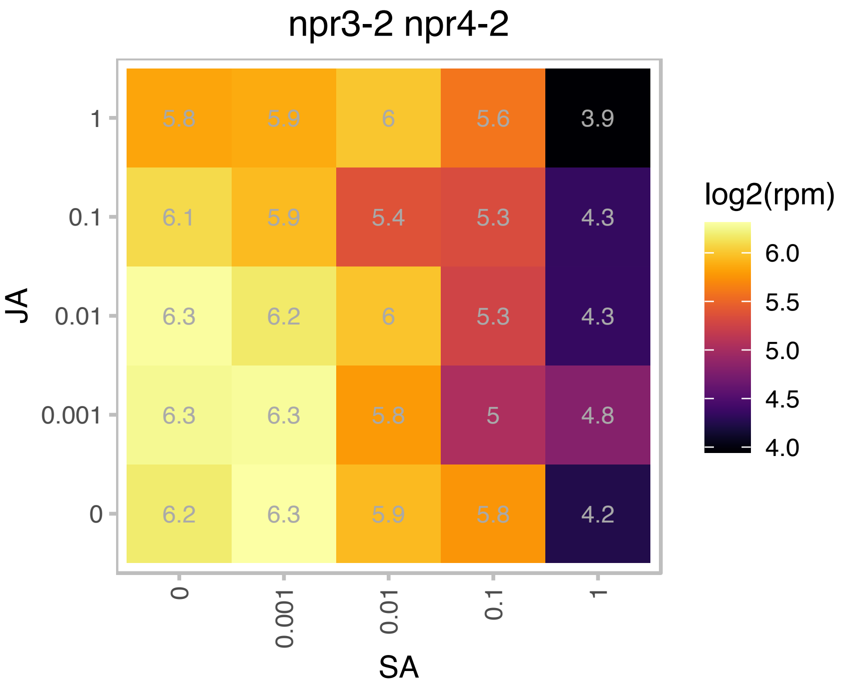


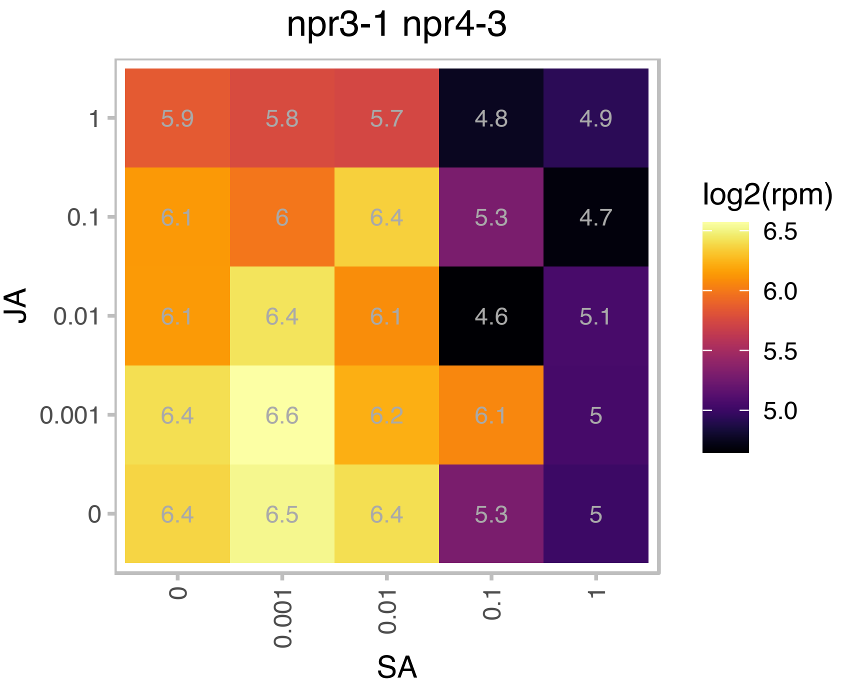

### AT4G25630.1 : ATFIB2, FIB2

fibrillarin 2

encodes a fibrillarin, a key nucleolar protein in eukaryotes which associates with box C/D small nucleolar RNAs (snoRNAs) directing 2'-O-ribose methylation of the rRNA. This gene also encodes a novel box C/D snoRNA, U60.2f in its fifth intron that accumulates in seedlings and that their targeted residue on the 25 S rRNA is methylated. The mRNA is cell-to-cell mobile.

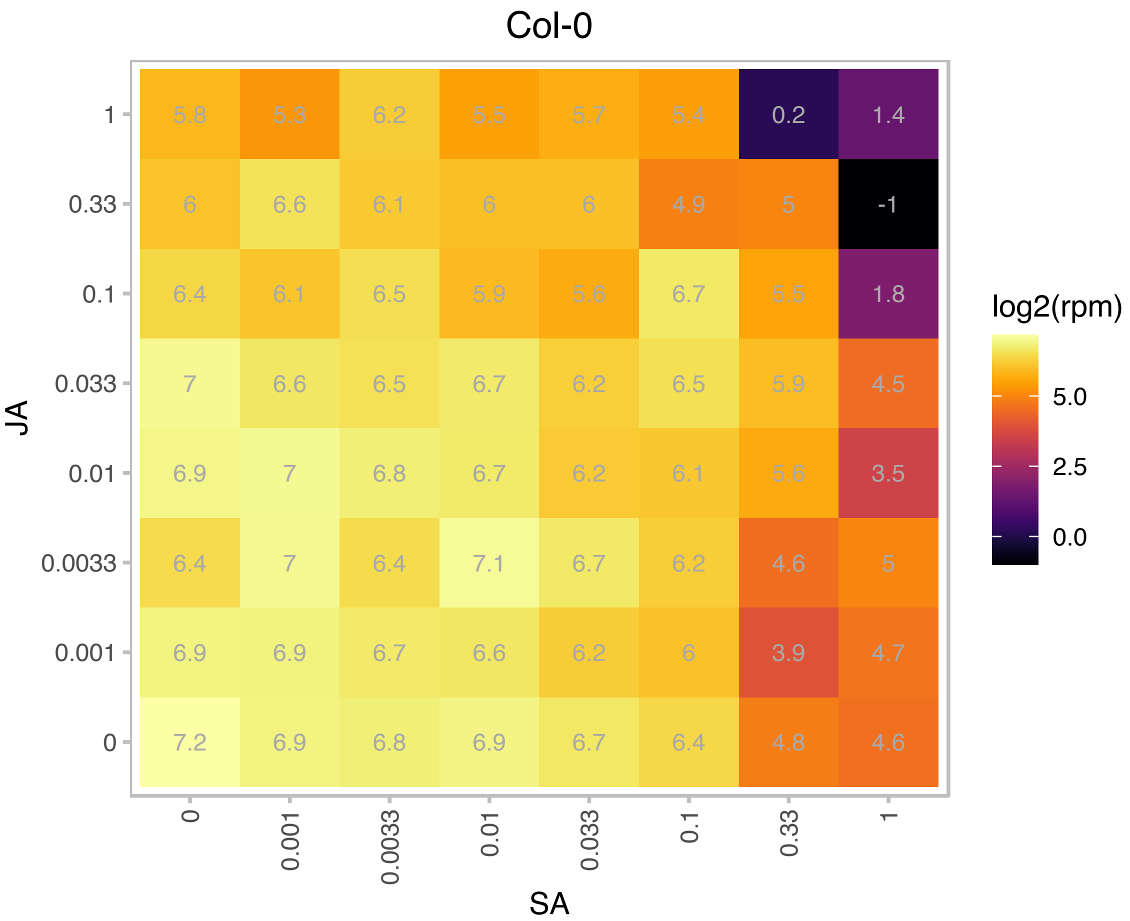


| | |
|---|---|
| Gene | AT4G25630 |
| is.expressed_genes | TRUE |
| is.responsive_genes | TRUE |
| is.apcluster | F-07 |

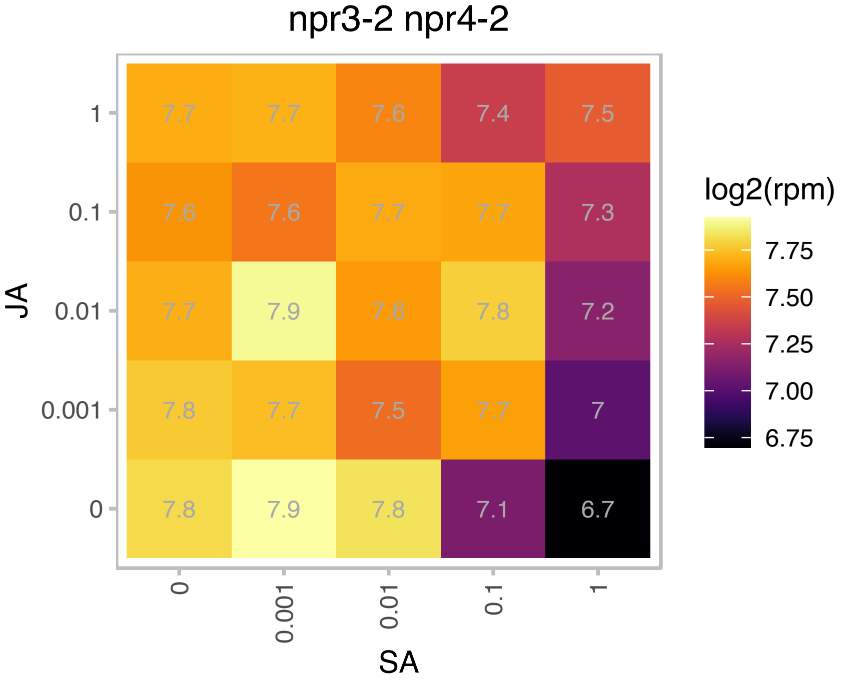


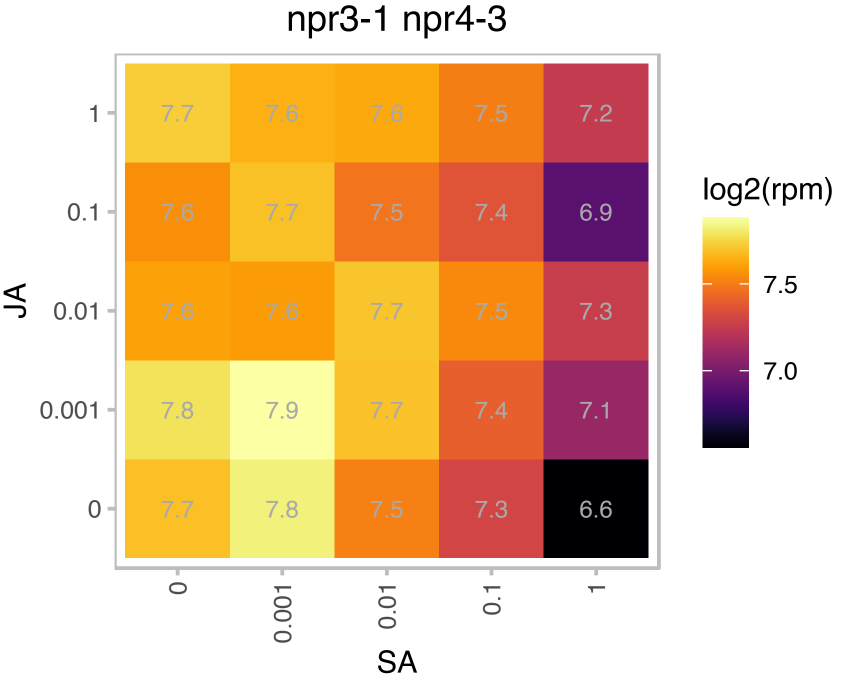

# AT4G25890.1 : NA

60S acidic ribosomal protein family

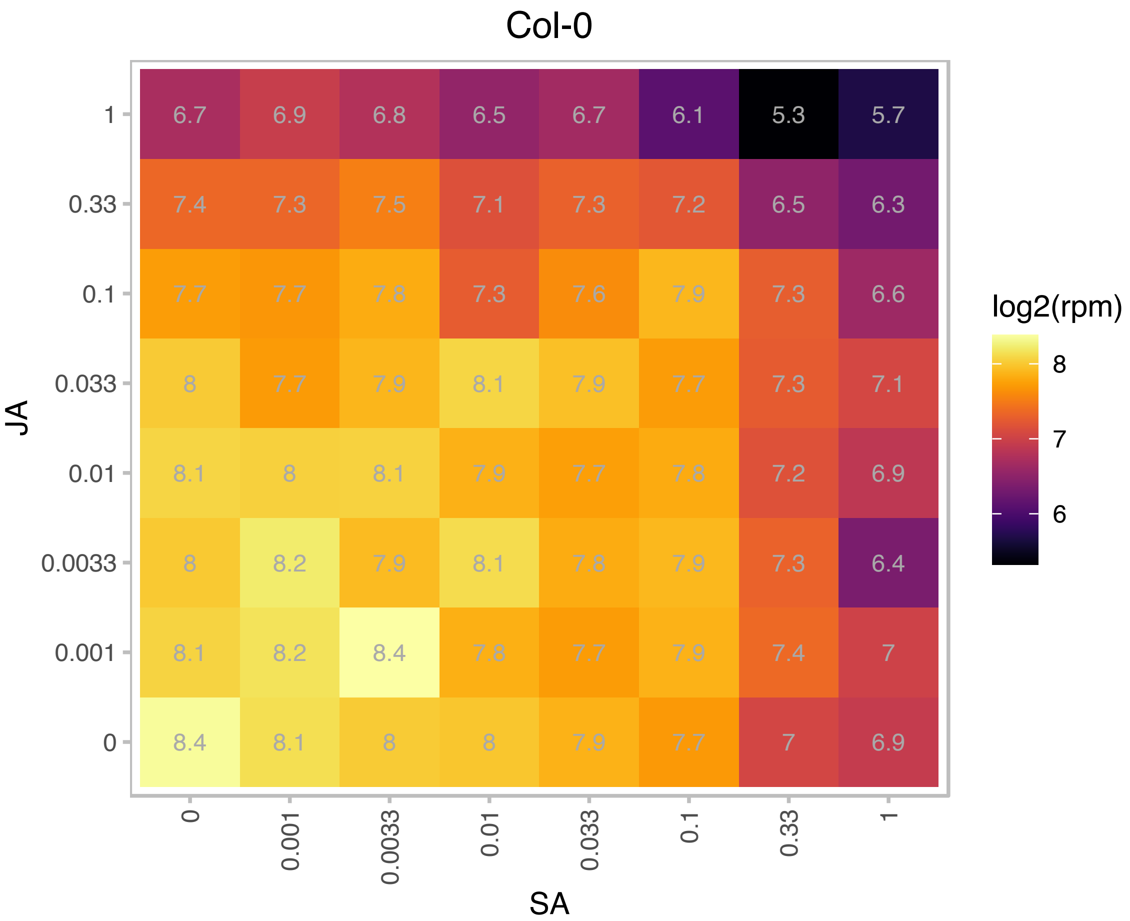


| | |
|---|---|
| Gene | AT4G25890 |
| is.expressed_genes | TRUE |
| is.responsive_genes | TRUE |
| is.apcluster | F-07 |

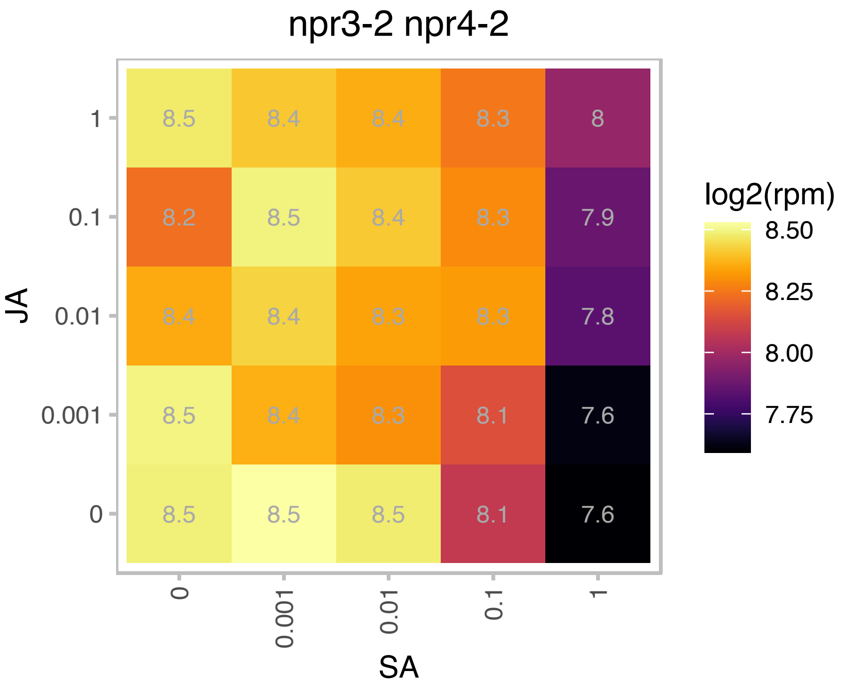


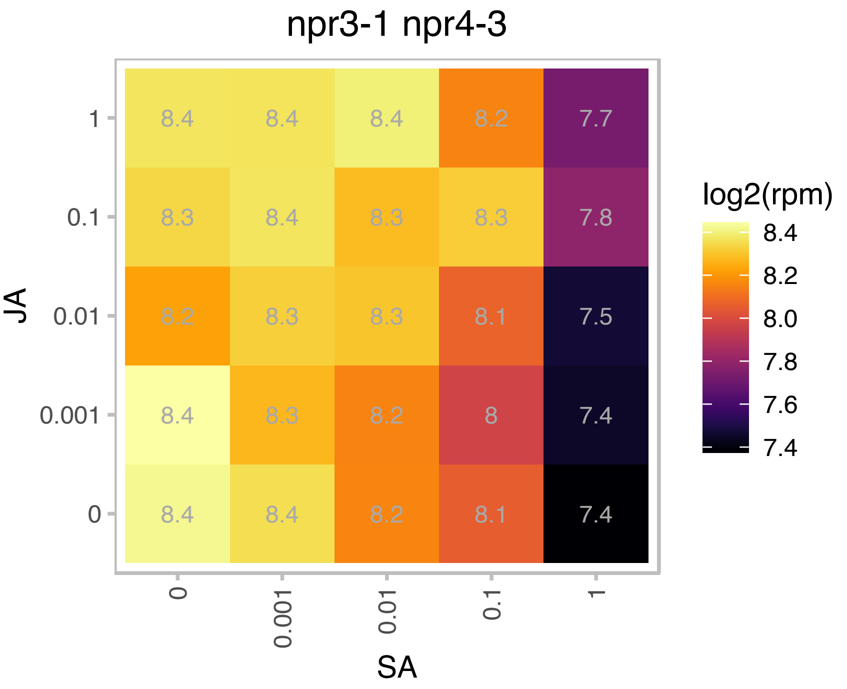

# AT4G26230.1 : NA

Ribosomal protein L31e family protein

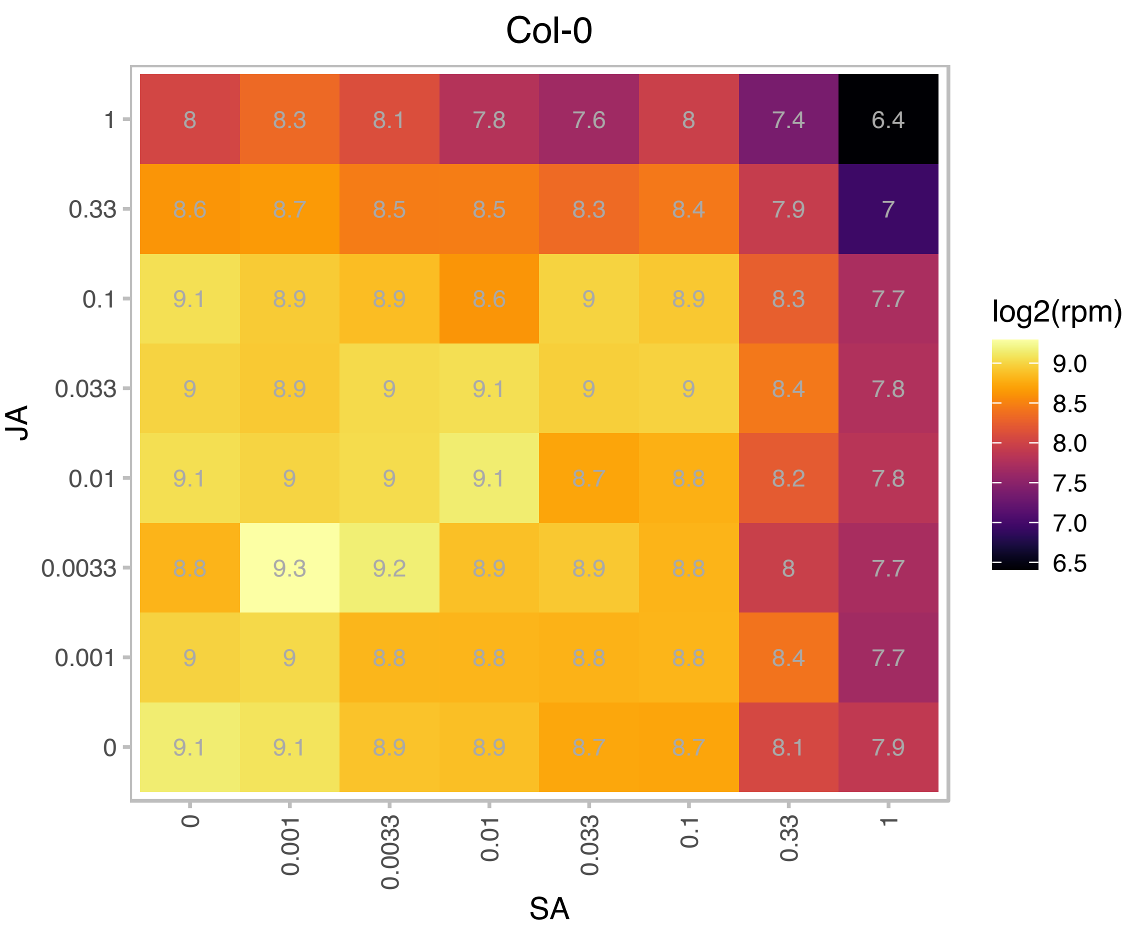


| | |
|---|---|
| Gene | AT4G26230 |
| is.expressed_genes | TRUE |
| is.responsive_genes | TRUE |
| is.apcluster | F-07 |

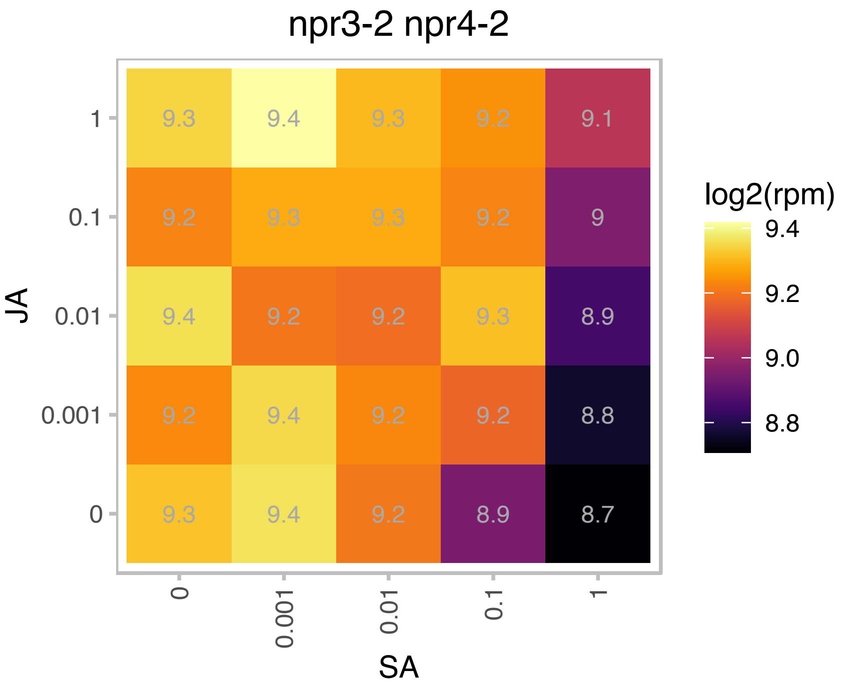


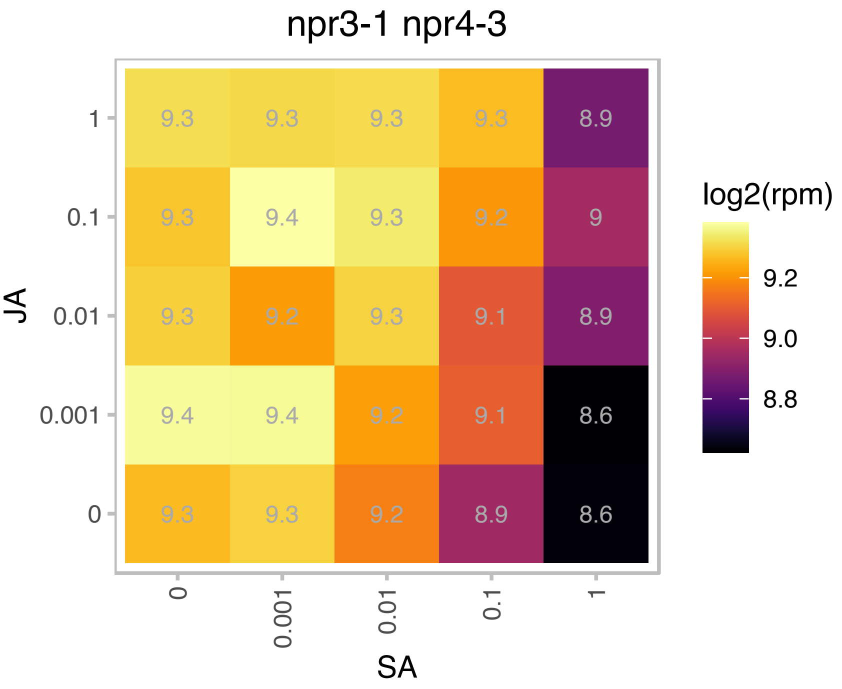

# AT4G28990.1 : NA

RNA-binding protein-related

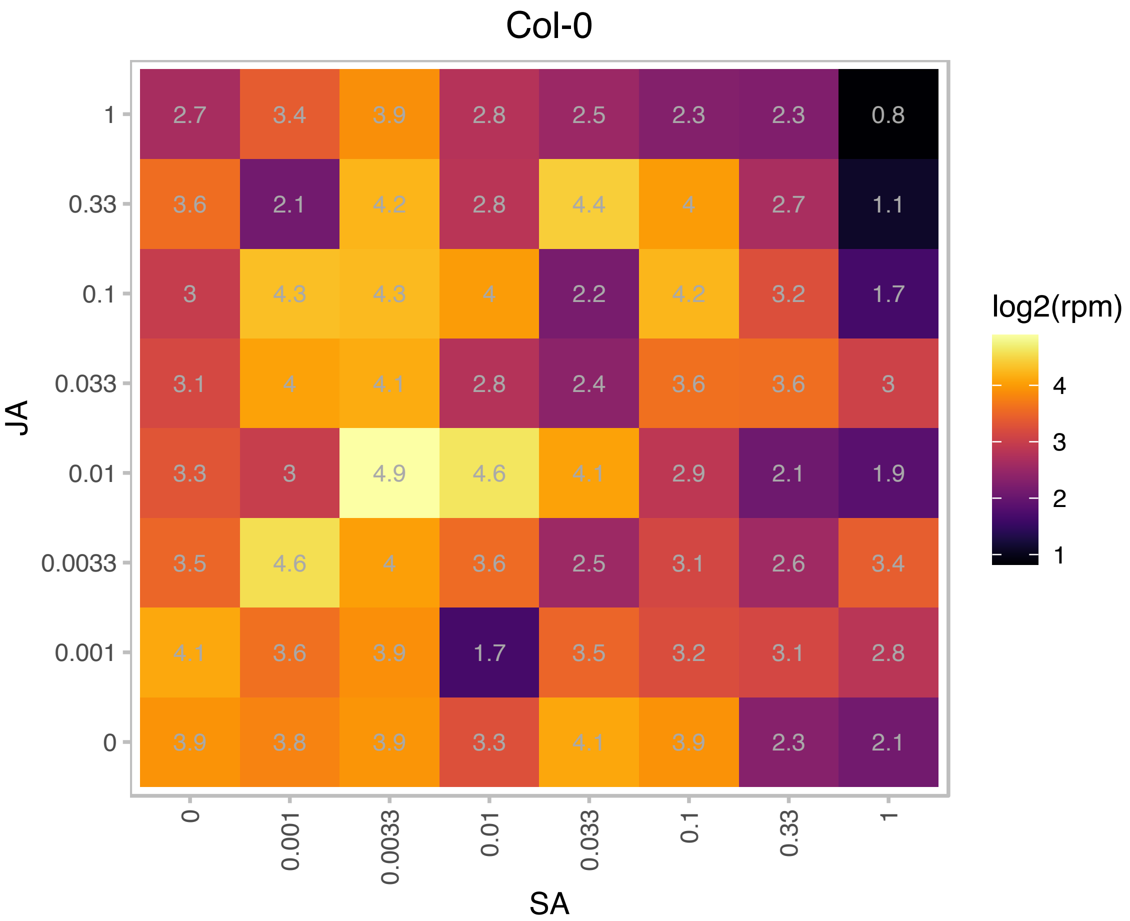


| | |
|---|---|
| Gene | AT4G28990 |
| is.expressed_genes | TRUE |
| is.responsive_genes | TRUE |
| is.apcluster | F-07 |

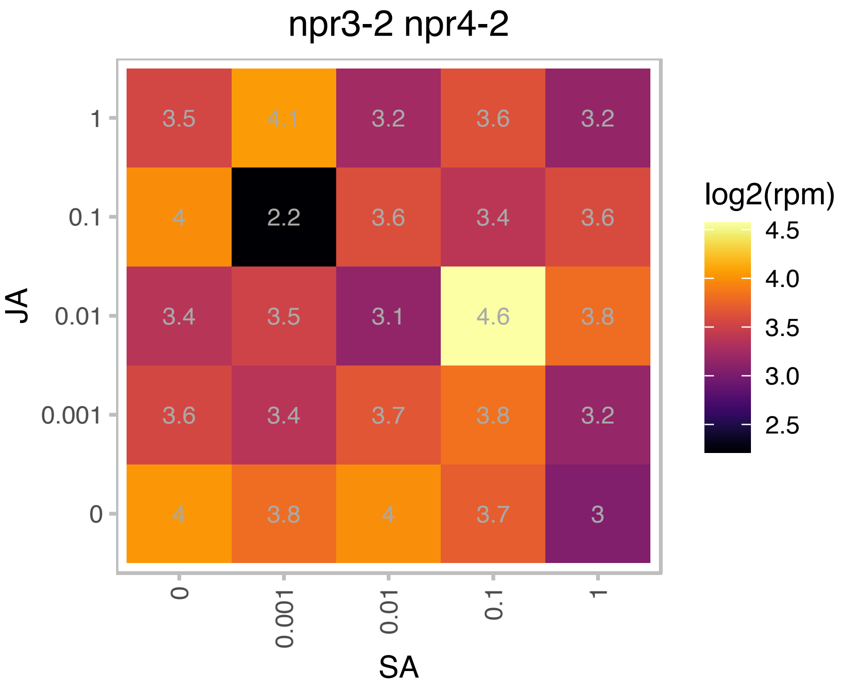


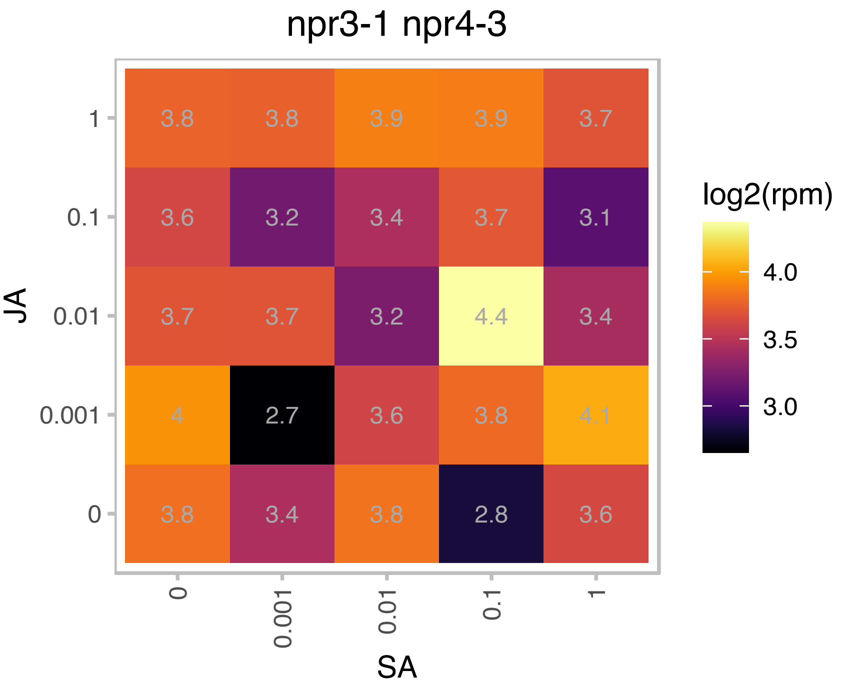

# AT4G29390.1 : NA

Ribosomal protein S30 family protein

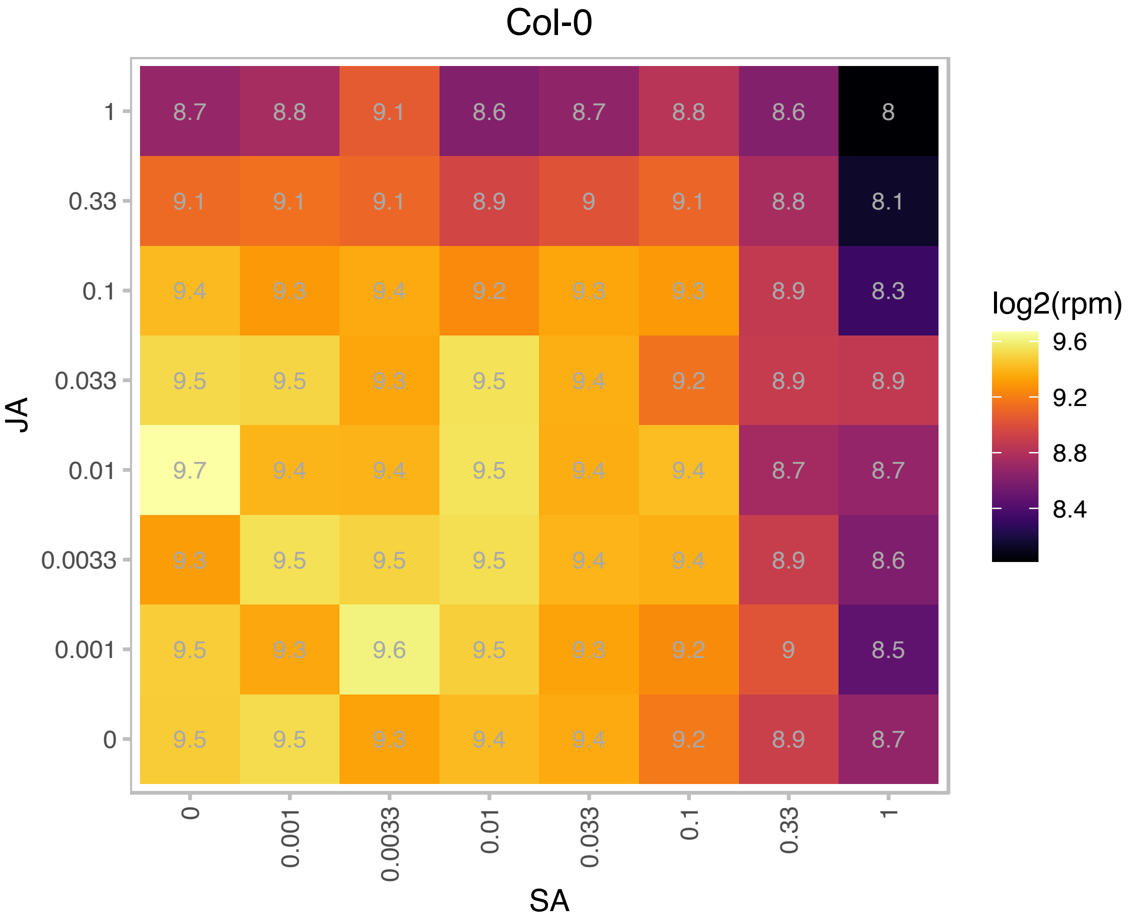


| | |
|---|---|
| Gene | AT4G29390 |
| is.expressed_genes | TRUE |
| is.responsive_genes | TRUE |
| is.apcluster | F-07 |

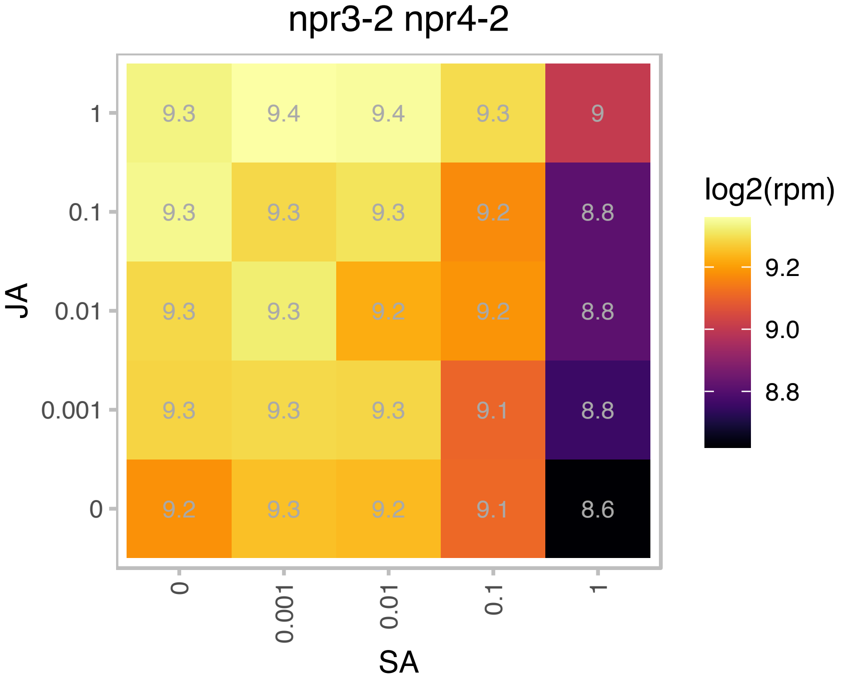


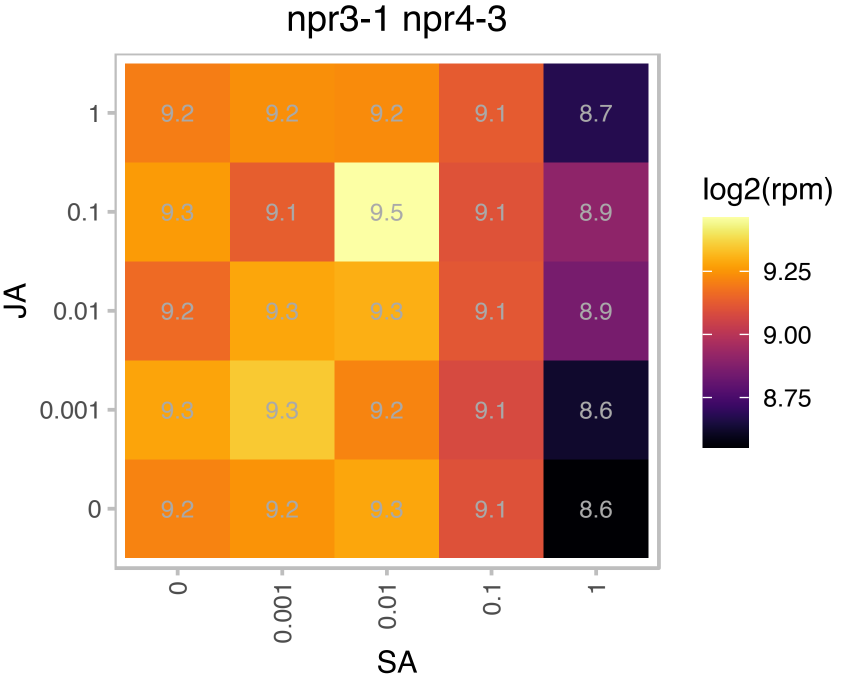

# AT4G29410.1 : NA

Ribosomal L28e protein family

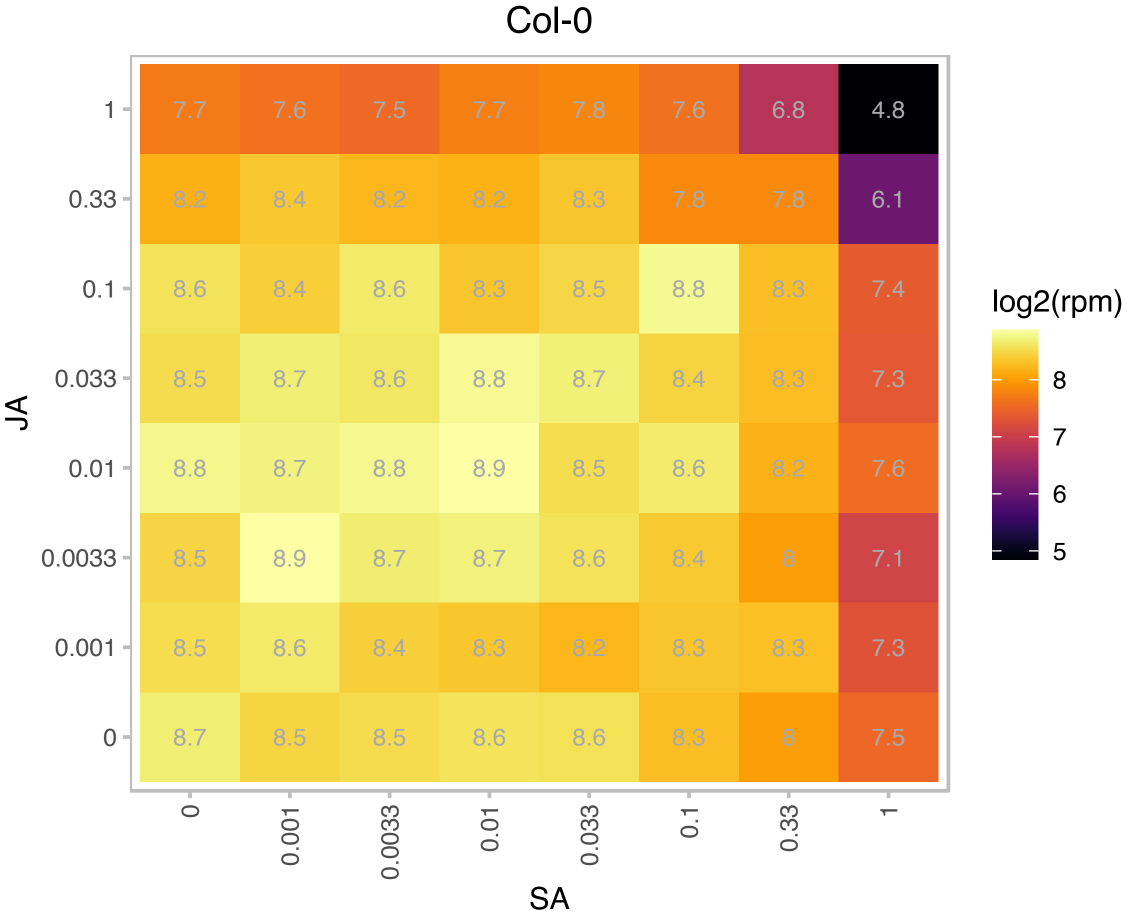


| | |
|---|---|
| Gene | AT4G29410 |
| is.expressed_genes | TRUE |
| is.responsive_genes | TRUE |
| is.apcluster | F-07 |

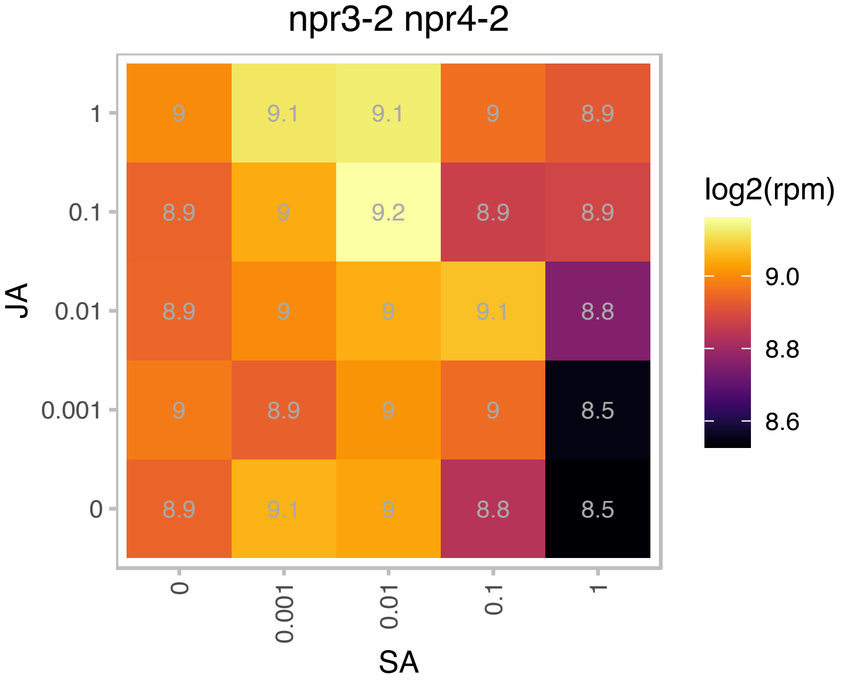


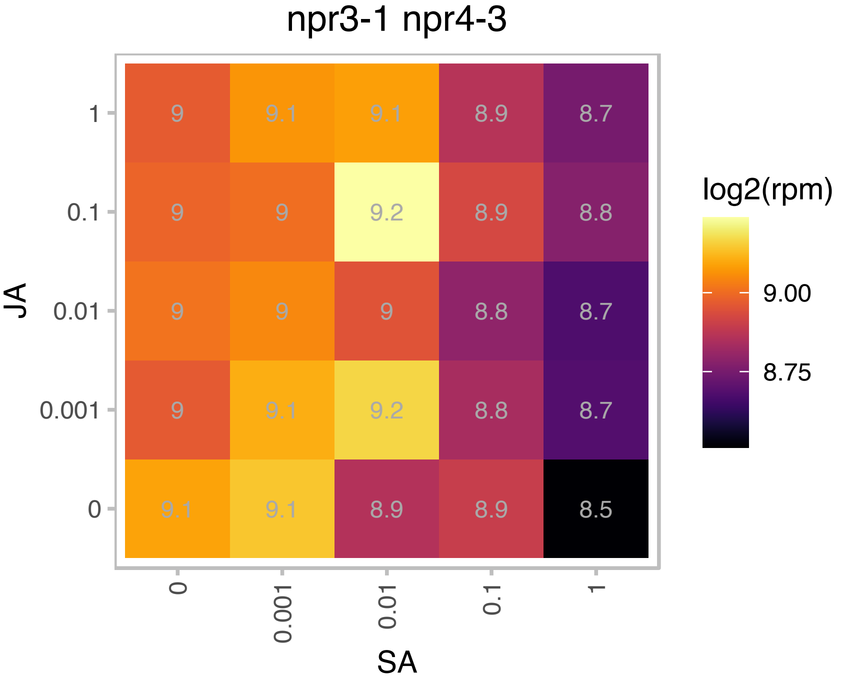

### AT4G29430.1 : rps15ae

ribosomal protein S15A E

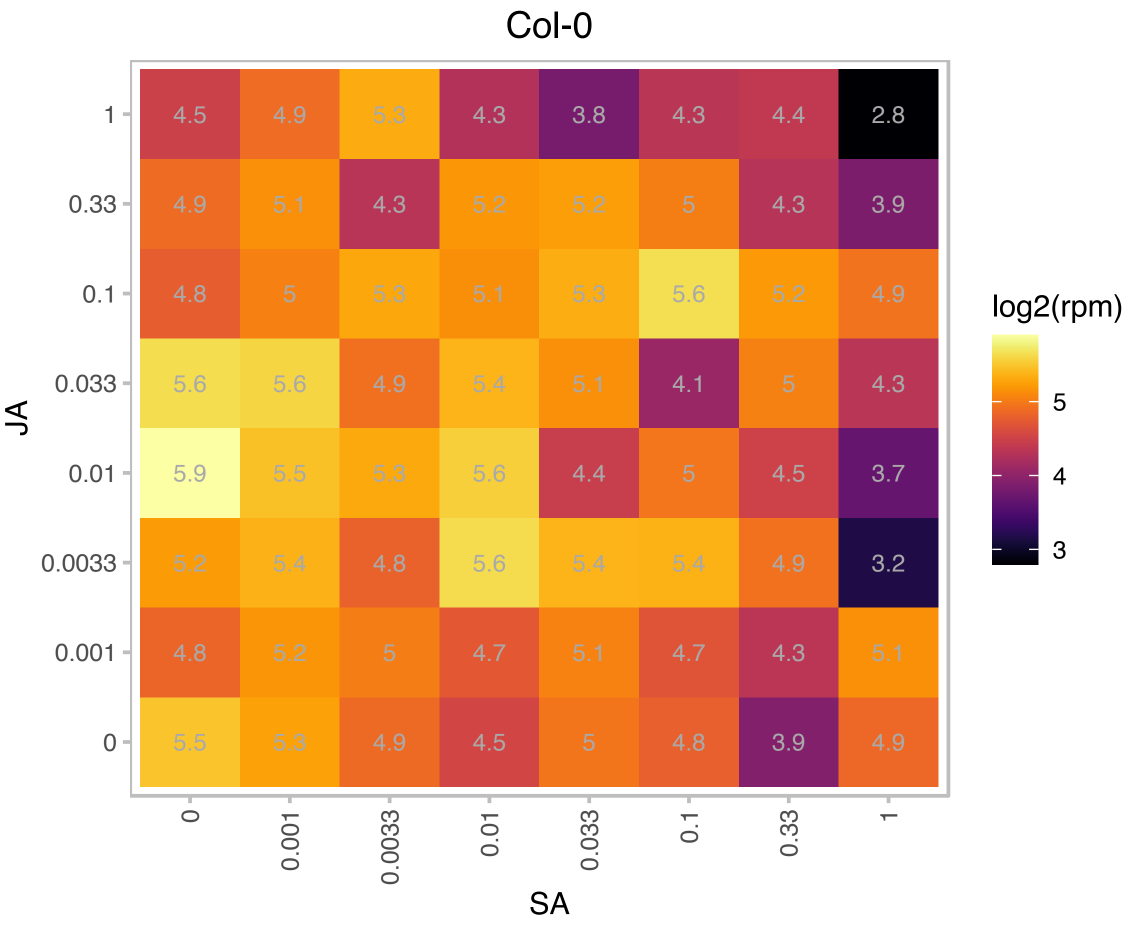


| | |
|---|---|
| Gene | AT4G29430 |
| is.expressed_genes | TRUE |
| is.responsive_genes | TRUE |
| is.apcluster | F-07 |

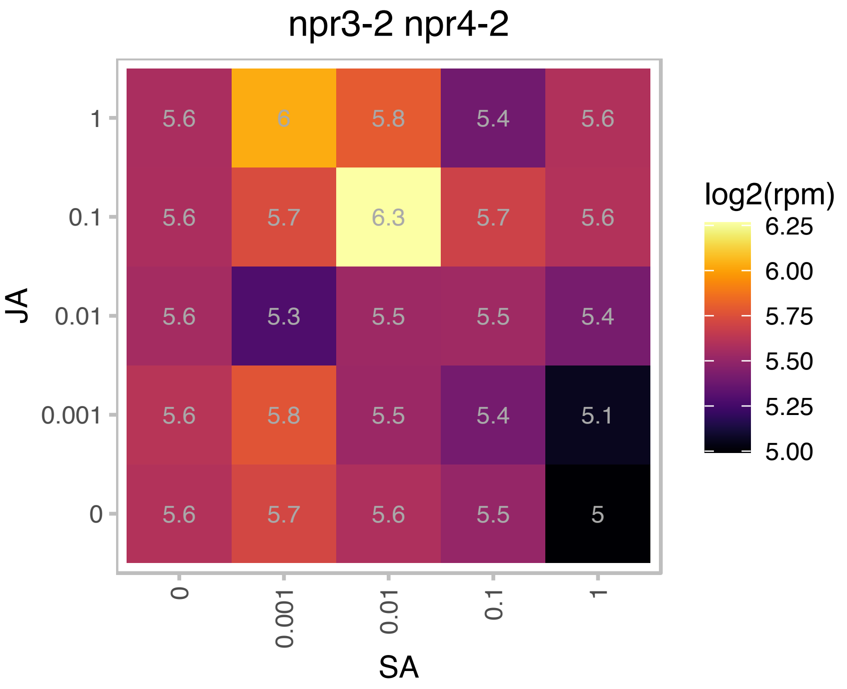


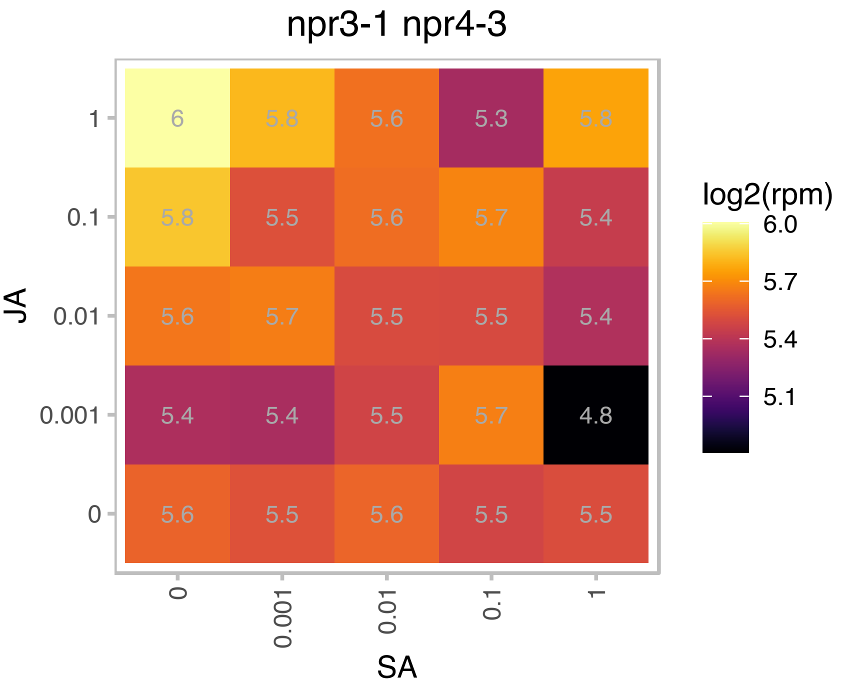

### AT4G31580.1 : At-RSZ22, AtRSZ22, RSZ22, RSZP22, SRZ

serine/arginine-rich 22

Encodes a Serine/arginine-rich (SR) protein RSZp22. SR proteins are splicing regulators that share a modular structure consisting of one or two N-terminal RNA recognition motif domains and a C-terminal RS-rich domain. RSZp22 is located in the nucleolus. It is a nucleocytoplasmic shuttling protein and an interacting partner to the Arabidopsis U1-70K. Barta et al (2010) have proposed a nomenclature for Serine/Arginine-Rich Protein Splicing Factors (SR proteins): Plant Cell. 2010, 22:2926.

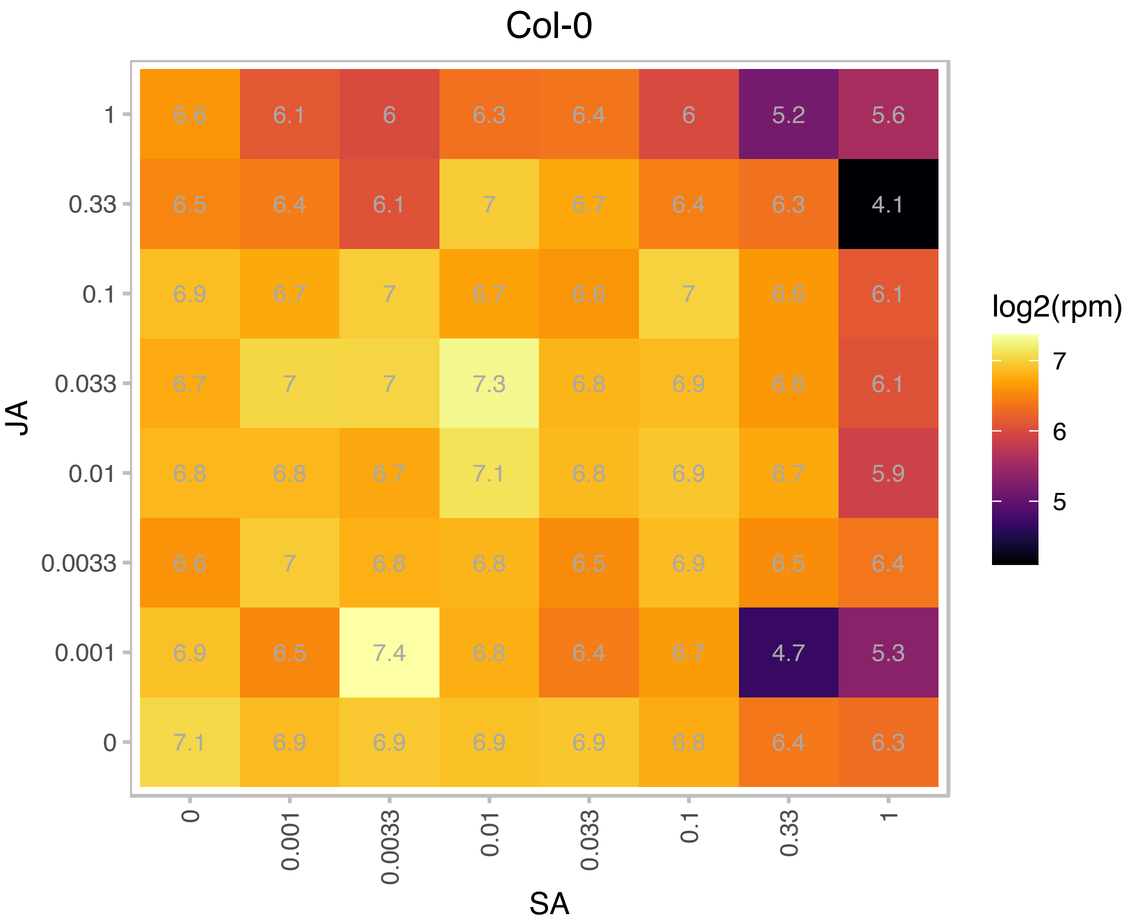


| | |
|---|---|
| Gene | AT4G31580 |
| is.expressed_genes | TRUE |
| is.responsive_genes | TRUE |
| is.apcluster | F-07 |

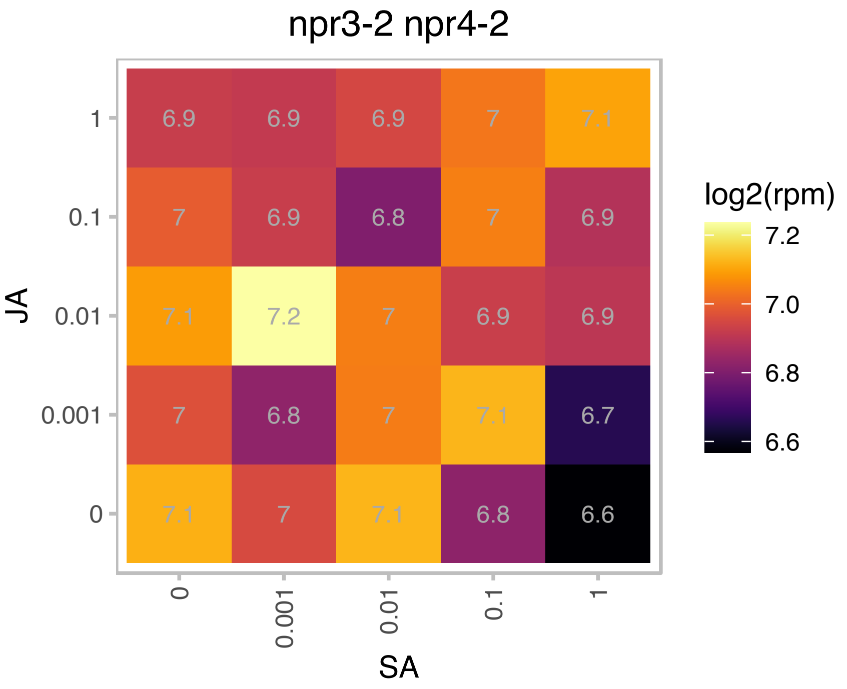


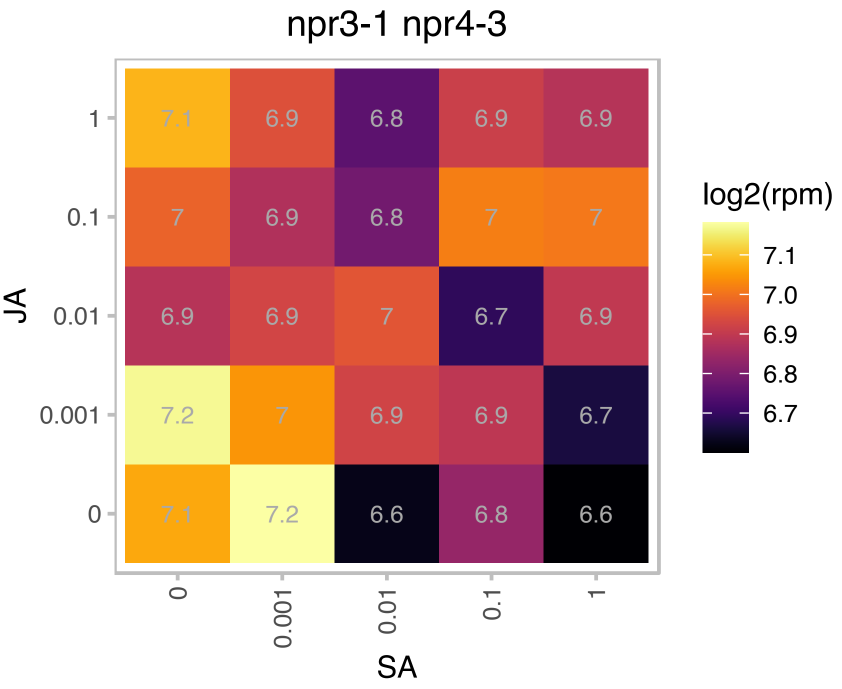

### AT4G31700.1 : AtRPS6, RPS6, RPS6A

ribosomal protein S6

Encodes a putative ribosomal protein S6 (rps6a). RPS6A and RPS6B are fully redundant and essential during gametogenesis.

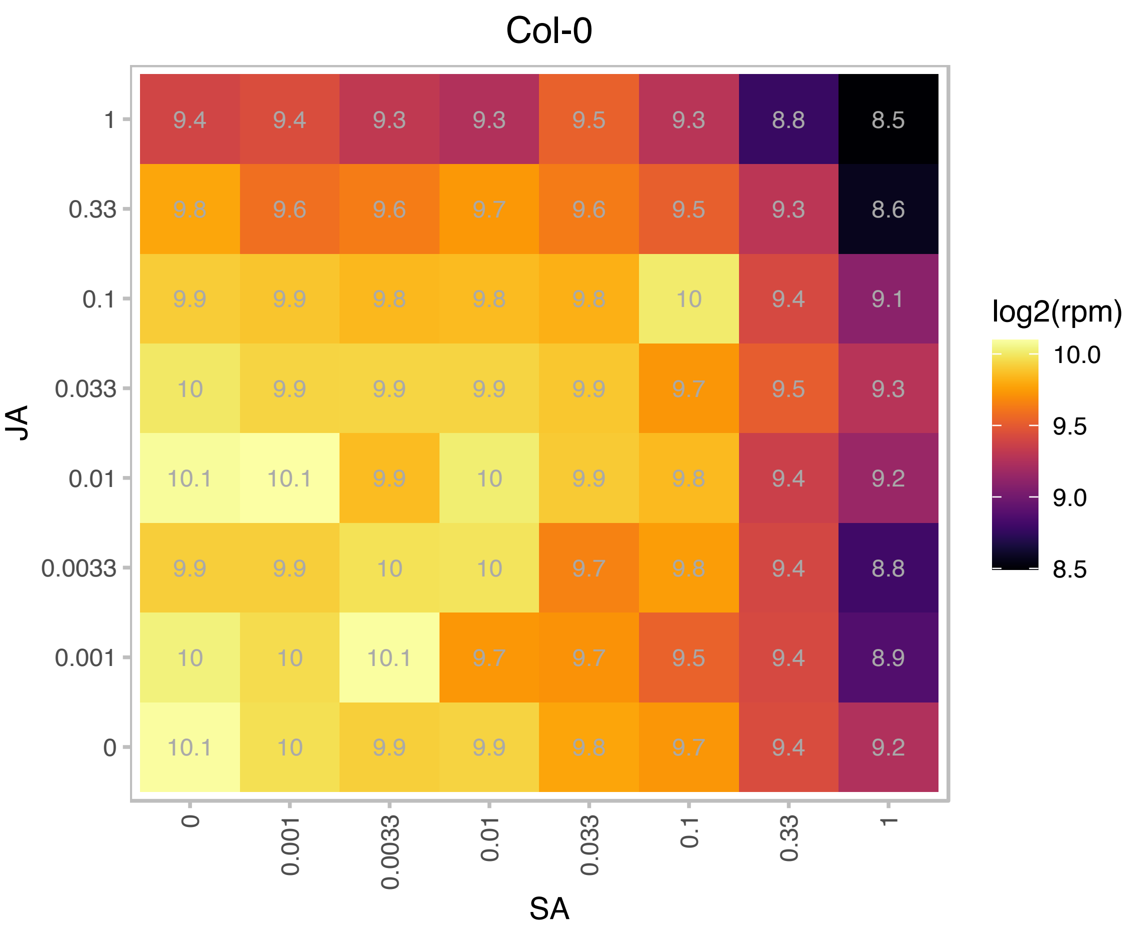


| | |
|---|---|
| Gene | AT4G31700 |
| is.expressed_genes | TRUE |
| is.responsive_genes | TRUE |
| is.apcluster | F-07 |

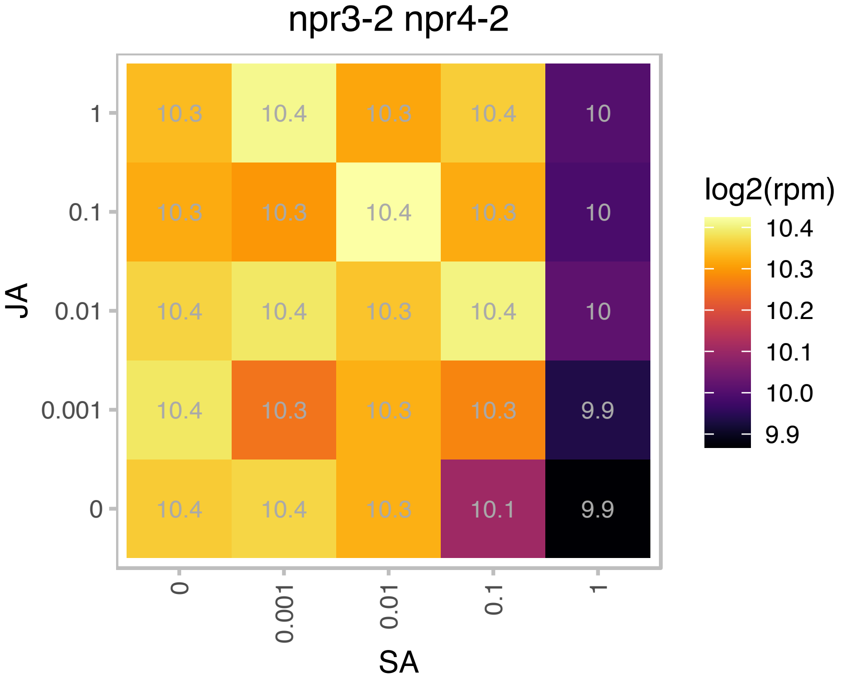


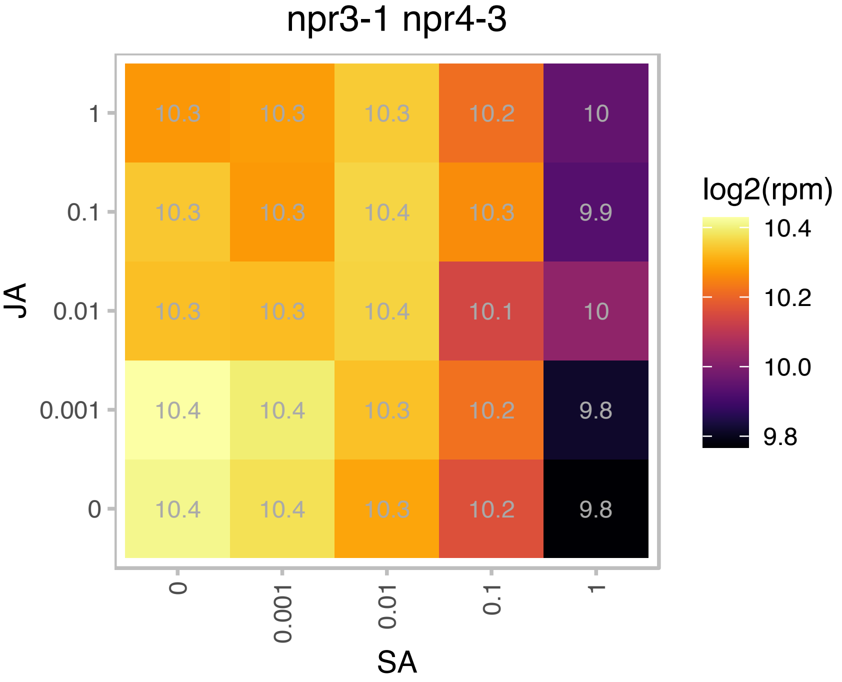

### AT4G36250.1 : ALDH3F1

aldehyde dehydrogenase 3F1

Encodes a putative aldehyde dehydrogenase. The gene is not responsive to osmotic stress and is expressed constitutively at a low level in plantlets and root cultures.

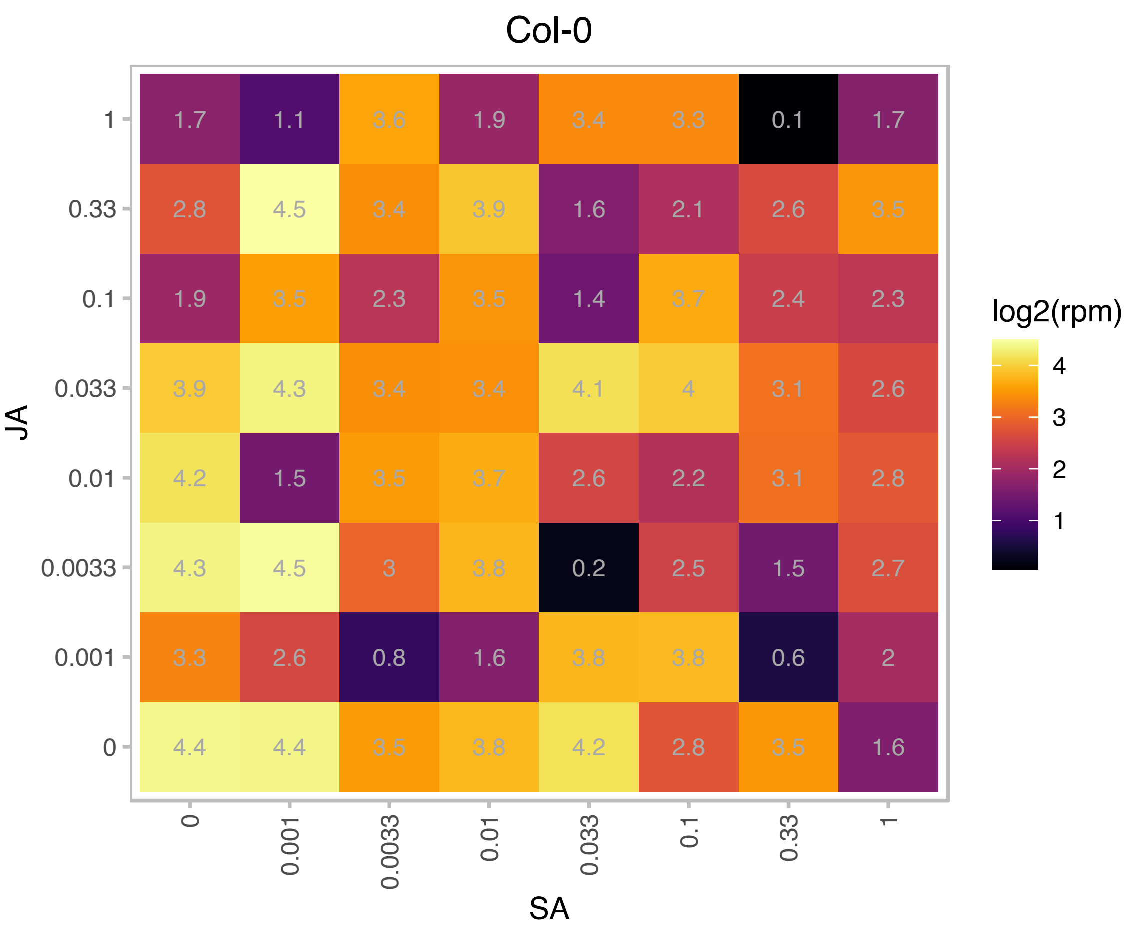


| | |
|---|---|
| Gene | AT4G36250 |
| is.expressed_genes | TRUE |
| is.responsive_genes | TRUE |
| is.apcluster | F-07 |

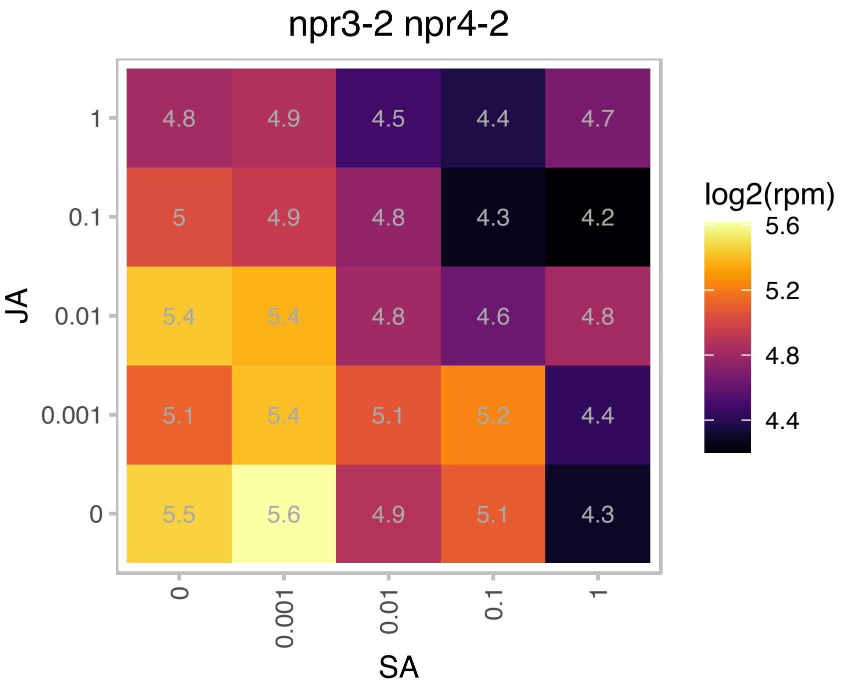


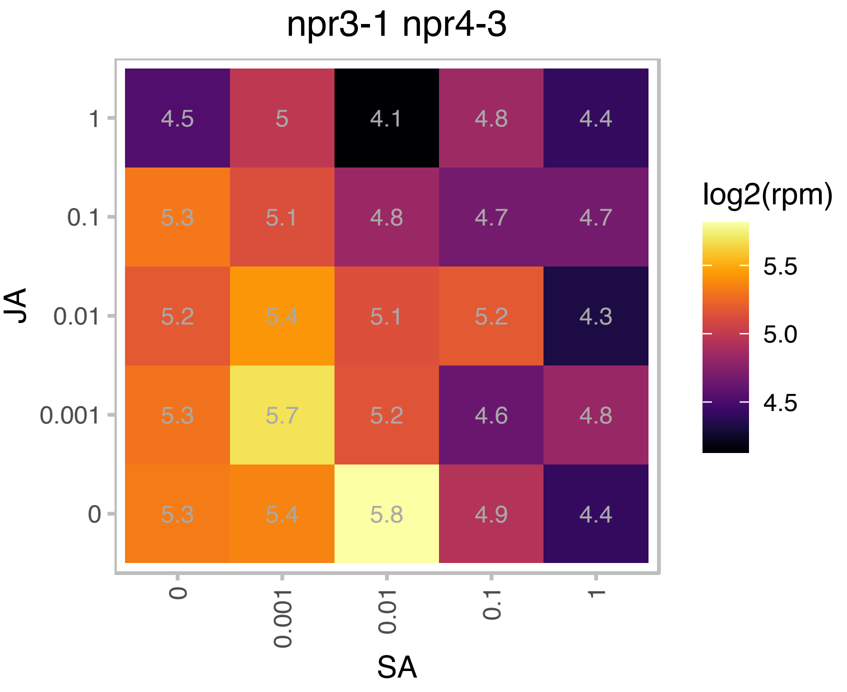

# AT4G36420.1 : NA

Ribosomal protein L12 family protein

Col-0

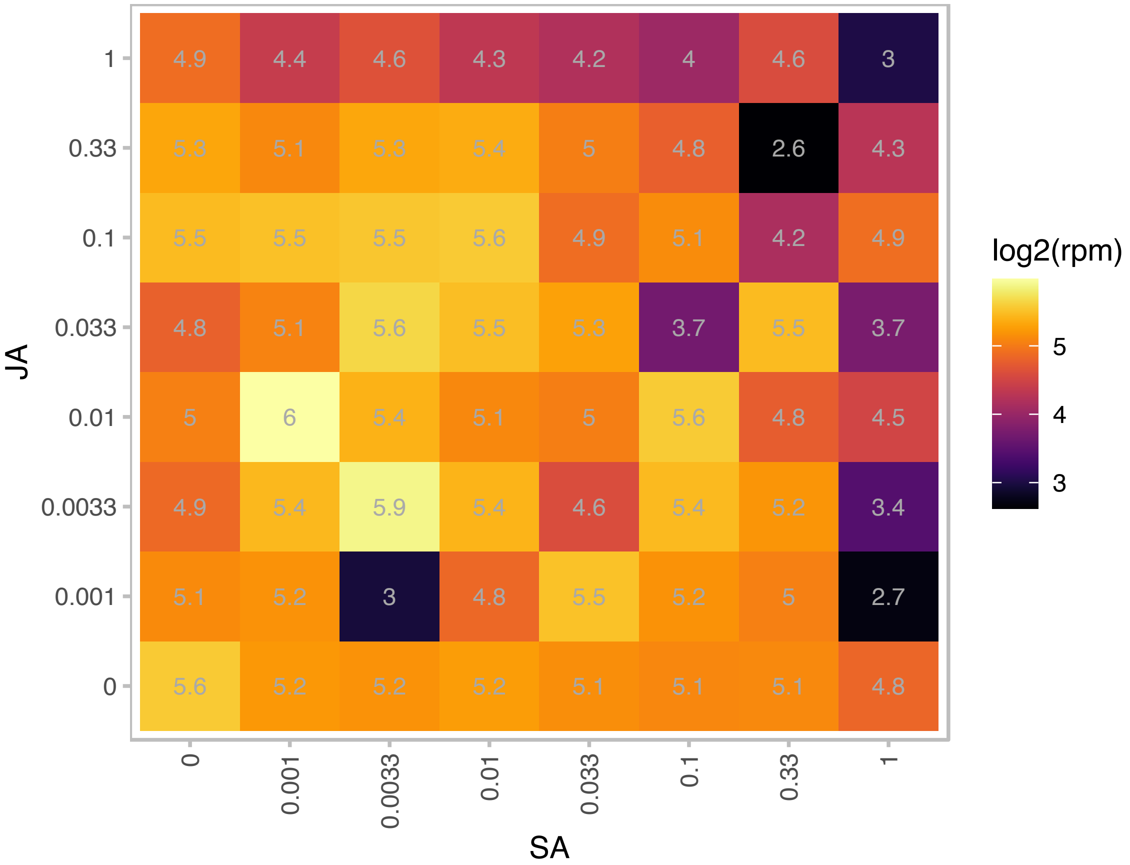


| | |
|---|---|
| Gene | AT4G36420 |
| is.expressed_genes | TRUE |
| is.responsive_genes | TRUE |
| is.apcluster | F-07 |

npr3-2 npr4-2

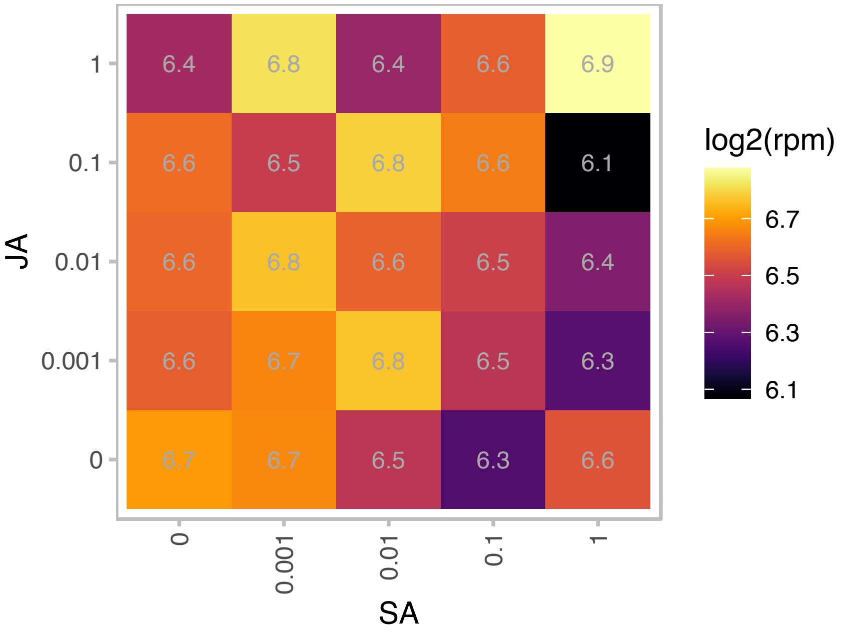


npr3-1 npr4-3

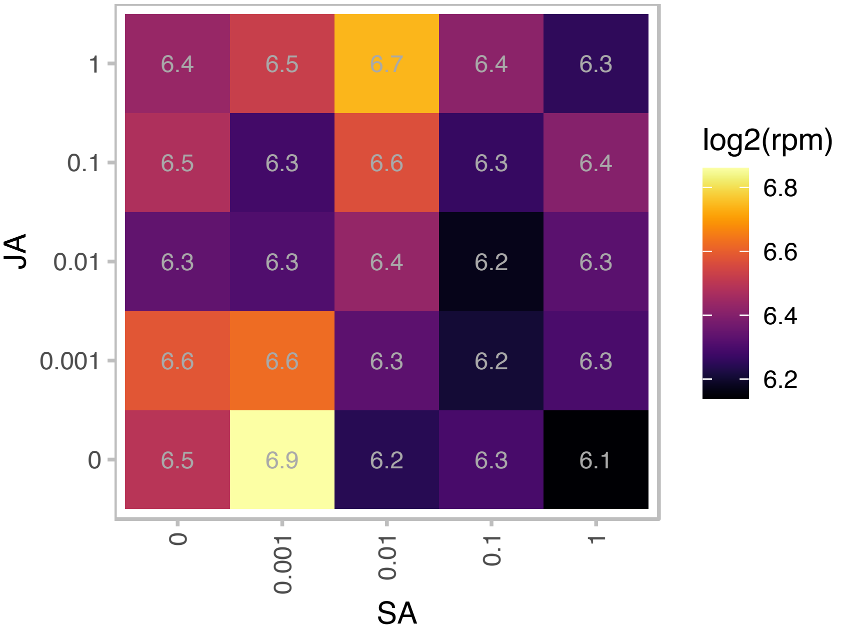

### AT4G37450.1 : AGP18, ATAGP18

arabinogalactan protein 18

AGP18 is a lysine-rich arabinogalactan-protein (AGP) and part of a multi-gene family of glycoproteins with approx. 50 members. It falls into one subclass with AGP17 and AGP19, other lysine-rich AGPs. It is expressed in young leaves, shoots, roots and flowers and is active in the regulation of the selection and survival of megaspores.

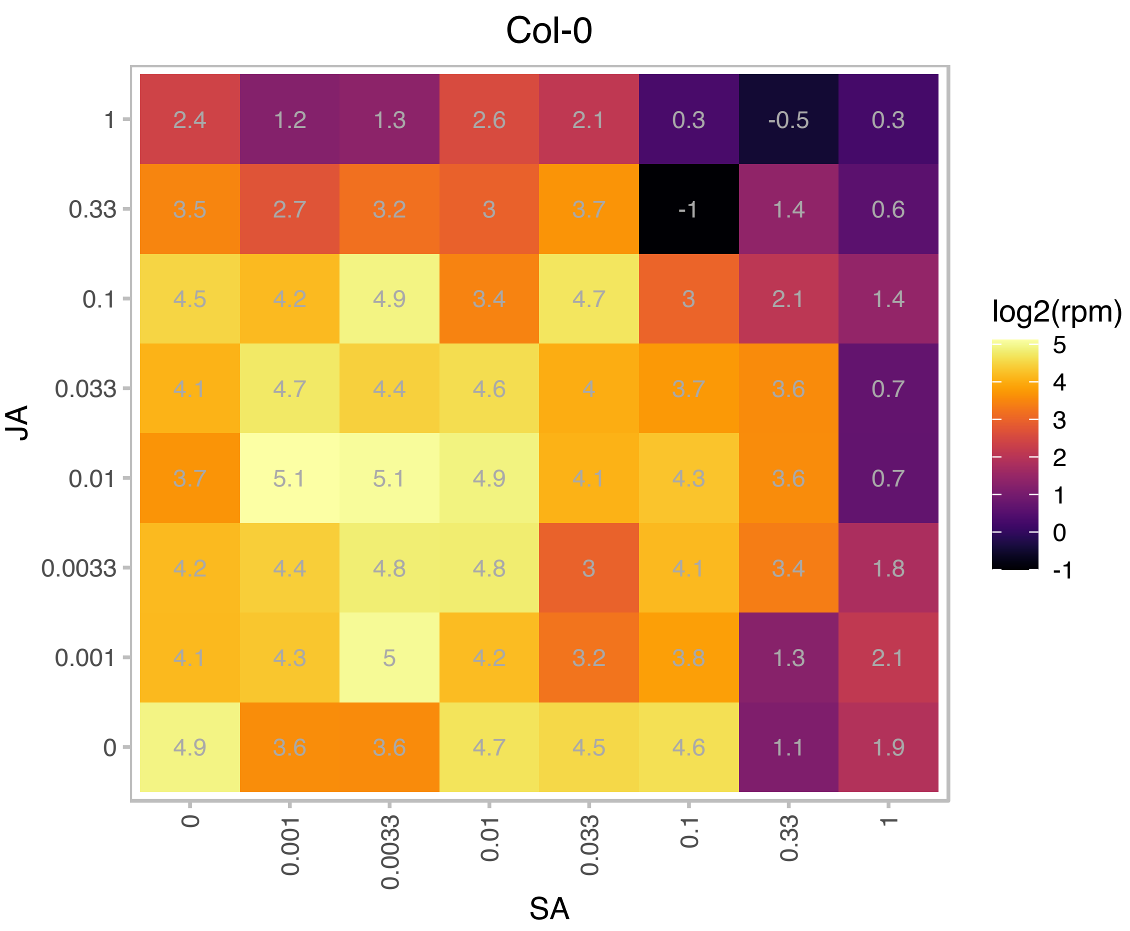


| | |
|---|---|
| Gene | AT4G37450 |
| is.expressed_genes | TRUE |
| is.responsive_genes | TRUE |
| is.apcluster | F-07 |

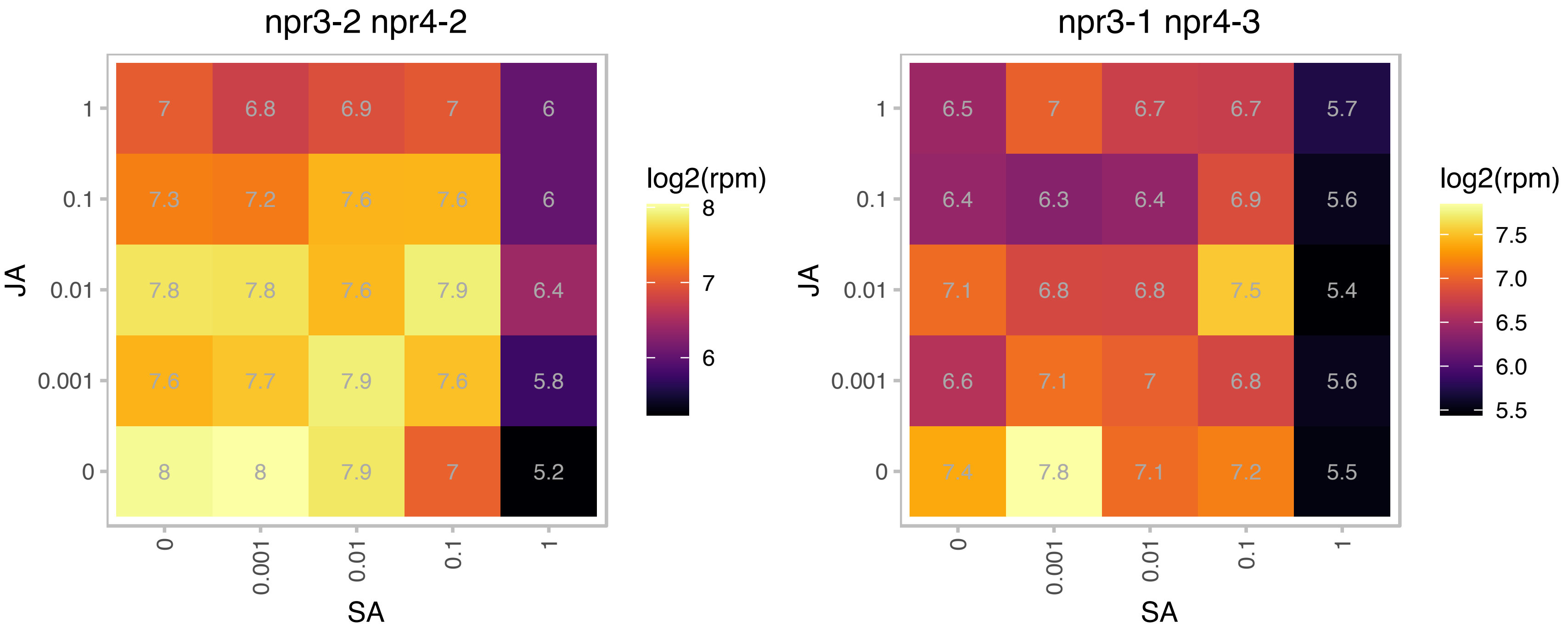

# AT4G39900.1 : NA

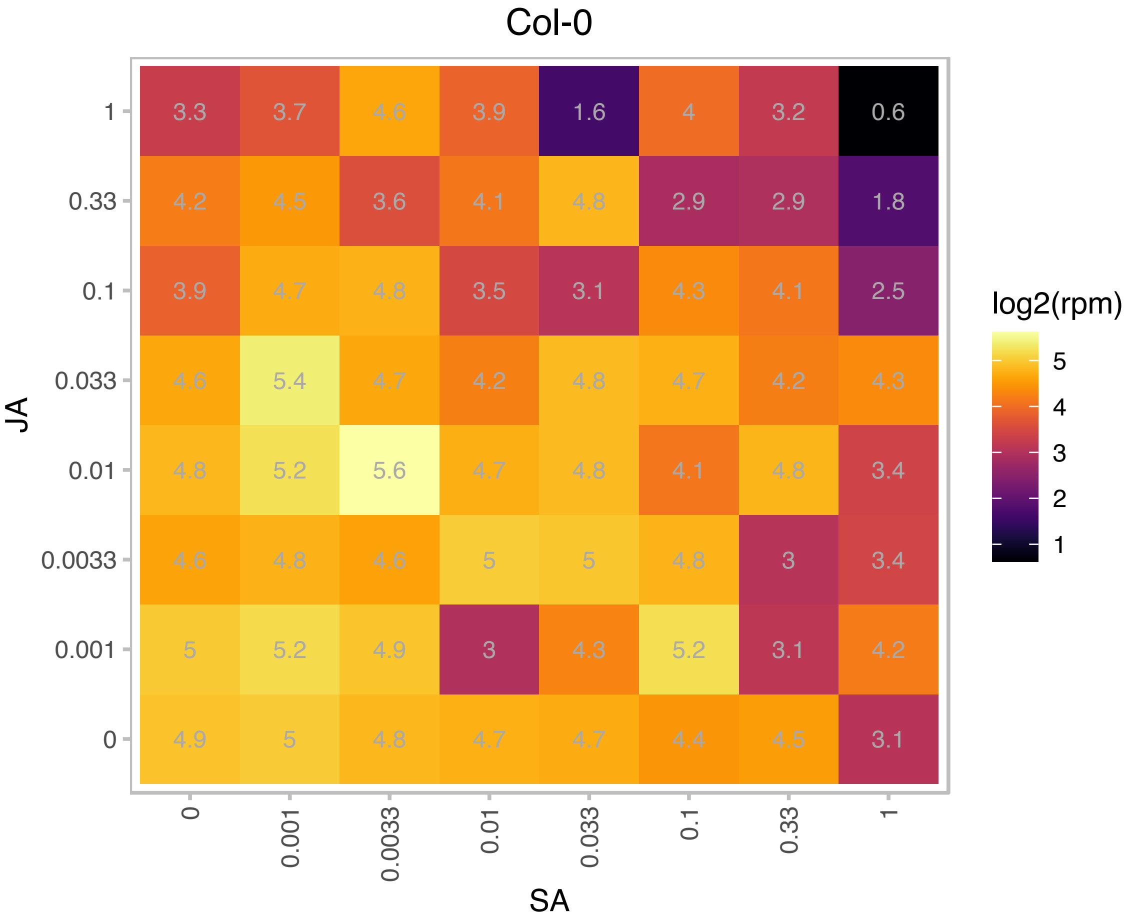


| | |
|---|---|
| Gene | AT4G39900 |
| is.expressed_genes | TRUE |
| is.responsive_genes | TRUE |
| is.apcluster | F-07 |

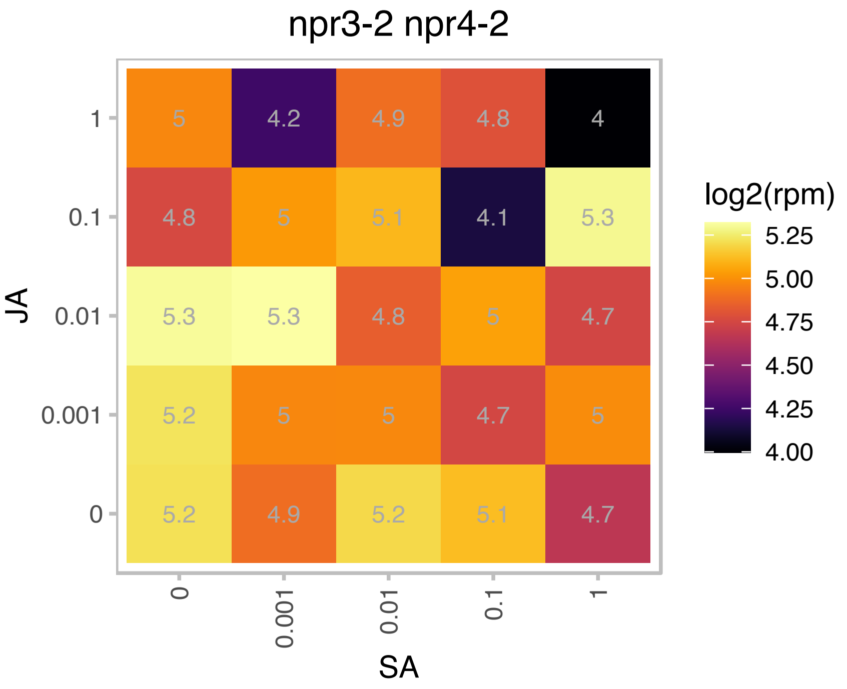


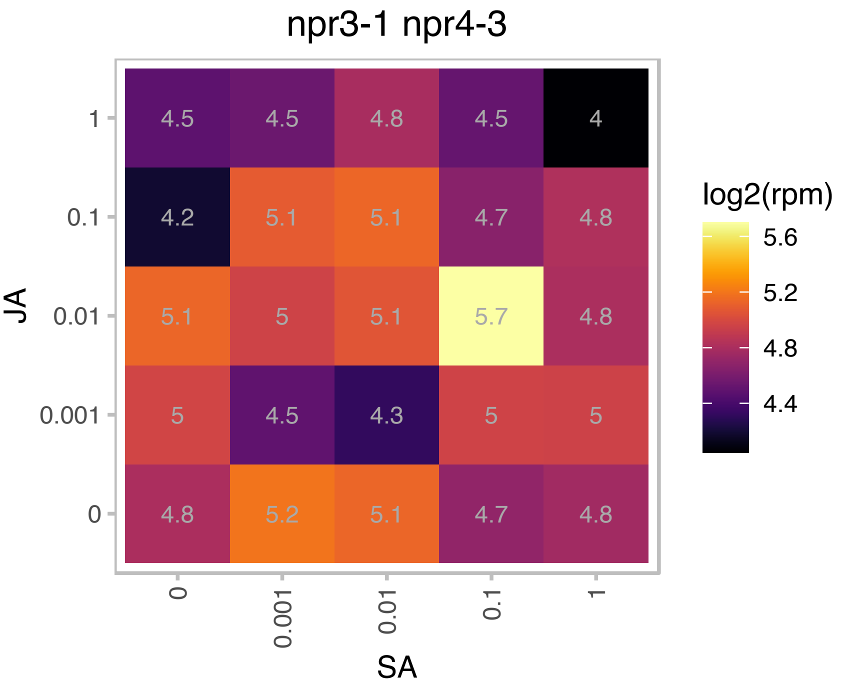

### AT4G39960.1 : DJA5

Molecular chaperone Hsp40/DnaJ family protein

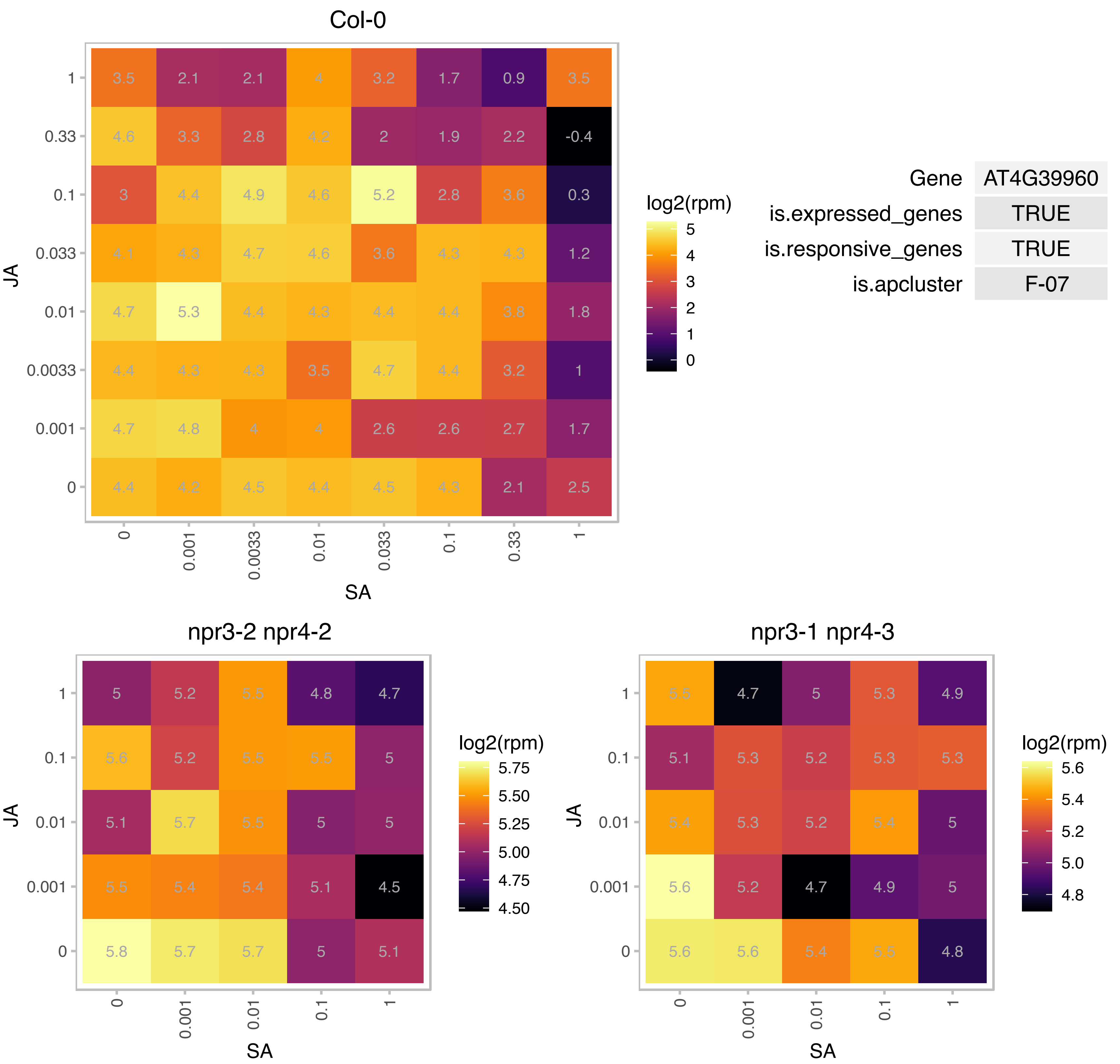


| | |
|---|---|
| Gene | AT4G39960 |
| is.expressed_genes | TRUE |
| is.responsive_genes | TRUE |
| is.apcluster | F-07 |

# AT5G02550.1 : NA

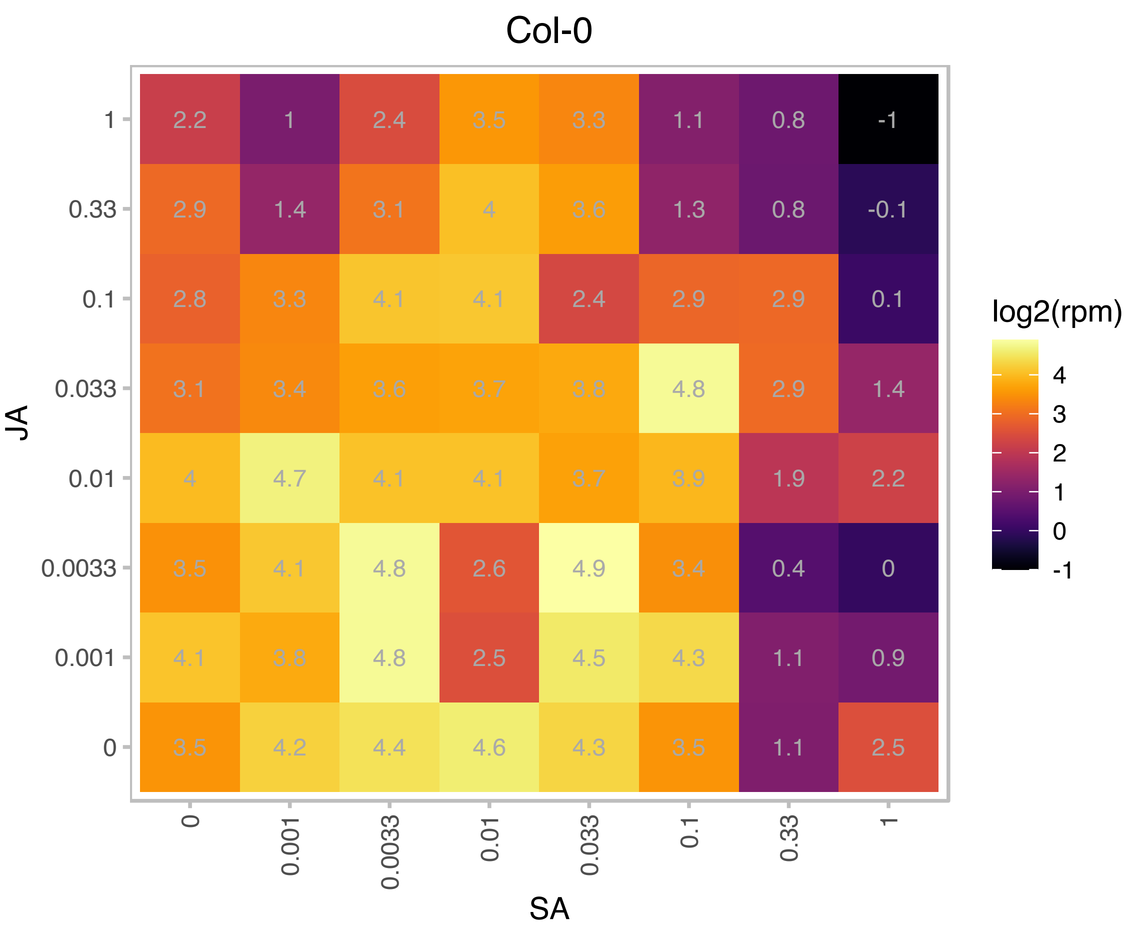


| | |
|---|---|
| Gene | AT5G02550 |
| is.expressed_genes | TRUE |
| is.responsive_genes | TRUE |
| is.apcluster | F-07 |

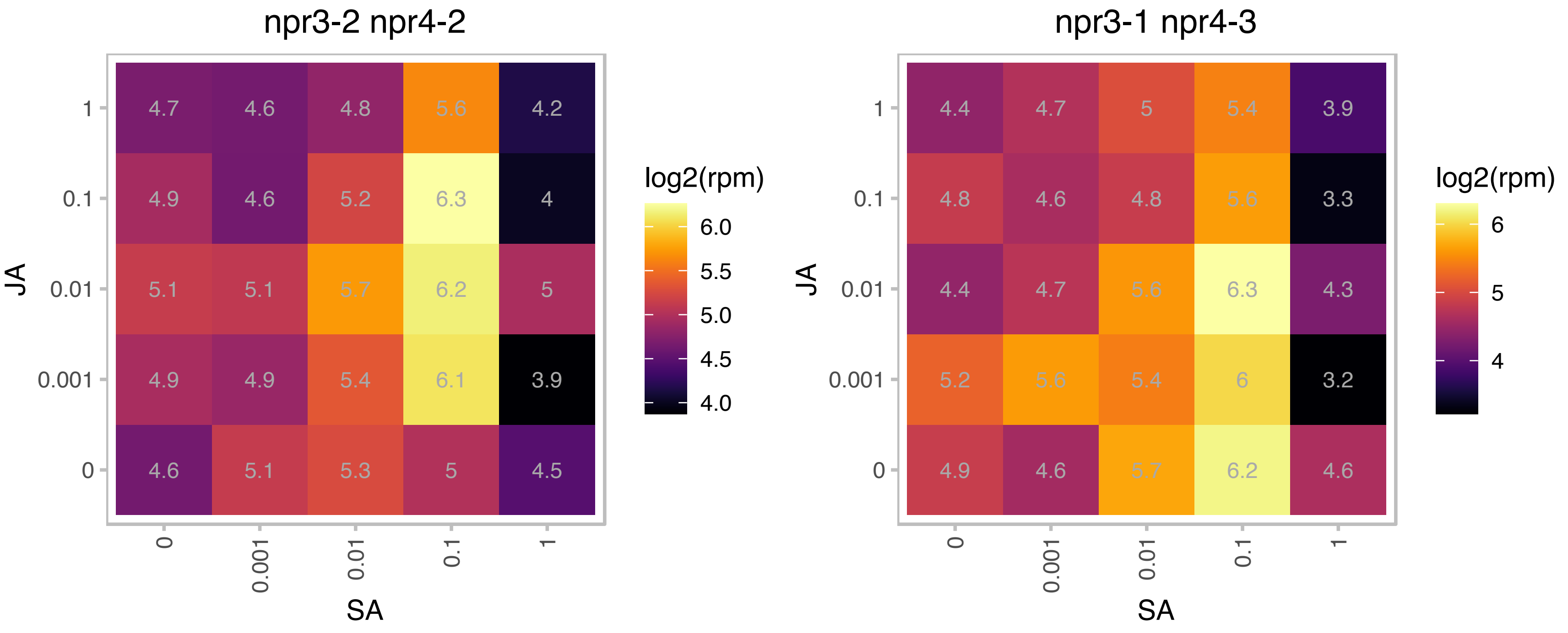

# AT5G02610.1 : NA

Ribosomal L29 family protein

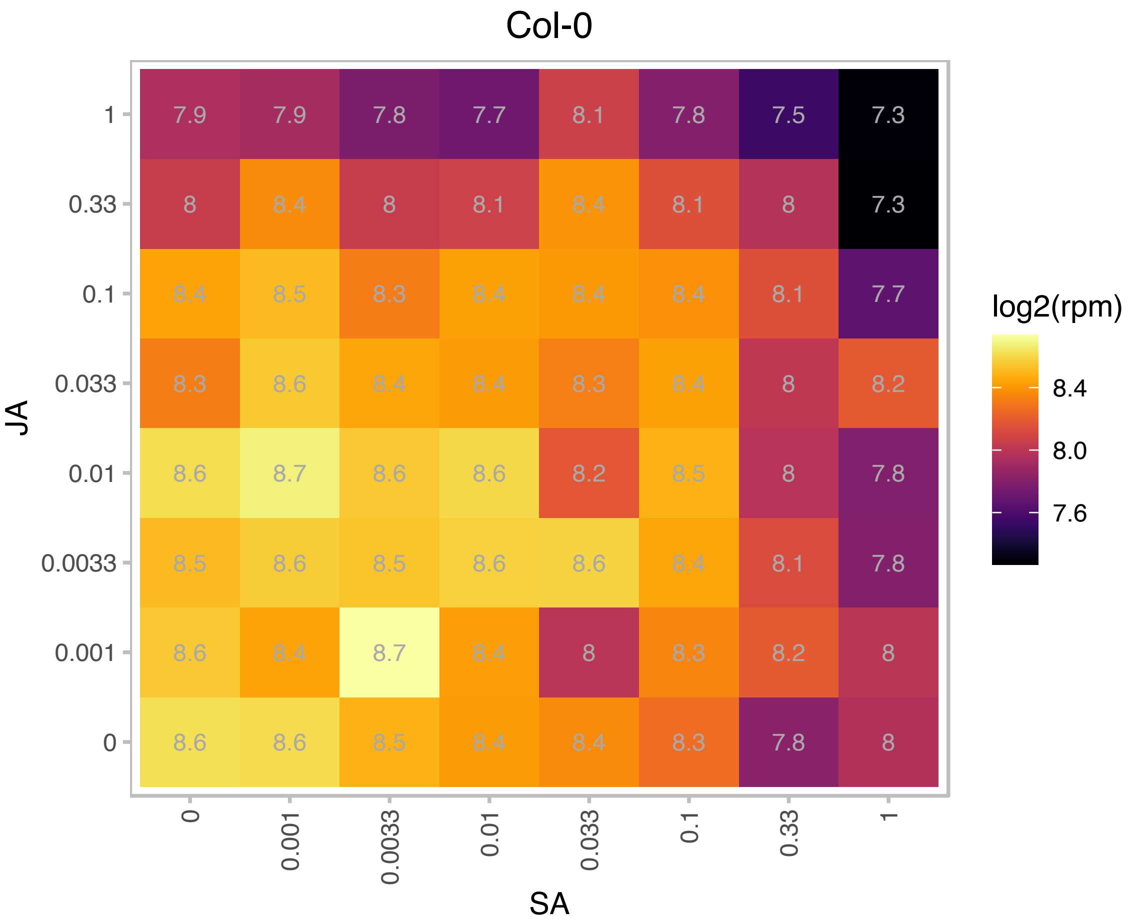


| | |
|---|---|
| Gene | AT5G02610 |
| is.expressed_genes | TRUE |
| is.responsive_genes | TRUE |
| is.apcluster | F-07 |

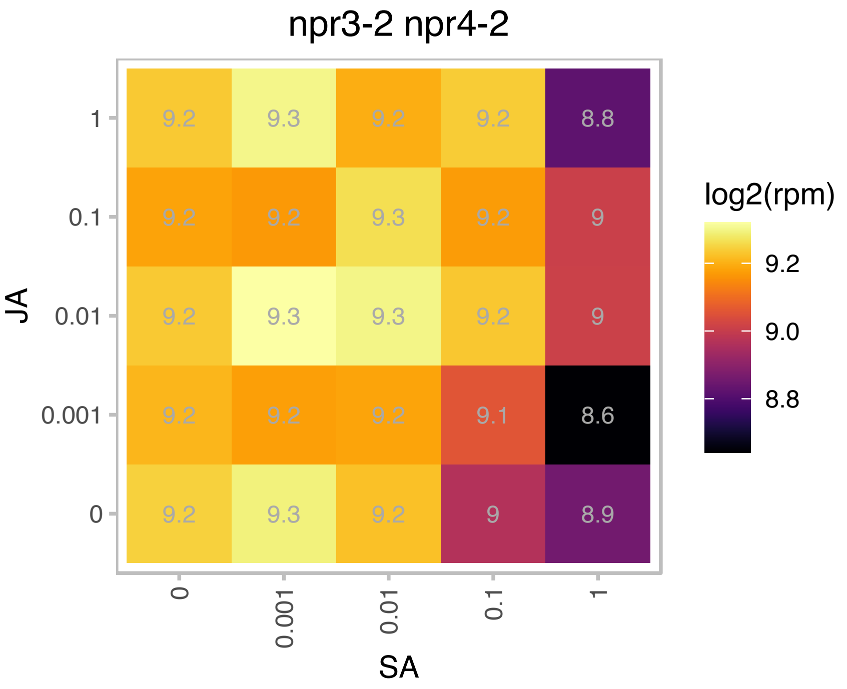


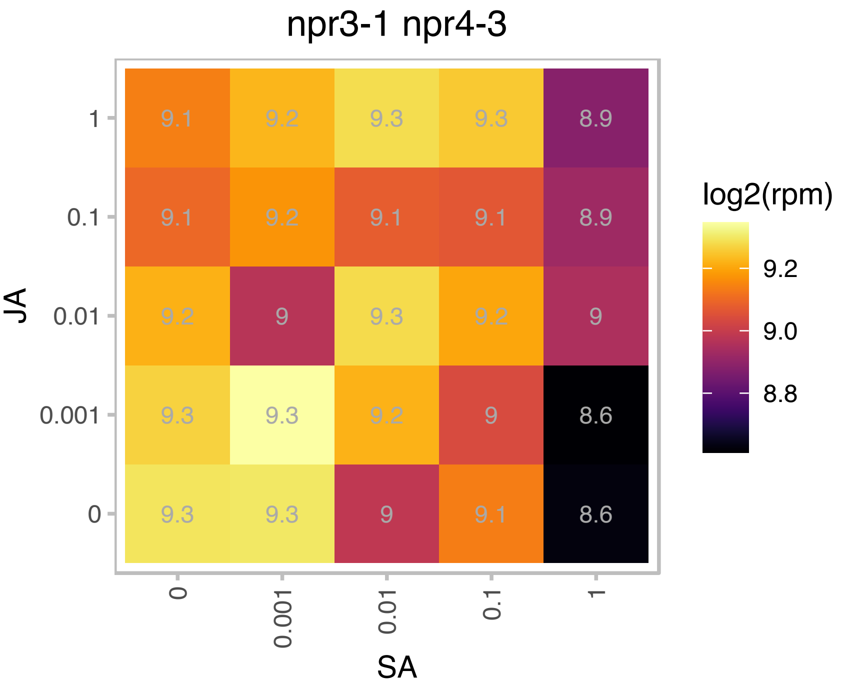

### AT5G02870.1 : RPL4

Ribosomal protein L4/L1 family

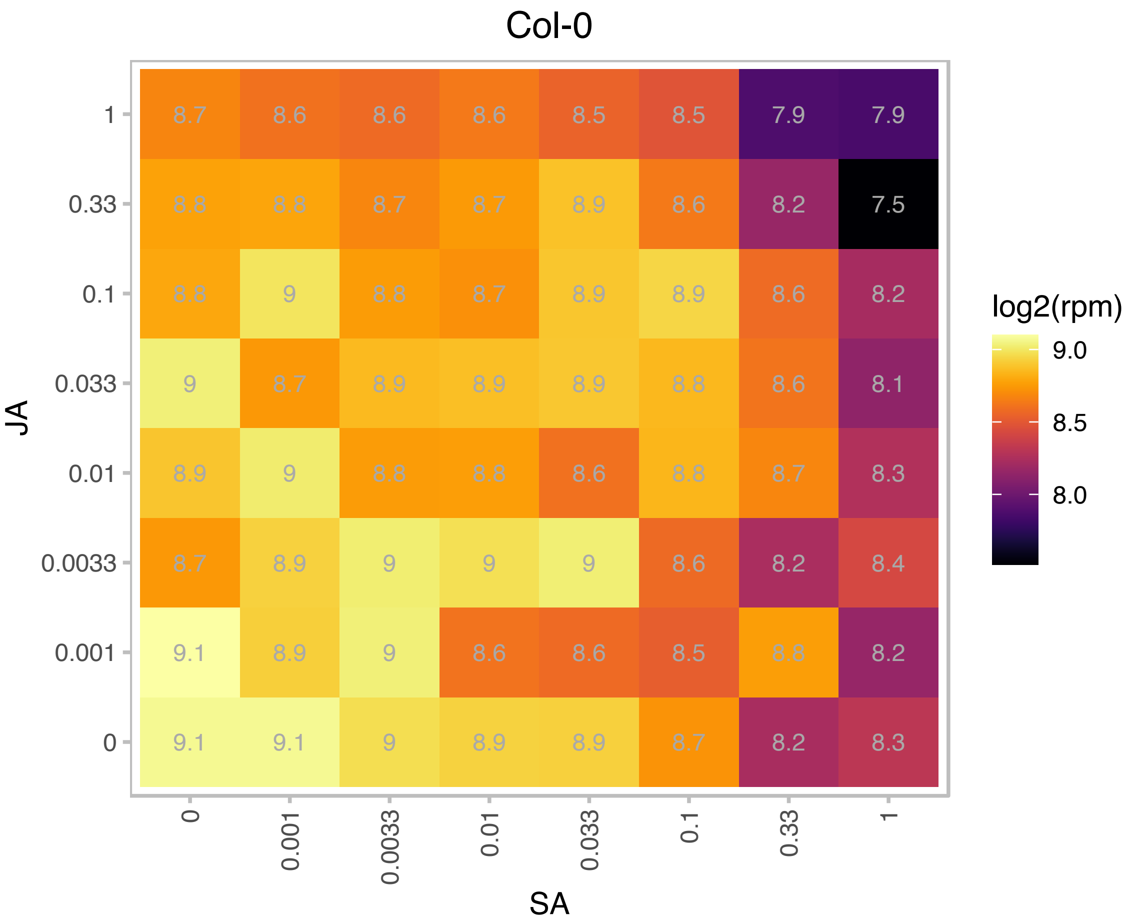


| | |
|---|---|
| Gene | AT5G02870 |
| is.expressed_genes | TRUE |
| is.responsive_genes | TRUE |
| is.apcluster | F-07 |

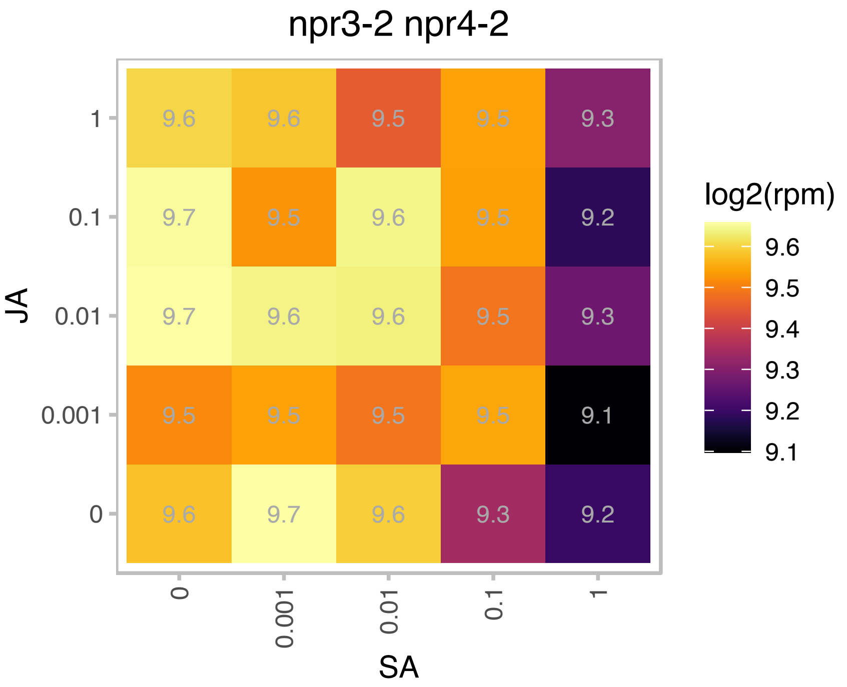


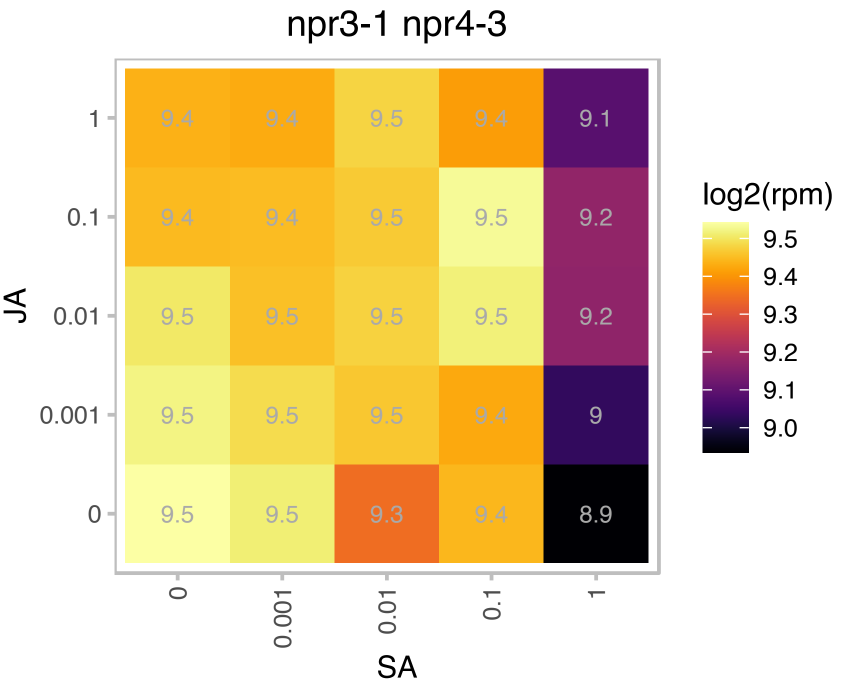

# AT5G02960.1 : NA

Ribosomal protein S12/S23 family protein

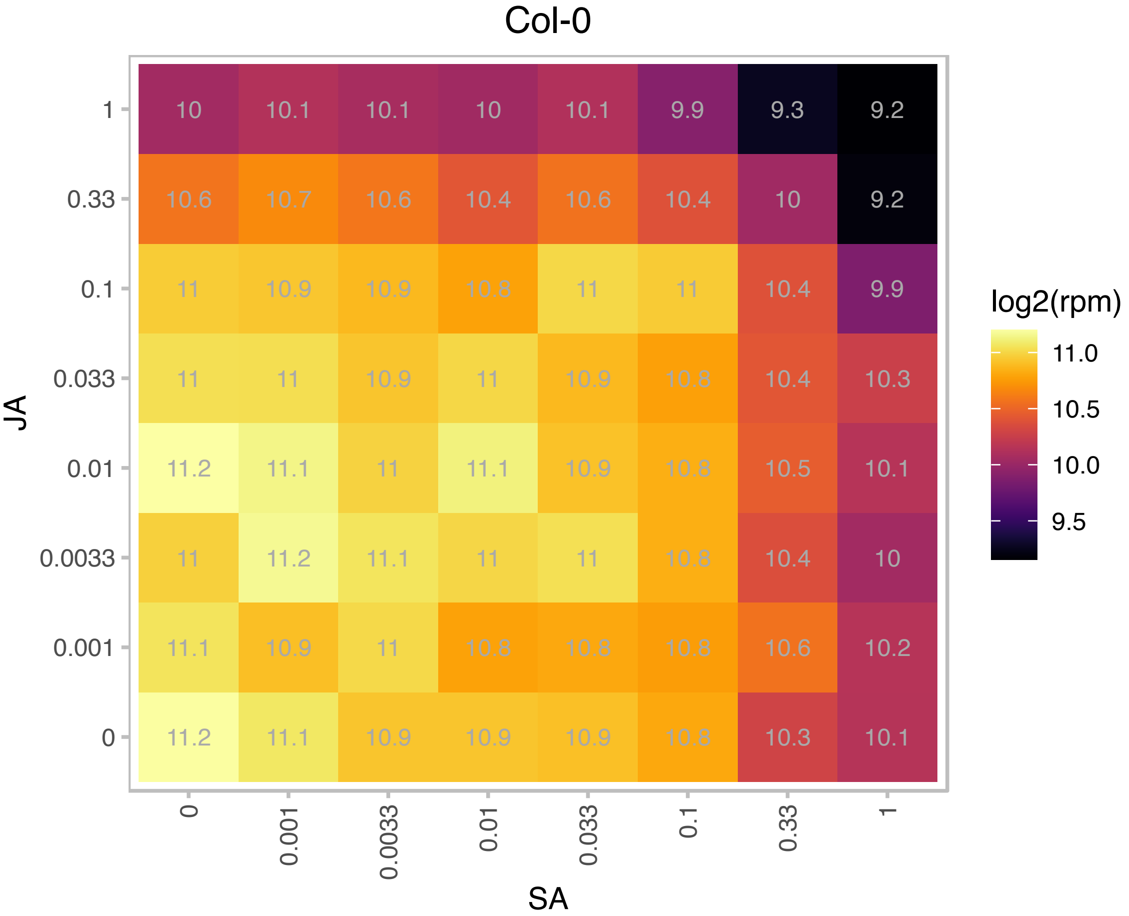


| | |
|---|---|
| Gene | AT5G02960 |
| is.expressed_genes | TRUE |
| is.responsive_genes | TRUE |
| is.apcluster | F-07 |

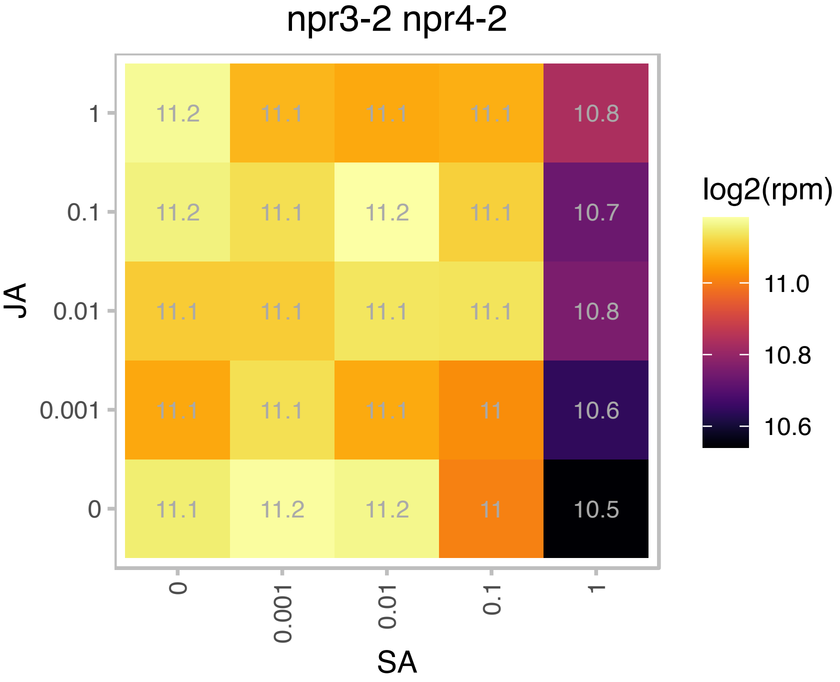


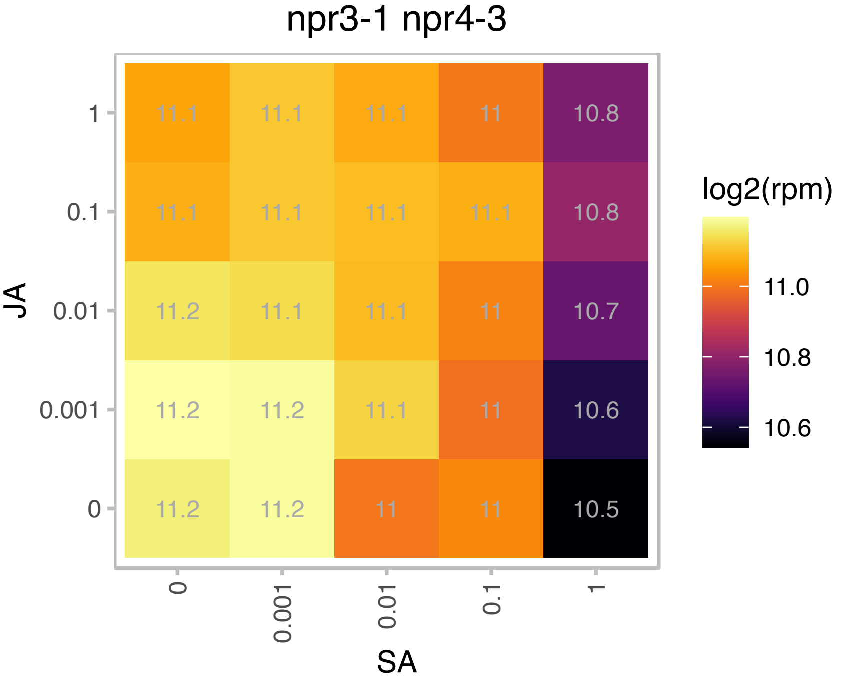

# AT5G04800.1 : NA

Ribosomal S17 family protein

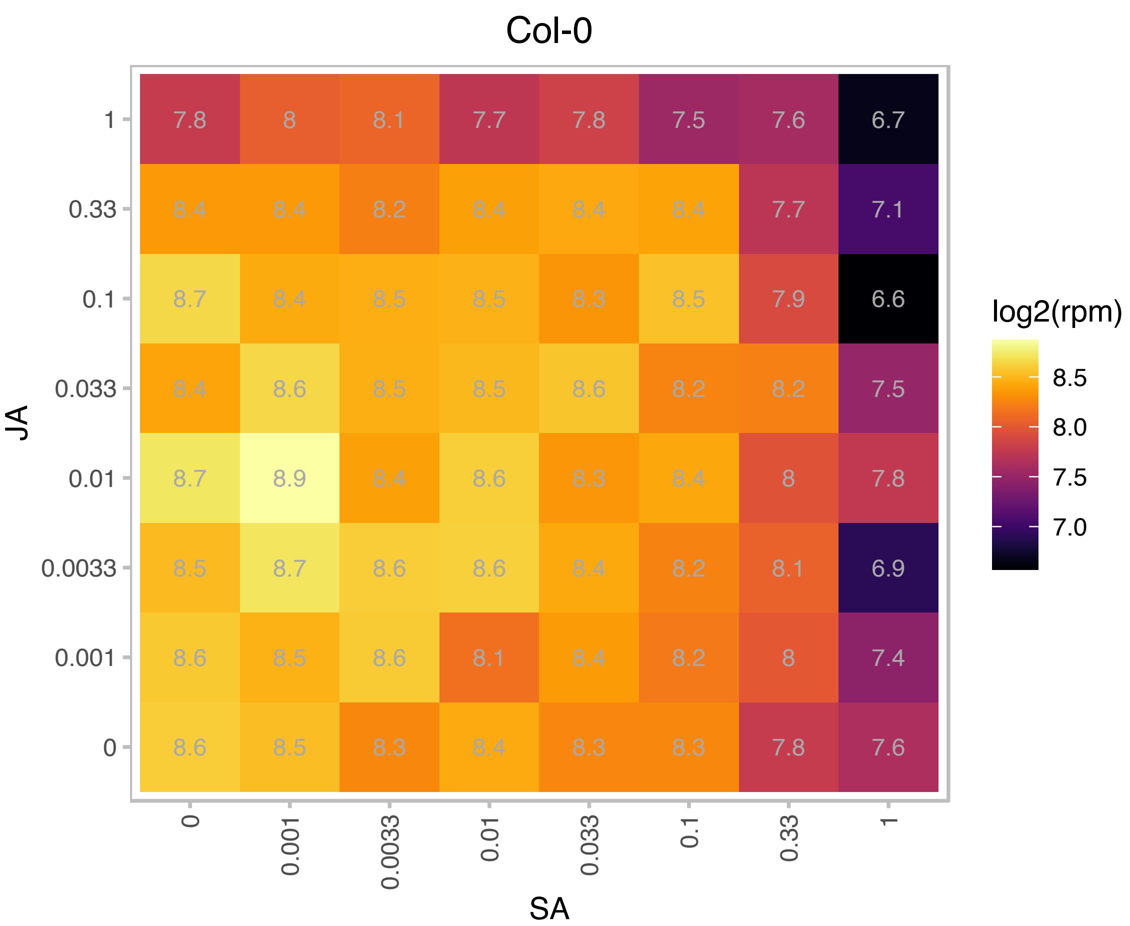


| | |
|---|---|
| Gene | AT5G04800 |
| is.expressed_genes | TRUE |
| is.responsive_genes | TRUE |
| is.apcluster | F-07 |

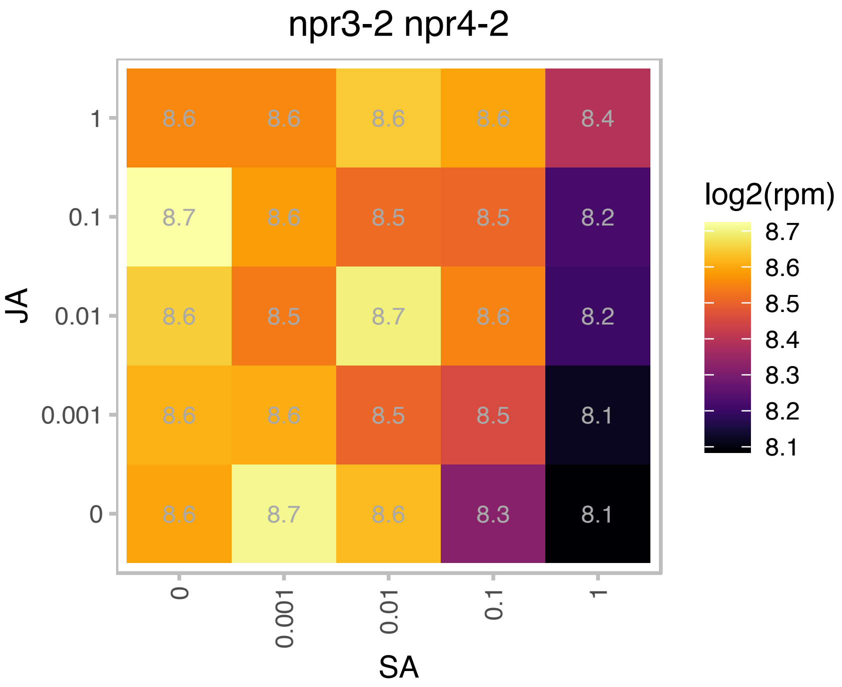


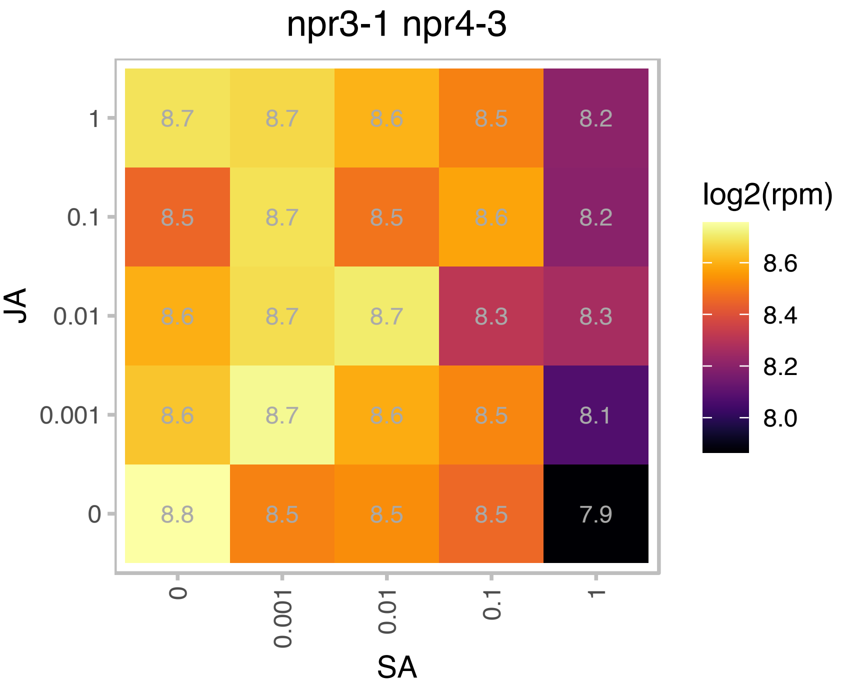

# AT5G05710.1 : NA

Pleckstrin homology (PH) domain superfamily protein

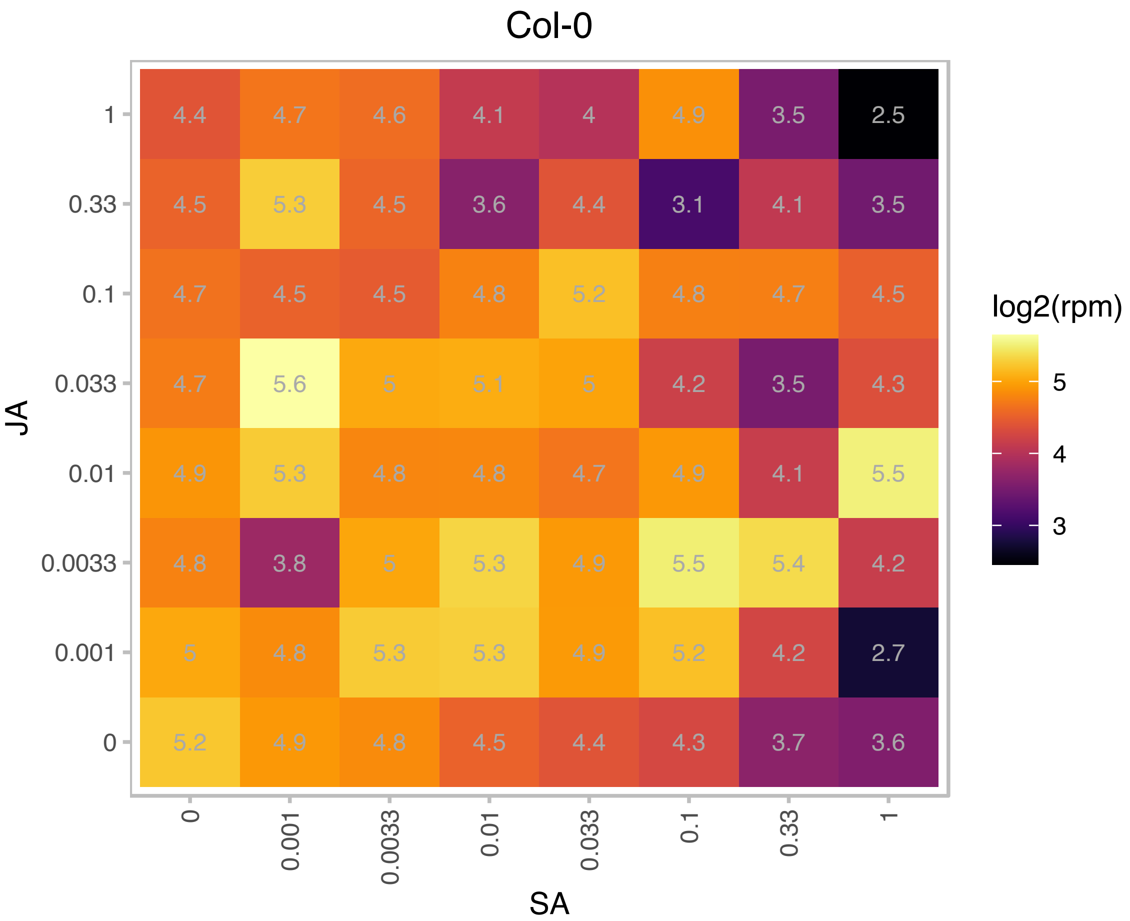


| | |
|---|---|
| Gene | AT5G05710 |
| is.expressed_genes | TRUE |
| is.responsive_genes | TRUE |
| is.apcluster | F-07 |

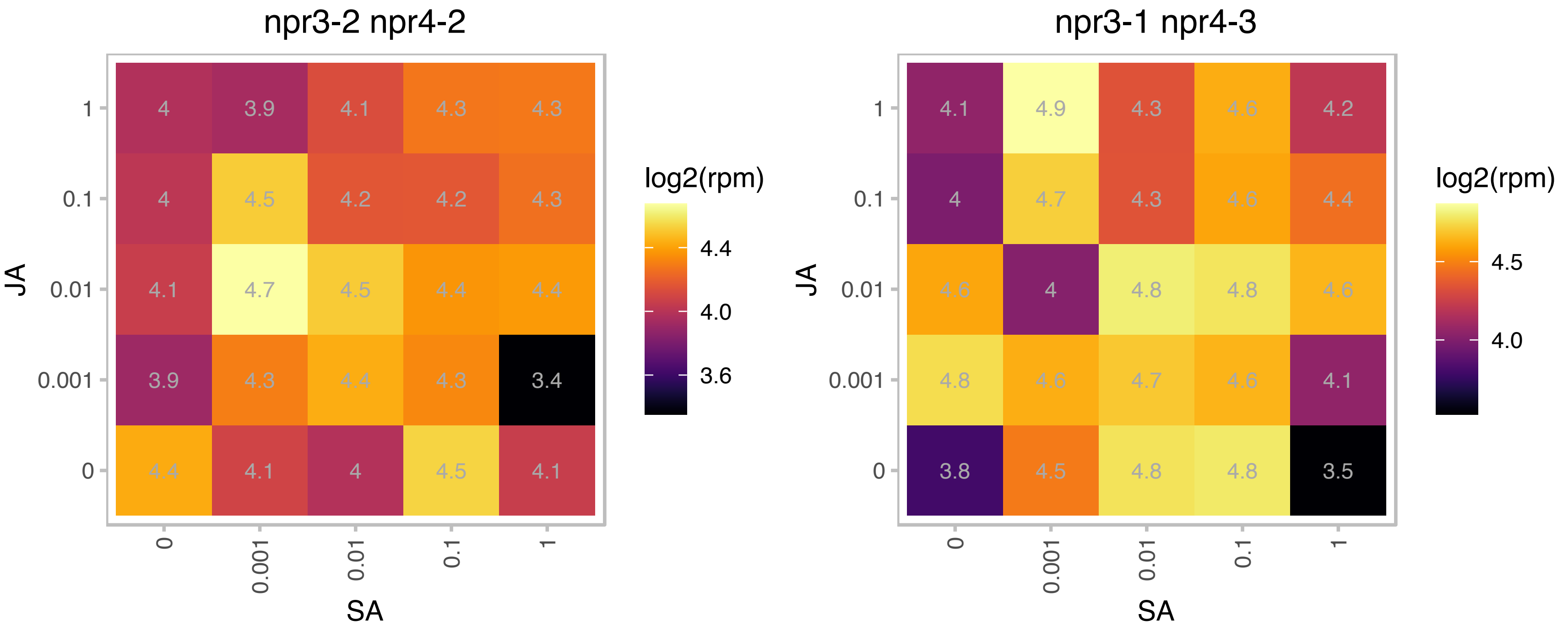

### AT5G06770.1 : KHZ2

KH domain-containing protein / zinc finger (CCCH type) family protein
KHZ2 is a CCCH zinc-finger and KH domain protein belonging to the VII subfamily. It is expressed throughout the plant. Highly similar to KHZ1.Double mutants with khz1 are late flowering. Overexpression leads to increased rates of leaf senescence.

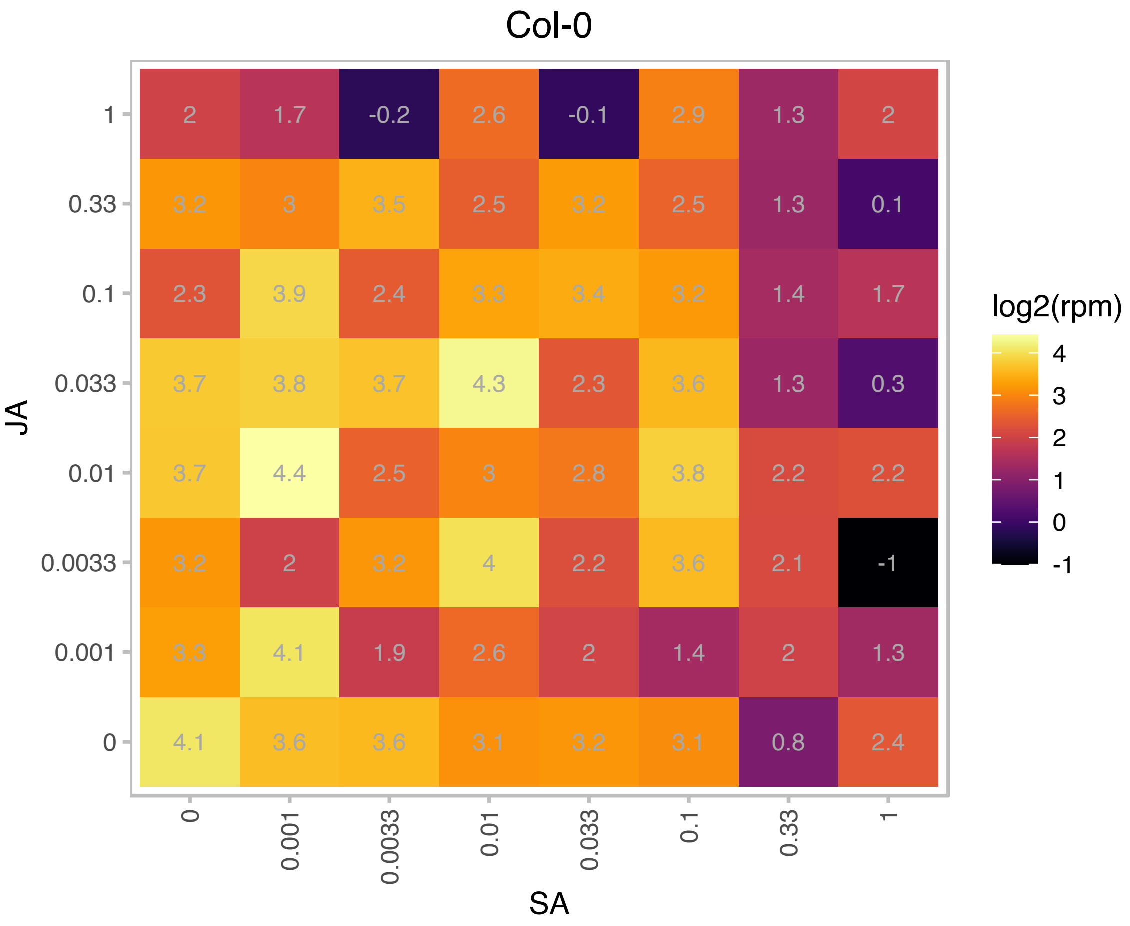


| | |
|---|---|
| Gene | AT5G06770 |
| is.expressed_genes | TRUE |
| is.responsive_genes | TRUE |
| is.apcluster | F-07 |

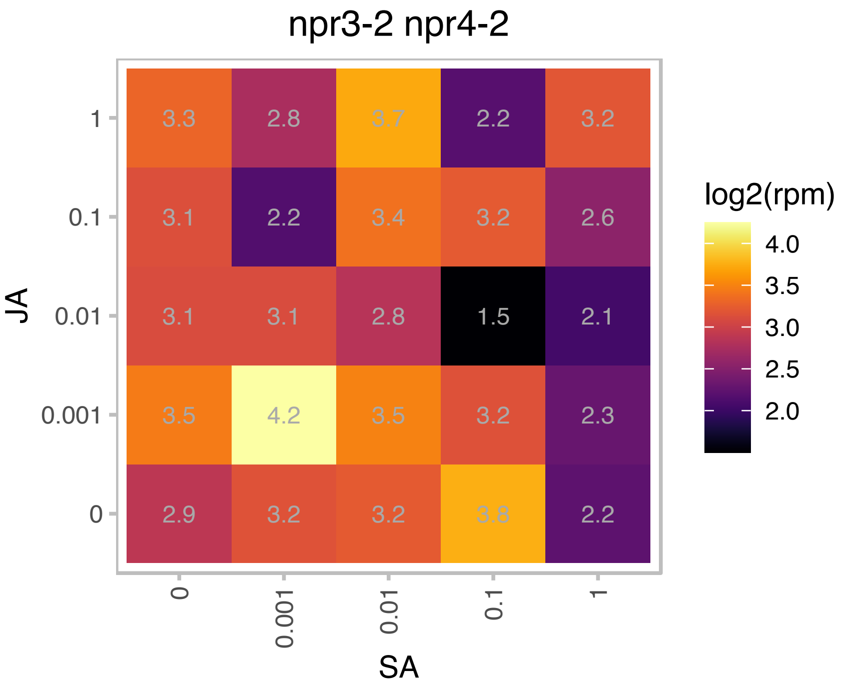


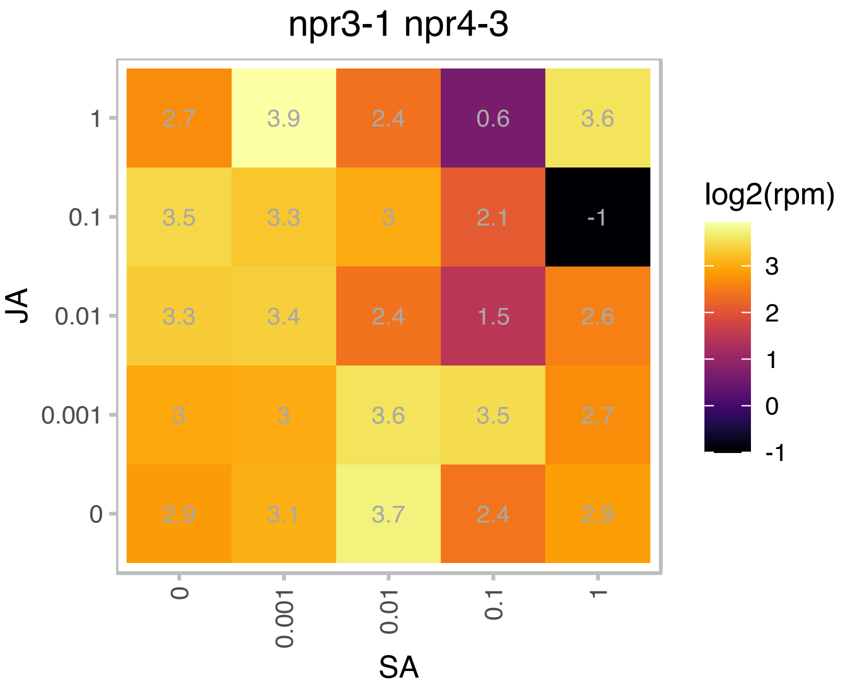

# AT5G07090.1 : NA

Ribosomal protein S4 (RPS4A) family protein

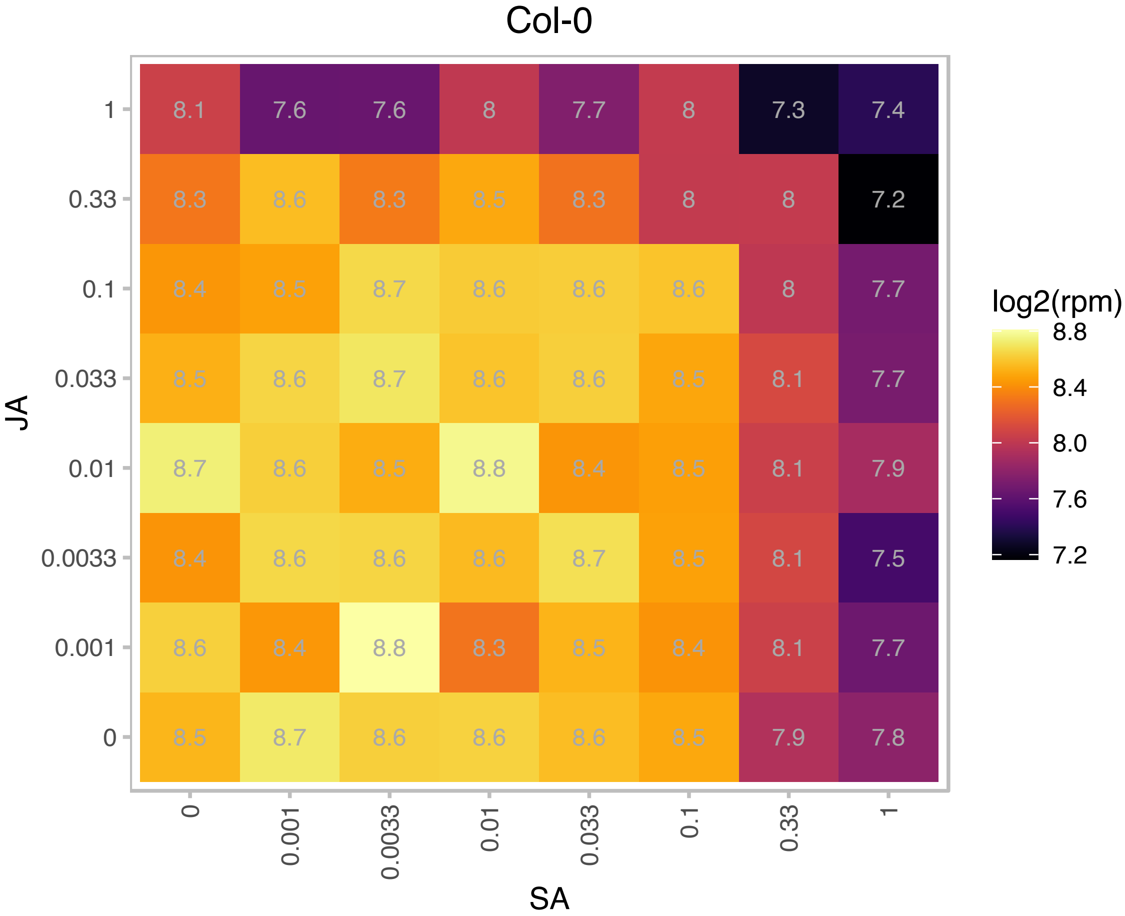


| | |
|---|---|
| Gene | AT5G07090 |
| is.expressed_genes | TRUE |
| is.responsive_genes | TRUE |
| is.apcluster | F-07 |

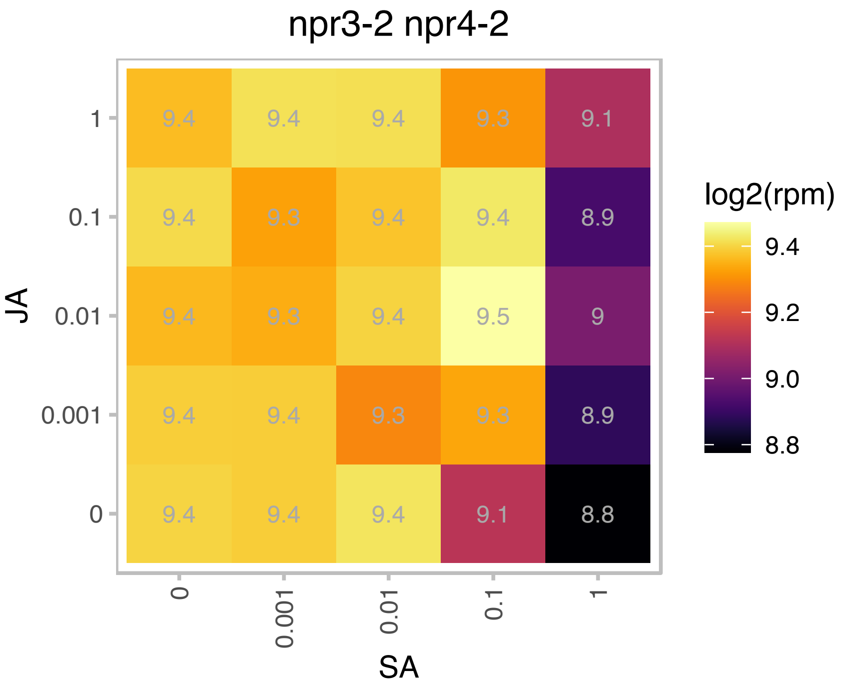


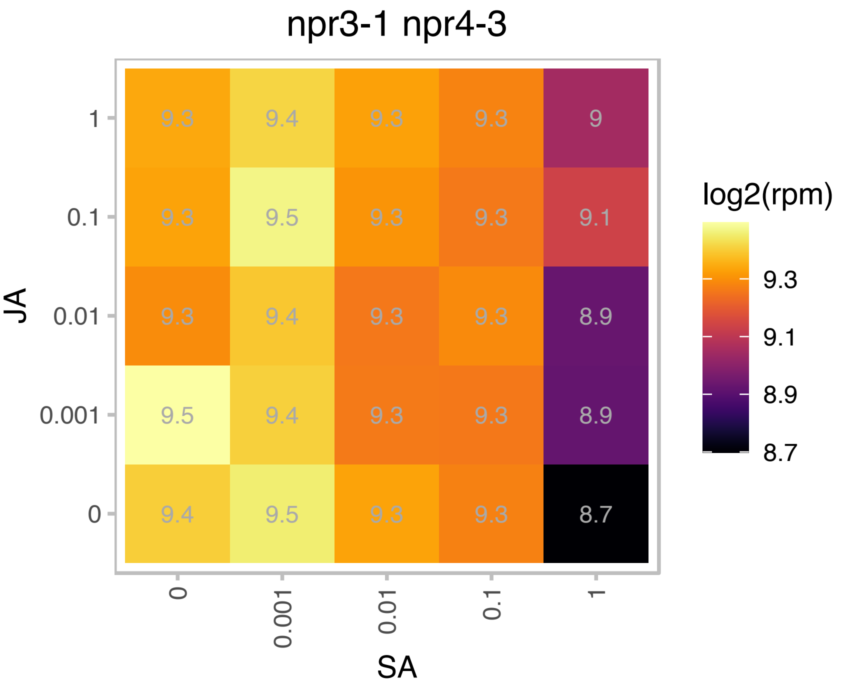

### AT5G07580.1 : DEWAX2, ERF106

Integrase-type DNA-binding superfamily protein
encodes a member of the ERF (ethylene response factor) subfamily B-3 of ERF/AP2 transcription factor family. The protein contains one AP2 domain. There are 18 members in this subfamily including ATERF-1, ATERF-2, AND ATERF-5.

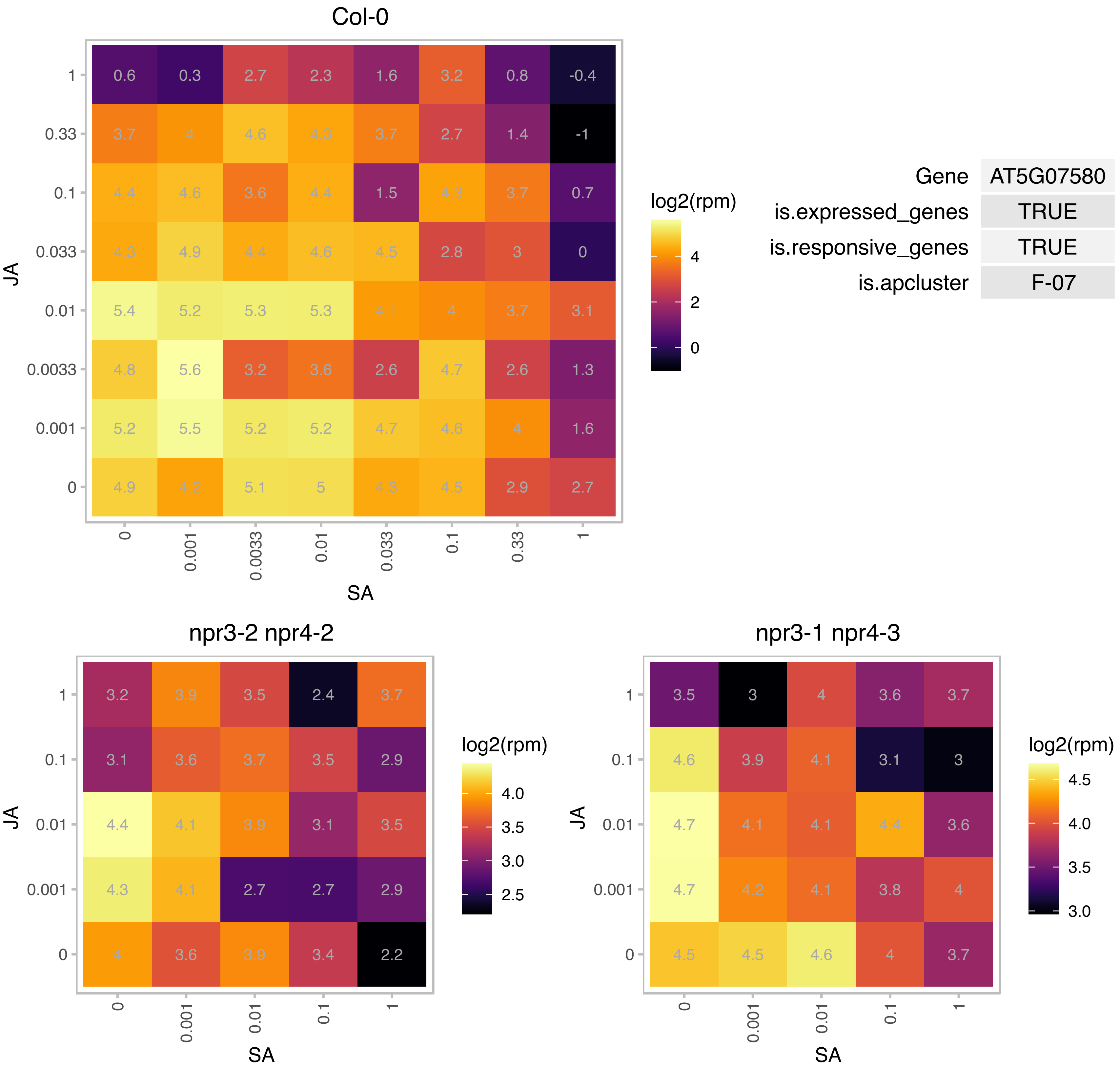


| | |
|---|---|
| Gene | AT5G07580 |
| is.expressed_genes | TRUE |
| is.responsive_genes | TRUE |
| is.apcluster | F-07 |

# AT5G09510.1 : NA

Ribosomal protein S19 family protein

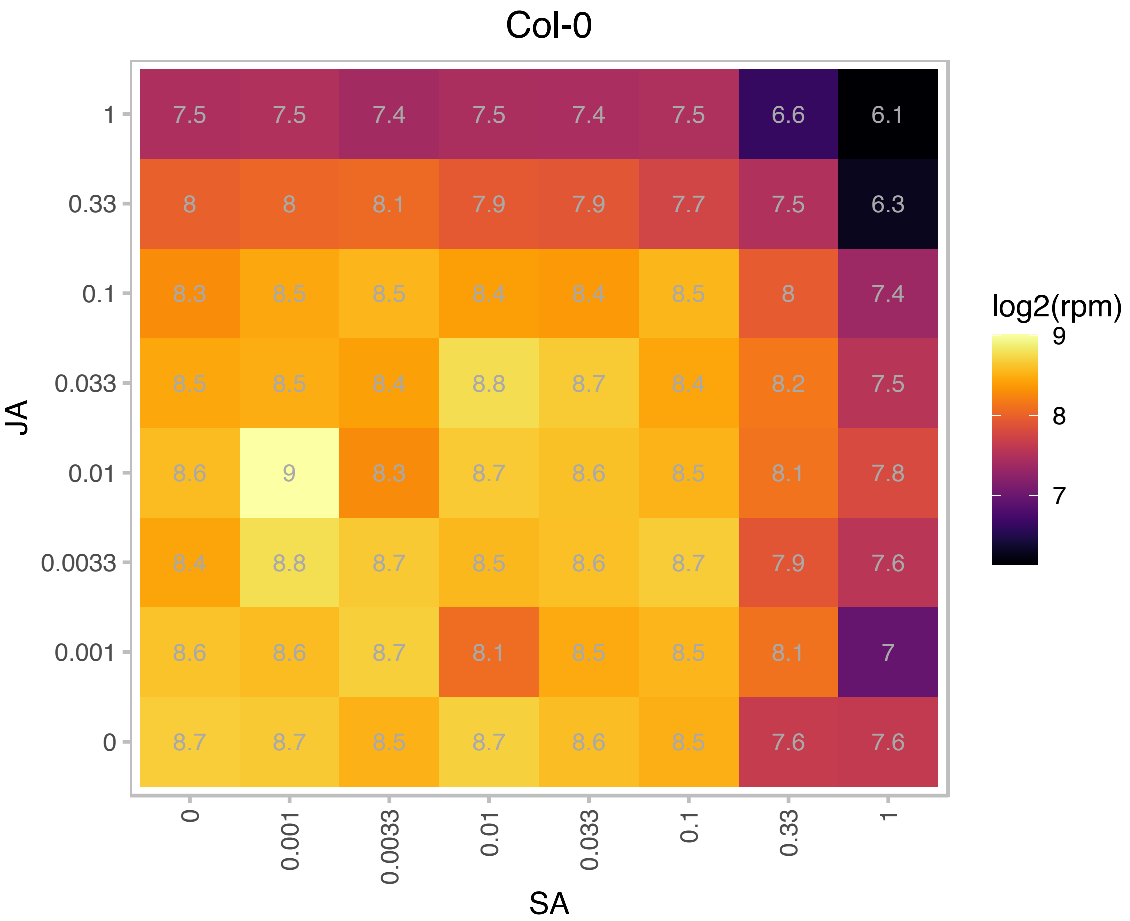


| | |
|---|---|
| Gene | AT5G09510 |
| is.expressed_genes | TRUE |
| is.responsive_genes | TRUE |
| is.apcluster | F-07 |

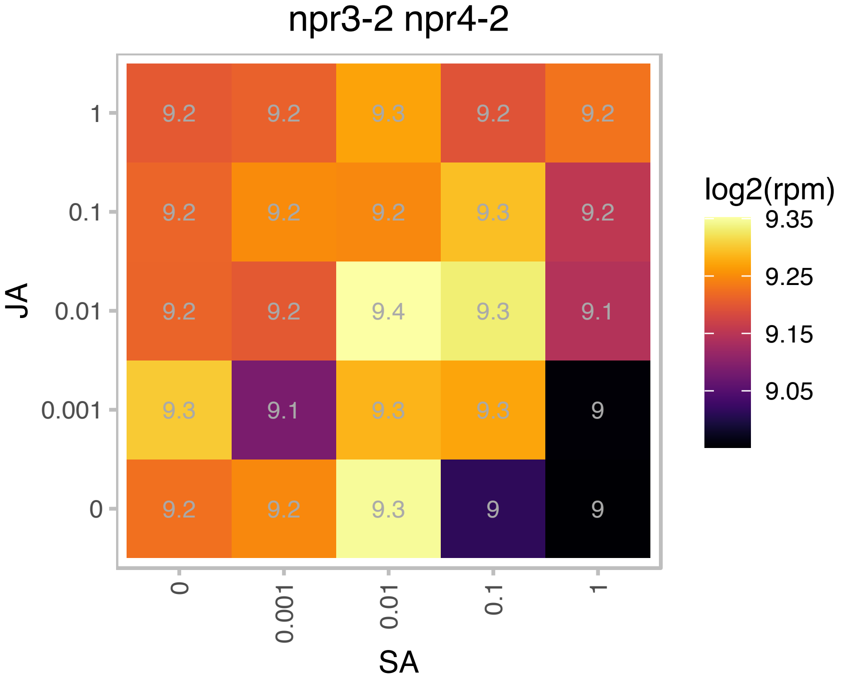


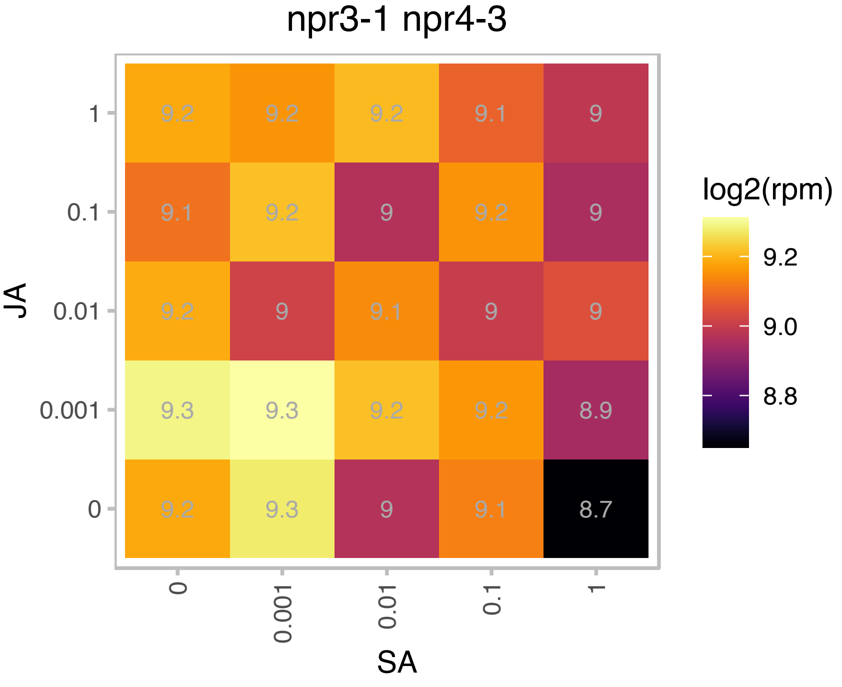

### AT5G12150.1 : PHGAP1

Rho GTPase activation protein (RhoGAP) with PH domain

Encodes a protein with similarity to REN1, a Rho GTPase activating protein.It is cytoplasmic and plasma membrane associated in interphase, but during mitosis localizes to the CDZ/CDS in a POK-dependent manner.

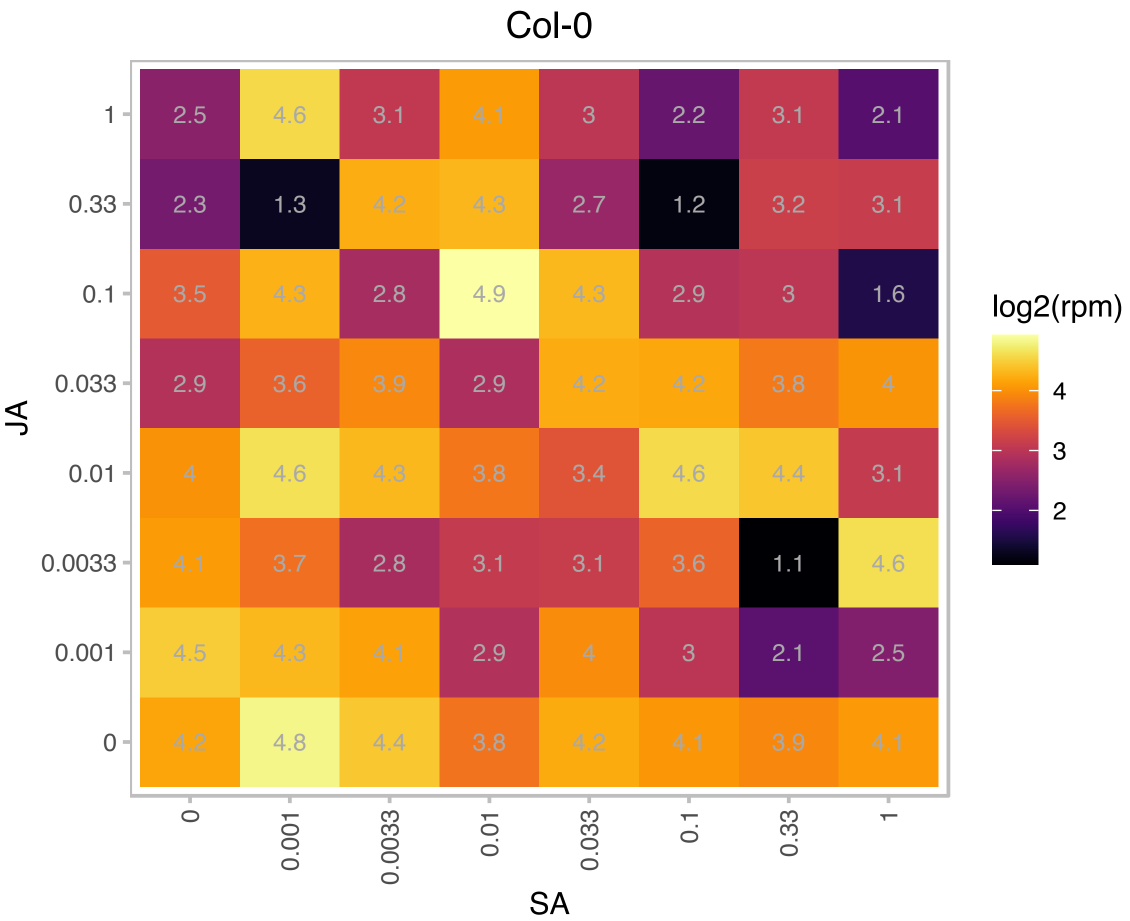


| | |
|---|---|
| Gene | AT5G12150 |
| is.expressed_genes | TRUE |
| is.responsive_genes | TRUE |
| is.apcluster | F-07 |

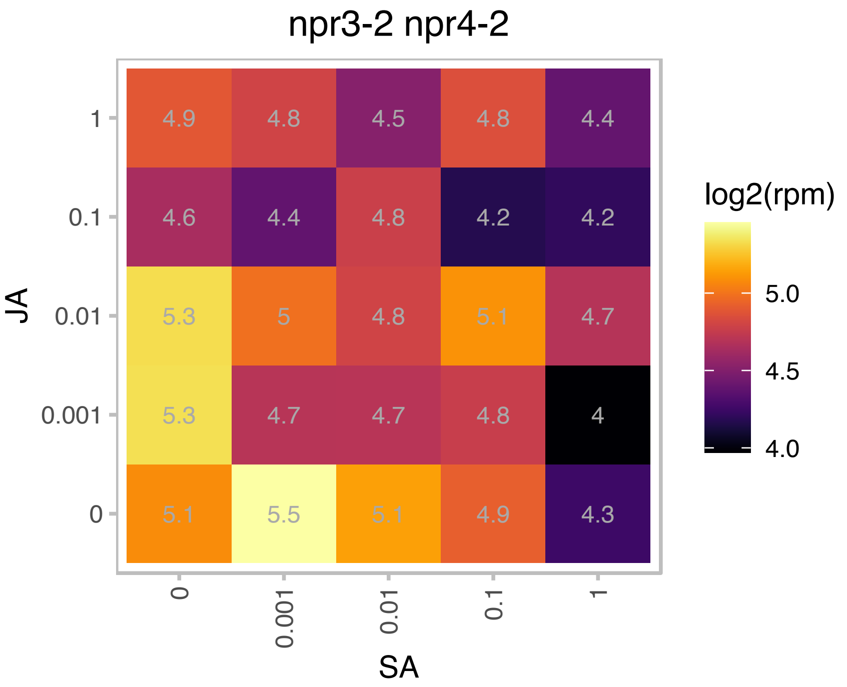


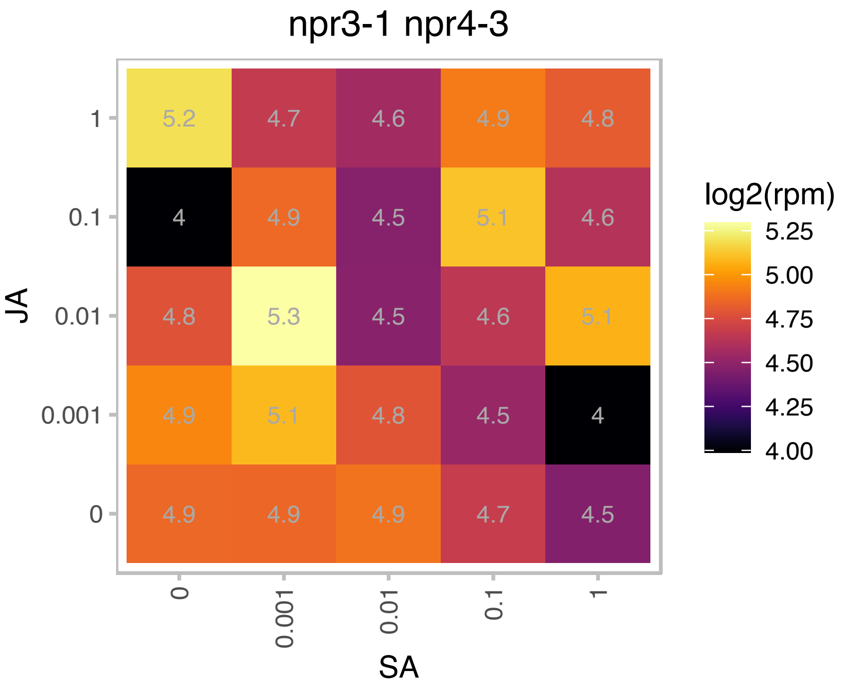

# AT5G13090.1 : NA

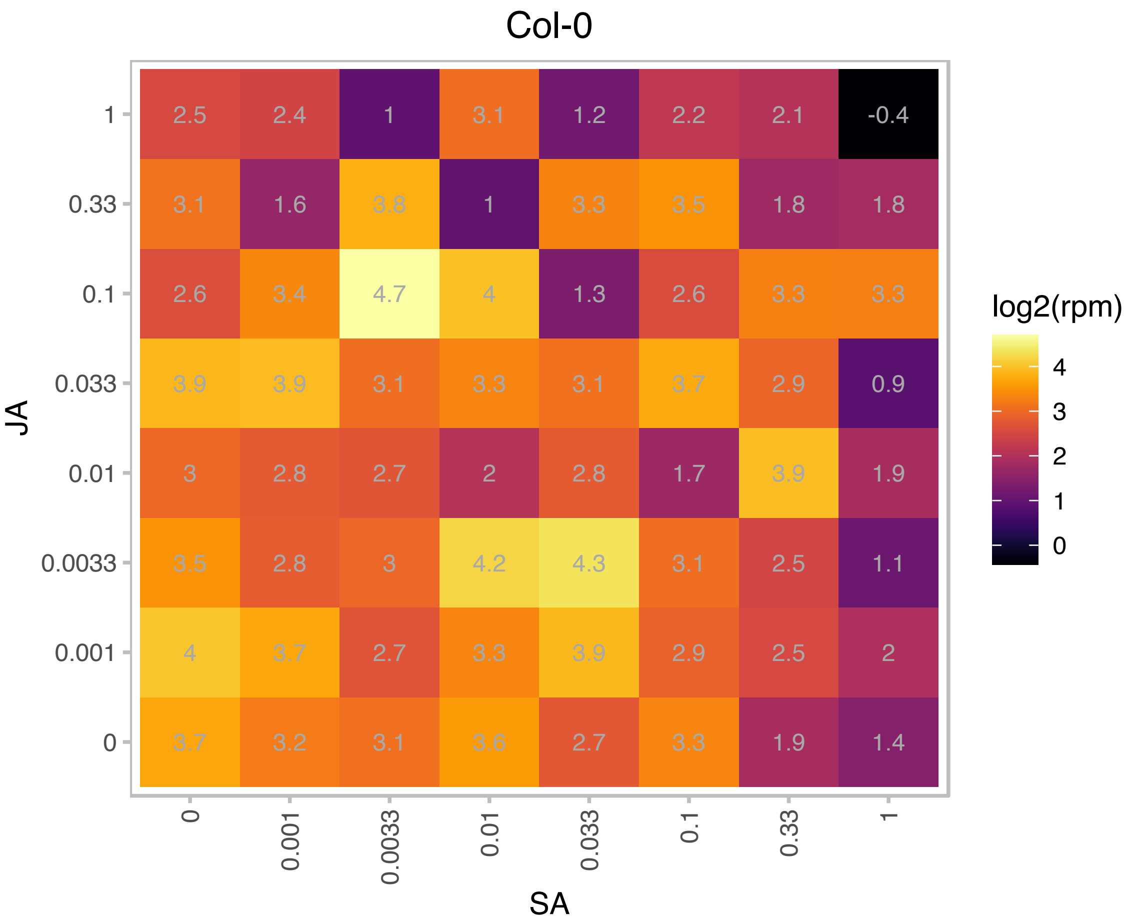


| | |
|---|---|
| Gene | AT5G13090 |
| is.expressed_genes | TRUE |
| is.responsive_genes | TRUE |
| is.apcluster | F-07 |

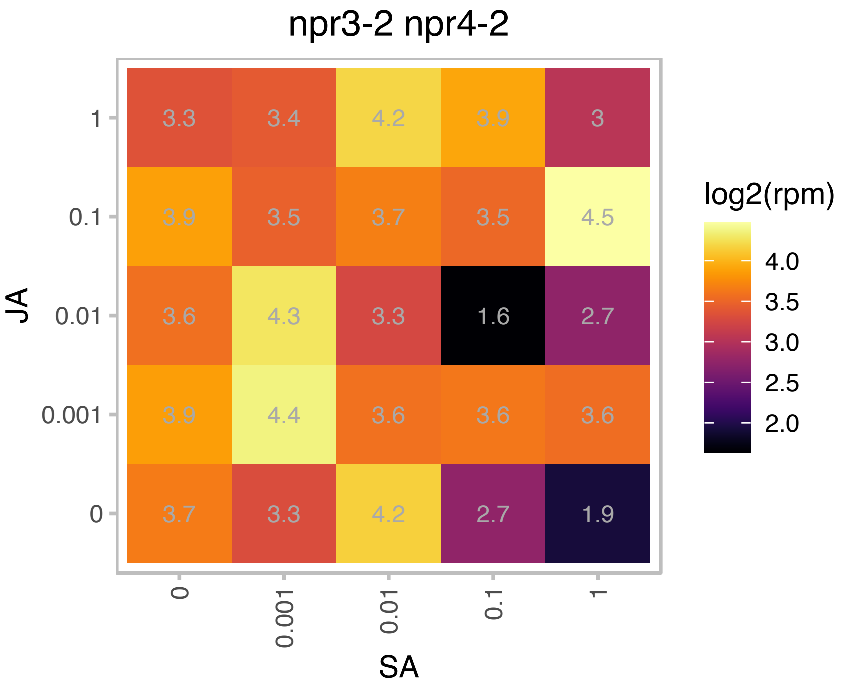


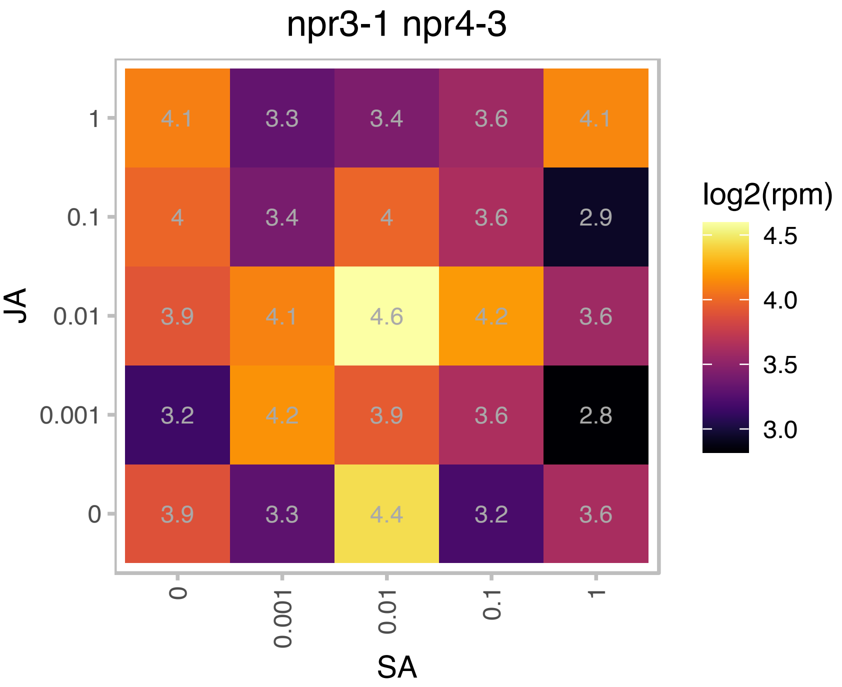

# AT5G14050.1 : NA

Transducin/WD40 repeat-like superfamily protein

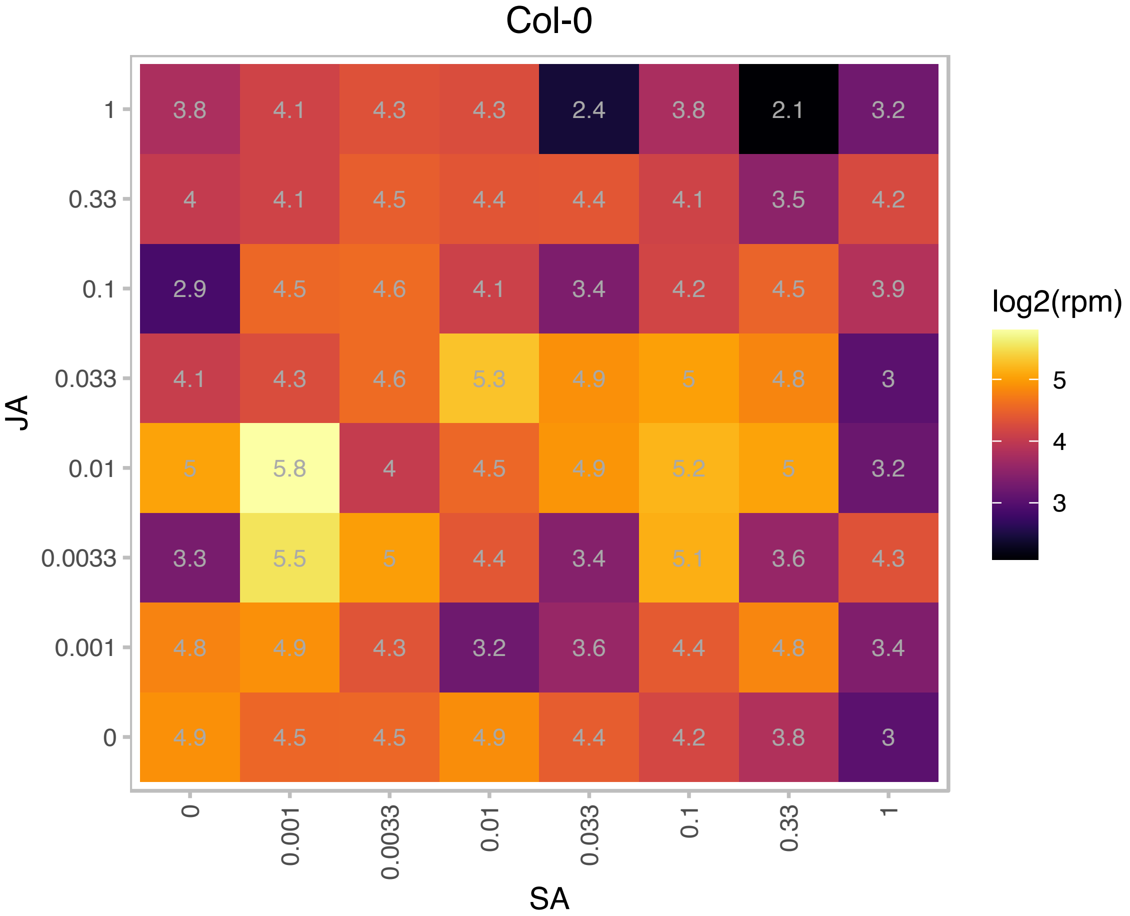


| | |
|---|---|
| Gene | AT5G14050 |
| is.expressed_genes | TRUE |
| is.responsive_genes | TRUE |
| is.apcluster | F-07 |

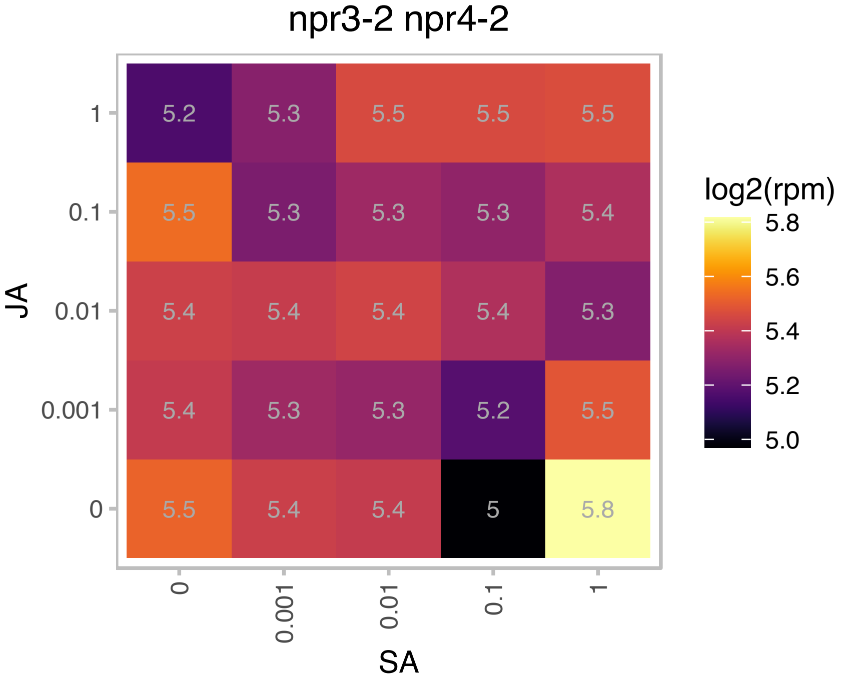


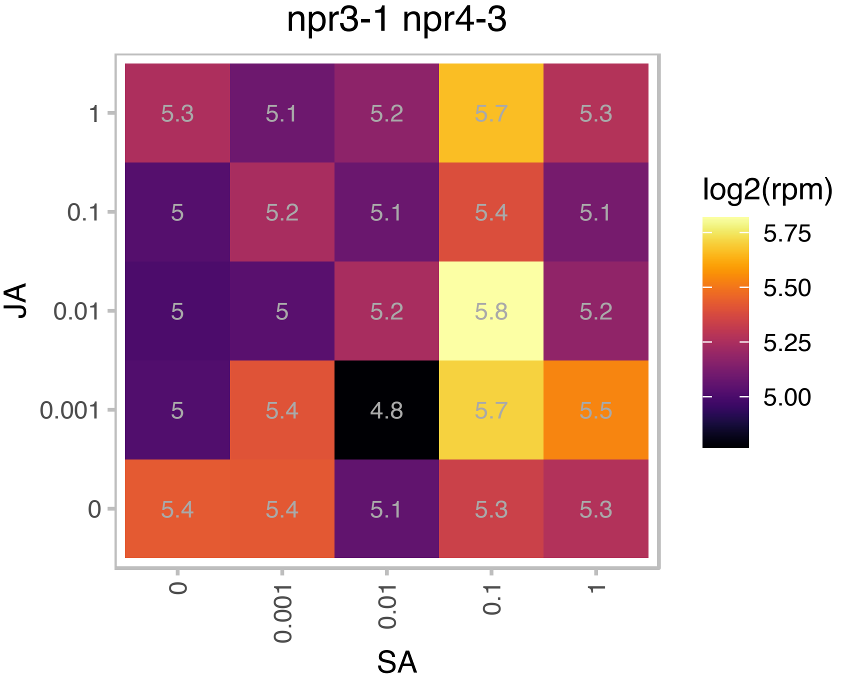

# AT5G14330.1 : NA

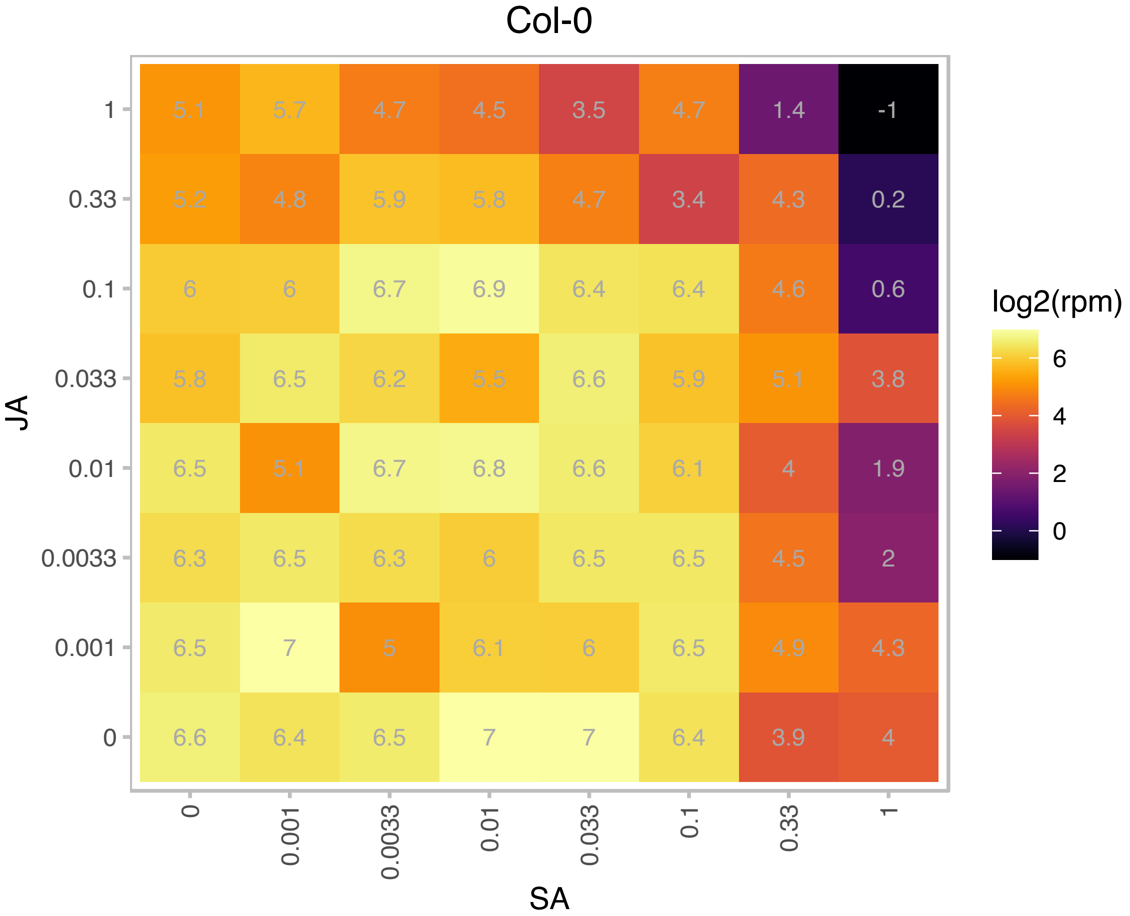


| | |
|---|---|
| Gene | AT5G14330 |
| is.expressed_genes | TRUE |
| is.responsive_genes | TRUE |
| is.apcluster | F-07 |

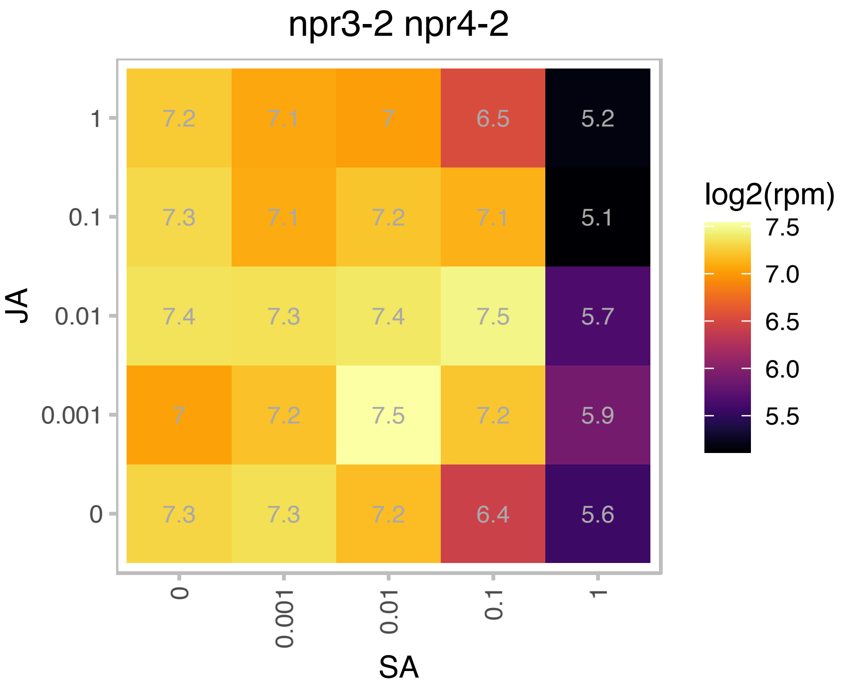


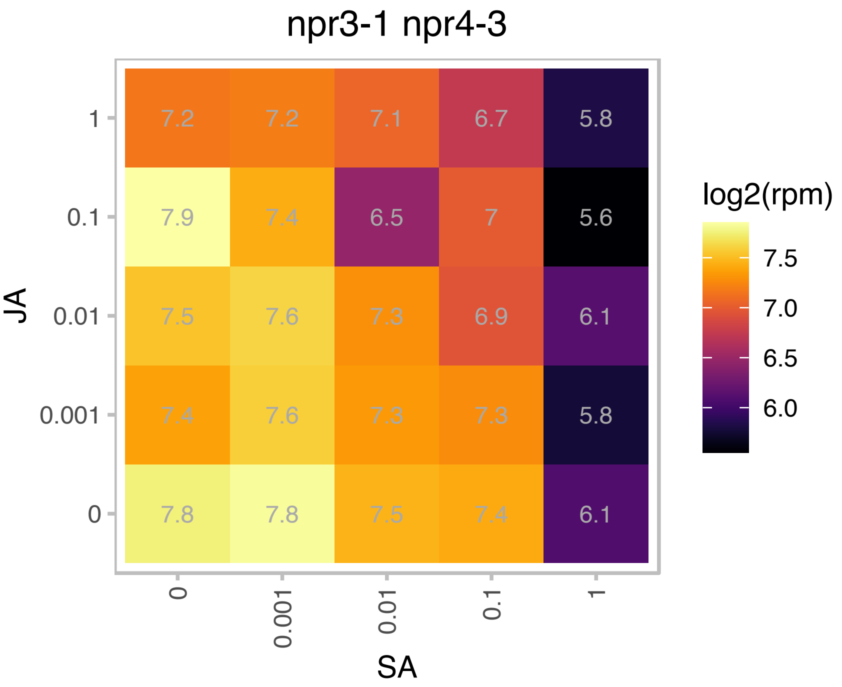

# AT5G15200.1 : NA

Ribosomal protein S4

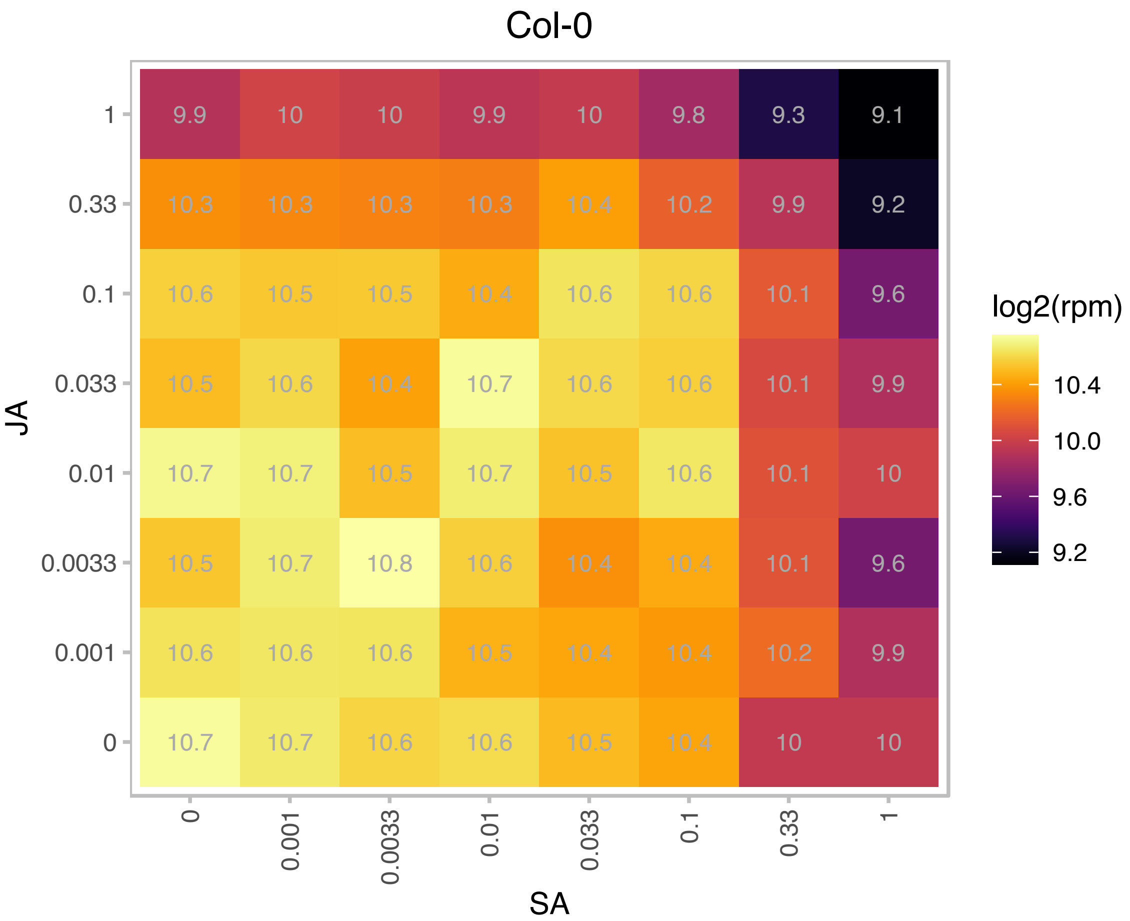


| | |
|---|---|
| Gene | AT5G15200 |
| is.expressed_genes | TRUE |
| is.responsive_genes | TRUE |
| is.apcluster | F-07 |

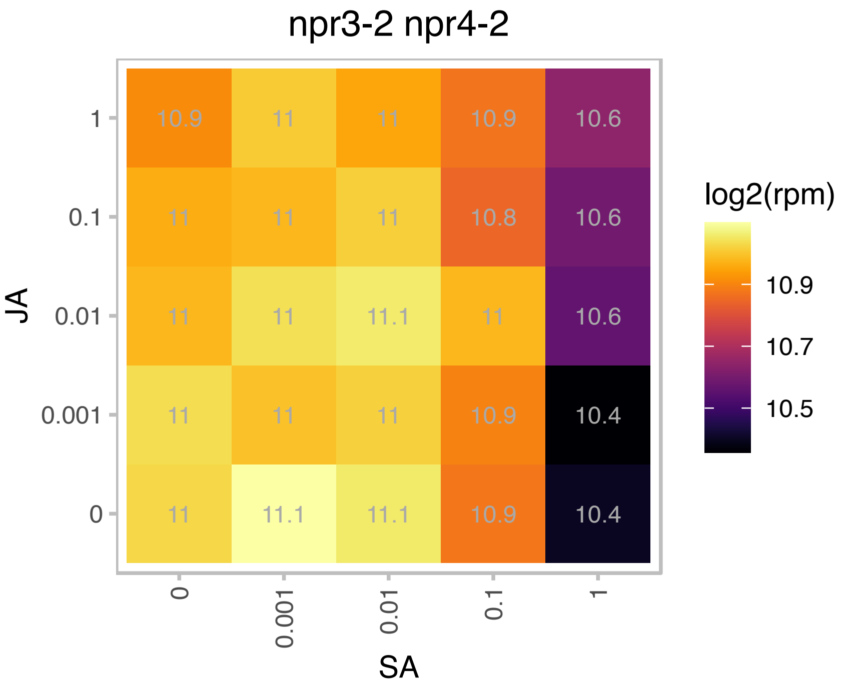


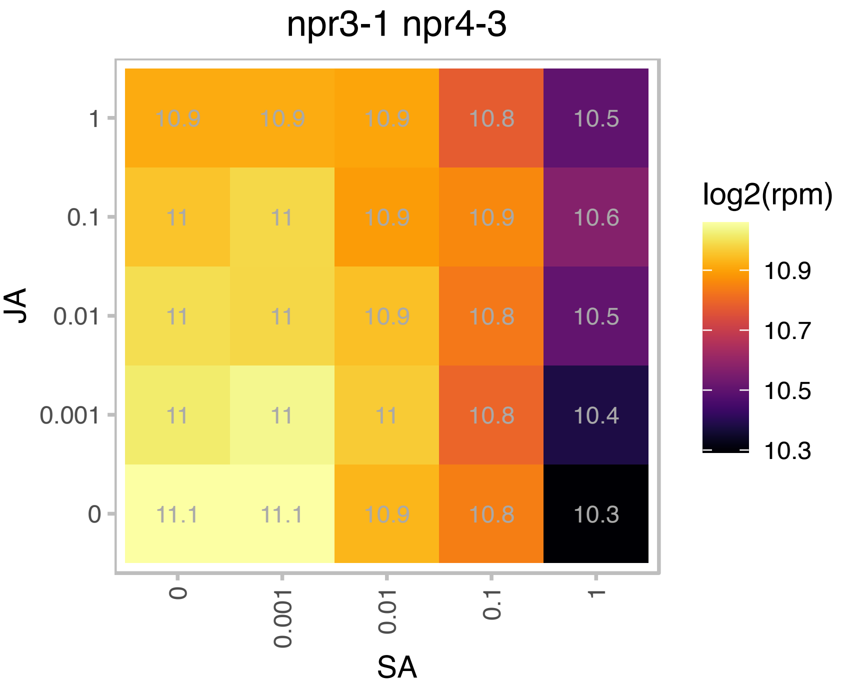

# AT5G15220.1 : NA

Ribosomal protein L27 family protein

| | |
|---|---|
| Gene | AT5G15220 |
| is.expressed_genes | TRUE |
| is.responsive_genes | TRUE |
| is.apcluster | F-07 |

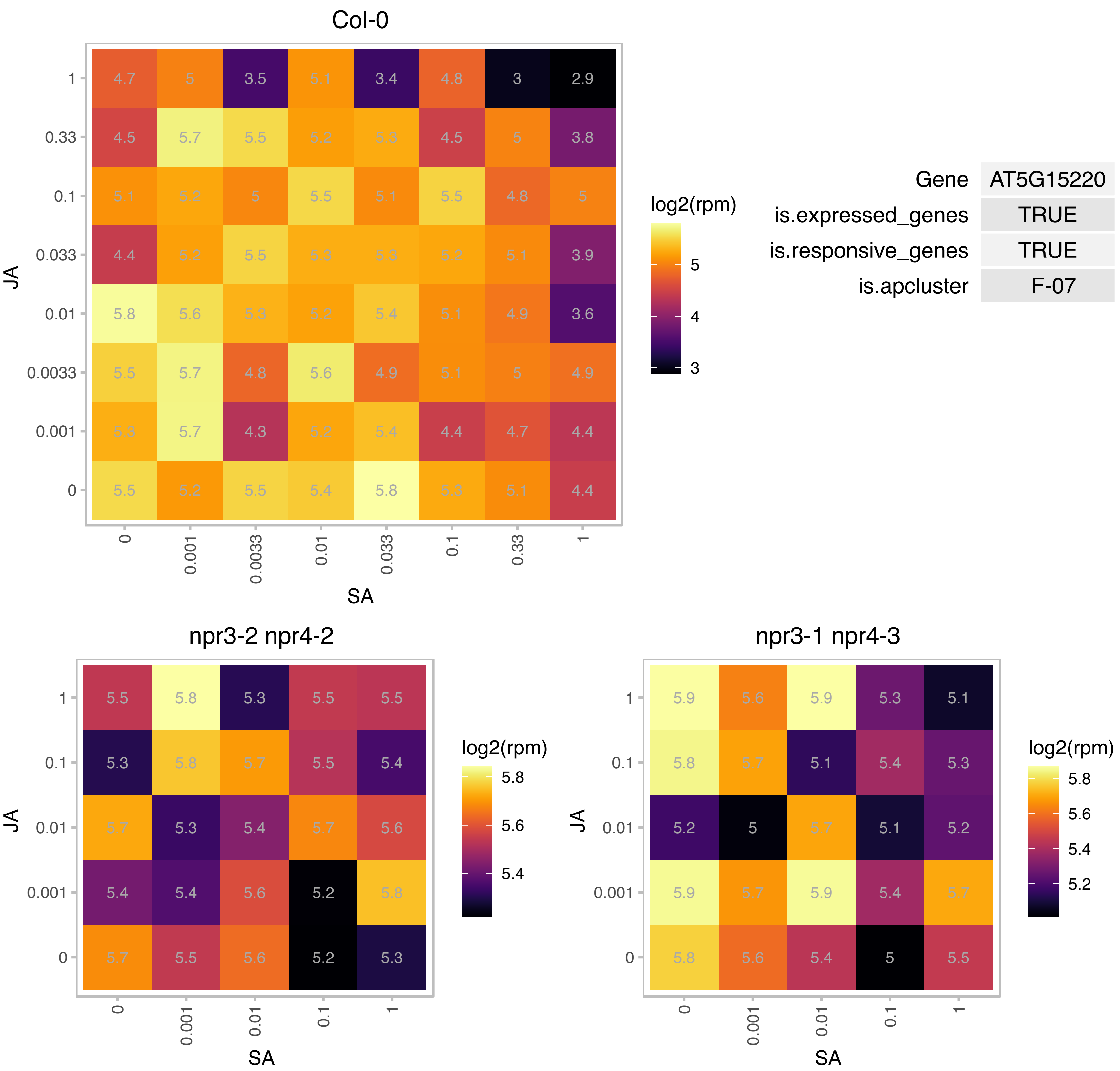

### AT5G15350.1 : AtENODL17, ENODL17

early nodulin-like protein 17

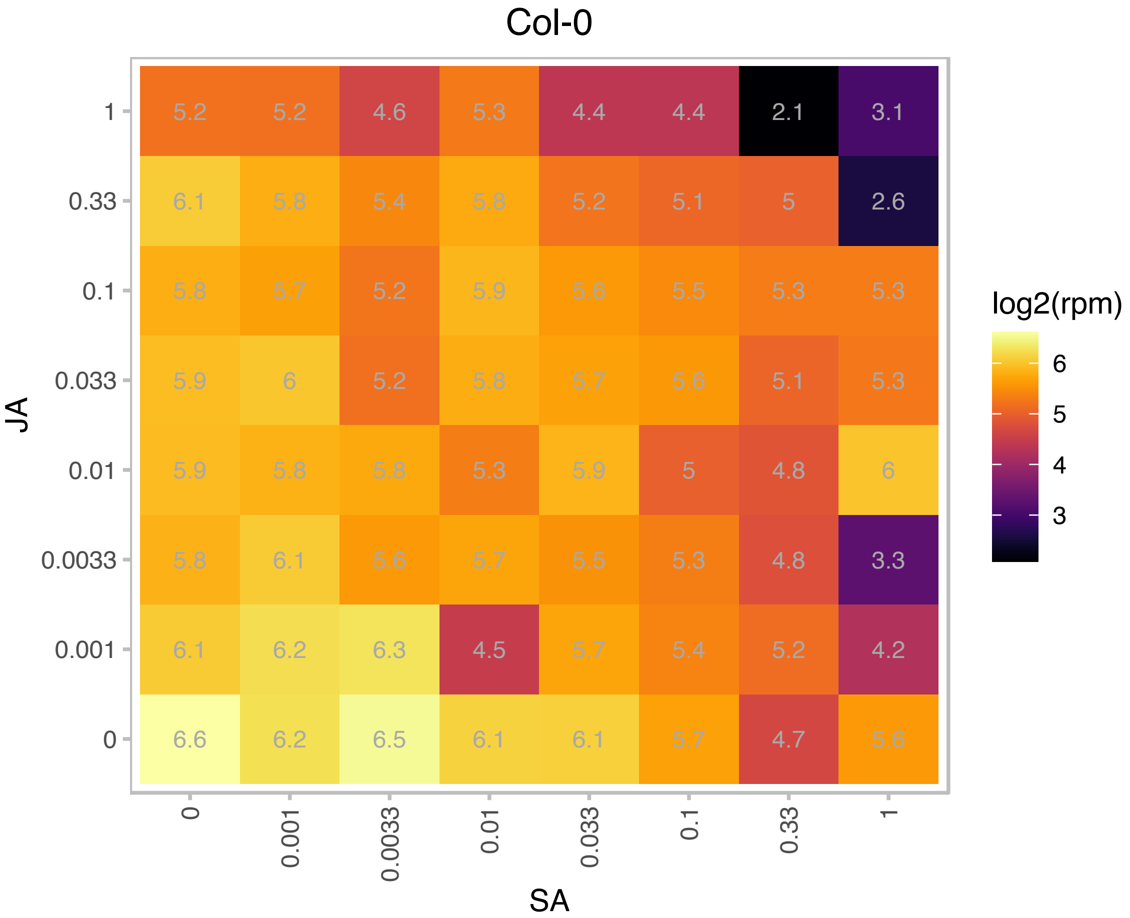


| | |
|---|---|
| Gene | AT5G15350 |
| is.expressed_genes | TRUE |
| is.responsive_genes | TRUE |
| is.apcluster | F-07 |

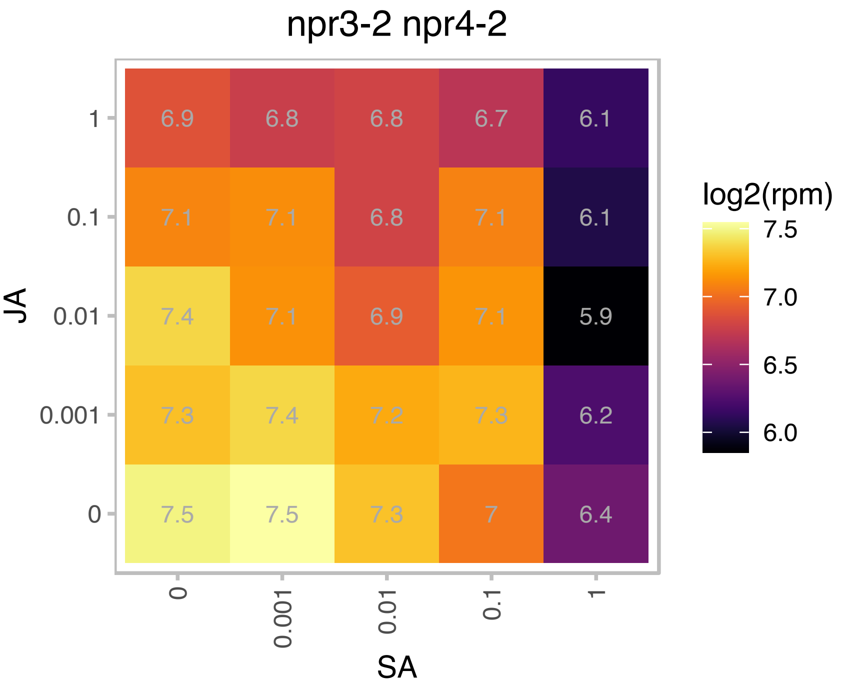


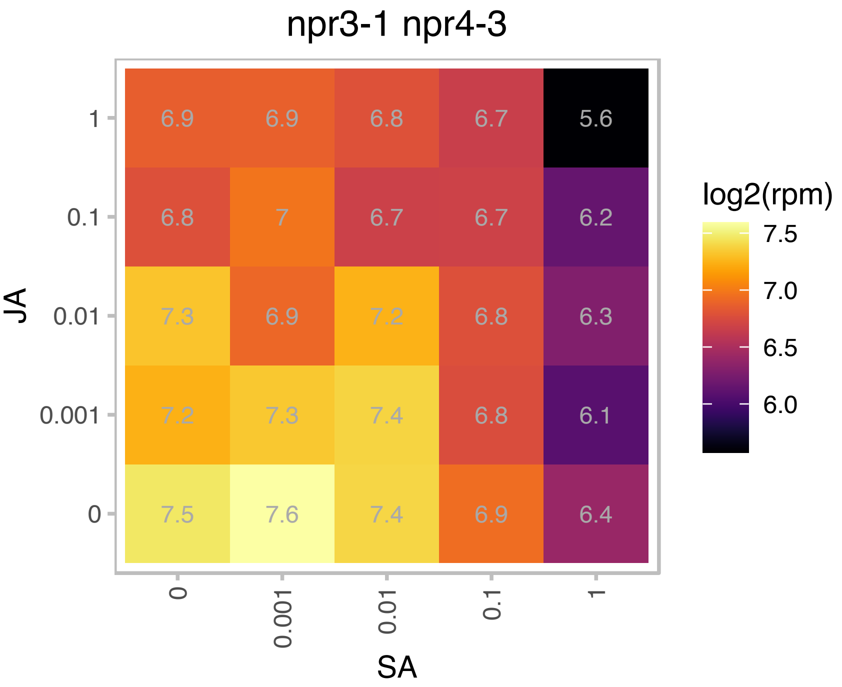

# AT5G15520.1 : NA

Ribosomal protein S19e family protein

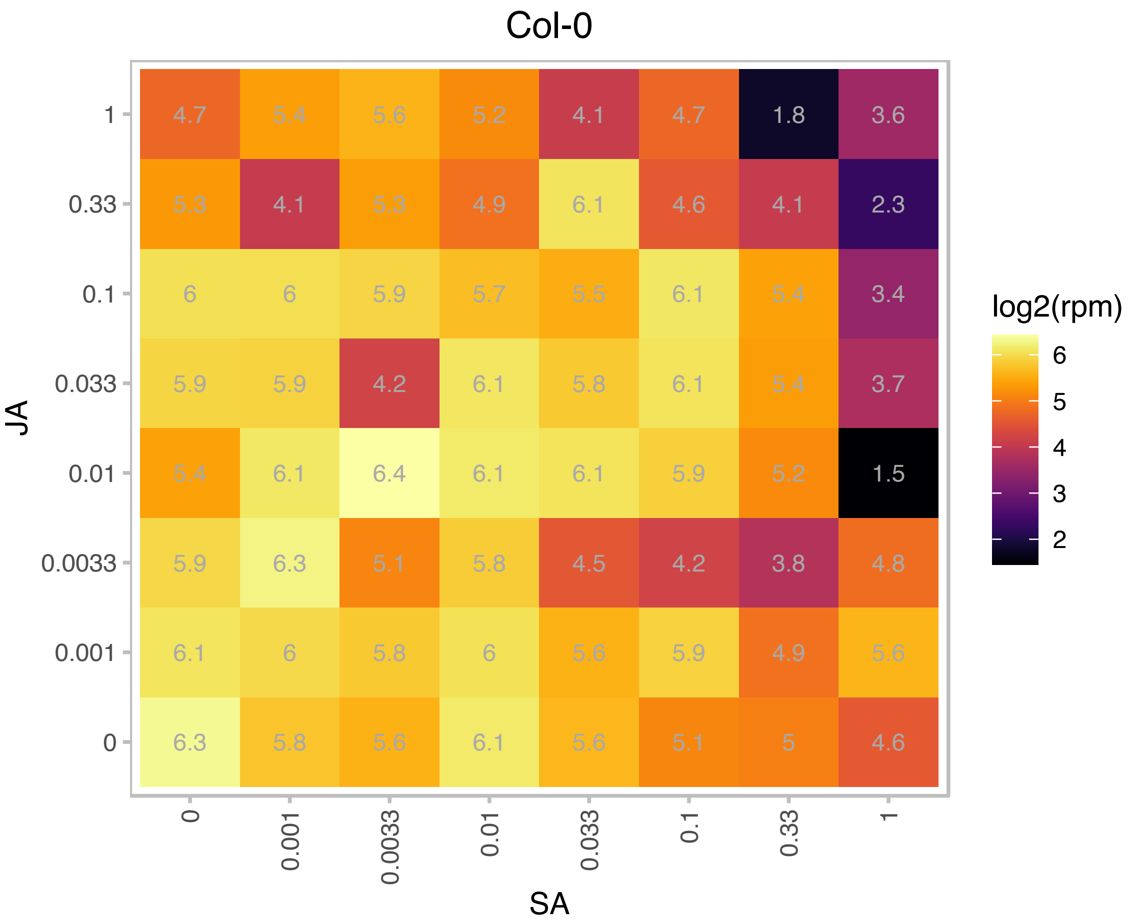


| | |
|---|---|
| Gene | AT5G15520 |
| is.expressed_genes | TRUE |
| is.responsive_genes | TRUE |
| is.apcluster | F-07 |

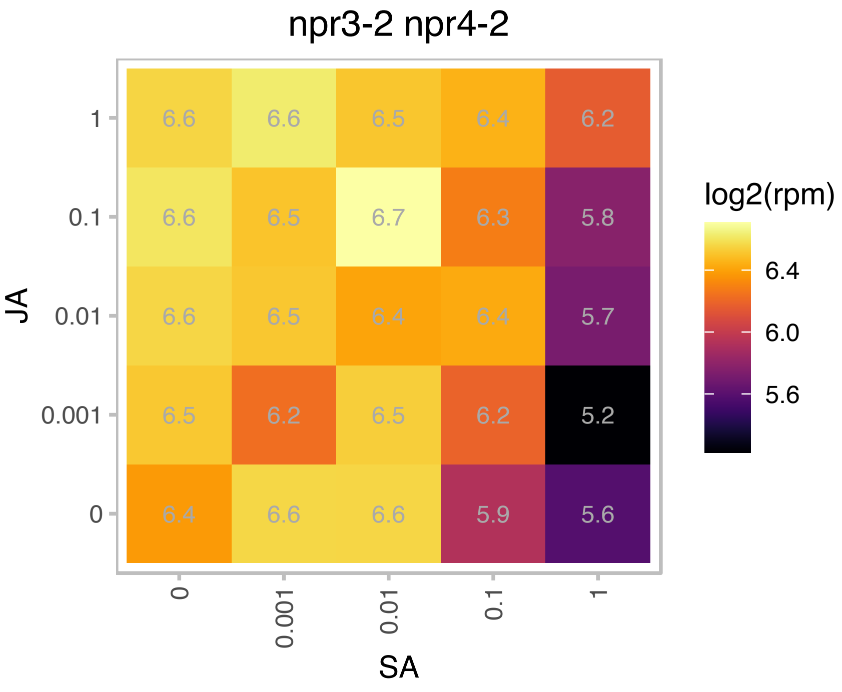


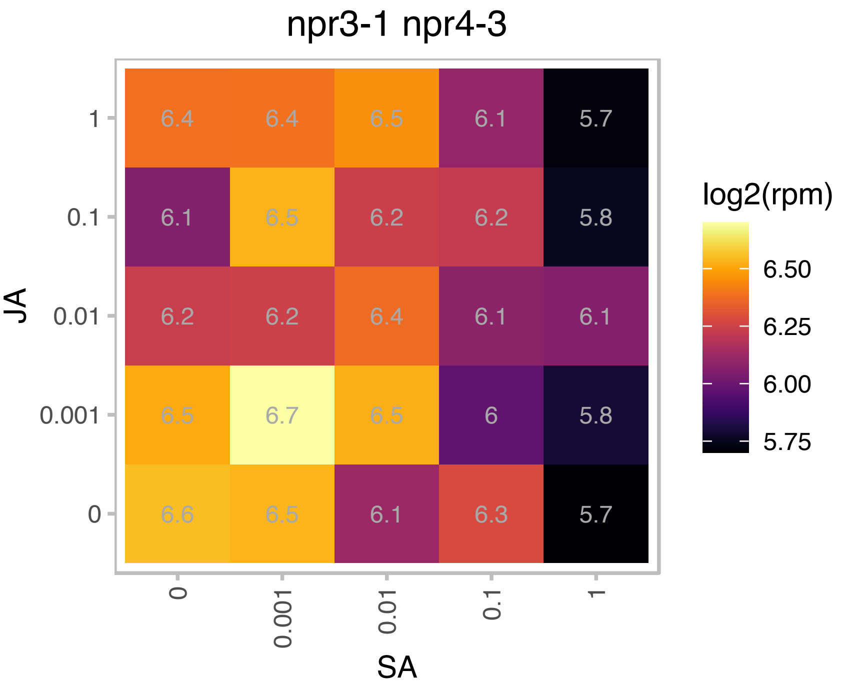

# AT5G16130.1 : NA

Ribosomal protein S7e family protein

| | |
|---|---|
| Gene | AT5G16130 |
| is.expressed_genes | TRUE |
| is.responsive_genes | TRUE |
| is.apcluster | F-07 |

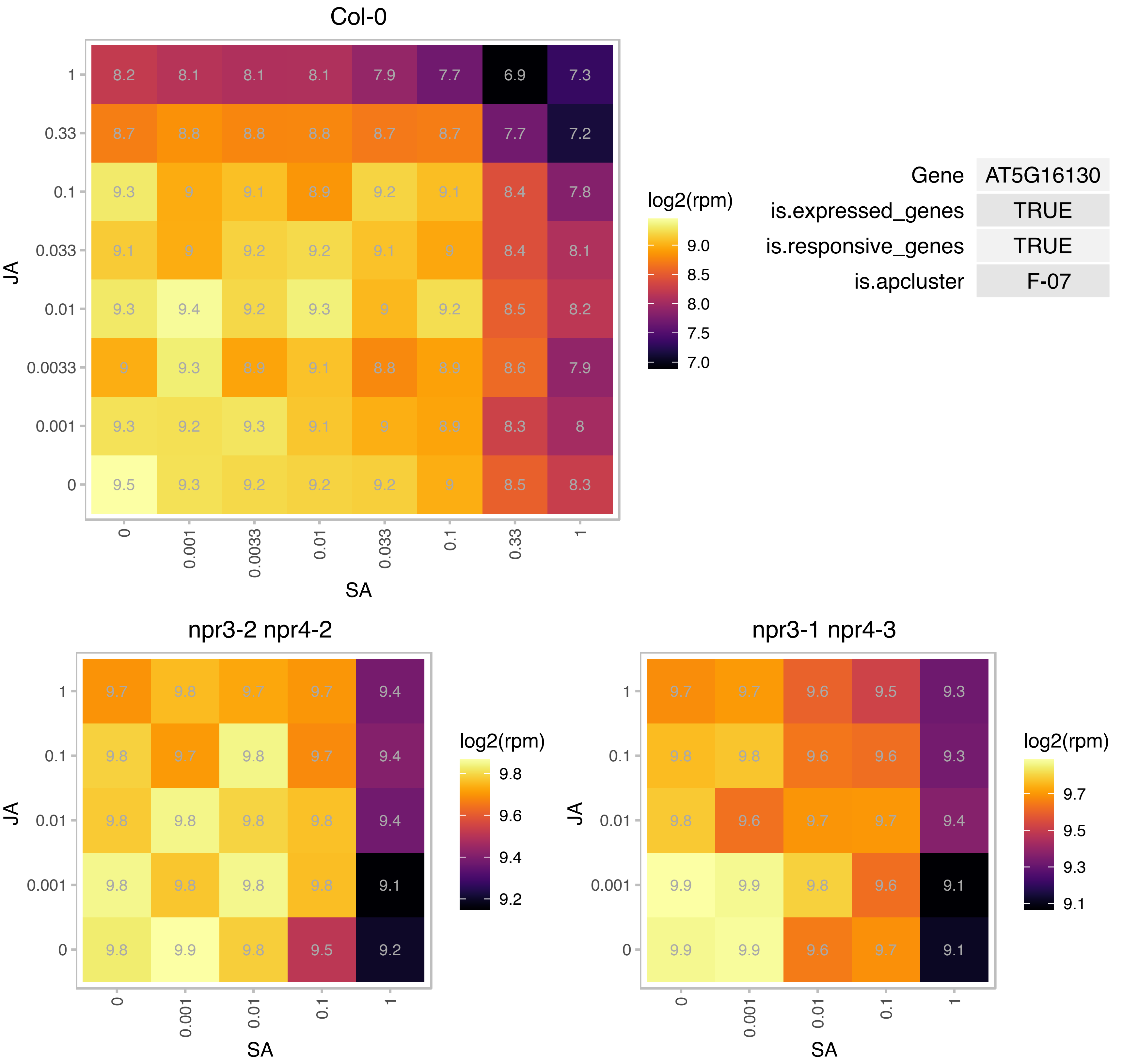

### AT5G16750.1 : TOZ

Transducin family protein / WD-40 repeat family protein

Encodes a nucleolar localized WD-40 repeat protein that is preferentially expressed in dividing cells and is required for regulated division planes and embryo development.

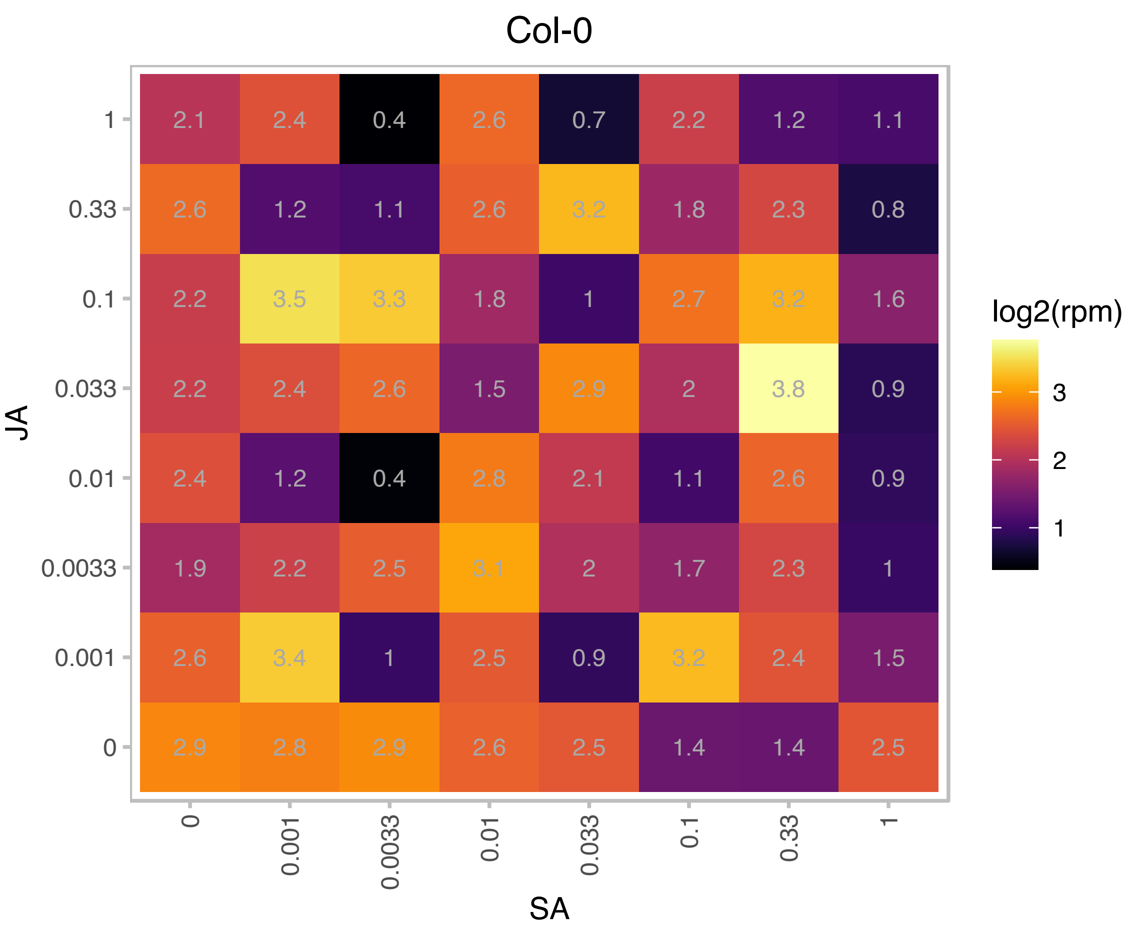


| | |
|---|---|
| Gene | AT5G16750 |
| is.expressed_genes | TRUE |
| is.responsive_genes | TRUE |
| is.apcluster | F-07 |

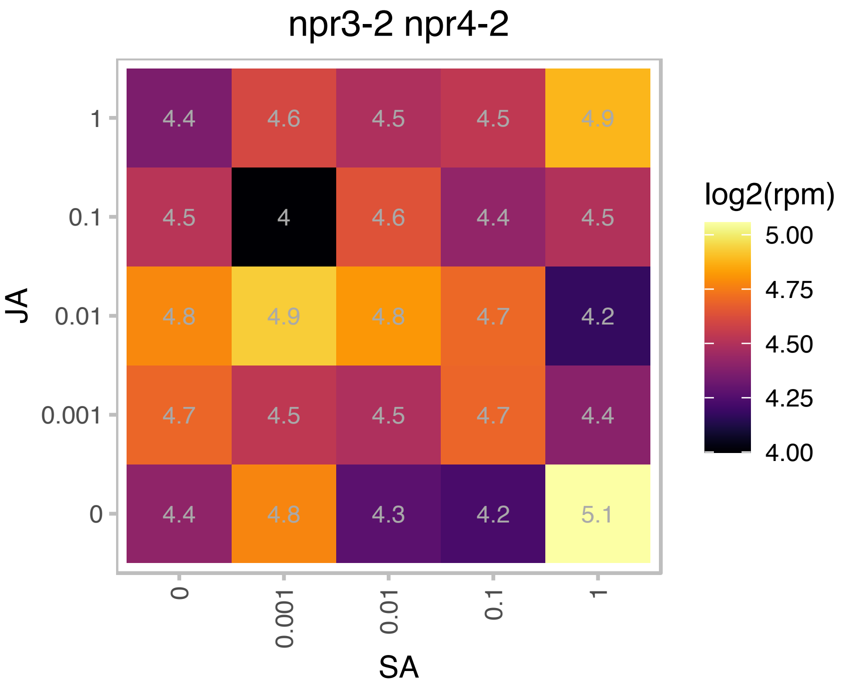


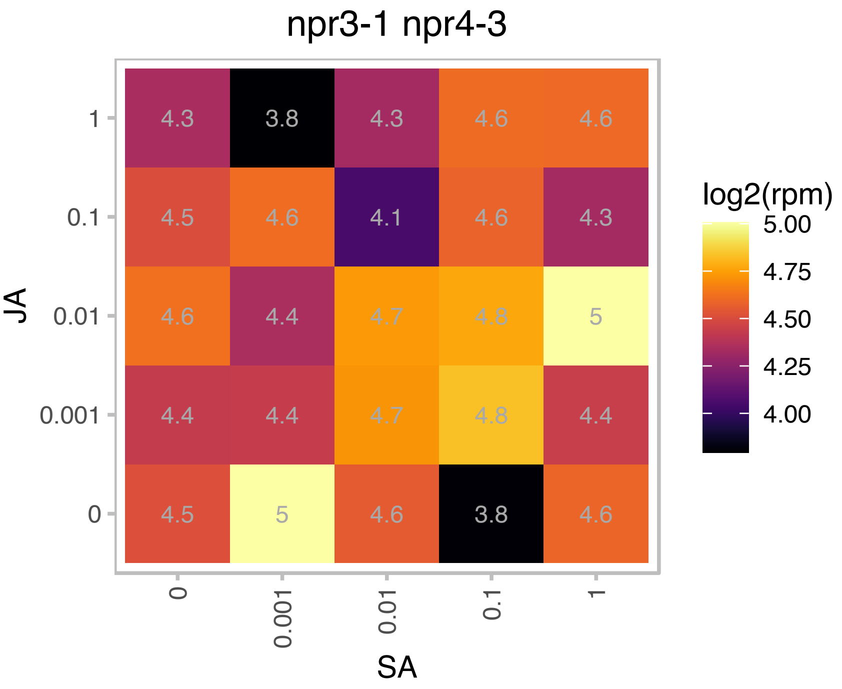

### AT5G18600.1 : ROXY10

Thioredoxin superfamily protein

Encodes a member of the CC-type glutaredoxin (ROXY) family that has been shown to interact with the transcription factor TGA2 and suppress ORA59 promoter activity.

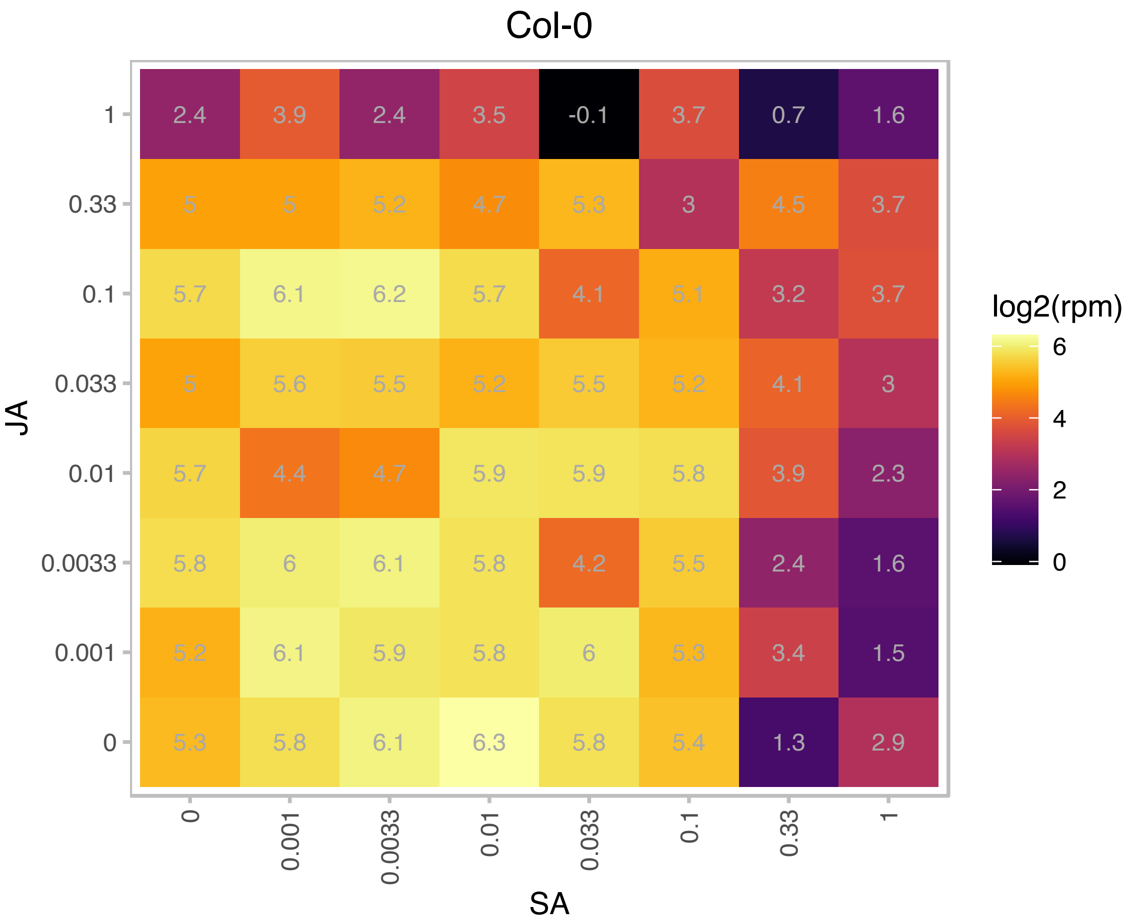


| | |
|---|---|
| Gene | AT5G18600 |
| is.expressed_genes | TRUE |
| is.responsive_genes | TRUE |
| is.apcluster | F-07 |

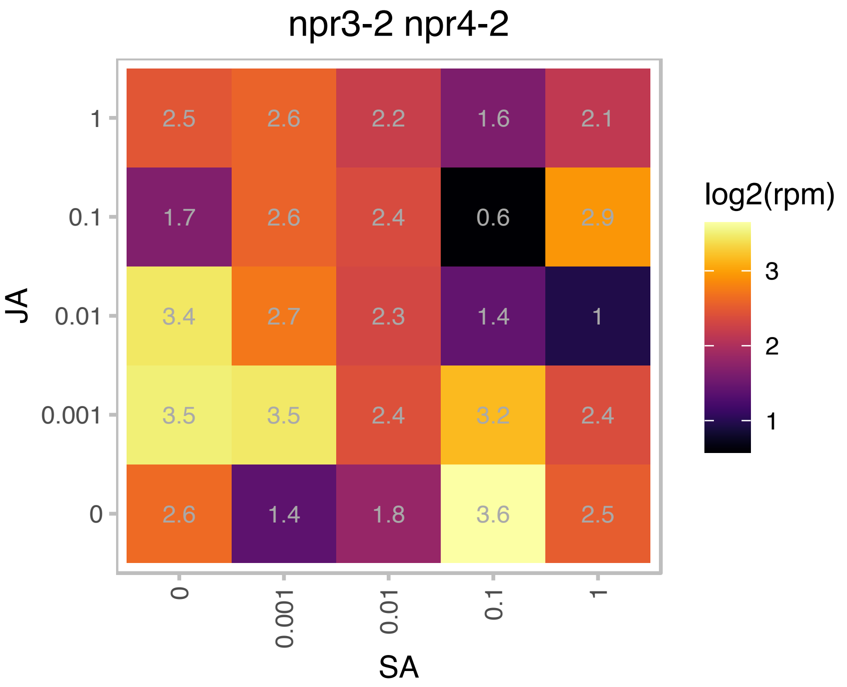


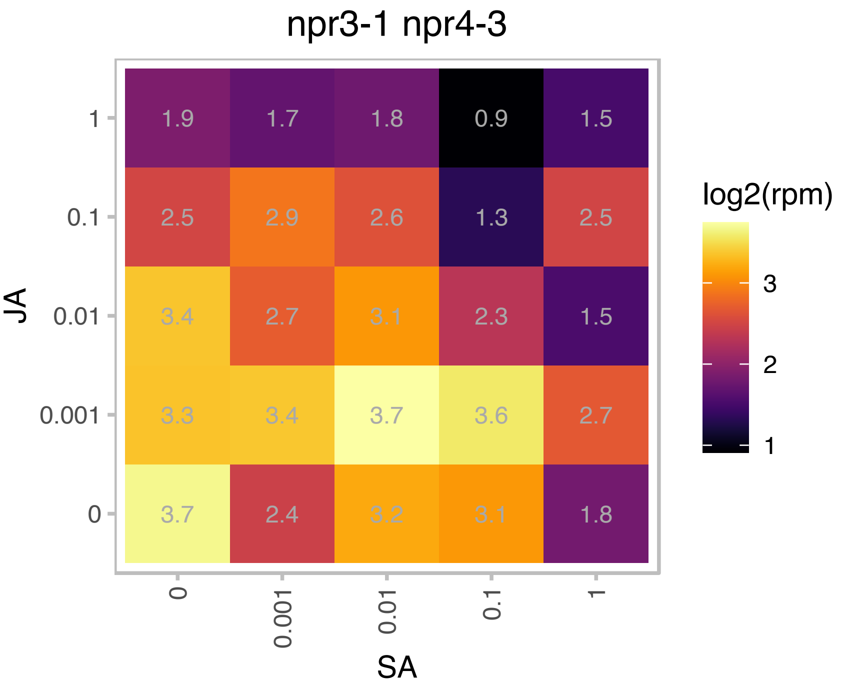

# AT5G18790.1 : NA

Ribosomal protein L33 family protein

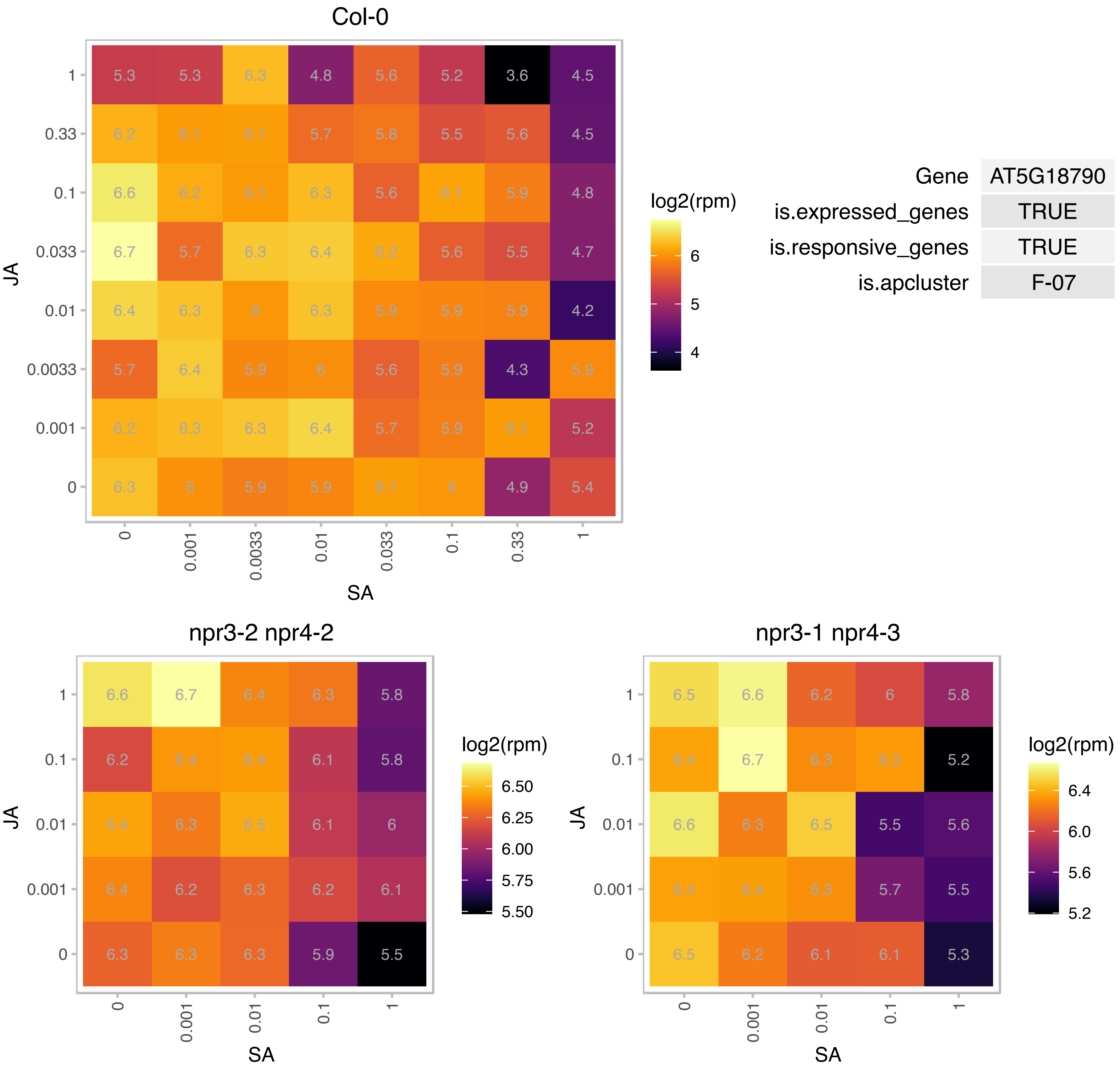


| | |
|---|---|
| Gene | AT5G18790 |
| is.expressed_genes | TRUE |
| is.responsive_genes | TRUE |
| is.apcluster | F-07 |

### AT5G19260.1 : FAF3

Protein of unknown function (DUF3049)

A member of the FAF family proteins encoded by the FANTASTIC FOUR (FAF) genes: AT4G02810 (FAF1), AT1G03 (FAF2), AT5G19260 (FAF3) and AT3G06020 (FAF4). FAFs have the potential to regulate shoot meristem size in Arabidopsis thaliana. FAFs can repress WUS, which ultimately leads to an arrest of meristem activity in FAF overexpressing lines.

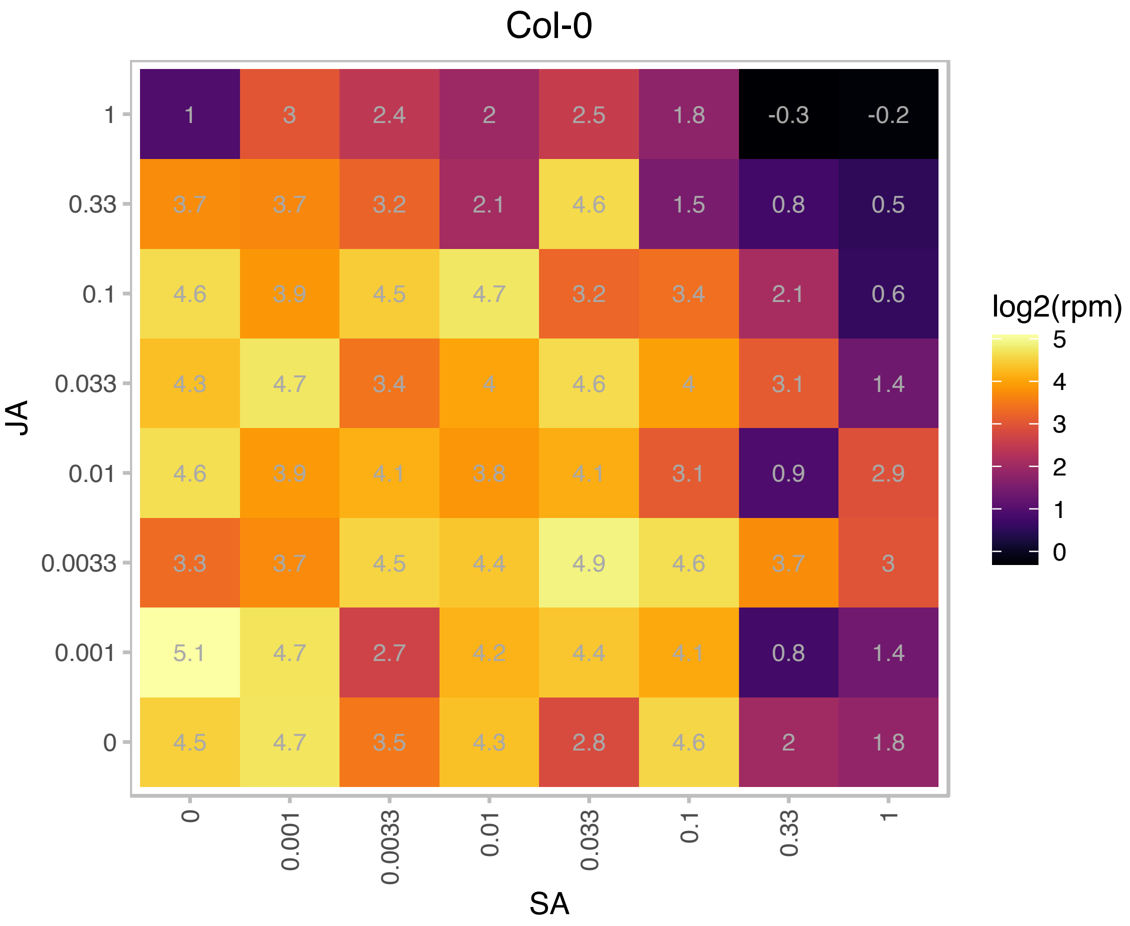


| | |
|---|---|
| Gene | AT5G19260 |
| is.expressed_genes | TRUE |
| is.responsive_genes | TRUE |
| is.apcluster | F-07 |

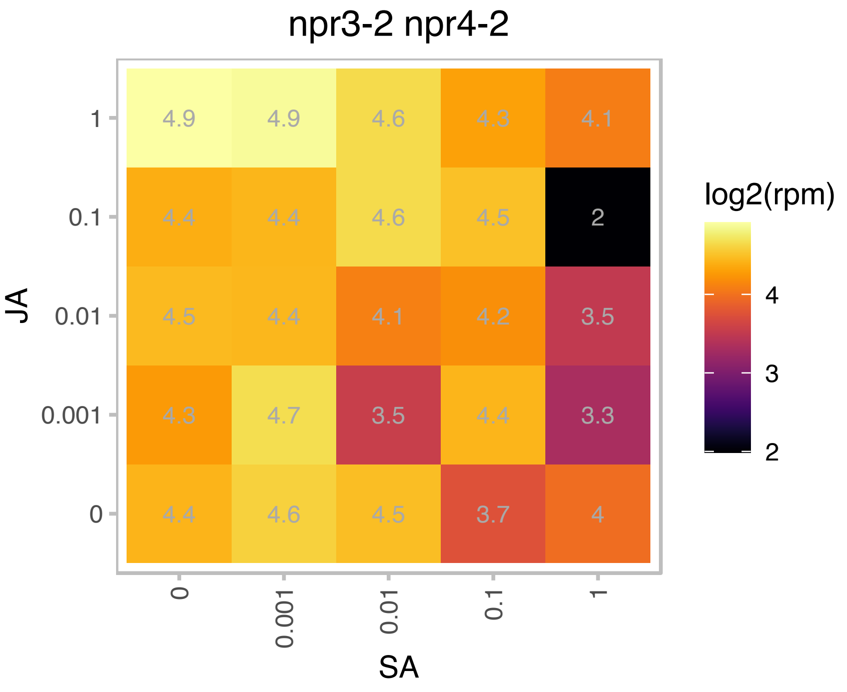


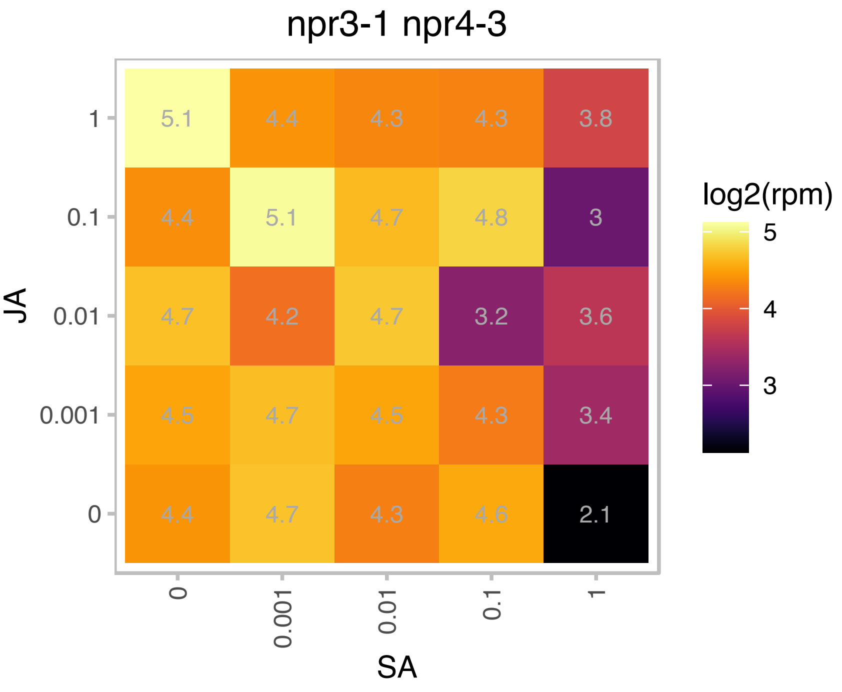

# AT5G20290.1 : NA

Ribosomal protein S8e family protein

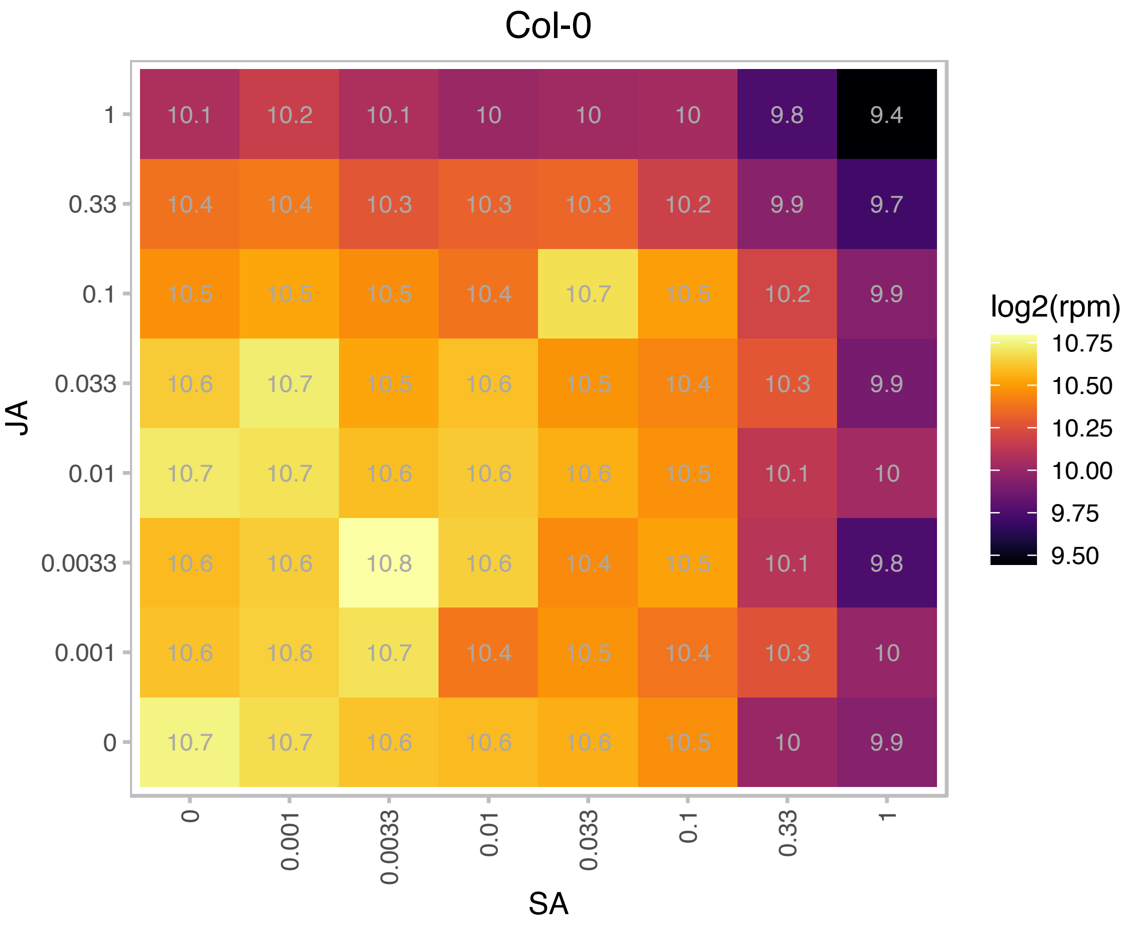


| | |
|---|---|
| Gene | AT5G20290 |
| is.expressed_genes | TRUE |
| is.responsive_genes | TRUE |
| is.apcluster | F-07 |

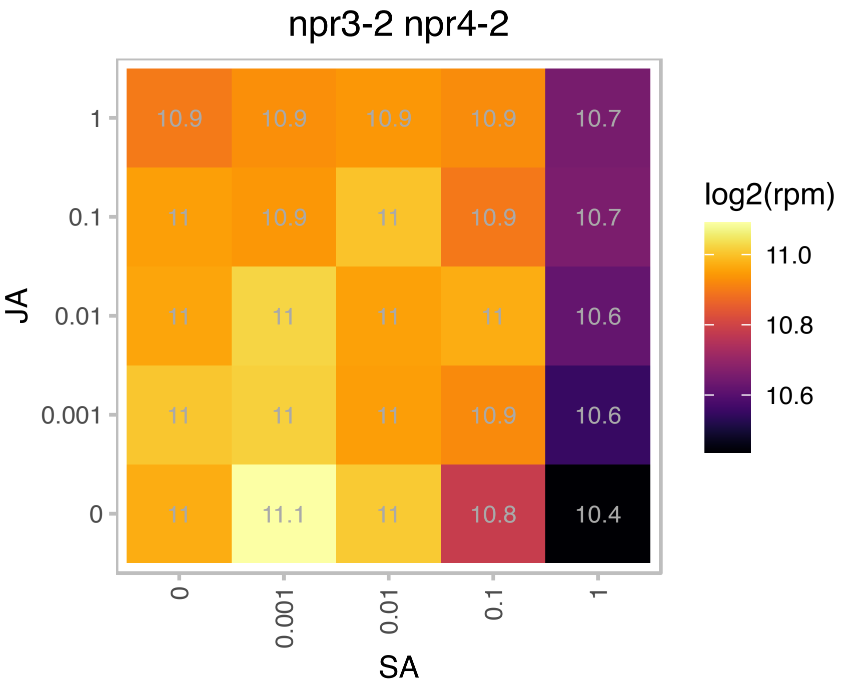


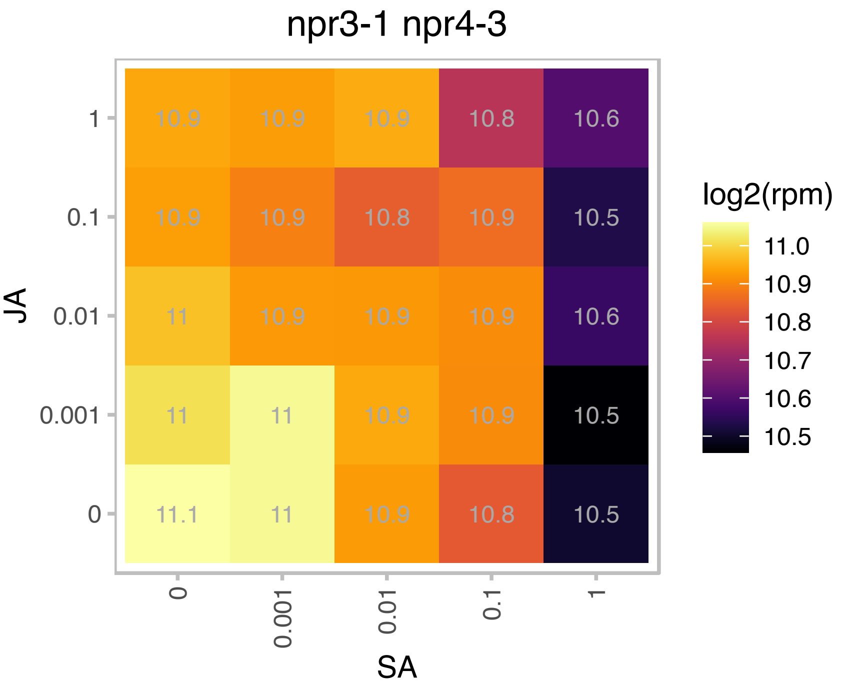

# AT5G22440.1 : NA

Ribosomal protein L1p/L10e family

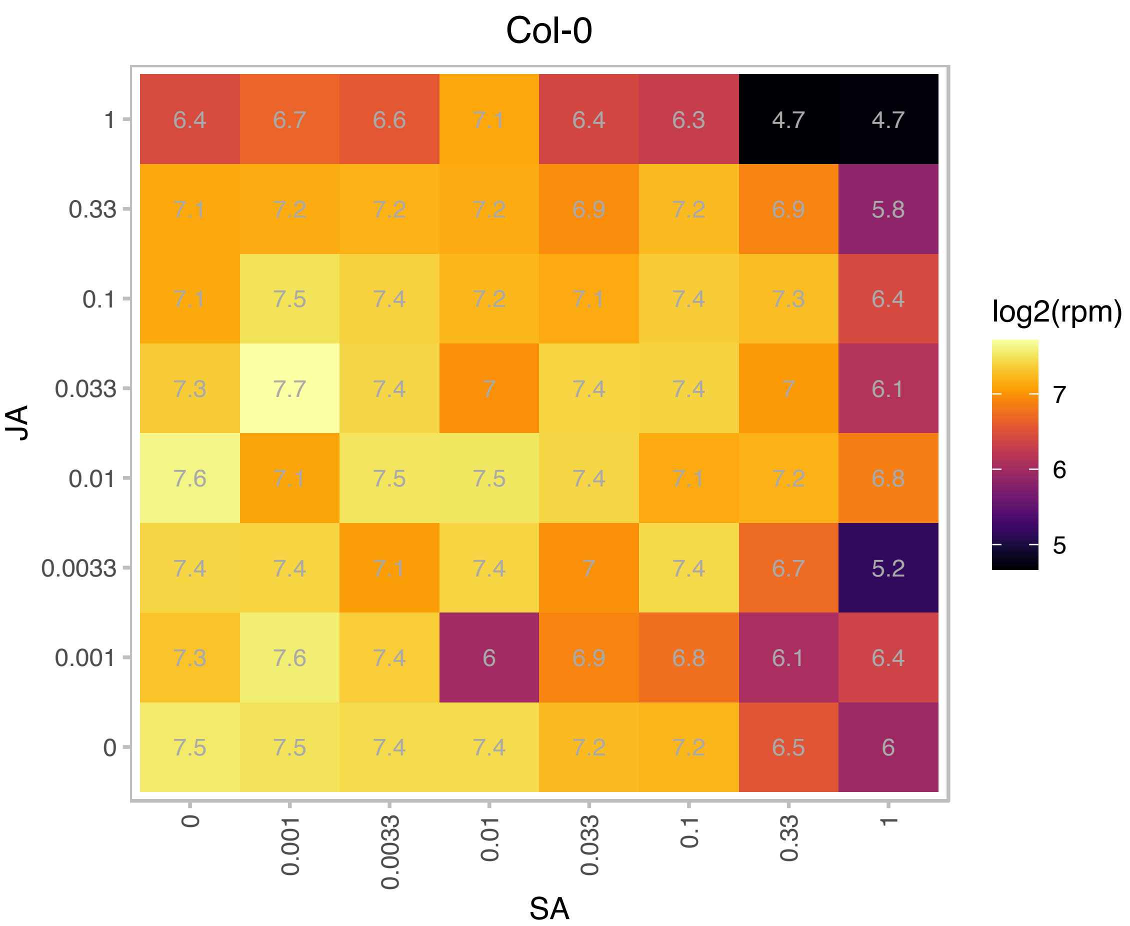


| | |
|---|---|
| Gene | AT5G22440 |
| is.expressed_genes | TRUE |
| is.responsive_genes | TRUE |
| is.apcluster | F-07 |

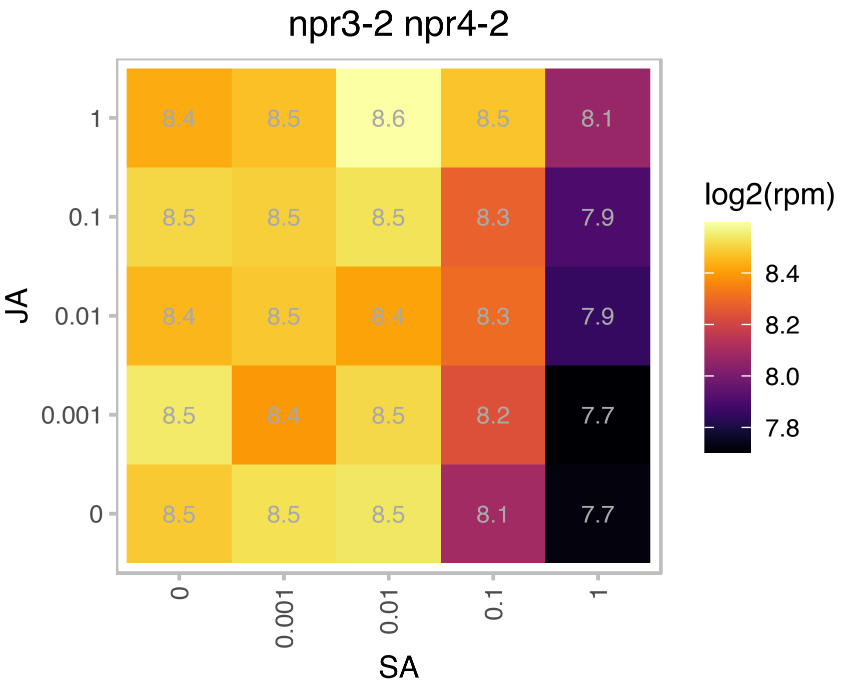


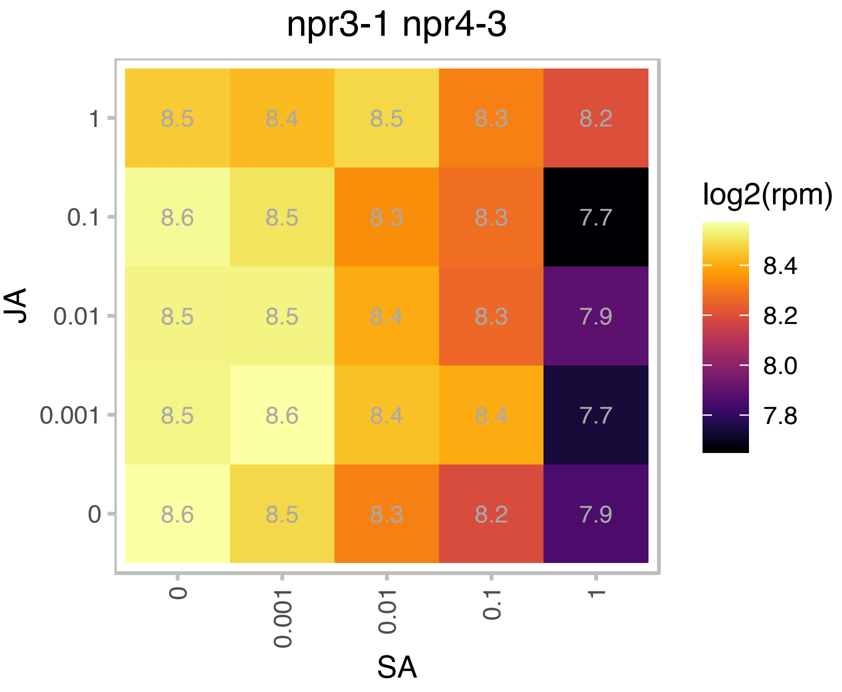

### AT5G22920.1 : AtRZPF34, CHYR1, RZPF34

CHY-type/CTCHY-type/RING-type Zinc finger protein

Encodes a protein with sequence similarity to RING, zinc finger proteins. Loss of function mutations show reduced (15%) stomatal aperture under non stress conditions.

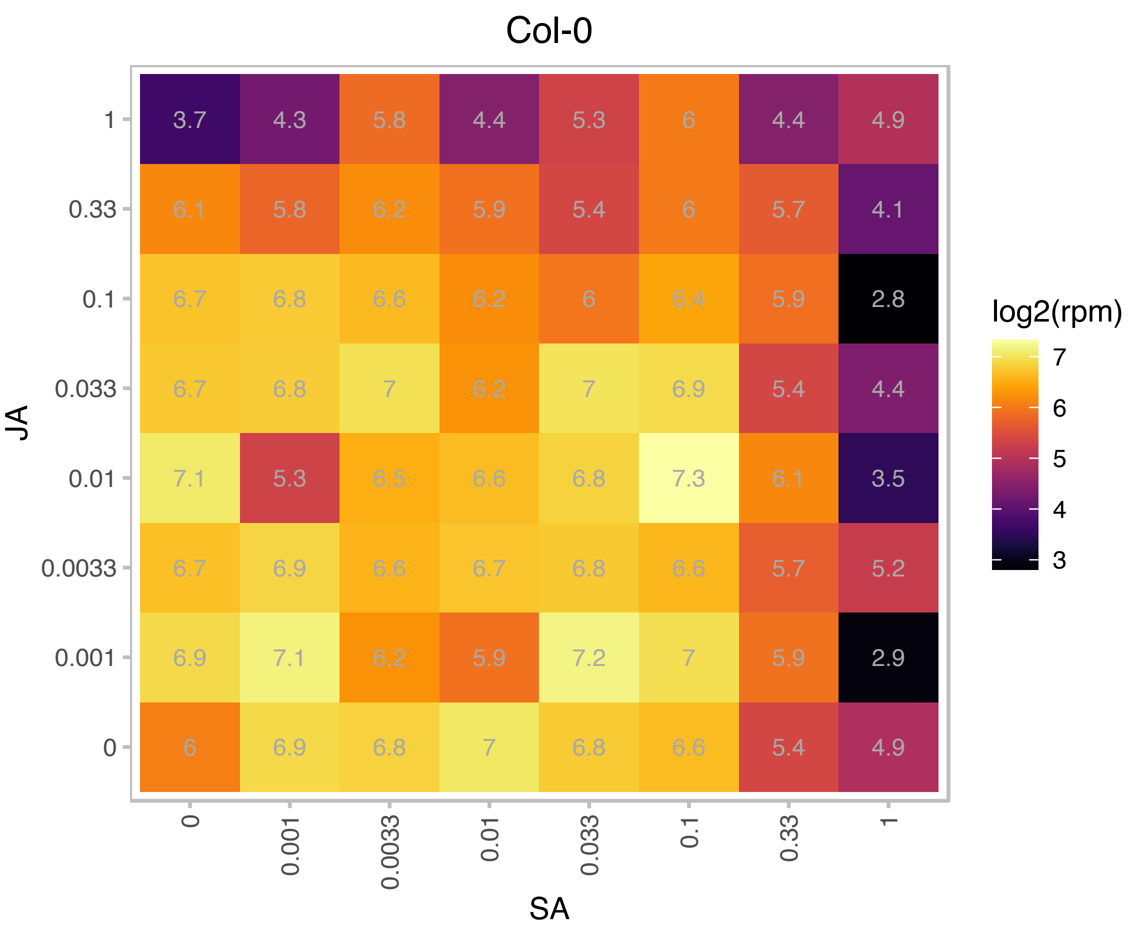


| | |
|---|---|
| Gene | AT5G22920 |
| is.expressed_genes | TRUE |
| is.responsive_genes | TRUE |
| is.apcluster | F-07 |

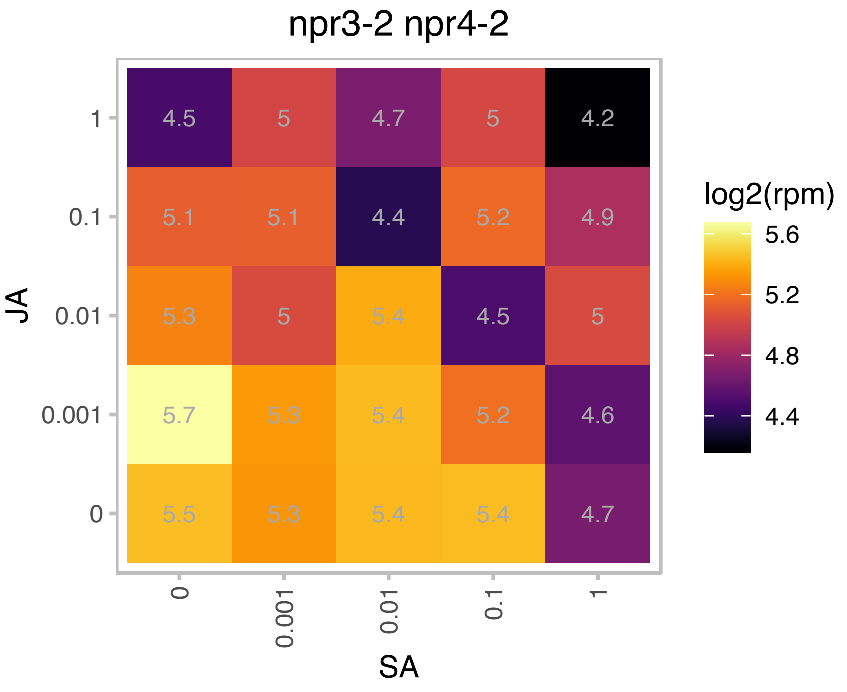


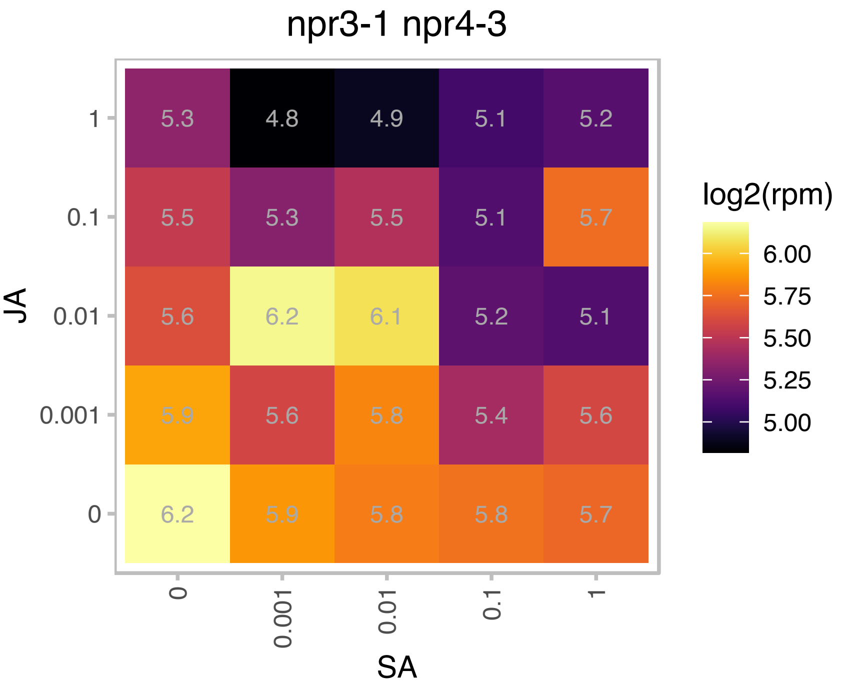

### AT5G23420.1 : HMGB6

high-mobility group box 6
Encodes HMGB6, a protein belonging to the subgroup of HMGB (high mobility group B) proteins.  Localized in the nucleus. Binds to supercoiled DNA in vitro. HMGB6 is phosphorylated by protein kinase CK2alpha within its acidic C-terminal domain.

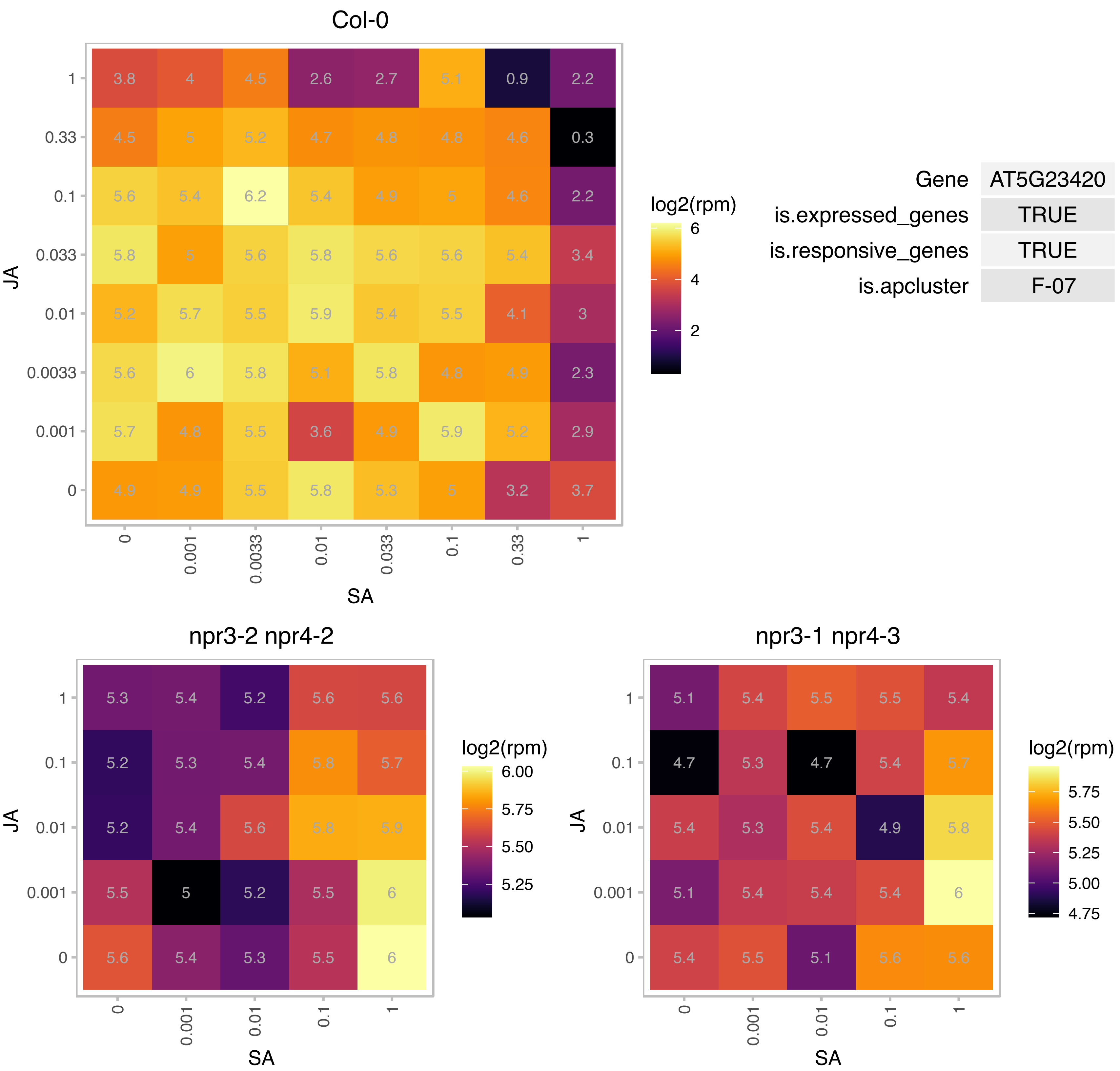

# AT5G23900.1 : NA

Ribosomal protein L13e family protein

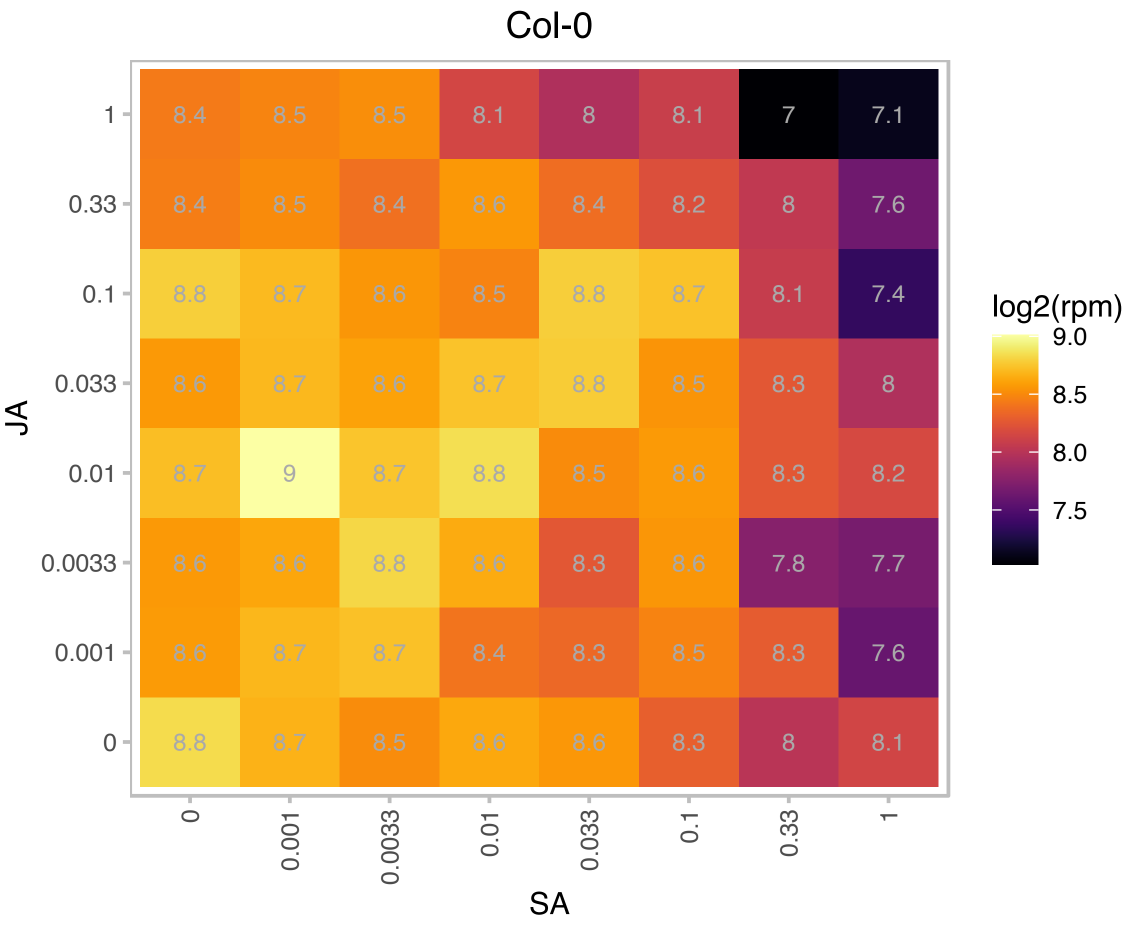


| | |
|---|---|
| Gene | AT5G23900 |
| is.expressed_genes | TRUE |
| is.responsive_genes | TRUE |
| is.apcluster | F-07 |

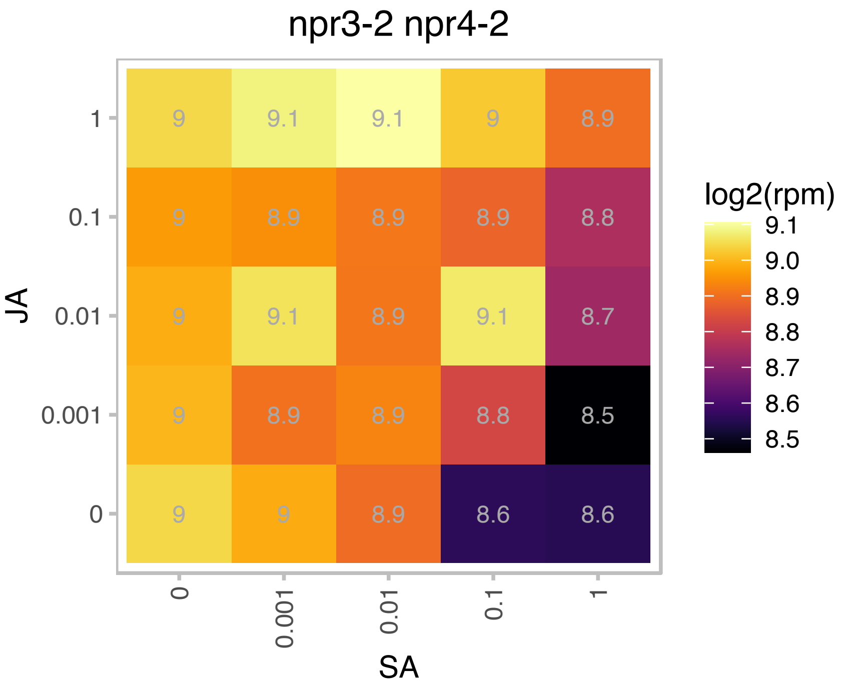


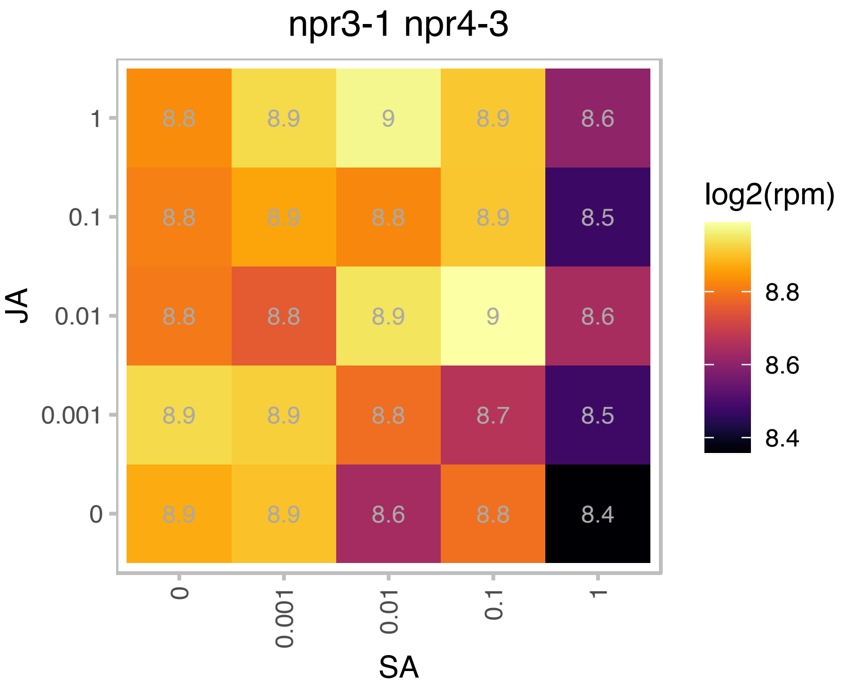

### AT5G27670.1 : h2a.w.7, HTA7

histone H2A 7

Encodes HTA7, a histone H2A protein.

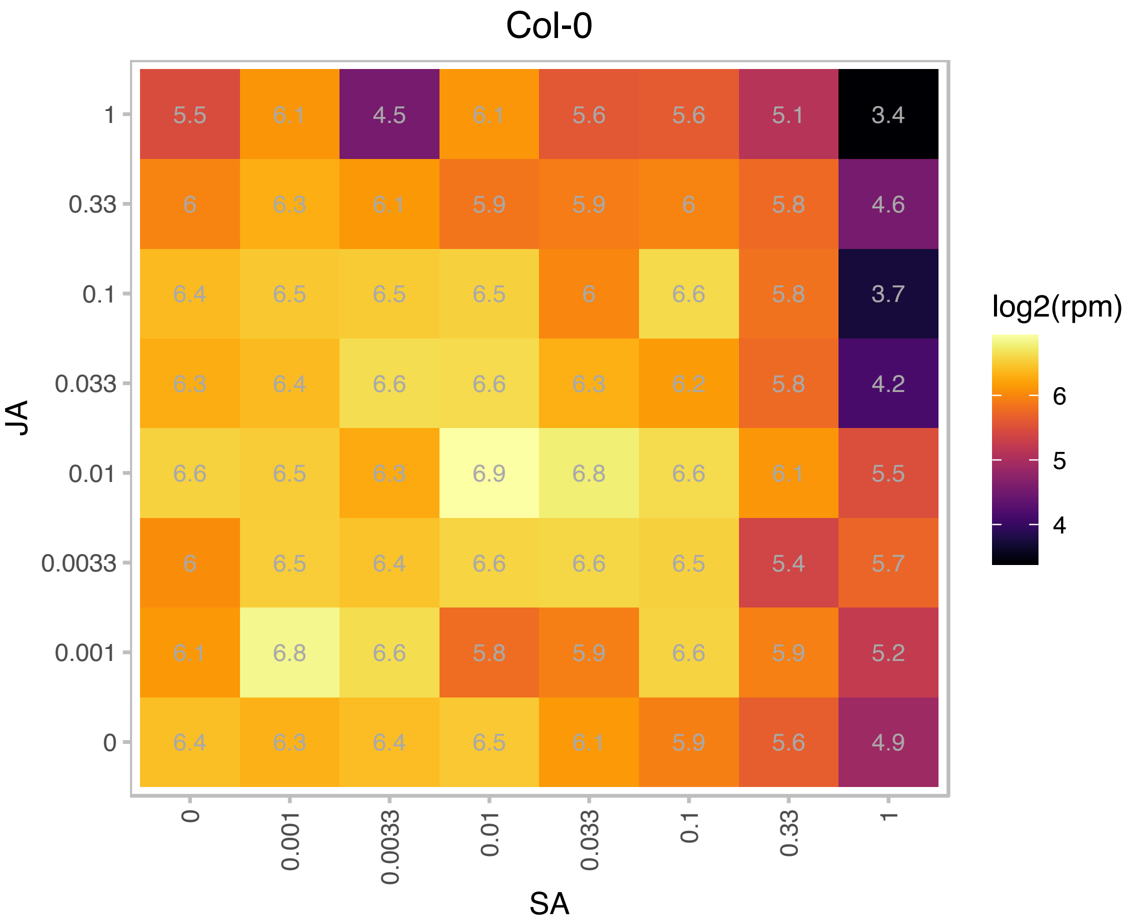


| | |
|---|---|
| Gene | AT5G27670 |
| is.expressed_genes | TRUE |
| is.responsive_genes | TRUE |
| is.apcluster | F-07 |

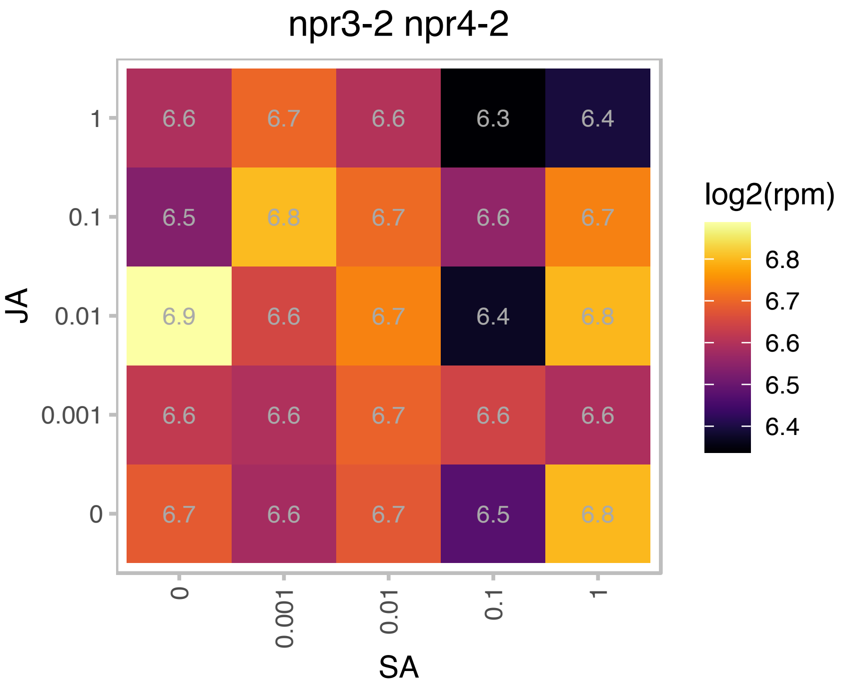


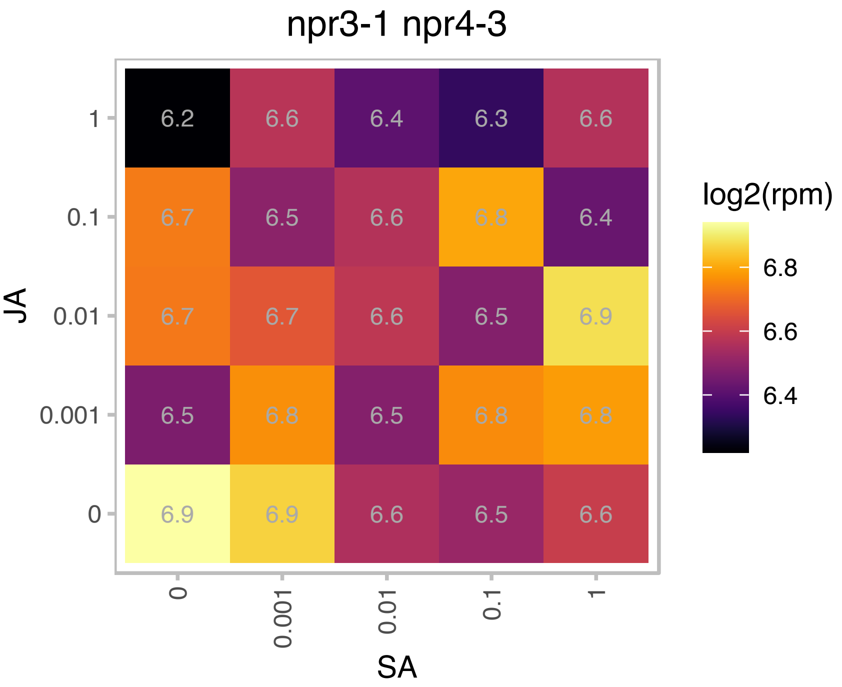

### AT5G27700.1 : EVR1L1, RPS21C

Ribosomal protein S21e

Cytosolic ribosomal protein. Similar to EVR1 and redundant with EVR1. Also enhances VAR2 mutant varigation, but to a lesser extent than evr1.

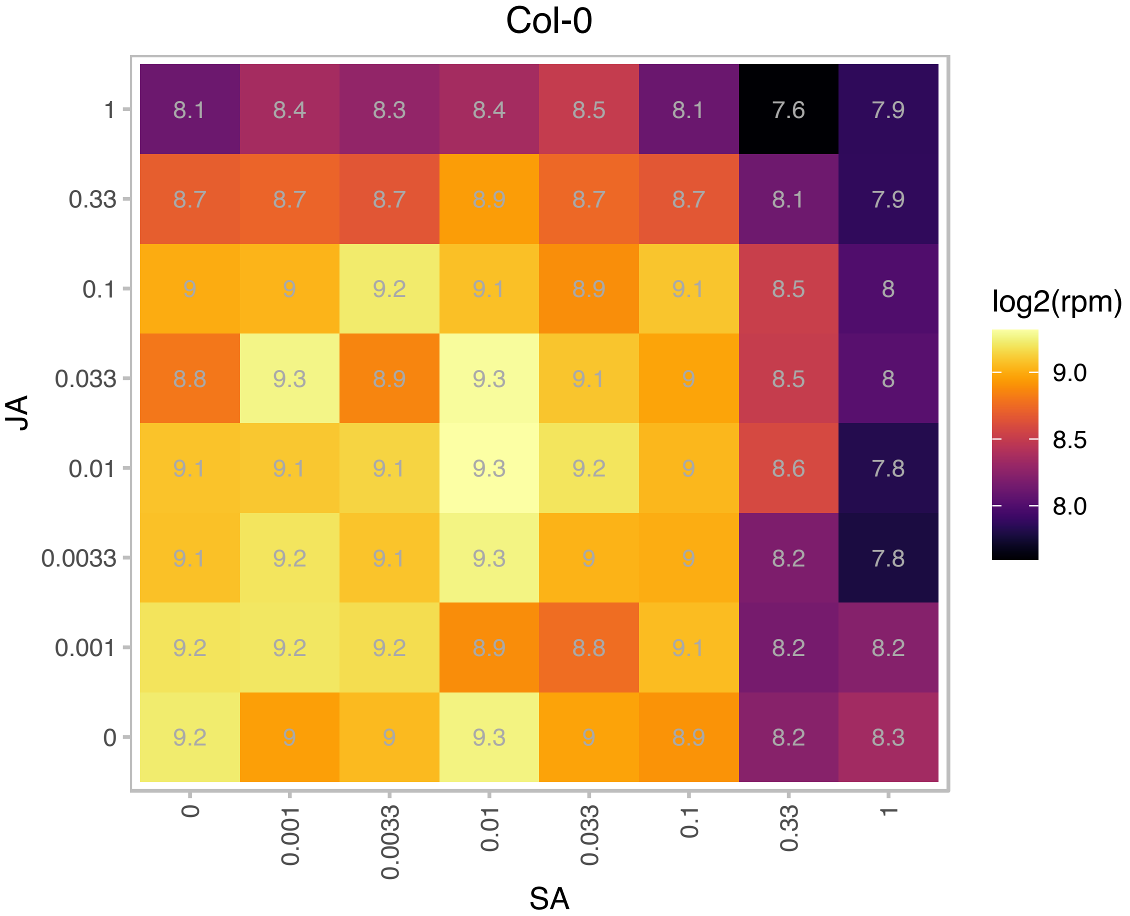


| | |
|---|---|
| Gene | AT5G27700 |
| is.expressed_genes | TRUE |
| is.responsive_genes | TRUE |
| is.apcluster | F-07 |

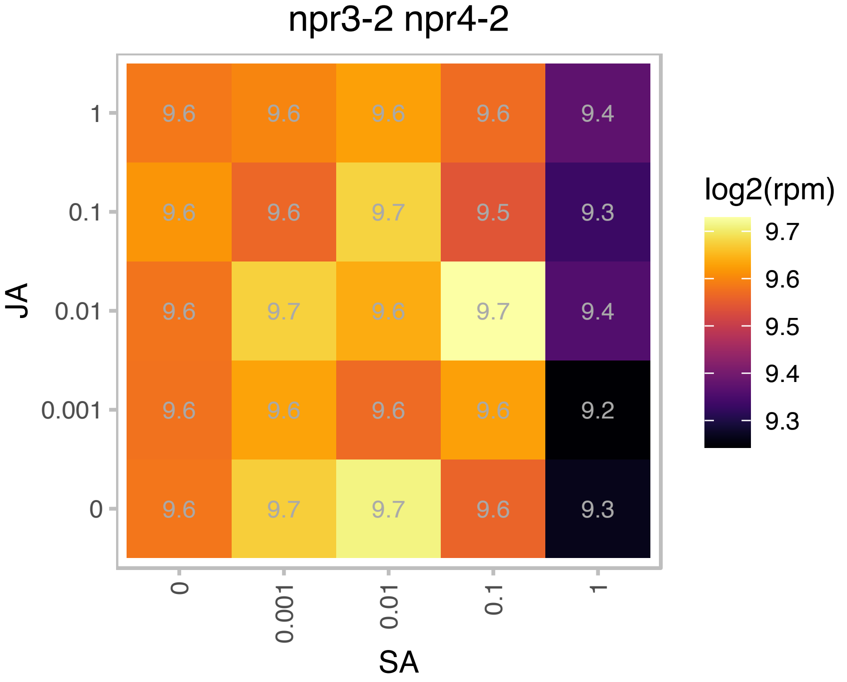


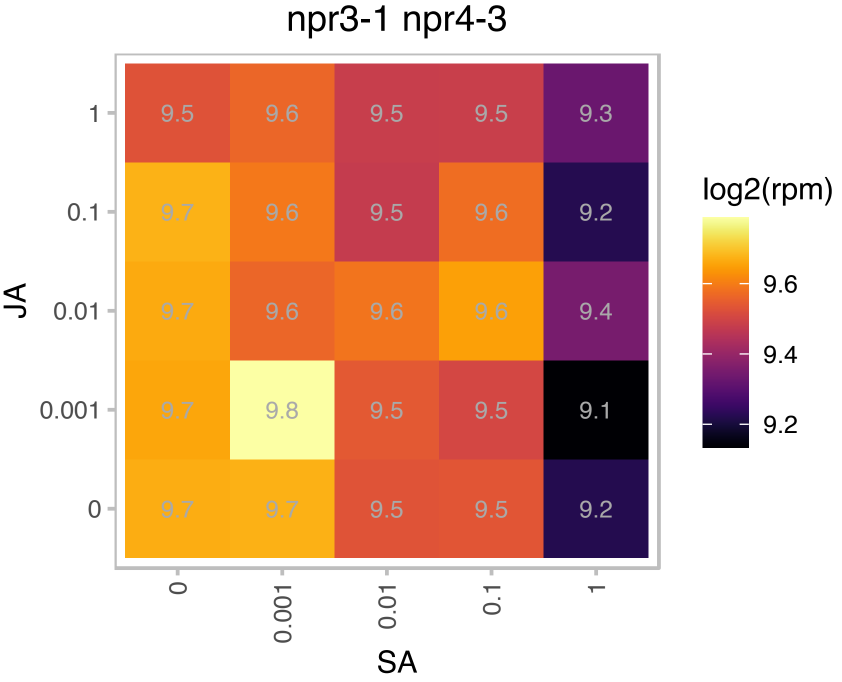

# AT5G27770.1 : NA

Ribosomal L22e protein family

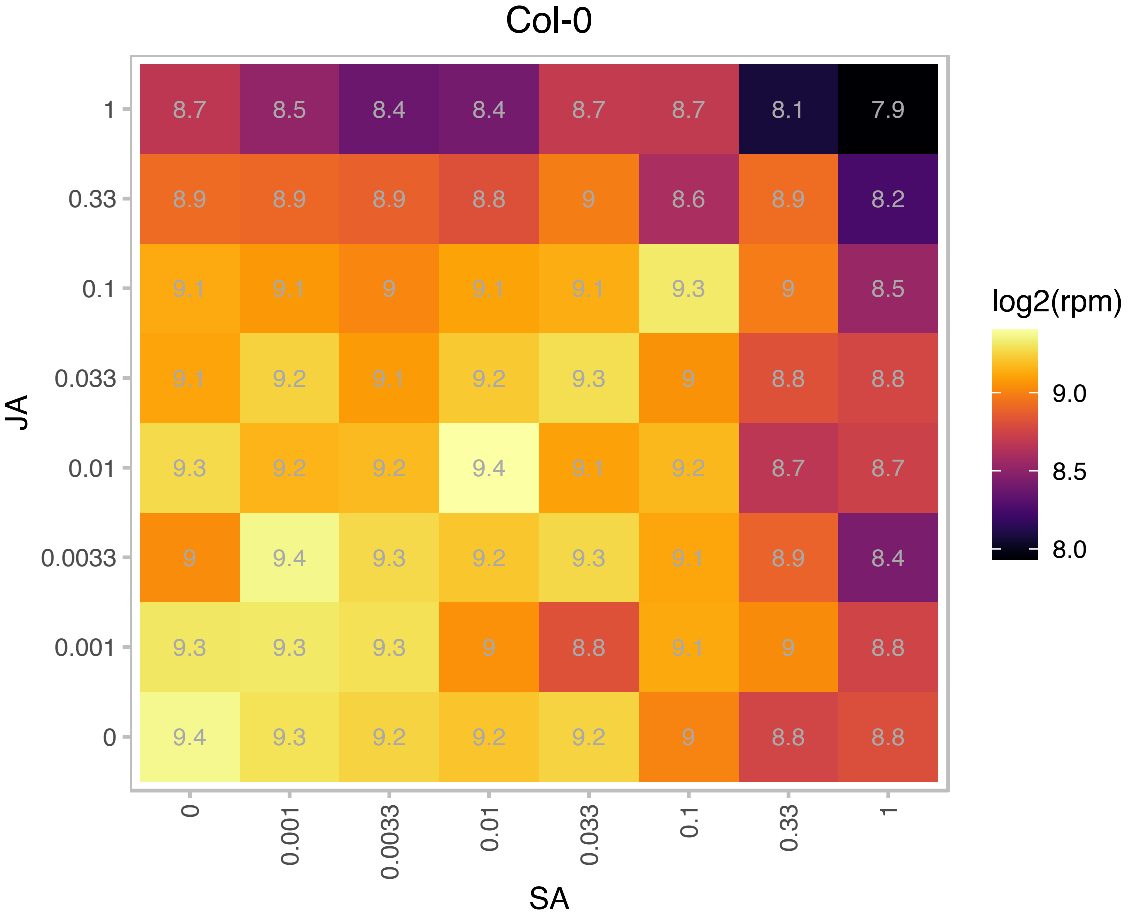


| | |
|---|---|
| Gene | AT5G27770 |
| is.expressed_genes | TRUE |
| is.responsive_genes | TRUE |
| is.apcluster | F-07 |

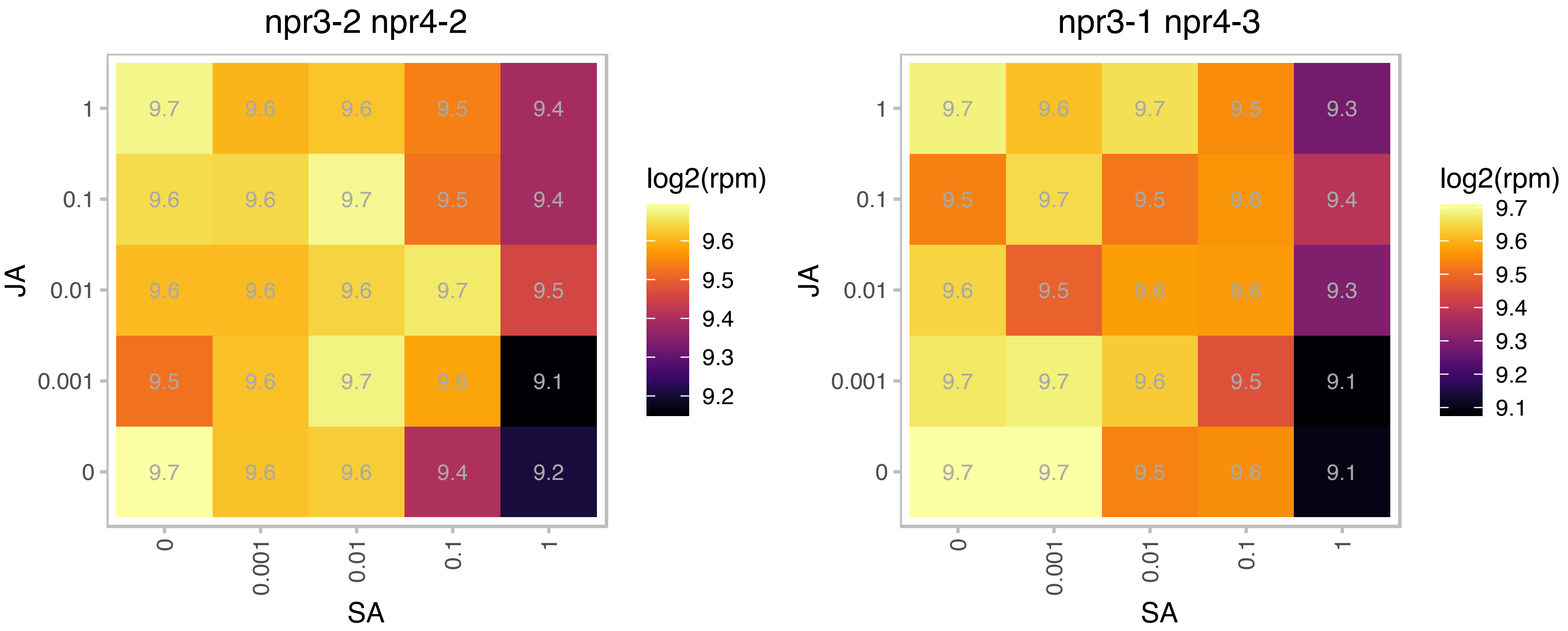

### AT5G37890.1 : AtSINAL7, SINAL7

Protein with RING/U-box and TRAF-like domains

Encodes an E3 protein ligase (based on a self-ubiquitination in vitro assay).

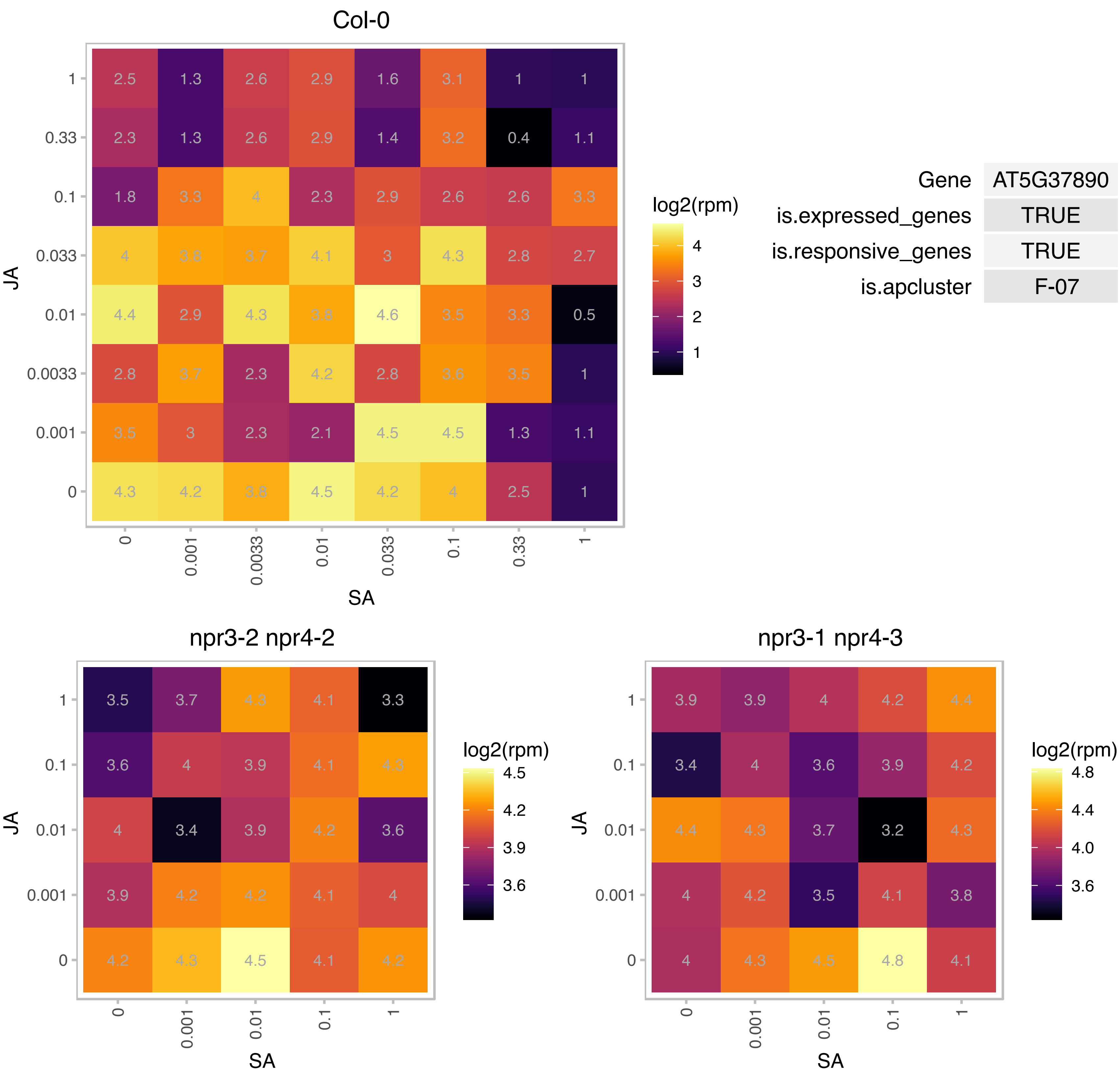


| | |
|---|---|
| Gene | AT5G37890 |
| is.expressed_genes | TRUE |
| is.responsive_genes | TRUE |
| is.apcluster | F-07 |

### AT5G39740.1 : OLI7, RPL5B

ribosomal protein L5 B

Encodes a ribosomal protein RPL5B that is involved in ribosome biogenesis and plays a role in organ size control by promoting cell proliferation and preventing compensation in normal leaf development.

#### Col-0

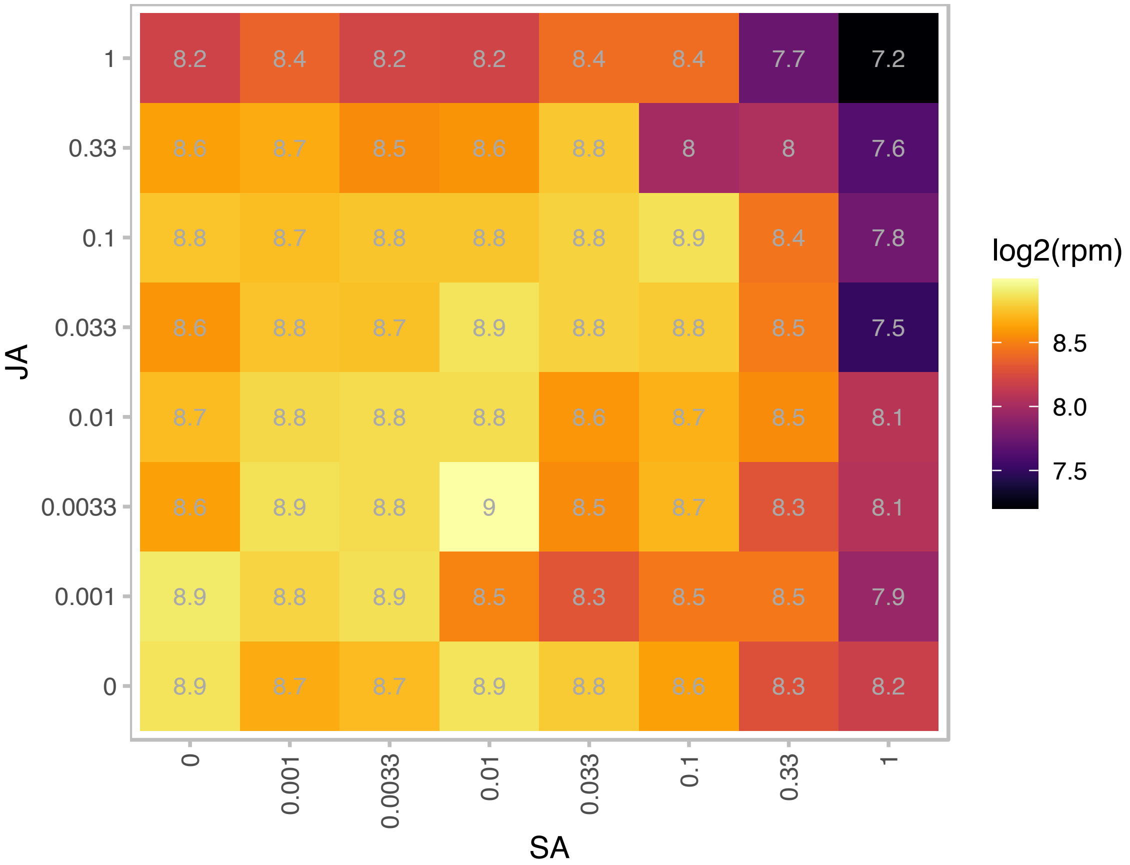


| | |
|---|---|
| Gene | AT5G39740 |
| is.expressed_genes | TRUE |
| is.responsive_genes | TRUE |
| is.apcluster | F-07 |

#### npr3-2 npr4-2

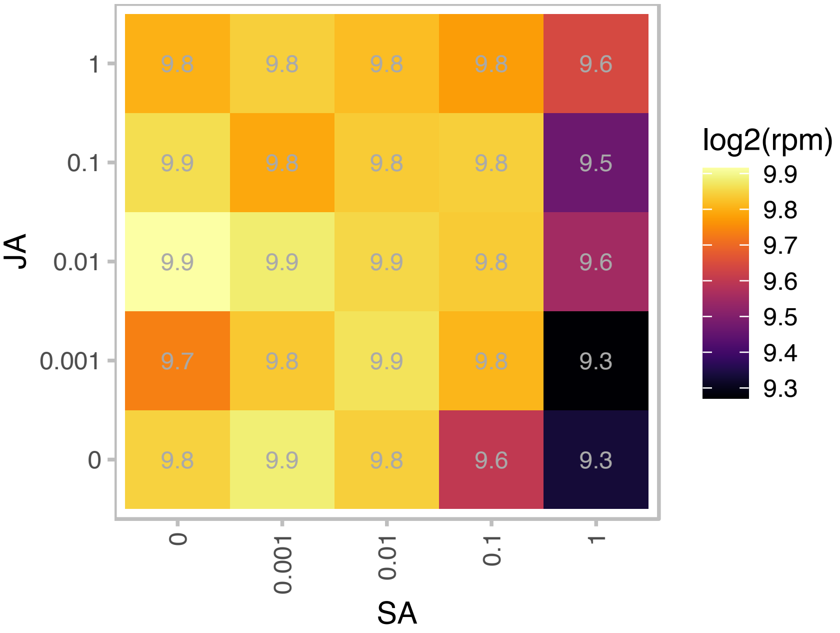


#### npr3-1 npr4-3

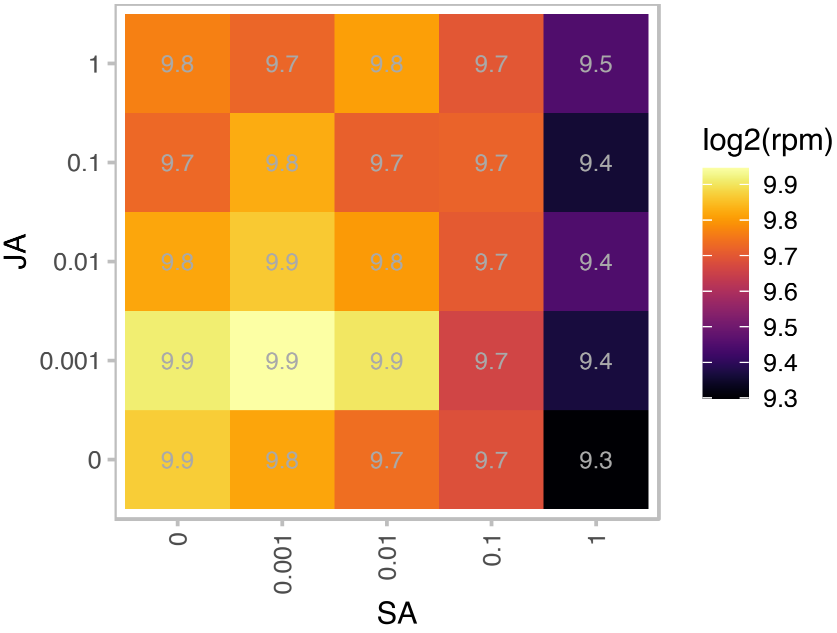

# AT5G39800.1 : NA

Mitochondrial ribosomal protein L27

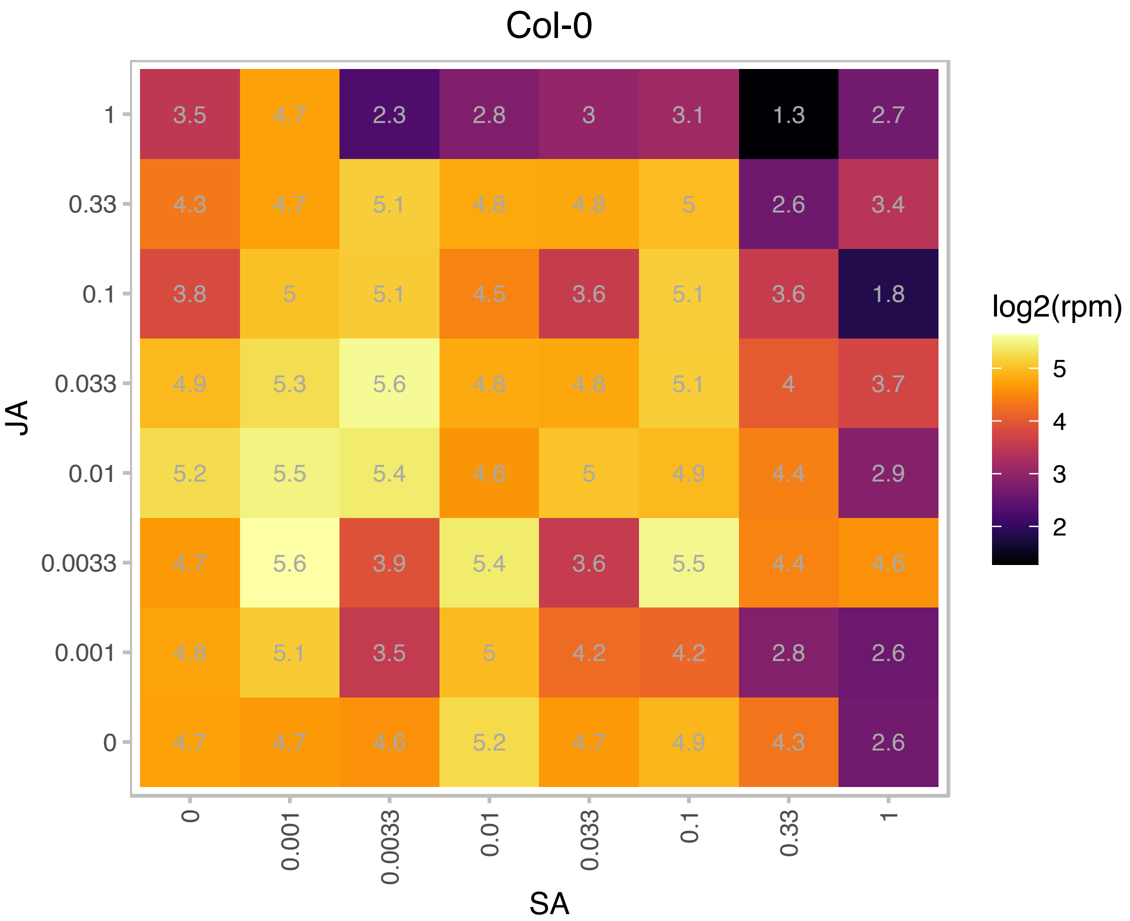


| | |
|---|---|
| Gene | AT5G39800 |
| is.expressed_genes | TRUE |
| is.responsive_genes | TRUE |
| is.apcluster | F-07 |

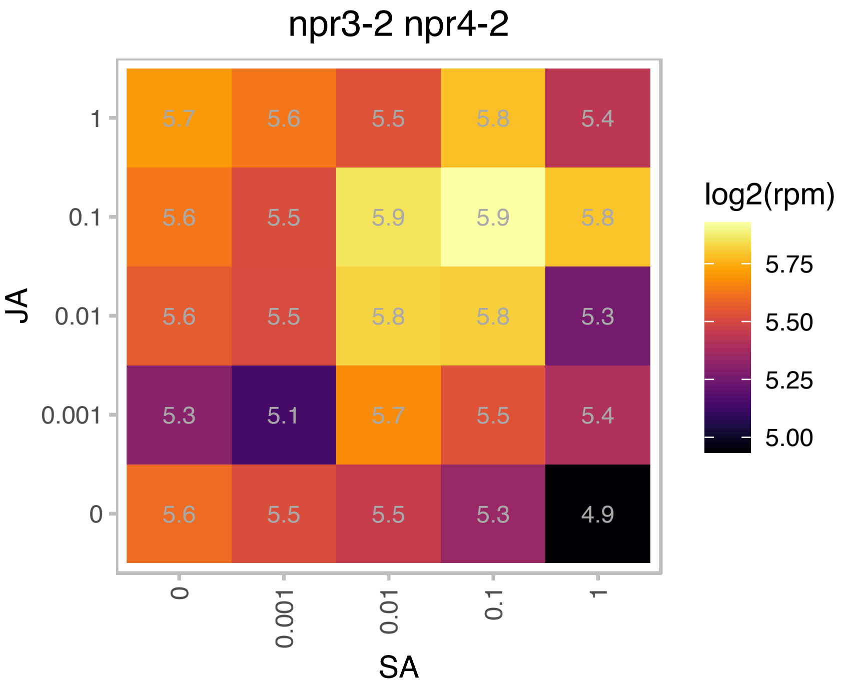


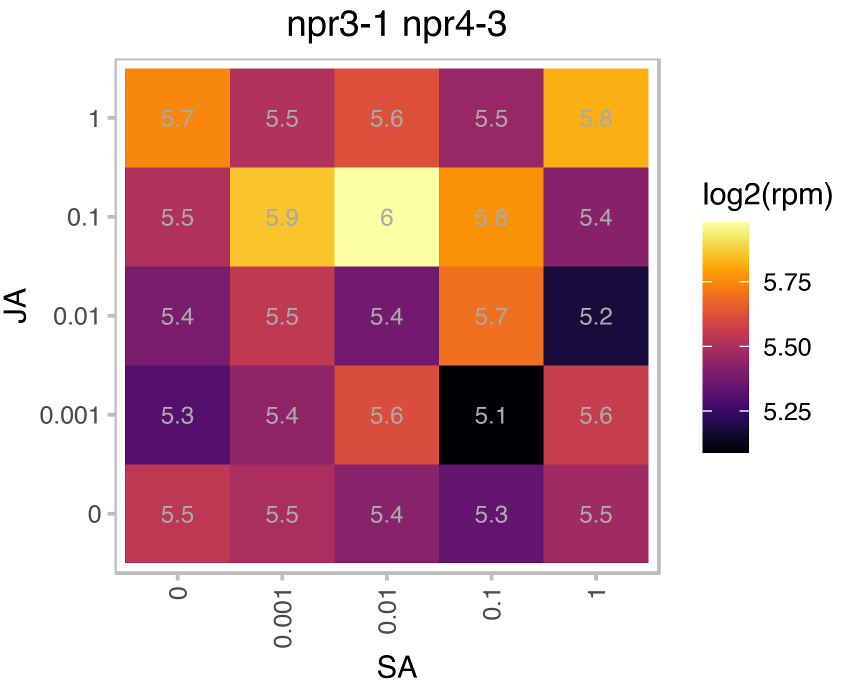

# AT5G40080.1 : NA

Mitochondrial ribosomal protein L27

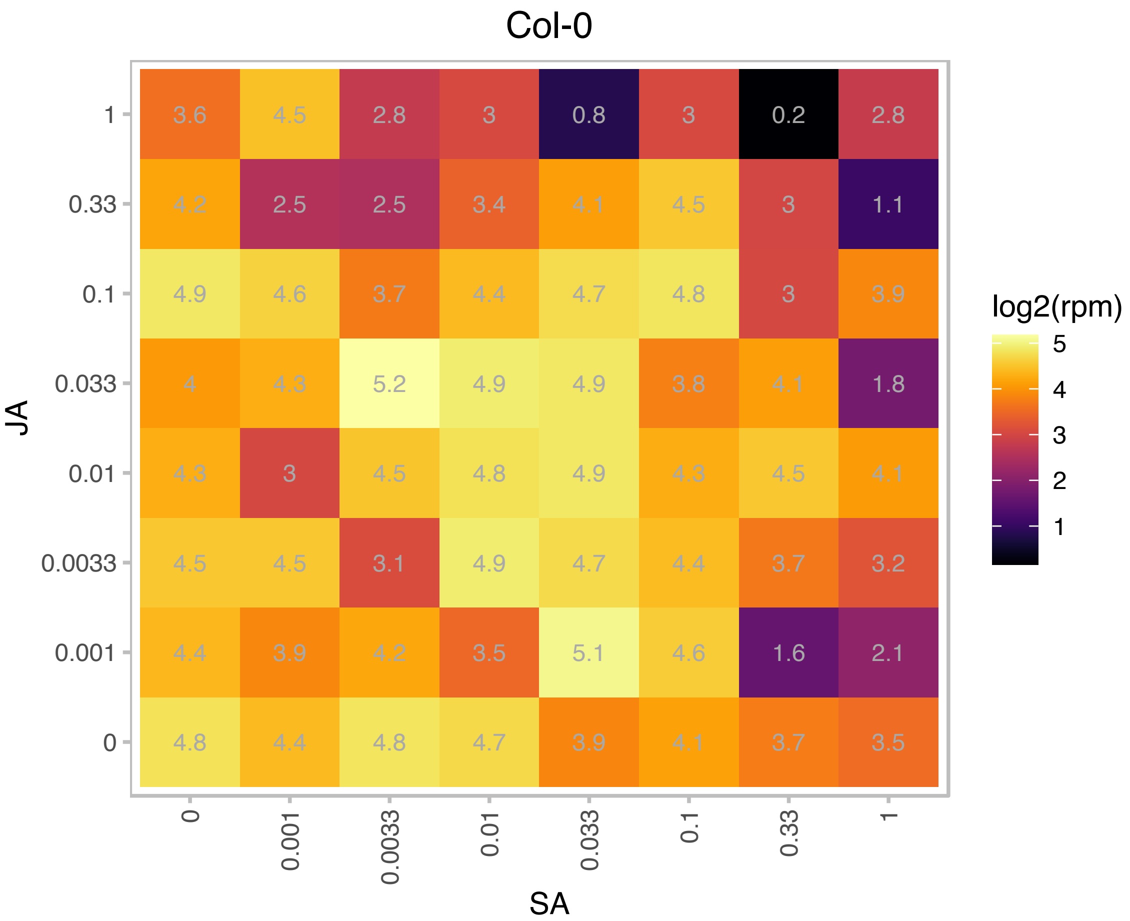


| | |
|---|---|
| Gene | AT5G40080 |
| is.expressed_genes | TRUE |
| is.responsive_genes | TRUE |
| is.apcluster | F-07 |

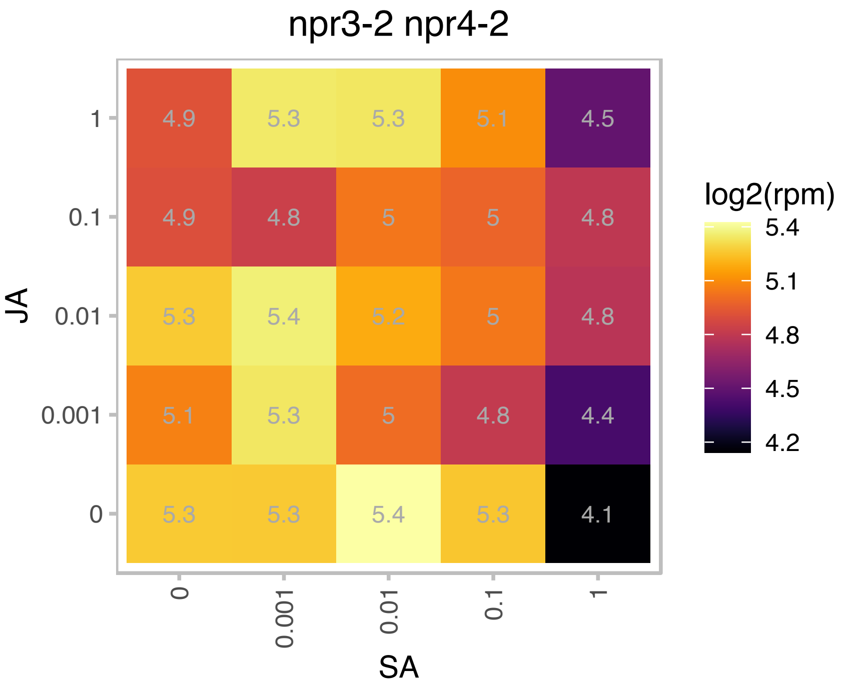


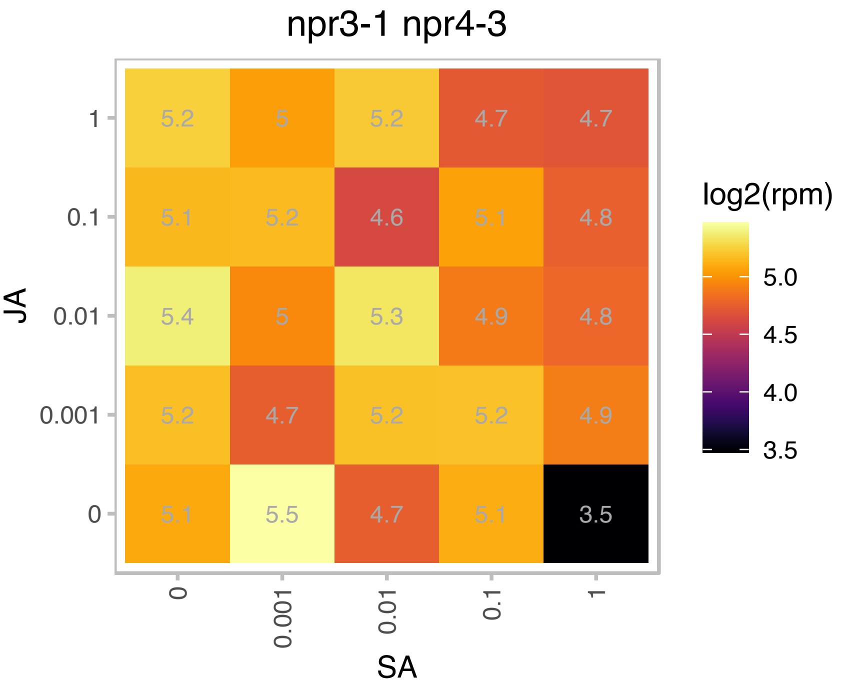

### AT5G43070.1 : WPP1

WPP domain protein 1
WPP family members contains an NE targeting domain. This domain, called the WPP domain after a highly conserved Trp-Pro-Pro motif, is necessary and sufficient for NE targeting of WPP1. RNAi suppression of WPP1 resulted in reduced mitotic activity.

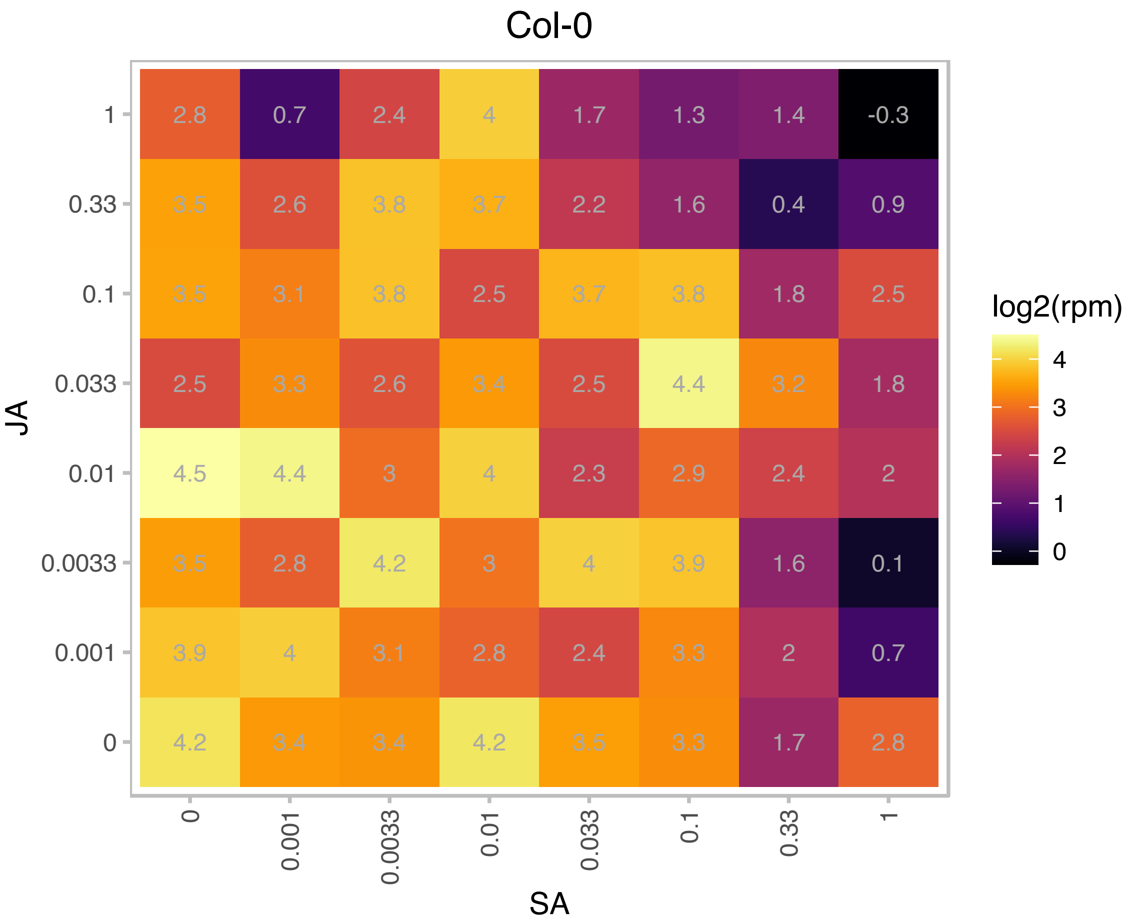


| | |
|---|---|
| Gene | AT5G43070 |
| is.expressed_genes | TRUE |
| is.responsive_genes | TRUE |
| is.apcluster | F-07 |

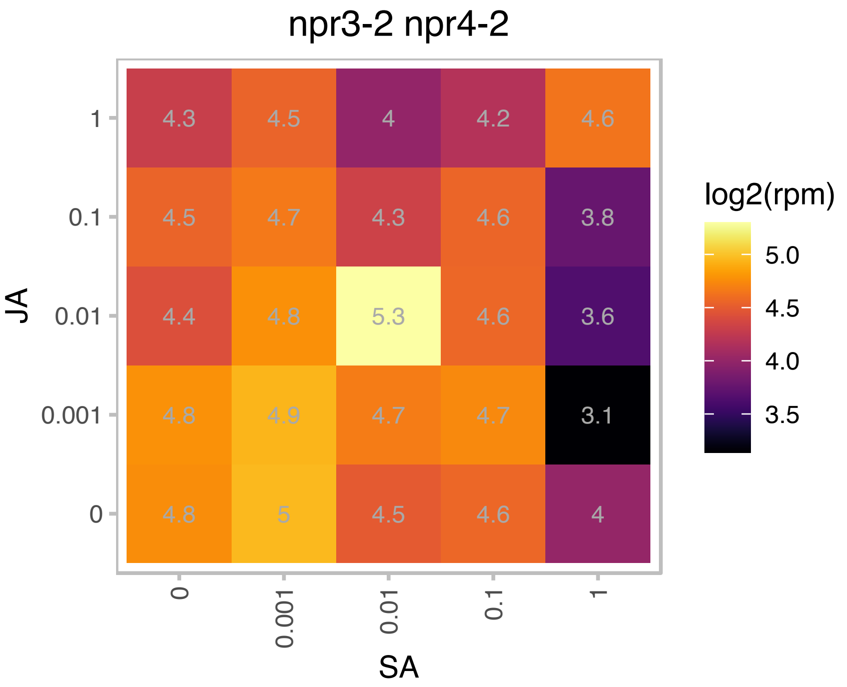


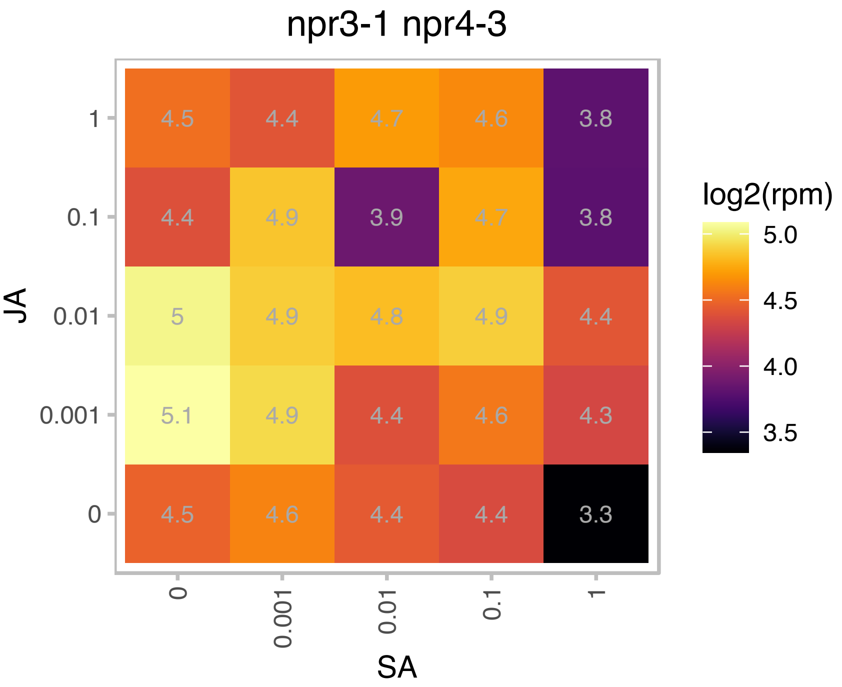

### AT5G43970.1 : ATTOM22-V, TOM22-V, TOM9-2

translocase of outer membrane 22-V

Subunit of the TOM complex, a translocase in the outer mitochondrial membrane that selectively allows proteins with a mitochondrial targeting sequence to enter the mitochondrion.

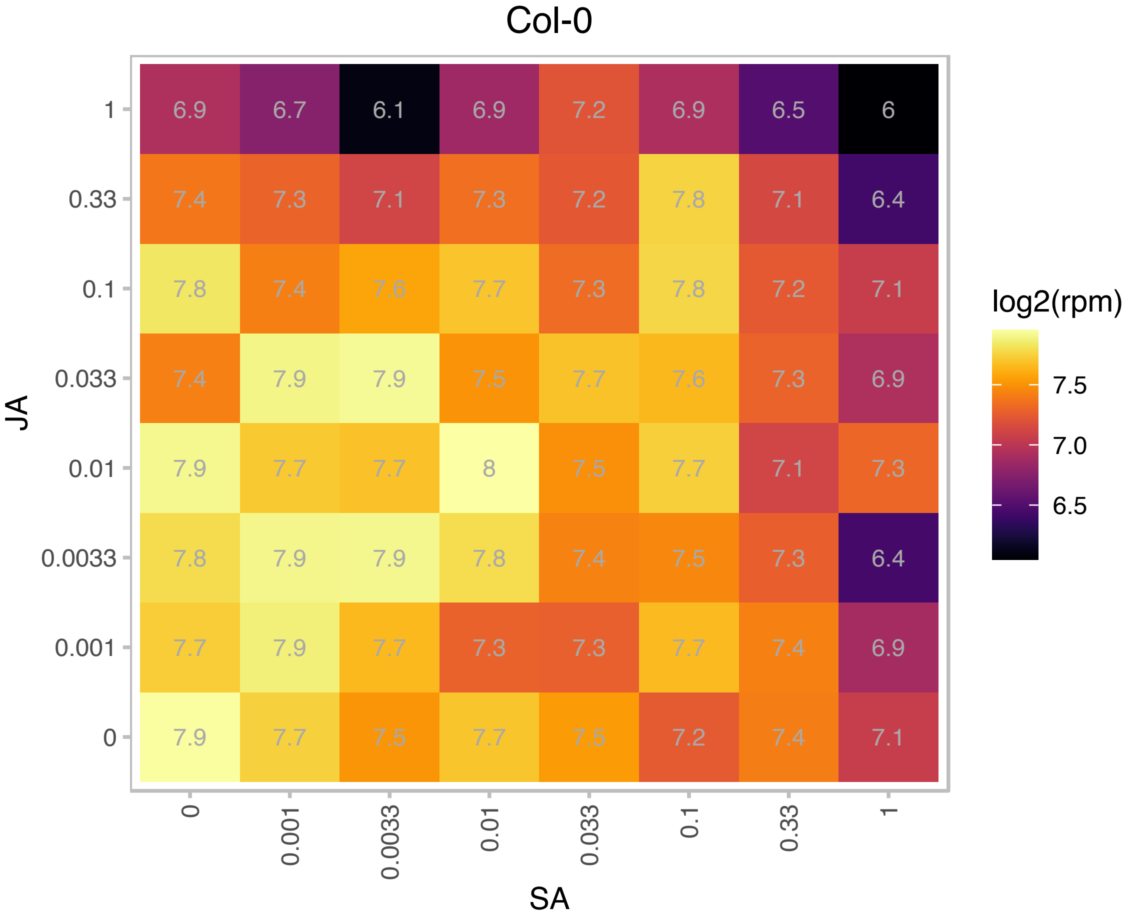


| | |
|---|---|
| Gene | AT5G43970 |
| is.expressed_genes | TRUE |
| is.responsive_genes | TRUE |
| is.apcluster | F-07 |

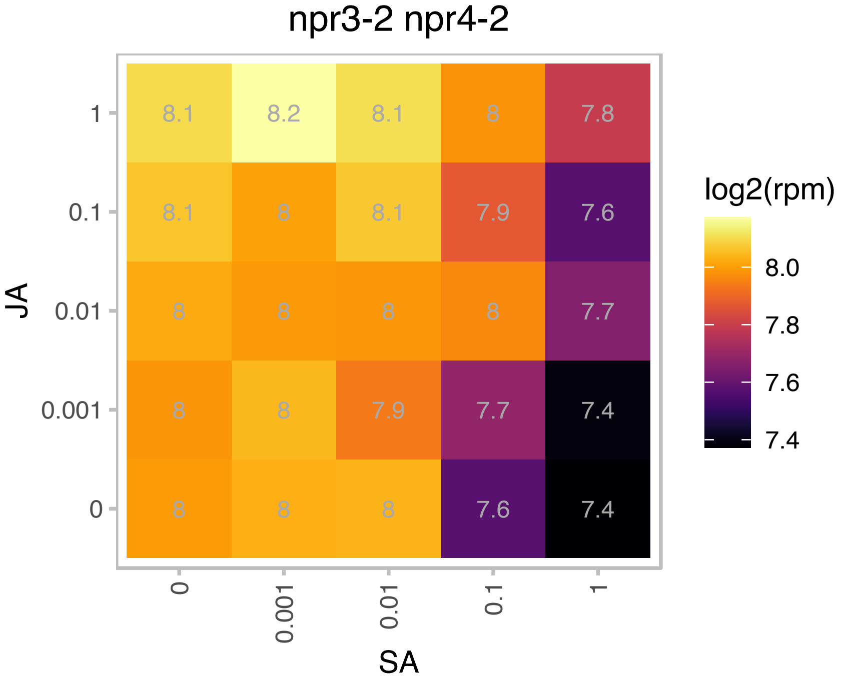


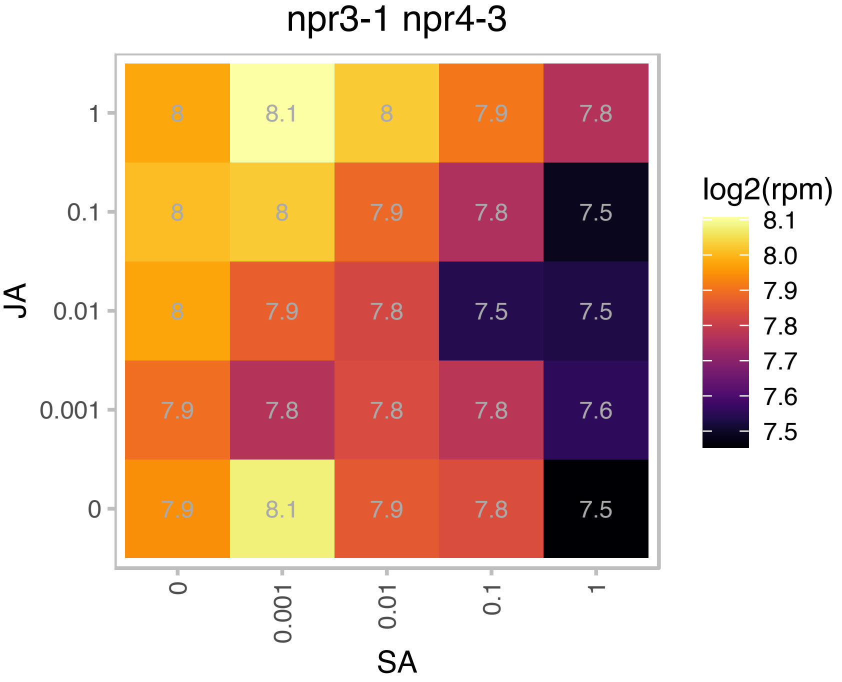

# AT5G45590.1 : NA

Ribosomal protein L35

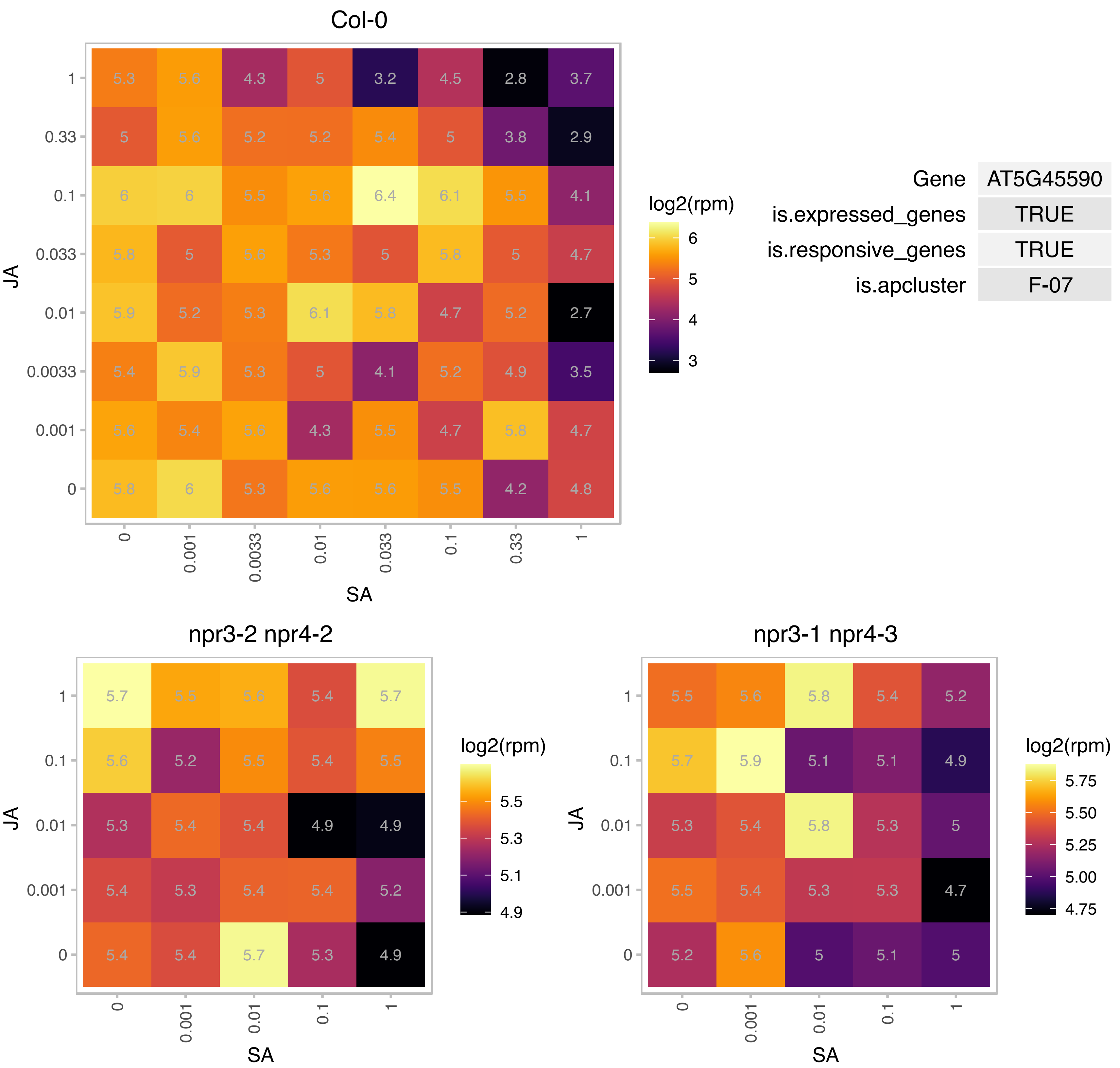


| | |
|---|---|
| Gene | AT5G45590 |
| is.expressed_genes | TRUE |
| is.responsive_genes | TRUE |
| is.apcluster | F-07 |

# AT5G45775.1 : NA

Ribosomal L5P family protein

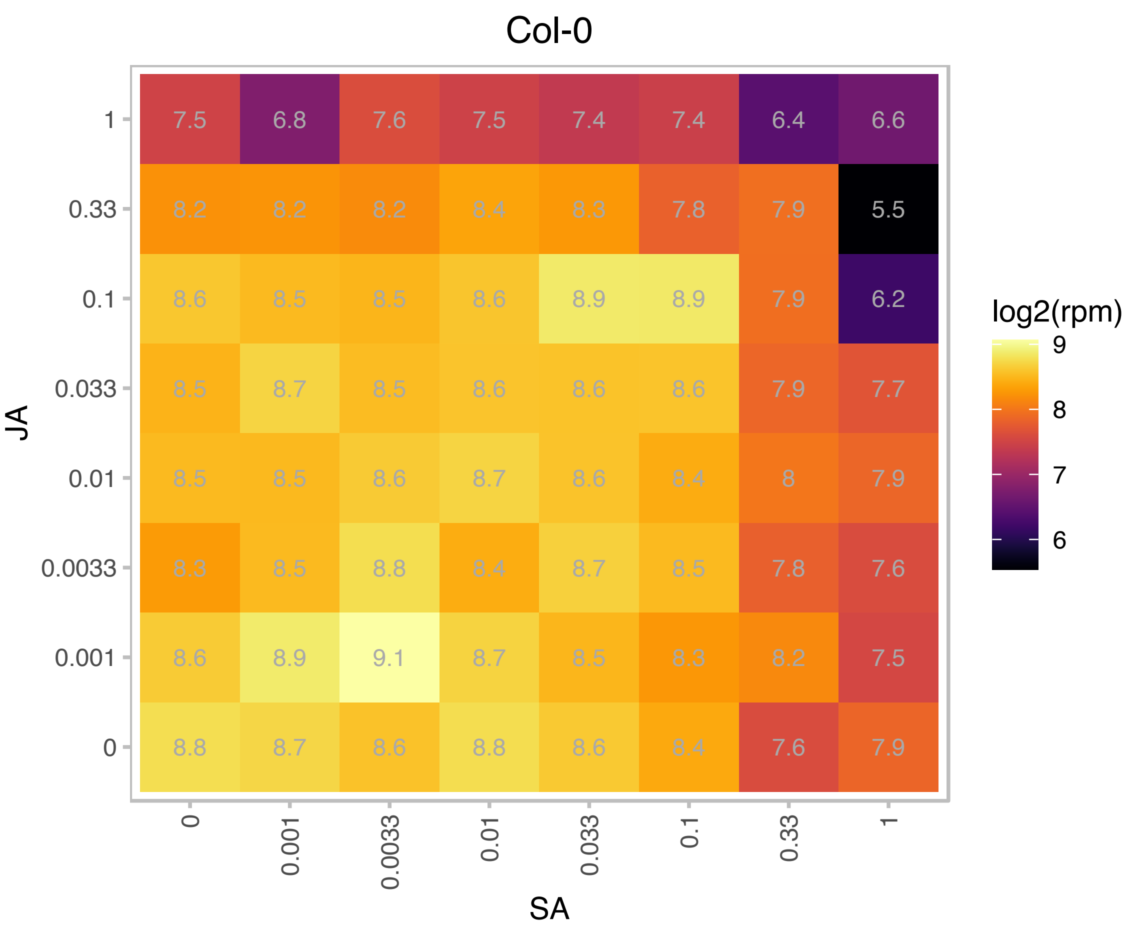


| | |
|---|---|
| Gene | AT5G45775 |
| is.expressed_genes | TRUE |
| is.responsive_genes | TRUE |
| is.apcluster | F-07 |

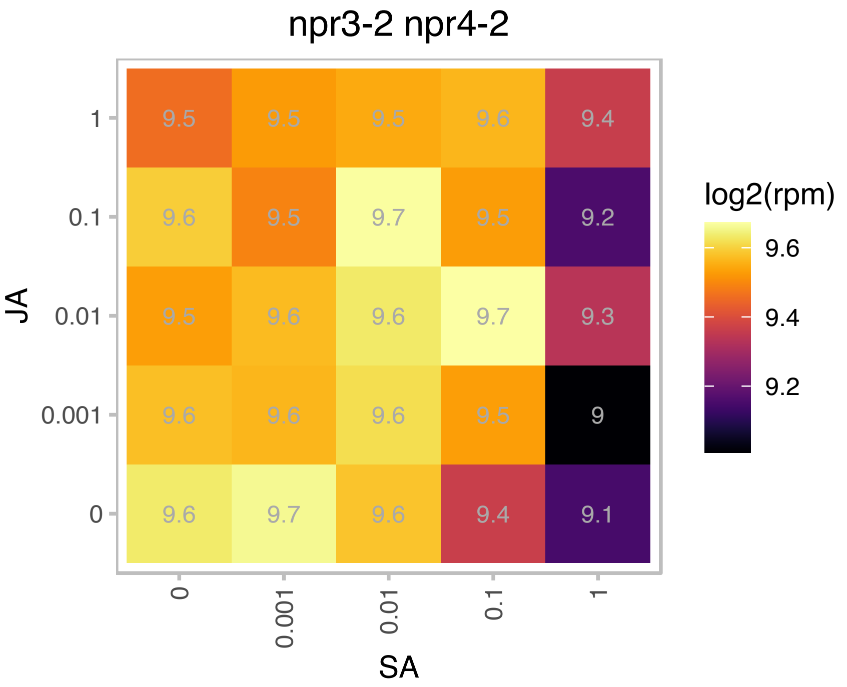


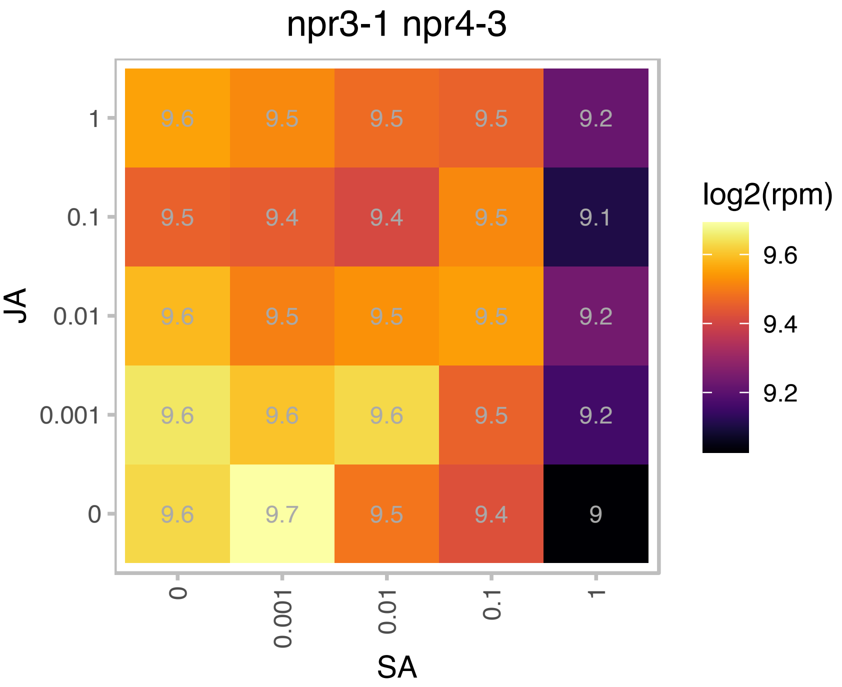

# AT5G46160.1 : NA

Ribosomal protein L14p/L23e family protein

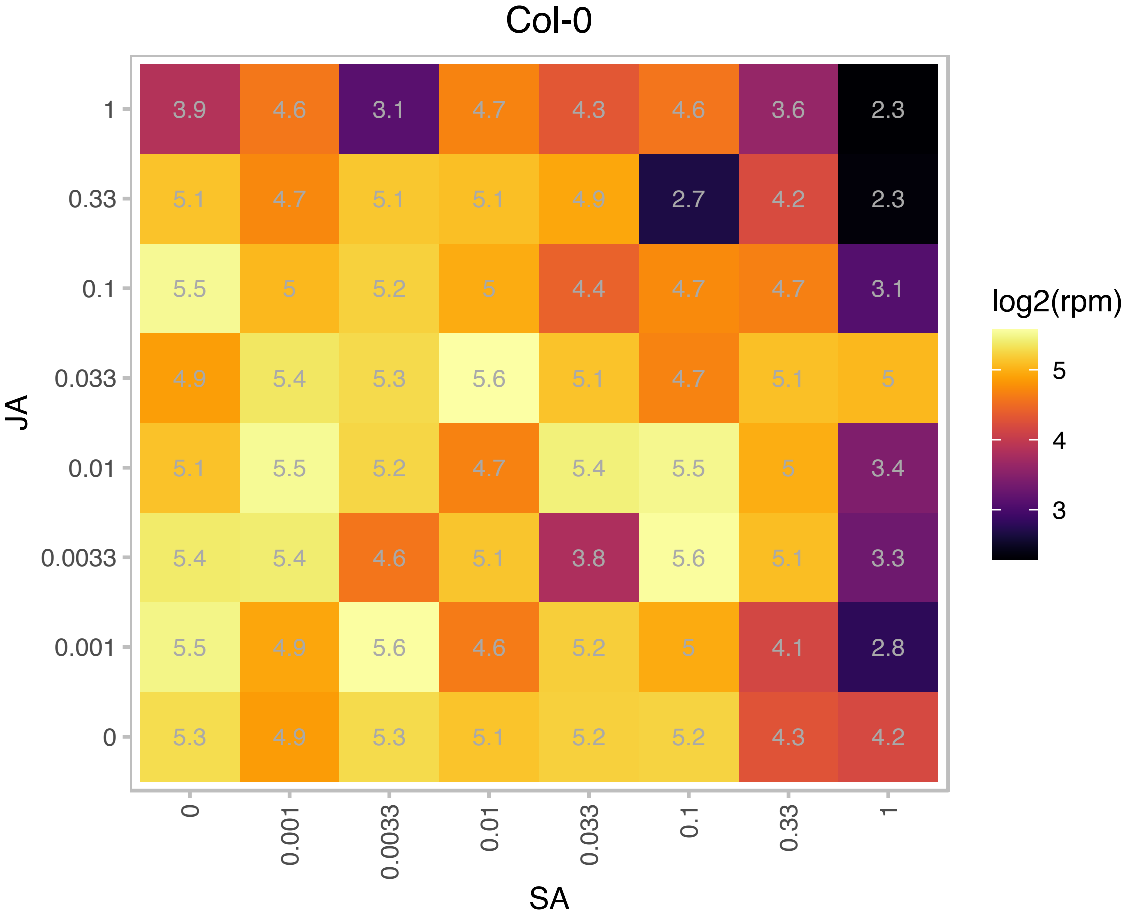


| | |
|---|---|
| Gene | AT5G46160 |
| is.expressed_genes | TRUE |
| is.responsive_genes | TRUE |
| is.apcluster | F-07 |

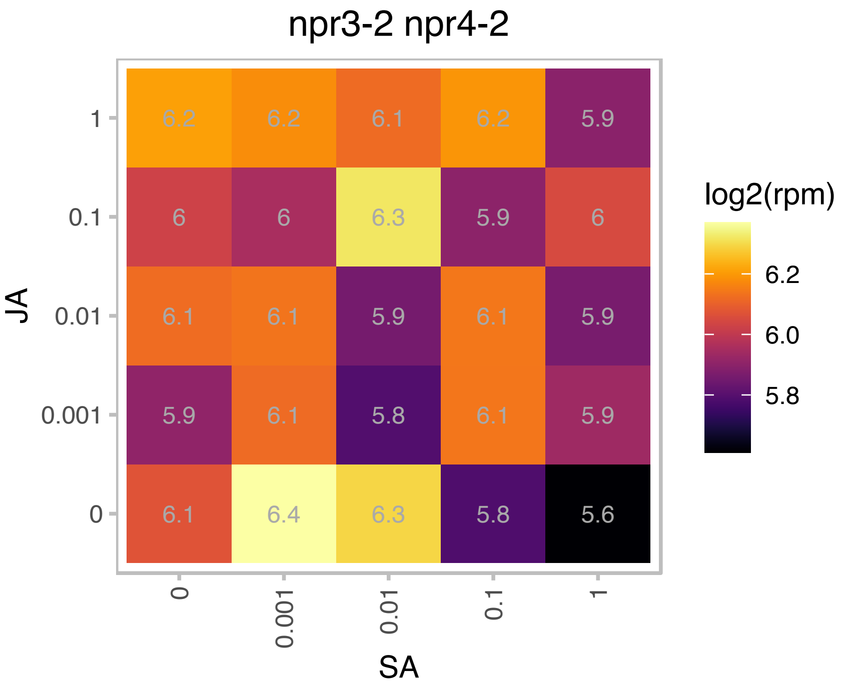


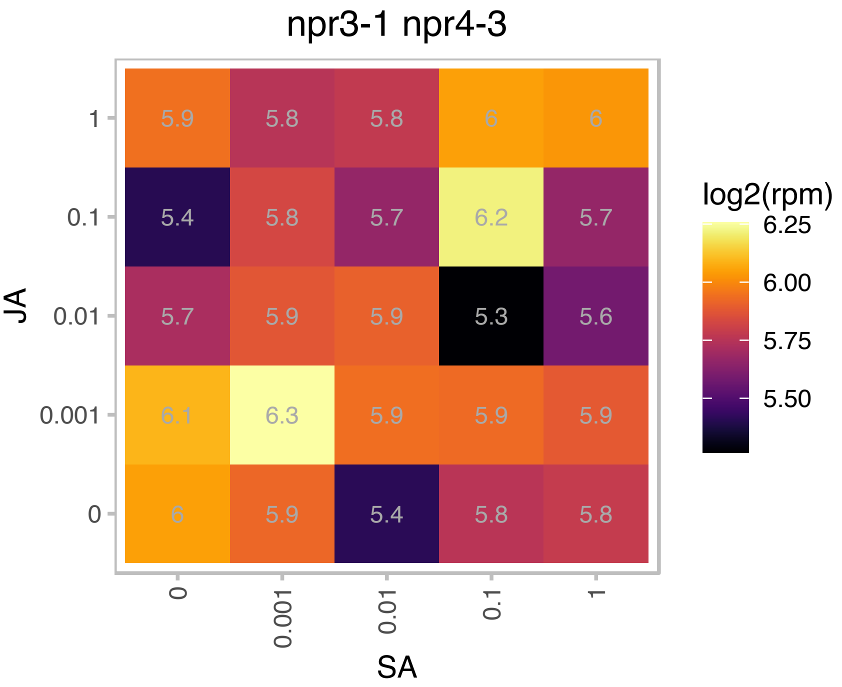

### AT5G46580.1 : SOT1

pentatricopeptide (PPR) repeat-containing protein

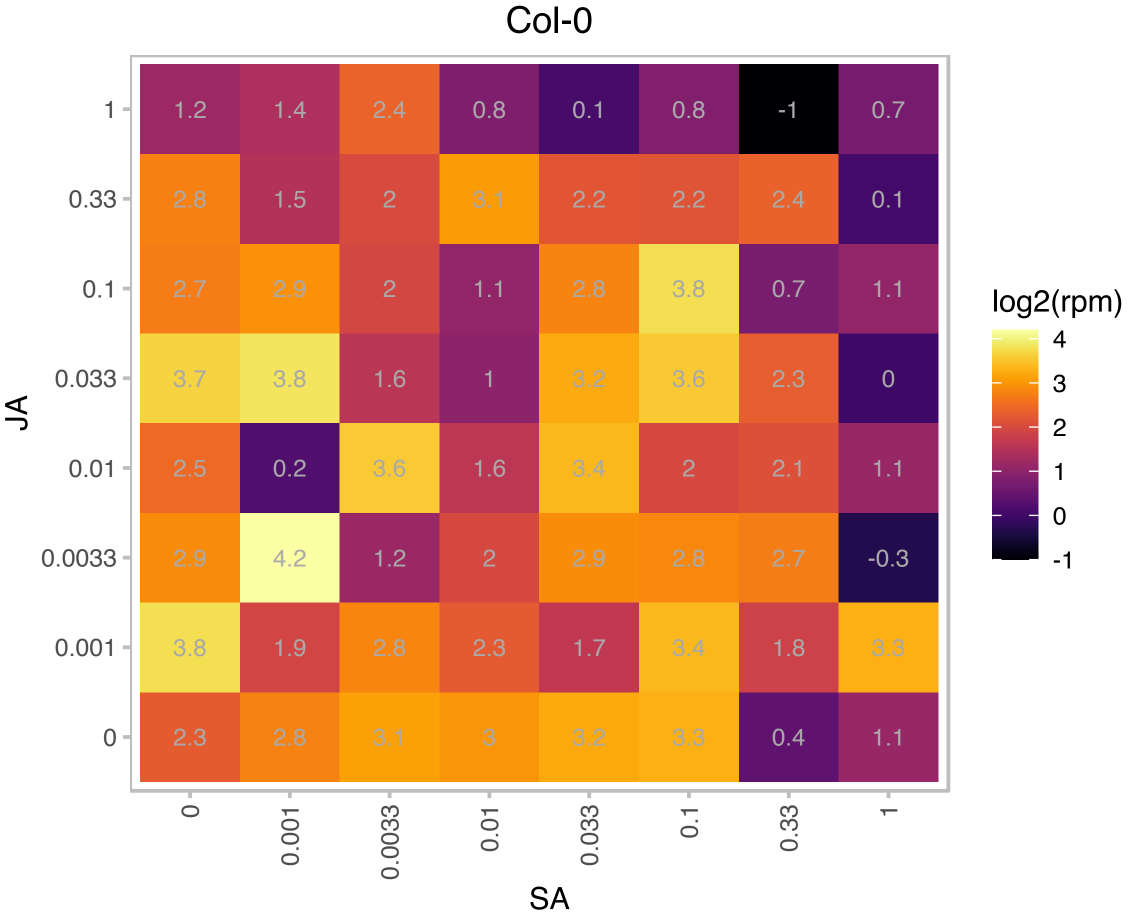


| | |
|---|---|
| Gene | AT5G46580 |
| is.expressed_genes | TRUE |
| is.responsive_genes | TRUE |
| is.apcluster | F-07 |

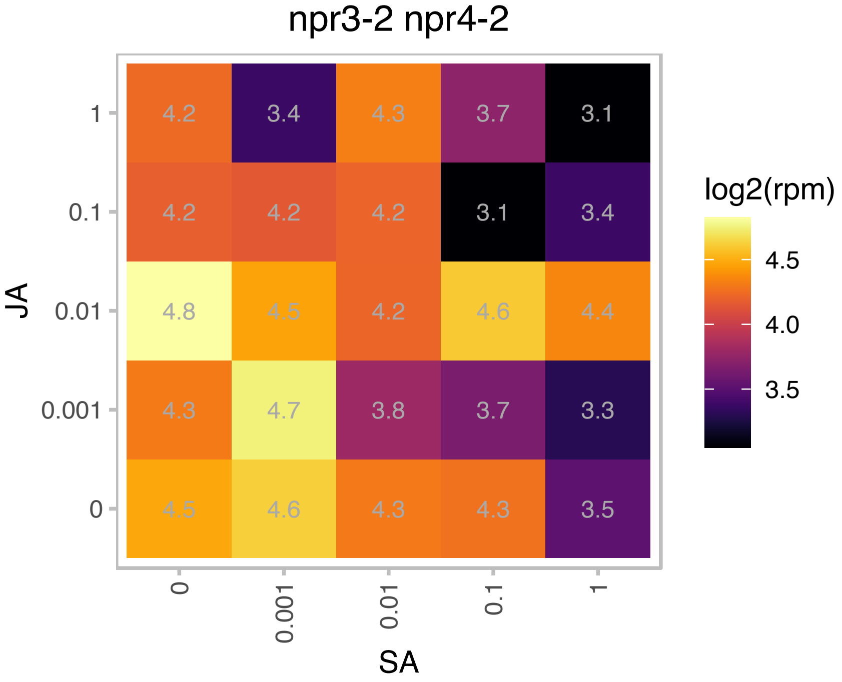


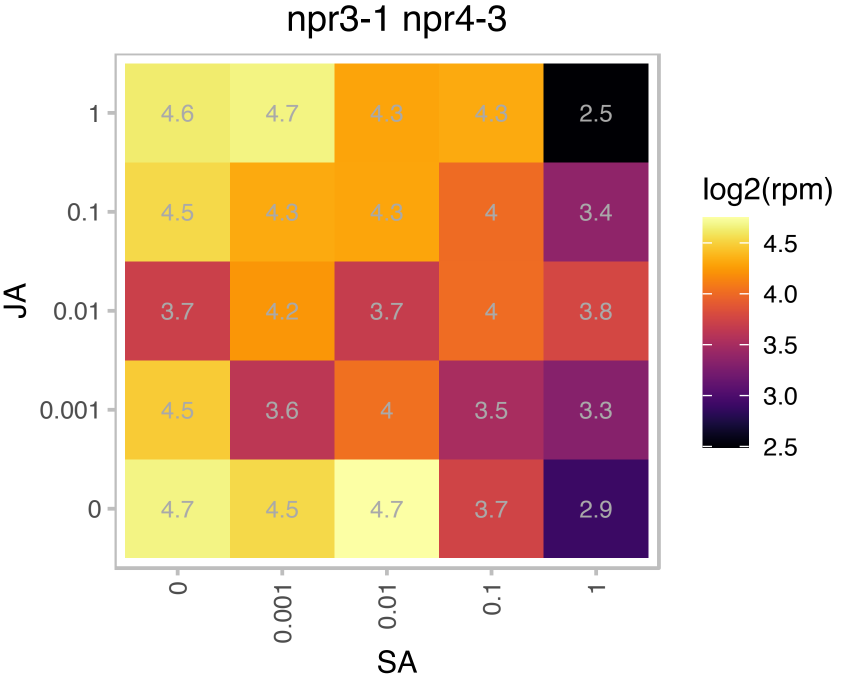

# AT5G47210.1 : NA

Hyaluronan / mRNA binding family

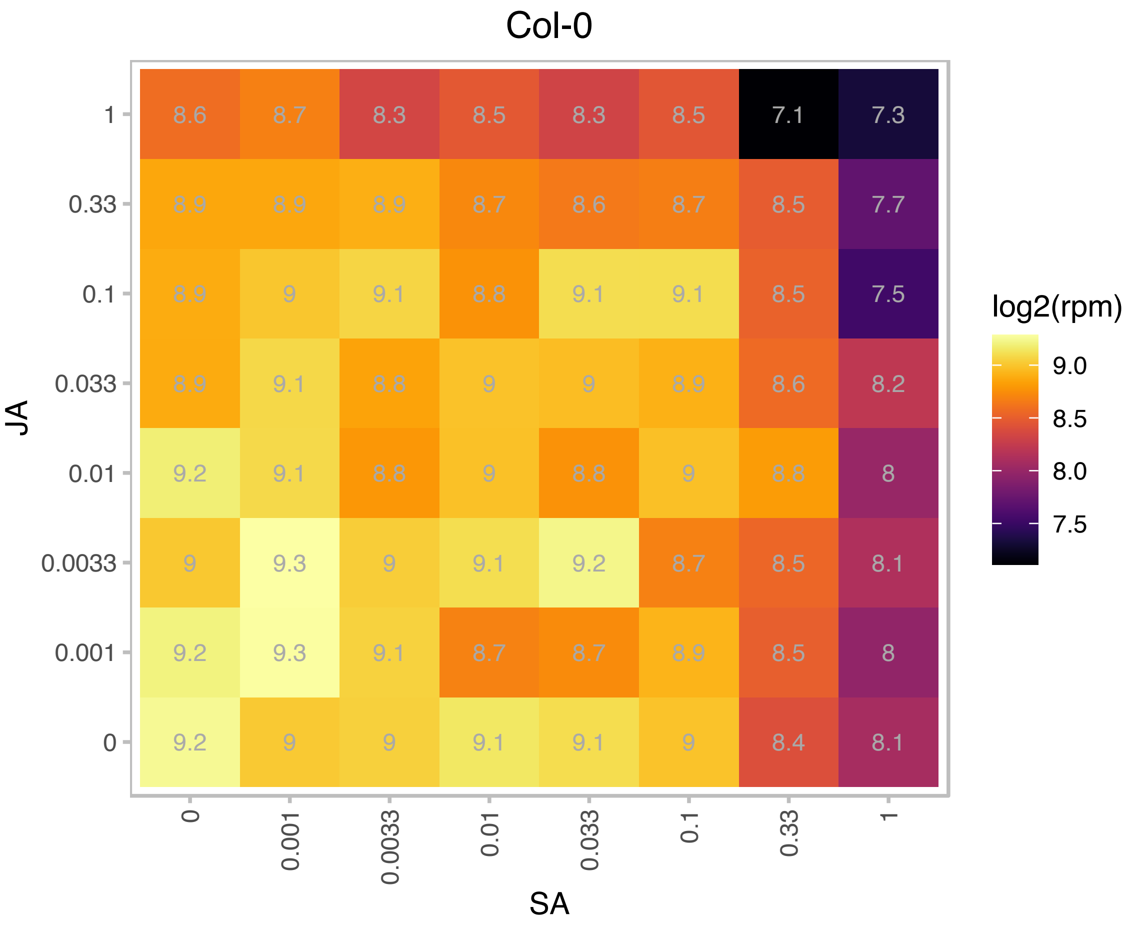


| | |
|---|---|
| Gene | AT5G47210 |
| is.expressed_genes | TRUE |
| is.responsive_genes | TRUE |
| is.apcluster | F-07 |

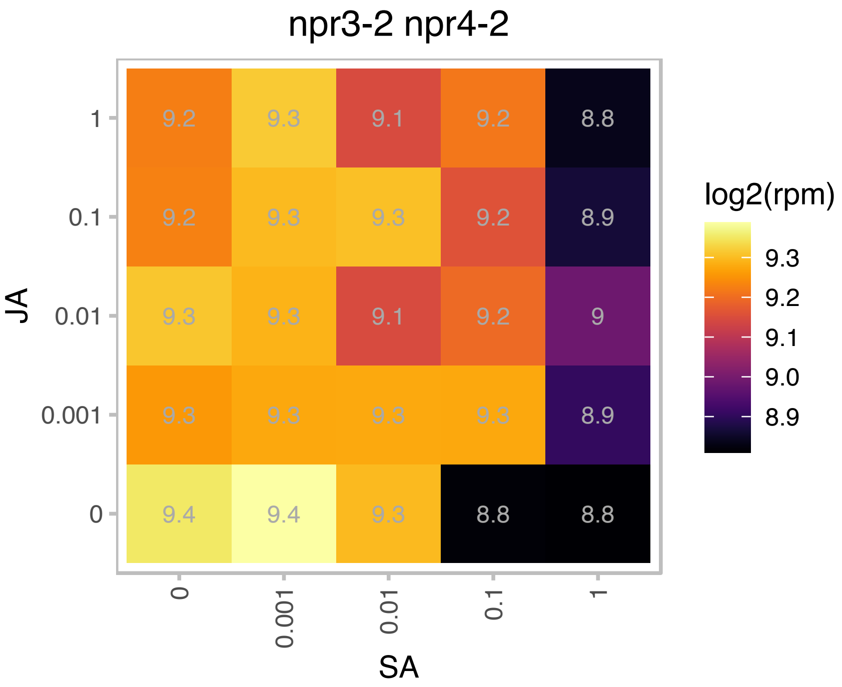


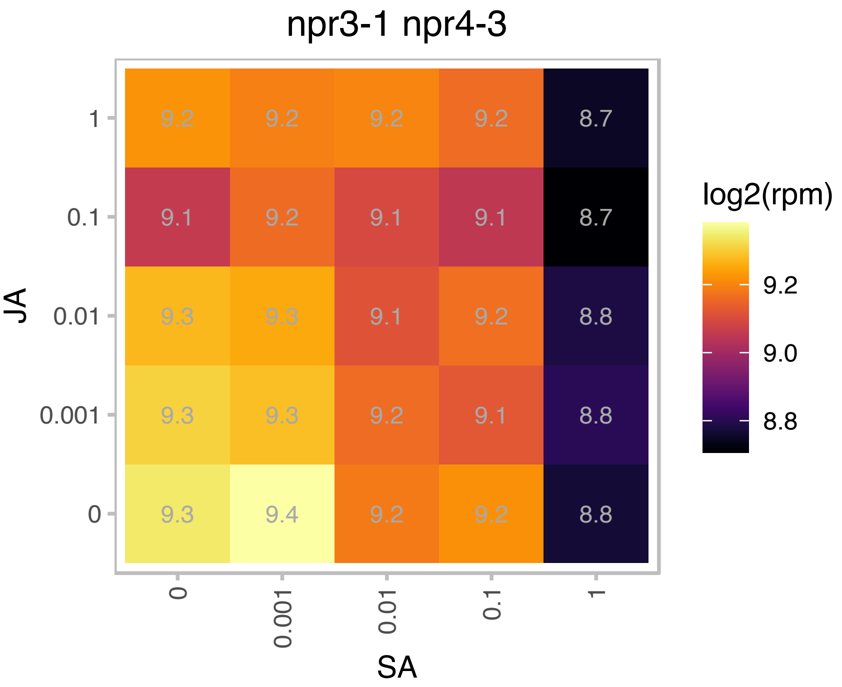

### AT5G49160.1 : DDM2, DMT01, DMT1, MET1, MET2, MET

methyltransferase 1
Encodes a cytosine methyltransferase MET1. Required for silencing of FWA paternal allele in endosperm. Two lines with RNAi constructs directed against DMT1 have reduced agrobacterium-mediated tumor formation. The mRNA is cell-to-cell mobile.

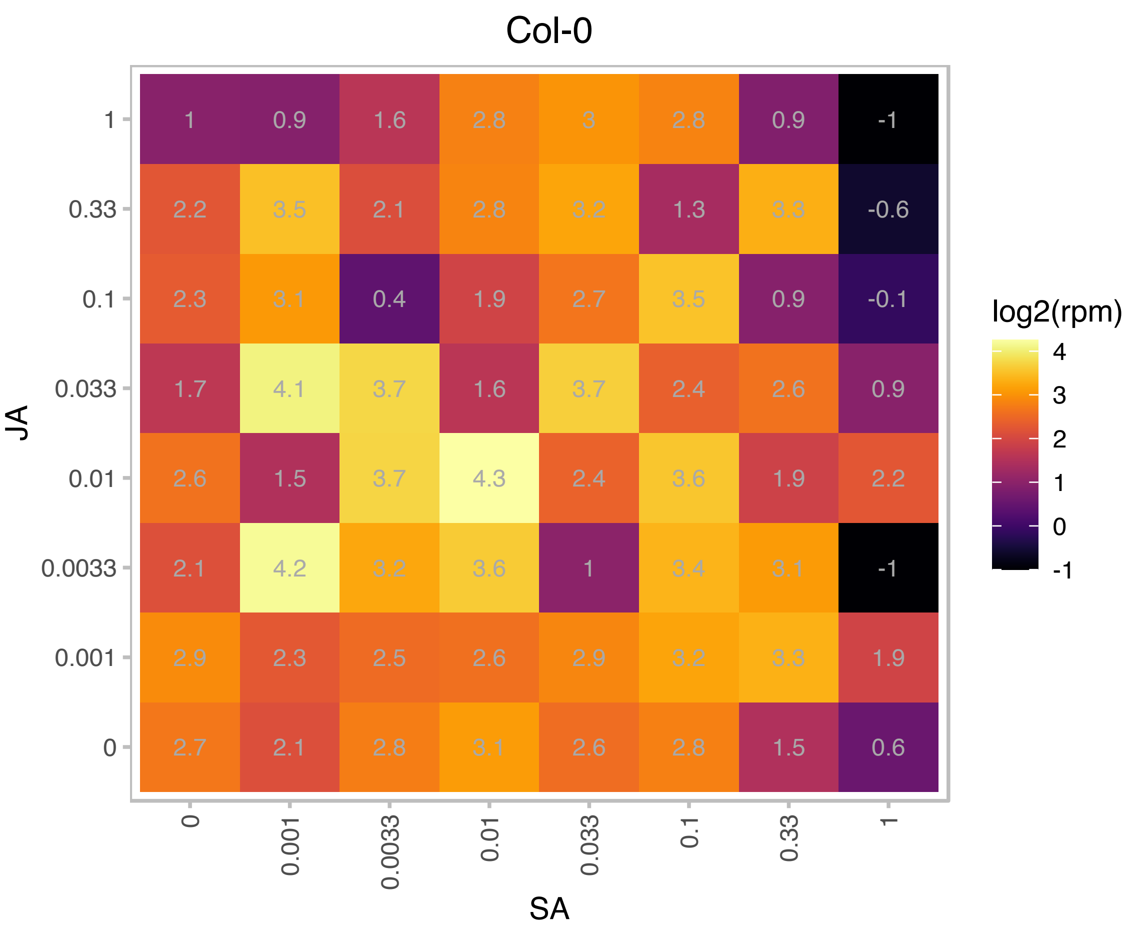


| | |
|---|---|
| Gene | AT5G49160 |
| is.expressed_genes | TRUE |
| is.responsive_genes | TRUE |
| is.apcluster | F-07 |

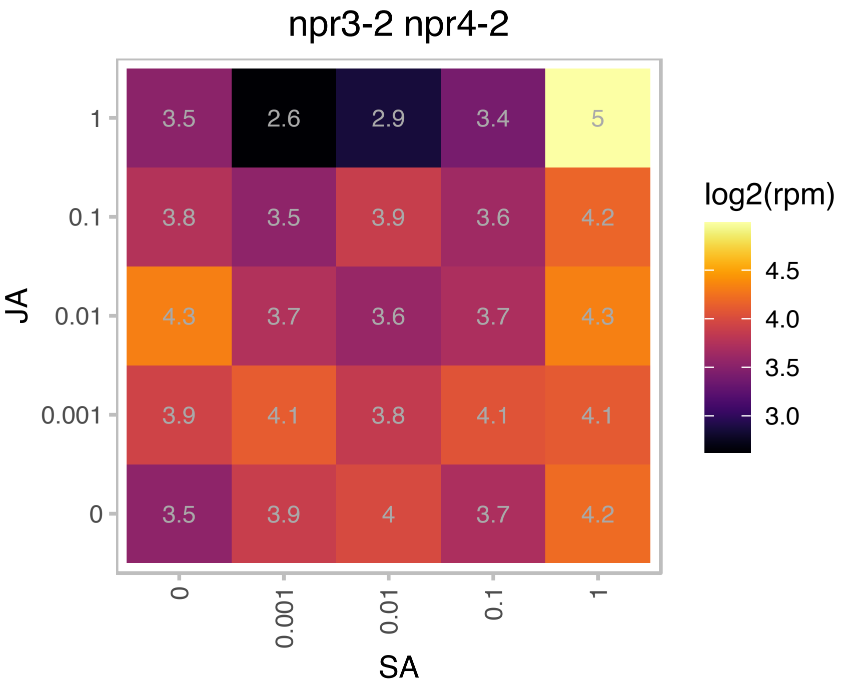


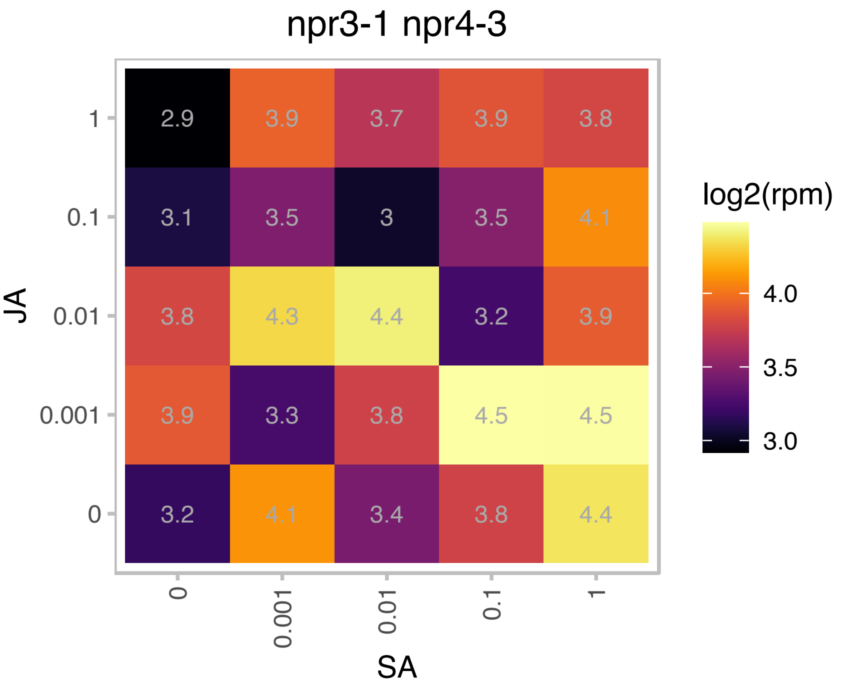

# AT5G50740.1 : NA

Heavy metal transport/detoxification superfamily protein

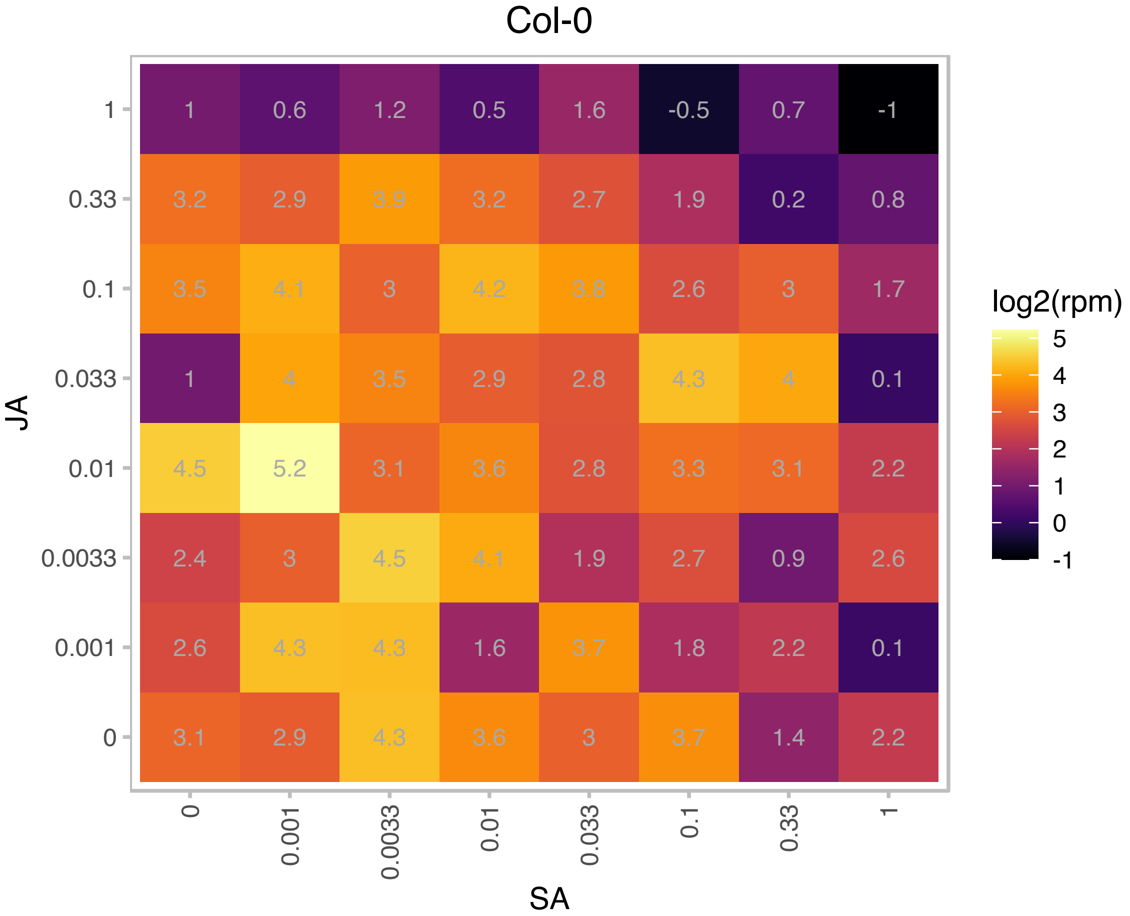


| | |
|---|---|
| Gene | AT5G50740 |
| is.expressed_genes | TRUE |
| is.responsive_genes | TRUE |
| is.apcluster | F-07 |

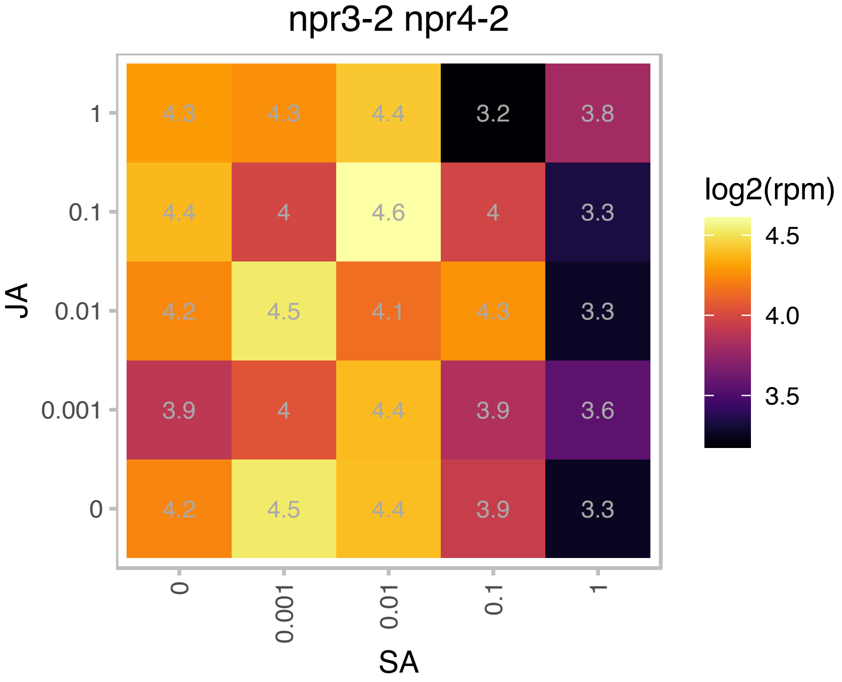


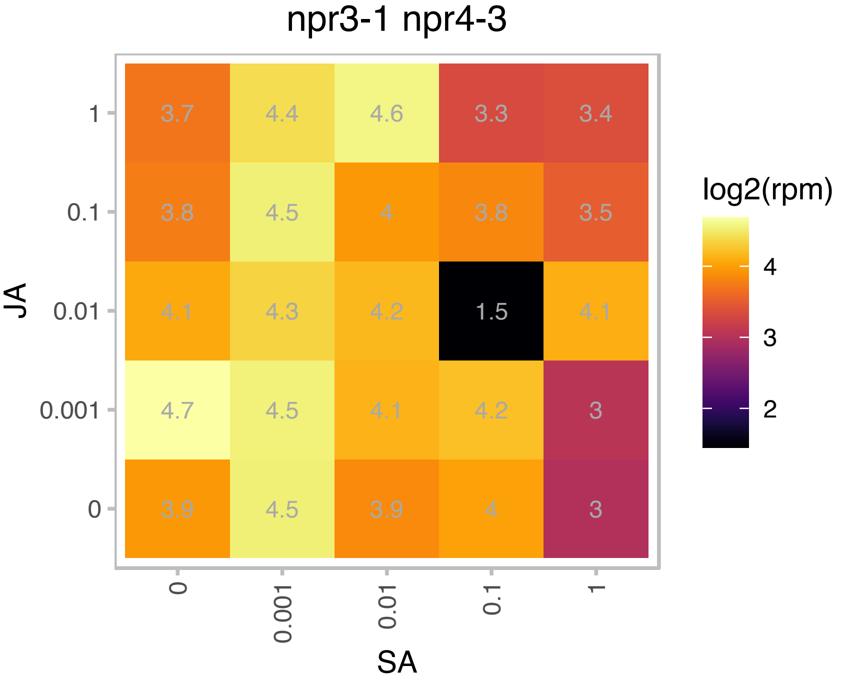

### AT5G50810.1 : TIM8

translocase inner membrane subunit 8

Encodes a small zinc finger-like protein that is a component of the mitochondrial protein import apparatus.

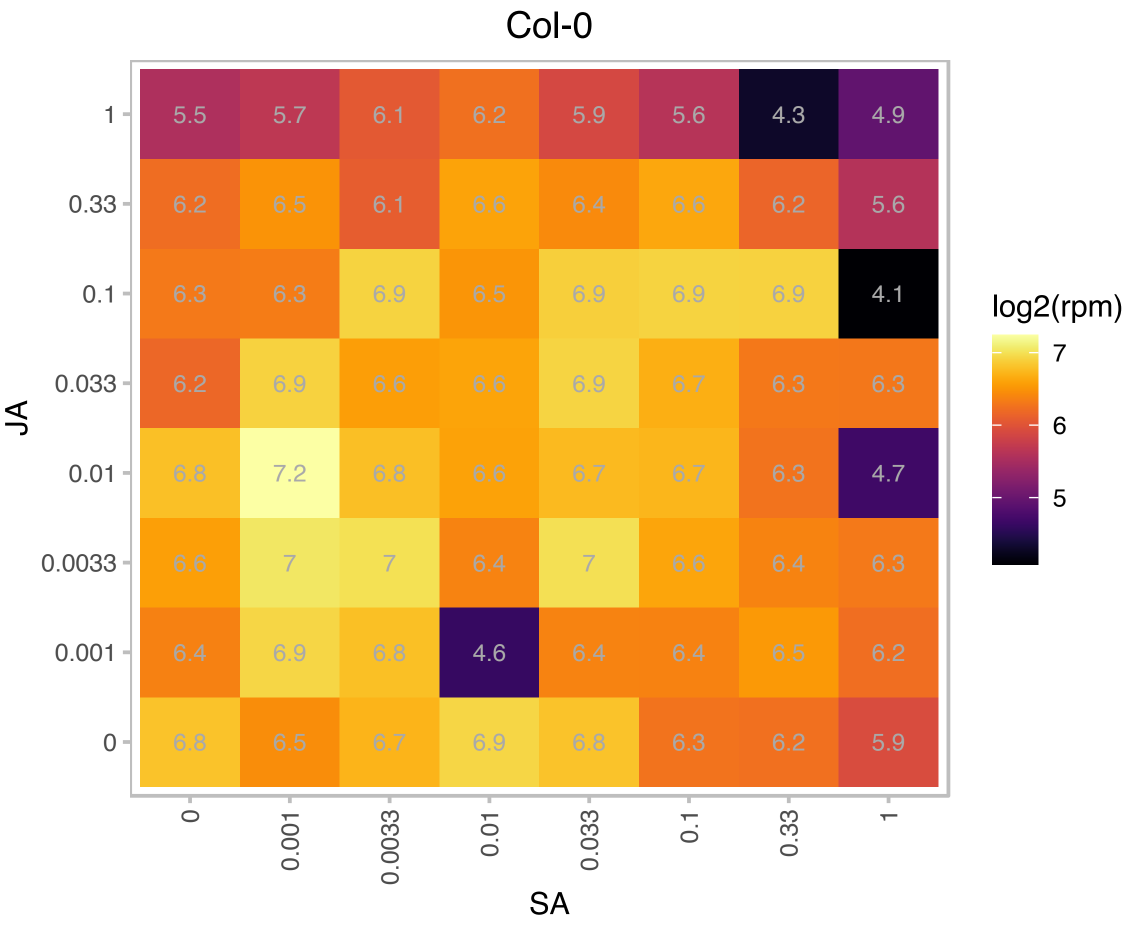


| | |
|---|---|
| Gene | AT5G50810 |
| is.expressed_genes | TRUE |
| is.responsive_genes | TRUE |
| is.apcluster | F-07 |

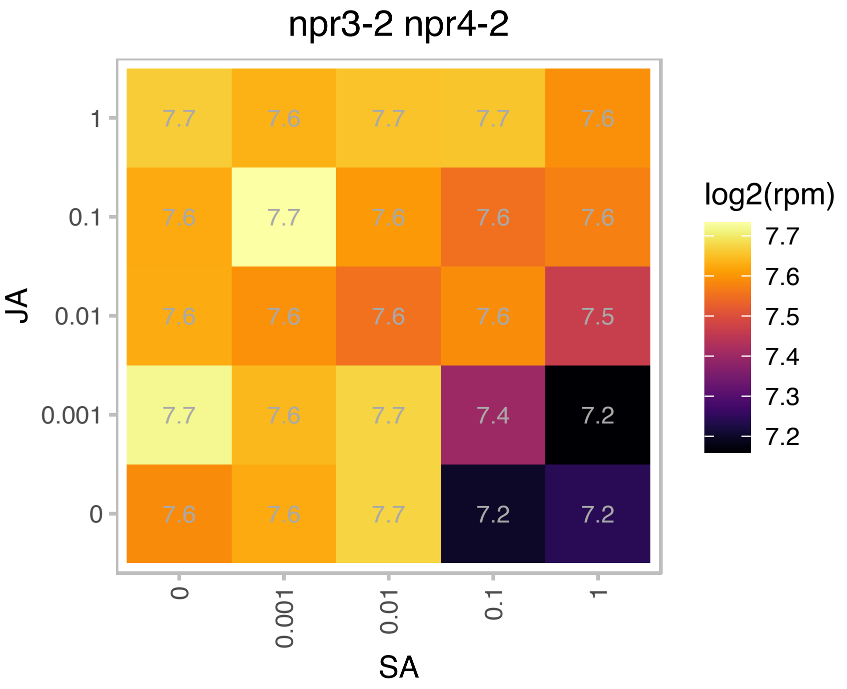


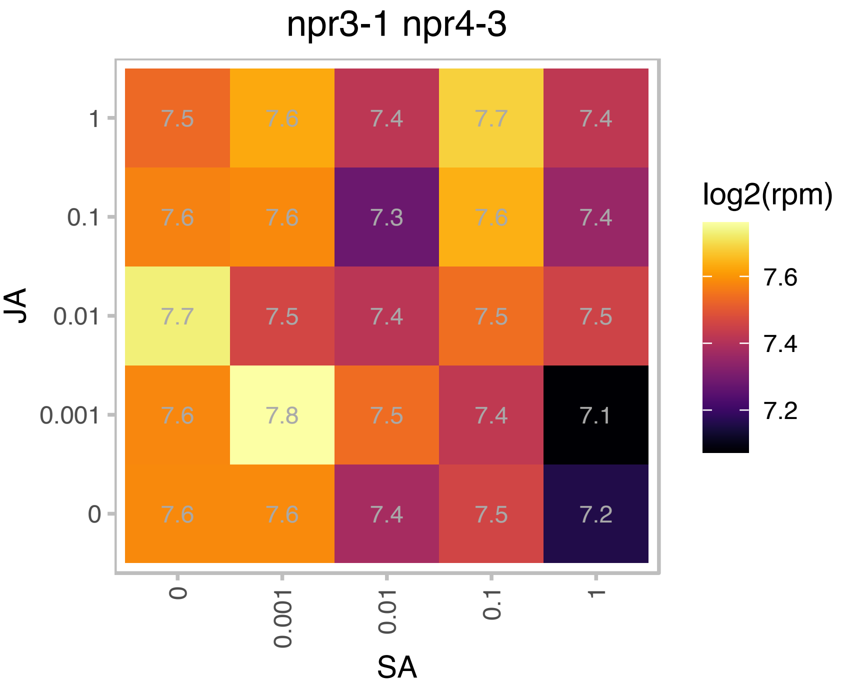

# AT5G52380.1 : NA

VASCULAR-RELATED NAC-DOMAIN 6

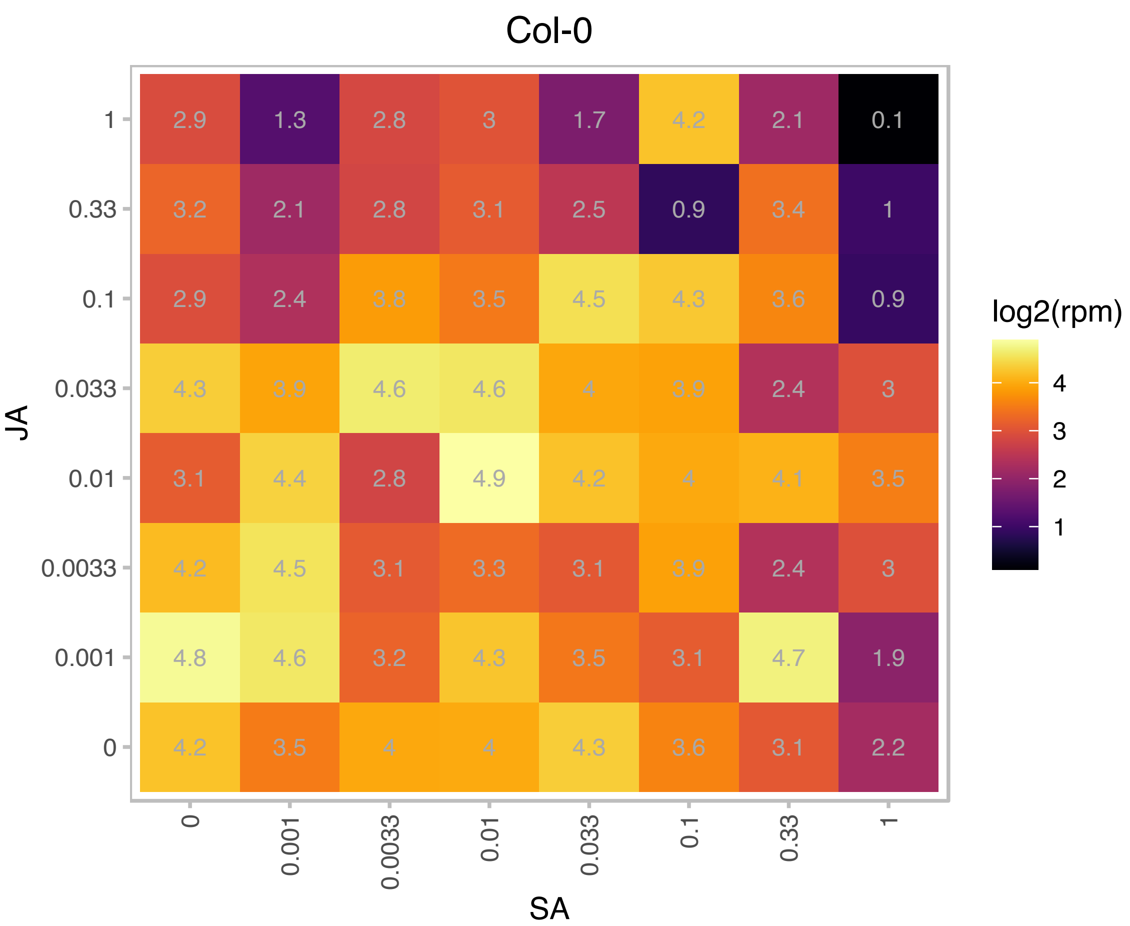


| | |
|---|---|
| Gene | AT5G52380 |
| is.expressed_genes | TRUE |
| is.responsive_genes | TRUE |
| is.apcluster | F-07 |

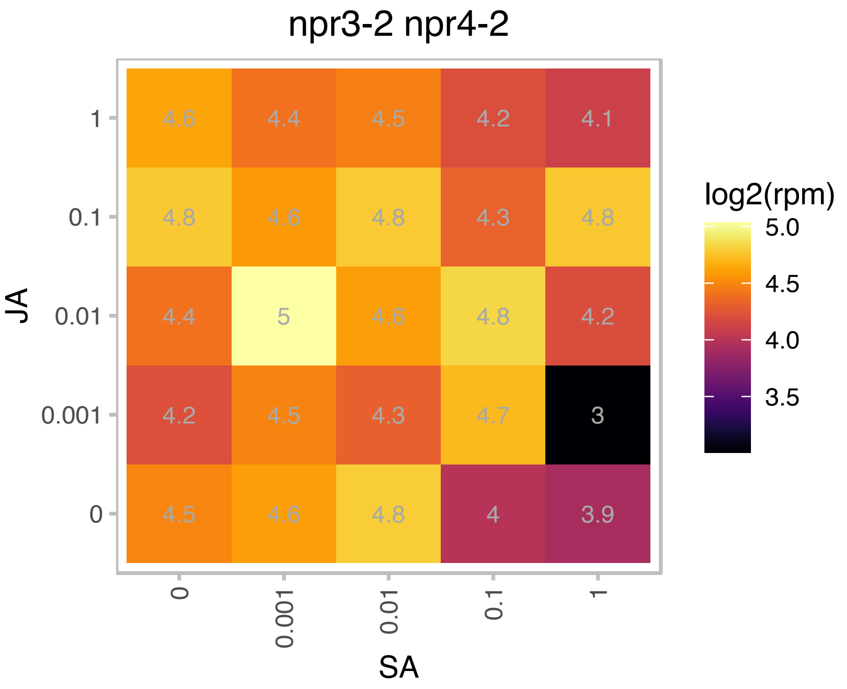


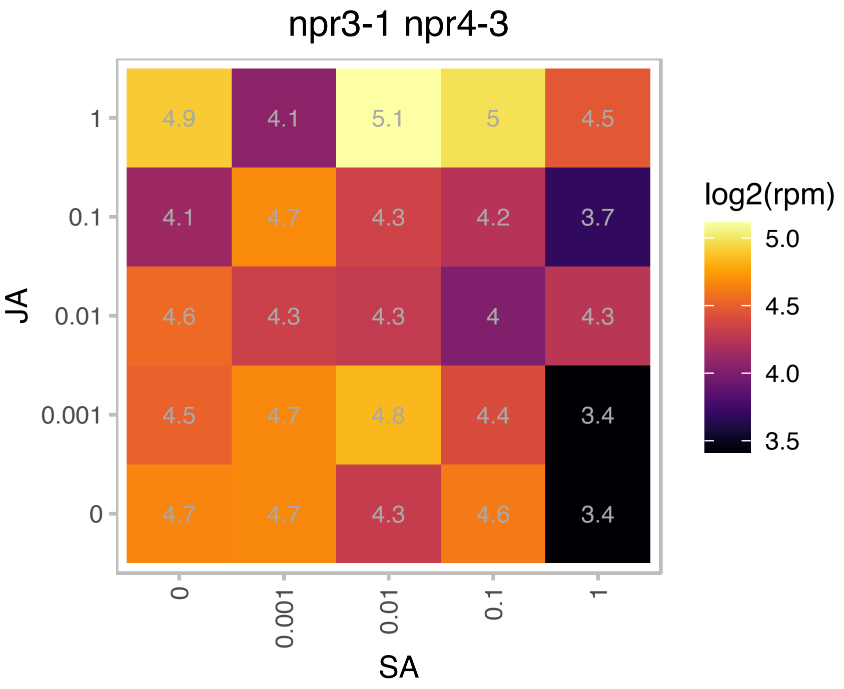

# AT5G52650.1 : NA

RNA binding Plectin/S10 domain-containing protein

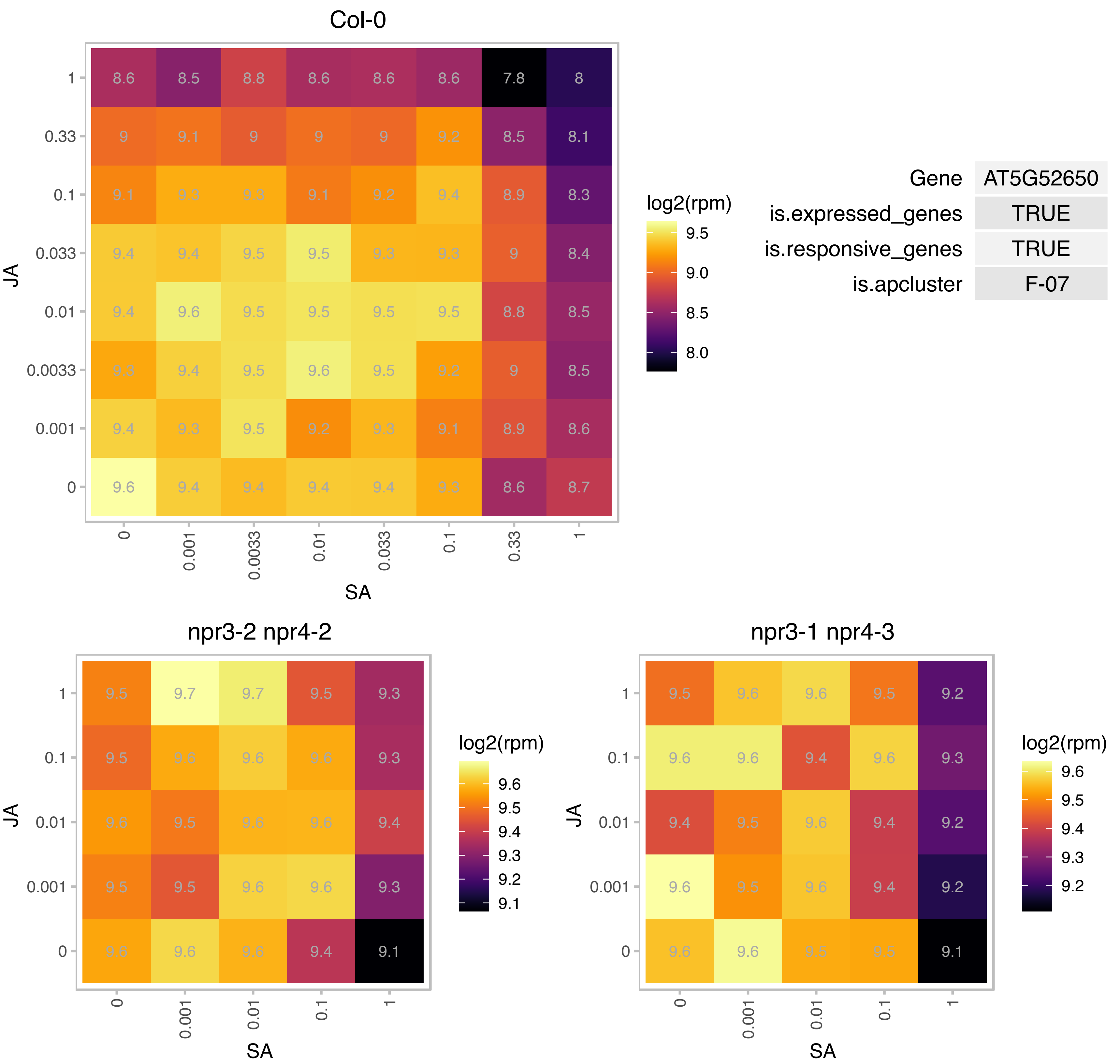


| | |
|---|---|
| Gene | AT5G52650 |
| is.expressed_genes | TRUE |
| is.responsive_genes | TRUE |
| is.apcluster | F-07 |

# AT5G53940.1 : NA

Yippee family putative zinc-binding protein

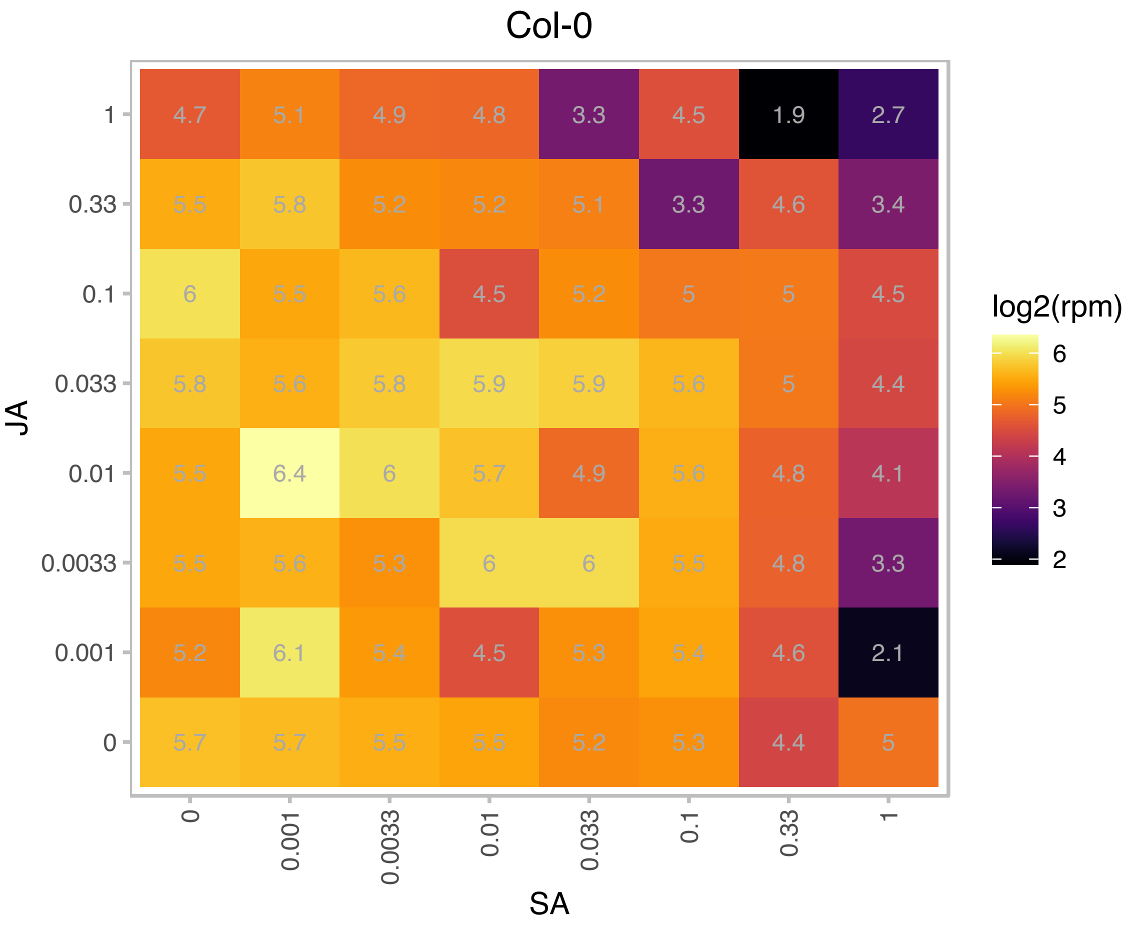


| | |
|---|---|
| Gene | AT5G53940 |
| is.expressed_genes | TRUE |
| is.responsive_genes | TRUE |
| is.apcluster | F-07 |

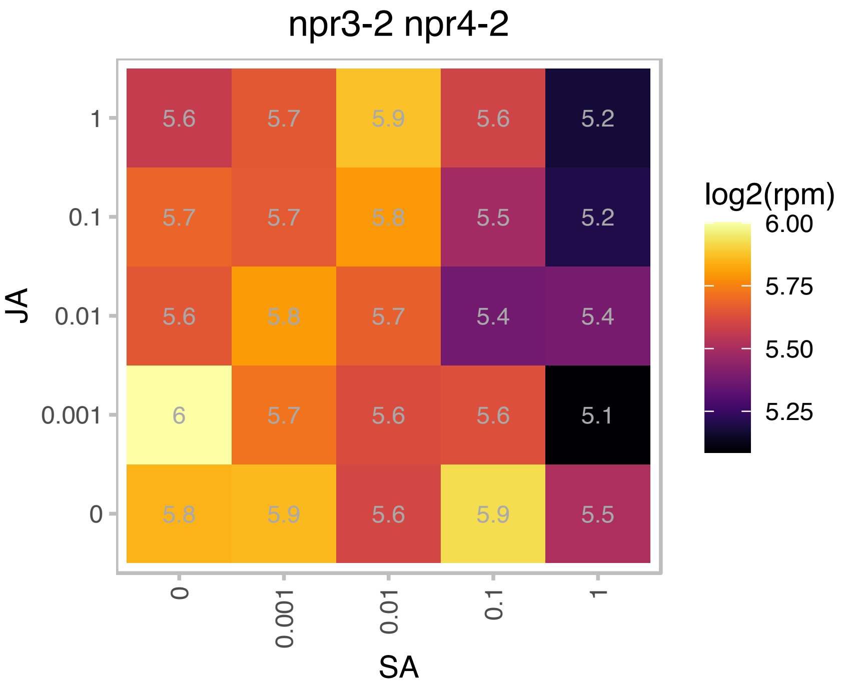


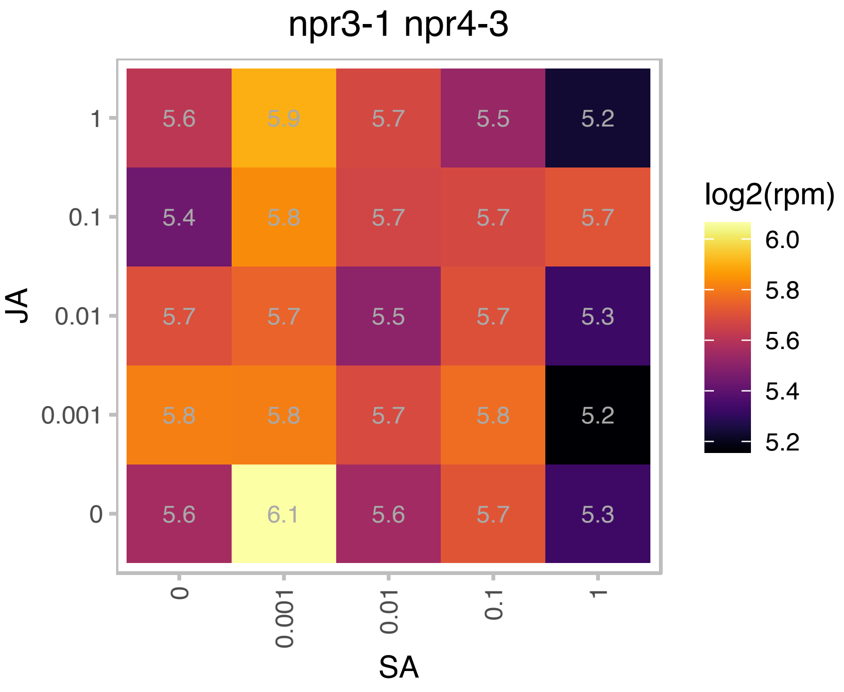

# AT5G55210.1 : NA

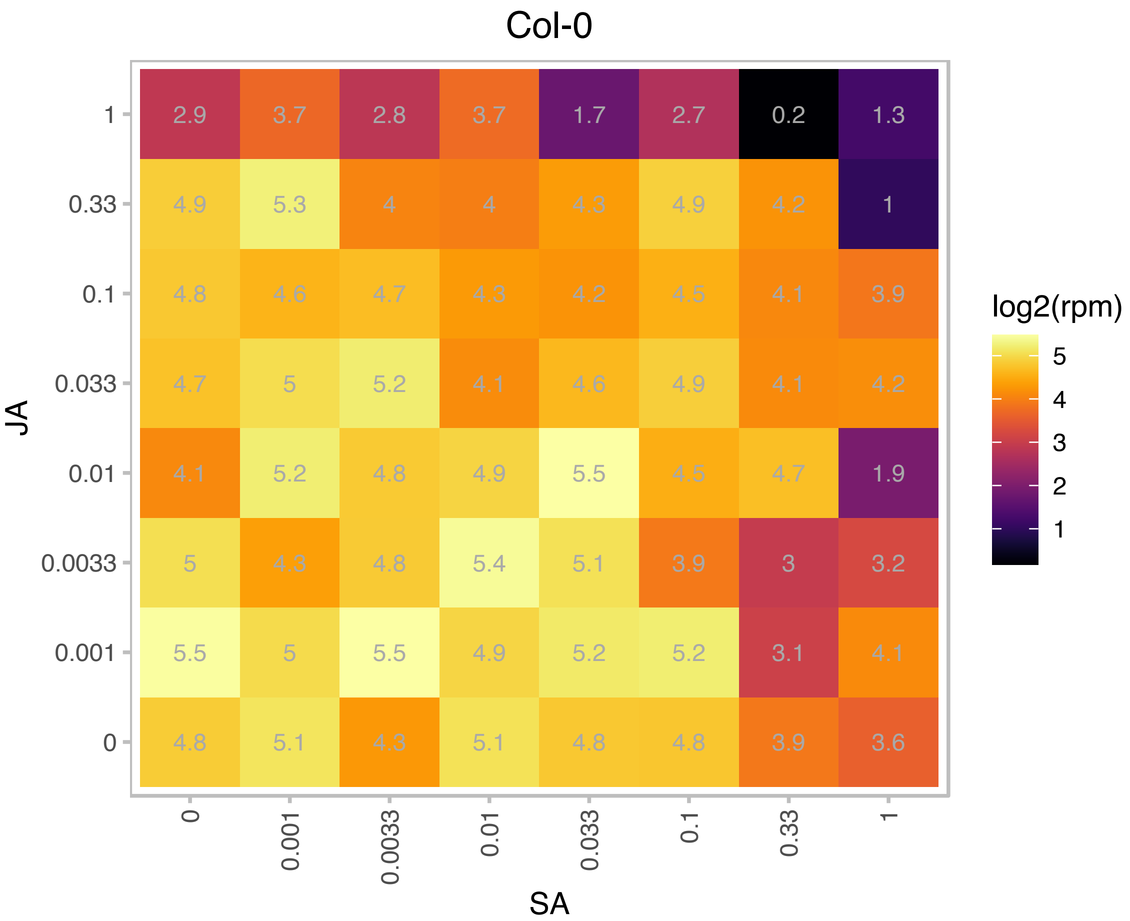


| | |
|---|---|
| Gene | AT5G55210 |
| is.expressed_genes | TRUE |
| is.responsive_genes | TRUE |
| is.apcluster | F-07 |

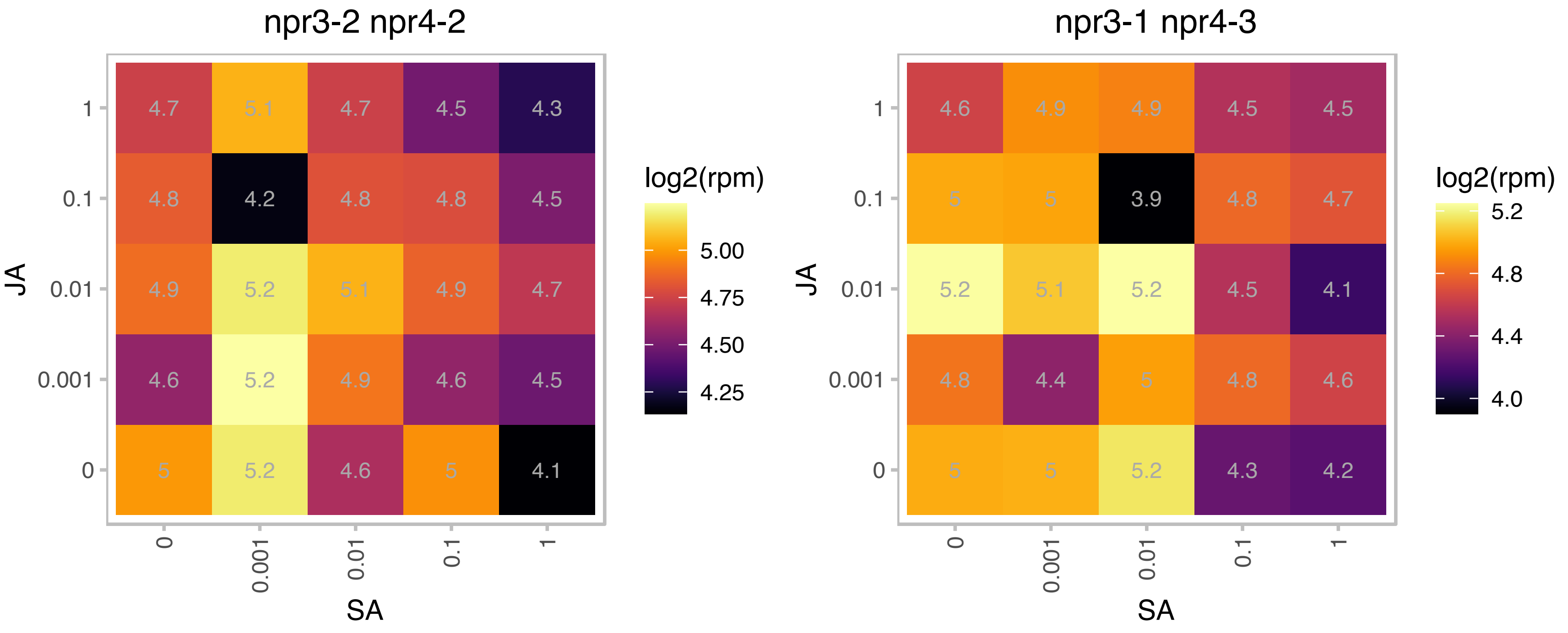

# AT5G55510.1 : HP22

Mitochondrial import inner membrane translocase subunit Tim17/Tim22/Tim23 family protein

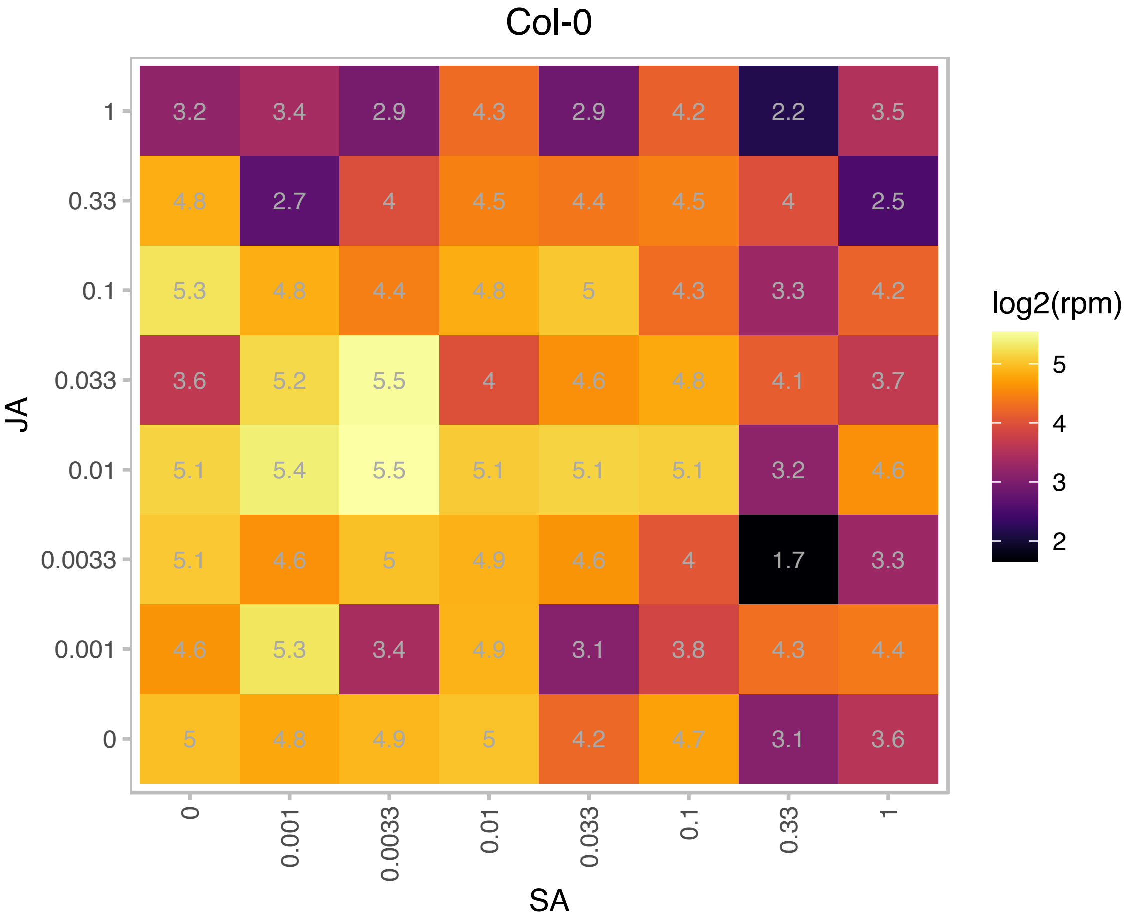


| | |
|---|---|
| Gene | AT5G55510 |
| is.expressed_genes | TRUE |
| is.responsive_genes | TRUE |
| is.apcluster | F-07 |

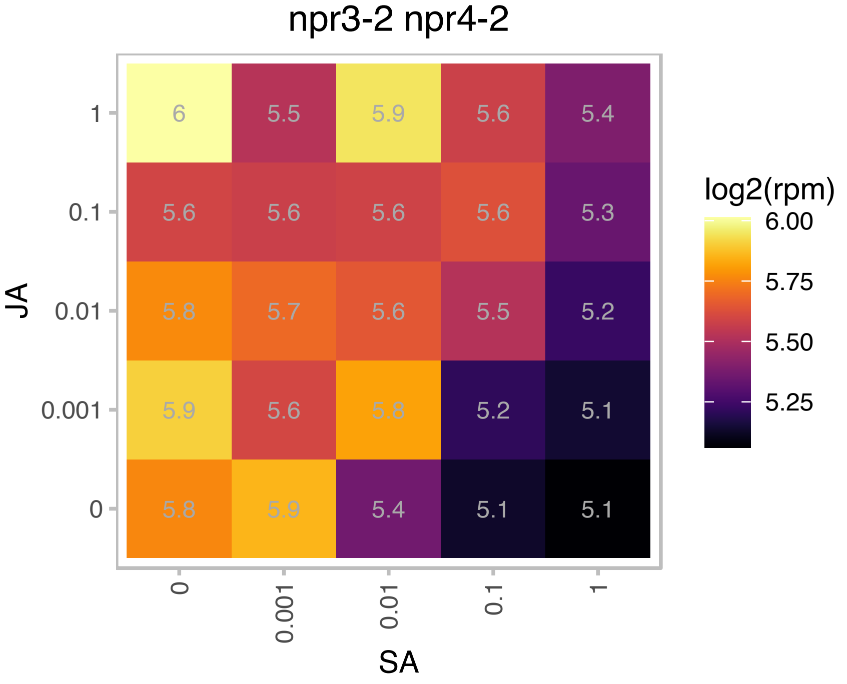


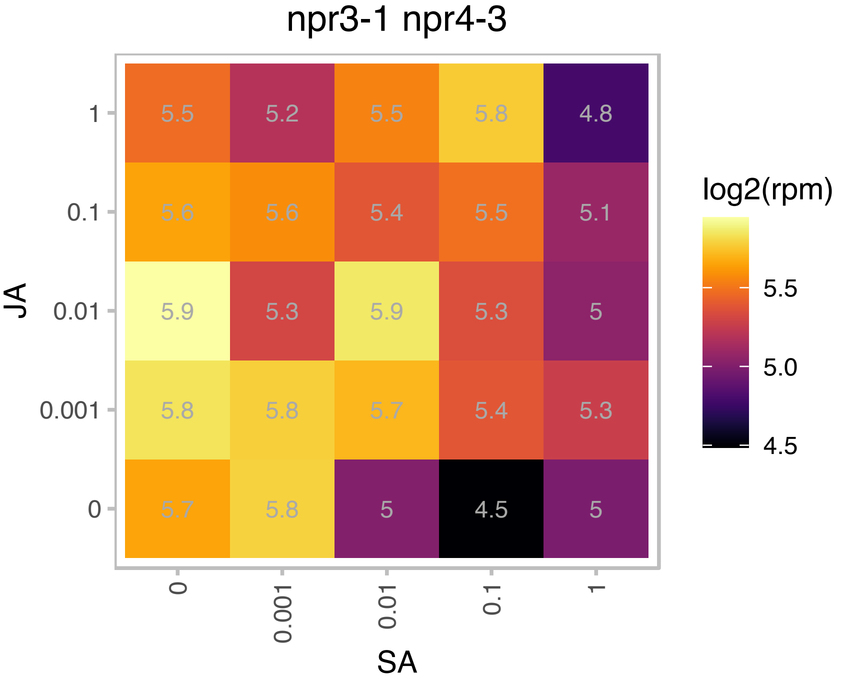

# AT5G56710.1 : NA

Ribosomal protein L31e family protein

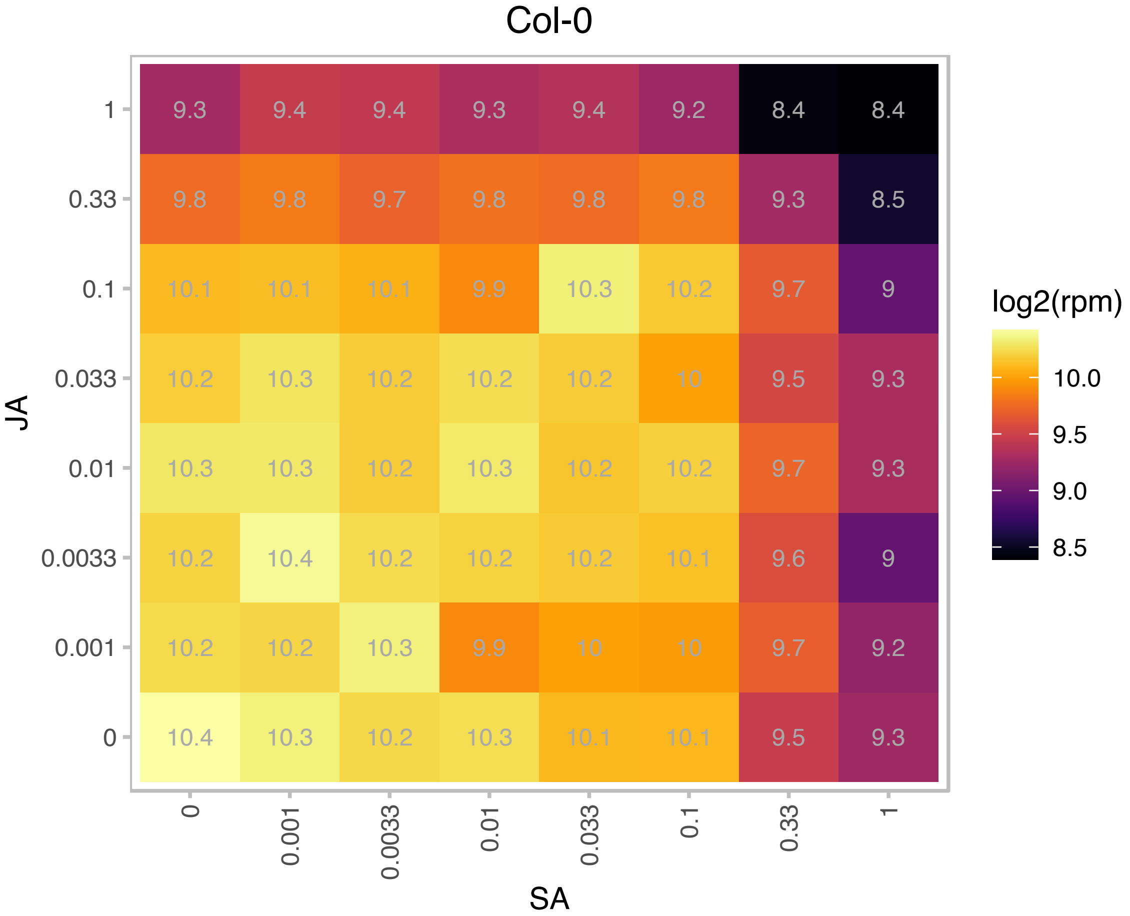


| | |
|---|---|
| Gene | AT5G56710 |
| is.expressed_genes | TRUE |
| is.responsive_genes | TRUE |
| is.apcluster | F-07 |

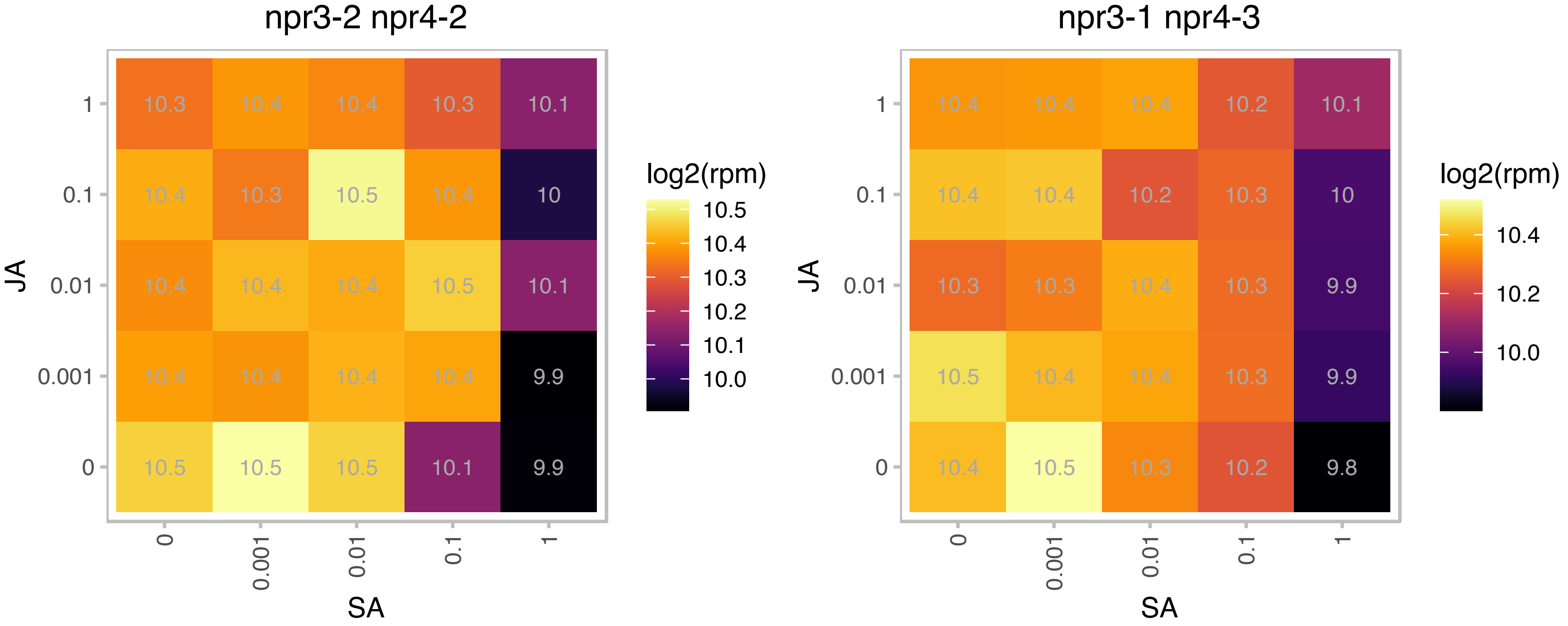

### AT5G57180.1 : CIA2

chloroplast import apparatus 2

Transcription regulator responsible for specific upregulation of the translocon genes atToc33 and atToc75 in leaves. Involved in protein import into chloroplast.

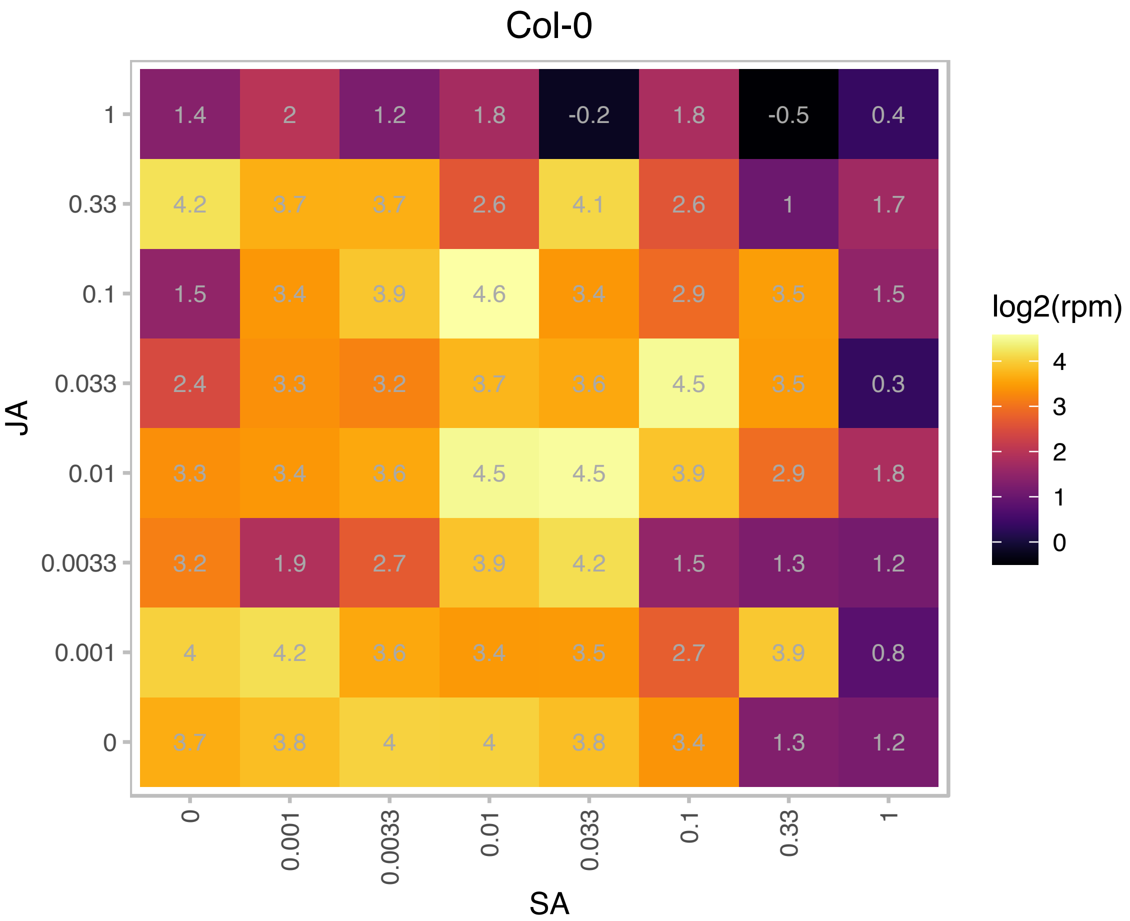


| | |
|---|---|
| Gene | AT5G57180 |
| is.expressed_genes | TRUE |
| is.responsive_genes | TRUE |
| is.apcluster | F-07 |

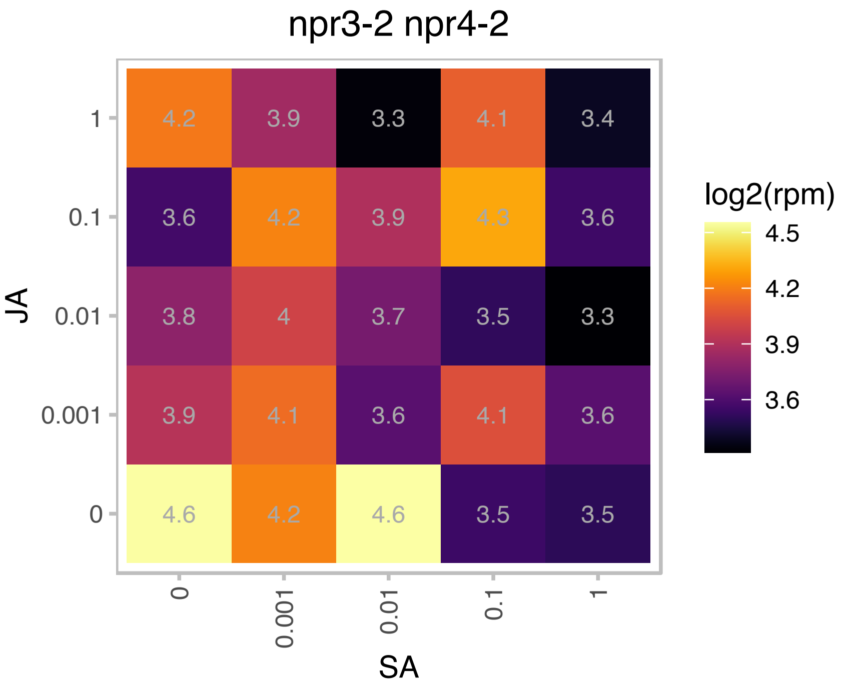


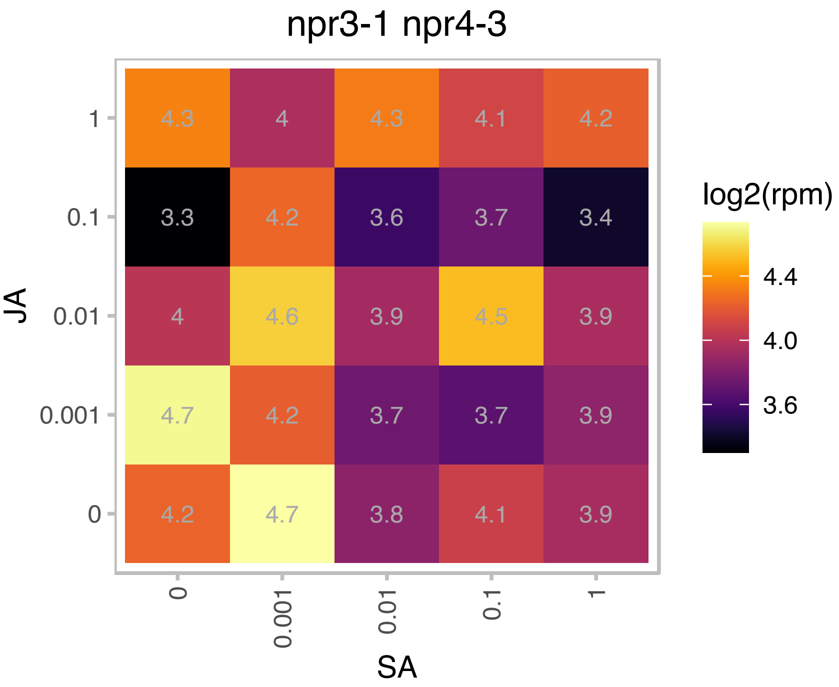

### AT5G57290.1 : AtP3B, P3B

60S acidic ribosomal protein family

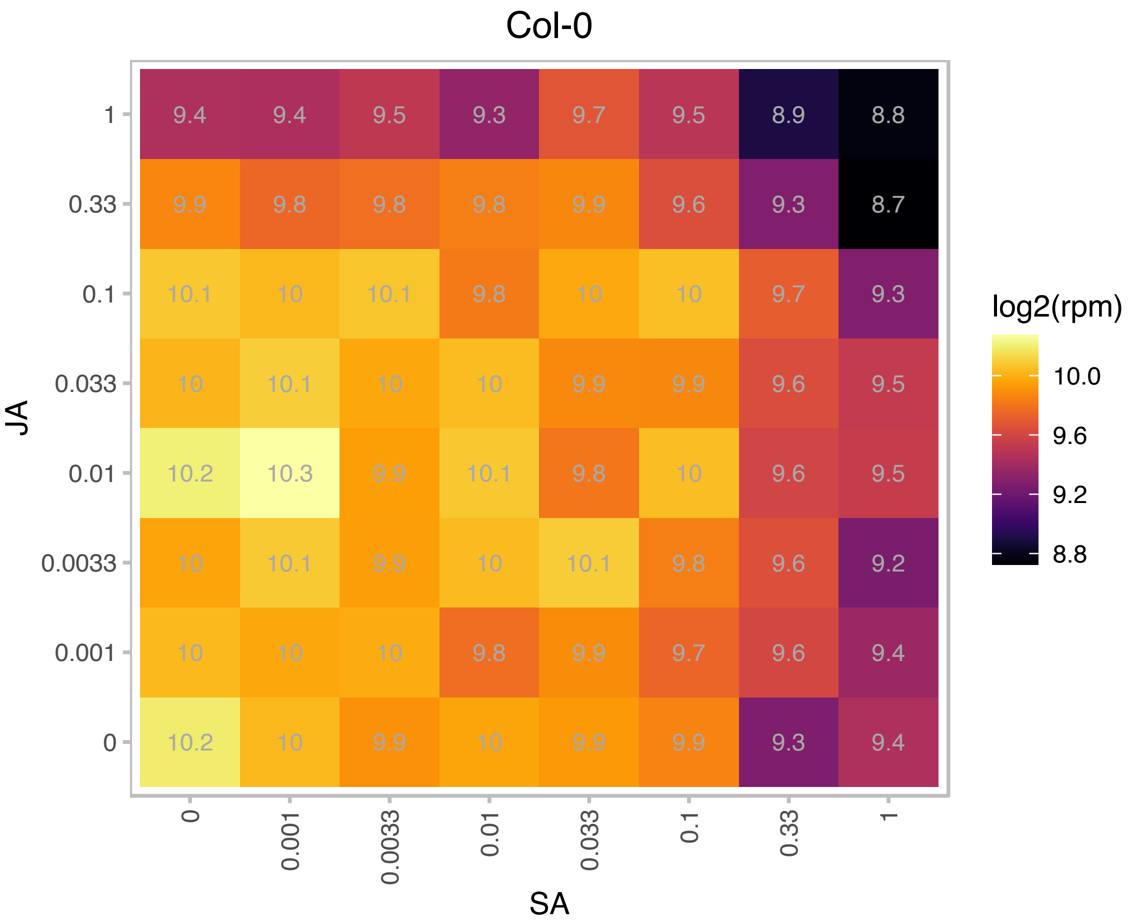


| | |
|---|---|
| Gene | AT5G57290 |
| is.expressed_genes | TRUE |
| is.responsive_genes | TRUE |
| is.apcluster | F-07 |

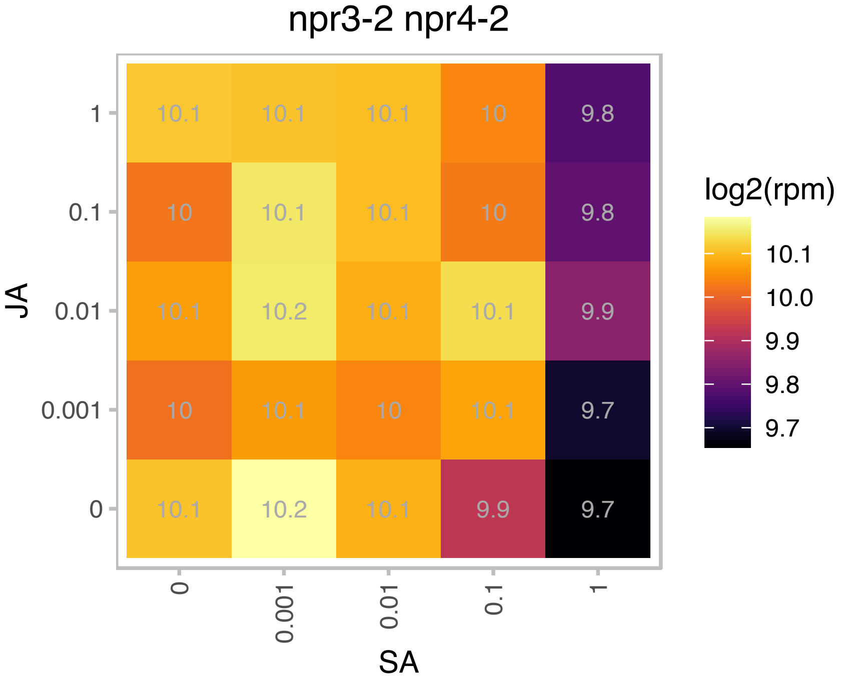


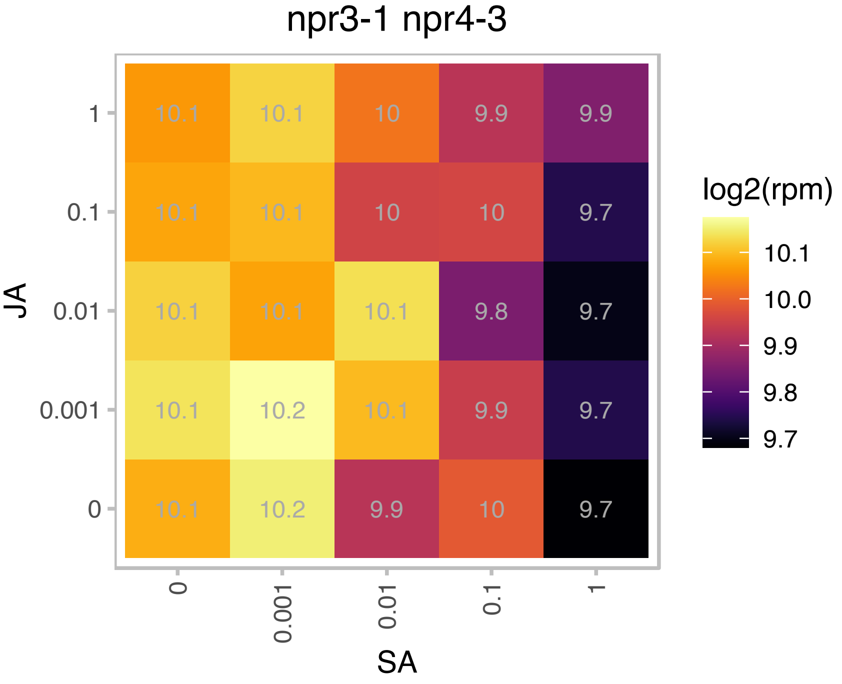

# AT5G57887.1 : NA

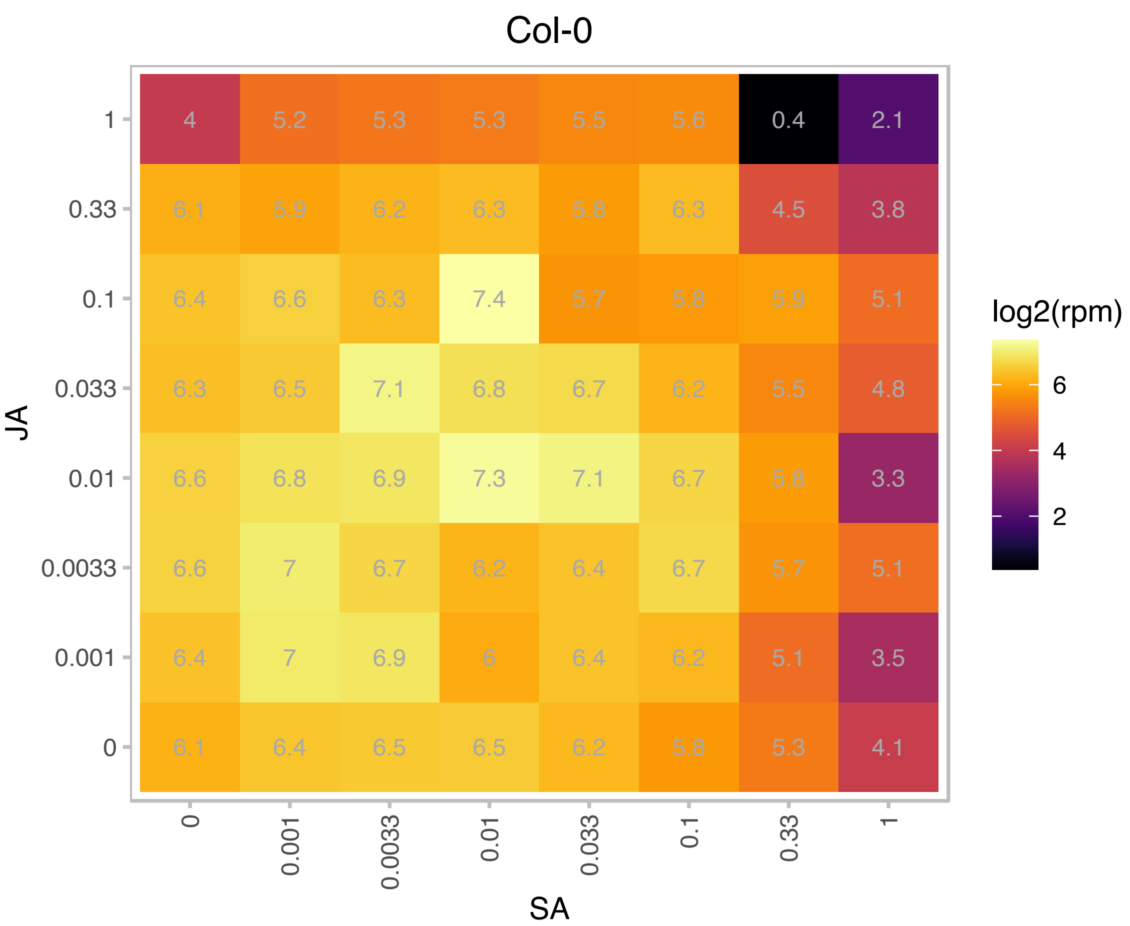


| | |
|---|---|
| Gene | AT5G57887 |
| is.expressed_genes | TRUE |
| is.responsive_genes | TRUE |
| is.apcluster | F-07 |

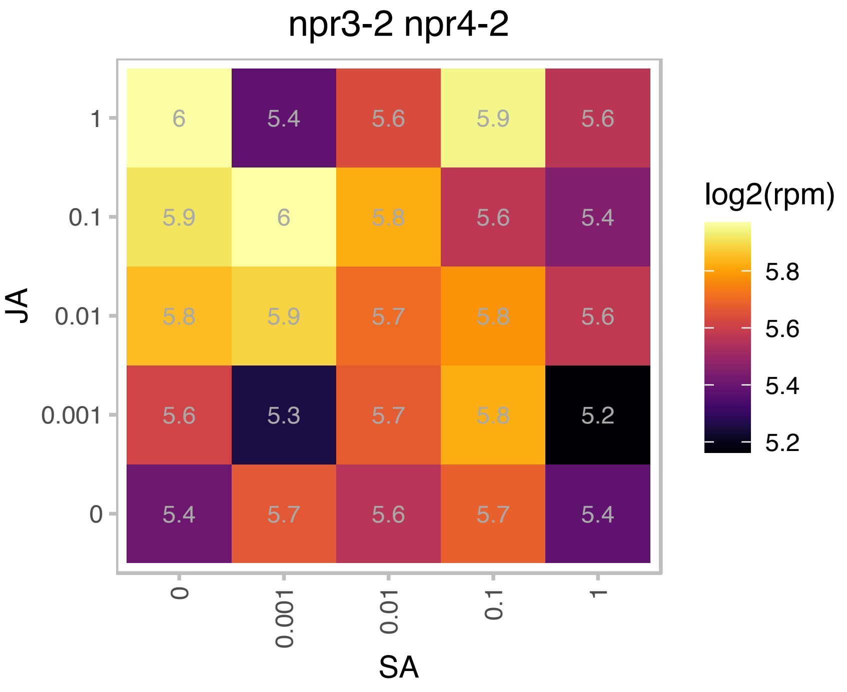


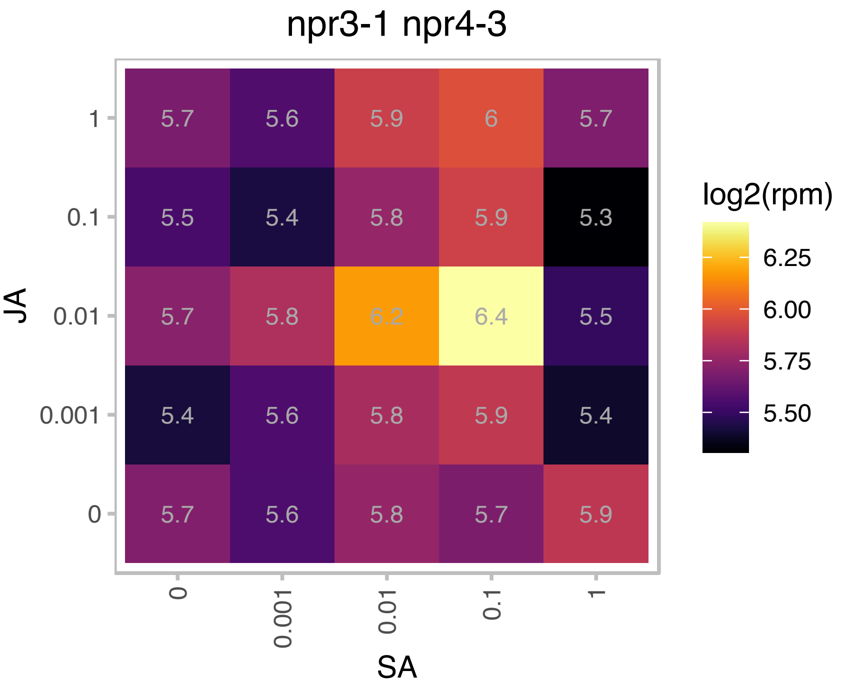

### AT5G57930.1 : APO2, emb1629

Arabidopsis thaliana protein of unknown function (DUF794)

ACCUMULATION OF PHOTOSYSTEM ONE 2

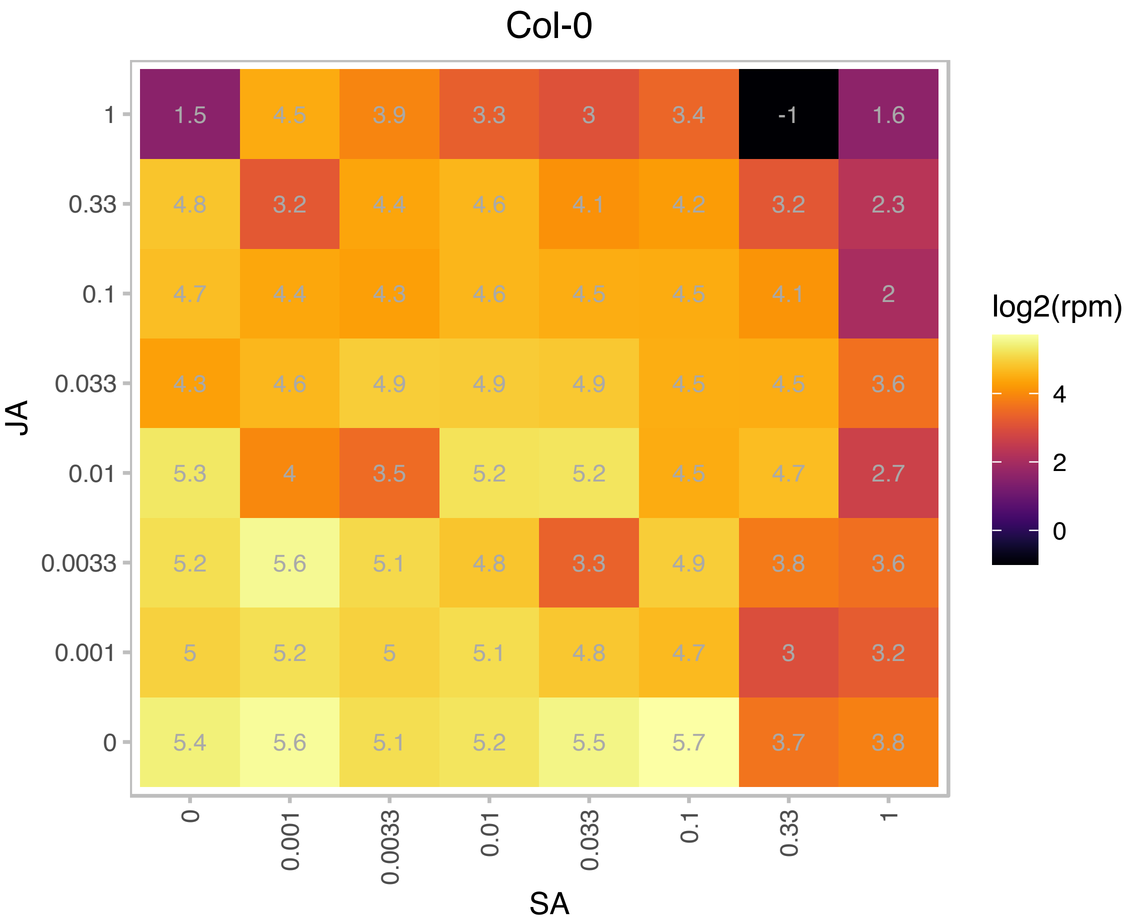


| | |
|---|---|
| Gene | AT5G57930 |
| is.expressed_genes | TRUE |
| is.responsive_genes | TRUE |
| is.apcluster | F-07 |

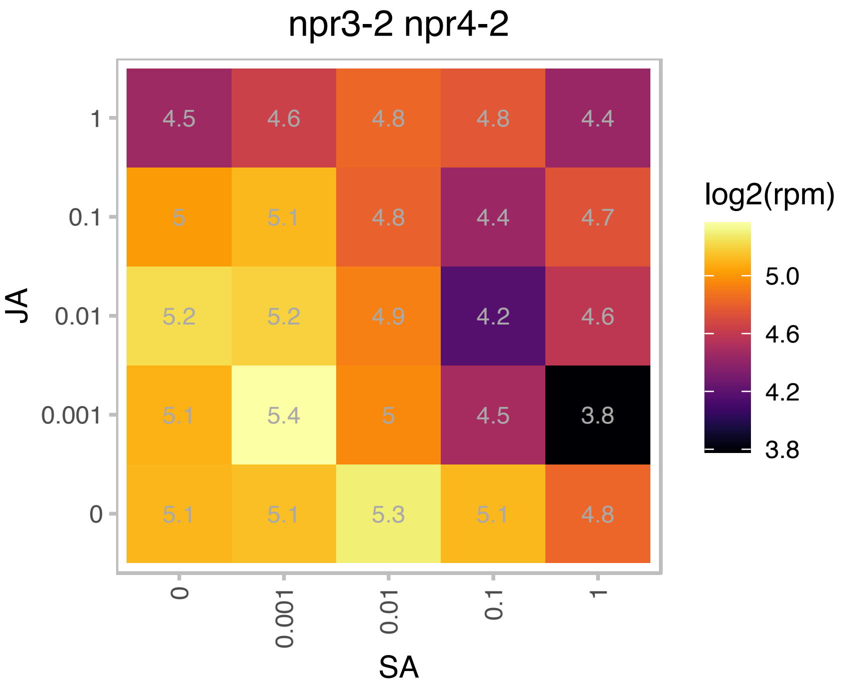


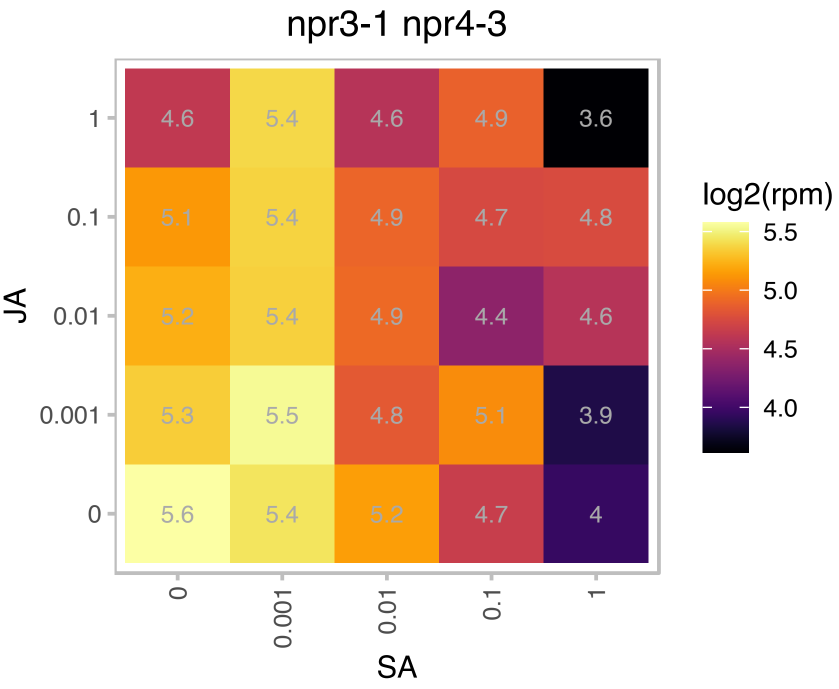

# AT5G58570.1 : NA

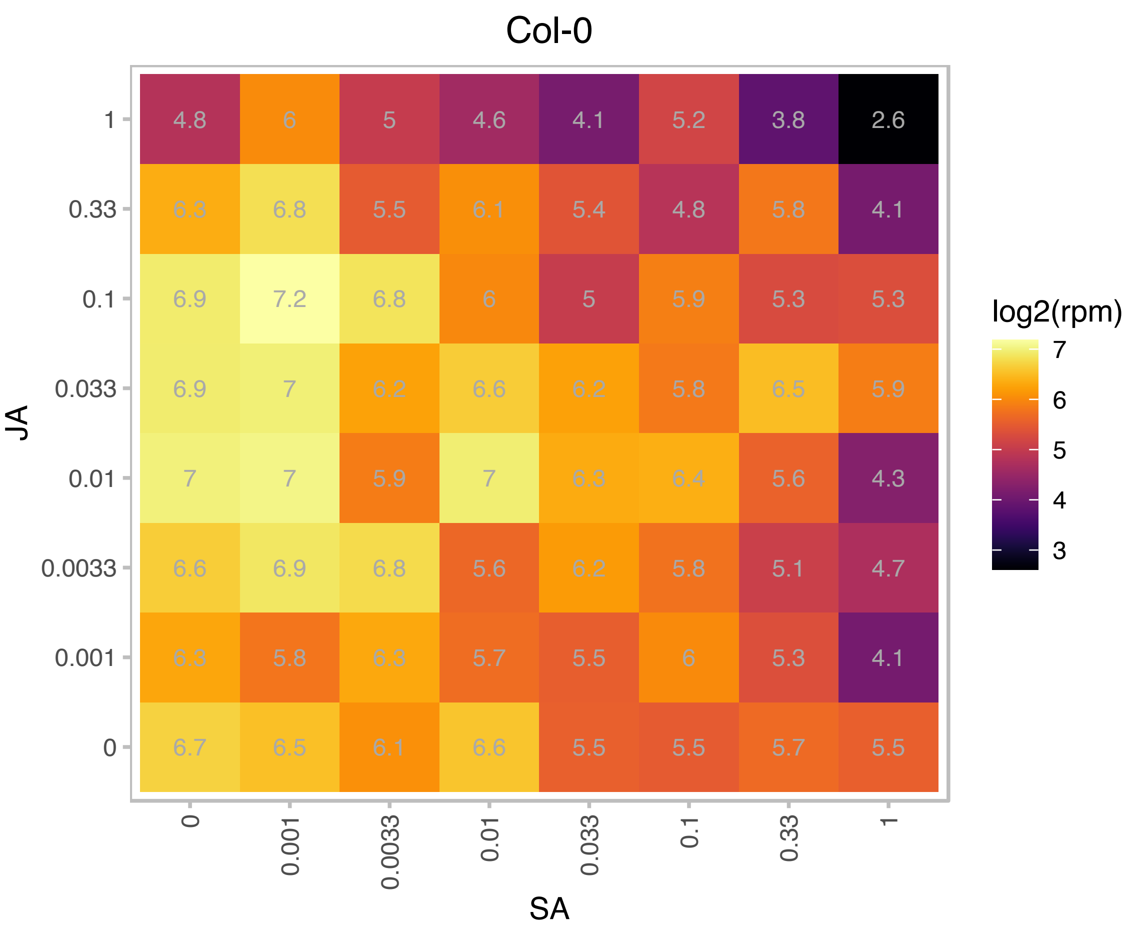


| | |
|---|---|
| Gene | AT5G58570 |
| is.expressed_genes | TRUE |
| is.responsive_genes | TRUE |
| is.apcluster | F-07 |

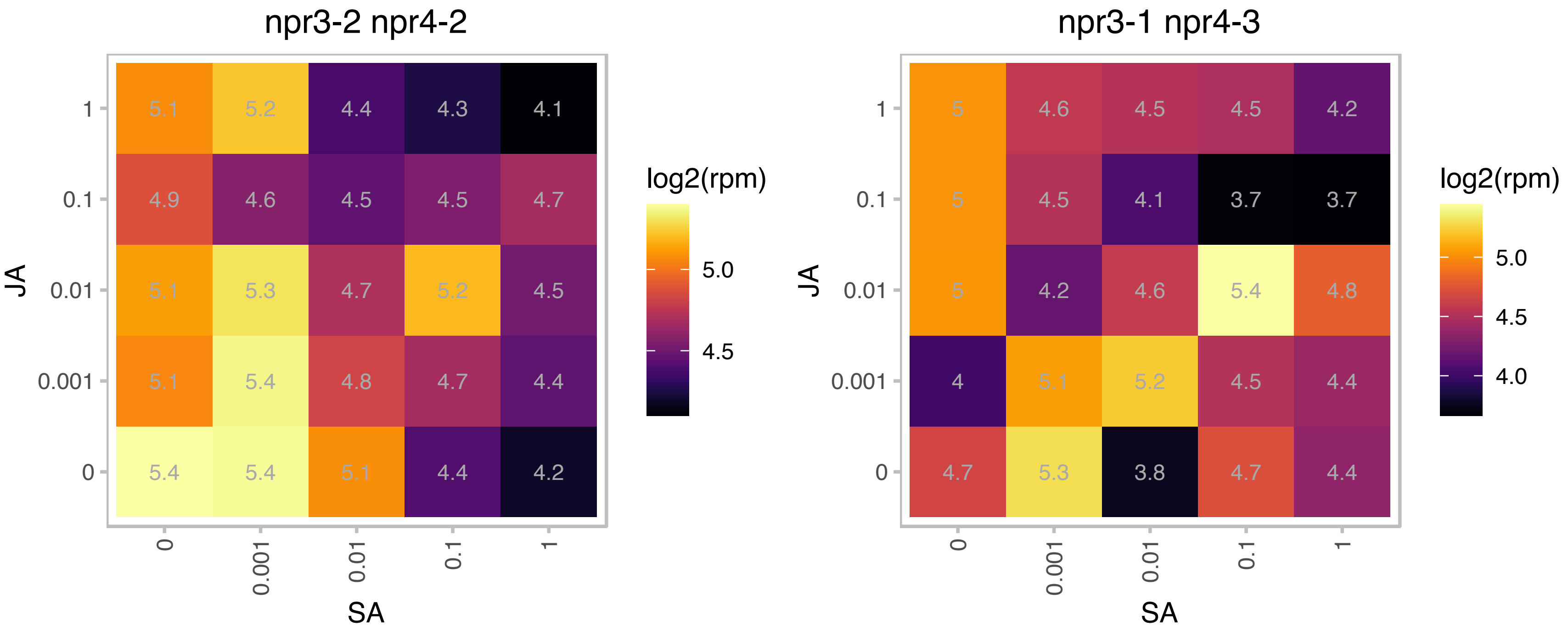

# AT5G59690.1 : NA

Histone superfamily protein

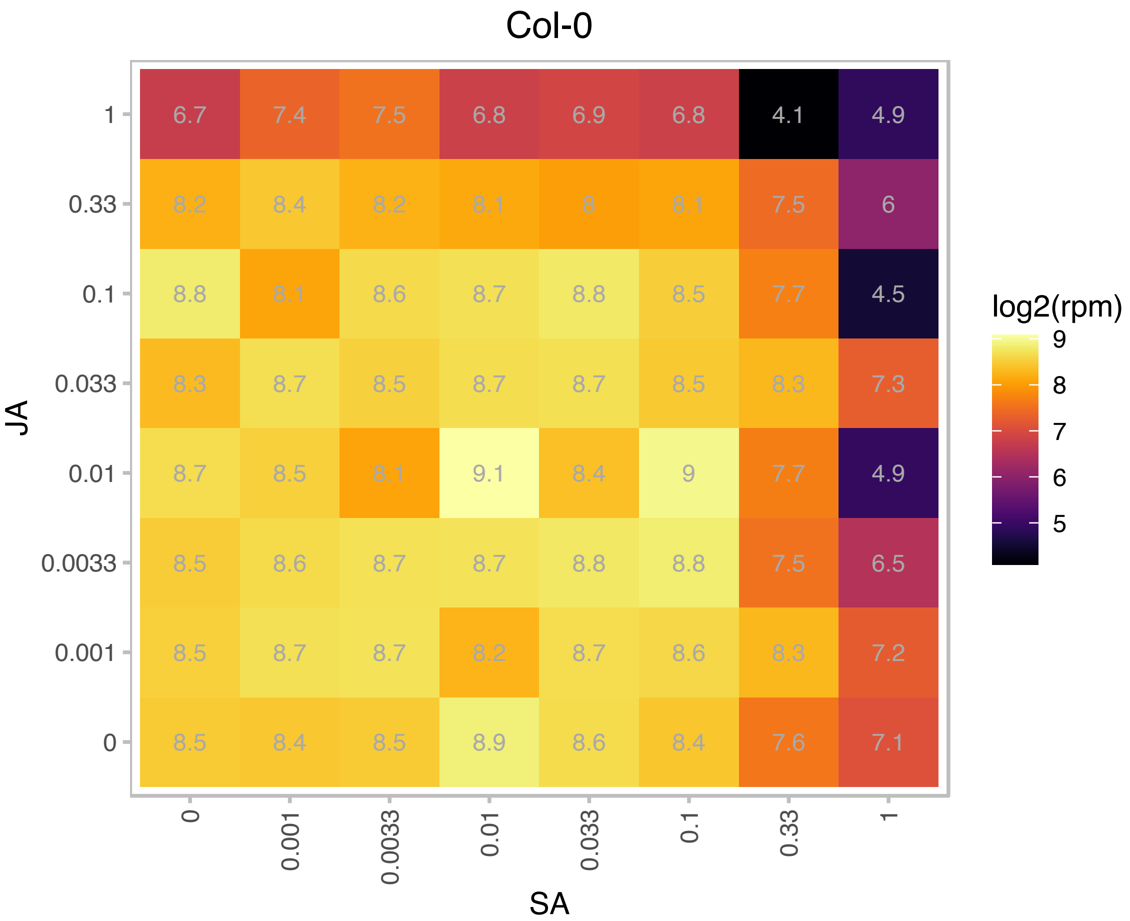


| | |
|---|---|
| Gene | AT5G59690 |
| is.expressed_genes | TRUE |
| is.responsive_genes | TRUE |
| is.apcluster | F-07 |

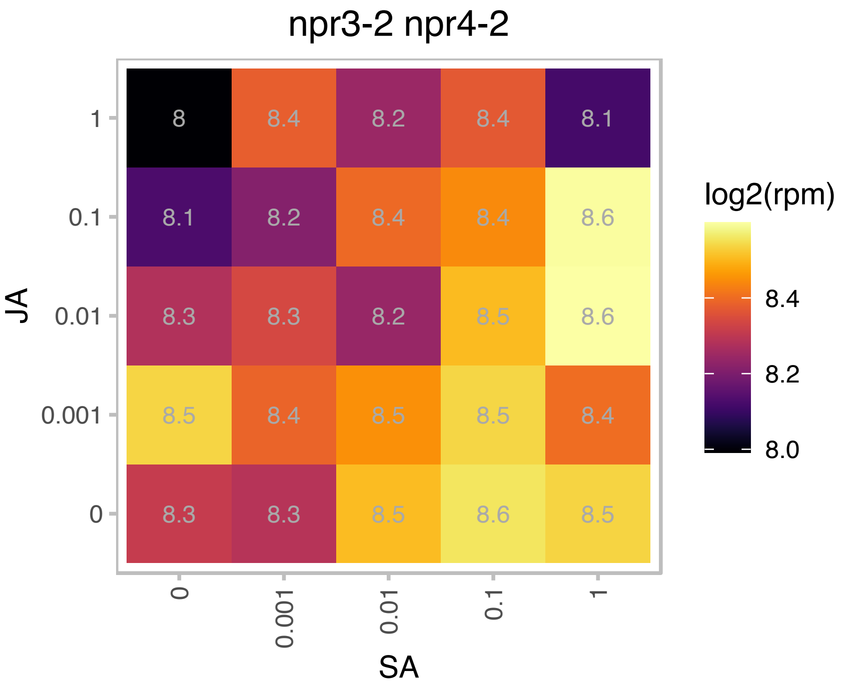


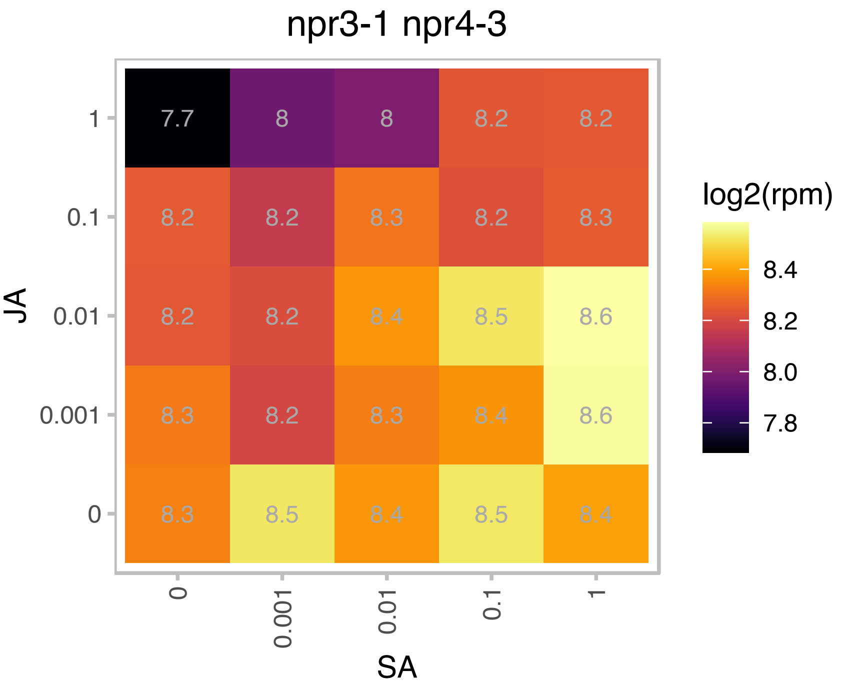

# AT5G59850.1 : NA

Ribosomal protein S8 family protein

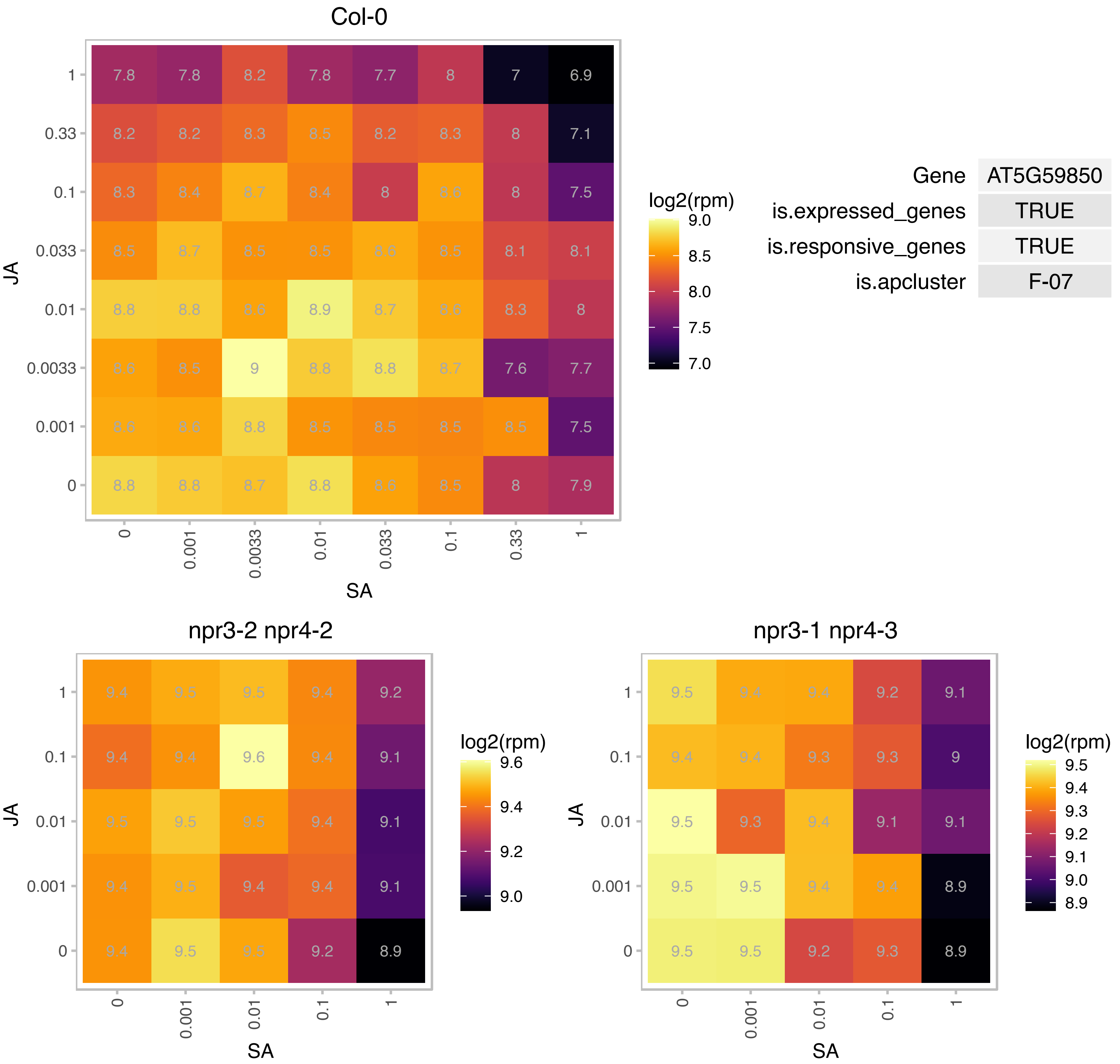


| | |
|---|---|
| Gene | AT5G59850 |
| is.expressed_genes | TRUE |
| is.responsive_genes | TRUE |
| is.apcluster | F-07 |

# AT5G59970.1 : NA

Histone superfamily protein

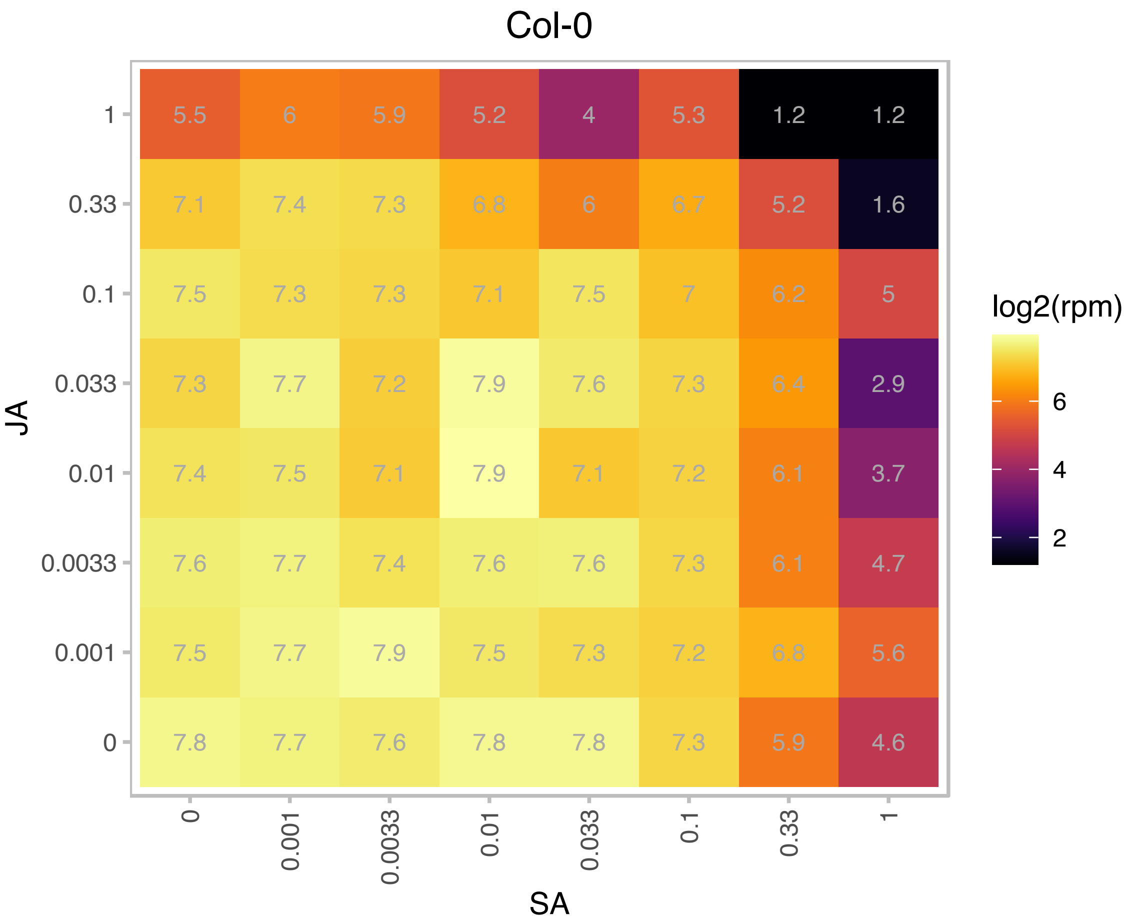

| | |
|---|---|
| Gene | AT5G59970 |
| is.expressed_genes | TRUE |
| is.responsive_genes | TRUE |
| is.apcluster | F-07 |

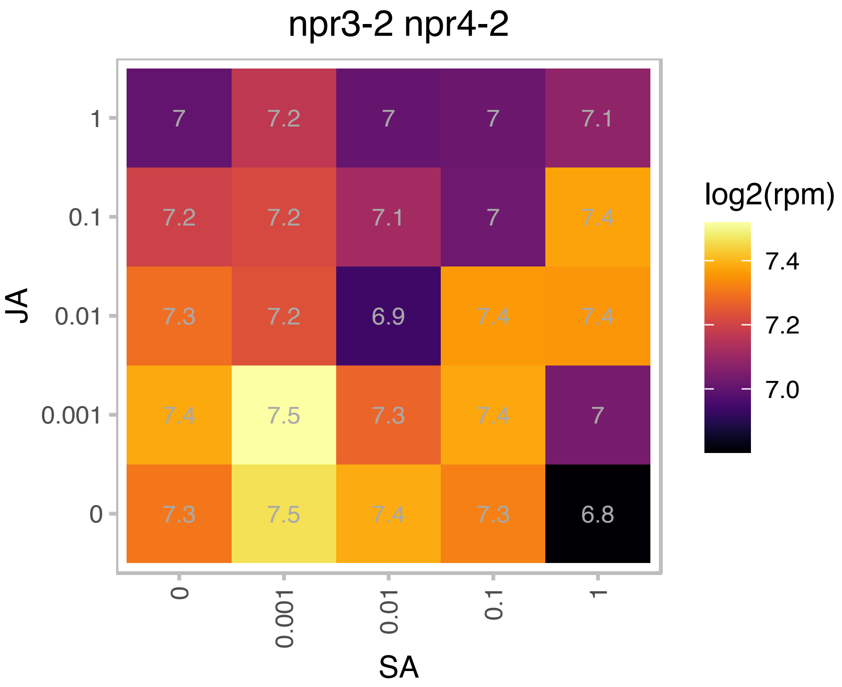

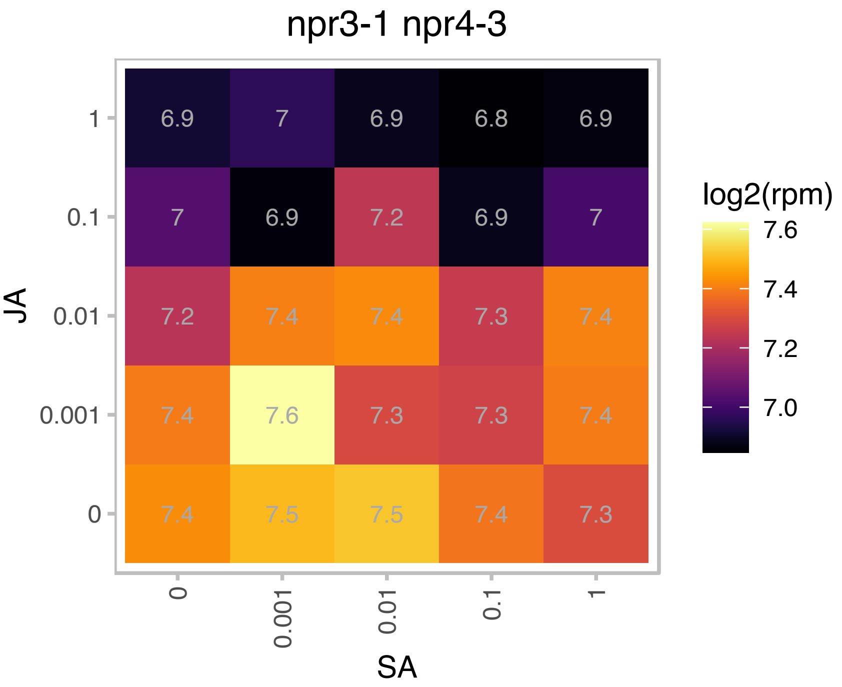

### AT5G60670.1 : RPL12C

Ribosomal protein L11 family protein

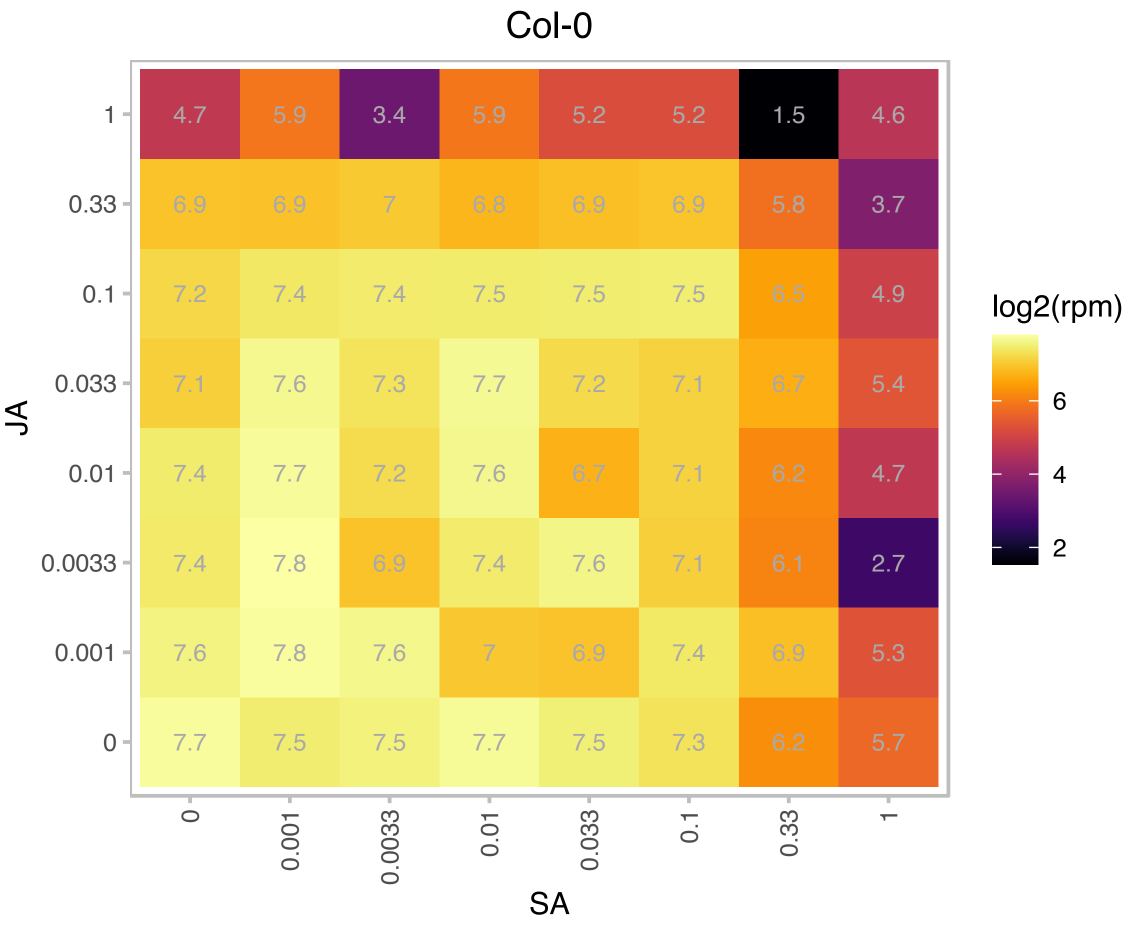


| | |
|---|---|
| Gene | AT5G60670 |
| is.expressed_genes | TRUE |
| is.responsive_genes | TRUE |
| is.apcluster | F-07 |

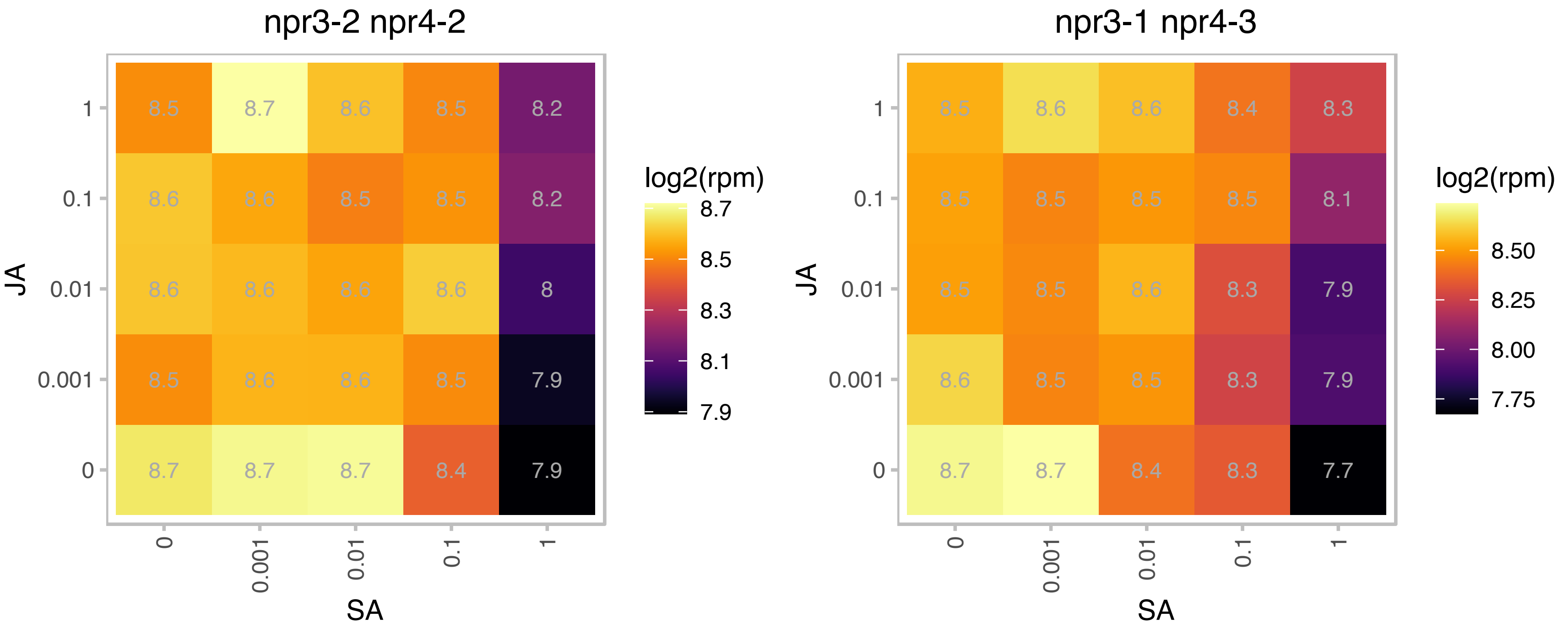

# AT5G61170.1 : NA

Ribosomal protein S19e family protein

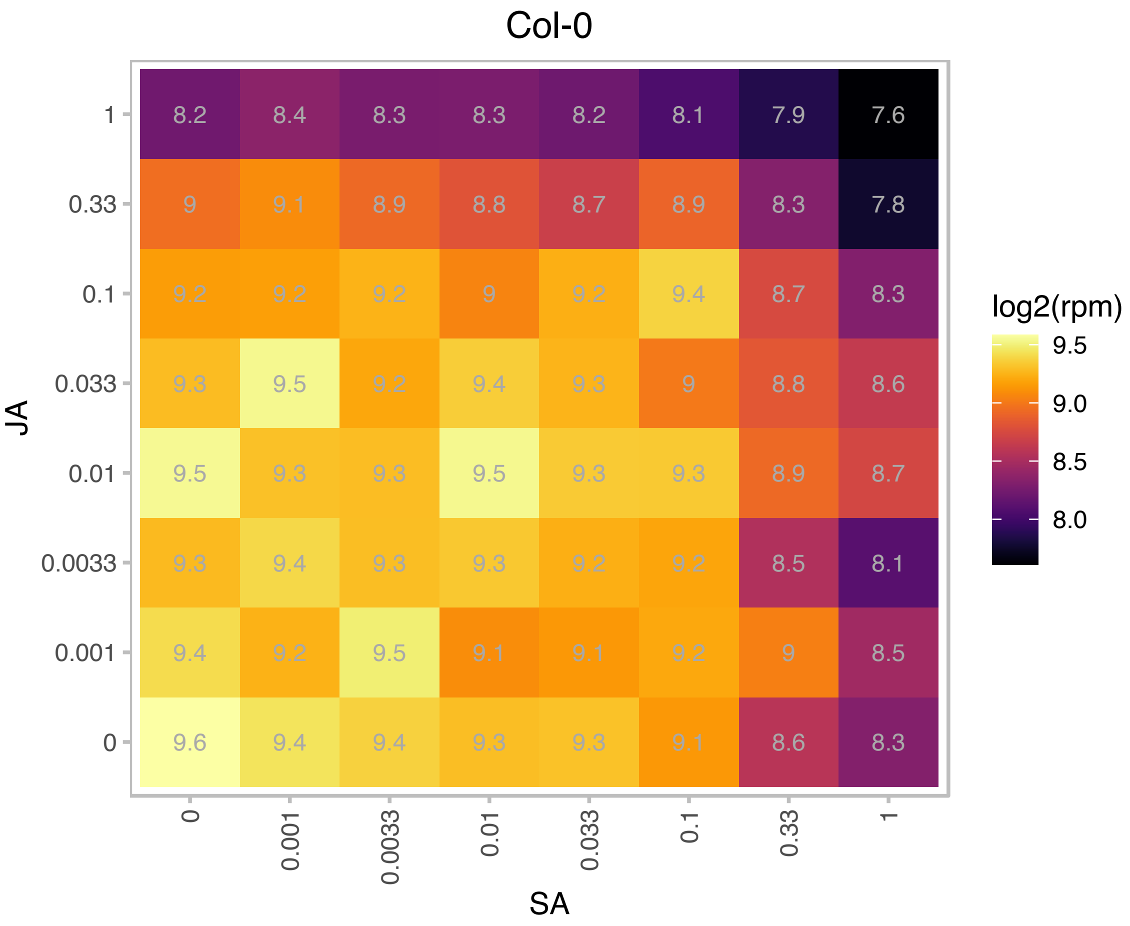


| | |
|---|---|
| Gene | AT5G61170 |
| is.expressed_genes | TRUE |
| is.responsive_genes | TRUE |
| is.apcluster | F-07 |

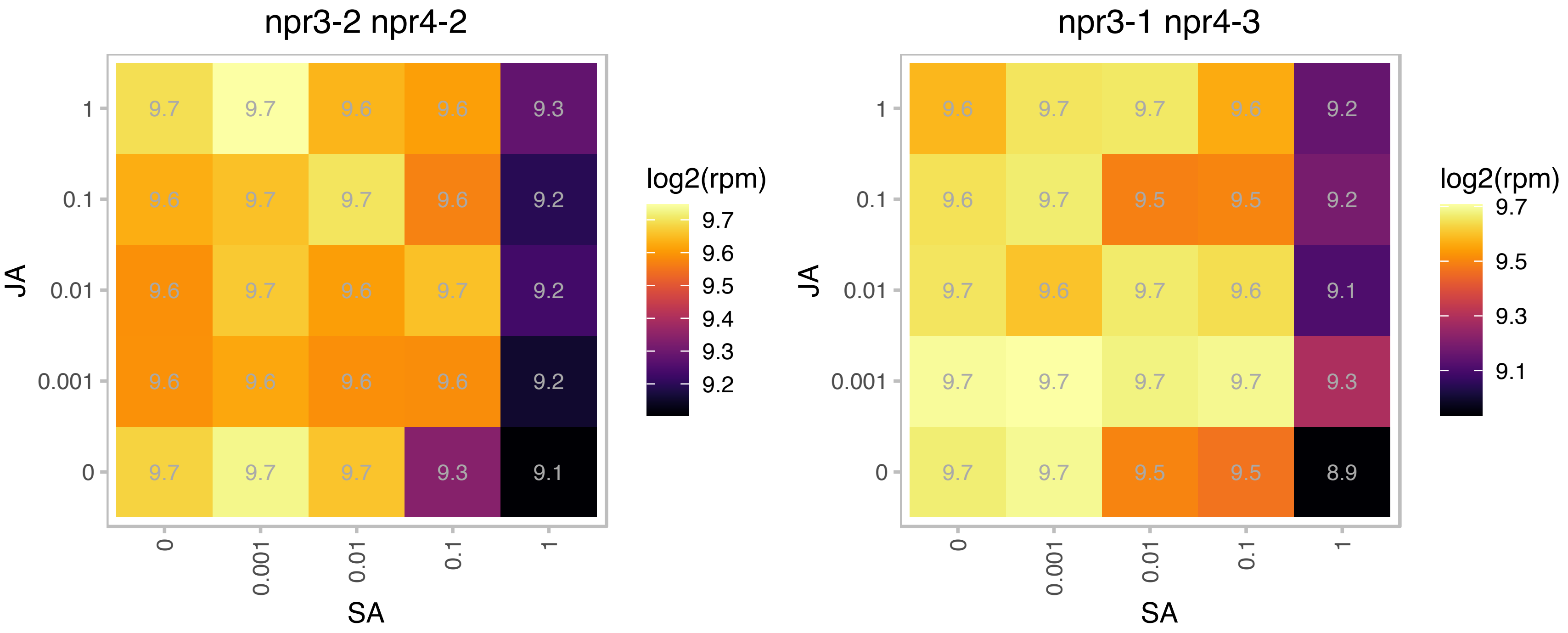

### AT5G61420.1 : AtMYB28, HAG1, MYB28, PMG1

myb domain protein 28

Encodes a nuclear localized member of the MYB transcription factor family. Involved in positive regulation of aliphatic glucosinolate biosynthesis.Expression is induced by touch, wounding and glucose.

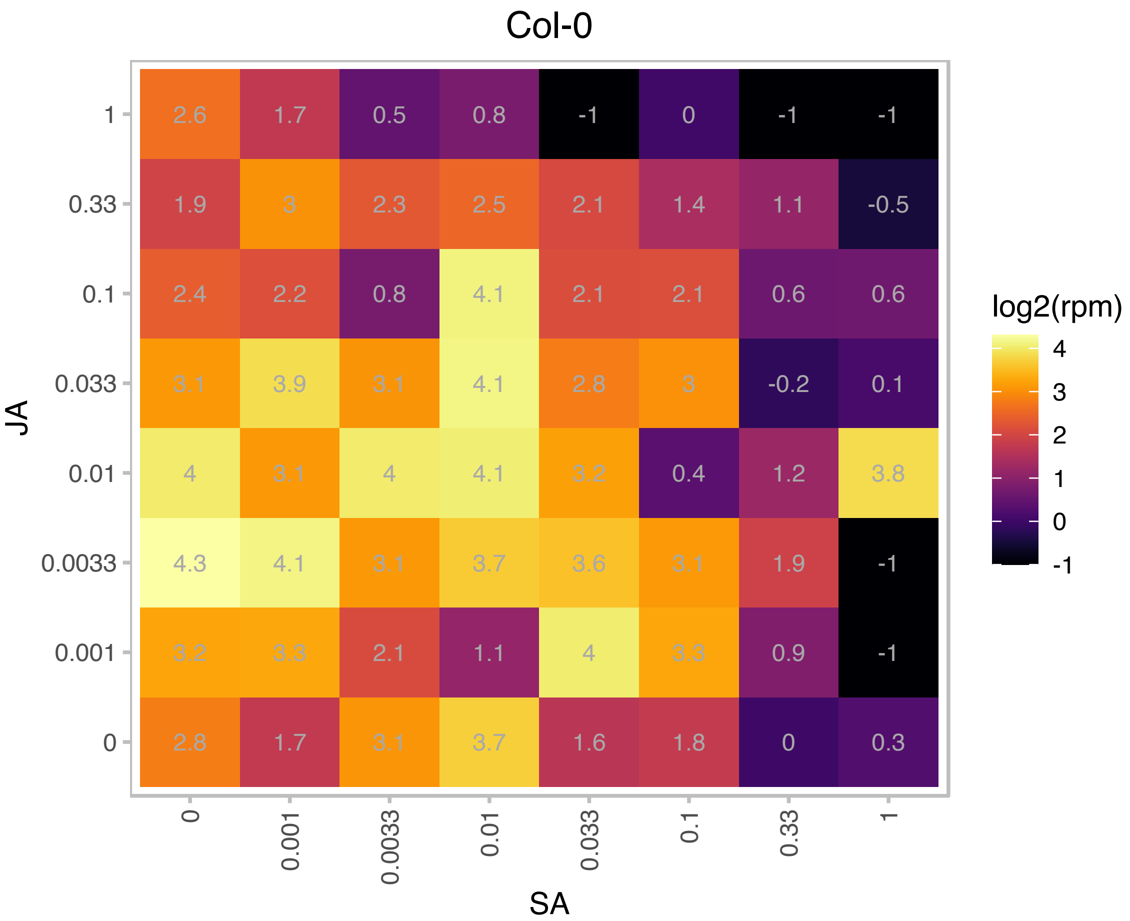


| | |
|---|---|
| Gene | AT5G61420 |
| is.expressed_genes | TRUE |
| is.responsive_genes | TRUE |
| is.apcluster | F-07 |

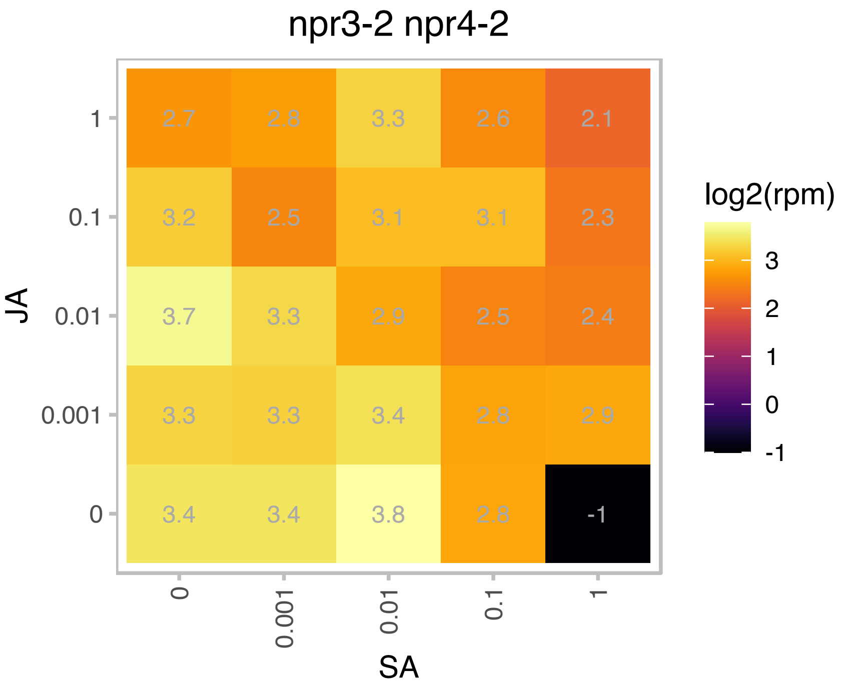


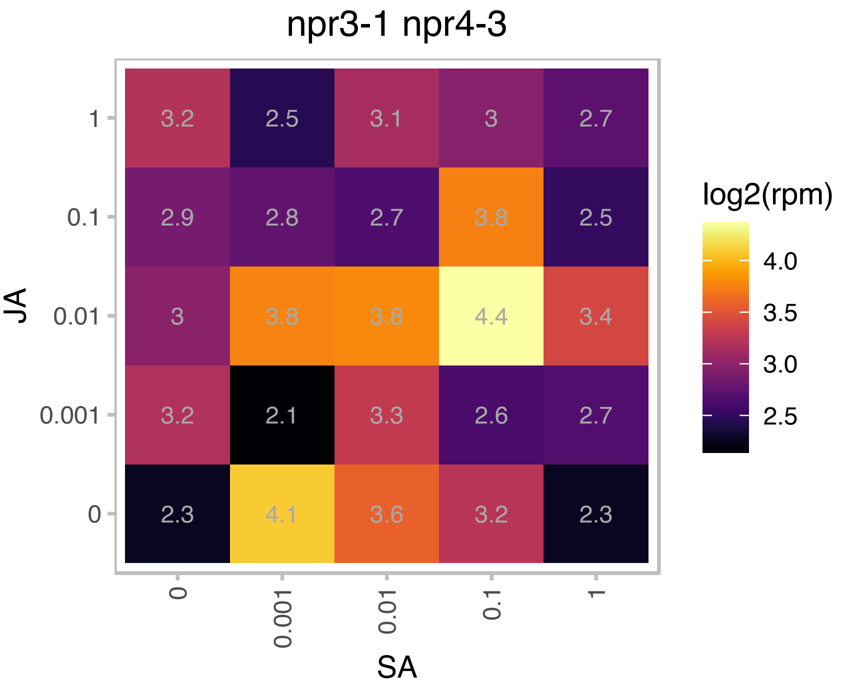

### AT5G61600.1 : ERF104

ethylene response factor 104
encodes a member of the ERF (ethylene response factor) subfamily B-3 of ERF/AP2 transcription factor family. The protein contains one AP2 domain. There are 18 members in this subfamily including ATERF-1, ATERF-2, AND ATERF-5.

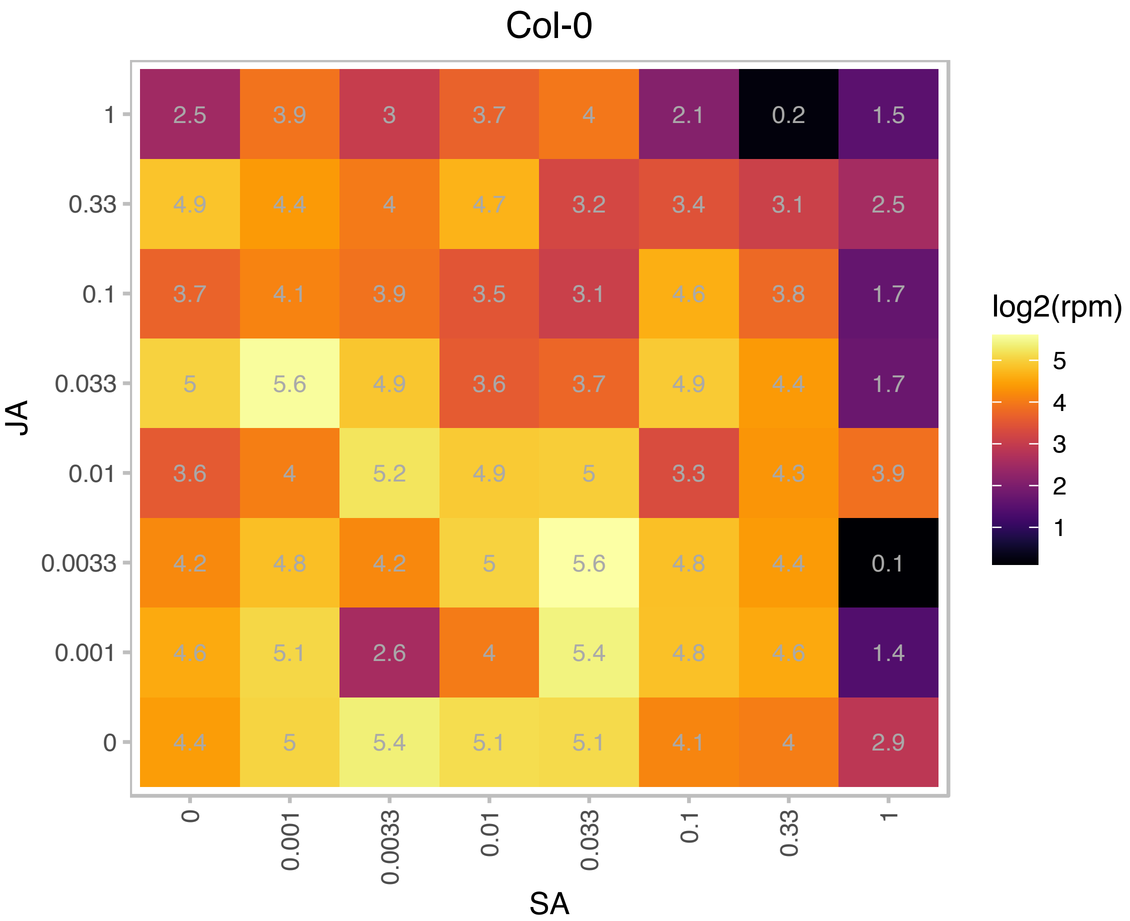


| | |
|---|---|
| Gene | AT5G61600 |
| is.expressed_genes | TRUE |
| is.responsive_genes | TRUE |
| is.apcluster | F-07 |

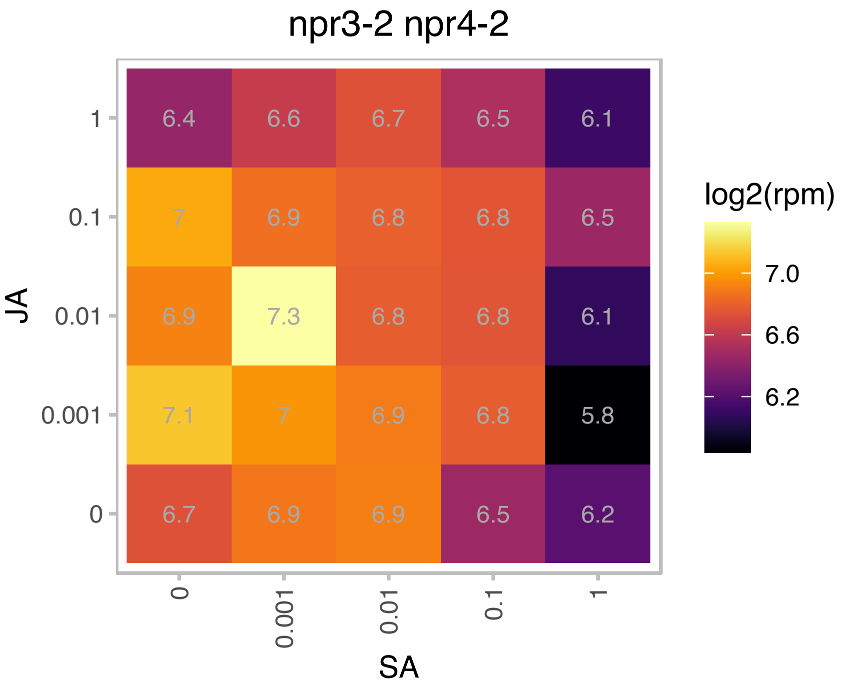


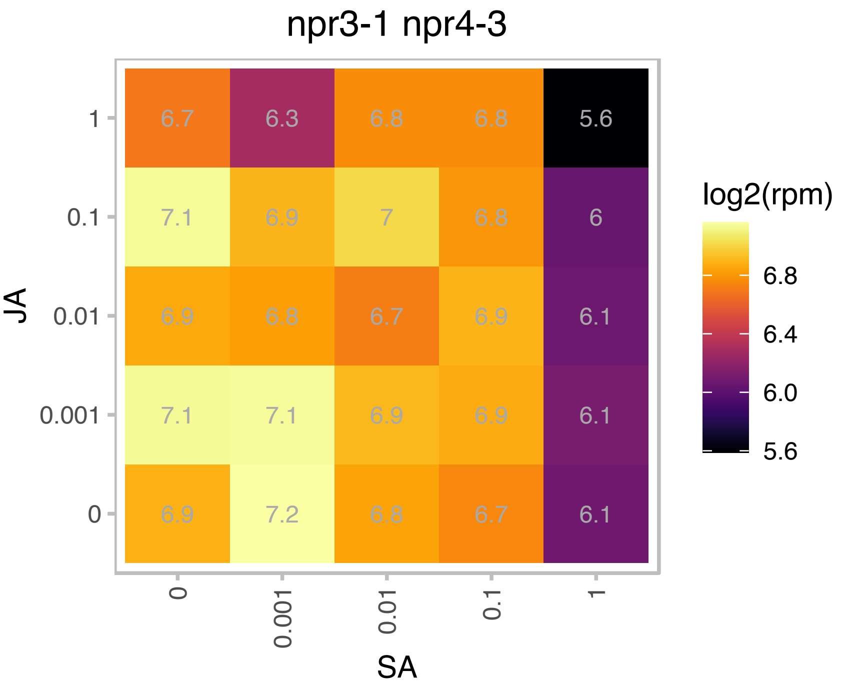

# AT5G62300.1 : NA

Ribosomal protein S10p/S20e family protein

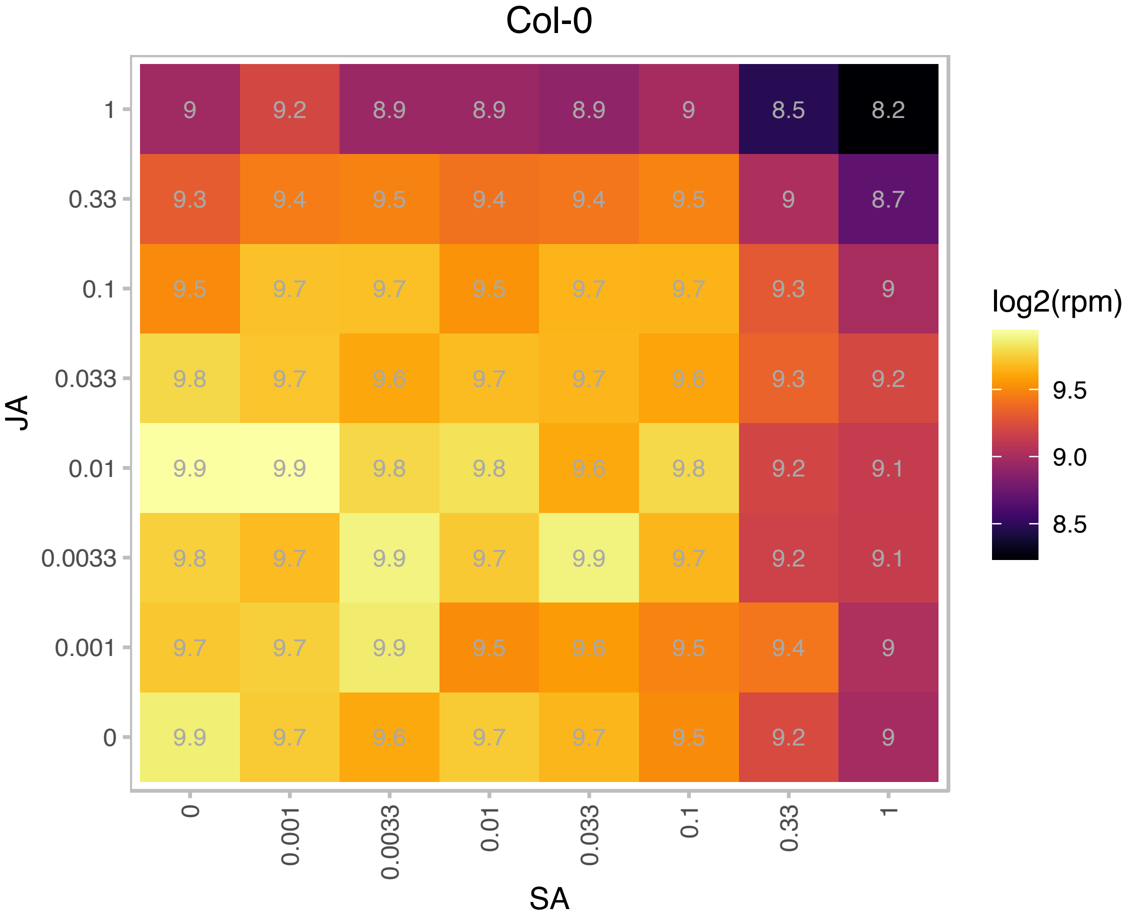


| | |
|---|---|
| Gene | AT5G62300 |
| is.expressed_genes | TRUE |
| is.responsive_genes | TRUE |
| is.apcluster | F-07 |

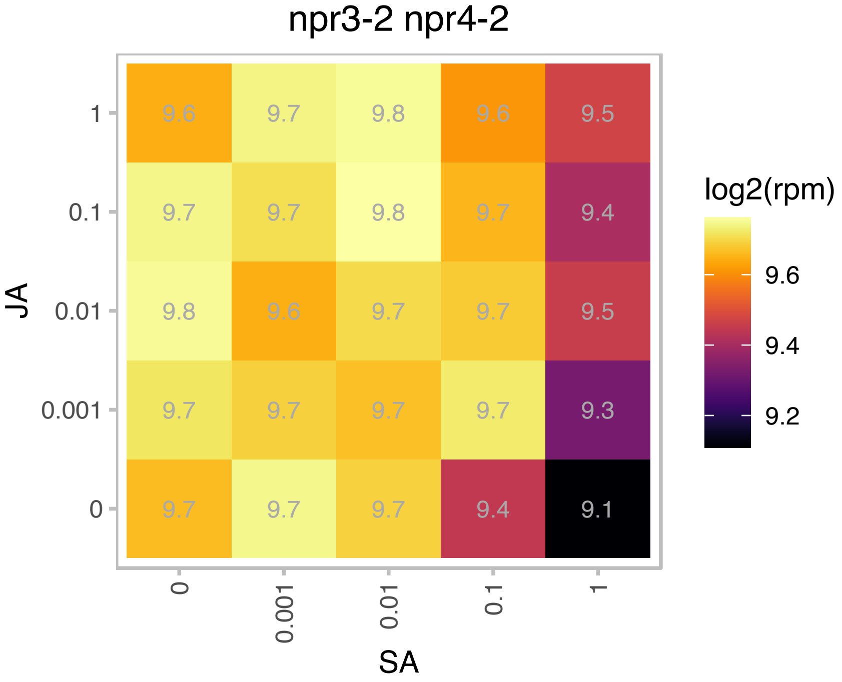


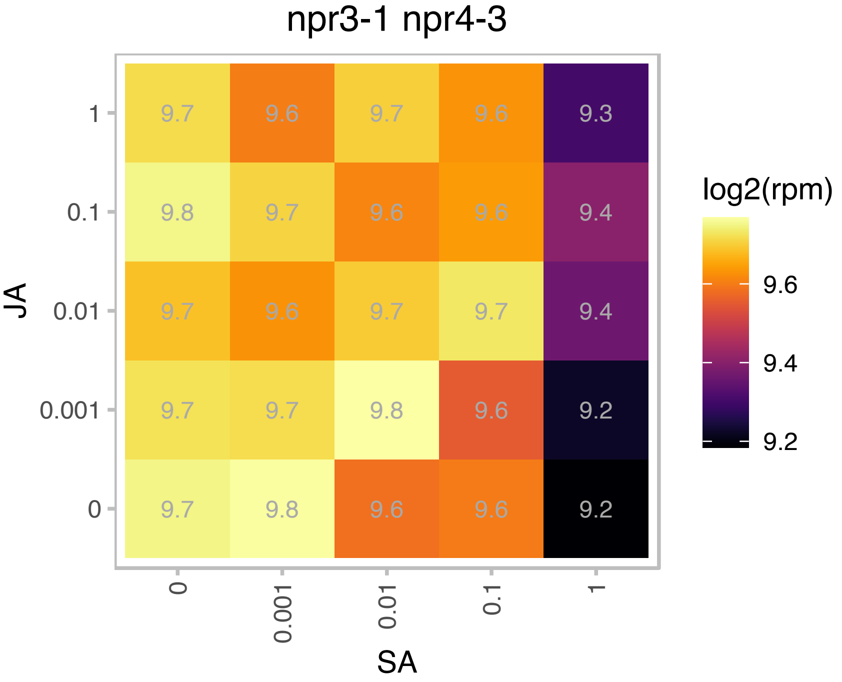

### AT5G62980.1 : FOLB2

Dihydroneopterin aldolase

Encodes an enzyme that can act as a aldolase or an epimerase for 7,8-dihydroneopterin and 7,8-dihydromonapterin in vitro. It is likely to act in folate biosynthesis as a homooctamer in vivo.

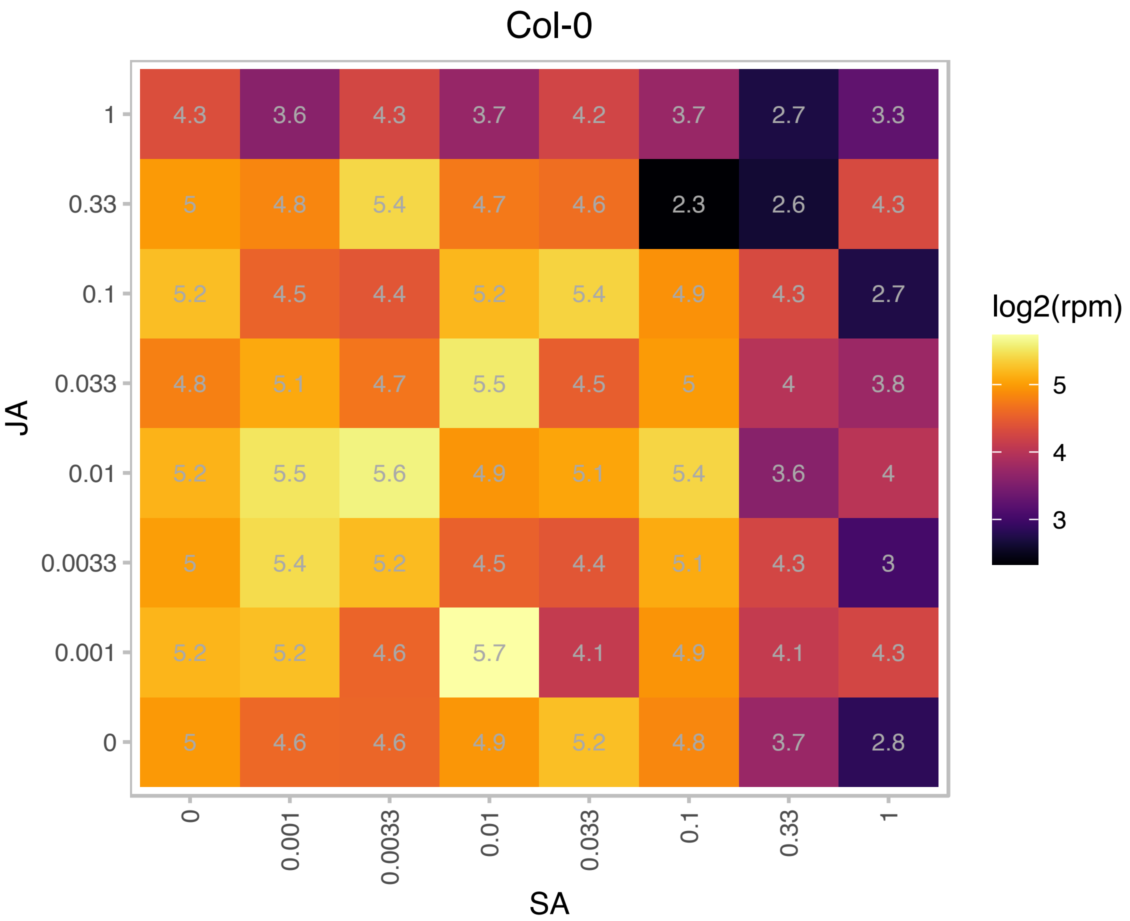


| | |
|---|---|
| Gene | AT5G62980 |
| is.expressed_genes | TRUE |
| is.responsive_genes | TRUE |
| is.apcluster | F-07 |

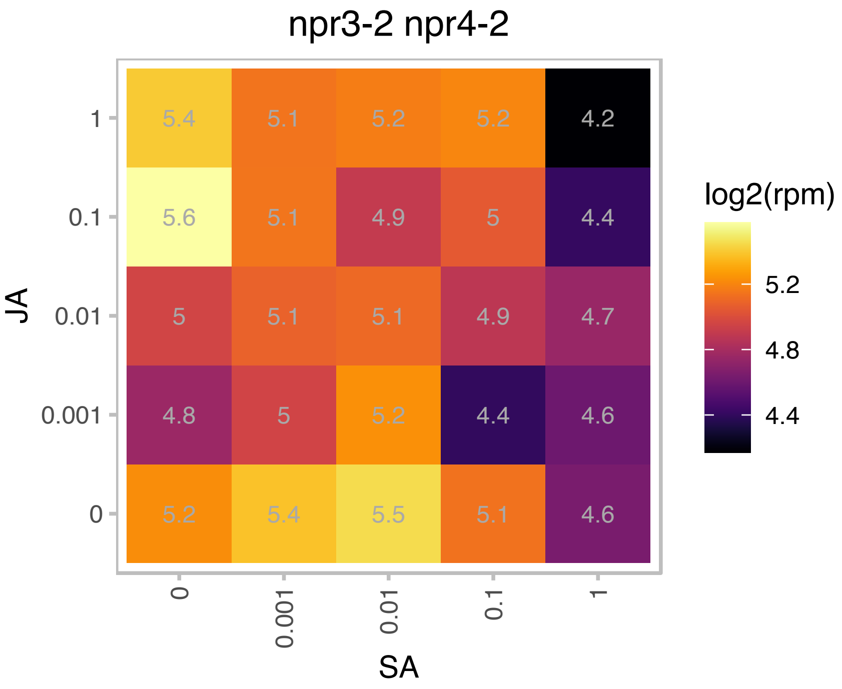


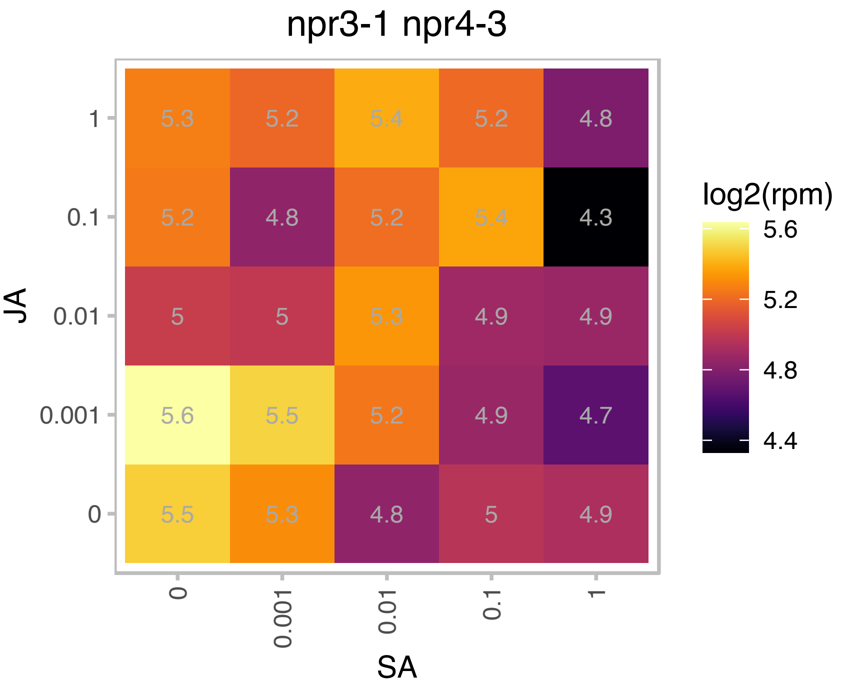

# AT5G63470.1 : NF-YC4

nuclear factor Y, subunit C4
Encodes a member of class of transcription regulators that his highly conserved across many plant species.In Arabidopsis and rice, NF-YC4 interacts with other members of this class and CO to regulate flowering. In Arabidopsis, it interacts with QQS to regulate C/N partitioning.

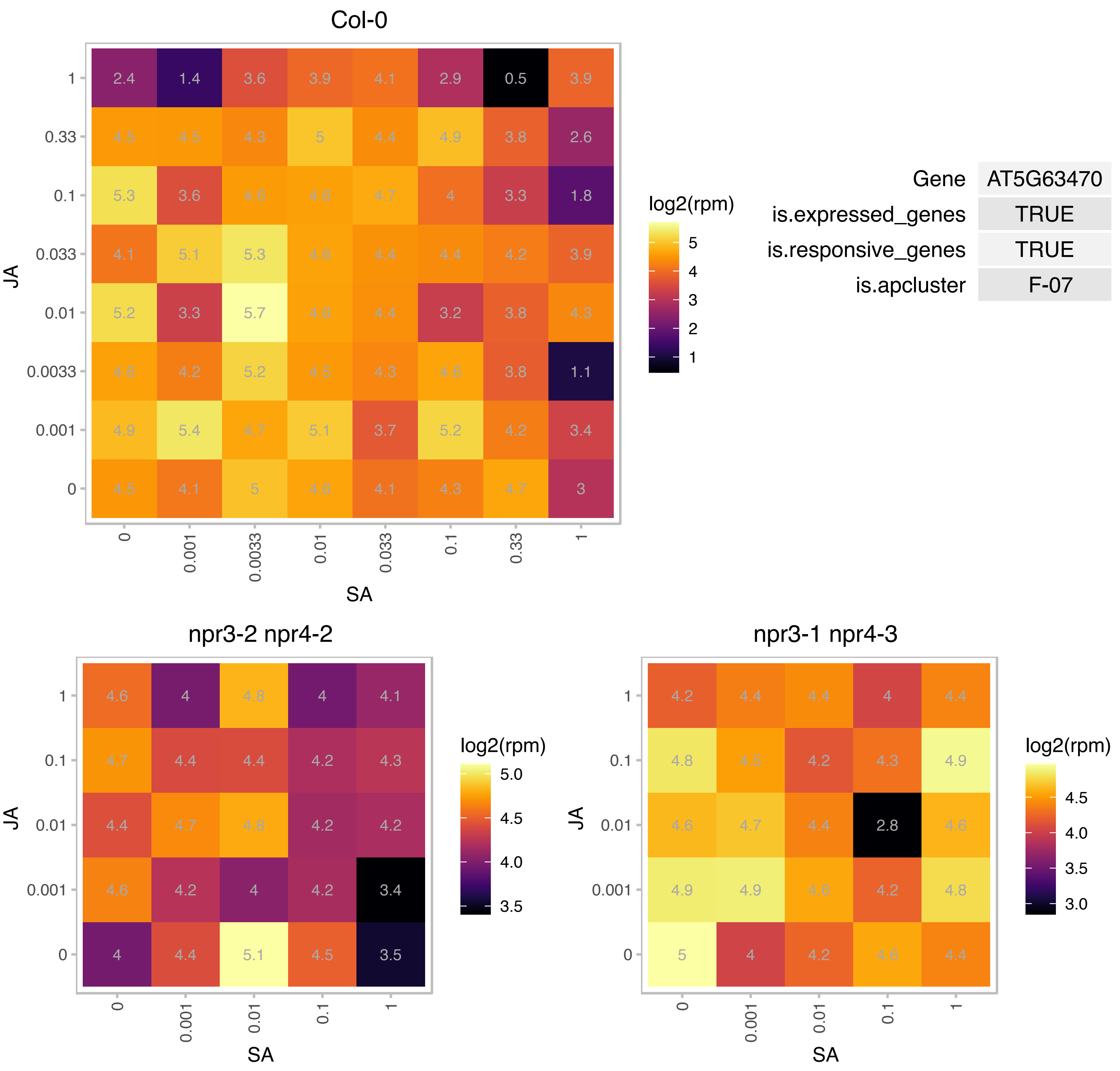


| | |
|---|---|
| Gene | AT5G63470 |
| is.expressed_genes | TRUE |
| is.responsive_genes | TRUE |
| is.apcluster | F-07 |

### AT5G64140.1 : RPS28

ribosomal protein S28

Encodes a putative ribosomal protein S28.

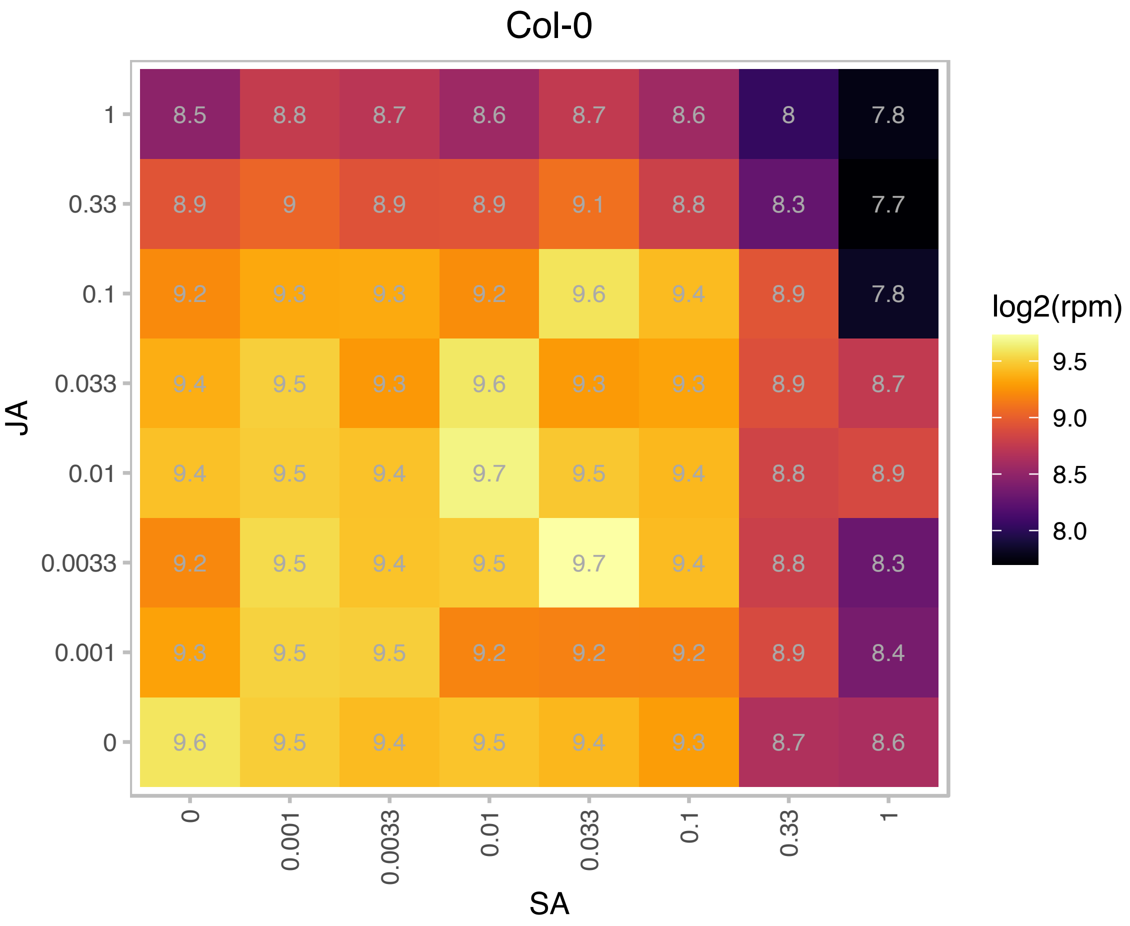


| Gene | AT5G64140 |
|---|---|
| is.expressed_genes | TRUE |
| is.responsive_genes | TRUE |
| is.apcluster | F-07 |

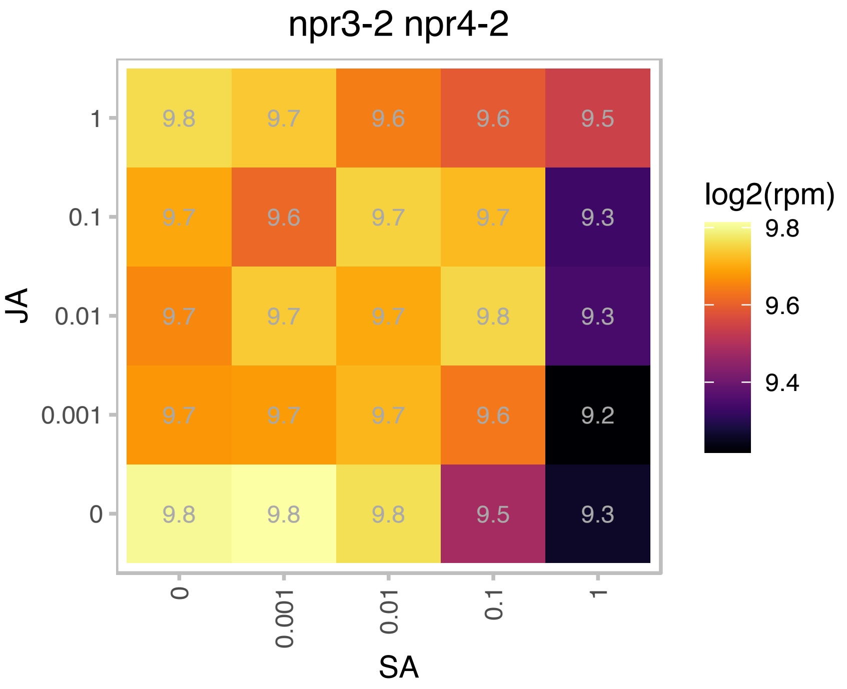


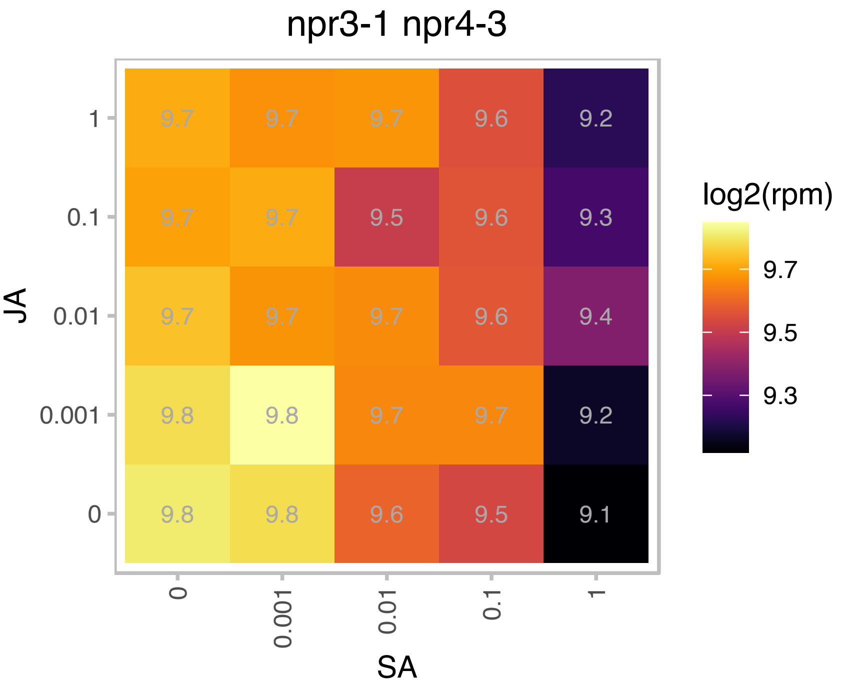

### AT5G64570.1 : ATBXL4, XYL4

beta-D-xylosidase 4

Encodes a beta-d-xylosidase that belongs to family 3 of glycoside hydrolases.

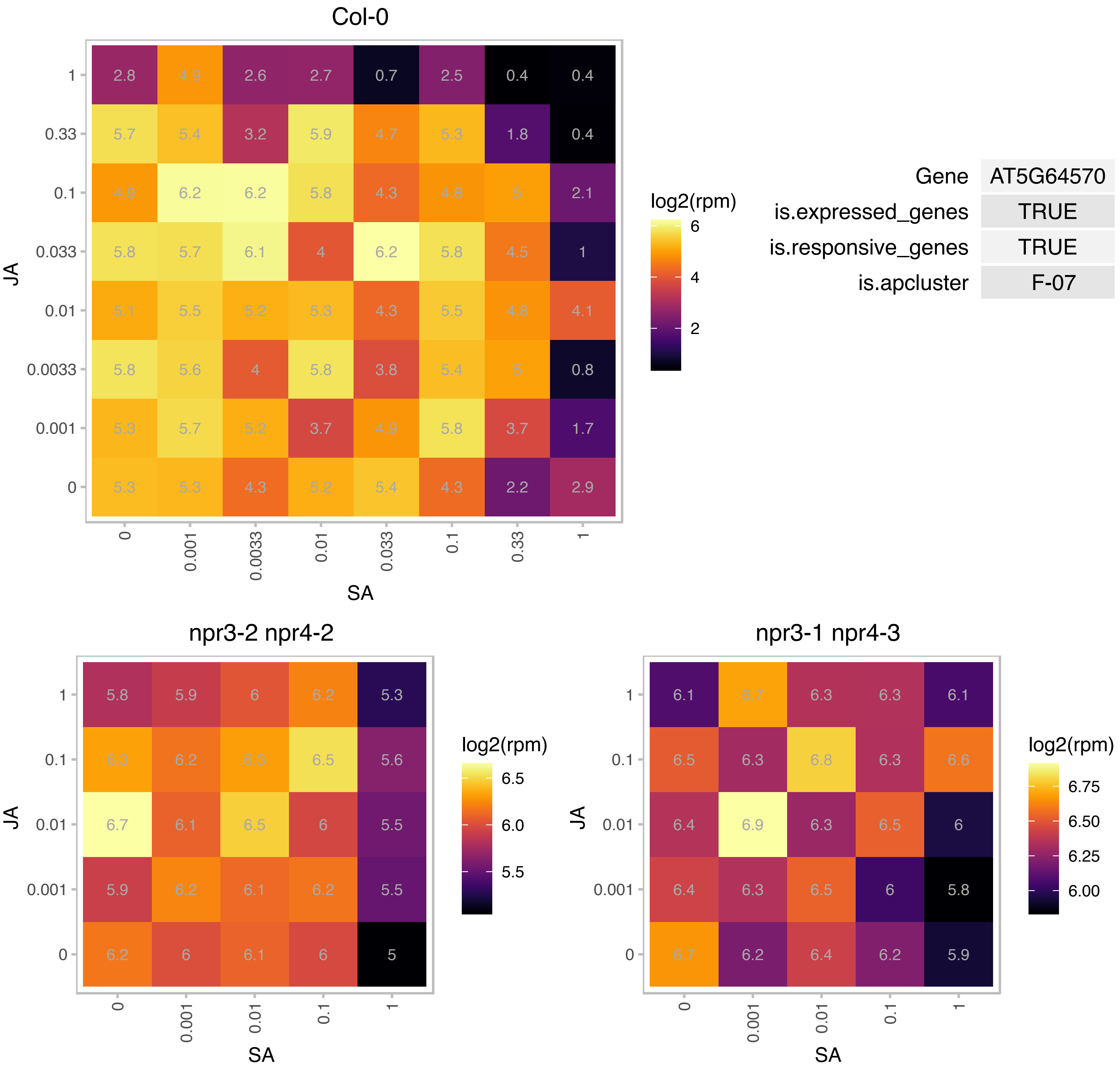


| | |
|---|---|
| Gene | AT5G64570 |
| is.expressed_genes | TRUE |
| is.responsive_genes | TRUE |
| is.apcluster | F-07 |

# AT5G64670.1 : NA

Ribosomal protein L18e/L15 superfamily protein

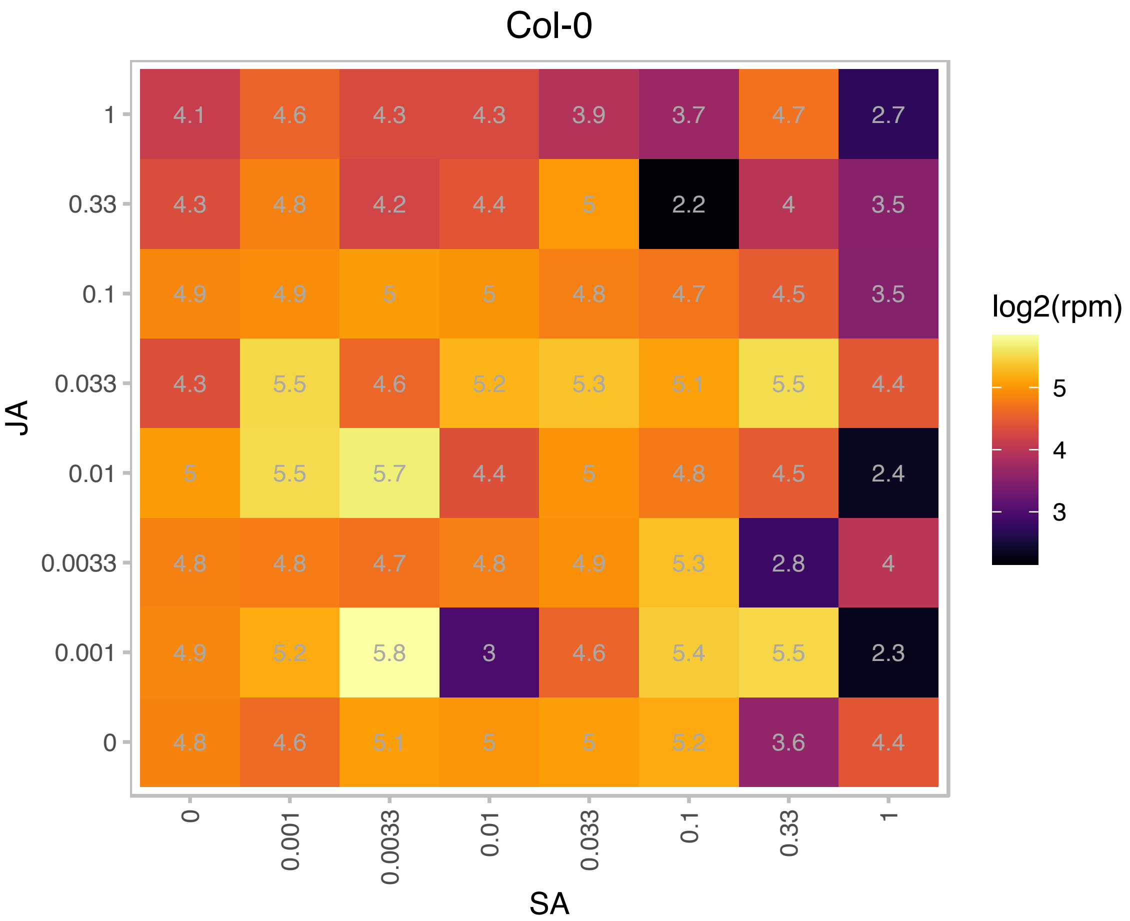


| | |
|---|---|
| Gene | AT5G64670 |
| is.expressed_genes | TRUE |
| is.responsive_genes | TRUE |
| is.apcluster | F-07 |

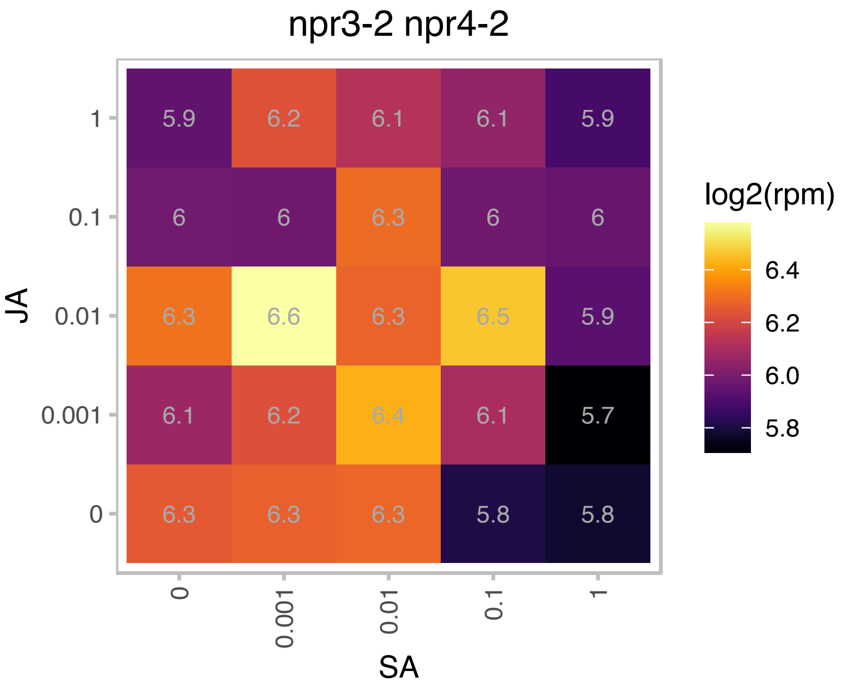


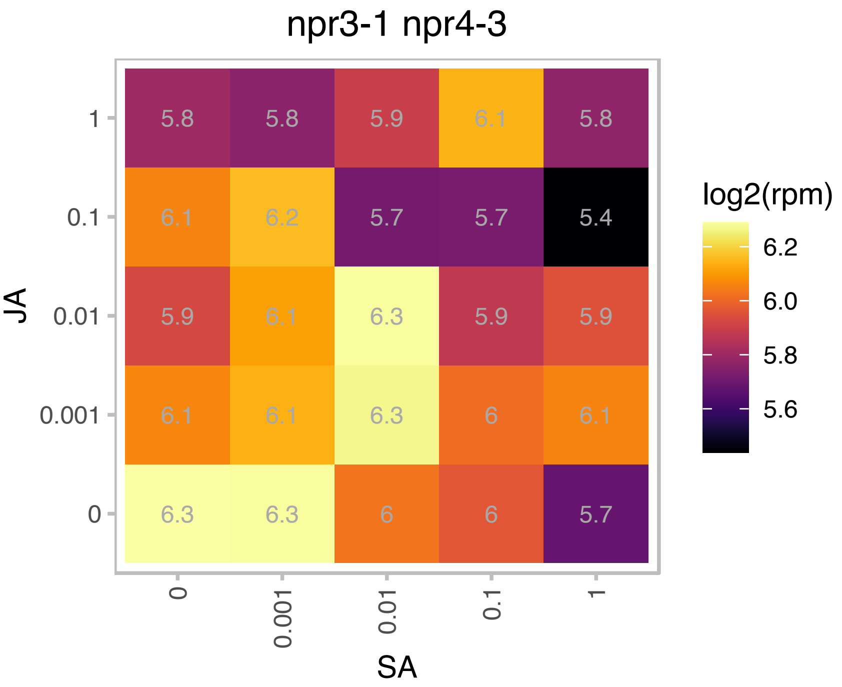

### AT5G65670.1 : IAA9

indole-3-acetic acid inducible 9

auxin (indole-3-acetic acid) induced gene The mRNA is cell-to-cell mobile.

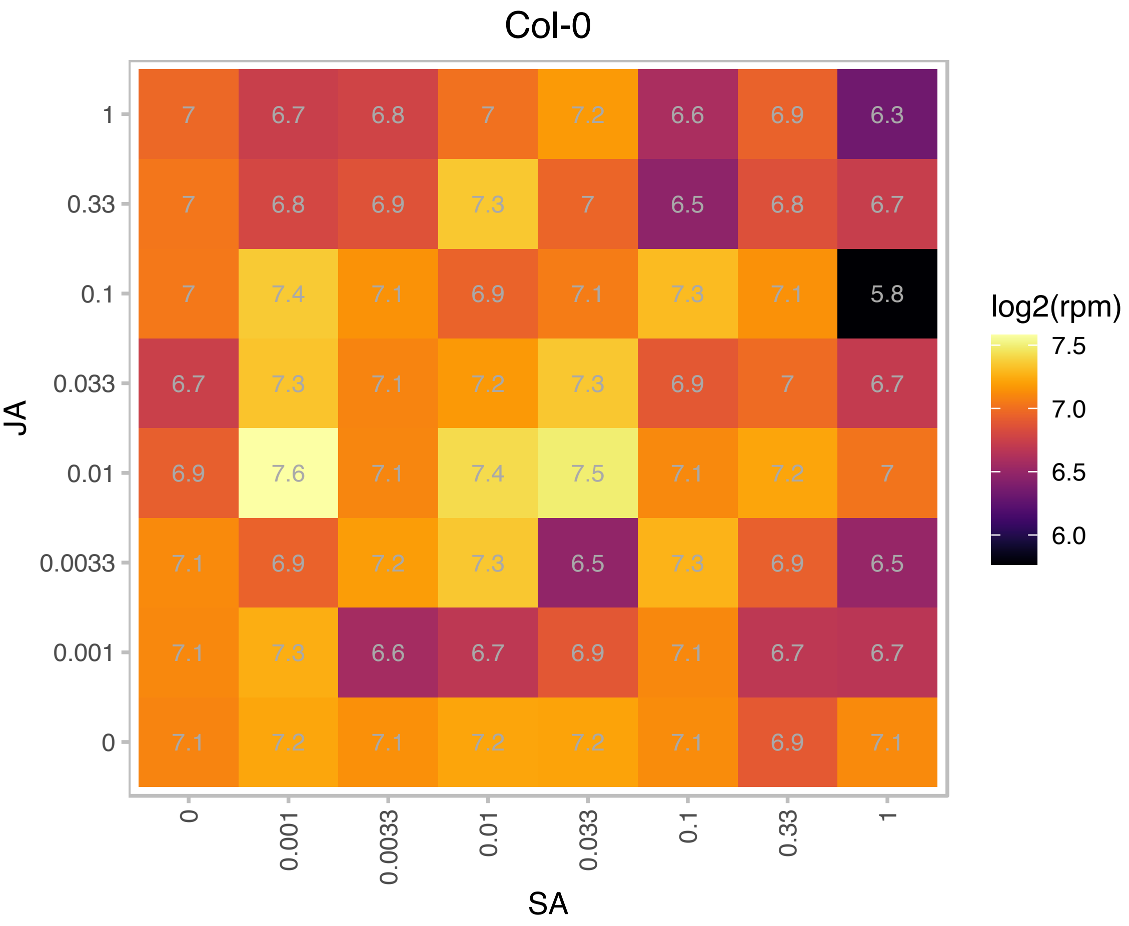


| | |
|---|---|
| Gene | AT5G65670 |
| is.expressed_genes | TRUE |
| is.responsive_genes | TRUE |
| is.apcluster | F-07 |

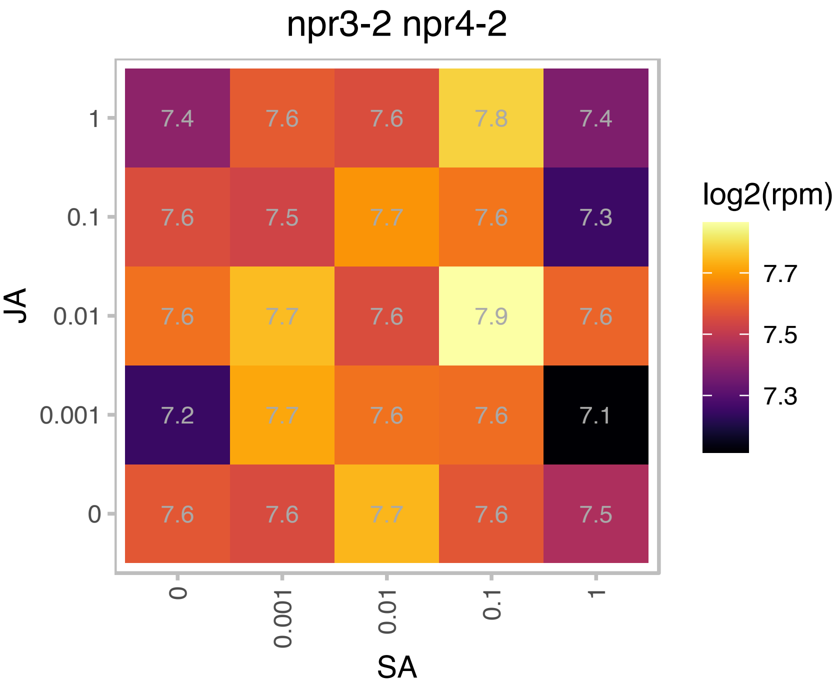


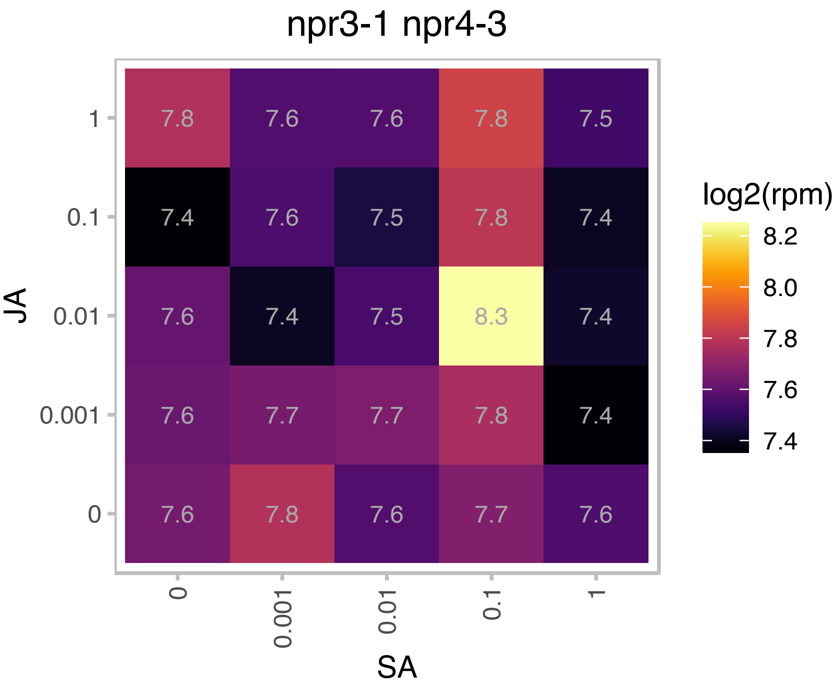

### AT5G67070.1 : RALF34, RALFL34

ralf-like 34

Member of a diversely expressed predicted peptide family showing sequence similarity to tobacco Rapid Alkalinization Factor (RALF), and is believed to play an essential role in the physiology of Arabidopsis. Consists of a single exon and is characterized by a conserved C-terminal motif and N-terminal signal peptide. The mRNA is cell-to-cell mobile.

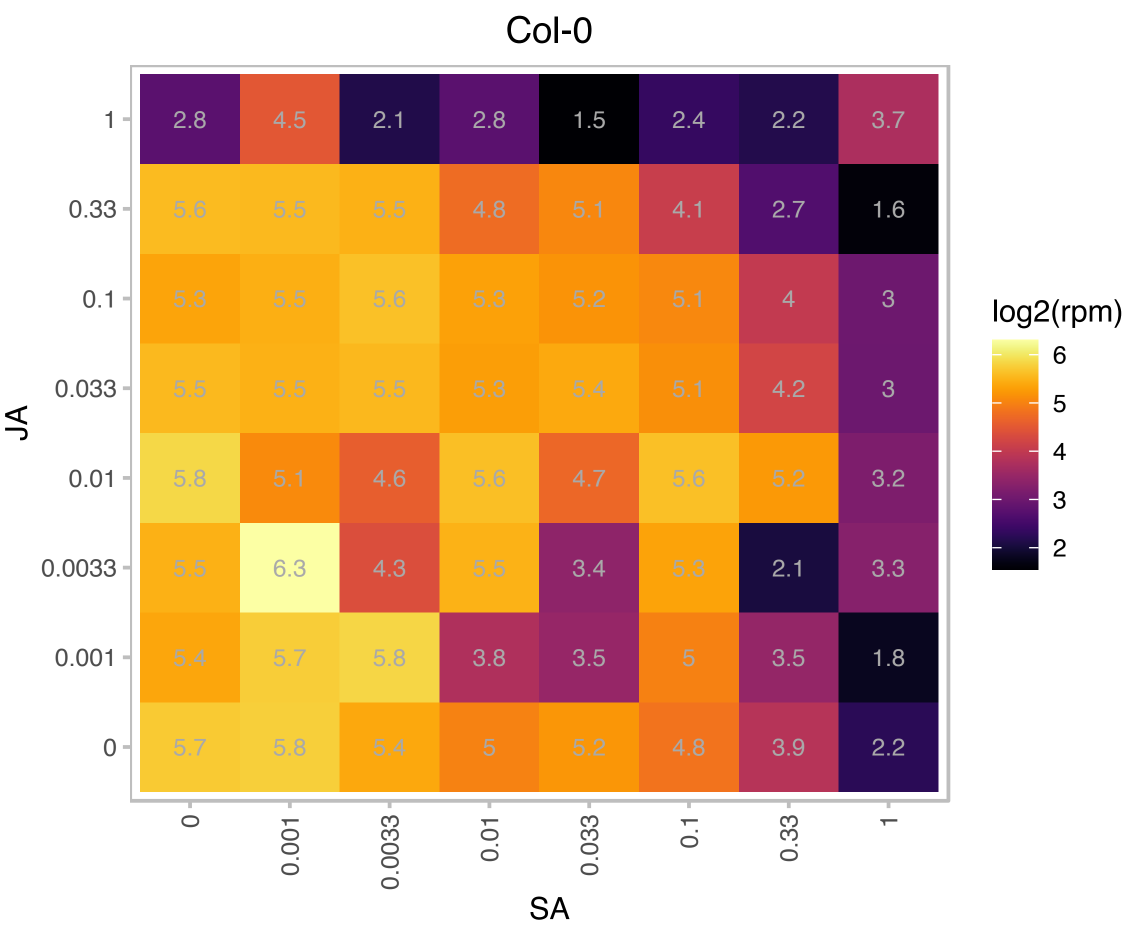


| | |
|---|---|
| Gene | AT5G67070 |
| is.expressed_genes | TRUE |
| is.responsive_genes | TRUE |
| is.apcluster | F-07 |

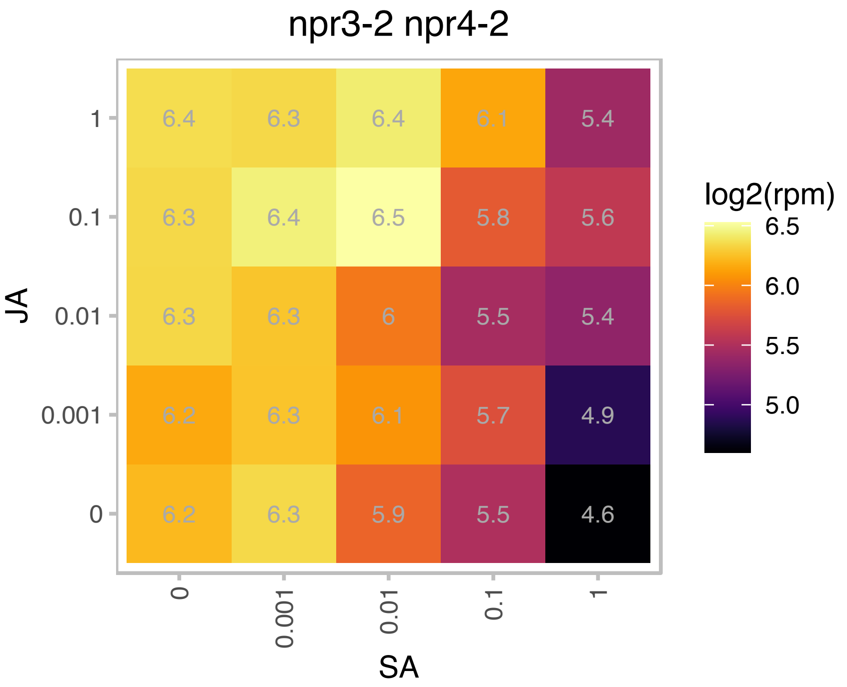


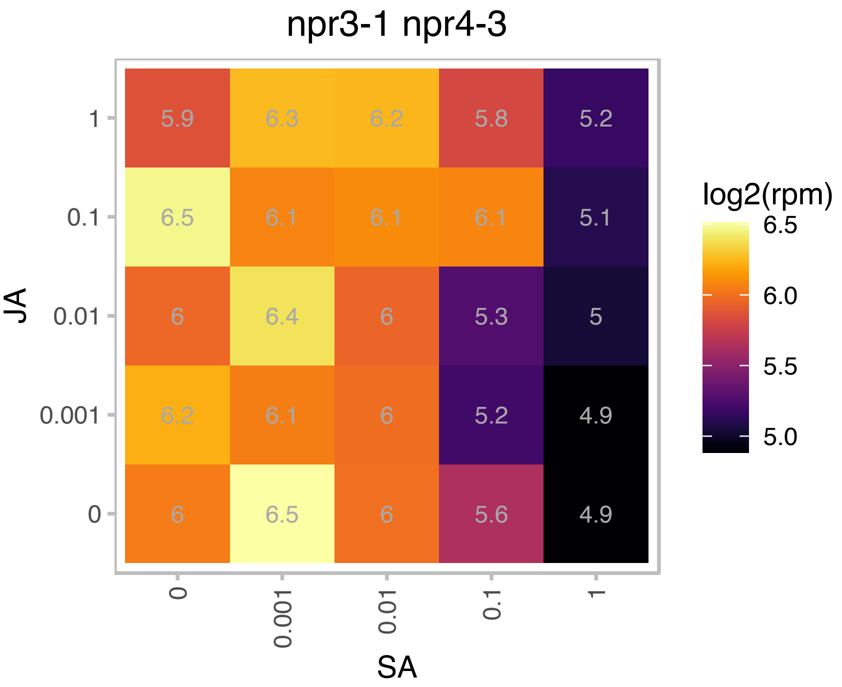

# AT1G02160.1 : NA

Cox19 family protein (CHCH motif)

Col-0

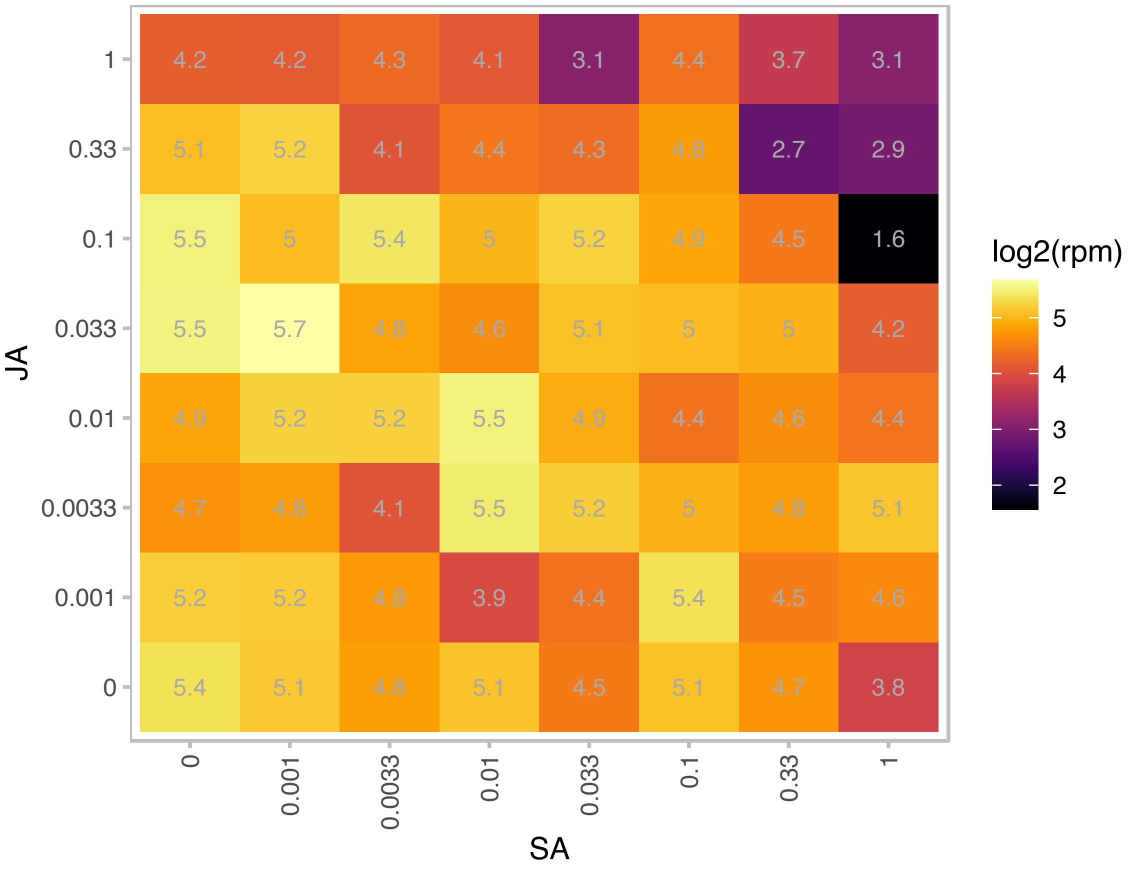


| | |
|---|---|
| Gene | AT1G02160 |
| is.expressed_genes | TRUE |
| is.responsive_genes | TRUE |
| is.apcluster | F-08 |

npr3-2 npr4-2

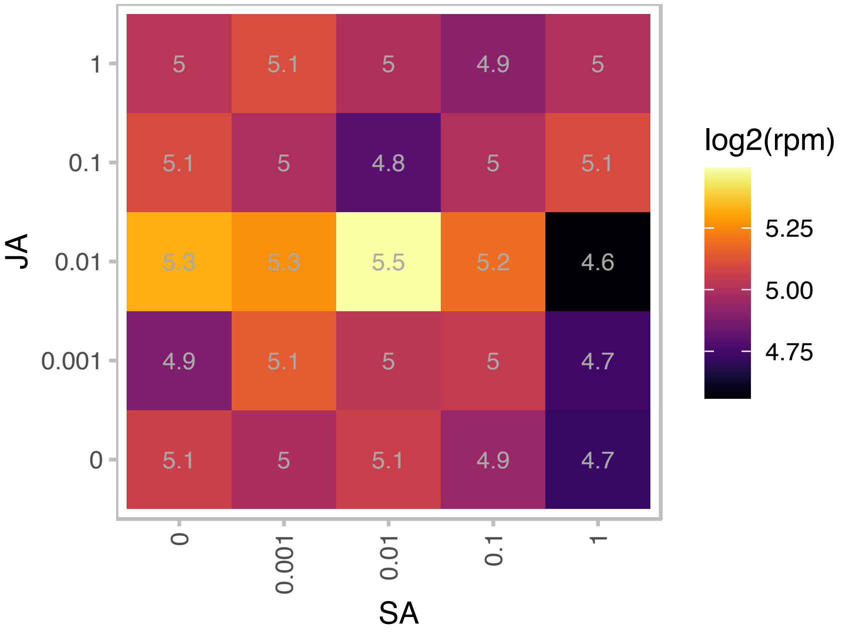


npr3-1 npr4-3

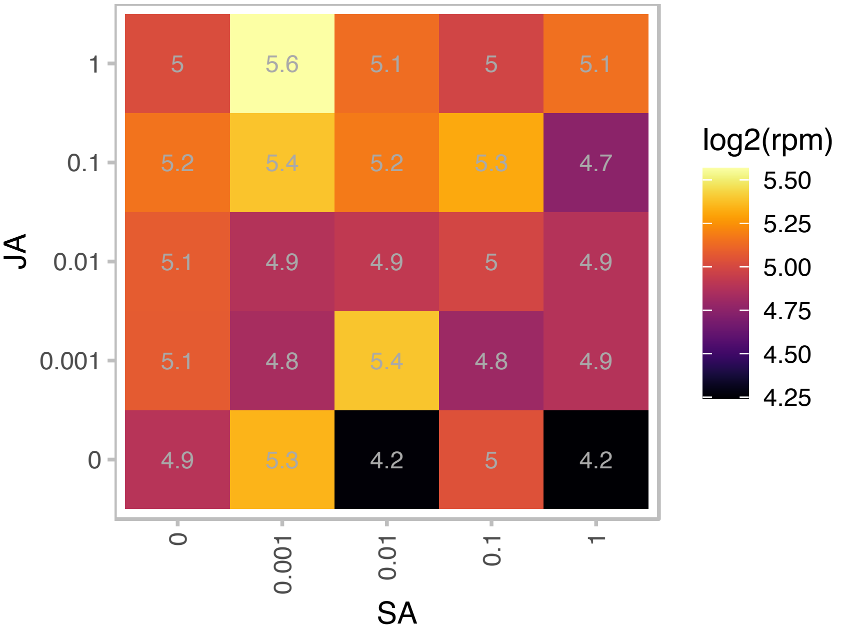

### AT1G02740.1 : MRG2

MRG family protein

| | |
|---|---|
| Gene | AT1G02740 |
| is.expressed_genes | TRUE |
| is.responsive_genes | TRUE |
| is.apcluster | F-08 |

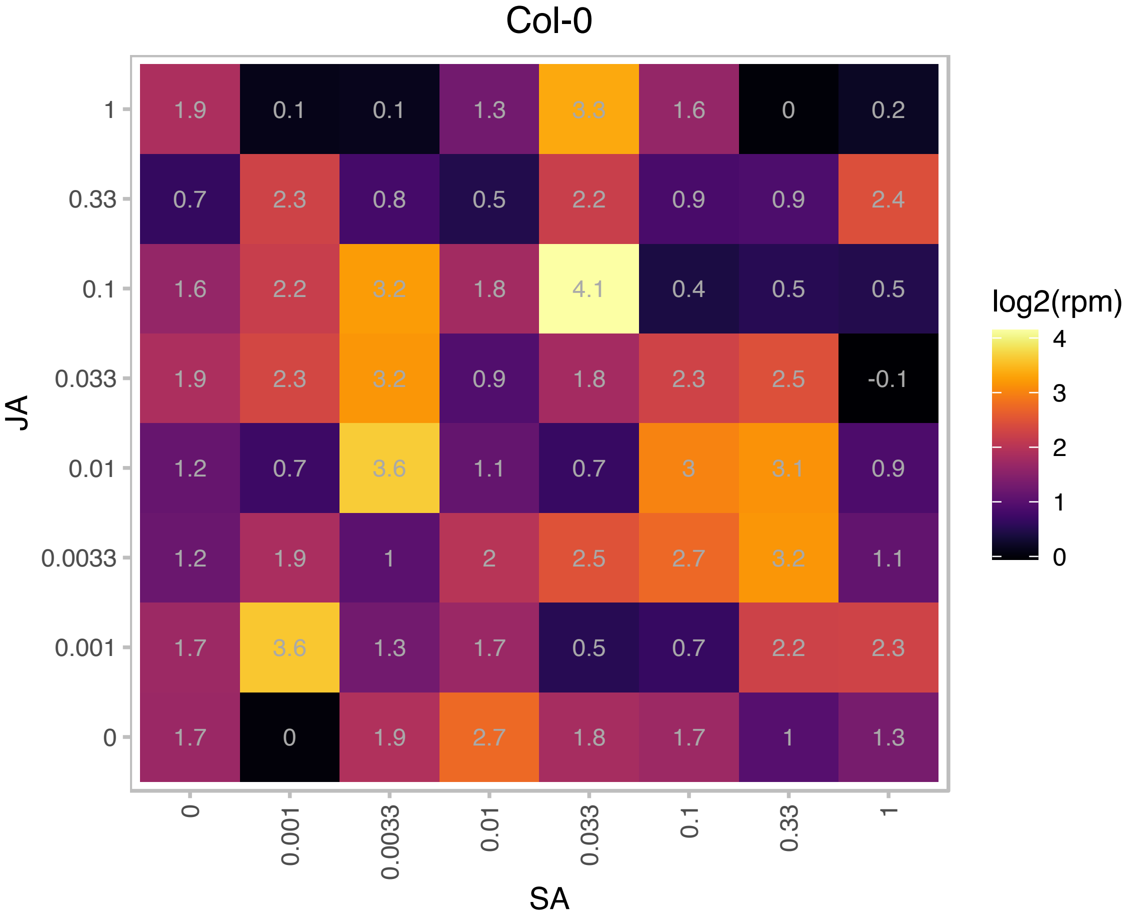


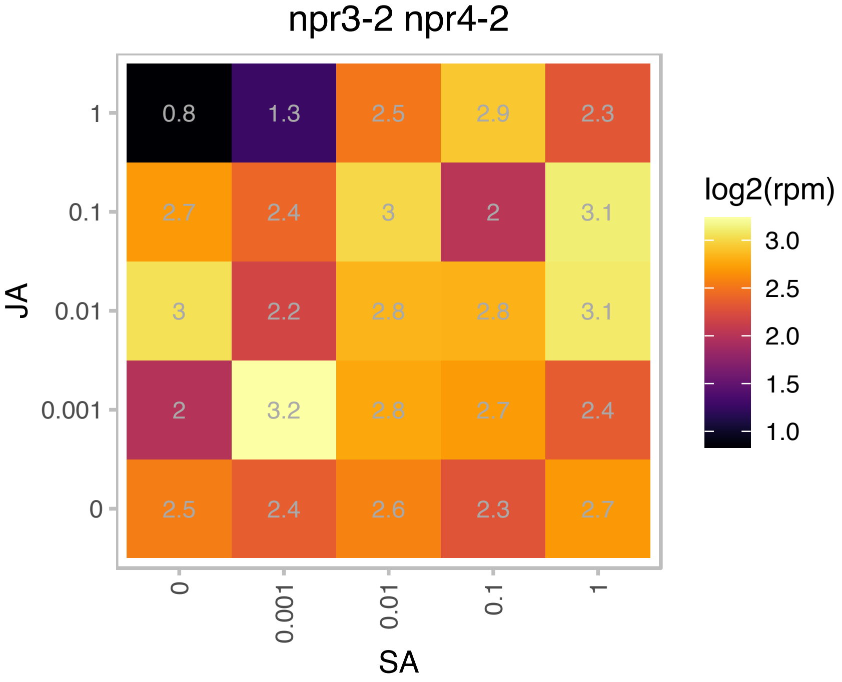


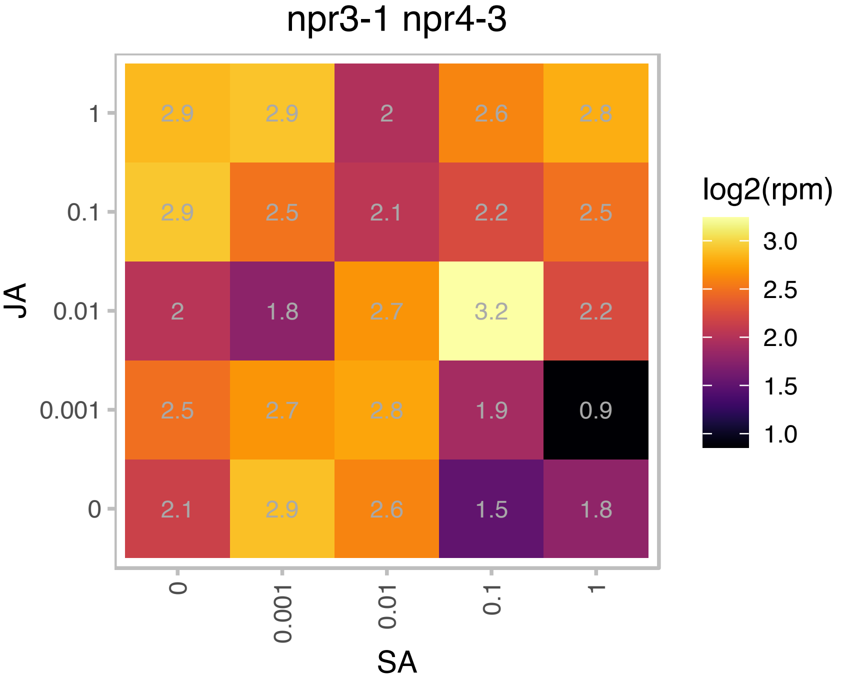

### AT1G03130.1 : PSAD-2

photosystem I subunit D-2

Encodes a protein predicted by sequence similarity with spinach PsaD to be photosystem I reaction center subunit II (PsaD2)

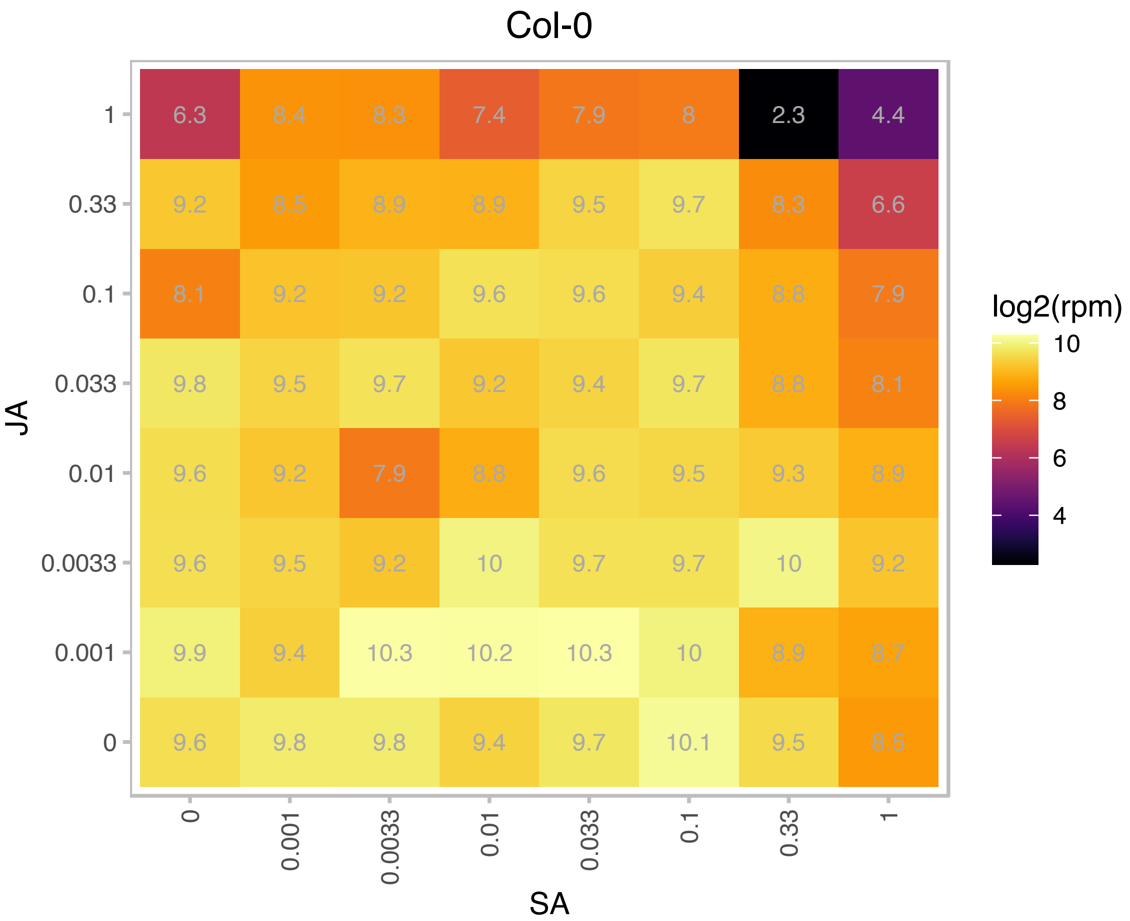


| | |
|---|---|
| Gene | AT1G03130 |
| is.expressed_genes | TRUE |
| is.responsive_genes | TRUE |
| is.apcluster | F-08 |

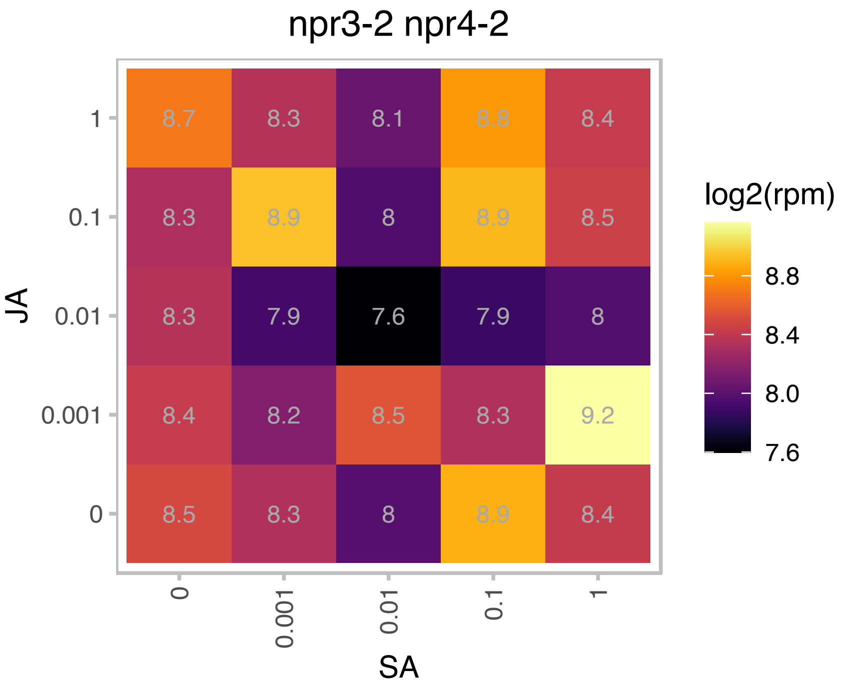


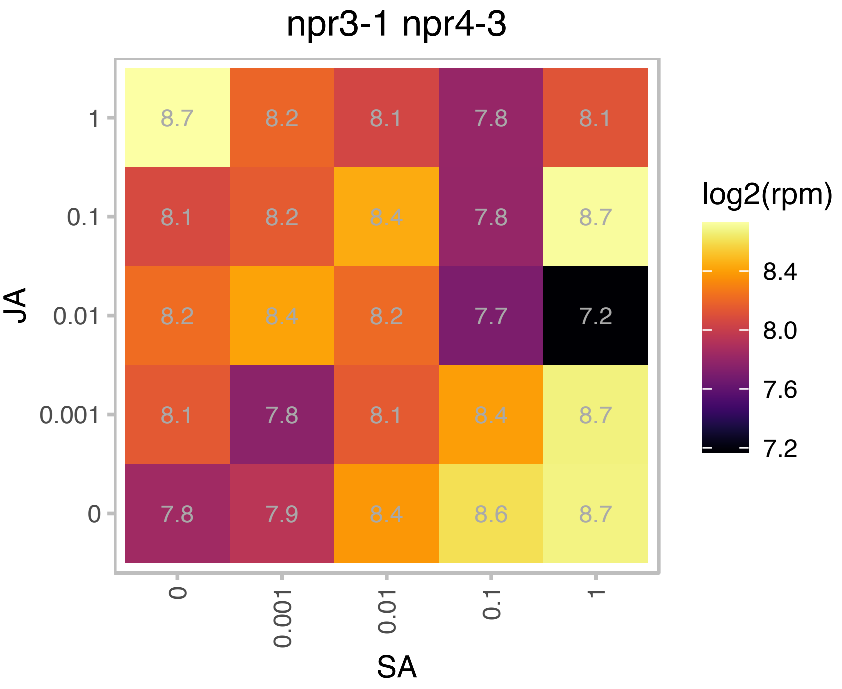

### AT1G04070.1 : ATTOM22-I, TOM22-I

translocase of outer membrane 22-I

Subunit of the TOM complex, a translocase in the outer mitochondrial membrane that selectively allows proteins with a mitochondrial targeting sequence to enter the mitochondrion.

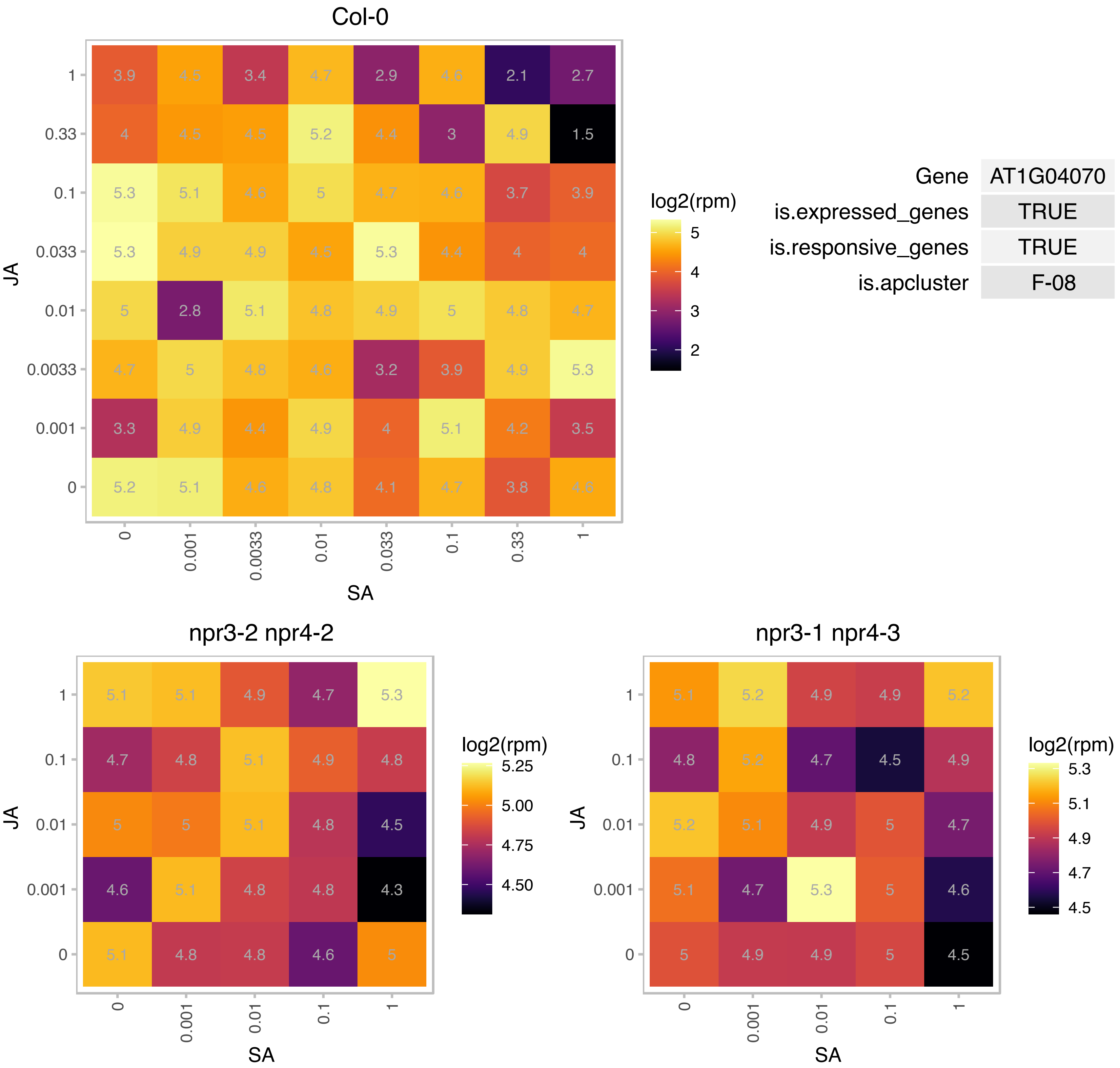


| | |
|---|---|
| Gene | AT1G04070 |
| is.expressed_genes | TRUE |
| is.responsive_genes | TRUE |
| is.apcluster | F-08 |

### AT1G04270.1 : RPS15

cytosolic ribosomal protein S15

Encodes cytosolic ribosomal protein S15.

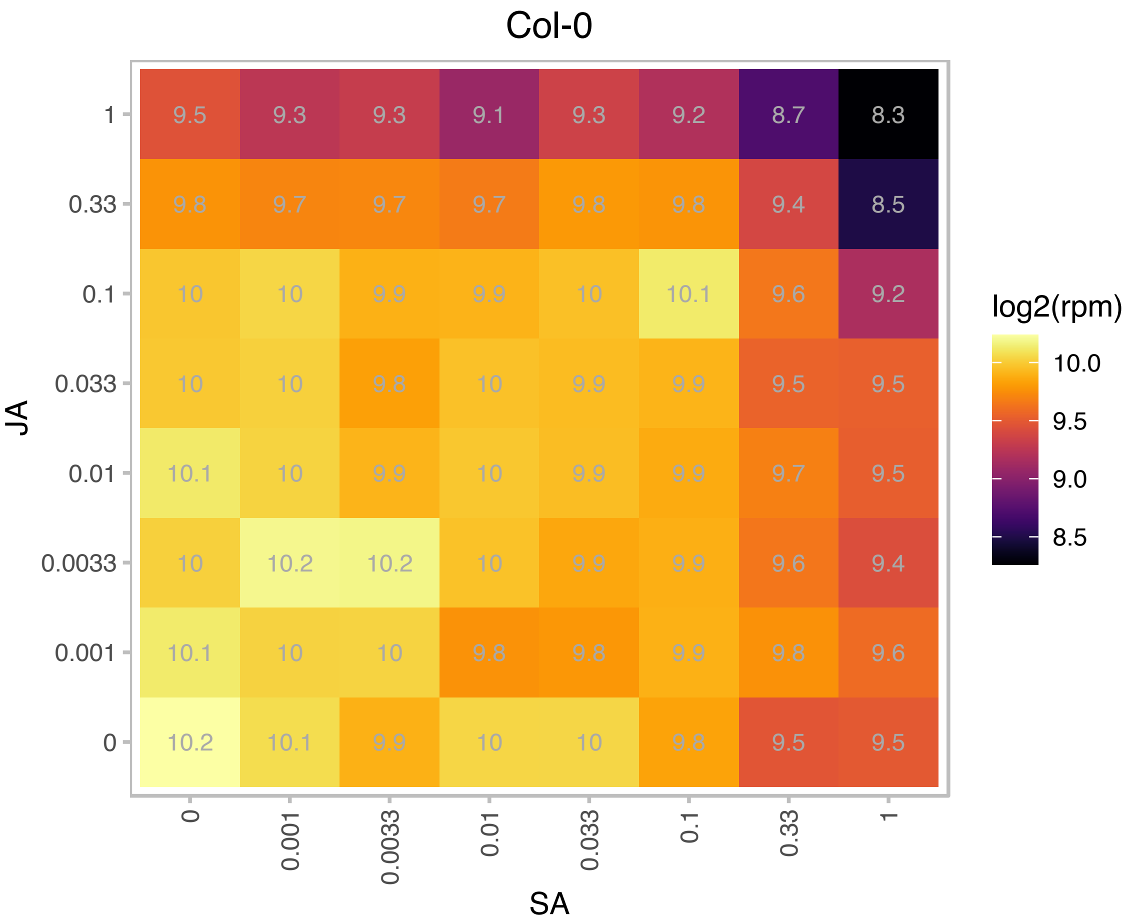


| | |
|---|---|
| Gene | AT1G04270 |
| is.expressed_genes | TRUE |
| is.responsive_genes | TRUE |
| is.apcluster | F-08 |

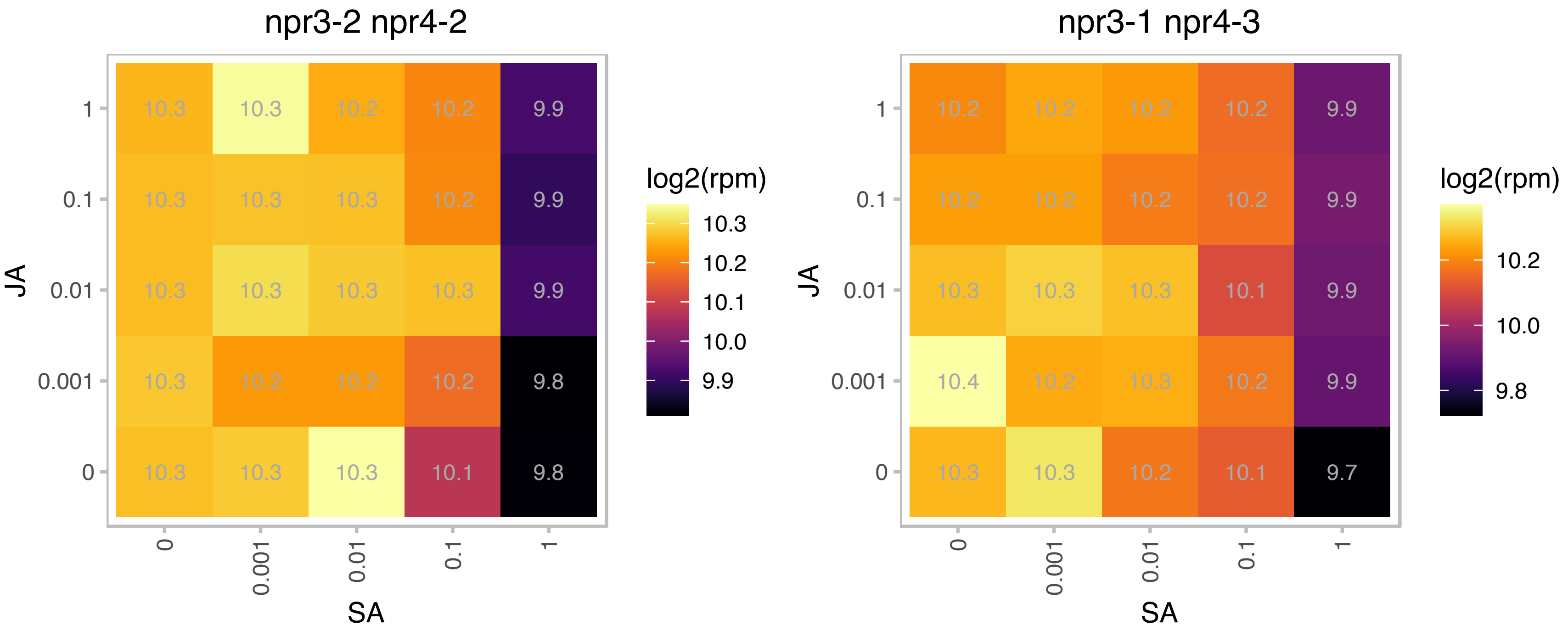

# AT1G04503.1 : NA

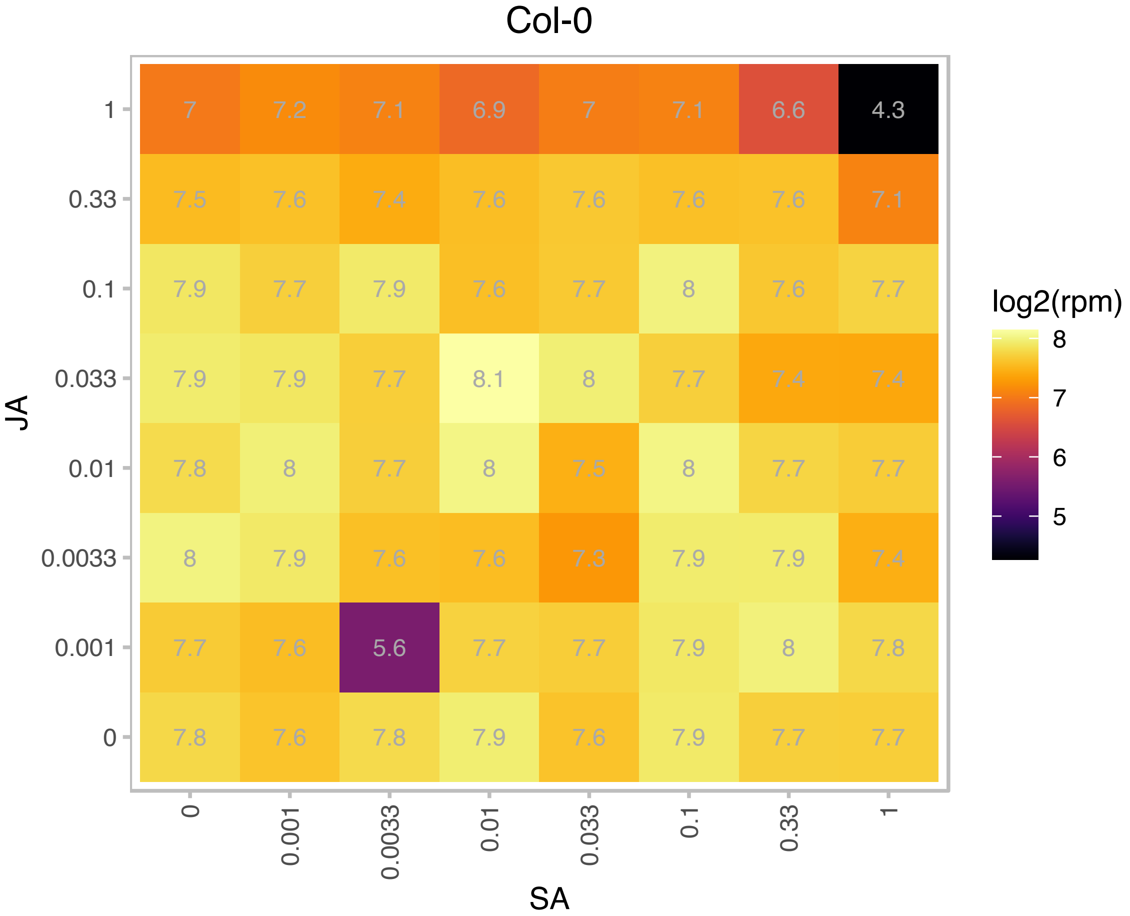


| | |
|---|---|
| Gene | AT1G04503 |
| is.expressed_genes | TRUE |
| is.responsive_genes | TRUE |
| is.apcluster | F-08 |

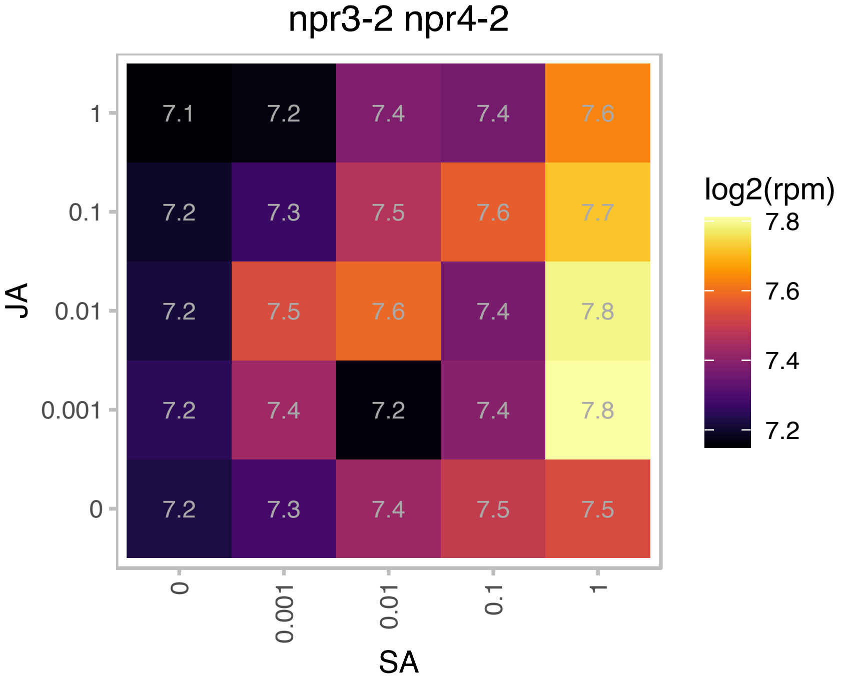


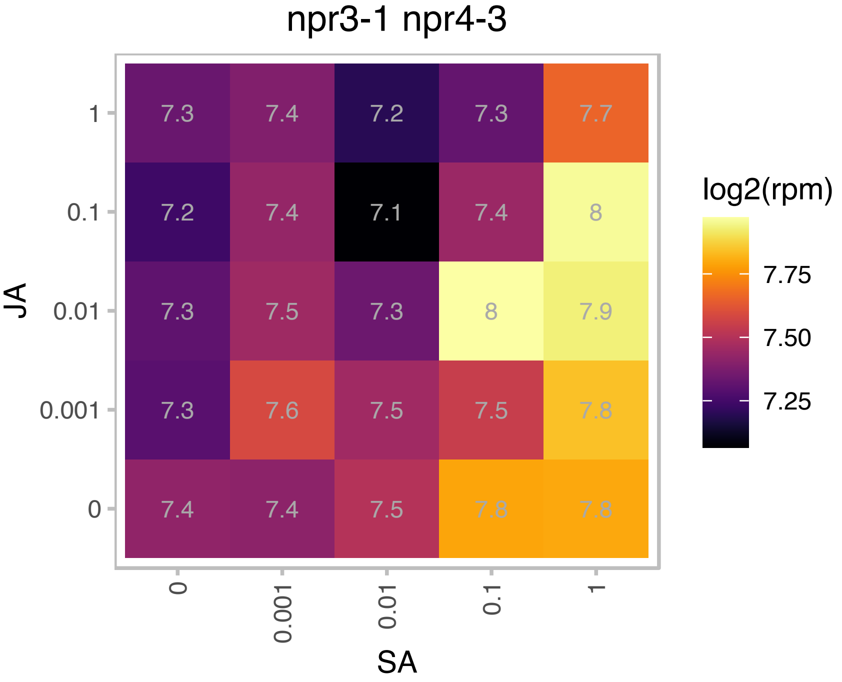

# AT1G06010.1 : NA

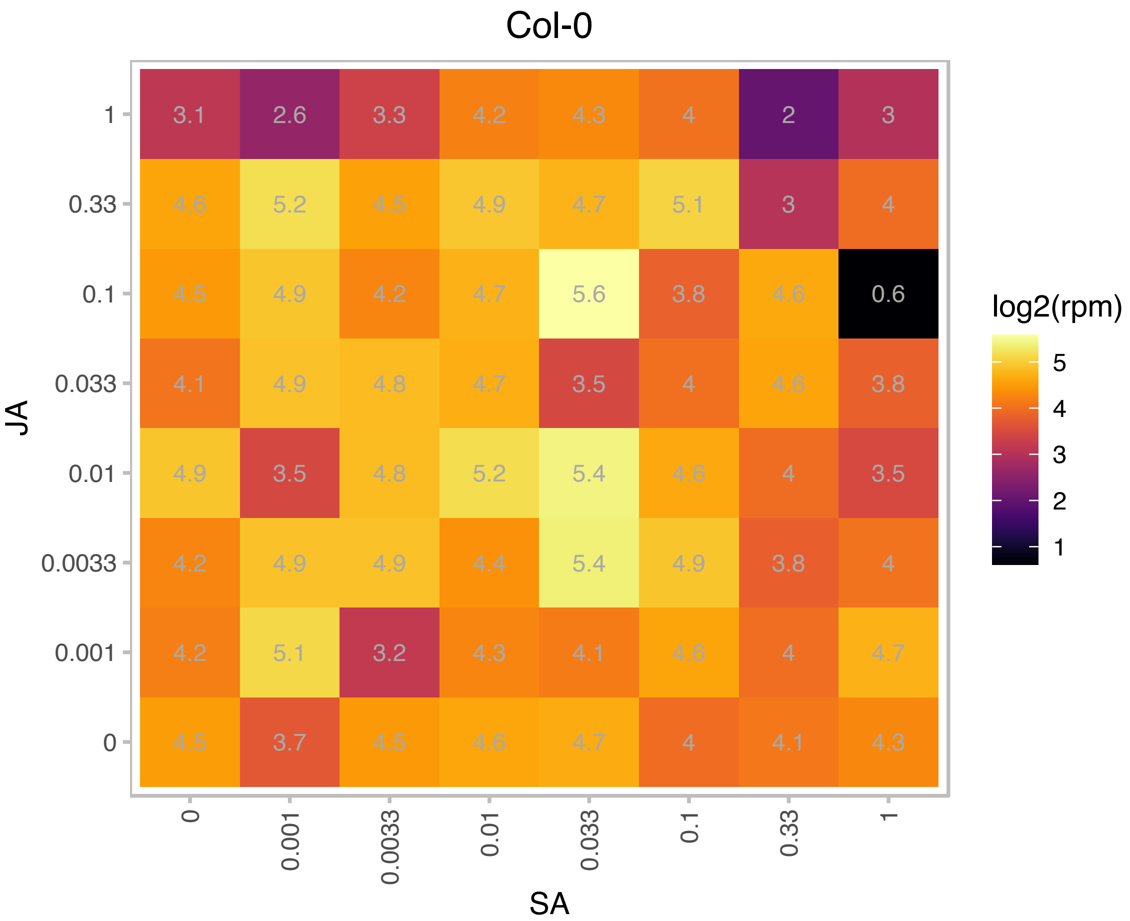


| | |
|---|---|
| Gene | AT1G06010 |
| is.expressed_genes | TRUE |
| is.responsive_genes | TRUE |
| is.apcluster | F-08 |

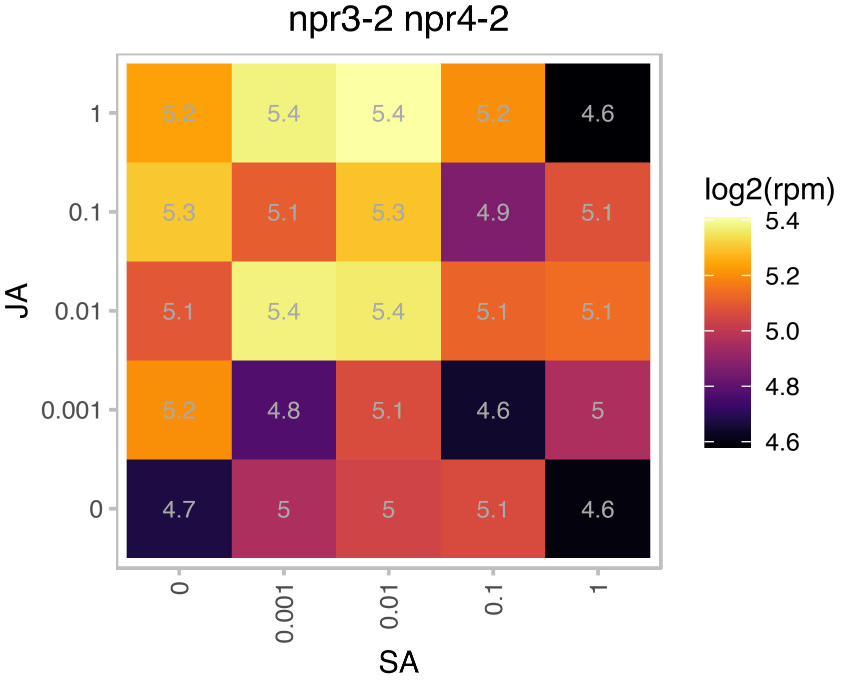


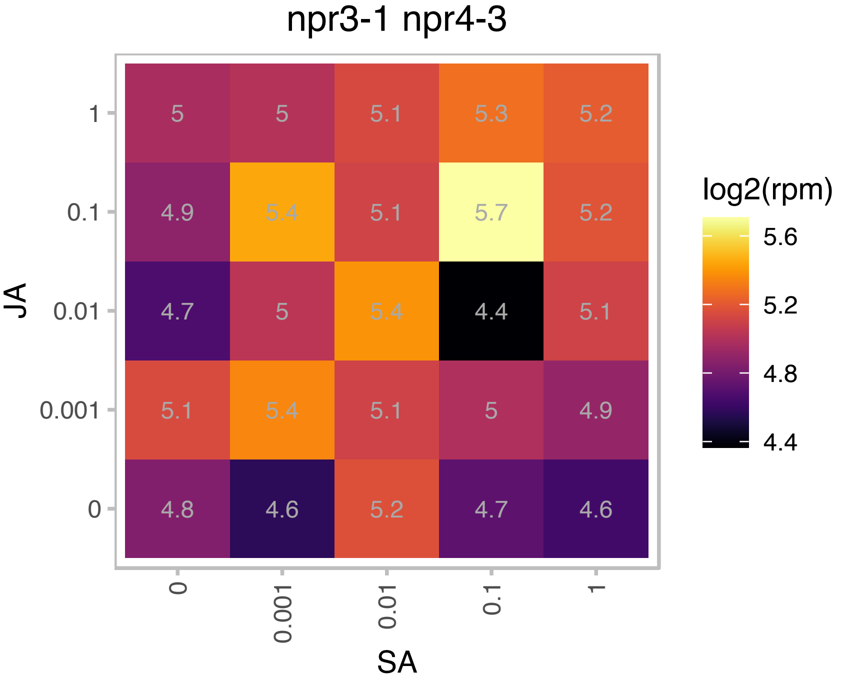

# AT1G06760.1 : H1.1

winged-helix DNA-binding transcription factor family protein

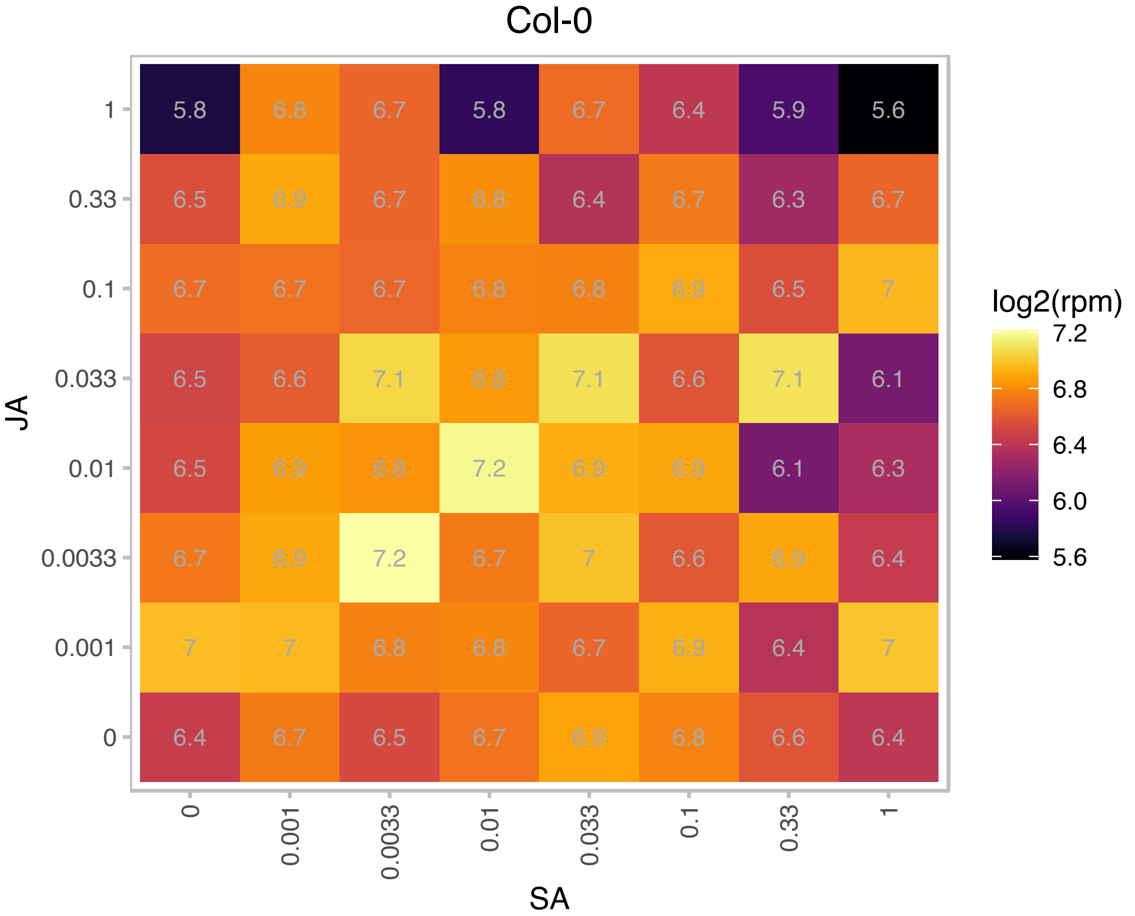


| | |
|---|---|
| Gene | AT1G06760 |
| is.expressed_genes | TRUE |
| is.responsive_genes | TRUE |
| is.apcluster | F-08 |

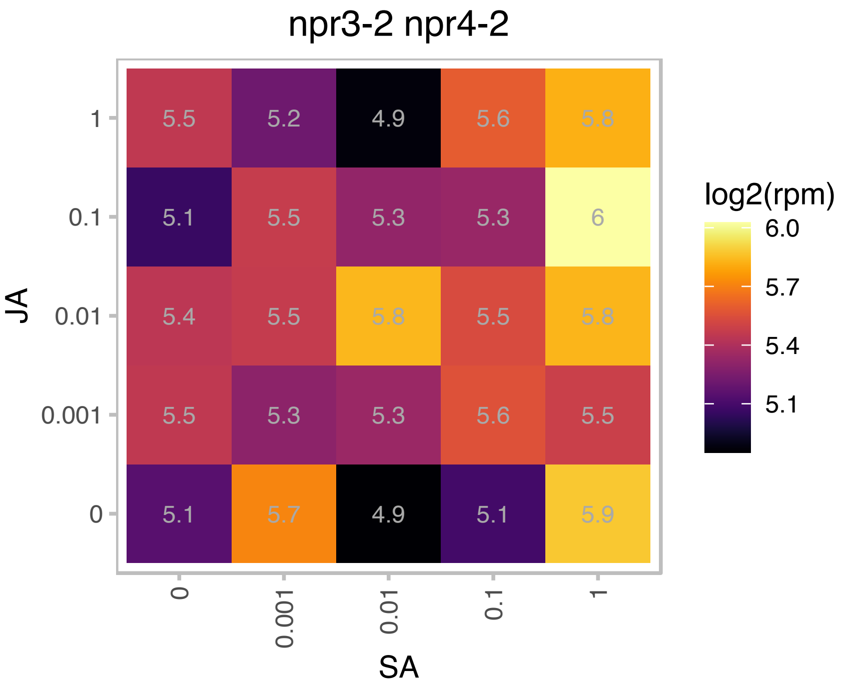


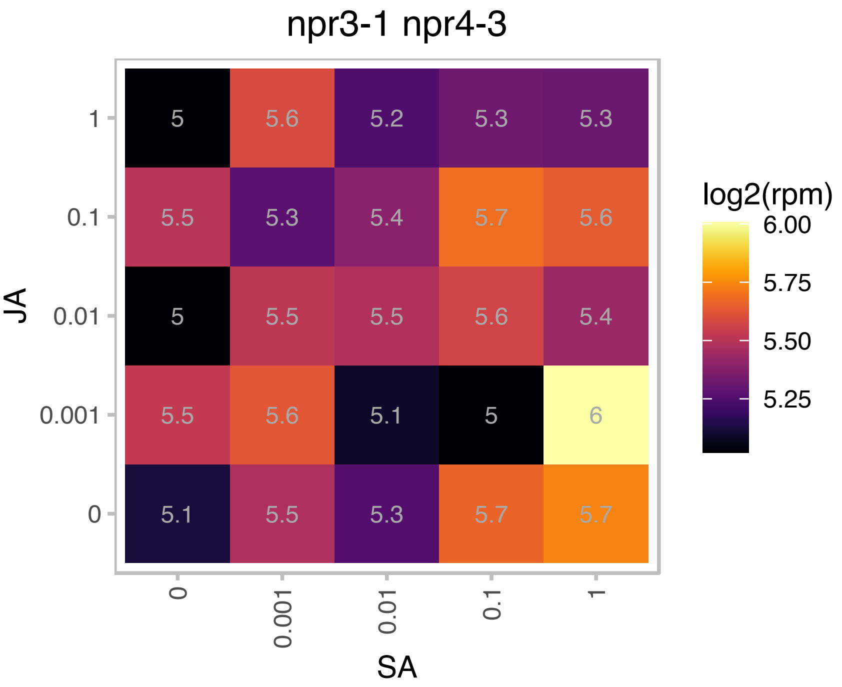

# AT1G07820.1 : NA

Histone superfamily protein

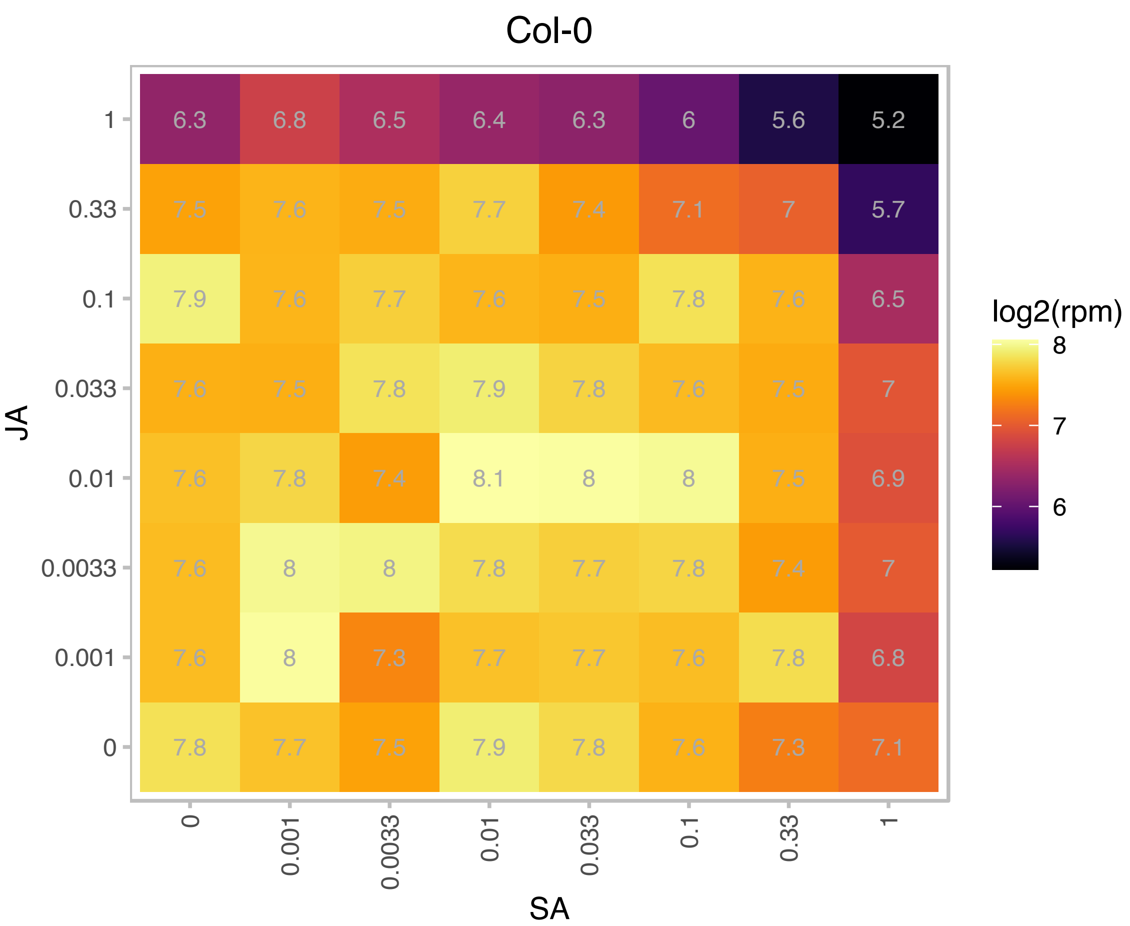


| | |
|---|---|
| Gene | AT1G07820 |
| is.expressed_genes | TRUE |
| is.responsive_genes | TRUE |
| is.apcluster | F-08 |

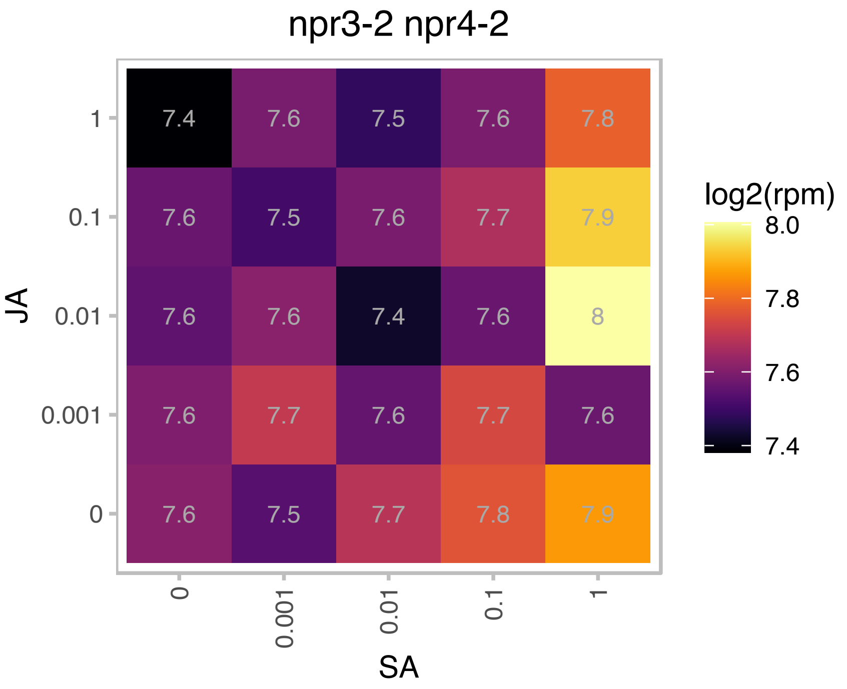


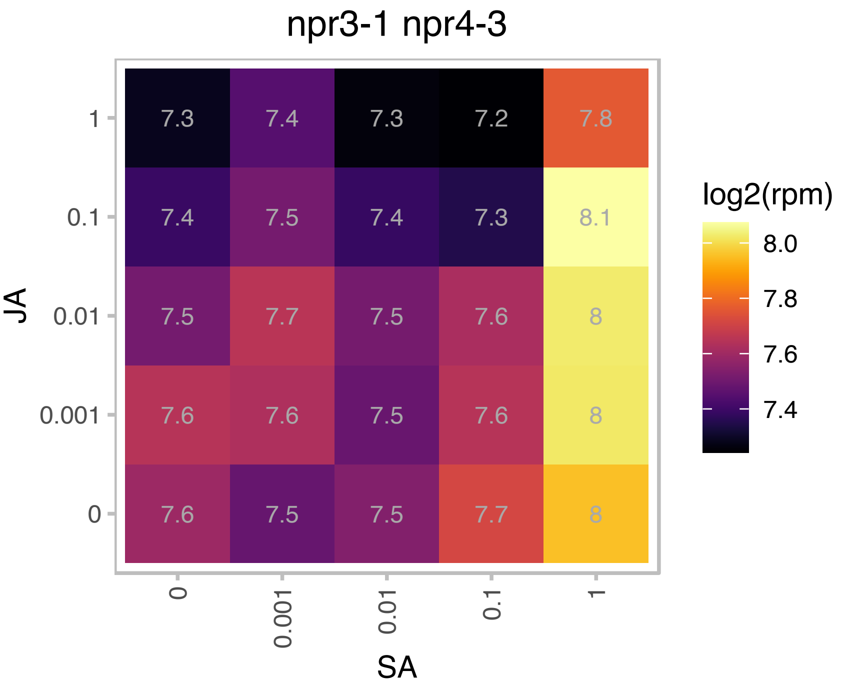

# AT1G07920.1 : NA

GTP binding Elongation factor Tu family protein

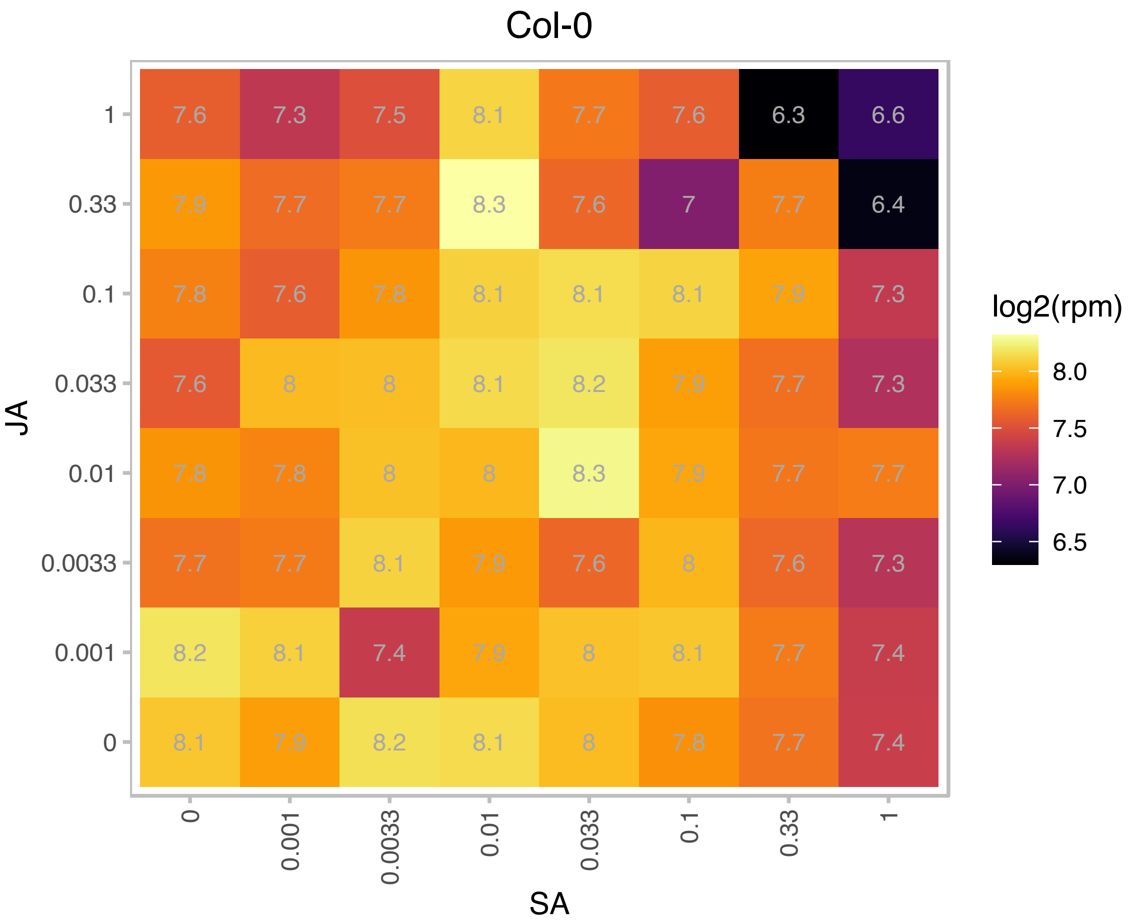


| Gene | AT1G07920 |
|---|---|
| is.expressed_genes | TRUE |
| is.responsive_genes | TRUE |
| is.apcluster | F-08 |

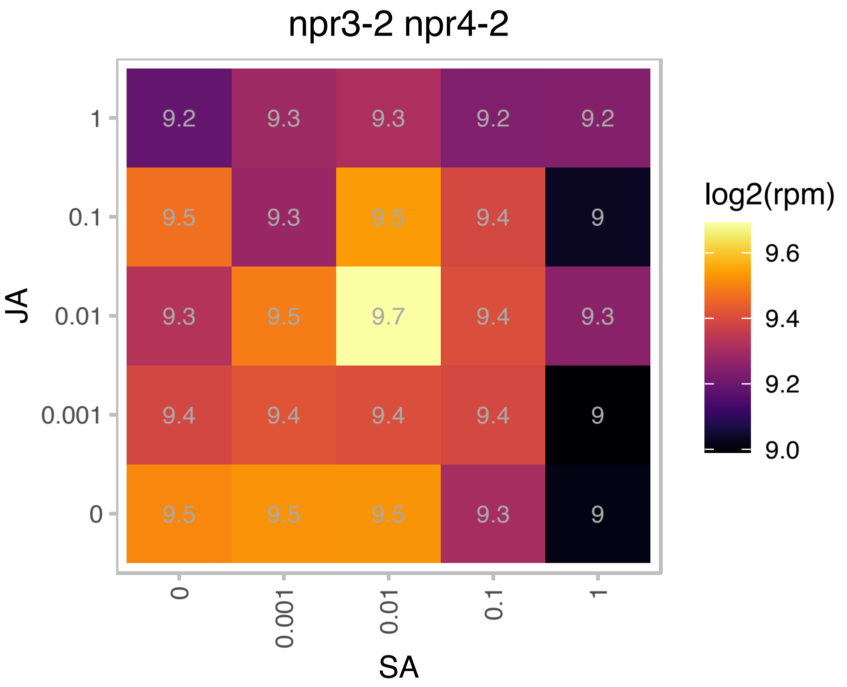


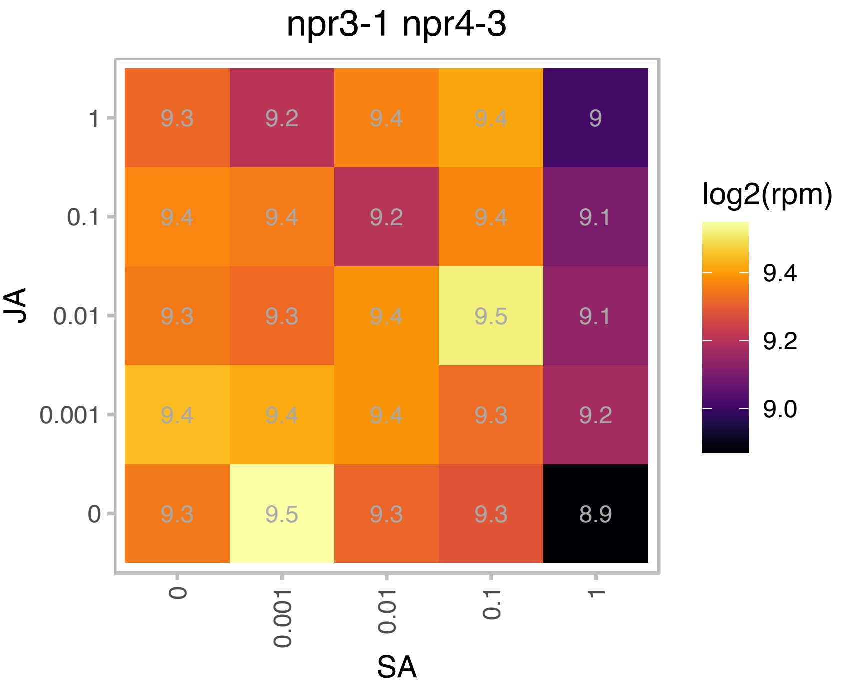

# AT1G08955.1 : NA

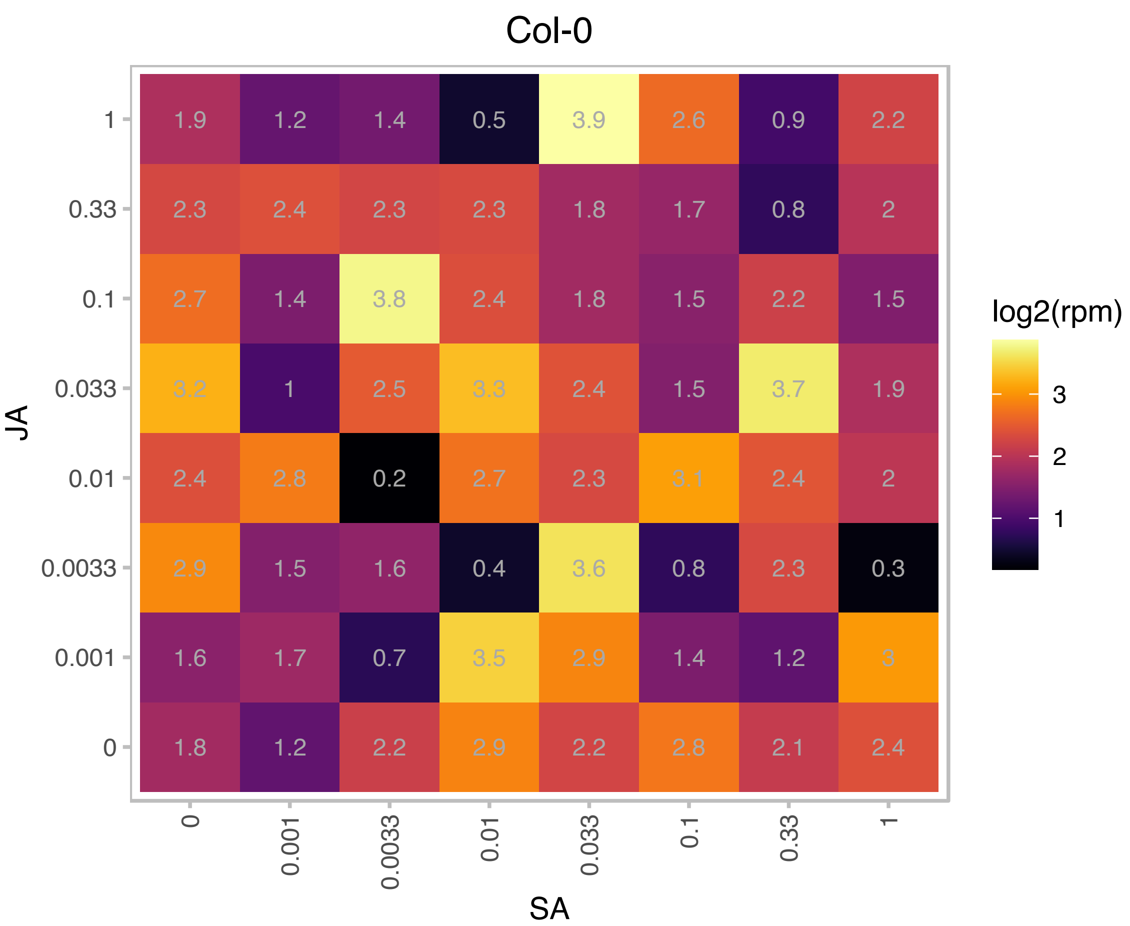


| | |
|---|---|
| Gene | AT1G08955 |
| is.expressed_genes | TRUE |
| is.responsive_genes | TRUE |
| is.apcluster | F-08 |

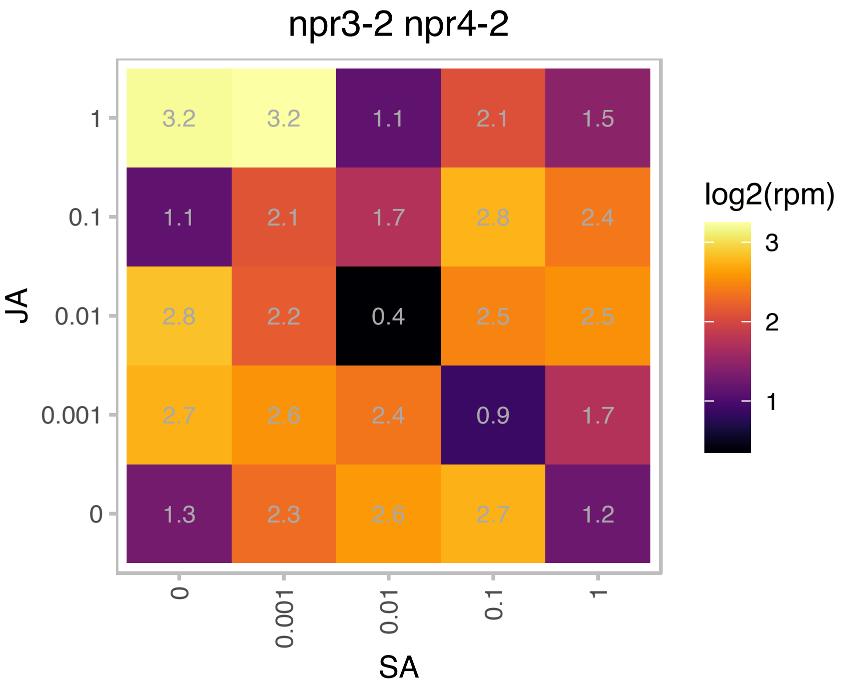


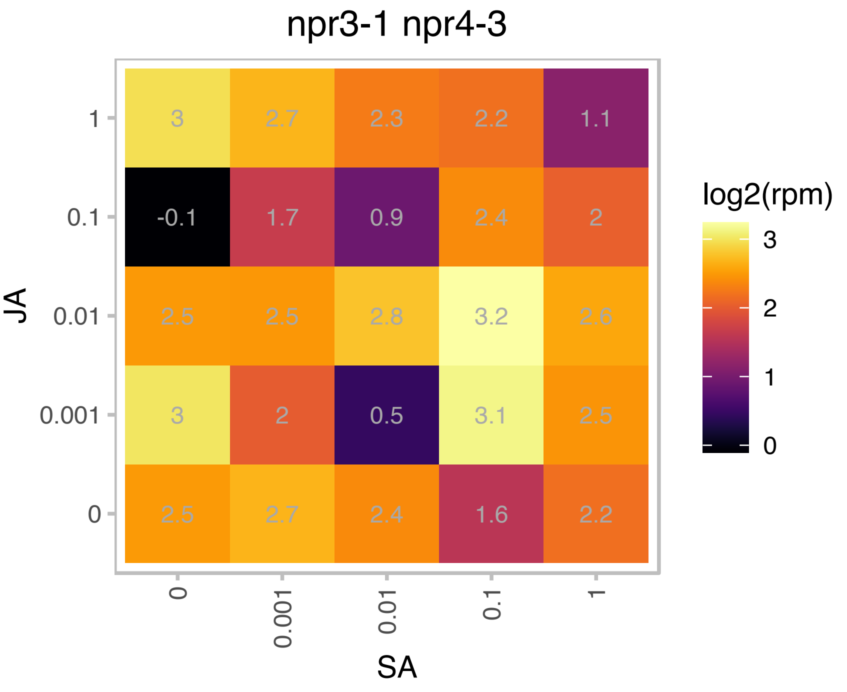

# AT1G09200.1 : H3.1

Histone superfamily protein

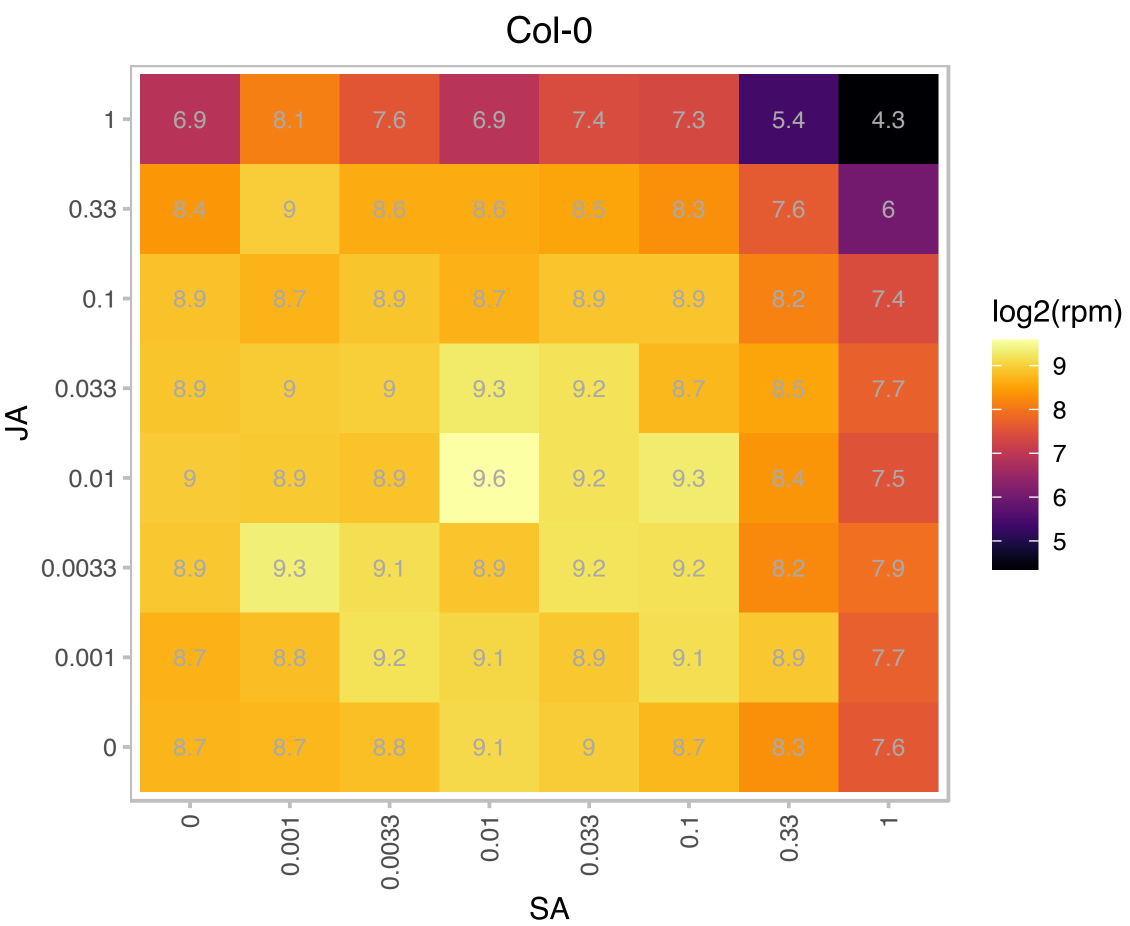


| | |
|---|---|
| Gene | AT1G09200 |
| is.expressed_genes | TRUE |
| is.responsive_genes | TRUE |
| is.apcluster | F-08 |

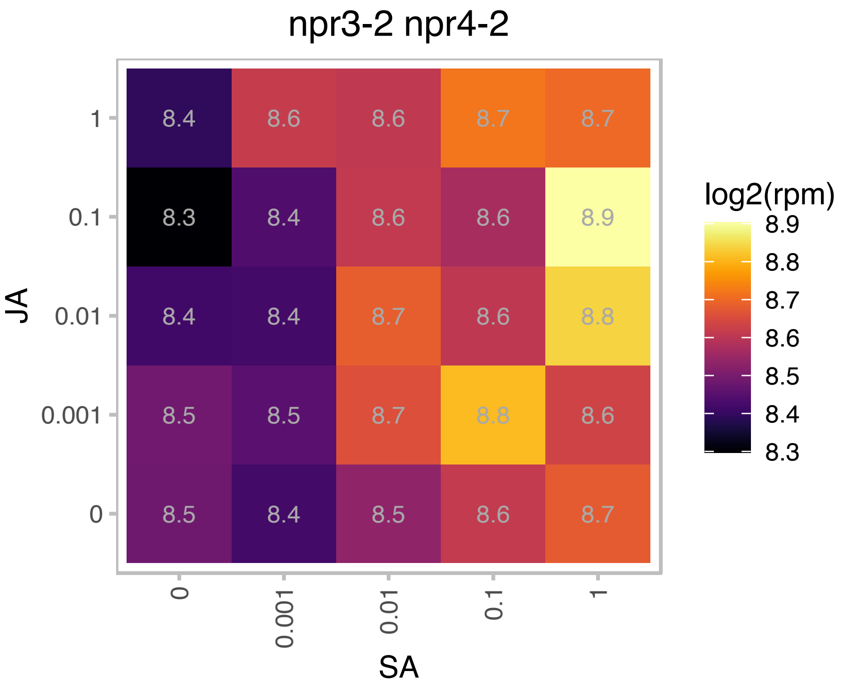


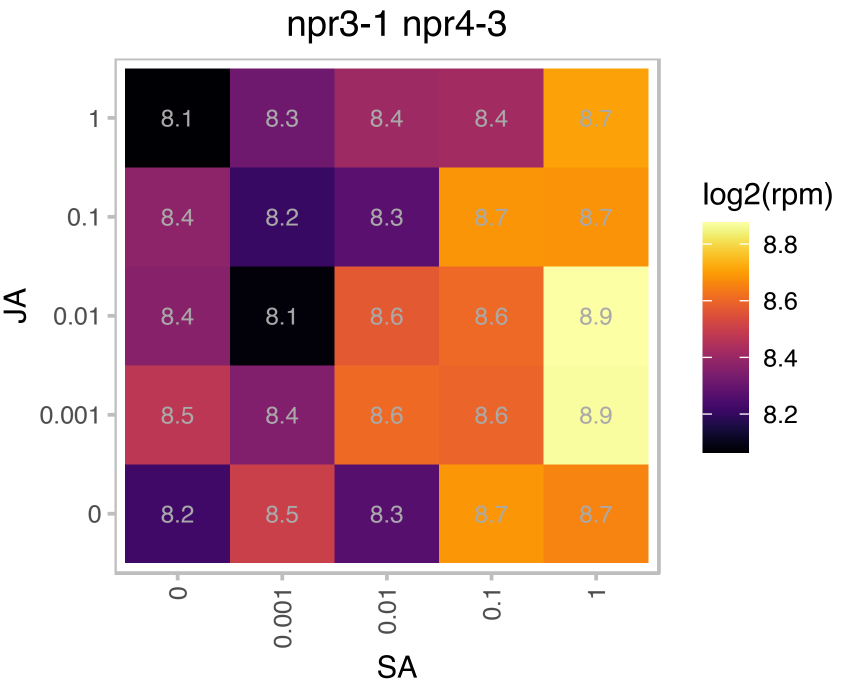

# AT1G11850.1 : NA

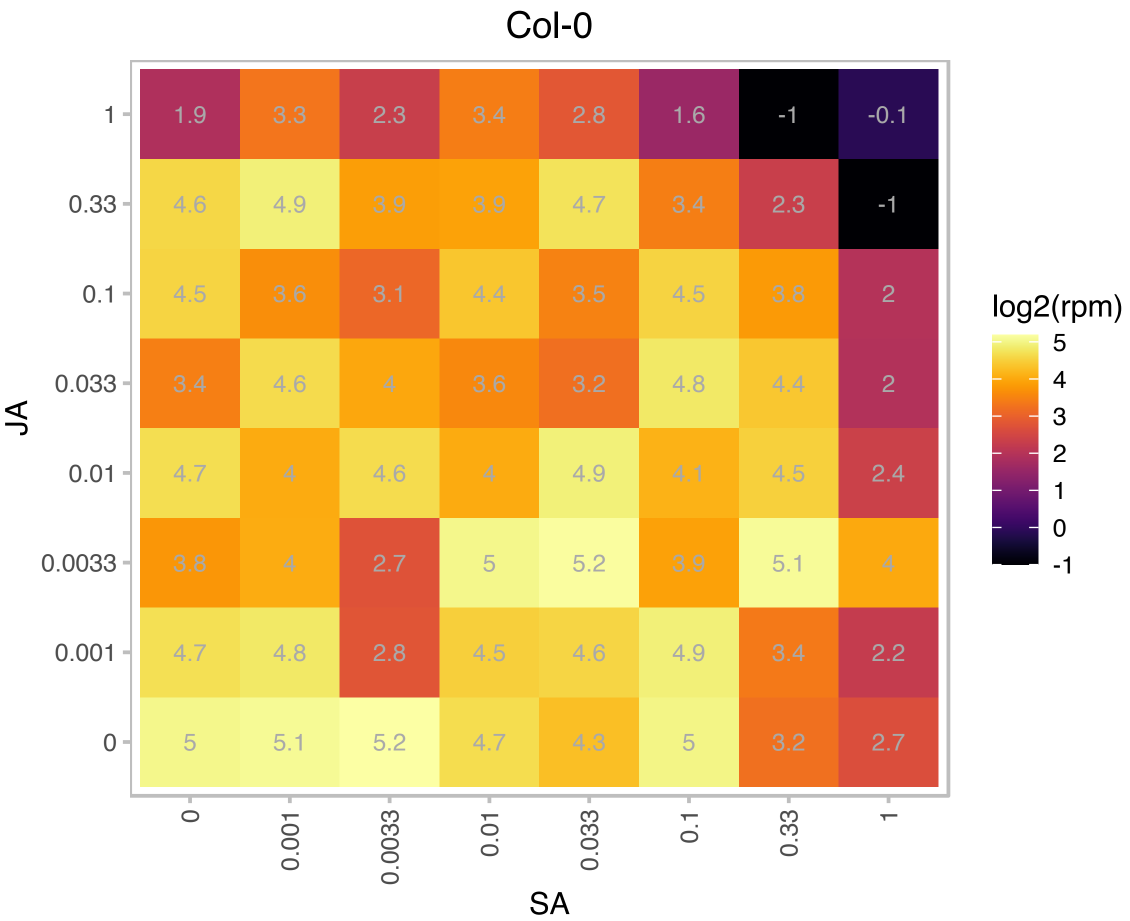


| | |
|---|---|
| Gene | AT1G11850 |
| is.expressed_genes | TRUE |
| is.responsive_genes | TRUE |
| is.apcluster | F-08 |

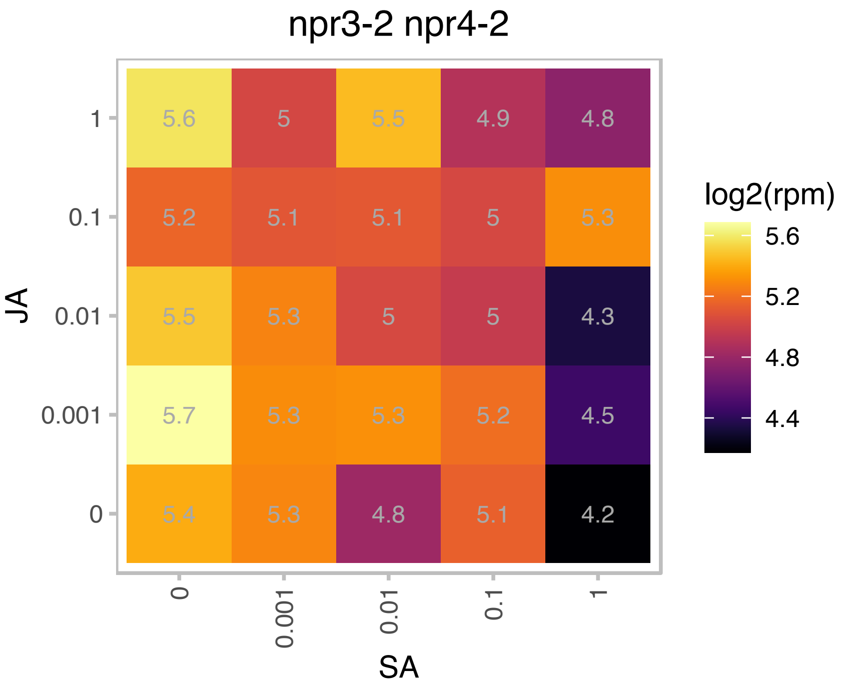


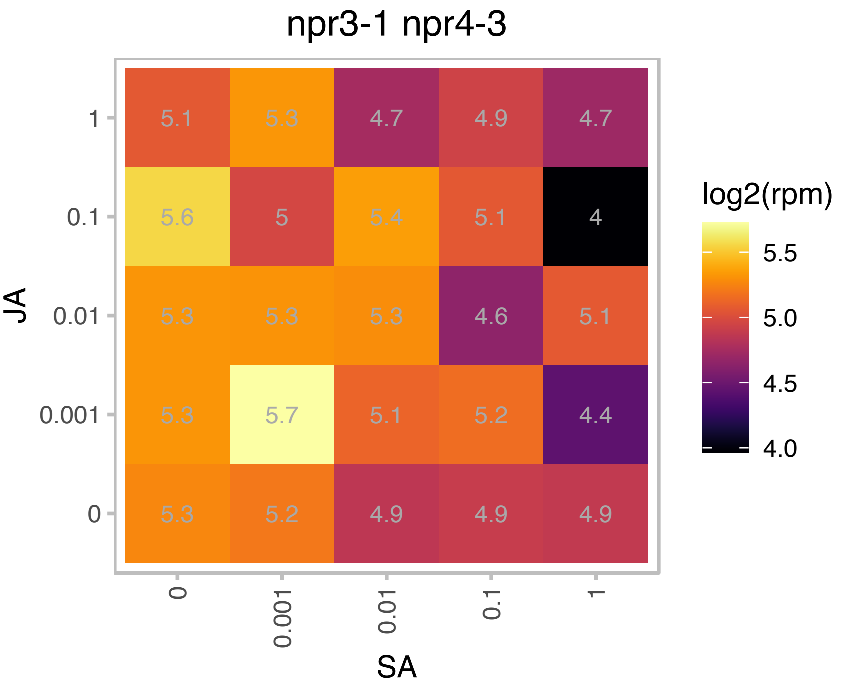

# AT1G12080.1 : NA

Vacuolar calcium-binding protein-related

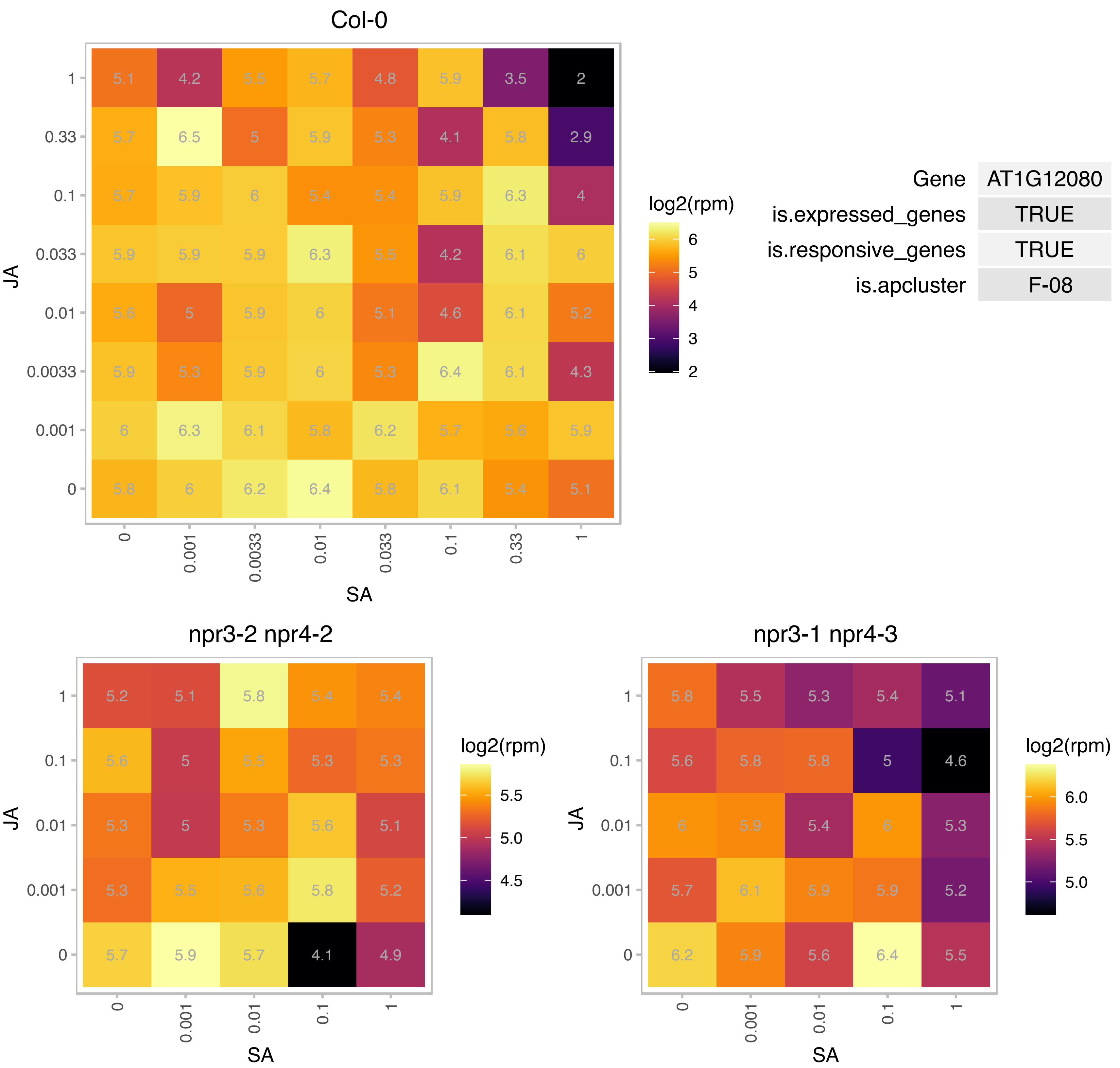


| | |
|---|---|
| Gene | AT1G12080 |
| is.expressed_genes | TRUE |
| is.responsive_genes | TRUE |
| is.apcluster | F-08 |

# AT1G12310.1 : NA

Calcium-binding EF-hand family protein

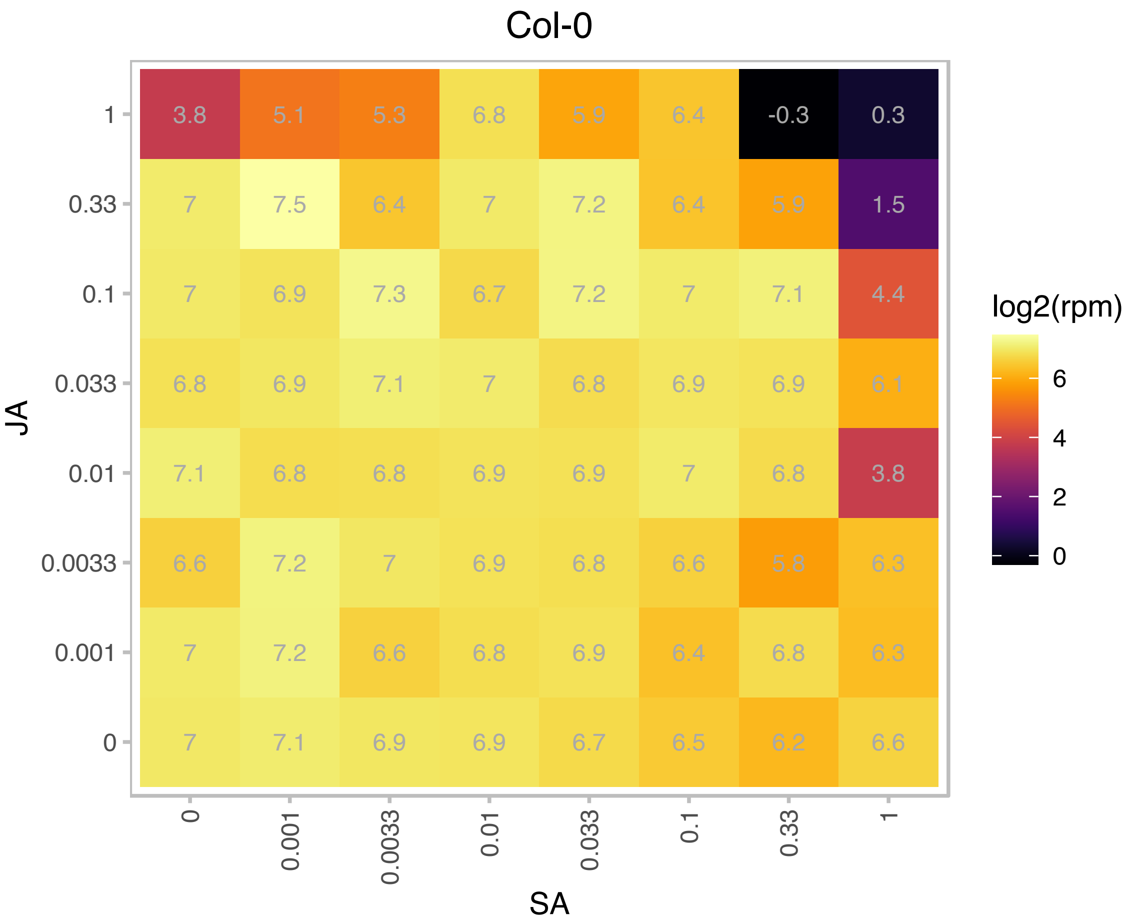


| | |
|---|---|
| Gene | AT1G12310 |
| is.expressed_genes | TRUE |
| is.responsive_genes | TRUE |
| is.apcluster | F-08 |

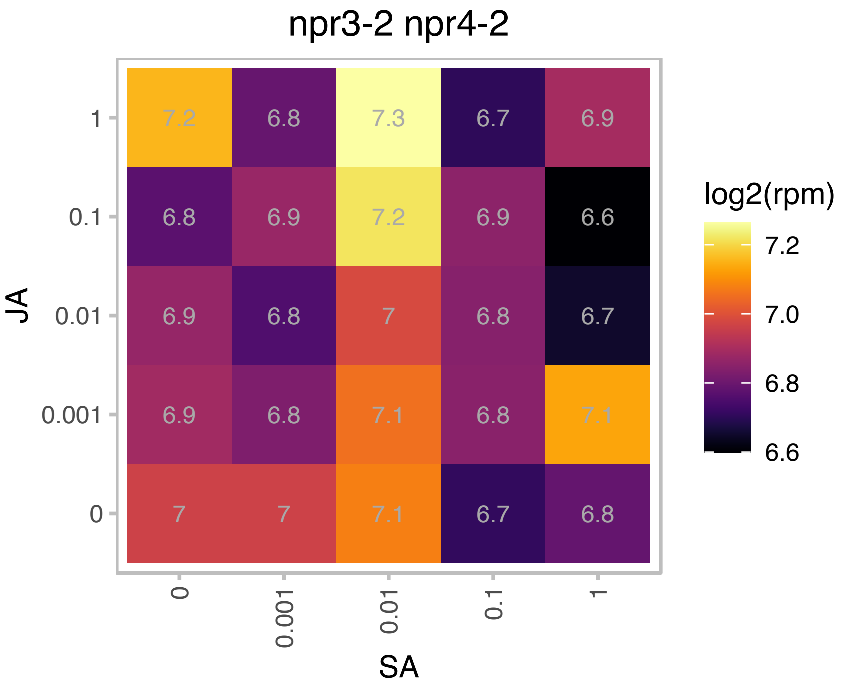


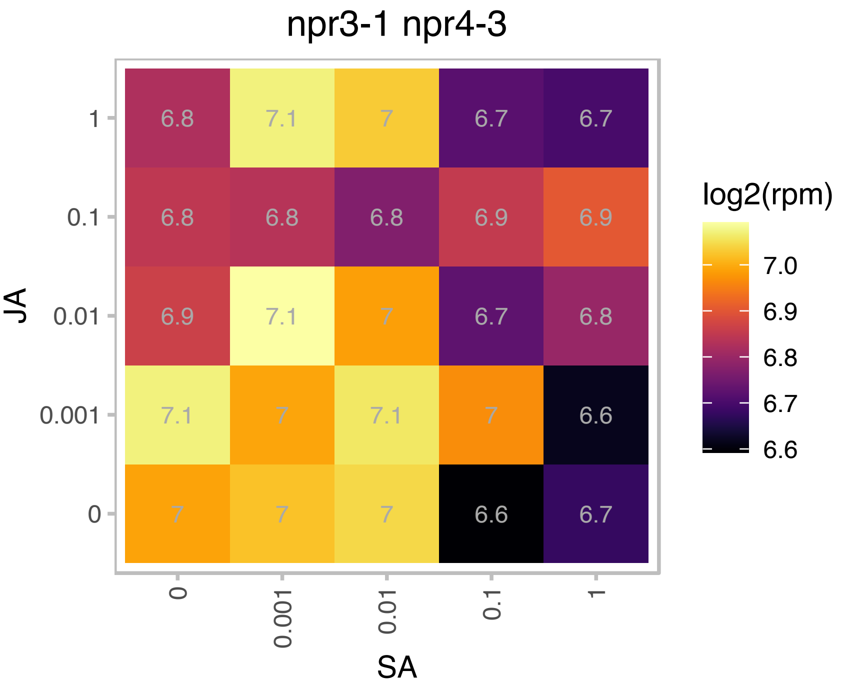

# AT1G14210.1 : NA

Ribonuclease T2 family protein

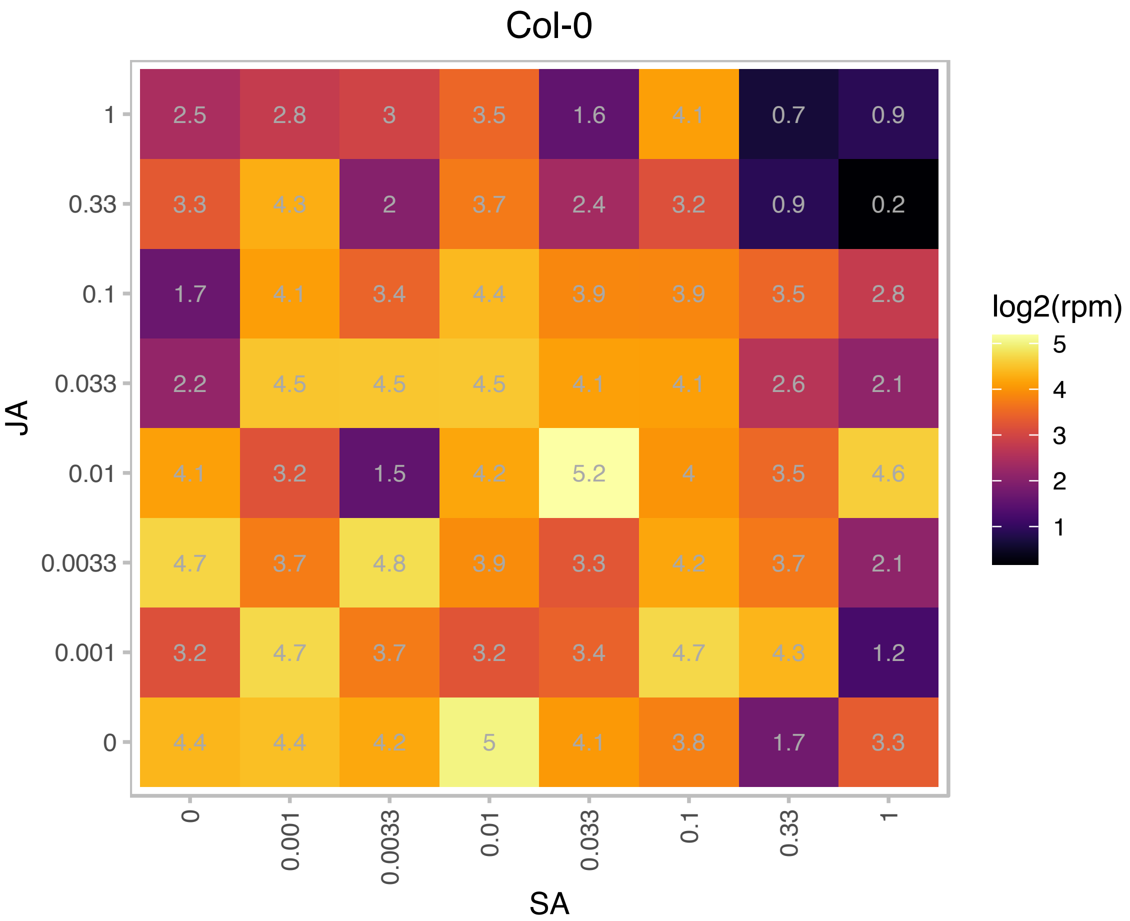


| | |
|---|---|
| Gene | AT1G14210 |
| is.expressed_genes | TRUE |
| is.responsive_genes | TRUE |
| is.apcluster | F-08 |

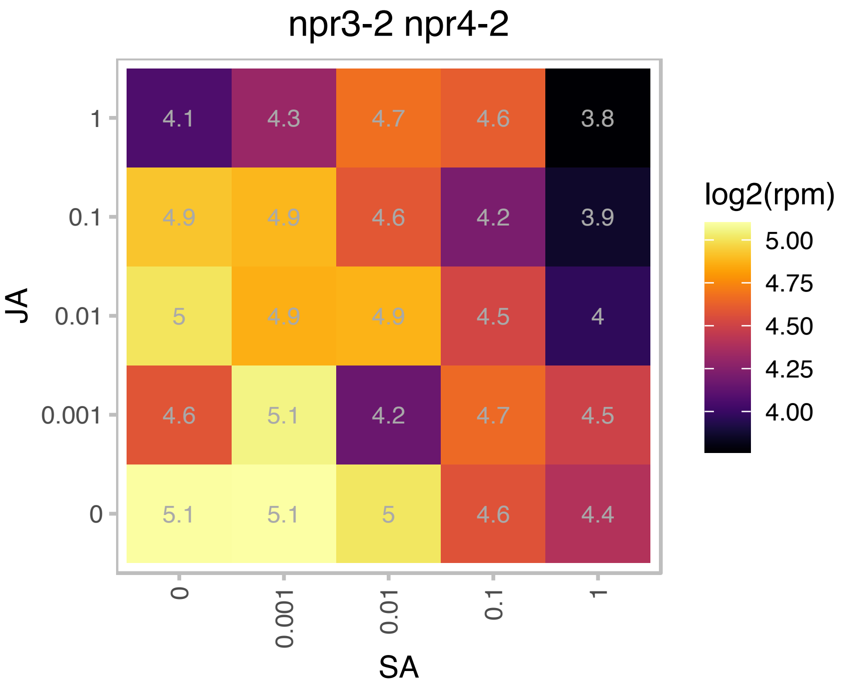


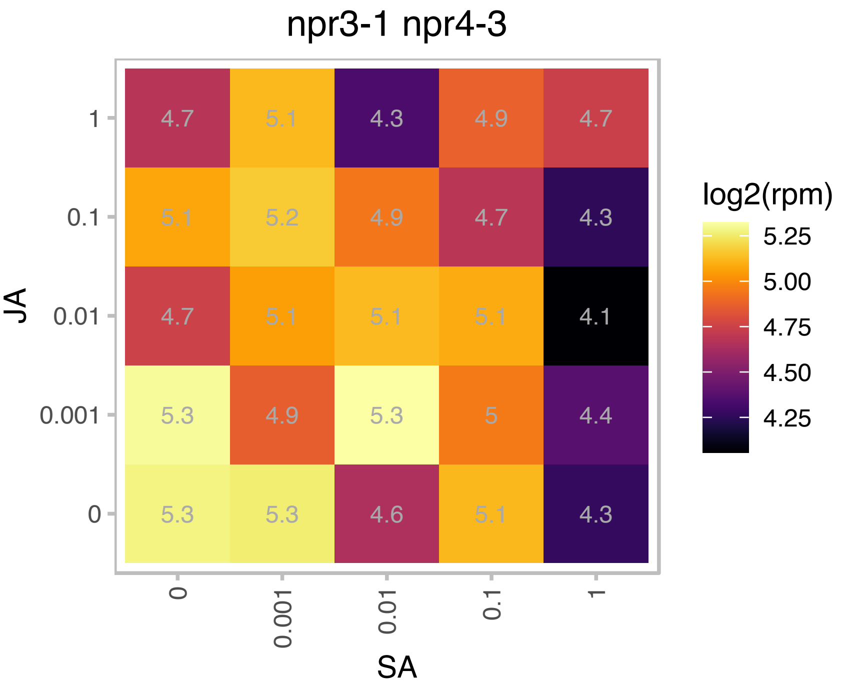

# AT1G15120.1 : NA

Ubiquinol-cytochrome C reductase hinge protein

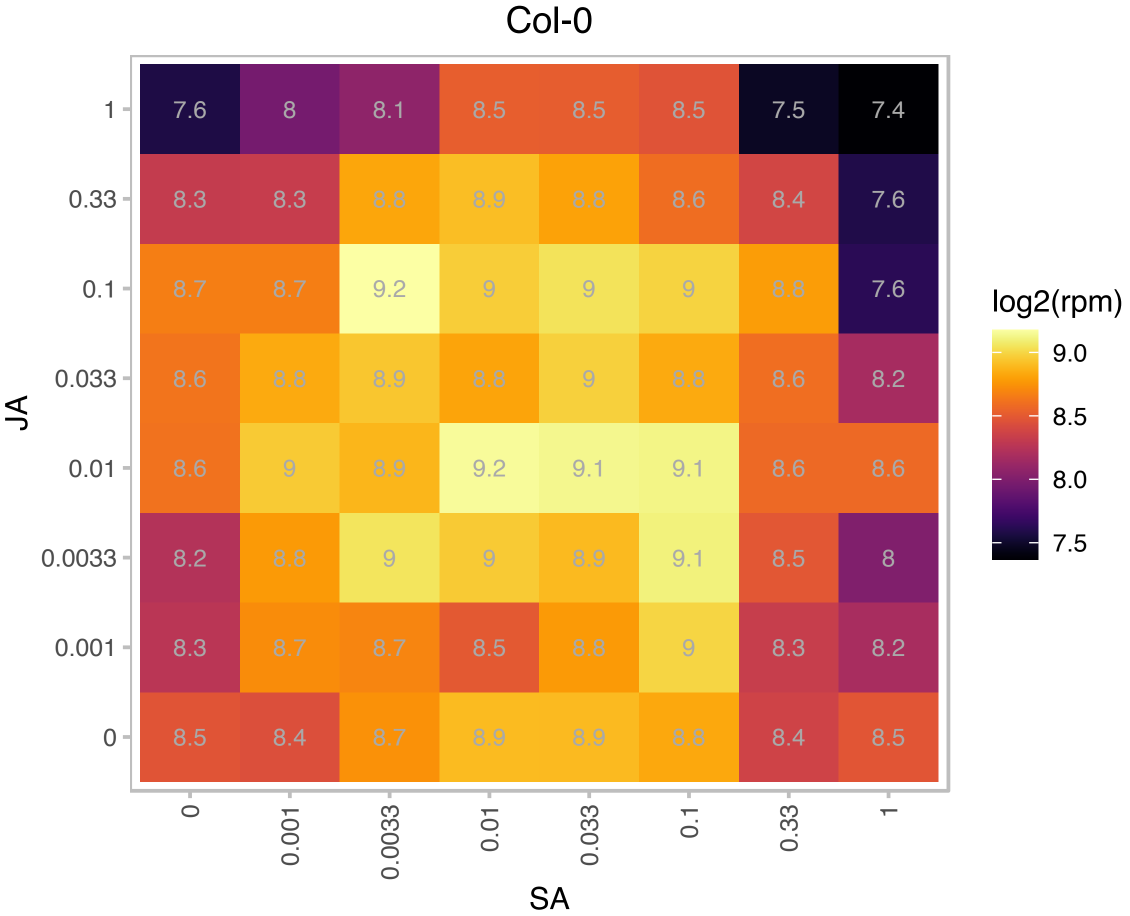


| Gene | AT1G15120 |
|---|---|
| is.expressed_genes | TRUE |
| is.responsive_genes | TRUE |
| is.apcluster | F-08 |

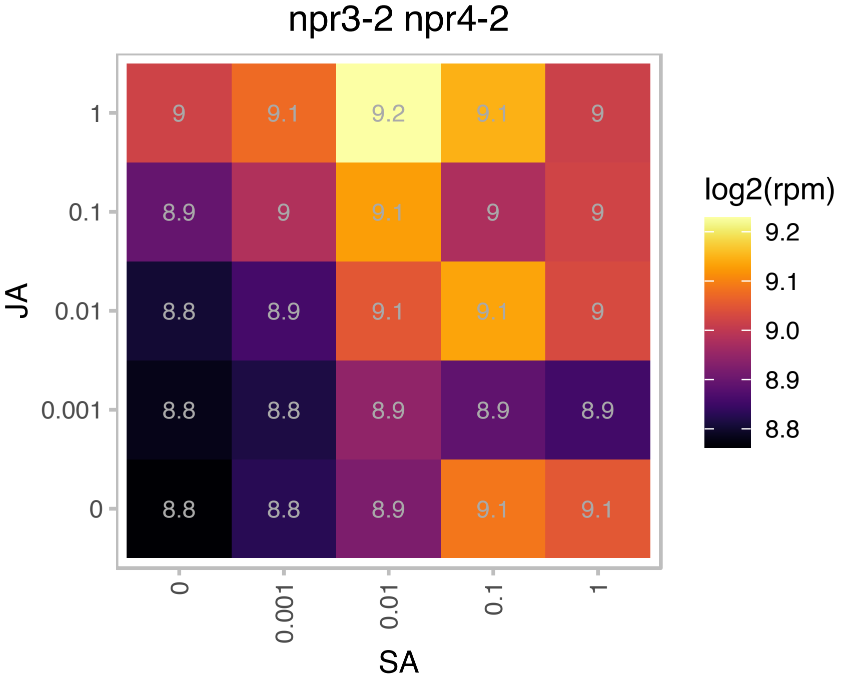


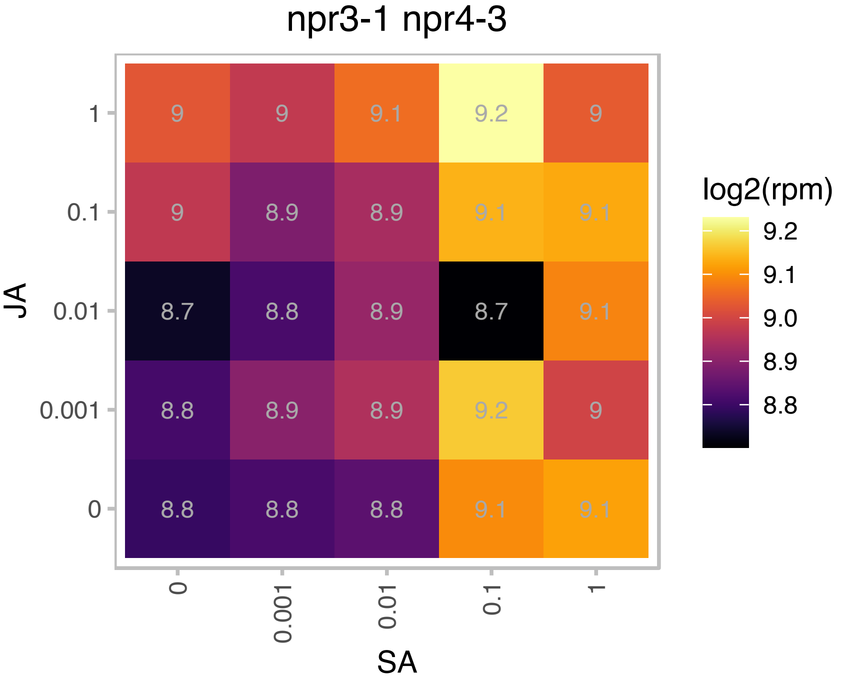

# AT1G16000.1 : NA

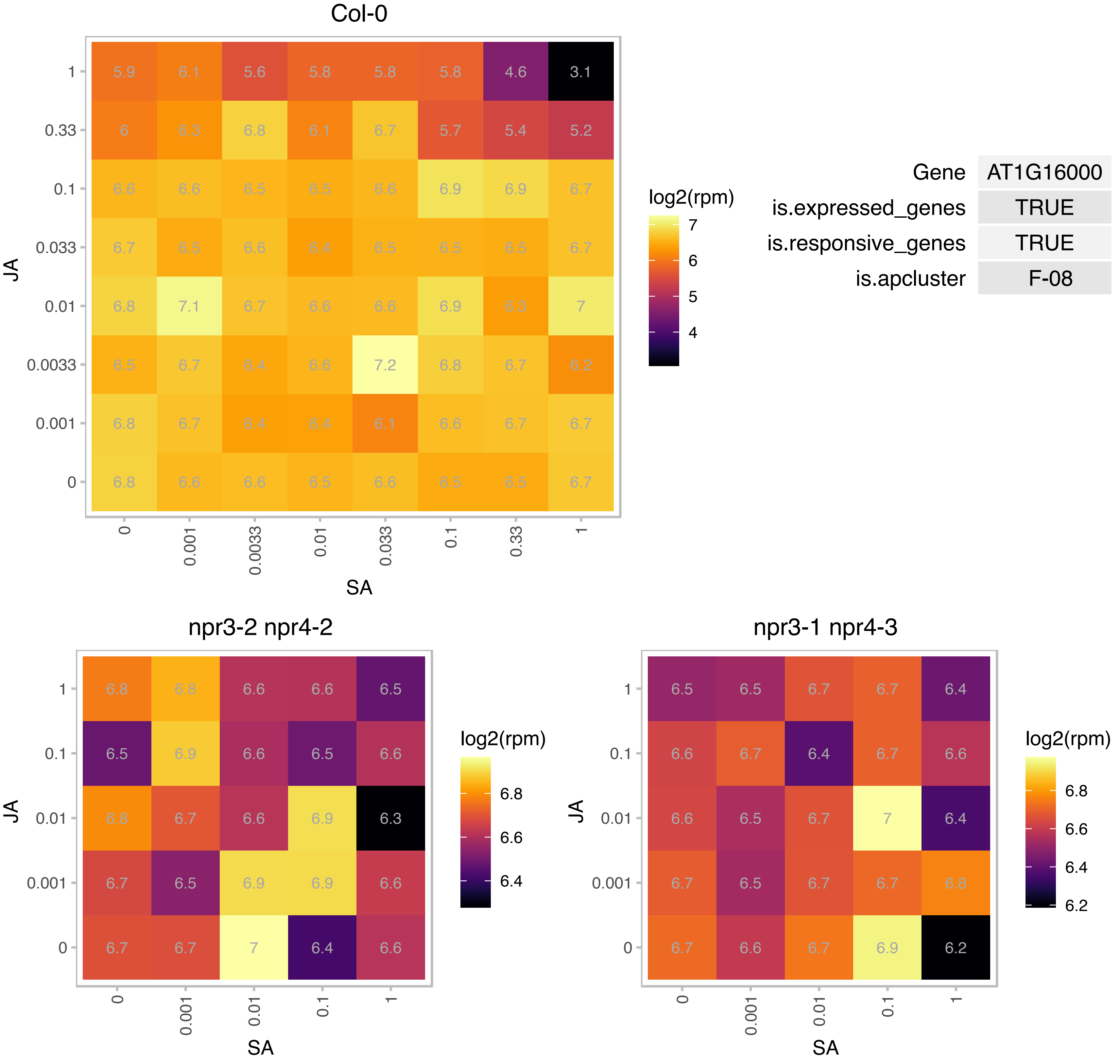


| | |
|---|---|
| Gene | AT1G16000 |
| is.expressed_genes | TRUE |
| is.responsive_genes | TRUE |
| is.apcluster | F-08 |

# AT1G16916.1 : NA

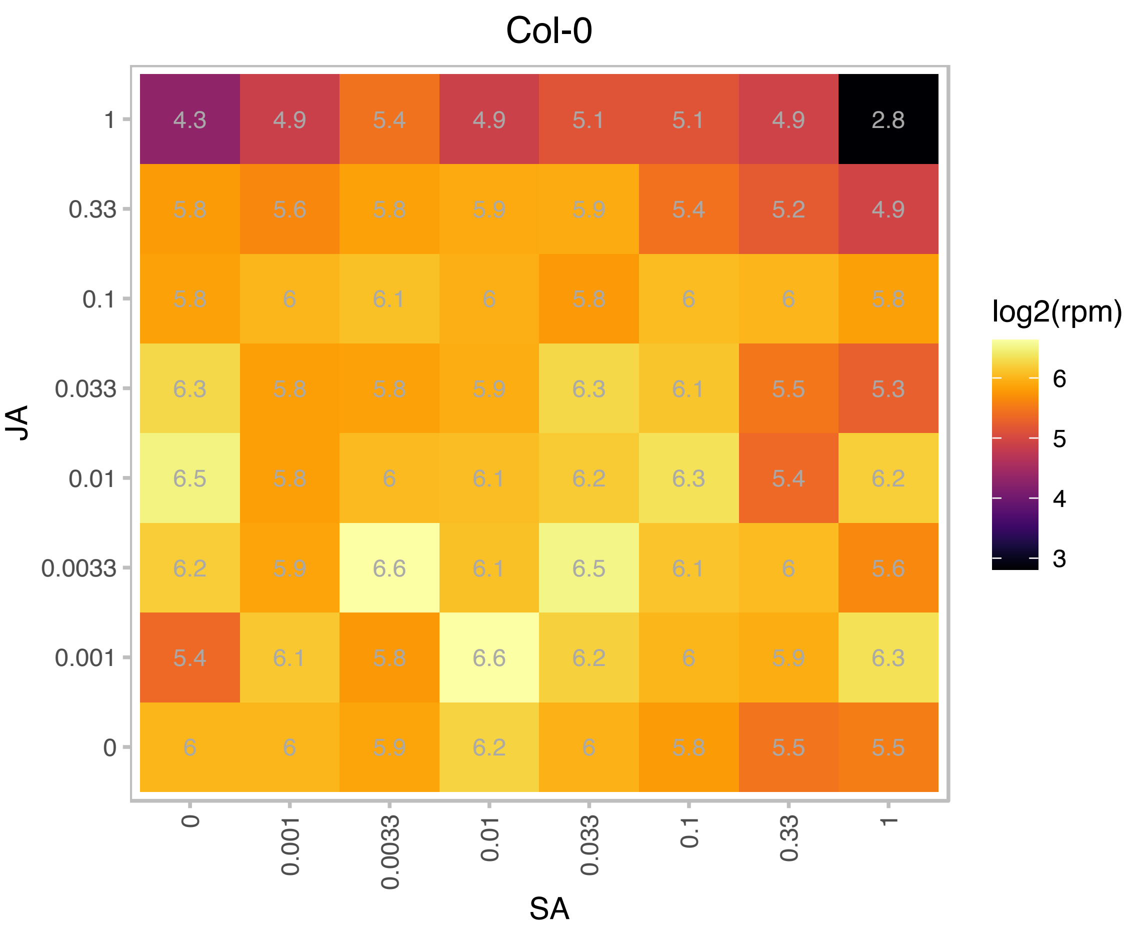


| | |
|---|---|
| Gene | AT1G16916 |
| is.expressed_genes | TRUE |
| is.responsive_genes | TRUE |
| is.apcluster | F-08 |

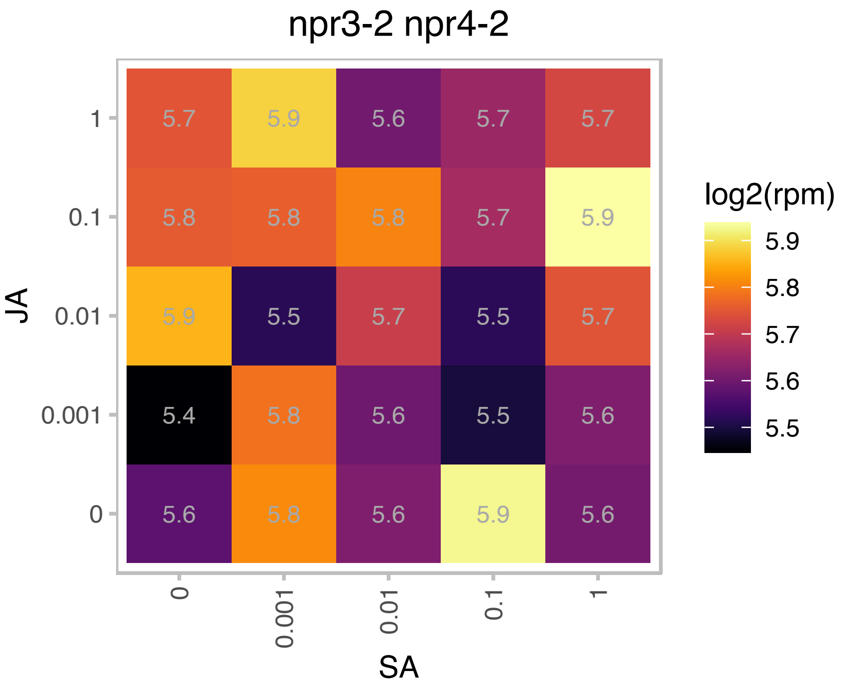


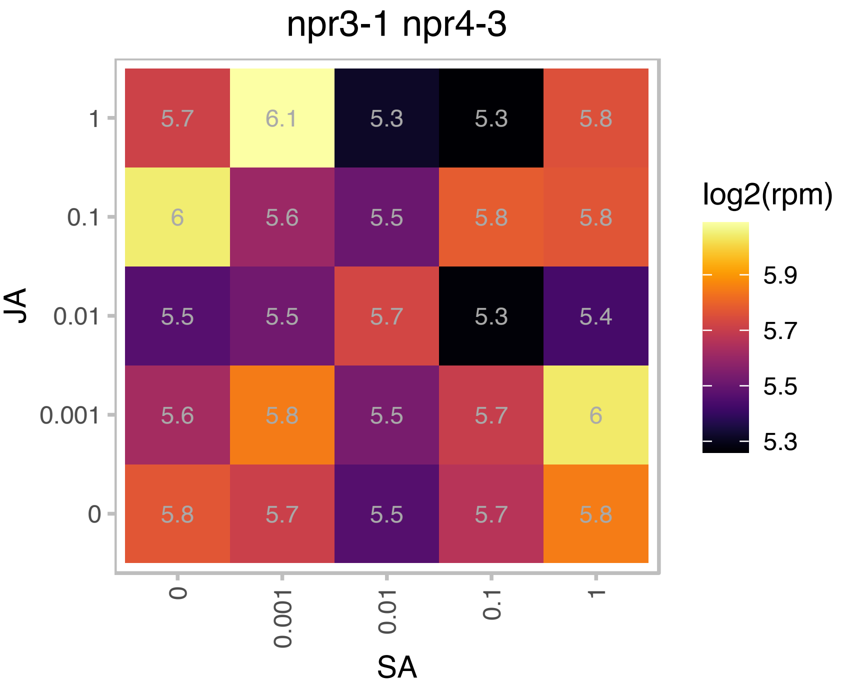

### AT1G19120.1 : LSM1A

Small nuclear ribonucleoprotein family protein

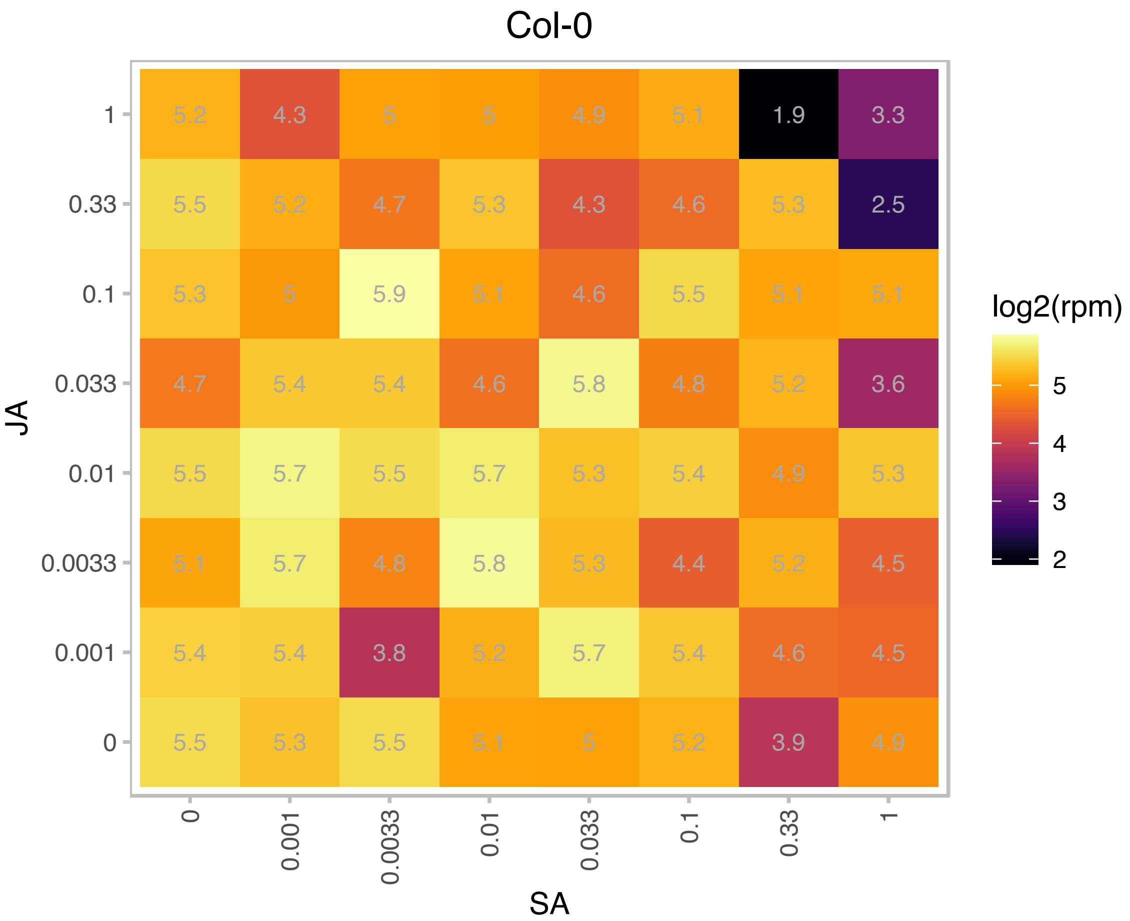


| | |
|---|---|
| Gene | AT1G19120 |
| is.expressed_genes | TRUE |
| is.responsive_genes | TRUE |
| is.apcluster | F-08 |

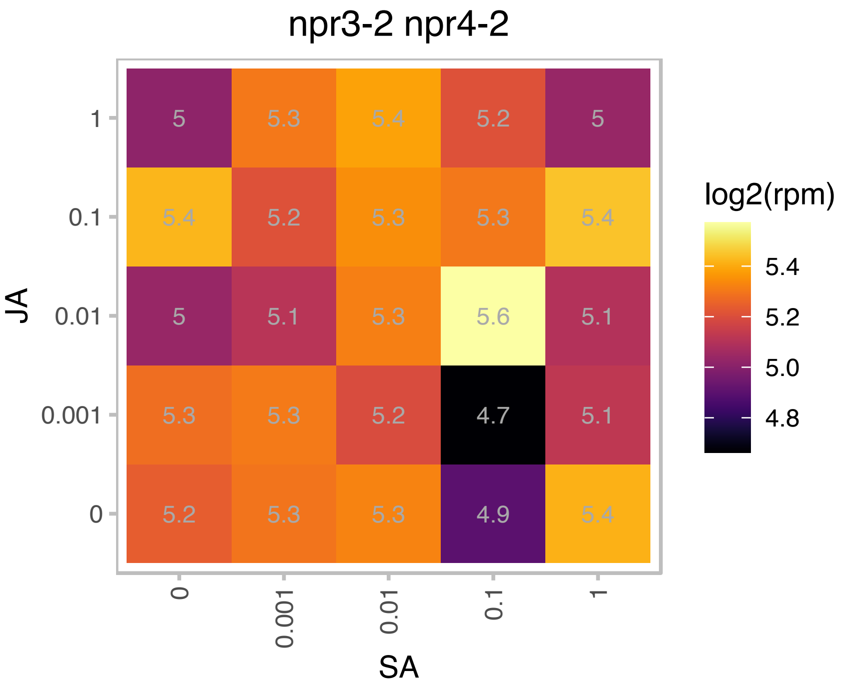


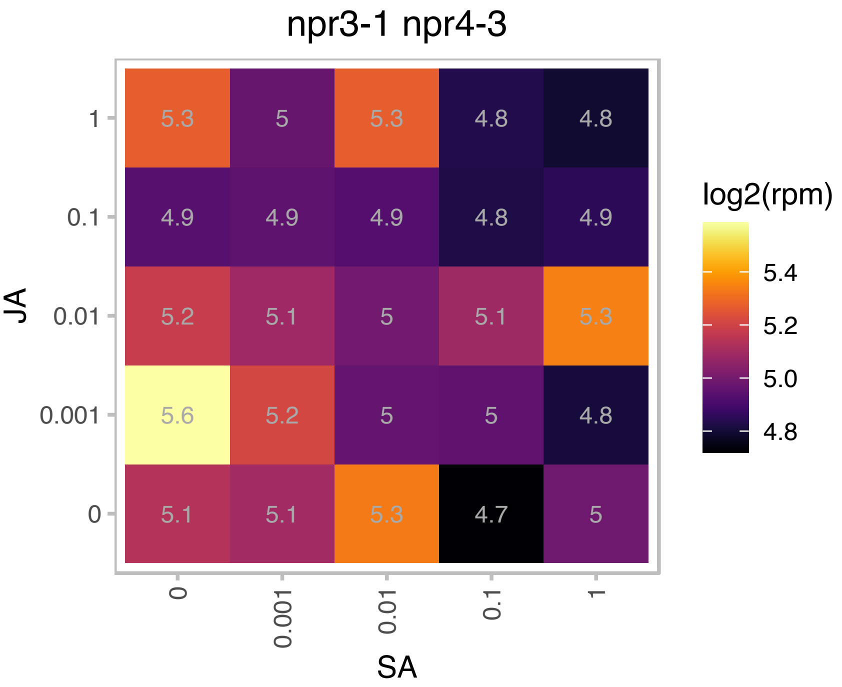

# AT1G20430.1 : NA

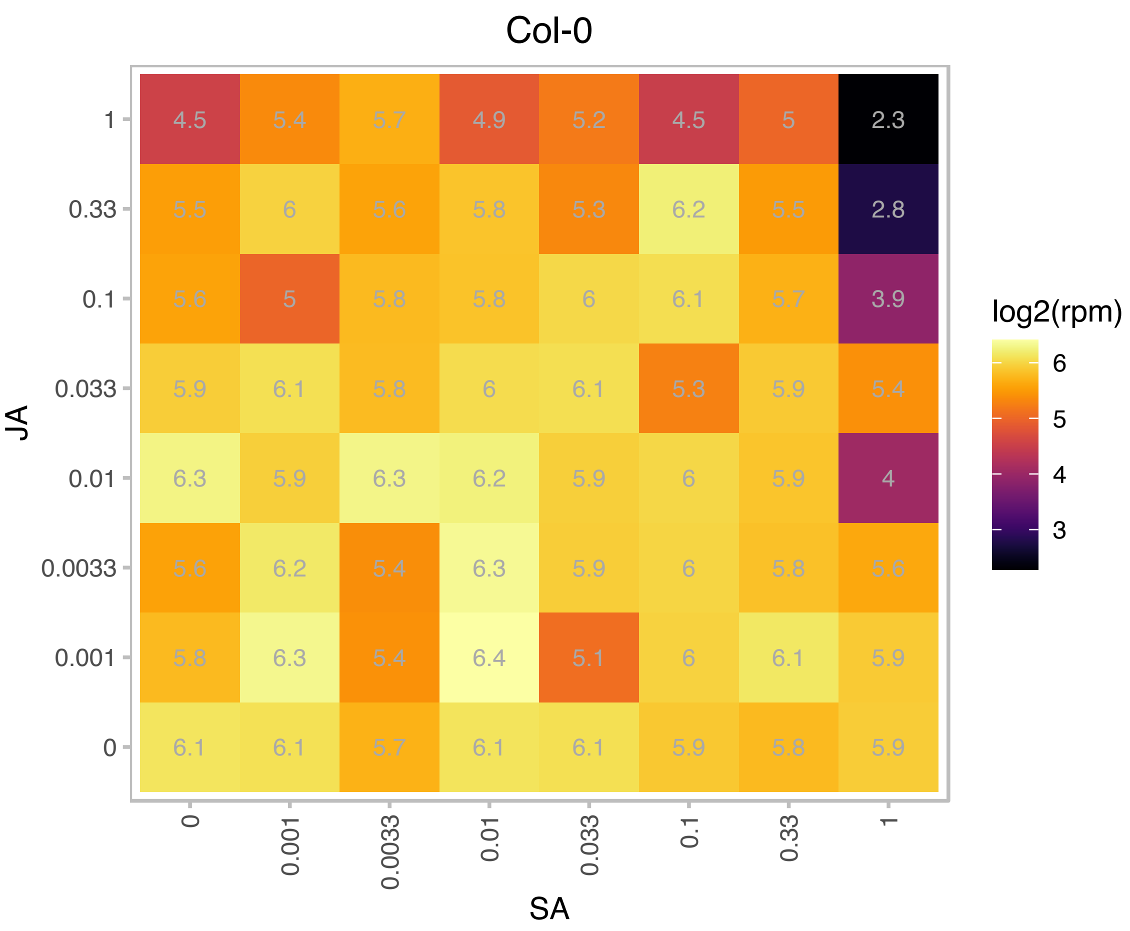


| | |
|---|---|
| Gene | AT1G20430 |
| is.expressed_genes | TRUE |
| is.responsive_genes | TRUE |
| is.apcluster | F-08 |

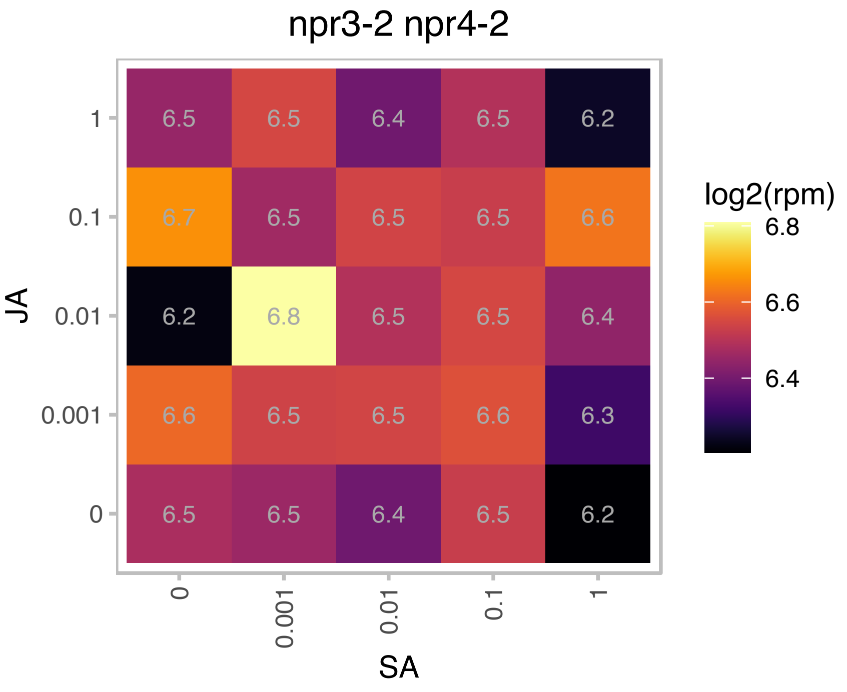


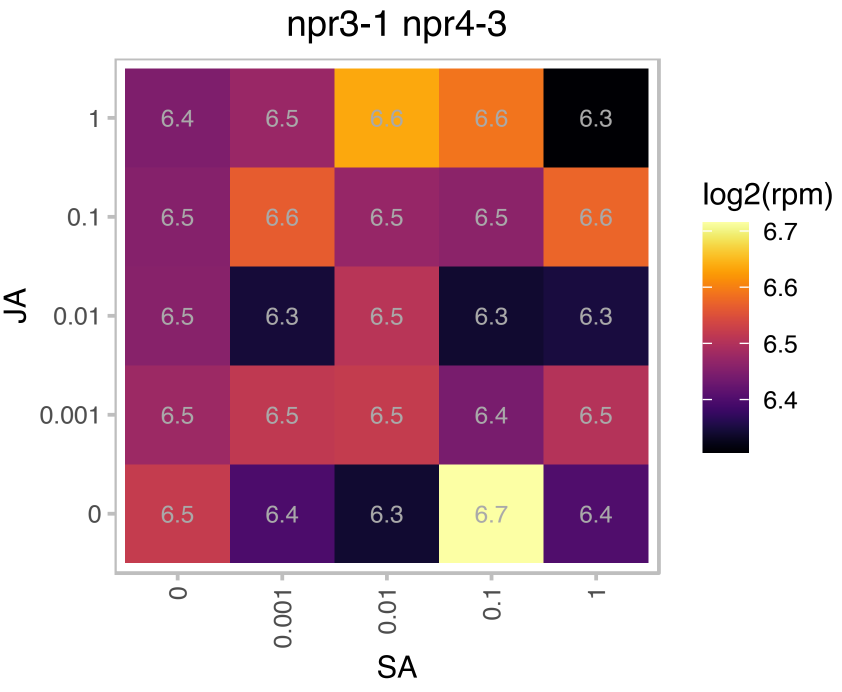

# AT1G20580.1 : NA

Small nuclear ribonucleoprotein family protein

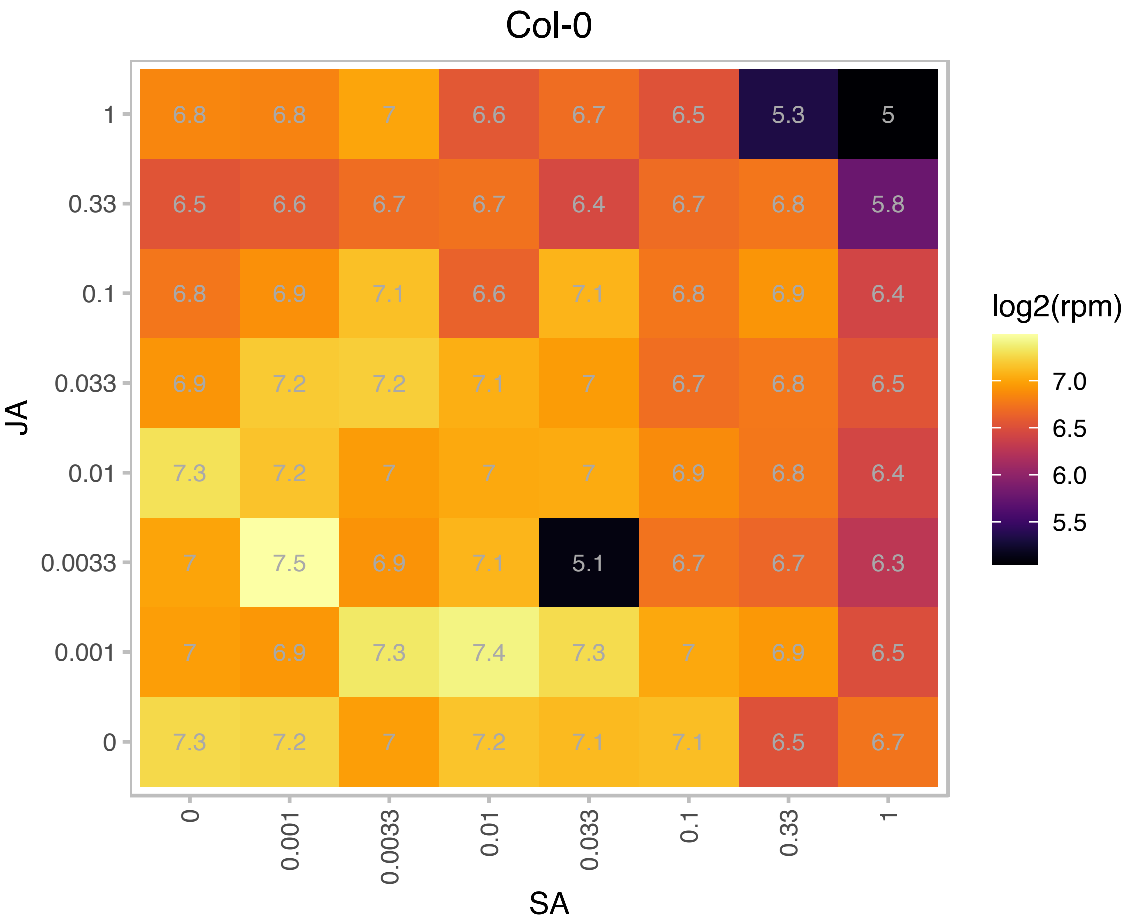


| | |
|---|---|
| Gene | AT1G20580 |
| is.expressed_genes | TRUE |
| is.responsive_genes | TRUE |
| is.apcluster | F-08 |

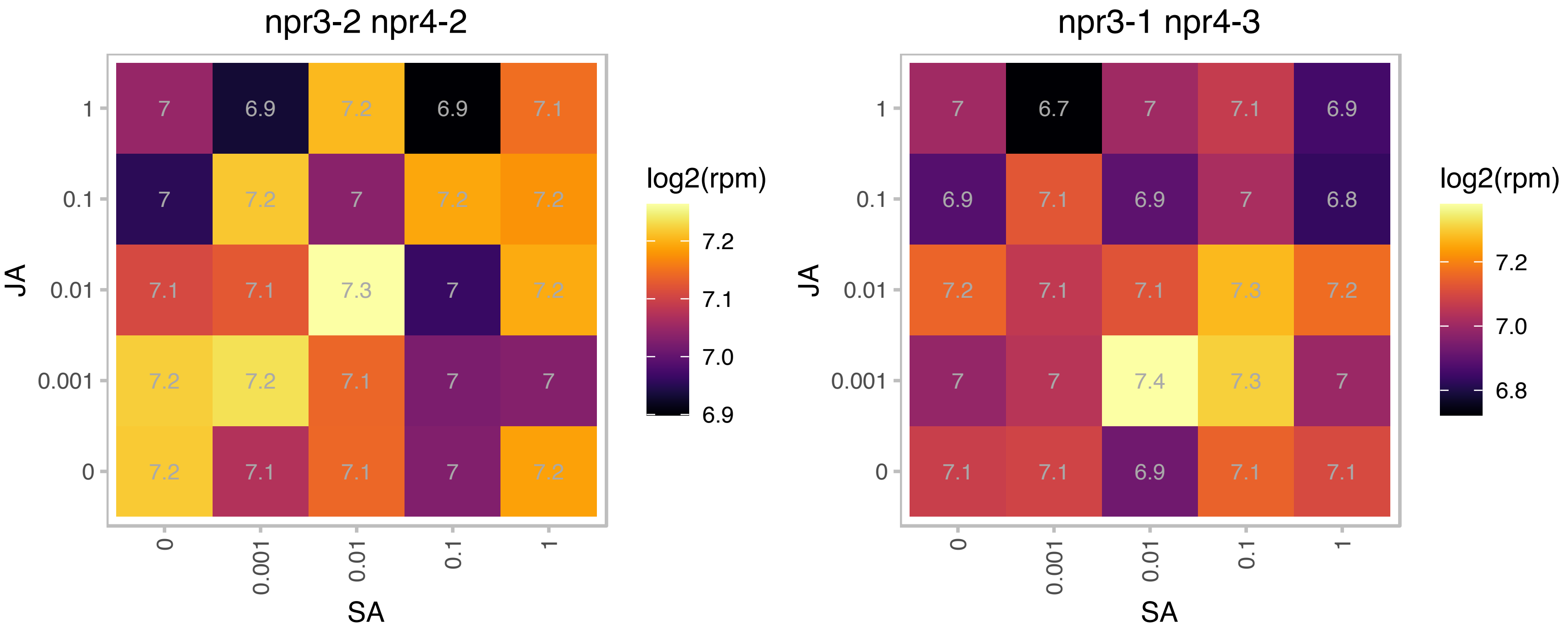

# AT1G21050.1 : NA

Protein of unknown function, DUF617

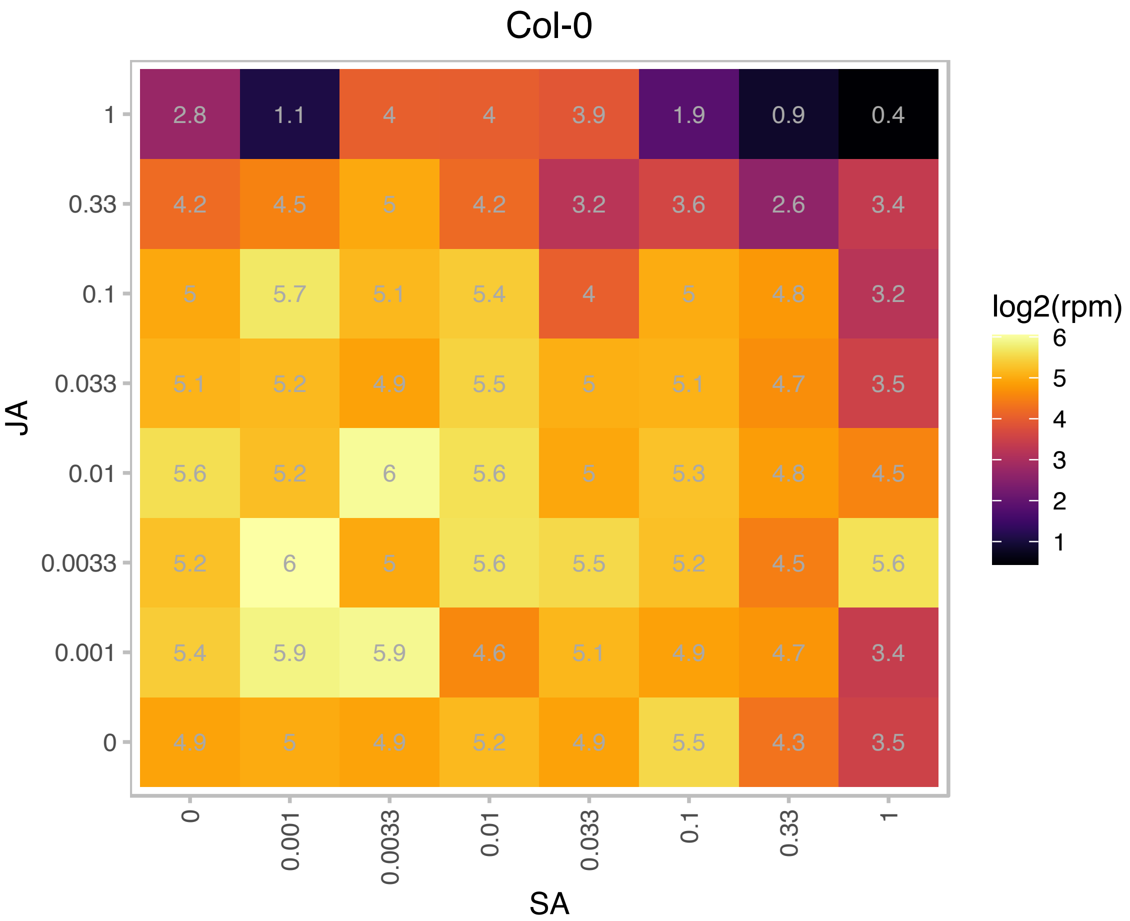


| | |
|---|---|
| Gene | AT1G21050 |
| is.expressed_genes | TRUE |
| is.responsive_genes | TRUE |
| is.apcluster | F-08 |

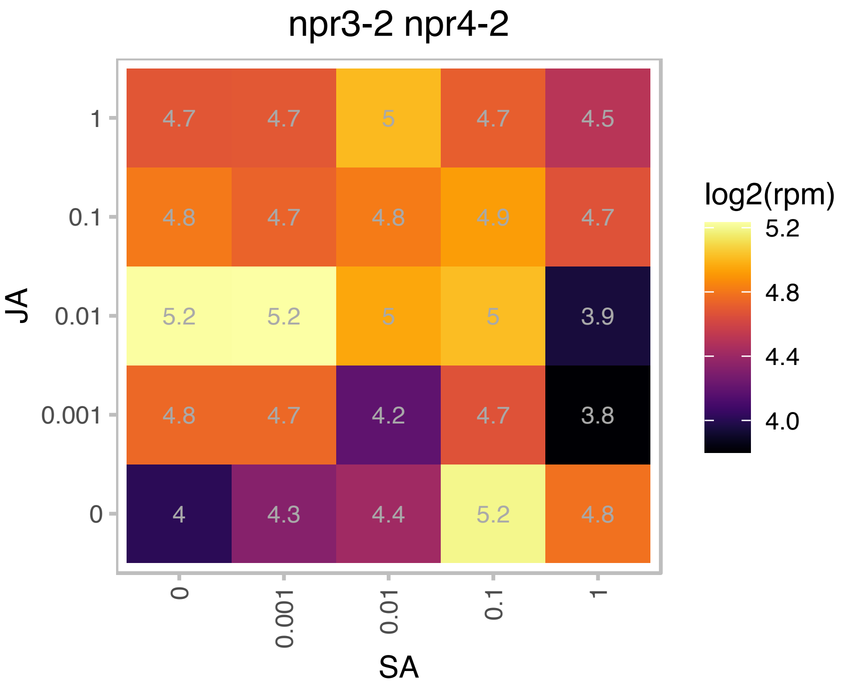


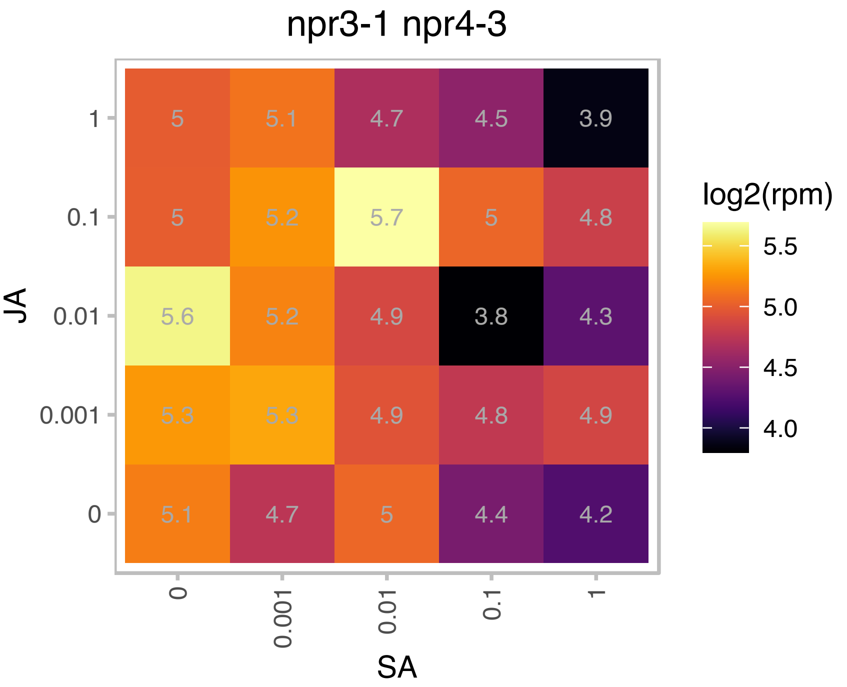

# AT1G25260.1 : NA

Ribosomal protein L10 family protein

| | |
|---|---|
| Gene | AT1G25260 |
| is.expressed_genes | TRUE |
| is.responsive_genes | TRUE |
| is.apcluster | F-08 |

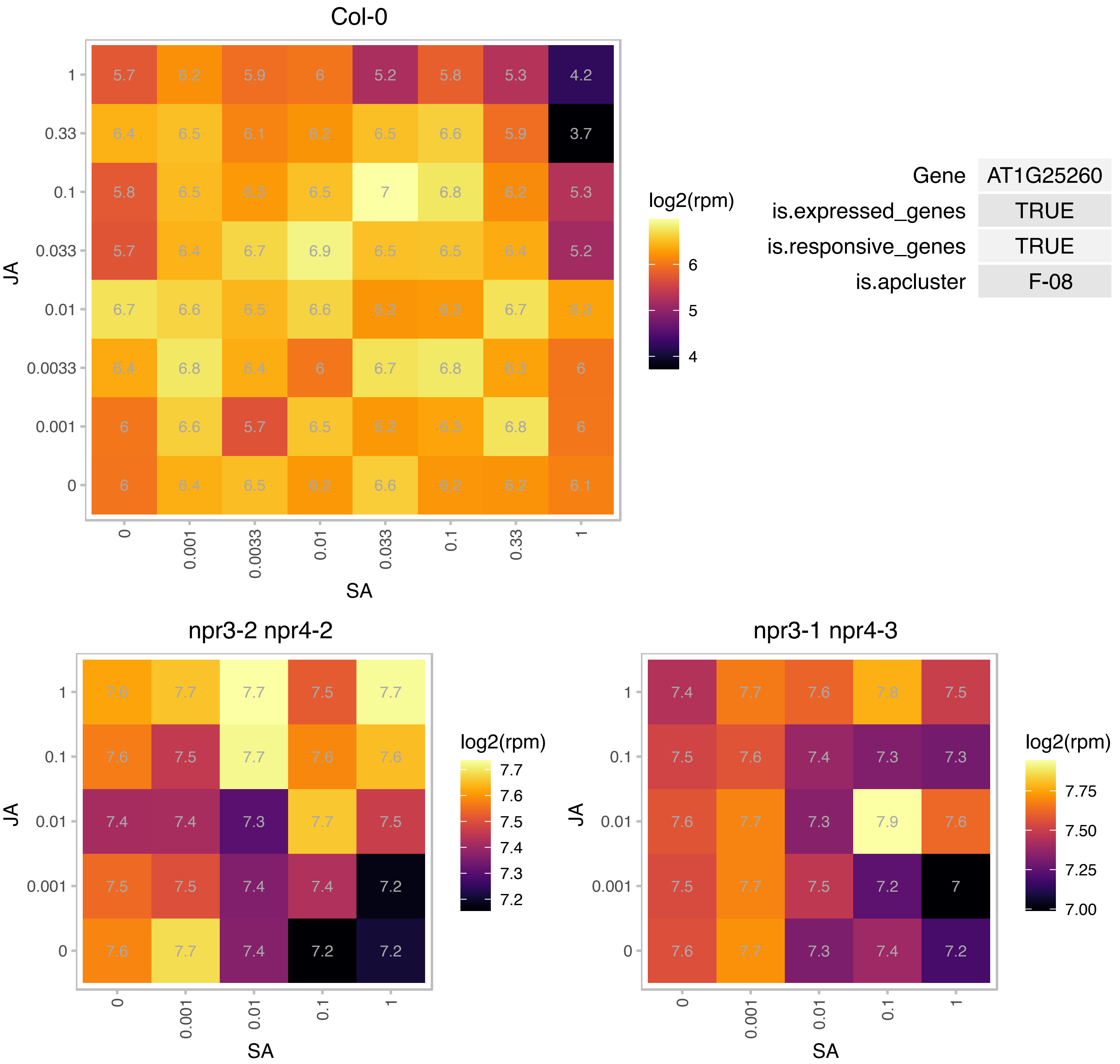

### AT1G25560.1 : AtTEM1, EDF1, TEM1

AP2/B3 transcription factor family protein

Encodes a member of the RAV transcription factor family that contains AP2 and B3 binding domains. Involved in the regulation of flowering under long days. Loss of function results in early flowering. Overexpression causes late flowering and repression of expression of FT. Novel transcriptional regulator involved in ethylene signaling. Promoter bound by EIN3. EDF1 in turn, binds to promoter elements in ethylene responsive genes.

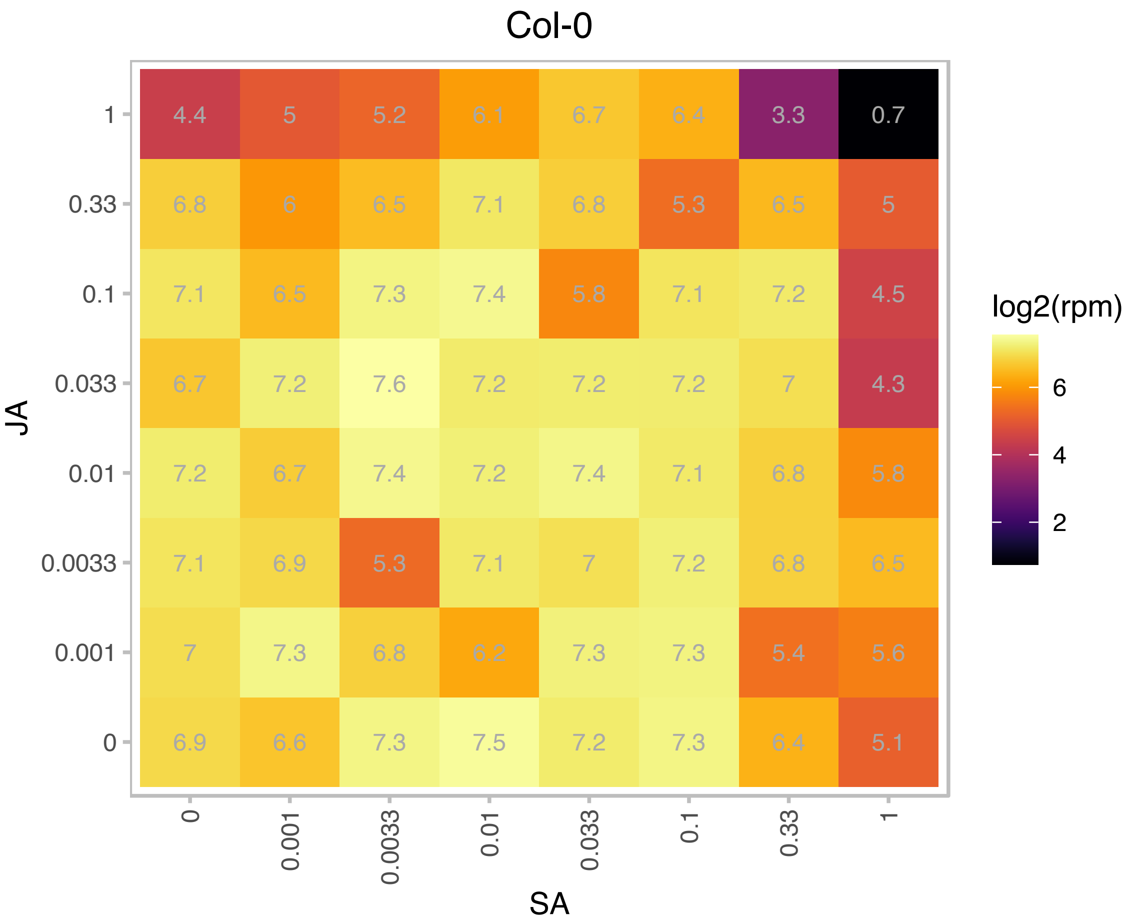


| | |
|---|---|
| Gene | AT1G25560 |
| is.expressed_genes | TRUE |
| is.responsive_genes | TRUE |
| is.apcluster | F-08 |

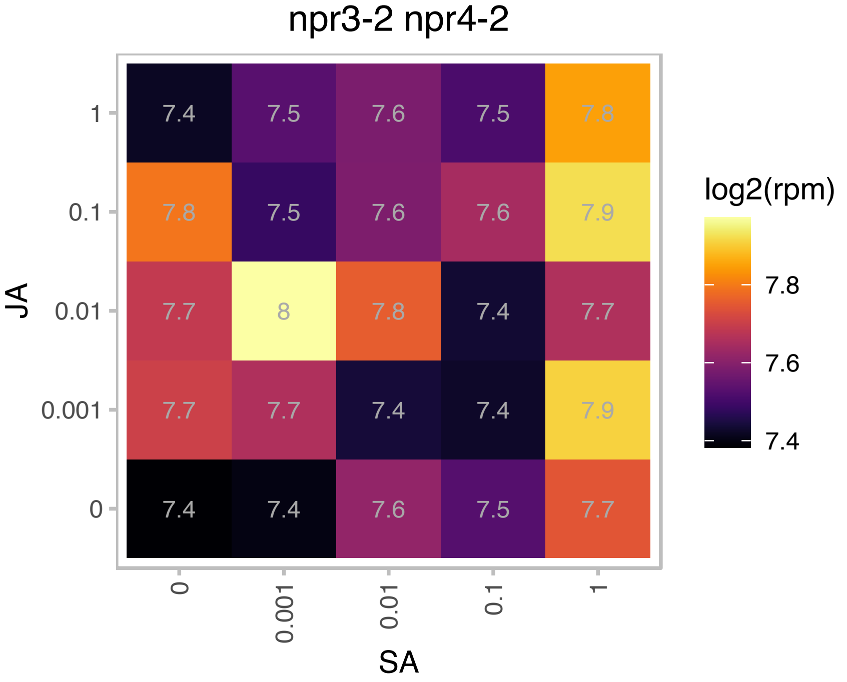


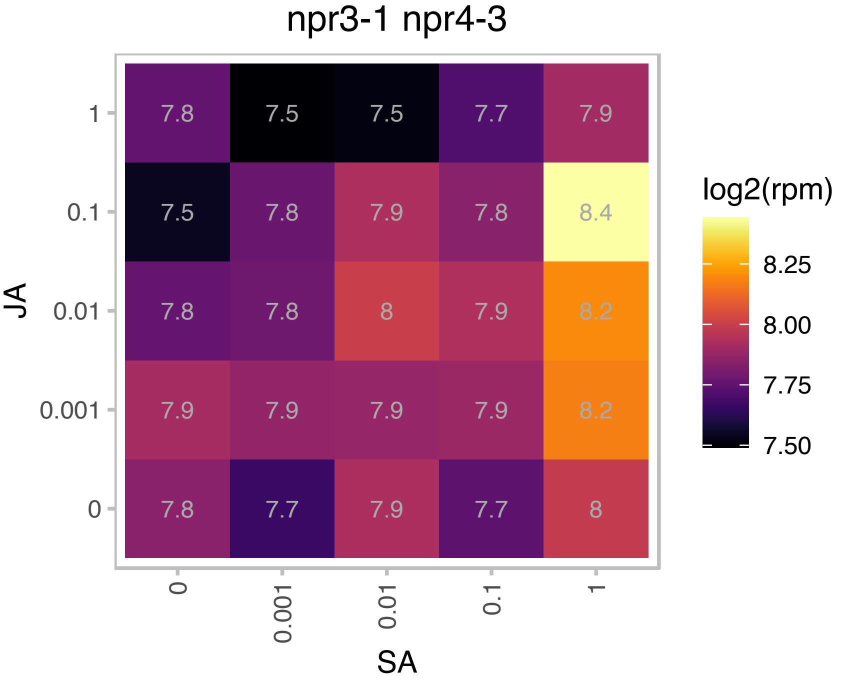

### AT1G26218.1 : sORF14

other RNA

Potential natural antisense gene, locus overlaps with AT1G26220. Has been identified as a translated small open reading frame by ribosome profiling.

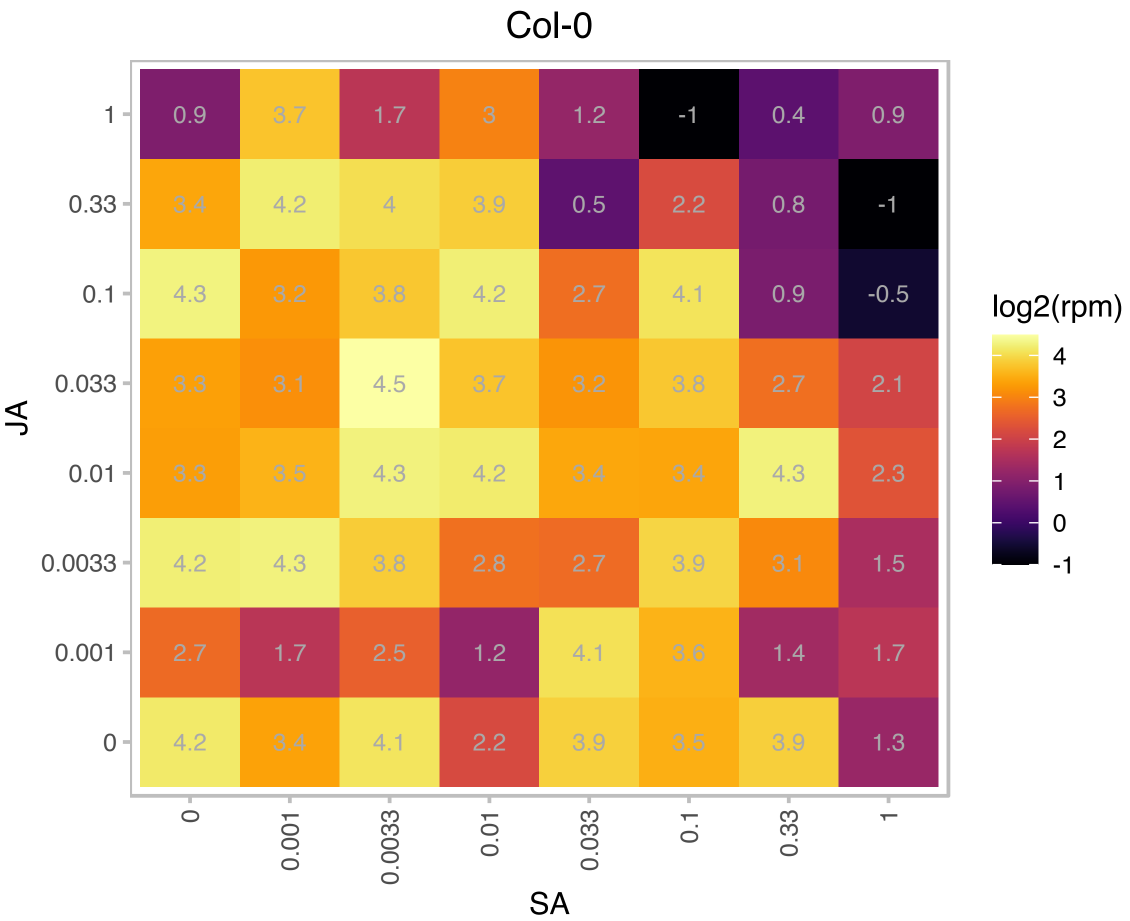


| | |
|---|---|
| Gene | AT1G26218 |
| is.expressed_genes | TRUE |
| is.responsive_genes | TRUE |
| is.apcluster | F-08 |

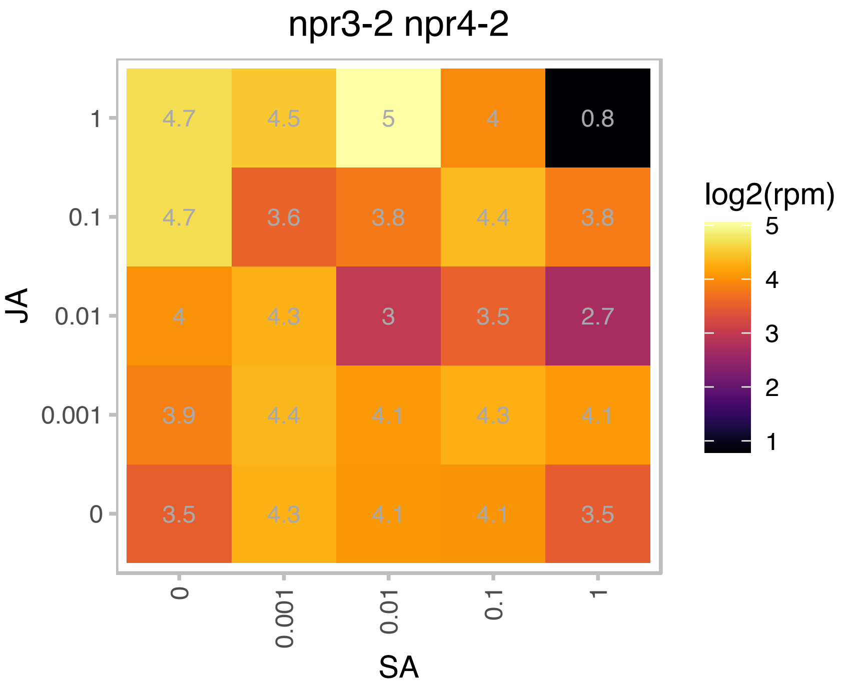


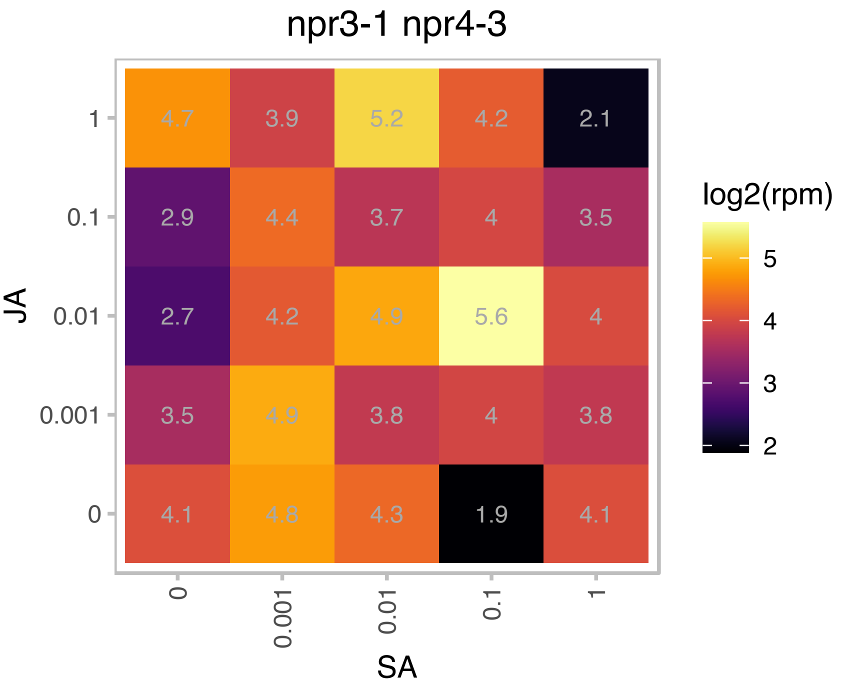

# AT1G27435.1 : NA

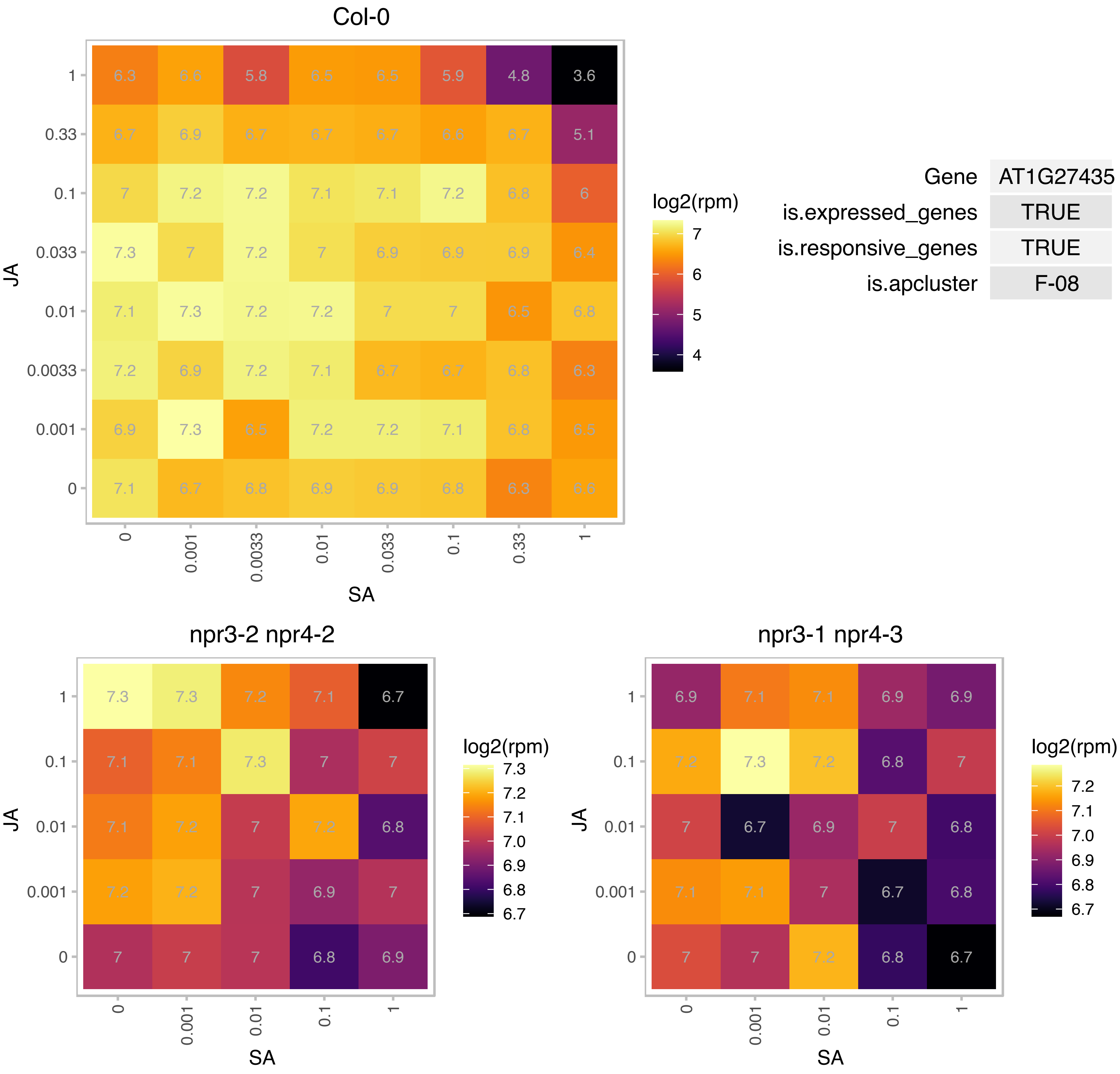

### AT1G27450.1 : APT1, ATAPT1

adenine phosphoribosyl transferase 1

Adenosine phosphoribosyl transferase(E.C:2.4.2.7), involved in the one-step salvage of adenine to AMP.

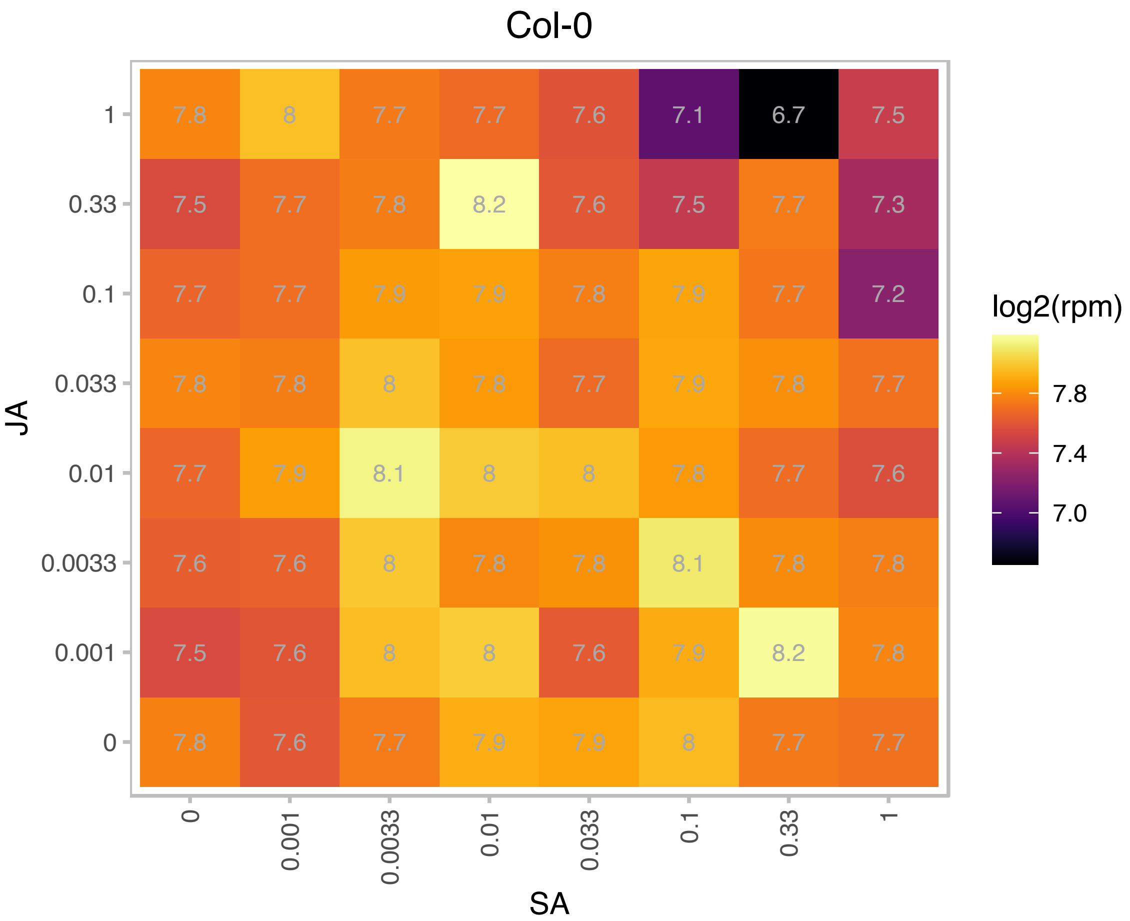


| | |
|---|---|
| Gene | AT1G27450 |
| is.expressed_genes | TRUE |
| is.responsive_genes | TRUE |
| is.apcluster | F-08 |

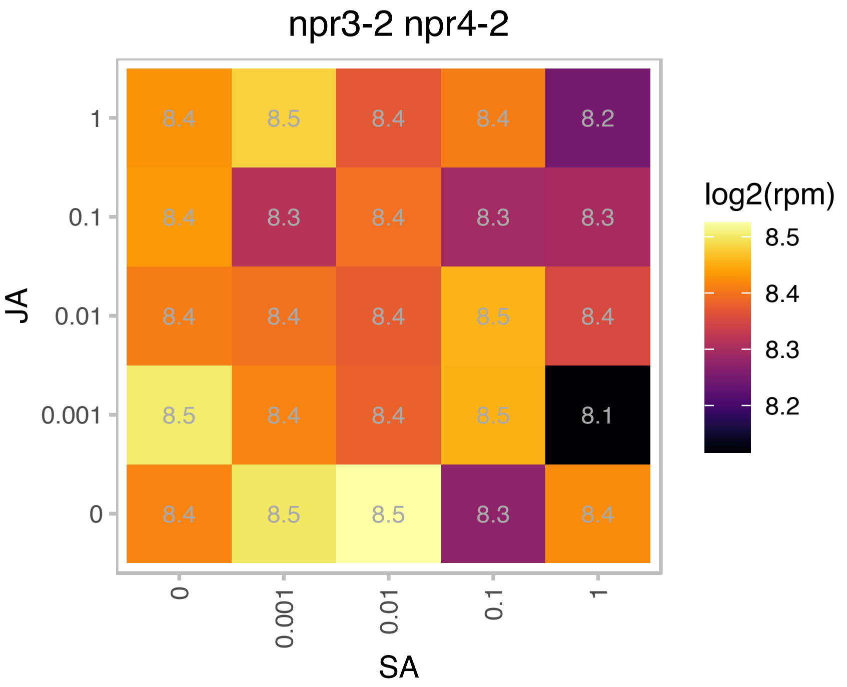


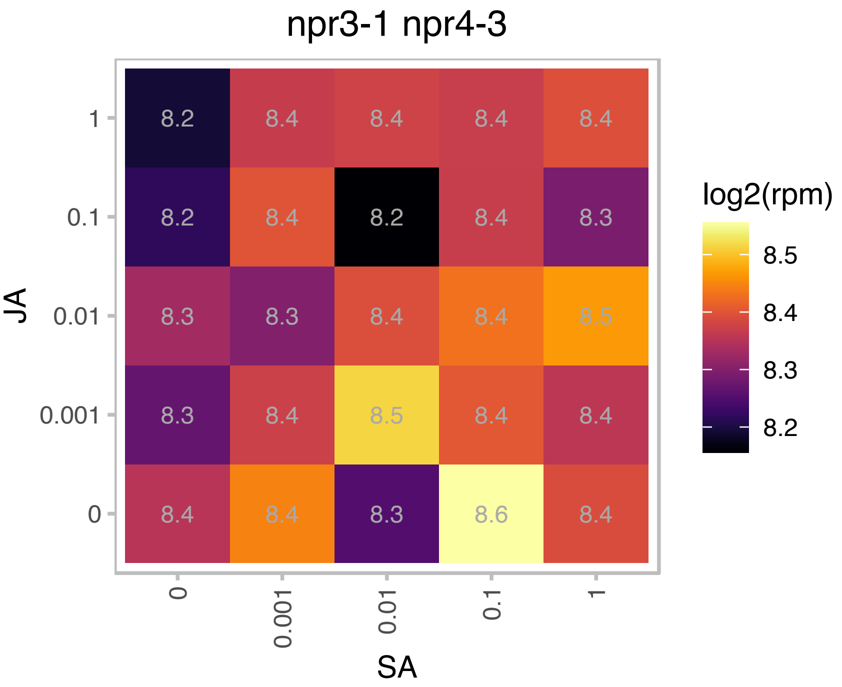

### AT1G33140.1 : PGY2

Ribosomal protein L6 family
Encodes ribosomal protein L9. Identified in a screen for enhancers of as1. as1/pgy double mutants show defects in leaf vascular patterning and adaxial cell fate. Double mutant analysis indicates pgy genes function in the same pathway as REV, KAN1 and KAN2. The mRNA is cell-to-cell mobile.

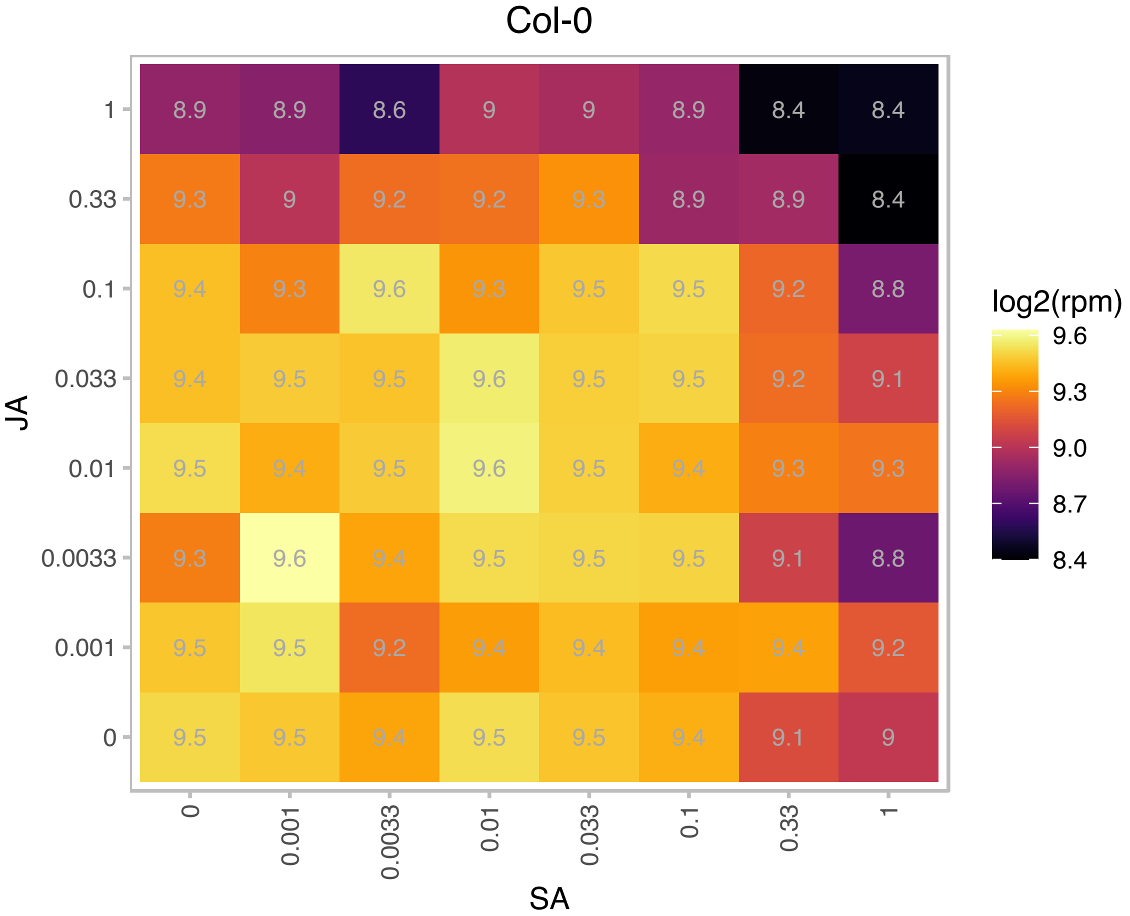


| | |
|---|---|
| Gene | AT1G33140 |
| is.expressed_genes | TRUE |
| is.responsive_genes | TRUE |
| is.apcluster | F-08 |

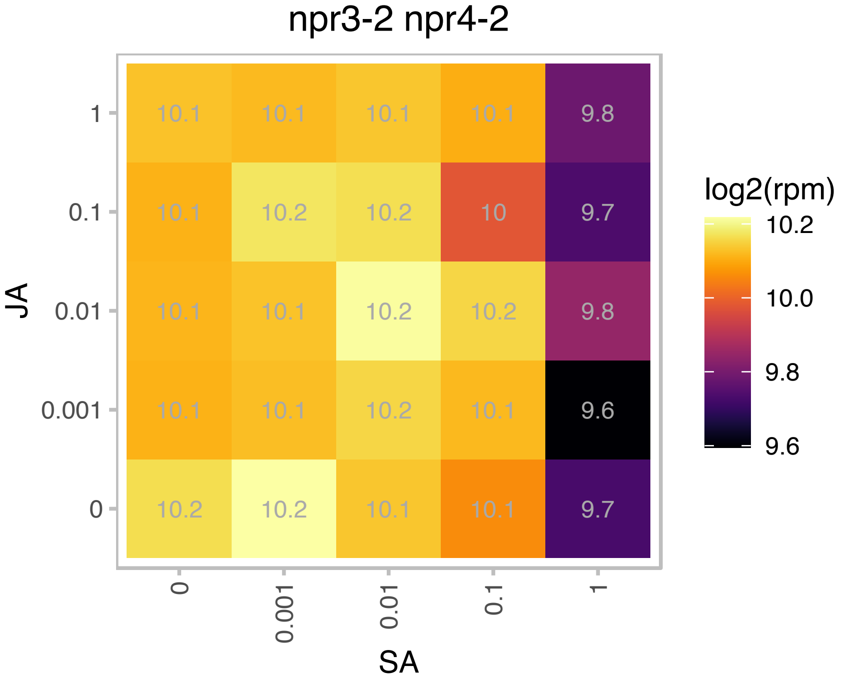


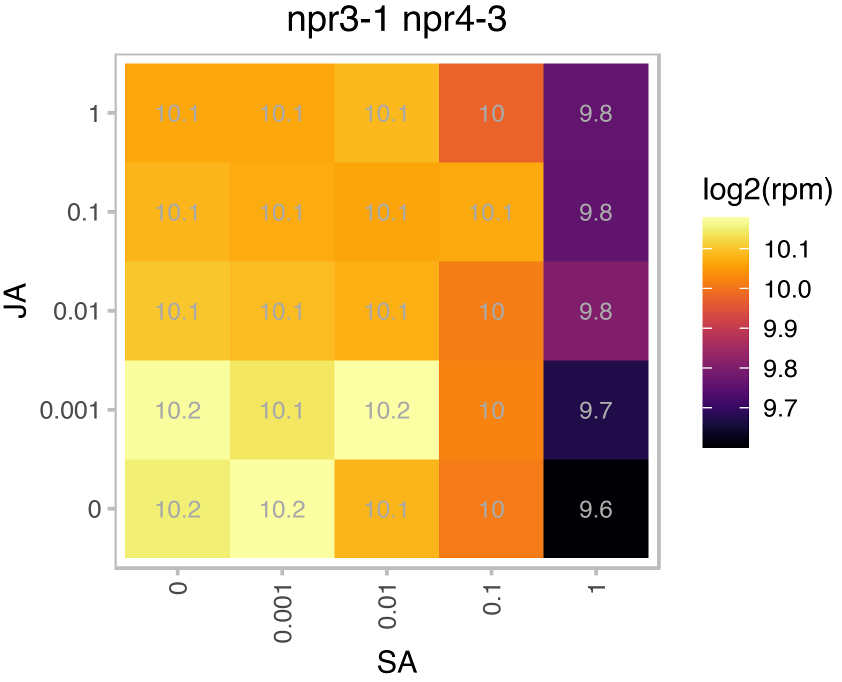

### AT1G36310.1 : AtTRM9, TRM9

S-adenosyl-L-methionine-dependent methyltransferases superfamily protein

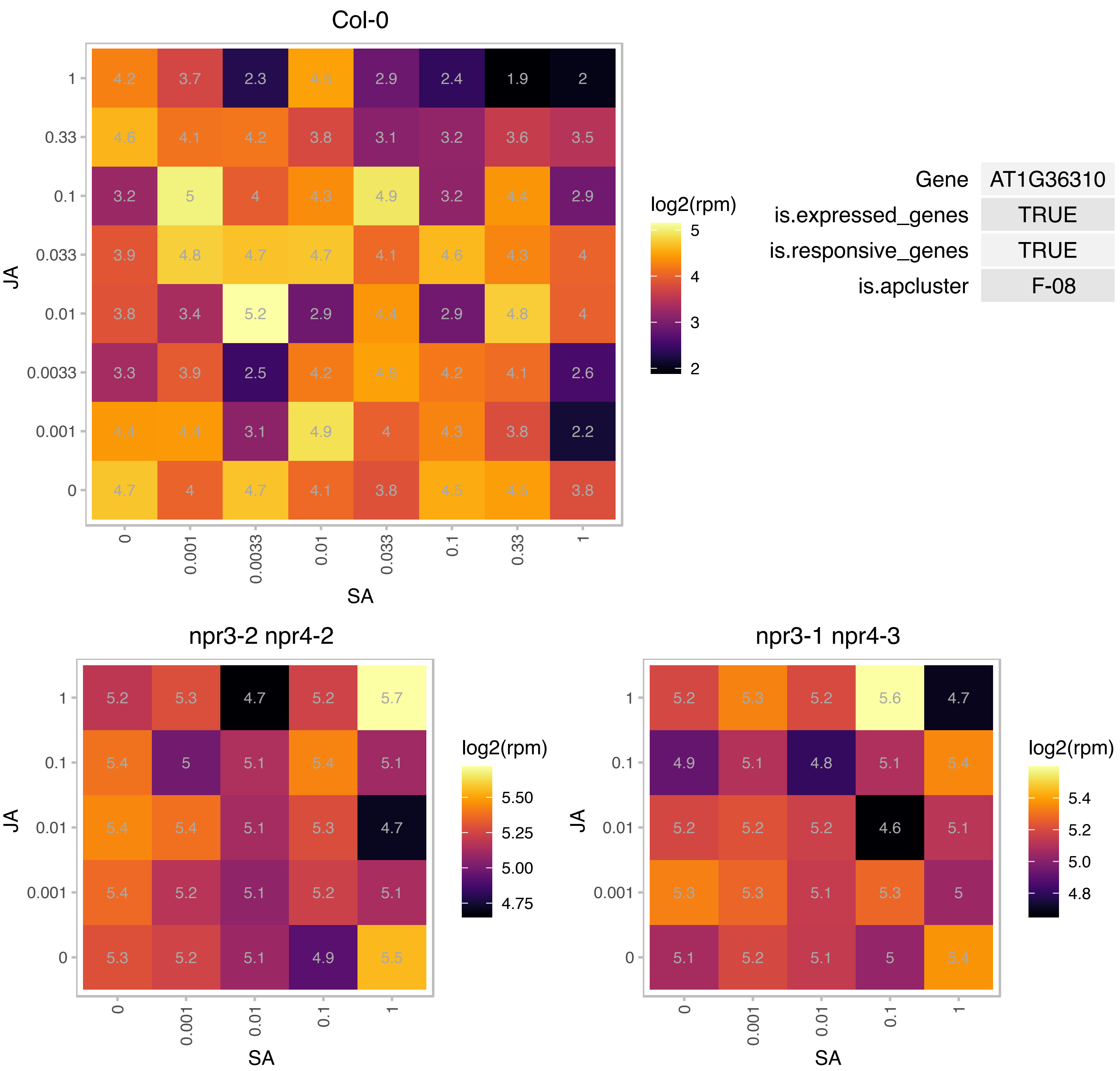

# AT1G41880.1 : NA

Ribosomal protein L35Ae family protein

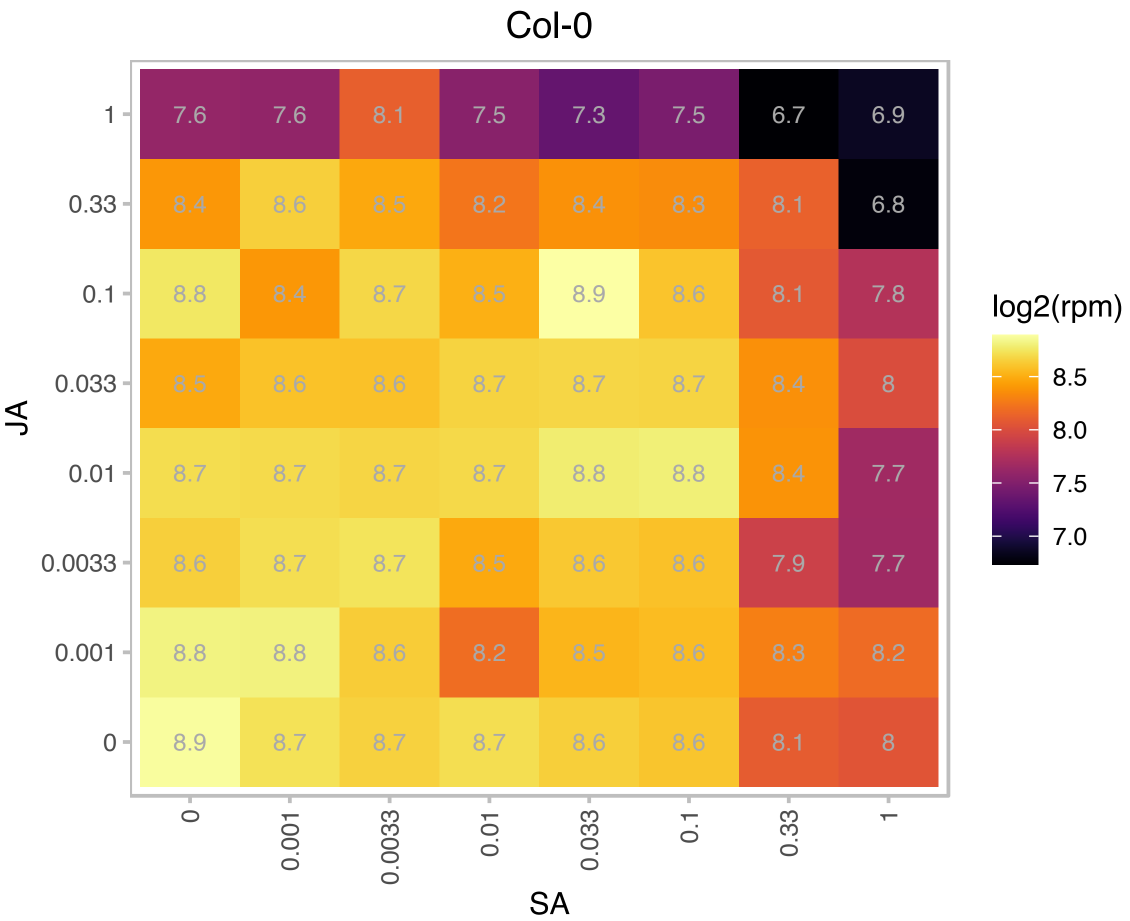


| | |
|---|---|
| Gene | AT1G41880 |
| is.expressed_genes | TRUE |
| is.responsive_genes | TRUE |
| is.apcluster | F-08 |

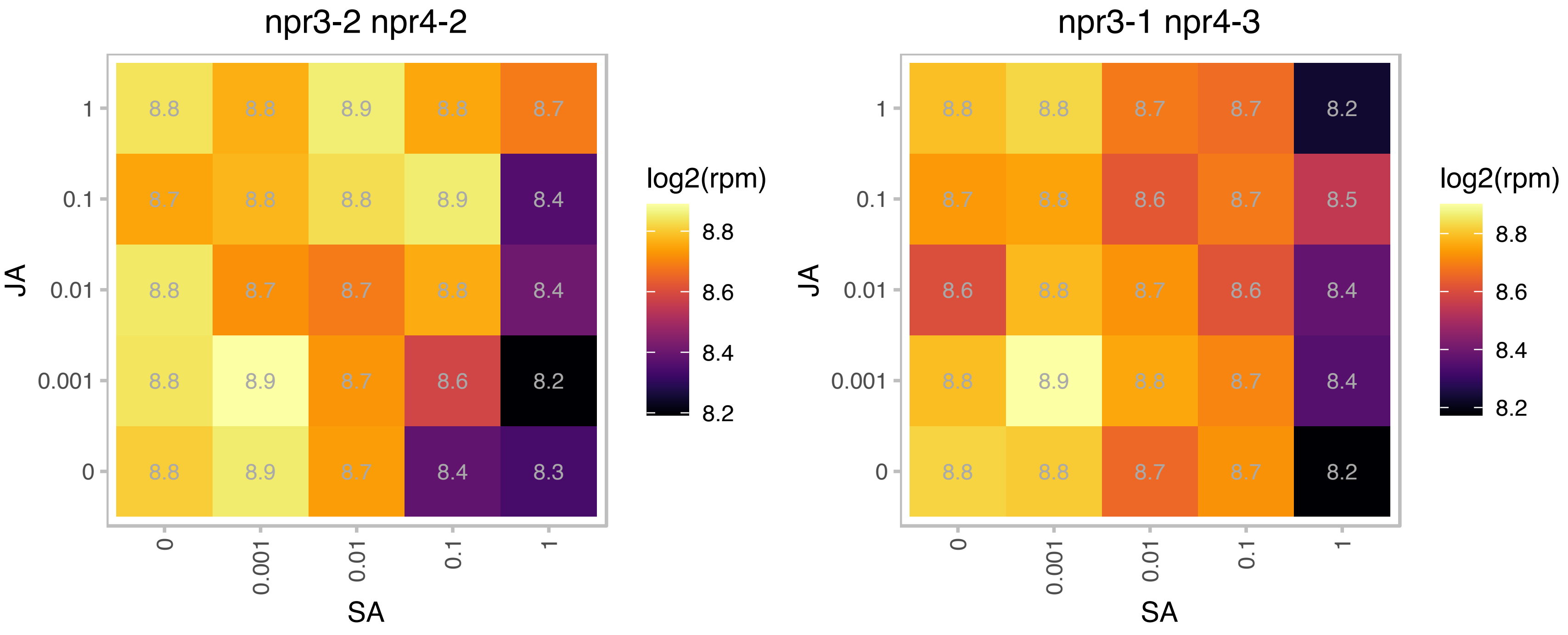

# AT1G42480.1 : NA

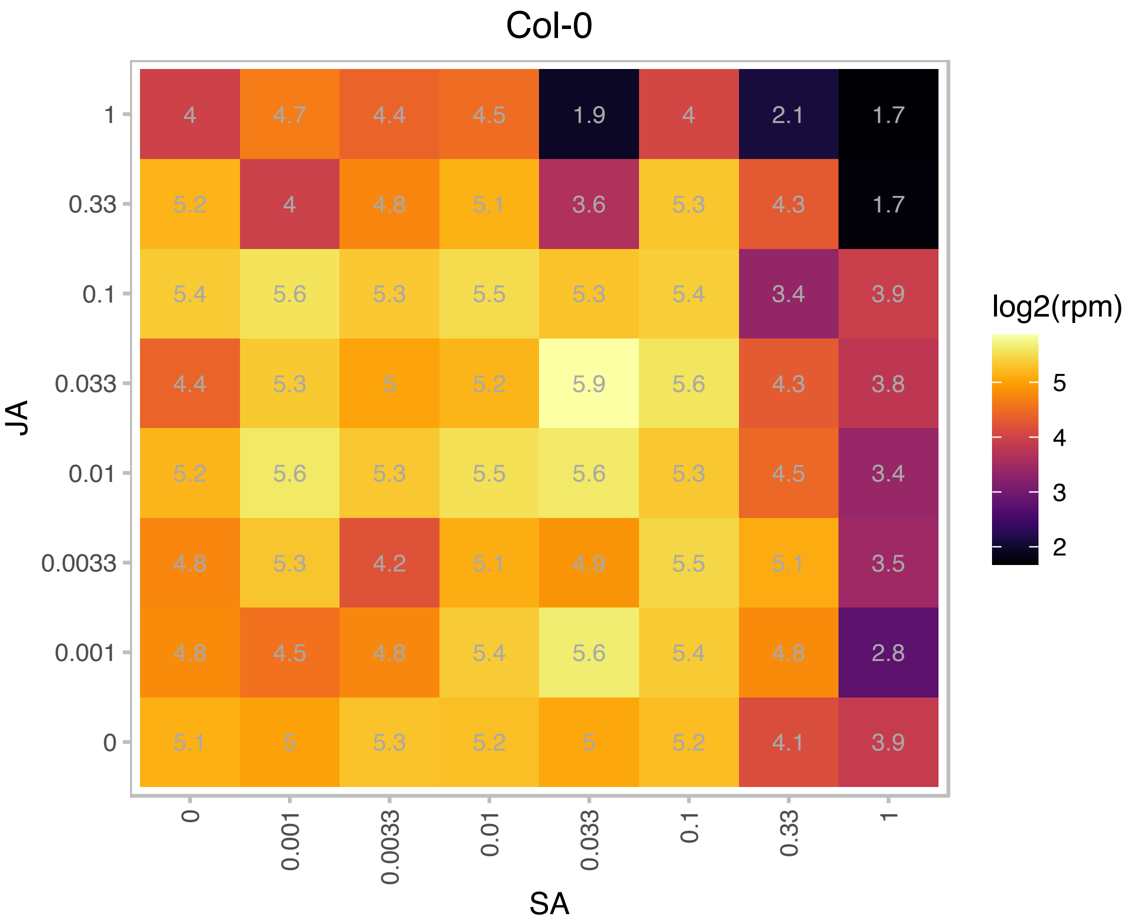


| | |
|---|---|
| Gene | AT1G42480 |
| is.expressed_genes | TRUE |
| is.responsive_genes | TRUE |
| is.apcluster | F-08 |

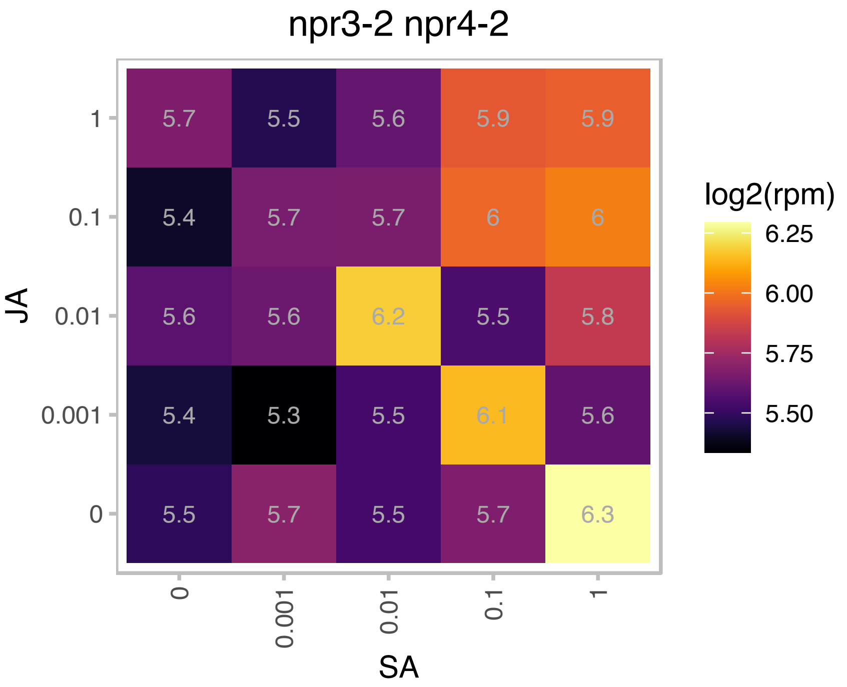


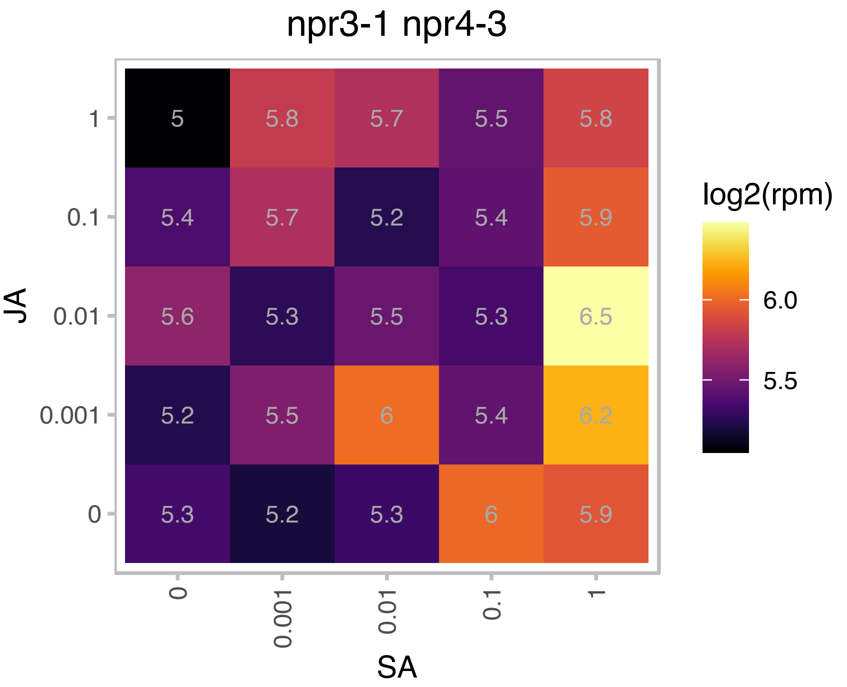

### AT1G48320.1 : DHNAT1

Thioesterase superfamily protein

Encodes one of the two functional DHNA-CoA (1,4-dihydroxy-2-naphthoyl-CoA) thioesterases found in Arabidopsis.

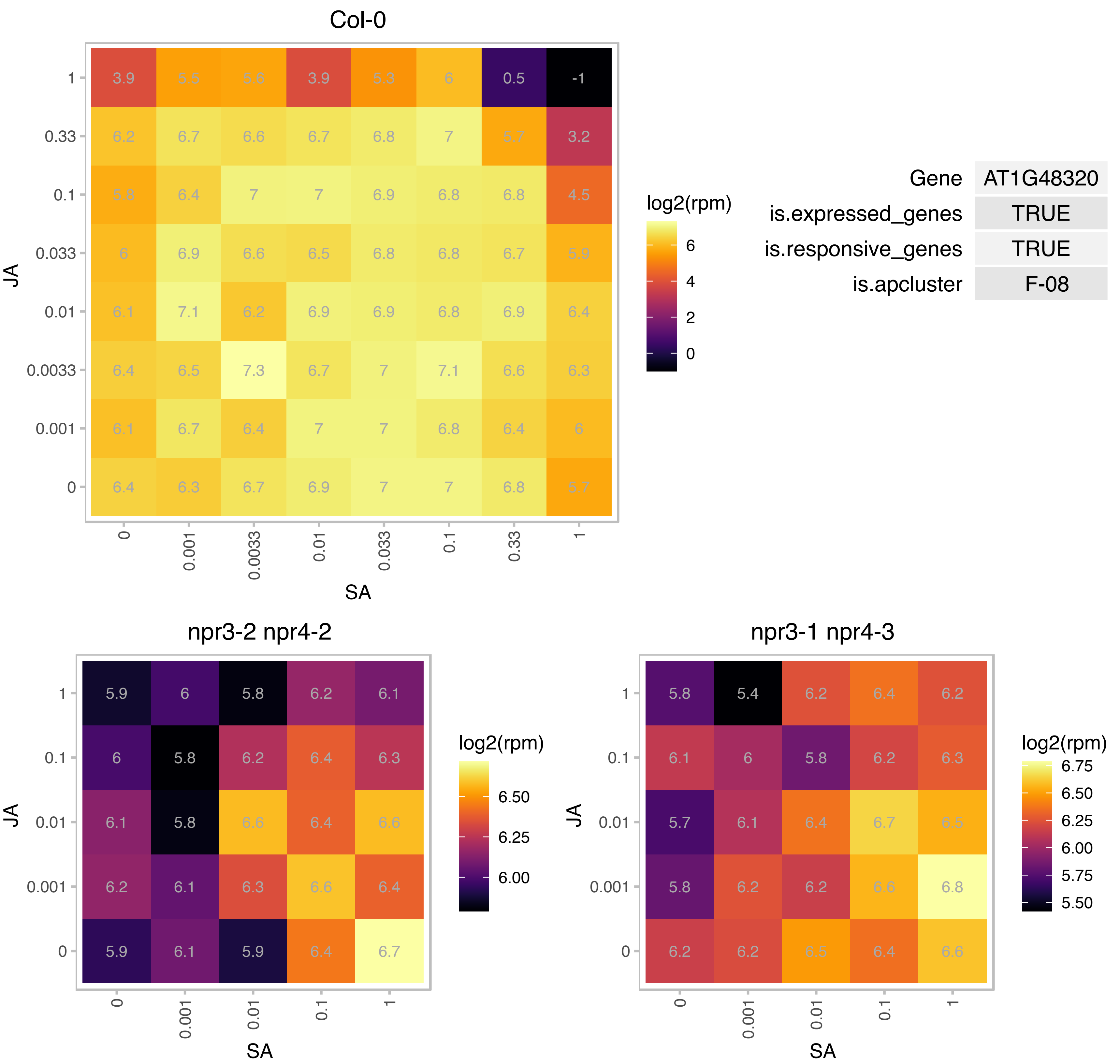


| | |
|---|---|
| Gene | AT1G48320 |
| is.expressed_genes | TRUE |
| is.responsive_genes | TRUE |
| is.apcluster | F-08 |

### AT1G48630.1 : RACK1B, RACK1B_AT

receptor for activated C kinase 1B
Encodes a protein with similarity to mammalian RACKs. RACKs function to shuttle activated protein kinase C to different subcellular sites and may also function as a scaffold through physical interactions with other proteins. RACK1B has no phenotype on its own and probably acts redundantly with RACK1A and RACK1C.

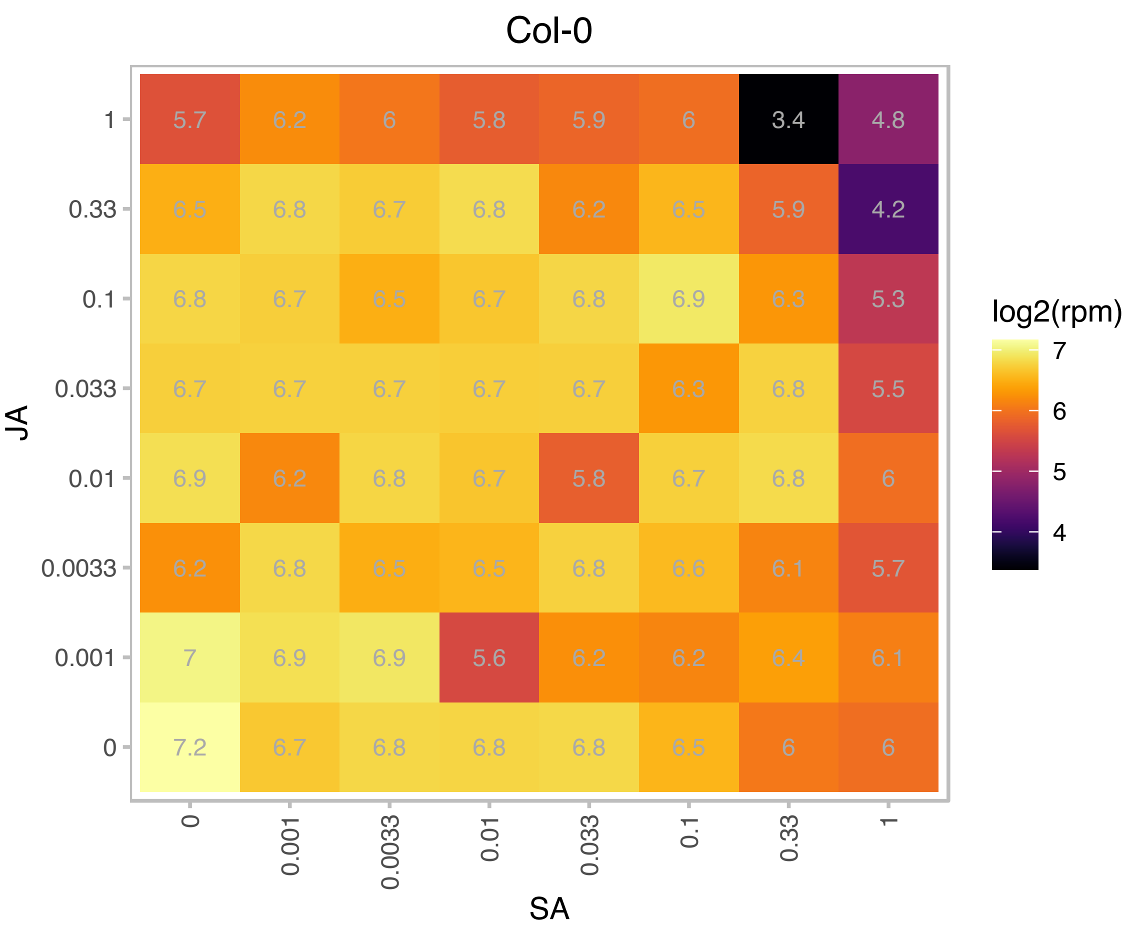


| | |
|---|---|
| Gene | AT1G48630 |
| is.expressed_genes | TRUE |
| is.responsive_genes | TRUE |
| is.apcluster | F-08 |

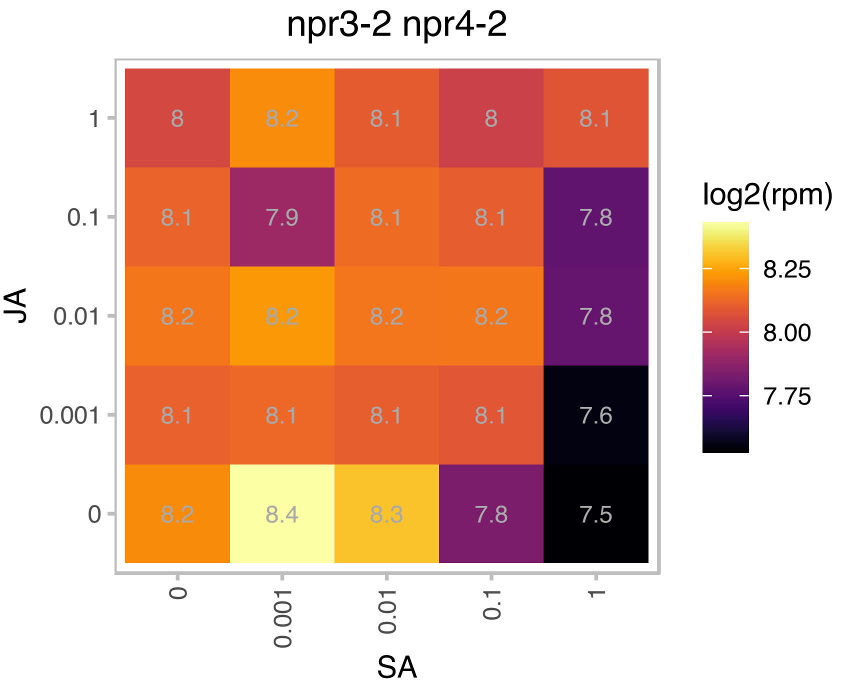


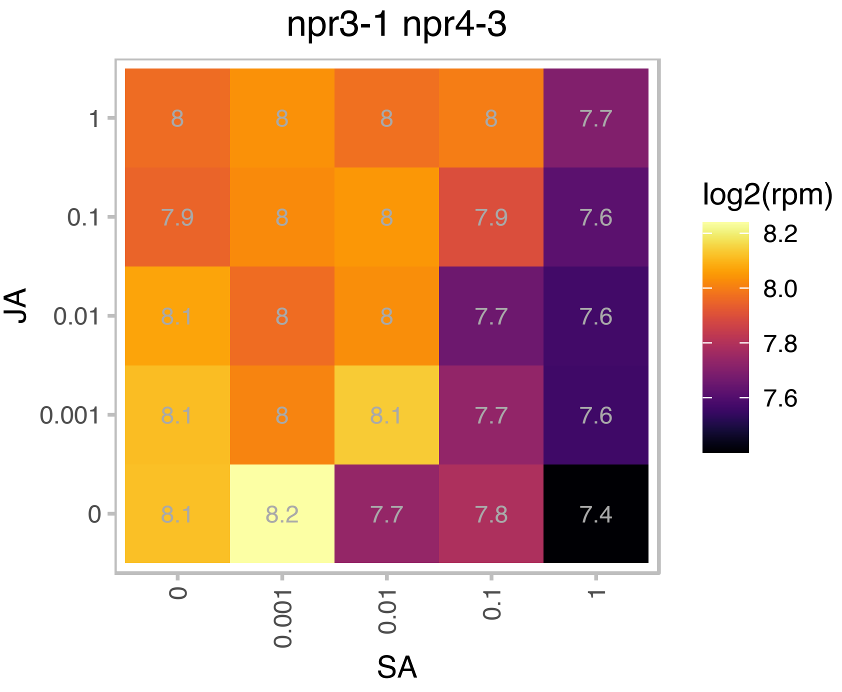

### AT1G49410.1 : TOM6

translocase of the outer mitochondrial membrane 6

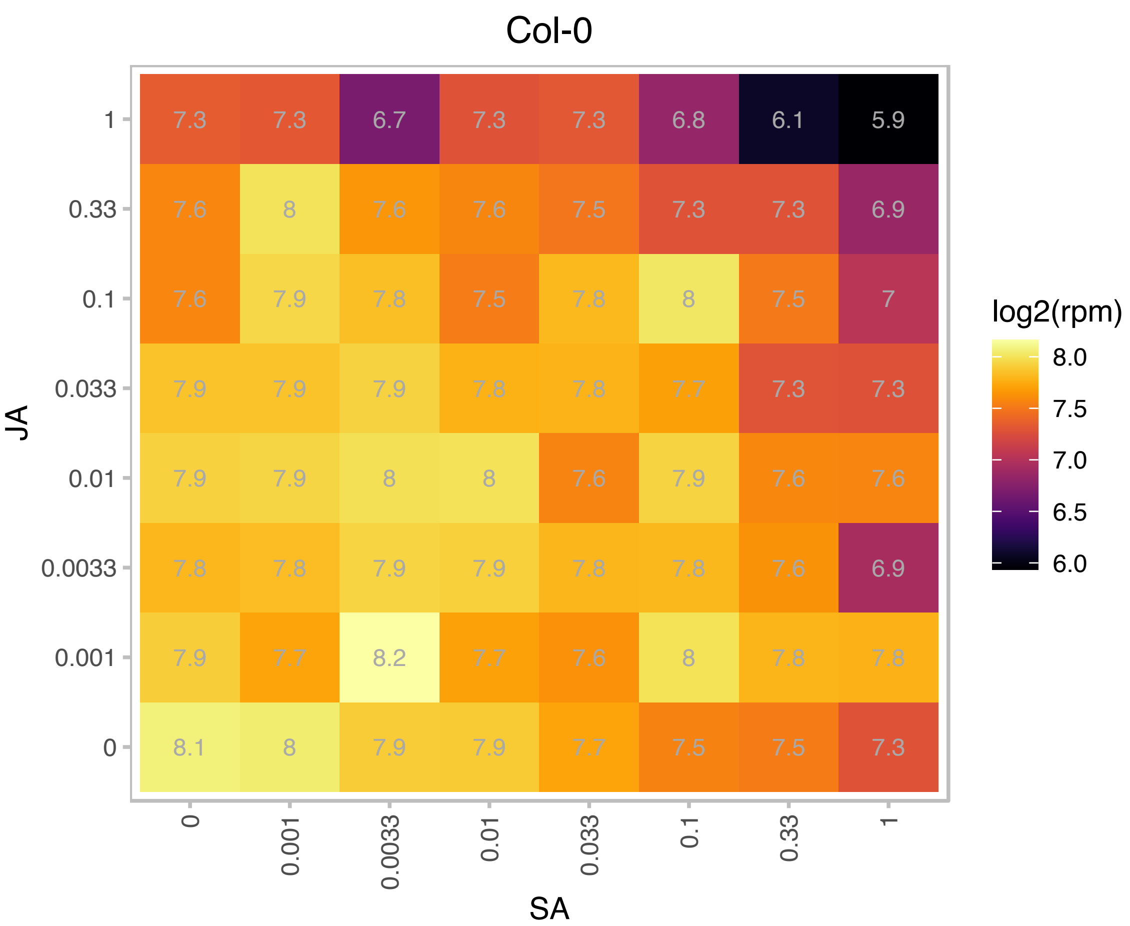


| | |
|---|---|
| Gene | AT1G49410 |
| is.expressed_genes | TRUE |
| is.responsive_genes | TRUE |
| is.apcluster | F-08 |

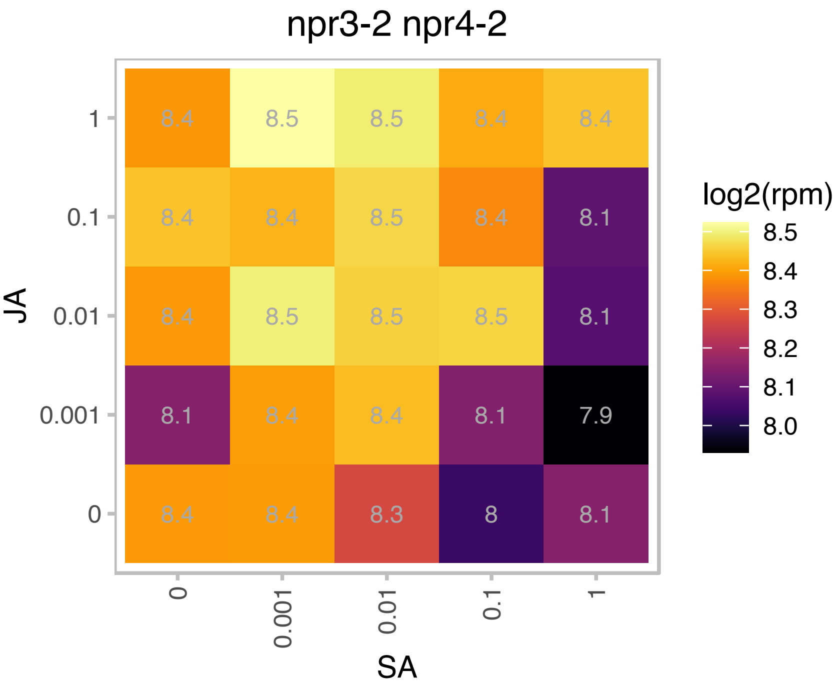


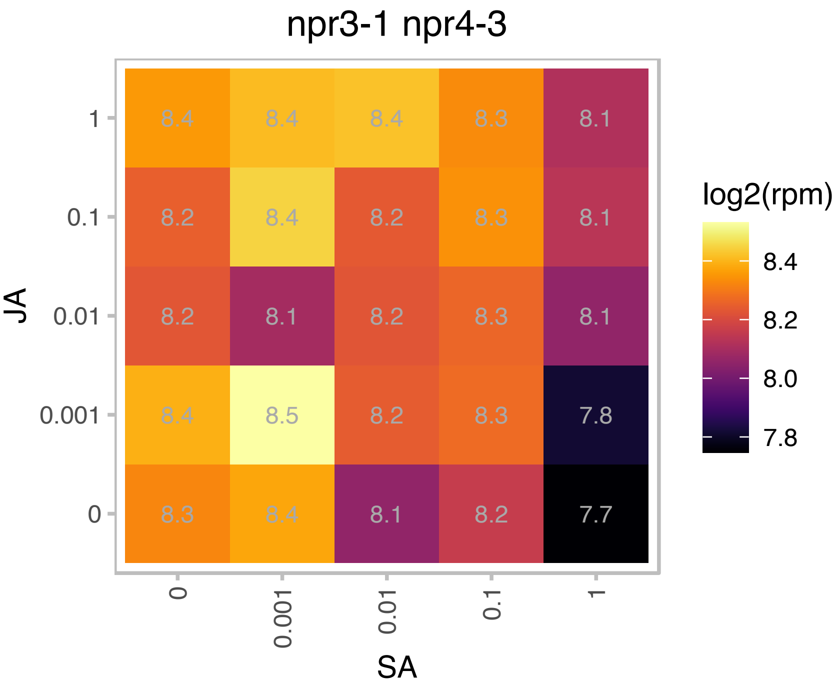

# AT1G49500.1 : NA

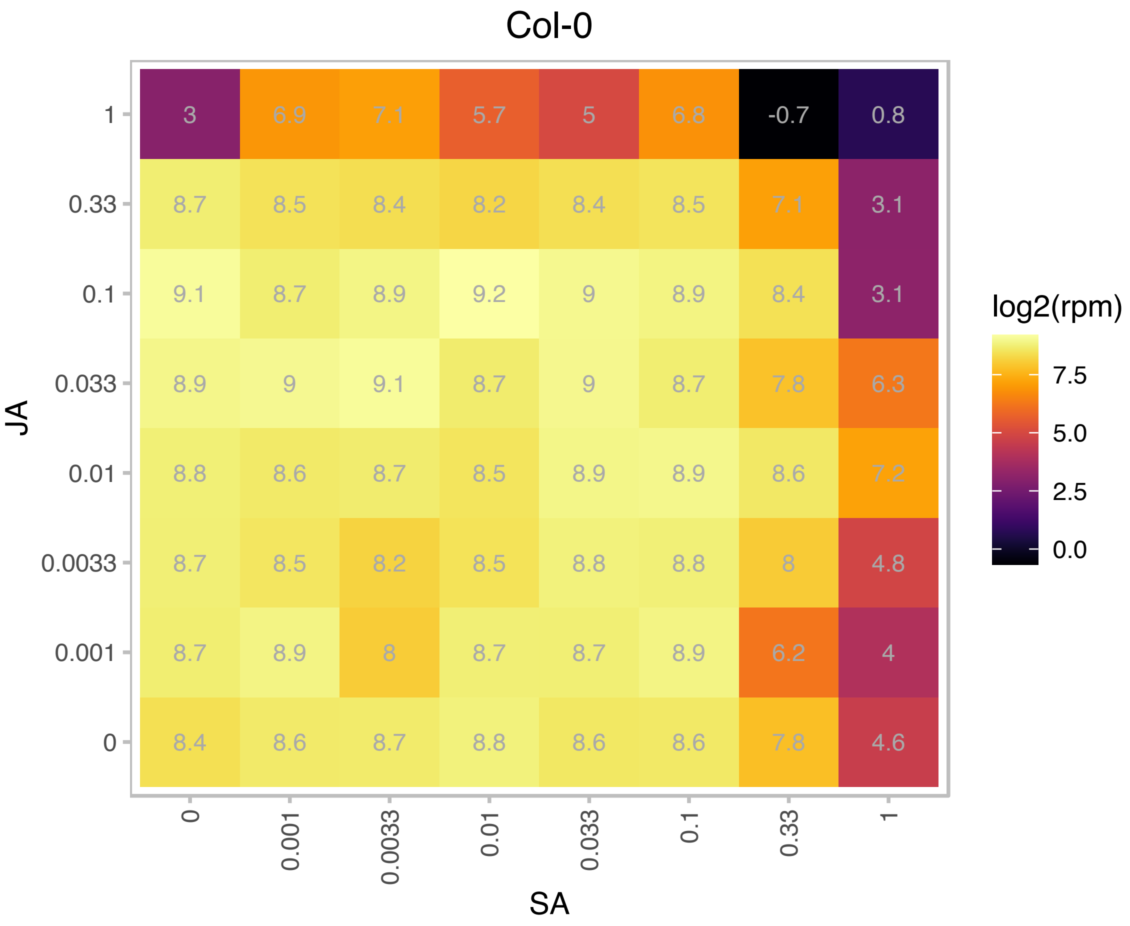


| | |
|---|---|
| Gene | AT1G49500 |
| is.expressed_genes | TRUE |
| is.responsive_genes | TRUE |
| is.apcluster | F-08 |

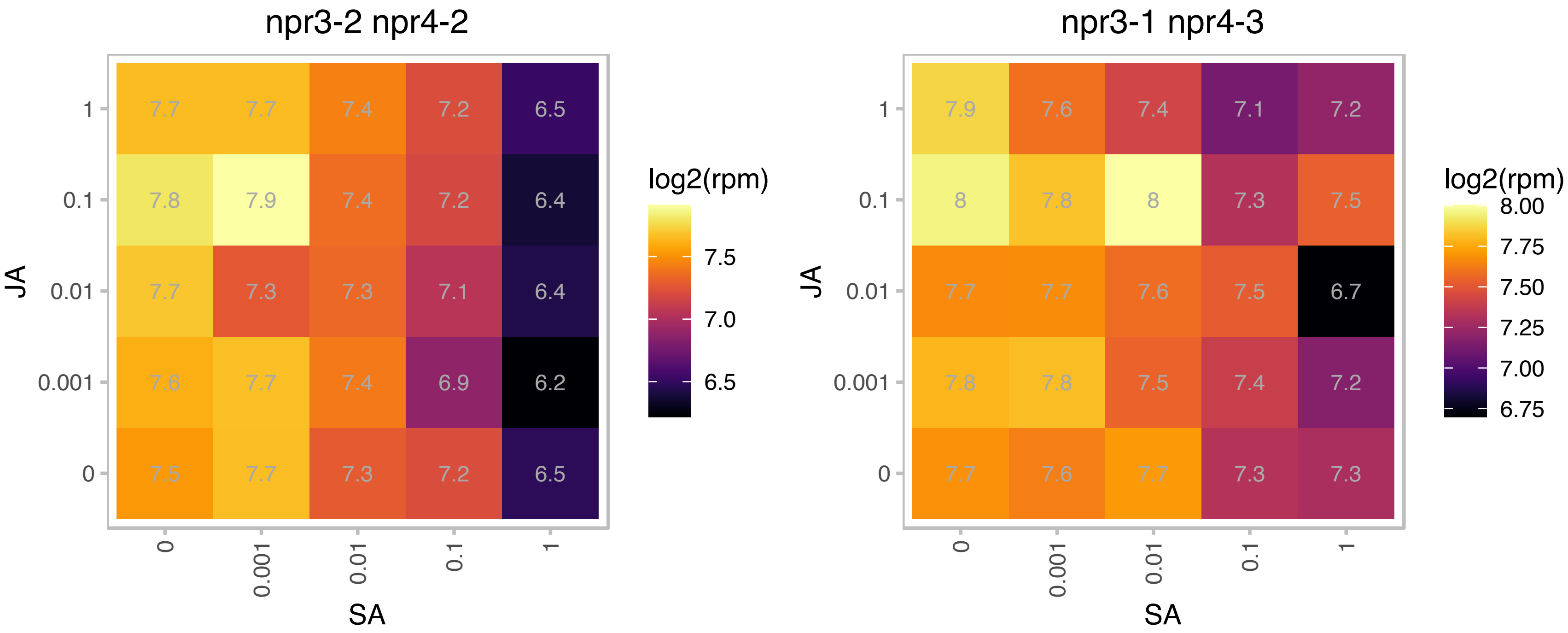

### AT1G49600.1 : ATRBP47A, RBP47A

RNA-binding protein 47A

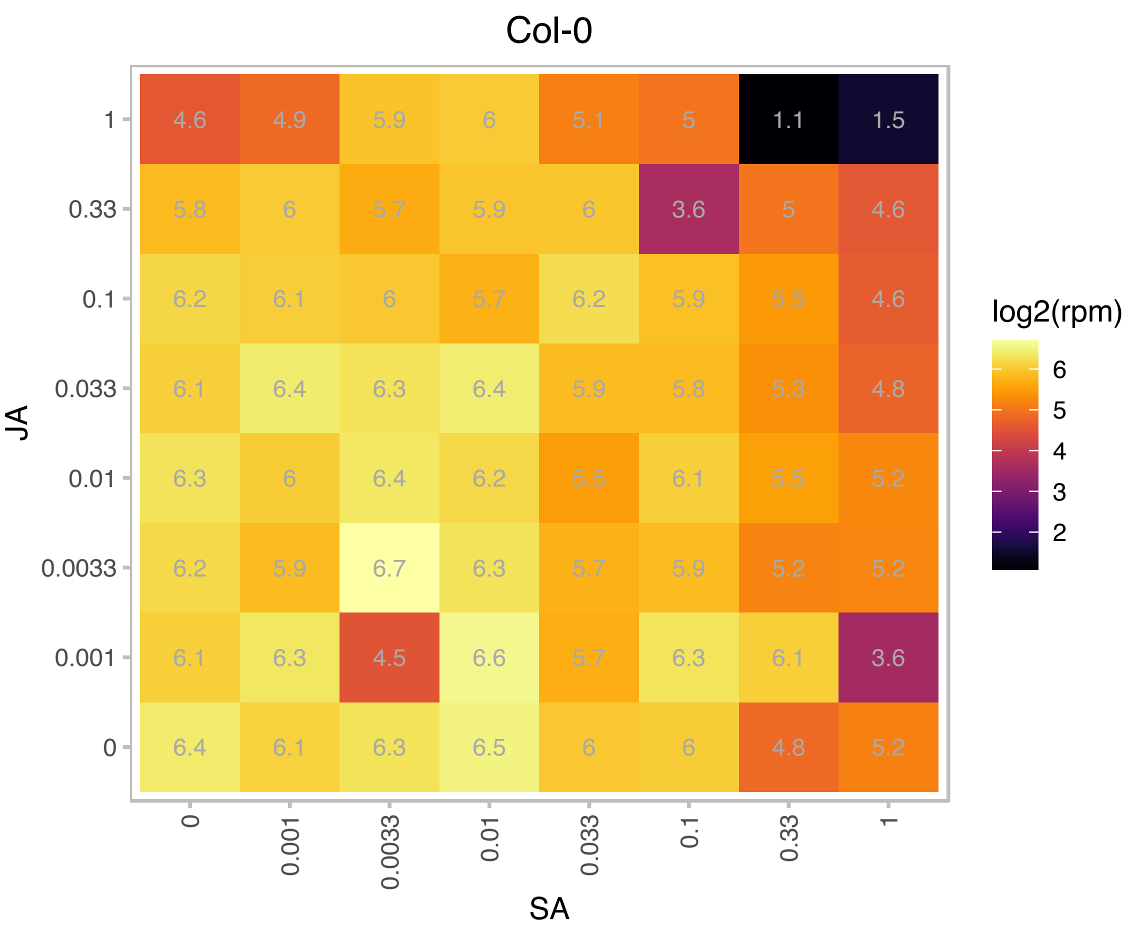


| Gene | AT1G49600 |
| --- | --- |
| is.expressed_genes | TRUE |
| is.responsive_genes | TRUE |
| is.apcluster | F-08 |

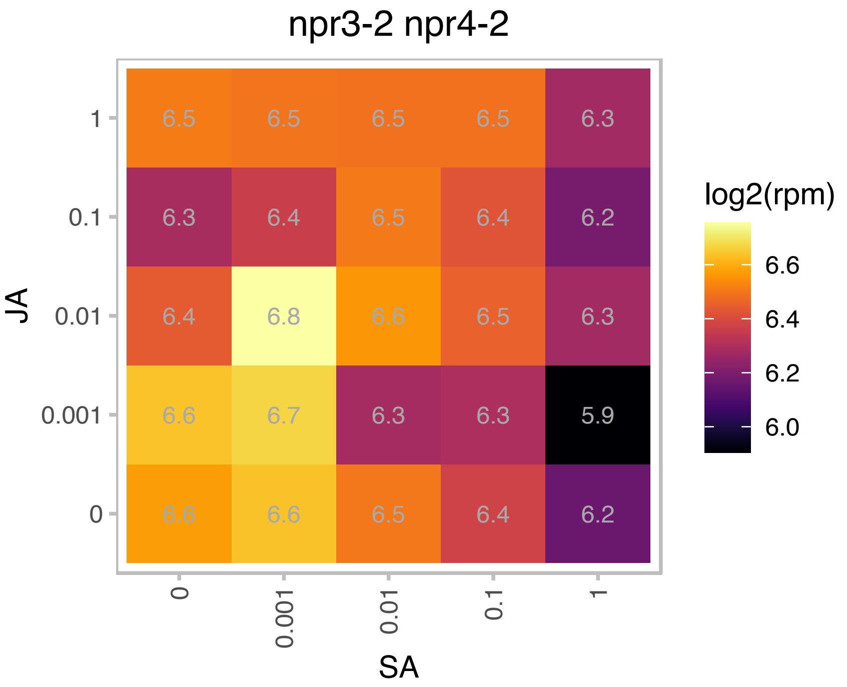


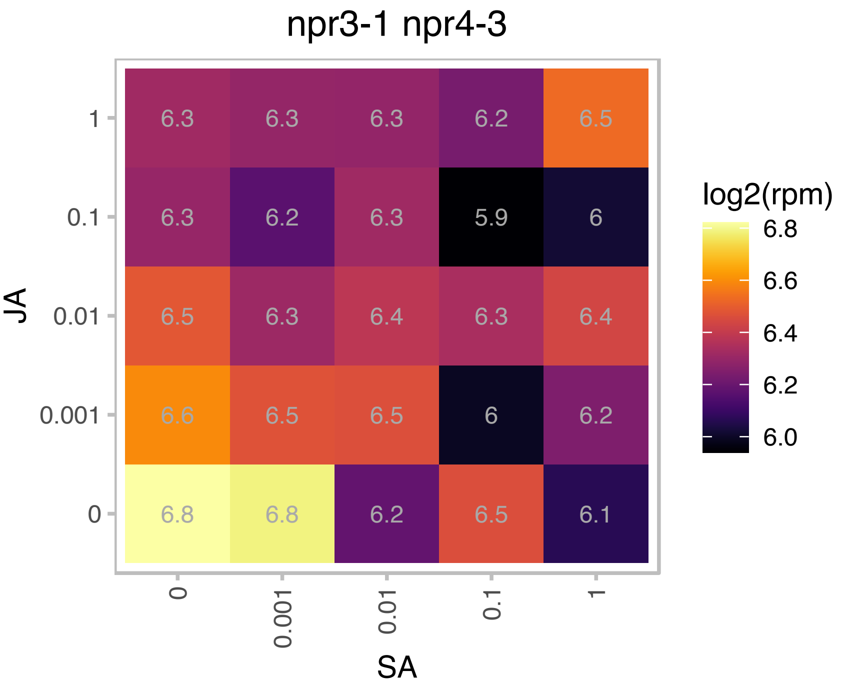

### AT1G49950.1 : ATTRB1, TRB1

telomere repeat binding factor 1

Encodes a telomeric DNA binding protein. In vitro, an N-terminal Myb domain enables it to interact with double-stranded telomeric repeats.Mutations in TRB1 and TRB3 enhance the lhp1 mutant phenotype. Triple mutants are more strongly affected than the respective double mutants with lhp1. TRB1 binds non-H3K27me3 target genes predominantly at the transcription start sites and its presence increases the expression of several of these targets, enriched for functions of primary metabolism. At H3K27me3 target genes, TRB1 binding is more distributed across gene bodies and at extended promoter regions, corresponding to an enrichment of putative binding sites "RMCCTAR". At these genes, TRB1 particiaptes in the repression because

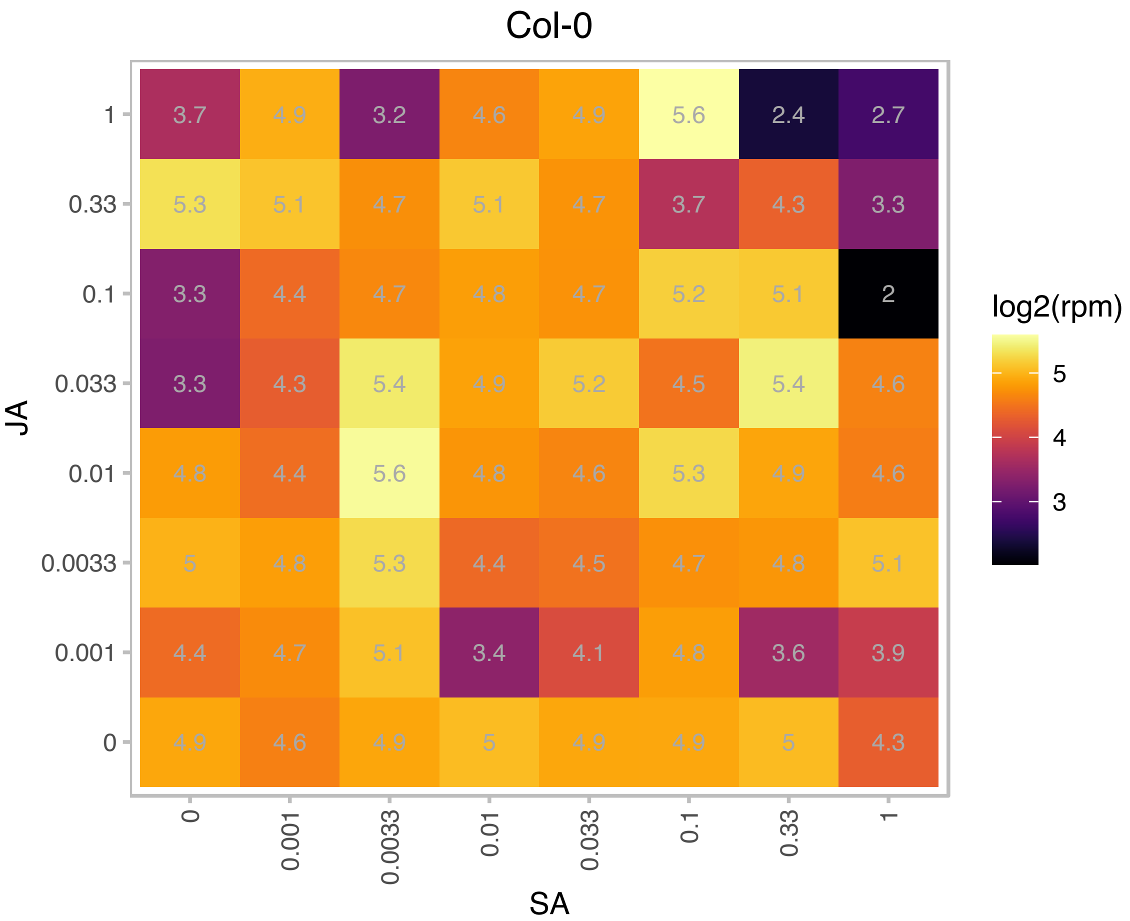


| | |
|---|---|
| Gene | AT1G49950 |
| is.expressed_genes | TRUE |
| is.responsive_genes | TRUE |
| is.apcluster | F-08 |

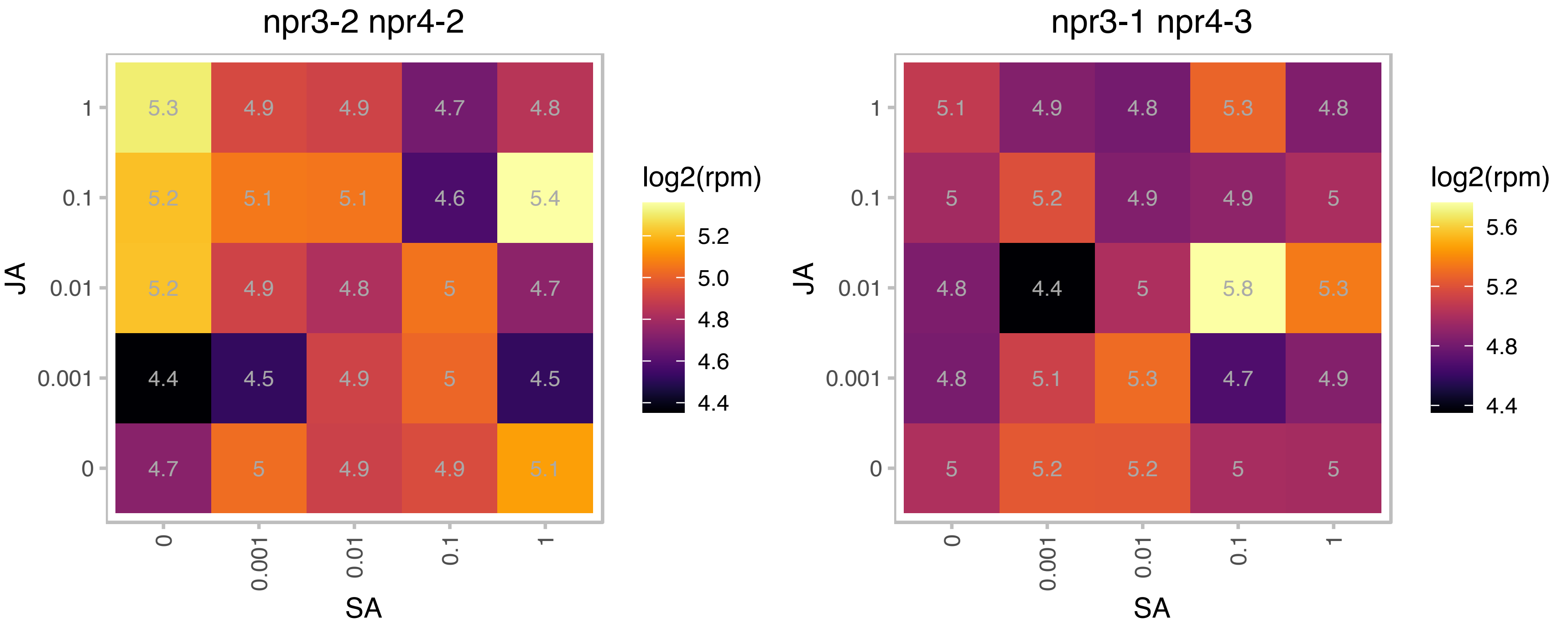

# AT1G51400.1 : NA

Photosystem II 5 kD protein

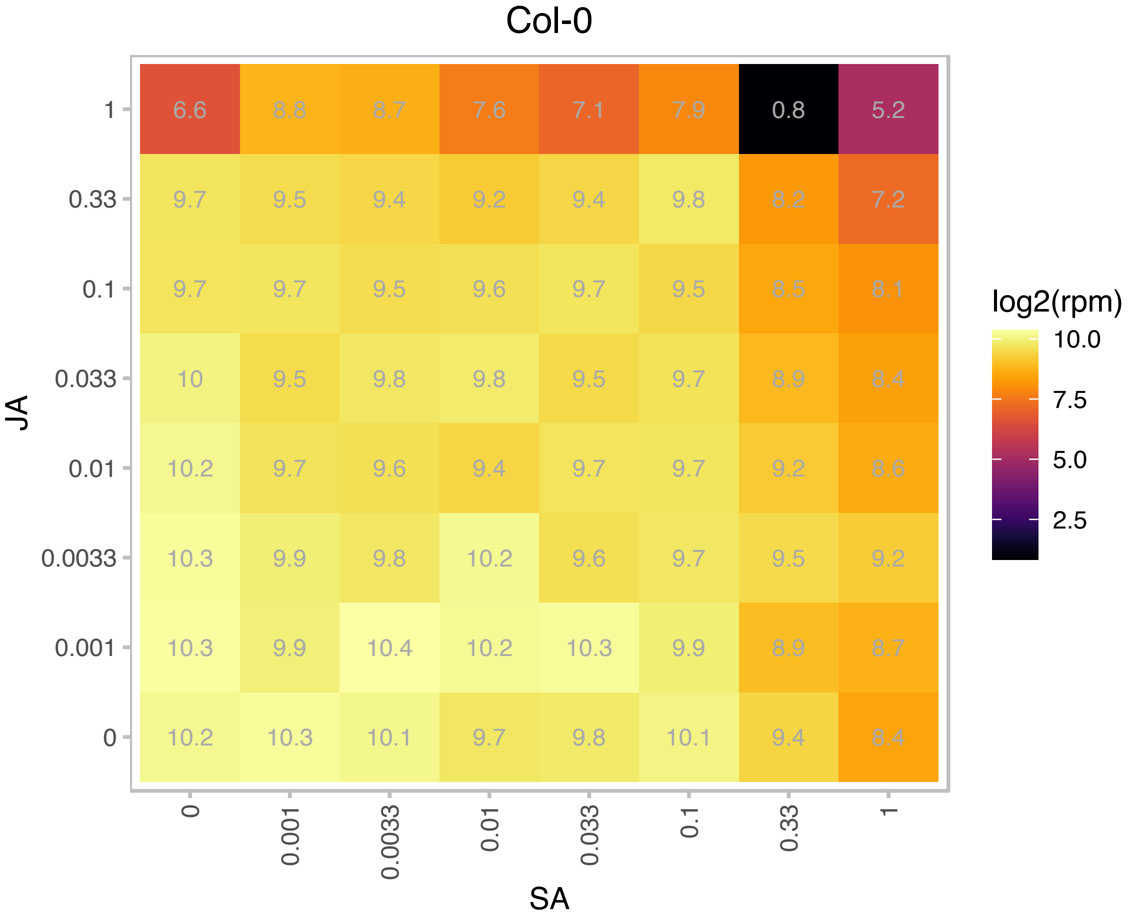


| | |
|---|---|
| Gene | AT1G51400 |
| is.expressed_genes | TRUE |
| is.responsive_genes | TRUE |
| is.apcluster | F-08 |

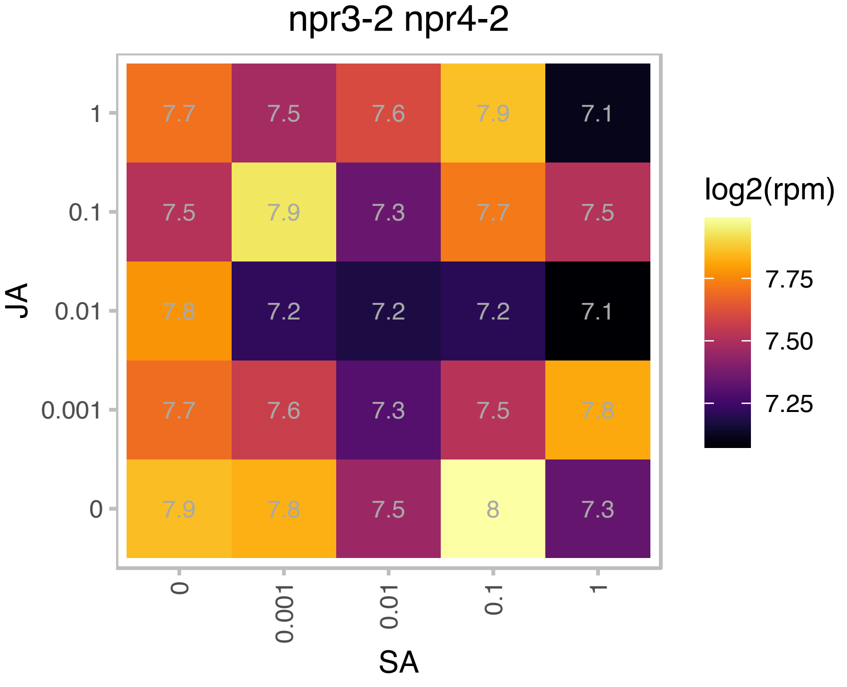


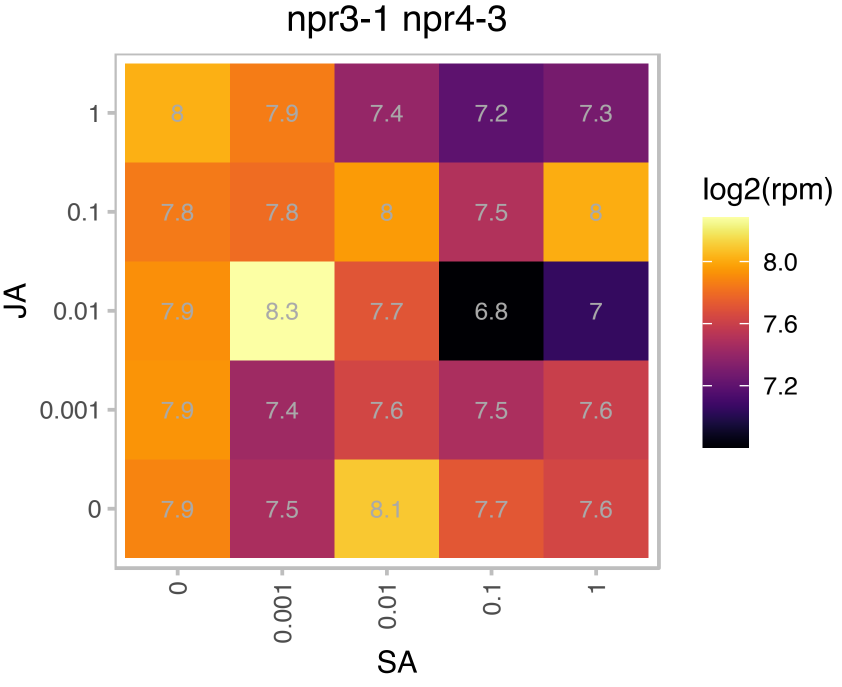

### AT1G52930.1 : atBRX1-2

Ribosomal RNA processing Brix domain protein
Encodes one of two Arabidopsis orthologs of yeast BRX1, a protein involved in maturation of the large ribosomal subunit.  The proteins are mainly localized in nucleolus.  Mutant plants are affected in pre-rRNA processing.

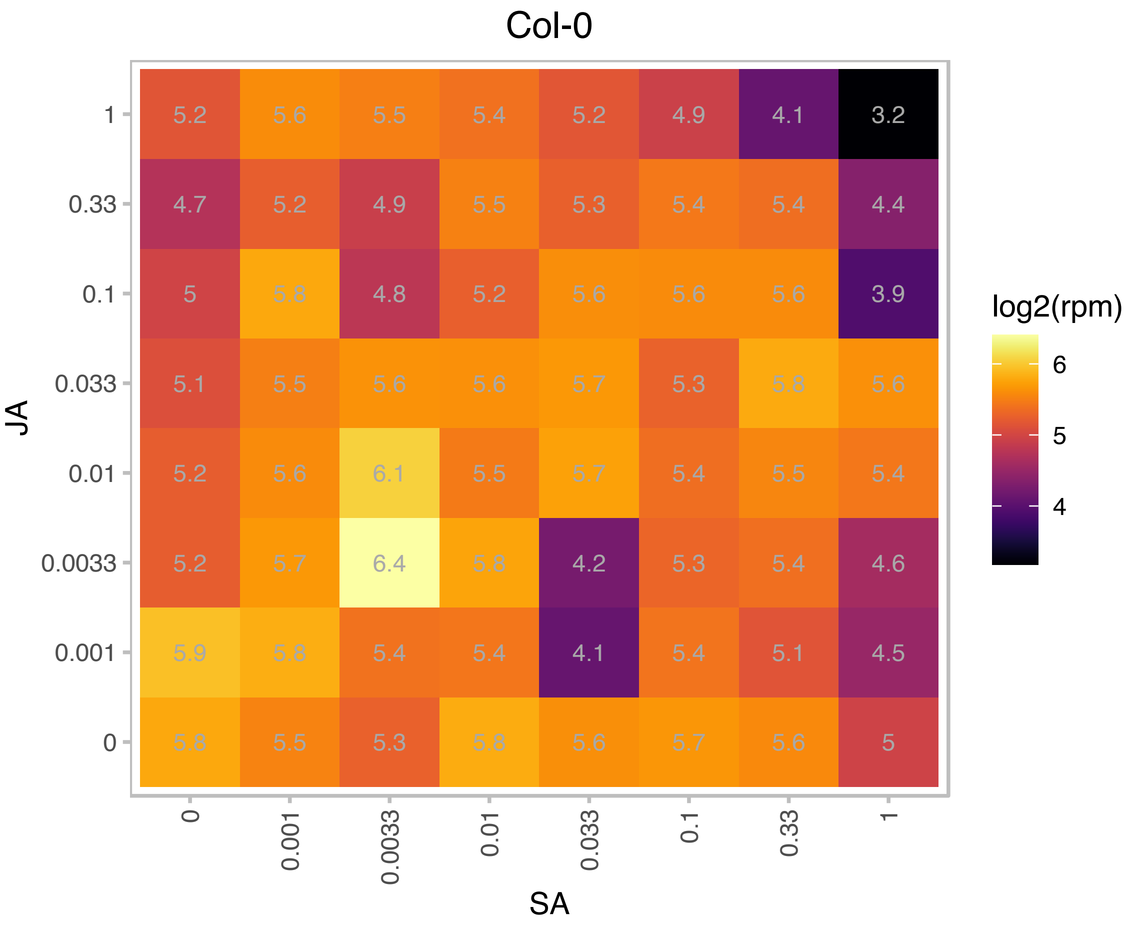


| | |
|---|---|
| Gene | AT1G52930 |
| is.expressed_genes | TRUE |
| is.responsive_genes | TRUE |
| is.apcluster | F-08 |

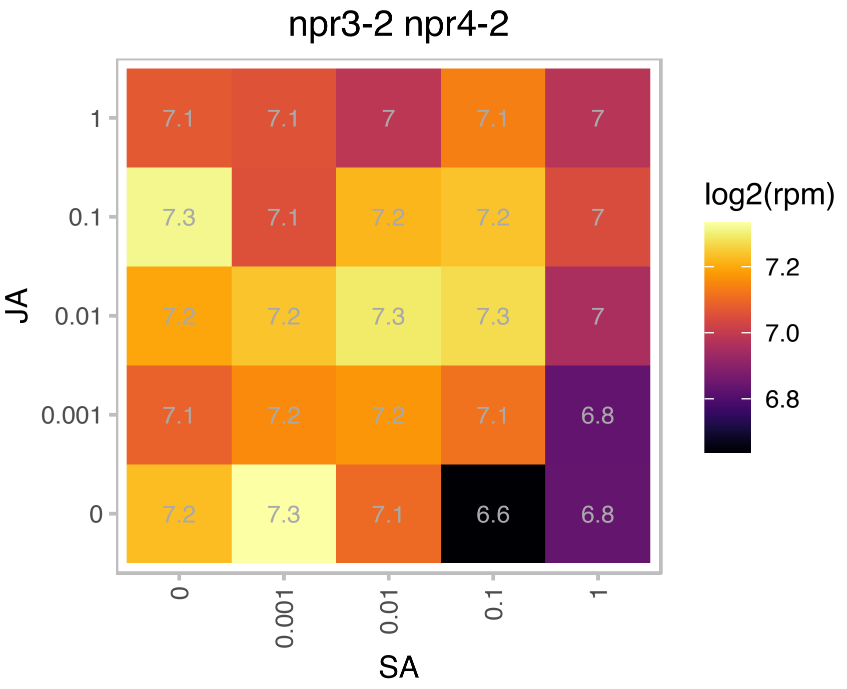


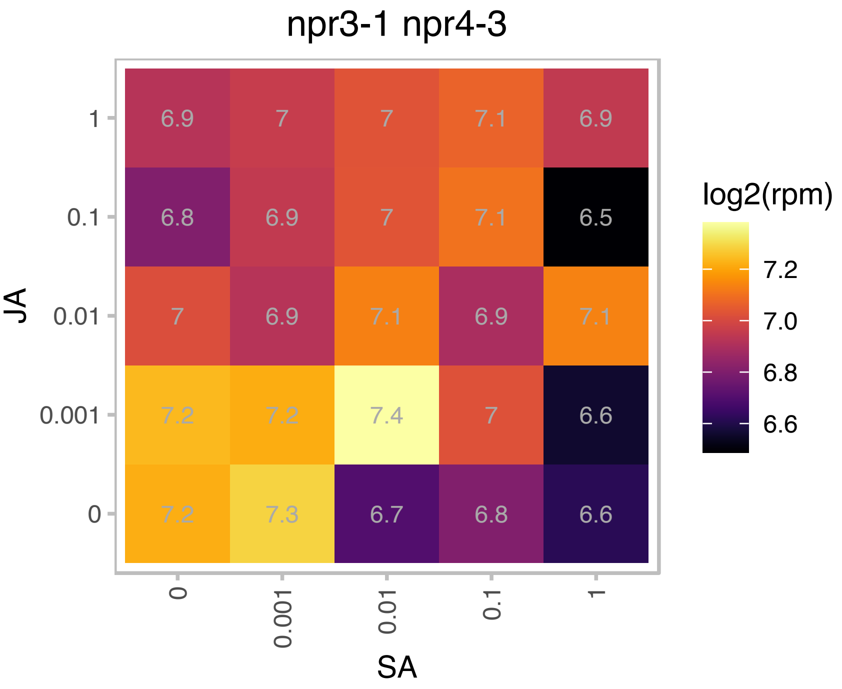

# AT1G55205.1 : NA

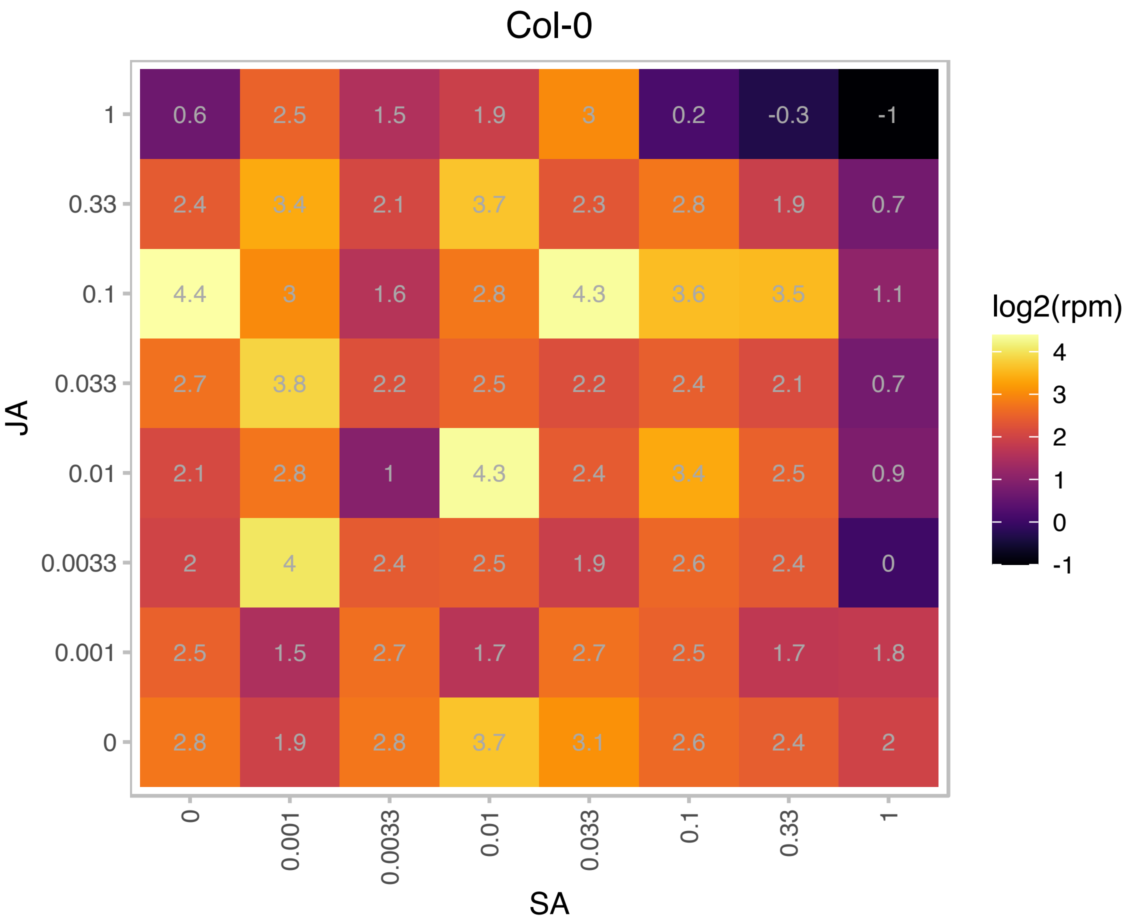


| | |
|---|---|
| Gene | AT1G55205 |
| is.expressed_genes | TRUE |
| is.responsive_genes | TRUE |
| is.apcluster | F-08 |

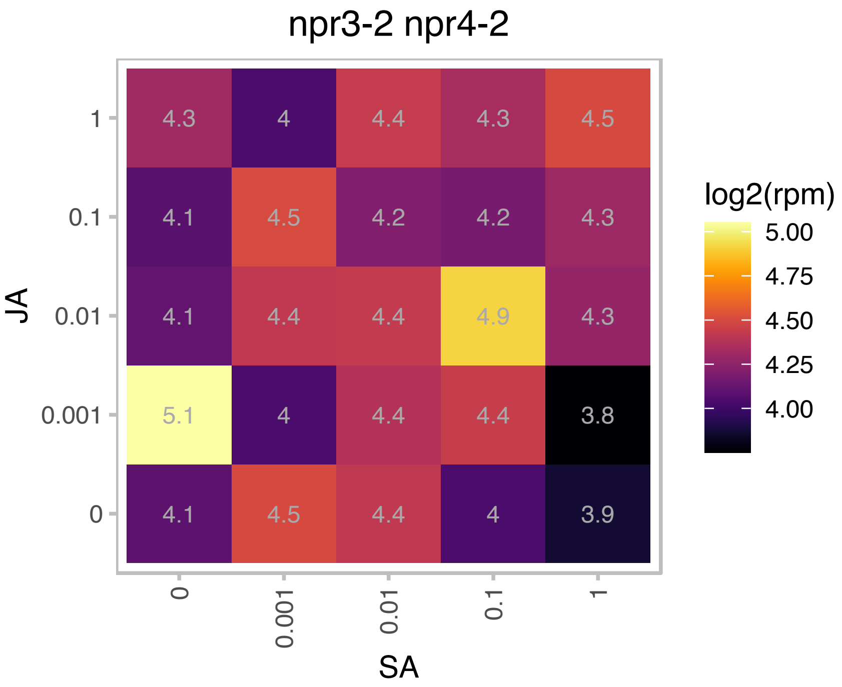


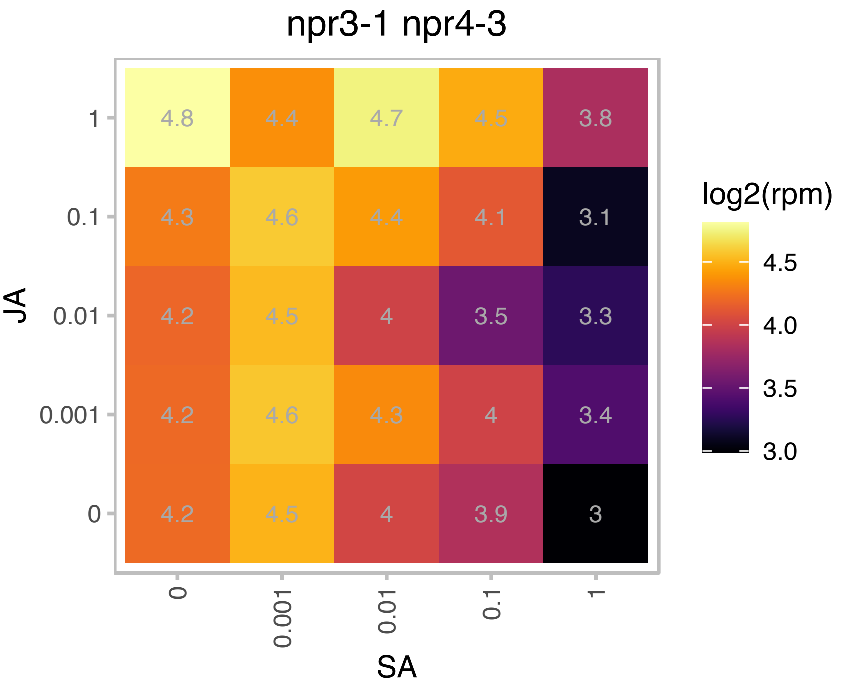

### AT1G55260.1 : LTPG6

Bifunctional inhibitor/lipid-transfer protein/seed storage 2S albumin superfamily protein

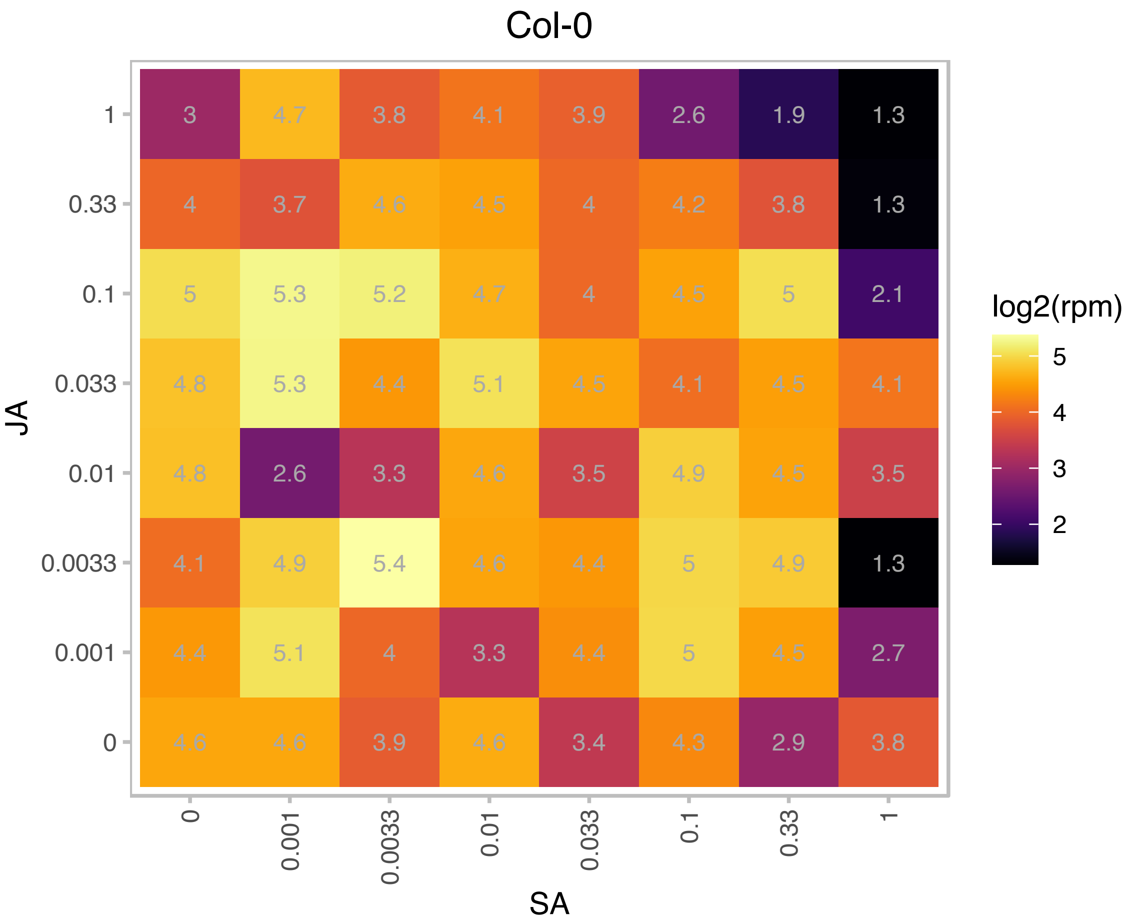


| | |
|---|---|
| Gene | AT1G55260 |
| is.expressed_genes | TRUE |
| is.responsive_genes | TRUE |
| is.apcluster | F-08 |

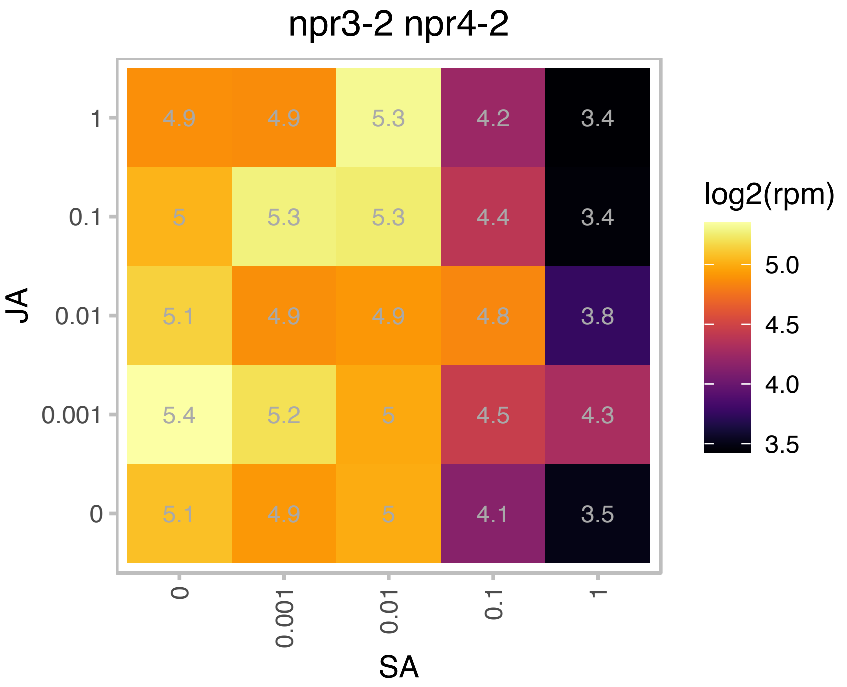


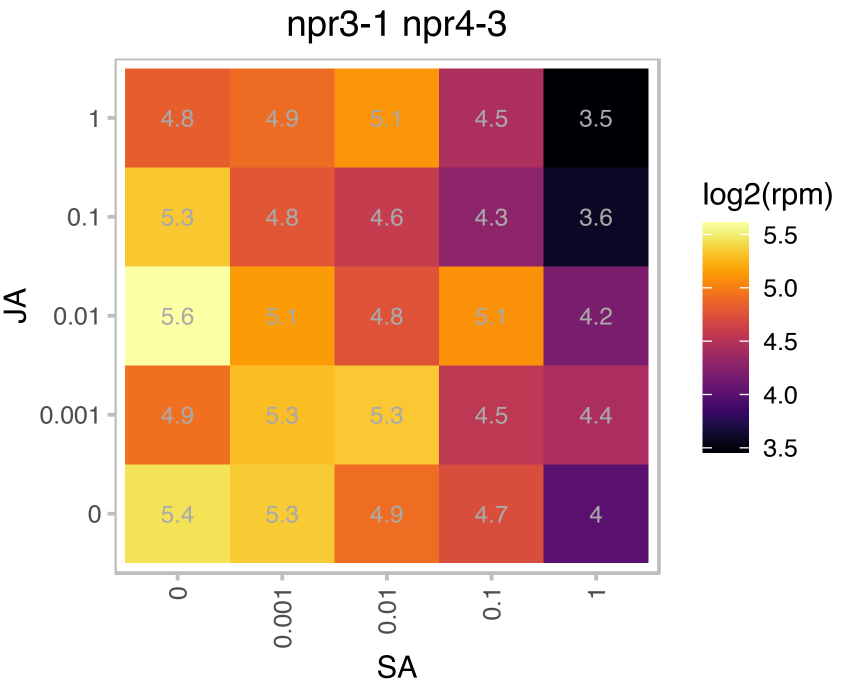

# AT1G56020.1 : NA

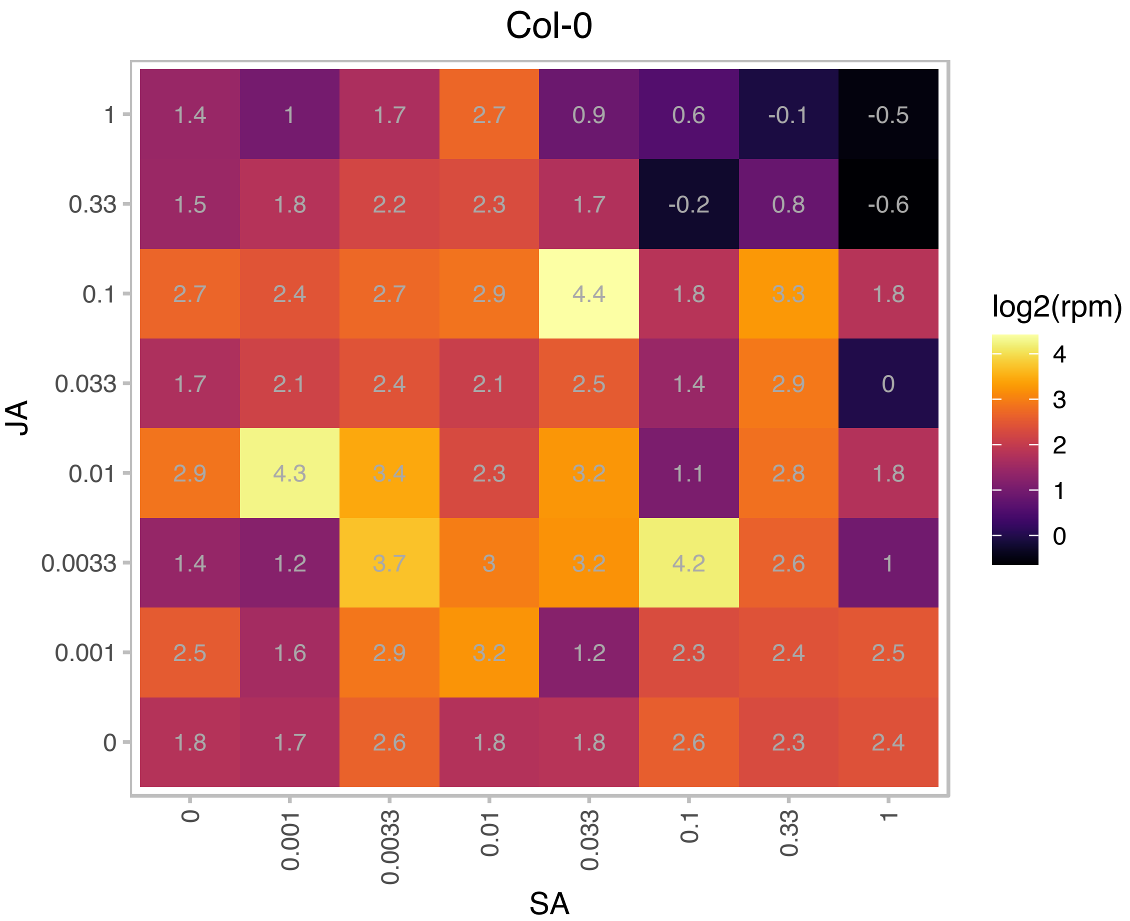


| | |
|---|---|
| Gene | AT1G56020 |
| is.expressed_genes | TRUE |
| is.responsive_genes | TRUE |
| is.apcluster | F-08 |

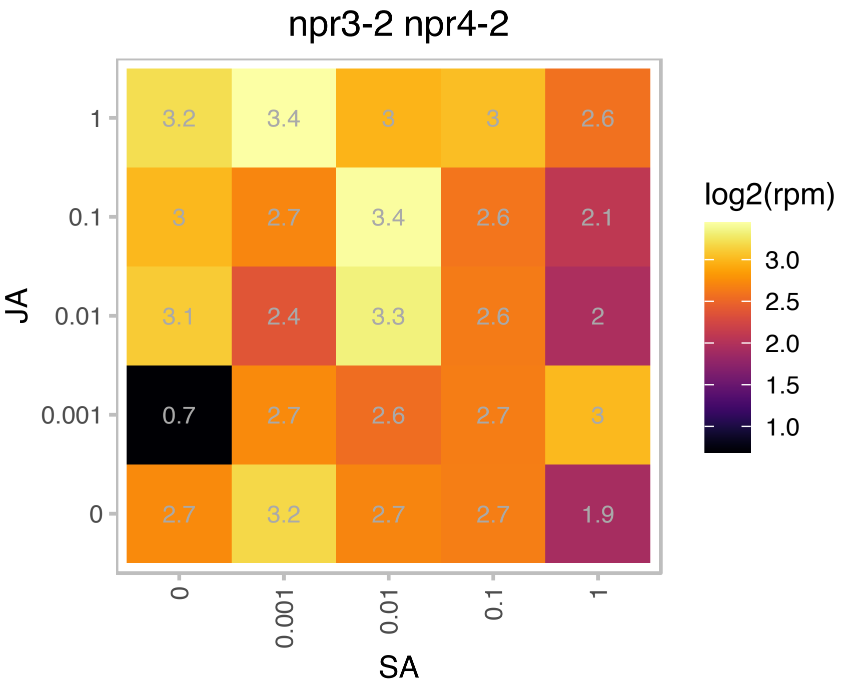


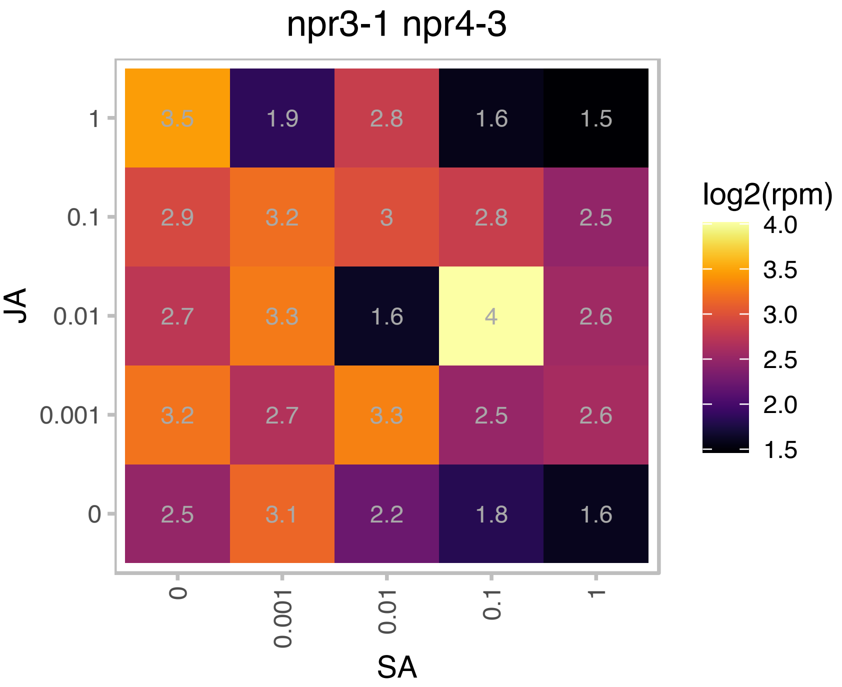

# AT1G59960.1 : NA

NAD(P)-linked oxidoreductase superfamily protein

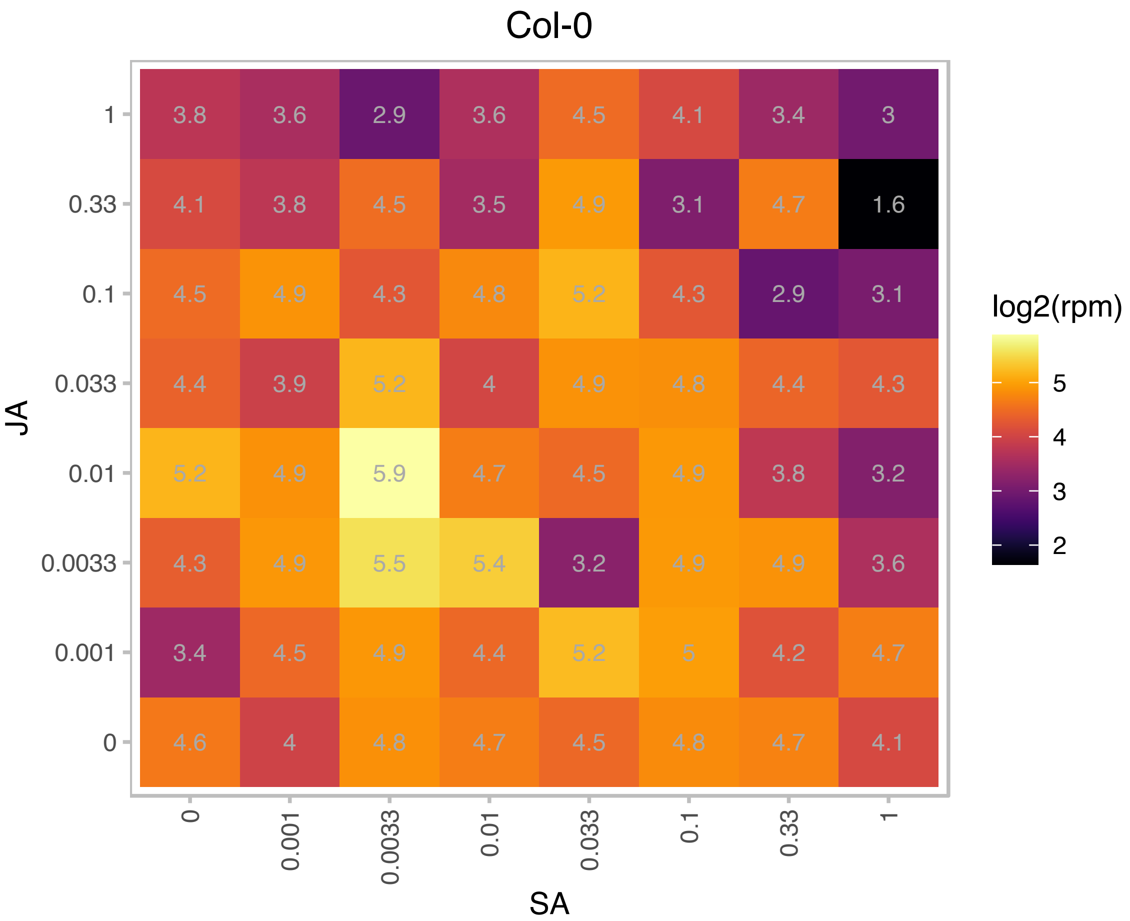


| | |
|---|---|
| Gene | AT1G59960 |
| is.expressed_genes | TRUE |
| is.responsive_genes | TRUE |
| is.apcluster | F-08 |

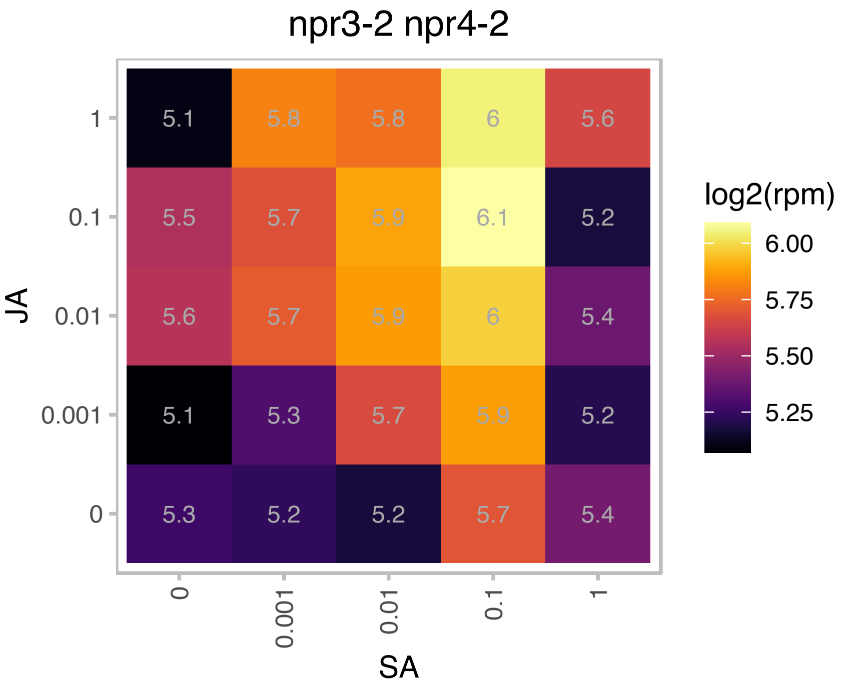


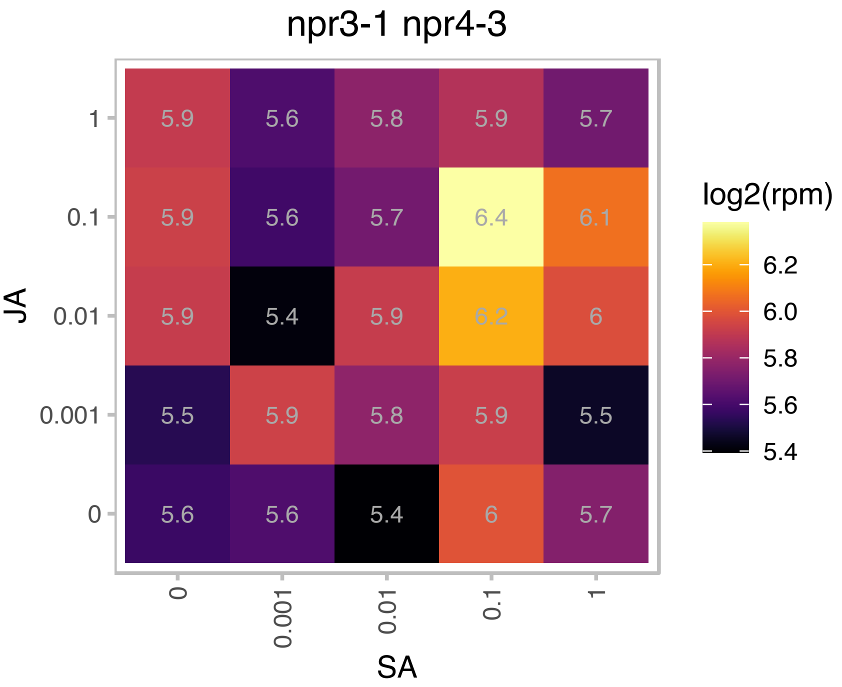

# AT1G60010.1 : NA

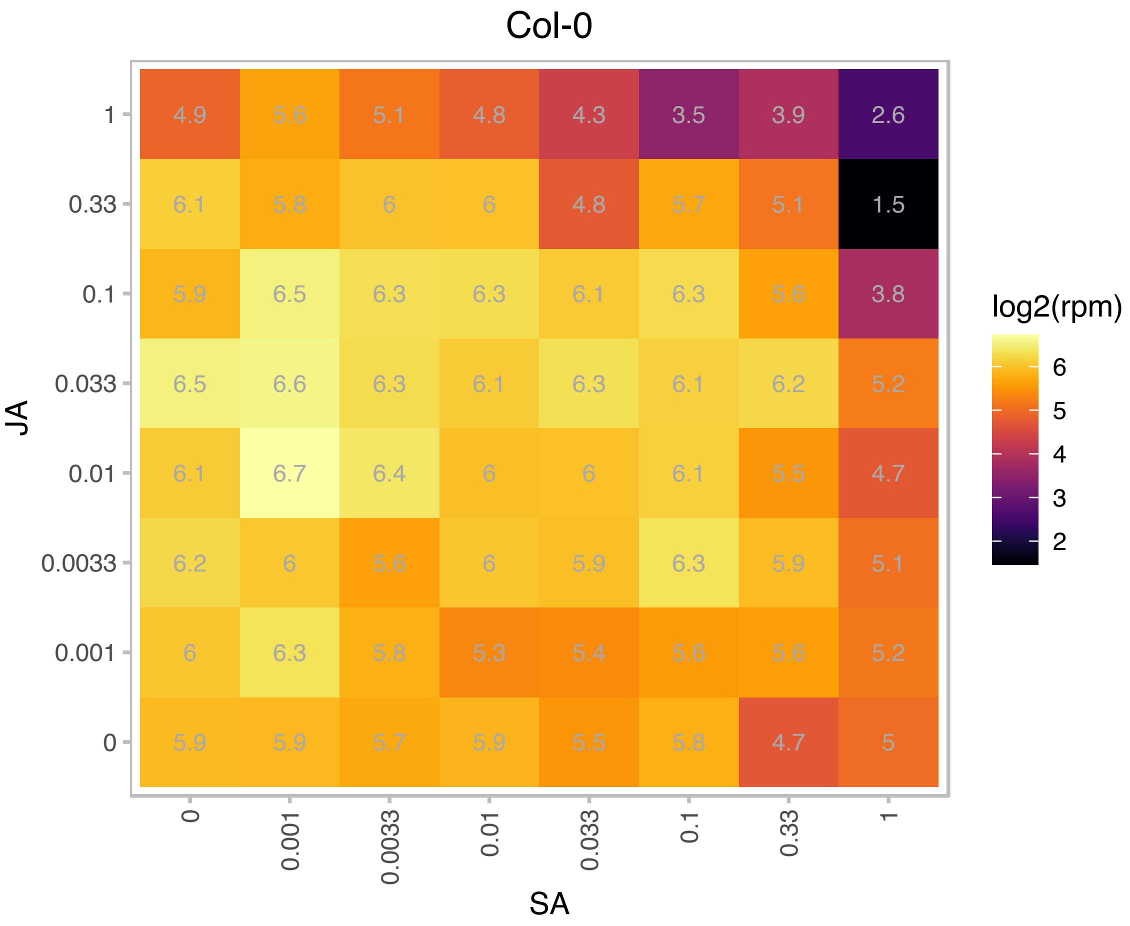


| | |
|---|---|
| Gene | AT1G60010 |
| is.expressed_genes | TRUE |
| is.responsive_genes | TRUE |
| is.apcluster | F-08 |

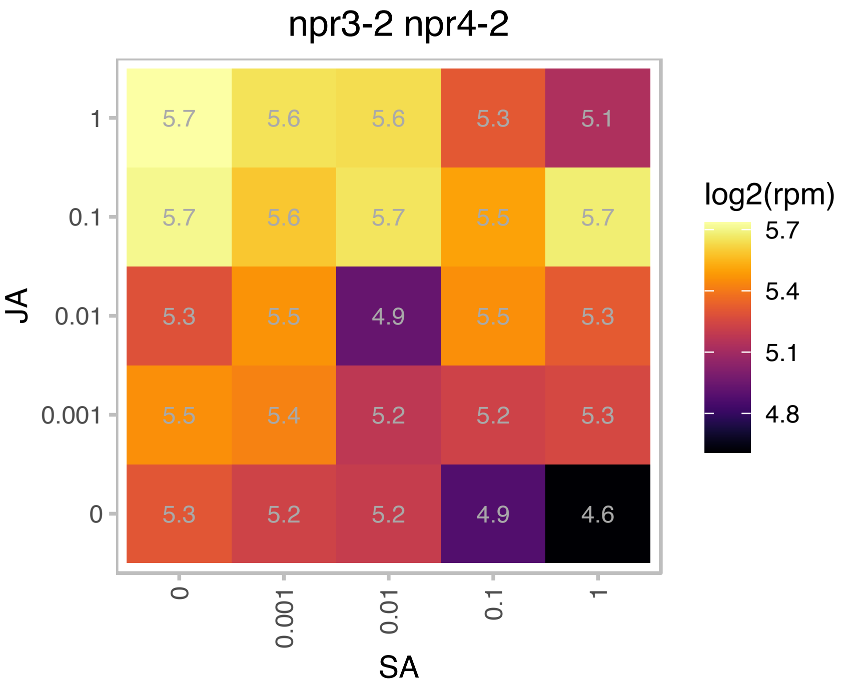


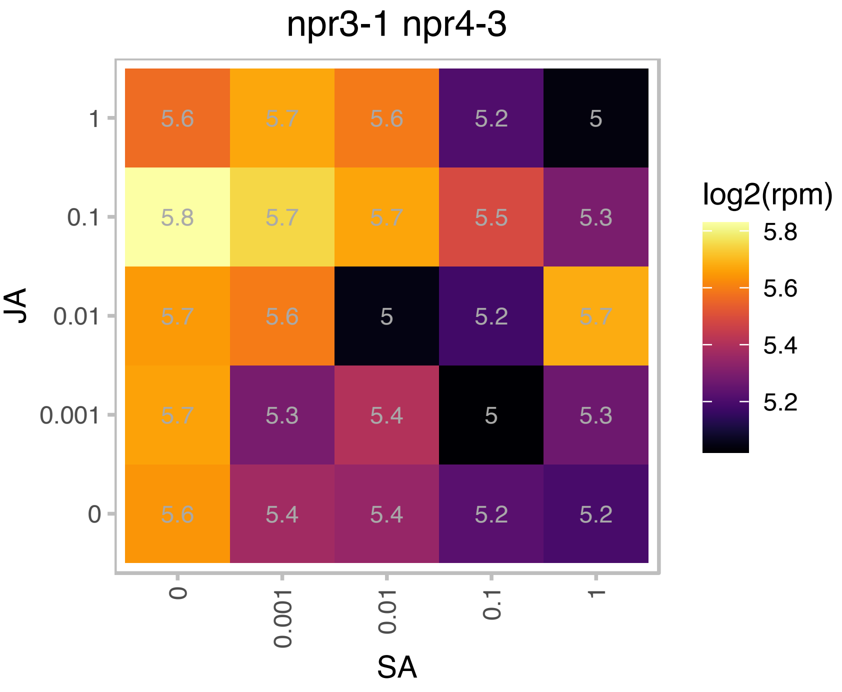

# AT1G64490.1 : NA

DEK, chromatin associated protein

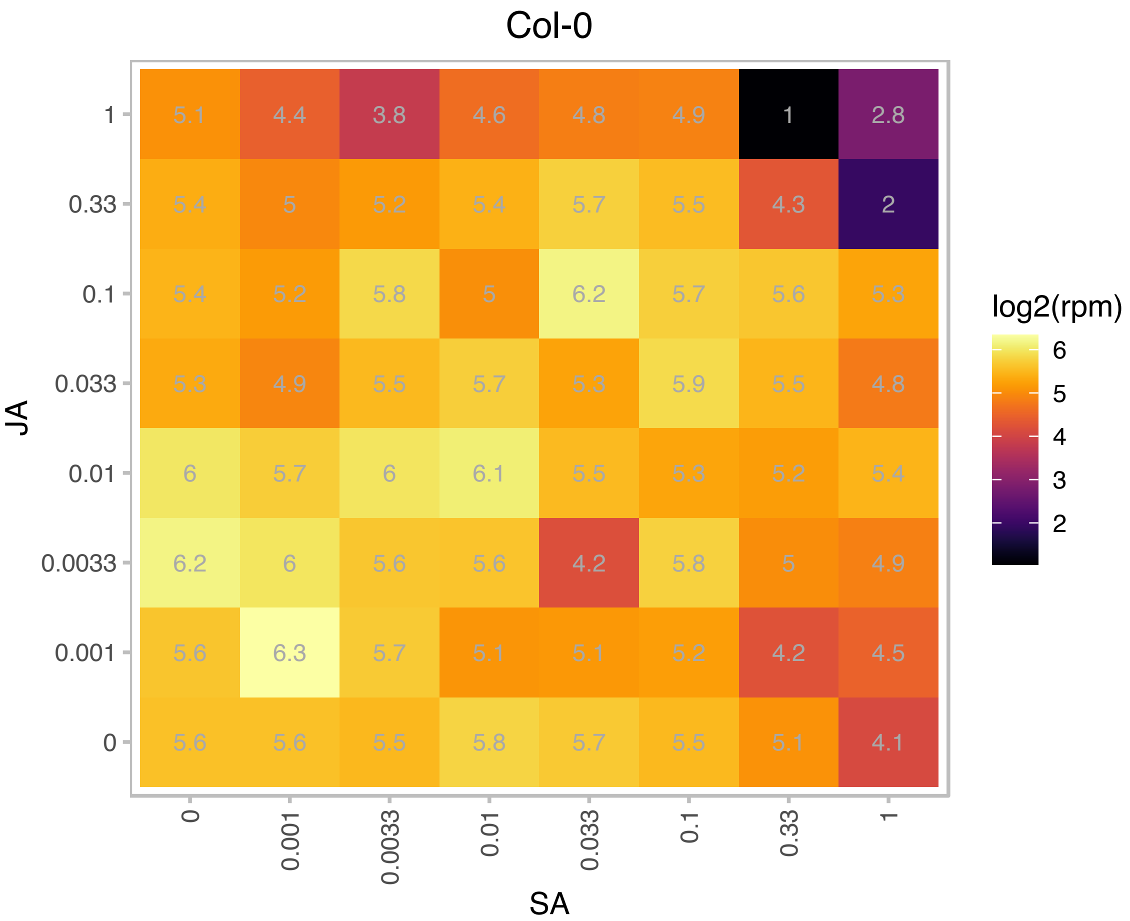


| | |
|---|---|
| Gene | AT1G64490 |
| is.expressed_genes | TRUE |
| is.responsive_genes | TRUE |
| is.apcluster | F-08 |

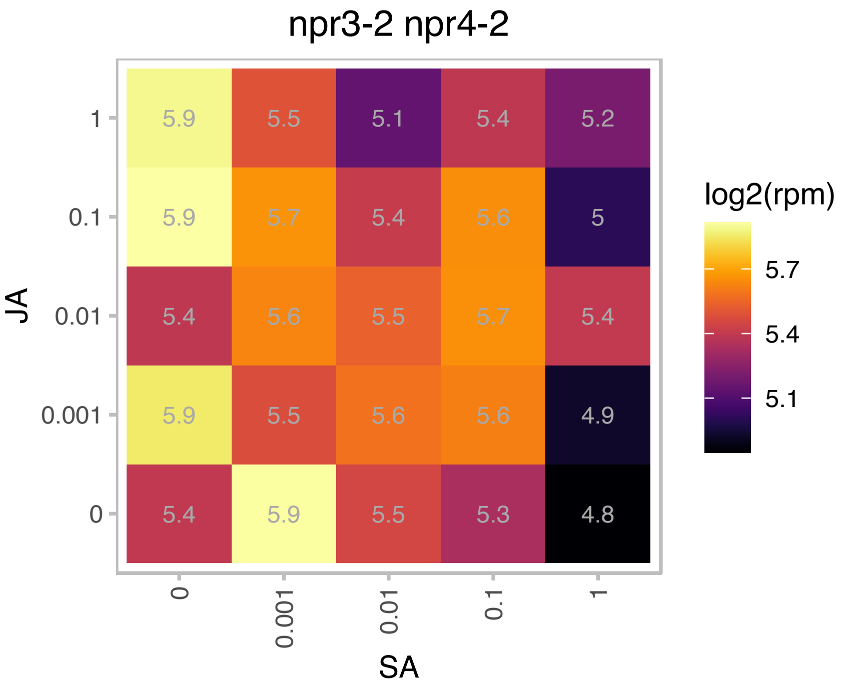


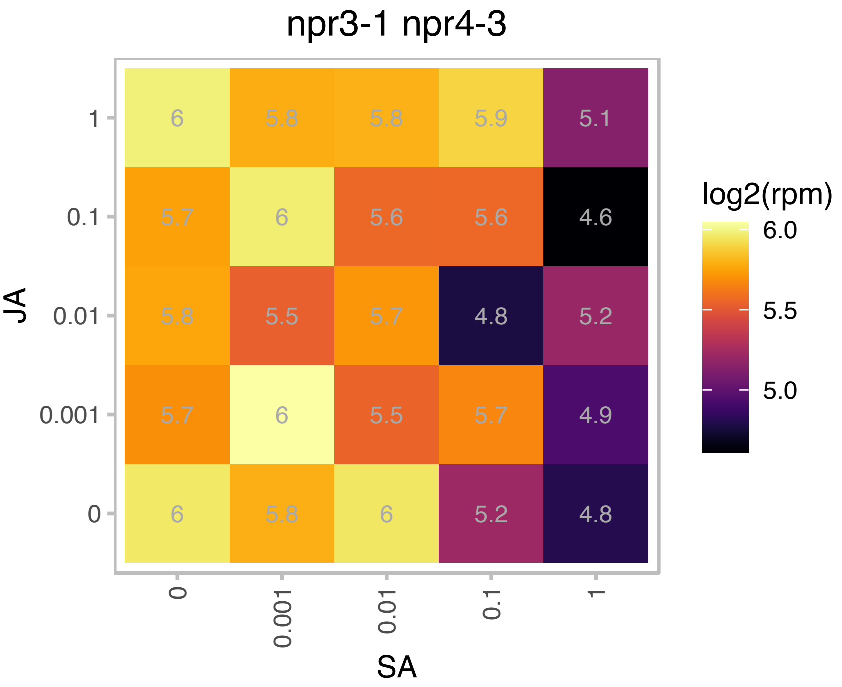

# AT1G65030.1 : NA

Transducin/WD40 repeat-like superfamily protein

This gene is predicted to encode a protein with a DWD motif. It can bind to DDB1a in Y2H assays, and DDB1b in co-IP assays, and may be involved in the formation of a CUL4-based E3 ubiquitin ligase

| | |
|---|---|
| Gene | AT1G65030 |
| is.expressed_genes | TRUE |
| is.responsive_genes | TRUE |
| is.apcluster | F-08 |

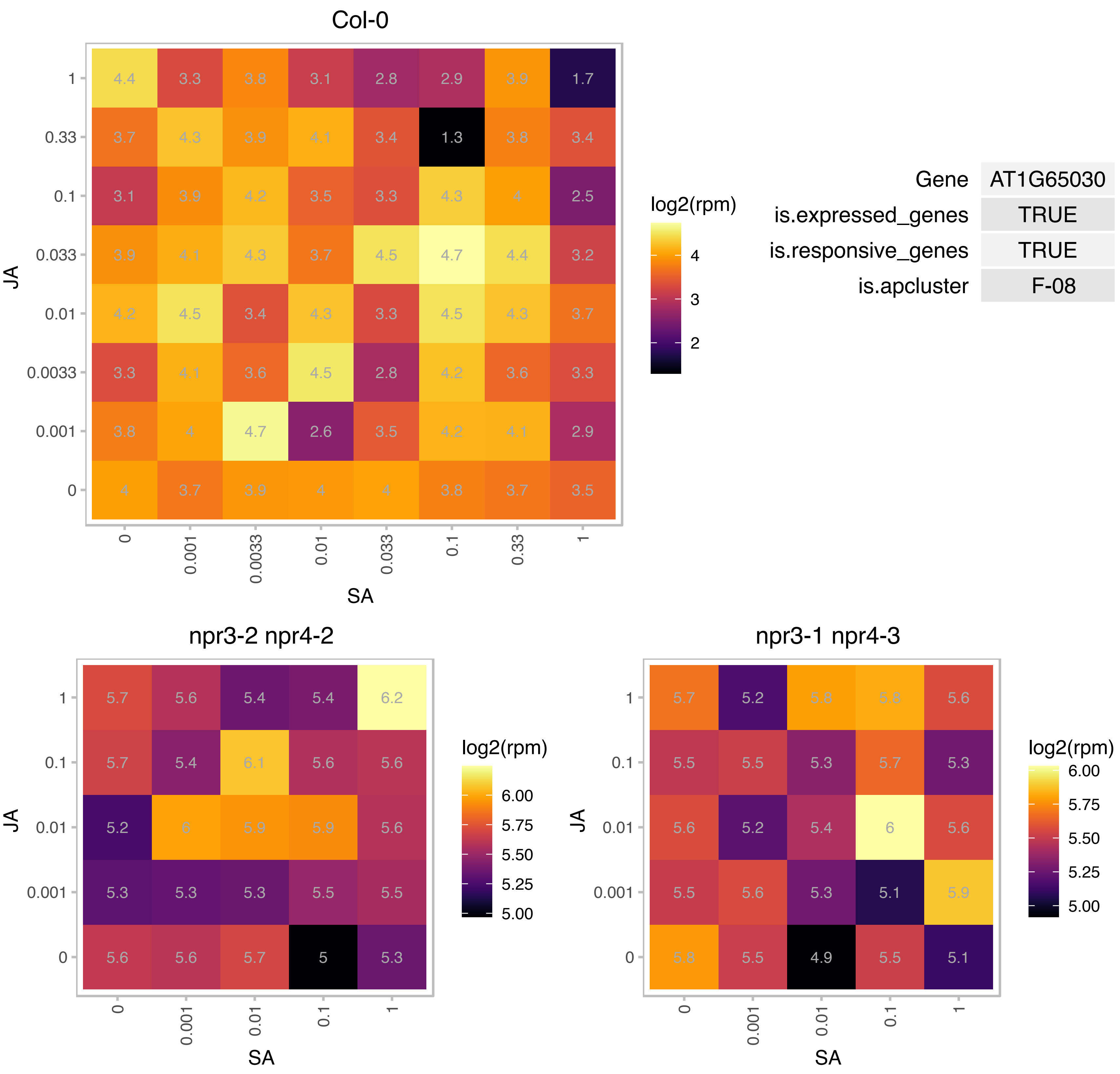

### AT1G67080.1 : ABA4

abscisic acid (aba)-deficient 4

Encodes a protein involved in the photoprotection of PSII. An aba4-1 mutant completely lacks neoxanthin,a component of the chromophore of the peripheral antenna system in PSII. ABA4 is required for neoxanthin biosynthesis, an intermediary step in abscisic acid biosynthesis, but no catalytic activity has been detected for the ABA4 protein.

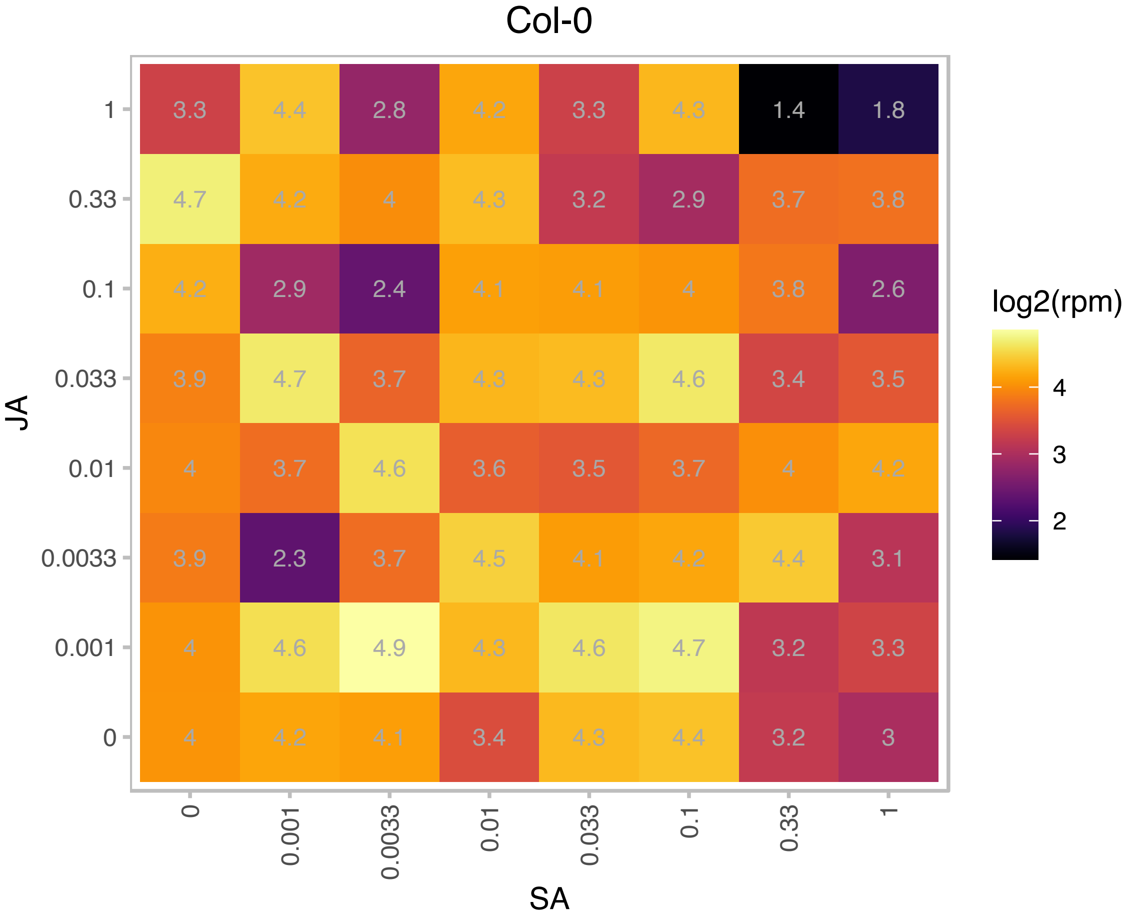


| JA \ SA | 0 | 0.001 | 0.0033 | 0.01 | 0.033 | 0.1 | 0.33 | 1 |
|---|---|---|---|---|---|---|---|---|
| 1 | 3.3 | 4.4 | 2.8 | 4.2 | 3.3 | 4.3 | 1.4 | 1.8 |
| 0.33 | 4.7 | 4.2 | 4 | 4.3 | 3.2 | 2.9 | 3.7 | 3.8 |
| 0.1 | 4.2 | 2.9 | 2.4 | 4.1 | 4.1 | 4 | 3.8 | 2.6 |
| 0.033 | 3.9 | 4.7 | 3.7 | 4.3 | 4.3 | 4.6 | 3.4 | 3.5 |
| 0.01 | 4 | 3.7 | 4.6 | 3.6 | 3.5 | 3.7 | 4 | 4.2 |
| 0.0033 | 3.9 | 2.3 | 3.7 | 4.5 | 4.1 | 4.2 | 4.4 | 3.1 |
| 0.001 | 4 | 4.6 | 4.9 | 4.3 | 4.6 | 4.7 | 3.2 | 3.3 |
| 0 | 4 | 4.2 | 4.1 | 3.4 | 4.3 | 4.4 | 3.2 | 3 |

| | |
|---|---|
| Gene | AT1G67080 |
| is.expressed_genes | TRUE |
| is.responsive_genes | TRUE |
| is.apcluster | F-08 |

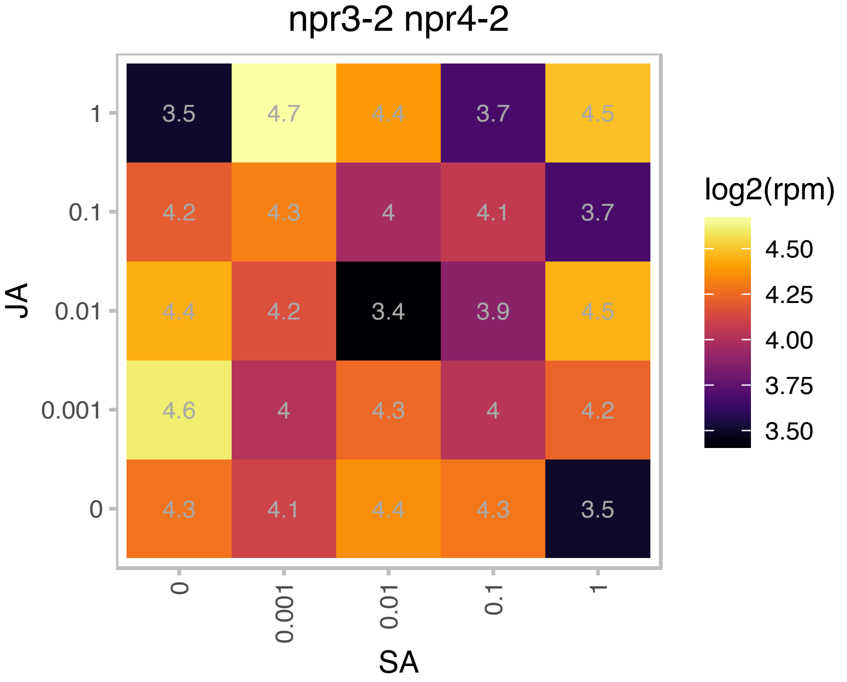


| JA \ SA | 0 | 0.001 | 0.01 | 0.1 | 1 |
|---|---|---|---|---|---|
| 1 | 3.5 | 4.7 | 4.4 | 3.7 | 4.5 |
| 0.1 | 4.2 | 4.3 | 4 | 4.1 | 3.7 |
| 0.01 | 4.4 | 4.2 | 3.4 | 3.9 | 4.5 |
| 0.001 | 4.6 | 4 | 4.3 | 4 | 4.2 |
| 0 | 4.3 | 4.1 | 4.4 | 4.3 | 3.5 |

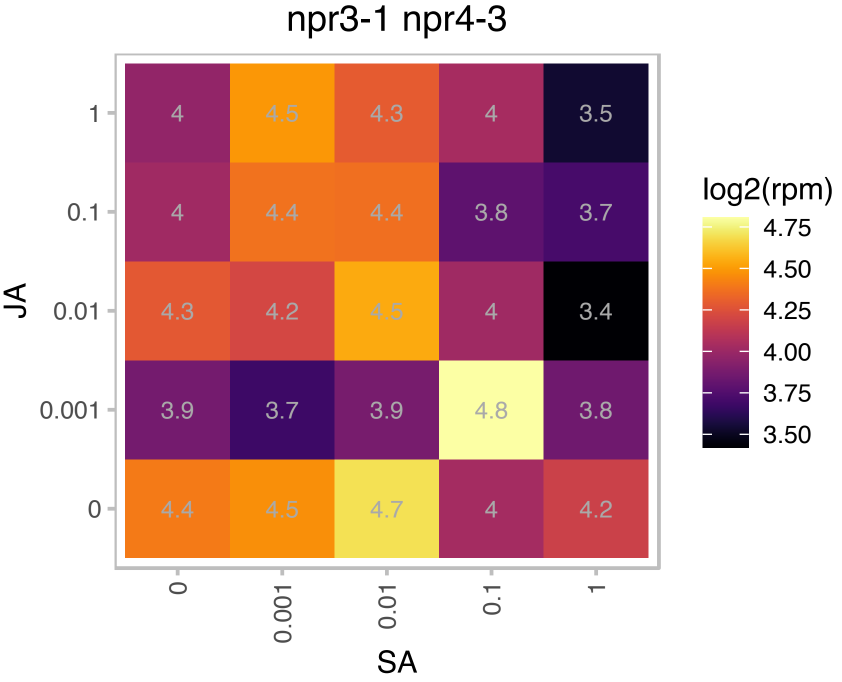


| JA \ SA | 0 | 0.001 | 0.01 | 0.1 | 1 |
|---|---|---|---|---|---|
| 1 | 4 | 4.5 | 4.3 | 4 | 3.5 |
| 0.1 | 4 | 4.4 | 4.4 | 3.8 | 3.7 |
| 0.01 | 4.3 | 4.2 | 4.5 | 4 | 3.4 |
| 0.001 | 3.9 | 3.7 | 3.9 | 4.8 | 3.8 |
| 0 | 4.4 | 4.5 | 4.7 | 4 | 4.2 |

# AT1G70600.1 : NA

Ribosomal protein L18e/L15 superfamily protein

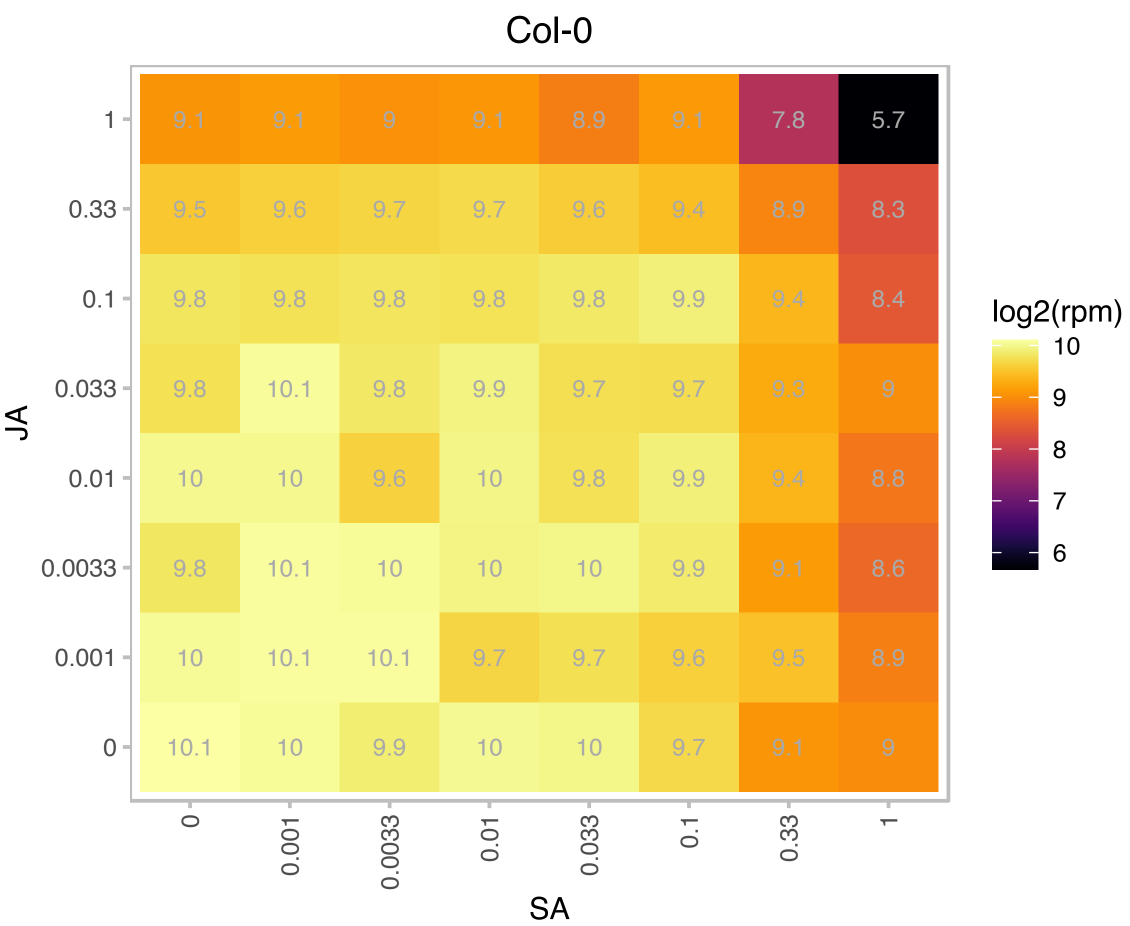


| | |
|---|---|
| Gene | AT1G70600 |
| is.expressed_genes | TRUE |
| is.responsive_genes | TRUE |
| is.apcluster | F-08 |

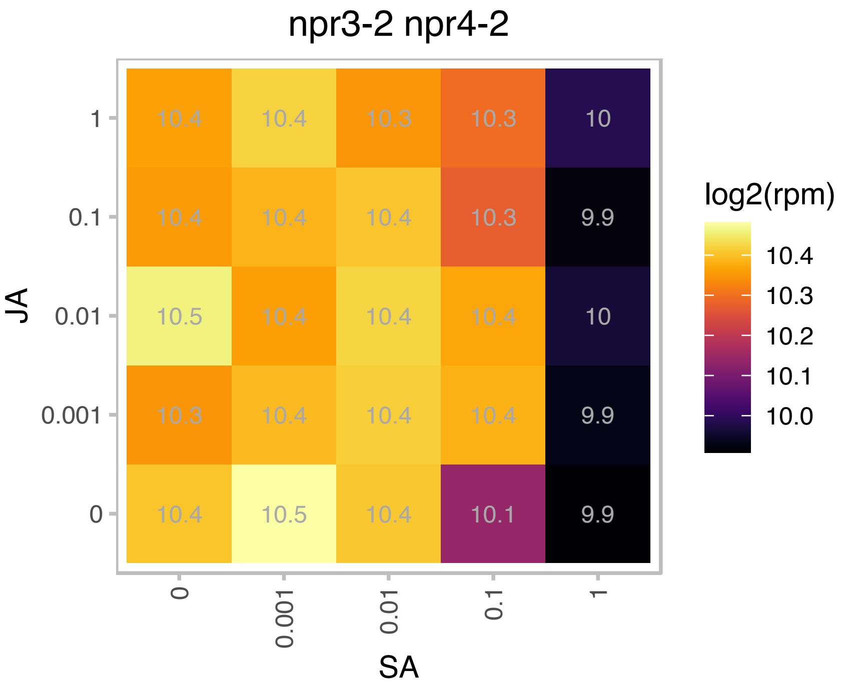


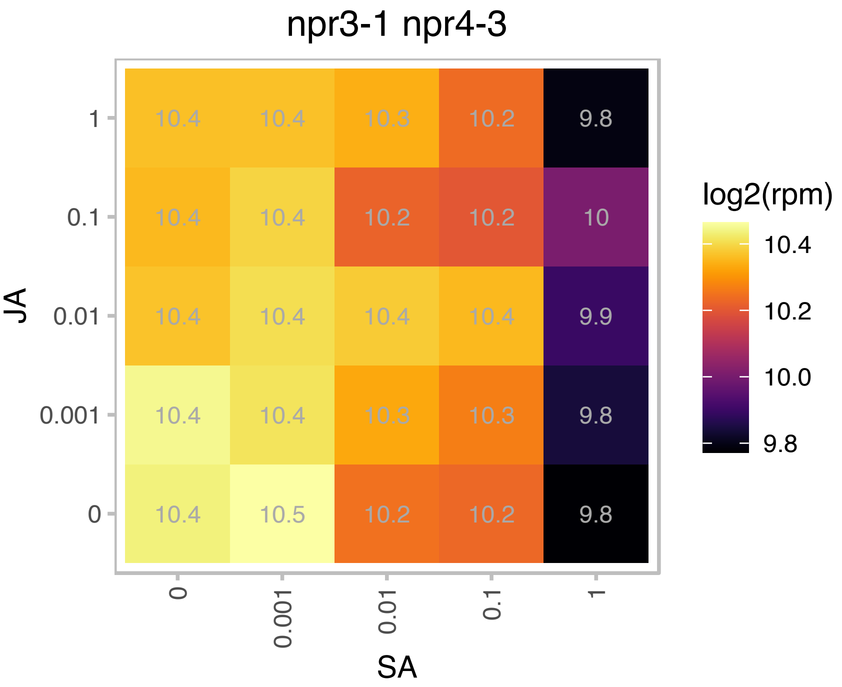

# AT1G71970.1 : NA

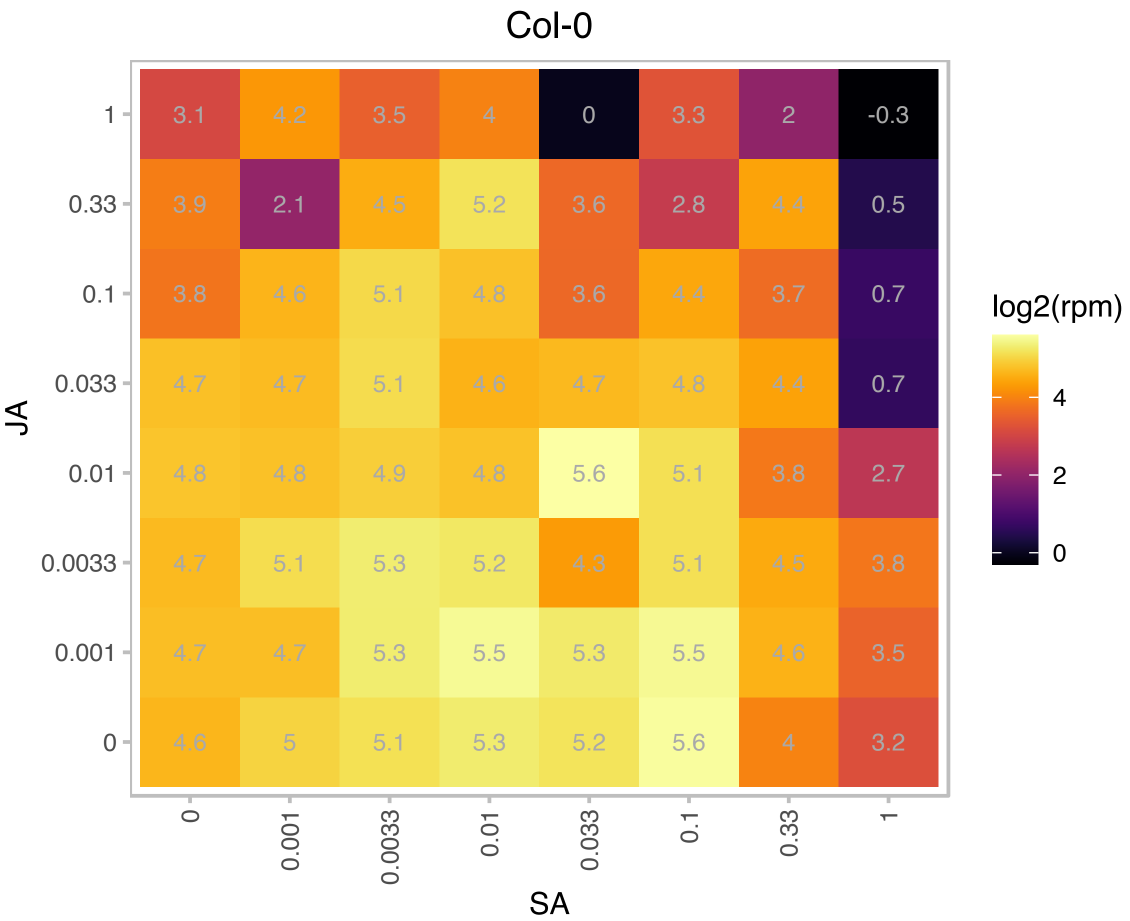


| | |
|---|---|
| Gene | AT1G71970 |
| is.expressed_genes | TRUE |
| is.responsive_genes | TRUE |
| is.apcluster | F-08 |

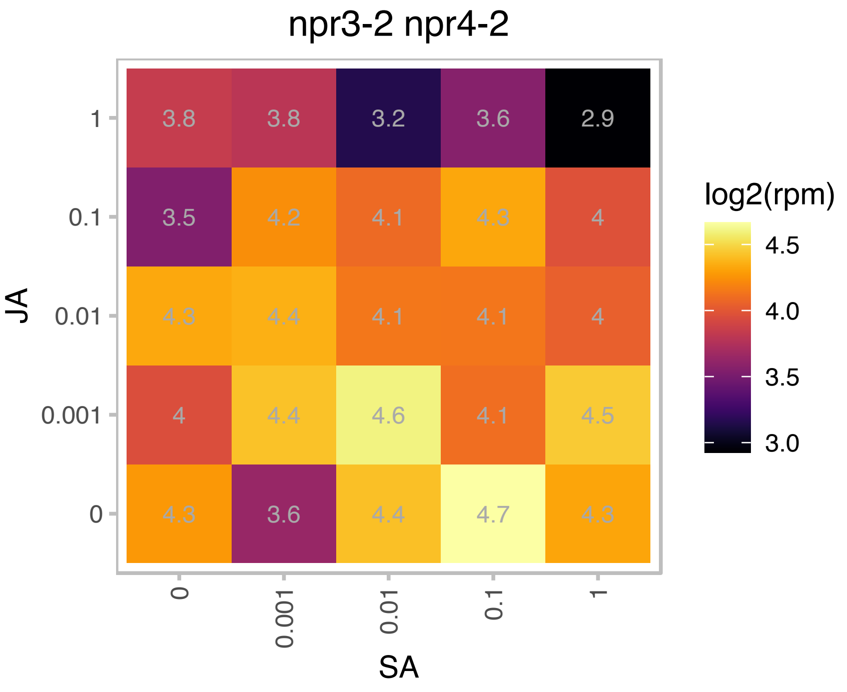


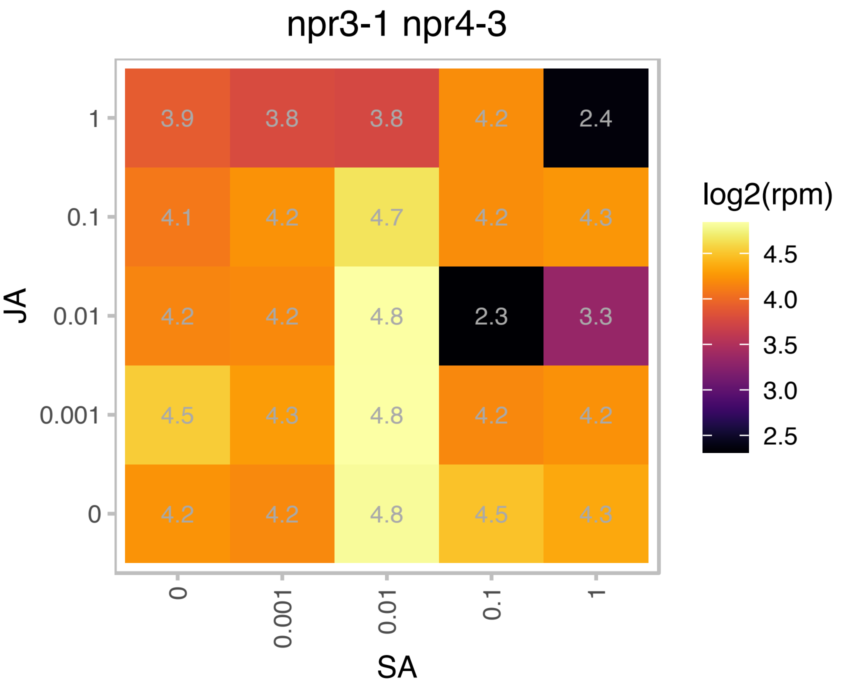

# AT1G73760.1 : NA

RING/U-box superfamily protein

| | |
|---|---|
| Gene | AT1G73760 |
| is.expressed_genes | TRUE |
| is.responsive_genes | TRUE |
| is.apcluster | F-08 |

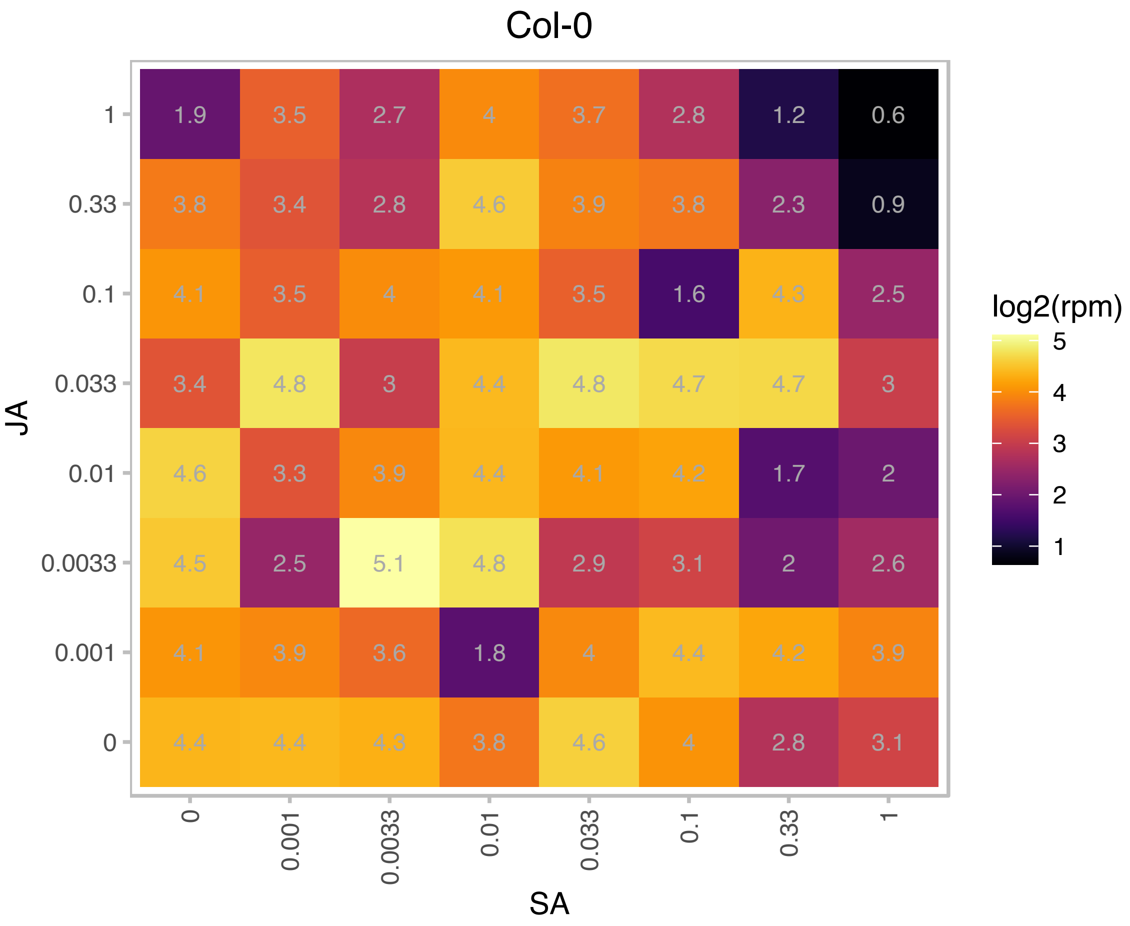


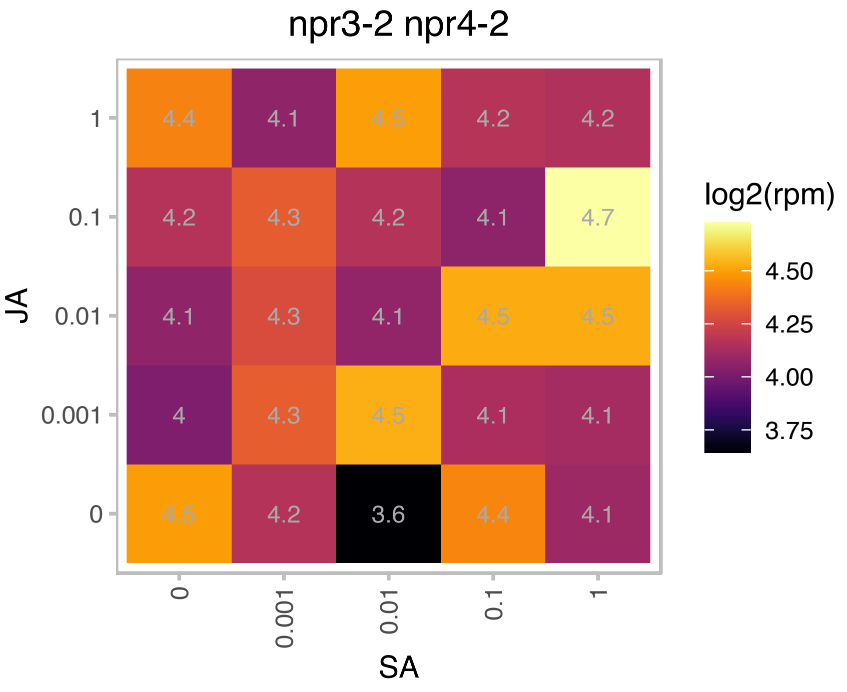


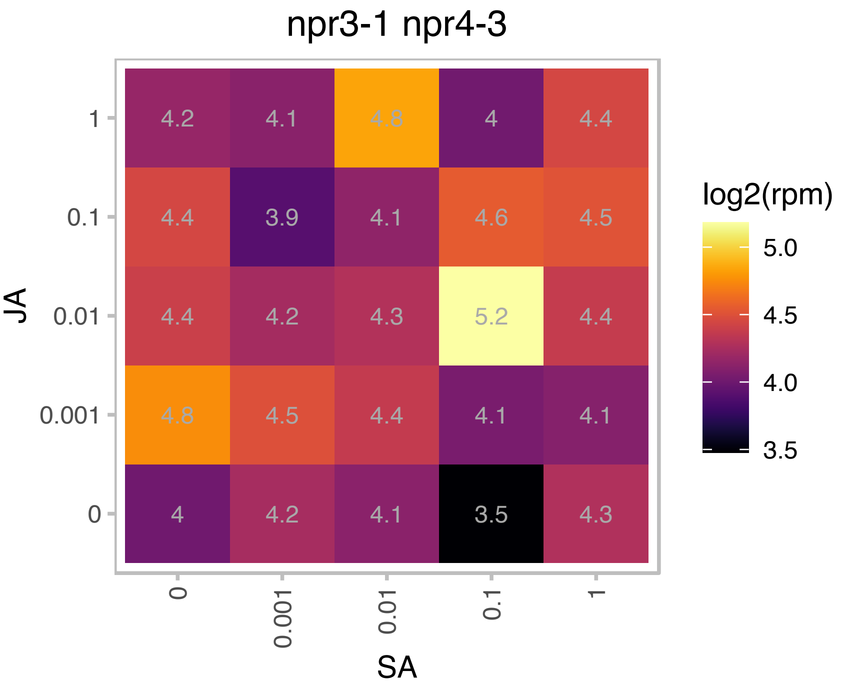

# AT1G75945.1 : NA

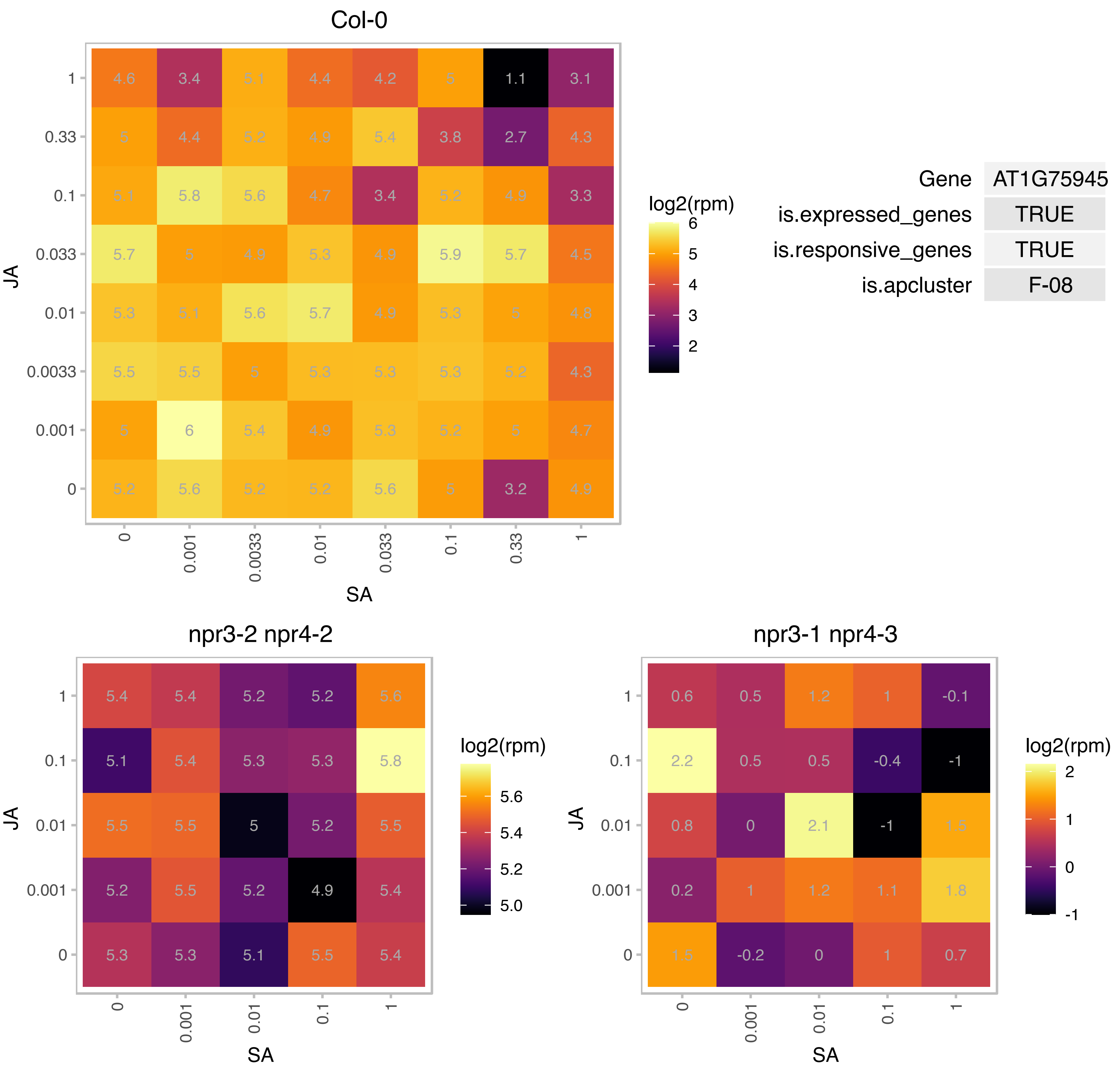


| | |
|---|---|
| Gene | AT1G75945 |
| is.expressed_genes | TRUE |
| is.responsive_genes | TRUE |
| is.apcluster | F-08 |

# AT1G75980.1 : NA

Single hybrid motif superfamily protein

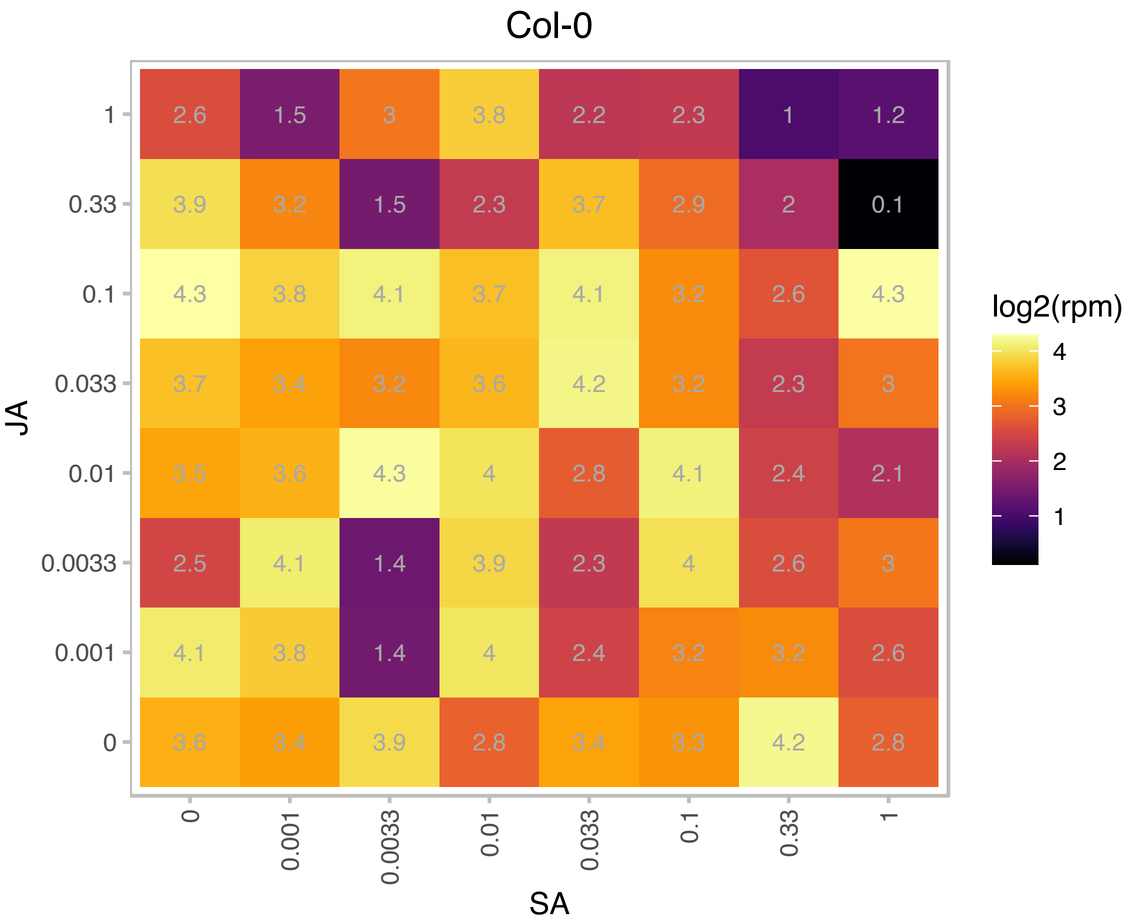


| | |
|---|---|
| Gene | AT1G75980 |
| is.expressed_genes | TRUE |
| is.responsive_genes | TRUE |
| is.apcluster | F-08 |

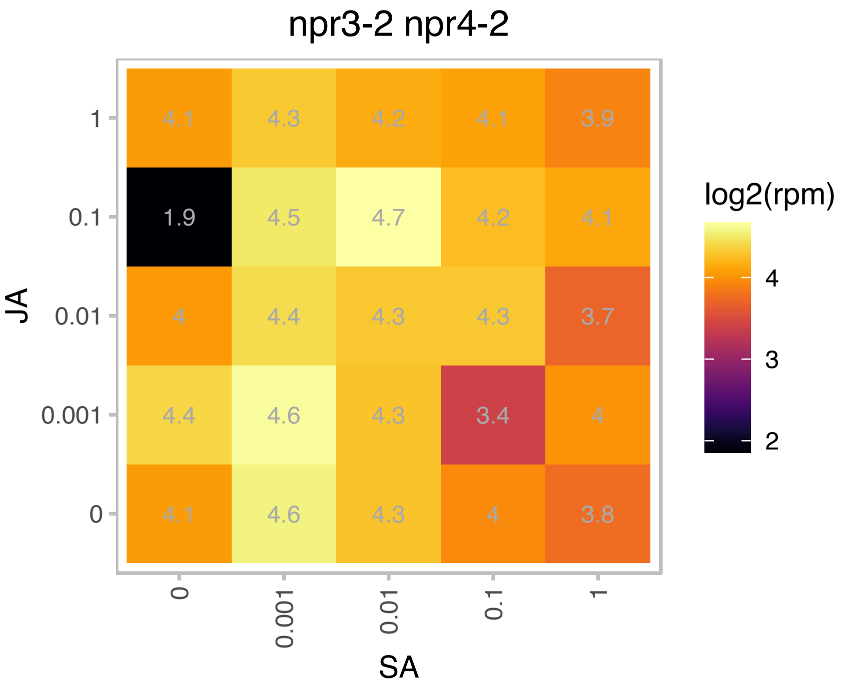


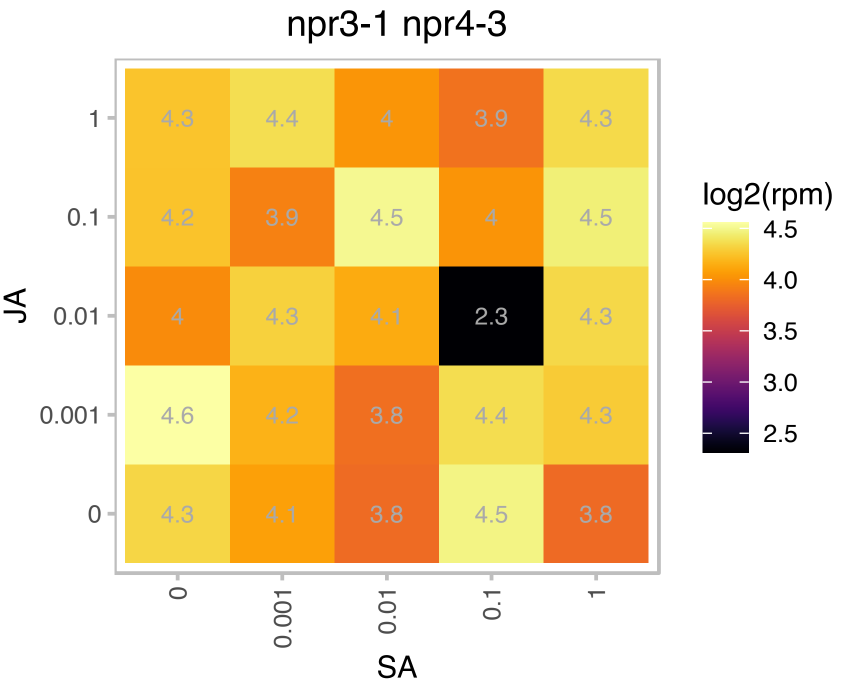

### AT1G76080.1 : ATCDSP32, CDSP32

chloroplastic drought-induced stress protein of 32 kD

Encodes a thioredoxin localized in chloroplast stroma.  Known as CDSP32 (CHLOROPLASTIC DROUGHT-INDUCE
PROTEIN OF 32 KD).

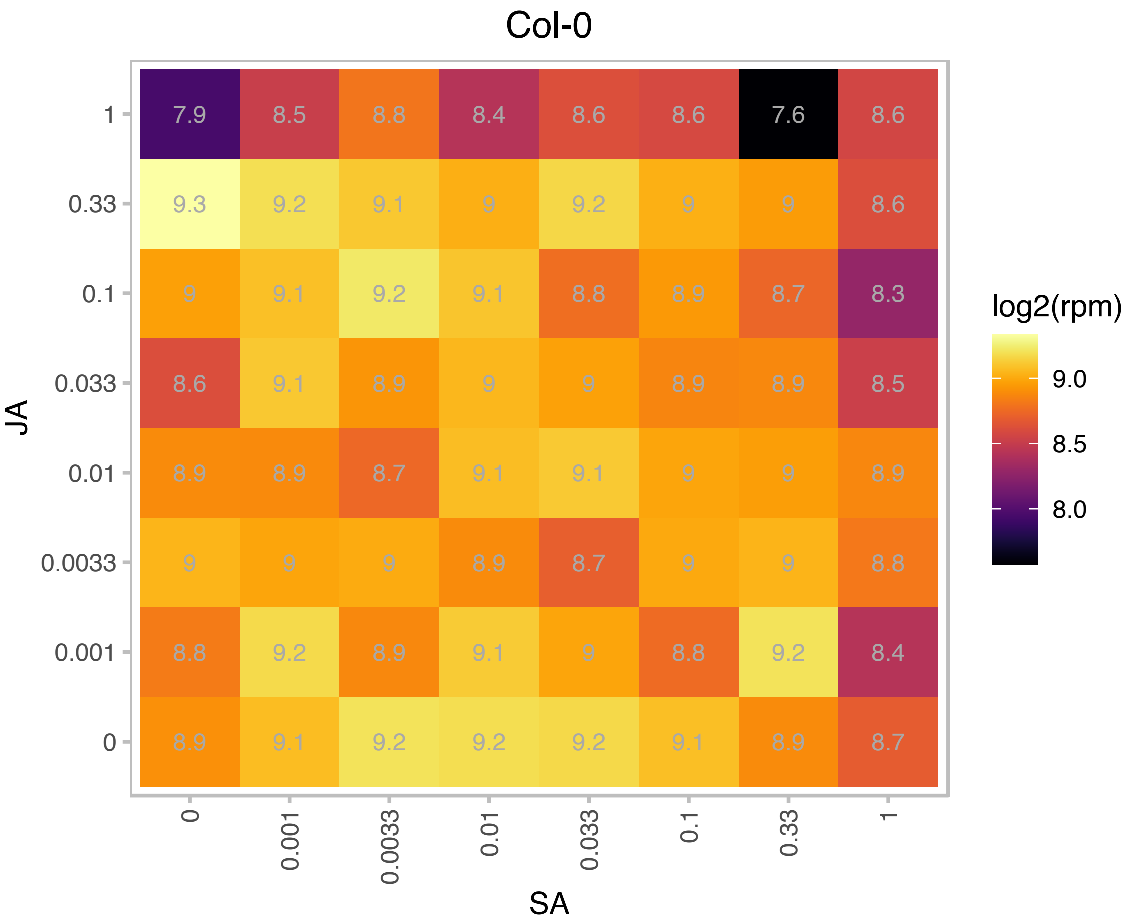


| | |
|---|---|
| Gene | AT1G76080 |
| is.expressed_genes | TRUE |
| is.responsive_genes | TRUE |
| is.apcluster | F-08 |

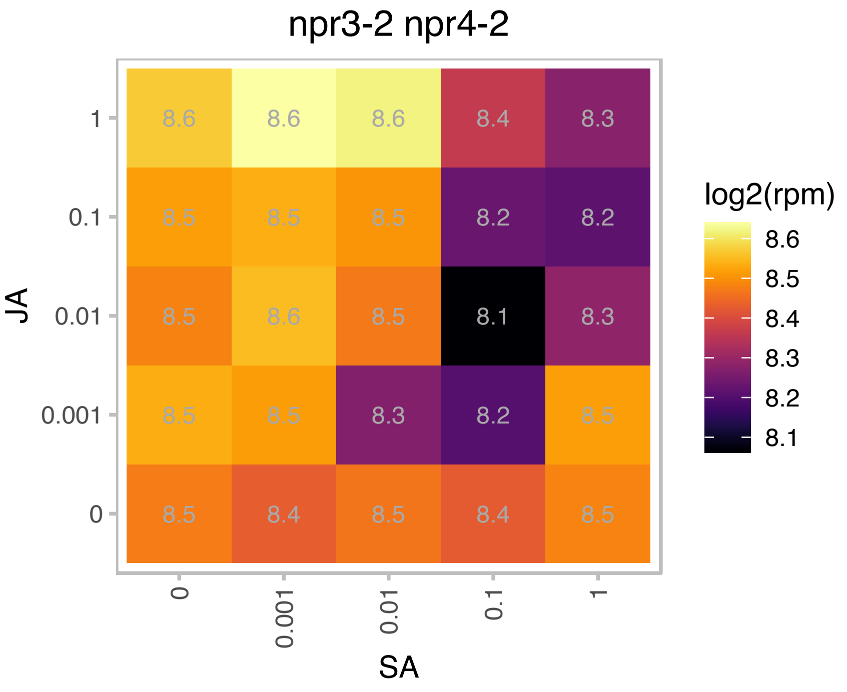


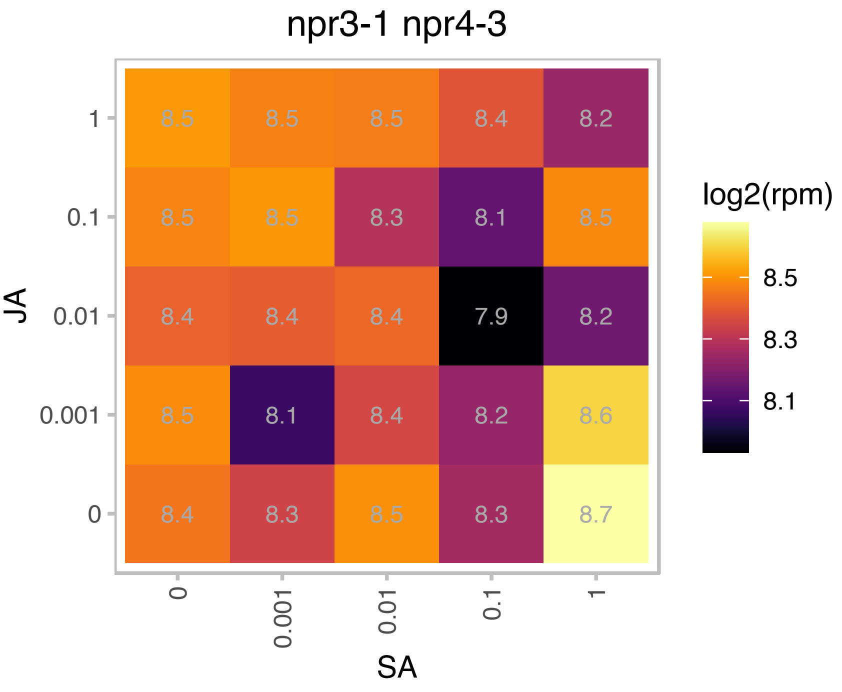

# AT1G77090.1 : NA

Mog1/PsbP/DUF1795-like photosystem II reaction center PsbP family protein

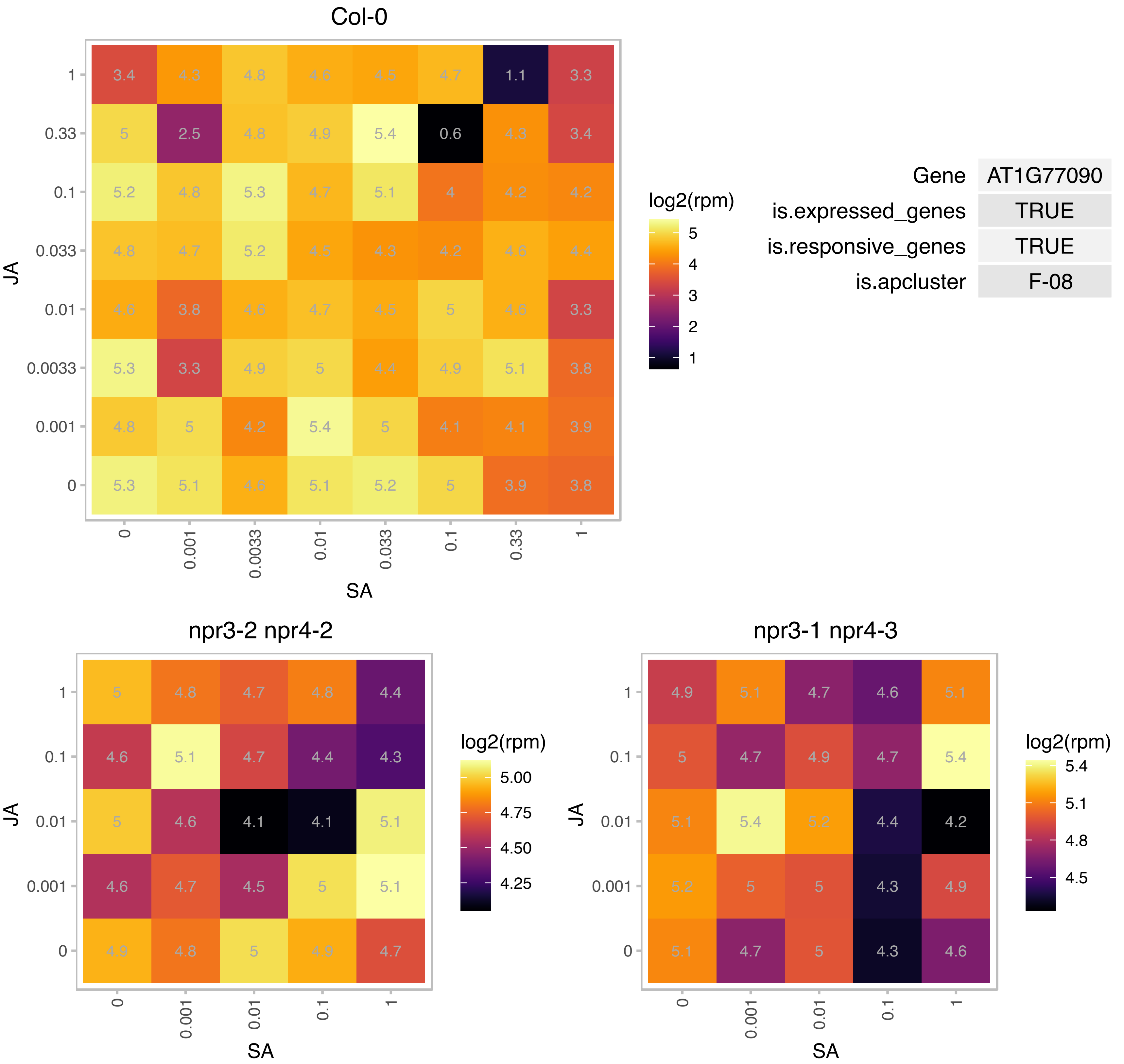


| | |
|---|---|
| Gene | AT1G77090 |
| is.expressed_genes | TRUE |
| is.responsive_genes | TRUE |
| is.apcluster | F-08 |

### AT1G77760.1 : GNR1, NIA1, NR1

nitrate reductase 1

Encodes the cytosolic minor isoform of nitrate reductase (NR). Involved in the first step of nitrate assimilation, it contributes about 15% of the nitrate reductase activity in shoots. Similar to molybdopterin oxidoreductases at the N-terminus, and to FAD/NAD-binding cytochrome reductases at the C-terminus. Cofactors: FAD, heme iron (cytochrome B-557), and molybdenum-pterin.

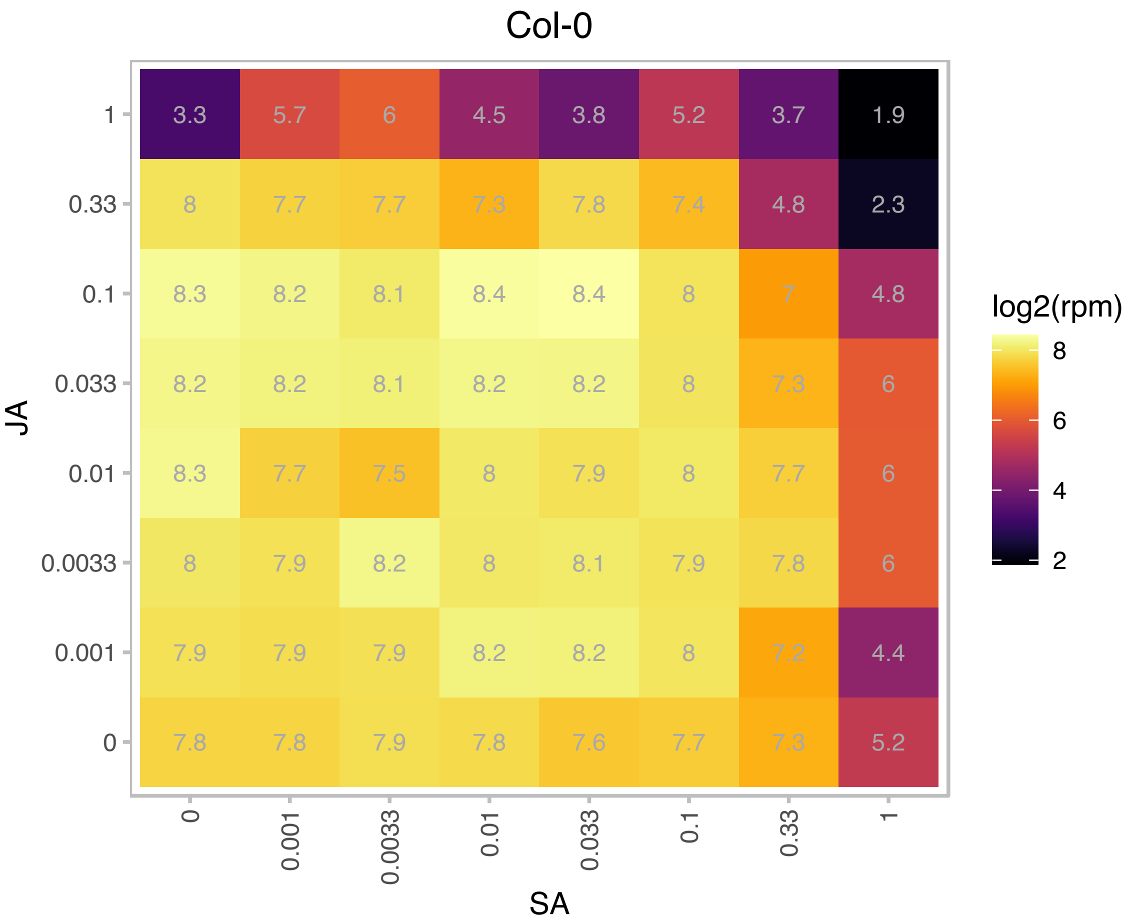


| | |
|---|---|
| Gene | AT1G77760 |
| is.expressed_genes | TRUE |
| is.responsive_genes | TRUE |
| is.apcluster | F-08 |

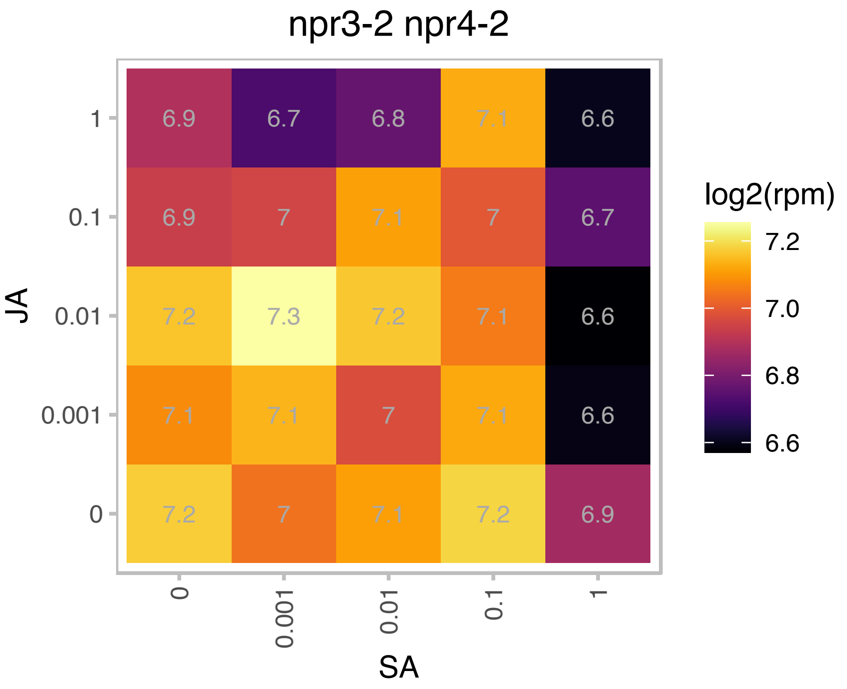


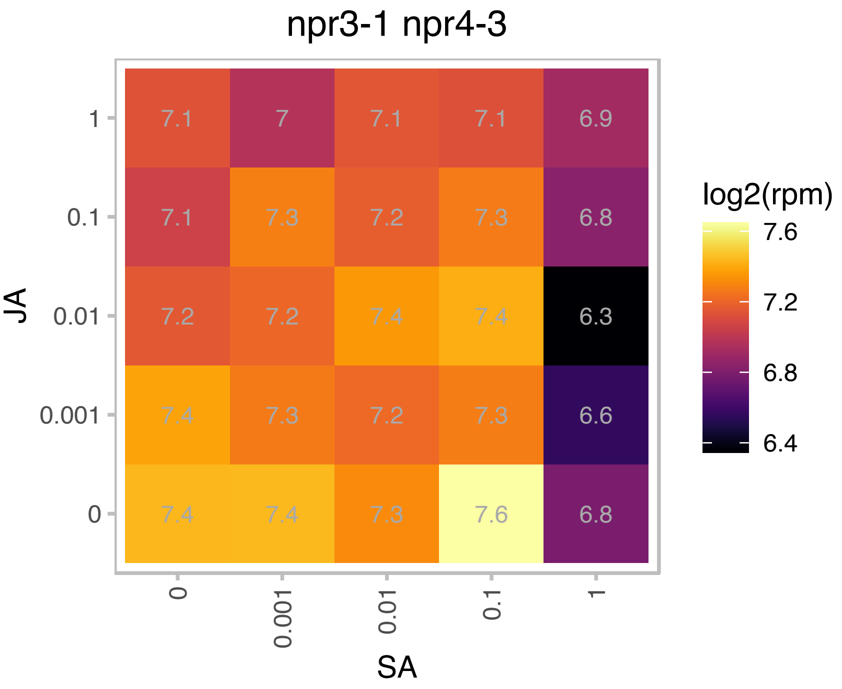

### AT1G80750.1 : RPL7A

Ribosomal protein L30/L7 family protein

Cytosolic ribosomal 60S subunit protein.

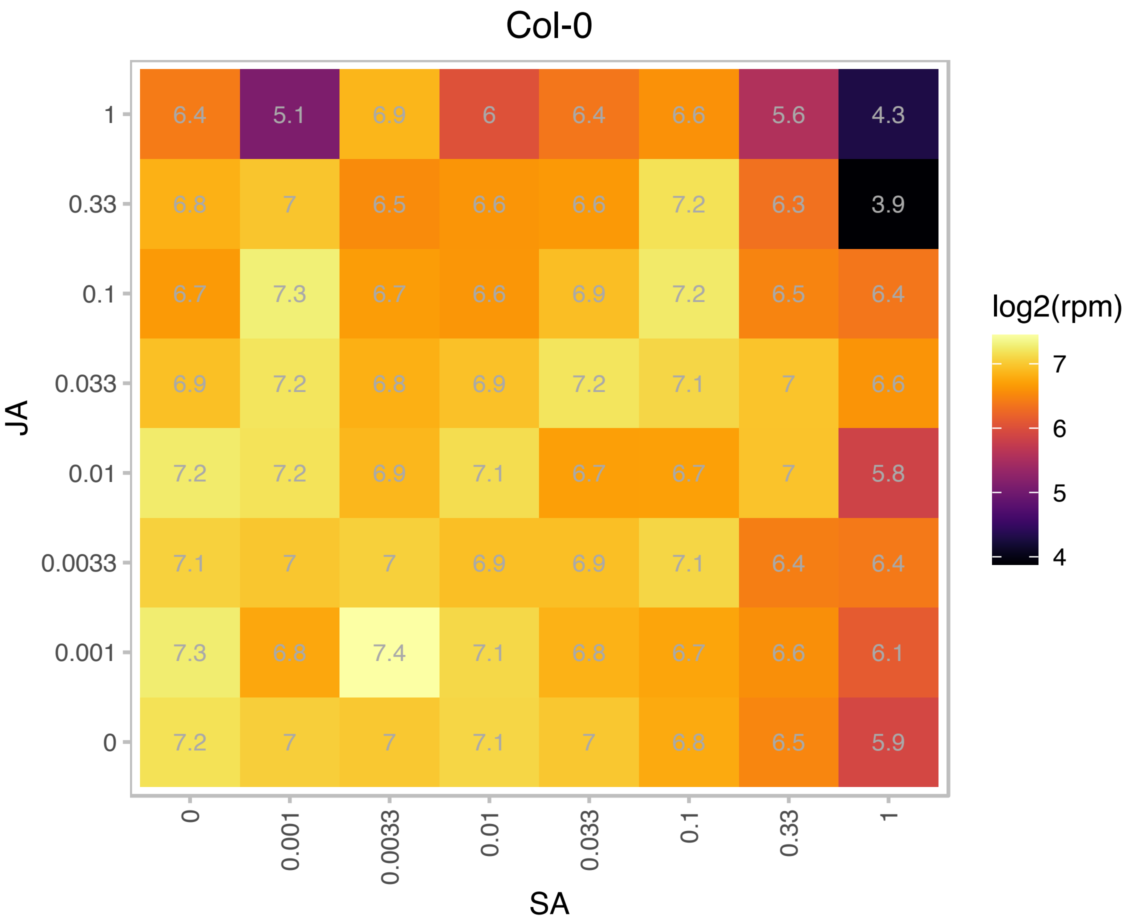


| | |
|---|---|
| Gene | AT1G80750 |
| is.expressed_genes | TRUE |
| is.responsive_genes | TRUE |
| is.apcluster | F-08 |

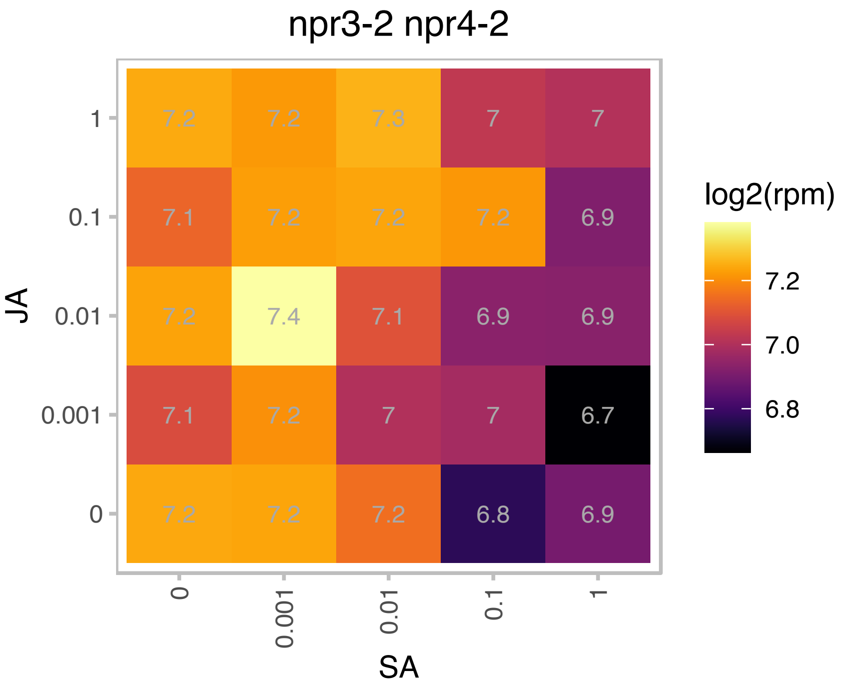


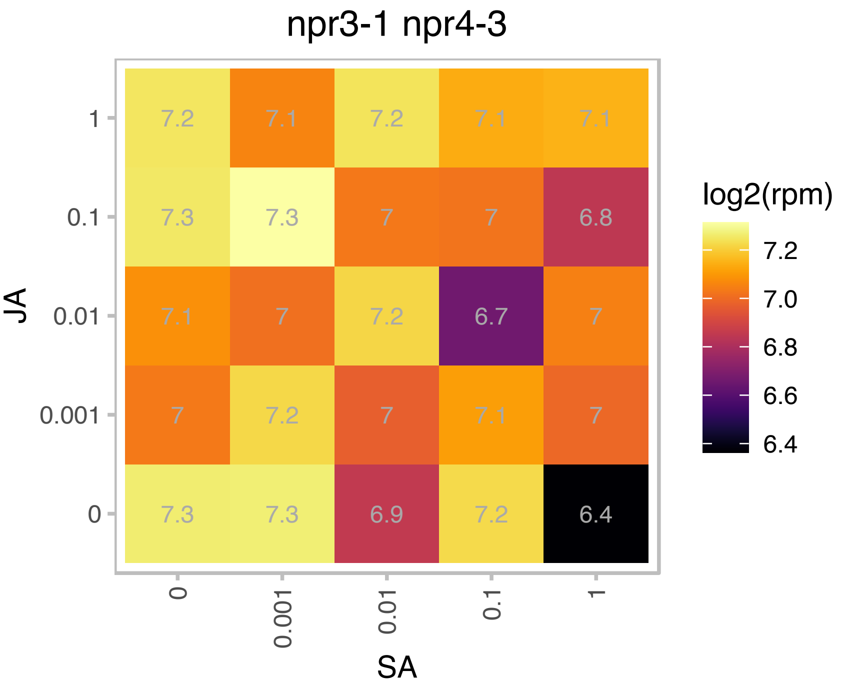

# AT2G01060.1 : NA

myb-like HTH transcriptional regulator family protein

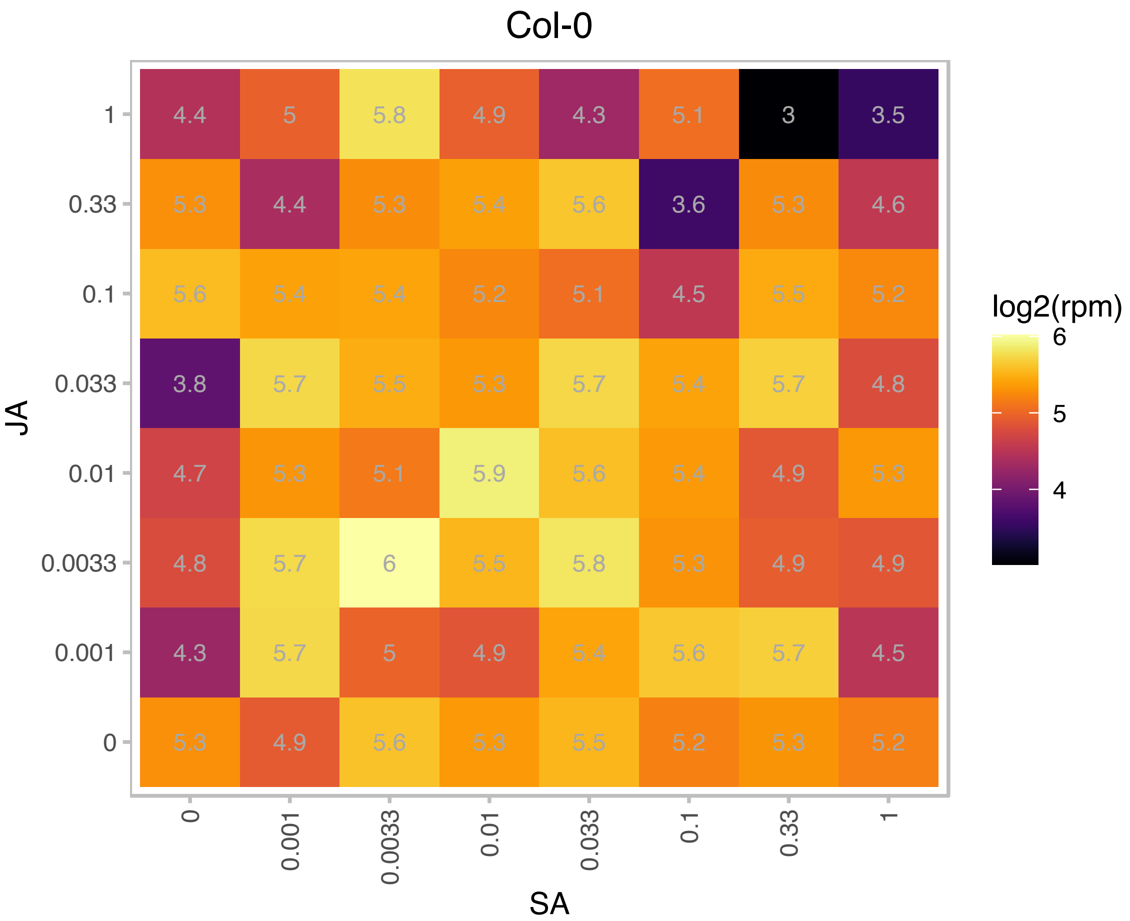


| | |
|---|---|
| Gene | AT2G01060 |
| is.expressed_genes | TRUE |
| is.responsive_genes | TRUE |
| is.apcluster | F-08 |

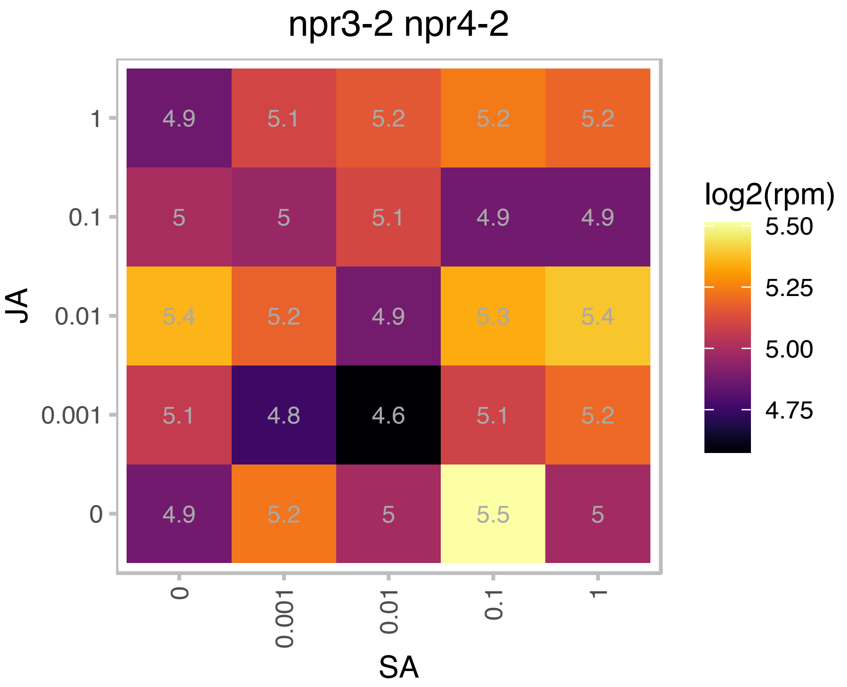


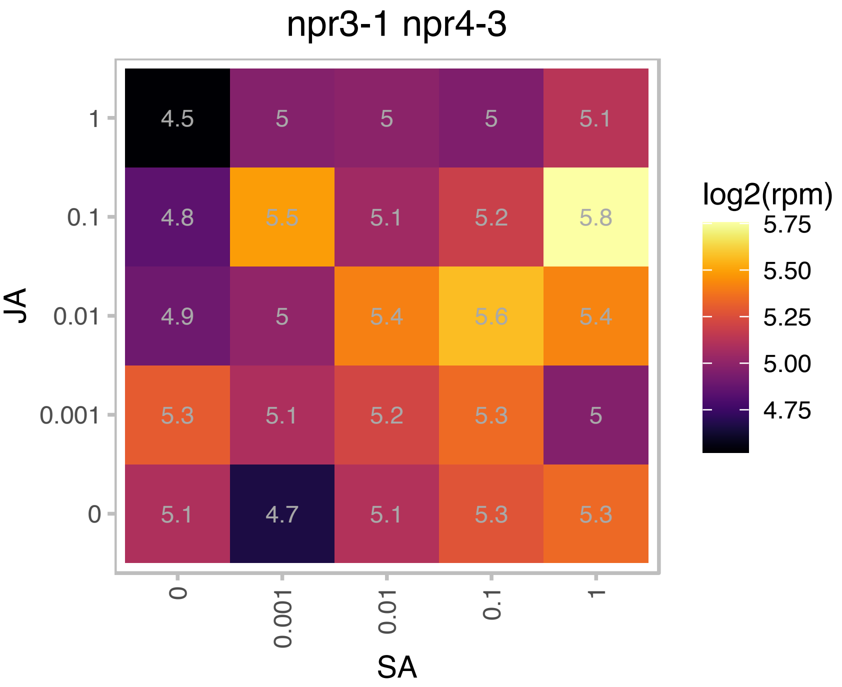

### AT2G01620.1 : MEE11

RNI-like superfamily protein

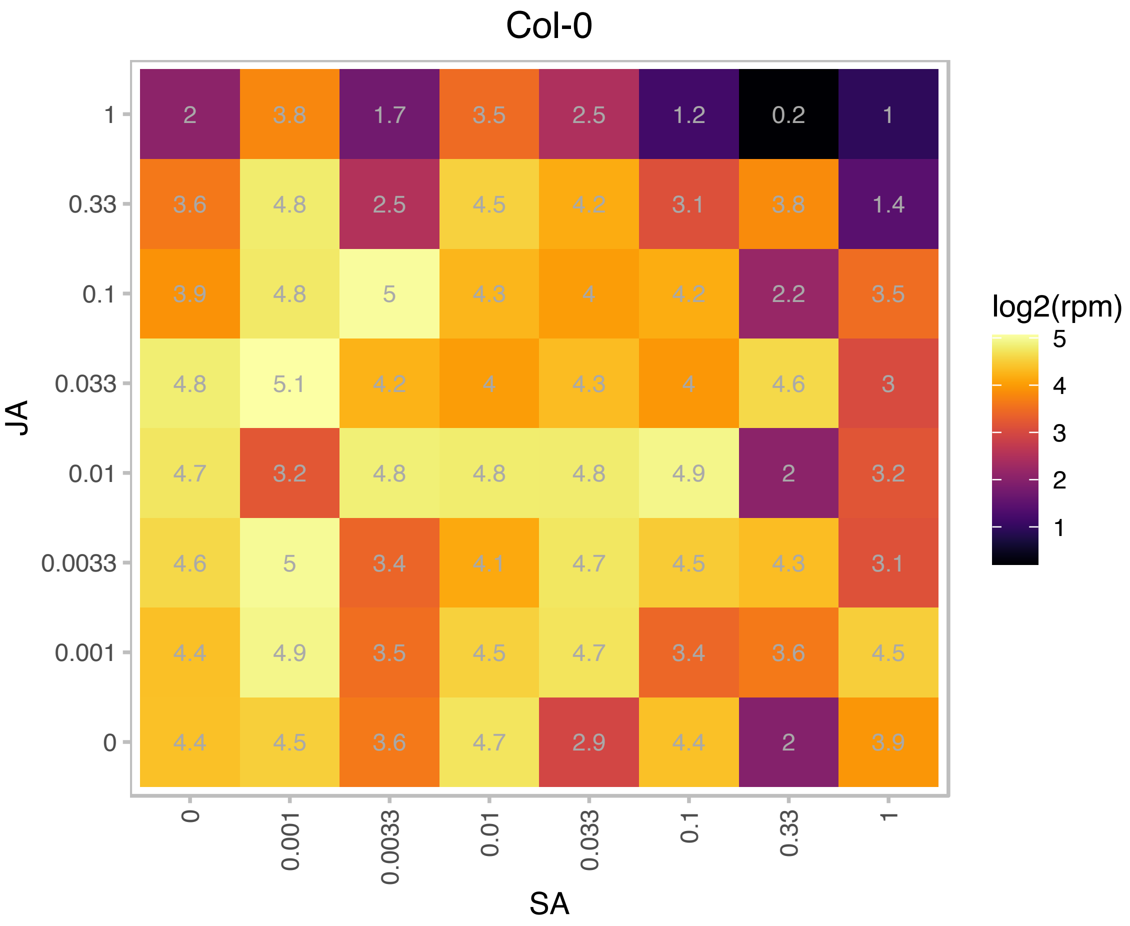


| | |
|---|---|
| Gene | AT2G01620 |
| is.expressed_genes | TRUE |
| is.responsive_genes | TRUE |
| is.apcluster | F-08 |

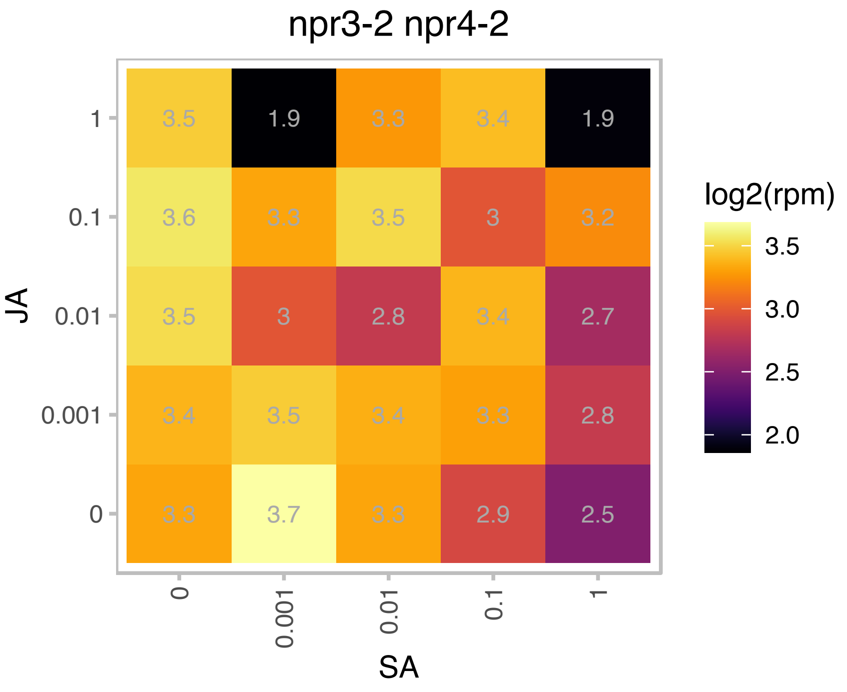


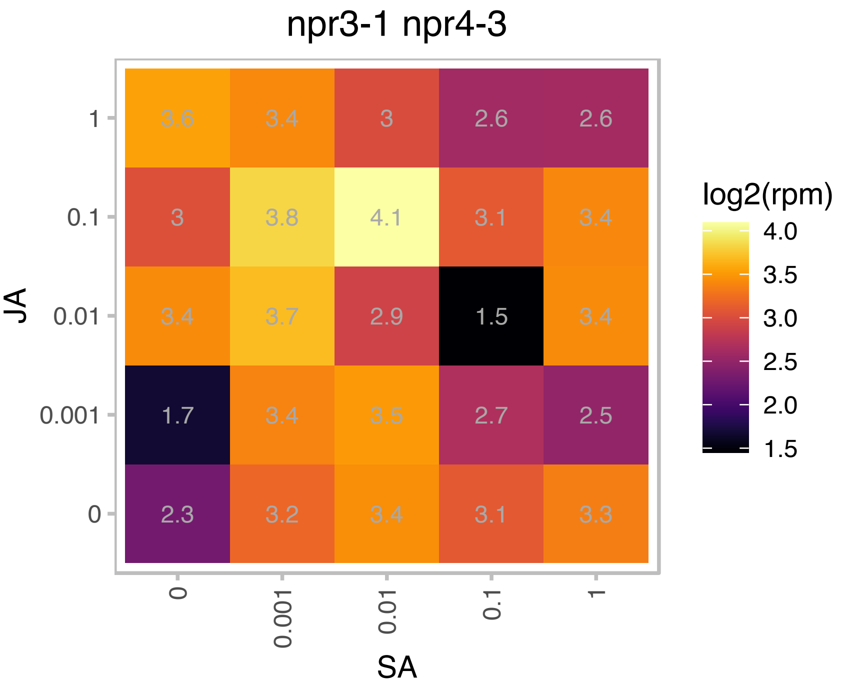

# AT2G11910.1 : NA

| | |
|---|---|
| Gene | AT2G11910 |
| is.expressed_genes | TRUE |
| is.responsive_genes | TRUE |
| is.apcluster | F-08 |

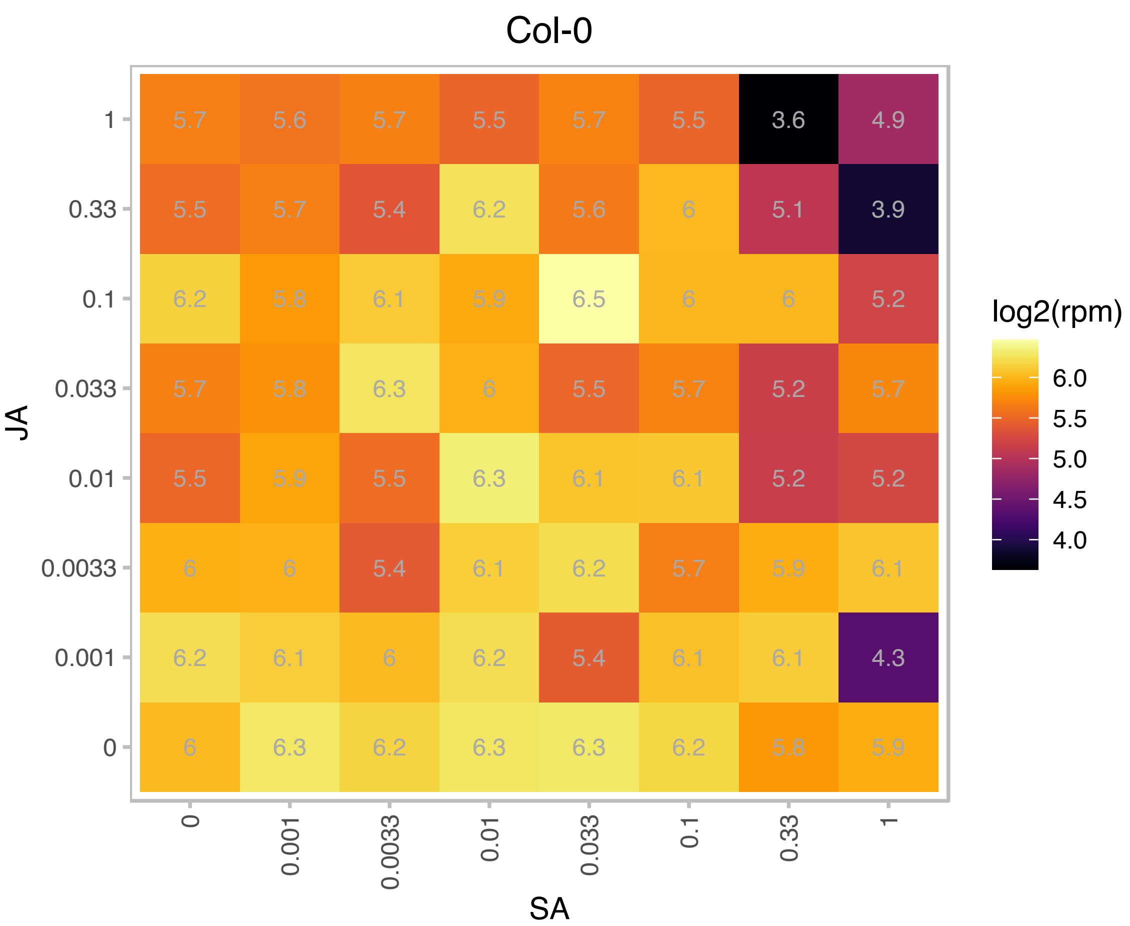


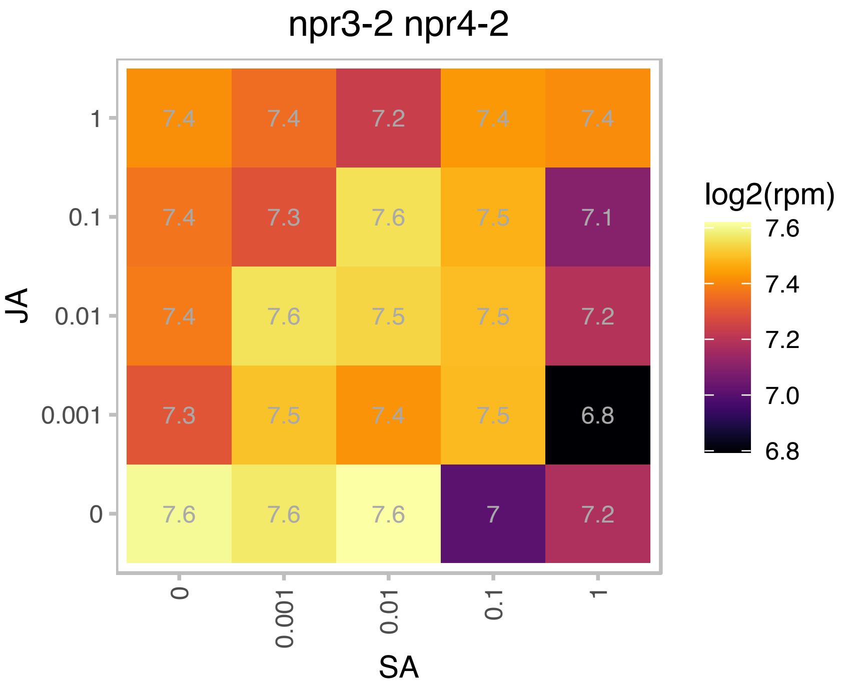


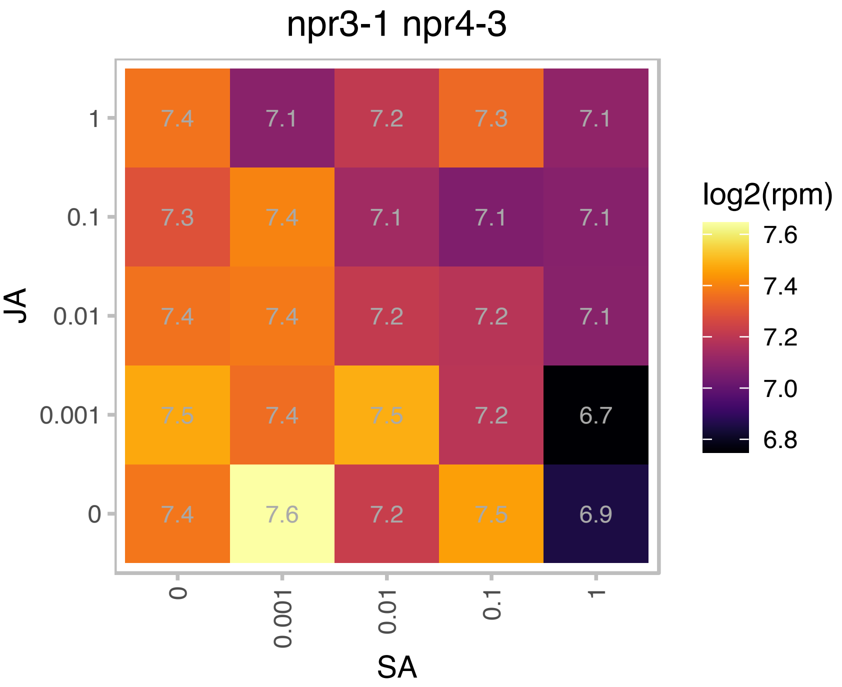

# AT2G18910.1 : NA

hydroxyproline-rich glycoprotein family protein

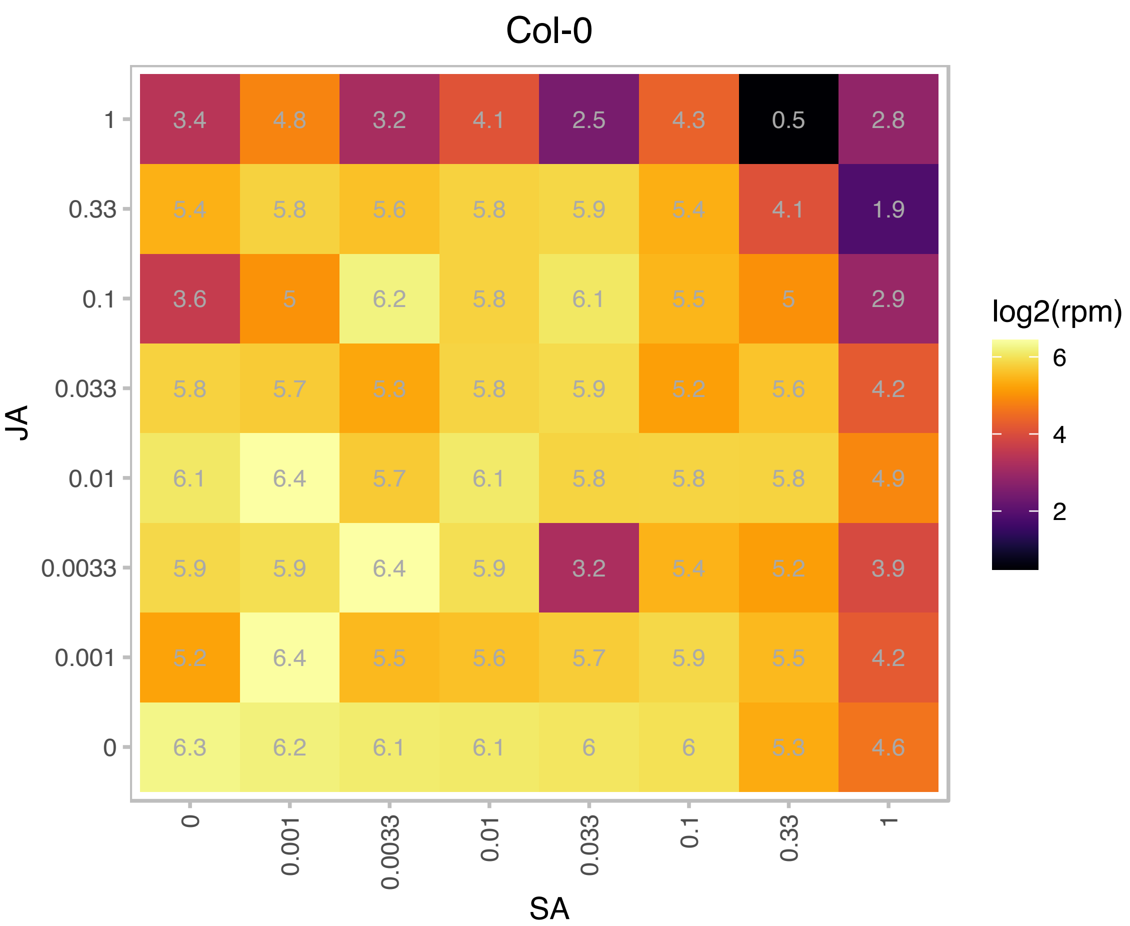


| | |
|---|---|
| Gene | AT2G18910 |
| is.expressed_genes | TRUE |
| is.responsive_genes | TRUE |
| is.apcluster | F-08 |

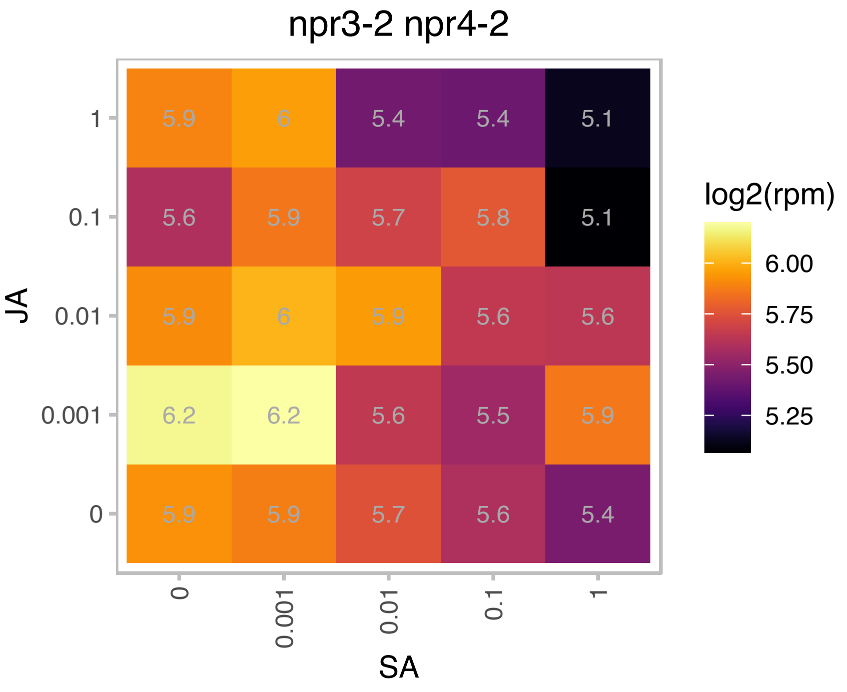


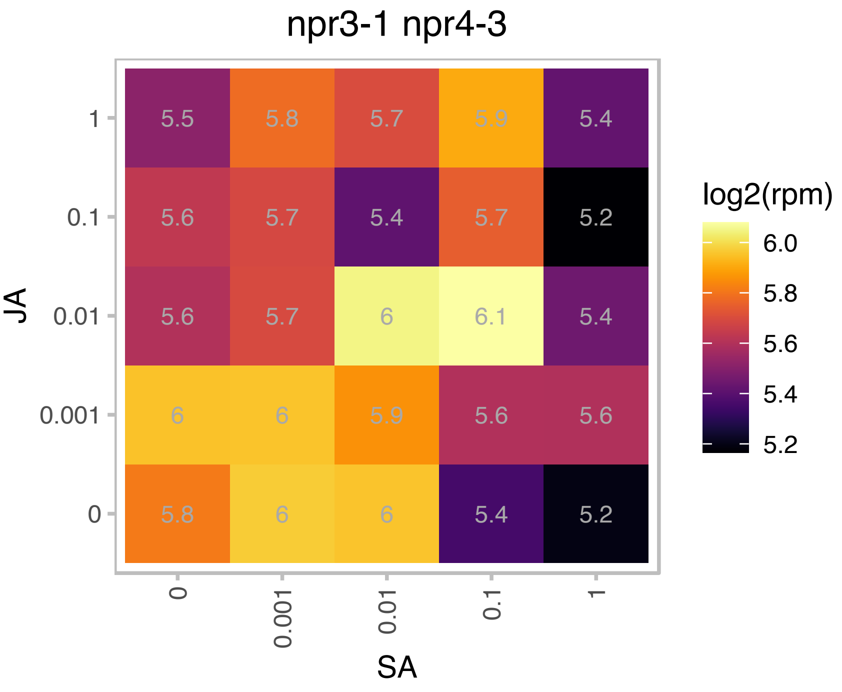

# AT2G20060.1 : NA

Ribosomal protein L4/L1 family

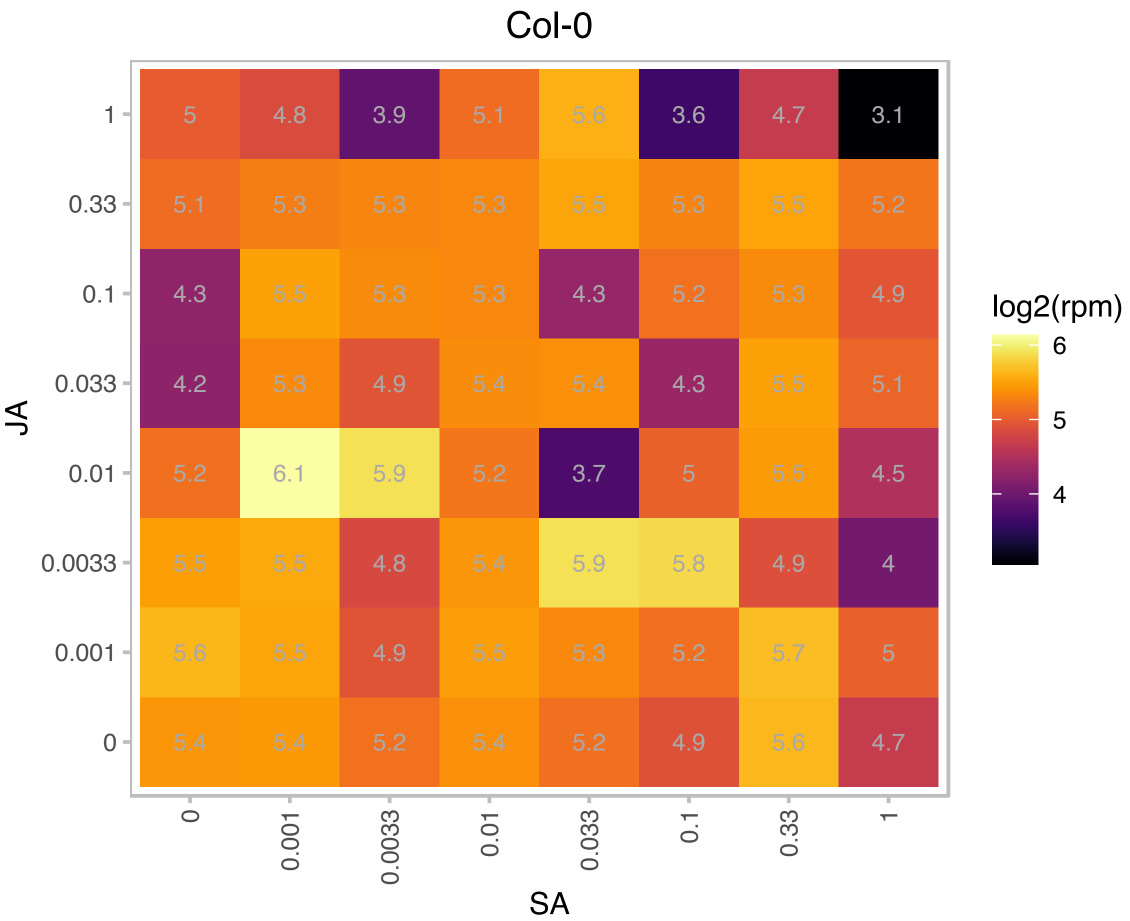


| | |
|---|---|
| Gene | AT2G20060 |
| is.expressed_genes | TRUE |
| is.responsive_genes | TRUE |
| is.apcluster | F-08 |

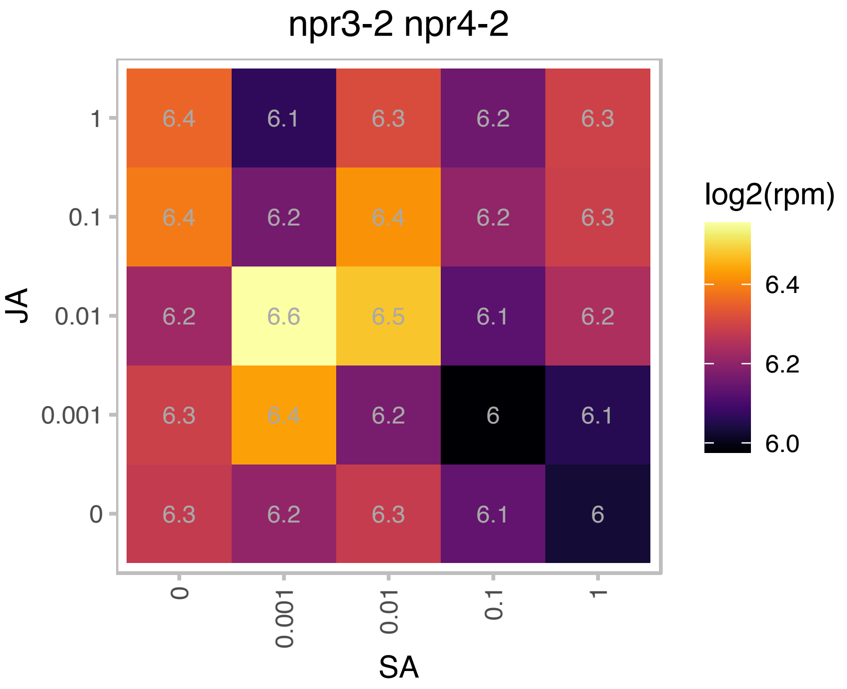


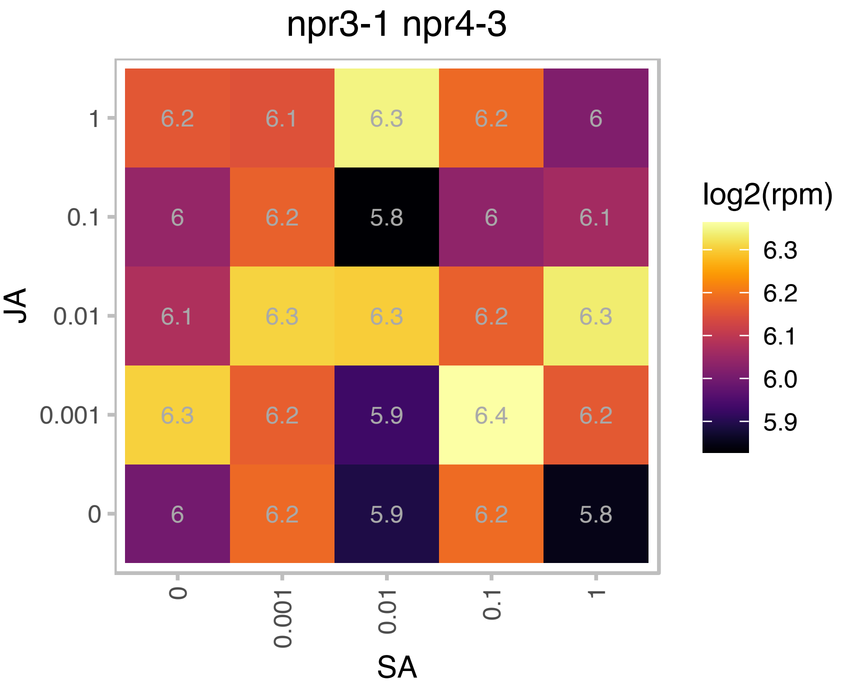

### AT2G22870.1 : EMB2001, EngB-1

P-loop containing nucleoside triphosphate hydrolases superfamily protein

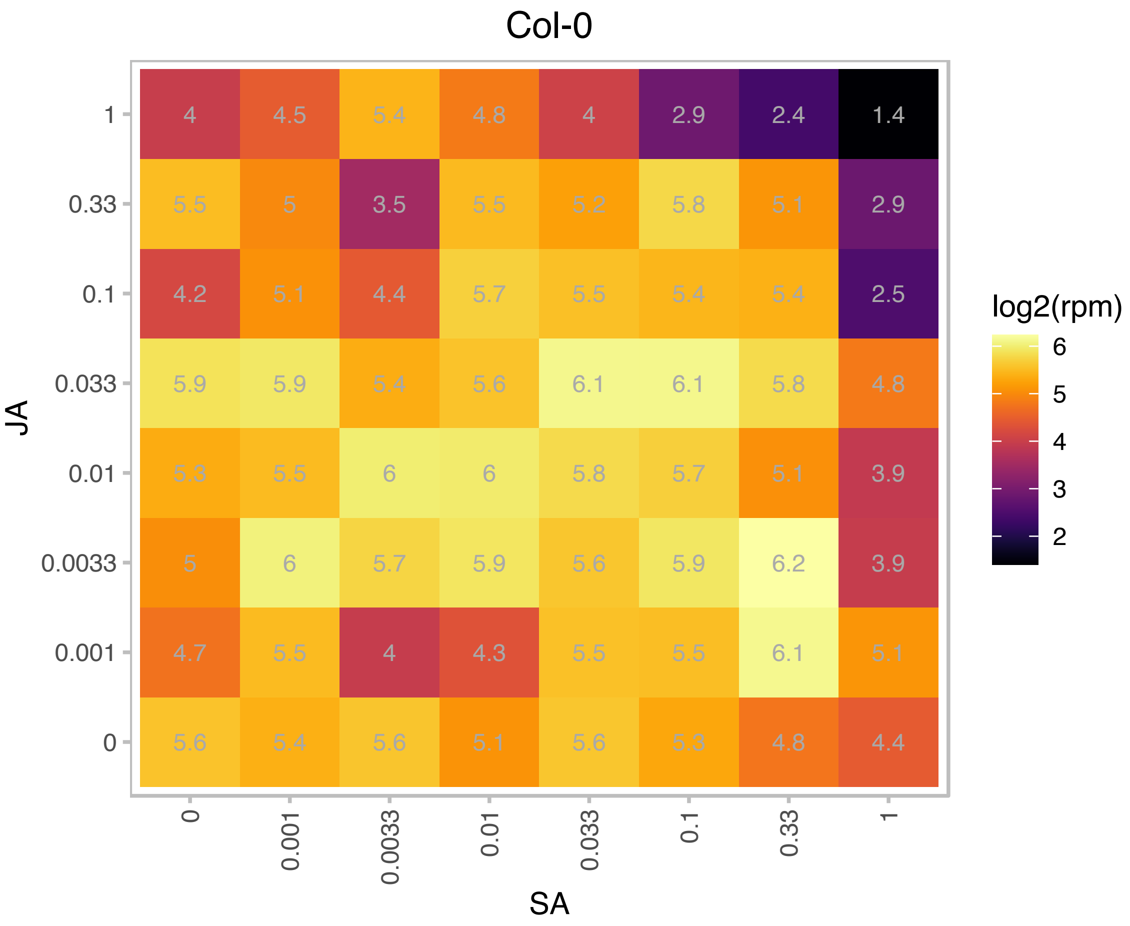


| | |
|---|---|
| Gene | AT2G22870 |
| is.expressed_genes | TRUE |
| is.responsive_genes | TRUE |
| is.apcluster | F-08 |

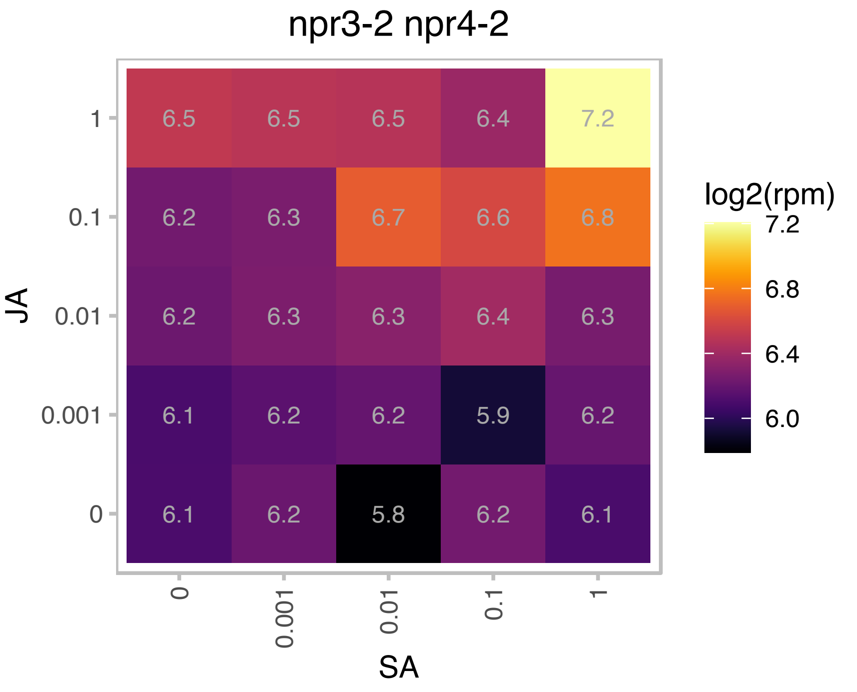


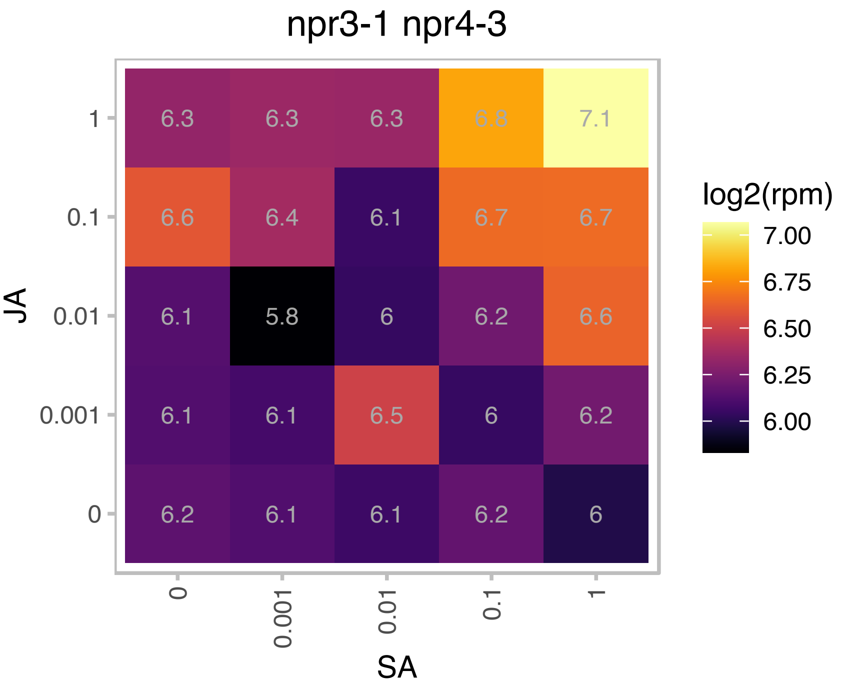

### AT2G24590.1 : At-RSZ22a, RSZ22a

RNA recognition motif and CCHC-type zinc finger domains containing protein

Barta et al (2010) have proposed a nomenclature for Serine/Arginine-Rich Protein Splicing Factors (SR proteins): Plant Cell. 2010, 22:2926.

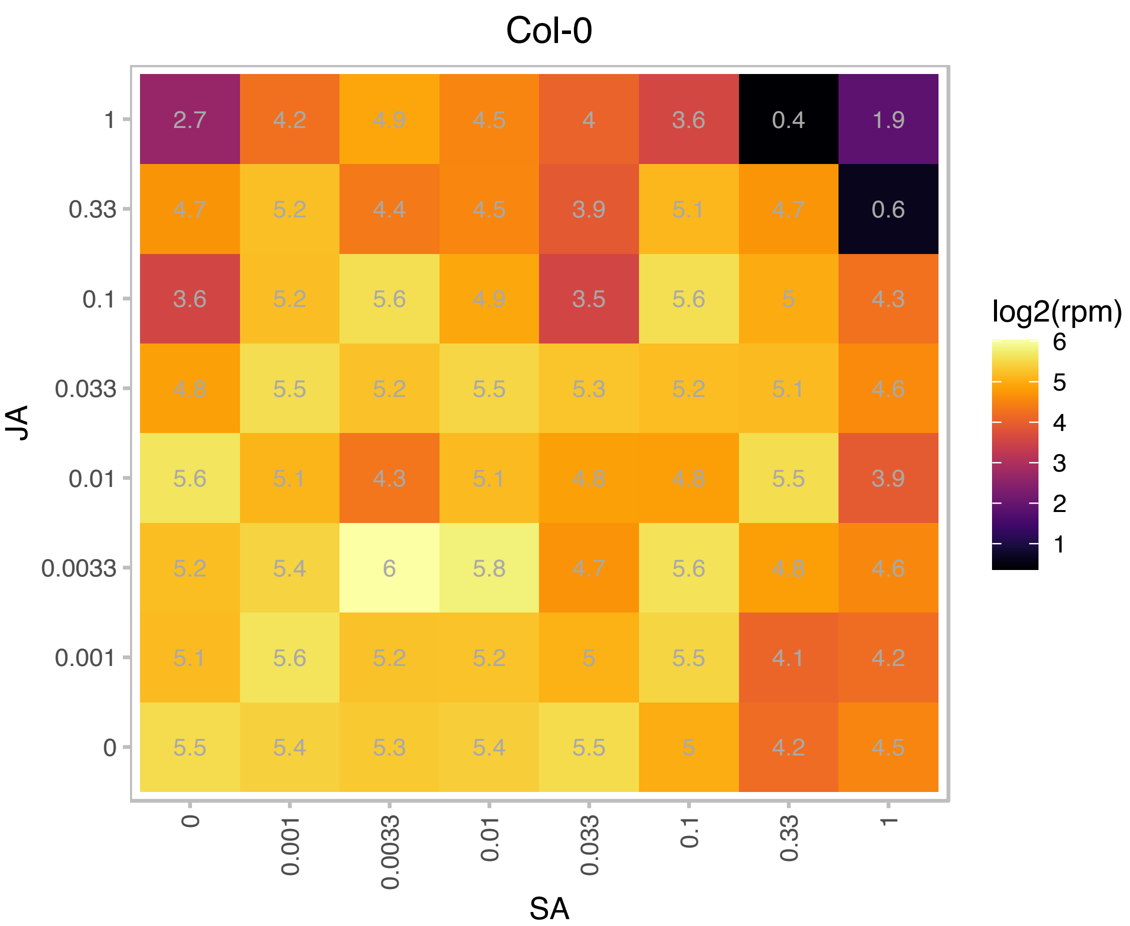


| | |
|---|---|
| Gene | AT2G24590 |
| is.expressed_genes | TRUE |
| is.responsive_genes | TRUE |
| is.apcluster | F-08 |

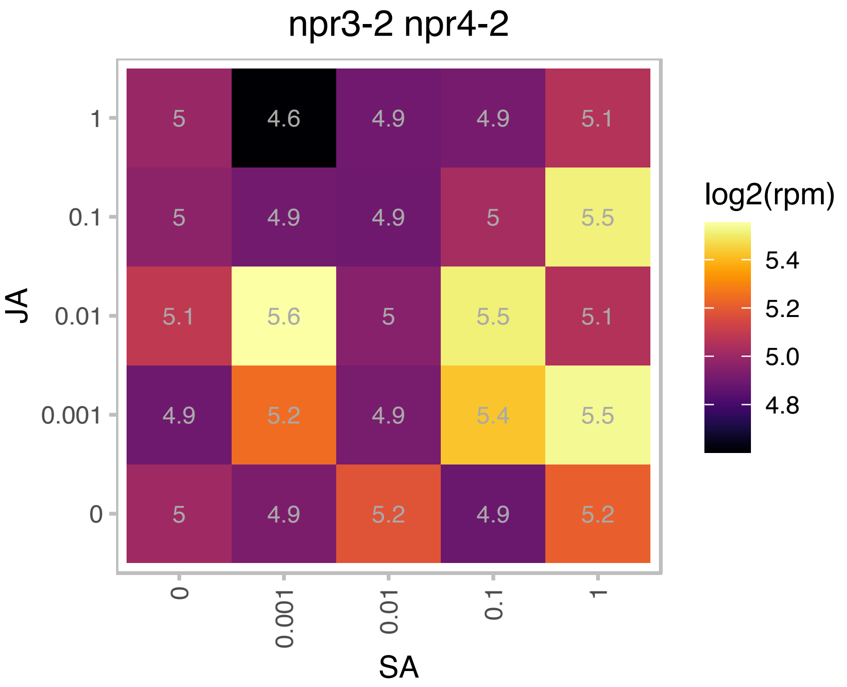


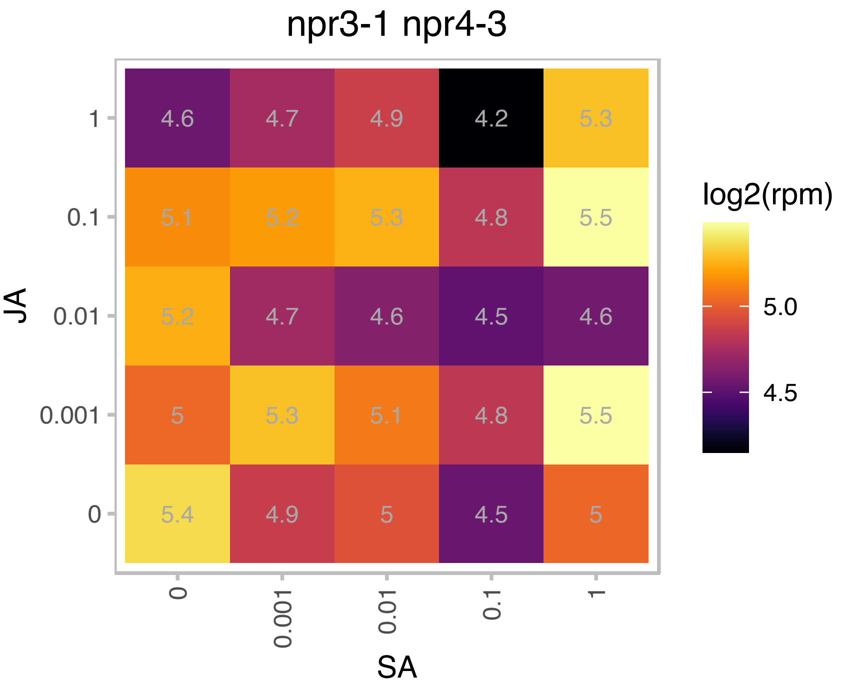

# AT2G25720.1 : NA

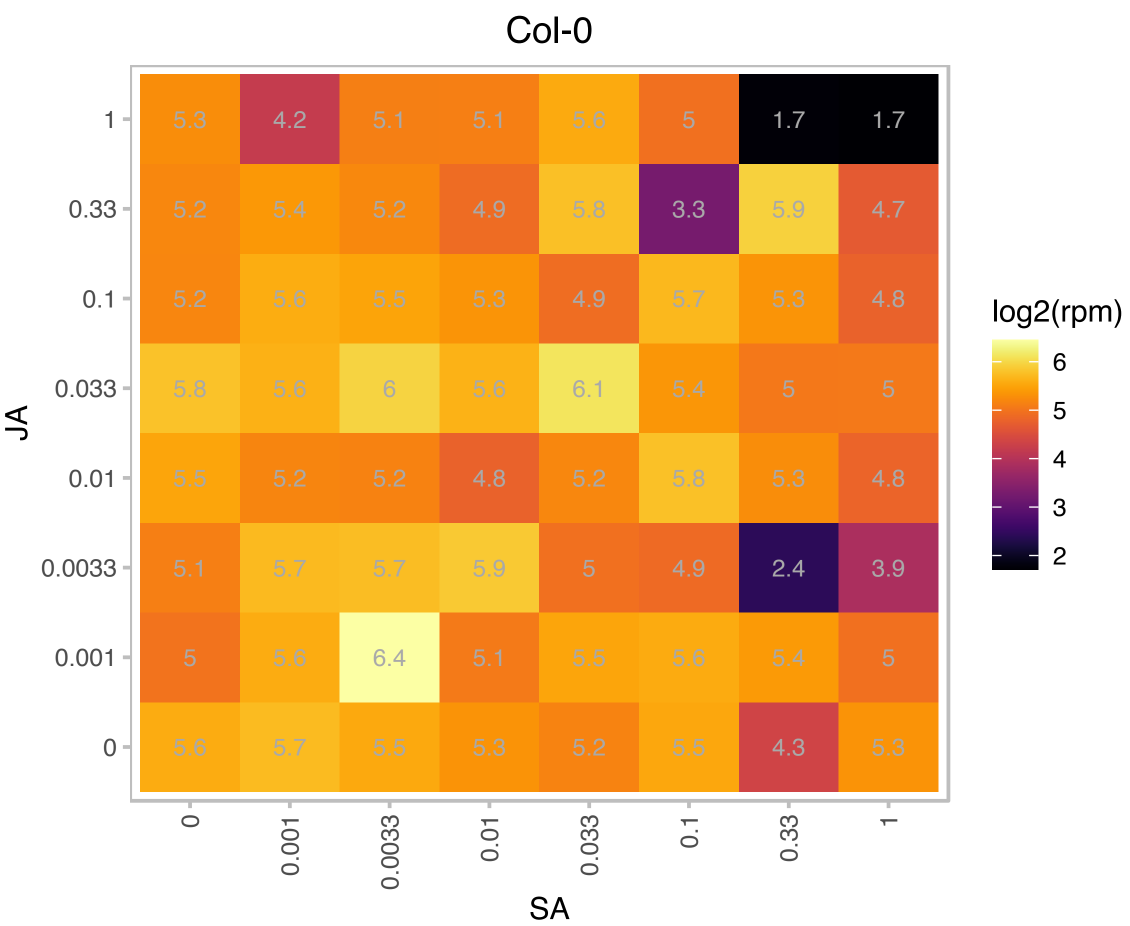


| Gene | AT2G25720 |
|---|---|
| is.expressed_genes | TRUE |
| is.responsive_genes | TRUE |
| is.apcluster | F-08 |

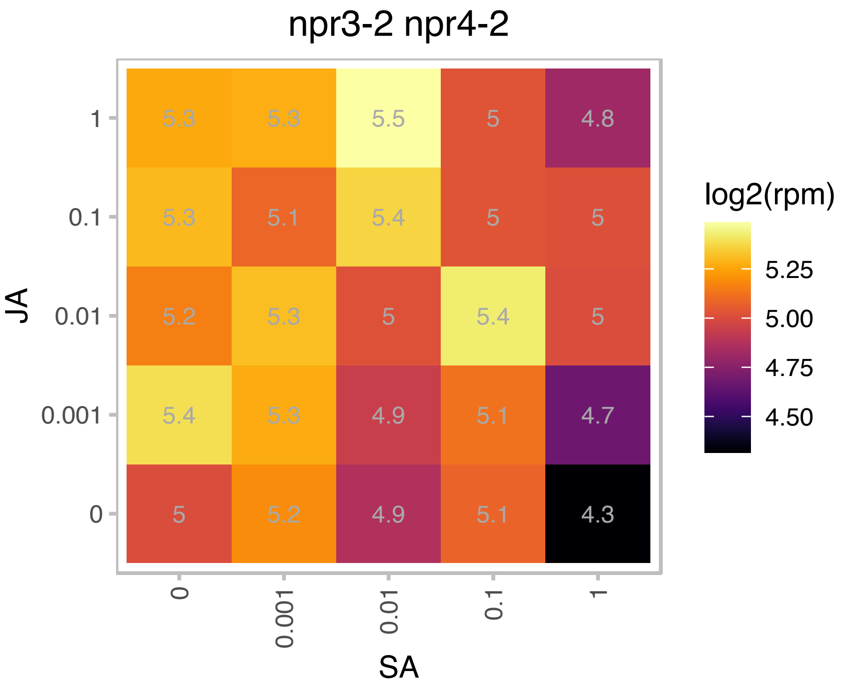


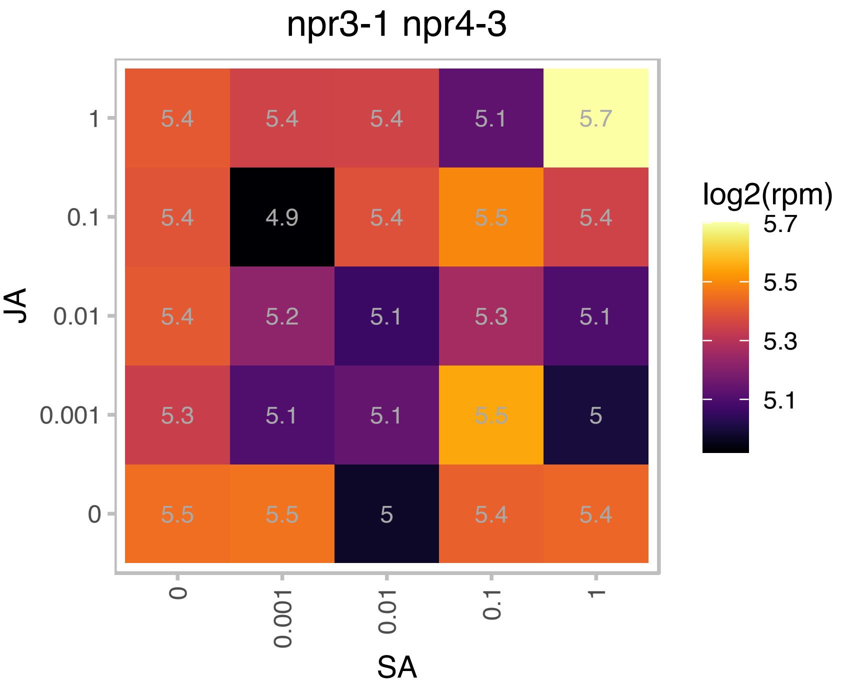

### AT2G26975.1 : COPT6

Ctr copper transporter family

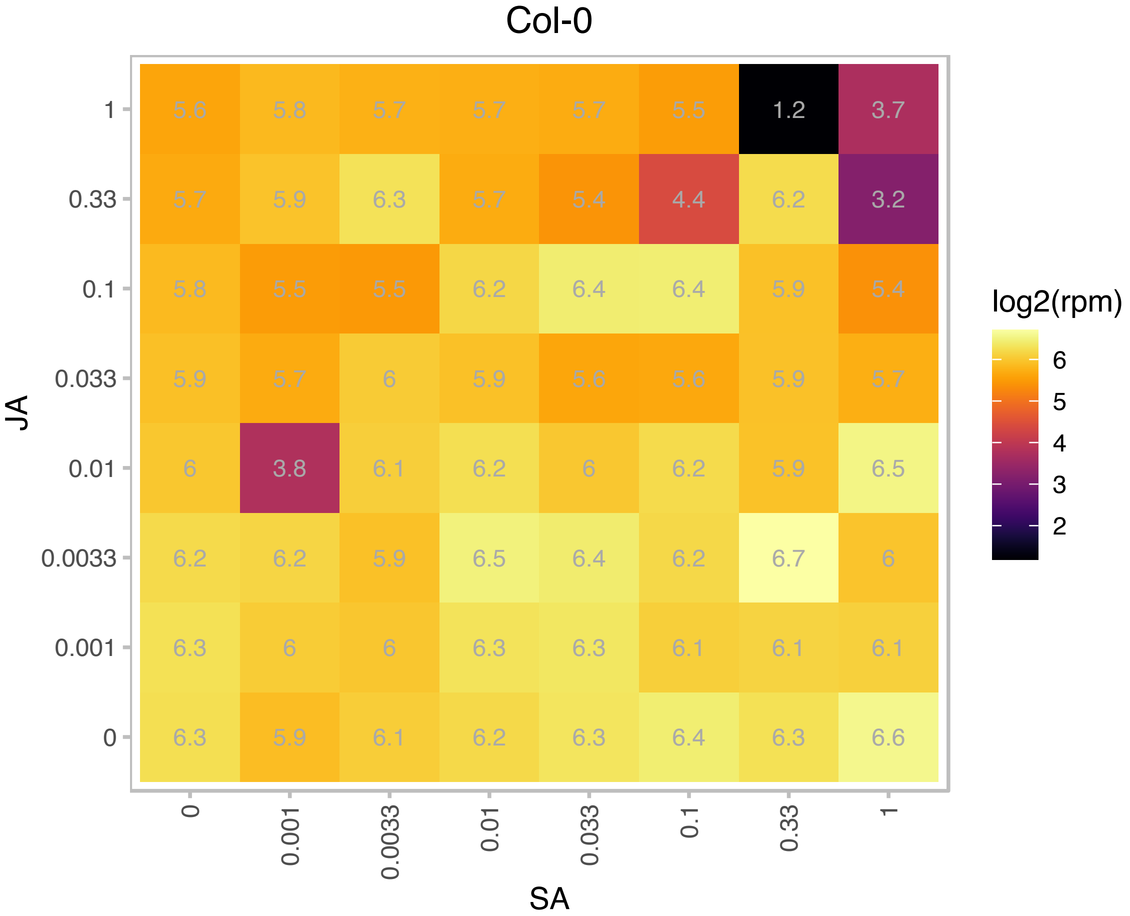


| | |
|---|---|
| Gene | AT2G26975 |
| is.expressed_genes | TRUE |
| is.responsive_genes | TRUE |
| is.apcluster | F-08 |

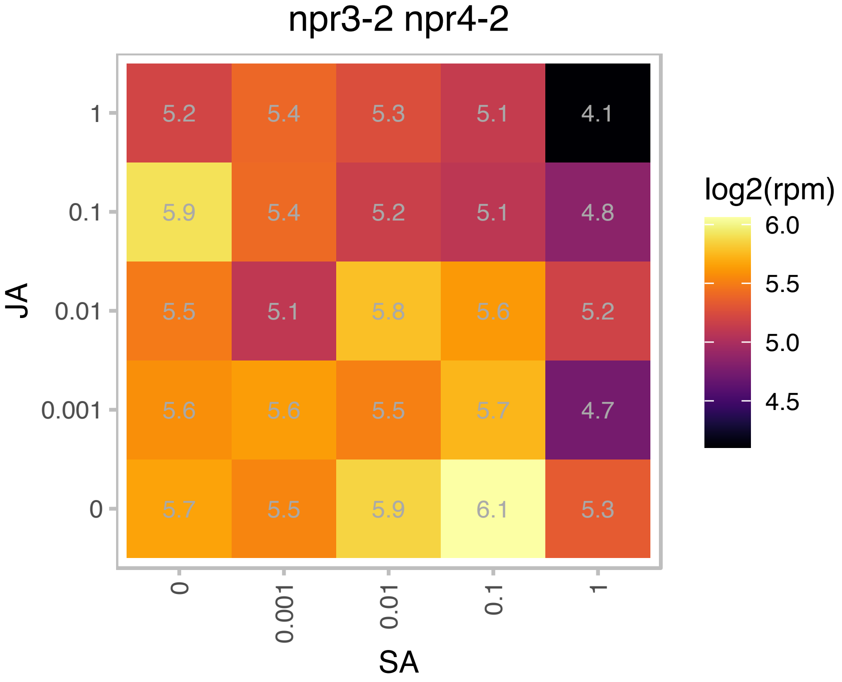


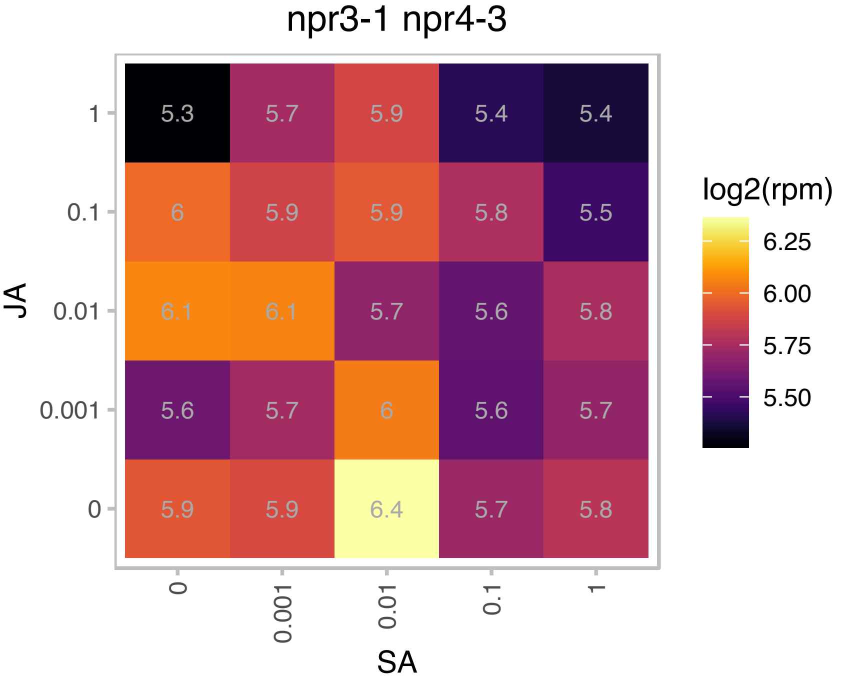

# AT2G27330.1 : NA

RNA-binding (RRM/RBD/RNP motifs) family protein

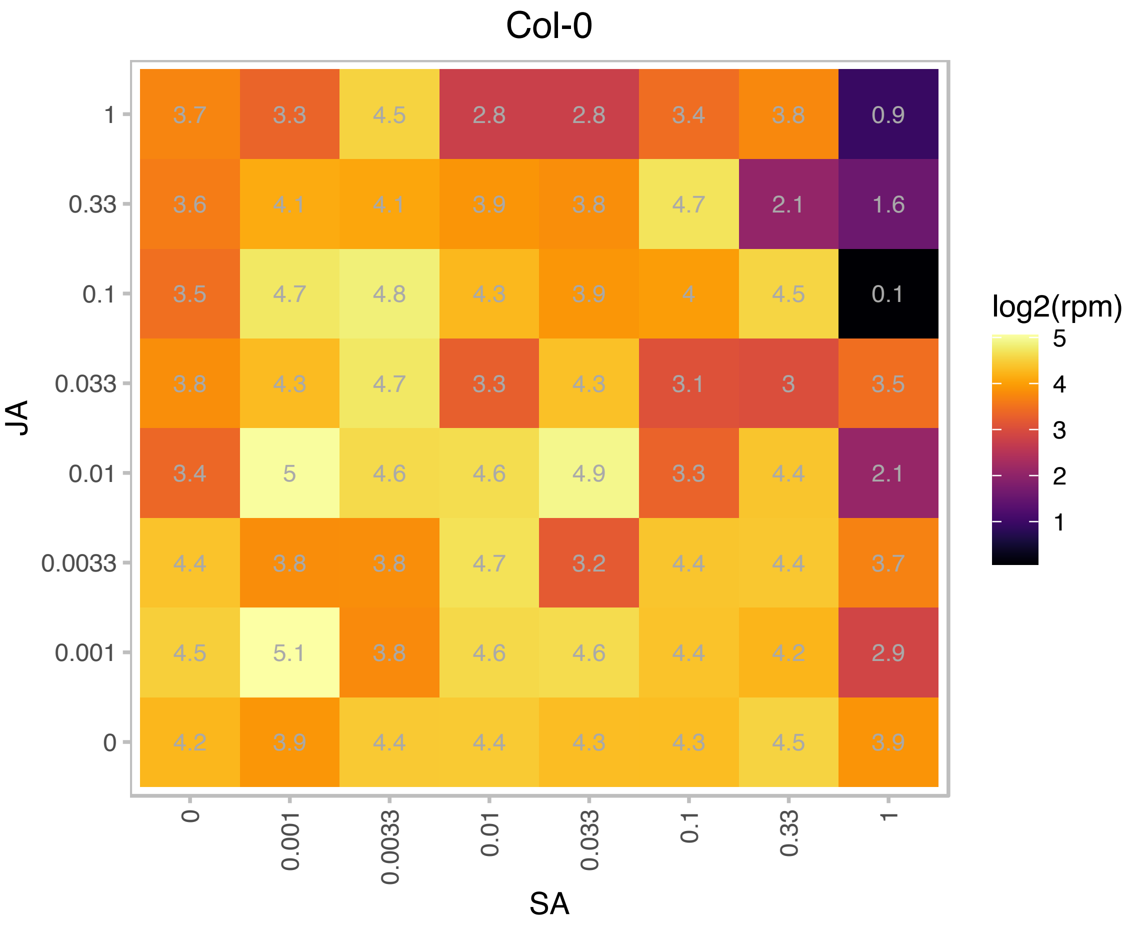


| | |
|---|---|
| Gene | AT2G27330 |
| is.expressed_genes | TRUE |
| is.responsive_genes | TRUE |
| is.apcluster | F-08 |

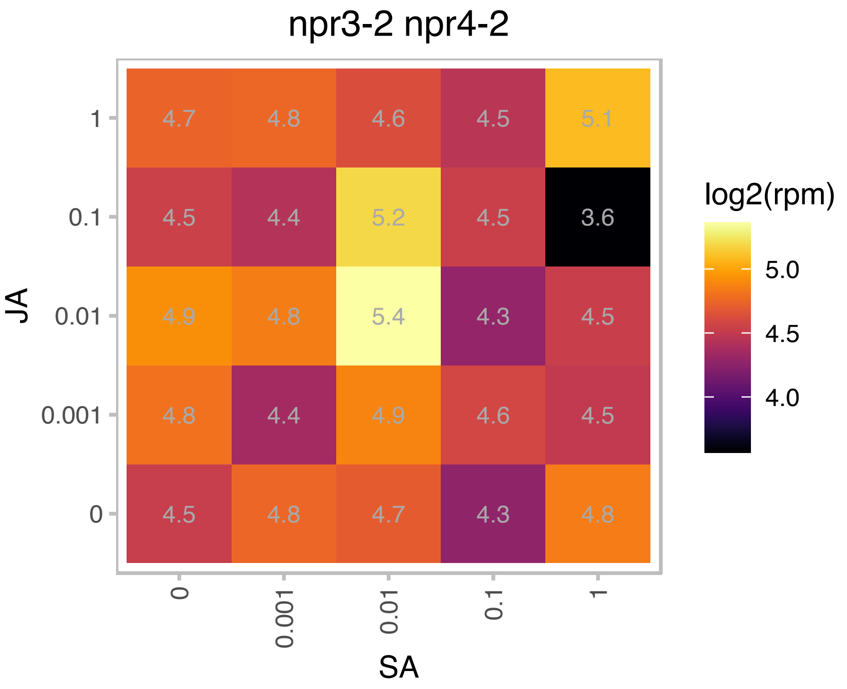


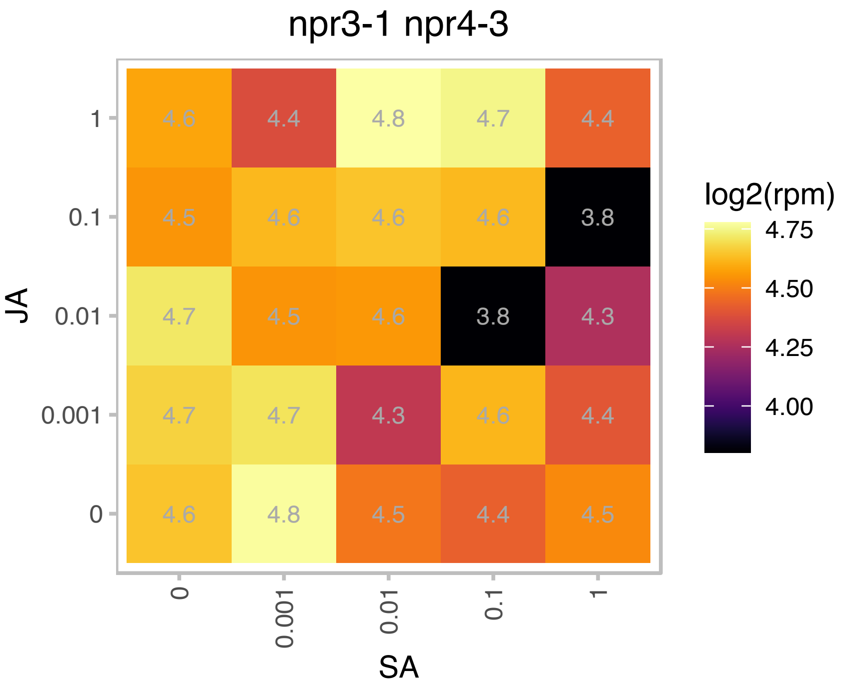

### AT2G28000.1 : CH-CPN60A, CPN60A, Cpn60alpha1, CPN

chaperonin-60alpha

Encodes chaperonin-60 alpha, a molecular chaperone involved in Rubisco folding.  Mutants display aberrant chloroplast and embryo development. The mRNA is cell-to-cell mobile.

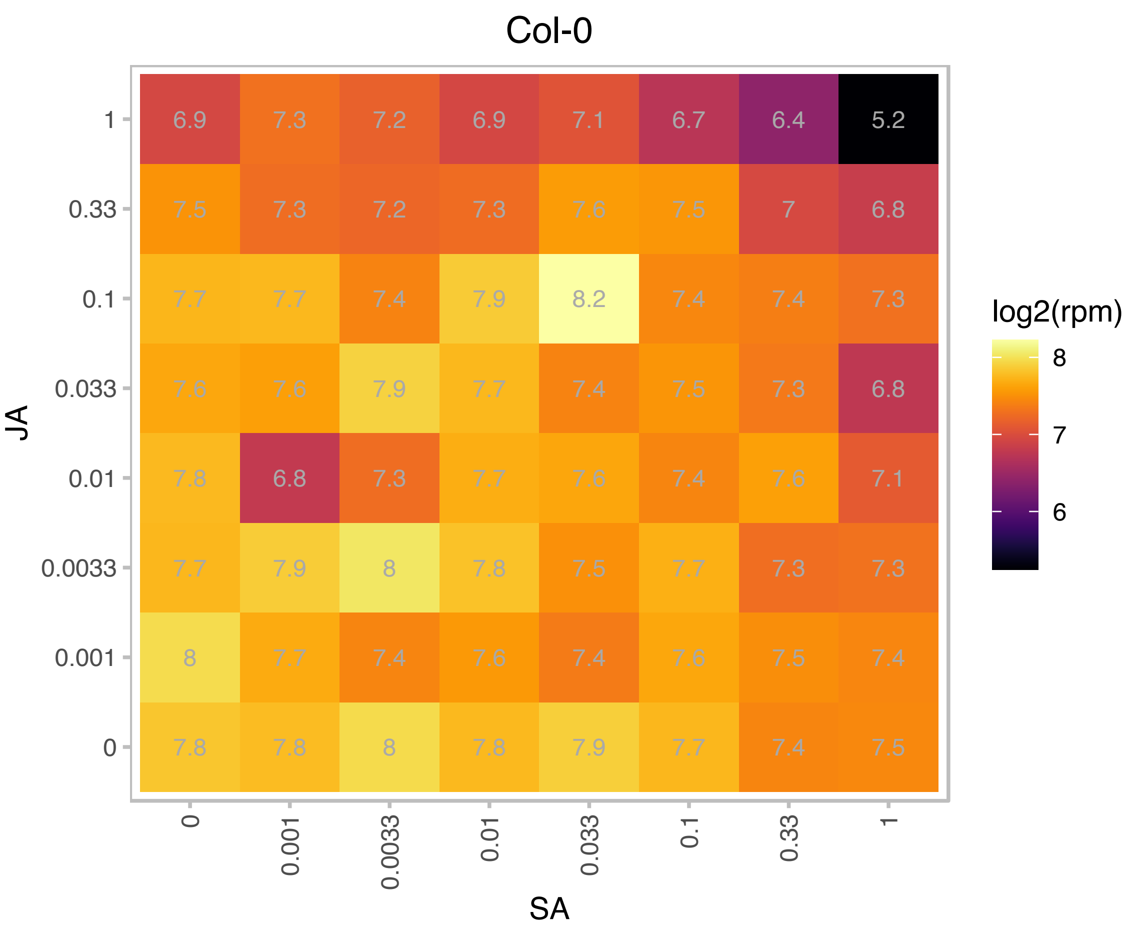


| | |
|---|---|
| Gene | AT2G28000 |
| is.expressed_genes | TRUE |
| is.responsive_genes | TRUE |
| is.apcluster | F-08 |

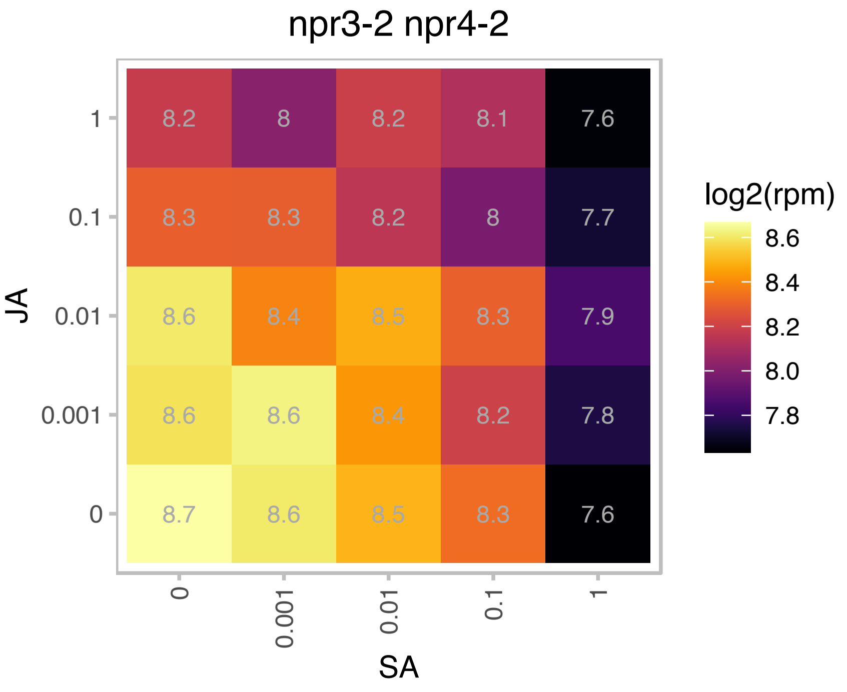


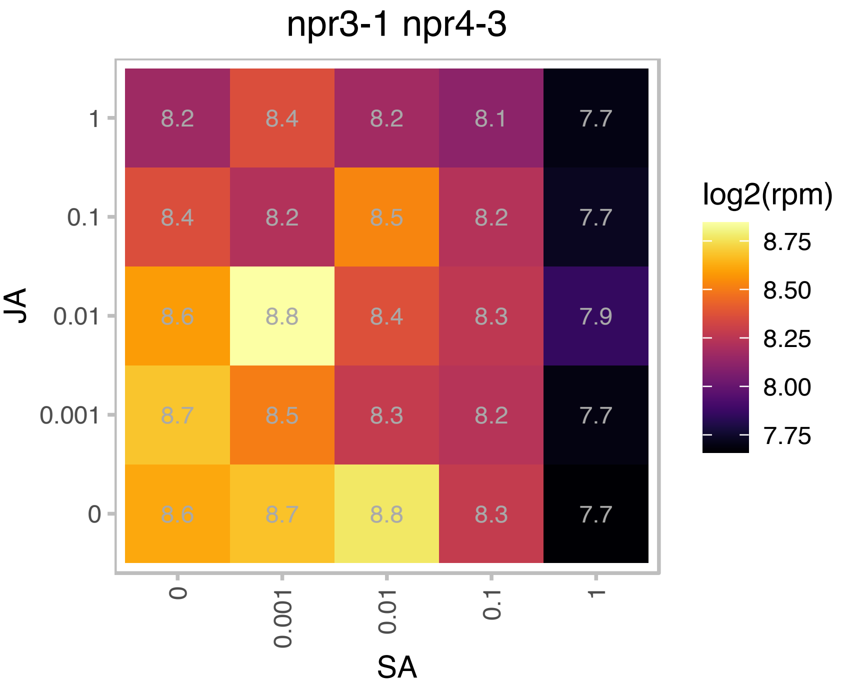

### AT2G28740.1 : HIS4

histone H4

histone 4

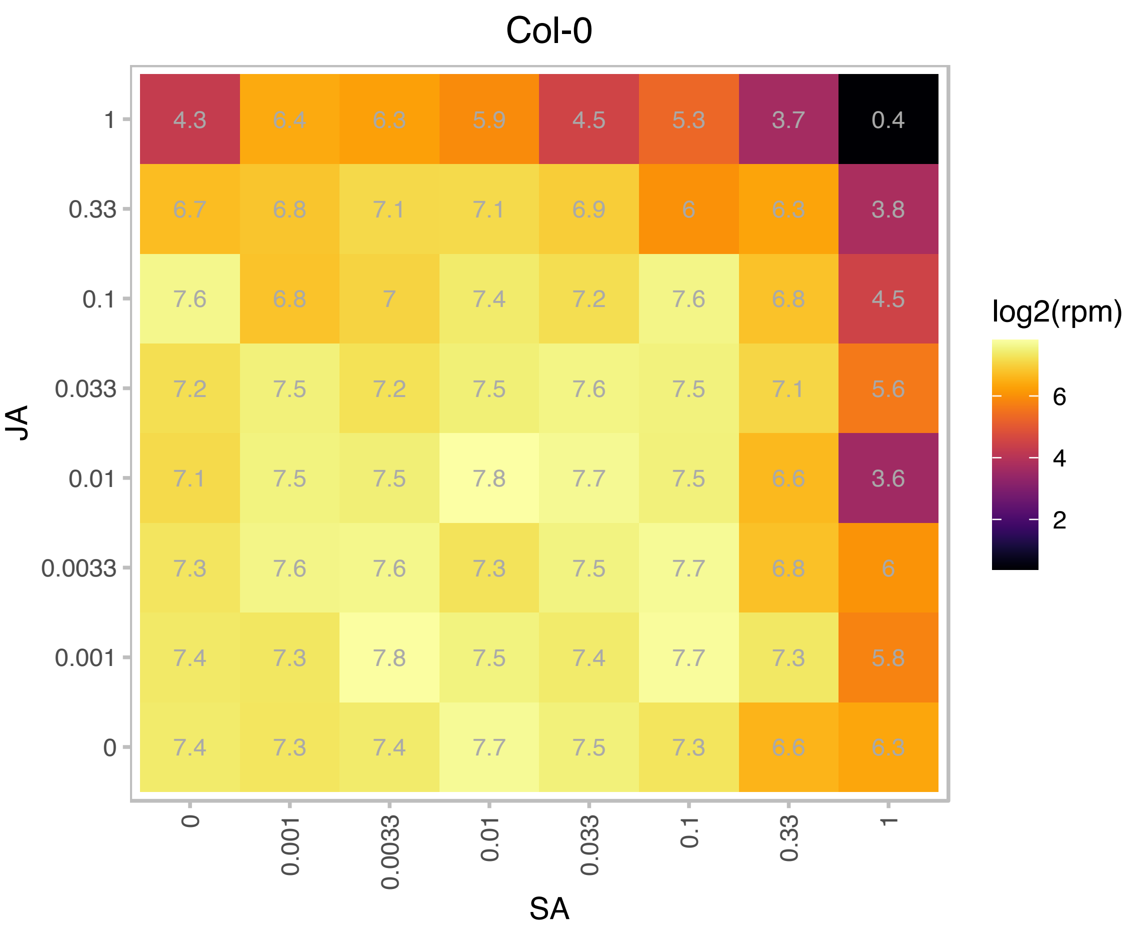


| | |
|---|---|
| Gene | AT2G28740 |
| is.expressed_genes | TRUE |
| is.responsive_genes | TRUE |
| is.apcluster | F-08 |

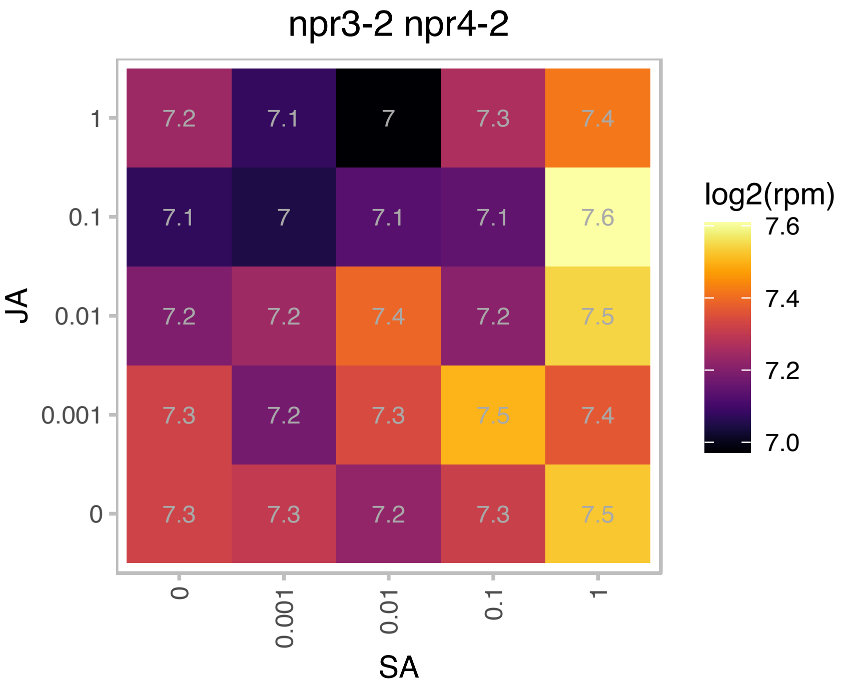


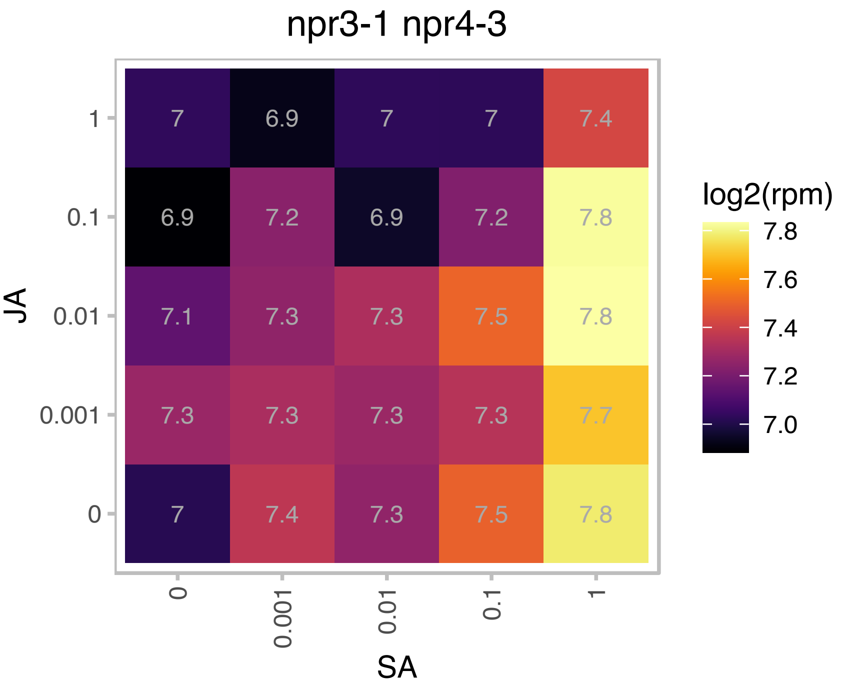

# AT2G30620.1 : H1.2

winged-helix DNA-binding transcription factor family protein

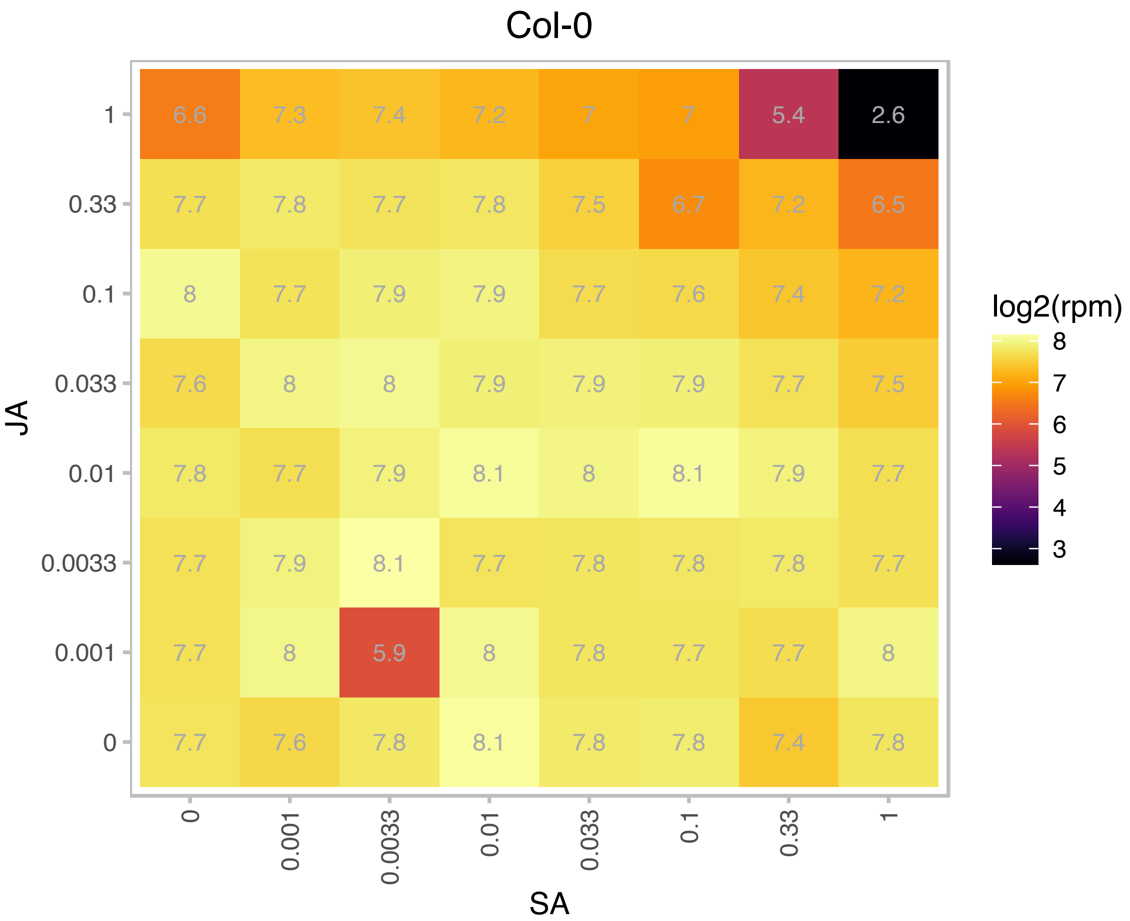


| | |
|---|---|
| Gene | AT2G30620 |
| is.expressed_genes | TRUE |
| is.responsive_genes | TRUE |
| is.apcluster | F-08 |

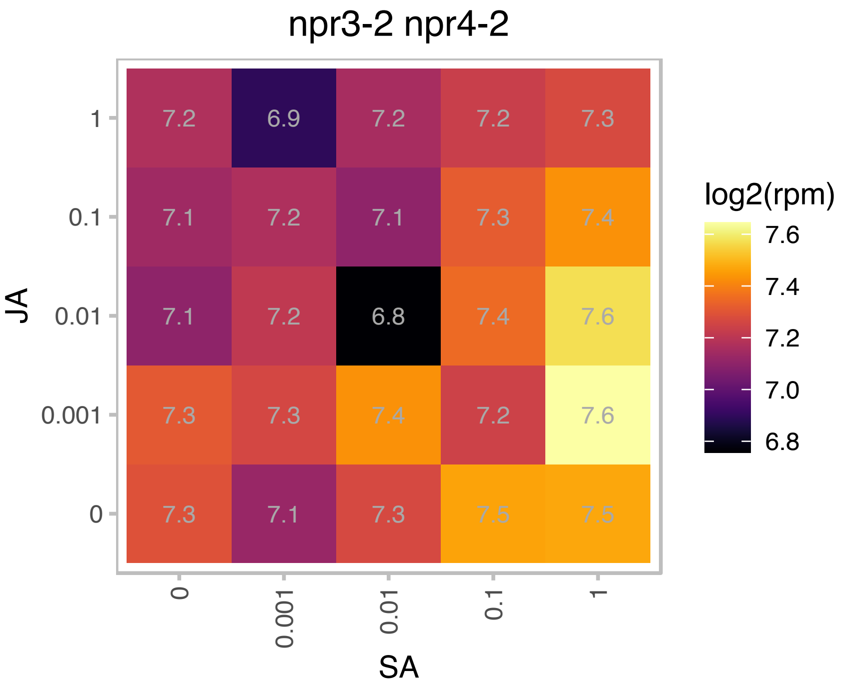


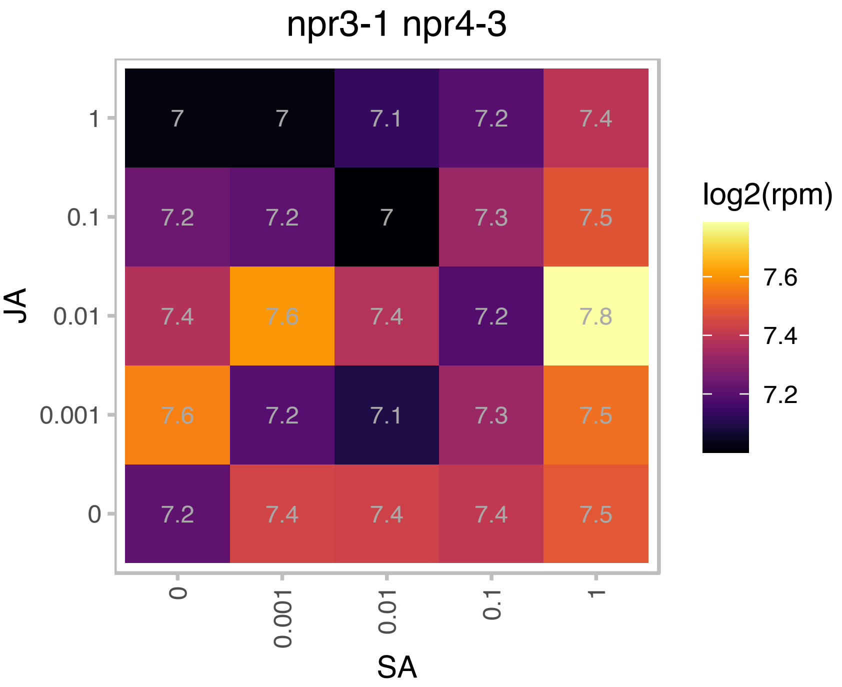

# AT2G31670.1 : UP3

Stress responsive alpha-beta barrel domain protein

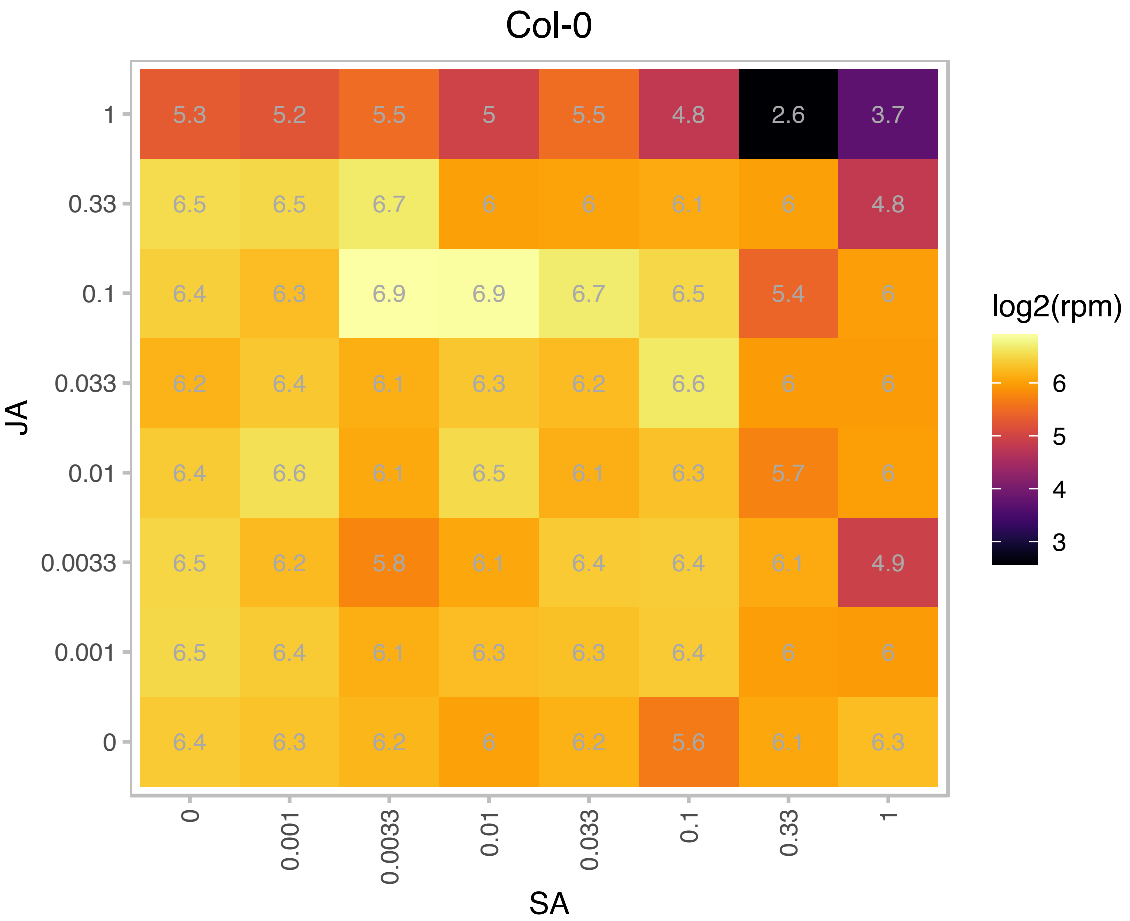


| | |
|---|---|
| Gene | AT2G31670 |
| is.expressed_genes | TRUE |
| is.responsive_genes | TRUE |
| is.apcluster | F-08 |

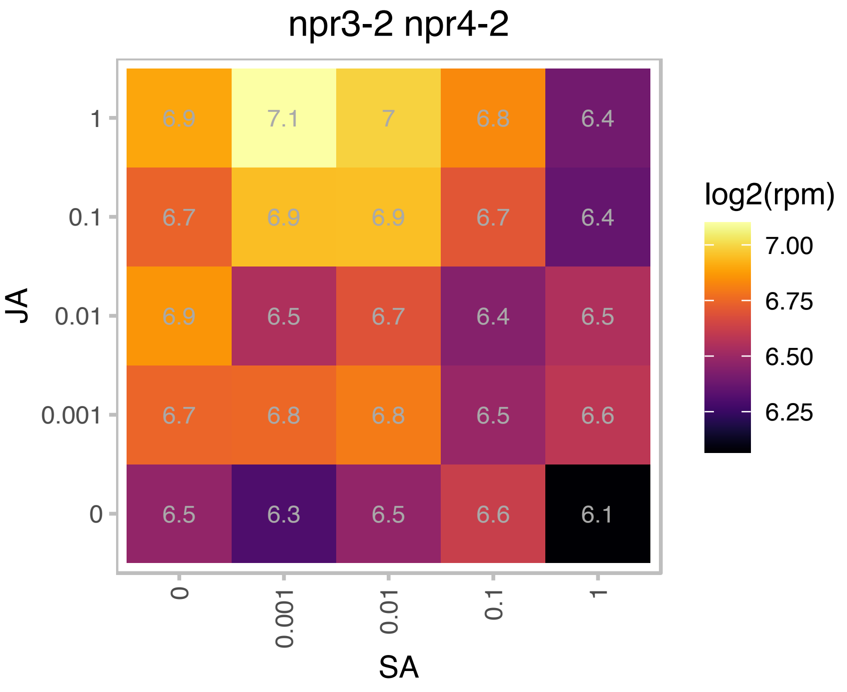


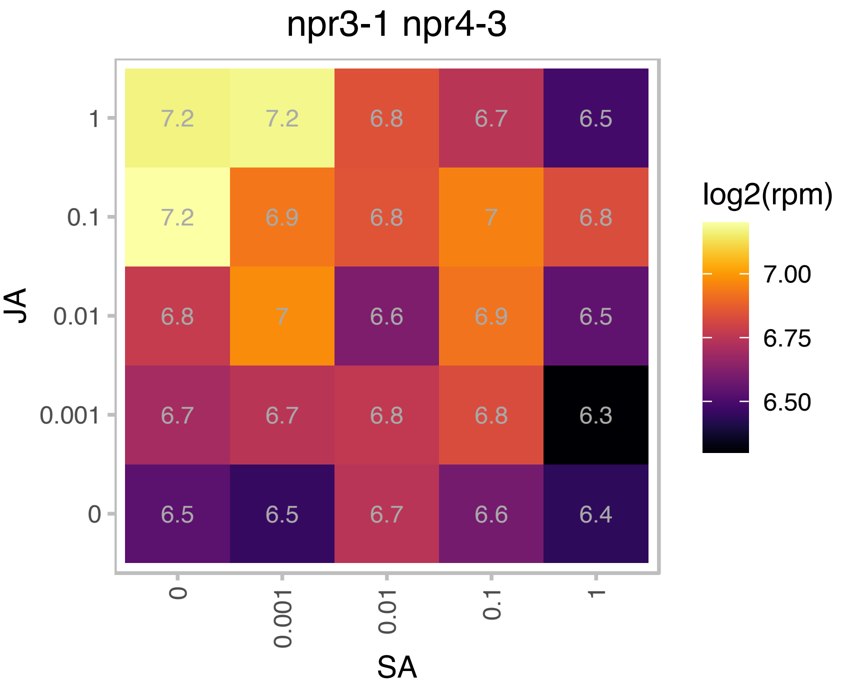

# AT2G31725.1 : NA

Eukaryotic protein of unknown function (DUF842)

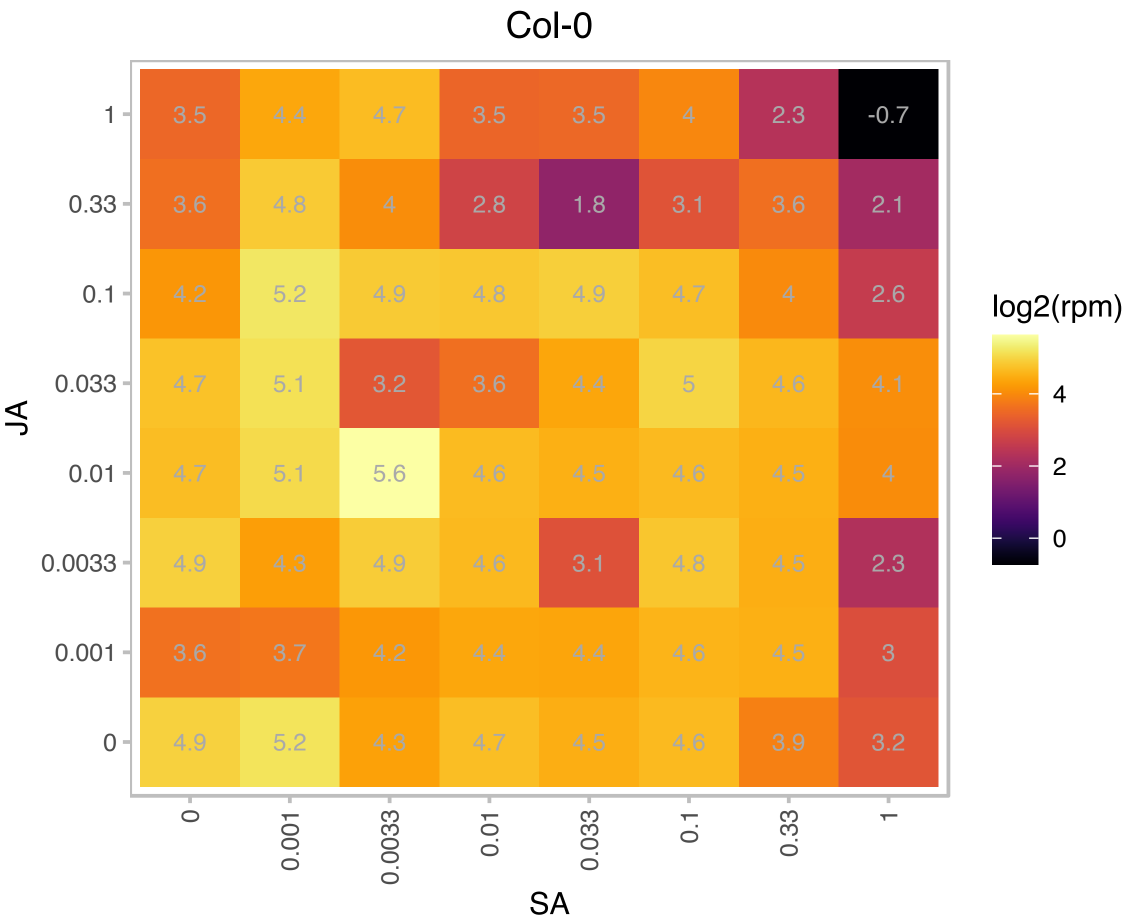


| | |
|---|---|
| Gene | AT2G31725 |
| is.expressed_genes | TRUE |
| is.responsive_genes | TRUE |
| is.apcluster | F-08 |

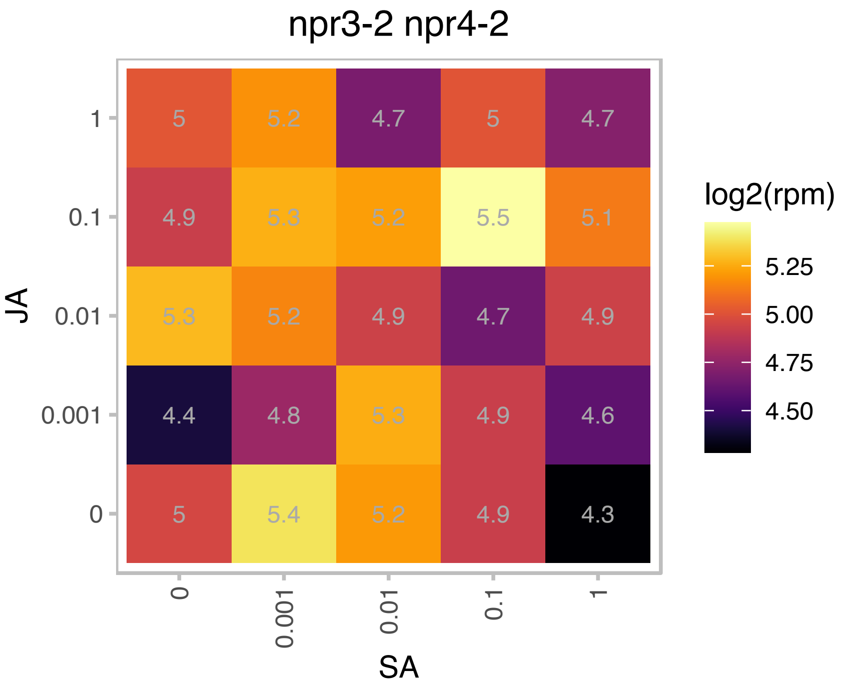


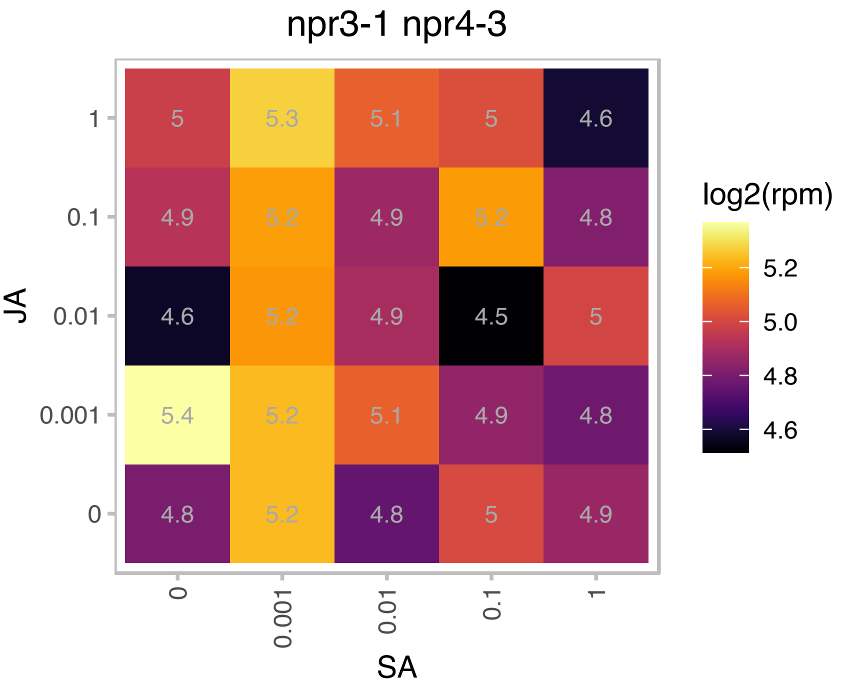

# AT2G32380.1 : NA

Transmembrane protein 97, predicted

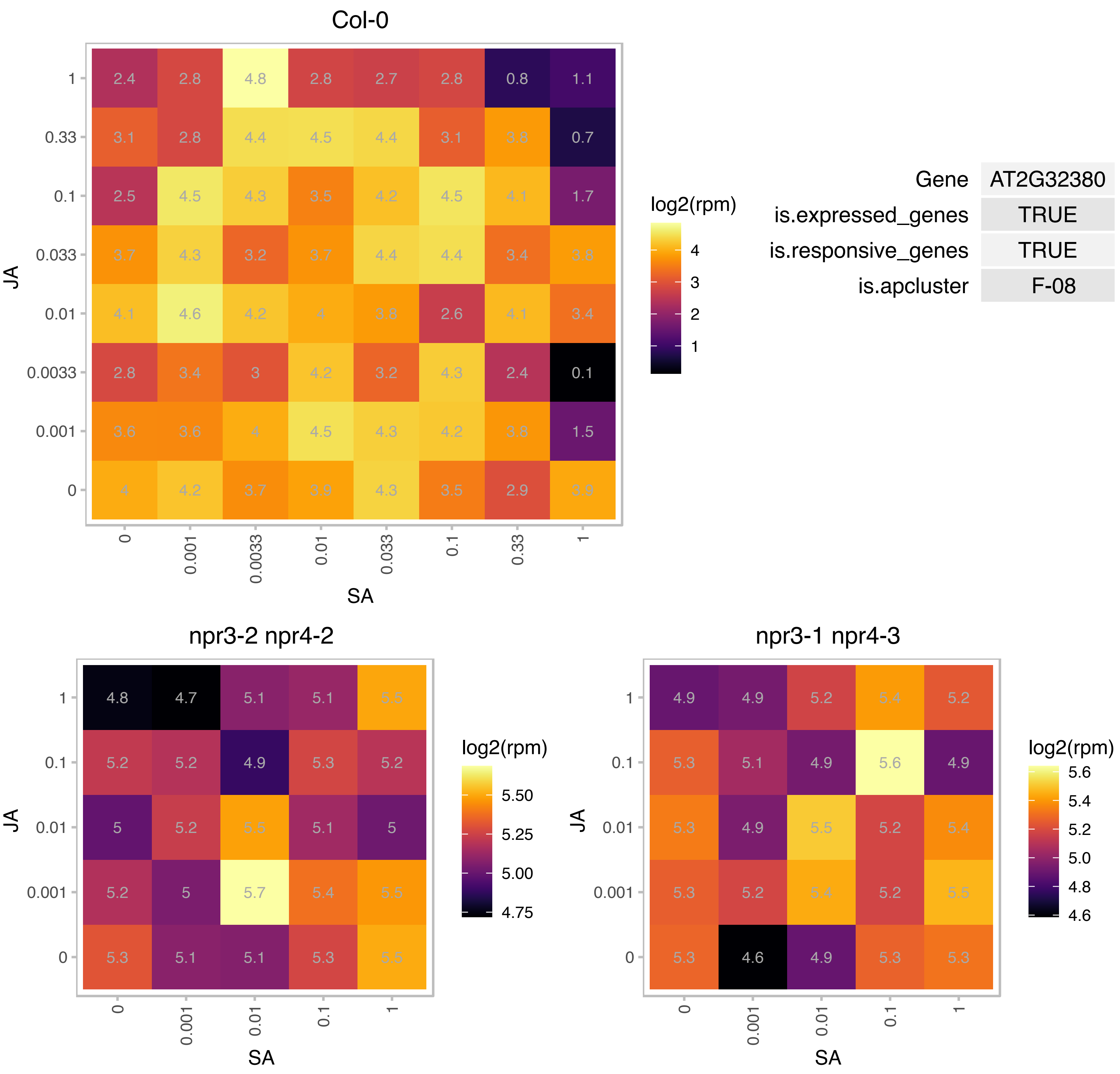

### AT2G32720.1 : ATCB5-B, B5 #4, CB5-B, CYTB5-D

cytochrome B5 isoform B

member of Cytochromes b5

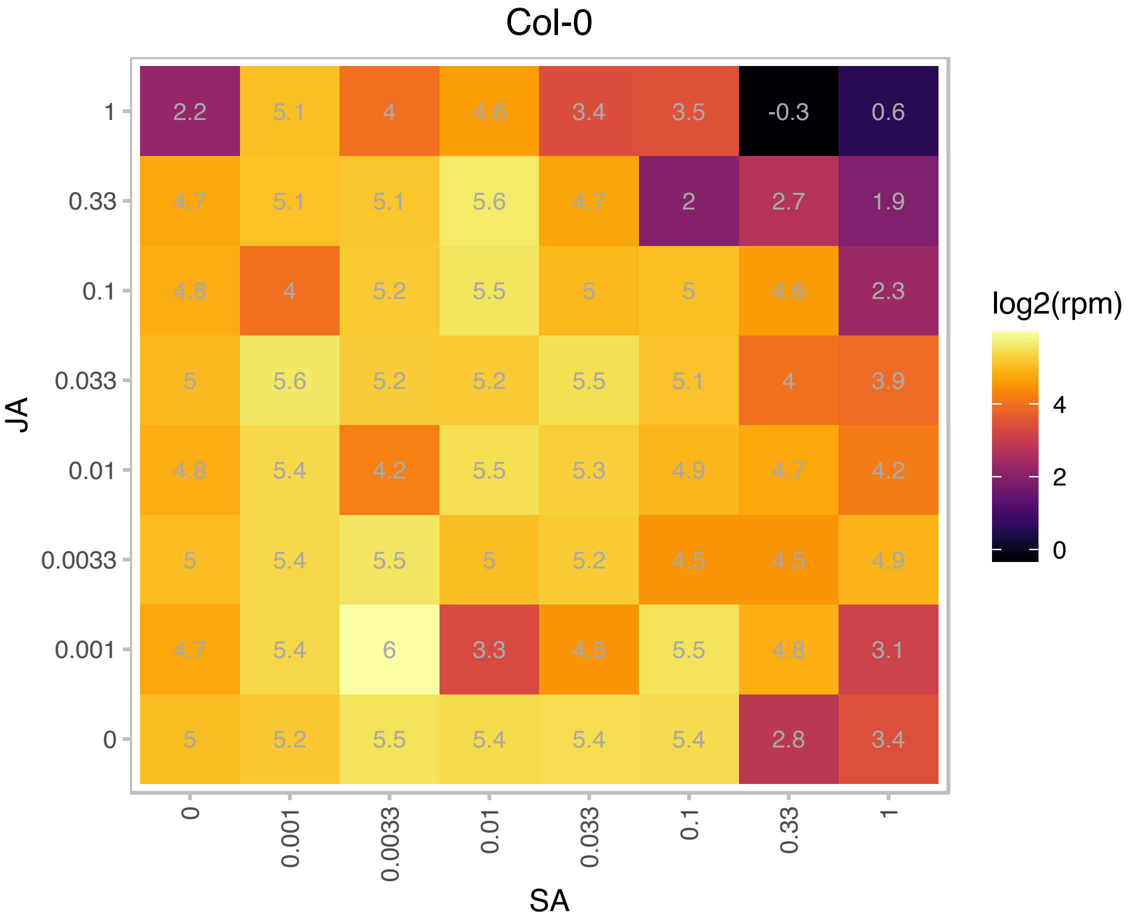


| | |
|---|---|
| Gene | AT2G32720 |
| is.expressed_genes | TRUE |
| is.responsive_genes | TRUE |
| is.apcluster | F-08 |

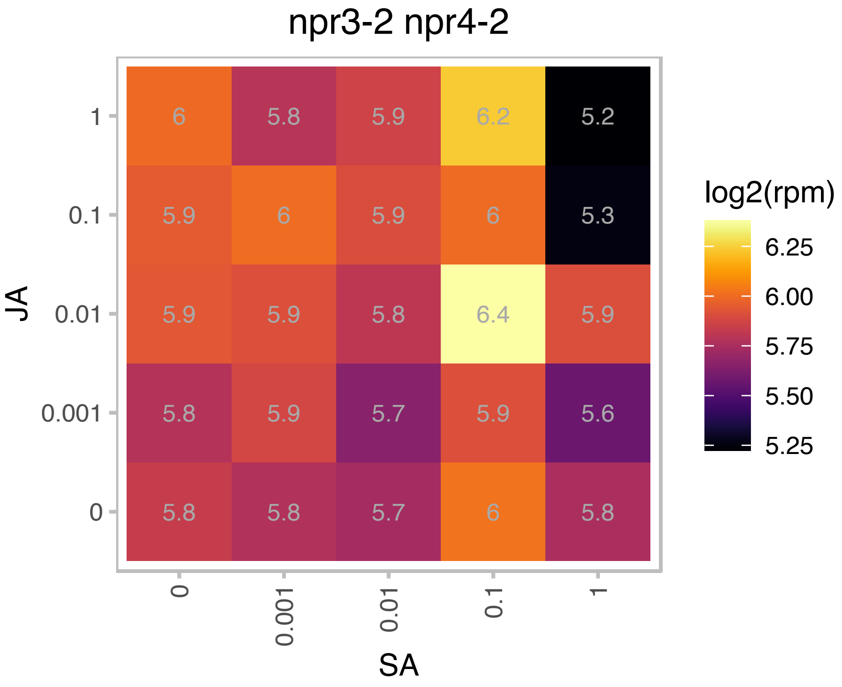


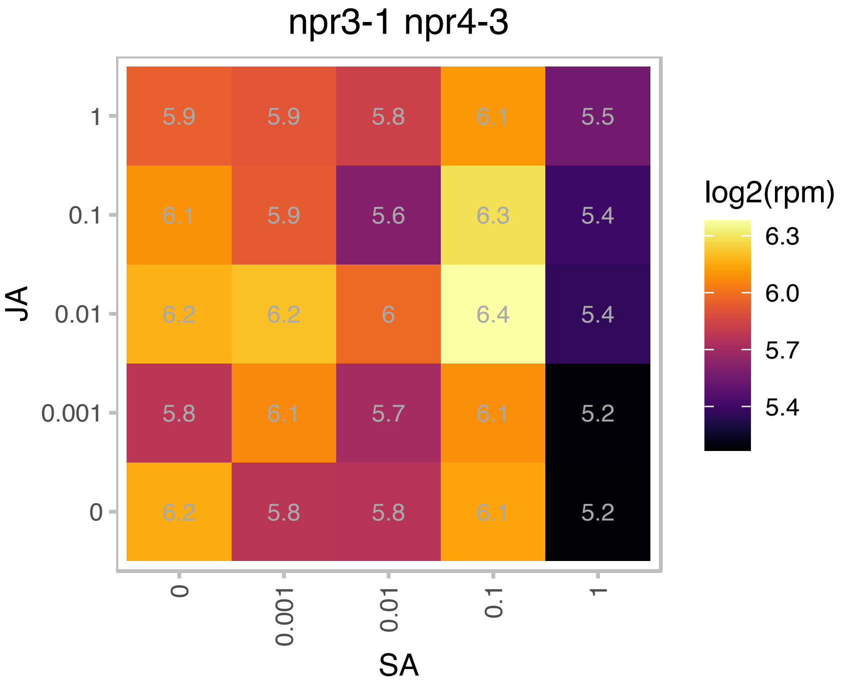

# AT2G33370.1 : NA

Ribosomal protein L14p/L23e family protein

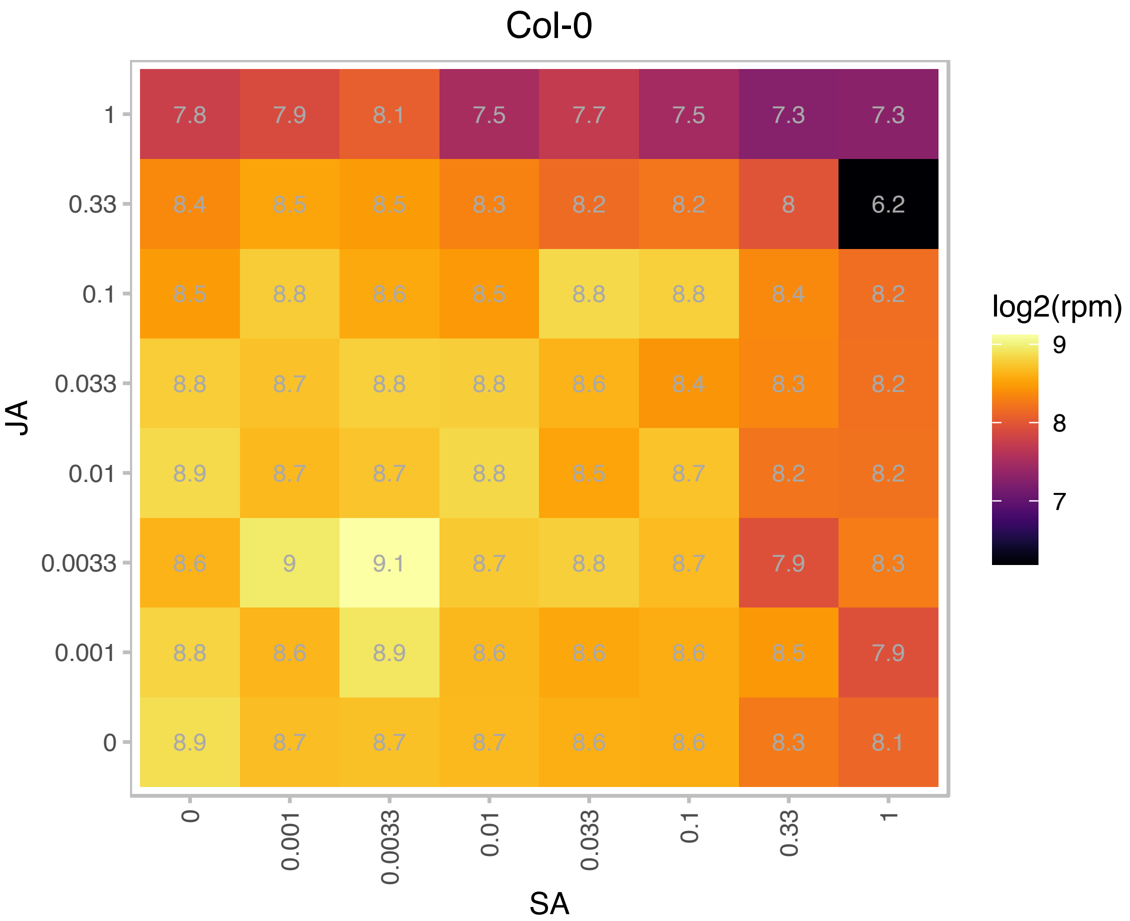


| | |
|---|---|
| Gene | AT2G33370 |
| is.expressed_genes | TRUE |
| is.responsive_genes | TRUE |
| is.apcluster | F-08 |

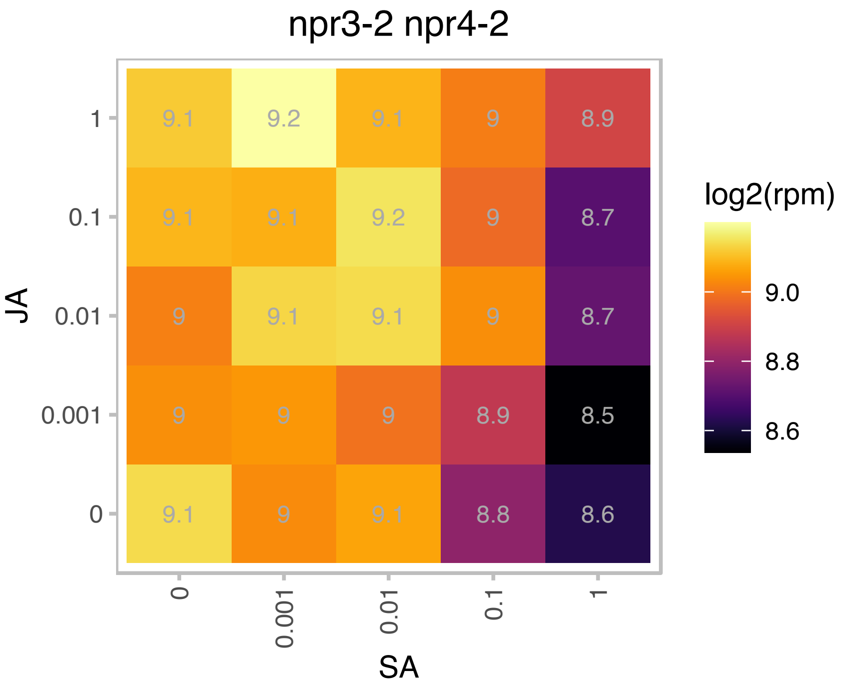


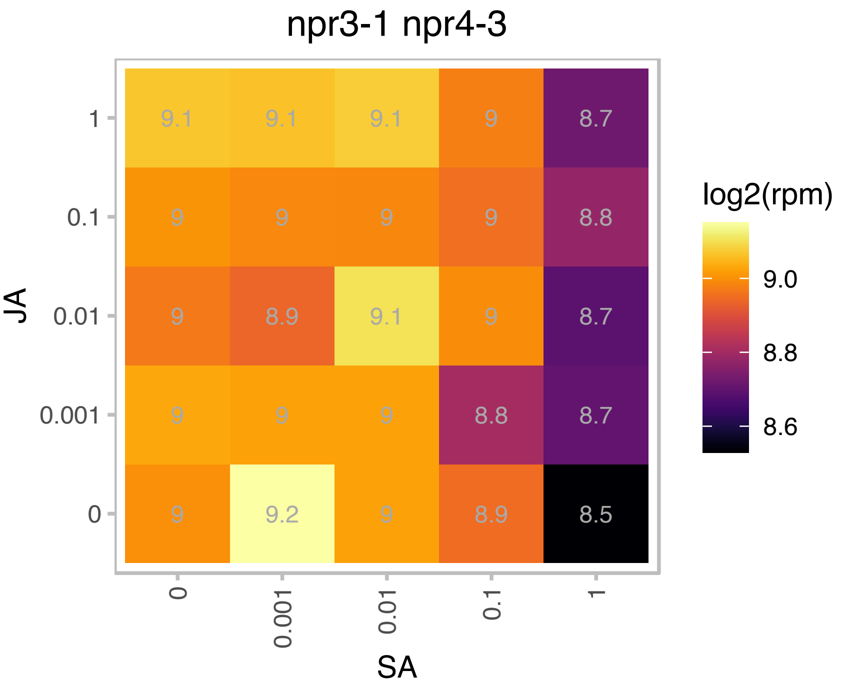

# AT2G33850.1 : NA

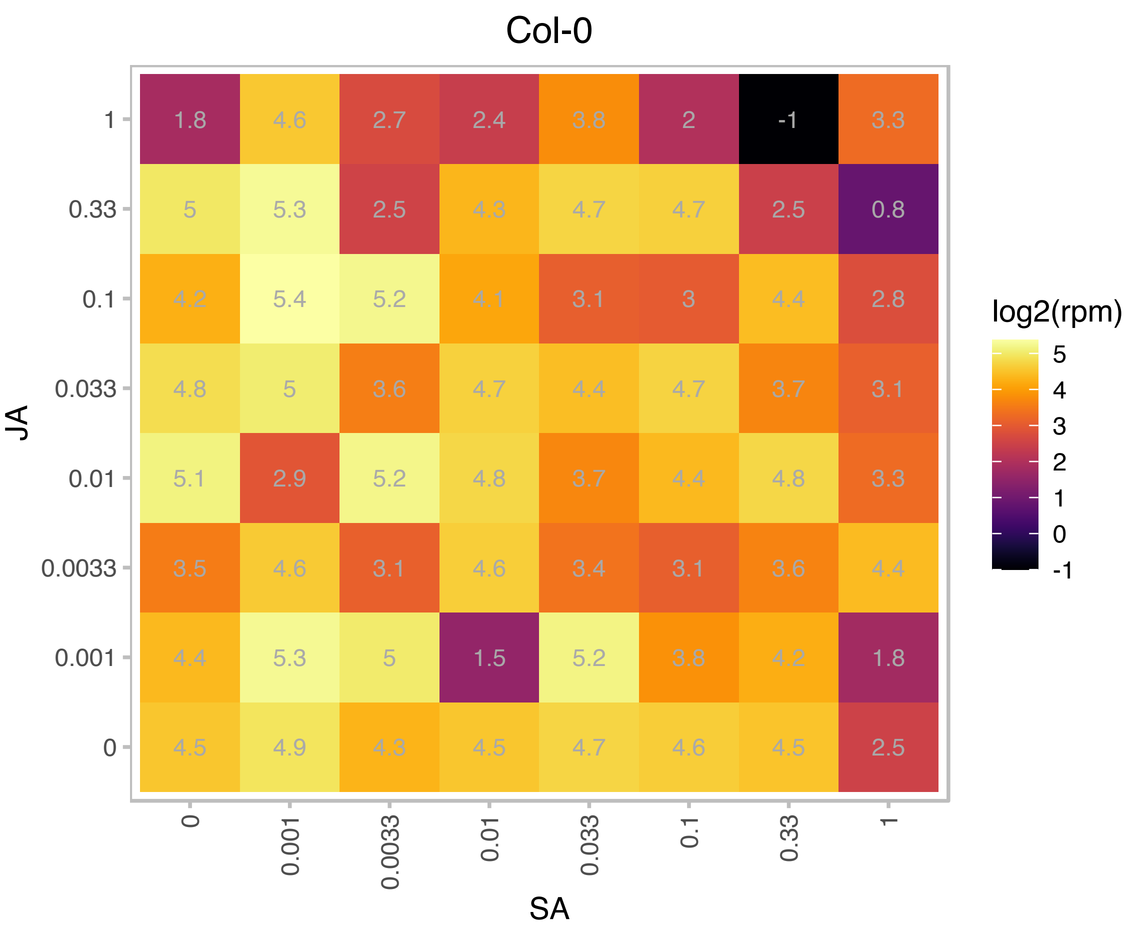


| Gene | AT2G33850 |
|---|---|
| is.expressed_genes | TRUE |
| is.responsive_genes | TRUE |
| is.apcluster | F-08 |

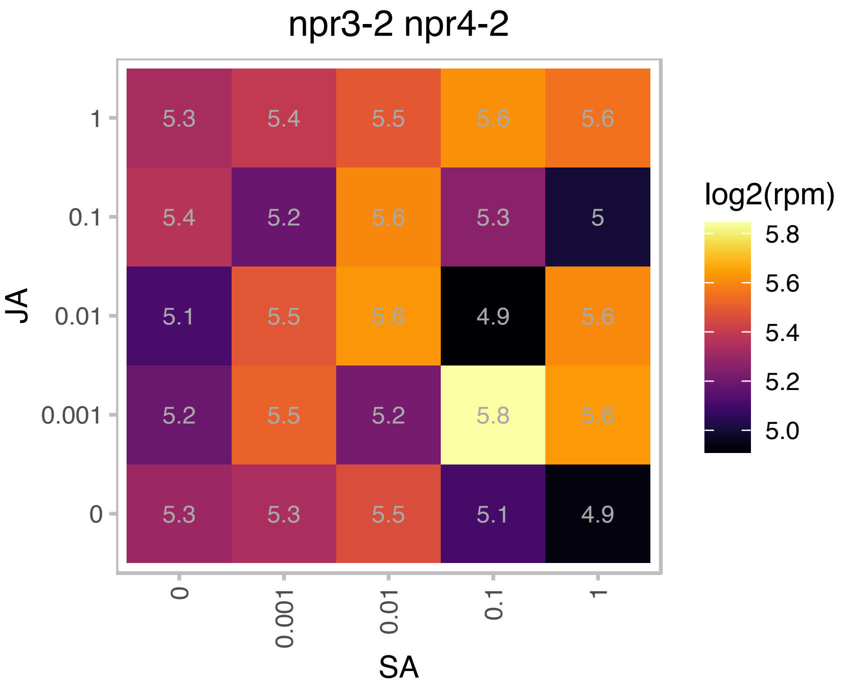


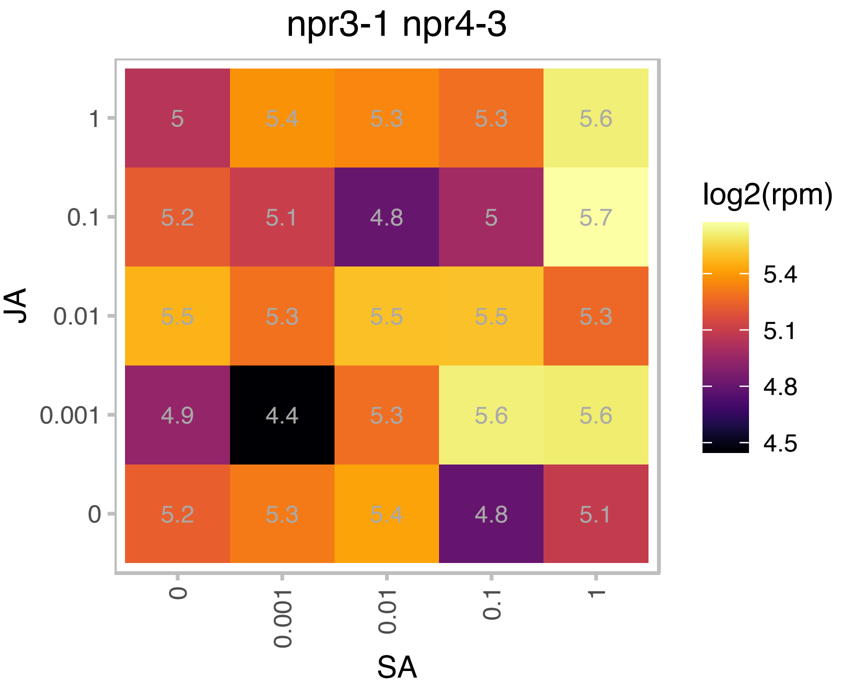

### AT2G34520.1 : RPS14

mitochondrial ribosomal protein S14

nuclear-encoded mitochondrial ribosomal protein S14

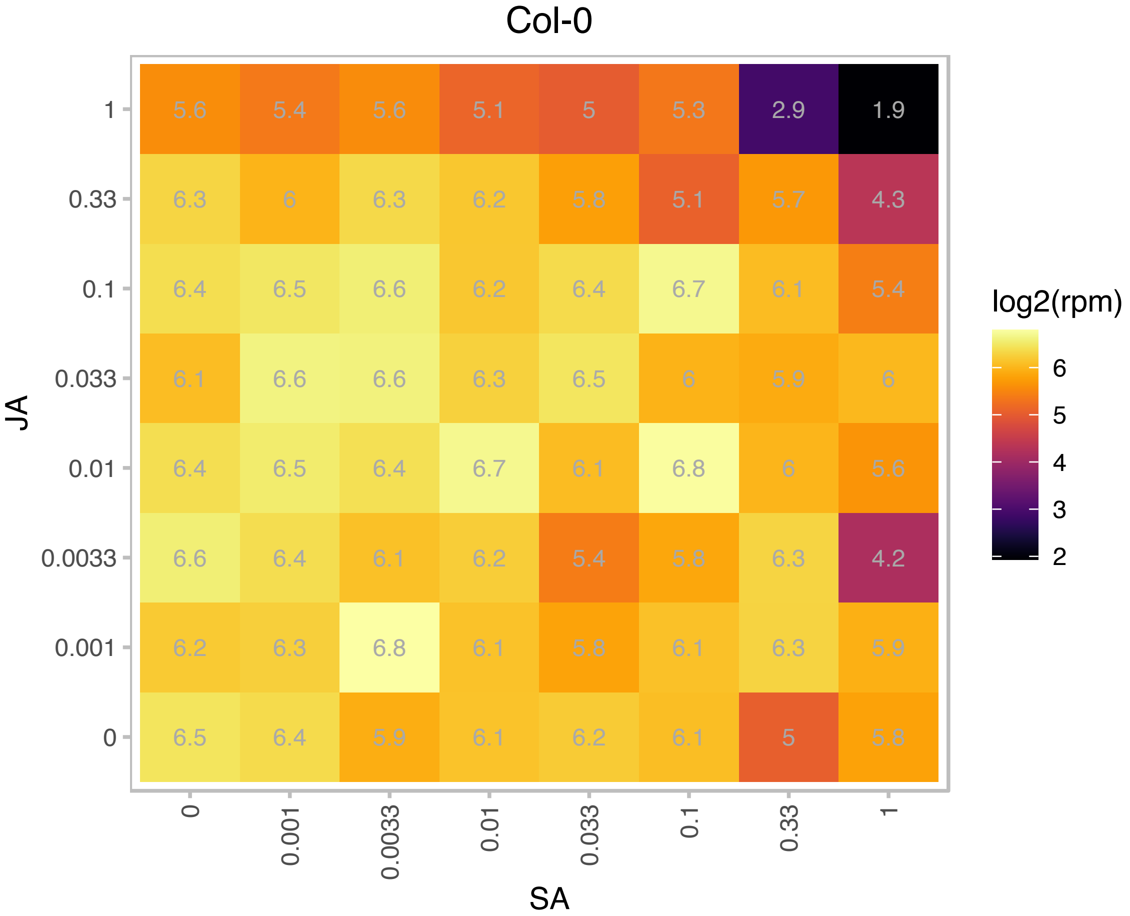


| | |
|---|---|
| Gene | AT2G34520 |
| is.expressed_genes | TRUE |
| is.responsive_genes | TRUE |
| is.apcluster | F-08 |

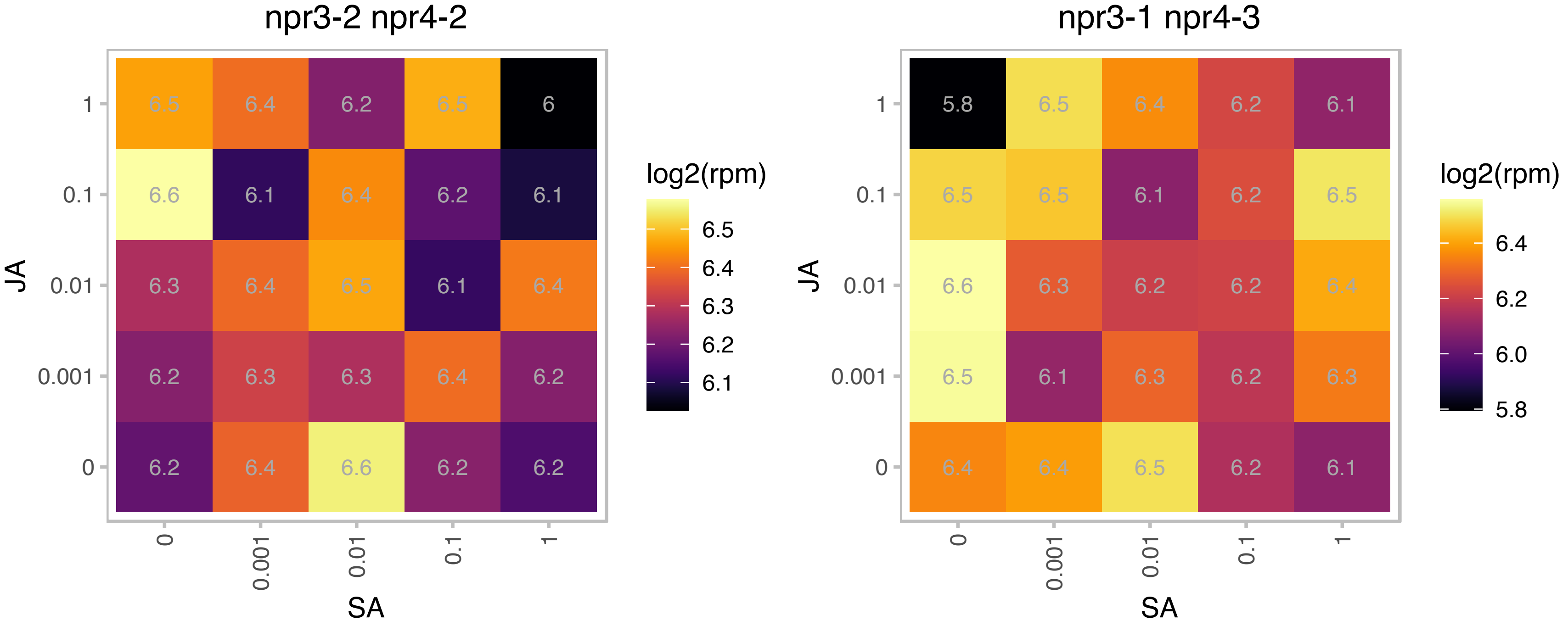

### AT2G38530.1 : AtLtpI-5, cdf3, LP2, LTP2

lipid transfer protein 2

Involved in lipid transfer between membranes and plays a role in maintaining the integrity of the cuticle-cell wall interface. Belongs to a family of Lipid transfer proteins. Sequence similarity to other plant/Arabidopsis LPT genes but highest similarity to LPT1. Stress and pathogen-inducible motifs found in the upstream region. Expressed in flower, leaves and siliques but absent in roots. Predicted to be a member of PR-14 pathogenesis-related protein family with the following members: At2g38540/LTP1, At2g38530/LTP2, At5g59320/LTP3, At5g59310/LTP4, At3g51600/LTP5, At3g08770/LTP6, At2g15050/LTP7, At2g18370/LTP8, At2g15325/LTP9, At5g01870/LTP10, At4g33355/LTP11, At3g51590/LTP12, At5g44265/LTP13, At5g62065/LTP14

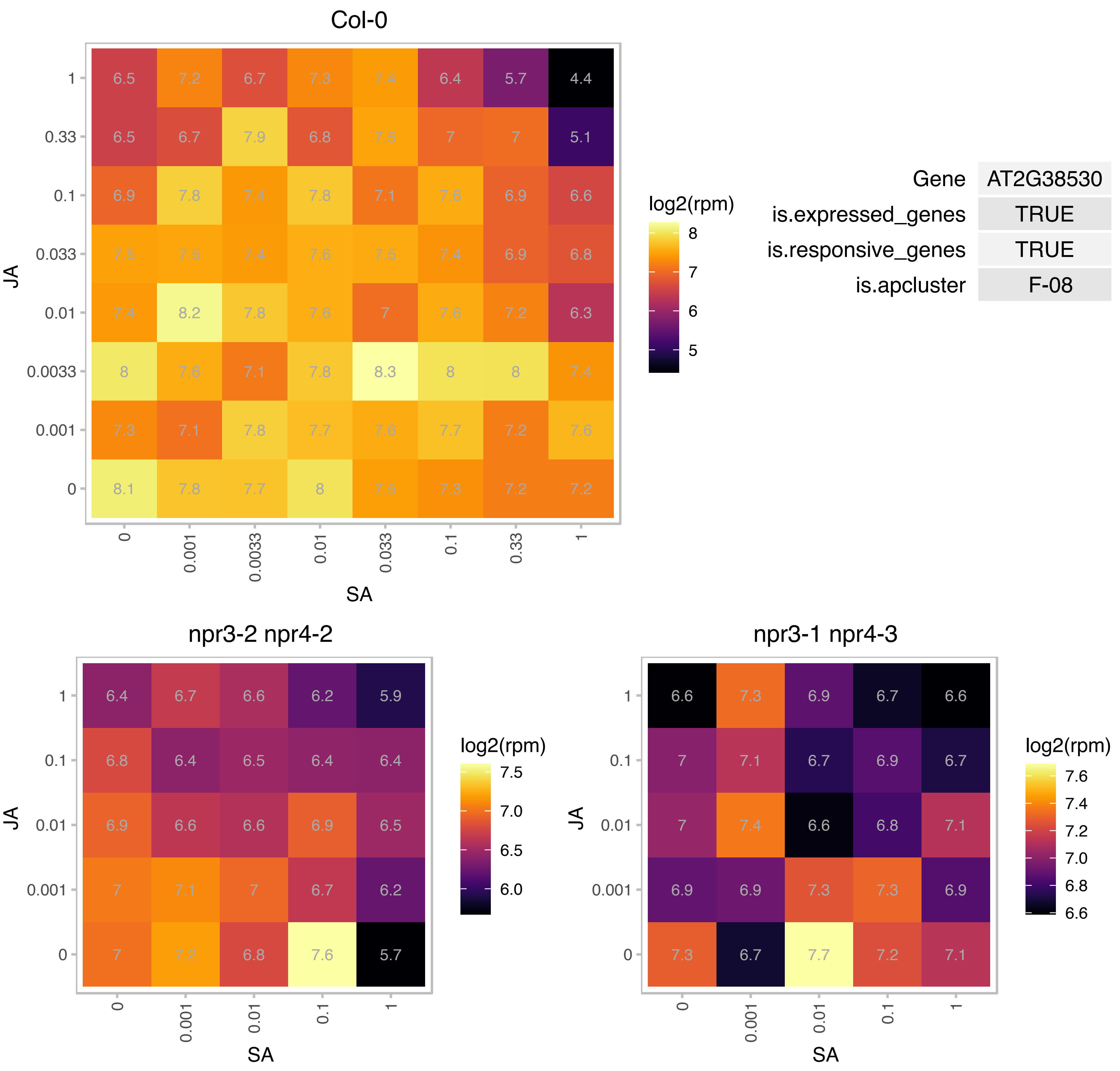

# AT2G39500.1 : NA

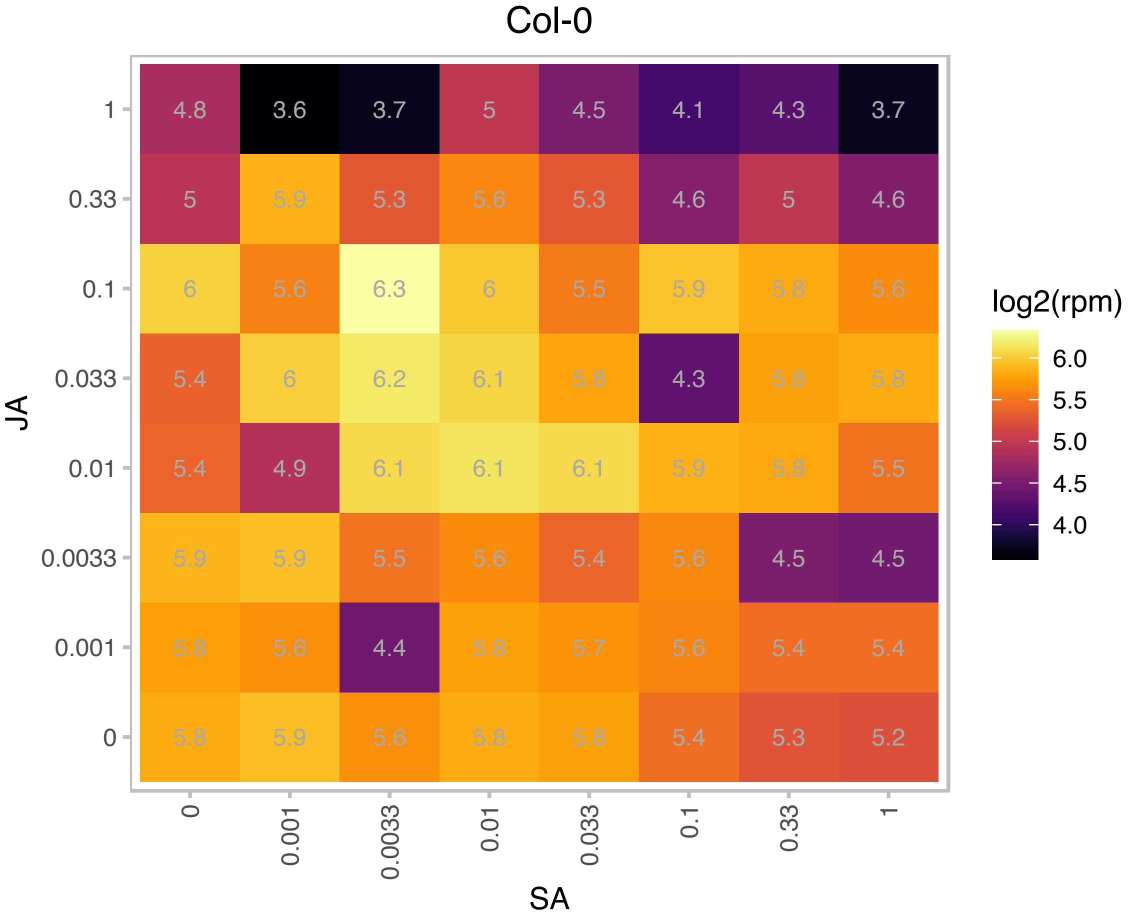


| | |
|---|---|
| Gene | AT2G39500 |
| is.expressed_genes | TRUE |
| is.responsive_genes | TRUE |
| is.apcluster | F-08 |

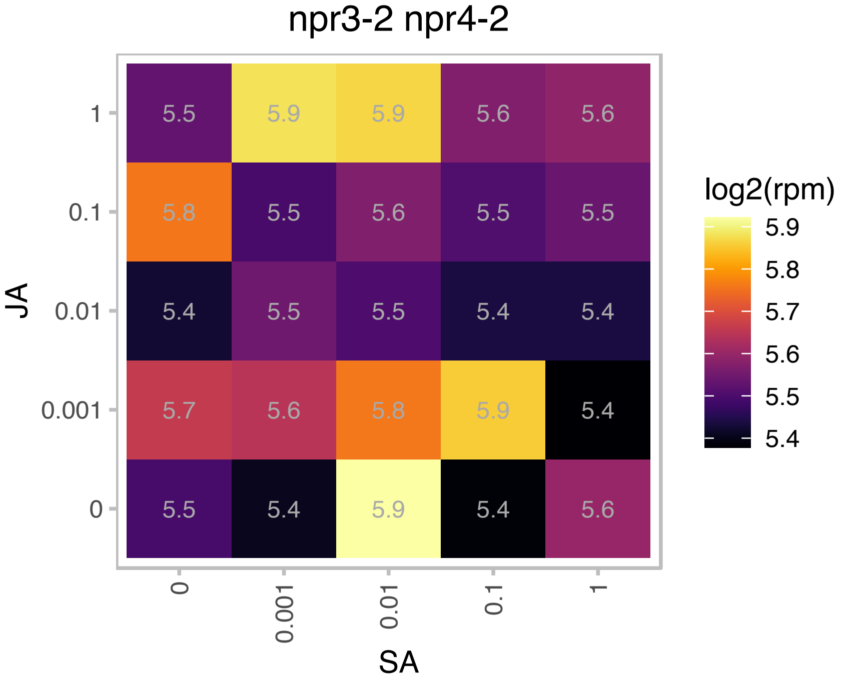


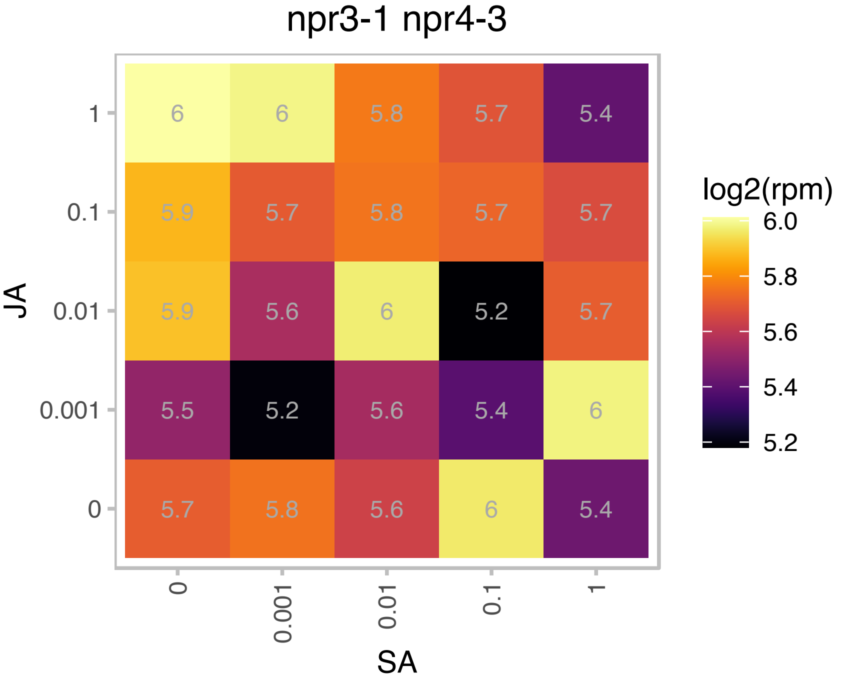

### AT2G40070.1 : BPP1

Encodes a microtubule-associated protein.

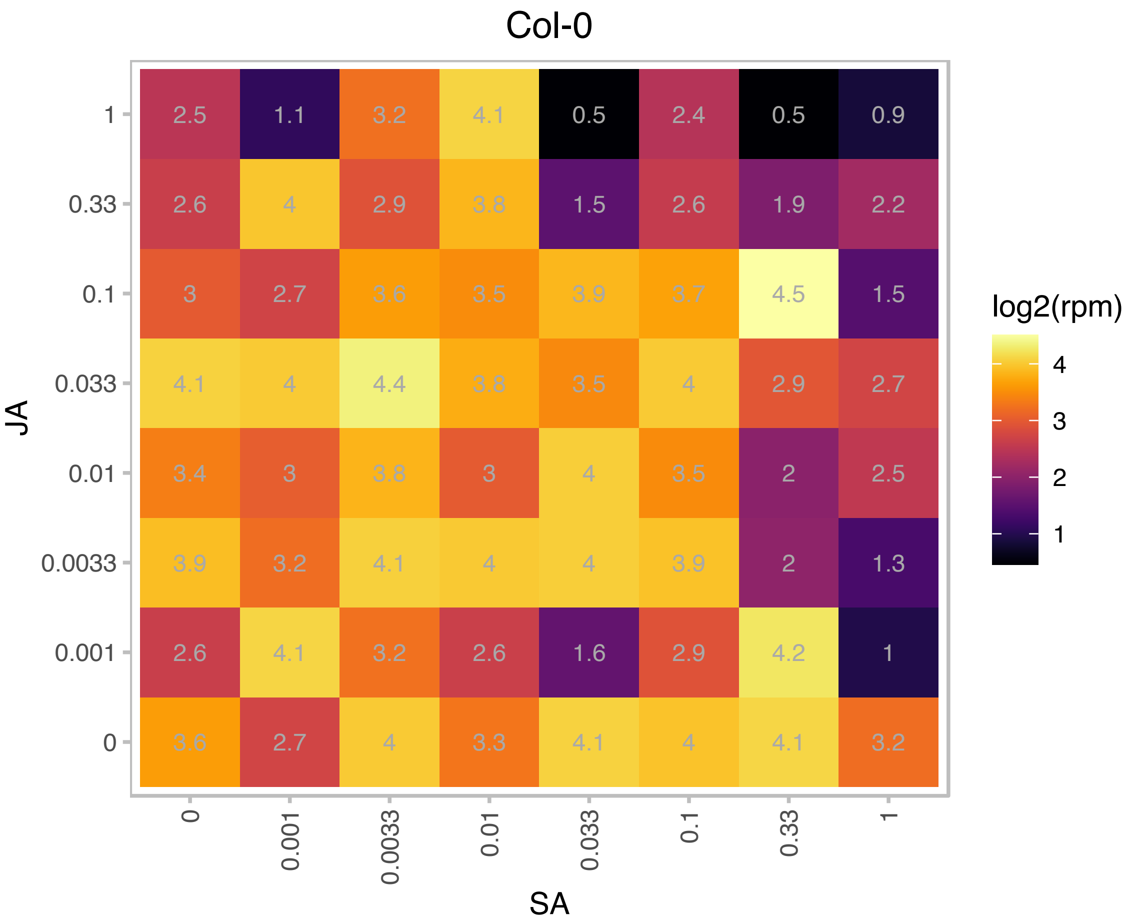


| | |
|---|---|
| Gene | AT2G40070 |
| is.expressed_genes | TRUE |
| is.responsive_genes | TRUE |
| is.apcluster | F-08 |

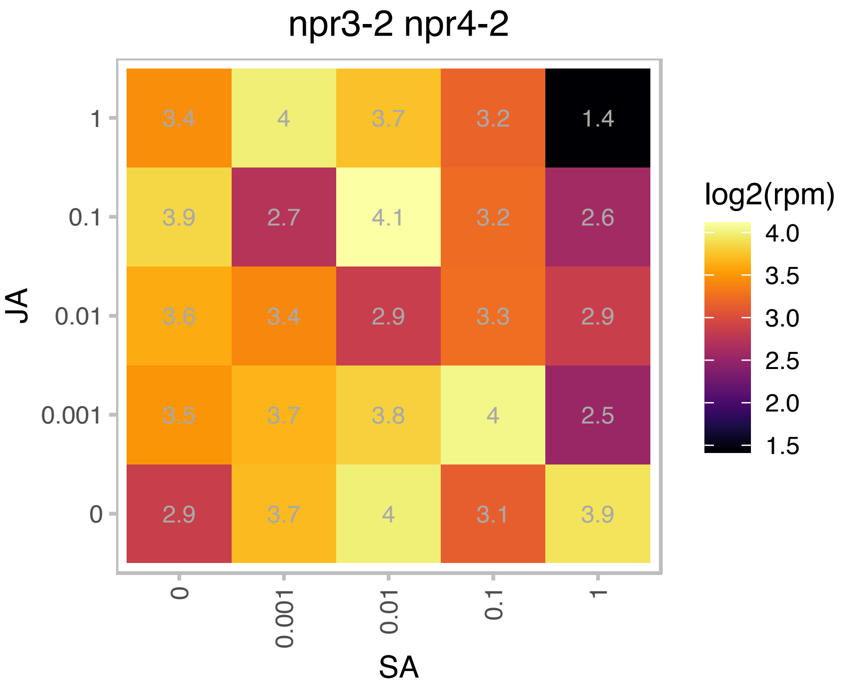


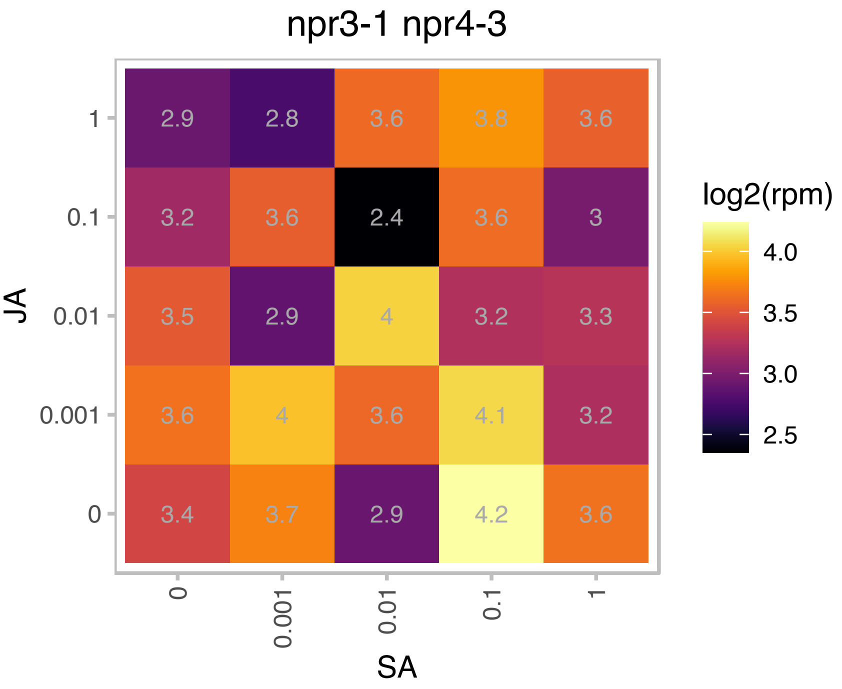

### AT2G40080.1 : ELF4

Encodes a novel nuclear 111 amino-acid phytochrome-regulated component of a negative feedback loop involving the circadian clock central oscillator components CCA1 and LHY. ELF4 is necessary for light-induced expression of both CCA1 and LHY, and conversely, CCA1 and LHY act negatively on light-induced ELF4 expression. ELF4 promotes clock accuracy and is required for sustained rhythms in the absence of daily light/dark cycles. It is involved in the phyB-mediated constant red light induced seedling de-etiolation process and may function to coregulate the expression of a subset of phyB-regulated genes.

Protein of unknown function (DUF1313)

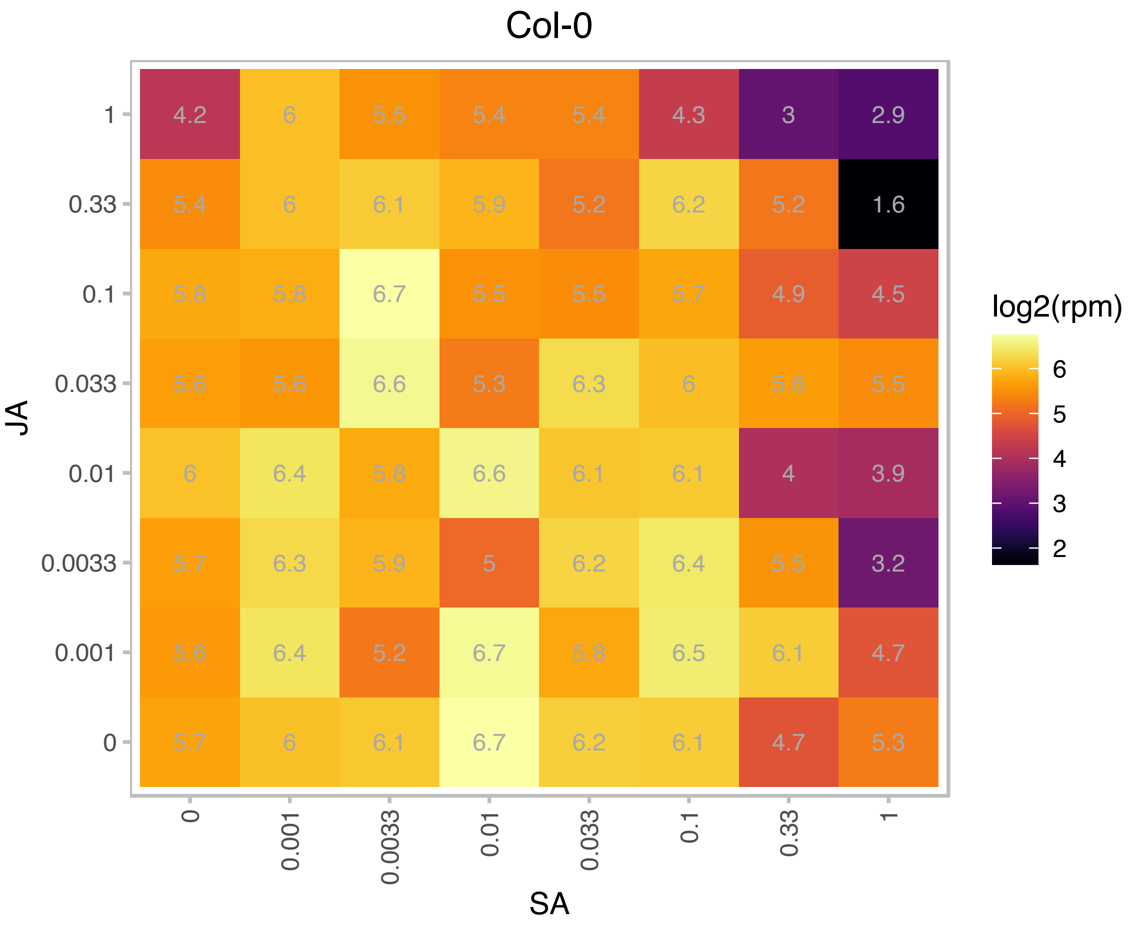


| | |
|---|---|
| Gene | AT2G40080 |
| is.expressed_genes | TRUE |
| is.responsive_genes | TRUE |
| is.apcluster | F-08 |

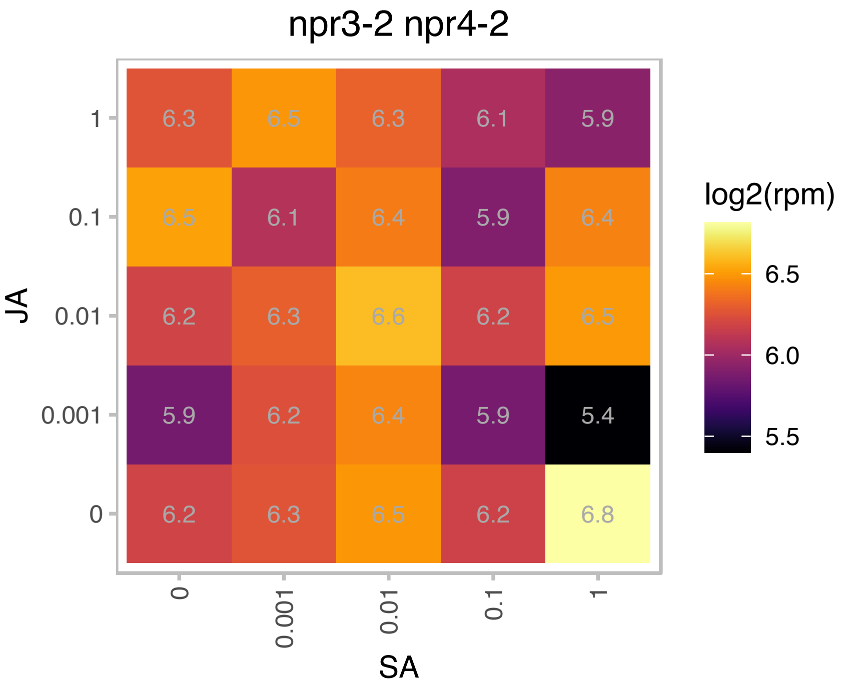


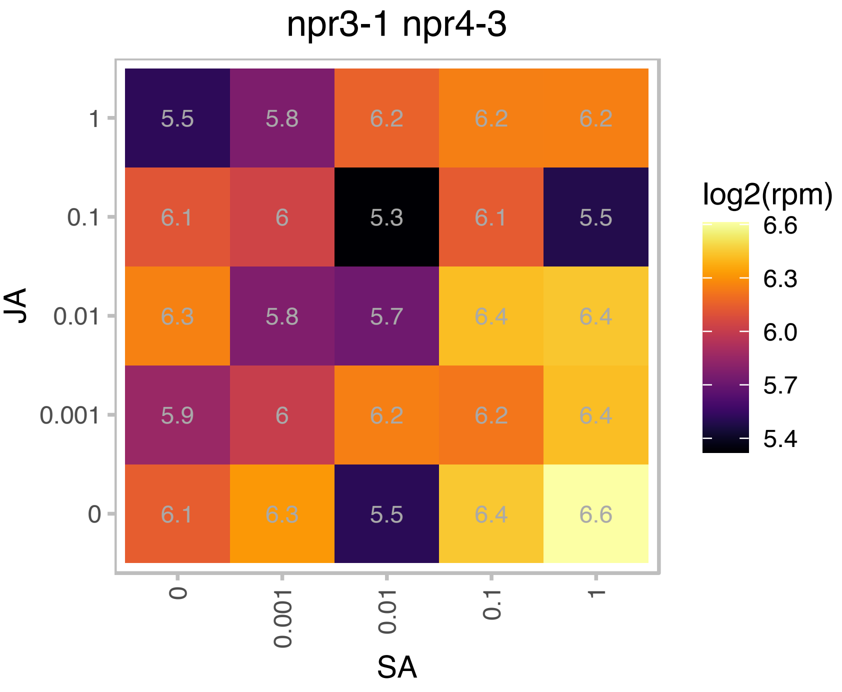

### AT2G40380.1 : PRA1.B2

prenylated RAB acceptor 1.B2

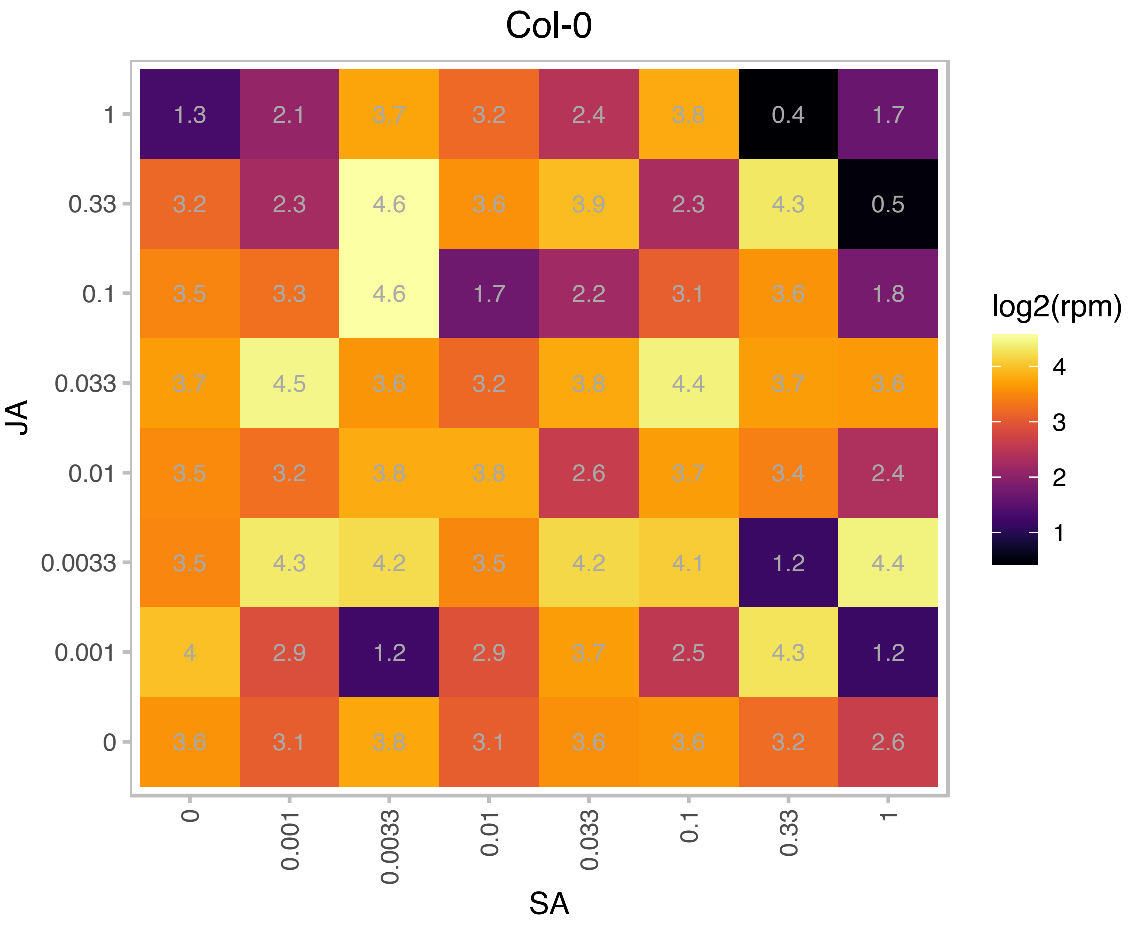


| | |
|---|---|
| Gene | AT2G40380 |
| is.expressed_genes | TRUE |
| is.responsive_genes | TRUE |
| is.apcluster | F-08 |

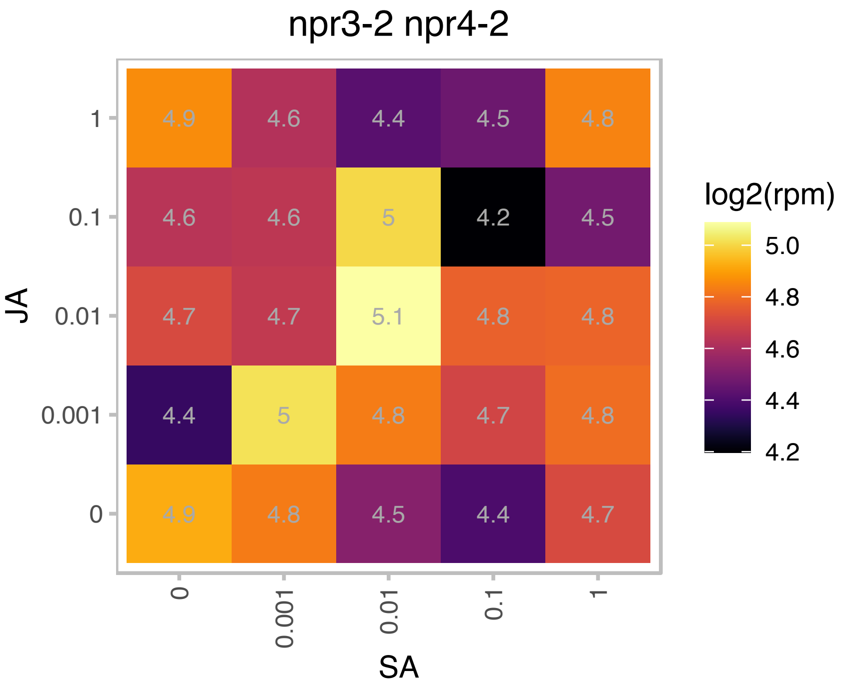


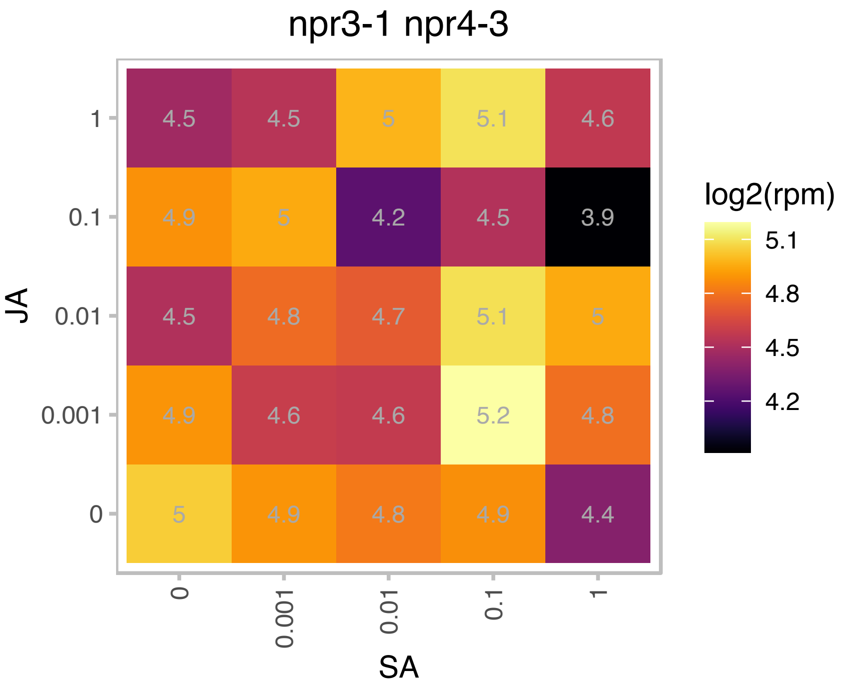

### AT2G42070.1 : ATNUDT23, ATNUDX23, NUDX23

nudix hydrolase homolog 23
Encodes a plastid-localized Nudix hydrolase that has FAD pyrophosphohydrolase activity. Negative feedback regulation of the metabolism of flavins through the hydrolysis of FAD by AtNUDX23 in plastids is involved in the flavin homeostasis in plant cells.

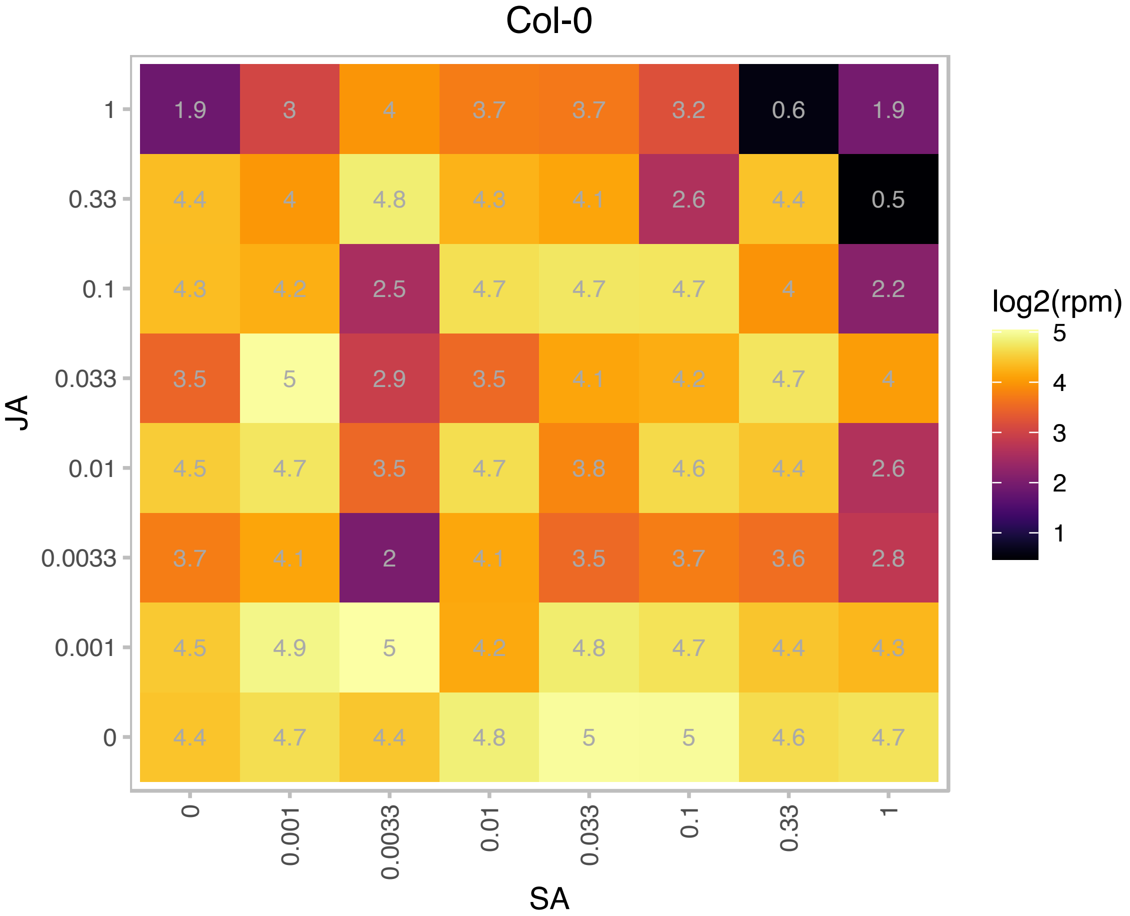


| | |
|---|---|
| Gene | AT2G42070 |
| is.expressed_genes | TRUE |
| is.responsive_genes | TRUE |
| is.apcluster | F-08 |

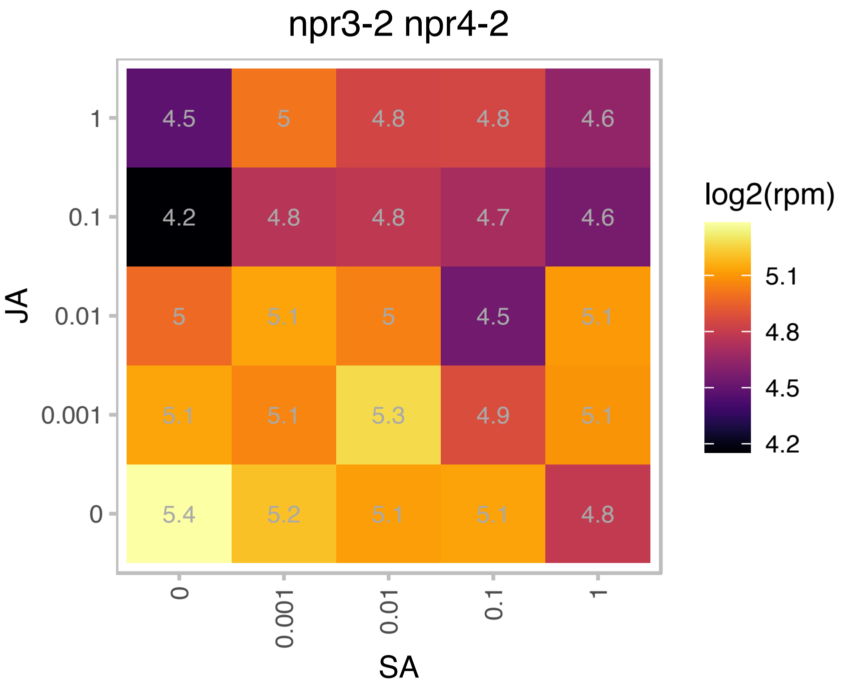


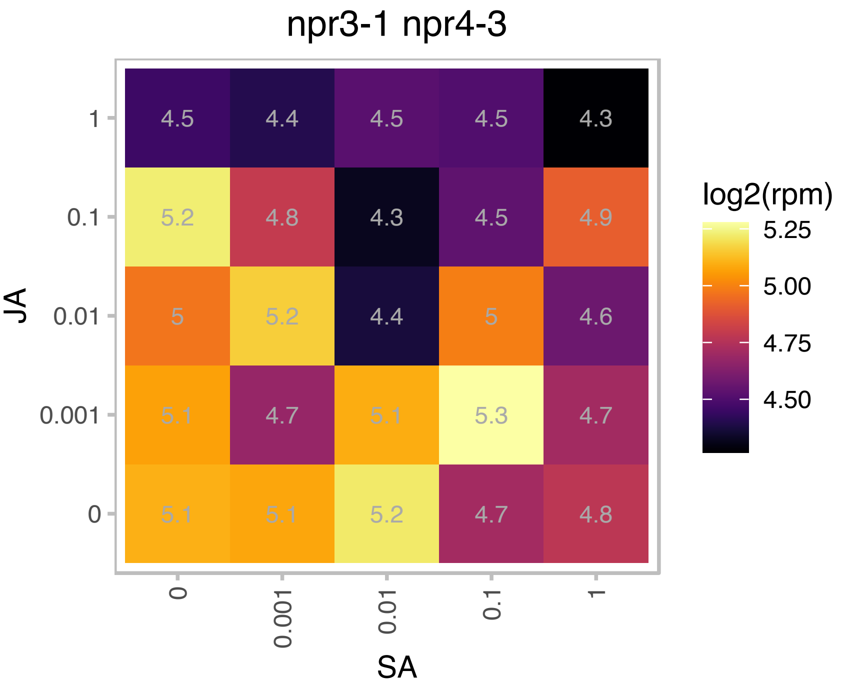

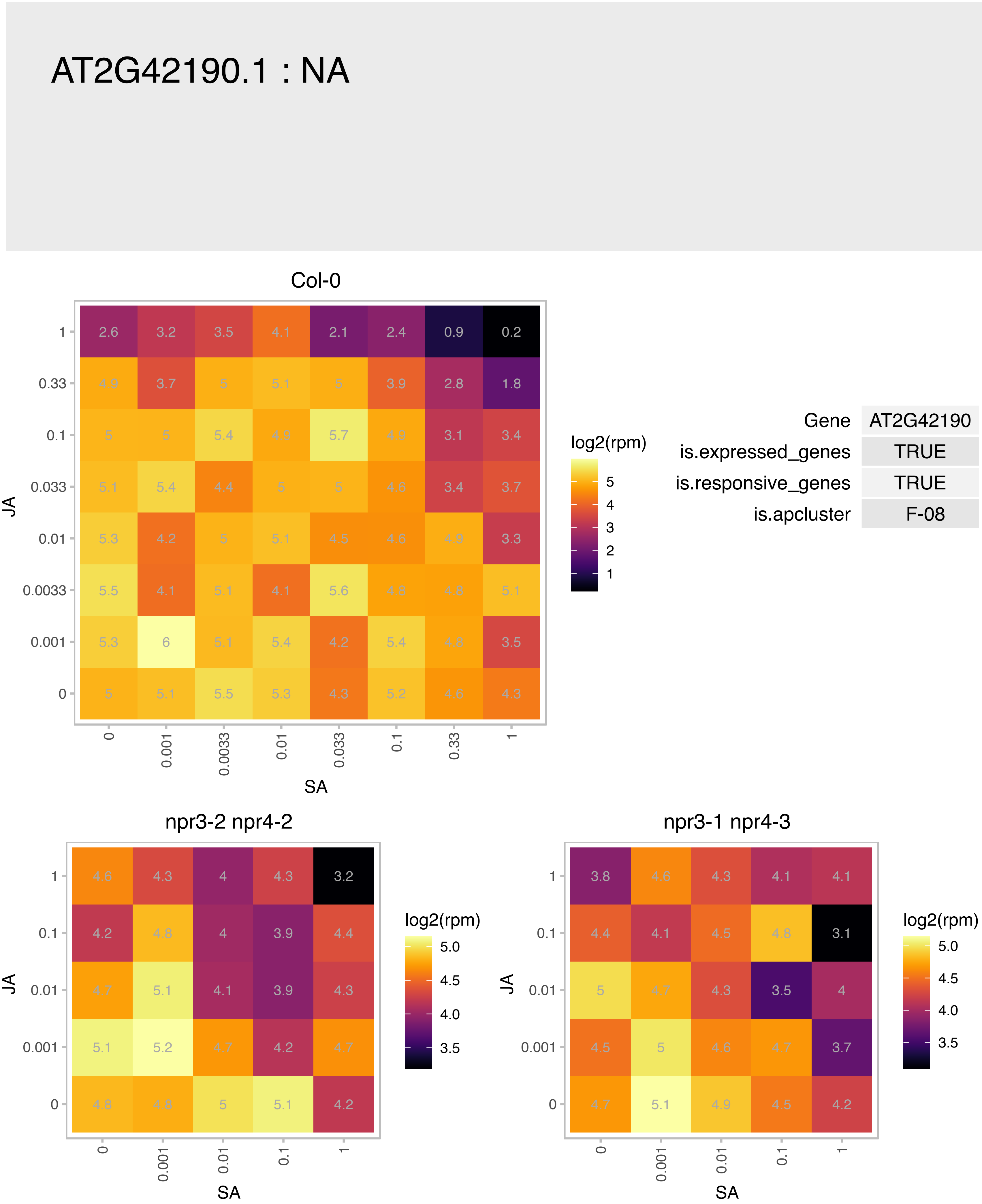
AT2G42190.1 : NA
Col-0
JA
SA
log2(rpm)
Gene AT2G42190
is.expressed_genes TRUE
is.responsive_genes TRUE
is.apcluster F-08
npr3-2 npr4-2
npr3-1 npr4-3

### AT2G42740.1 : RPL16A

ribosomal protein large subunit 16A

encodes a cytosolic ribosomal protein L16, which is a constituent of 60S large ribosomal complex. Gene is expressed in root stele and anthers and expression is induced by auxin treatment.

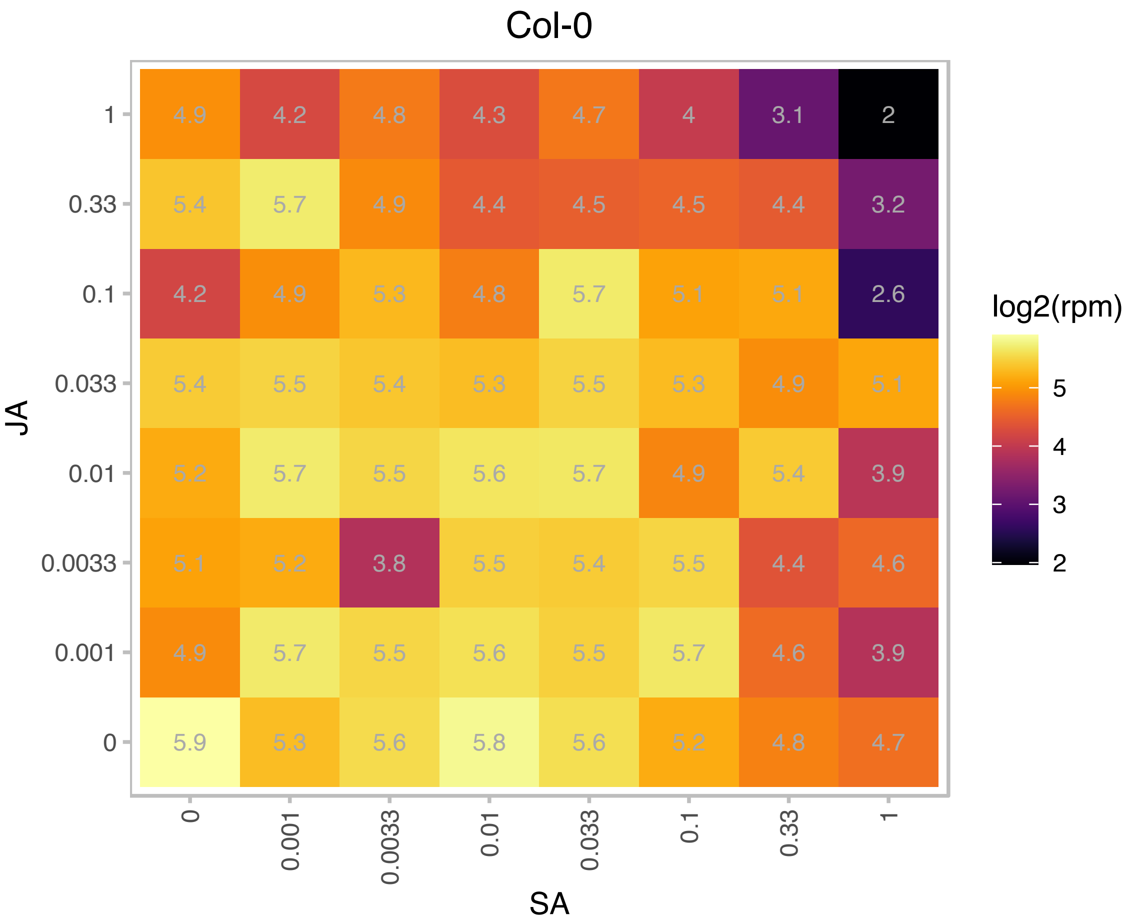


| | |
|---|---|
| Gene | AT2G42740 |
| is.expressed_genes | TRUE |
| is.responsive_genes | TRUE |
| is.apcluster | F-08 |

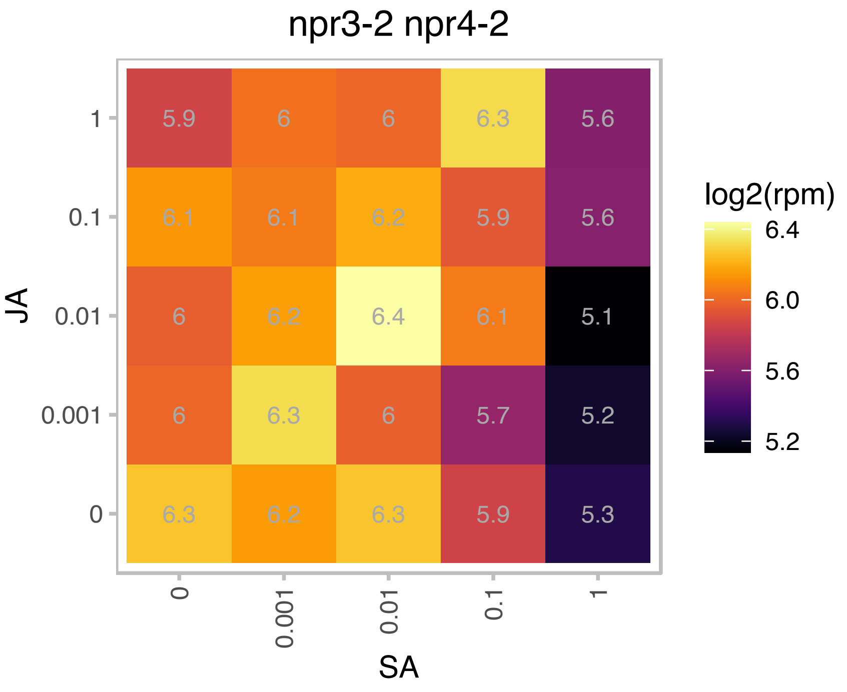


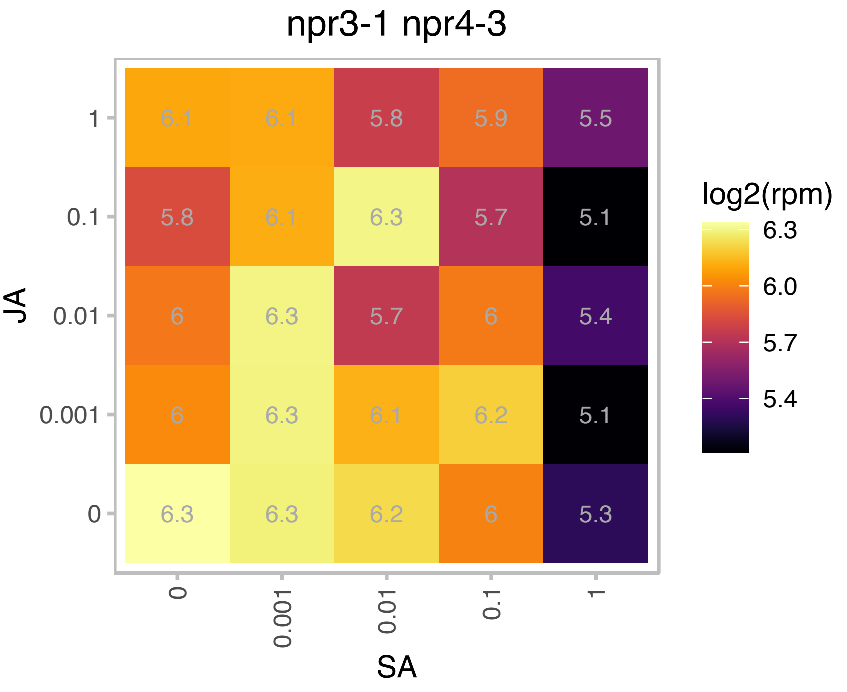

# AT2G45250.1 : NA

Integral membrane protein hemolysin-III homolog

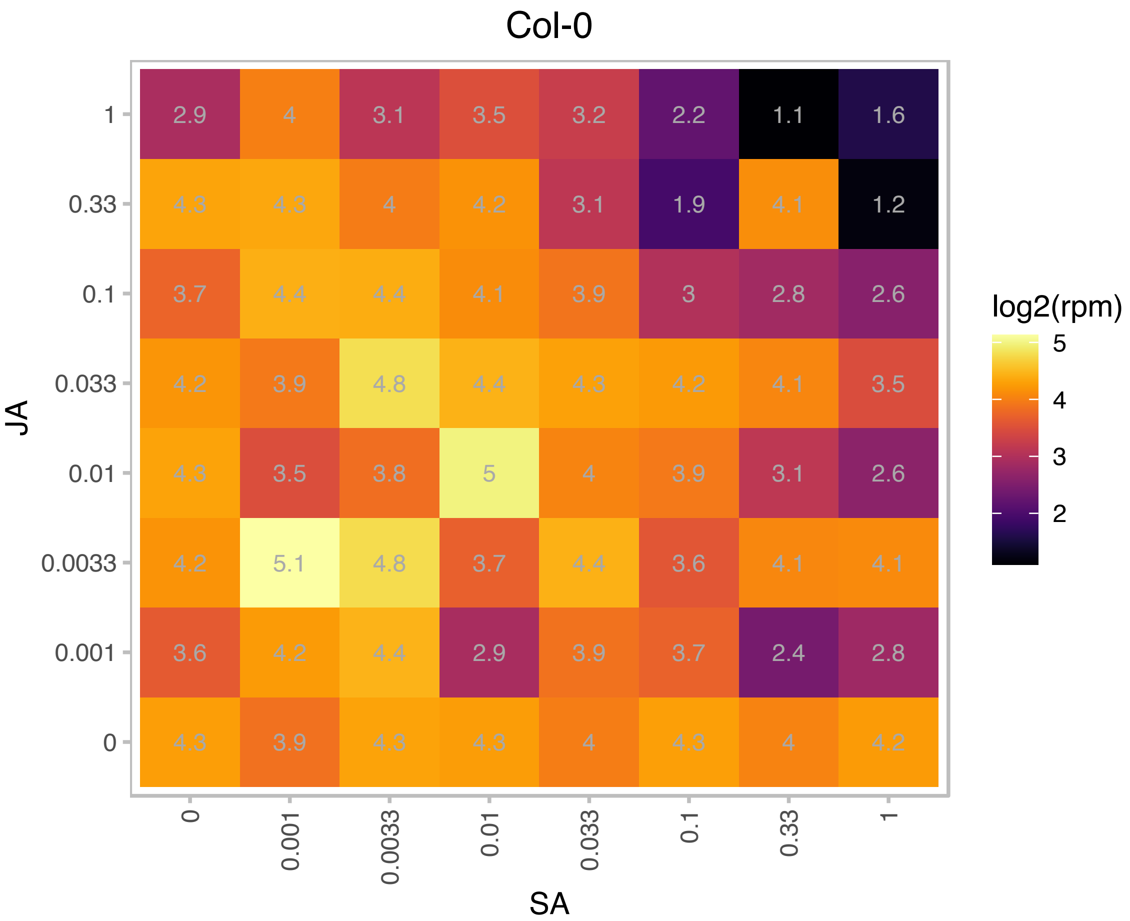


| | |
|---|---|
| Gene | AT2G45250 |
| is.expressed_genes | TRUE |
| is.responsive_genes | TRUE |
| is.apcluster | F-08 |

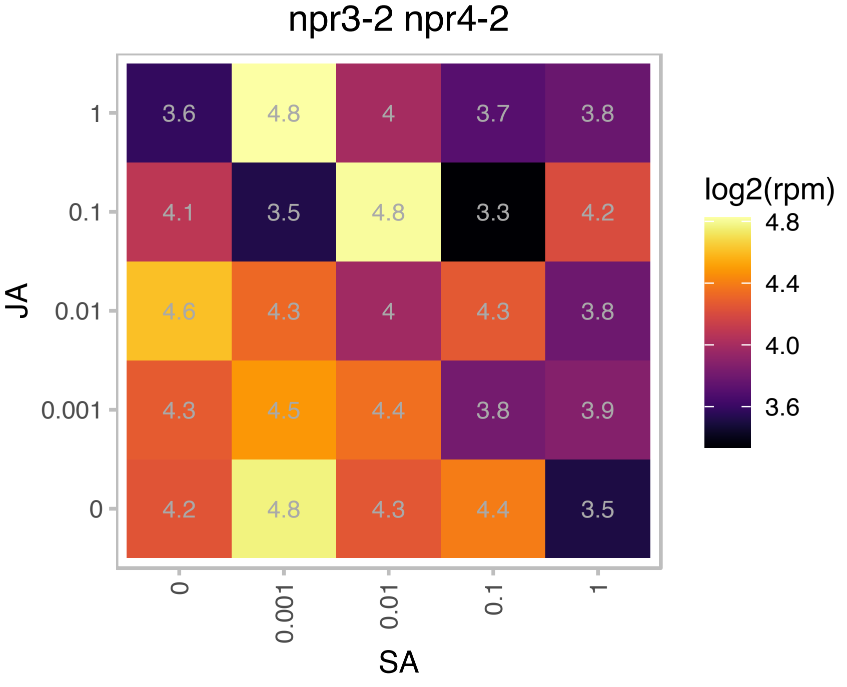


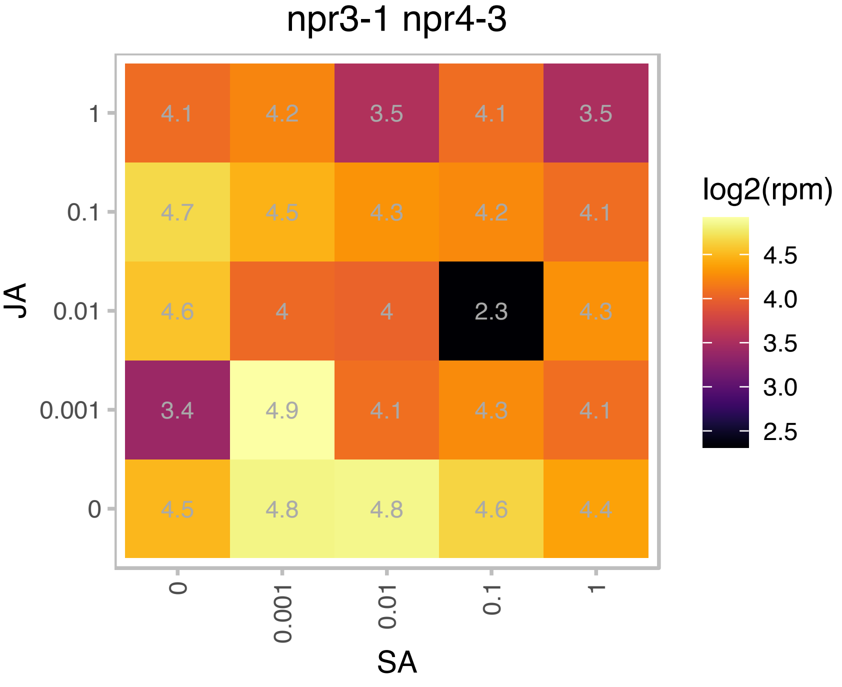

# AT2G45860.1 : NA

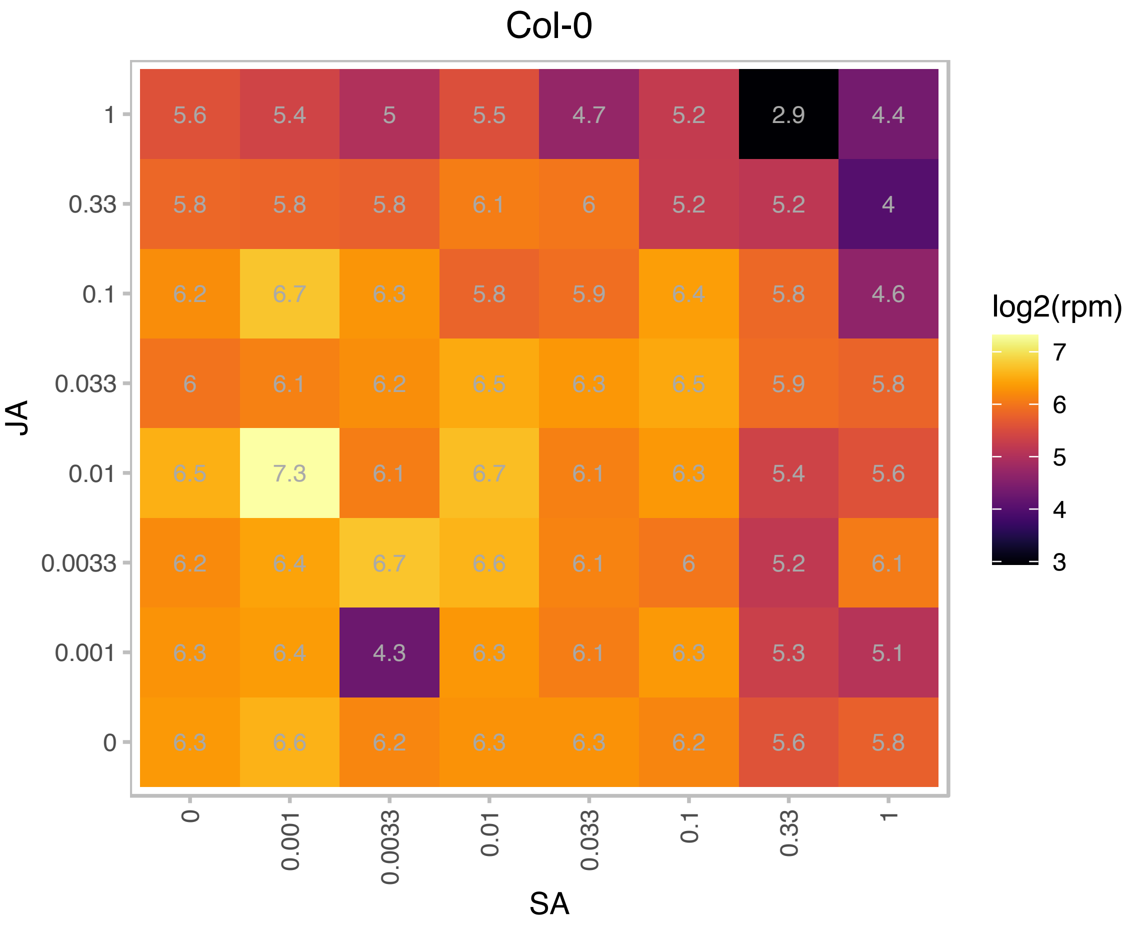


| | |
|---|---|
| Gene | AT2G45860 |
| is.expressed_genes | TRUE |
| is.responsive_genes | TRUE |
| is.apcluster | F-08 |

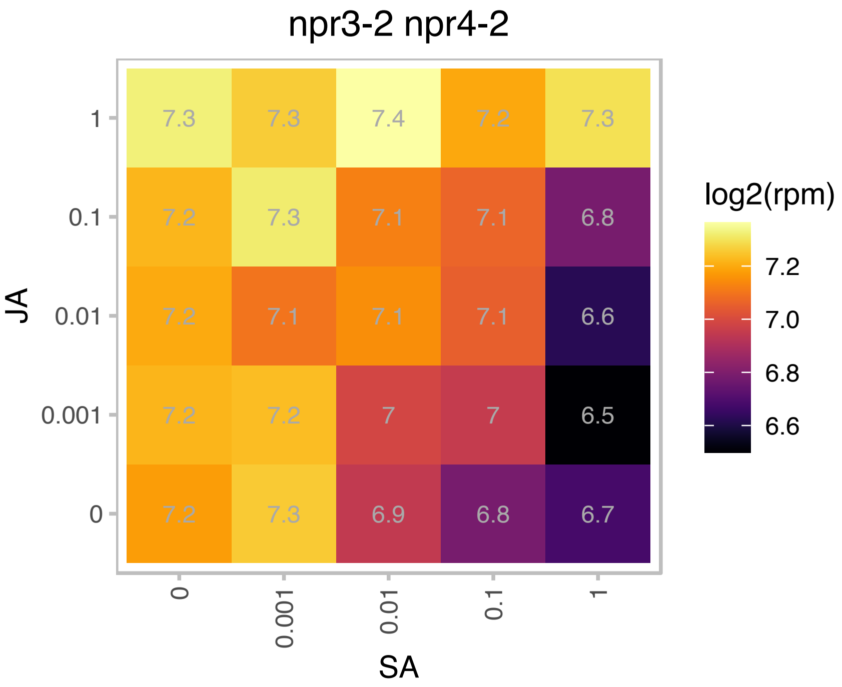


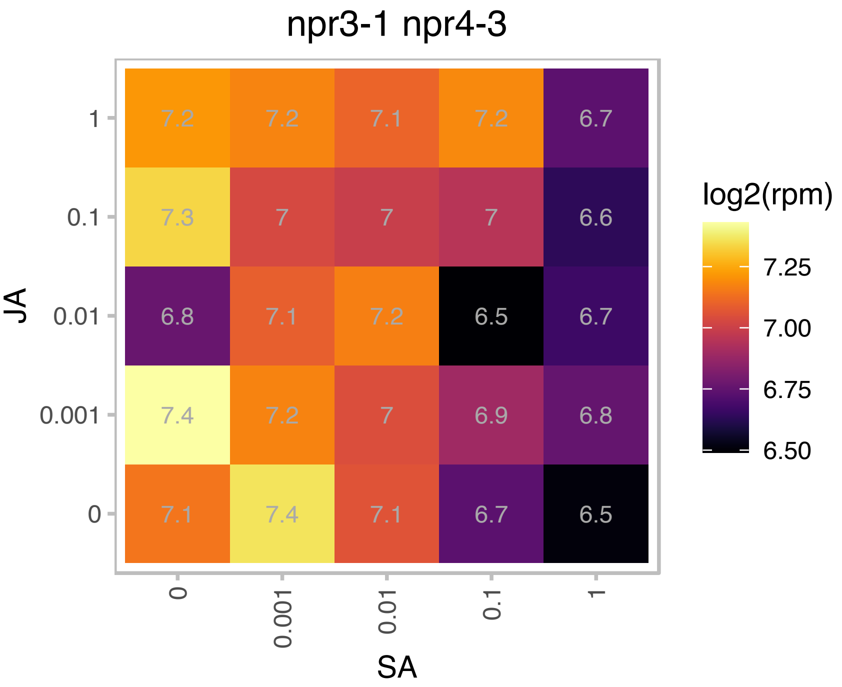

# AT2G46230.1 : NA

PIN domain-like family protein

| | |
|---|---|
| Gene | AT2G46230 |
| is.expressed_genes | TRUE |
| is.responsive_genes | TRUE |
| is.apcluster | F-08 |

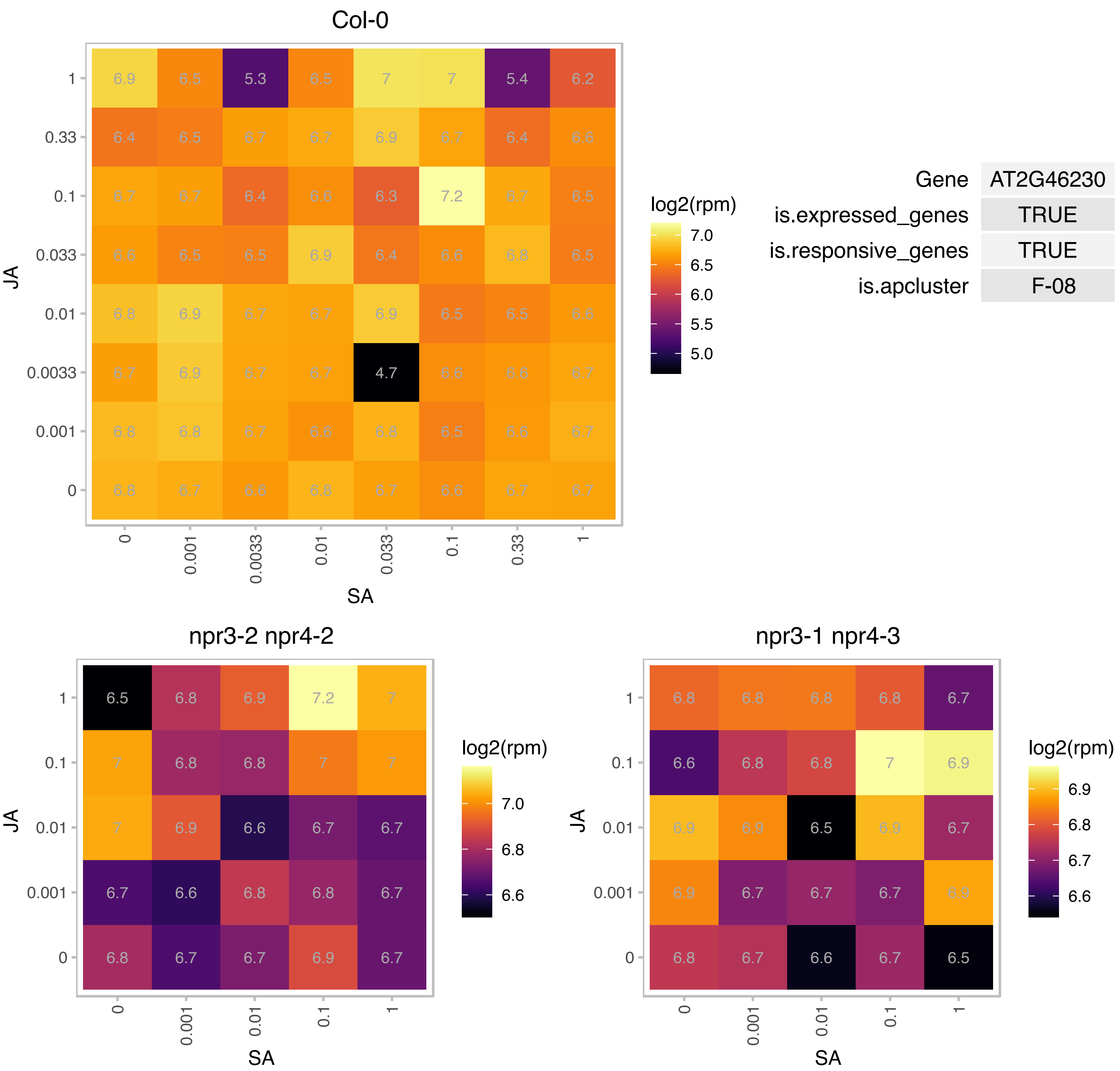

### AT2G46870.1 : NGA1

AP2/B3-like transcriptional factor family protein

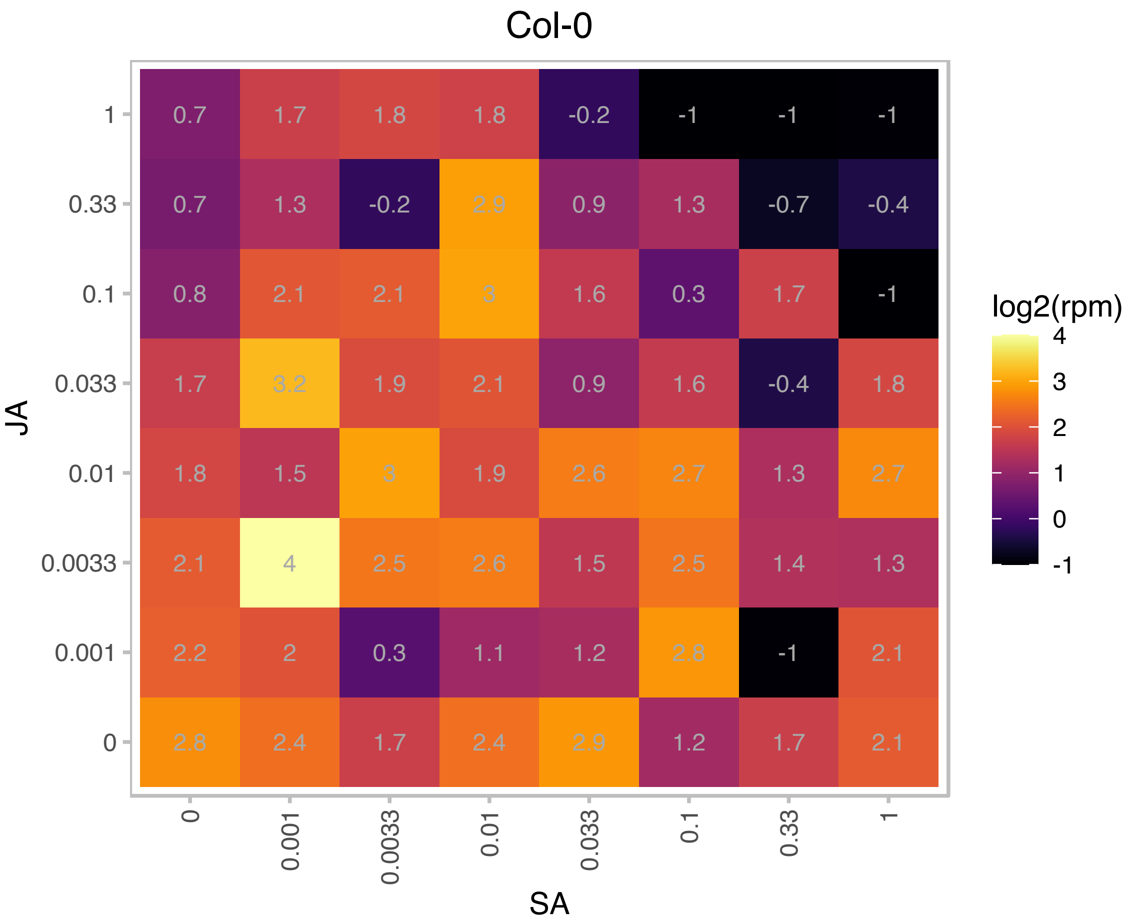


| | |
|---|---|
| Gene | AT2G46870 |
| is.expressed_genes | TRUE |
| is.responsive_genes | TRUE |
| is.apcluster | F-08 |

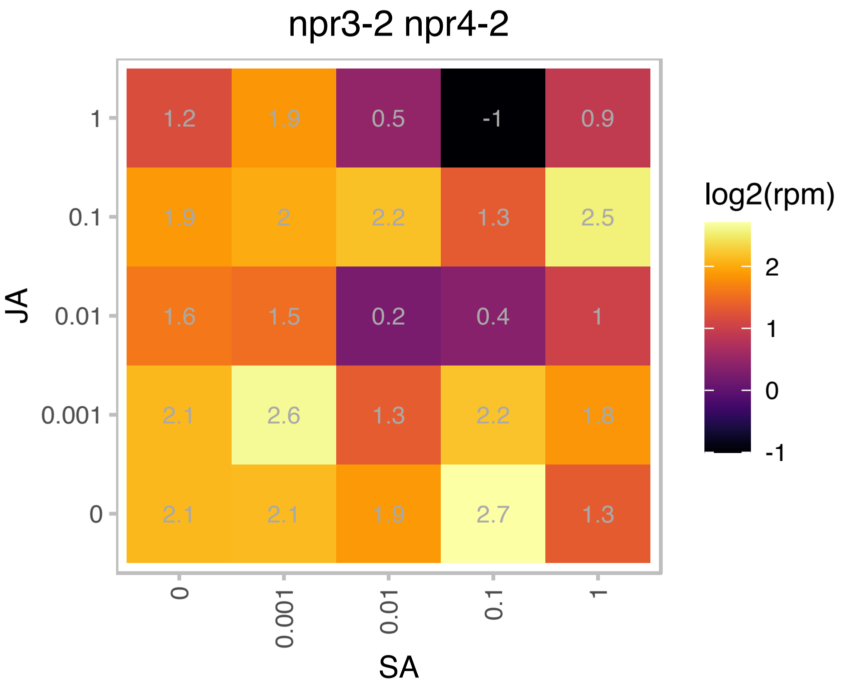


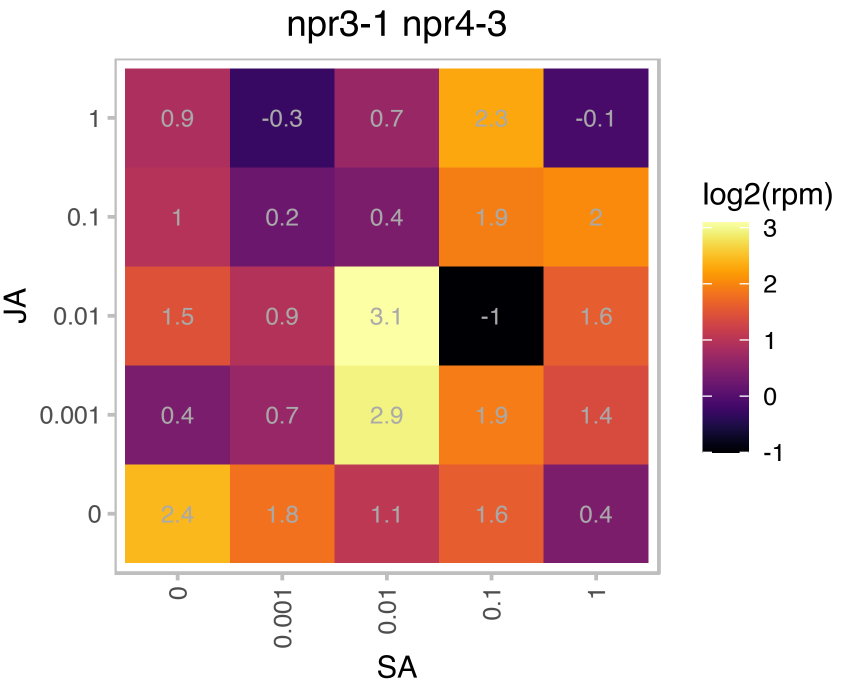

# AT3G01130.1 : NA

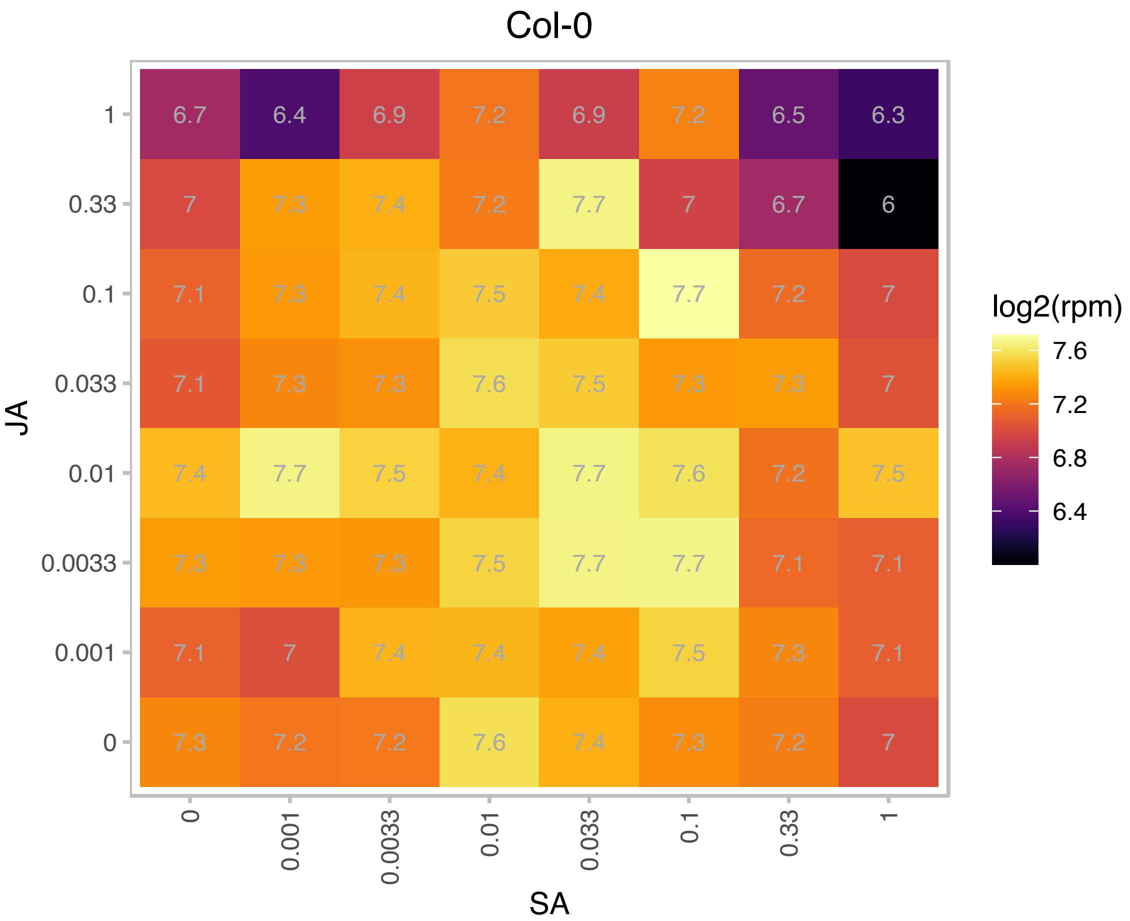


| | |
|---|---|
| Gene | AT3G01130 |
| is.expressed_genes | TRUE |
| is.responsive_genes | TRUE |
| is.apcluster | F-08 |

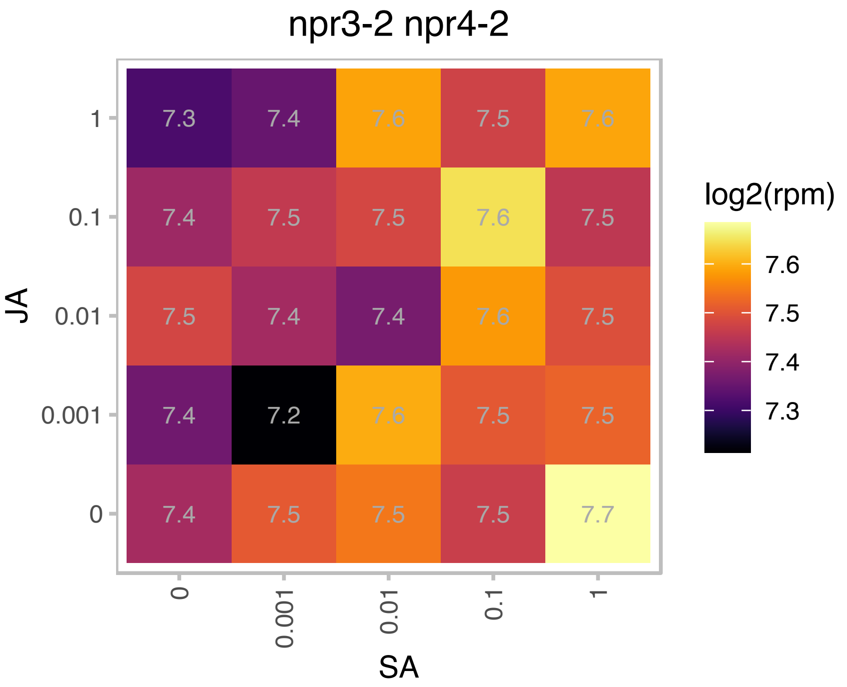


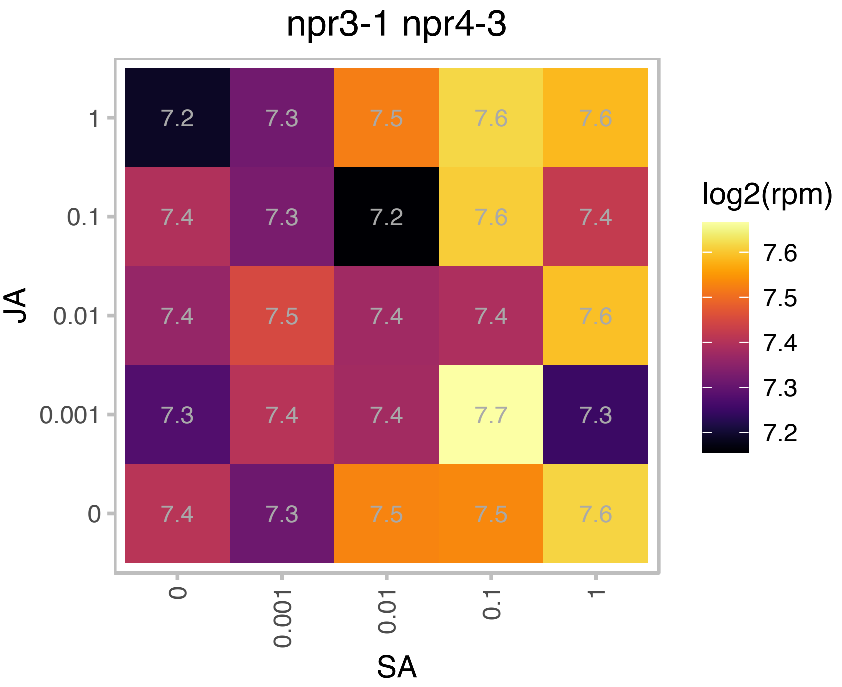

# AT3G01740.1 : NA

Mitochondrial ribosomal protein L37

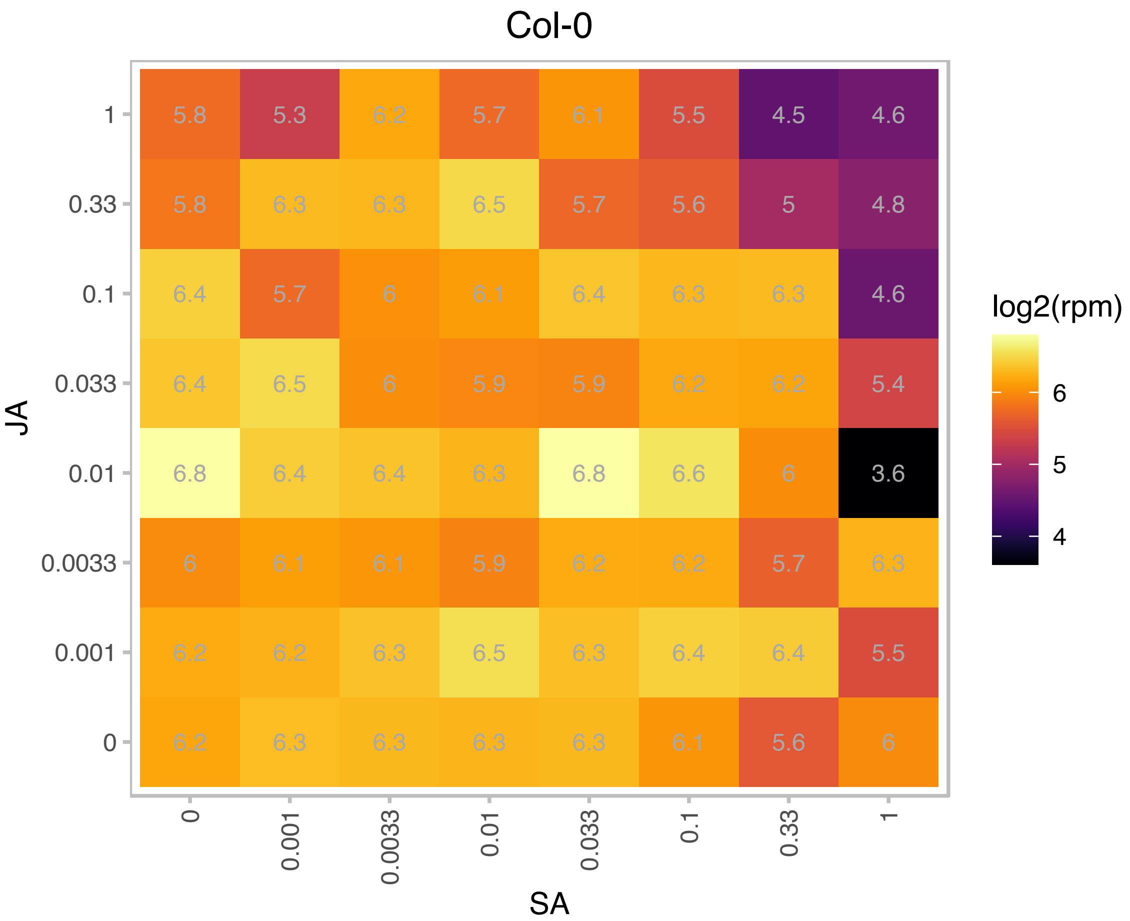


| | |
|---|---|
| Gene | AT3G01740 |
| is.expressed_genes | TRUE |
| is.responsive_genes | TRUE |
| is.apcluster | F-08 |

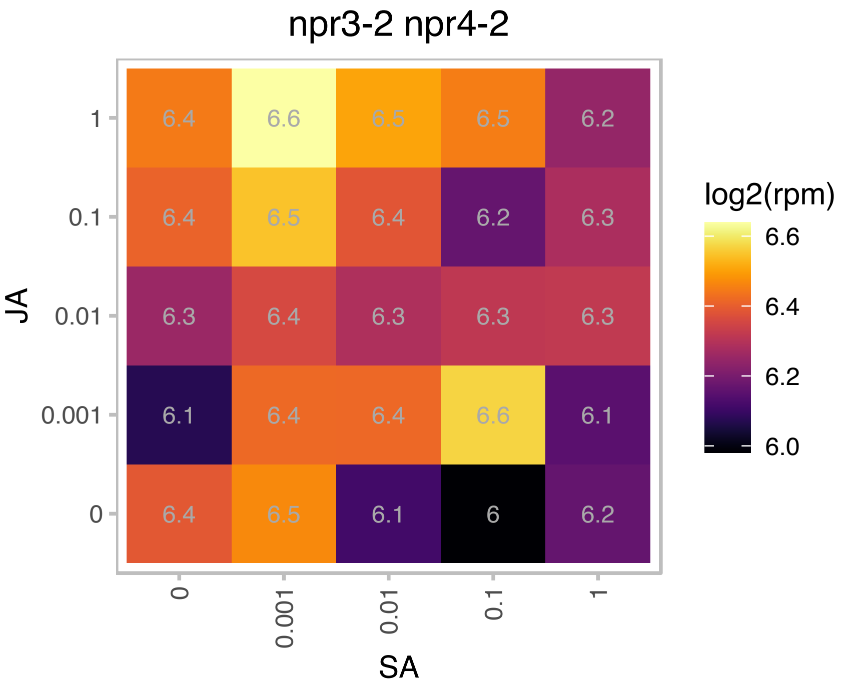


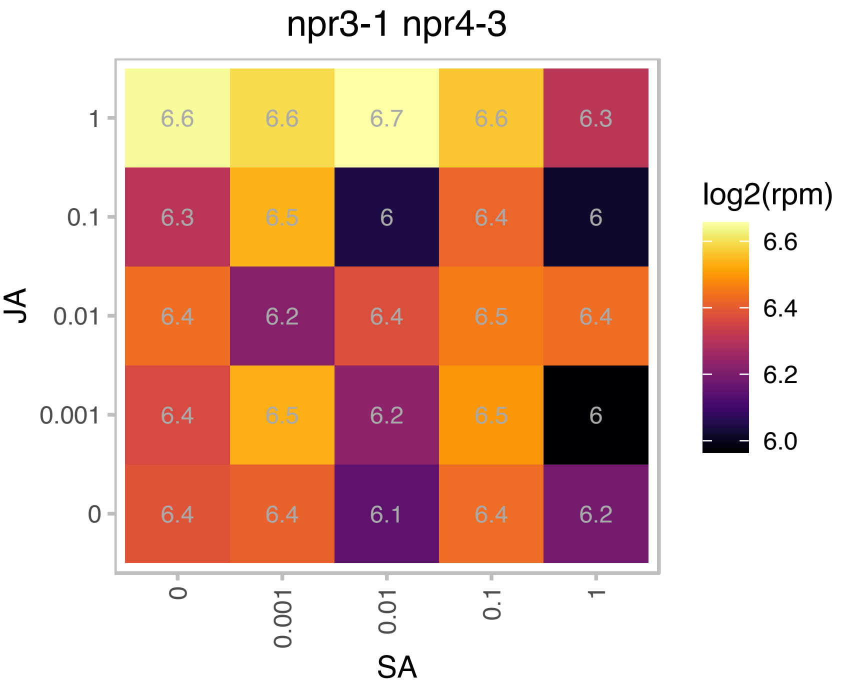

# AT3G02120.1 : NA

hydroxyproline-rich glycoprotein family protein

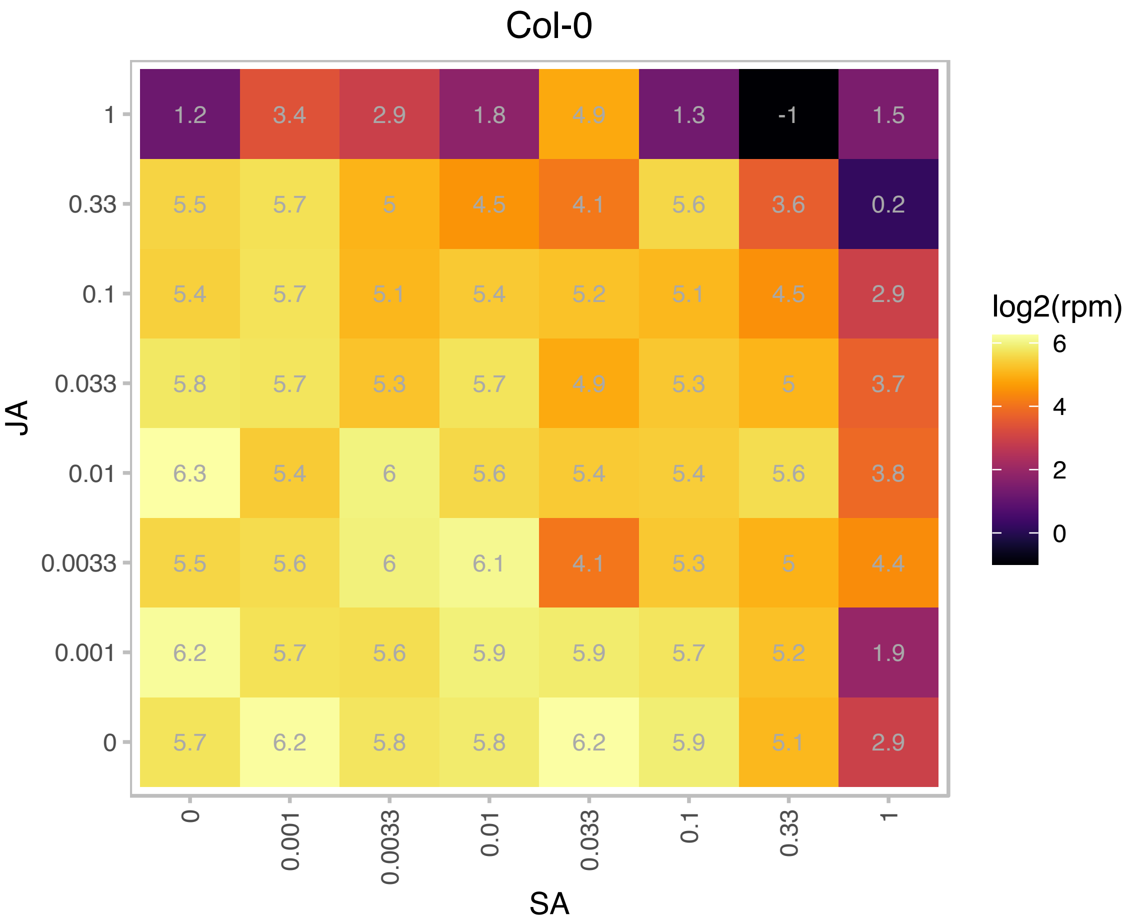


| | |
|---|---|
| Gene | AT3G02120 |
| is.expressed_genes | TRUE |
| is.responsive_genes | TRUE |
| is.apcluster | F-08 |

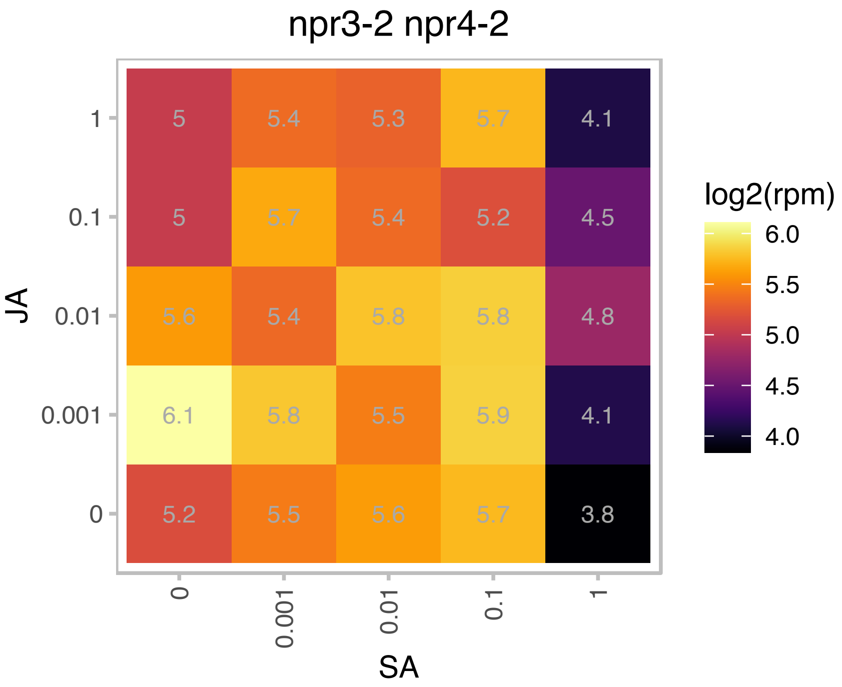


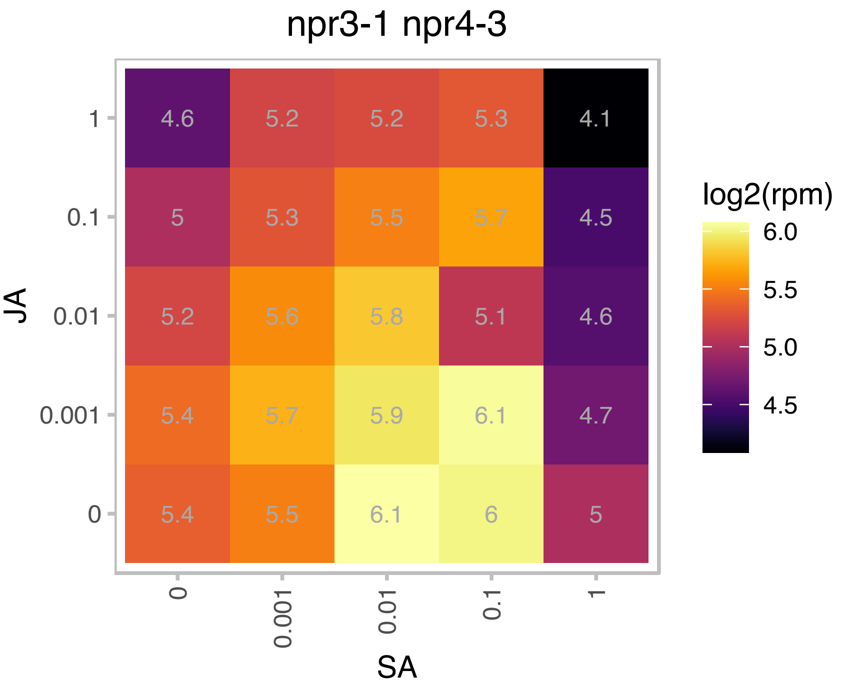

# AT3G02180.1 : SP1L3

SPIRAL1-like3
SPIRAL1-LIKE3 belongs to a six-member gene family in Arabidopsis; all members share high sequence similarity in amino- and carboxy-terminal regions. Regulates cortical microtubule organization. Mutant plants exhibit altered patterns of root, leaf and petal growth as a result of defective anisotropic cell expansion.

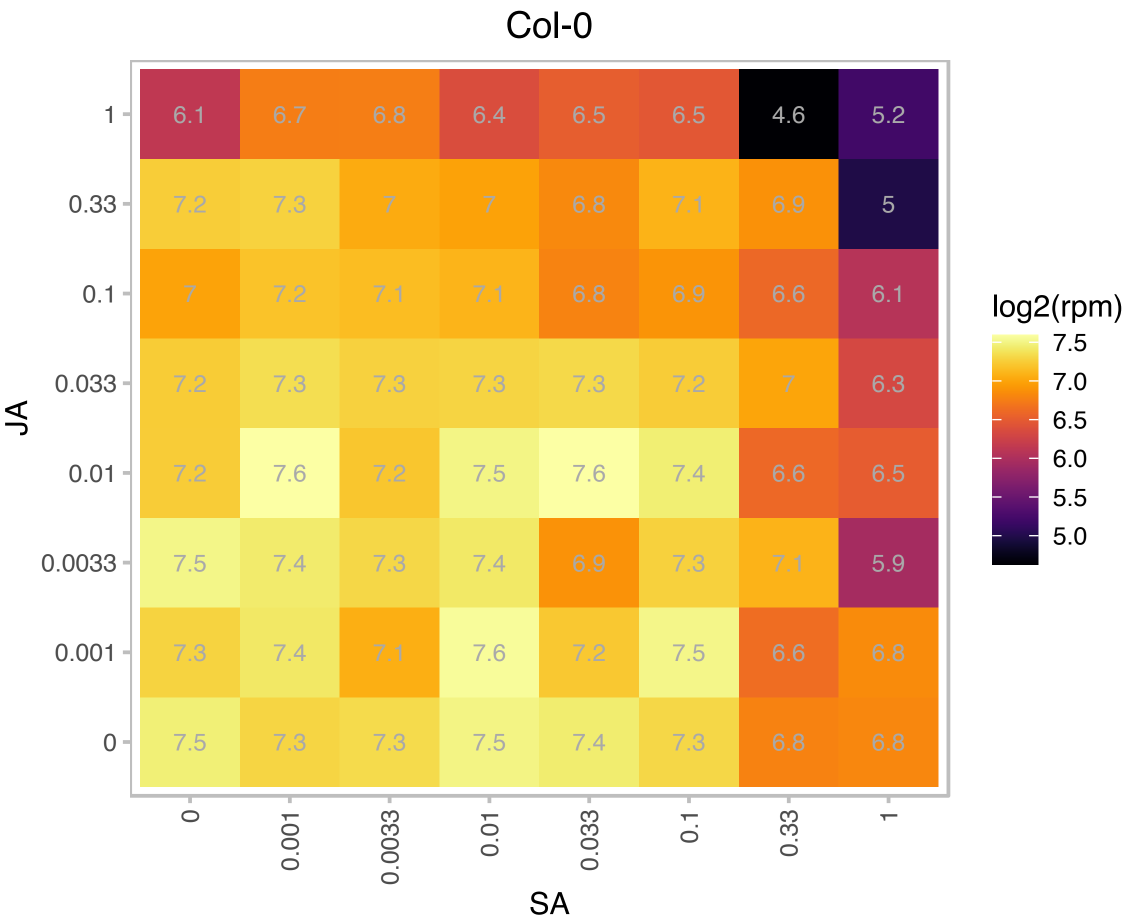


| | |
|---|---|
| Gene | AT3G02180 |
| is.expressed_genes | TRUE |
| is.responsive_genes | TRUE |
| is.apcluster | F-08 |

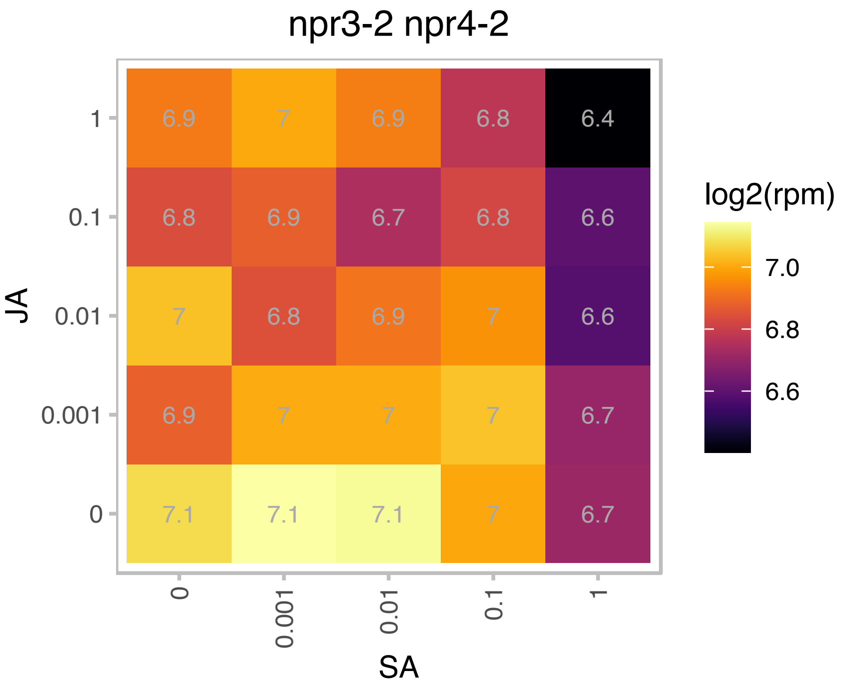


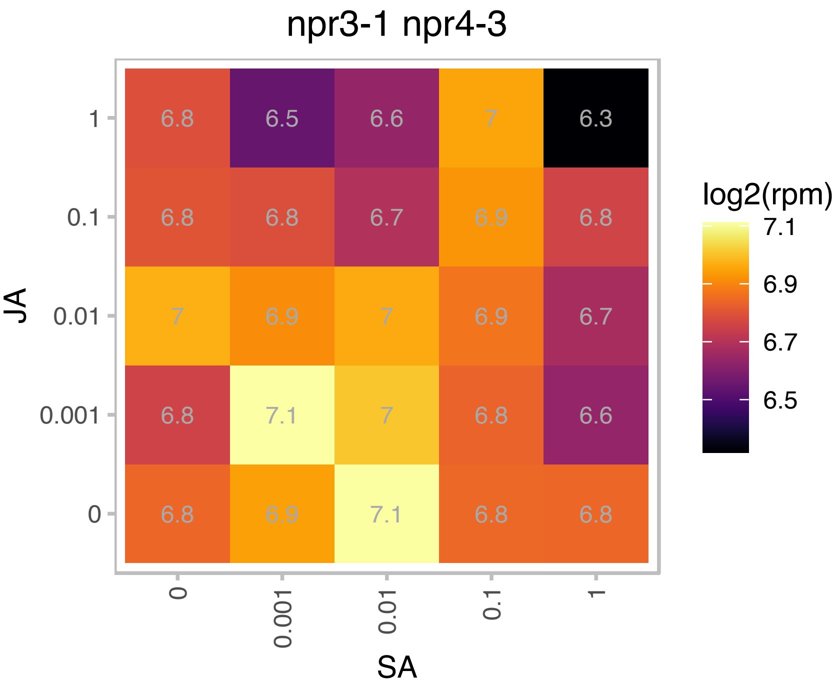

# AT3G02220.1 : NA

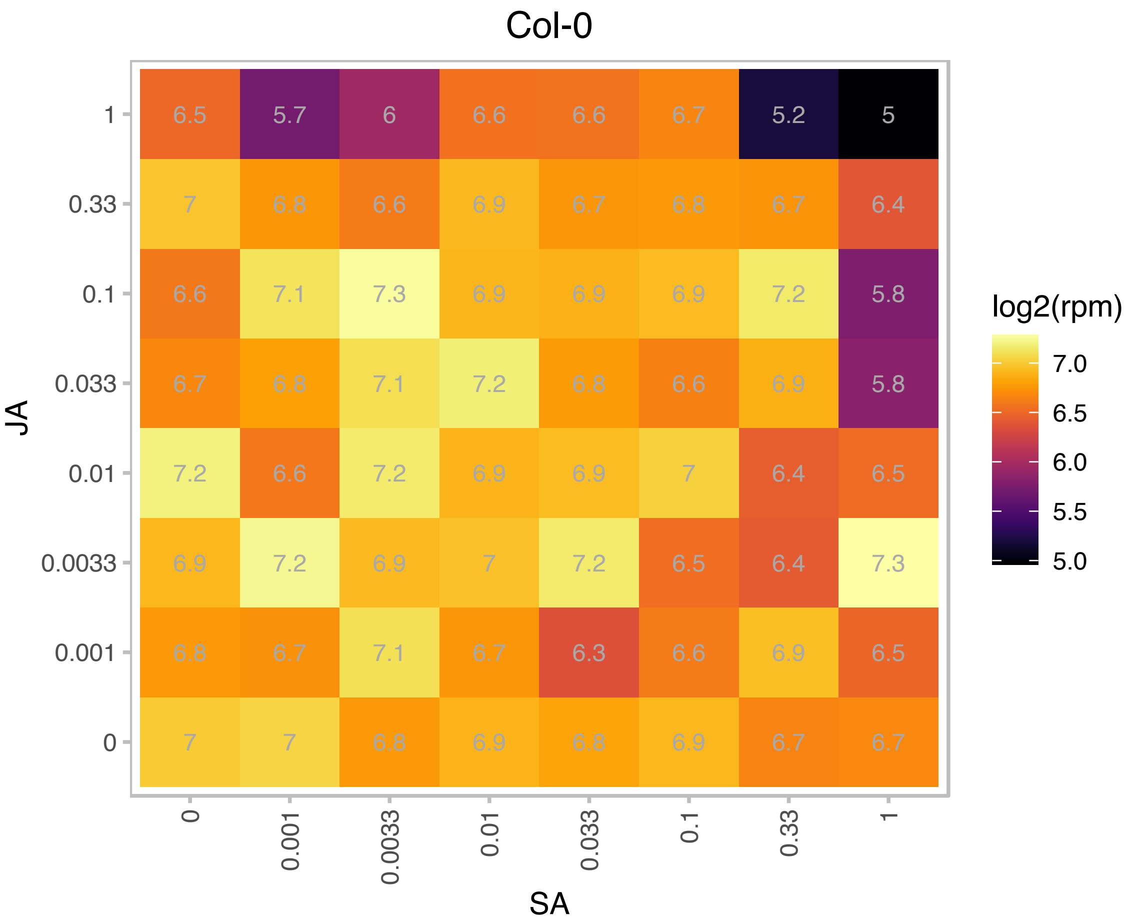


| | |
|---|---|
| Gene | AT3G02220 |
| is.expressed_genes | TRUE |
| is.responsive_genes | TRUE |
| is.apcluster | F-08 |

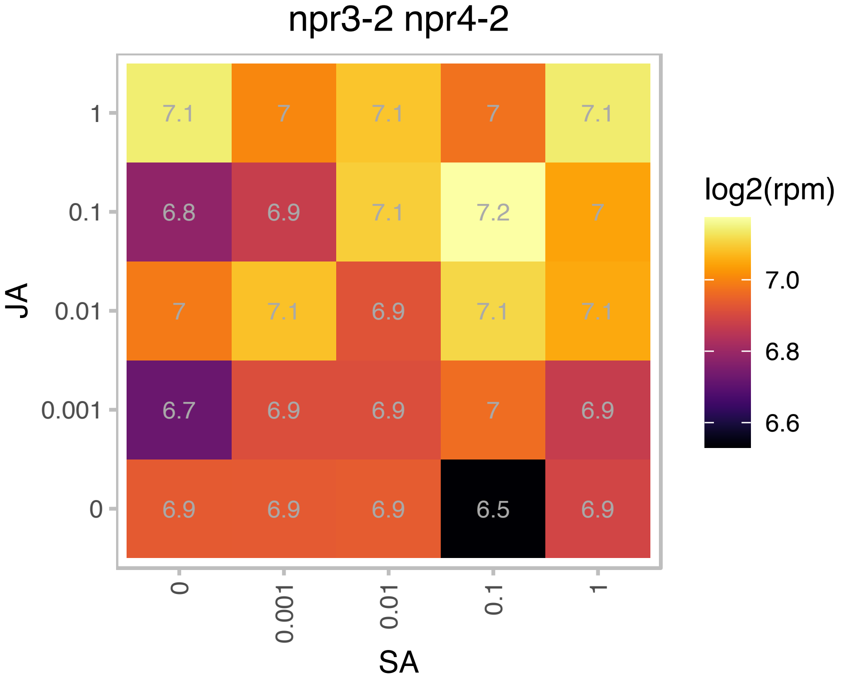


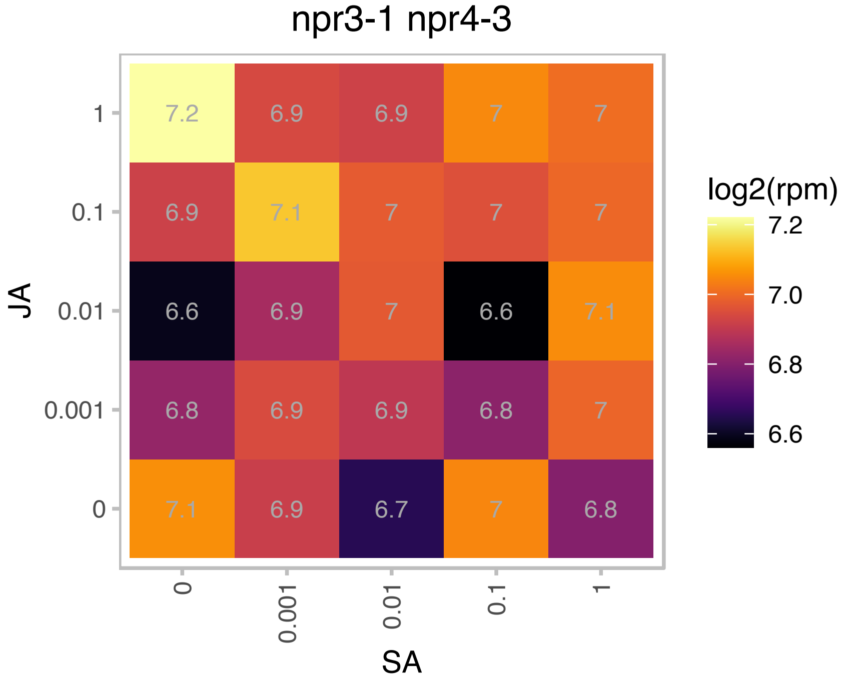

# AT3G05625.1 : NA

Tetratricopeptide repeat (TPR)-like superfamily protein

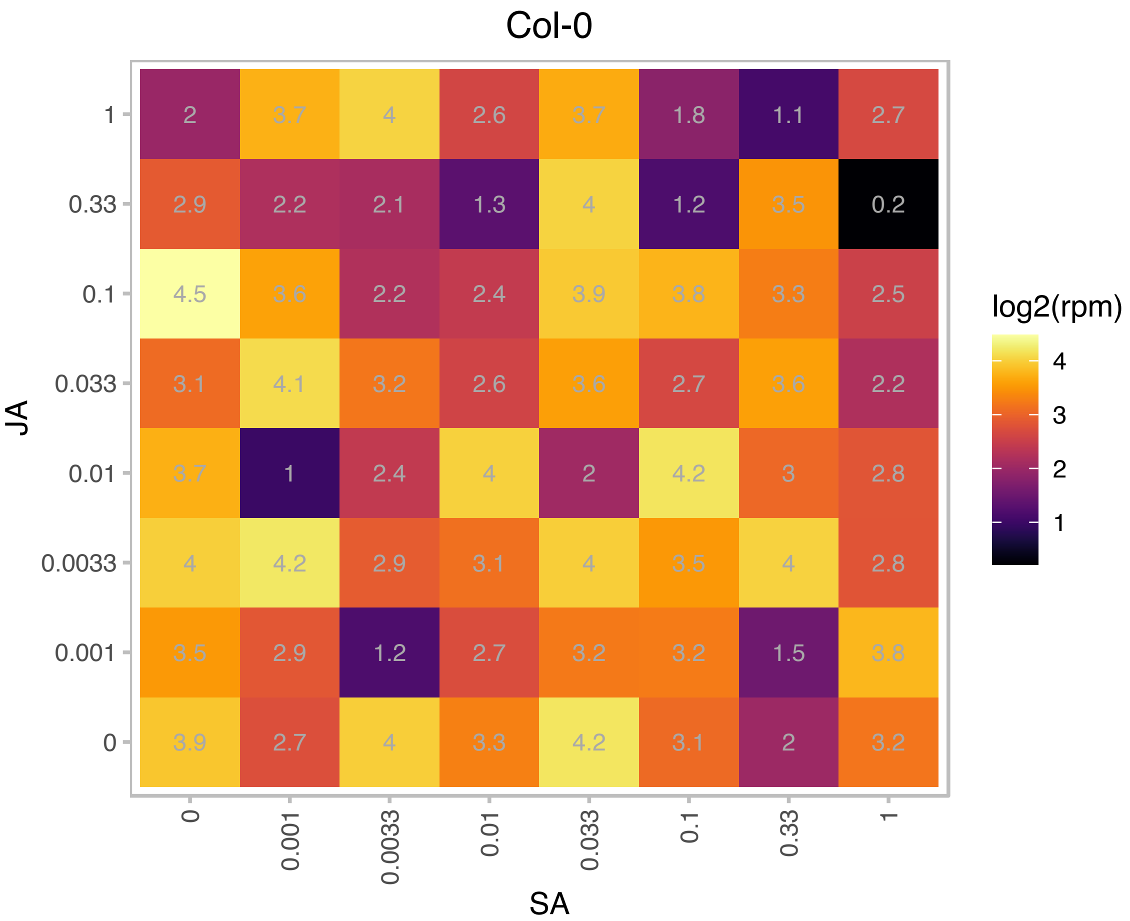


| | |
|---|---|
| Gene | AT3G05625 |
| is.expressed_genes | TRUE |
| is.responsive_genes | TRUE |
| is.apcluster | F-08 |

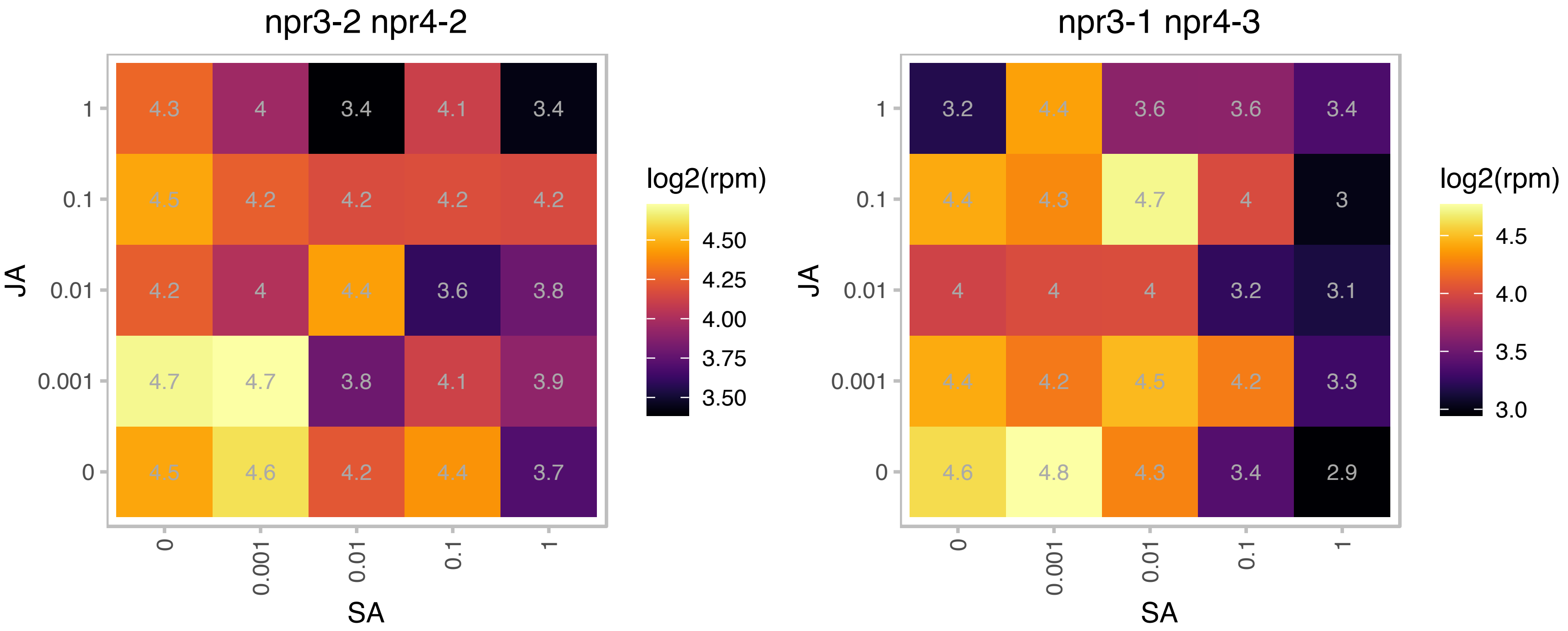

# AT3G06610.1 : NA

DNA-binding enhancer protein-related

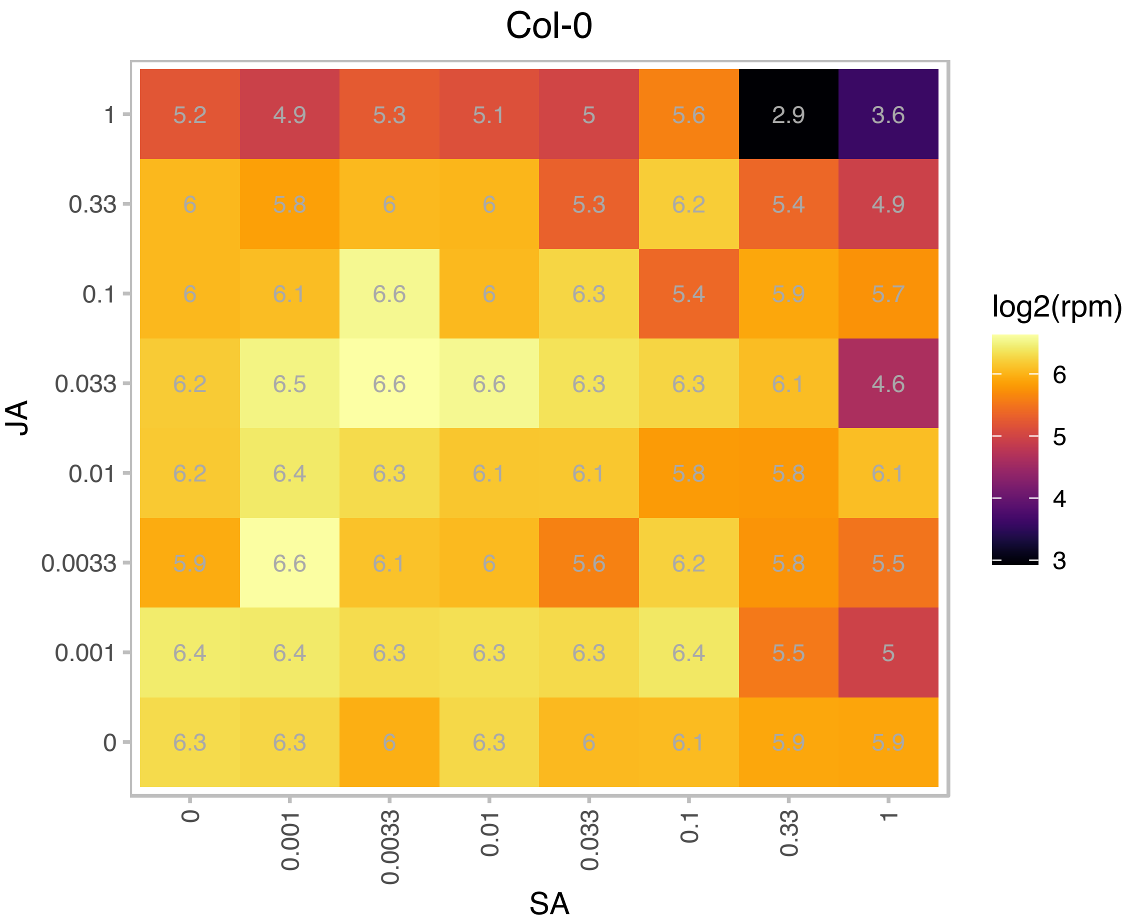


| | |
|---|---|
| Gene | AT3G06610 |
| is.expressed_genes | TRUE |
| is.responsive_genes | TRUE |
| is.apcluster | F-08 |

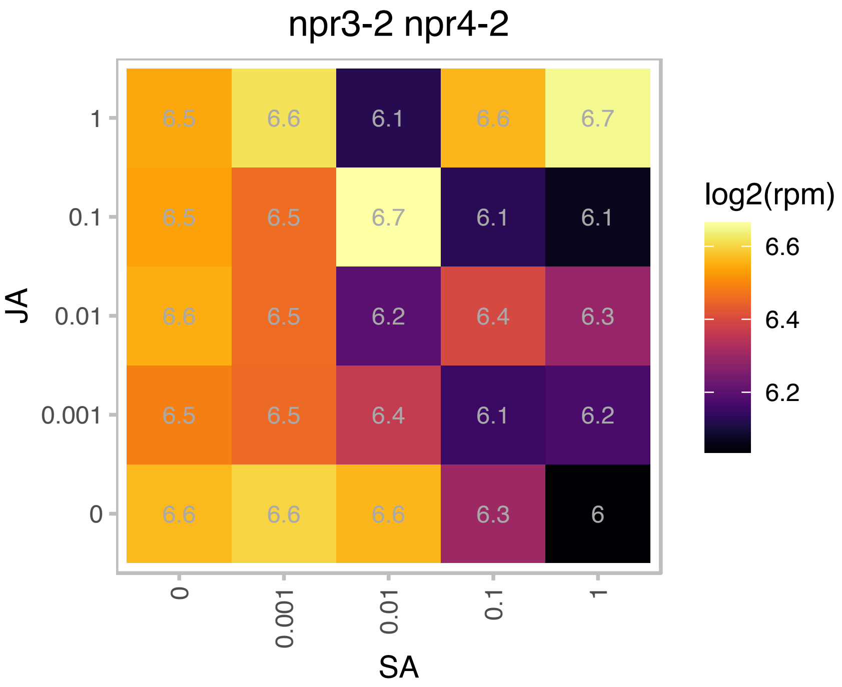


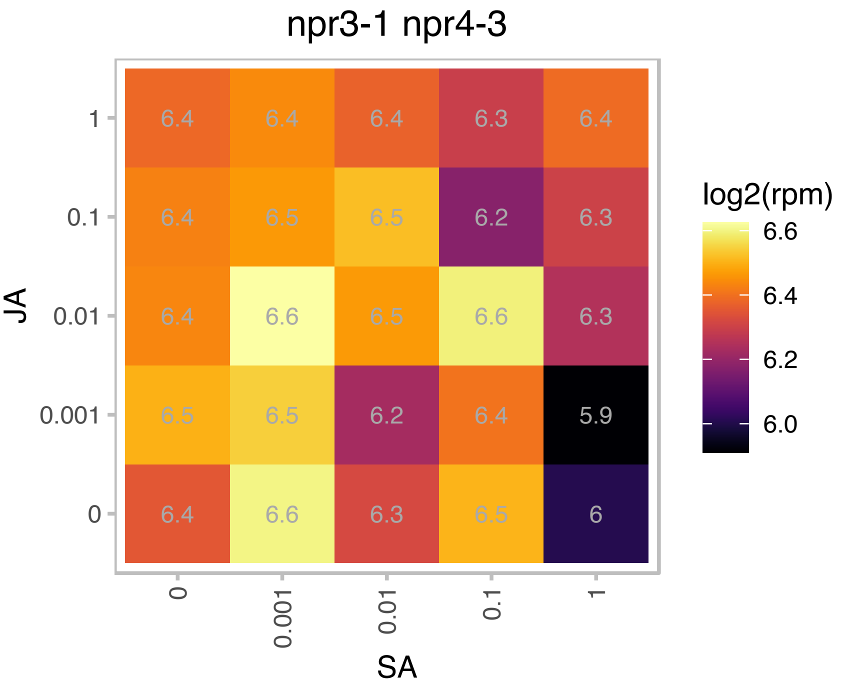

# AT3G10090.1 : NA

Nucleic acid-binding, OB-fold-like protein

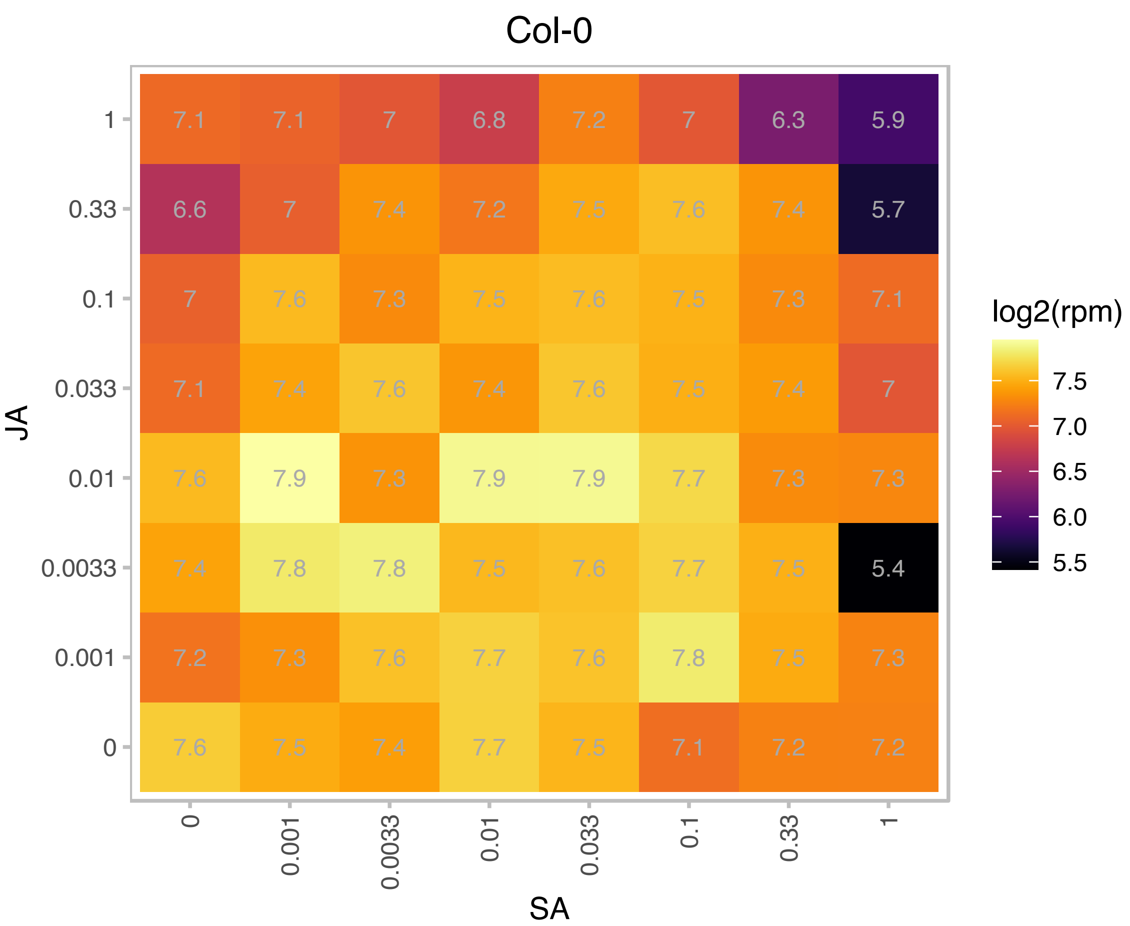


| | |
|---|---|
| Gene | AT3G10090 |
| is.expressed_genes | TRUE |
| is.responsive_genes | TRUE |
| is.apcluster | F-08 |

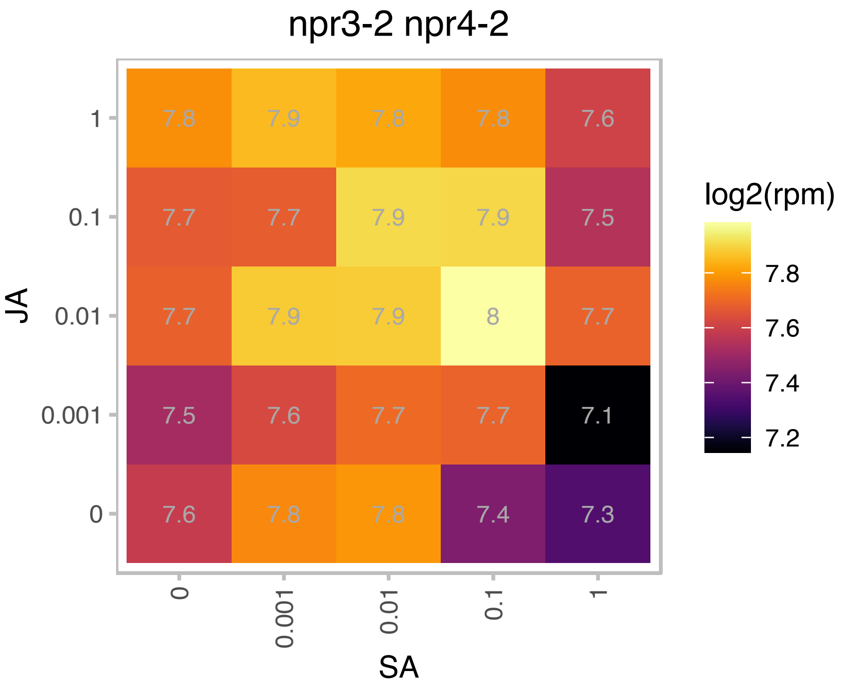


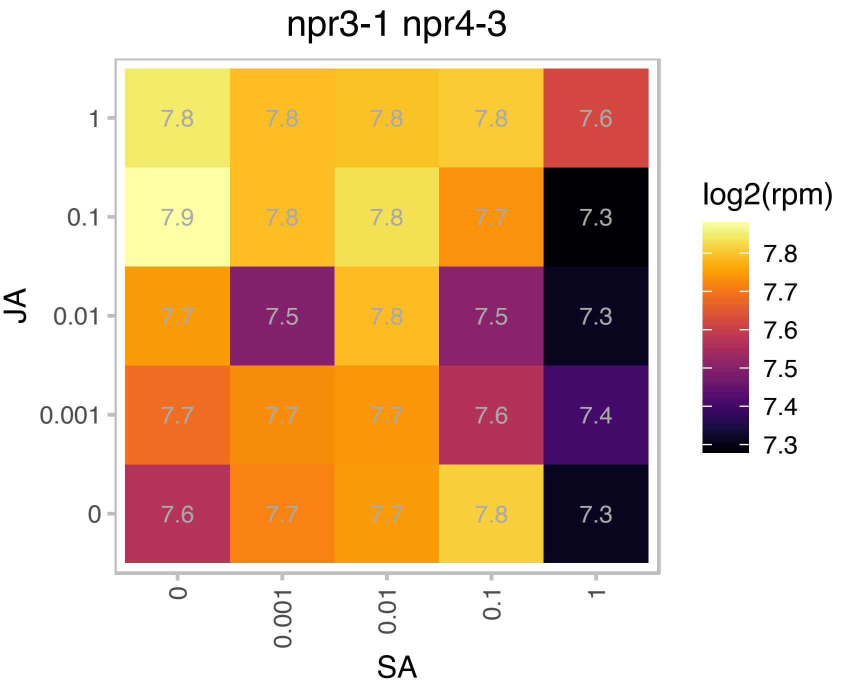

### AT3G11940.1 : AML1, ATRPS5A, RPS5A

ribosomal protein 5A
One of two genes encoding the ribosomal protein S5. Mutants have semi-dominant developmental phenotypes. Most cell-division processes are delayed or disturbed in the heterozygous mutant, and development is completely arrested at an early embryonic stage in the homozygous mutant.

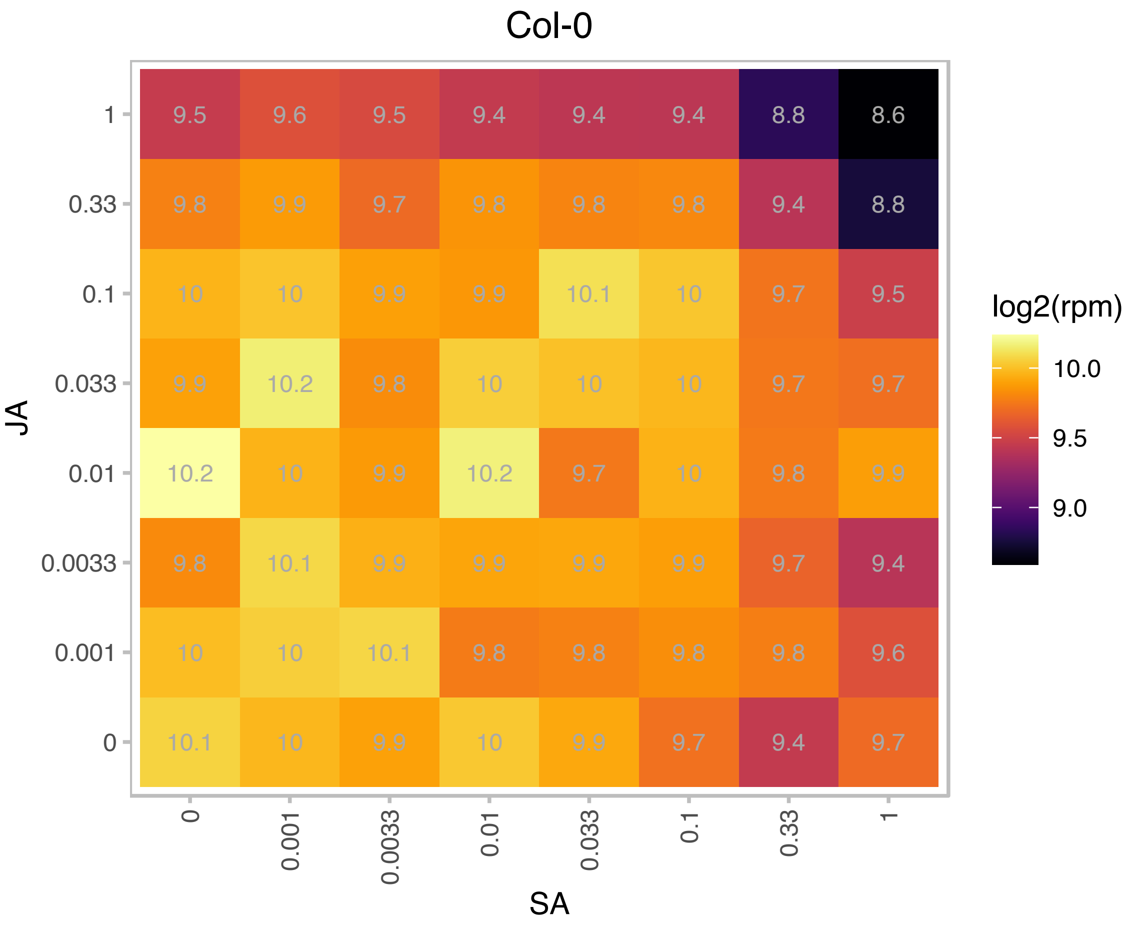


| | |
|---|---|
| Gene | AT3G11940 |
| is.expressed_genes | TRUE |
| is.responsive_genes | TRUE |
| is.apcluster | F-08 |

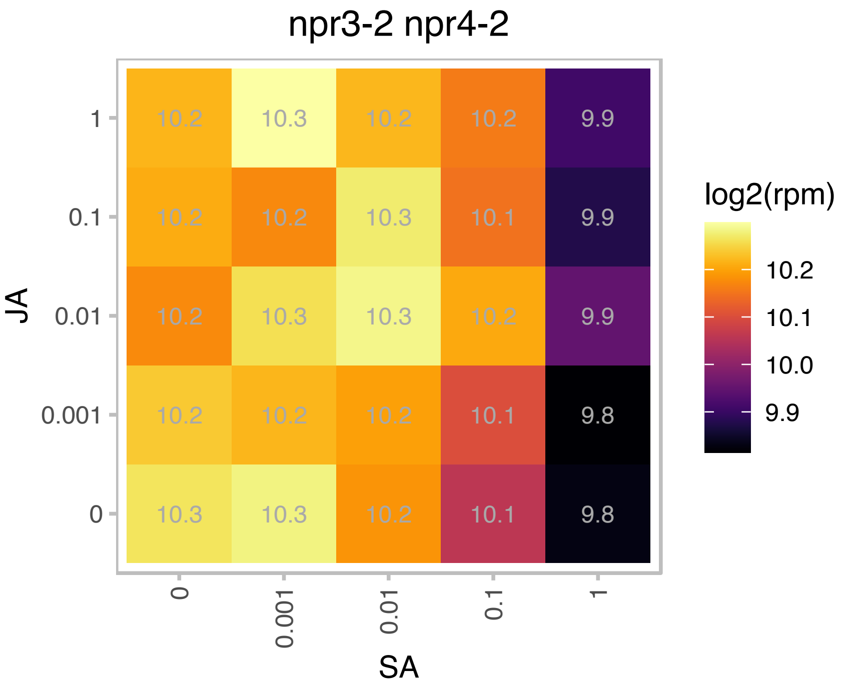


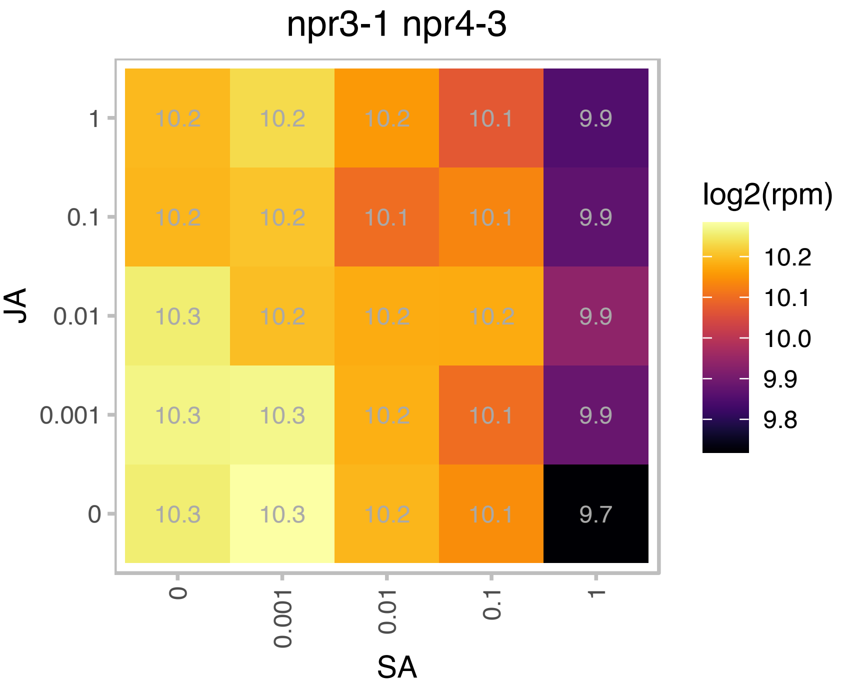

# AT3G13882.1 : NA

Ribosomal protein L34

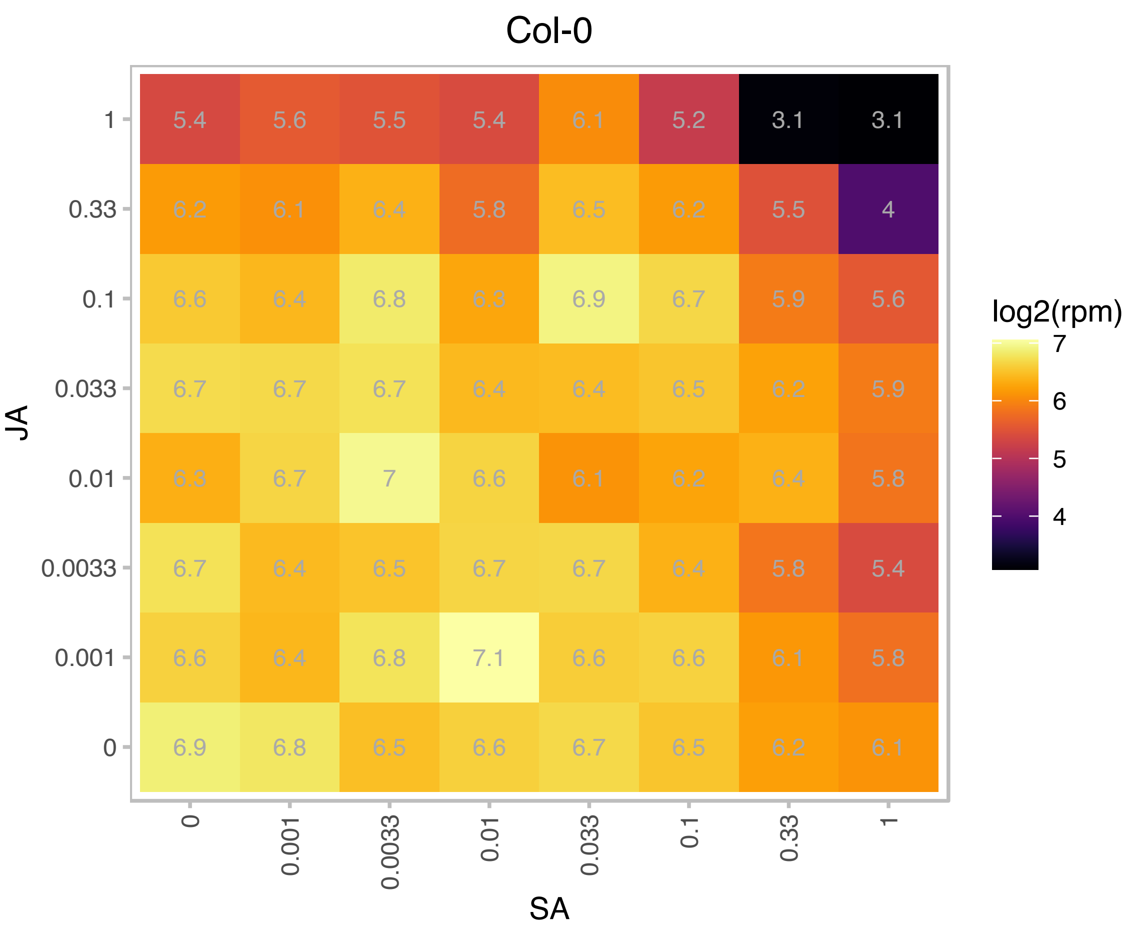


| | |
|---|---|
| Gene | AT3G13882 |
| is.expressed_genes | TRUE |
| is.responsive_genes | TRUE |
| is.apcluster | F-08 |

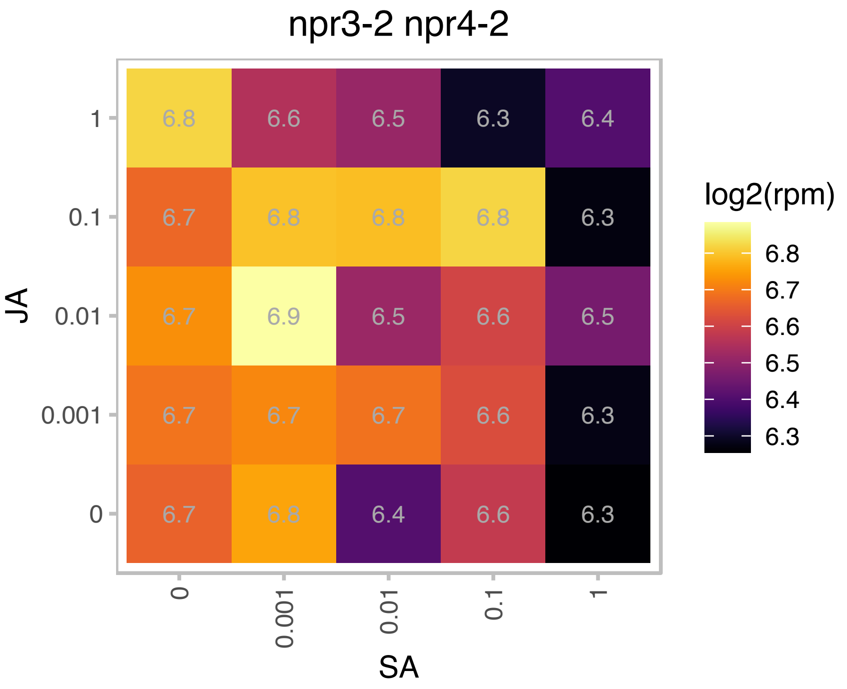


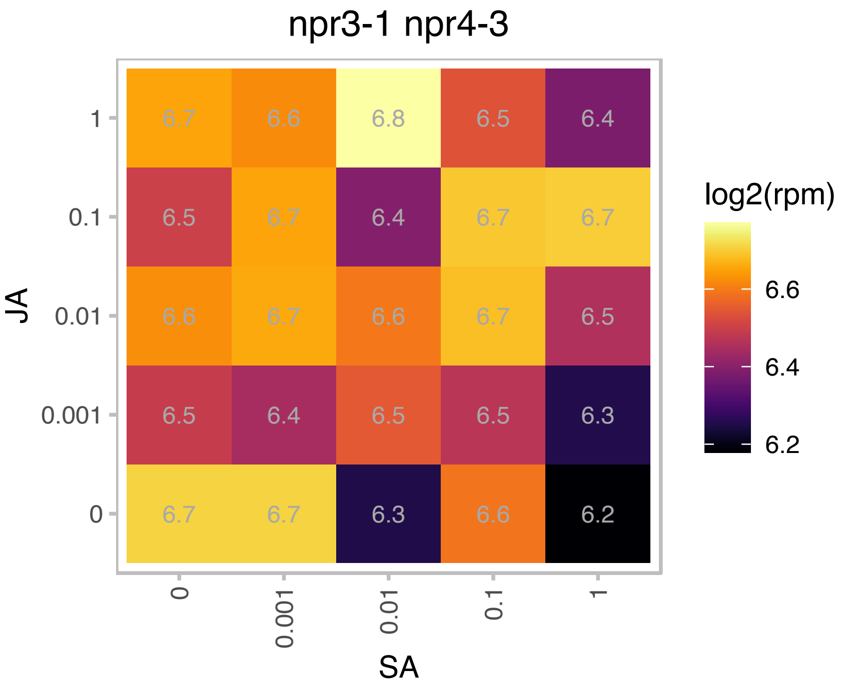

### AT3G16000.1 : MFP1

MAR binding filament-like protein 1

Encodes a DNA-binding protein that binds to plastid DNA non-specifically and is associated with nucleoids and thylakoid membranes. The expression of the gene is correlated with the development of thylakoid membranes.MFP1 is required for the normal initiation of starch granules in Arabidopsis chloroplasts. Mutants lacking MFP1 have fewer starch granules per chloroplast relative to wild-type plants. MFP1 interacts with PTST2 (At1g27070), which is also involved in starch granule initiation. MFP1 is required for proper PTST2 location within the chloroplast.

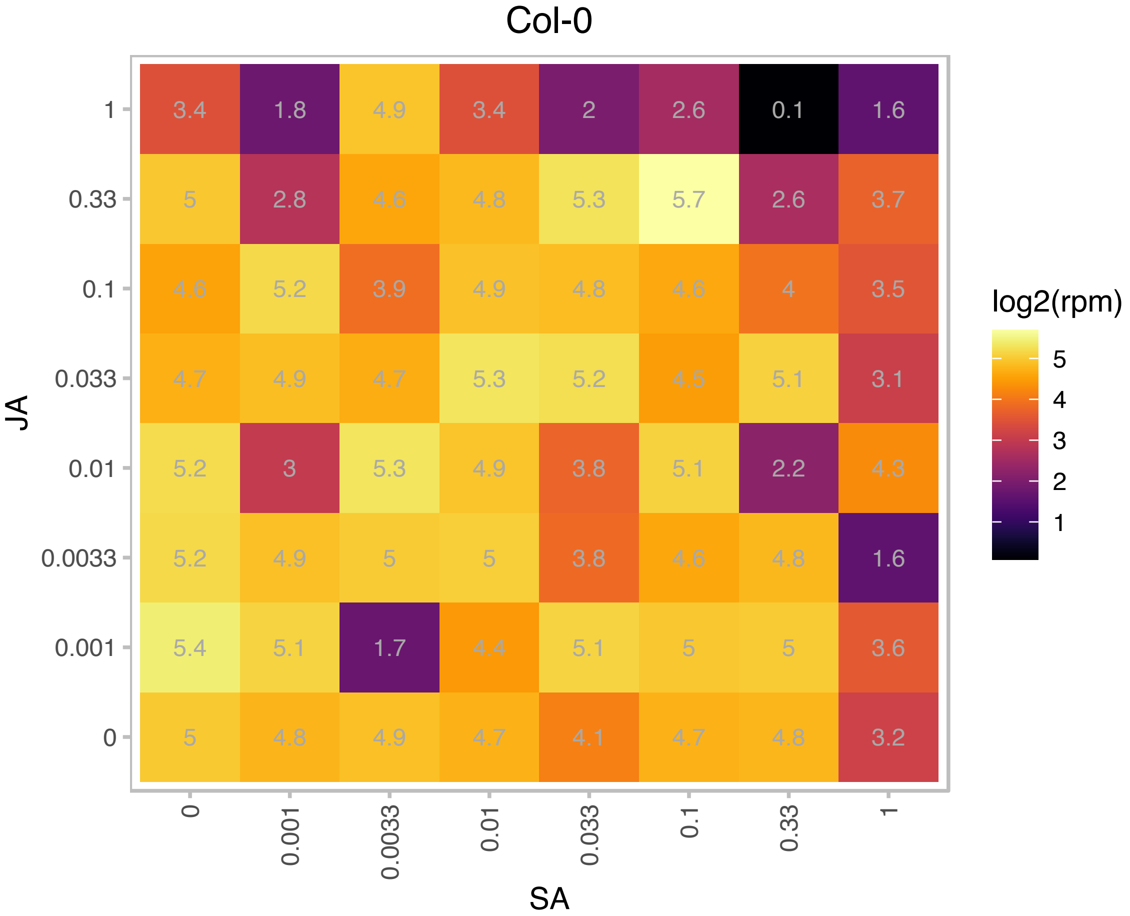


| | |
|---|---|
| Gene | AT3G16000 |
| is.expressed_genes | TRUE |
| is.responsive_genes | TRUE |
| is.apcluster | F-08 |

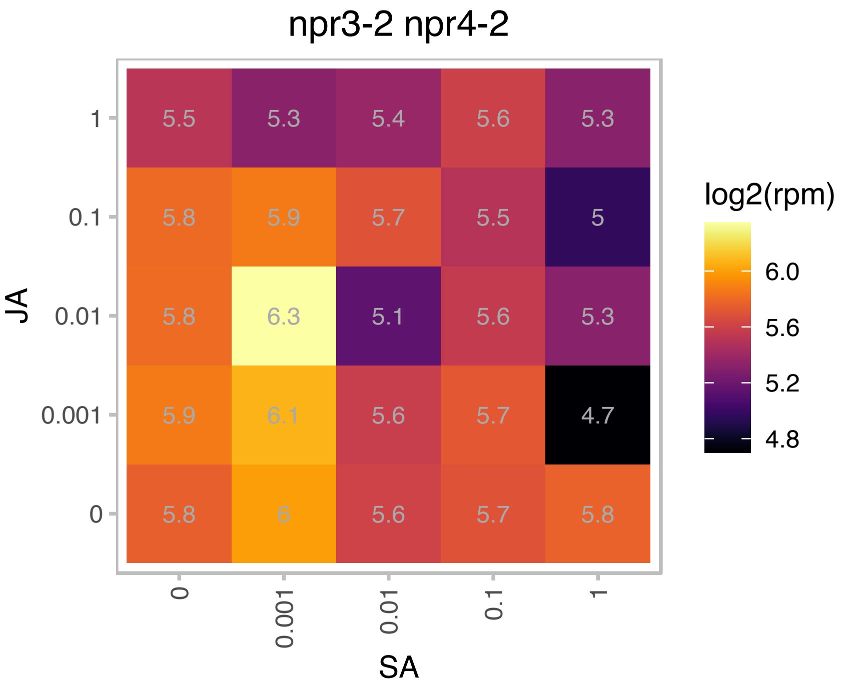


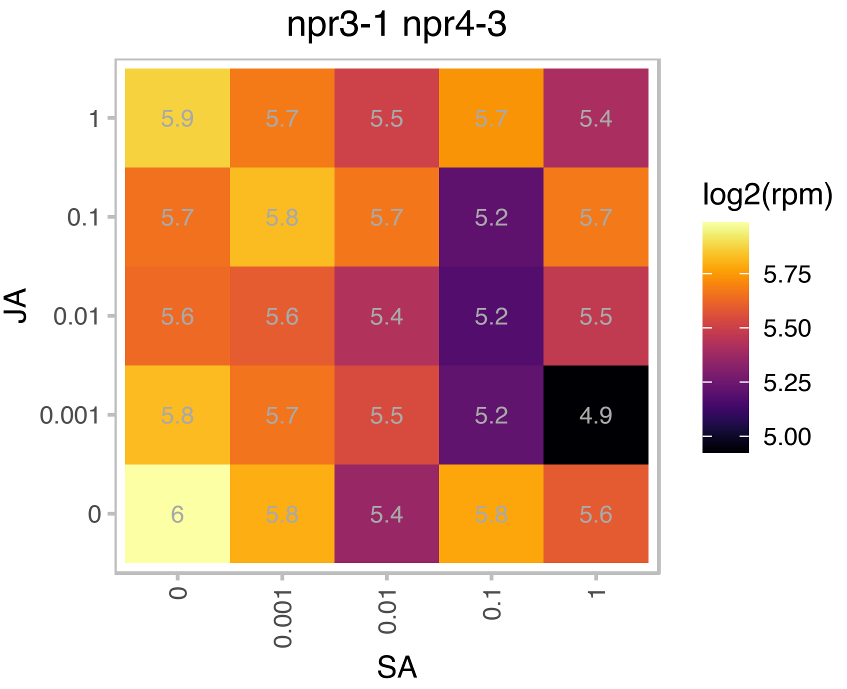

### AT3G18035.1 : HON4

winged-helix DNA-binding transcription factor family protein

A linker histone like protein

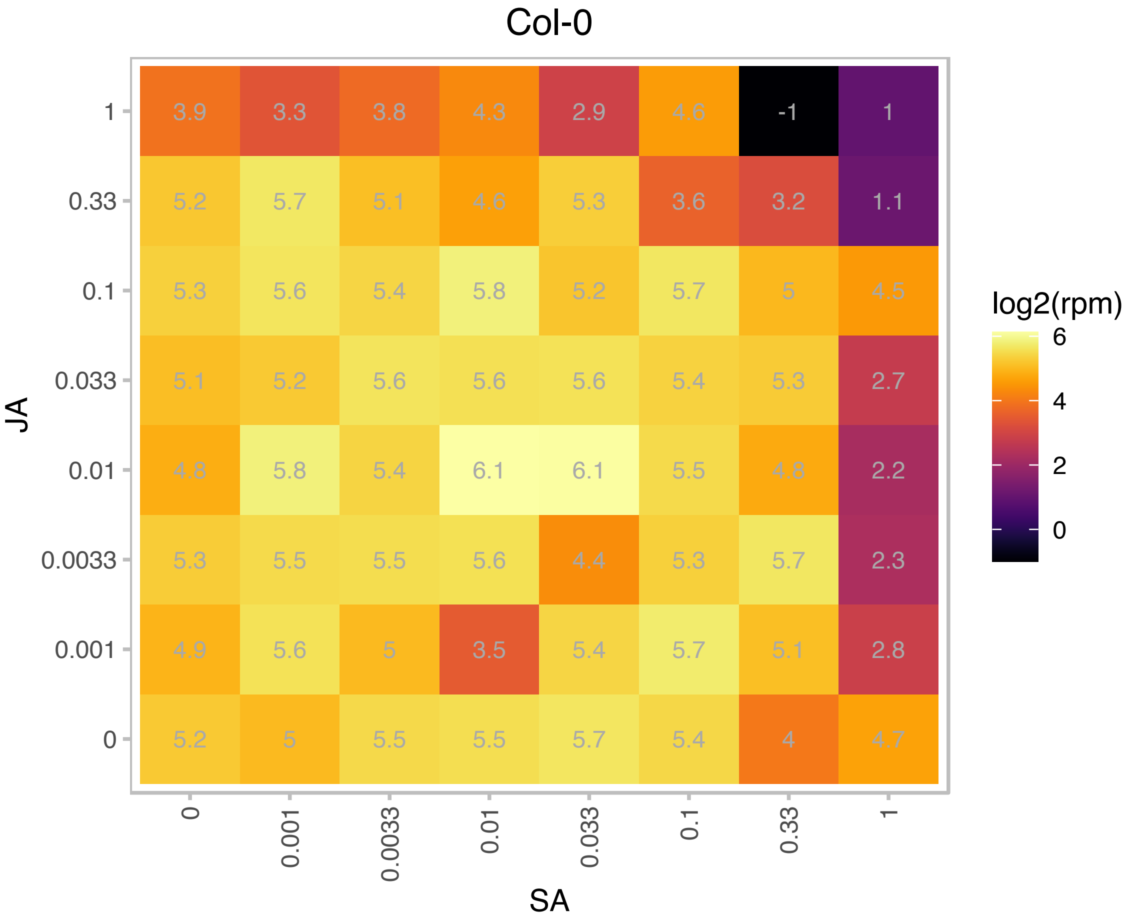


| | |
|---|---|
| Gene | AT3G18035 |
| is.expressed_genes | TRUE |
| is.responsive_genes | TRUE |
| is.apcluster | F-08 |

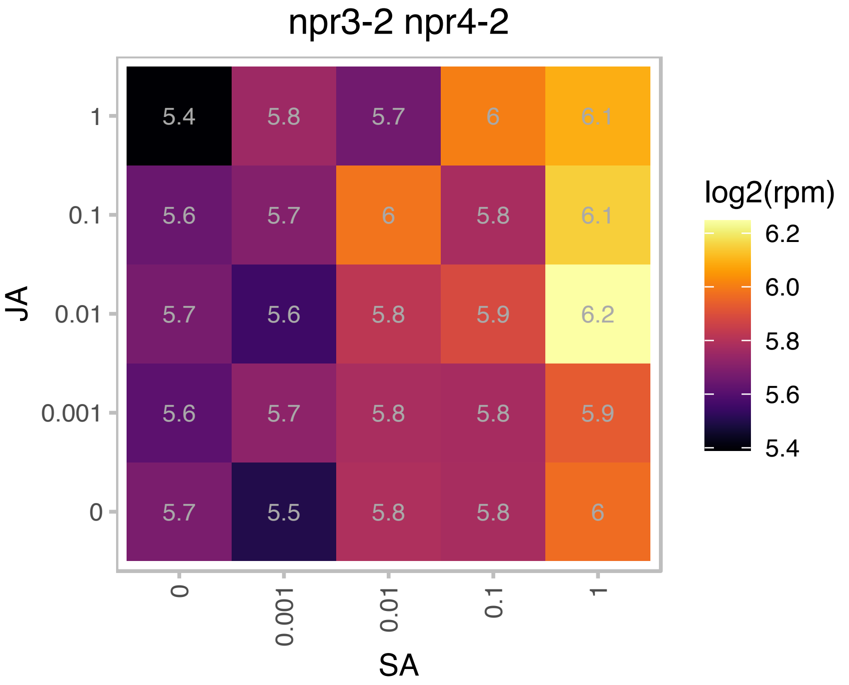


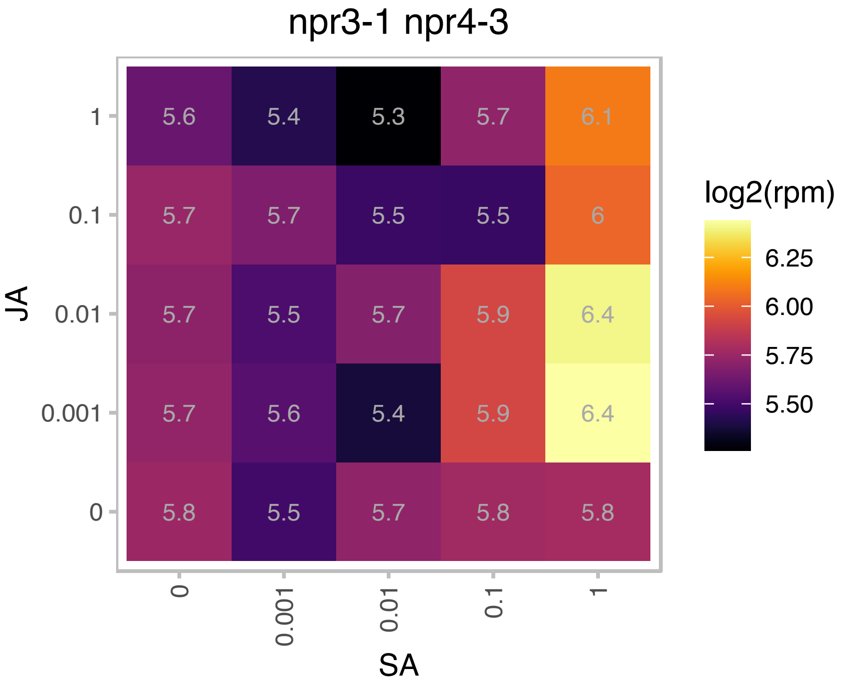

### AT3G18165.1 : MOS4

modifier of snc1,4

Encodes MOS4 (Modifier of snc1, 4), a nuclear protein homologous to human Breast Cancer-Amplified Sequence (BCAS2). MOS4 interacts with AtCDC5 and PRL1. All three proteins are essential for plant innate immunity.

Col-0

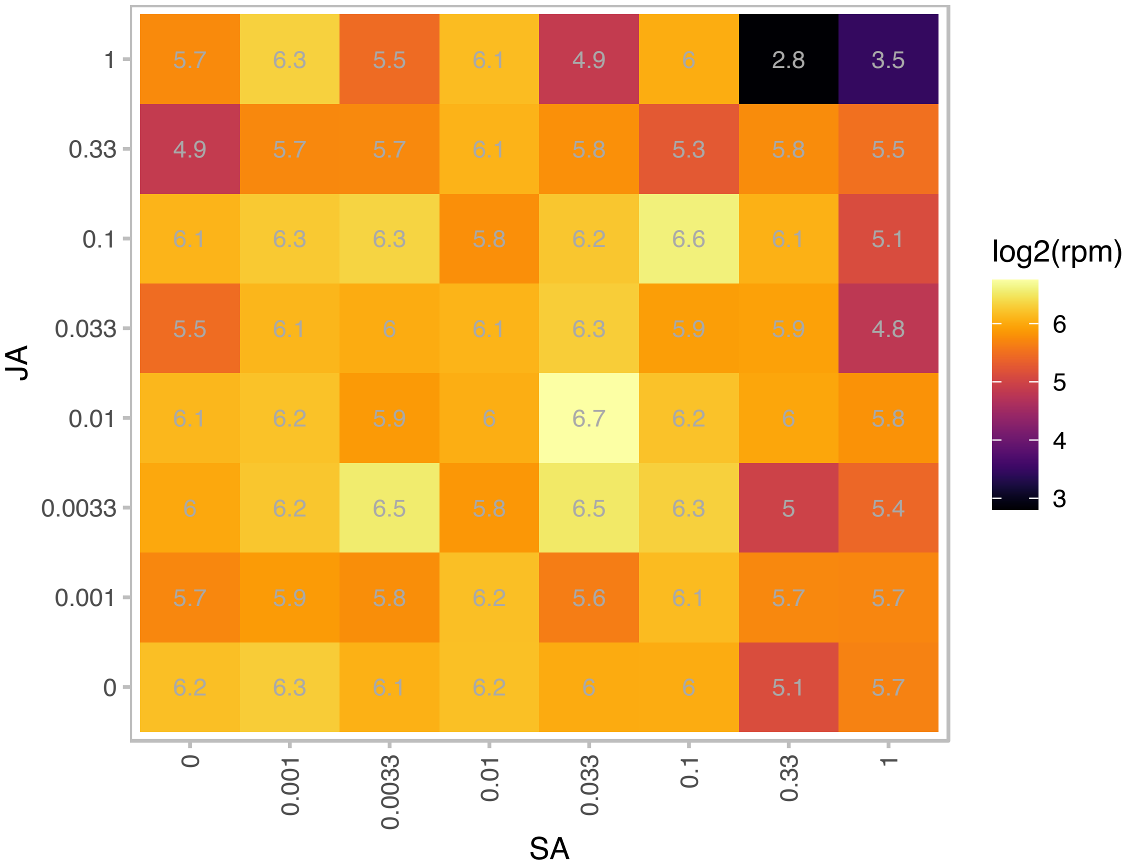


| | |
|---|---|
| Gene | AT3G18165 |
| is.expressed_genes | TRUE |
| is.responsive_genes | TRUE |
| is.apcluster | F-08 |

npr3-2 npr4-2

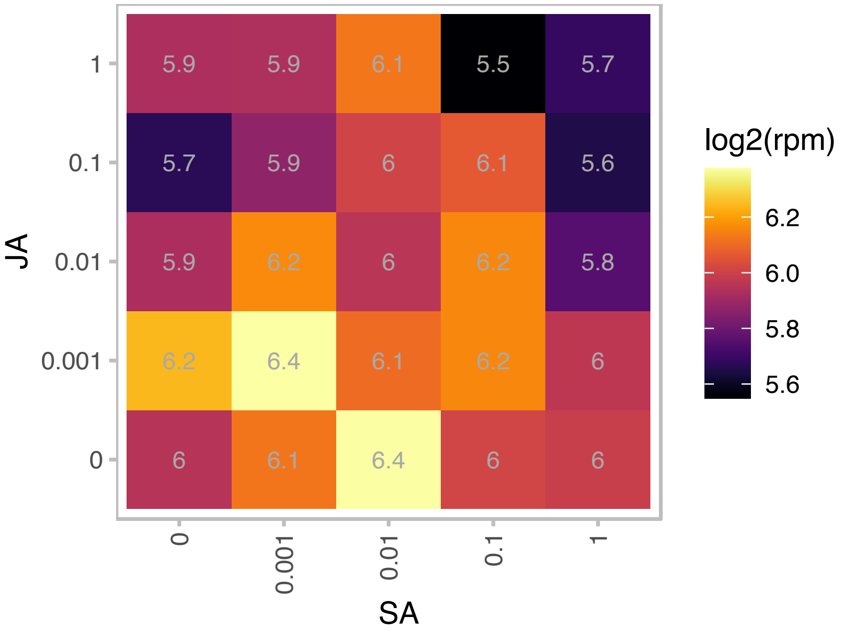


npr3-1 npr4-3

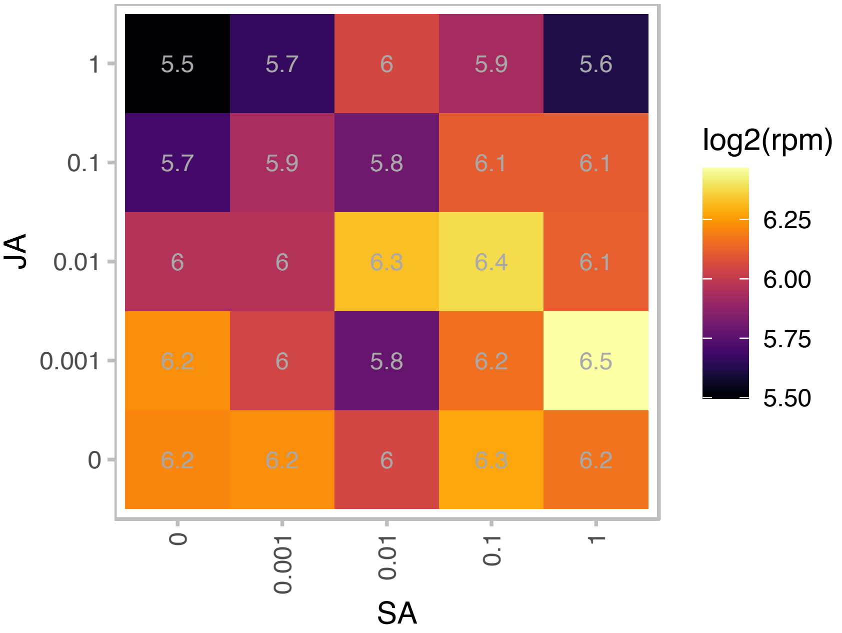

# AT3G19030.1 : NA

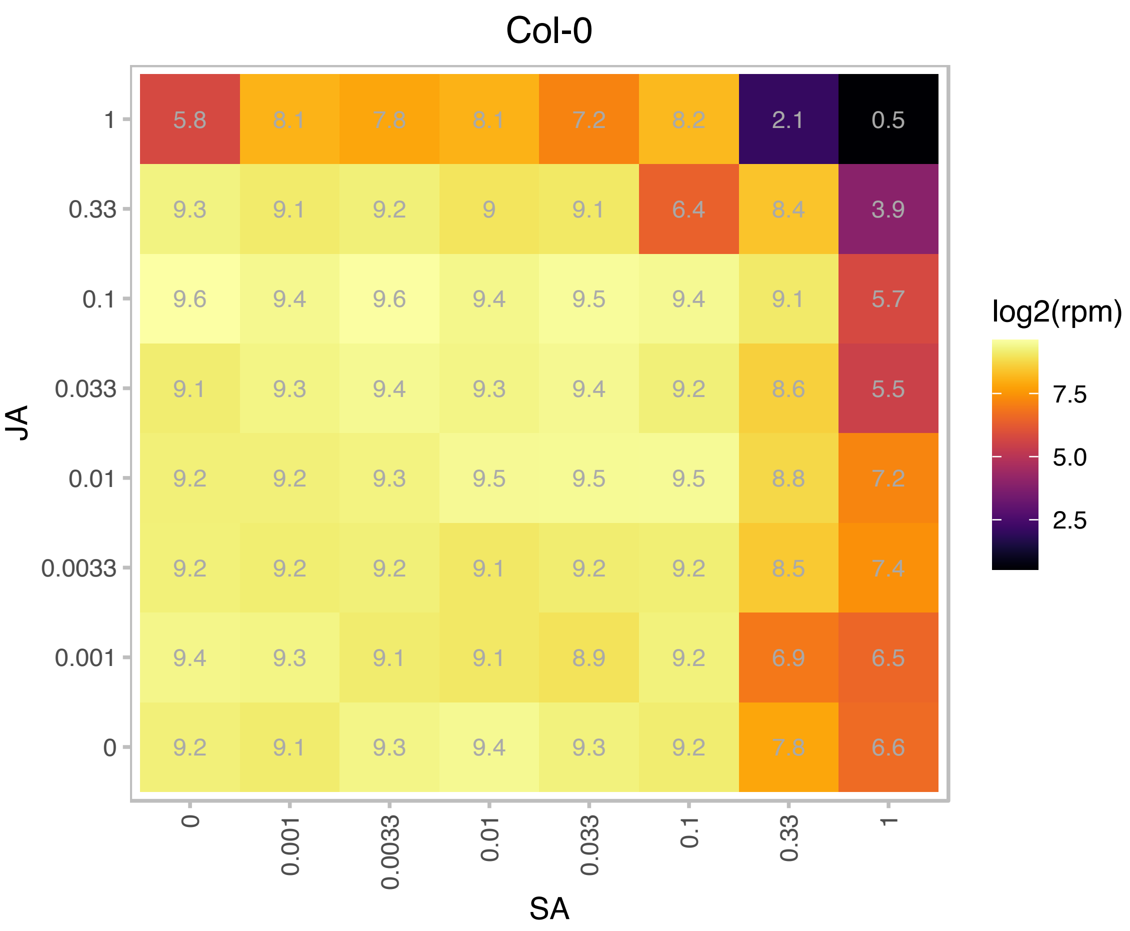


| | |
|---|---|
| Gene | AT3G19030 |
| is.expressed_genes | TRUE |
| is.responsive_genes | TRUE |
| is.apcluster | F-08 |

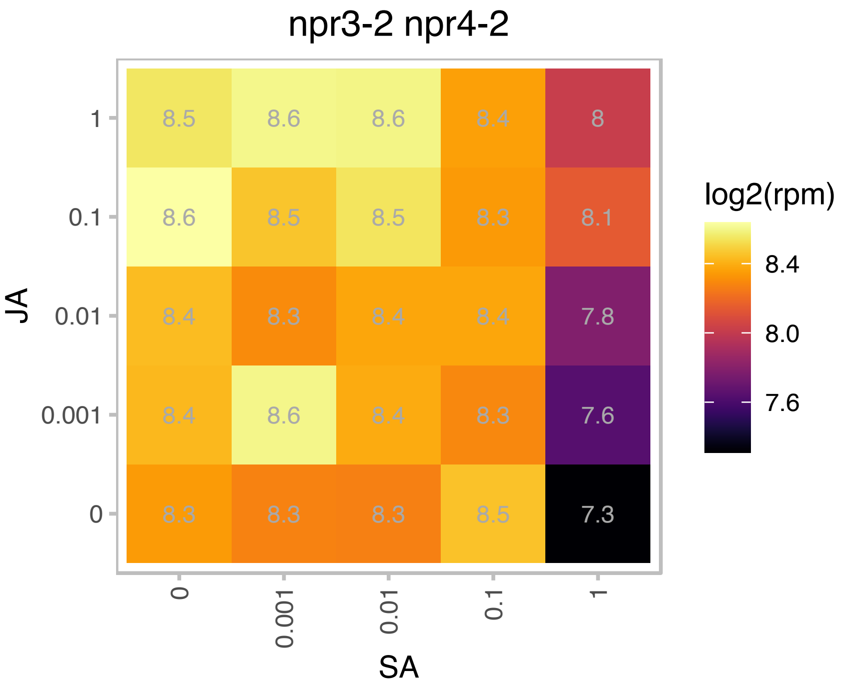


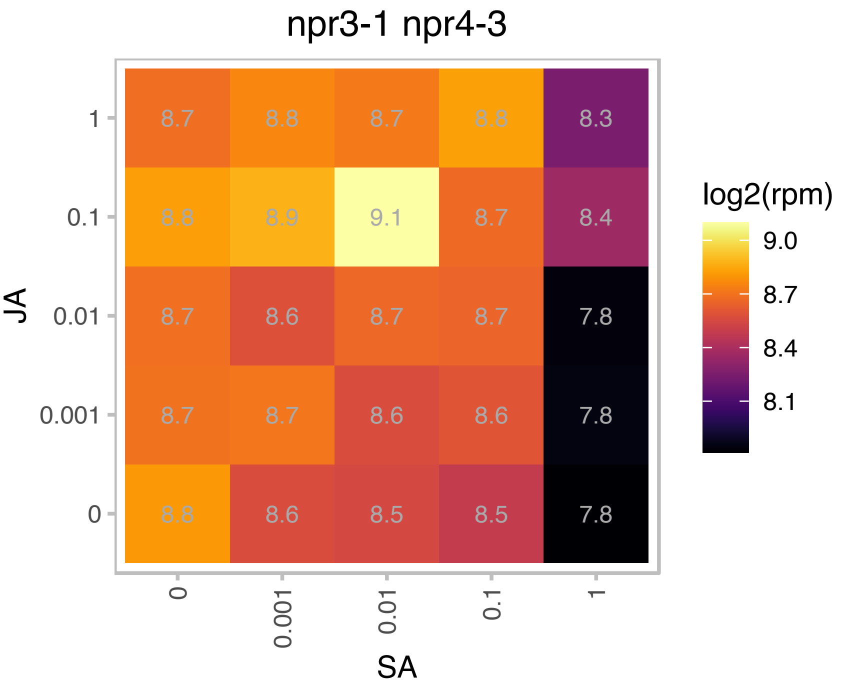

# AT3G19650.1 : NA

cyclin-related

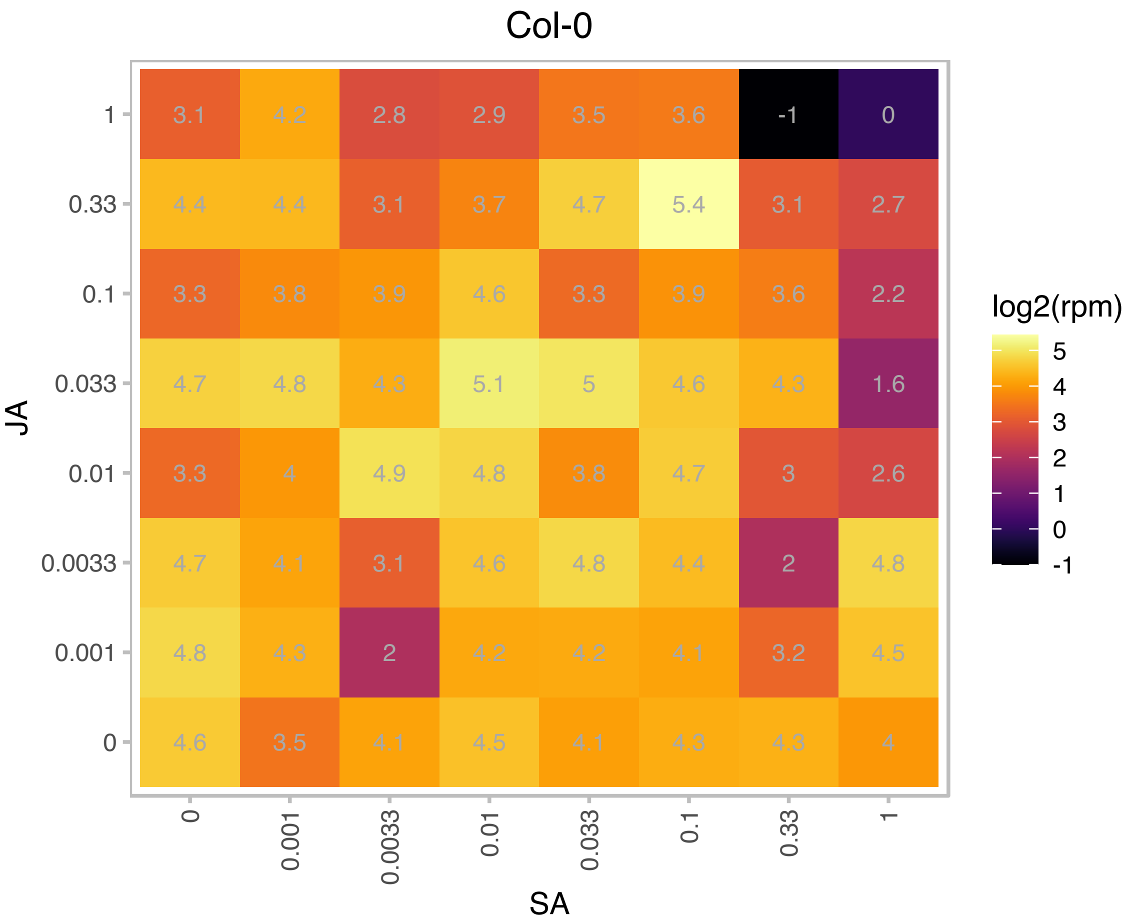


| | |
|---|---|
| Gene | AT3G19650 |
| is.expressed_genes | TRUE |
| is.responsive_genes | TRUE |
| is.apcluster | F-08 |

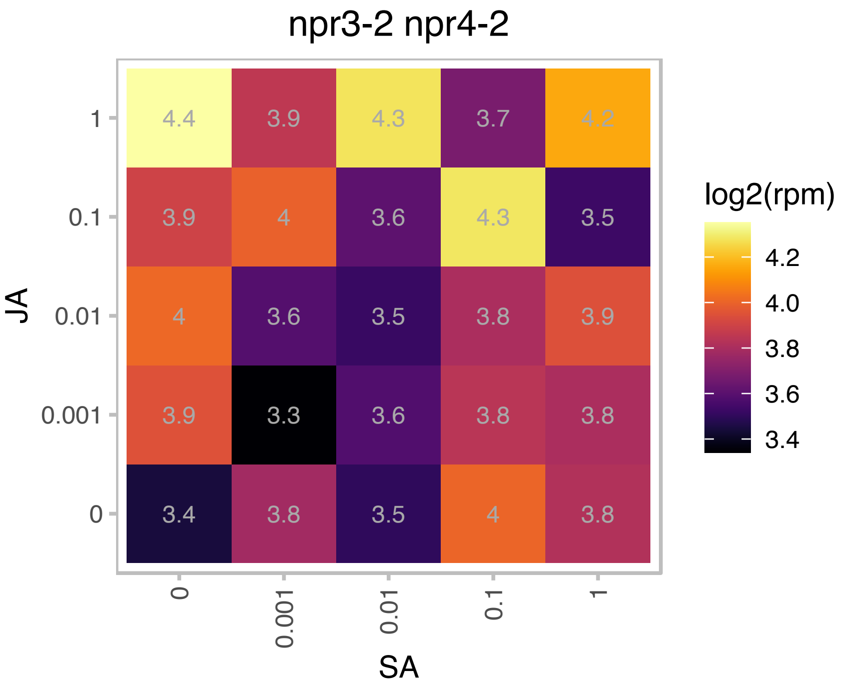


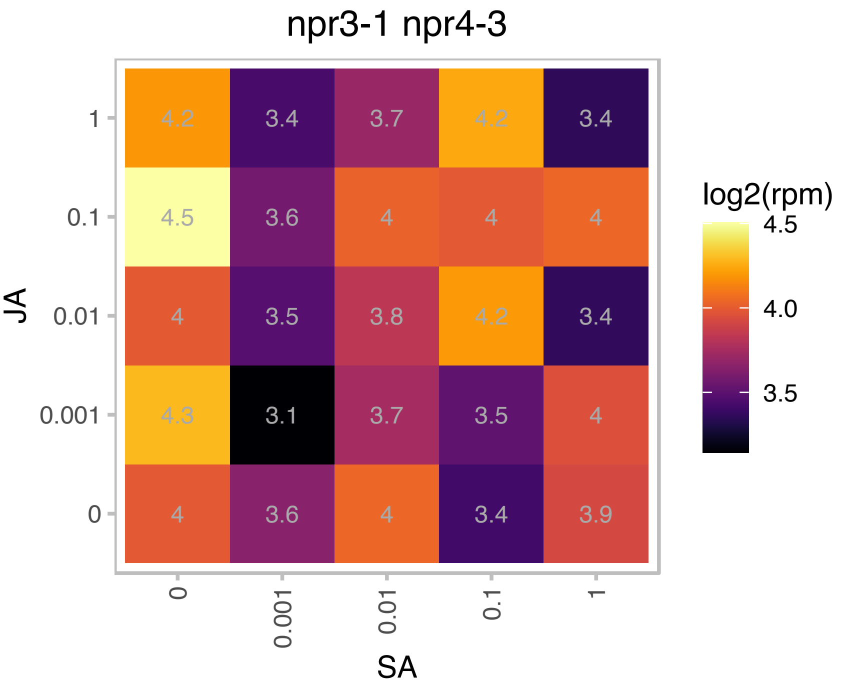

### AT3G21750.1 : UGT71B1

UDP-glucosyl transferase 71B1

Encodes a glucosyltransferase that can attach glucose to a number of hydroxylated phenolic compounds as well as quercetins in vitro

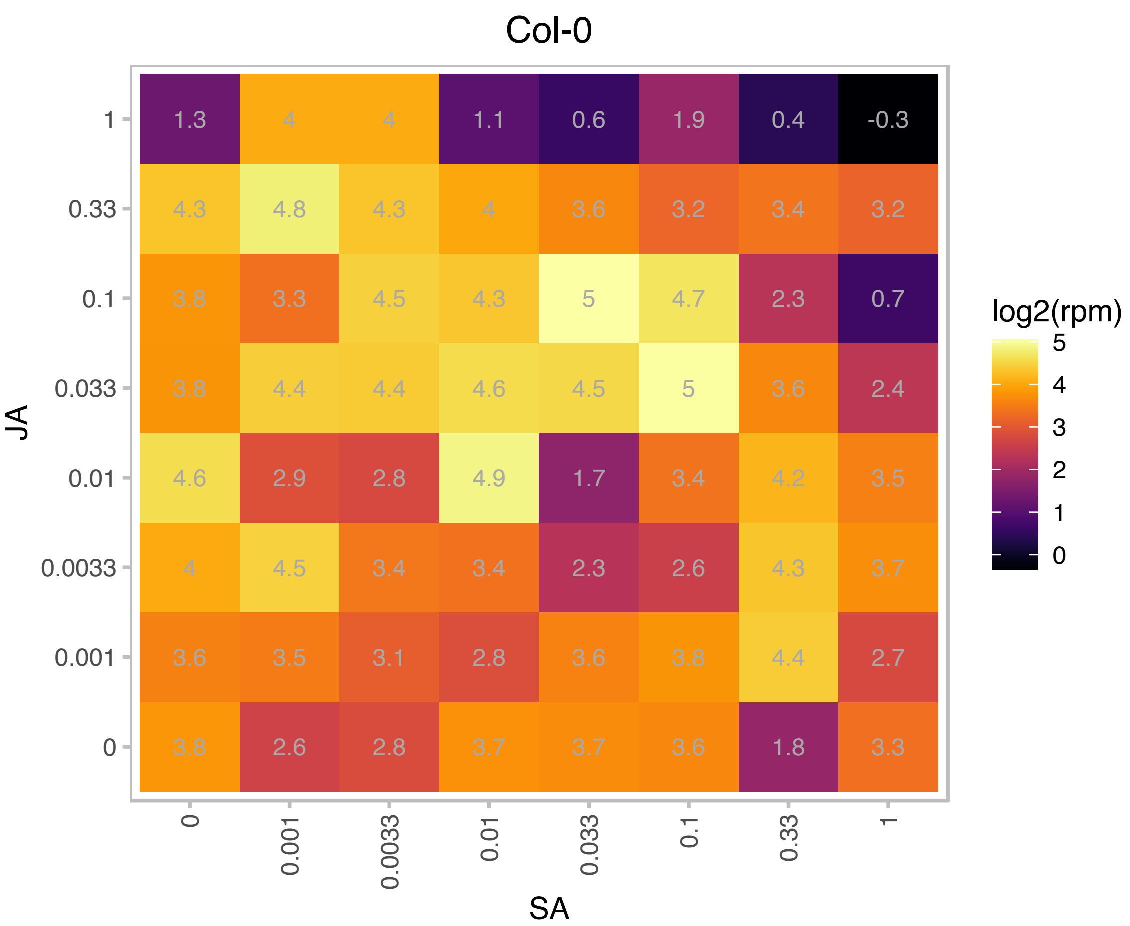


| | |
|---|---|
| Gene | AT3G21750 |
| is.expressed_genes | TRUE |
| is.responsive_genes | TRUE |
| is.apcluster | F-08 |

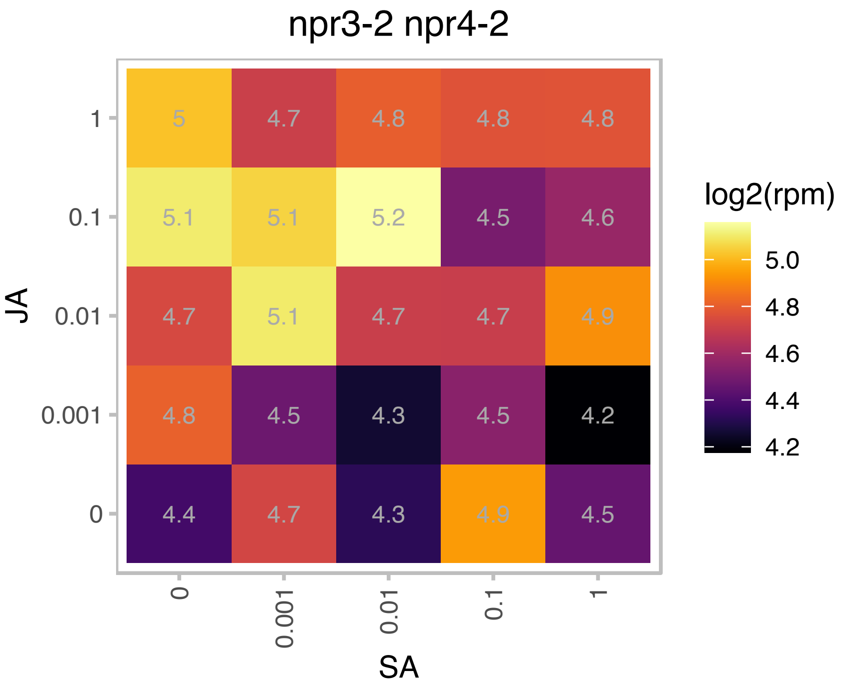


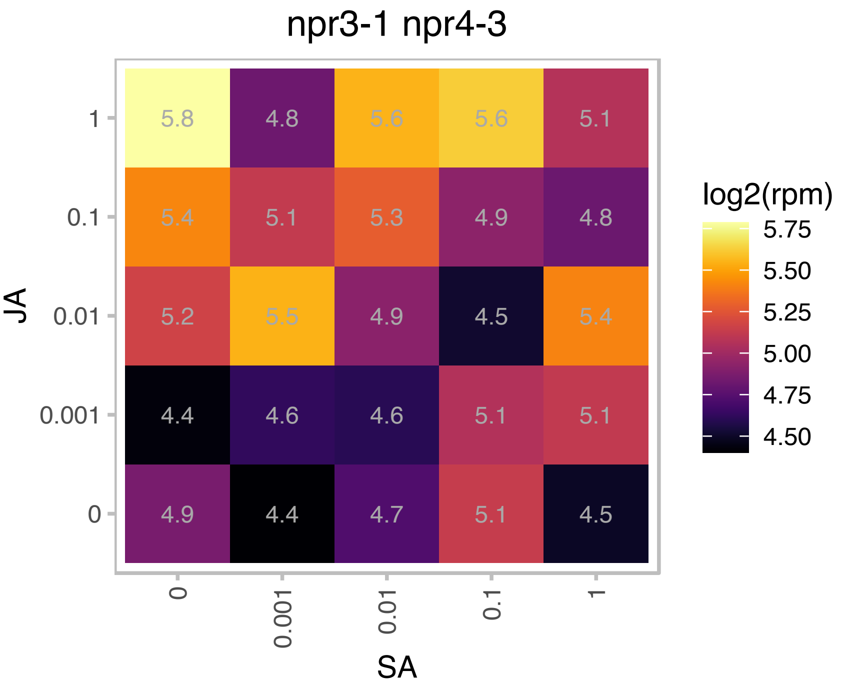

# AT3G22121.1 : NA

other RNA

Potential natural antisense gene, locus overlaps with AT3G22120 The mRNA is cell-to-cell mobile.

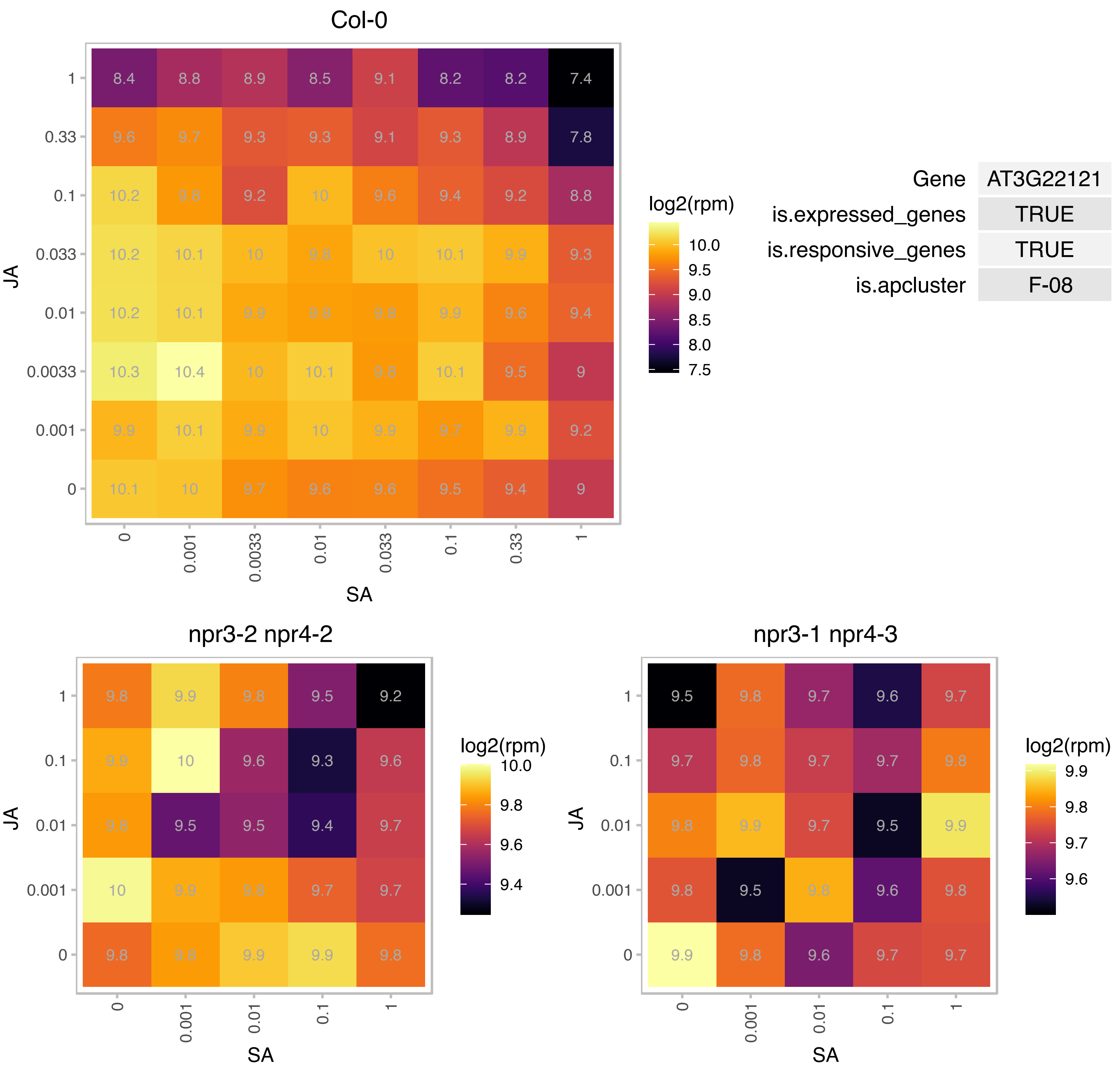


| | |
|---|---|
| Gene | AT3G22121 |
| is.expressed_genes | TRUE |
| is.responsive_genes | TRUE |
| is.apcluster | F-08 |

### AT3G22780.1 : ATTSO1, TSO1

Tesmin/TSO1-like CXC domain-containing protein

putative DNA binding protein (tso1) mRNA, tso1-3 allele,

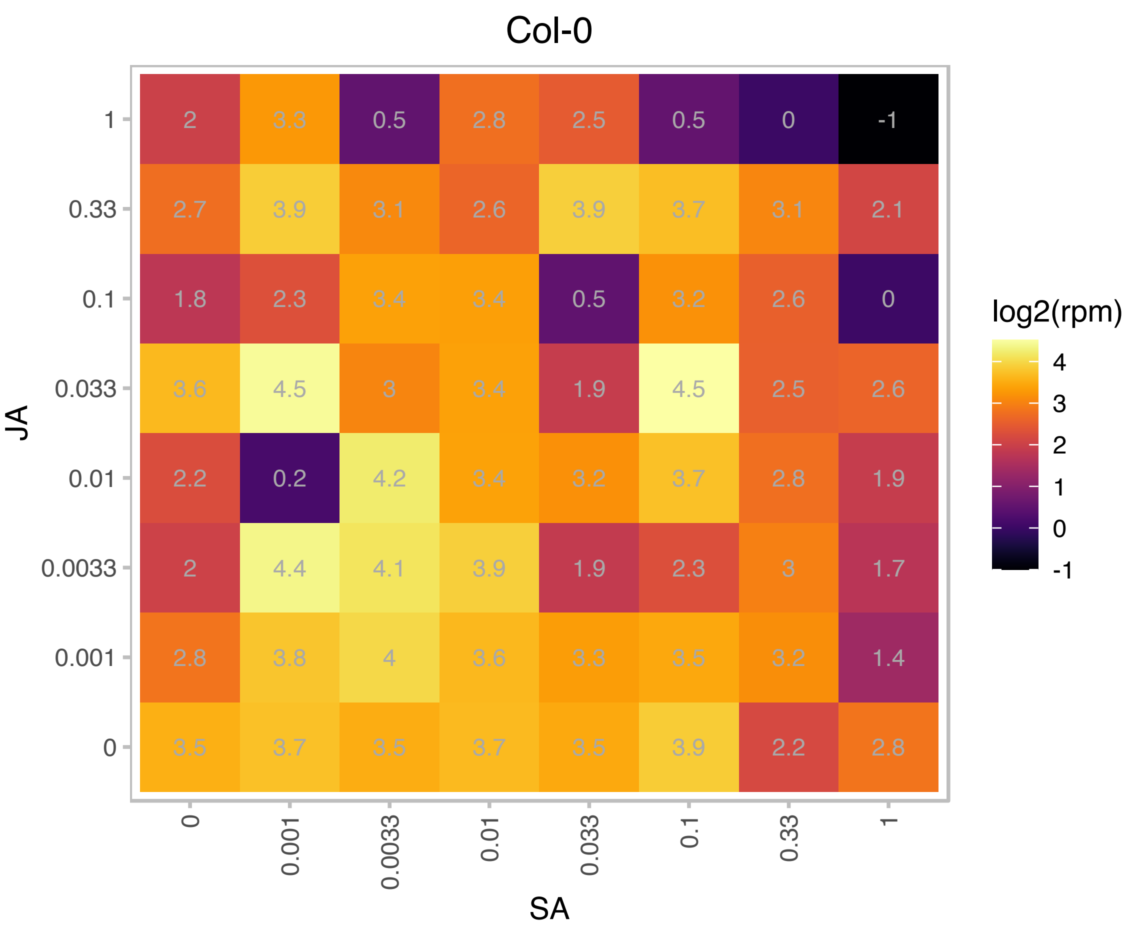


| | |
|---|---|
| Gene | AT3G22780 |
| is.expressed_genes | TRUE |
| is.responsive_genes | TRUE |
| is.apcluster | F-08 |

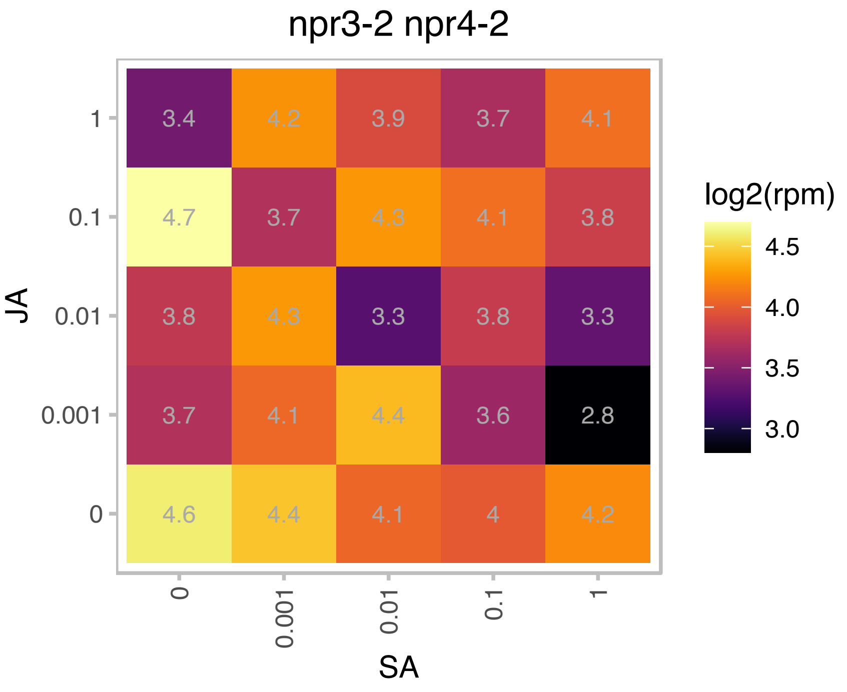


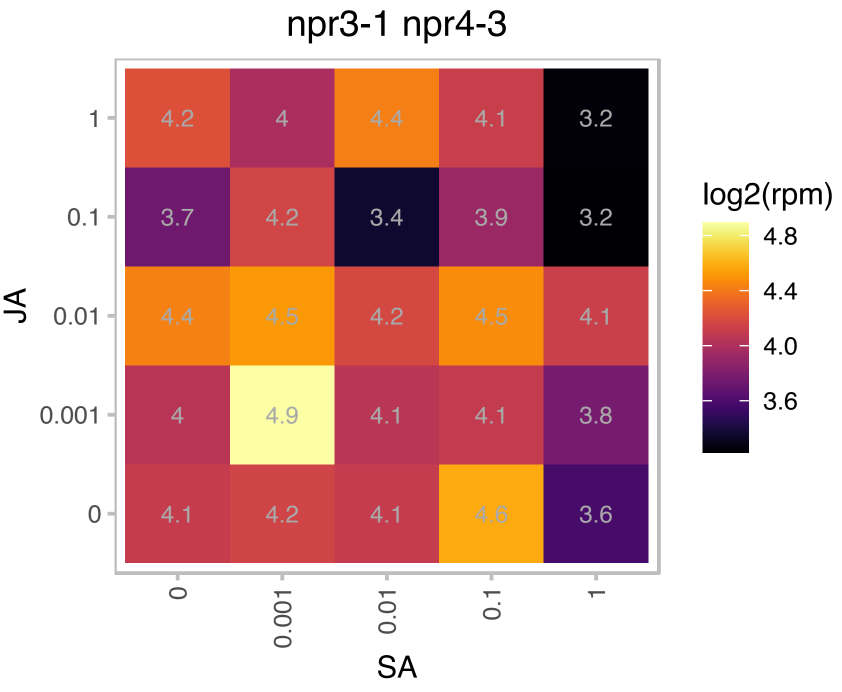

### AT3G25520.1 : ATL5, OLI5, PGY3, RPL5A

ribosomal protein L5

Encodes ribosomal protein L5 that binds to 5S ribosomal RNA and in involved in its export from the nucleus to the cytoplasm. Identified in a screen for enhancers of as1. as1/pgy double mutants show defects in leaf vascular patterning and adaxial cell fate. Double mutant analysis indicates pgy genes function in the same pathway as REV, KAN1 and KAN2.

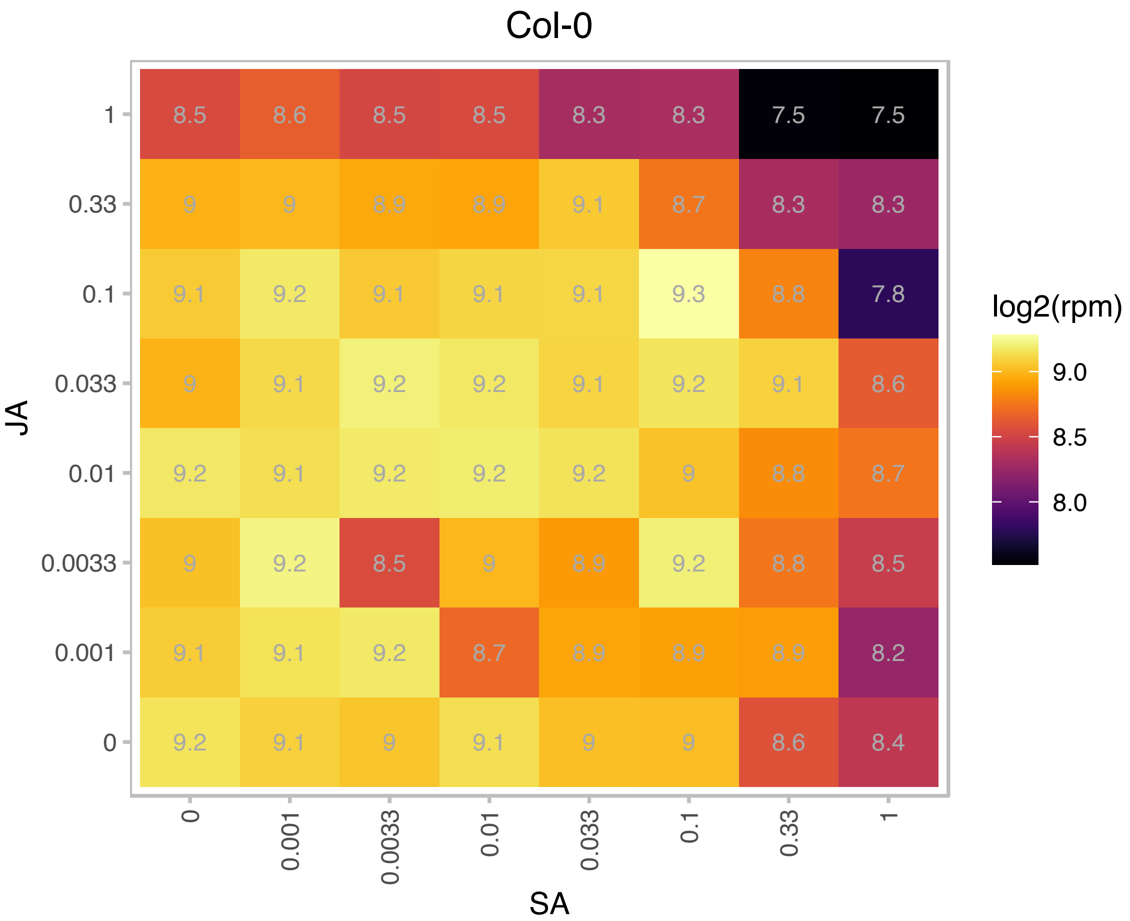


| | |
|---|---|
| Gene | AT3G25520 |
| is.expressed_genes | TRUE |
| is.responsive_genes | TRUE |
| is.apcluster | F-08 |

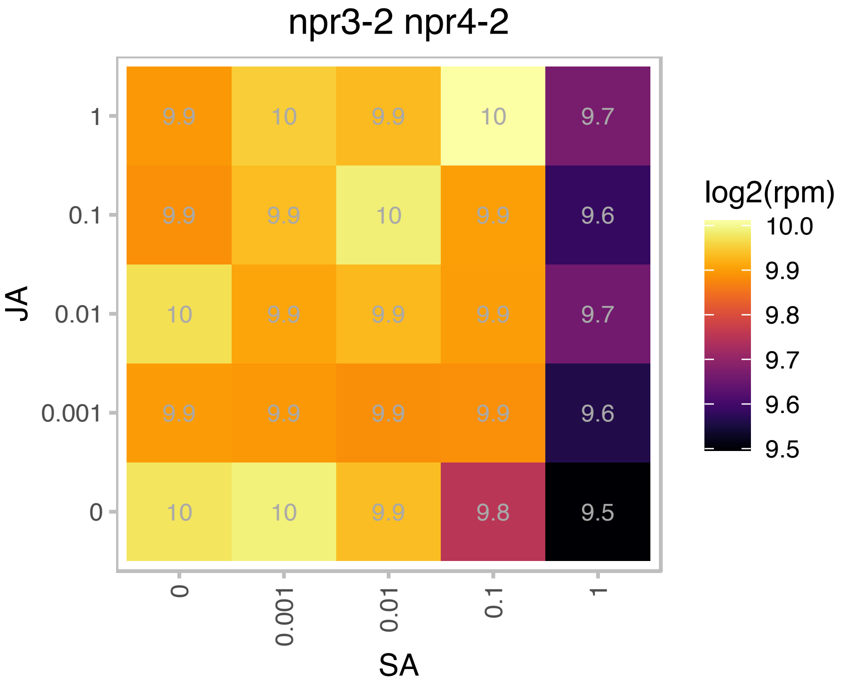


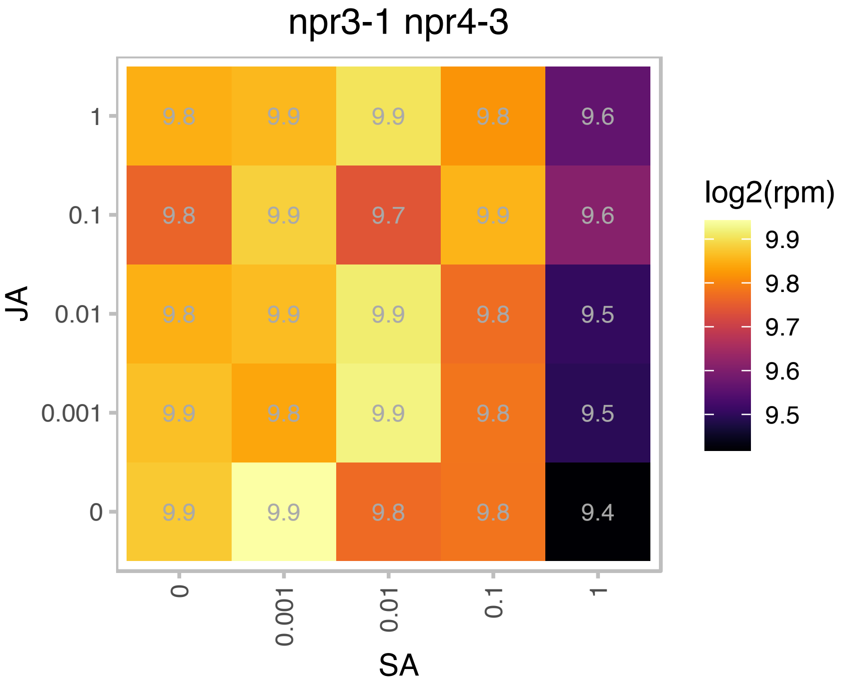

# AT3G27050.1 : NA

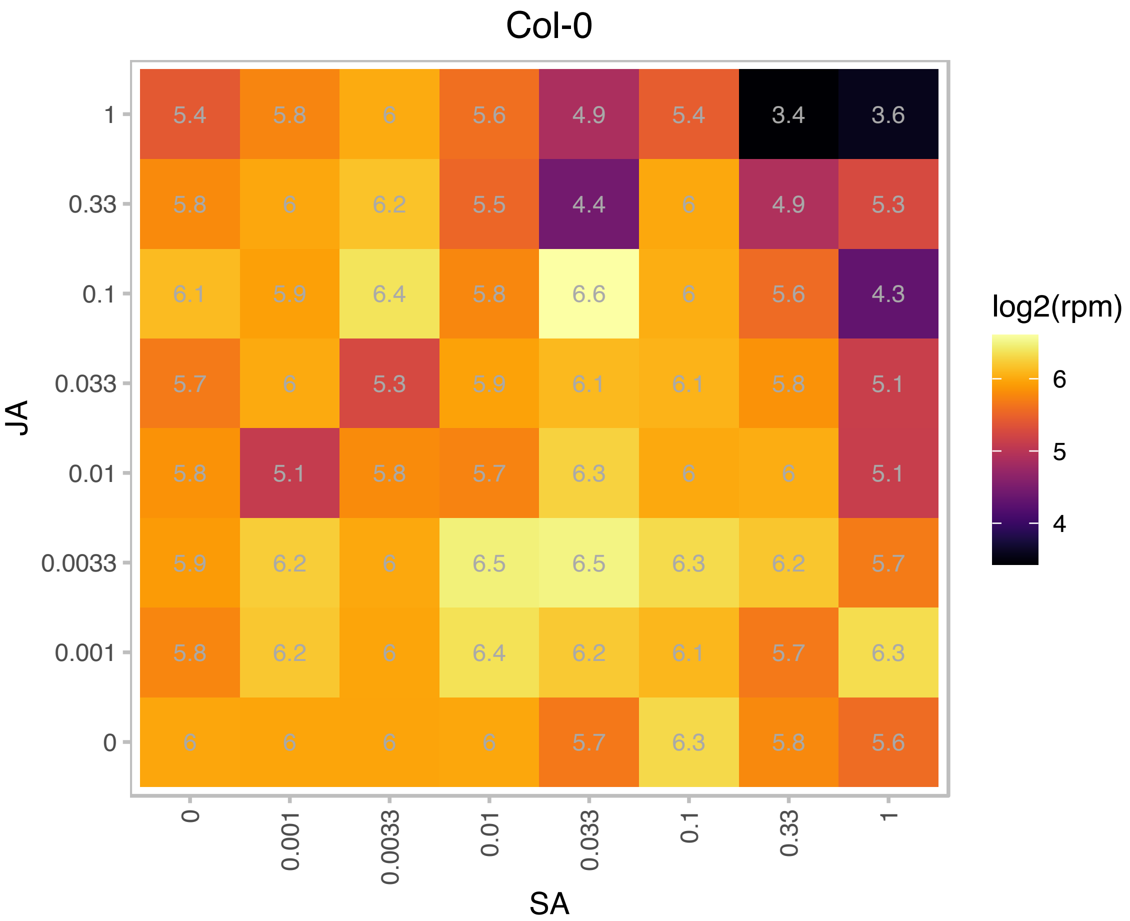


| | |
|---|---|
| Gene | AT3G27050 |
| is.expressed_genes | TRUE |
| is.responsive_genes | TRUE |
| is.apcluster | F-08 |

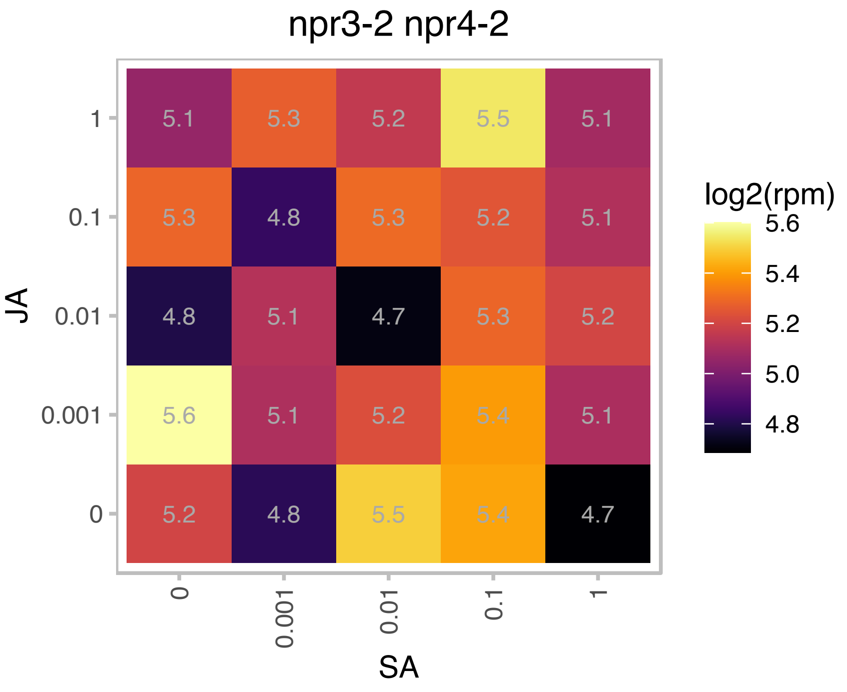


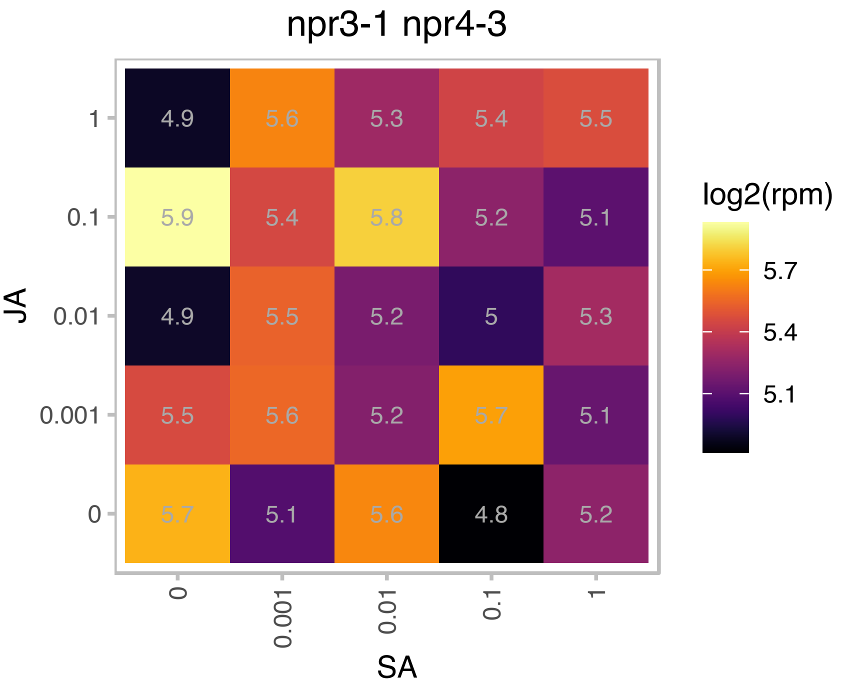

### AT3G27360.1 : H3.1, HTR3

Histone superfamily protein

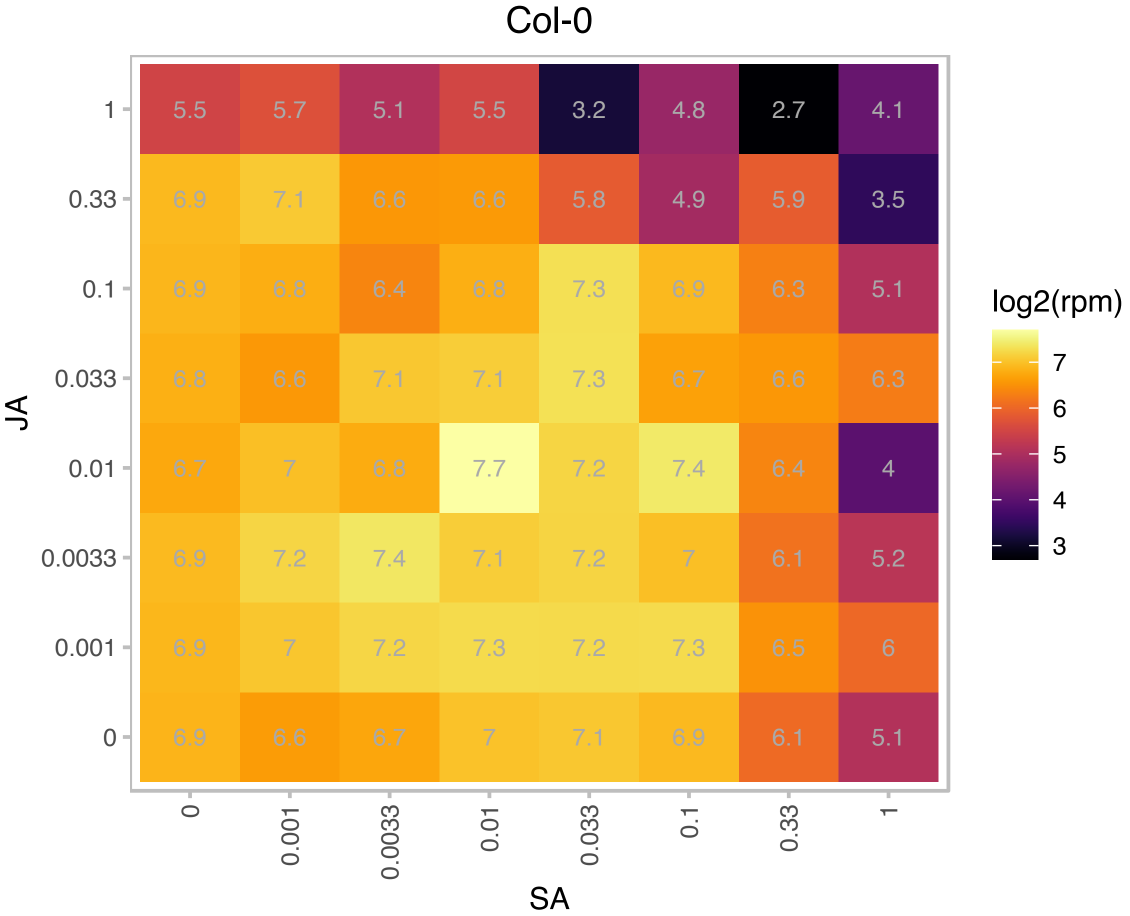


| | |
|---|---|
| Gene | AT3G27360 |
| is.expressed_genes | TRUE |
| is.responsive_genes | TRUE |
| is.apcluster | F-08 |

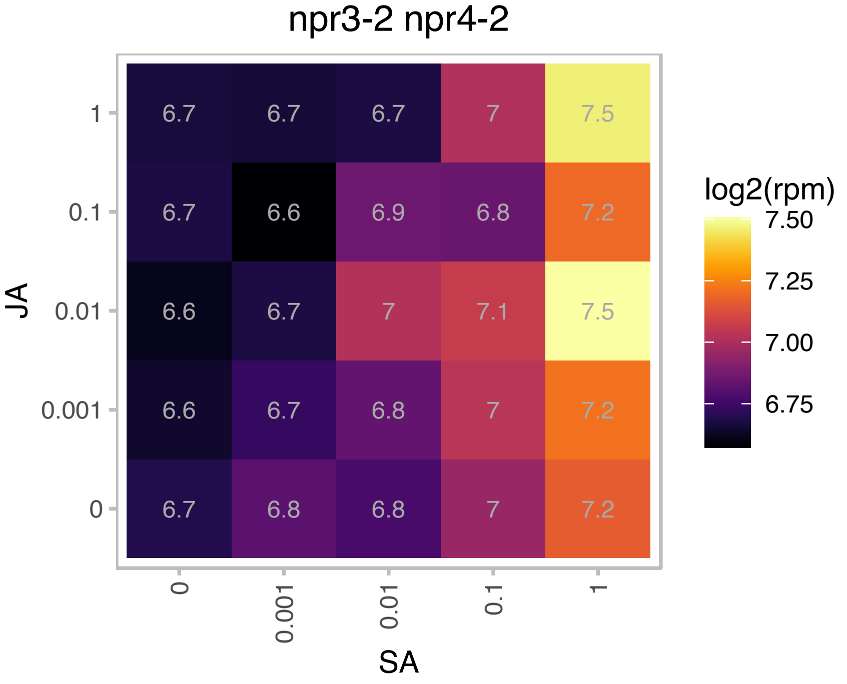


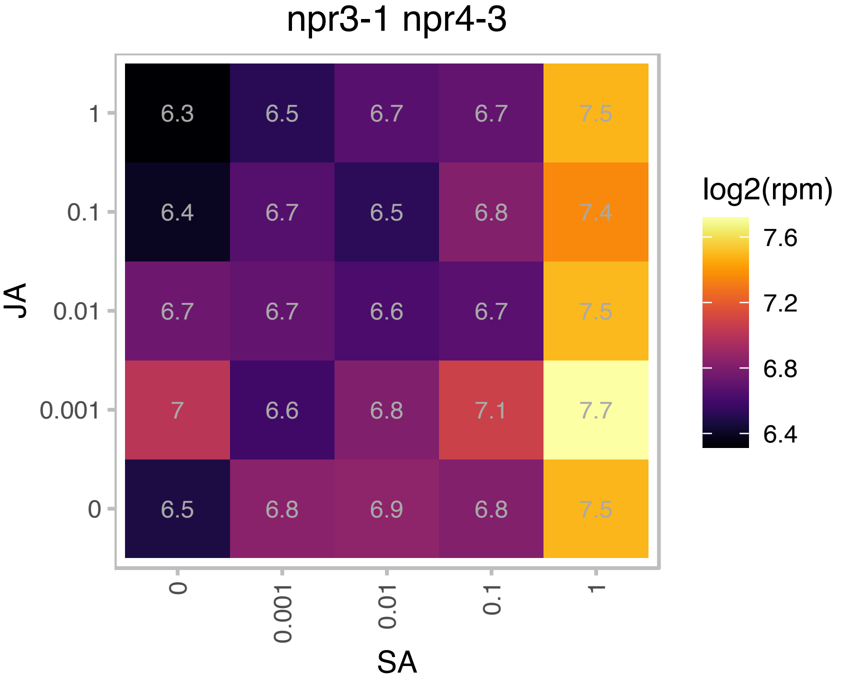

# AT3G27906.1 : NA

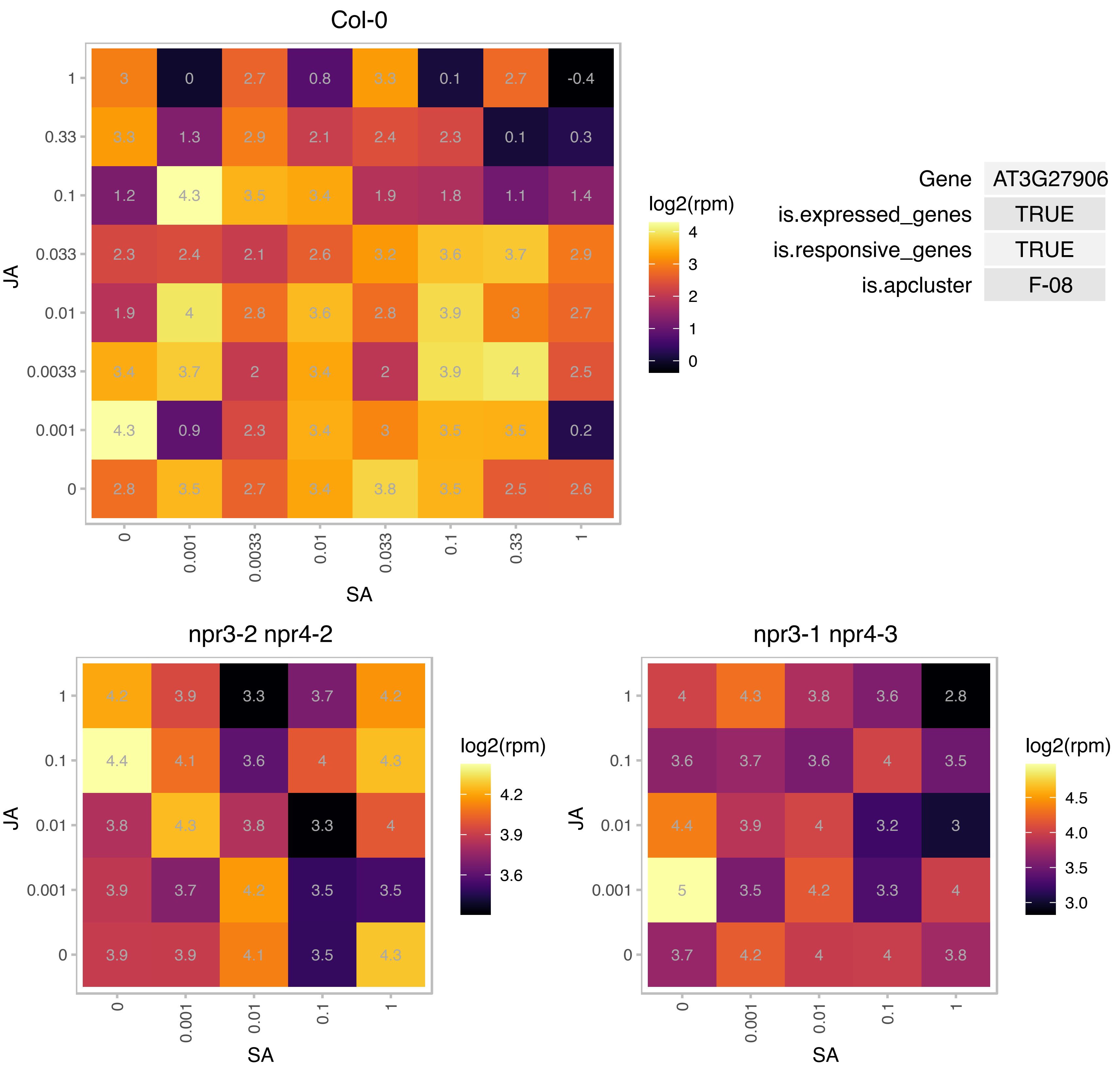


| | |
|---|---|
| Gene | AT3G27906 |
| is.expressed_genes | TRUE |
| is.responsive_genes | TRUE |
| is.apcluster | F-08 |

### AT3G44750.1 : ATHD2A, HD2A, HDA3, HDT1

histone deacetylase 3

Encodes a histone deacetylase. Controls the development of adaxial/abaxial leaf polarity. Two lines with RNAi-directed against this gene show reduced Agrobacterium-mediated DNA transformation of the roots.

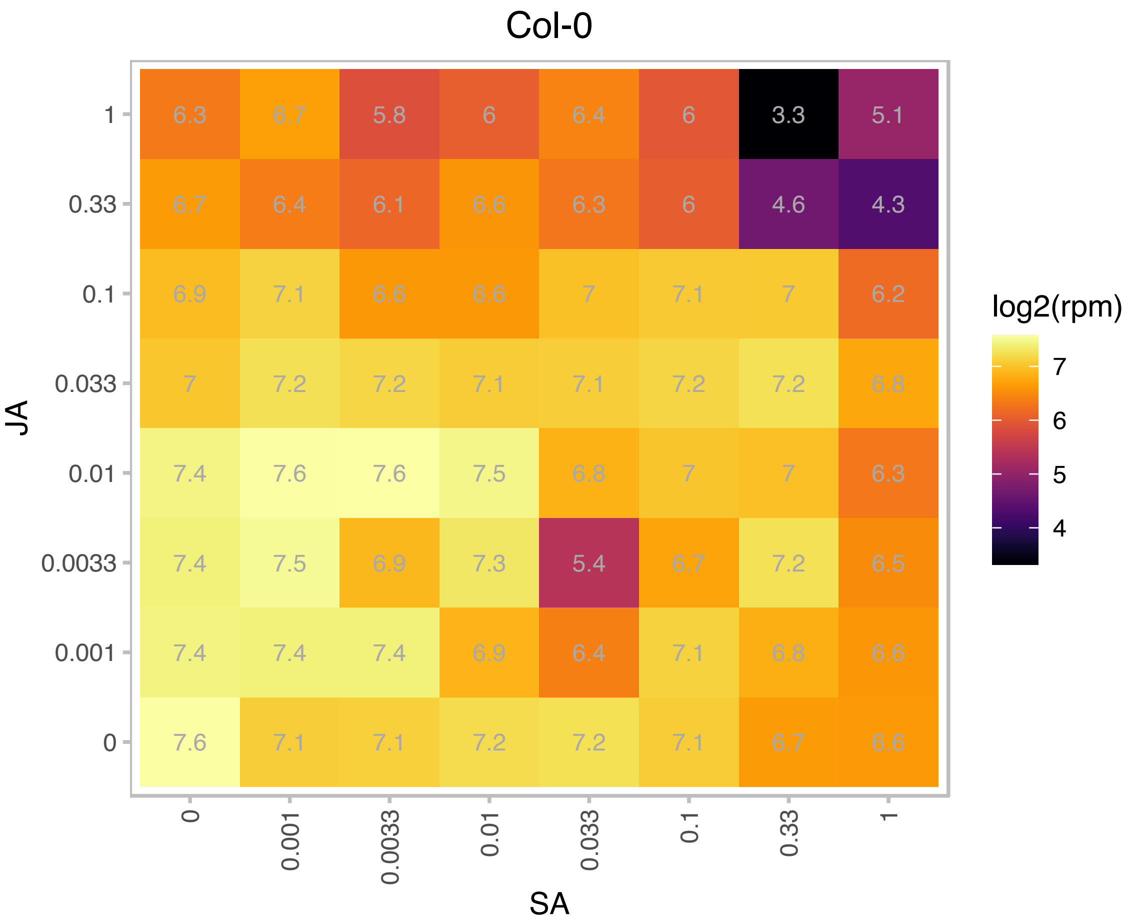


| | |
|---|---|
| Gene | AT3G44750 |
| is.expressed_genes | TRUE |
| is.responsive_genes | TRUE |
| is.apcluster | F-08 |

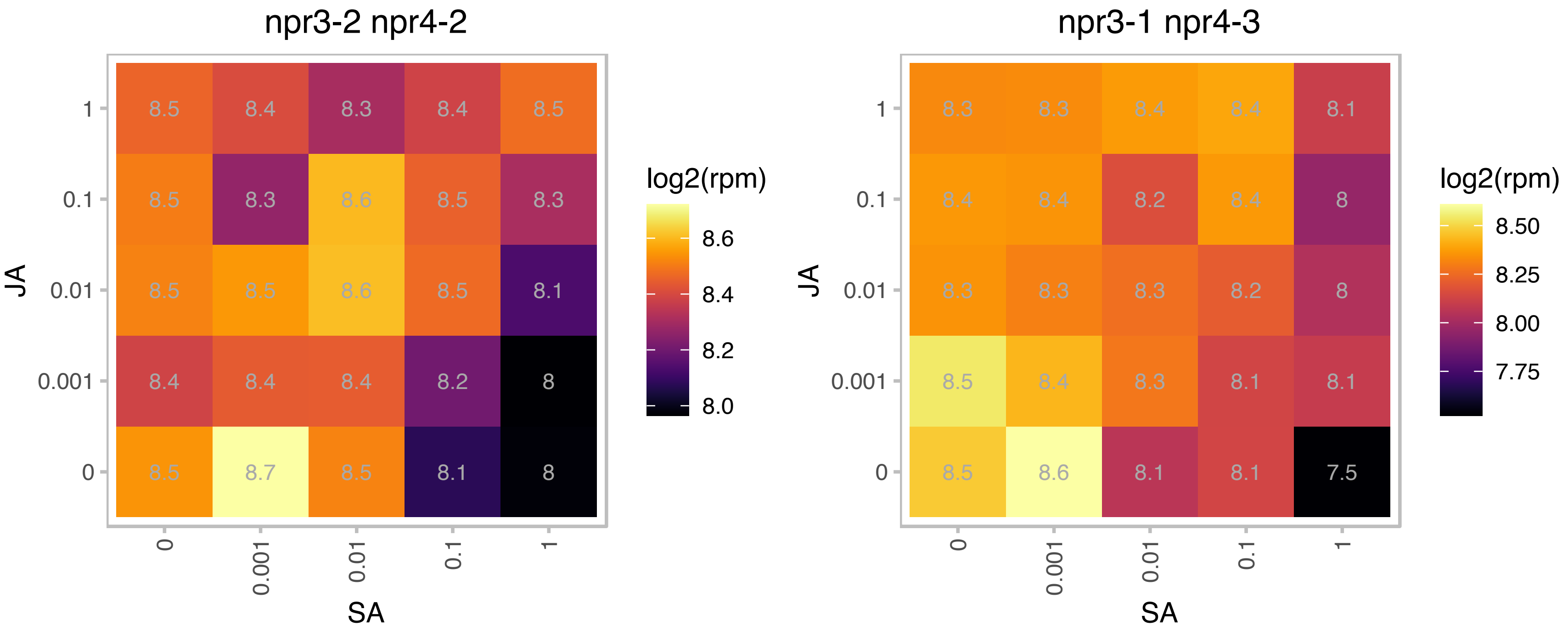

# AT3G45930.1 : NA

Histone superfamily protein

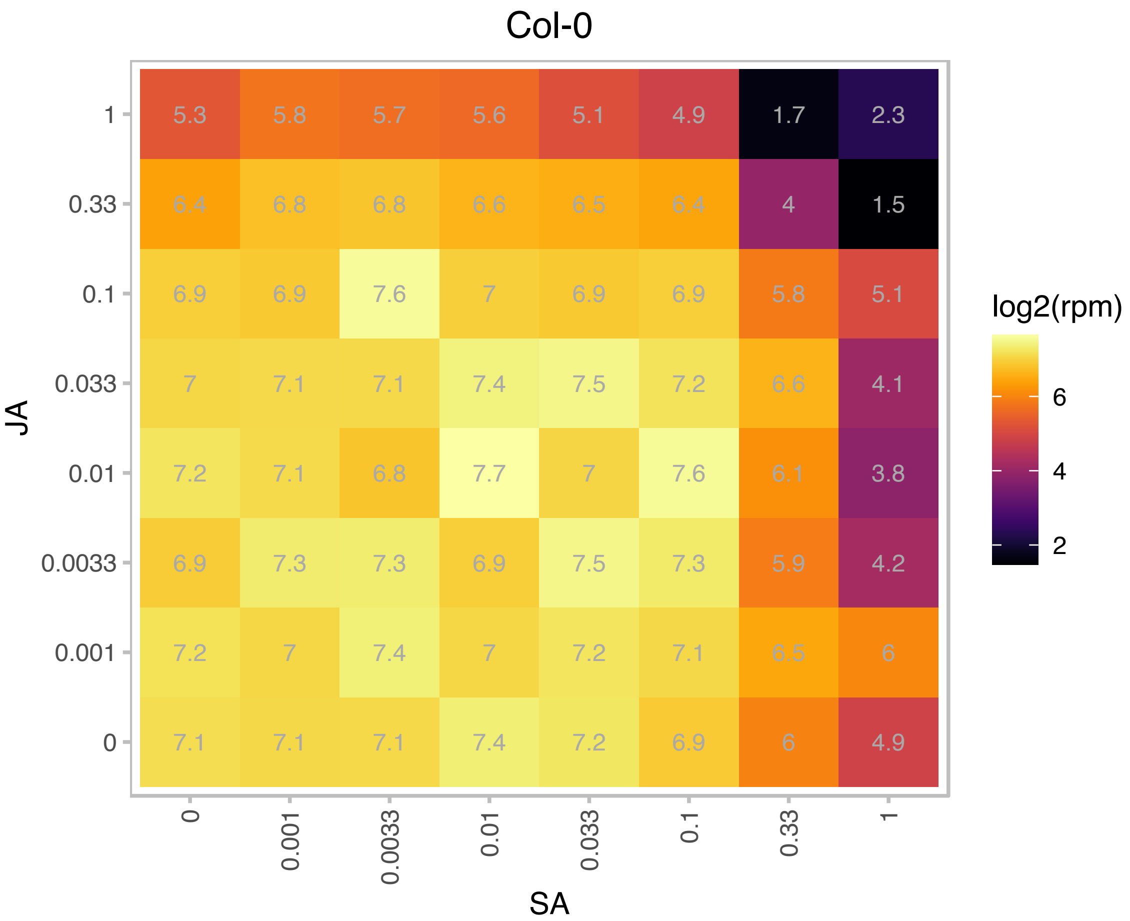


| | |
|---|---|
| Gene | AT3G45930 |
| is.expressed_genes | TRUE |
| is.responsive_genes | TRUE |
| is.apcluster | F-08 |

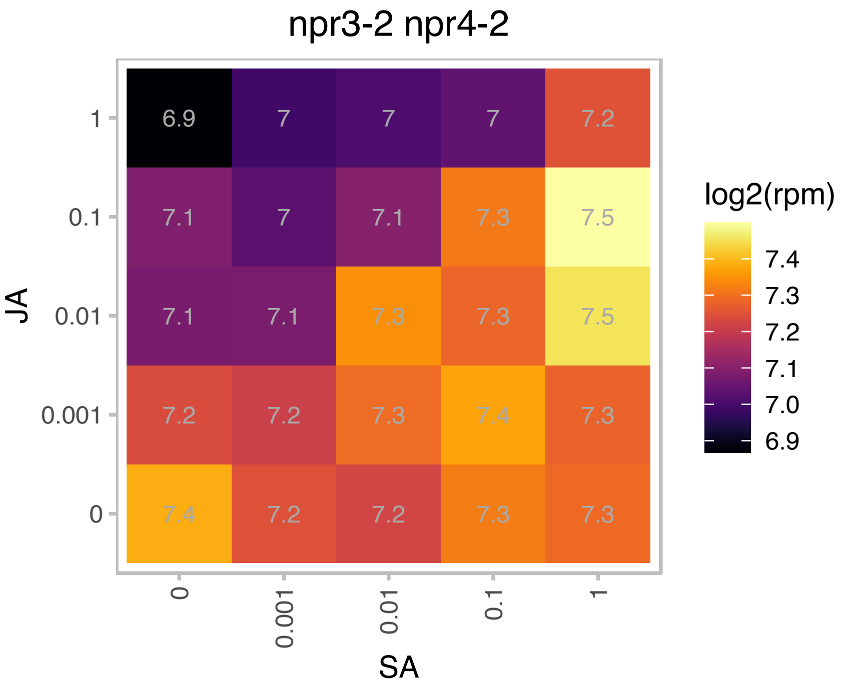


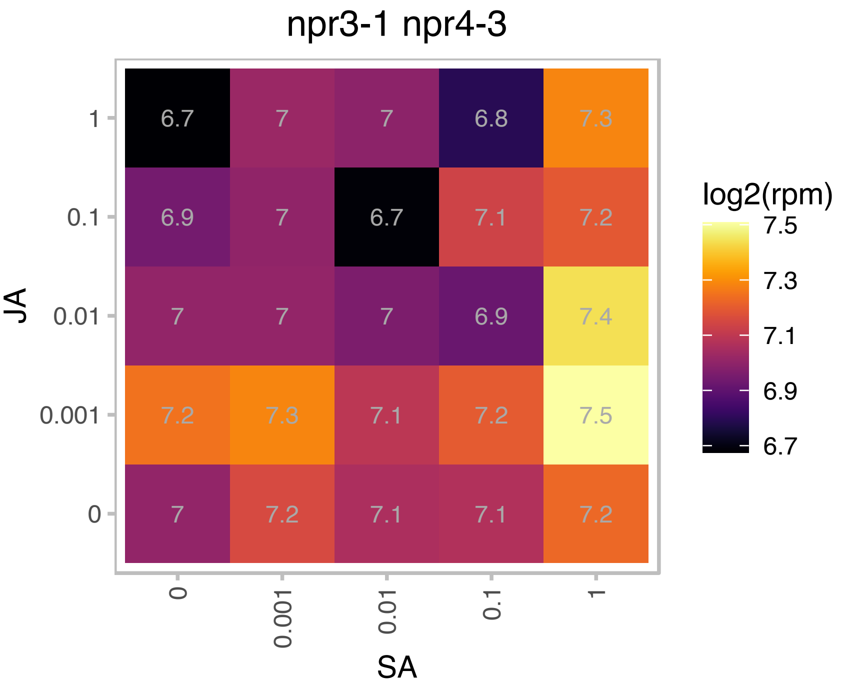

### AT3G45980.1 : H2B, HTB9

Histone superfamily protein
Encodes a histone 2B (H2B) protein. This protein can be ubiquitinated in planta, and this modification depends on the HUB1 and HUB2 E3 ubiquitin ligases as well as the UBC1 and UBC2 E2 ubiquitin conjugating enzymes. Lysine 146 appears to be the site of the ubiquitin addition.

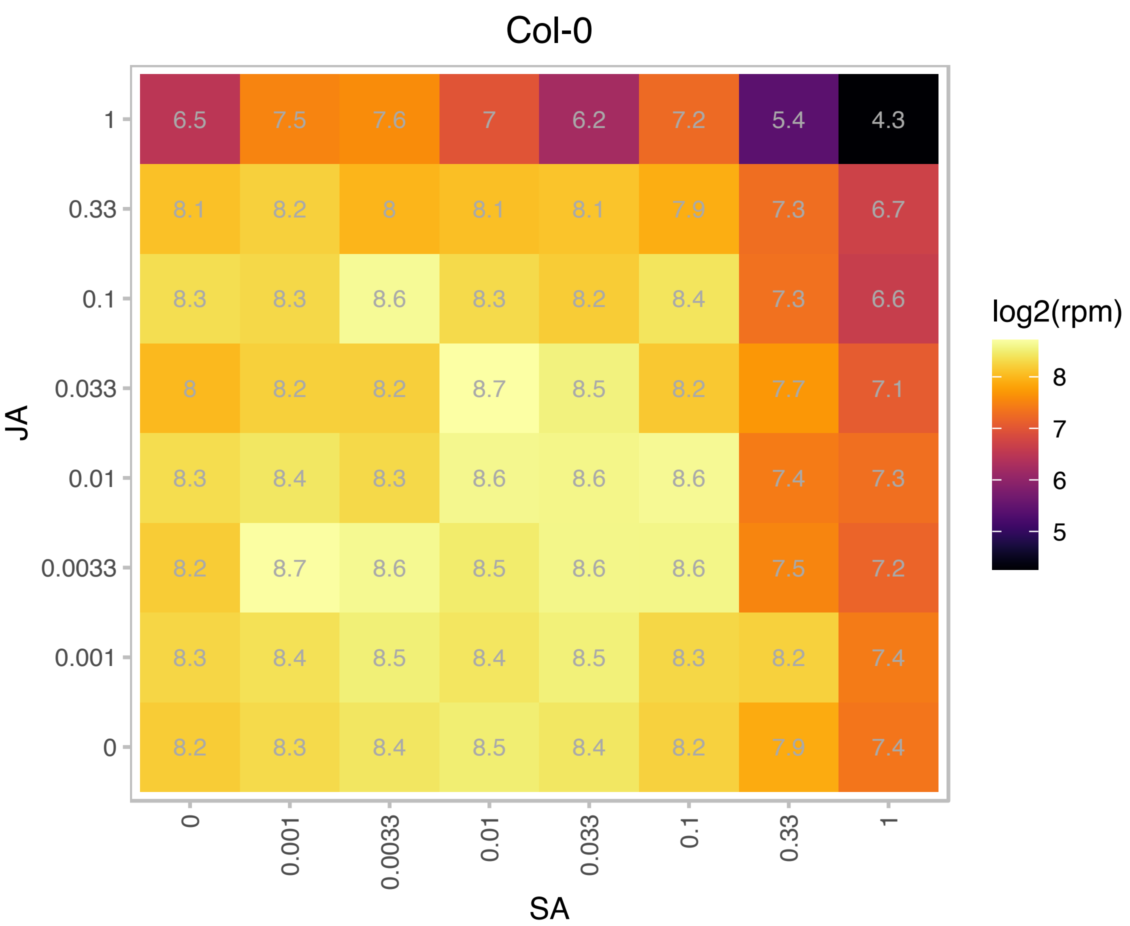


| | |
|---|---|
| Gene | AT3G45980 |
| is.expressed_genes | TRUE |
| is.responsive_genes | TRUE |
| is.apcluster | F-08 |

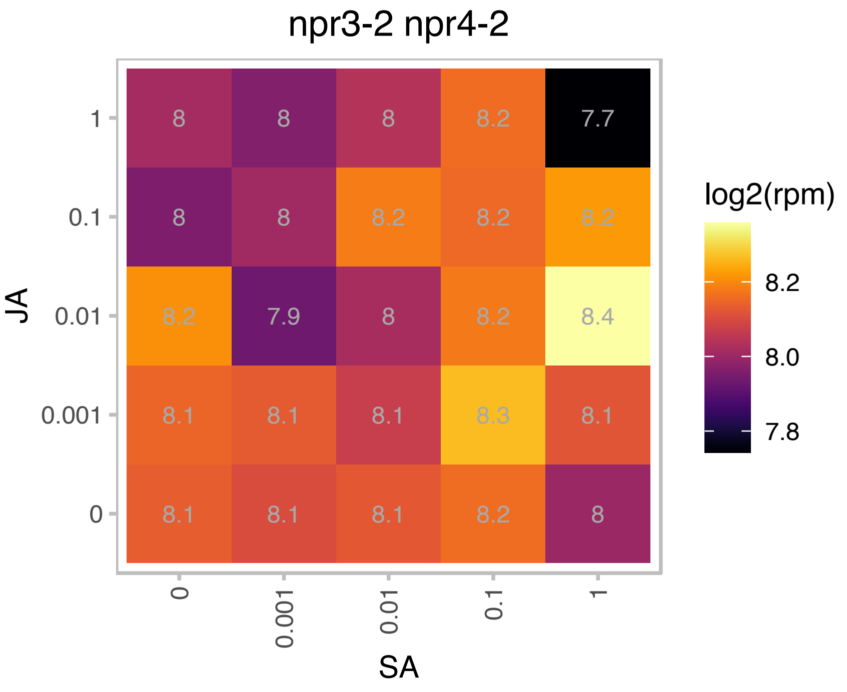


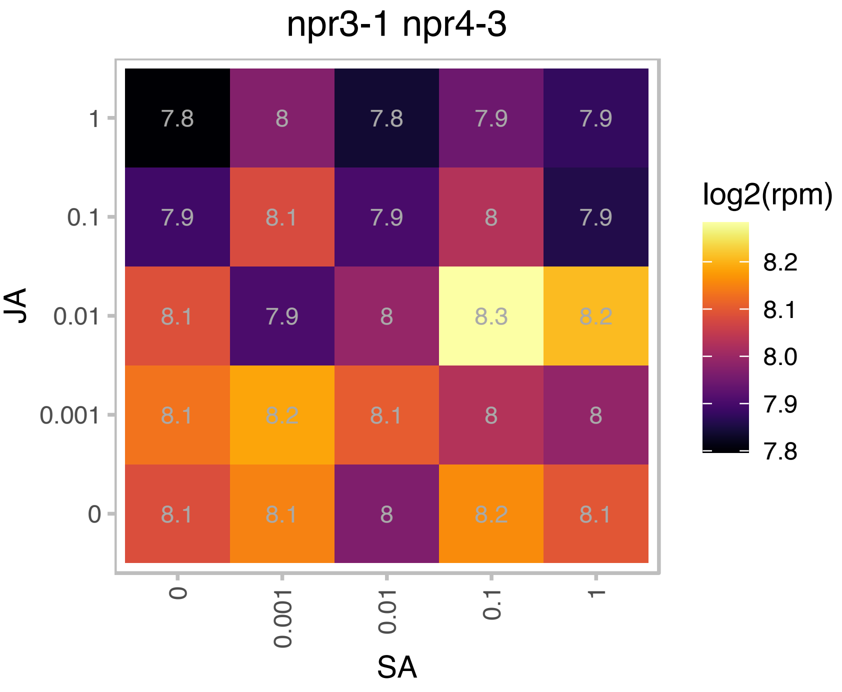

### AT3G46030.1 : HTB11

Histone superfamily protein

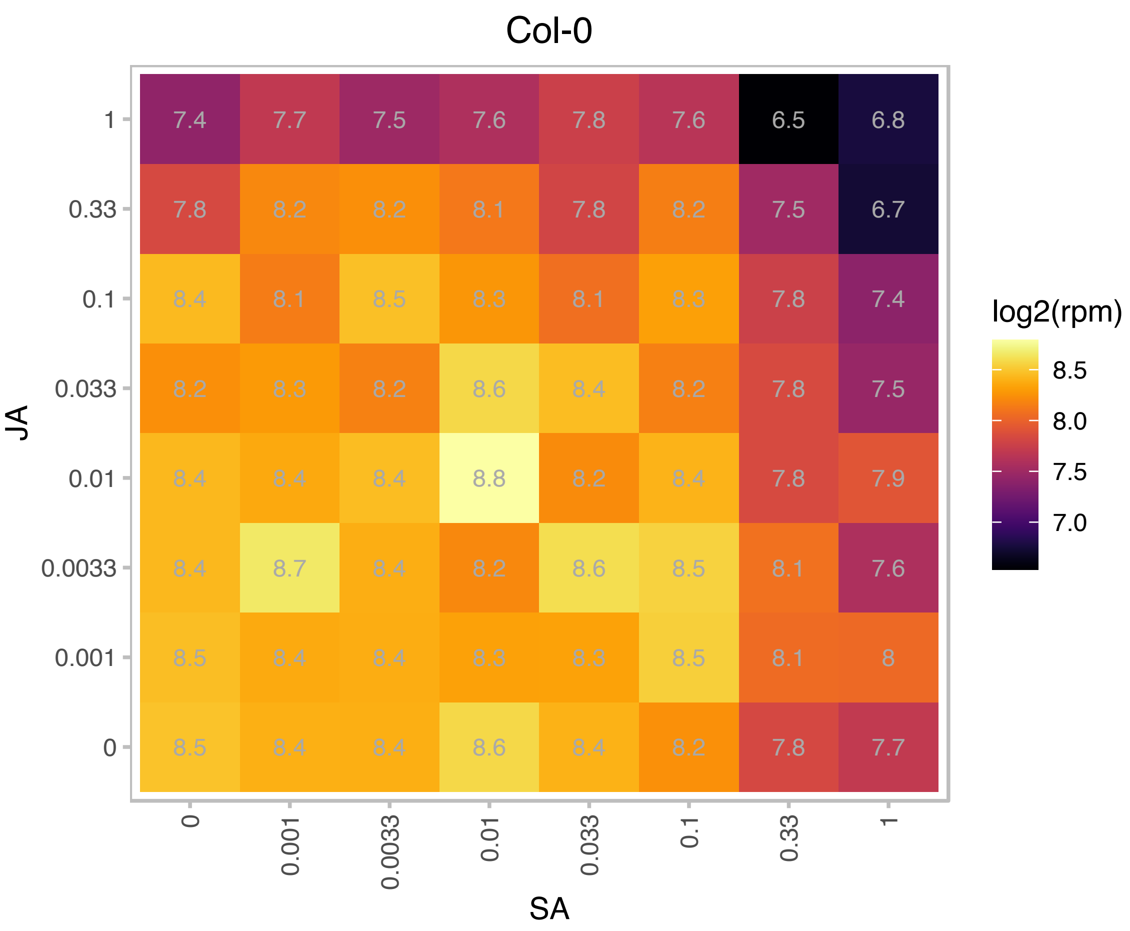


| | |
|---|---|
| Gene | AT3G46030 |
| is.expressed_genes | TRUE |
| is.responsive_genes | TRUE |
| is.apcluster | F-08 |

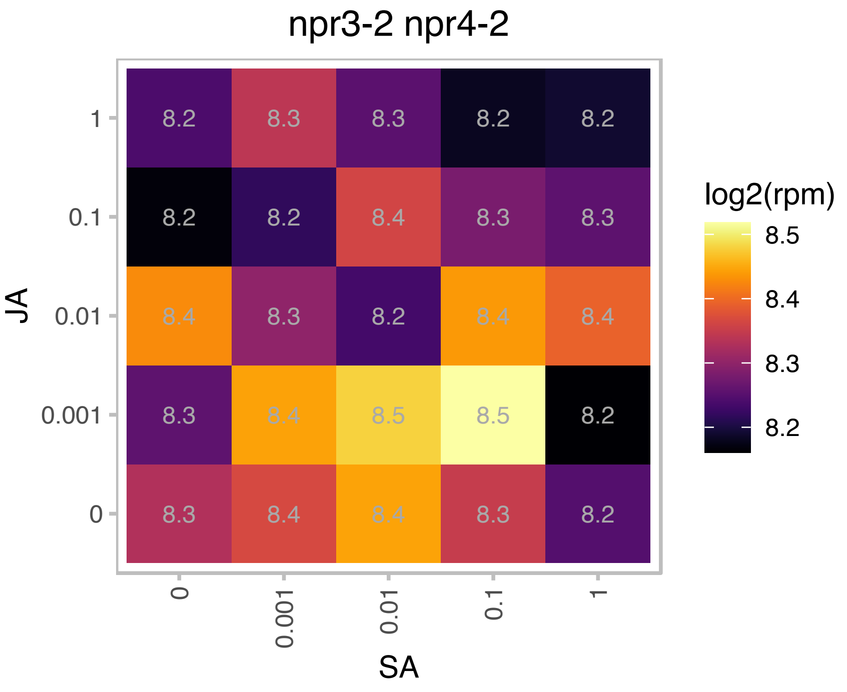


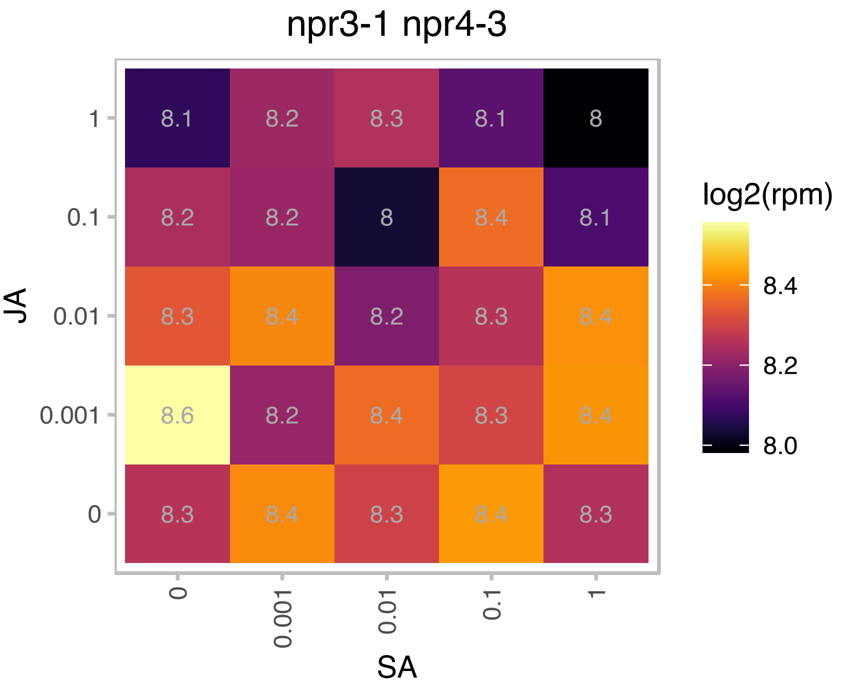

# AT3G47370.1 : NA

Ribosomal protein S10p/S20e family protein

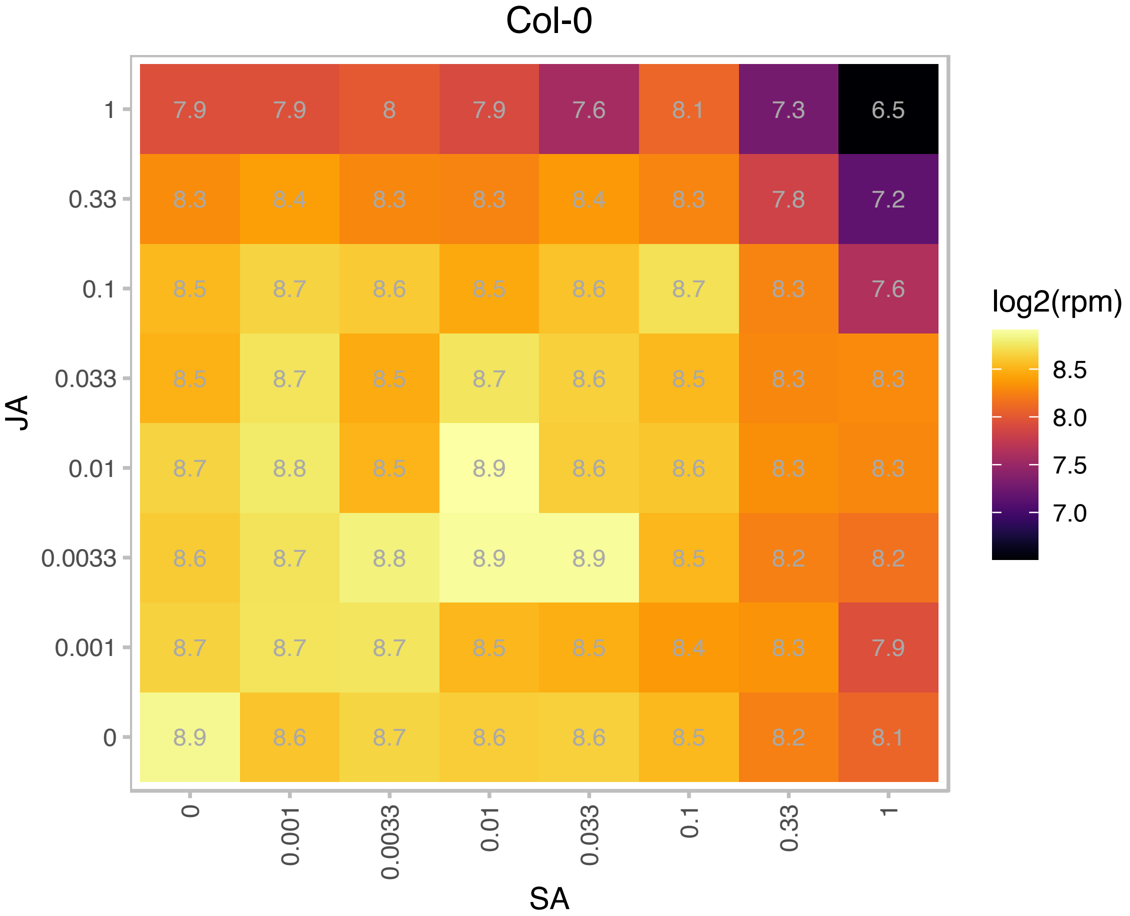


| | |
|---|---|
| Gene | AT3G47370 |
| is.expressed_genes | TRUE |
| is.responsive_genes | TRUE |
| is.apcluster | F-08 |

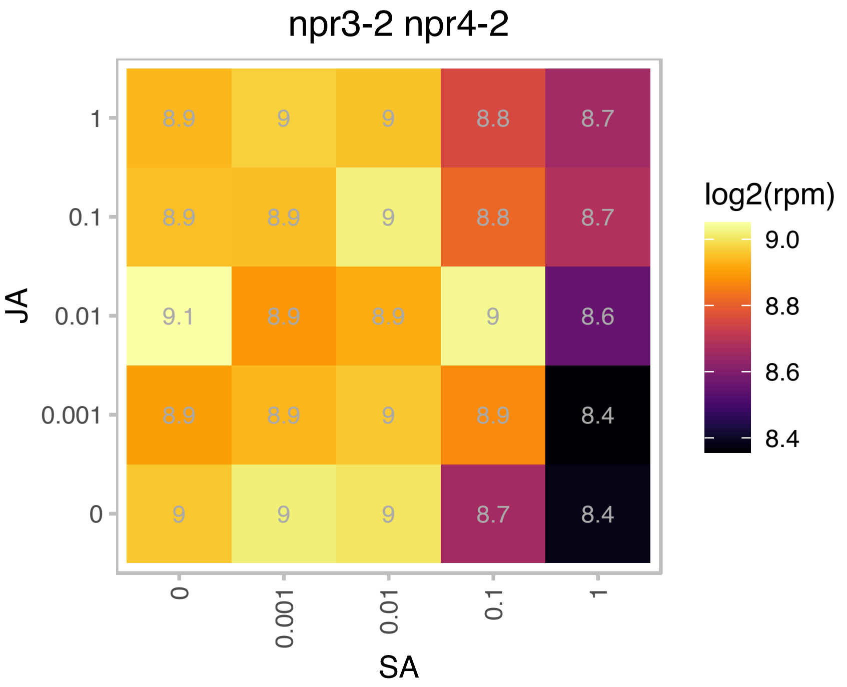


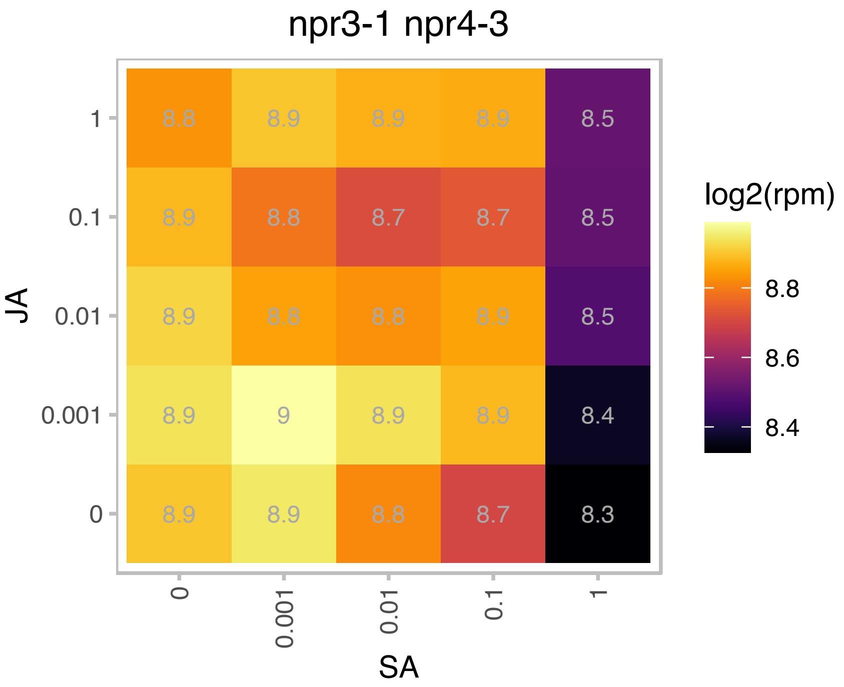

# AT3G47490.1 : NA

HNH endonuclease

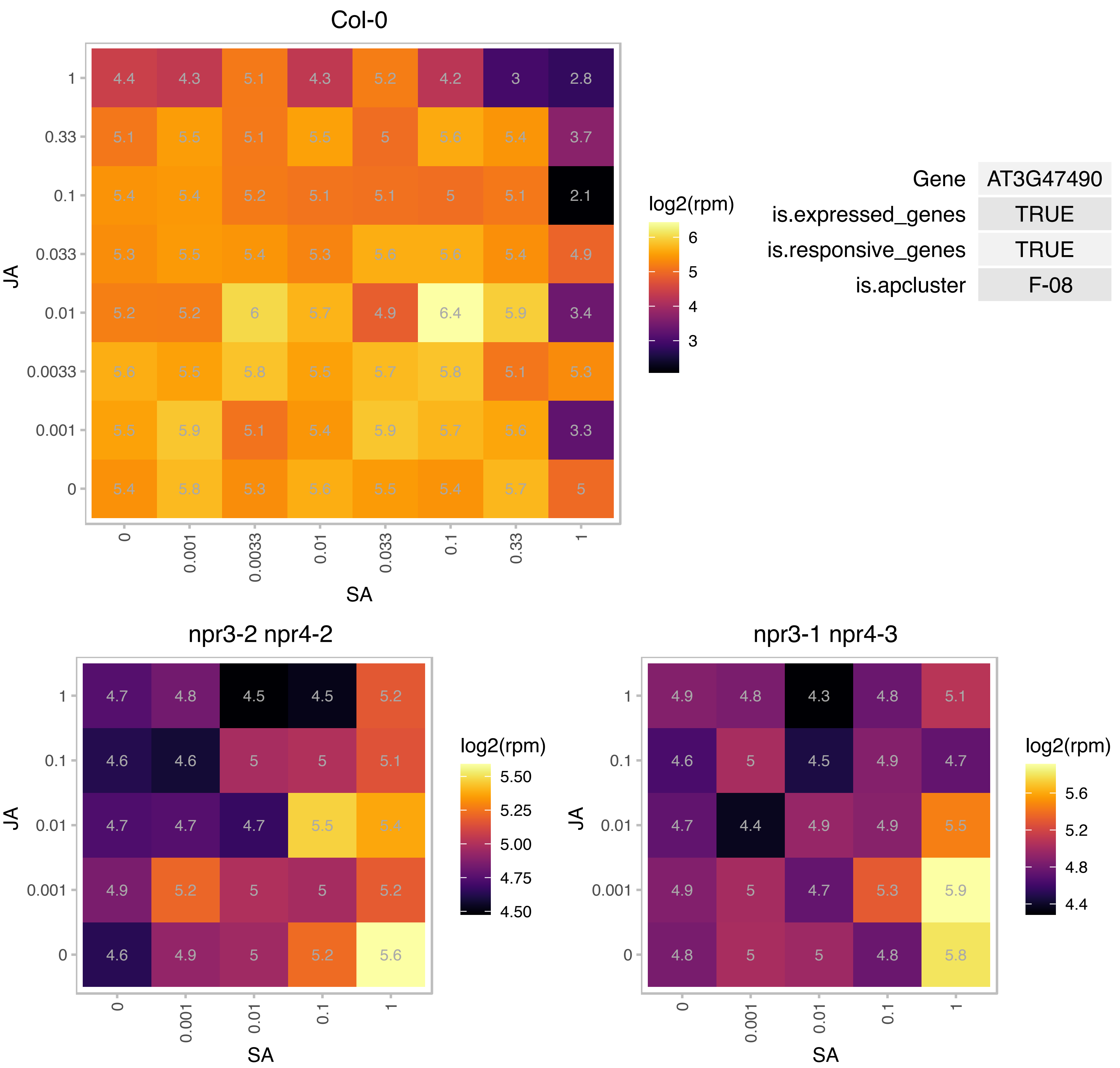


| | |
|---|---|
| Gene | AT3G47490 |
| is.expressed_genes | TRUE |
| is.responsive_genes | TRUE |
| is.apcluster | F-08 |

### AT3G51420.1 : ATSSL4, SSL4

strictosidine synthase-like 4

Although this enzyme is predicted to encode a strictosidine synthase (SS), it lacks a conserved catalytic glutamate residue found in active SS enzymes and it is not expected to have SS activity.

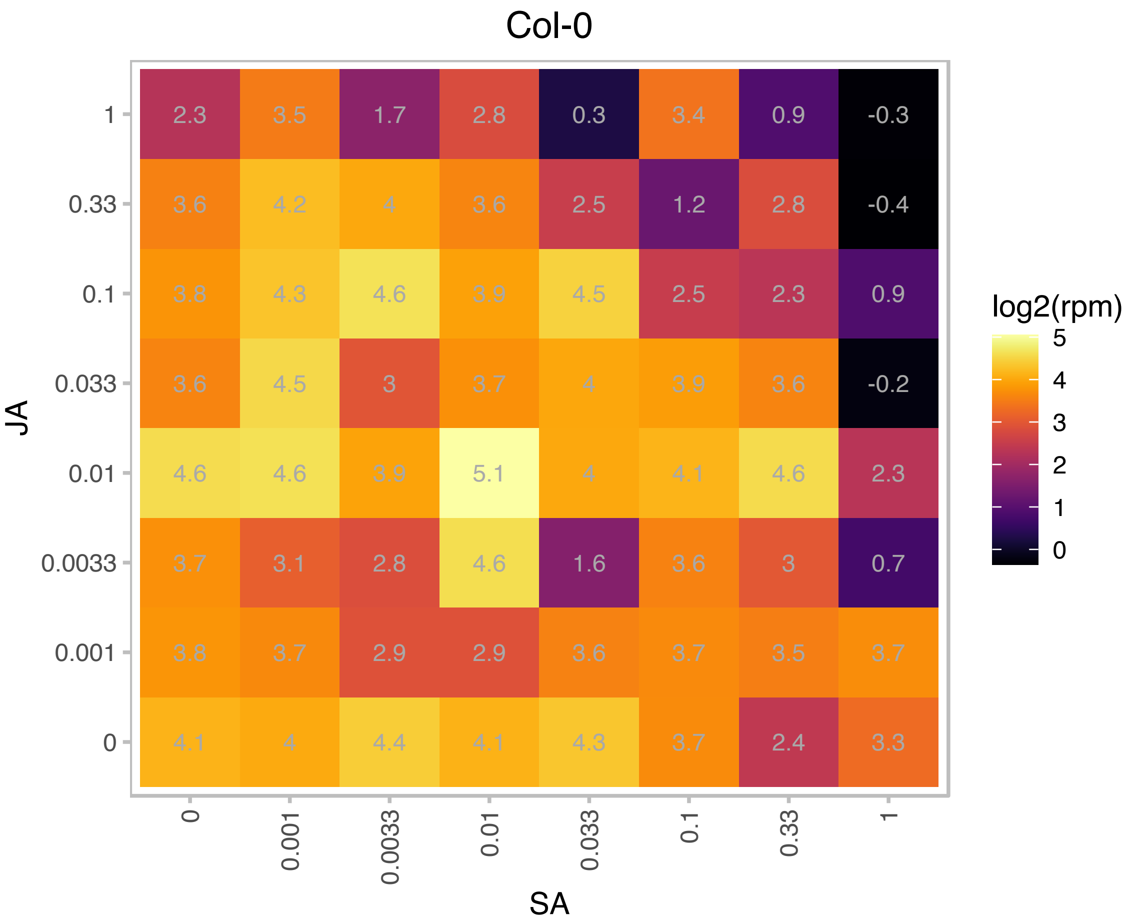


| | |
|---|---|
| Gene | AT3G51420 |
| is.expressed_genes | TRUE |
| is.responsive_genes | TRUE |
| is.apcluster | F-08 |

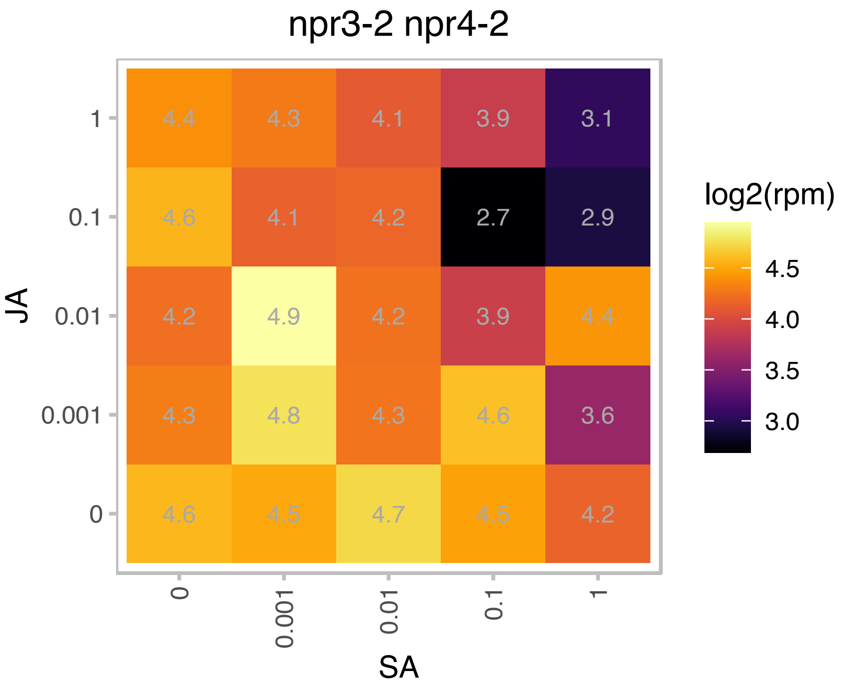


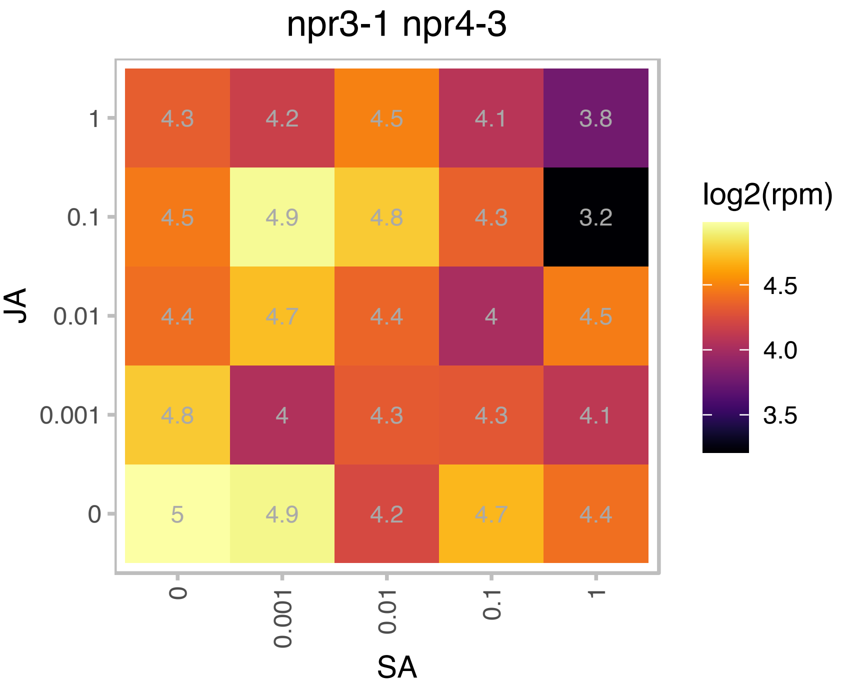

### AT3G51600.1 : LTP5

lipid transfer protein 5

Predicted to encode a PR (pathogenesis-related) protein. Belongs to the lipid transfer protein (PR-14) family with the following members: At2g38540/LTP1, At2g38530/LTP2, At5g59320/LTP3, At5g59310/LTP4, At3g51600/LTP5, At3g08770/LTP6, At2g15050/LTP7, At2g18370/LTP8, At2g15325/LTP9, At5g01870/LTP10, At4g33355/LTP11, At3g51590/LTP12, At5g44265/LTP13, At5g62065/LTP14, At4g08530/LTP15.

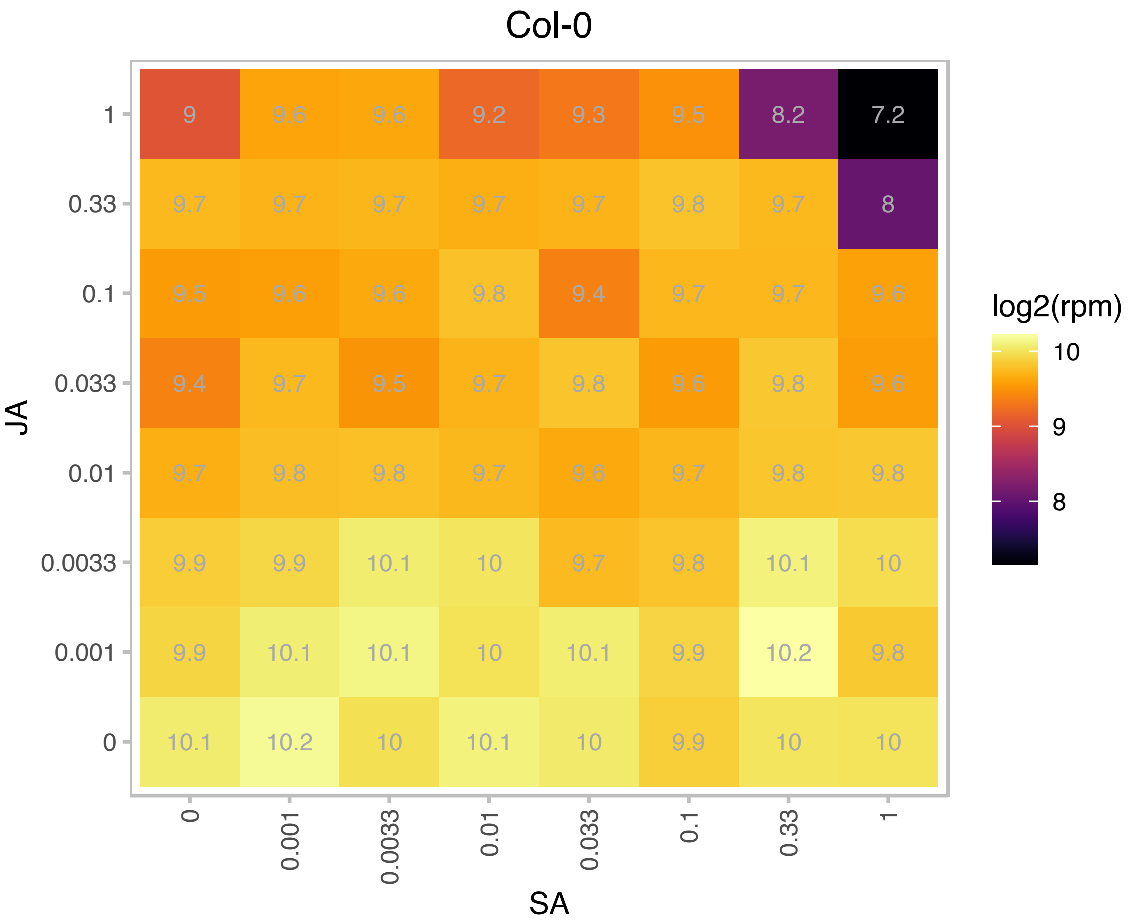


| | |
|---|---|
| Gene | AT3G51600 |
| is.expressed_genes | TRUE |
| is.responsive_genes | TRUE |
| is.apcluster | F-08 |

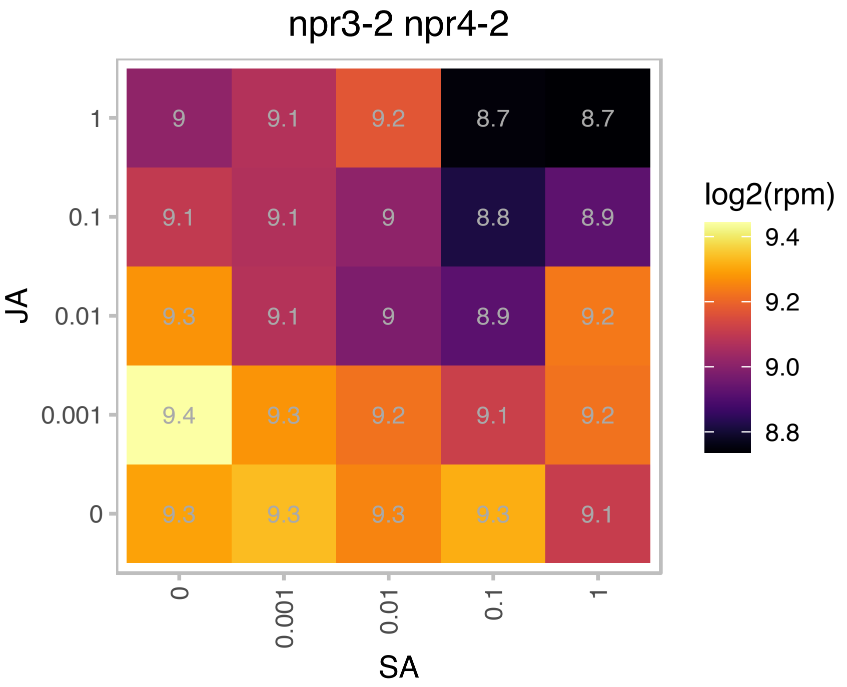


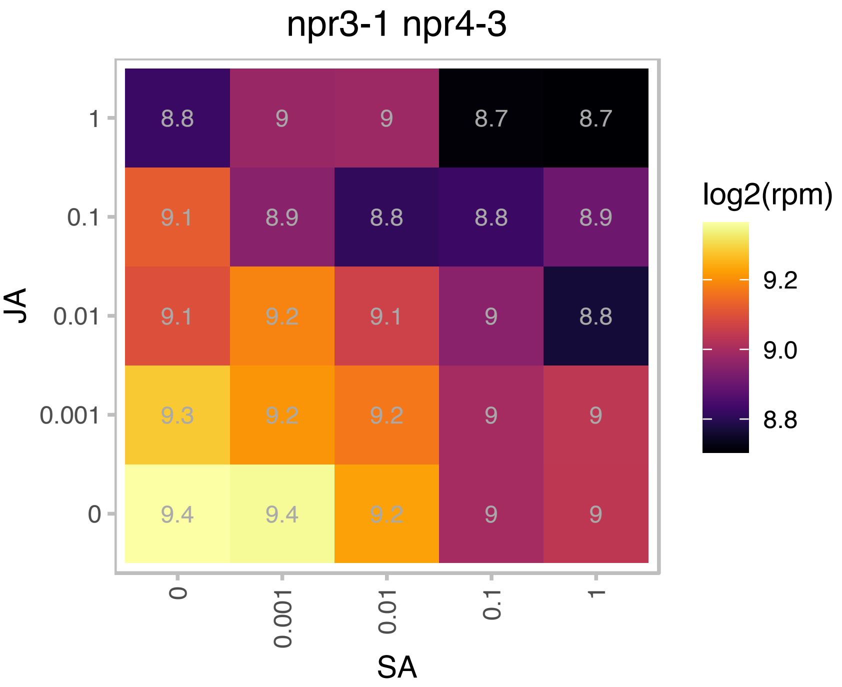

# AT3G52040.1 : NA

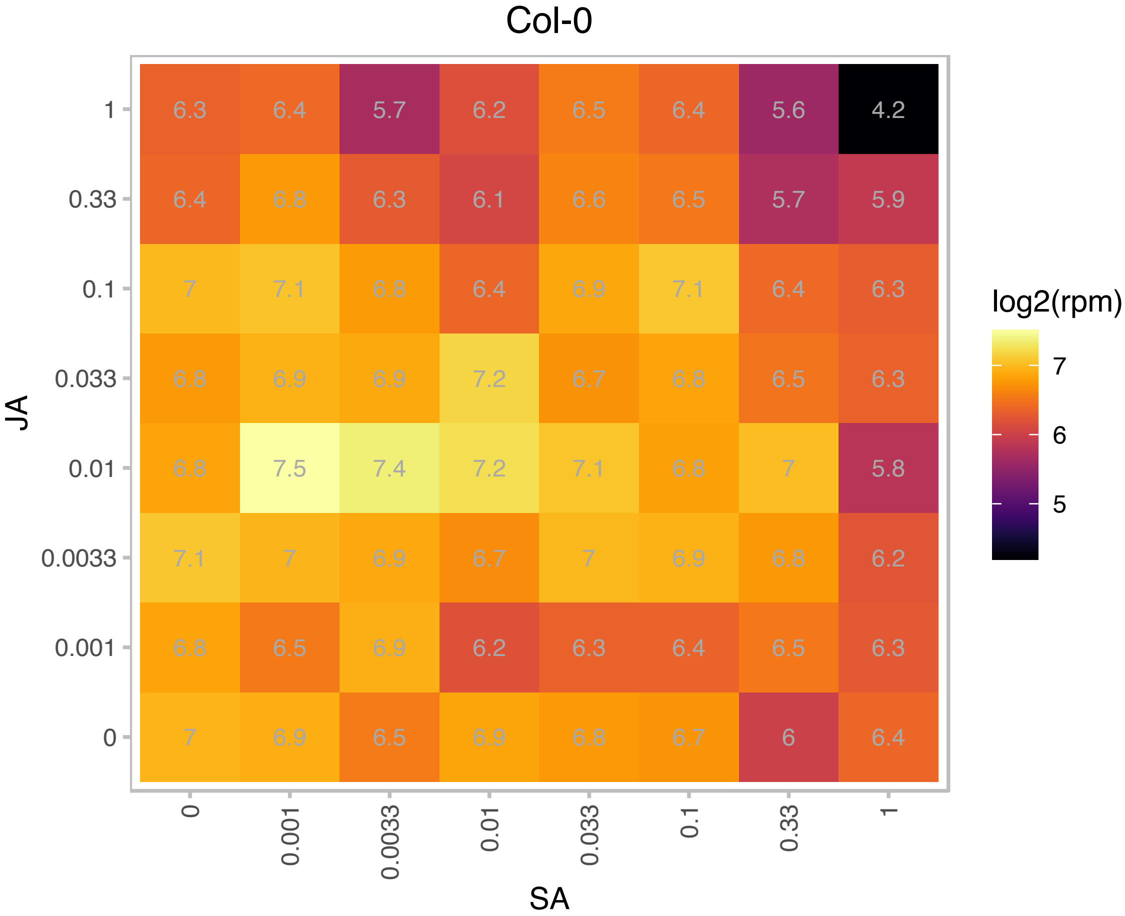


| | |
|---|---|
| Gene | AT3G52040 |
| is.expressed_genes | TRUE |
| is.responsive_genes | TRUE |
| is.apcluster | F-08 |

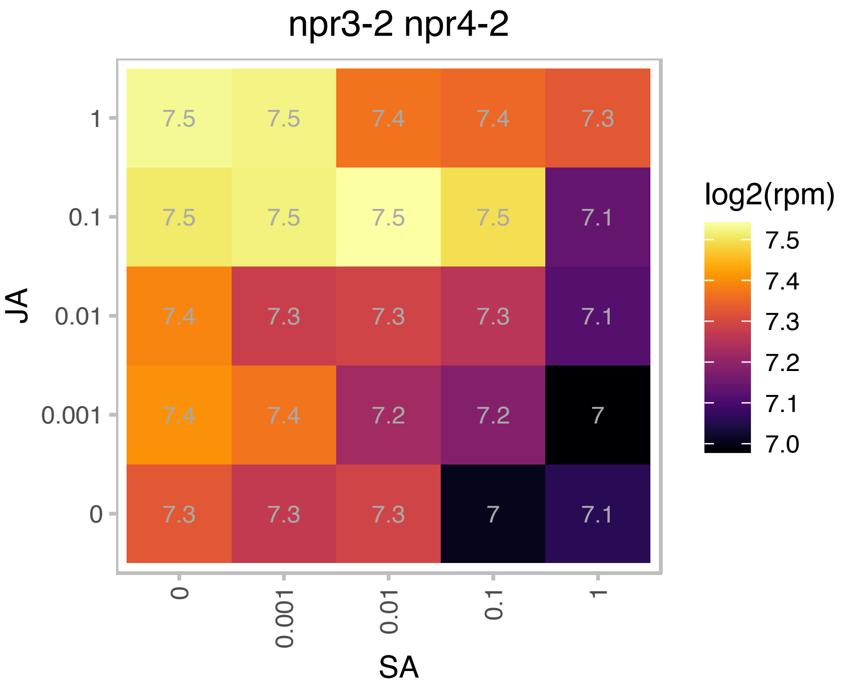


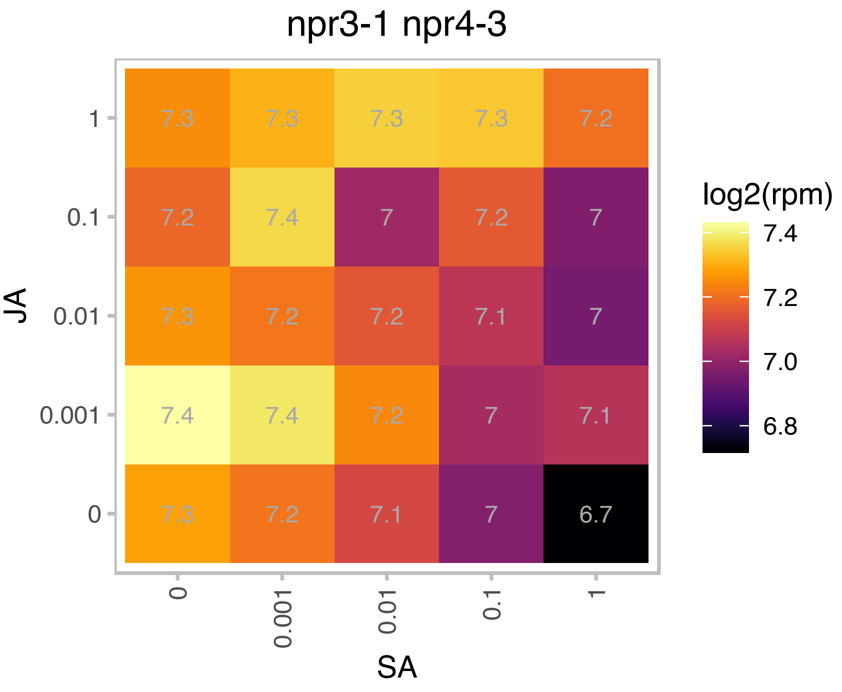

# AT3G53870.1 : NA

Ribosomal protein S3 family protein

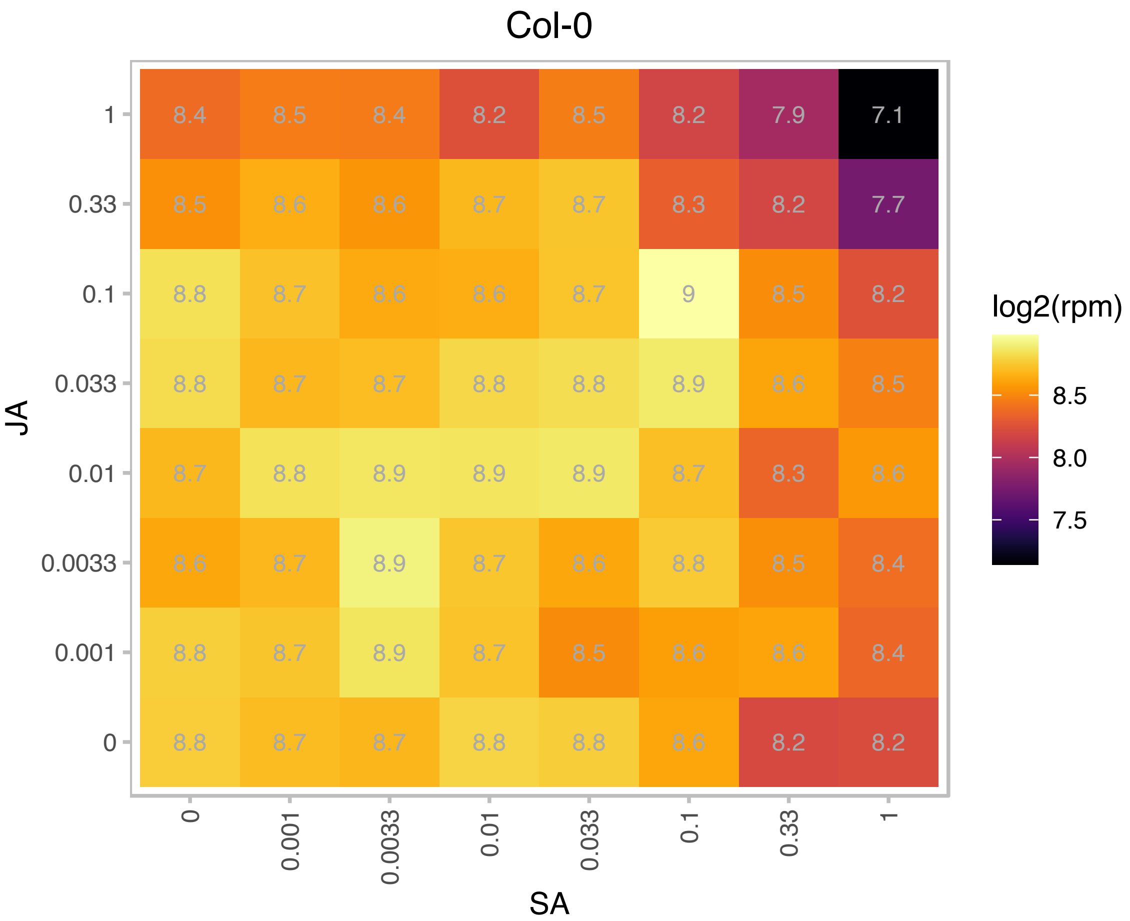


| | |
|---|---|
| Gene | AT3G53870 |
| is.expressed_genes | TRUE |
| is.responsive_genes | TRUE |
| is.apcluster | F-08 |

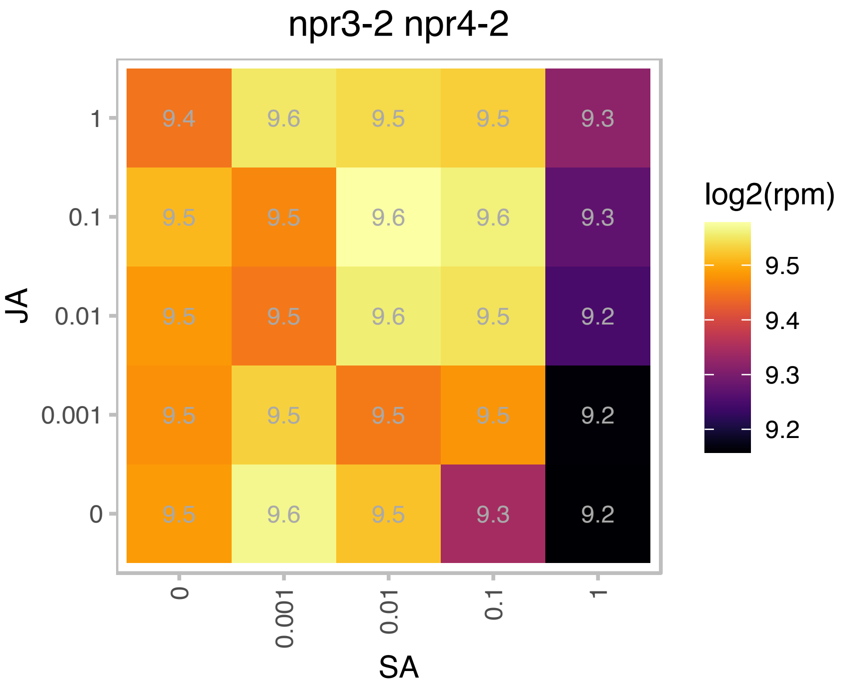


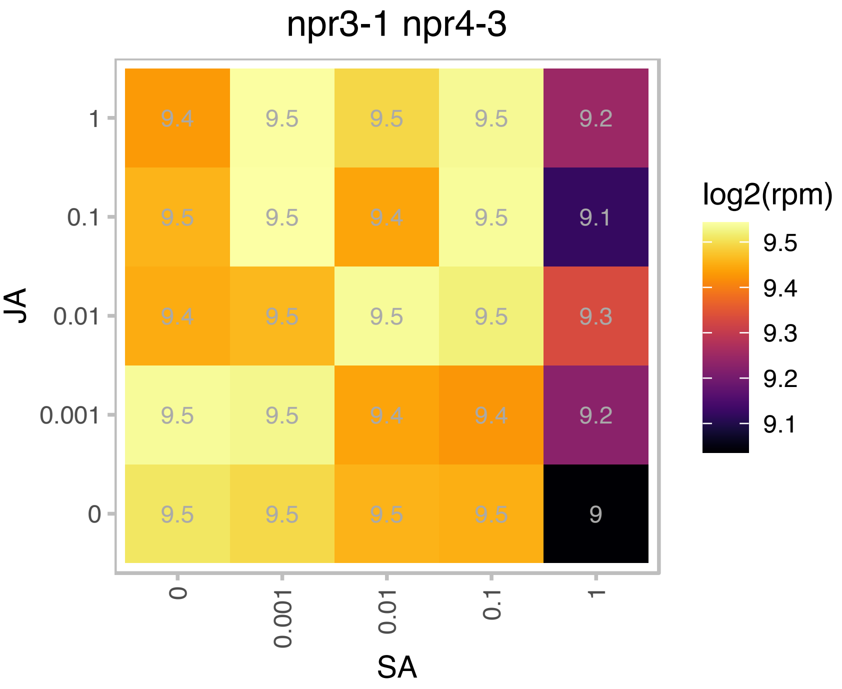

### AT3G54810.1 : BME3, BME3-ZF, GATA8

Plant-specific GATA-type zinc finger transcription factor family protein

Encodes a protein containing a GATA type zinc finger domain that is expressed in the embryo axis and involved in germination. Mutants have a reduced rate of germination even when stratified.

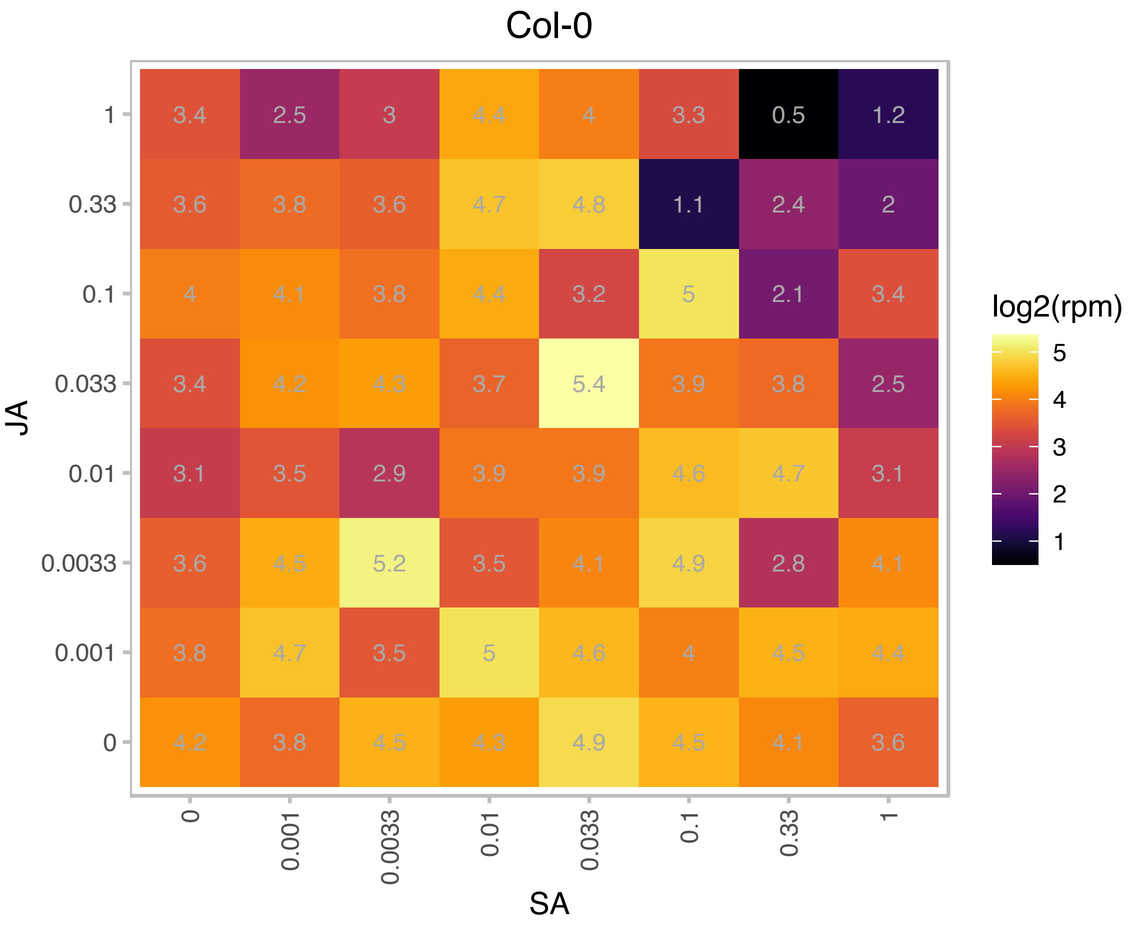


| | |
|---|---|
| Gene | AT3G54810 |
| is.expressed_genes | TRUE |
| is.responsive_genes | TRUE |
| is.apcluster | F-08 |

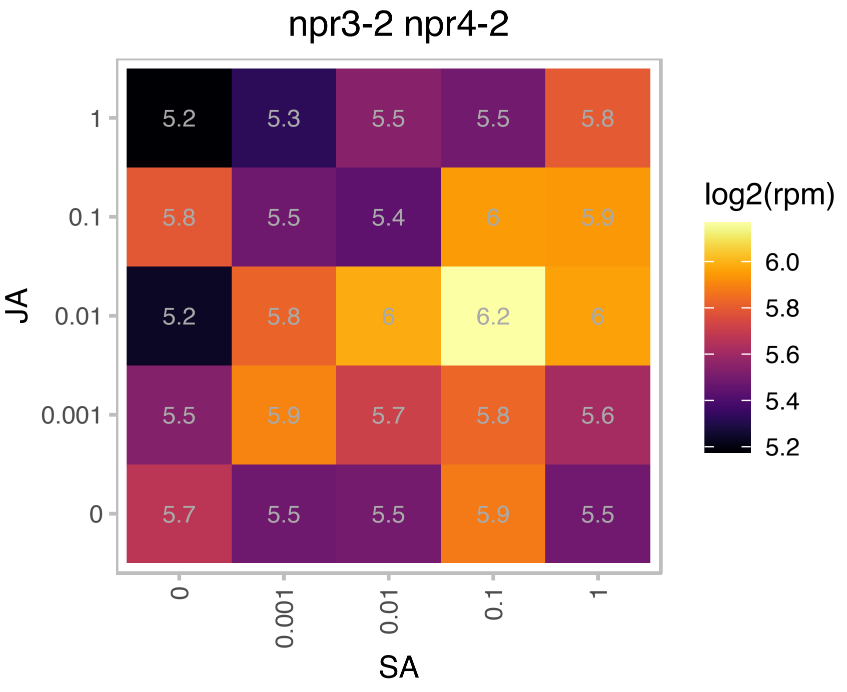


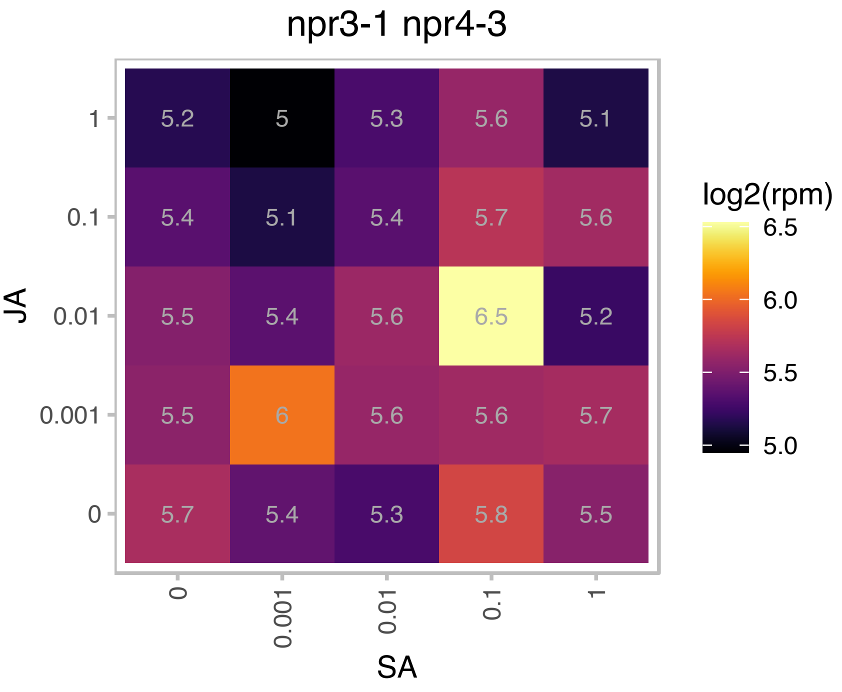

### AT3G56070.1 : ROC2

rotamase cyclophilin 2

rotamase cyclophilin 2 (ROC2) exhibiting peptidyl-prolyl cis-trans isomerase activity involved in signal transduction.

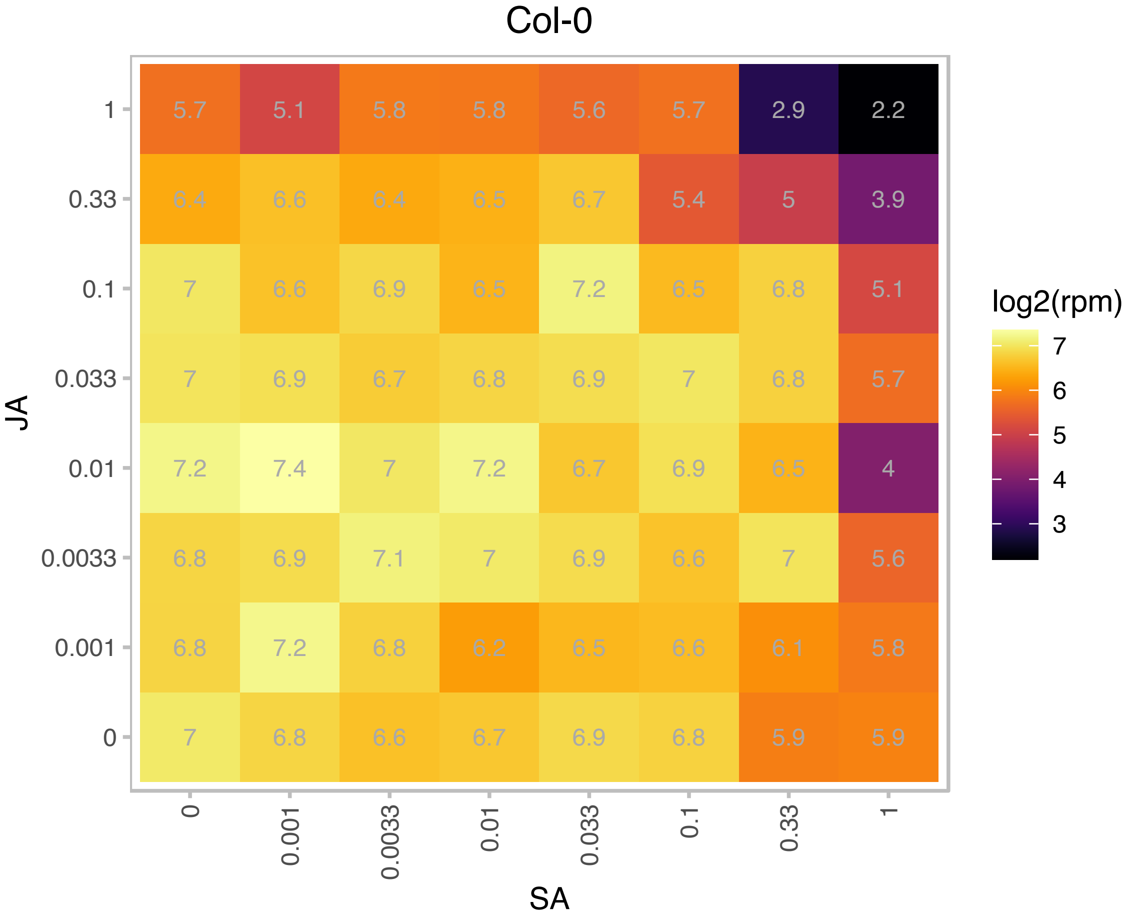


| | |
|---|---|
| Gene | AT3G56070 |
| is.expressed_genes | TRUE |
| is.responsive_genes | TRUE |
| is.apcluster | F-08 |

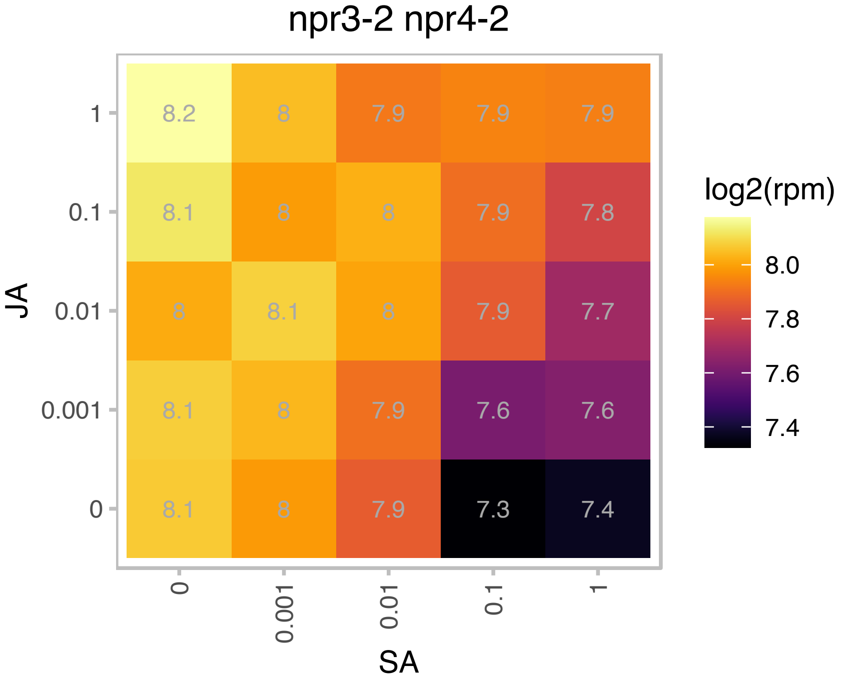


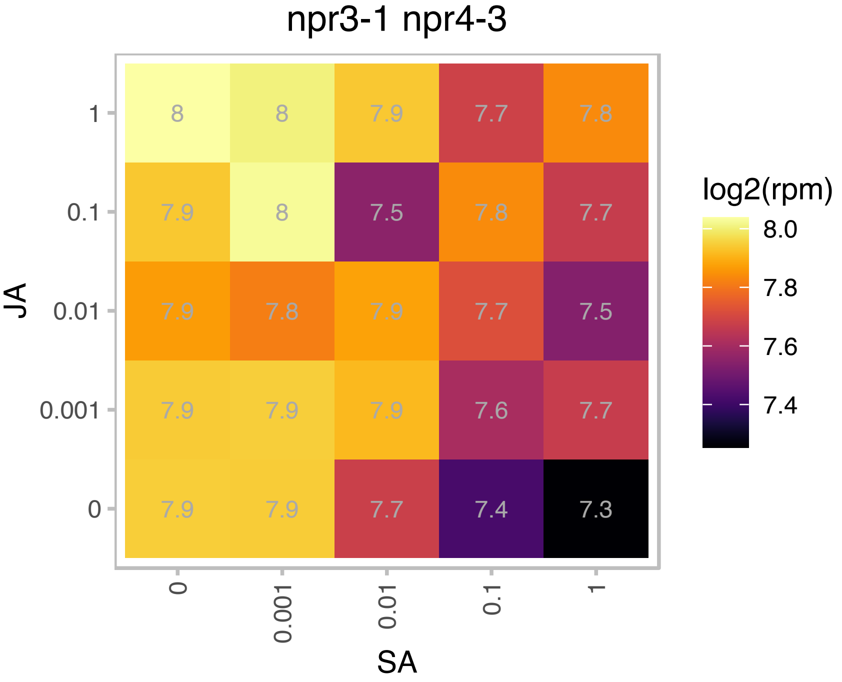

### AT3G57150.1 : AtCBF5, AtNAP57, CBF5, NAP57

homologue of NAP57

Encodes a putative pseudouridine synthase (NAP57).

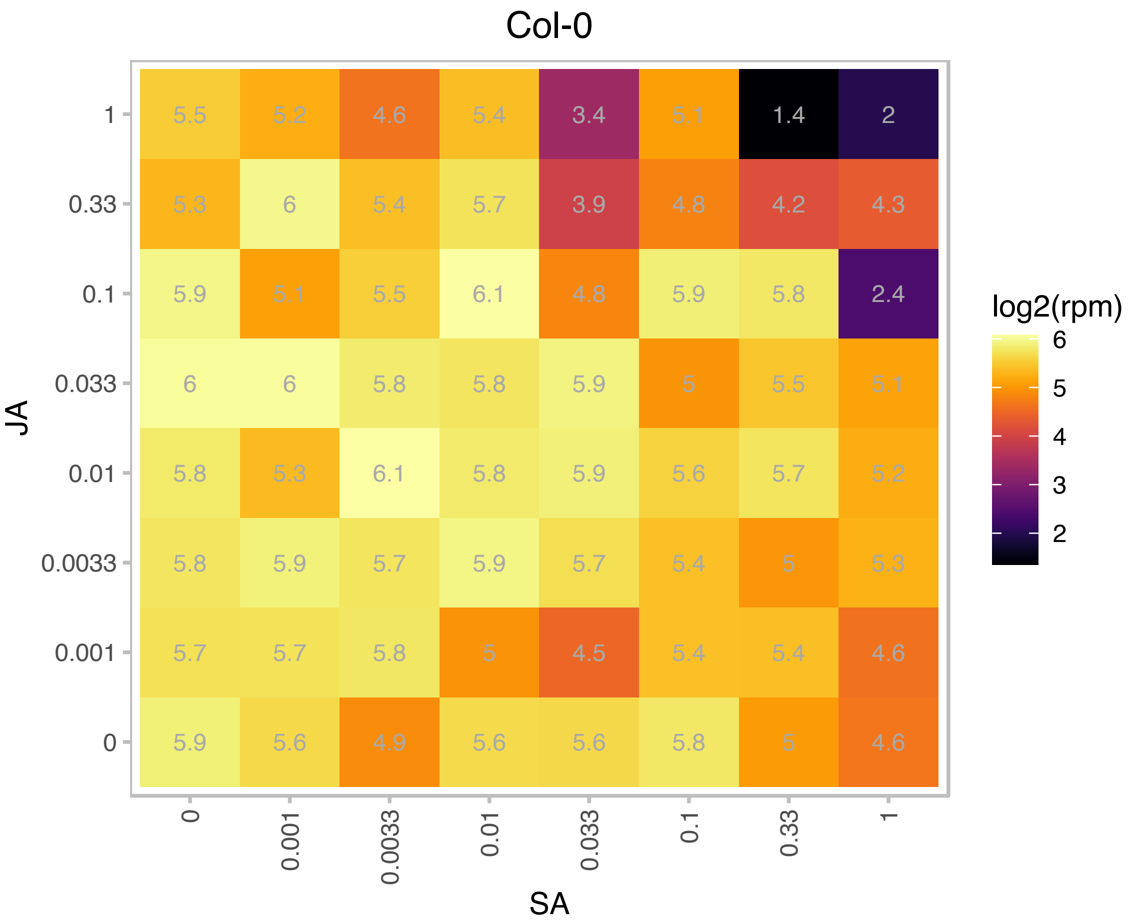


| | |
|---|---|
| Gene | AT3G57150 |
| is.expressed_genes | TRUE |
| is.responsive_genes | TRUE |
| is.apcluster | F-08 |

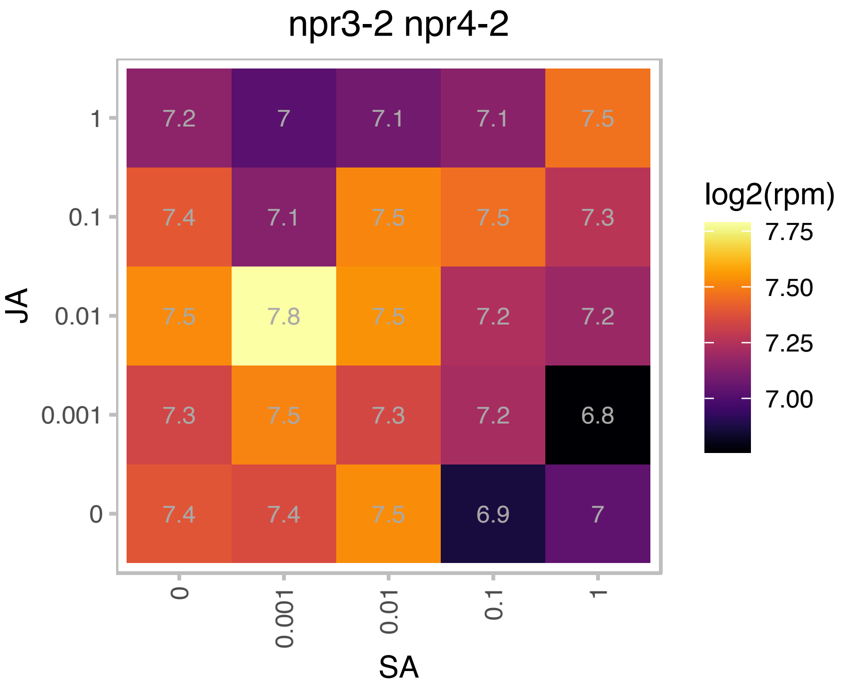


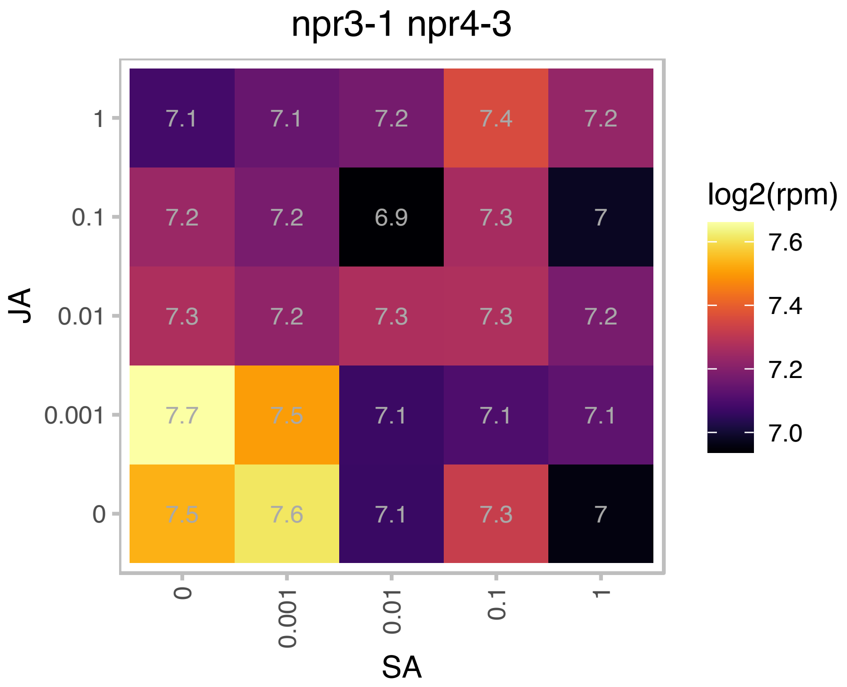

### AT3G59400.1 : GUN4

enzyme binding;tetrapyrrole binding

GUN, genomes uncoupled, is necessary for coupling the expression of some nuclear genes to the functional state of the chloroplast. Binds to the magnesium chelatase complex and promotes formation of the substrate,a tetrapyrrole signaling molecule. Porphyrin-binding protein that enhances the activity of Mg-chelatase. Although required for chlorophyll accumulation under normal growth conditions, GUN4 is not essential for chlorophyll synthesis.

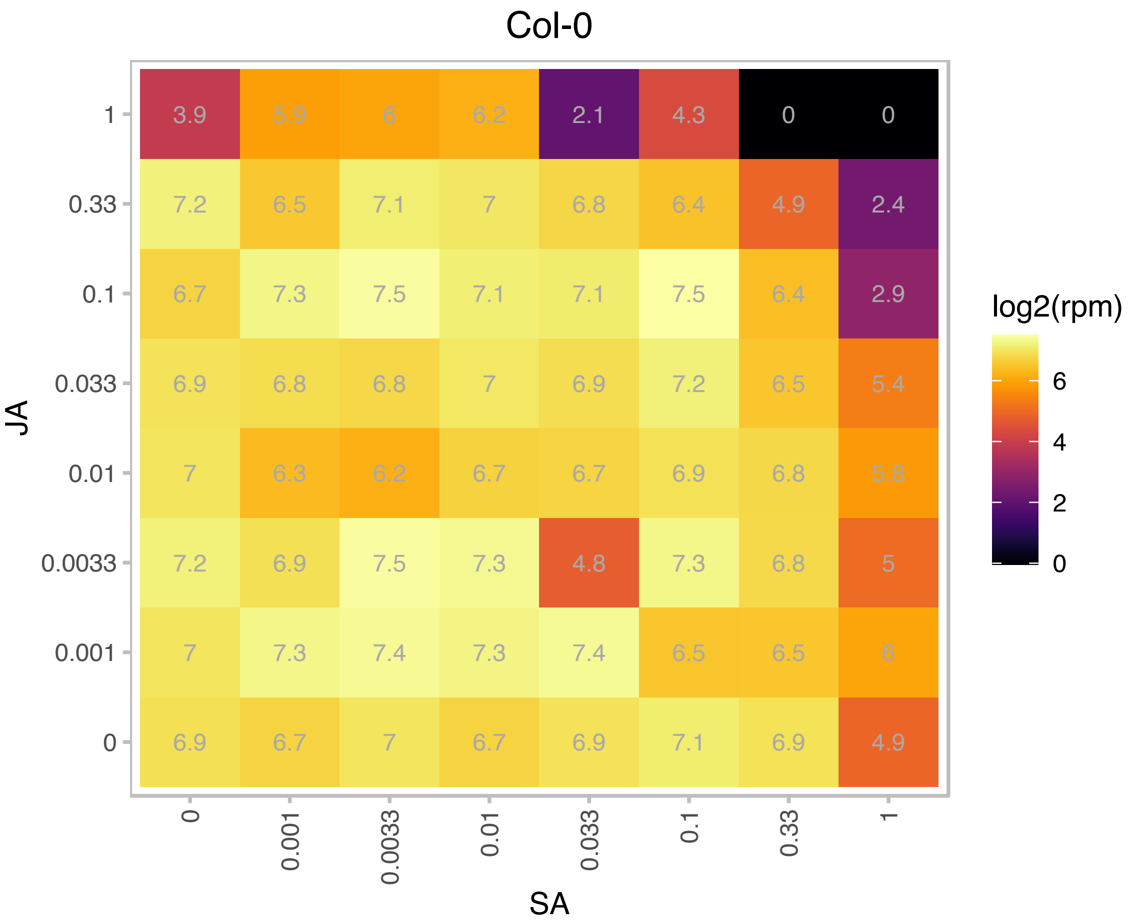


| | |
|---|---|
| Gene | AT3G59400 |
| is.expressed_genes | TRUE |
| is.responsive_genes | TRUE |
| is.apcluster | F-08 |

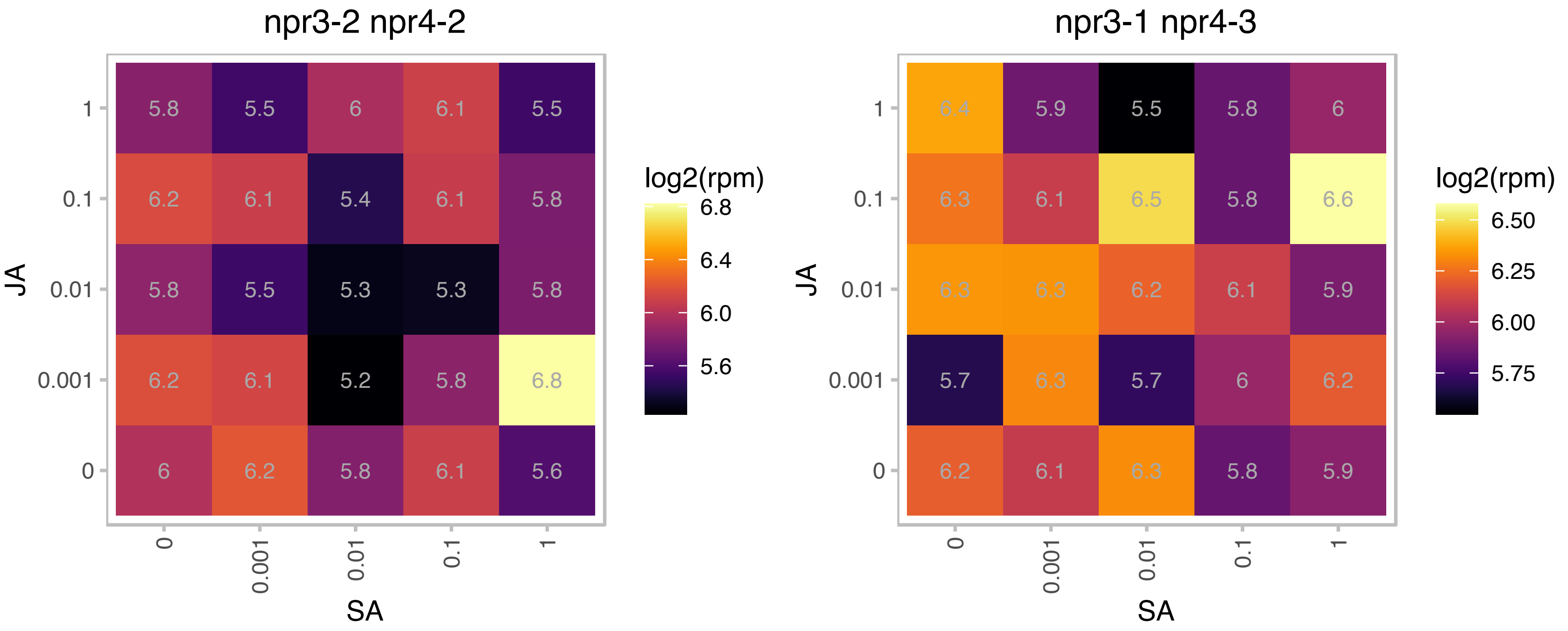

# AT3G59840.1 : NA

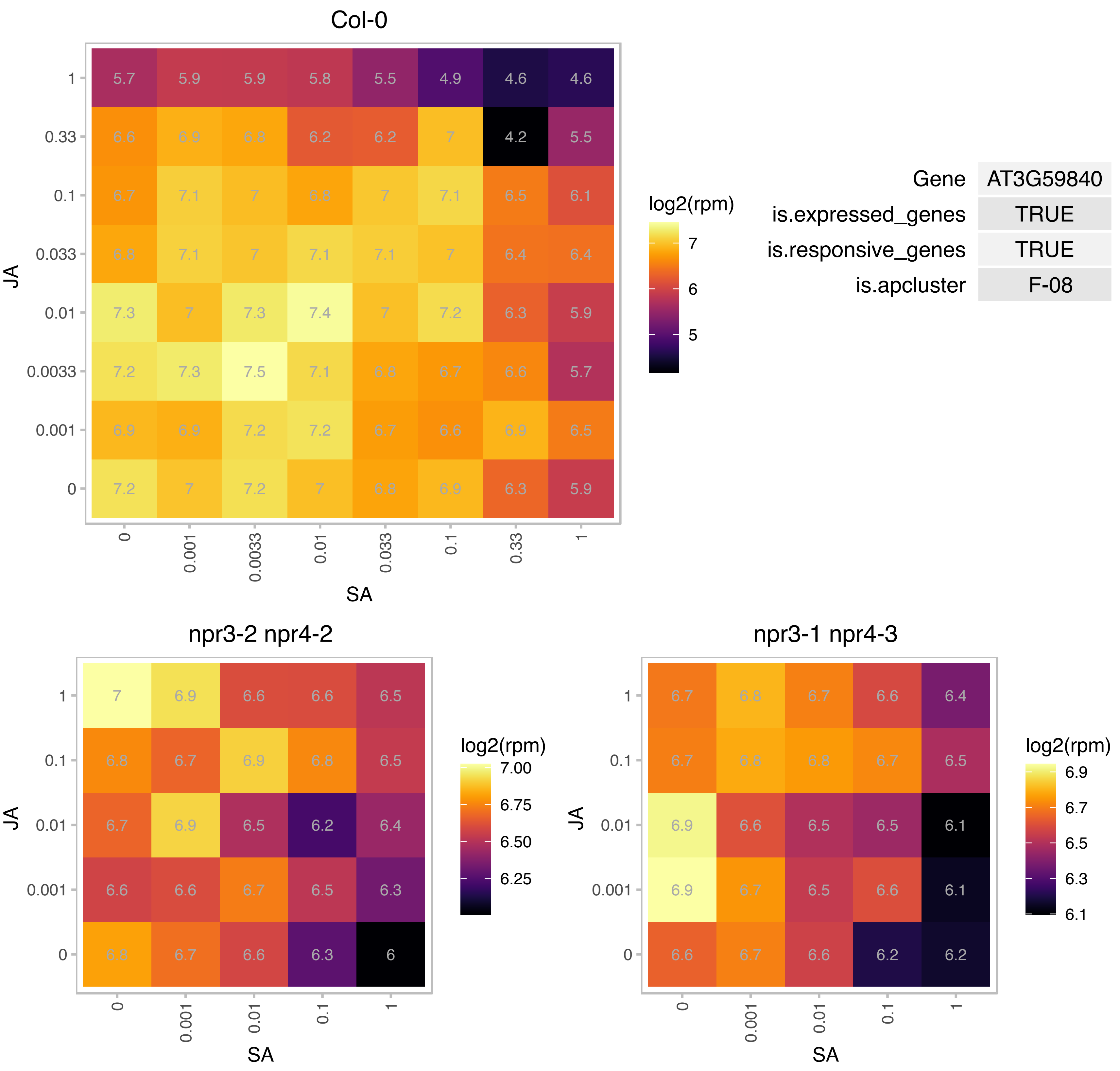

### AT3G61620.1 : RRP41

3'-5'-exoribonuclease family protein

exonuclease RRP41 (RRP41)

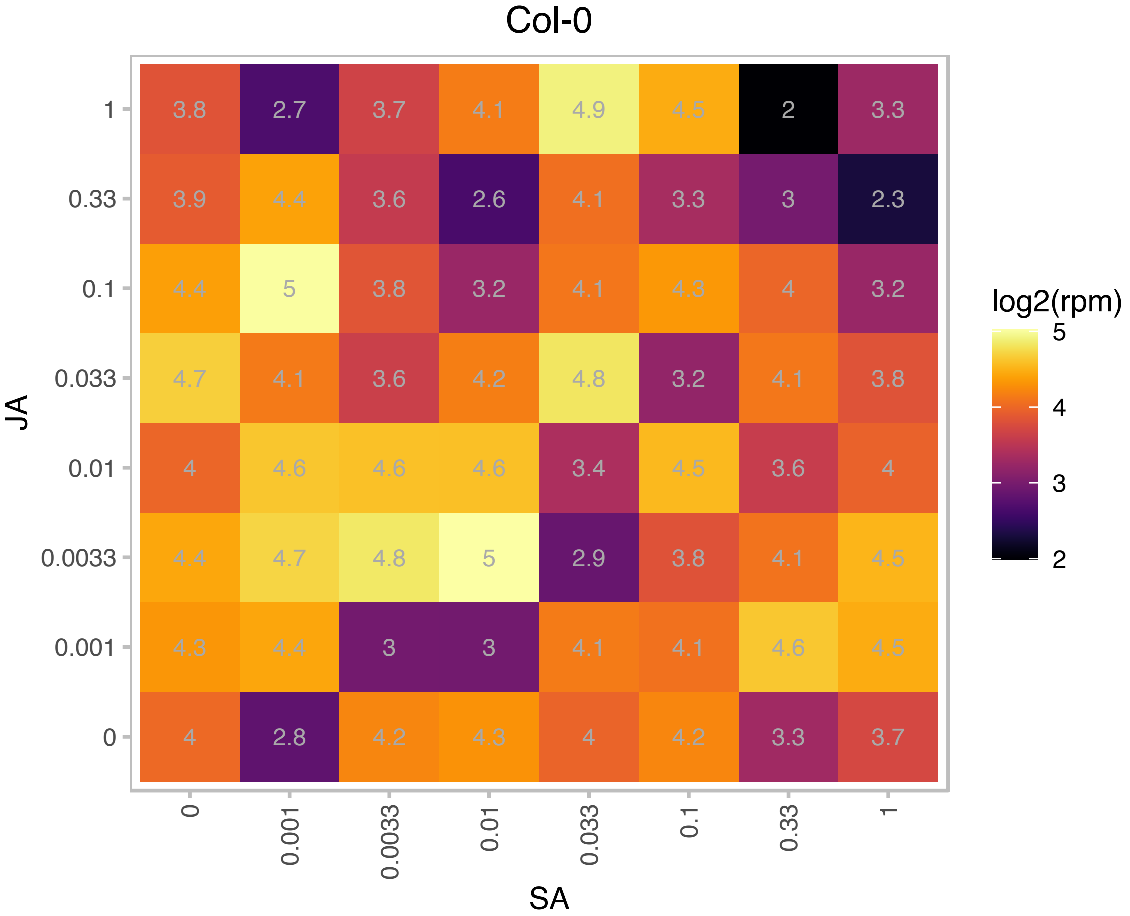


| | |
|---|---|
| Gene | AT3G61620 |
| is.expressed_genes | TRUE |
| is.responsive_genes | TRUE |
| is.apcluster | F-08 |

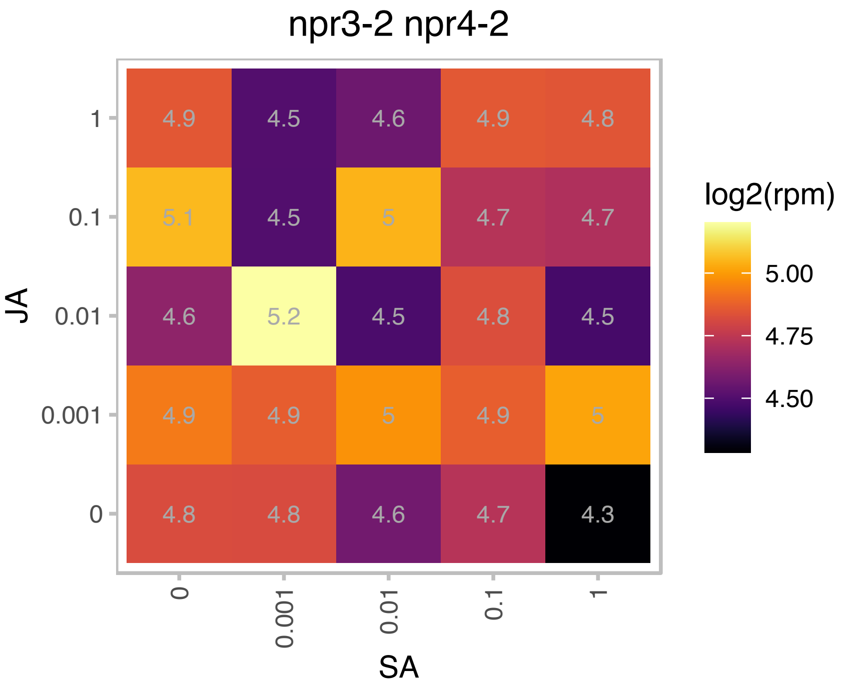


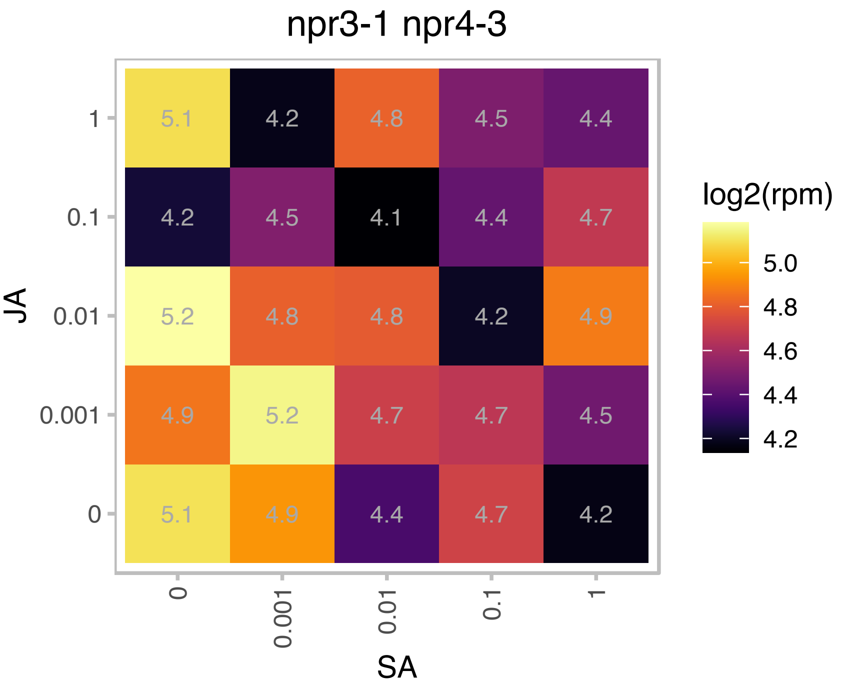

# AT3G62460.1 : NA

Putative endonuclease or glycosyl hydrolase

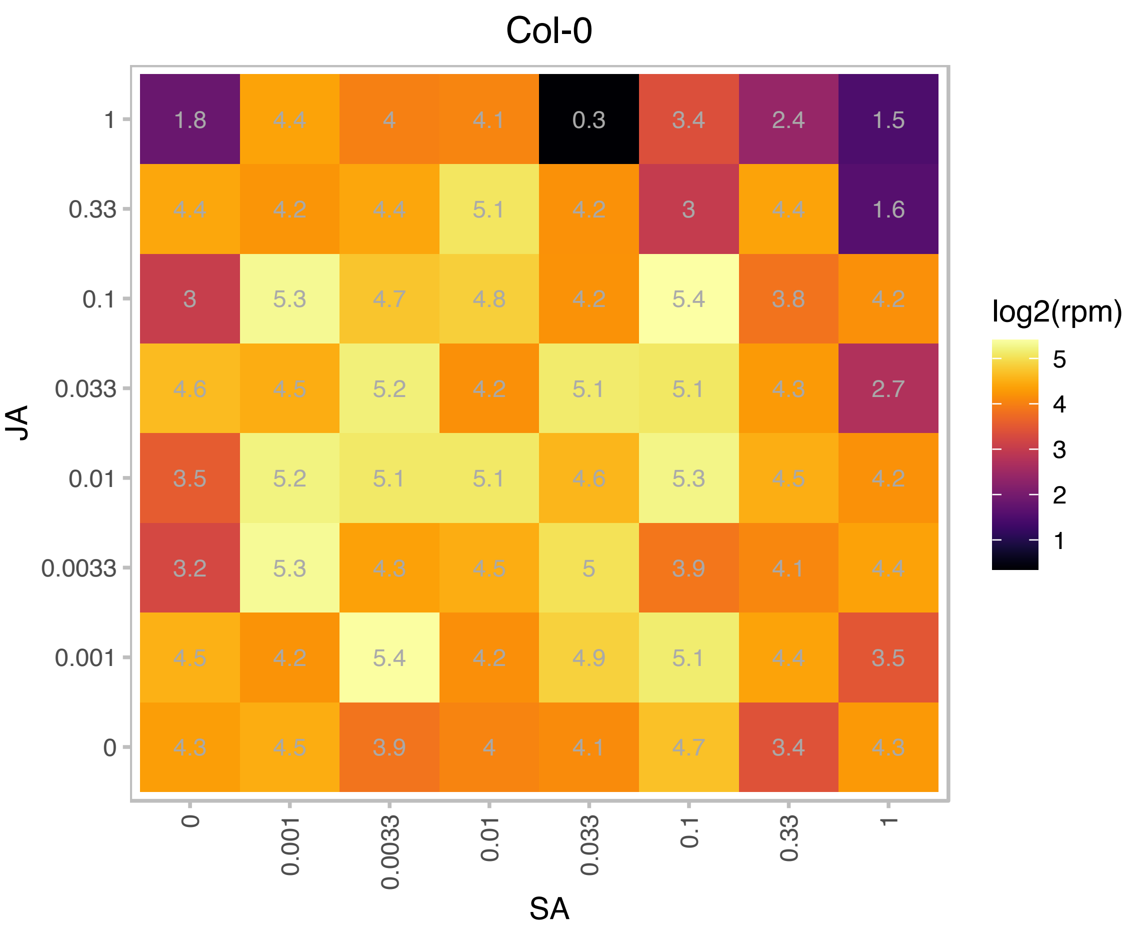


| | |
|---|---|
| Gene | AT3G62460 |
| is.expressed_genes | TRUE |
| is.responsive_genes | TRUE |
| is.apcluster | F-08 |

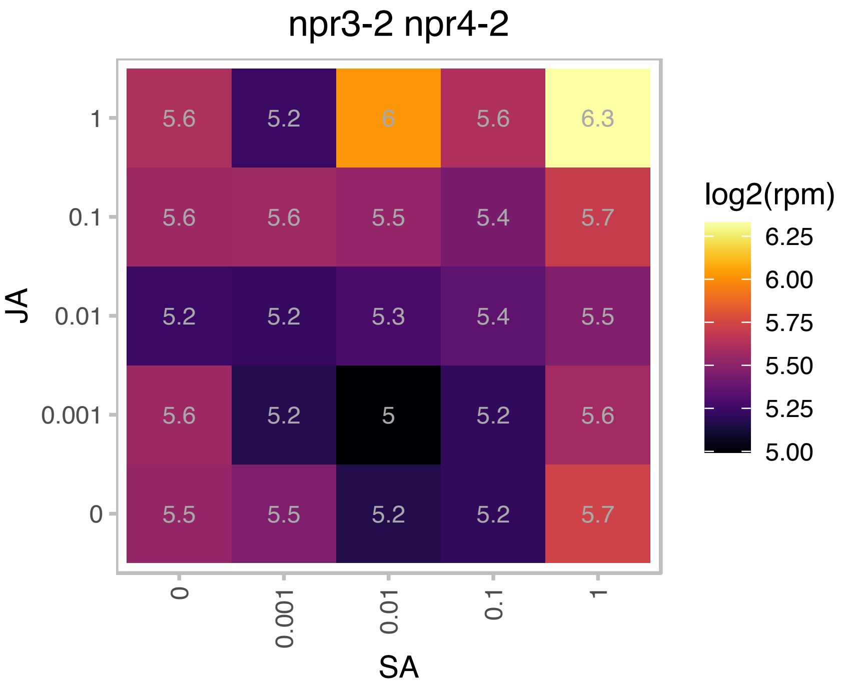


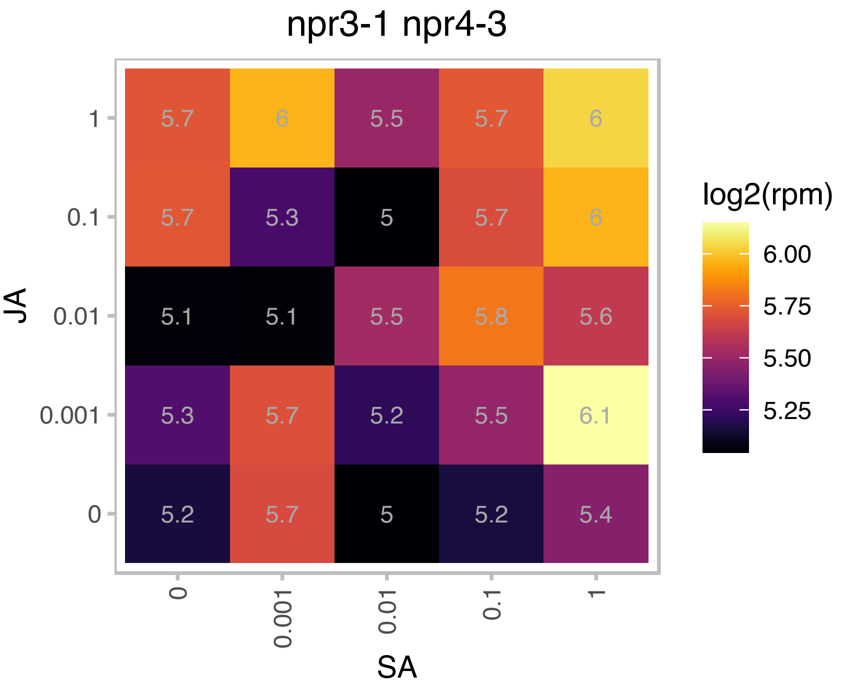

### AT3G63250.1 : ATHMT-2, HMT-2, HMT2

homocysteine methyltransferase 2
Encodes a homocysteine methyltransferase (HMT). Among the three HMT coding genes in the genome, HMT2 is responsible for a significant proportion of HMT activity in the flower stalks and silique hulls. However, HMT2 does not significantly contribute to the total HMT activity in seeds.

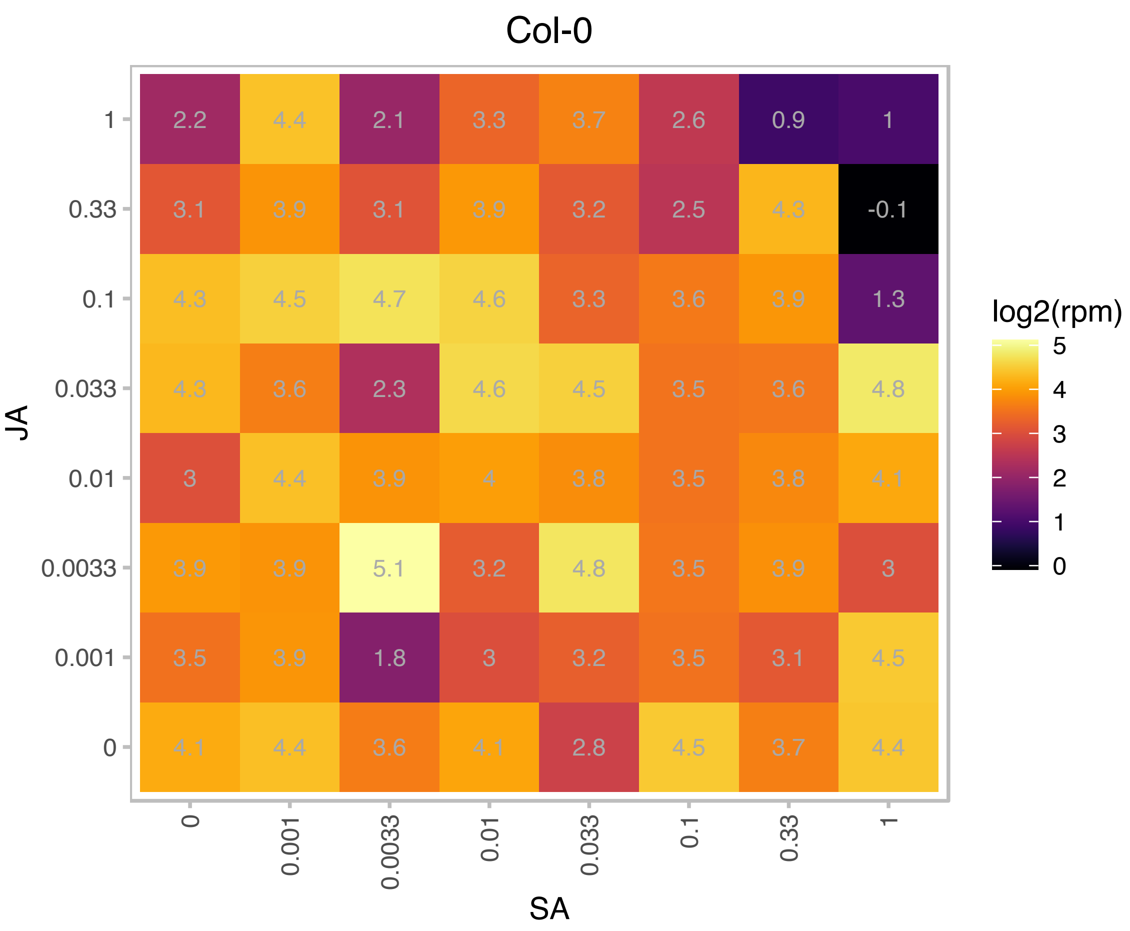


| | |
|---|---|
| Gene | AT3G63250 |
| is.expressed_genes | TRUE |
| is.responsive_genes | TRUE |
| is.apcluster | F-08 |

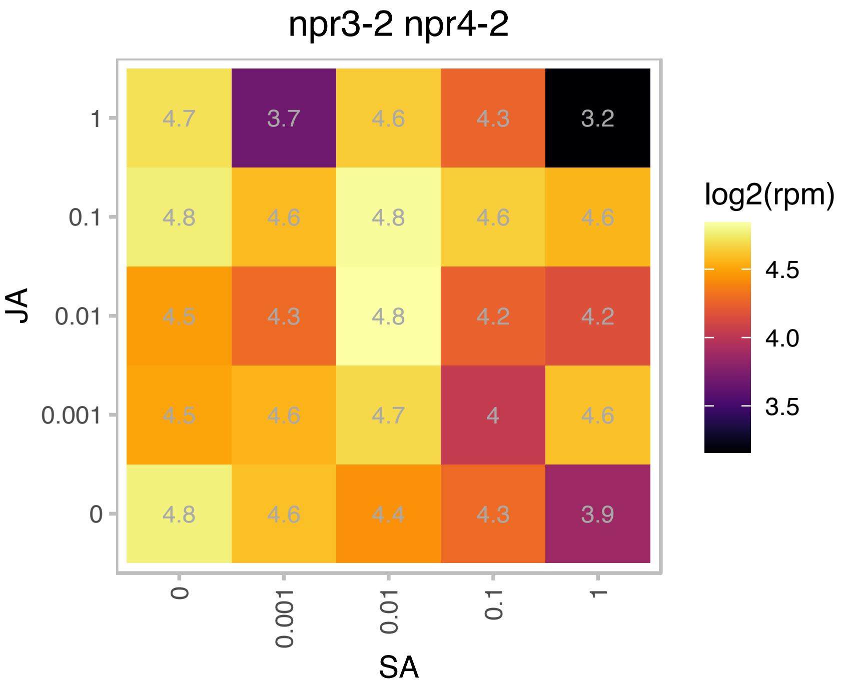


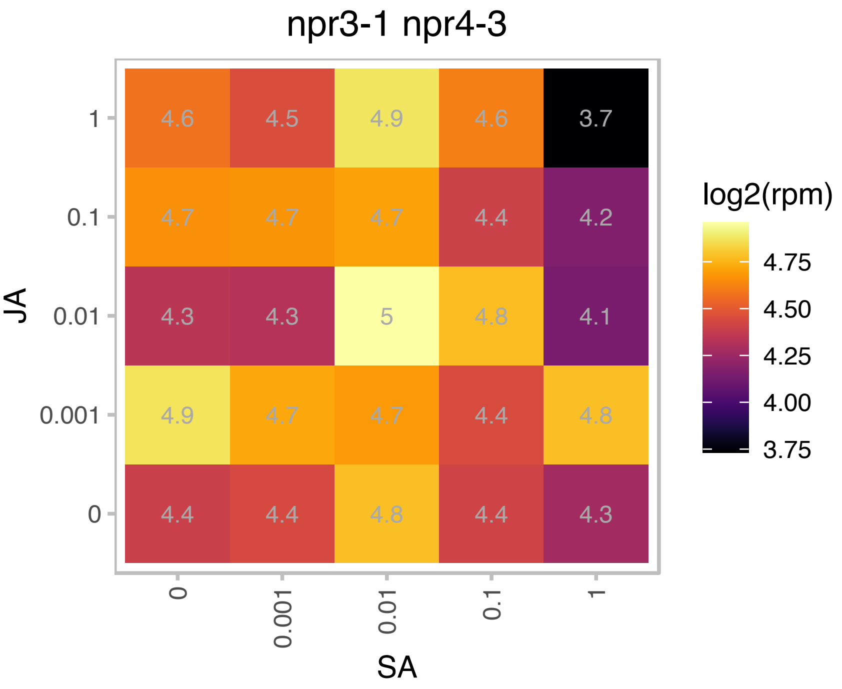

# AT4G01790.1 : NA

Ribosomal protein L7Ae/L30e/S12e/Gadd45 family protein

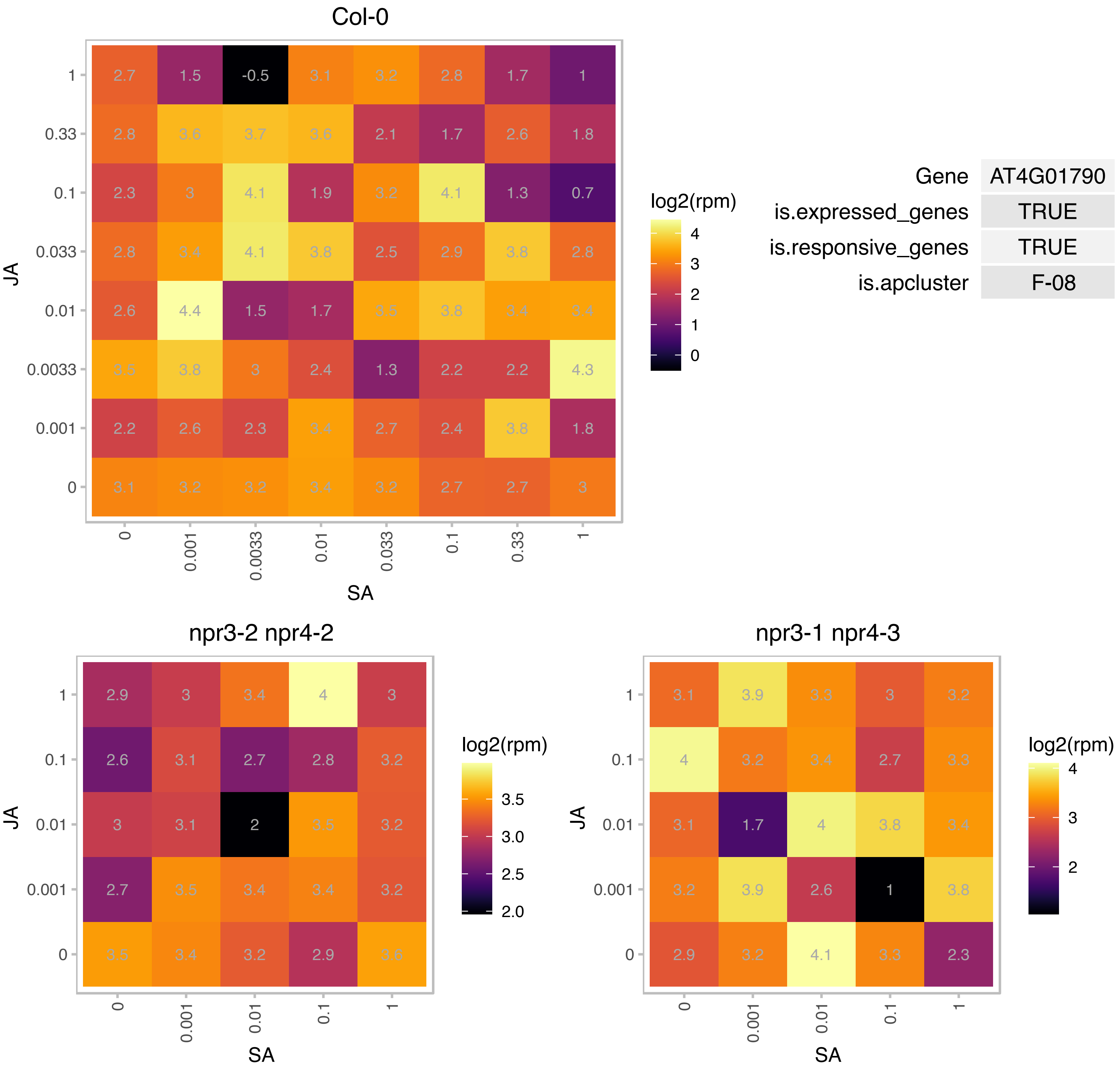


| | |
|---|---|
| Gene | AT4G01790 |
| is.expressed_genes | TRUE |
| is.responsive_genes | TRUE |
| is.apcluster | F-08 |

# AT4G04614.1 : NA

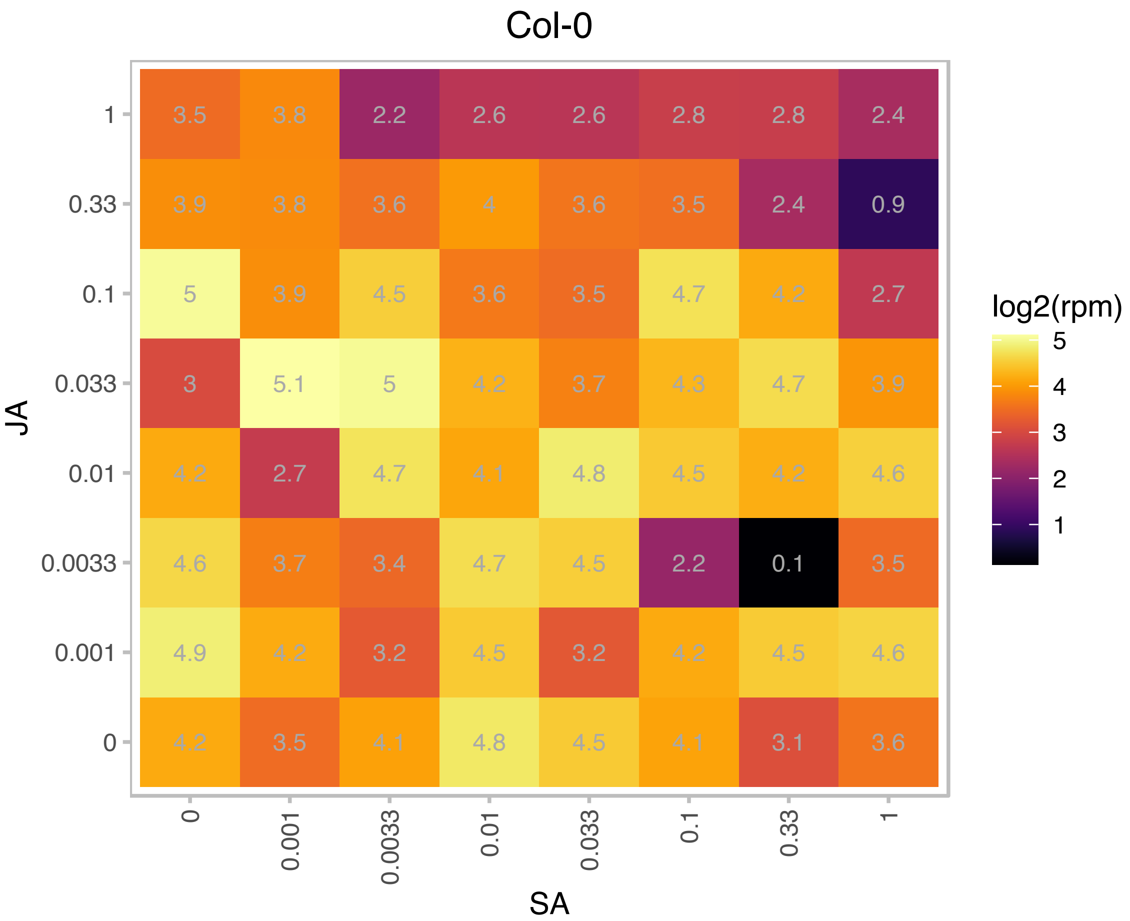


| | |
|---|---|
| Gene | AT4G04614 |
| is.expressed_genes | TRUE |
| is.responsive_genes | TRUE |
| is.apcluster | F-08 |

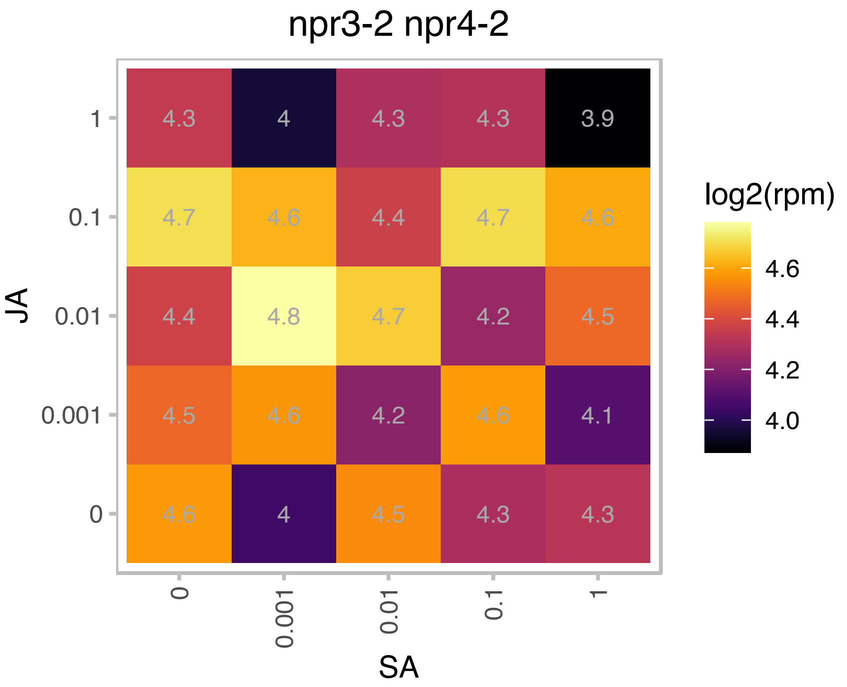


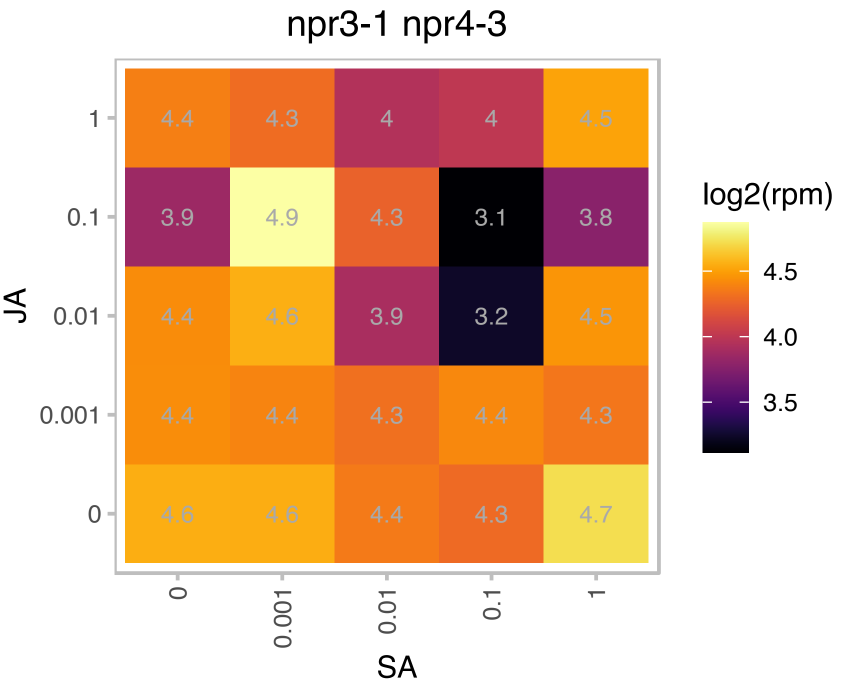

### AT4G09800.1 : RPS18C

S18 ribosomal protein

encodes a ribosomal protein S18C, a constituent of the small subunit of the ribosomal complex

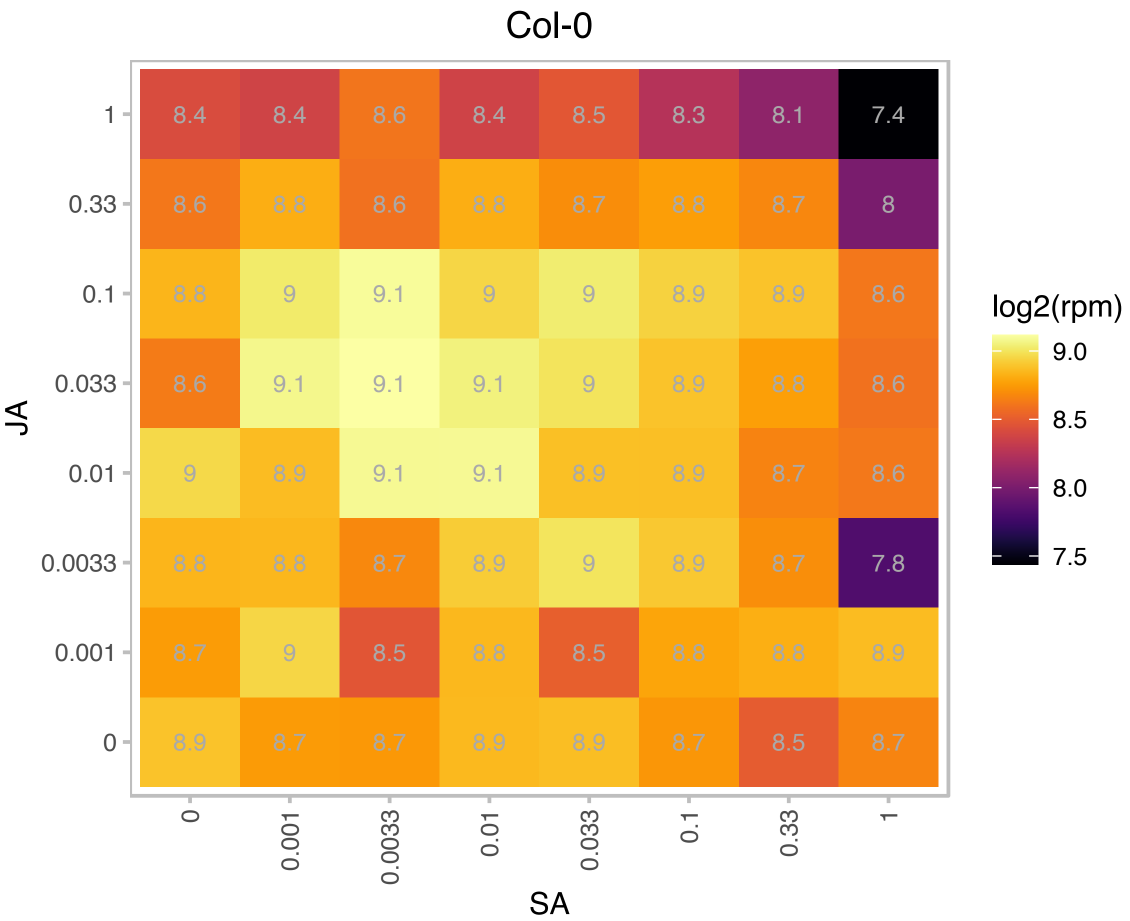


| | |
|---|---|
| Gene | AT4G09800 |
| is.expressed_genes | TRUE |
| is.responsive_genes | TRUE |
| is.apcluster | F-08 |

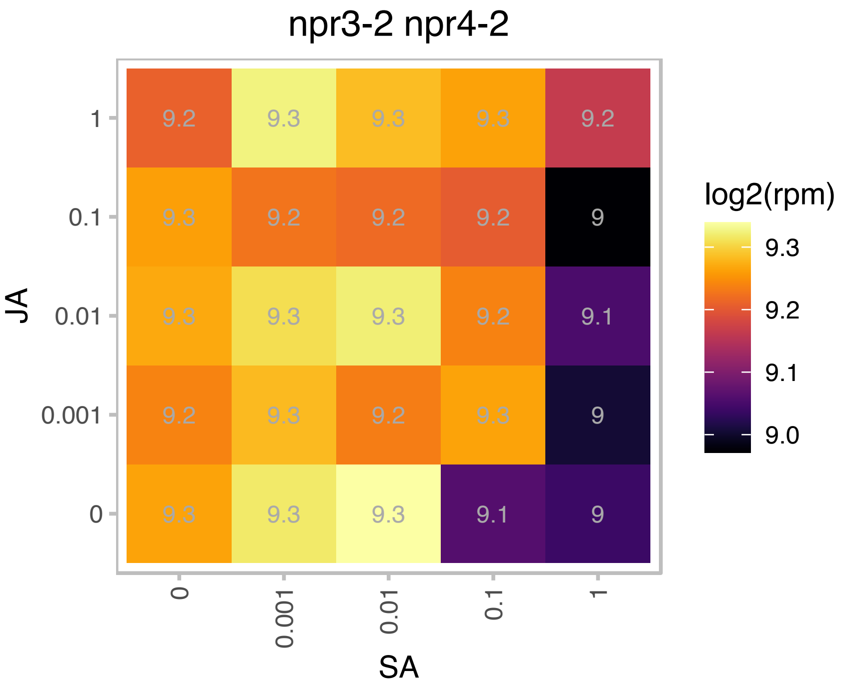


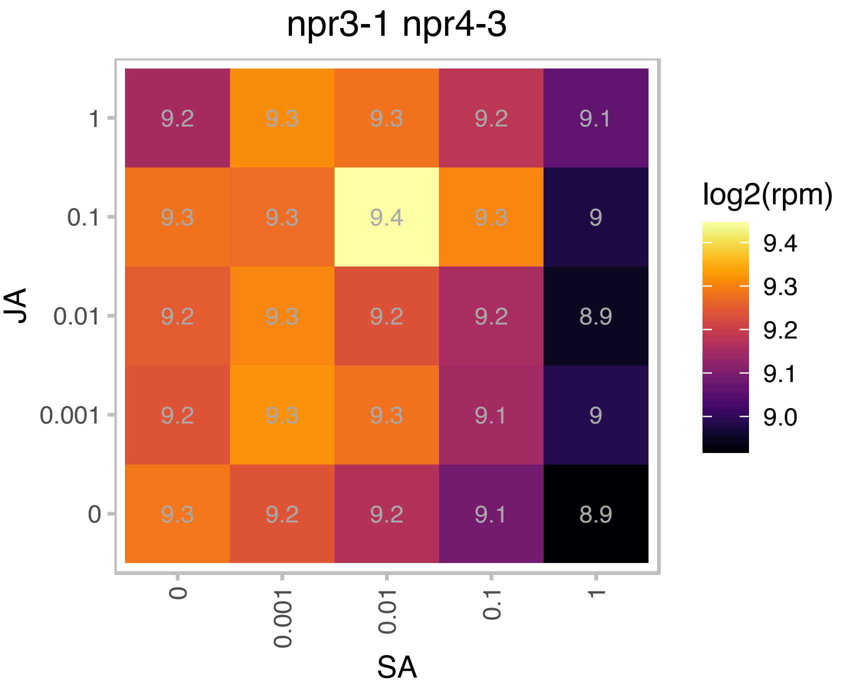

# AT4G10450.1 : NA

Ribosomal protein L6 family

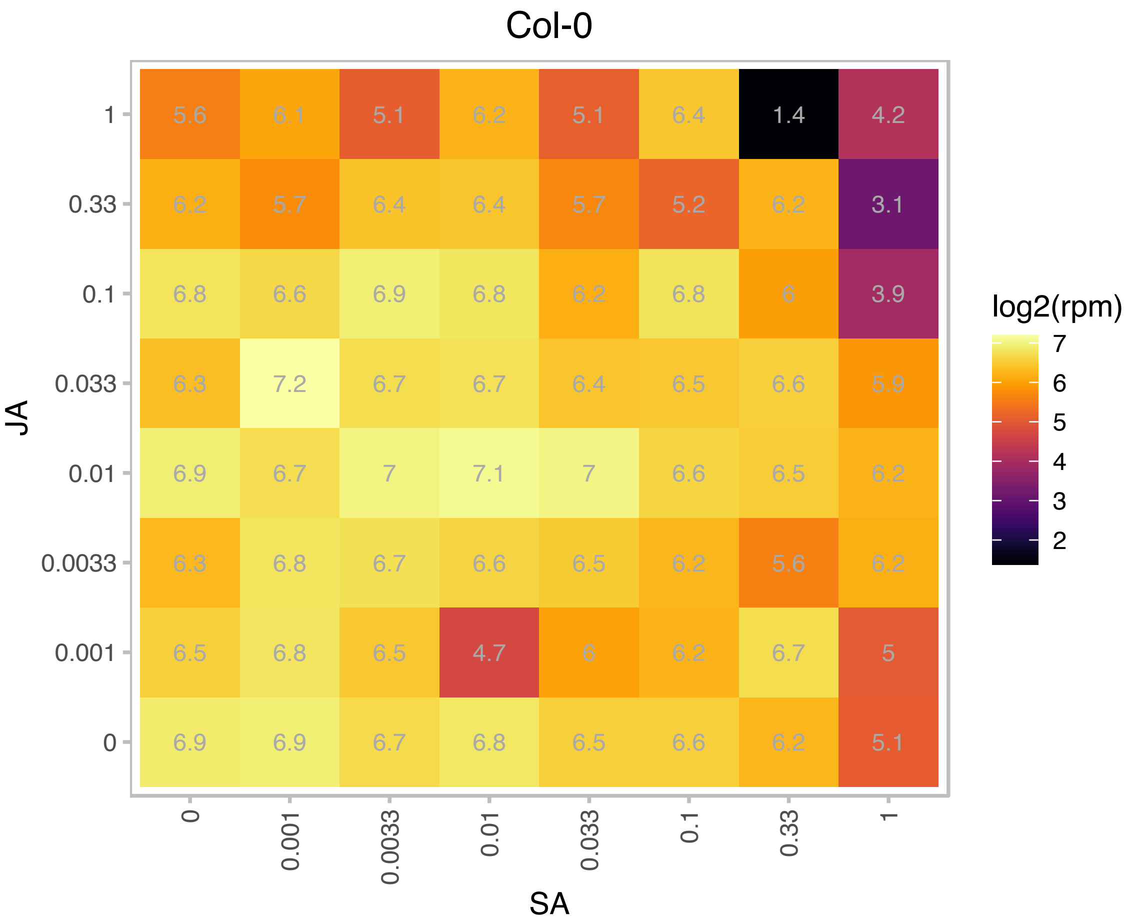


| | |
|---|---|
| Gene | AT4G10450 |
| is.expressed_genes | TRUE |
| is.responsive_genes | TRUE |
| is.apcluster | F-08 |

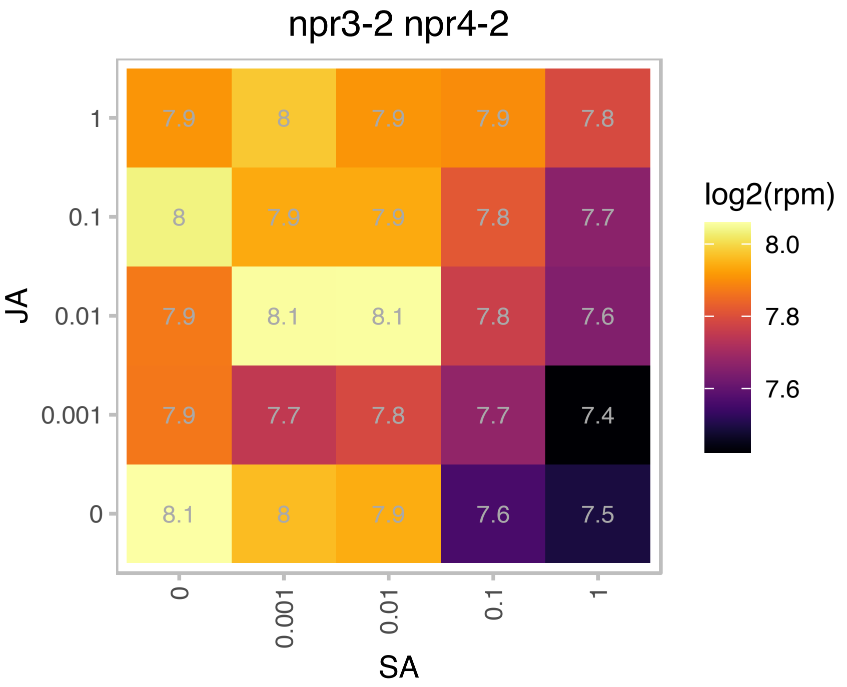


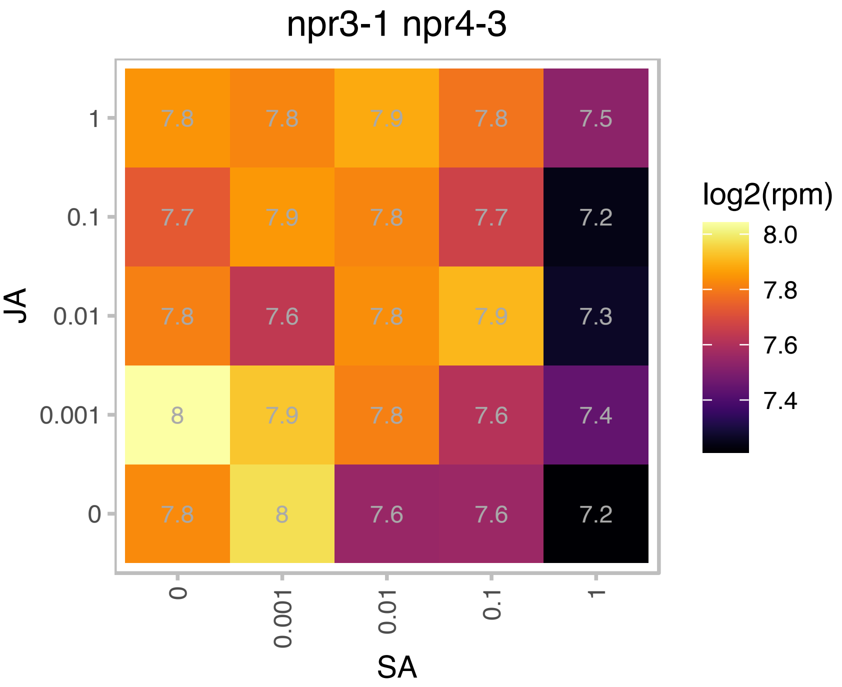

# AT4G13220.1 : NA

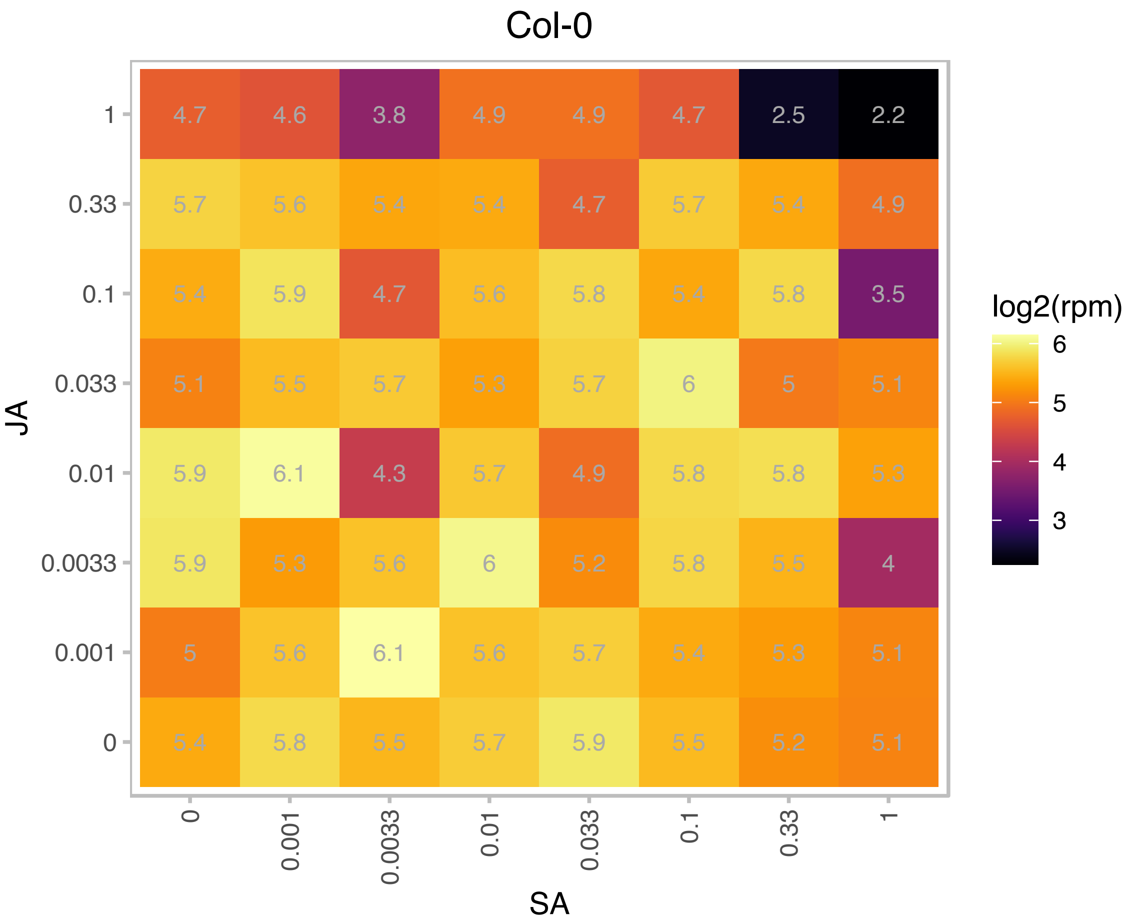


| | |
|---|---|
| Gene | AT4G13220 |
| is.expressed_genes | TRUE |
| is.responsive_genes | TRUE |
| is.apcluster | F-08 |

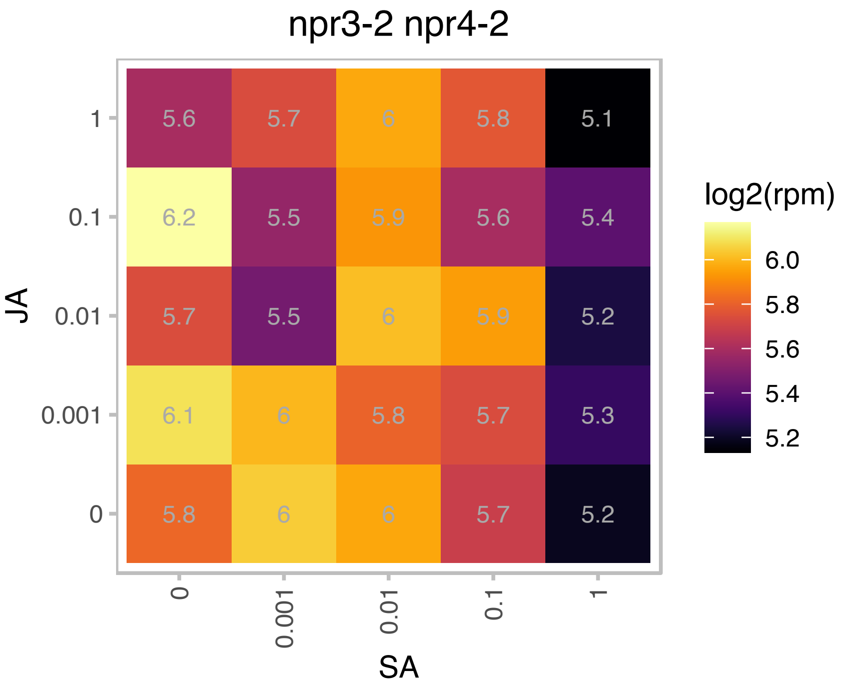


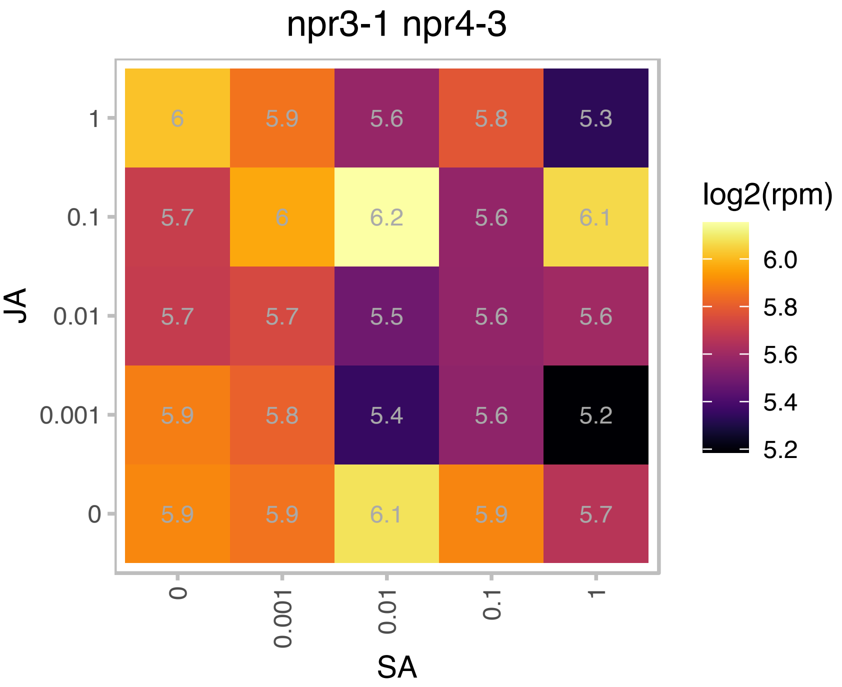

# AT4G13495.1 : NA

other RNA

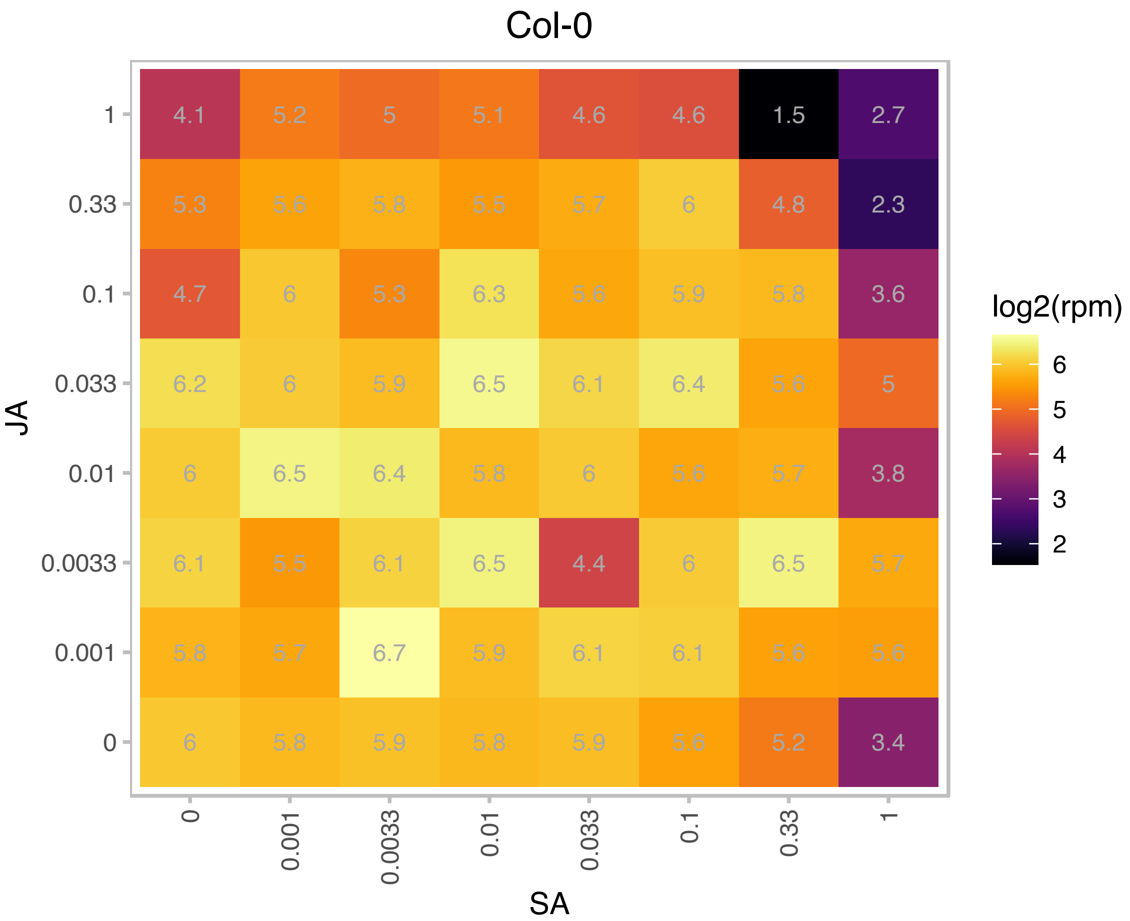


| Gene | AT4G13495 |
|---|---|
| is.expressed_genes | TRUE |
| is.responsive_genes | TRUE |
| is.apcluster | F-08 |

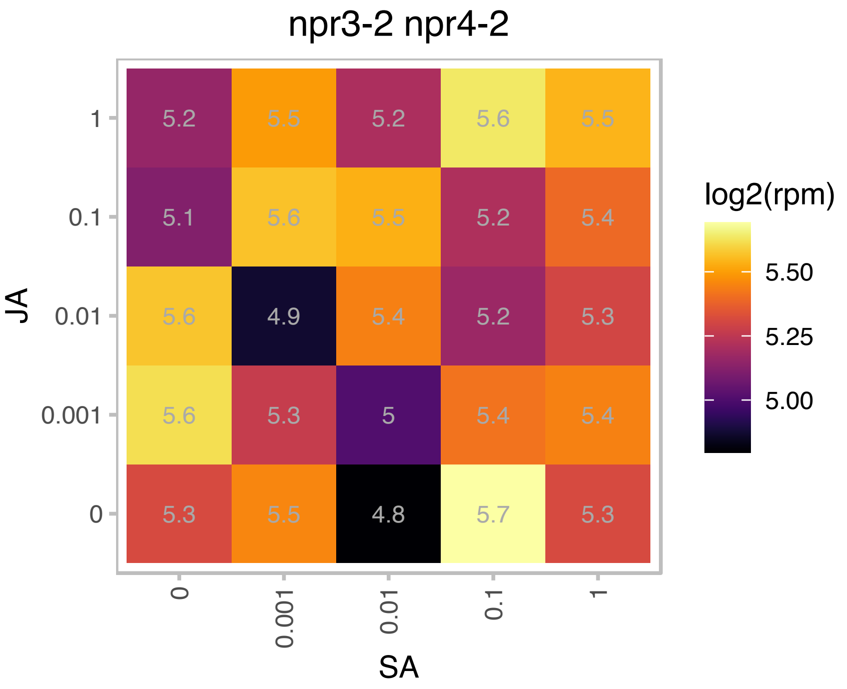


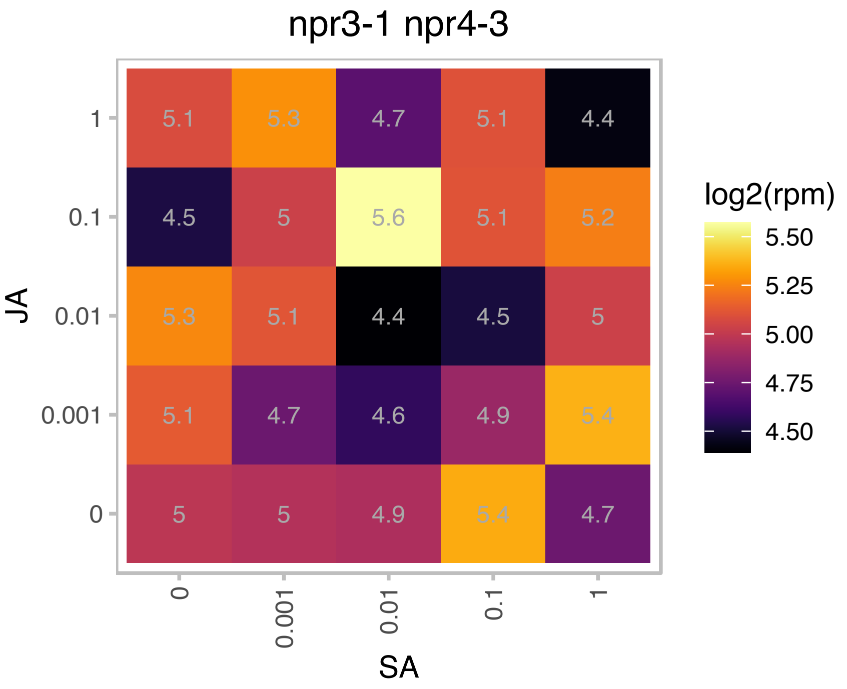

### AT4G13830.1 : DJC26, J20

DNAJ-like 20

DnaJ-like protein (J20); nuclear gene

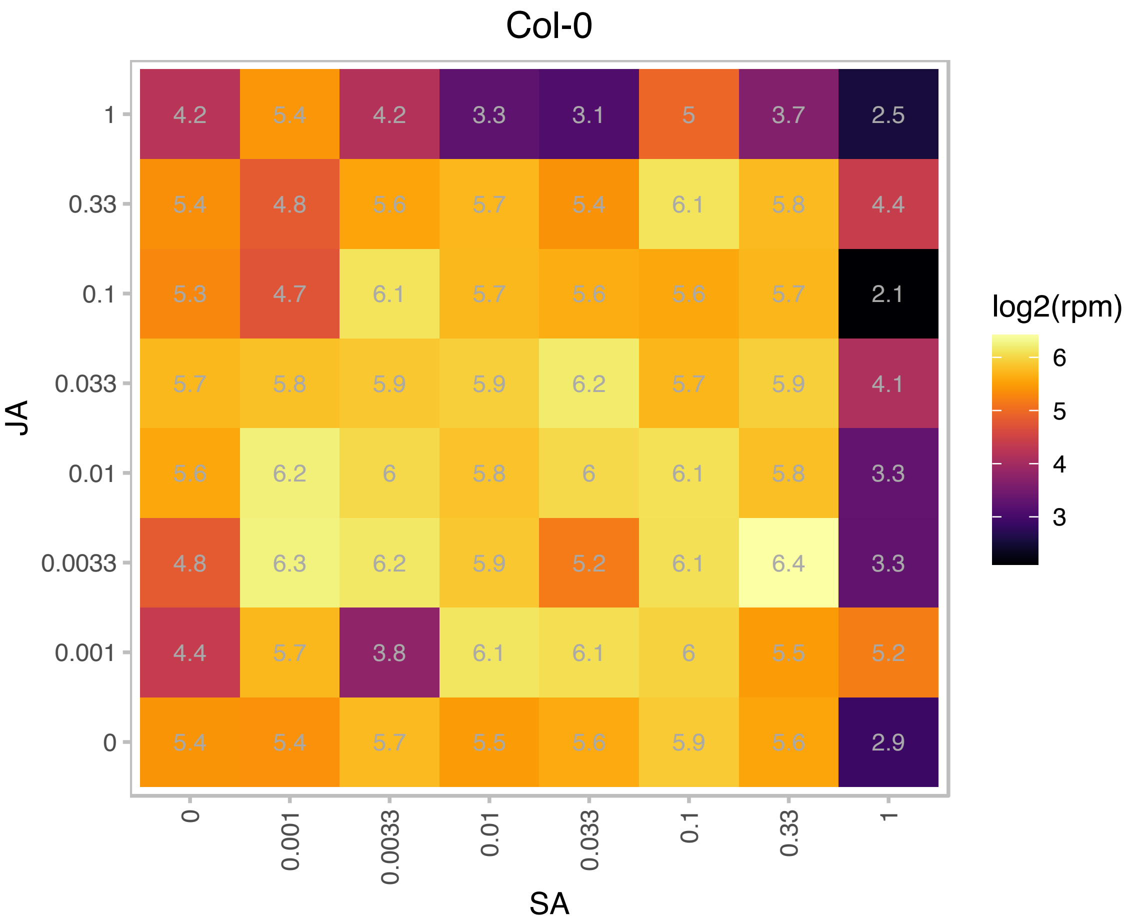


| | |
|---|---|
| Gene | AT4G13830 |
| is.expressed_genes | TRUE |
| is.responsive_genes | TRUE |
| is.apcluster | F-08 |

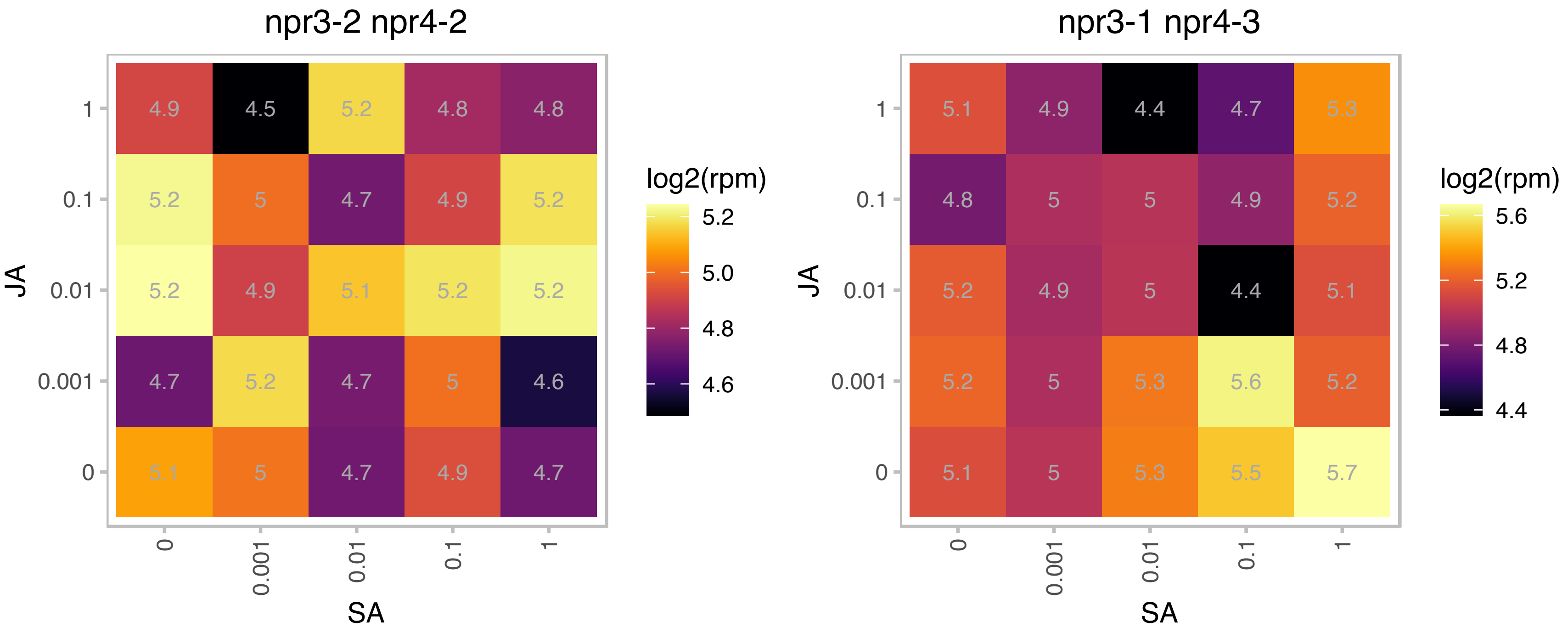

### AT4G14455.1 : ATBET12, ATBS14B, BET12, BS14B

Target SNARE coiled-coil domain protein

Encodes a Bet1/Sft1-like SNARE protein, which can only partially suppresses the temperature-sensitive growth defect in <i>sft1-1</i> yeast cells; however, it cannot support the deletion of the yeast BET1 gene (<i>bet1Δ</i>). In yeast, Bet1p is the v-SNARE (soluble N-ethylmaleimide-sensitive factor adaptor protein receptor, V-type) involved in trafficking between the ER and Golgi.

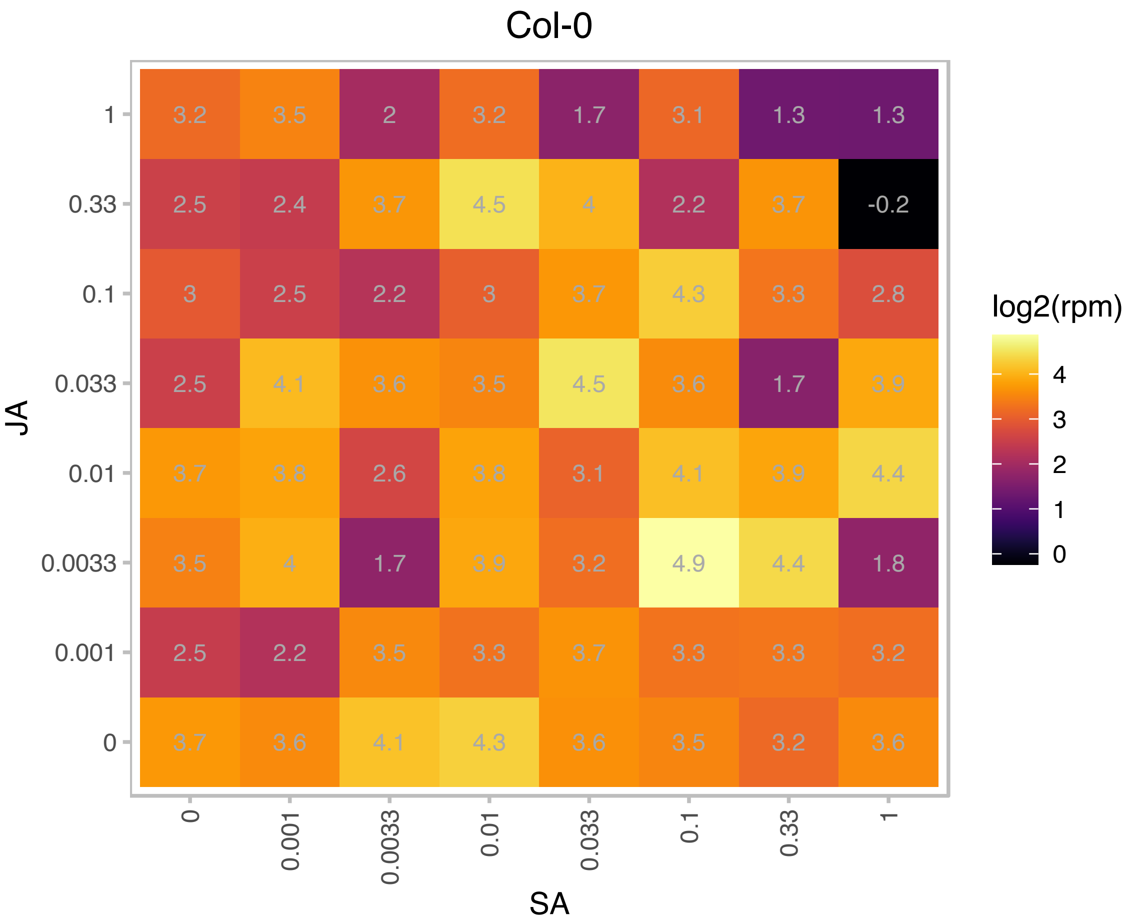


| | |
|---|---|
| Gene | AT4G14455 |
| is.expressed_genes | TRUE |
| is.responsive_genes | TRUE |
| is.apcluster | F-08 |

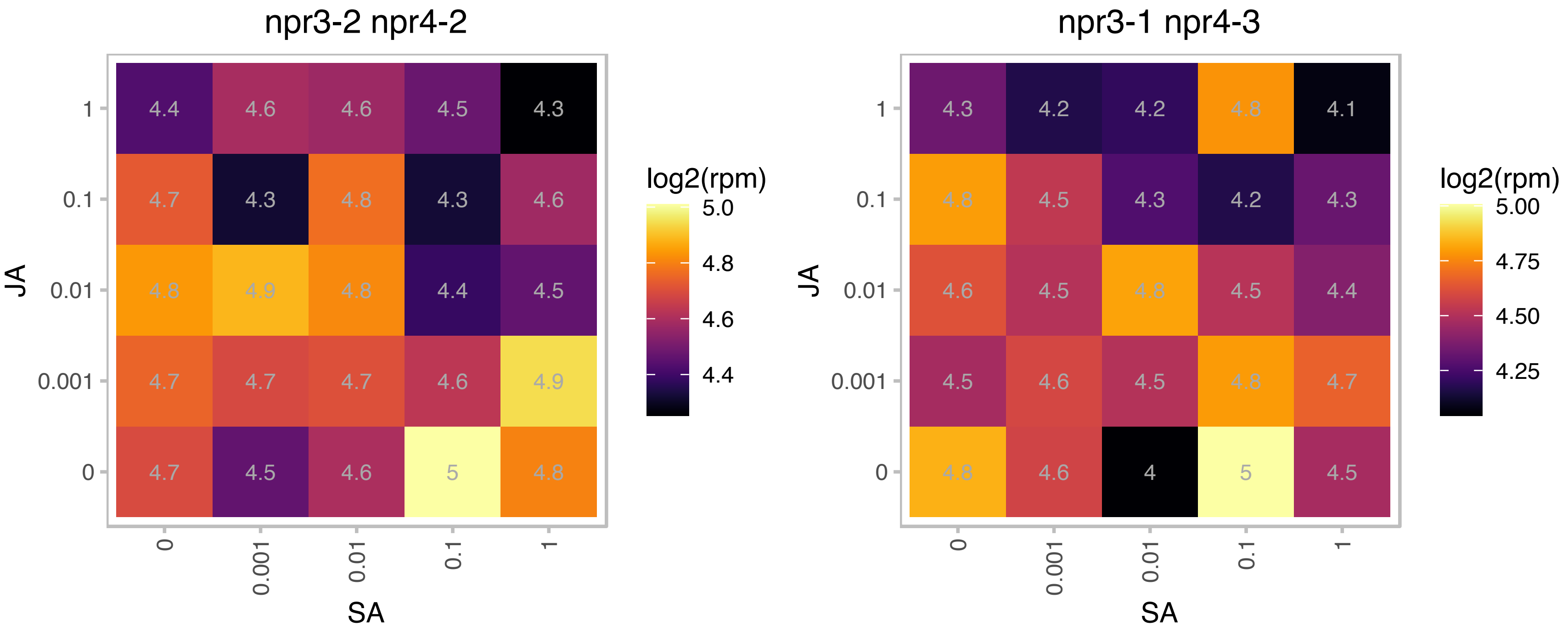

### AT4G17050.1 : UGLYAH

ureidoglycine aminohydrolase

Encodes a protein with ureidoglycine aminohydrolase activity.

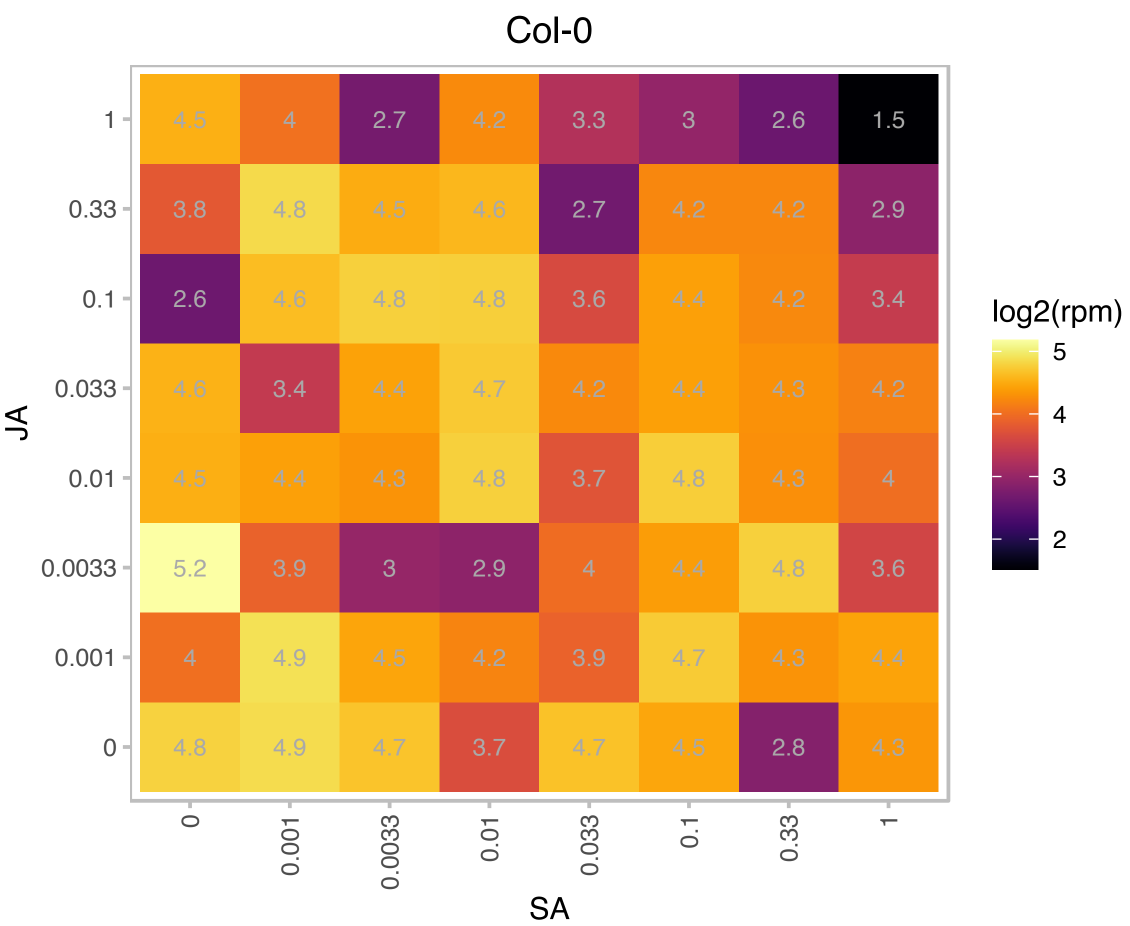


| | |
|---|---|
| Gene | AT4G17050 |
| is.expressed_genes | TRUE |
| is.responsive_genes | TRUE |
| is.apcluster | F-08 |

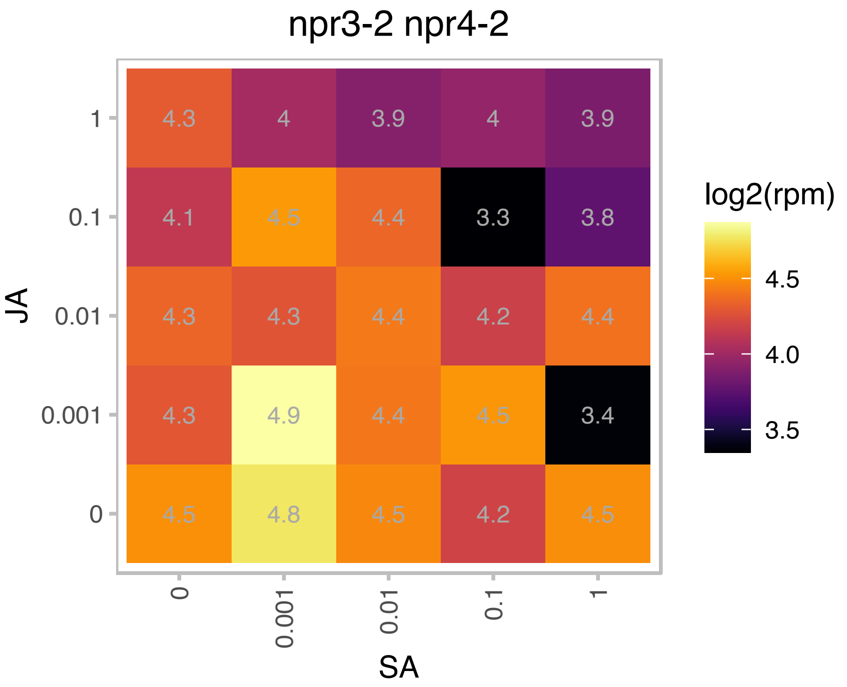


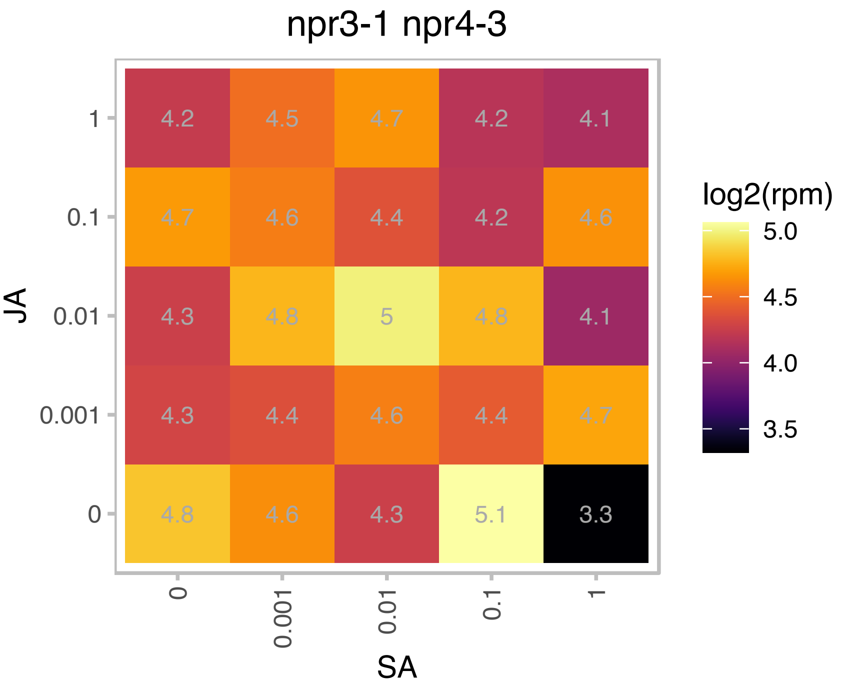

# AT4G18593.1 : NA

dual specificity protein phosphatase-related

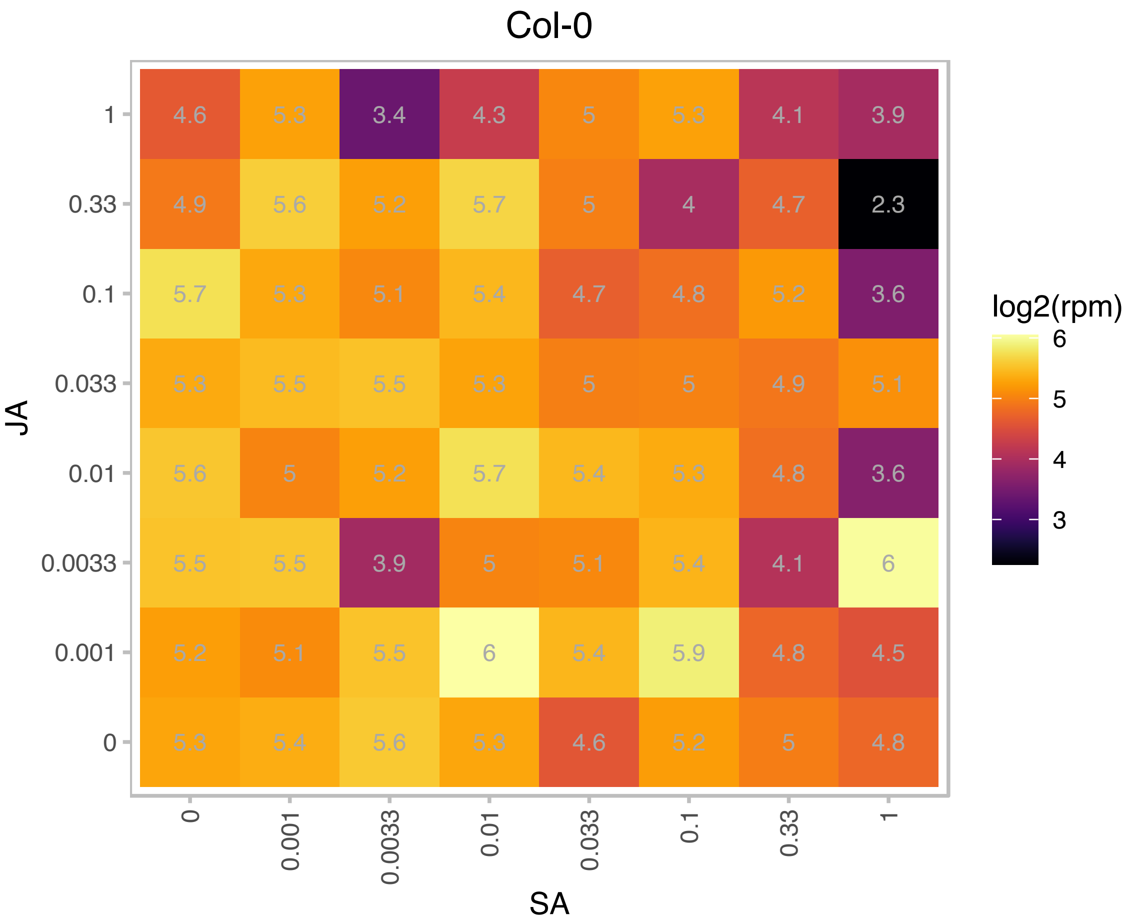


| | |
|---|---|
| Gene | AT4G18593 |
| is.expressed_genes | TRUE |
| is.responsive_genes | TRUE |
| is.apcluster | F-08 |

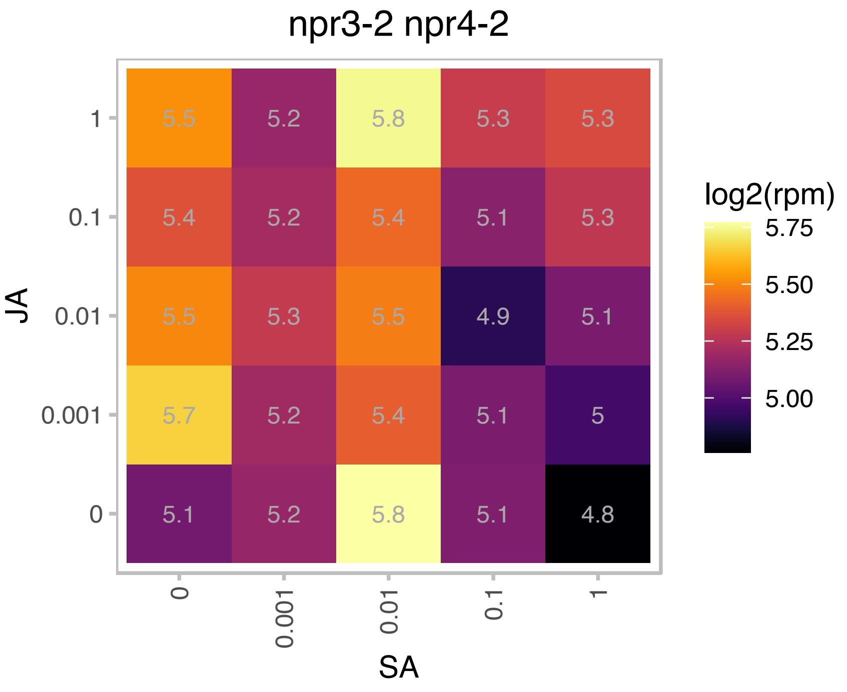


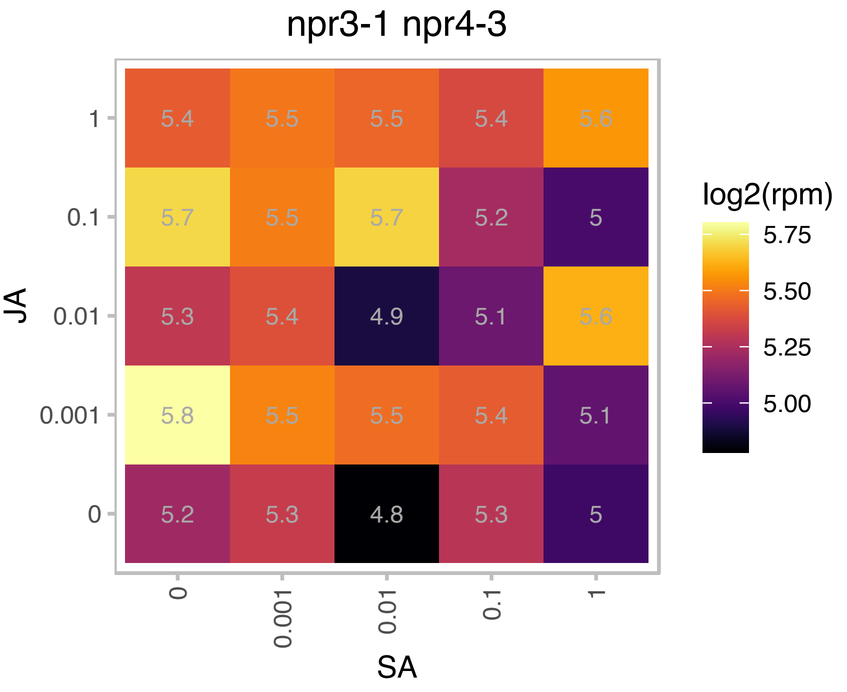

# AT4G22380.1 : NA

Ribosomal protein L7Ae/L30e/S12e/Gadd45 family protein

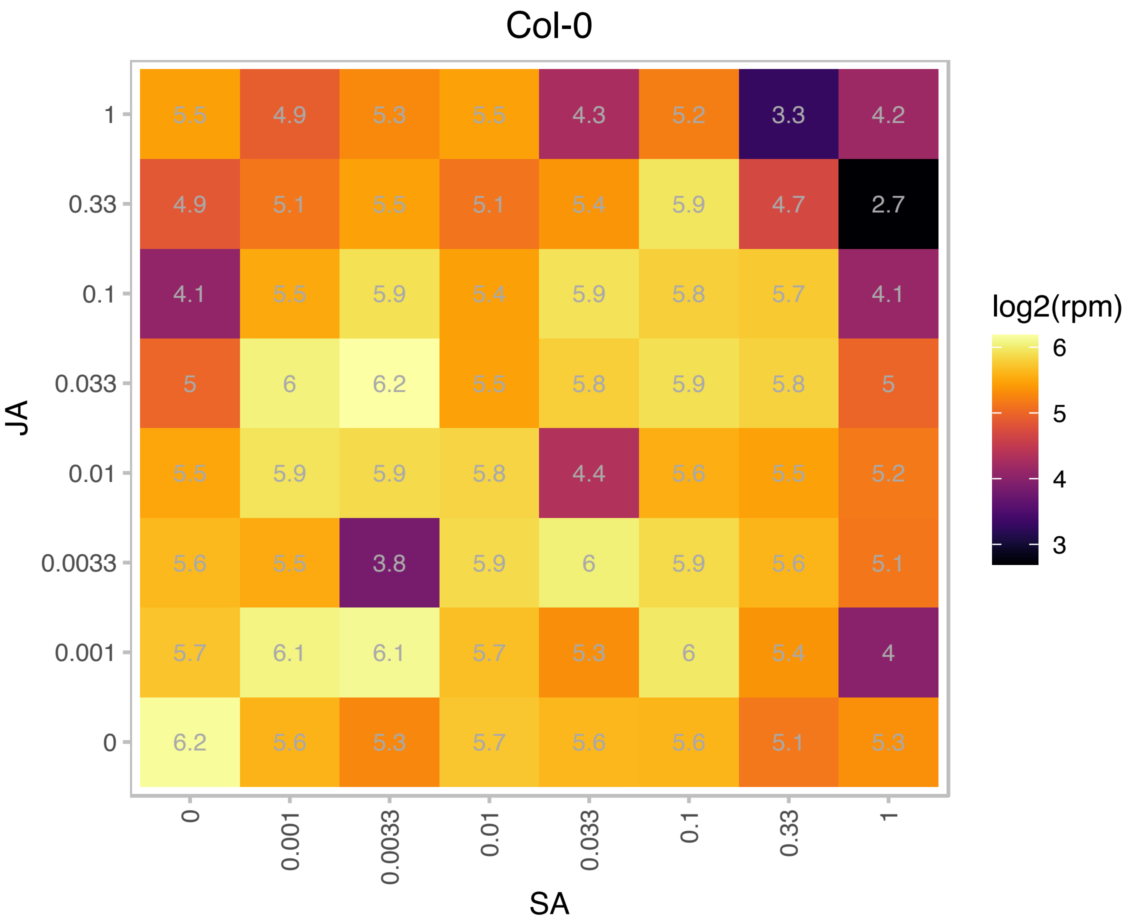


| | |
|---|---|
| Gene | AT4G22380 |
| is.expressed_genes | TRUE |
| is.responsive_genes | TRUE |
| is.apcluster | F-08 |

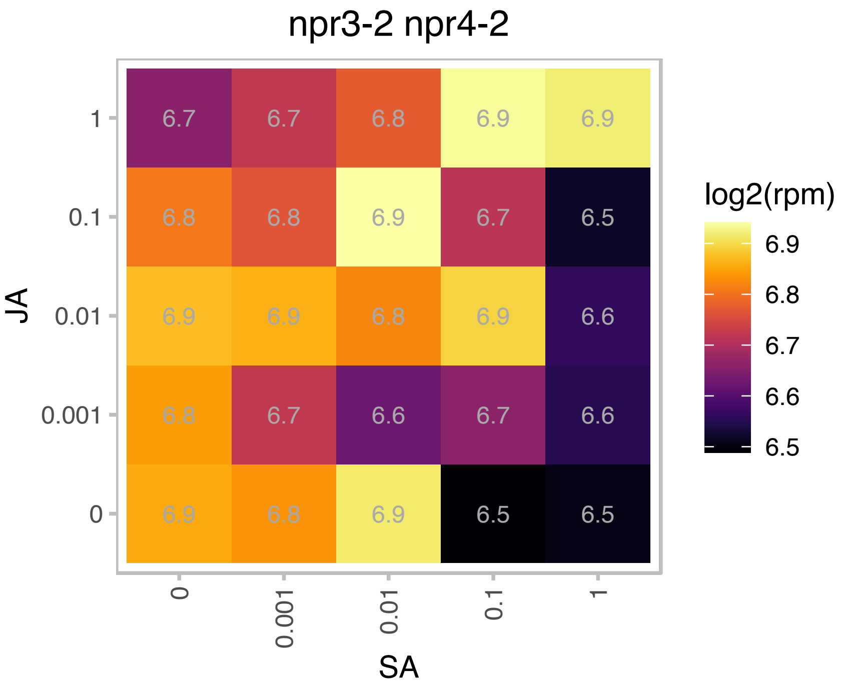


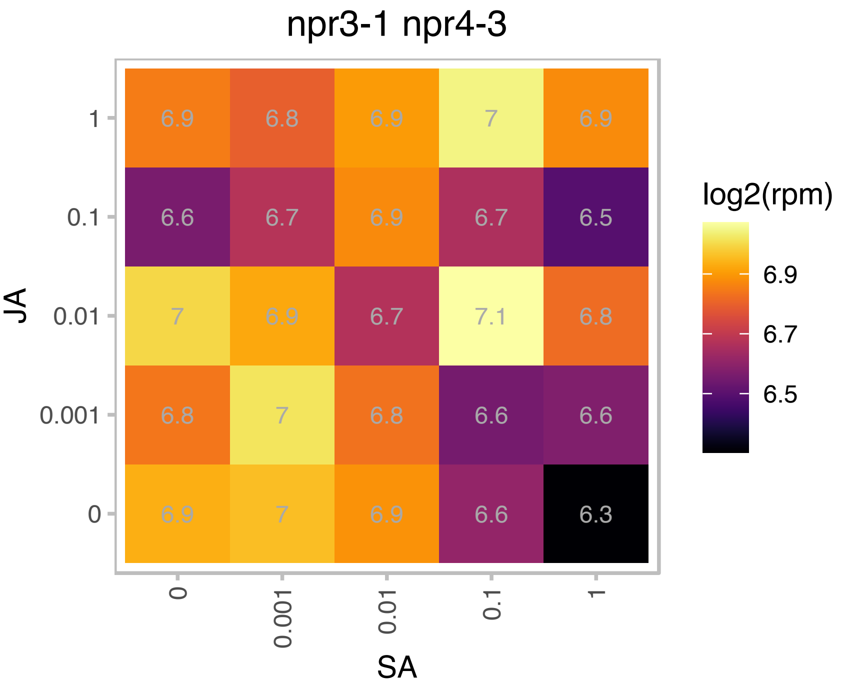

# AT4G22475.1 : NA

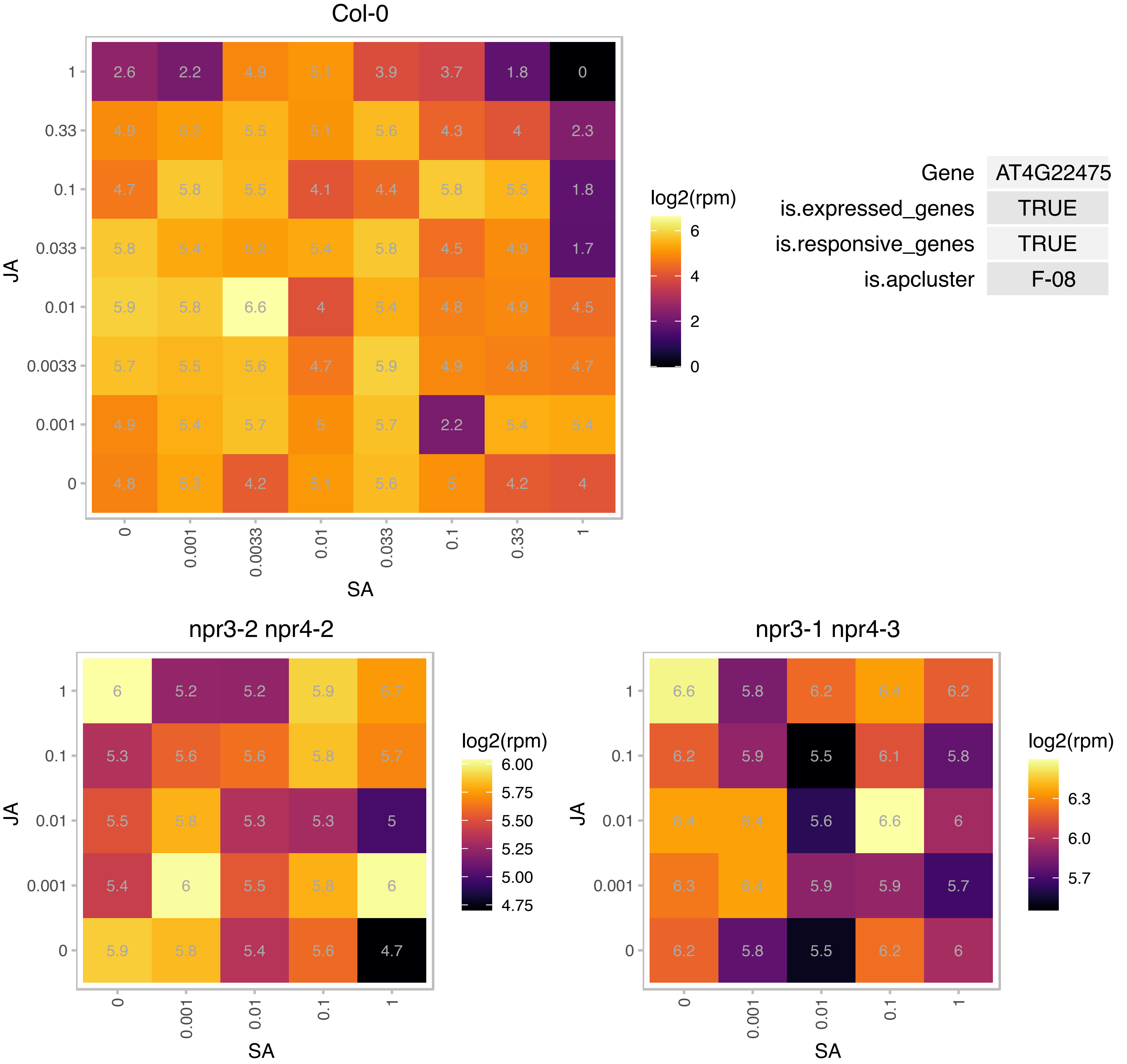


| Gene | AT4G22475 |
|---|---|
| is.expressed_genes | TRUE |
| is.responsive_genes | TRUE |
| is.apcluster | F-08 |

# AT4G24275.1 : NA

Identified as a screen for stress-responsive genes.

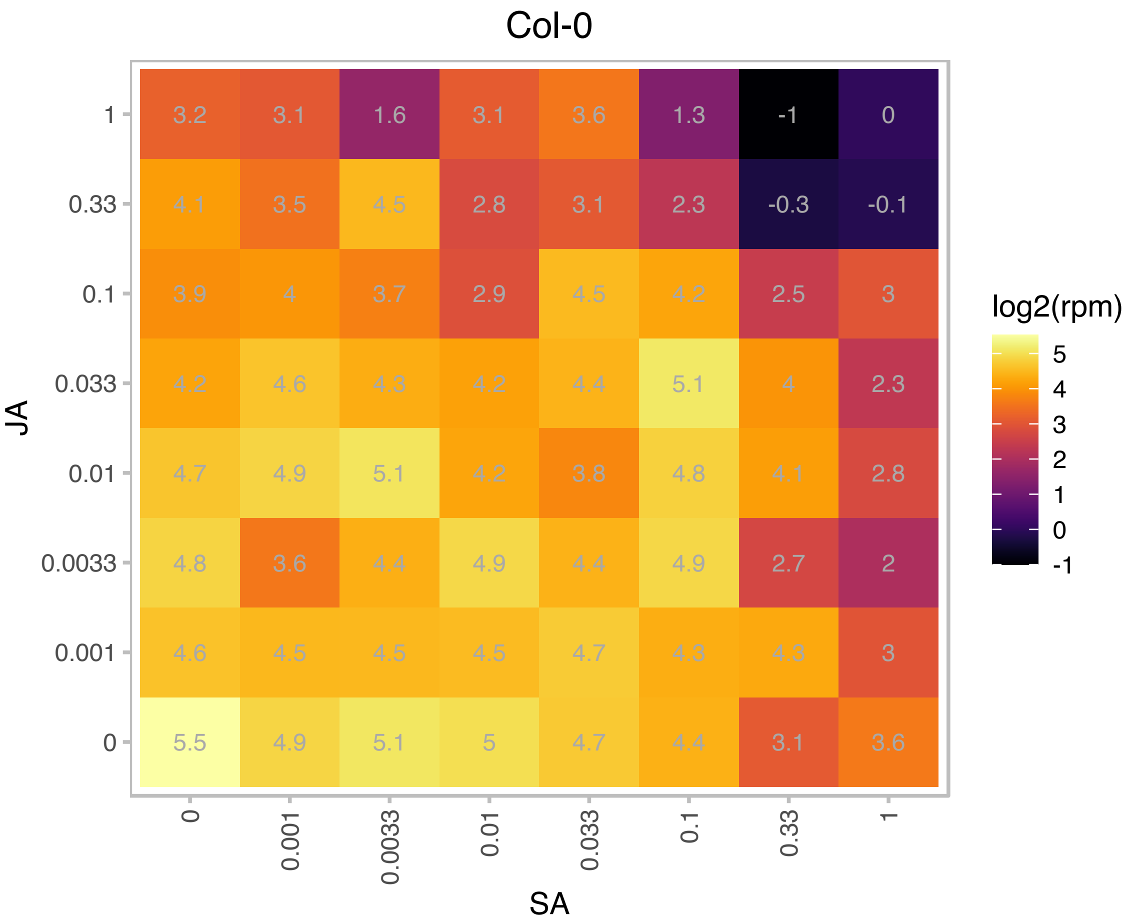


| | |
|---|---|
| Gene | AT4G24275 |
| is.expressed_genes | TRUE |
| is.responsive_genes | TRUE |
| is.apcluster | F-08 |

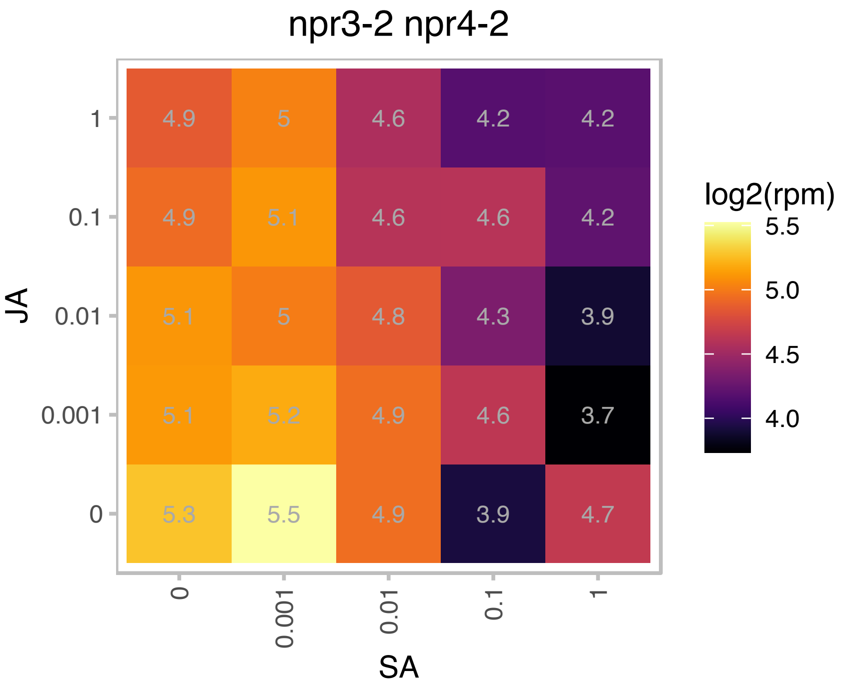


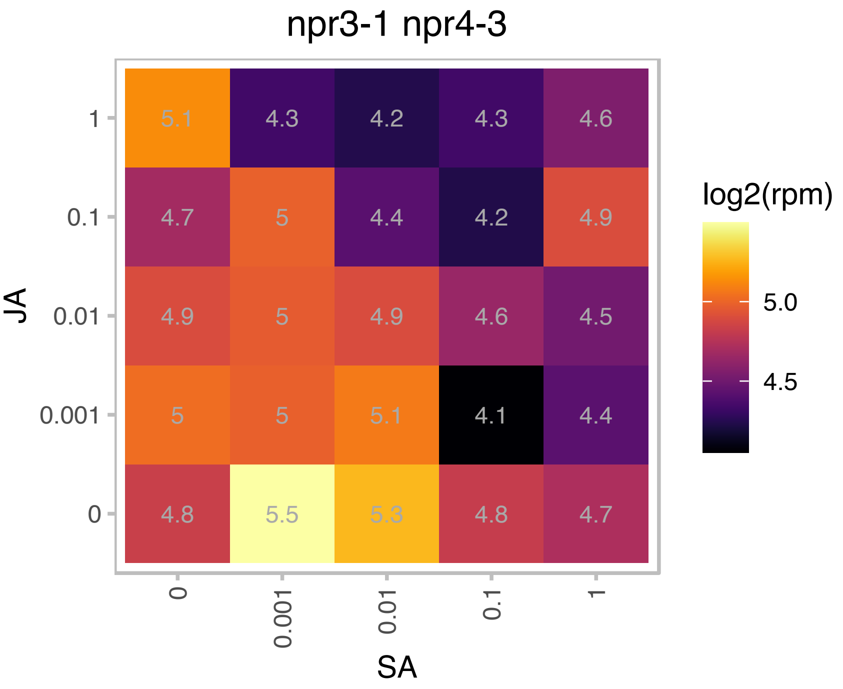

### AT4G25340.1 : ATFKBP53, FKBP53

FK506 BINDING PROTEIN 53

Encodes a member of the FKBP-type immunophilin family that functions as a histone chaparone. Binds to 18S rDNA and represses its expression.

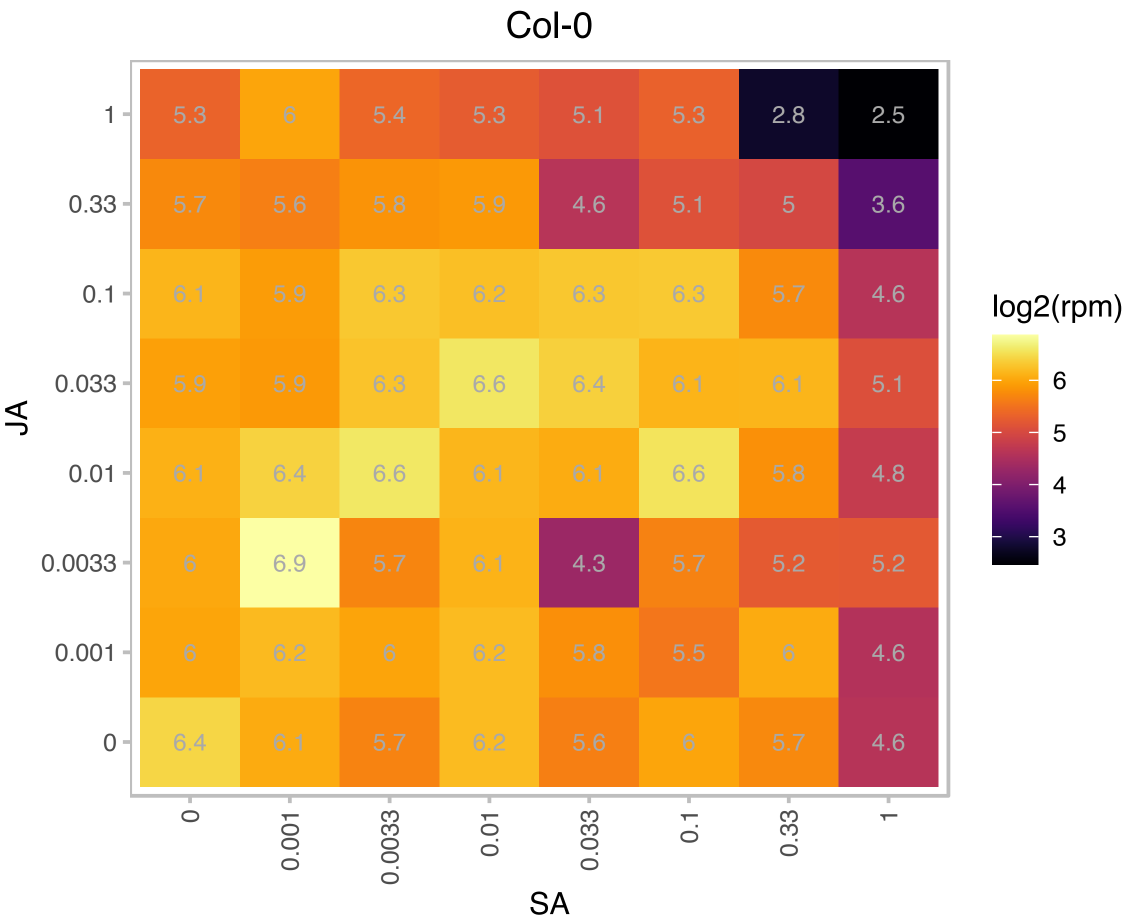


| | |
|---|---|
| Gene | AT4G25340 |
| is.expressed_genes | TRUE |
| is.responsive_genes | TRUE |
| is.apcluster | F-08 |

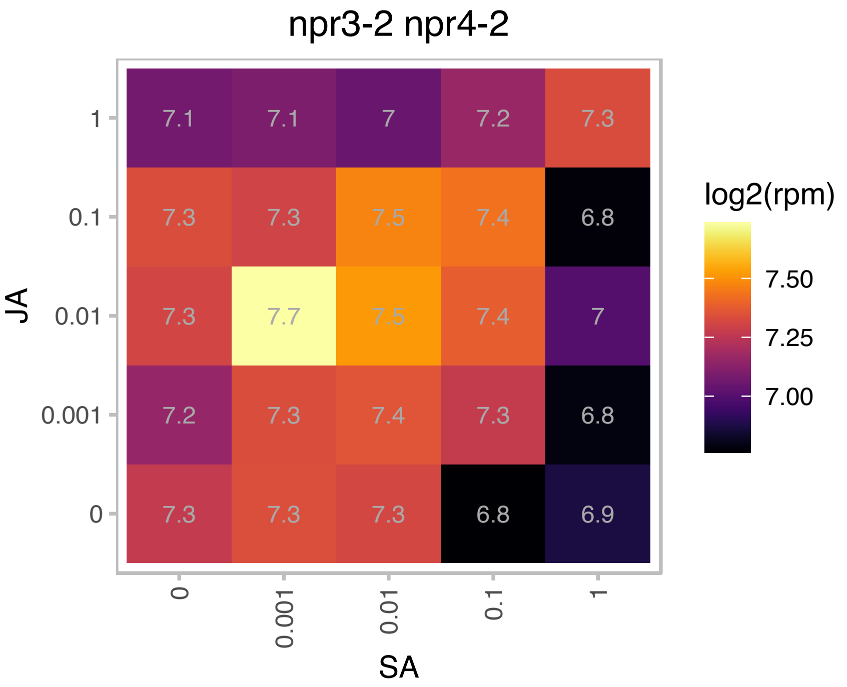


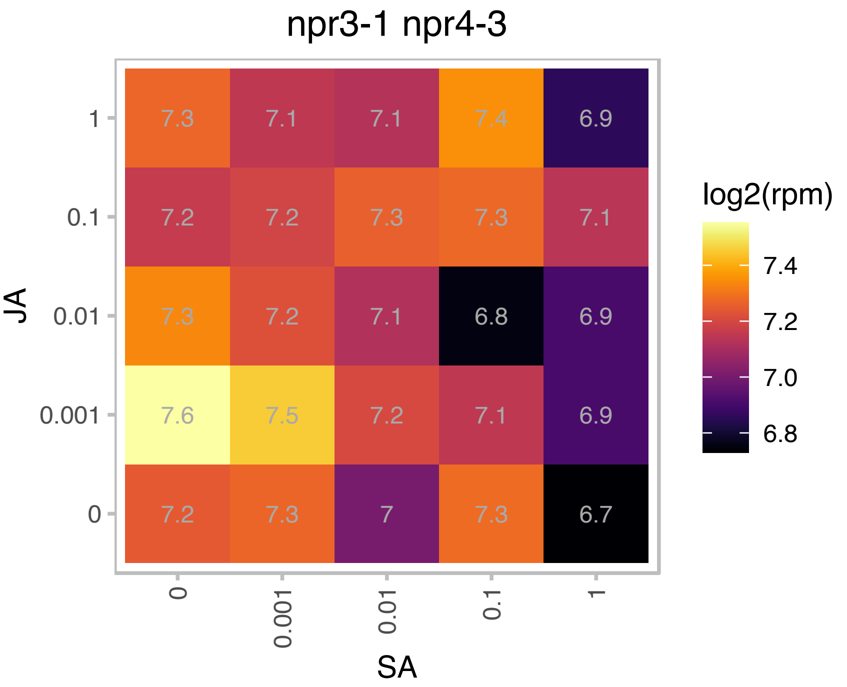

### AT4G26320.1 : AGP13

arabinogalactan protein 13

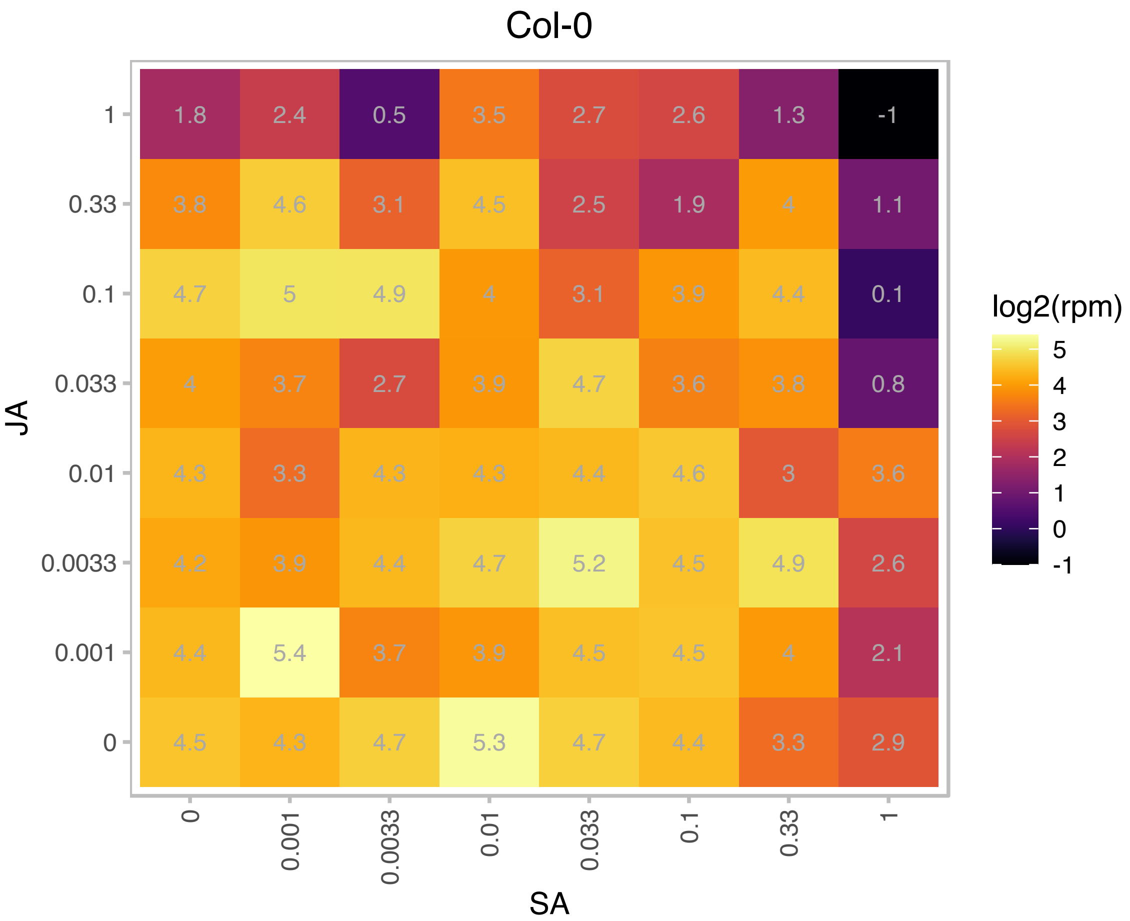


| | |
|---|---|
| Gene | AT4G26320 |
| is.expressed_genes | TRUE |
| is.responsive_genes | TRUE |
| is.apcluster | F-08 |

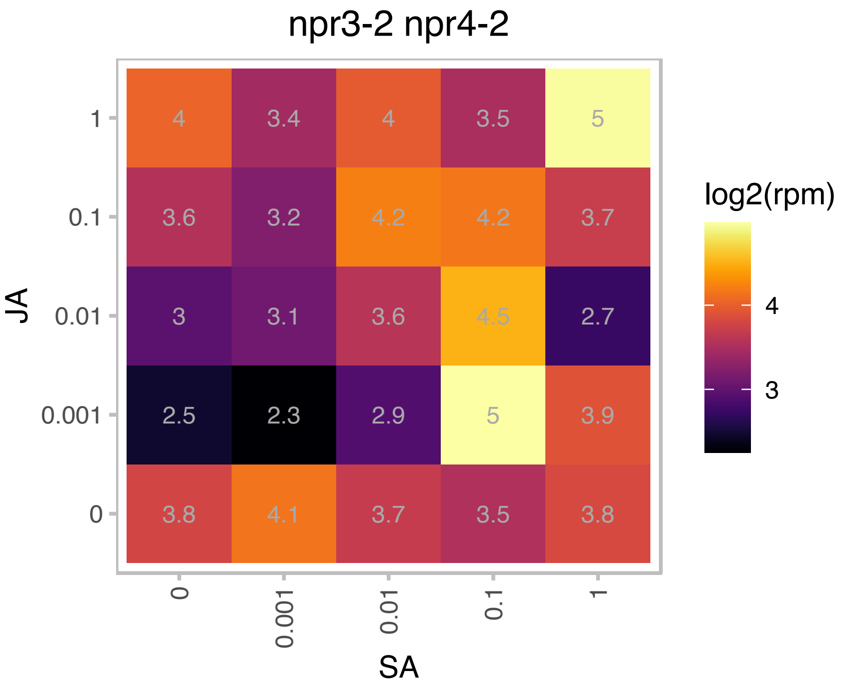


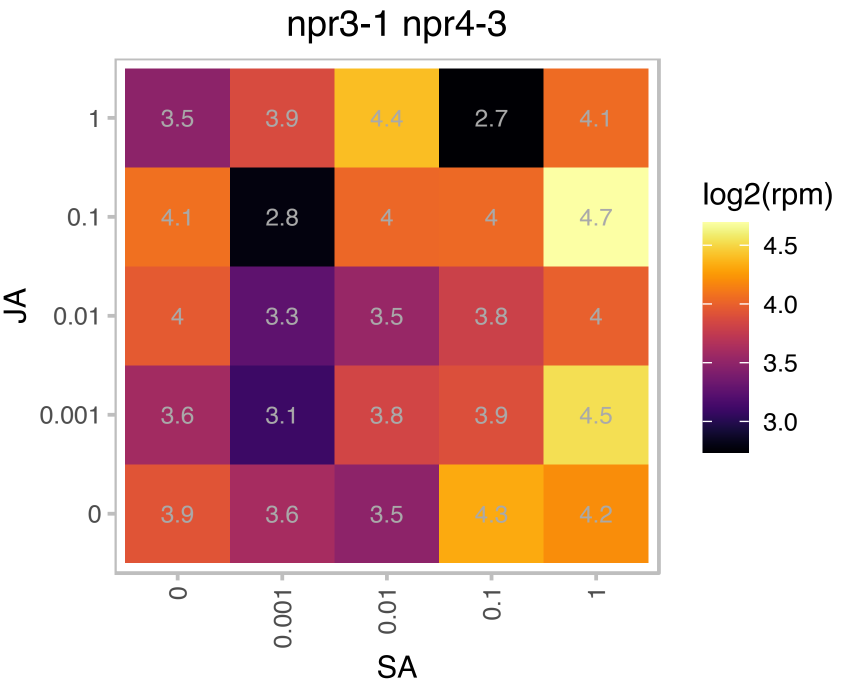

### AT4G26840.1 : ATSUMO1, SUM1, SUMO 1, SUMO1

small ubiquitin-like modifier 1
Encodes a small ubiquitin-like modifier (SUMO) polypeptide that becomes covalently attached to various intracellular protein targets, much like ubiquitination, leading to post-translational modification of those targets.

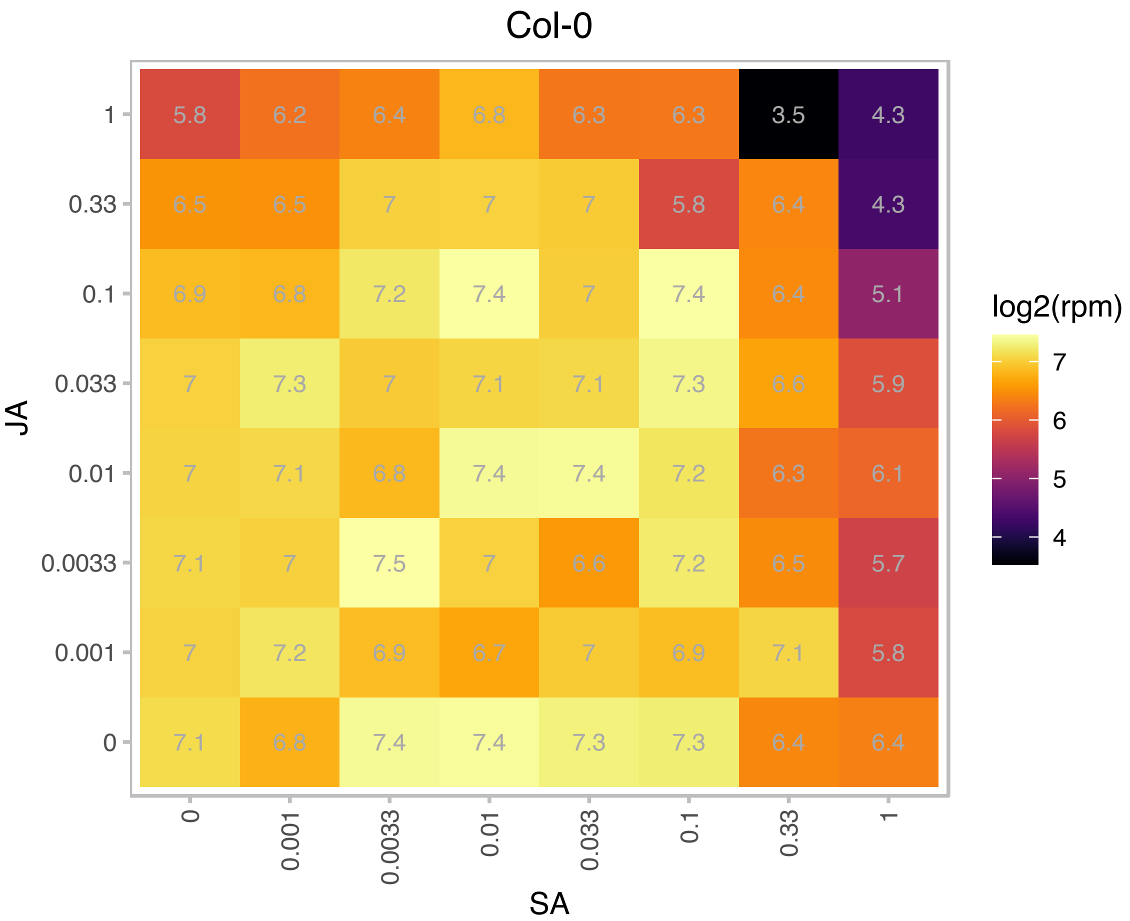


| | |
|---|---|
| Gene | AT4G26840 |
| is.expressed_genes | TRUE |
| is.responsive_genes | TRUE |
| is.apcluster | F-08 |

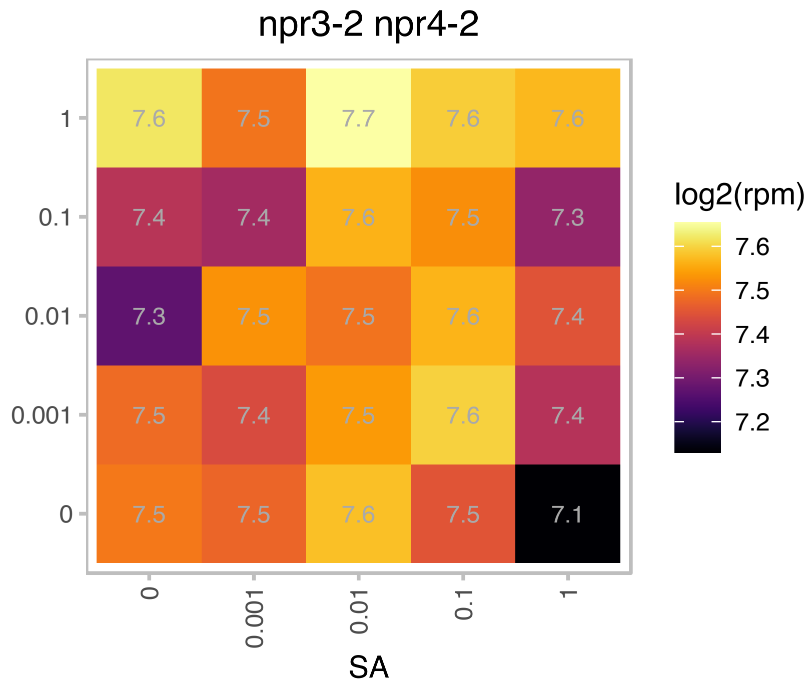


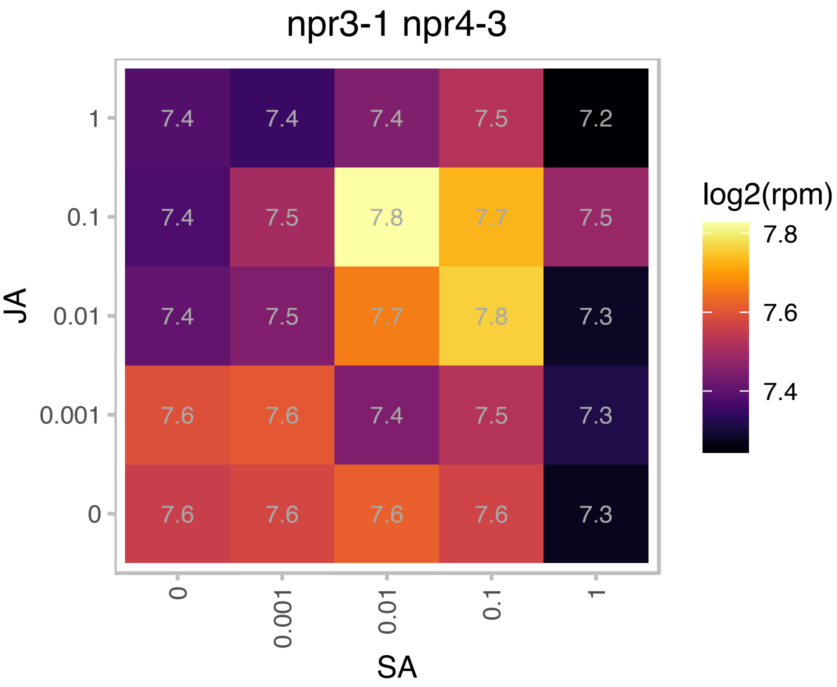

# AT4G26960.1 : NA

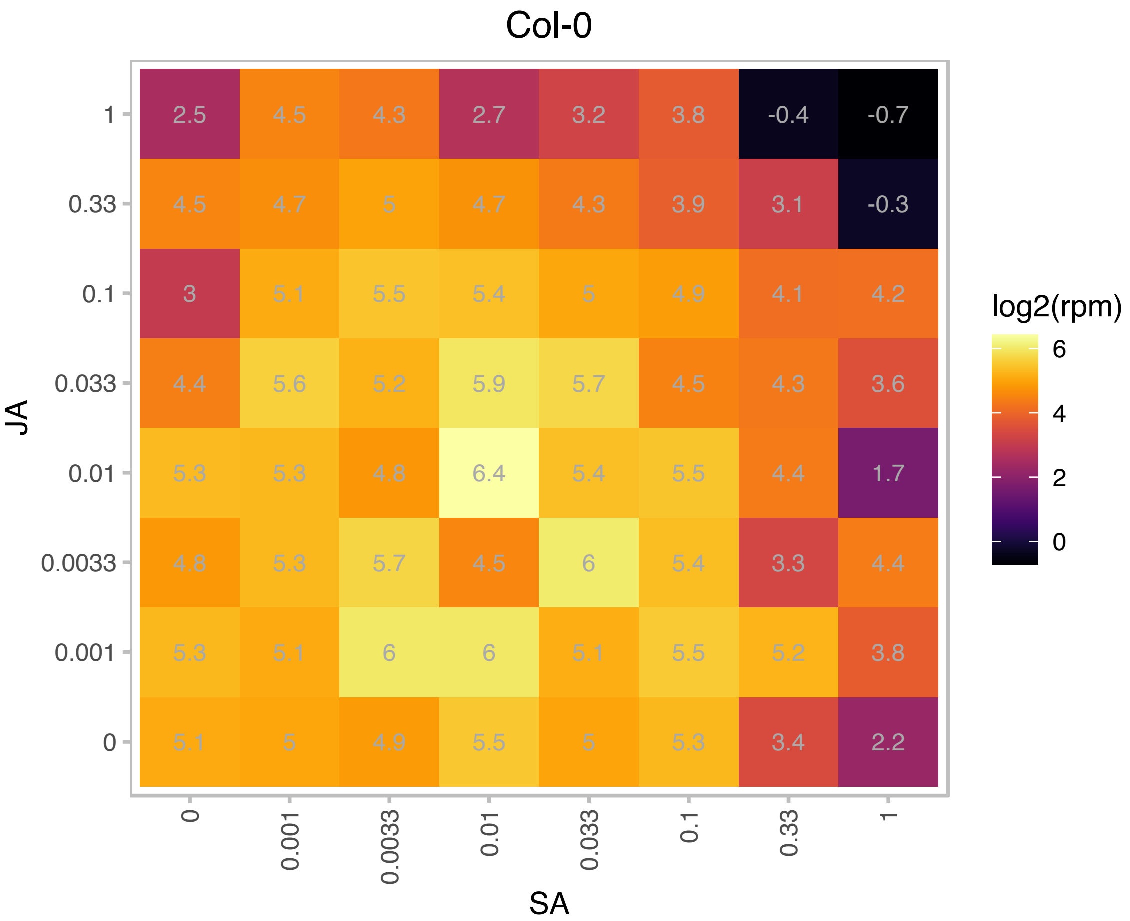


| | |
|---|---|
| Gene | AT4G26960 |
| is.expressed_genes | TRUE |
| is.responsive_genes | TRUE |
| is.apcluster | F-08 |

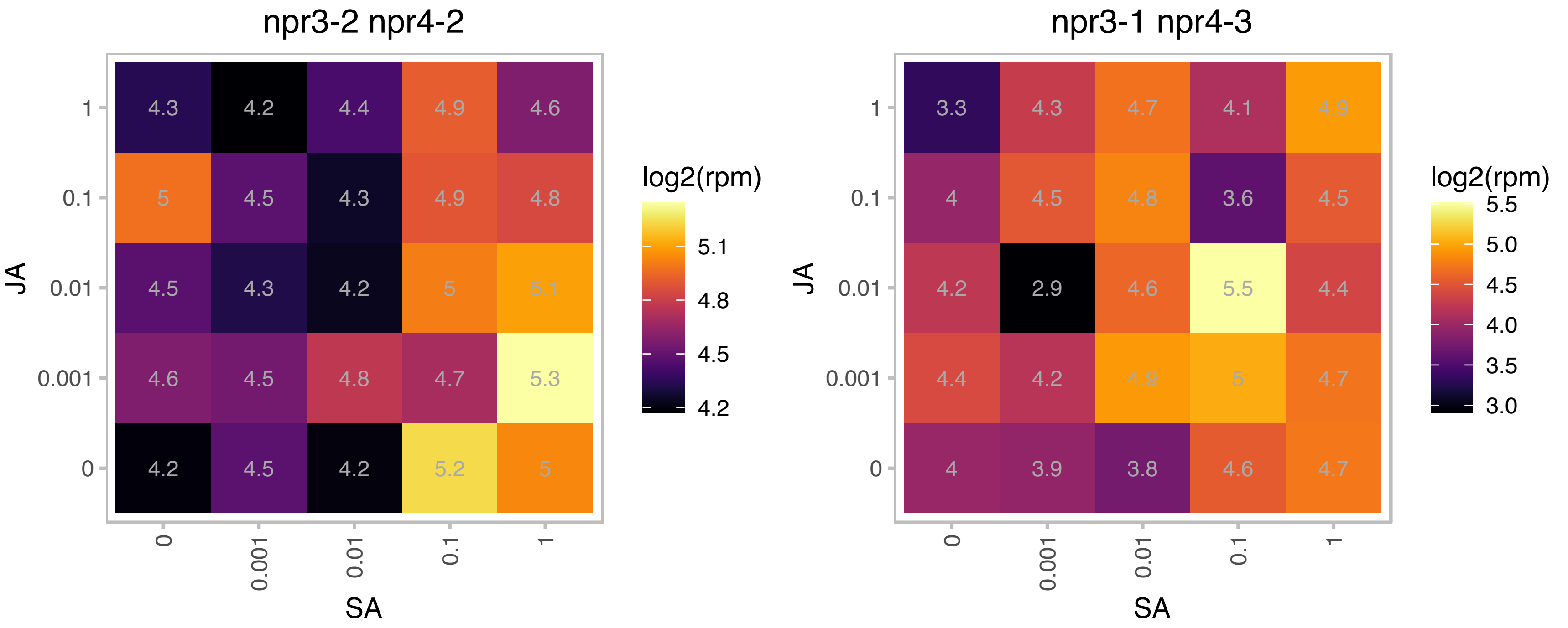

### AT4G27230.1 : HTA2

histone H2A 2

Encodes HTA2, a histone H2A protein.

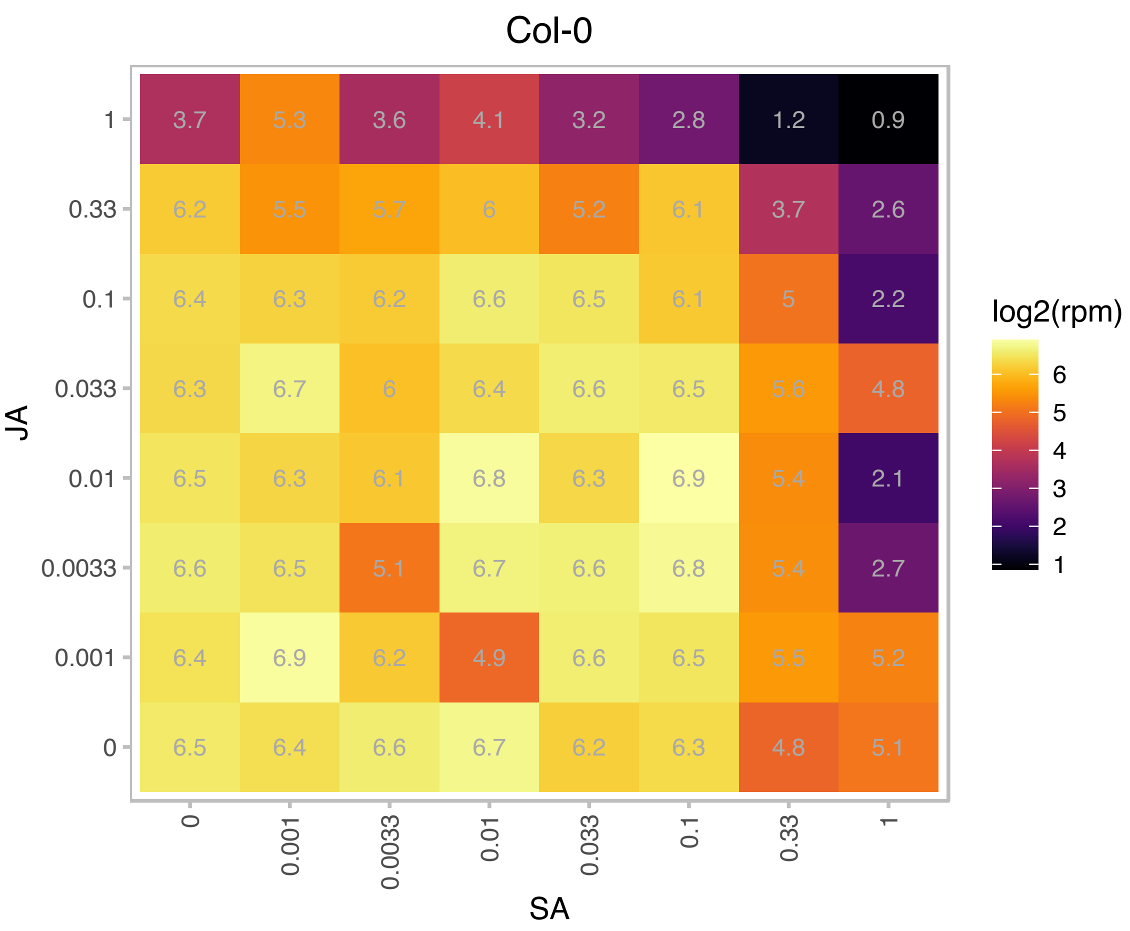


| | |
|---|---|
| Gene | AT4G27230 |
| is.expressed_genes | TRUE |
| is.responsive_genes | TRUE |
| is.apcluster | F-08 |

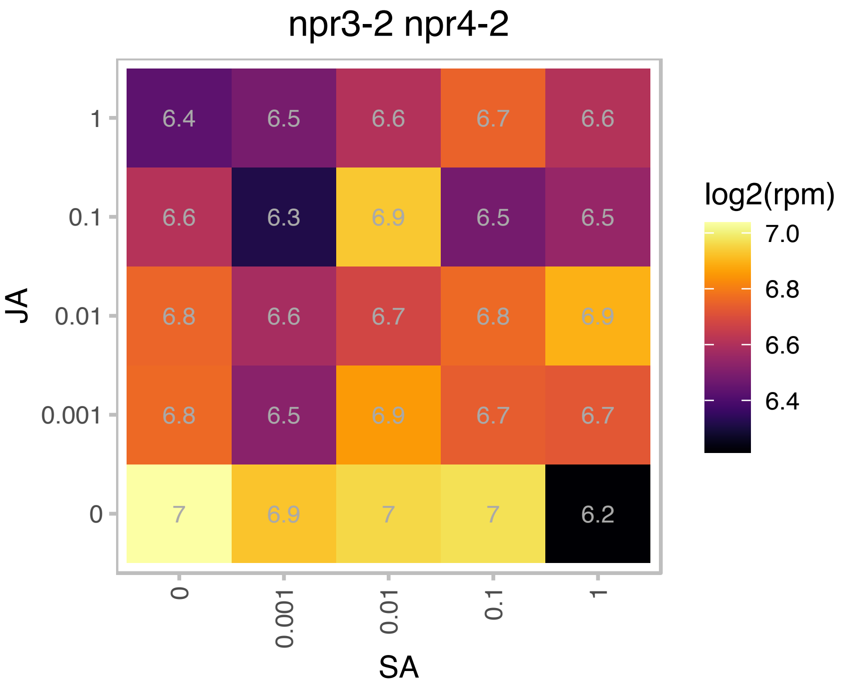


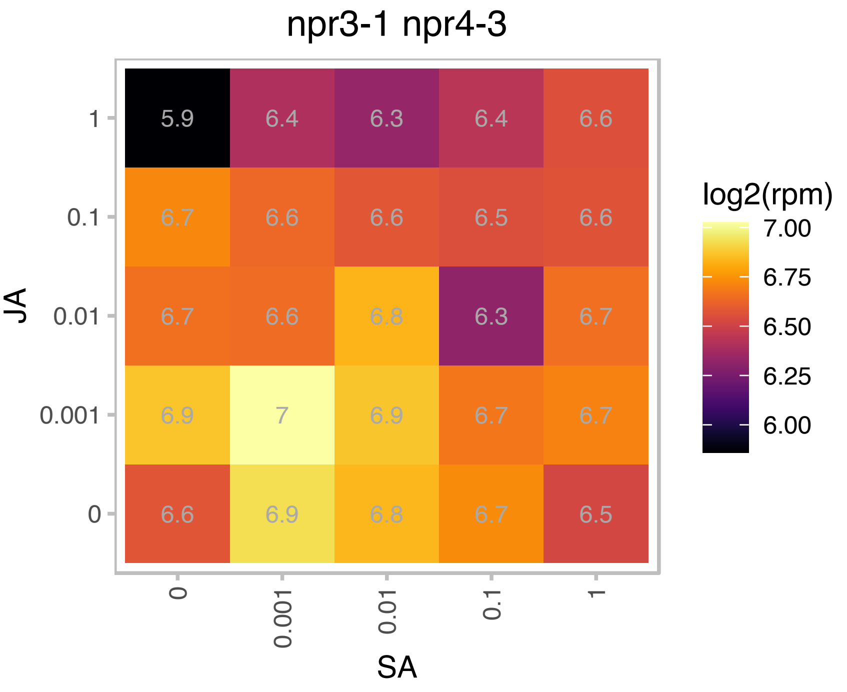

### AT4G29510.1 : ATPRMT11, ATPRMT1B, PRMT11, PRMT1

arginine methyltransferase 11

Has arginine N-methyltransferase activity. Modifies AtMBD7.

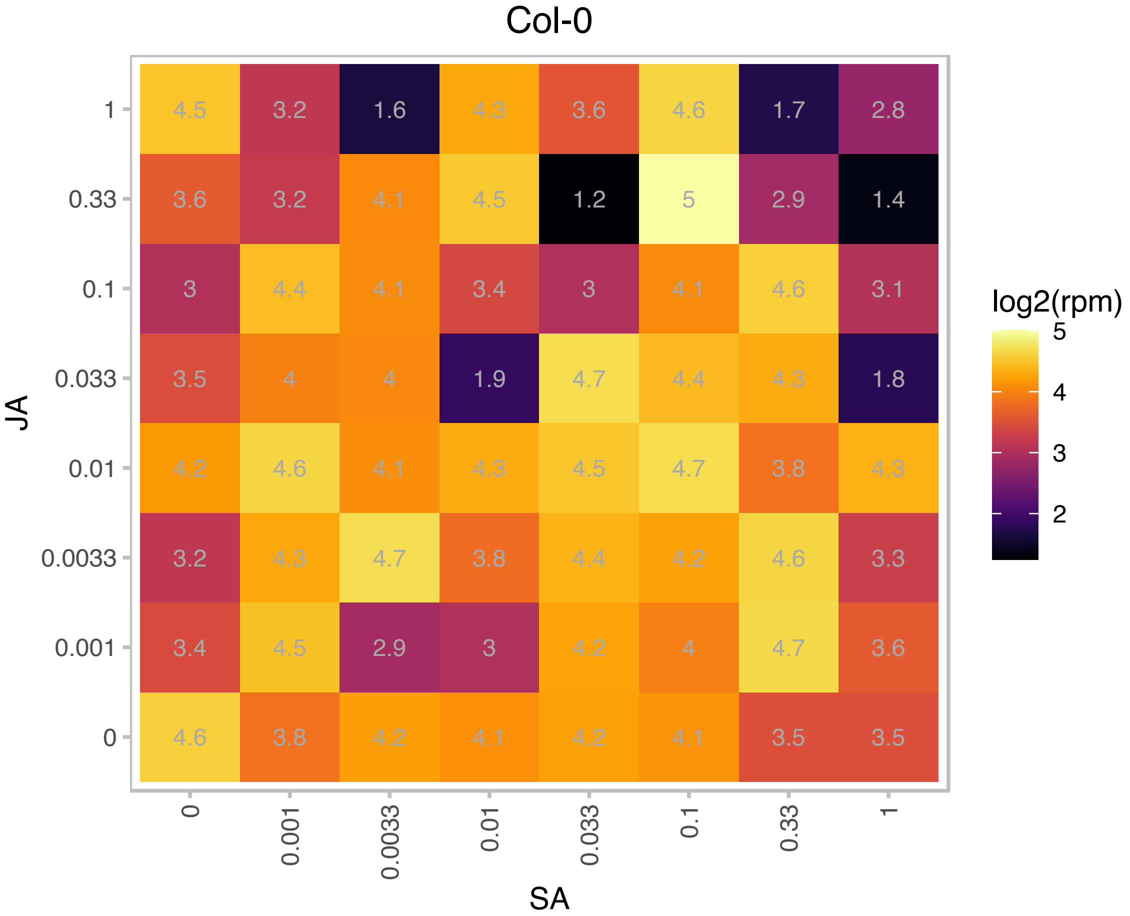


| | |
|---|---|
| Gene | AT4G29510 |
| is.expressed_genes | TRUE |
| is.responsive_genes | TRUE |
| is.apcluster | F-08 |

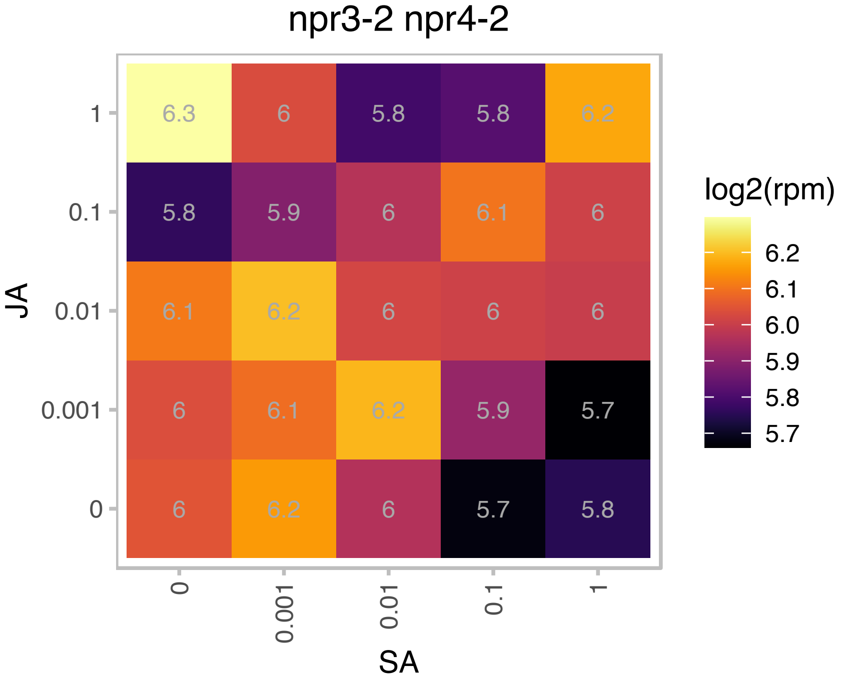


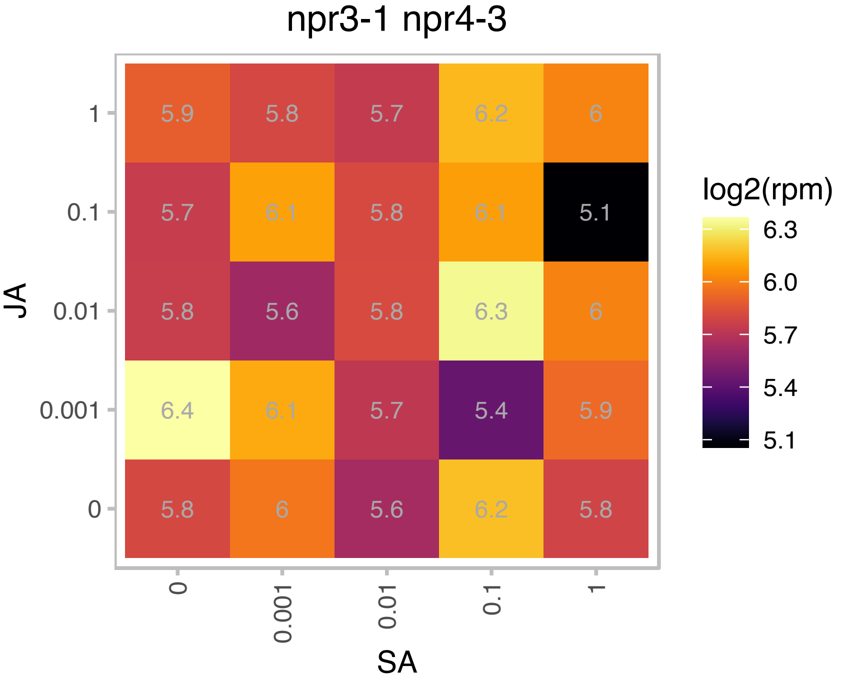

### AT4G34050.1 : CCoAOMT1

S-adenosyl-L-methionine-dependent methyltransferases superfamily protein

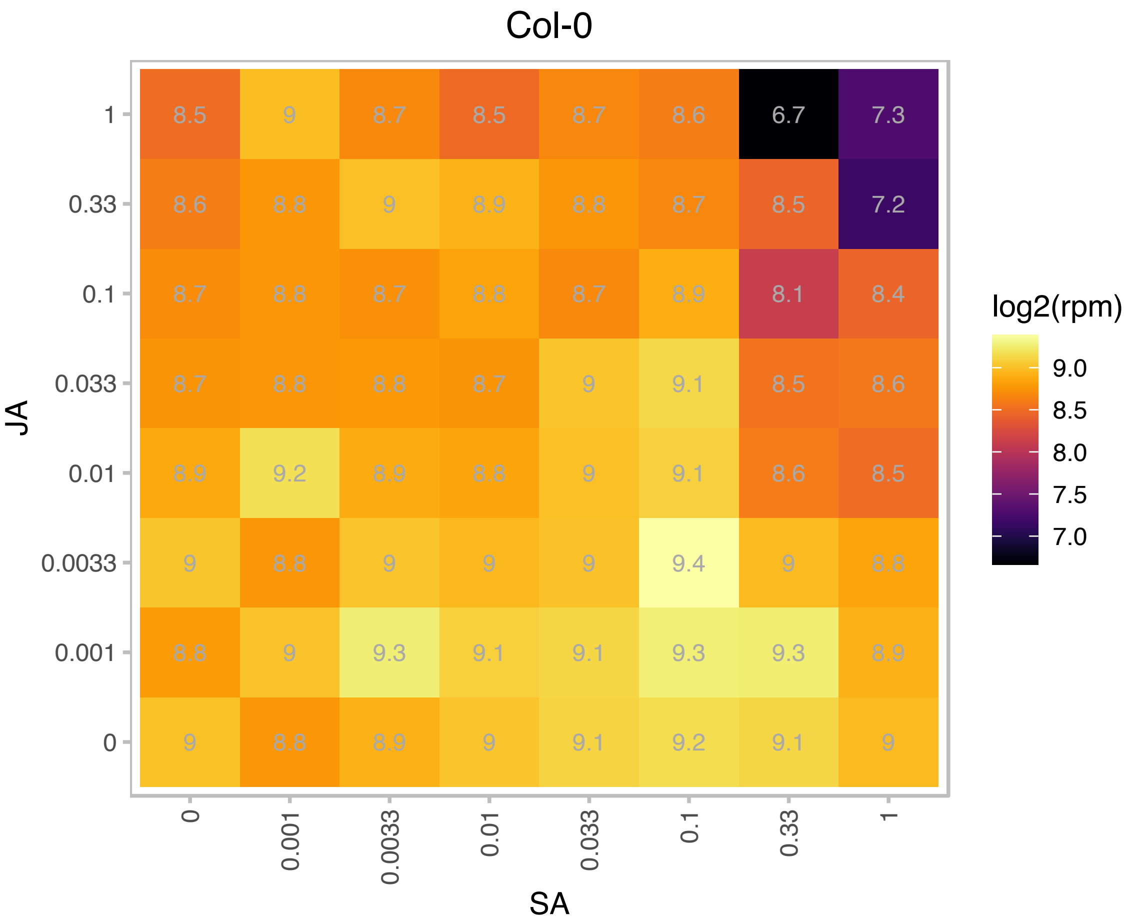


| | |
|---|---|
| Gene | AT4G34050 |
| is.expressed_genes | TRUE |
| is.responsive_genes | TRUE |
| is.apcluster | F-08 |

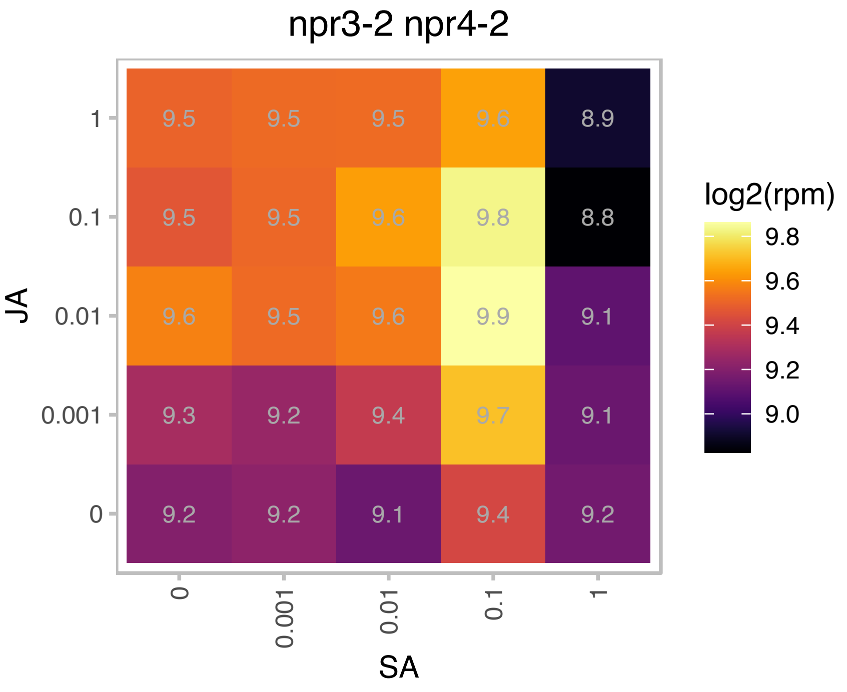


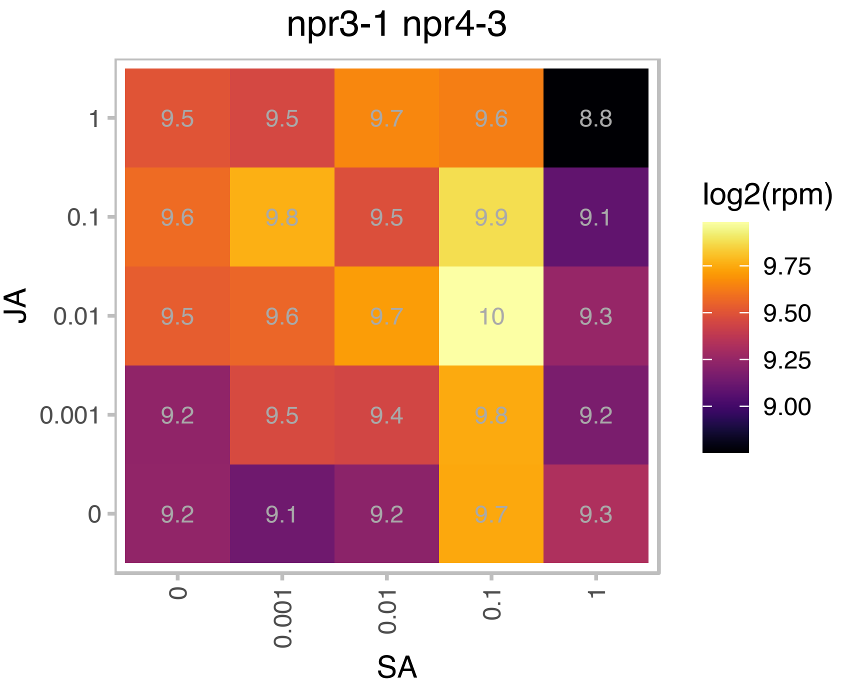

### AT4G35480.1 : RHA3B

RING-H2 finger A3B

Encodes a putative RING-H2 finger protein RHA3b.

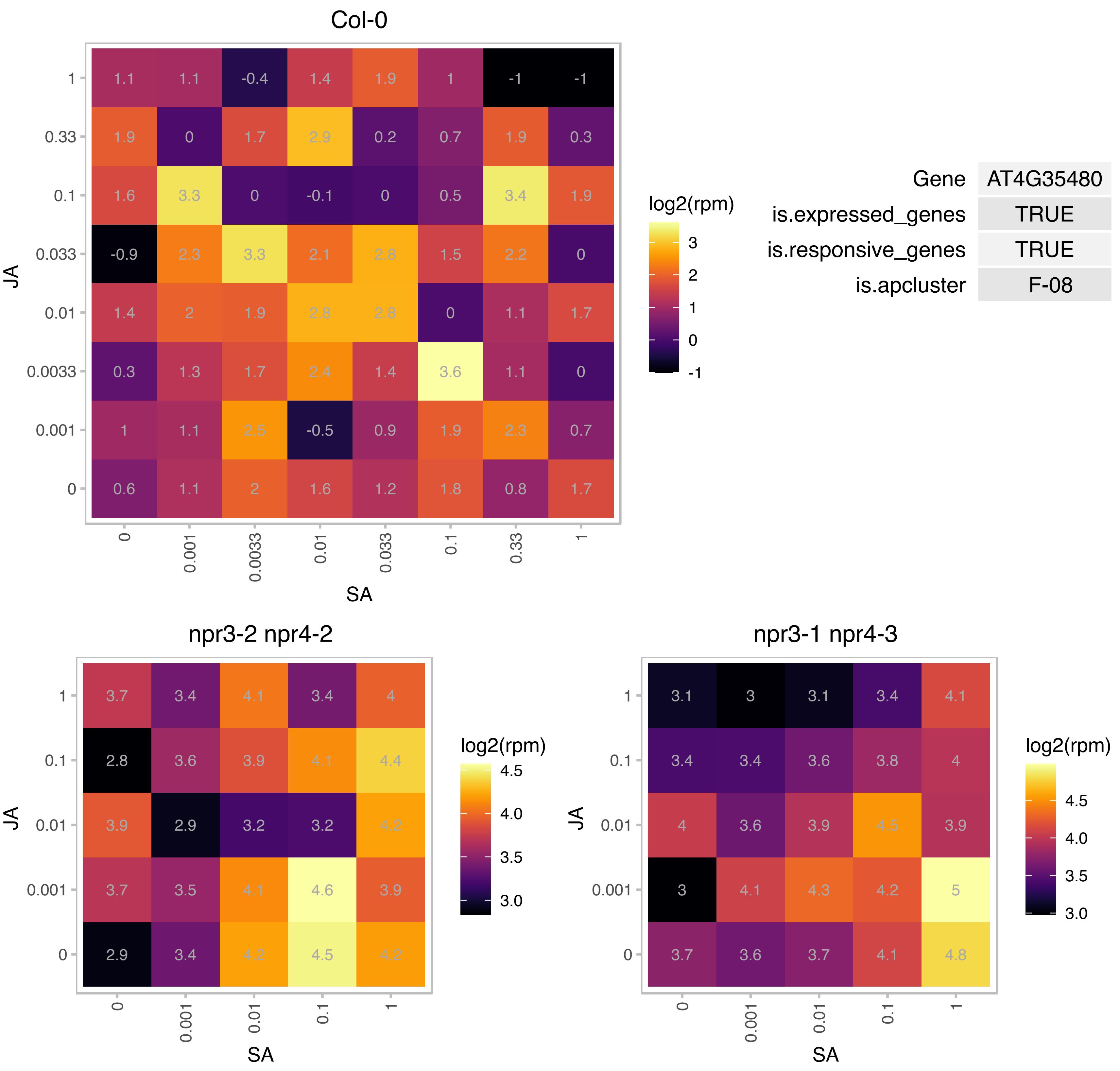


| | |
|---|---|
| Gene | AT4G35480 |
| is.expressed_genes | TRUE |
| is.responsive_genes | TRUE |
| is.apcluster | F-08 |

# AT4G39300.1 : NA

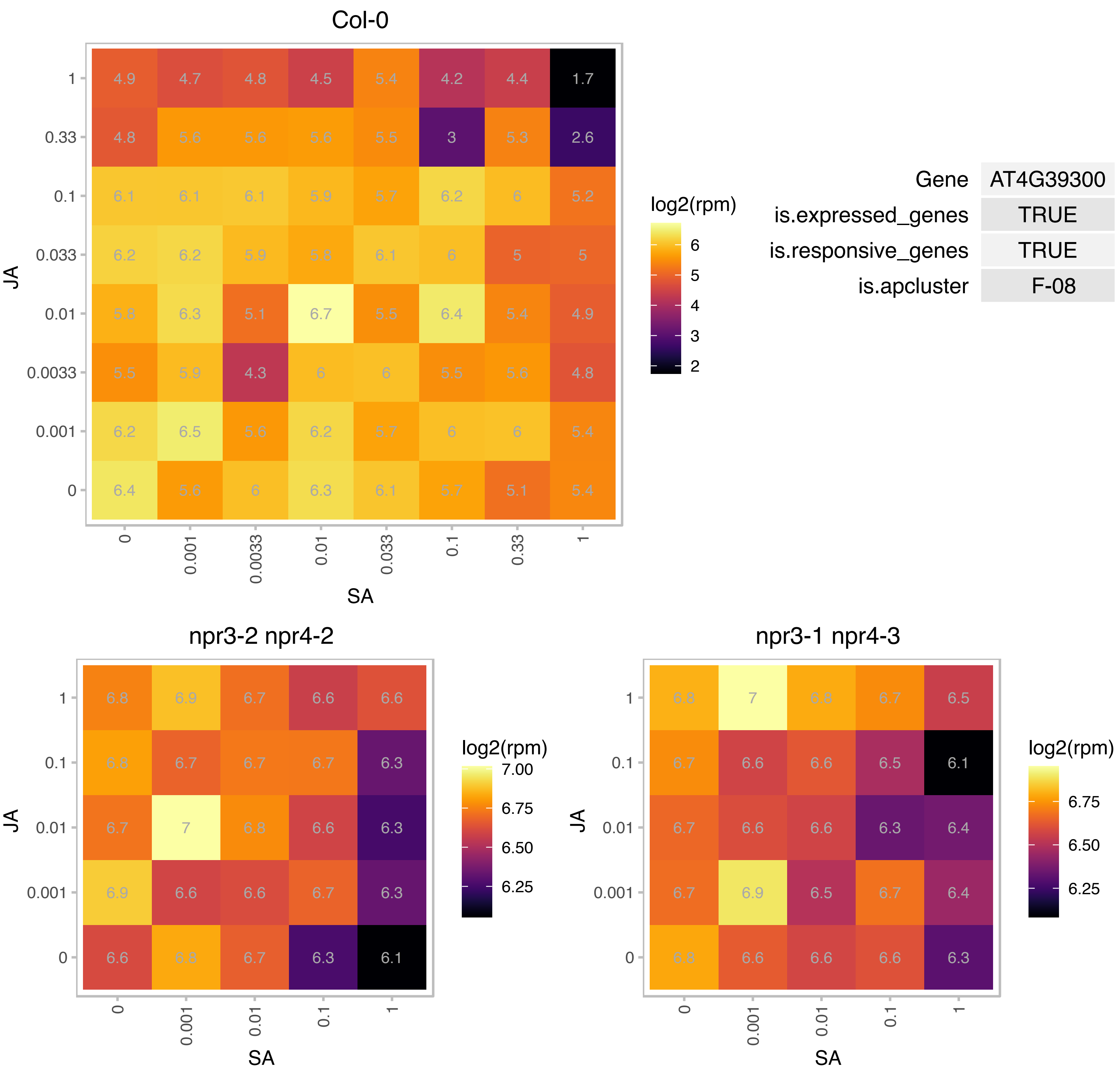

# AT5G01020.1 : NA

Protein kinase superfamily protein

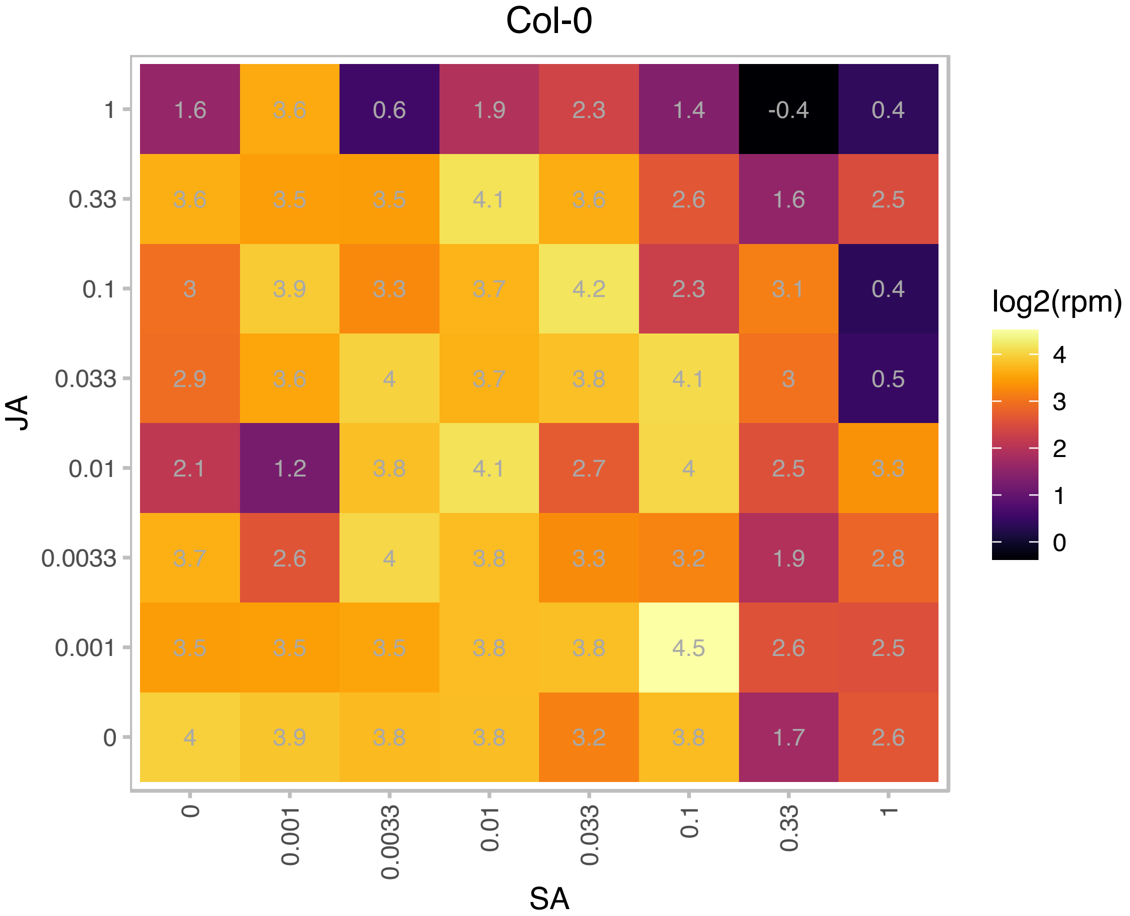


| | |
|---|---|
| Gene | AT5G01020 |
| is.expressed_genes | TRUE |
| is.responsive_genes | TRUE |
| is.apcluster | F-08 |

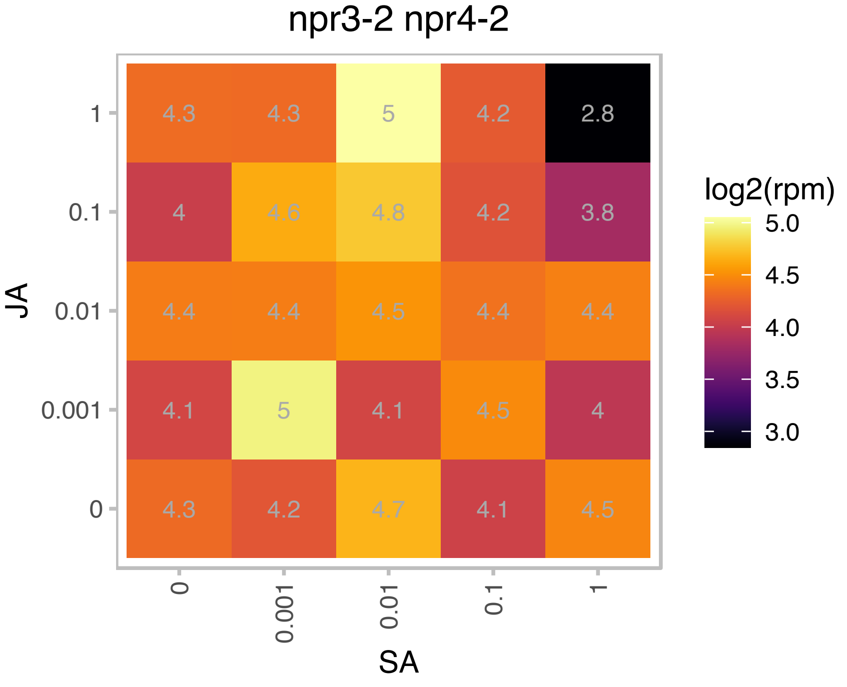


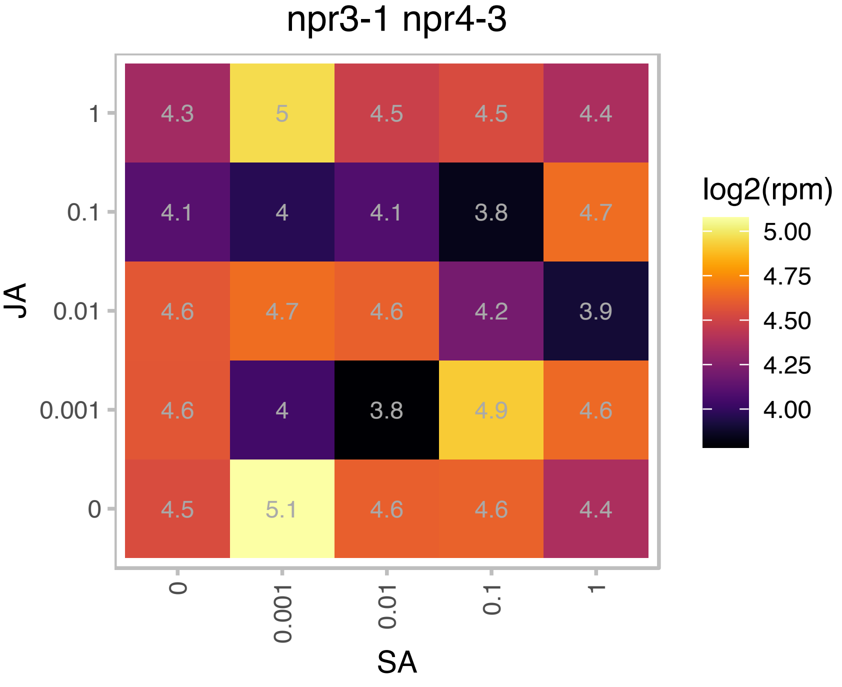

### AT5G02530.1 : ALY2

RNA-binding (RRM/RBD/RNP motifs) family protein

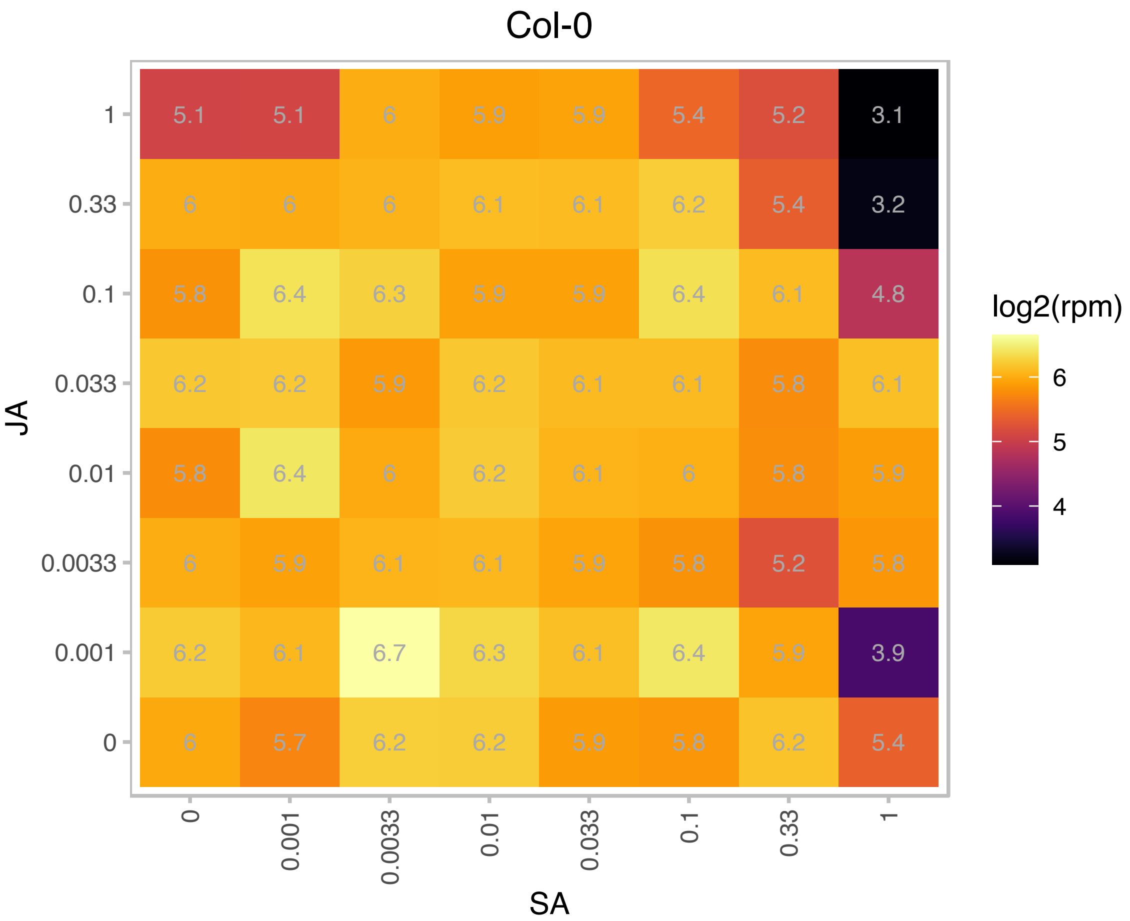


| | |
|---|---|
| Gene | AT5G02530 |
| is.expressed_genes | TRUE |
| is.responsive_genes | TRUE |
| is.apcluster | F-08 |

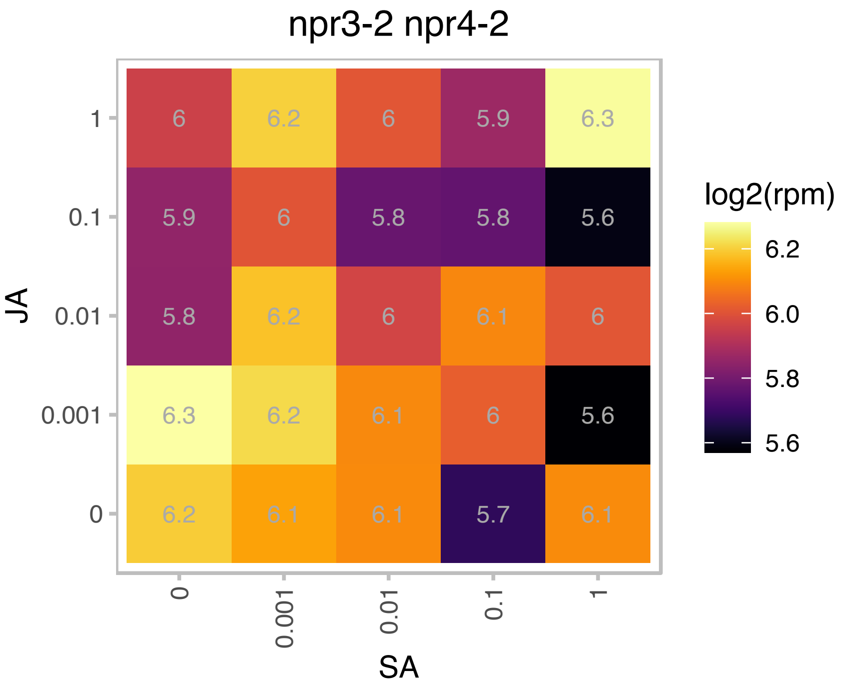


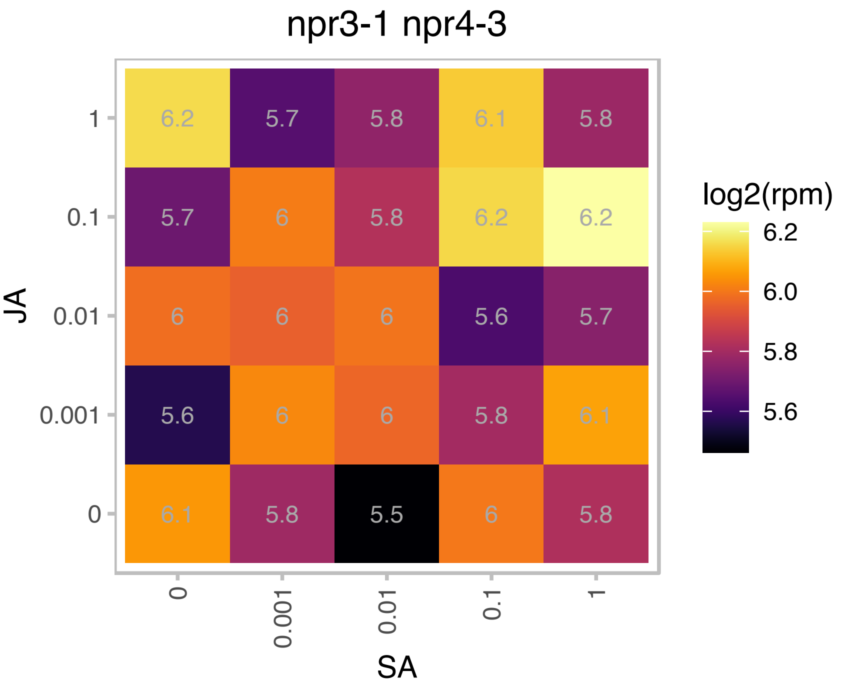

# AT5G05990.1 : NA

Mitochondrial glycoprotein family protein

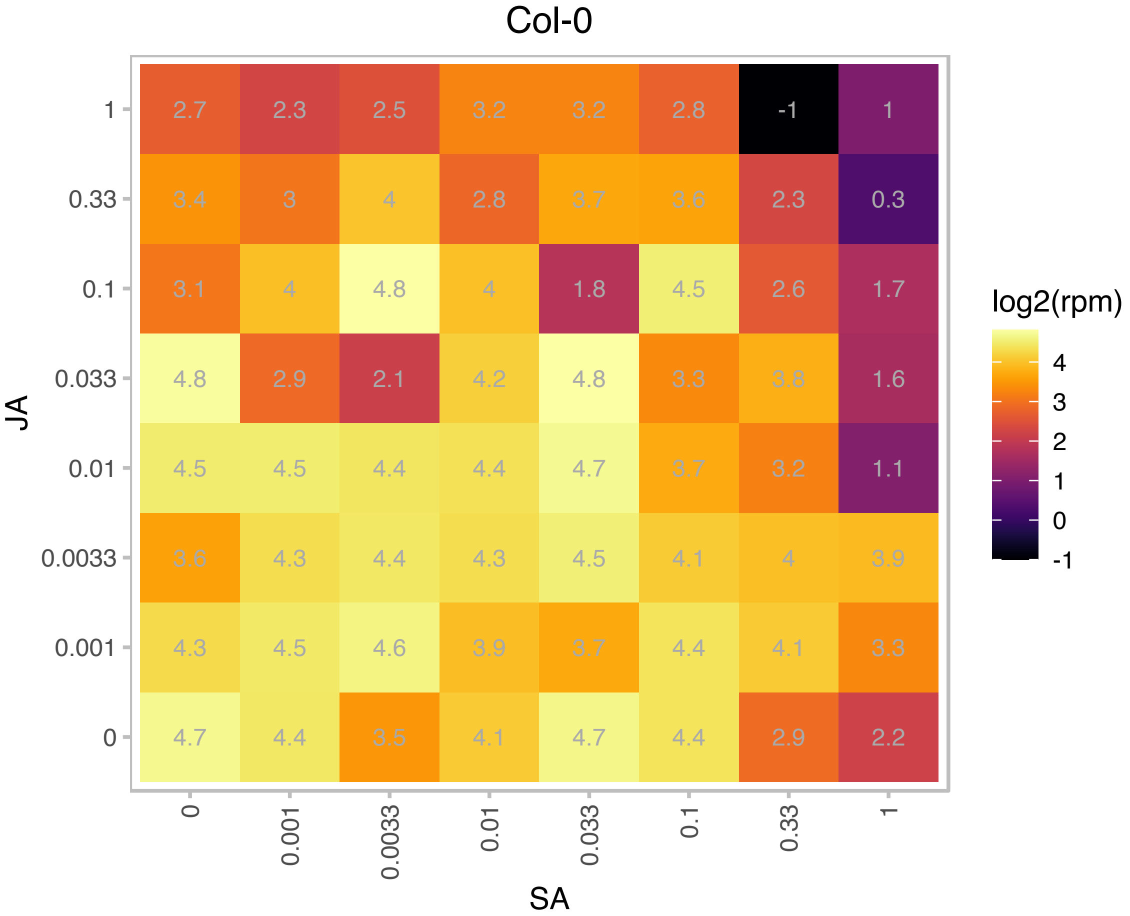


| | |
|---|---|
| Gene | AT5G05990 |
| is.expressed_genes | TRUE |
| is.responsive_genes | TRUE |
| is.apcluster | F-08 |

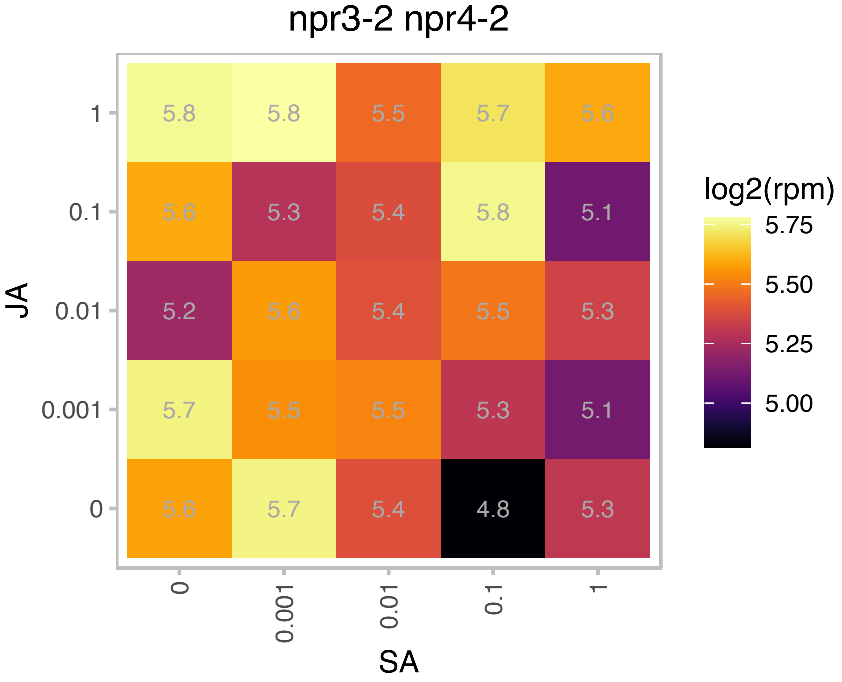


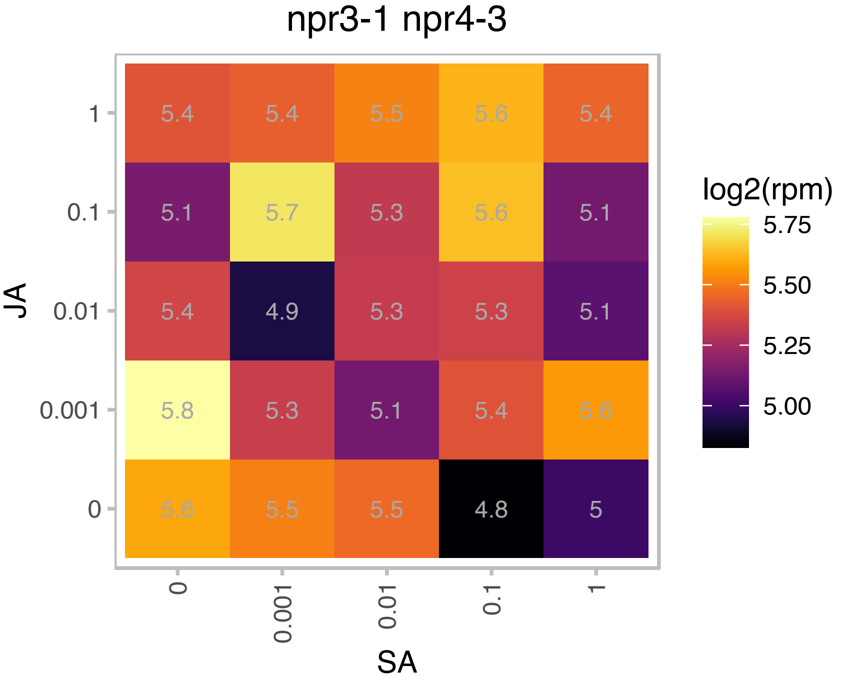

### AT5G07830.1 : AtGUS2, GUS2

glucuronidase 2

Belongs to the plant glycoside hydrolase family 79. Encodes a protein with several posttranslational modification sites including O-β-GlcNAc attachment sites and serine-, threonine- and tyrosine-phosphorylation sites, suggesting that this protein is extensively modified posttranslationally. The protein is predicted (WoLF PSORT program) to be membrane-associated.  It is involved in cell elongation. The mRNA is cell-to-cell mobile.

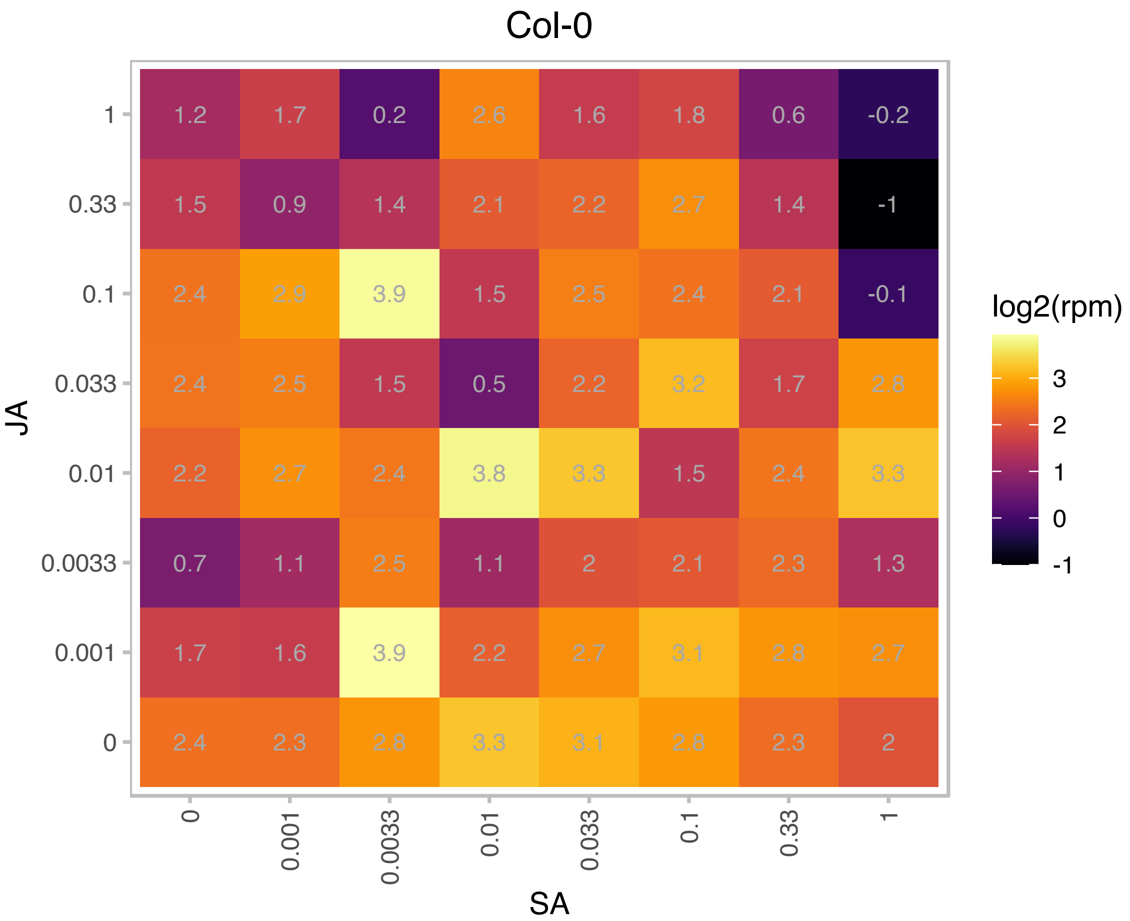


| | |
|---|---|
| Gene | AT5G07830 |
| is.expressed_genes | TRUE |
| is.responsive_genes | TRUE |
| is.apcluster | F-08 |

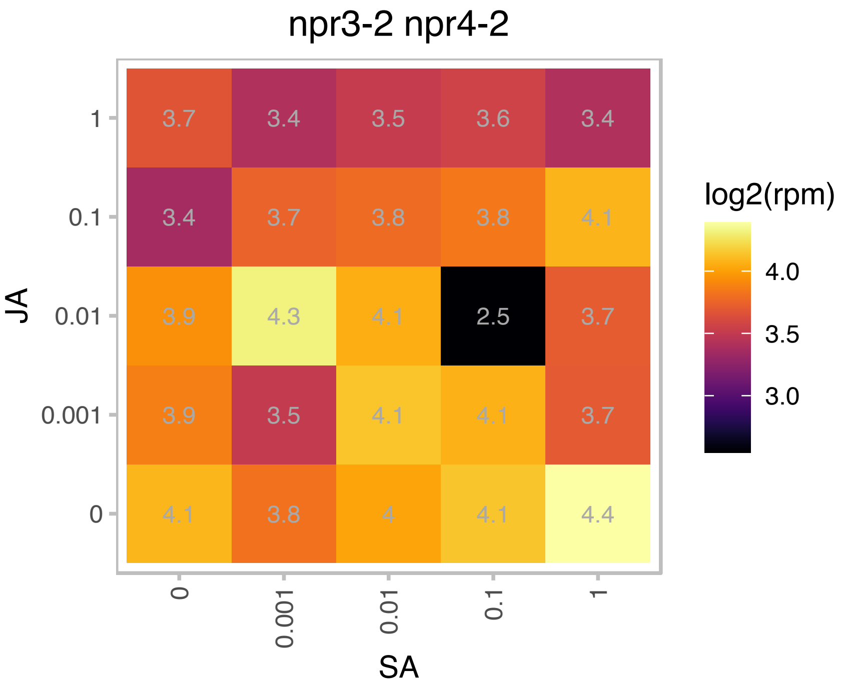


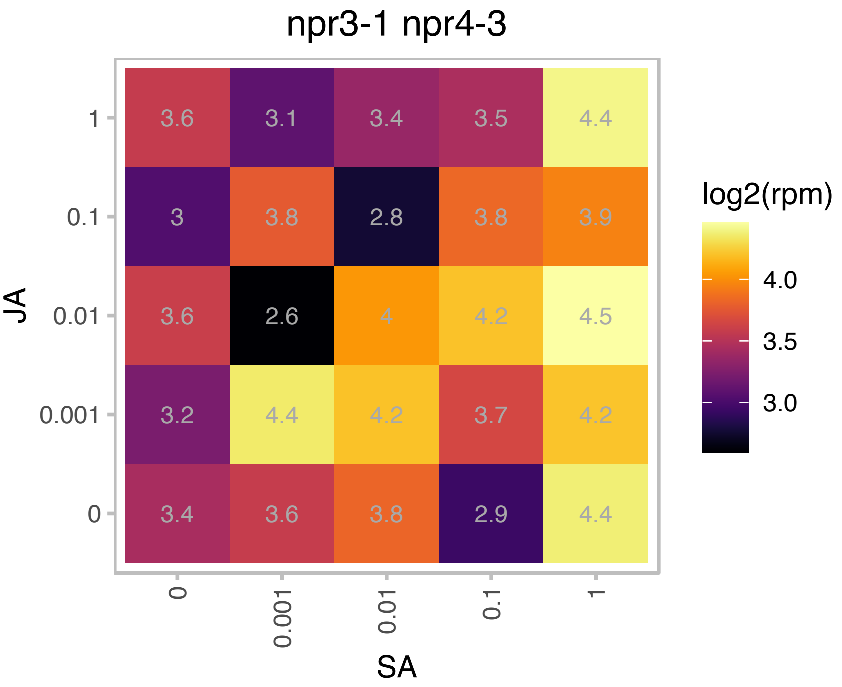

# AT5G08180.1 : NA

Ribosomal protein L7Ae/L30e/S12e/Gadd45 family protein

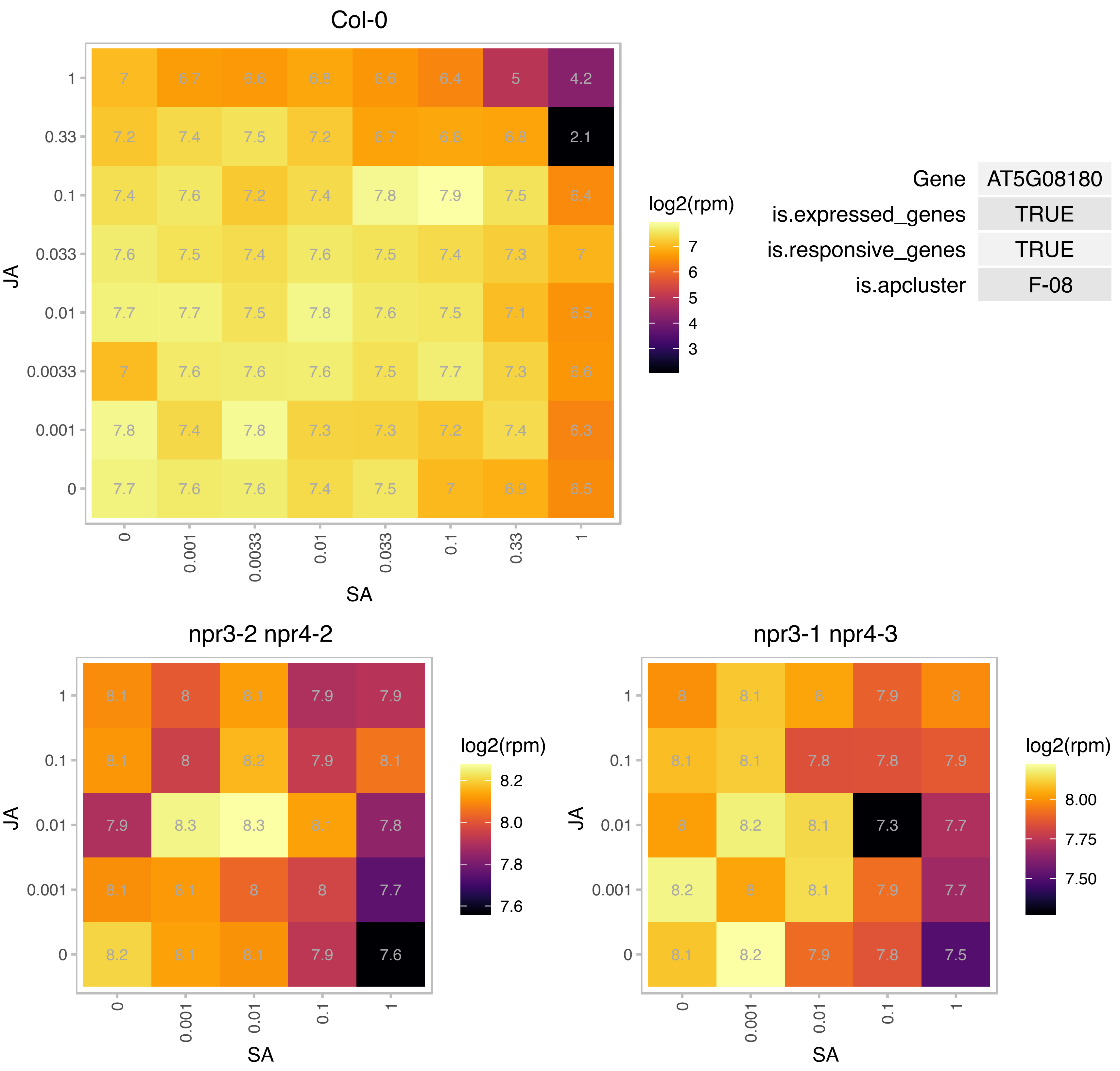

### AT5G09920.1 : ATRPB15.9, NRPB4, RPB15.9, RPB15.9.9

RNA polymerase II, Rpb4, core protein

Non-catalytic subunit specific to DNA-dependent RNA polymerase II; the ortholog of budding yeast RPB4)

| | |
|---|---|
| Gene | AT5G09920 |
| is.expressed_genes | TRUE |
| is.responsive_genes | TRUE |
| is.apcluster | F-08 |

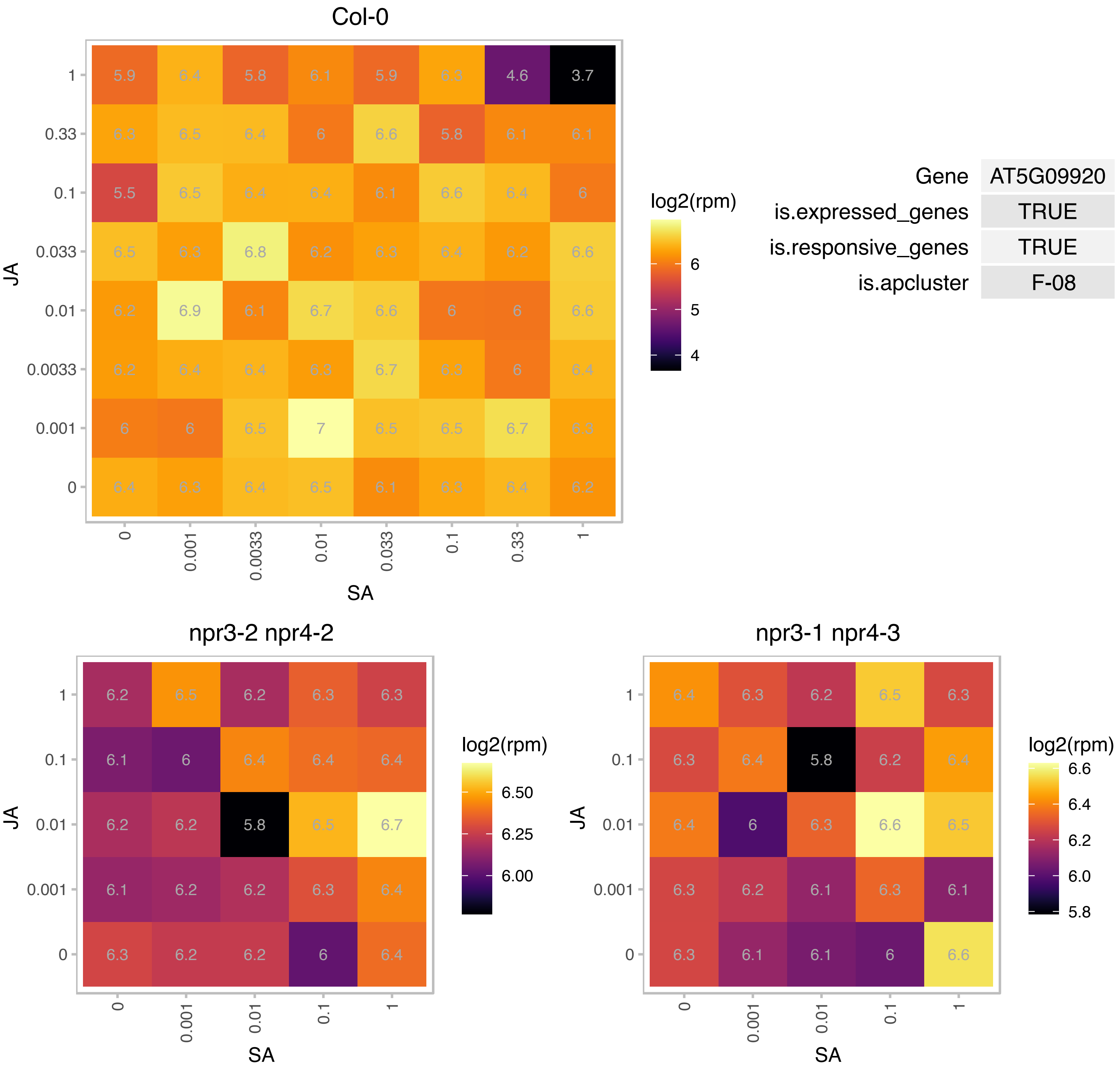

### AT5G10390.1 : H3.1, HTR13

Histone superfamily protein

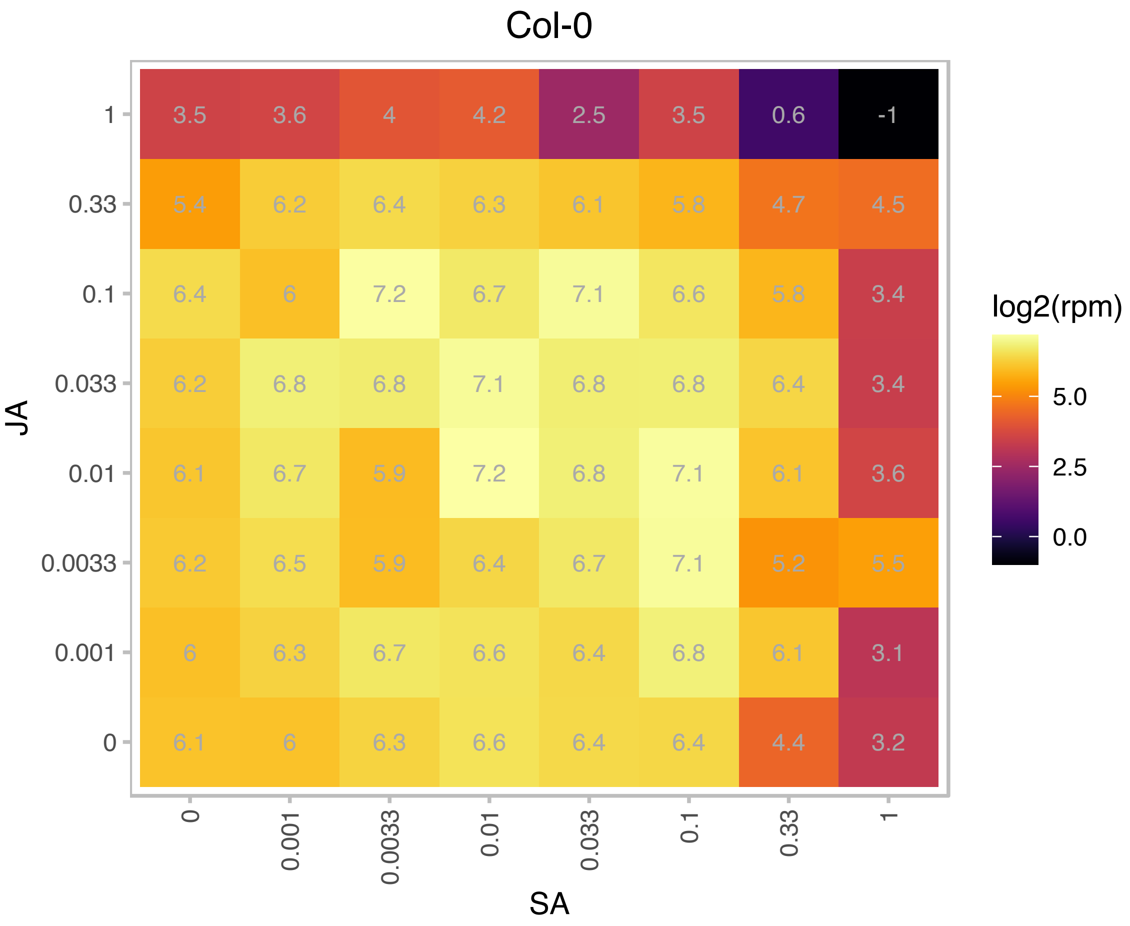


| | |
|---|---|
| Gene | AT5G10390 |
| is.expressed_genes | TRUE |
| is.responsive_genes | TRUE |
| is.apcluster | F-08 |

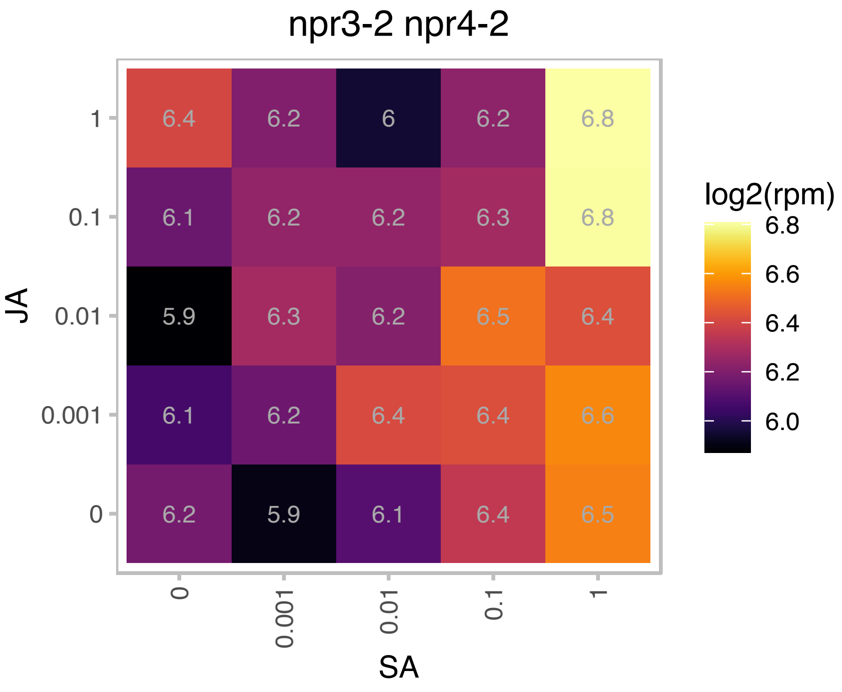


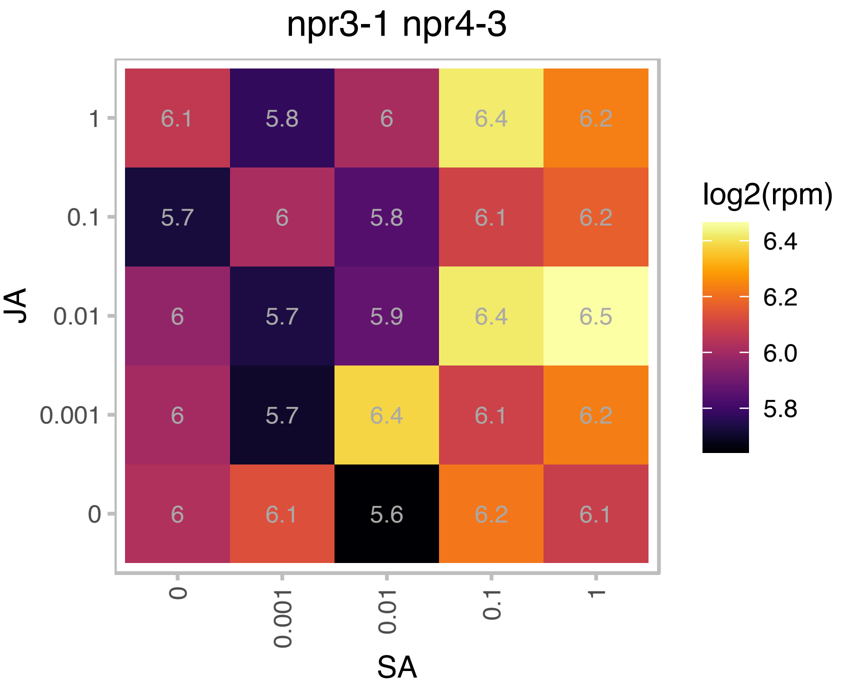

### AT5G10400.1 : H3.1, HTR9

Histone superfamily protein

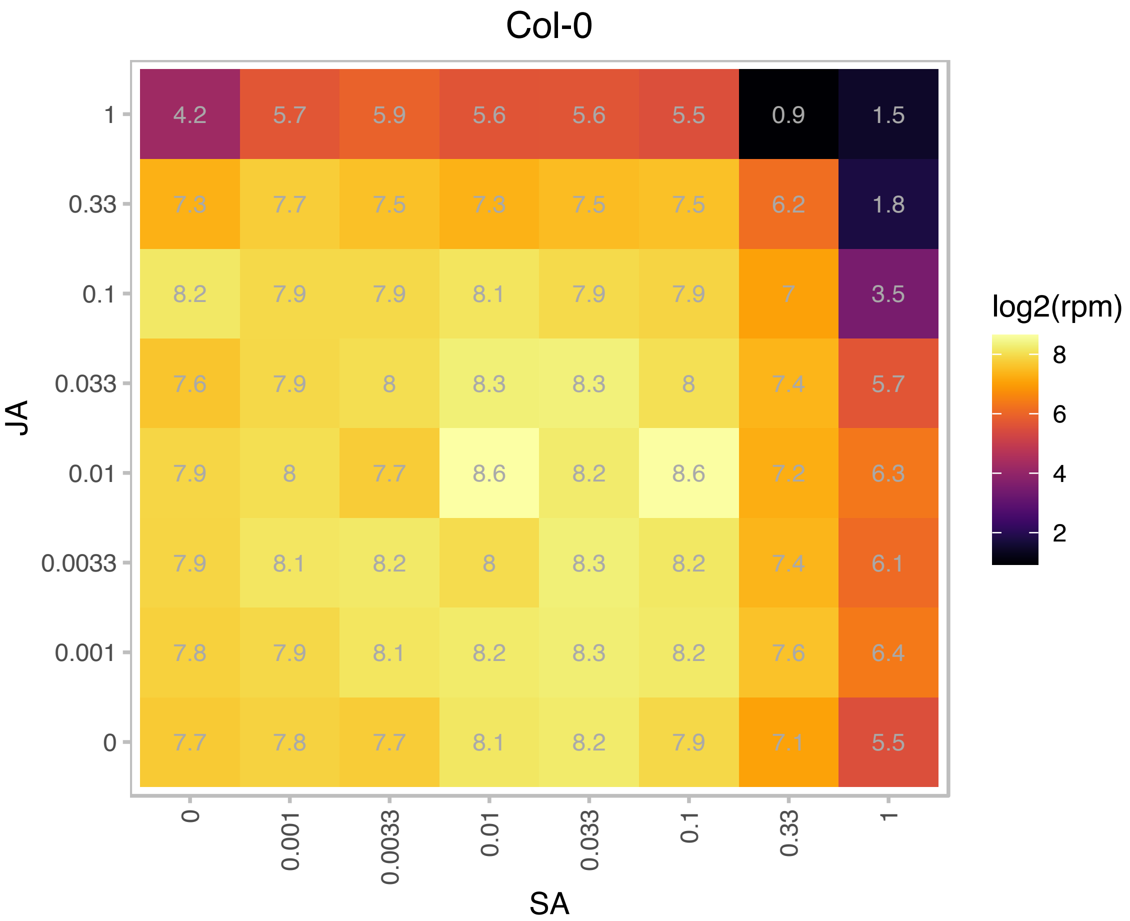


| | |
|---|---|
| Gene | AT5G10400 |
| is.expressed_genes | TRUE |
| is.responsive_genes | TRUE |
| is.apcluster | F-08 |

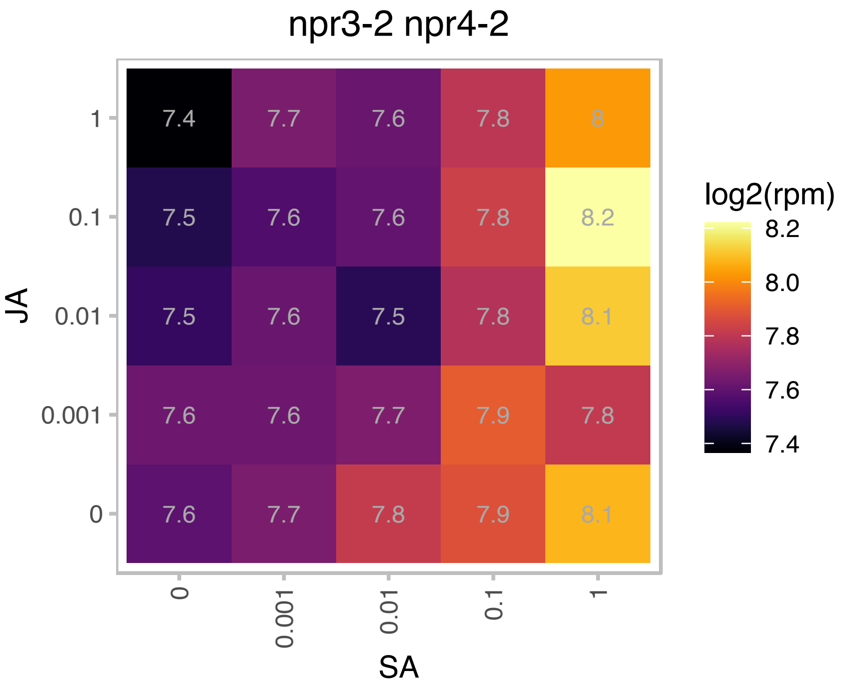


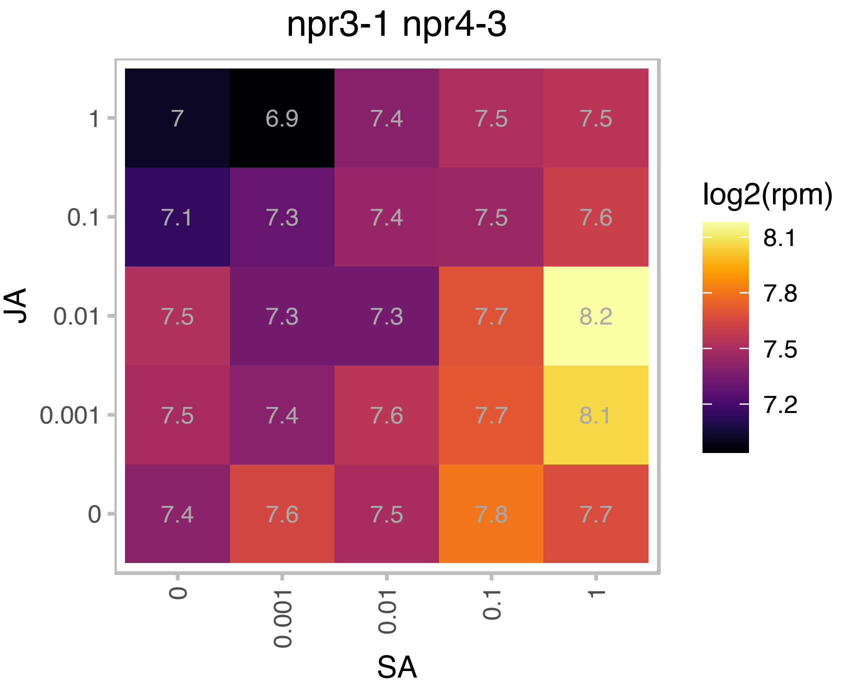

# AT5G10910.1 : NA

mraW methylase family protein

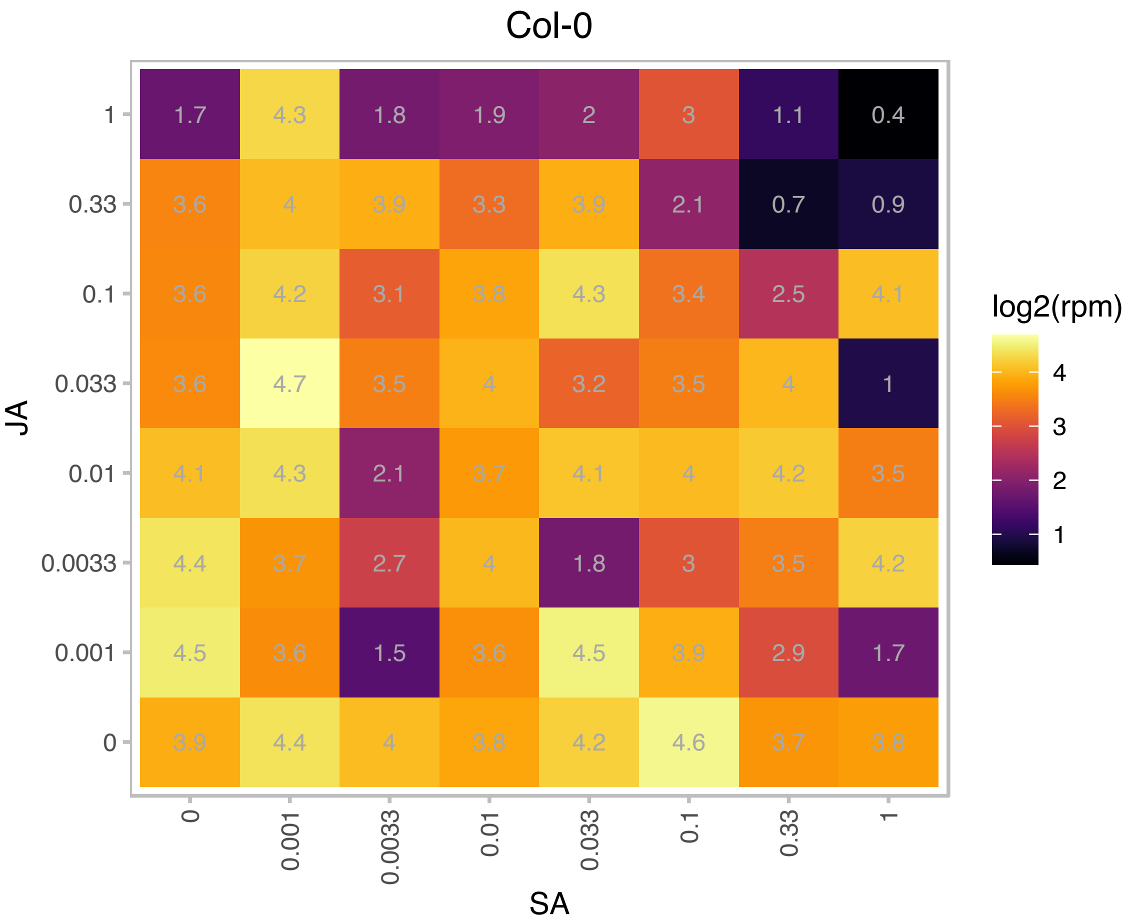


| | |
|---|---|
| Gene | AT5G10910 |
| is.expressed_genes | TRUE |
| is.responsive_genes | TRUE |
| is.apcluster | F-08 |

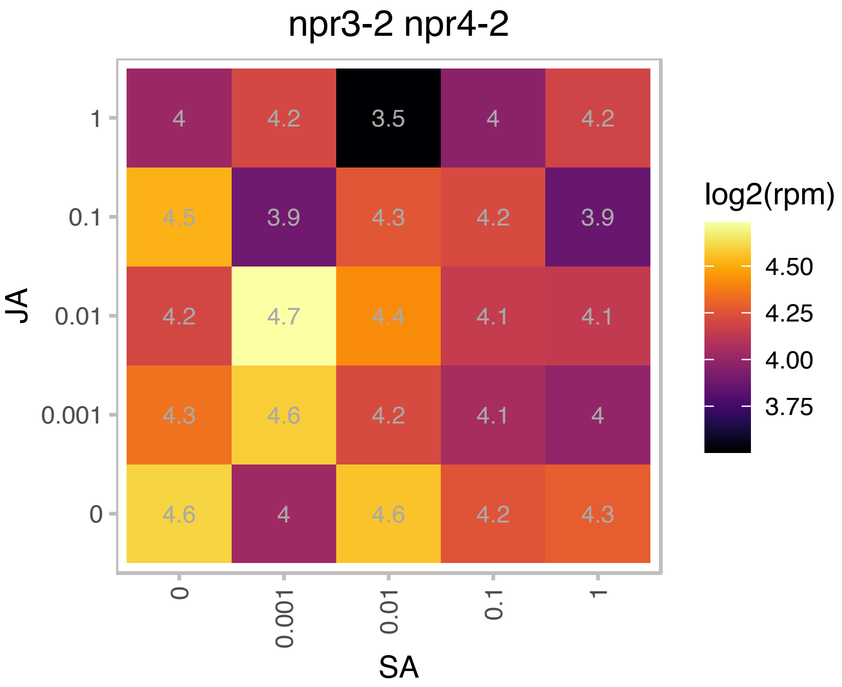


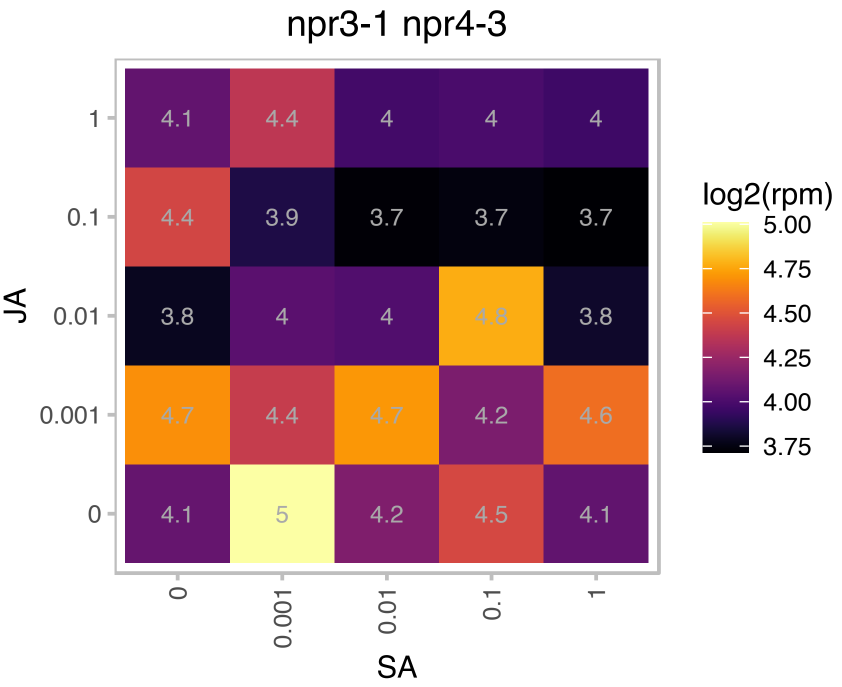

# AT5G11070.1 : NA

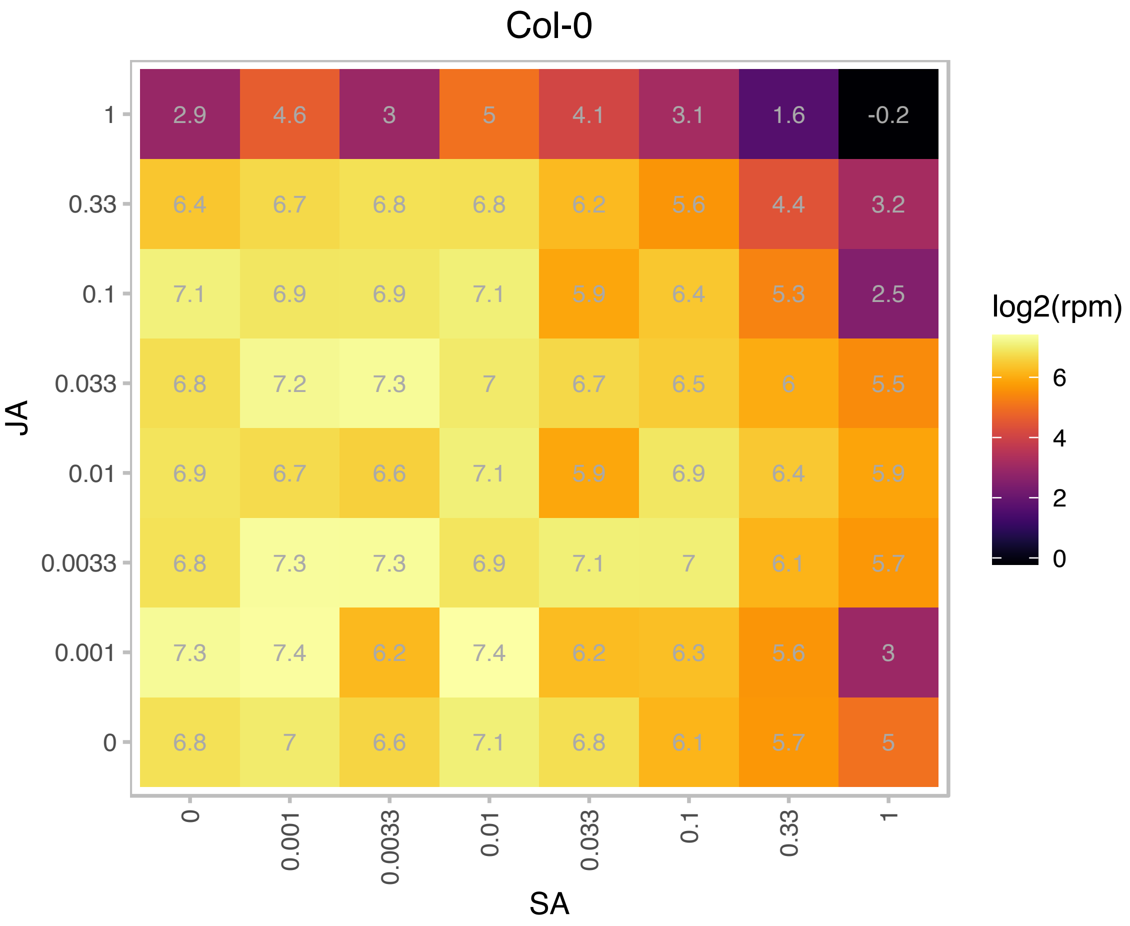


| | |
|---|---|
| Gene | AT5G11070 |
| is.expressed_genes | TRUE |
| is.responsive_genes | TRUE |
| is.apcluster | F-08 |

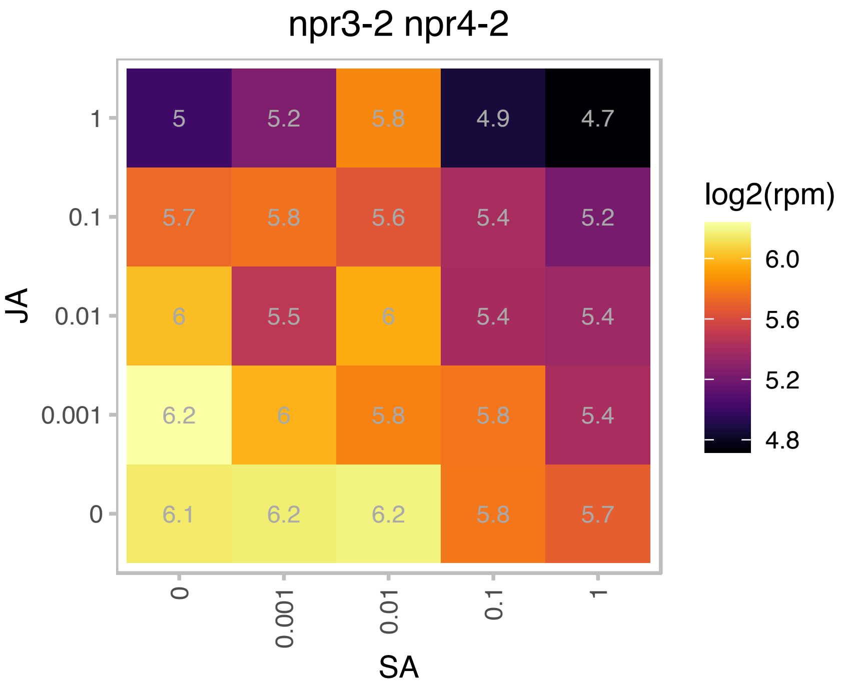


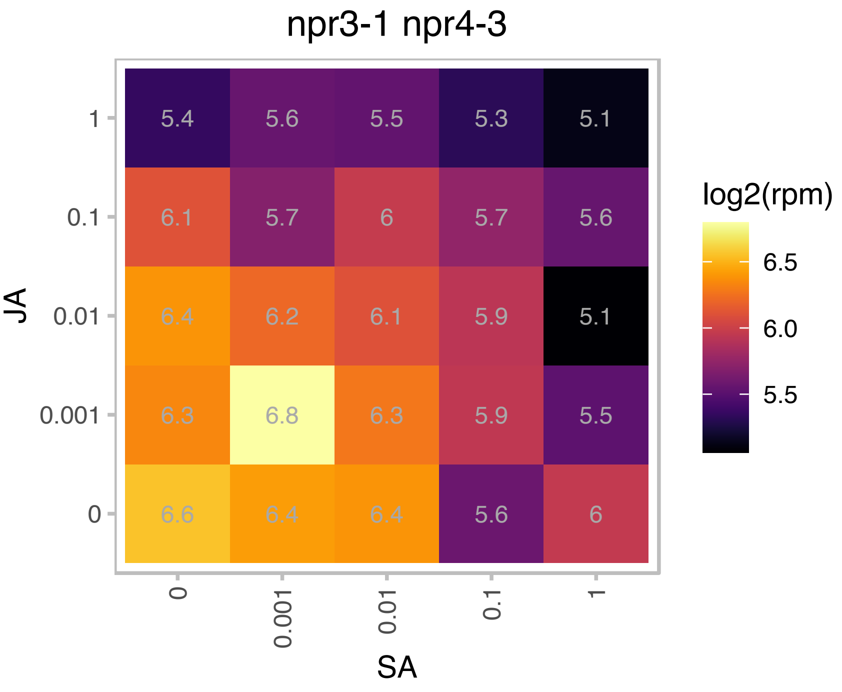

# AT5G16950.1 : NA

| | |
|---|---|
| Gene | AT5G16950 |
| is.expressed_genes | TRUE |
| is.responsive_genes | TRUE |
| is.apcluster | F-08 |

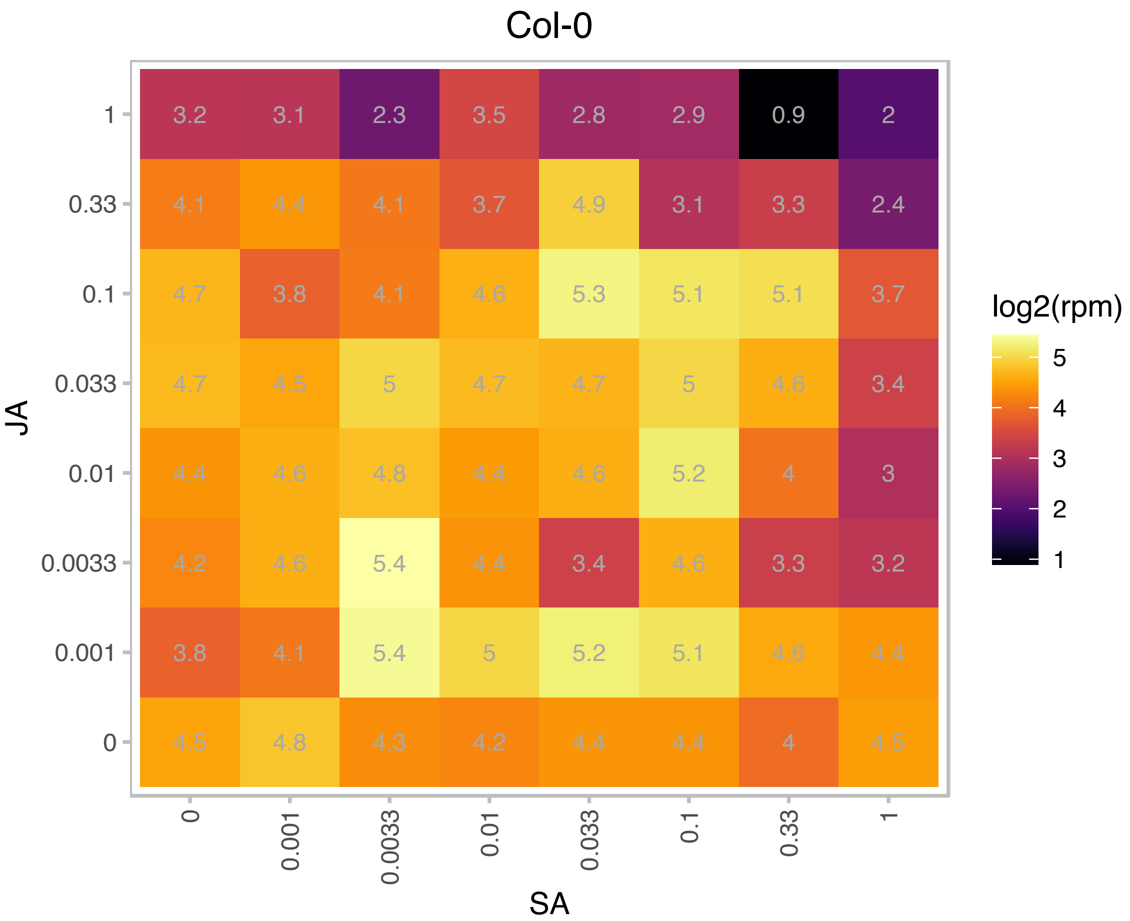


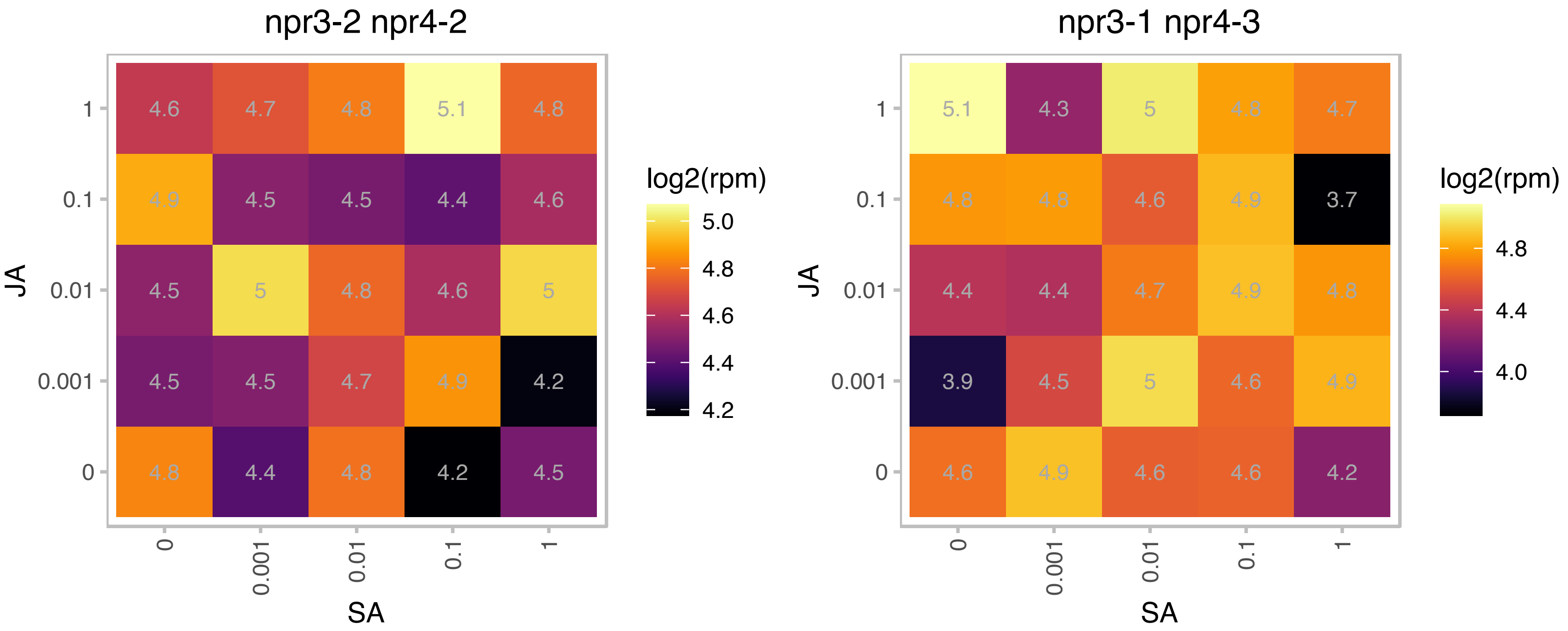

# AT5G19151.1 : NA

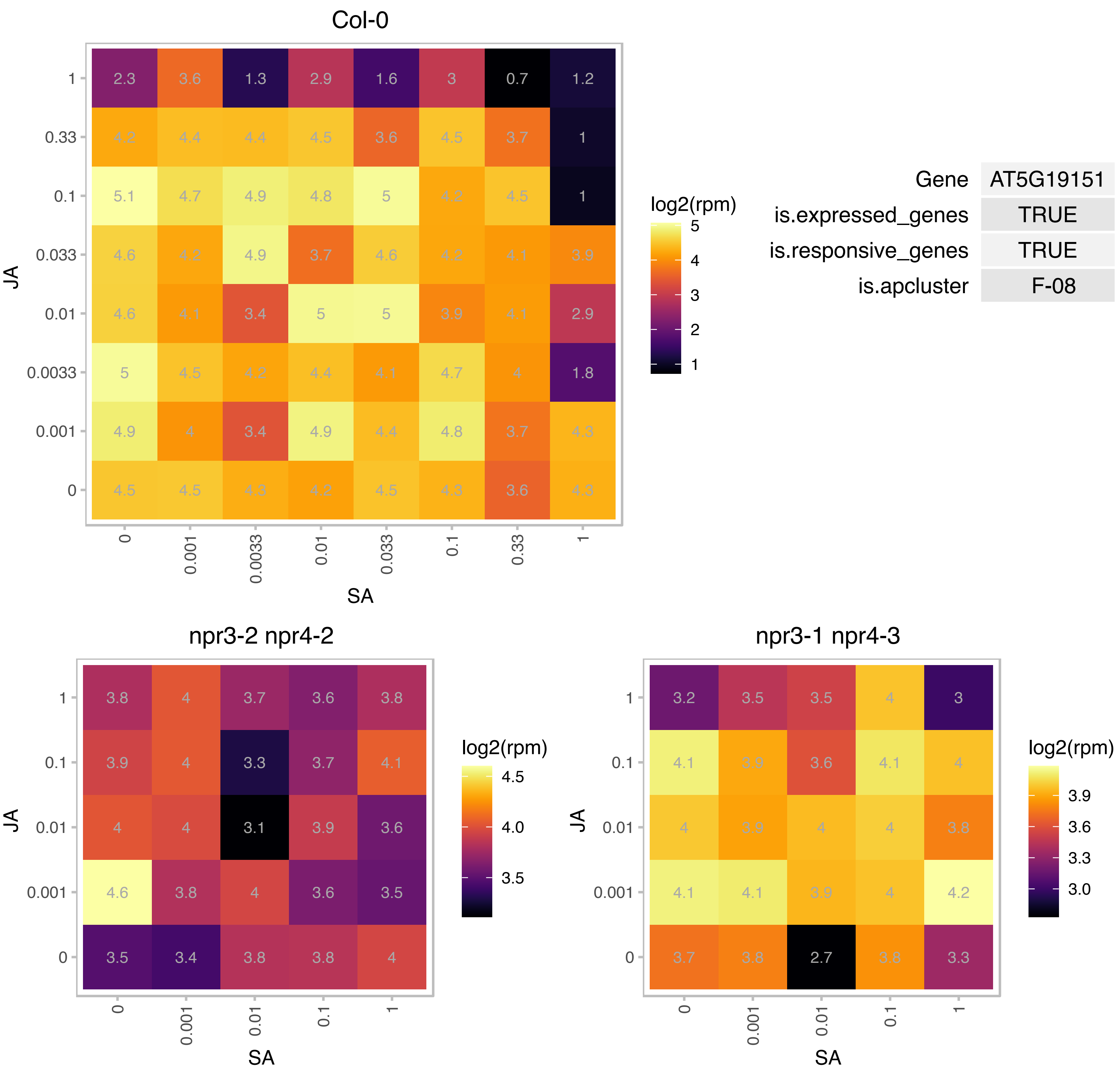

# AT5G20160.1 : NA

Ribosomal protein L7Ae/L30e/S12e/Gadd45 family protein

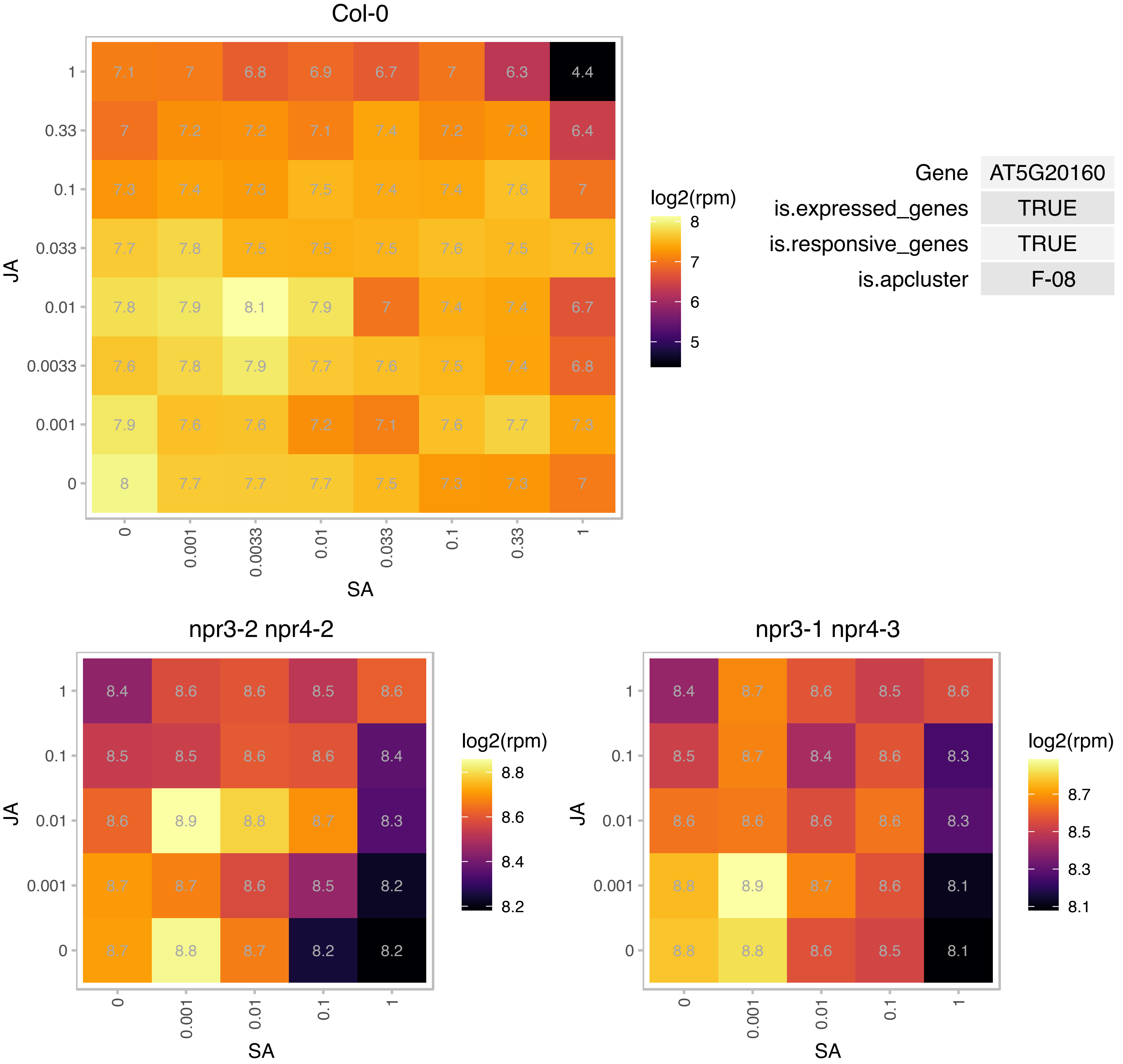


| | |
|---|---|
| Gene | AT5G20160 |
| is.expressed_genes | TRUE |
| is.responsive_genes | TRUE |
| is.apcluster | F-08 |

# AT5G20700.1 : NA

Protein of unknown function (DUF581)

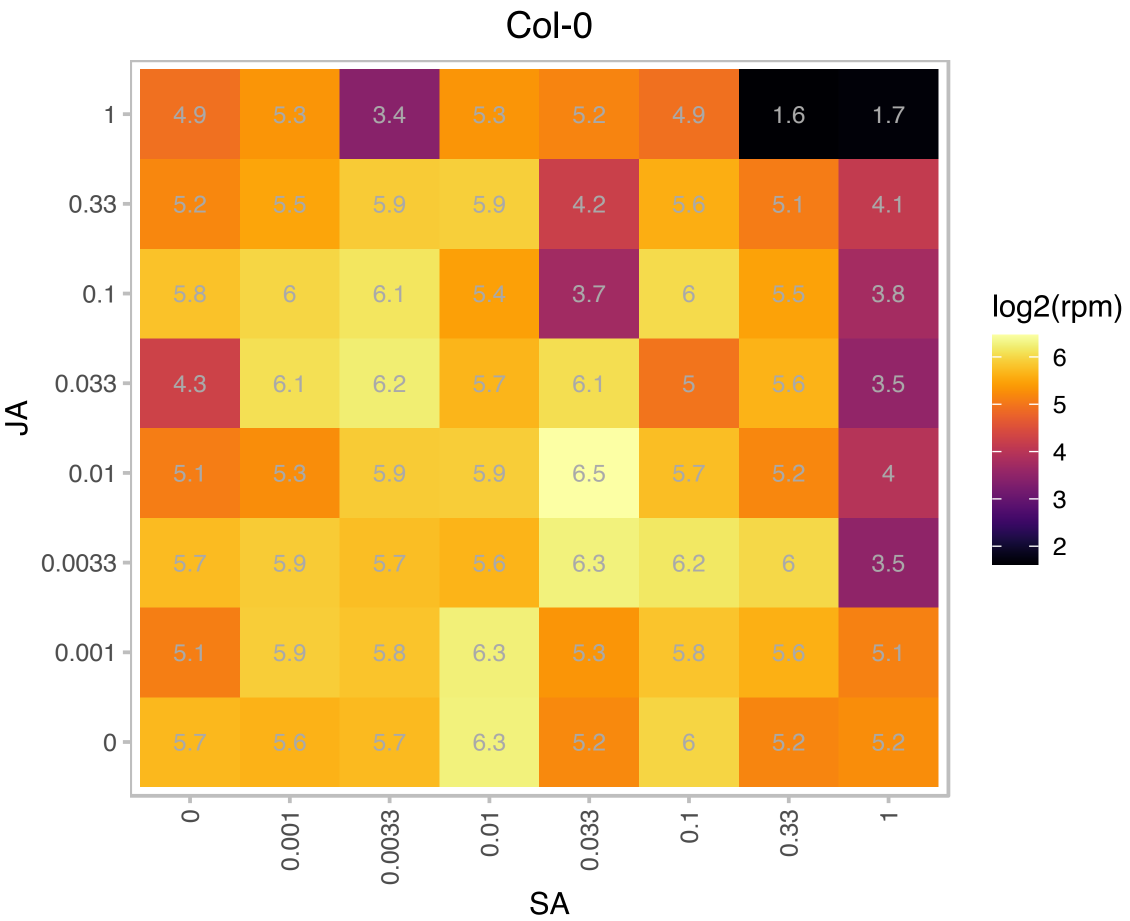


| | |
|---|---|
| Gene | AT5G20700 |
| is.expressed_genes | TRUE |
| is.responsive_genes | TRUE |
| is.apcluster | F-08 |

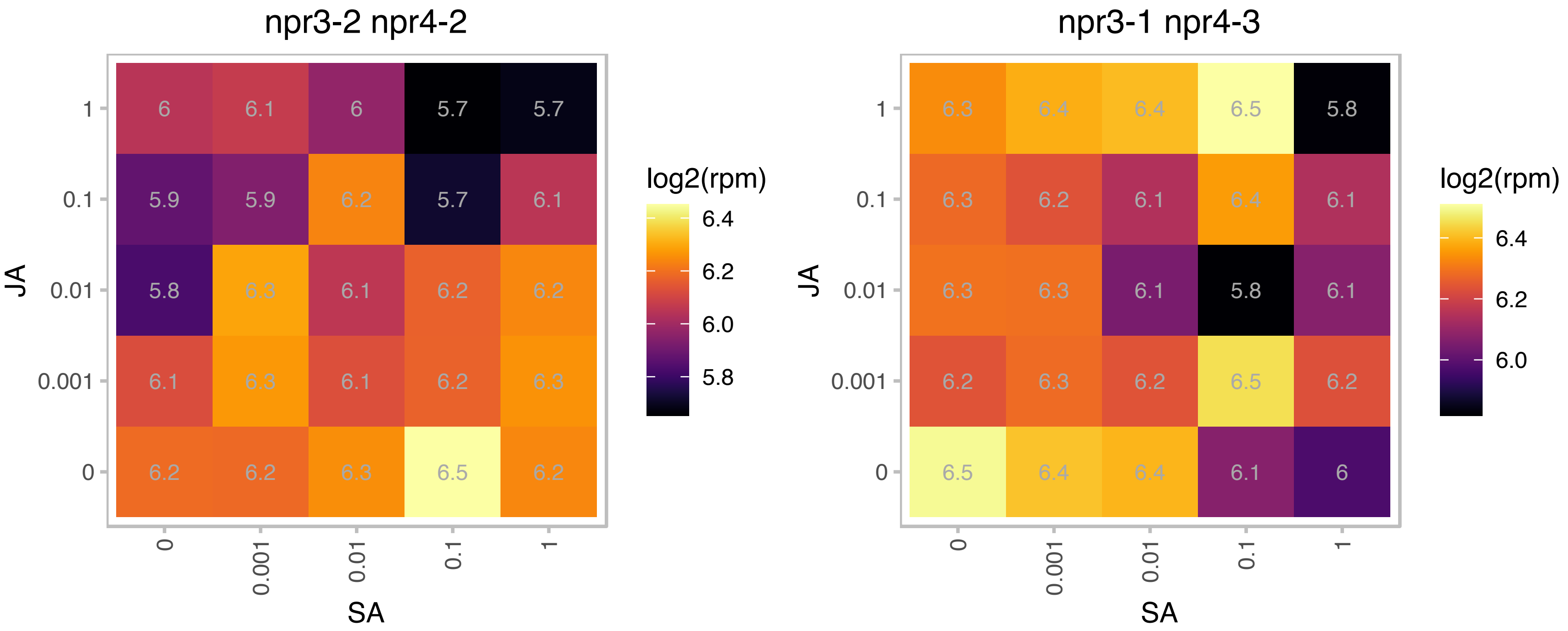

### AT5G22880.1 : H2B, HTB2

histone B2

Encodes a histone 2B (H2B) protein. This protein can be ubiquitinated in planta, and this modification depends on the HUB1 and HUB2 E3 ubiquitin ligases.

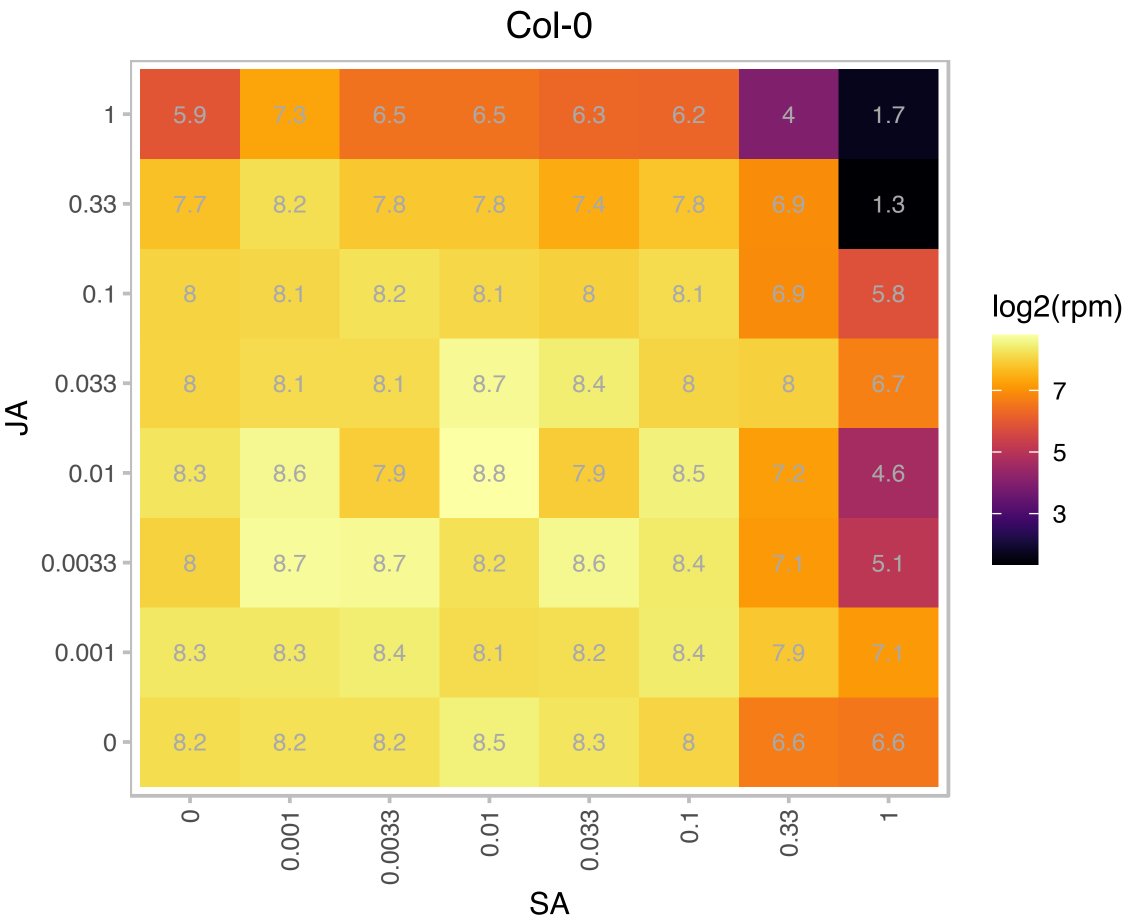


| | |
|---|---|
| Gene | AT5G22880 |
| is.expressed_genes | TRUE |
| is.responsive_genes | TRUE |
| is.apcluster | F-08 |

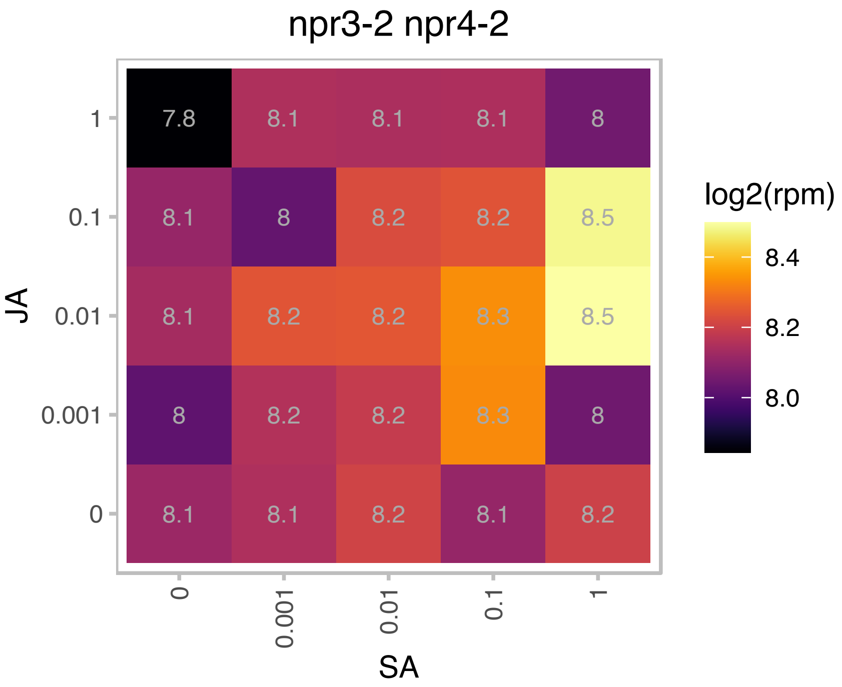


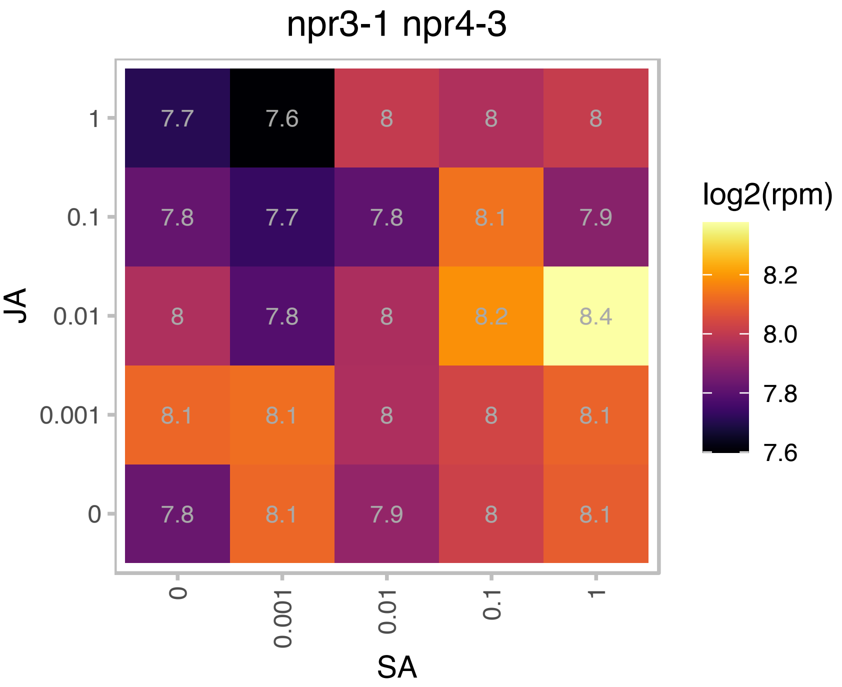

### AT5G23060.1 : CaS

calcium sensing receptor
Encodes a chloroplast-localized protein that modulates cytoplasmic Ca2+ concentration and is crucial for proper stomatal regulation in response to elevations of external Ca2+. Phosphorylation of this protein is dependent on calcium.

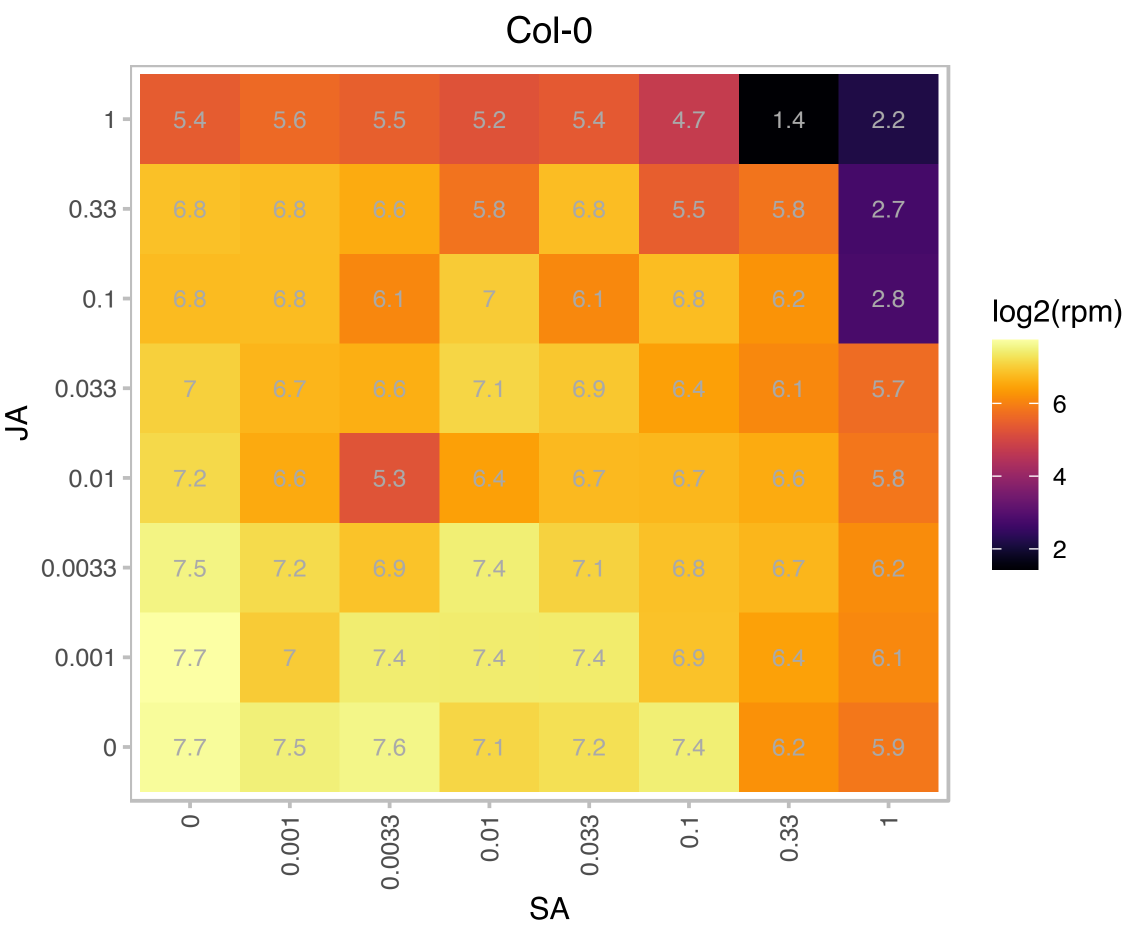


| | |
|---|---|
| Gene | AT5G23060 |
| is.expressed_genes | TRUE |
| is.responsive_genes | TRUE |
| is.apcluster | F-08 |

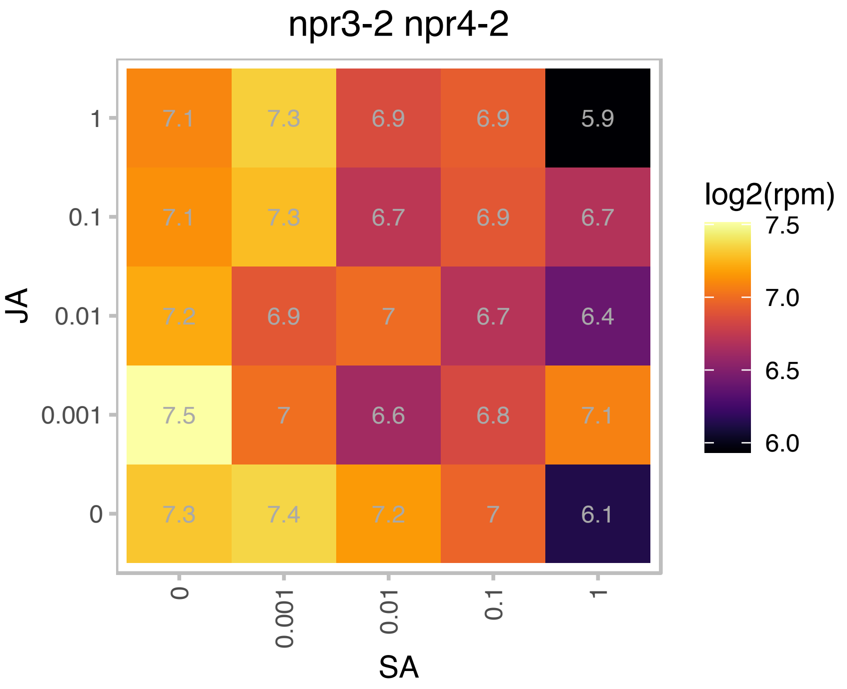


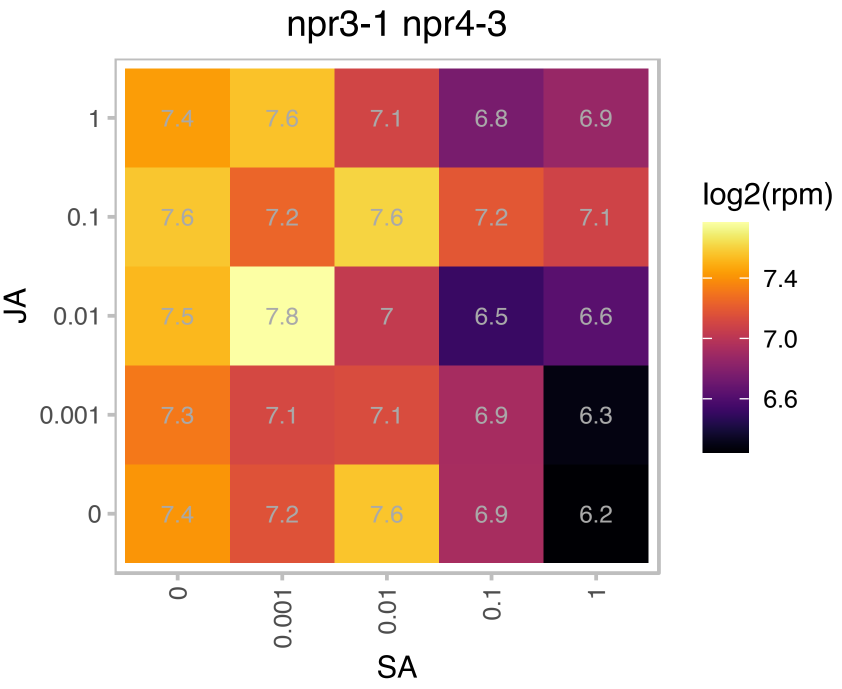

### AT5G23440.1 : FTRA1

ferredoxin/thioredoxin reductase subunit A (variable subunit) 1

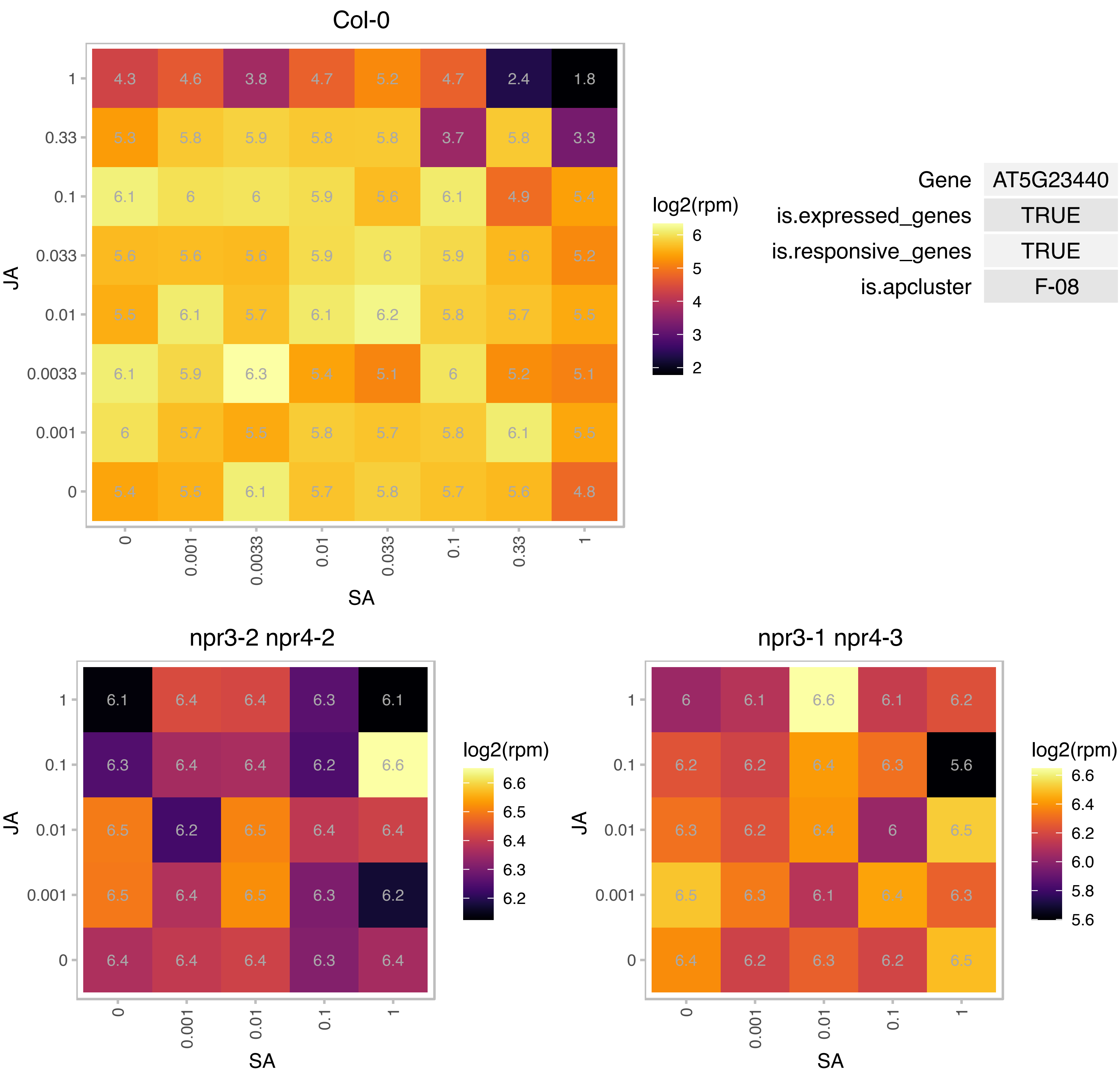


| | |
|---|---|
| Gene | AT5G23440 |
| is.expressed_genes | TRUE |
| is.responsive_genes | TRUE |
| is.apcluster | F-08 |

# AT5G23760.1 : NA

Copper transport protein family

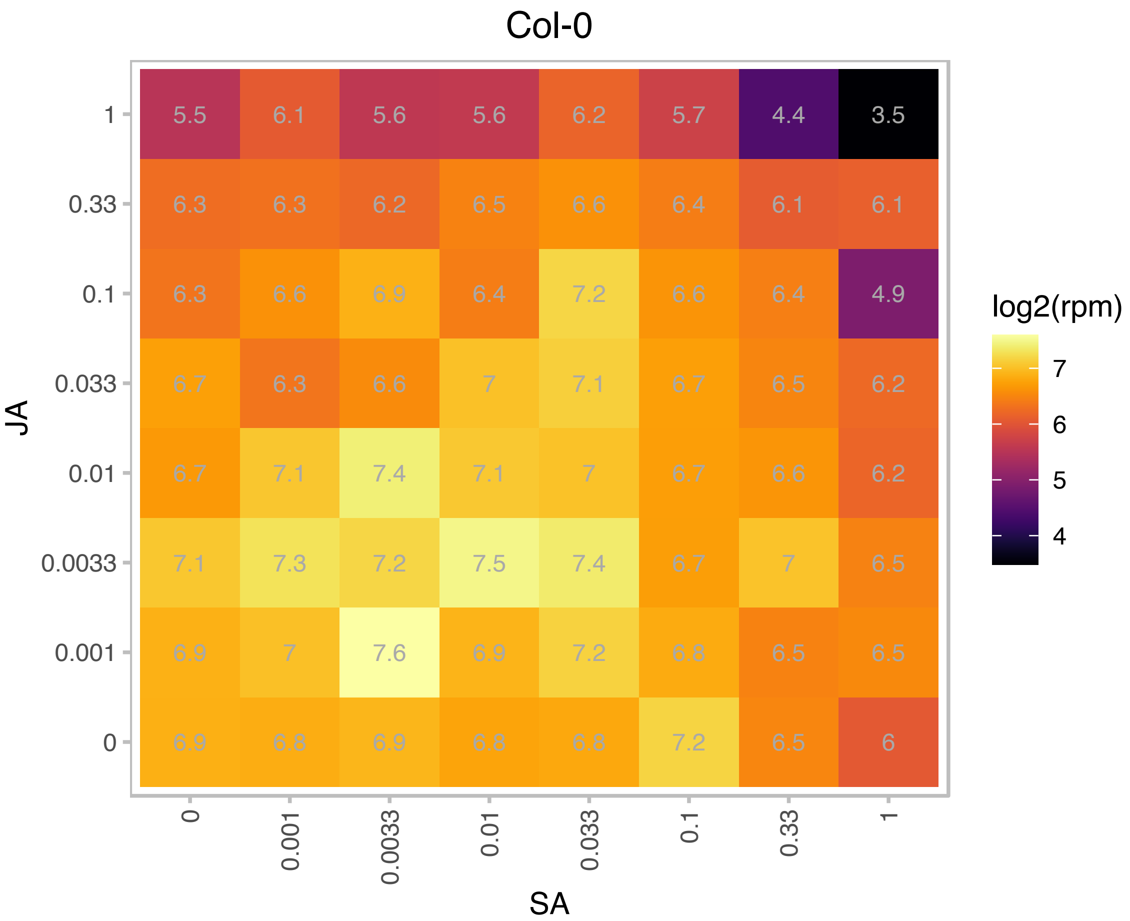


| | |
|---|---|
| Gene | AT5G23760 |
| is.expressed_genes | TRUE |
| is.responsive_genes | TRUE |
| is.apcluster | F-08 |

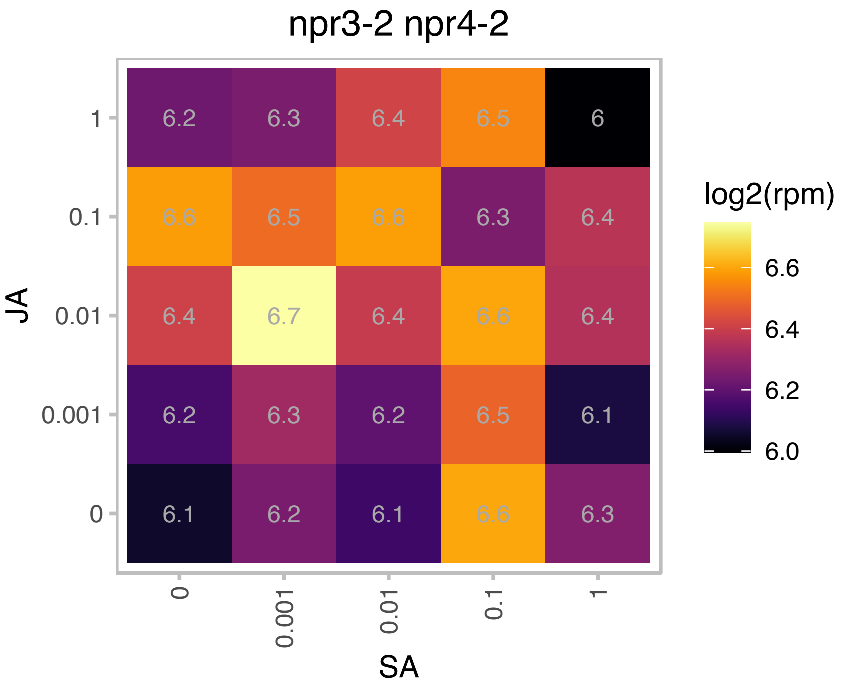


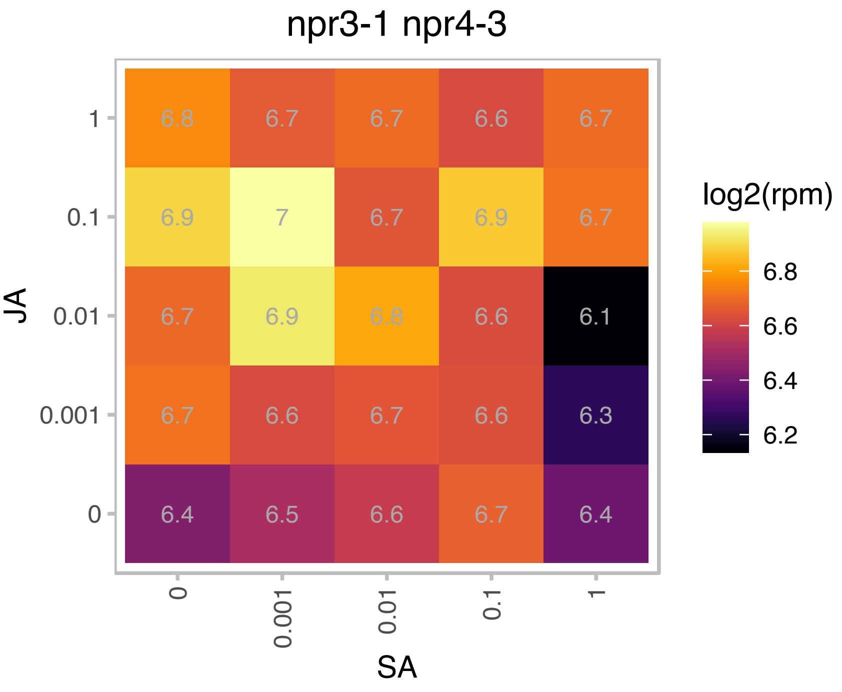

### AT5G24840.1 : AtTRM8a, TRM8a

tRNA (guanine-N-7) methyltransferase

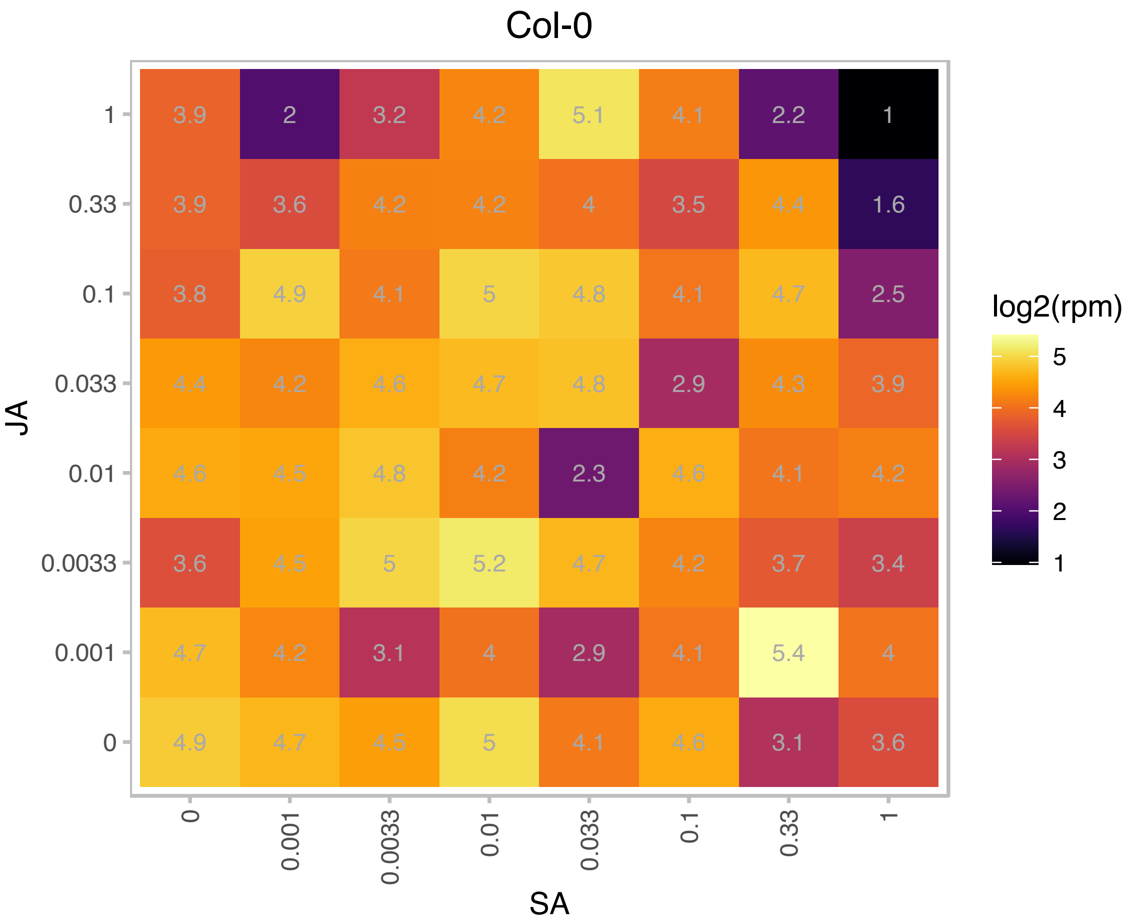


| | |
|---|---|
| Gene | AT5G24840 |
| is.expressed_genes | TRUE |
| is.responsive_genes | TRUE |
| is.apcluster | F-08 |

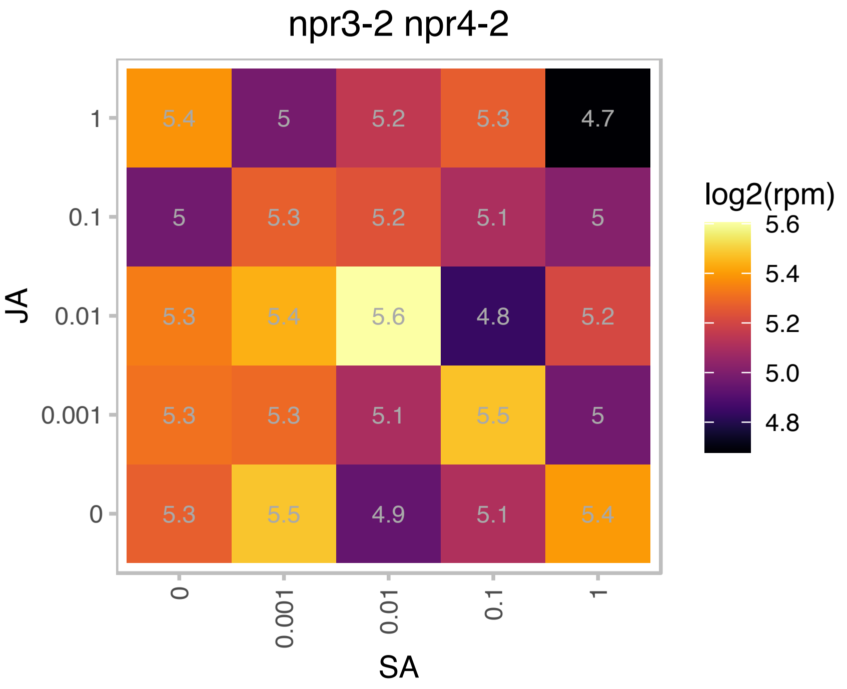


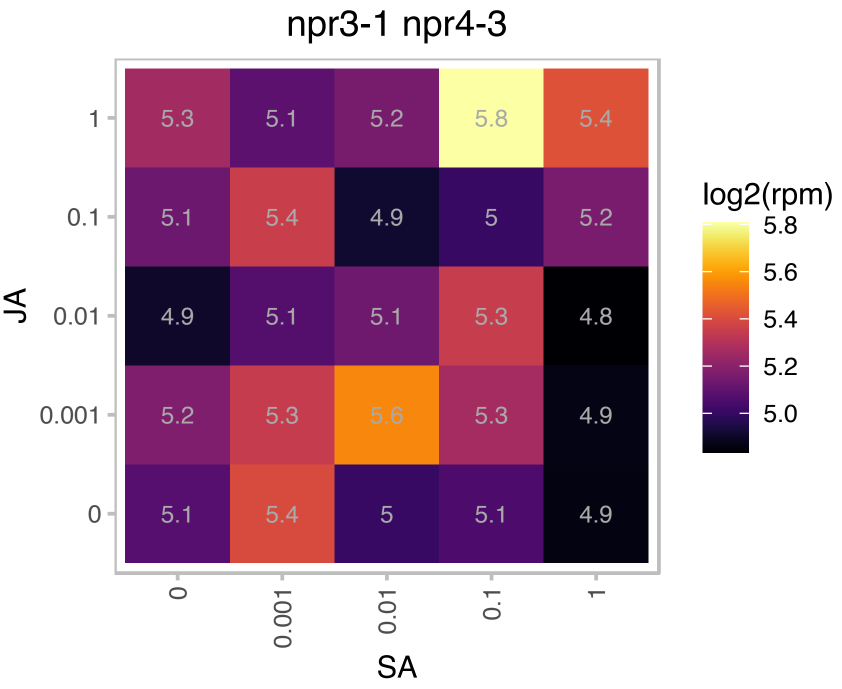

# AT5G30495.1 : NA

Fcf2 pre-rRNA processing protein

| | |
|---|---|
| Gene | AT5G30495 |
| is.expressed_genes | TRUE |
| is.responsive_genes | TRUE |
| is.apcluster | F-08 |

Col-0

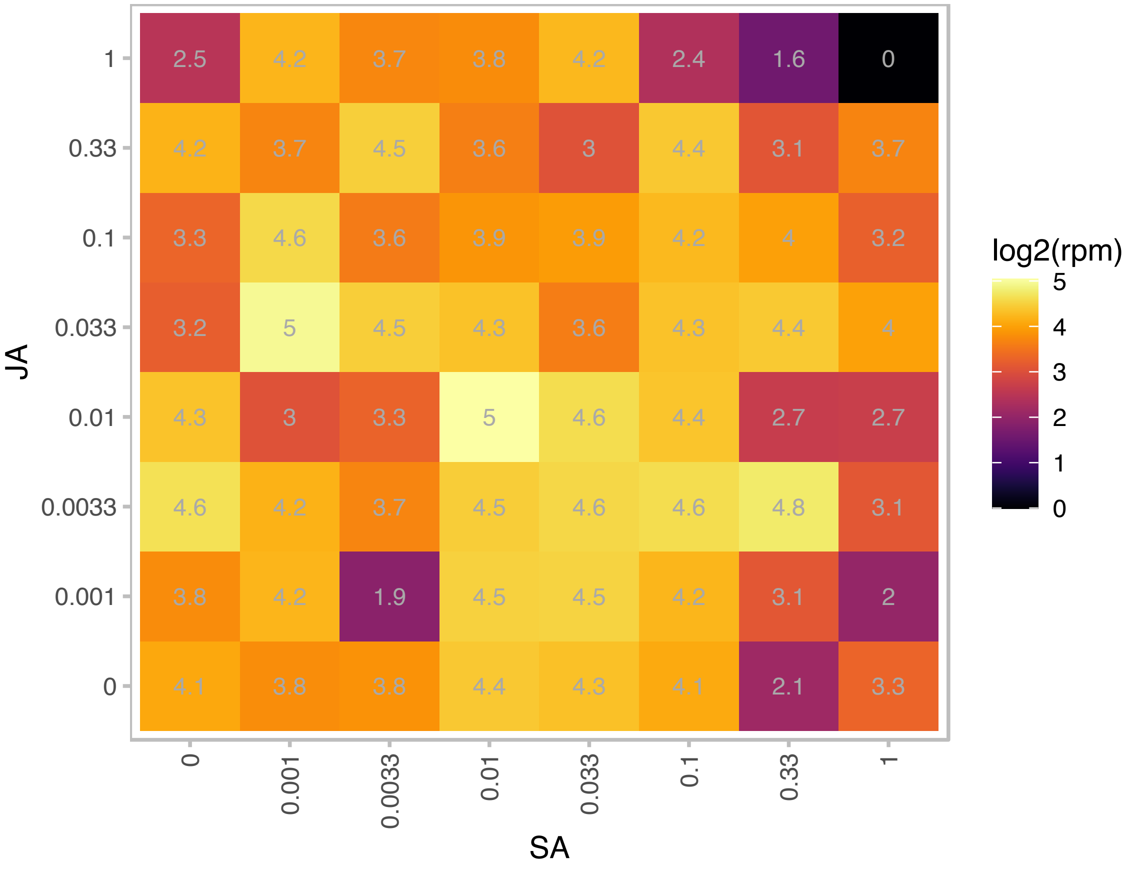


npr3-2 npr4-2

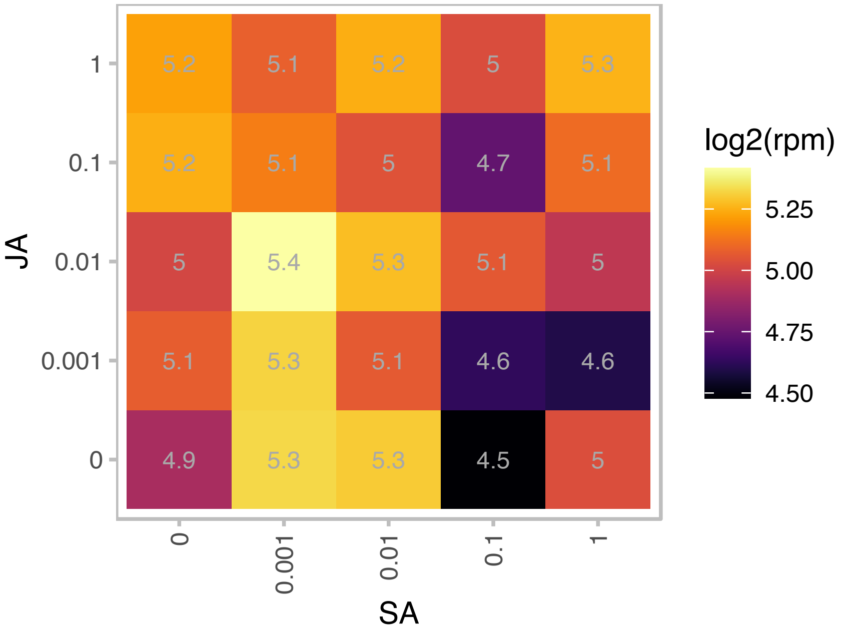


npr3-1 npr4-3

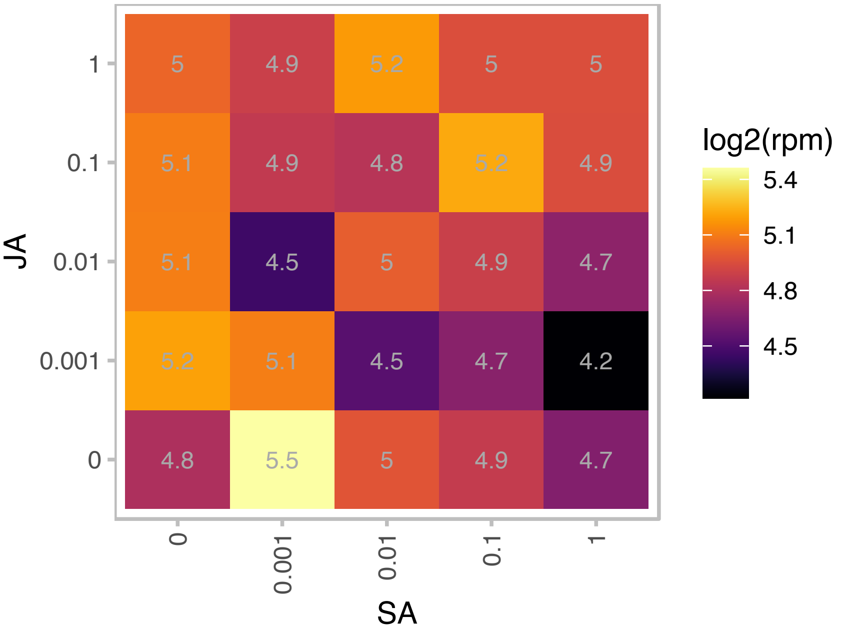

### AT5G37770.1 : AtCML24, CML24, TCH2

EF-hand calcium-binding protein family

Encodes a protein with 40% similarity to calmodulin. Binds Ca(2+) and, as a consequence, undergoes conformational changes. CML24 expression occurs in all major organs, and transcript levels are increased from 2- to 15-fold in plants subjected to touch, darkness, heat, cold, hydrogen peroxide, abscisic acid (ABA), and indole-3-acetic acid. However, CML24 protein accumulation changes were not detectable. The putative CML24 regulatory region confers reporter expression at sites of predicted mechanical stress; in regions undergoing growth; in vascular tissues and various floral organs; and in stomata, trichomes, and hydathodes. CML24-underexpressing transgenics are resistant to ABA inhibition of germination and seedling growth, are [illegible]

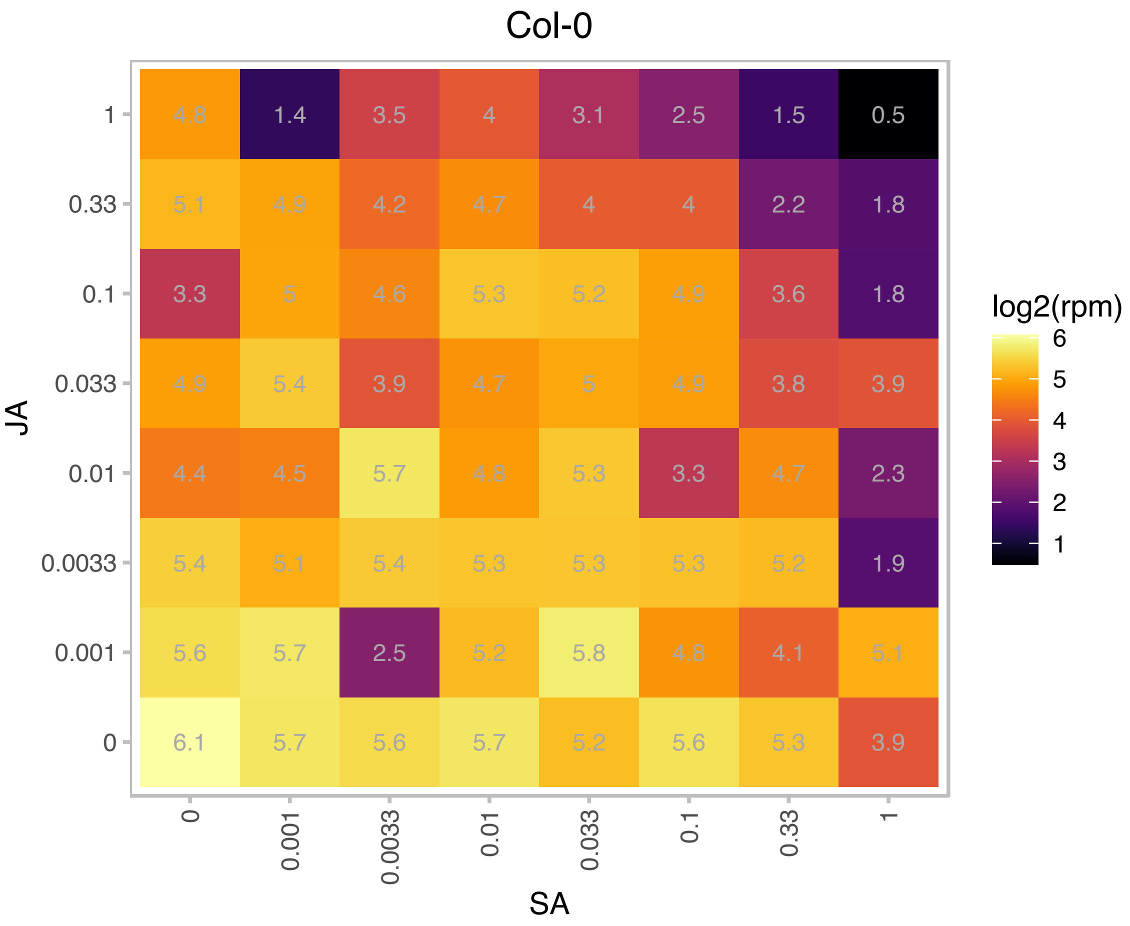


| | |
|---|---|
| Gene | AT5G37770 |
| is.expressed_genes | TRUE |
| is.responsive_genes | TRUE |
| is.apcluster | F-08 |

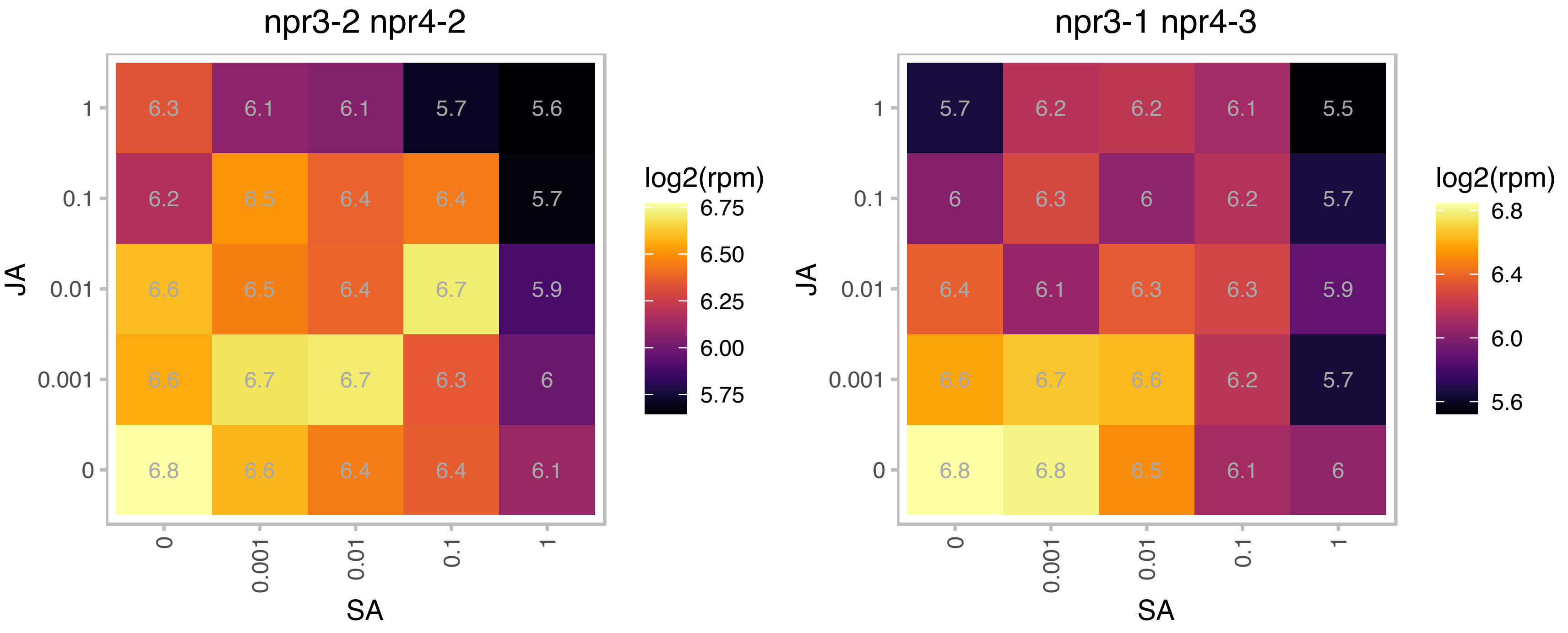

### AT5G39760.1 : AtHB23, HB23, ZHD10

homeobox protein 23

Functions together with TZP in co-regulation of the expression of blue-light dependent transcriptional regulators. Coassociates with and regulates the expression of light-regulated loci as well as transcriptional regulators to shape plant development in response to environmental stimuli with targets in RNA processing factors as well as proteins involved in salt stress and ABA signaling, in addition to embryo development. Acts downstream of TZP action with regard to blue-light-regulated hypocotyl elongation.

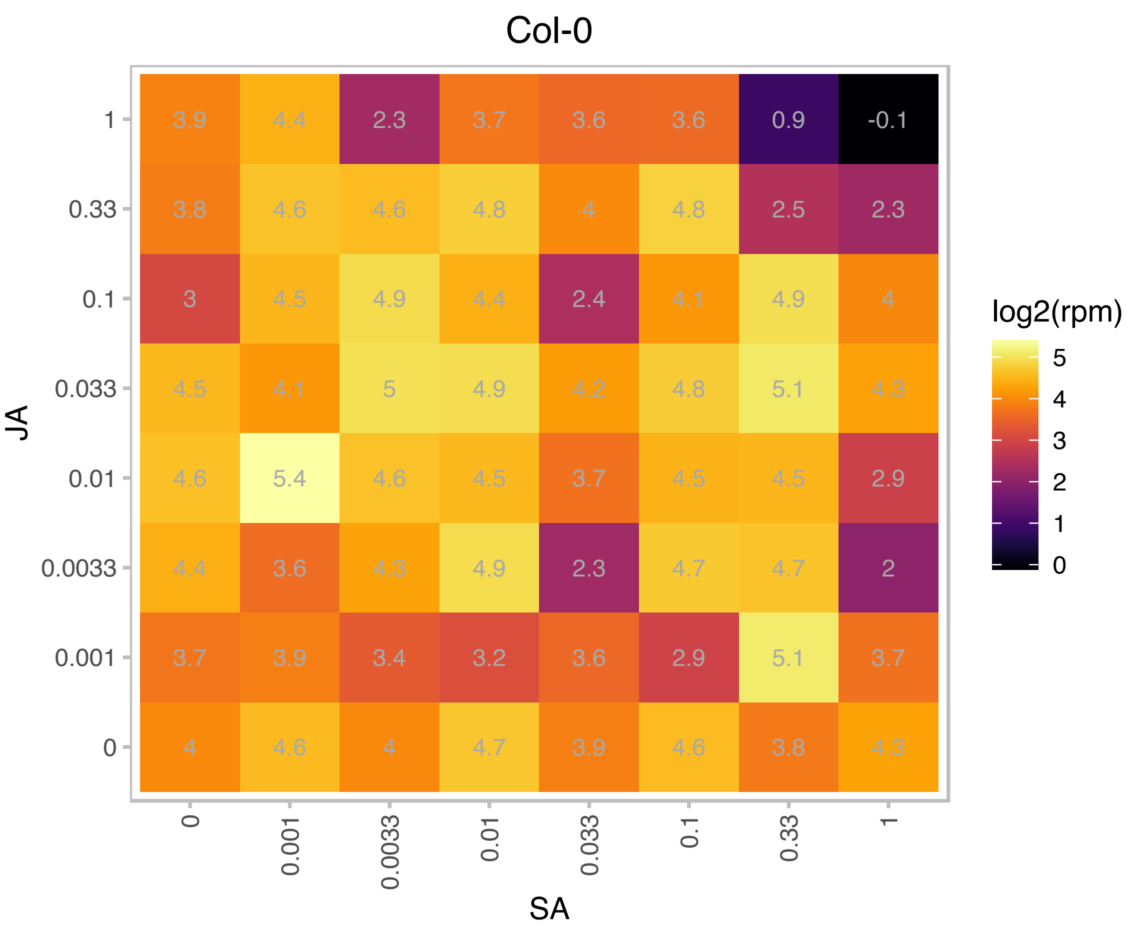


| | |
|---|---|
| Gene | AT5G39760 |
| is.expressed_genes | TRUE |
| is.responsive_genes | TRUE |
| is.apcluster | F-08 |

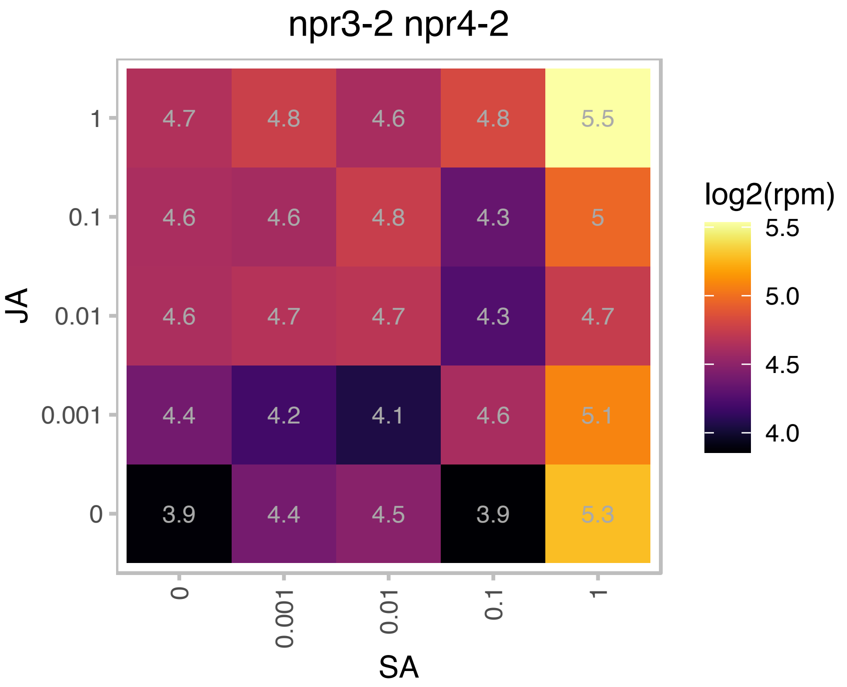


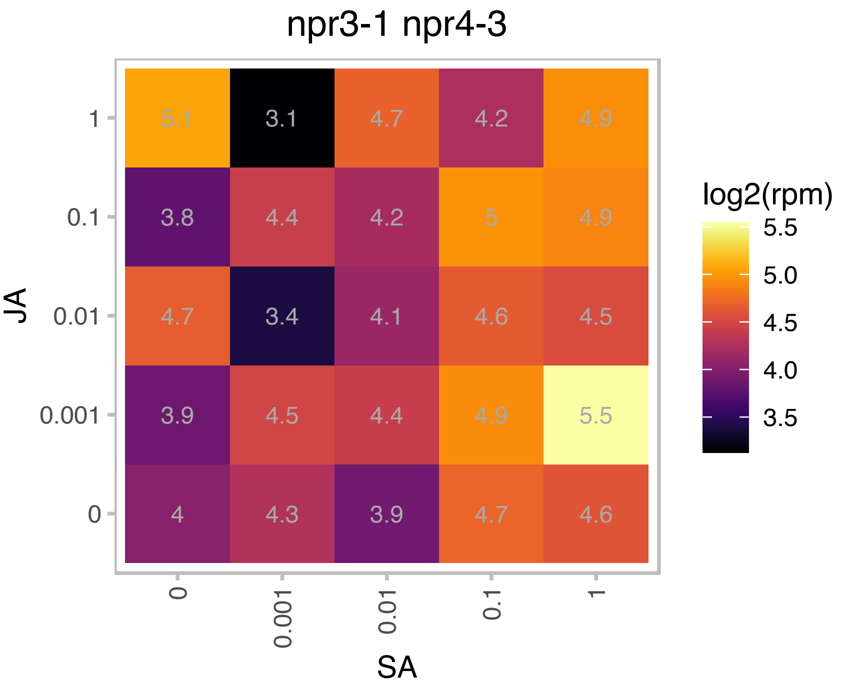

### AT5G48870.1 : AtLSM5, AtSAD1, LSM5, SAD1

Small nuclear ribonucleoprotein family protein
SAD1 encodes a polypeptide similar to multifunctional Sm-like snRNP proteins that are required for mRNA splicing, export, and degradation. Mutation in this gene increases plant sensitivity to drought stress and ABA in seed germination, root growth, and the expression of some stress-responsive genes.

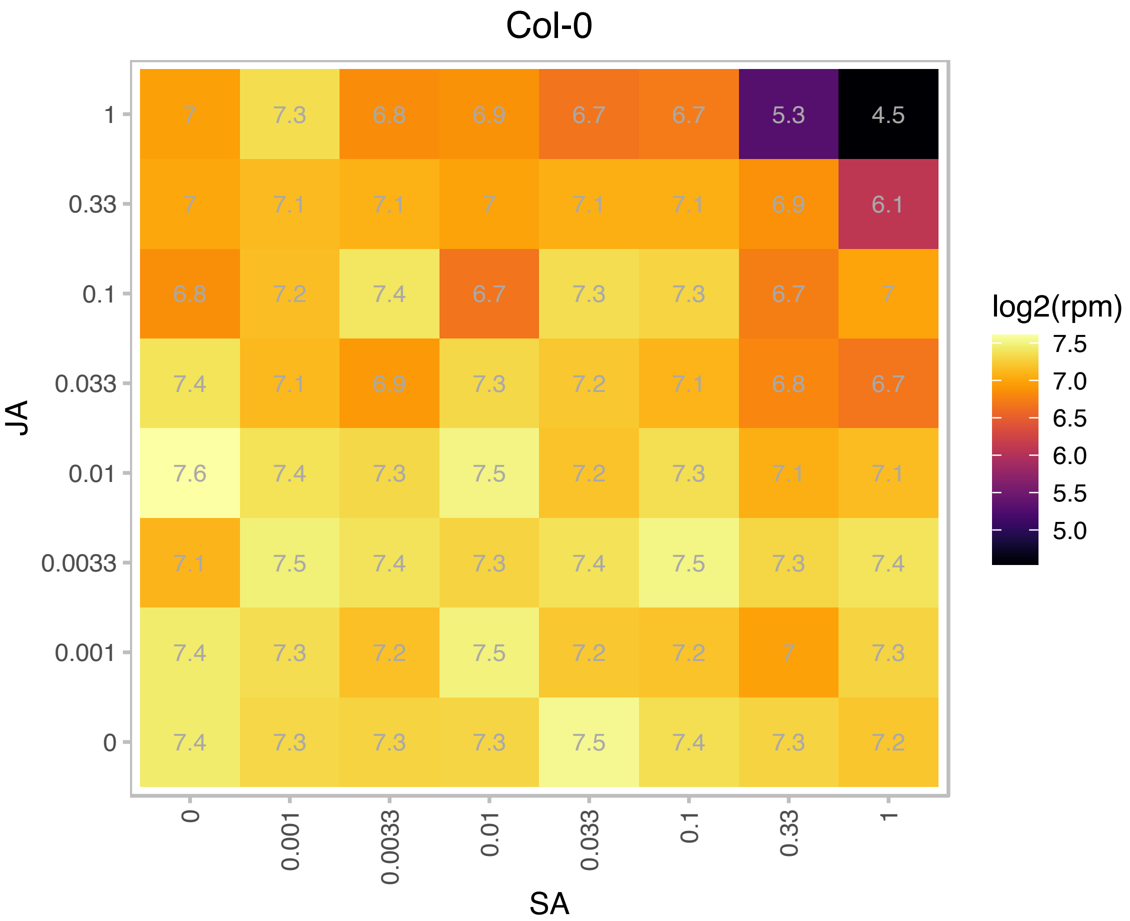


| | |
|---|---|
| Gene | AT5G48870 |
| is.expressed_genes | TRUE |
| is.responsive_genes | TRUE |
| is.apcluster | F-08 |

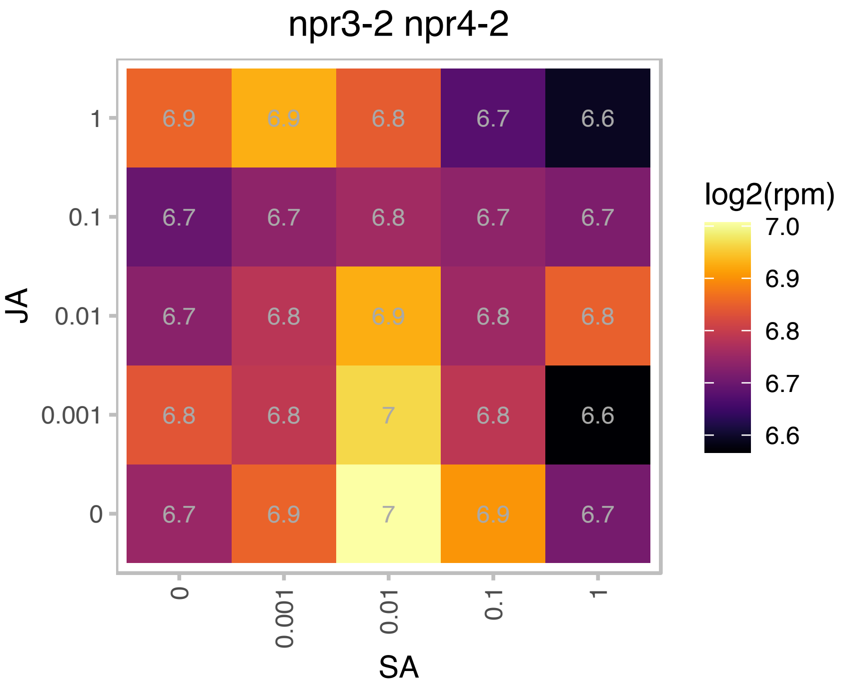


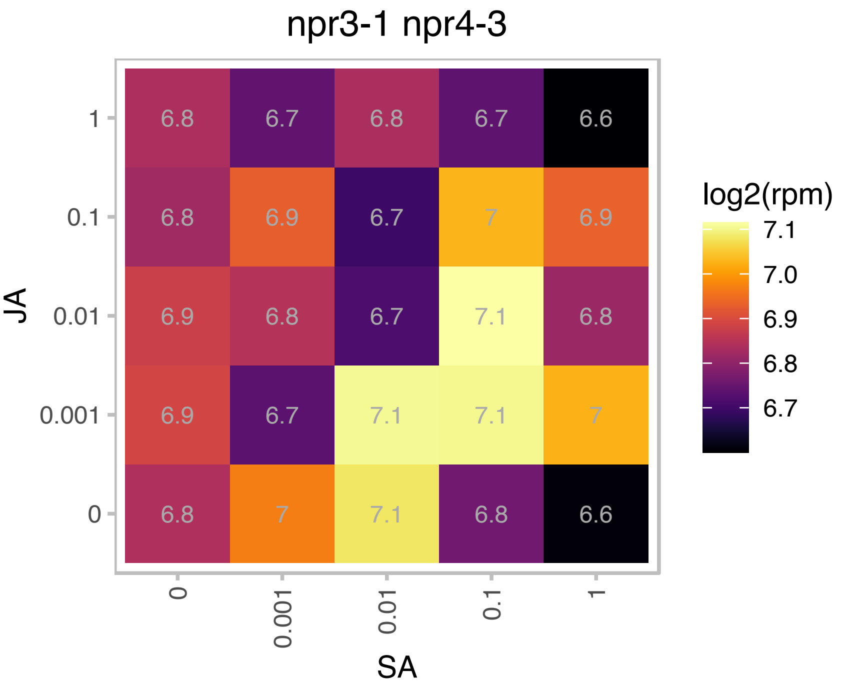

### AT5G54580.1 : ORRM2

RNA-binding (RRM/RBD/RNP motifs) family protein

Encodes an RNA recognition motif (RRM) and is involved in C-> U RNA editing in mitochondria.

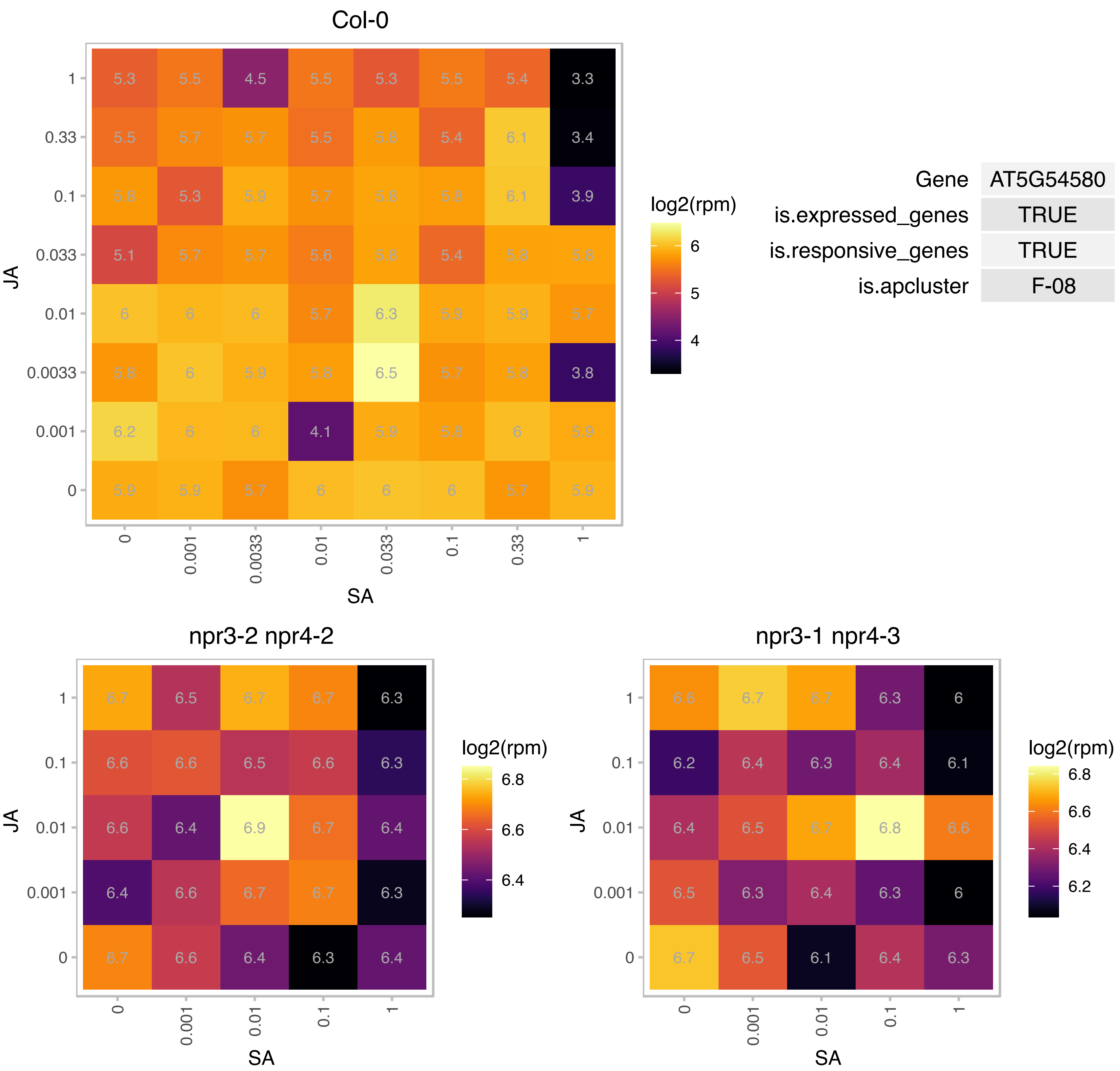


| | |
|---|---|
| Gene | AT5G54580 |
| is.expressed_genes | TRUE |
| is.responsive_genes | TRUE |
| is.apcluster | F-08 |

# AT5G58375.1 : NA

Methyltransferase-related protein

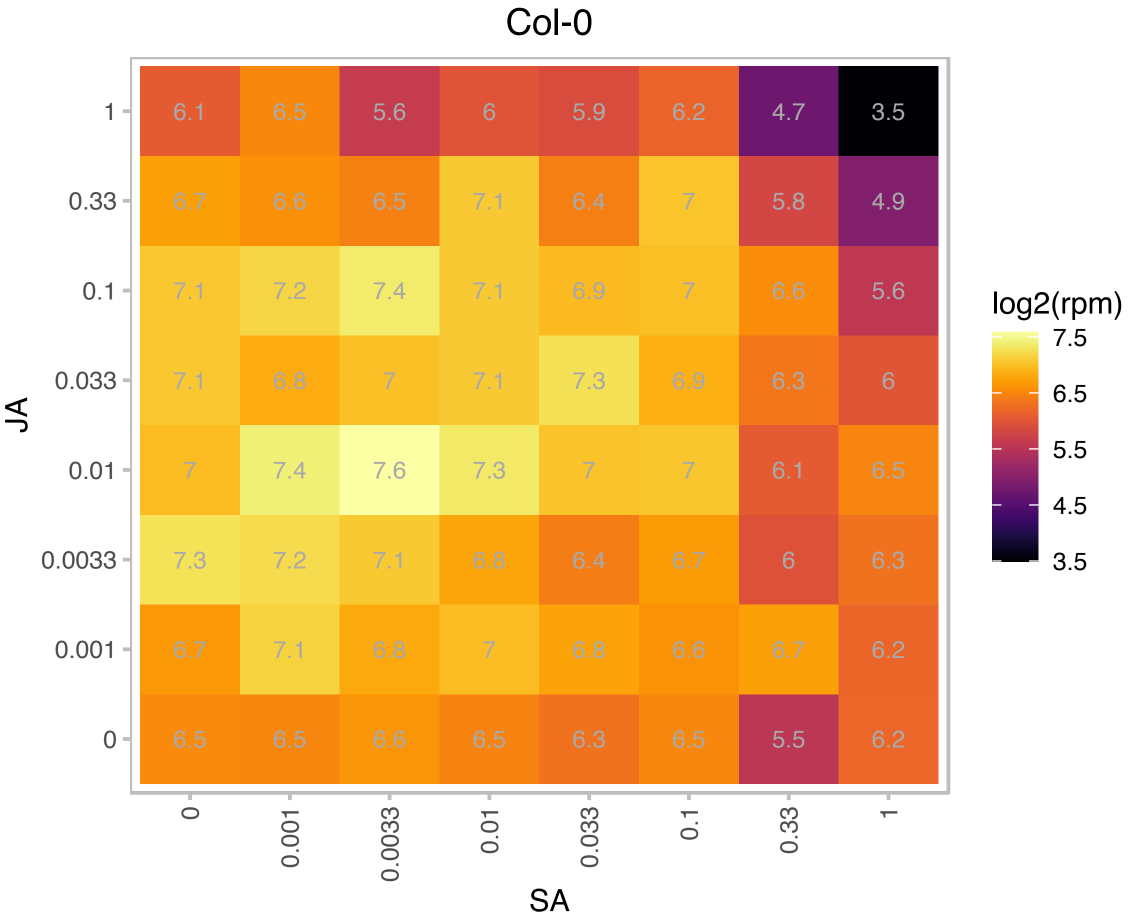


| | |
|---|---|
| Gene | AT5G58375 |
| is.expressed_genes | TRUE |
| is.responsive_genes | TRUE |
| is.apcluster | F-08 |

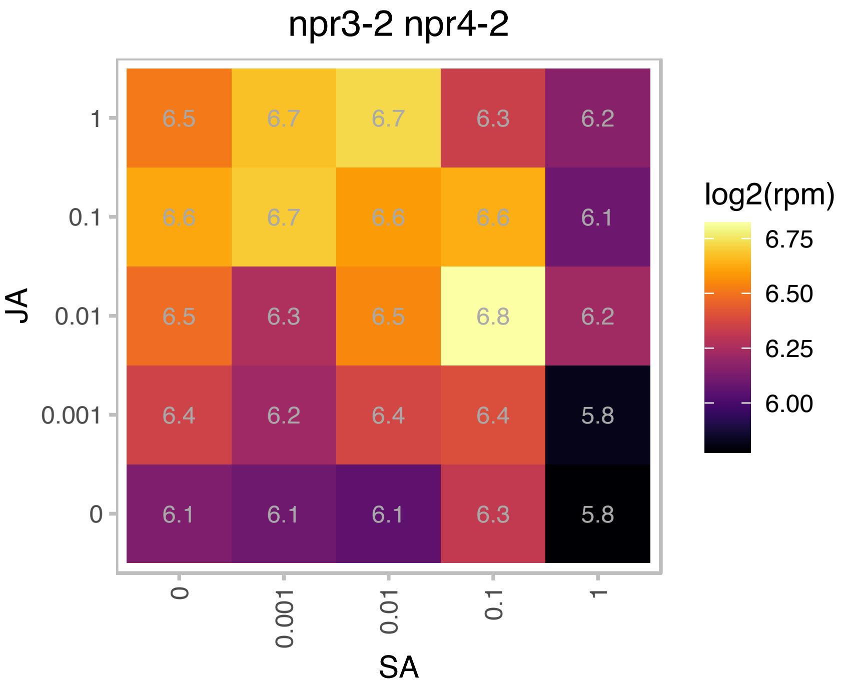


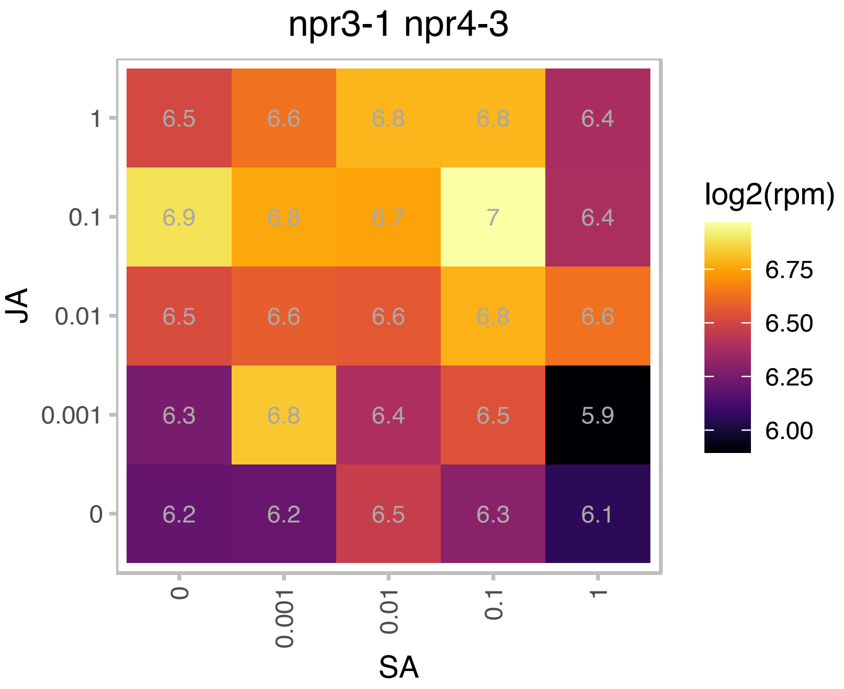

### AT5G59870.1 : h2a.w.6, HTA6

histone H2A 6

Encodes HTA6, a histone H2A protein.

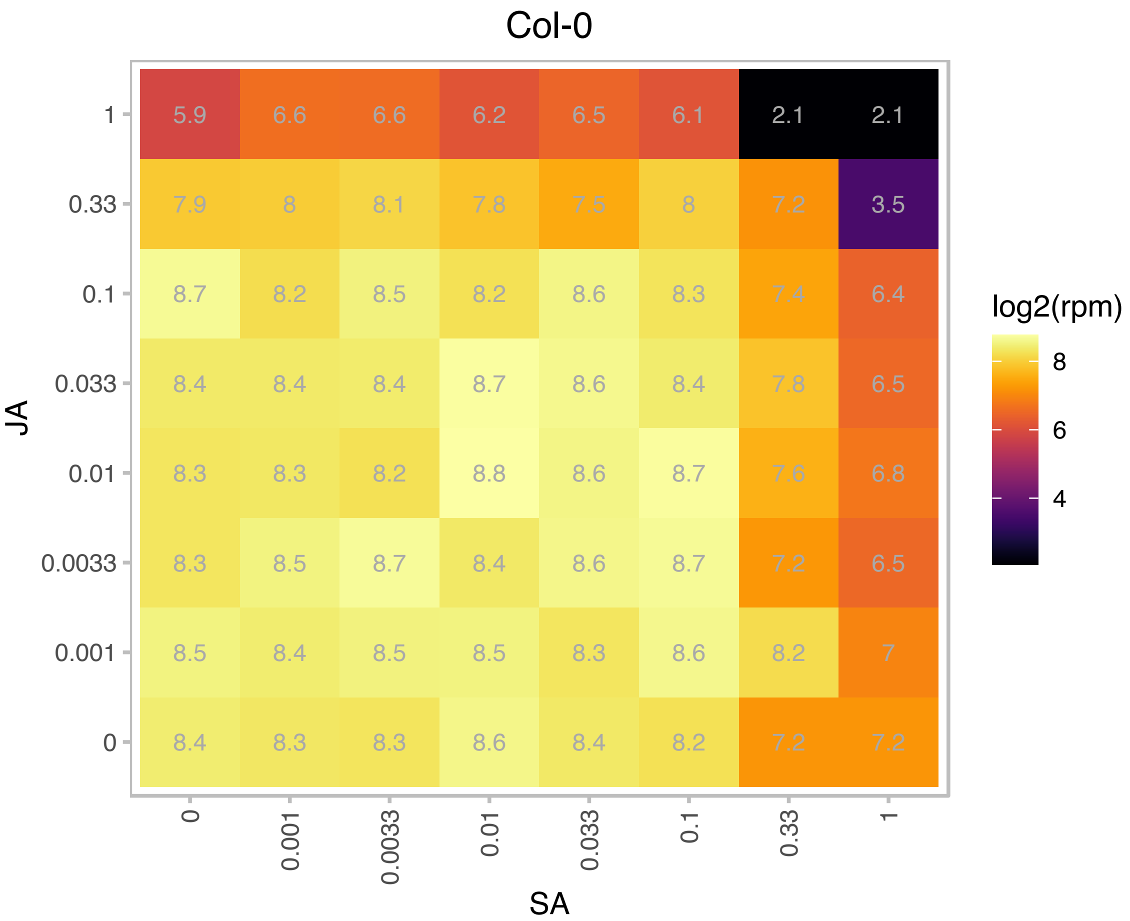


| | |
|---|---|
| Gene | AT5G59870 |
| is.expressed_genes | TRUE |
| is.responsive_genes | TRUE |
| is.apcluster | F-08 |

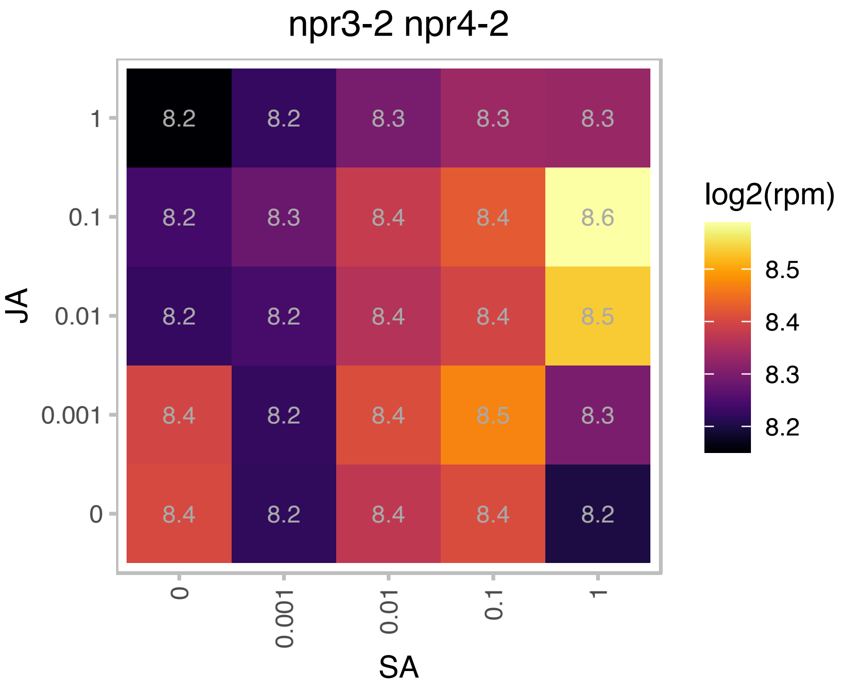


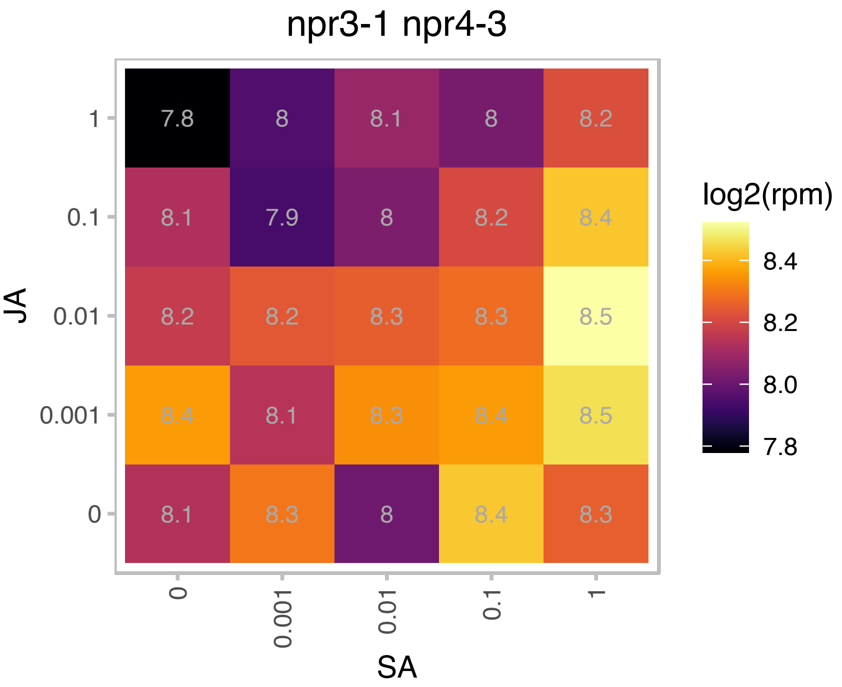

### AT5G59950.1 : ALY1

RNA-binding (RRM/RBD/RNP motifs) family protein

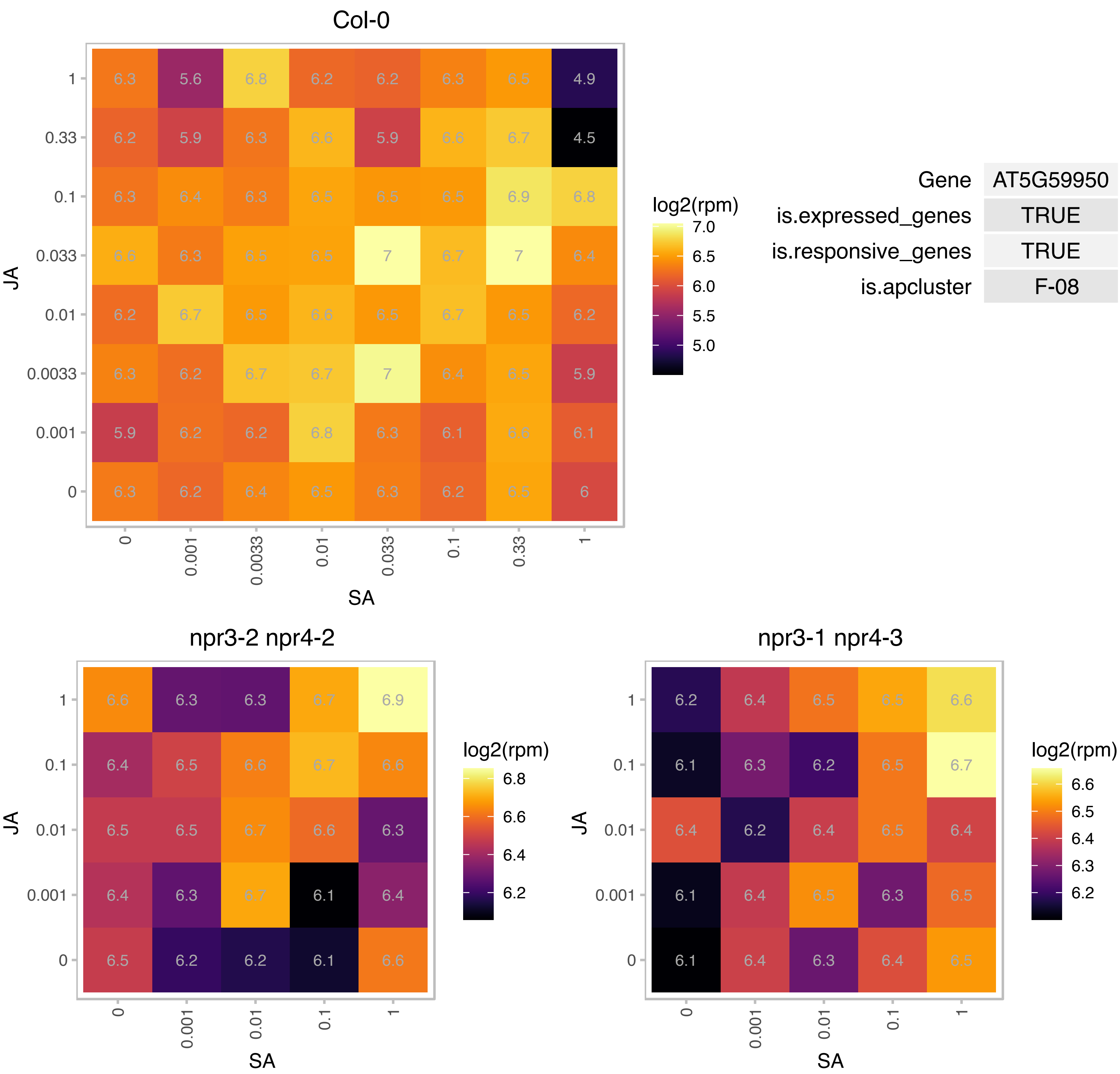


| | |
|---|---|
| Gene | AT5G59950 |
| is.expressed_genes | TRUE |
| is.responsive_genes | TRUE |
| is.apcluster | F-08 |

# AT5G65360.1 : H3.1

Histone superfamily protein

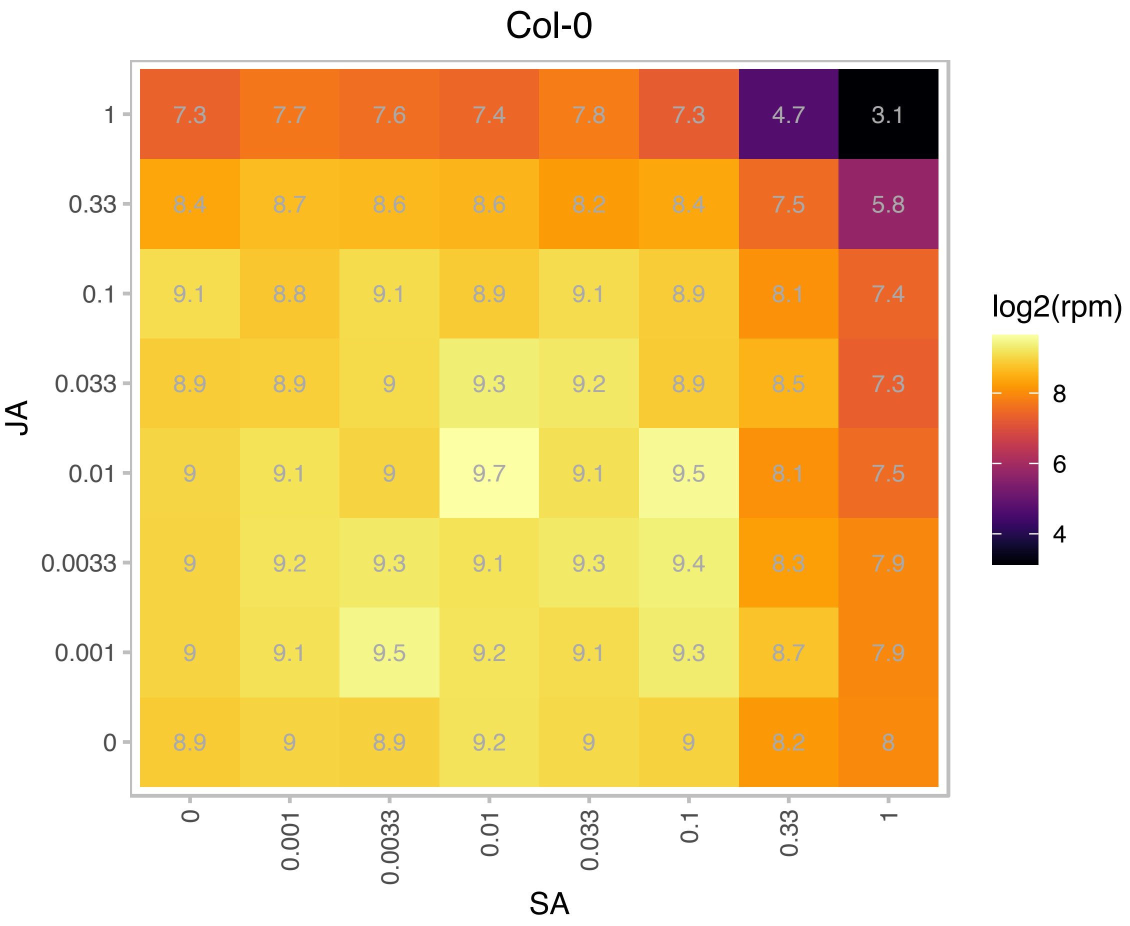


| | |
|---|---|
| Gene | AT5G65360 |
| is.expressed_genes | TRUE |
| is.responsive_genes | TRUE |
| is.apcluster | F-08 |

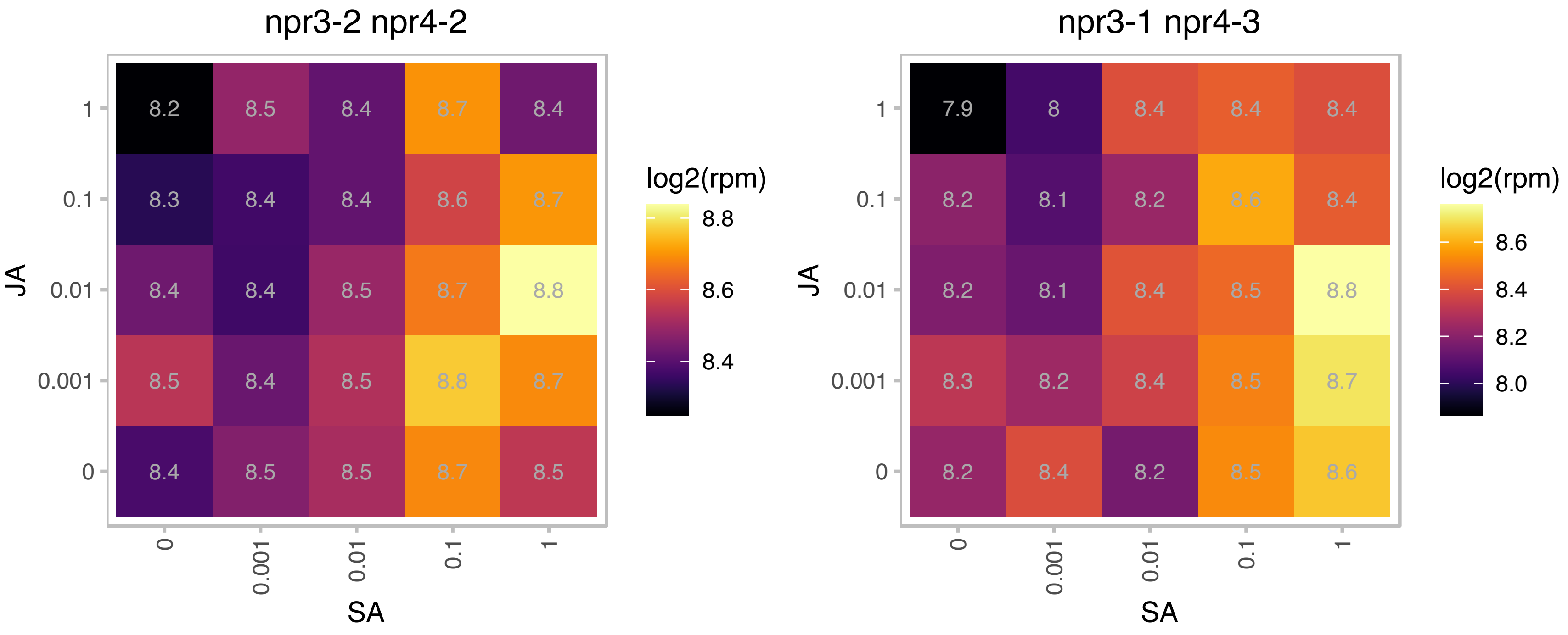

# AT5G67390.1 : NA

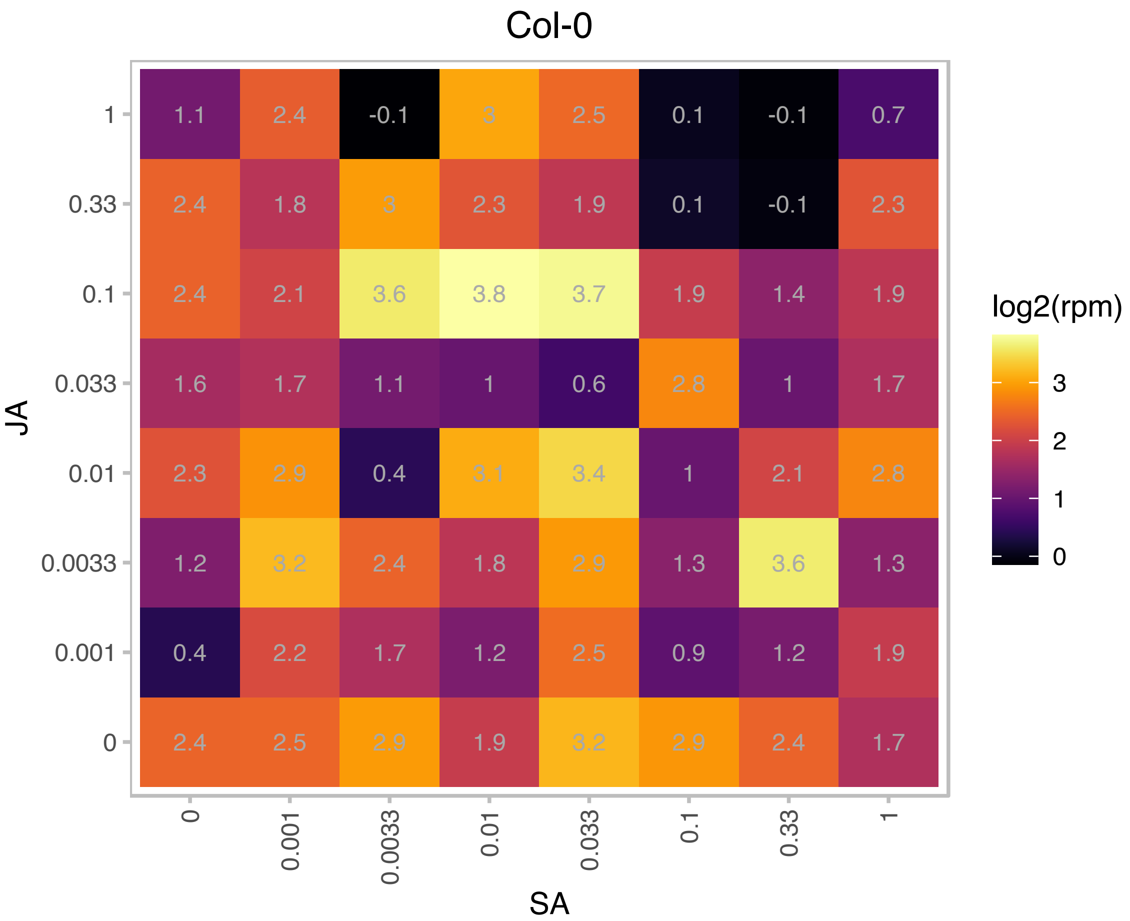


| | |
|---|---|
| Gene | AT5G67390 |
| is.expressed_genes | TRUE |
| is.responsive_genes | TRUE |
| is.apcluster | F-08 |

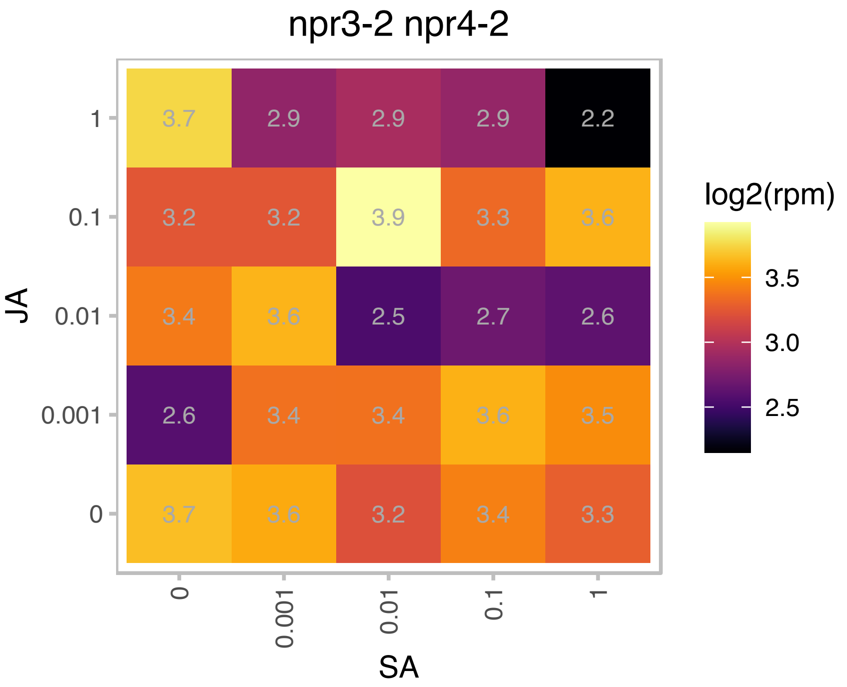


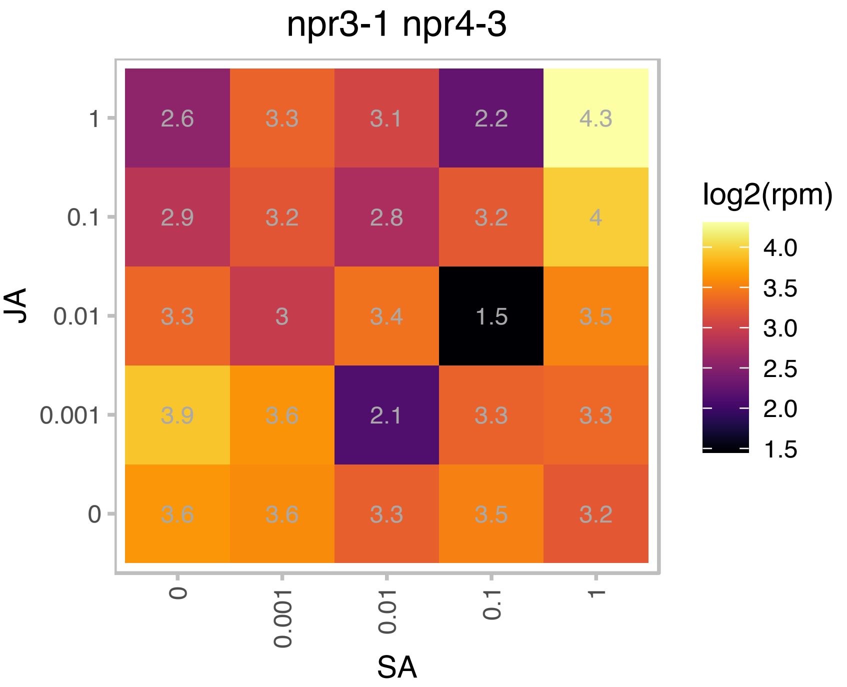

### AT5G67420.1 : ASL39, LBD37

LOB domain-containing protein 37

Encodes a LOB-domain protein involved in nitrogen metabolism and affecting leaf morphogenesis.

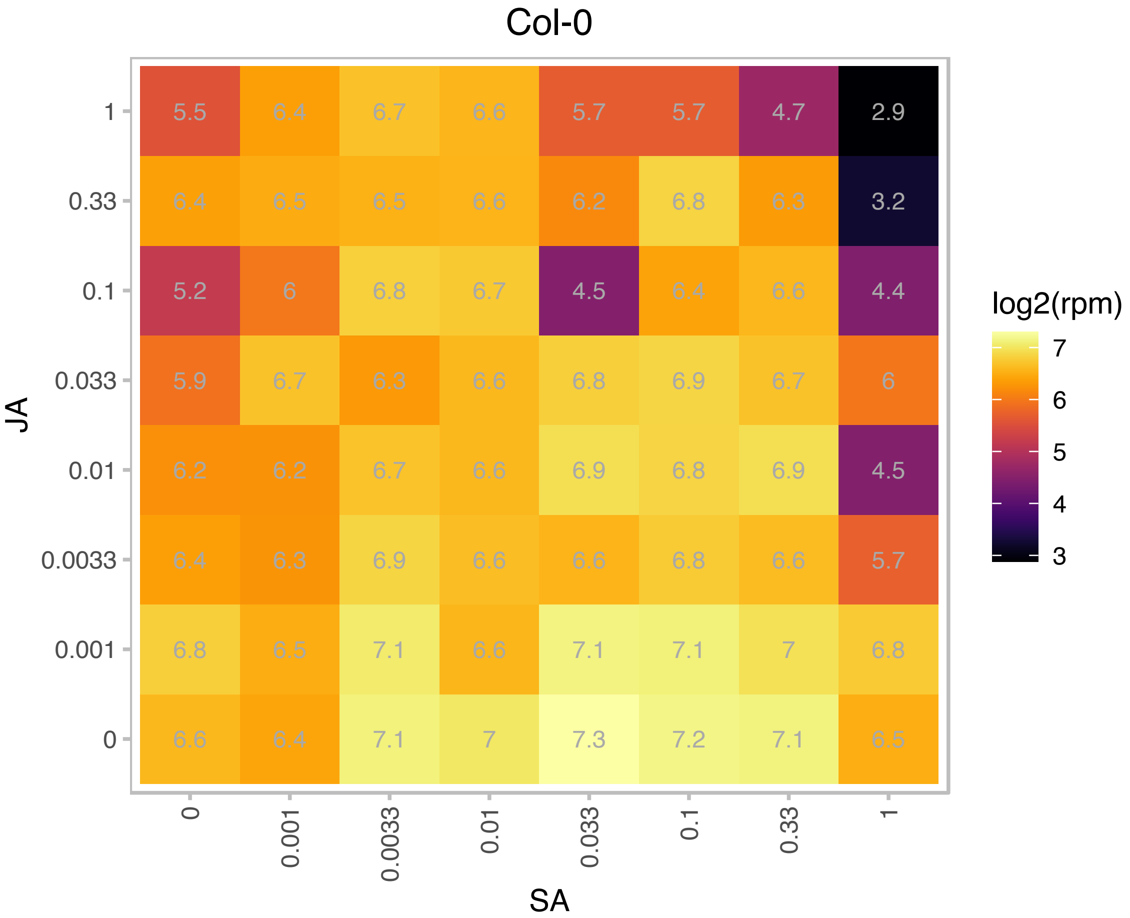


| Gene | AT5G67420 |
|---|---|
| is.expressed_genes | TRUE |
| is.responsive_genes | TRUE |
| is.apcluster | F-08 |

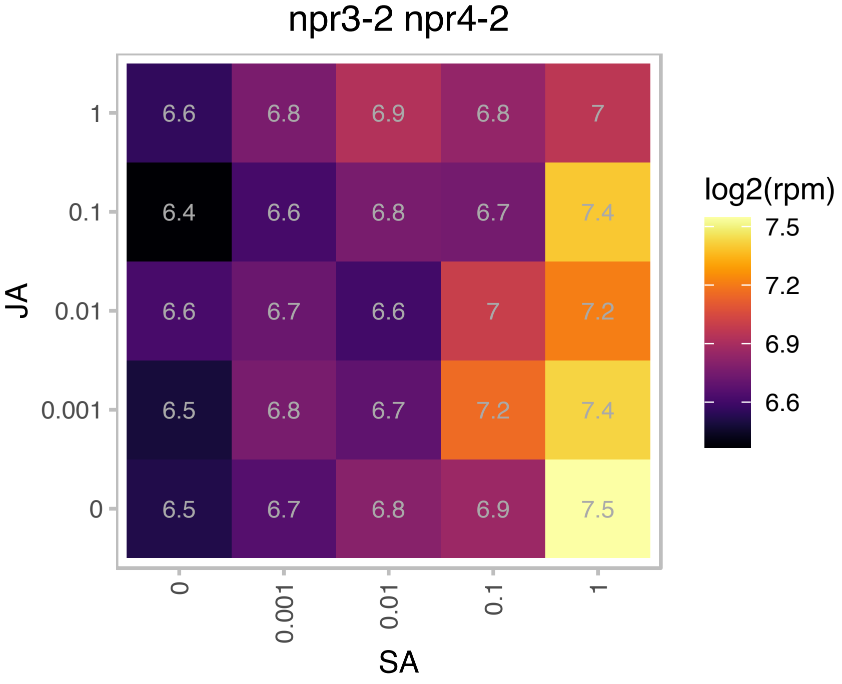


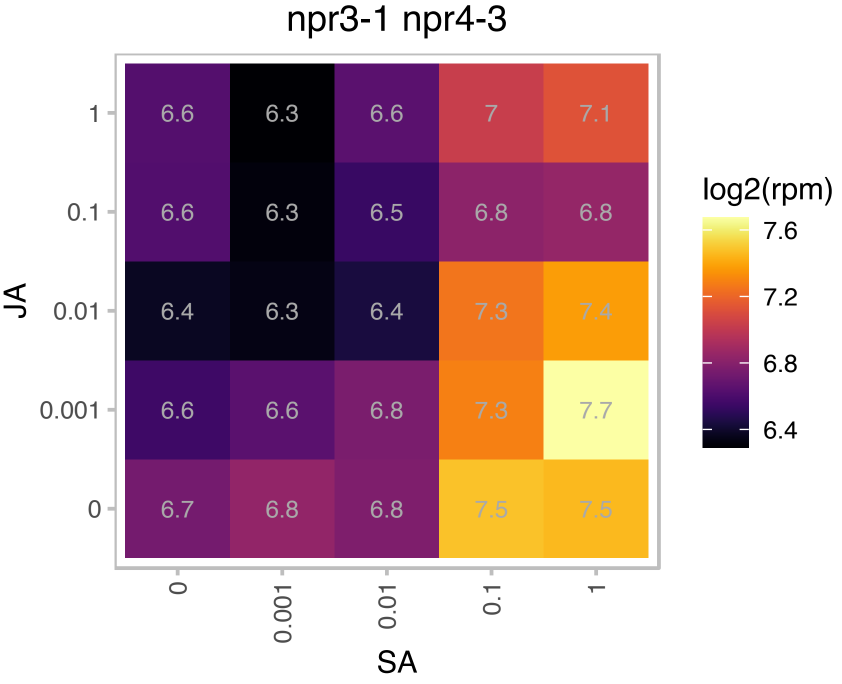

# AT5G67510.1 : NA

Translation protein SH3-like family protein

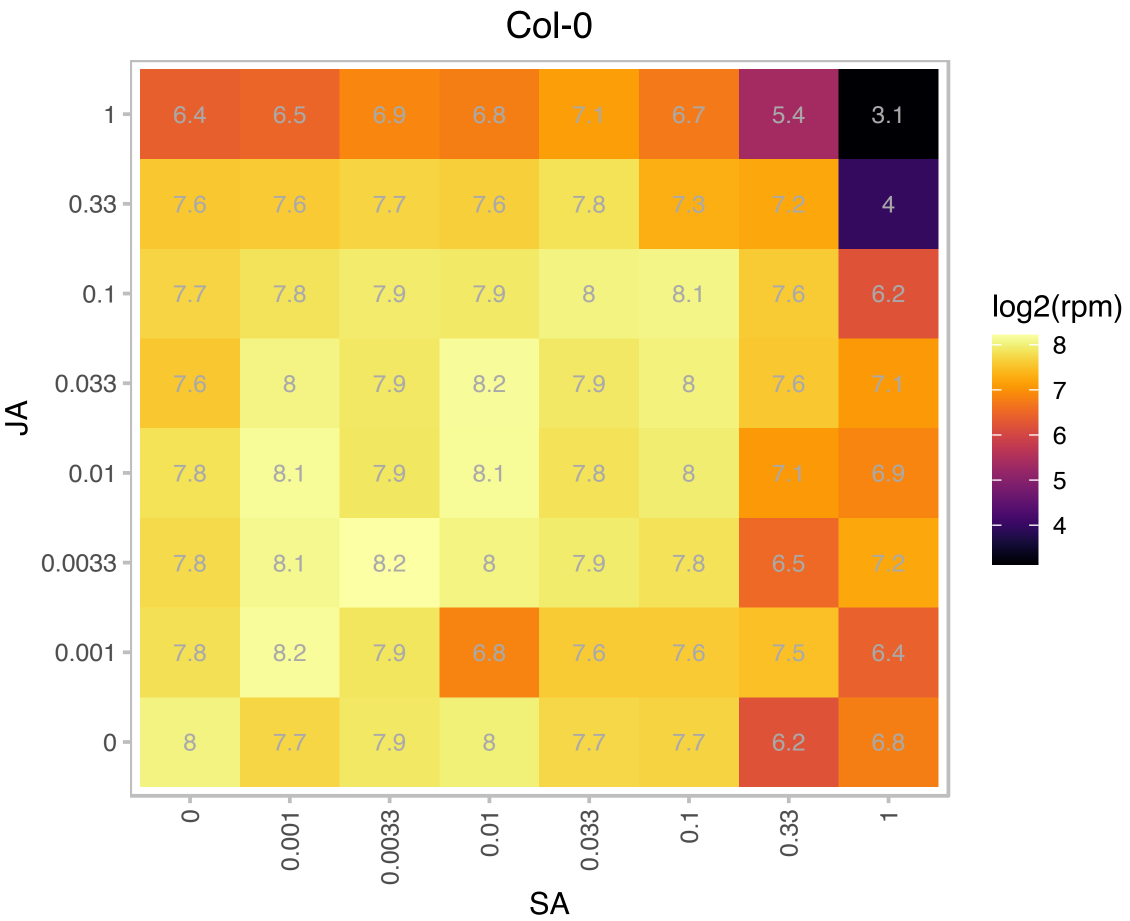


| | |
|---|---|
| Gene | AT5G67510 |
| is.expressed_genes | TRUE |
| is.responsive_genes | TRUE |
| is.apcluster | F-08 |

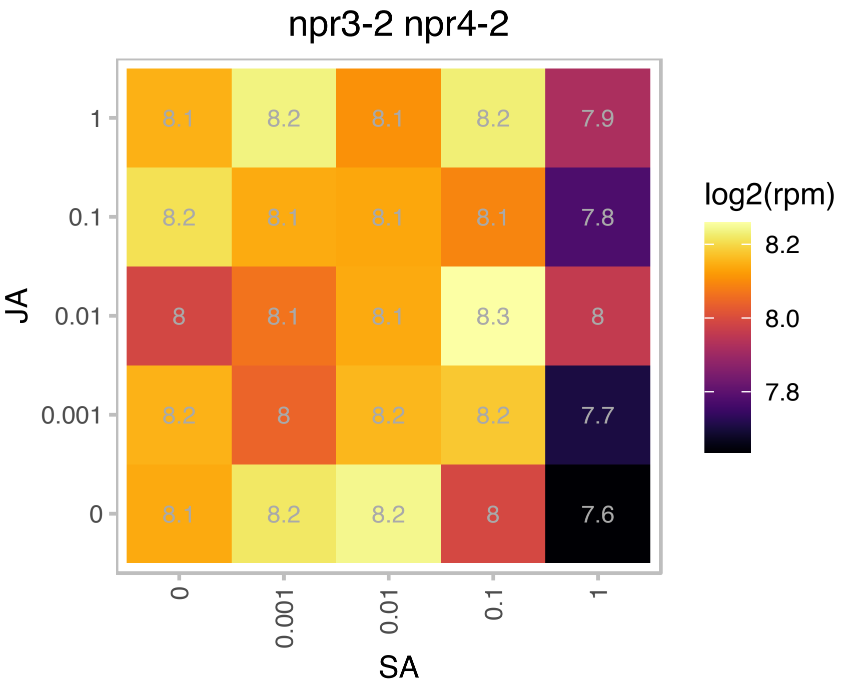


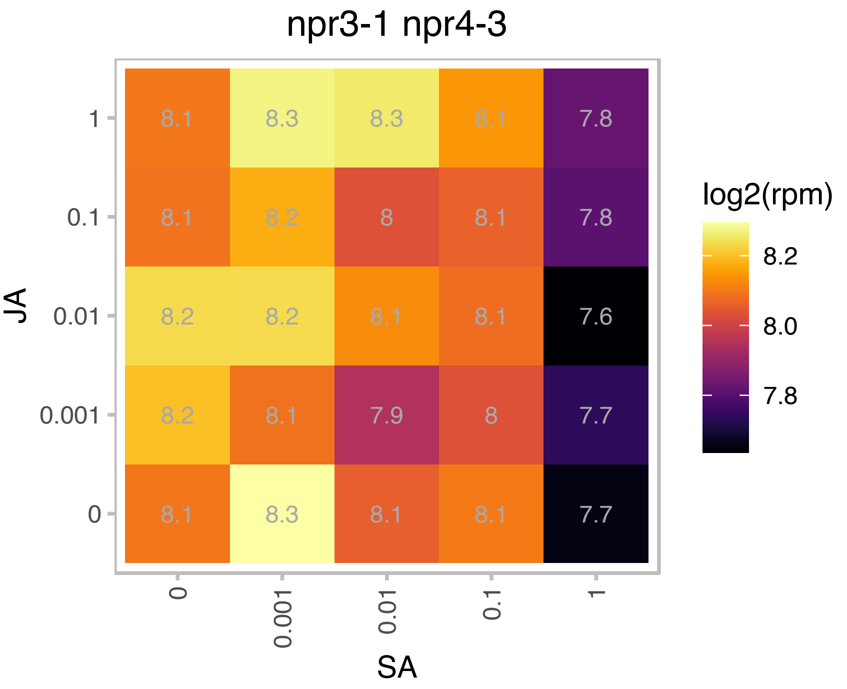

### AT1G04640.1 : LIP2

lipoyltransferase 2

Lipoyltransferase, located in mitochondria but not found in chloroplasts

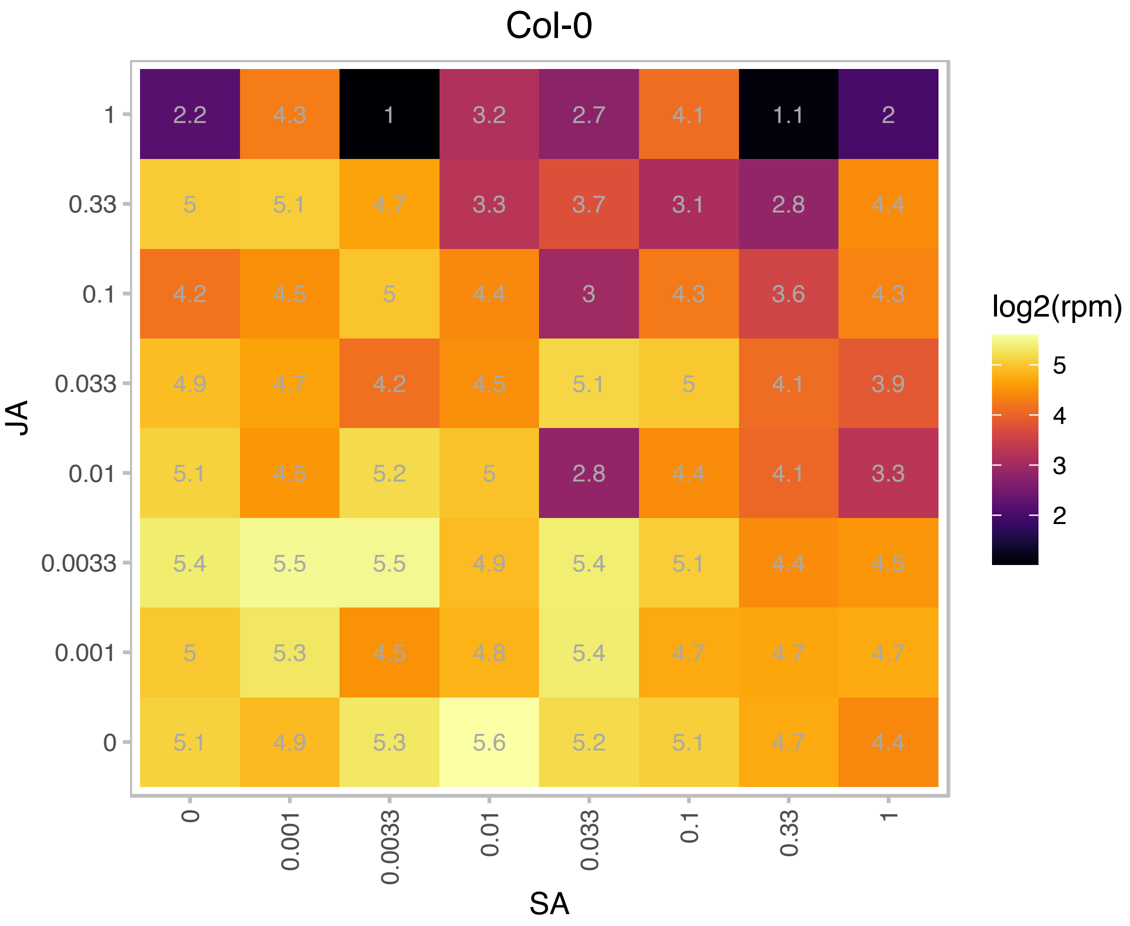


| | |
|---|---|
| Gene | AT1G04640 |
| is.expressed_genes | TRUE |
| is.responsive_genes | TRUE |
| is.apcluster | F-09 |

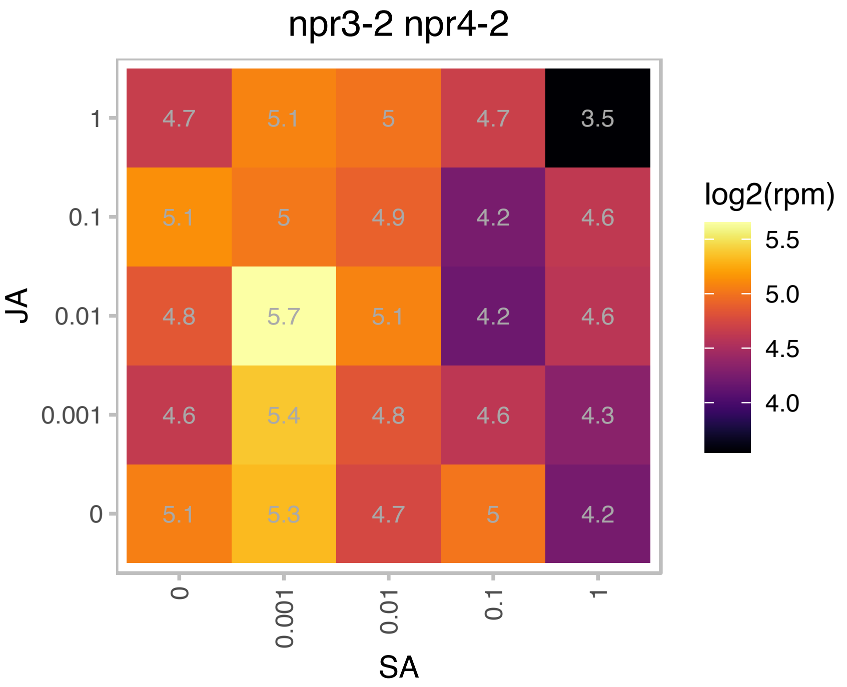


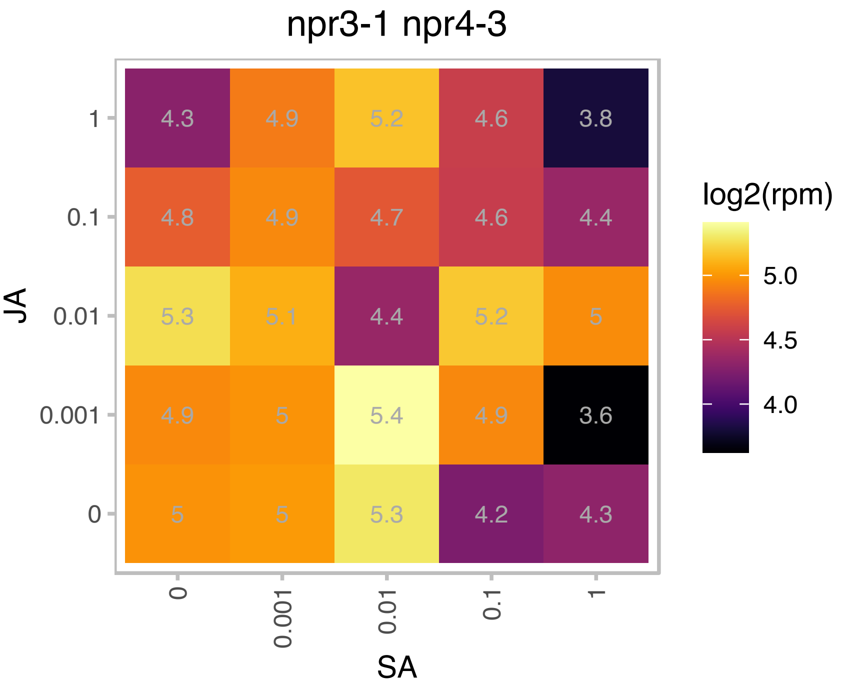

# AT1G05835.1 : NA

PHD finger protein

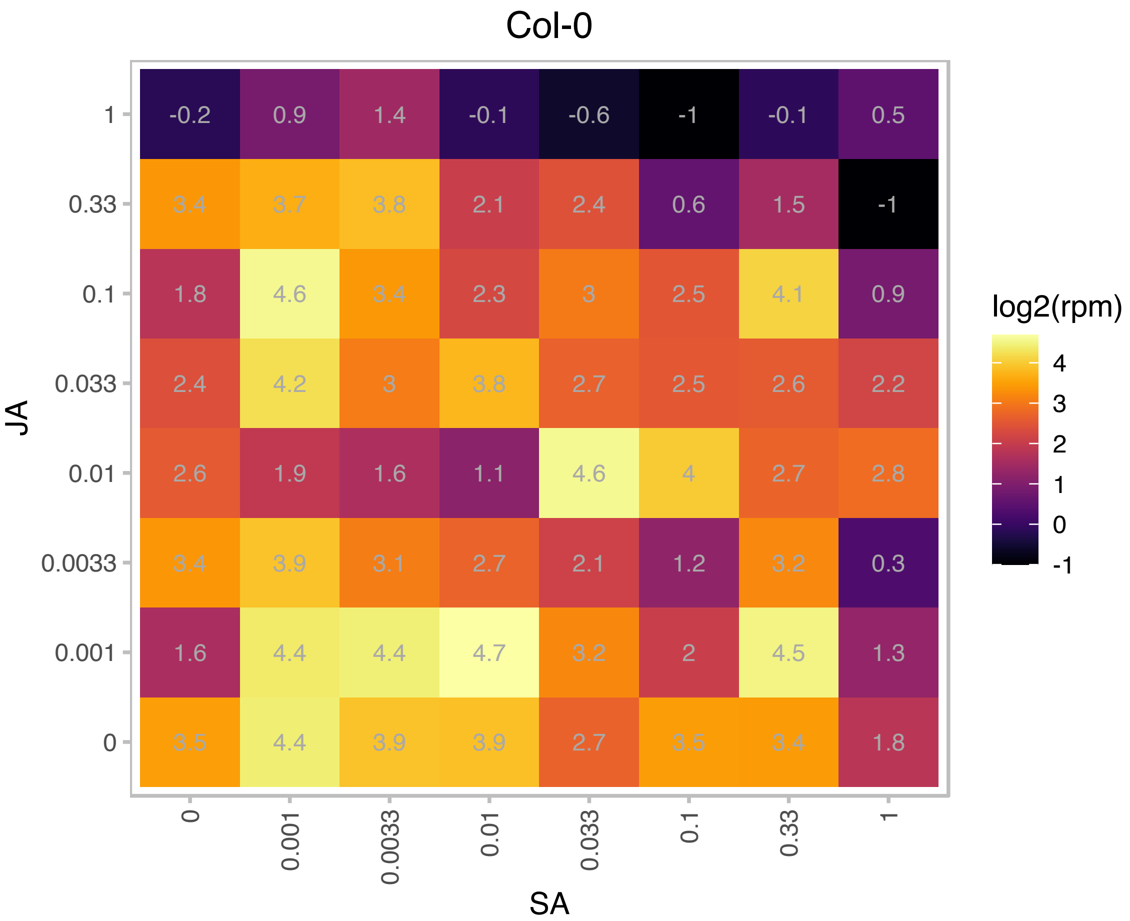


| | |
|---|---|
| Gene | AT1G05835 |
| is.expressed_genes | TRUE |
| is.responsive_genes | TRUE |
| is.apcluster | F-09 |

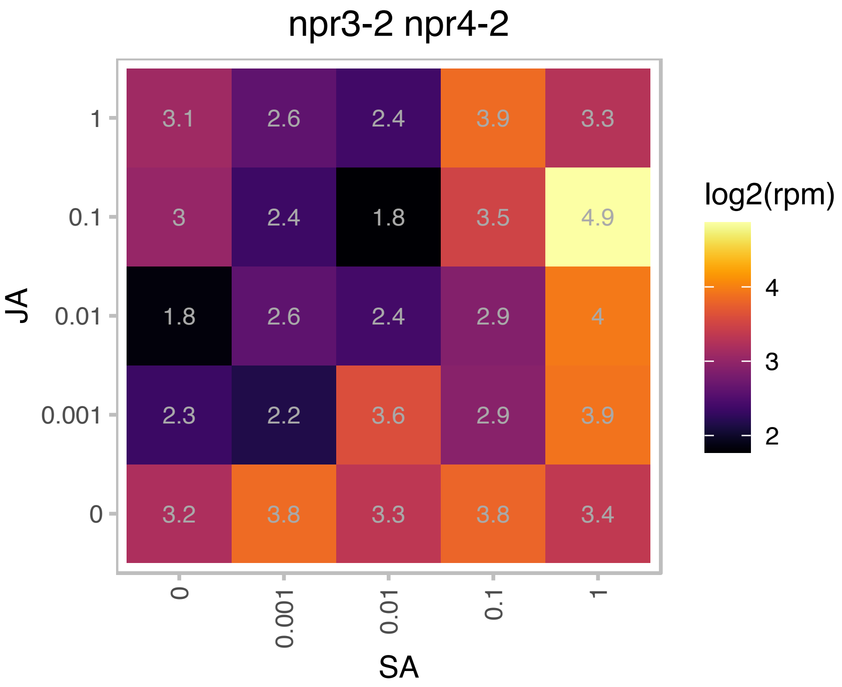


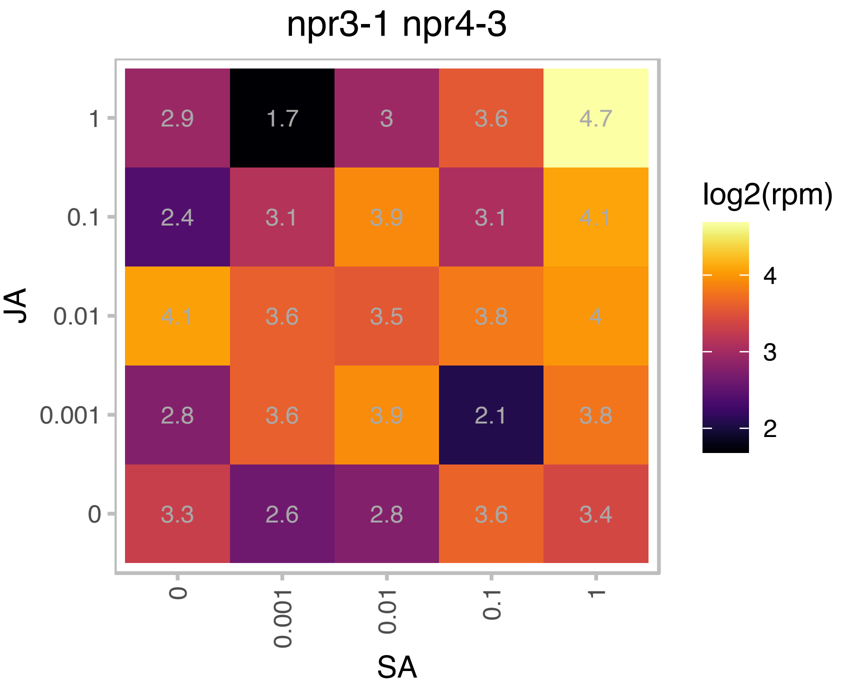

### AT1G11530.1 : ATCXXS1, CXXS1

C-terminal cysteine residue is changed to a serine 1
Encodes a monocysteinic thioredoxin, thioredoxin in which the second cysteine of the redox site is replaced by a serine, with low disulfide reductase but efficient disulfide isomerase activity. The mRNA is cell-to-cell mobile.

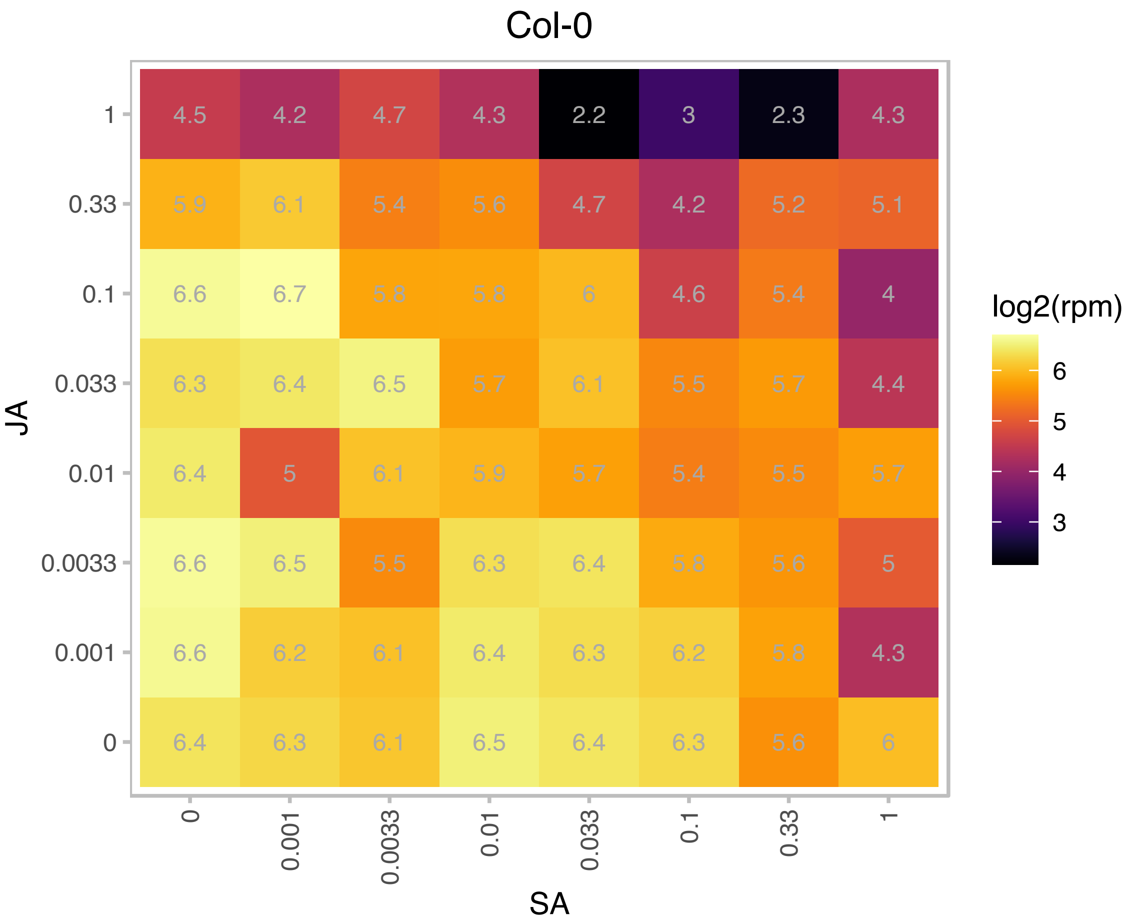


| | |
|---|---|
| Gene | AT1G11530 |
| is.expressed_genes | TRUE |
| is.responsive_genes | TRUE |
| is.apcluster | F-09 |

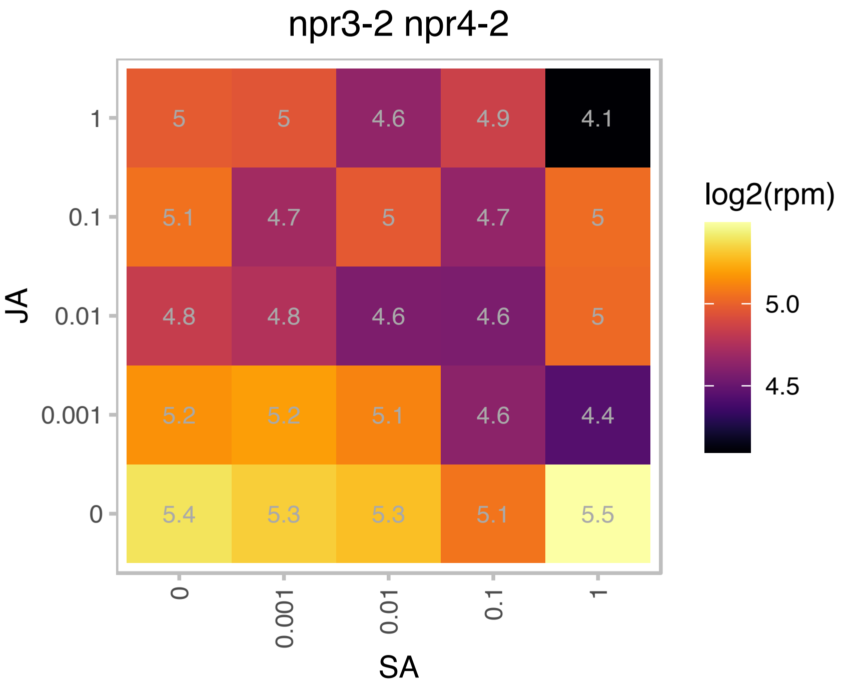


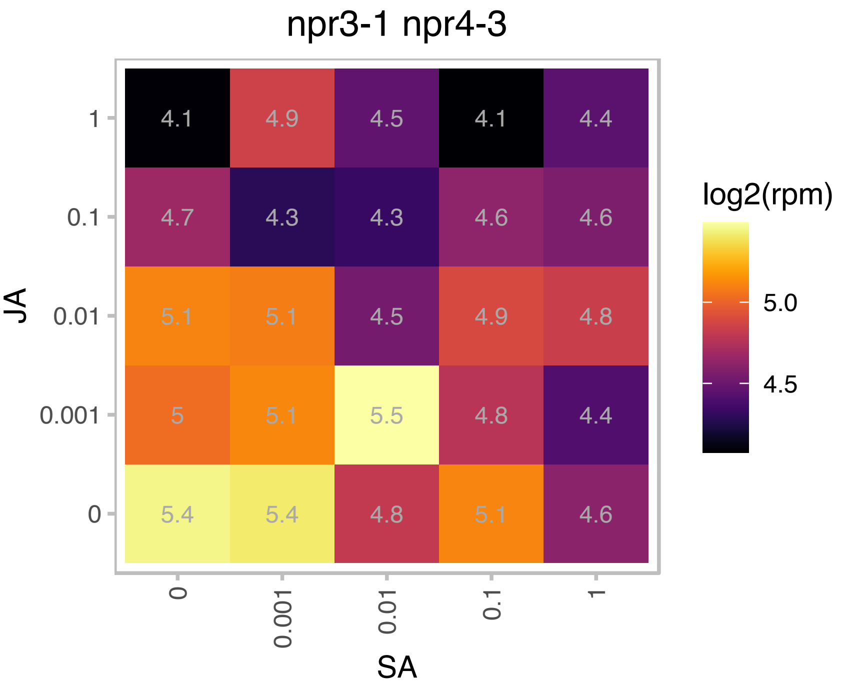

### AT1G13950.1 : ATELF5A-1, EIF-5A, EIF5A, ELF5A-1

eukaryotic elongation factor 5A-1
Encodes eukaryotic translation initiation factor 5A (EIF-5A).In mammalian cells it functions as a shuttle protein that translocates mRNA from the nucleus to cytoplasmic ribosomes. Overexpression results in an increase in both primary and secondary xylem formation. In RNAi suppressed lines, xylem formation is reduced.

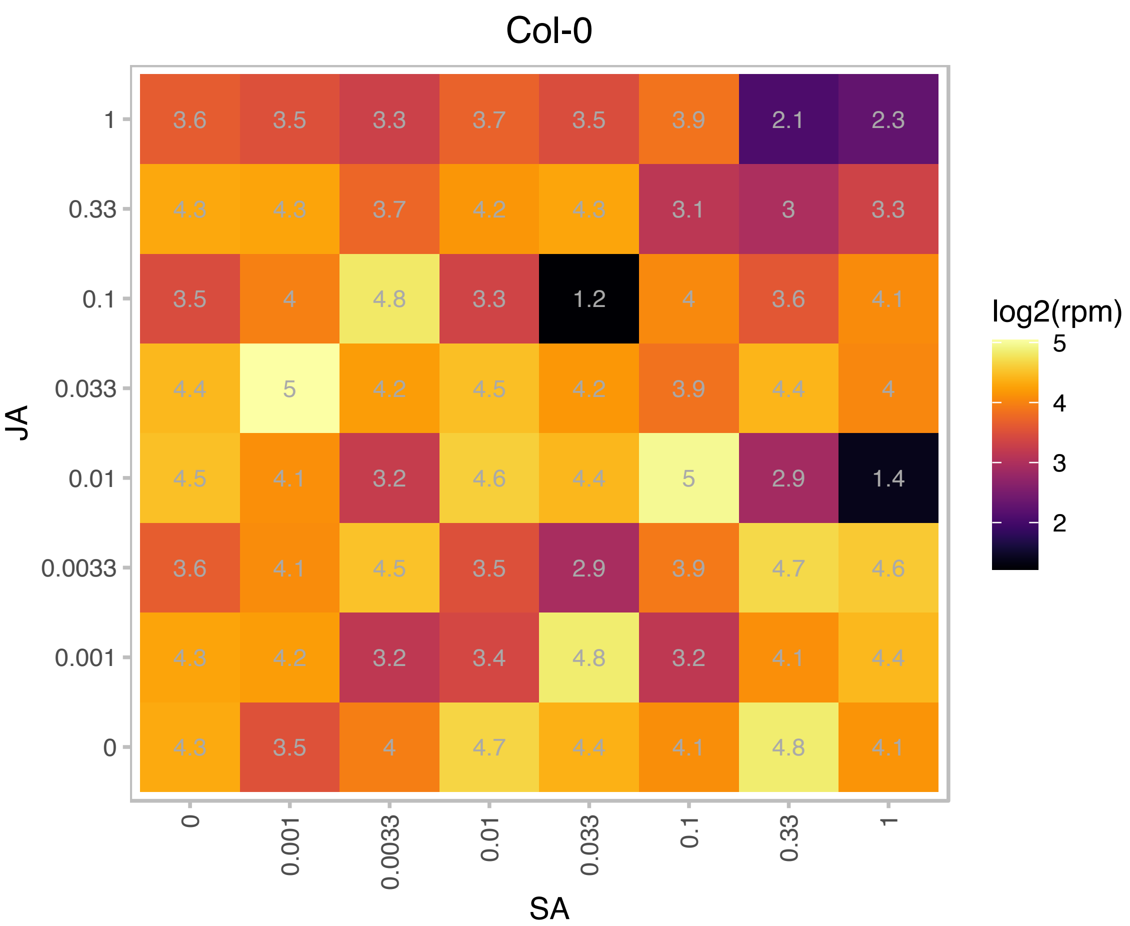


| | |
|---|---|
| Gene | AT1G13950 |
| is.expressed_genes | TRUE |
| is.responsive_genes | TRUE |
| is.apcluster | F-09 |

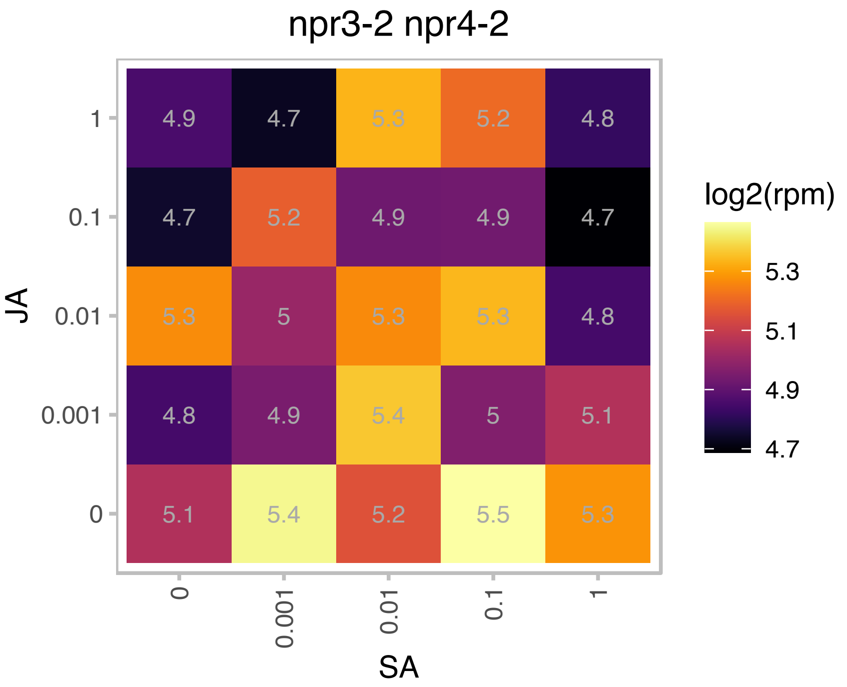


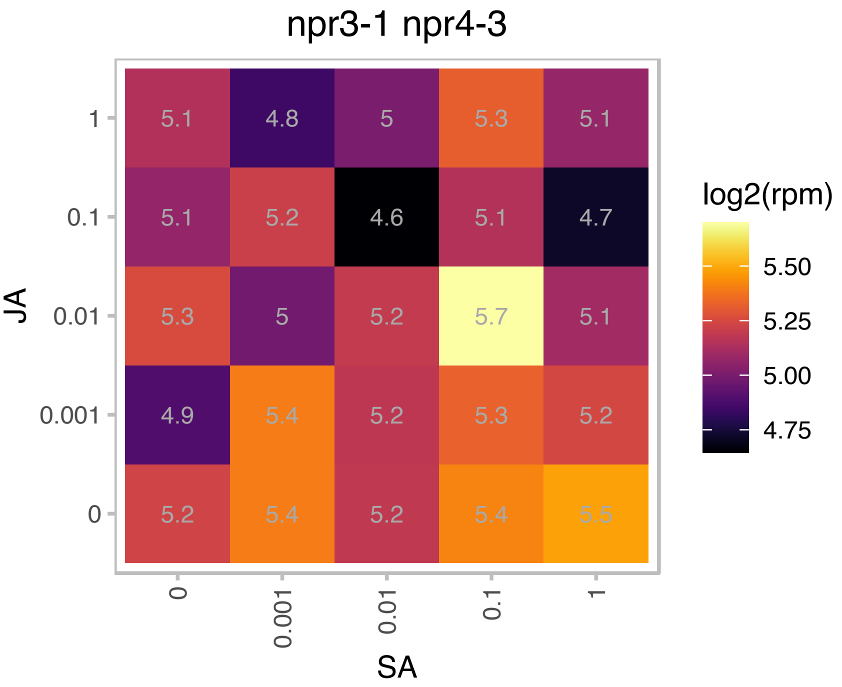

# AT1G16350.1 : NA

Aldolase-type TIM barrel family protein

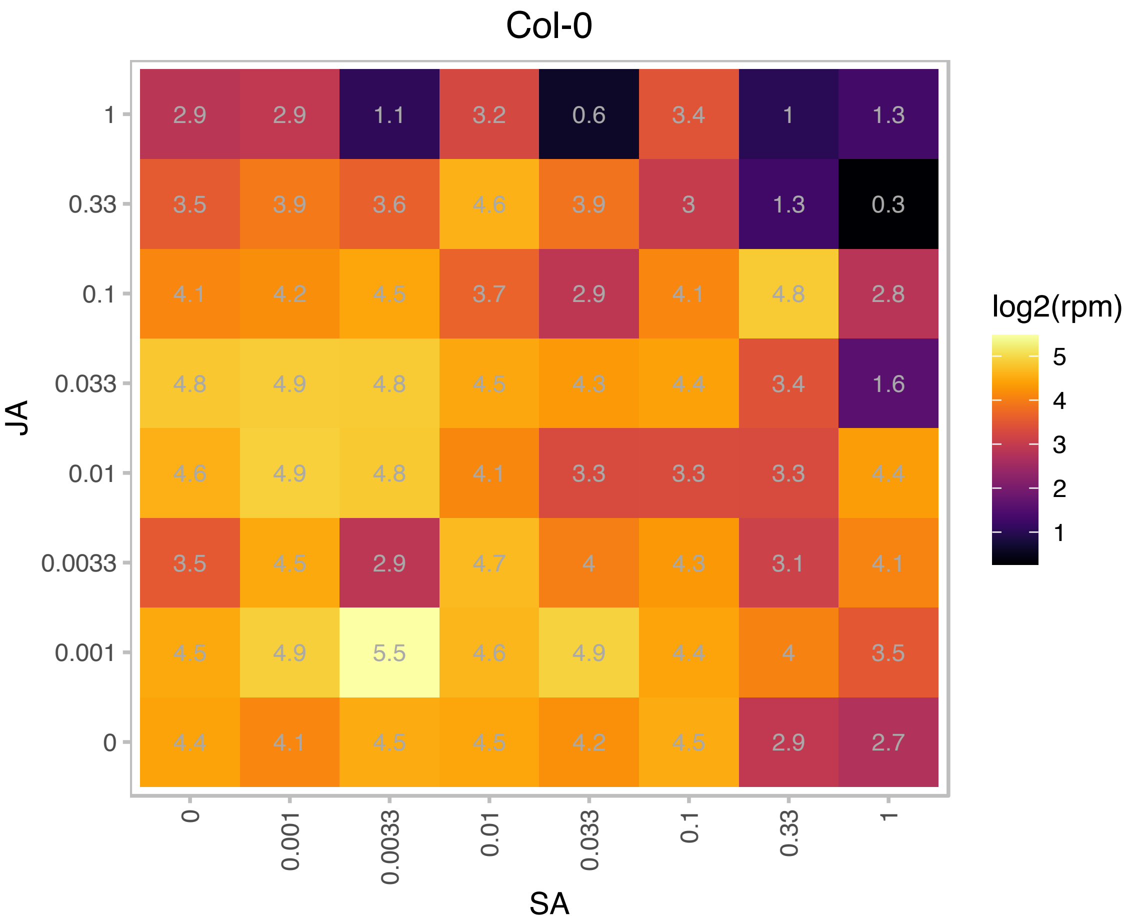


| | |
|---|---|
| Gene | AT1G16350 |
| is.expressed_genes | TRUE |
| is.responsive_genes | TRUE |
| is.apcluster | F-09 |

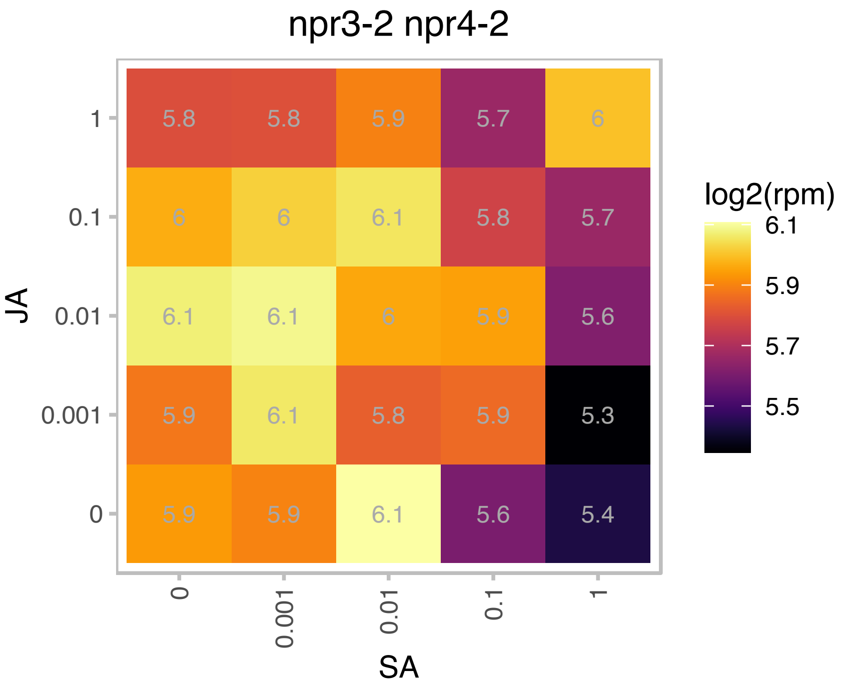


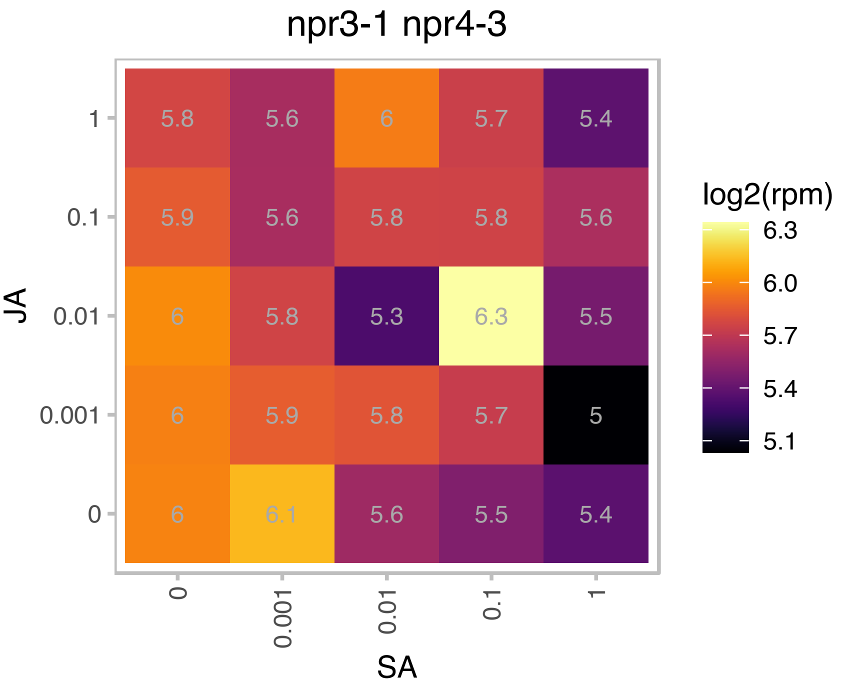

### AT1G17200.1 : CASPL2A1

Uncharacterised protein family (UPF0497)

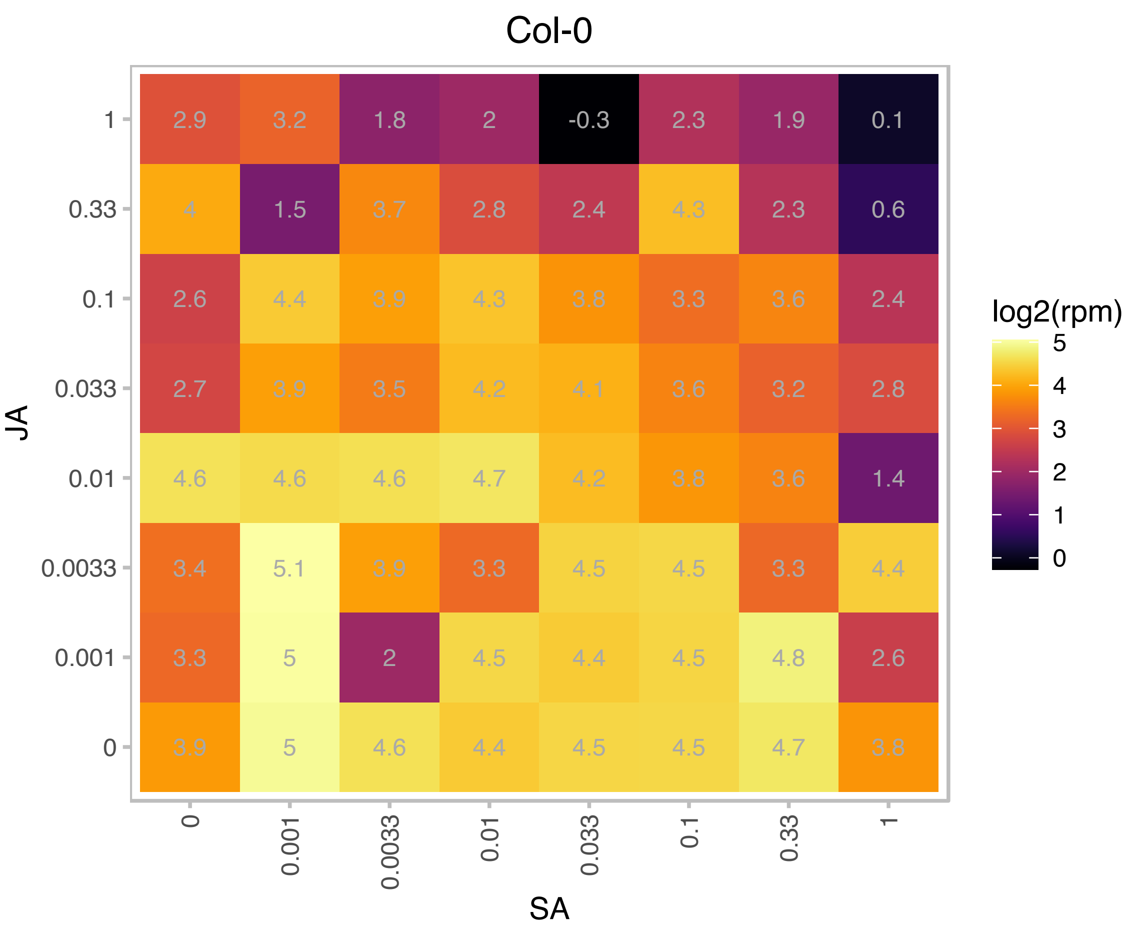


| Gene | AT1G17200 |
|---|---|
| is.expressed_genes | TRUE |
| is.responsive_genes | TRUE |
| is.apcluster | F-09 |

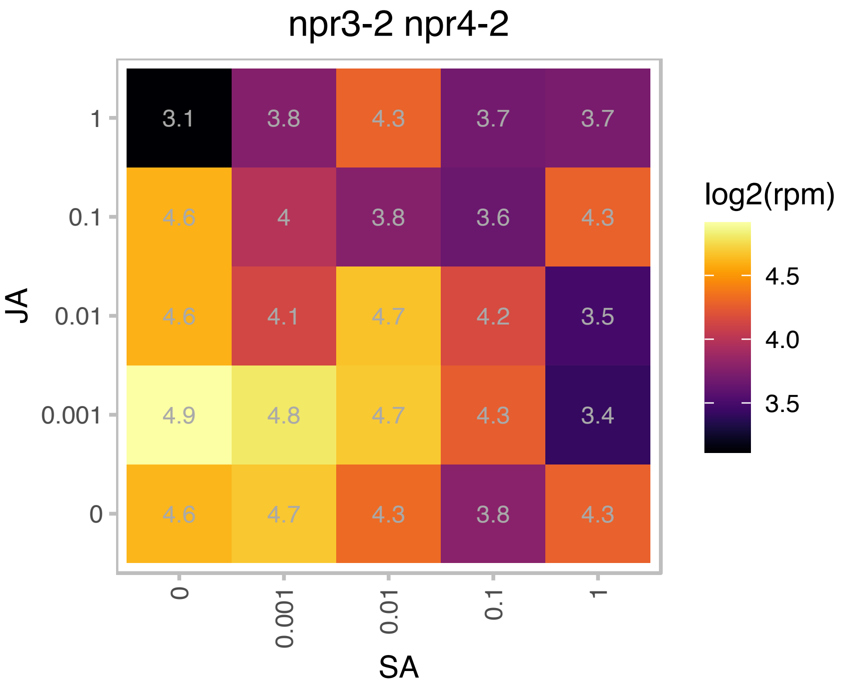


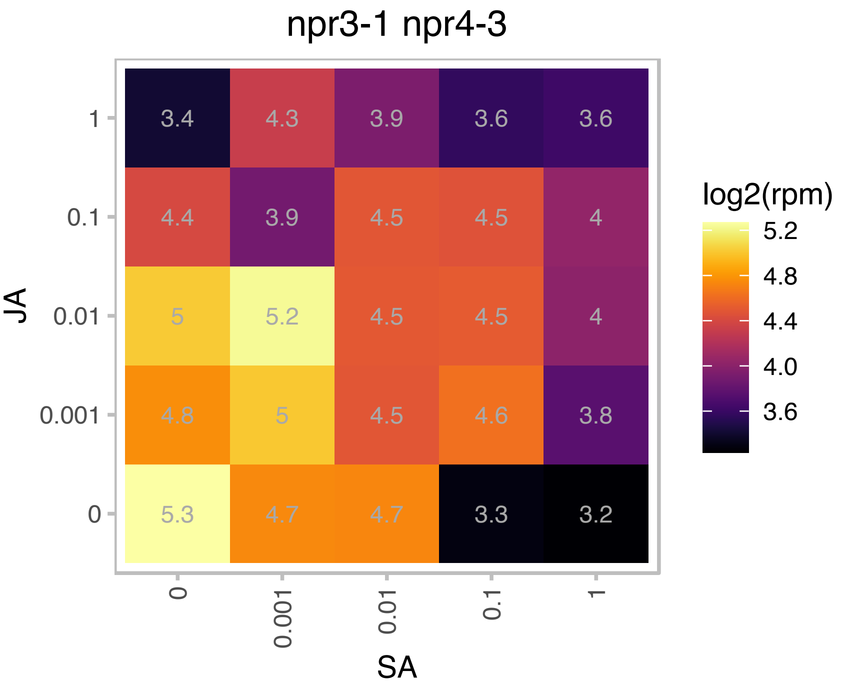

### AT1G17560.1 : HLL

Ribosomal protein L14p/L23e family protein

Encodes HUELLENLOS (HLL), a mitochondrial ribosome protein, similar to L14 ribosomal protein of eubacteria. HLL is essential for normal ovule development.

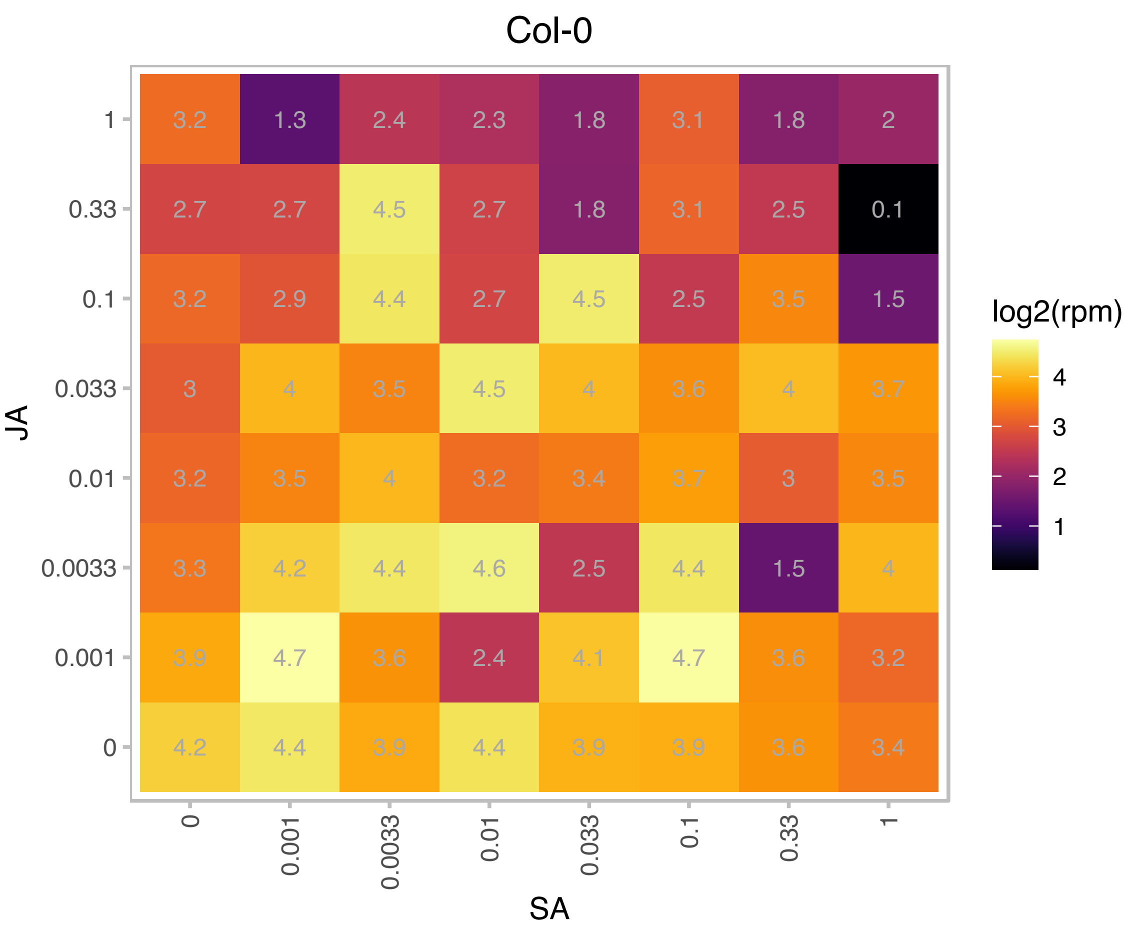


| | |
|---|---|
| Gene | AT1G17560 |
| is.expressed_genes | TRUE |
| is.responsive_genes | TRUE |
| is.apcluster | F-09 |

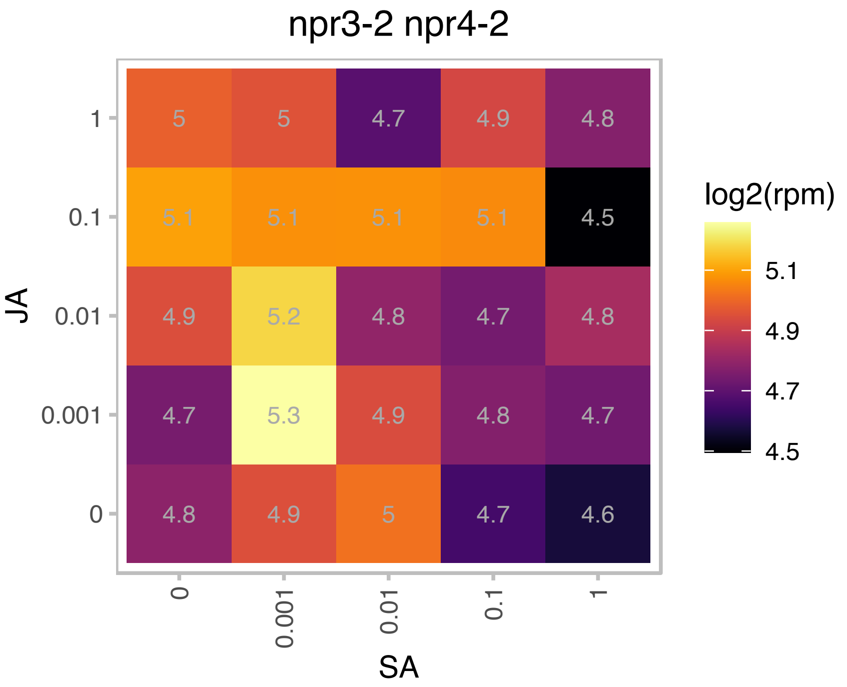


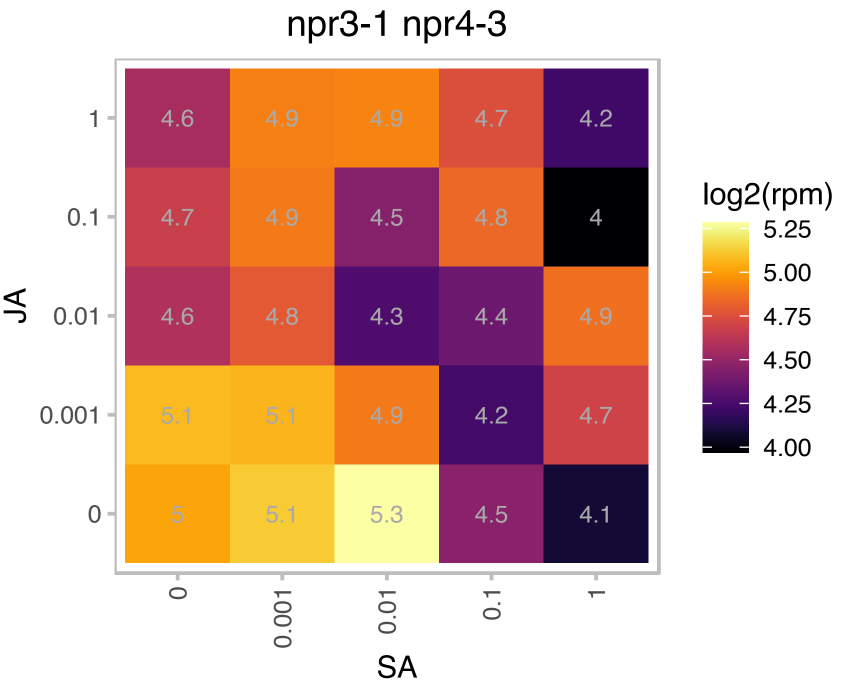

### AT1G18080.1 : ATARCA, AtRACK1, RACK1A, RACK1A AT

Transducin/WD40 repeat-like superfamily protein
Encodes the Arabidopsis thaliana homolog of the tobacco WD-40 repeat ArcA gene. It is a scaffold protein that acts downstream of and is phosphorylated by WNK8 in the glucose responsiveness and flowering pathways . Knockout mutants show insensitivity to gibberellin in GA-induced seed germination.

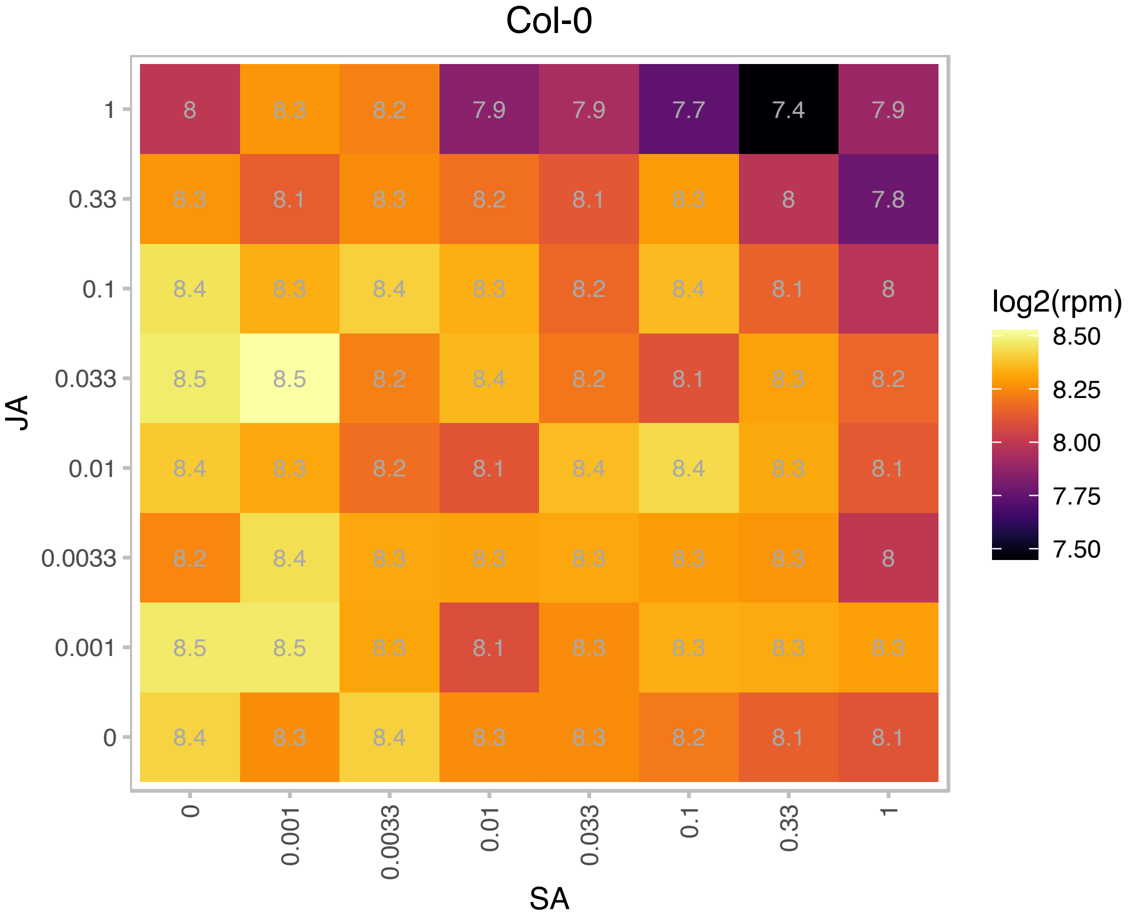


| | |
|---|---|
| Gene | AT1G18080 |
| is.expressed_genes | TRUE |
| is.responsive_genes | TRUE |
| is.apcluster | F-09 |

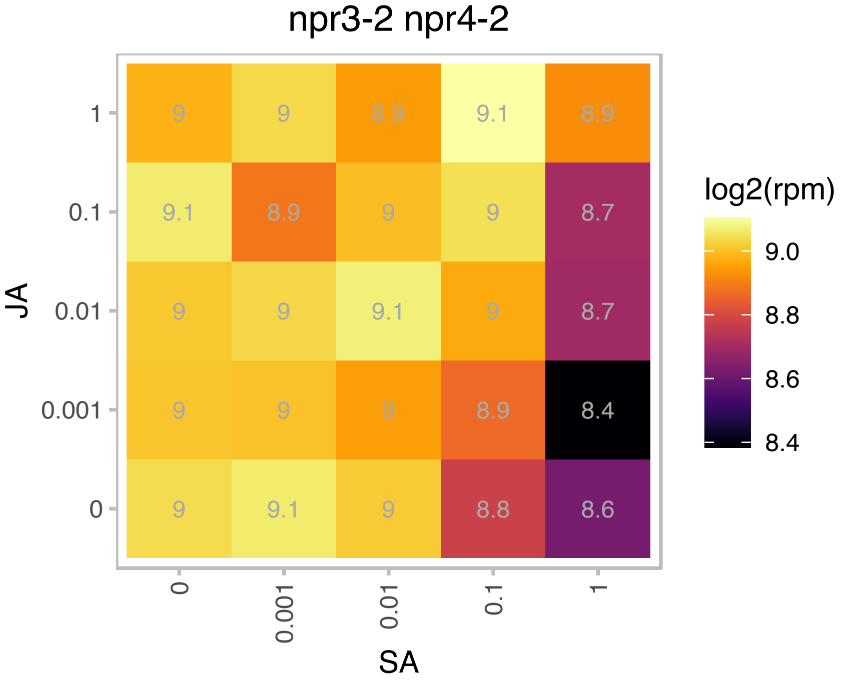


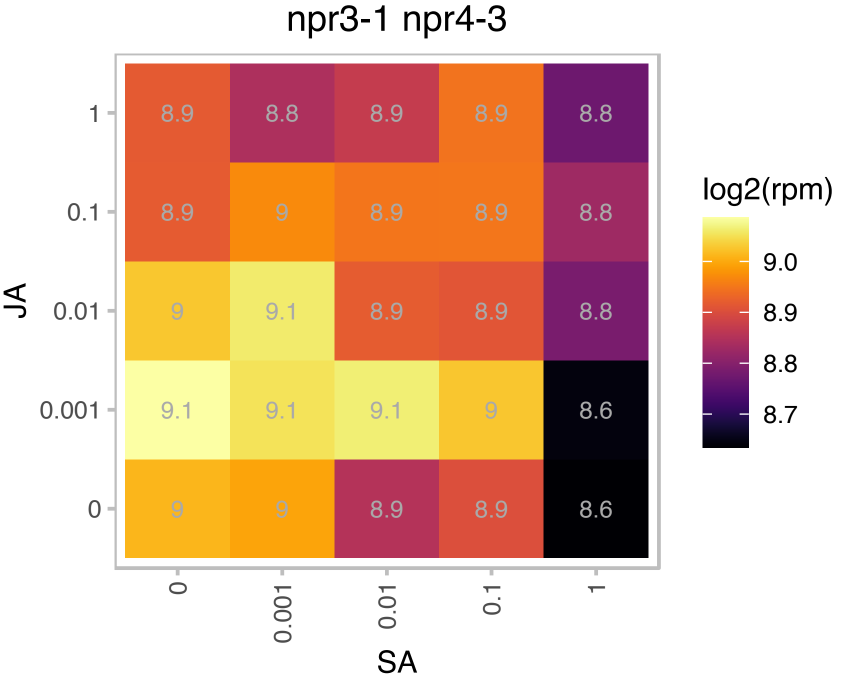

# AT1G22885.1 : NA

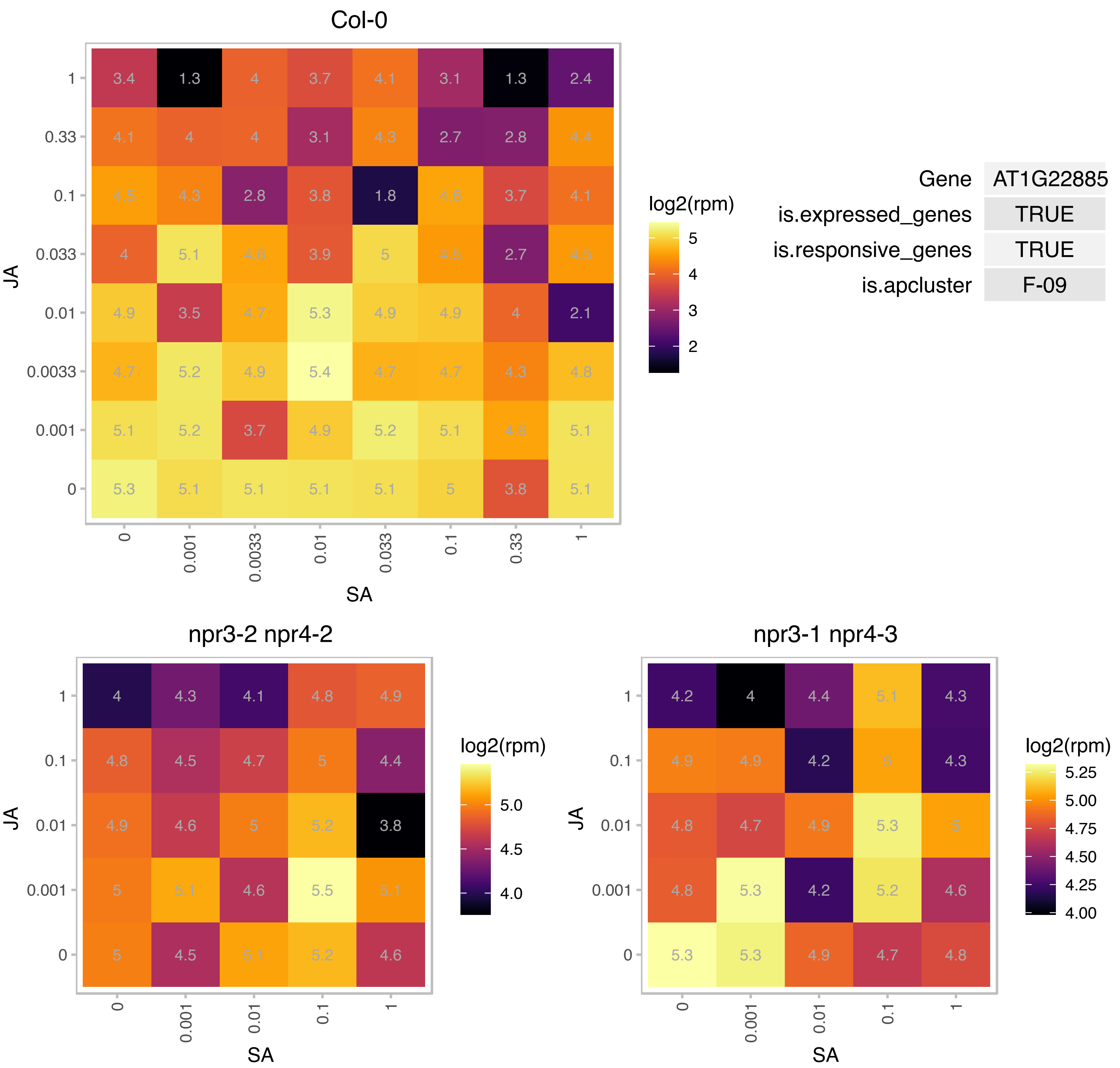


| | |
|---|---|
| Gene | AT1G22885 |
| is.expressed_genes | TRUE |
| is.responsive_genes | TRUE |
| is.apcluster | F-09 |

### AT1G25550.1 : HHO3, NIGT1.1

myb-like transcription factor family protein

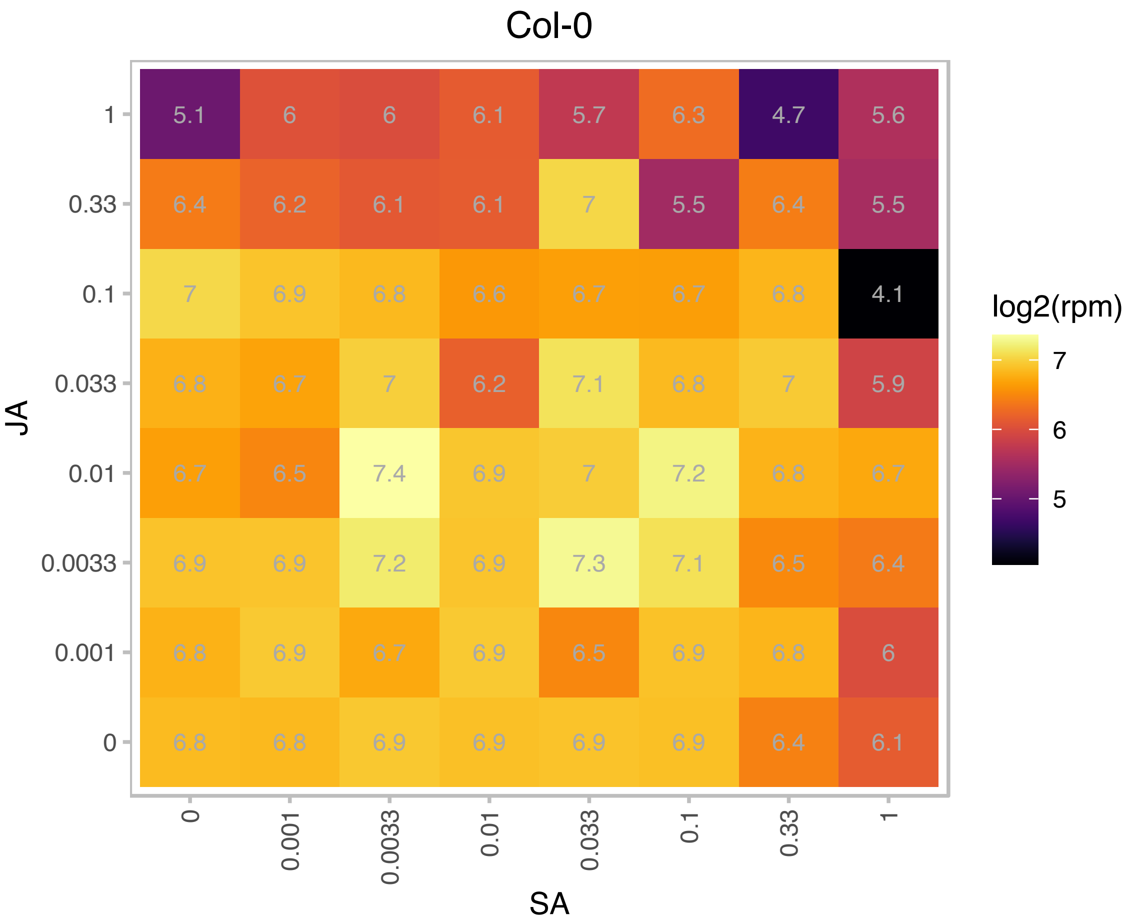


| | |
|---|---|
| Gene | AT1G25550 |
| is.expressed_genes | TRUE |
| is.responsive_genes | TRUE |
| is.apcluster | F-09 |

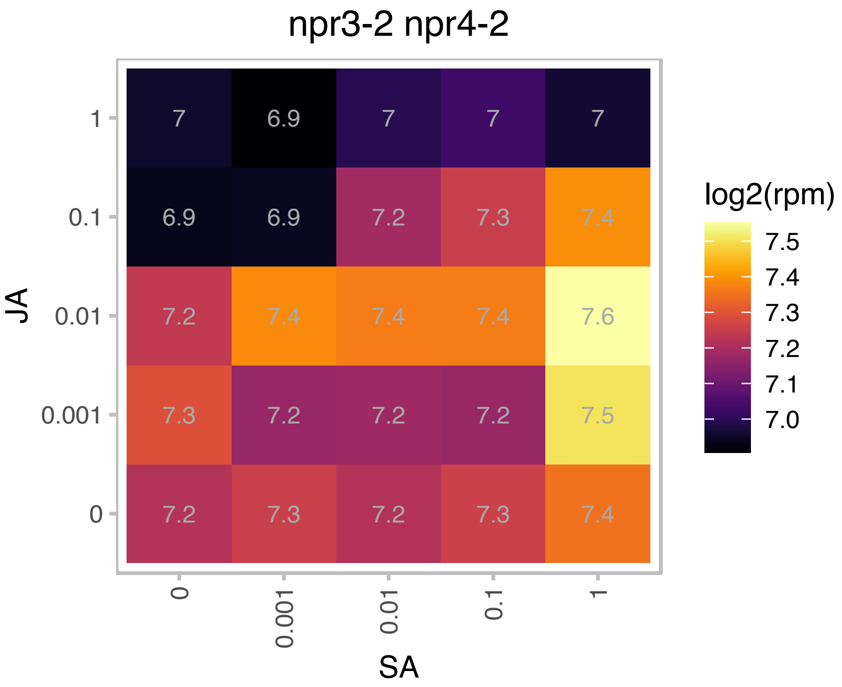


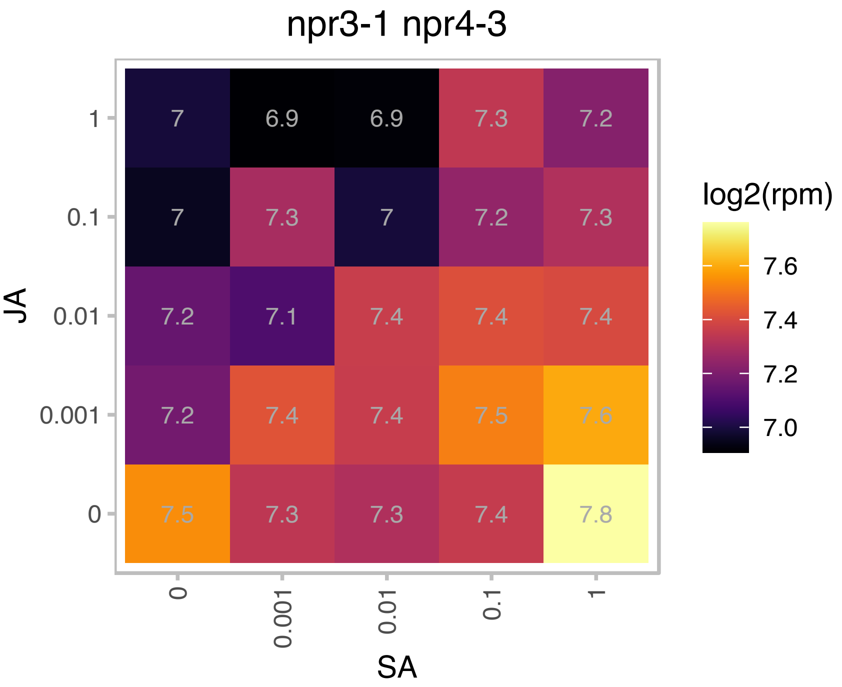

# AT1G26740.1 : NA

Ribosomal L32p protein family

| | |
|---|---|
| Gene | AT1G26740 |
| is.expressed_genes | TRUE |
| is.responsive_genes | TRUE |
| is.apcluster | F-09 |

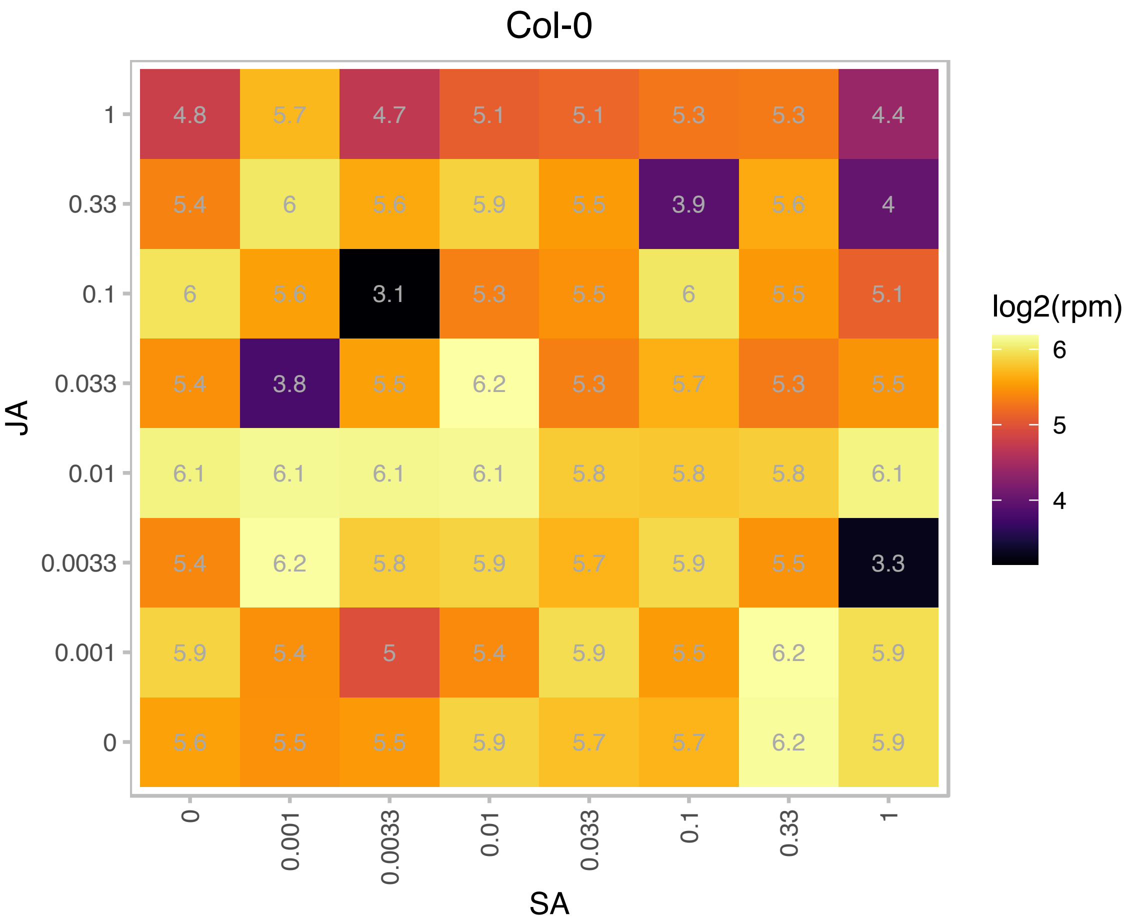


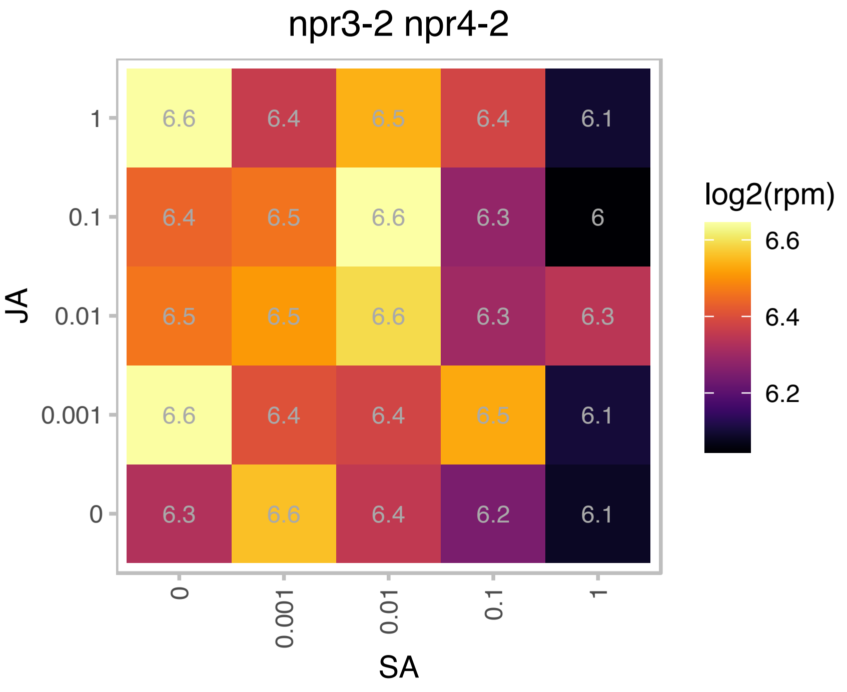


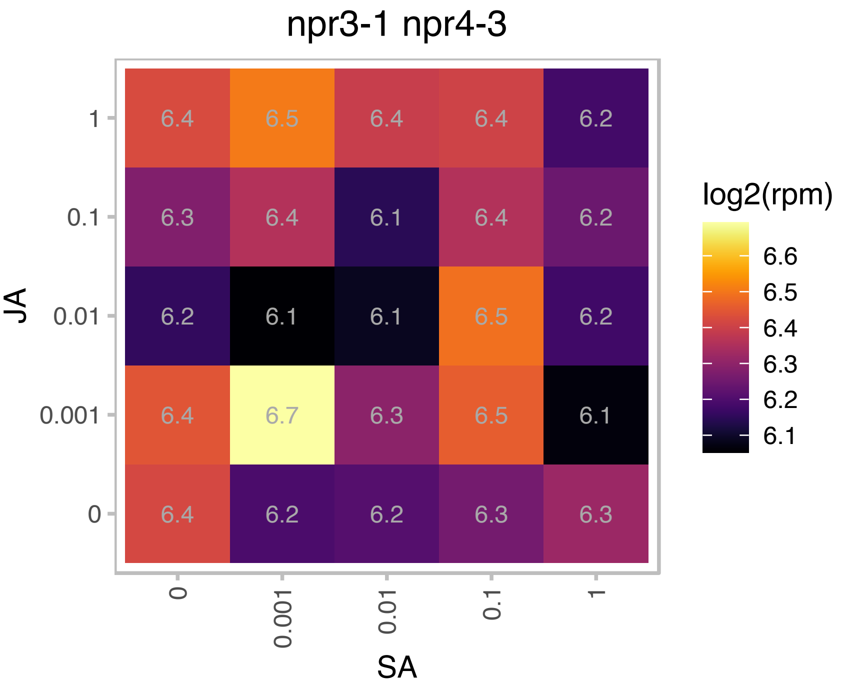

### AT1G31760.1 : SWIB5

SWIB/MDM2 domain superfamily protein

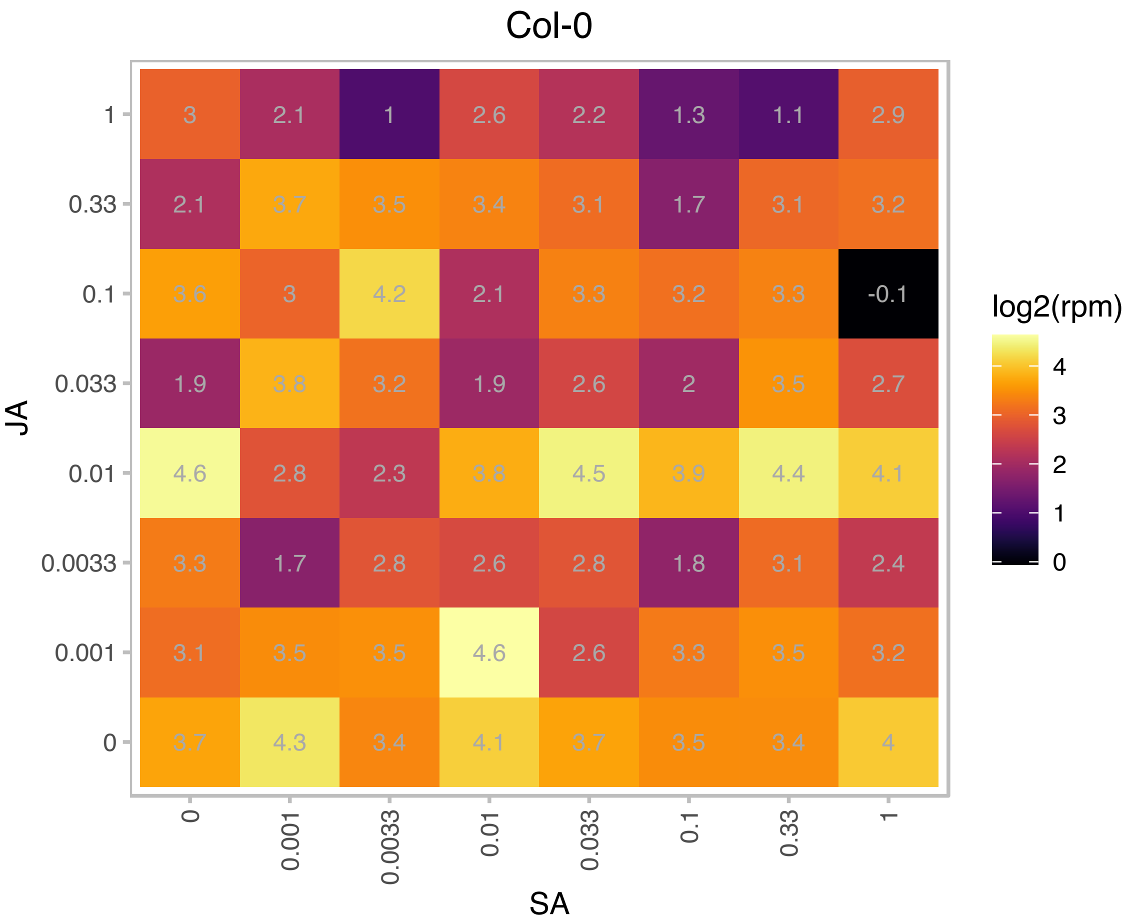


| | |
|---|---|
| Gene | AT1G31760 |
| is.expressed_genes | TRUE |
| is.responsive_genes | TRUE |
| is.apcluster | F-09 |

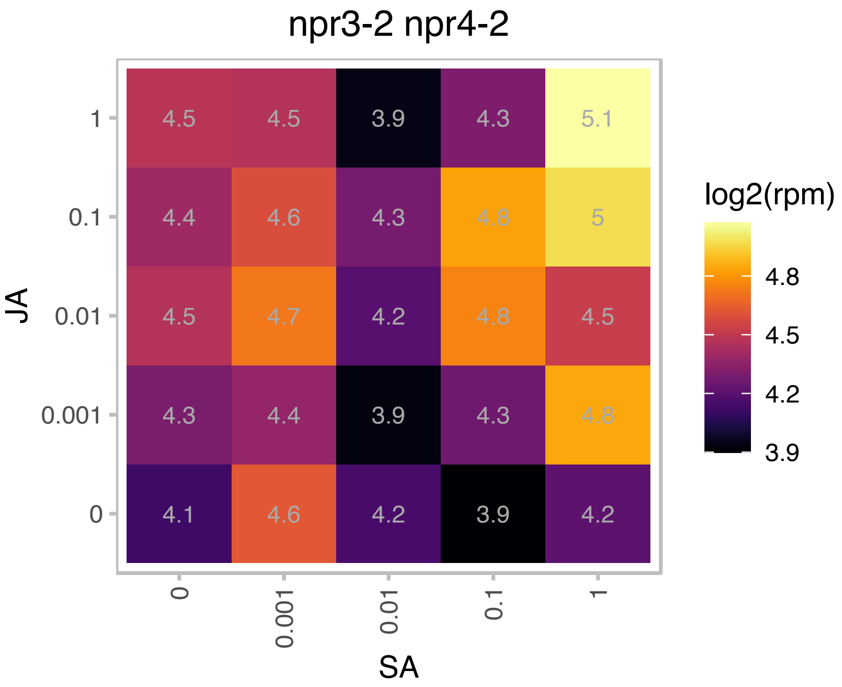


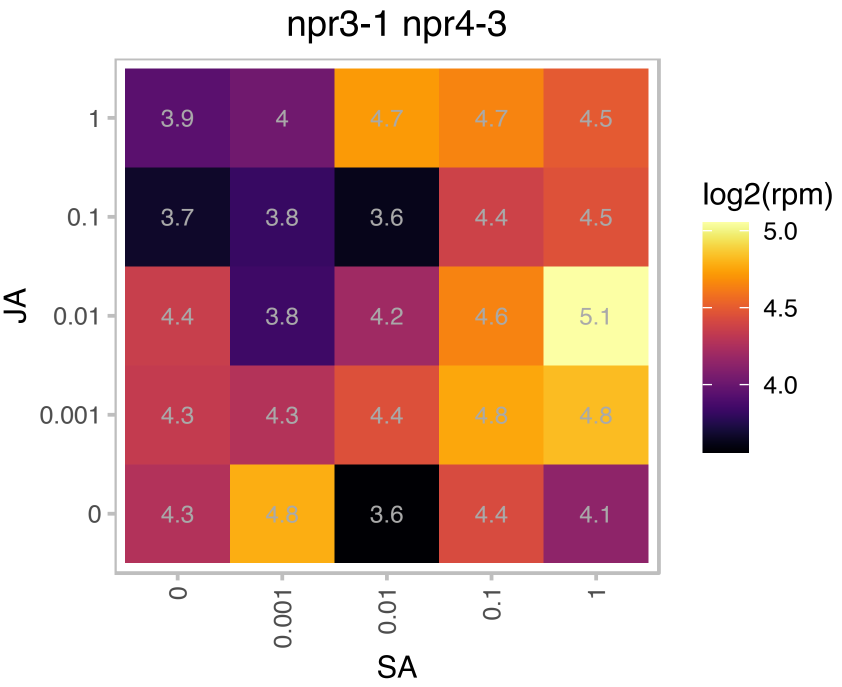

# AT1G34570.1 : NA

Essential protein Yae1, N-terminal

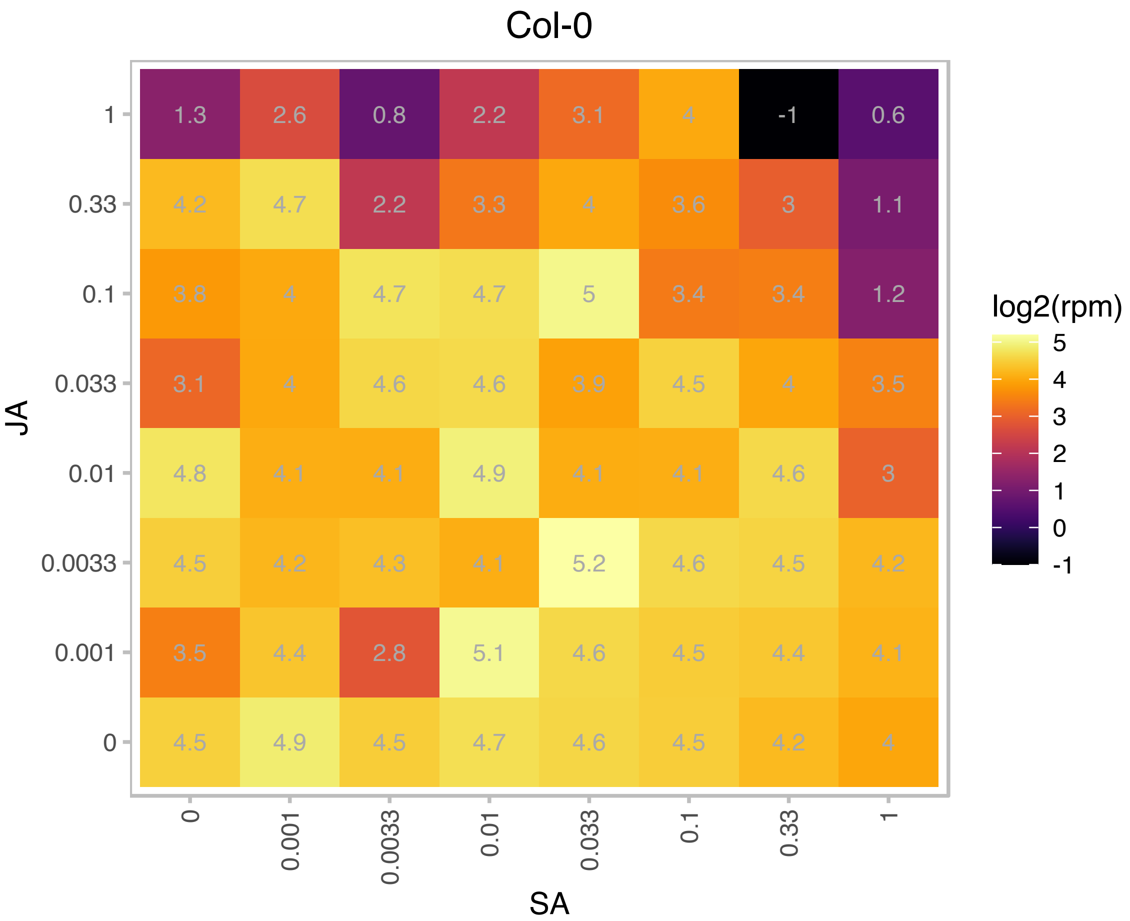


| | |
|---|---|
| Gene | AT1G34570 |
| is.expressed_genes | TRUE |
| is.responsive_genes | TRUE |
| is.apcluster | F-09 |

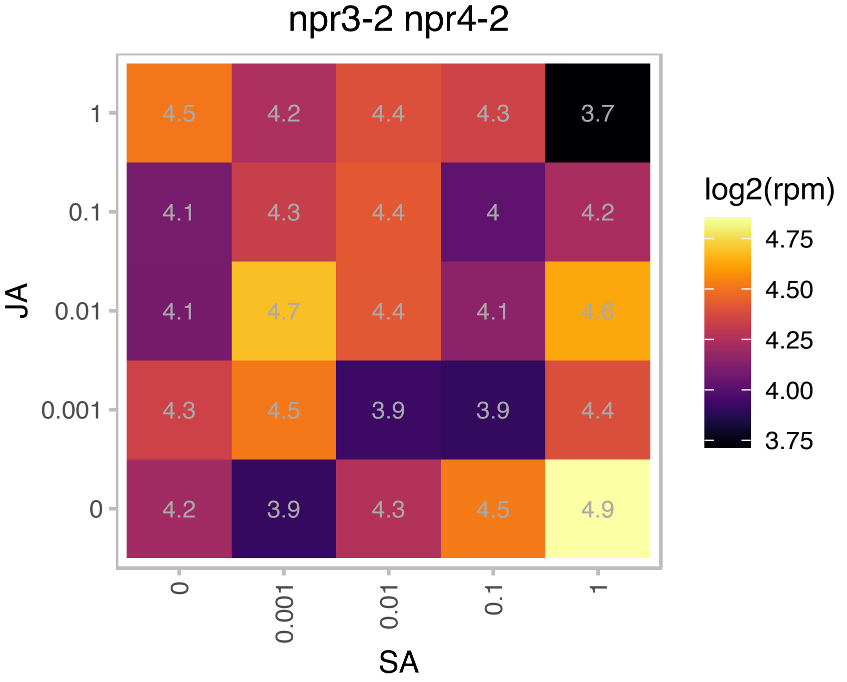


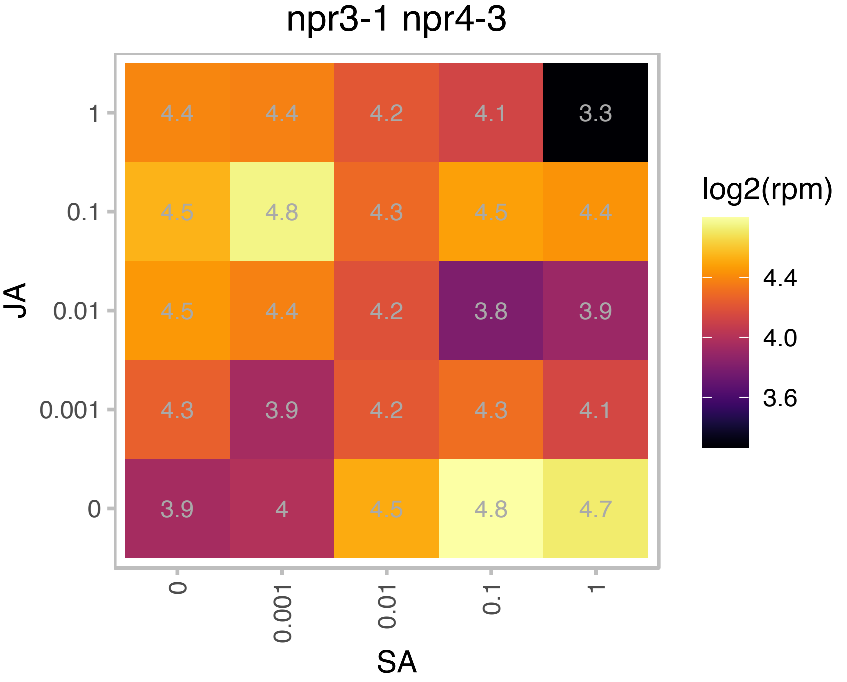

### AT1G37130.1 : ATNR2, B29, CHL3, NIA2, NIA2-1, NR, NR[illegible]

nitrate reductase 2
Identified as a mutant resistant to chlorate. Encodes nitrate reductase structural gene. Involved in nitrate assimilation. Has nitrate reductase activity. Up-regulated by the fungus P. indica. Binds transcription factor At2g35940. The mRNA is cell-to-cell mobile.

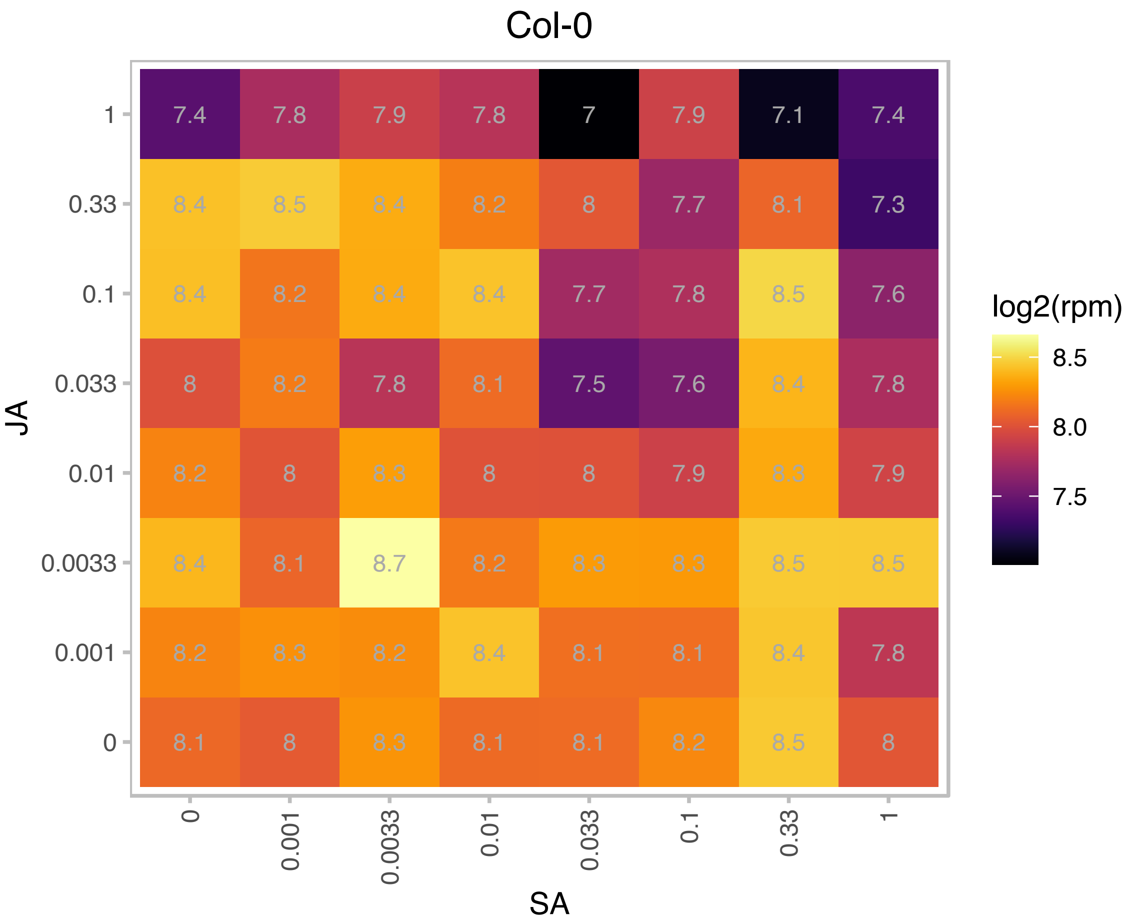


| | |
|---|---|
| Gene | AT1G37130 |
| is.expressed_genes | TRUE |
| is.responsive_genes | TRUE |
| is.apcluster | F-09 |

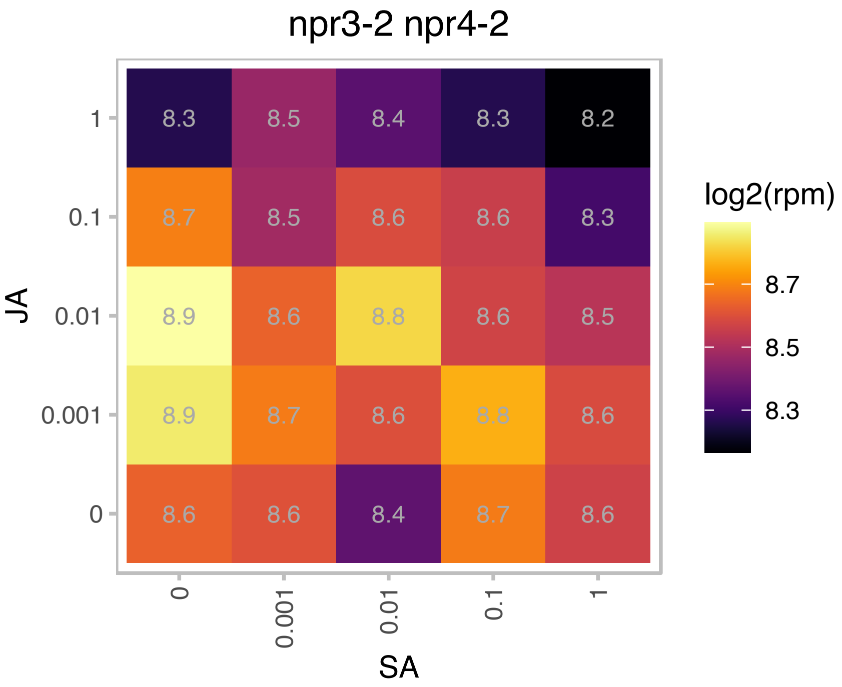


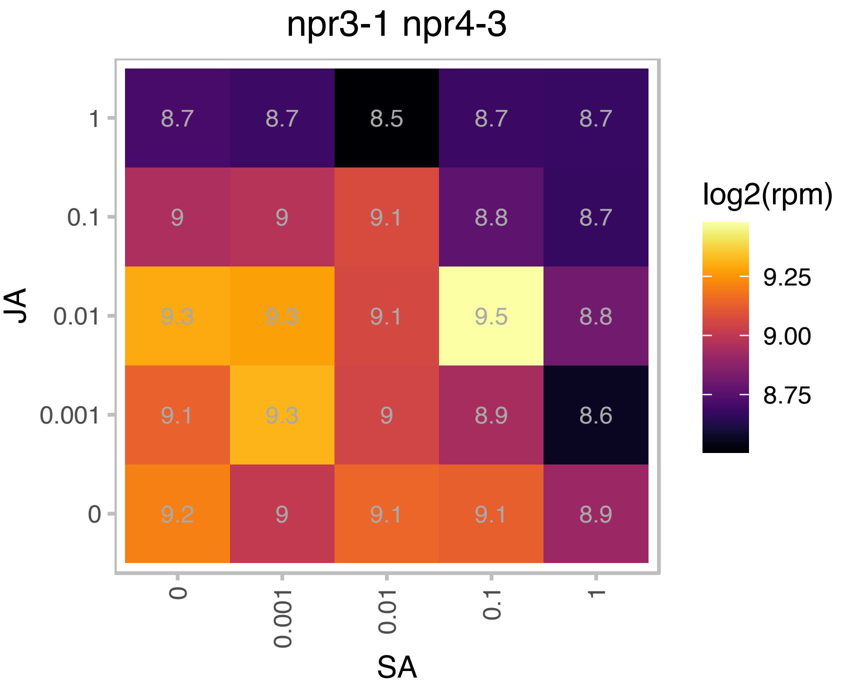

# AT1G49245.1 : NA

Prefoldin chaperone subunit family protein

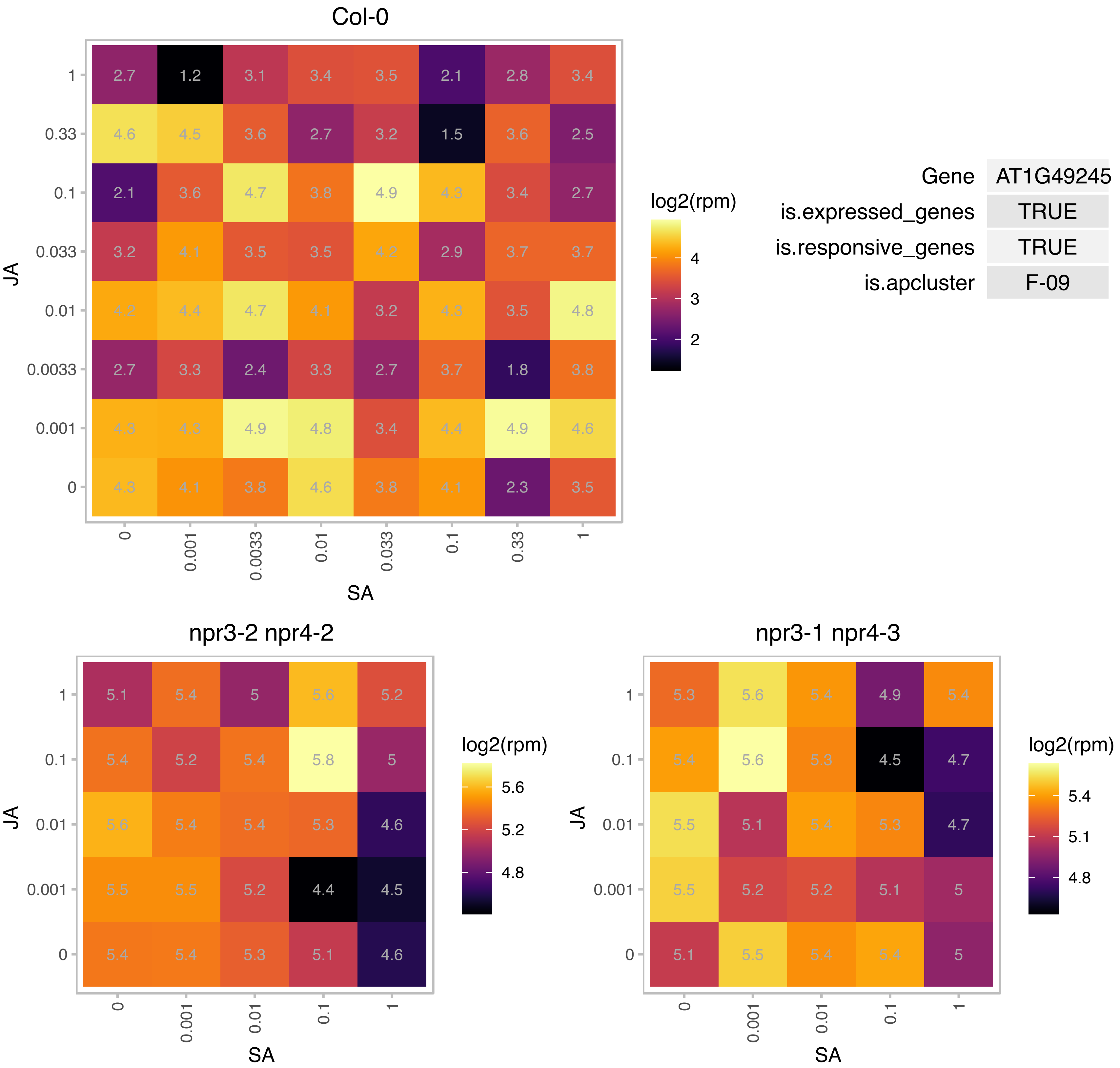


| | |
|---|---|
| Gene | AT1G49245 |
| is.expressed_genes | TRUE |
| is.responsive_genes | TRUE |
| is.apcluster | F-09 |

### AT1G50170.1 : ATSIRB, SIRB

sirohydrochlorin ferrochelatase B

encodes sirohydrochlorin ferrochelatase catalyzing the last step of the siroheme biosynthesis

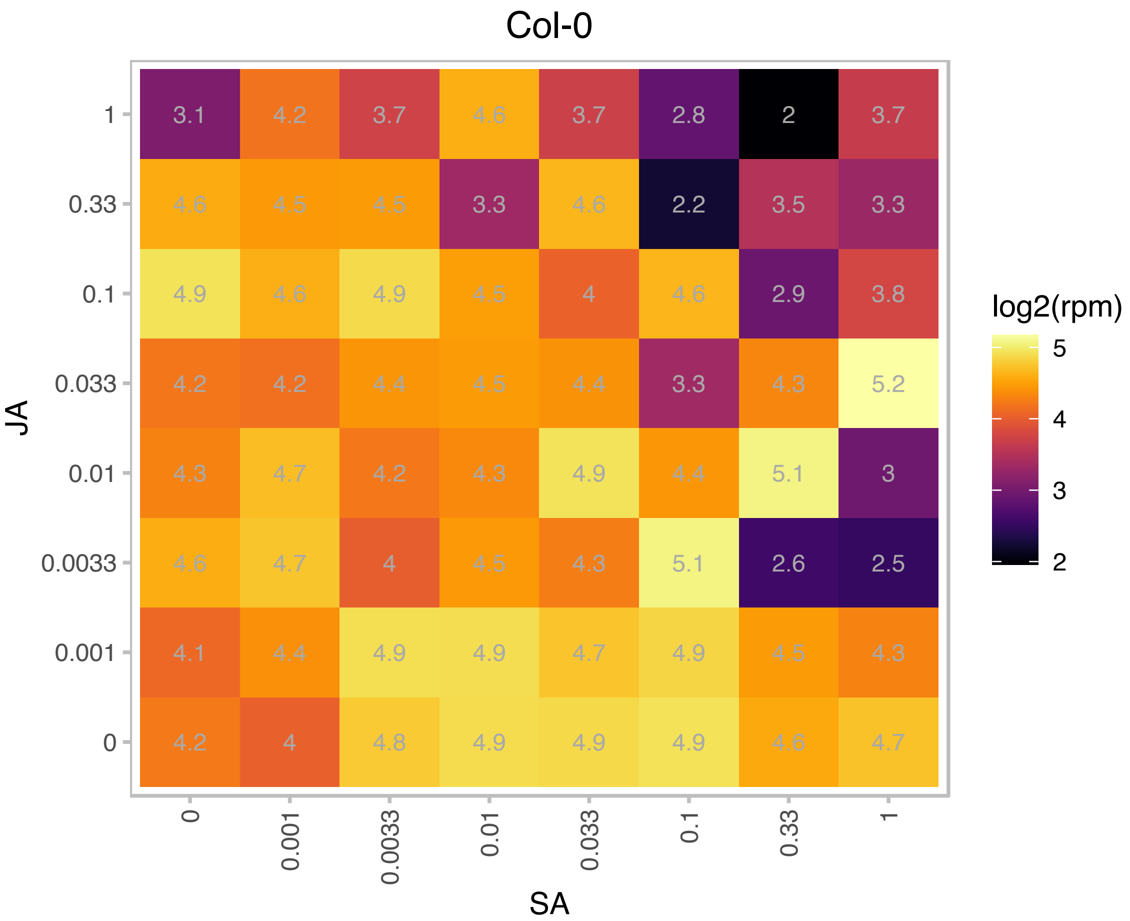


| | |
|---|---|
| Gene | AT1G50170 |
| is.expressed_genes | TRUE |
| is.responsive_genes | TRUE |
| is.apcluster | F-09 |

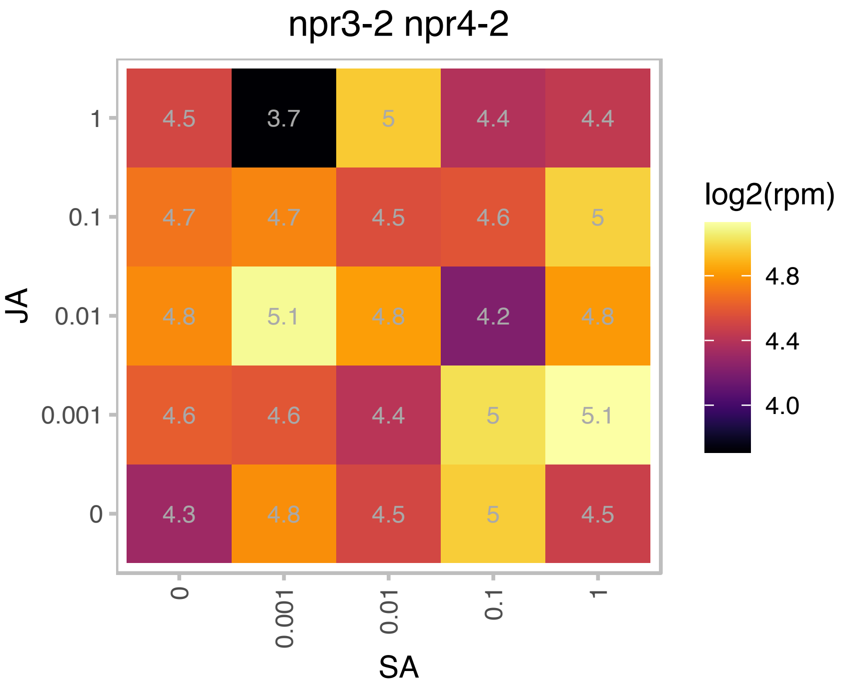


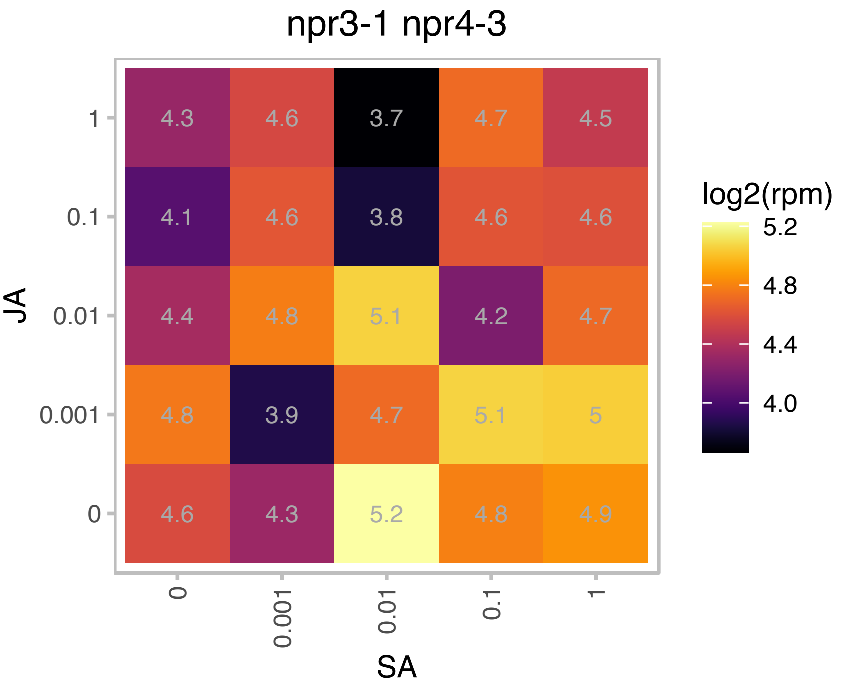

### AT1G51570.1 : FTIP4, MCTP4

Calcium-dependent lipid-binding (CaLB domain) plant phosphoribosyltransferase family protein
Required for maintenance of inflorescence and shoot SAMs and normal development of the derived vascular cambium, functions in the SAM to promote continuous organogenesis, affects SAM development through STM, whe it affects intracellular localization of STM in SAM cells in the peripheral region and prevents STM localization toward the cell wall of SAM cells in the peripheral region.

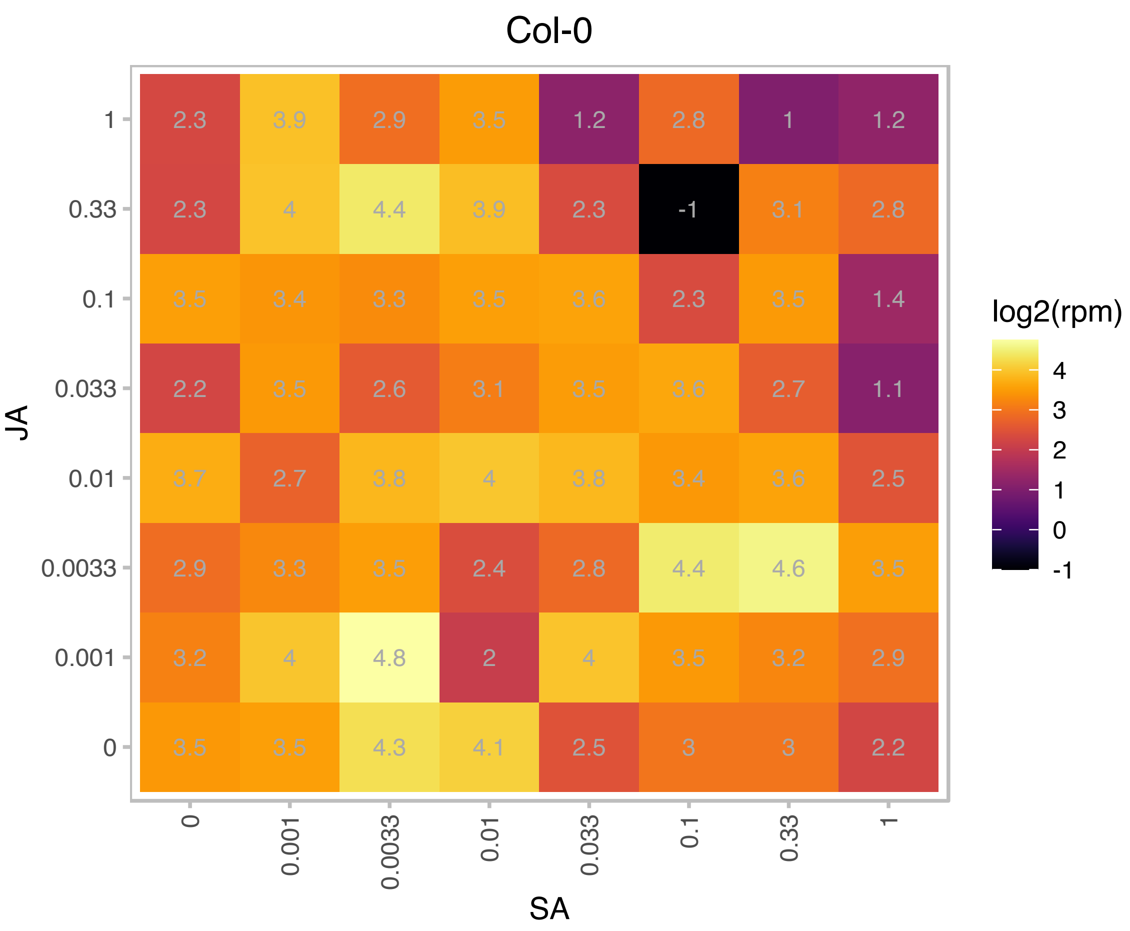


| | |
|---|---|
| Gene | AT1G51570 |
| is.expressed_genes | TRUE |
| is.responsive_genes | TRUE |
| is.apcluster | F-09 |

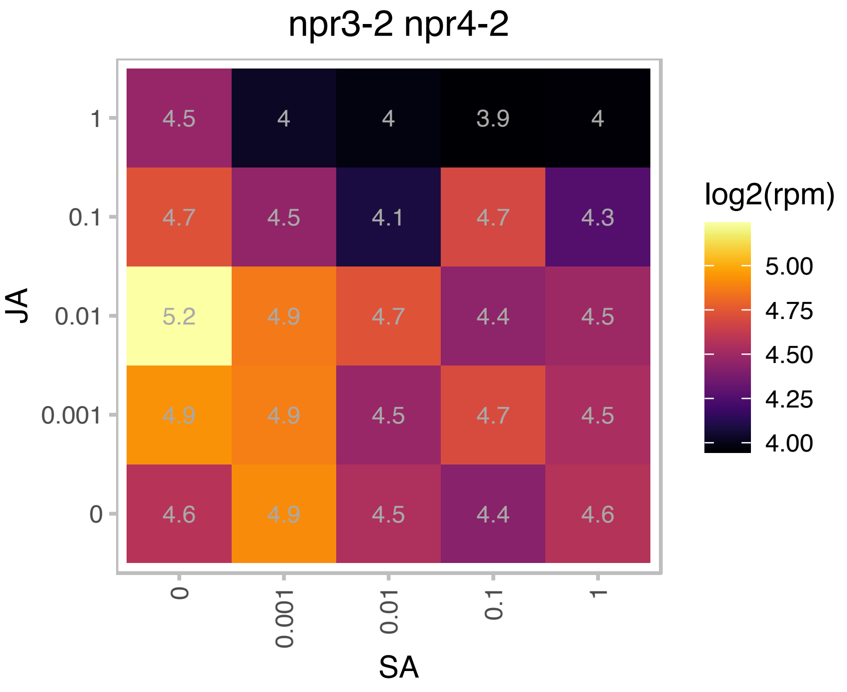


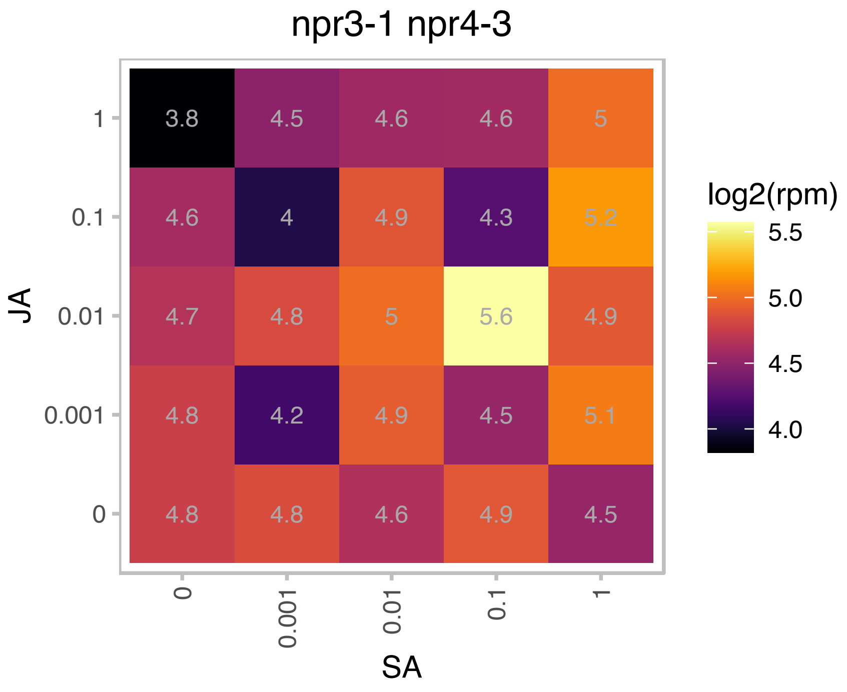

# AT1G55360.1 : NA

Protein of Unknown Function (DUF239)

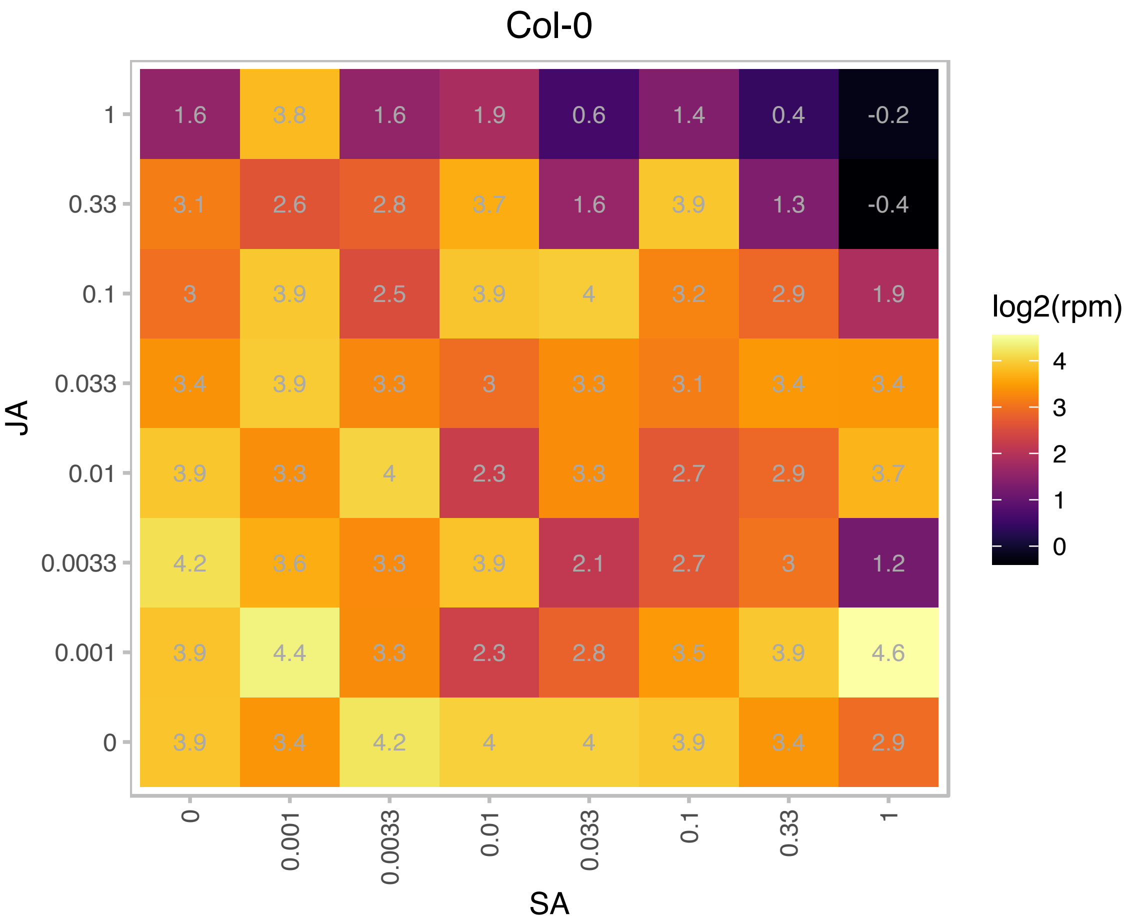


| | |
|---|---|
| Gene | AT1G55360 |
| is.expressed_genes | TRUE |
| is.responsive_genes | TRUE |
| is.apcluster | F-09 |

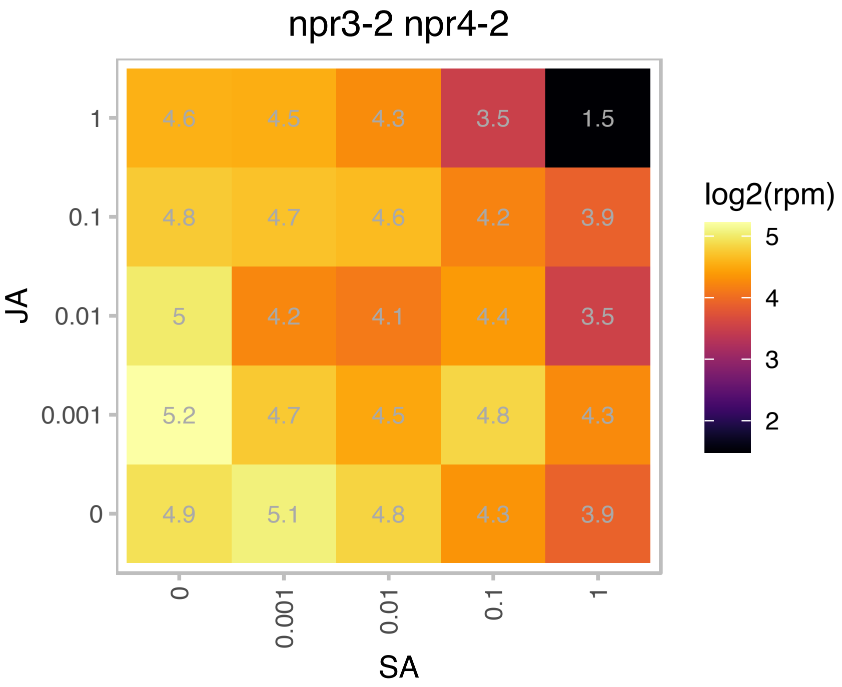


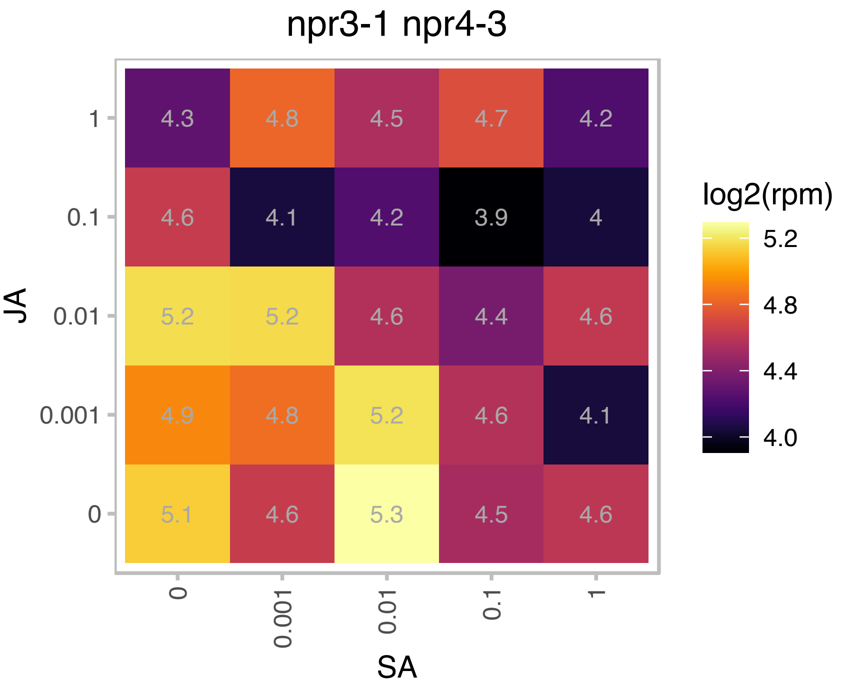

### AT1G60660.1 : ATCB5LP, B5 #5, CB5LP

cytochrome B5-like protein

member of Cytochromes b5

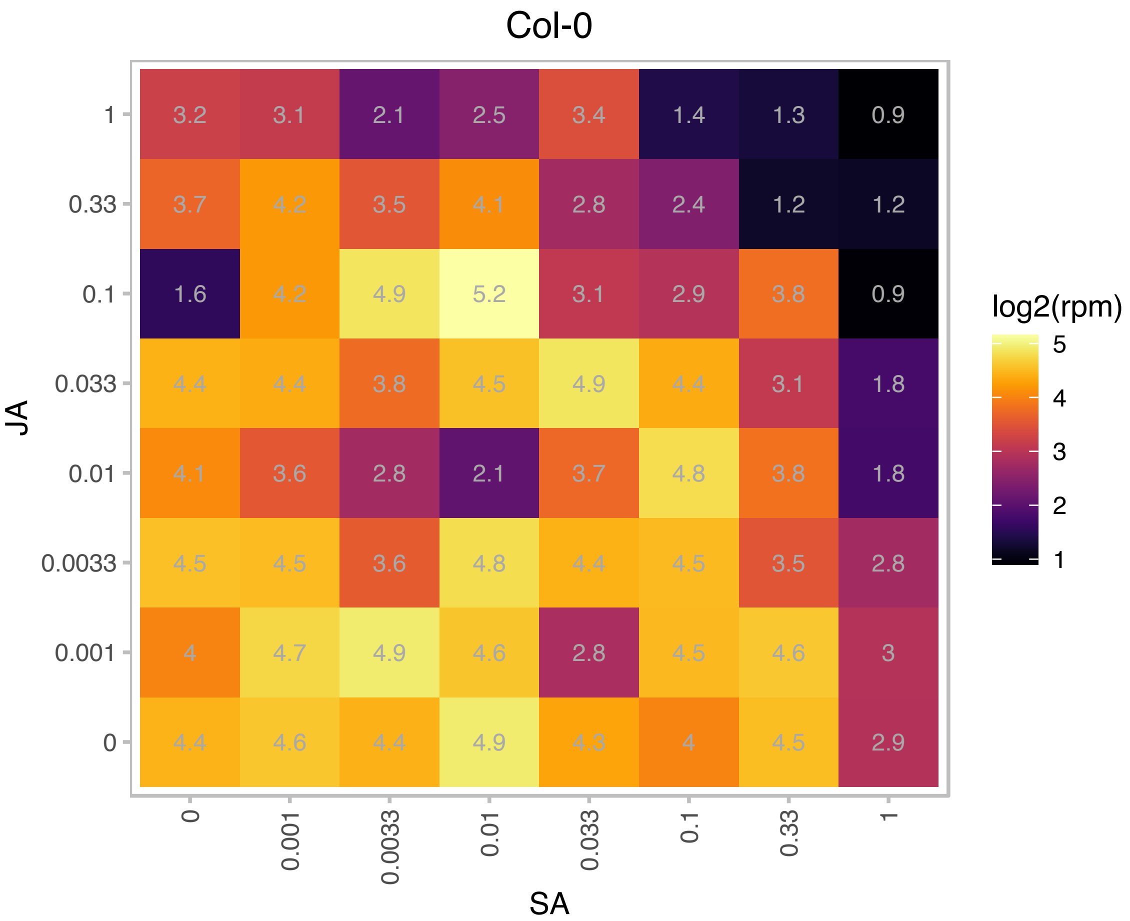


| | |
|---|---|
| Gene | AT1G60660 |
| is.expressed_genes | TRUE |
| is.responsive_genes | TRUE |
| is.apcluster | F-09 |

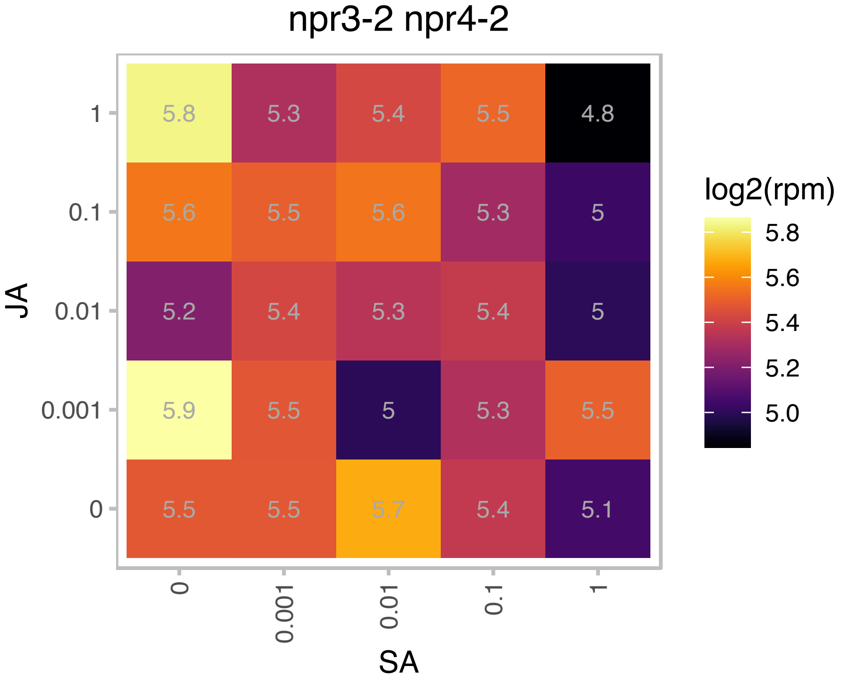


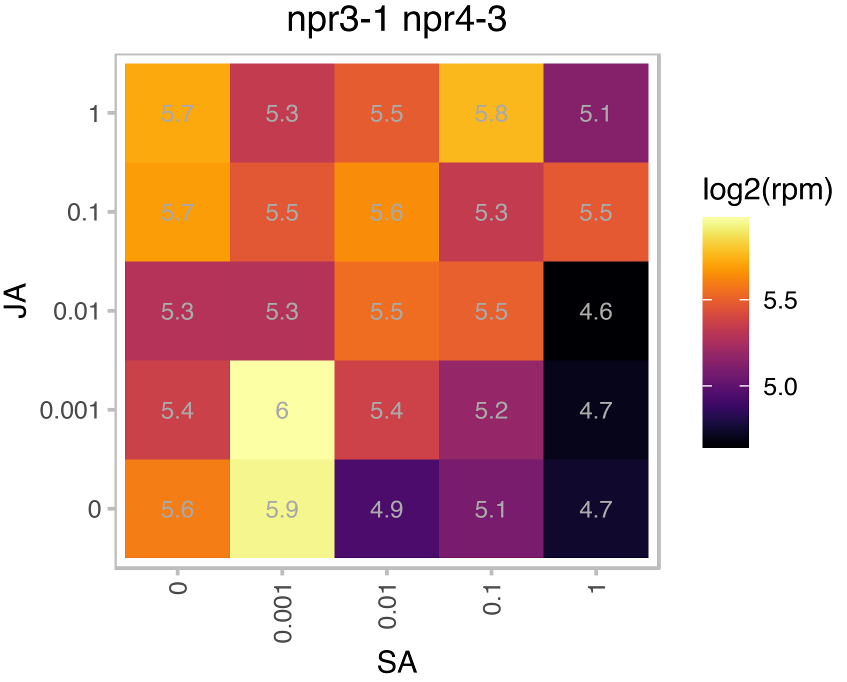

# AT1G61990.1 : NA

Mitochondrial transcription termination factor family protein

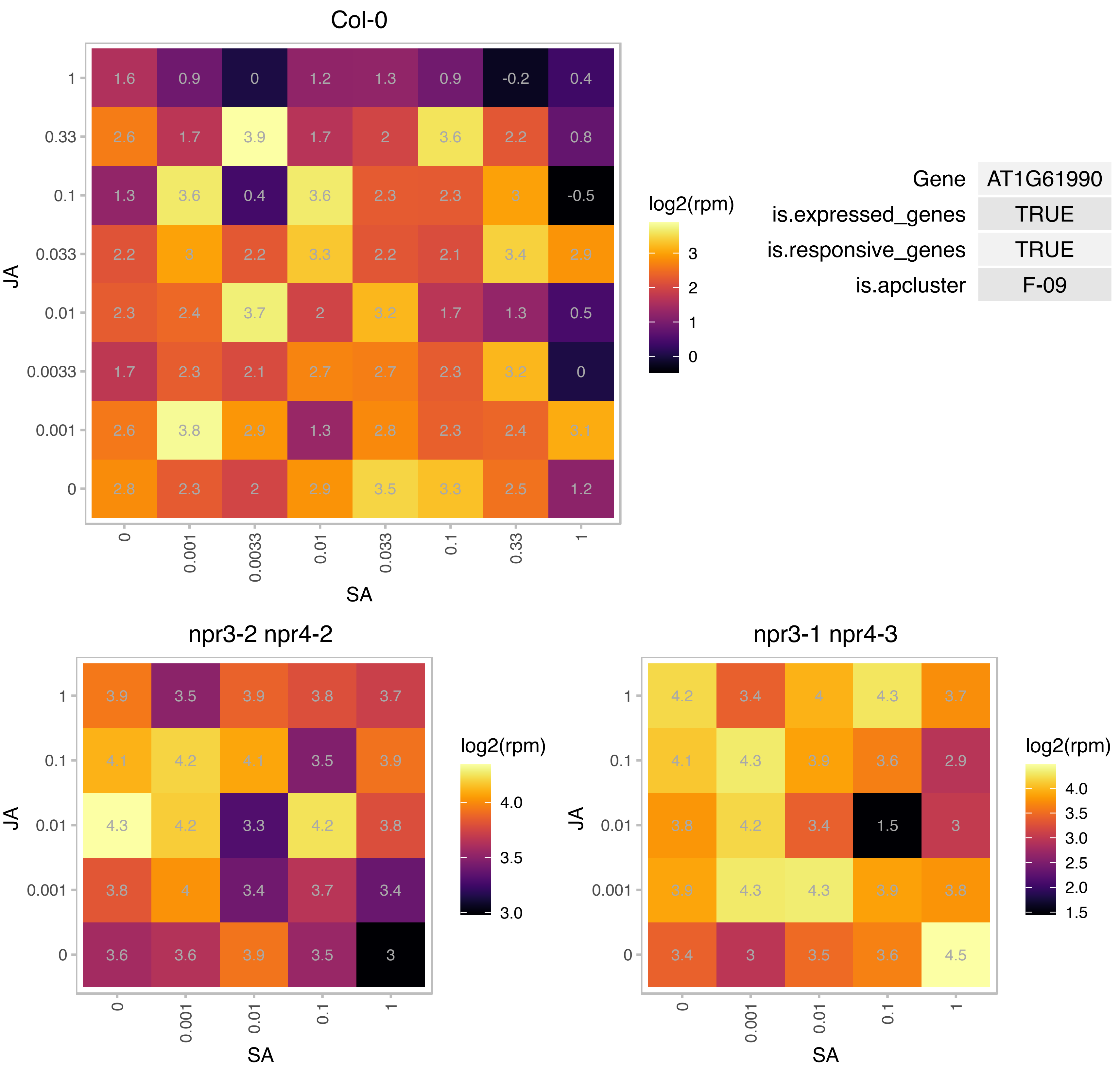


| | |
|---|---|
| Gene | AT1G61990 |
| is.expressed_genes | TRUE |
| is.responsive_genes | TRUE |
| is.apcluster | F-09 |

# AT1G65295.1 : NA

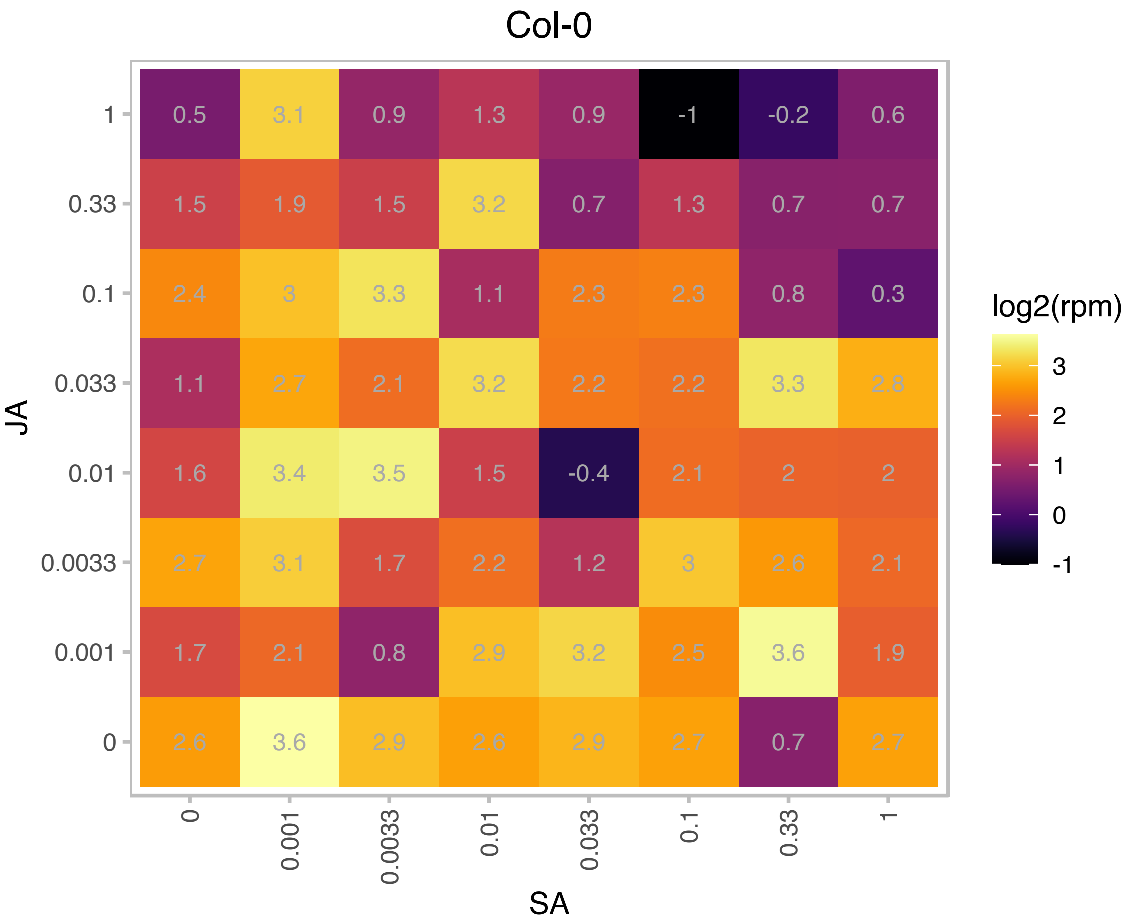


| | |
|---|---|
| Gene | AT1G65295 |
| is.expressed_genes | TRUE |
| is.responsive_genes | TRUE |
| is.apcluster | F-09 |

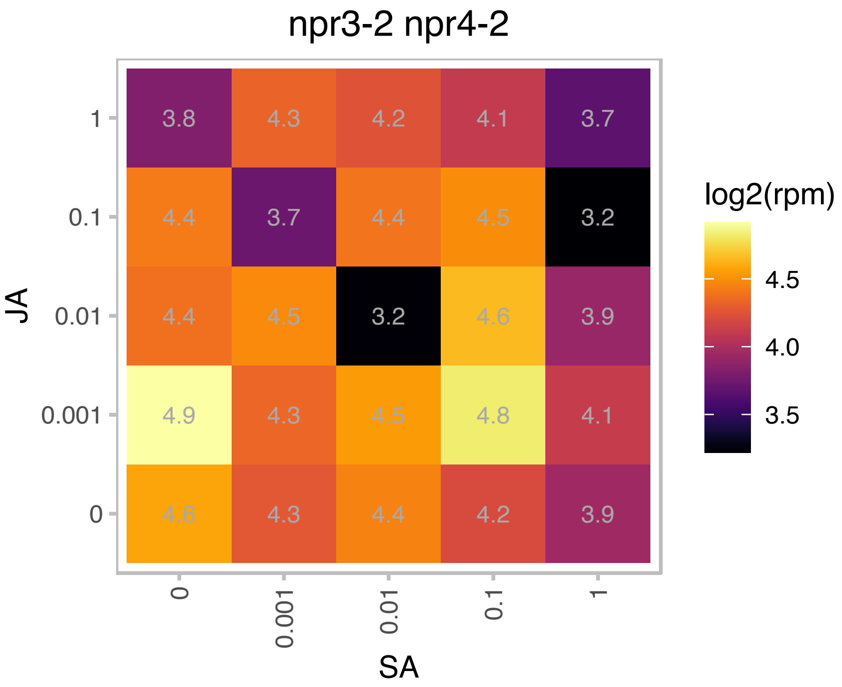


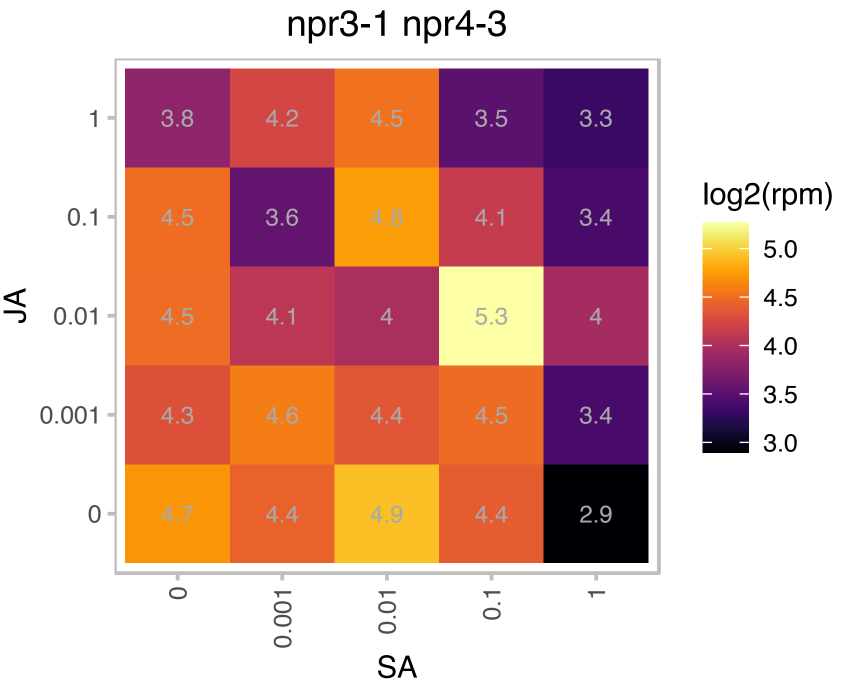

# AT1G65490.1 : NA

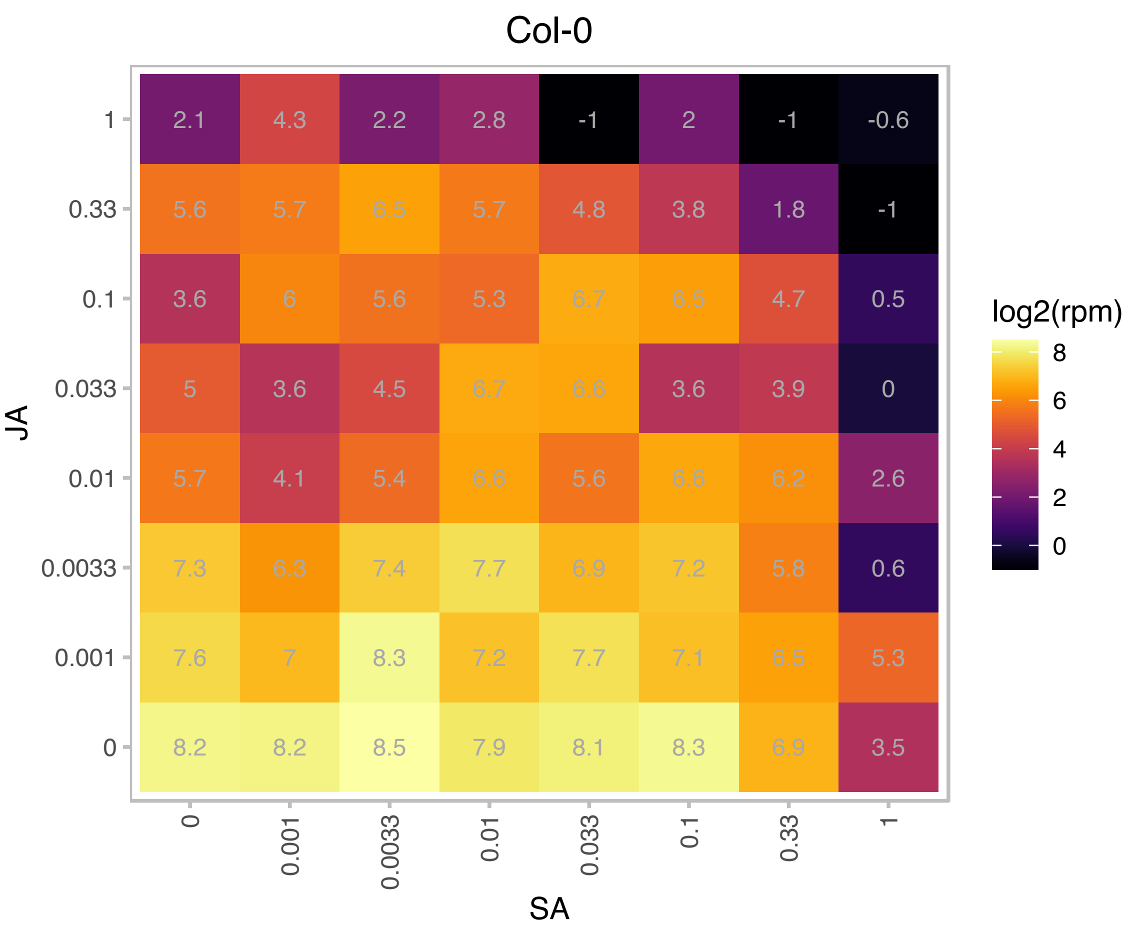


| | |
|---|---|
| Gene | AT1G65490 |
| is.expressed_genes | TRUE |
| is.responsive_genes | TRUE |
| is.apcluster | F-09 |

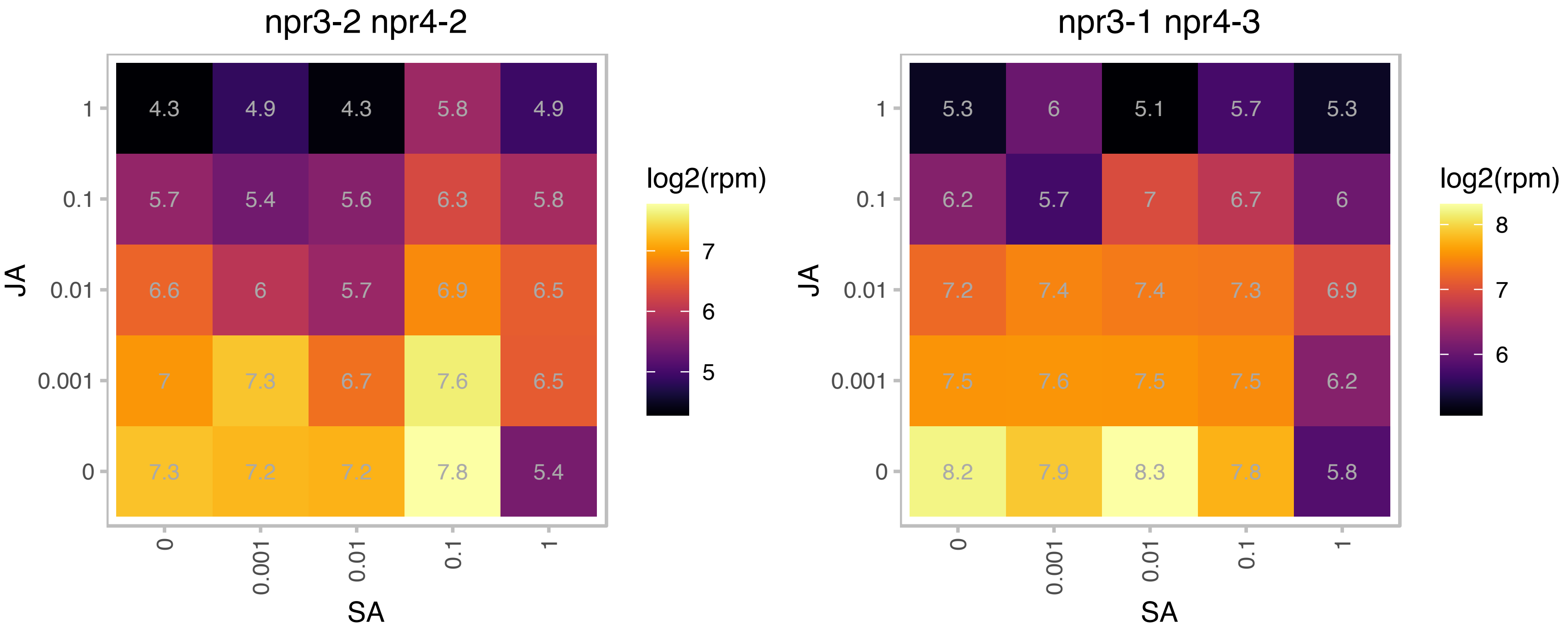

# AT1G66080.1 : NA

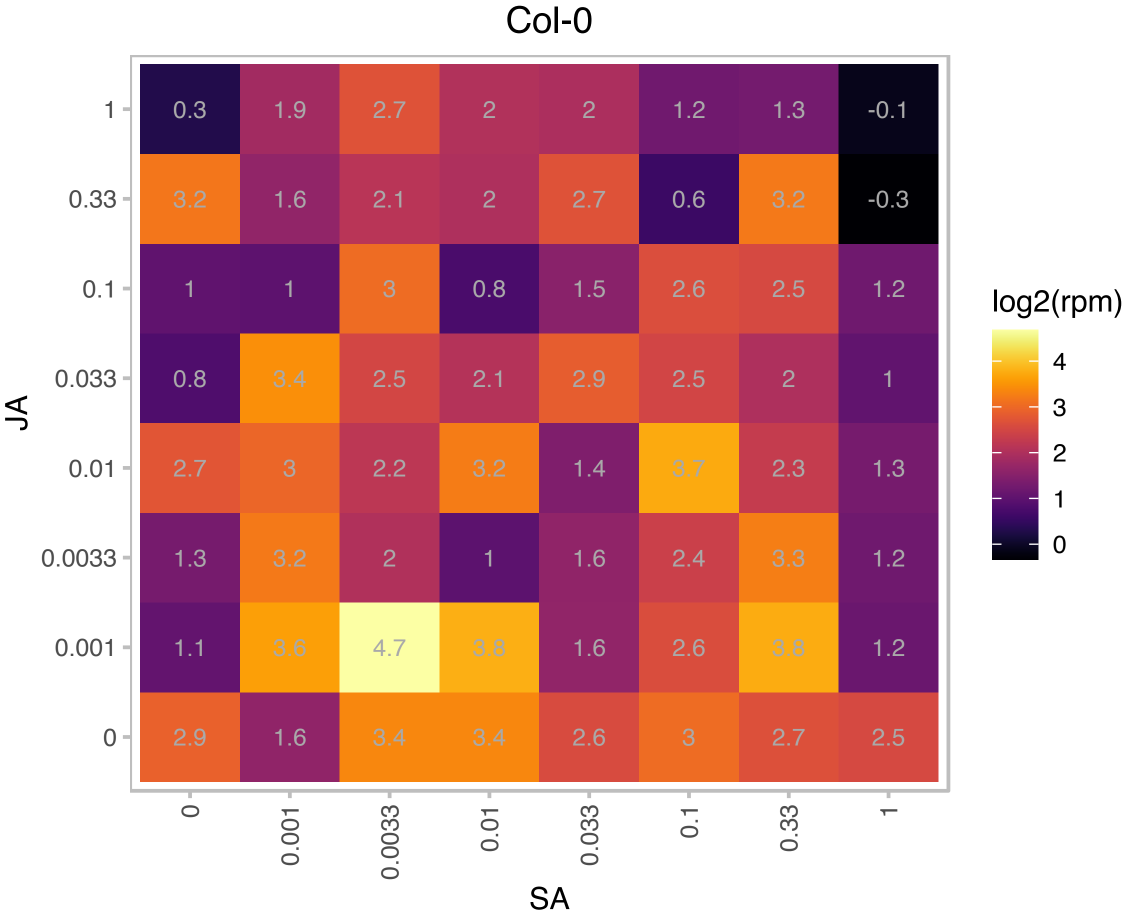


| | |
|---|---|
| Gene | AT1G66080 |
| is.expressed_genes | TRUE |
| is.responsive_genes | TRUE |
| is.apcluster | F-09 |

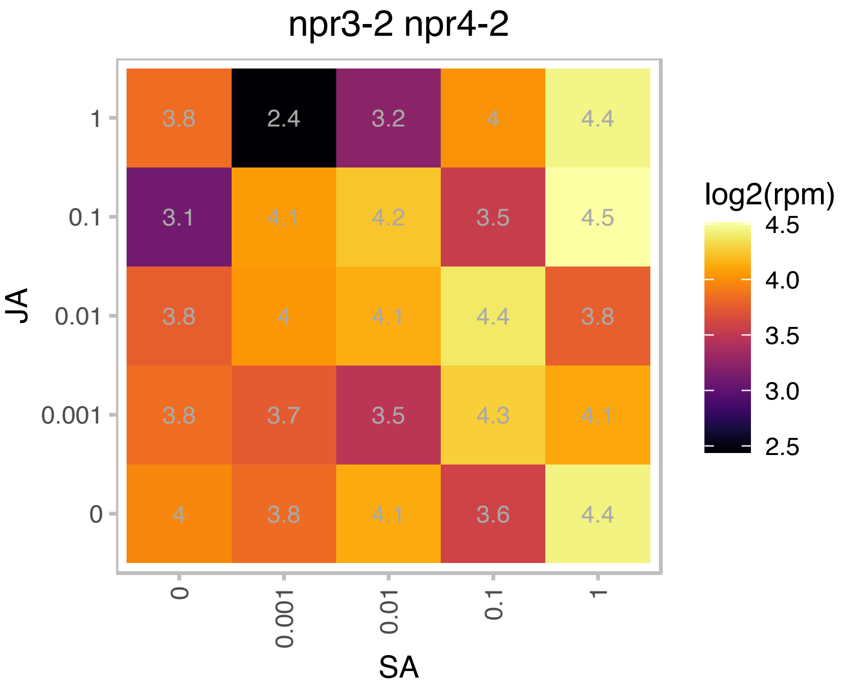


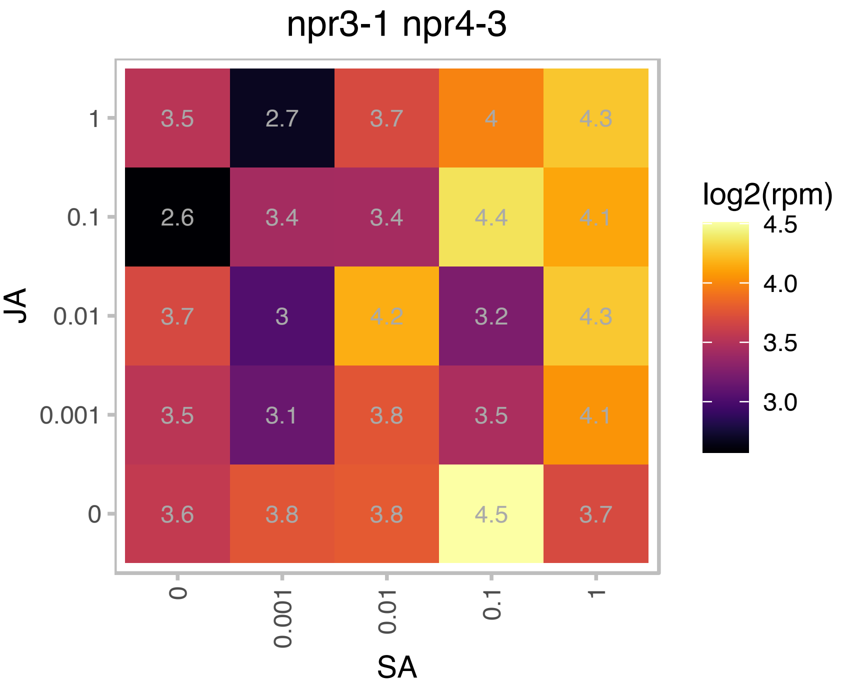

### AT1G69530.1 : AT-EXP1, ATEXP1, ATEXPA1, ATHEXP ALI

expansin A1

Member of Alpha-Expansin Gene Family. Naming convention from the Expansin Working Group (Kende et al, Plant Mol Bio).  Involved in the formation of nematode-induced syncytia in roots of Arabidopsis thaliana.

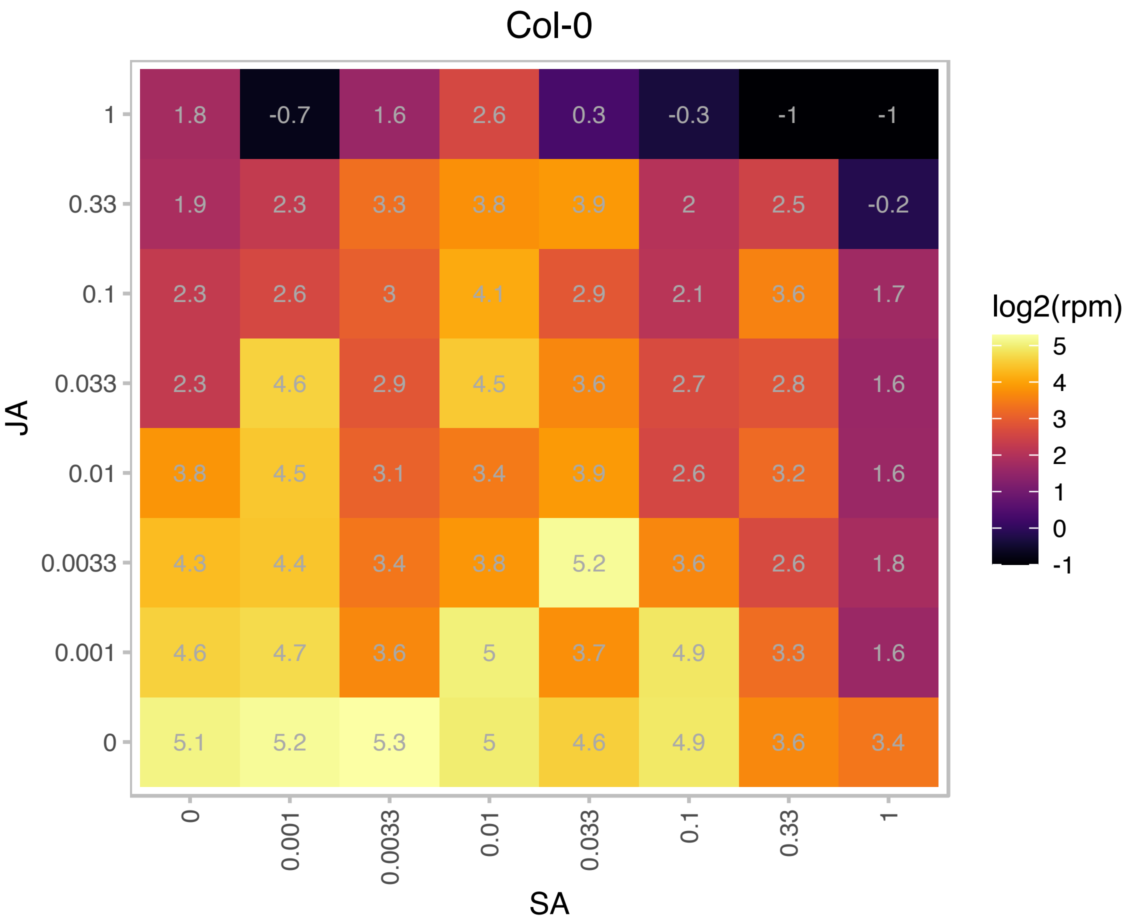


| | |
|---|---|
| Gene | AT1G69530 |
| is.expressed_genes | TRUE |
| is.responsive_genes | TRUE |
| is.apcluster | F-09 |

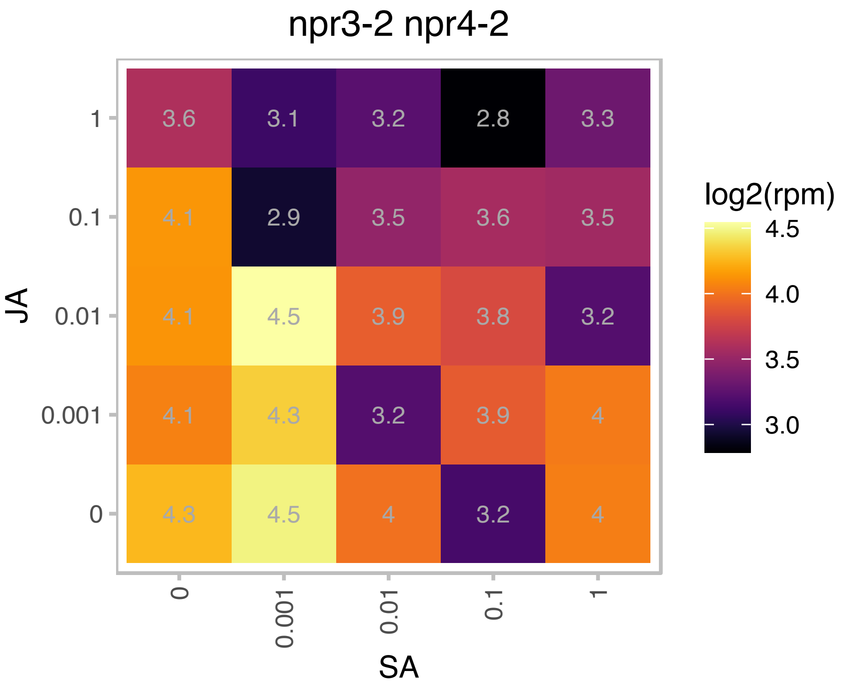


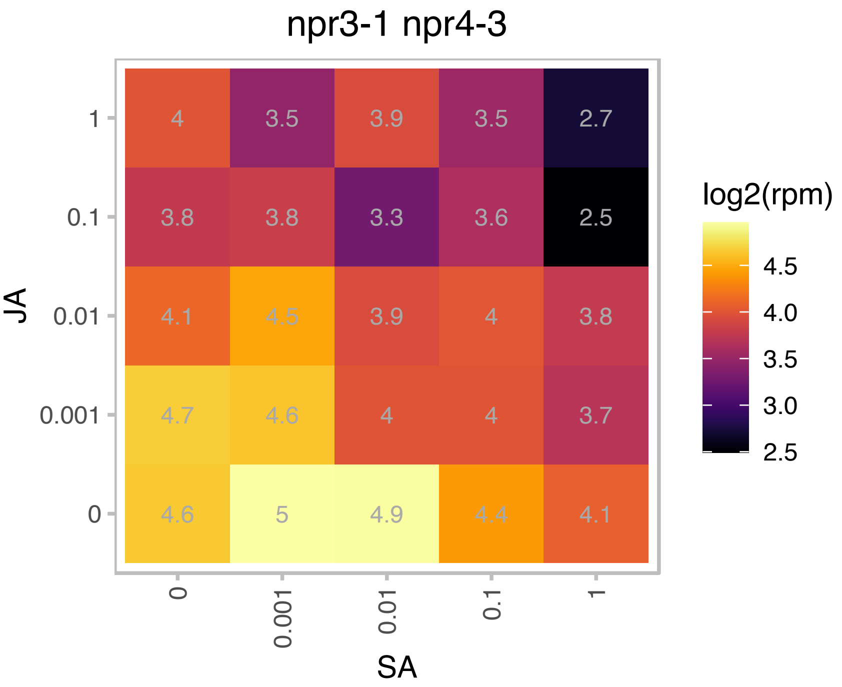

# AT1G73620.1 : NA

Pathogenesis-related thaumatin superfamily protein

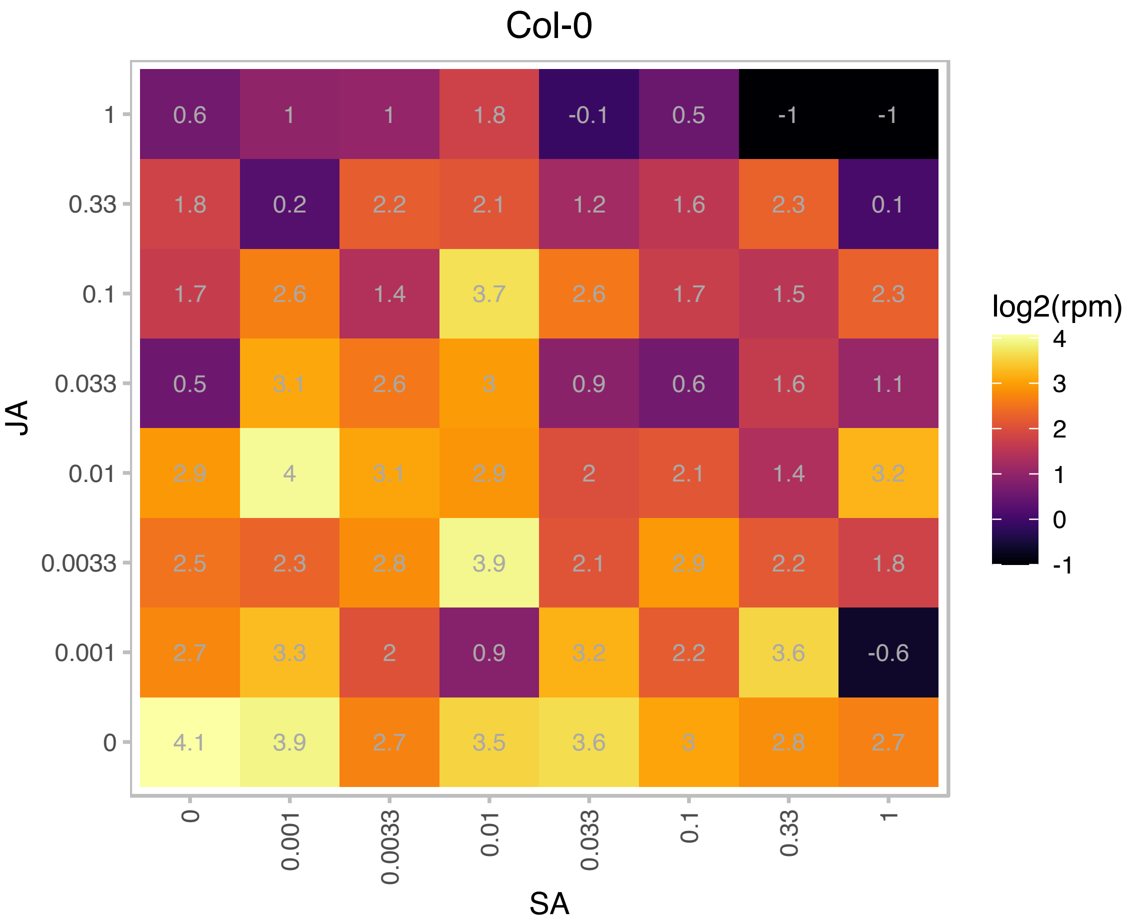


| | |
|---|---|
| Gene | AT1G73620 |
| is.expressed_genes | TRUE |
| is.responsive_genes | TRUE |
| is.apcluster | F-09 |

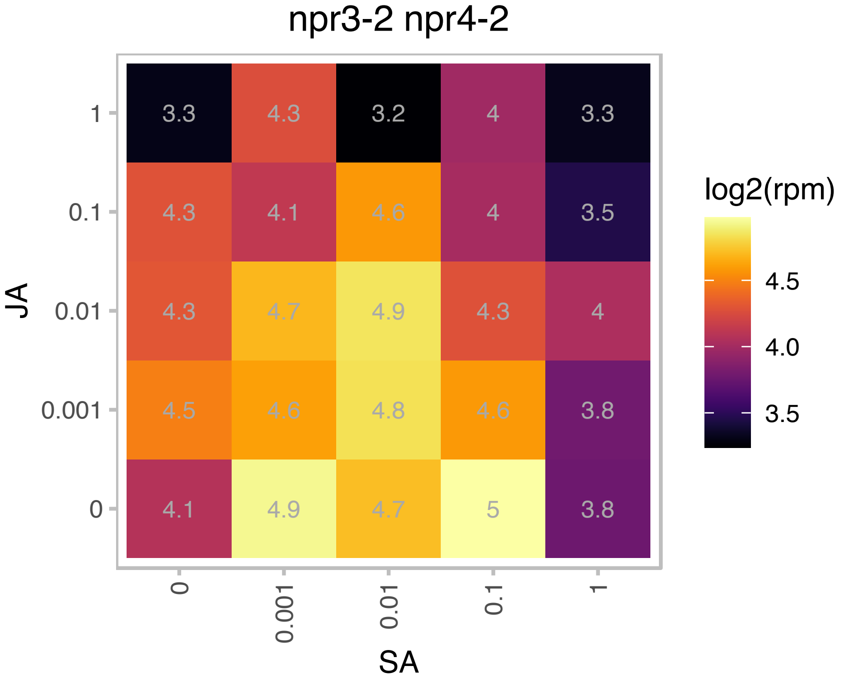


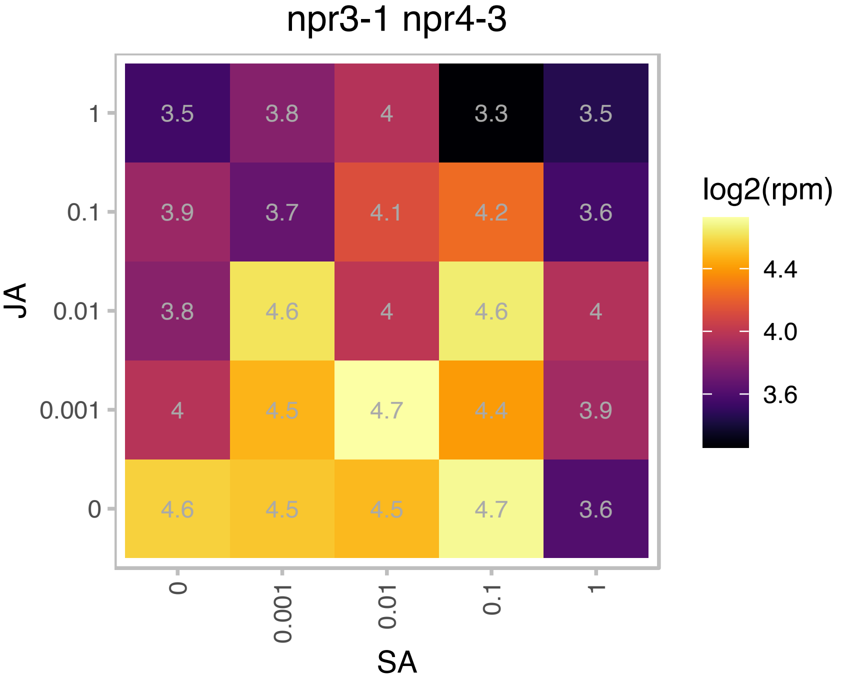

# AT1G74470.1 : NA

Pyridine nucleotide-disulphide oxidoreductase family protein
Encodes for a multifunctional protein with geranylgeranyl reductase activity shown to catalyze the reduction of prenylated geranylgeranyl-chlorophyll a to phytyl-chlorophyll a (chlorophyll a) and free geranylgeranyl pyrophosphate to phytyl pyrophosphate. The mRNA is cell-to-cell mobile.

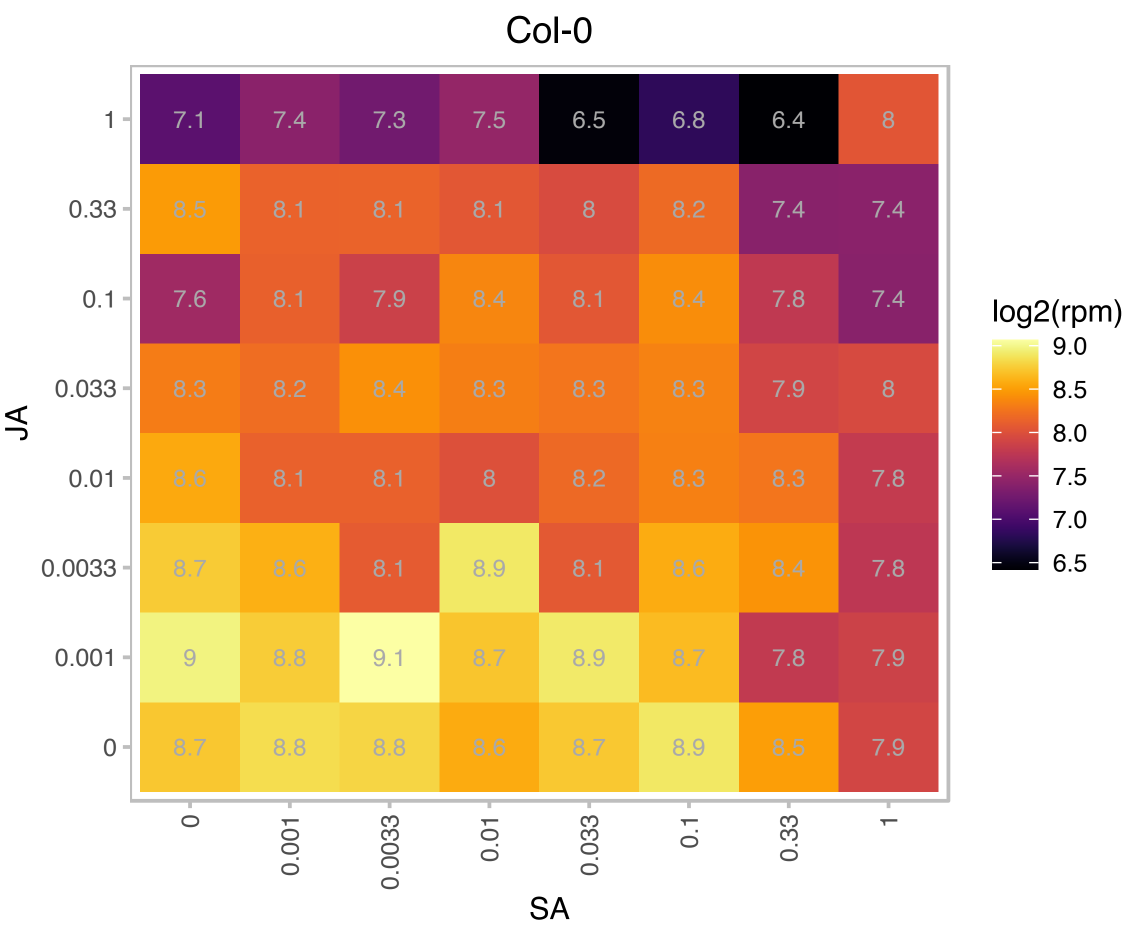


| | |
|---|---|
| Gene | AT1G74470 |
| is.expressed_genes | TRUE |
| is.responsive_genes | TRUE |
| is.apcluster | F-09 |

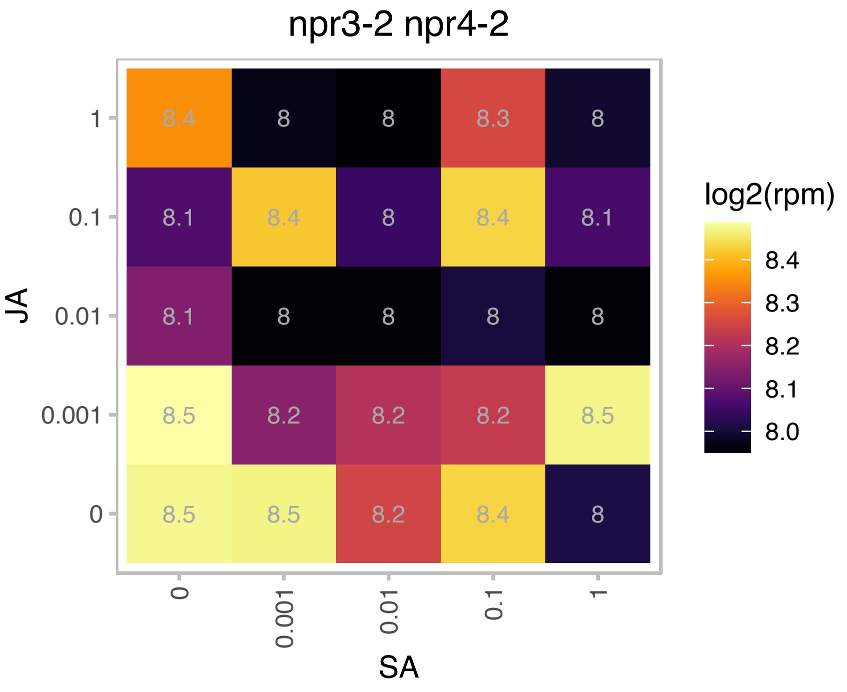


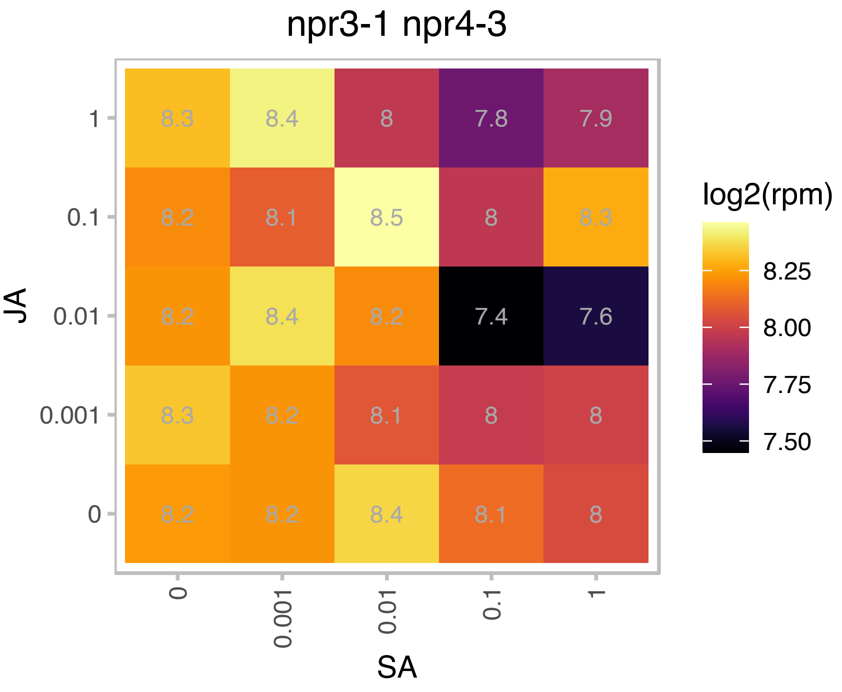

# AT1G76405.1 : NA

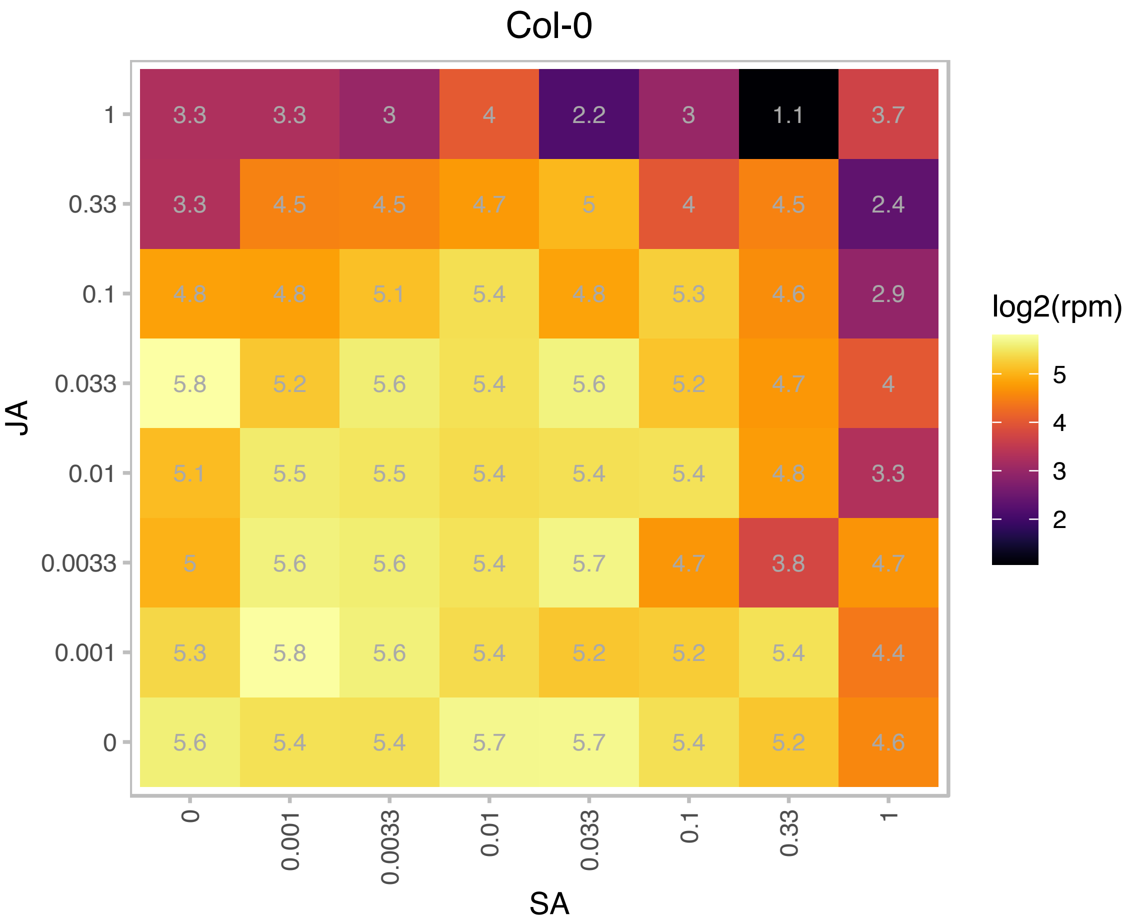


| | |
|---|---|
| Gene | AT1G76405 |
| is.expressed_genes | TRUE |
| is.responsive_genes | TRUE |
| is.apcluster | F-09 |

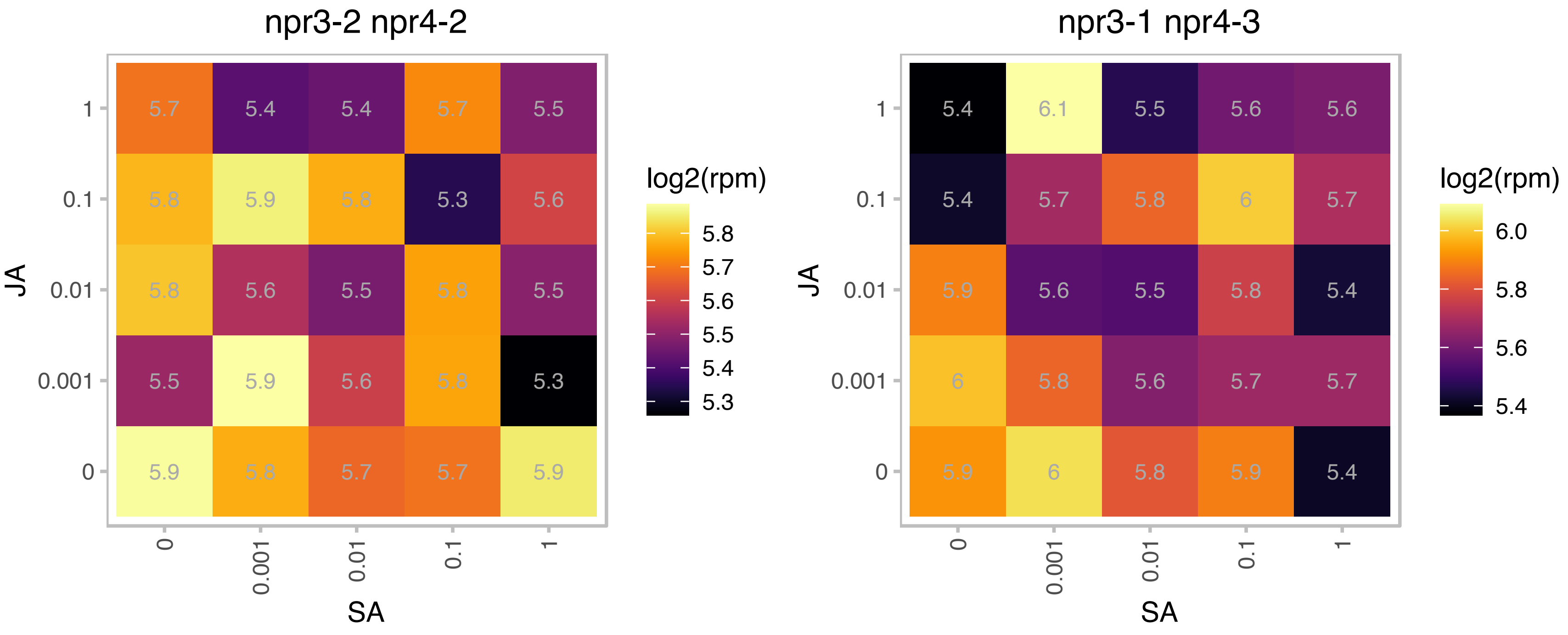

# AT1G77122.1 : NA

Uncharacterised protein family UPF0090

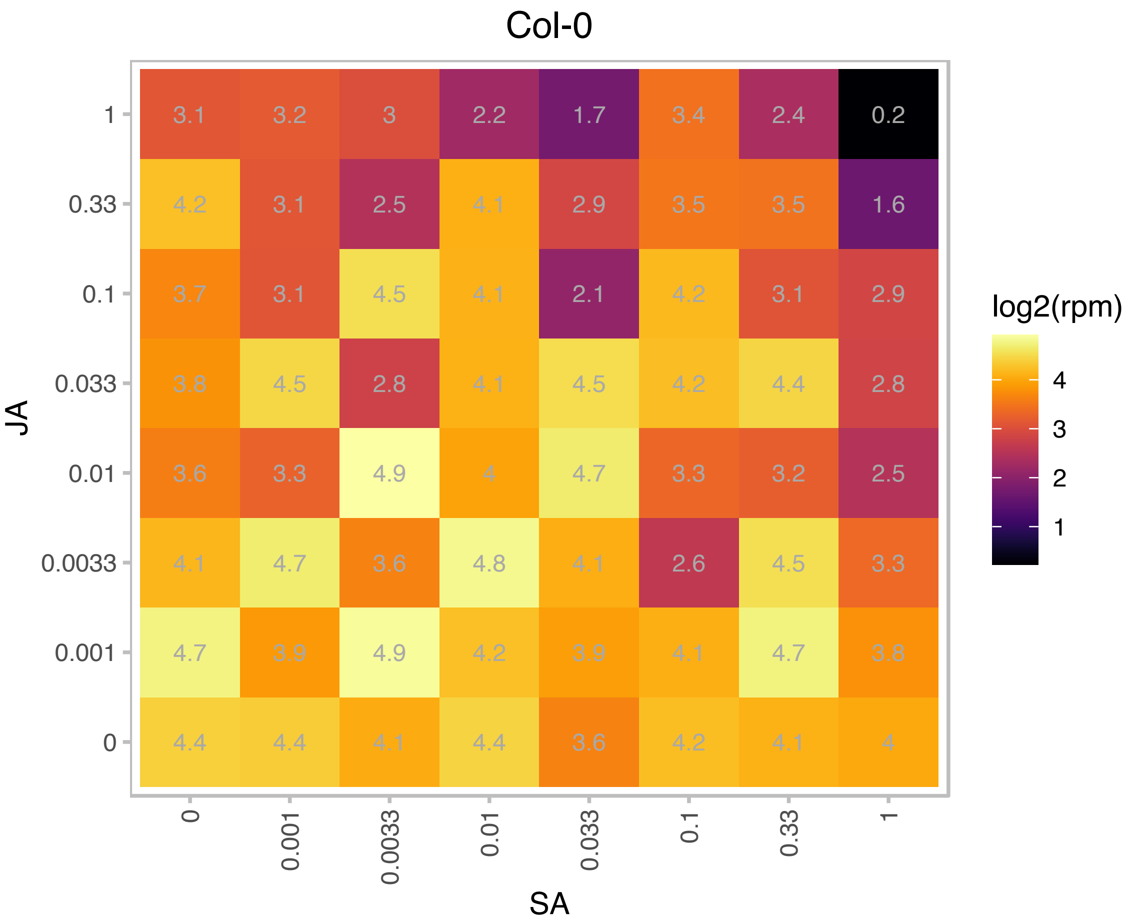


| | |
|---|---|
| Gene | AT1G77122 |
| is.expressed_genes | TRUE |
| is.responsive_genes | TRUE |
| is.apcluster | F-09 |

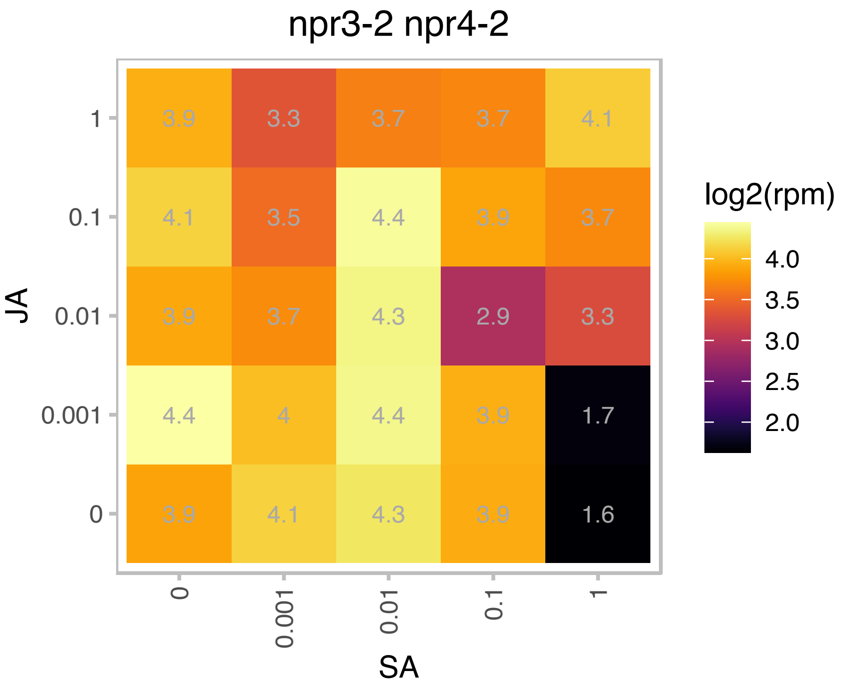


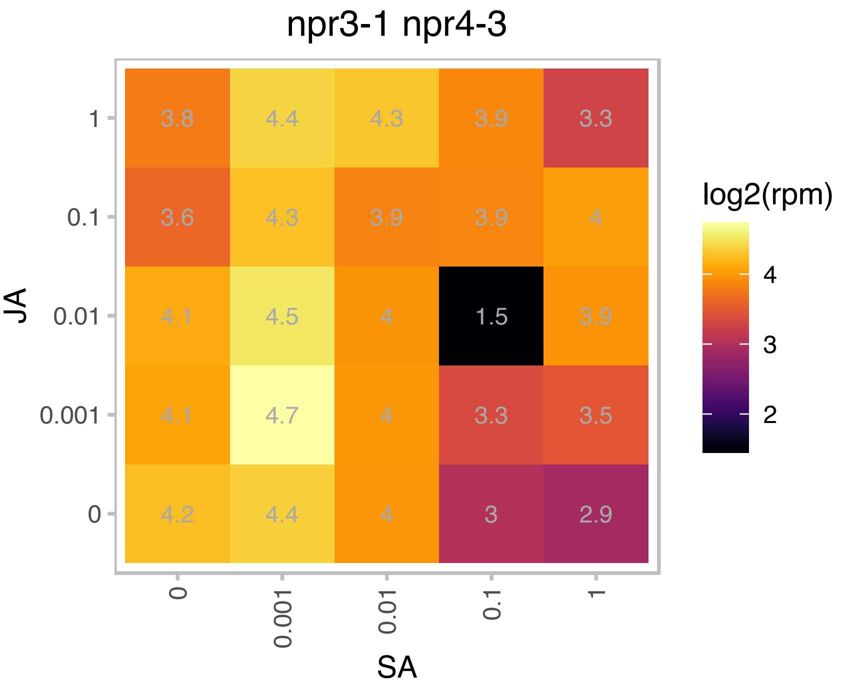

# AT1G77750.1 : NA

Ribosomal protein S13/S18 family

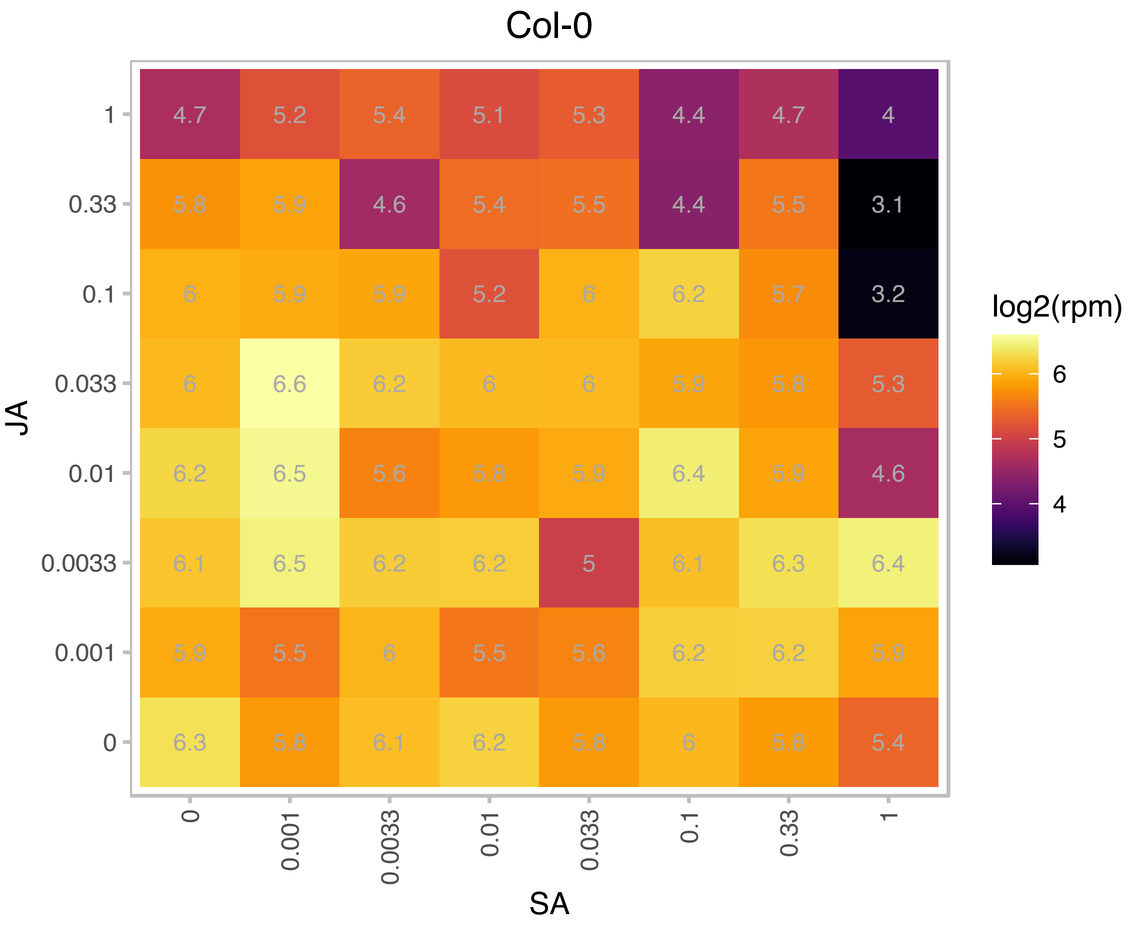


| | |
|---|---|
| Gene | AT1G77750 |
| is.expressed_genes | TRUE |
| is.responsive_genes | TRUE |
| is.apcluster | F-09 |

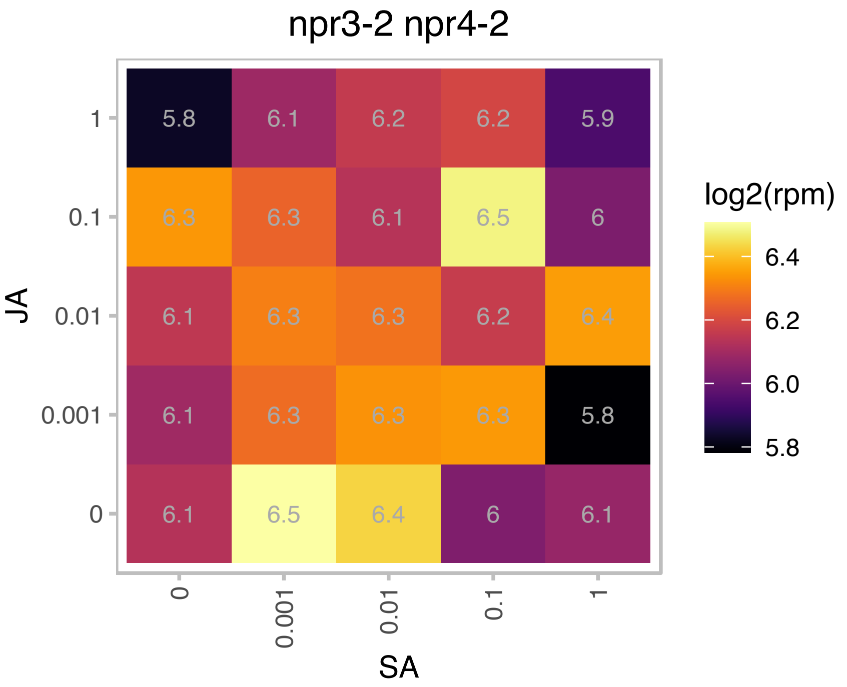


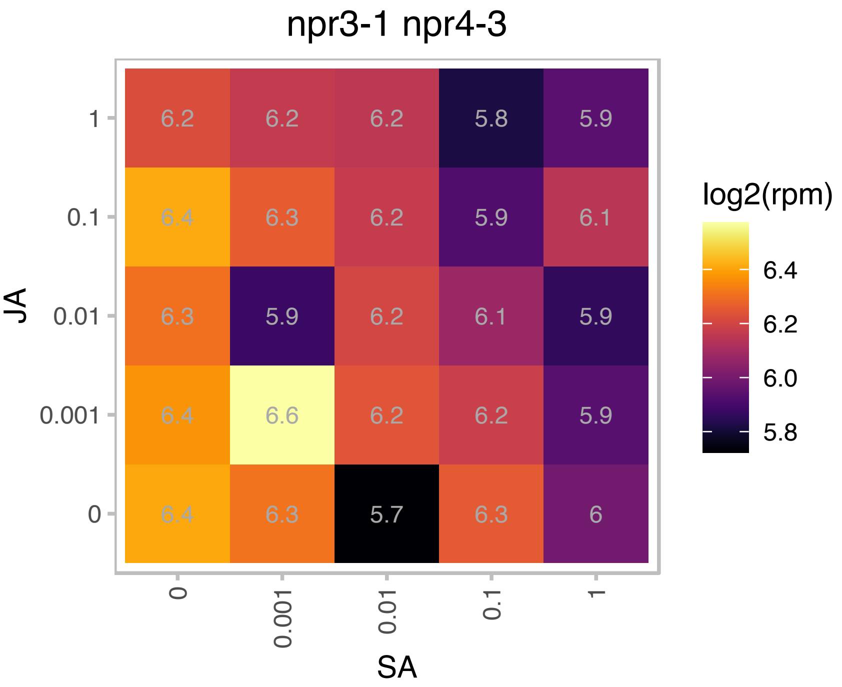

# AT2G01090.1 : NA

Ubiquinol-cytochrome C reductase hinge protein

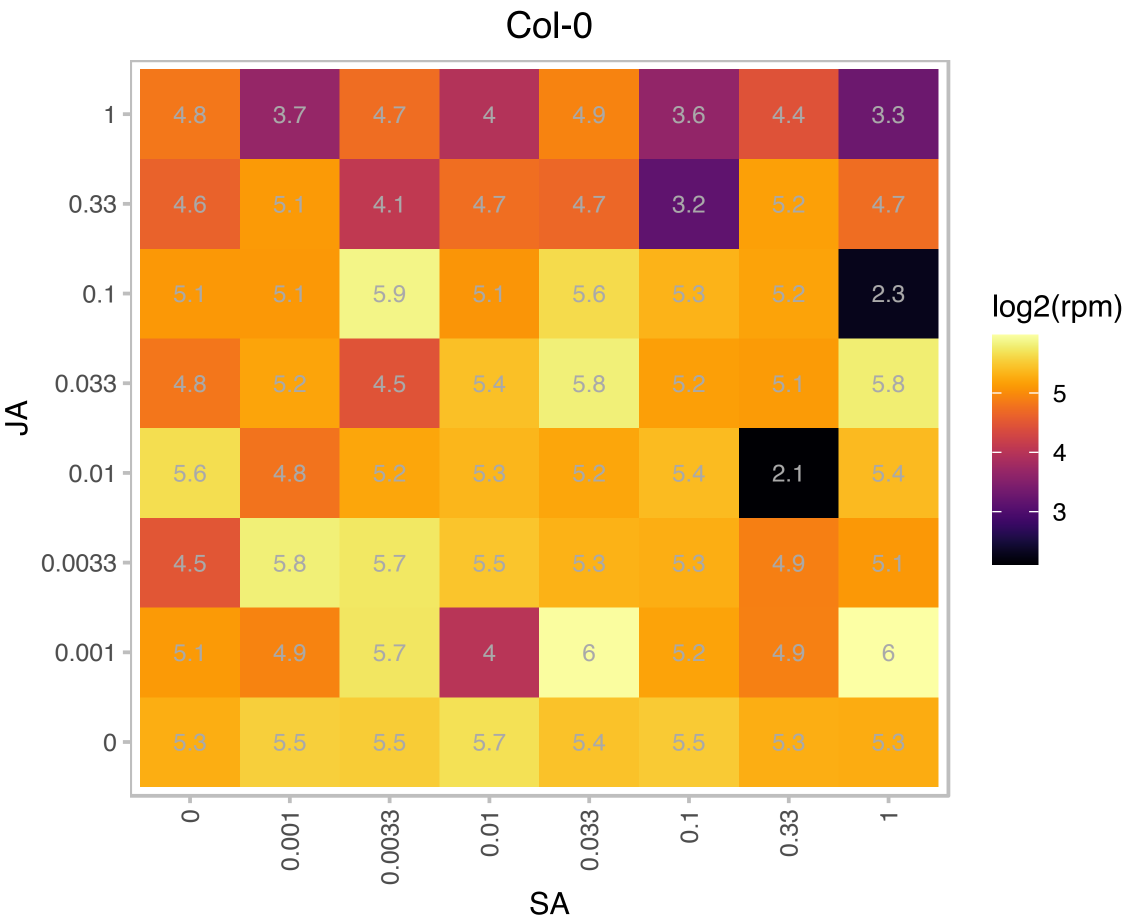


| | |
|---|---|
| Gene | AT2G01090 |
| is.expressed_genes | TRUE |
| is.responsive_genes | TRUE |
| is.apcluster | F-09 |

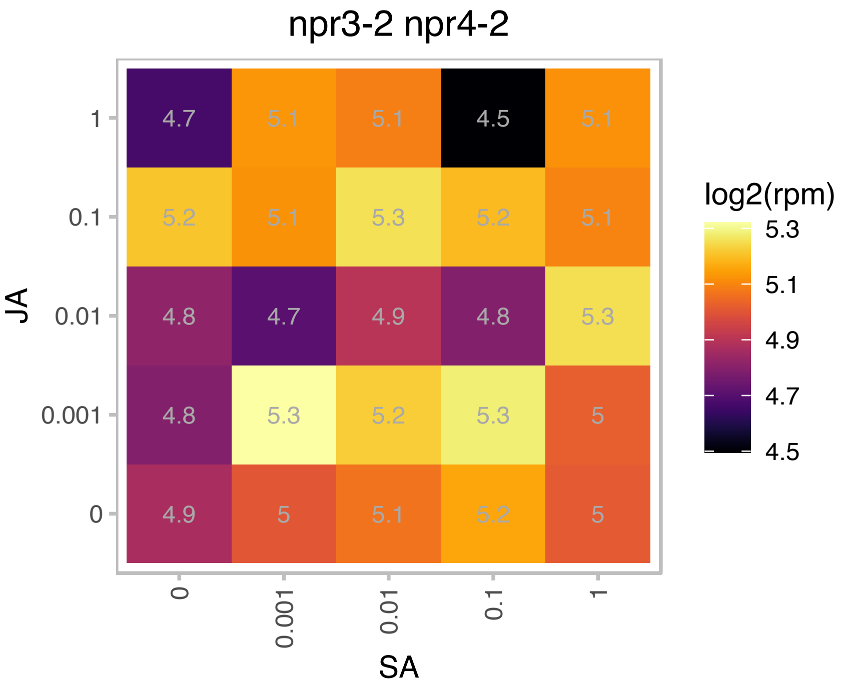


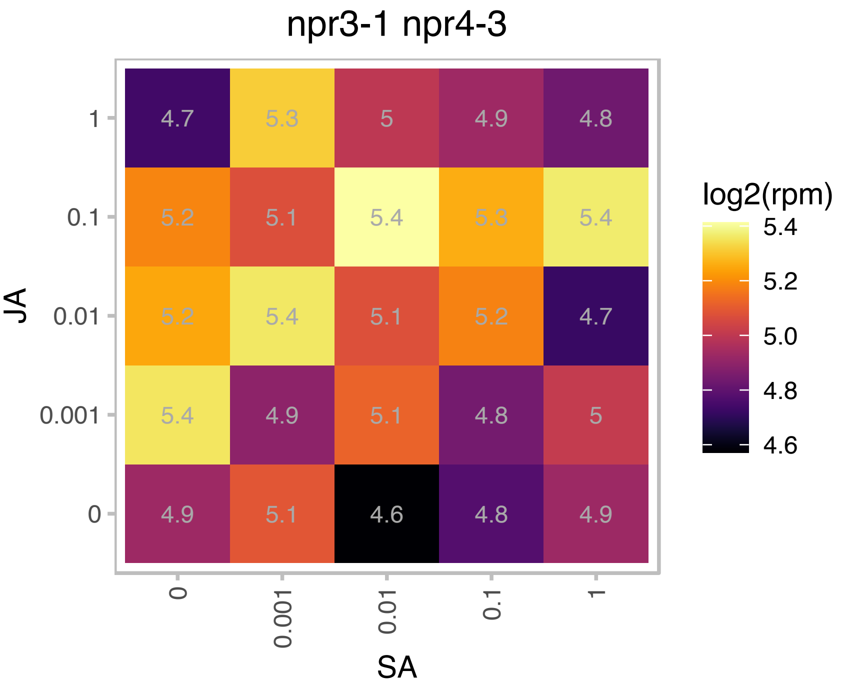

### AT2G03440.1 : ATNRP1, NRP1

nodulin-related protein 1

Induced at the transcriptional level by Pseudomonas syringae pv. tomato infection.

#### Col-0

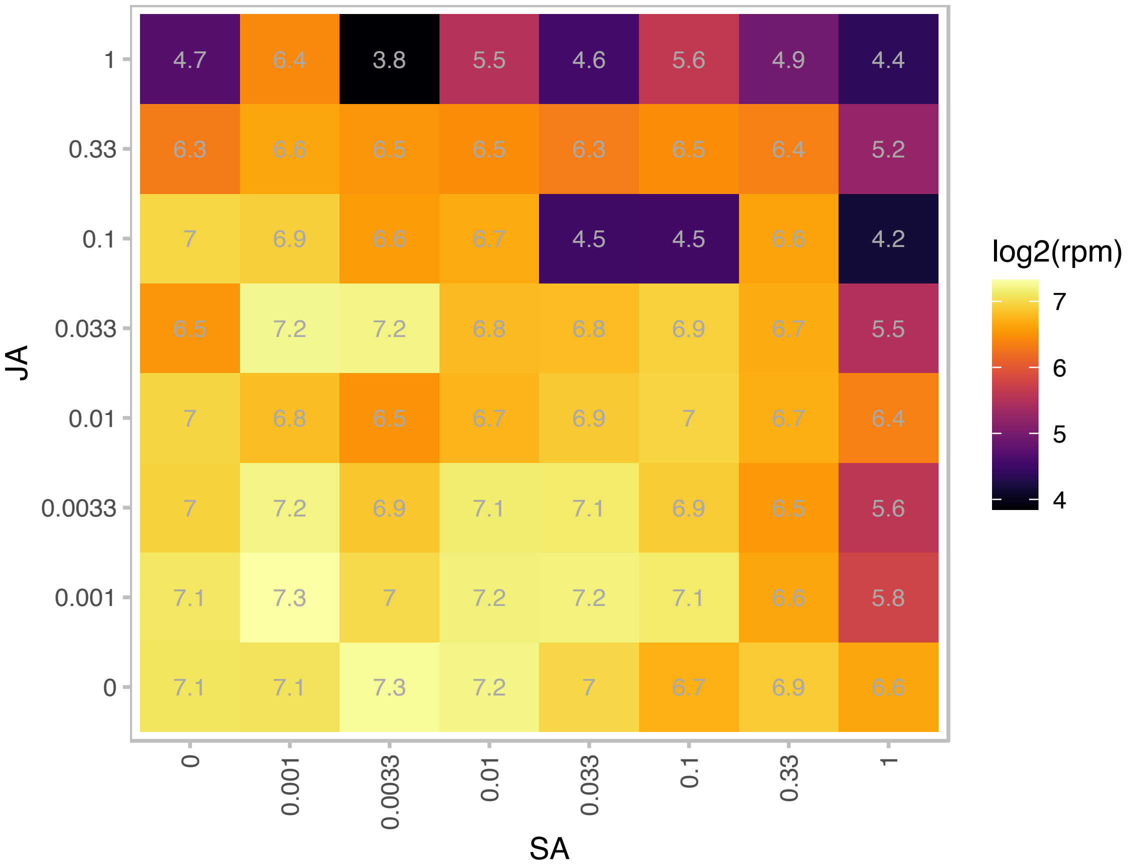


| Gene | AT2G03440 |
|---|---|
| is.expressed_genes | TRUE |
| is.responsive_genes | TRUE |
| is.apcluster | F-09 |

#### npr3-2 npr4-2

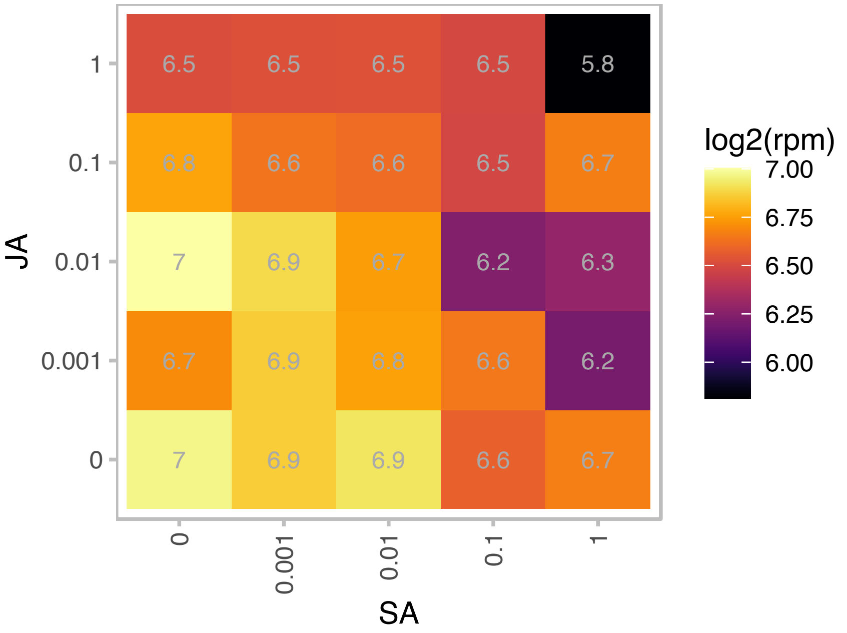


#### npr3-1 npr4-3

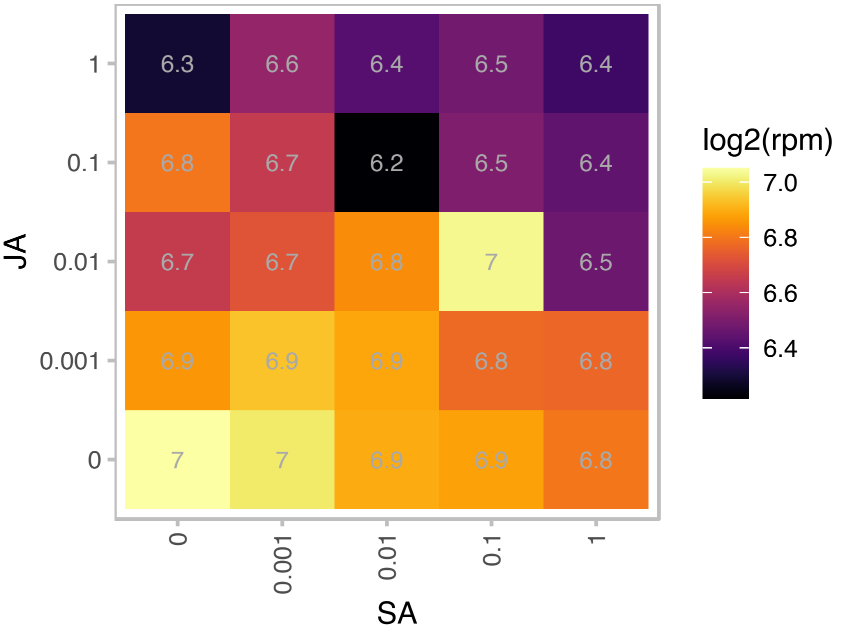

### AT2G05070.1 : LHCB2, LHCB2.2

photosystem II light harvesting complex gene 2.2

Encodes Lhcb2.2. Belongs to the Lhc super-gene family encodes the light-harvesting chlorophyll a/b-binding (LHC) proteins that constitute the antenna system of the photosynthetic apparatus.

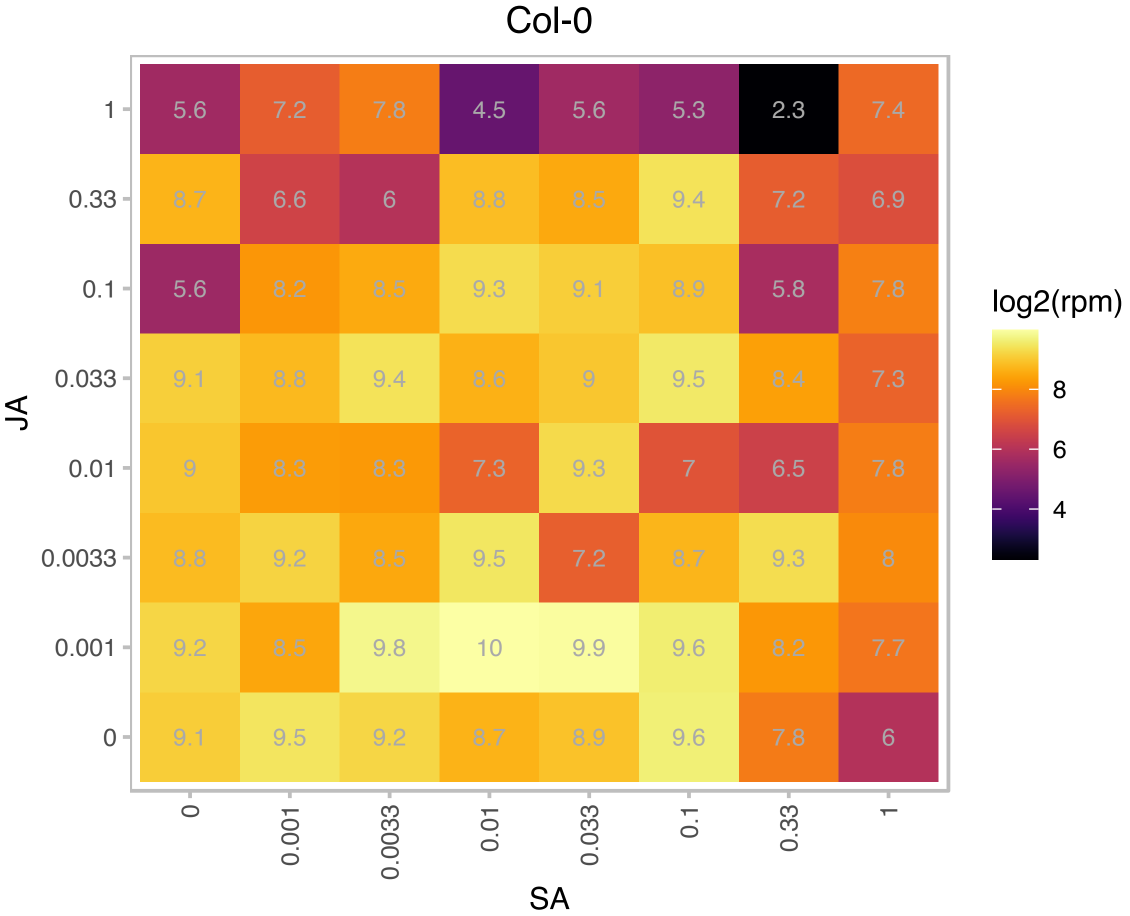


| | |
|---|---|
| Gene | AT2G05070 |
| is.expressed_genes | TRUE |
| is.responsive_genes | TRUE |
| is.apcluster | F-09 |

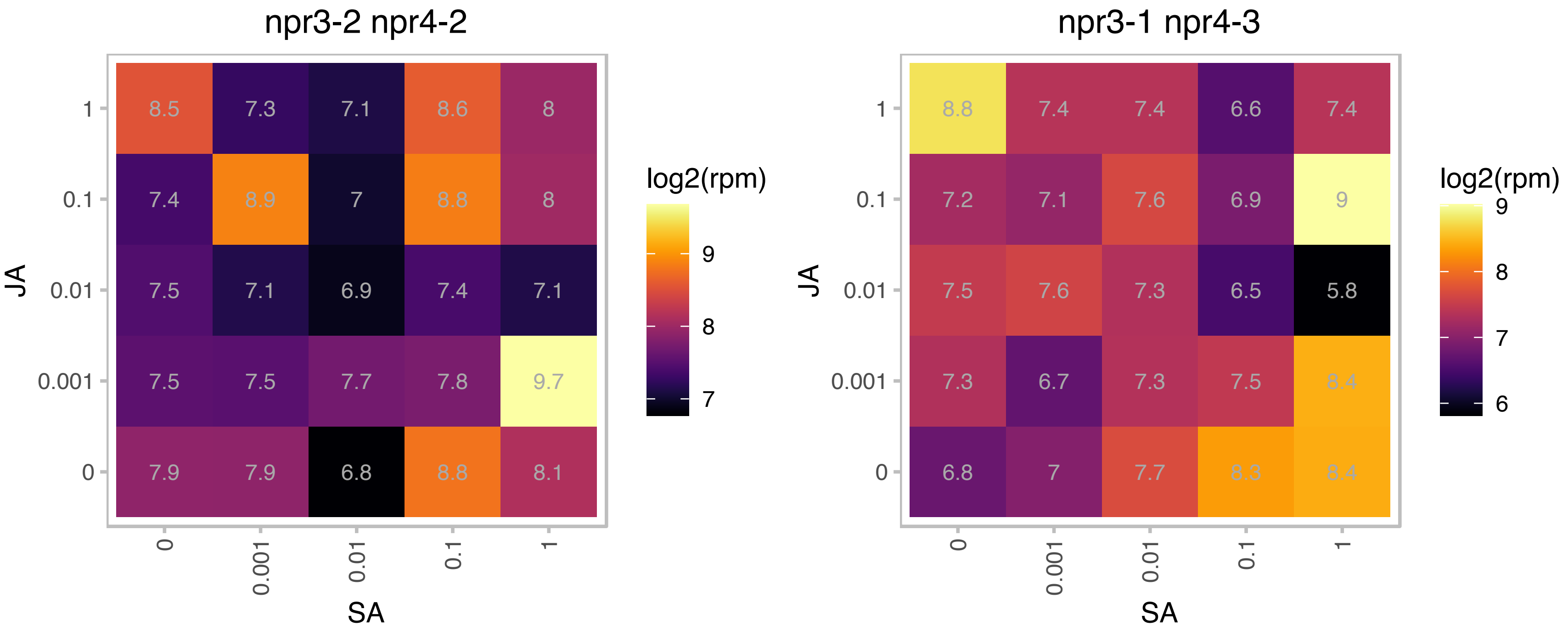

### AT2G06010.1 : ORG4

OBP3-responsive gene 4
encodes a novel protein whose expression level is induced in lines overexpressing salicylic-acid (SA)-inducible Arabidopsis DNA binding with one finger (Dof) transcription factor, called OBF-binding protein 3.

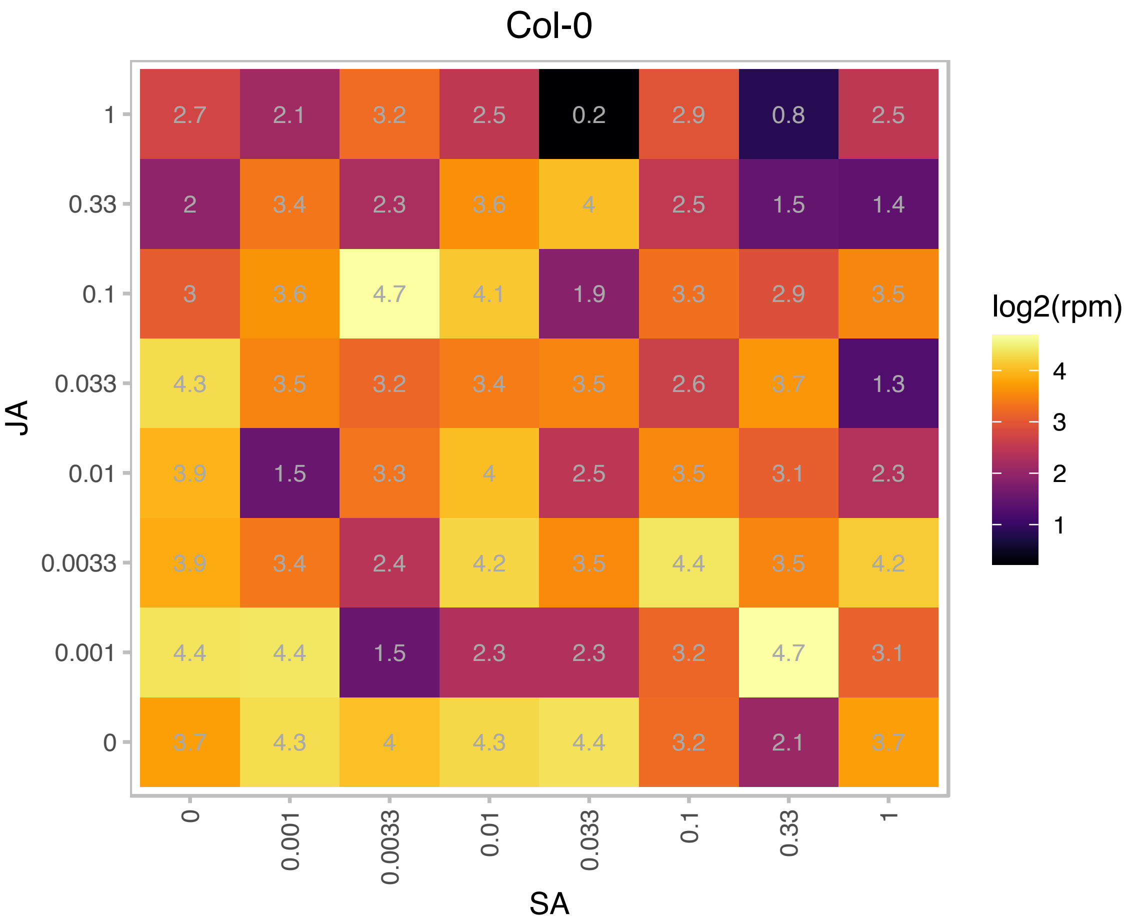


| | |
|---|---|
| Gene | AT2G06010 |
| is.expressed_genes | TRUE |
| is.responsive_genes | TRUE |
| is.apcluster | F-09 |

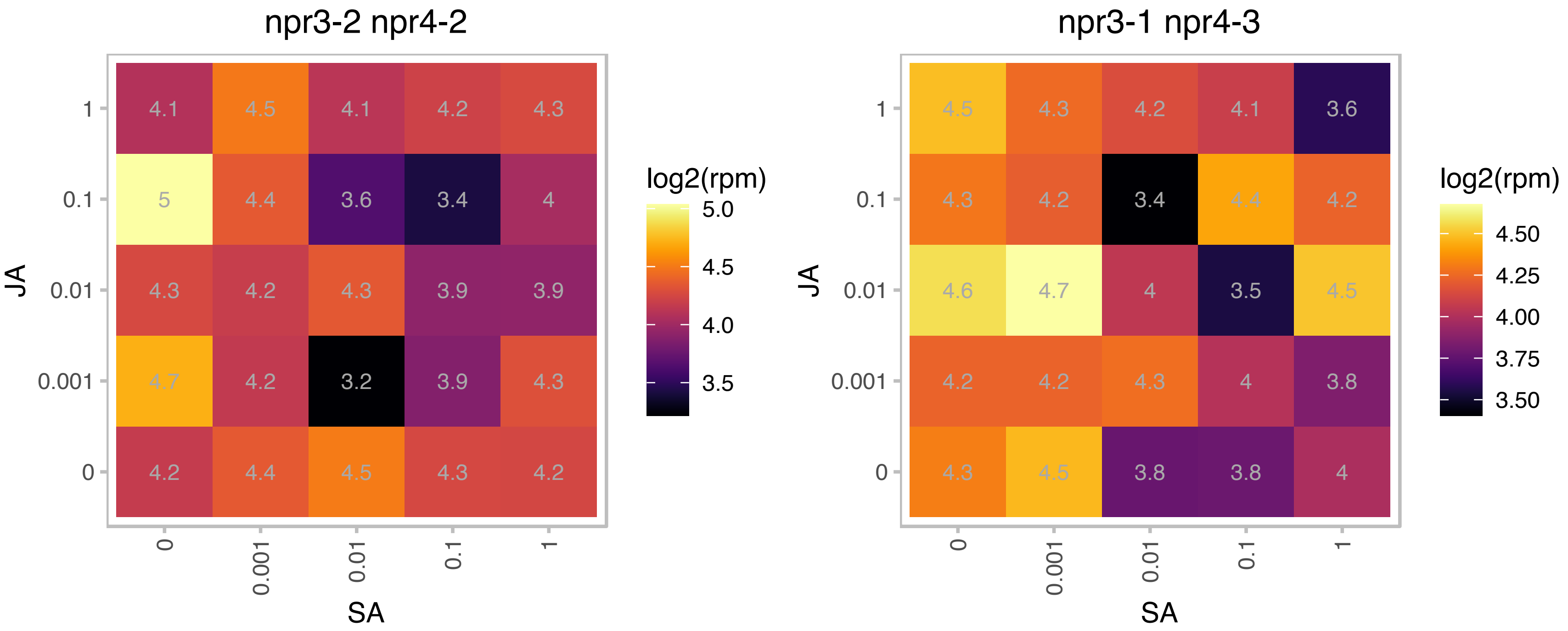

### AT2G19720.1 : rps15ab

ribosomal protein S15A B

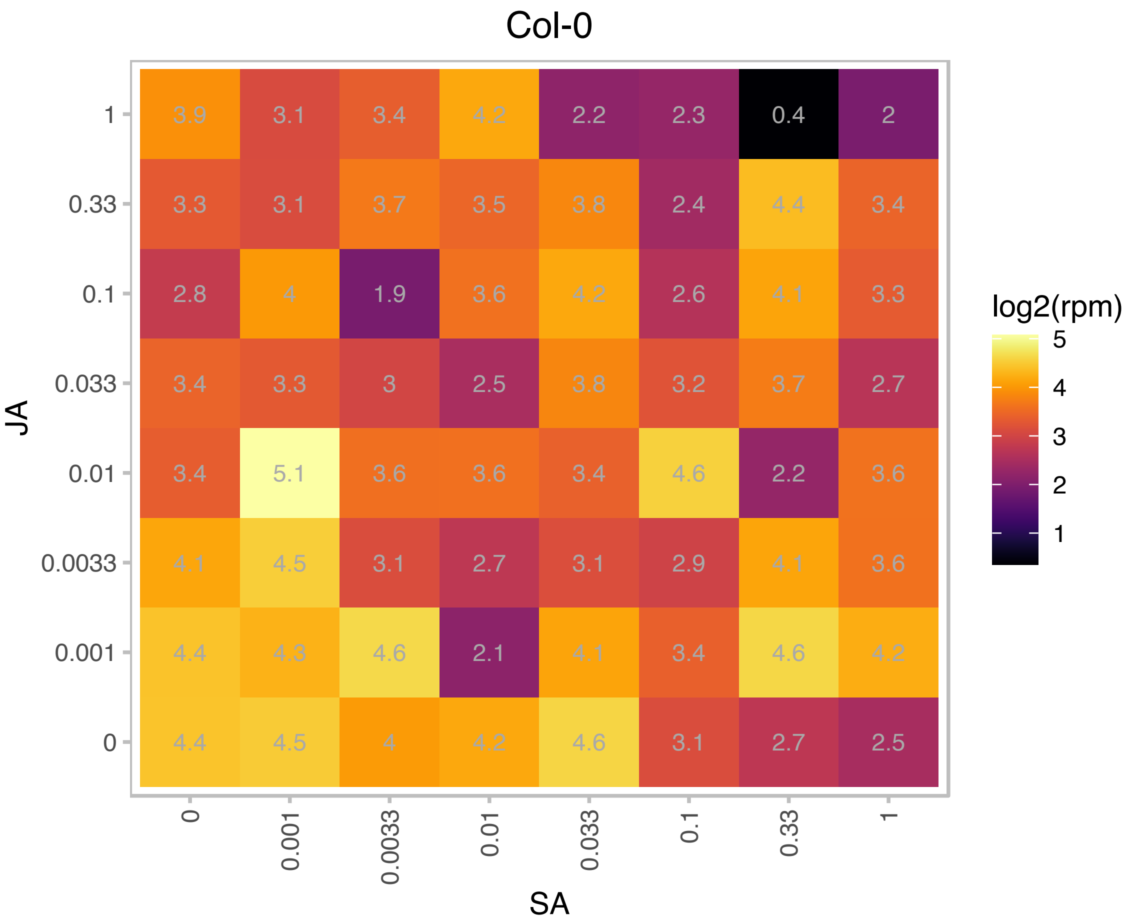


| | |
|---|---|
| Gene | AT2G19720 |
| is.expressed_genes | TRUE |
| is.responsive_genes | TRUE |
| is.apcluster | F-09 |

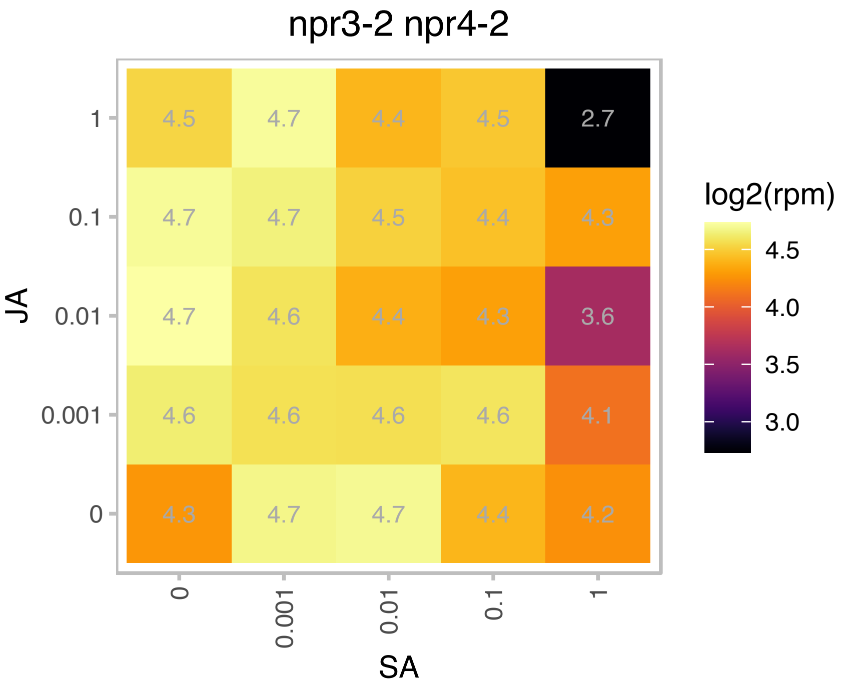


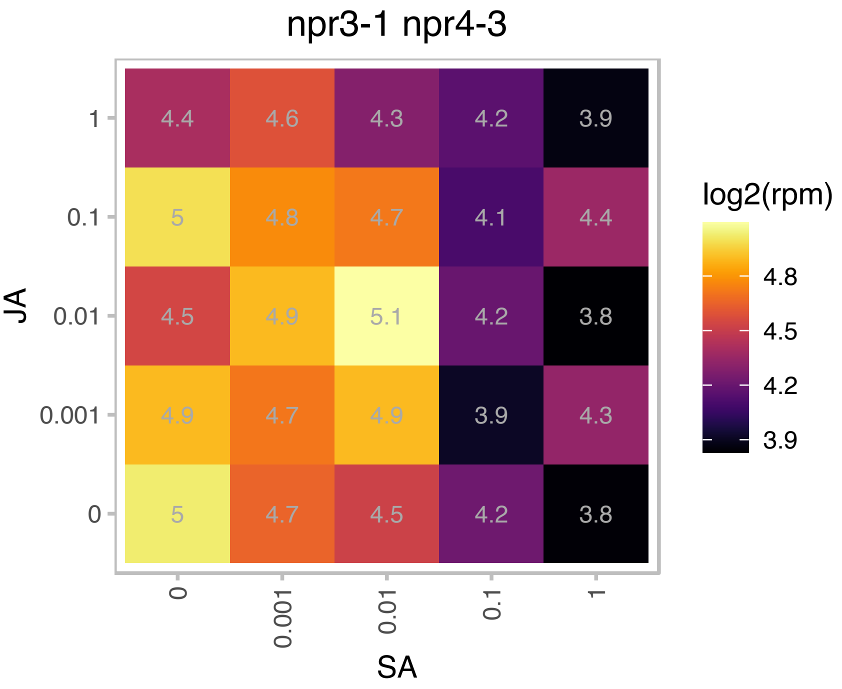

# AT2G19870.1 : NA

tRNA/rRNA methyltransferase (SpoU) family protein

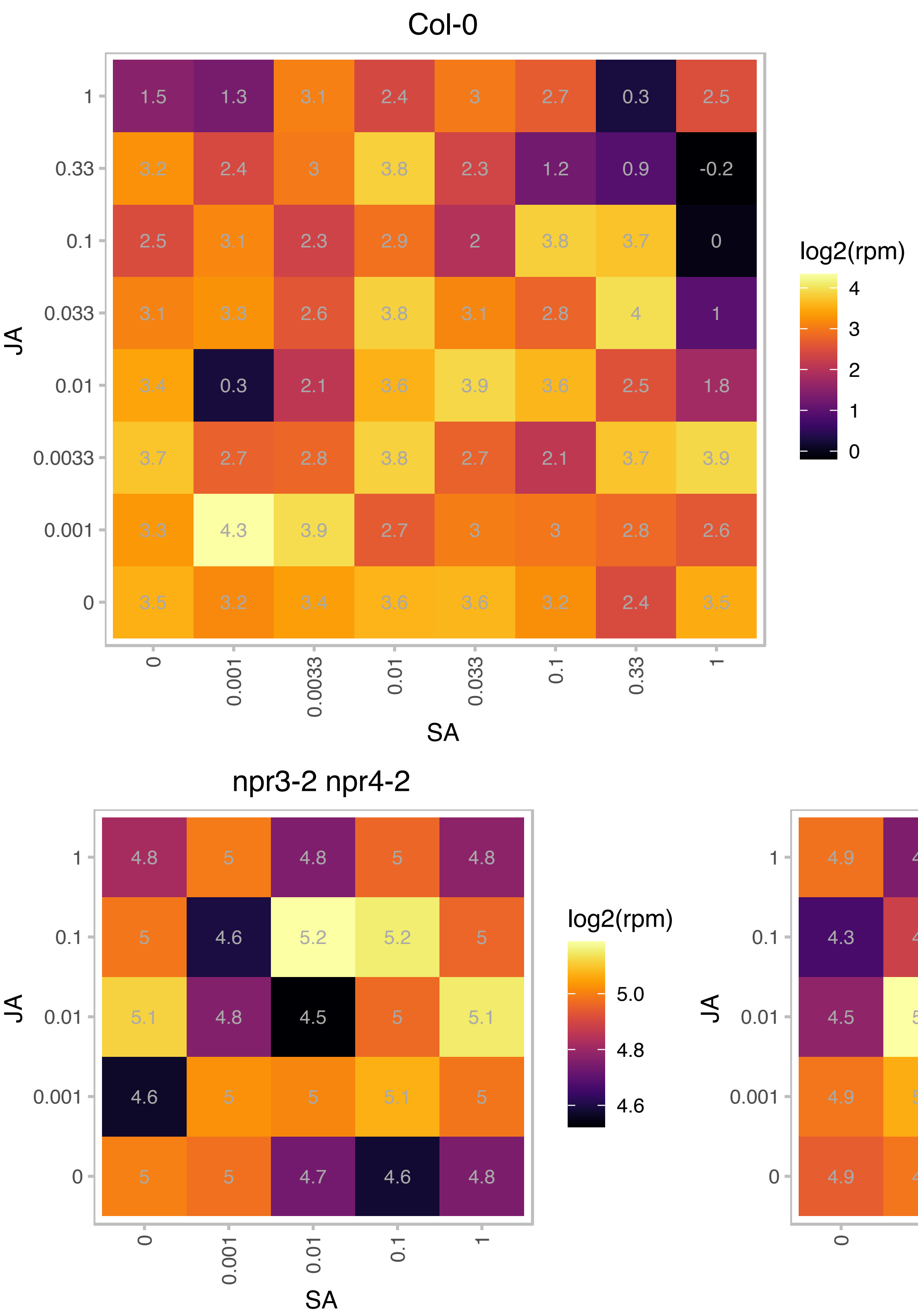


| | |
|---|---|
| Gene | AT2G19870 |
| is.expressed_genes | TRUE |
| is.responsive_genes | TRUE |
| is.apcluster | F-09 |

### AT2G21210.1 : SAUR6

SAUR-like auxin-responsive protein family

Putative auxin-regulated protein whose expression is downregulated in response to chitin oligomers.

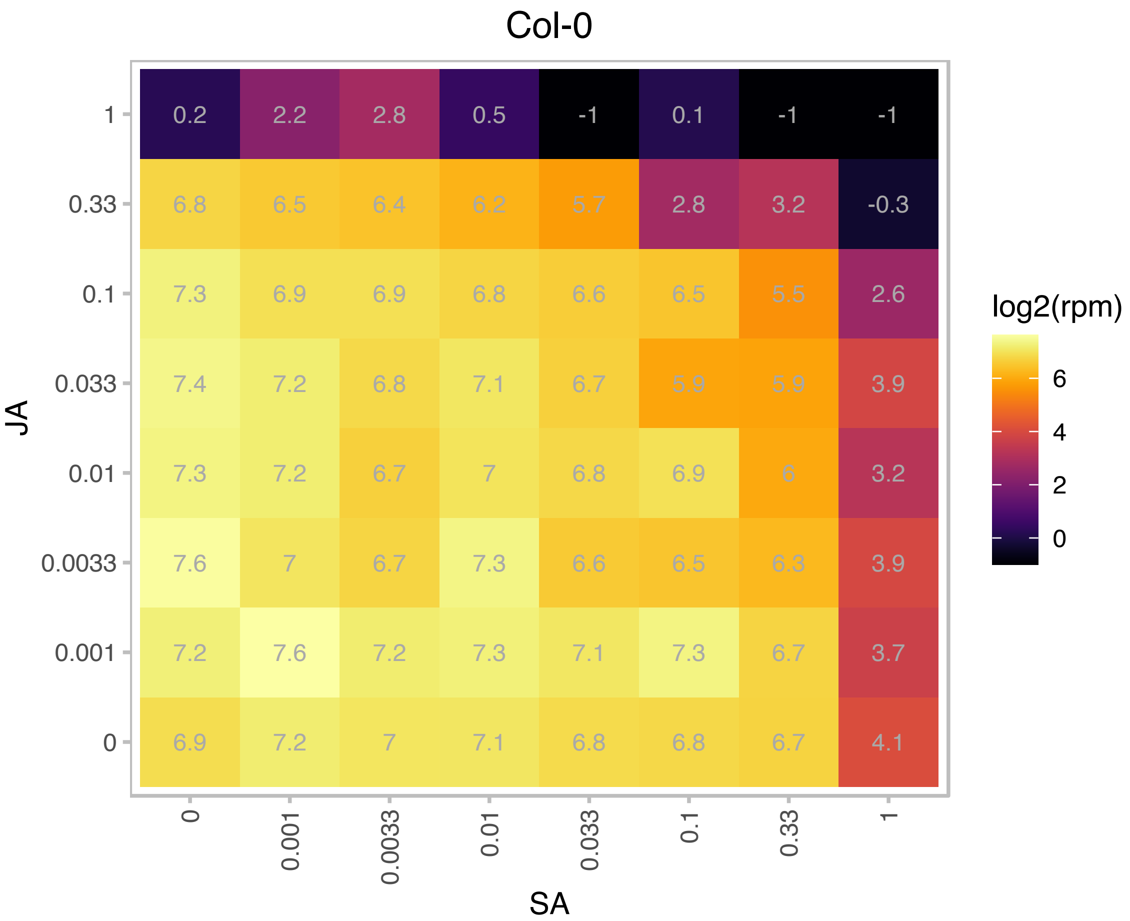


| | |
|---|---|
| Gene | AT2G21210 |
| is.expressed_genes | TRUE |
| is.responsive_genes | TRUE |
| is.apcluster | F-09 |

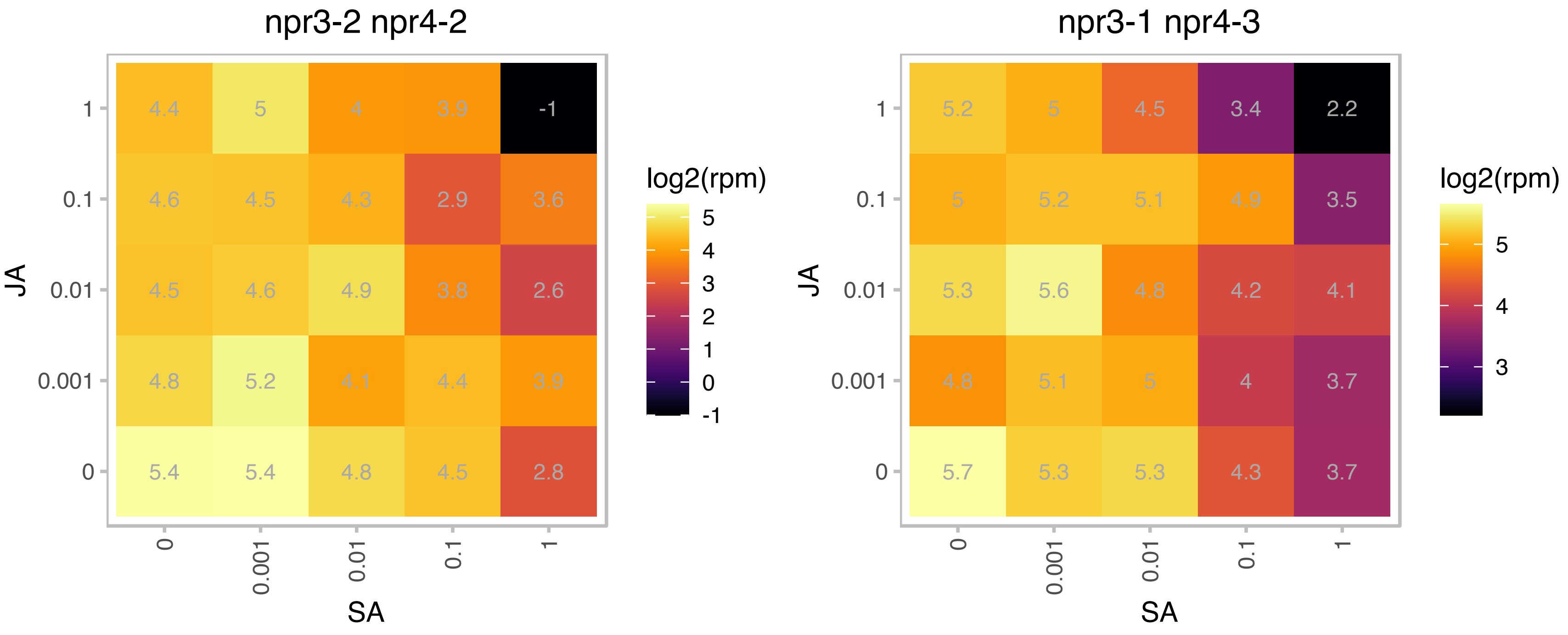

### AT2G21970.1 : SEP2

stress enhanced protein 2

stress enhanced protein 2 (SEP2) chlorophyll a/b-binding protein

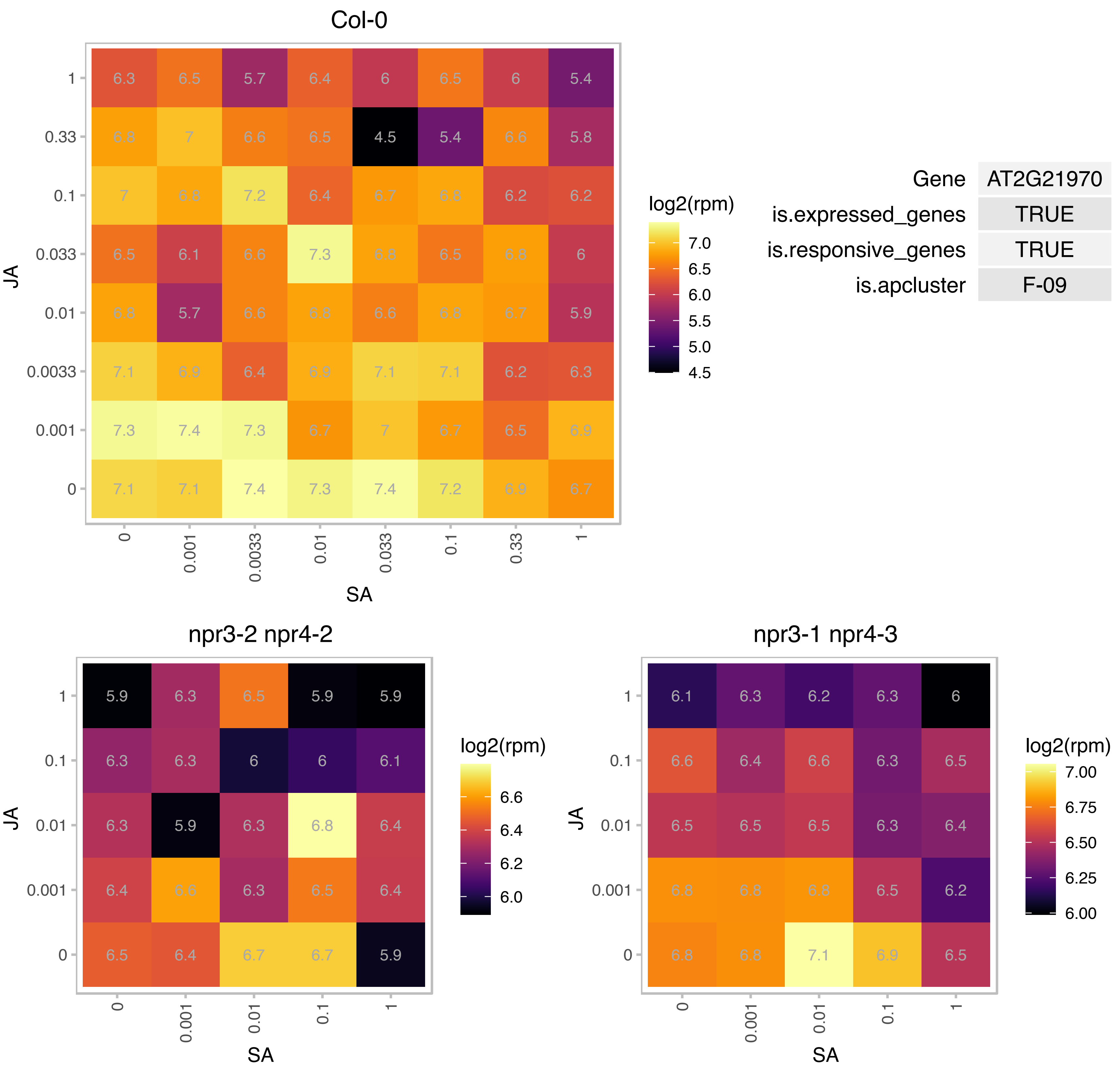


| | |
|---|---|
| Gene | AT2G21970 |
| is.expressed_genes | TRUE |
| is.responsive_genes | TRUE |
| is.apcluster | F-09 |

# AT2G22080.1 : NA

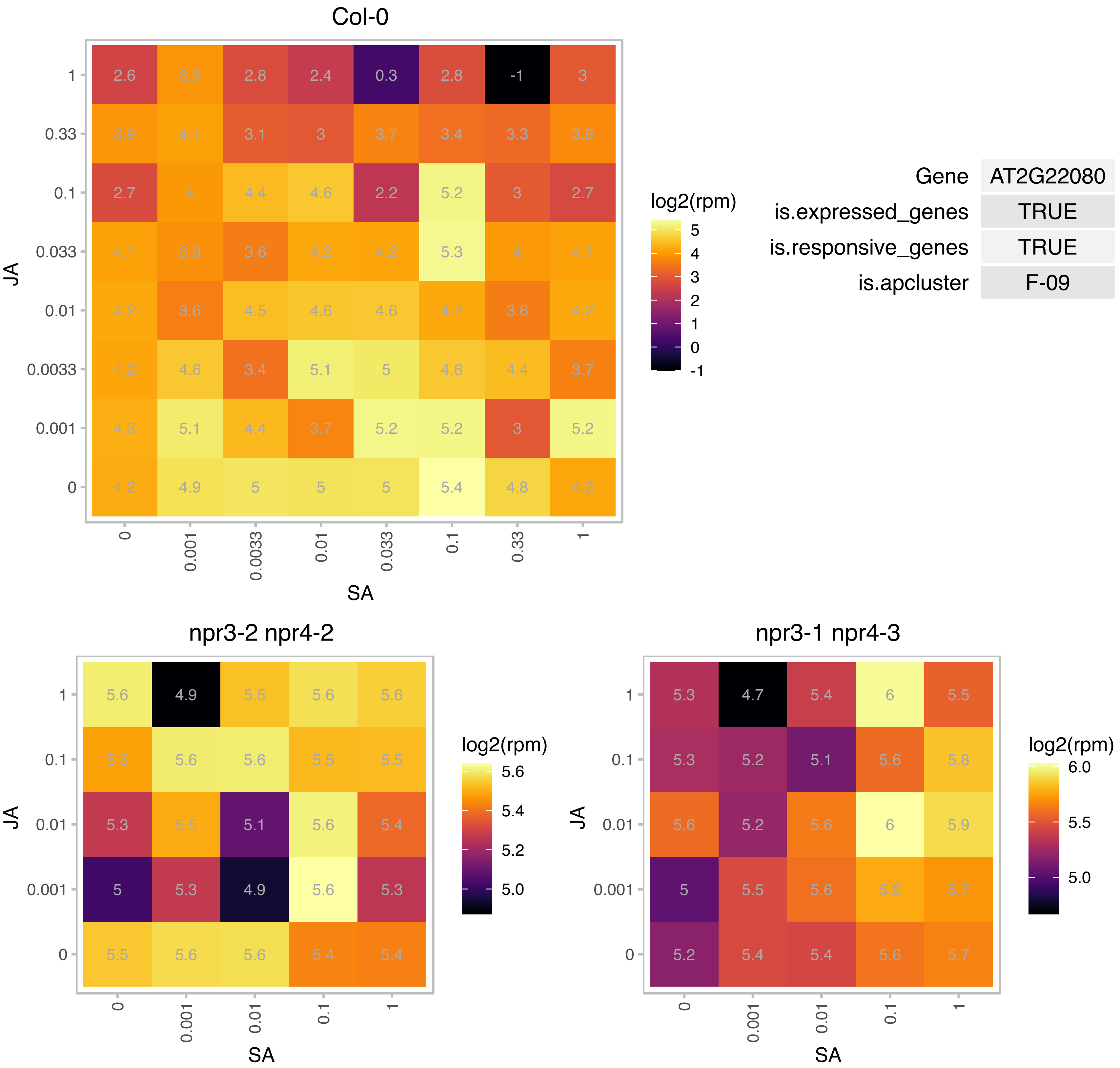


| | |
|---|---|
| Gene | AT2G22080 |
| is.expressed_genes | TRUE |
| is.responsive_genes | TRUE |
| is.apcluster | F-09 |

### AT2G26250.1 : FDH, KCS10

3-ketoacyl-CoA synthase 10

epidermis-specific, encodes KCS10, a putative 3-ketoacyl-CoA synthase. probably involved in the synthesis of long-chain lipids found in the cuticle.

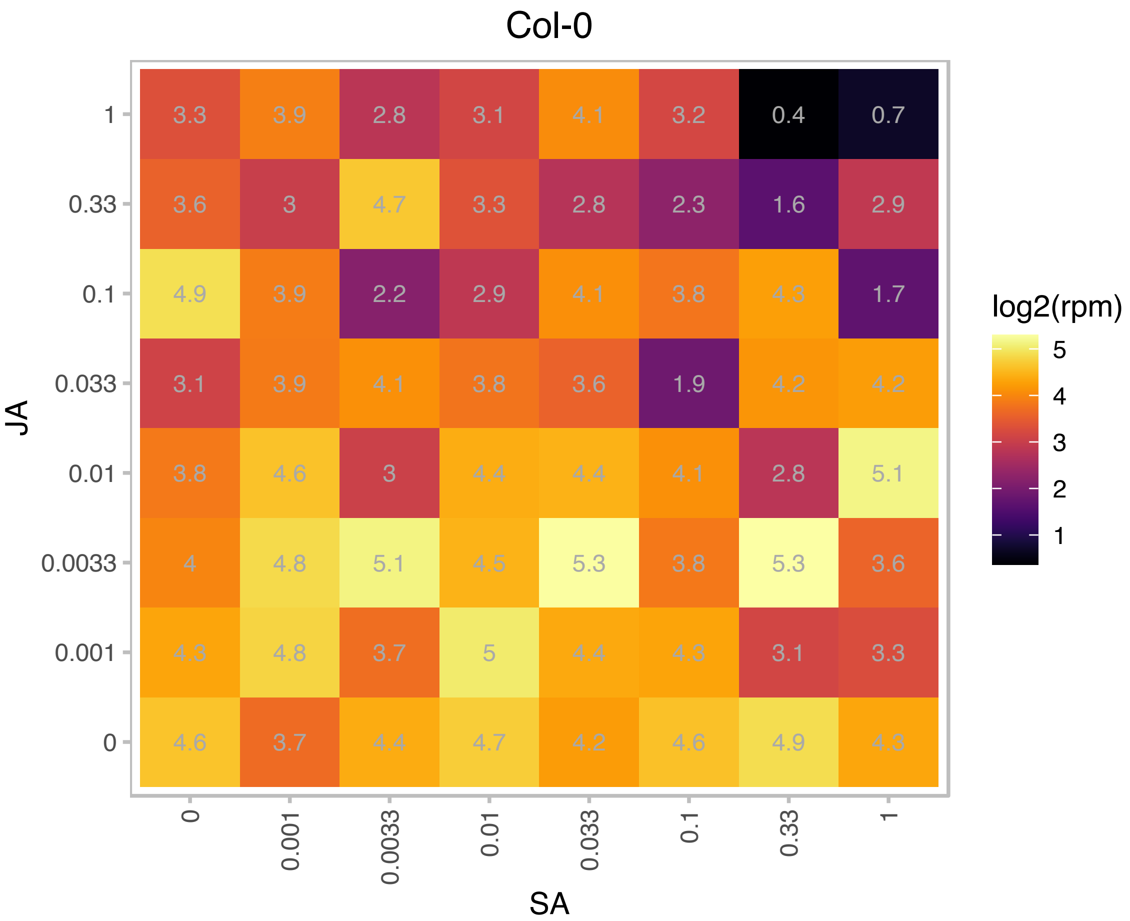


| | |
|---|---|
| Gene | AT2G26250 |
| is.expressed_genes | TRUE |
| is.responsive_genes | TRUE |
| is.apcluster | F-09 |

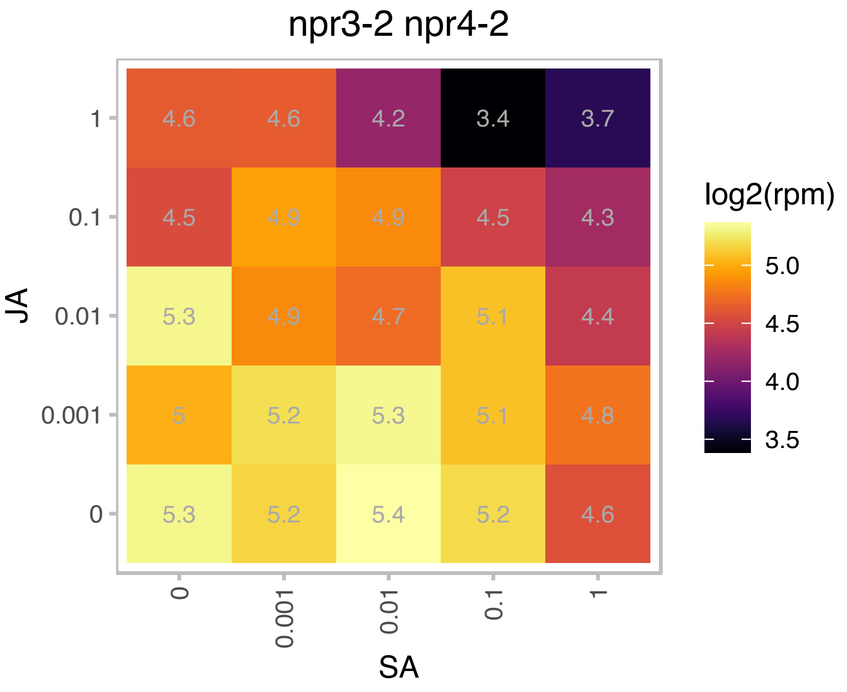


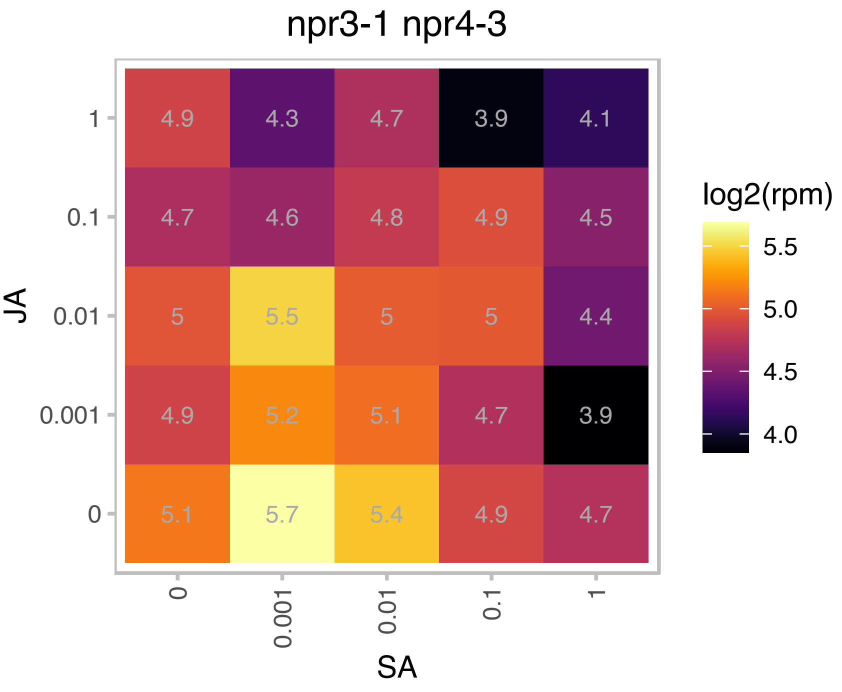

### AT2G31840.1 : MRL7-L

Thioredoxin superfamily protein

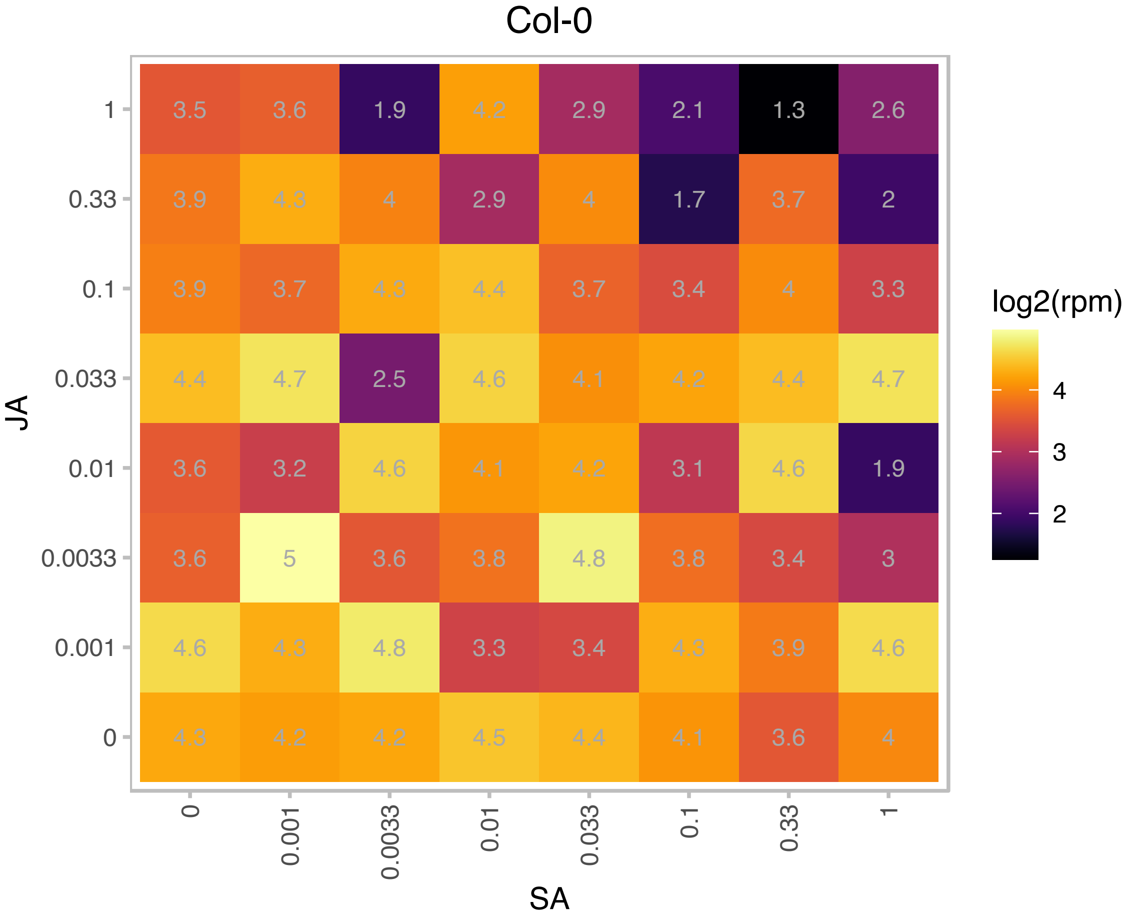


| | |
|---|---|
| Gene | AT2G31840 |
| is.expressed_genes | TRUE |
| is.responsive_genes | TRUE |
| is.apcluster | F-09 |

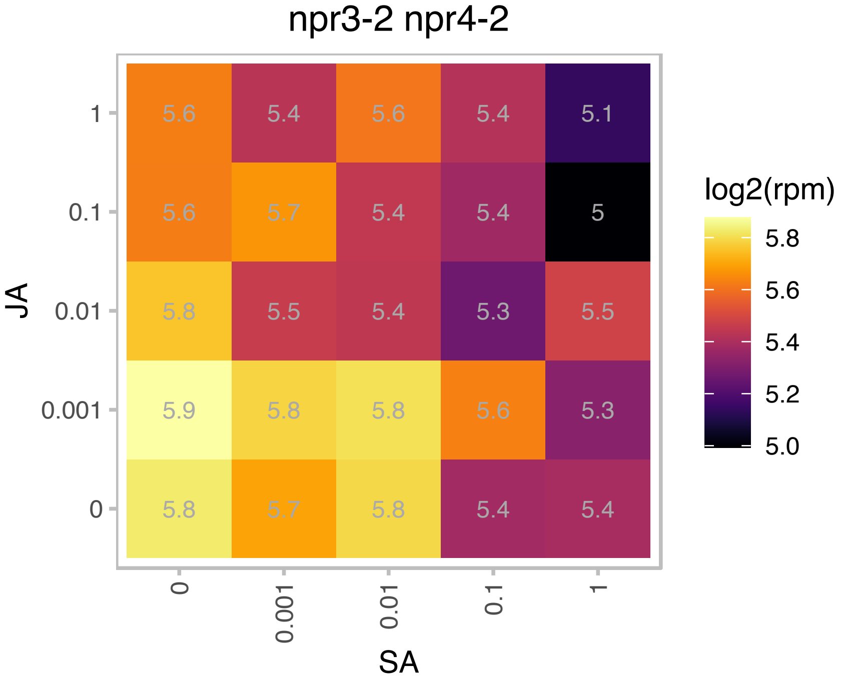


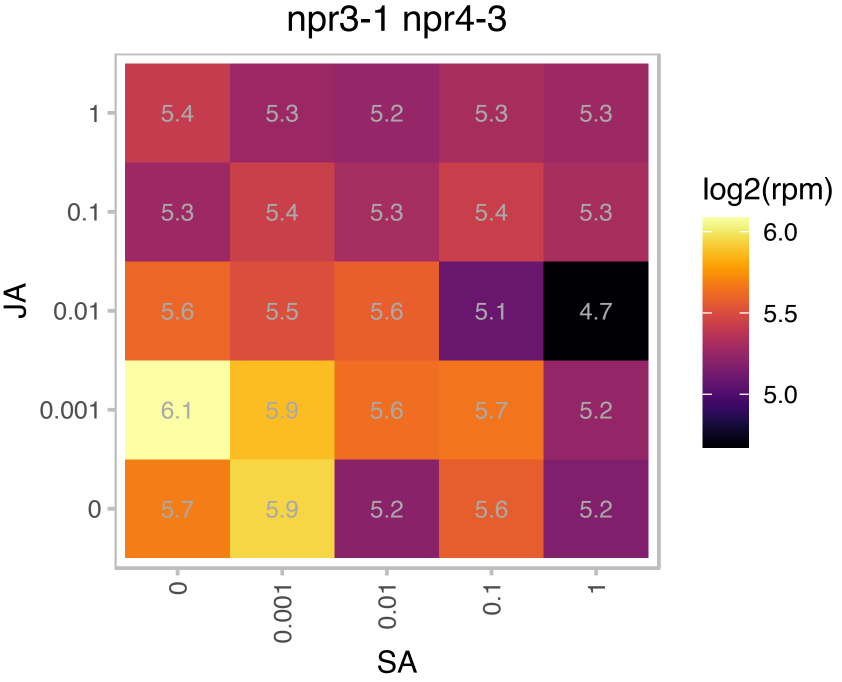

# AT2G37120.1 : NA

S1FA-like DNA-binding protein

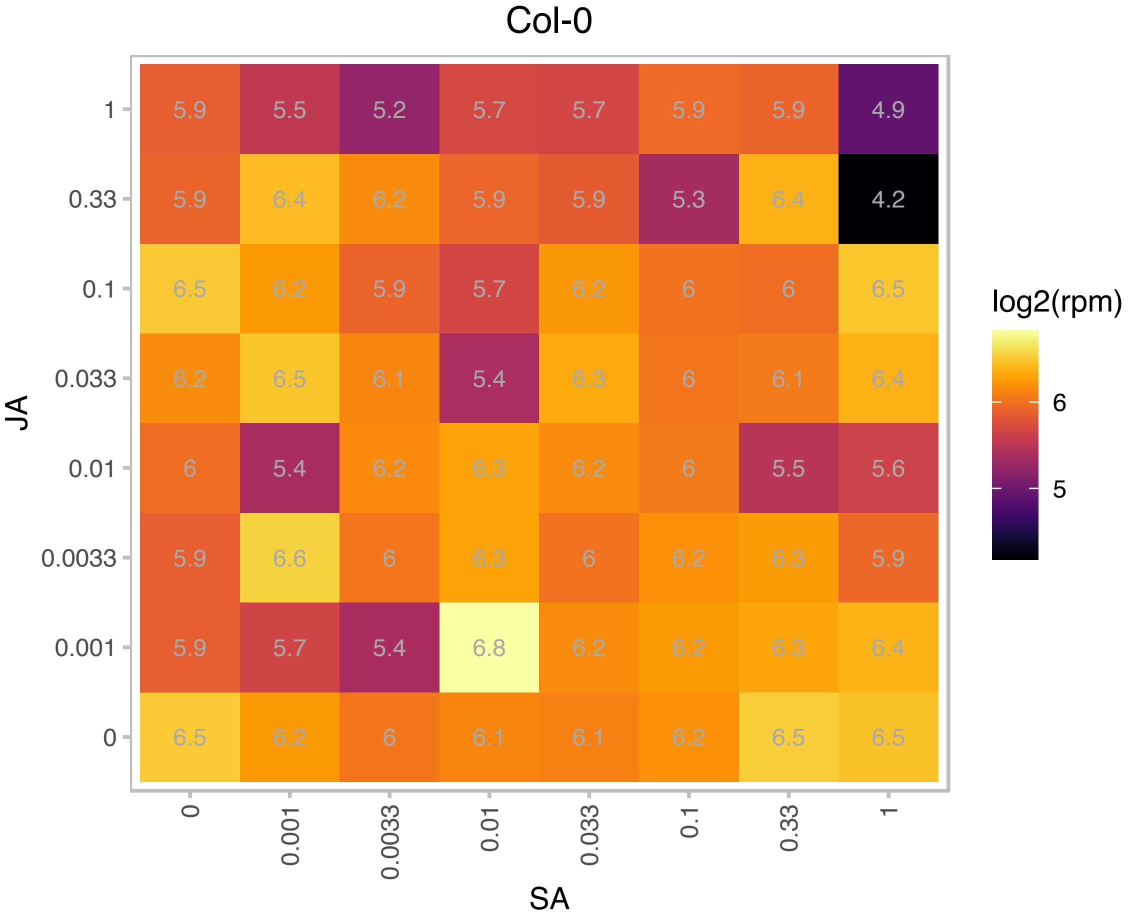


| | |
|---|---|
| Gene | AT2G37120 |
| is.expressed_genes | TRUE |
| is.responsive_genes | TRUE |
| is.apcluster | F-09 |

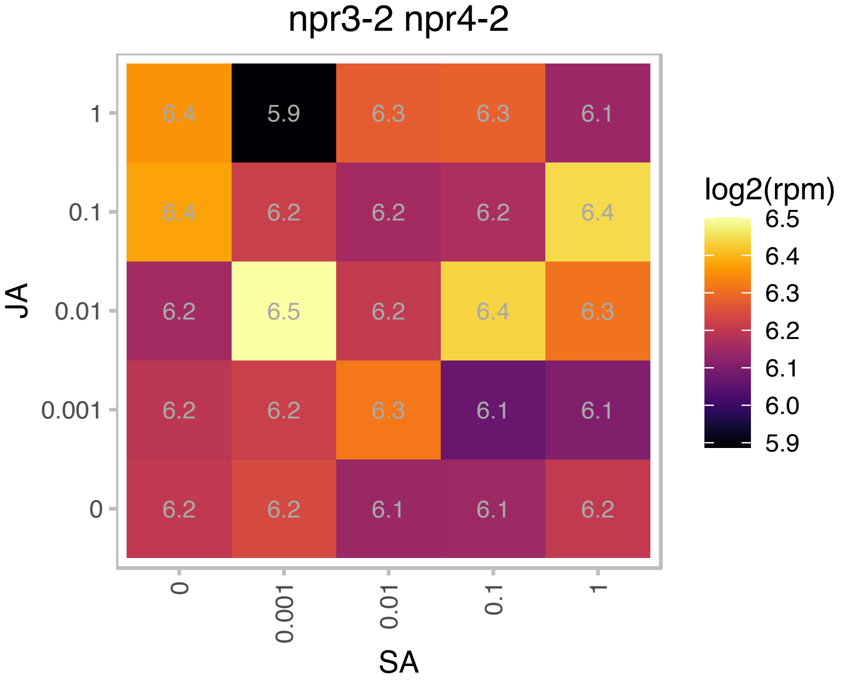


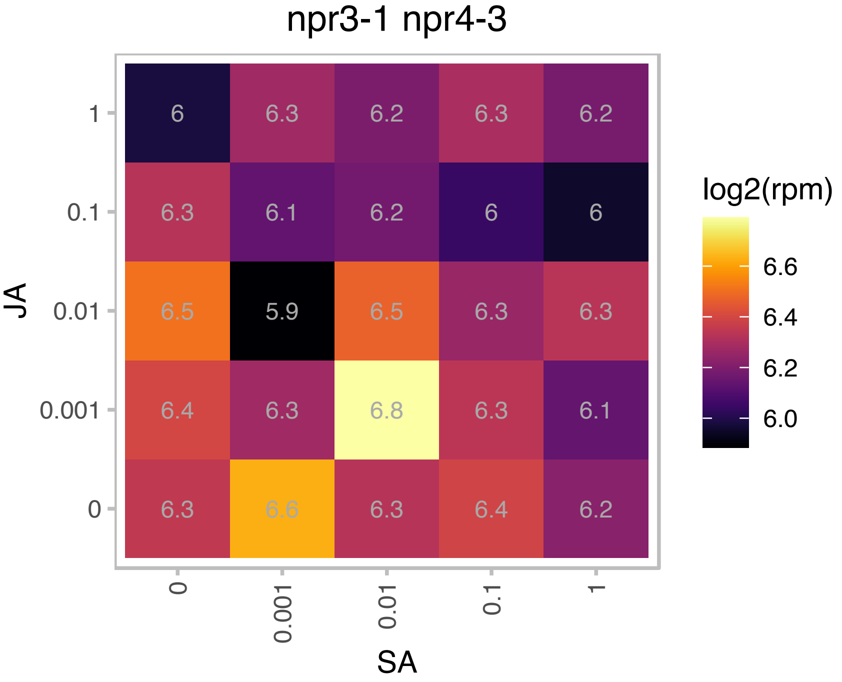

### AT2G38540.1 : ATLTP1, AtLtpI-4, LP1, LTP1

lipid transfer protein 1

Non-specific lipid transfer protein. Binds calmodulin in a Ca2+-independent manner. Localized to the cell wall. Specifically expressed in L1 epidermal layer. Predicted to be a member of PR-14 pathogenesis-related protein family with the following members: At2g38540/LTP1, At2g38530/LTP2, At5g59320/LTP3, At5g59310/LTP4, At3g51600/LTP5, At3g08770/LTP6, At2g15050/LTP7, At2g18370/LTP8, At2g15325/LTP9, At5g01870/LTP10, At4g33355/LTP11, At3g51590/LTP12, At5g44265/LTP13, At5g62065/LTP14, At4g08530/LTP15. The mRNA is cell-to mobile.

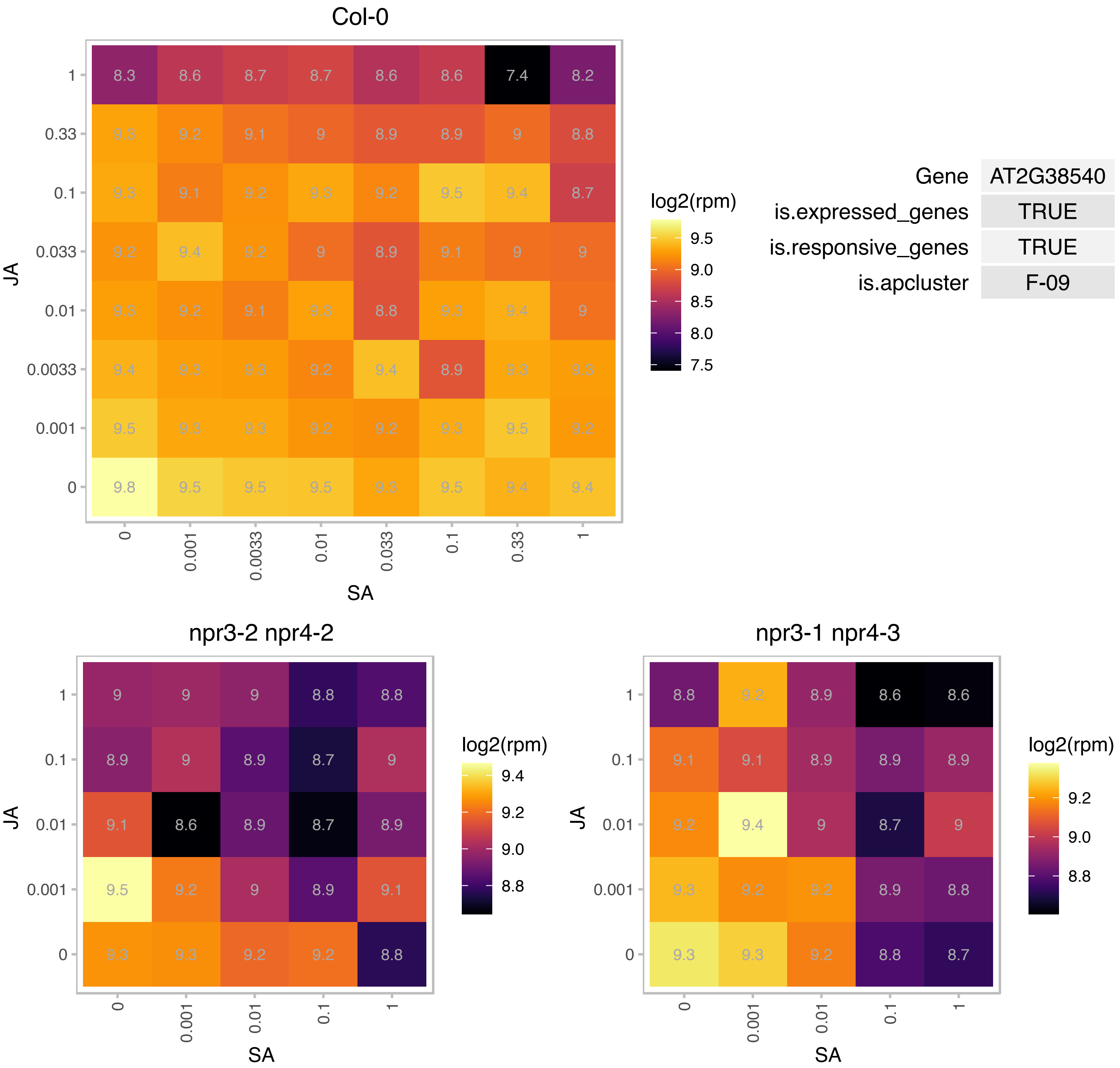


| | |
|---|---|
| Gene | AT2G38540 |
| is.expressed_genes | TRUE |
| is.responsive_genes | TRUE |
| is.apcluster | F-09 |

# AT2G40510.1 : NA

Ribosomal protein S26e family protein

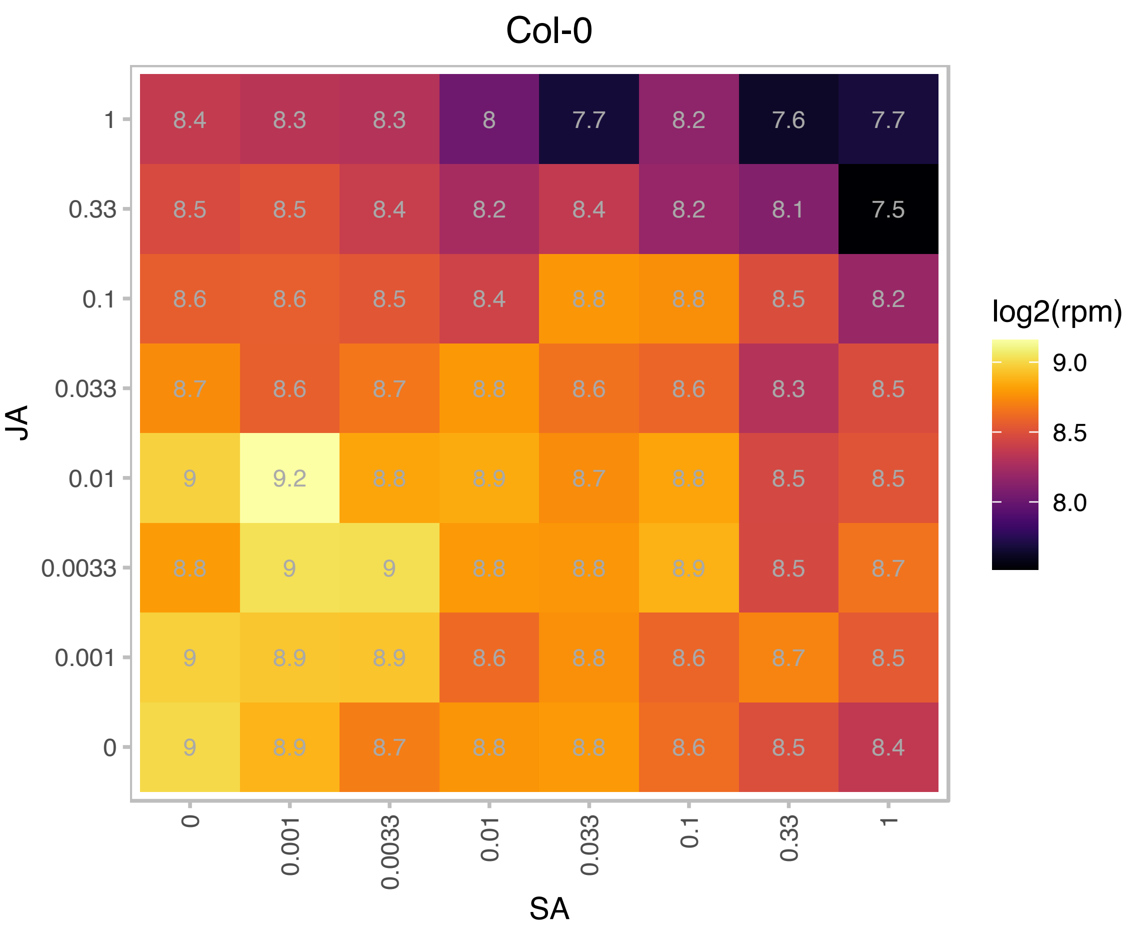


| | |
|---|---|
| Gene | AT2G40510 |
| is.expressed_genes | TRUE |
| is.responsive_genes | TRUE |
| is.apcluster | F-09 |

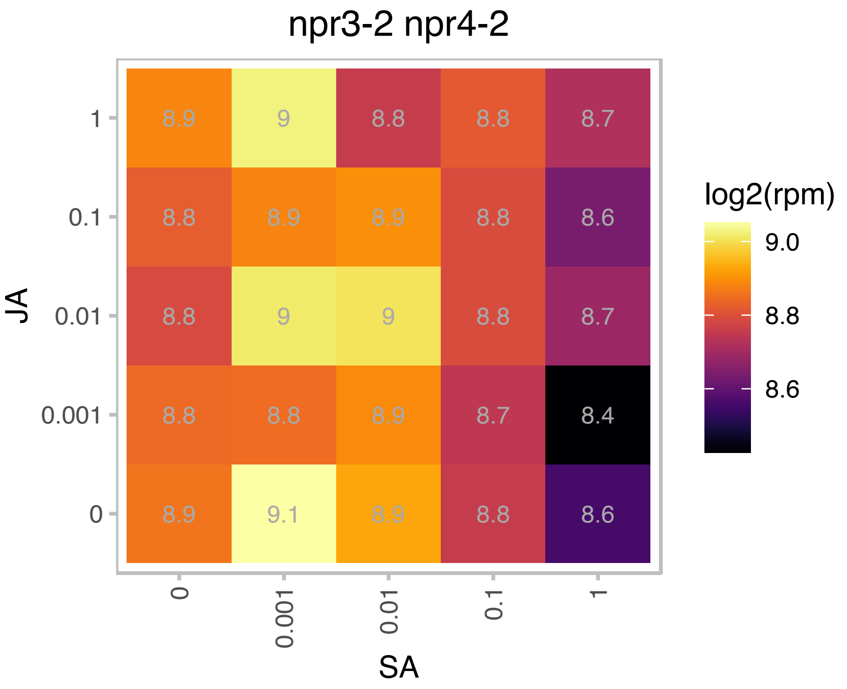


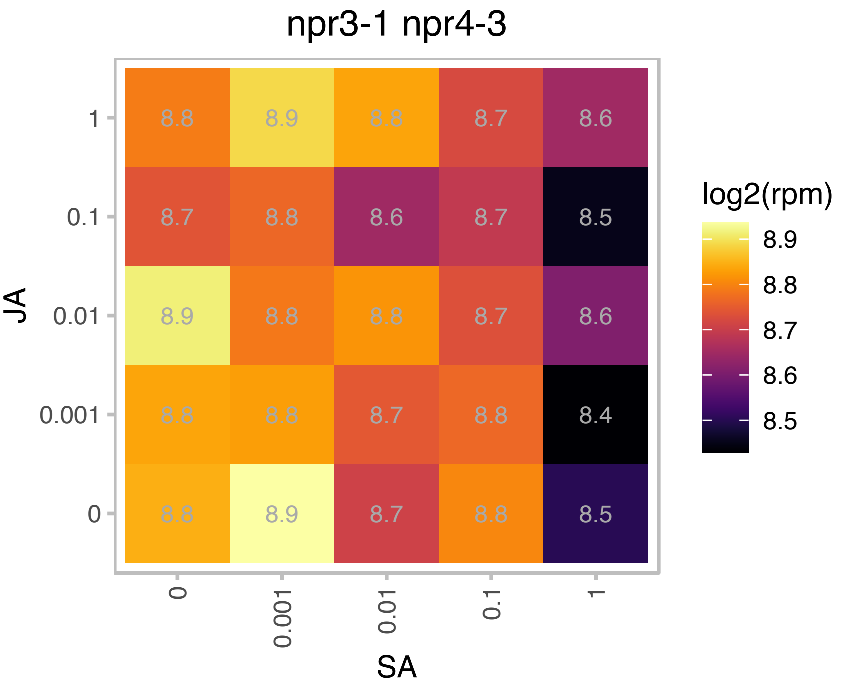

# AT2G43770.1 : NA

Transducin/WD40 repeat-like superfamily protein

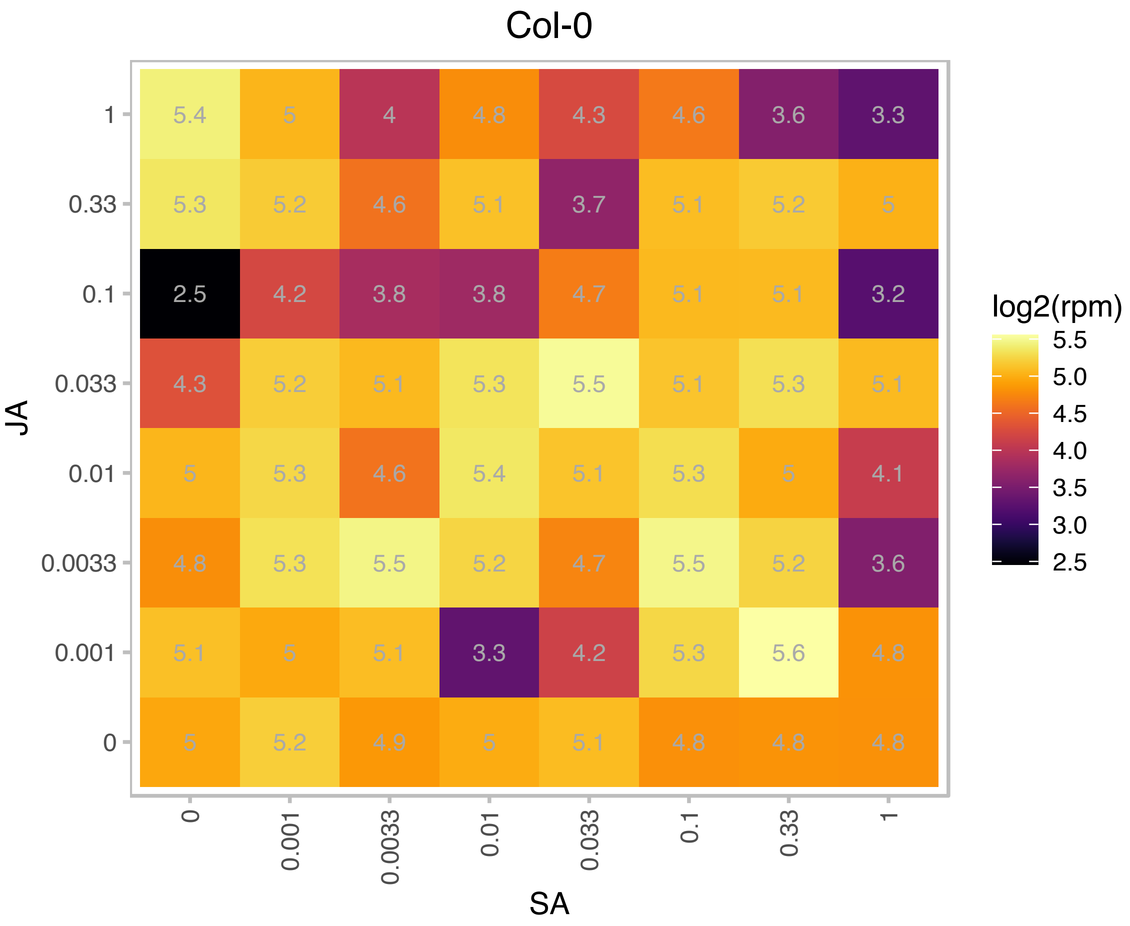


| | |
|---|---|
| Gene | AT2G43770 |
| is.expressed_genes | TRUE |
| is.responsive_genes | TRUE |
| is.apcluster | F-09 |

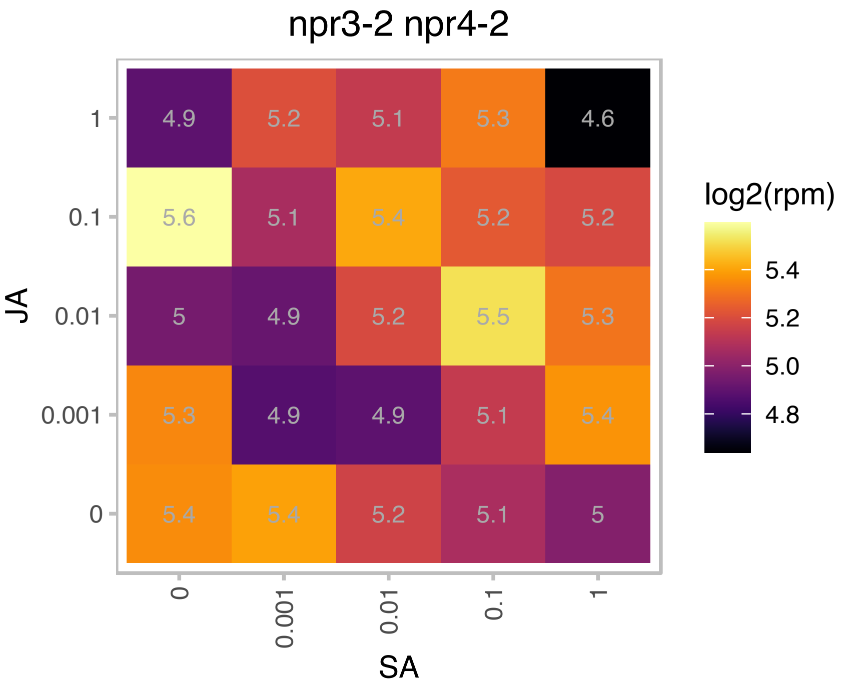


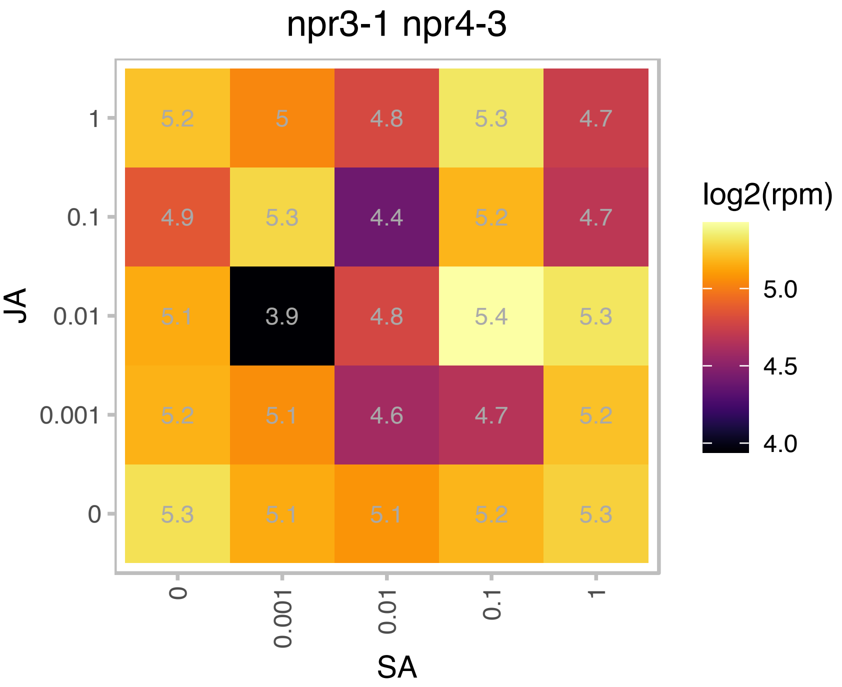

### AT2G47840.1 : AtTic20-II, Tic20-II

Uncharacterised conserved protein ycf60

Encodes a component of the TIC (translocon at the inner envelope membrane of chloroplasts) protein translocation machinery mediating the protein translocation across the inner envelope of plastids. The Arabidopsis genome encodes four Tic20 homologous proteins, AT1G04940(Tic20-I), AT2G47840(Tic20-II), AT4G03320(Tic20-IV) and AT5G55710(Tic20-V).

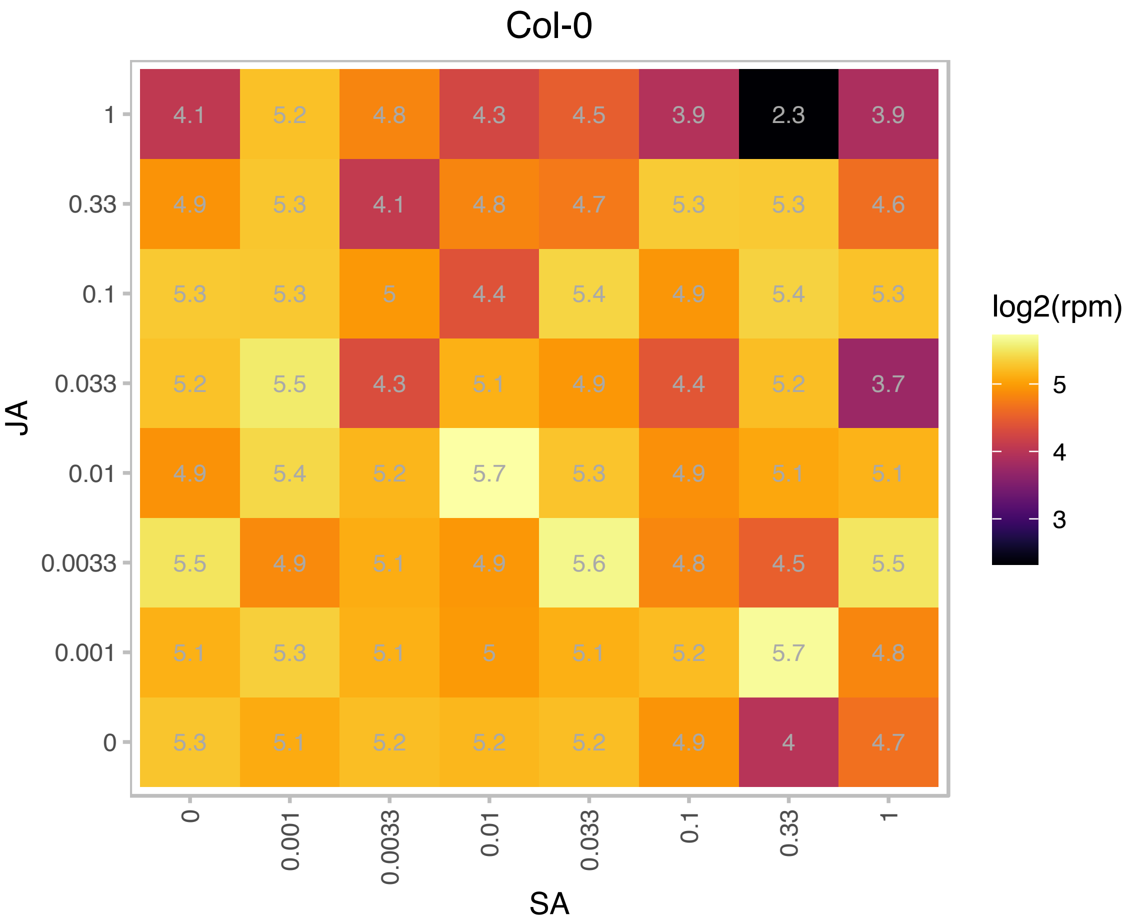


| | |
|---|---|
| Gene | AT2G47840 |
| is.expressed_genes | TRUE |
| is.responsive_genes | TRUE |
| is.apcluster | F-09 |

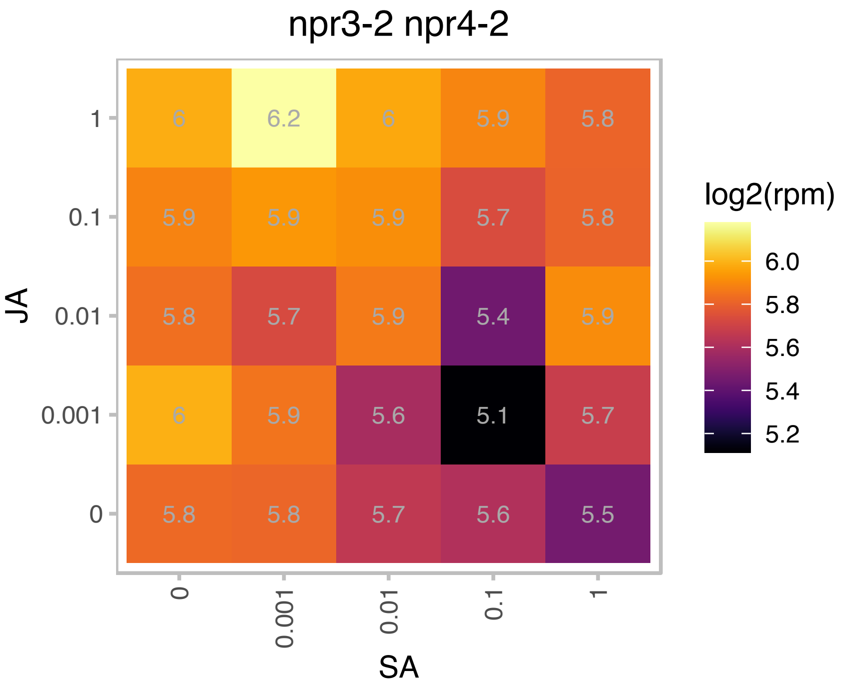


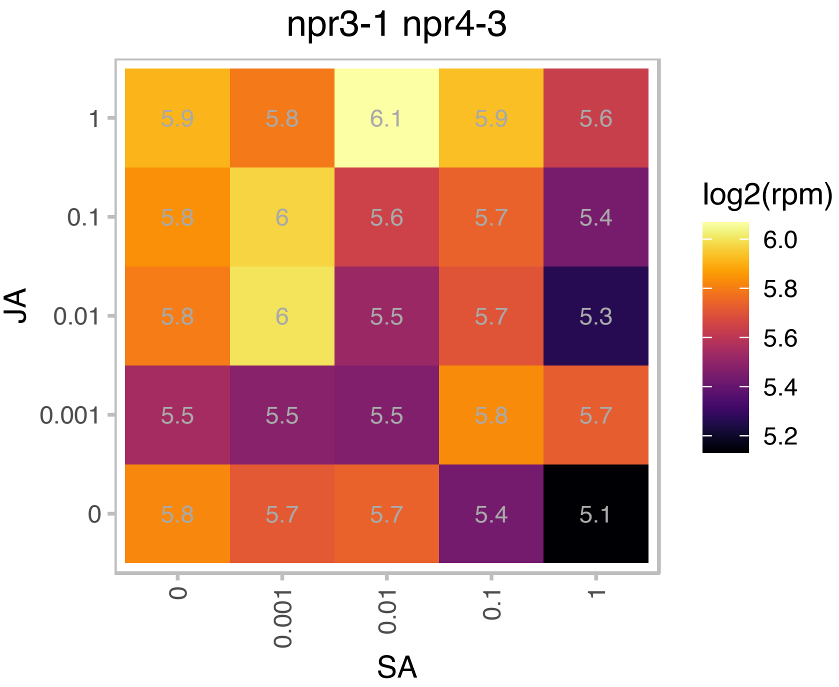

### AT3G02350.1 : GAUT9

galacturonosyltransferase 9

Encodes a protein with putative galacturonosyltransferase activity. The mRNA is cell-to-cell mobile.

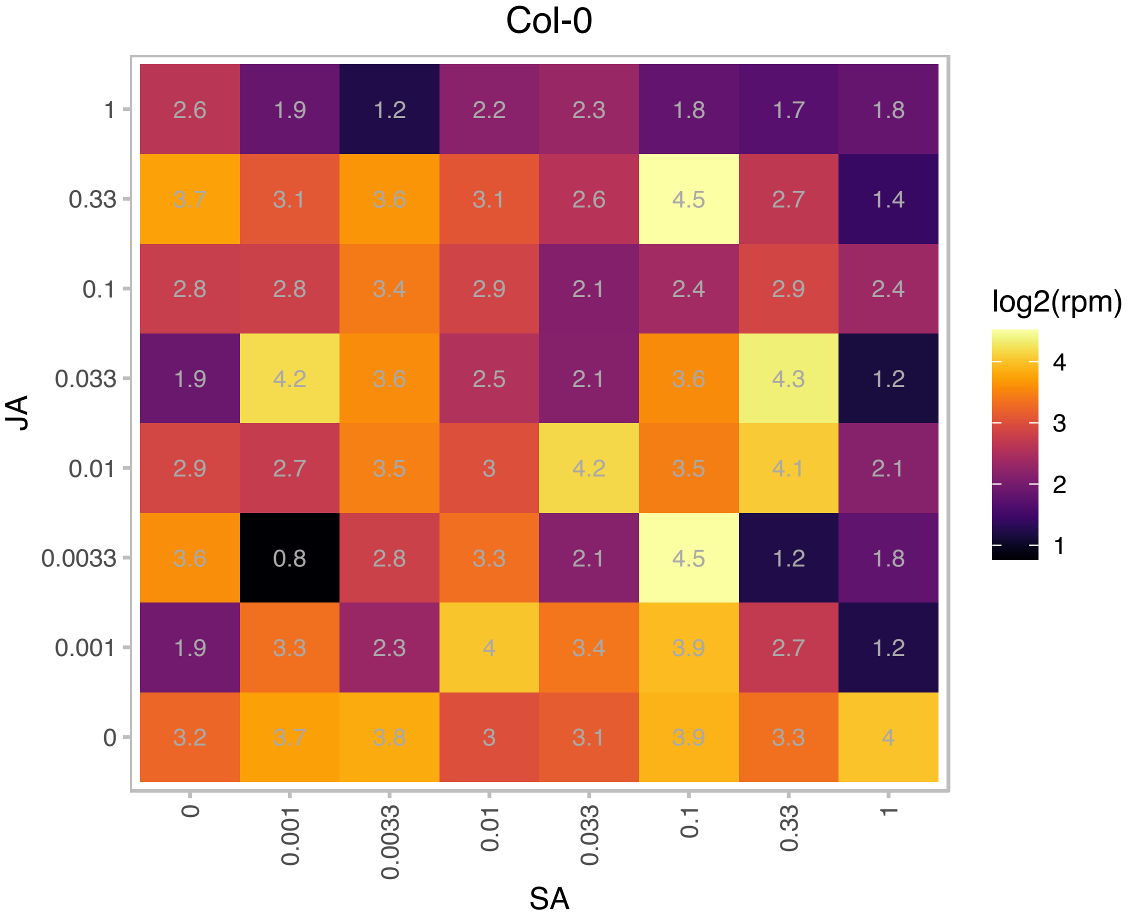


| | |
|---|---|
| Gene | AT3G02350 |
| is.expressed_genes | TRUE |
| is.responsive_genes | TRUE |
| is.apcluster | F-09 |

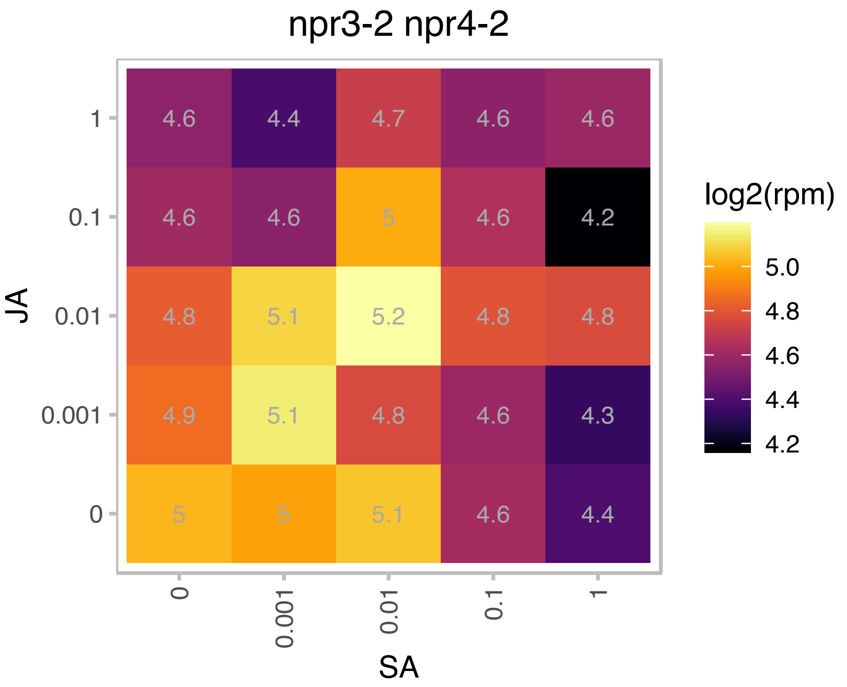


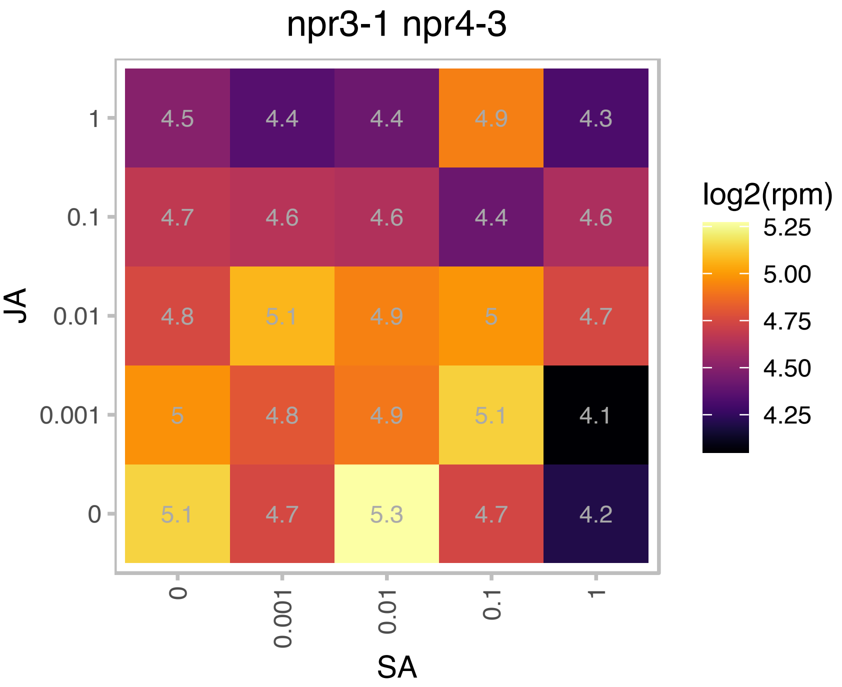

# AT3G07440.1 : NA

| | |
|---|---|
| Gene | AT3G07440 |
| is.expressed_genes | TRUE |
| is.responsive_genes | TRUE |
| is.apcluster | F-09 |

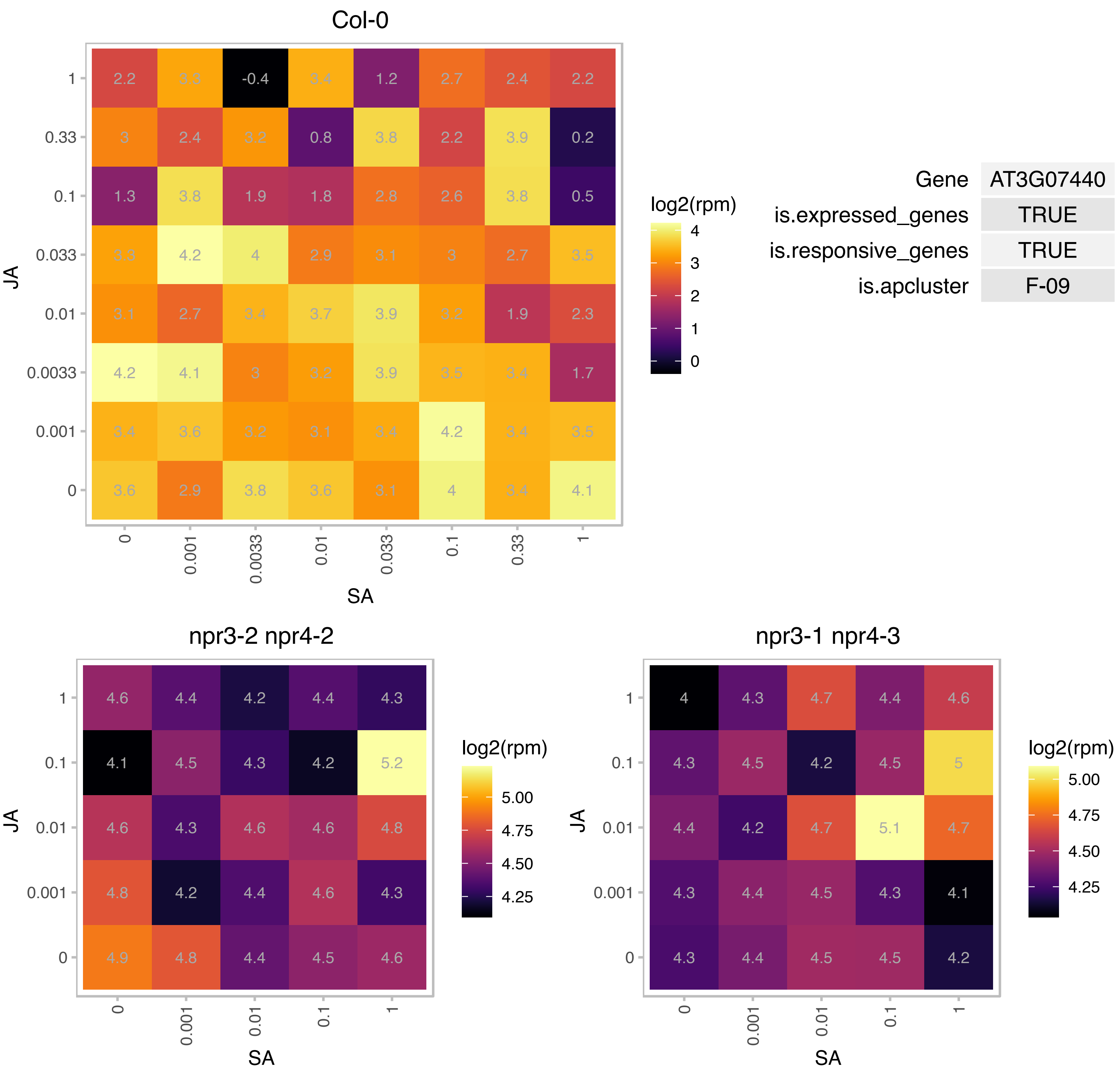

### AT3G08940.1 : LHCB4.2

light harvesting complex photosystem II

Lhcb4.2 protein (Lhcb4.2, protein involved in the light harvesting complex of photosystem II The mRNA is cell-to-cell mobile.

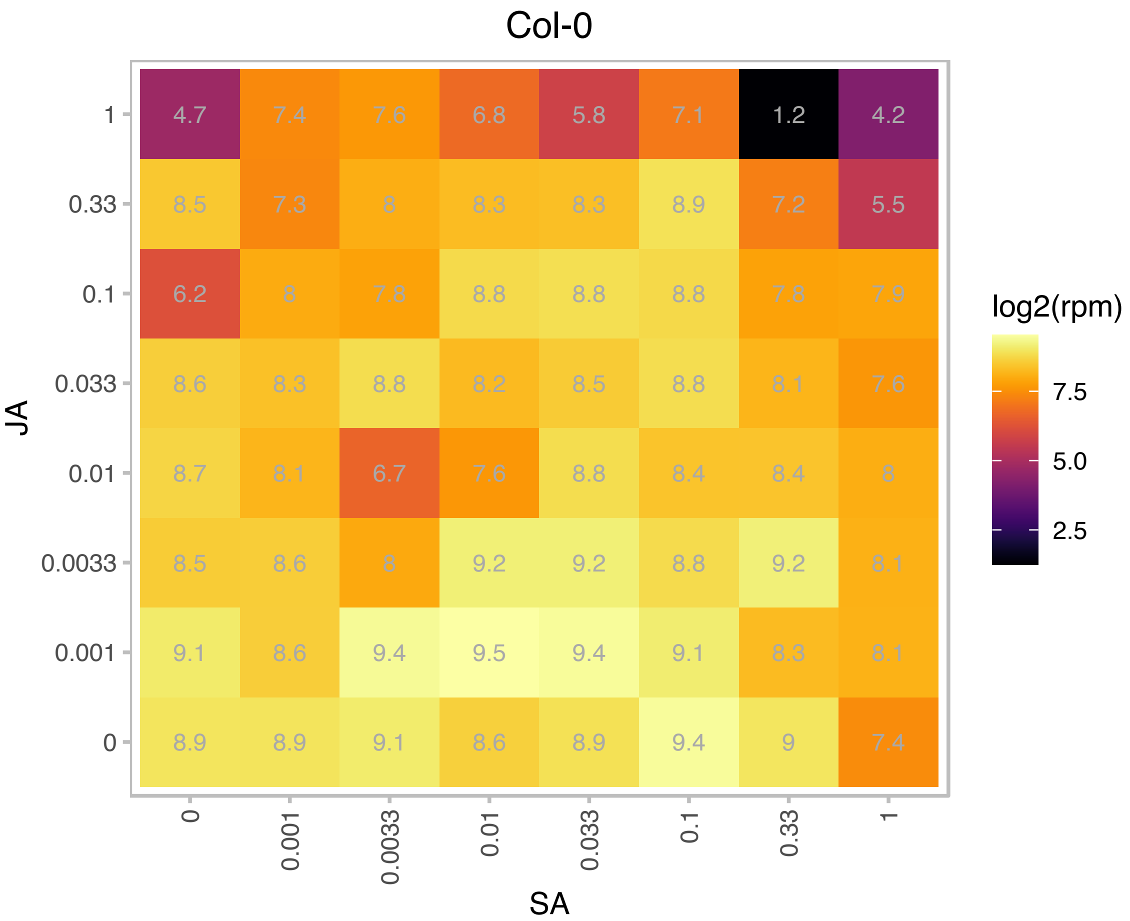


| | |
|---|---|
| Gene | AT3G08940 |
| is.expressed_genes | TRUE |
| is.responsive_genes | TRUE |
| is.apcluster | F-09 |

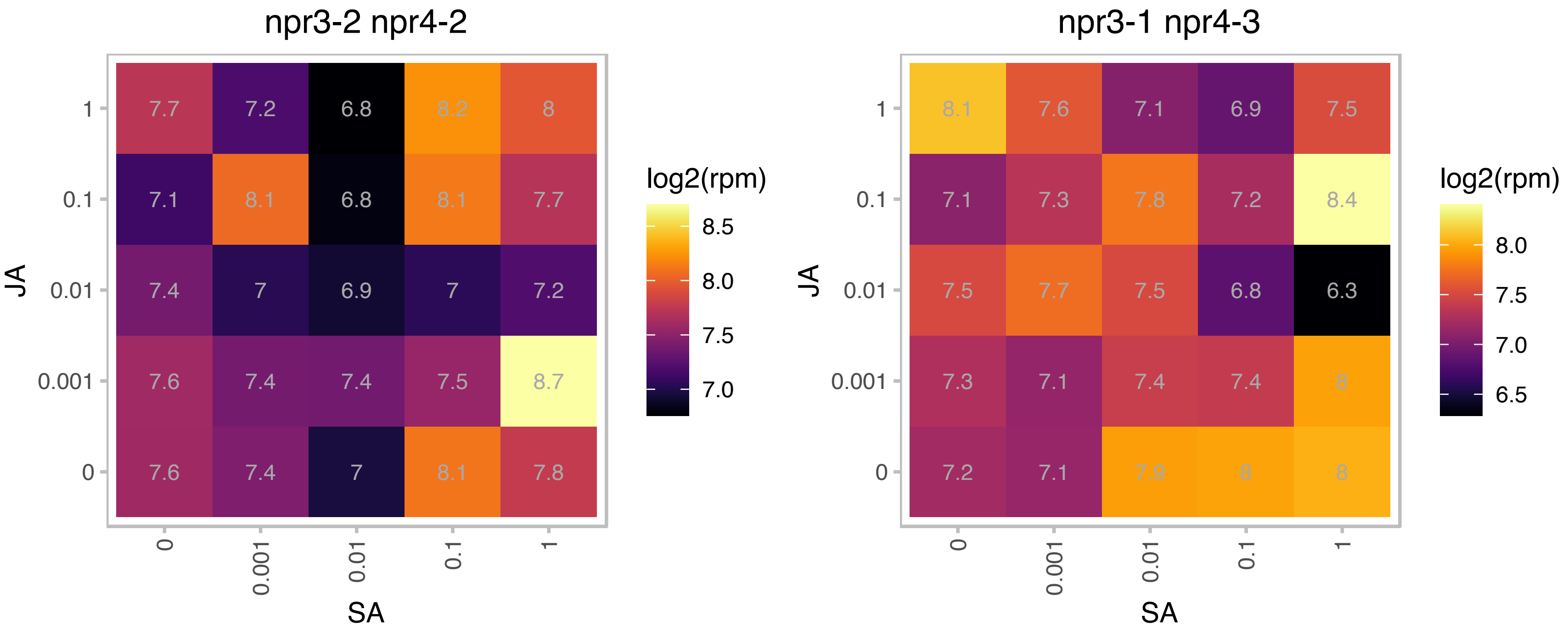

### AT3G14110.1 : FLU

Tetratricopeptide repeat (TPR)-like superfamily protein

Encodes a novel coiled-coil, TPR domain containing protein that is localized to the chloroplast membrane and is involved in chlorophyll biosynthesis. Mutants accumulate protochlorophyllide, an intermediate in the chlorophyll biosynthesis pathway, in dark and release singlet oxygen in plastids in a controlled and non-invasive manner upon a dark/light shift.

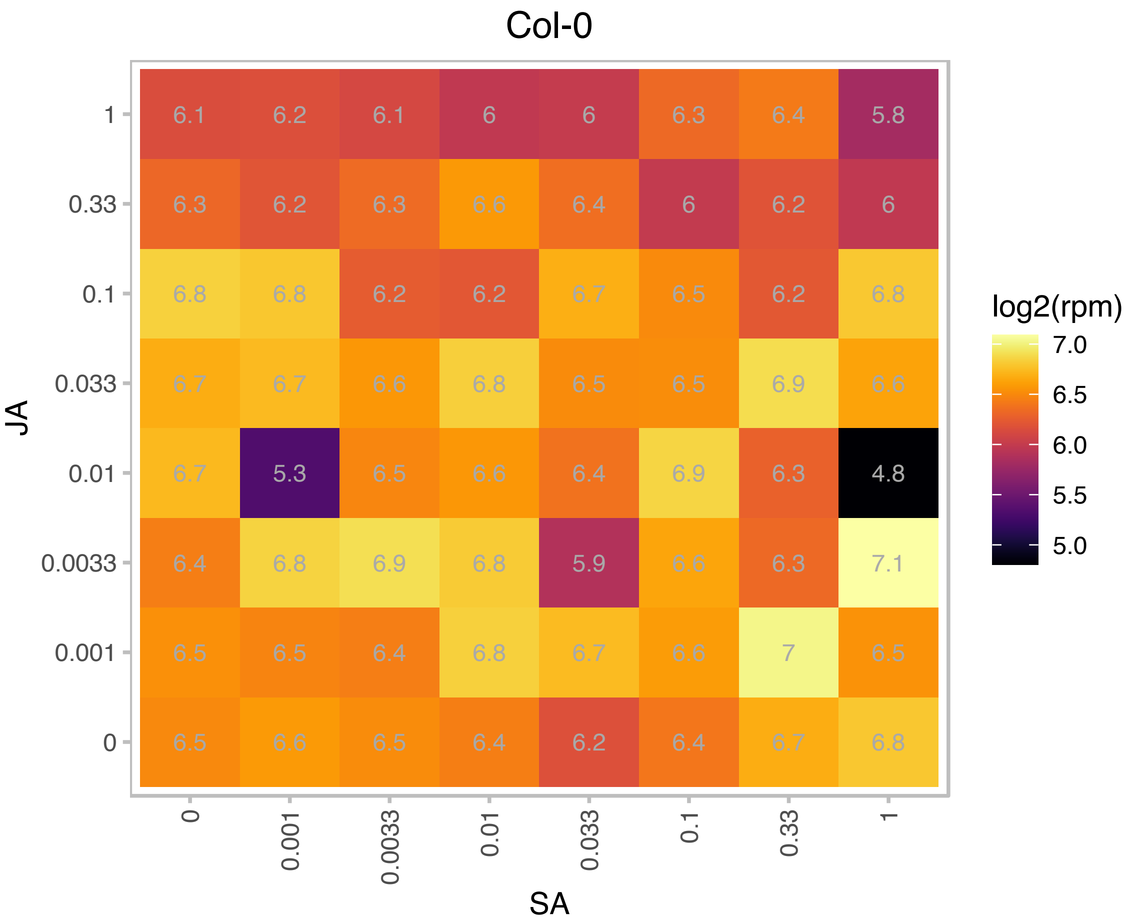


| | |
|---|---|
| Gene | AT3G14110 |
| is.expressed_genes | TRUE |
| is.responsive_genes | TRUE |
| is.apcluster | F-09 |

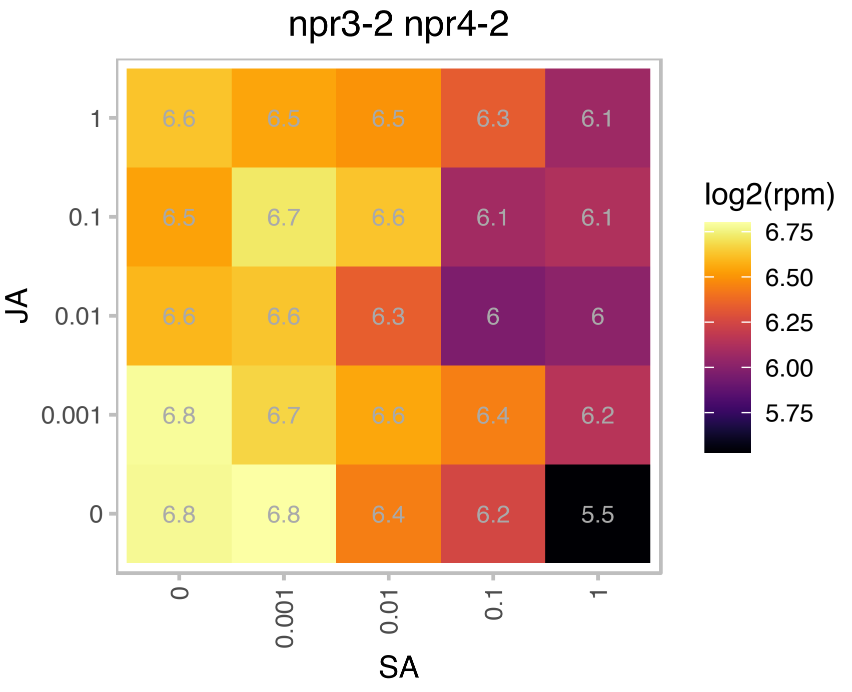


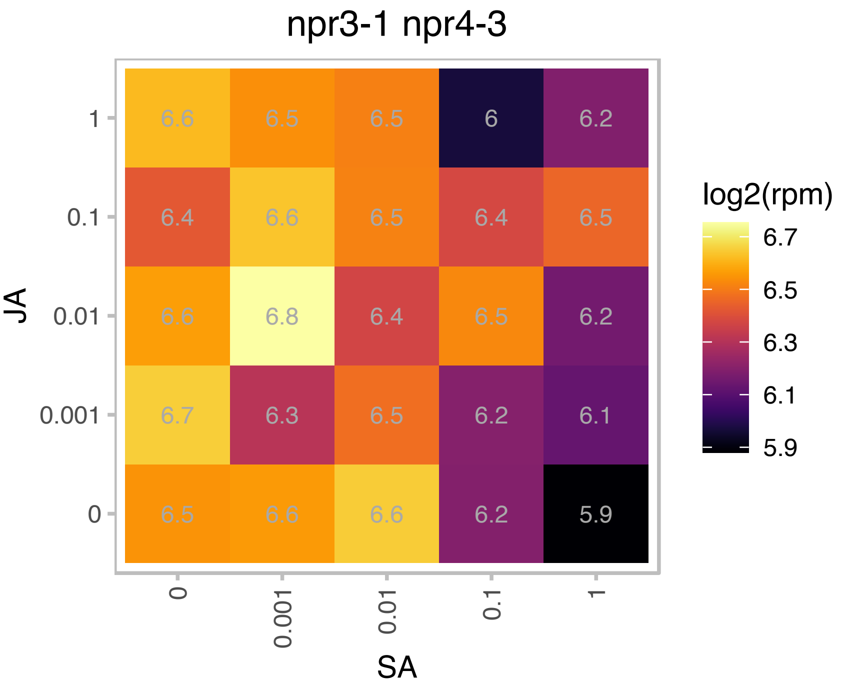

### AT3G14210.1 : ESM1

epithiospecifier modifier 1
A semidominant QTL which has an epistatic effect on the Epithiospecifier gene. Represses nitrile formation and favors isothiocyanate production during glucosinolate hydrolysis. The functional allele deters the insect herbivory T. ni.

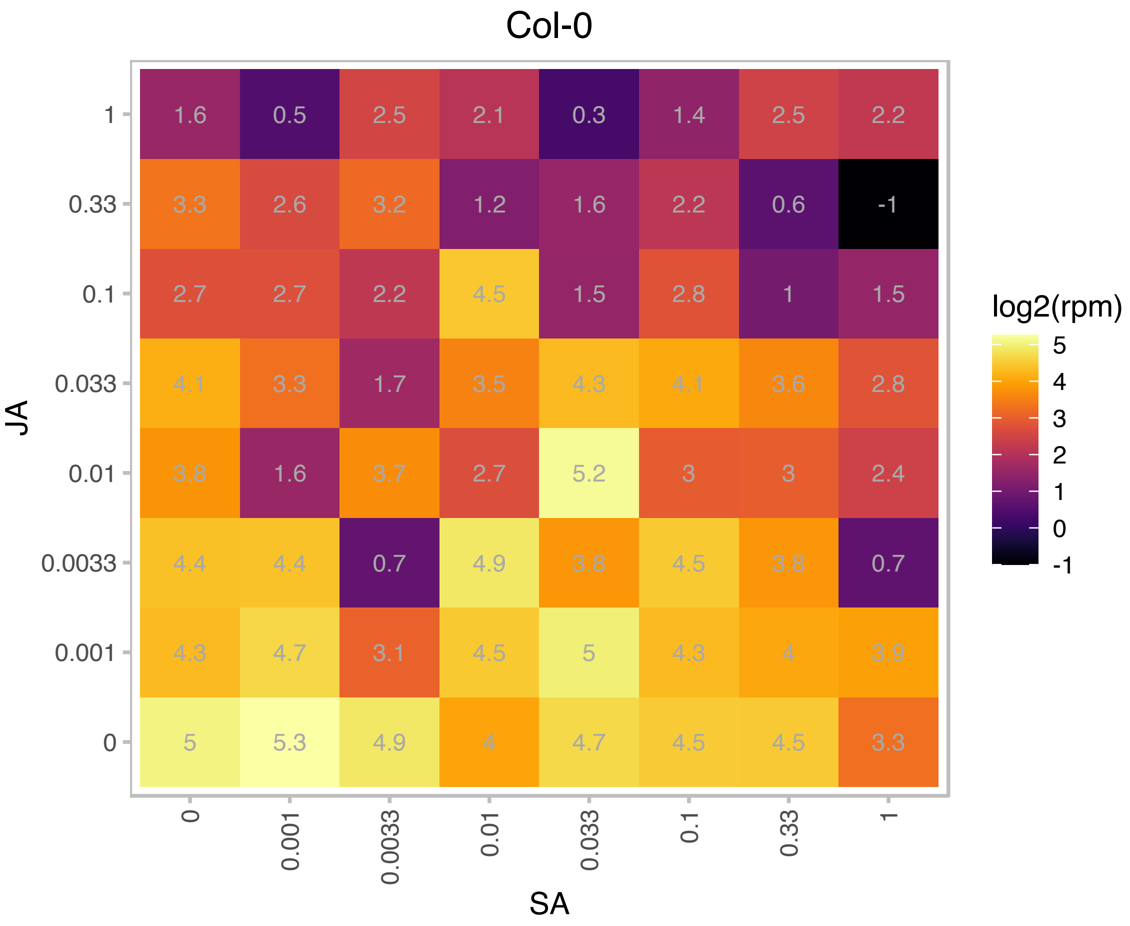


| | |
|---|---|
| Gene | AT3G14210 |
| is.expressed_genes | TRUE |
| is.responsive_genes | TRUE |
| is.apcluster | F-09 |

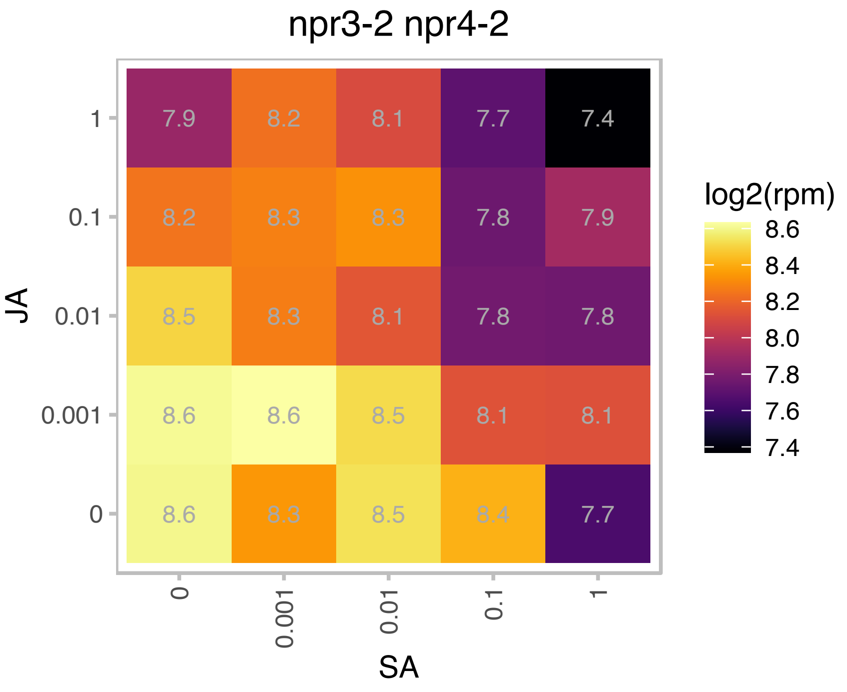


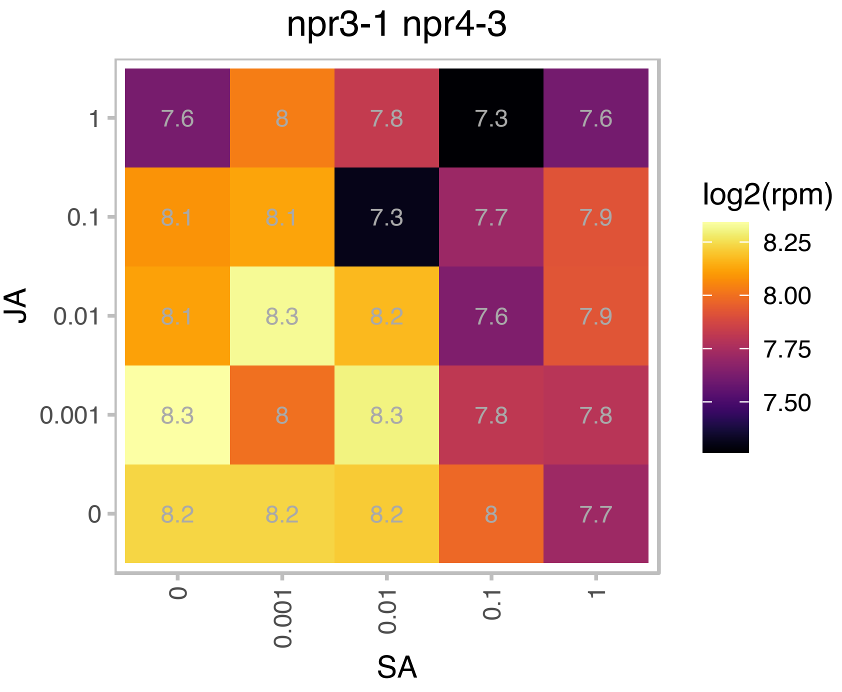

### AT3G17210.1 : ATHS1, HS1

heat stable protein 1

Encodes a heat stable protein with antimicrobial and antifungal activity.

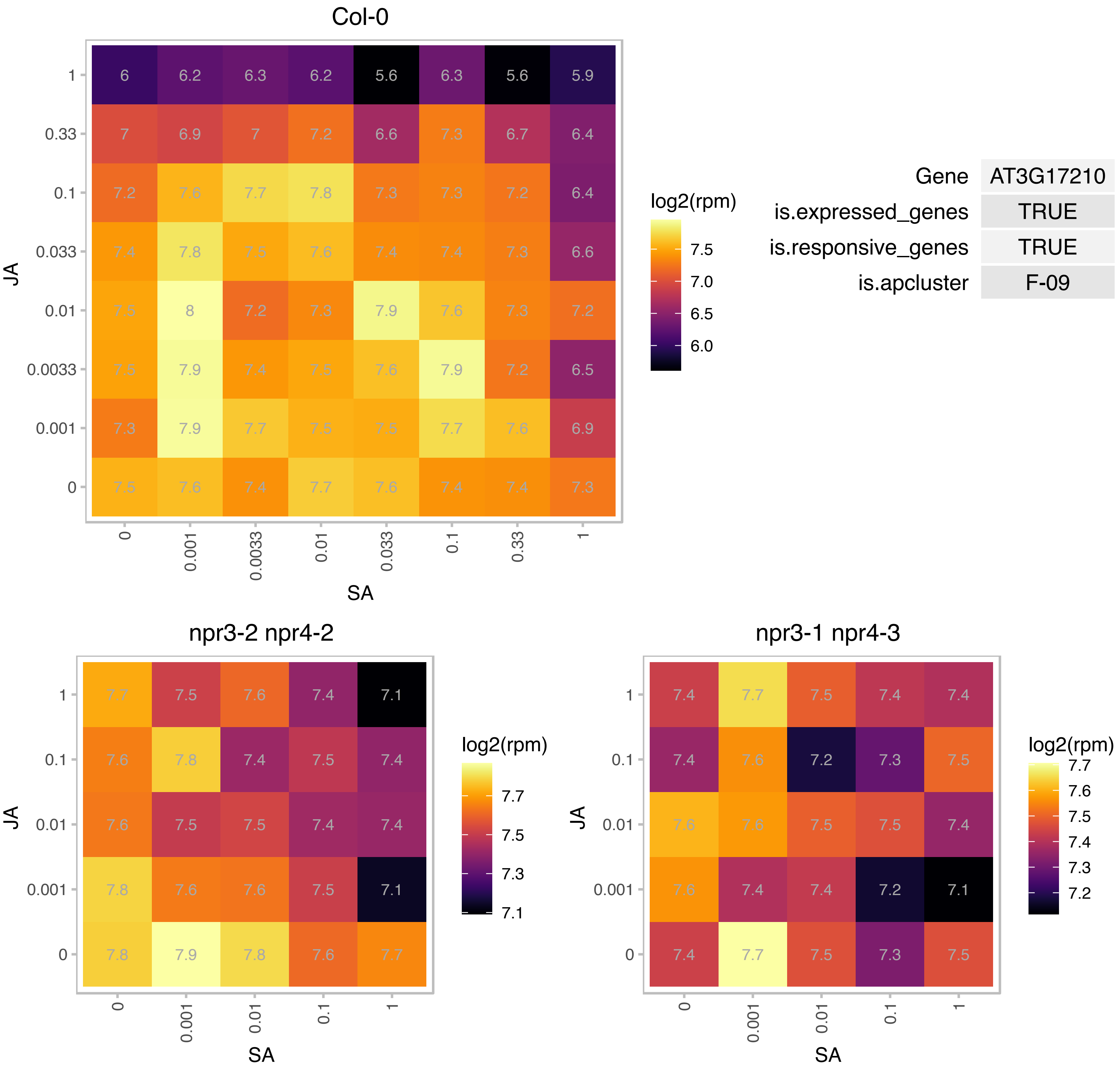


| | |
|---|---|
| Gene | AT3G17210 |
| is.expressed_genes | TRUE |
| is.responsive_genes | TRUE |
| is.apcluster | F-09 |

# AT3G18050.1 : NA

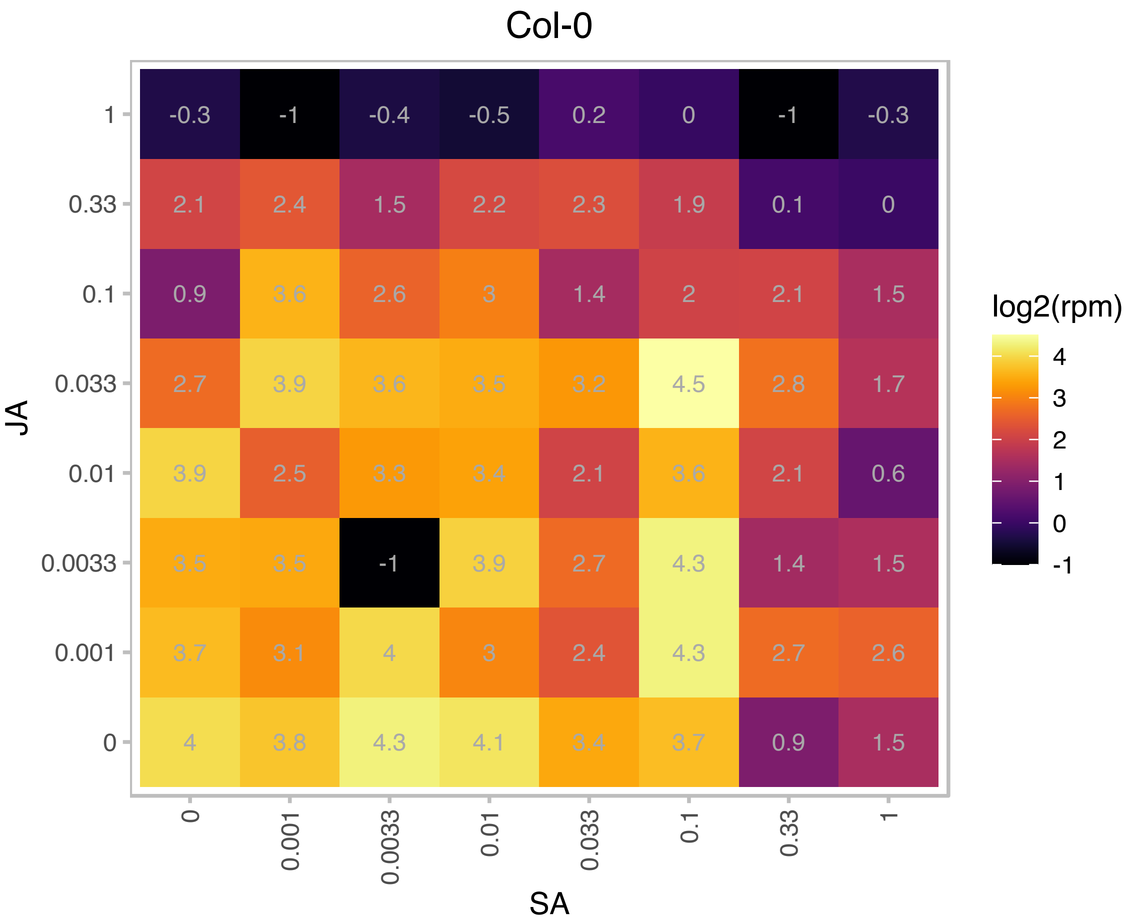


| | |
|---|---|
| Gene | AT3G18050 |
| is.expressed_genes | TRUE |
| is.responsive_genes | TRUE |
| is.apcluster | F-09 |

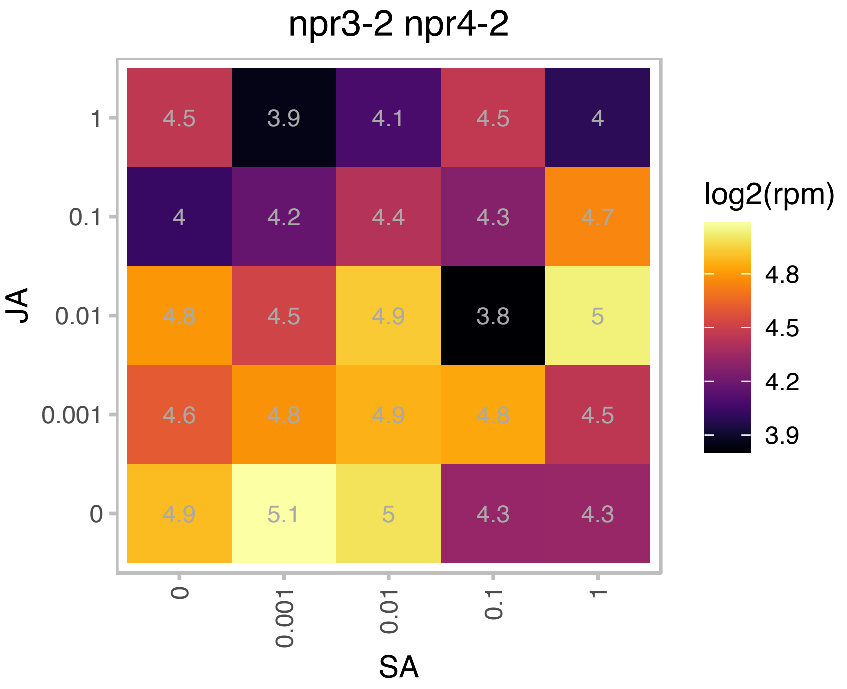


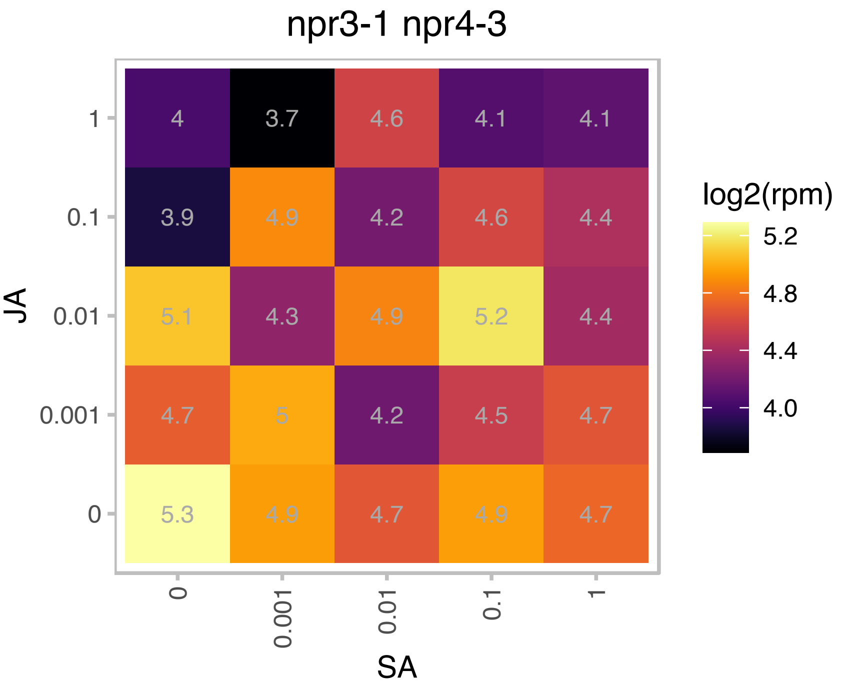

### AT3G21055.1 : PSBTN

photosystem II subunit T

Encodes photosystem II 5 kD protein subunit PSII-T. This is a nuclear-encoded gene (PsbTn) which also has a plastid-encoded paralog (PsbTc).

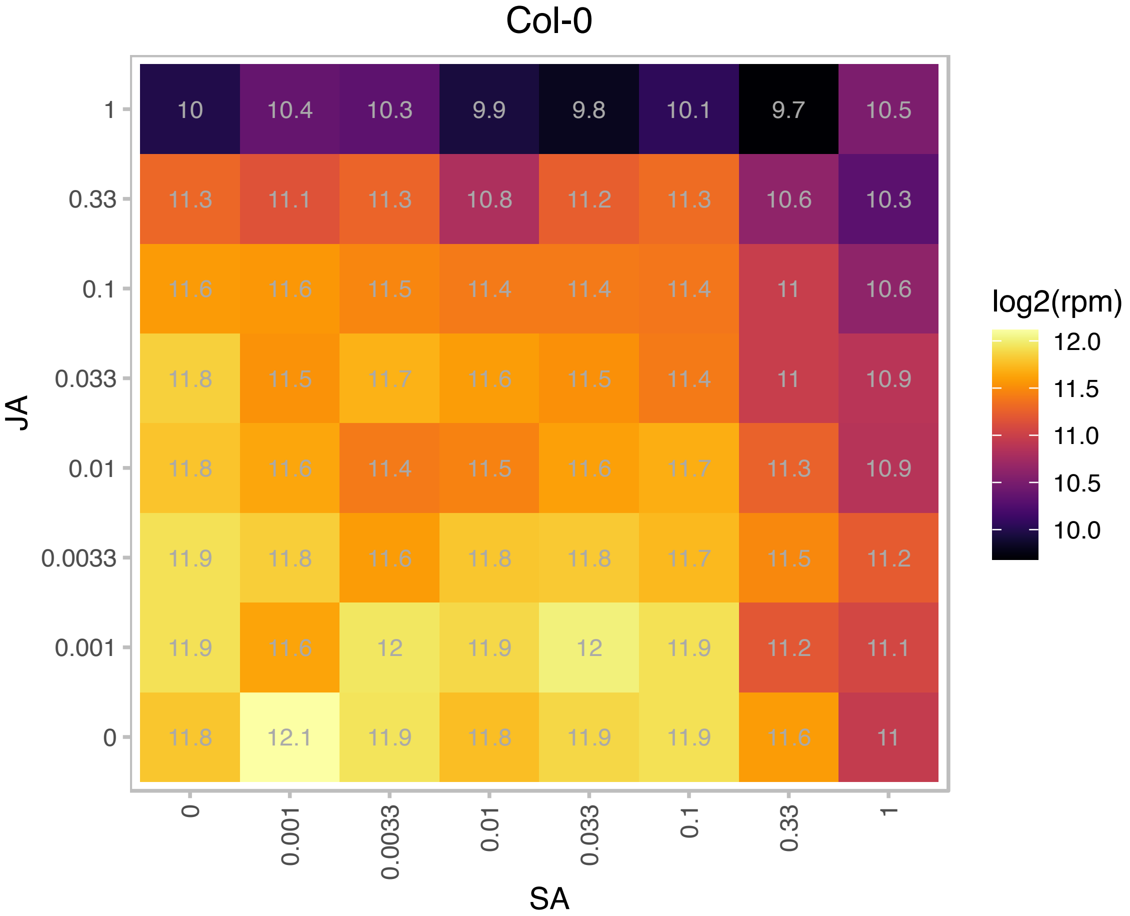


| | |
|---|---|
| Gene | AT3G21055 |
| is.expressed_genes | TRUE |
| is.responsive_genes | TRUE |
| is.apcluster | F-09 |

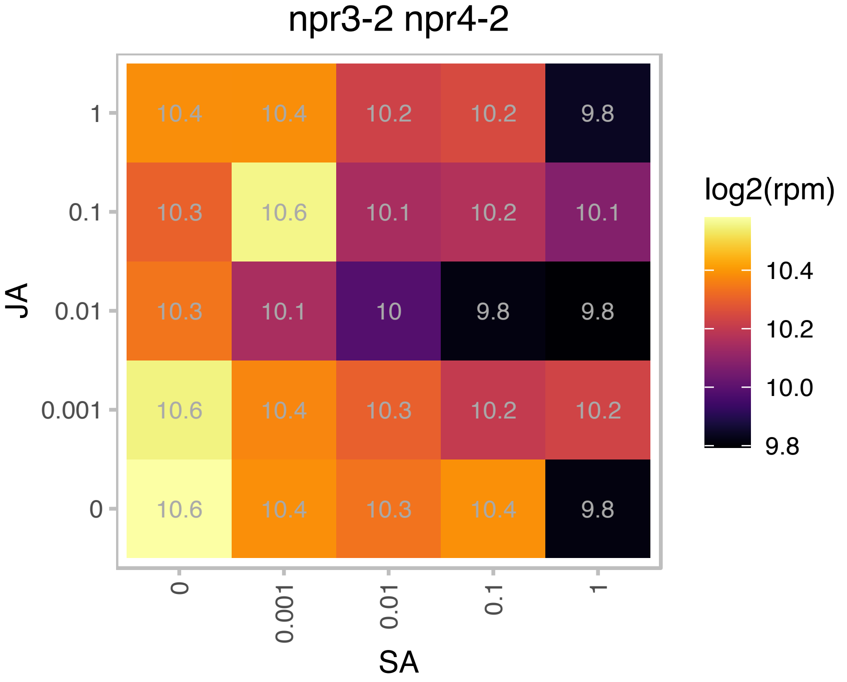


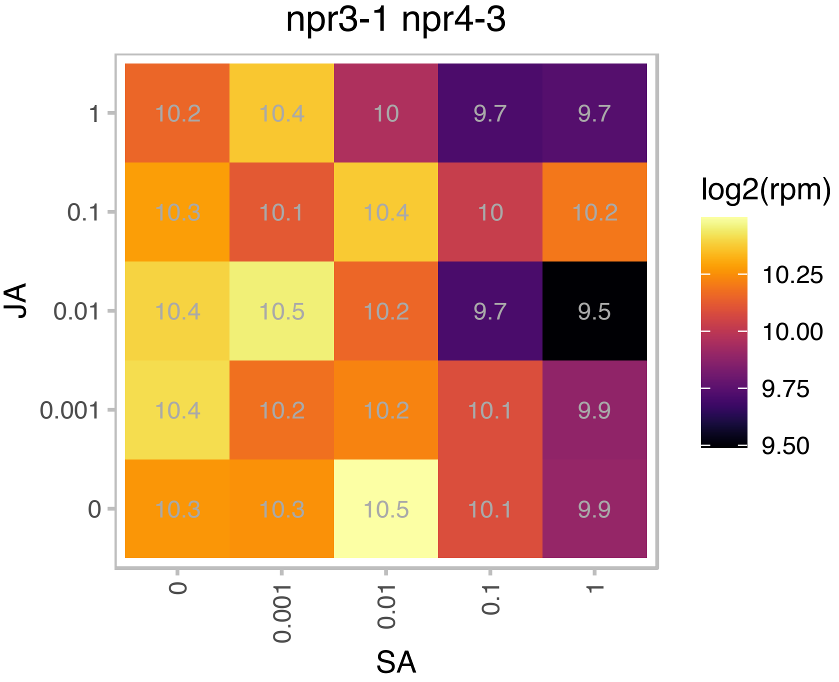

# AT3G26510.1 : NA

Octicosapeptide/Phox/Bem1p family protein

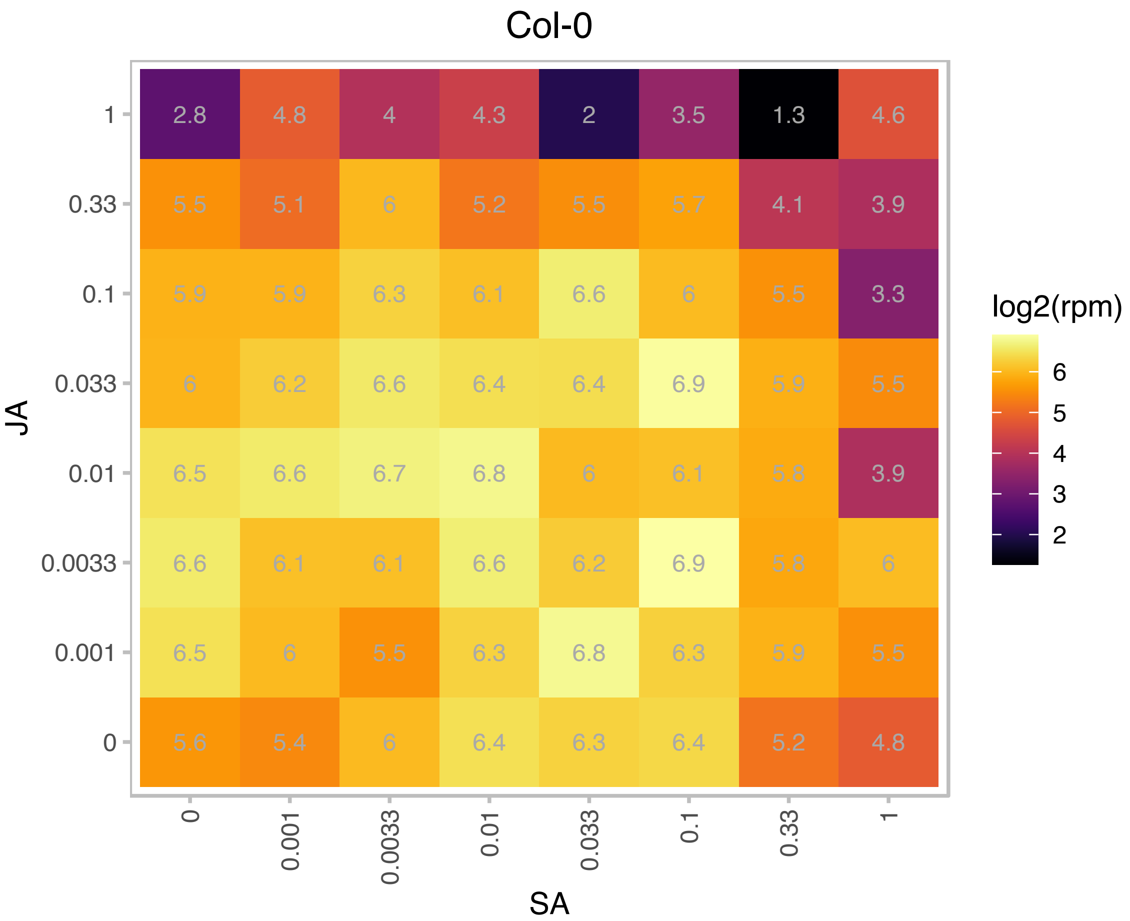


| | |
|---|---|
| Gene | AT3G26510 |
| is.expressed_genes | TRUE |
| is.responsive_genes | TRUE |
| is.apcluster | F-09 |

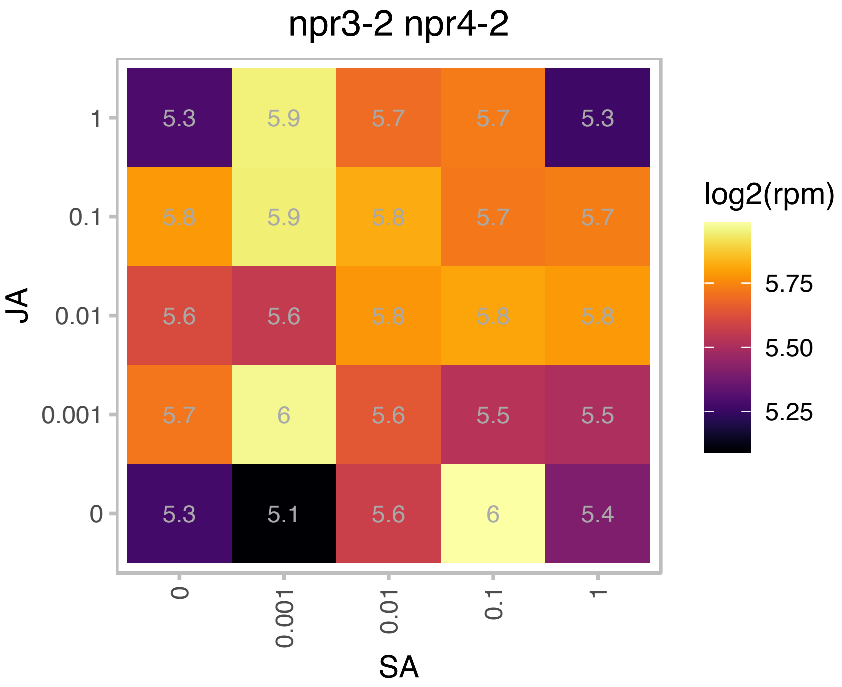


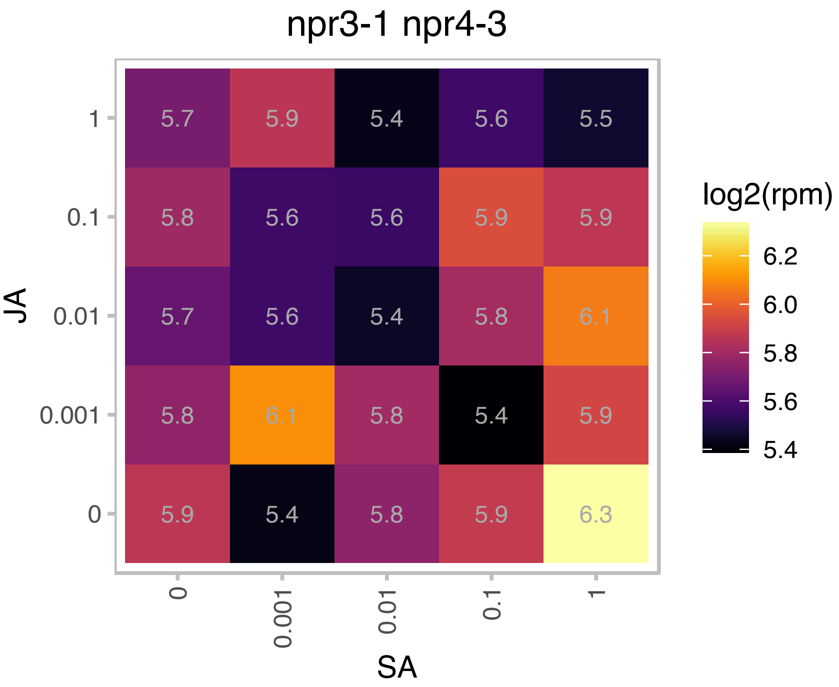

# AT3G45050.1 : NA

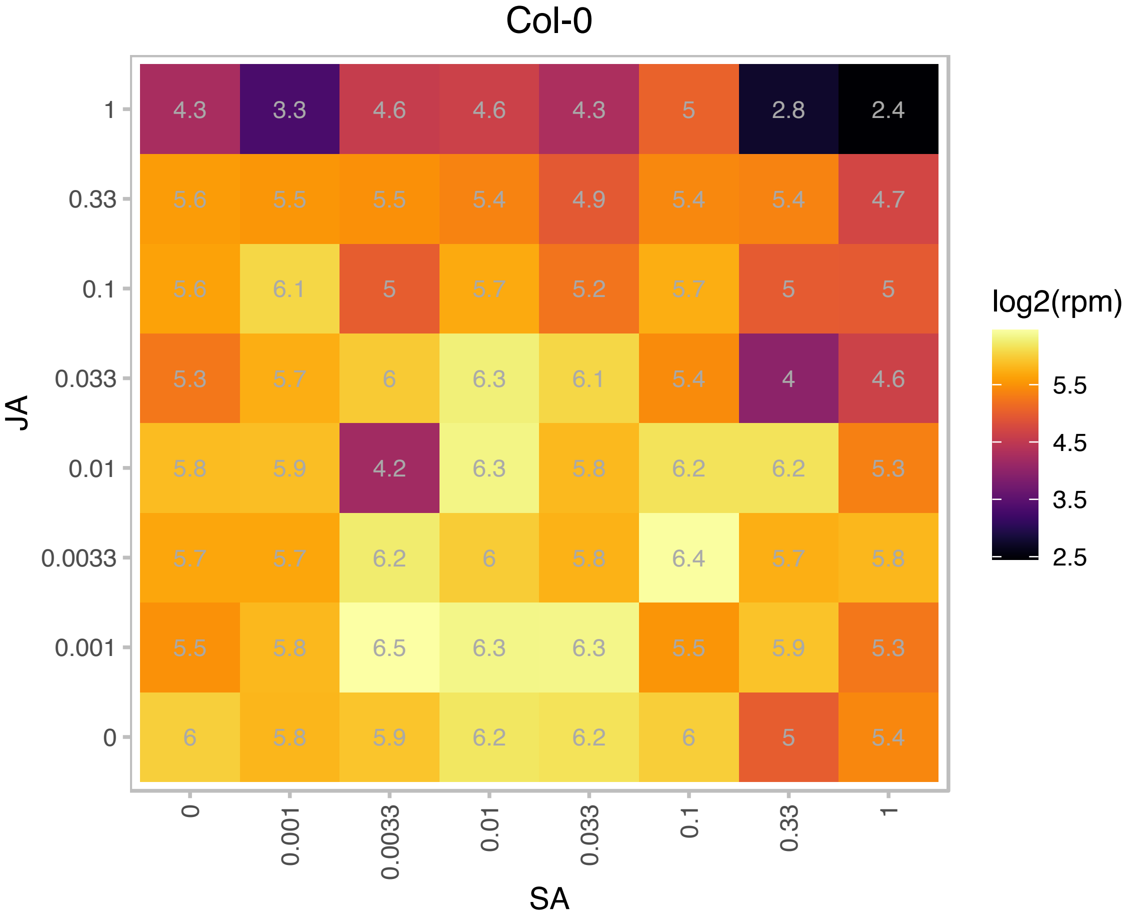


| | |
|---|---|
| Gene | AT3G45050 |
| is.expressed_genes | TRUE |
| is.responsive_genes | TRUE |
| is.apcluster | F-09 |

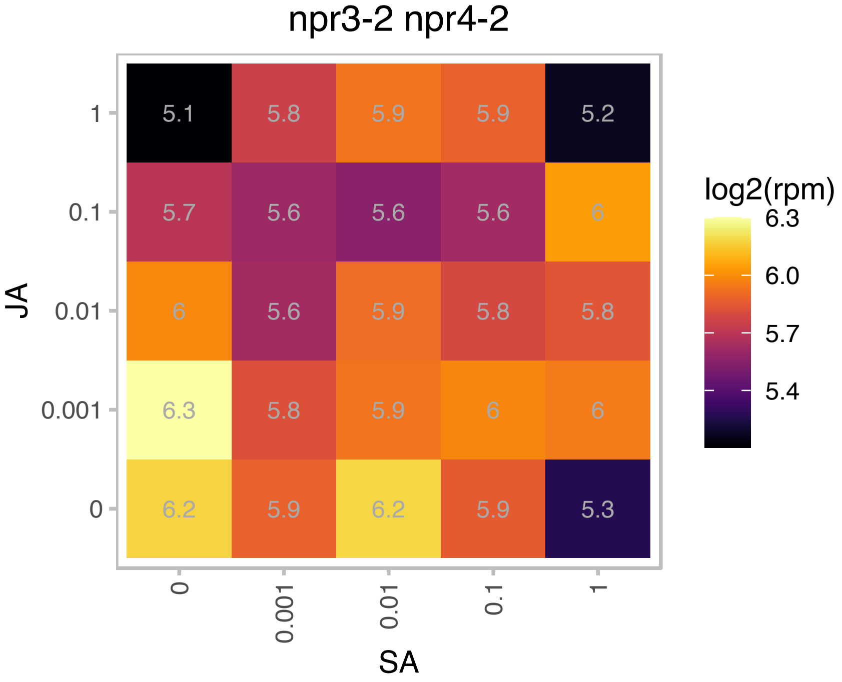


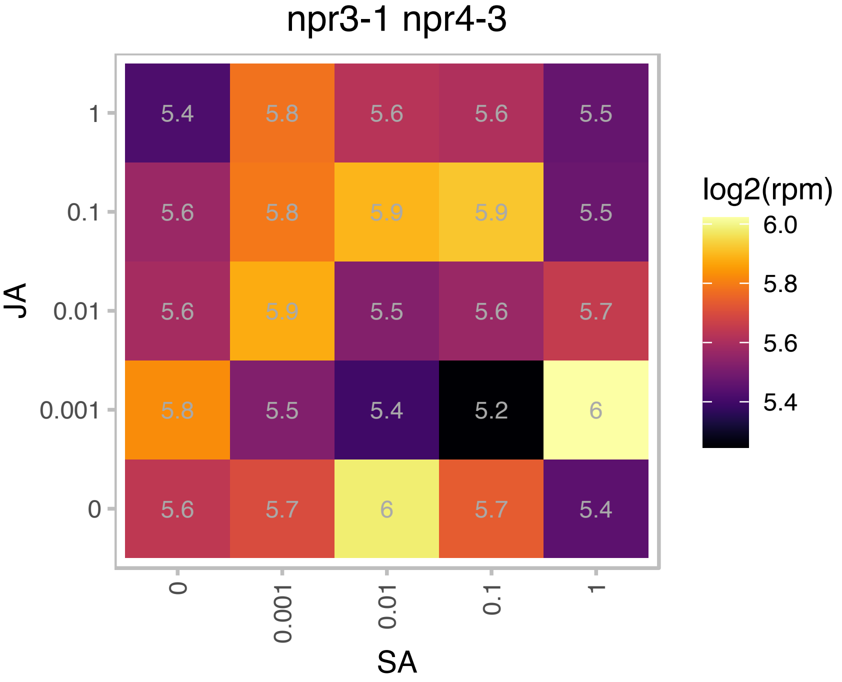

# AT3G46630.1 : NA

Protein of unknown function (DUF3223)

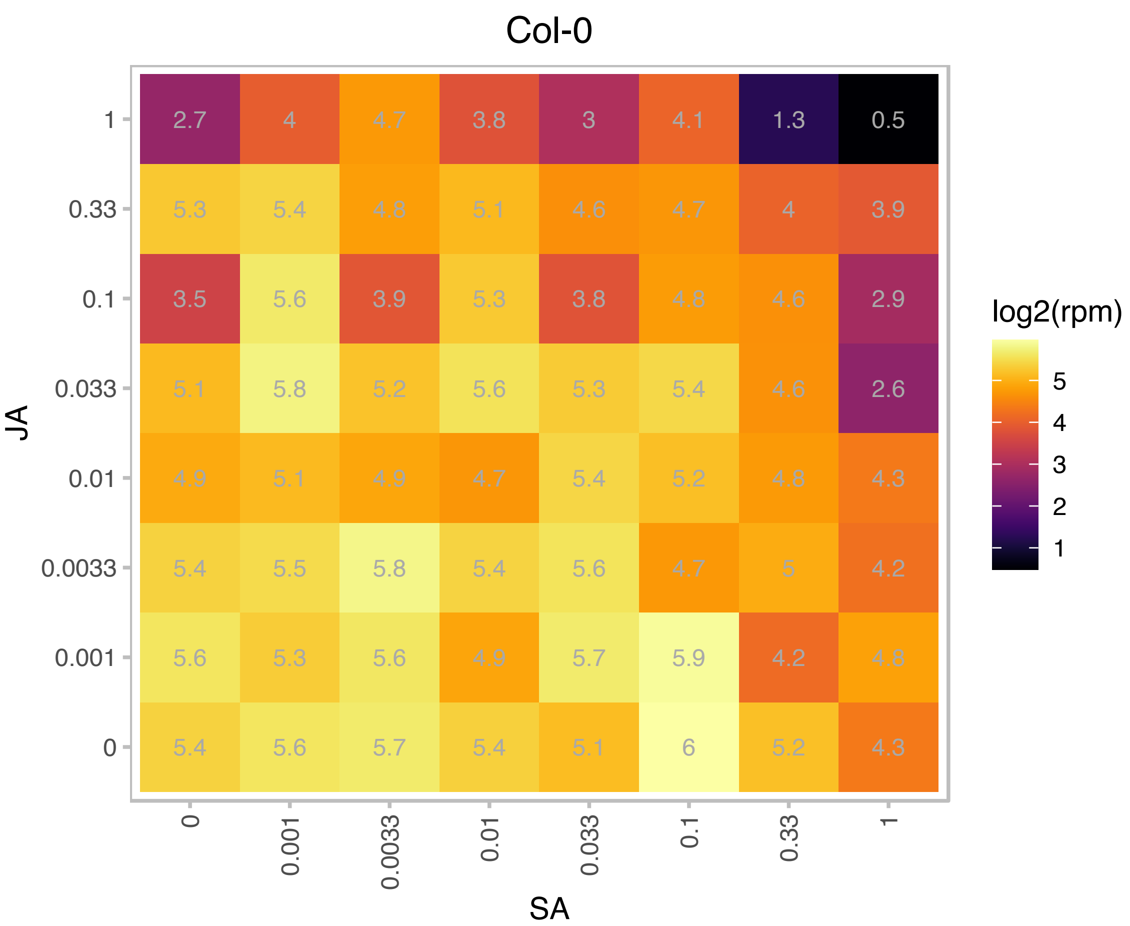


| | |
|---|---|
| Gene | AT3G46630 |
| is.expressed_genes | TRUE |
| is.responsive_genes | TRUE |
| is.apcluster | F-09 |

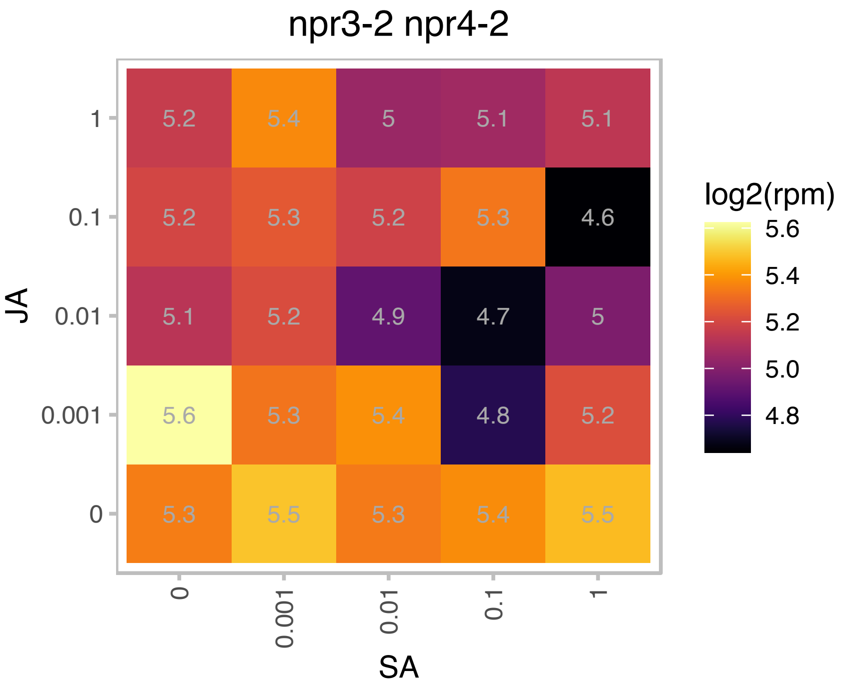


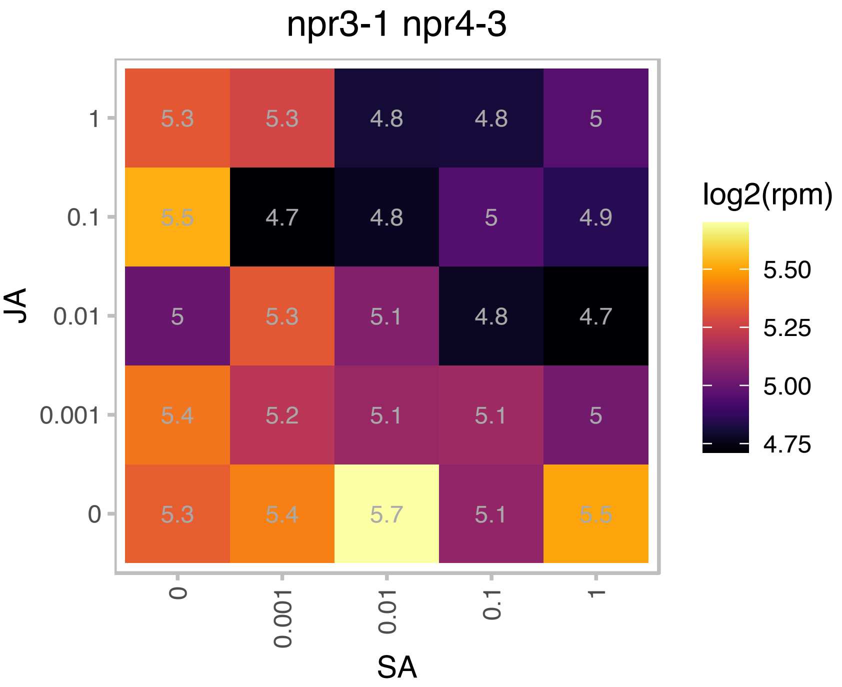

### AT3G53850.1 : CASPL5B2

Uncharacterised protein family (UPF0497)

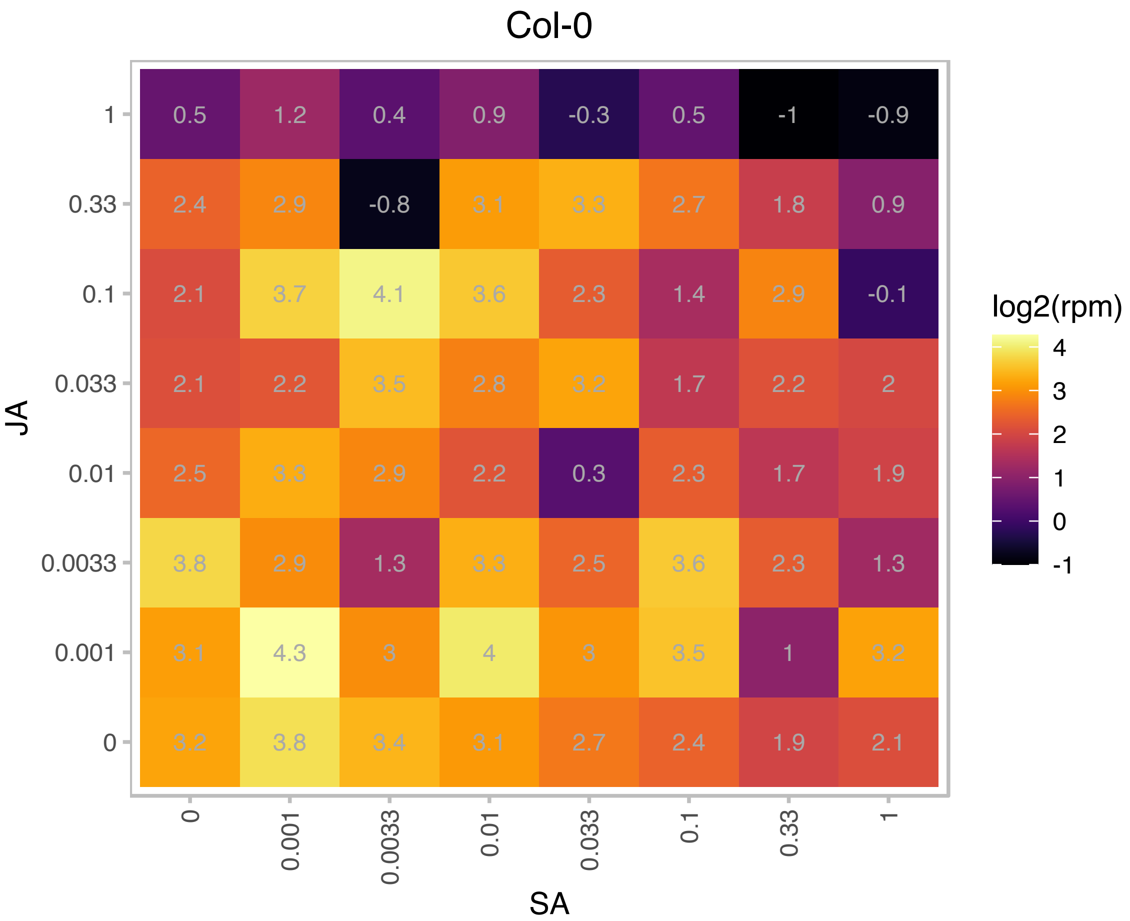


| | |
|---|---|
| Gene | AT3G53850 |
| is.expressed_genes | TRUE |
| is.responsive_genes | TRUE |
| is.apcluster | F-09 |

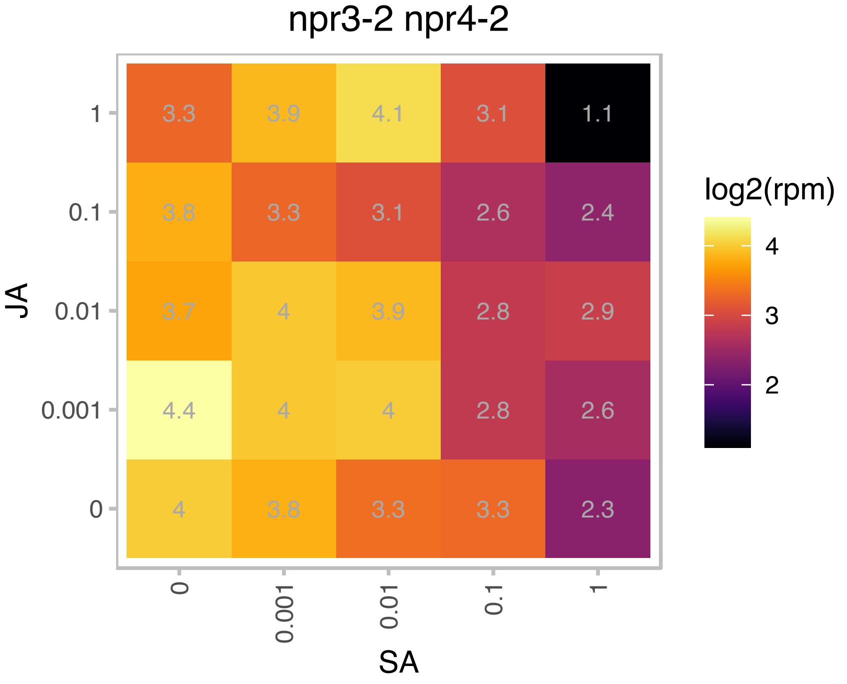


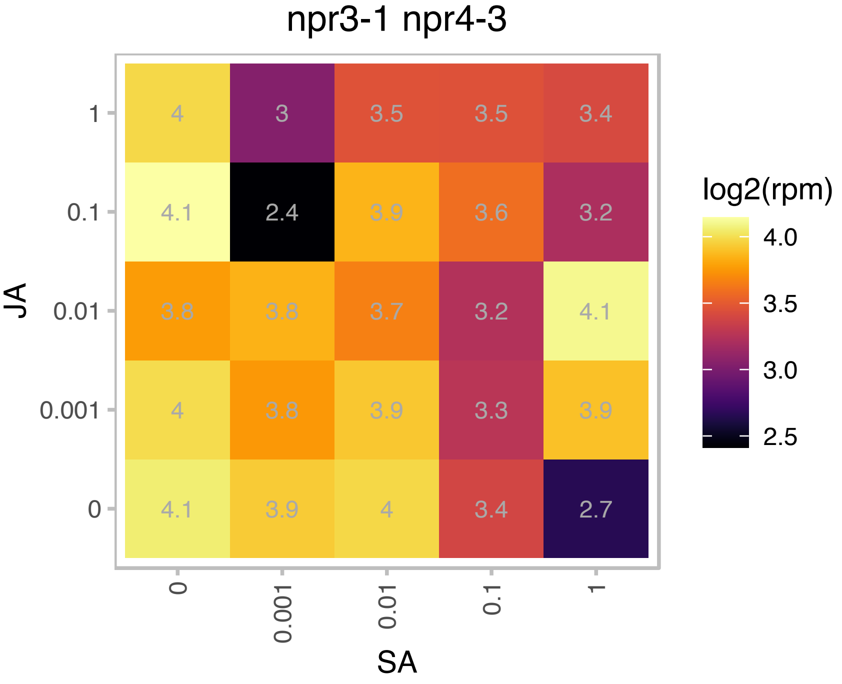

### AT3G54560.1 : H2A.Z, HTA11

histone H2A 11

Encodes HTA11, a histone H2A protein. Loss of all H2A.Z (triple mutant with HTA8 and HTA9) results in a reduction in DNA methylation of transposons but not that of genes.  Loss of H2A.Z causes misregulation of many genes involved in the response to developmental and environmental cues, and that these genes tend to have high levels of gene-body H2A.Z.

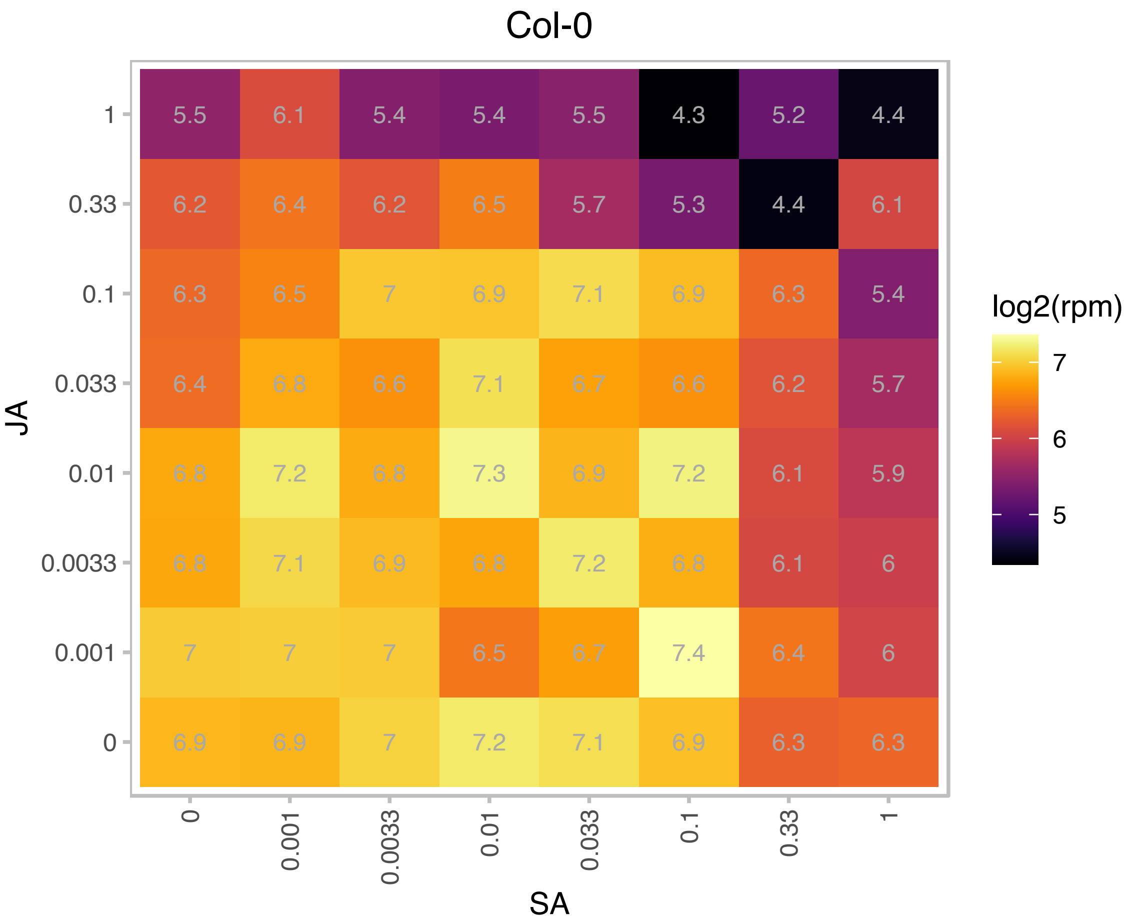


| | |
|---|---|
| Gene | AT3G54560 |
| is.expressed_genes | TRUE |
| is.responsive_genes | TRUE |
| is.apcluster | F-09 |

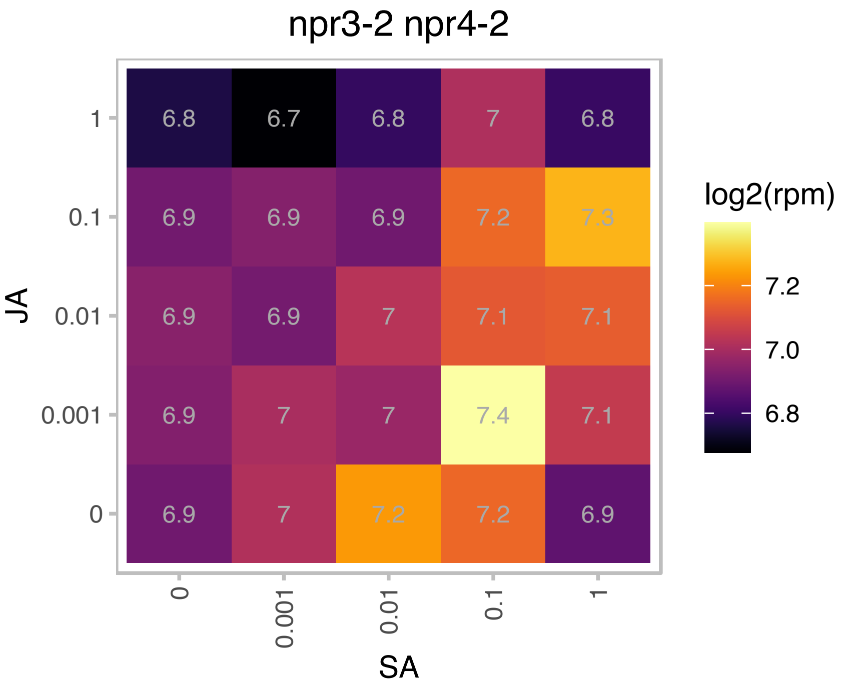


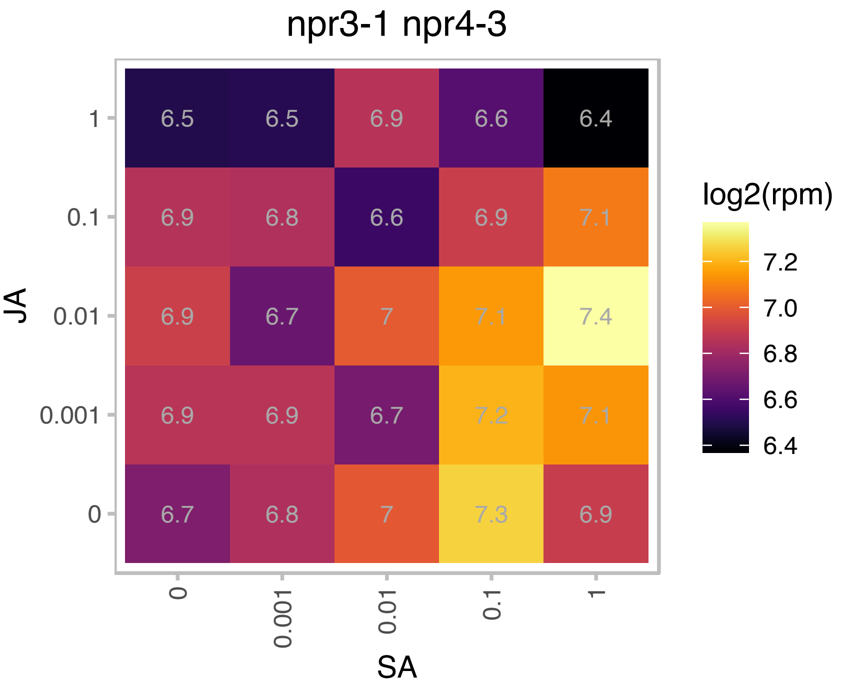

# AT3G59980.1 : NA

Nucleic acid-binding, OB-fold-like protein

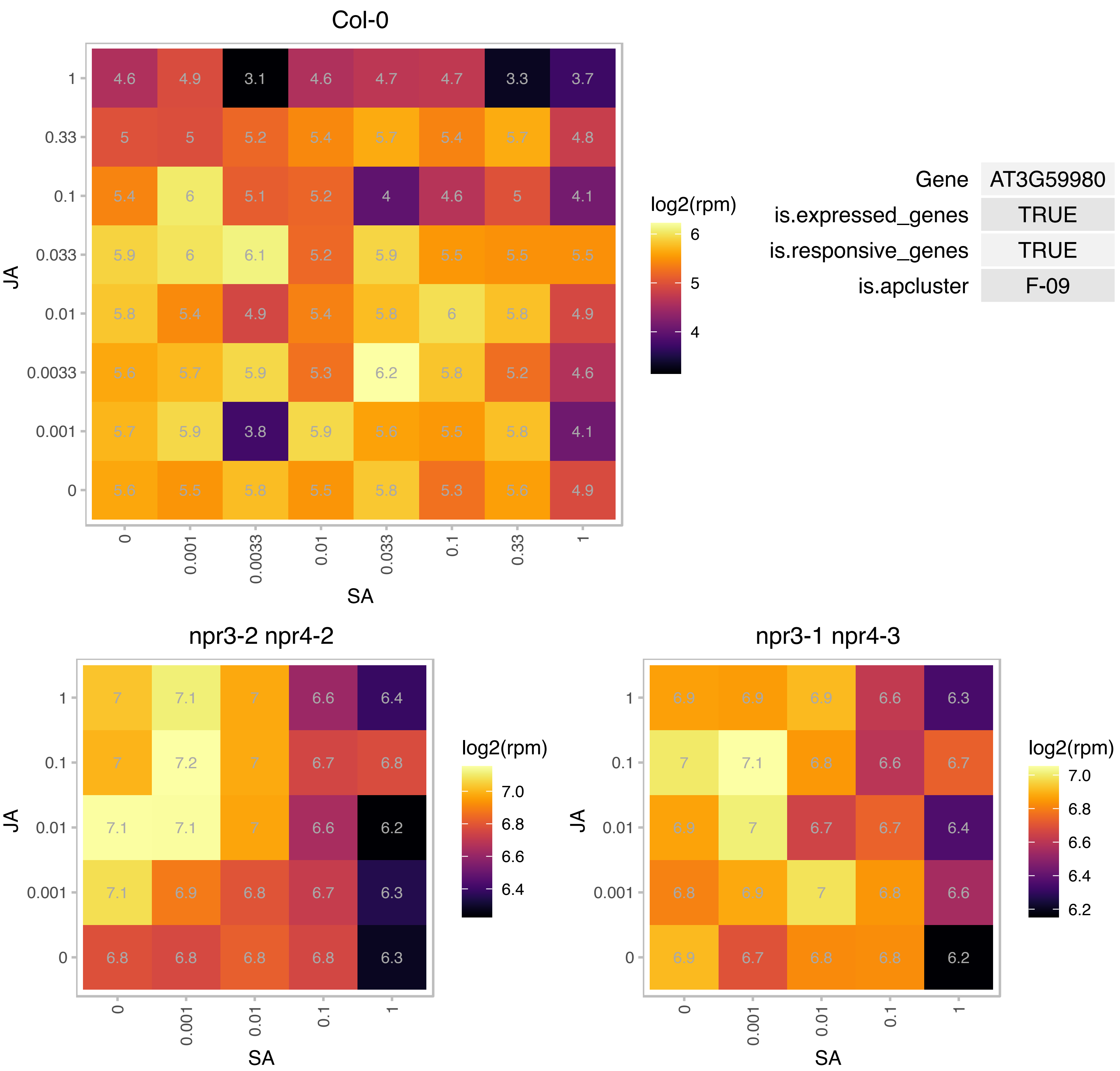


| | |
|---|---|
| Gene | AT3G59980 |
| is.expressed_genes | TRUE |
| is.responsive_genes | TRUE |
| is.apcluster | F-09 |

### AT3G60530.1 : GATA4

GATA transcription factor 4

Encodes a member of the GATA factor family of zinc finger transcription factors.

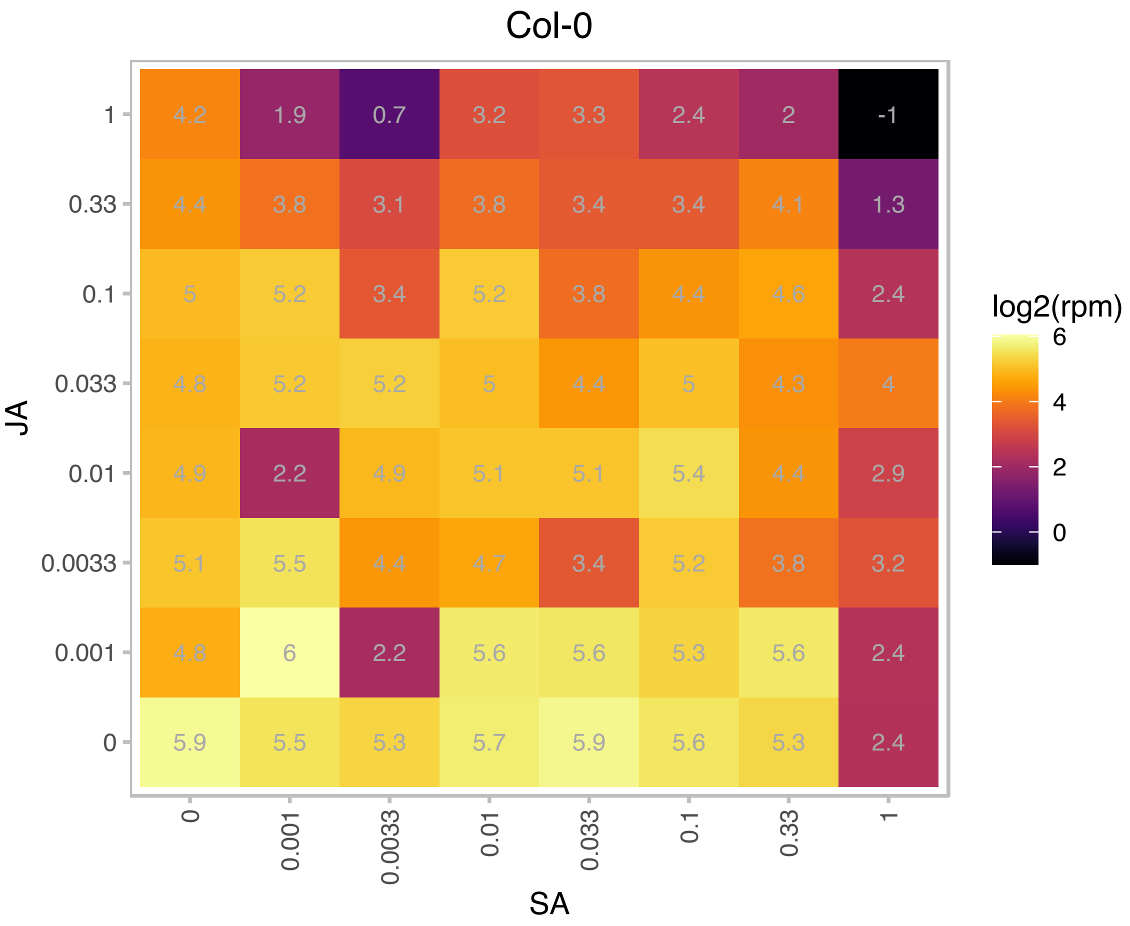


| | |
|---|---|
| Gene | AT3G60530 |
| is.expressed_genes | TRUE |
| is.responsive_genes | TRUE |
| is.apcluster | F-09 |

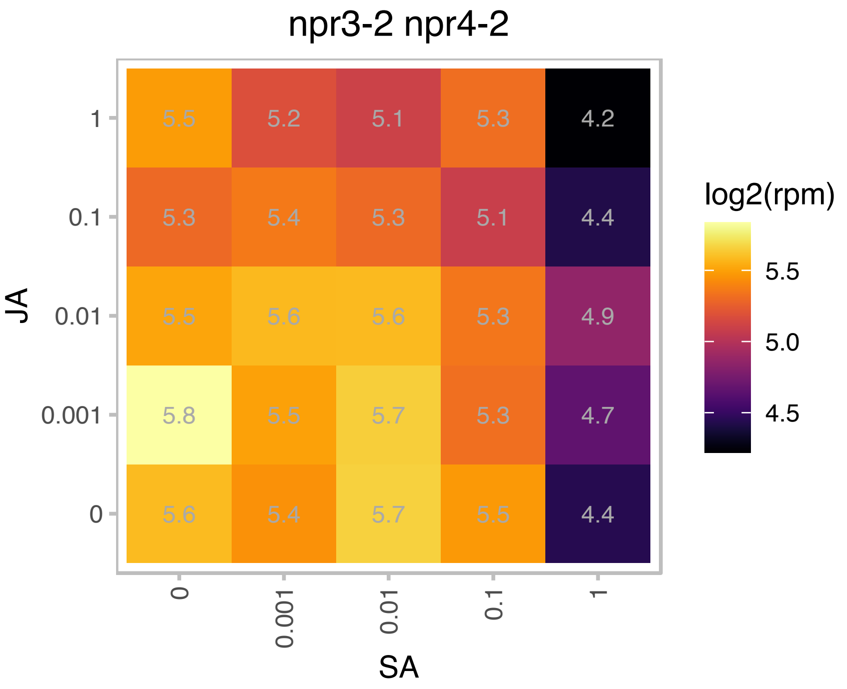


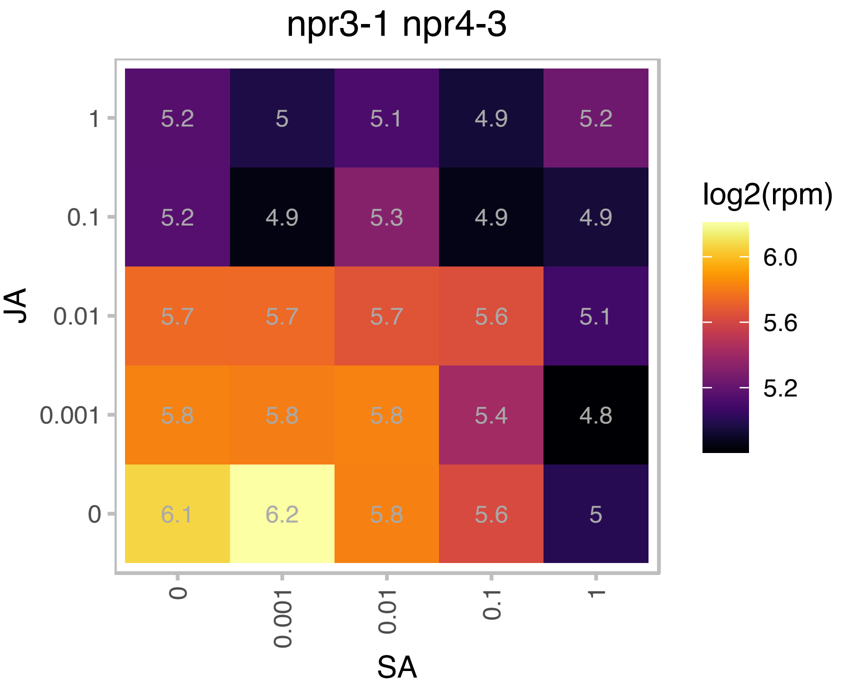

### AT3G60620.1 : CDS5

cytidinediphosphate diacylglycerol synthase 5

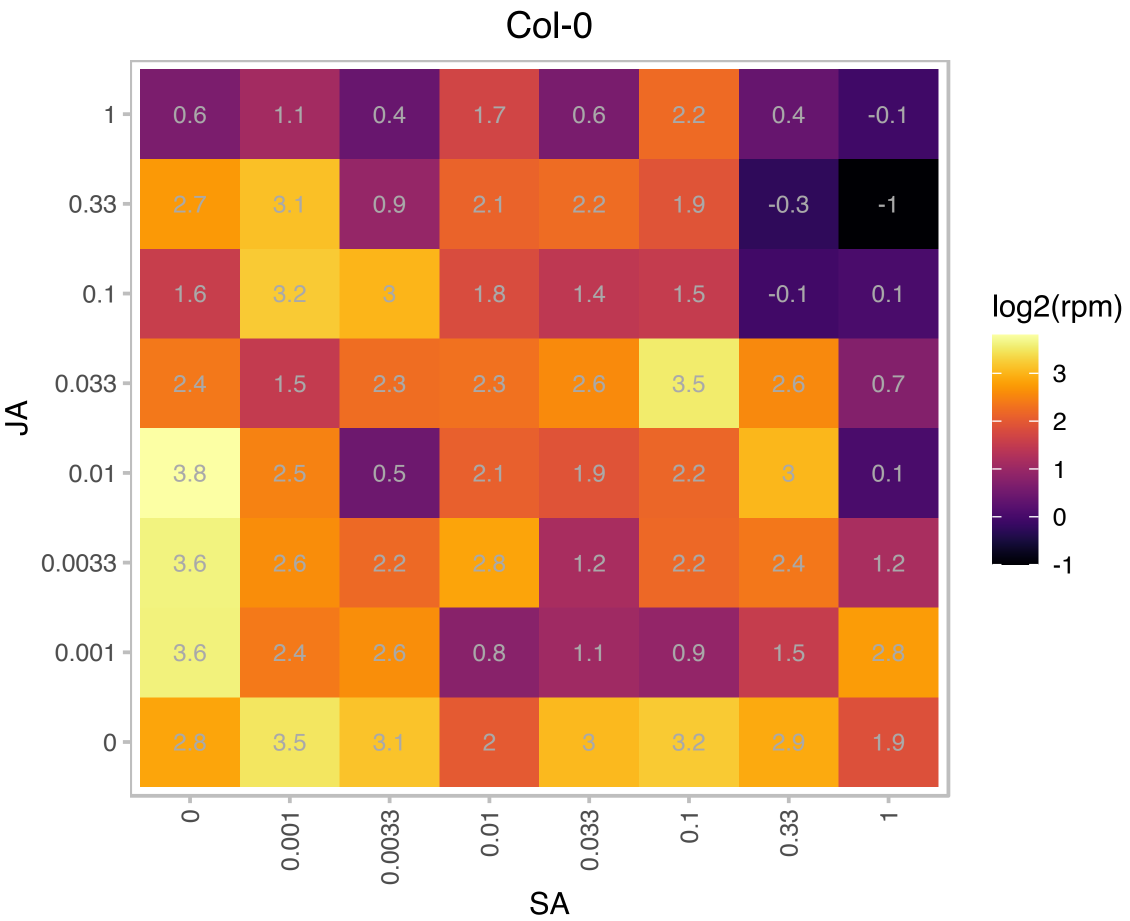


| | |
|---|---|
| Gene | AT3G60620 |
| is.expressed_genes | TRUE |
| is.responsive_genes | TRUE |
| is.apcluster | F-09 |

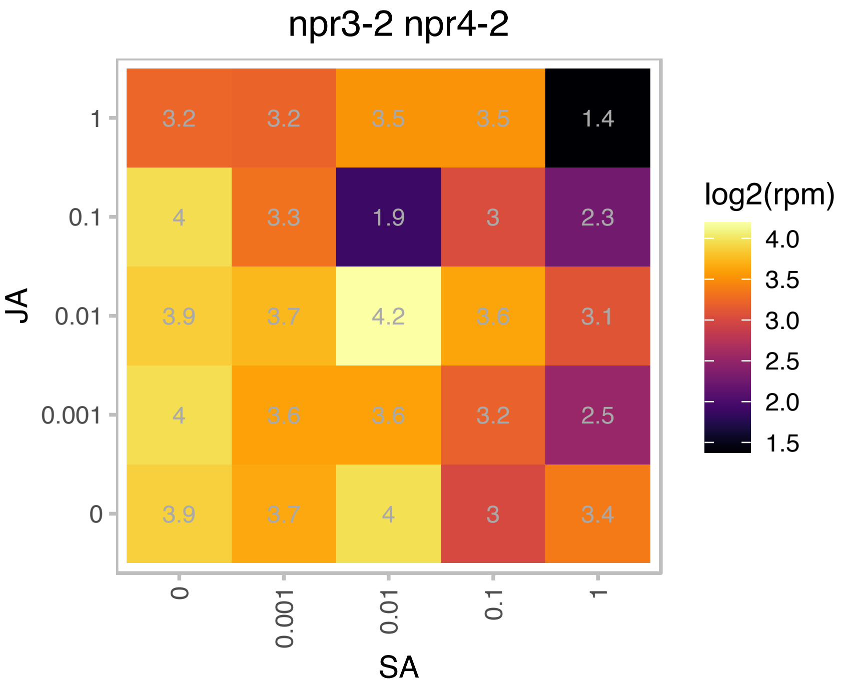


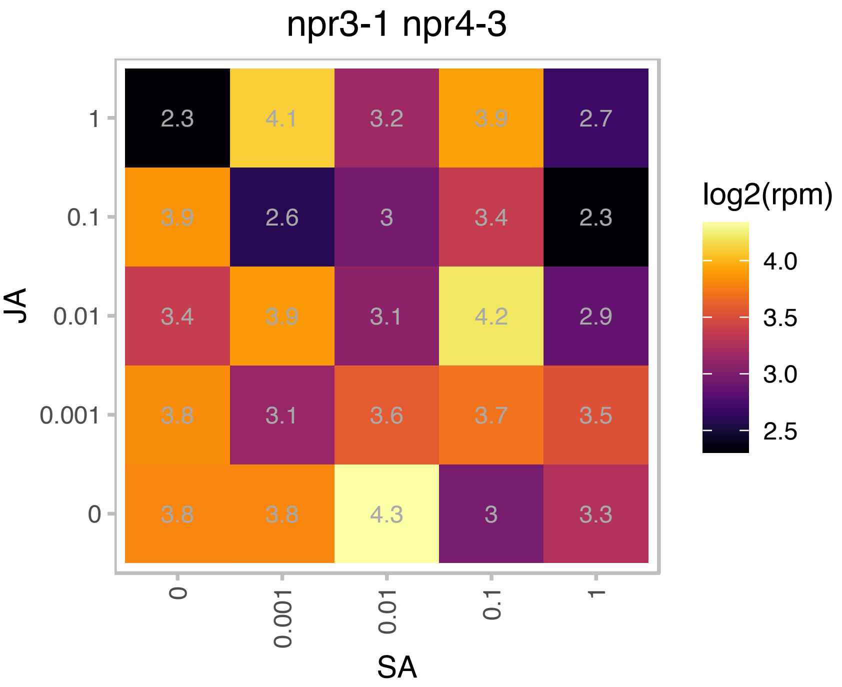

# AT4G00165.1 : NA

Bifunctional inhibitor/lipid-transfer protein/seed storage 2S albumin superfamily protein

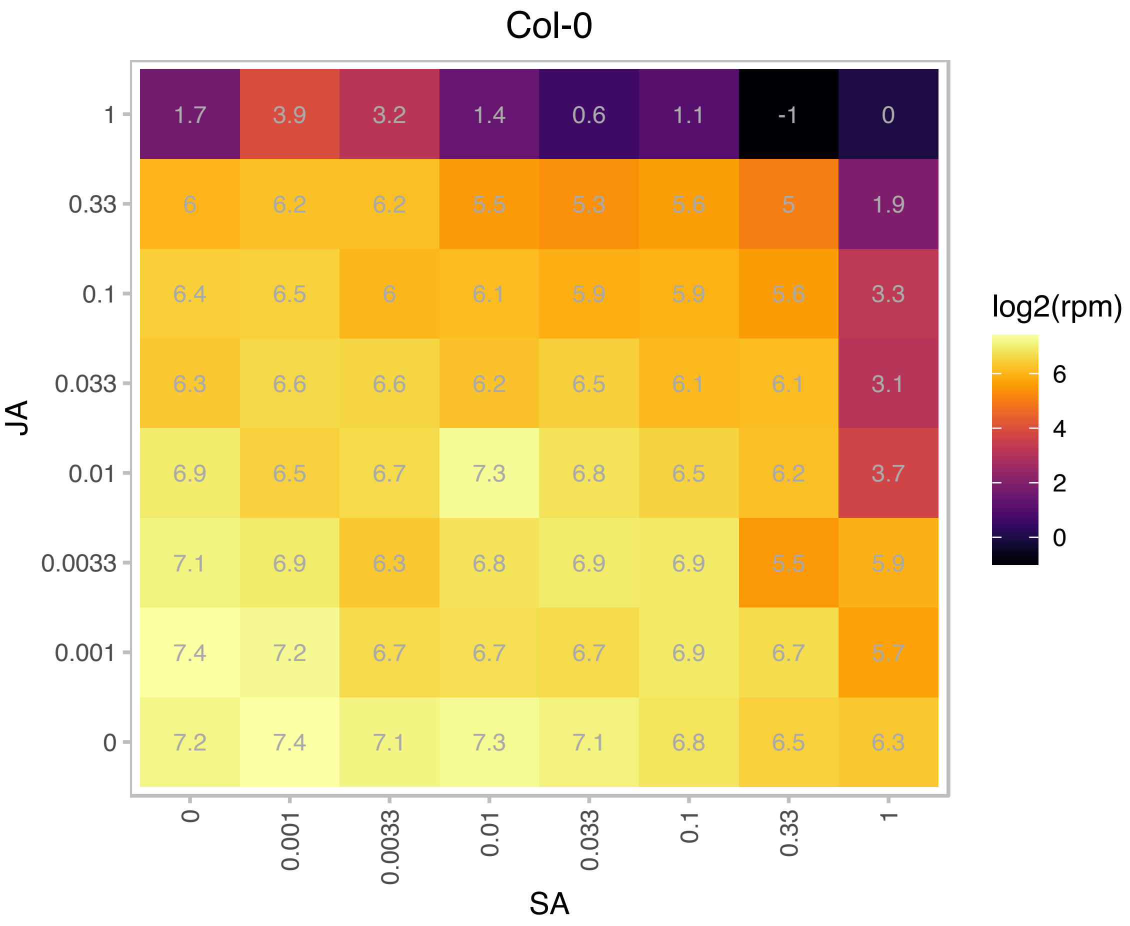


| | |
|---|---|
| Gene | AT4G00165 |
| is.expressed_genes | TRUE |
| is.responsive_genes | TRUE |
| is.apcluster | F-09 |

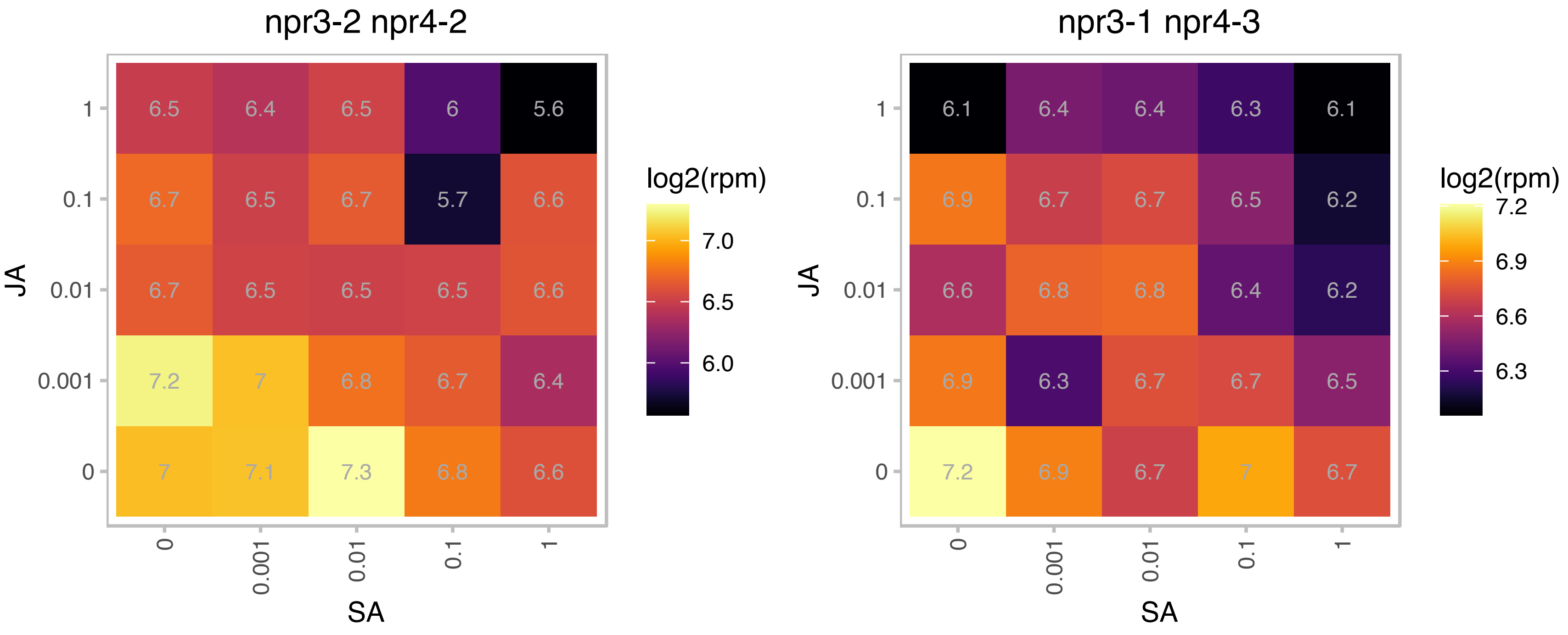

### AT4G02990.1 : BSM, RUG2

Mitochondrial transcription termination factor family protein
Encodes BELAYA SMERT (BSM), a plastid-localized protein homologous to mitochondrial transcription termination factors (mTERF) found in animal. Mutant bsm cells are albino, are compromised in growth, and suffer defects in global plastidic gene expression. The mRNA is cell-to-cell mobile.

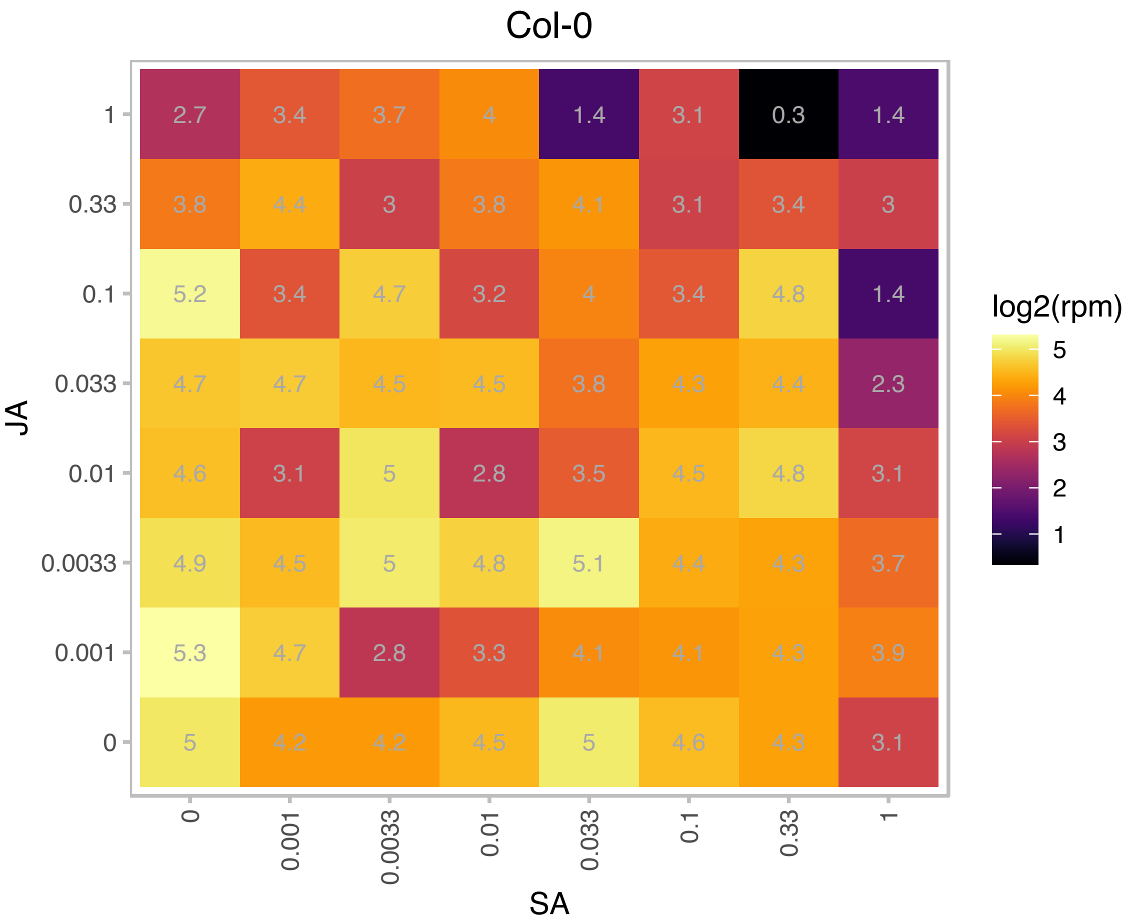


| | |
|---|---|
| Gene | AT4G02990 |
| is.expressed_genes | TRUE |
| is.responsive_genes | TRUE |
| is.apcluster | F-09 |

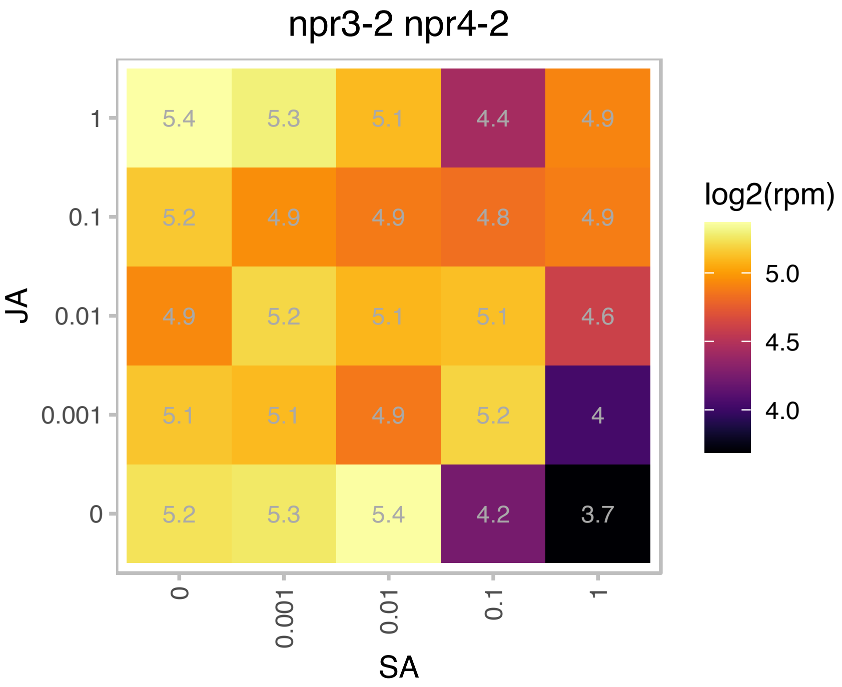


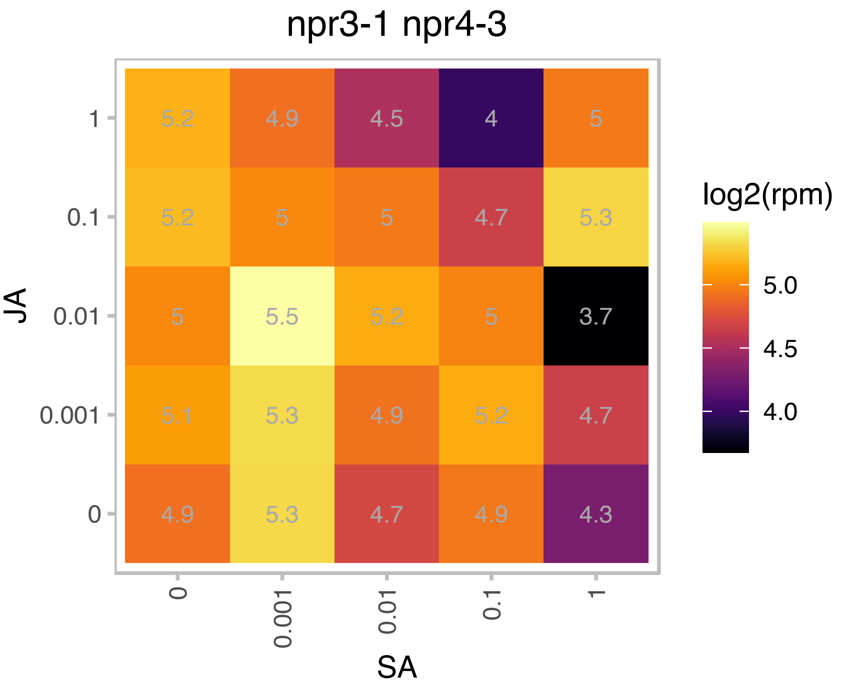

### AT4G09320.1 : ATNDK1, NDK1, NDPK1

Nucleoside diphosphate kinase family protein

nucleoside diphosphate kinase type 1 (NDPK1) gene, complete The mRNA is cell-to-cell mobile.

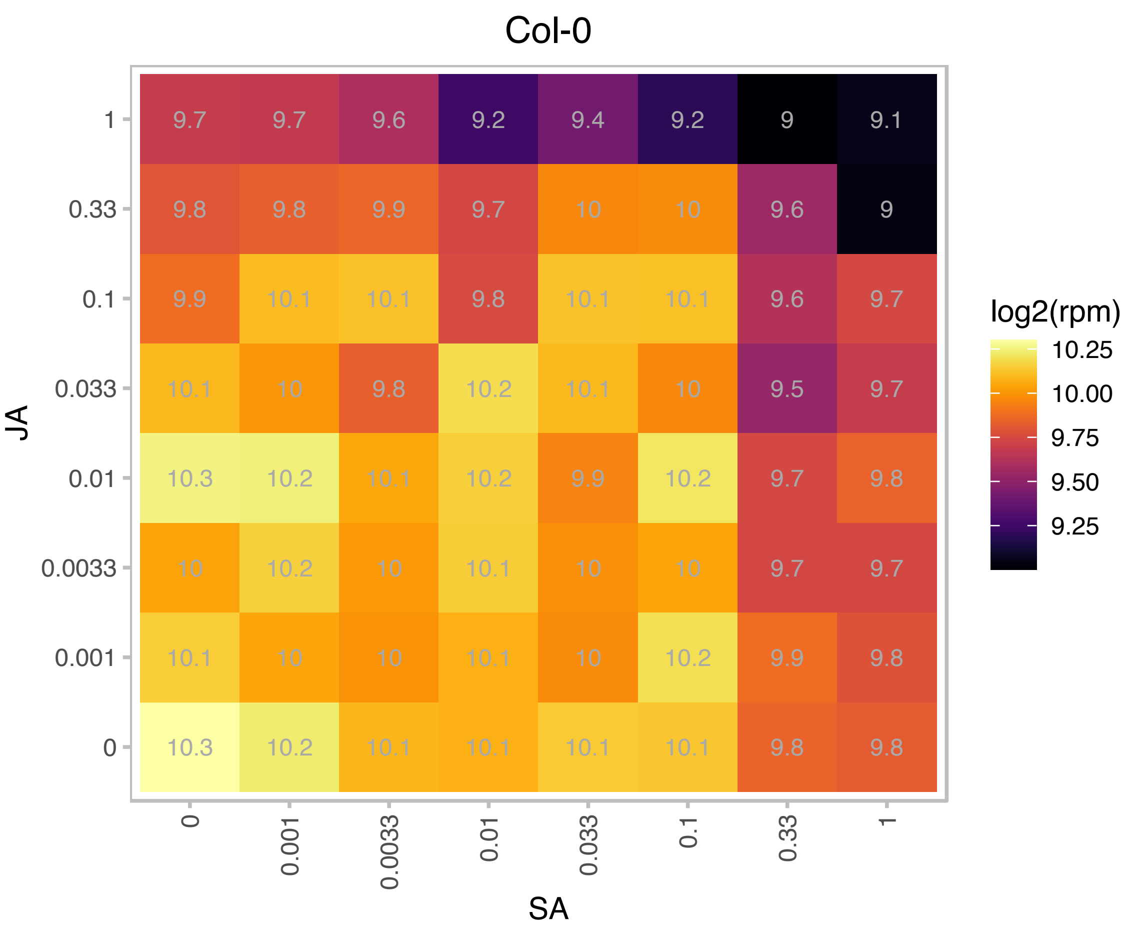


| | |
|---|---|
| Gene | AT4G09320 |
| is.expressed_genes | TRUE |
| is.responsive_genes | TRUE |
| is.apcluster | F-09 |

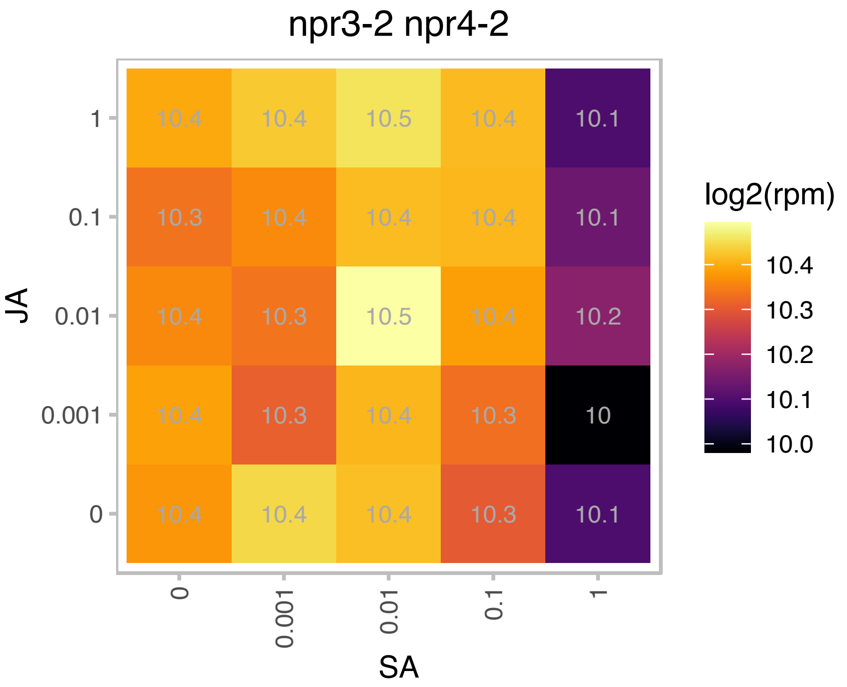


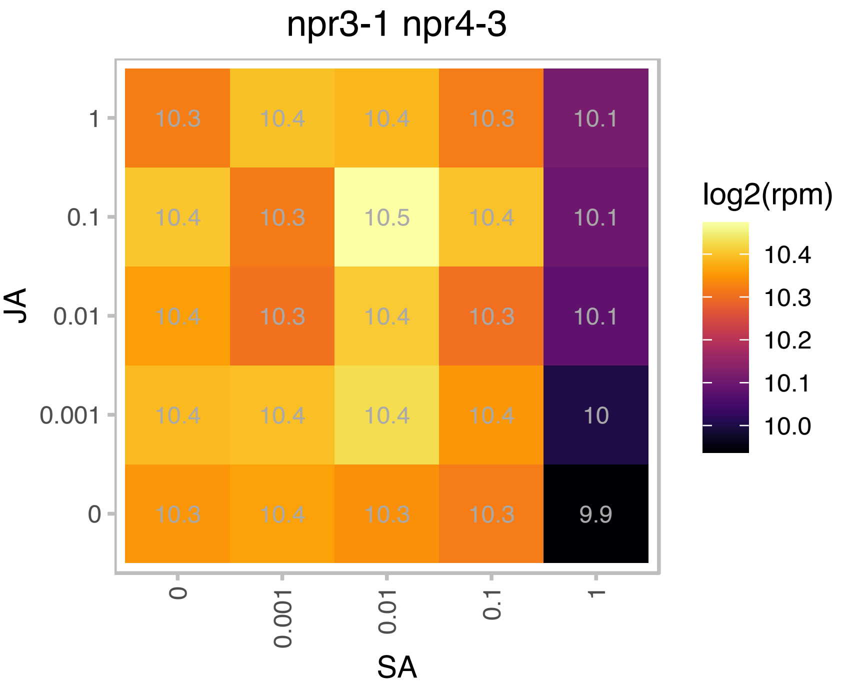

### AT4G09550.1 : ATGIP1, GIP1, GIP1a

AtGCP3 interacting protein 1

Encodes a gamma-tubulin complex protein that plays a role in gamma-tubulin complex localization, spindle stability and chromosomal segregation.

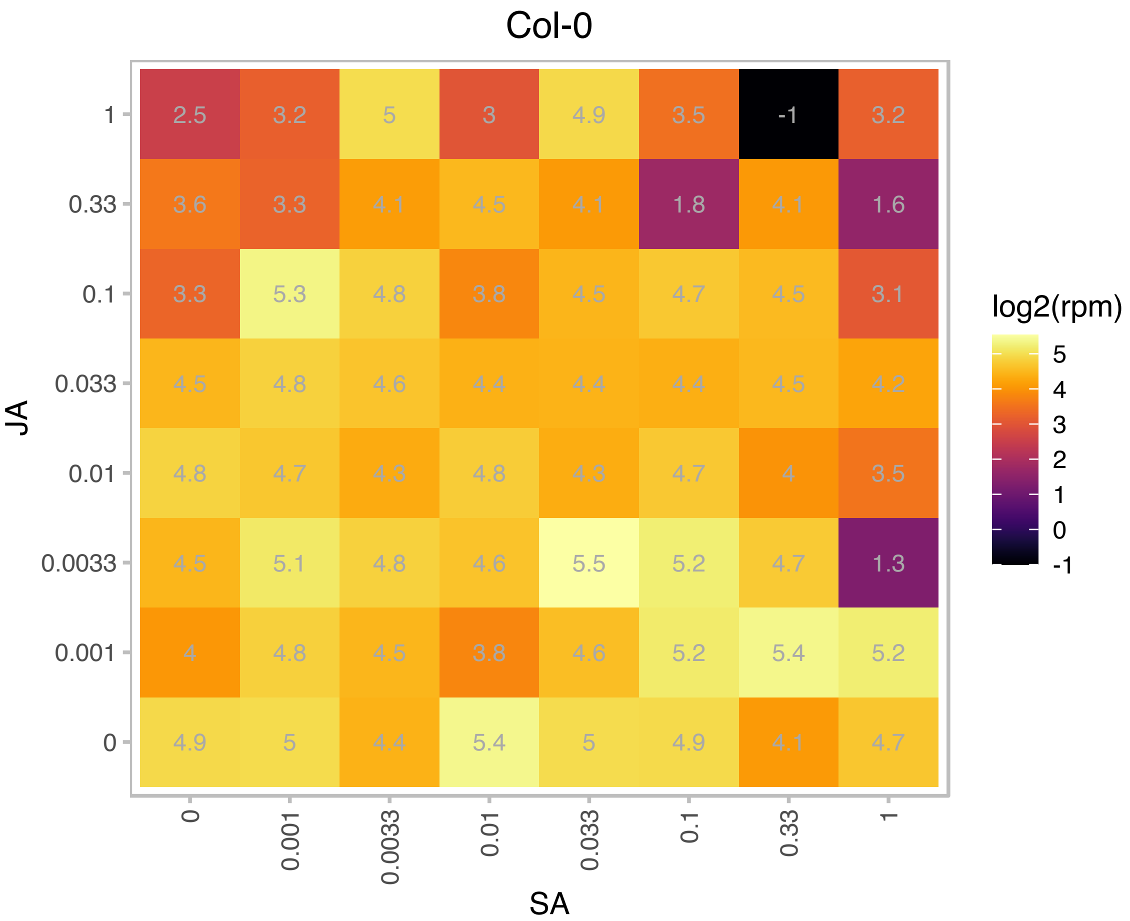


| | |
|---|---|
| Gene | AT4G09550 |
| is.expressed_genes | TRUE |
| is.responsive_genes | TRUE |
| is.apcluster | F-09 |

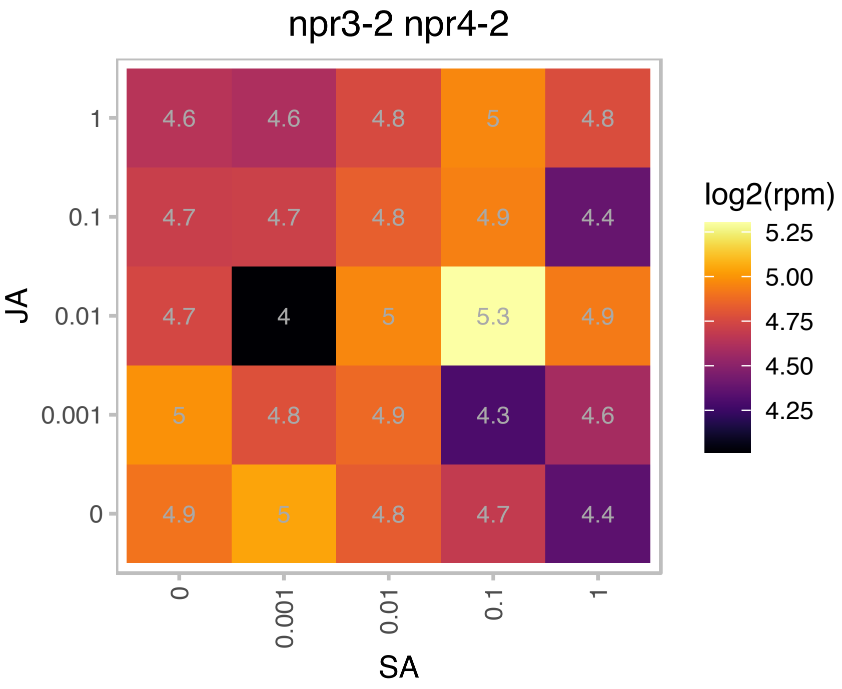


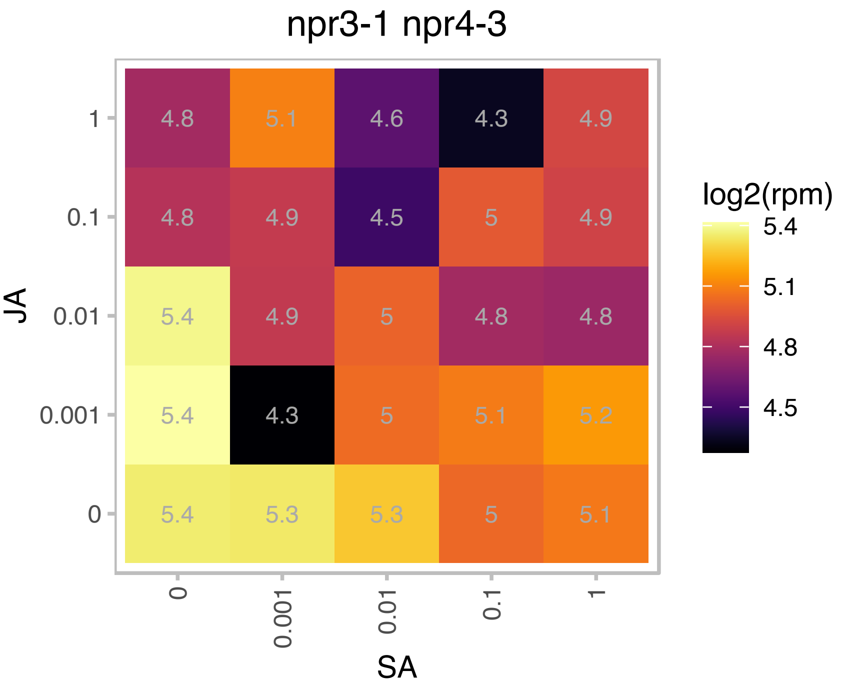

# AT4G12600.1 : NA

Ribosomal protein L7Ae/L30e/S12e/Gadd45 family protein

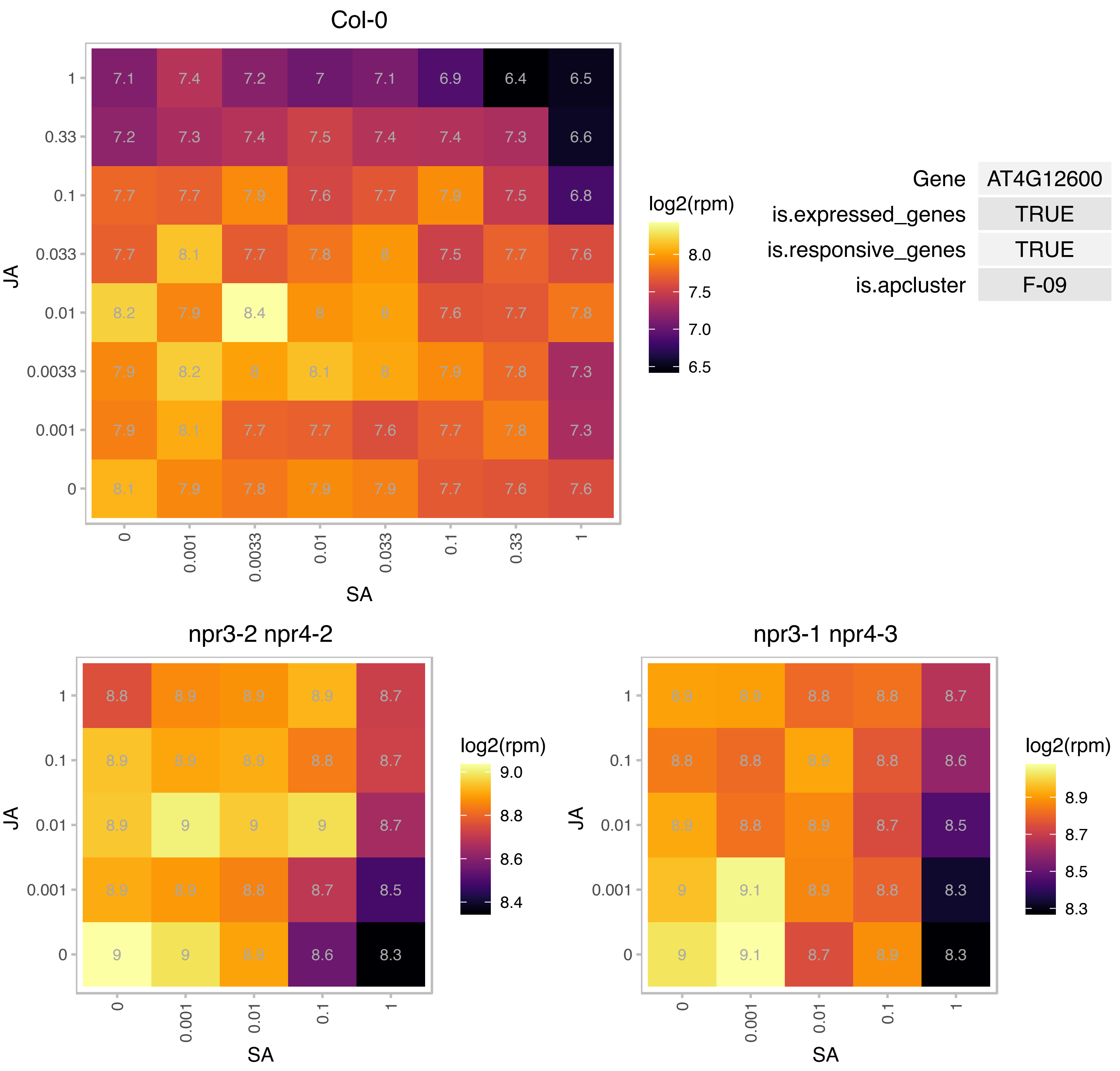

# AT4G16880.1 : NA

Leucine-rich repeat (LRR) family protein

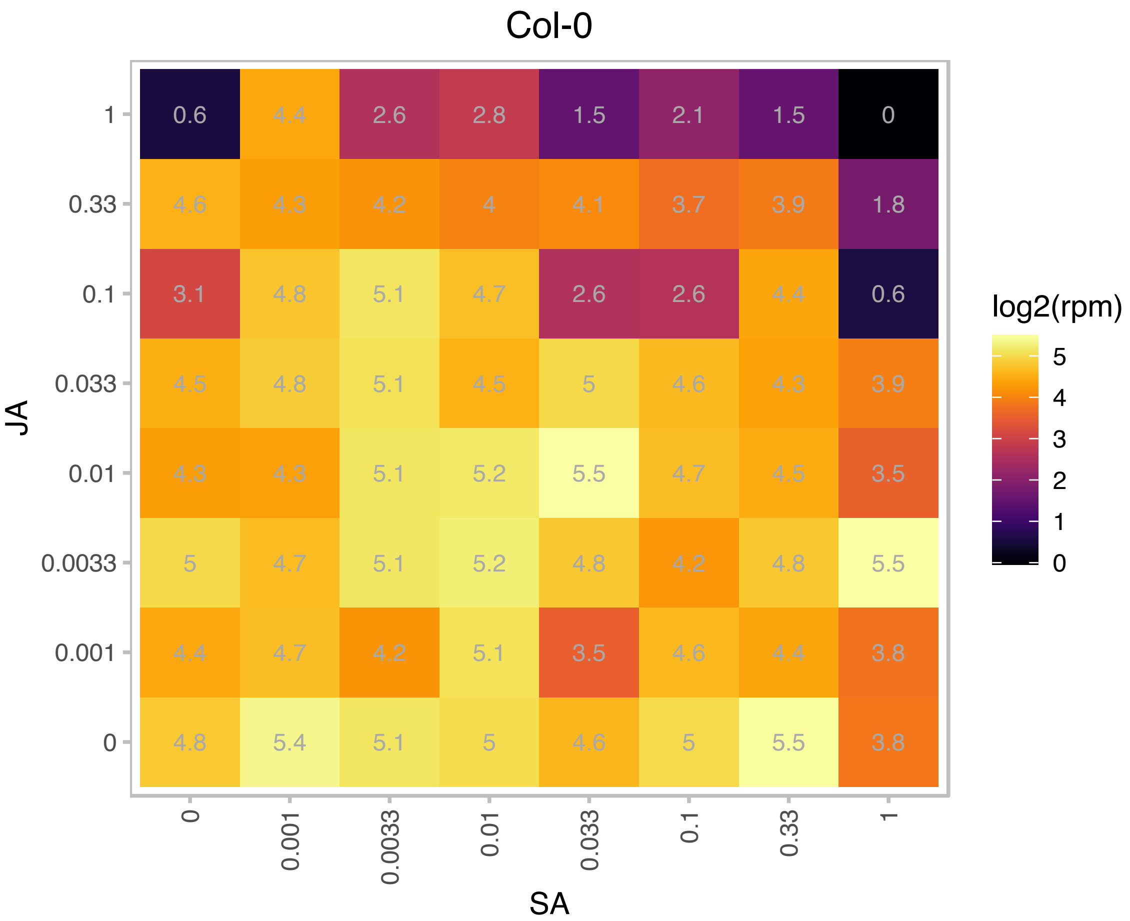


| | |
|---|---|
| Gene | AT4G16880 |
| is.expressed_genes | TRUE |
| is.responsive_genes | TRUE |
| is.apcluster | F-09 |

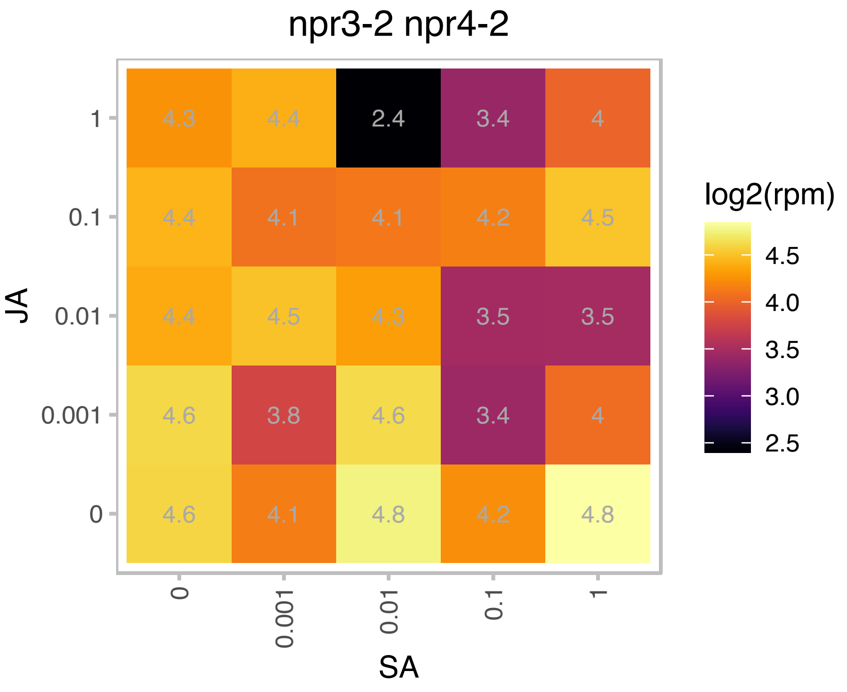


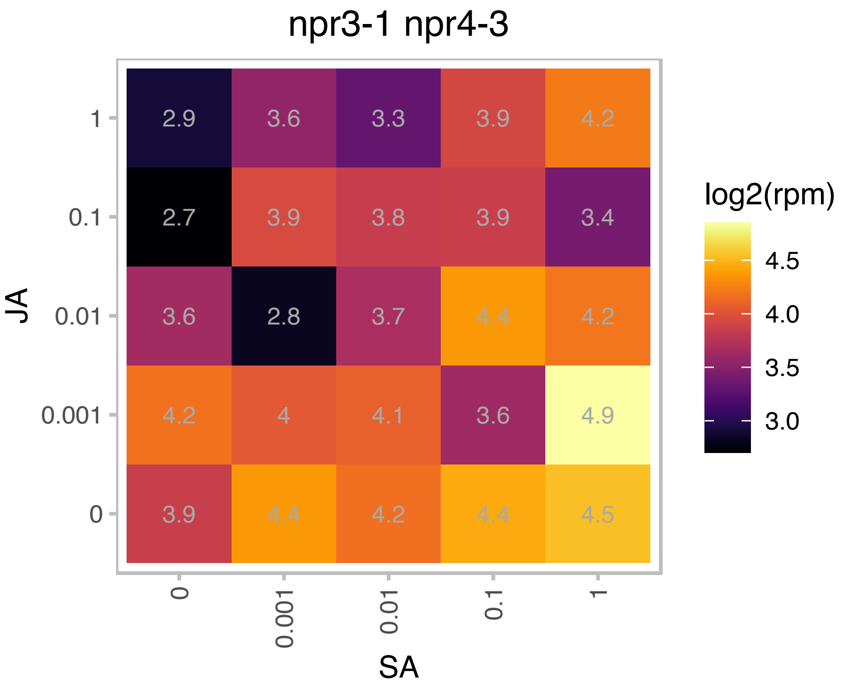

# AT4G16983.1 : NA

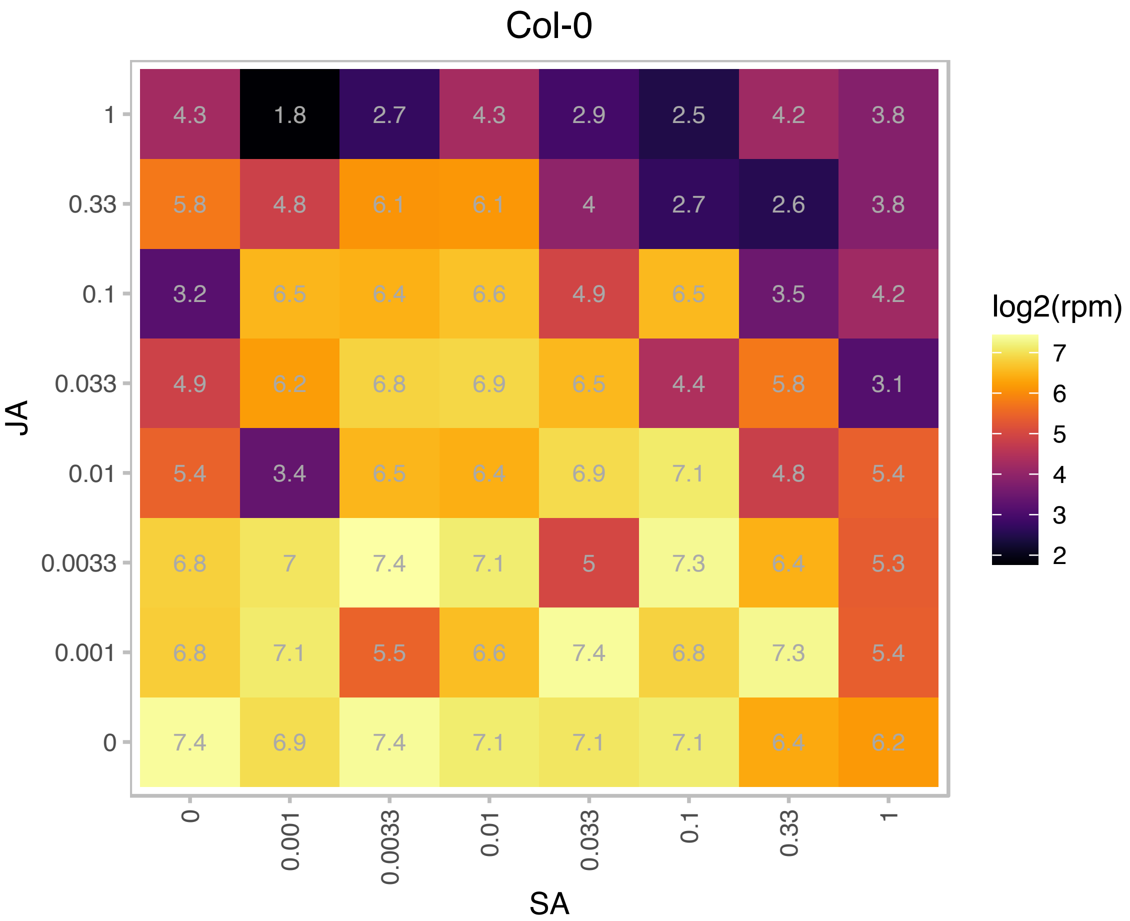


| | |
|---|---|
| Gene | AT4G16983 |
| is.expressed_genes | TRUE |
| is.responsive_genes | TRUE |
| is.apcluster | F-09 |

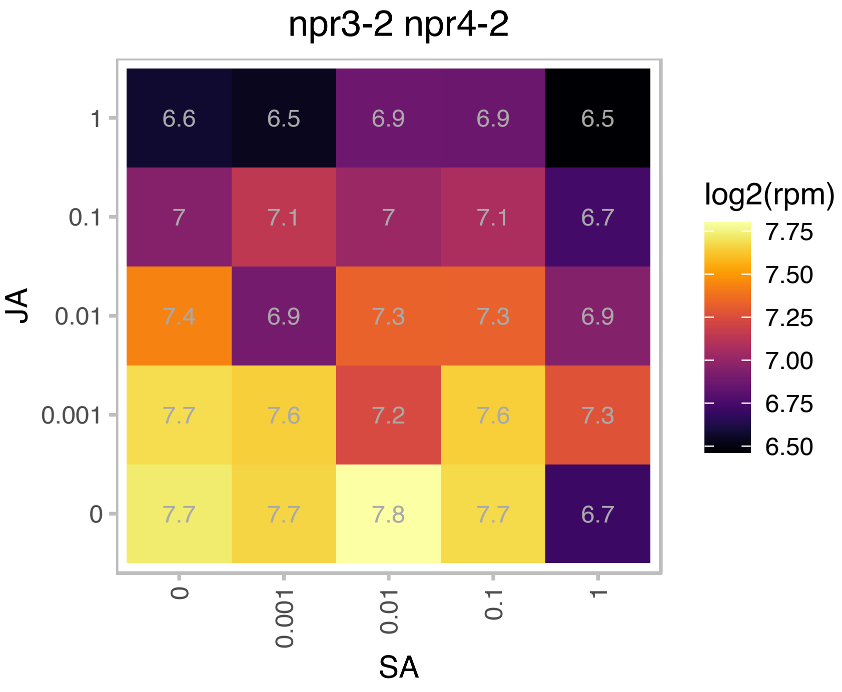


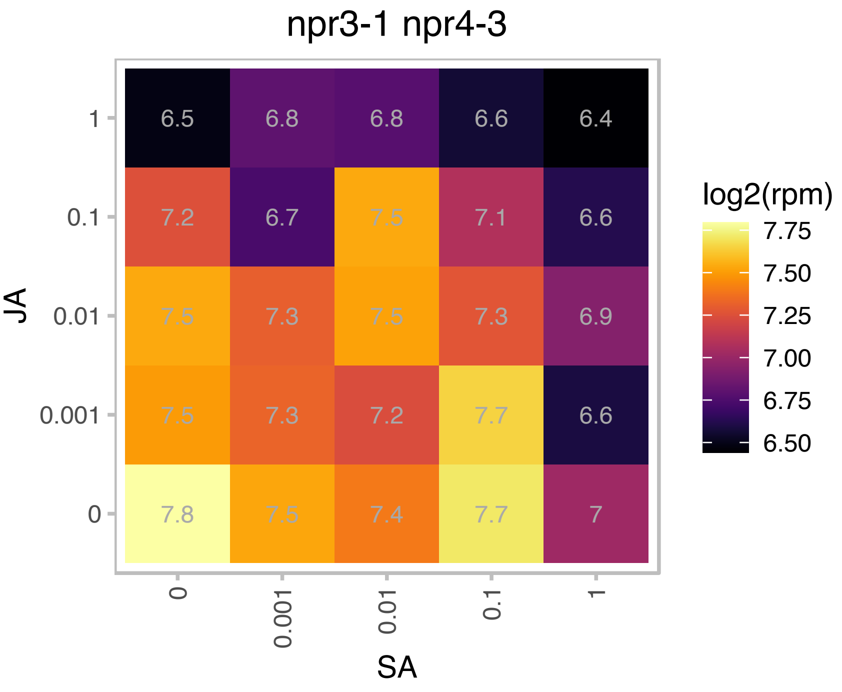

### AT4G18370.1 : DEG5, DEGP5, HHOA

DEGP protease 5

Encodes DEG5.  Forms a hexamer with DEG8 in the thylakoid lumen.  Involved in the cleavage of photodamaged D
protein of photosystem II (PSII).

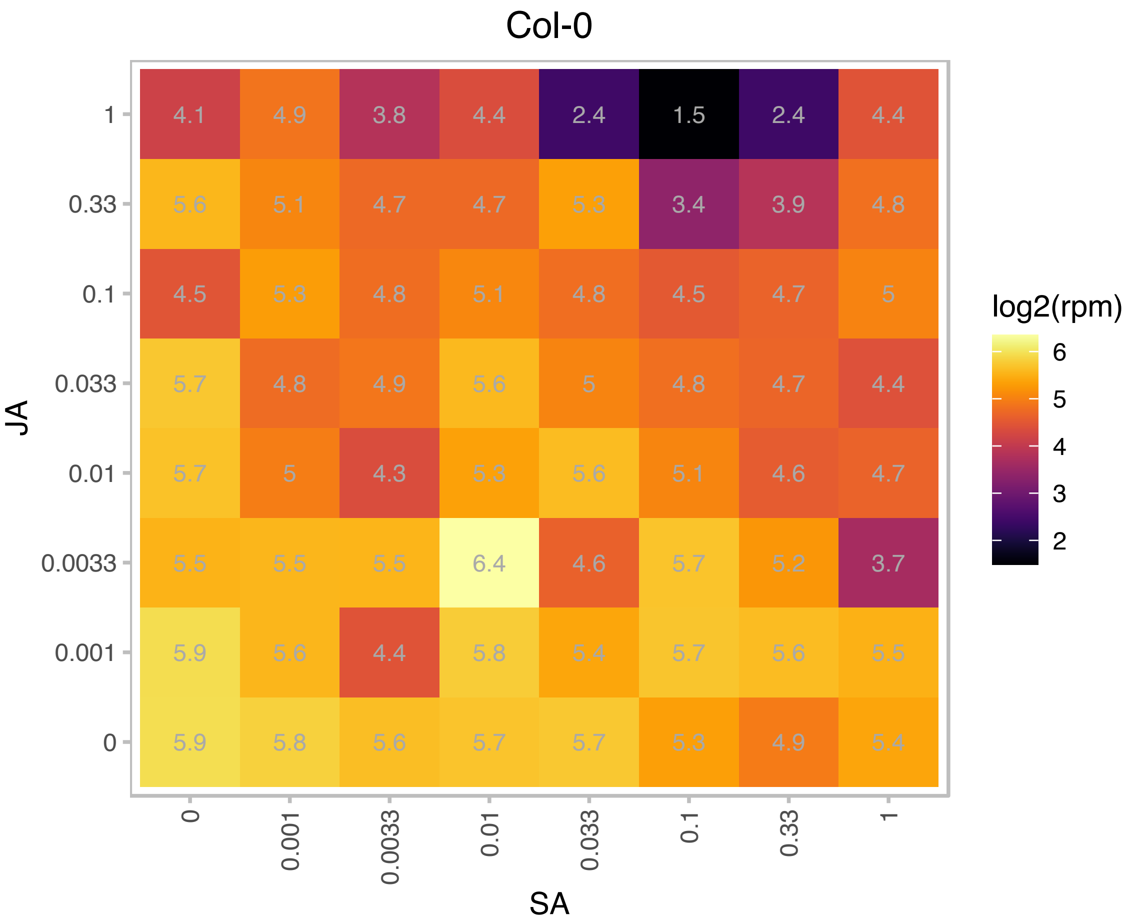


| | |
|---|---|
| Gene | AT4G18370 |
| is.expressed_genes | TRUE |
| is.responsive_genes | TRUE |
| is.apcluster | F-09 |

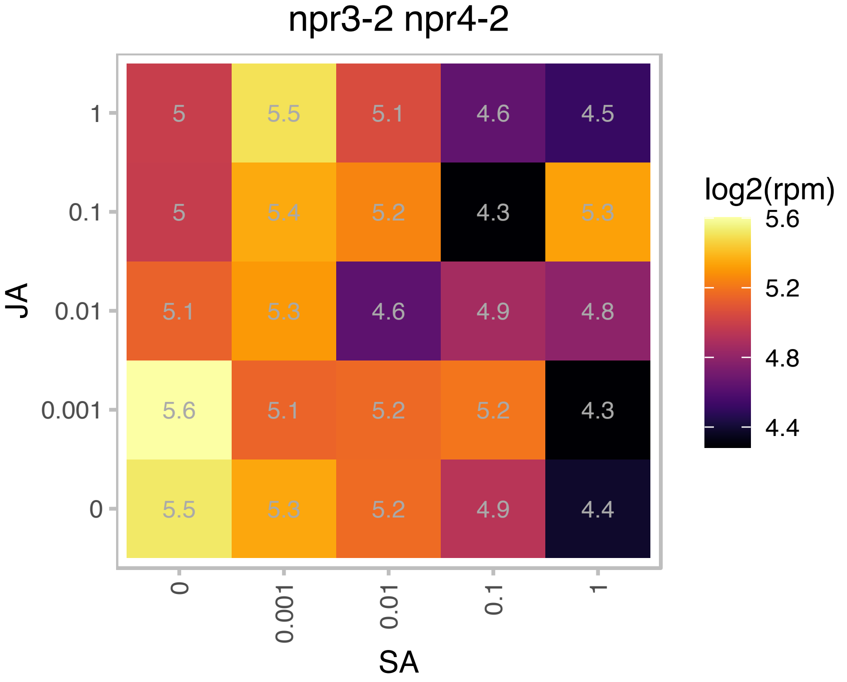


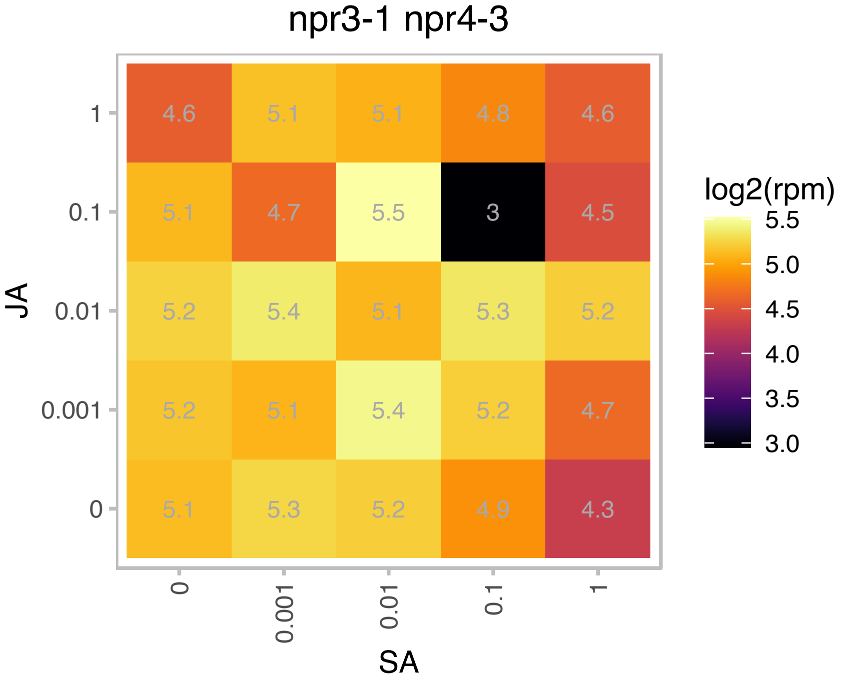

### AT4G18590.1 : RPA3B

Nucleic acid-binding, OB-fold-like protein

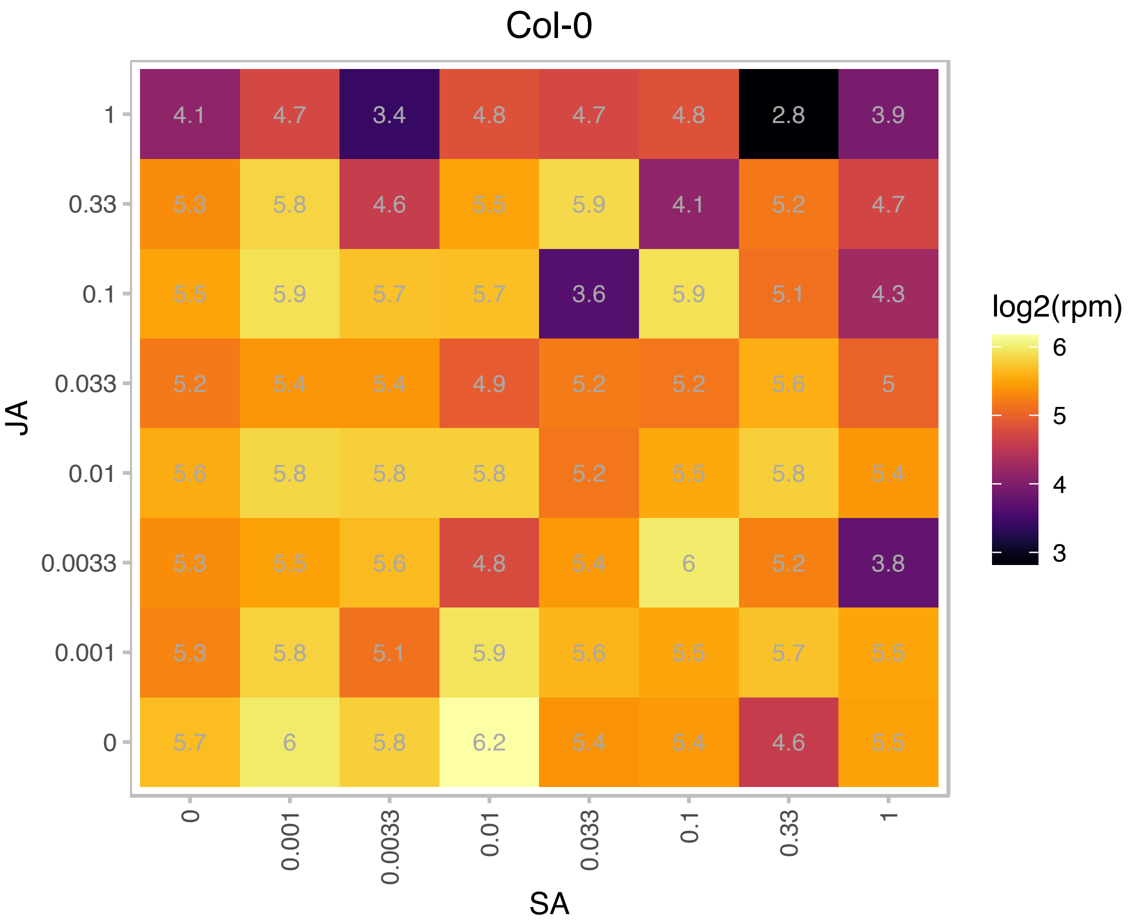


| | |
|---|---|
| Gene | AT4G18590 |
| is.expressed_genes | TRUE |
| is.responsive_genes | TRUE |
| is.apcluster | F-09 |

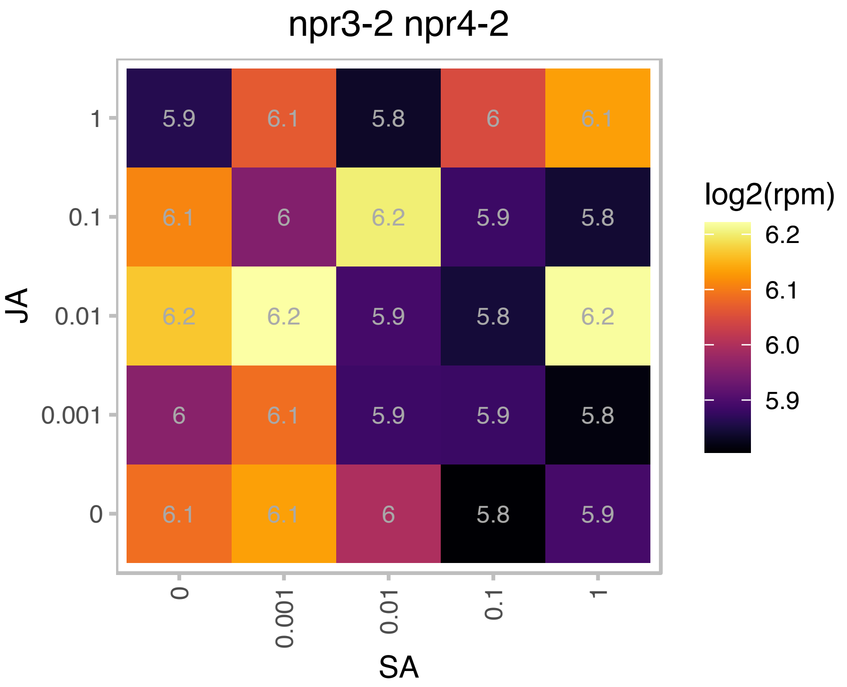


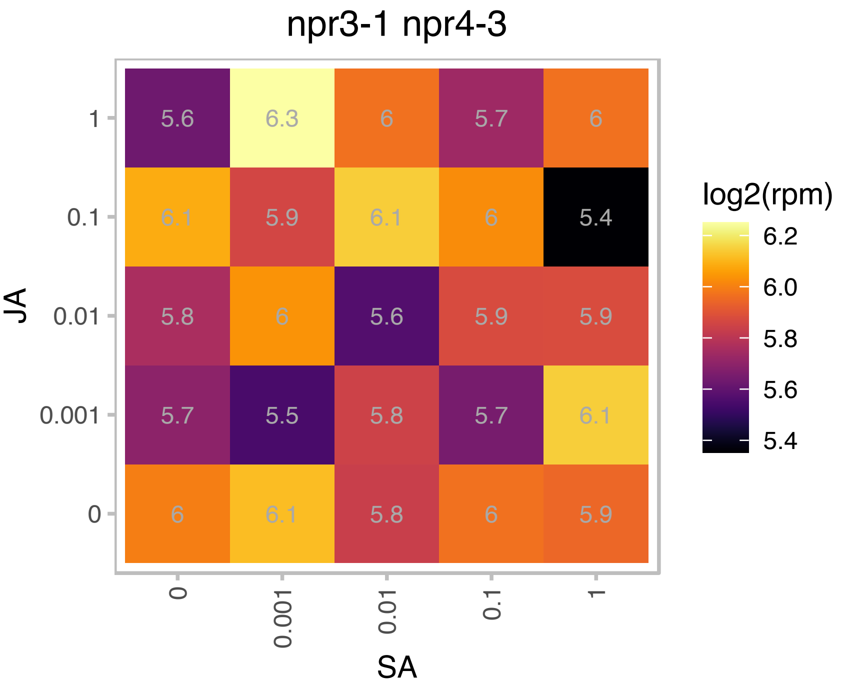

# AT4G22190.1 : NA

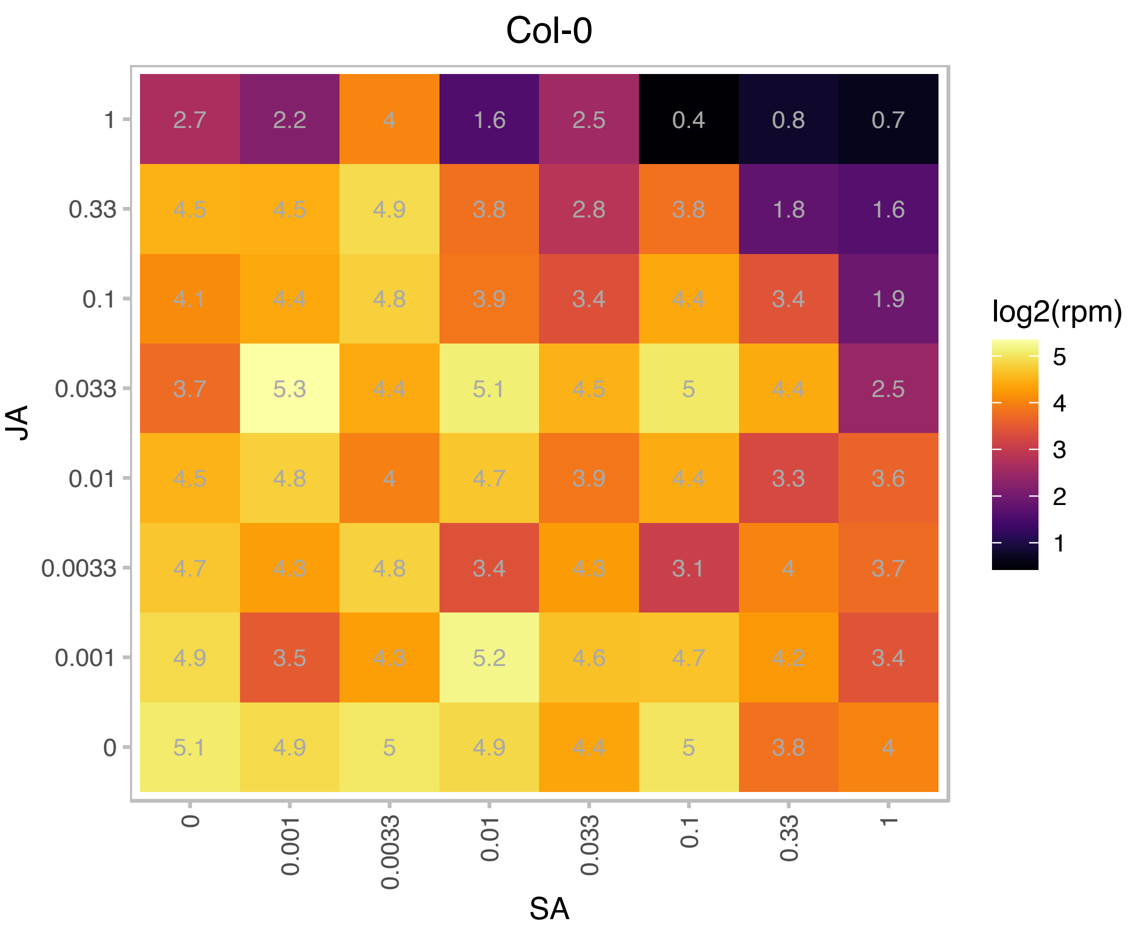


| | |
|---|---|
| Gene | AT4G22190 |
| is.expressed_genes | TRUE |
| is.responsive_genes | TRUE |
| is.apcluster | F-09 |

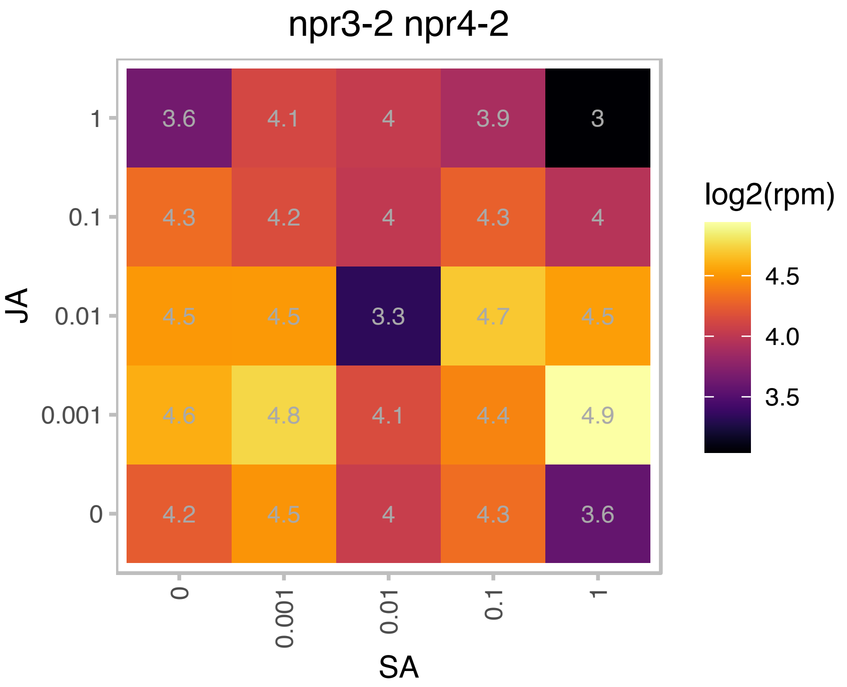


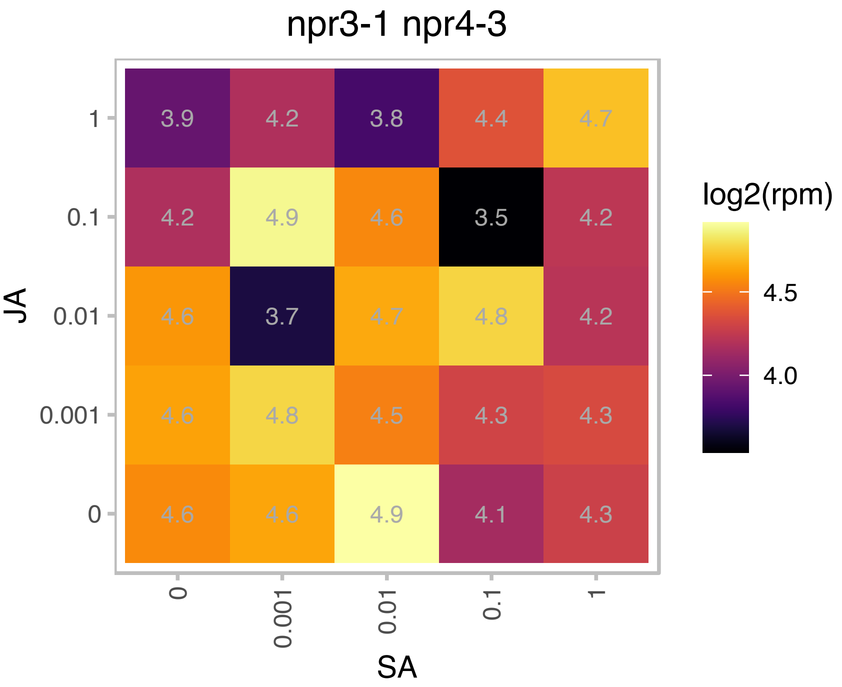

### AT4G23890.1 : CRR31, NdhS

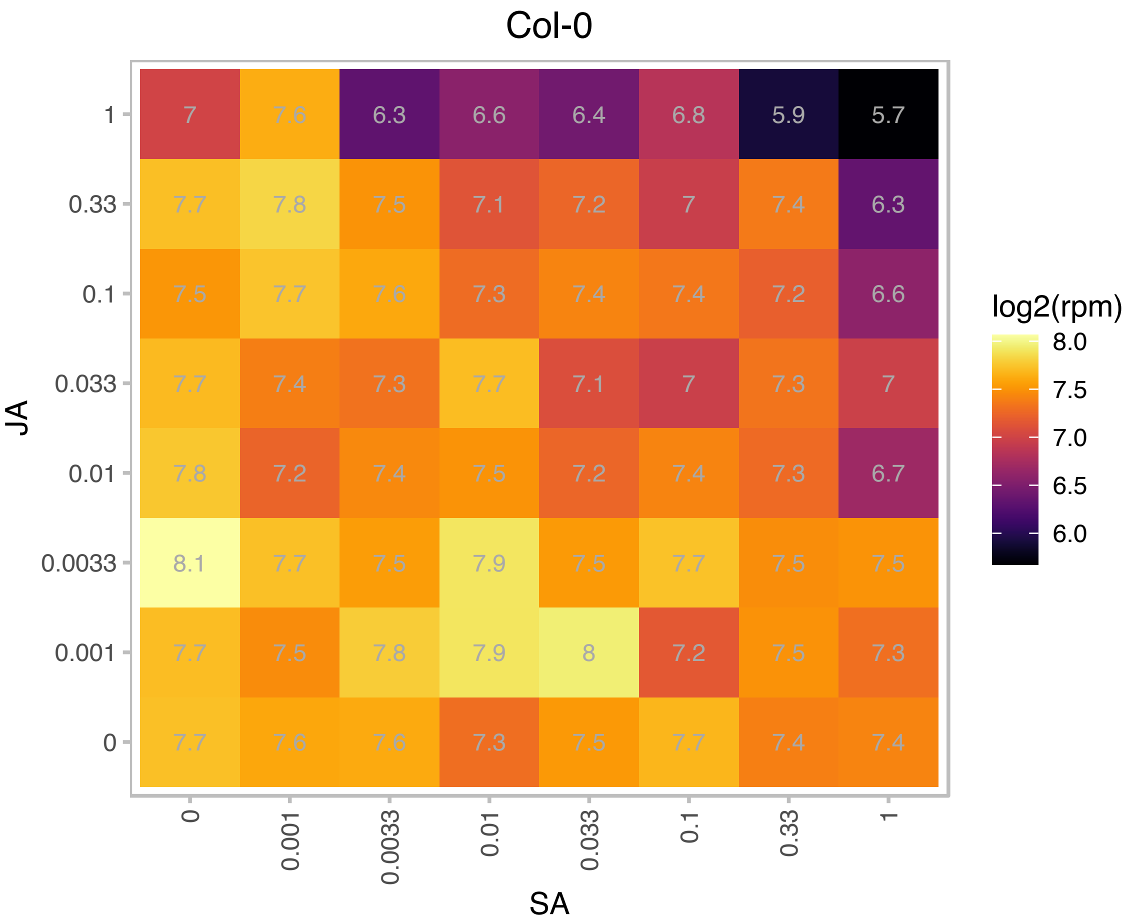


| | |
|---|---|
| Gene | AT4G23890 |
| is.expressed_genes | TRUE |
| is.responsive_genes | TRUE |
| is.apcluster | F-09 |

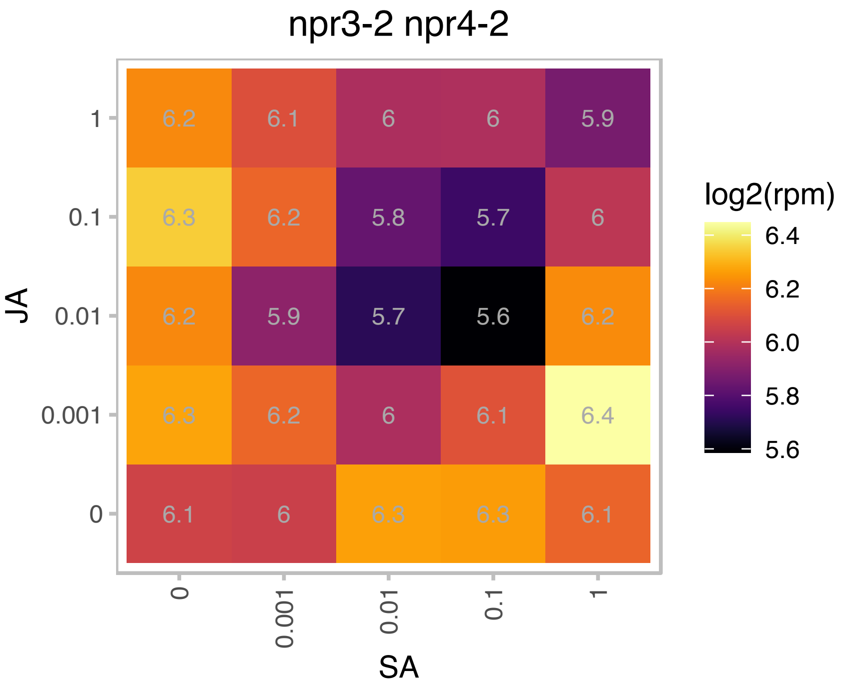


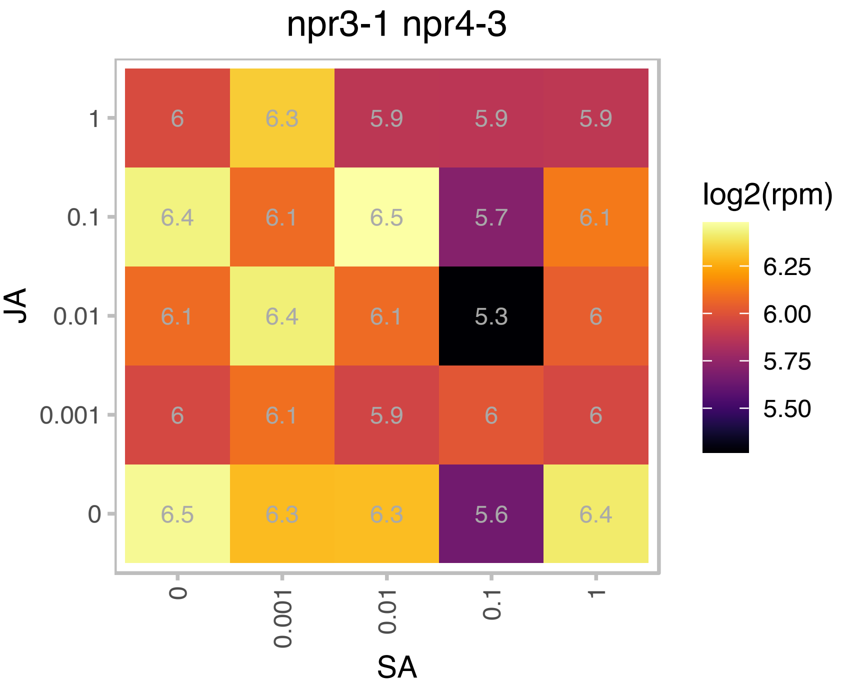

# AT4G24700.1 : NA

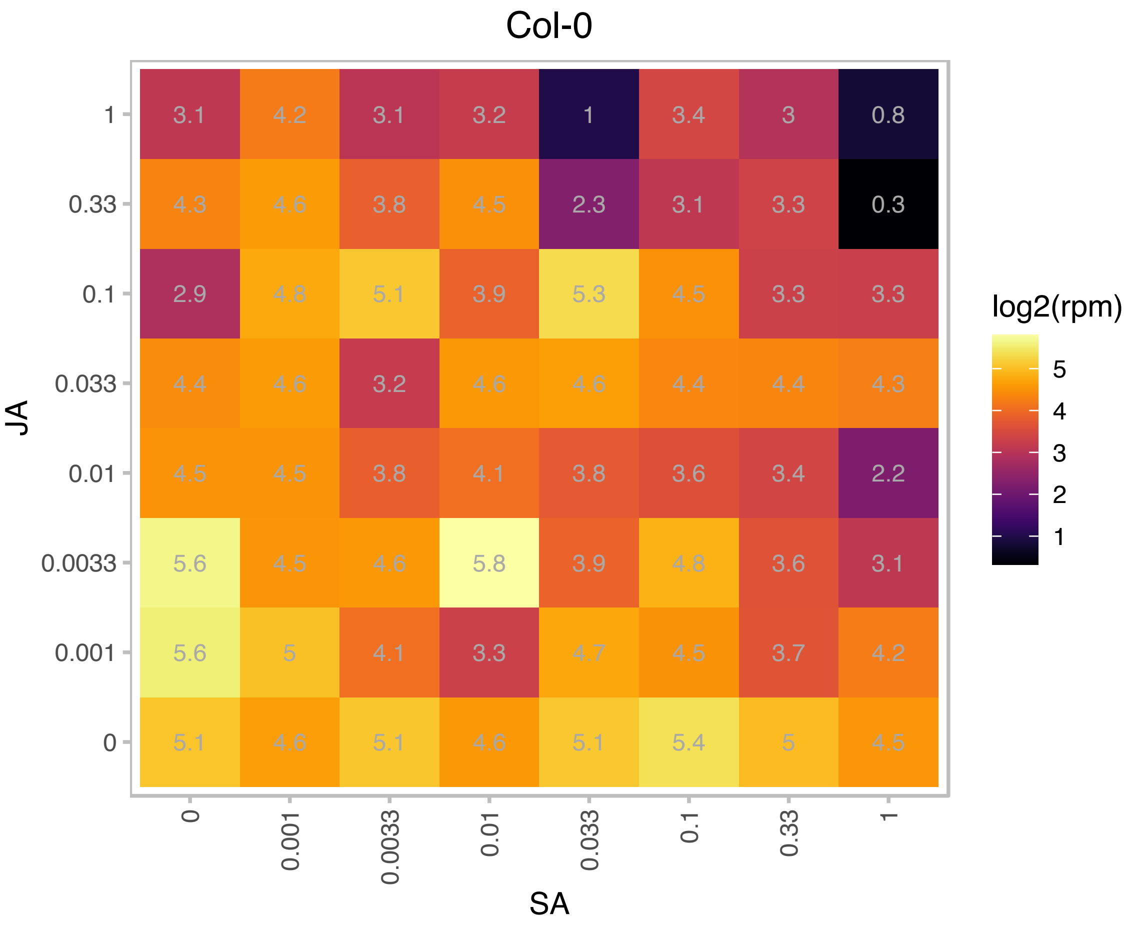


| | |
|---|---|
| Gene | AT4G24700 |
| is.expressed_genes | TRUE |
| is.responsive_genes | TRUE |
| is.apcluster | F-09 |

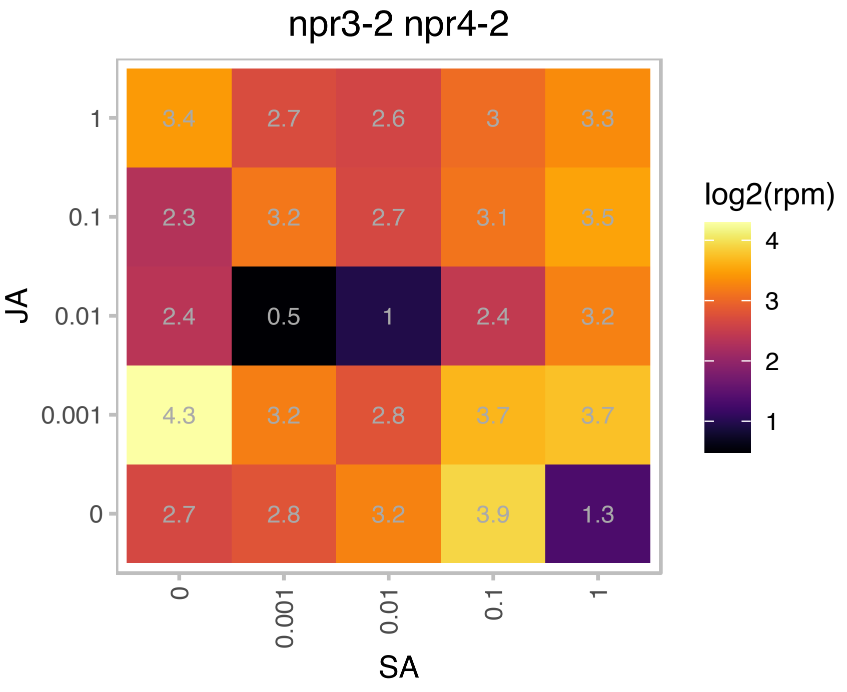


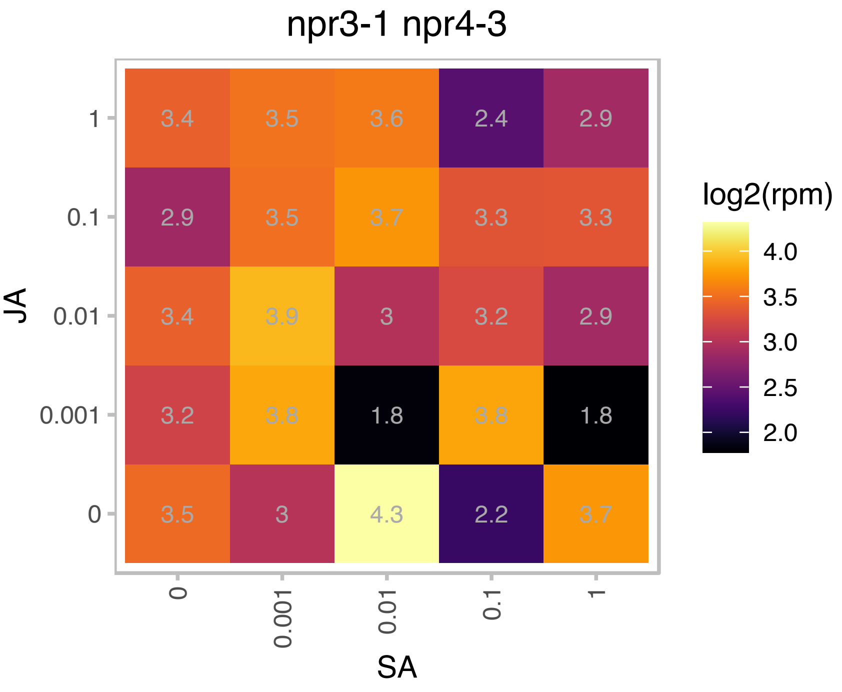

# AT4G32605.1 : NA

Mitochondrial glycoprotein family protein

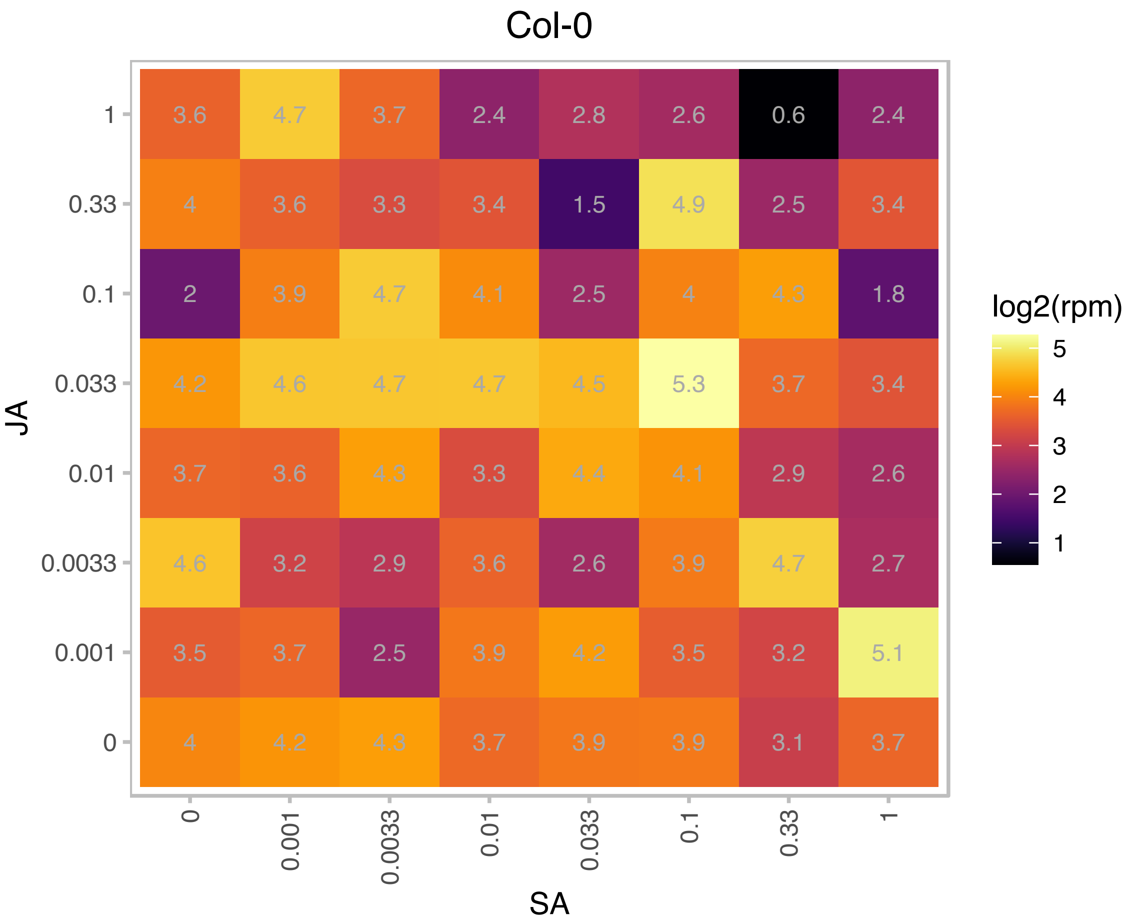


| | |
|---|---|
| Gene | AT4G32605 |
| is.expressed_genes | TRUE |
| is.responsive_genes | TRUE |
| is.apcluster | F-09 |

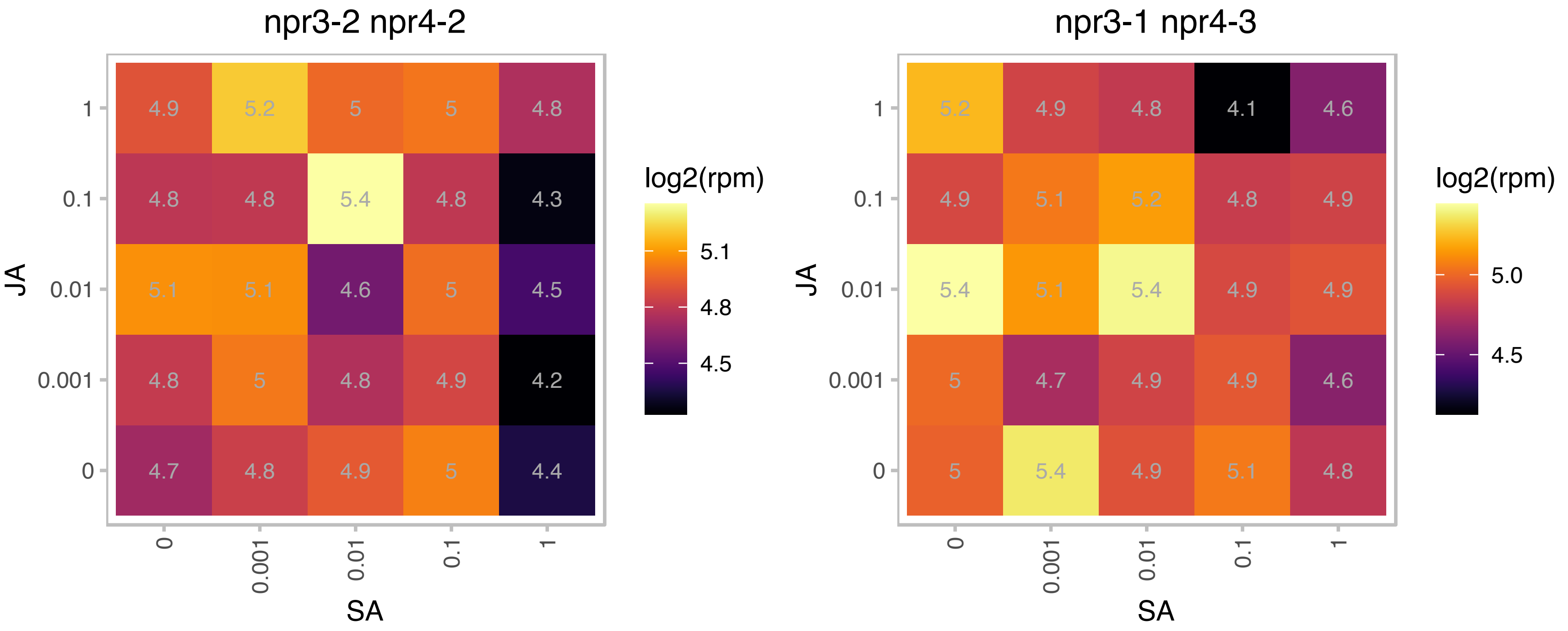

# AT4G33640.1 : NA

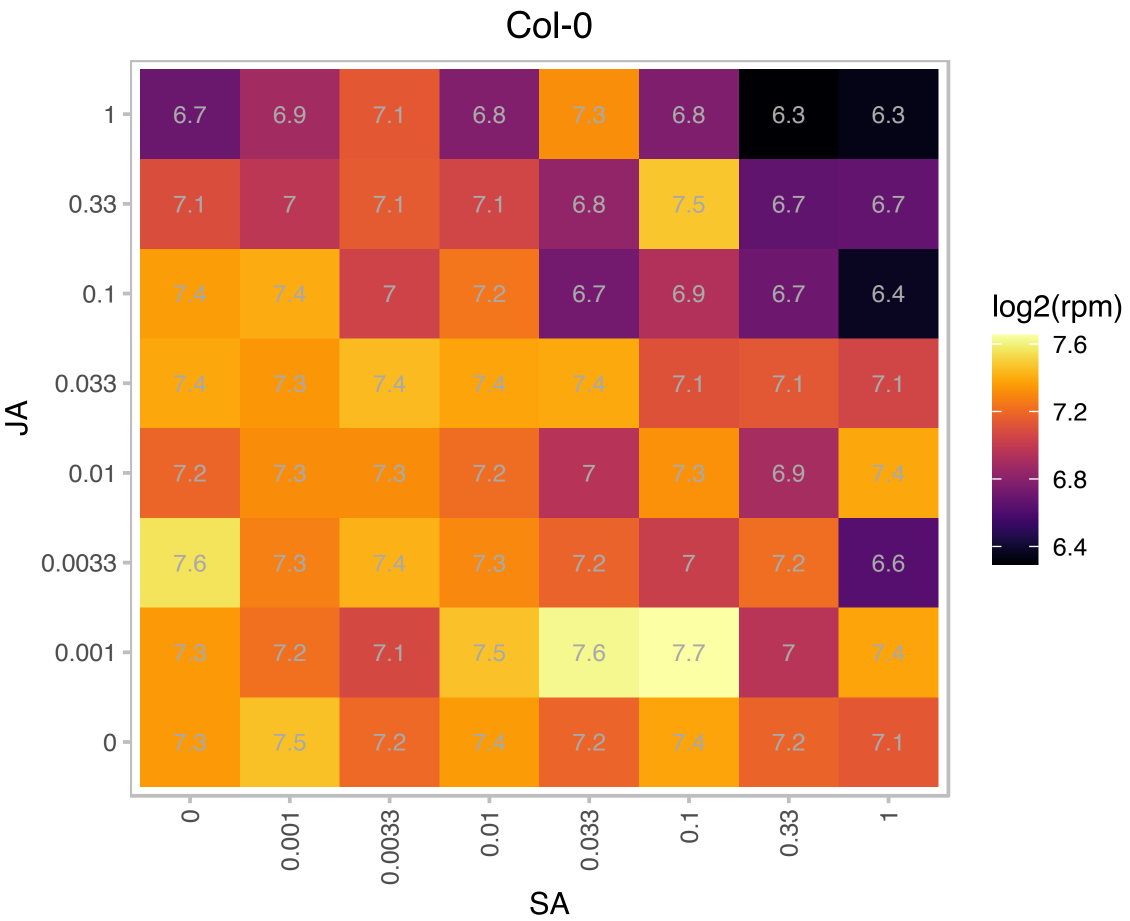


| | |
|---|---|
| Gene | AT4G33640 |
| is.expressed_genes | TRUE |
| is.responsive_genes | TRUE |
| is.apcluster | F-09 |

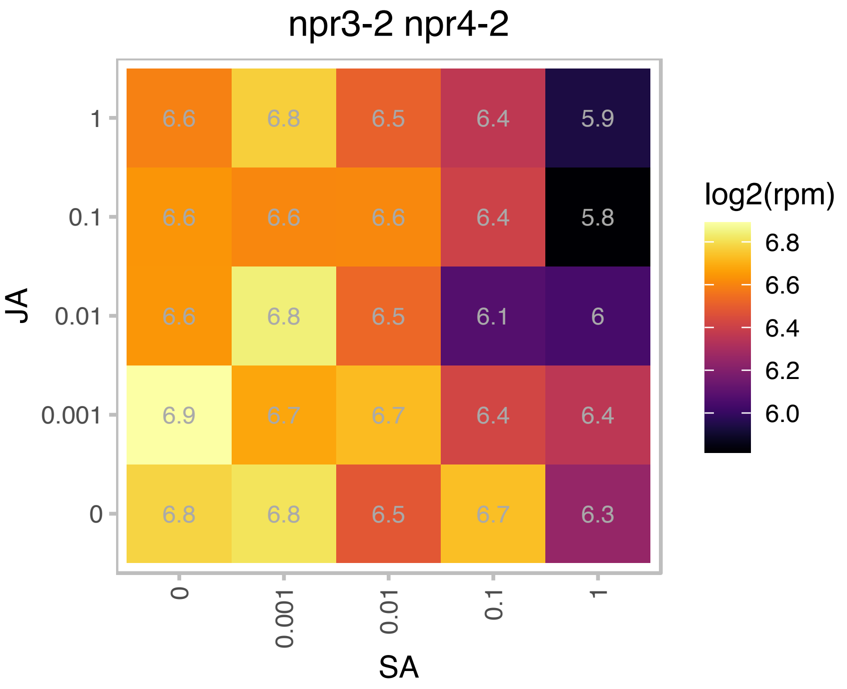


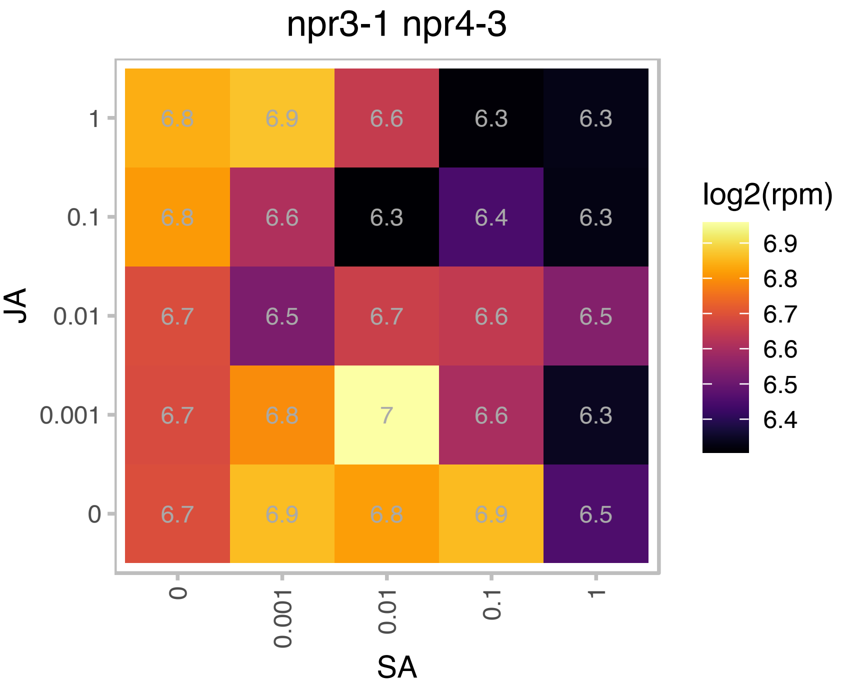

# AT4G38370.1 : NA

Phosphoglycerate mutase family protein

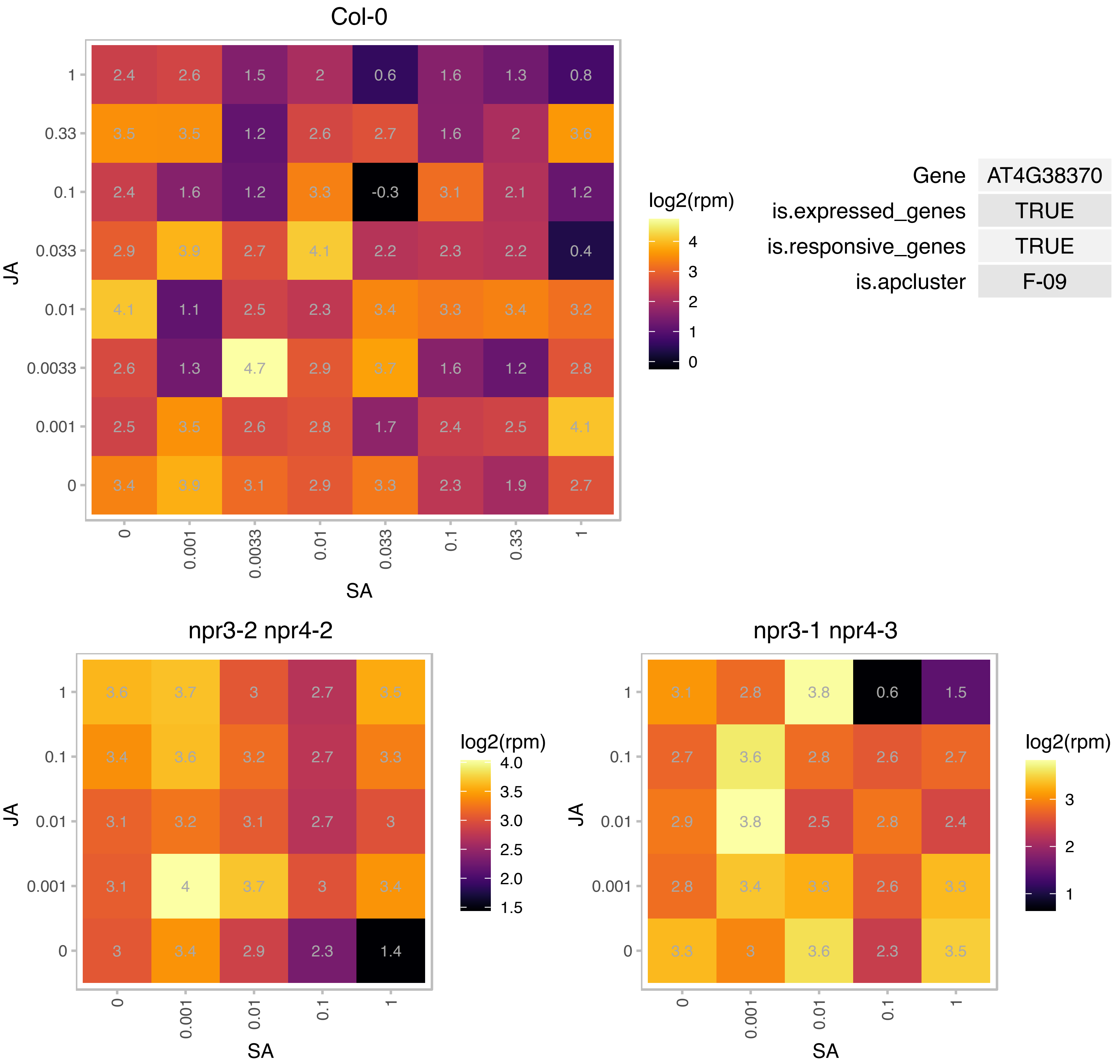


| | |
|---|---|
| Gene | AT4G38370 |
| is.expressed_genes | TRUE |
| is.responsive_genes | TRUE |
| is.apcluster | F-09 |

### AT5G07240.1 : IQD24

IQ-domain 24

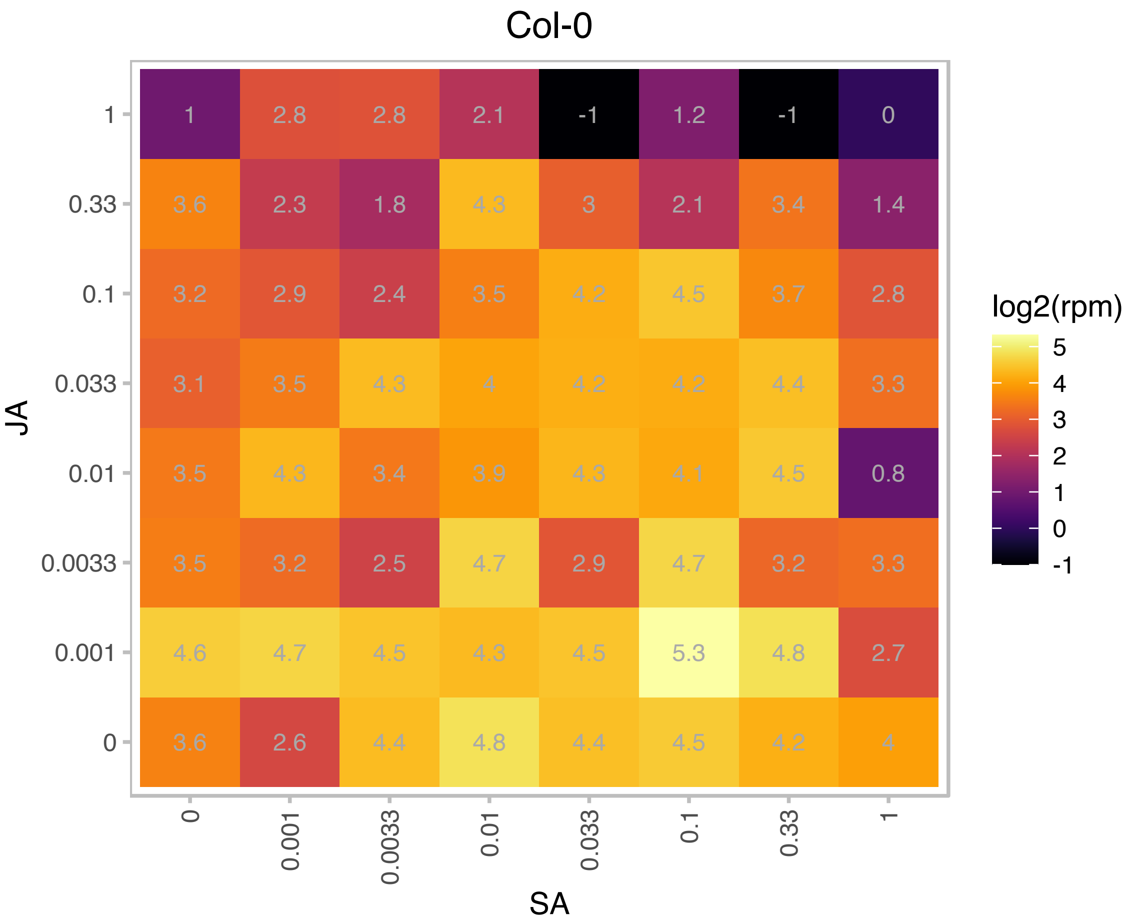


| | |
|---|---|
| Gene | AT5G07240 |
| is.expressed_genes | TRUE |
| is.responsive_genes | TRUE |
| is.apcluster | F-09 |

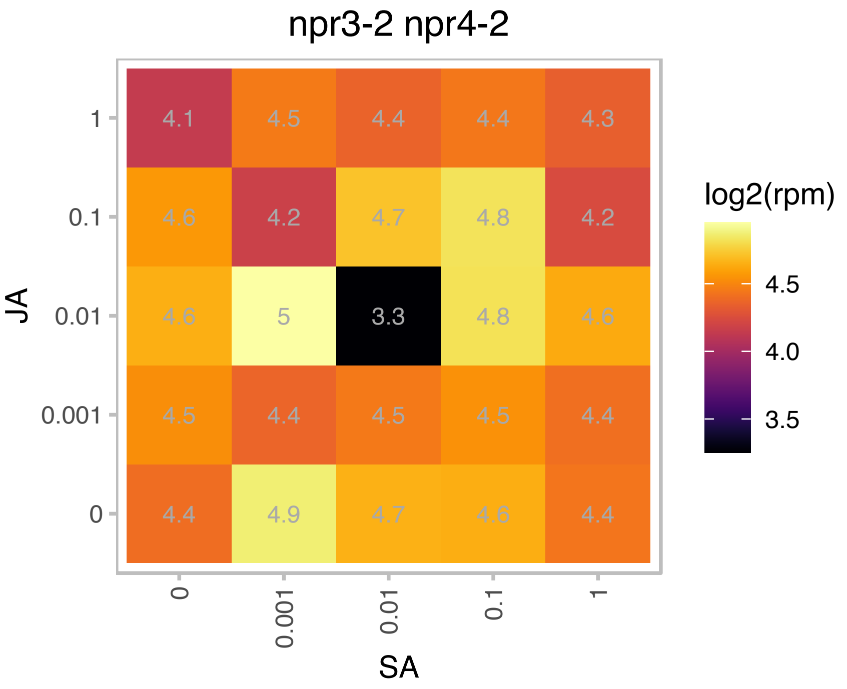


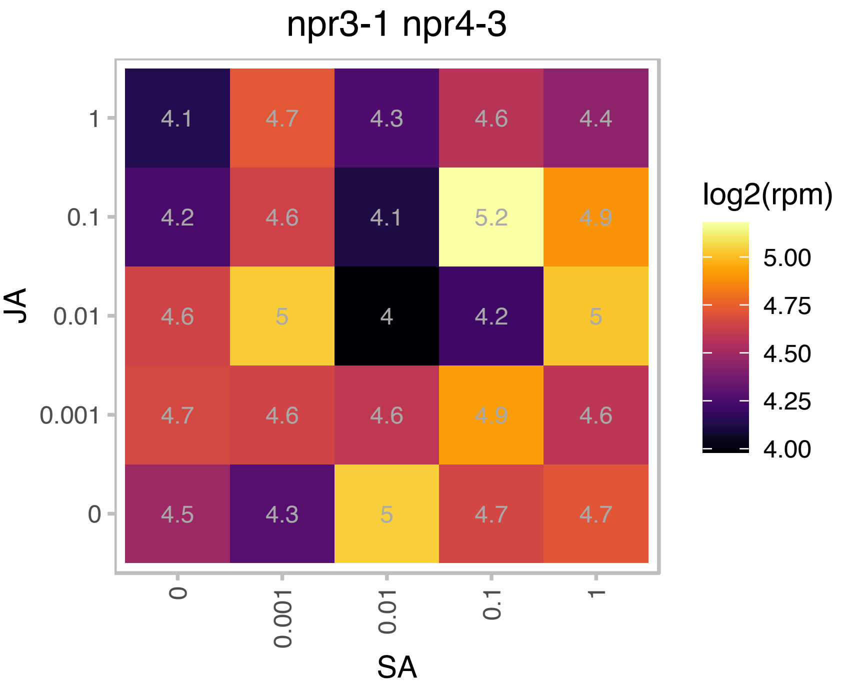

# AT5G13140.1 : NA

Pollen Ole e 1 allergen and extensin family protein

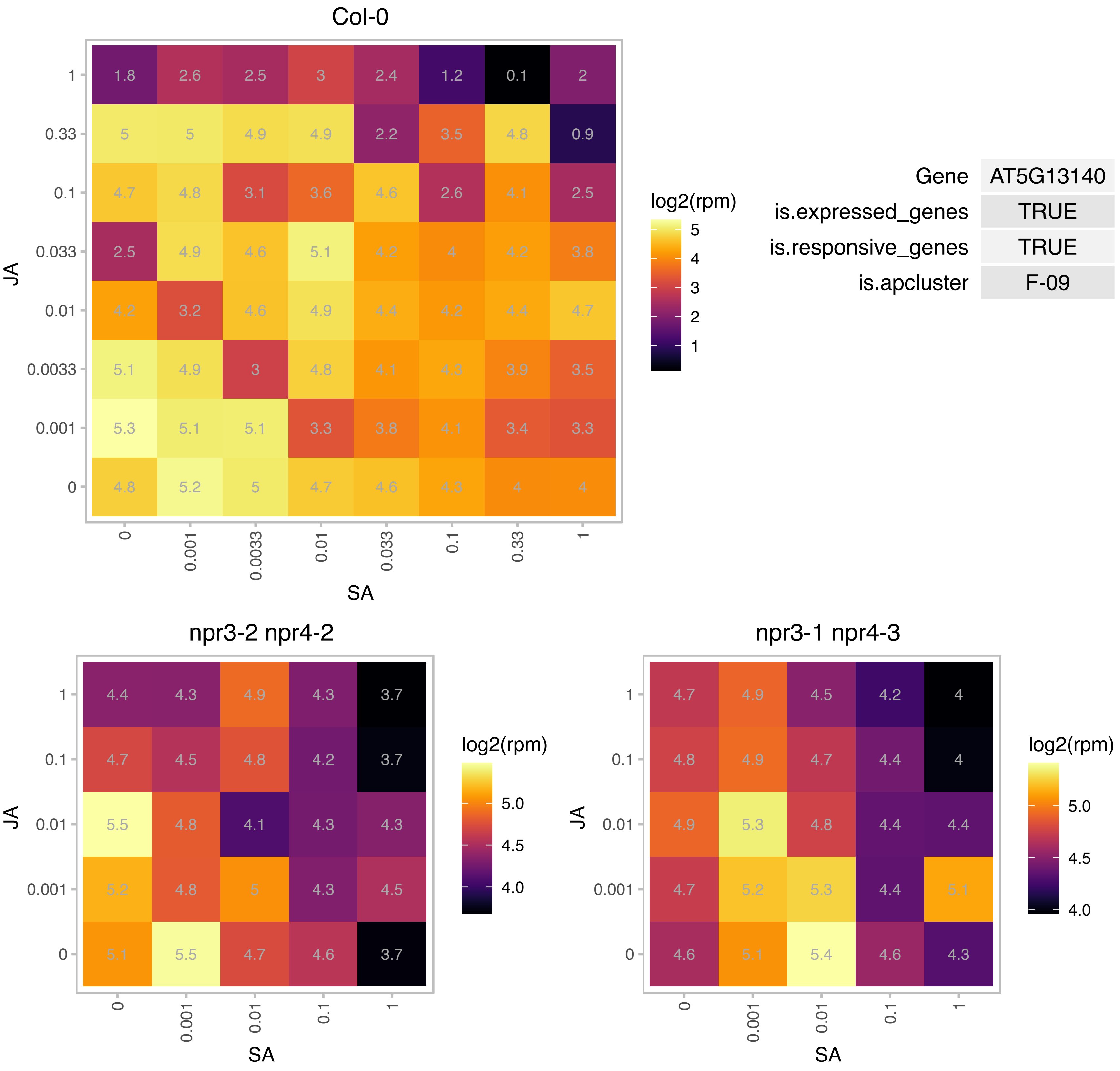


| | |
|---|---|
| Gene | AT5G13140 |
| is.expressed_genes | TRUE |
| is.responsive_genes | TRUE |
| is.apcluster | F-09 |

# AT5G14410.1 : NA

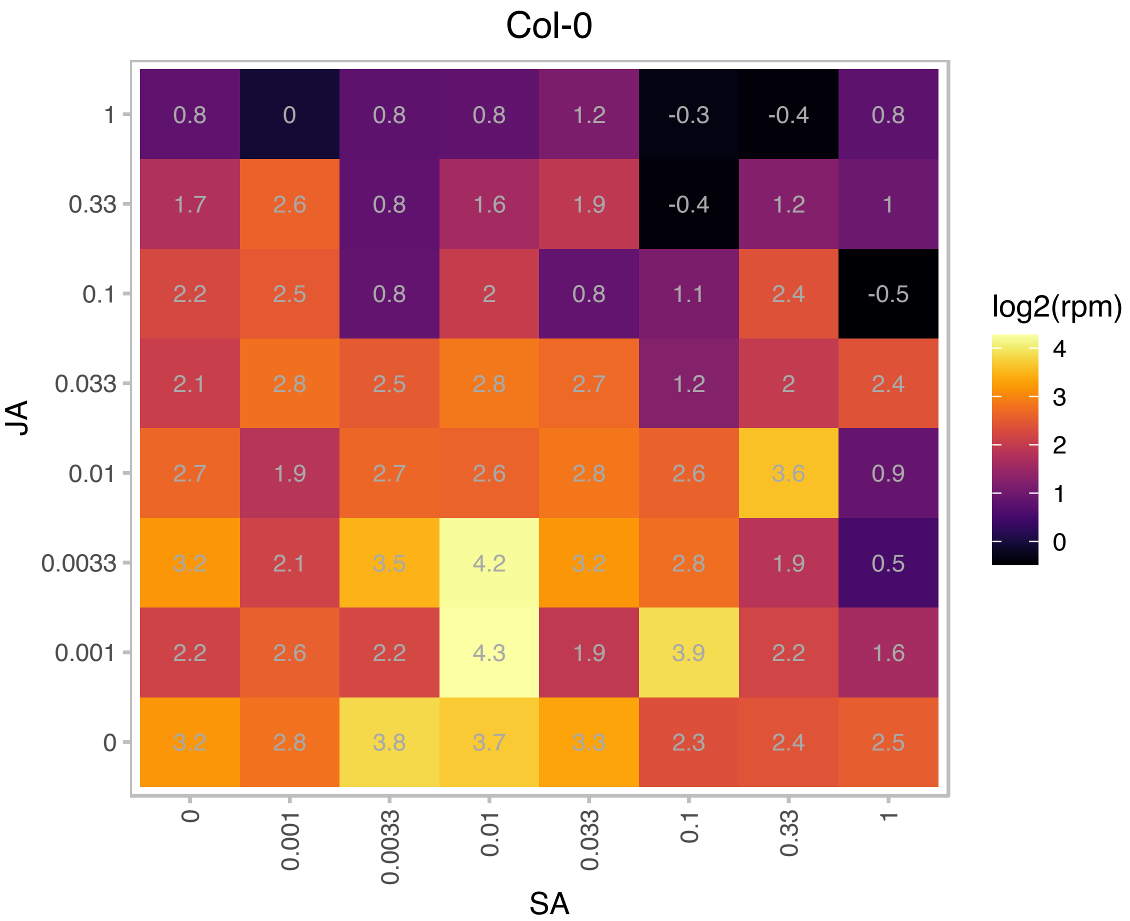


| | |
|---|---|
| Gene | AT5G14410 |
| is.expressed_genes | TRUE |
| is.responsive_genes | TRUE |
| is.apcluster | F-09 |

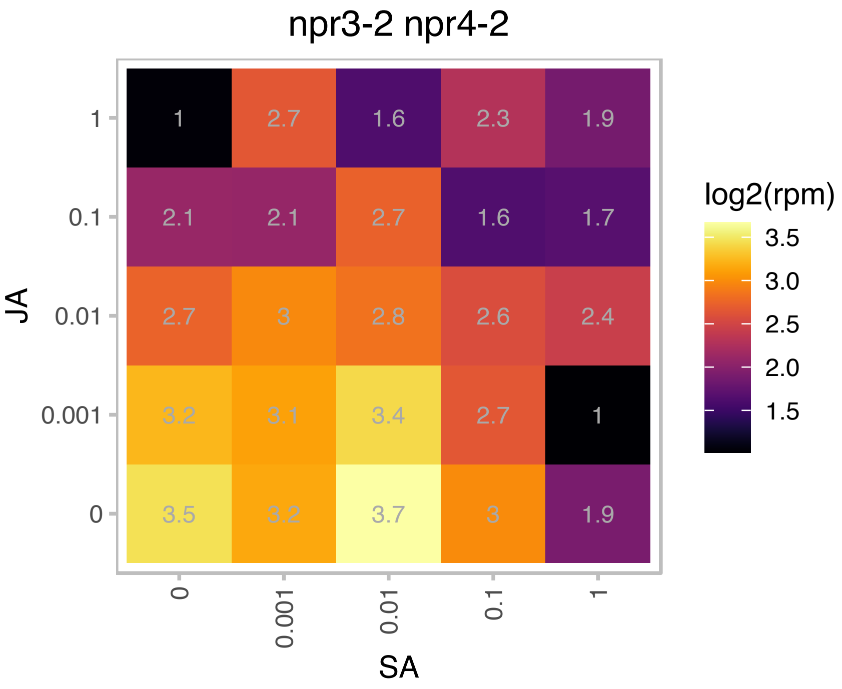


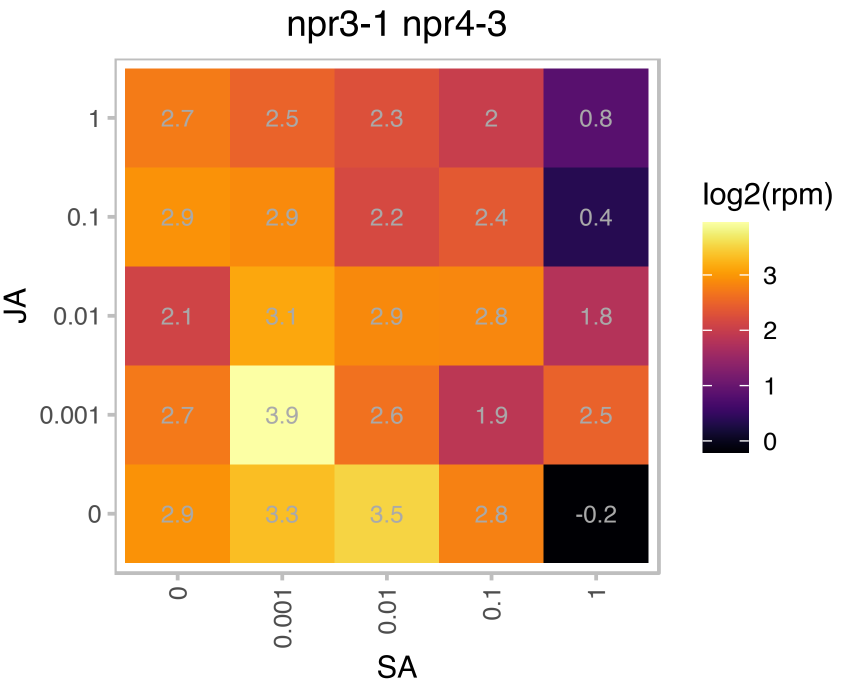

# AT5G16030.1 : NA

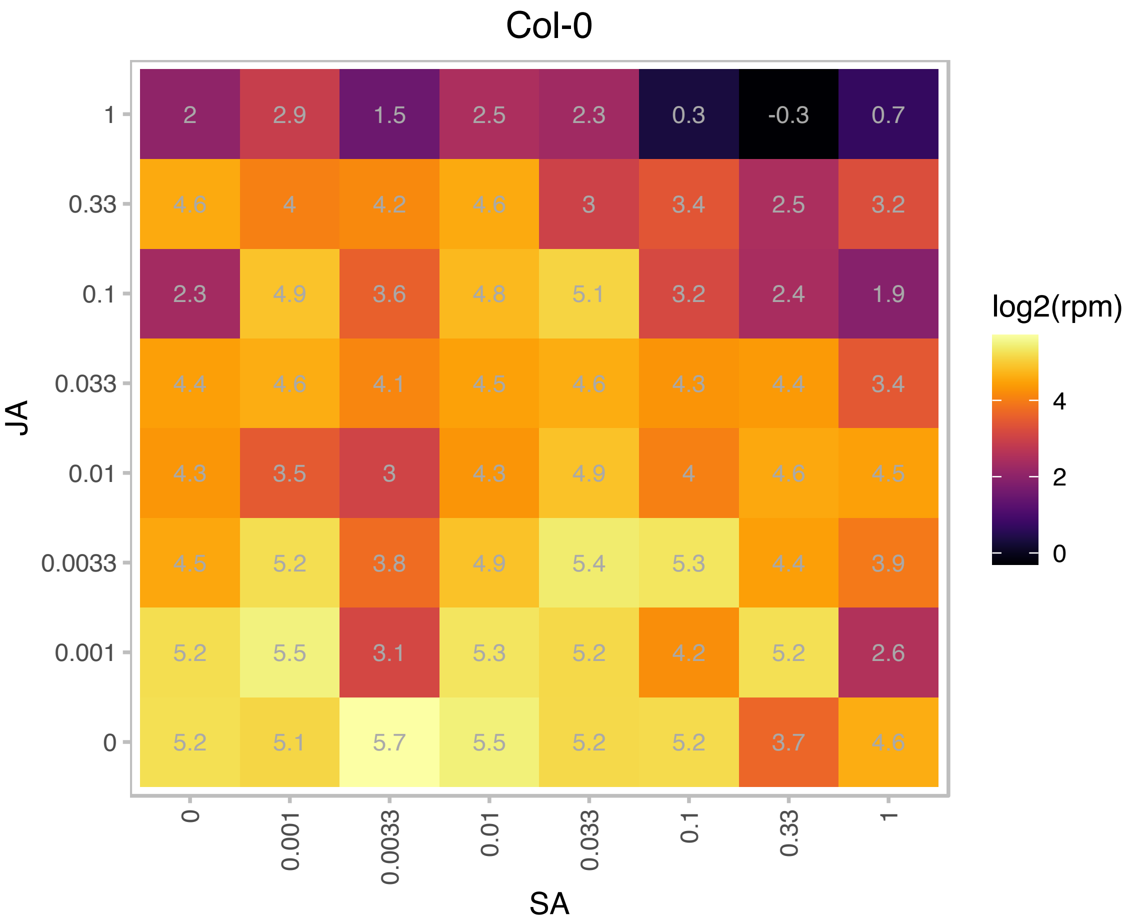


| | |
|---|---|
| Gene | AT5G16030 |
| is.expressed_genes | TRUE |
| is.responsive_genes | TRUE |
| is.apcluster | F-09 |

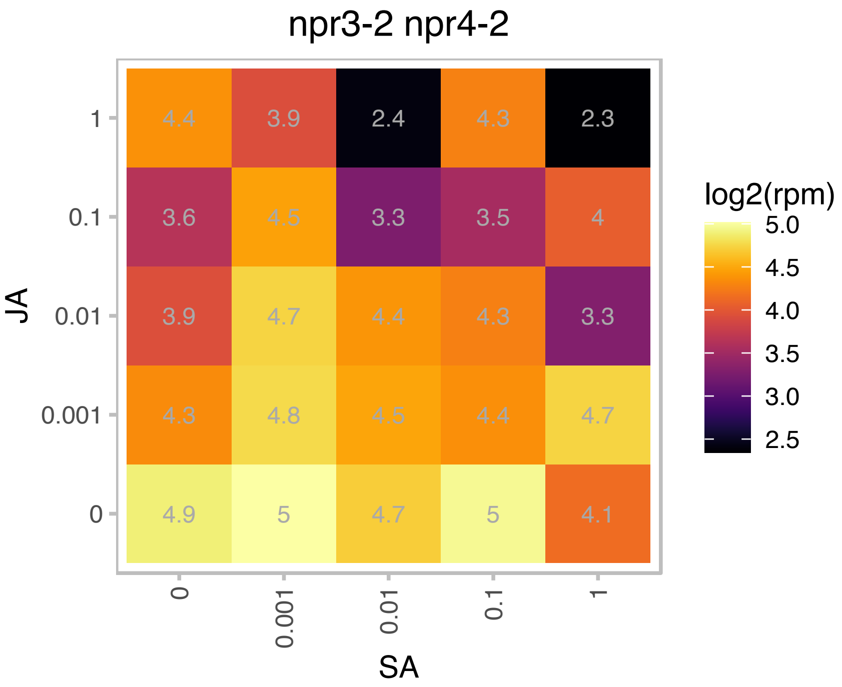


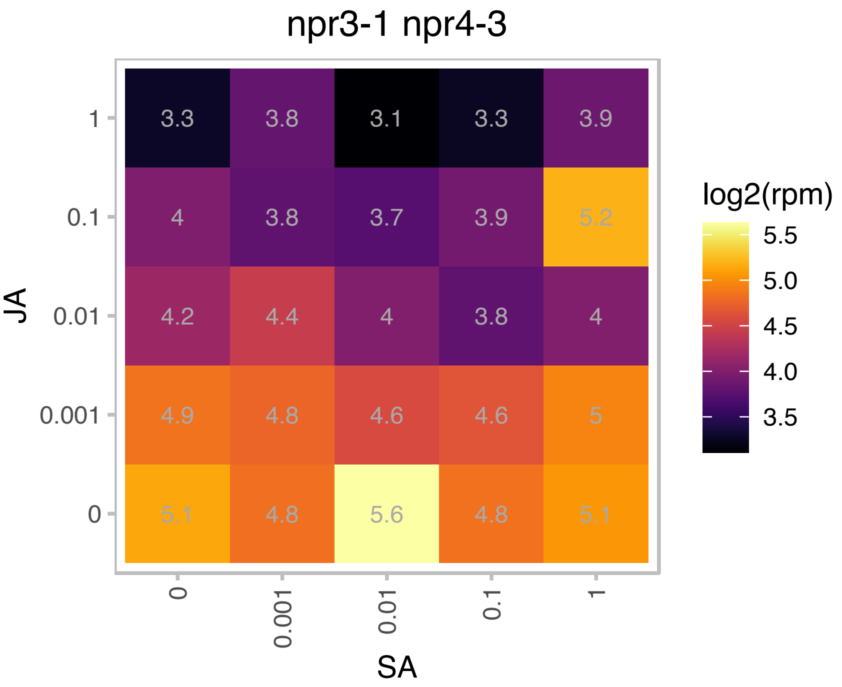

# AT5G16250.1 : NA

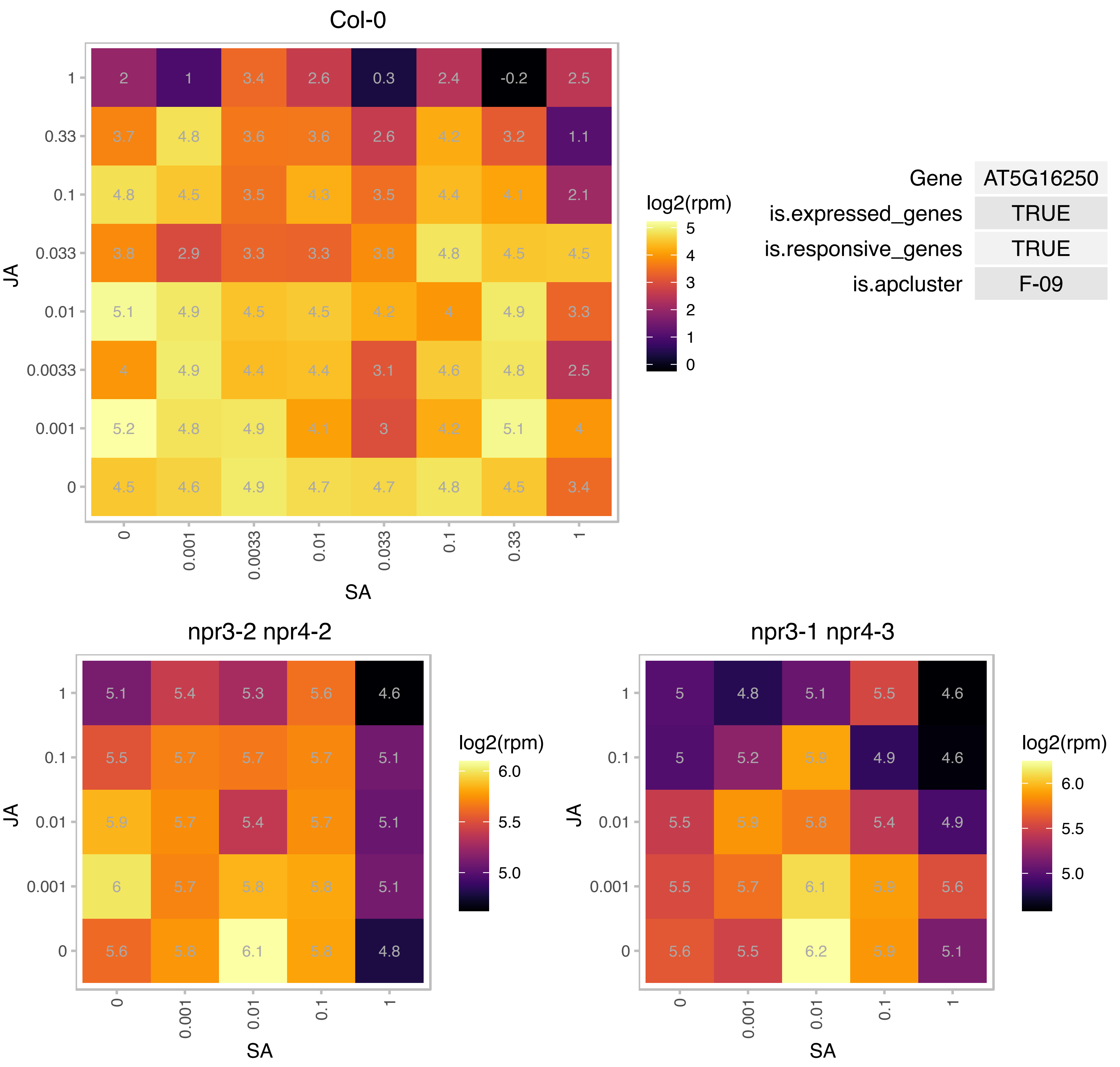


| | |
|---|---|
| Gene | AT5G16250 |
| is.expressed_genes | TRUE |
| is.responsive_genes | TRUE |
| is.apcluster | F-09 |

### AT5G18140.1 : DJC69

Chaperone DnaJ-domain superfamily protein

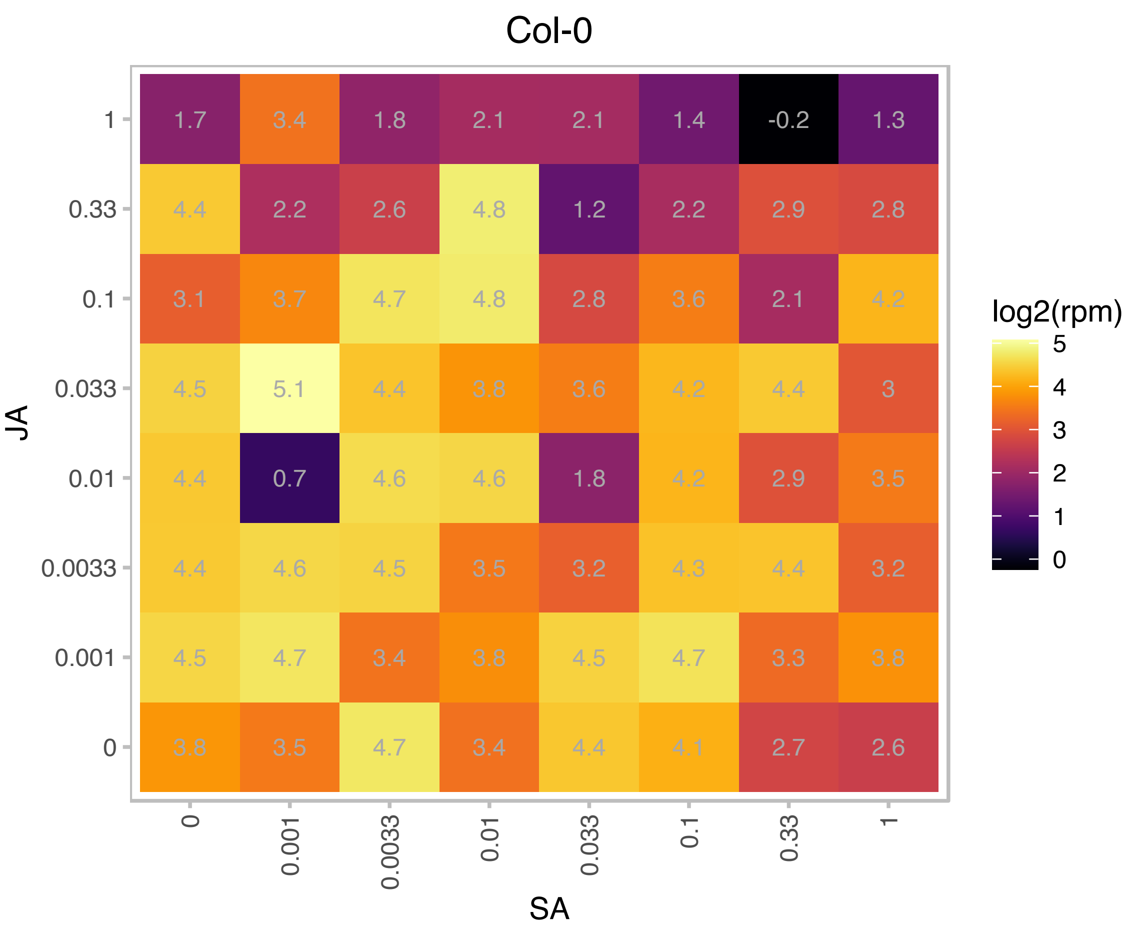


| | |
|---|---|
| Gene | AT5G18140 |
| is.expressed_genes | TRUE |
| is.responsive_genes | TRUE |
| is.apcluster | F-09 |

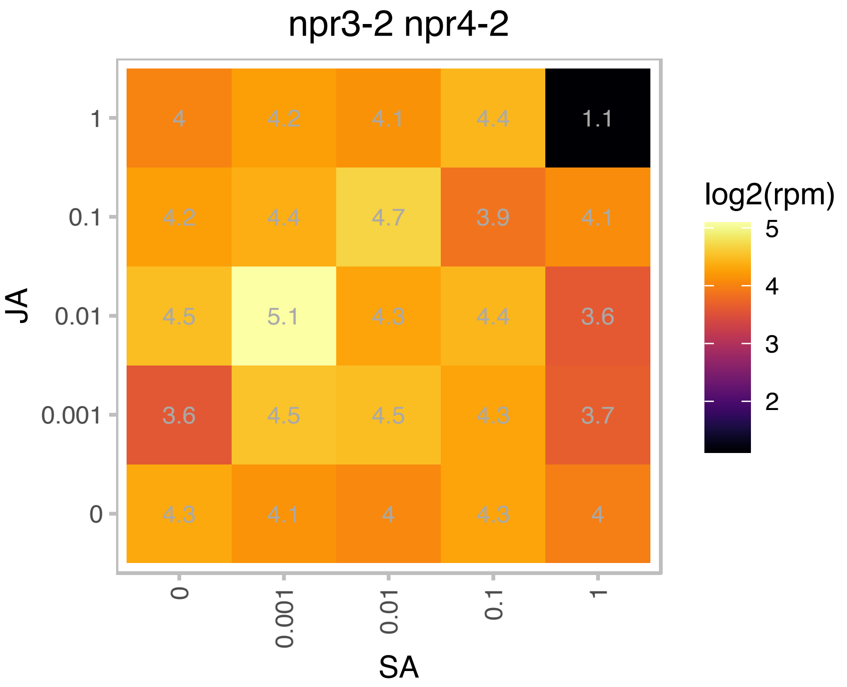


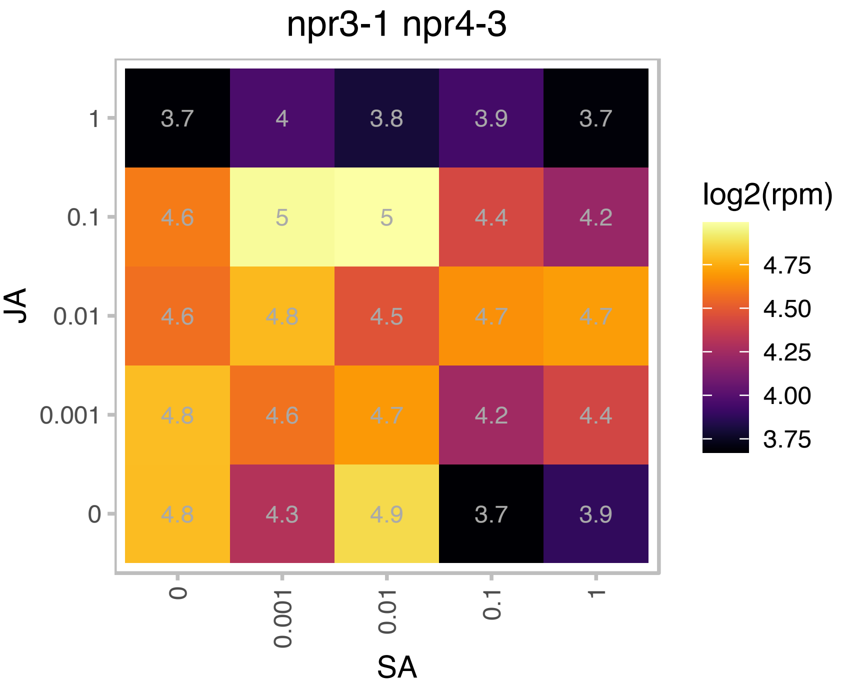

### AT5G19140.1 : AILP1, ATAILP1

Aluminium induced protein with YGL and LRDR motifs

Col-0

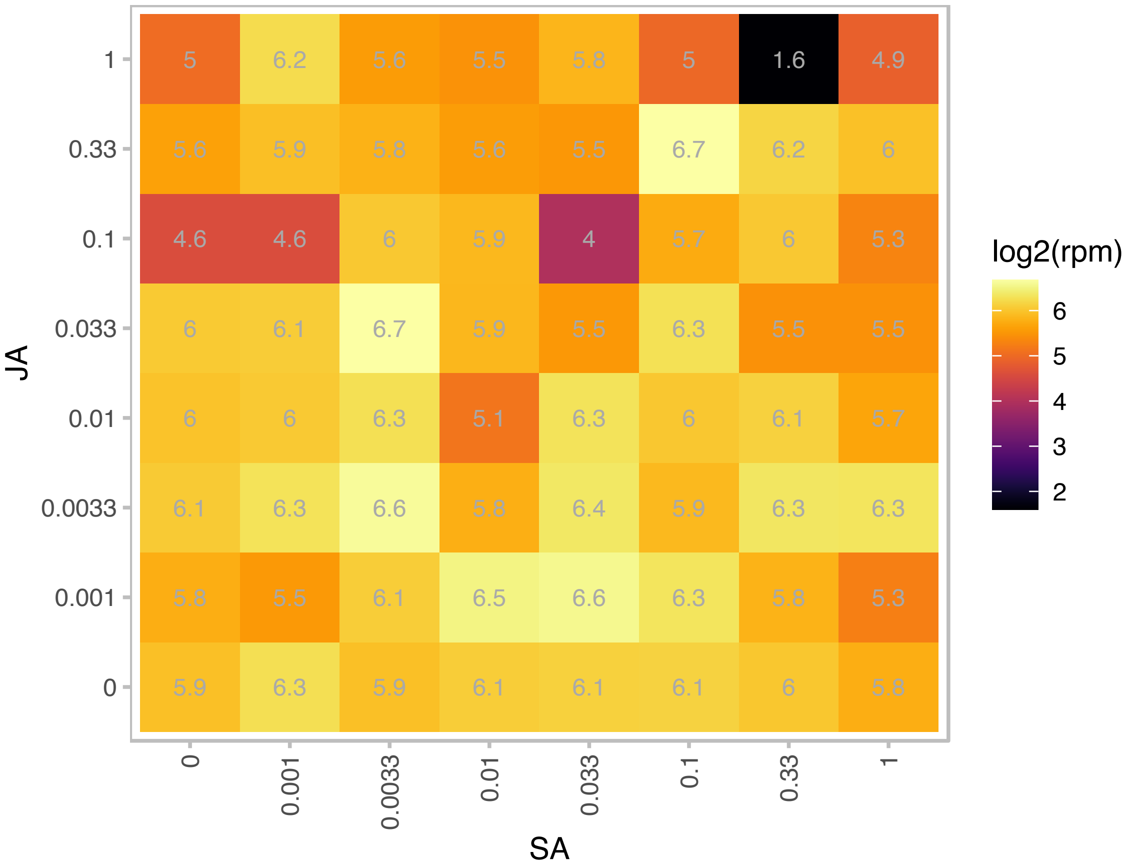


| | |
|---|---|
| Gene | AT5G19140 |
| is.expressed_genes | TRUE |
| is.responsive_genes | TRUE |
| is.apcluster | F-09 |

npr3-2 npr4-2

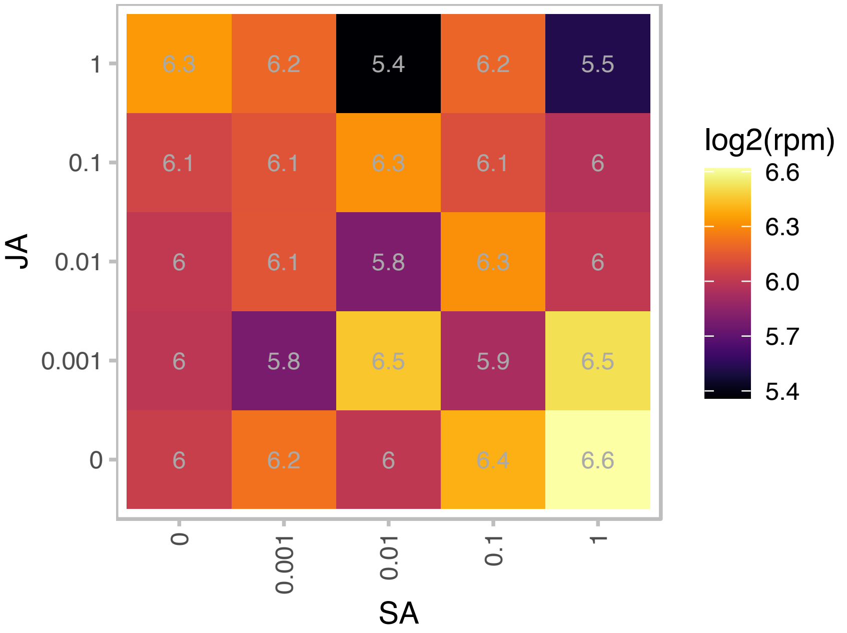


npr3-1 npr4-3

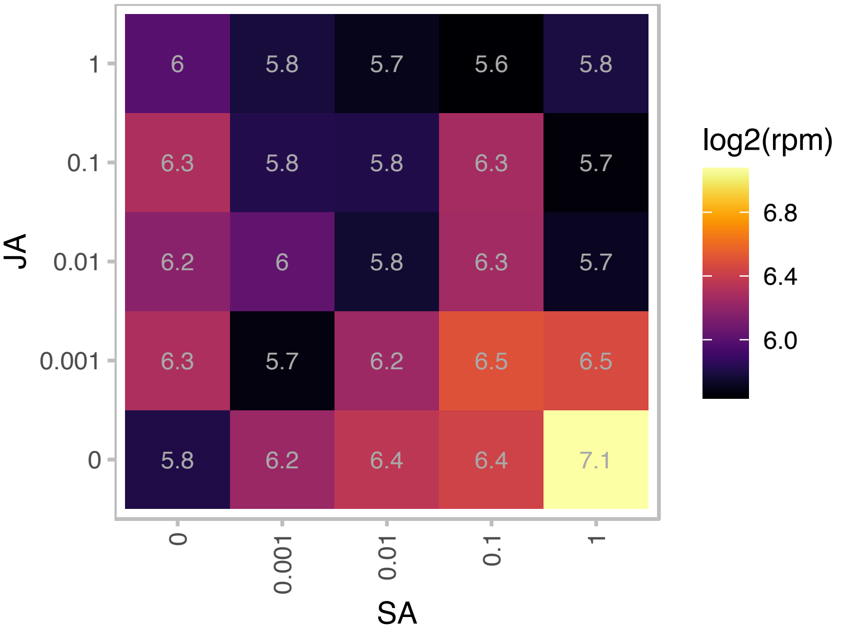

### AT5G20720.1 : ATCPN21, CHCPN10, CPN10, CPN20, CP

chaperonin 20

Encodes a chloroplast co-chaperonin with similarity to CPN21 from spinach, E.coli GroES.

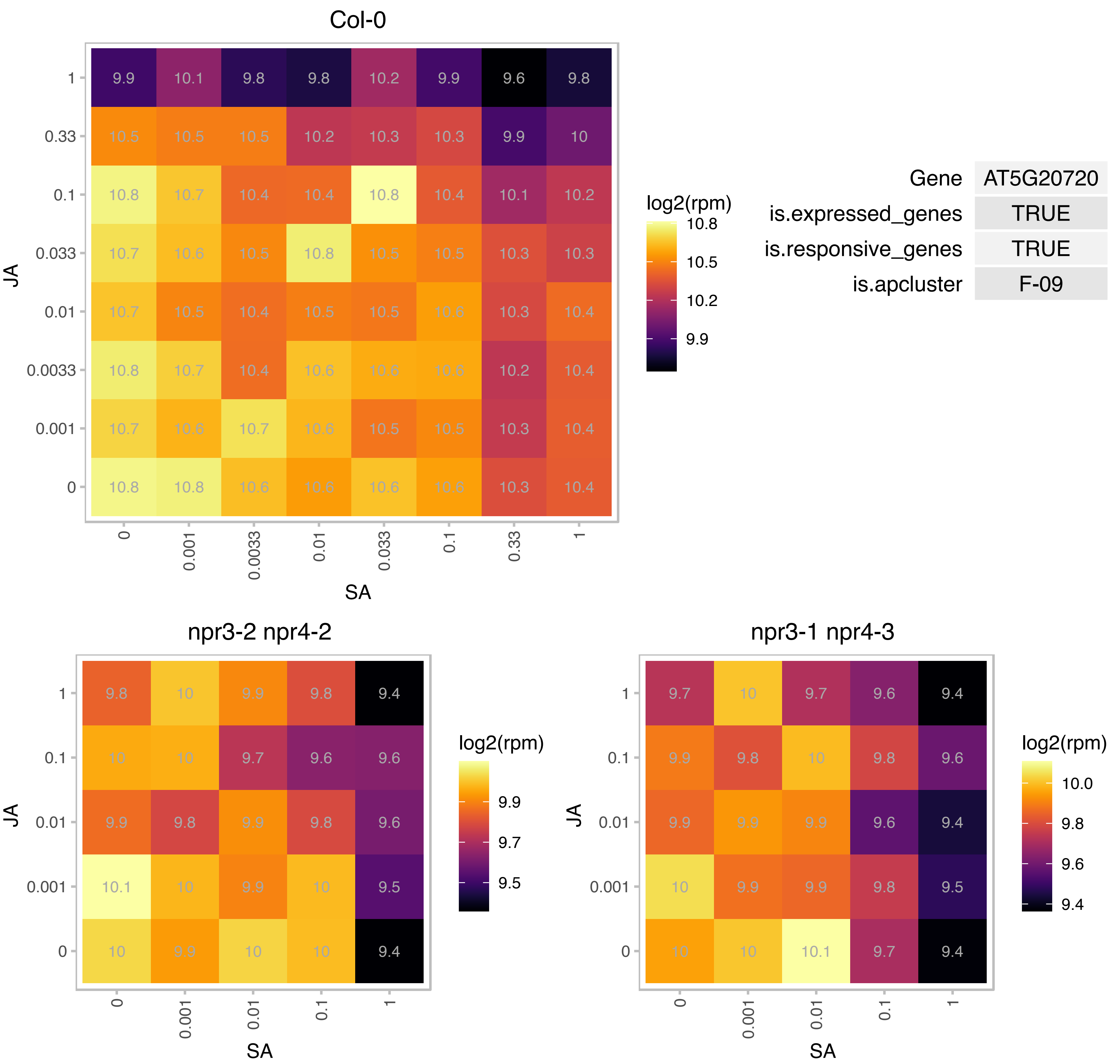


| | |
|---|---|
| Gene | AT5G20720 |
| is.expressed_genes | TRUE |
| is.responsive_genes | TRUE |
| is.apcluster | F-09 |

### AT5G20950.1 : BGLC1

Glycosyl hydrolase family protein

Encodes a beta-glucosidase involved in xyloglucan metabolism.

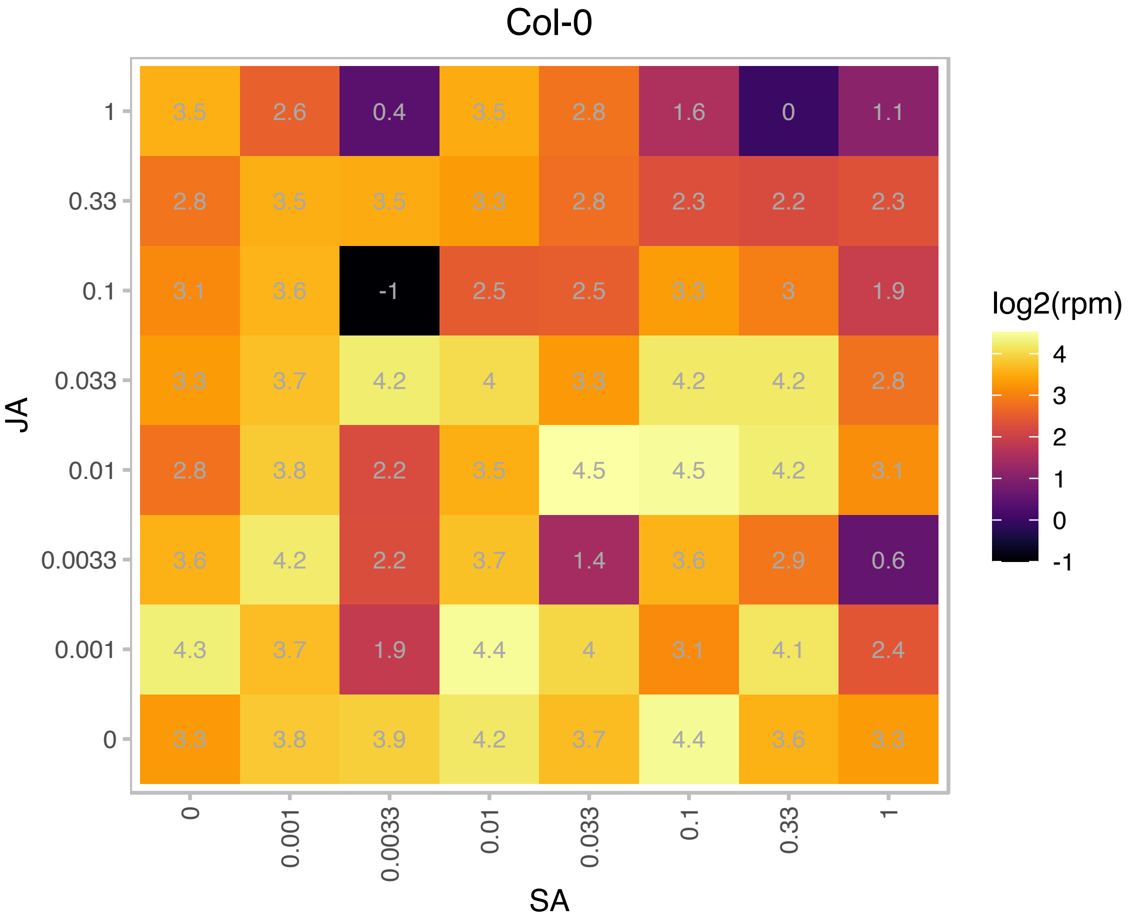


| Gene | AT5G20950 |
|---|---|
| is.expressed_genes | TRUE |
| is.responsive_genes | TRUE |
| is.apcluster | F-09 |

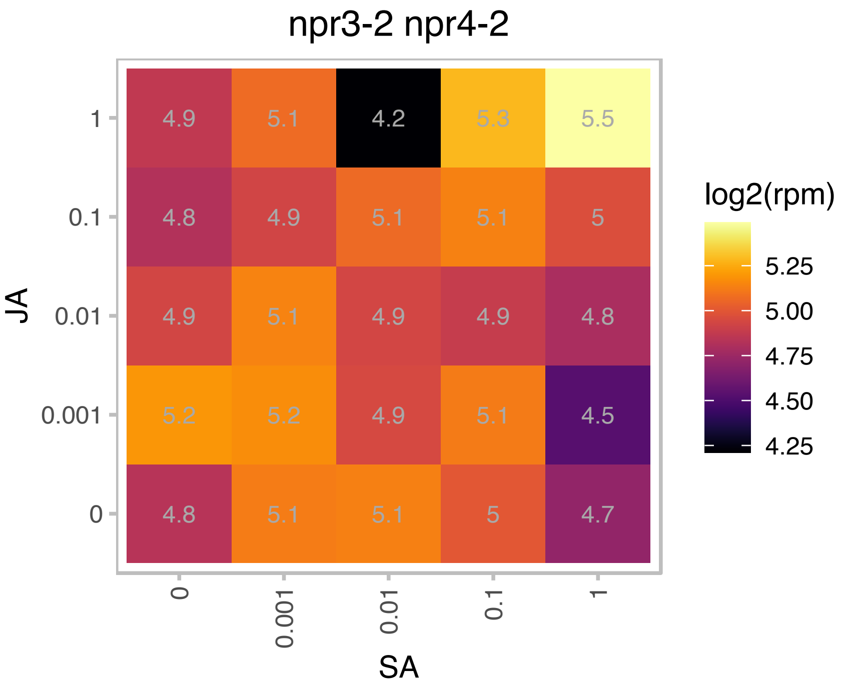


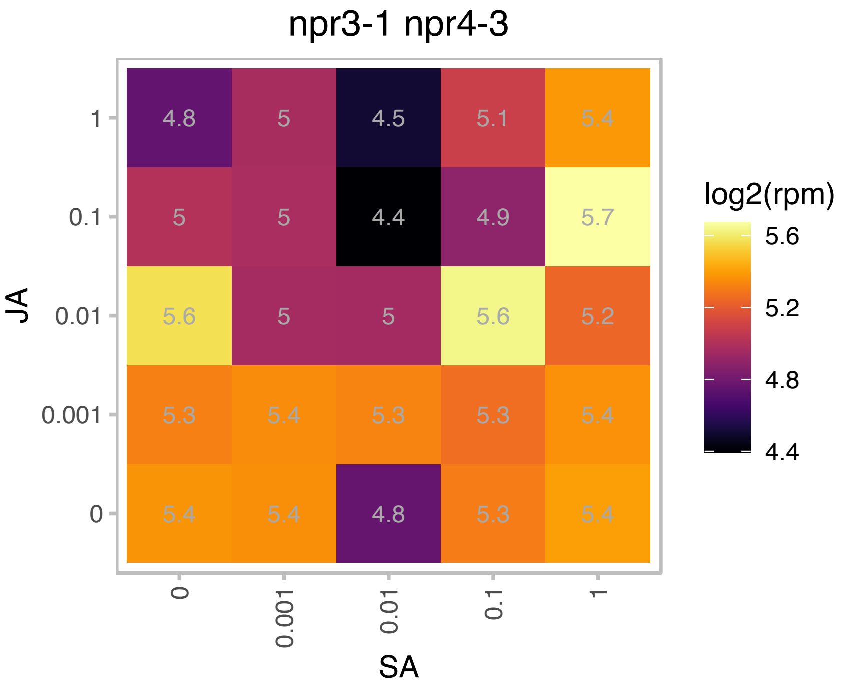

# AT5G22020.1 : NA

Calcium-dependent phosphotriesterase superfamily protein

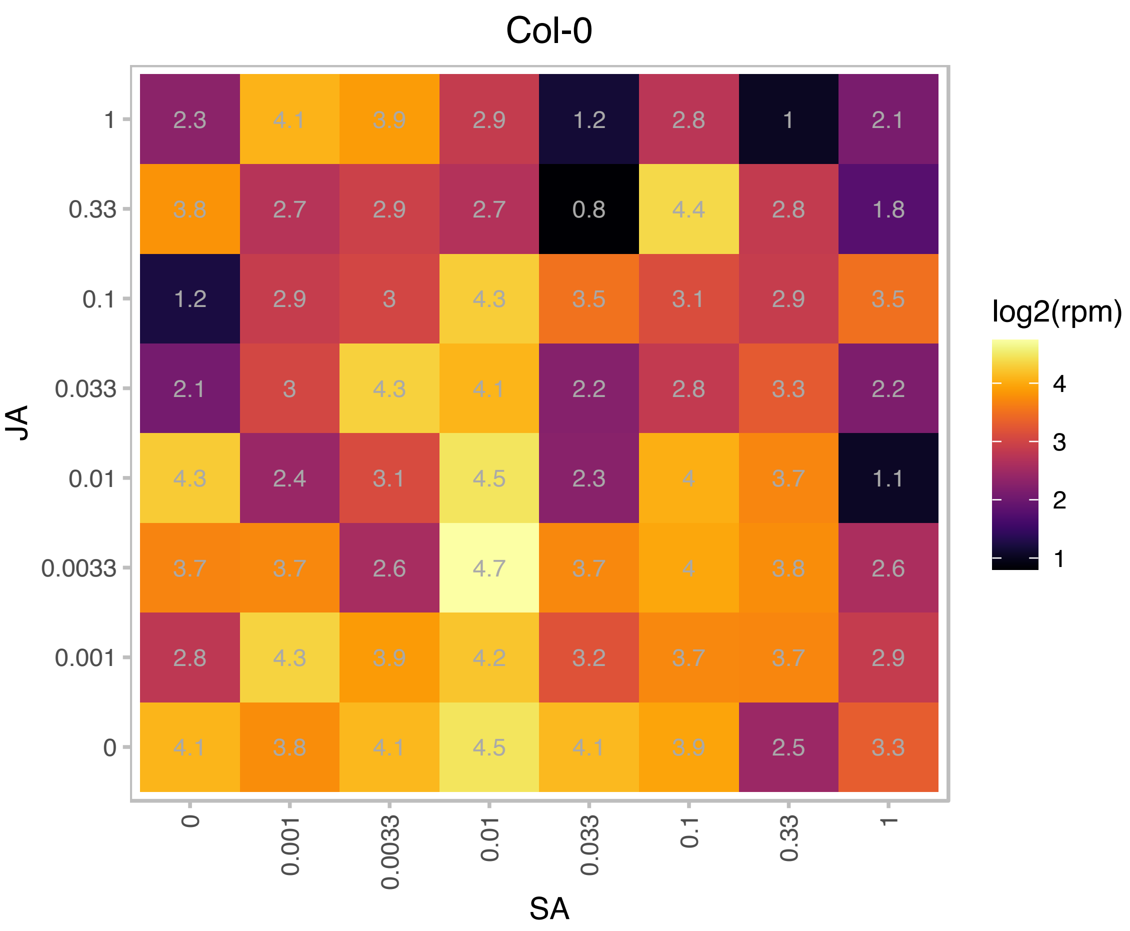


| | |
|---|---|
| Gene | AT5G22020 |
| is.expressed_genes | TRUE |
| is.responsive_genes | TRUE |
| is.apcluster | F-09 |

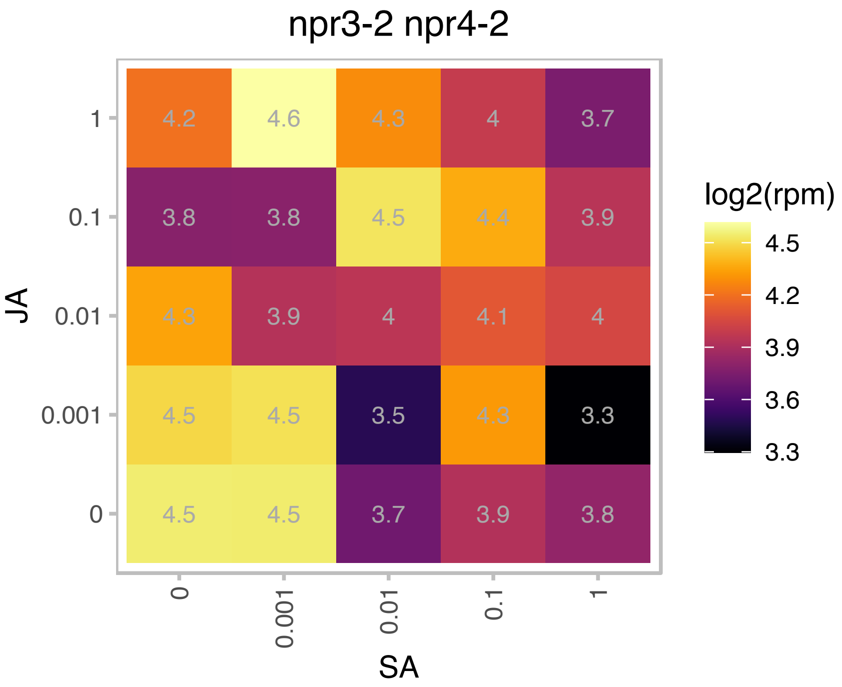


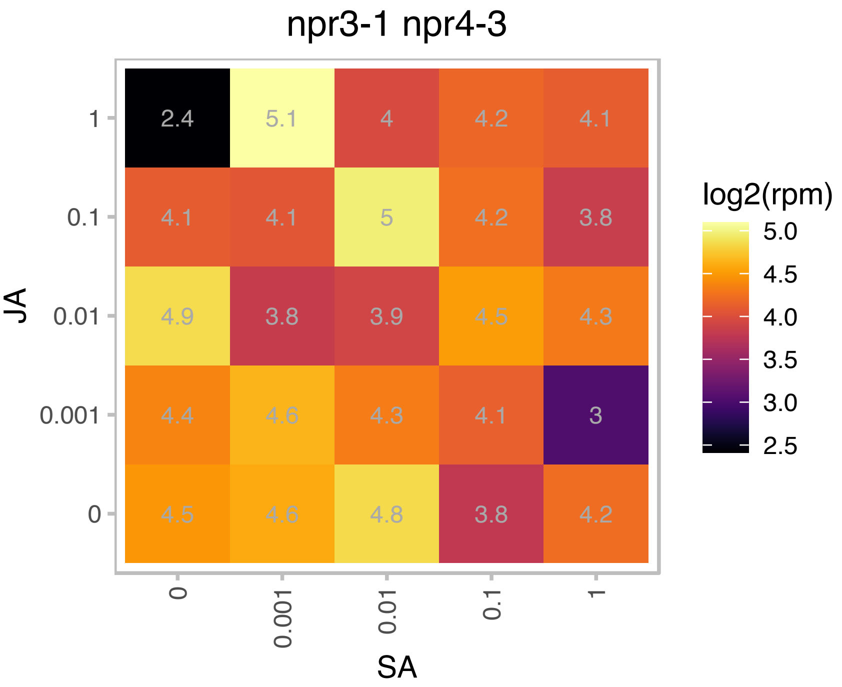

# AT5G37290.1 : NA

ARM repeat superfamily protein

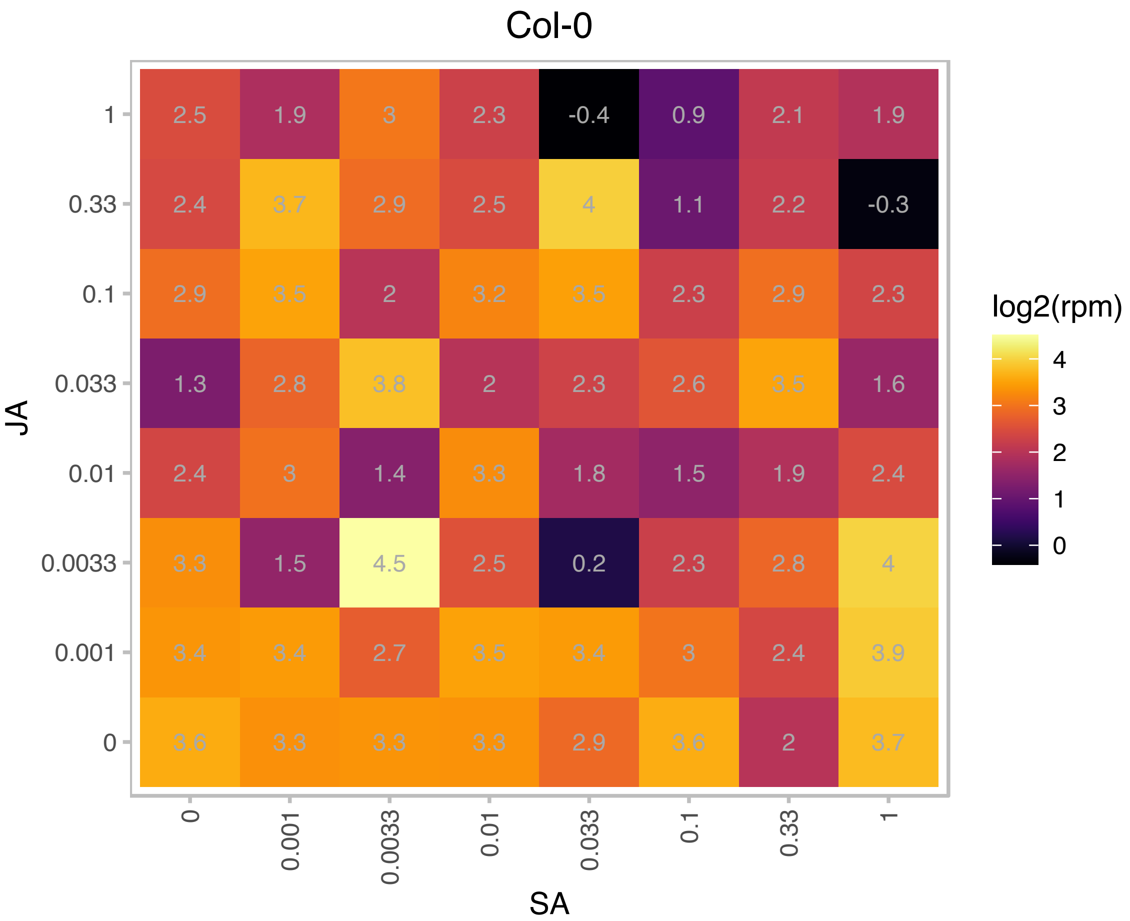


| | |
|---|---|
| Gene | AT5G37290 |
| is.expressed_genes | TRUE |
| is.responsive_genes | TRUE |
| is.apcluster | F-09 |

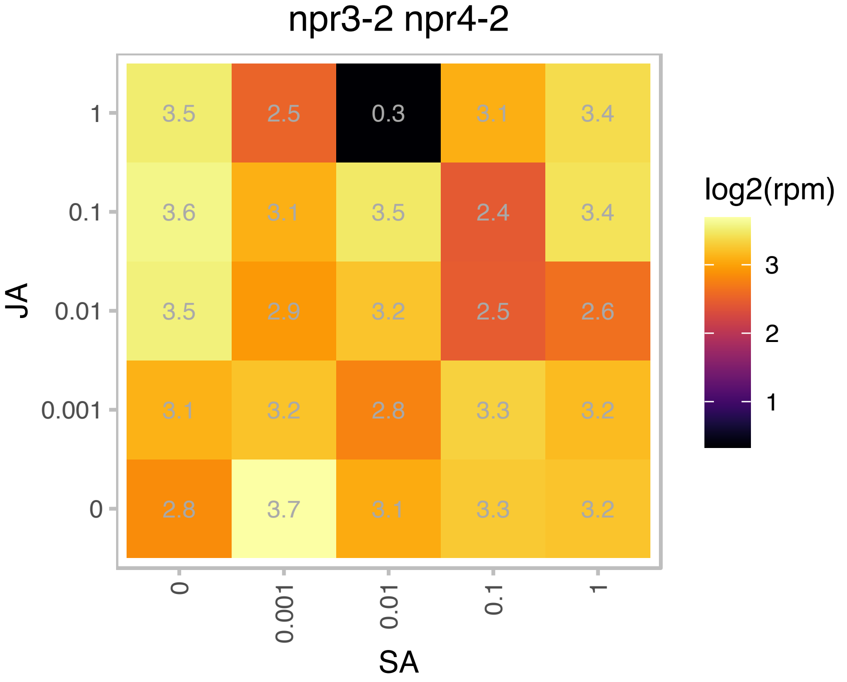


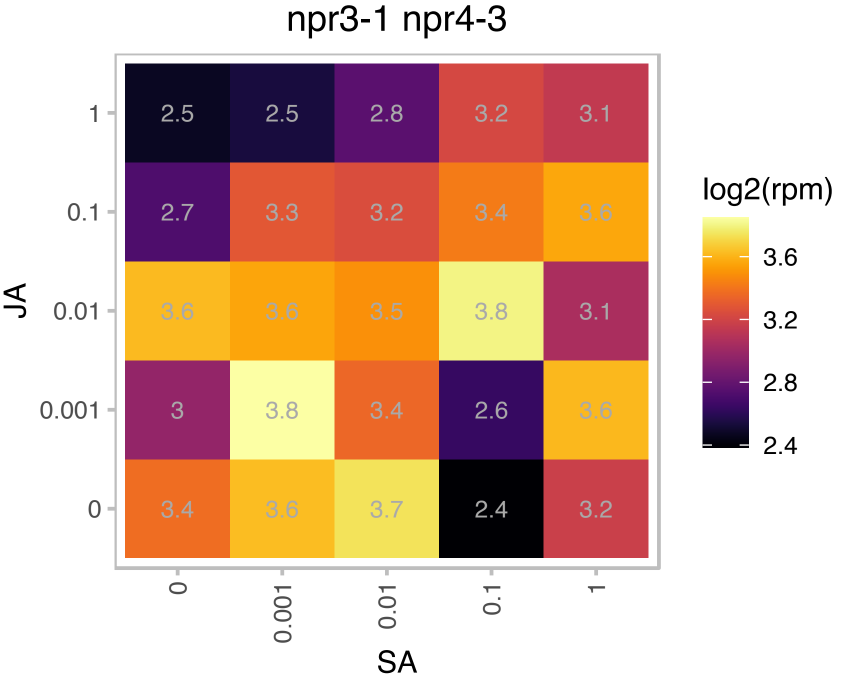

# AT5G38980.1 : NA

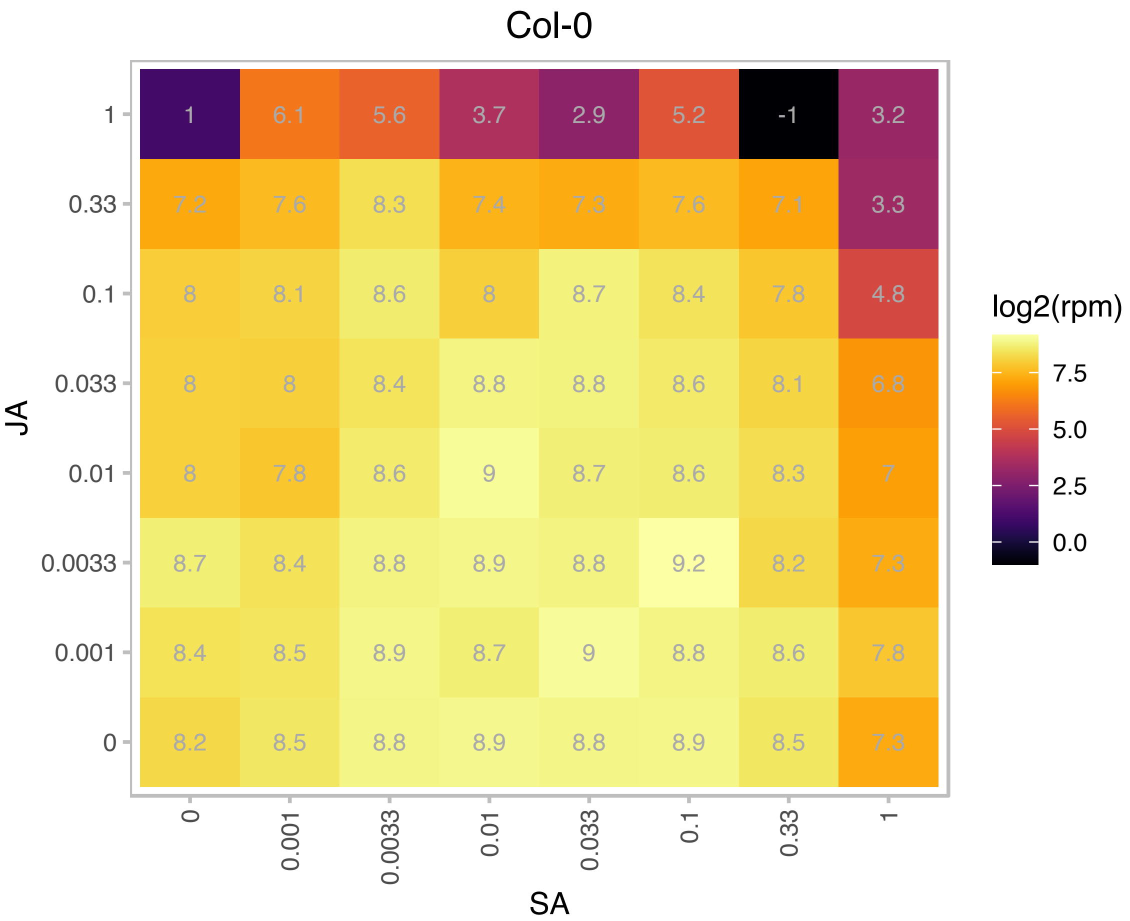


| | |
|---|---|
| Gene | AT5G38980 |
| is.expressed_genes | TRUE |
| is.responsive_genes | TRUE |
| is.apcluster | F-09 |

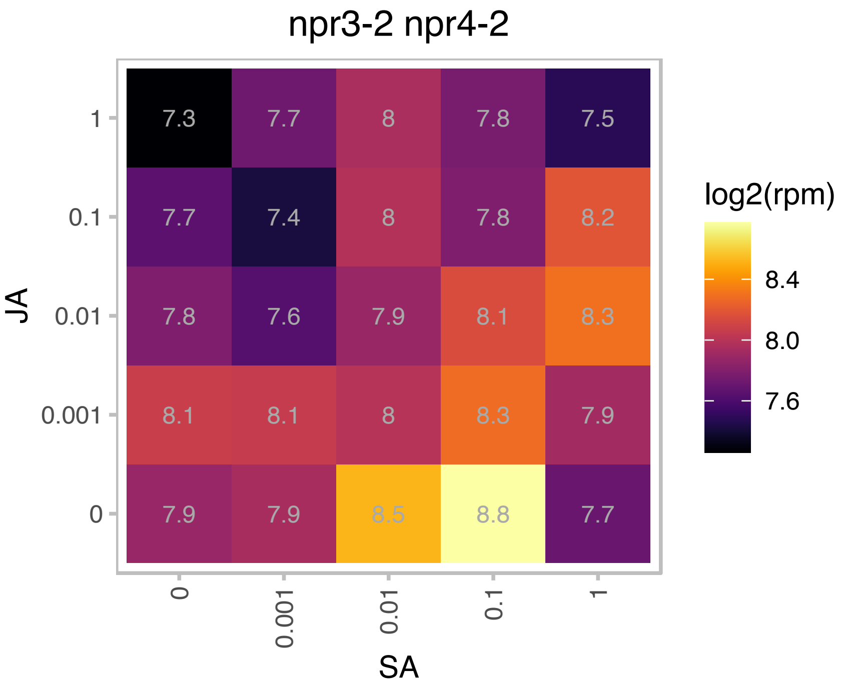


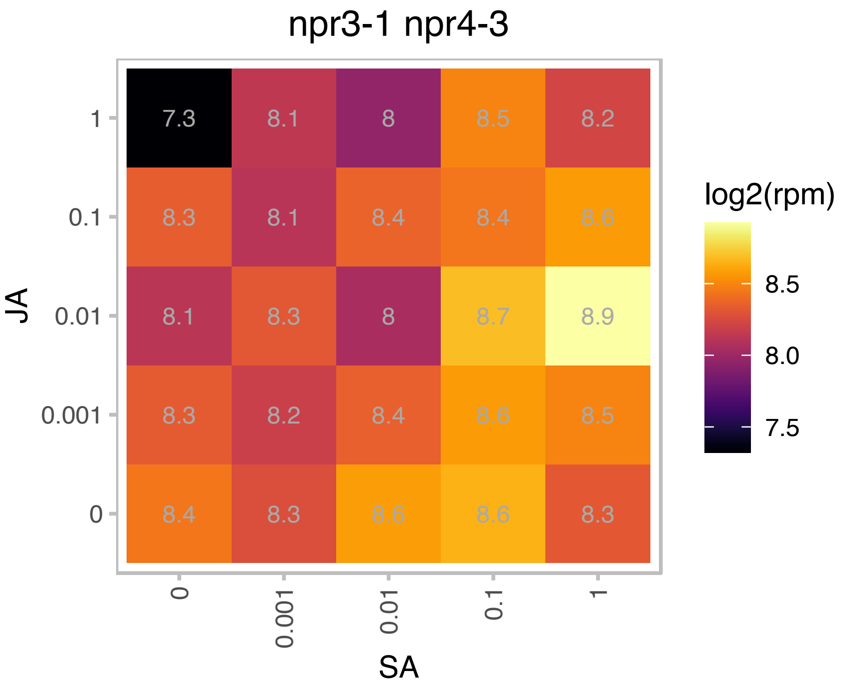

### AT5G41600.1 : BTI3, RTNLB4

VIRB2-interacting protein 3

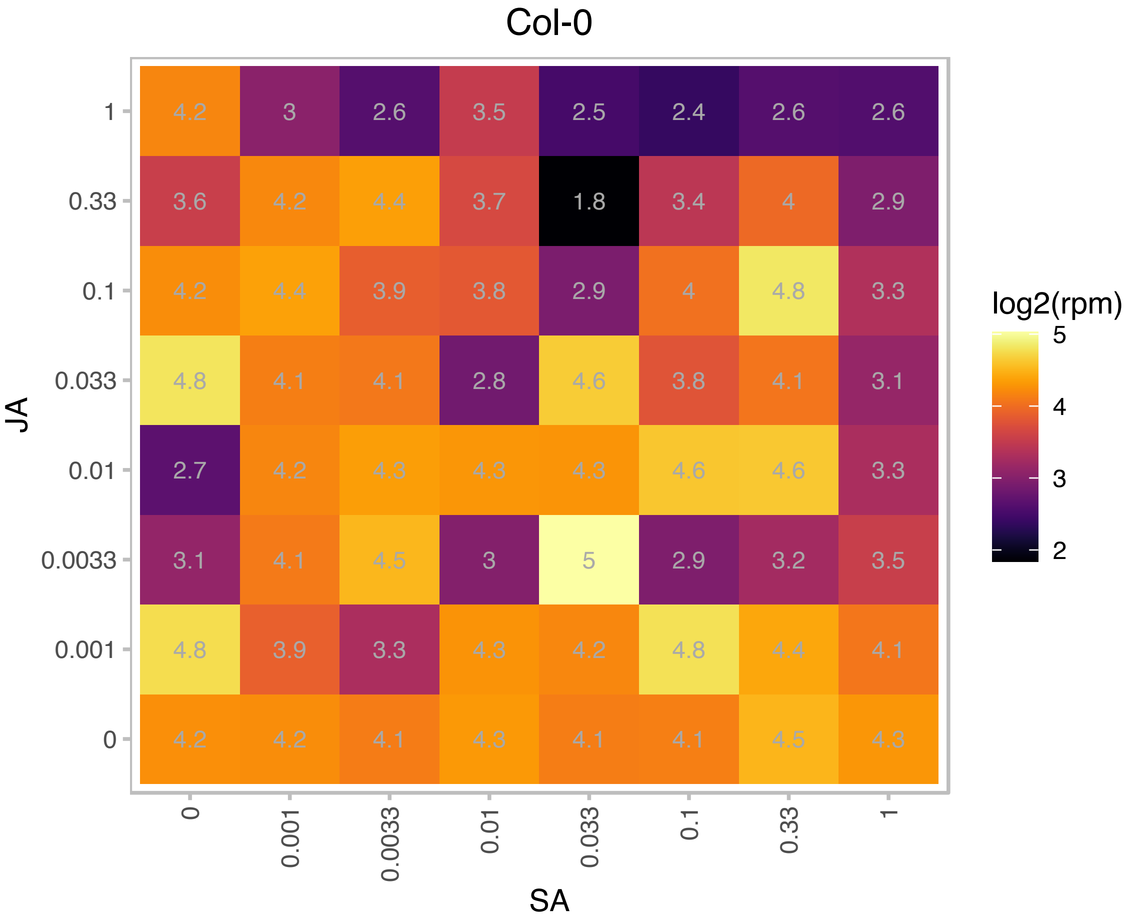


| | |
|---|---|
| Gene | AT5G41600 |
| is.expressed_genes | TRUE |
| is.responsive_genes | TRUE |
| is.apcluster | F-09 |

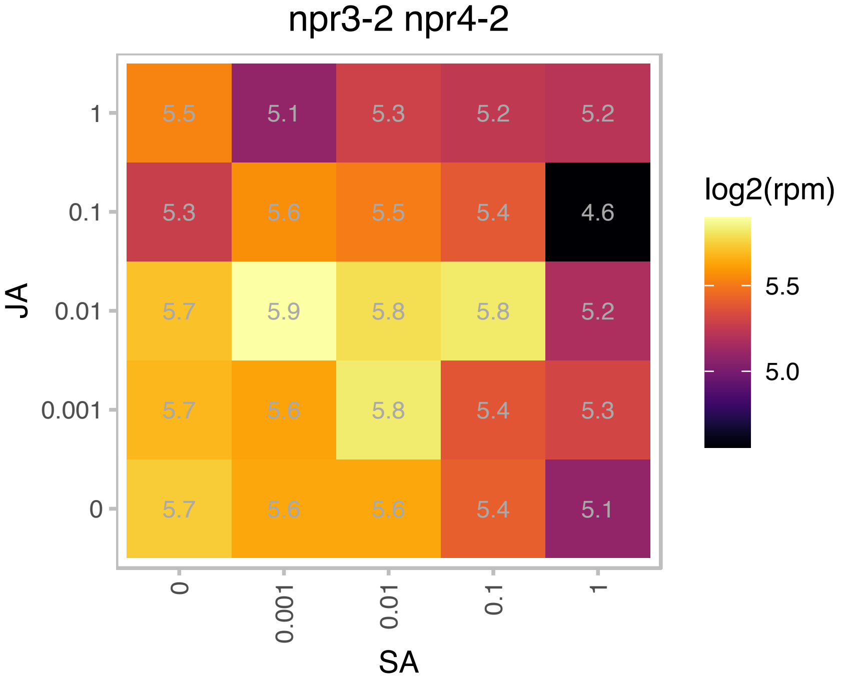


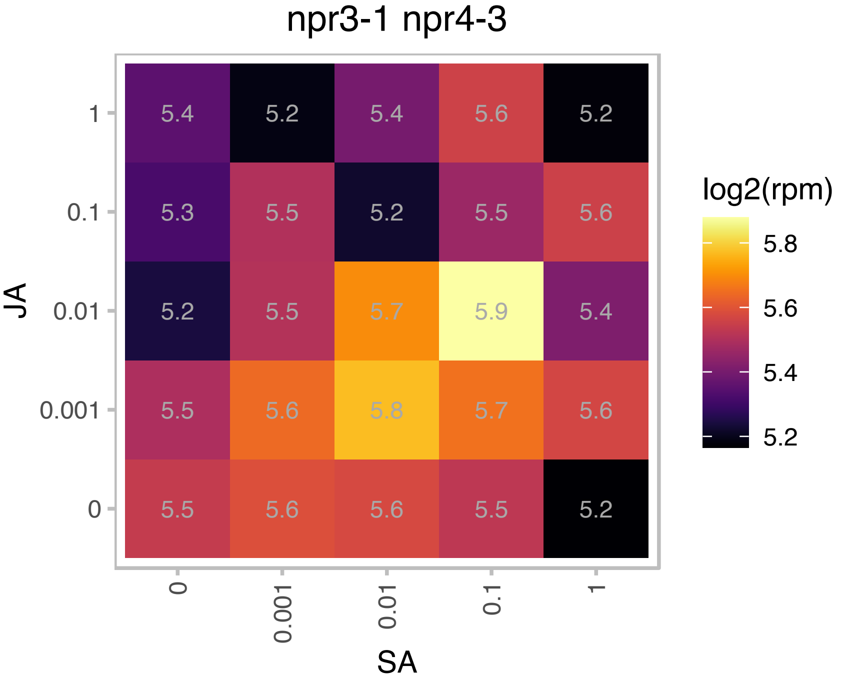

# AT5G44580.1 : NA

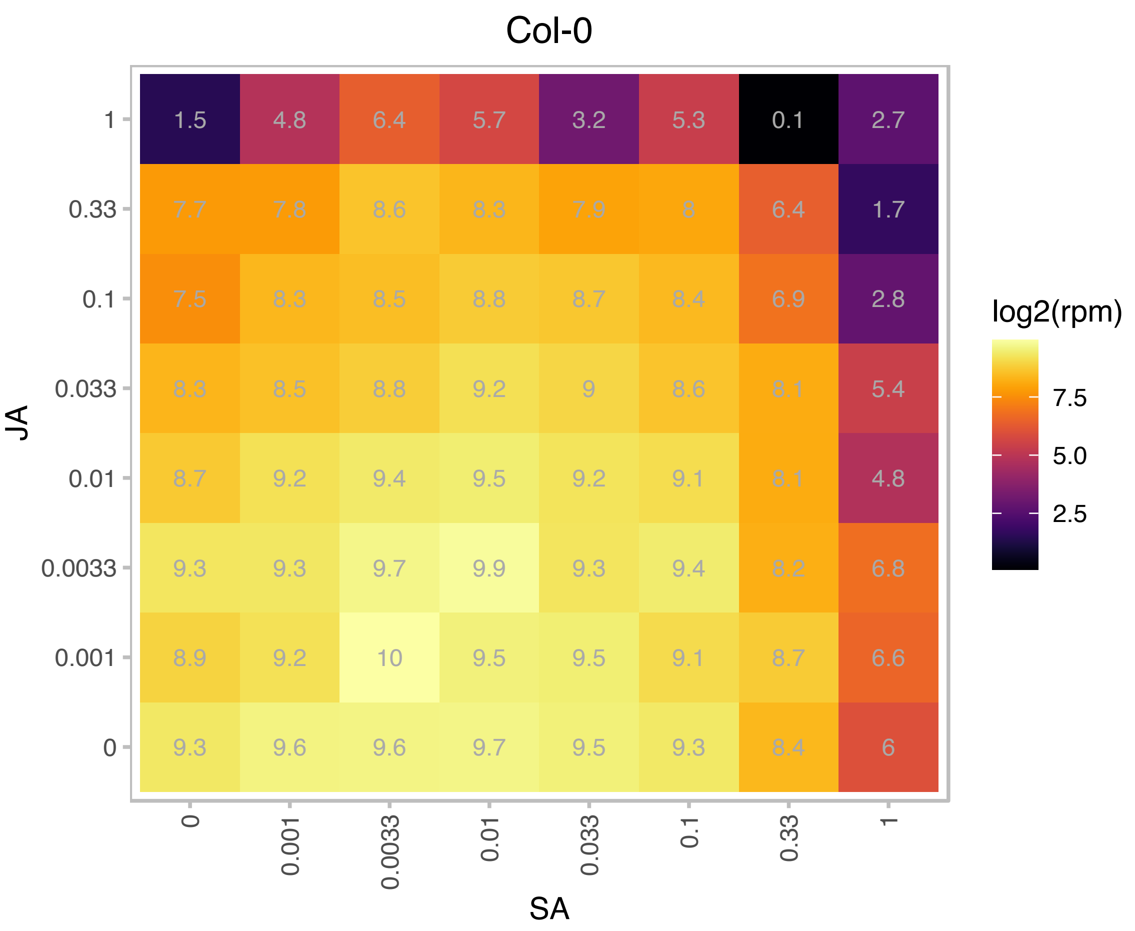


| | |
|---|---|
| Gene | AT5G44580 |
| is.expressed_genes | TRUE |
| is.responsive_genes | TRUE |
| is.apcluster | F-09 |

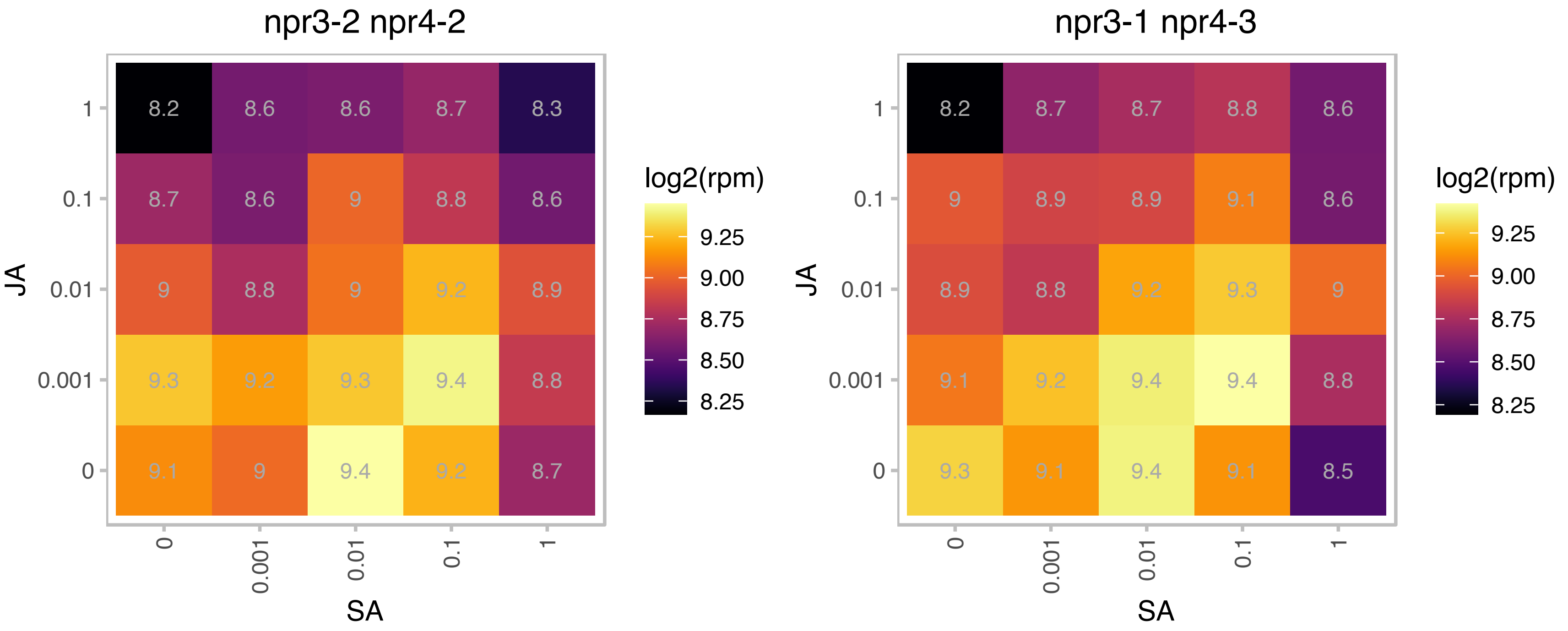

### AT5G46800.1 : BOU

Mitochondrial substrate carrier family protein

Seedling lethal mutation; Mitochondrial Carnitine Acyl Carrier-Like Protein

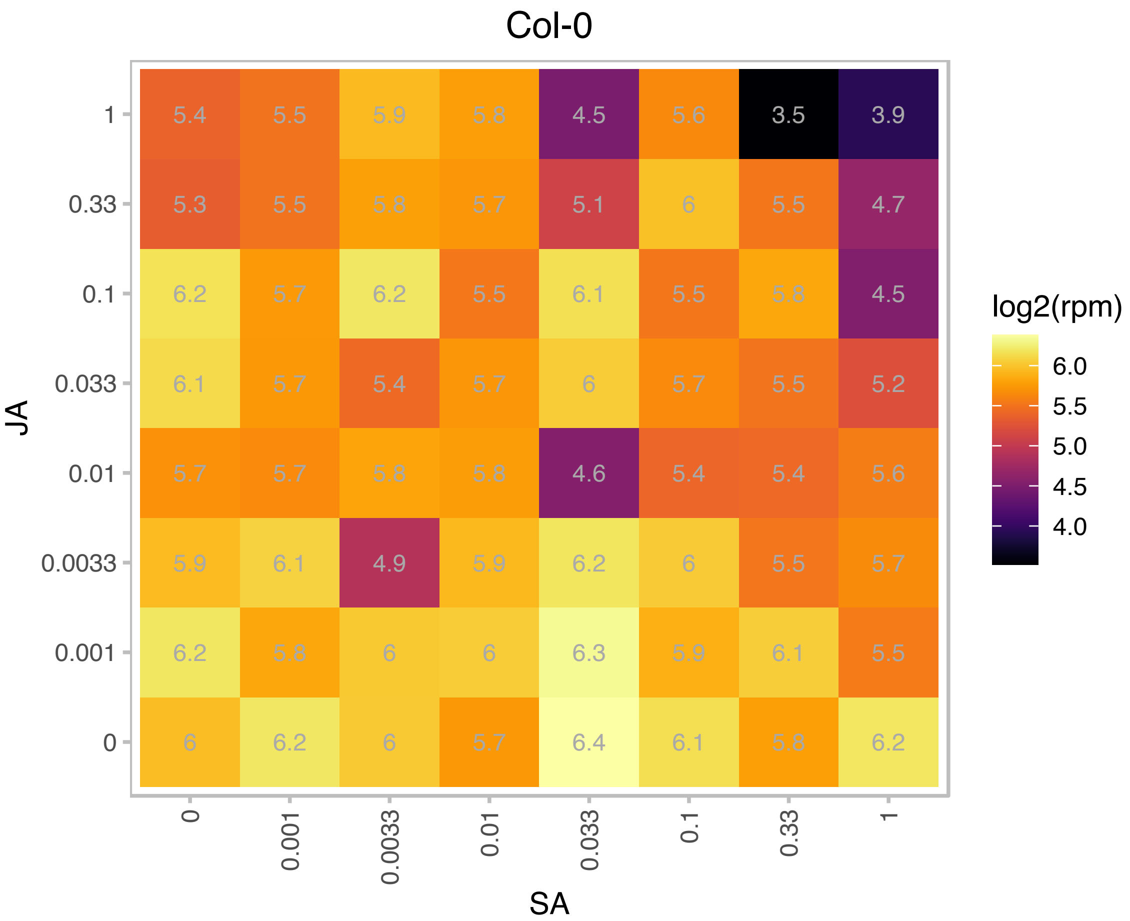


| | |
|---|---|
| Gene | AT5G46800 |
| is.expressed_genes | TRUE |
| is.responsive_genes | TRUE |
| is.apcluster | F-09 |

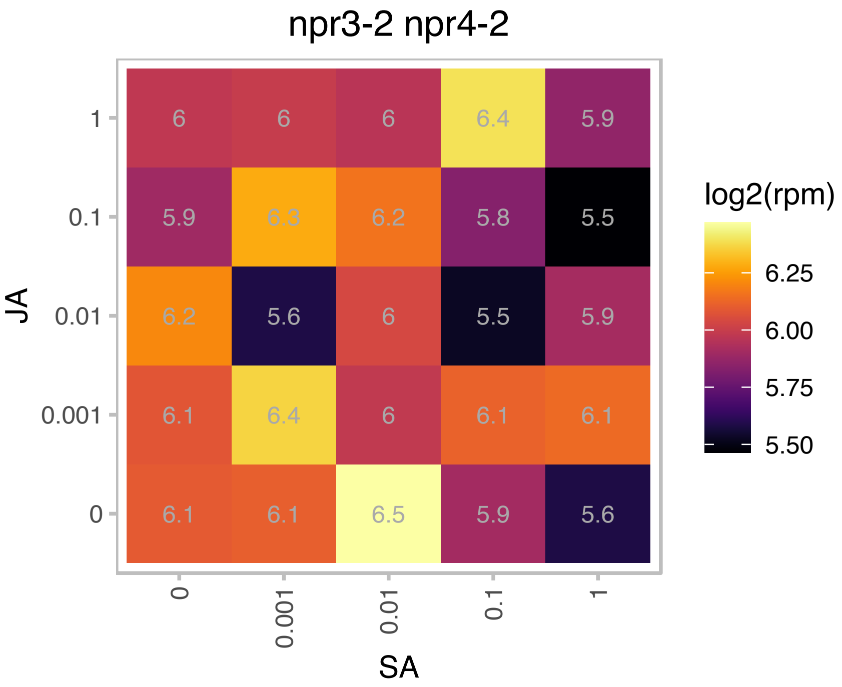


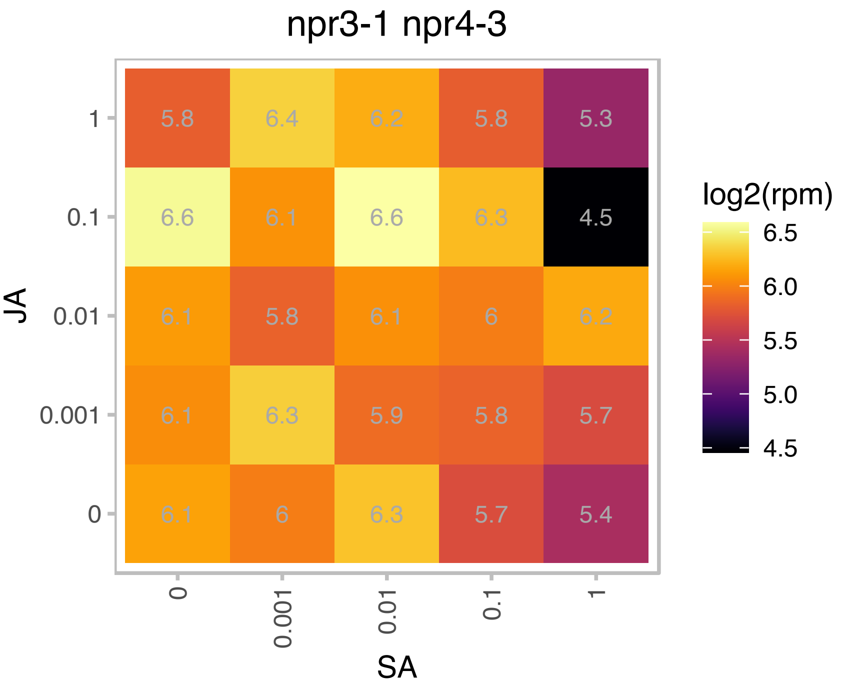

# AT5G47640.1 : NF-YB2

nuclear factor Y, subunit B2

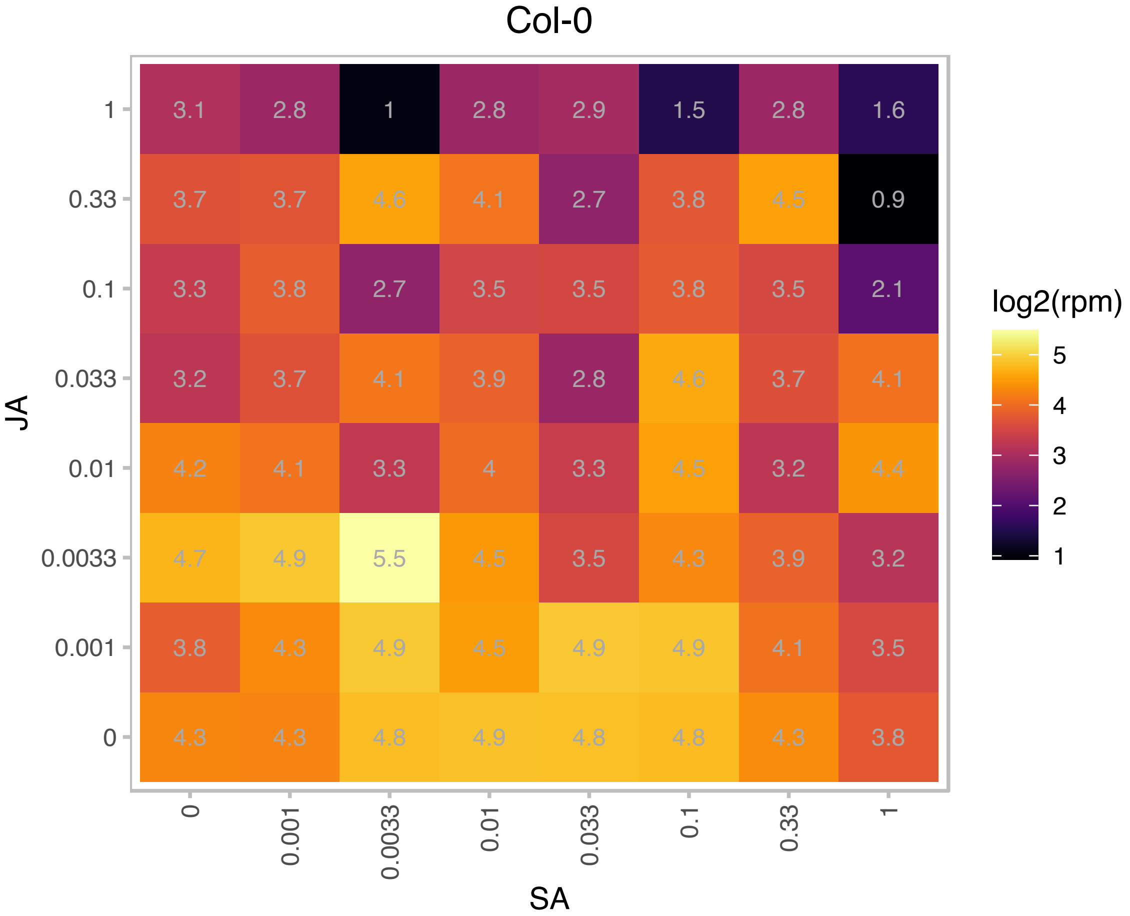


| | |
|---|---|
| Gene | AT5G47640 |
| is.expressed_genes | TRUE |
| is.responsive_genes | TRUE |
| is.apcluster | F-09 |

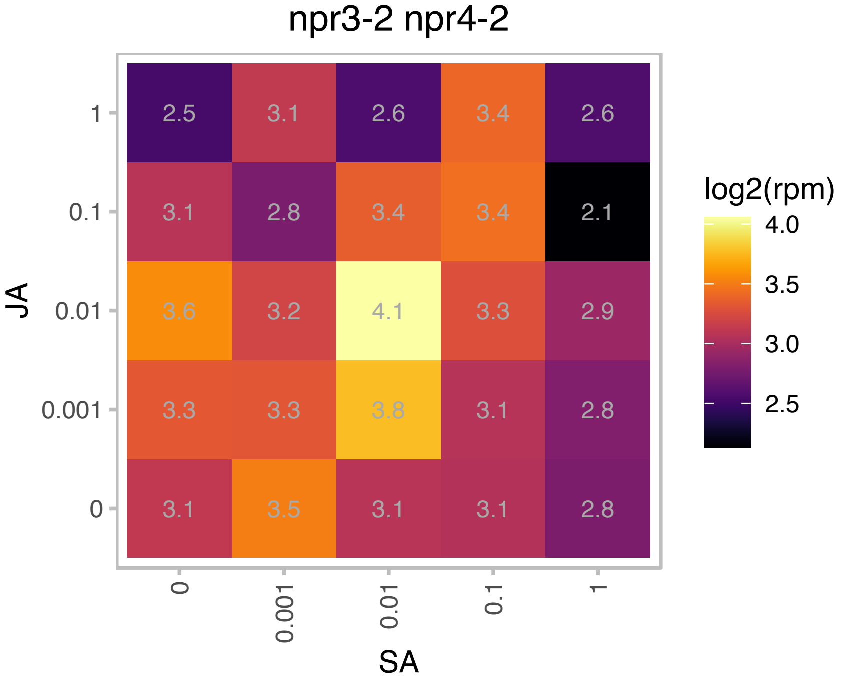


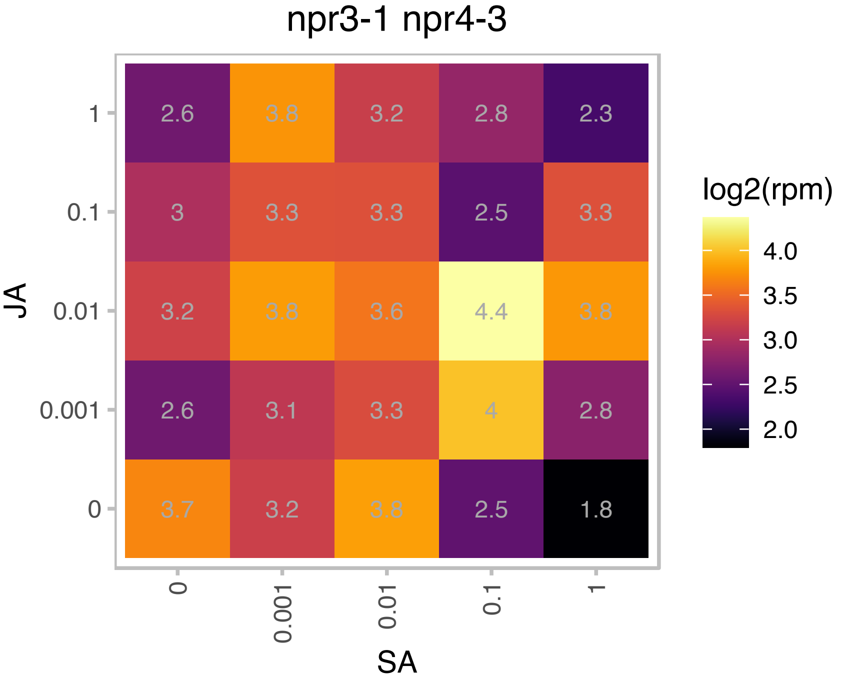

### AT5G53250.1 : AGP22, ATAGP22

arabinogalactan protein 22

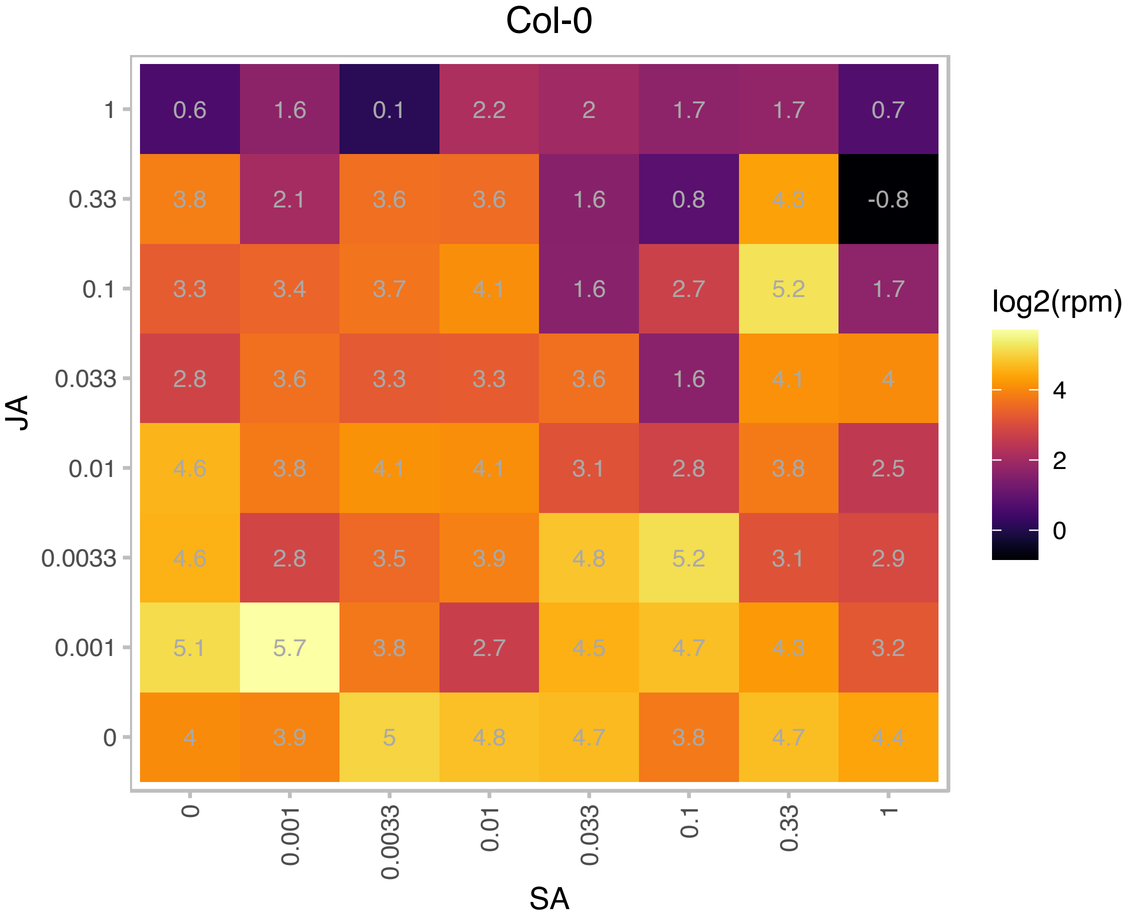


| | |
|---|---|
| Gene | AT5G53250 |
| is.expressed_genes | TRUE |
| is.responsive_genes | TRUE |
| is.apcluster | F-09 |

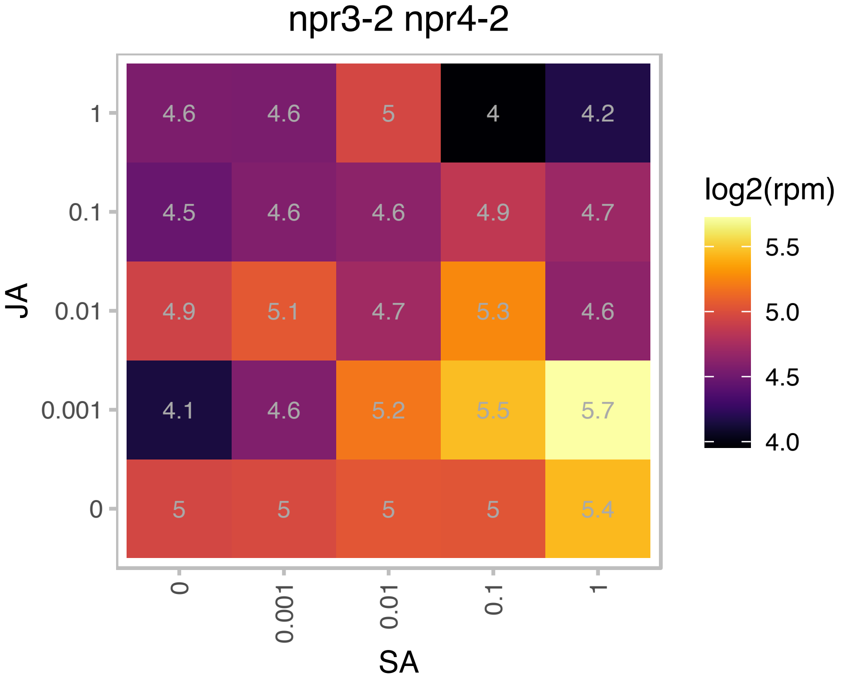


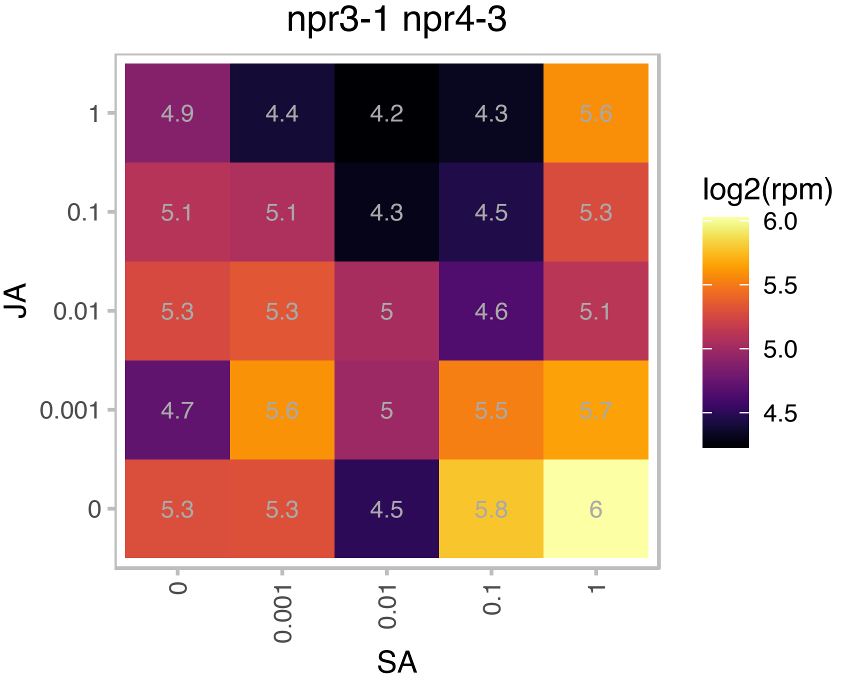

# AT5G55125.1 : NA

Ribosomal protein L31

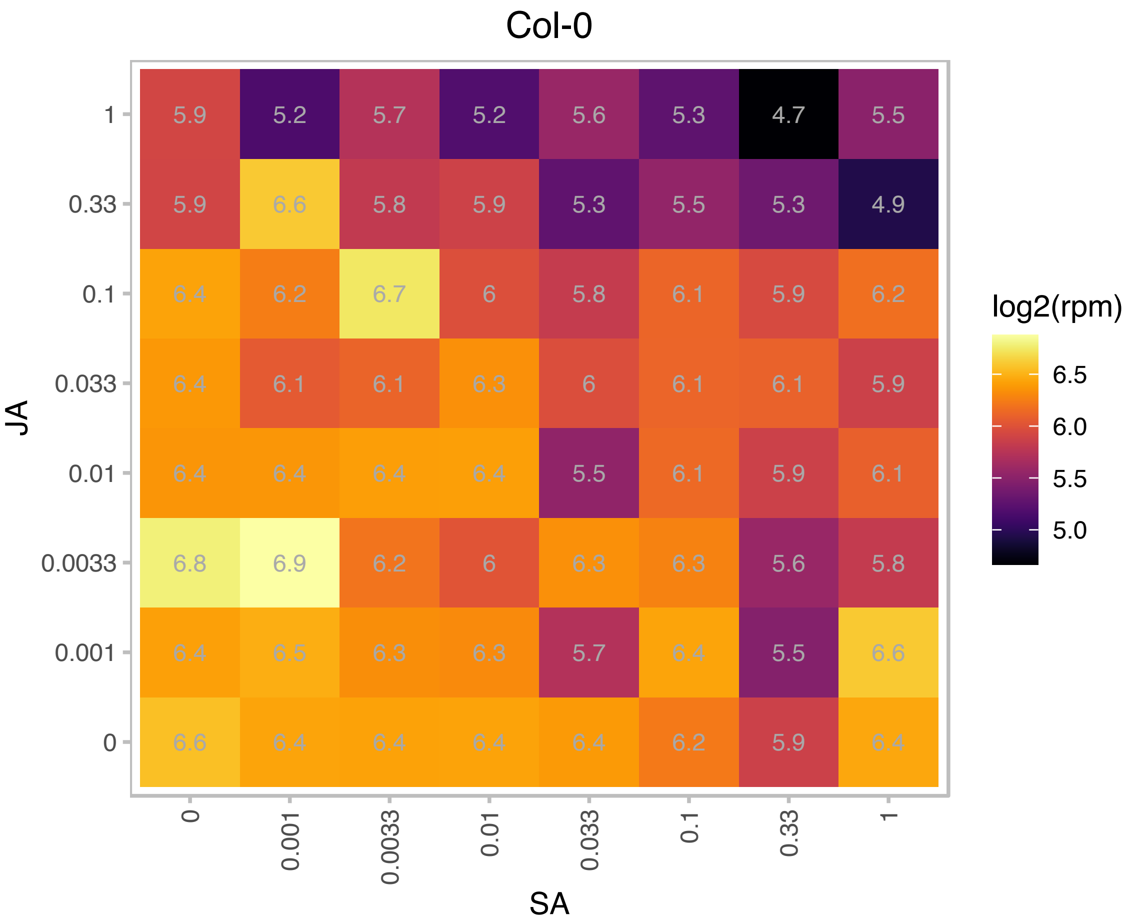


| | |
|---|---|
| Gene | AT5G55125 |
| is.expressed_genes | TRUE |
| is.responsive_genes | TRUE |
| is.apcluster | F-09 |

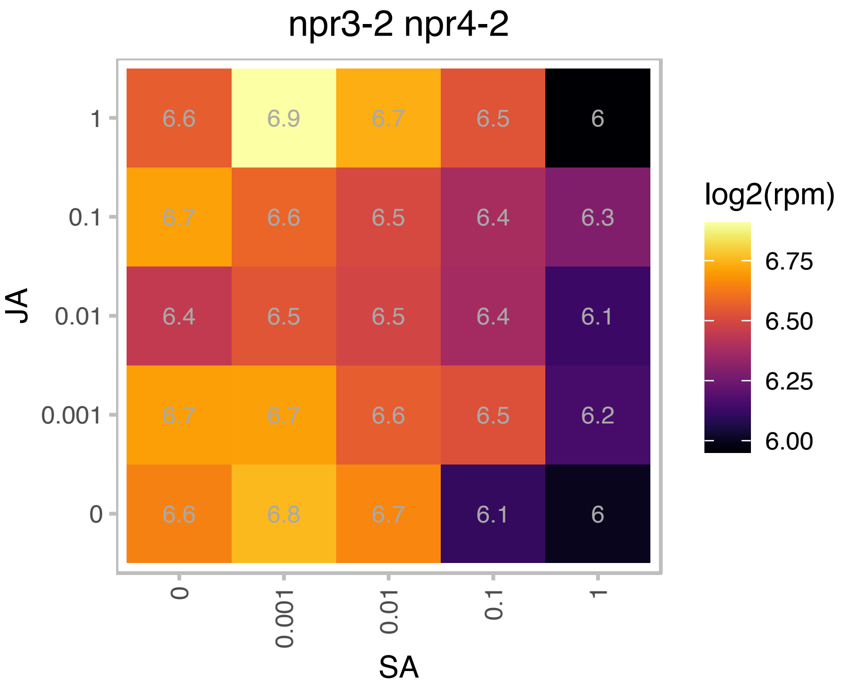


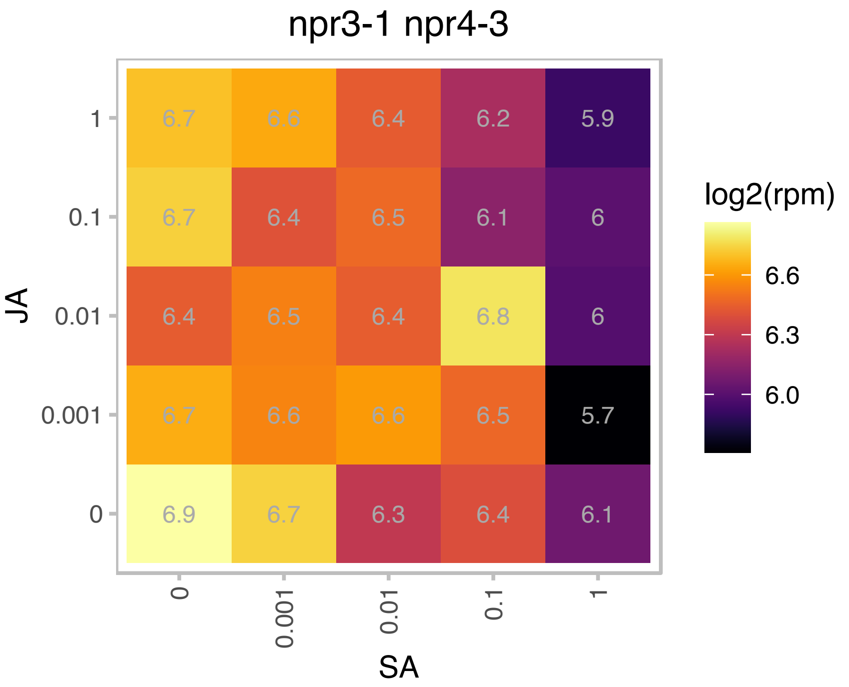

# AT5G60400.1 : NA

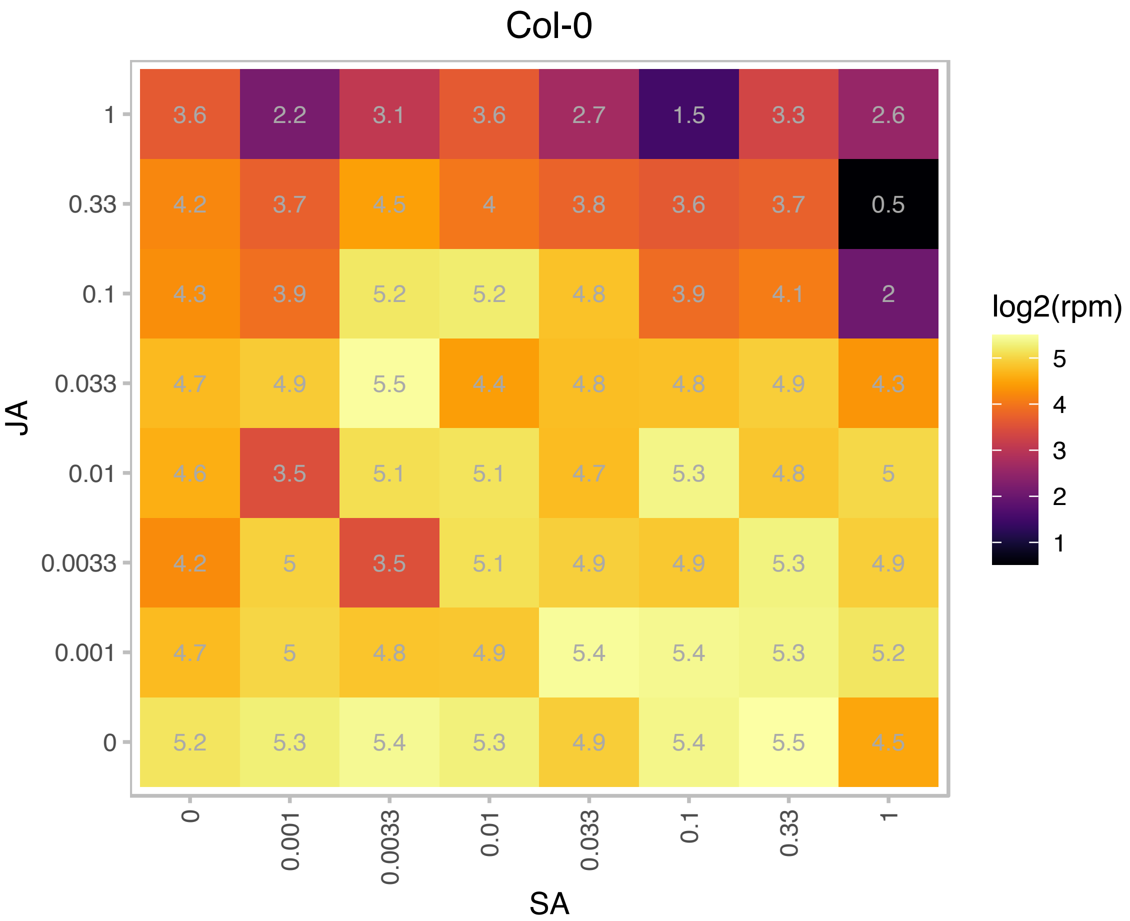


| | |
|---|---|
| Gene | AT5G60400 |
| is.expressed_genes | TRUE |
| is.responsive_genes | TRUE |
| is.apcluster | F-09 |

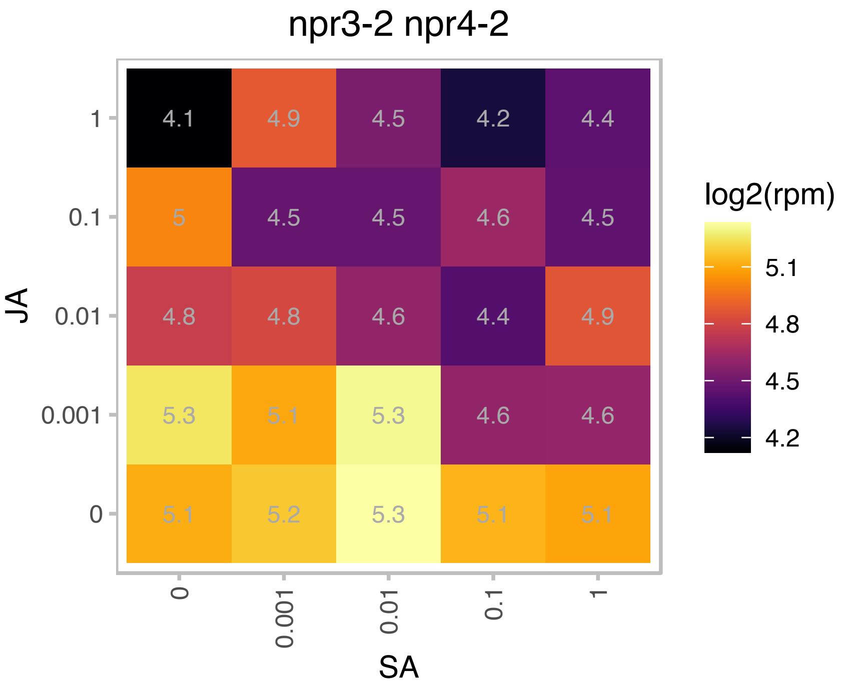


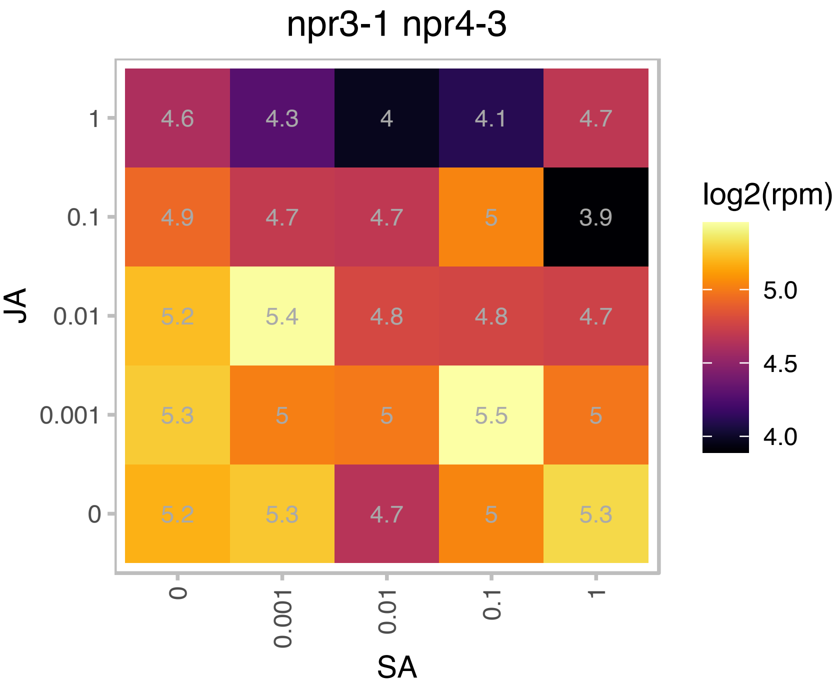

# AT5G65880.1 : NA

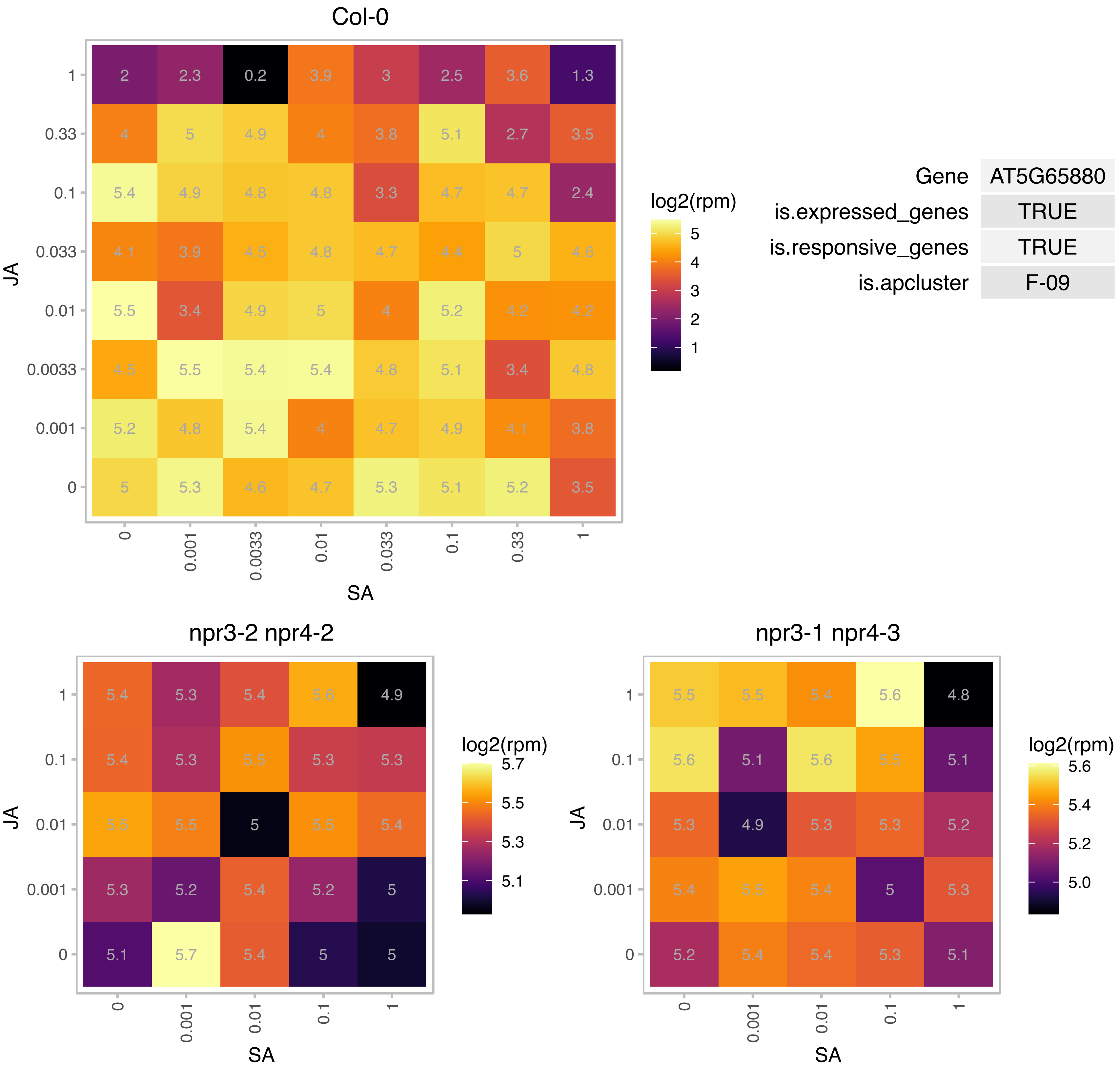


| | |
|---|---|
| Gene | AT5G65880 |
| is.expressed_genes | TRUE |
| is.responsive_genes | TRUE |
| is.apcluster | F-09 |

# AT5G67640.1 : NA

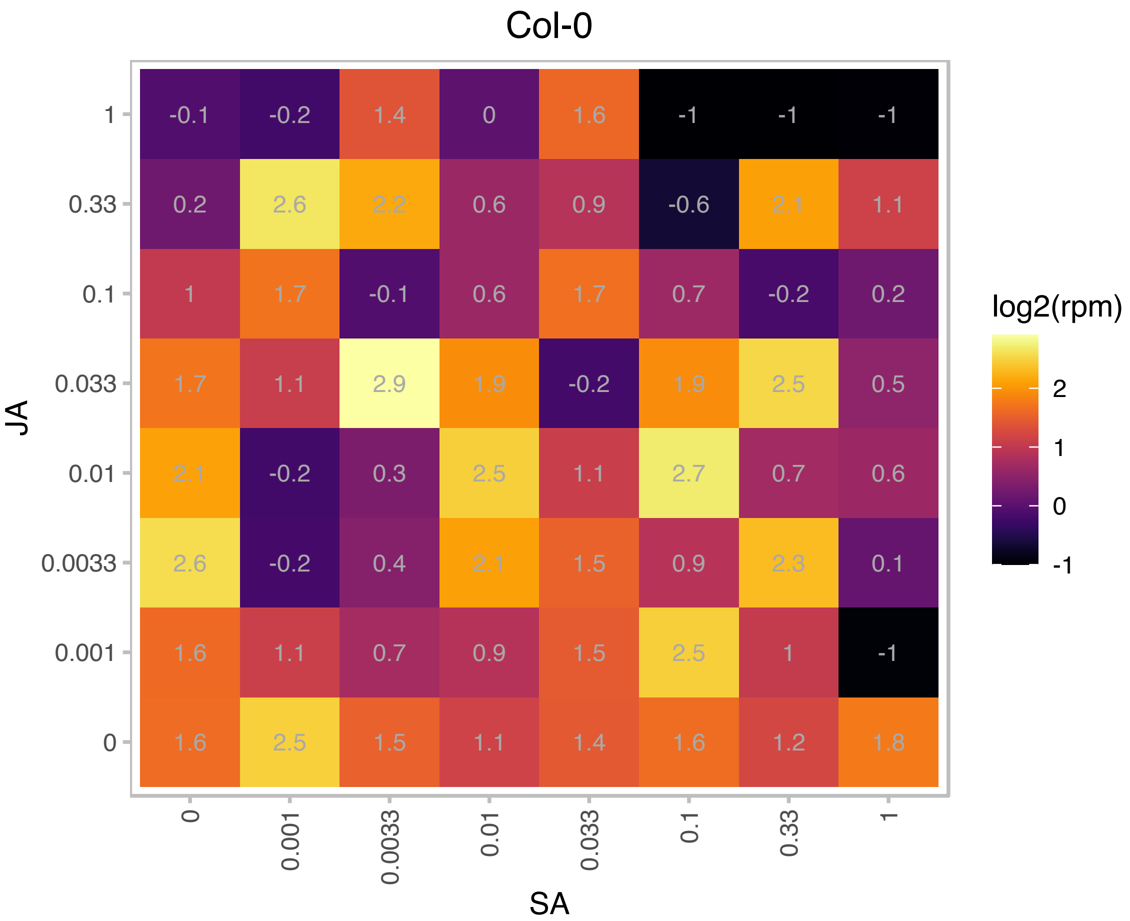


| | |
|---|---|
| Gene | AT5G67640 |
| is.expressed_genes | TRUE |
| is.responsive_genes | TRUE |
| is.apcluster | F-09 |

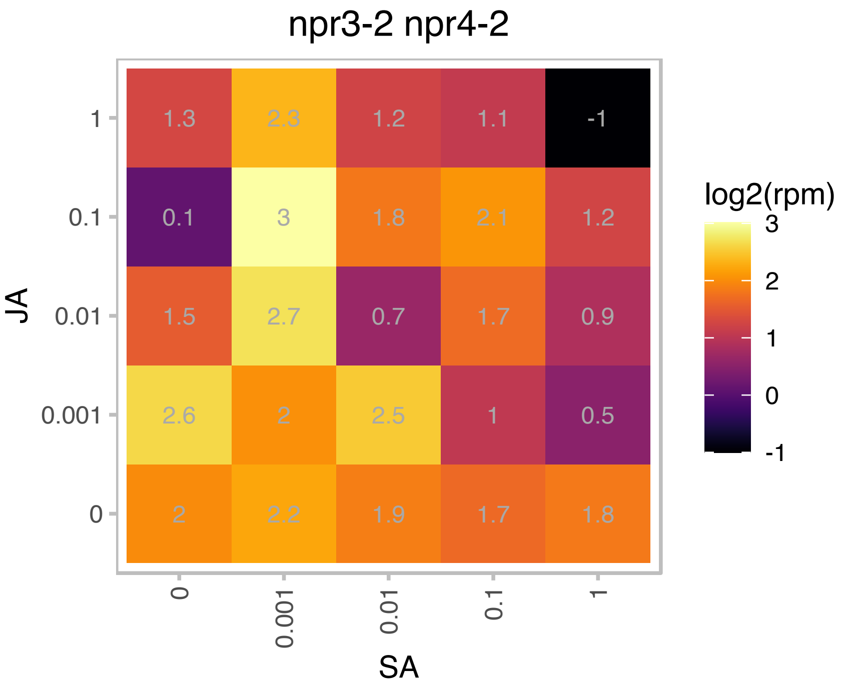


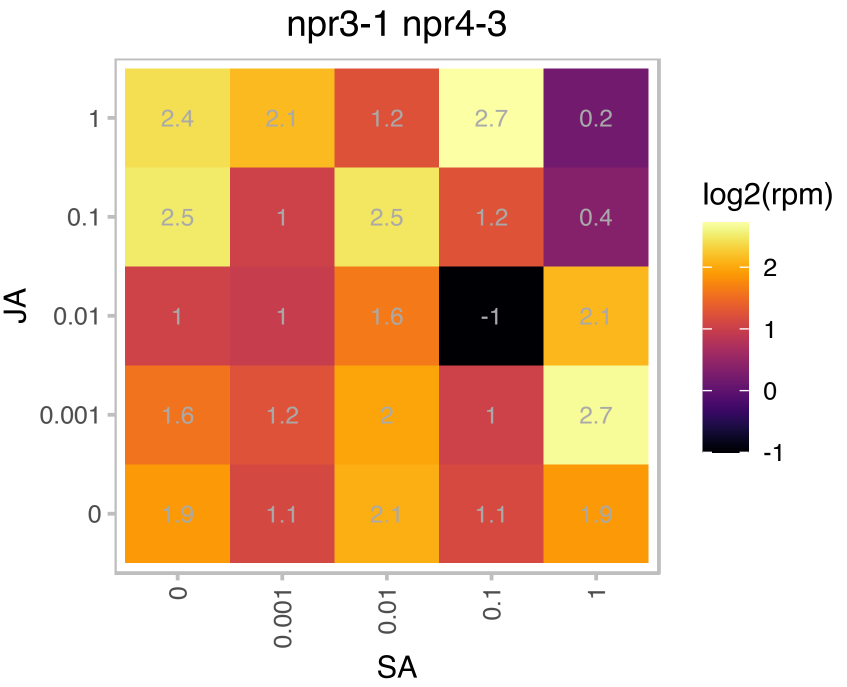

# AT1G01080.1 : NA

RNA-binding (RRM/RBD/RNP motifs) family protein

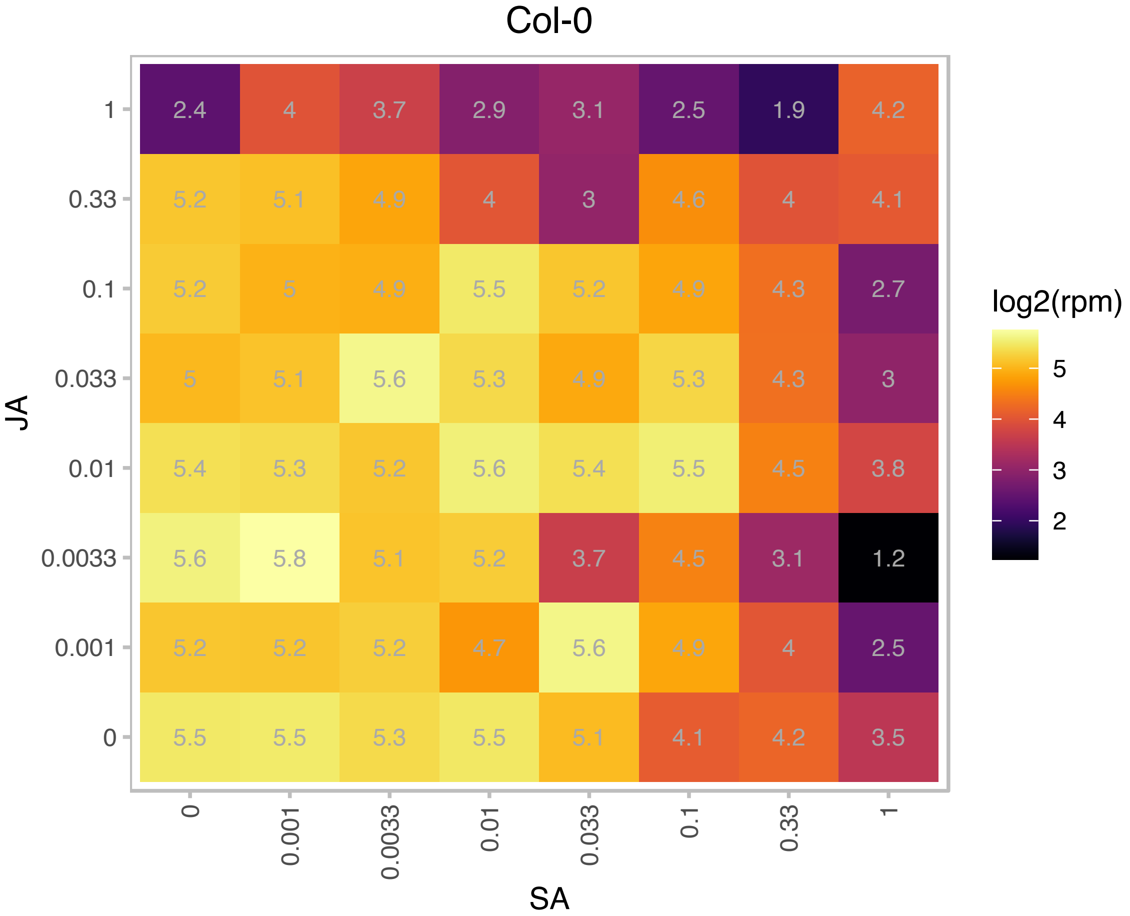


| | |
|---|---|
| Gene | AT1G01080 |
| is.expressed_genes | TRUE |
| is.responsive_genes | TRUE |
| is.apcluster | F-10 |

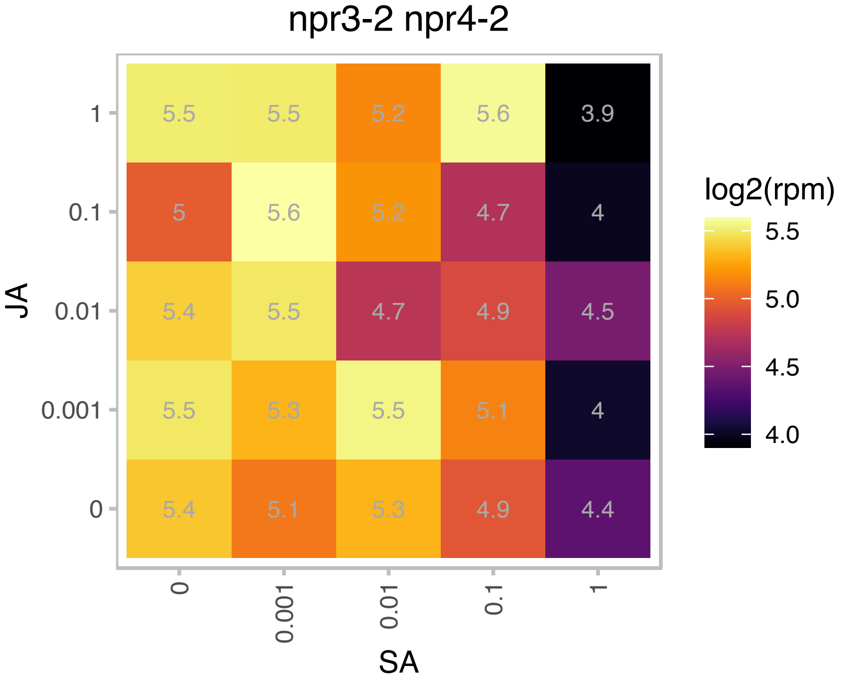


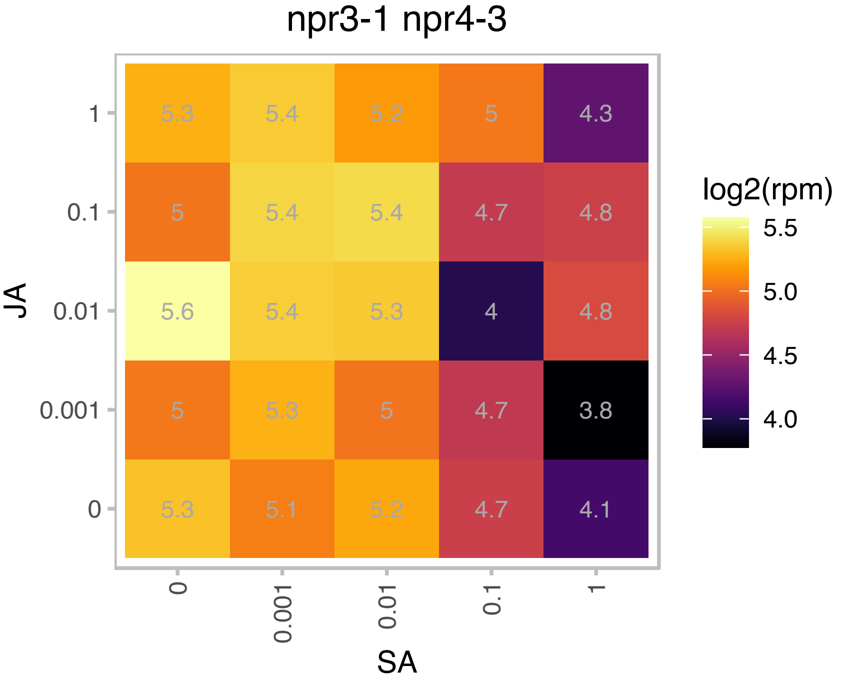

### AT1G04940.1 : atTIC20, AtTic20-I, TIC20, Tic20-I

translocon at the inner envelope membrane of chloroplasts 20

Tic20 is believed to function as a component of the protein-conducting channel at the inner envelope membrane. Genes AT1G04940 and AT1G04945 were switched for the TAIR7 genome release to give consistency w MIPs annotation. The Arabidopsis genome encodes four Tic20 homologous proteins, AT1G04940(Tic20-I), AT2G47840(Tic20-II), AT4G03320(Tic20-IV) and AT5G55710(Tic20-V).

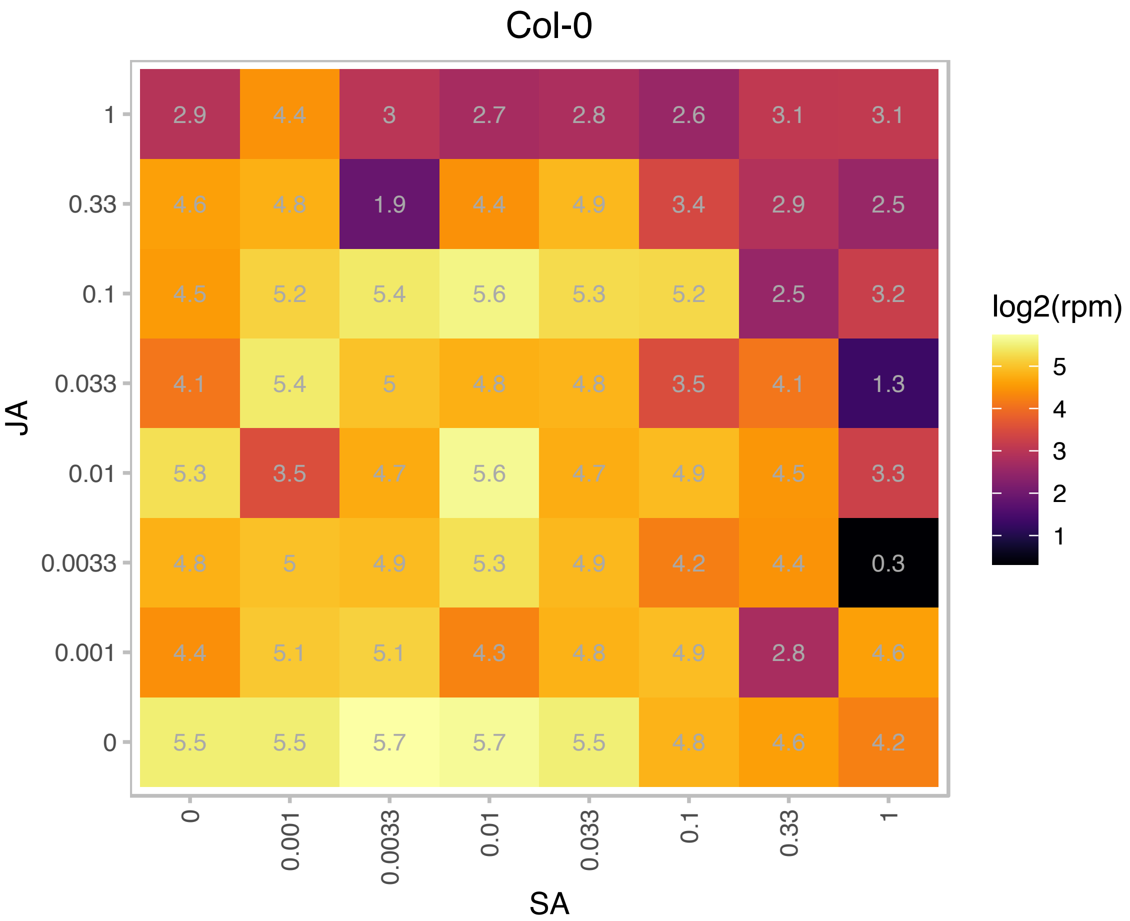


| | |
|---|---|
| Gene | AT1G04940 |
| is.expressed_genes | TRUE |
| is.responsive_genes | TRUE |
| is.apcluster | F-10 |

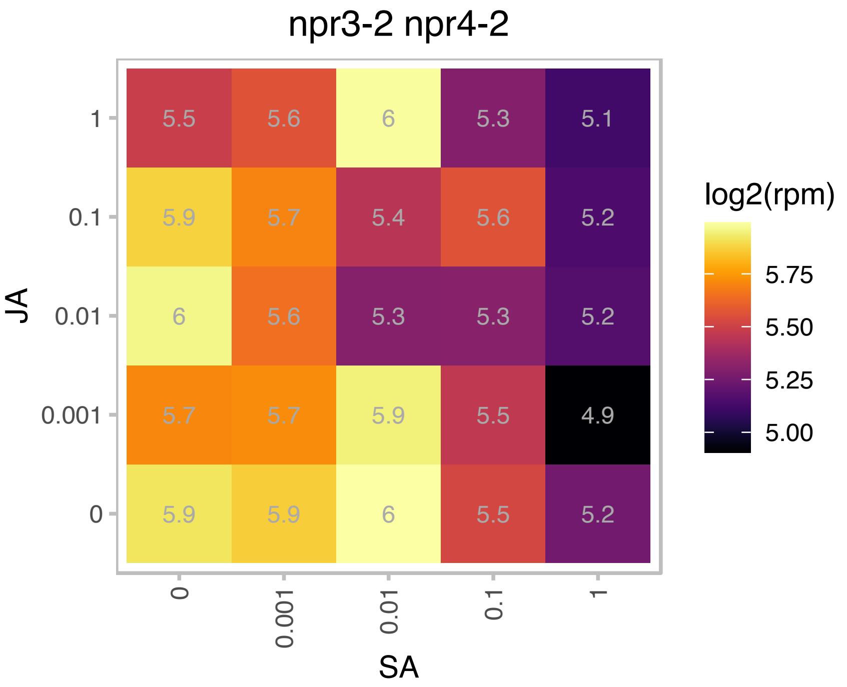


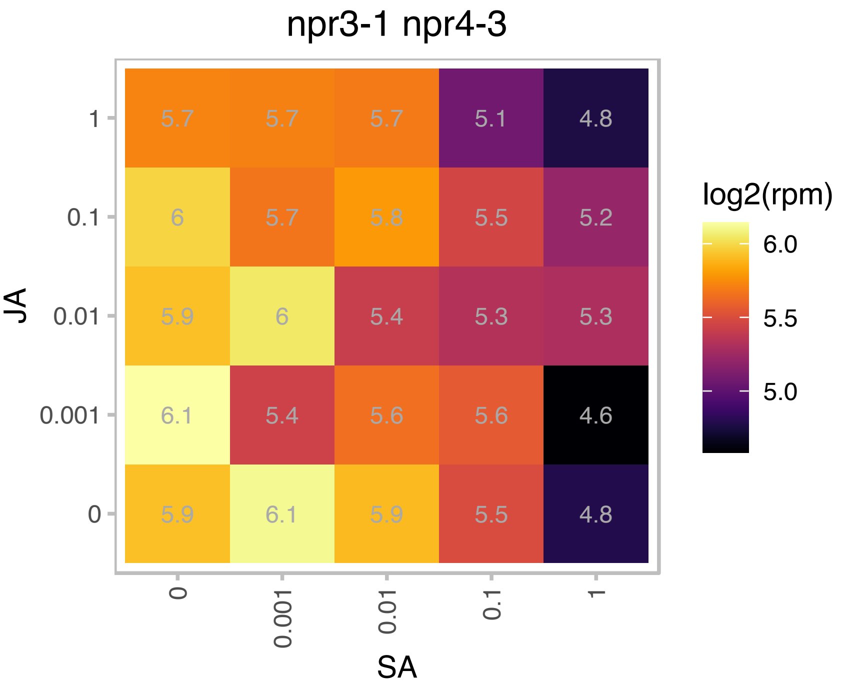

### AT1G06460.1 : ACD31.2, ACD32.1

alpha-crystallin domain 32.1

ACD32.1 encodes an alpha-crystallin domain containing protein with homology to small heat shock proteins.

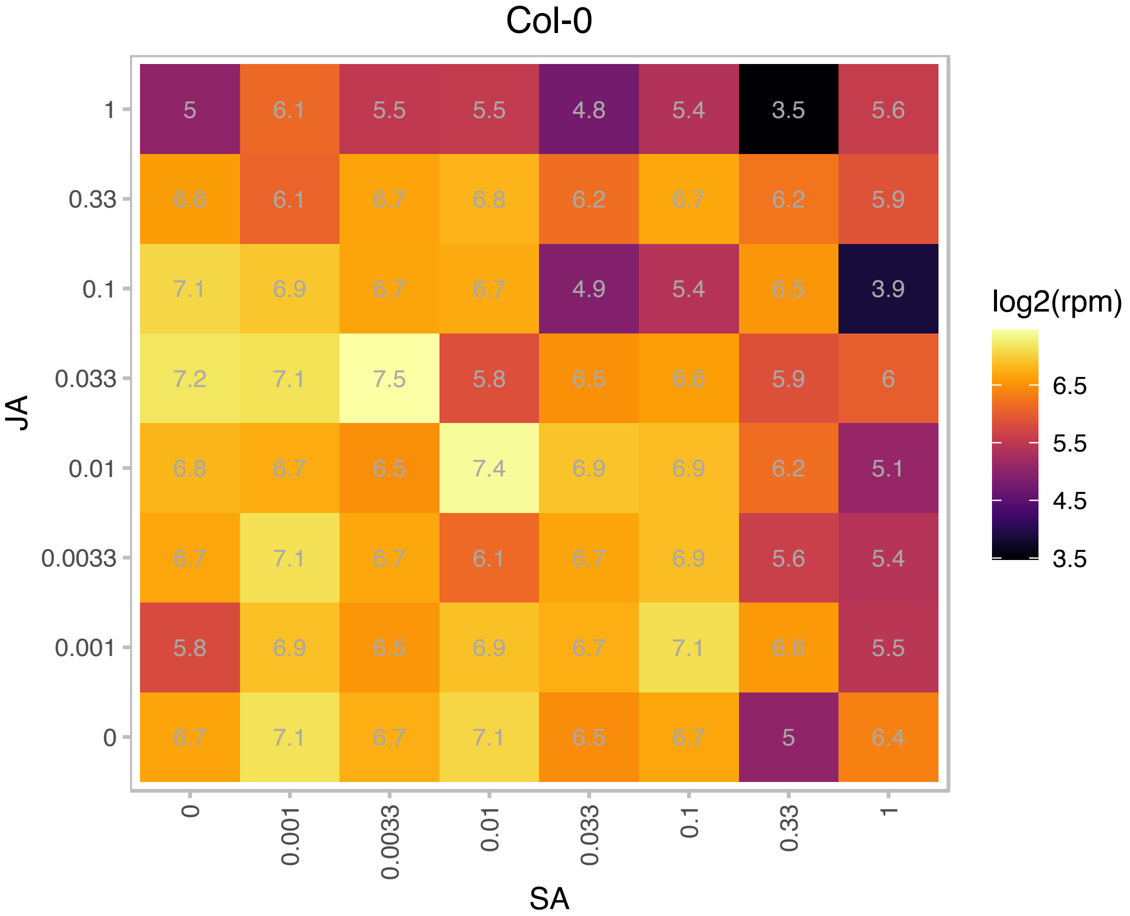


| | |
|---|---|
| Gene | AT1G06460 |
| is.expressed_genes | TRUE |
| is.responsive_genes | TRUE |
| is.apcluster | F-10 |

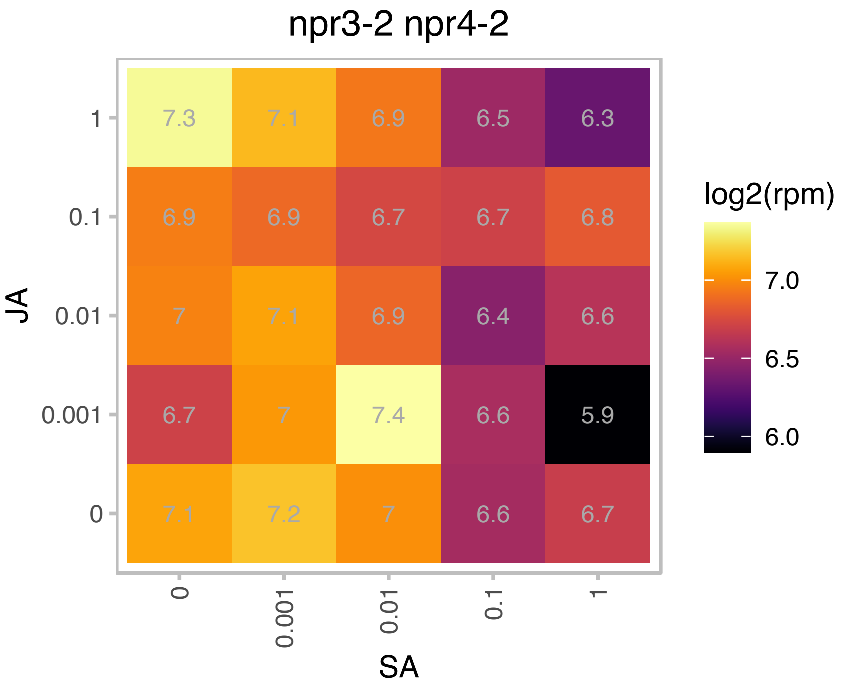


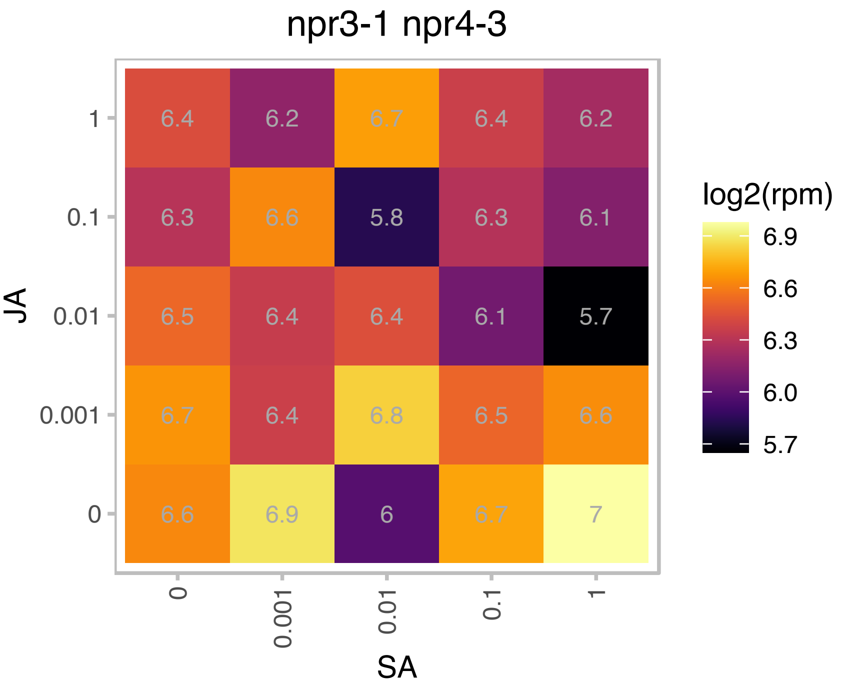

# AT1G08757.1 : NA

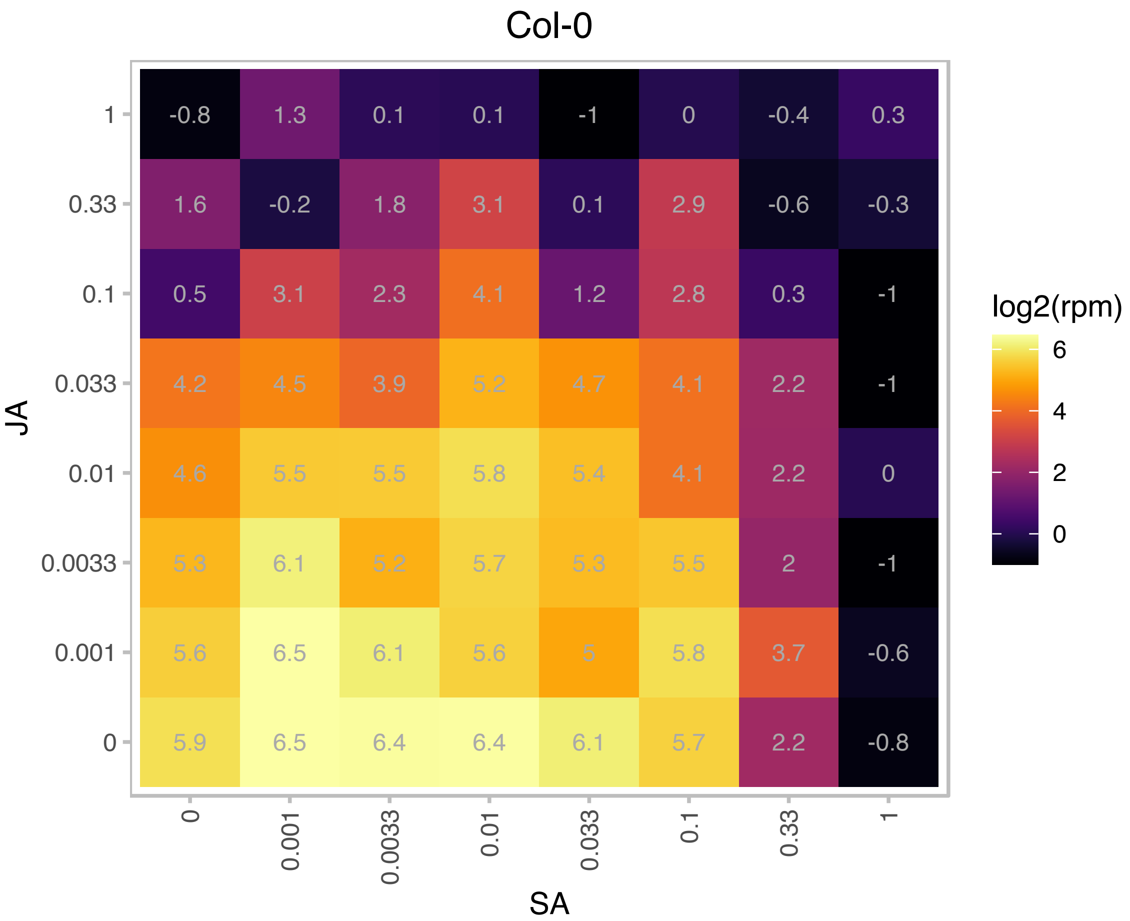


| | |
|---|---|
| Gene | AT1G08757 |
| is.expressed_genes | TRUE |
| is.responsive_genes | TRUE |
| is.apcluster | F-10 |

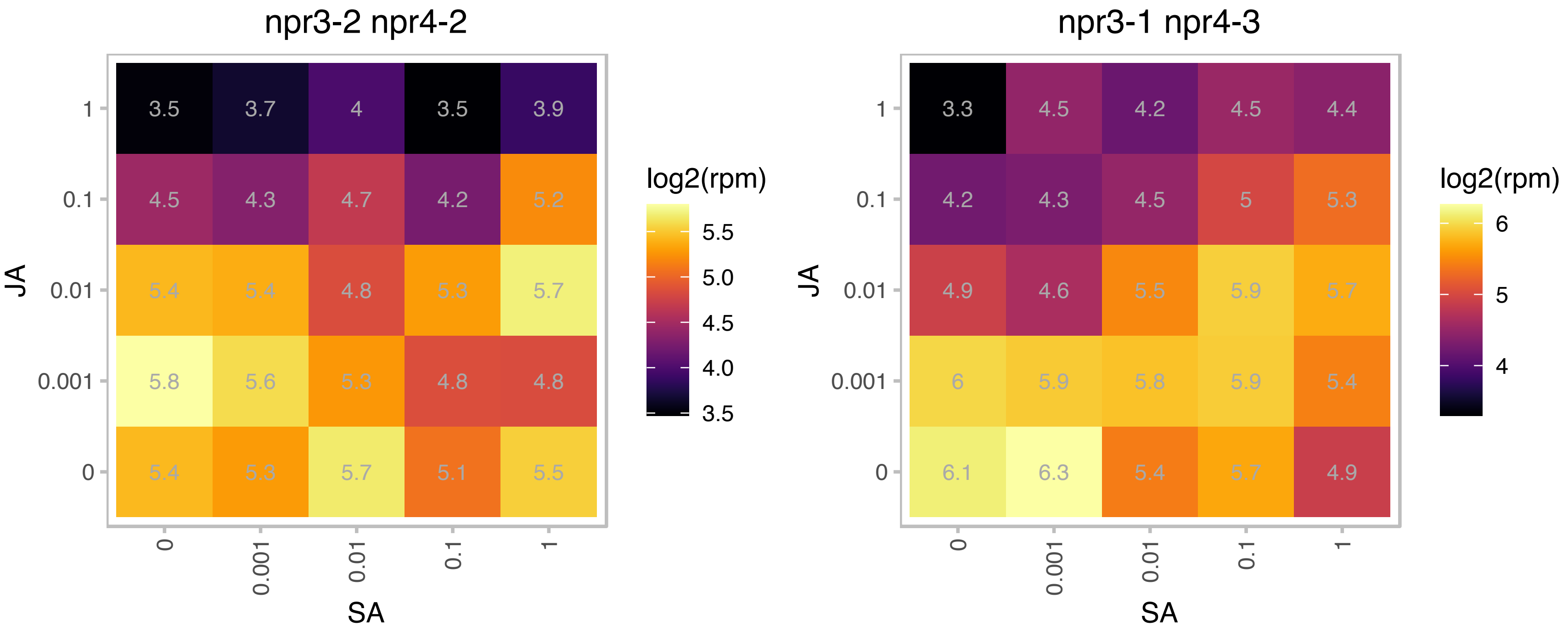

### AT1G09530.1 : PAP3, PIF3, POC1

phytochrome interacting factor 3

Transcription factor interacting with photoreceptors phyA and phyB. Forms a ternary complex in vitro with G-box element of the promoters of LHY, CCA1. Acts as a negative regulator of phyB signalling. It degrades rapidly after irradiation of dark grown seedlings in a process controlled by phytochromes. Does not play a significant role in controlling light input and function of the circadian clockwork. Binds to G- and E-boxes, but not to other ACEs. Binds to anthocyanin biosynthetic genes in a light- and HY5-independent fashion. PIF3 function as a transcriptional activator can be functionally and mechanistically separated from its role in [illegible]

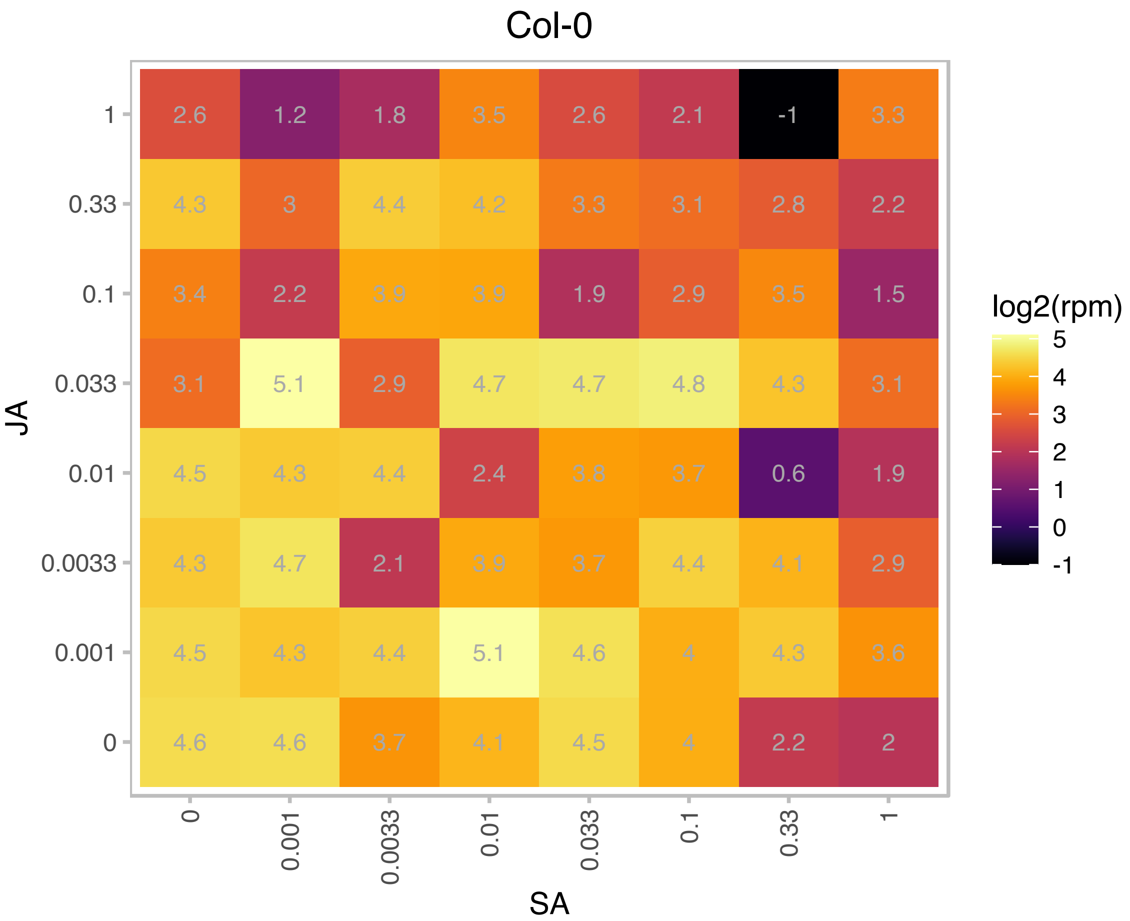


| | |
|---|---|
| Gene | AT1G09530 |
| is.expressed_genes | TRUE |
| is.responsive_genes | TRUE |
| is.apcluster | F-10 |

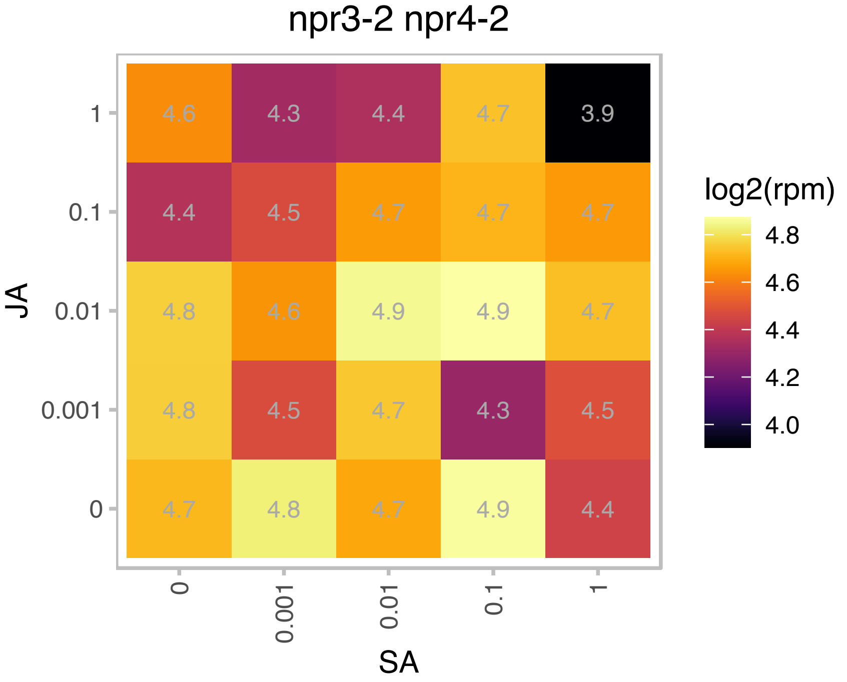


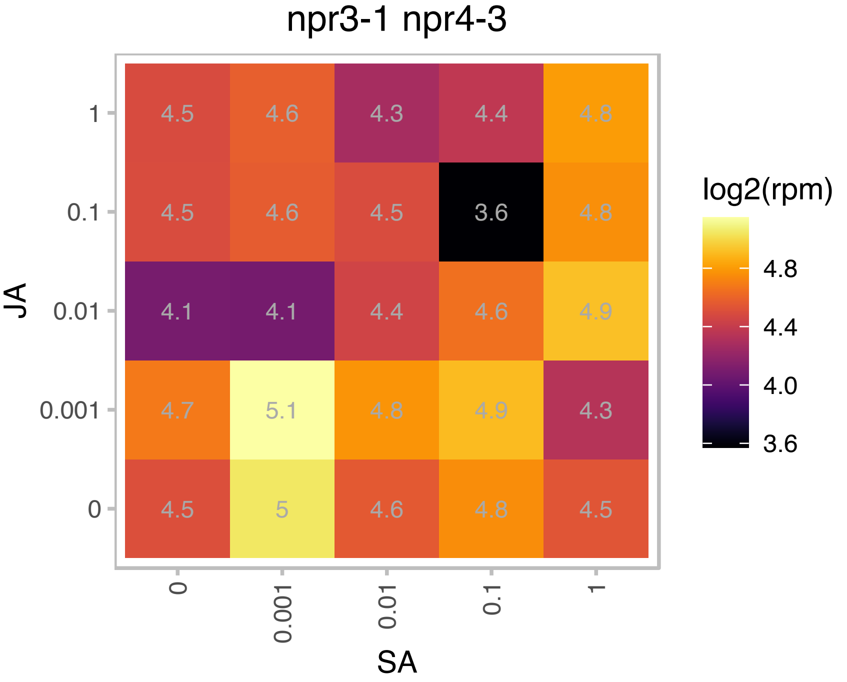

# AT1G14345.1 : NA

NAD(P)-linked oxidoreductase superfamily protein

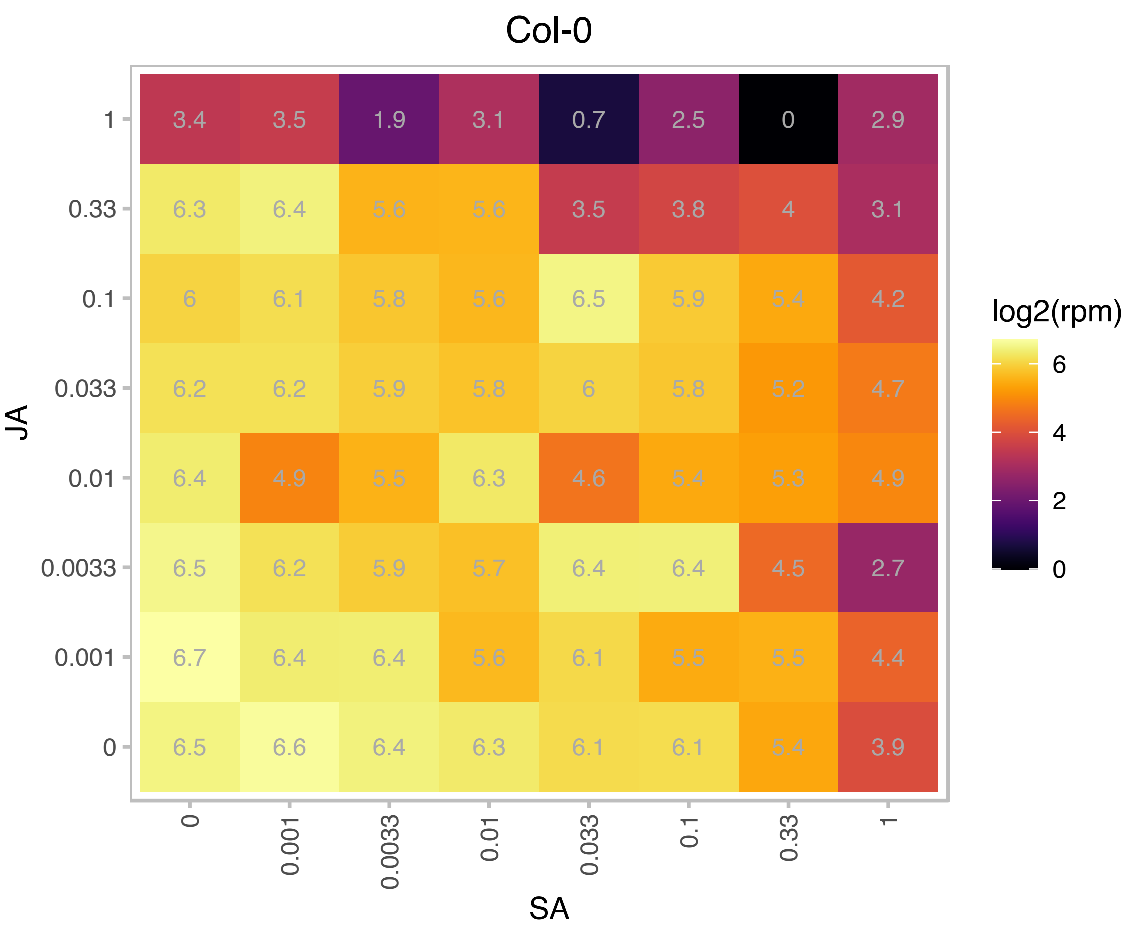


| | |
|---|---|
| Gene | AT1G14345 |
| is.expressed_genes | TRUE |
| is.responsive_genes | TRUE |
| is.apcluster | F-10 |

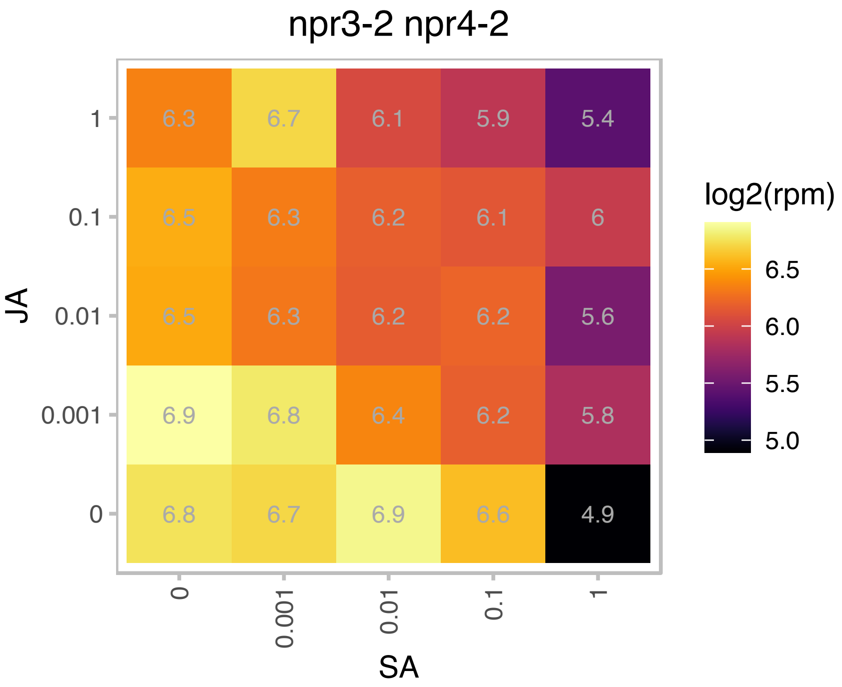


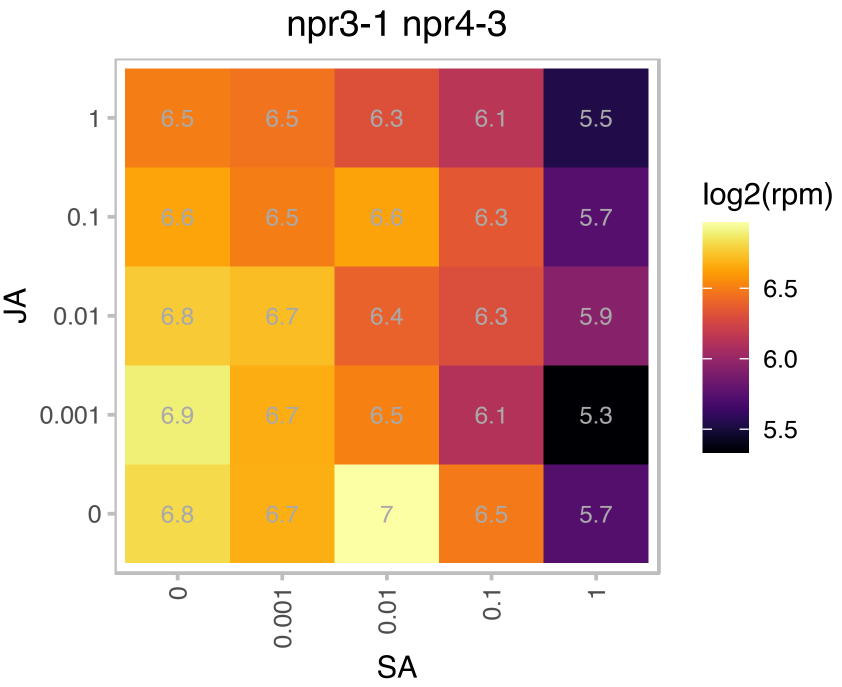

### AT1G15390.1 : ATDEF1, PDF1A

peptide deformylase 1A

encodes a peptide deformylase-like protein. Removes N-formyl groups, a prerequisite for the action of methionine aminopeptidase during protein synthesis. Targeted to mitochondria. Requires Zn for catalysis.

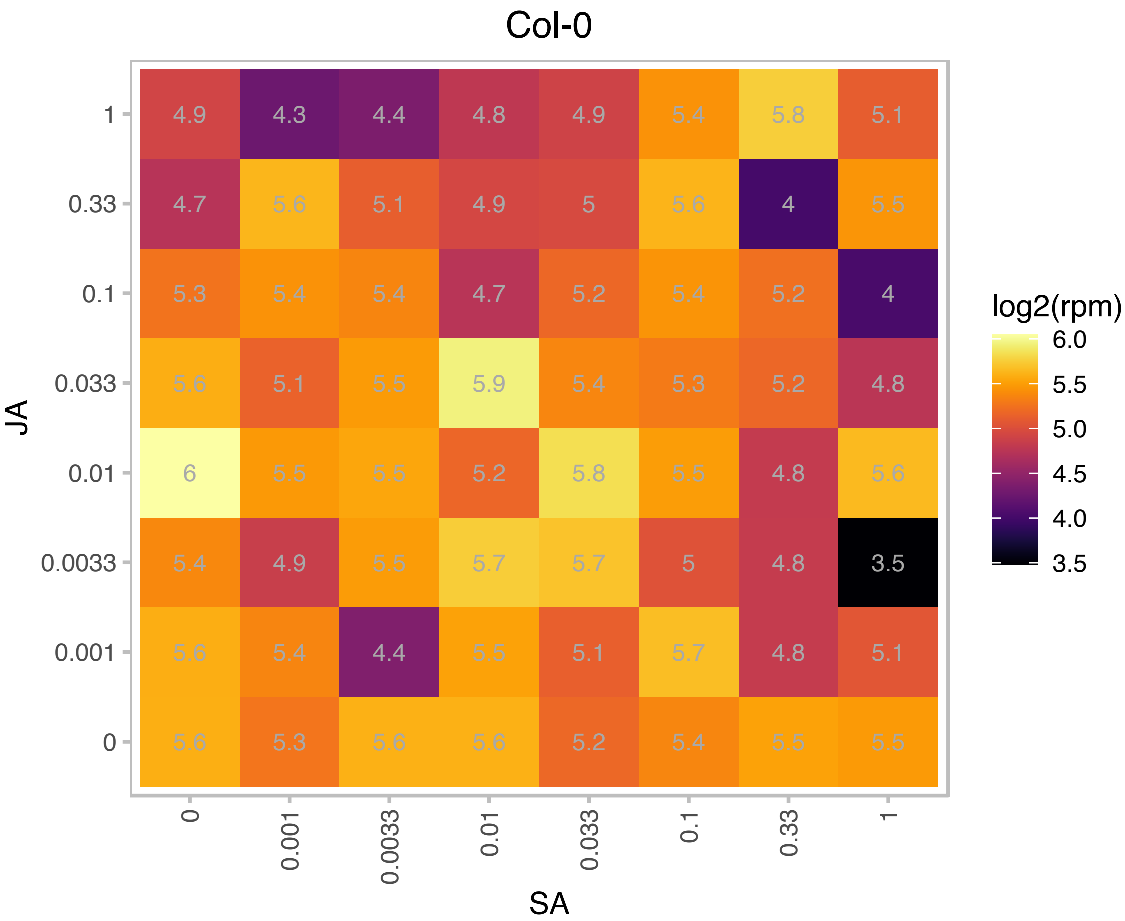


| | |
|---|---|
| Gene | AT1G15390 |
| is.expressed_genes | TRUE |
| is.responsive_genes | TRUE |
| is.apcluster | F-10 |

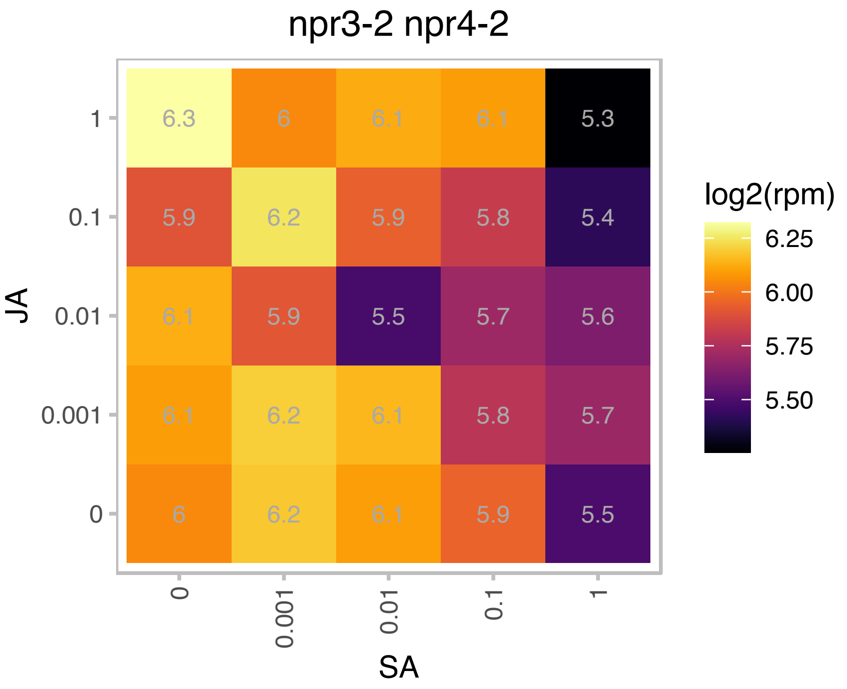


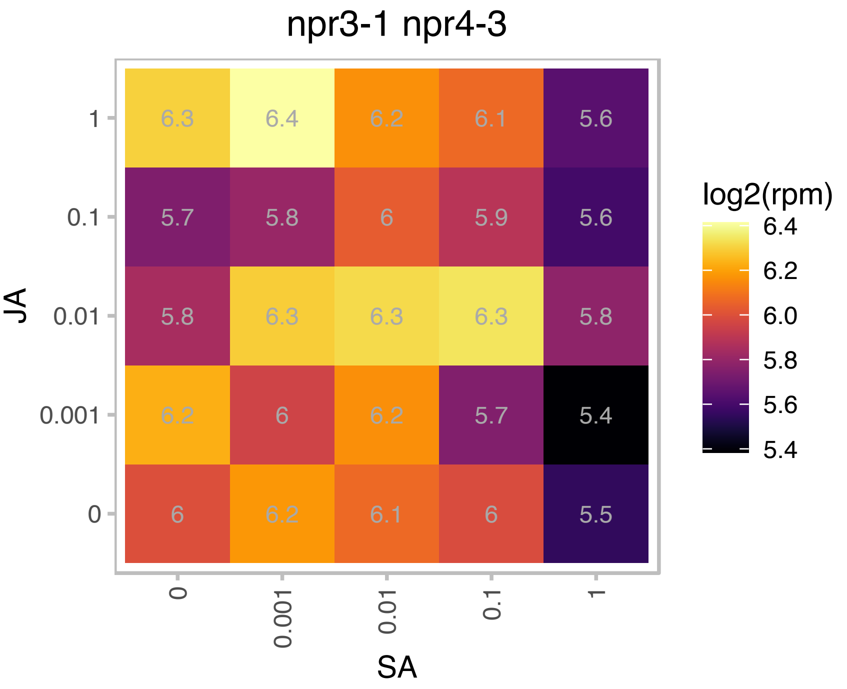

# AT1G19380.1 : NA

Protein of unknown function (DUF1195)

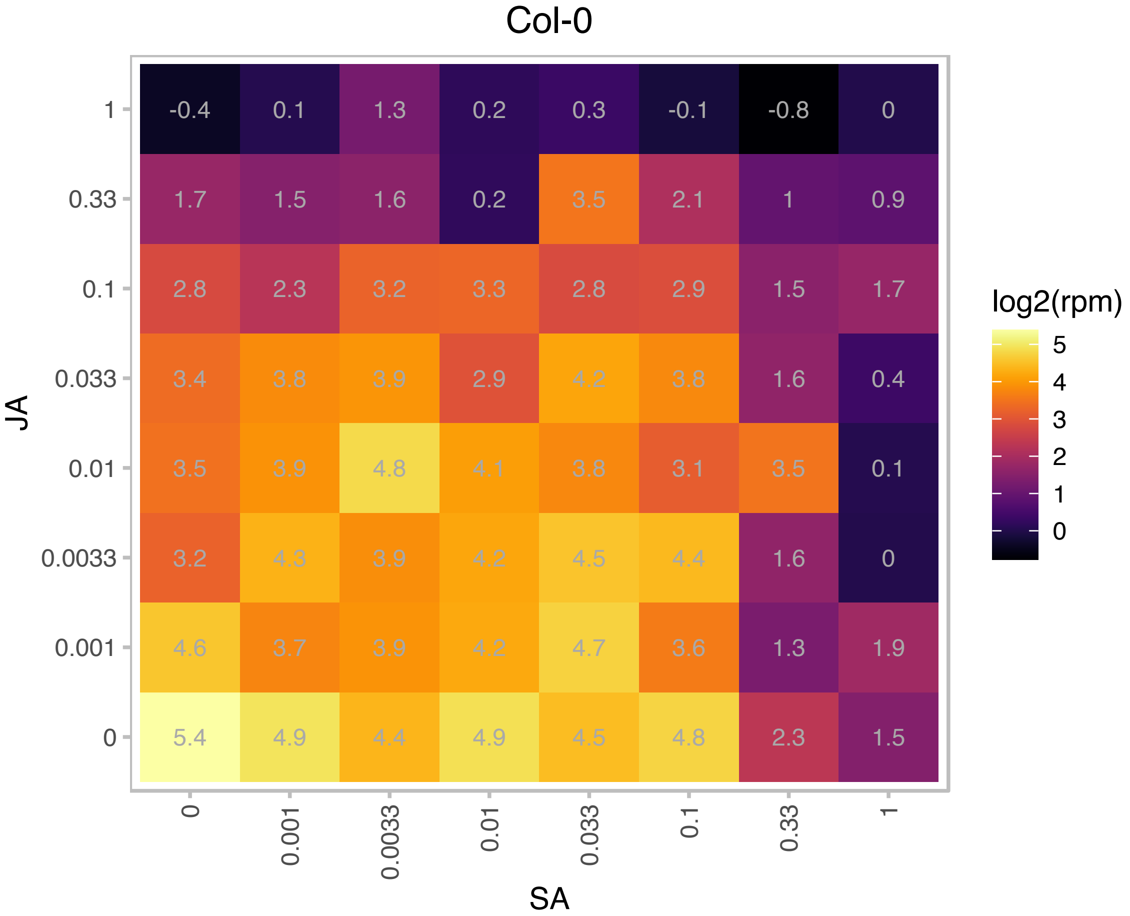


| | |
|---|---|
| Gene | AT1G19380 |
| is.expressed_genes | TRUE |
| is.responsive_genes | TRUE |
| is.apcluster | F-10 |

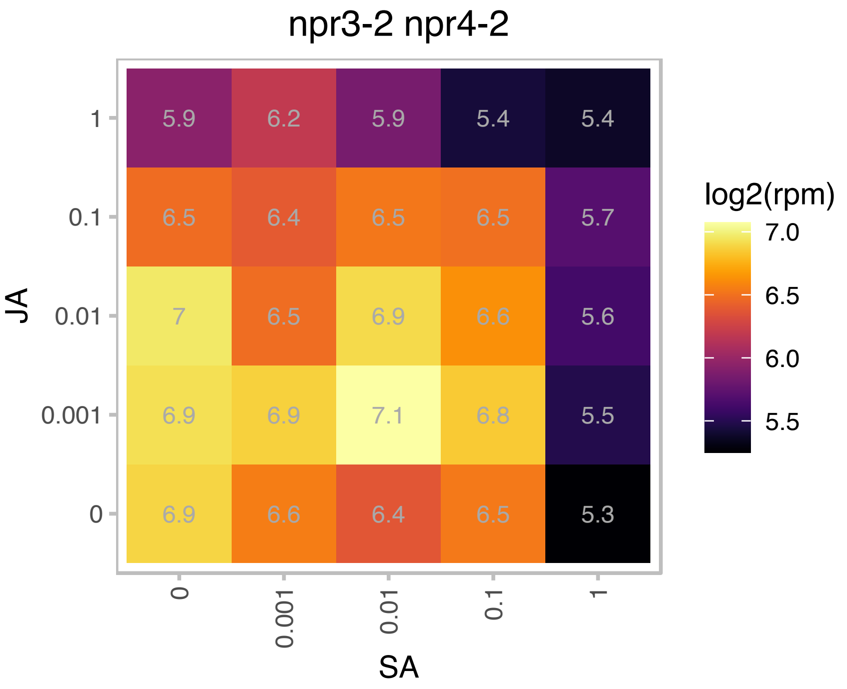


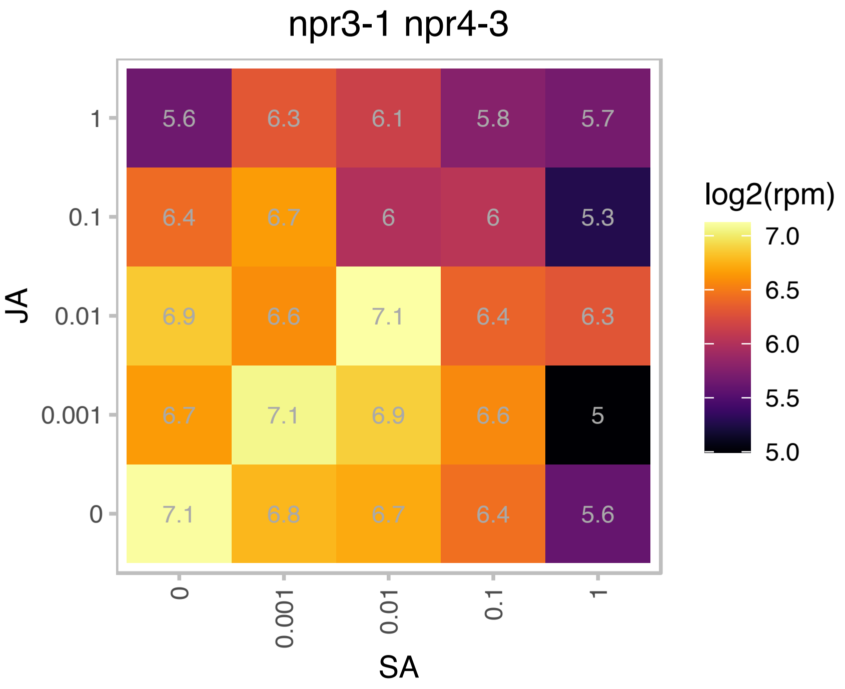

# AT1G22630.1 : NA

| | |
|---|---|
| Gene | AT1G22630 |
| is.expressed_genes | TRUE |
| is.responsive_genes | TRUE |
| is.apcluster | F-10 |

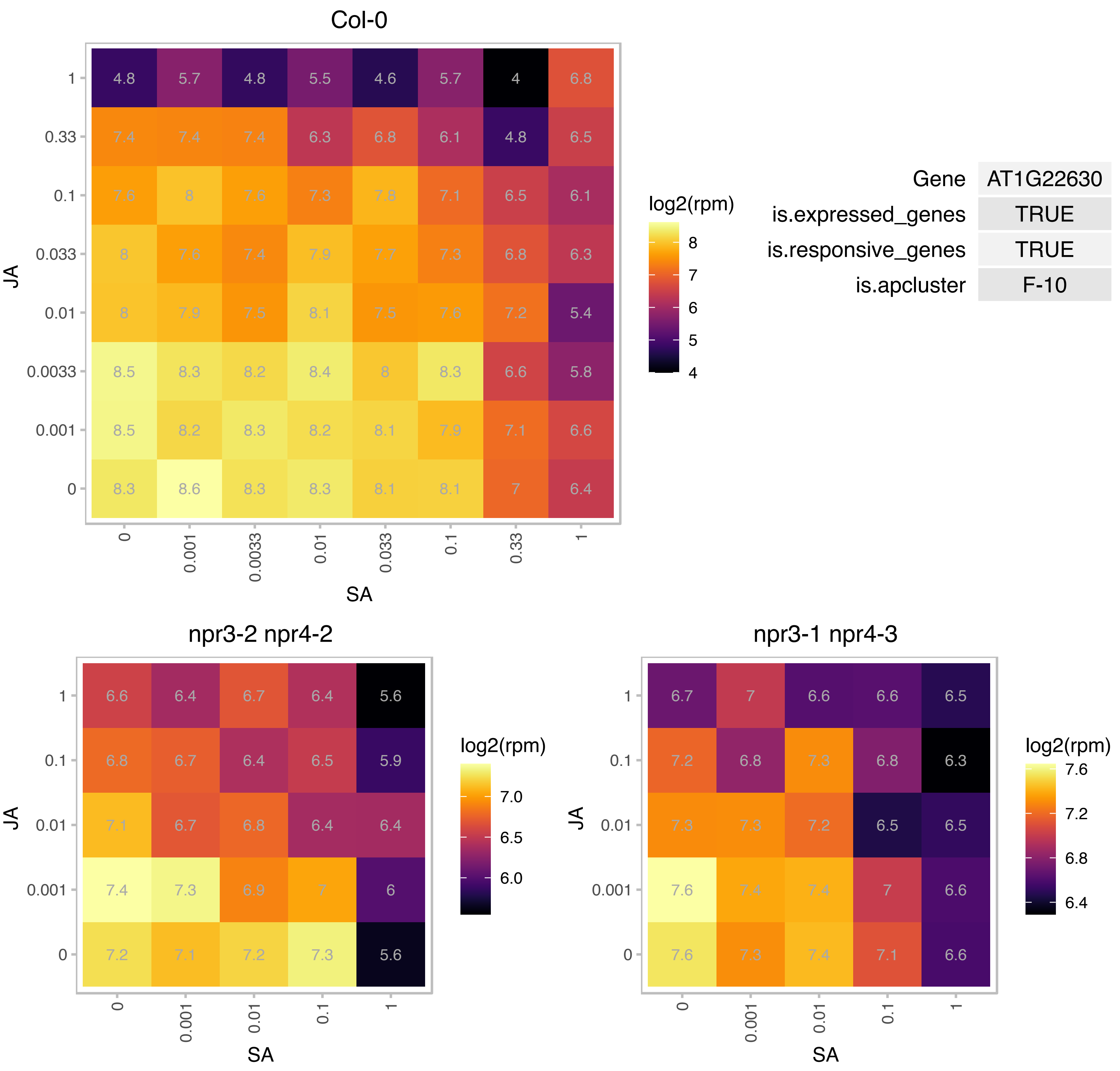

# AT1G27020.1 : NA

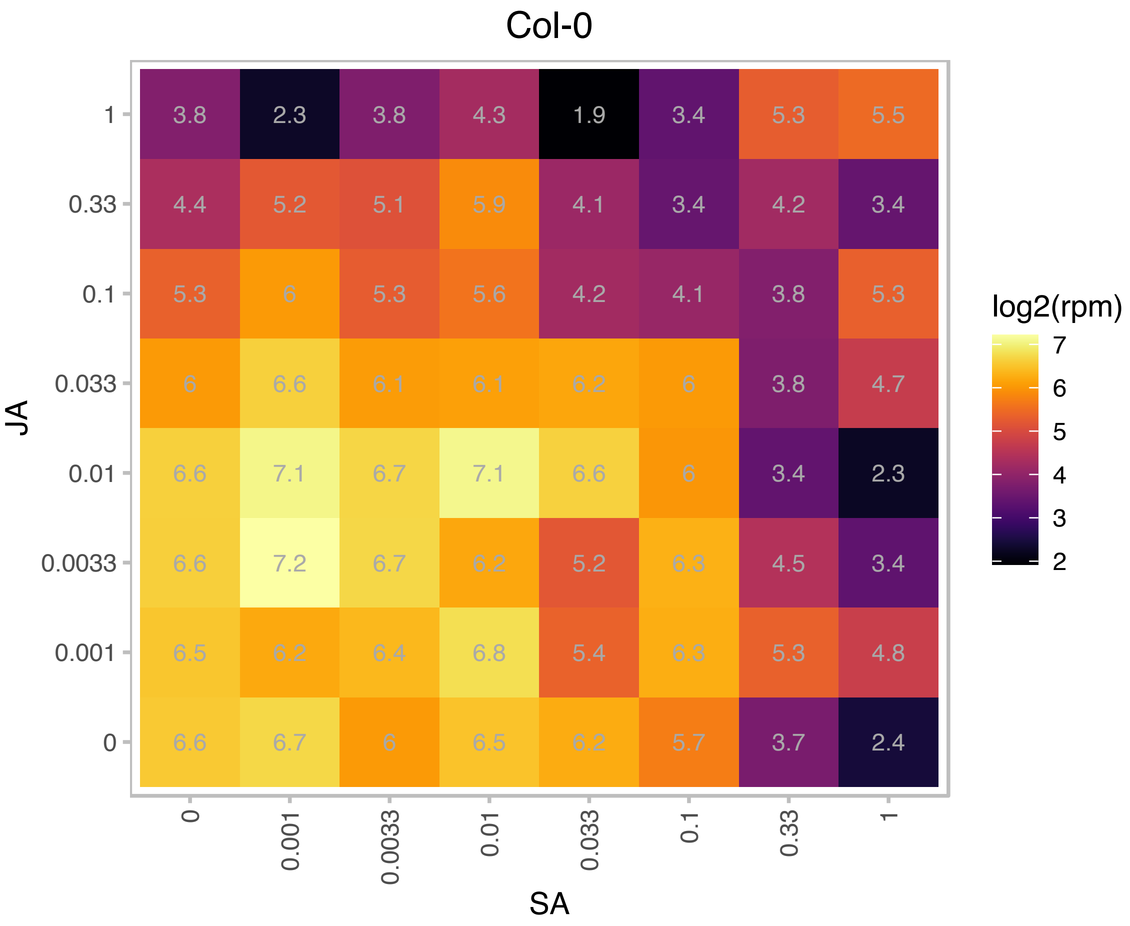


| | |
|---|---|
| Gene | AT1G27020 |
| is.expressed_genes | TRUE |
| is.responsive_genes | TRUE |
| is.apcluster | F-10 |

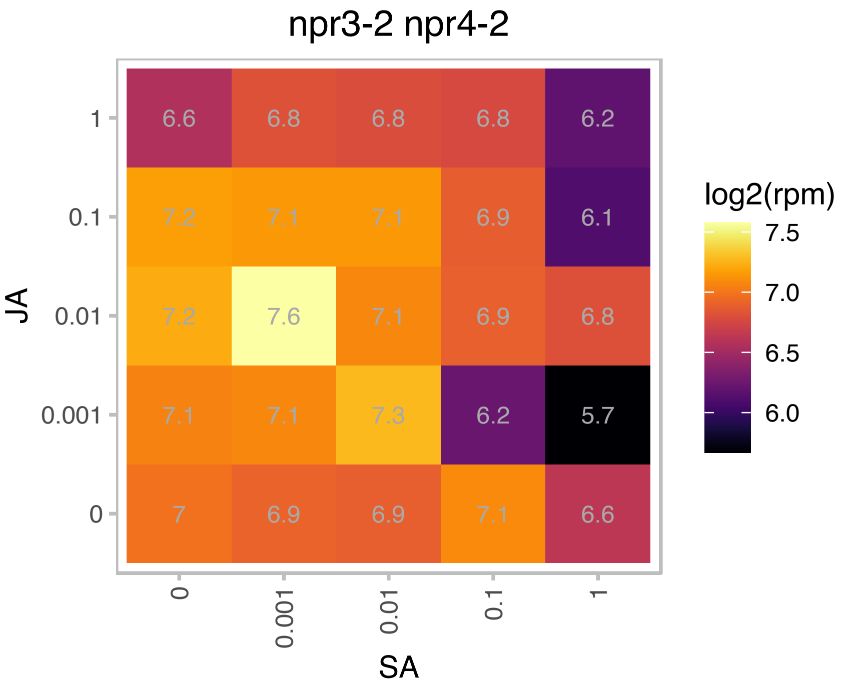


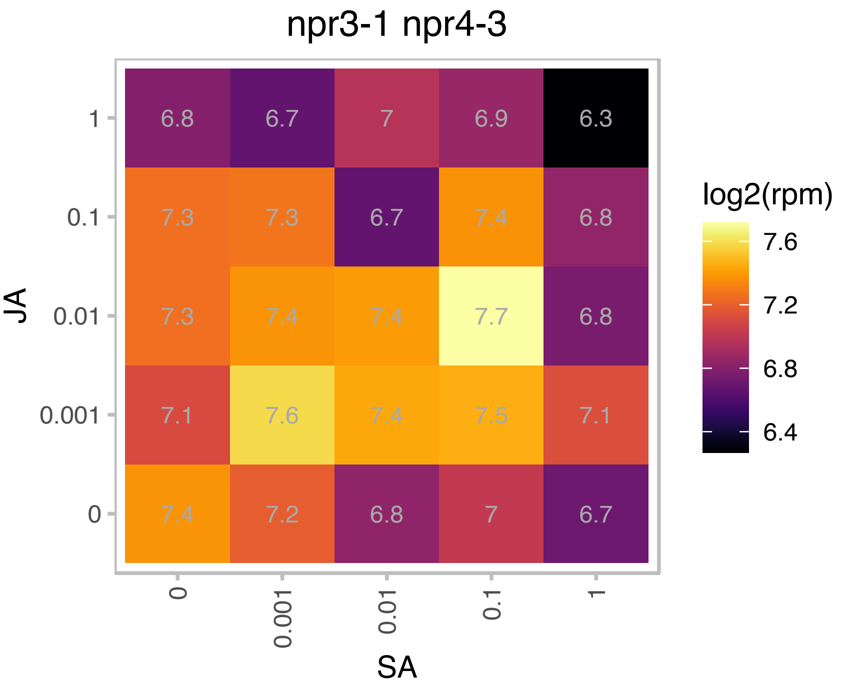

### AT1G30510.1 : ATRFNR2, RFNR2

root FNR 2

Encodes a root-type ferredoxin:NADP(H) oxidoreductase.

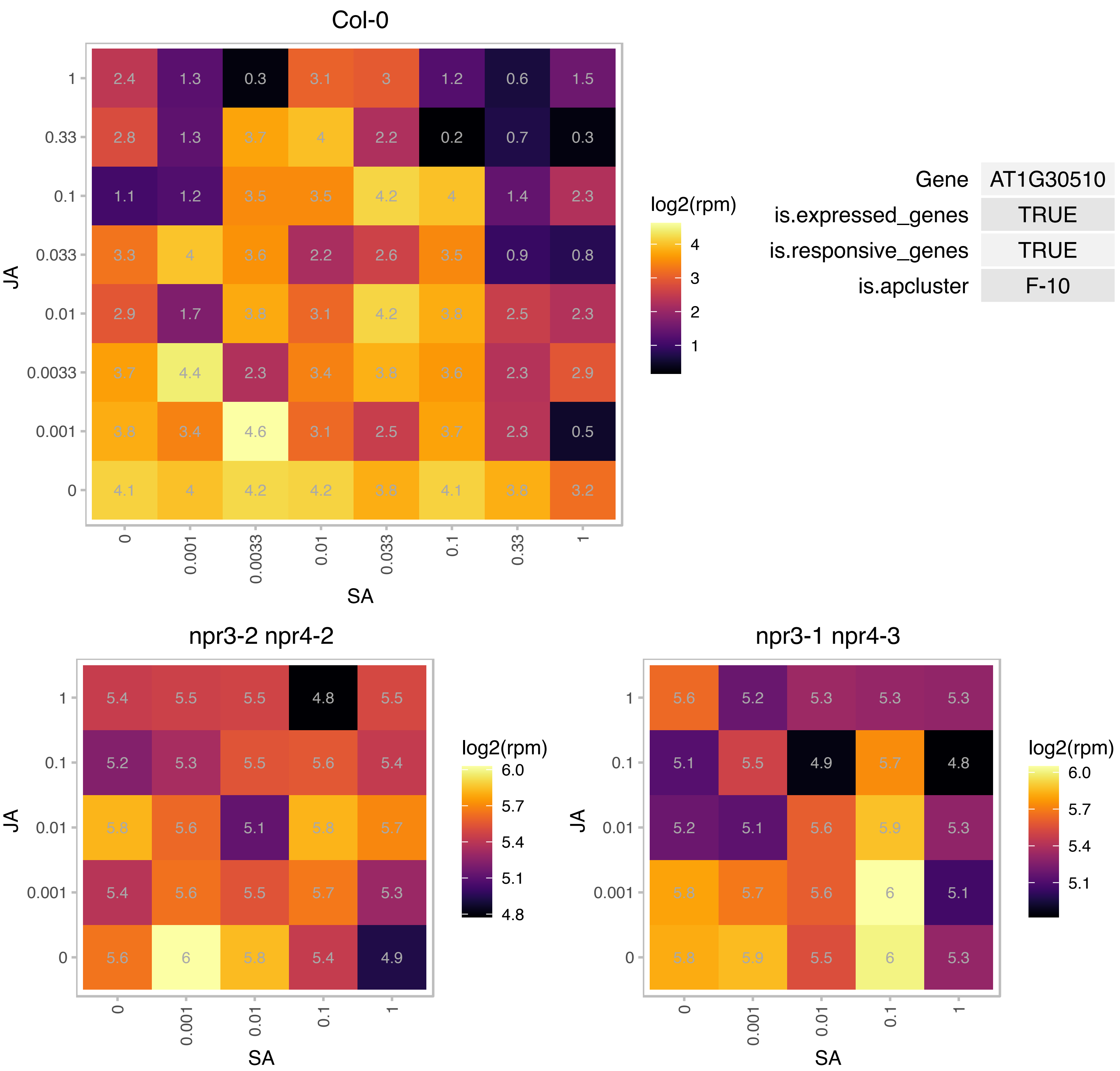

# AT1G47520.1 : NA

transposable element gene

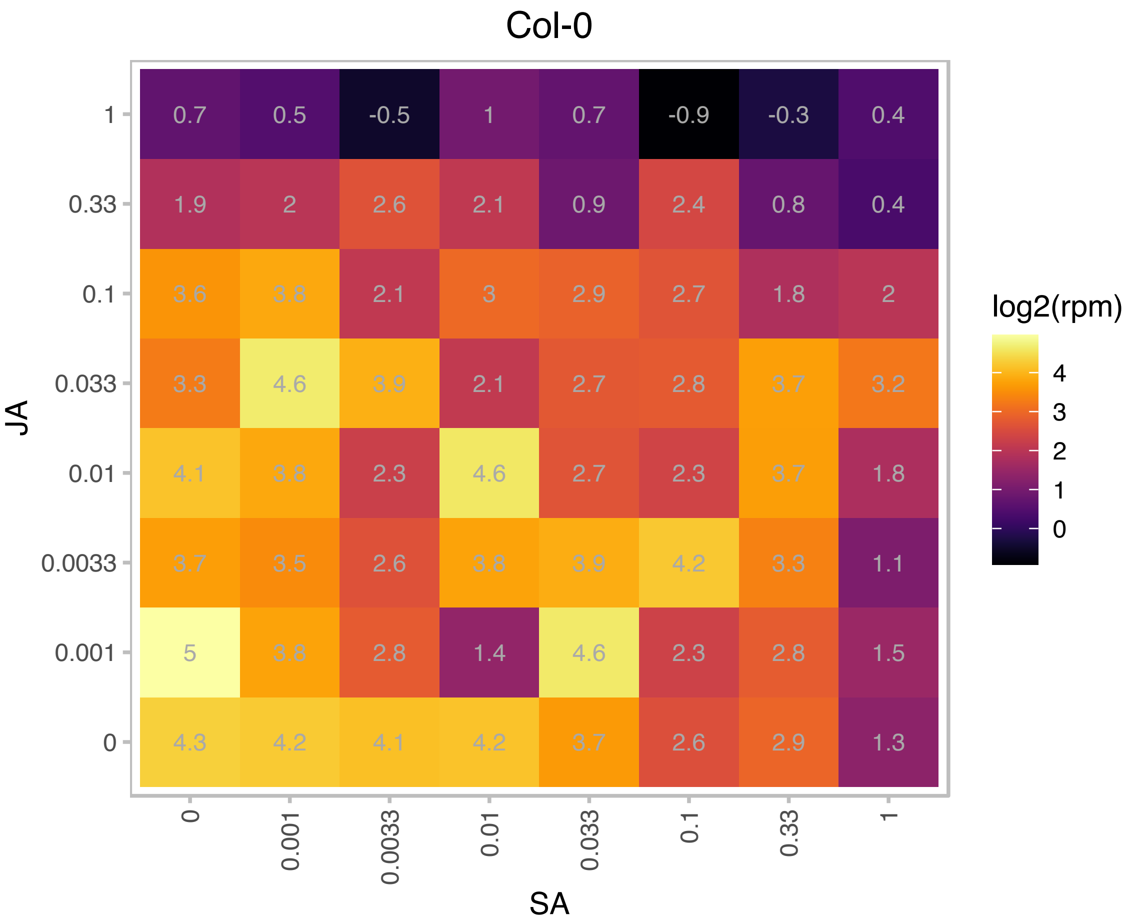


| | |
|---|---|
| Gene | AT1G47520 |
| is.expressed_genes | TRUE |
| is.responsive_genes | TRUE |
| is.apcluster | F-10 |

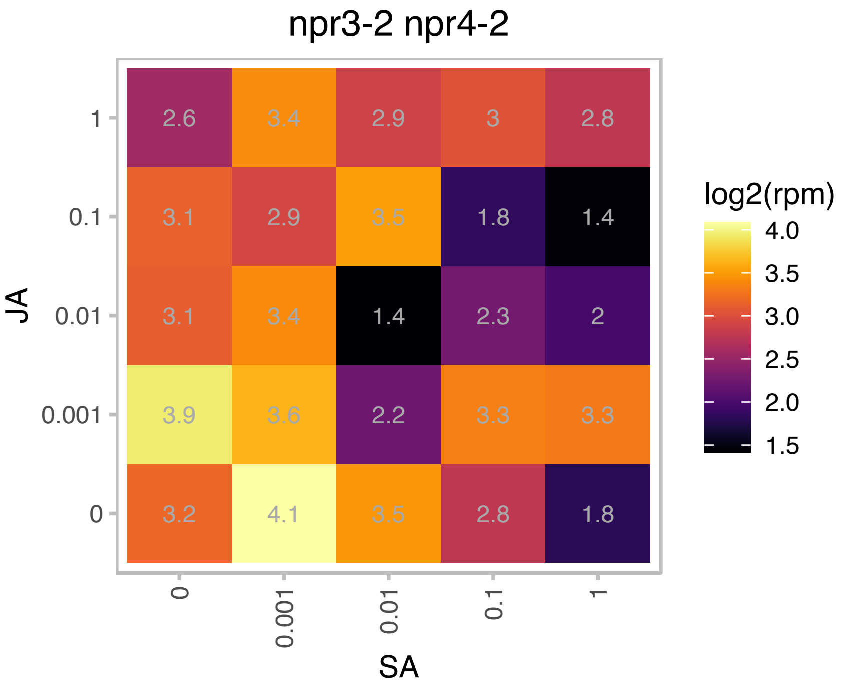


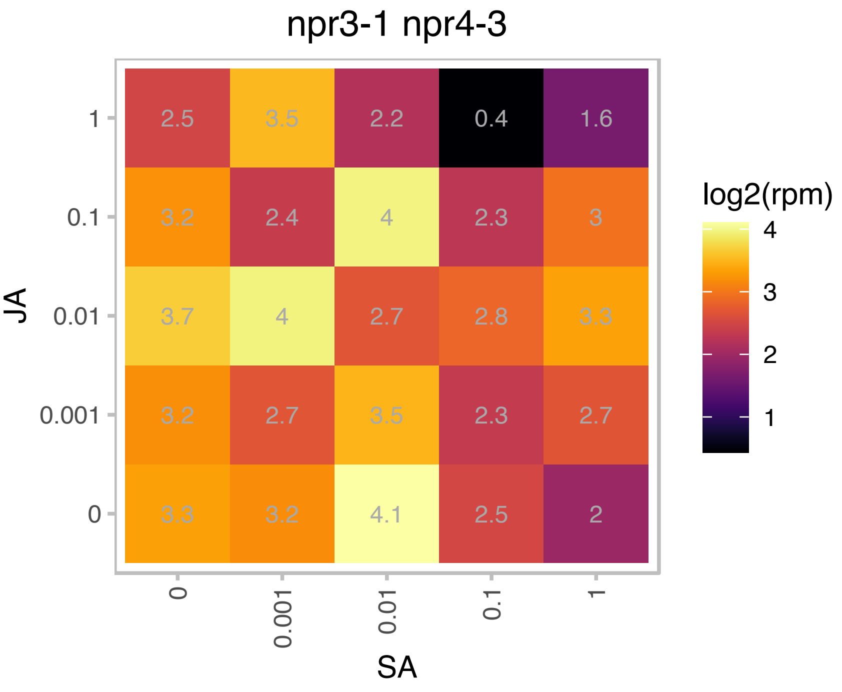

# AT1G49975.1 : NA

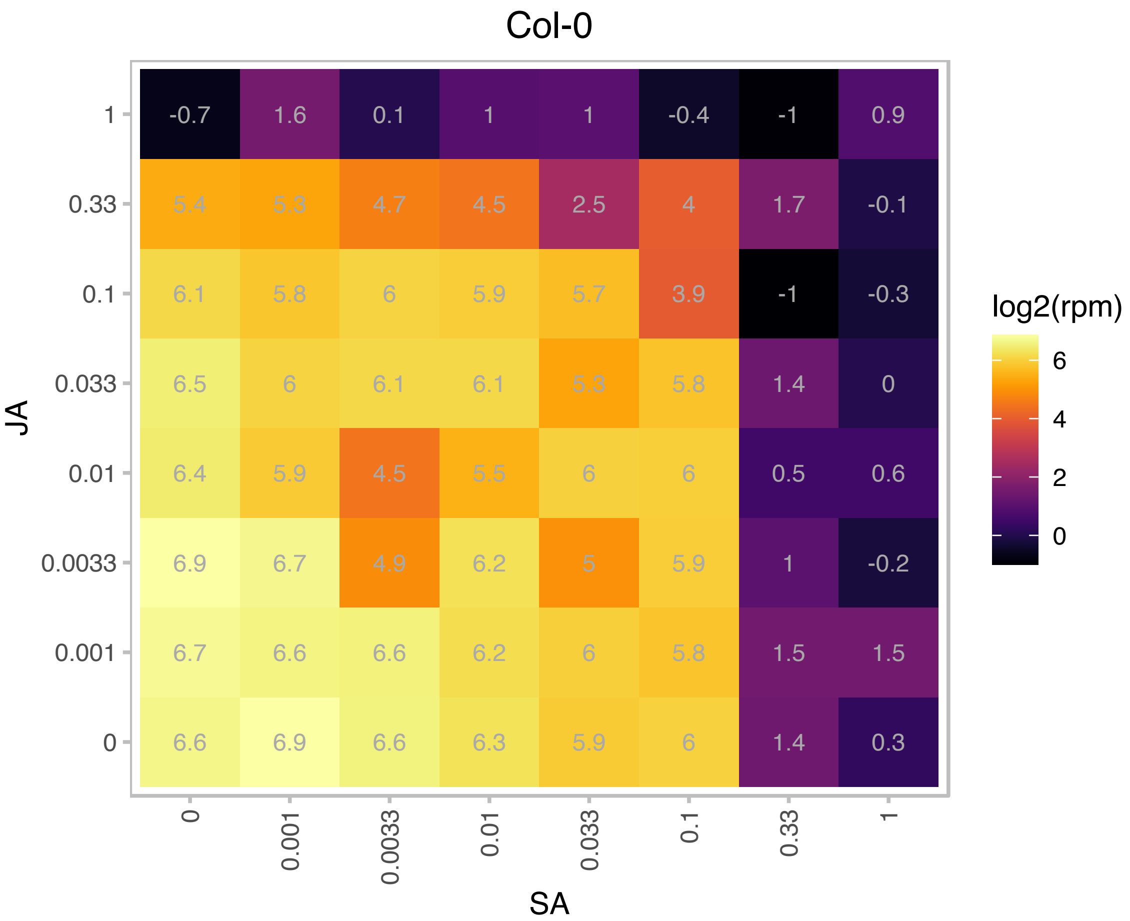


| | |
|---|---|
| Gene | AT1G49975 |
| is.expressed_genes | TRUE |
| is.responsive_genes | TRUE |
| is.apcluster | F-10 |

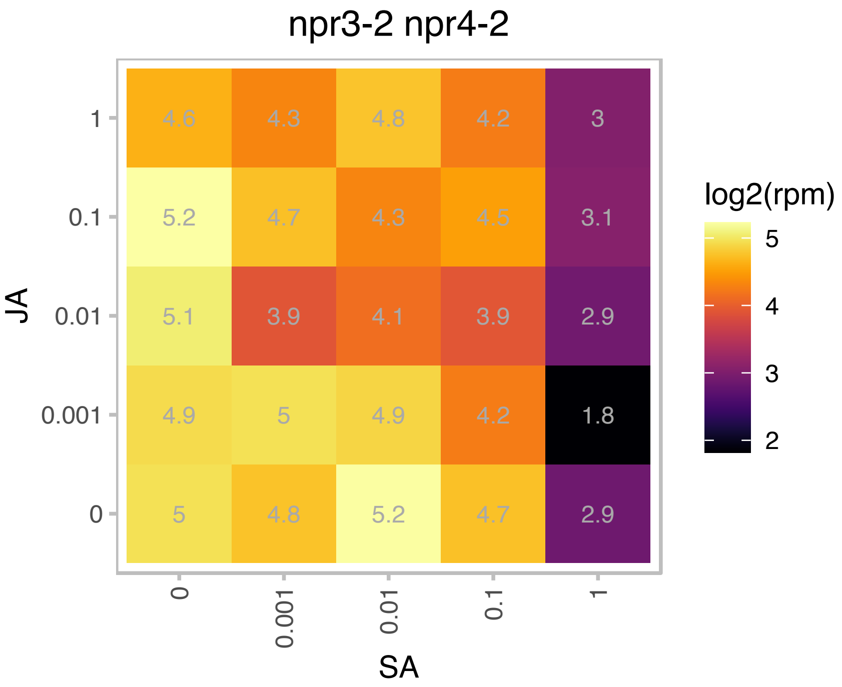


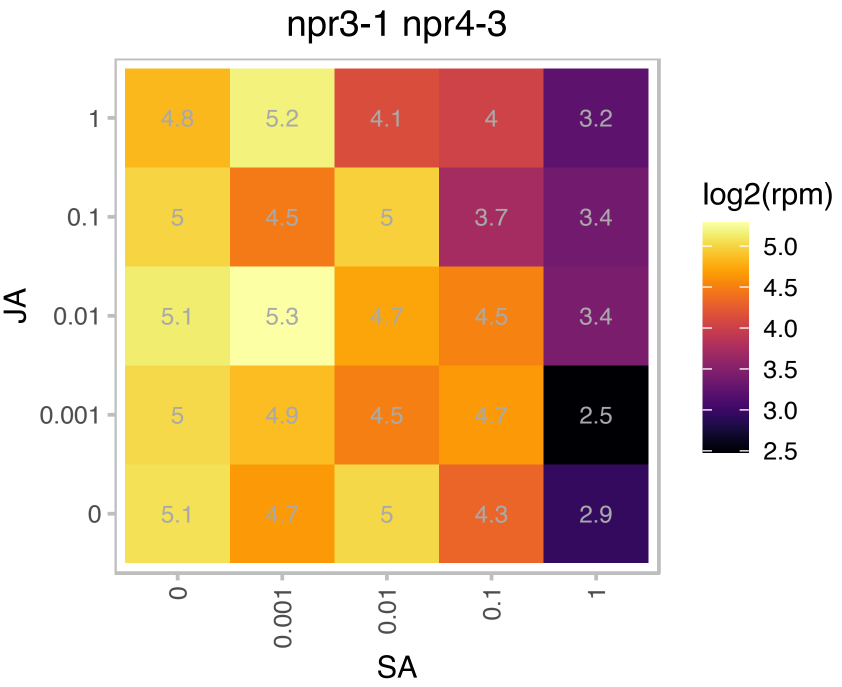

### AT1G50900.1 : GDC1, LTD

Ankyrin repeat family protein

Encodes GDC1 (Grana Deficient Chloroplast 1), an ankyrin domain containing protein required fro chloroplast thylakoid grana formation. The mRNA is cell-to-cell mobile.

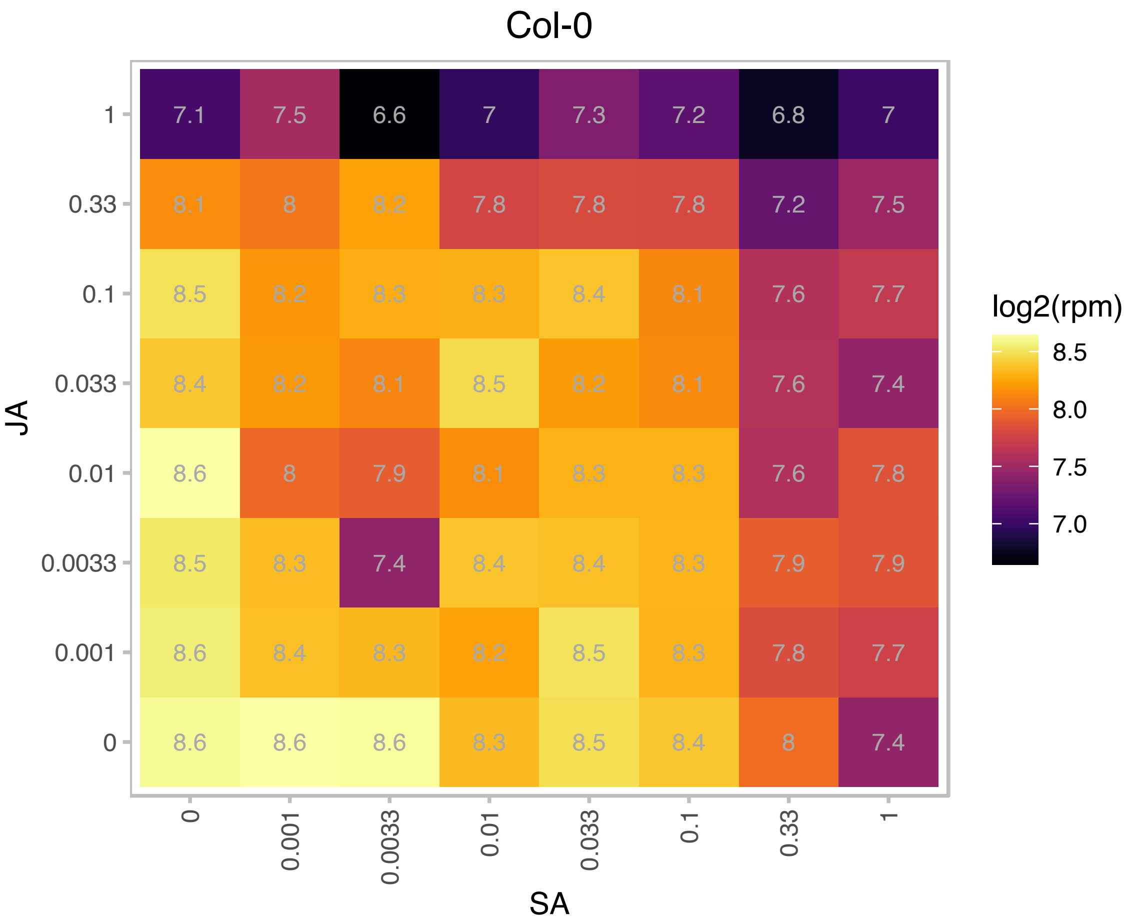


| | |
|---|---|
| Gene | AT1G50900 |
| is.expressed_genes | TRUE |
| is.responsive_genes | TRUE |
| is.apcluster | F-10 |

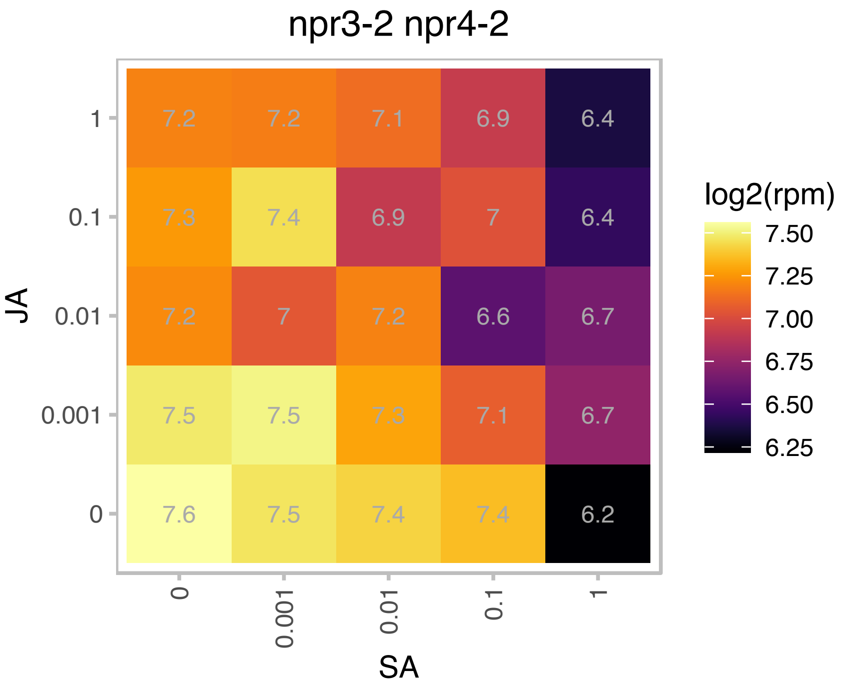


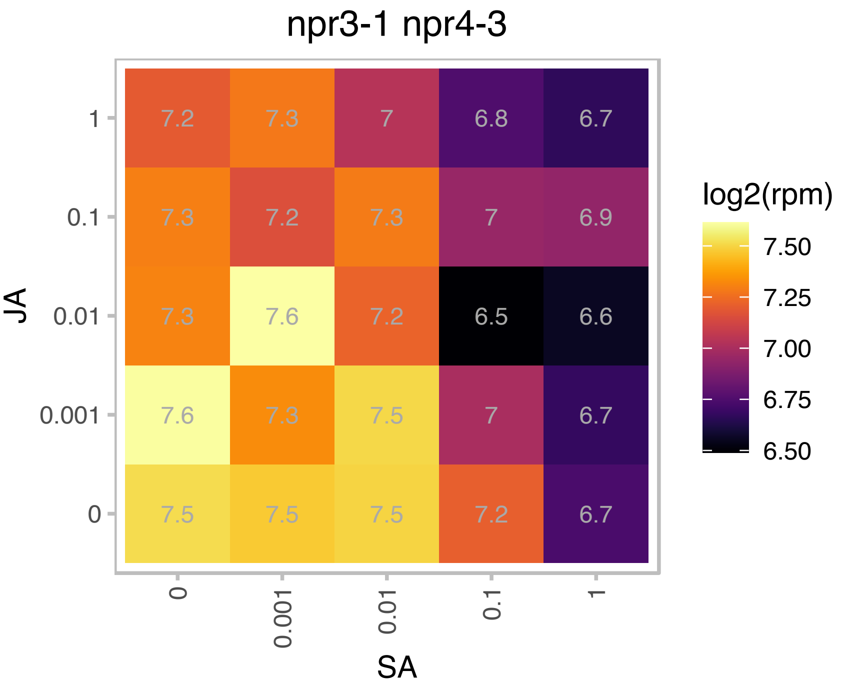

### AT1G52230.1 : PSAH-2, PSAH2, PSI-H

photosystem I subunit H2

Phosphorylation of this protein is dependent on calcium. The mRNA is cell-to-cell mobile.

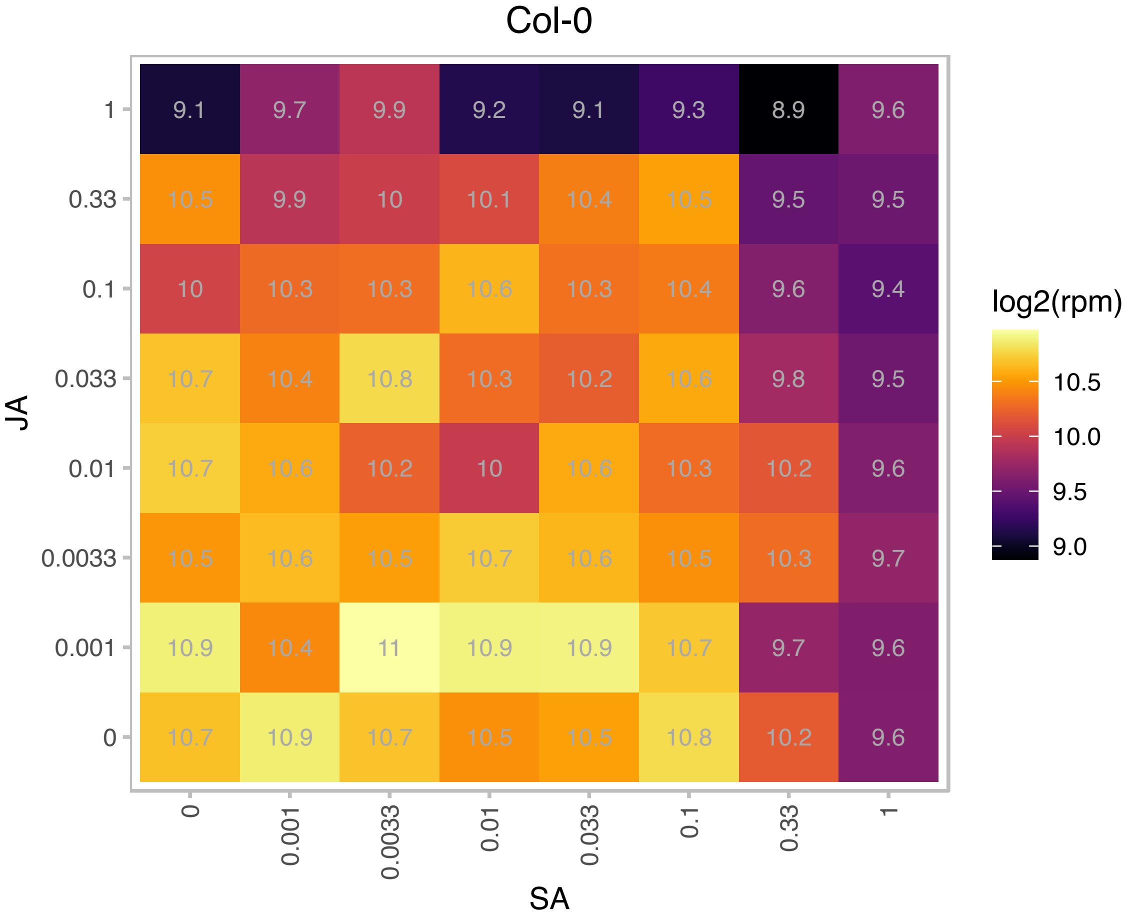


| | |
|---|---|
| Gene | AT1G52230 |
| is.expressed_genes | TRUE |
| is.responsive_genes | TRUE |
| is.apcluster | F-10 |

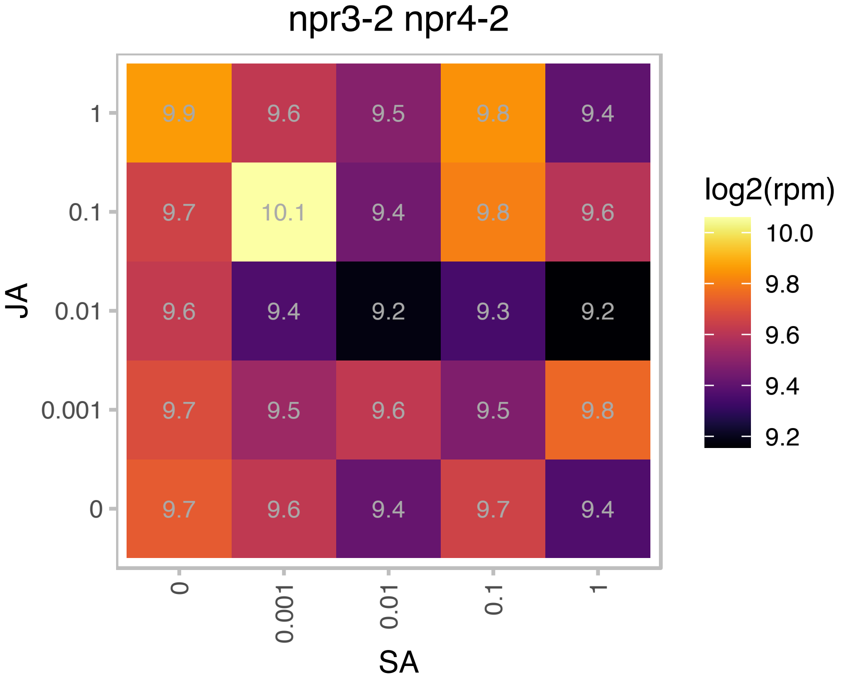


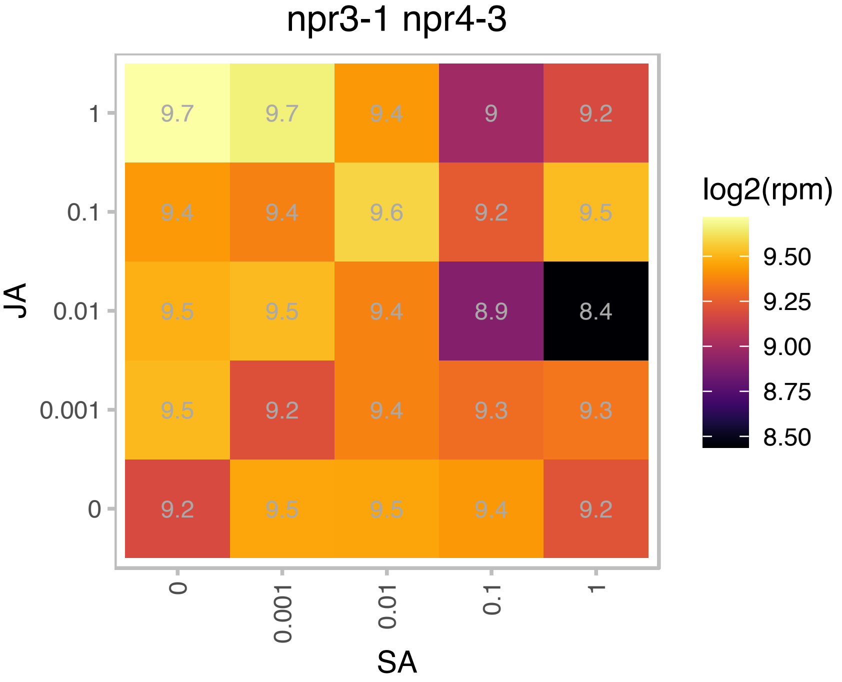

### AT1G60600.1 : ABC4

UbiA prenyltransferase family protein
Encodes a protein similar to 1,4-dihydroxy-2-naphthoic acid phytyltransferase involved in phylloquinone and plastoquinone biosynthesis. Mutants are pale green and heterotrophic with defects in photosynthetic electron transport.

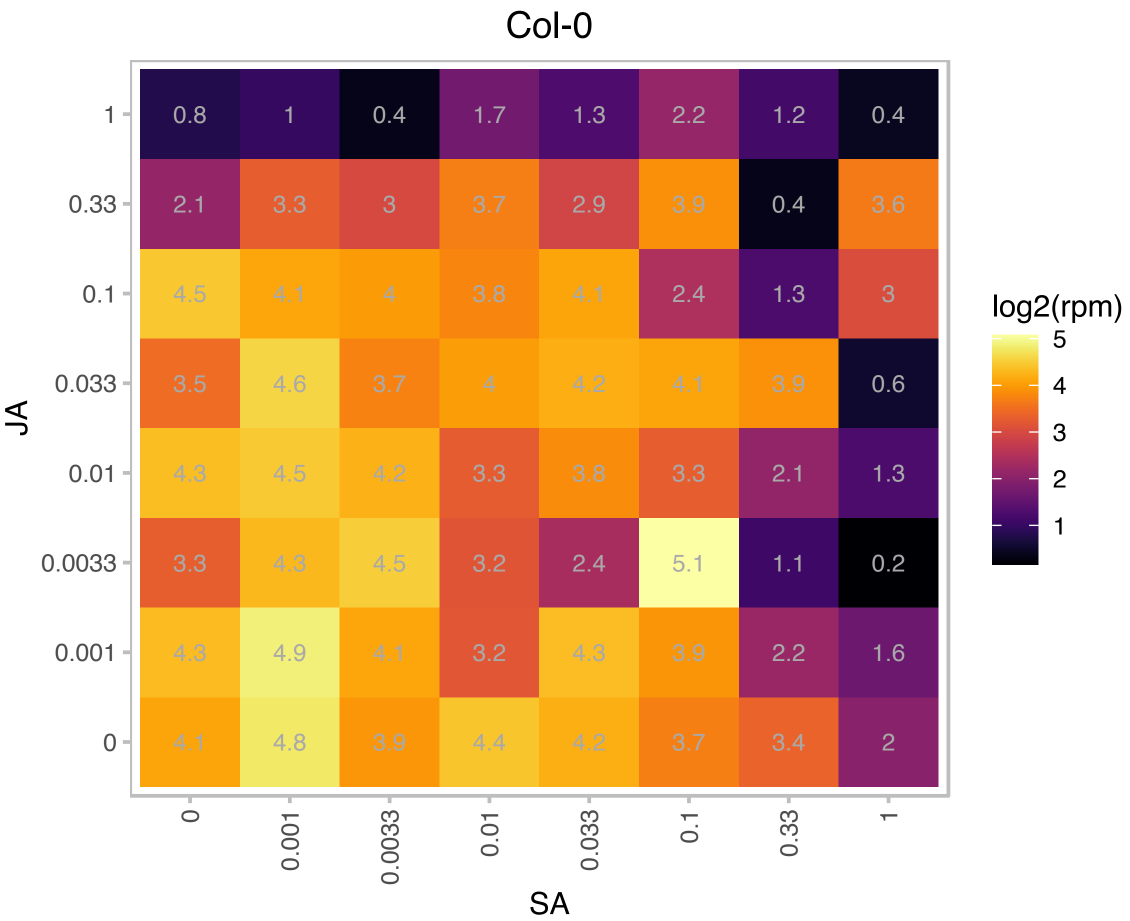


| | |
|---|---|
| Gene | AT1G60600 |
| is.expressed_genes | TRUE |
| is.responsive_genes | TRUE |
| is.apcluster | F-10 |

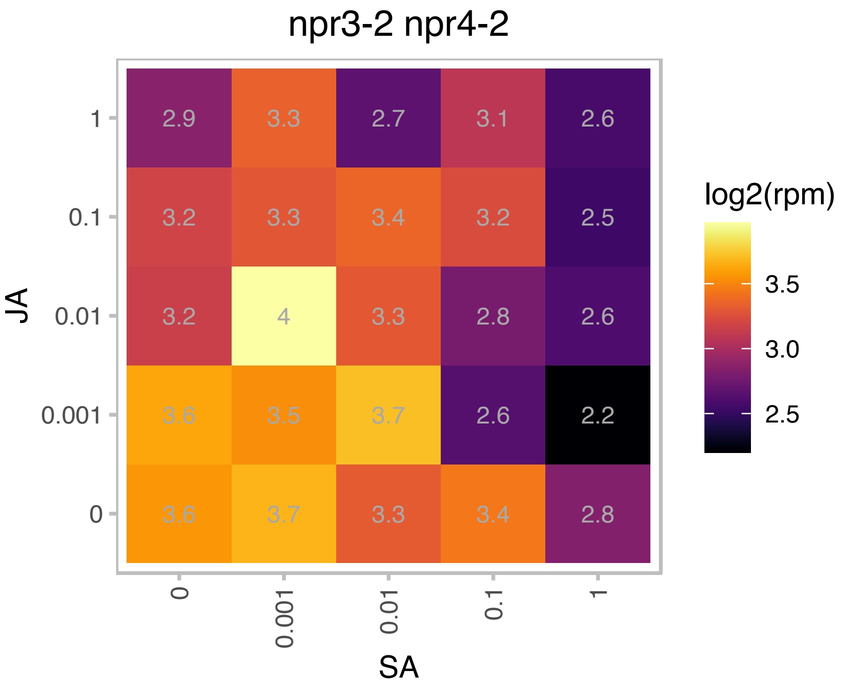


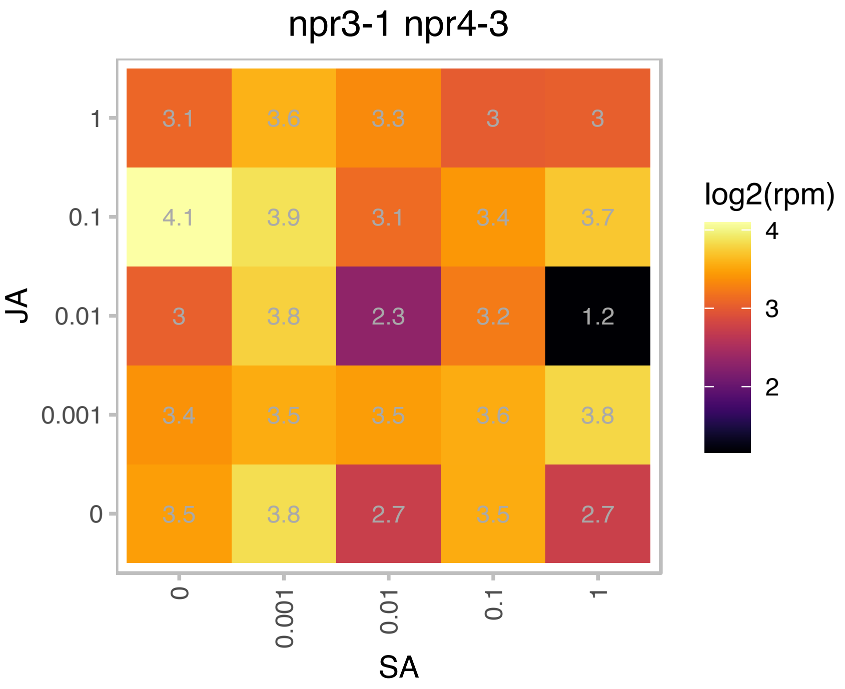

# AT1G61450.1 : NA

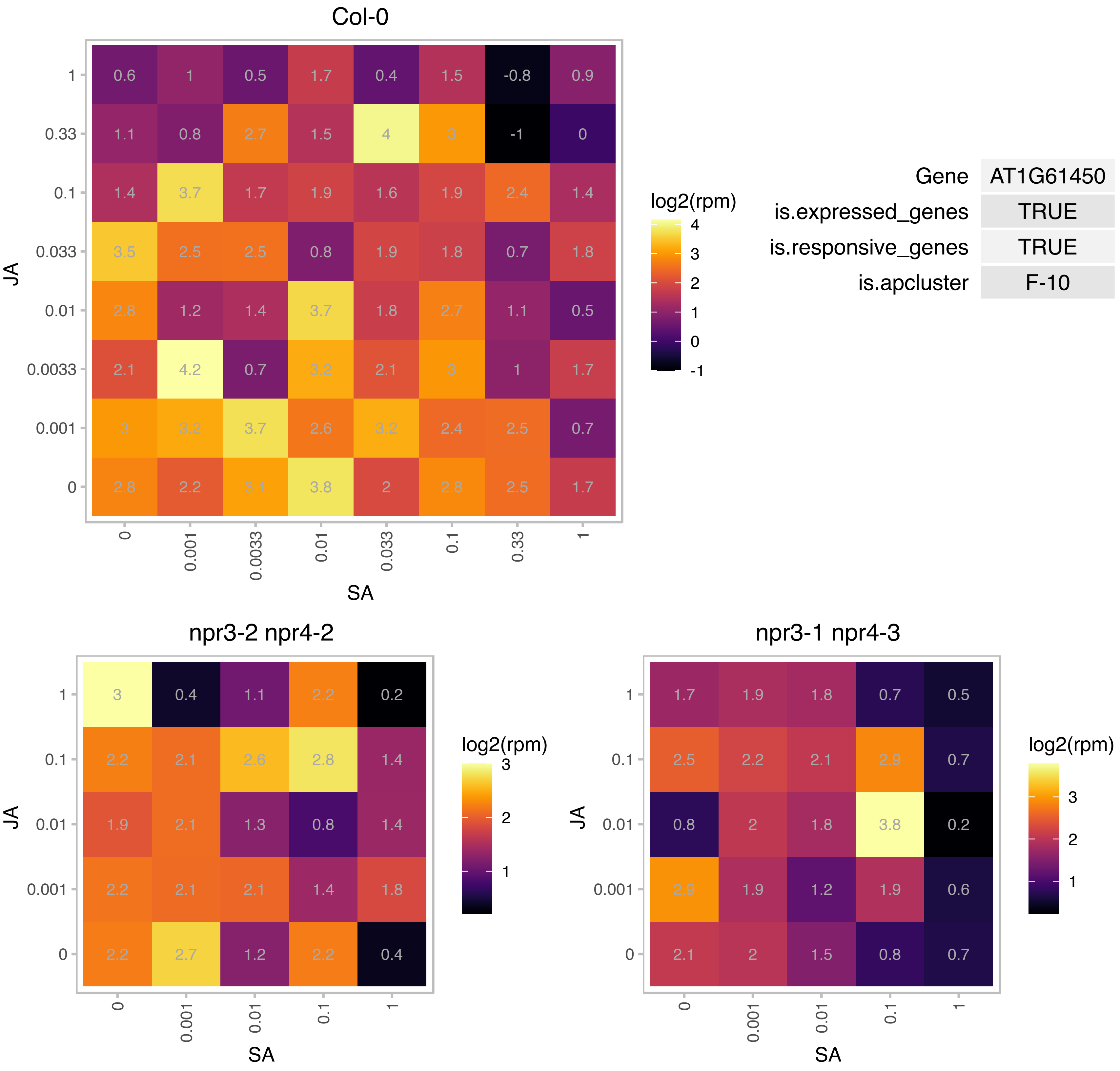

# AT1G64355.1 : NA

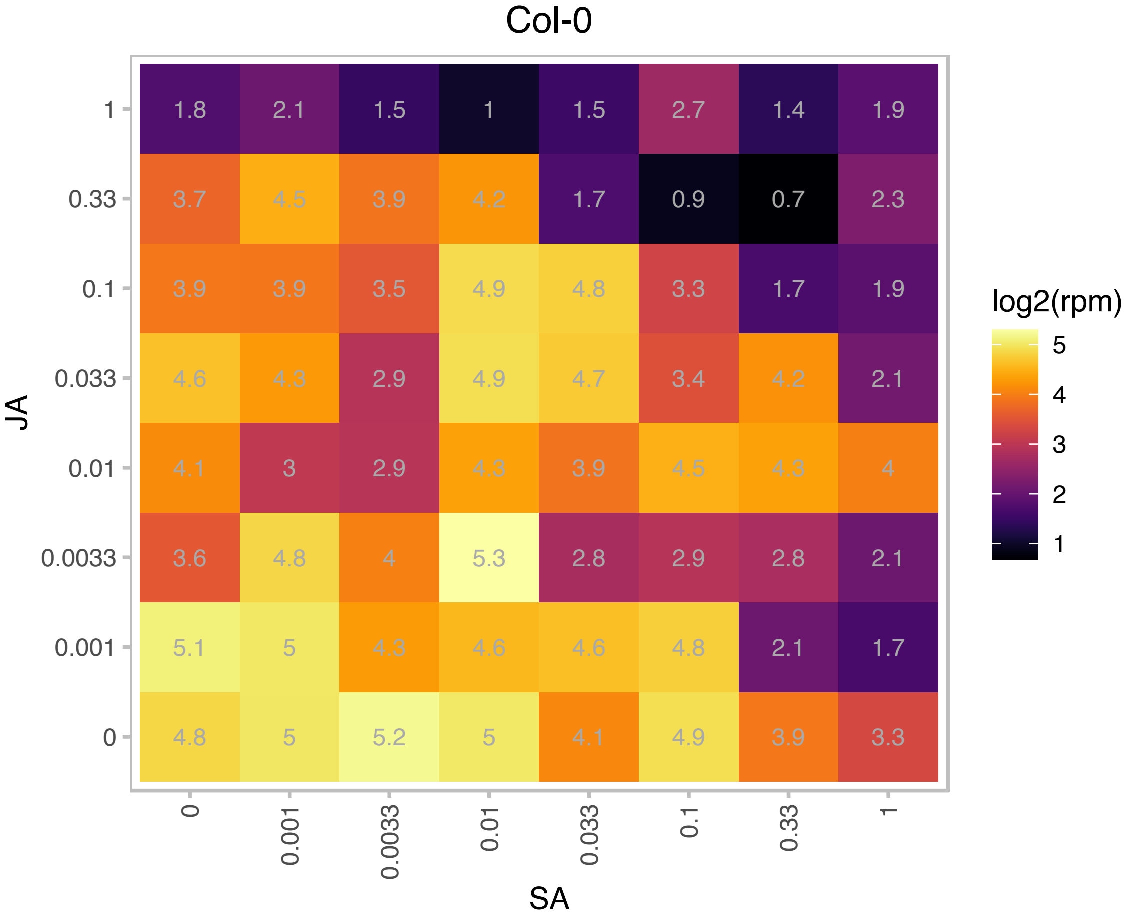


| | |
|---|---|
| Gene | AT1G64355 |
| is.expressed_genes | TRUE |
| is.responsive_genes | TRUE |
| is.apcluster | F-10 |

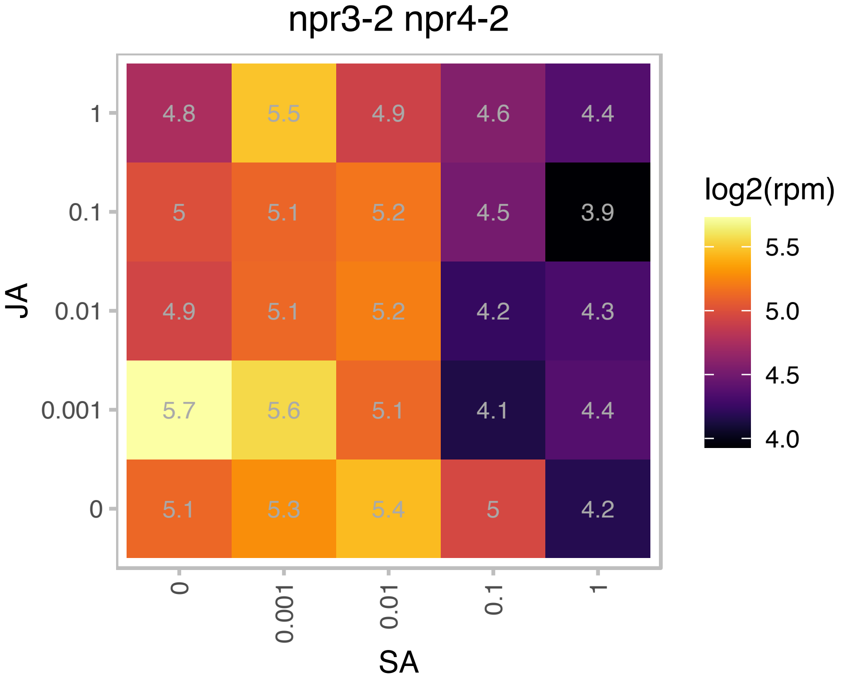


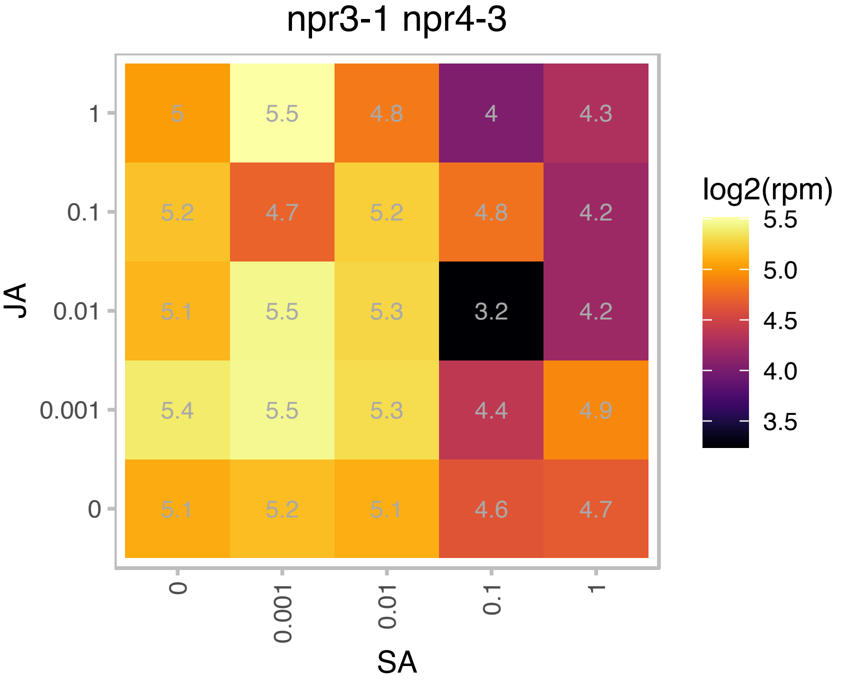

# AT1G68238.1 : NA

Col-0

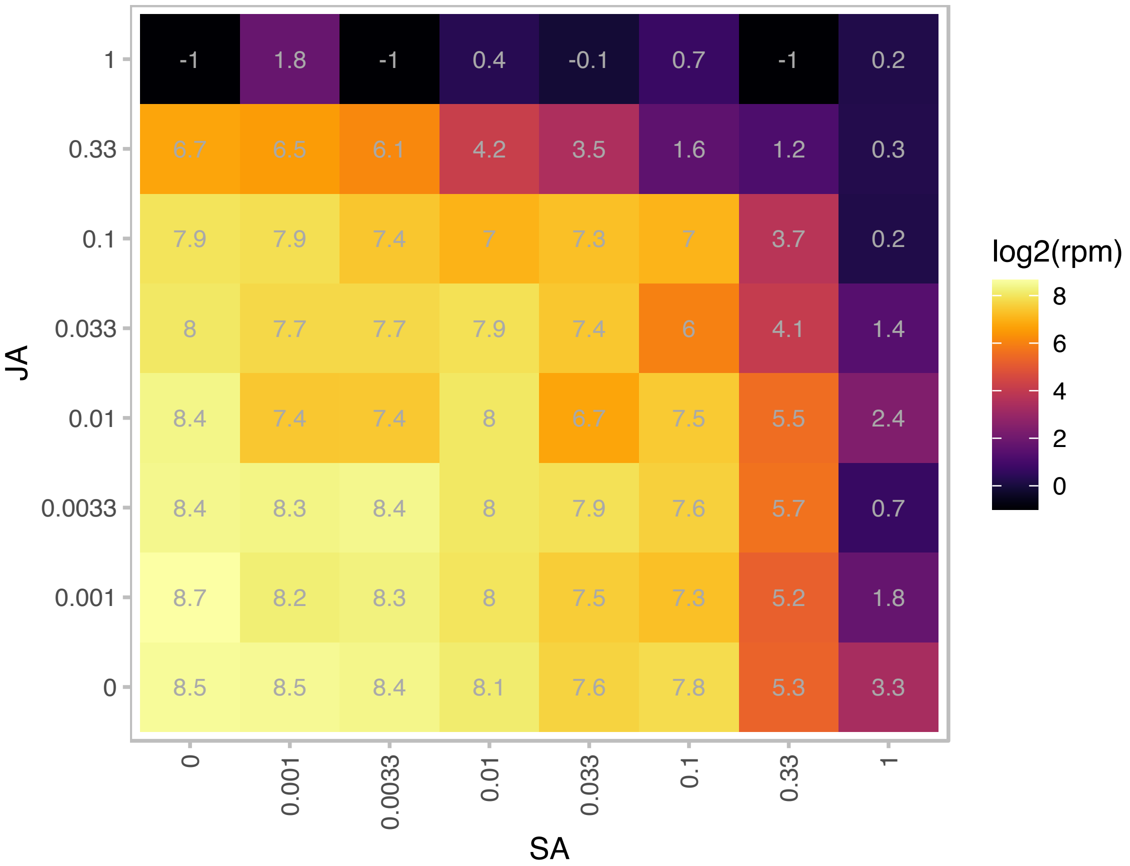


| | |
|---|---|
| Gene | AT1G68238 |
| is.expressed_genes | TRUE |
| is.responsive_genes | TRUE |
| is.apcluster | F-10 |

npr3-2 npr4-2

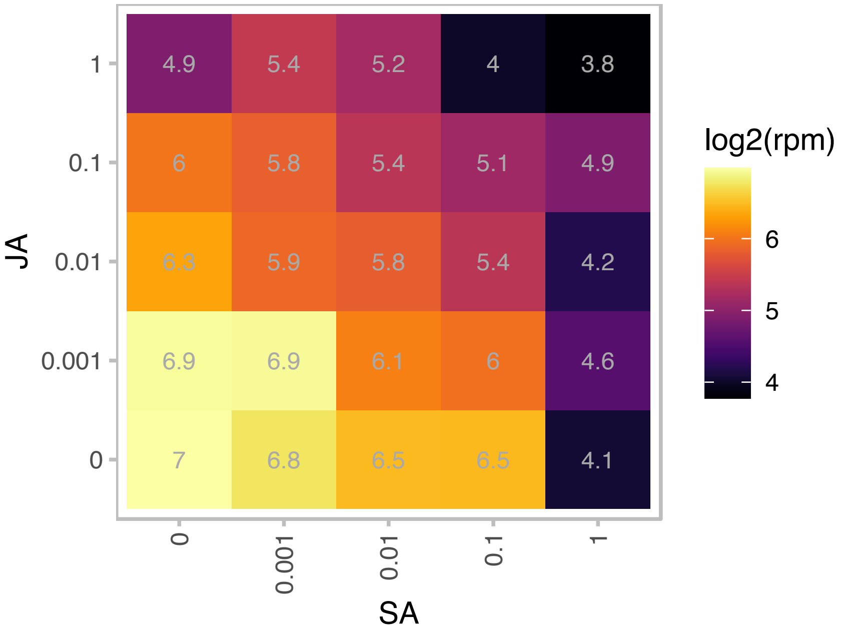


npr3-1 npr4-3

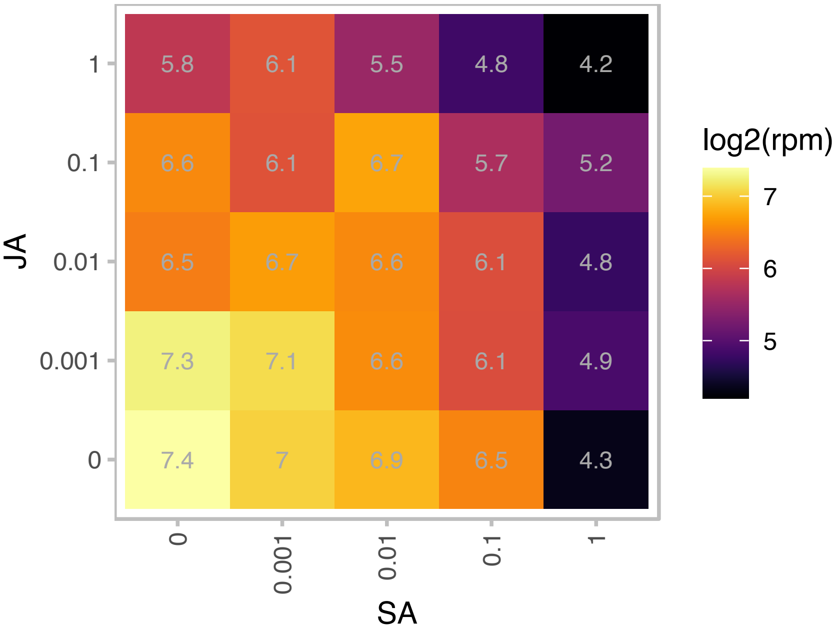

# AT1G75190.1 : NA

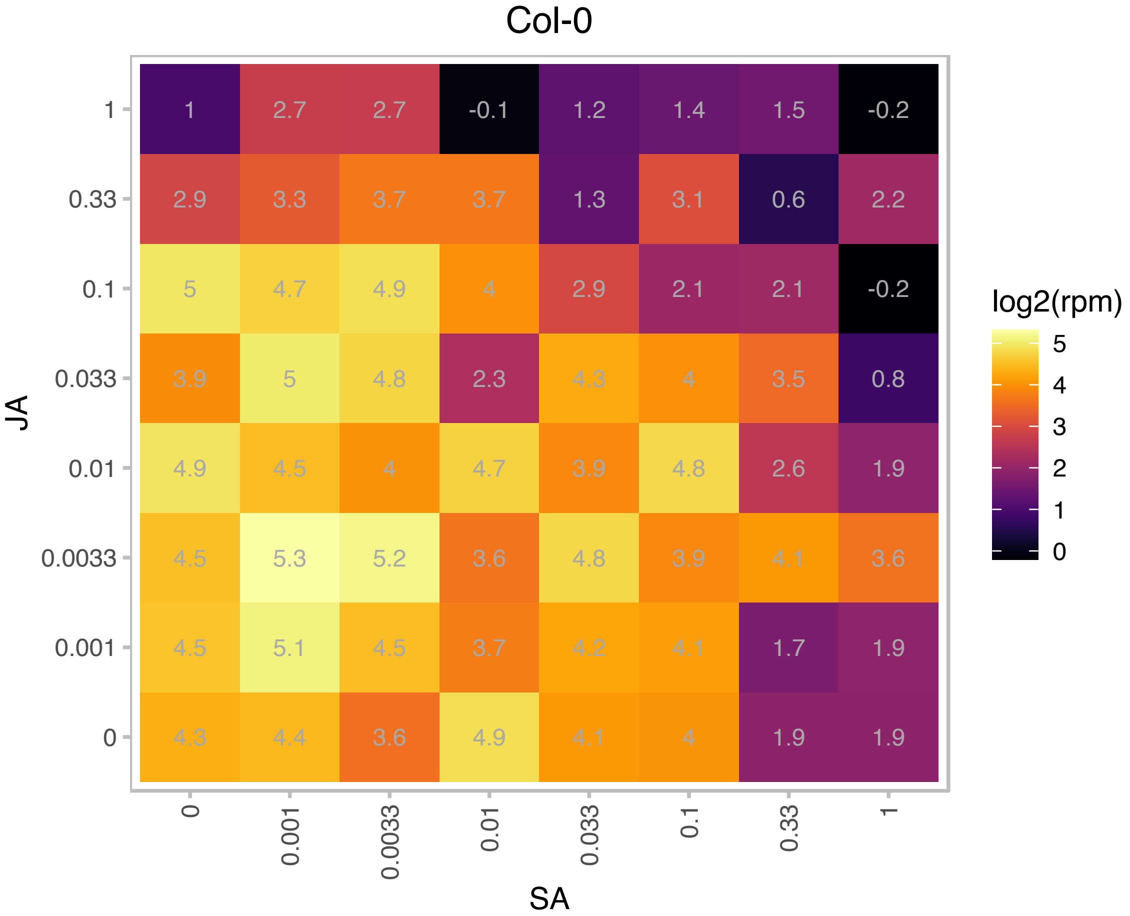


| | |
|---|---|
| Gene | AT1G75190 |
| is.expressed_genes | TRUE |
| is.responsive_genes | TRUE |
| is.apcluster | F-10 |

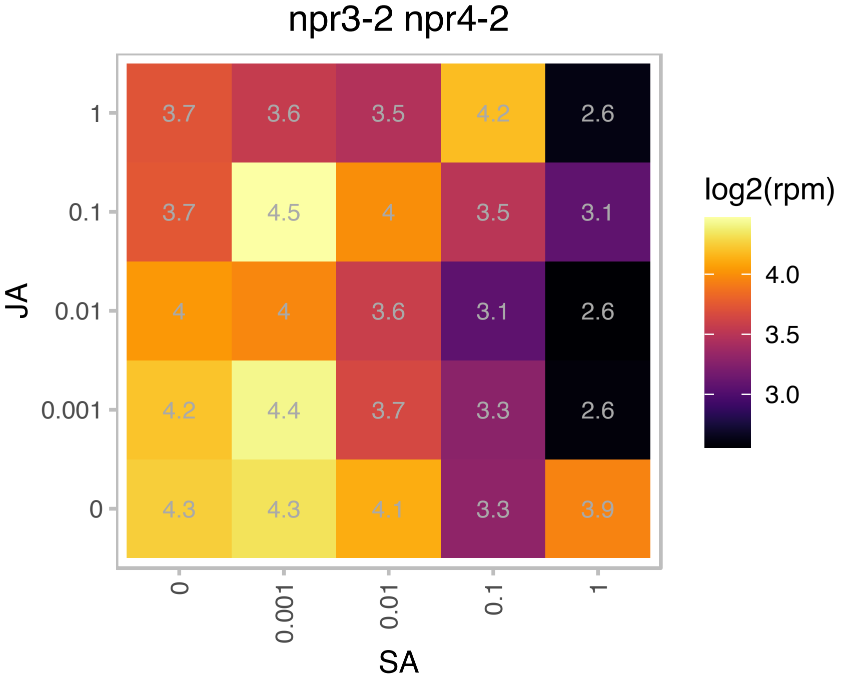


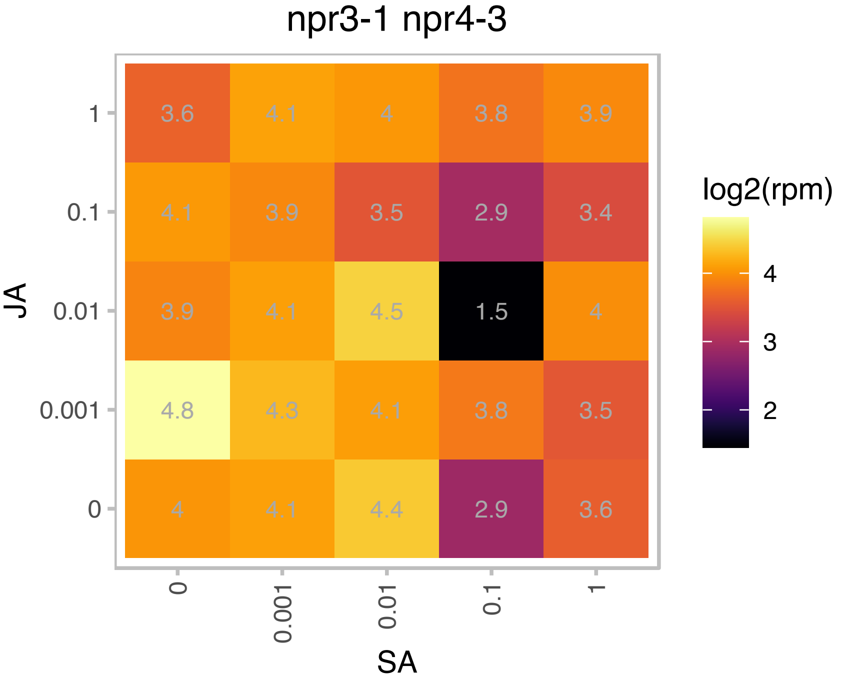

# AT1G78995.1 : NA

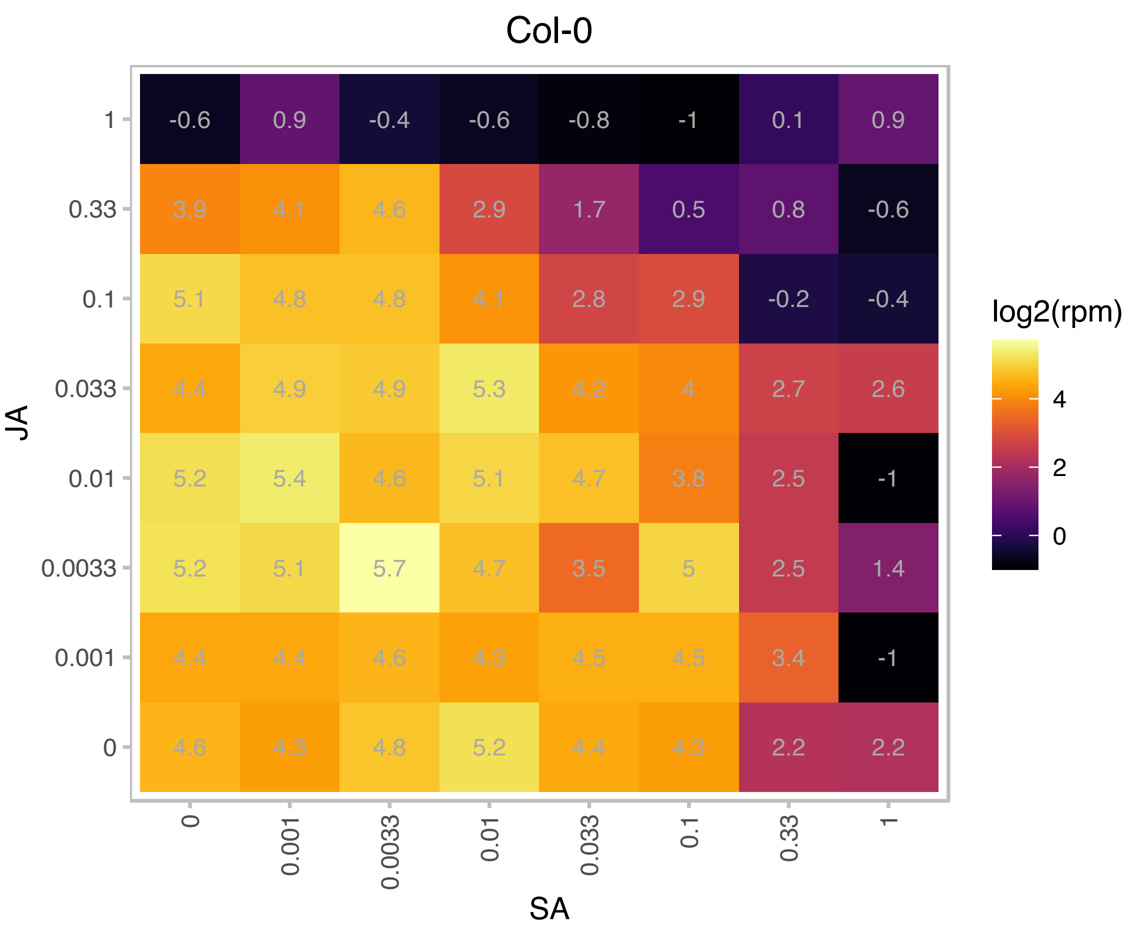


| | |
|---|---|
| Gene | AT1G78995 |
| is.expressed_genes | TRUE |
| is.responsive_genes | TRUE |
| is.apcluster | F-10 |

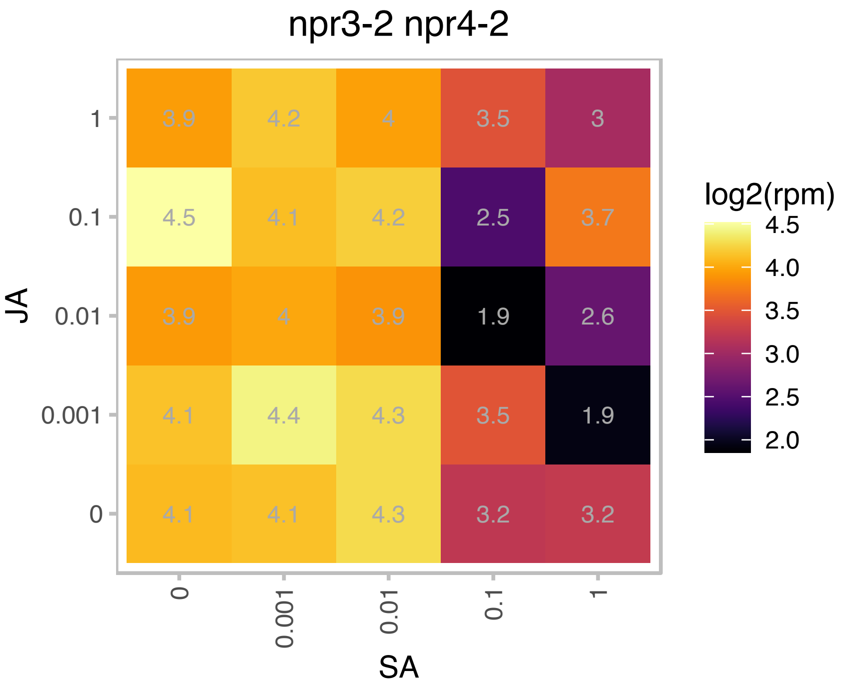


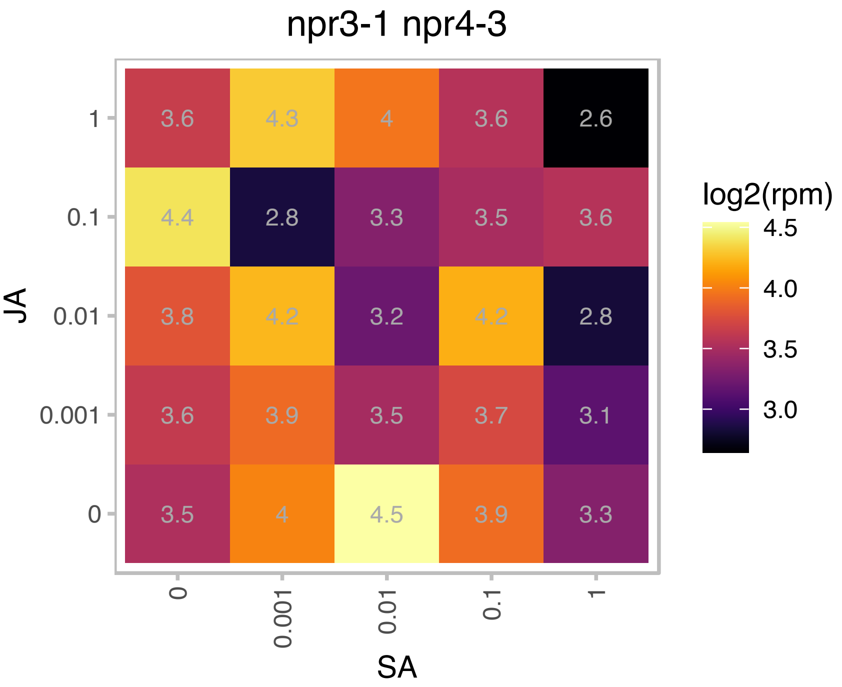

# AT2G01755.1 : NA

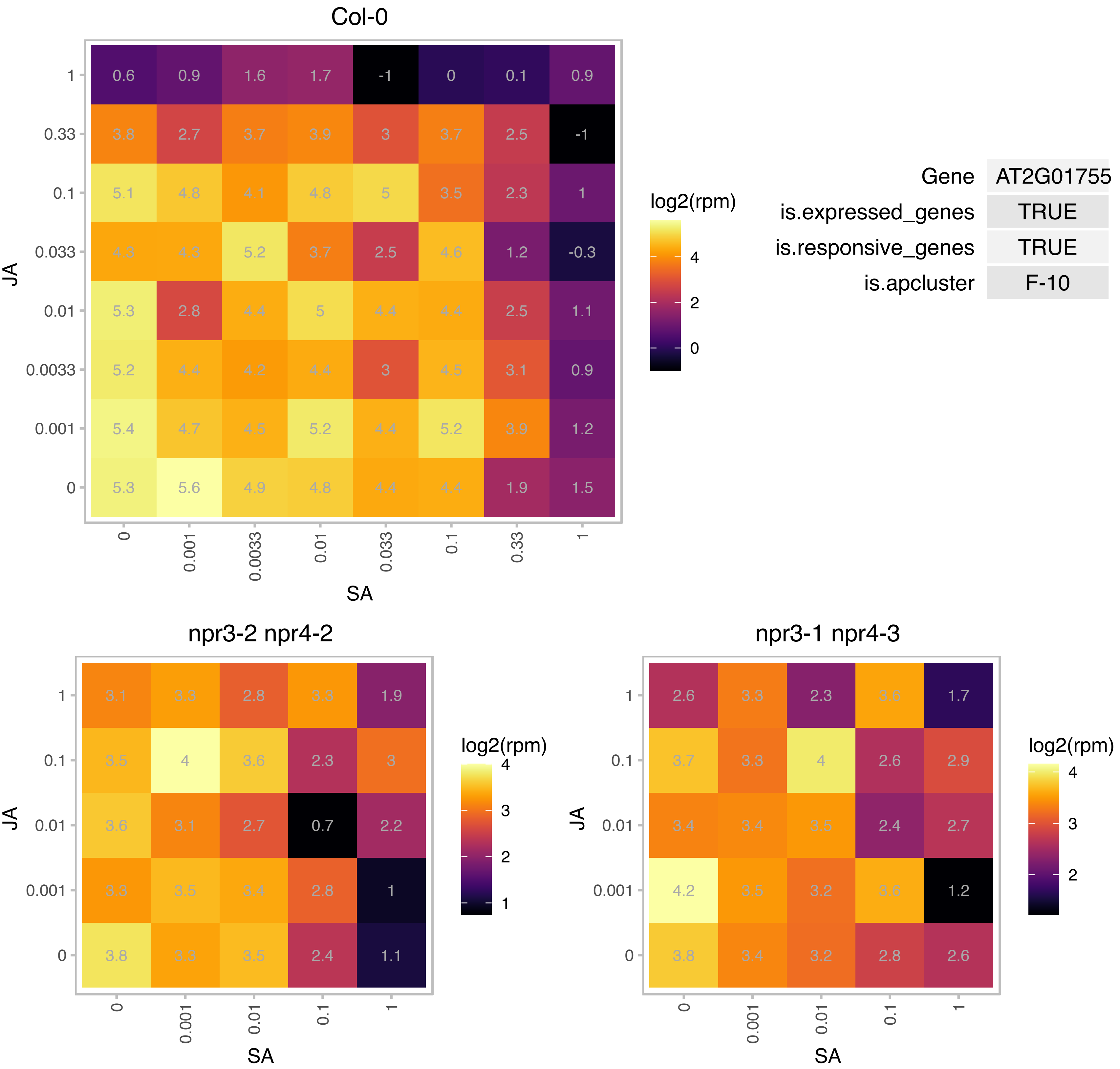

### AT2G06520.1 : PSBX

photosystem II subunit X

Encodes a protein with sequence similarity to the spinach photosystem II subunit PsbX.

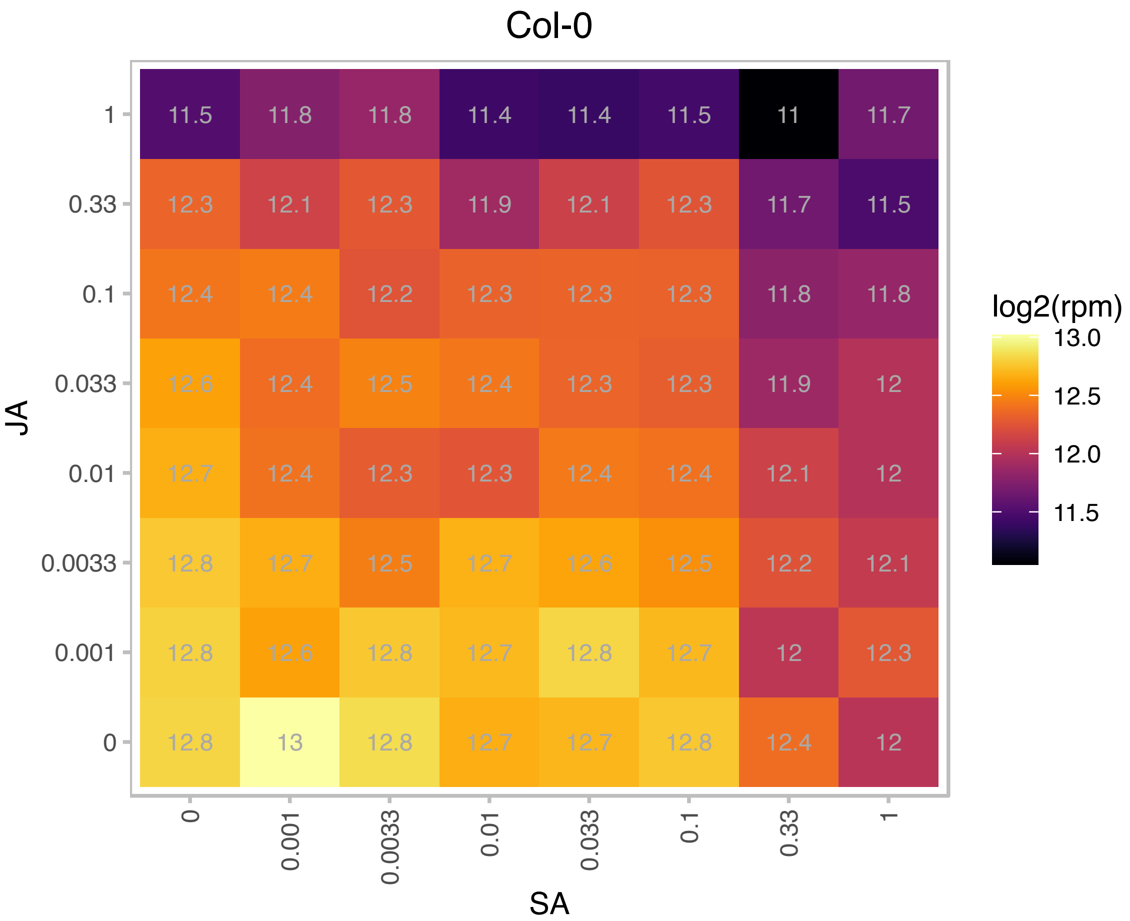


| | |
|---|---|
| Gene | AT2G06520 |
| is.expressed_genes | TRUE |
| is.responsive_genes | TRUE |
| is.apcluster | F-10 |

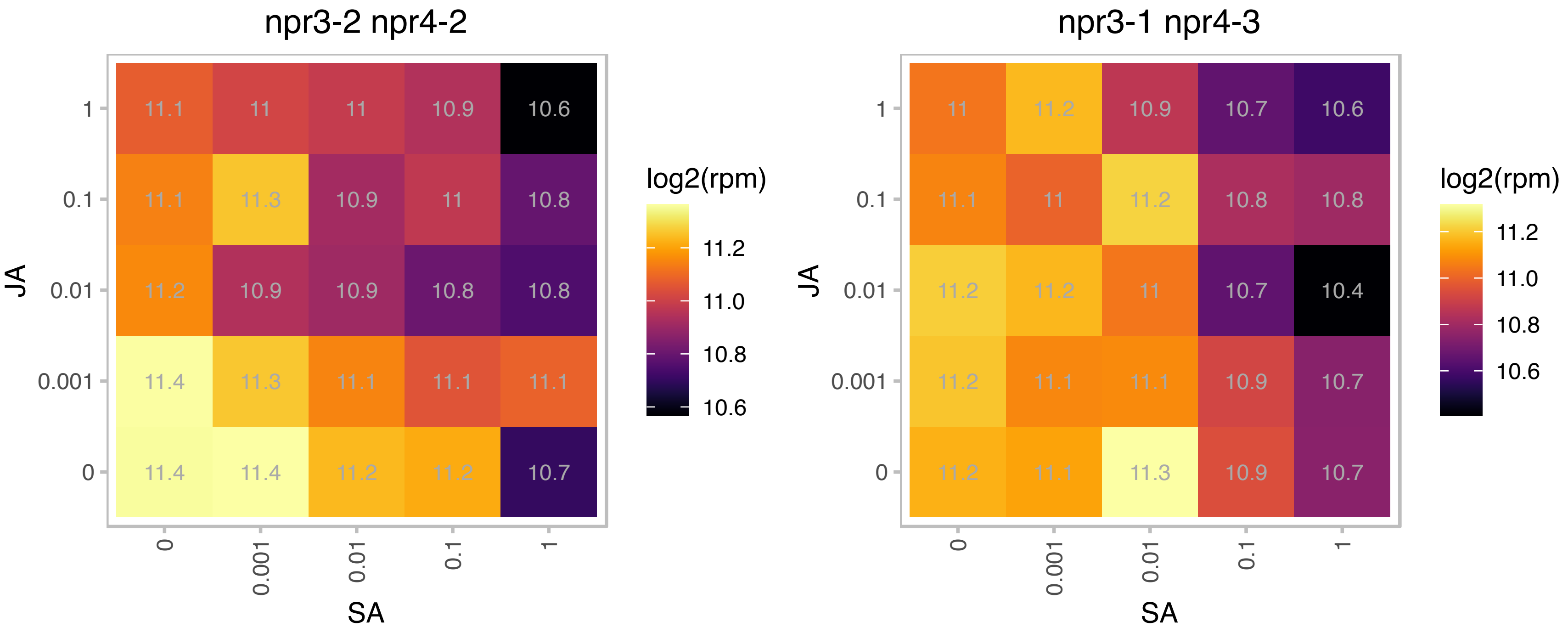

### AT2G15050.1 : LTP, LTP7

lipid transfer protein

Predicted to encode a PR (pathogenesis-related) protein. Belongs to the lipid transfer protein (PR-14) family with the following members: At2g38540/LTP1, At2g38530/LTP2, At5g59320/LTP3, At5g59310/LTP4, At3g51600/LTP5, At3g08770/LTP6, At2g15050/LTP7, At2g18370/LTP8, At2g15325/LTP9, At5g01870/LTP10, At4g33355/LTP11, At3g51590/LTP12, At5g44265/LTP13, At5g62065/LTP14, At4g08530/LTP15.

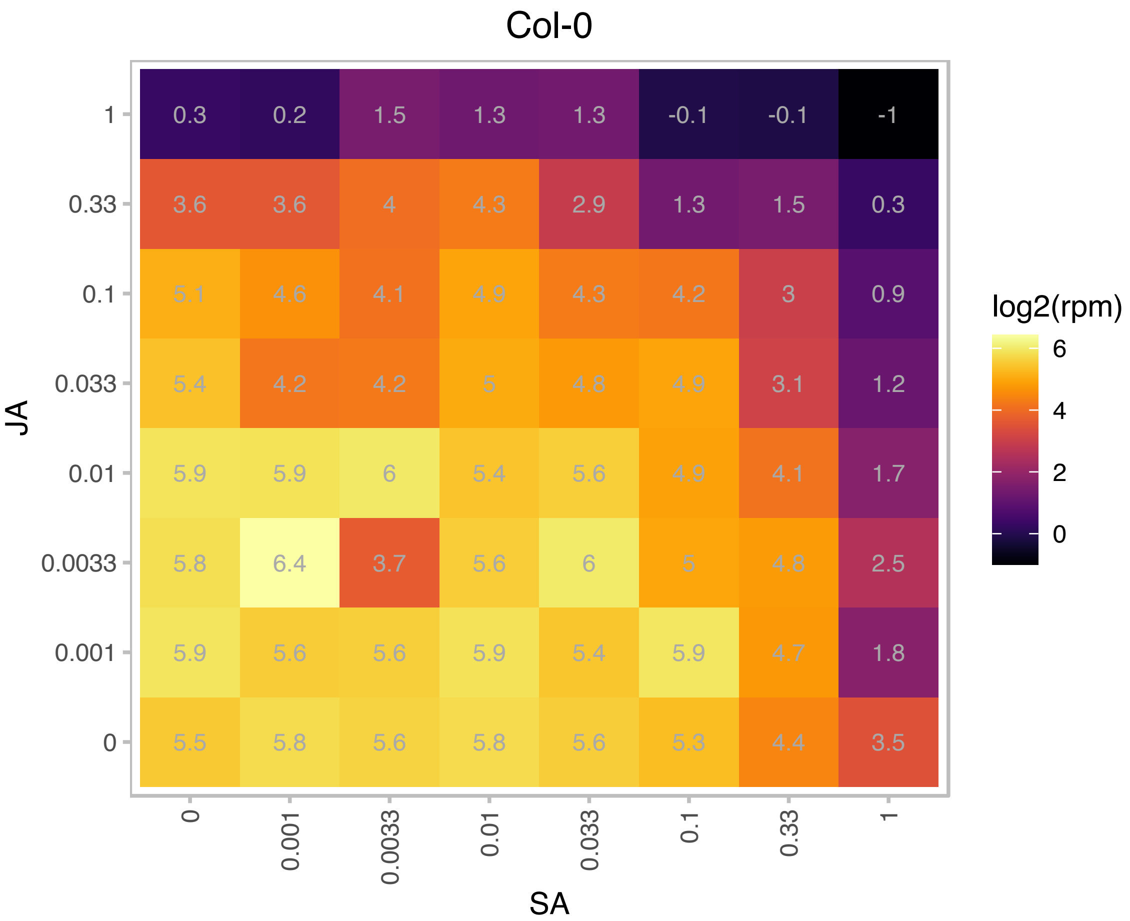


| | |
|---|---|
| Gene | AT2G15050 |
| is.expressed_genes | TRUE |
| is.responsive_genes | TRUE |
| is.apcluster | F-10 |

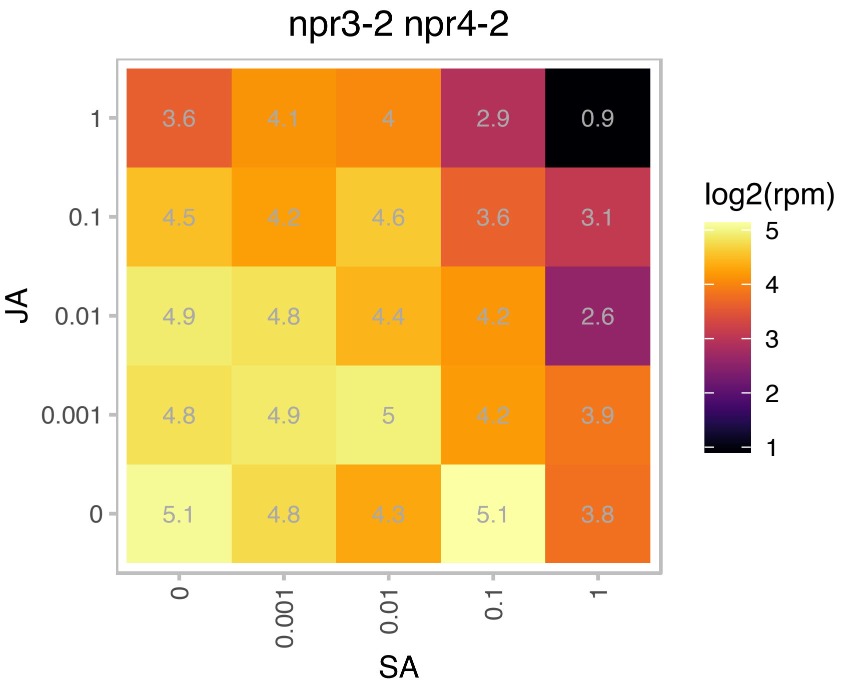


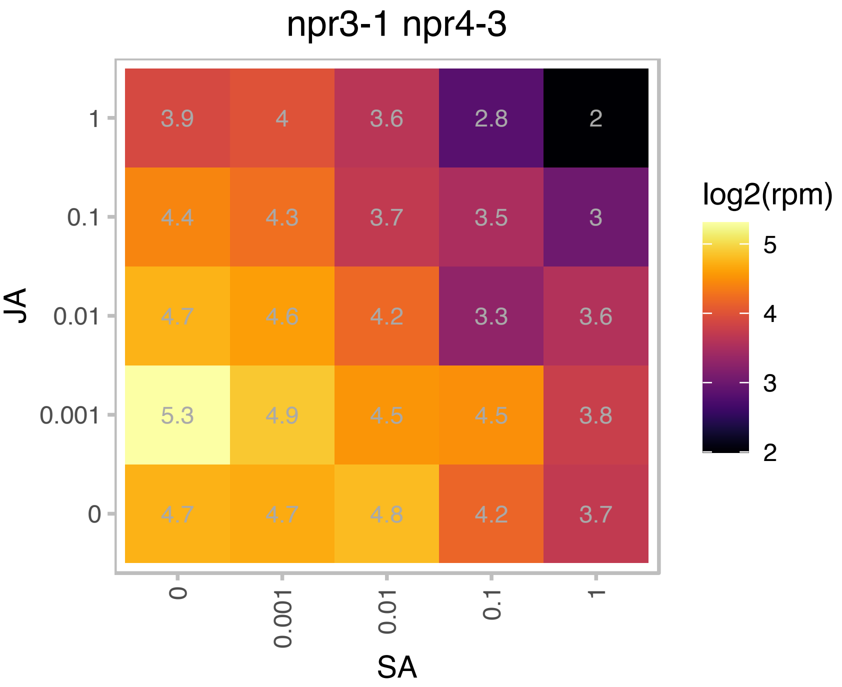

### AT2G15620.1 : ATHNIR, NIR, NIR1

nitrite reductase 1

Involved in the second step of nitrate assimilation.  Its expression is induced by nitrate. The mRNA is cell-to-cell mobile.

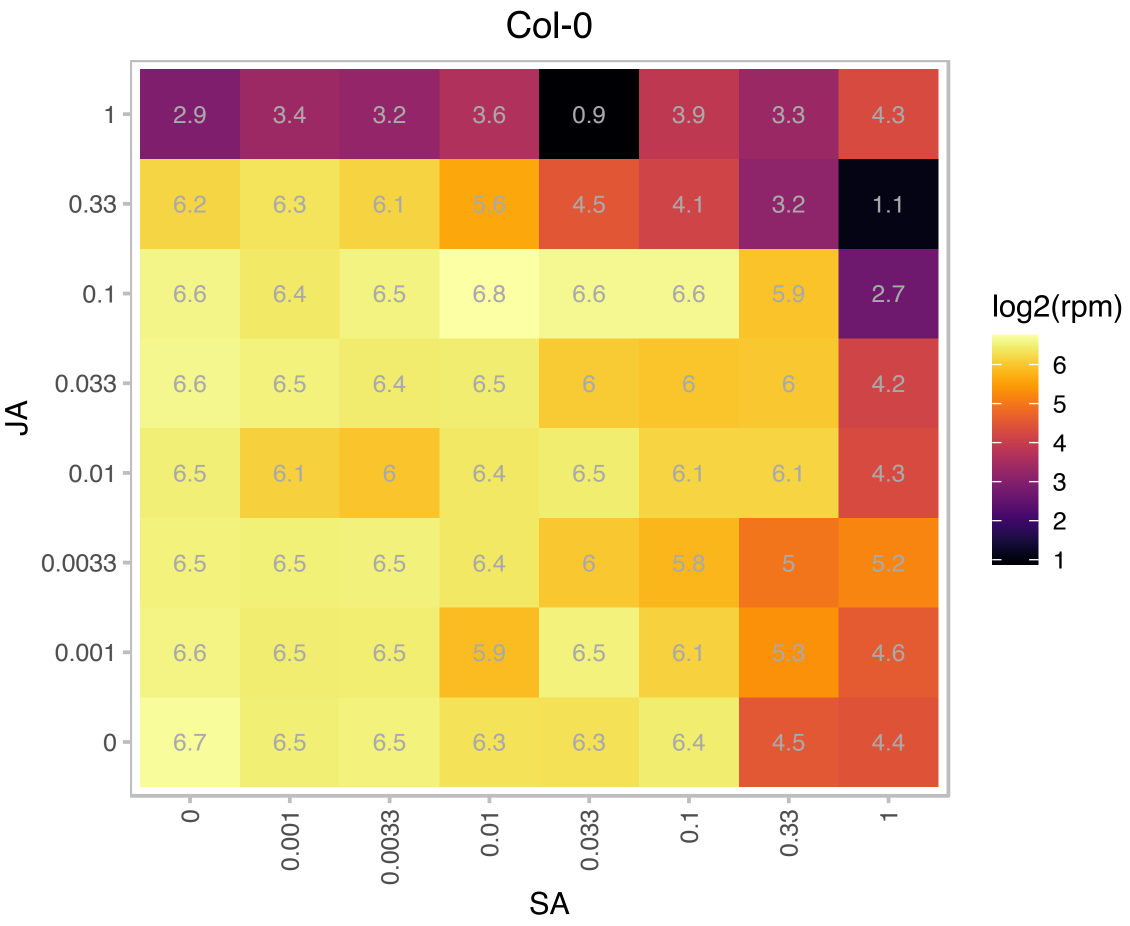


| | |
|---|---|
| Gene | AT2G15620 |
| is.expressed_genes | TRUE |
| is.responsive_genes | TRUE |
| is.apcluster | F-10 |

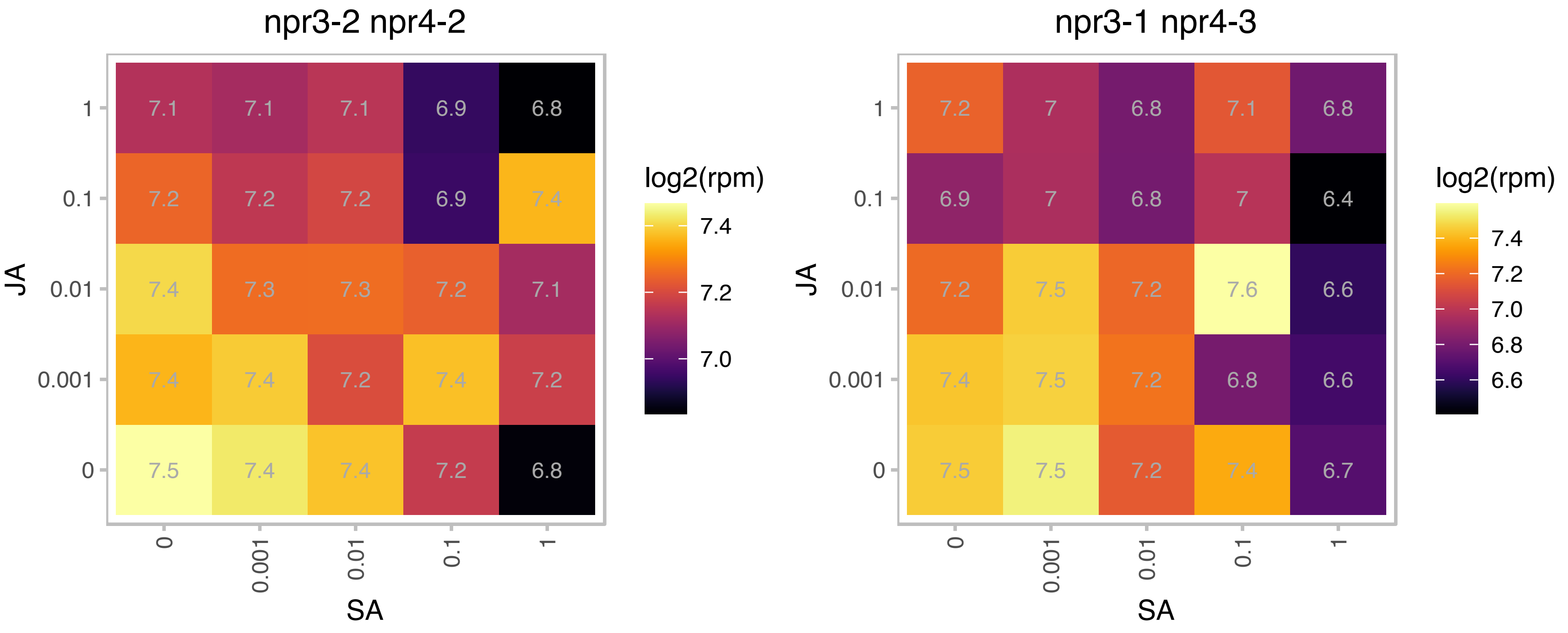

# AT2G17695.1 : NA

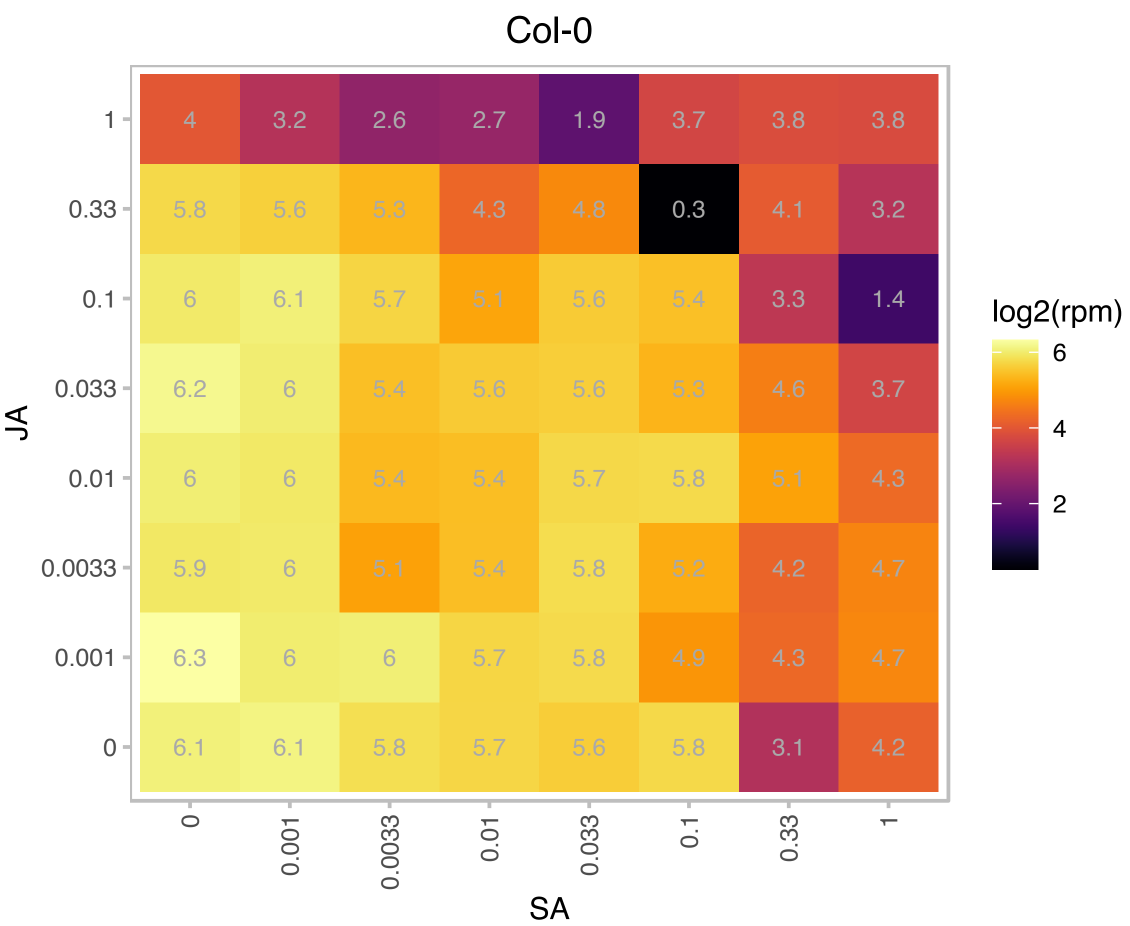


| | |
|---|---|
| Gene | AT2G17695 |
| is.expressed_genes | TRUE |
| is.responsive_genes | TRUE |
| is.apcluster | F-10 |

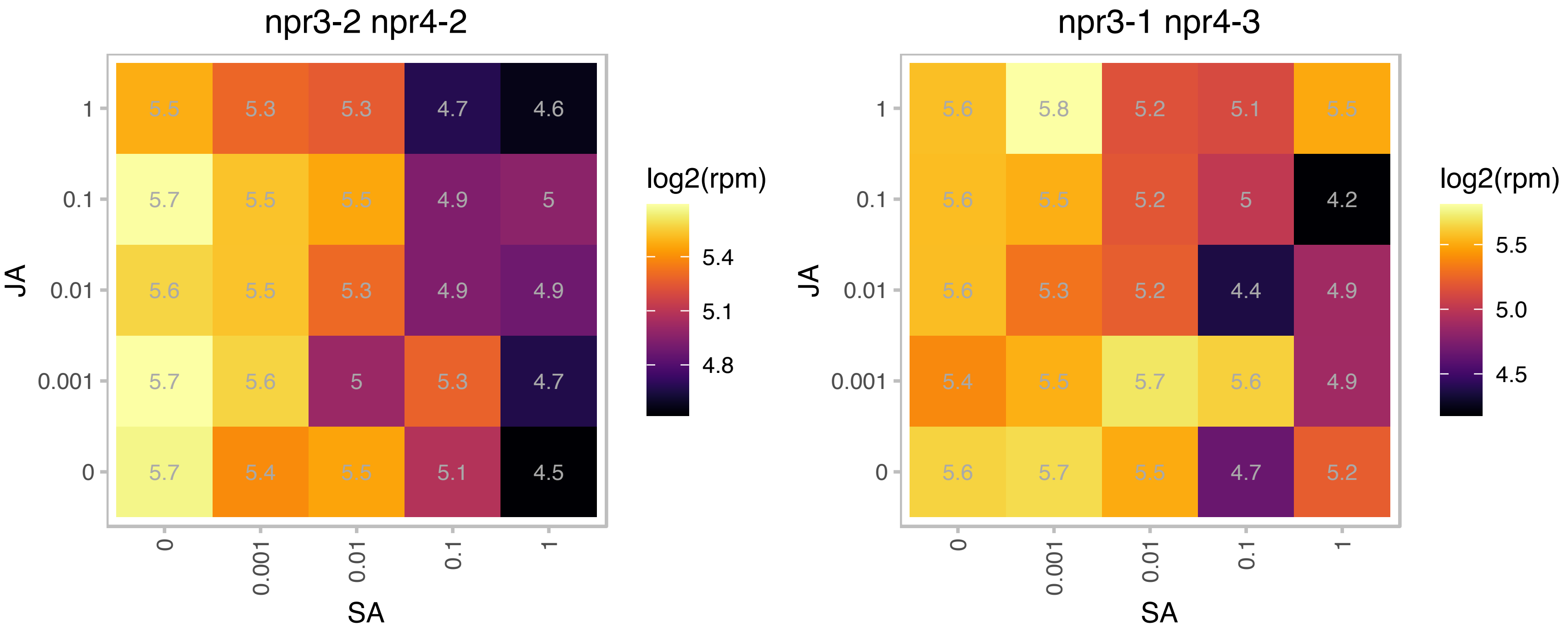

### AT2G20490.1 : EDA27, NOP10

nucleolar RNA-binding Nop10p family protein

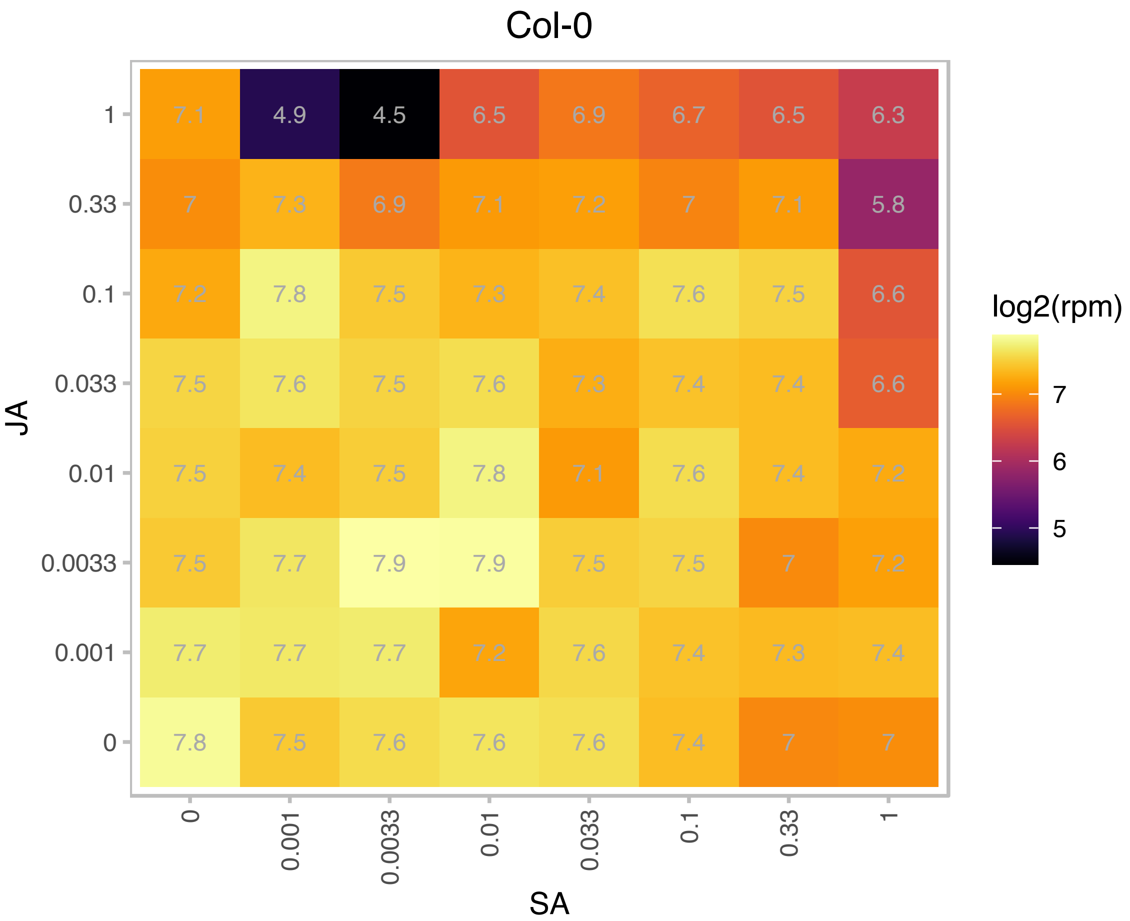


| | |
|---|---|
| Gene | AT2G20490 |
| is.expressed_genes | TRUE |
| is.responsive_genes | TRUE |
| is.apcluster | F-10 |

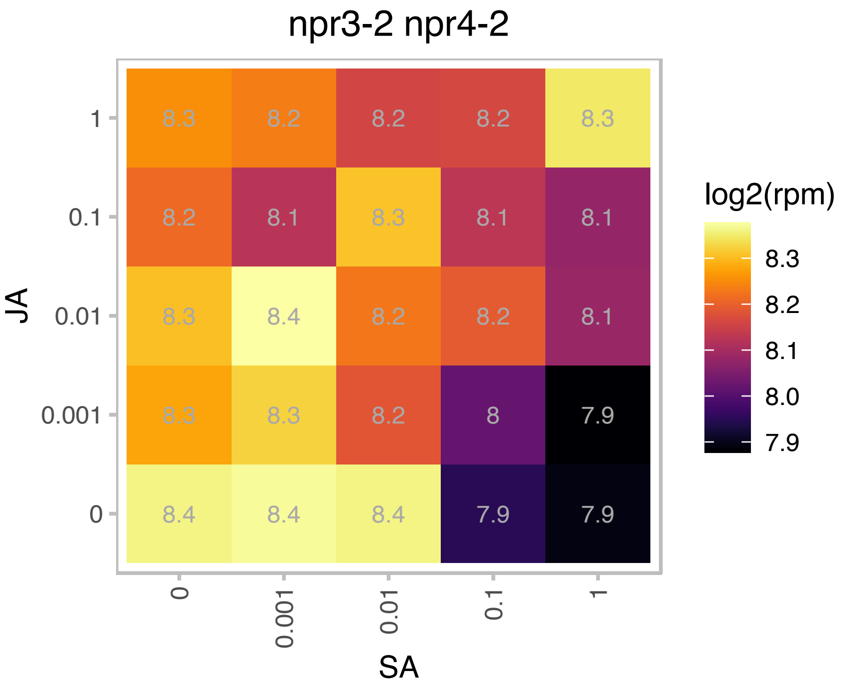


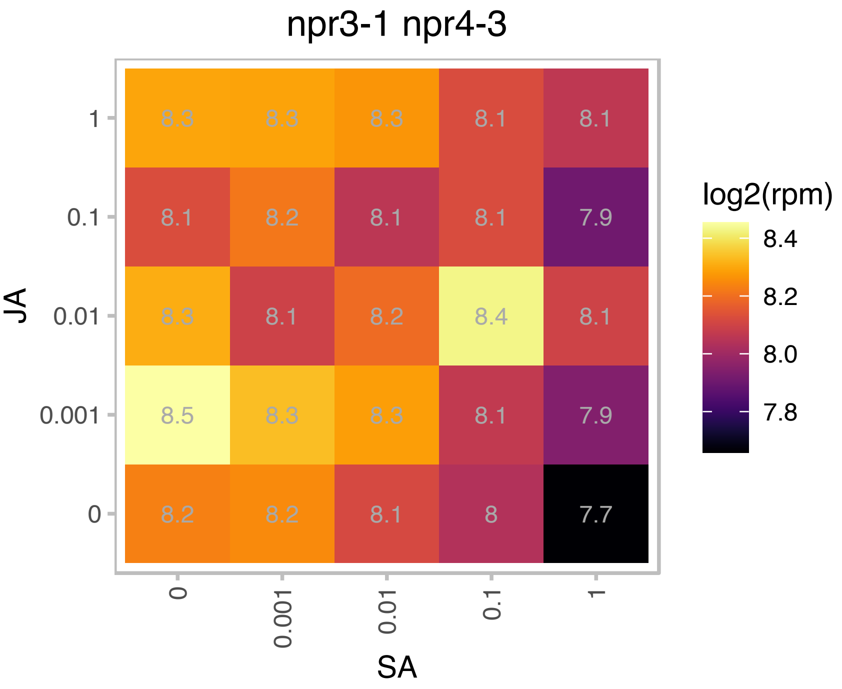

# AT2G22122.1 : NA

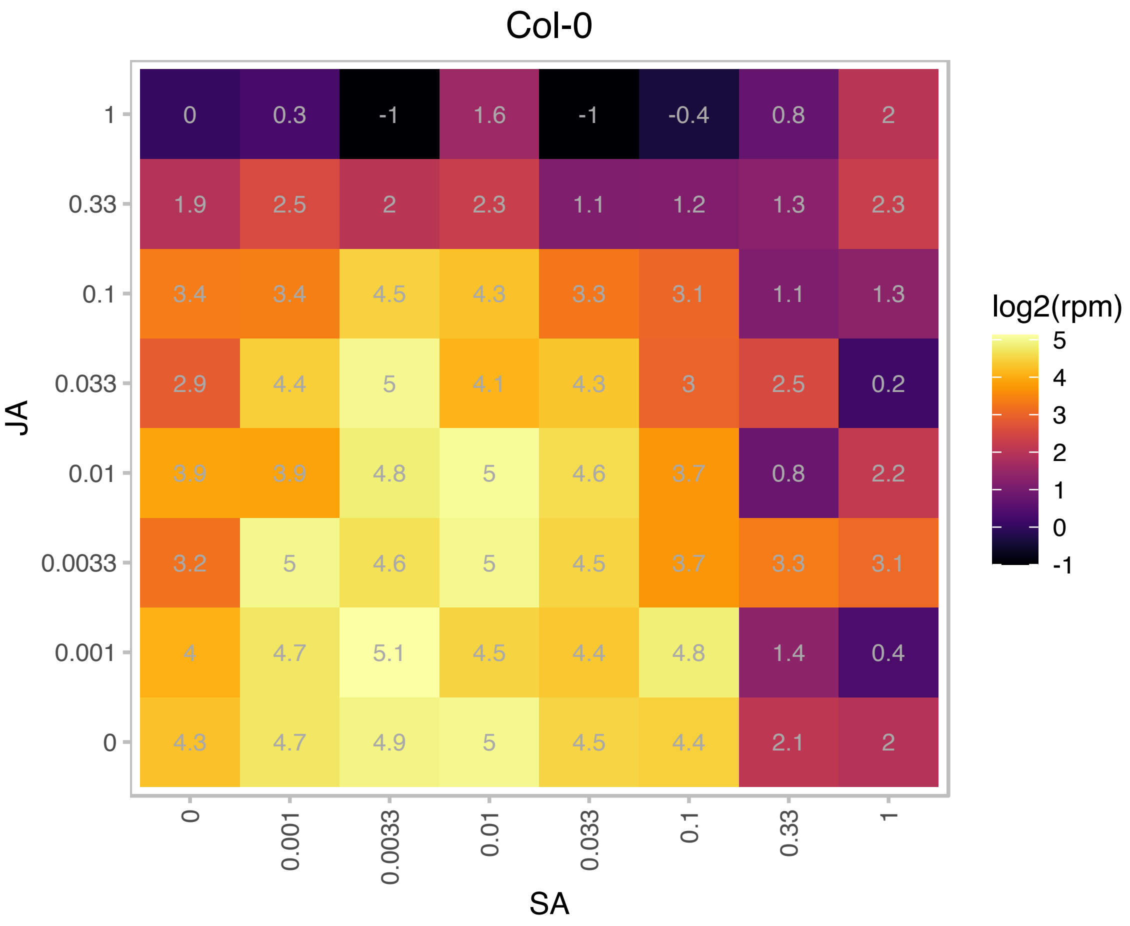


| | |
|---|---|
| Gene | AT2G22122 |
| is.expressed_genes | TRUE |
| is.responsive_genes | TRUE |
| is.apcluster | F-10 |

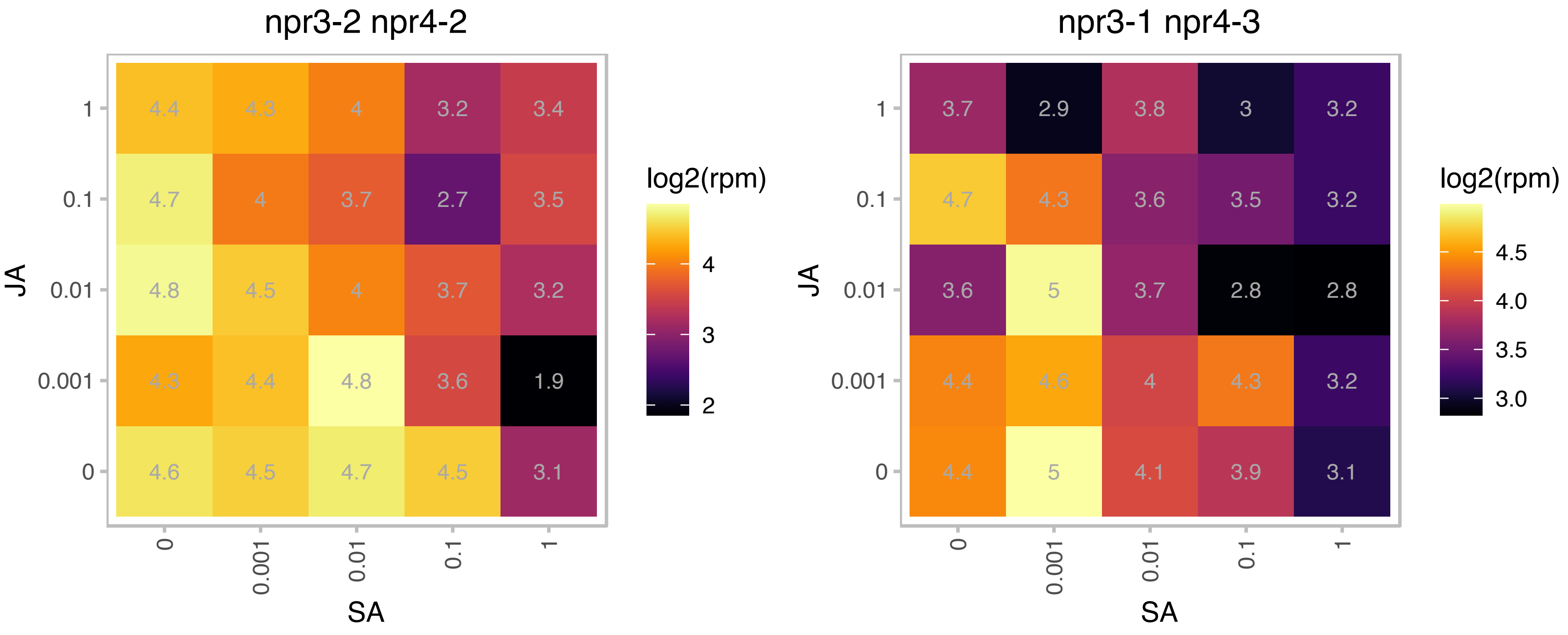

### AT2G27400.1 : TAS1A

TAS1A; other RNA

Trans-acting siRNA1a primary transcript (TAS1a). Regulated by miR173.

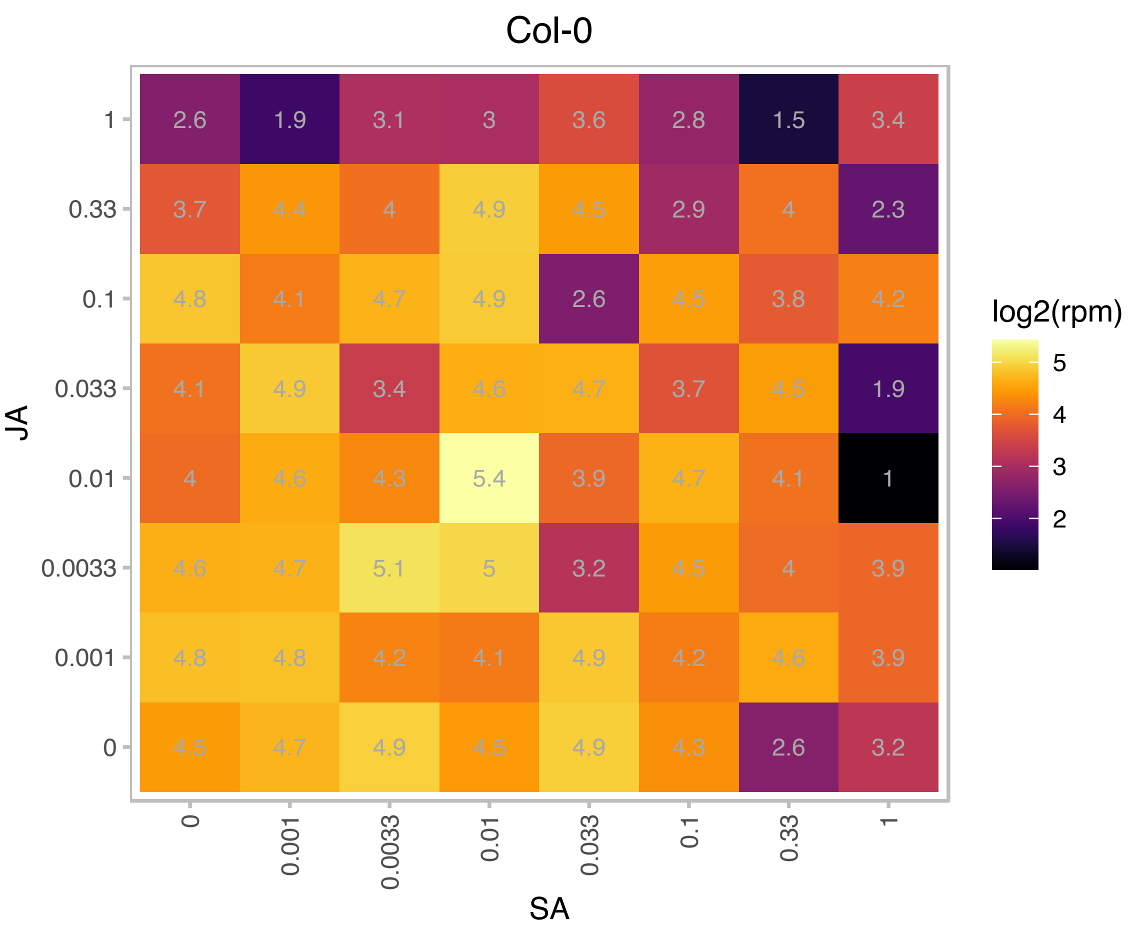


| | |
|---|---|
| Gene | AT2G27400 |
| is.expressed_genes | TRUE |
| is.responsive_genes | TRUE |
| is.apcluster | F-10 |

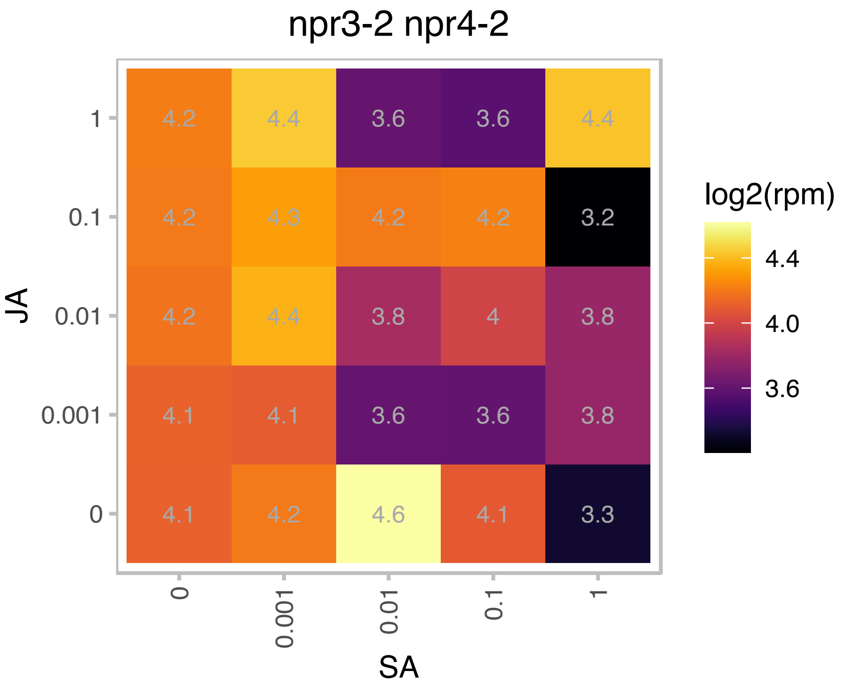


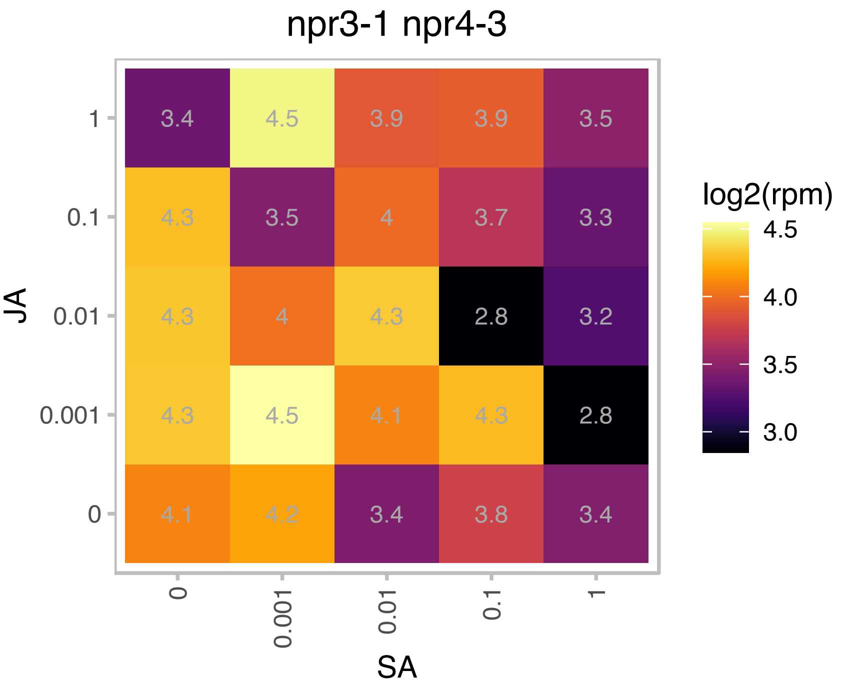

# AT2G28720.1 : NA

Histone superfamily protein

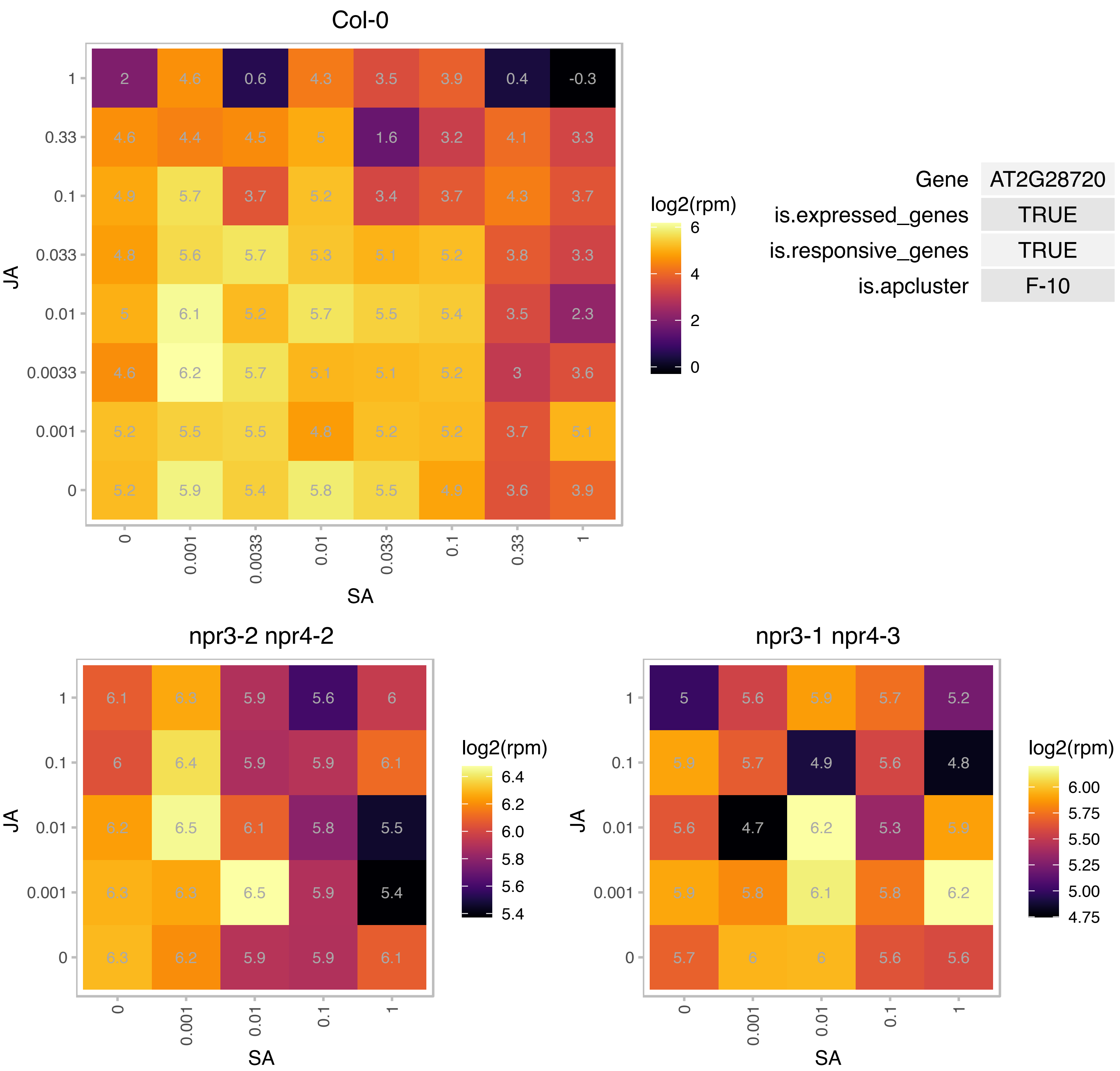


| | |
|---|---|
| Gene | AT2G28720 |
| is.expressed_genes | TRUE |
| is.responsive_genes | TRUE |
| is.apcluster | F-10 |

### AT2G28950.1 : ATEXP6, ATEXPA6, ATHEXP ALPHA 1.8, E

expansin A6

Encodes an expansin. Naming convention from the Expansin Working Group (Kende et al, 2004. Plant Mol Bio). Involved in the formation of nematode-induced syncytia in roots of Arabidopsis thaliana.

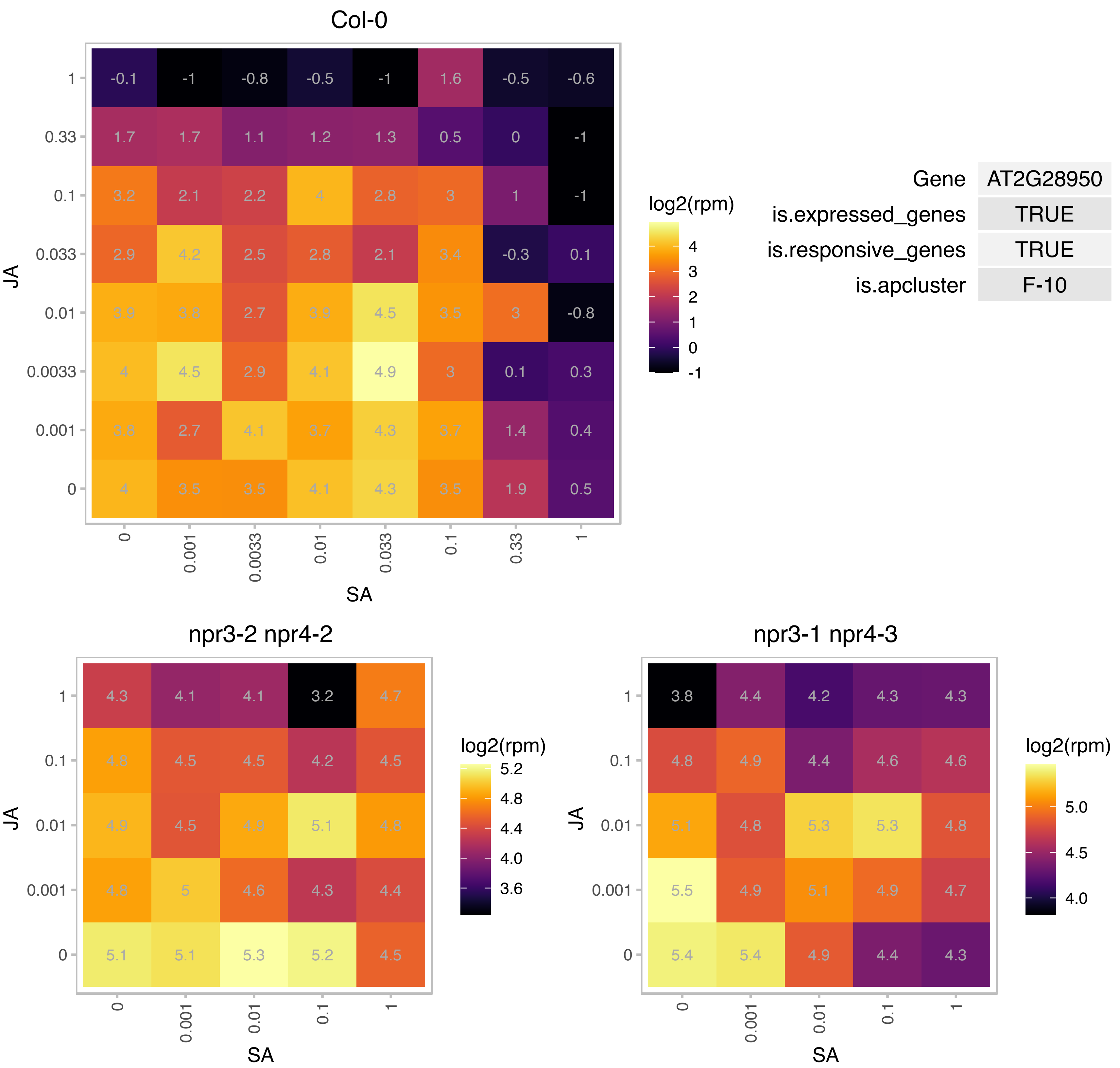


| | |
|---|---|
| Gene | AT2G28950 |
| is.expressed_genes | TRUE |
| is.responsive_genes | TRUE |
| is.apcluster | F-10 |

### AT2G32180.1 : PTAC18

plastid transcriptionally active 18

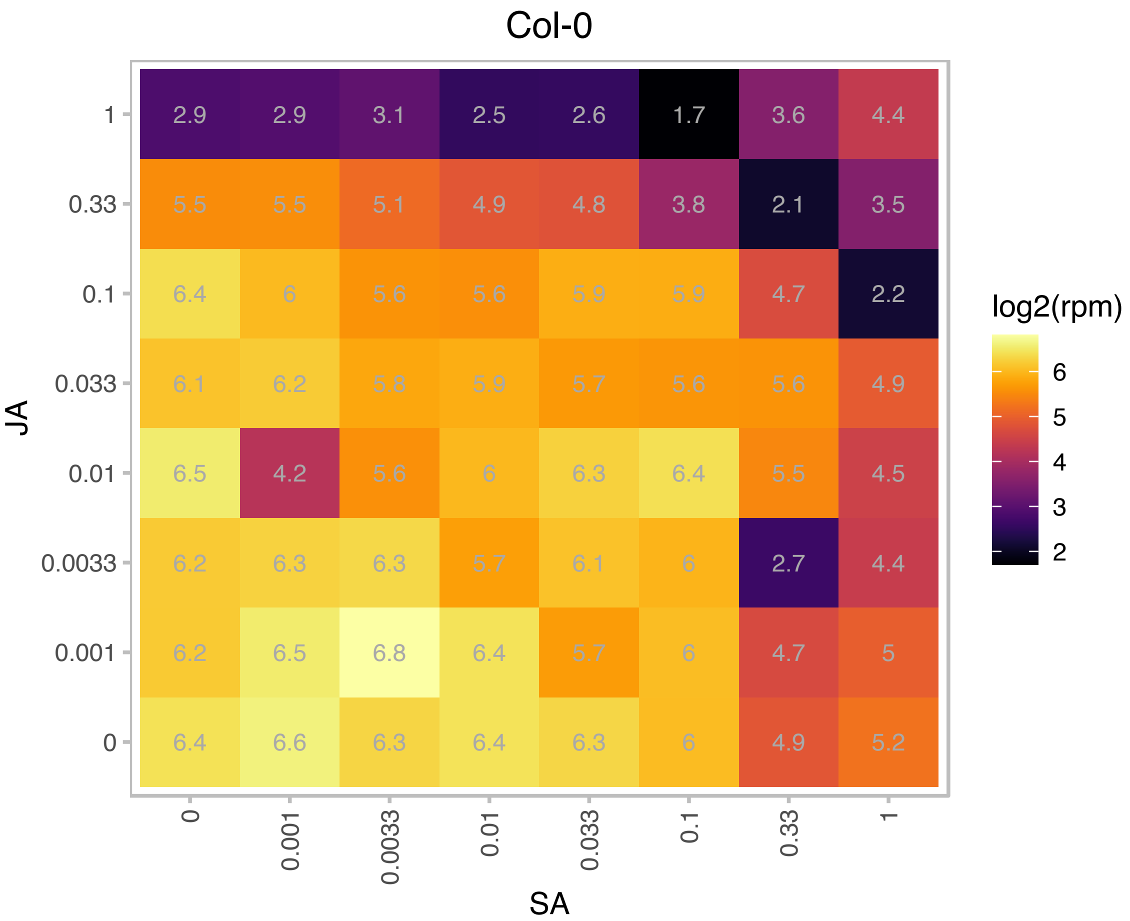


| | |
|---|---|
| Gene | AT2G32180 |
| is.expressed_genes | TRUE |
| is.responsive_genes | TRUE |
| is.apcluster | F-10 |

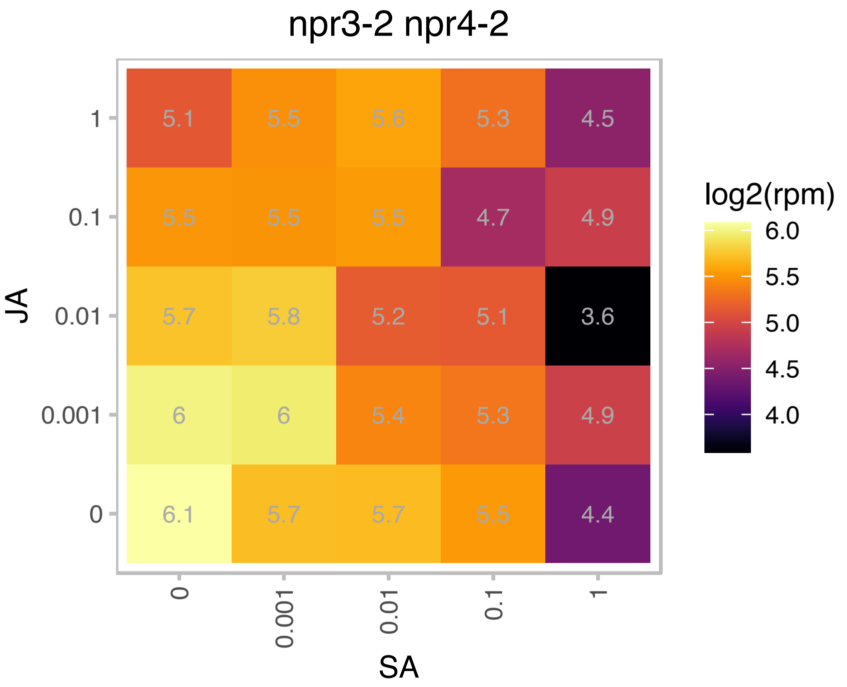


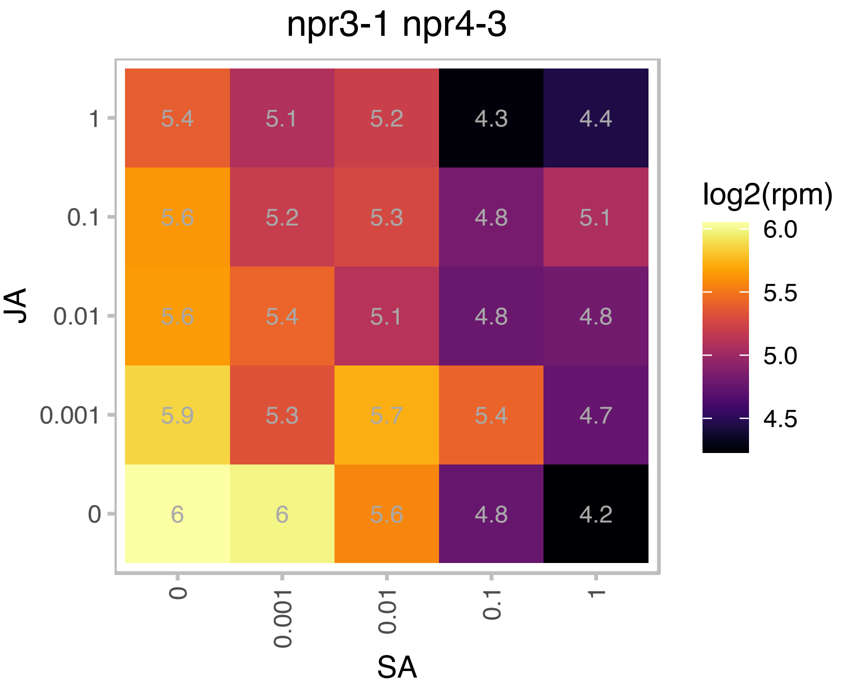

# AT2G32650.1 : NA

RmlC-like cupins superfamily protein

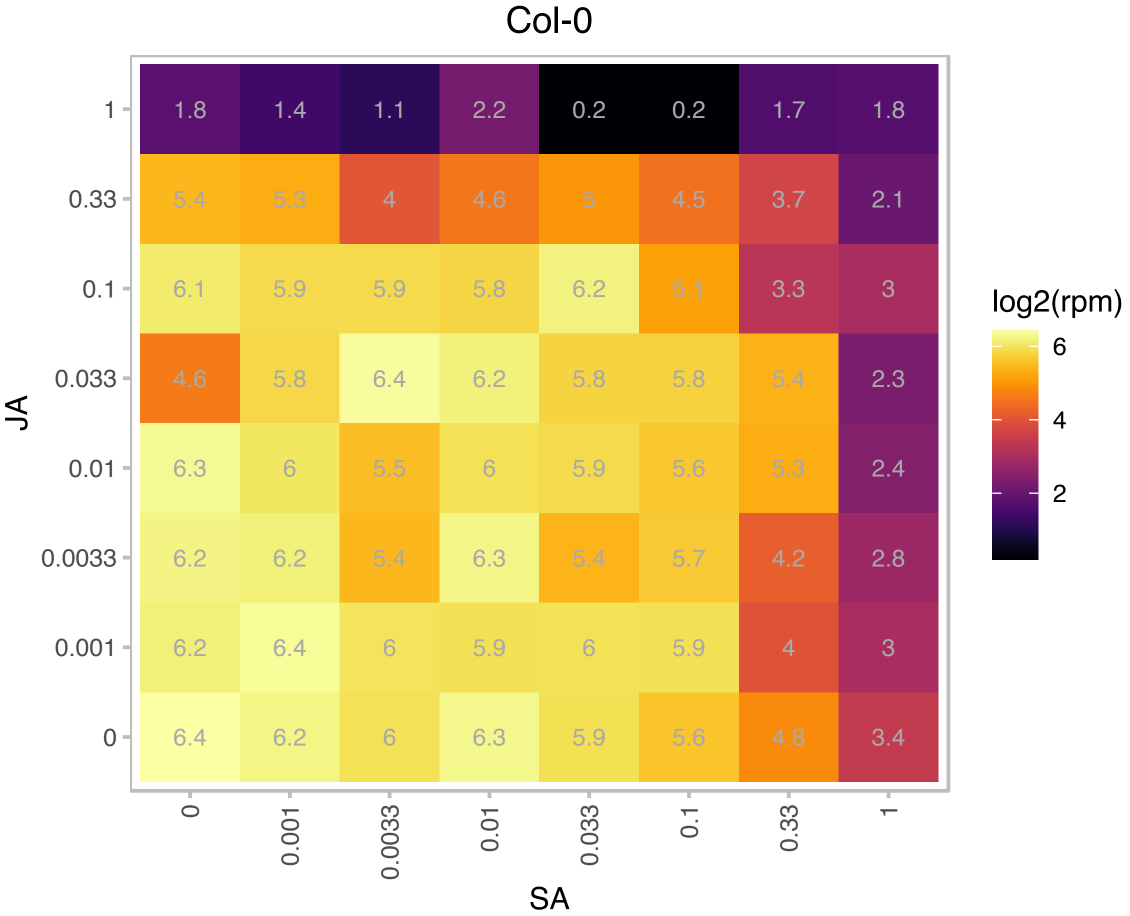


| | |
|---|---|
| Gene | AT2G32650 |
| is.expressed_genes | TRUE |
| is.responsive_genes | TRUE |
| is.apcluster | F-10 |

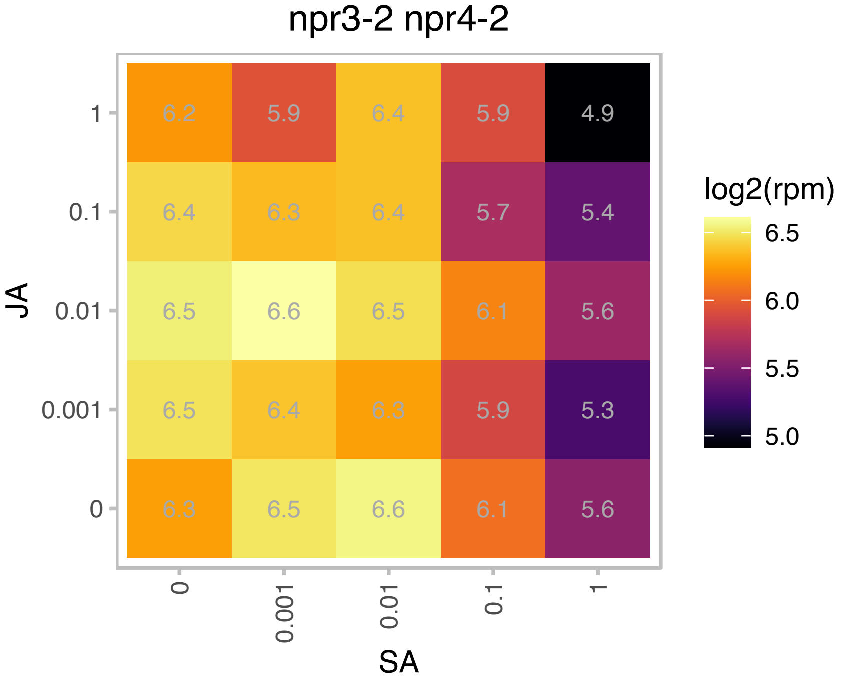


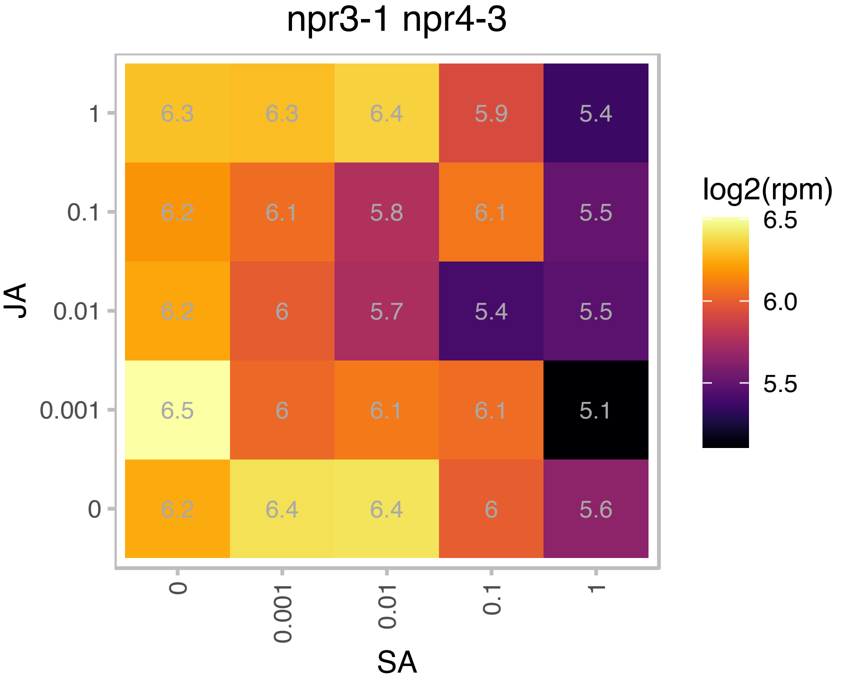

# AT2G33180.1 : NA

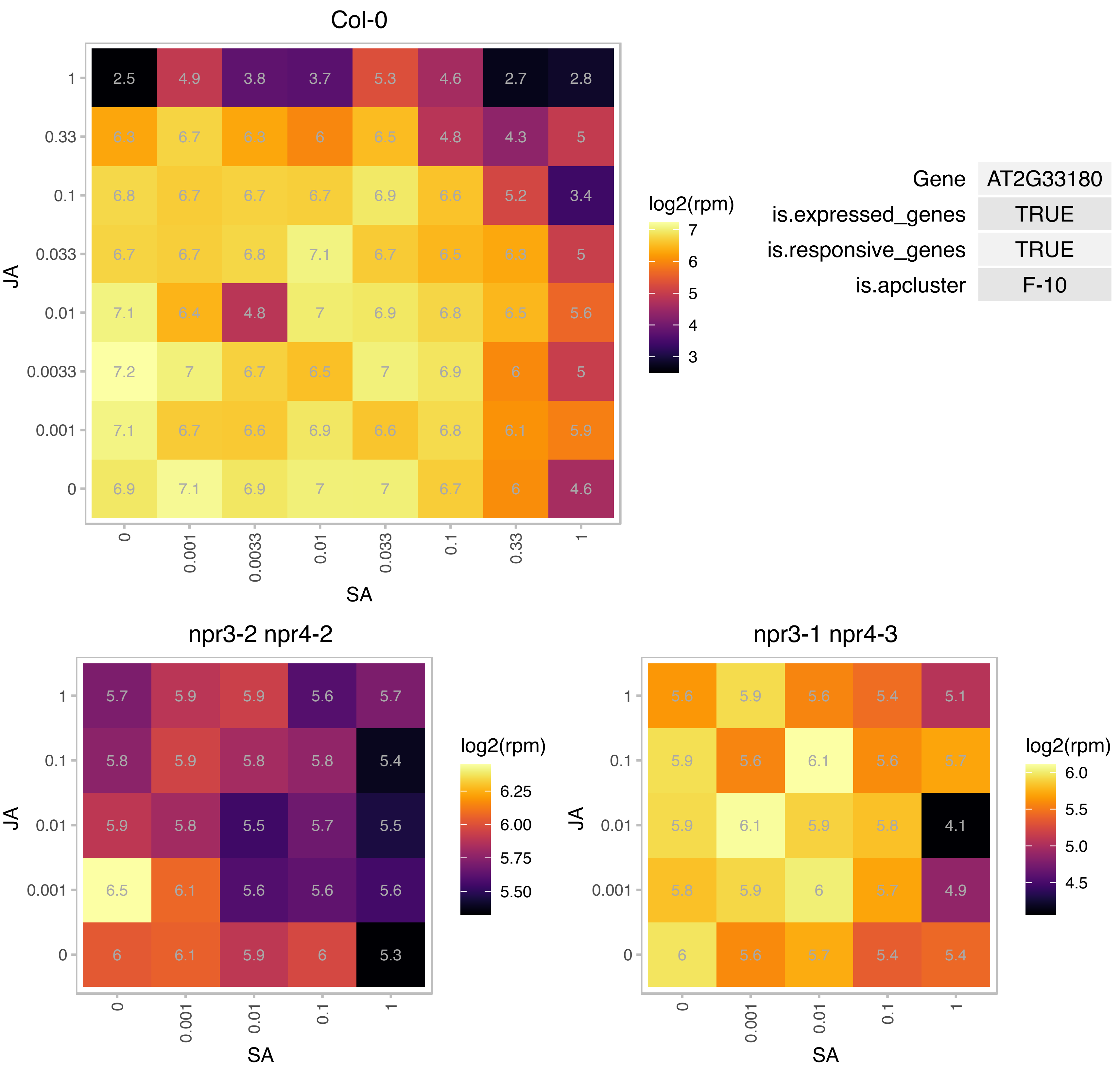


| | |
|---|---|
| Gene | AT2G33180 |
| is.expressed_genes | TRUE |
| is.responsive_genes | TRUE |
| is.apcluster | F-10 |

# AT2G33400.1 : NA

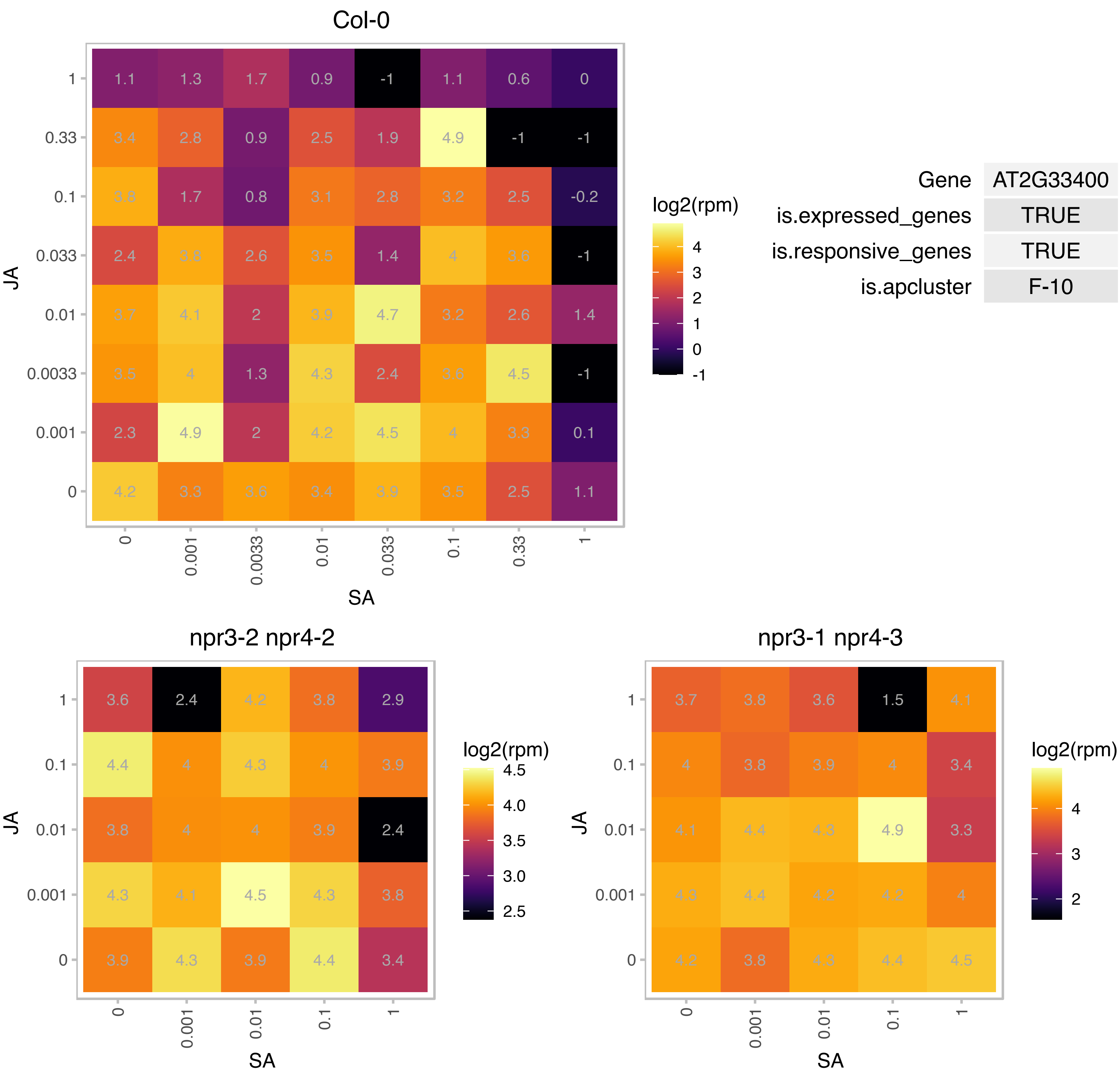


| | |
|---|---|
| Gene | AT2G33400 |
| is.expressed_genes | TRUE |
| is.responsive_genes | TRUE |
| is.apcluster | F-10 |

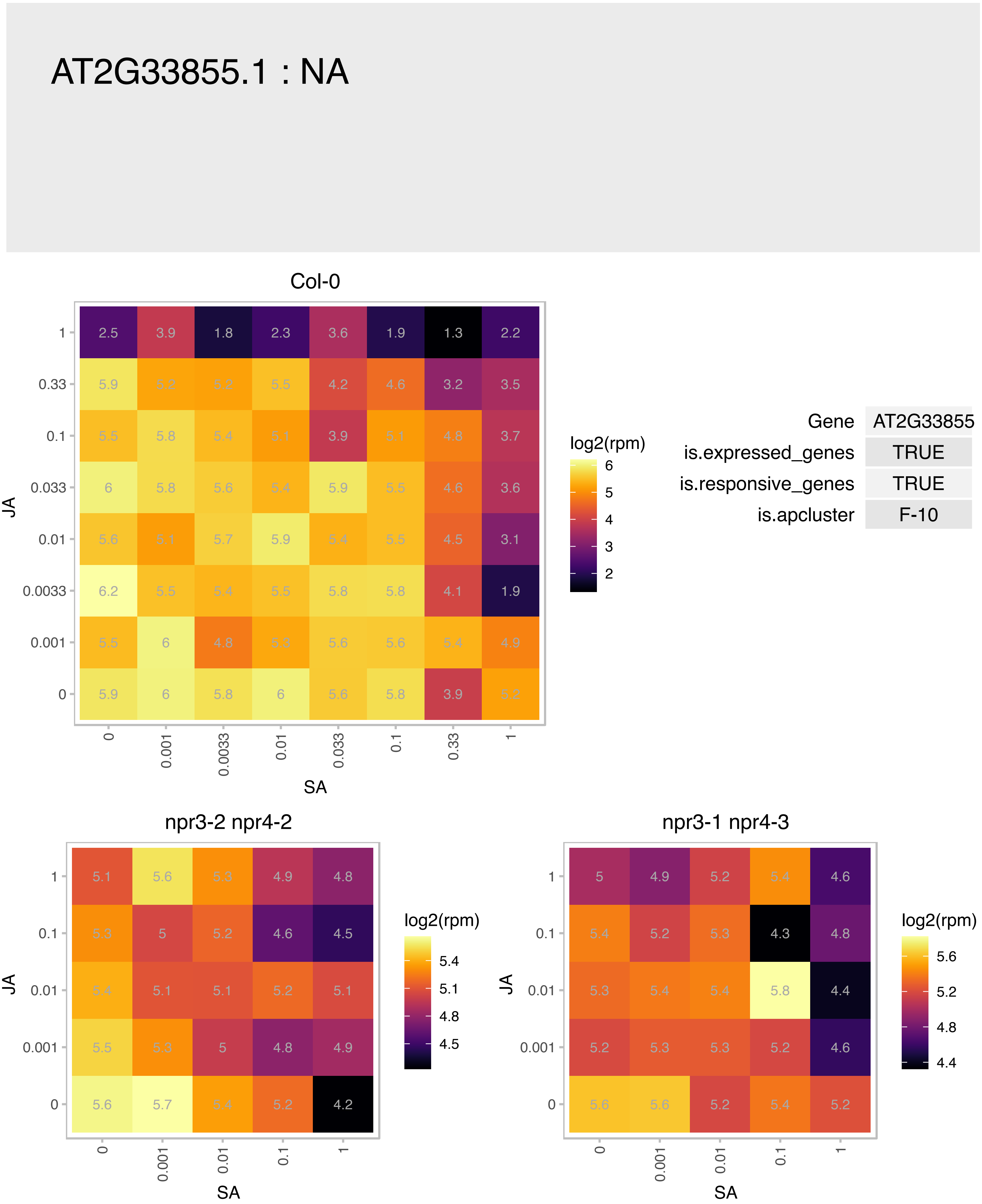

AT2G33855.1 : NA
Col-0
JA
SA
log2(rpm)
Gene AT2G33855
is.expressed_genes TRUE
is.responsive_genes TRUE
is.apcluster F-10
npr3-2 npr4-2
npr3-1 npr4-3

### AT2G34640.1 : HMR, PTAC12, TAC12

plastid transcriptionally active 12

Present in transcriptionally active plastid chromosomes. Involved in plastid gene expression.

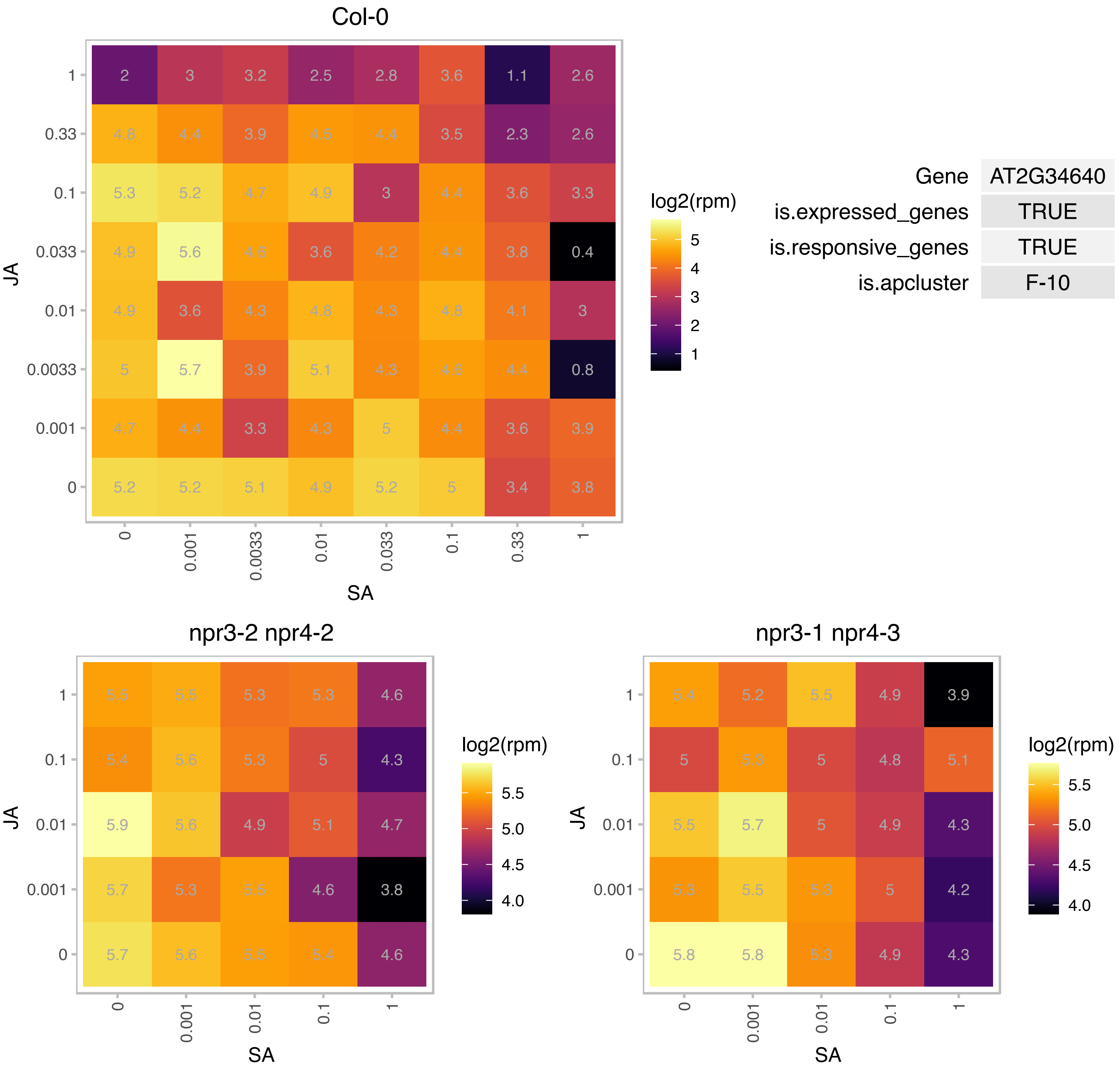


| | |
|---|---|
| Gene | AT2G34640 |
| is.expressed_genes | TRUE |
| is.responsive_genes | TRUE |
| is.apcluster | F-10 |

# AT2G35820.1 : NA

ureidoglycolate hydrolases

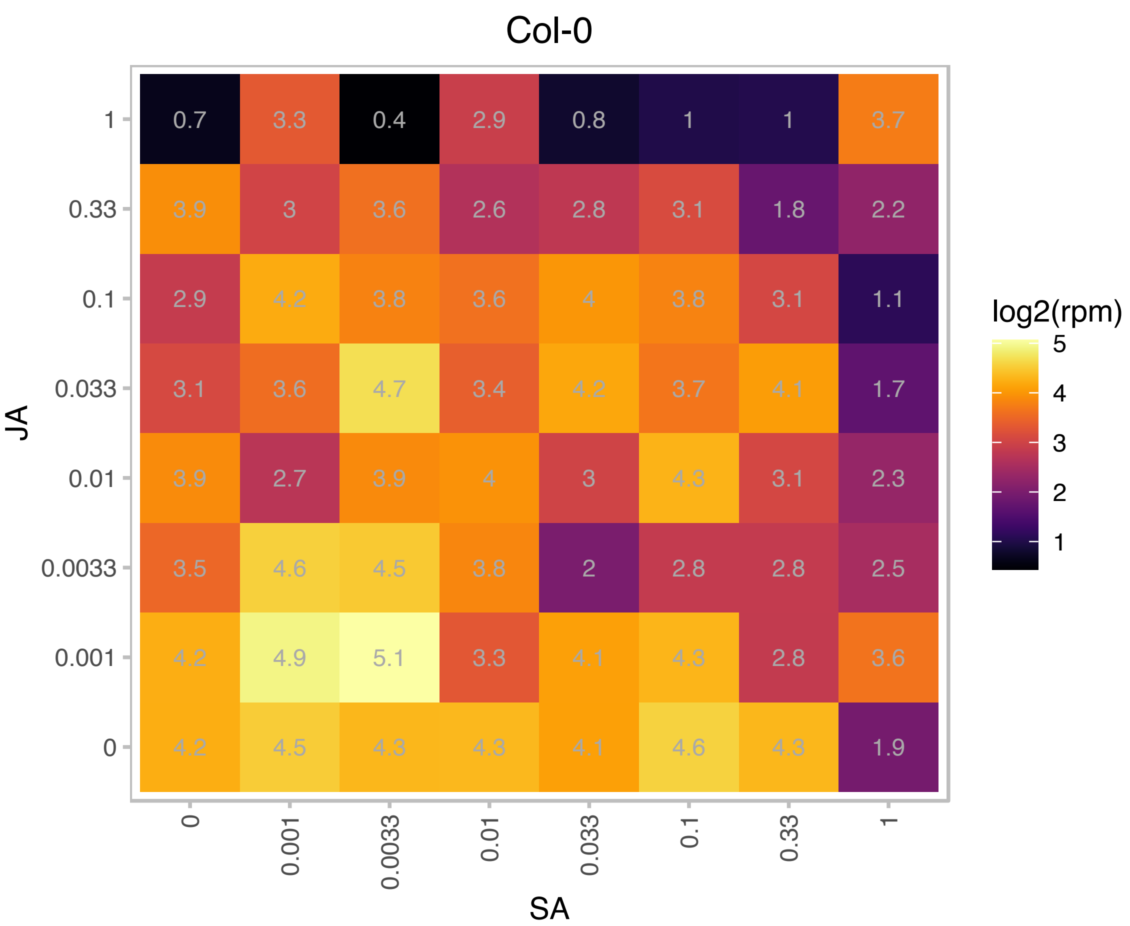


| | |
|---|---|
| Gene | AT2G35820 |
| is.expressed_genes | TRUE |
| is.responsive_genes | TRUE |
| is.apcluster | F-10 |

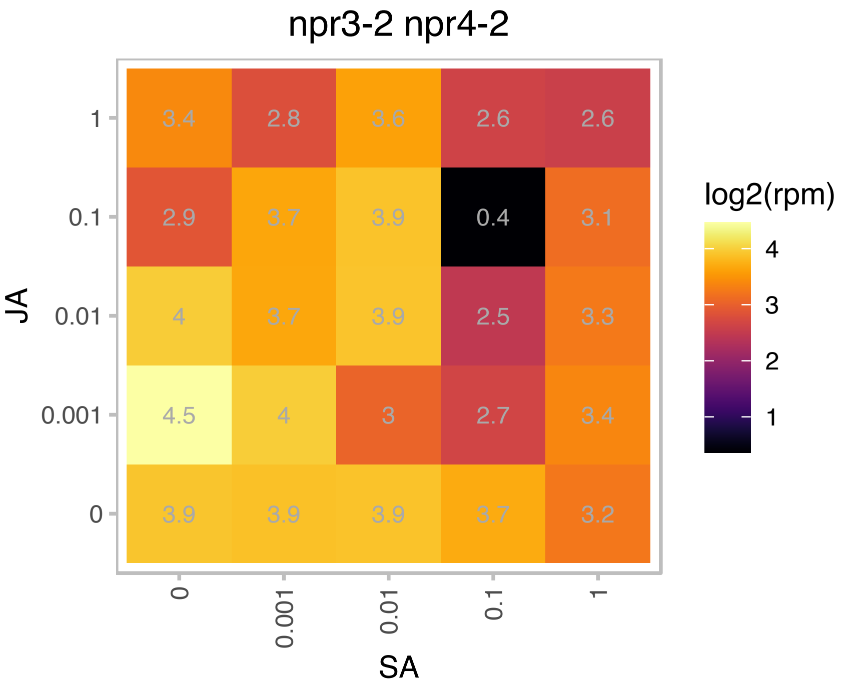


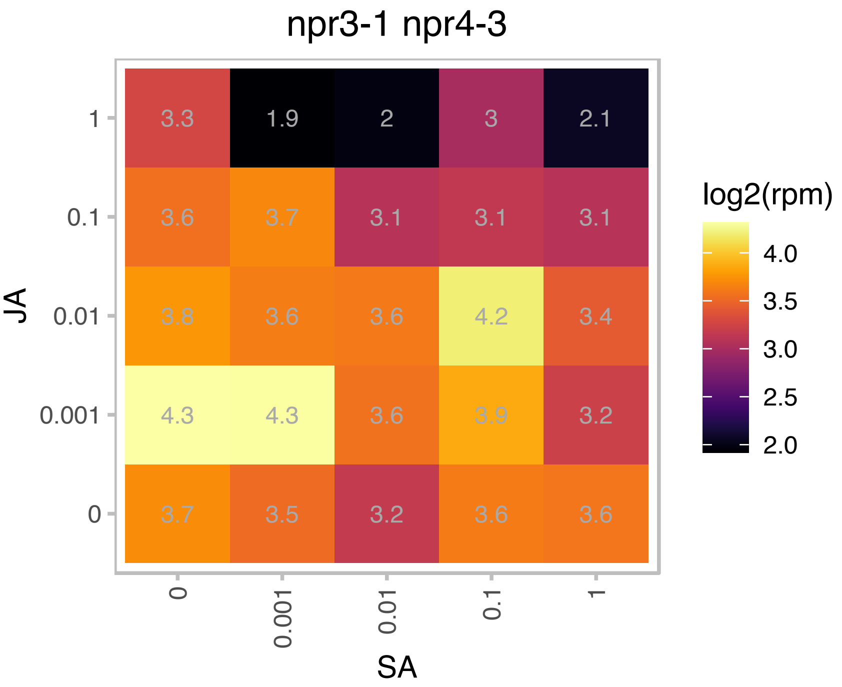

### AT2G36000.1 : EMB3114

Mitochondrial transcription termination factor family protein

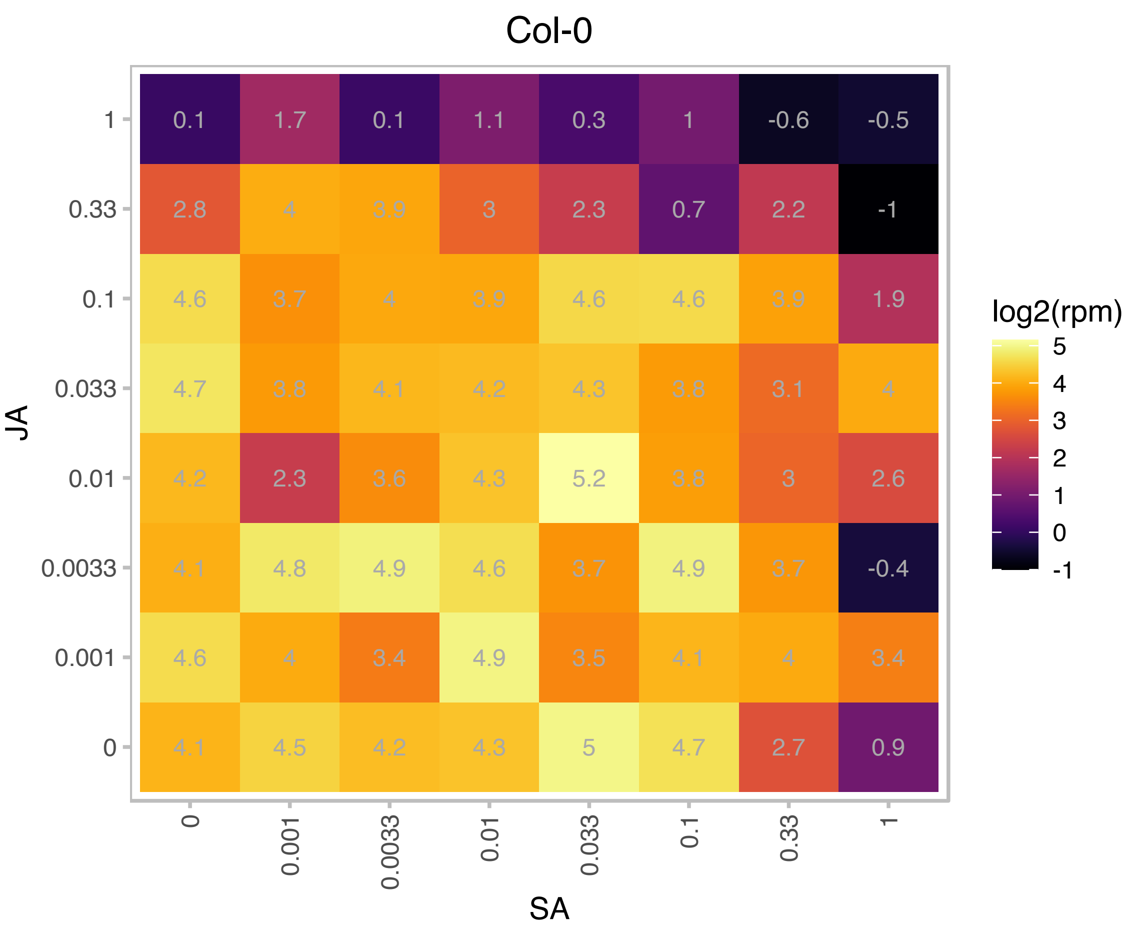


| | |
|---|---|
| Gene | AT2G36000 |
| is.expressed_genes | TRUE |
| is.responsive_genes | TRUE |
| is.apcluster | F-10 |

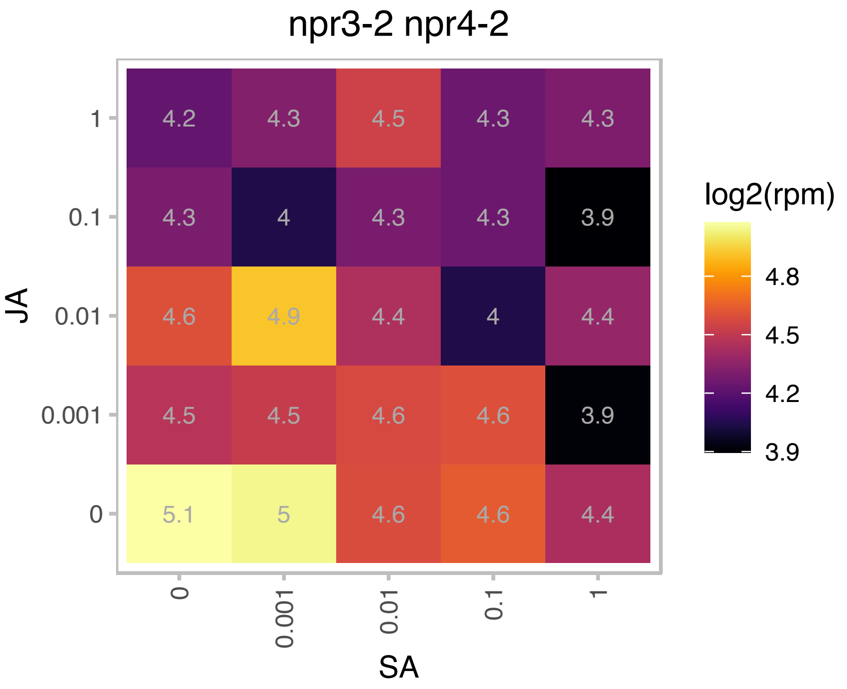


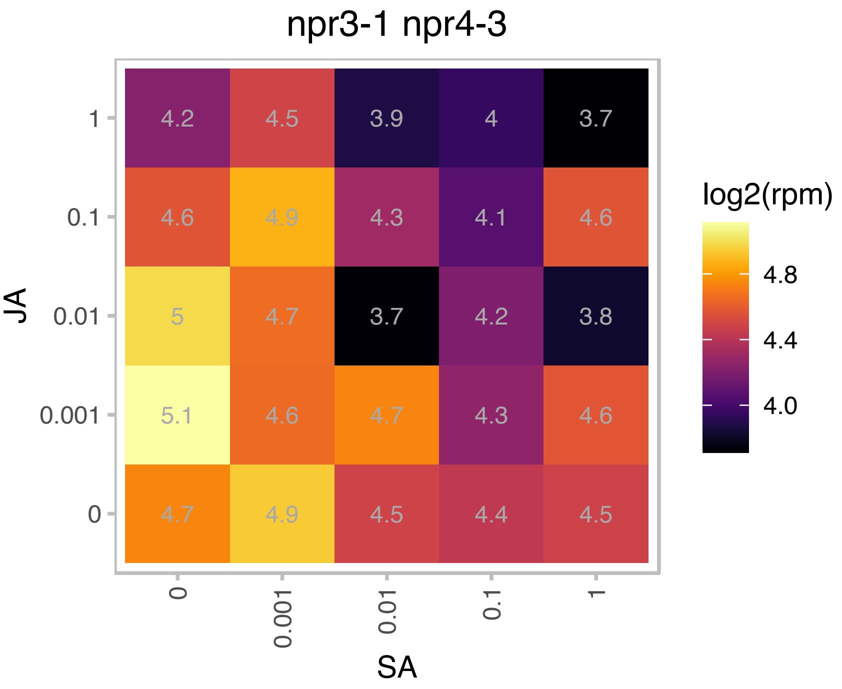

# AT2G37470.1 : NA

Histone superfamily protein

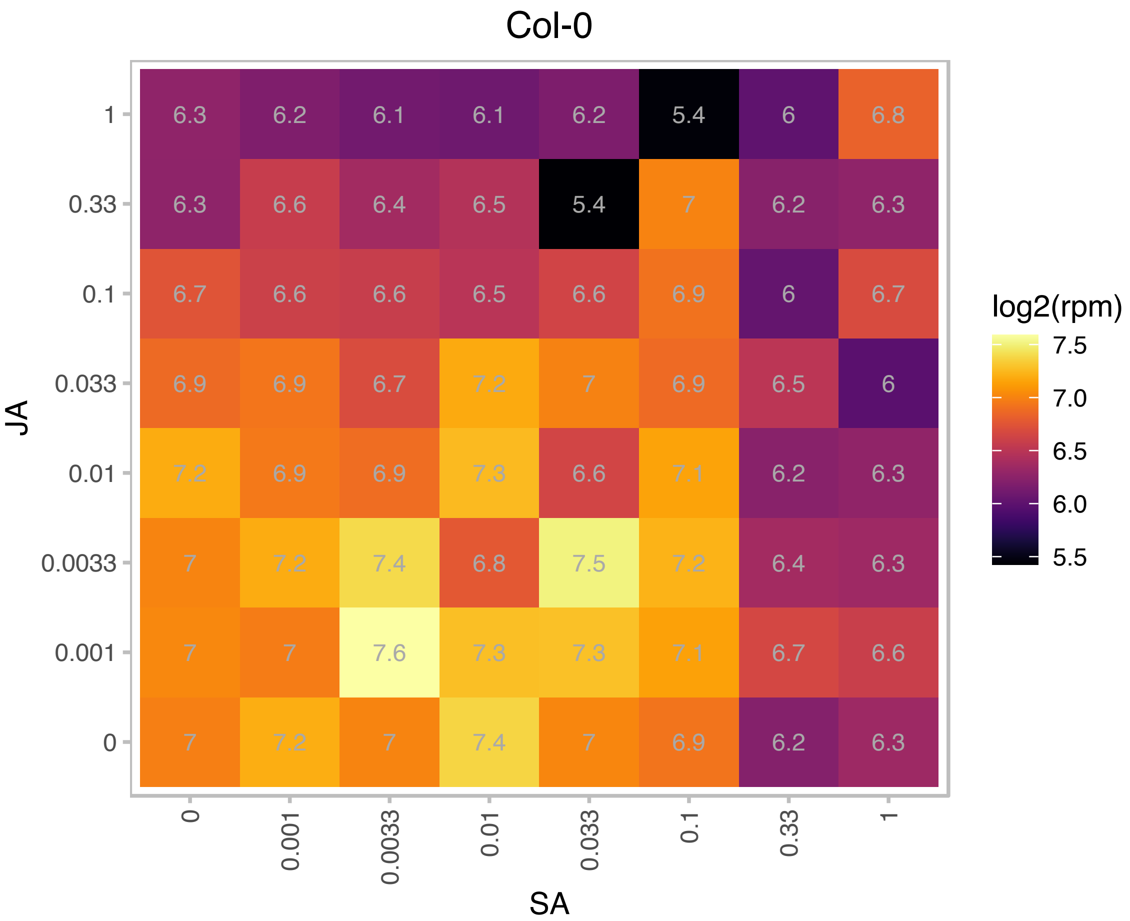


| | |
|---|---|
| Gene | AT2G37470 |
| is.expressed_genes | TRUE |
| is.responsive_genes | TRUE |
| is.apcluster | F-10 |

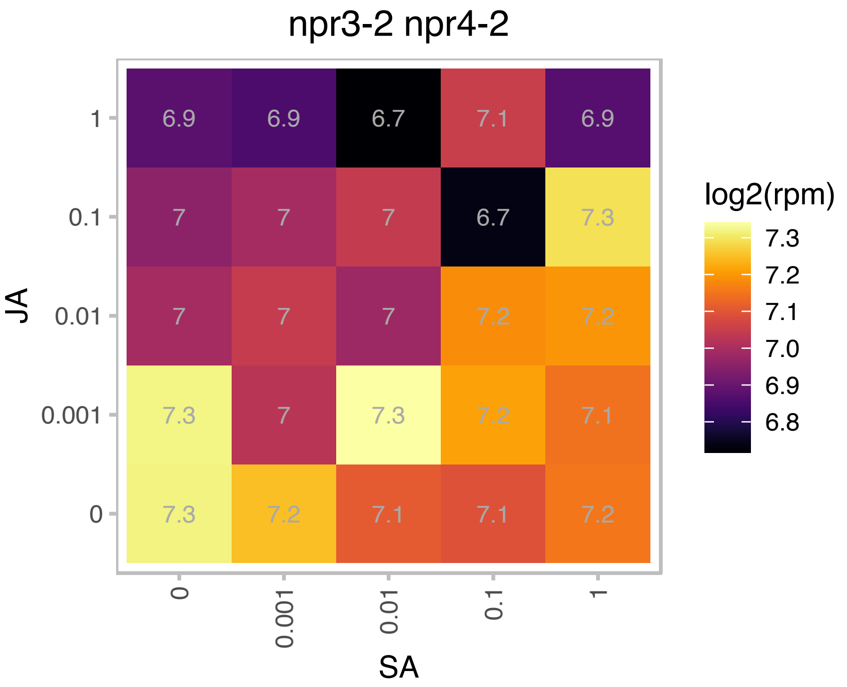


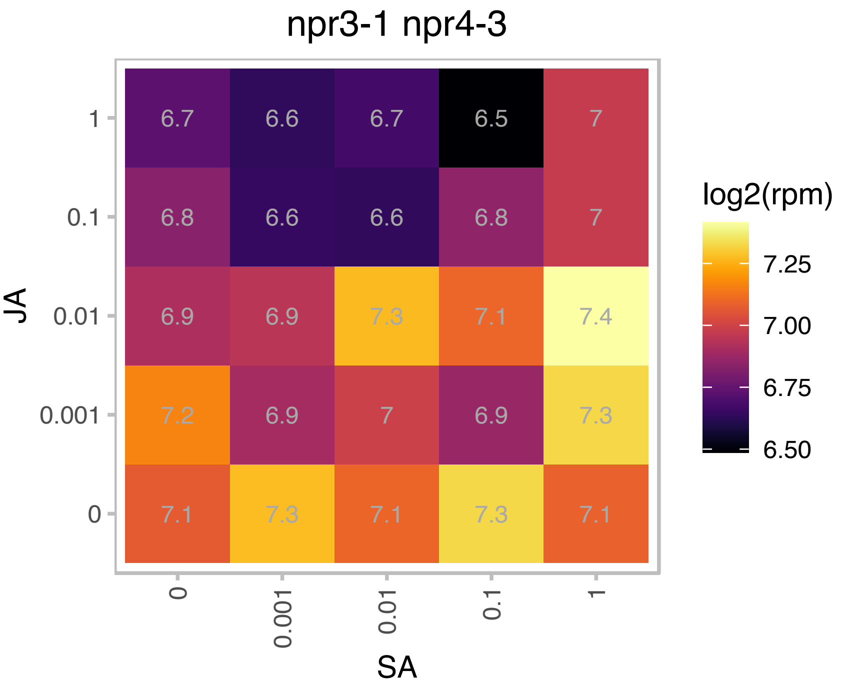

# AT2G37660.1 : NA

NAD(P)-binding Rossmann-fold superfamily protein

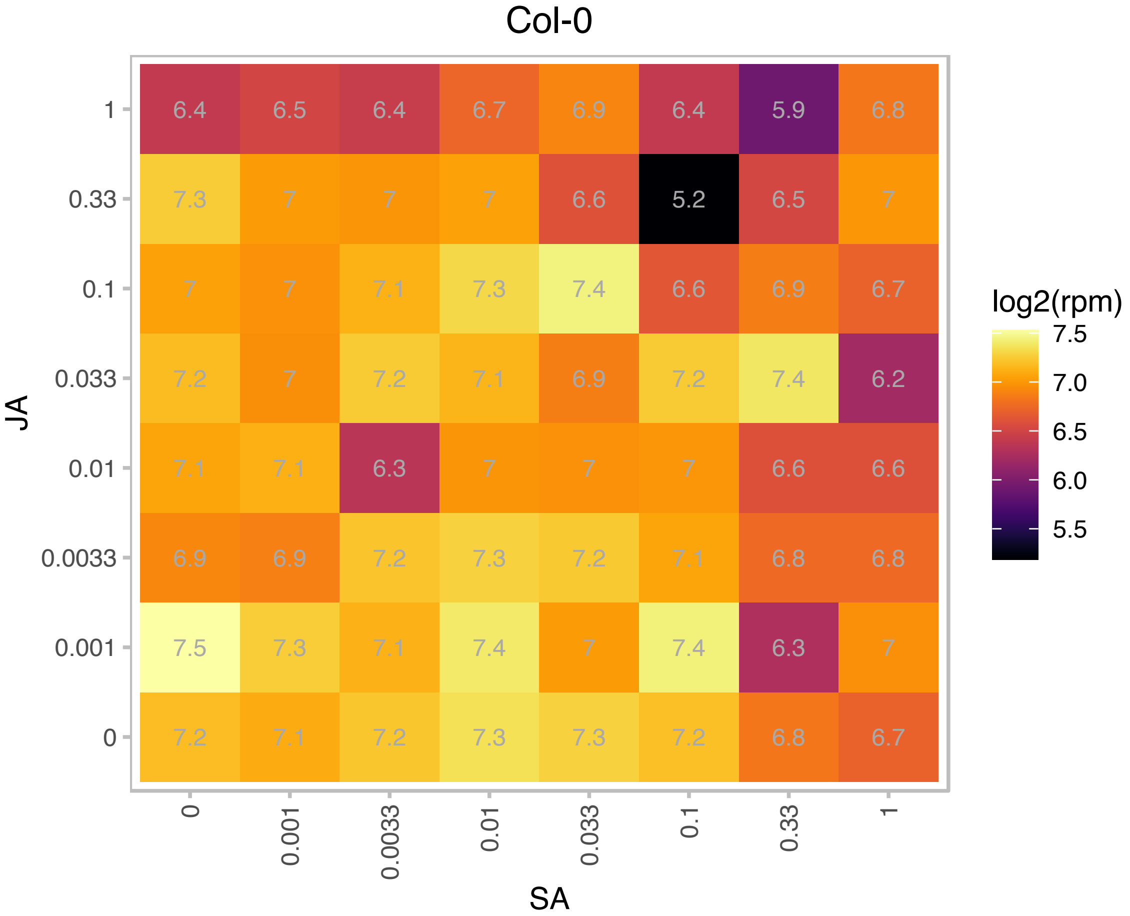


| | |
|---|---|
| Gene | AT2G37660 |
| is.expressed_genes | TRUE |
| is.responsive_genes | TRUE |
| is.apcluster | F-10 |

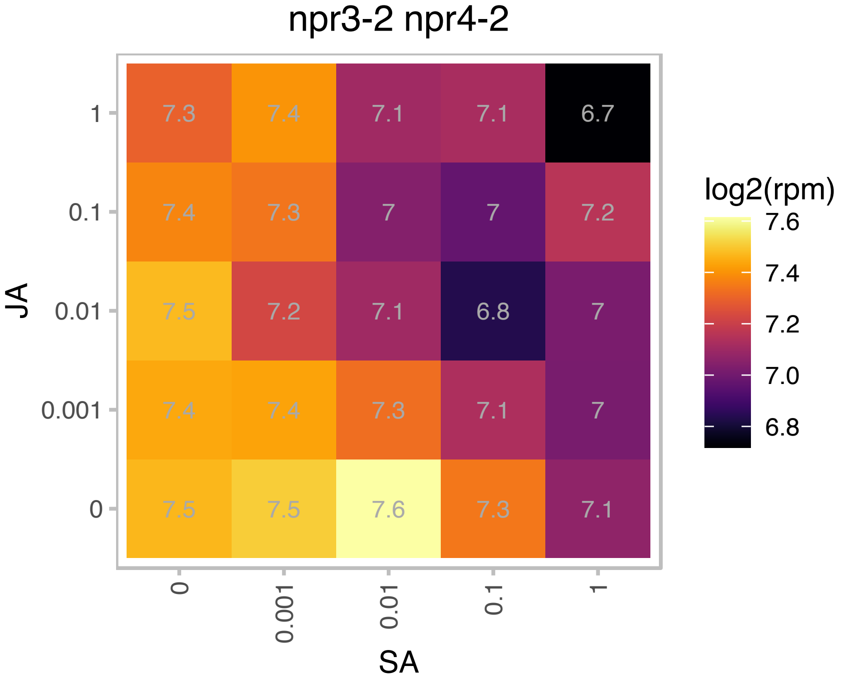


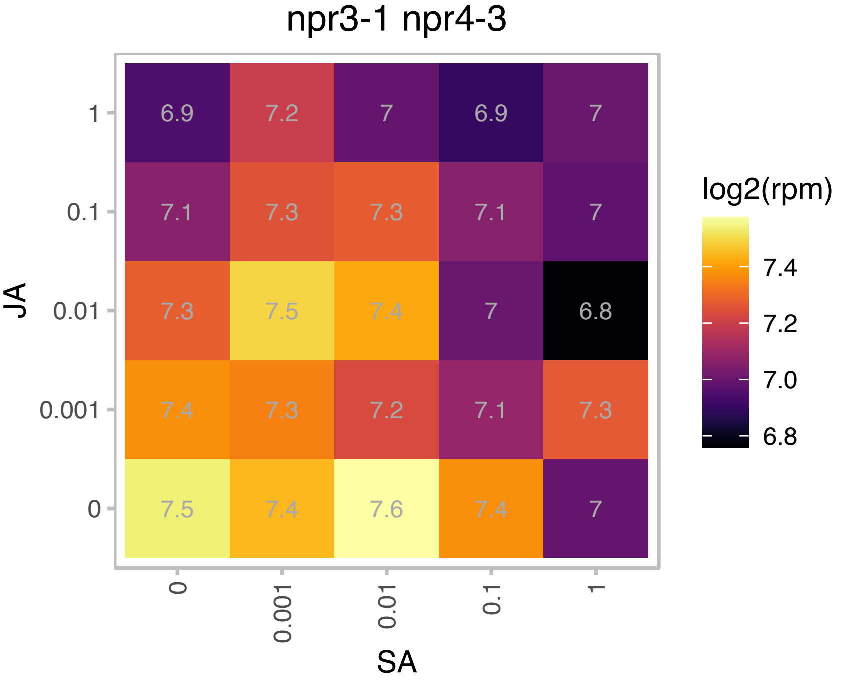

### AT2G39705.1 : DVL11, RTFL8

ROTUNDIFOLIA like 8

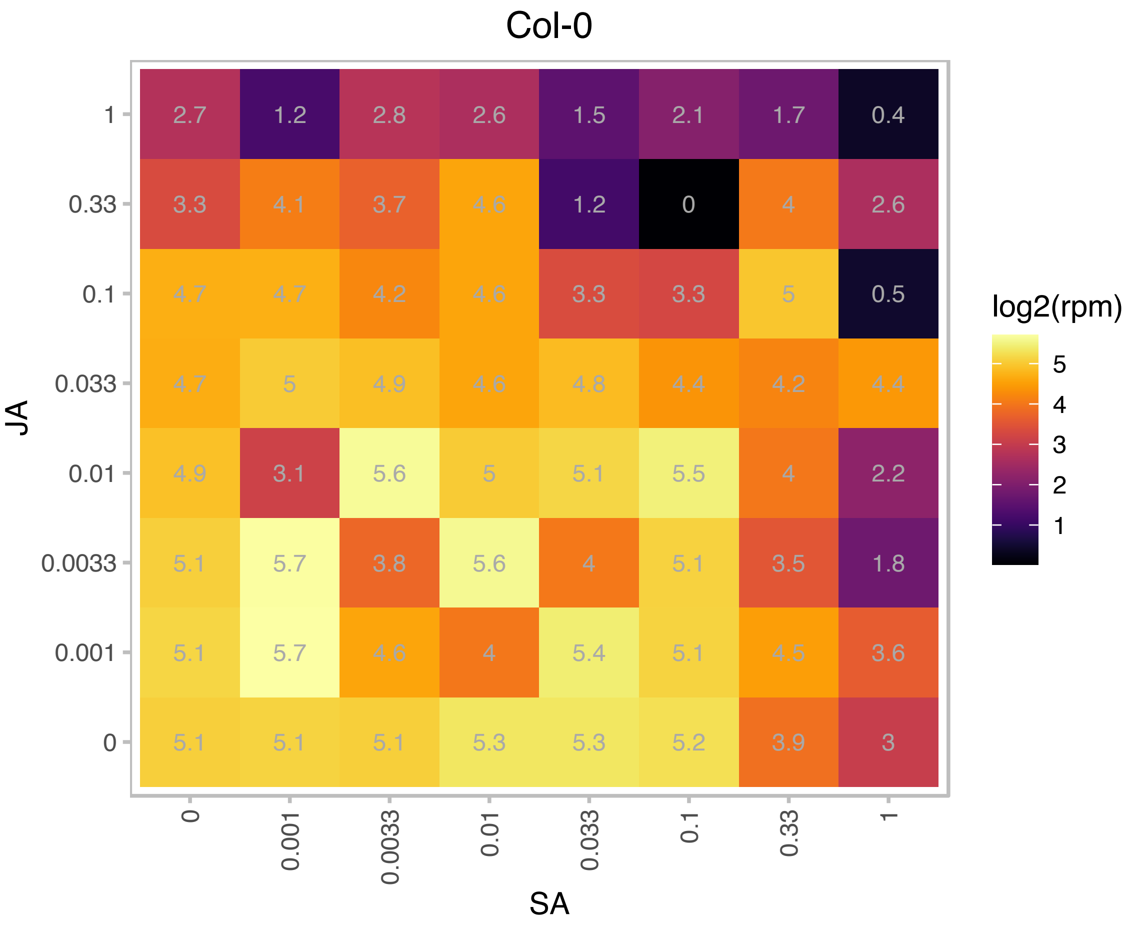


| | |
|---|---|
| Gene | AT2G39705 |
| is.expressed_genes | TRUE |
| is.responsive_genes | TRUE |
| is.apcluster | F-10 |

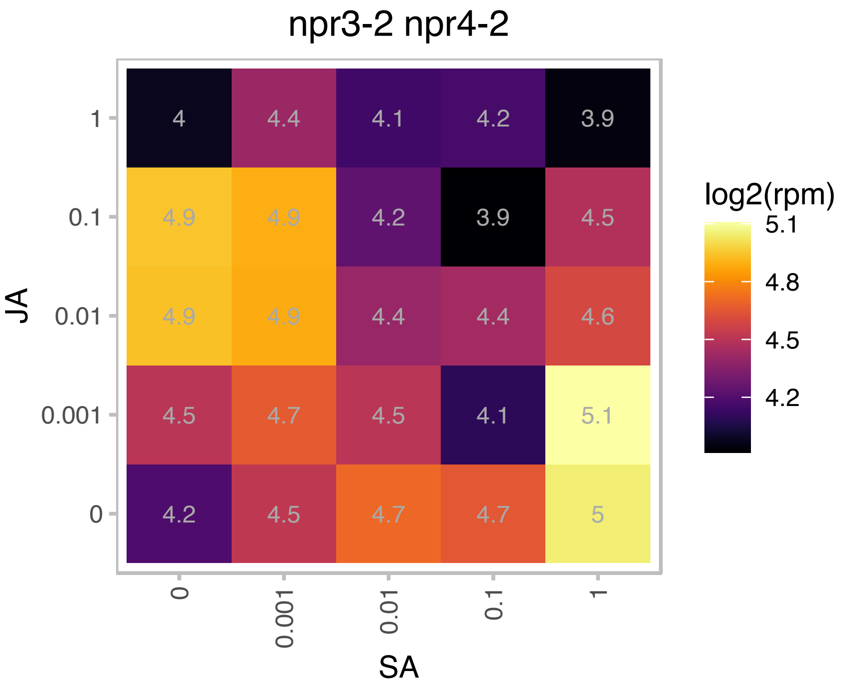


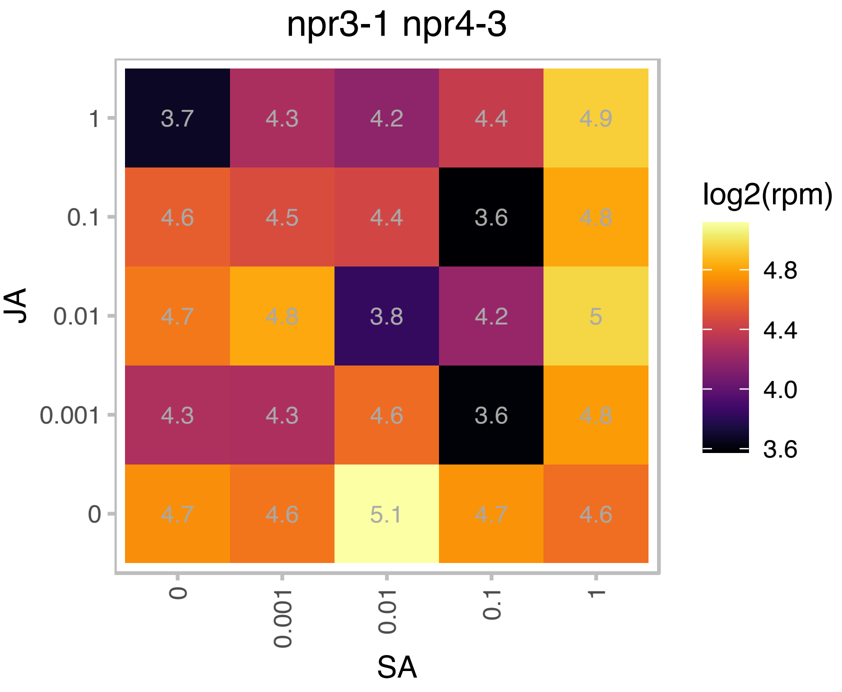

# AT2G41950.1 : NA

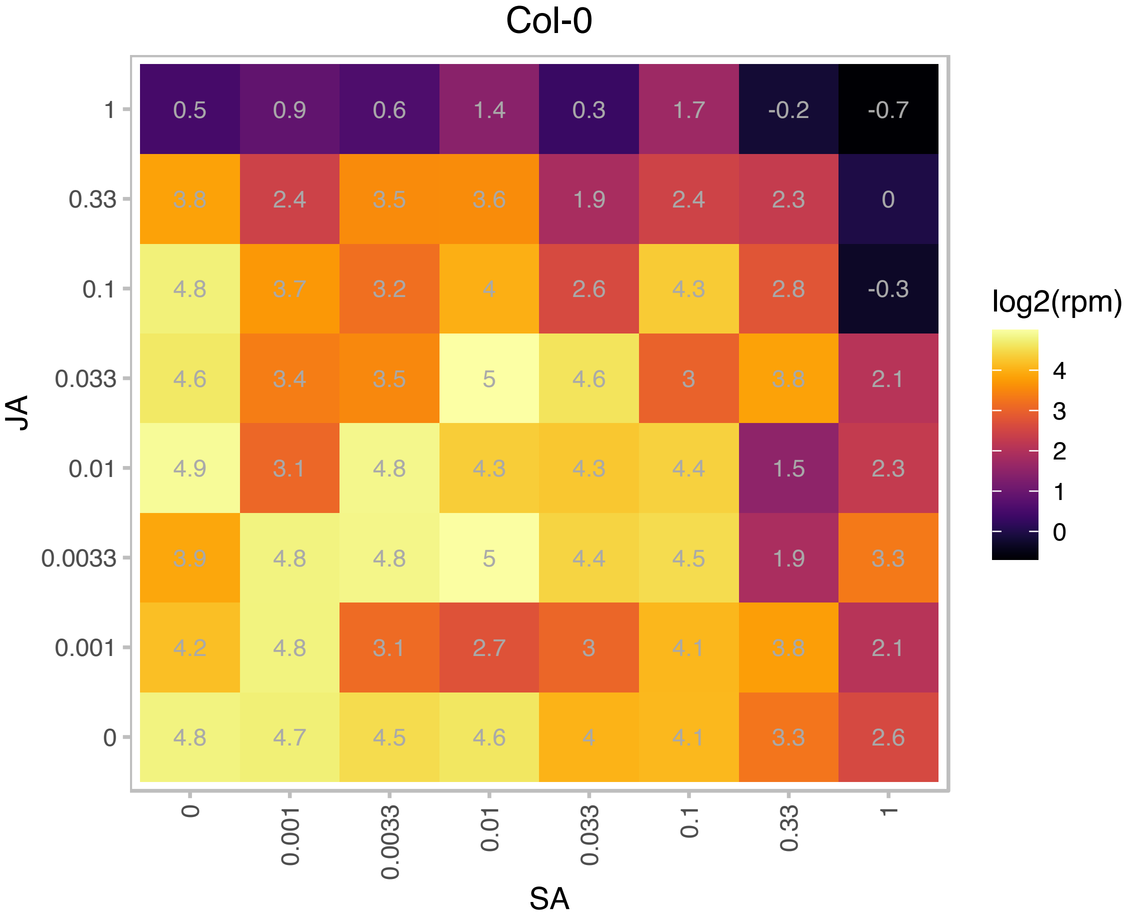


| | |
|---|---|
| Gene | AT2G41950 |
| is.expressed_genes | TRUE |
| is.responsive_genes | TRUE |
| is.apcluster | F-10 |

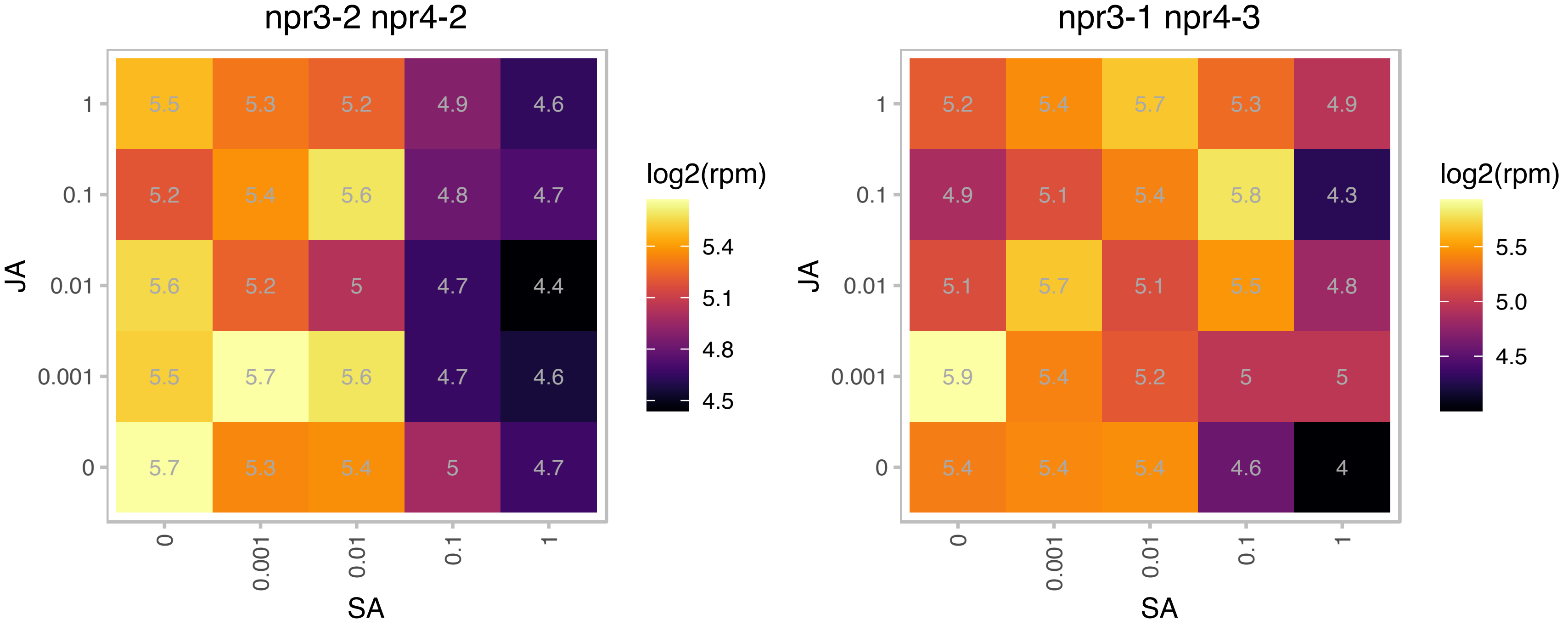

# AT2G42110.1 : NA

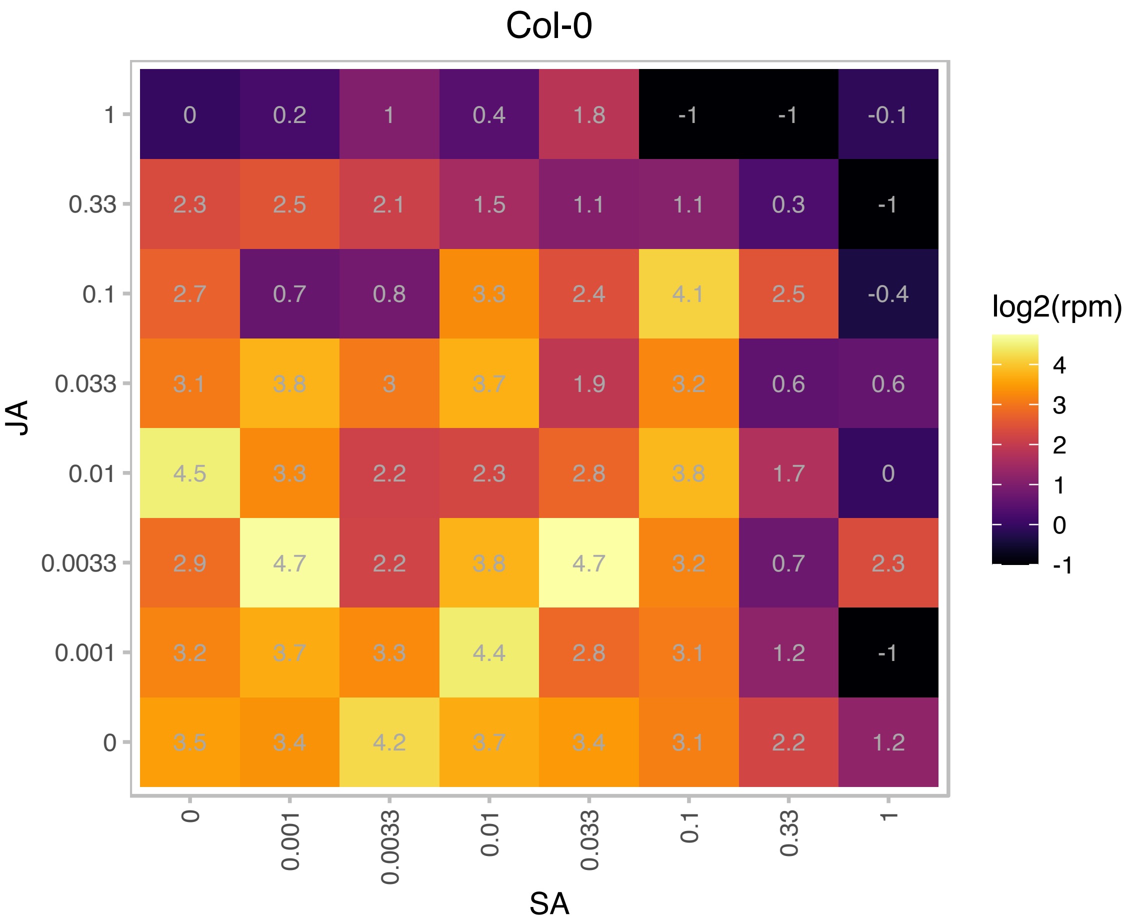


| | |
|---|---|
| Gene | AT2G42110 |
| is.expressed_genes | TRUE |
| is.responsive_genes | TRUE |
| is.apcluster | F-10 |

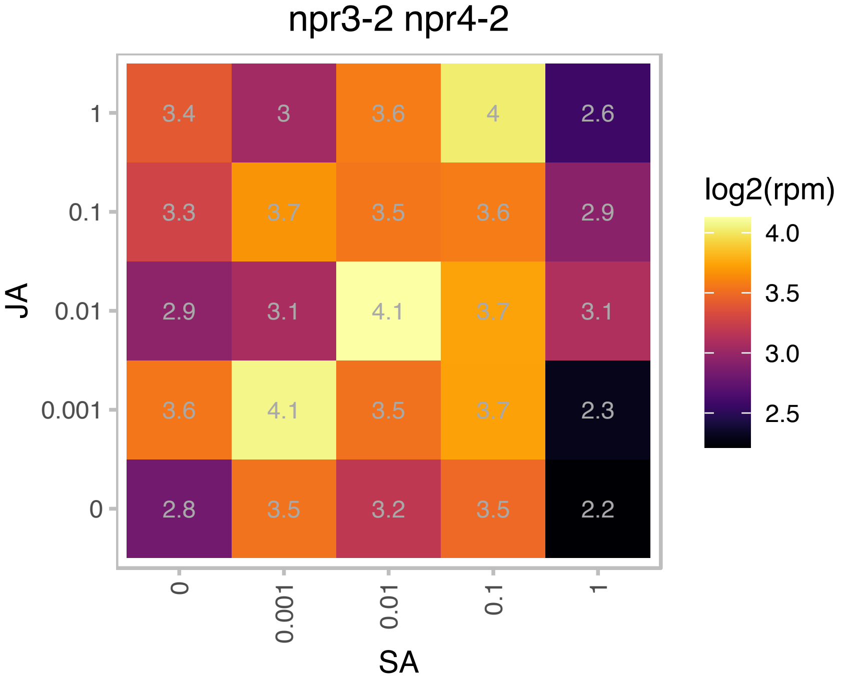


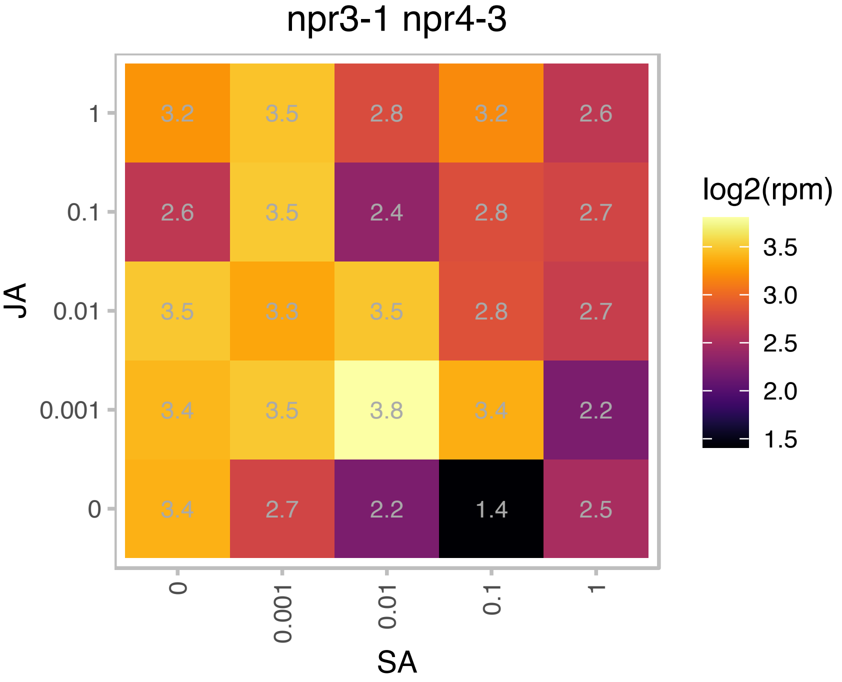

# AT2G42220.1 : NA

Rhodanese/Cell cycle control phosphatase superfamily protein

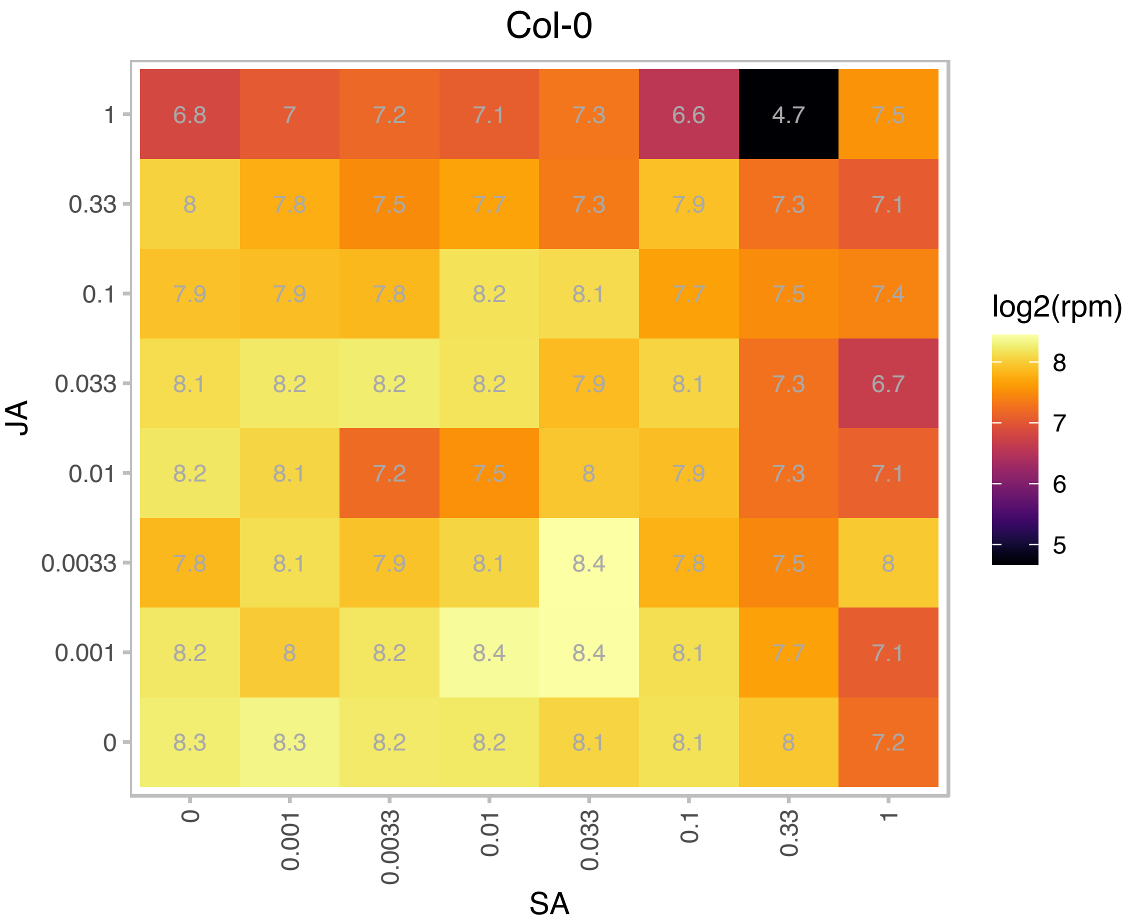


| | |
|---|---|
| Gene | AT2G42220 |
| is.expressed_genes | TRUE |
| is.responsive_genes | TRUE |
| is.apcluster | F-10 |

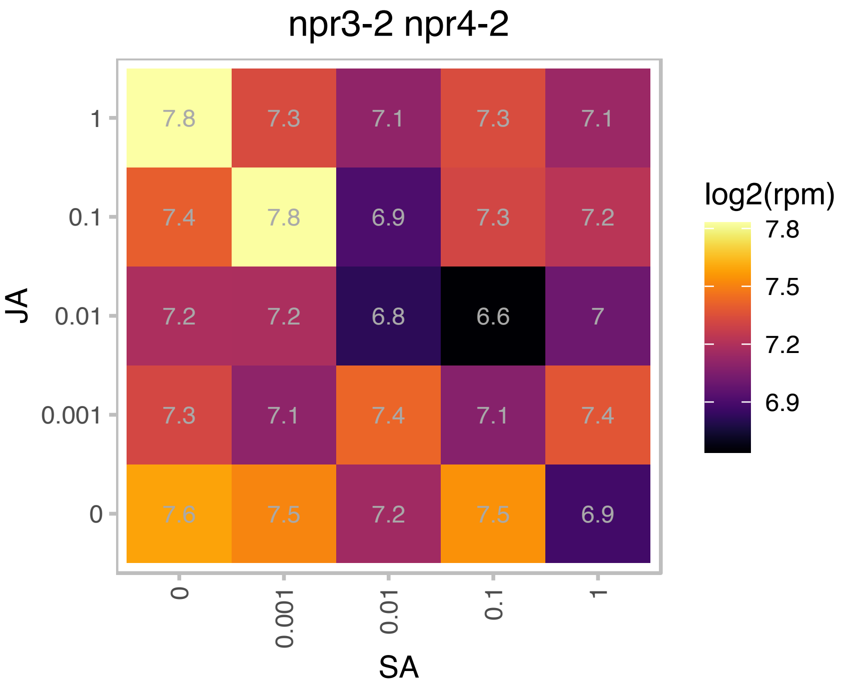


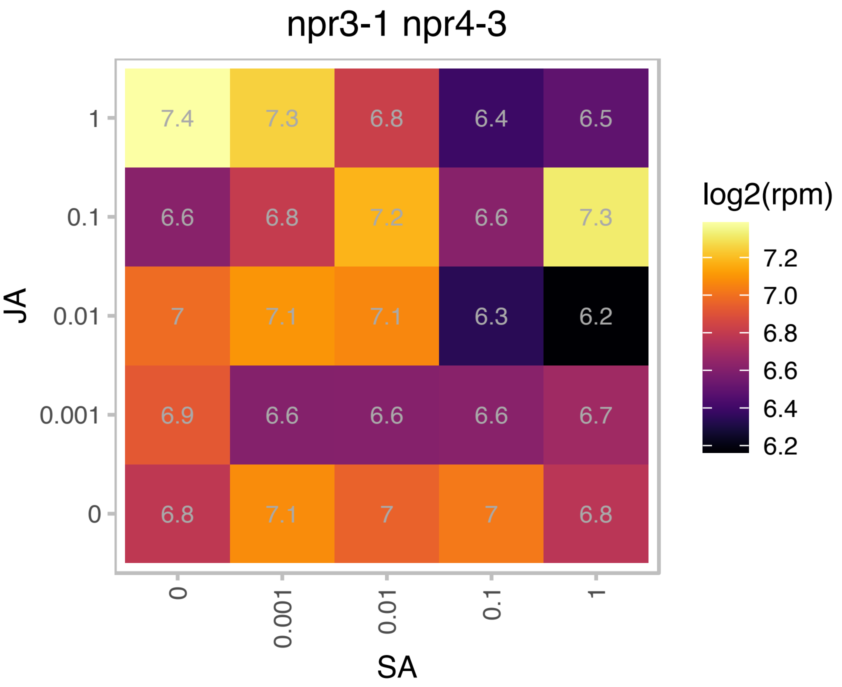

# AT2G42690.1 : NA

alpha/beta-Hydrolases superfamily protein

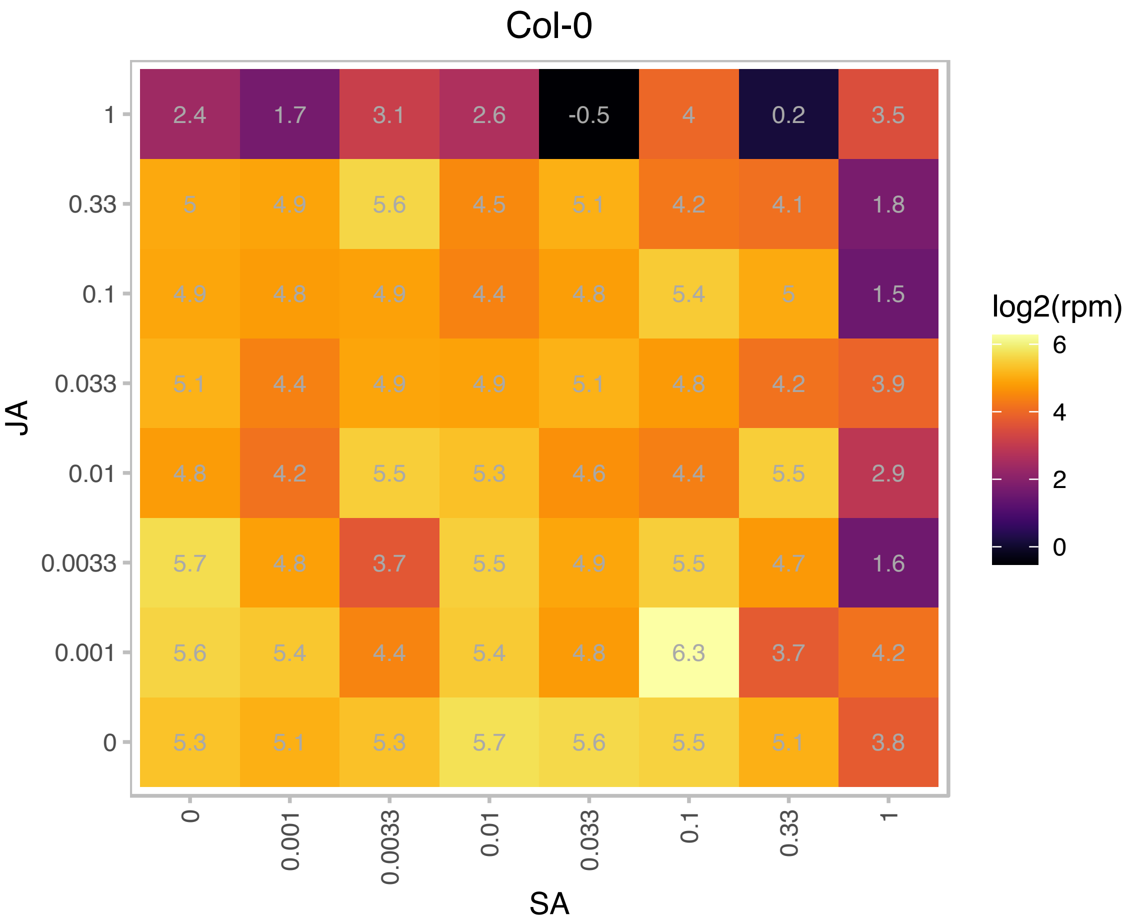


| | |
|---|---|
| Gene | AT2G42690 |
| is.expressed_genes | TRUE |
| is.responsive_genes | TRUE |
| is.apcluster | F-10 |

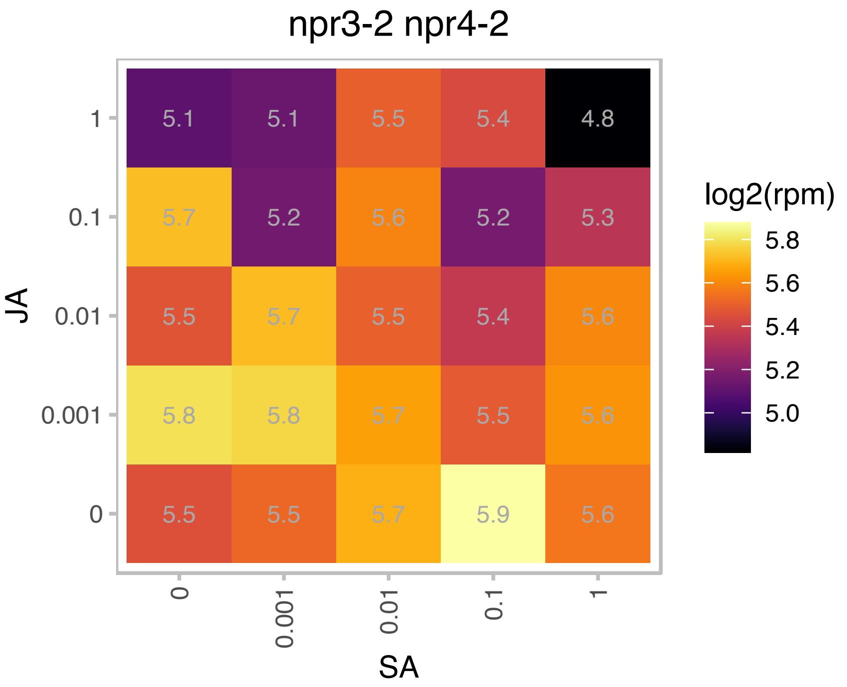


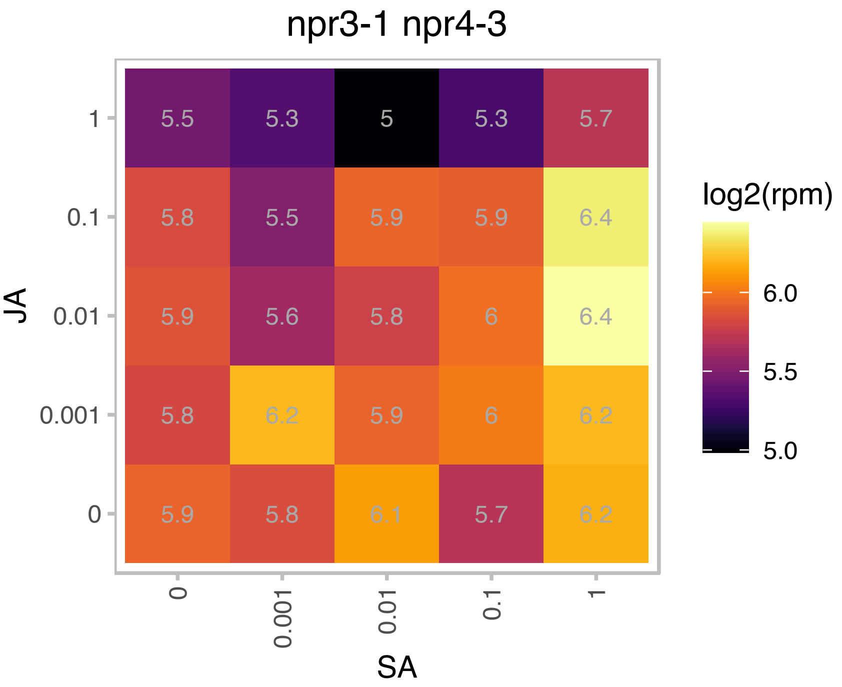

# AT2G45180.1 : NA

Bifunctional inhibitor/lipid-transfer protein/seed storage 2S albumin superfamily protein

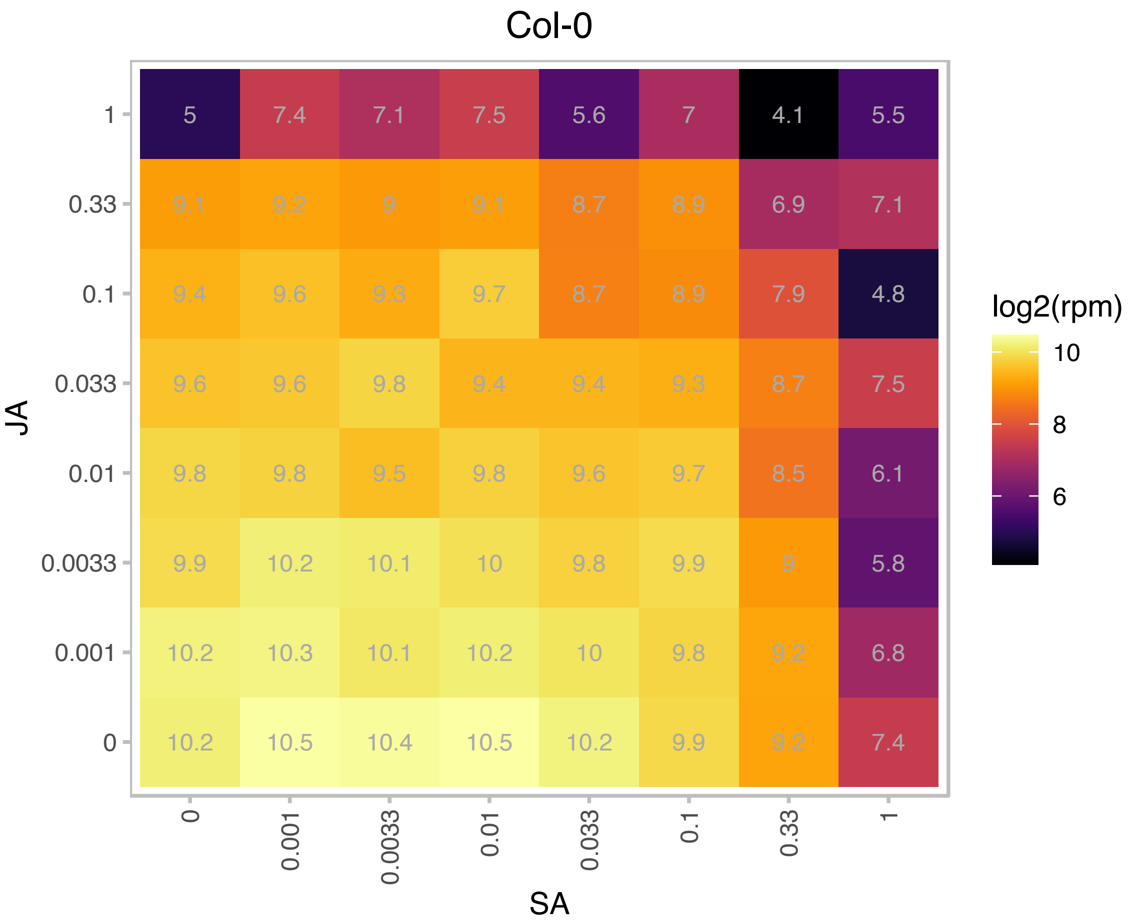


| | |
|---|---|
| Gene | AT2G45180 |
| is.expressed_genes | TRUE |
| is.responsive_genes | TRUE |
| is.apcluster | F-10 |

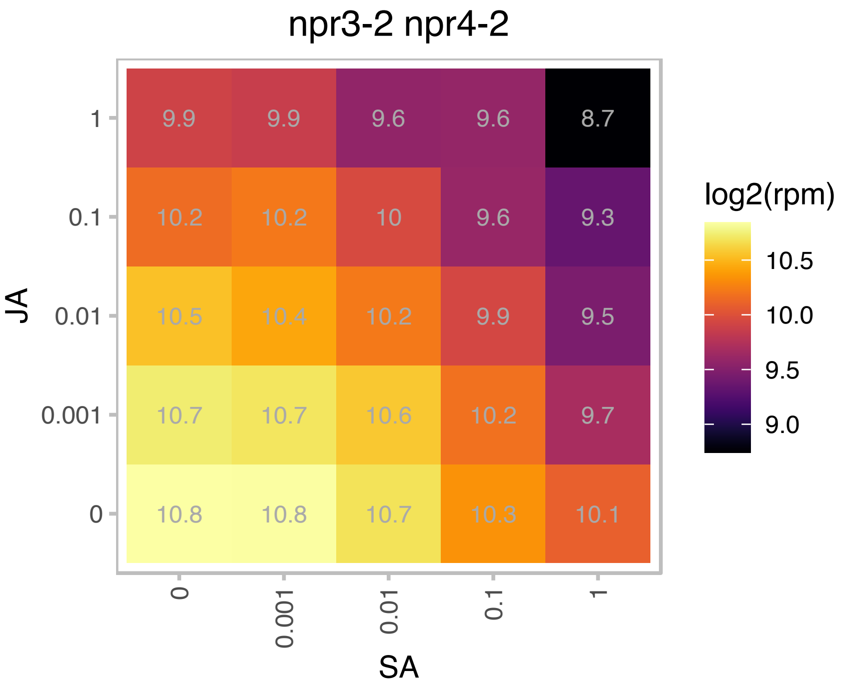


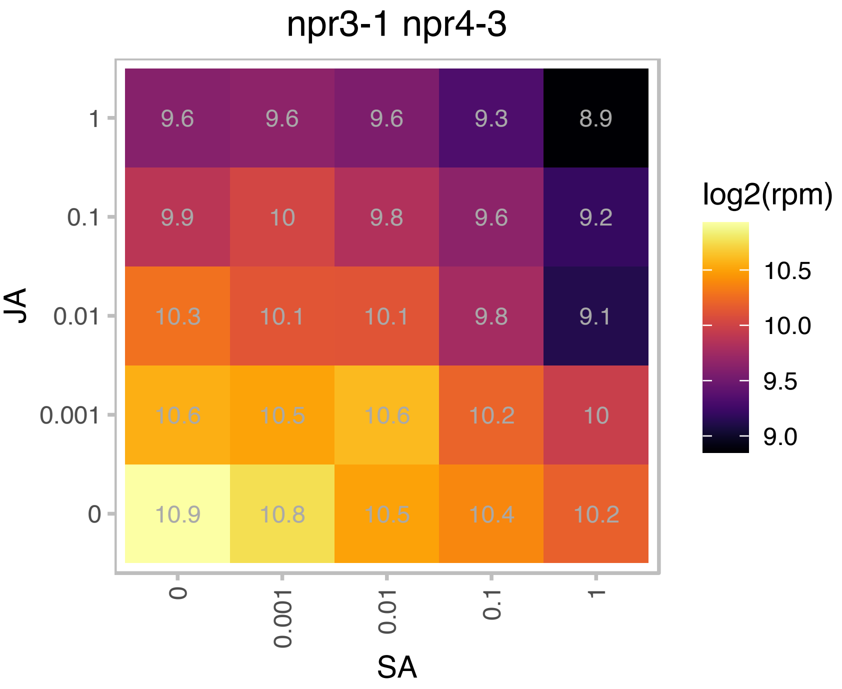

### AT2G47910.1 : CRR6

chlororespiratory reduction 6

Encodes a chloroplast thylakoid membrane protein. Required for the assembly/accumulation of the NAD(P)H dehydrogenase complex of the photosynthetic electron transport chain.

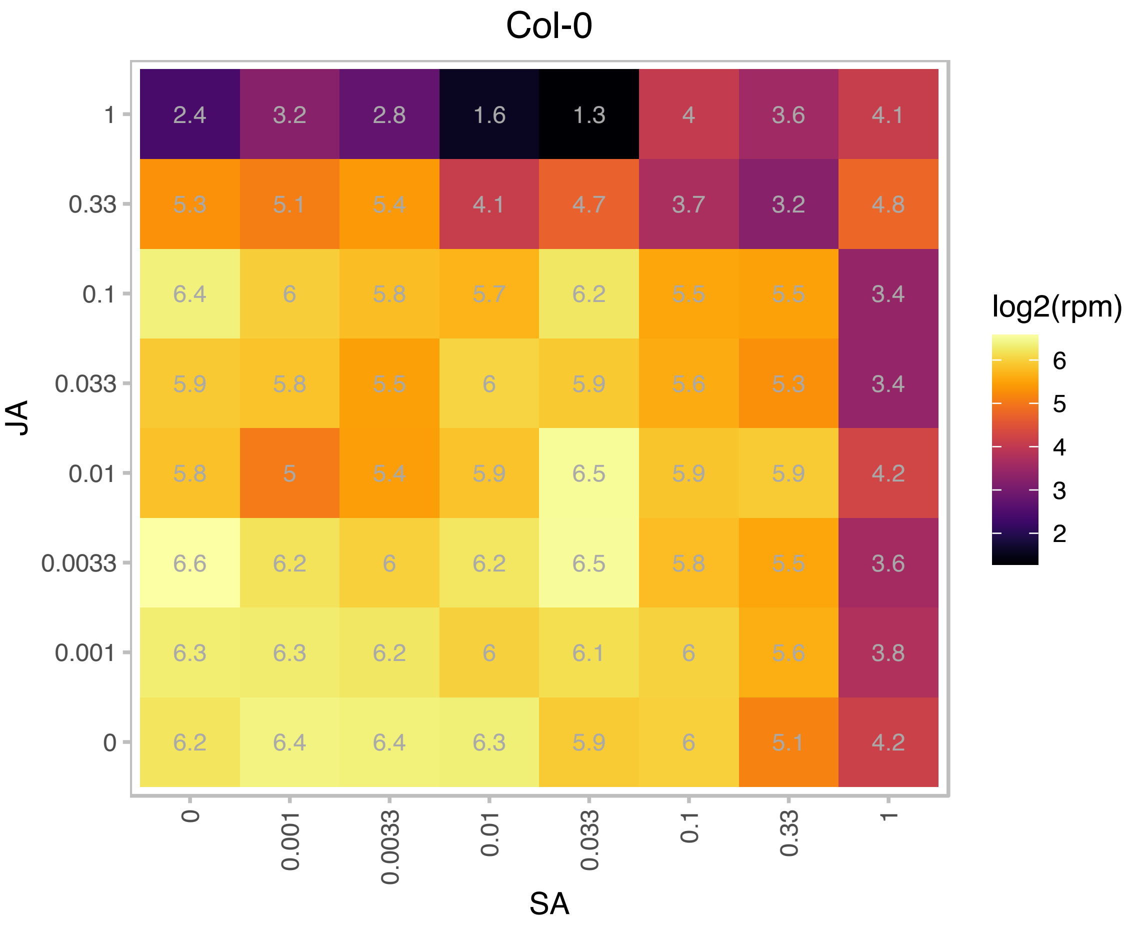


| | |
|---|---|
| Gene | AT2G47910 |
| is.expressed_genes | TRUE |
| is.responsive_genes | TRUE |
| is.apcluster | F-10 |

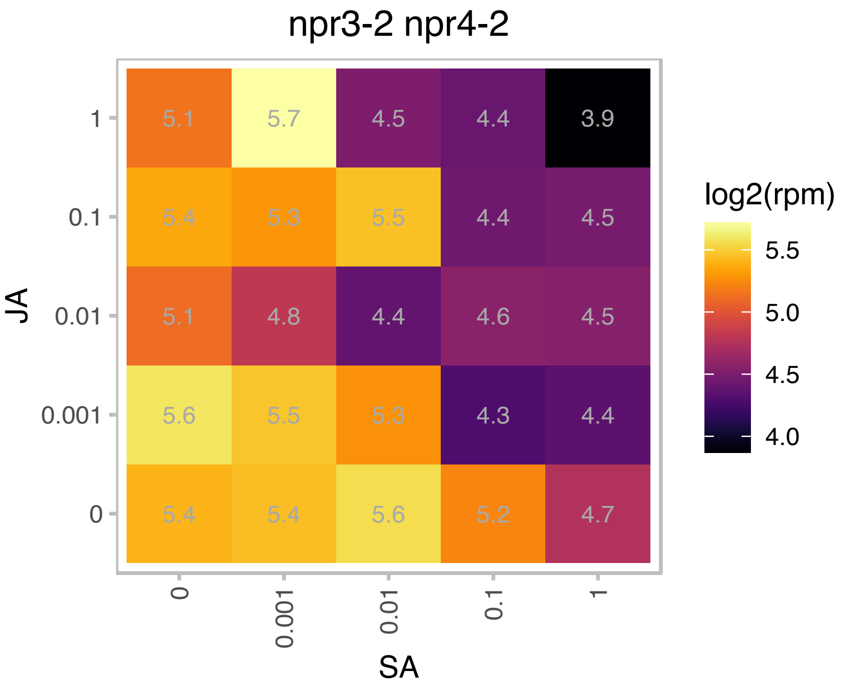


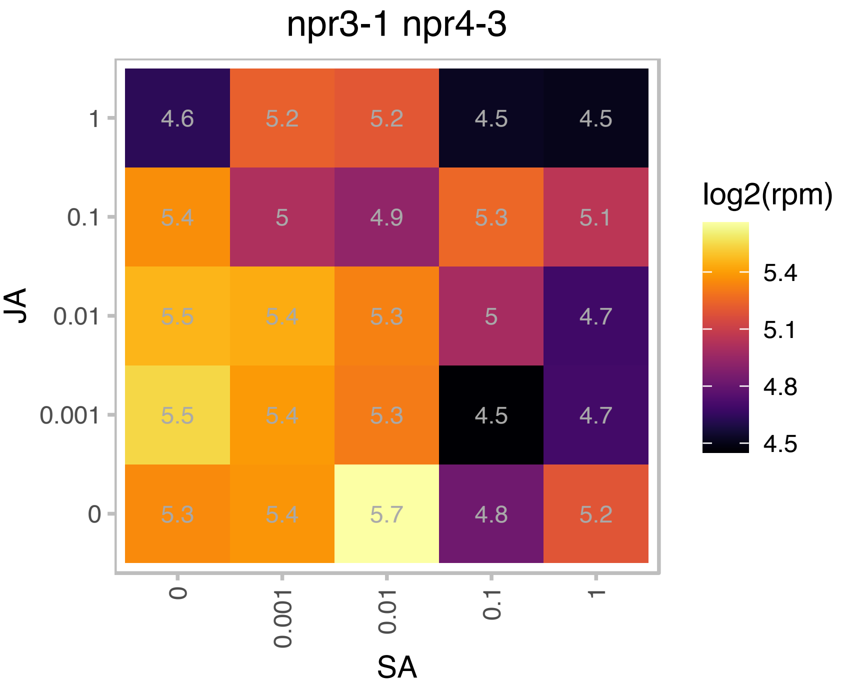

# AT3G01690.1 : NA

alpha/beta-Hydrolases superfamily protein

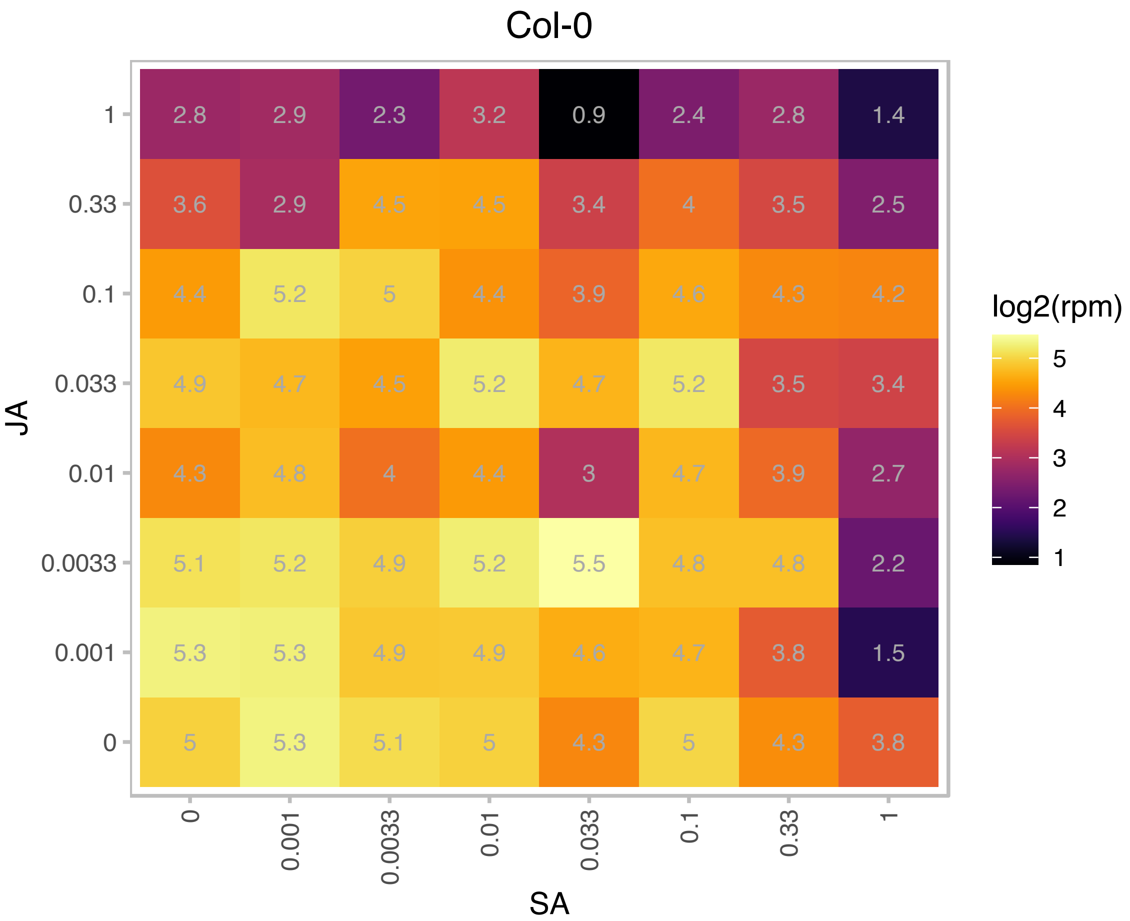


| | |
|---|---|
| Gene | AT3G01690 |
| is.expressed_genes | TRUE |
| is.responsive_genes | TRUE |
| is.apcluster | F-10 |

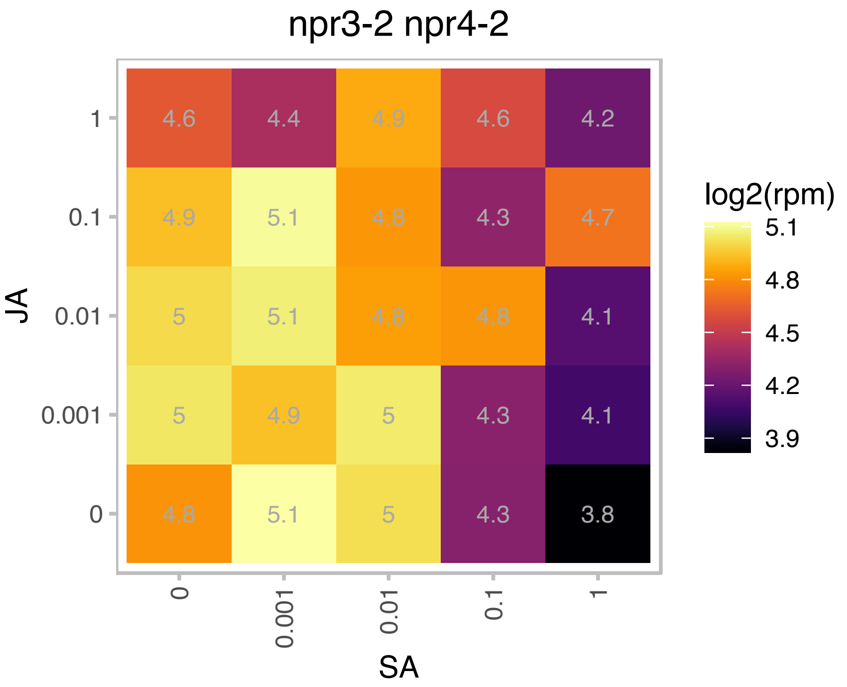


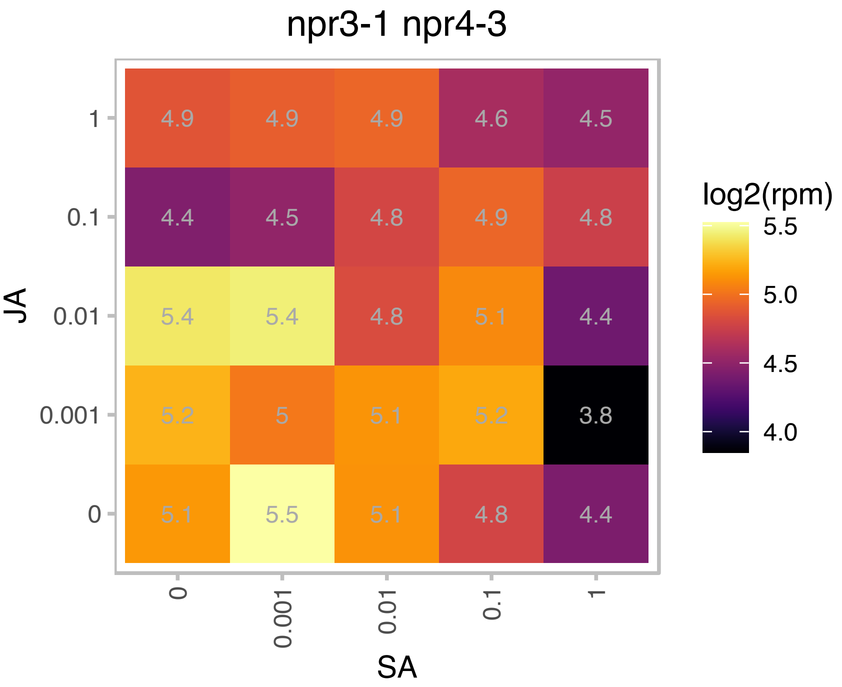

# AT3G10405.1 : NA

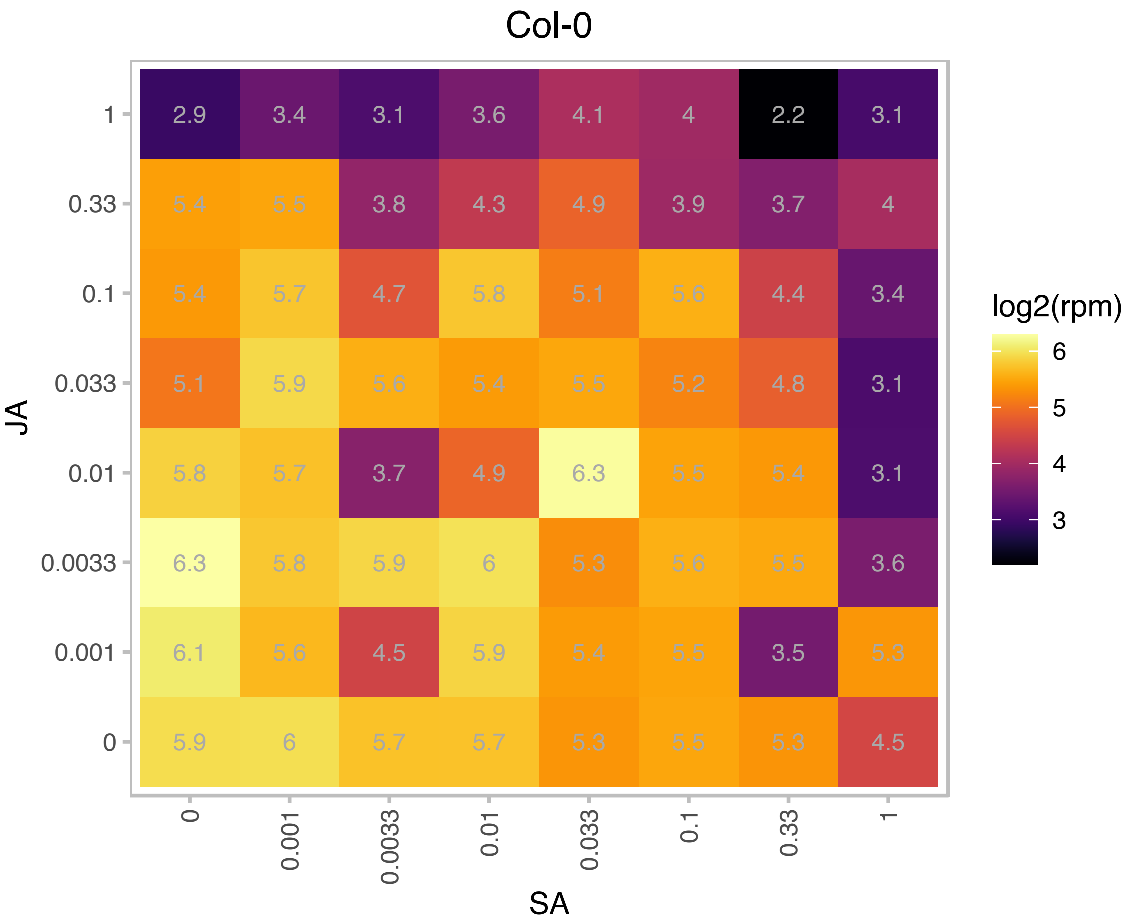


| | |
|---|---|
| Gene | AT3G10405 |
| is.expressed_genes | TRUE |
| is.responsive_genes | TRUE |
| is.apcluster | F-10 |

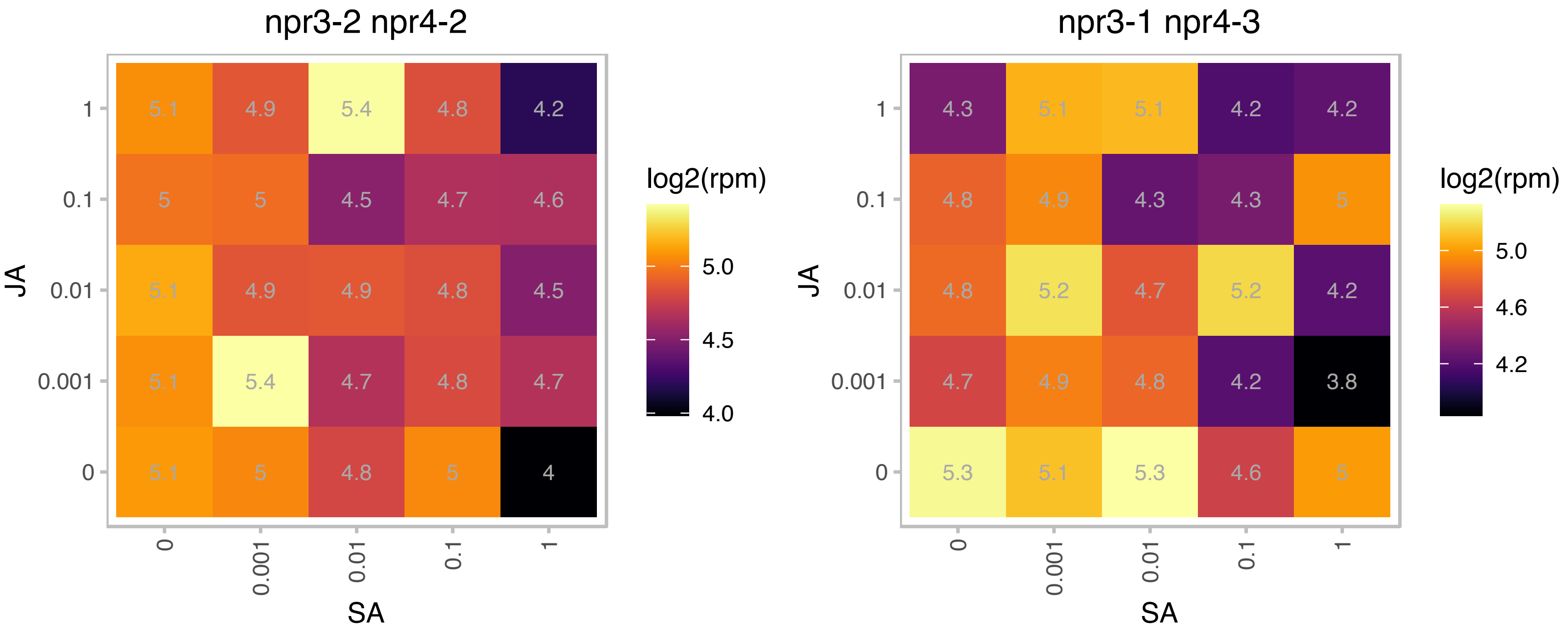

# AT3G12210.1 : NA

DNA binding

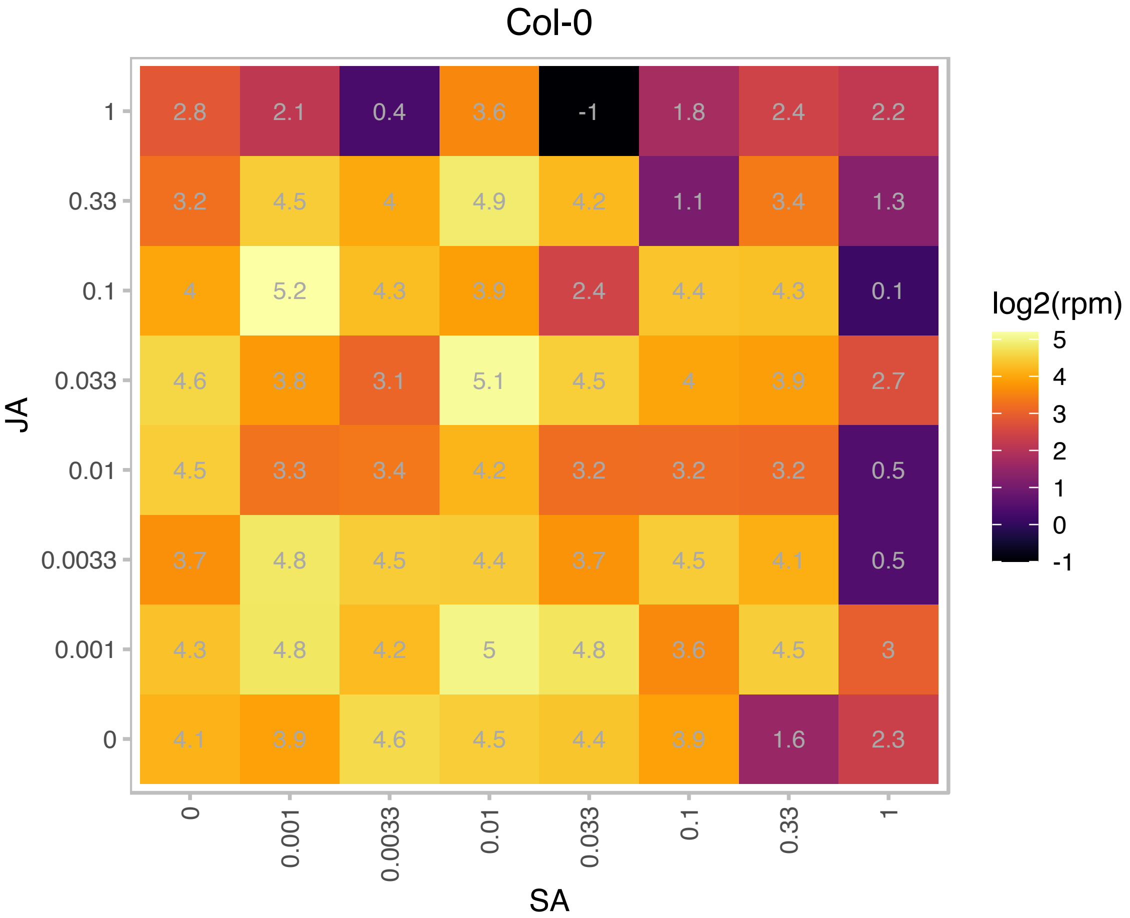


| | |
|---|---|
| Gene | AT3G12210 |
| is.expressed_genes | TRUE |
| is.responsive_genes | TRUE |
| is.apcluster | F-10 |

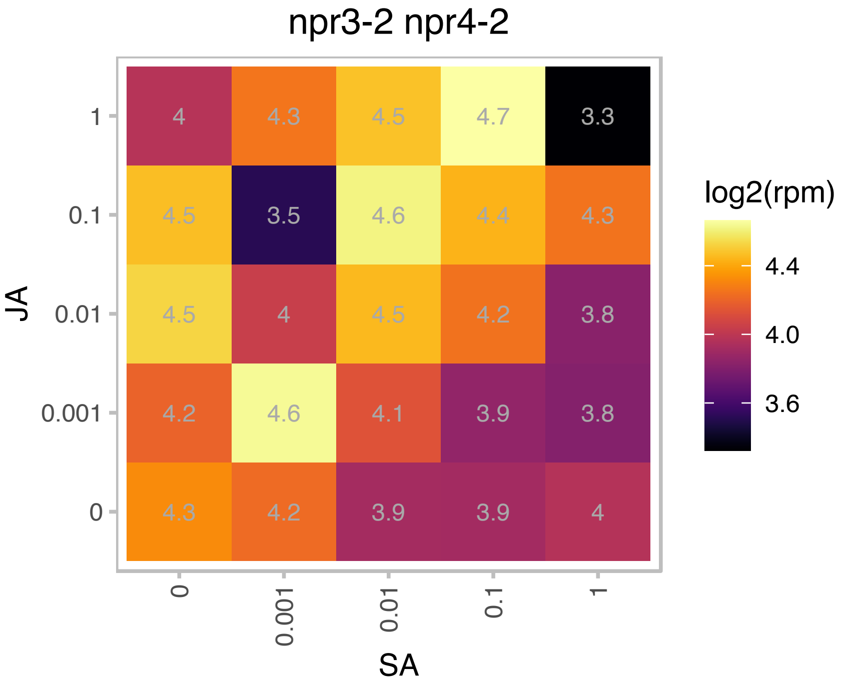


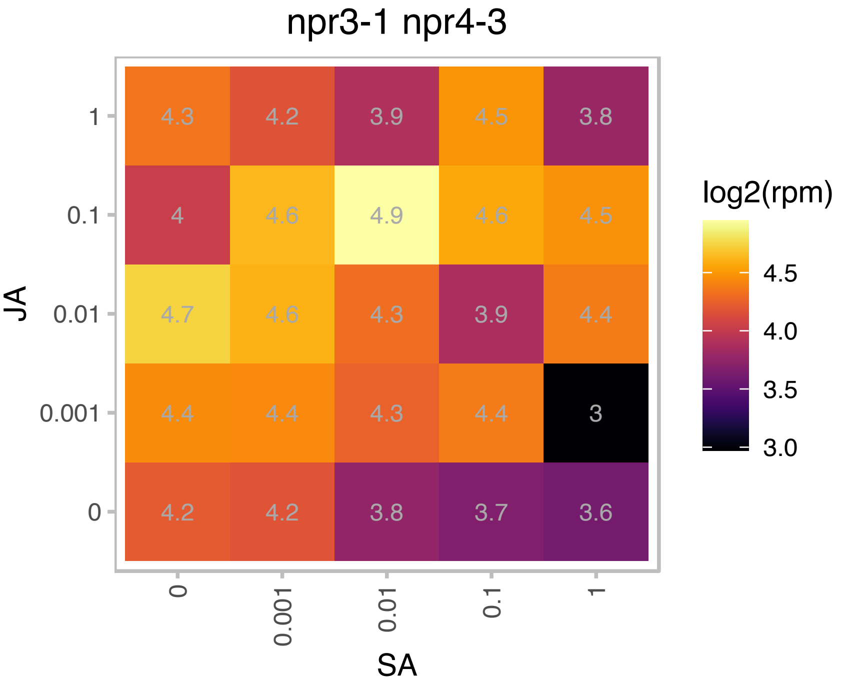

### AT3G13120.1 : PRPS10

Ribosomal protein S10p/S20e family protein

#### Col-0

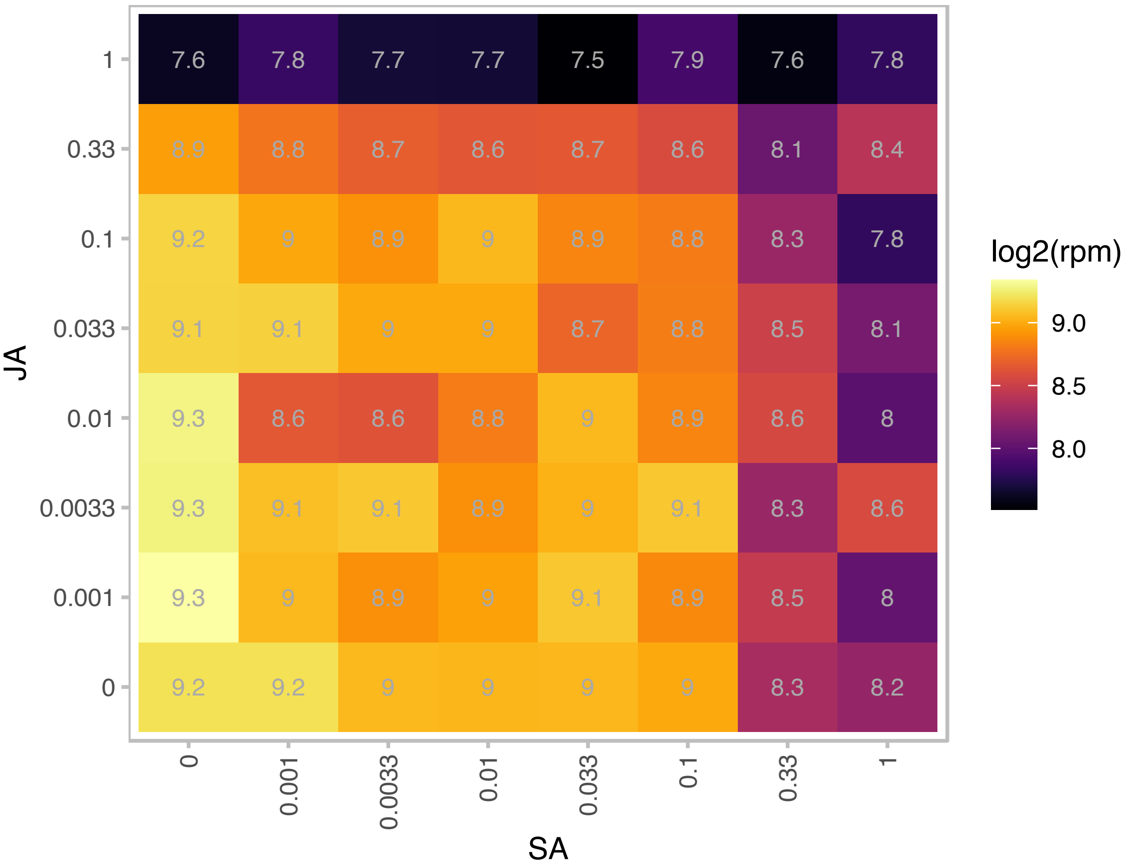


| | |
|---|---|
| Gene | AT3G13120 |
| is.expressed_genes | TRUE |
| is.responsive_genes | TRUE |
| is.apcluster | F-10 |

#### npr3-2 npr4-2

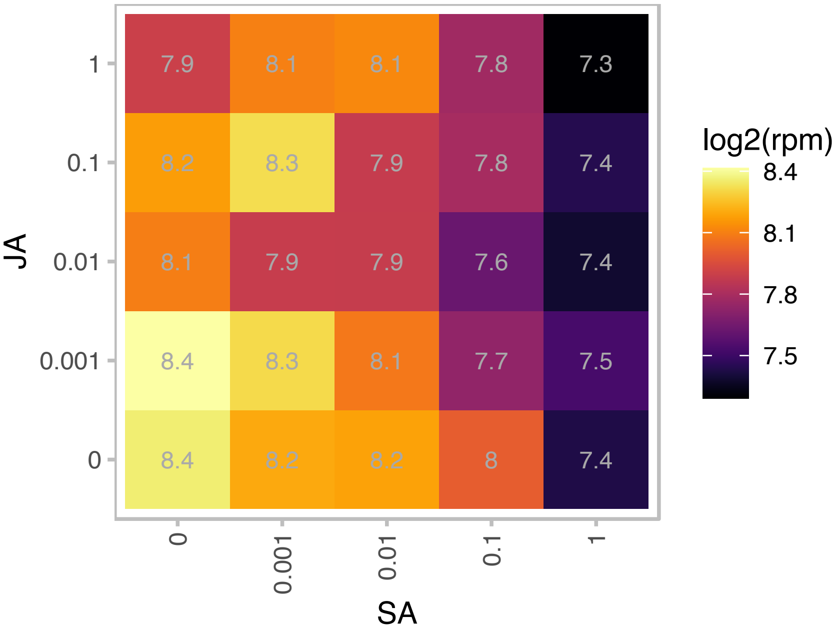


#### npr3-1 npr4-3

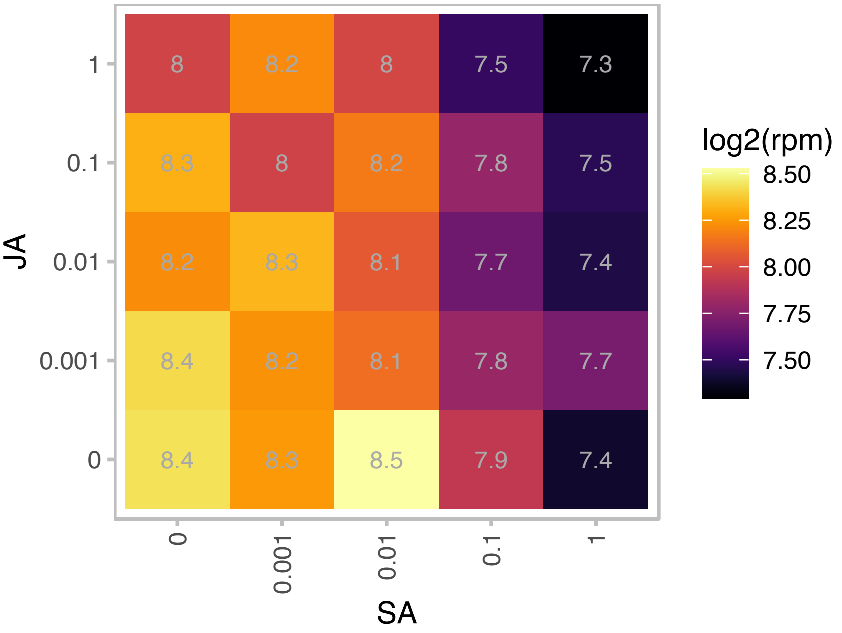

# AT3G13677.1 : NA

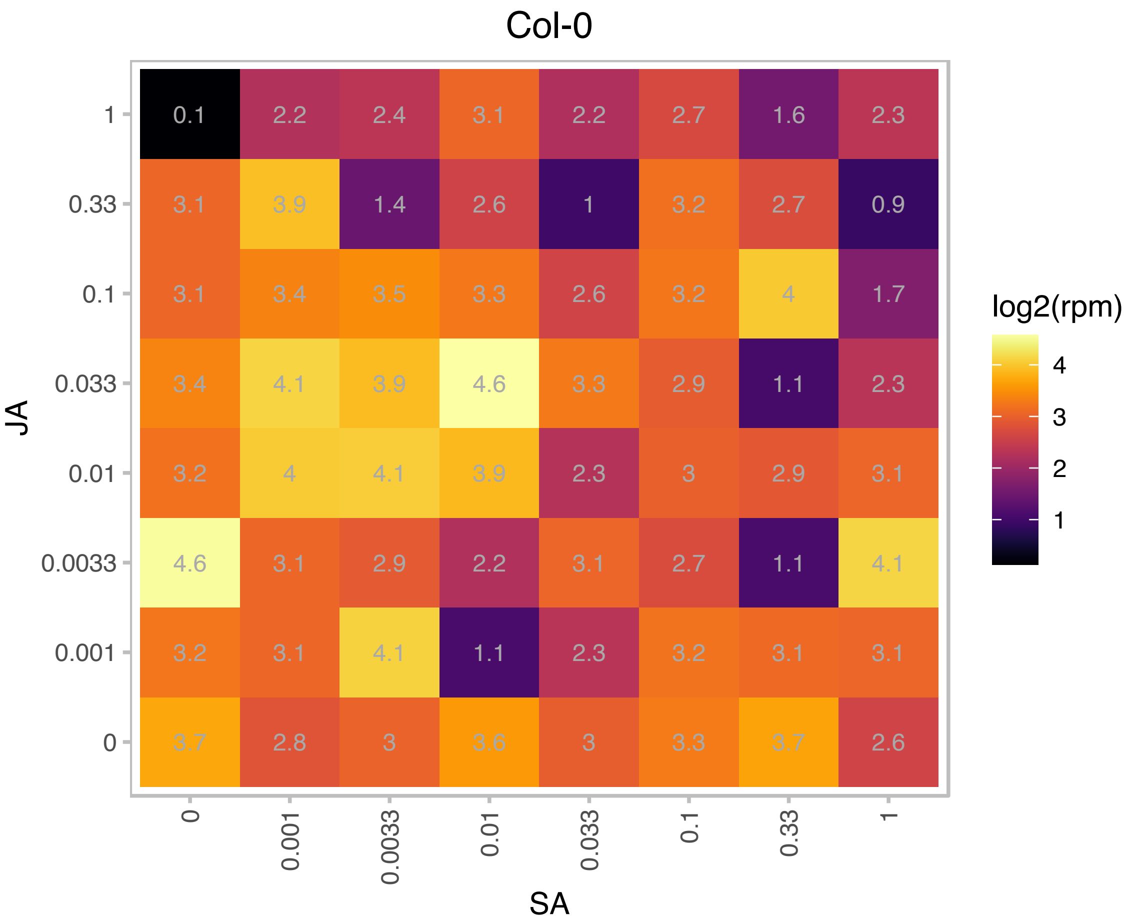


| | |
|---|---|
| Gene | AT3G13677 |
| is.expressed_genes | TRUE |
| is.responsive_genes | TRUE |
| is.apcluster | F-10 |

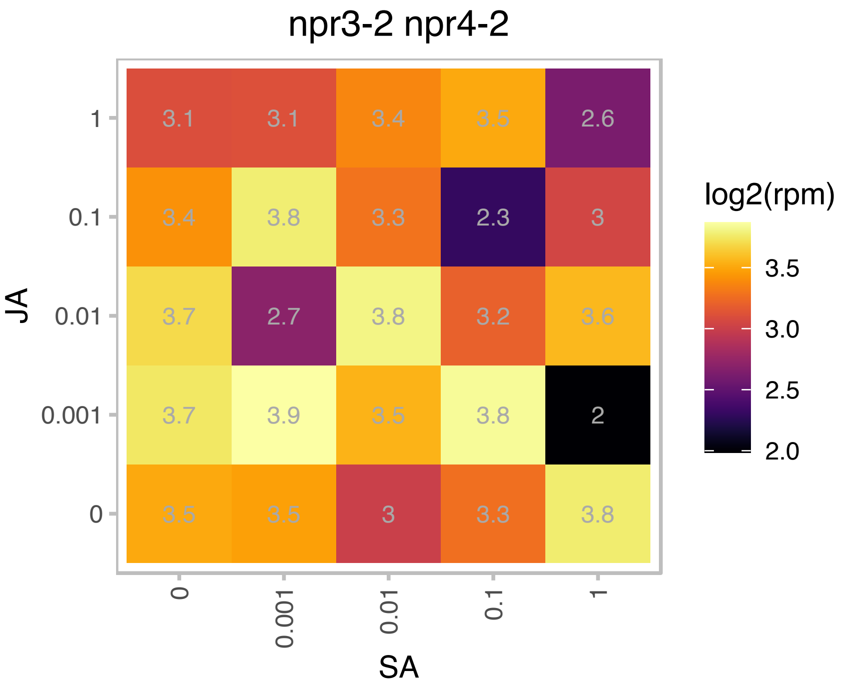


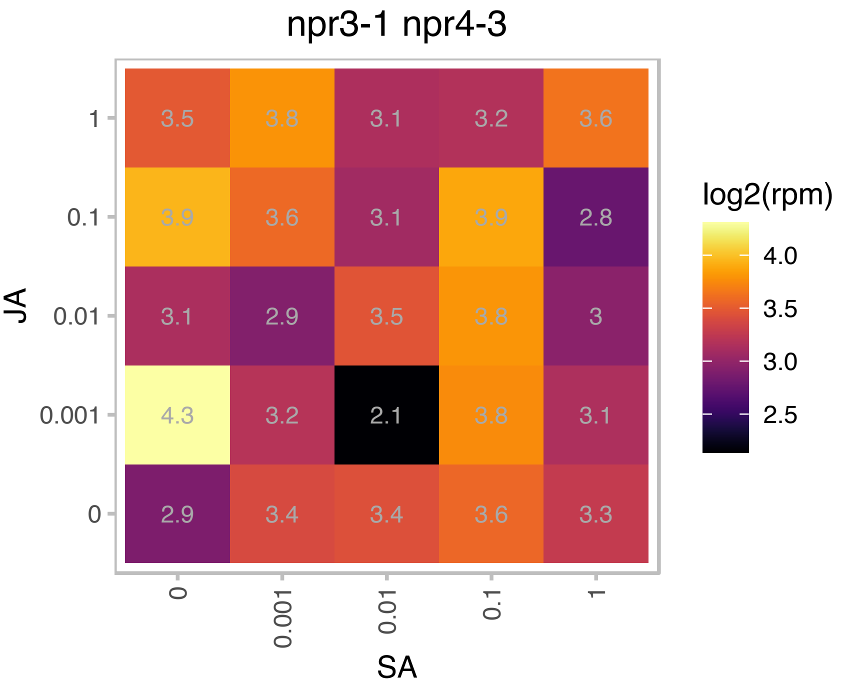

# AT3G15630.1 : NA

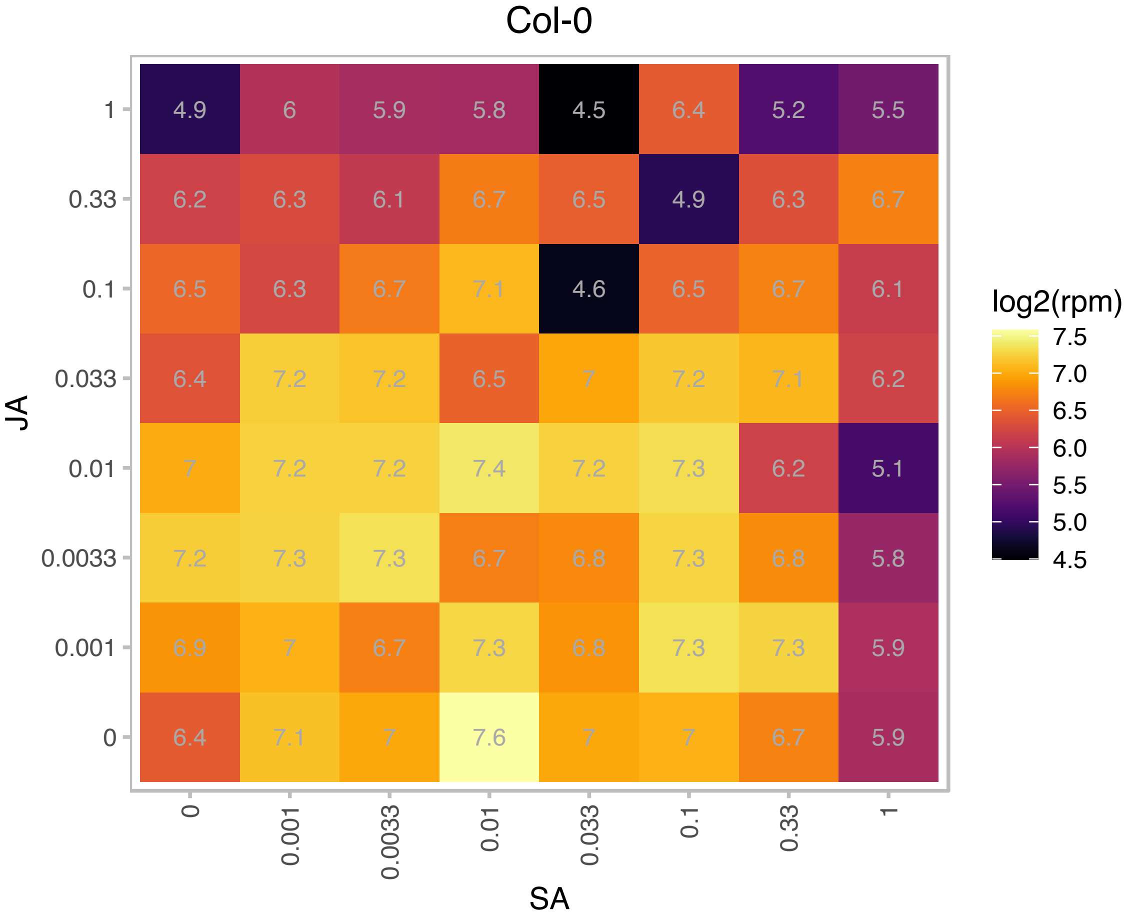


| | |
|---|---|
| Gene | AT3G15630 |
| is.expressed_genes | TRUE |
| is.responsive_genes | TRUE |
| is.apcluster | F-10 |

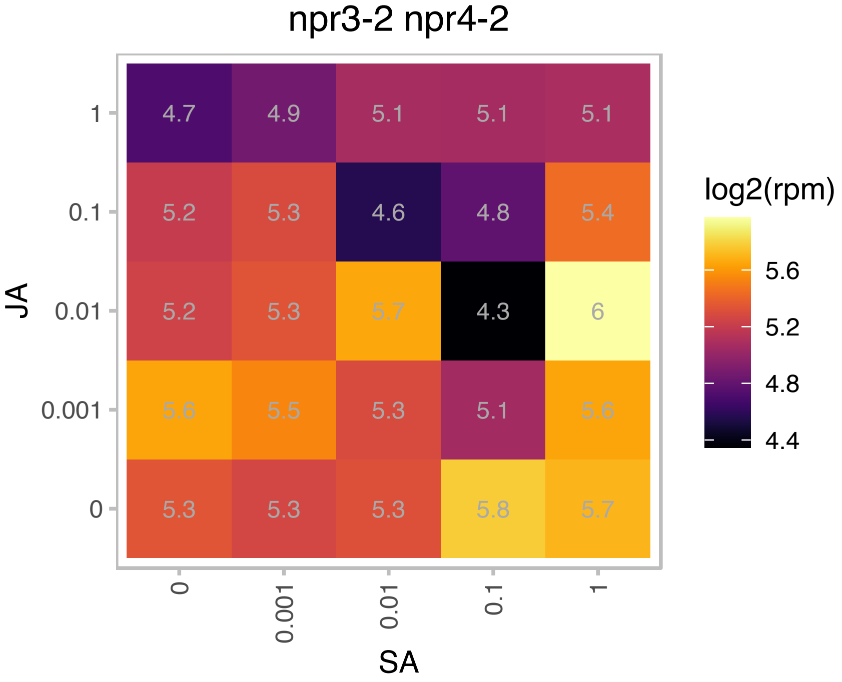


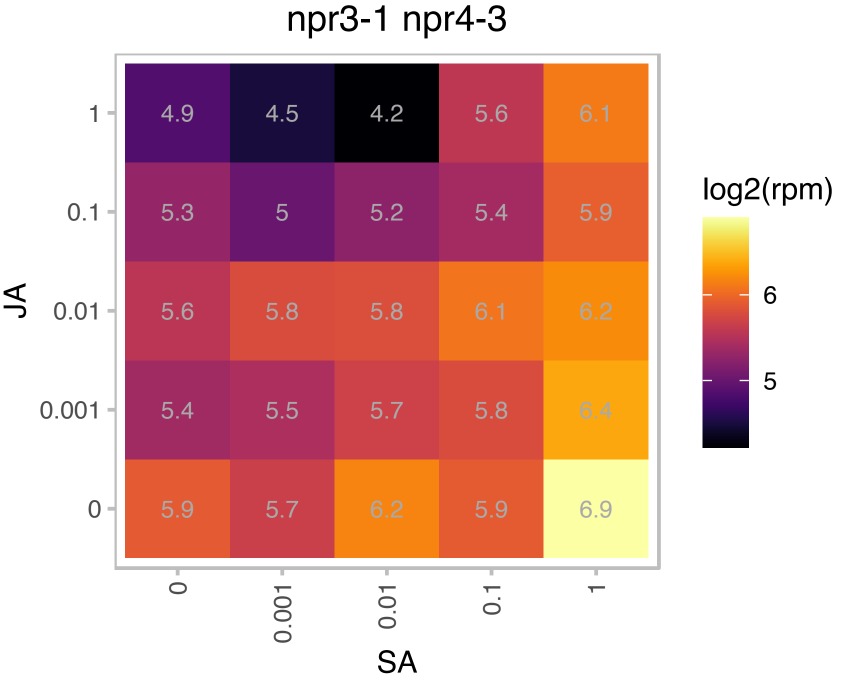

### AT3G17170.1 : RFC3

Translation elongation factor EF1B/ribosomal protein S6 family protein

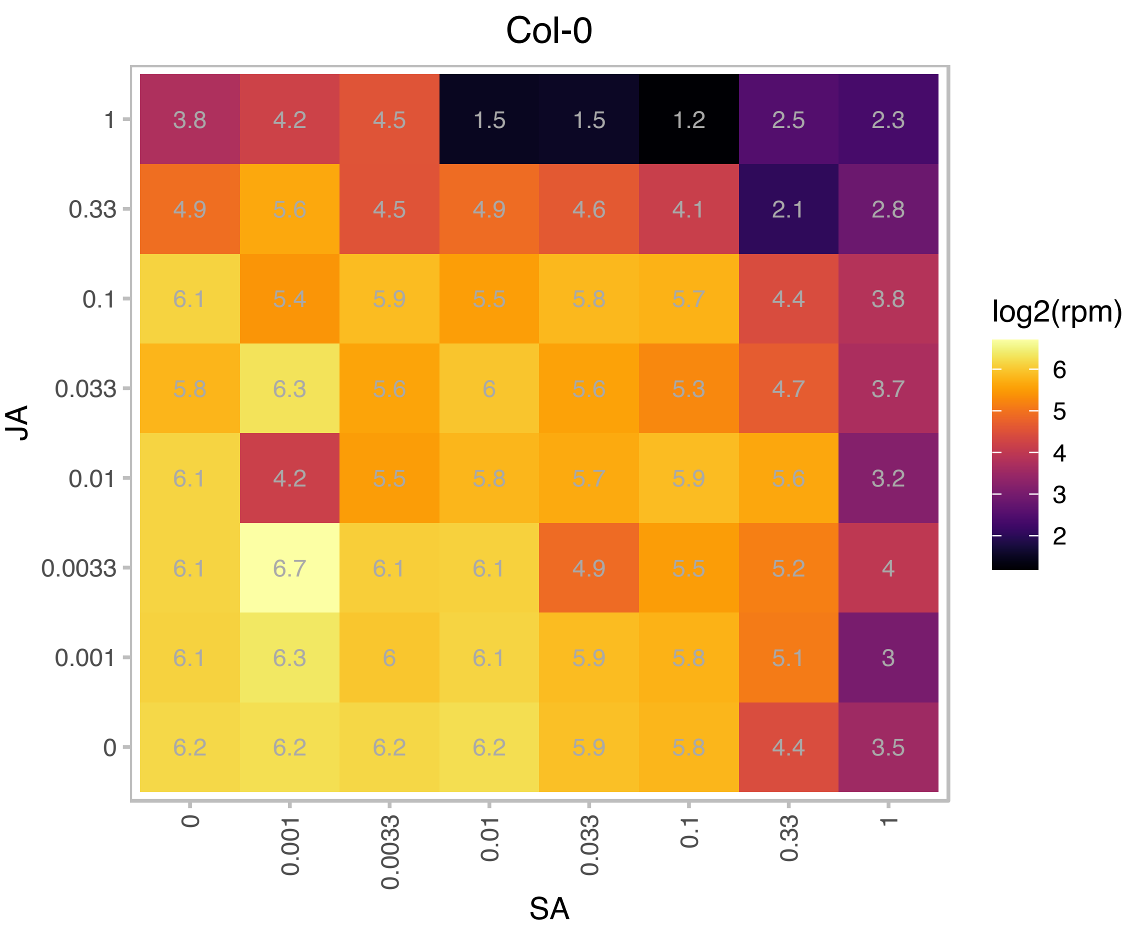


| | |
|---|---|
| Gene | AT3G17170 |
| is.expressed_genes | TRUE |
| is.responsive_genes | TRUE |
| is.apcluster | F-10 |

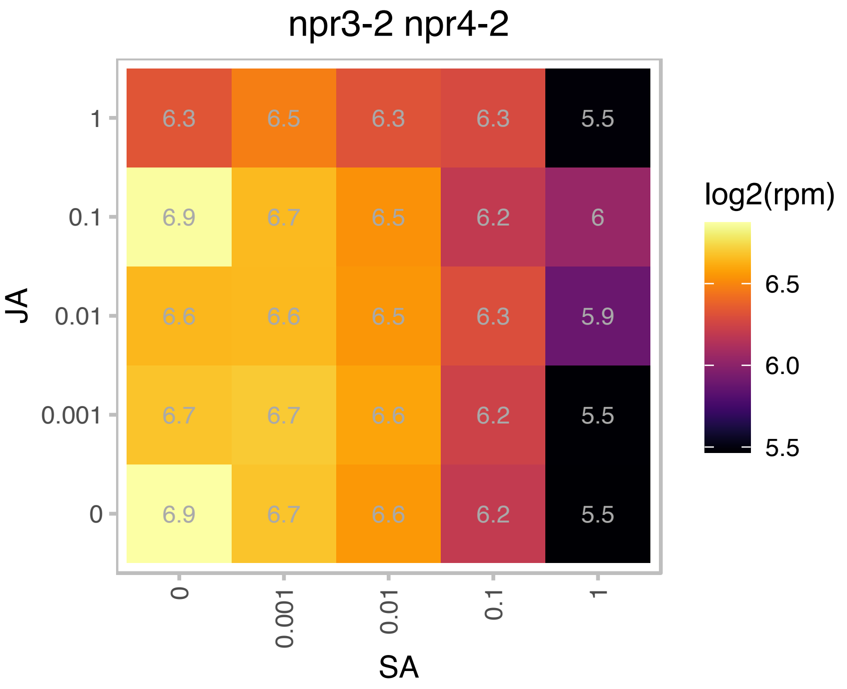


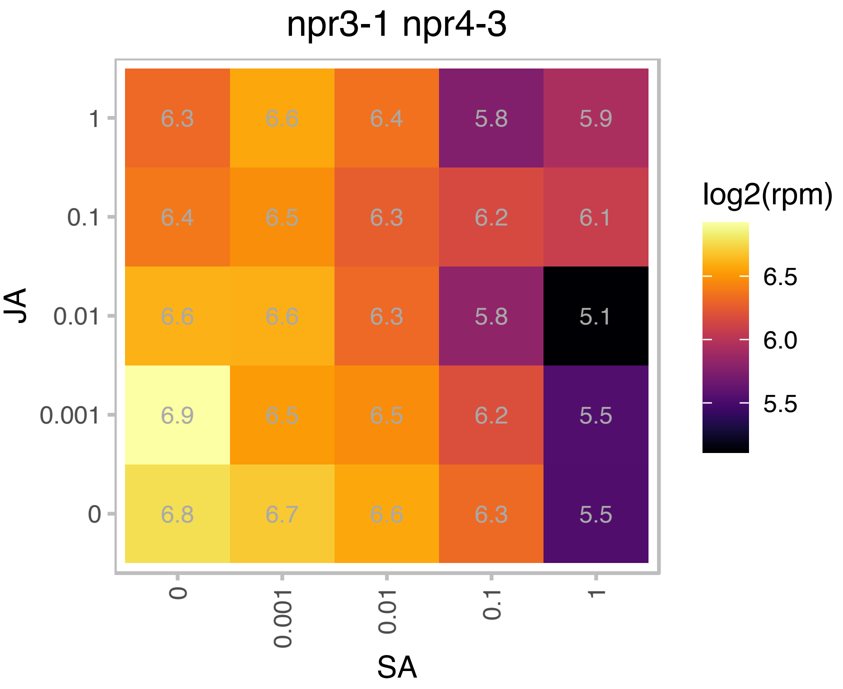

### AT3G18000.1 : AtPMT1, NMT1, PEAMT, XPL1

S-adenosyl-L-methionine-dependent methyltransferases superfamily protein

Arabidopsis thaliana N-methyltransferase-like protein mRNA. Reduce transmission through pollen.

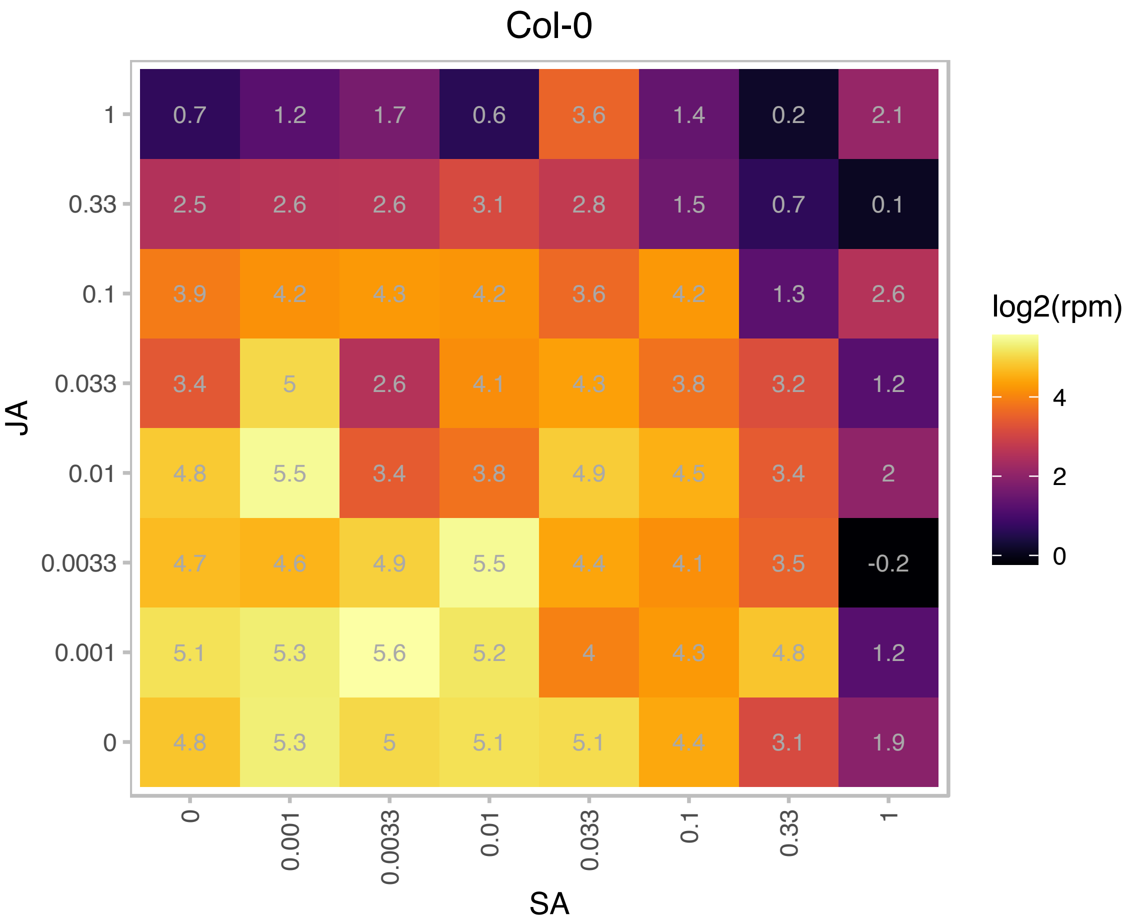


| | |
|---|---|
| Gene | AT3G18000 |
| is.expressed_genes | TRUE |
| is.responsive_genes | TRUE |
| is.apcluster | F-10 |

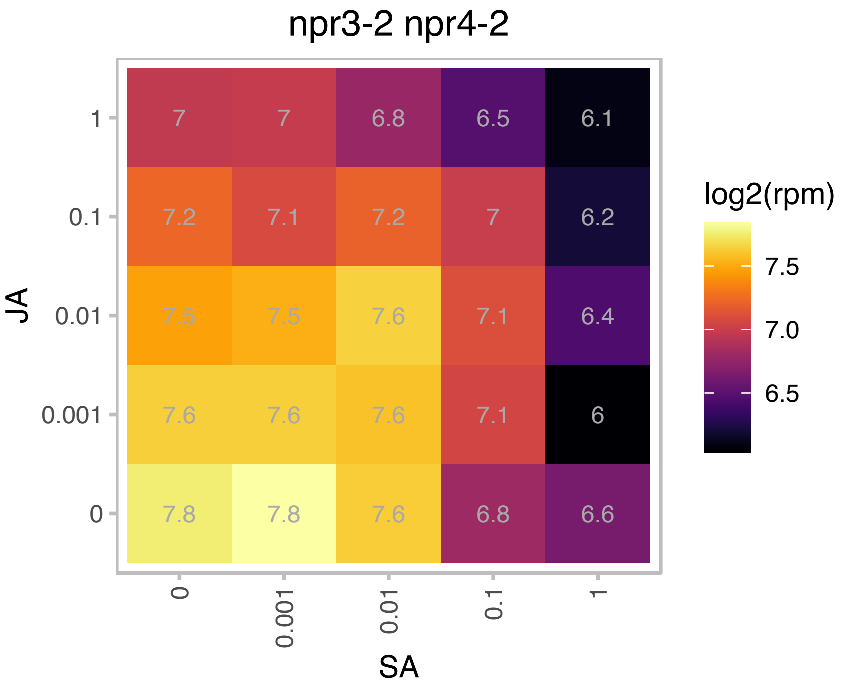


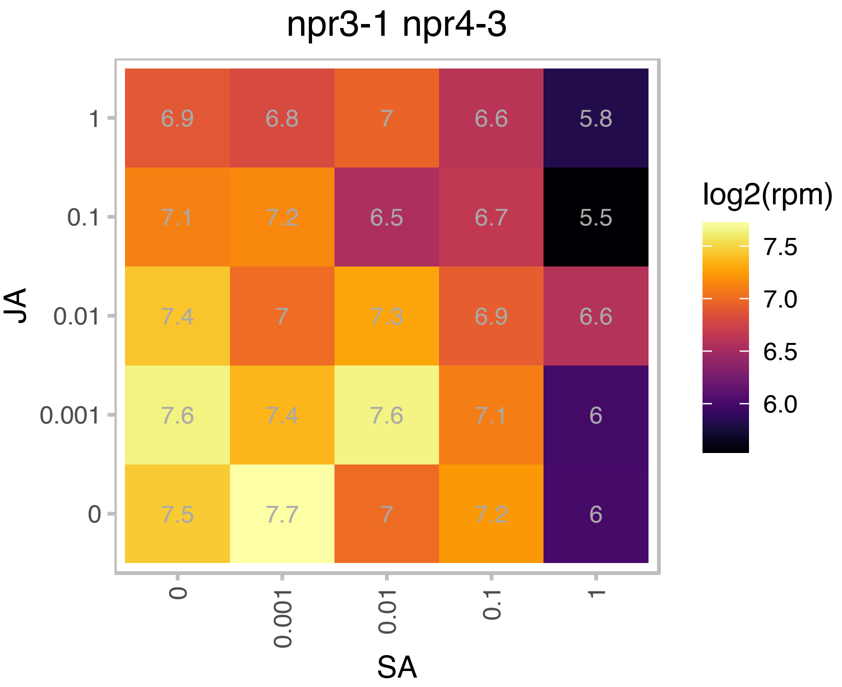

### AT3G19450.1 : ATCAD4, CAD, CAD-C, CAD4

GroES-like zinc-binding alcohol dehydrogenase family protein
Encodes a catalytically active cinnamyl alcohol dehydrogenase which uses p-coumaryl aldehyde as a preferred substrate. It can also use caffeyl, coniferyl and d-hydroxyconiferyl aldehydes as substrates. The mRNA is cell-to-cell mobile.

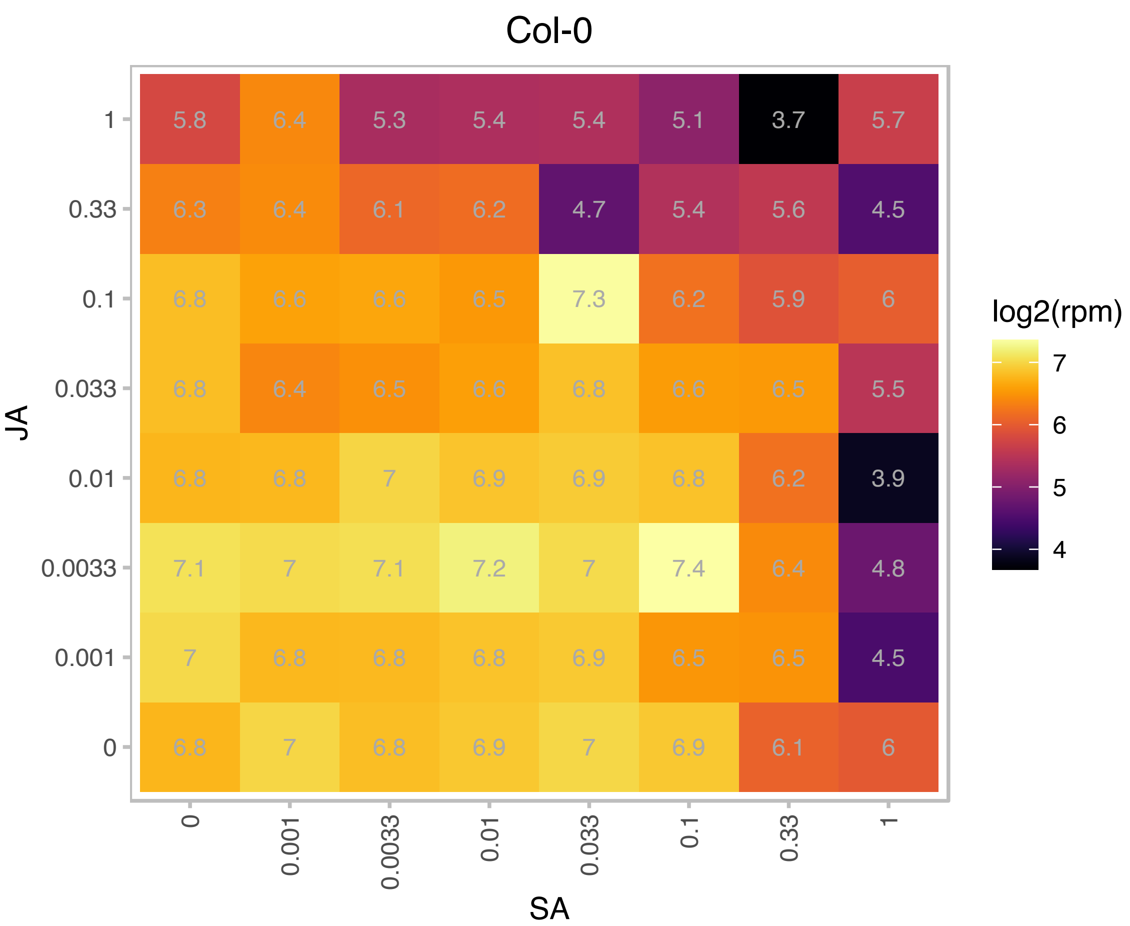


| | |
|---|---|
| Gene | AT3G19450 |
| is.expressed_genes | TRUE |
| is.responsive_genes | TRUE |
| is.apcluster | F-10 |

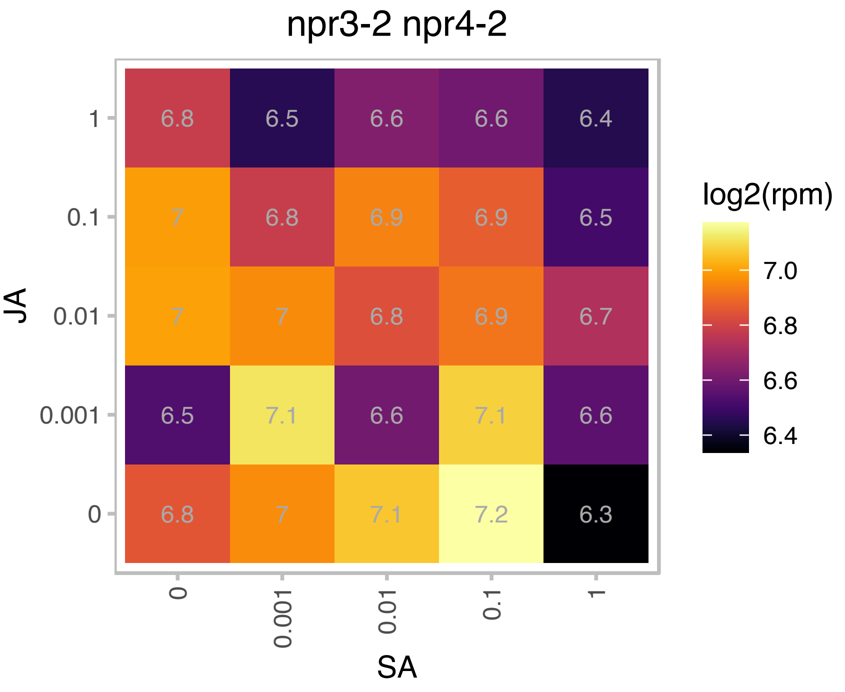


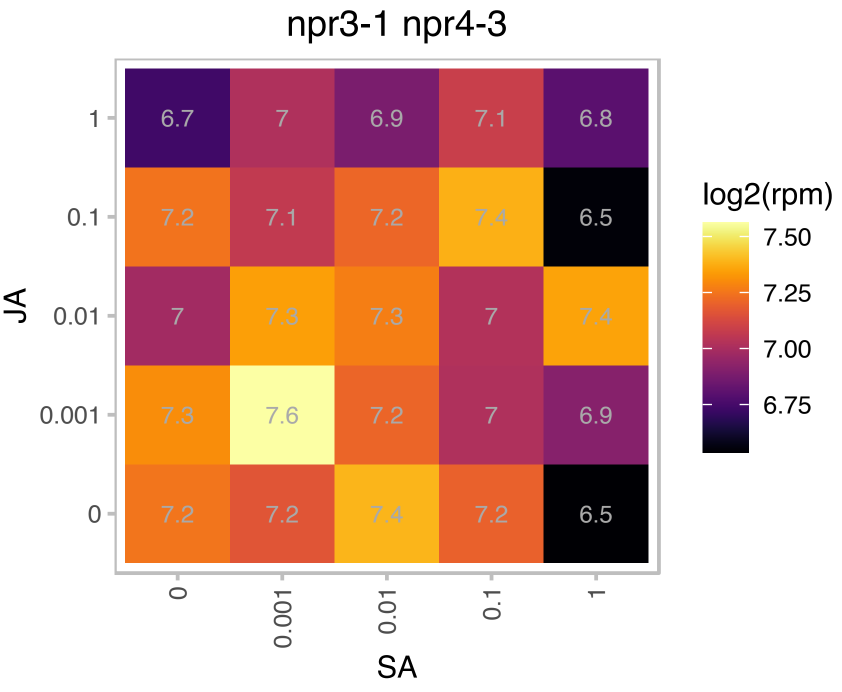

### AT3G20670.1 : HTA13

histone H2A 13

Encodes HTA13, a histone H2A protein.

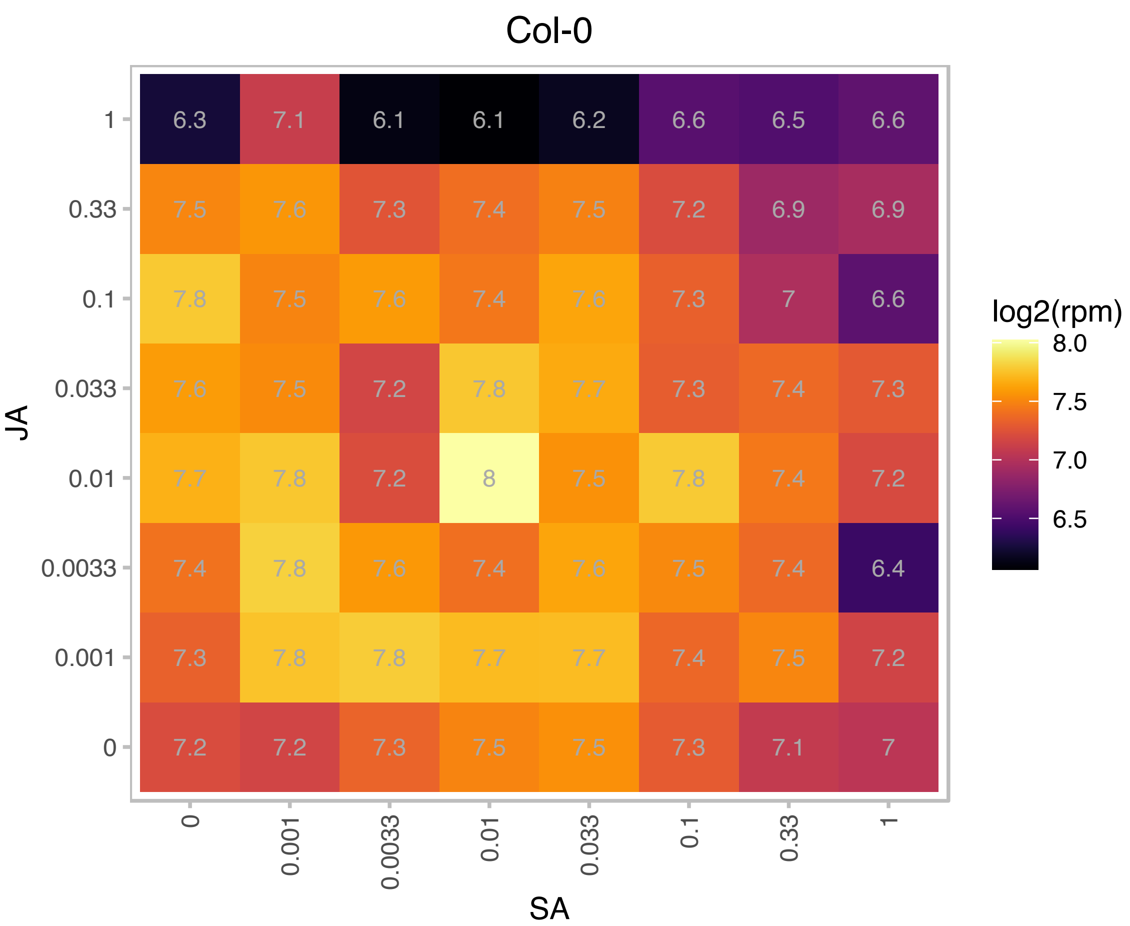


| | |
|---|---|
| Gene | AT3G20670 |
| is.expressed_genes | TRUE |
| is.responsive_genes | TRUE |
| is.apcluster | F-10 |

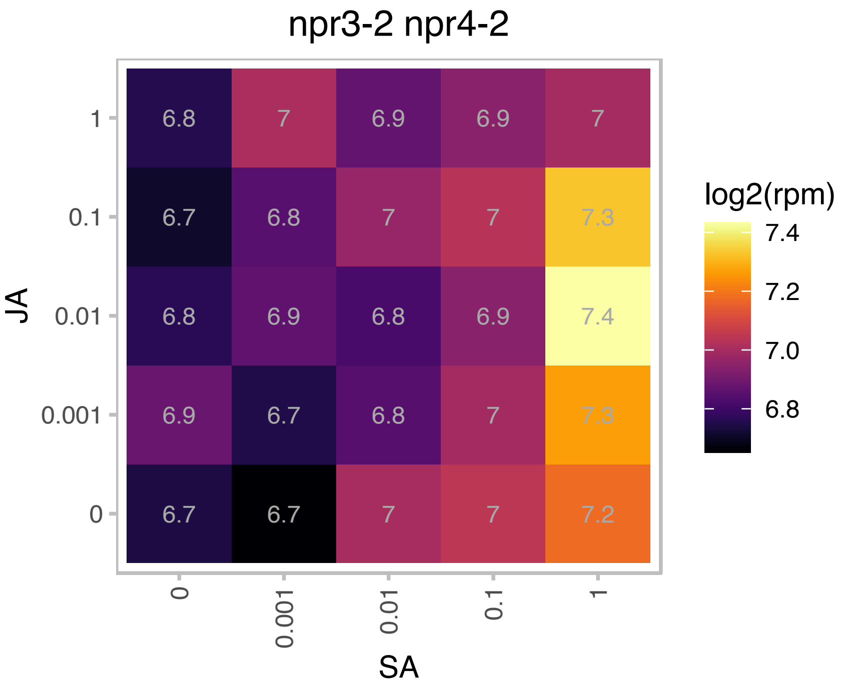


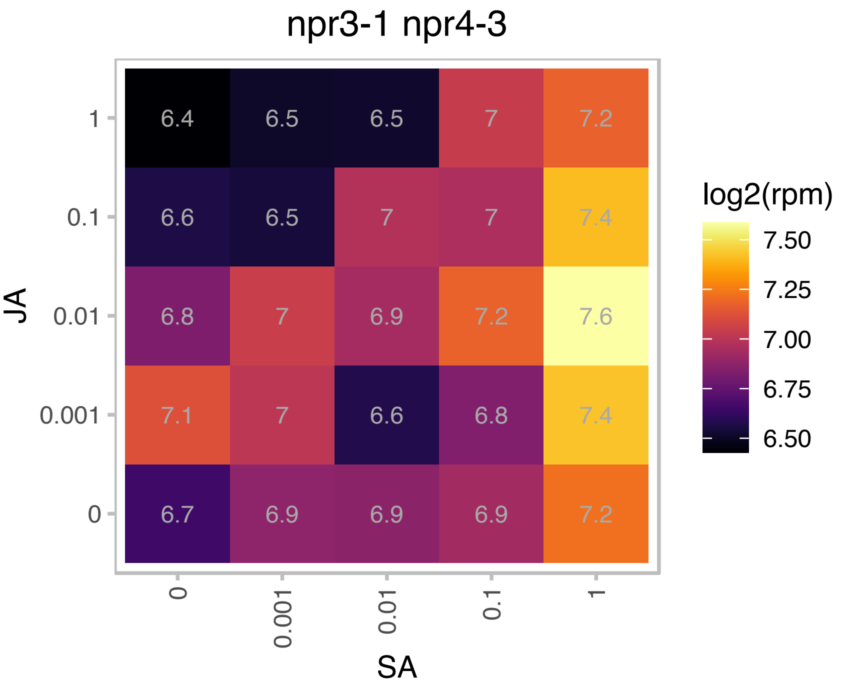

### AT3G27750.1 : EMB3123, THA8

Encodes a pentatricopeptide repeat (PPR) protein required for the splicing of specific group II introns. Null alleles are embryo lethal.

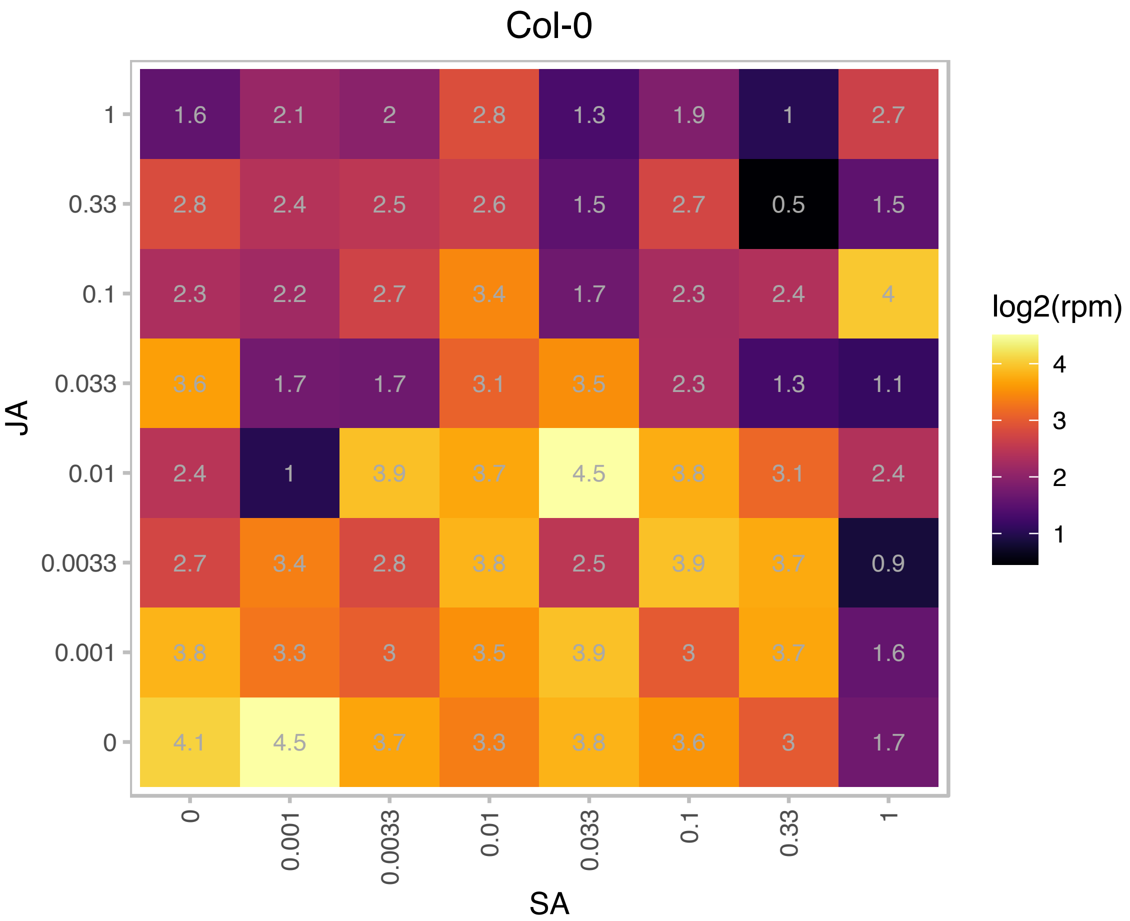


| | |
|---|---|
| Gene | AT3G27750 |
| is.expressed_genes | TRUE |
| is.responsive_genes | TRUE |
| is.apcluster | F-10 |

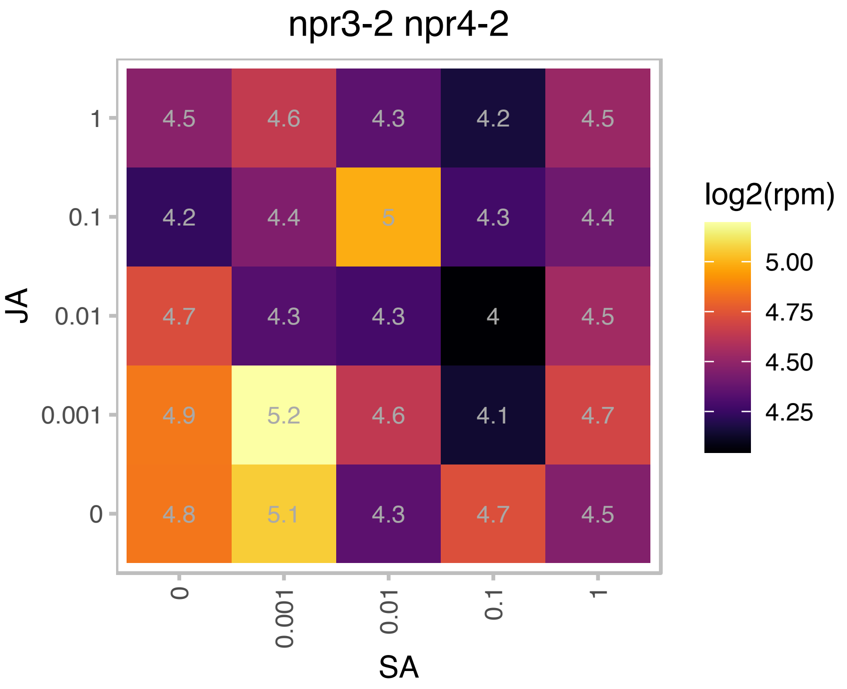


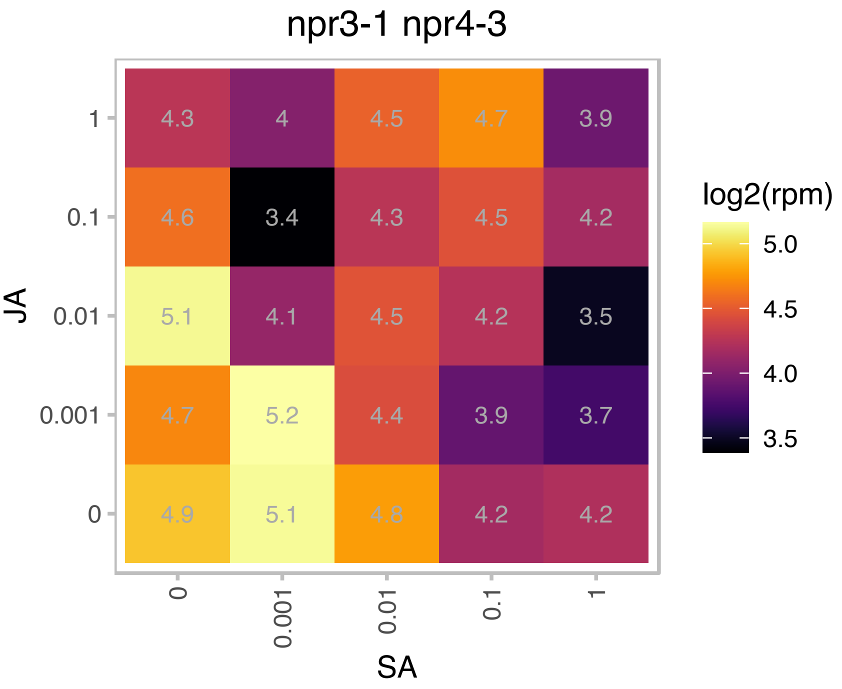

# AT3G43980.1 : NA

Ribosomal protein S14p/S29e family protein

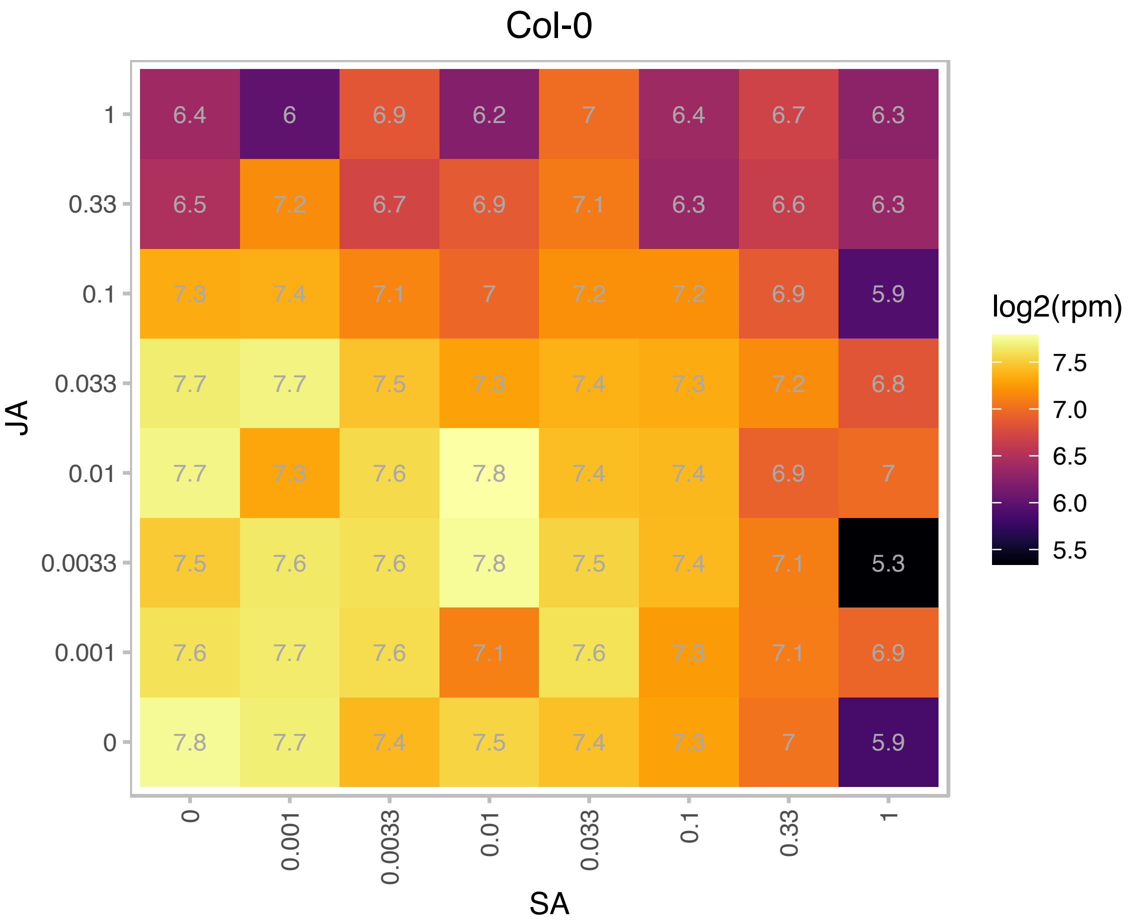


| | |
|---|---|
| Gene | AT3G43980 |
| is.expressed_genes | TRUE |
| is.responsive_genes | TRUE |
| is.apcluster | F-10 |

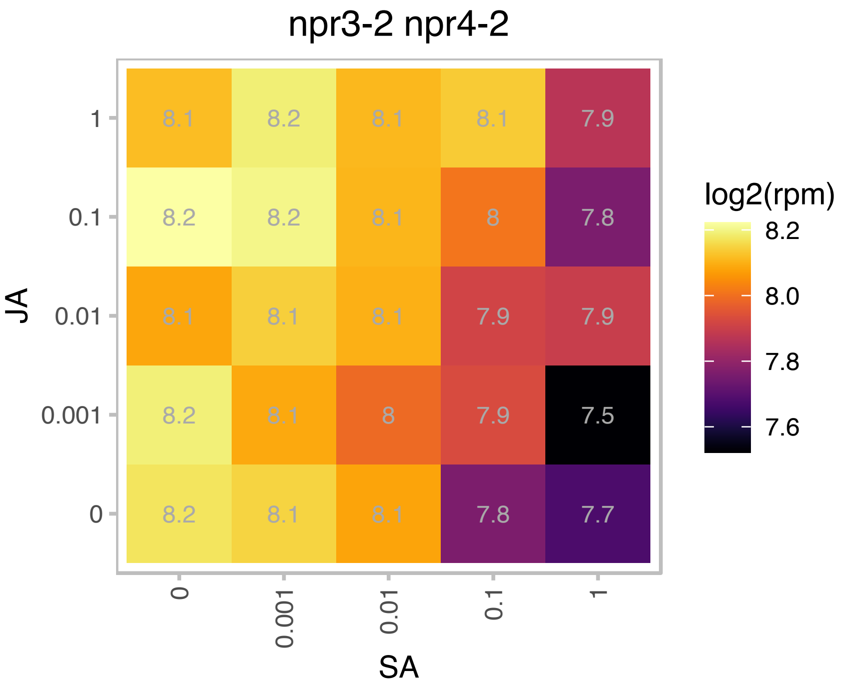


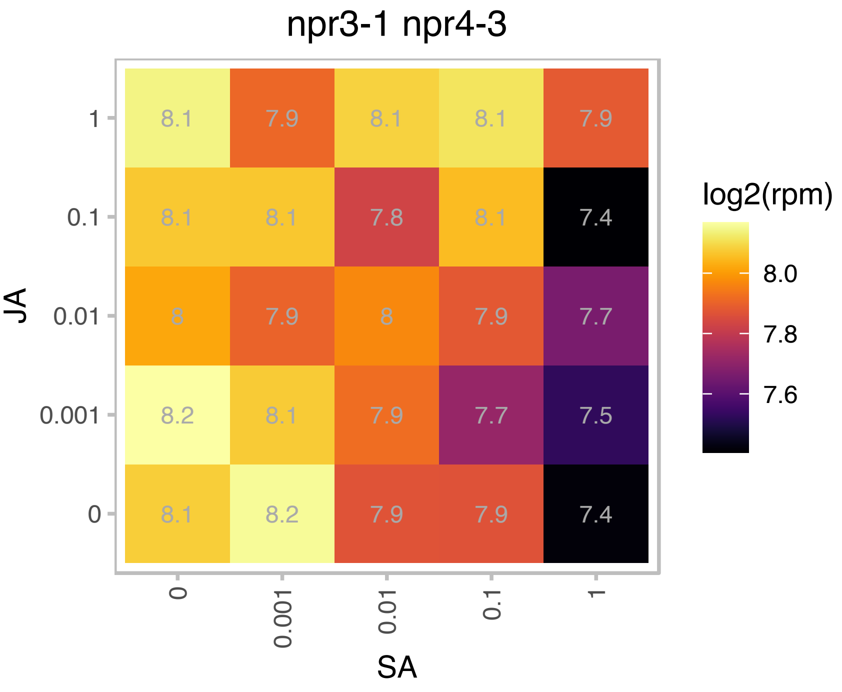

# AT3G44970.1 : NA

Cytochrome P450 superfamily protein

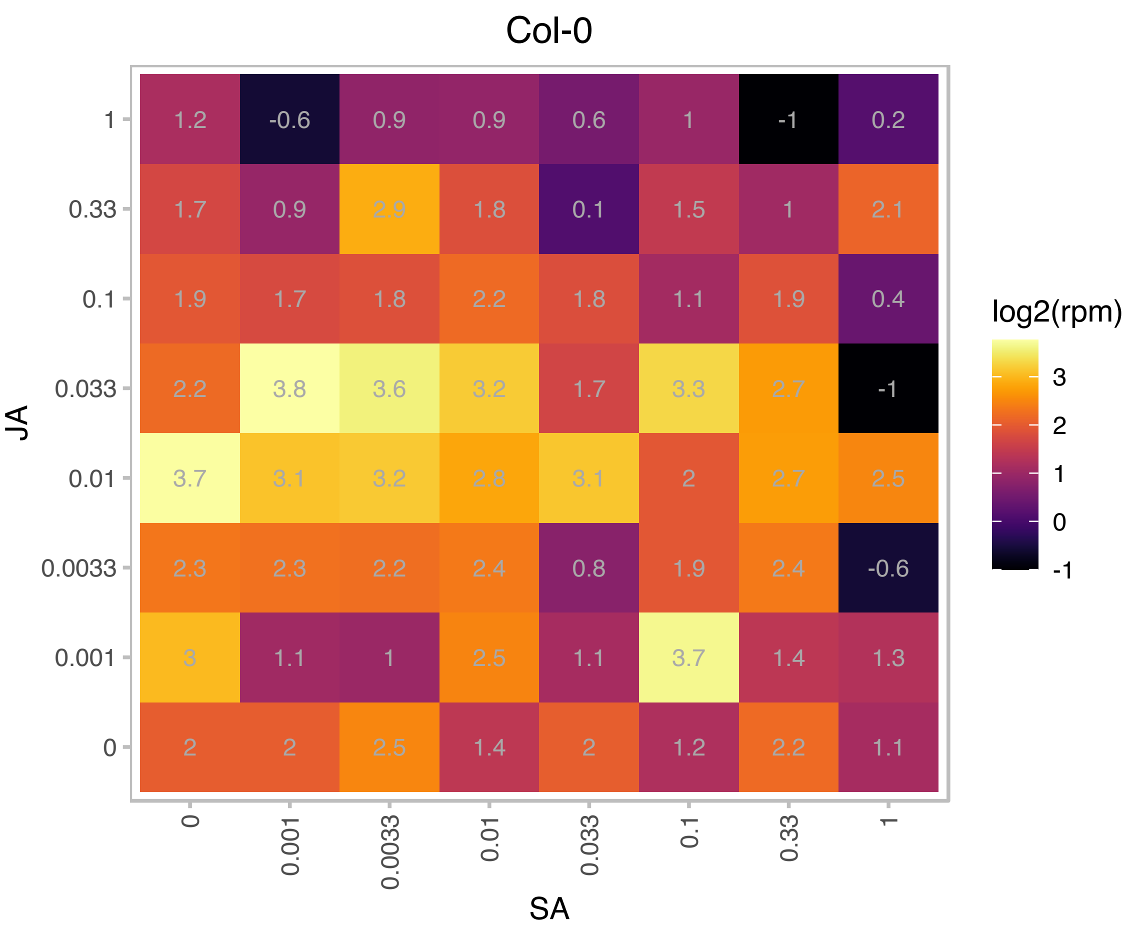


| | |
|---|---|
| Gene | AT3G44970 |
| is.expressed_genes | TRUE |
| is.responsive_genes | TRUE |
| is.apcluster | F-10 |

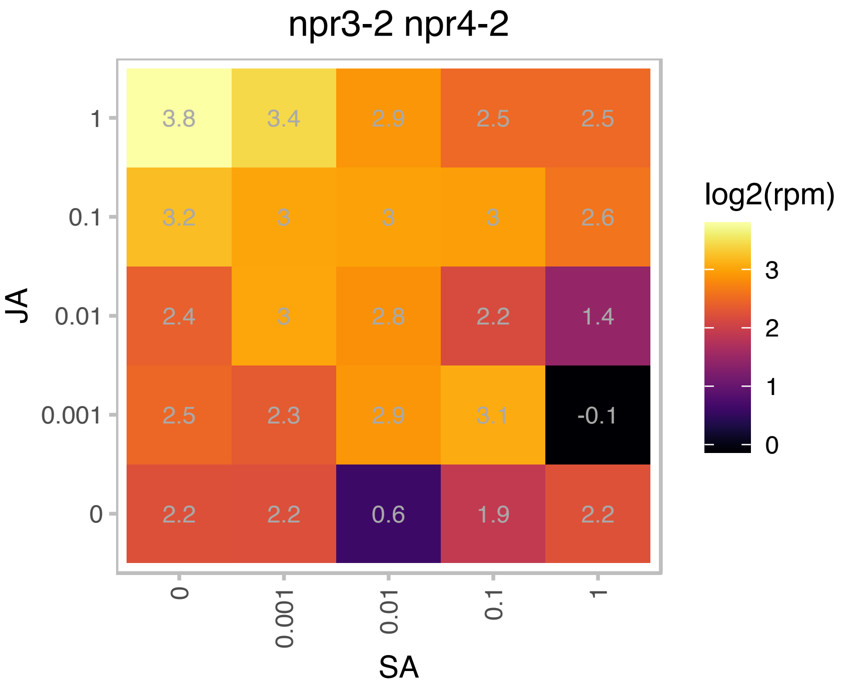


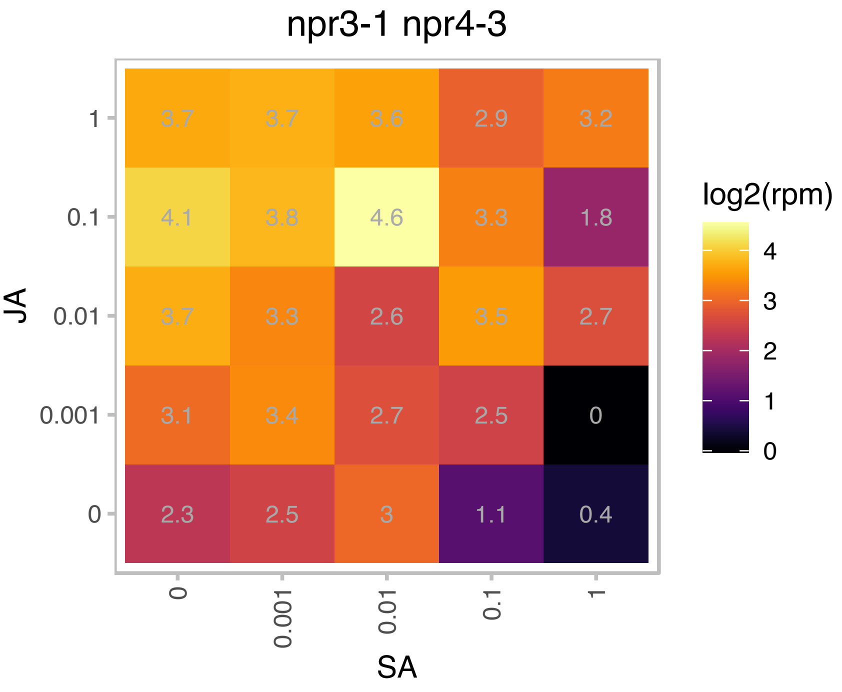

### AT3G48360.1 : ATBT2, BT2

BTB and TAZ domain protein 2
Encodes a protein (BT2) that is an essential component of the TAC1-mediated telomerase activation pathway. Acts redundantly with BT3 and BT1 during female gametophyte development and with BT3 during male gametophyt development. BT2 also mediates multiple responses to nutrients, stresses, and hormones.

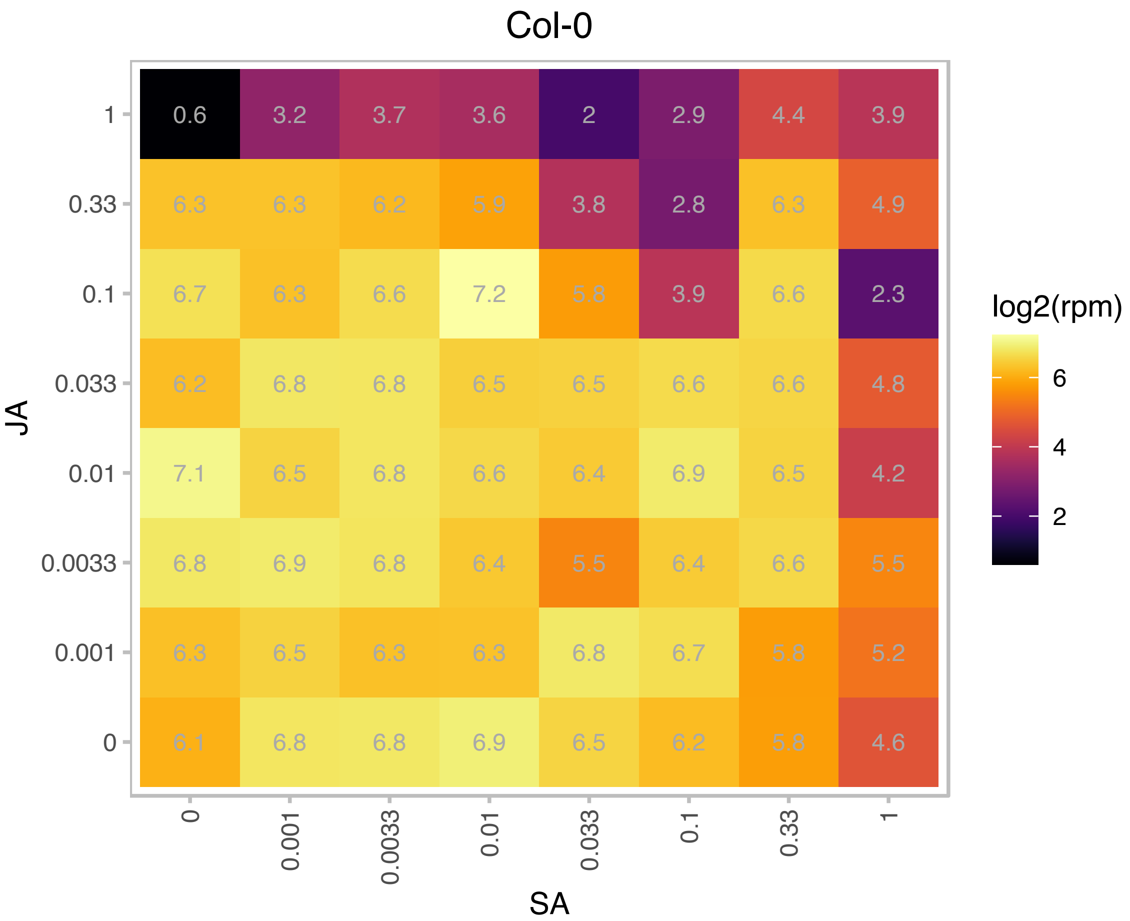


| | |
|---|---|
| Gene | AT3G48360 |
| is.expressed_genes | TRUE |
| is.responsive_genes | TRUE |
| is.apcluster | F-10 |

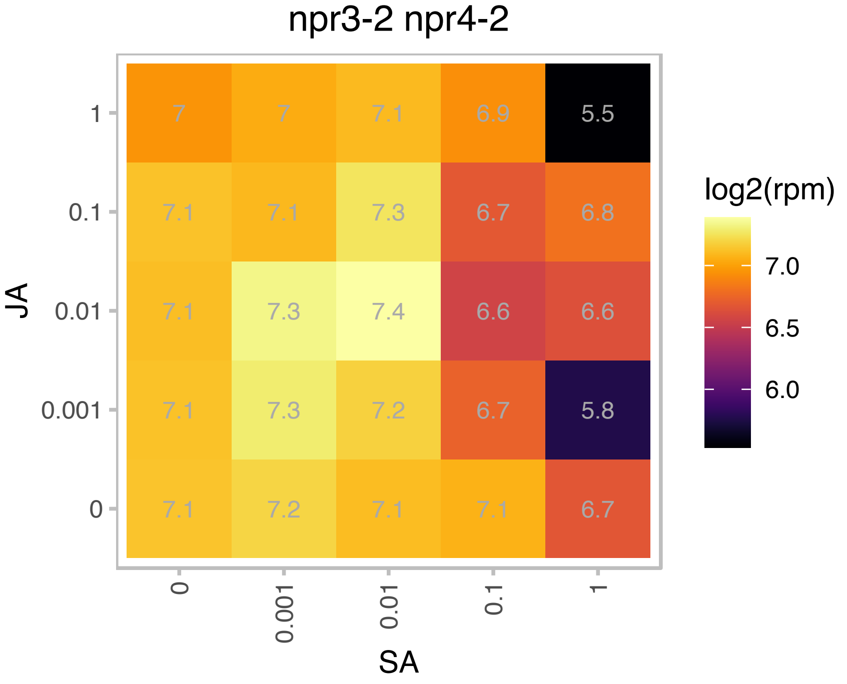


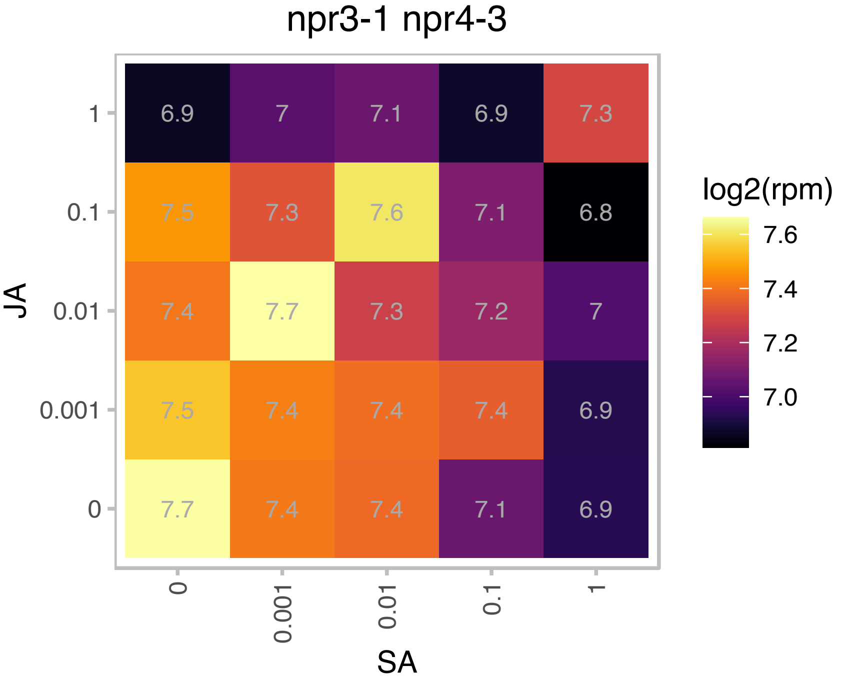

### AT3G49470.1 : NACA2

nascent polypeptide-associated complex subunit alpha-like protein 2

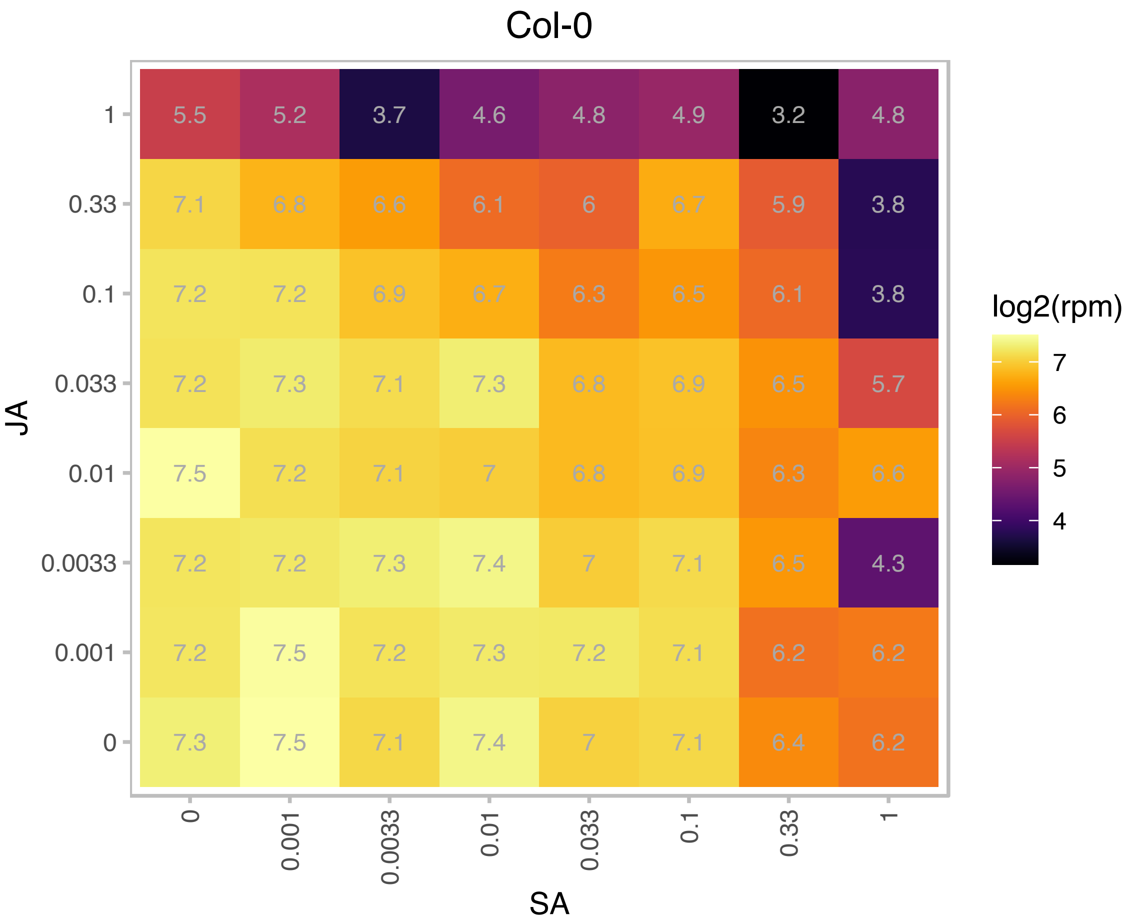


| | |
|---|---|
| Gene | AT3G49470 |
| is.expressed_genes | TRUE |
| is.responsive_genes | TRUE |
| is.apcluster | F-10 |

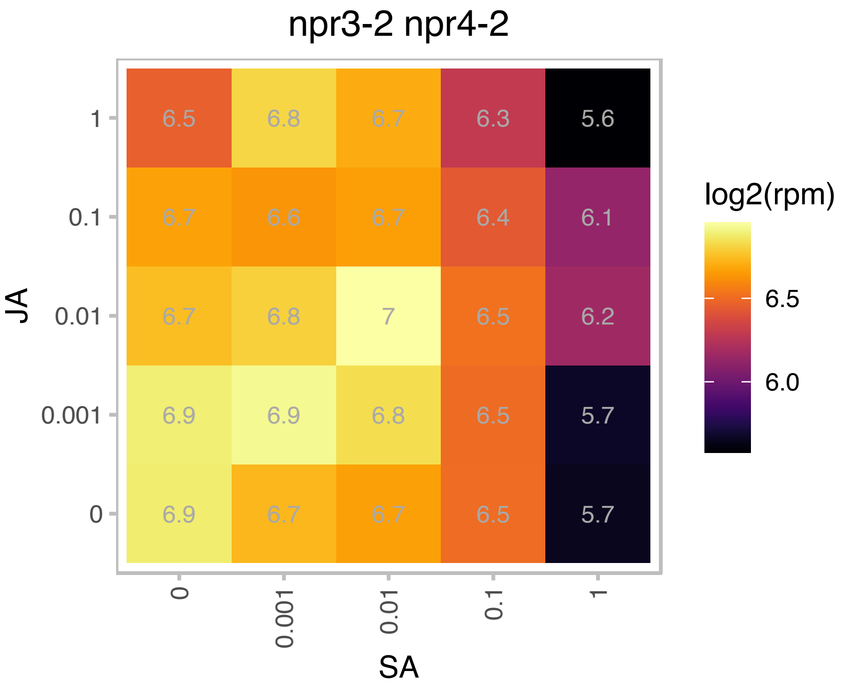


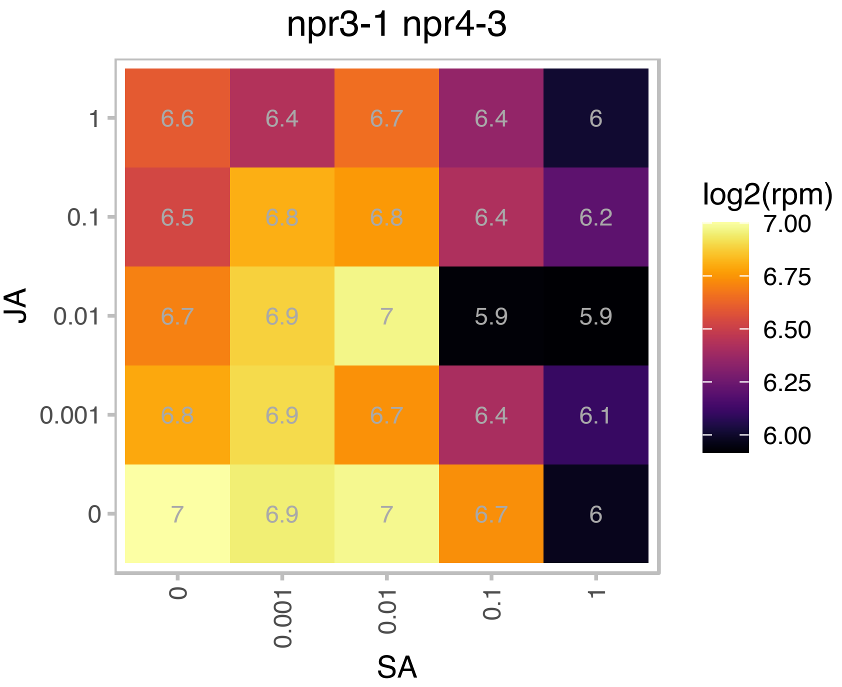

### AT3G50740.1 : UGT72E1

UDP-glucosyl transferase 72E1

UGT72E1 is an UDPG:coniferyl alcohol glucosyltransferase which specifically glucosylates sinapyl- and coniferyl aldehydes. The enzyme is thought to be involved in lignin metabolism.

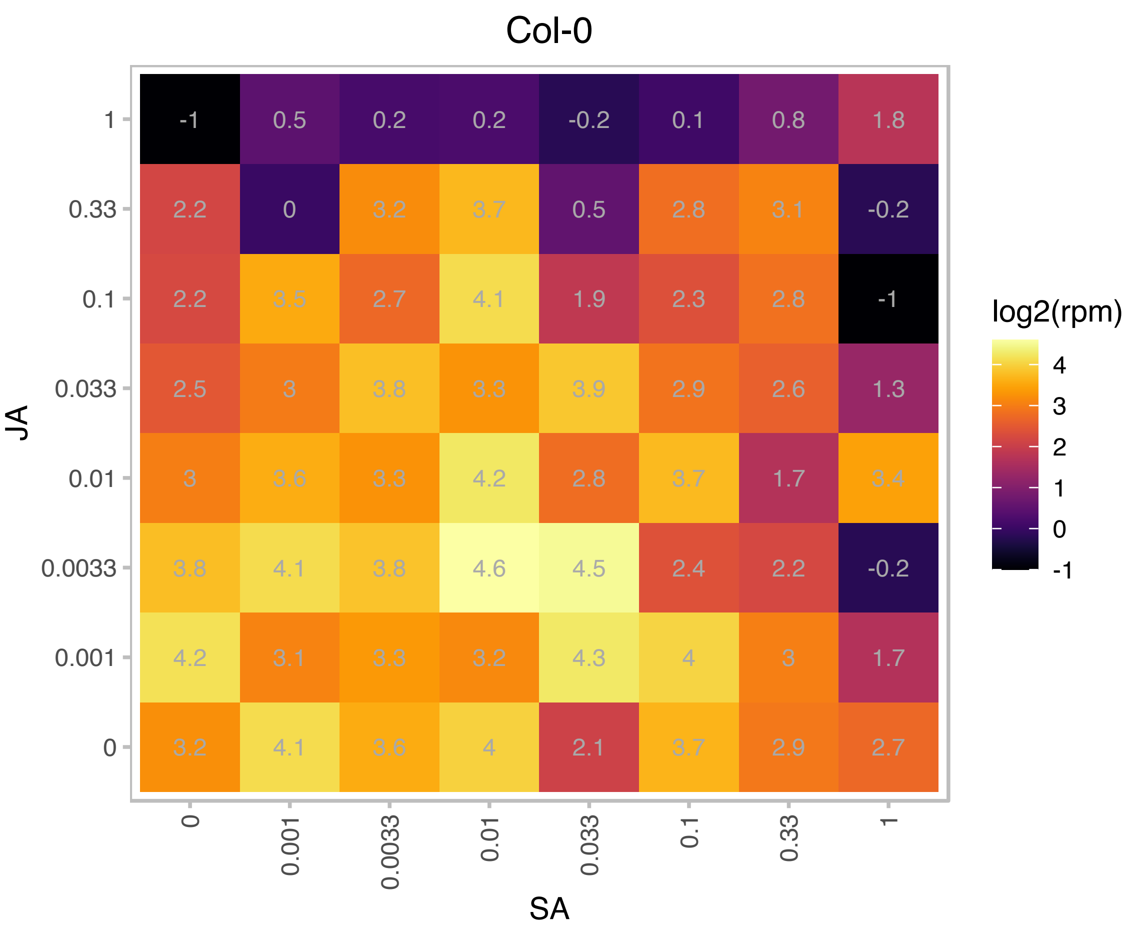


| | |
|---|---|
| Gene | AT3G50740 |
| is.expressed_genes | TRUE |
| is.responsive_genes | TRUE |
| is.apcluster | F-10 |

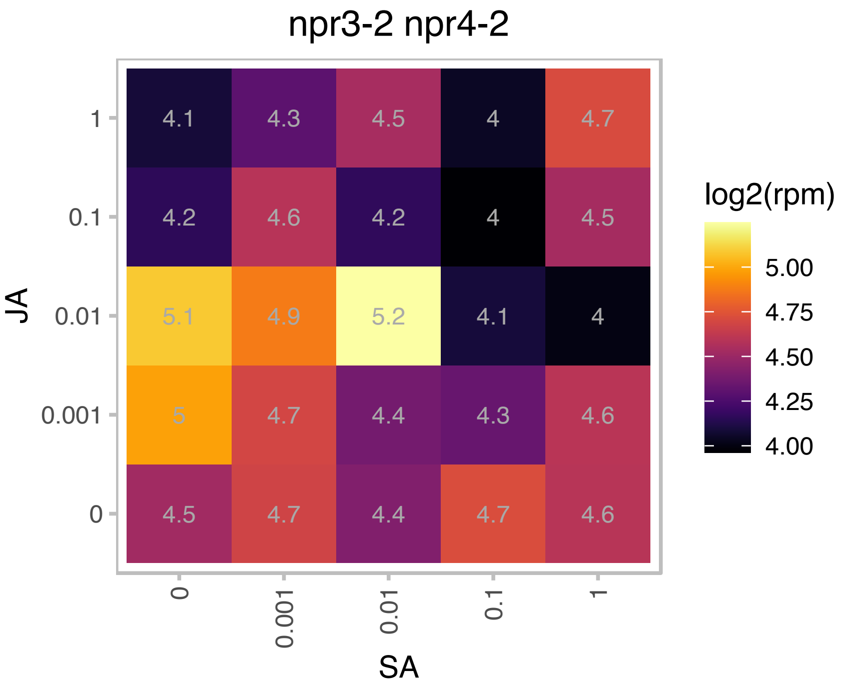


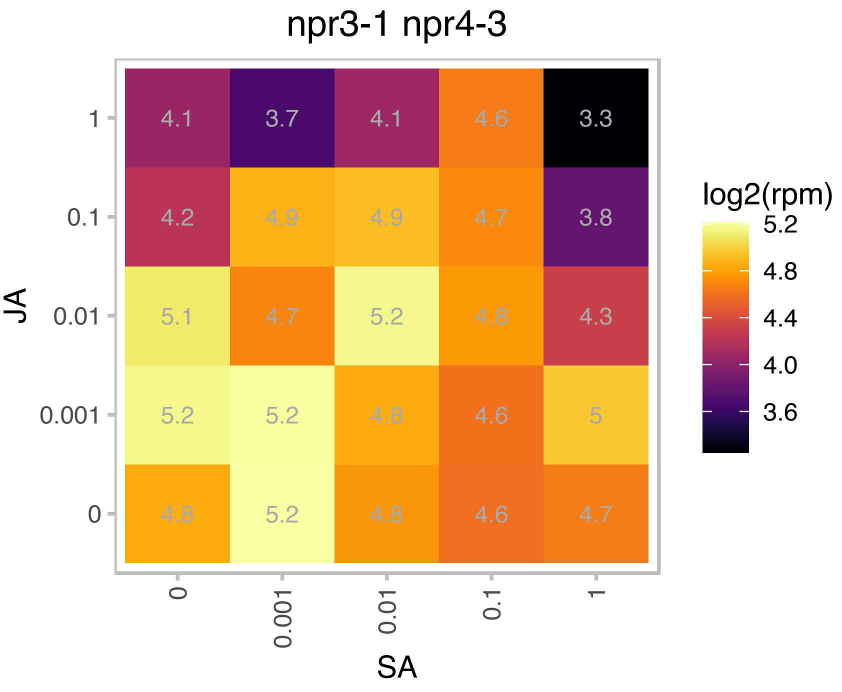

# AT3G53460.1 : CP29

chloroplast RNA-binding protein 29

Encodes a nuclear gene with a consensus RNA-binding domain that is localized to the chloroplast.

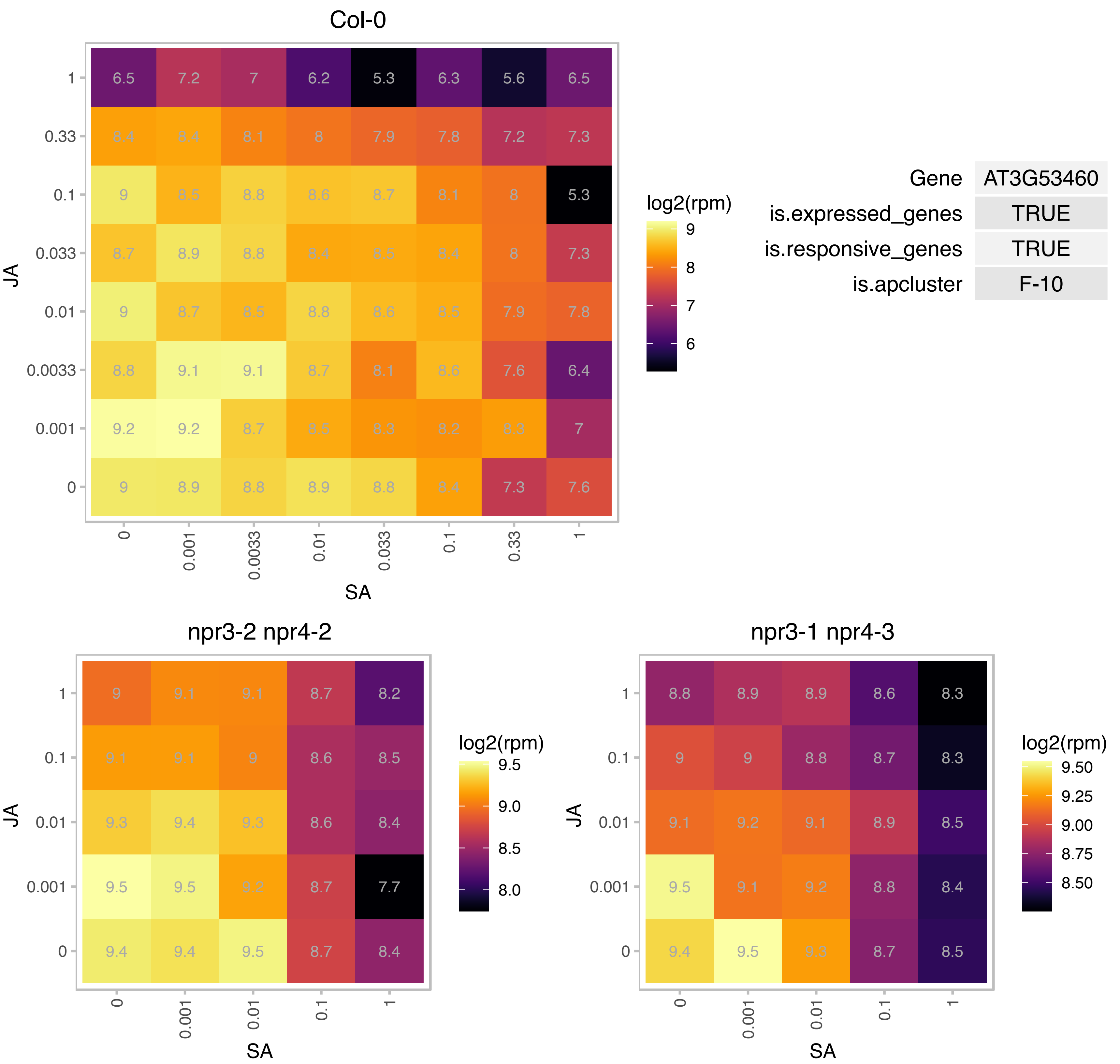


| | |
|---|---|
| Gene | AT3G53460 |
| is.expressed_genes | TRUE |
| is.responsive_genes | TRUE |
| is.apcluster | F-10 |

# AT3G55240.1 : NA

Plant protein 1589 of unknown function

Overexpression leads to PEL (Pseudo-Etiolation in Light) phenotype.

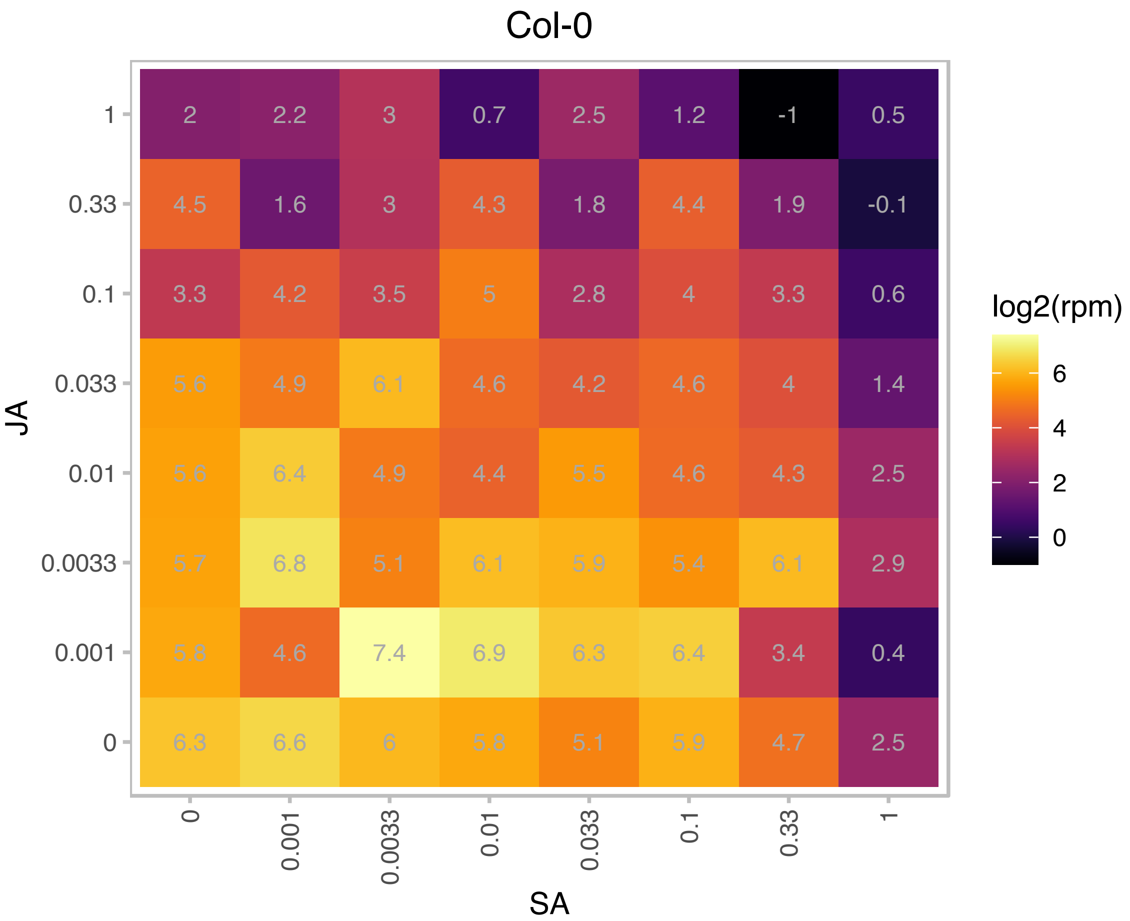


| | |
|---|---|
| Gene | AT3G55240 |
| is.expressed_genes | TRUE |
| is.responsive_genes | TRUE |
| is.apcluster | F-10 |

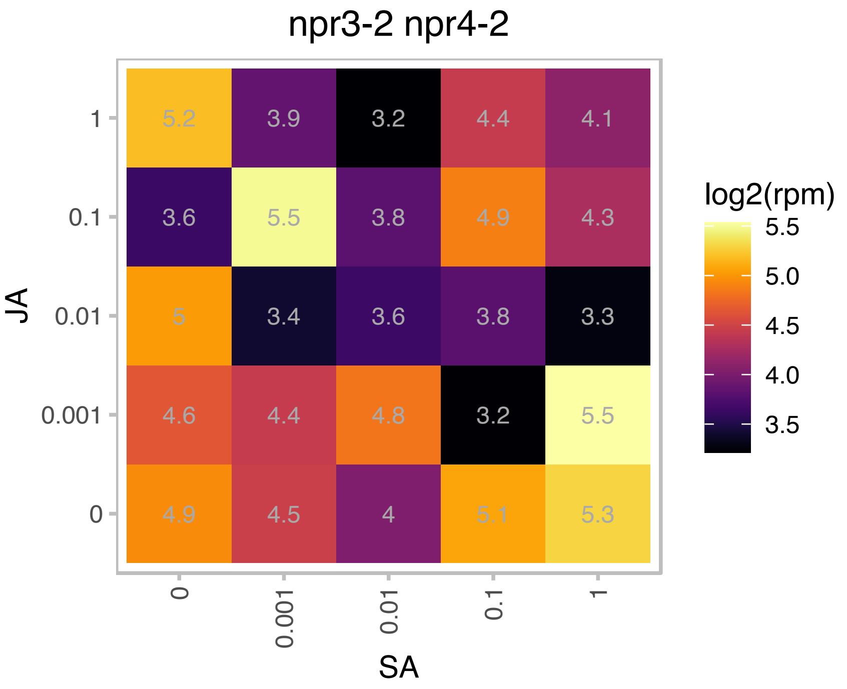


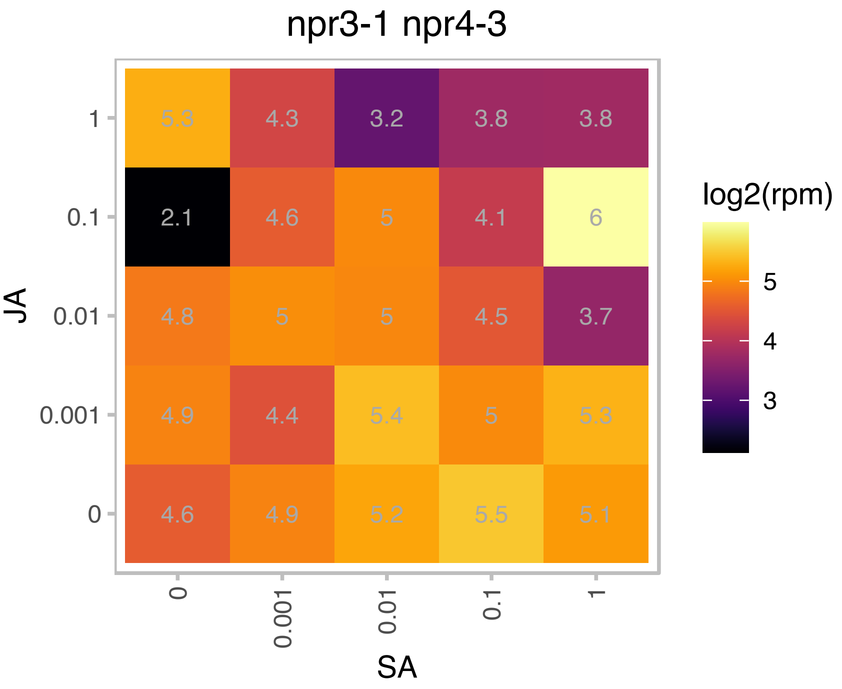

### AT3G61440.1 : ARATH;BSAS3;1, ATCYSC1, CAS-C1, CYS

cysteine synthase C1
Encodes a cysteine synthase isomer CysC1. The isomer is however less effective in cysteine biosynthesis. It is involved in beta-cyanoalanine biosynthesis, an intermediate of cyanide detoxification pathway. The mRNA is cell-to-cell mobile.

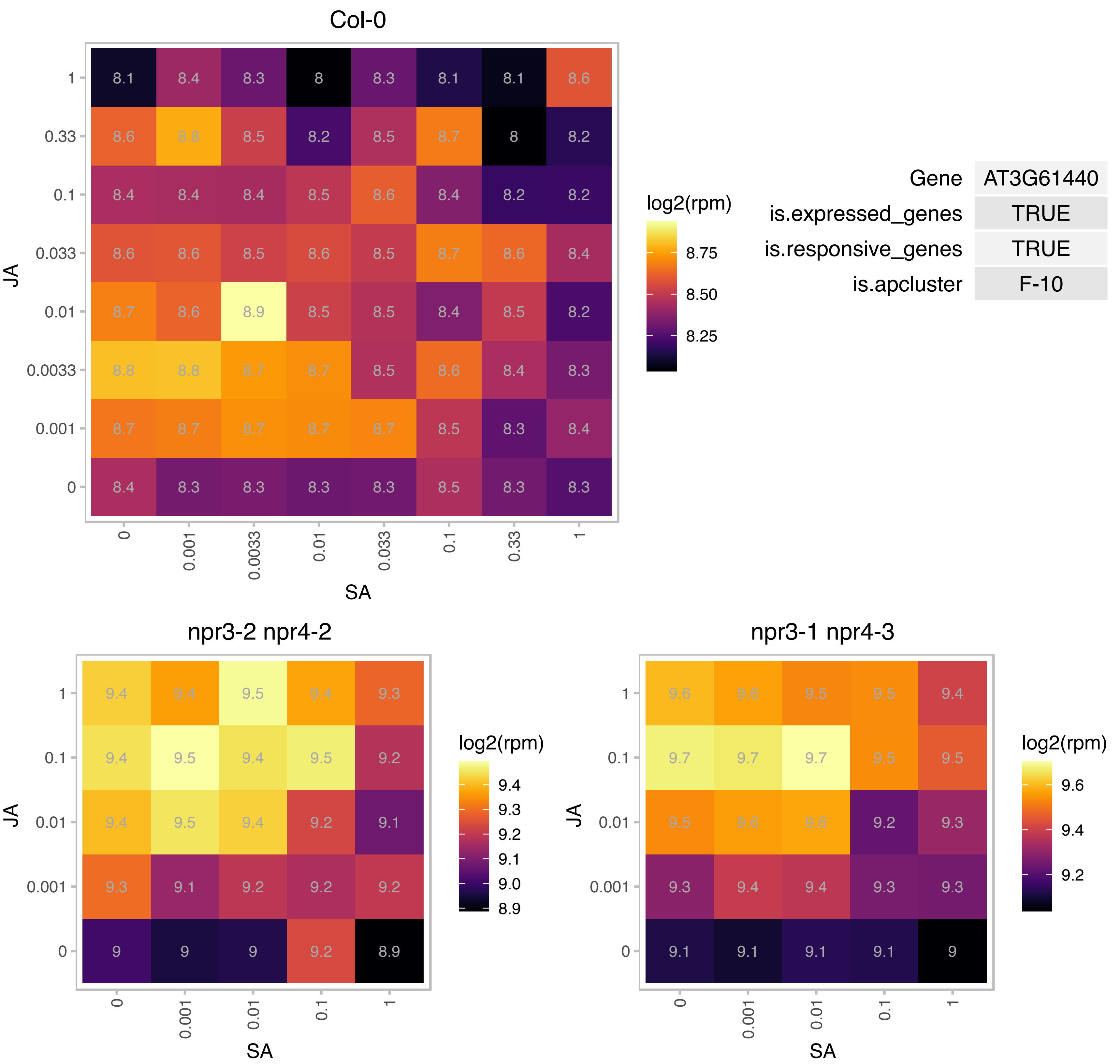


| | |
|---|---|
| Gene | AT3G61440 |
| is.expressed_genes | TRUE |
| is.responsive_genes | TRUE |
| is.apcluster | F-10 |

# AT3G62410.1 : CP12, CP12-2

CP12 domain-containing protein 2

CP12-2 encodes a small peptide found in the chloroplast stroma. It belongs to the CP12 gene family thought to be involved in the formation of a supramolecular complex with glyceraldehyde-3-phosphate dehydrogenase (GAPDH) and phosphoribulokinase (PRK) embedded in the Calvin cycle. CP12-2 is coordinately regulated by light with the photosynthetic GAPDH and PRK. The annotation of this gene is based on article 32494. The mRNA is cell-to-cell mobile.

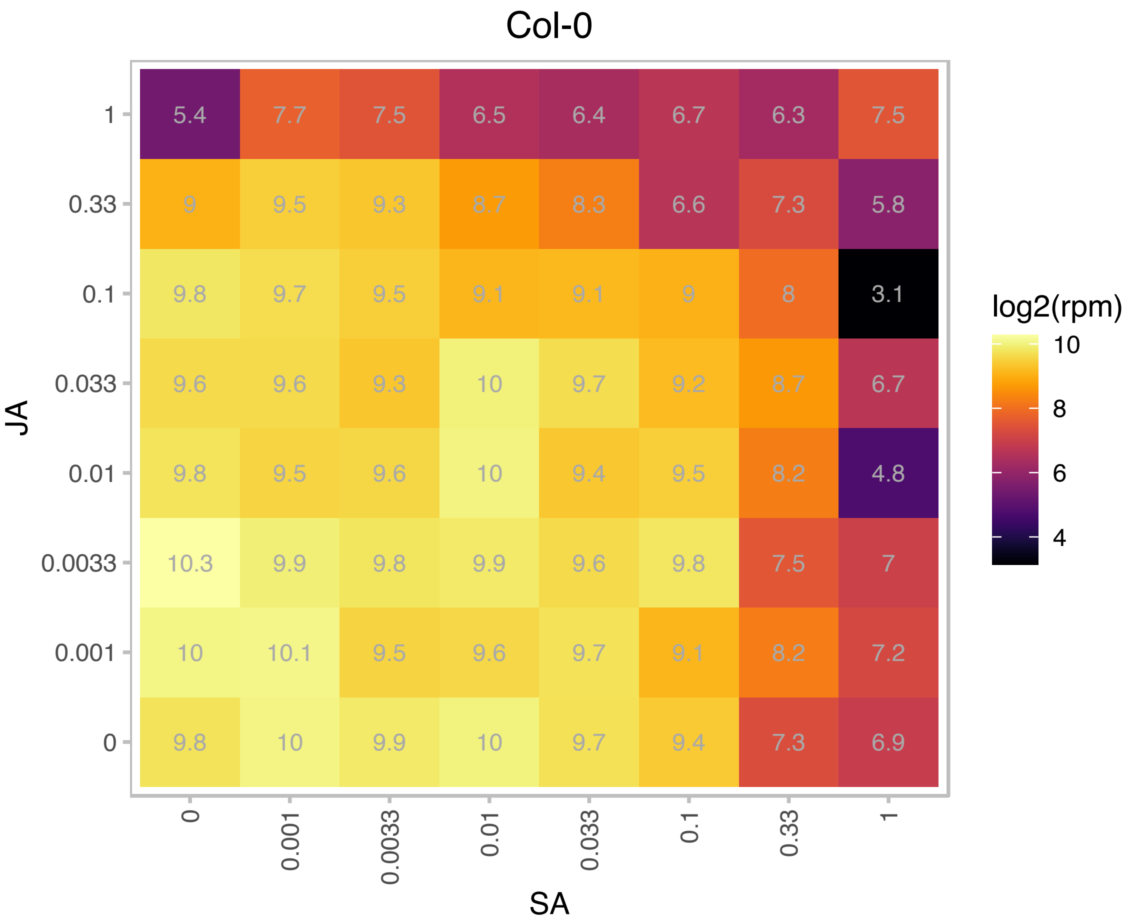


| | |
|---|---|
| Gene | AT3G62410 |
| is.expressed_genes | TRUE |
| is.responsive_genes | TRUE |
| is.apcluster | F-10 |

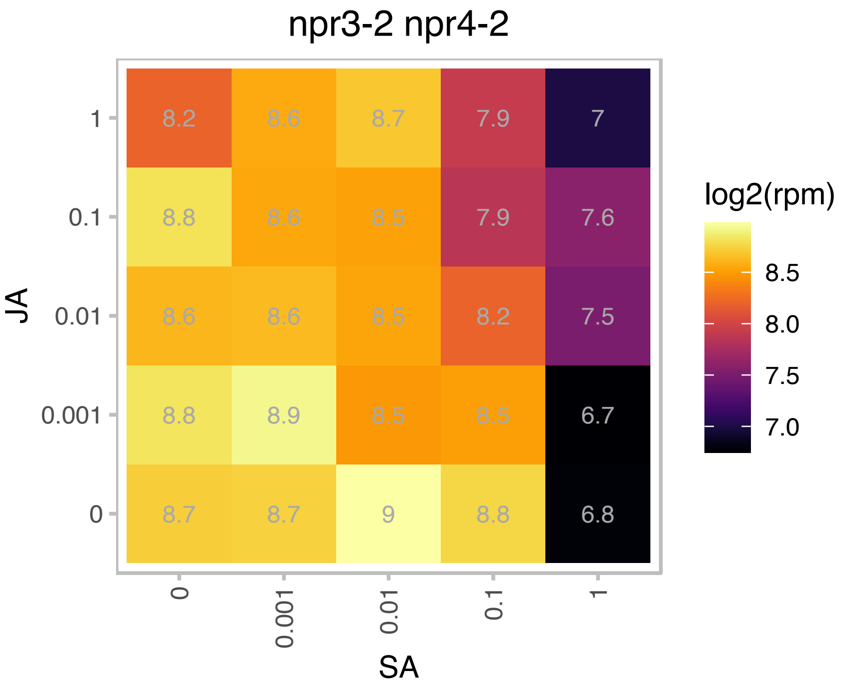


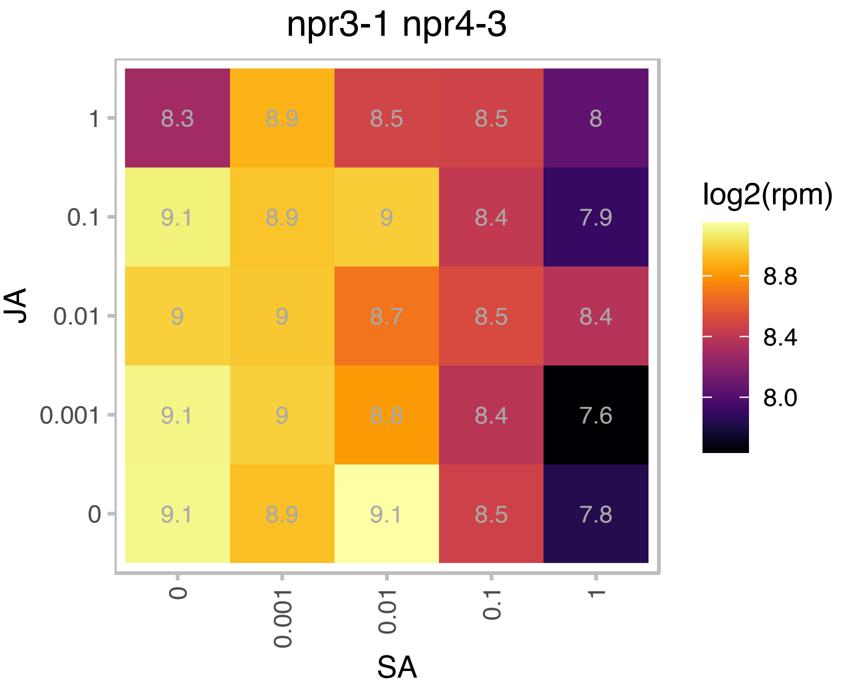

# AT4G01580.1 : NA

AP2/B3-like transcriptional factor family protein

| | |
|---|---|
| Gene | AT4G01580 |
| is.expressed_genes | TRUE |
| is.responsive_genes | TRUE |
| is.apcluster | F-10 |

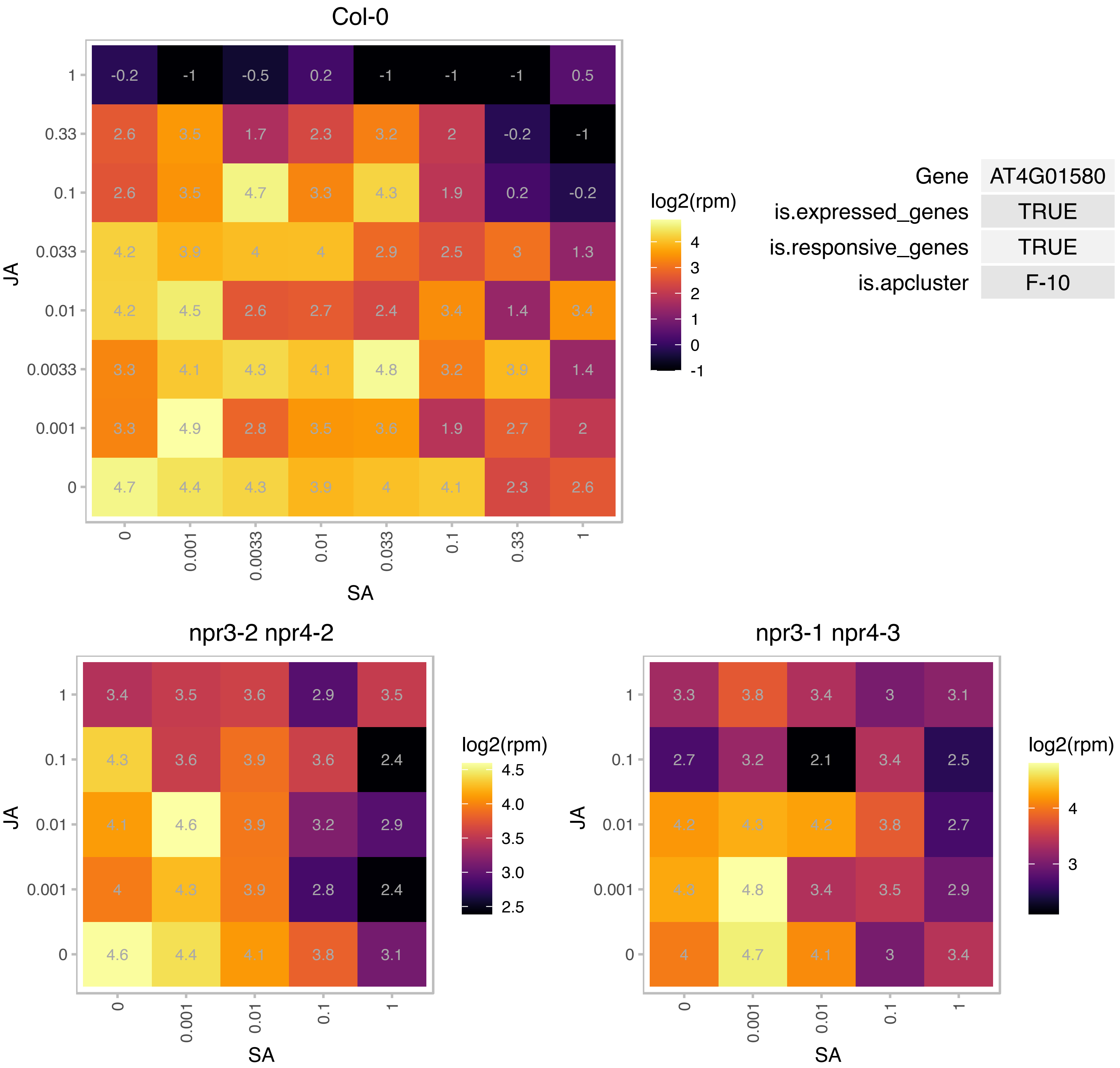

# AT4G03150.1 : NA

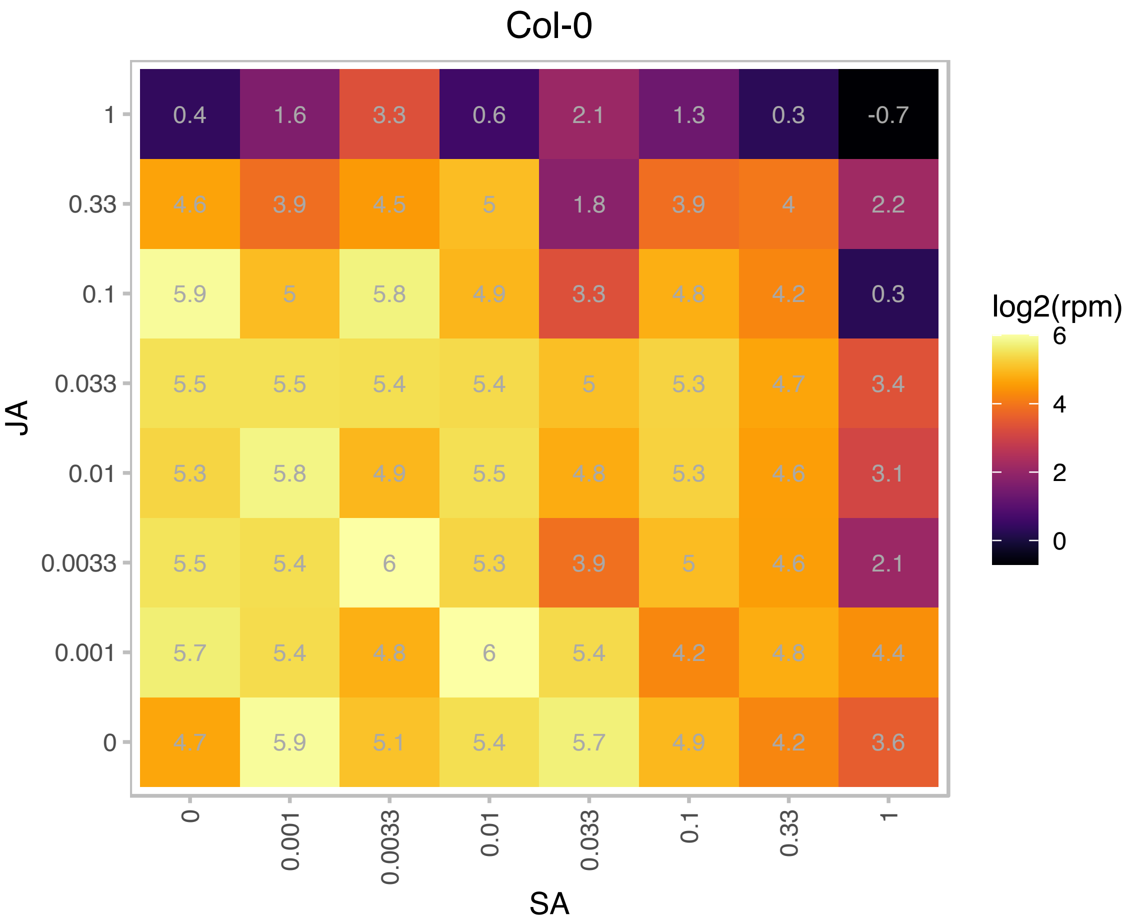


| | |
|---|---|
| Gene | AT4G03150 |
| is.expressed_genes | TRUE |
| is.responsive_genes | TRUE |
| is.apcluster | F-10 |

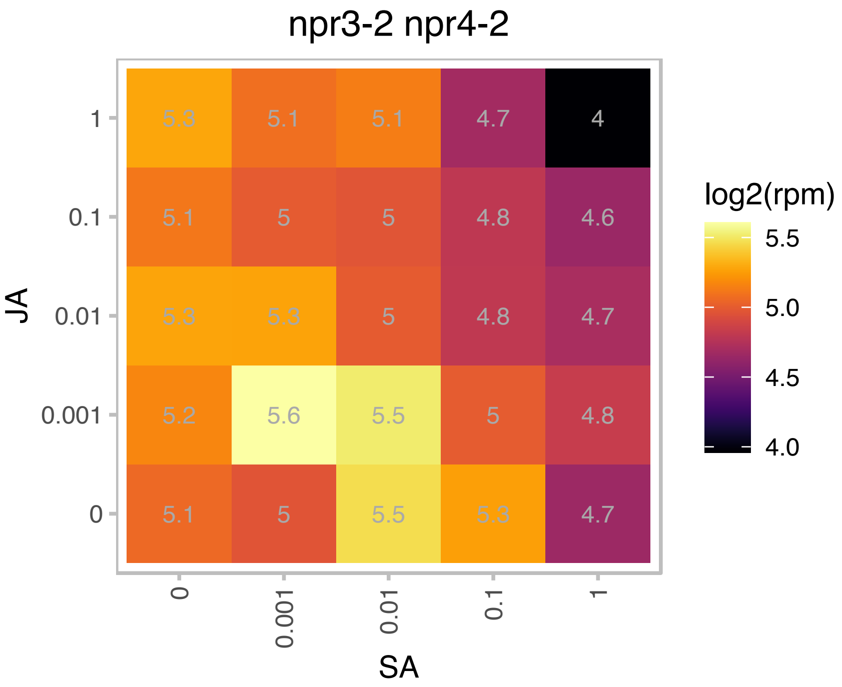


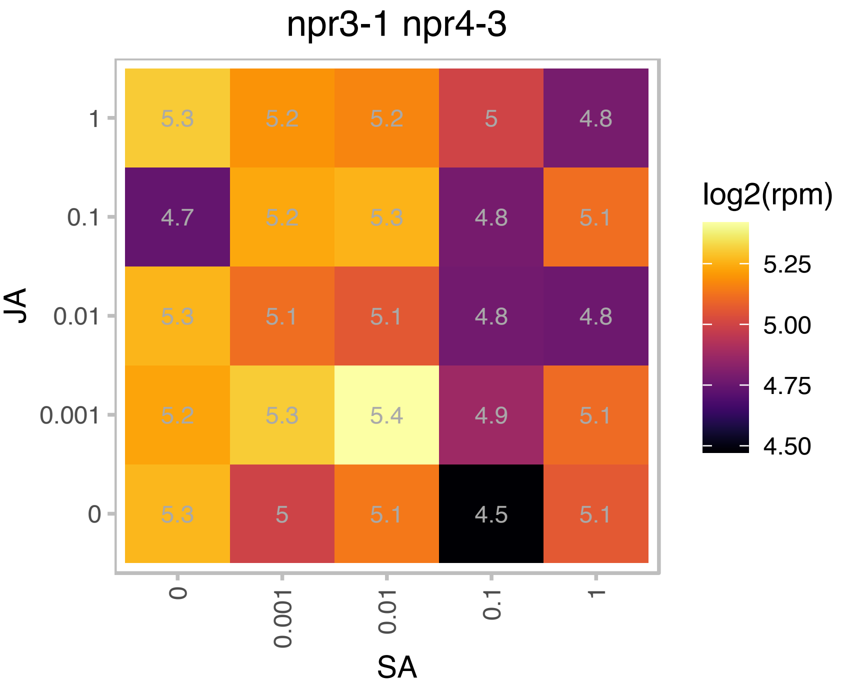

### AT4G09650.1 : ATPD, PDE332

ATP synthase delta-subunit gene

Encodes the chloroplast ATPase delta-subunit. The mRNA is cell-to-cell mobile.

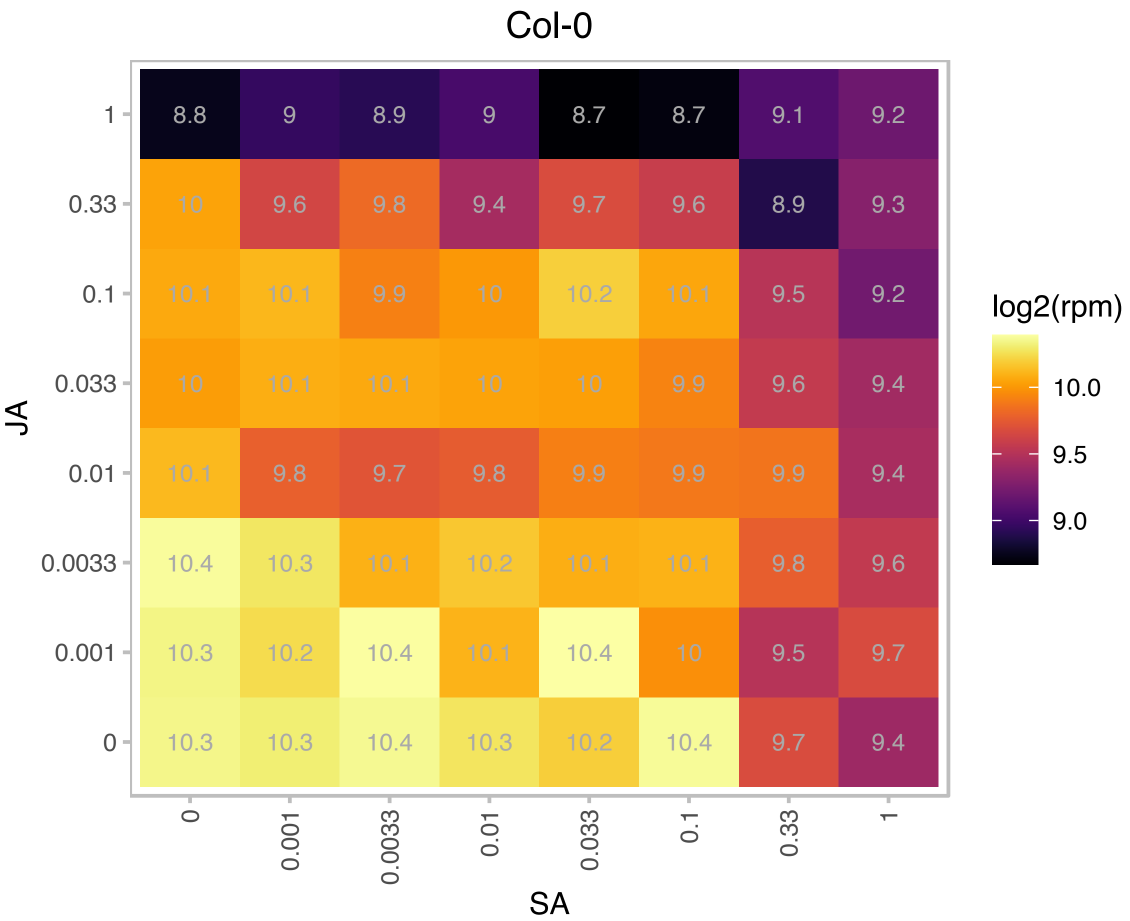


| | |
|---|---|
| Gene | AT4G09650 |
| is.expressed_genes | TRUE |
| is.responsive_genes | TRUE |
| is.apcluster | F-10 |

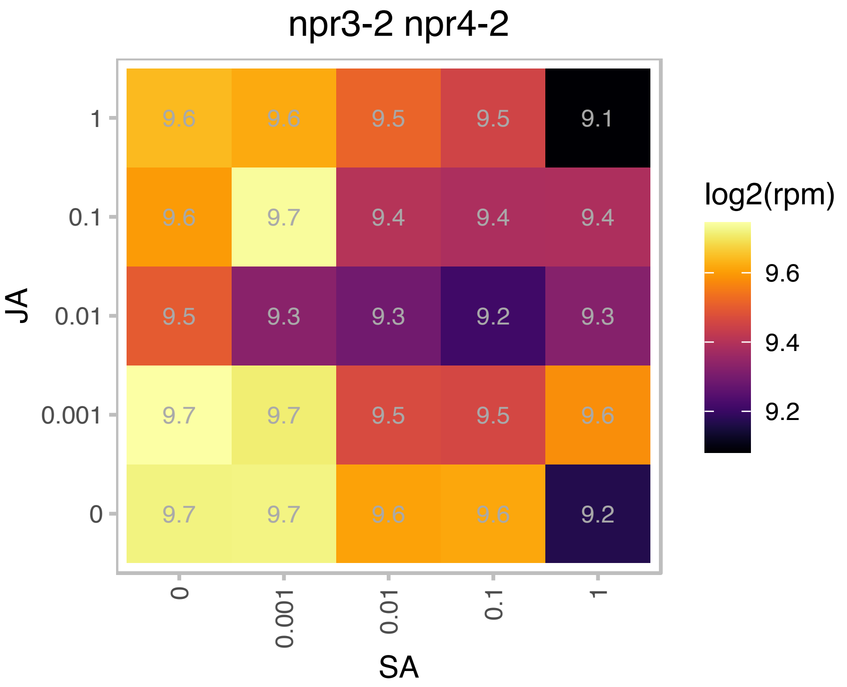


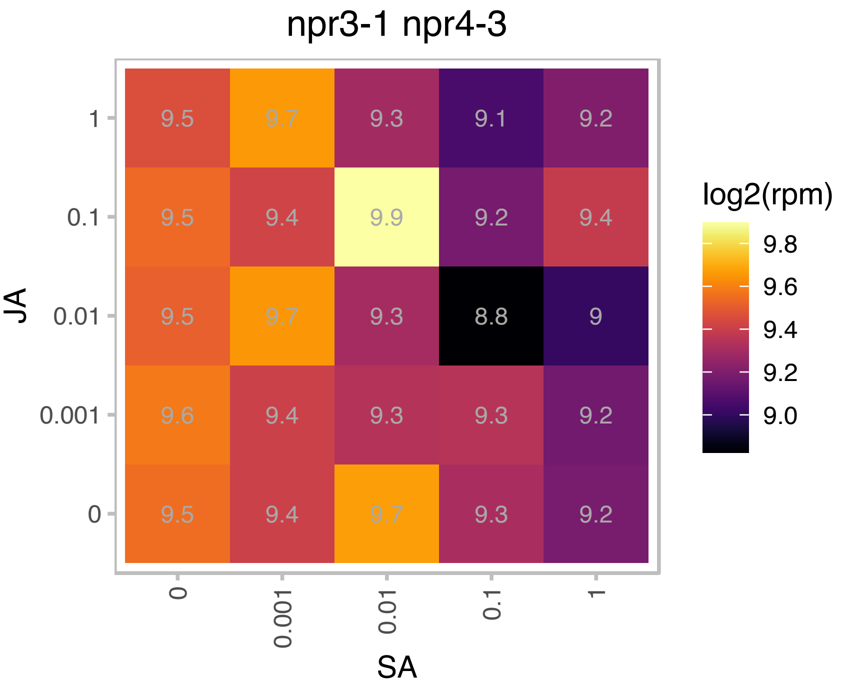

### AT4G14890.1 : FdC1

2Fe-2S ferredoxin-like superfamily protein

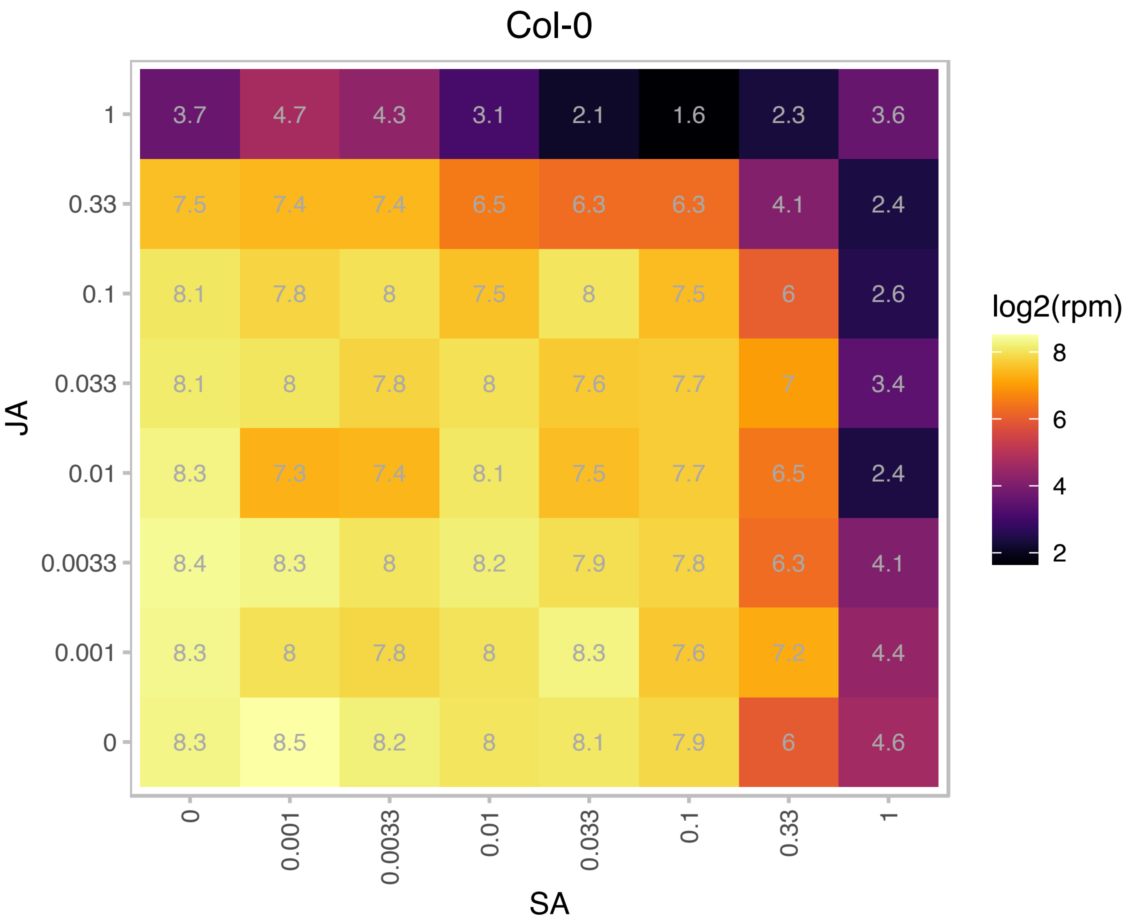


| | |
|---|---|
| Gene | AT4G14890 |
| is.expressed_genes | TRUE |
| is.responsive_genes | TRUE |
| is.apcluster | F-10 |

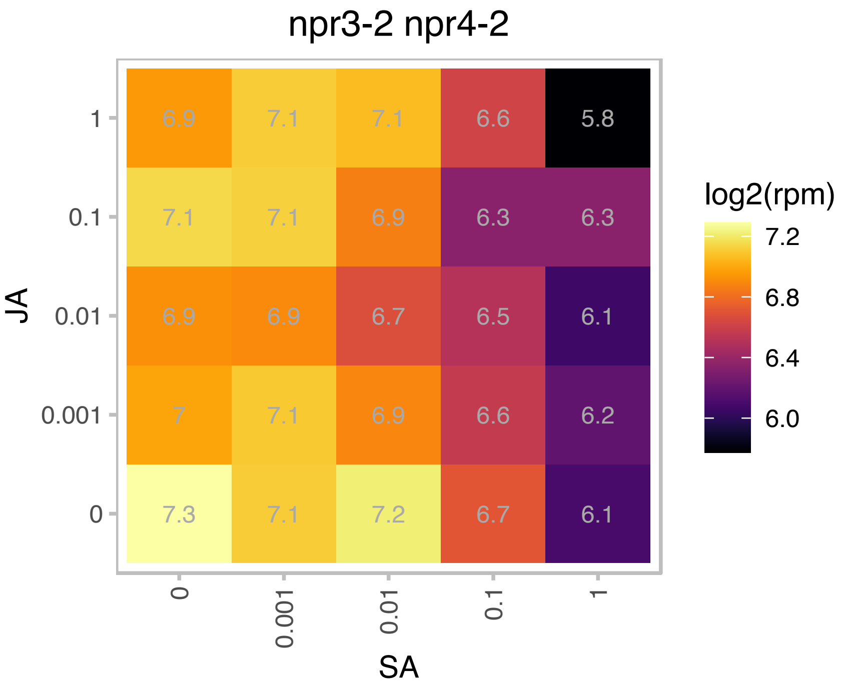


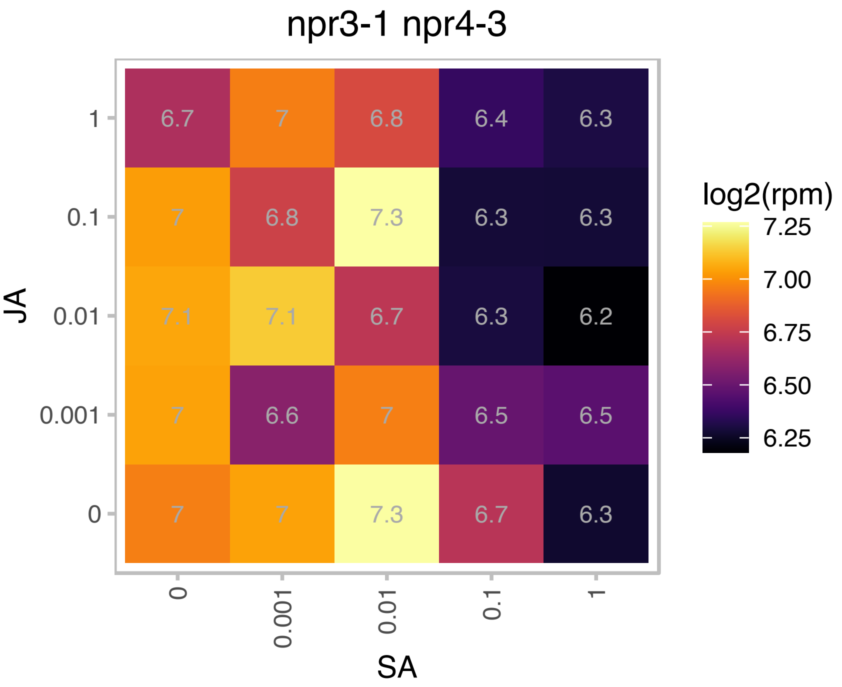

### AT4G15130.1 : ATCCT2, CCT2

phosphorylcholine cytidylyltransferase2

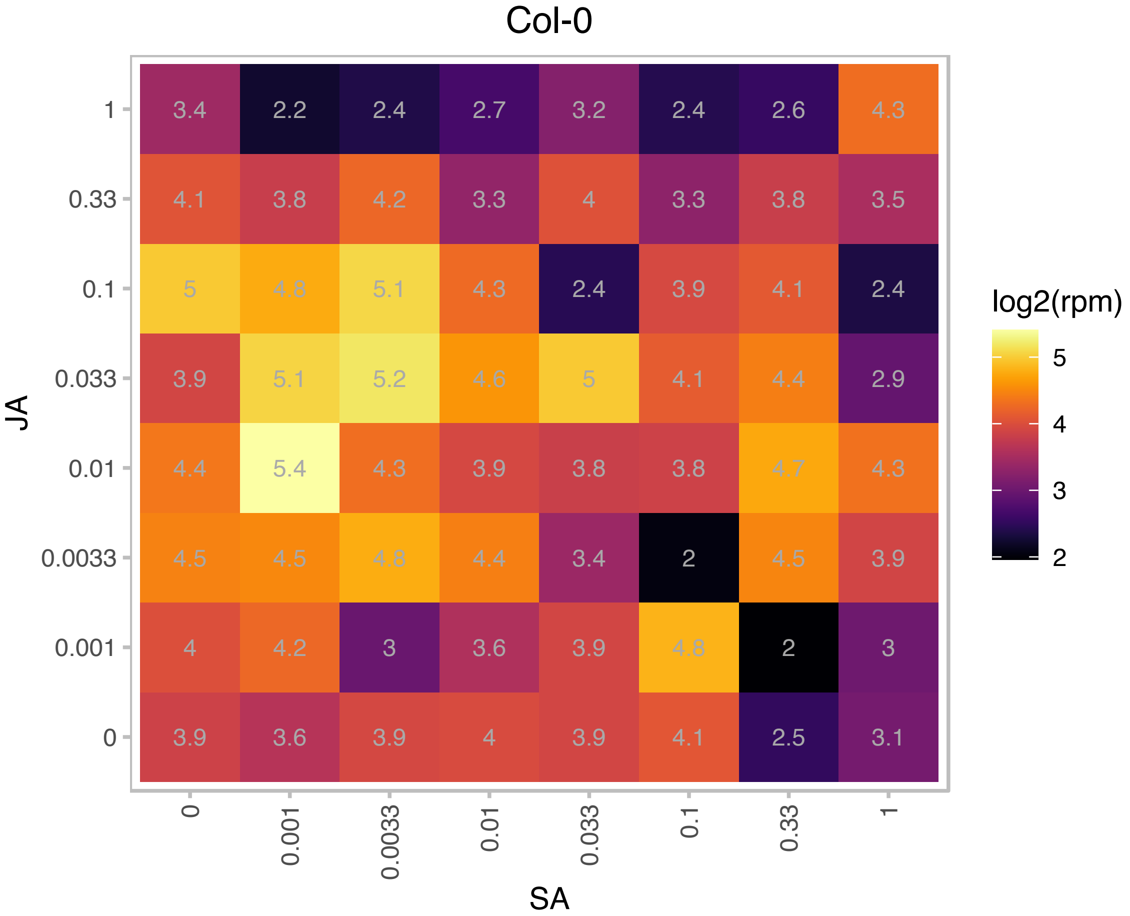


| | |
|---|---|
| Gene | AT4G15130 |
| is.expressed_genes | TRUE |
| is.responsive_genes | TRUE |
| is.apcluster | F-10 |

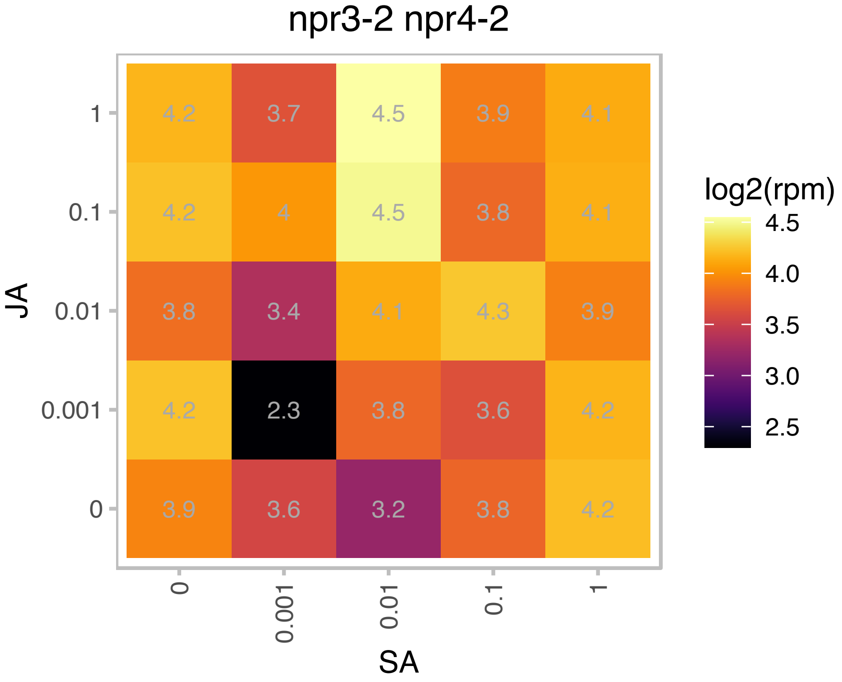


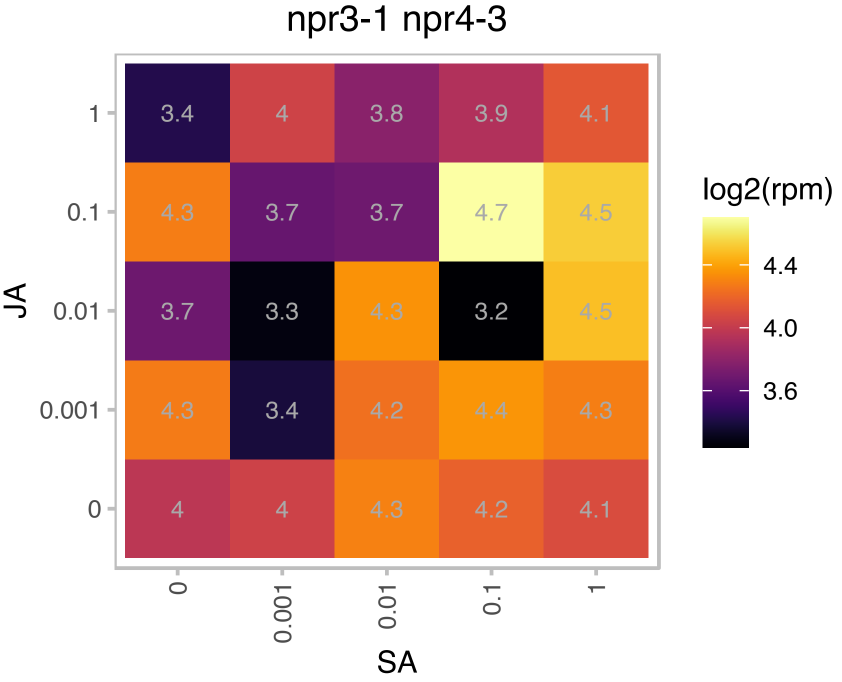

# AT4G15545.1 : NA

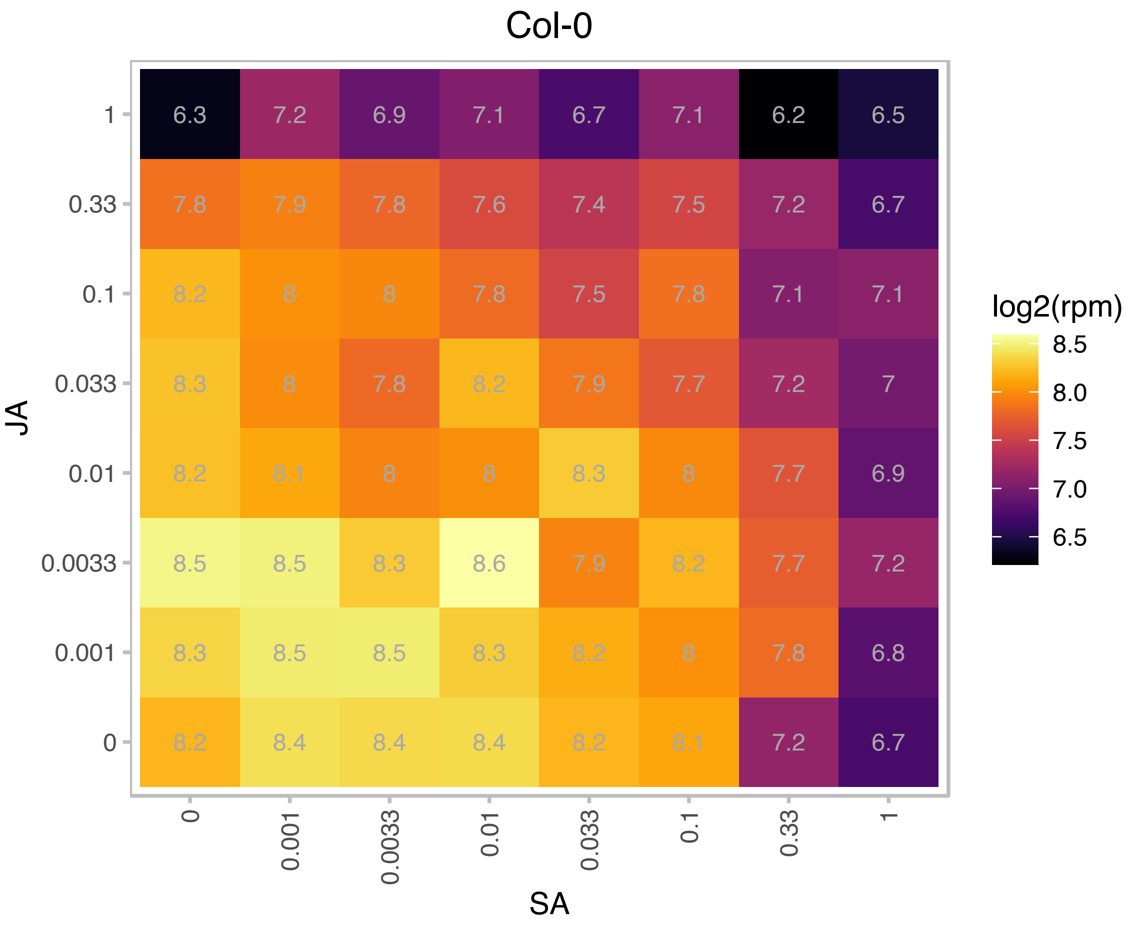


| | |
|---|---|
| Gene | AT4G15545 |
| is.expressed_genes | TRUE |
| is.responsive_genes | TRUE |
| is.apcluster | F-10 |

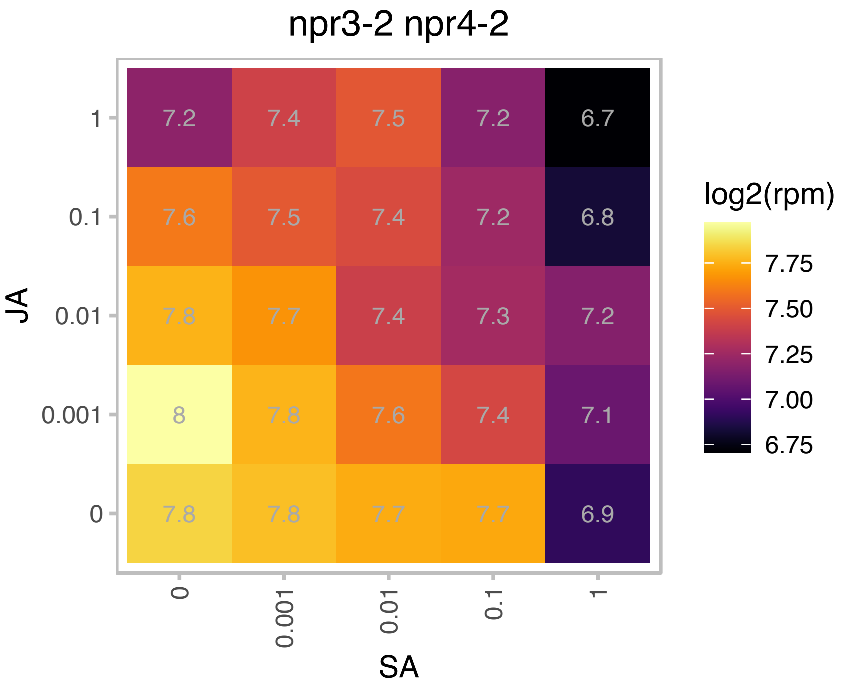


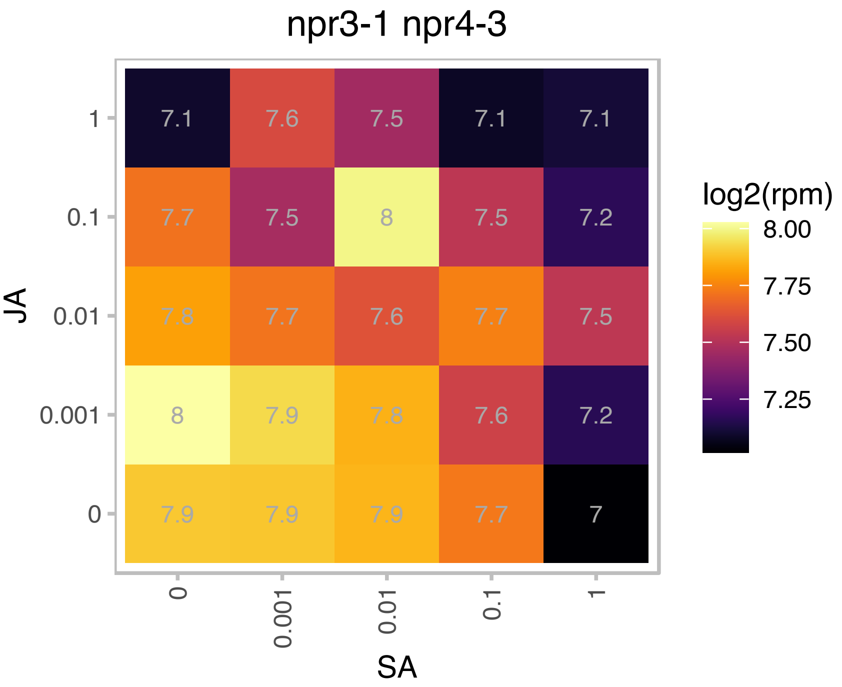

# AT4G16146.1 : NA

cAMP-regulated phosphoprotein 19-related protein

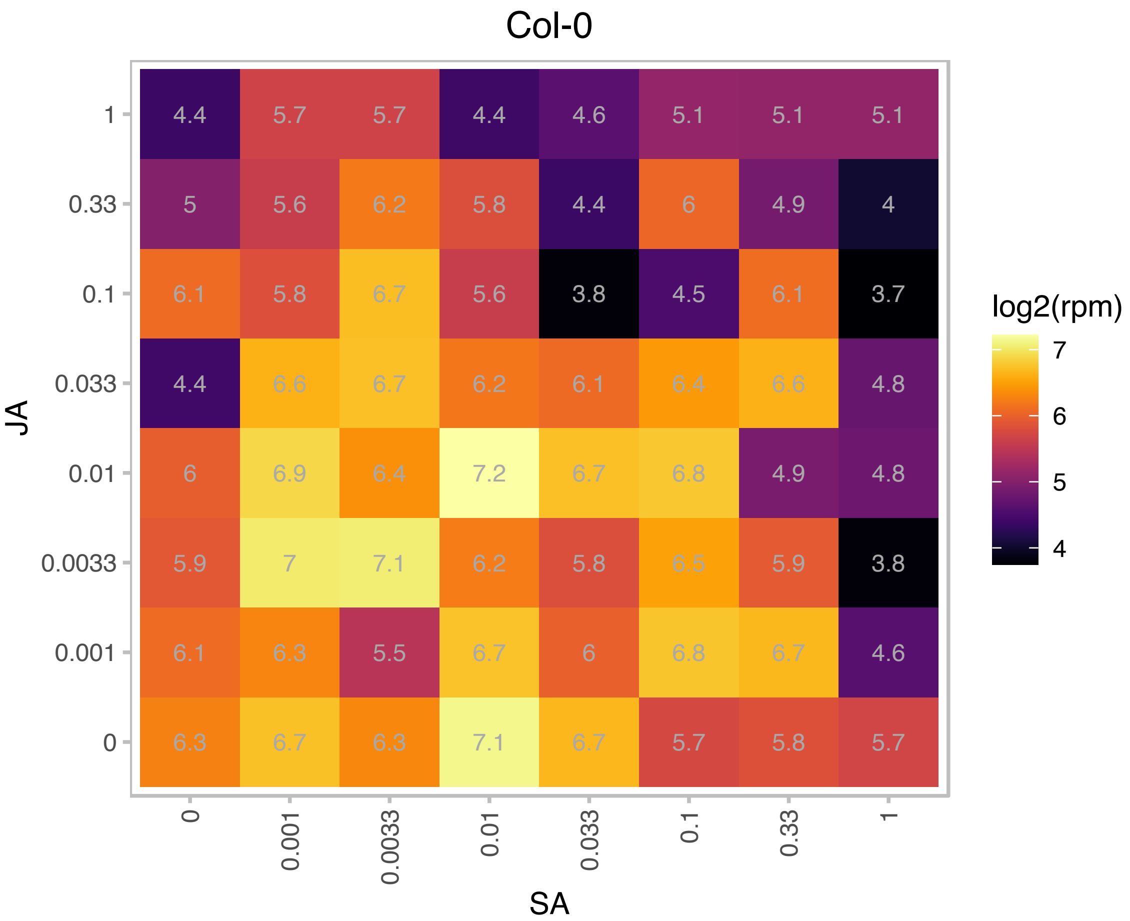


| | |
|---|---|
| Gene | AT4G16146 |
| is.expressed_genes | TRUE |
| is.responsive_genes | TRUE |
| is.apcluster | F-10 |

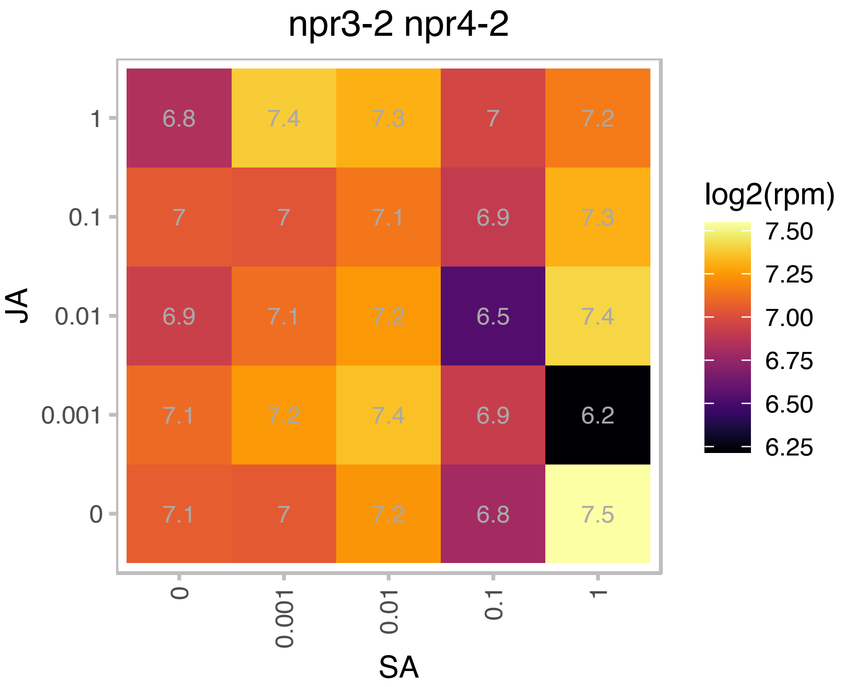


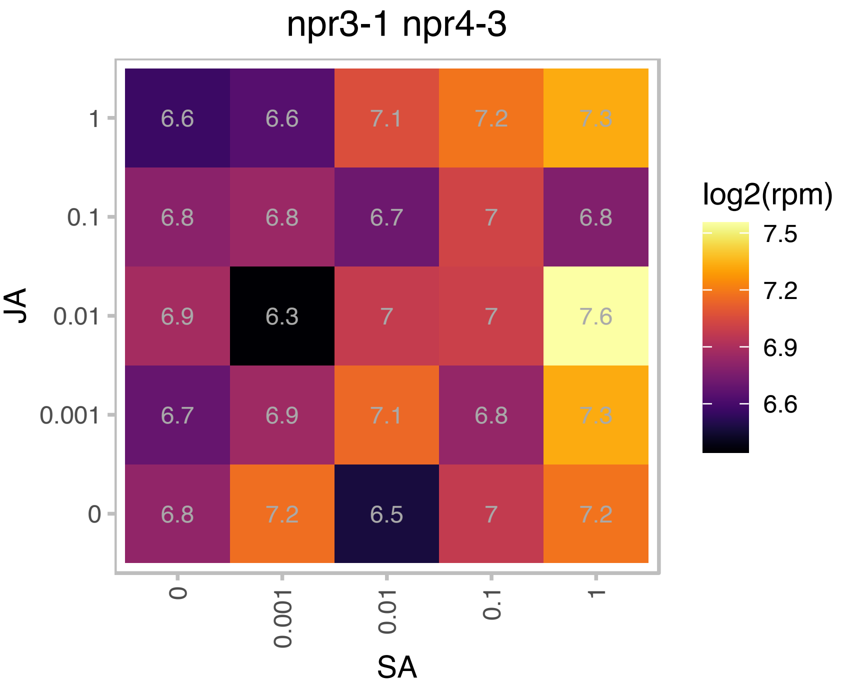

# AT4G16980.1 : NA

arabinogalactan-protein family

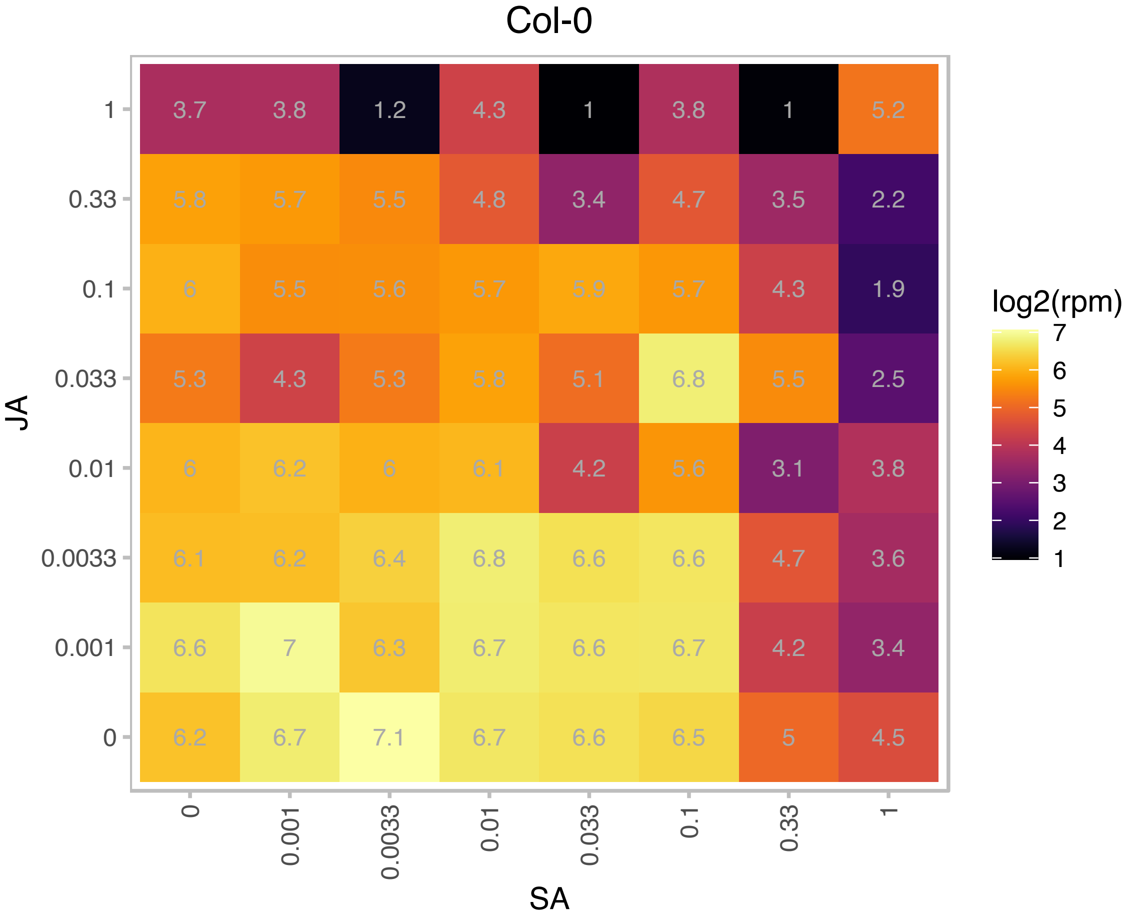


| | |
|---|---|
| Gene | AT4G16980 |
| is.expressed_genes | TRUE |
| is.responsive_genes | TRUE |
| is.apcluster | F-10 |

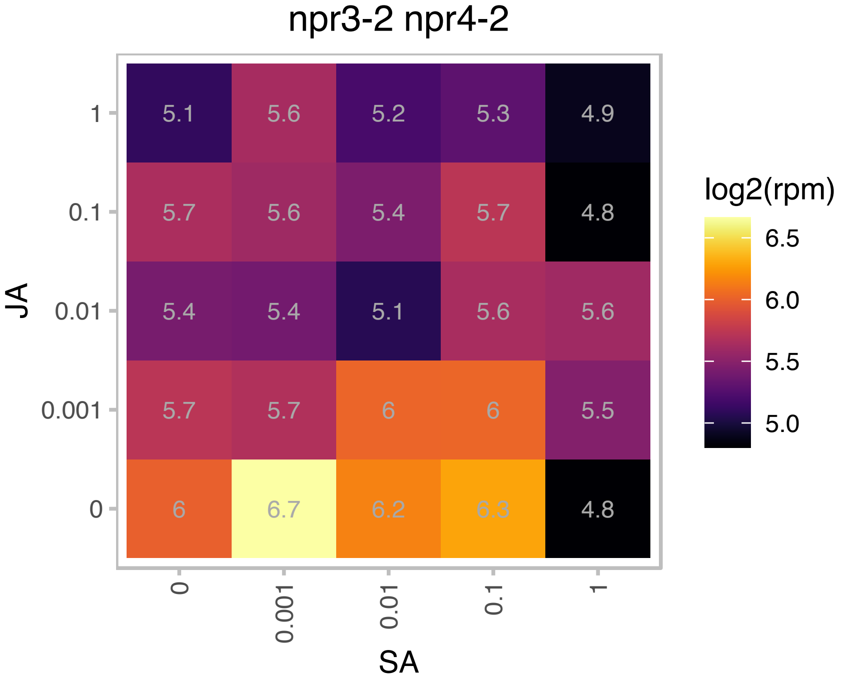


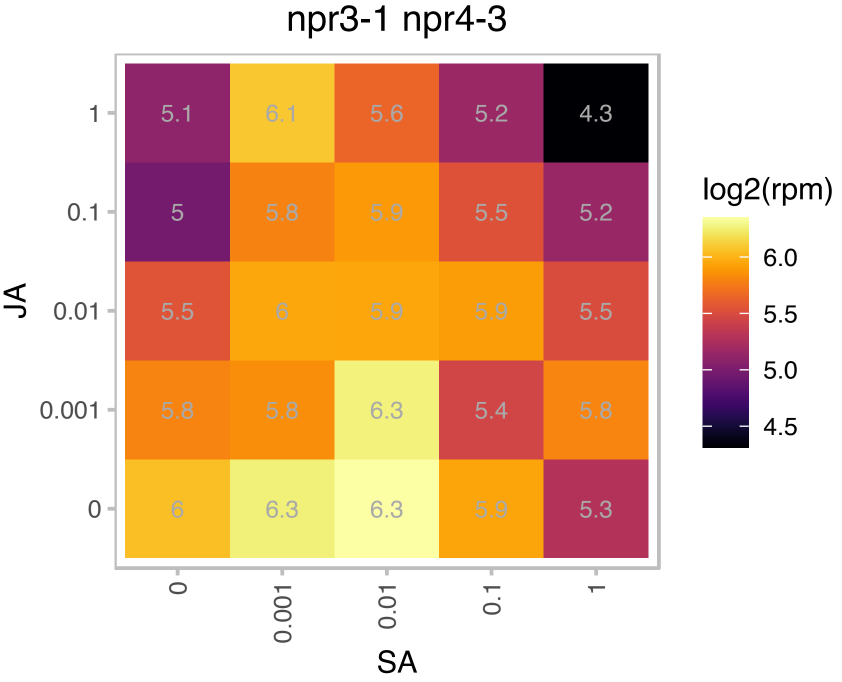

# AT4G17085.1 : NA

Putative membrane lipoprotein

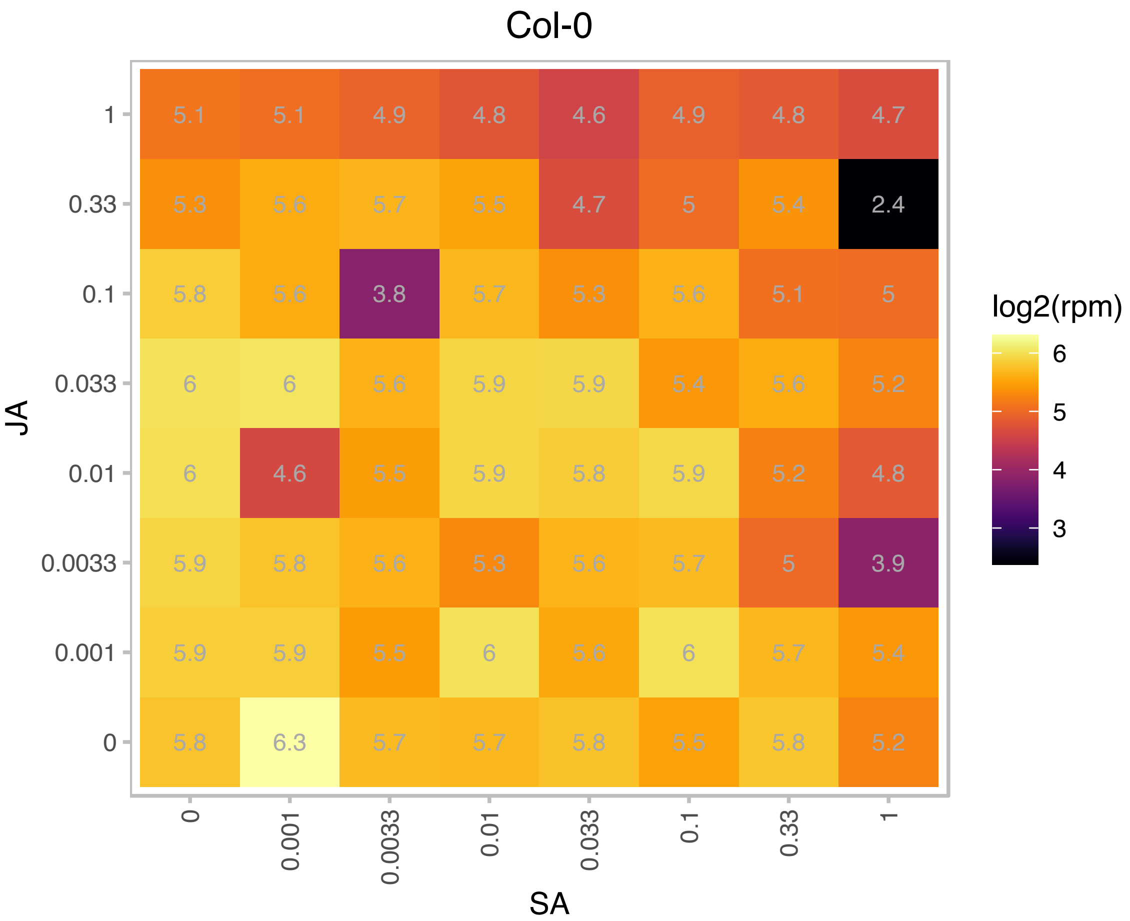


| | |
|---|---|
| Gene | AT4G17085 |
| is.expressed_genes | TRUE |
| is.responsive_genes | TRUE |
| is.apcluster | F-10 |

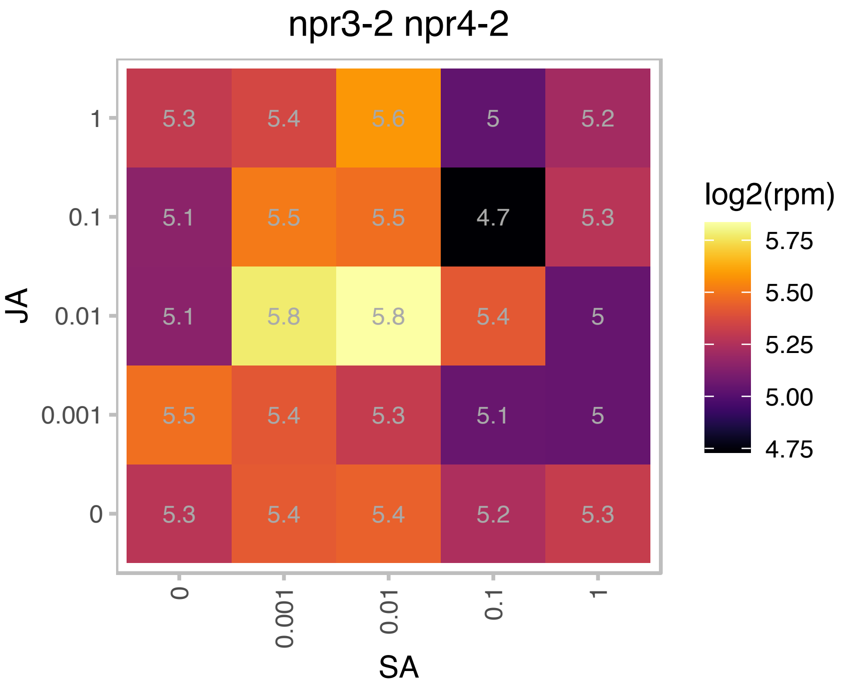


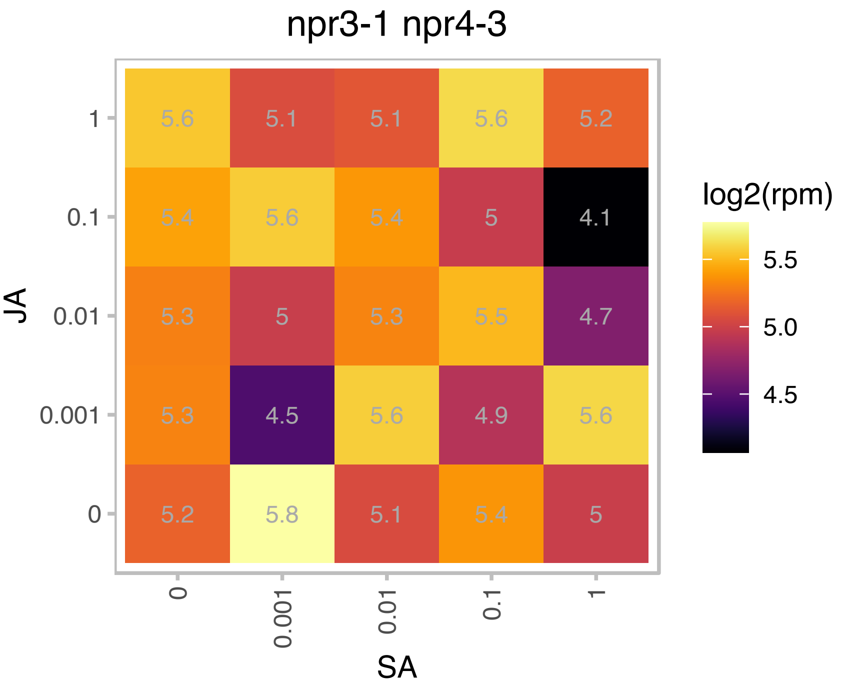

### AT4G19100.1 : PAM68

Protein of unknown function (DUF3464)

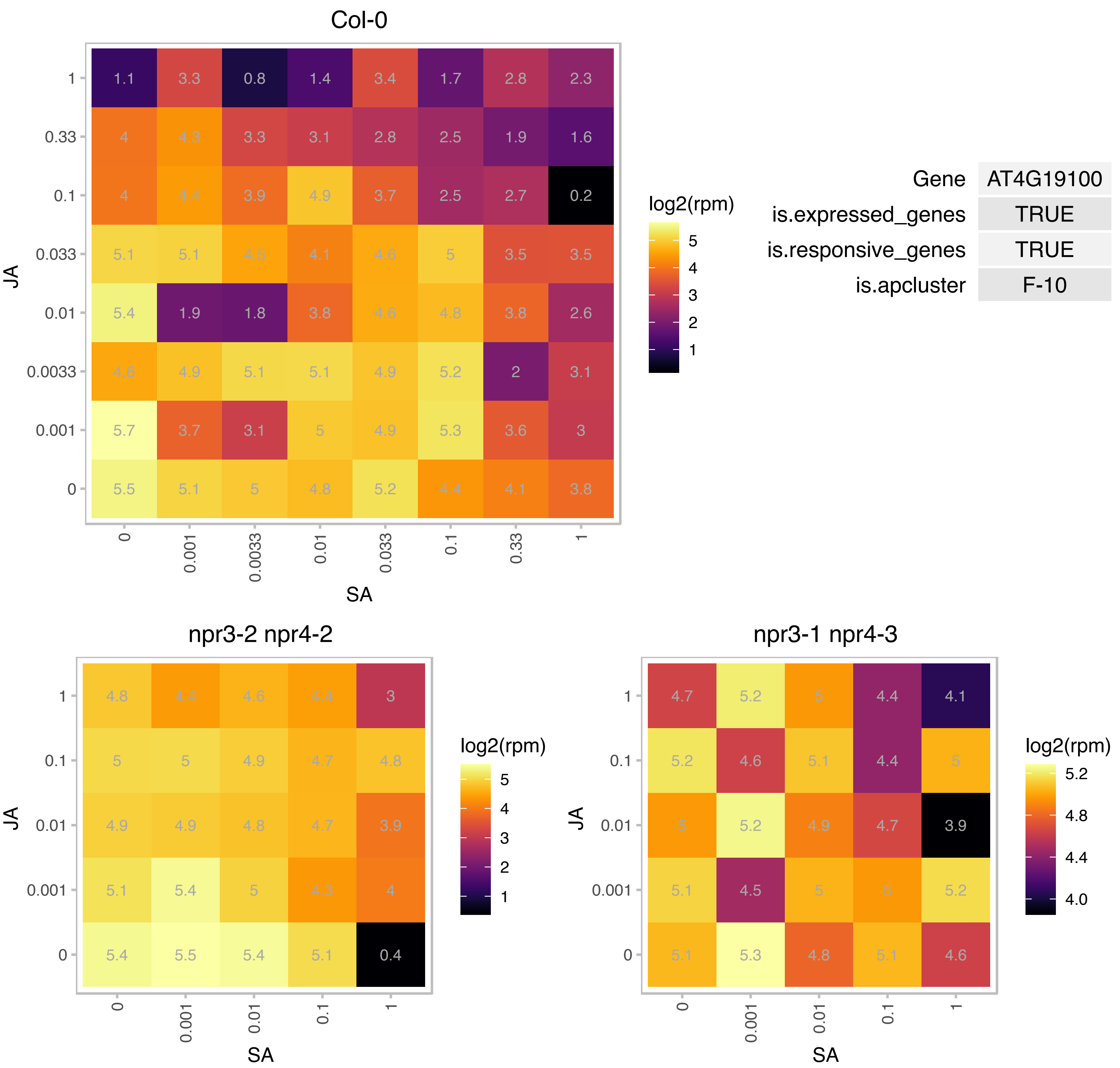


| | |
|---|---|
| Gene | AT4G19100 |
| is.expressed_genes | TRUE |
| is.responsive_genes | TRUE |
| is.apcluster | F-10 |

### AT4G21445.1 : CRR9

CRR9 gene encodes a novel stromal protein without any known functional domains or motifs. It is highly conserved in cyanobacteria and land plants but not in green algae.

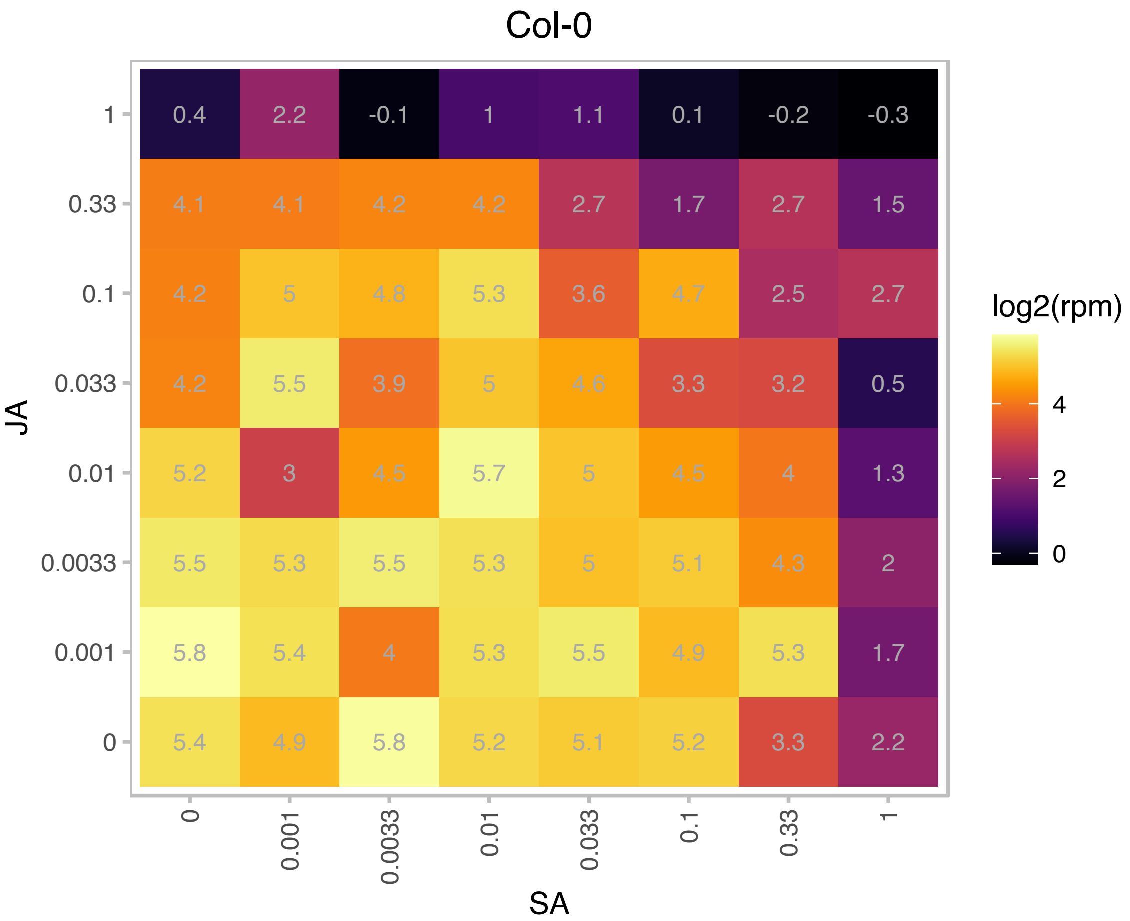


| | |
|---|---|
| Gene | AT4G21445 |
| is.expressed_genes | TRUE |
| is.responsive_genes | TRUE |
| is.apcluster | F-10 |

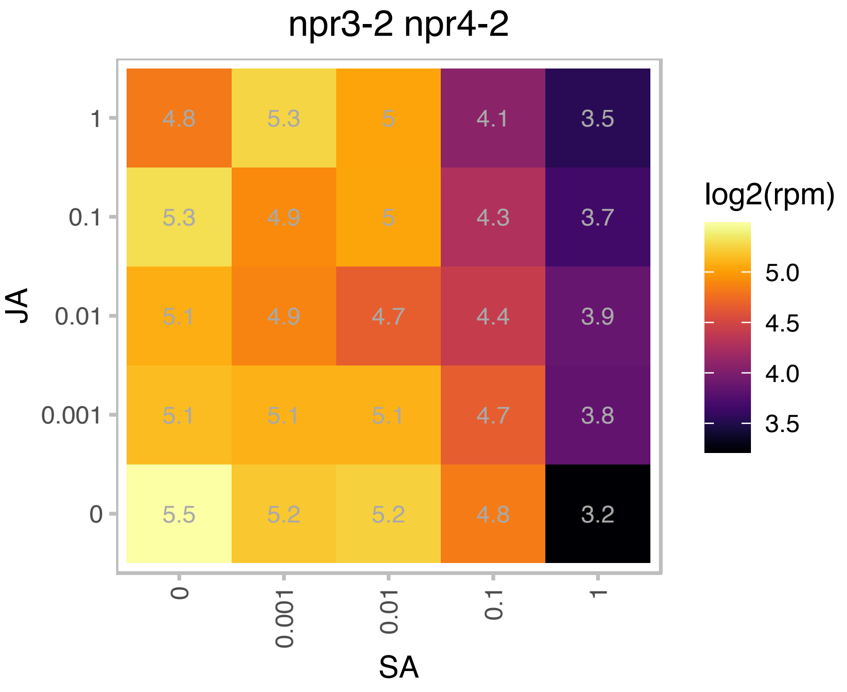


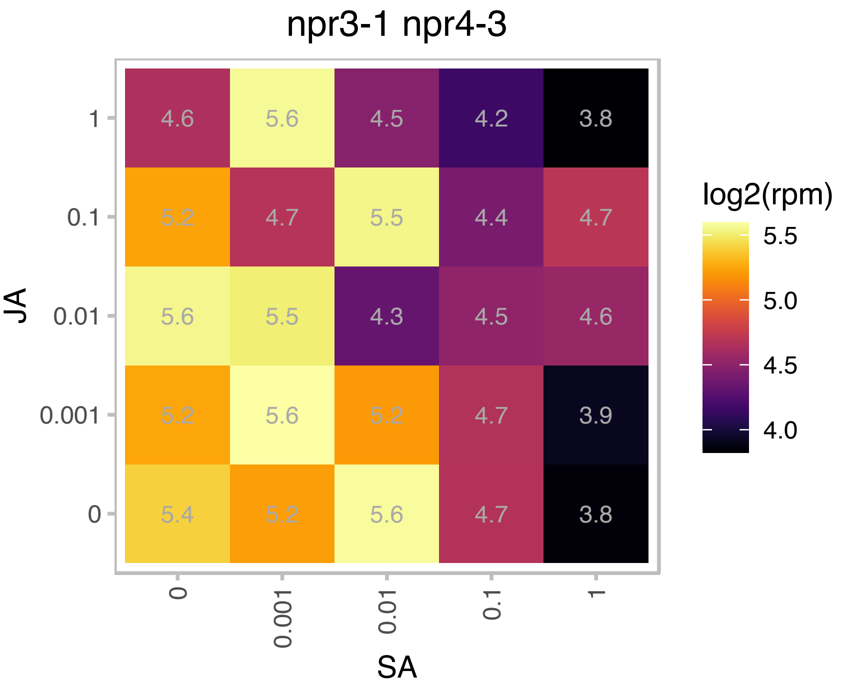

# AT4G22490.1 : NA

Bifunctional inhibitor/lipid-transfer protein/seed storage 2S albumin superfamily protein

Col-0

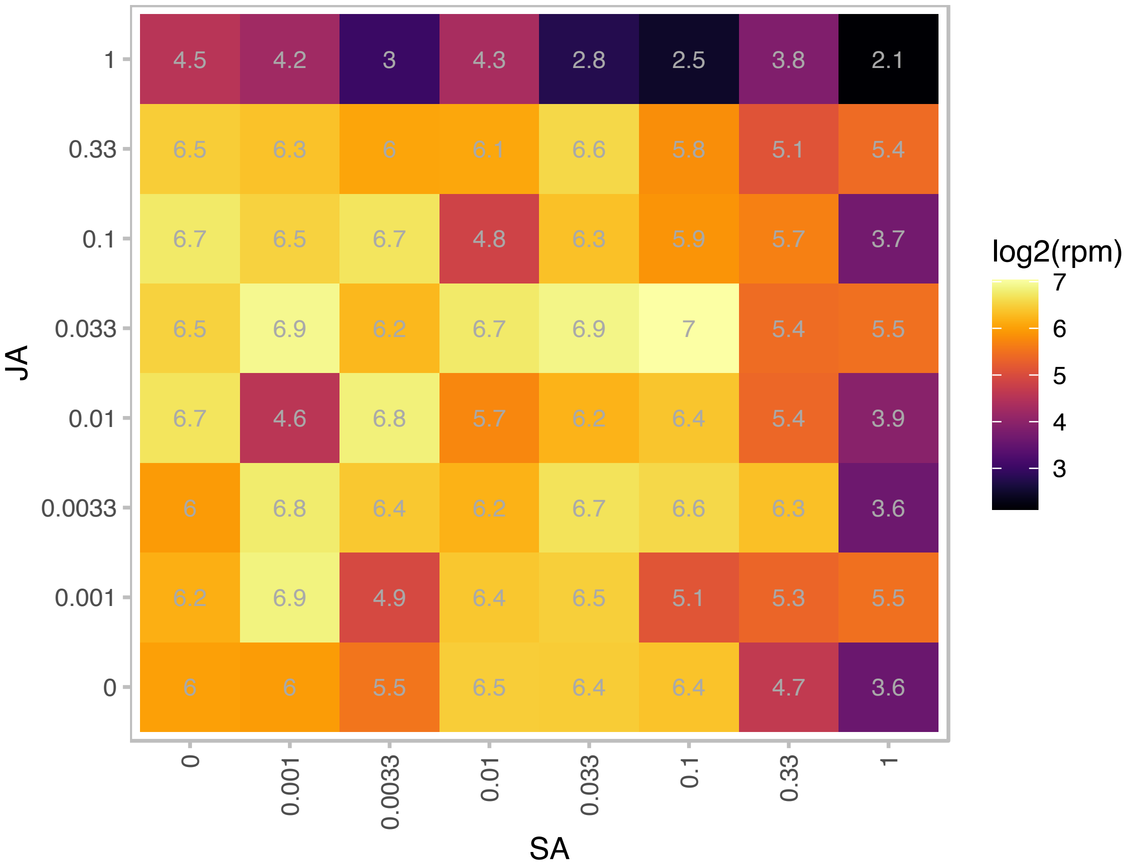


| | |
|---|---|
| Gene | AT4G22490 |
| is.expressed_genes | TRUE |
| is.responsive_genes | TRUE |
| is.apcluster | F-10 |

npr3-2 npr4-2

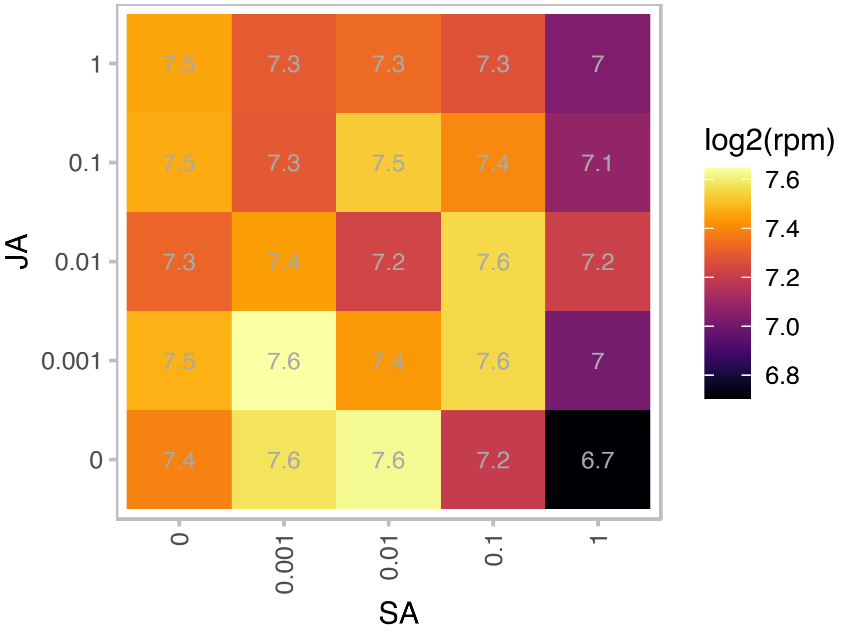


npr3-1 npr4-3

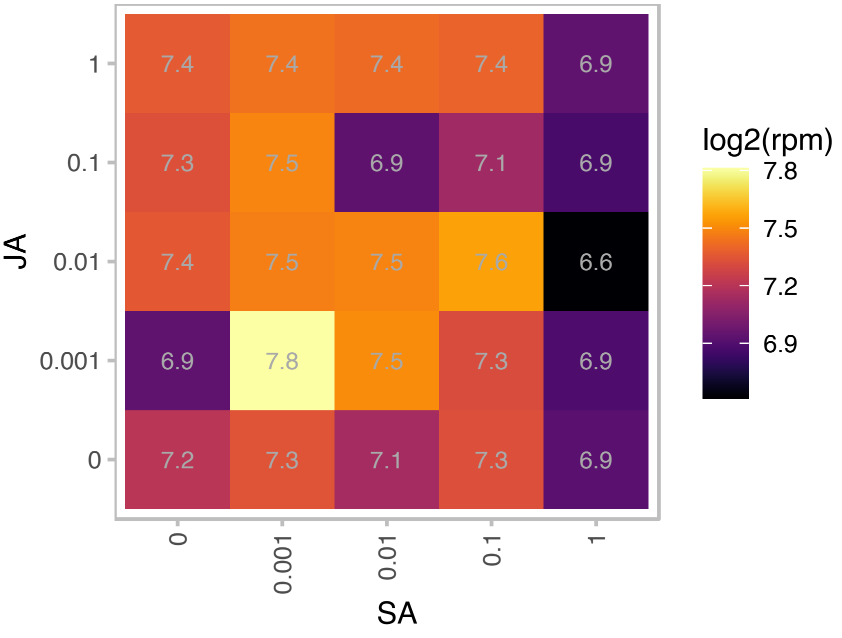

# AT4G23770.1 : NA

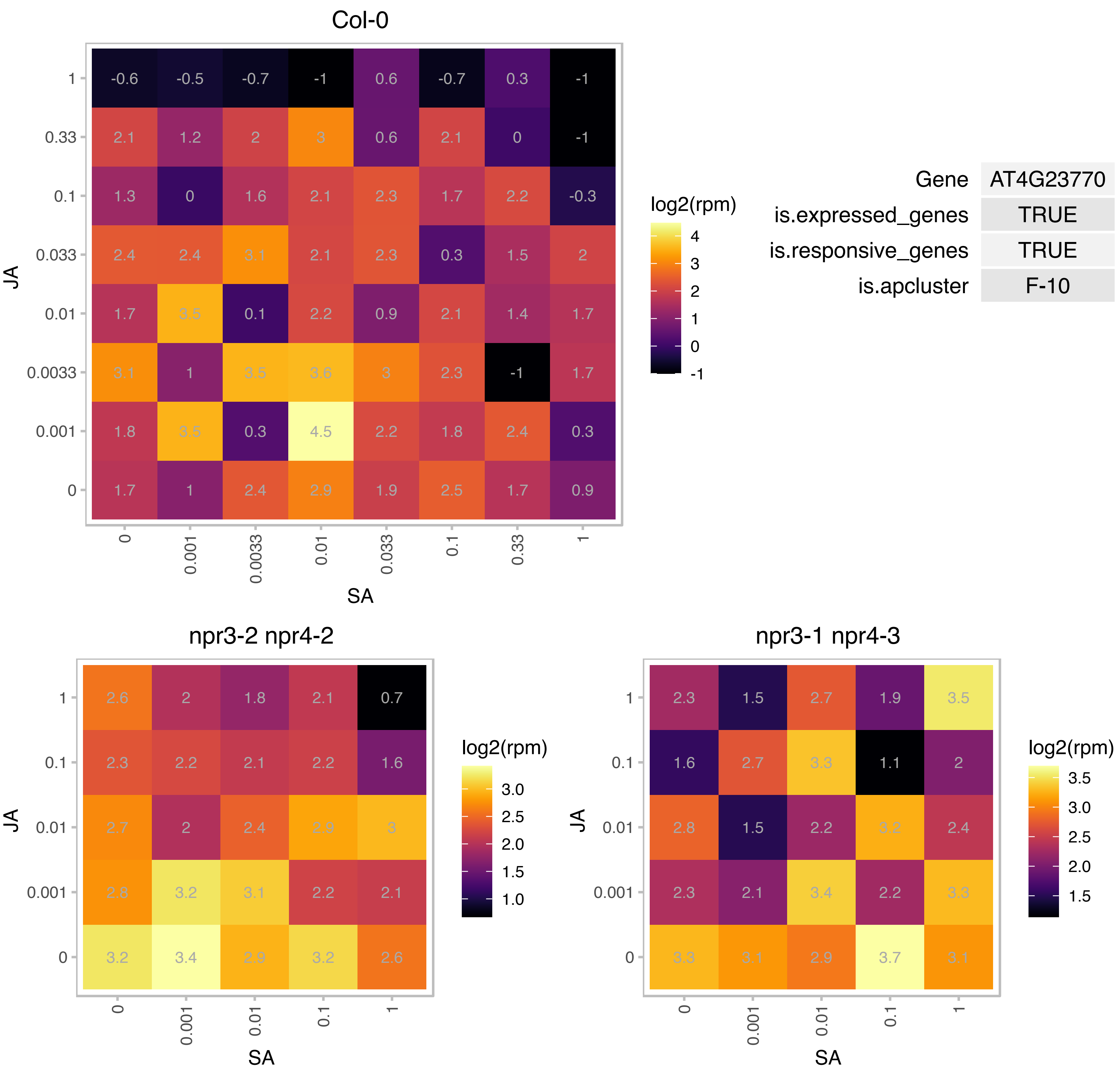

### AT4G25500.1 : At-RS40, AT-SRP40, ATRSP35, ATRSP40,

arginine/serine-rich splicing factor35

Encodes an arginine/serine-rich splicing factor. The transcript is alternatively spliced and is differentially expressed in different tissues (flowers, roots, stems, and leaves) examined. Barta et al (2010) have proposed a nomenclature for Serine/Arginine-Rich Protein Splicing Factors (SR proteins): Plant Cell. 2010, 22:2926. RS40 binds to HYL1 and co-localizes to the nuclear dicing body. Along with RS41, it appears to be involved in pri-miRNA processing and miRNA biogenesis (DOI:10.1093/nar/gkv751).

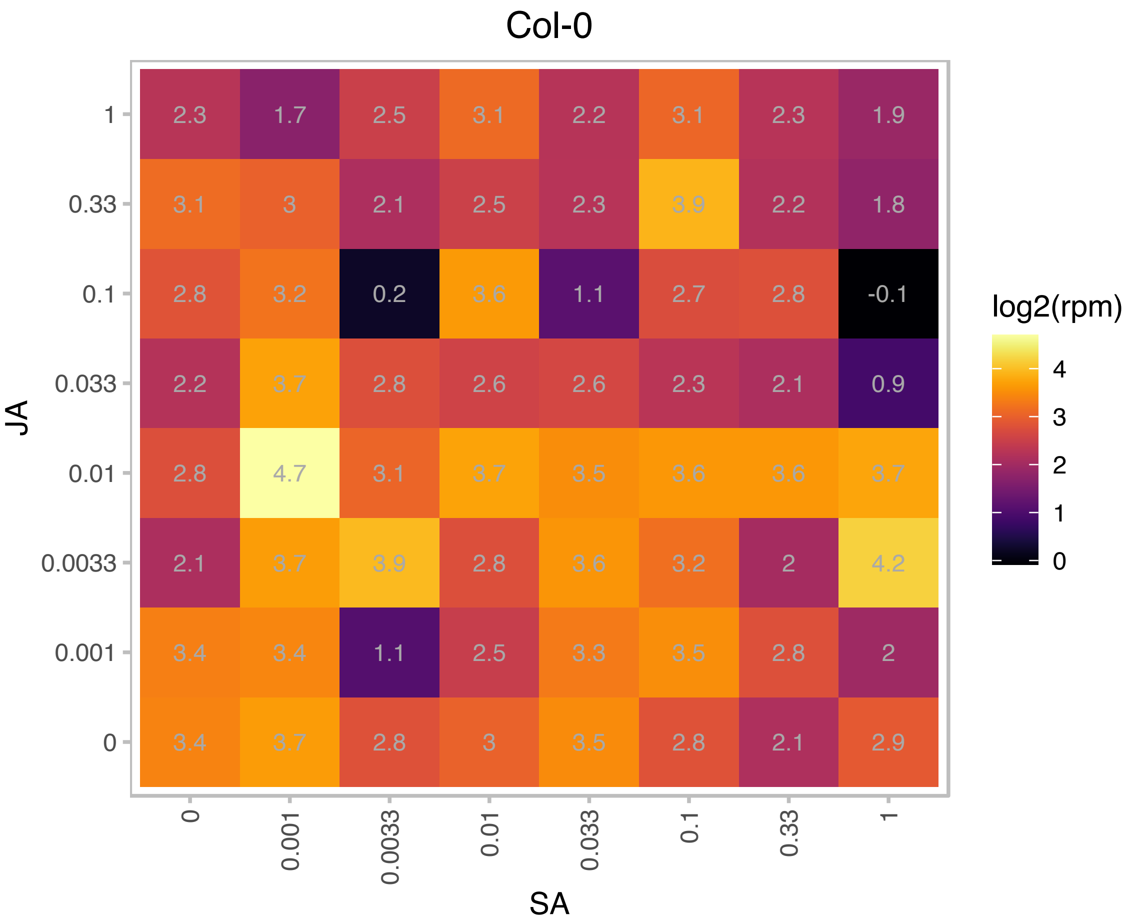


| | |
|---|---|
| Gene | AT4G25500 |
| is.expressed_genes | TRUE |
| is.responsive_genes | TRUE |
| is.apcluster | F-10 |

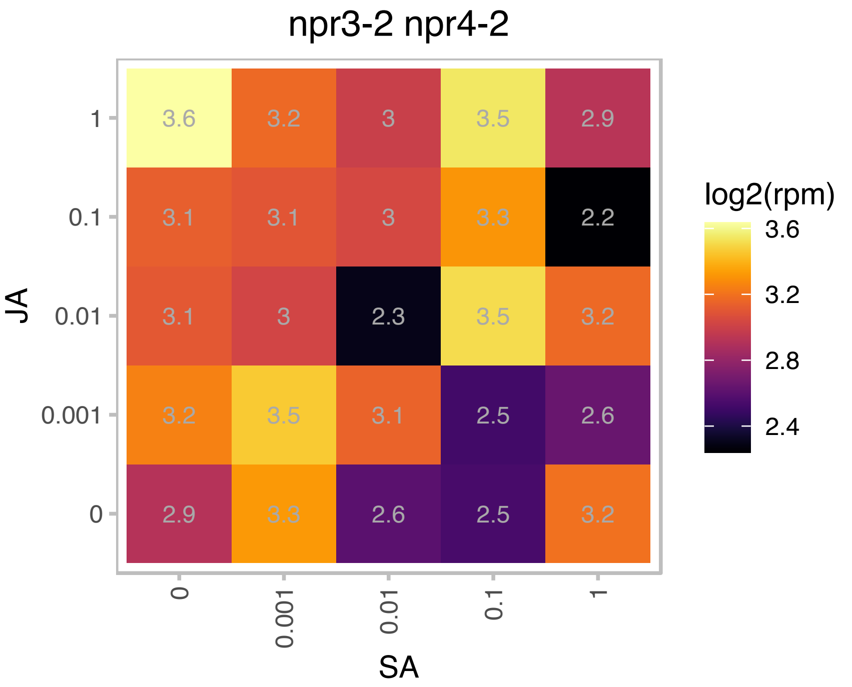


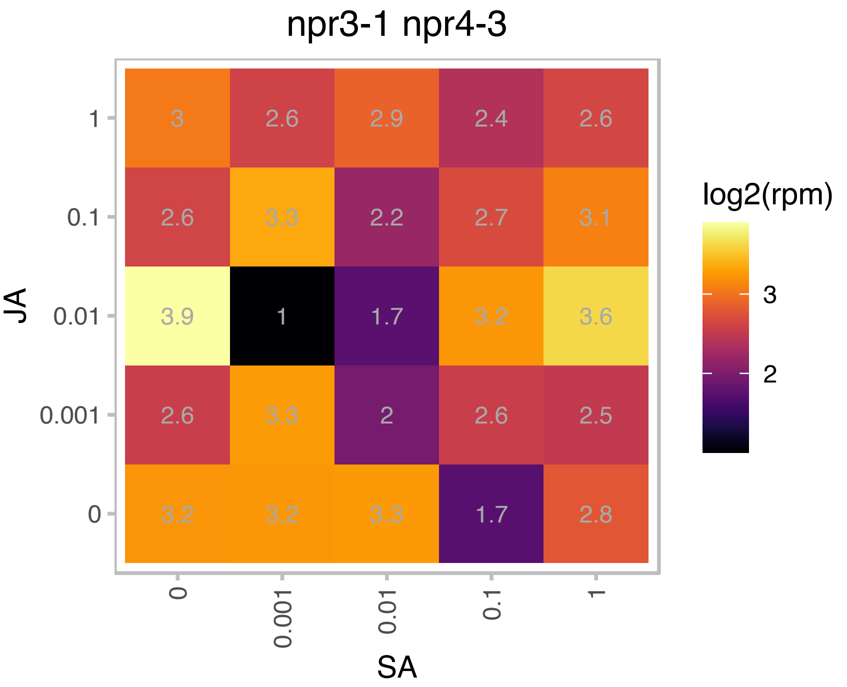

# AT4G25740.1 : NA

RNA binding Plectin/S10 domain-containing protein

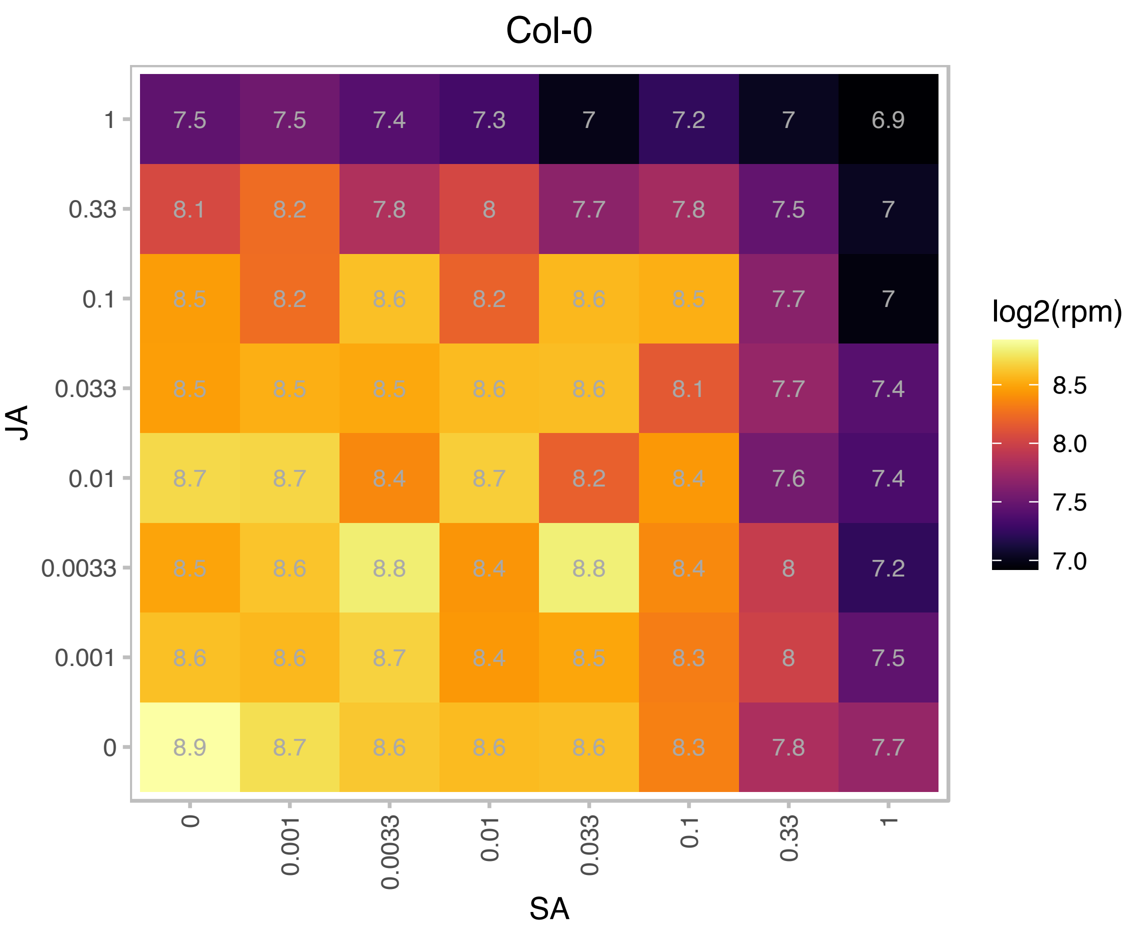


| | |
|---|---|
| Gene | AT4G25740 |
| is.expressed_genes | TRUE |
| is.responsive_genes | TRUE |
| is.apcluster | F-10 |

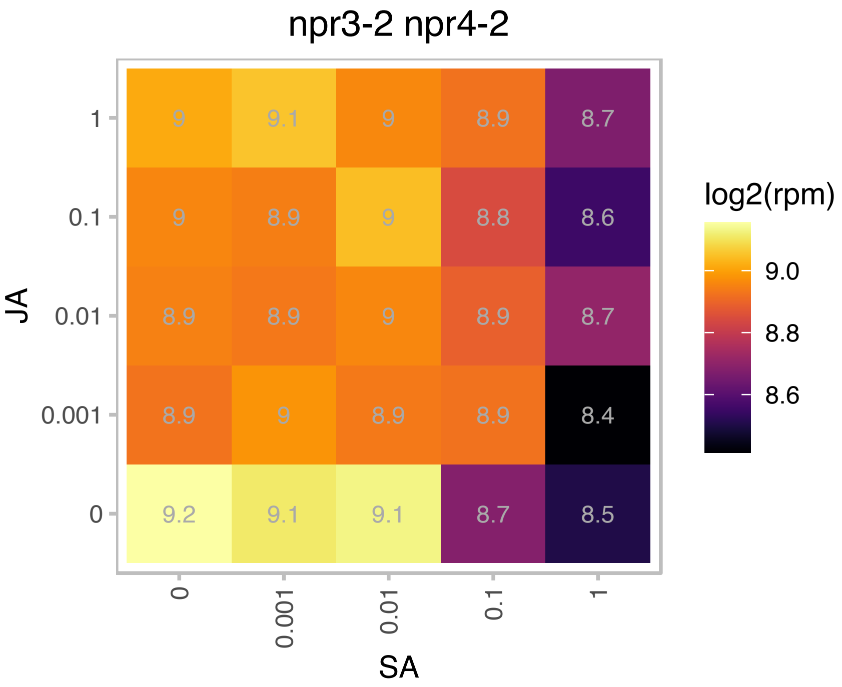


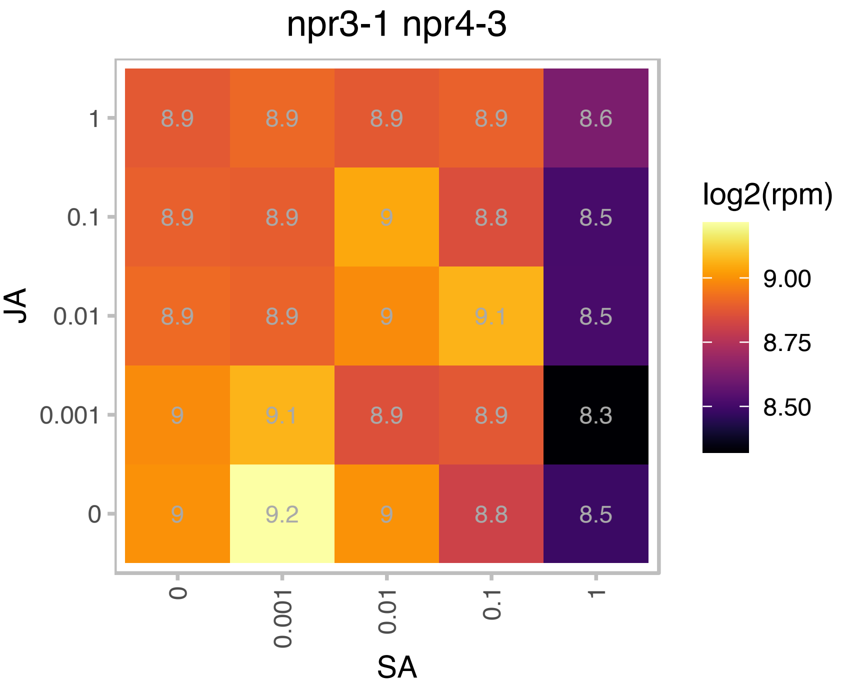

### AT4G30220.1 : RUXF, SMFA

small nuclear ribonucleoprotein F

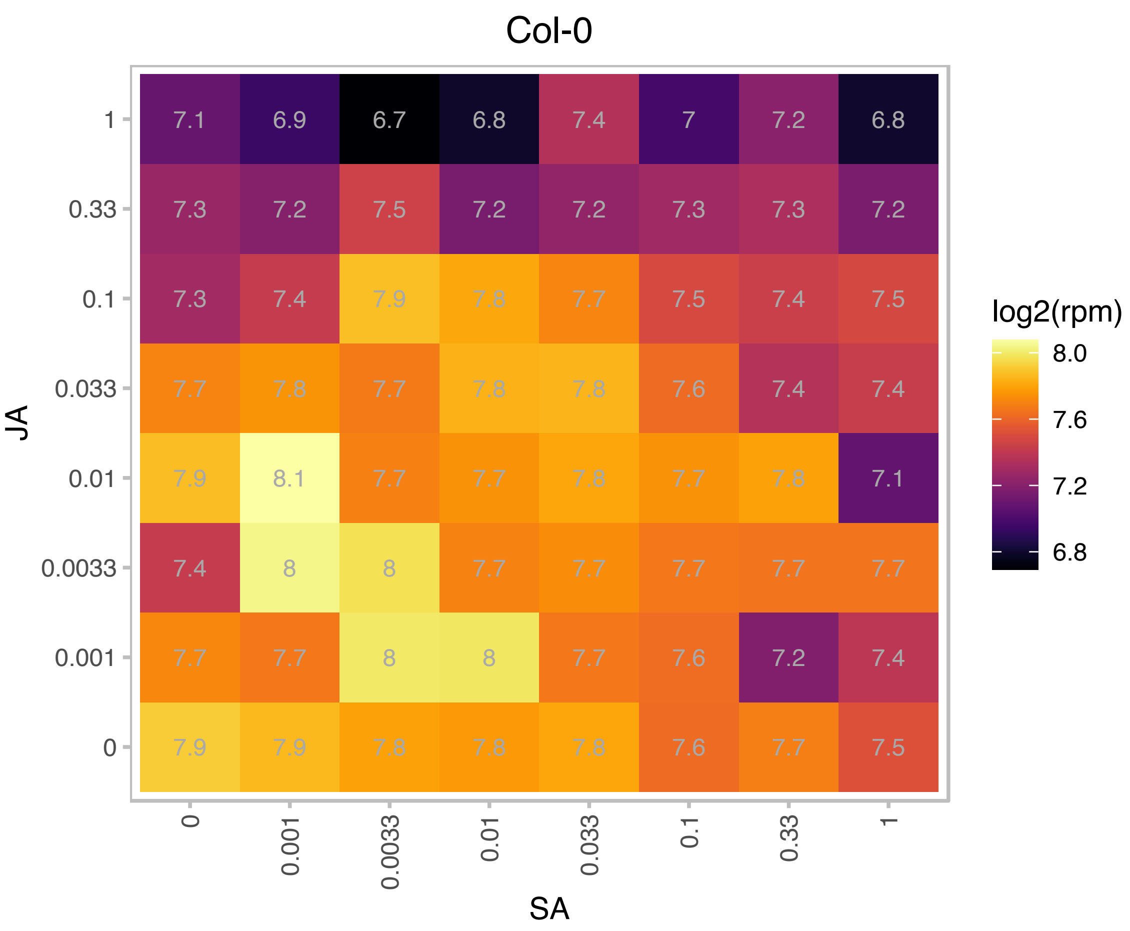


| | |
|---|---|
| Gene | AT4G30220 |
| is.expressed_genes | TRUE |
| is.responsive_genes | TRUE |
| is.apcluster | F-10 |

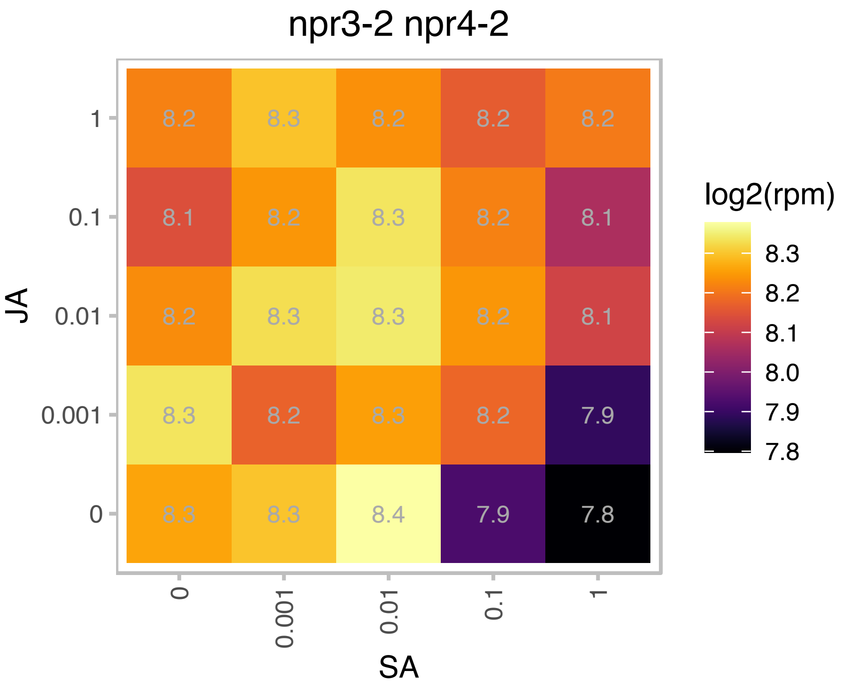


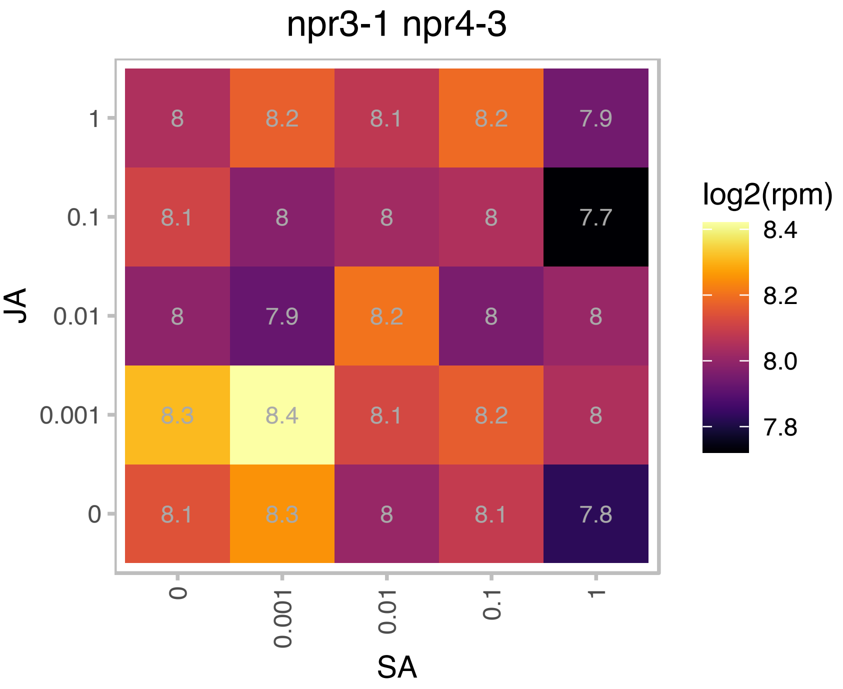

# AT4G30330.1 : NA

Small nuclear ribonucleoprotein family protein

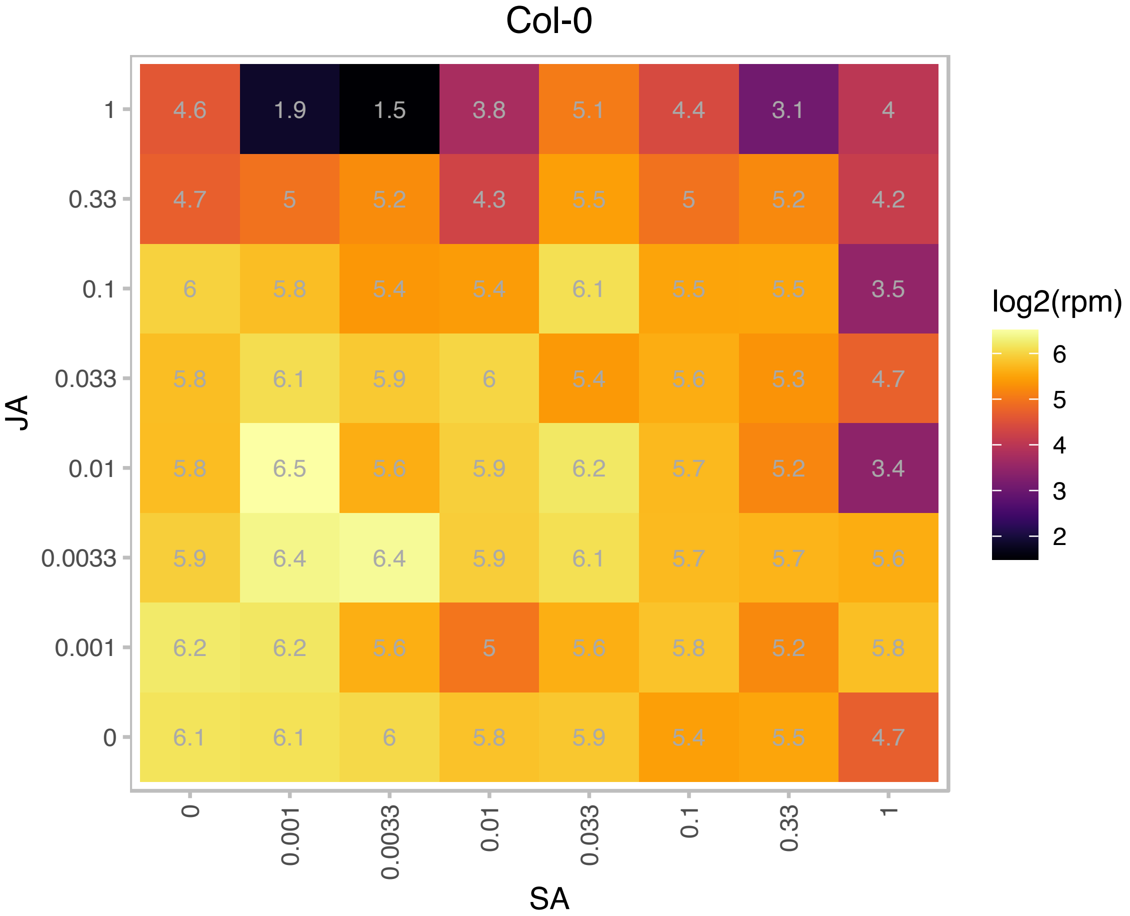


| | |
|---|---|
| Gene | AT4G30330 |
| is.expressed_genes | TRUE |
| is.responsive_genes | TRUE |
| is.apcluster | F-10 |

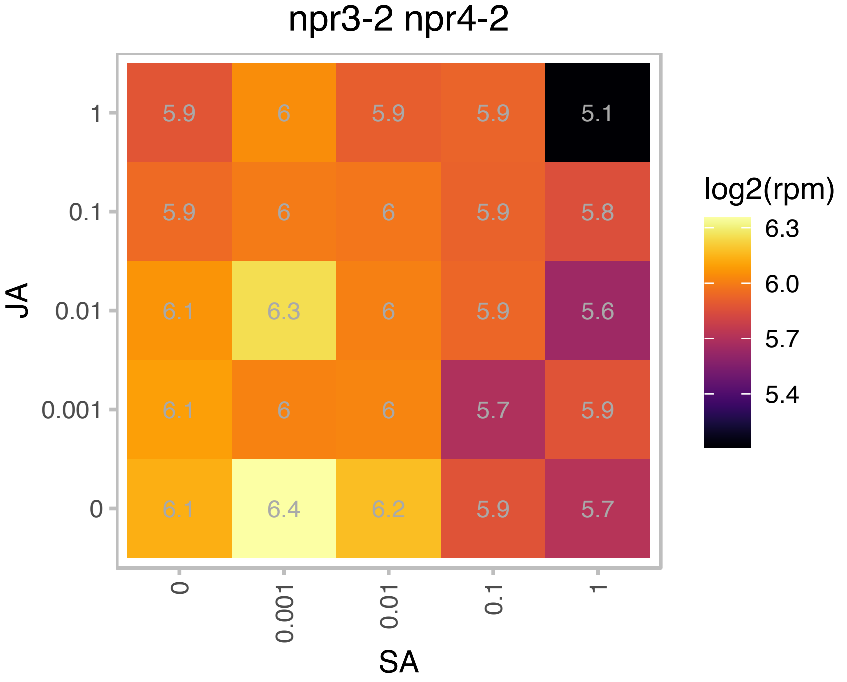


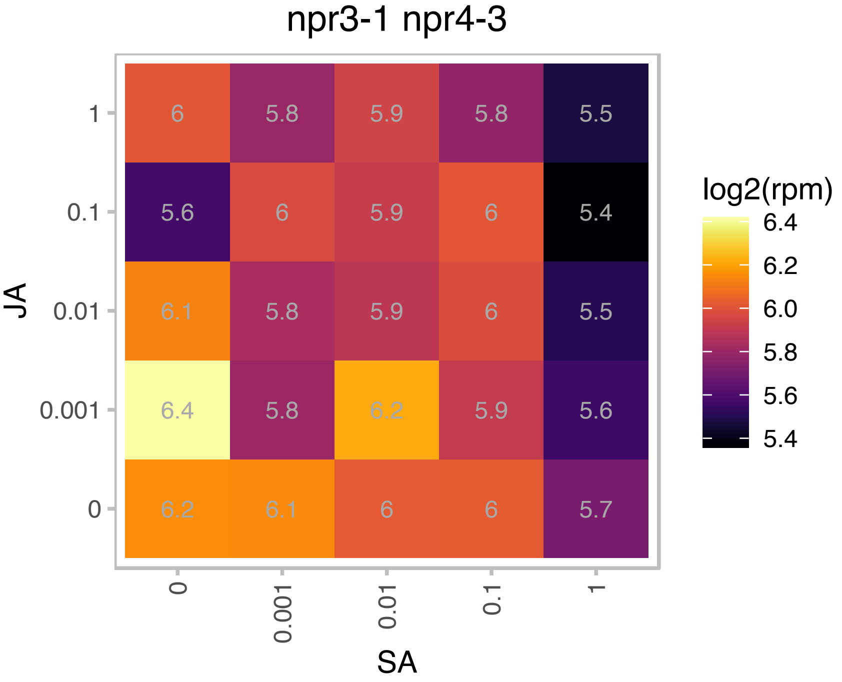

### AT4G30620.1 : STCL

Uncharacterised BCR, YbaB family COG0718

Homolog of STIC2, recent duplication.

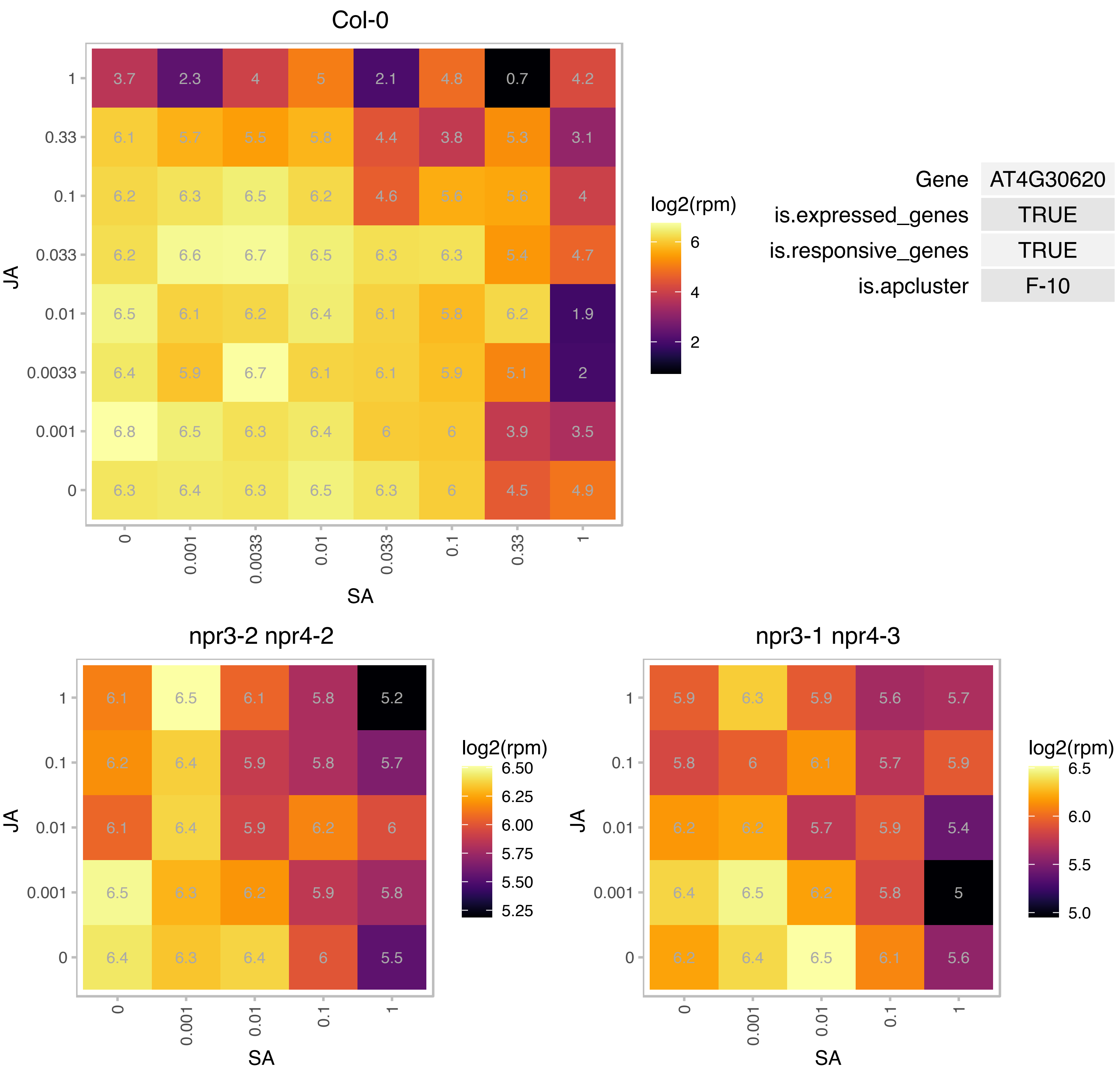


| | |
|---|---|
| Gene | AT4G30620 |
| is.expressed_genes | TRUE |
| is.responsive_genes | TRUE |
| is.apcluster | F-10 |

# AT4G32240.1 : NA

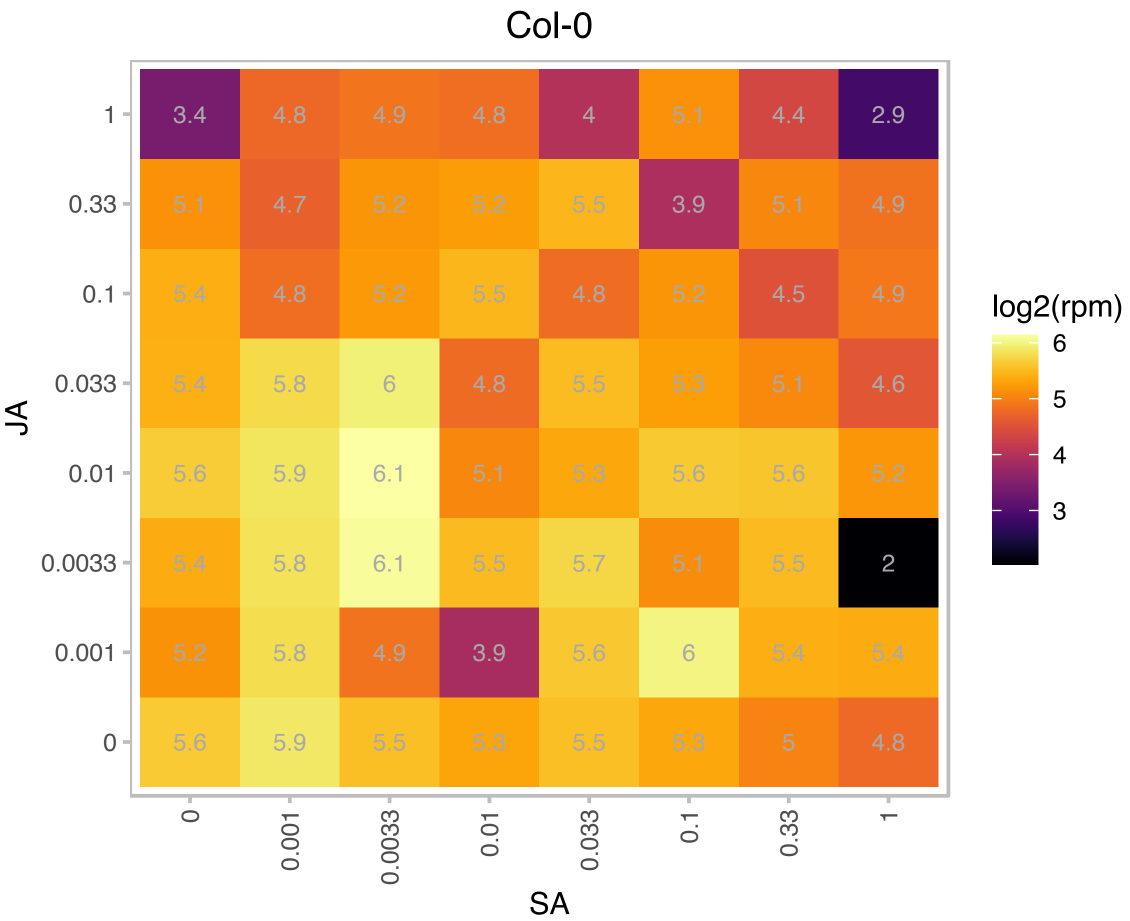


| | |
|---|---|
| Gene | AT4G32240 |
| is.expressed_genes | TRUE |
| is.responsive_genes | TRUE |
| is.apcluster | F-10 |

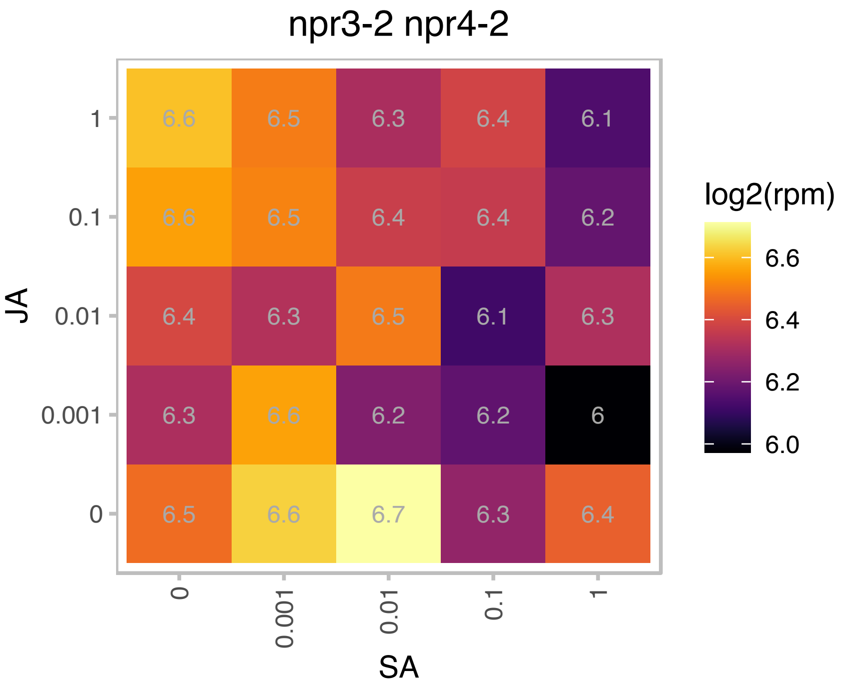


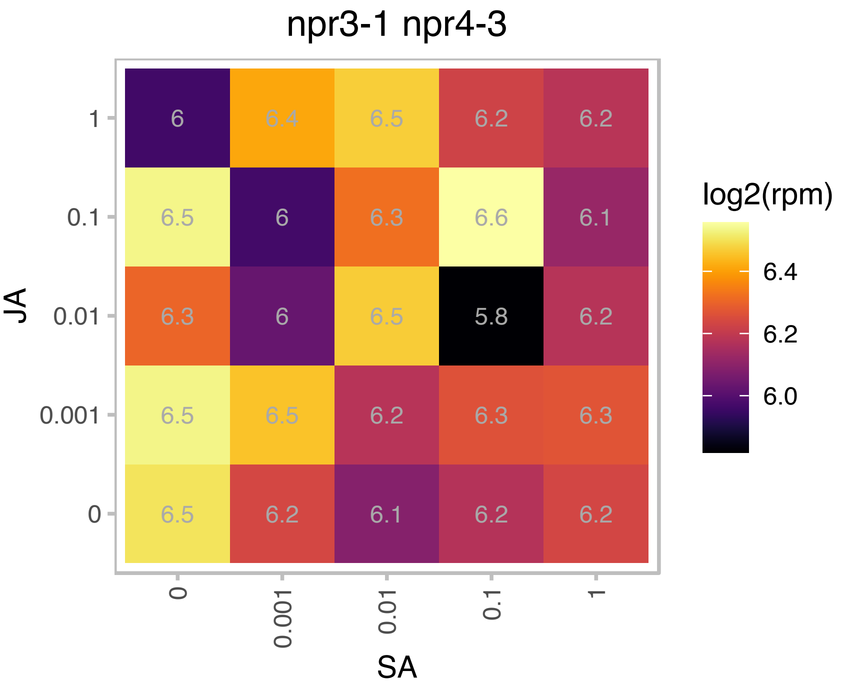

# AT4G32340.1 : NA

Tetratricopeptide repeat (TPR)-like superfamily protein

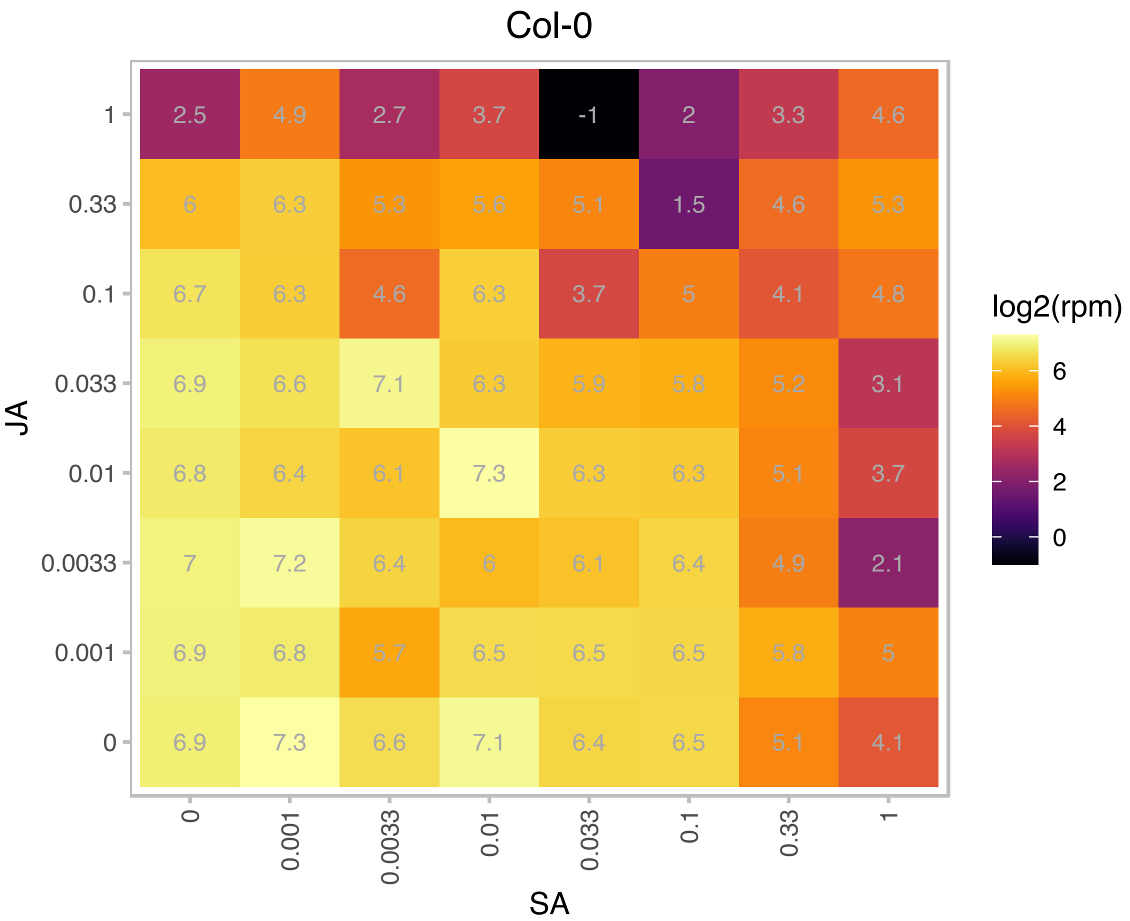


| | |
|---|---|
| Gene | AT4G32340 |
| is.expressed_genes | TRUE |
| is.responsive_genes | TRUE |
| is.apcluster | F-10 |

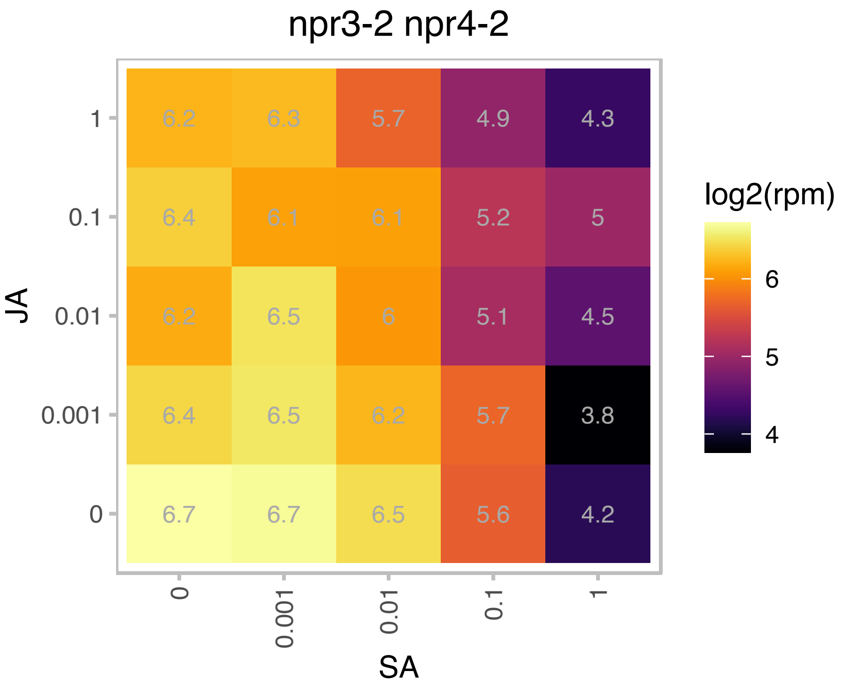


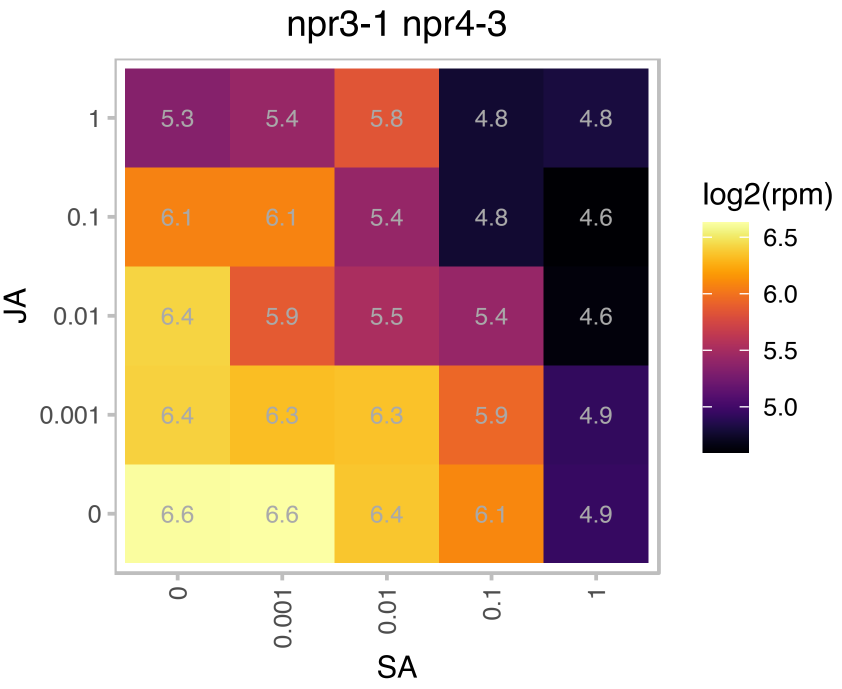

# AT4G32915.1 : NA

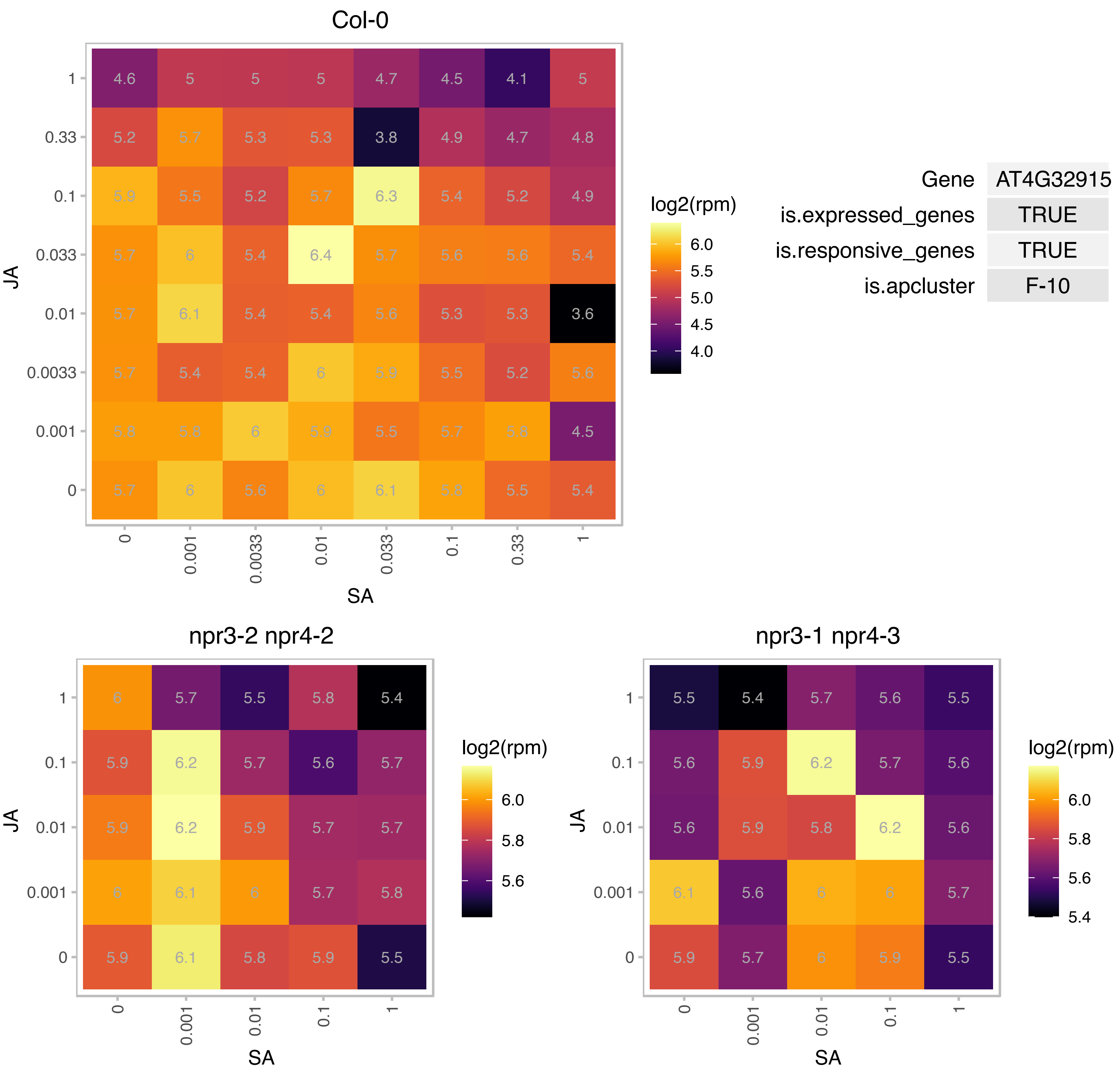

### AT4G33270.1 : AtCDC20.1, CDC20.1

Transducin family protein / WD-40 repeat family protein

Encodes a CDC20 protein that interacts with APC subunits, components of the mitochondrial checkpoint complex and mitotic cyclin substrates and is indispensable for normal plant development and fertility.

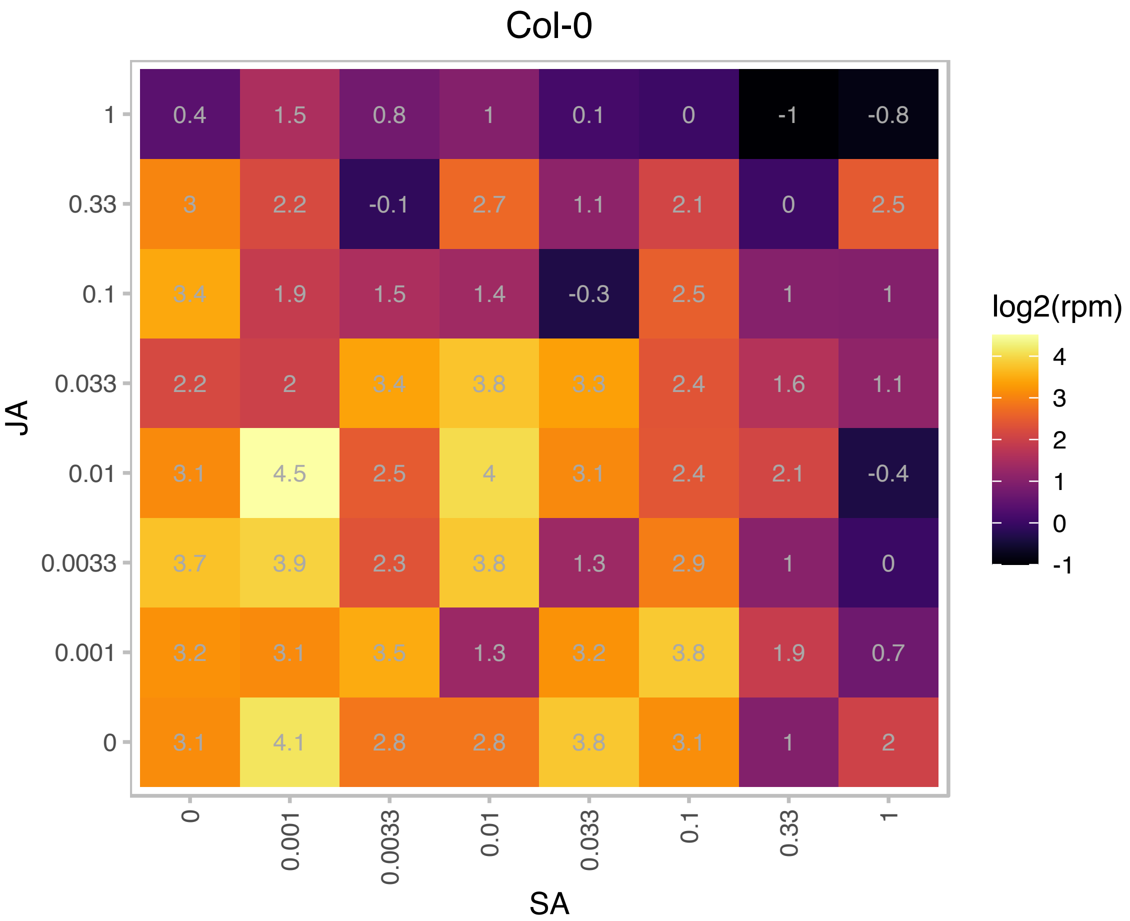


| | |
|---|---|
| Gene | AT4G33270 |
| is.expressed_genes | TRUE |
| is.responsive_genes | TRUE |
| is.apcluster | F-10 |

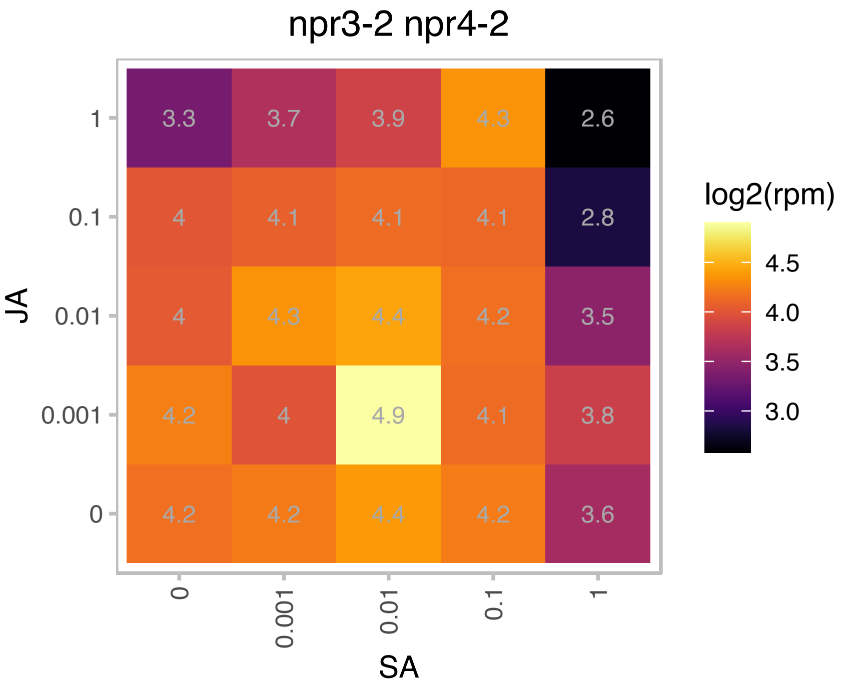


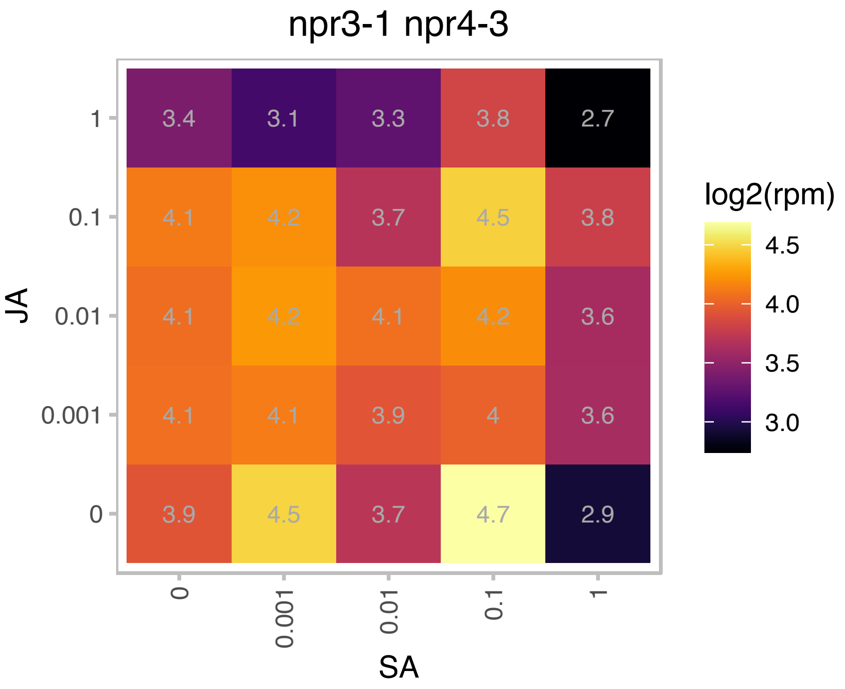

### AT4G34290.1 : SWIB3

SWIB/MDM2 domain superfamily protein

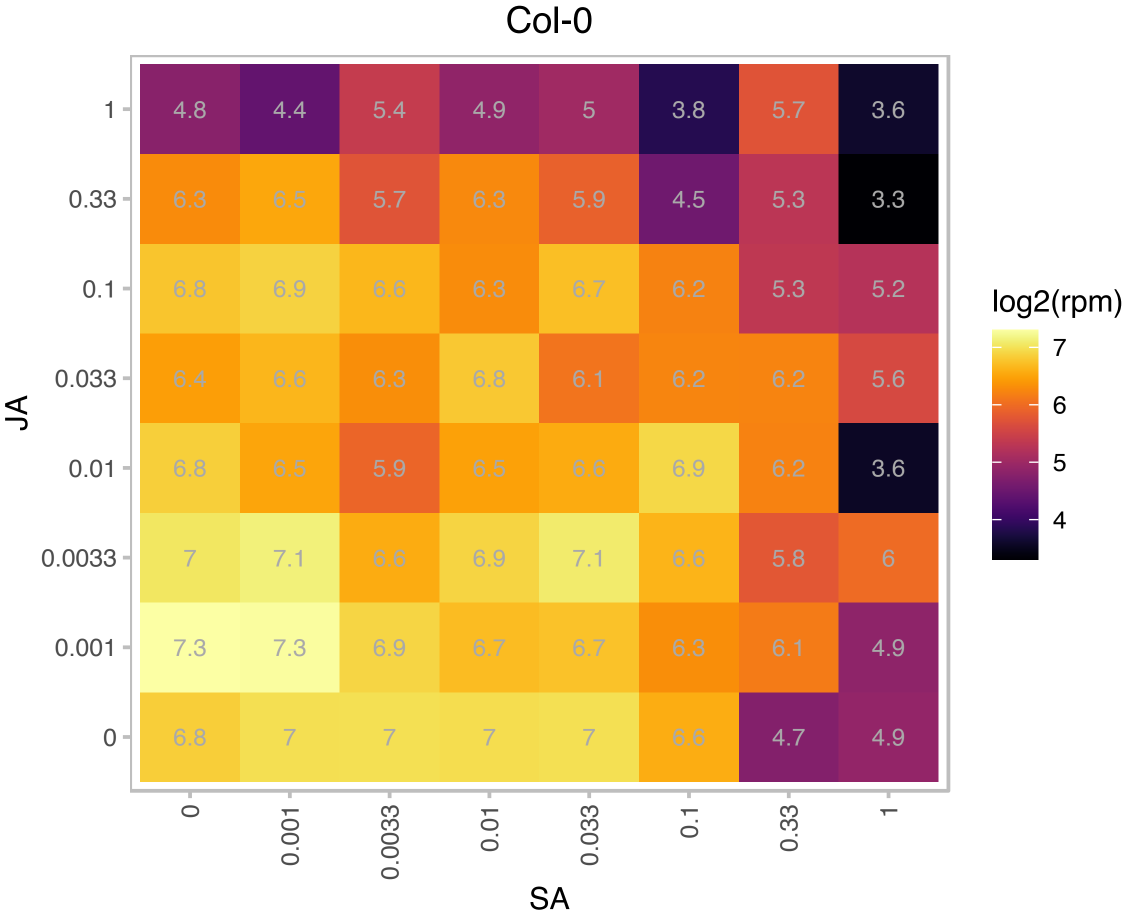


| | |
|---|---|
| Gene | AT4G34290 |
| is.expressed_genes | TRUE |
| is.responsive_genes | TRUE |
| is.apcluster | F-10 |

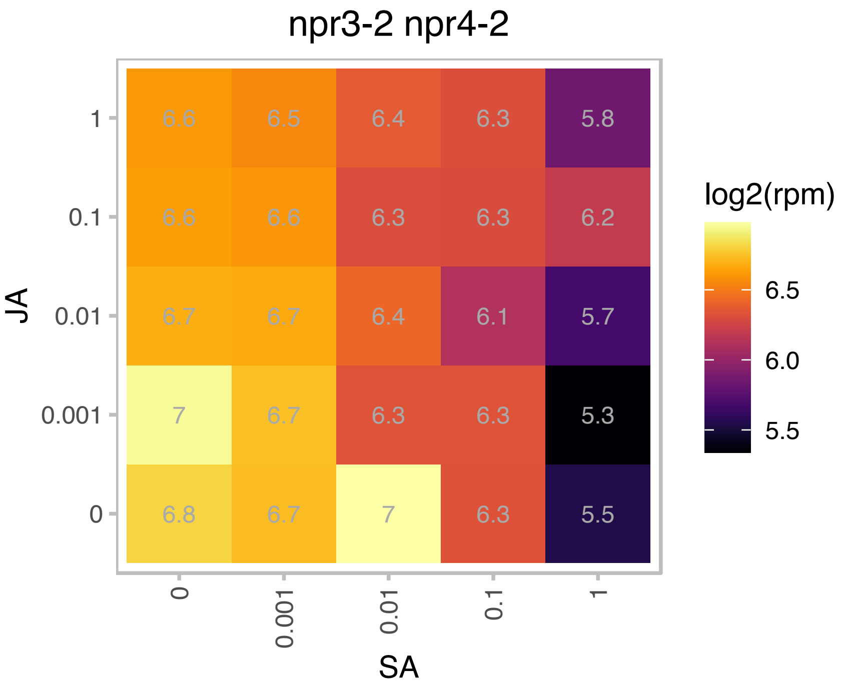


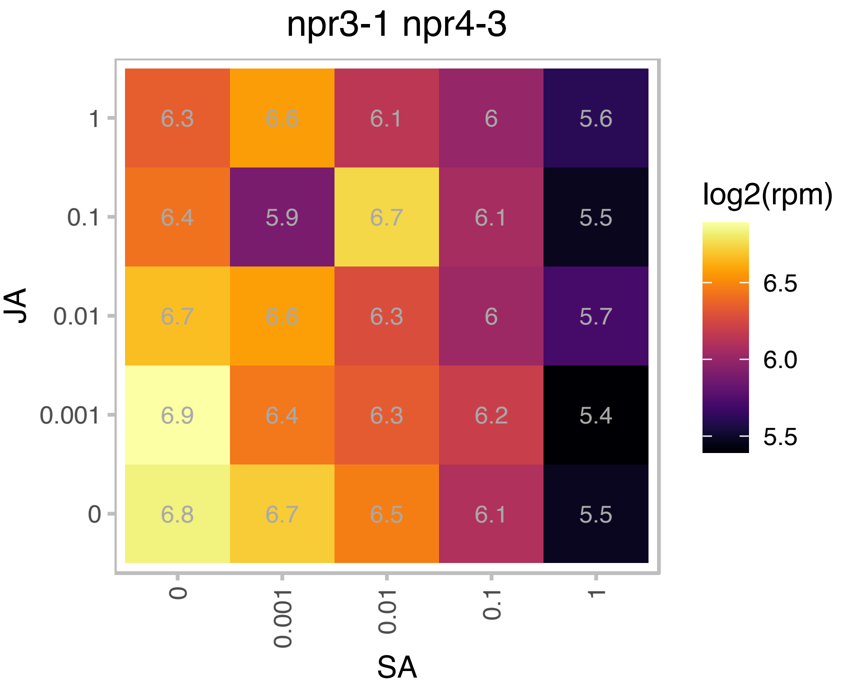

### AT4G36670.1 : AtPLT6, AtPMT6, PLT6, PMT6

Major facilitator superfamily protein

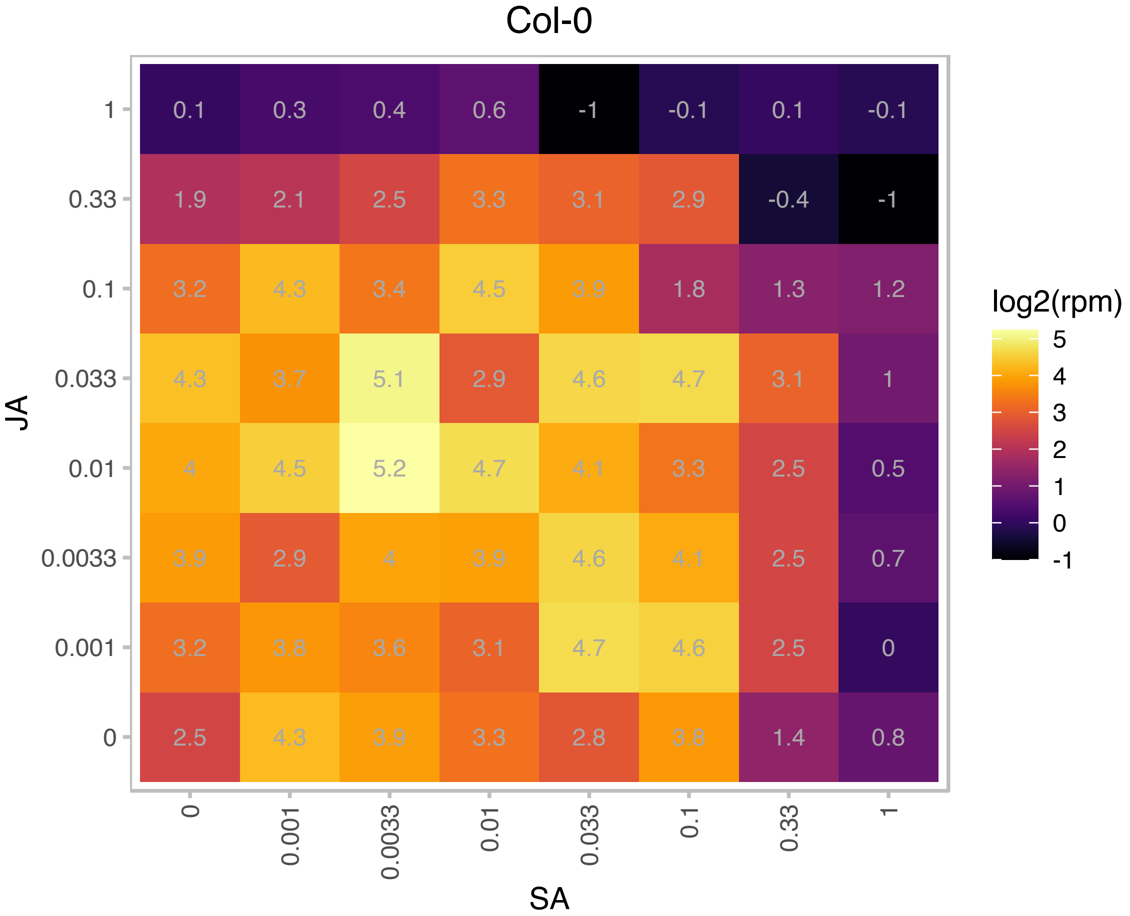


| | |
|---|---|
| Gene | AT4G36670 |
| is.expressed_genes | TRUE |
| is.responsive_genes | TRUE |
| is.apcluster | F-10 |

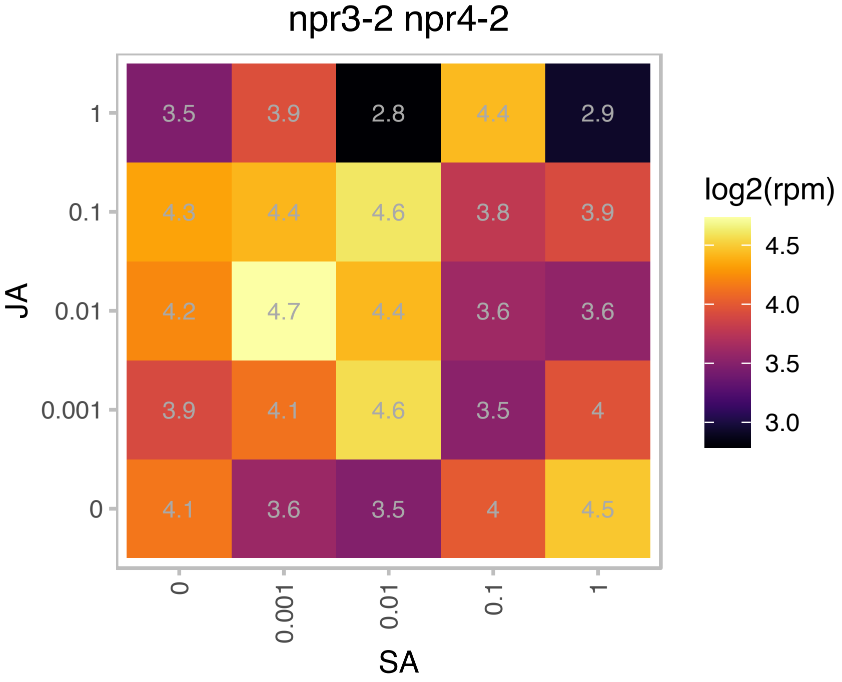


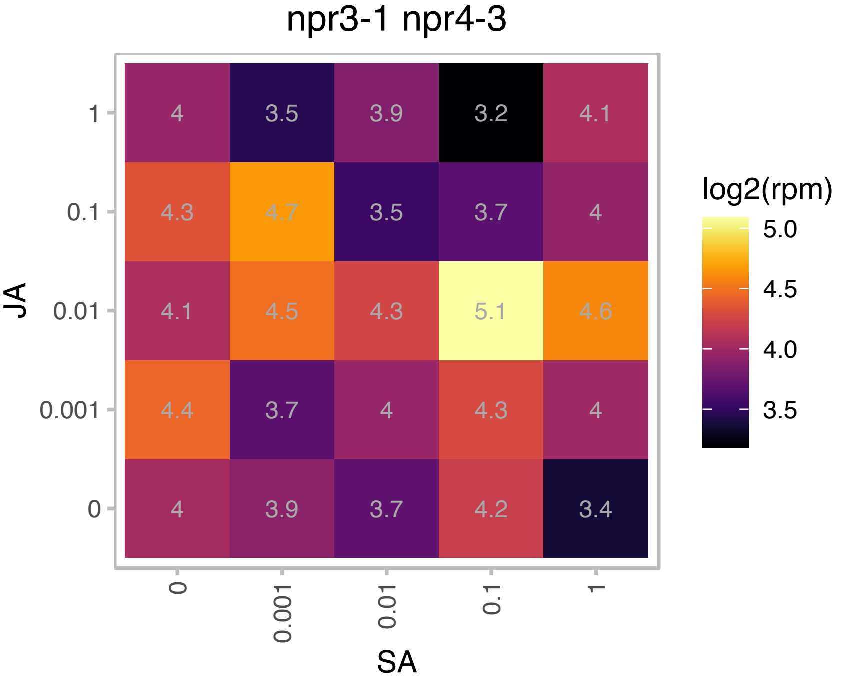

### AT4G37300.1 : MEE59

maternal effect embryo arrest 59

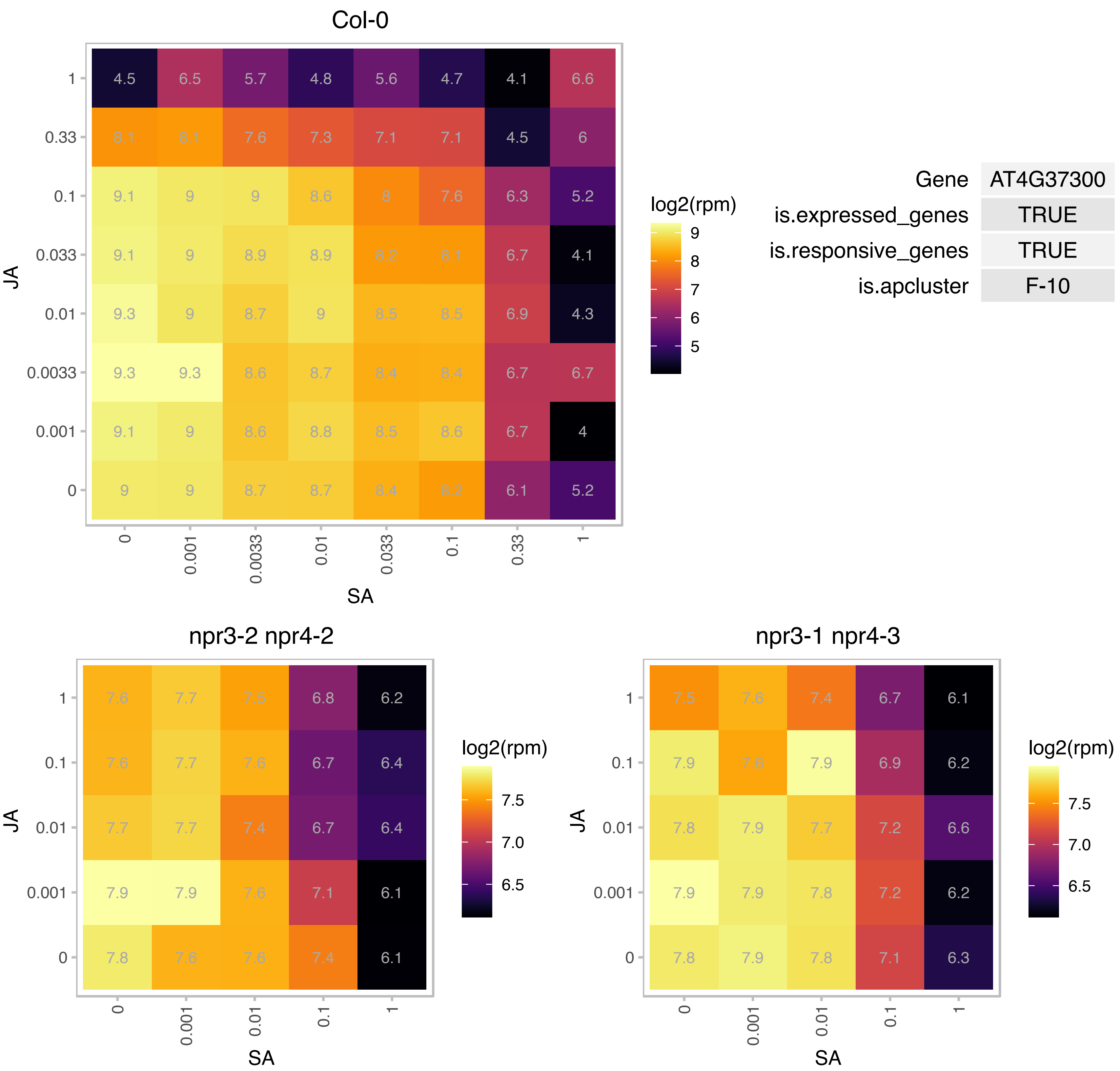


| | |
|---|---|
| Gene | AT4G37300 |
| is.expressed_genes | TRUE |
| is.responsive_genes | TRUE |
| is.apcluster | F-10 |

### AT4G38160.1 : mTERF6, pde191

Mitochondrial transcription termination factor family protein

Encodes a member of the mitochondrial transcription termination factor family. MTERF6 is localized to both the mitochondria and chloroplast. It is required for ILE tRNA maturation during chloroplast development. Loss of function of MTERF6 results in plants with defective chloroplasts or reduced rate of photosynthesis when MTERF6 expression is low.

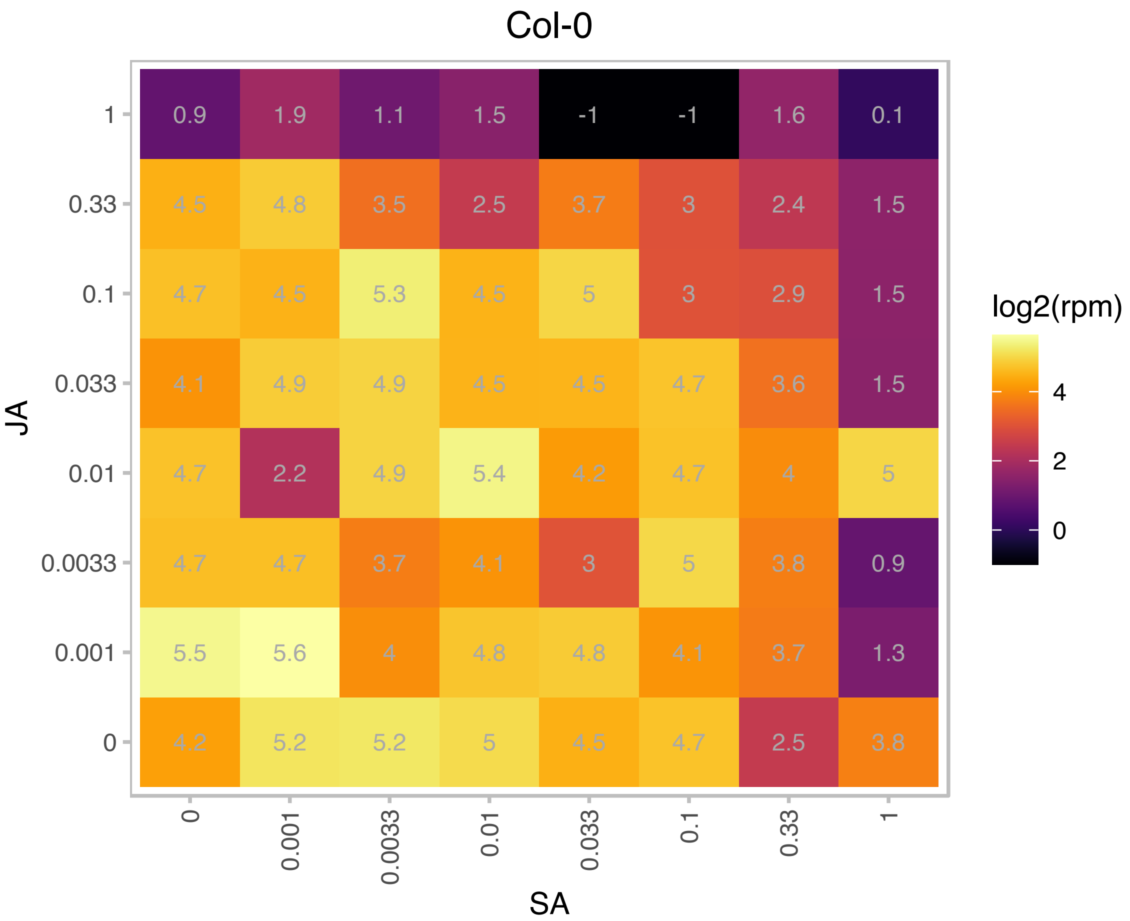


| | |
|---|---|
| Gene | AT4G38160 |
| is.expressed_genes | TRUE |
| is.responsive_genes | TRUE |
| is.apcluster | F-10 |

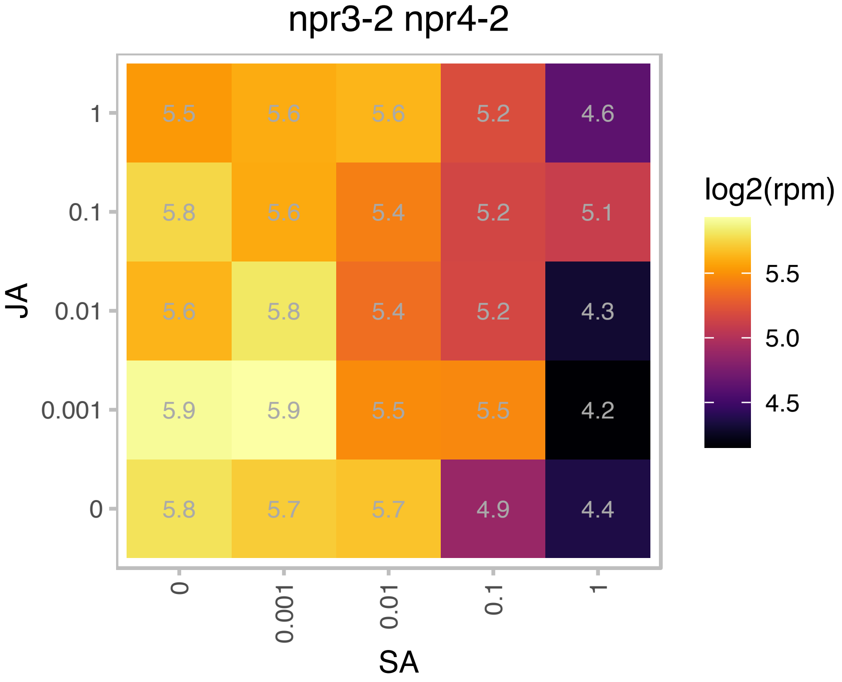


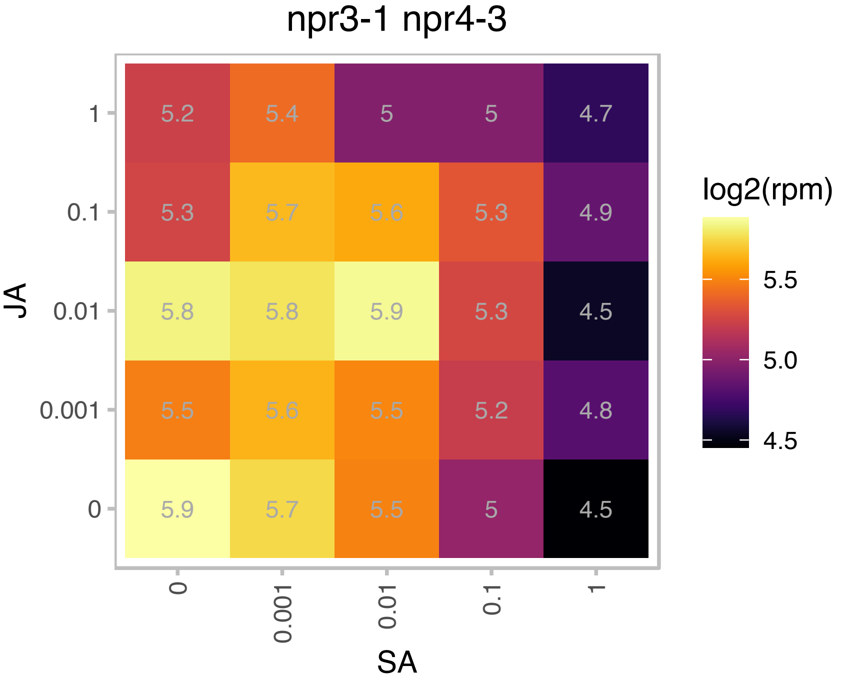

# AT4G38280.1 : NA

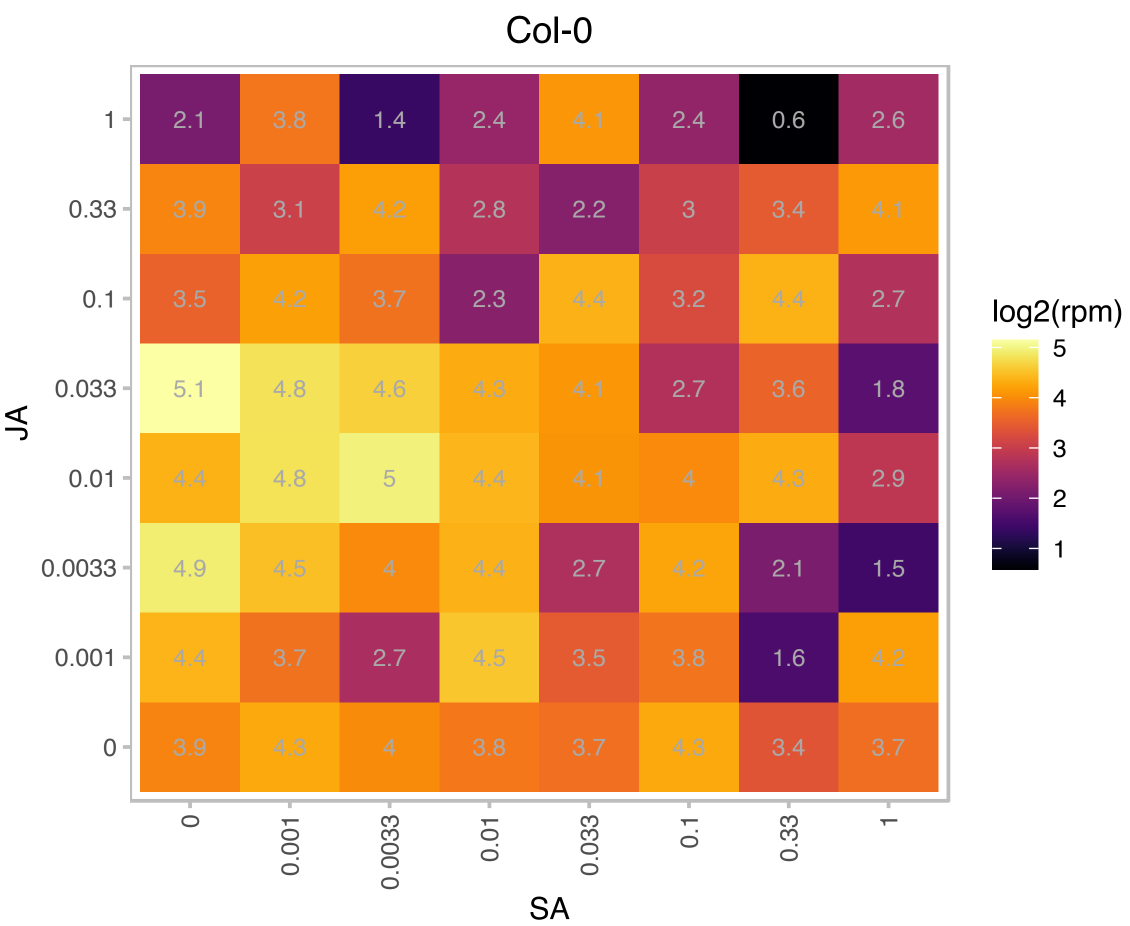


| | |
|---|---|
| Gene | AT4G38280 |
| is.expressed_genes | TRUE |
| is.responsive_genes | TRUE |
| is.apcluster | F-10 |

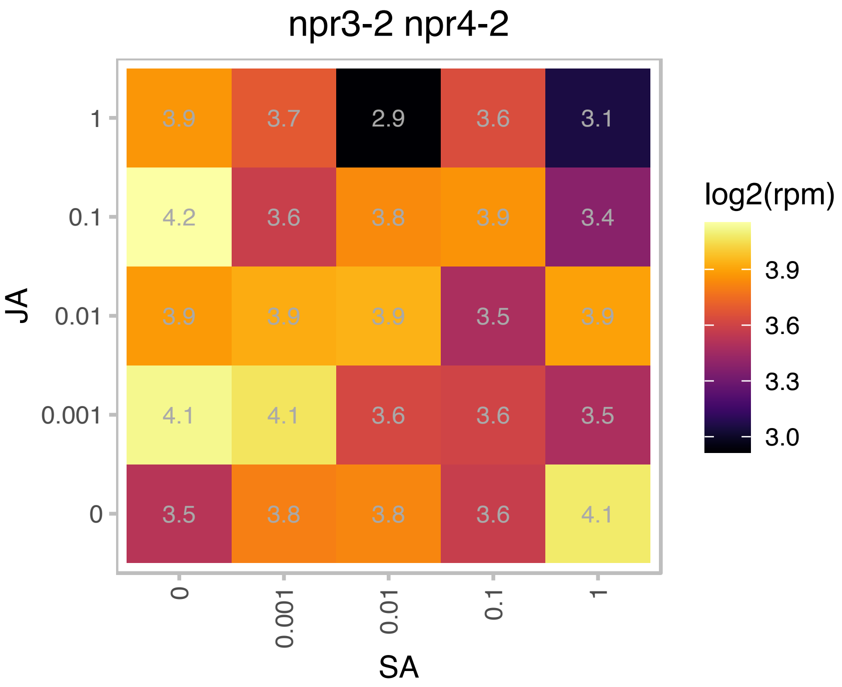


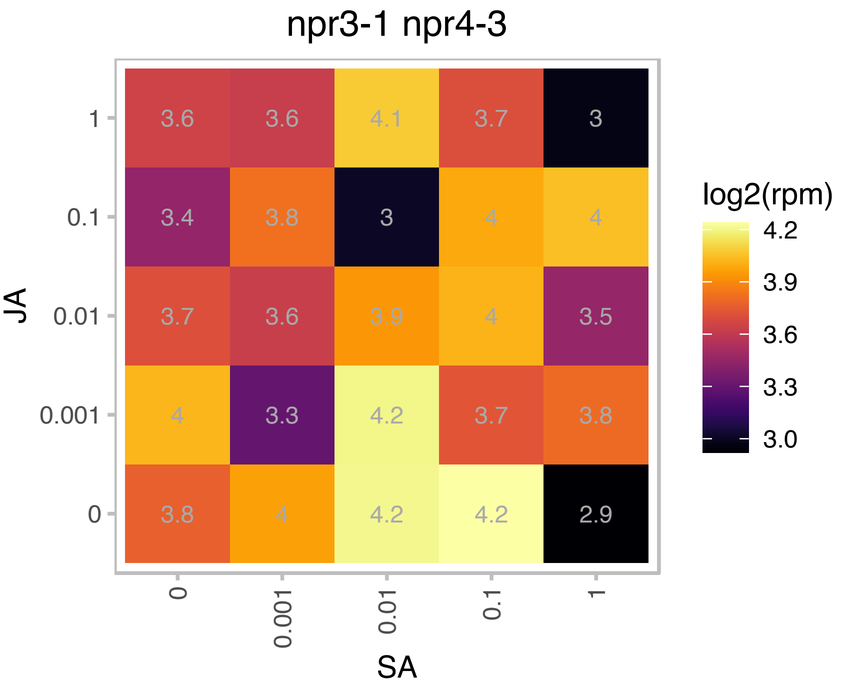

### AT4G38490.1 : SECE2

| | |
|---|---|
| Gene | AT4G38490 |
| is.expressed_genes | TRUE |
| is.responsive_genes | TRUE |
| is.apcluster | F-10 |

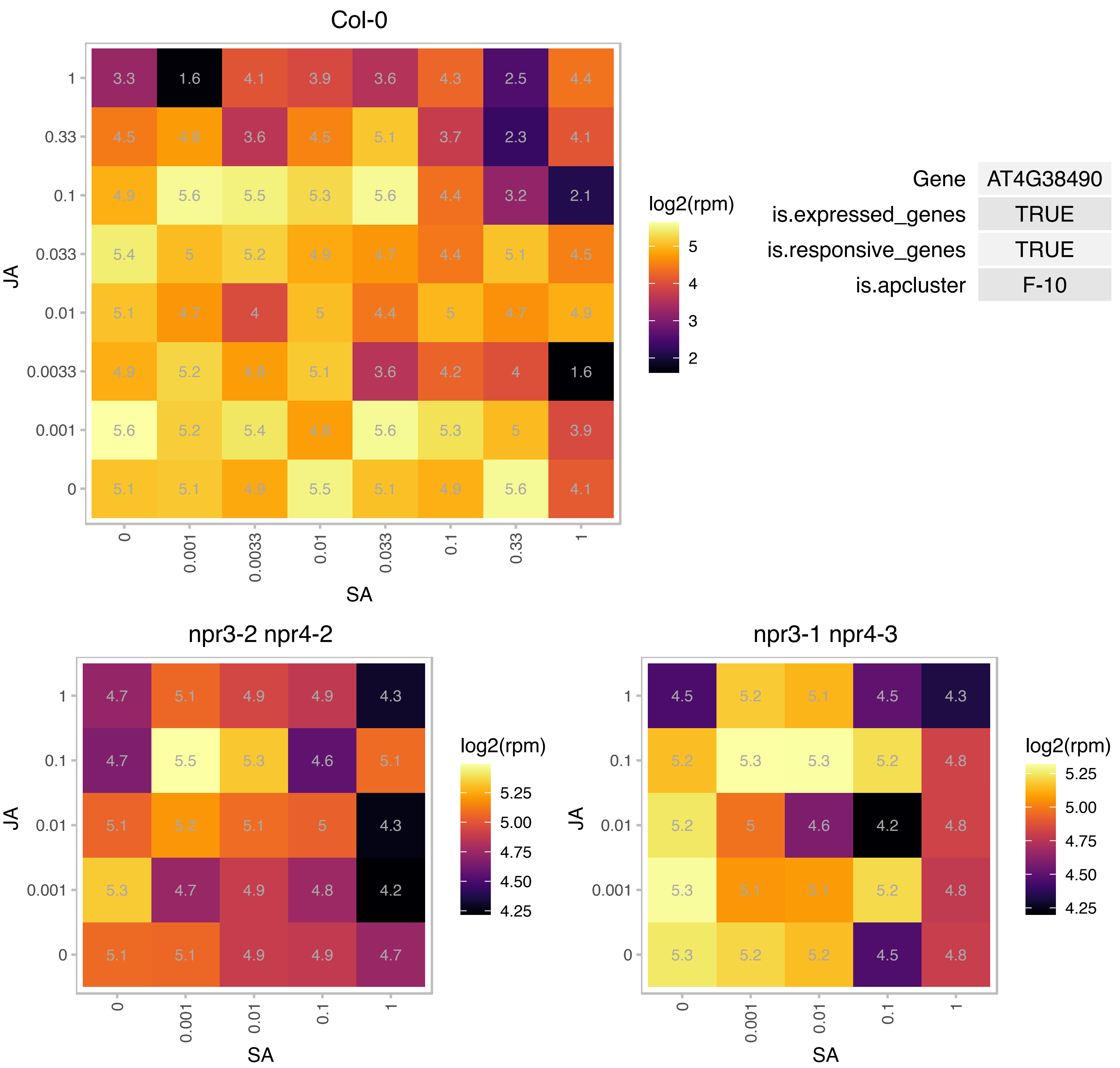

# AT5G02710.1 : NA

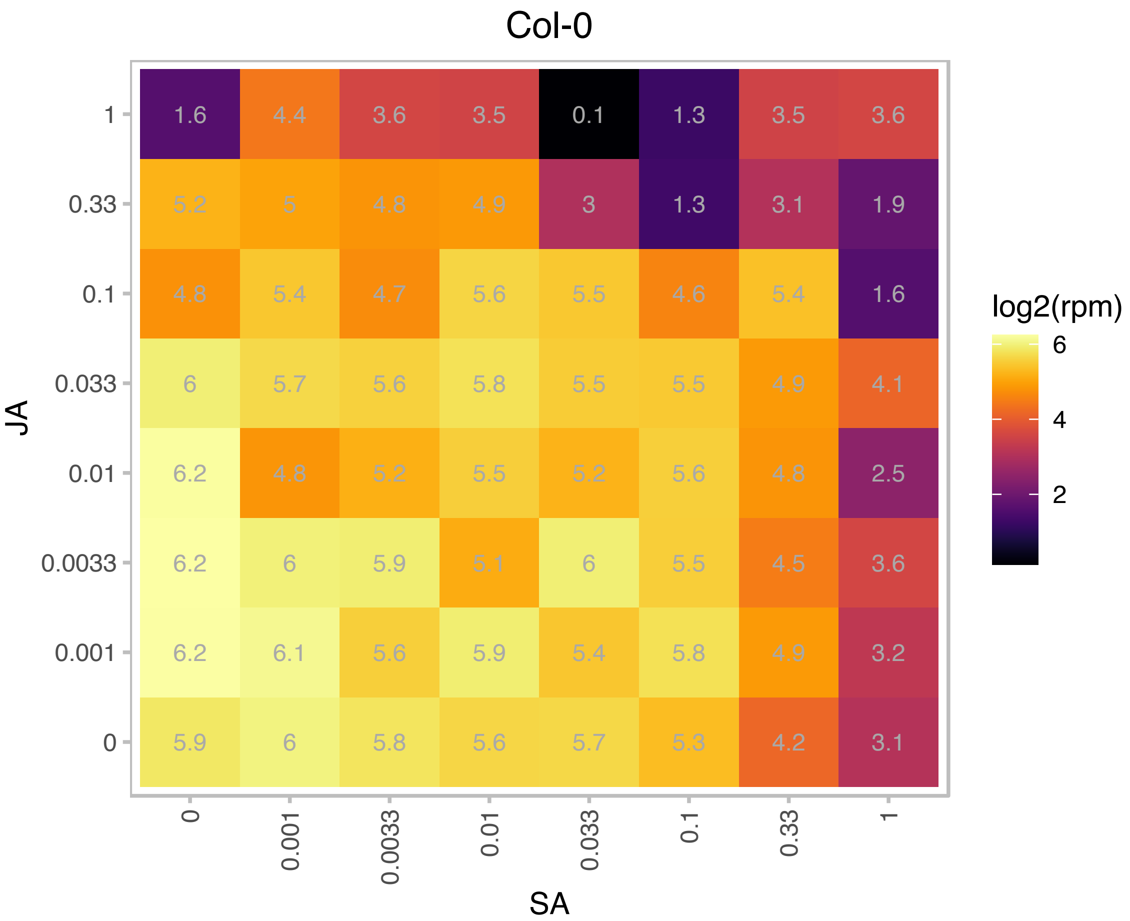


| | |
|---|---|
| Gene | AT5G02710 |
| is.expressed_genes | TRUE |
| is.responsive_genes | TRUE |
| is.apcluster | F-10 |

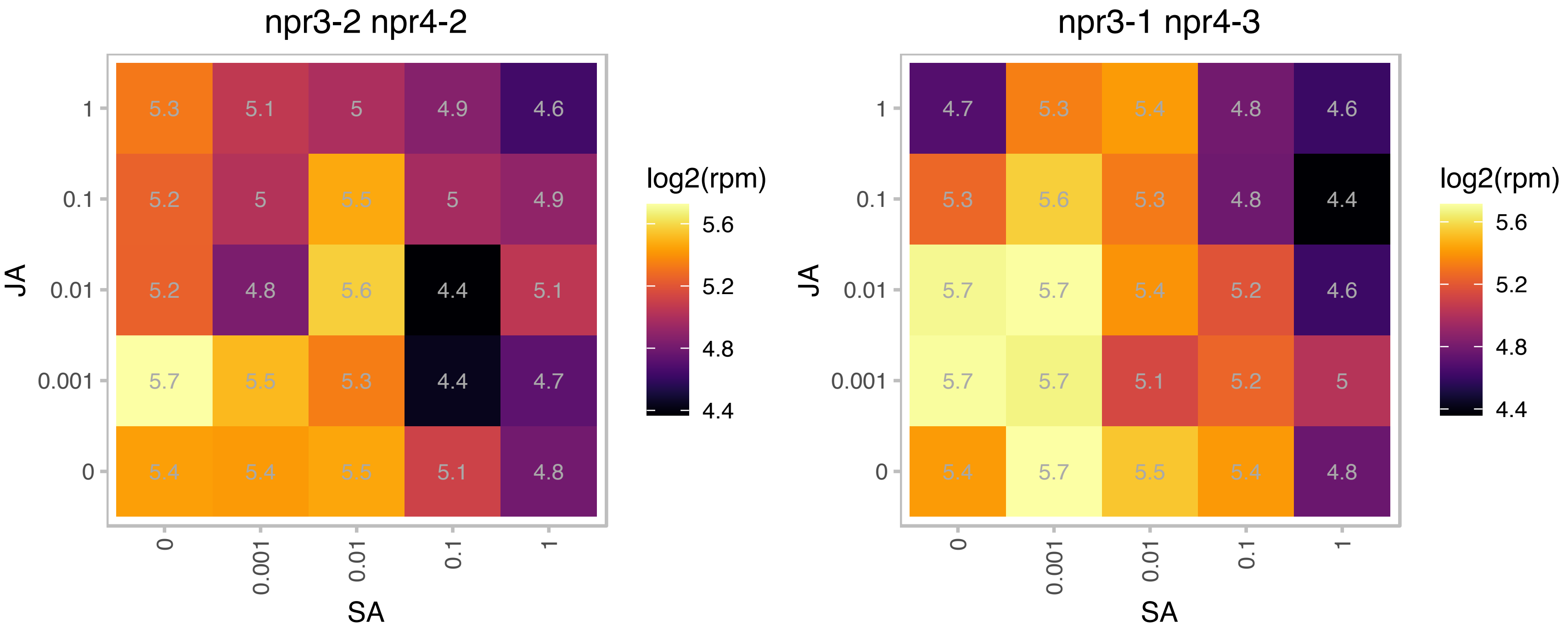

### AT5G06690.1 : WCRKC1

WCRKC thioredoxin 1

Encodes a thioredoxin (WCRKC1) localized in chloroplast stroma.  Contains a WCRKC motif.

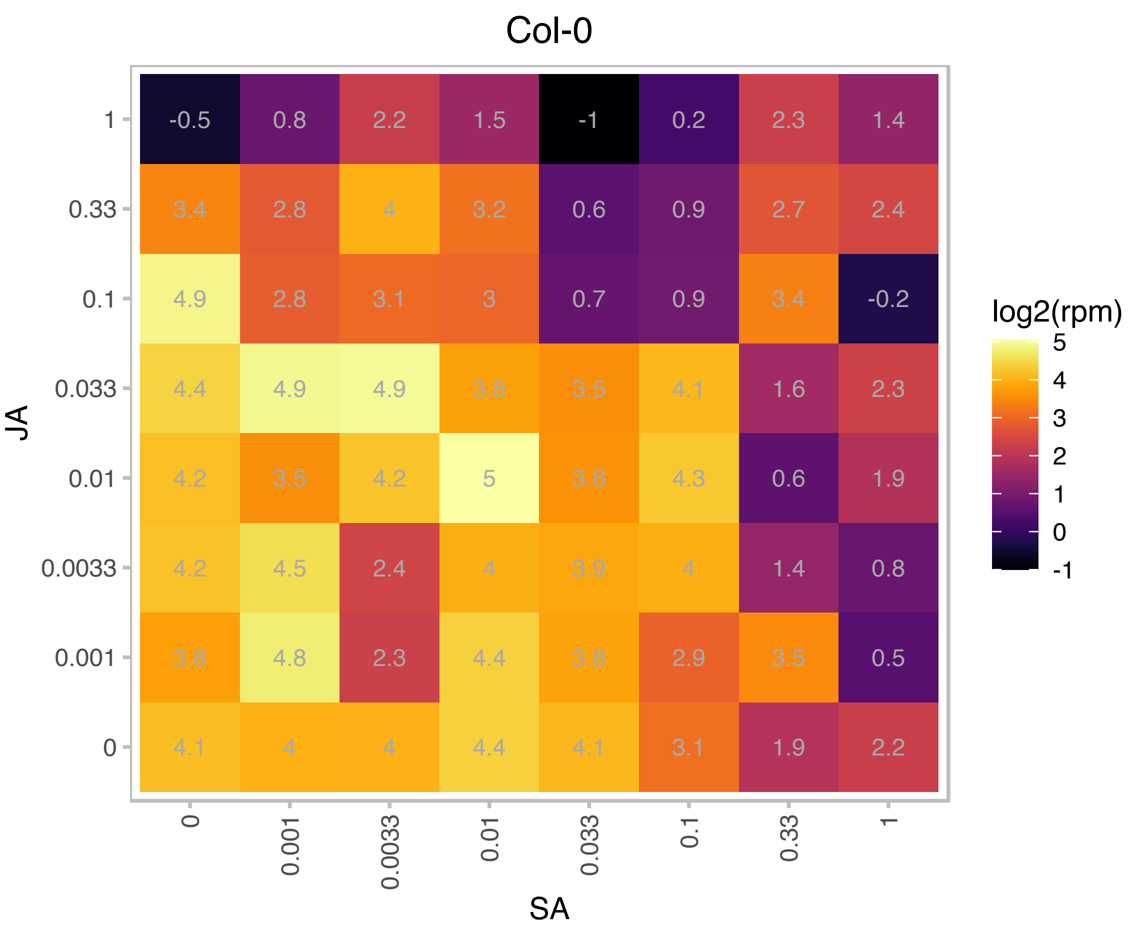


| | |
|---|---|
| Gene | AT5G06690 |
| is.expressed_genes | TRUE |
| is.responsive_genes | TRUE |
| is.apcluster | F-10 |

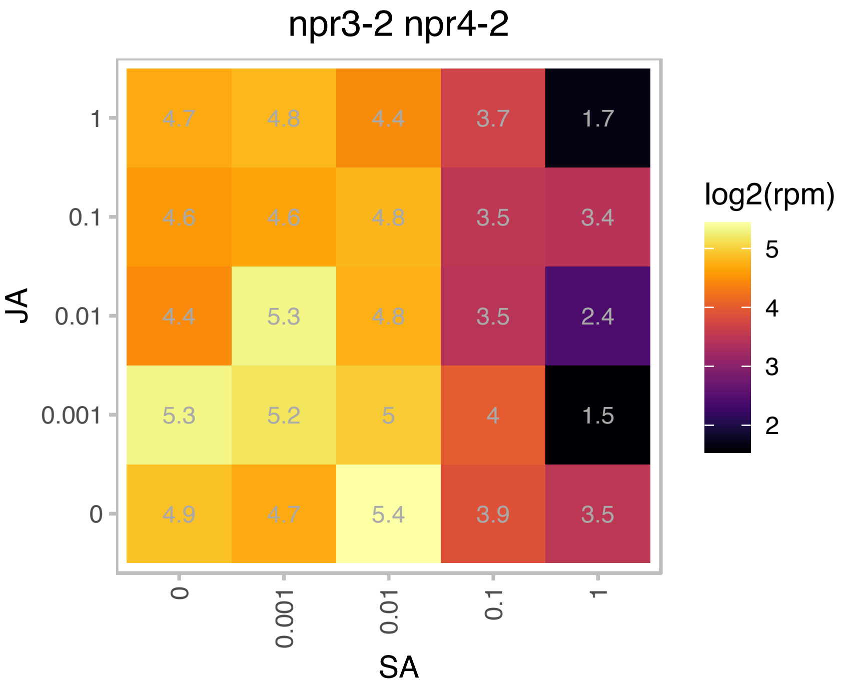


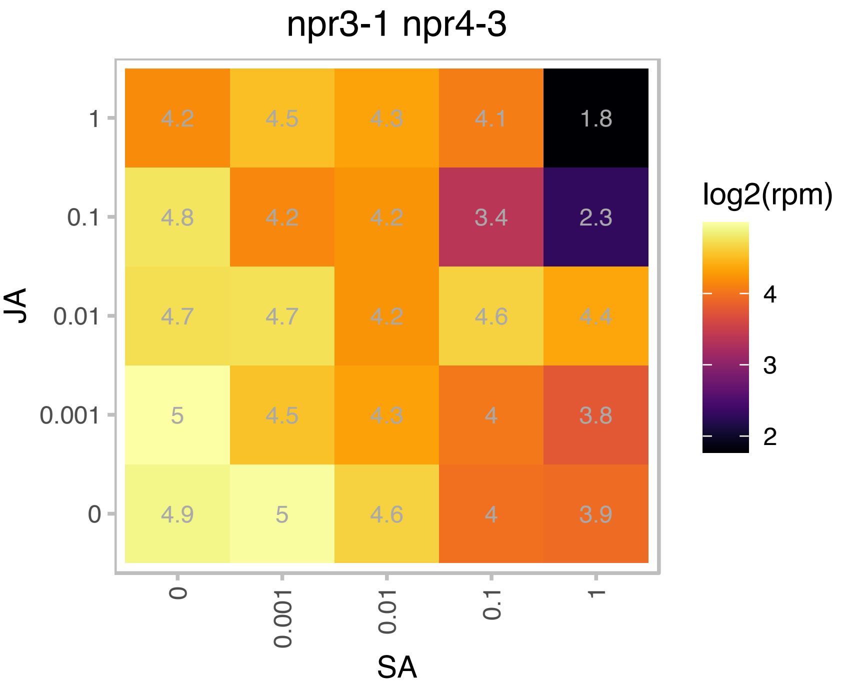

### AT5G08280.1 : HEMC, RUG1

hydroxymethylbilane synthase

Encodes a protein with porphobilinogen deaminase activity. This protein is targeted to the chloroplast. Mutants spontaneously develop chlorotic leaf lesions in the absence of pathogen attack, resembling the phenotype of lesion-mimic mutants. It has been shown to interact with the PPR protein AtECB2 for chloroplast RNA editing.

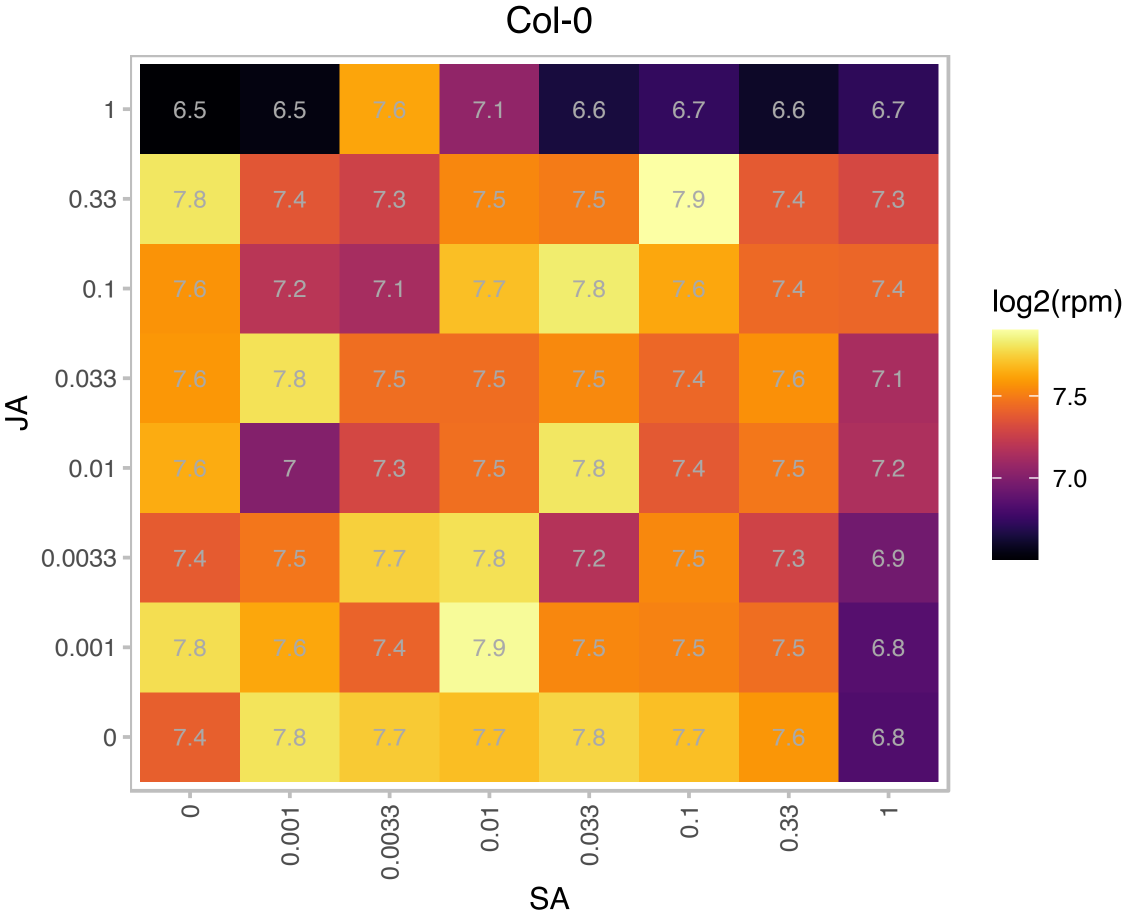


| | |
|---|---|
| Gene | AT5G08280 |
| is.expressed_genes | TRUE |
| is.responsive_genes | TRUE |
| is.apcluster | F-10 |

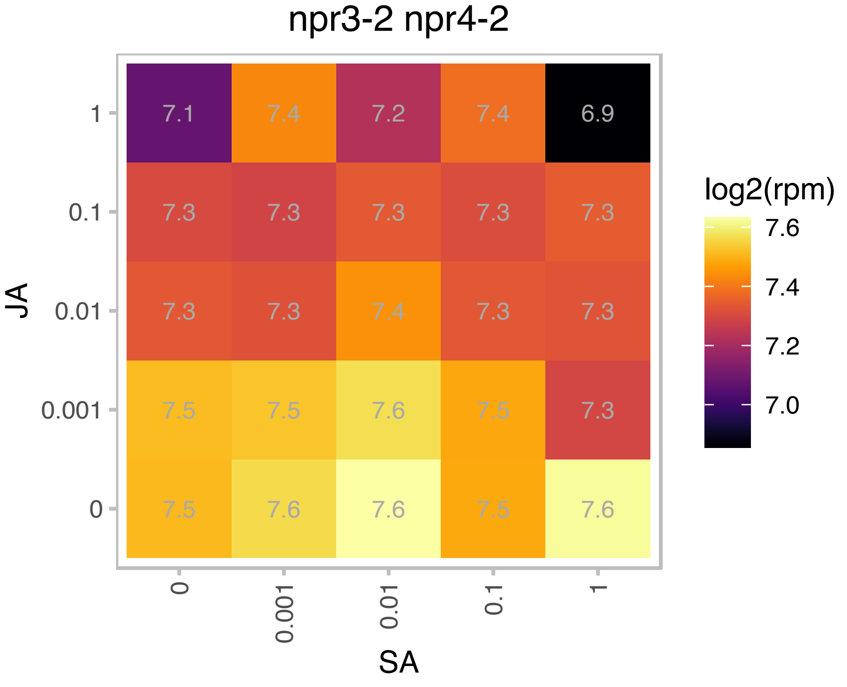


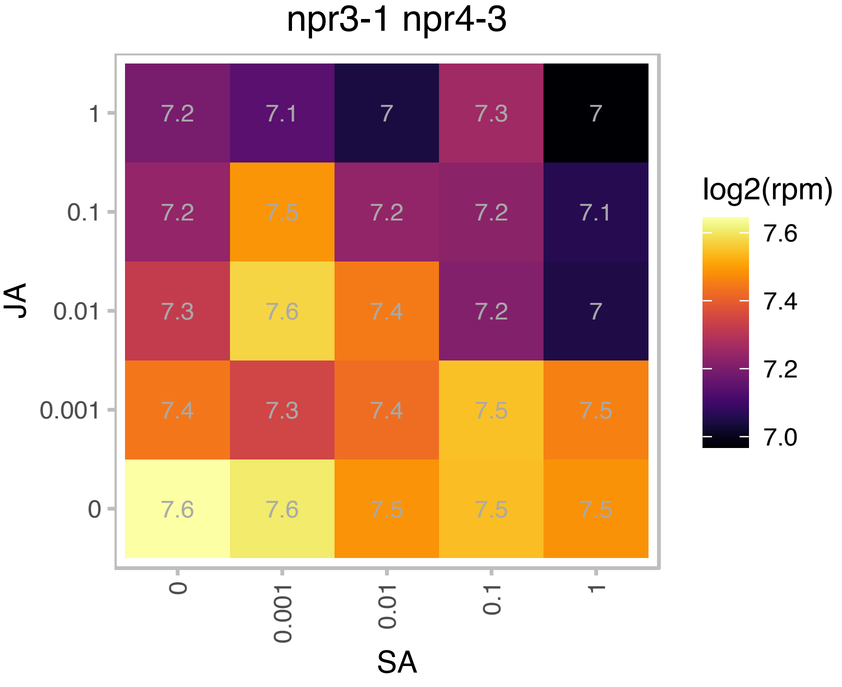

# AT5G09770.1 : NA

Ribosomal protein L17 family protein

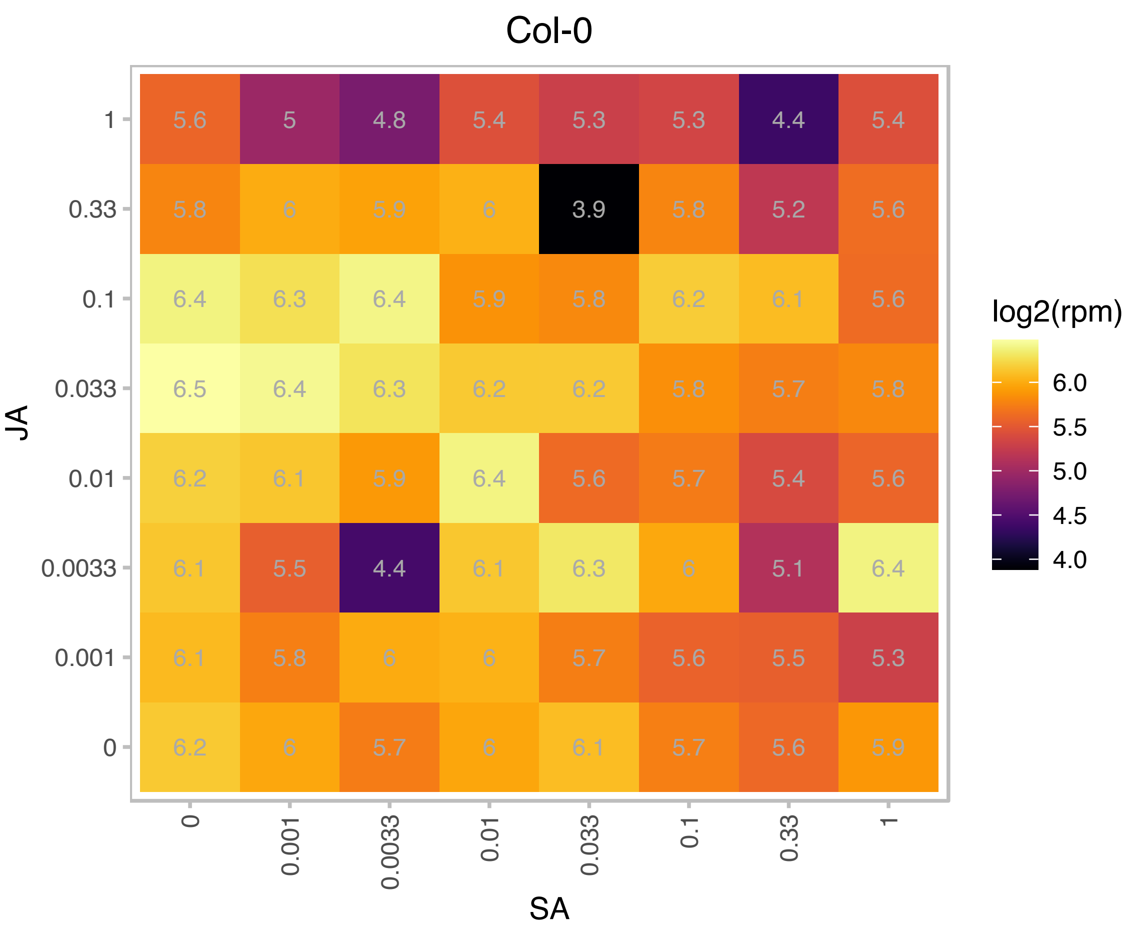


| | |
|---|---|
| Gene | AT5G09770 |
| is.expressed_genes | TRUE |
| is.responsive_genes | TRUE |
| is.apcluster | F-10 |

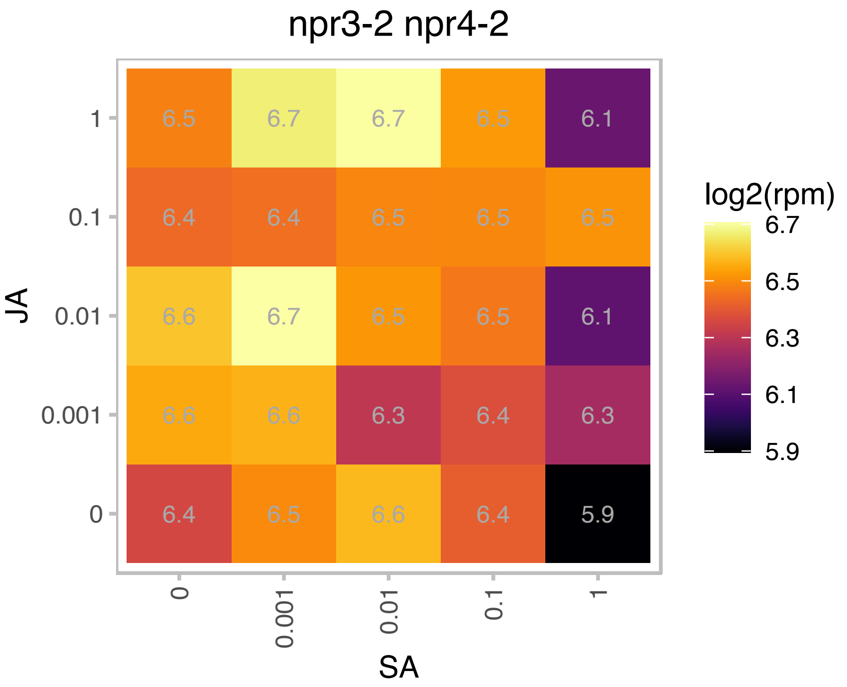


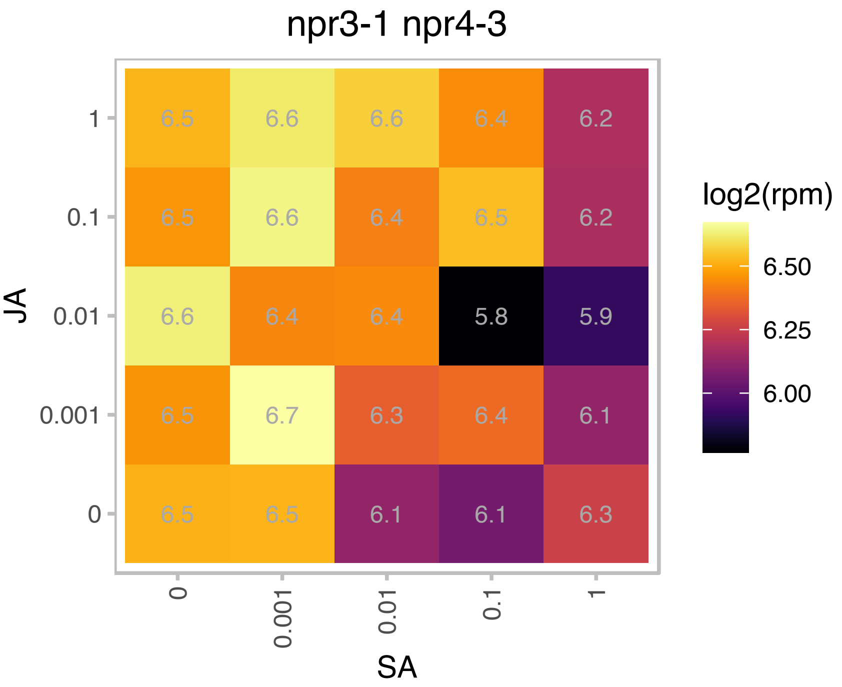

# AT5G19510.1 : NA

Translation elongation factor EF1B/ribosomal protein S6 family protein

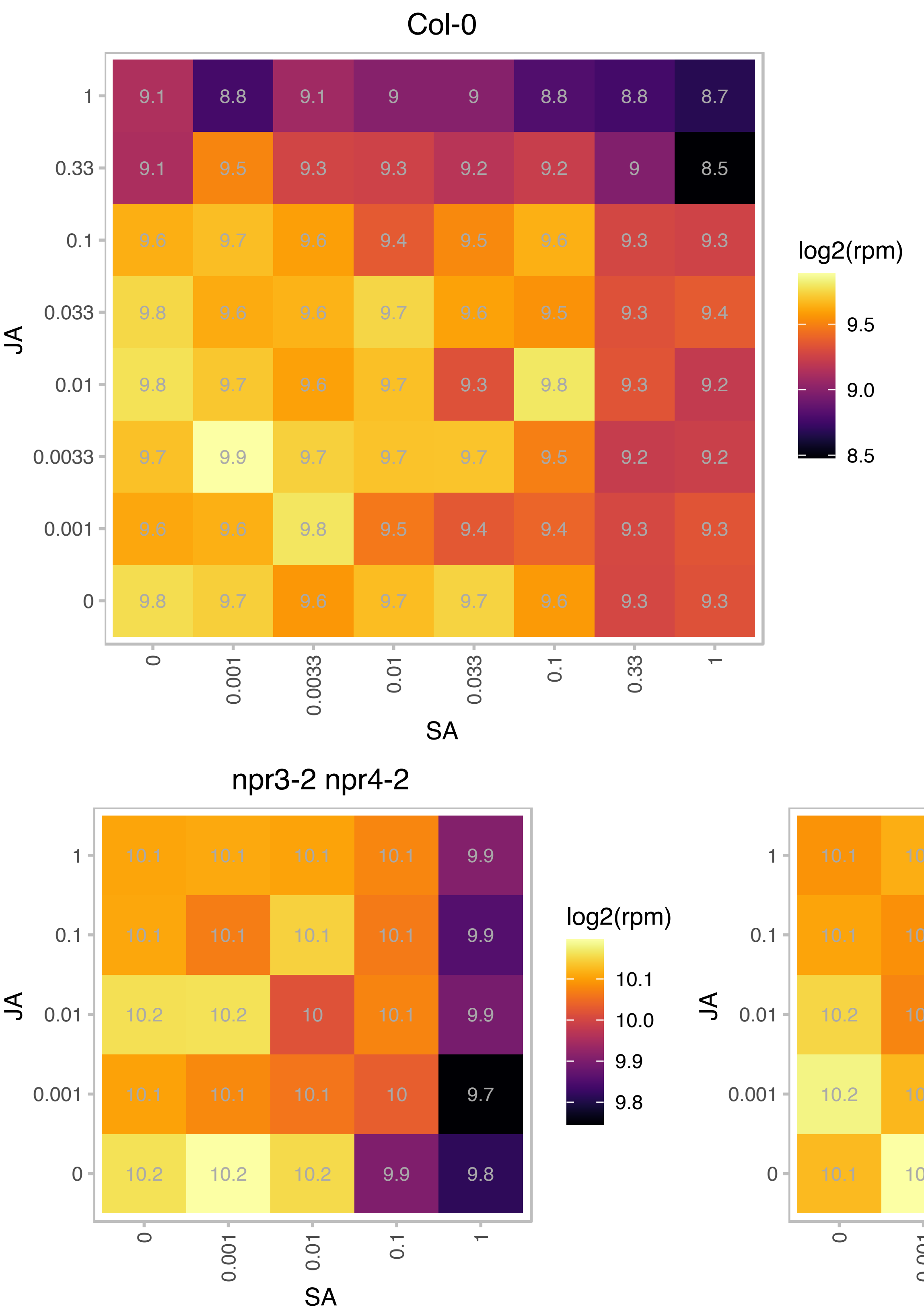


| | |
|---|---|
| Gene | AT5G19510 |
| is.expressed_genes | TRUE |
| is.responsive_genes | TRUE |
| is.apcluster | F-10 |

# AT5G19570.1 : NA

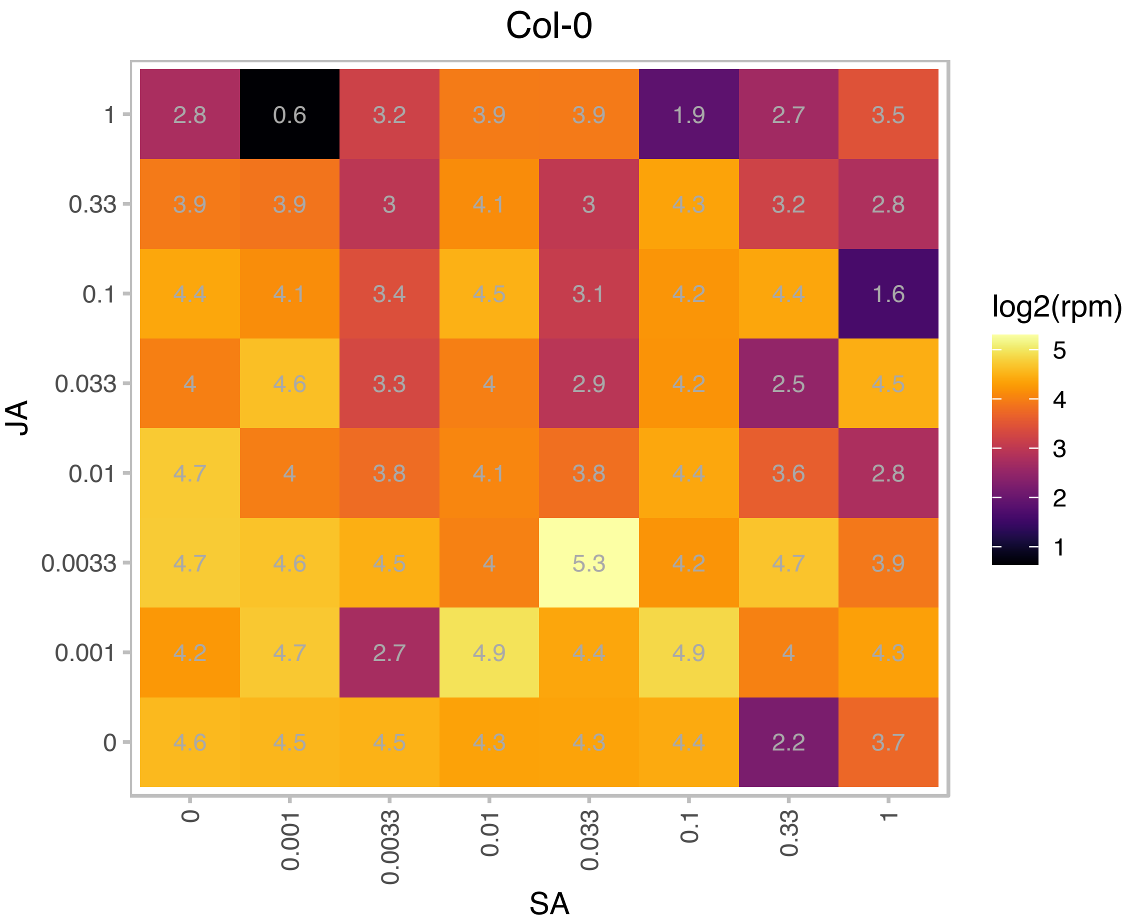


| | |
|---|---|
| Gene | AT5G19570 |
| is.expressed_genes | TRUE |
| is.responsive_genes | TRUE |
| is.apcluster | F-10 |

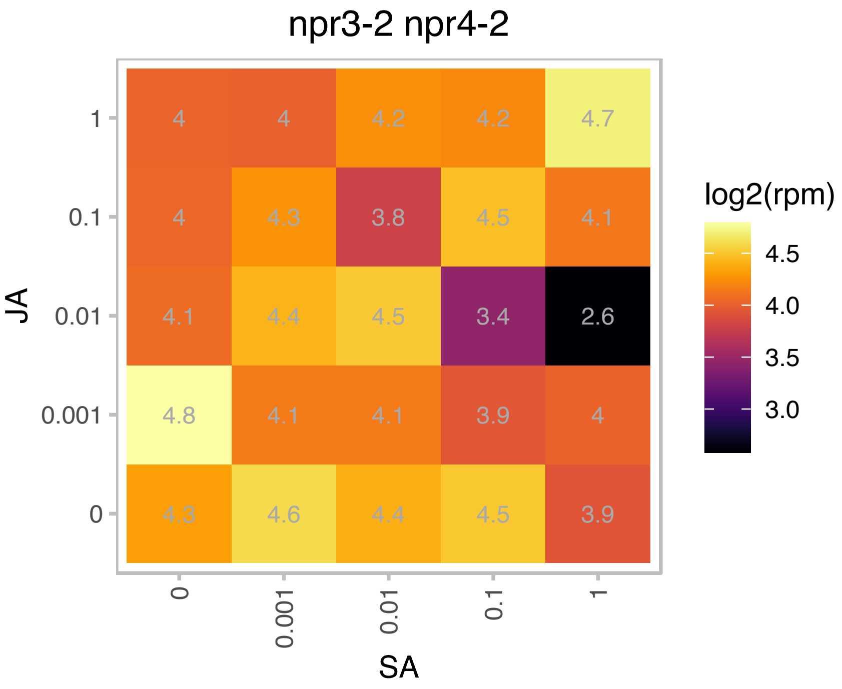


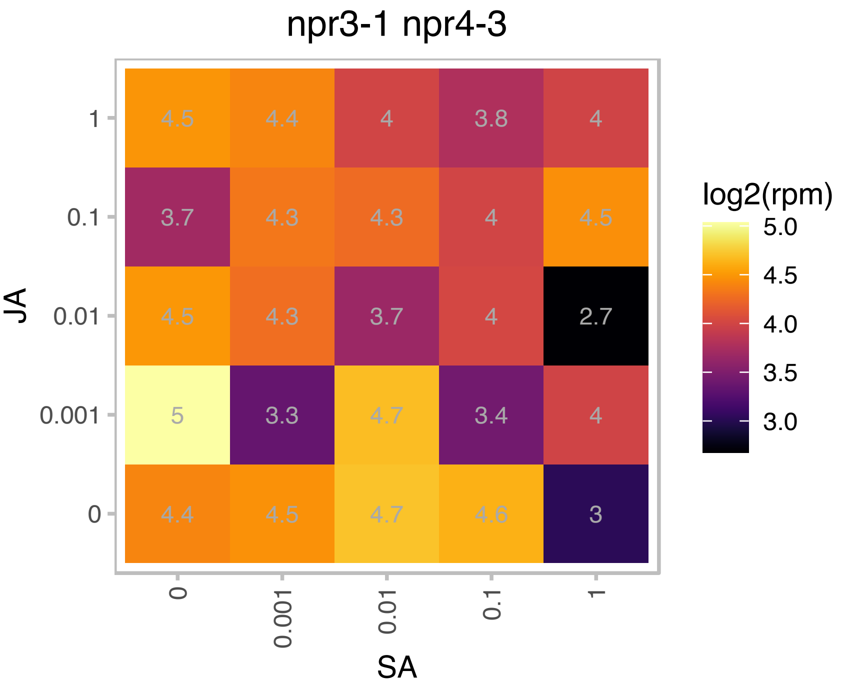

### AT5G20935.1 : CRR42

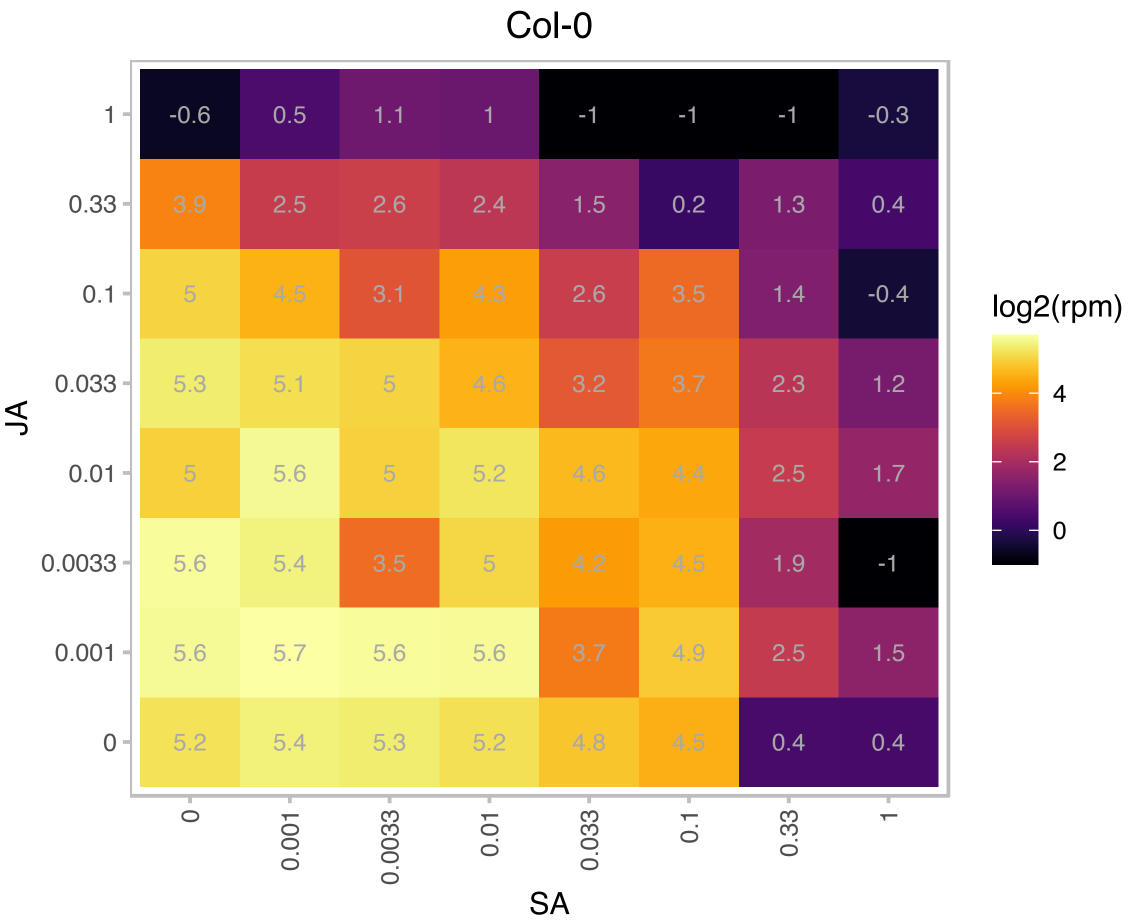


| | |
|---|---|
| Gene | AT5G20935 |
| is.expressed_genes | TRUE |
| is.responsive_genes | TRUE |
| is.apcluster | F-10 |

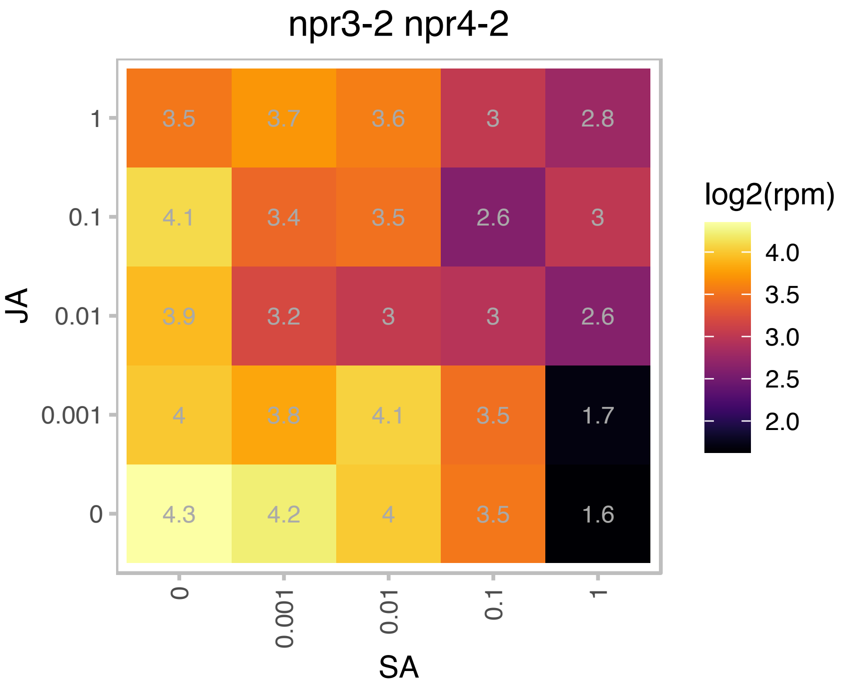


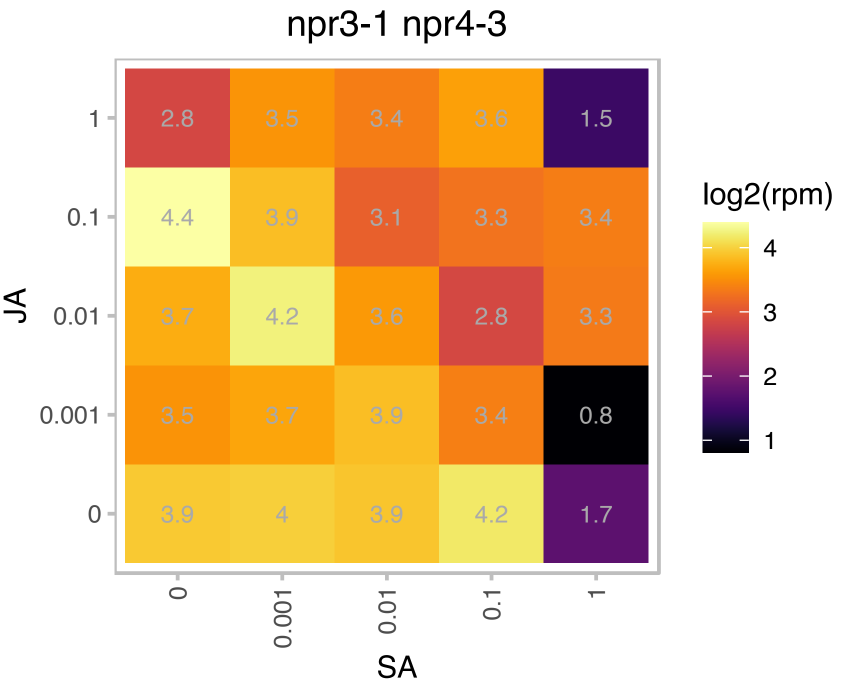

### AT5G23740.1 : RPS11-BETA

ribosomal protein S11-beta

Encodes a putative ribosomal protein S11 (RPS11-beta).

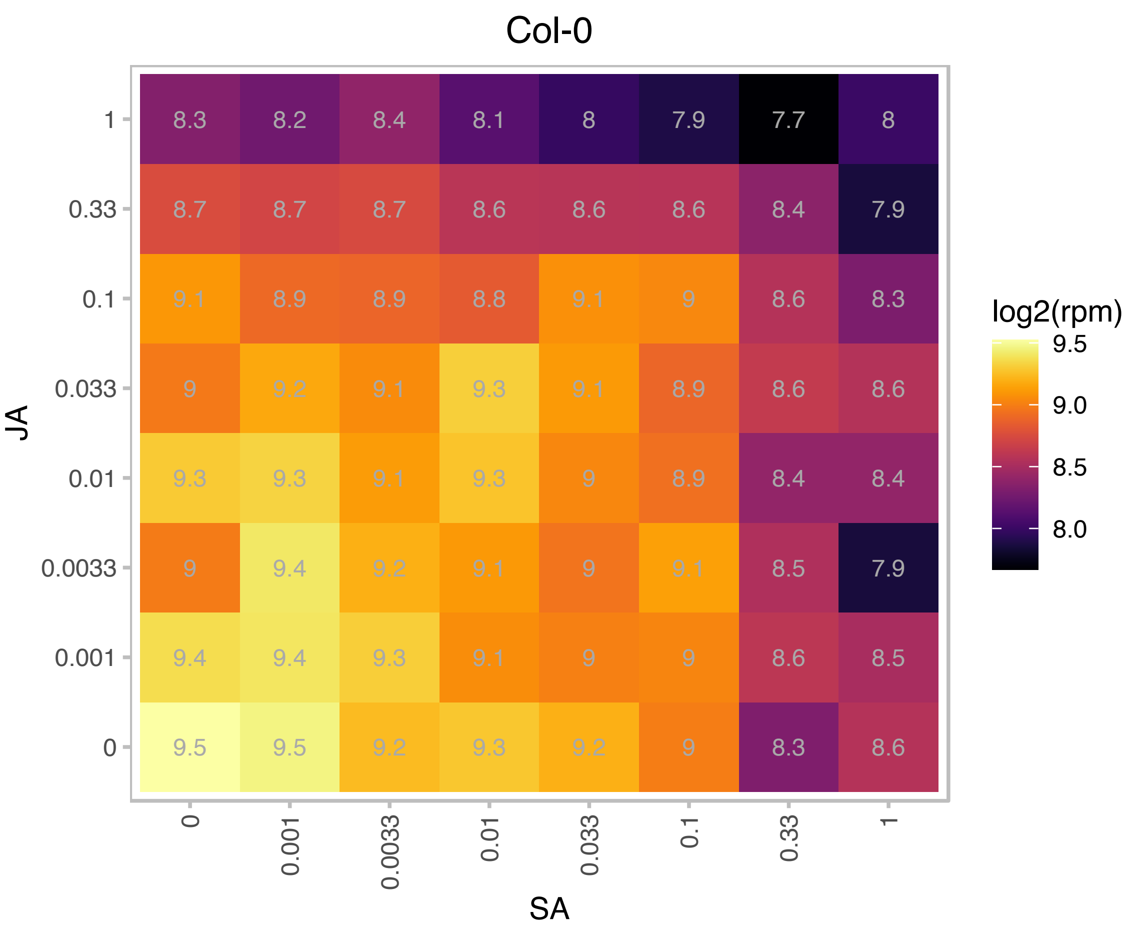


| | |
|---|---|
| Gene | AT5G23740 |
| is.expressed_genes | TRUE |
| is.responsive_genes | TRUE |
| is.apcluster | F-10 |

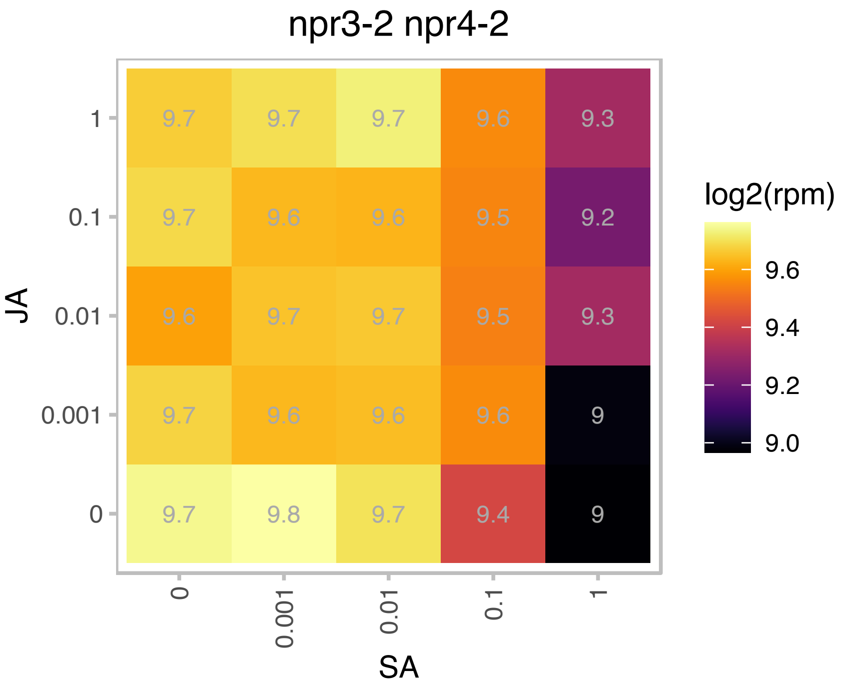


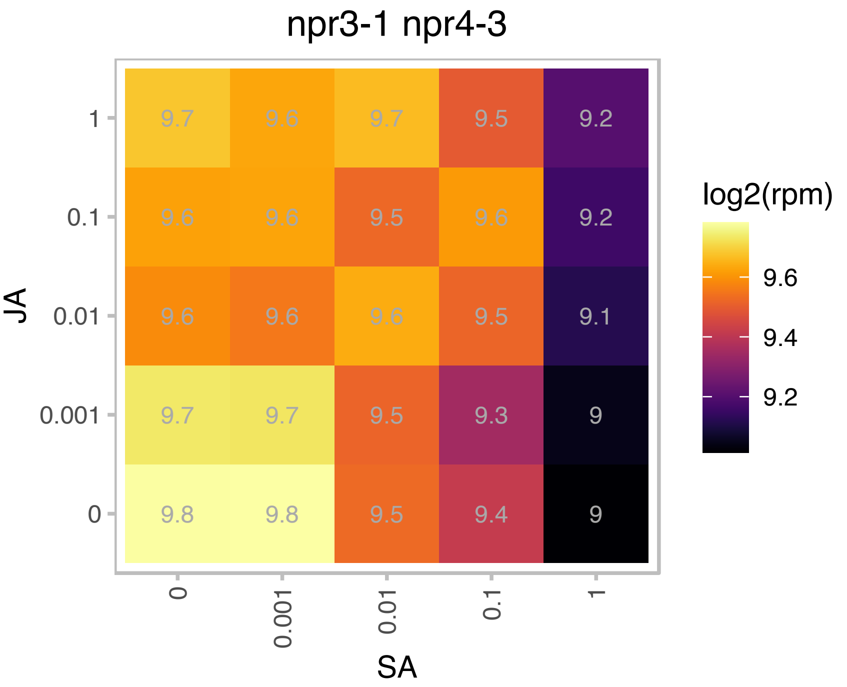

### AT5G24314.1 : PDE225, PTAC7, TAC7

plastid transcriptionally active7

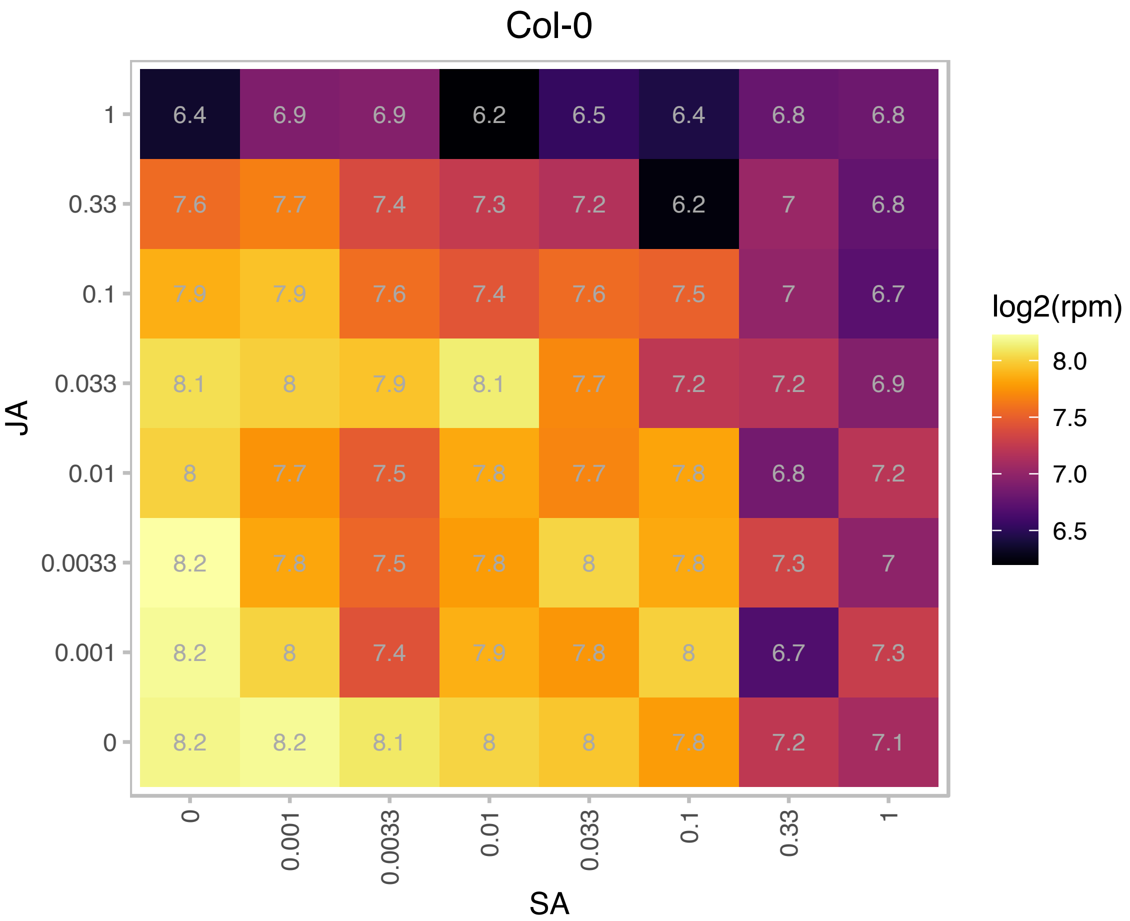


| | |
|---|---|
| Gene | AT5G24314 |
| is.expressed_genes | TRUE |
| is.responsive_genes | TRUE |
| is.apcluster | F-10 |

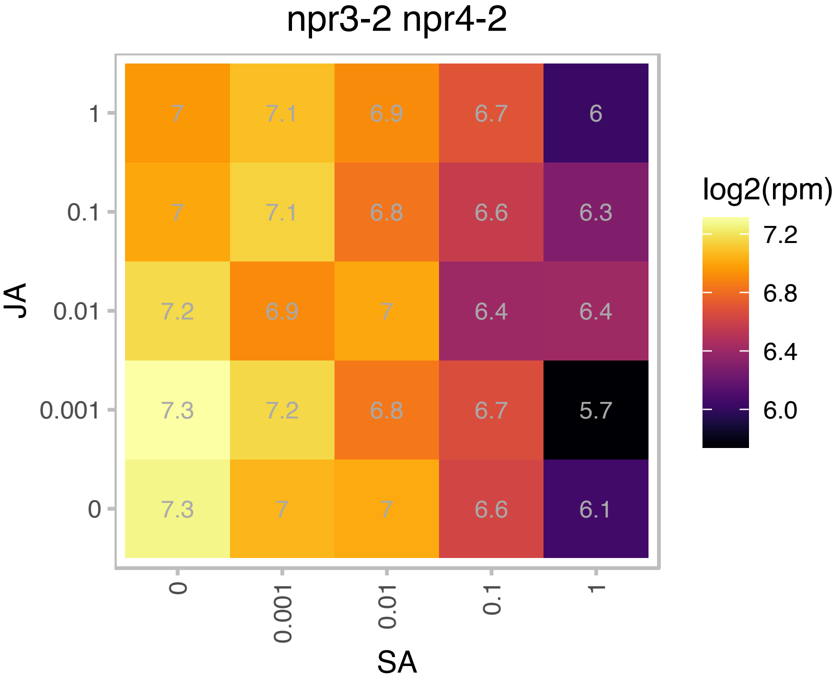


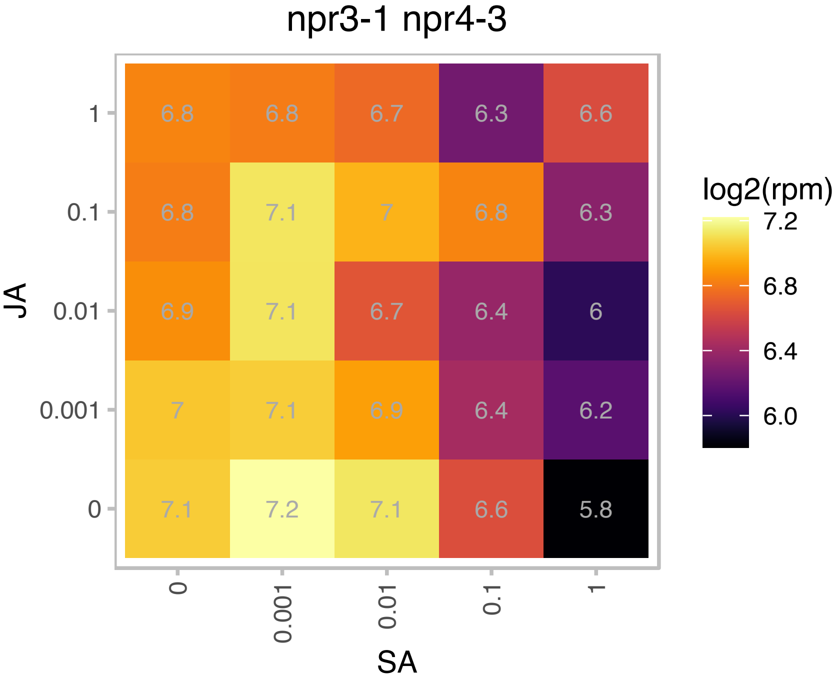

### AT5G26742.1 : AtRH3, emb1138, RH3

DEAD box RNA helicase (RH3)

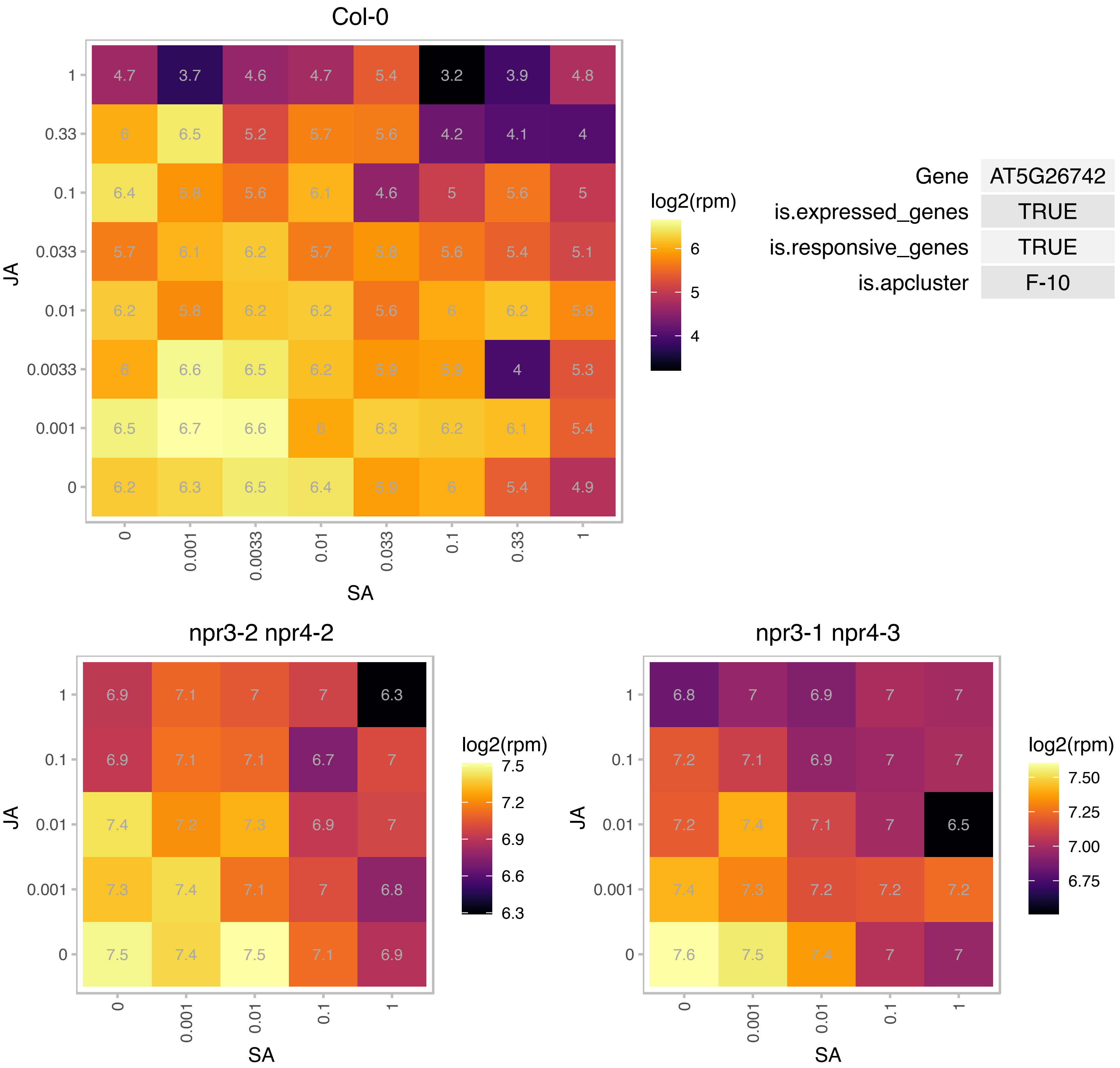


| | |
|---|---|
| Gene | AT5G26742 |
| is.expressed_genes | TRUE |
| is.responsive_genes | TRUE |
| is.apcluster | F-10 |

### AT5G36120.1 : atylmg3, CCB3, YLMG3

cofactor assembly, complex C (B6F)

| Gene | AT5G36120 |
|---|---|
| is.expressed_genes | TRUE |
| is.responsive_genes | TRUE |
| is.apcluster | F-10 |

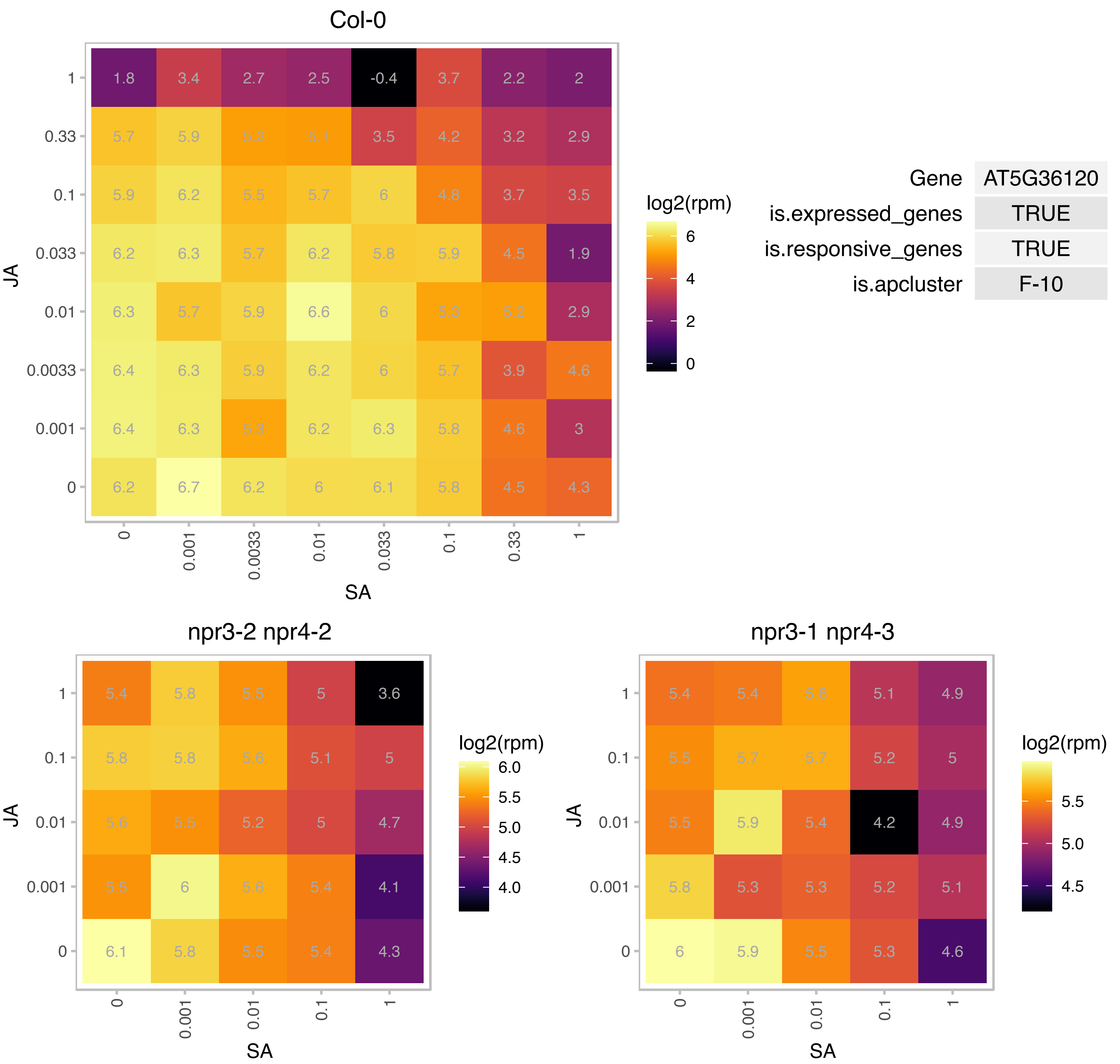

### AT5G38430.1 : RBCS1B

Ribulose bisphosphate carboxylase (small chain) family protein

Encodes a member of the Rubisco small subunit (RBCS) multigene family: RBCS1A (At1g67090), RBCS1B (At5g38430), RBCS2B (At5g38420), and RBCS3B (At5g38410).

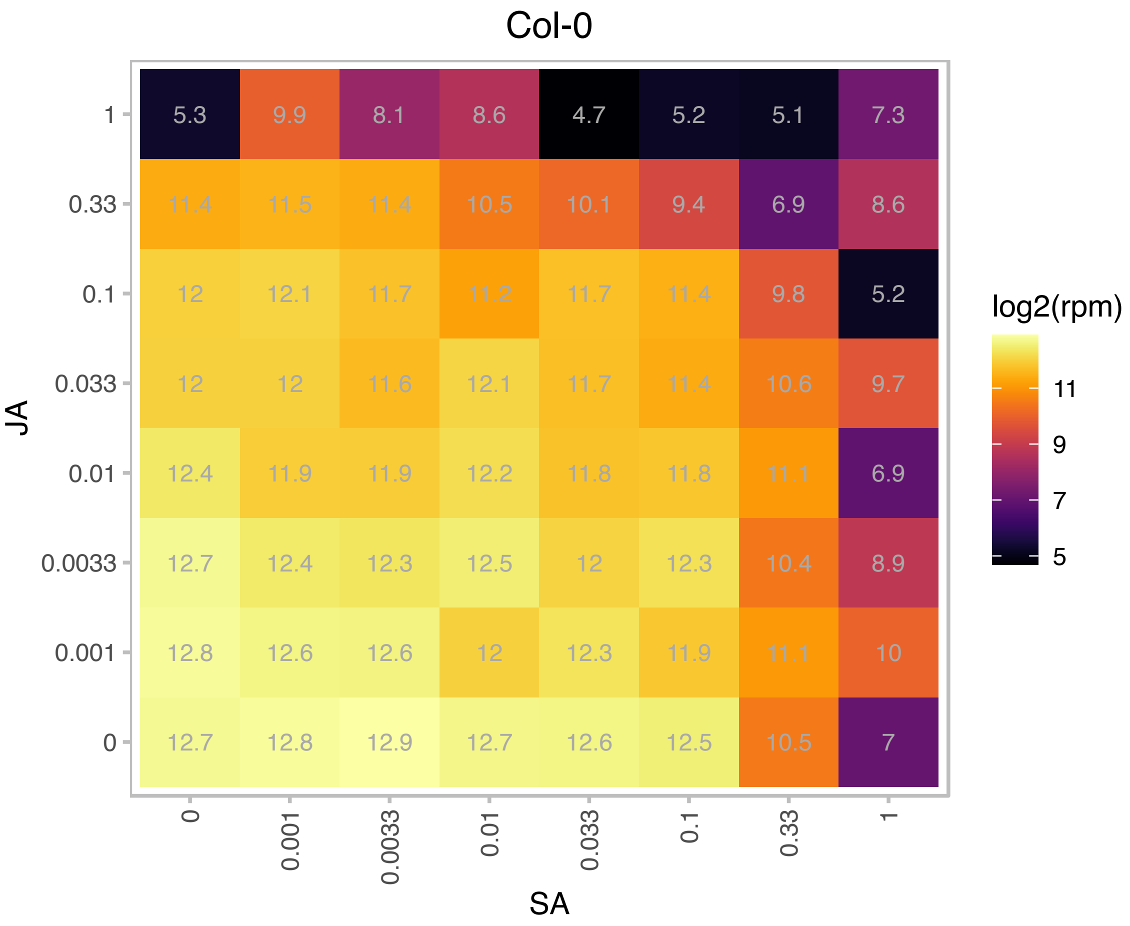


| | |
|---|---|
| Gene | AT5G38430 |
| is.expressed_genes | TRUE |
| is.responsive_genes | TRUE |
| is.apcluster | F-10 |

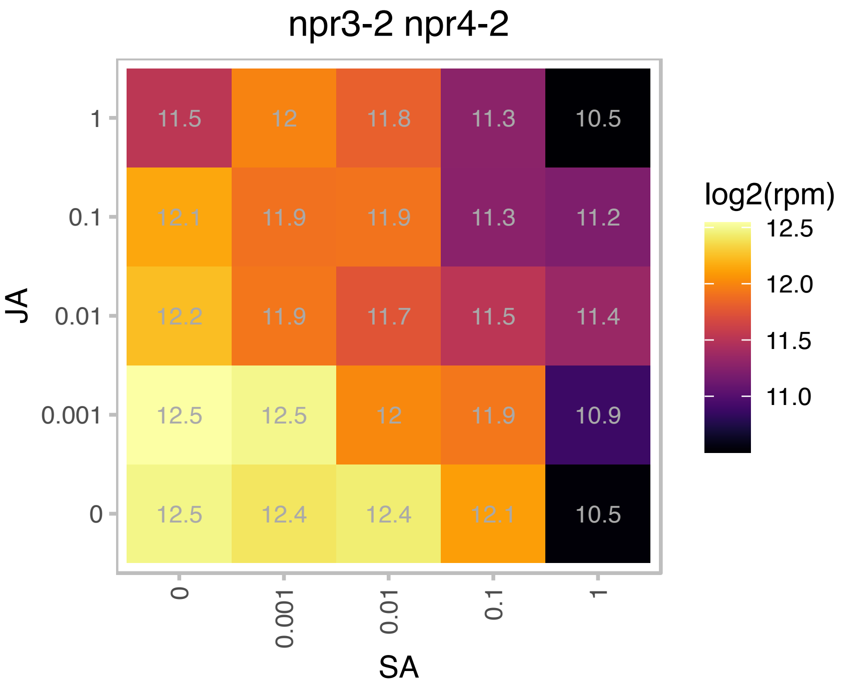


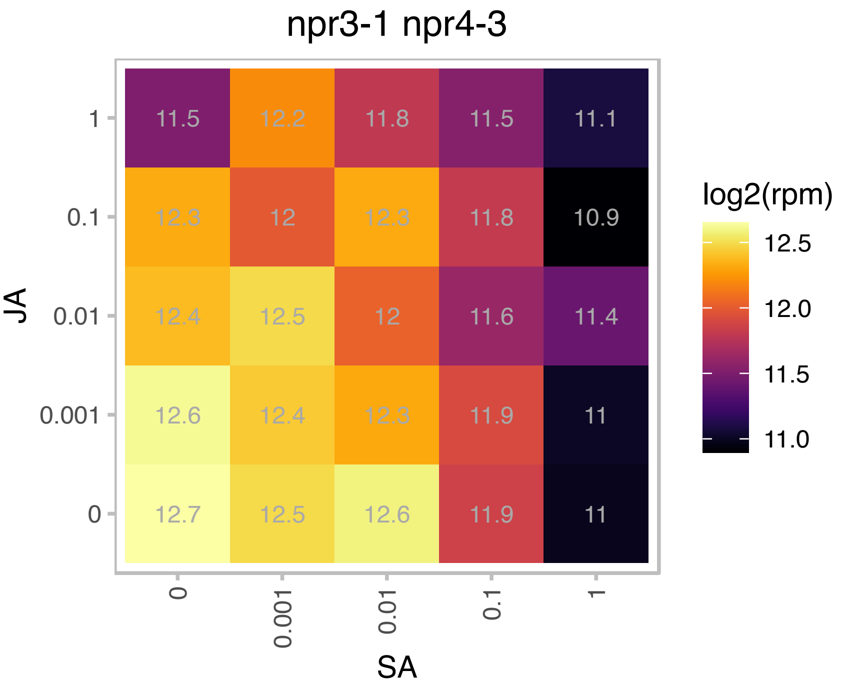

# AT5G39530.1 : NA

Protein of unknown function (DUF1997)

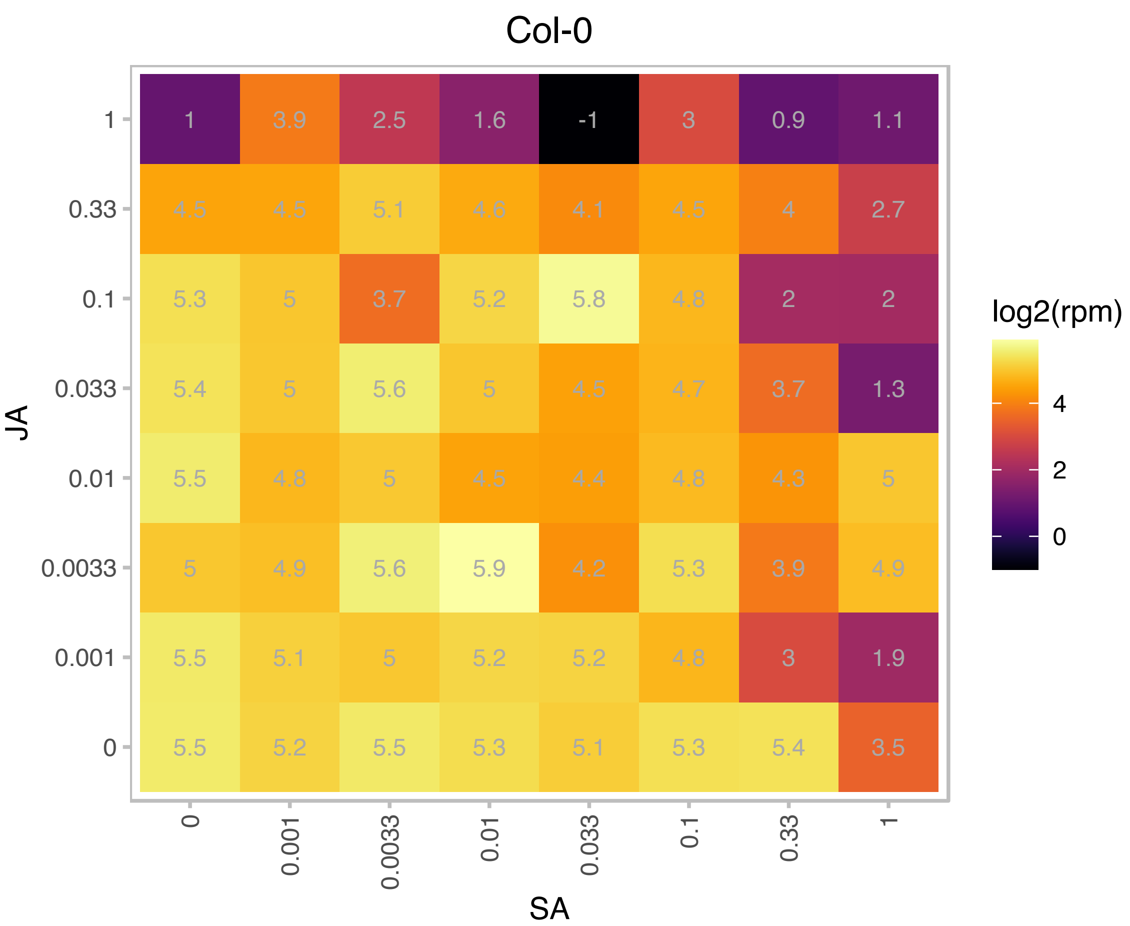


| | |
|---|---|
| Gene | AT5G39530 |
| is.expressed_genes | TRUE |
| is.responsive_genes | TRUE |
| is.apcluster | F-10 |

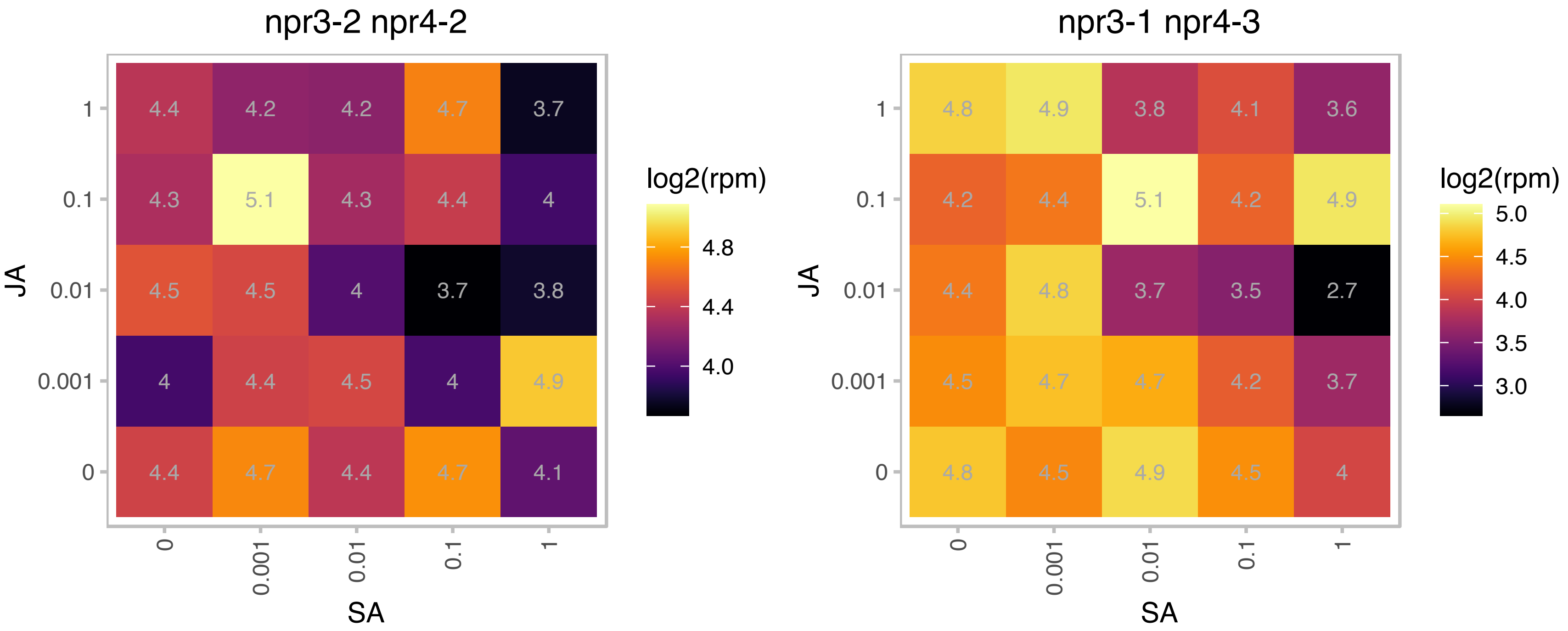

# AT5G42070.1 : NA

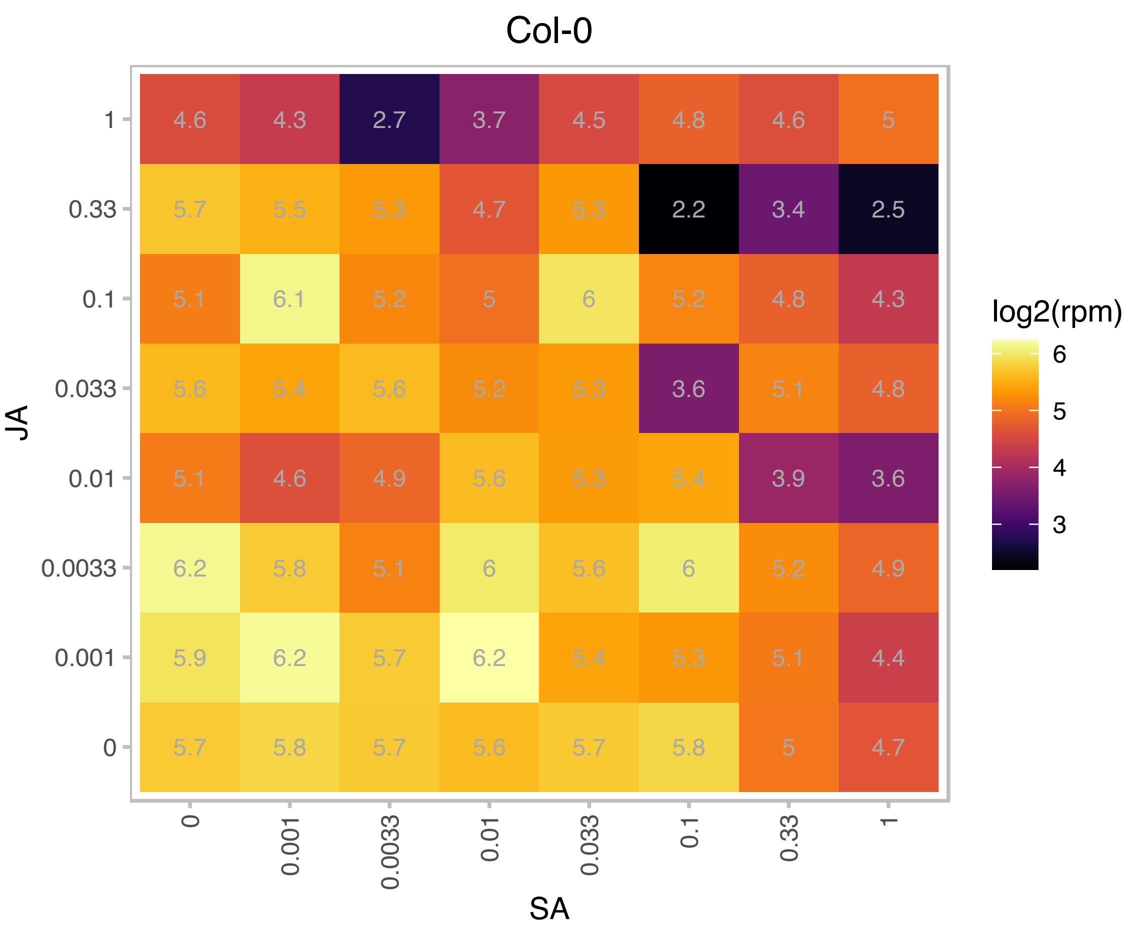


| | |
|---|---|
| Gene | AT5G42070 |
| is.expressed_genes | TRUE |
| is.responsive_genes | TRUE |
| is.apcluster | F-10 |

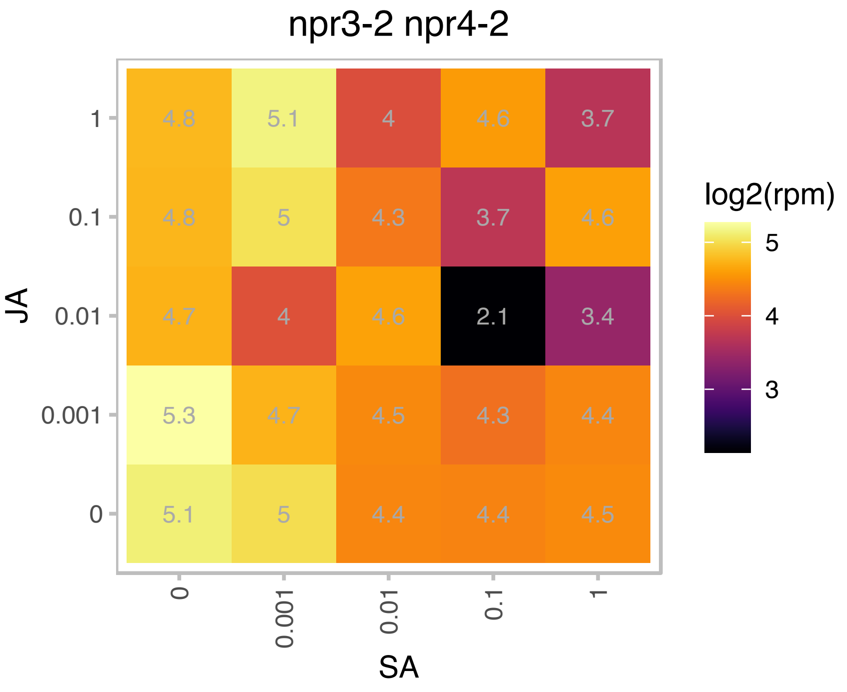


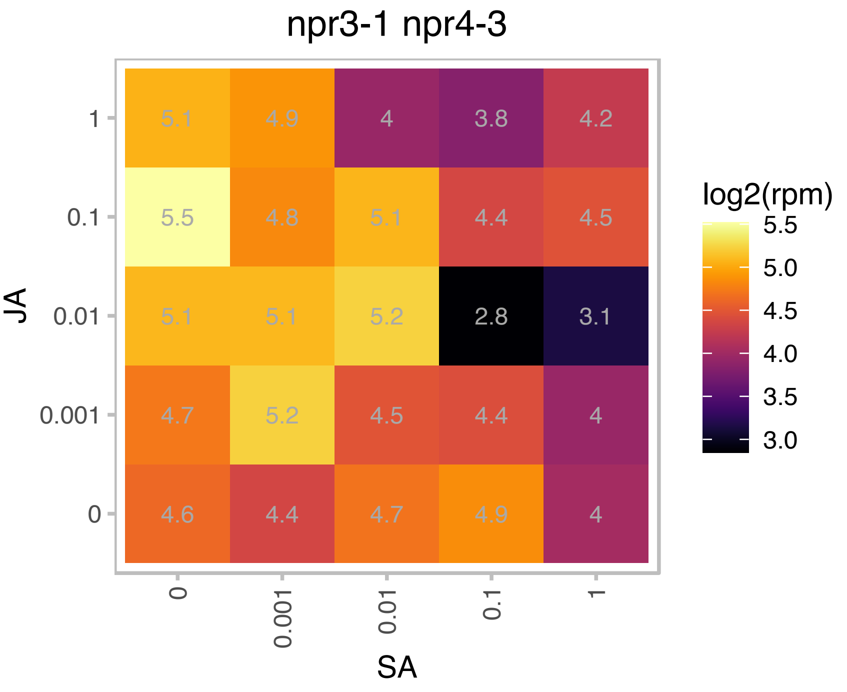

### AT5G45930.1 : CHL I2, CHLI-2, CHLI2

magnesium chelatase i2

encodes a second Chl I gene (CHLI2), a subunit of magnesium chelatase which is required for chlorophyll biosynthesis.  Has ATPase activity, regulated by TRX-f. Involved in the assembly of the Mg chelatase complex.

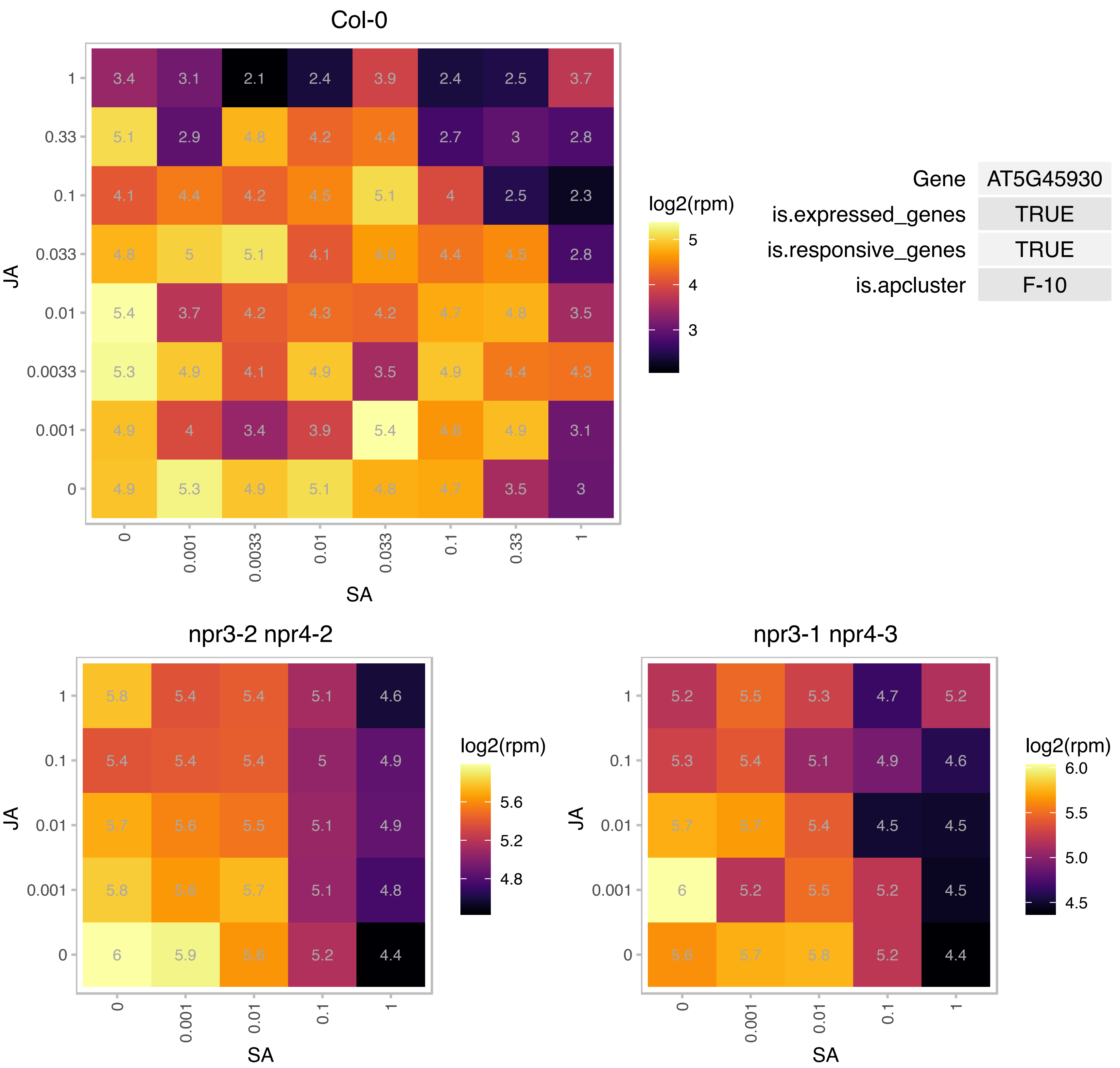


| | |
|---|---|
| Gene | AT5G45930 |
| is.expressed_genes | TRUE |
| is.responsive_genes | TRUE |
| is.apcluster | F-10 |

### AT5G47630.1 : mtACP3

mitochondrial acyl carrier protein 3

Encodes a member of the mitochondrial acyl carrier protein (ACP) family.  Its role is currently obscure as it is weakly expressed and has not yet been identified by mitochondrial proteome analysis.

| | |
|---|---|
| Gene | AT5G47630 |
| is.expressed_genes | TRUE |
| is.responsive_genes | TRUE |
| is.apcluster | F-10 |

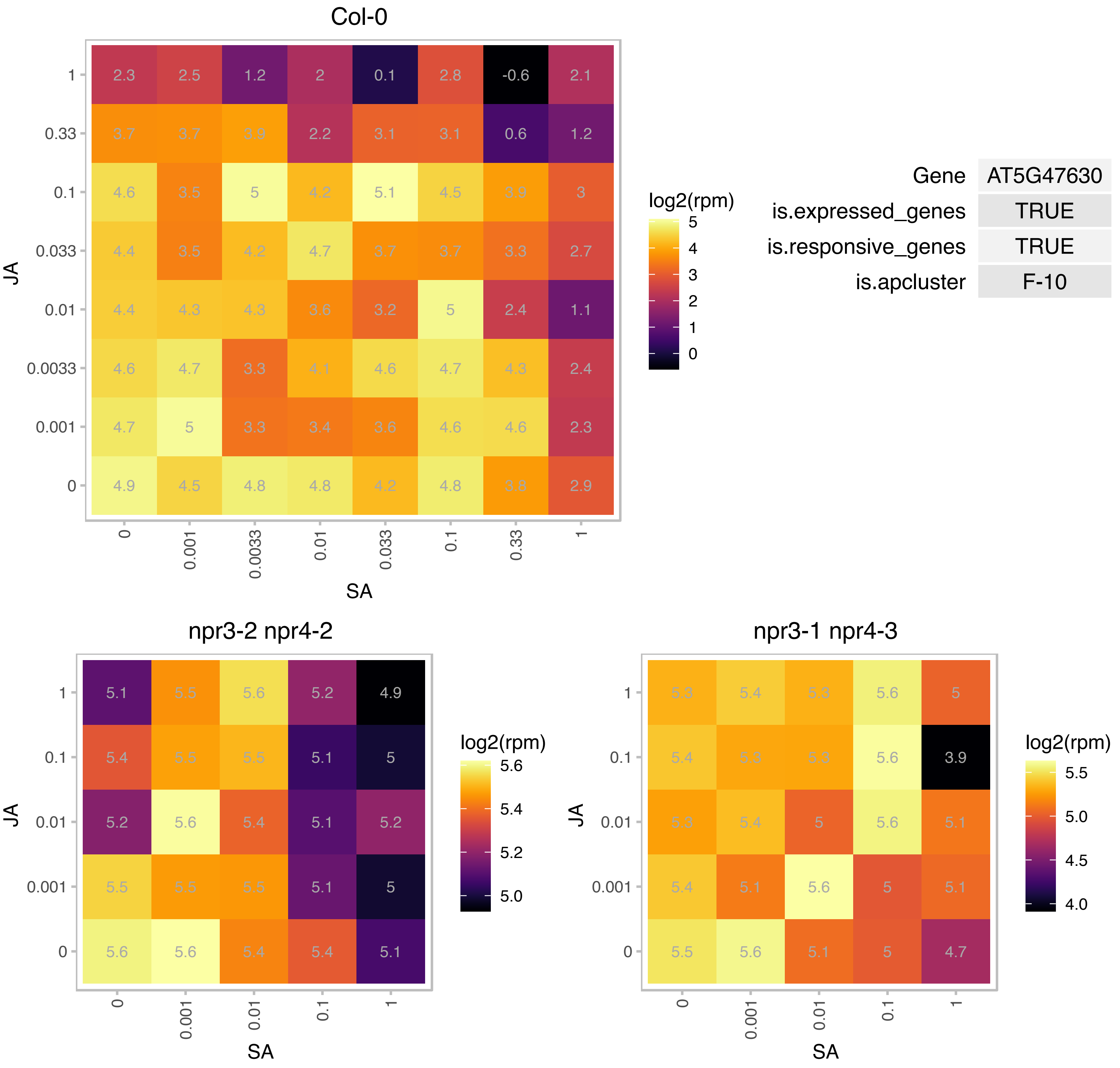

### AT5G48490.1 : DIR1-like

Bifunctional inhibitor/lipid-transfer protein/seed storage 2S albumin superfamily protein

Encodes a protein with similarity to a lipid transfer protein that may contribute to systemic acquired resistance (SAR).

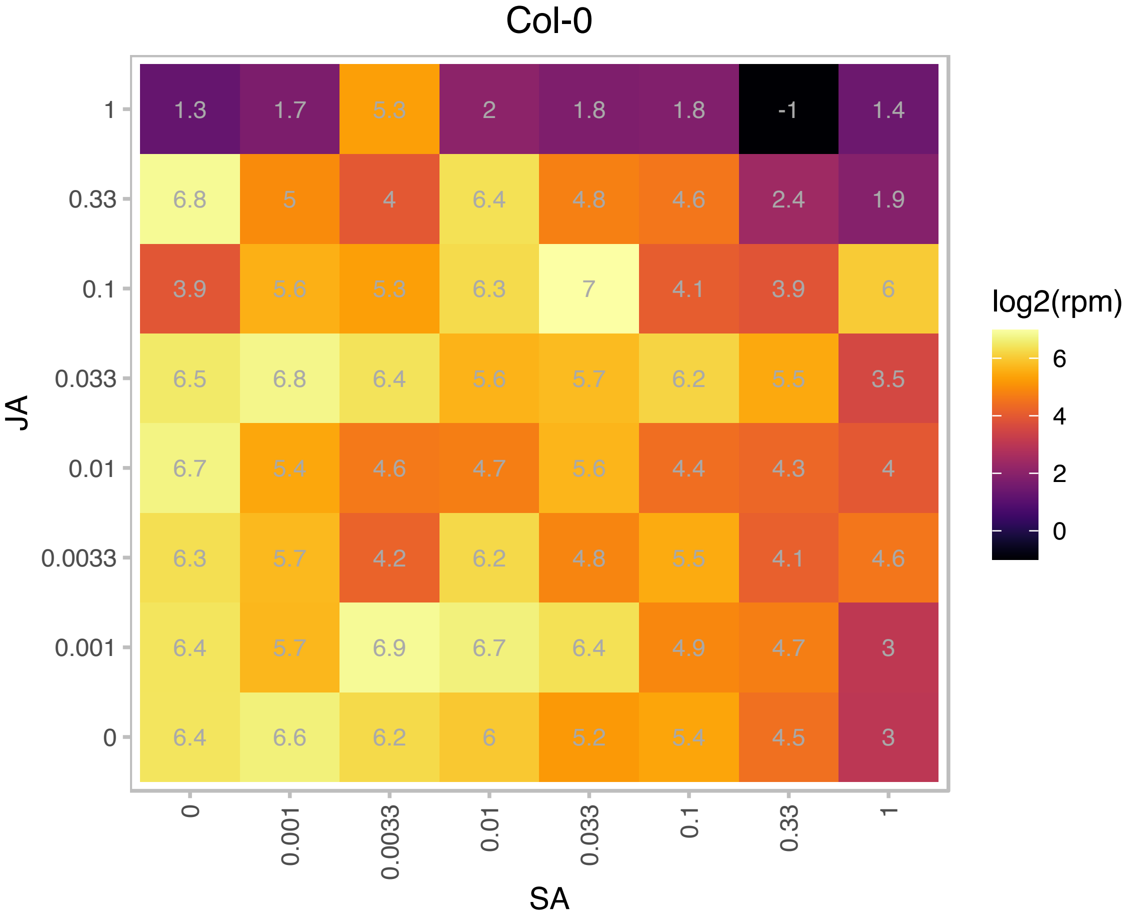


| | |
|---|---|
| Gene | AT5G48490 |
| is.expressed_genes | TRUE |
| is.responsive_genes | TRUE |
| is.apcluster | F-10 |

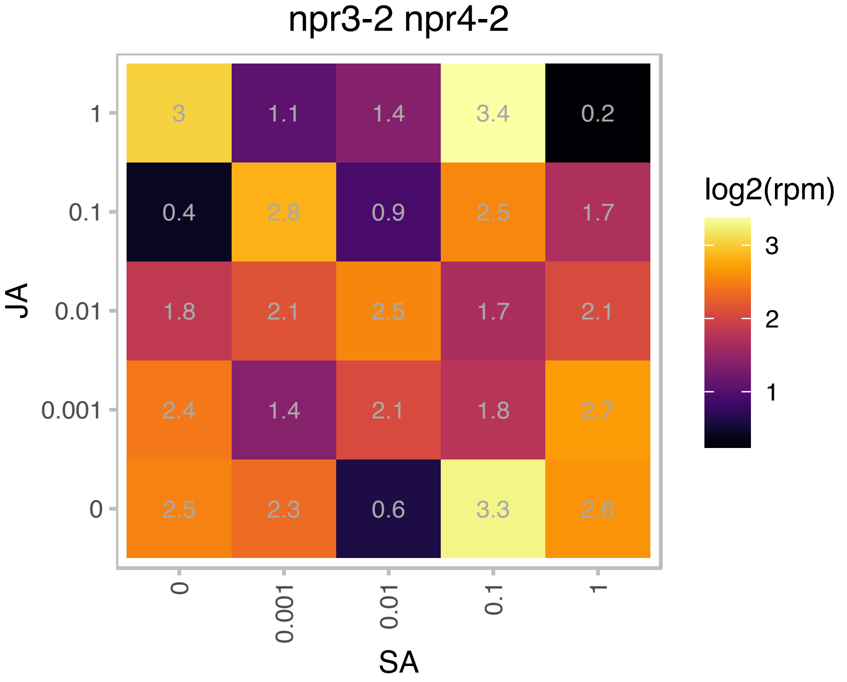


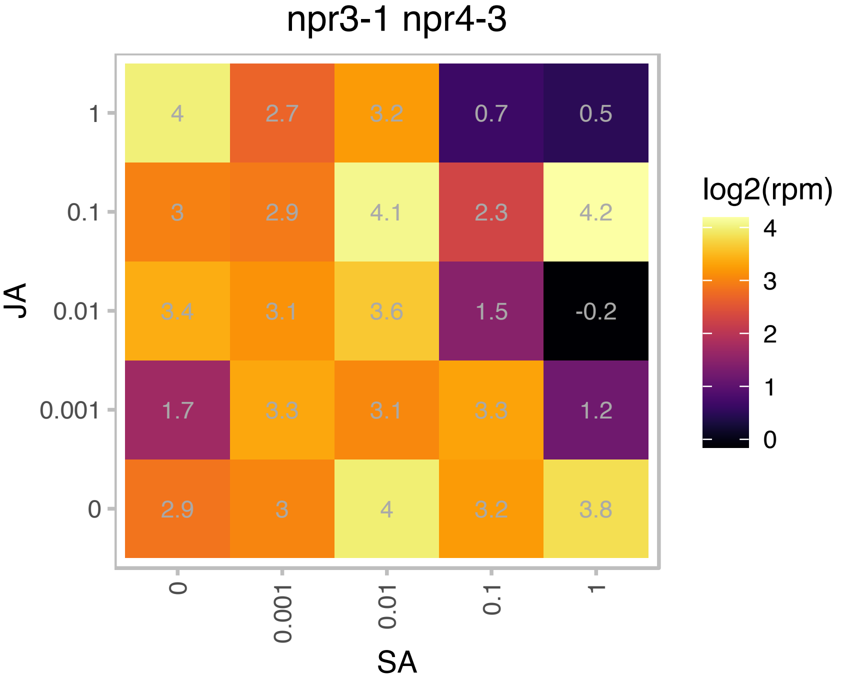

### AT5G51190.1 : ERF105

Integrase-type DNA-binding superfamily protein
encodes a member of the ERF (ethylene response factor) subfamily B-3 of ERF/AP2 transcription factor family. The protein contains one AP2 domain. There are 18 members in this subfamily including ATERF-1, ATERF-2, AND ATERF-5.

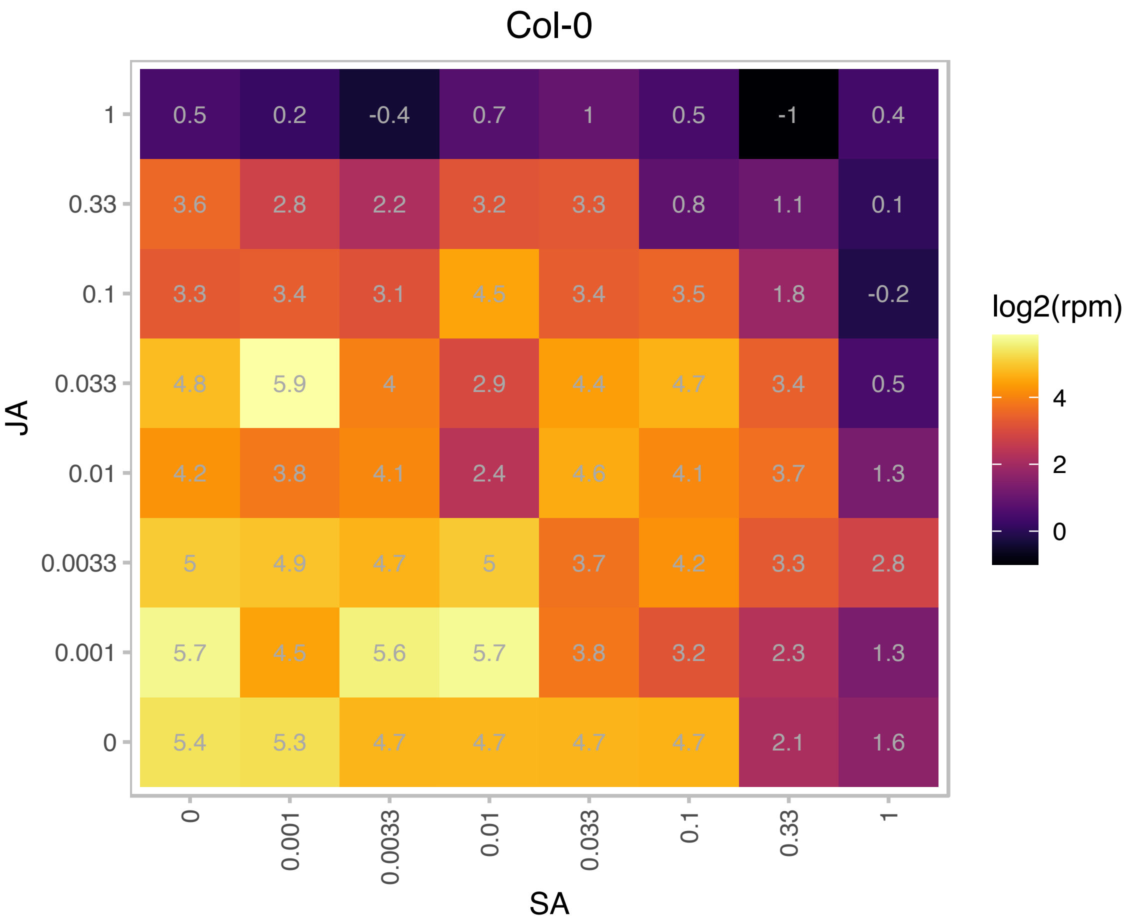


| | |
|---|---|
| Gene | AT5G51190 |
| is.expressed_genes | TRUE |
| is.responsive_genes | TRUE |
| is.apcluster | F-10 |

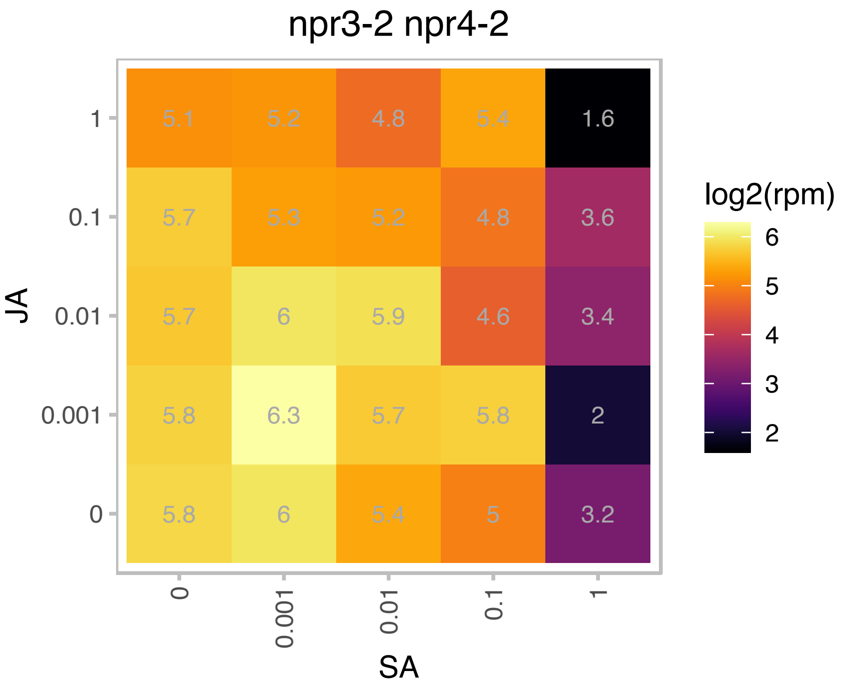


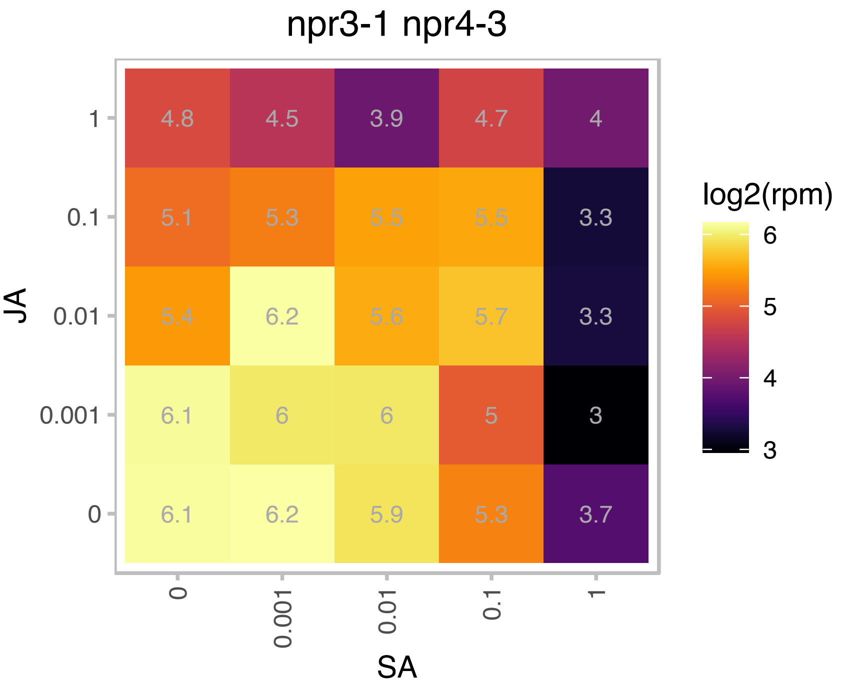

# AT5G51960.1 : NA

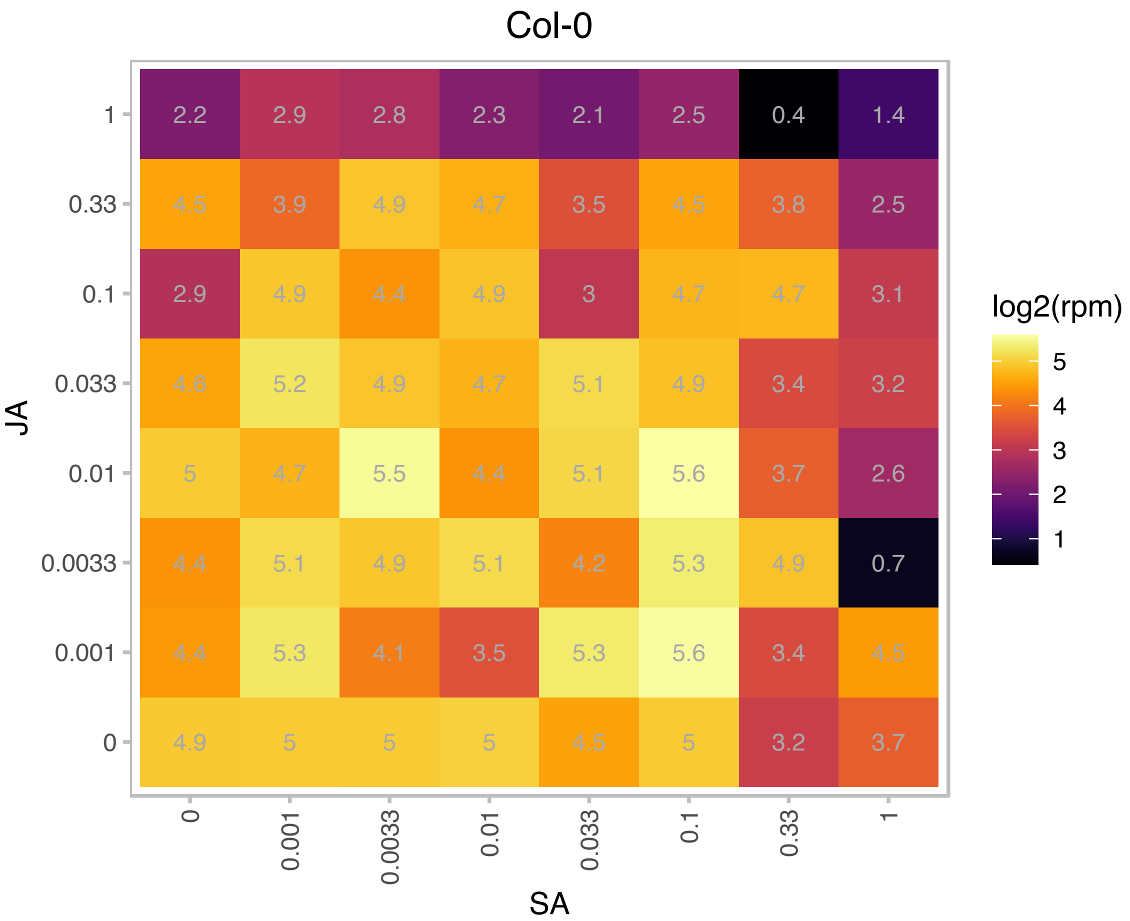


| | |
|---|---|
| Gene | AT5G51960 |
| is.expressed_genes | TRUE |
| is.responsive_genes | TRUE |
| is.apcluster | F-10 |

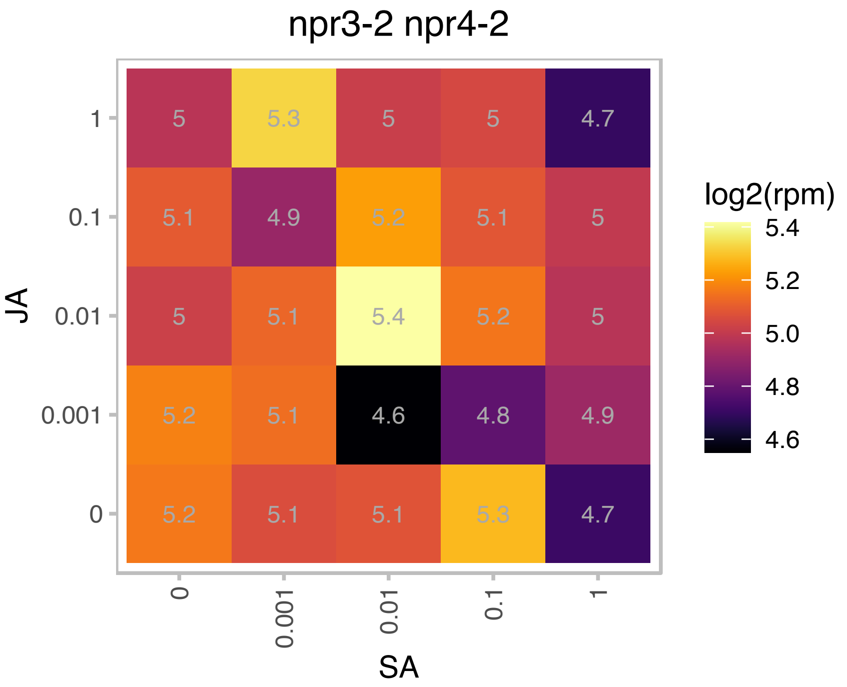


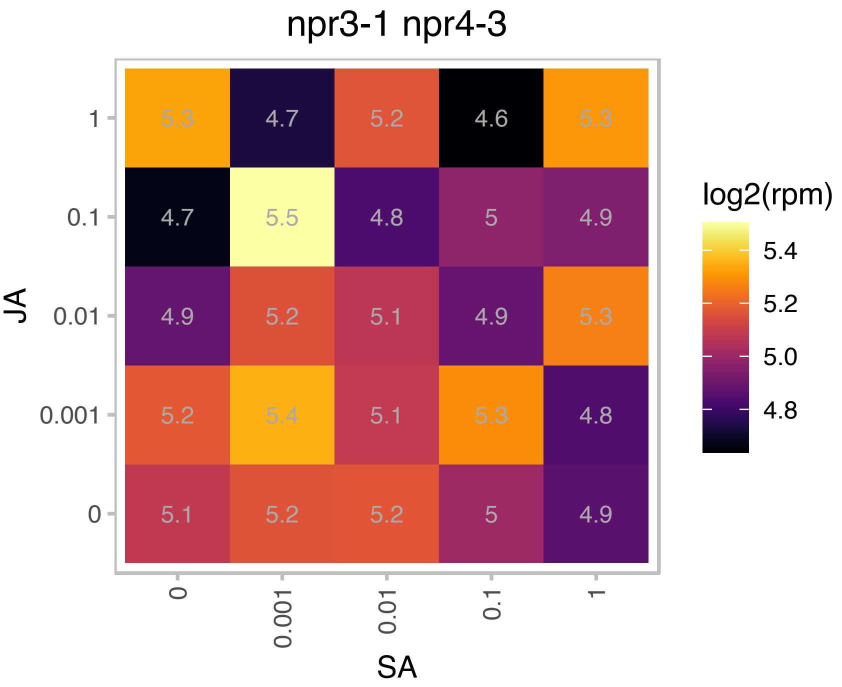

### AT5G52100.1 : CRR1

Dihydrodipicolinate reductase, bacterial/plant
Is essential for chloroplast NAD(P)H dehydrogenase activity, which is involved in electron transfer between PSII and PSI. Likely functions in biogenesis or stabilization of the NAD(P)H dehydrogenase complex. The mRNA is cell-to-cell mobile.

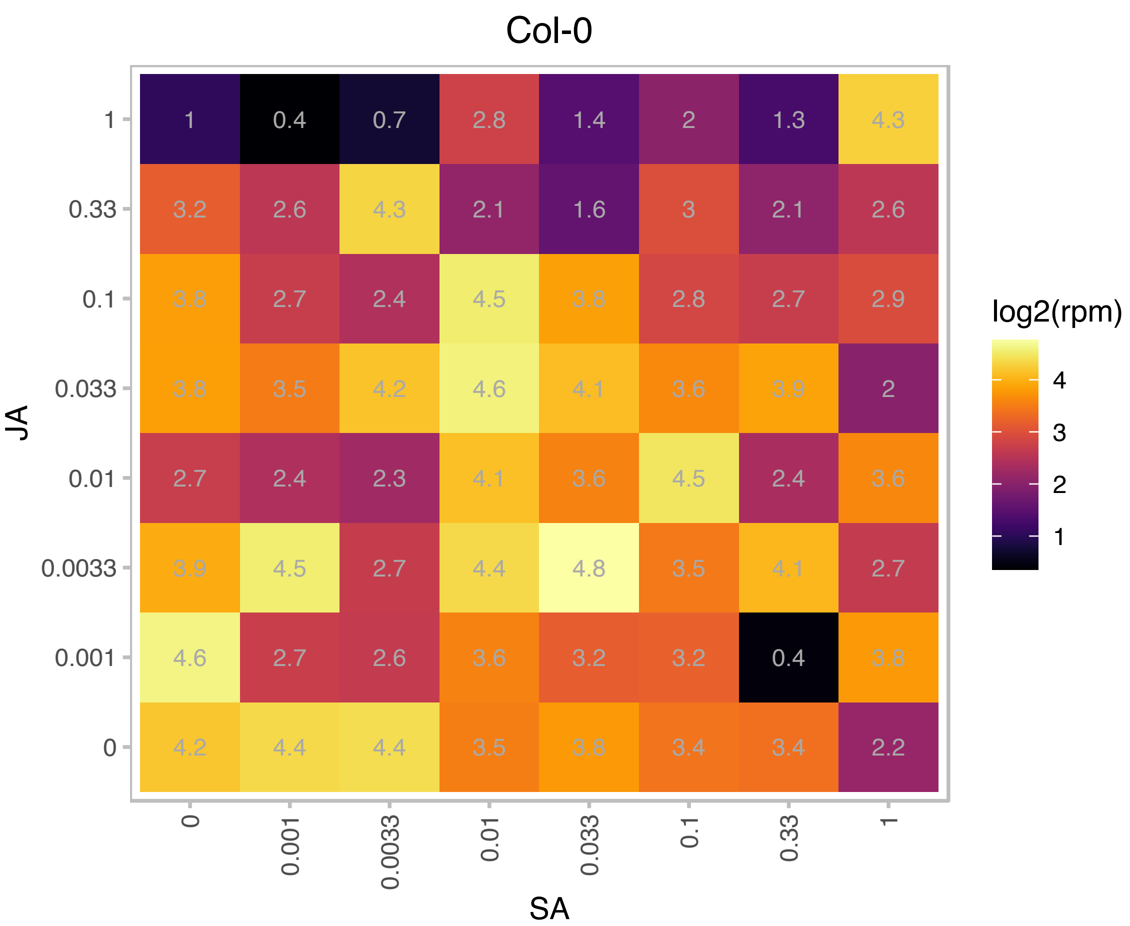


| | |
|---|---|
| Gene | AT5G52100 |
| is.expressed_genes | TRUE |
| is.responsive_genes | TRUE |
| is.apcluster | F-10 |

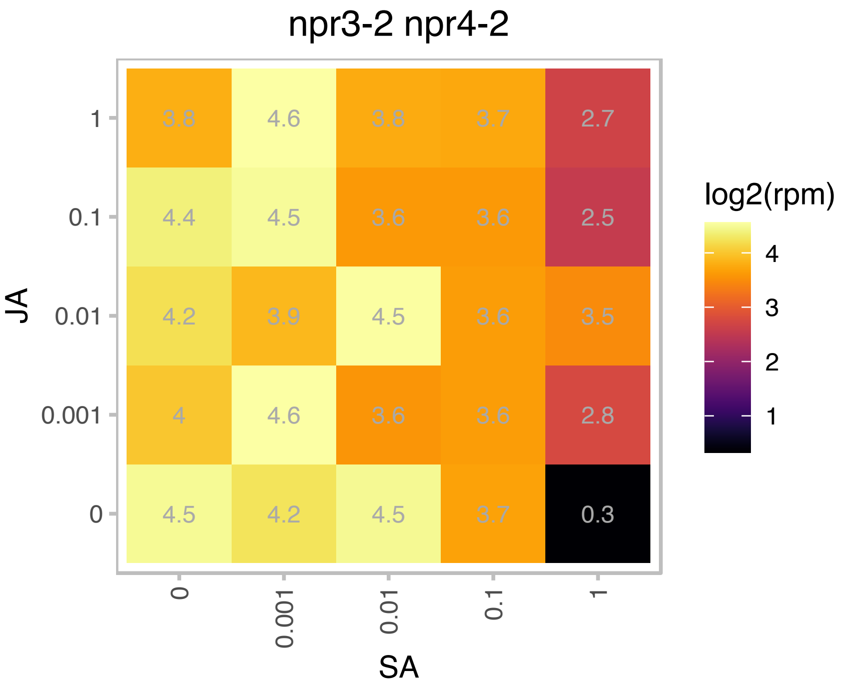


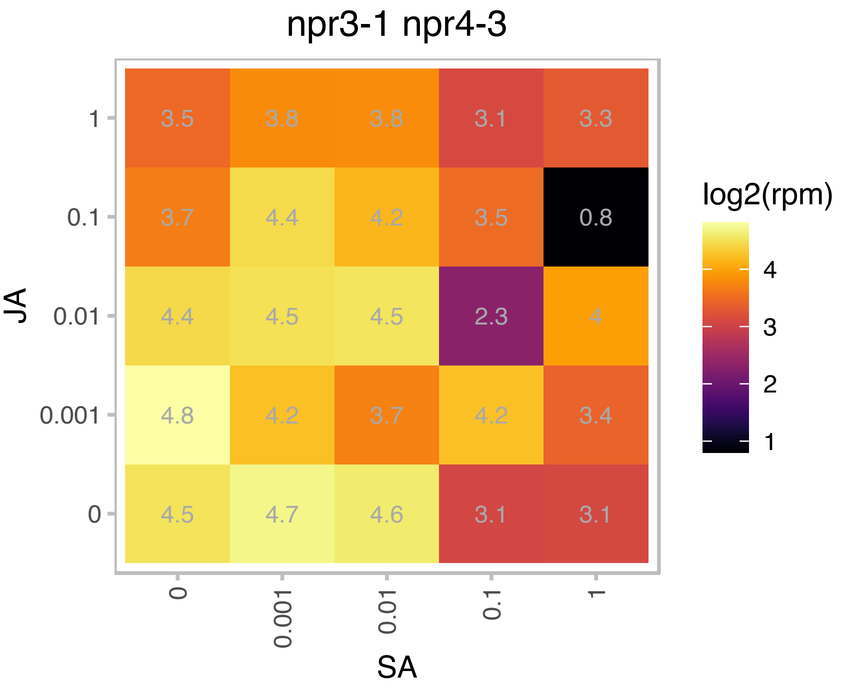

# AT5G52780.1 : NA

Protein of unknown function (DUF3464)

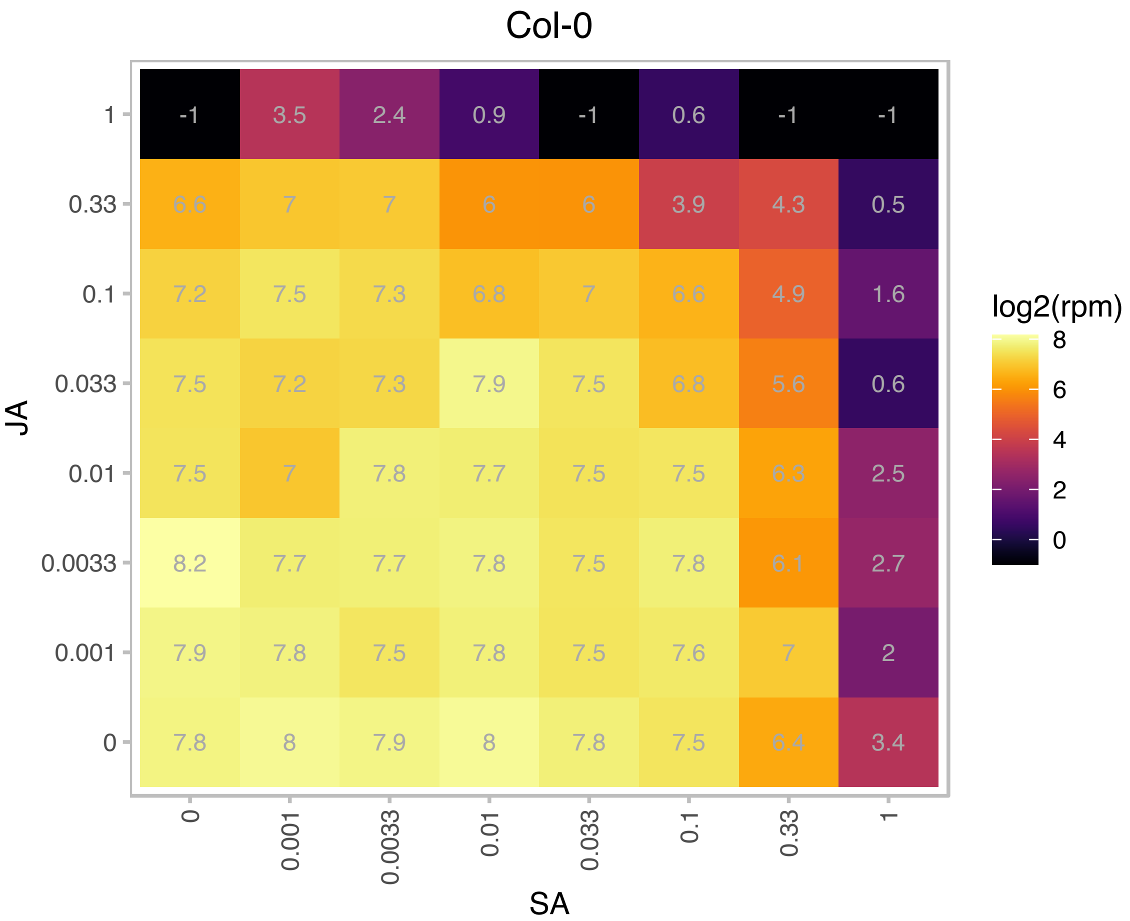


| | |
|---|---|
| Gene | AT5G52780 |
| is.expressed_genes | TRUE |
| is.responsive_genes | TRUE |
| is.apcluster | F-10 |

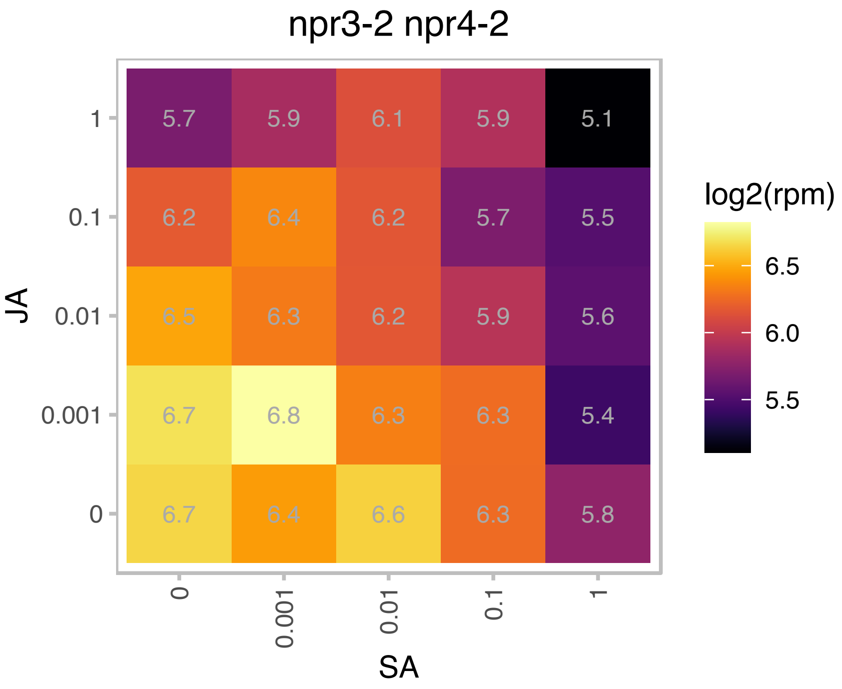


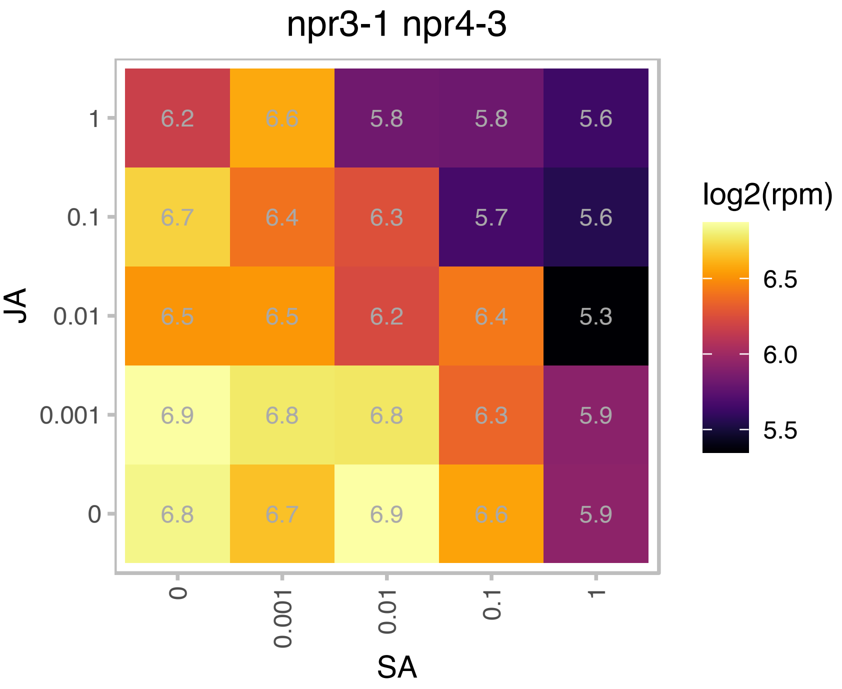

### AT5G53160.1 : PYL8, RCAR3

regulatory components of ABA receptor 3

Encodes RCAR3, a regulatory component of ABA receptor. Interacts with protein phosphatase 2Cs ABI1 and ABI2. Stimulates ABA signaling. The mRNA is cell-to-cell mobile.

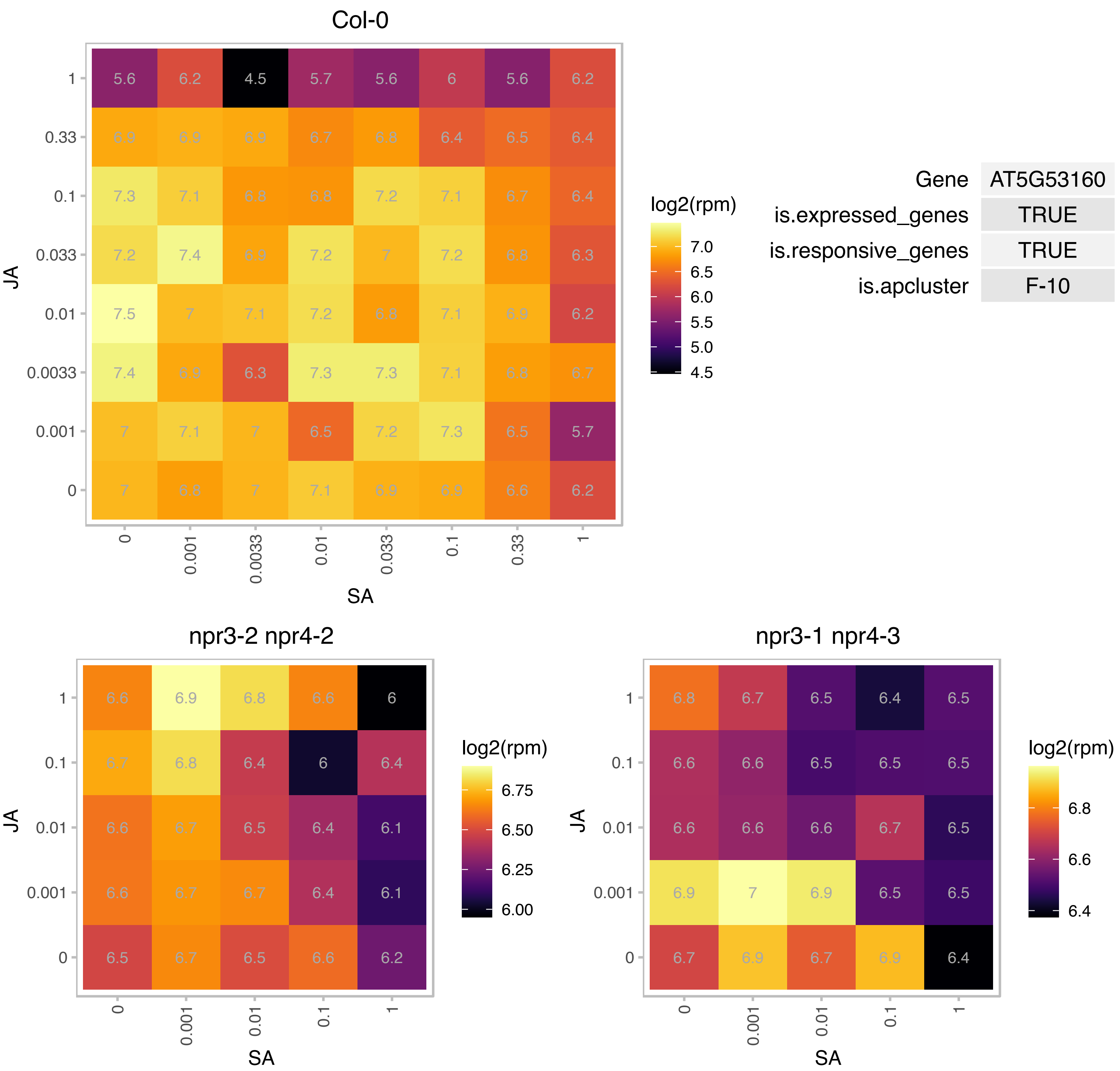


| | |
|---|---|
| Gene | AT5G53160 |
| is.expressed_genes | TRUE |
| is.responsive_genes | TRUE |
| is.apcluster | F-10 |

### AT5G54160.1 : AtCOMT, ATOMT1, COMT1, OMT1, OMT3

O-methyltransferase 1

A caffeic acid/5-hydroxyferulic acid O-methyltransferase. Interacts with 14-4-3 proteins in yeast 2 hybrid assay. AtOMT1 (At5g54160) encodes a flavonol 3?-O-methyltransferase that is highly active towards quercetin and myricetin. The substrate specificity identifies the enzyme as flavonol 3?-methyltransferase which replaces the former annotation of the gene to encode a caffeic acid/5-hydroxyferulic acid O-methyltransferase The mRNA is cell-to-cell mobile.

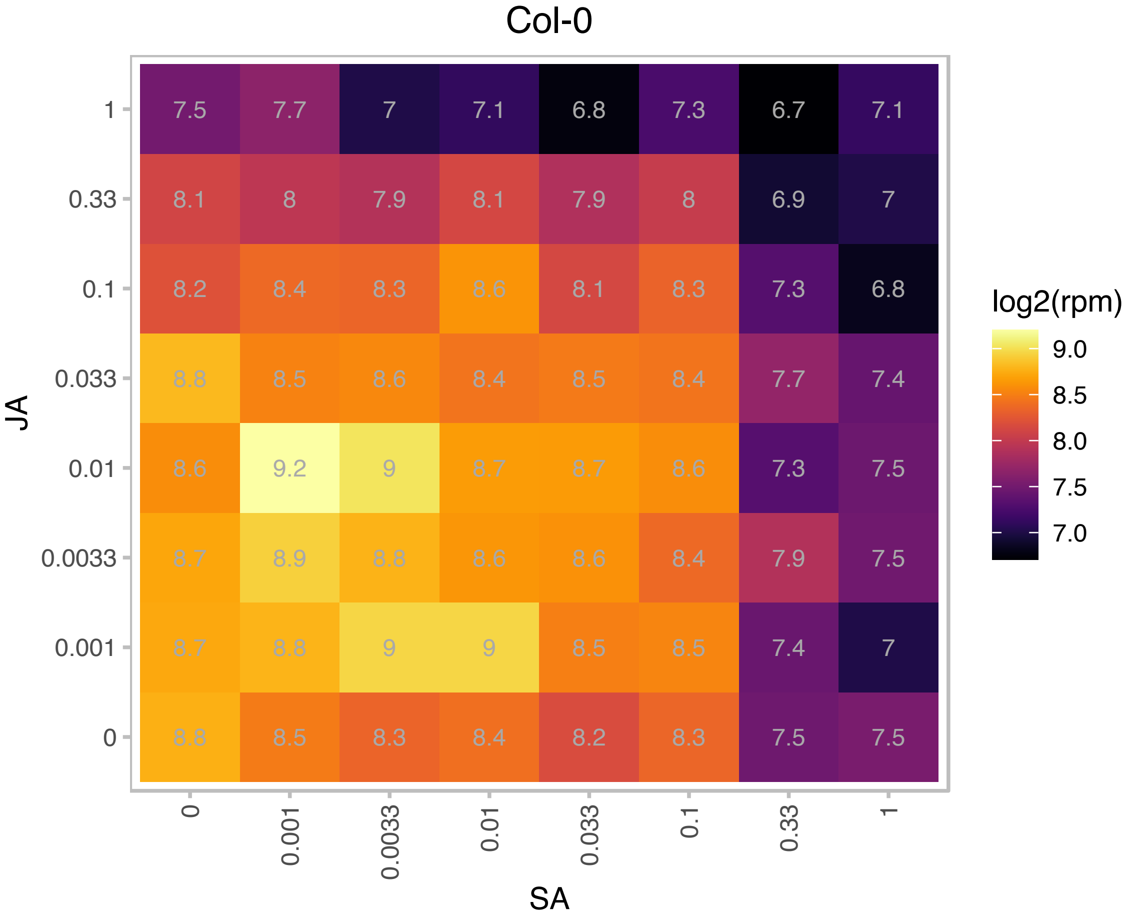


| | |
|---|---|
| Gene | AT5G54160 |
| is.expressed_genes | TRUE |
| is.responsive_genes | TRUE |
| is.apcluster | F-10 |

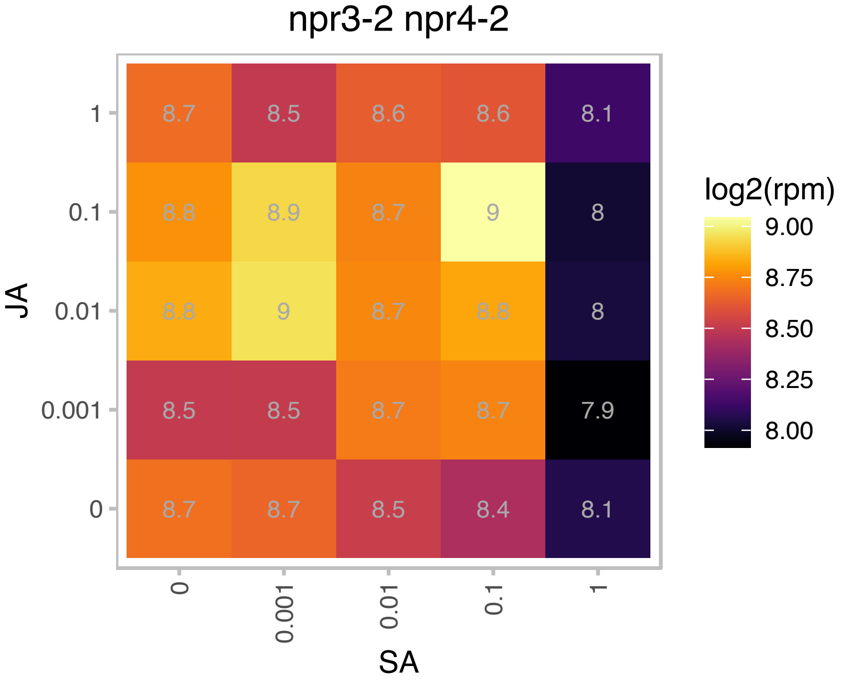


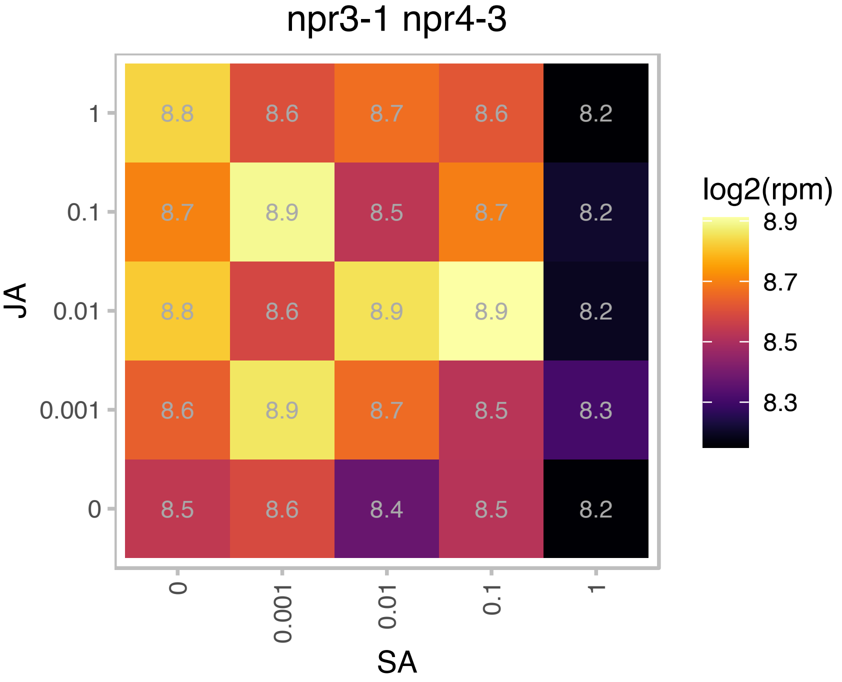

# AT5G56120.1 : NA

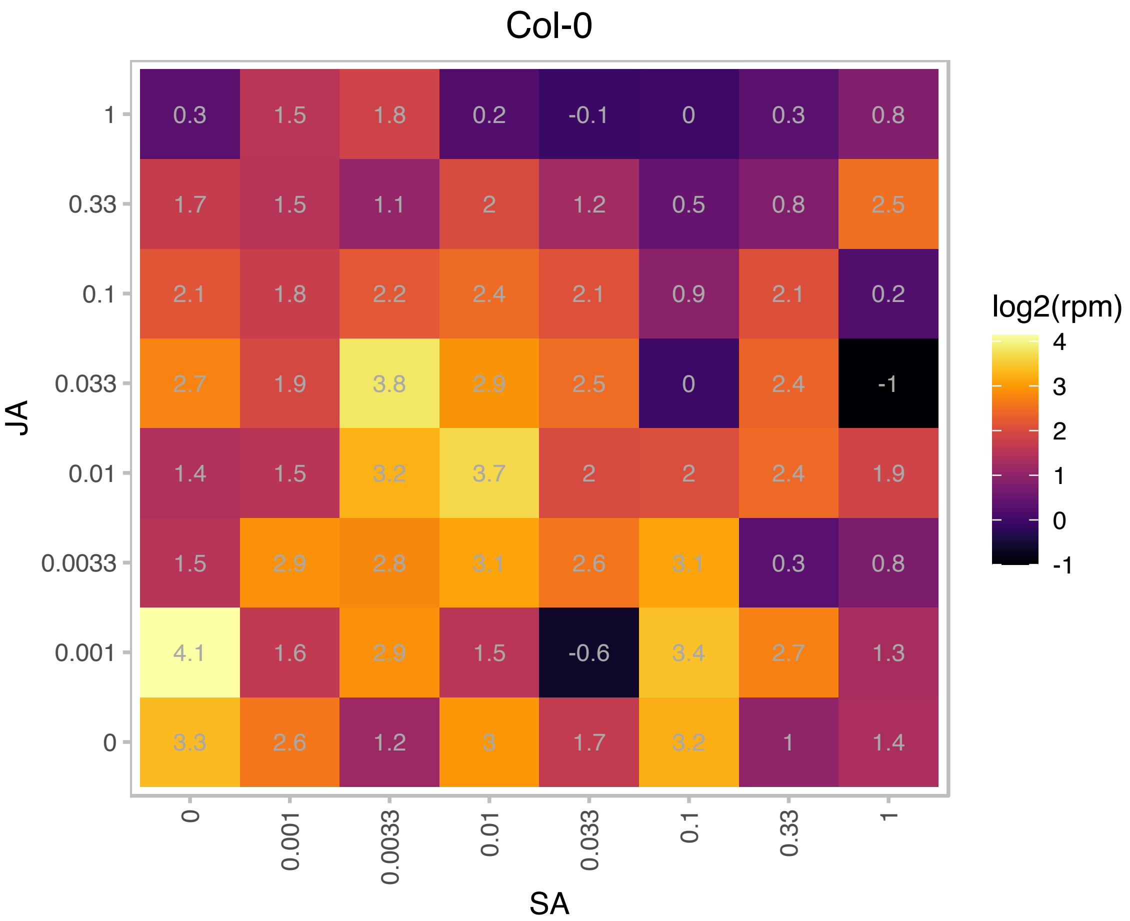


| | |
|---|---|
| Gene | AT5G56120 |
| is.expressed_genes | TRUE |
| is.responsive_genes | TRUE |
| is.apcluster | F-10 |

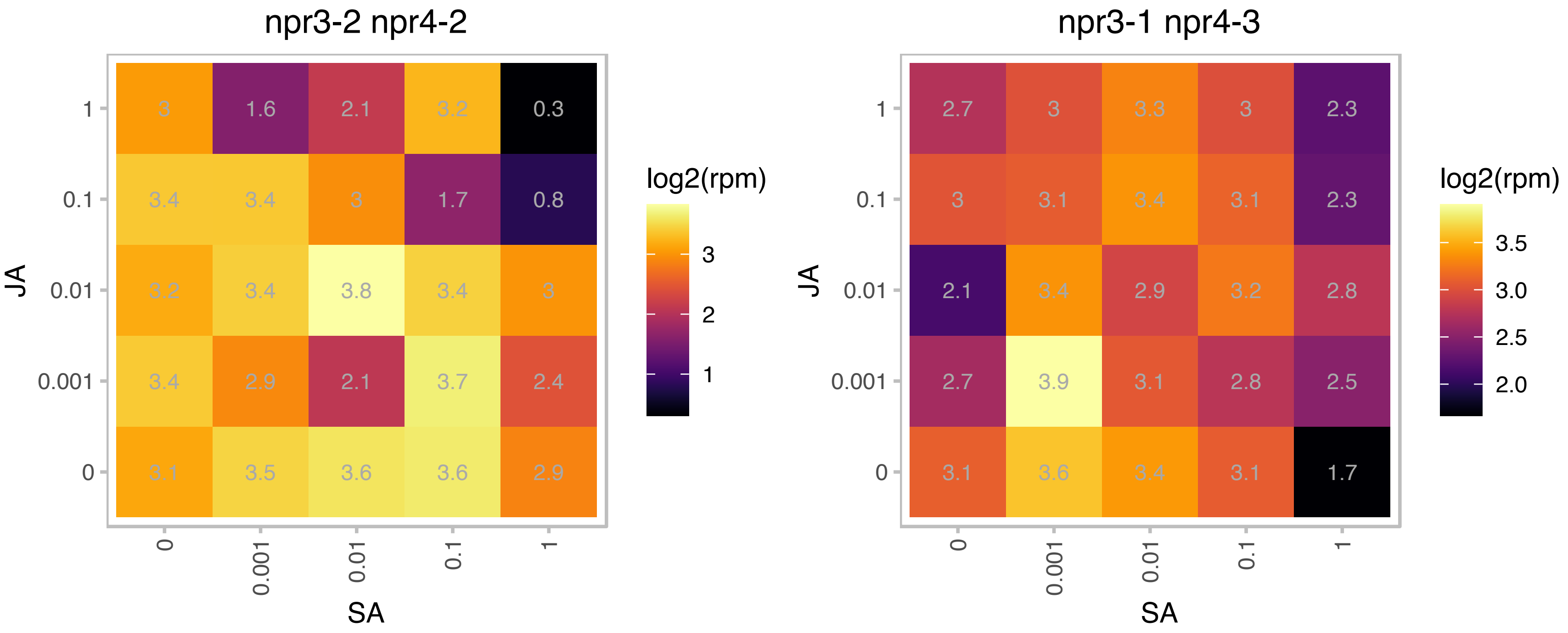

### AT5G64620.1 : ATC/VIF2, C/VIF2

cell wall / vacuolar inhibitor of fructosidase 2

Plant cell wall (CWI) and vacuolar invertases (VI) play important roles in carbohydrate metabolism, stress responses and sugar signaling.

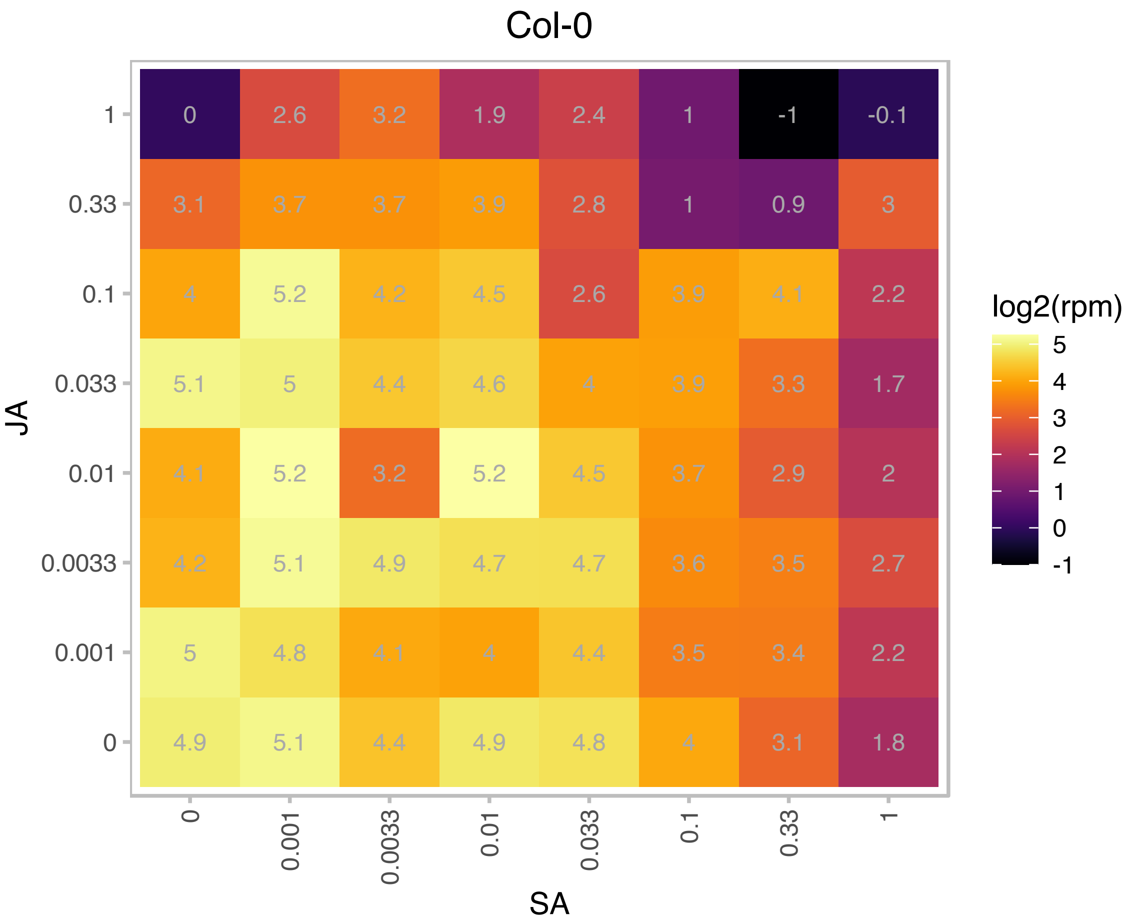


| | |
|---|---|
| Gene | AT5G64620 |
| is.expressed_genes | TRUE |
| is.responsive_genes | TRUE |
| is.apcluster | F-10 |

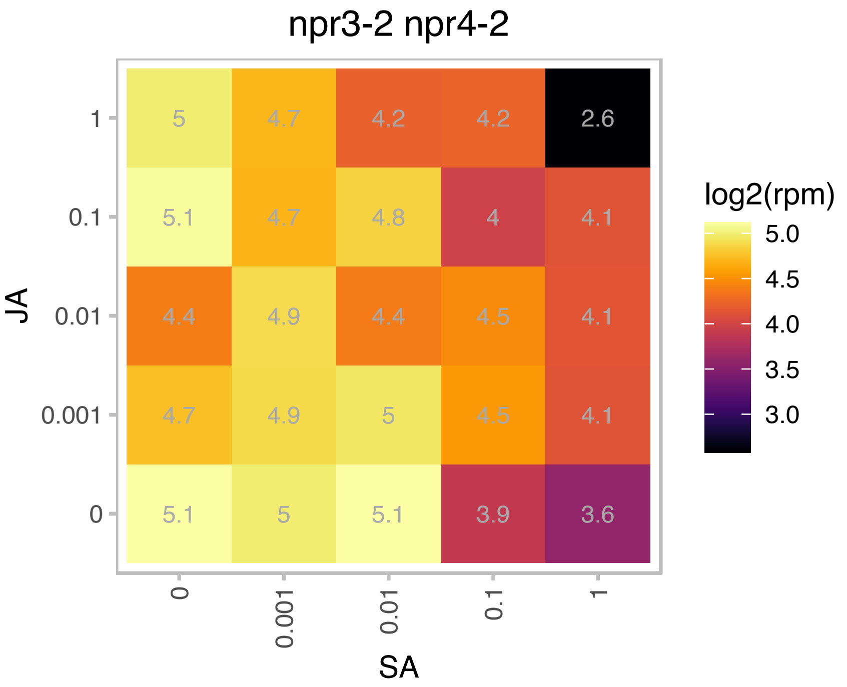


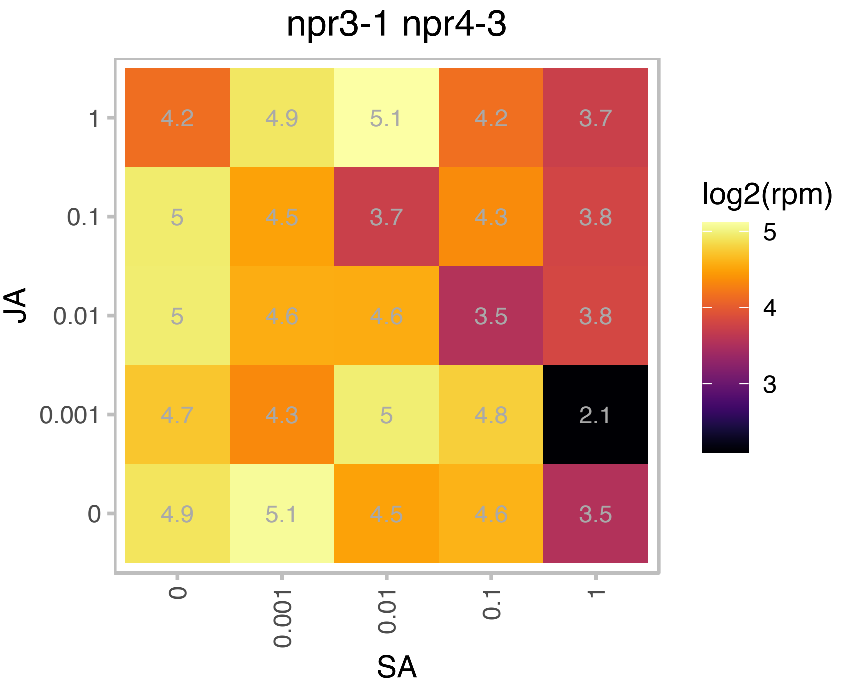

# AT5G66000.1 : NA

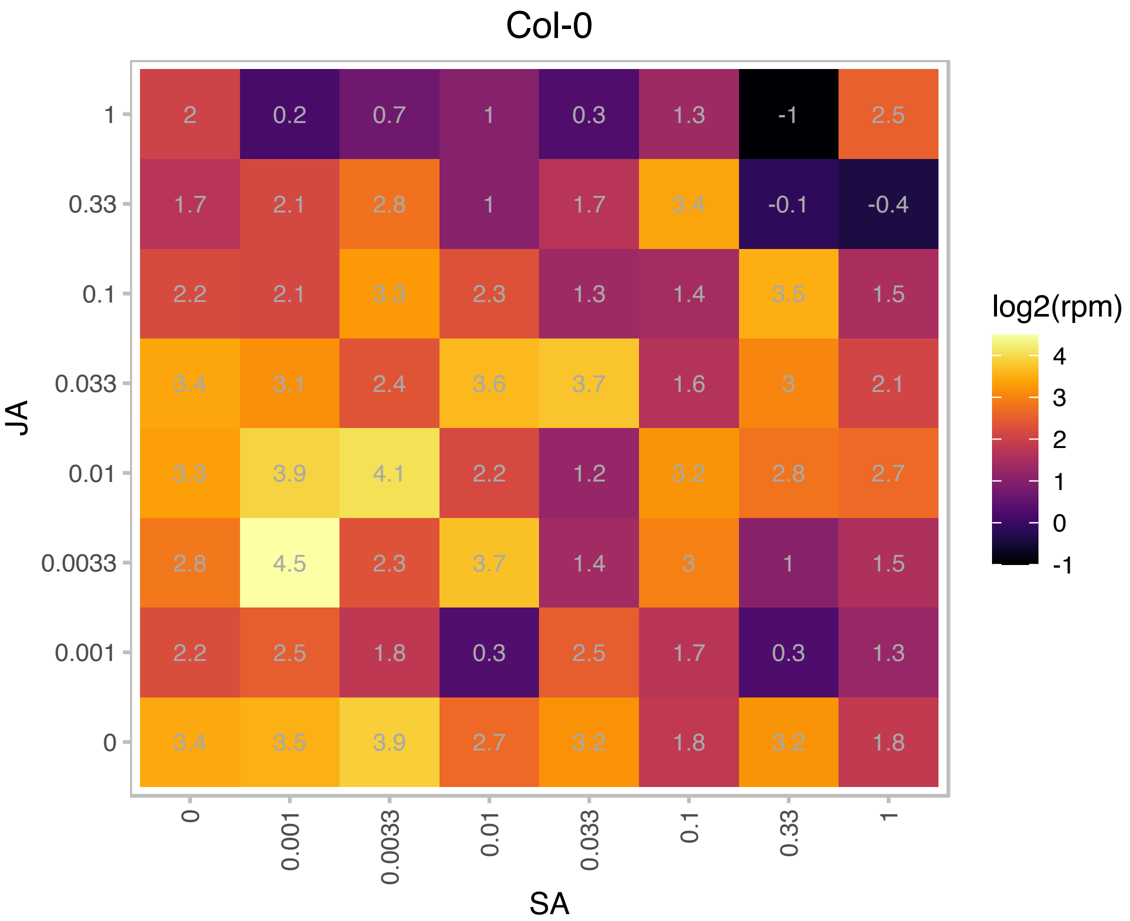


| | |
|---|---|
| Gene | AT5G66000 |
| is.expressed_genes | TRUE |
| is.responsive_genes | TRUE |
| is.apcluster | F-10 |

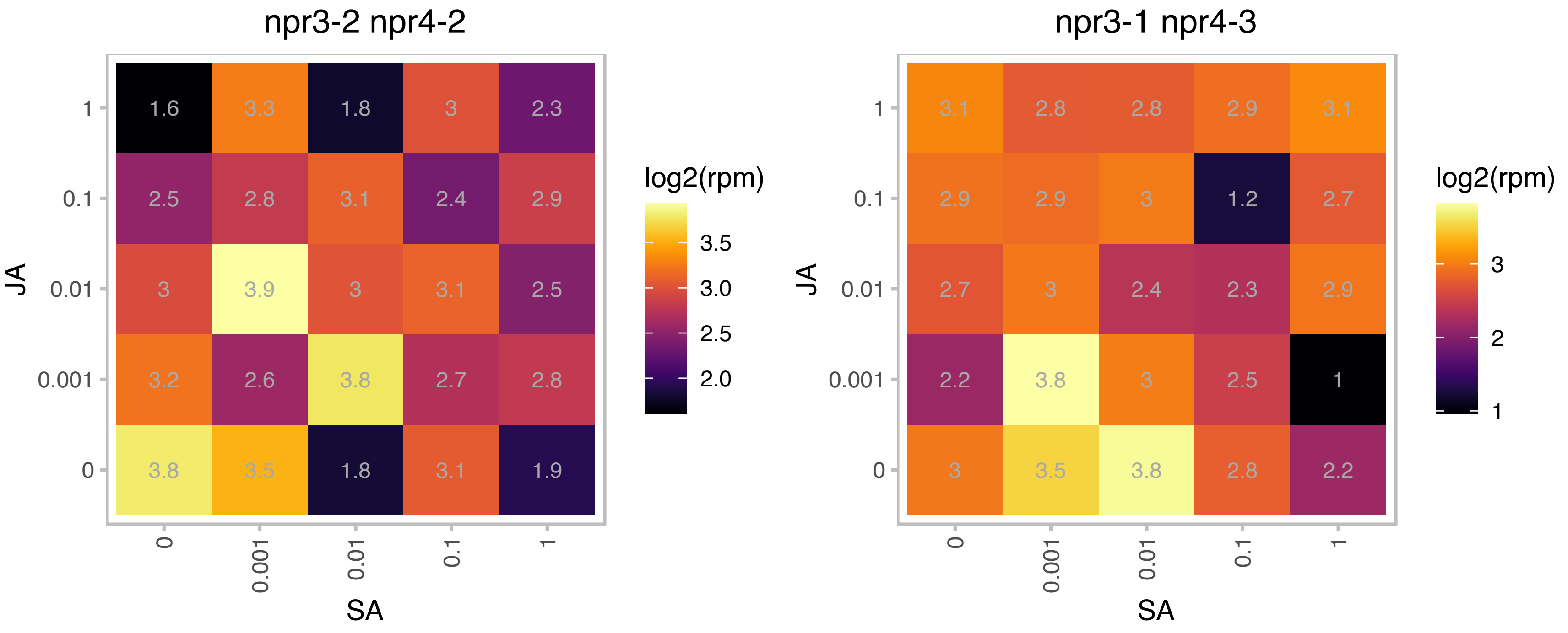

# AT5G66490.1 : NA

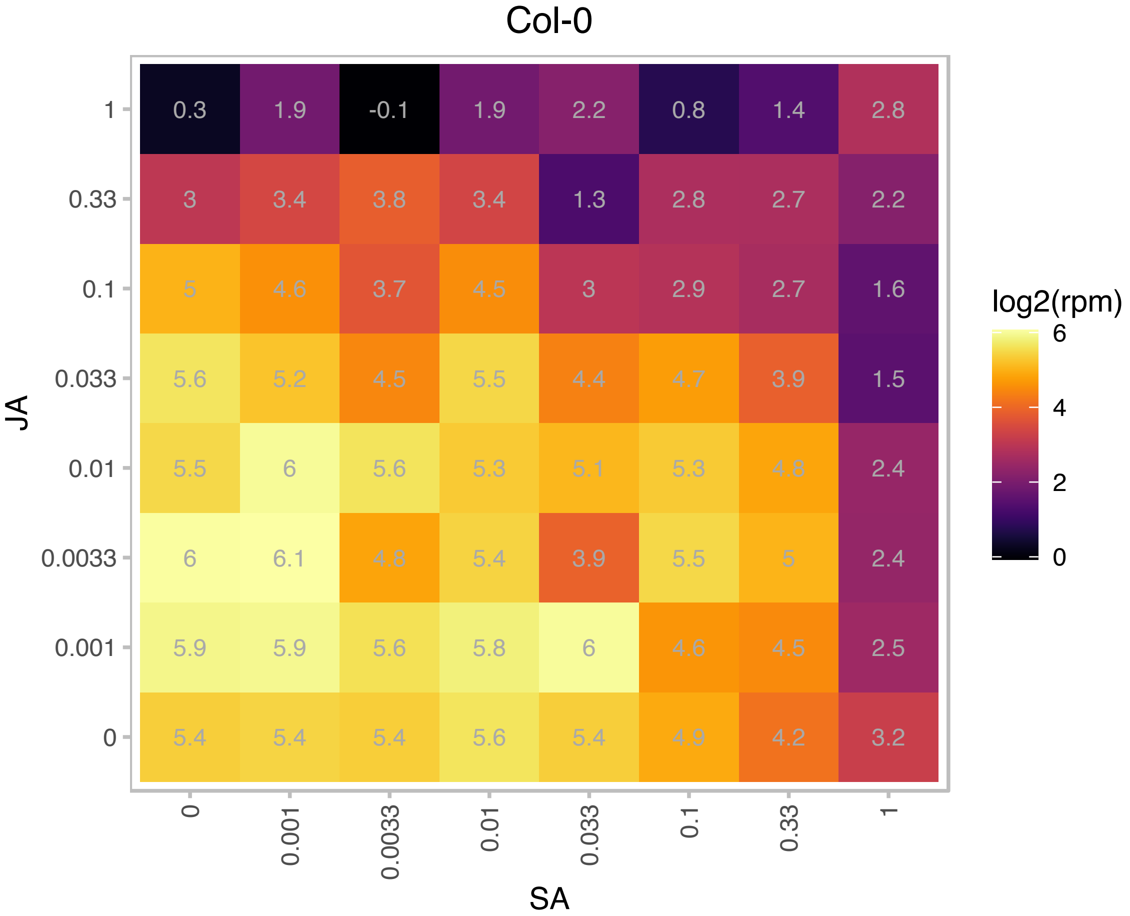


| | |
|---|---|
| Gene | AT5G66490 |
| is.expressed_genes | TRUE |
| is.responsive_genes | TRUE |
| is.apcluster | F-10 |

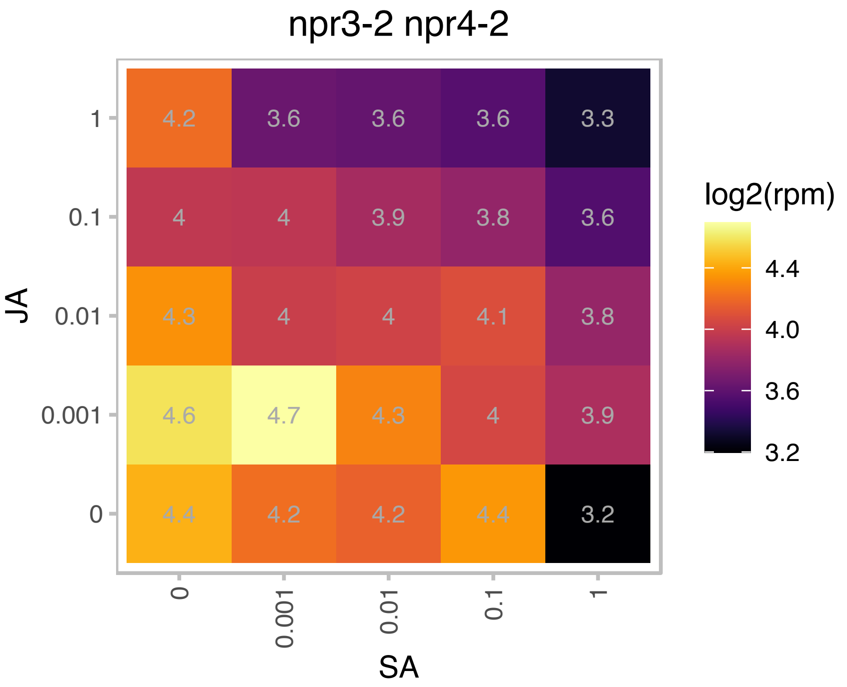


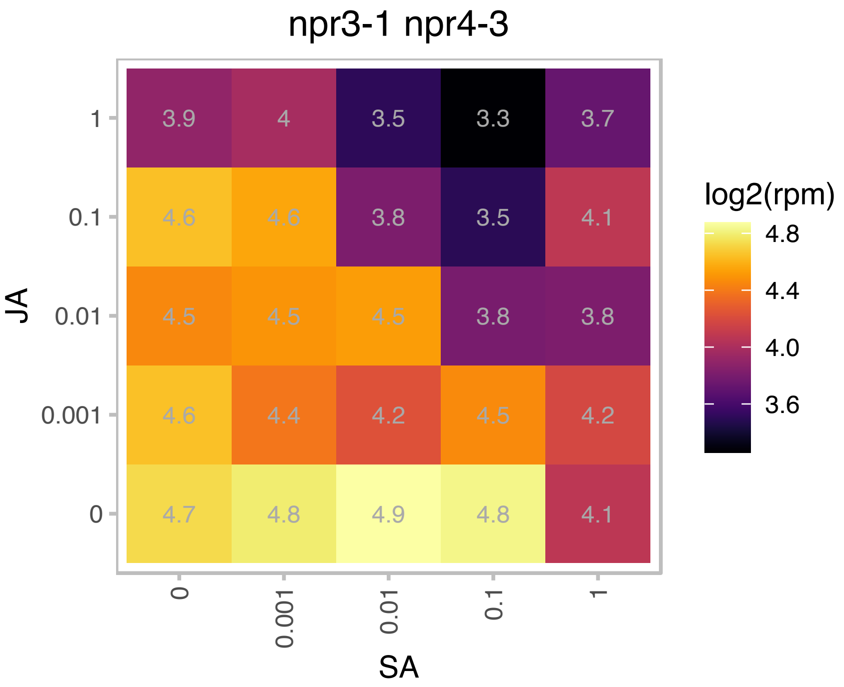

### AT1G04820.1 : TOR2, TUA4

tubulin alpha-4 chain

Encodes an alpha tubulin isoform that is expressed in roots, leaves and flowers.

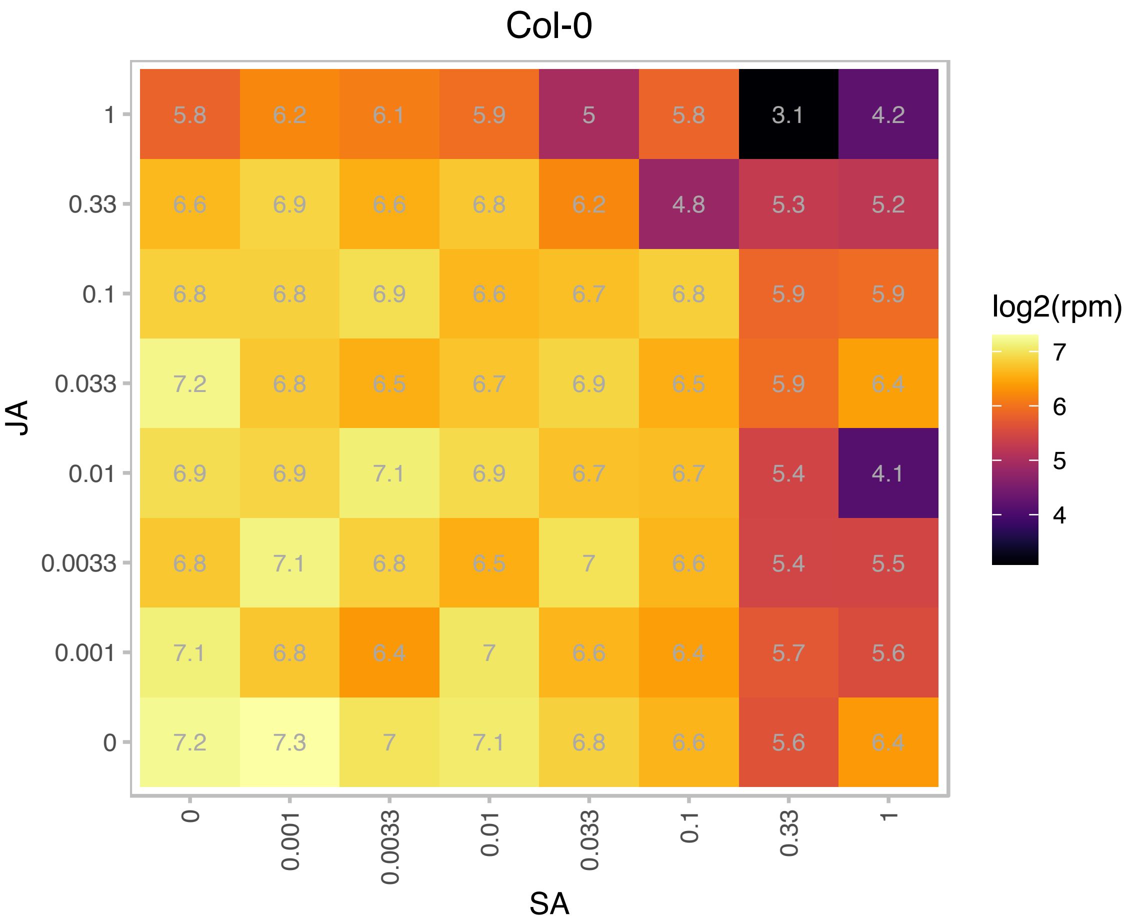


| | |
|---|---|
| Gene | AT1G04820 |
| is.expressed_genes | TRUE |
| is.responsive_genes | TRUE |
| is.apcluster | F-11 |

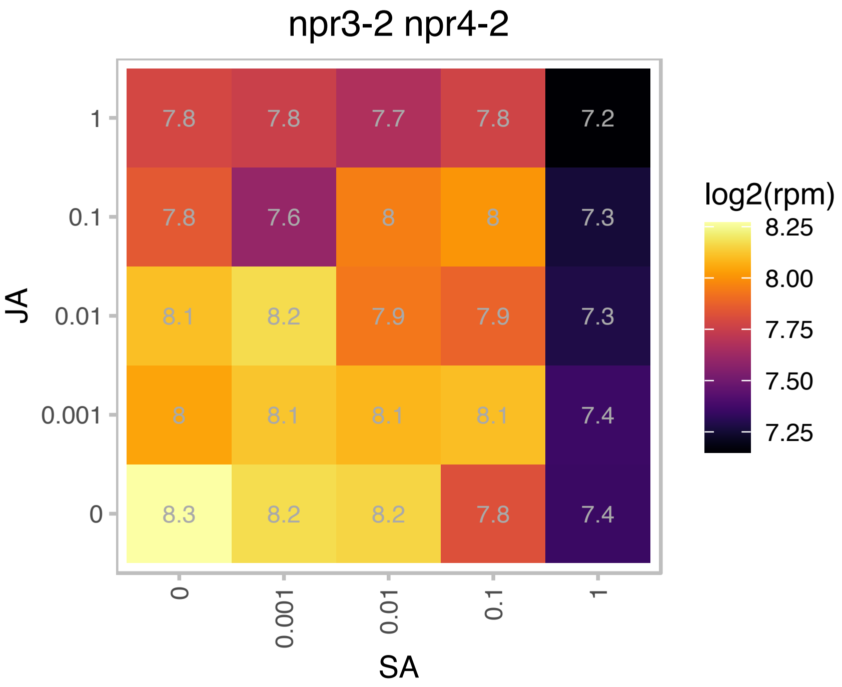


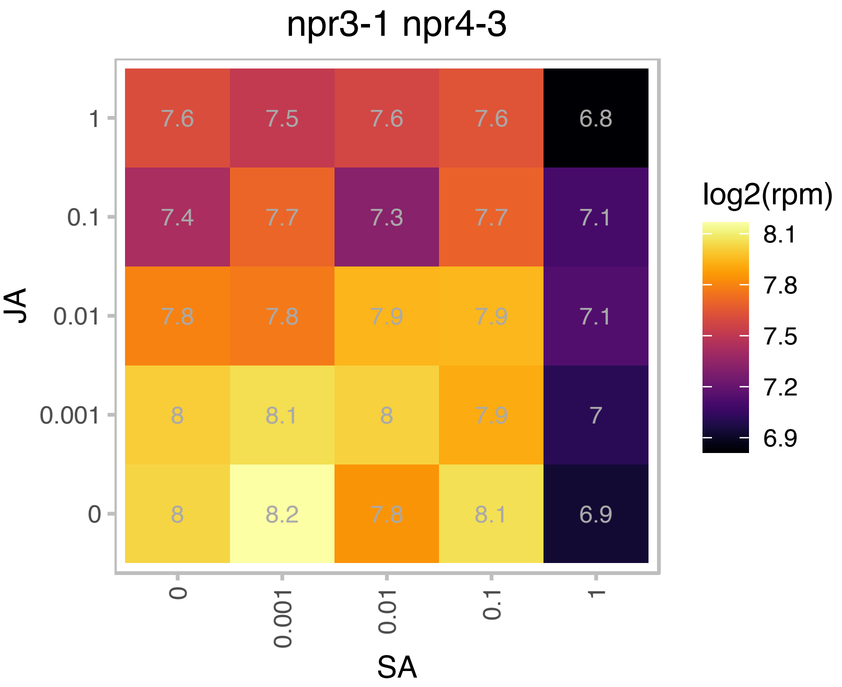

# AT1G06500.1 : NA

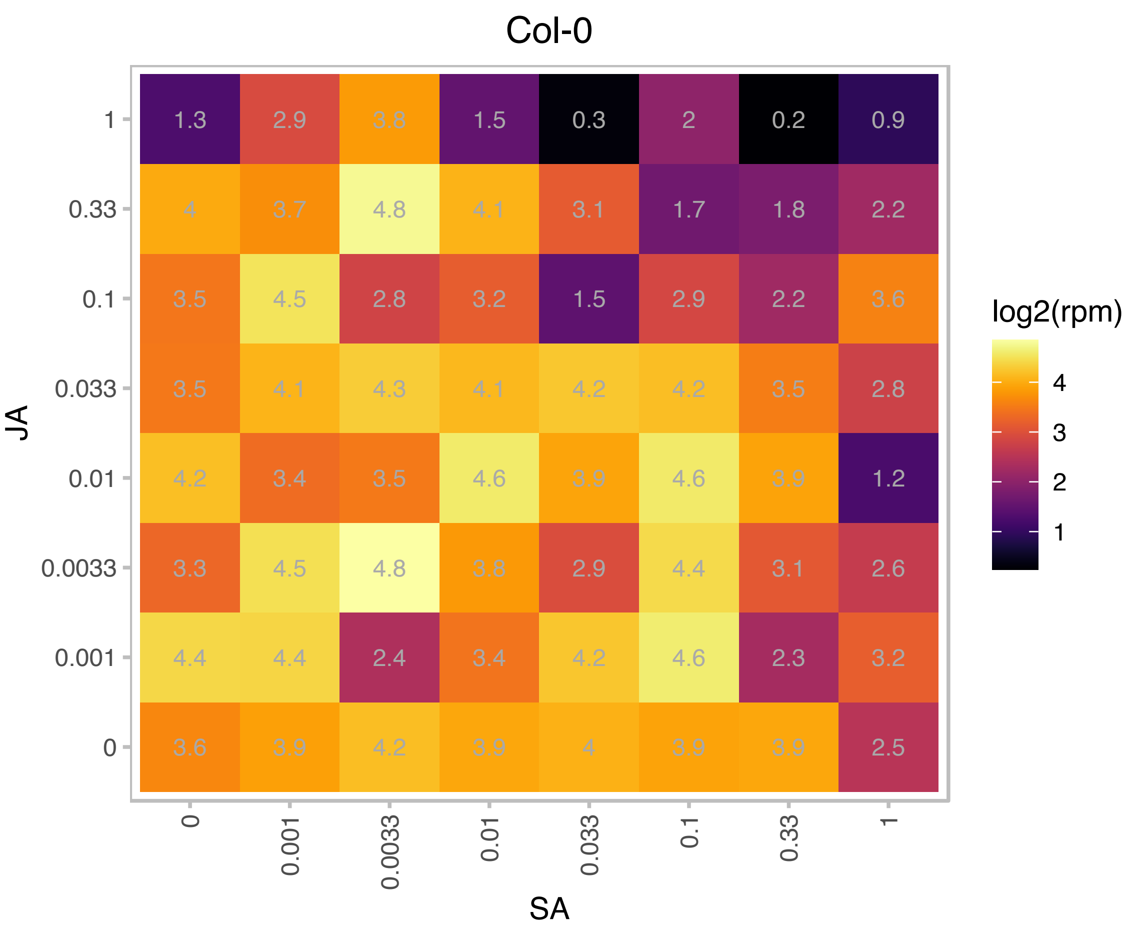


| | |
|---|---|
| Gene | AT1G06500 |
| is.expressed_genes | TRUE |
| is.responsive_genes | TRUE |
| is.apcluster | F-11 |

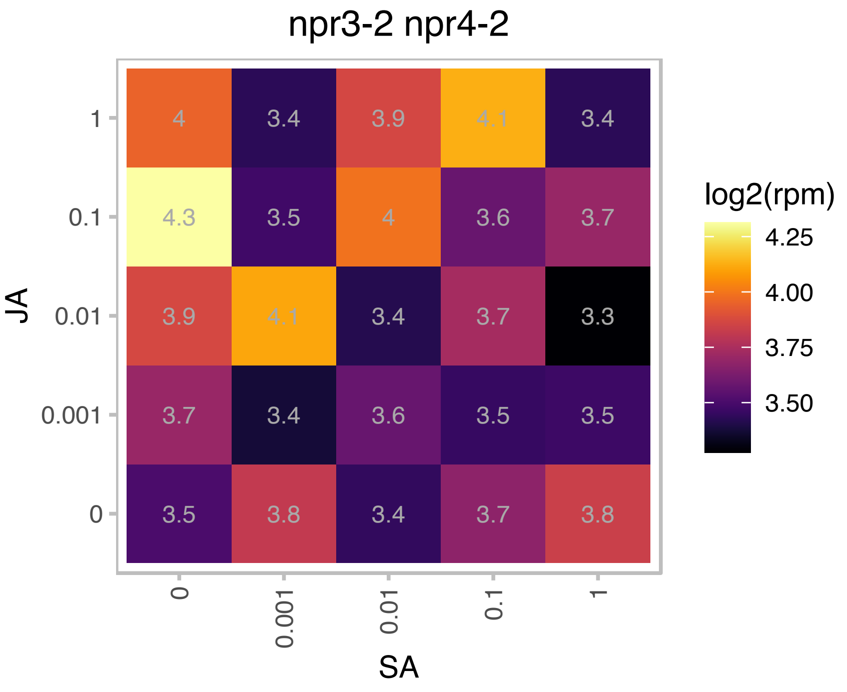


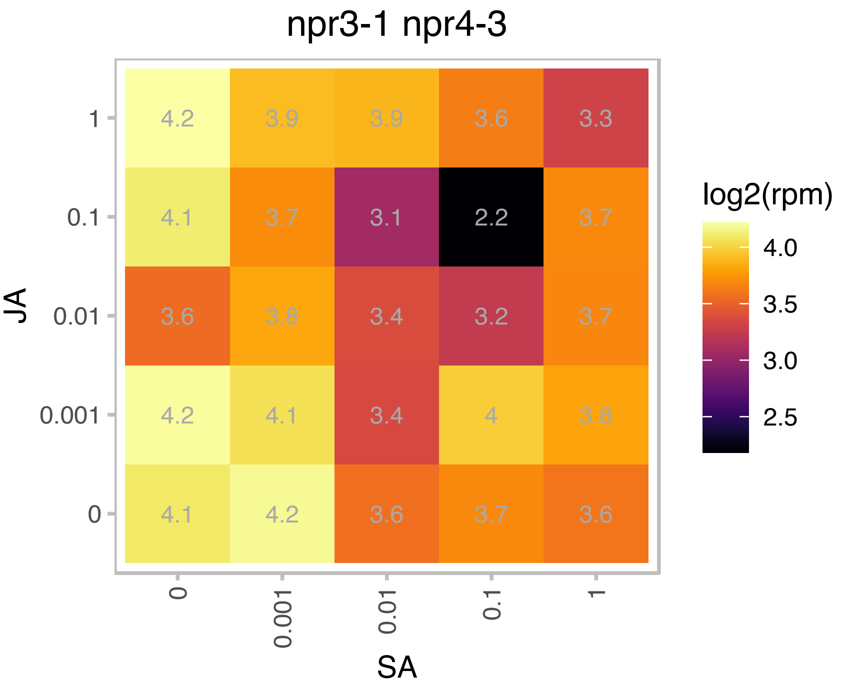

### AT1G10960.1 : ATFD1, FD1

ferredoxin 1

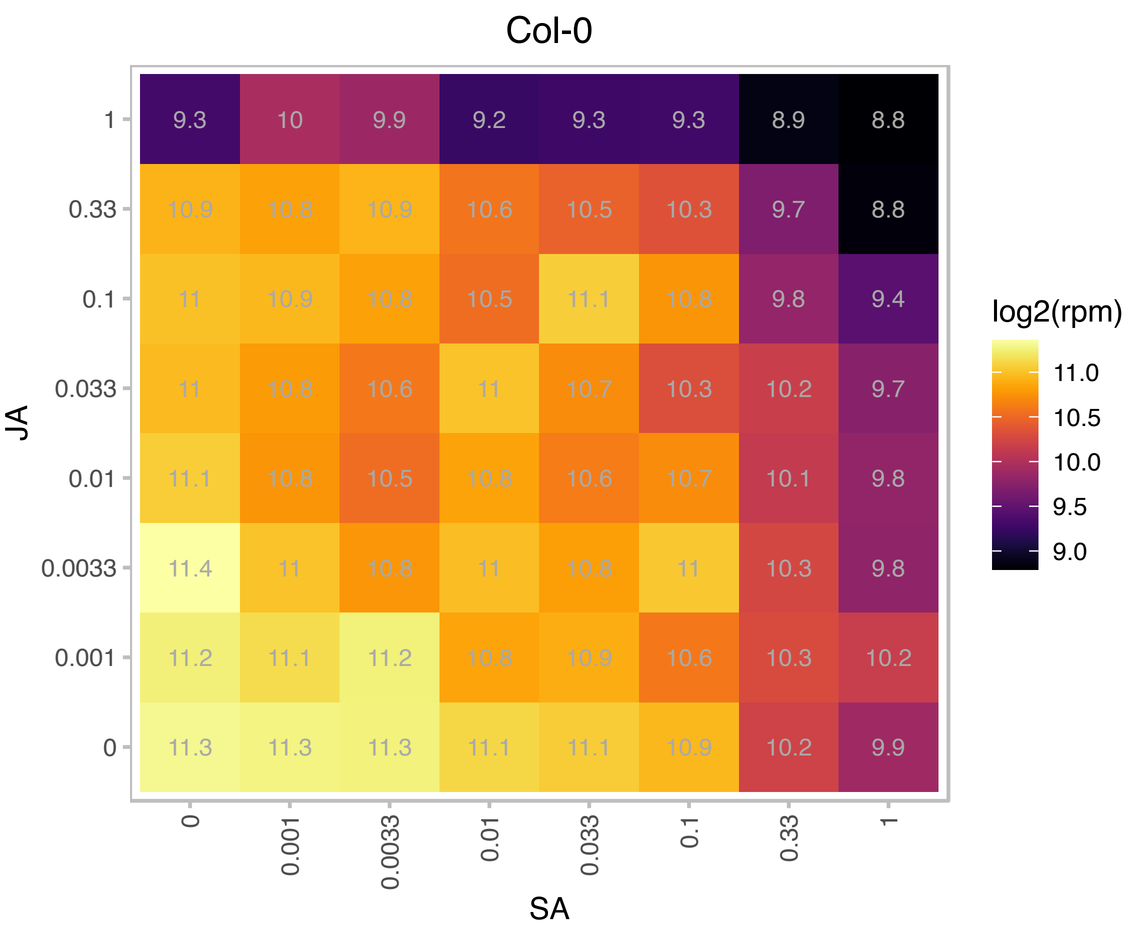


| | |
|---|---|
| Gene | AT1G10960 |
| is.expressed_genes | TRUE |
| is.responsive_genes | TRUE |
| is.apcluster | F-11 |

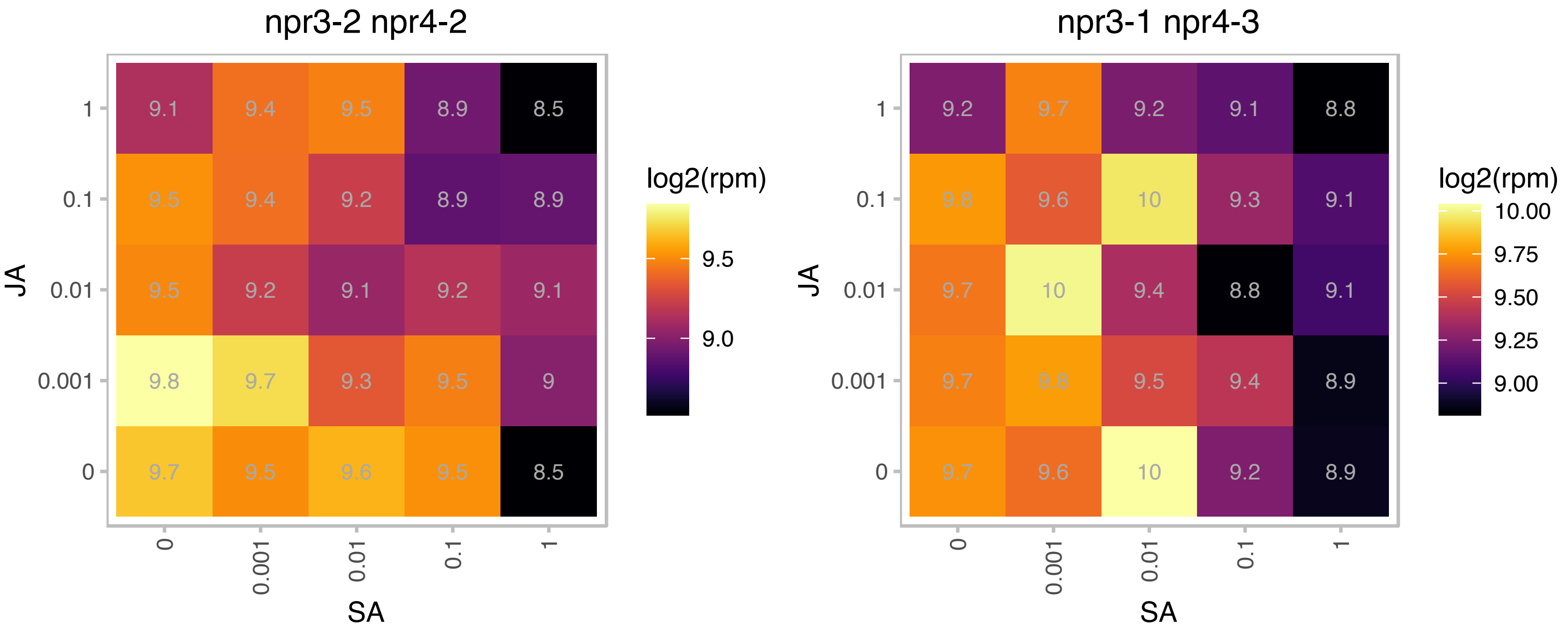

### AT1G13690.1 : ATE1

ATPase E1

AtE1 - stimulates the ATPase activity of DnaK/DnaJ

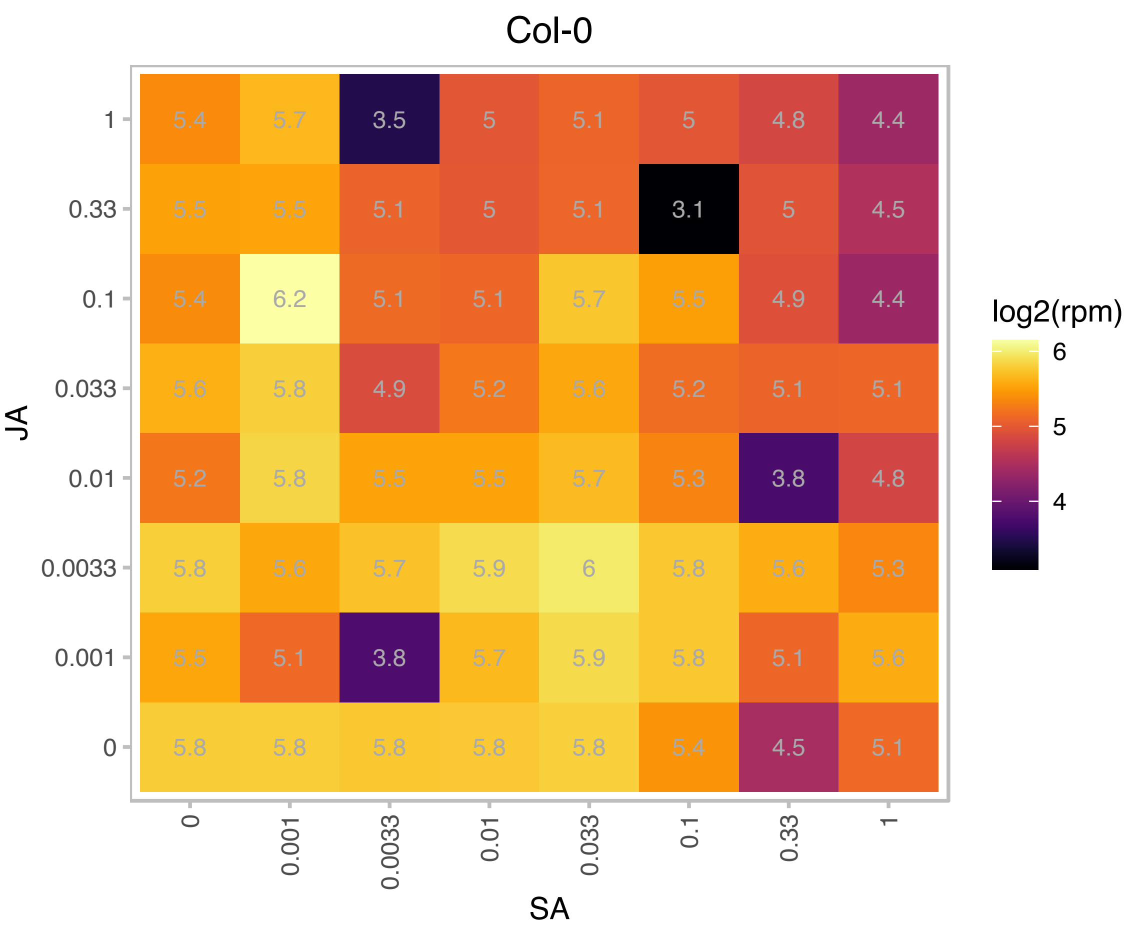


| | |
|---|---|
| Gene | AT1G13690 |
| is.expressed_genes | TRUE |
| is.responsive_genes | TRUE |
| is.apcluster | F-11 |

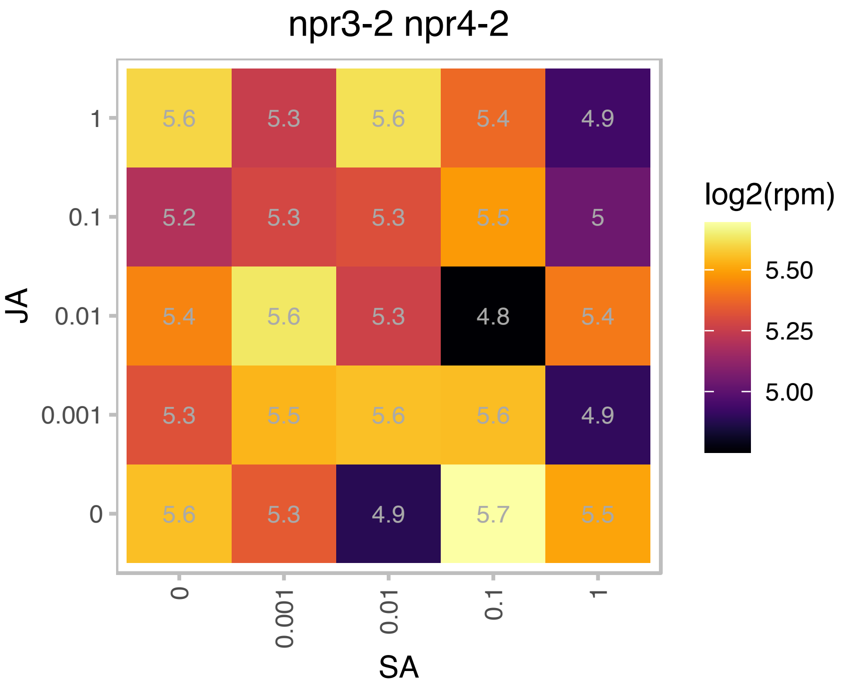


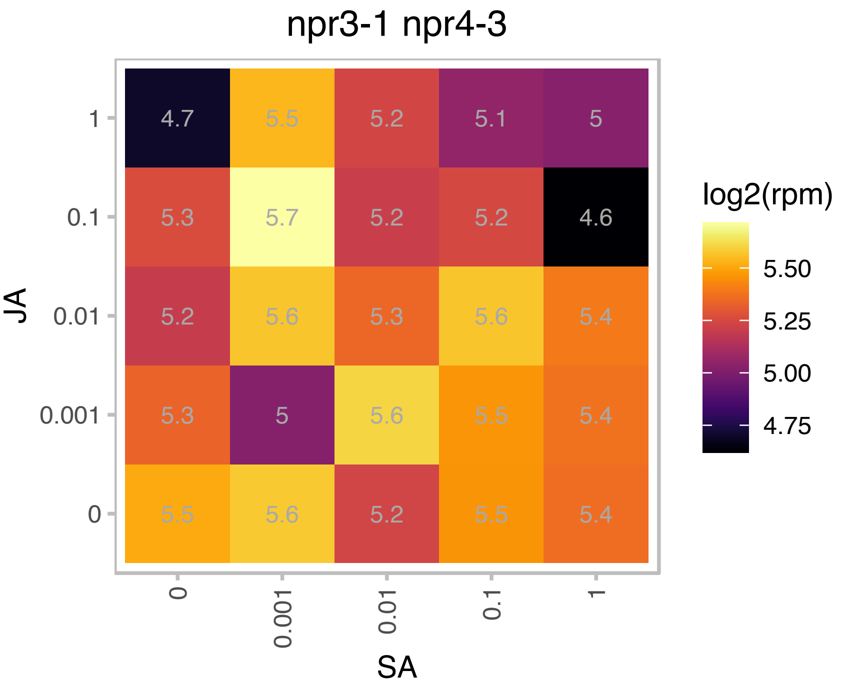

# AT1G14960.1 : NA

Polyketide cyclase/dehydrase and lipid transport superfamily protein

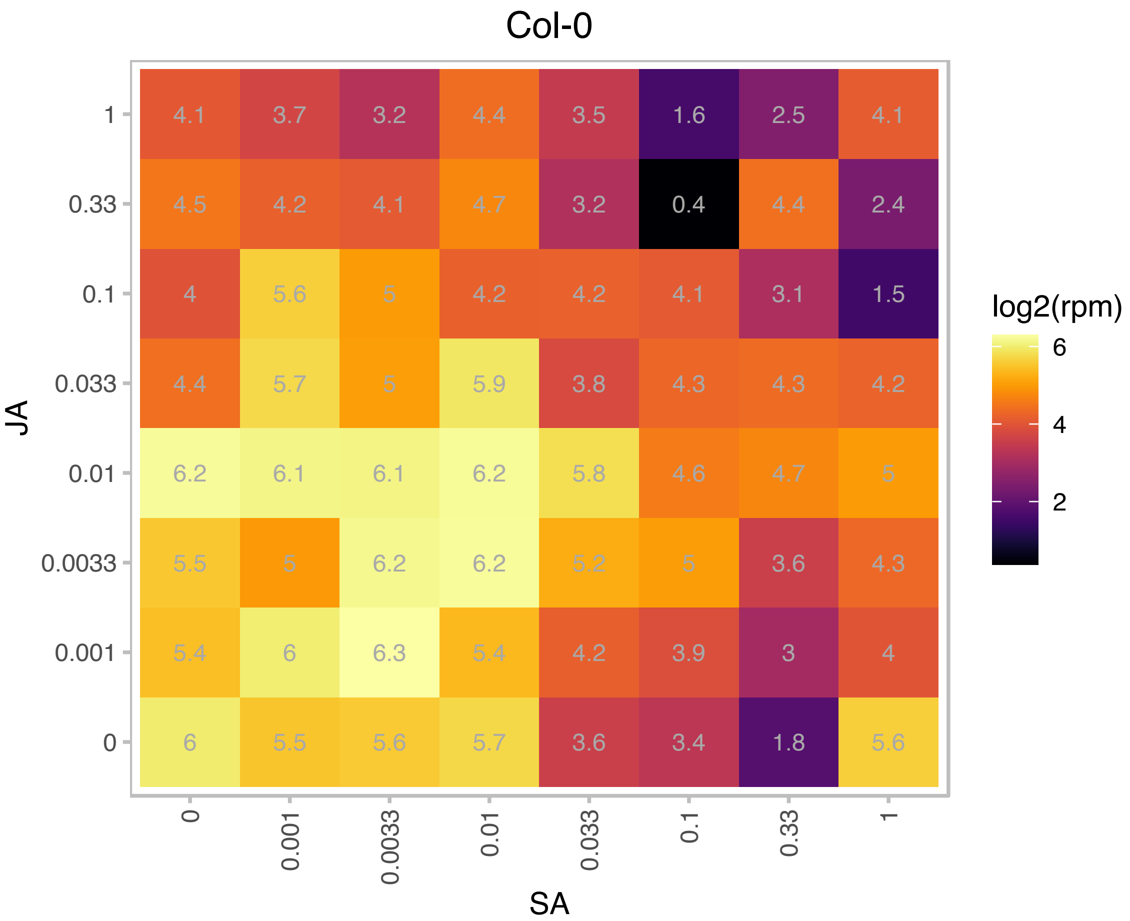


| | |
|---|---|
| Gene | AT1G14960 |
| is.expressed_genes | TRUE |
| is.responsive_genes | TRUE |
| is.apcluster | F-11 |

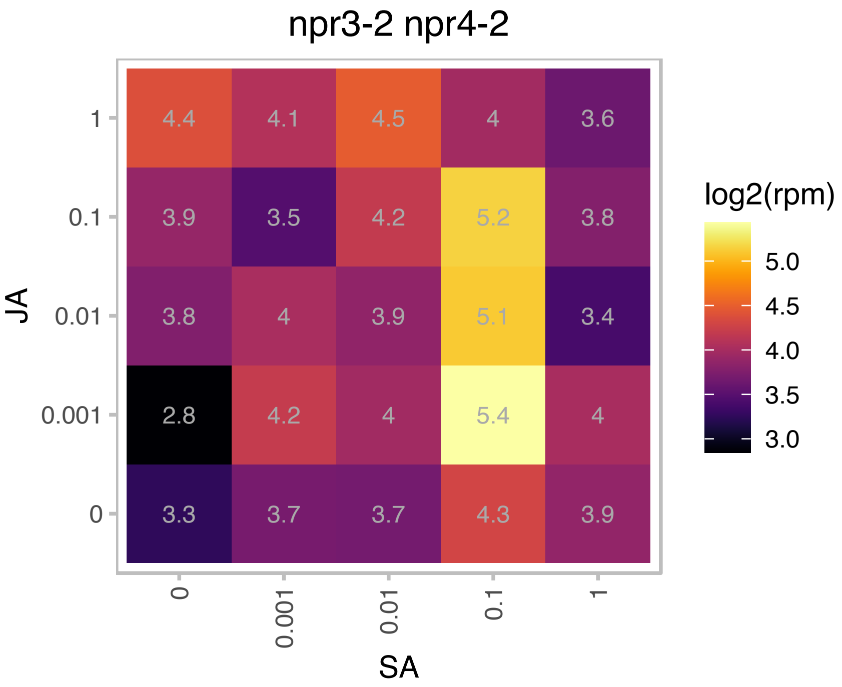


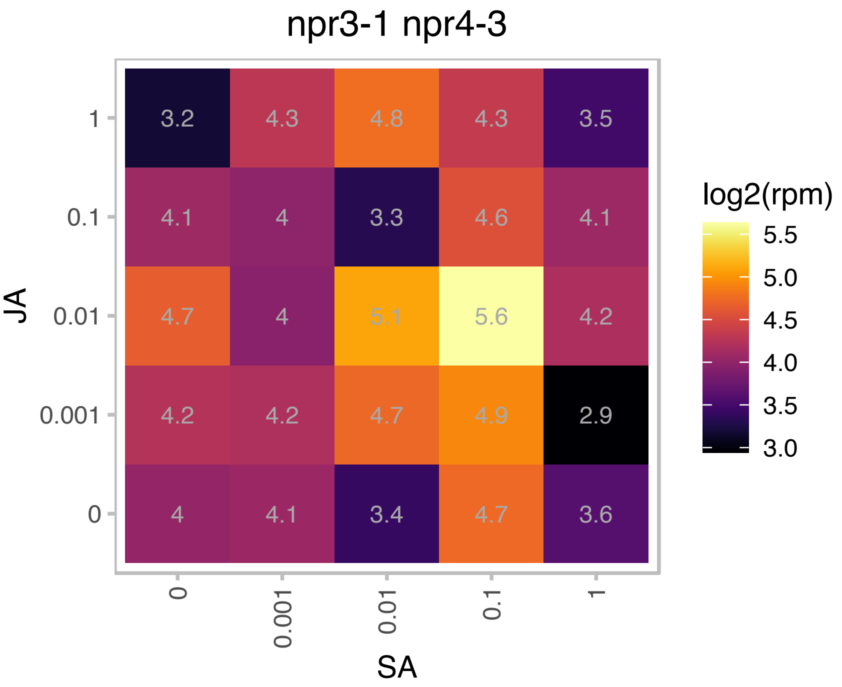

# AT1G15260.1 : NA

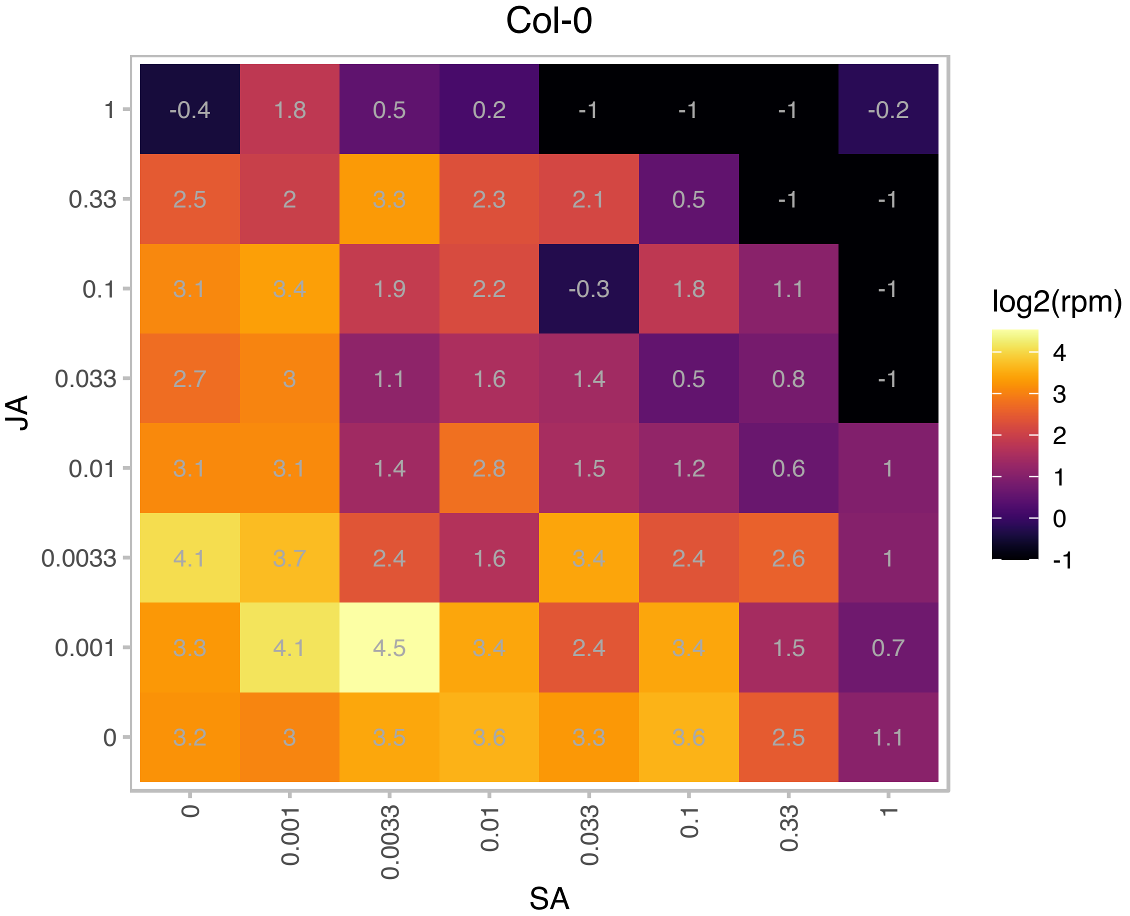


| | |
|---|---|
| Gene | AT1G15260 |
| is.expressed_genes | TRUE |
| is.responsive_genes | TRUE |
| is.apcluster | F-11 |

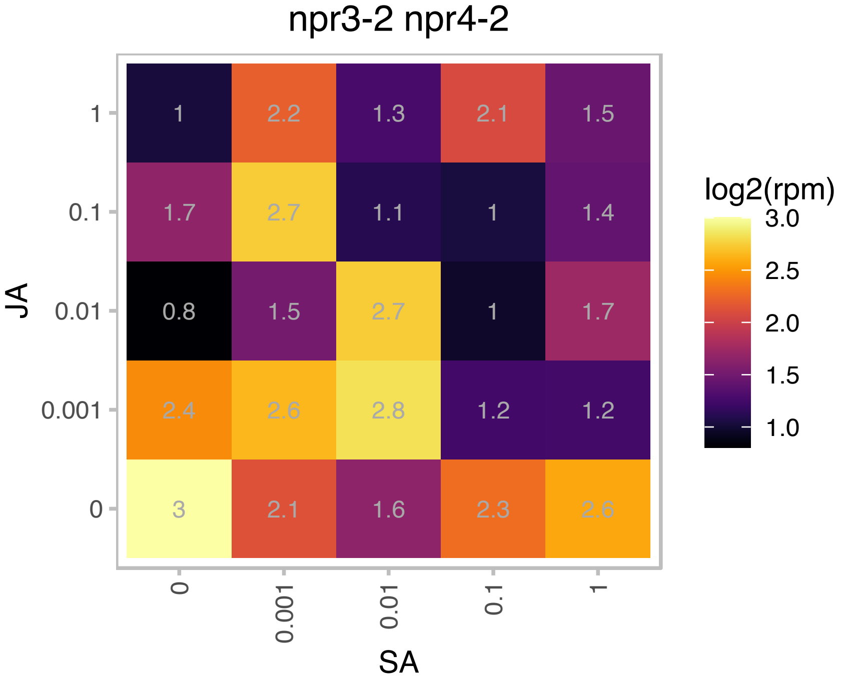


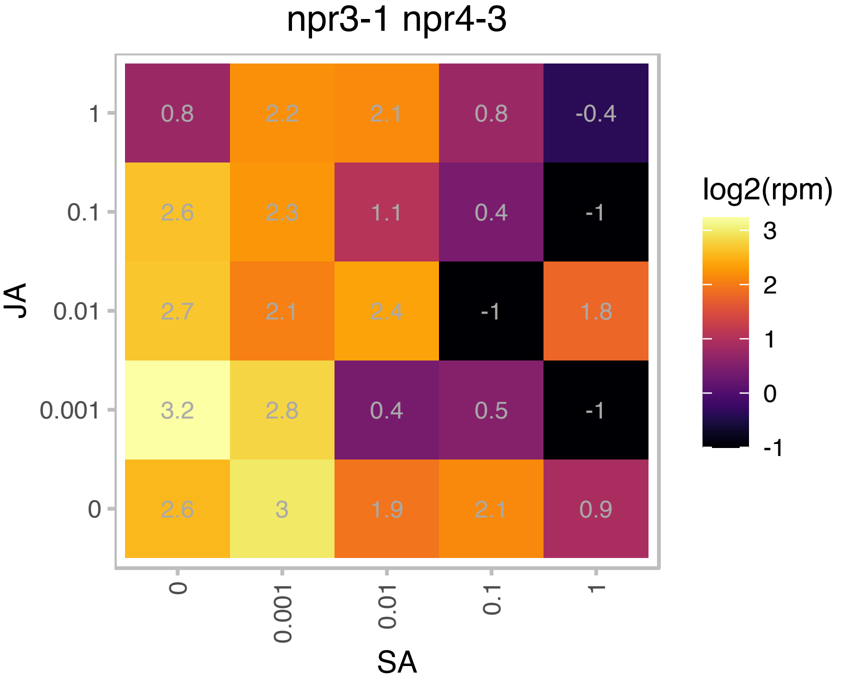

### AT1G16920.1 : ATRABA1B, BEX5, RAB11, RABA1b

RAB GTPase homolog A1B

small GTP-binding protein (Rab11)similar to YPT3/RAB11 proteins in yeast and mammals, respectively. YPT3/RAB11 is involved in intracellular protein trafficking.

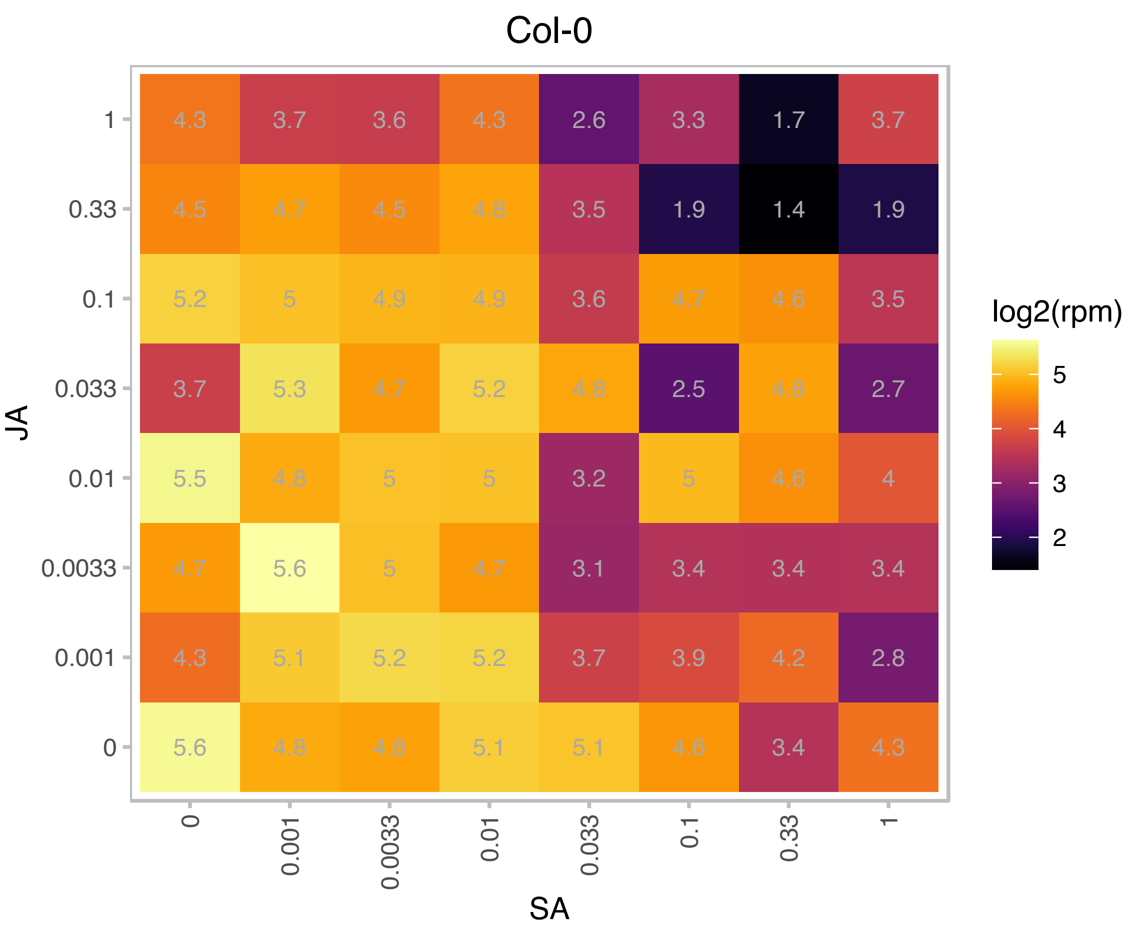


| | |
|---|---|
| Gene | AT1G16920 |
| is.expressed_genes | TRUE |
| is.responsive_genes | TRUE |
| is.apcluster | F-11 |

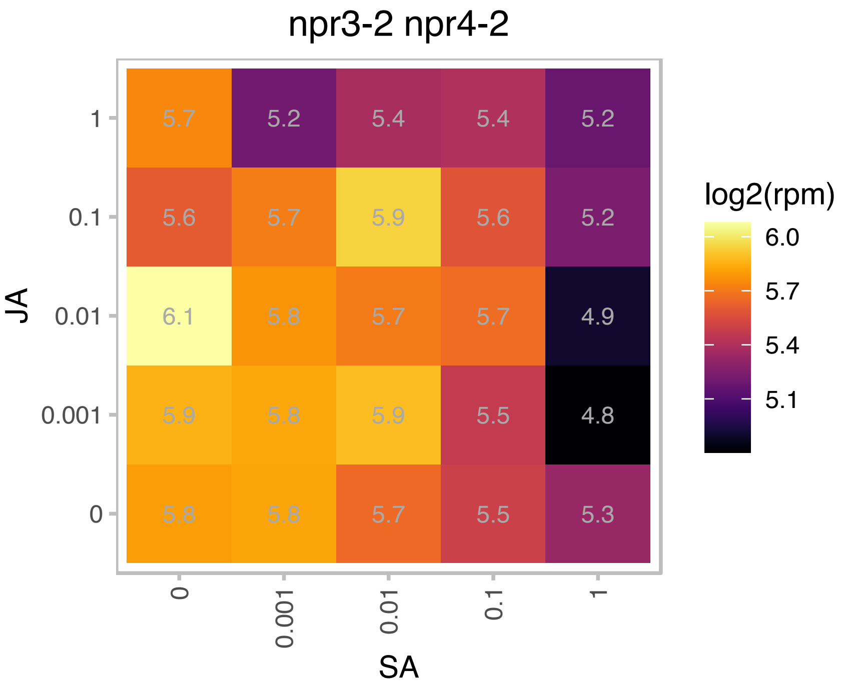


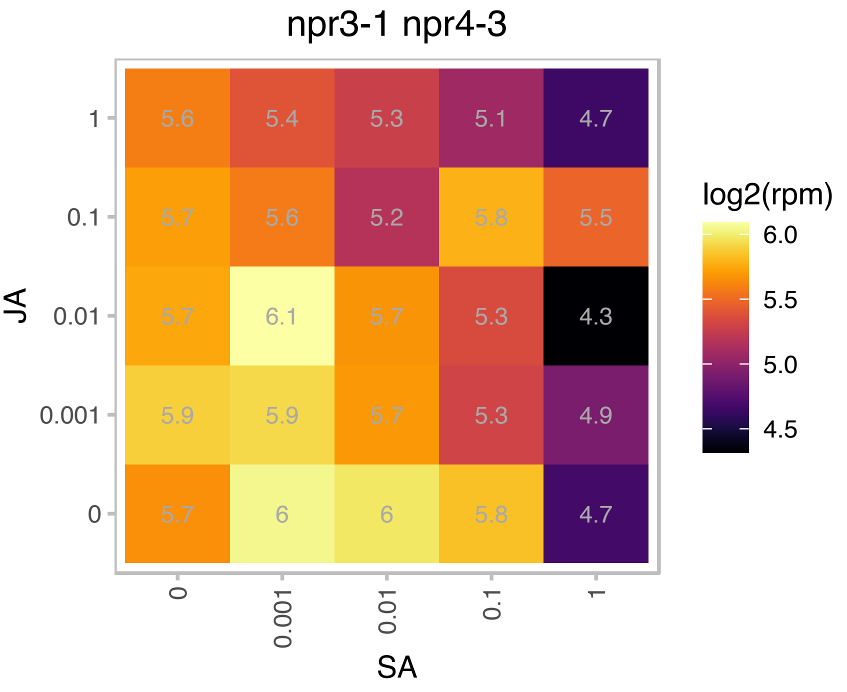

### AT1G17100.1 : AtHBP1, HBP1

SOUL heme-binding family protein

| | |
|---|---|
| Gene | AT1G17100 |
| is.expressed_genes | TRUE |
| is.responsive_genes | TRUE |
| is.apcluster | F-11 |

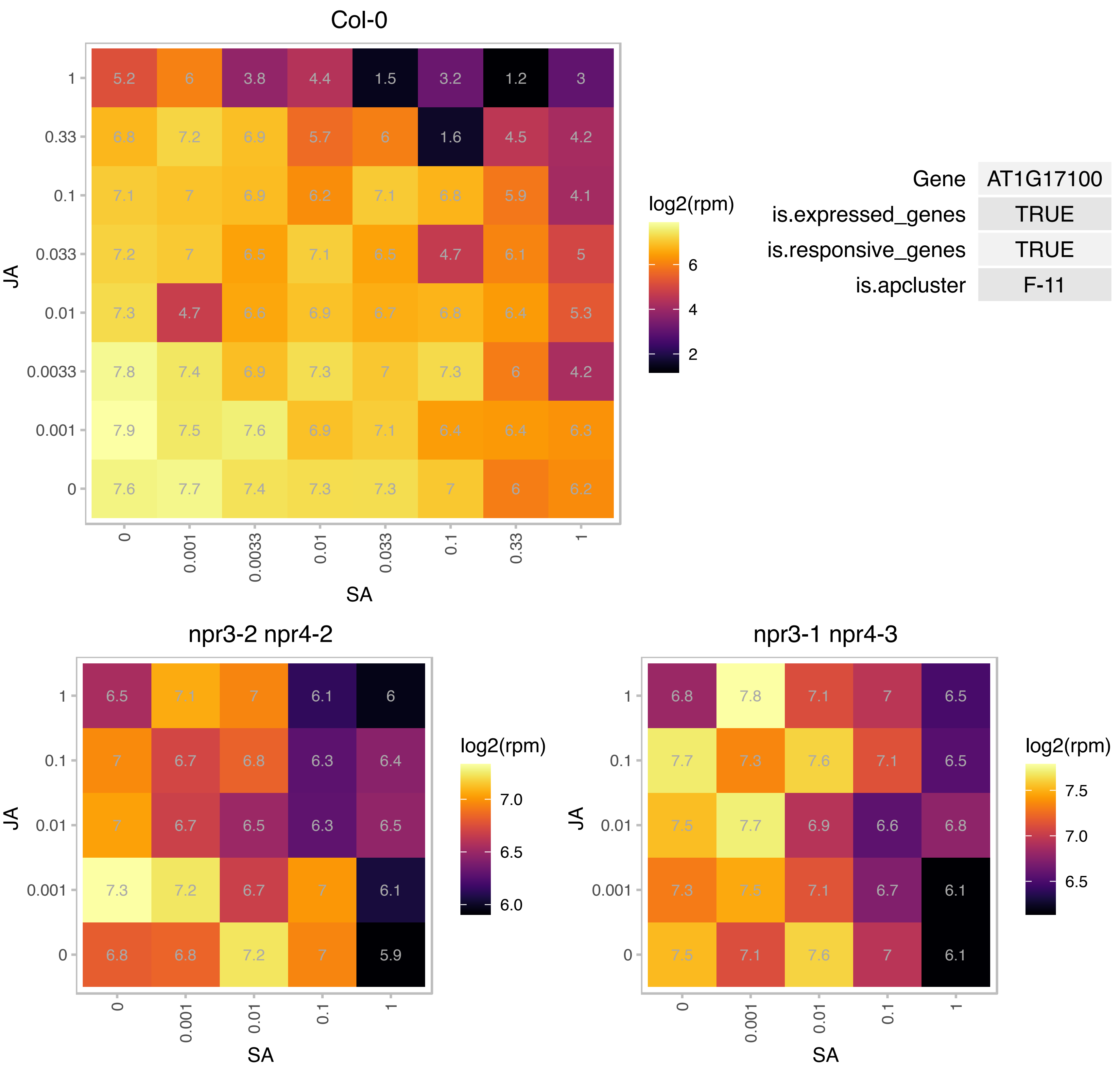

### AT1G17880.1 : ATBTF3, BTF3

basic transcription factor 3

Col-0

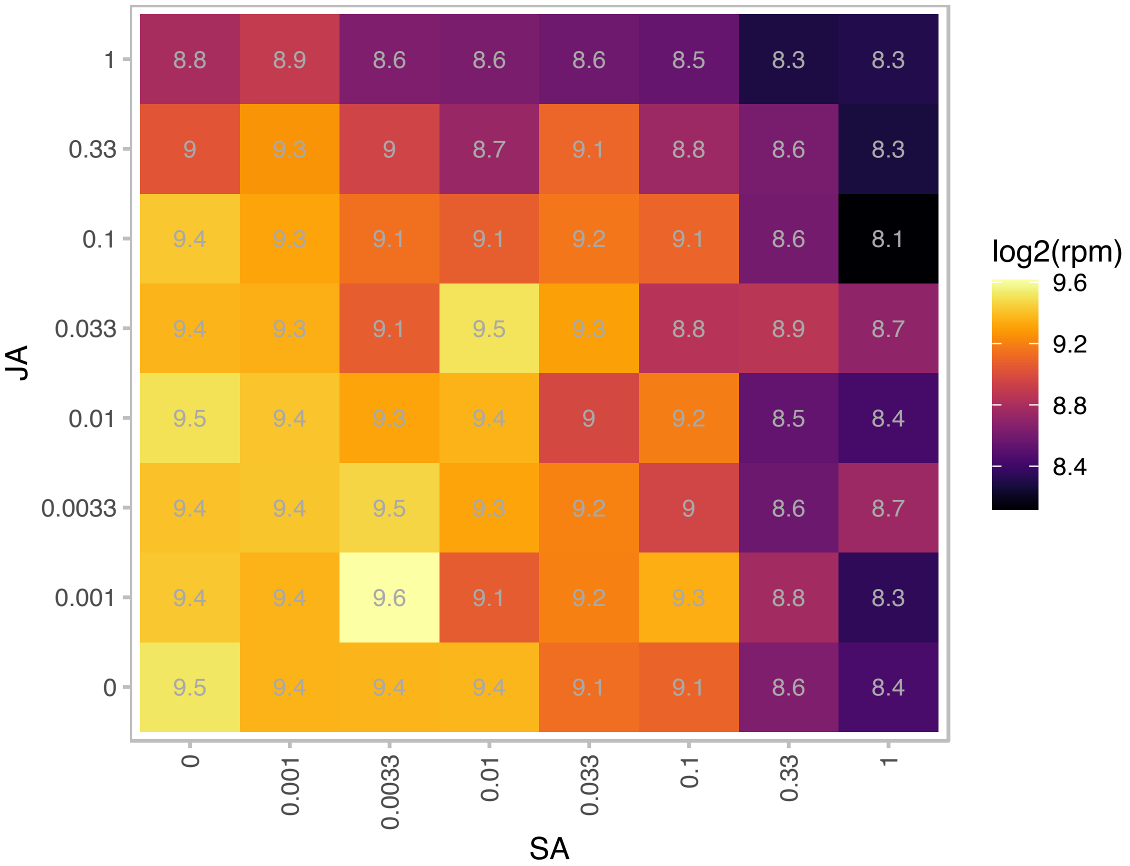


| | |
|---|---|
| Gene | AT1G17880 |
| is.expressed_genes | TRUE |
| is.responsive_genes | TRUE |
| is.apcluster | F-11 |

npr3-2 npr4-2

npr3-1 npr4-3

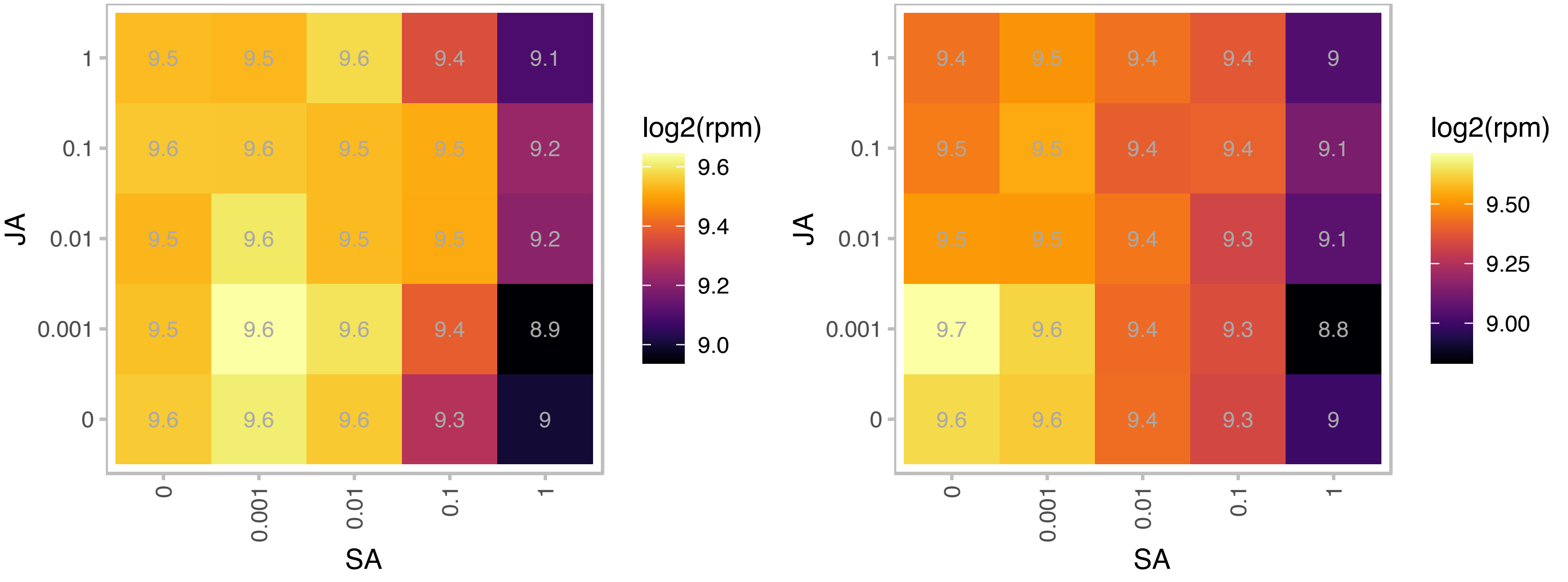

### AT1G21460.1 : AtSWEET1, SWEET1

Nodulin MtN3 family protein

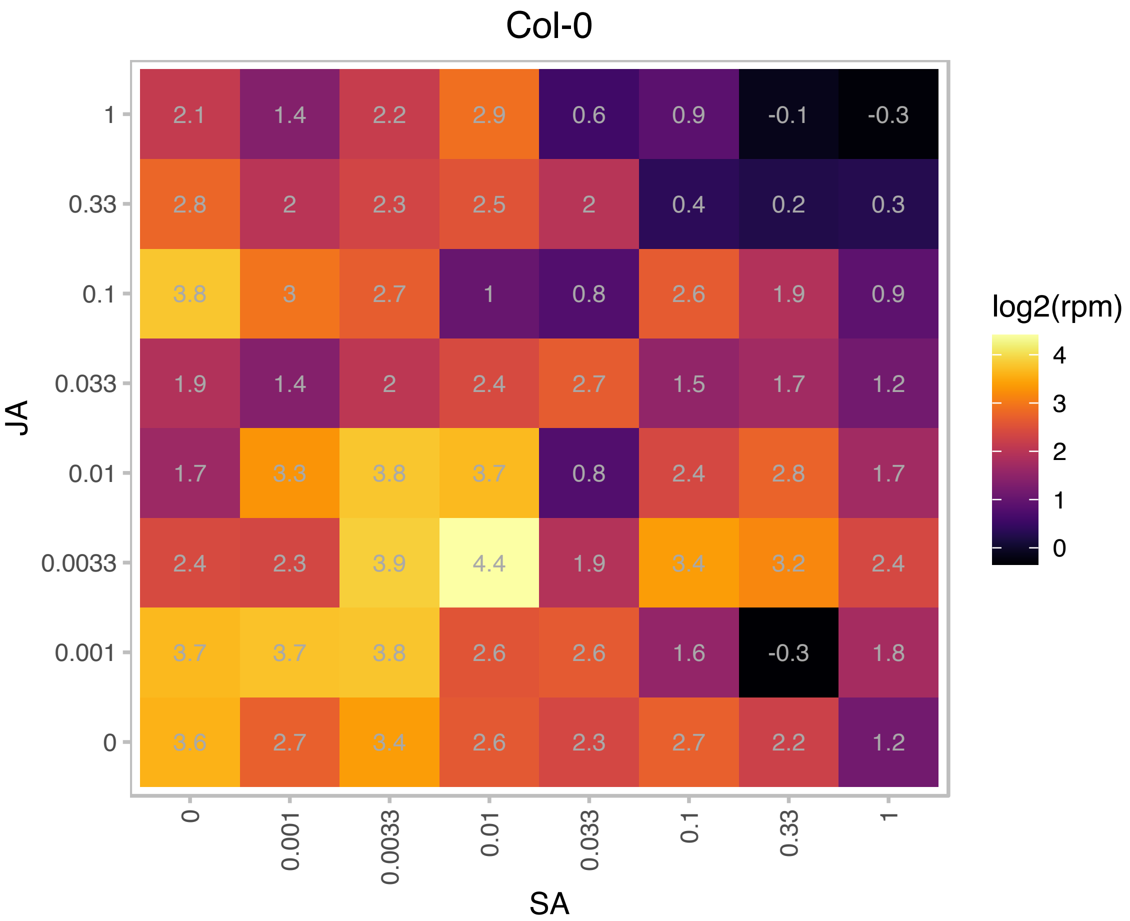


| | |
|---|---|
| Gene | AT1G21460 |
| is.expressed_genes | TRUE |
| is.responsive_genes | TRUE |
| is.apcluster | F-11 |

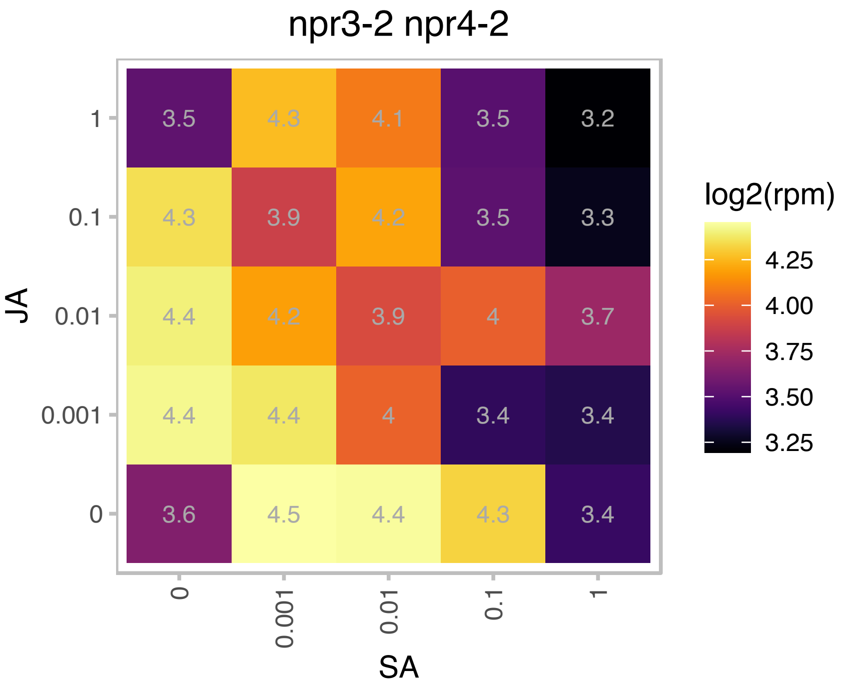


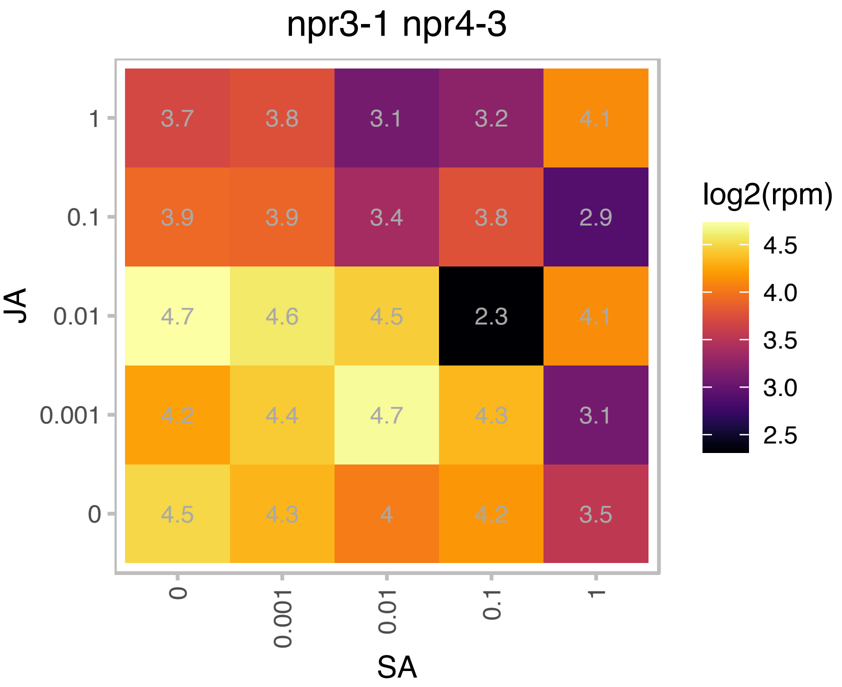

### AT1G21600.1 : PTAC6

plastid transcriptionally active 6

Present in transcriptionally active plastid chromosomes. Involved in plastid gene expression.

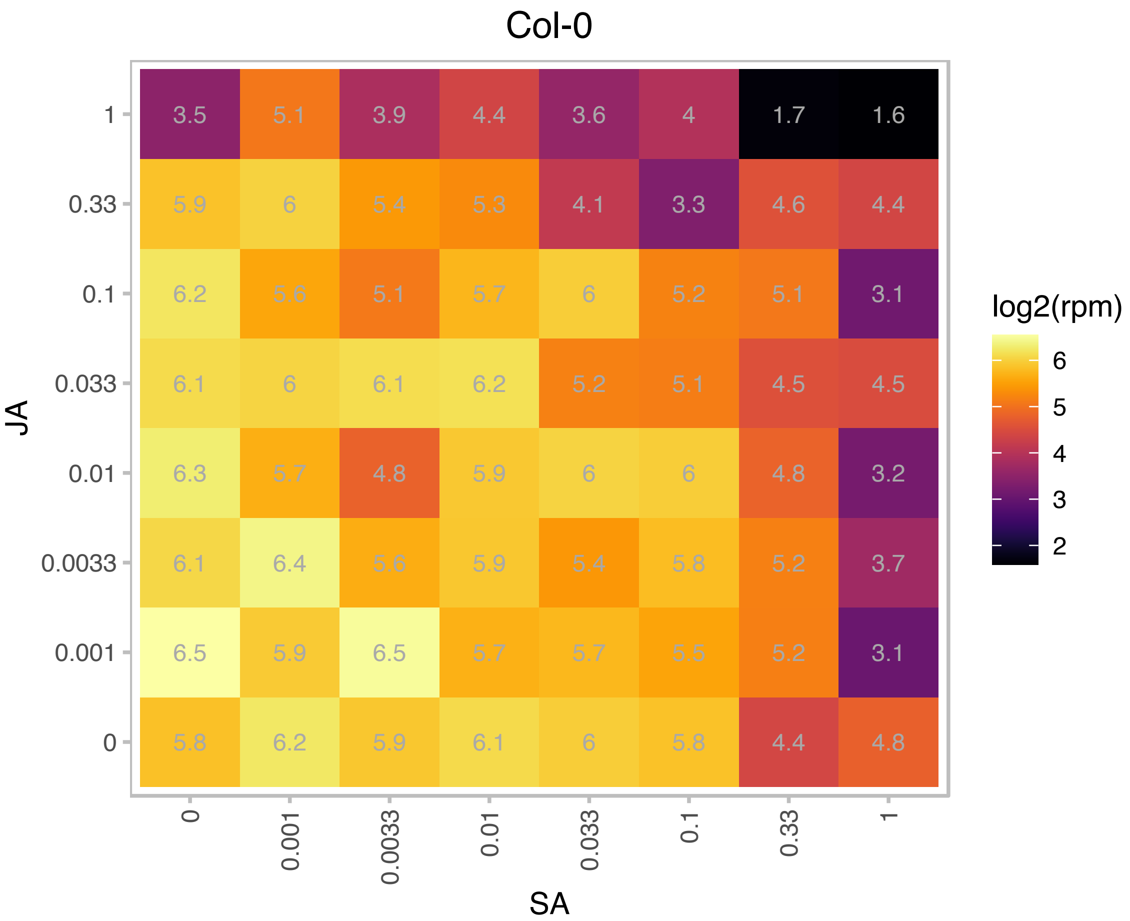


| | |
|---|---|
| Gene | AT1G21600 |
| is.expressed_genes | TRUE |
| is.responsive_genes | TRUE |
| is.apcluster | F-11 |

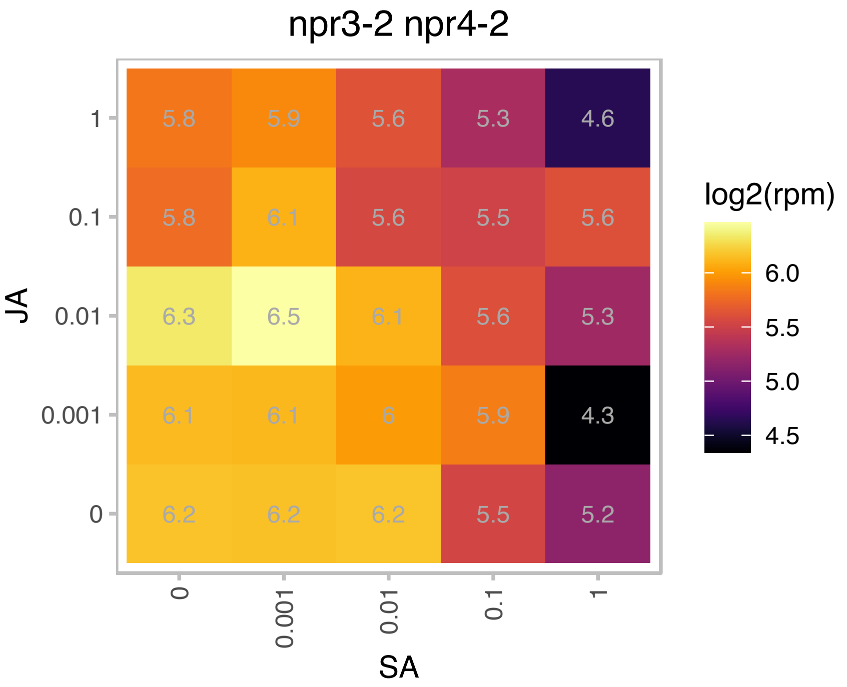


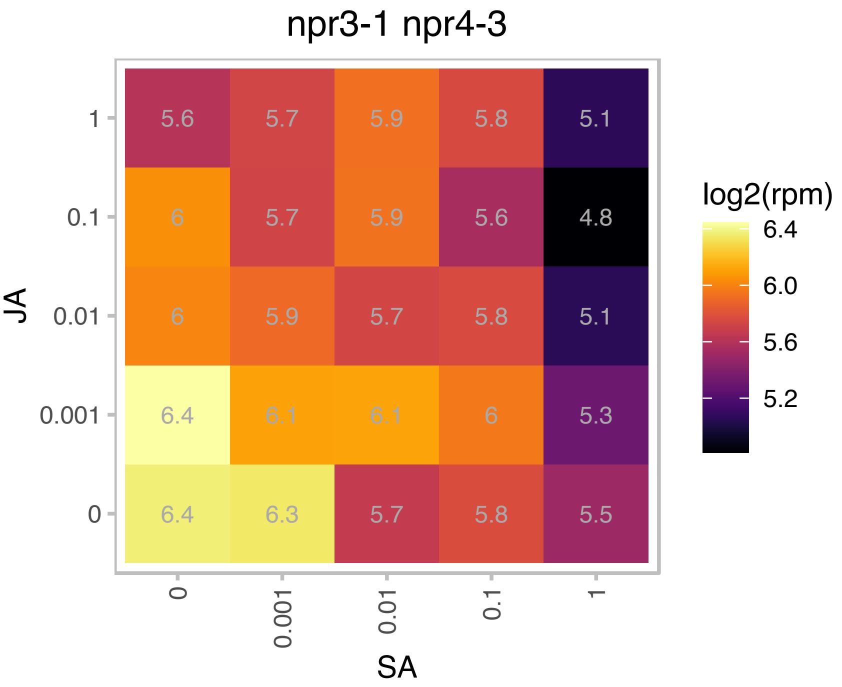

### AT1G22270.1 : AtTRM112b, TRM112b

Trm112p-like protein

Encodes SMO2 (Small Organ 2). Modulates progression of cell division during organ growth.

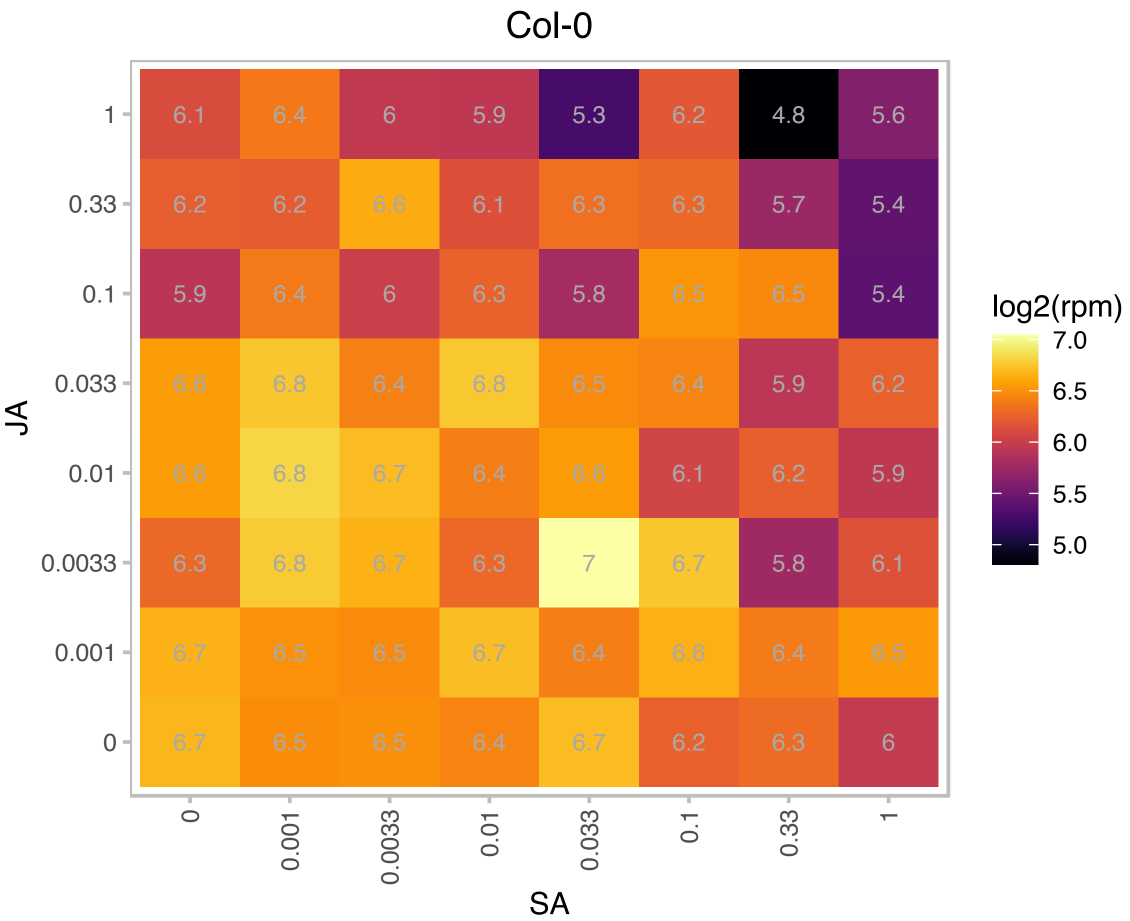


| | |
|---|---|
| Gene | AT1G22270 |
| is.expressed_genes | TRUE |
| is.responsive_genes | TRUE |
| is.apcluster | F-11 |

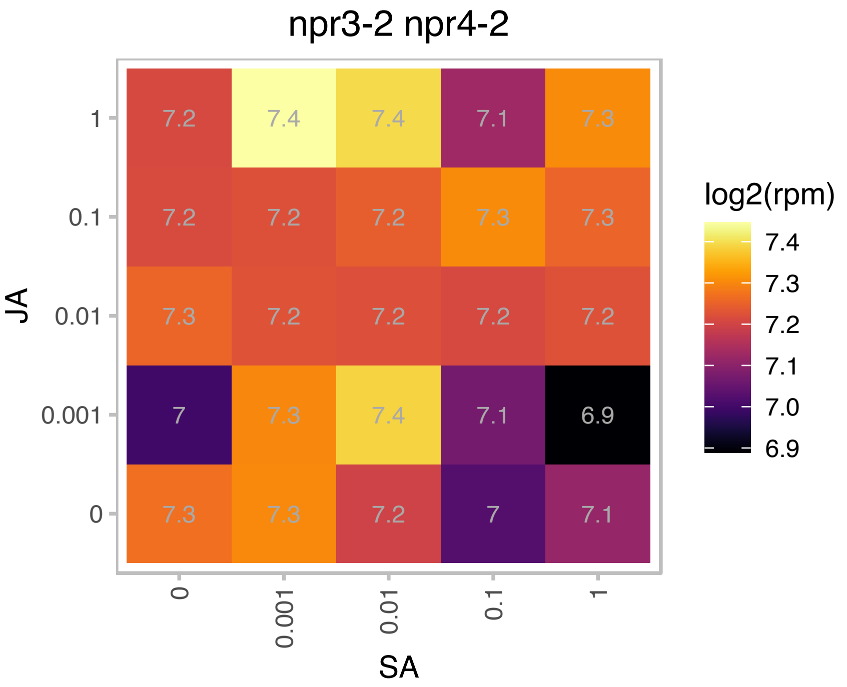


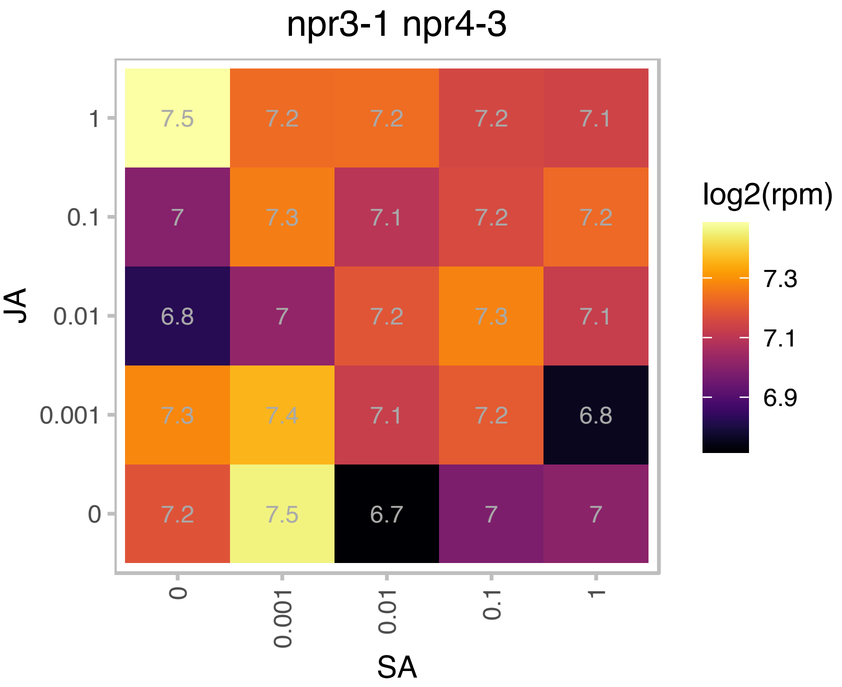

### AT1G22590.1 : AGL87

AGAMOUS-like 87

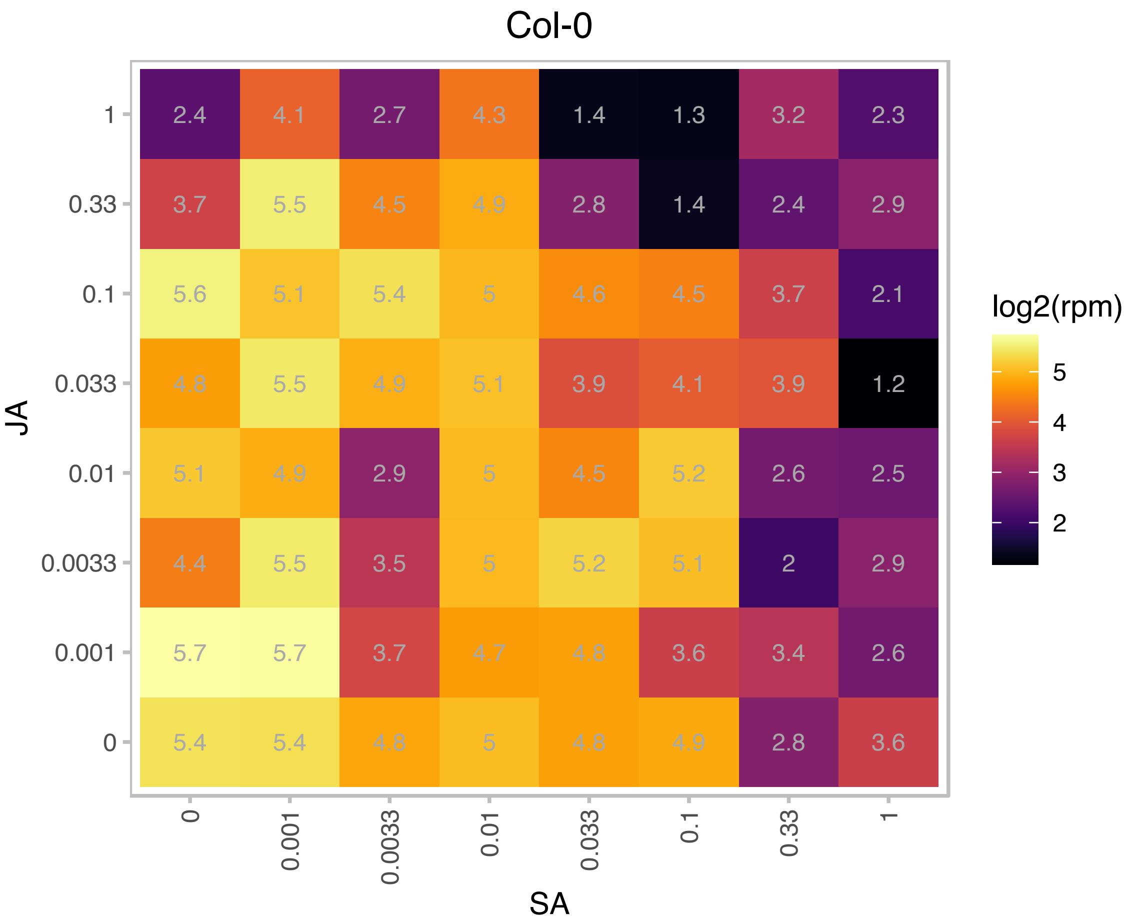


| | |
|---|---|
| Gene | AT1G22590 |
| is.expressed_genes | TRUE |
| is.responsive_genes | TRUE |
| is.apcluster | F-11 |

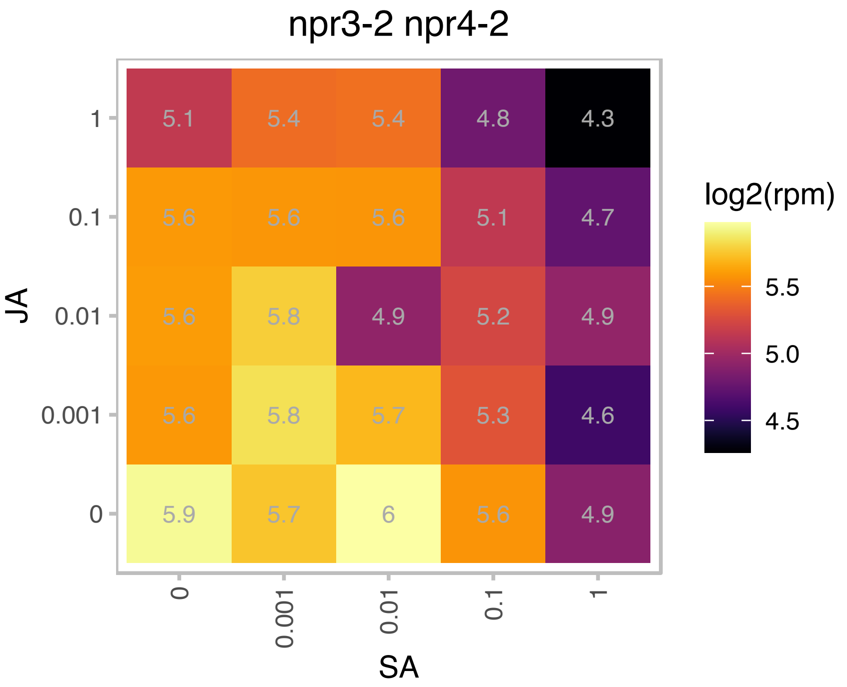


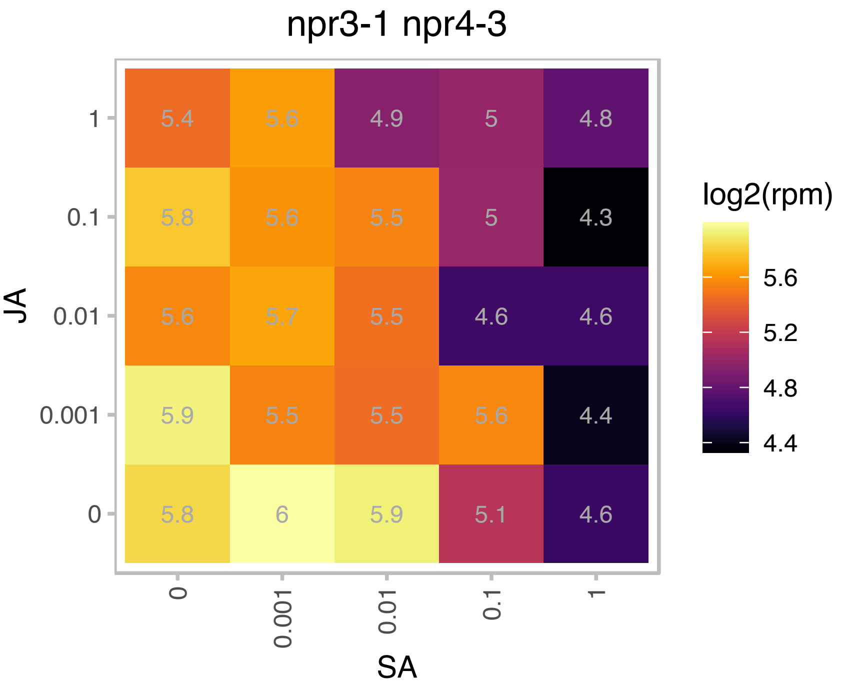

# AT1G26220.1 : NA

Acyl-CoA N-acyltransferases (NAT) superfamily protein

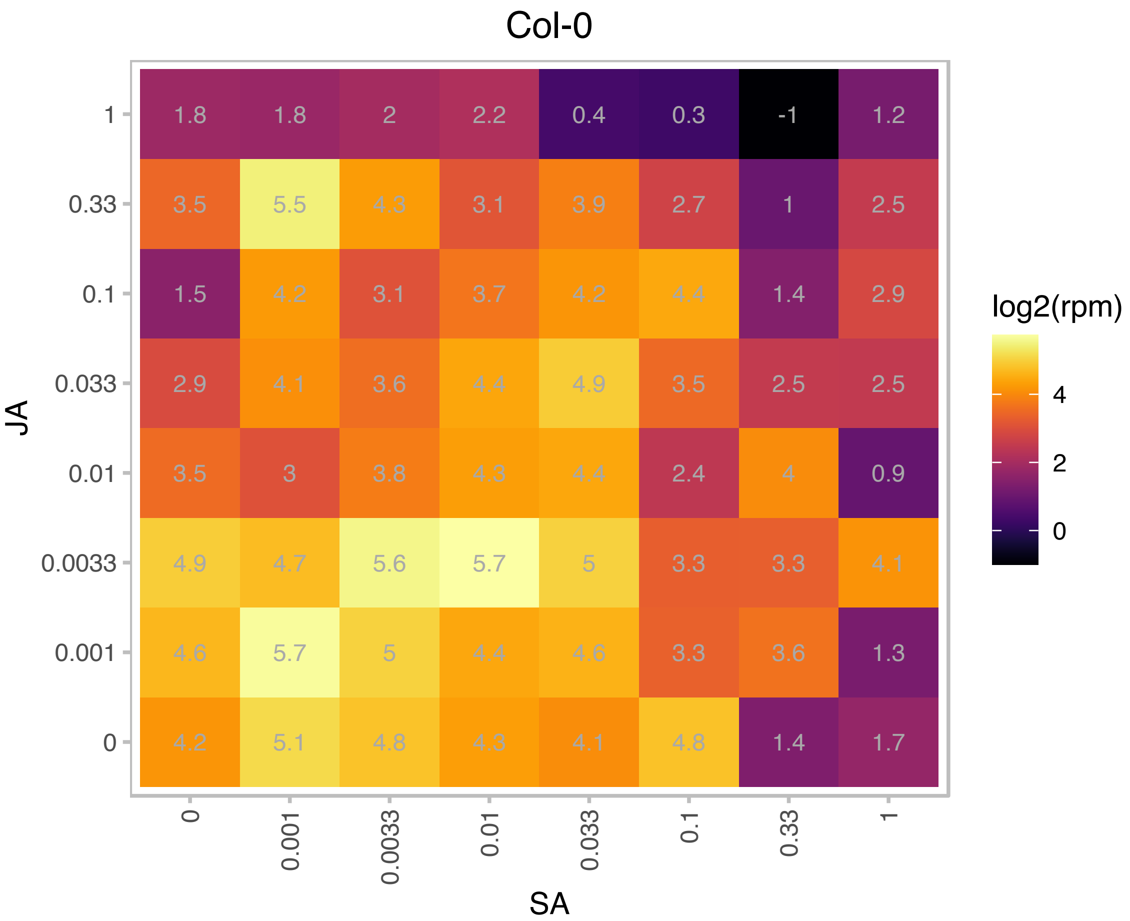


| | |
|---|---|
| Gene | AT1G26220 |
| is.expressed_genes | TRUE |
| is.responsive_genes | TRUE |
| is.apcluster | F-11 |

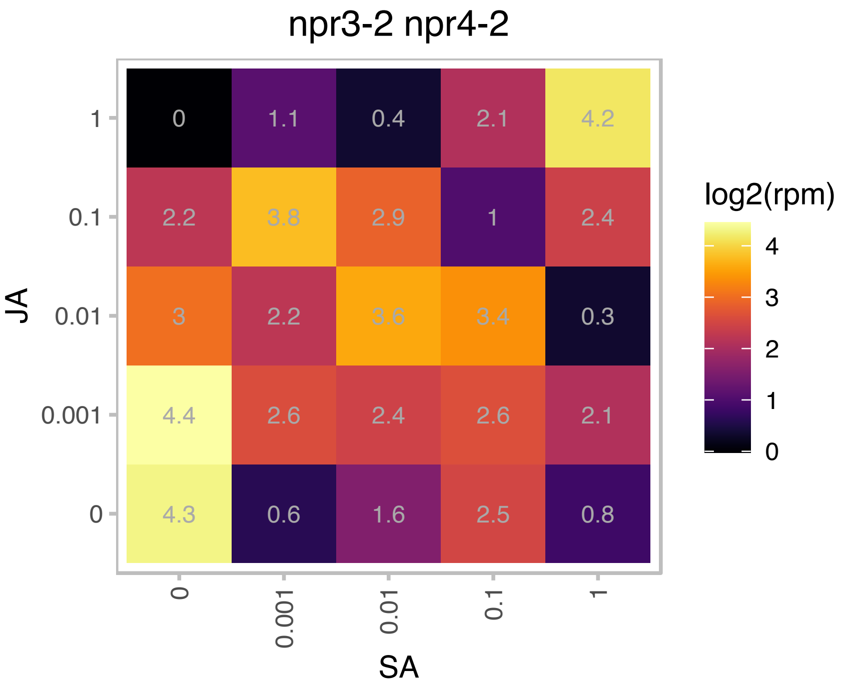


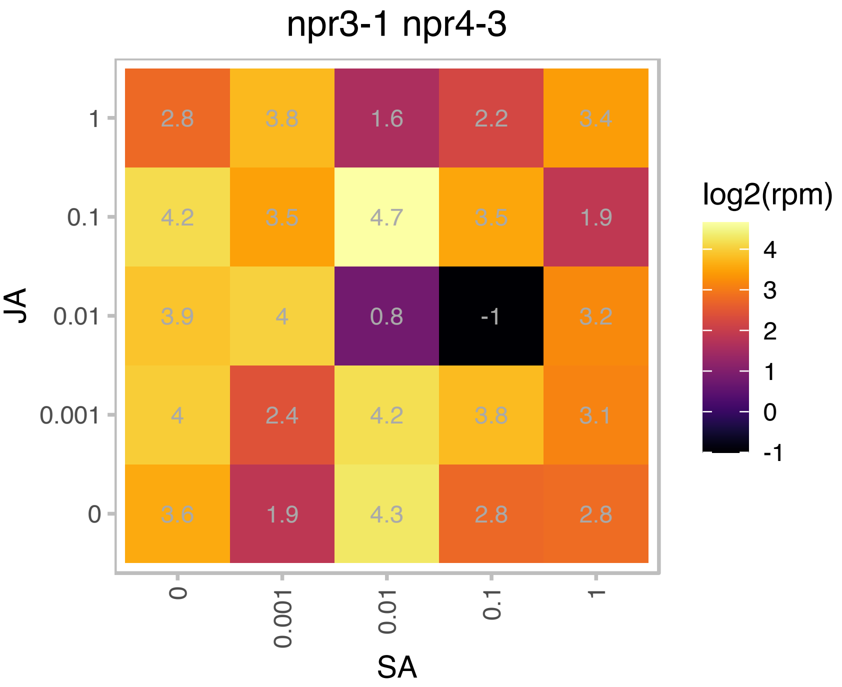

# AT1G26761.1 : NA

Arabinanase/levansucrase/invertase

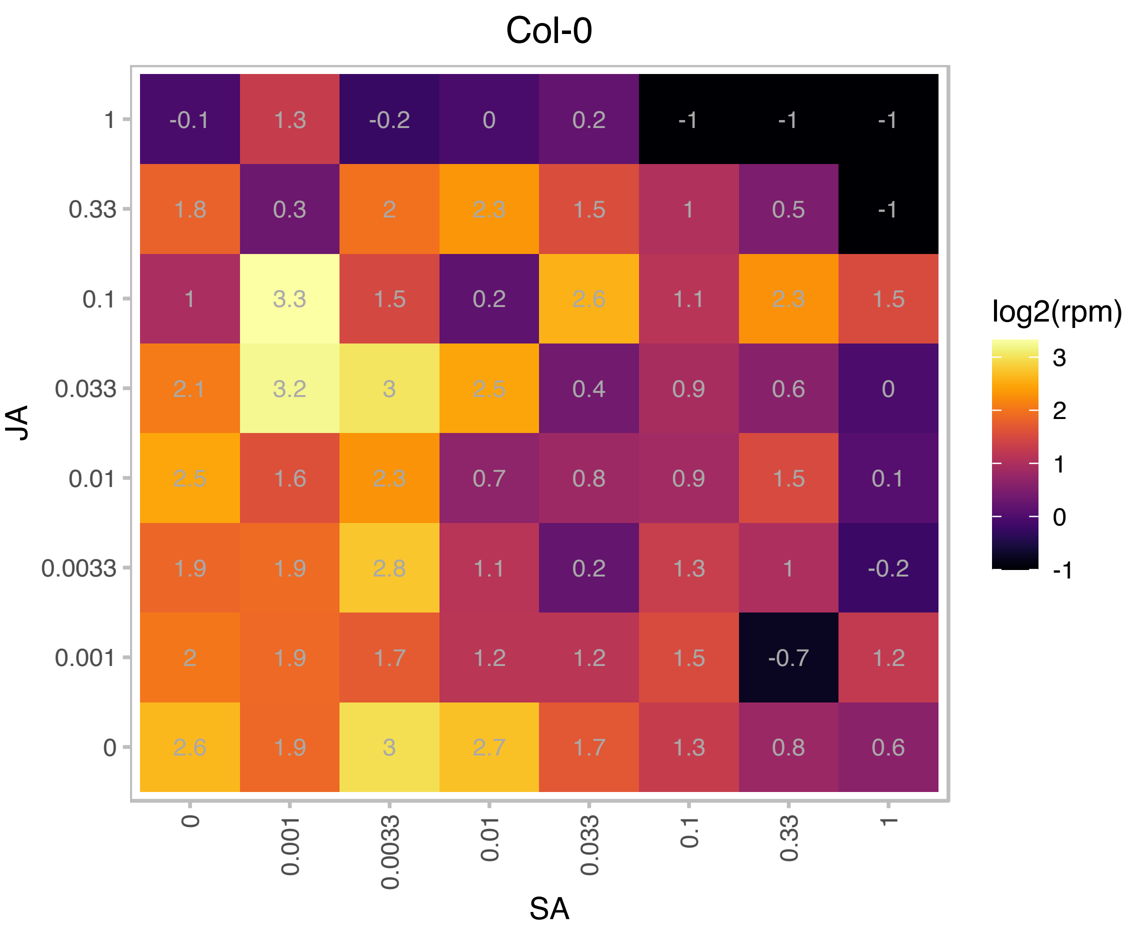


| | |
|---|---|
| Gene | AT1G26761 |
| is.expressed_genes | TRUE |
| is.responsive_genes | TRUE |
| is.apcluster | F-11 |

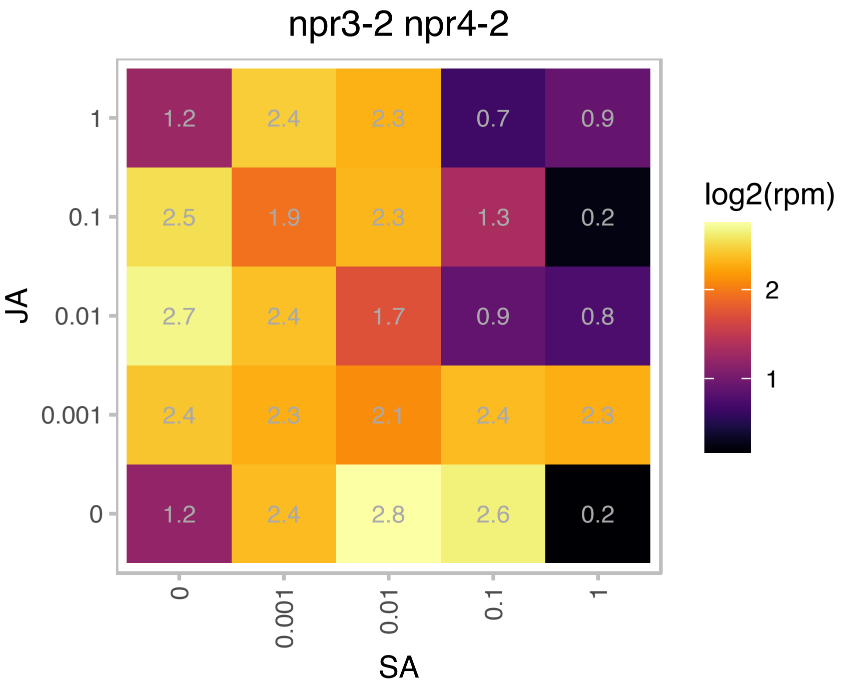


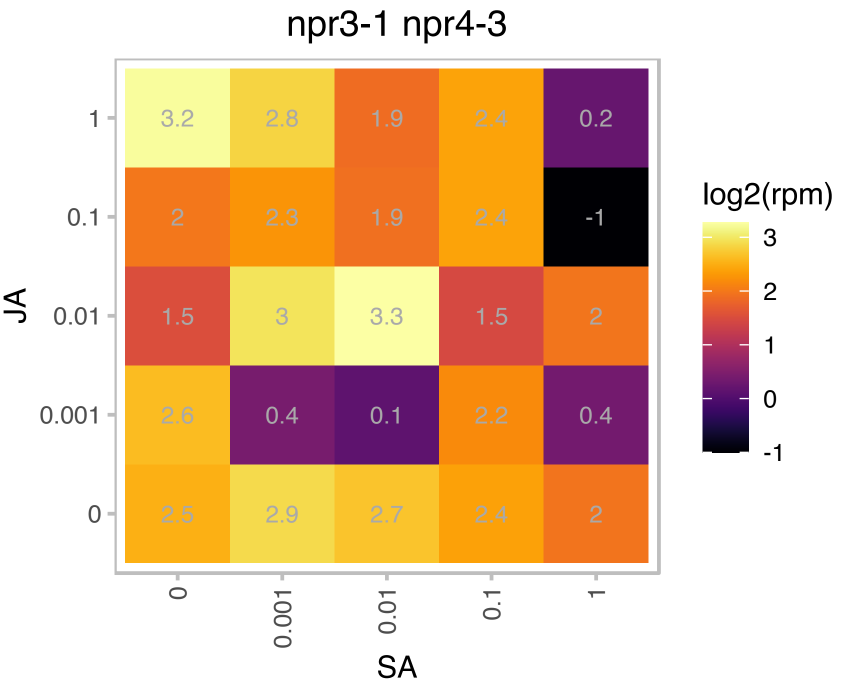

### AT1G28670.1 : ARAB-1

GDSL-like Lipase/Acylhydrolase superfamily protein

Arabidopsis thaliana lipase

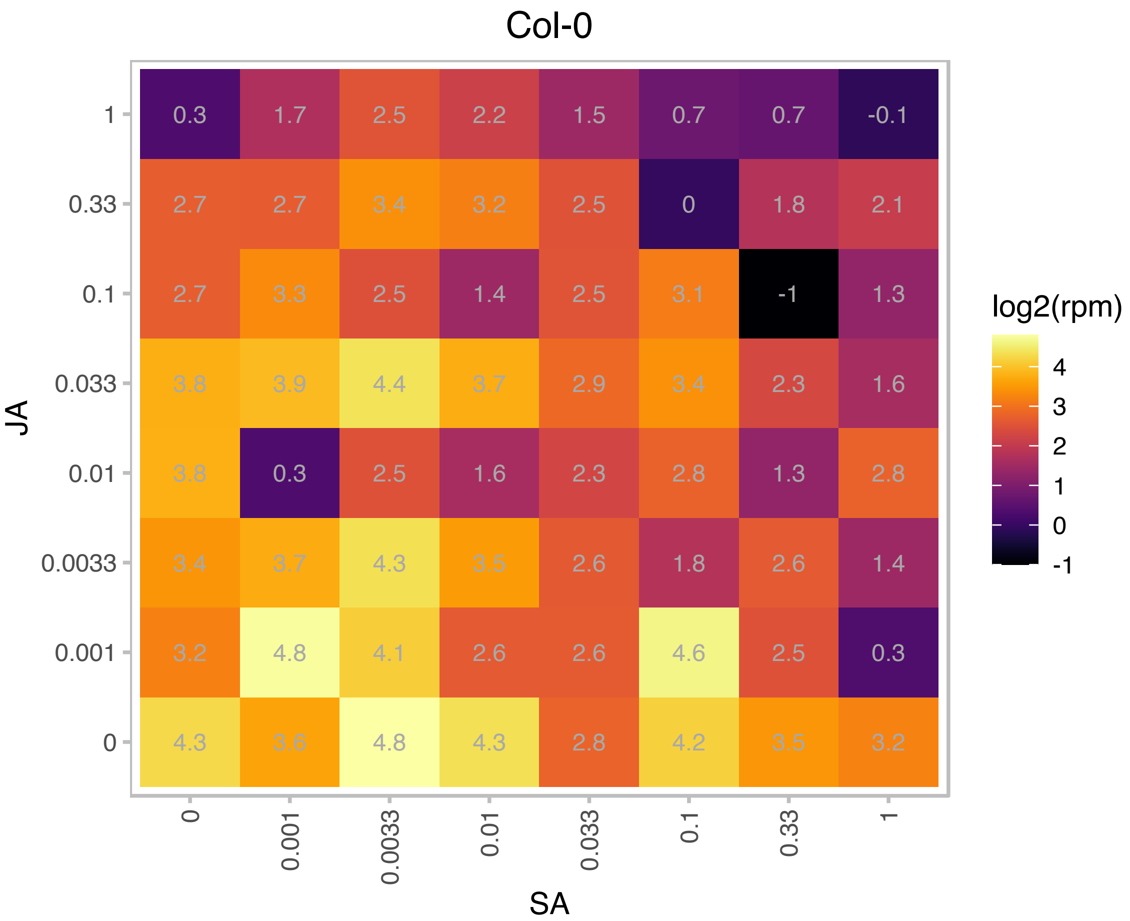


| | |
|---|---|
| Gene | AT1G28670 |
| is.expressed_genes | TRUE |
| is.responsive_genes | TRUE |
| is.apcluster | F-11 |

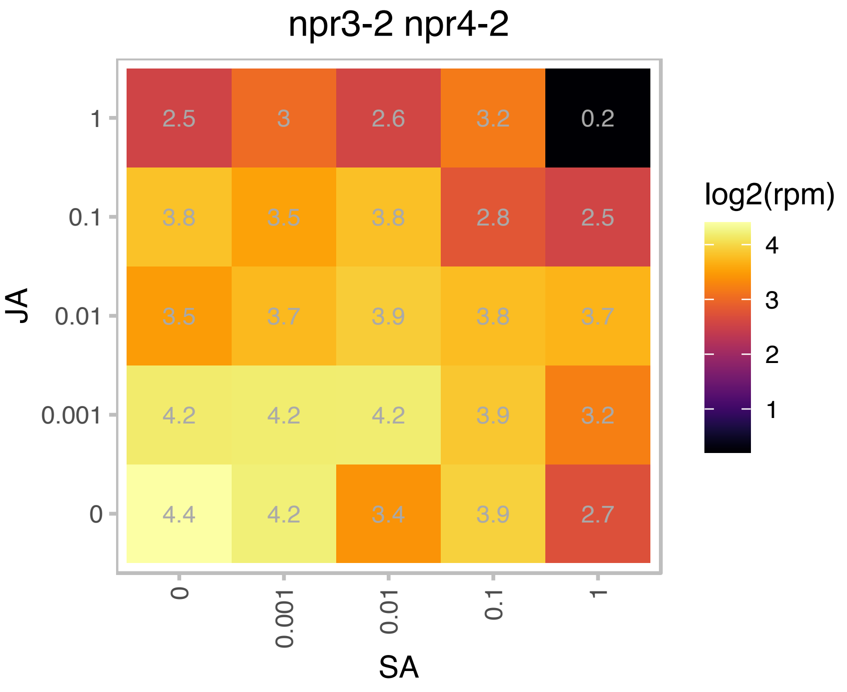


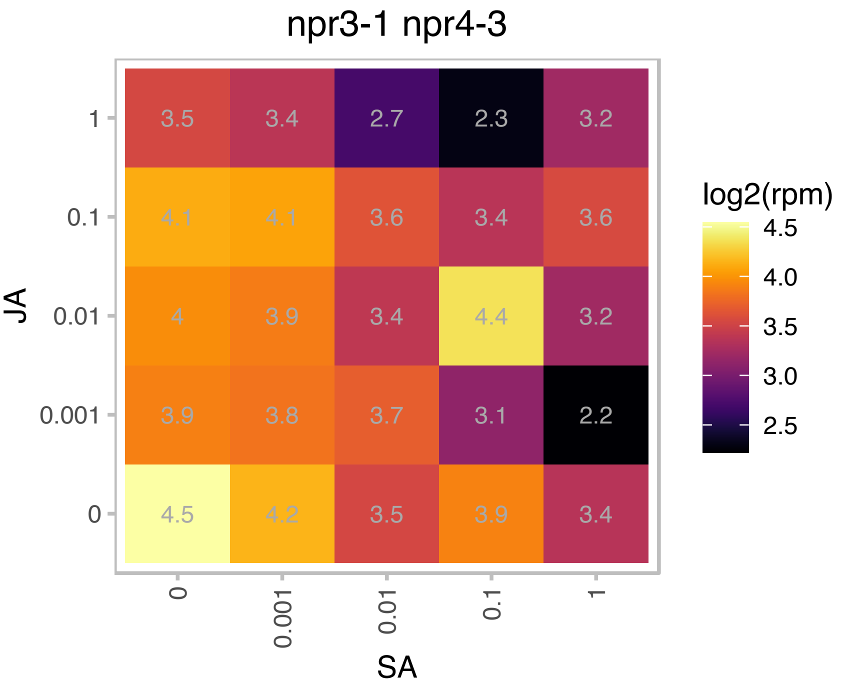

# AT1G44920.1 : NA

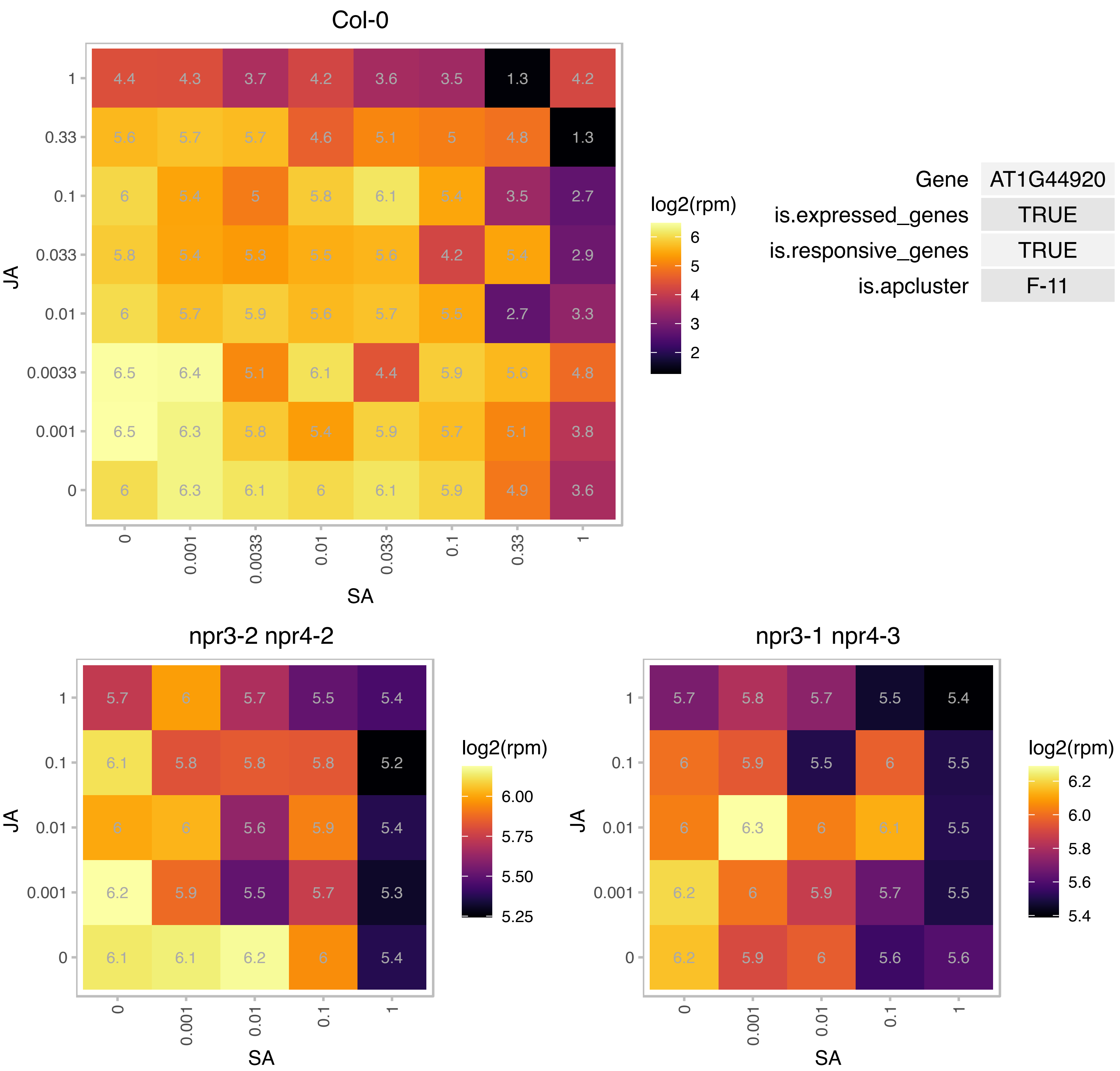


| | |
|---|---|
| Gene | AT1G44920 |
| is.expressed_genes | TRUE |
| is.responsive_genes | TRUE |
| is.apcluster | F-11 |

# AT1G45230.1 : NA

Protein of unknown function (DUF3223)

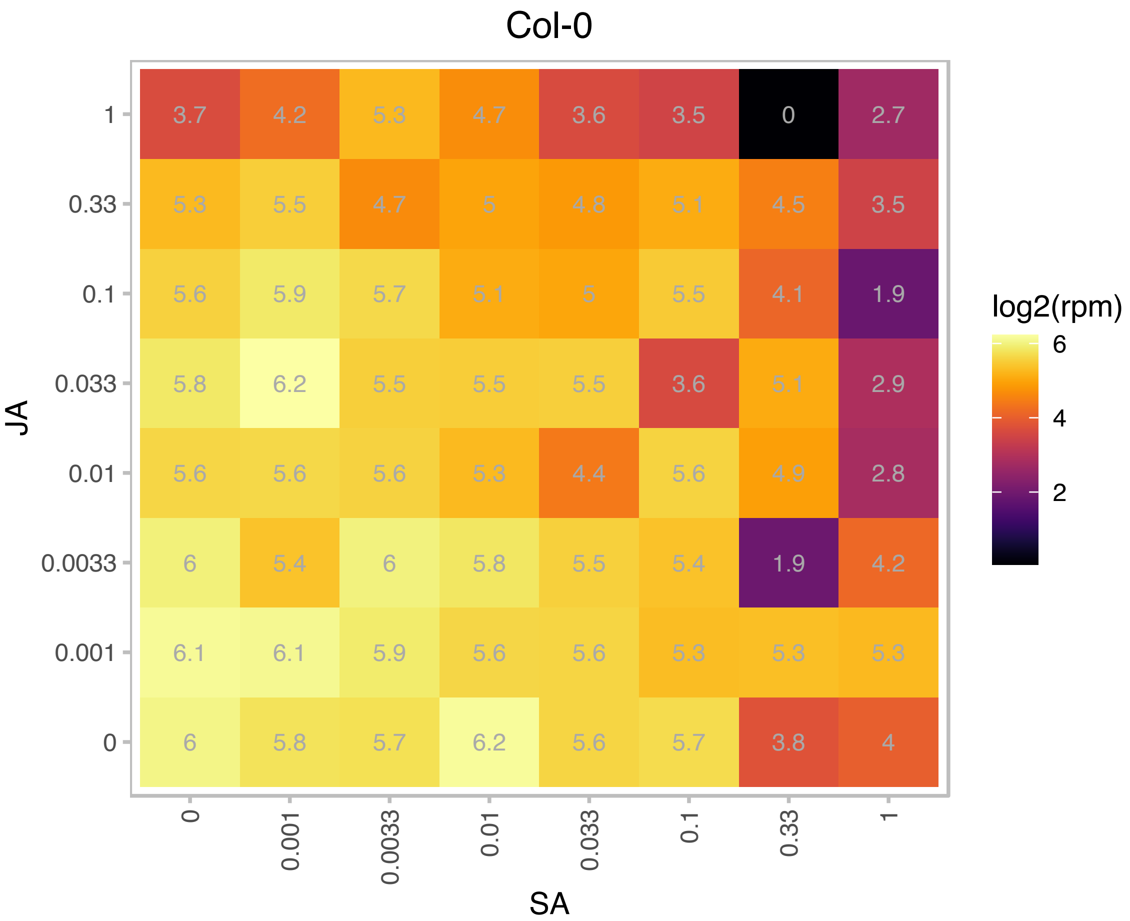


| | |
|---|---|
| Gene | AT1G45230 |
| is.expressed_genes | TRUE |
| is.responsive_genes | TRUE |
| is.apcluster | F-11 |

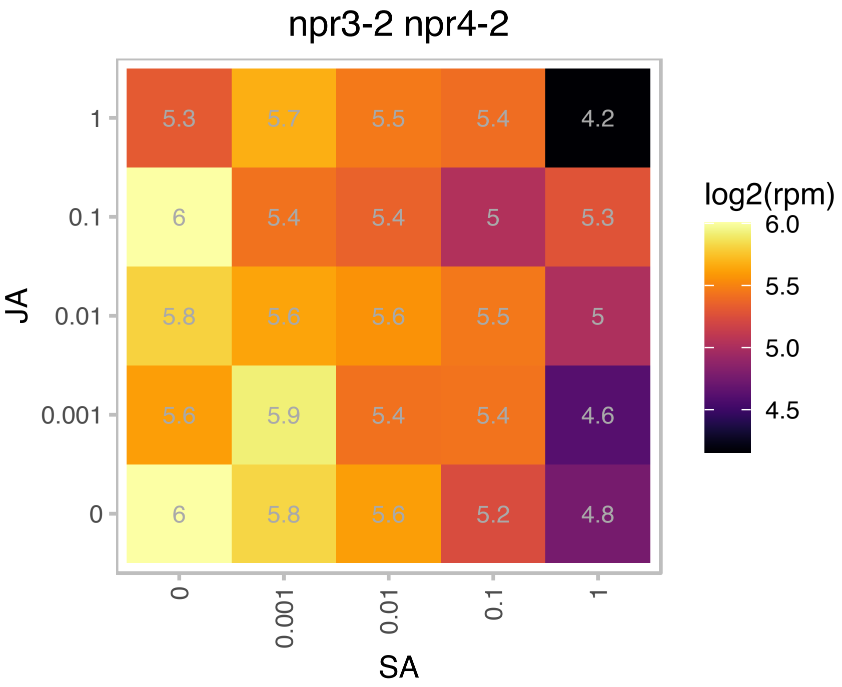


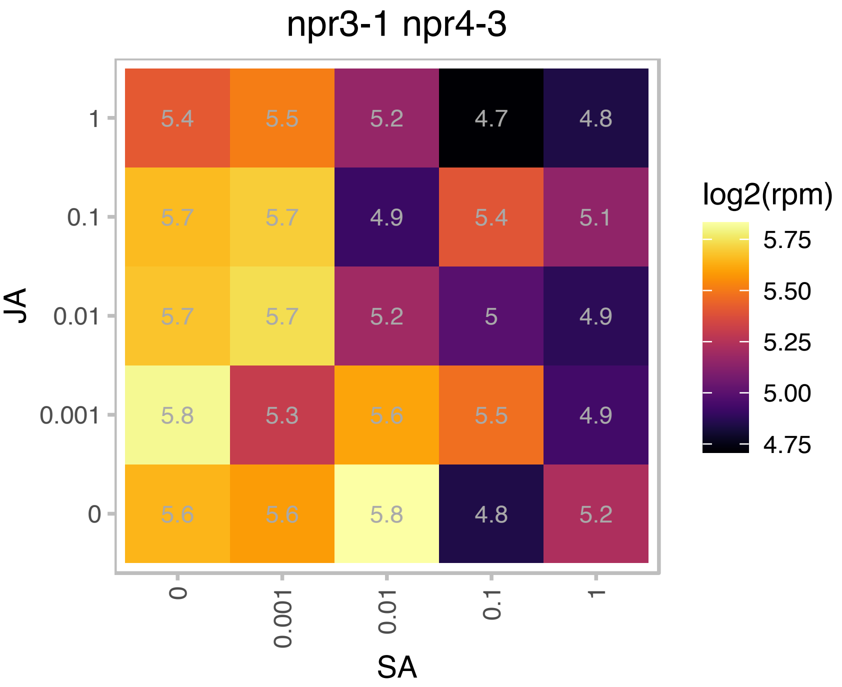

### AT1G47580.1 : DYW1

Pentatricopeptide repeat (PPR) superfamily protein

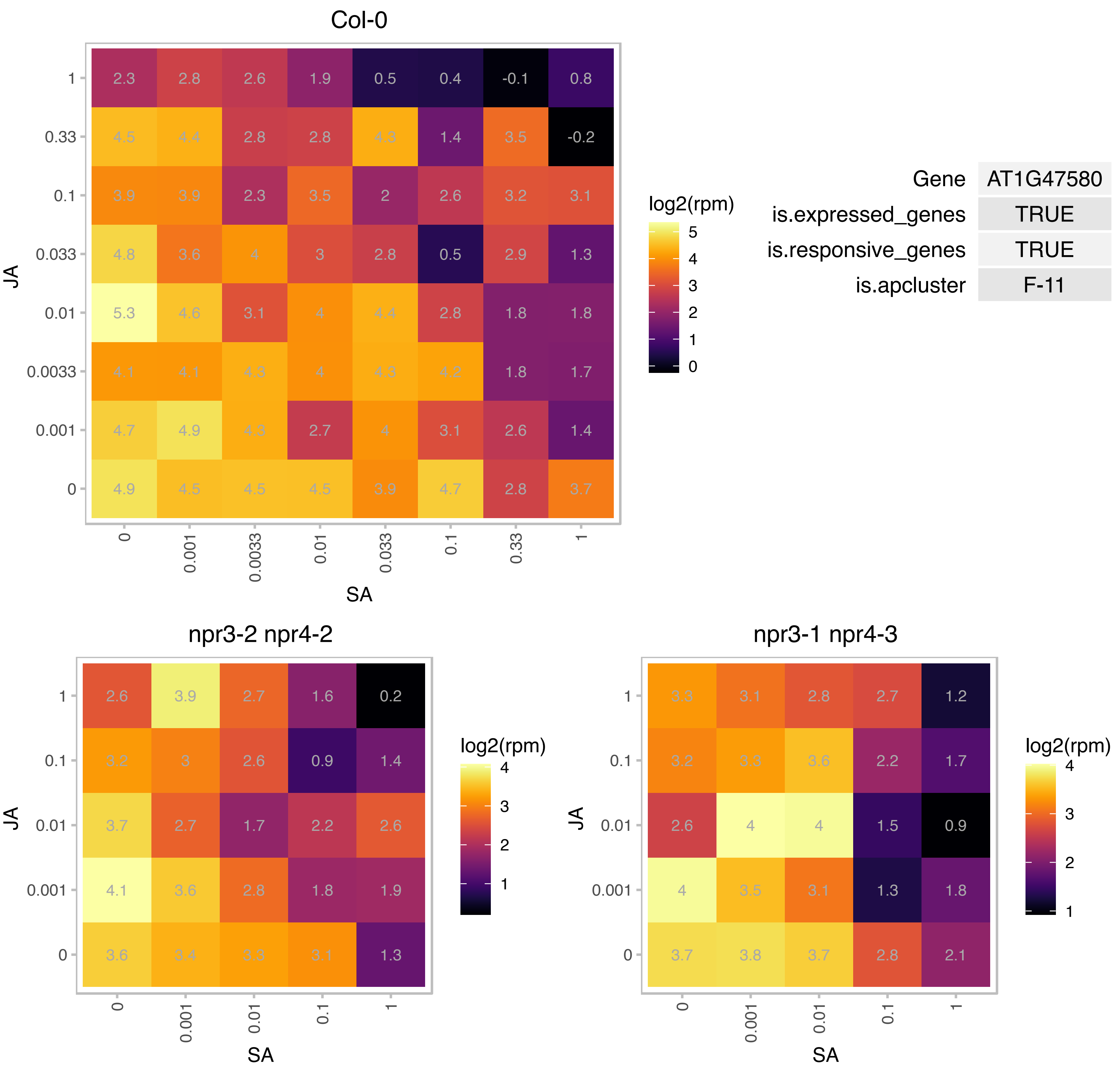


| | |
|---|---|
| Gene | AT1G47580 |
| is.expressed_genes | TRUE |
| is.responsive_genes | TRUE |
| is.apcluster | F-11 |

### AT1G49010.1 : MYBS1

Duplicated homeodomain-like superfamily protein

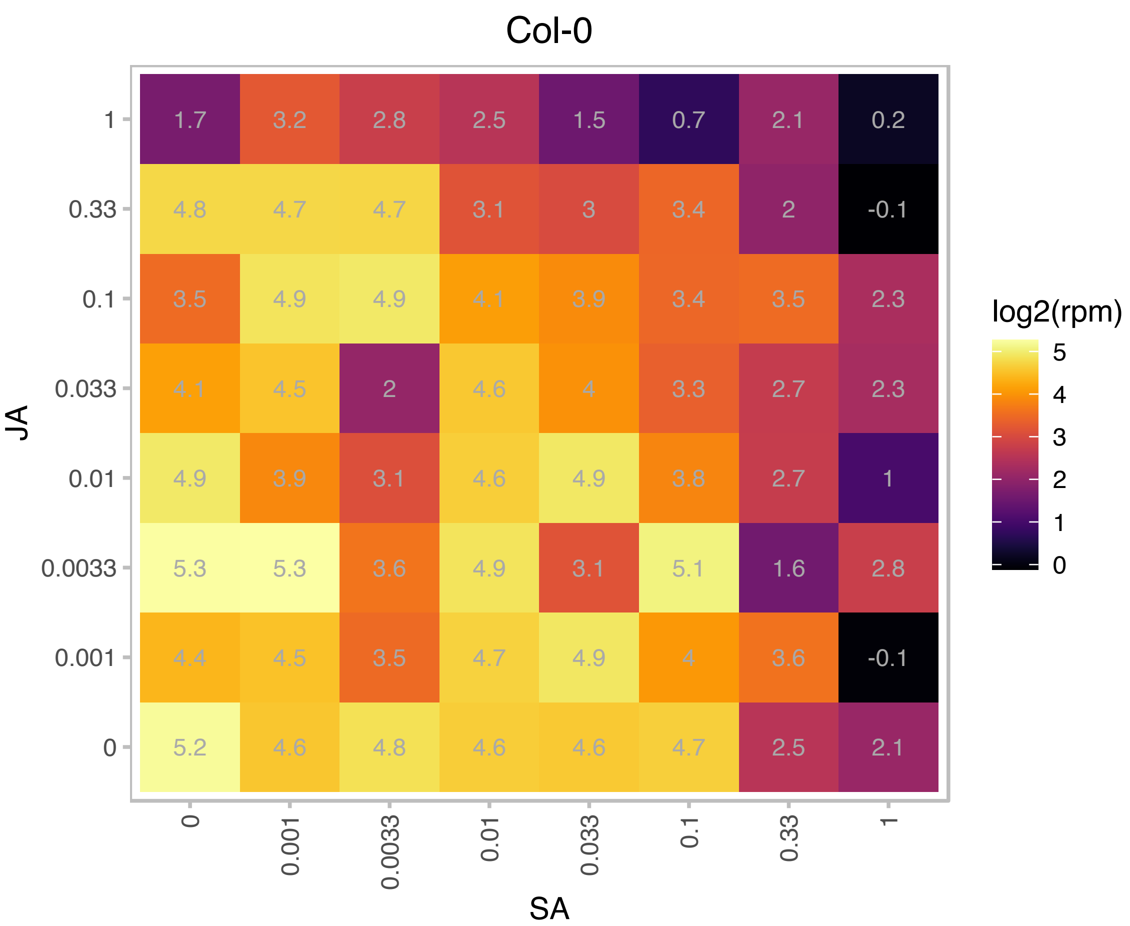


| | |
|---|---|
| Gene | AT1G49010 |
| is.expressed_genes | TRUE |
| is.responsive_genes | TRUE |
| is.apcluster | F-11 |

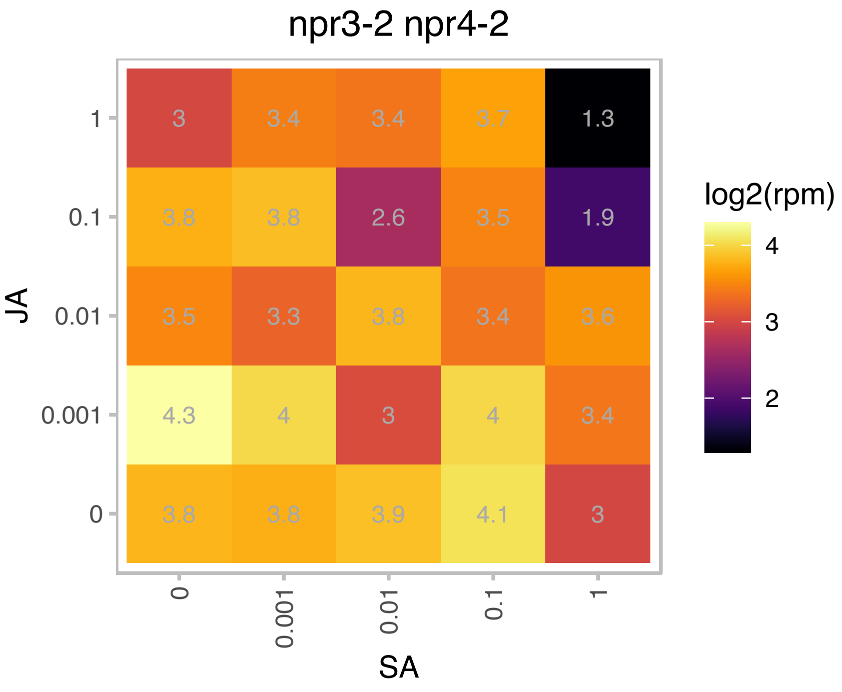


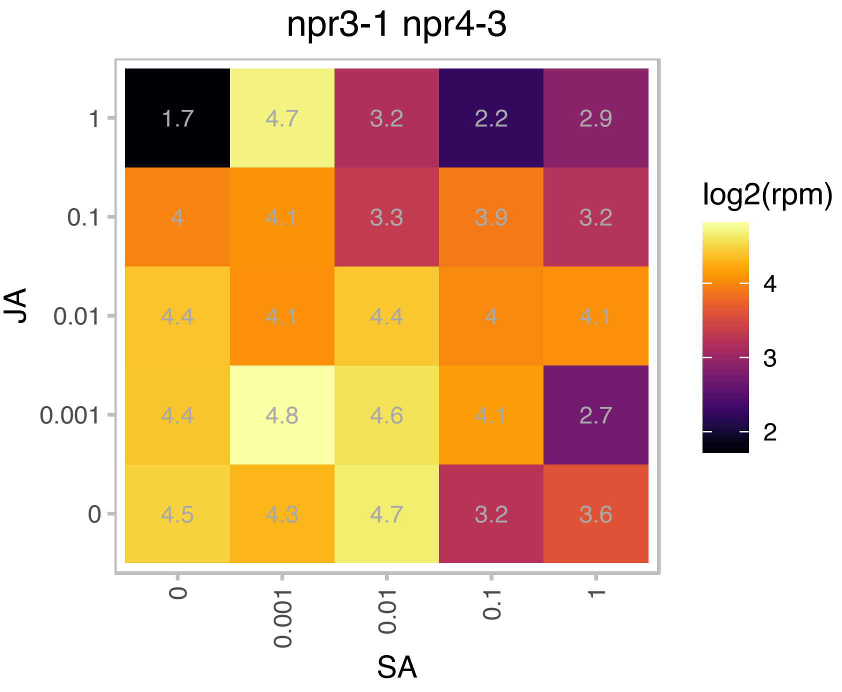

### AT1G50490.1 : UBC20

ubiquitin-conjugating enzyme 20

Encodes one of two ubiquitin-conjugating enzymes belonging to the E2-C gene family (the other being UBC19). Transcript is always found in diving cells, but also in other non-dividing cells.

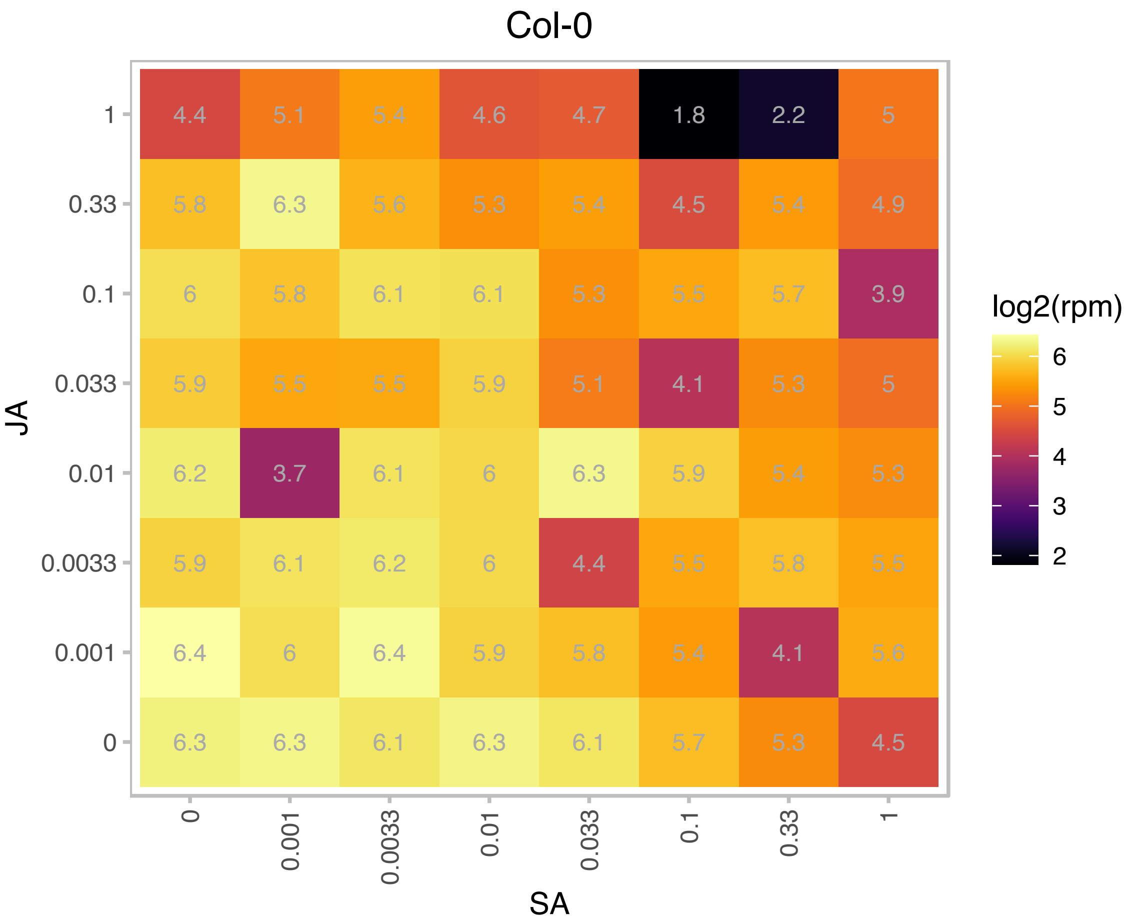


| | |
|---|---|
| Gene | AT1G50490 |
| is.expressed_genes | TRUE |
| is.responsive_genes | TRUE |
| is.apcluster | F-11 |

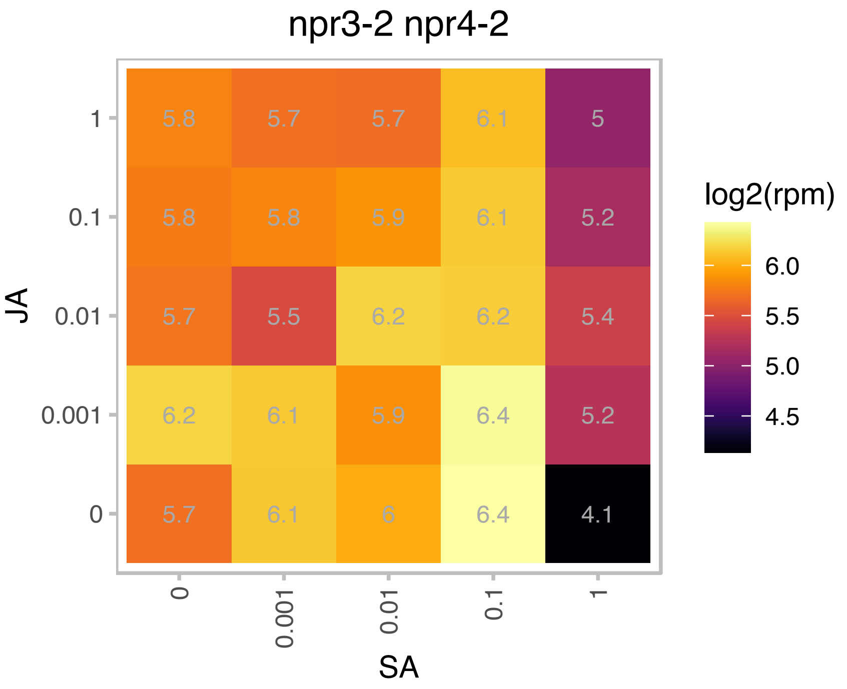


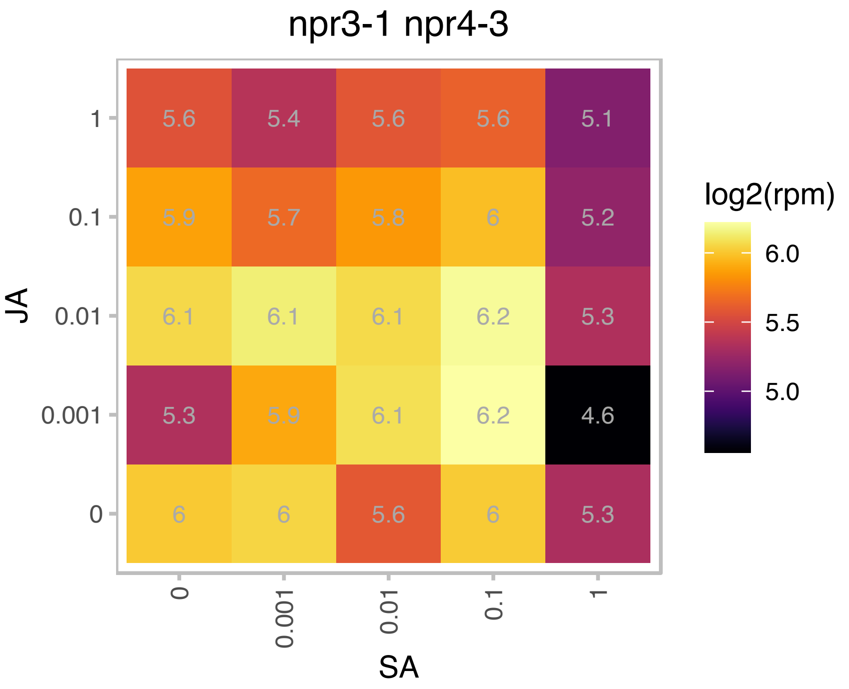

# AT1G54500.1 : NA

Rubredoxin-like superfamily protein
RBD1 is a thylakoid membrane-bound iron-binding protein that is required for the proper assembly of photosystem II in Arabidopsis. It is found in all oxygenic photoautotrophic organisms (plants, algae and cyanobacteria).

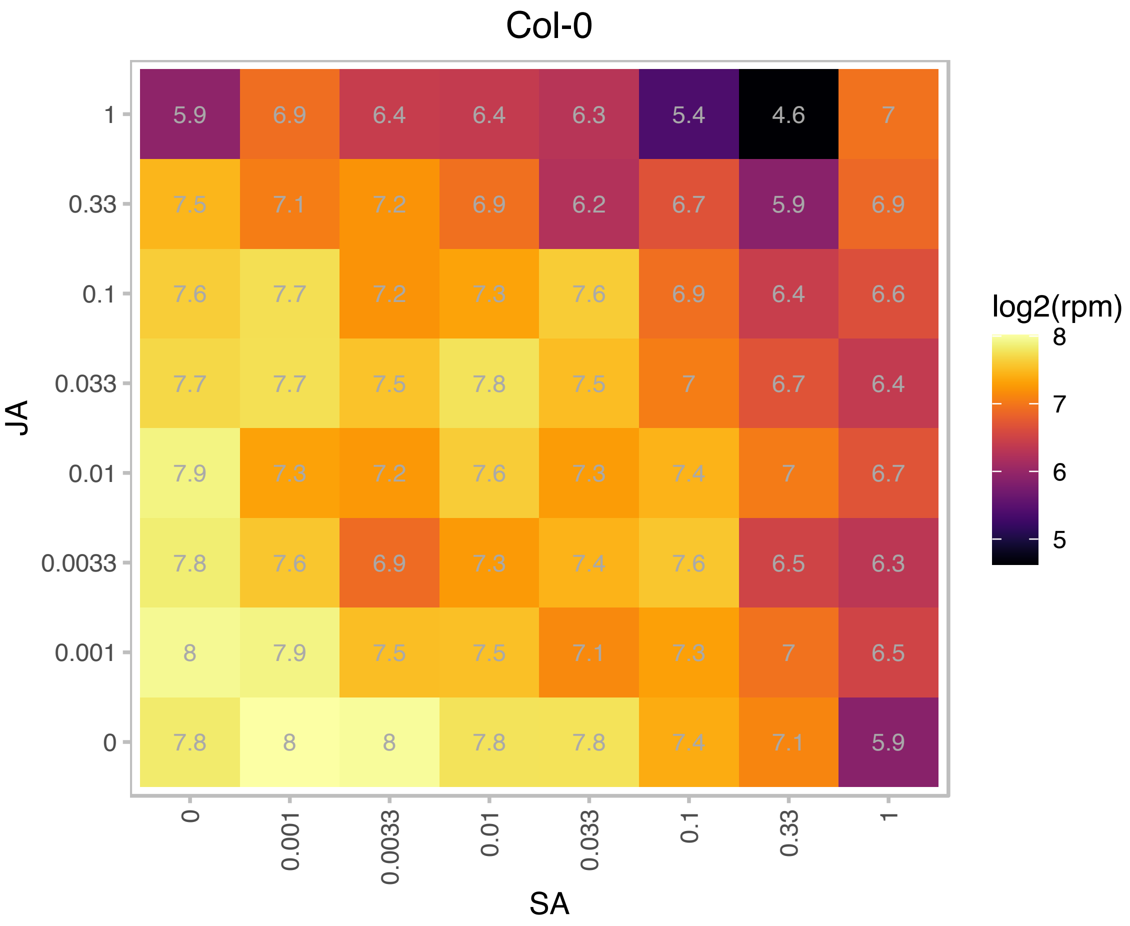


| | |
|---|---|
| Gene | AT1G54500 |
| is.expressed_genes | TRUE |
| is.responsive_genes | TRUE |
| is.apcluster | F-11 |

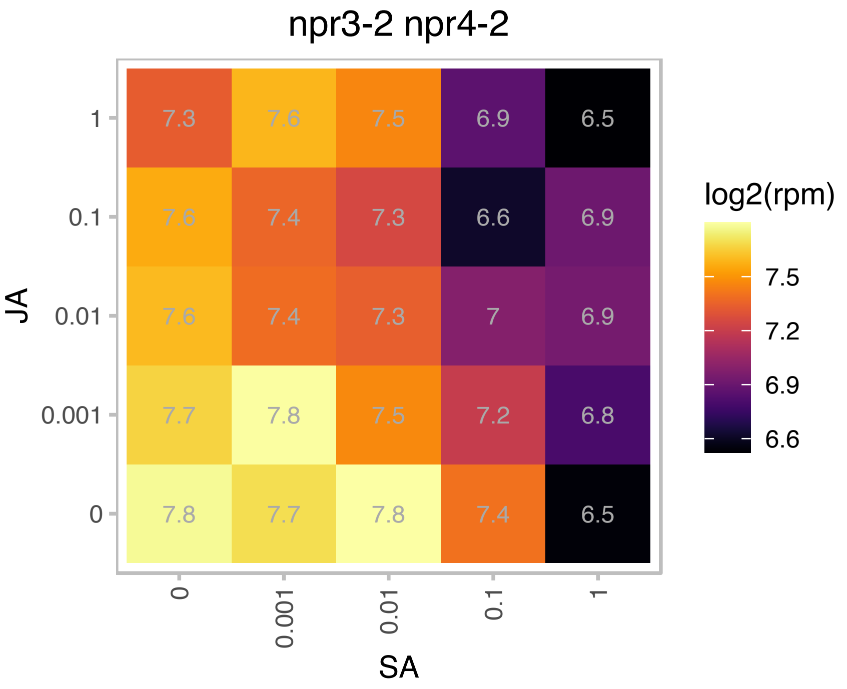


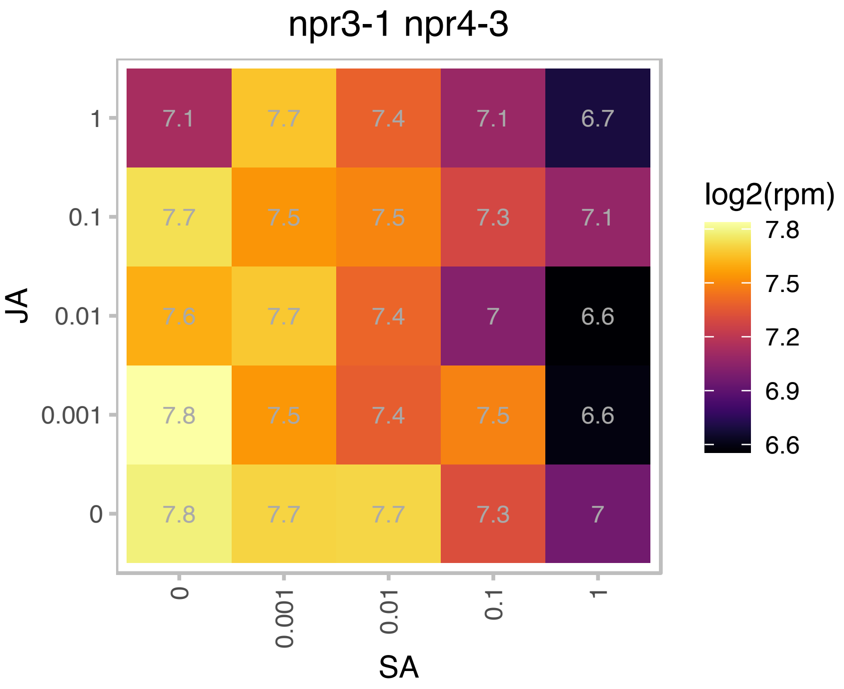

# AT1G56220.1 : NA

Dormancy/auxin associated family protein

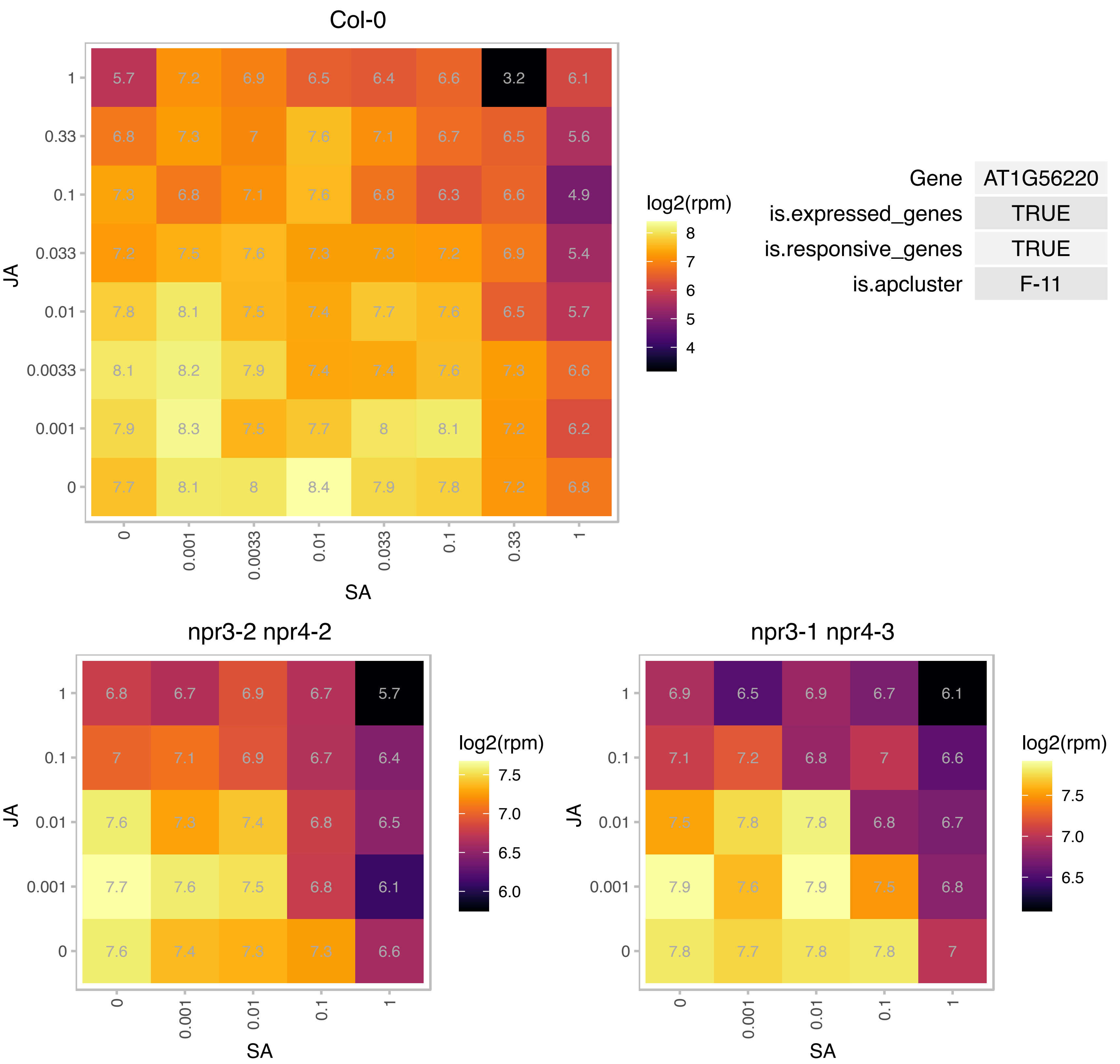


| | |
|---|---|
| Gene | AT1G56220 |
| is.expressed_genes | TRUE |
| is.responsive_genes | TRUE |
| is.apcluster | F-11 |

### AT1G64105.1 : ANAC027, NAC027

NAC domain containing protein 27

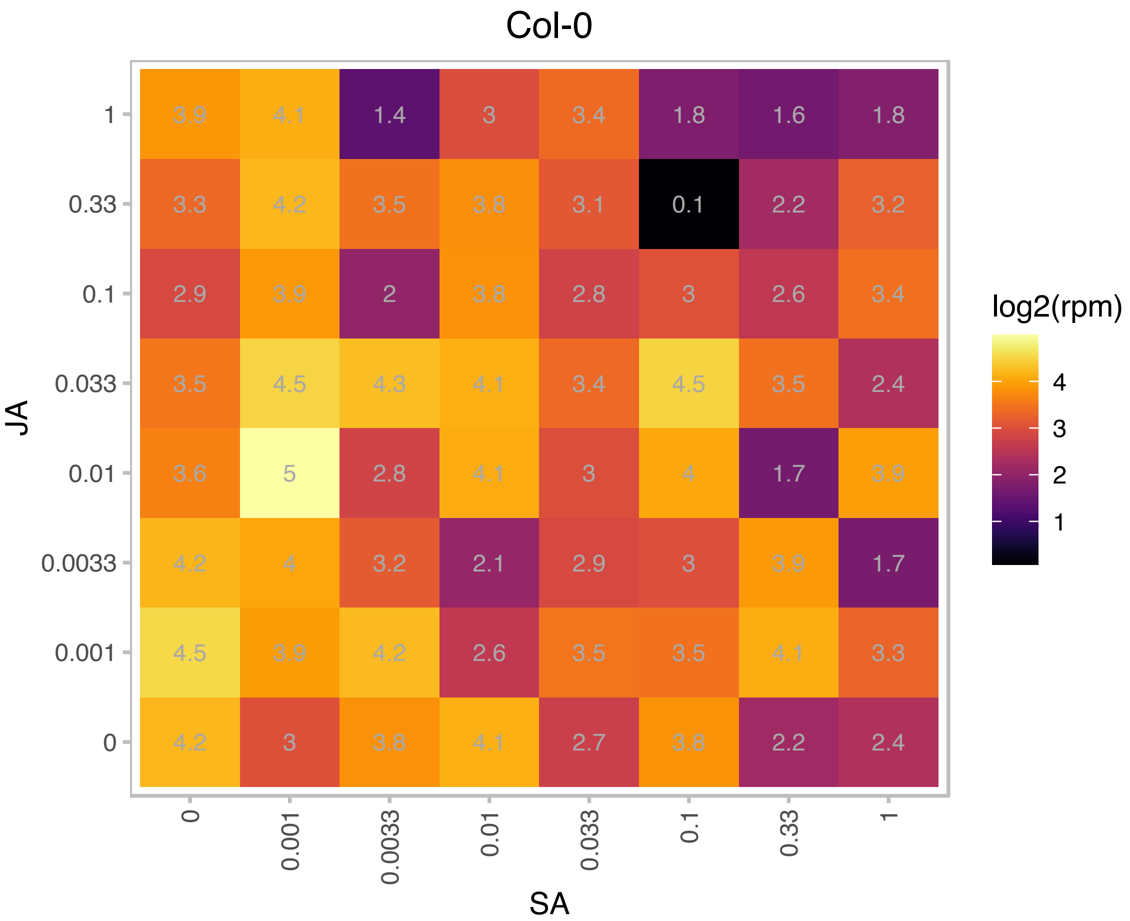


| | |
|---|---|
| Gene | AT1G64105 |
| is.expressed_genes | TRUE |
| is.responsive_genes | TRUE |
| is.apcluster | F-11 |

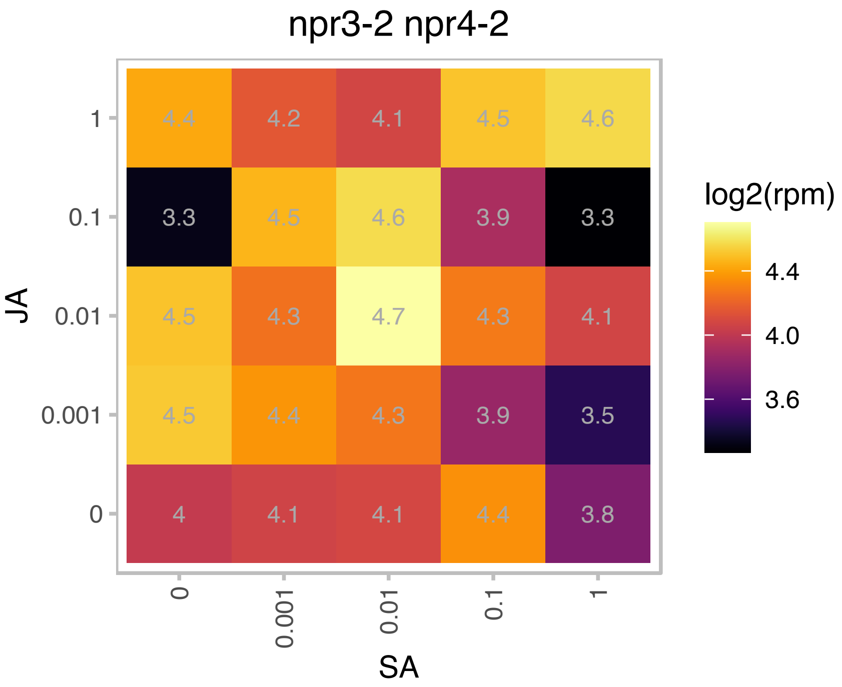


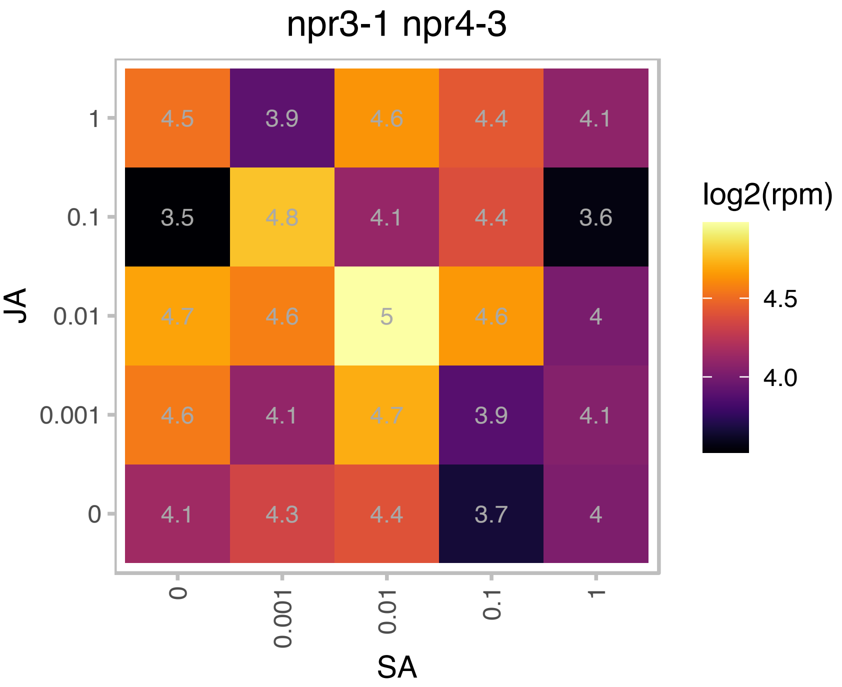

# AT1G66130.1 : NA

NAD(P)-binding Rossmann-fold superfamily protein

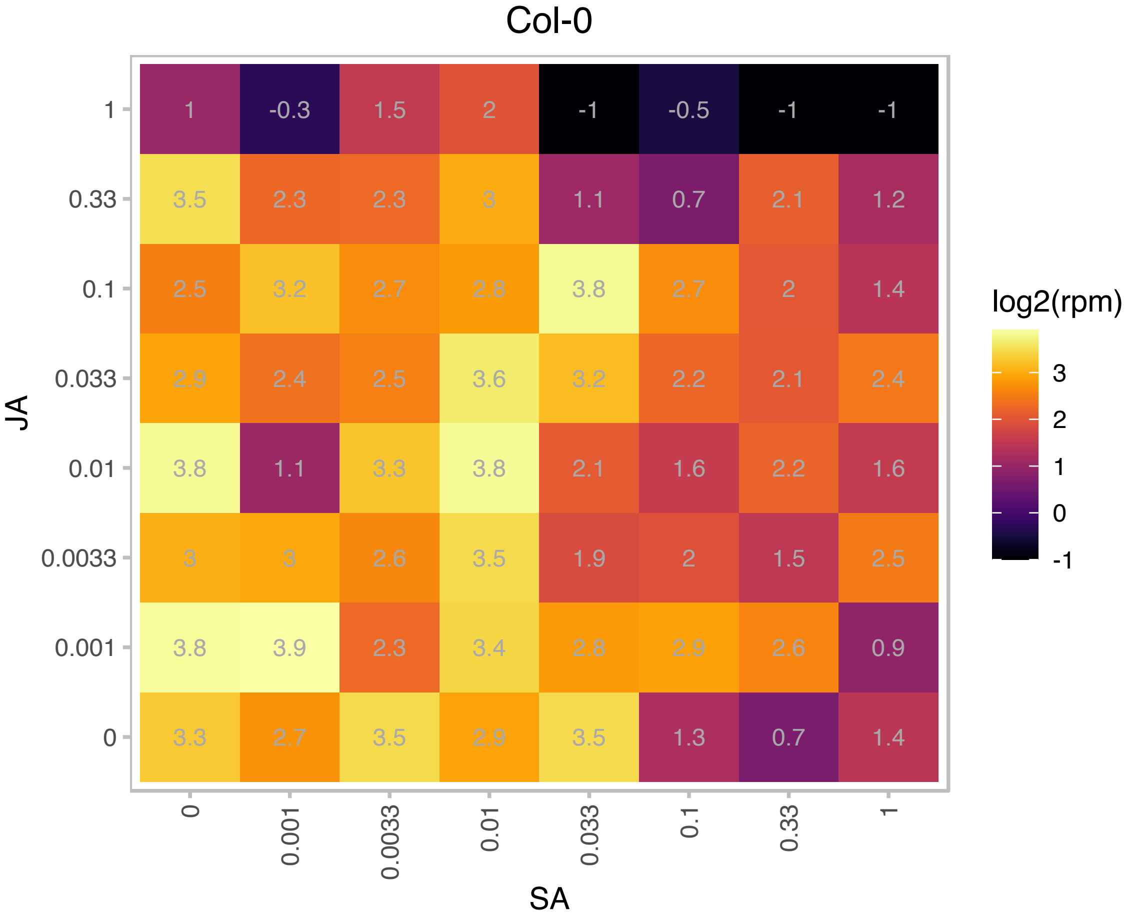


| | |
|---|---|
| Gene | AT1G66130 |
| is.expressed_genes | TRUE |
| is.responsive_genes | TRUE |
| is.apcluster | F-11 |

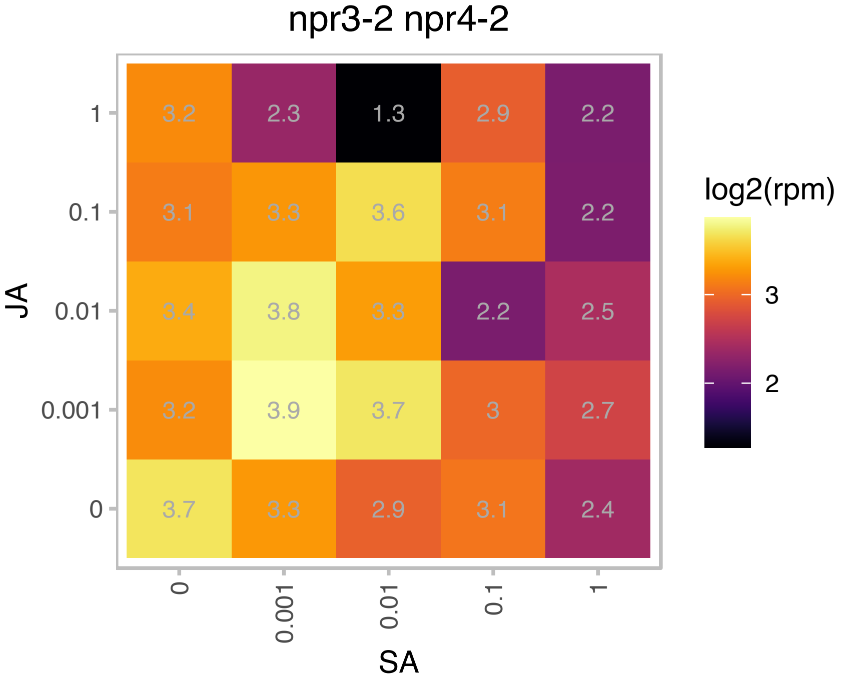


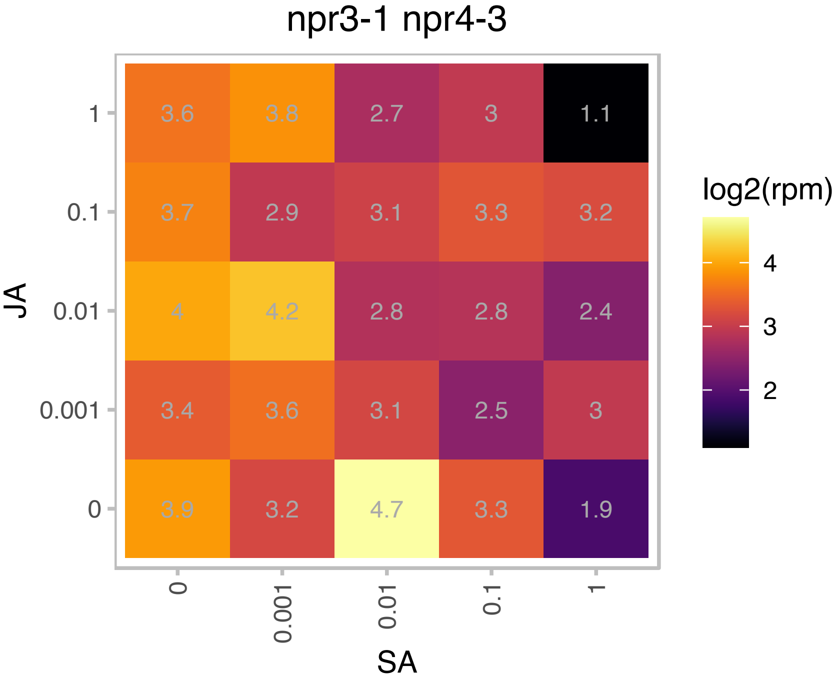

# AT1G66820.1 : NA

glycine-rich protein

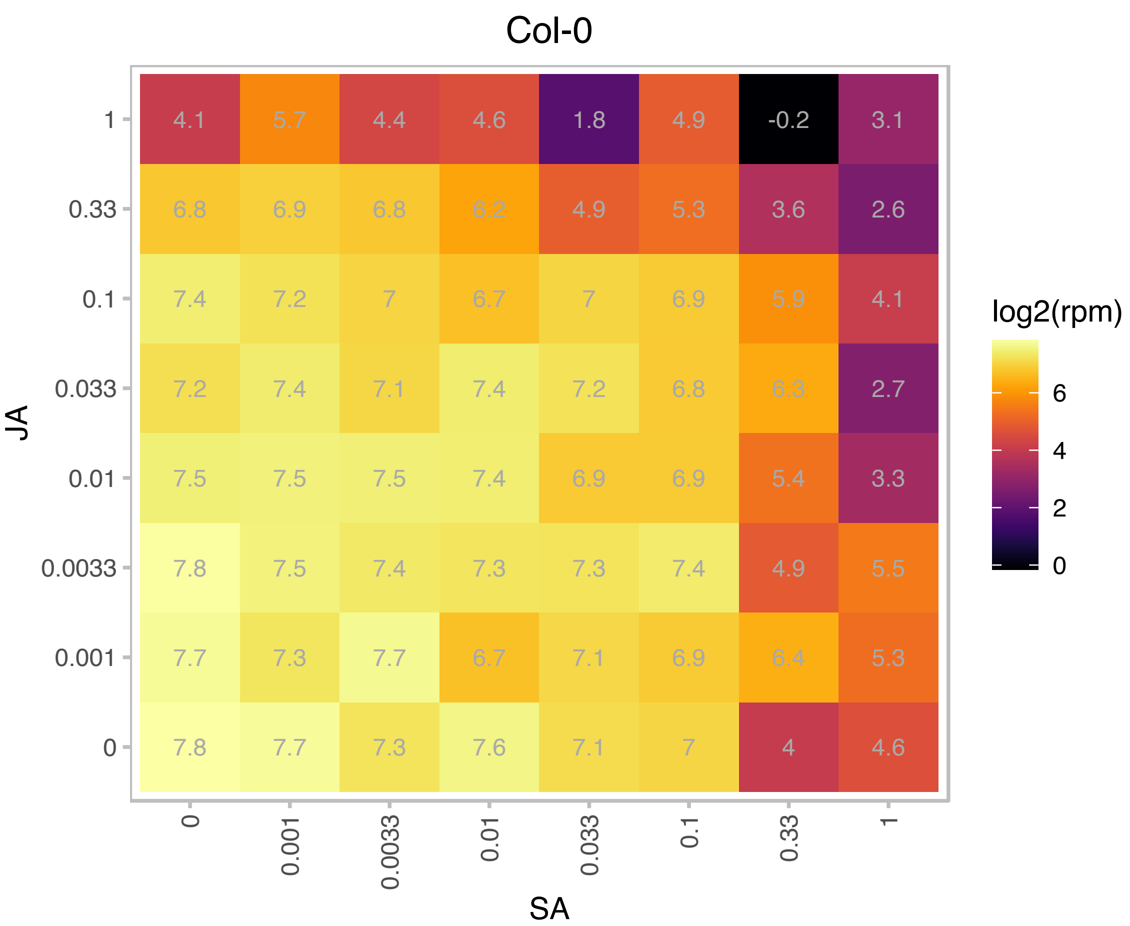


| | |
|---|---|
| Gene | AT1G66820 |
| is.expressed_genes | TRUE |
| is.responsive_genes | TRUE |
| is.apcluster | F-11 |

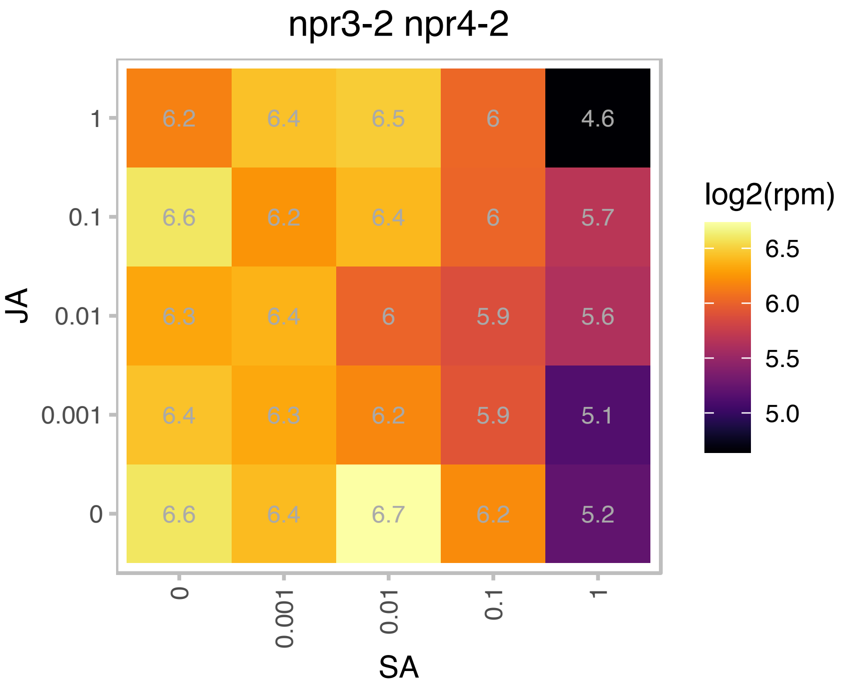


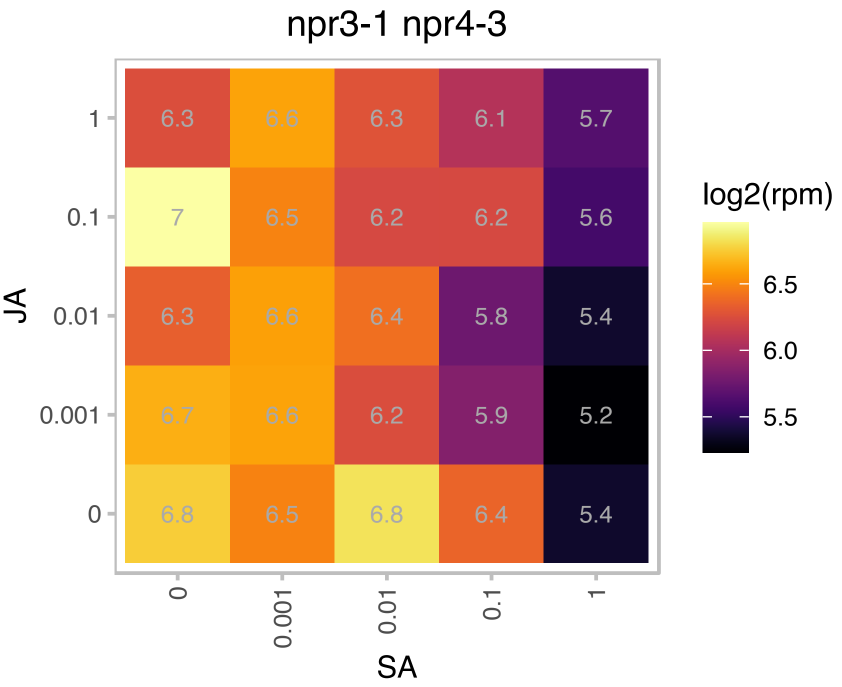

# AT1G72030.1 : NA

Acyl-CoA N-acyltransferases (NAT) superfamily protein

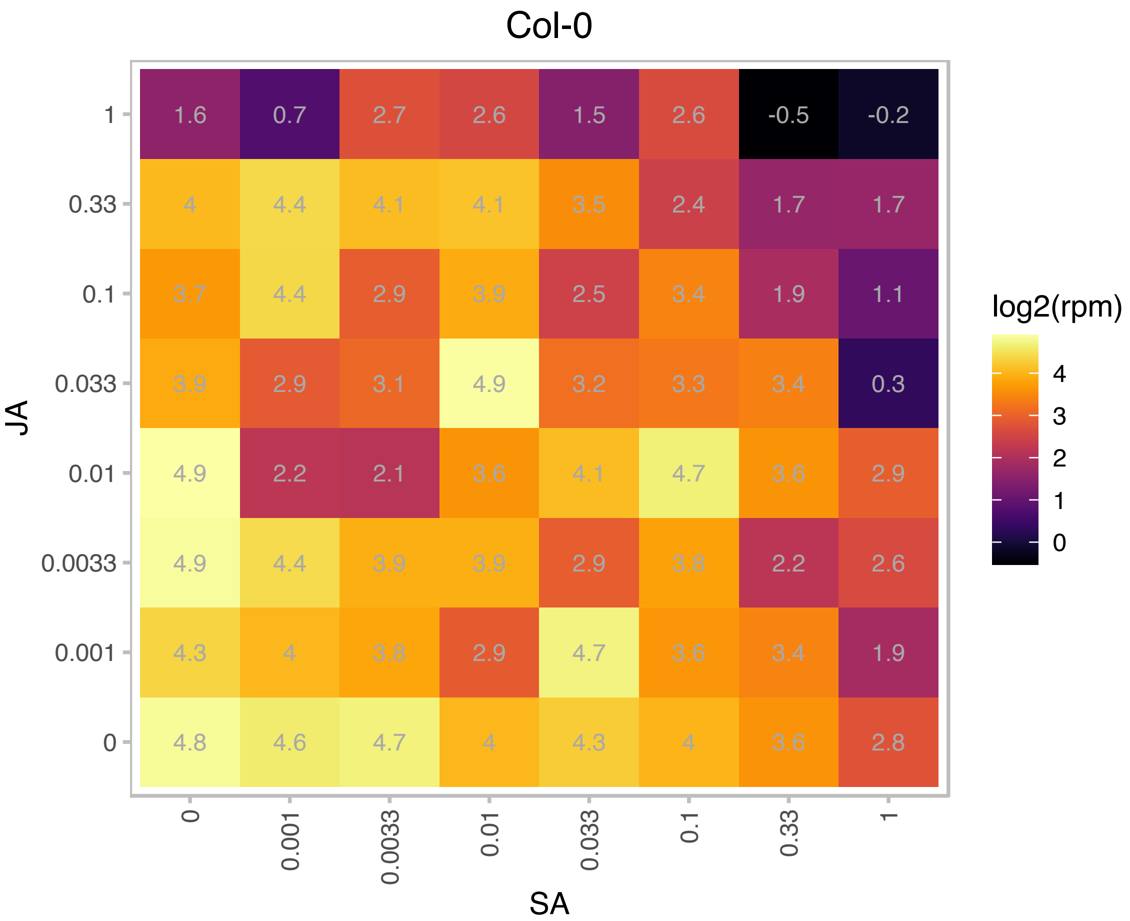


| | |
|---|---|
| Gene | AT1G72030 |
| is.expressed_genes | TRUE |
| is.responsive_genes | TRUE |
| is.apcluster | F-11 |

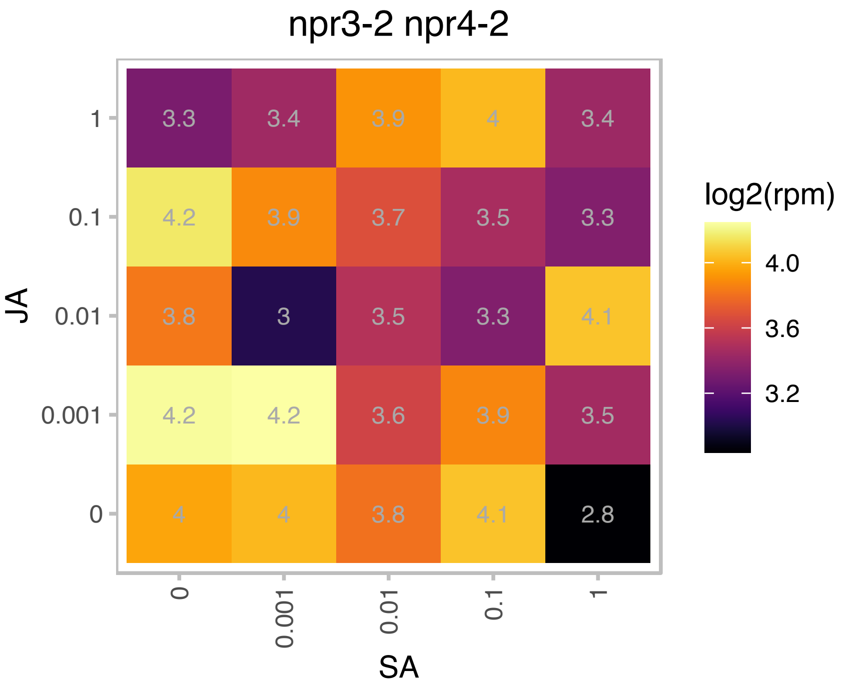


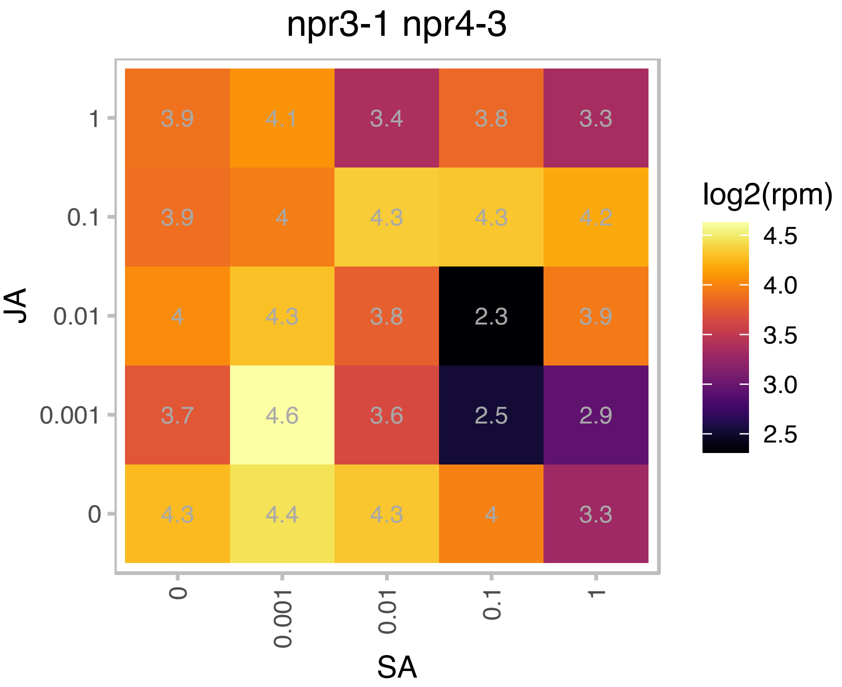

# AT1G72645.1 : NA

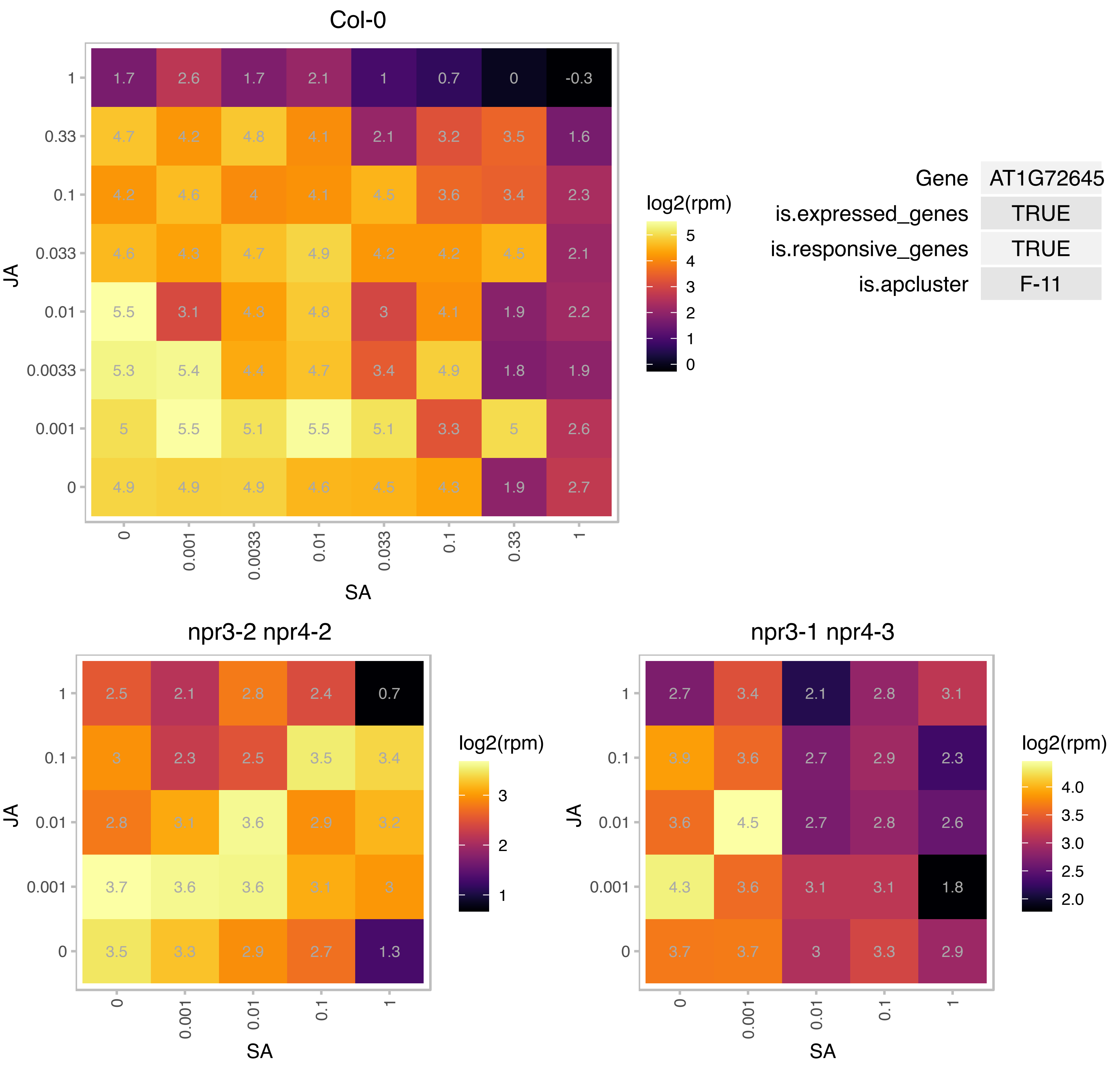

# AT1G74070.1 : NA

Cyclophilin-like peptidyl-prolyl cis-trans isomerase family protein

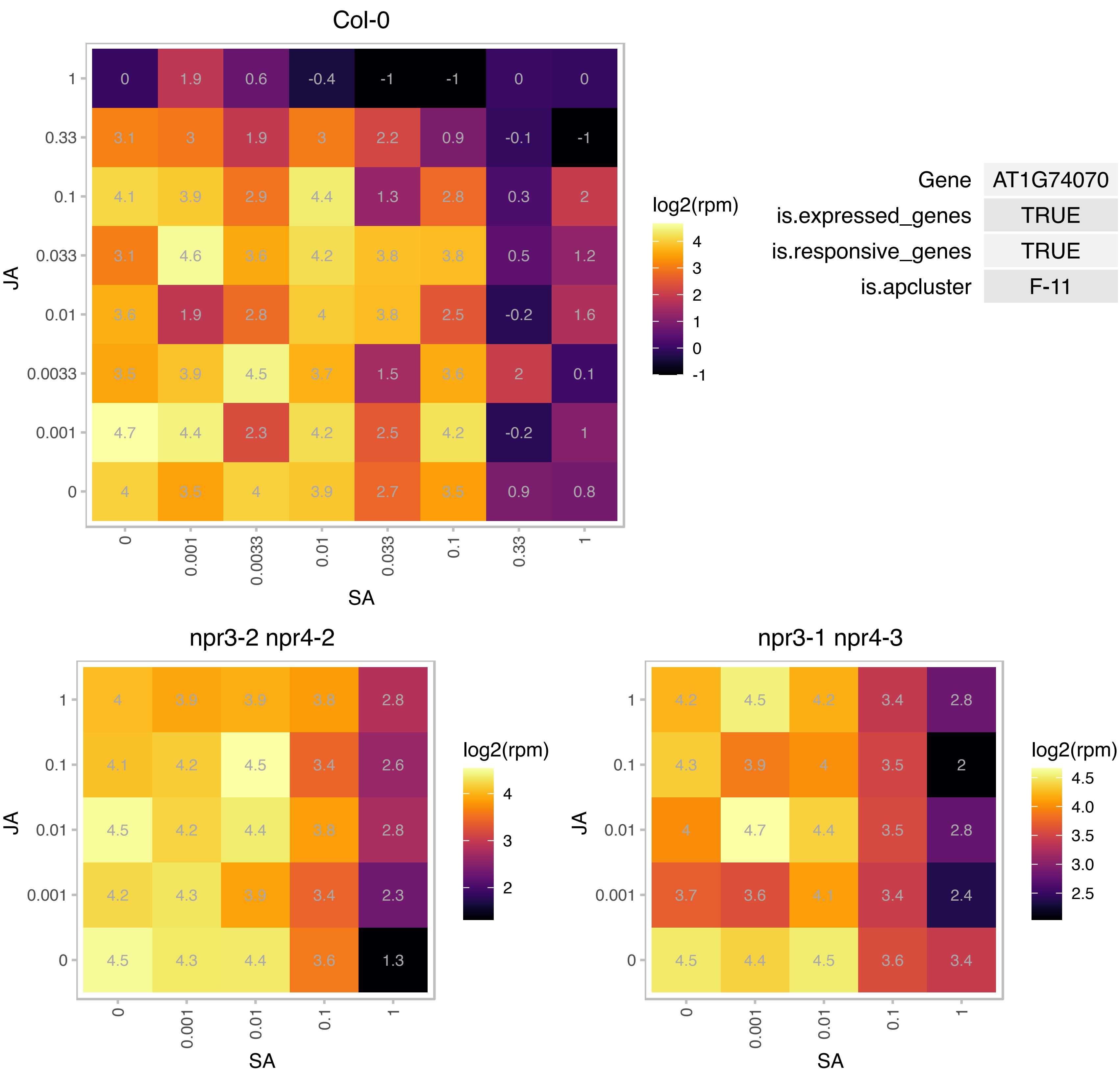


| | |
|---|---|
| Gene | AT1G74070 |
| is.expressed_genes | TRUE |
| is.responsive_genes | TRUE |
| is.apcluster | F-11 |

### AT1G75840.1 : ARAC5, ATGP3, ATROP4, ROP4

RAC-like GTP binding protein 5

Belongs to the plant-specific ROP group of Rho GTPases; localized to the plasma membrane of tips of root hairs; involved in polar growth control. The mRNA is cell-to-cell mobile.

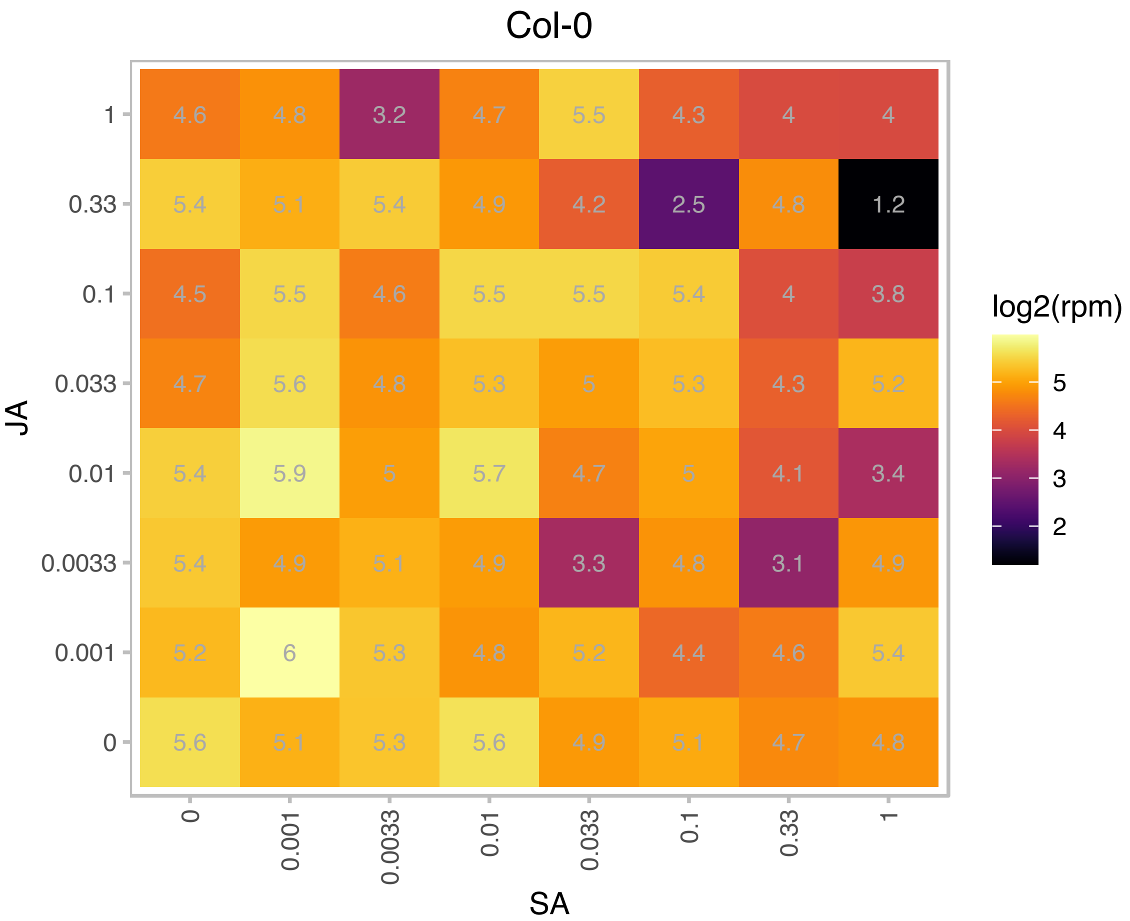


| | |
|---|---|
| Gene | AT1G75840 |
| is.expressed_genes | TRUE |
| is.responsive_genes | TRUE |
| is.apcluster | F-11 |

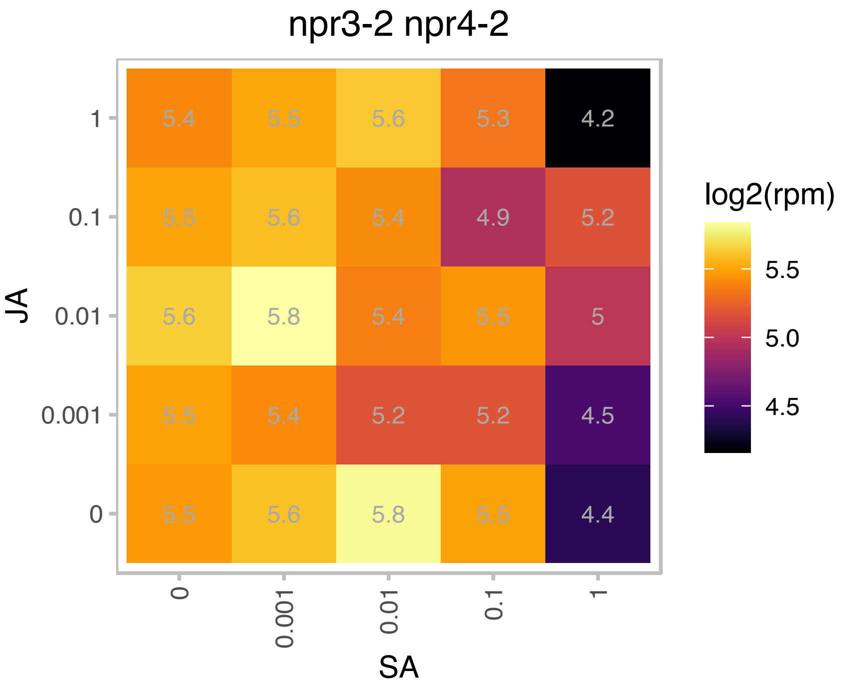


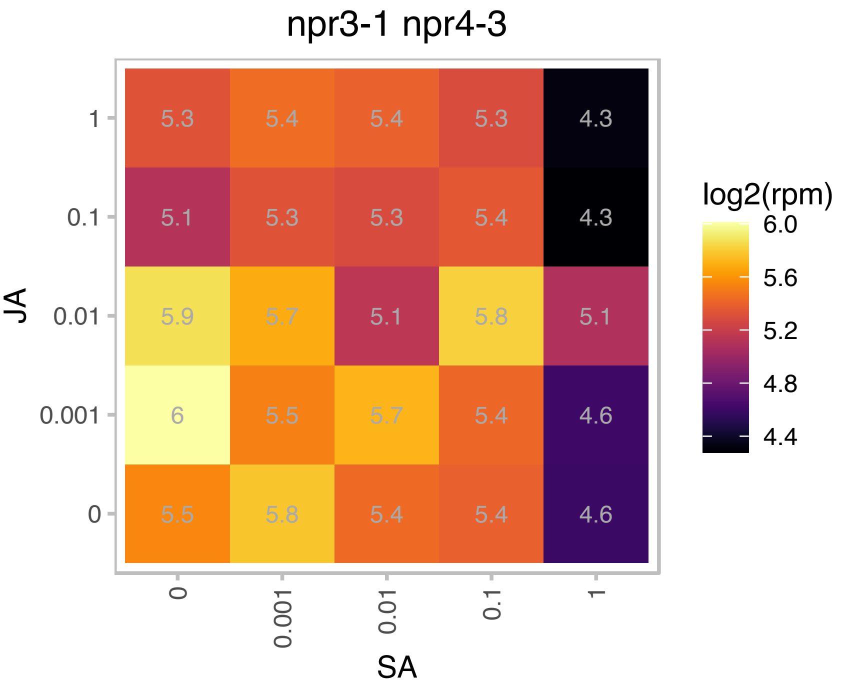

### AT1G76540.1 : CDKB2;1

cyclin-dependent kinase B2;1

Encodes a cyclin-dependent protein kinase involved in regulation of the G2/M transition of the mitotic cell cycle.  Specifically binds to the cyclin CYCD4;1, expressed in shoot meristem, young leaves and vascular tissue during the G2/M phase. Required for proper organization of the shoot apical meristem and for hormone signaling.

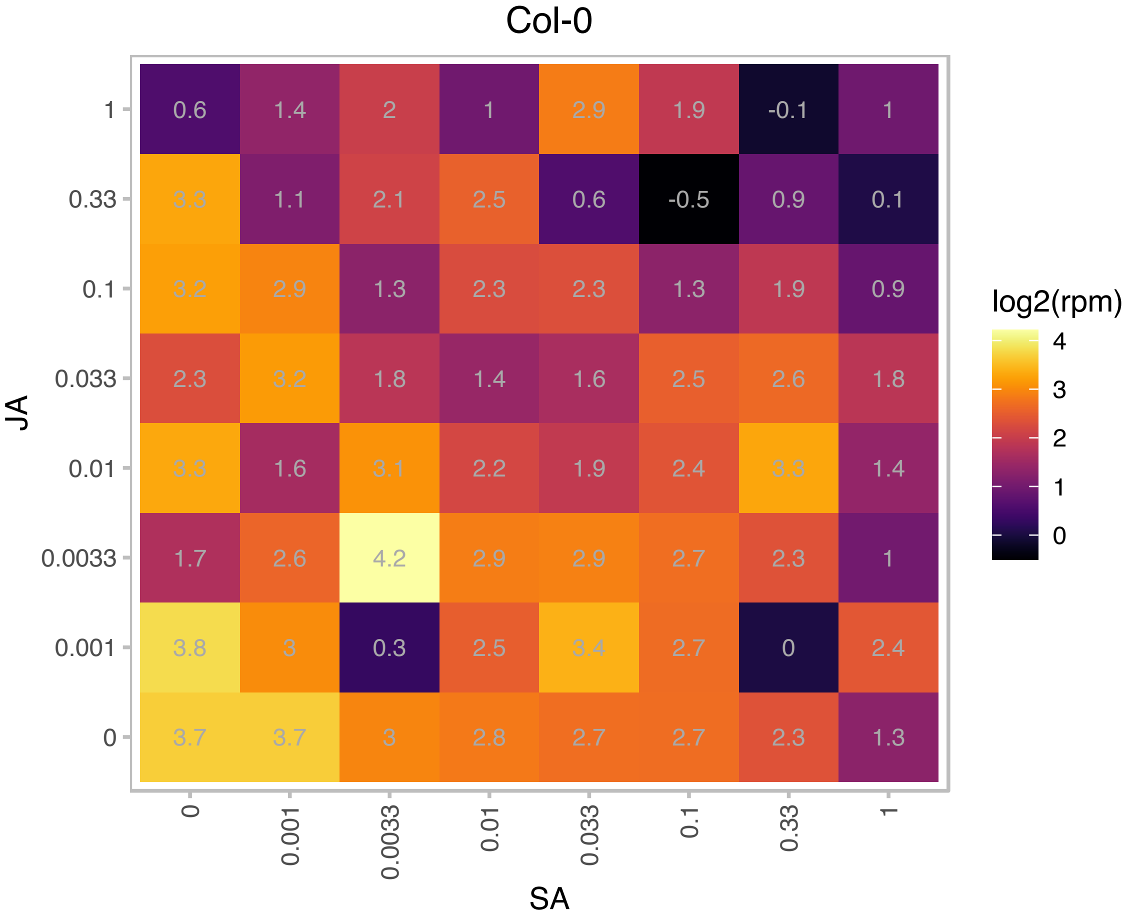


| | |
|---|---|
| Gene | AT1G76540 |
| is.expressed_genes | TRUE |
| is.responsive_genes | TRUE |
| is.apcluster | F-11 |

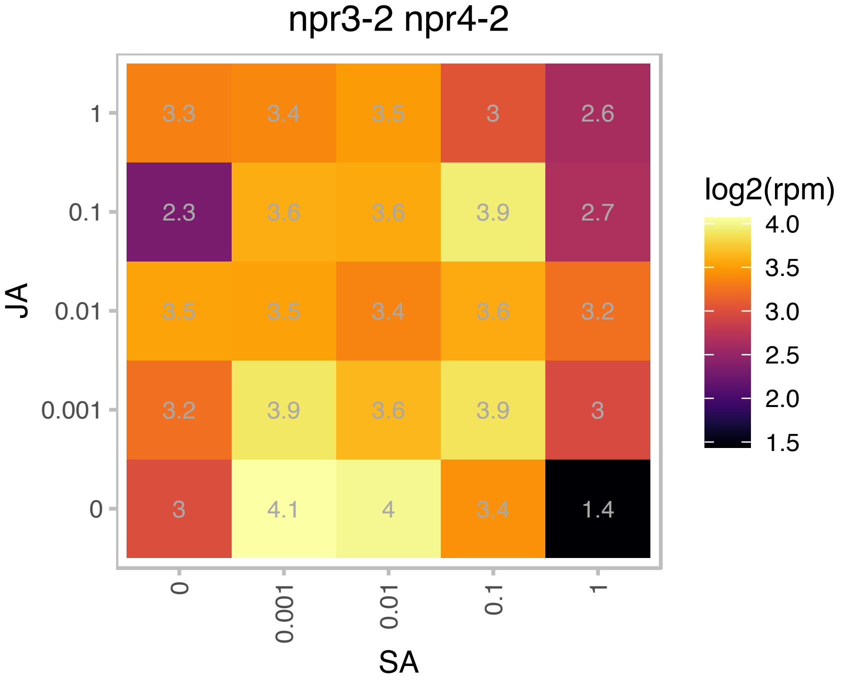


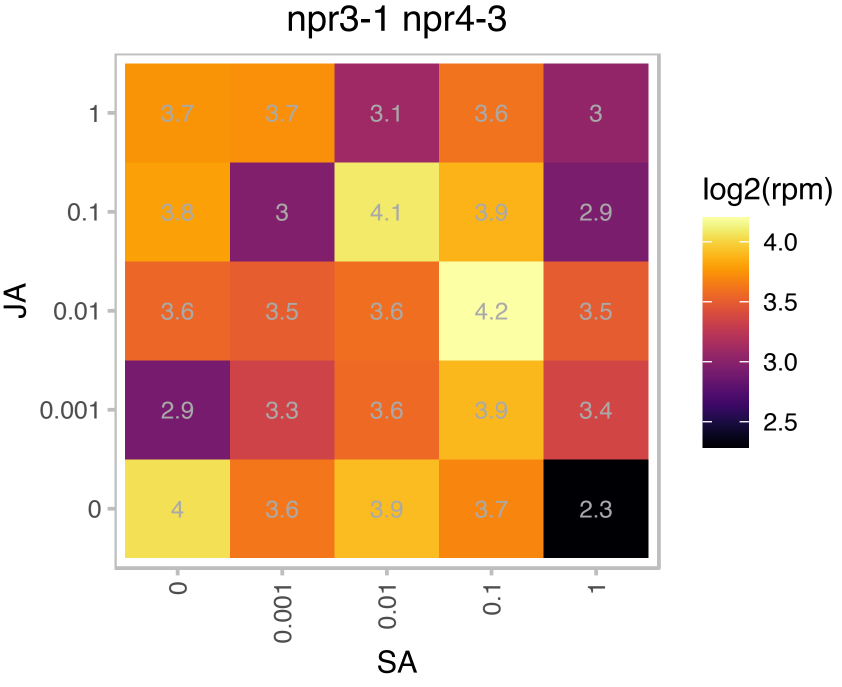

# AT1G79660.1 : NA

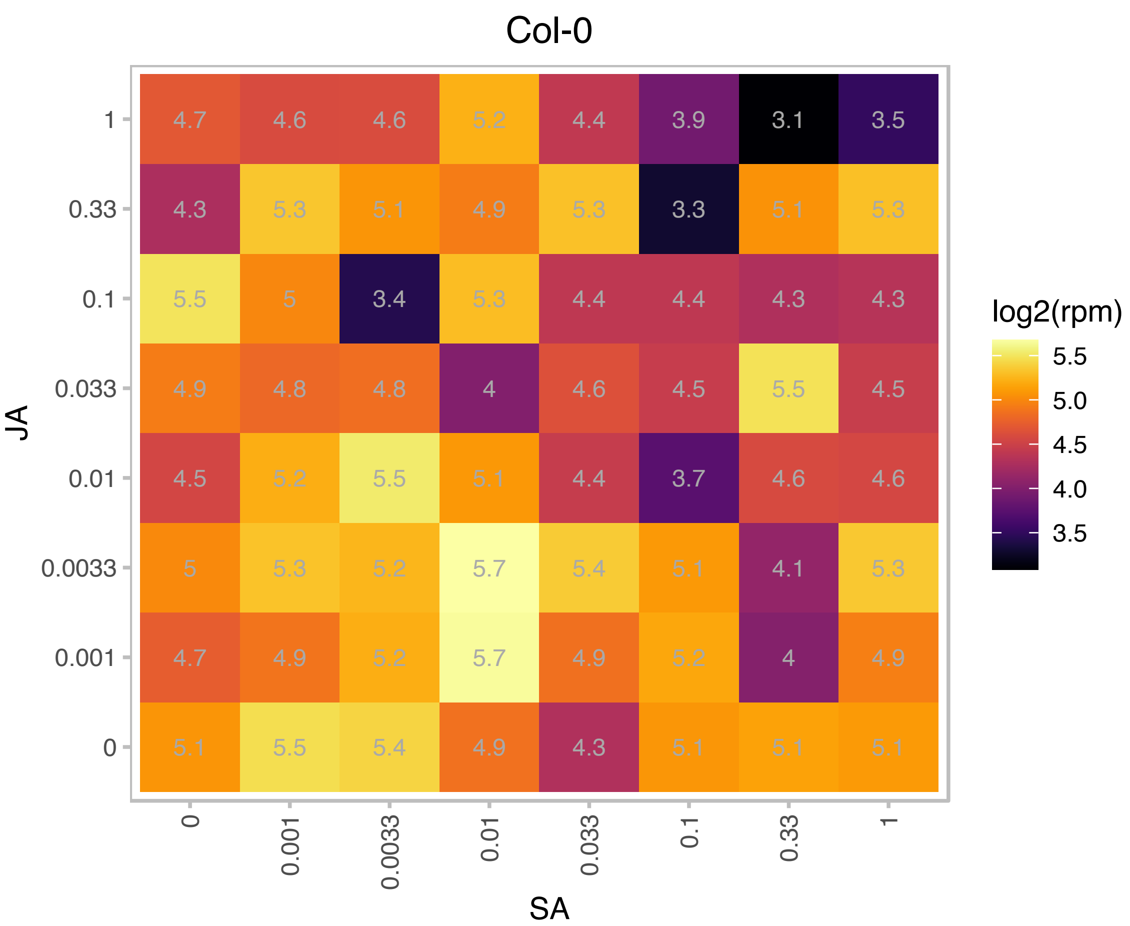


| | |
|---|---|
| Gene | AT1G79660 |
| is.expressed_genes | TRUE |
| is.responsive_genes | TRUE |
| is.apcluster | F-11 |

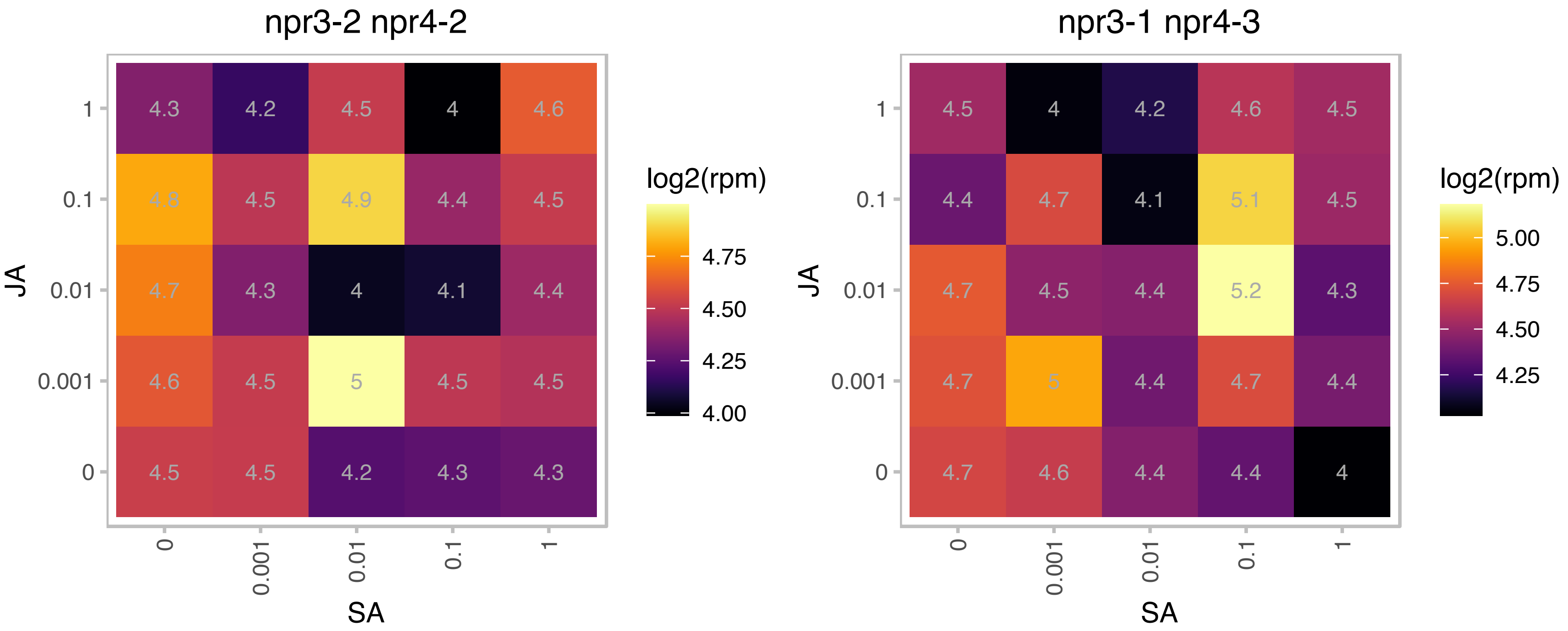

# AT2G03420.1 : NA

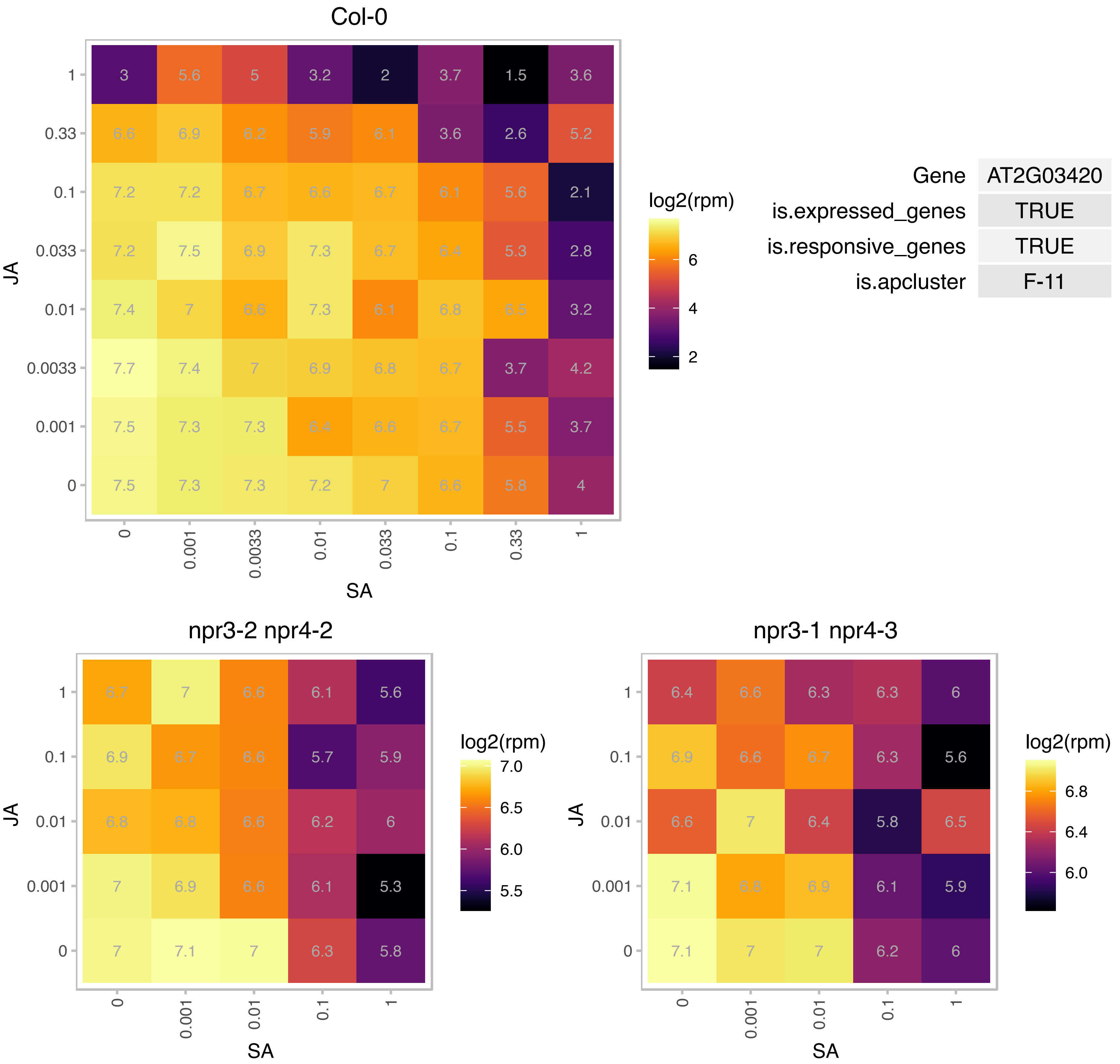


| | |
|---|---|
| Gene | AT2G03420 |
| is.expressed_genes | TRUE |
| is.responsive_genes | TRUE |
| is.apcluster | F-11 |

# AT2G04378.1 : NA

Beta-galactosidase related protein

Col-0

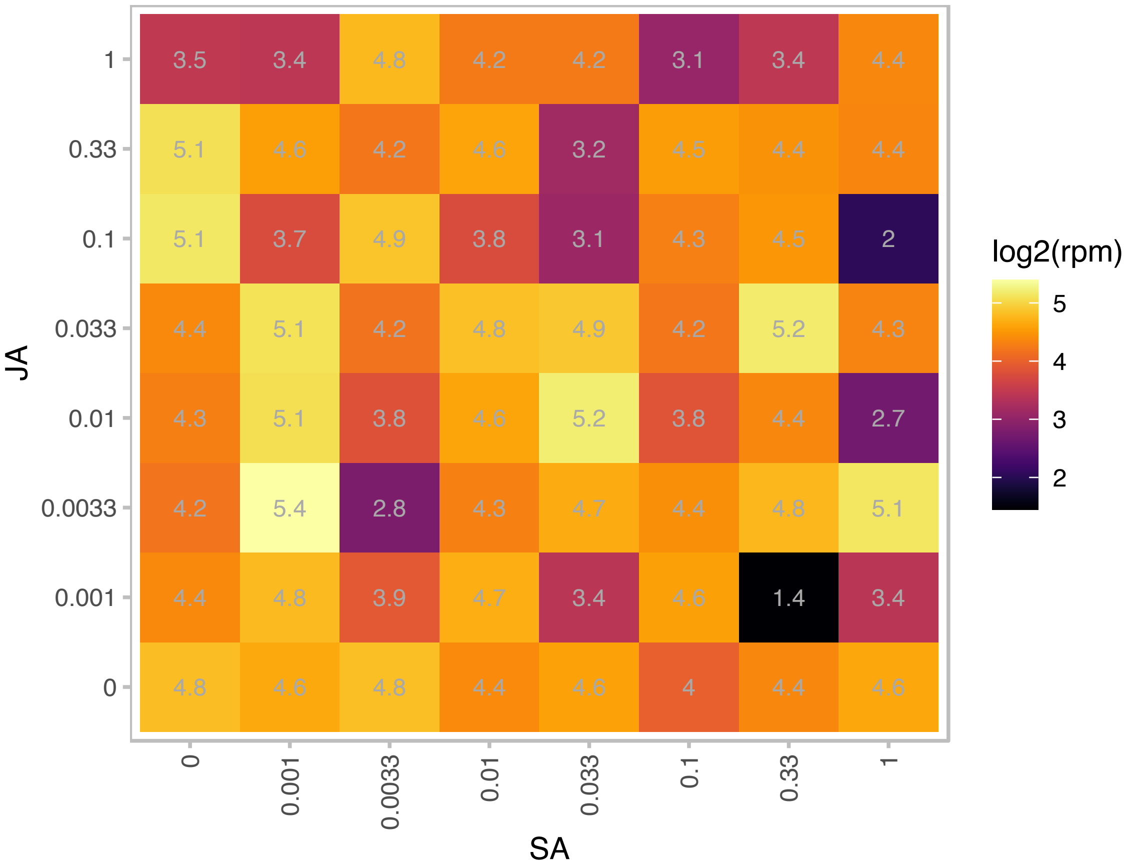


| | |
|---|---|
| Gene | AT2G04378 |
| is.expressed_genes | TRUE |
| is.responsive_genes | TRUE |
| is.apcluster | F-11 |

npr3-2 npr4-2

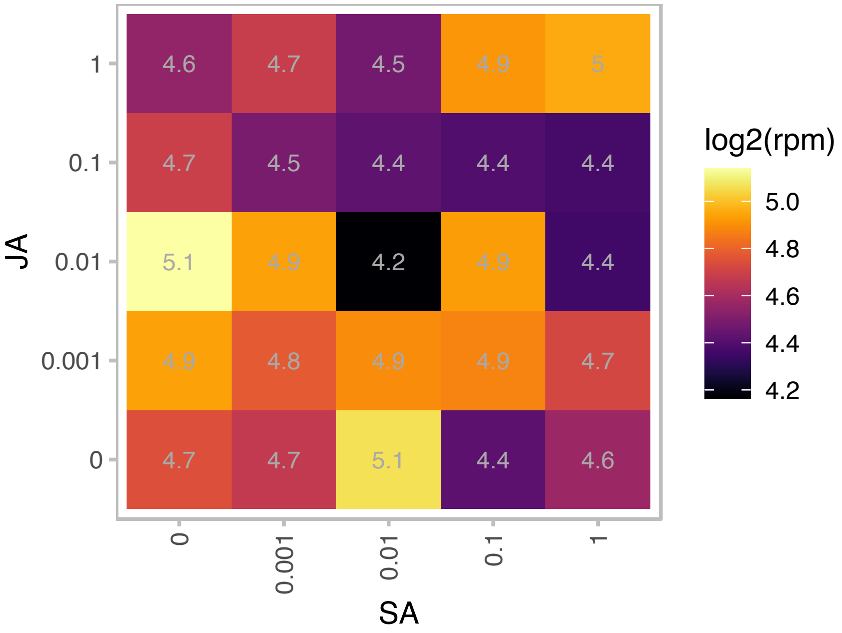


npr3-1 npr4-3

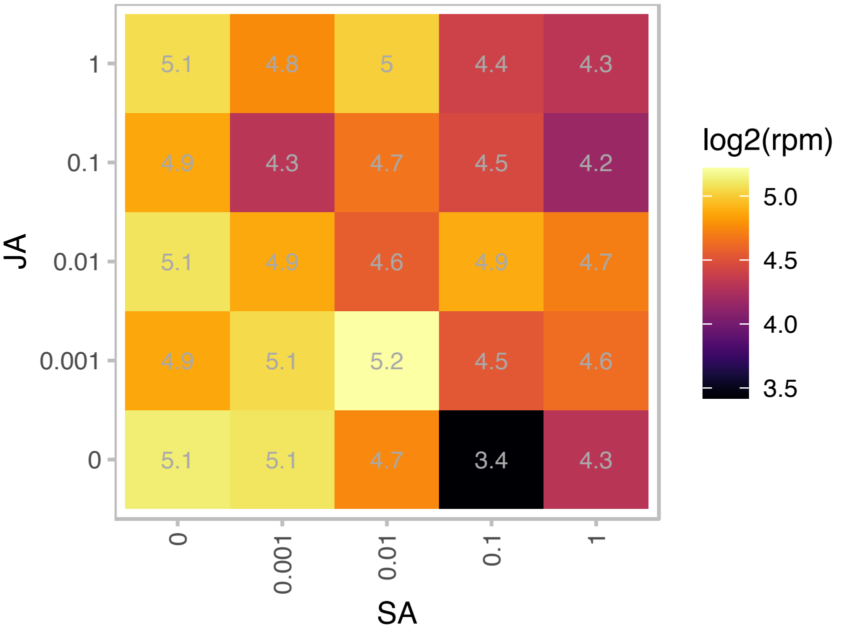

### AT2G15890.1 : CBP1, MEE14

maternal effect embryo arrest 14

Encodes CBP1, a regulator of transcription initiation in central cell-mediated pollen tube guidance.

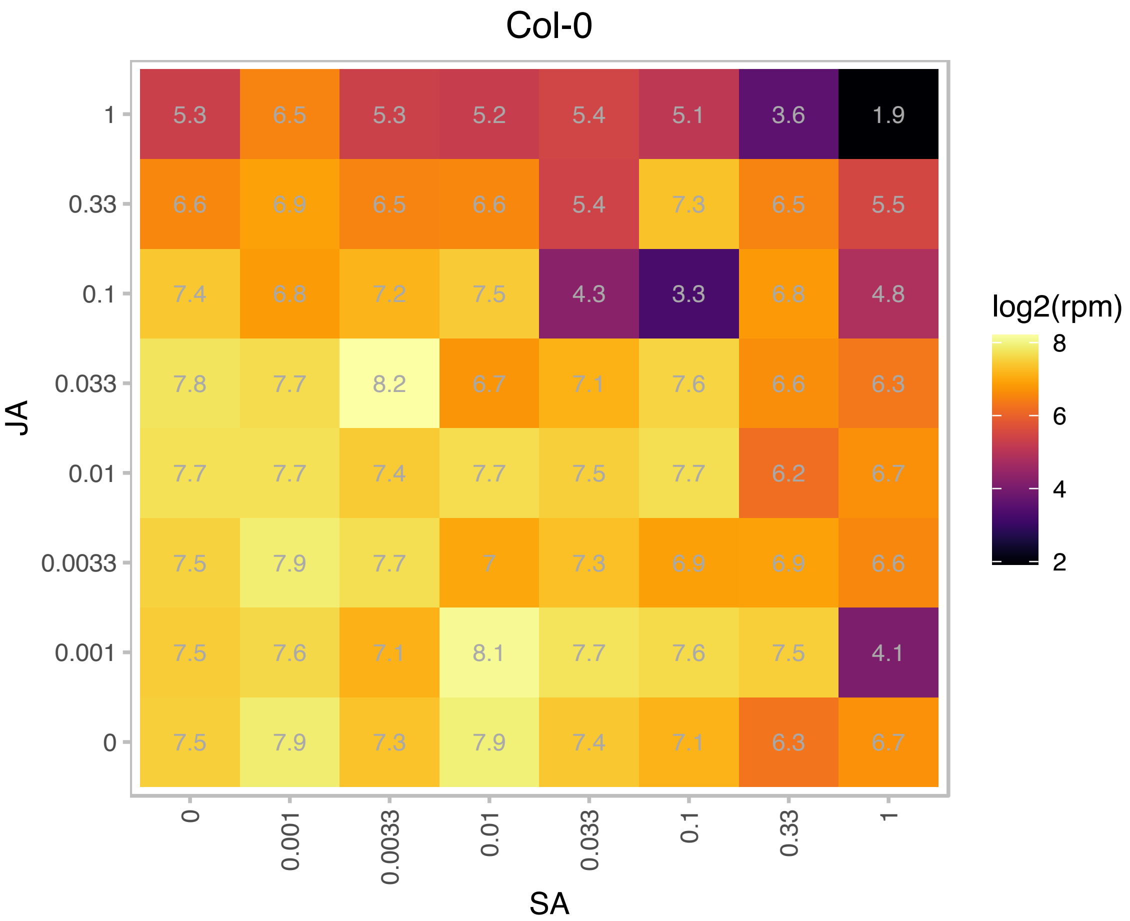


| | |
|---|---|
| Gene | AT2G15890 |
| is.expressed_genes | TRUE |
| is.responsive_genes | TRUE |
| is.apcluster | F-11 |

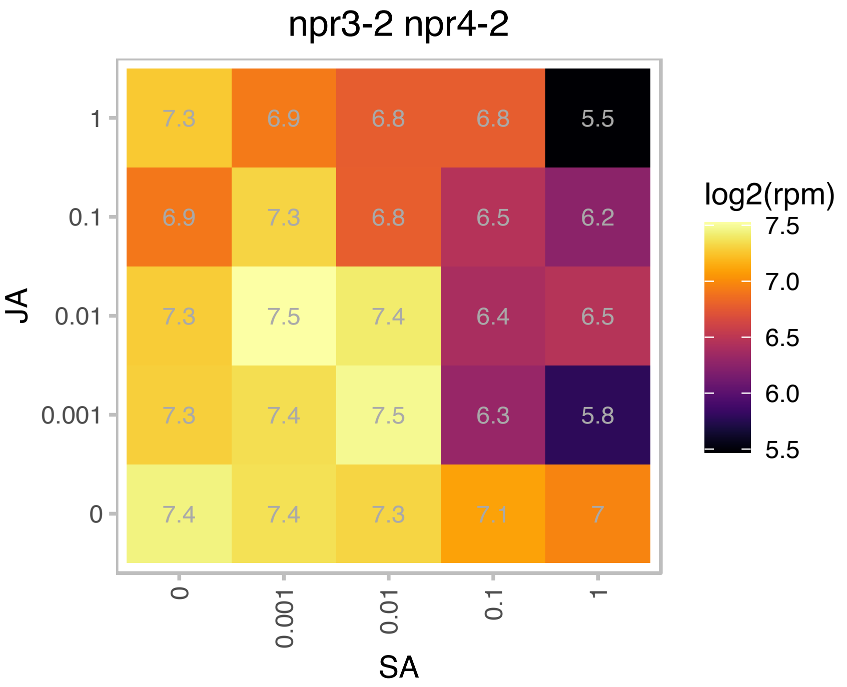


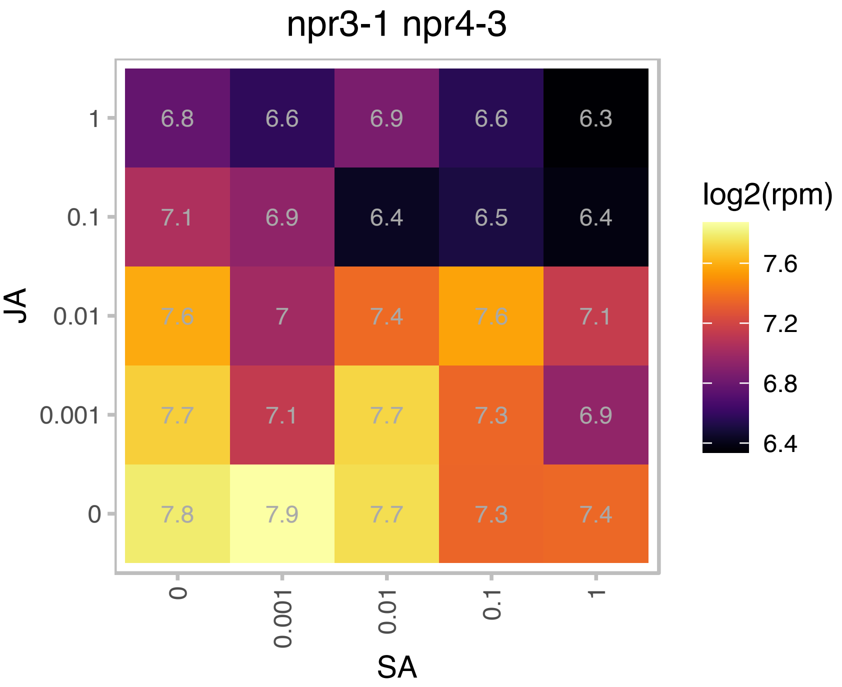

### AT2G19670.1 : ATPRMT1A, PRMT1A

protein arginine methyltransferase 1A

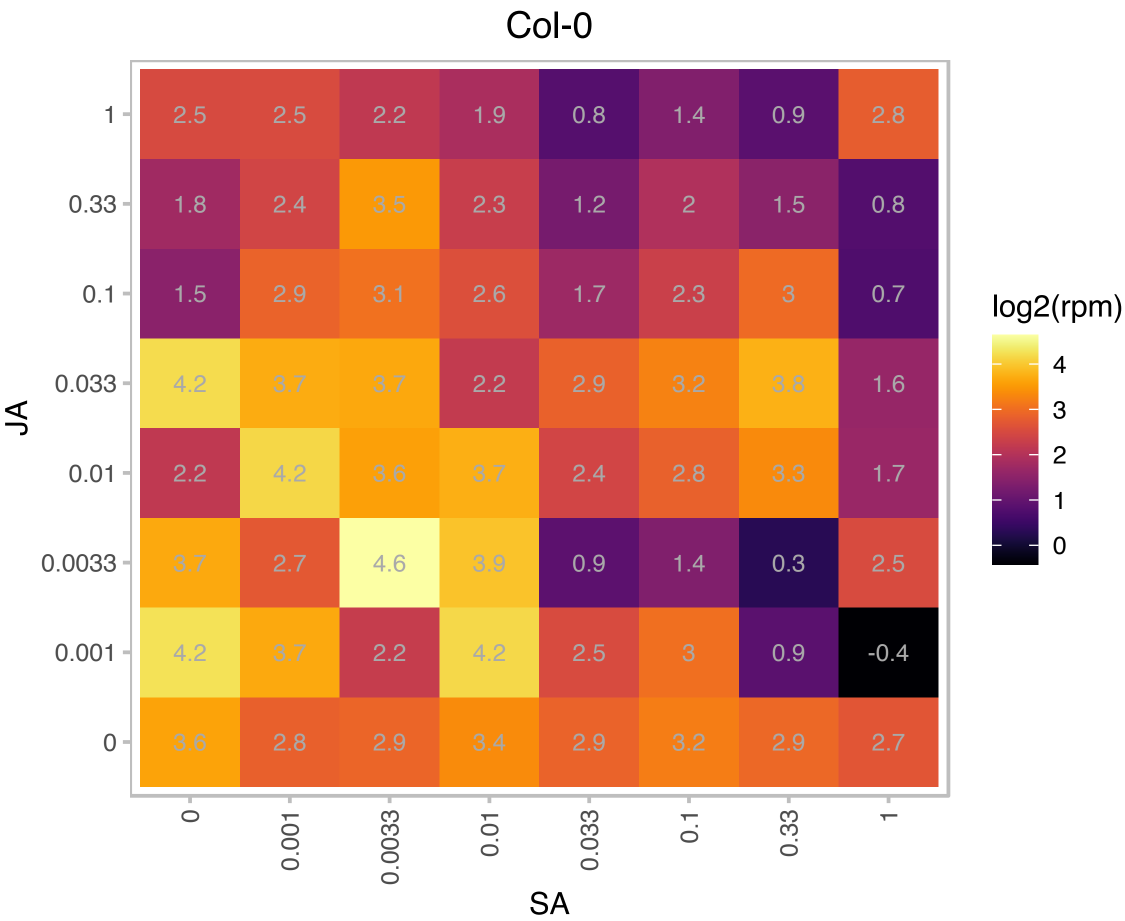


| | |
|---|---|
| Gene | AT2G19670 |
| is.expressed_genes | TRUE |
| is.responsive_genes | TRUE |
| is.apcluster | F-11 |

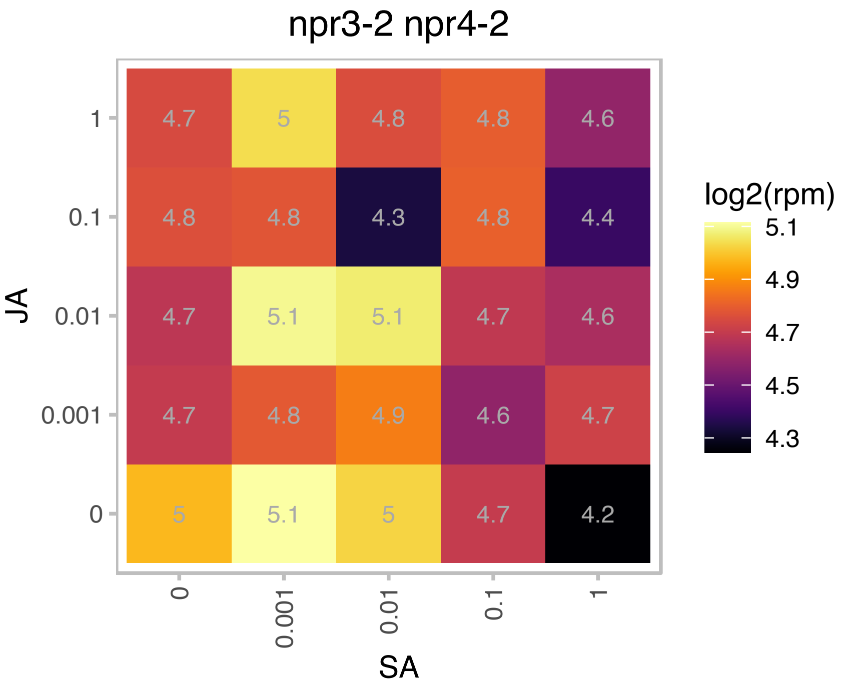


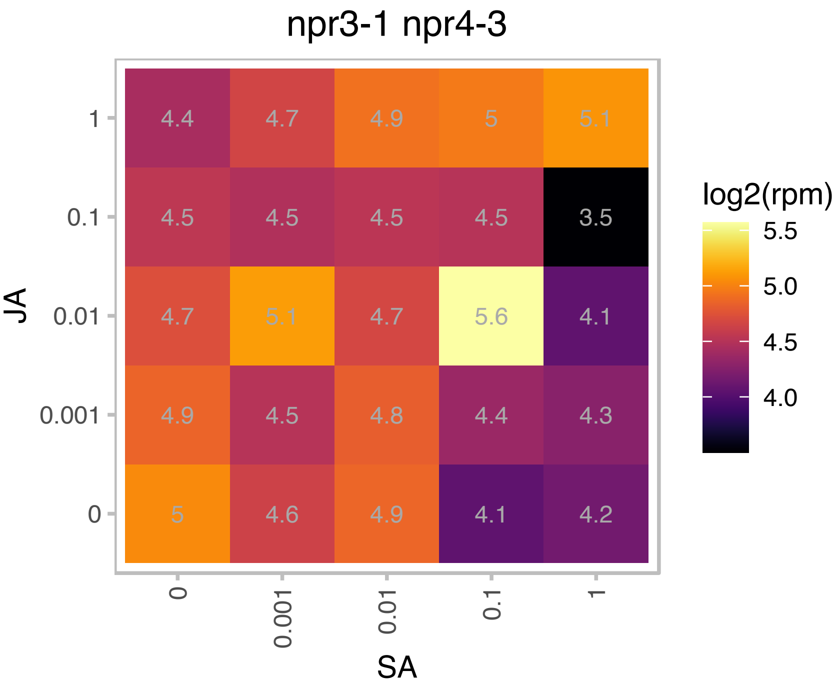

### AT2G22670.1 : IAA8

indoleacetic acid-induced protein 8

Encodes a transcriptional repressor of the auxin response that is auxin inducible and is involved in lateral root formation. The mRNA is cell-to-cell mobile.

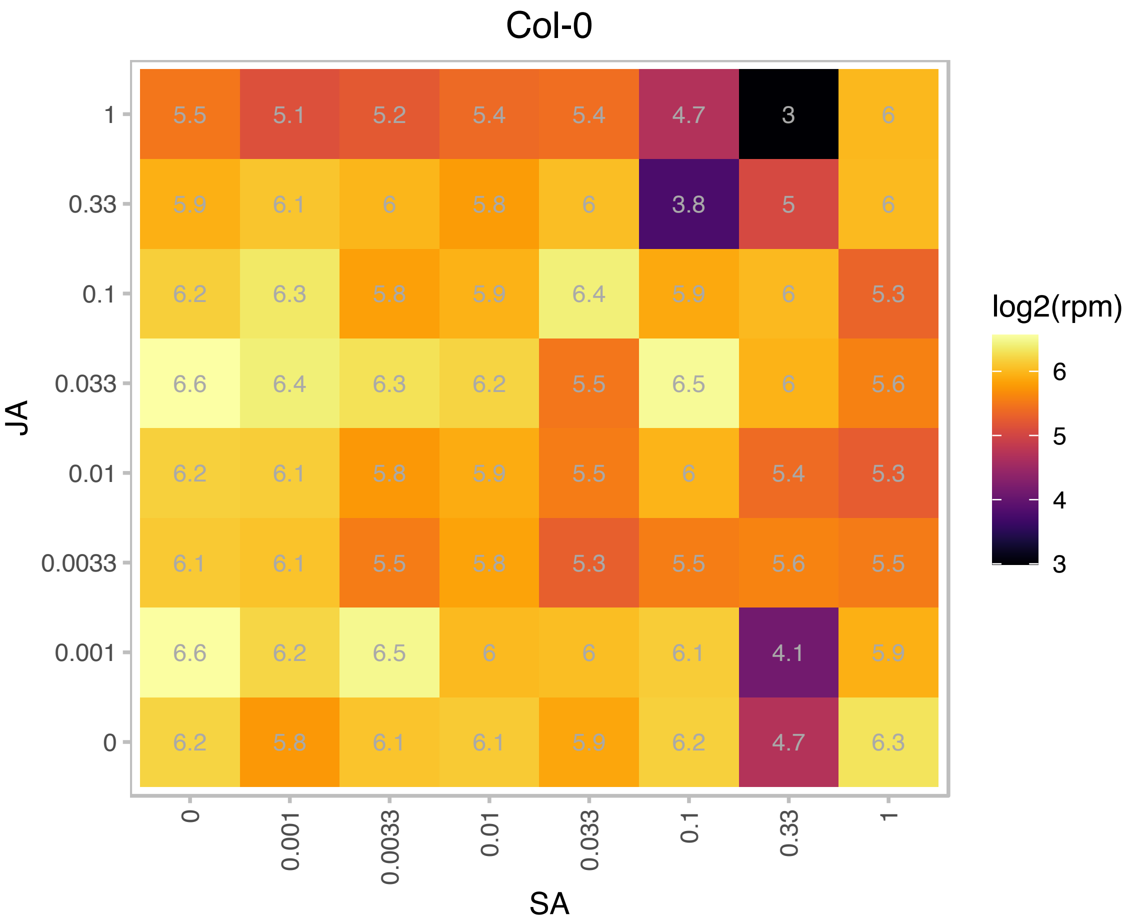


| | |
|---|---|
| Gene | AT2G22670 |
| is.expressed_genes | TRUE |
| is.responsive_genes | TRUE |
| is.apcluster | F-11 |

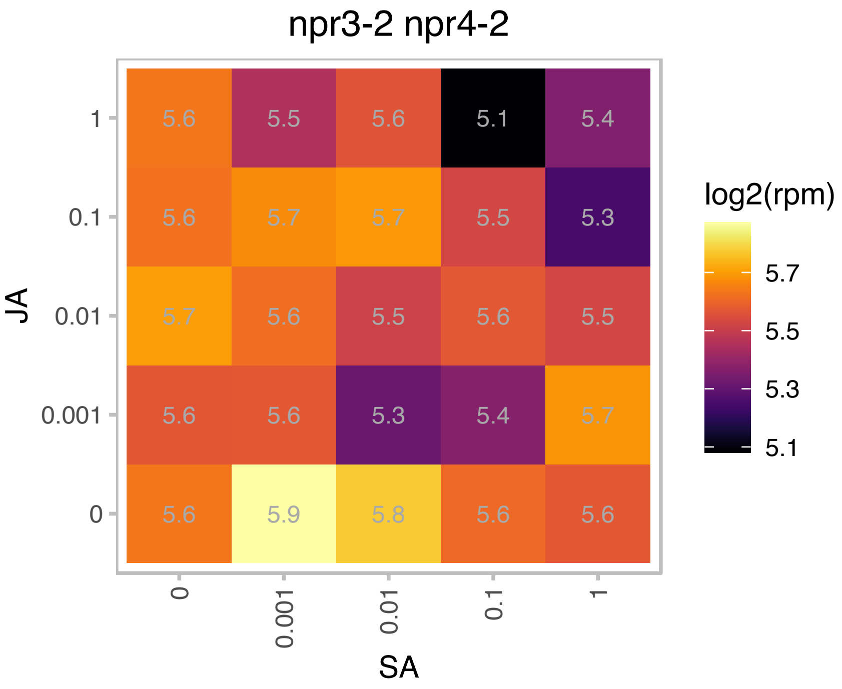


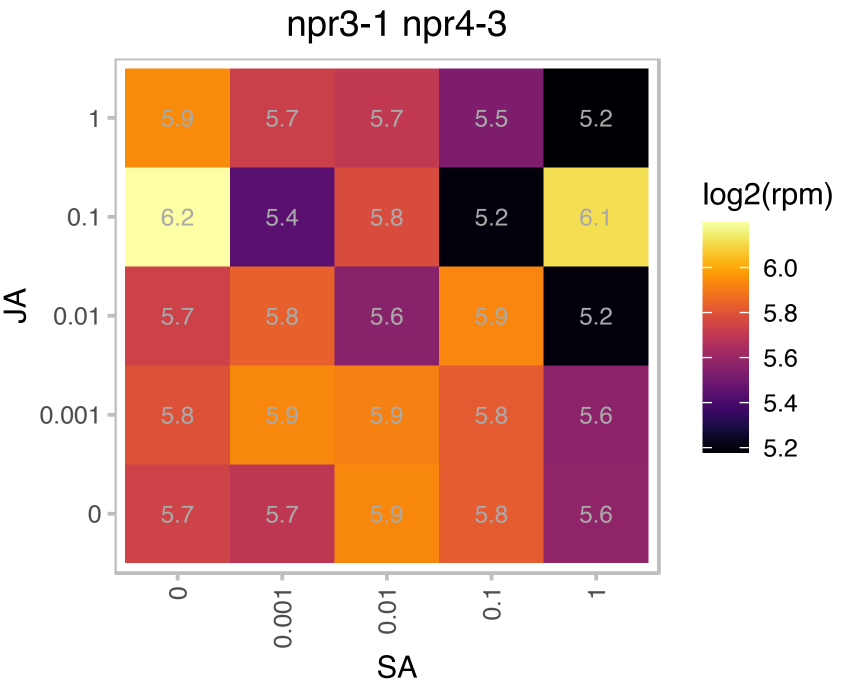

### AT2G23600.1 : ACL, ATME8, ATMES2, ME8, MES2

acetone-cyanohydrin lyase
Encodes a protein shown to have carboxylesterase activity, methyl salicylate esterase activity, methyl jasmonate esterase activity, and methyl IAA esterase activity in vitro. MES2 appears to be involved in MeSA hydrolysis in planta. This protein does not act on MeGA4, or MEGA9 in vitro.

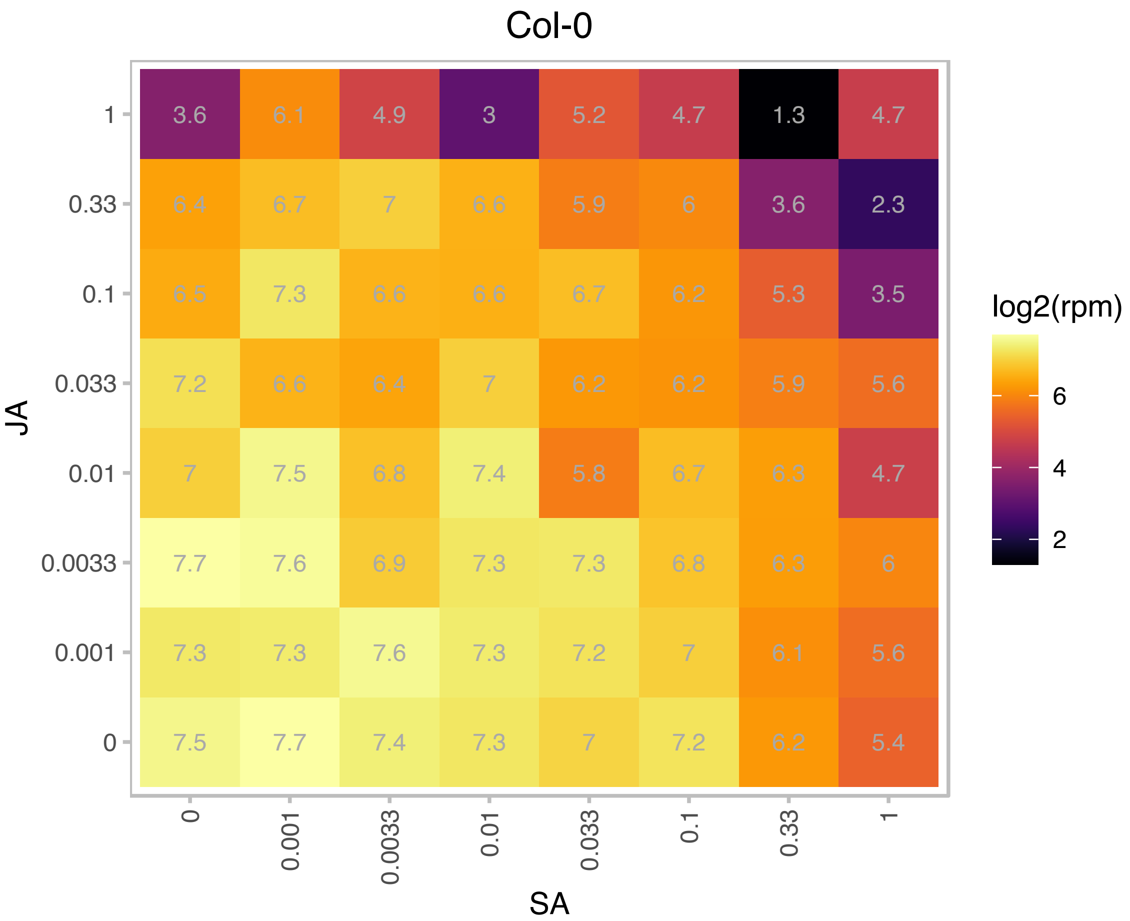


| | |
|---|---|
| Gene | AT2G23600 |
| is.expressed_genes | TRUE |
| is.responsive_genes | TRUE |
| is.apcluster | F-11 |

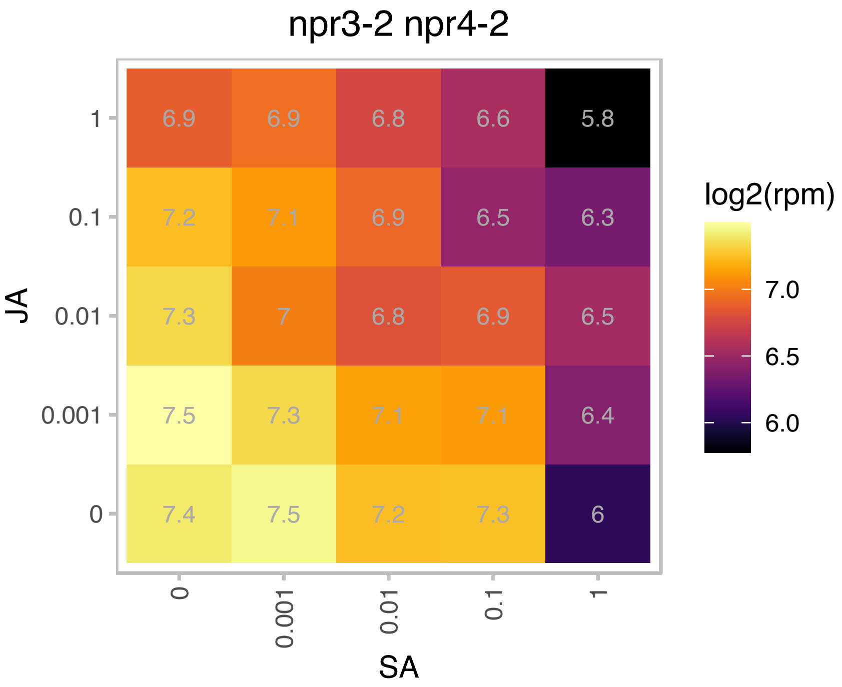


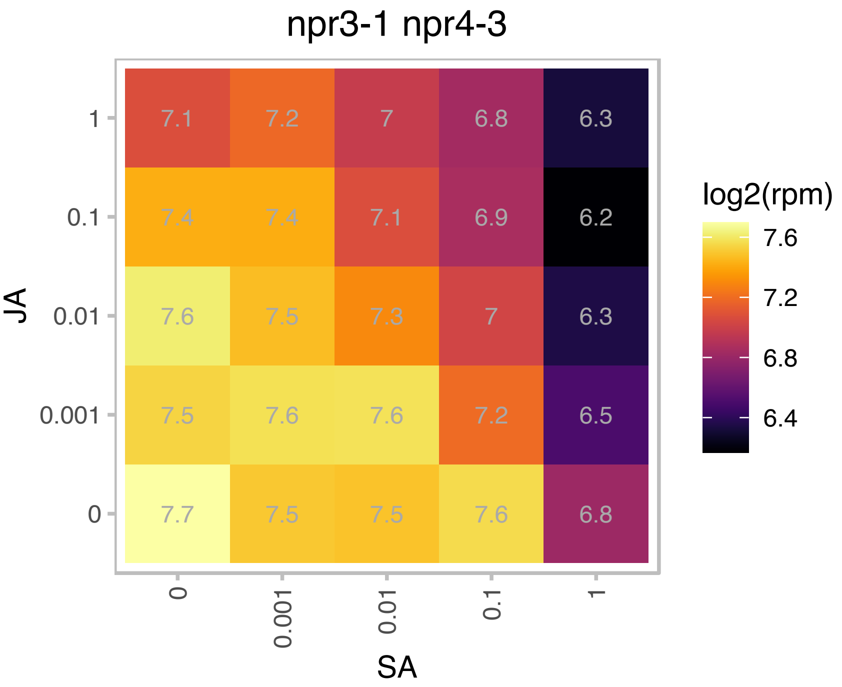

### AT2G23670.1 : YCF37

homolog of Synechocystis YCF37

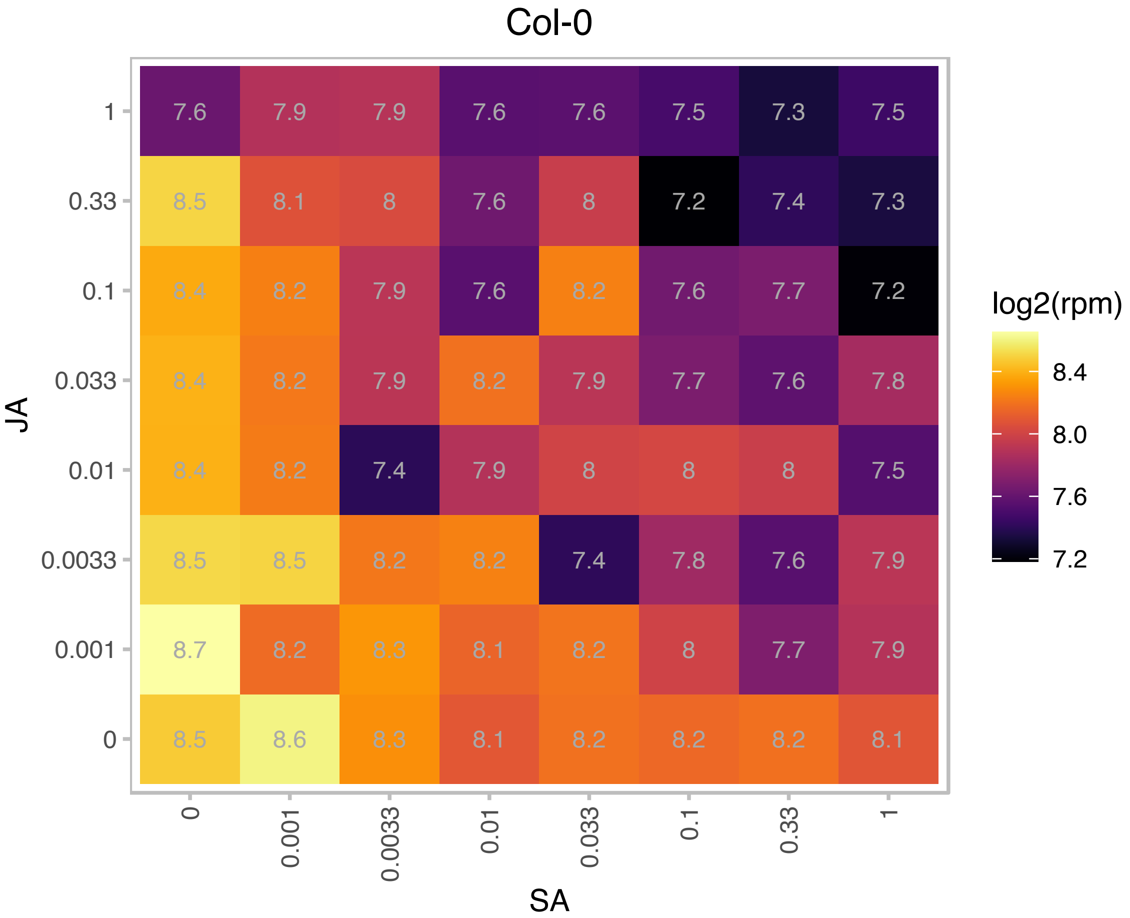


| Gene | AT2G23670 |
|---|---|
| is.expressed_genes | TRUE |
| is.responsive_genes | TRUE |
| is.apcluster | F-11 |

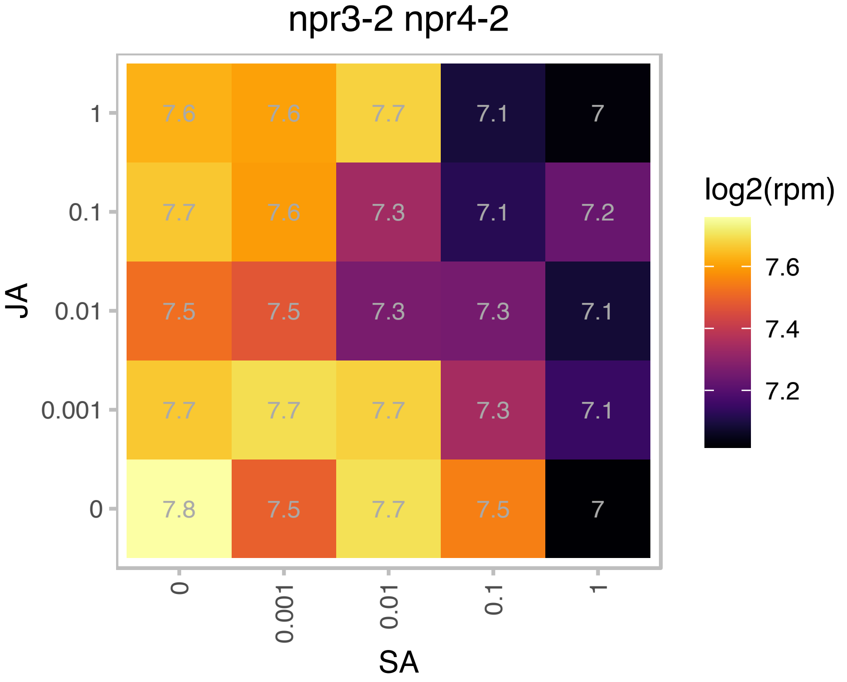


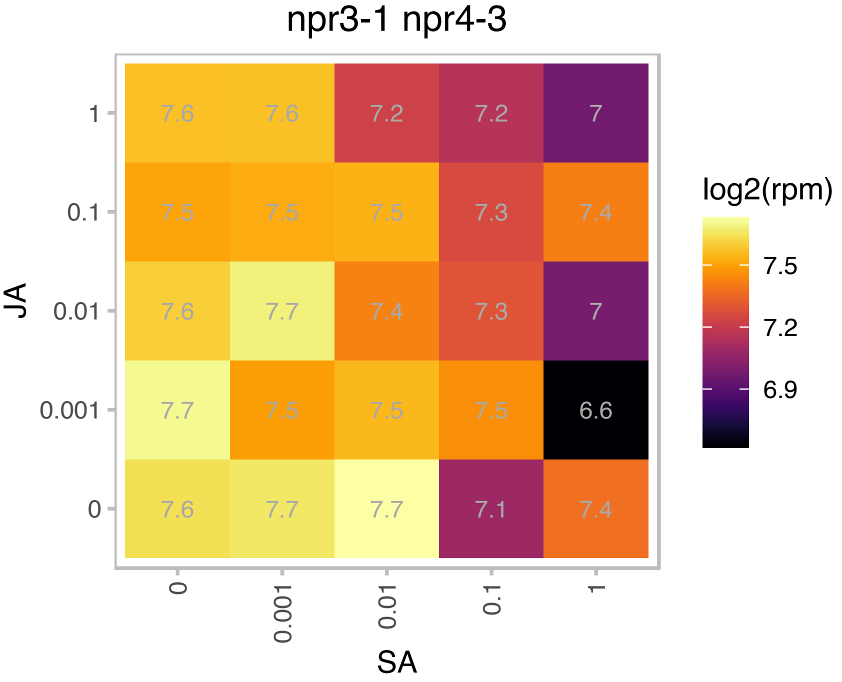

### AT2G25060.1 : AtENODL14, ENODL14

early nodulin-like protein 14

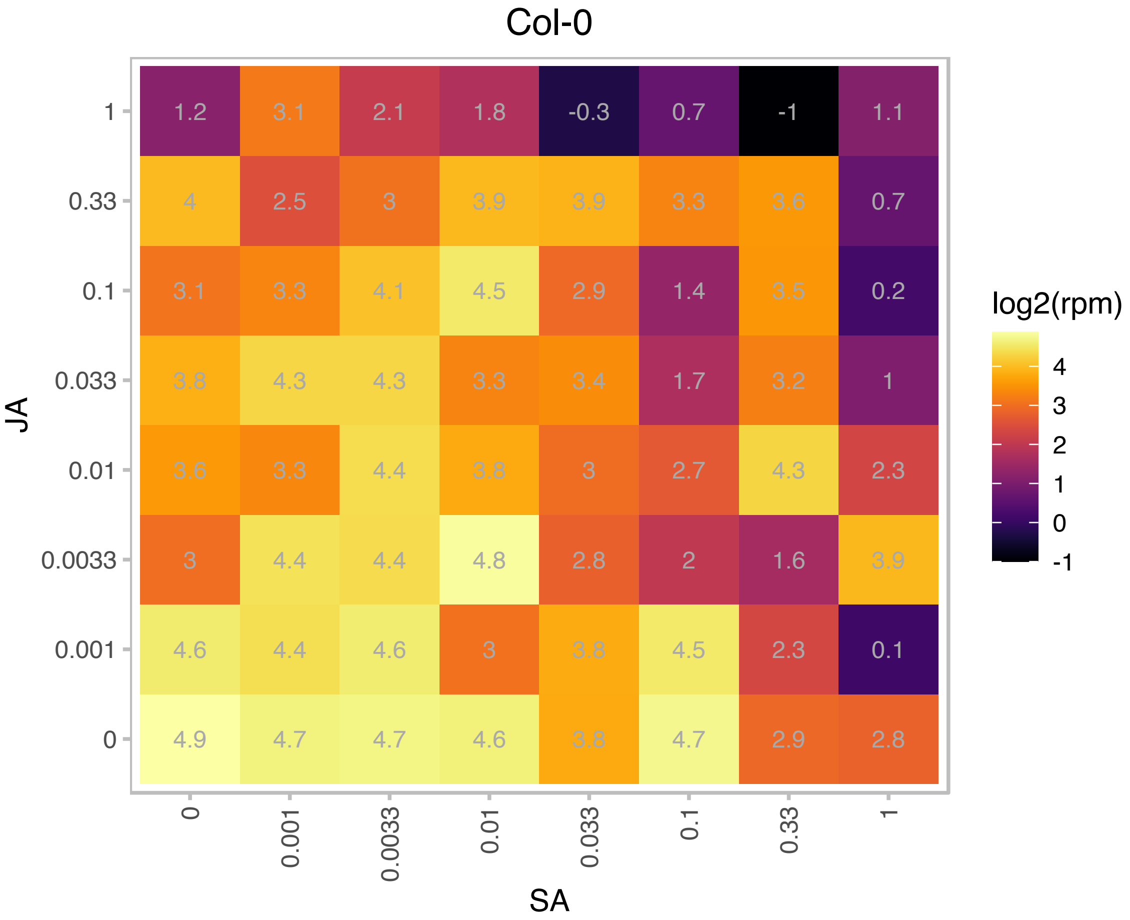


| | |
|---|---|
| Gene | AT2G25060 |
| is.expressed_genes | TRUE |
| is.responsive_genes | TRUE |
| is.apcluster | F-11 |

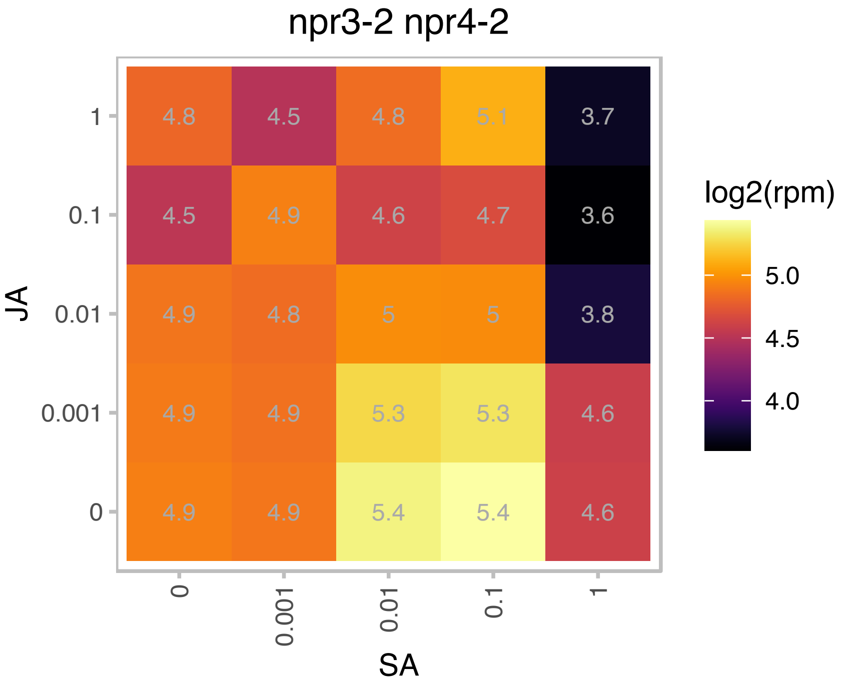


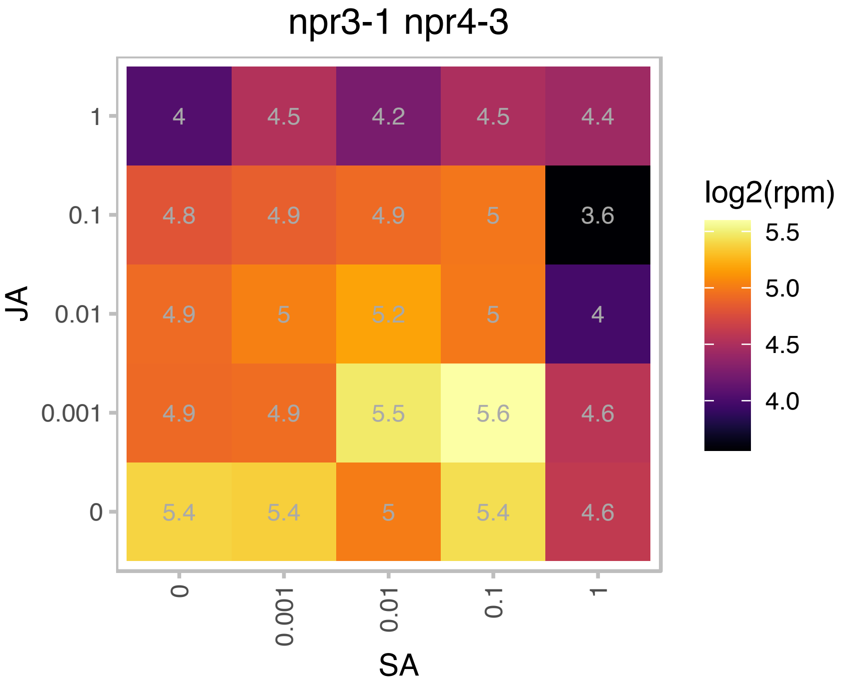

# AT2G28605.1 : NA

Photosystem II reaction center PsbP family protein

Encodes a PsbP domain-OEC23 like protein localized in thylakoid (peripheral-lumenal side).

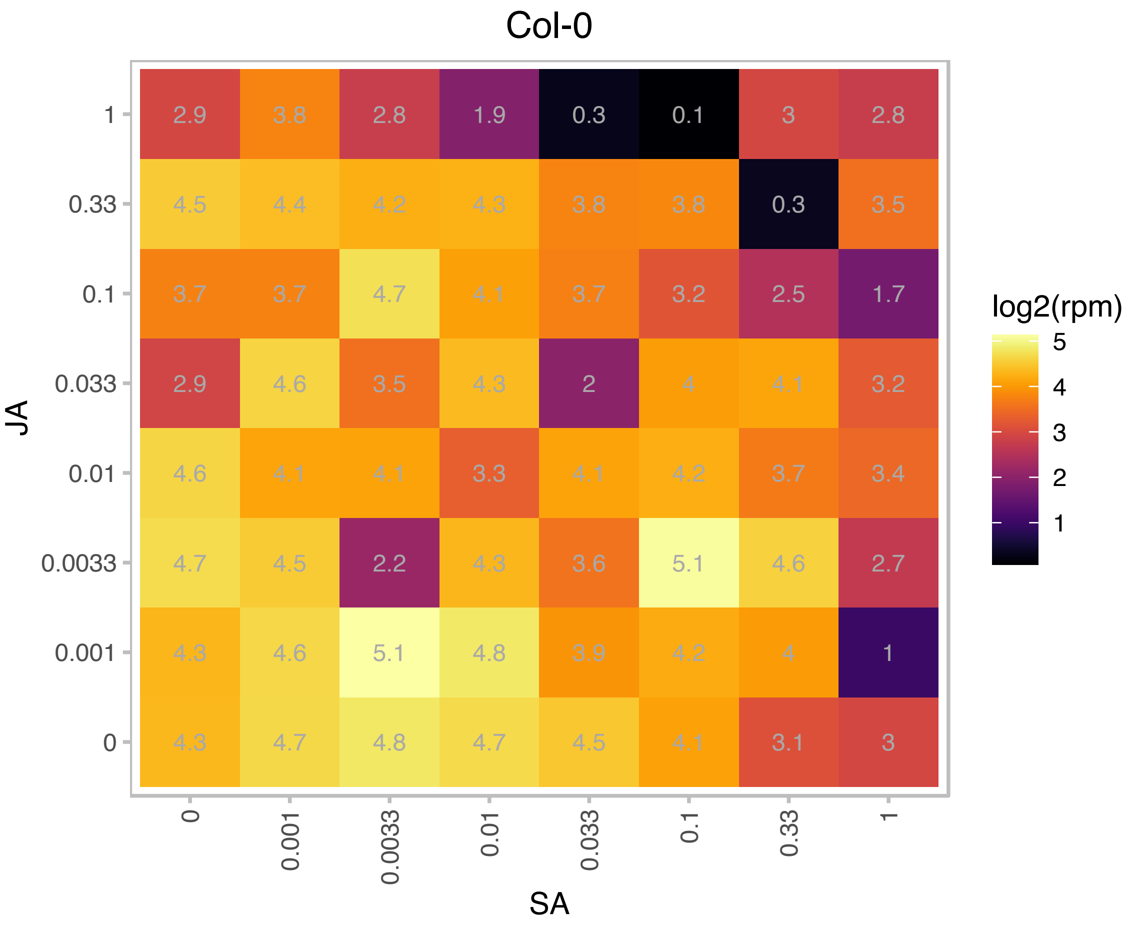


| | |
|---|---|
| Gene | AT2G28605 |
| is.expressed_genes | TRUE |
| is.responsive_genes | TRUE |
| is.apcluster | F-11 |

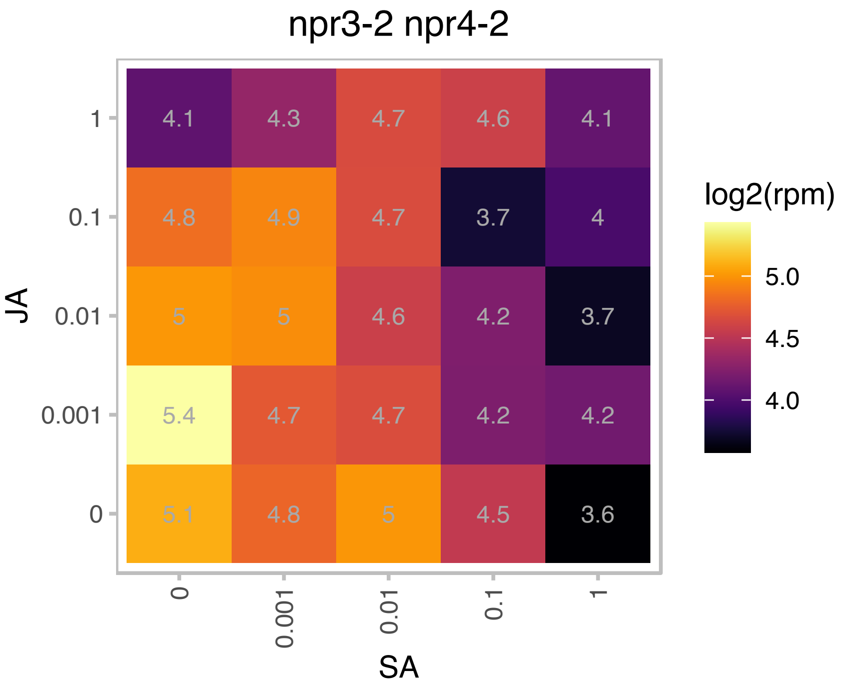


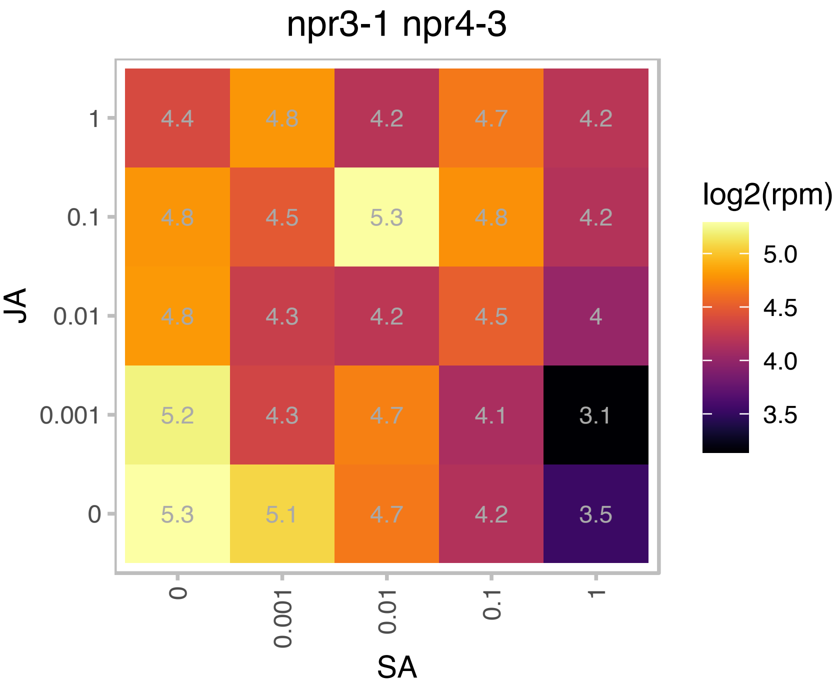

# AT2G29320.1 : NA

NAD(P)-binding Rossmann-fold superfamily protein

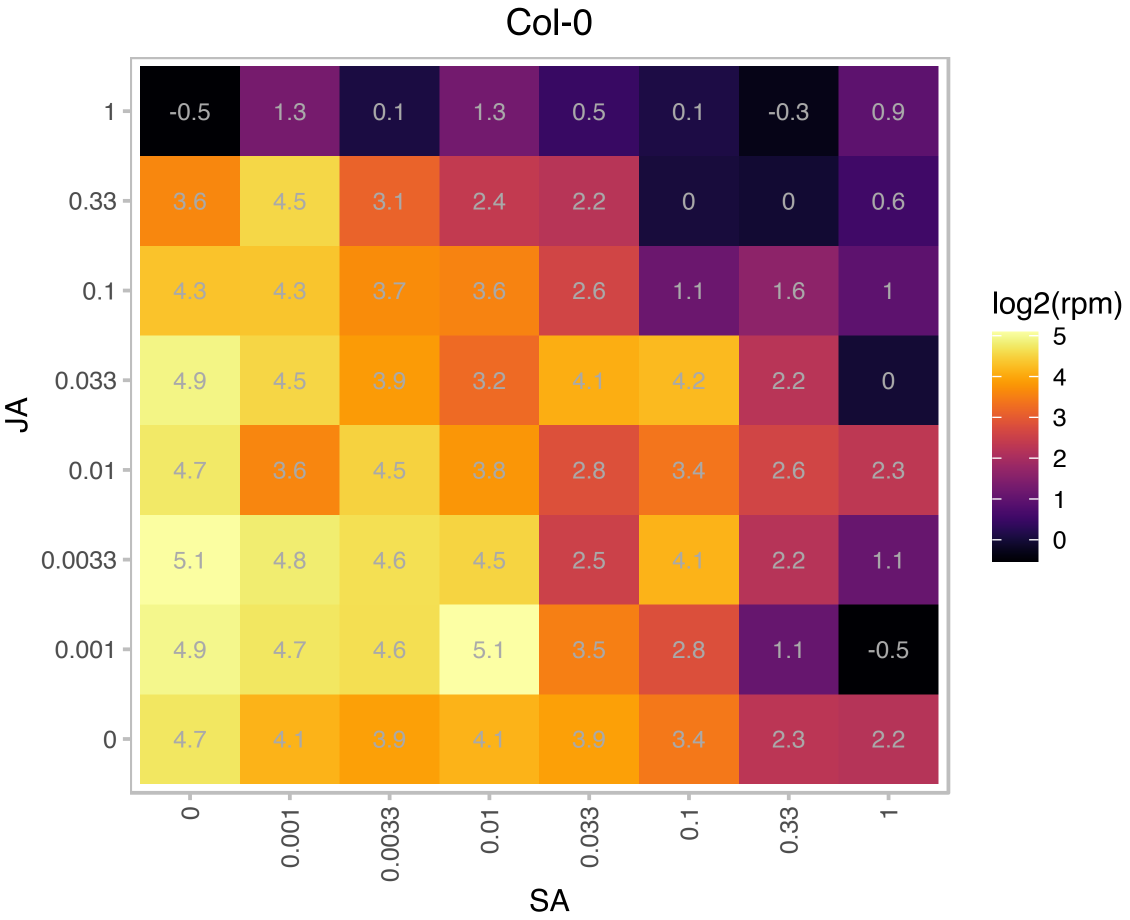


| | |
|---|---|
| Gene | AT2G29320 |
| is.expressed_genes | TRUE |
| is.responsive_genes | TRUE |
| is.apcluster | F-11 |

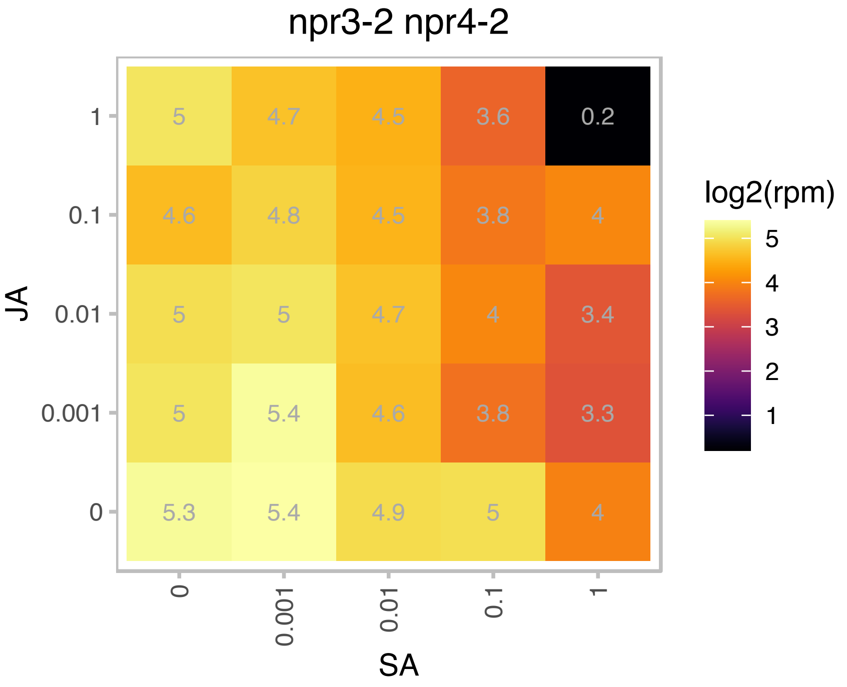


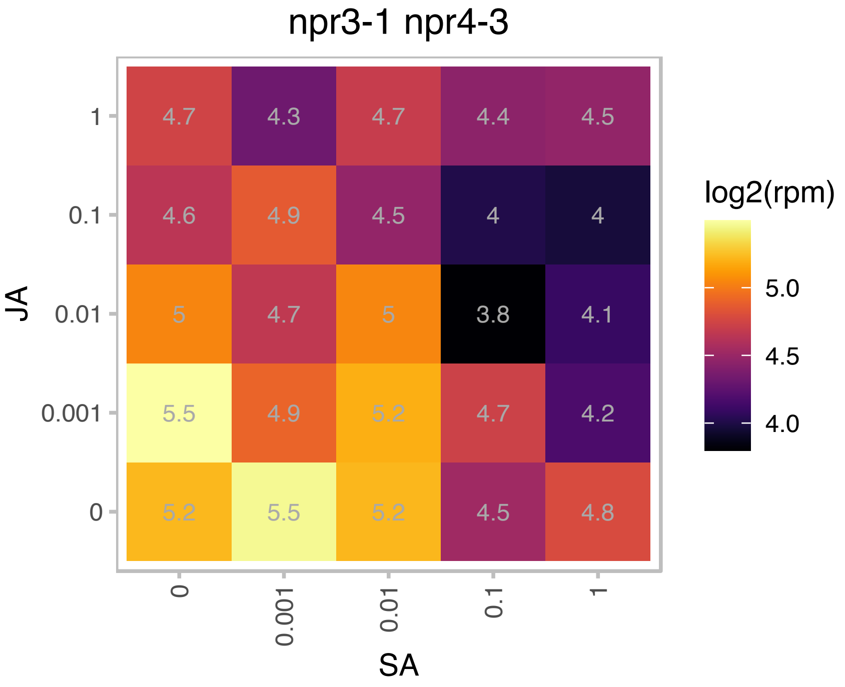

### AT2G31270.1 : ATCDT1A, CDT1, CDT1A

homolog of yeast CDT1 A

Encodes a cyclin-dependent protein kinase. Involved in nuclear DNA replication and plastid division. Located in nucleus and chloroplast.

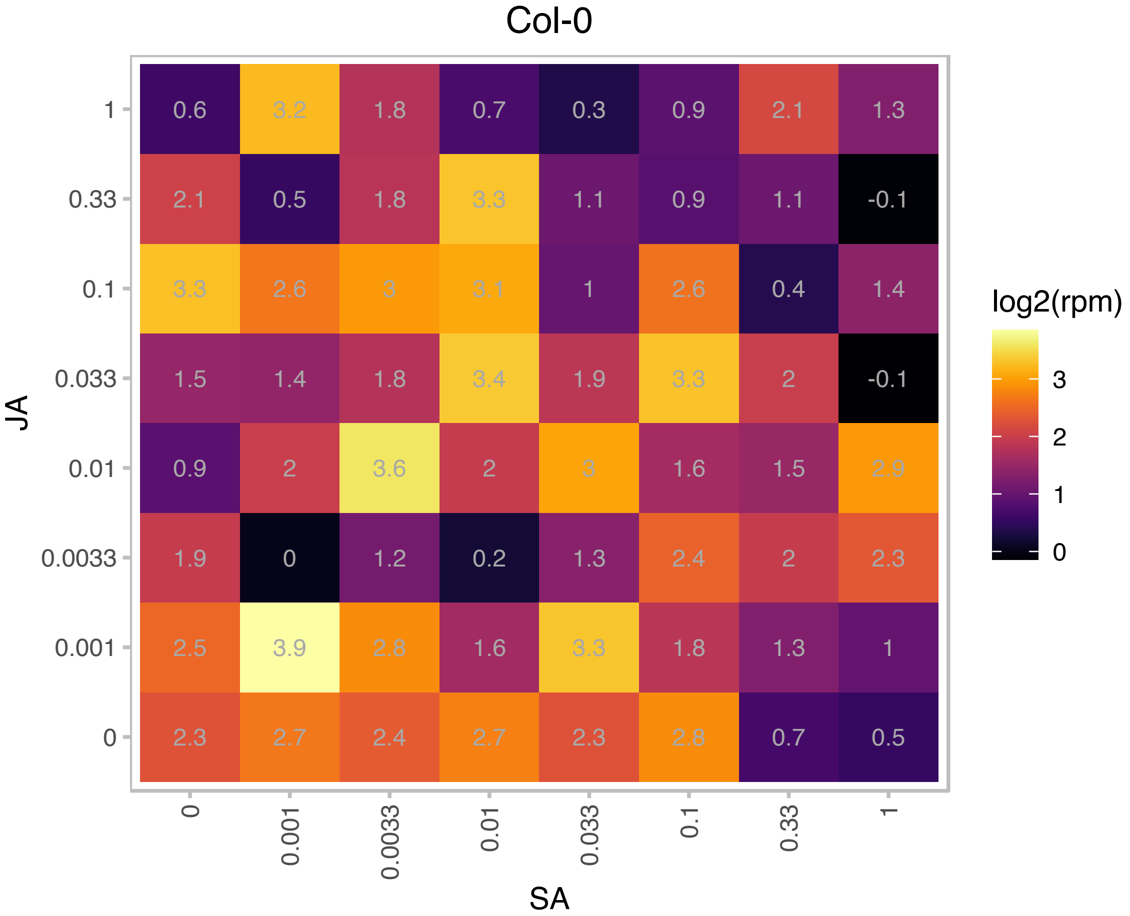


| | |
|---|---|
| Gene | AT2G31270 |
| is.expressed_genes | TRUE |
| is.responsive_genes | TRUE |
| is.apcluster | F-11 |

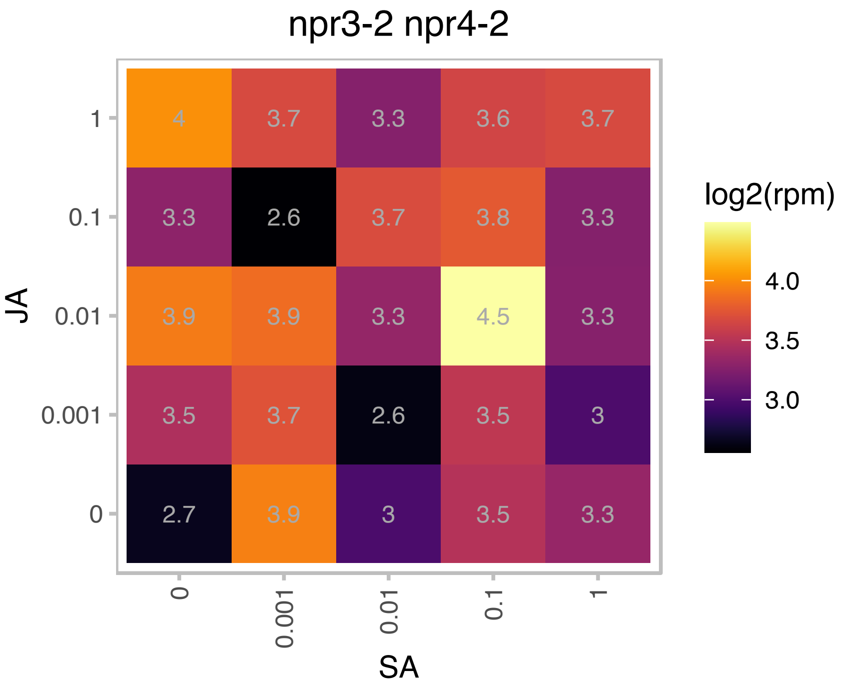


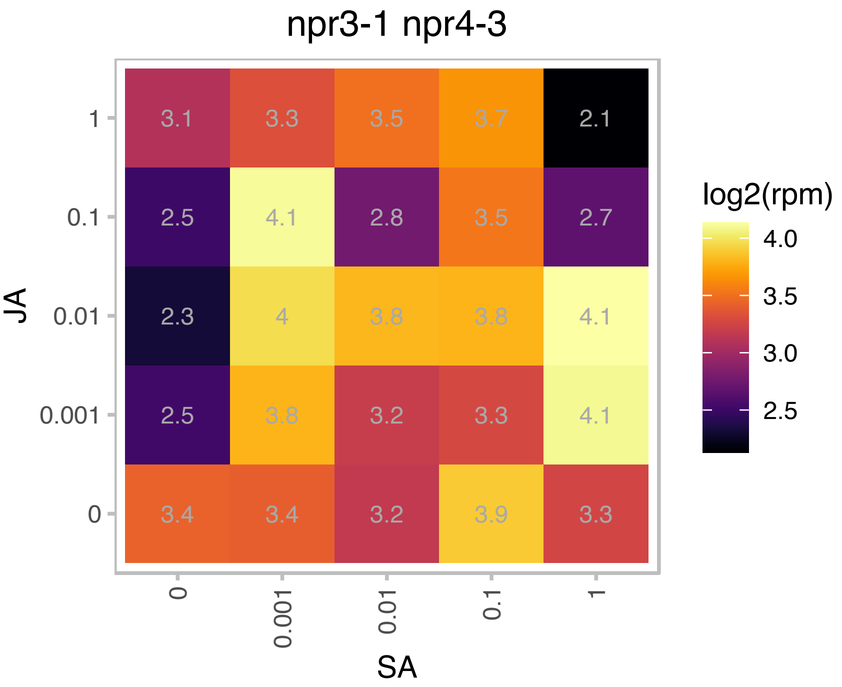

### AT2G32765.1 : ATSUMO5, SUM5, SUMO5

small ubiquitinrelated modifier 5
Encodes a small ubiquitin-like modifier (SUMO) protein that becomes covalently attached to various intracellular protein targets through an isopeptide bond. SUMOylation typically has a post-translational effect on the behavior of the target protein.

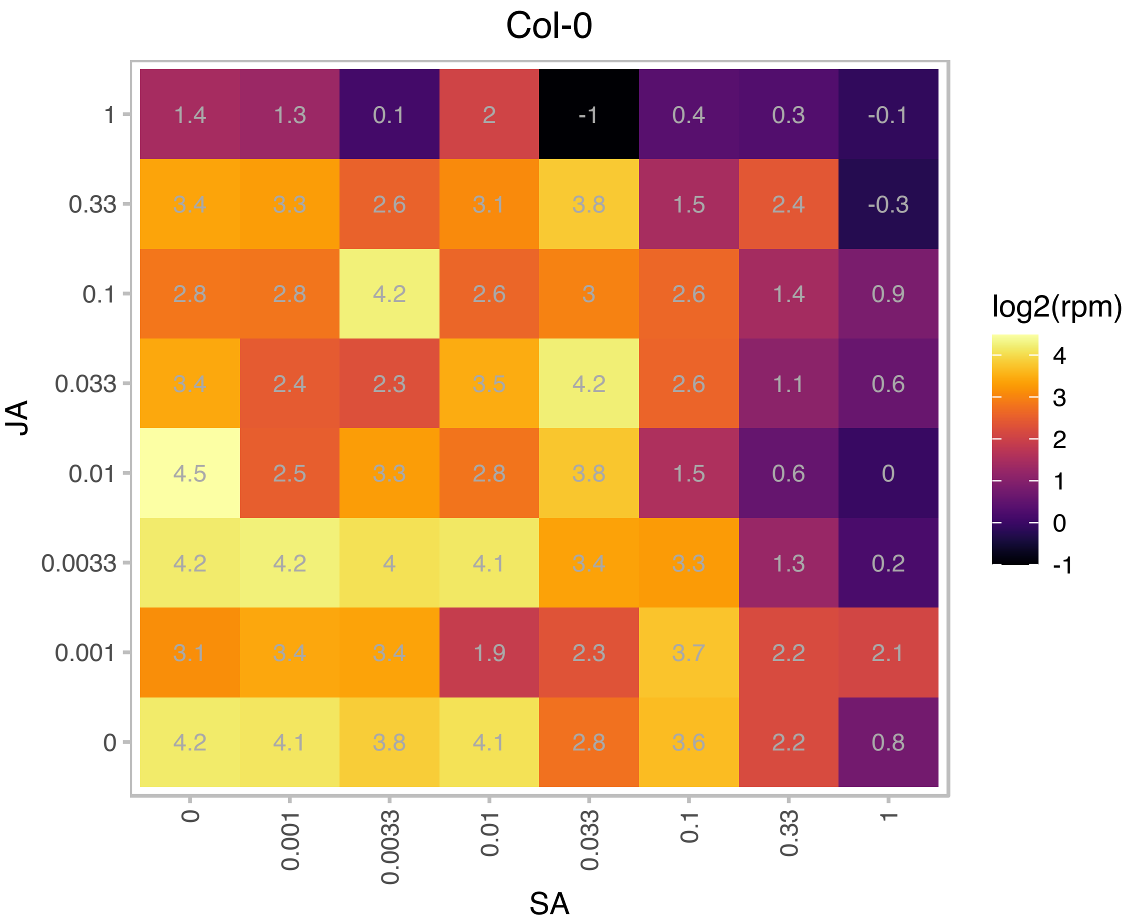


| | |
|---|---|
| Gene | AT2G32765 |
| is.expressed_genes | TRUE |
| is.responsive_genes | TRUE |
| is.apcluster | F-11 |

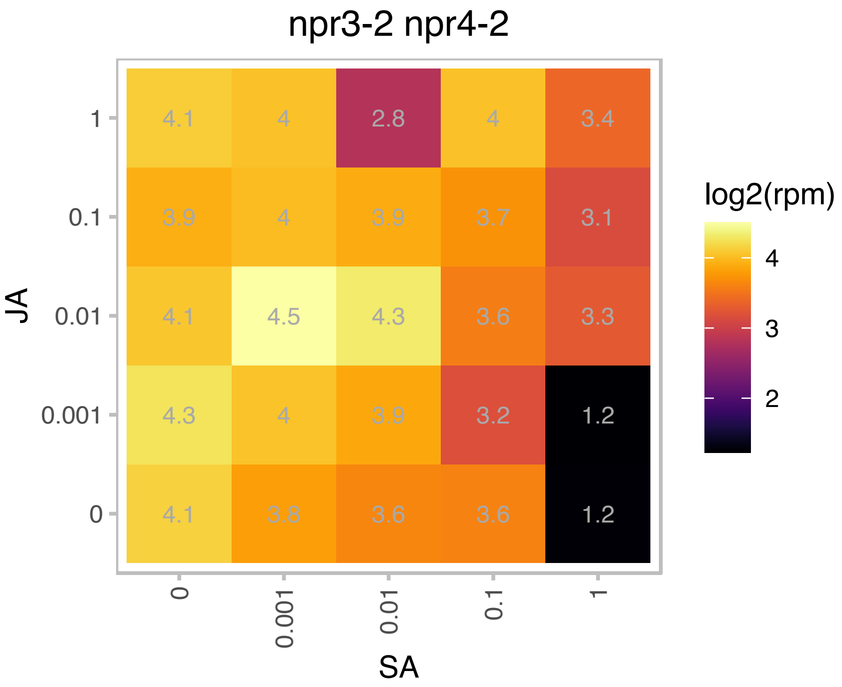


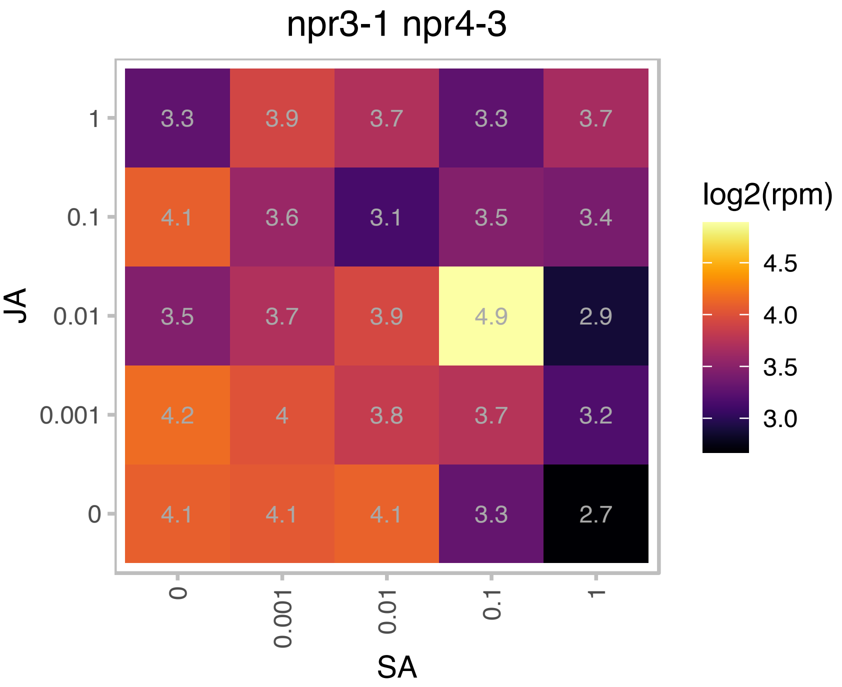

### AT2G36130.1 : Cyclophilin

Cyclophilin-like peptidyl-prolyl cis-trans isomerase family protein

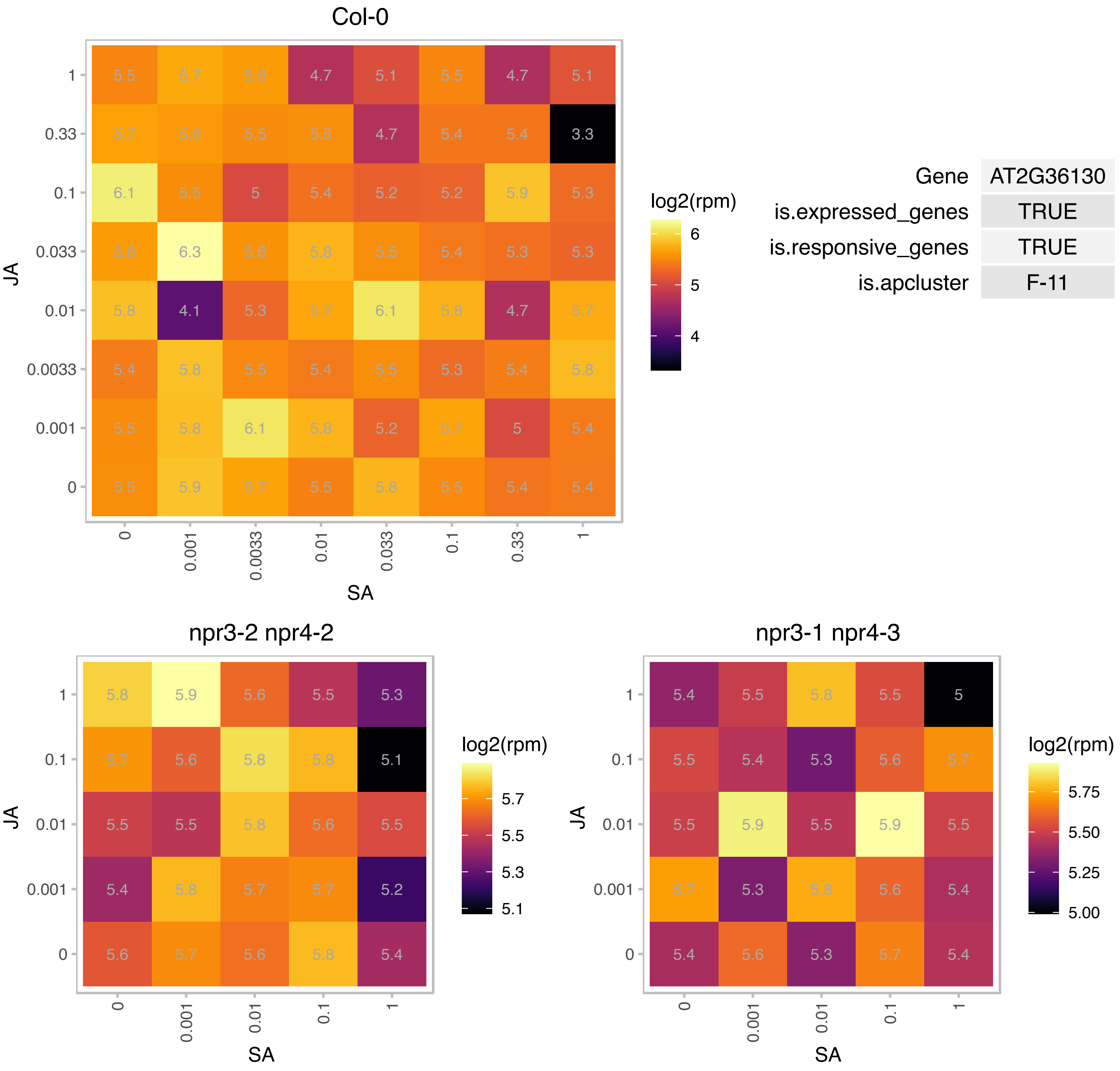

### AT2G36990.1 : ATSIG6, SIG6, SIGF, SOLDAT8

RNApolymerase sigma-subunit F

Encodes a general sigma factor in chloroplasts and is probably responsible for the recognition of sigma 70 type standard bacteria-type multi-subunit RNA polymerase (PEP) promoters in young cotyledons. It is a substrate for regulatory phosphorylation by cpCK2, a nuclear-coded plastid-targeted casein kinase 2, that has been implicated as a key component in plant sigma factor phosphorylation and transcriptional regulation.

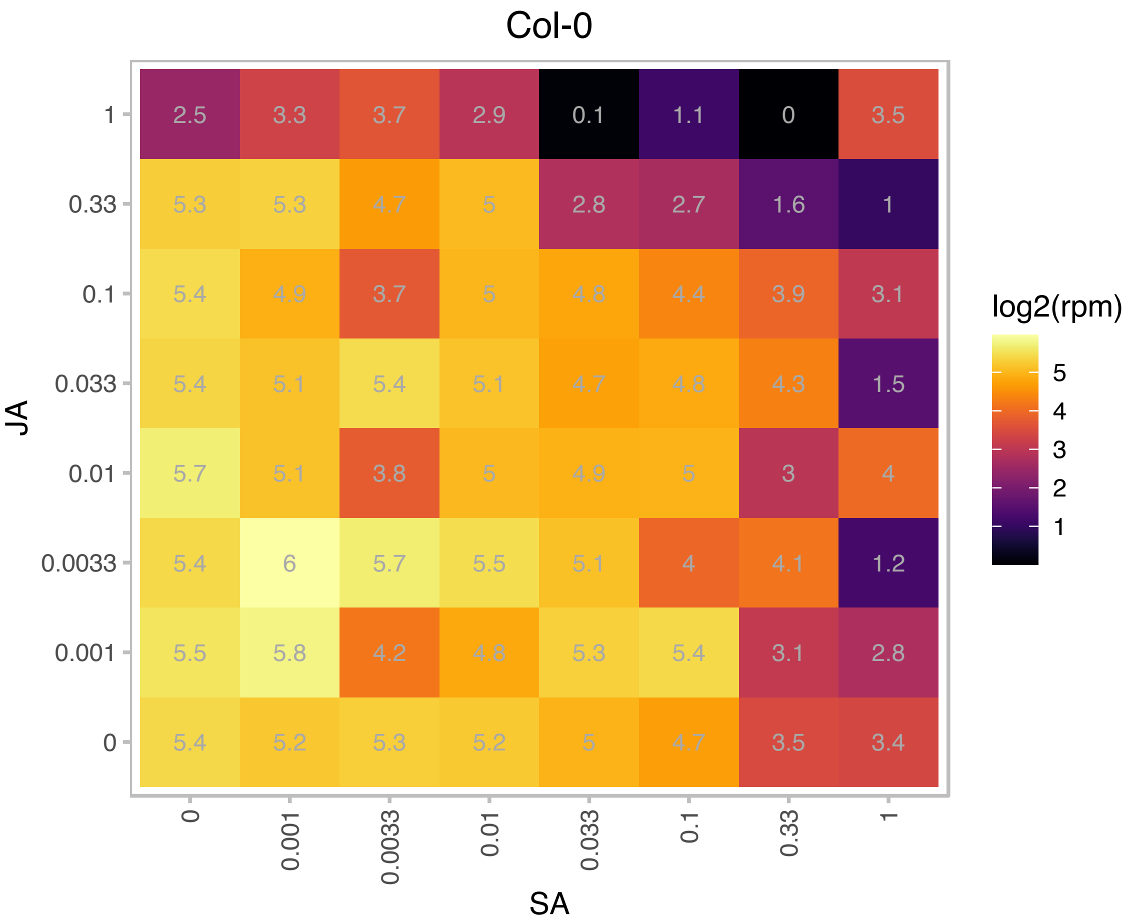


| | |
|---|---|
| Gene | AT2G36990 |
| is.expressed_genes | TRUE |
| is.responsive_genes | TRUE |
| is.apcluster | F-11 |

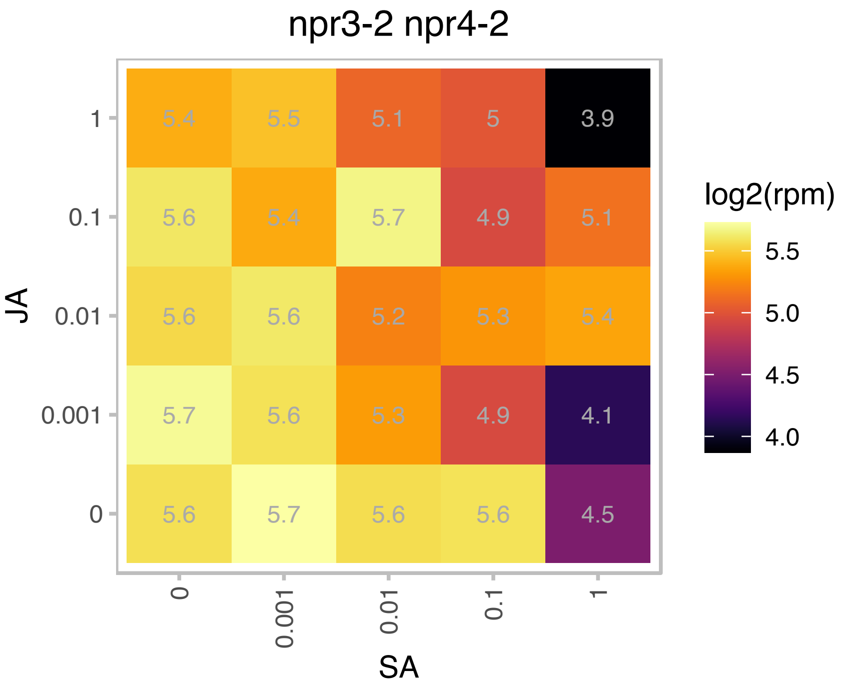


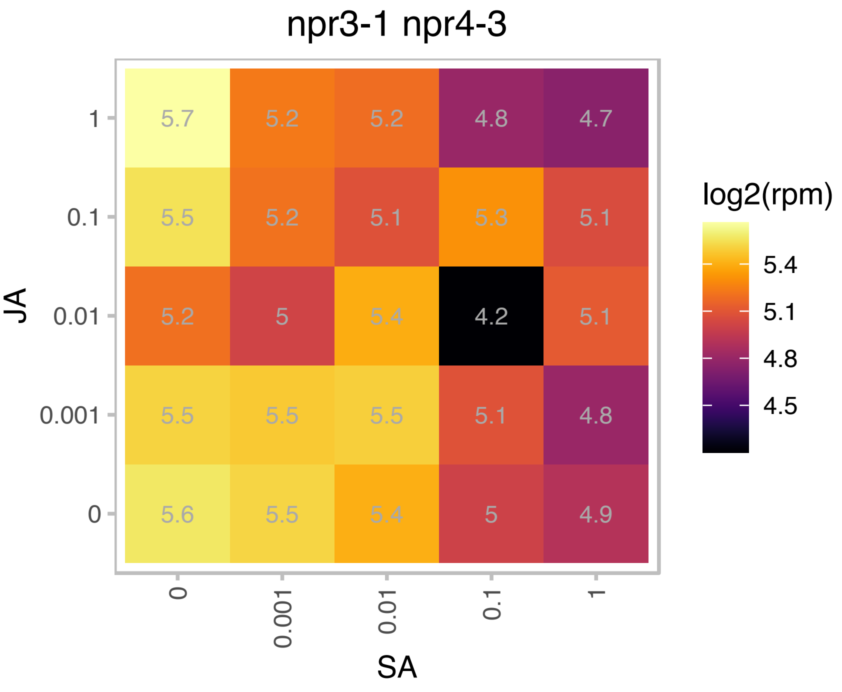

### AT2G38170.1 : ATCAX1, CAX1, RCI4

cation exchanger 1
Encodes a high affinity vacuolar calcium antiporter.  The residue His 338 is critical to Ca2+ transport activity. Disruption of CAX1 reduces manganese and zinc of shoot tissue and results in a decrease in the activity of vacuolar V-type proton ATPase.

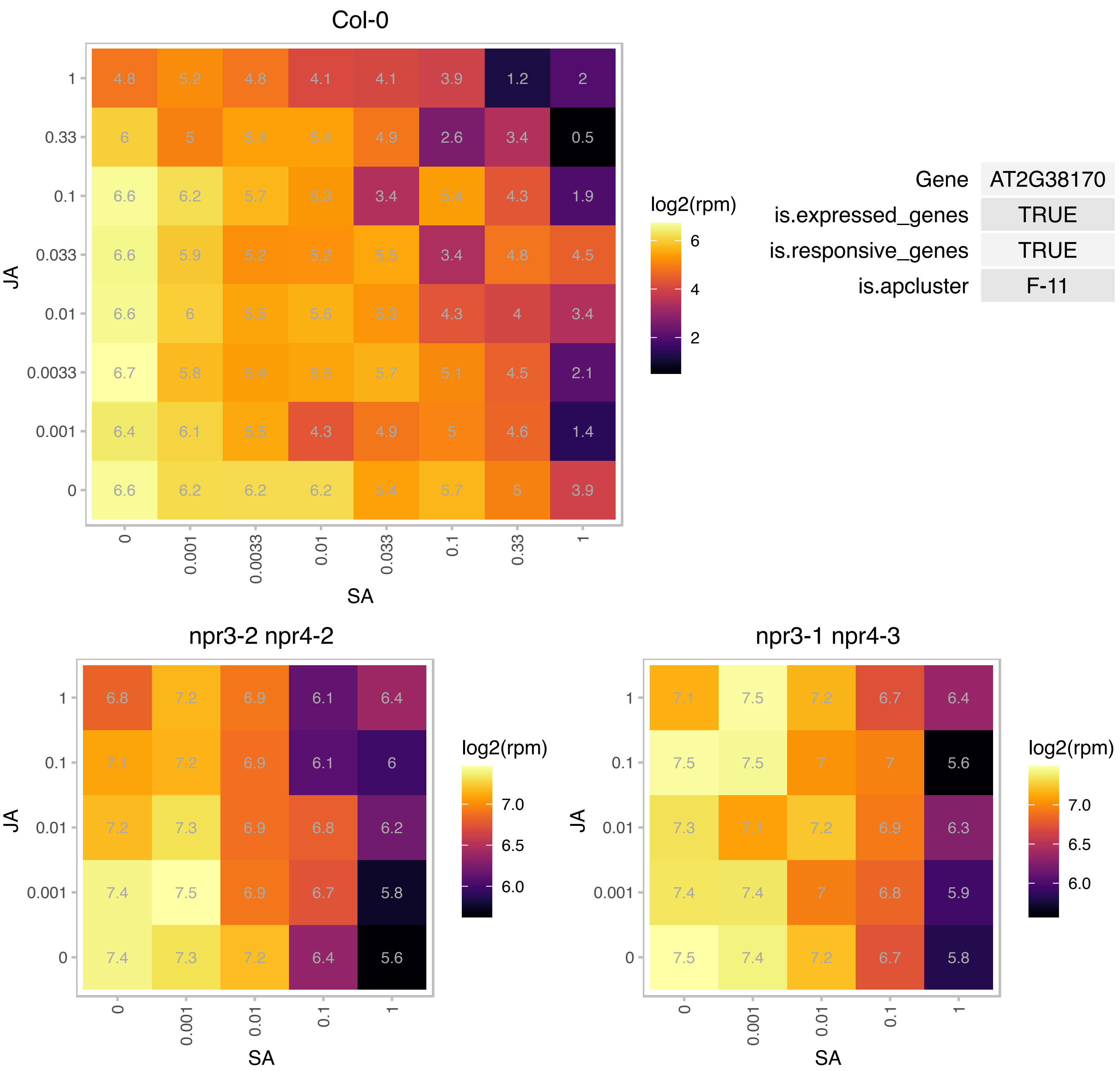


| | |
|---|---|
| Gene | AT2G38170 |
| is.expressed_genes | TRUE |
| is.responsive_genes | TRUE |
| is.apcluster | F-11 |

### AT2G38210.1 : PDX1L4

putative PDX1-like protein 4

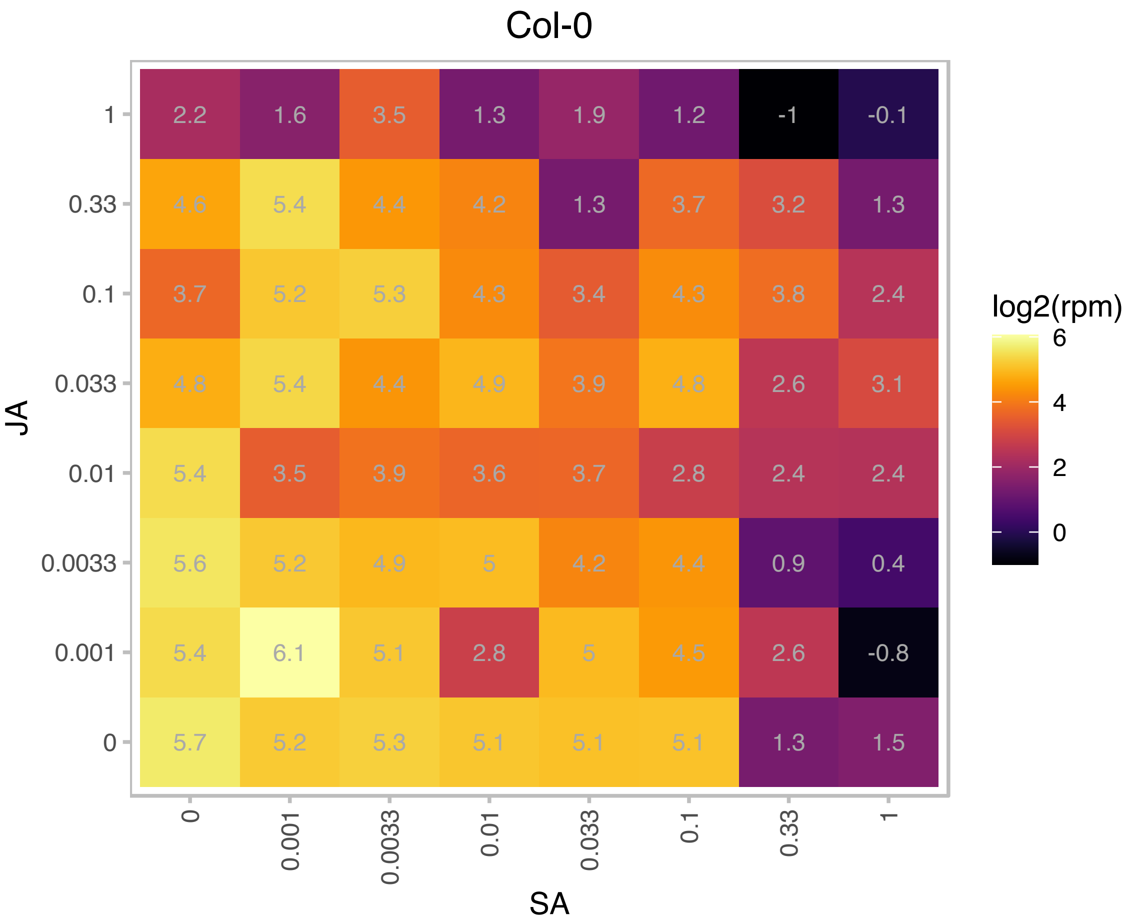


| | |
|---|---|
| Gene | AT2G38210 |
| is.expressed_genes | TRUE |
| is.responsive_genes | TRUE |
| is.apcluster | F-11 |

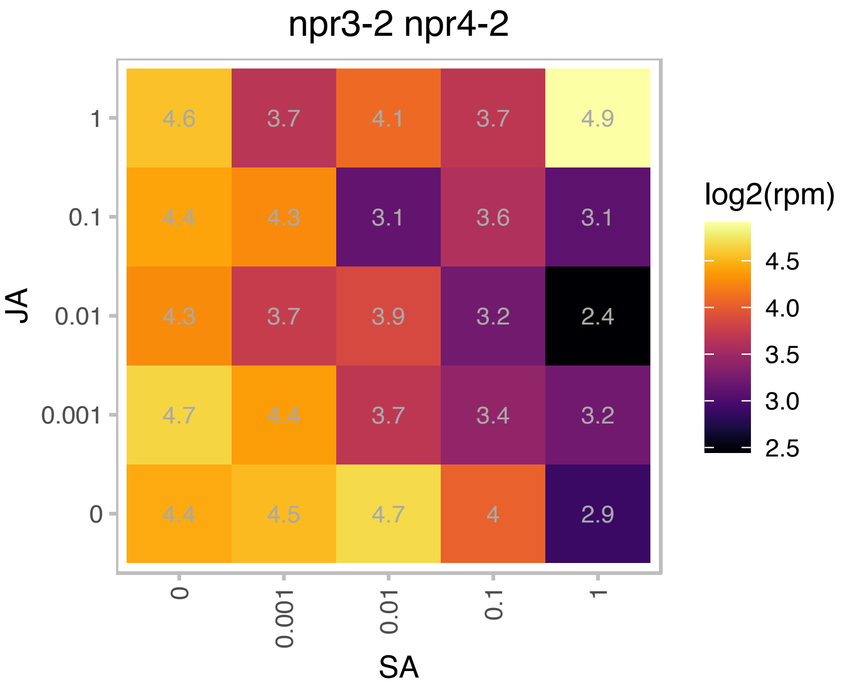


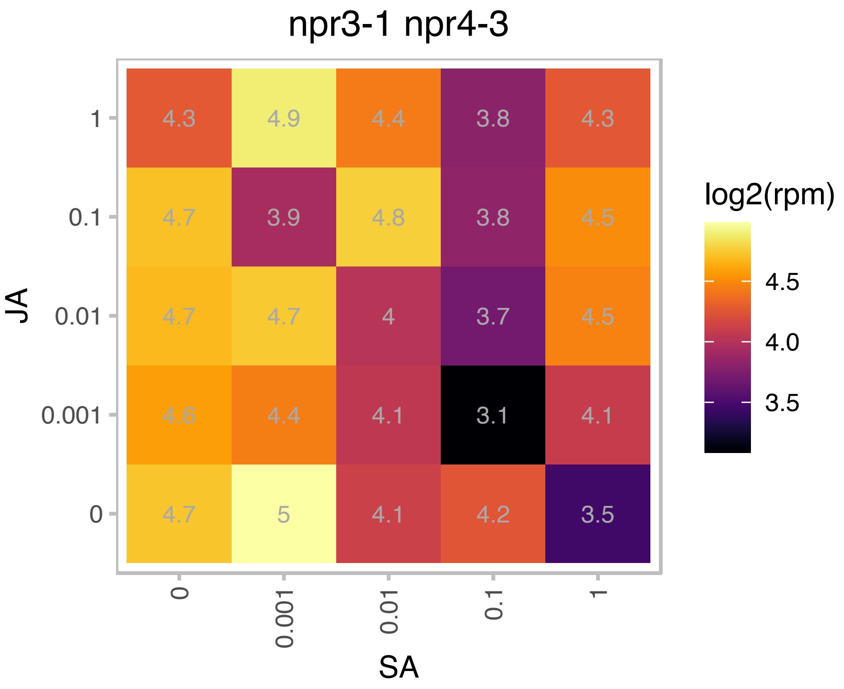

# AT3G02060.1 : NA

DEAD/DEAH box helicase, putative

| | |
|---|---|
| Gene | AT3G02060 |
| is.expressed_genes | TRUE |
| is.responsive_genes | TRUE |
| is.apcluster | F-11 |

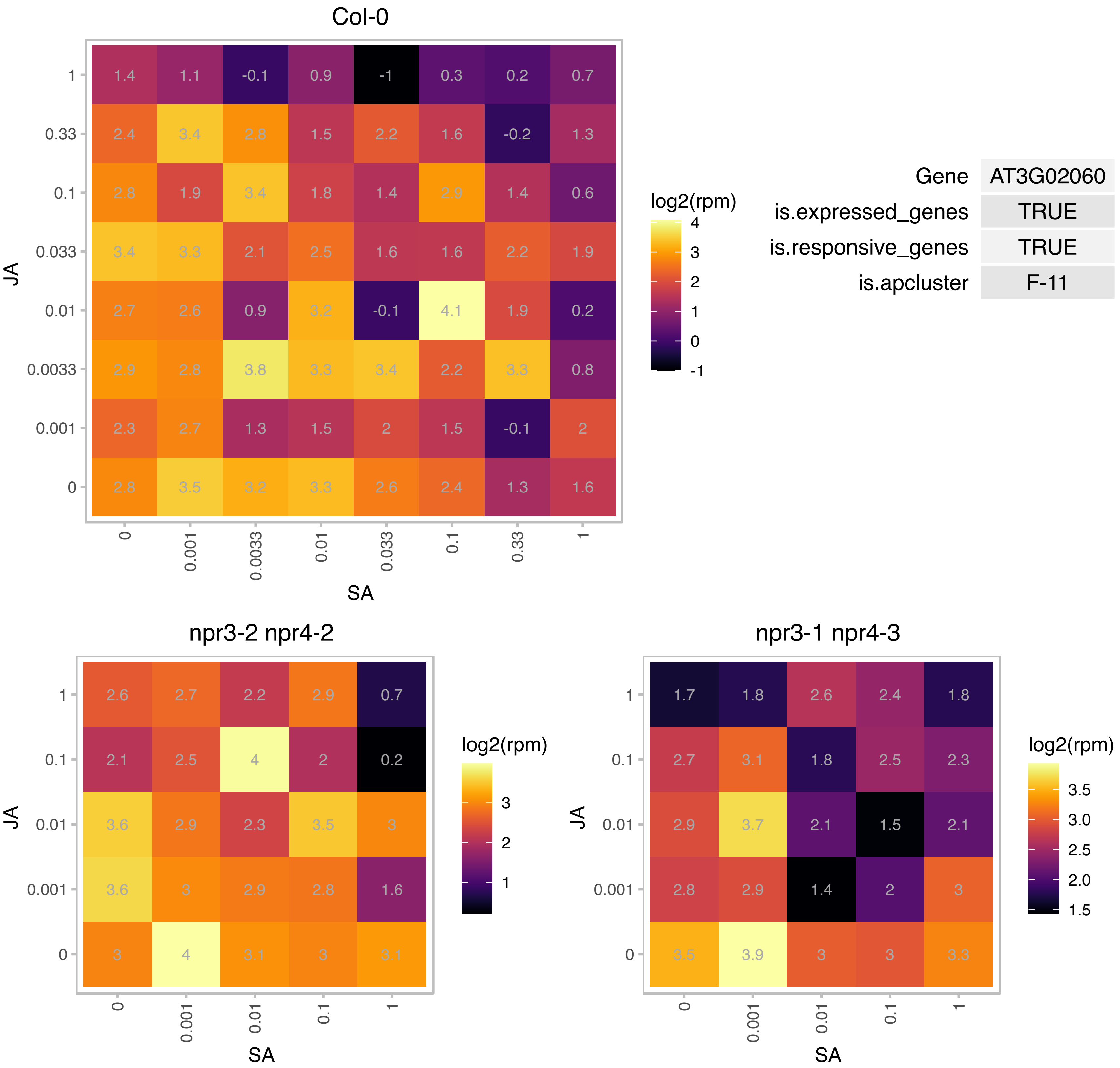

### AT3G02380.1 : ATCOL2, BBX3, COL2

CONSTANS-like 2

homologous to the flowering-time gene CONSTANS (CO) encoding zinc-finger proteins

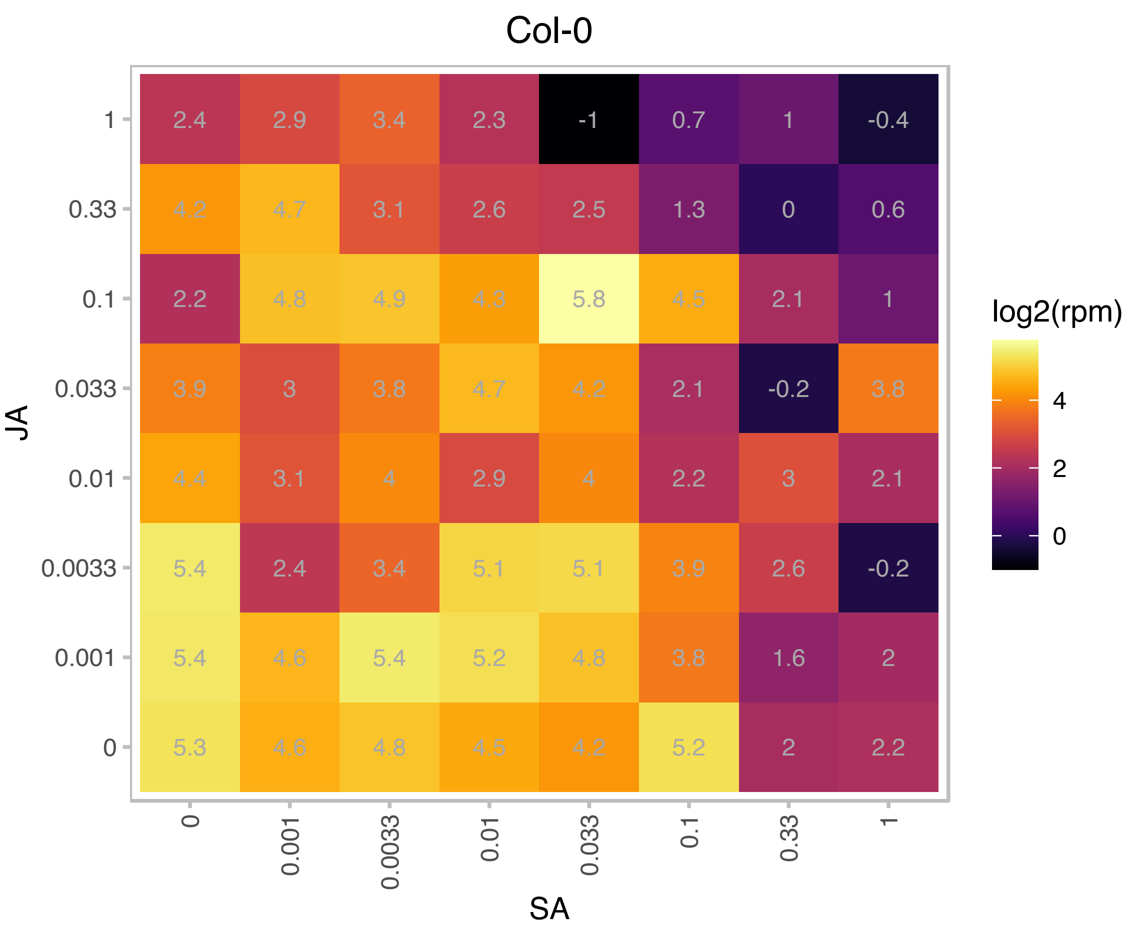


| | |
|---|---|
| Gene | AT3G02380 |
| is.expressed_genes | TRUE |
| is.responsive_genes | TRUE |
| is.apcluster | F-11 |

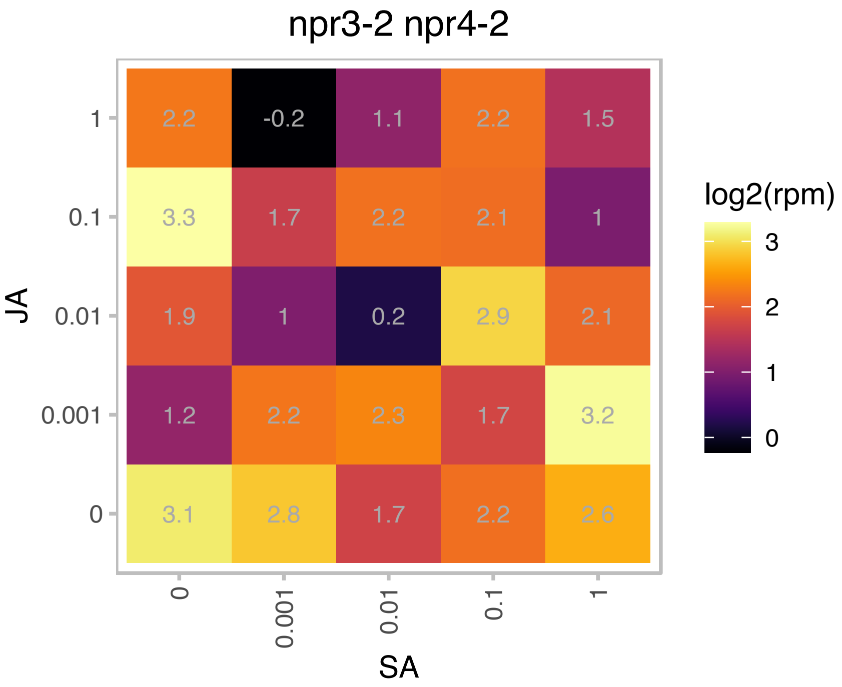


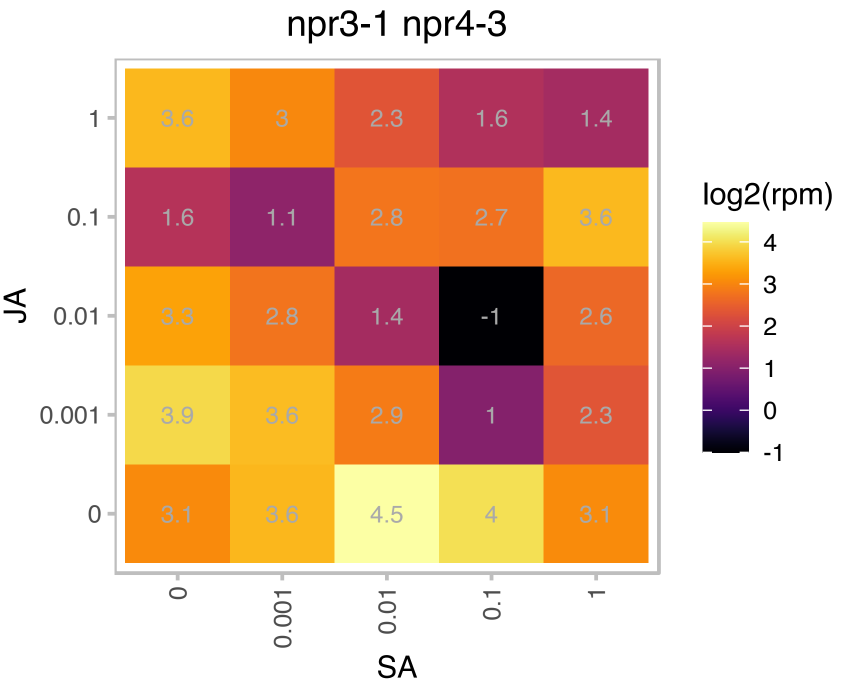

# AT3G06130.1 : NA

Heavy metal transport/detoxification superfamily protein

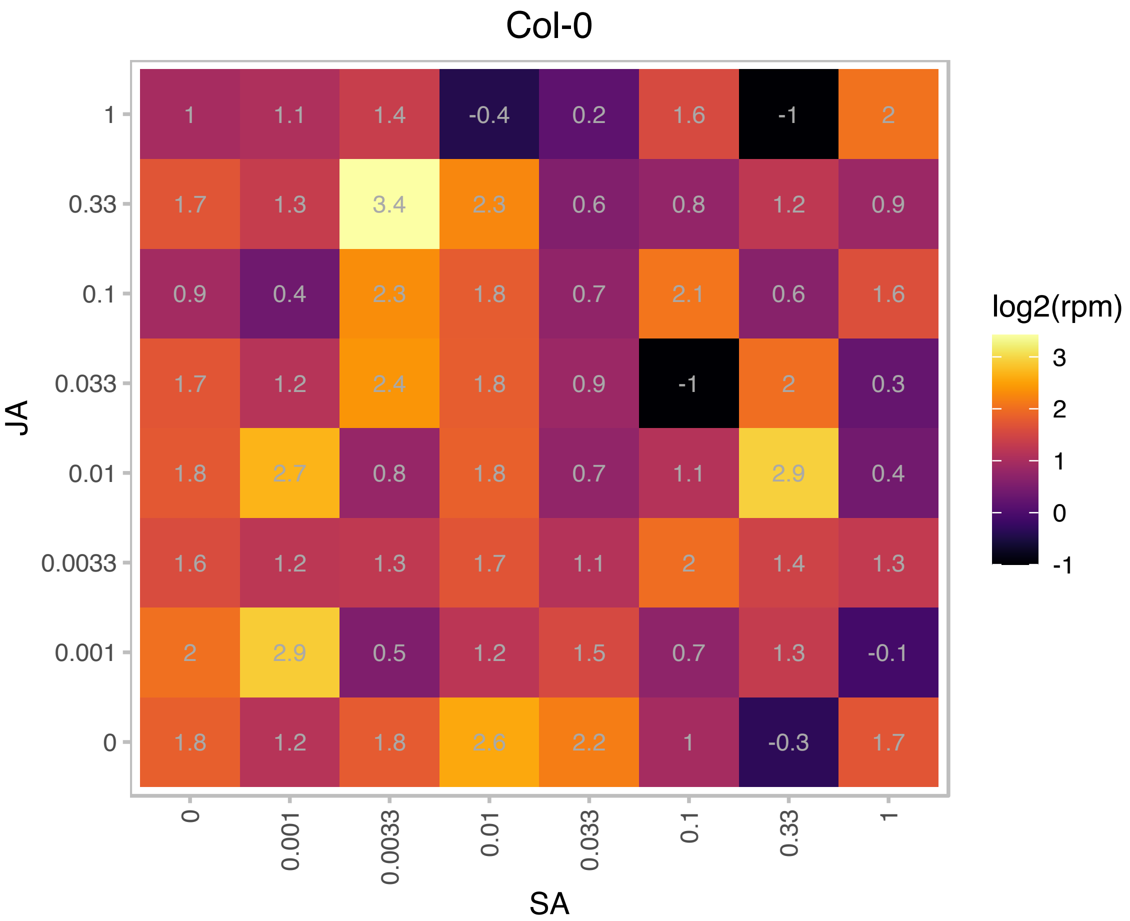


| | |
|---|---|
| Gene | AT3G06130 |
| is.expressed_genes | TRUE |
| is.responsive_genes | TRUE |
| is.apcluster | F-11 |

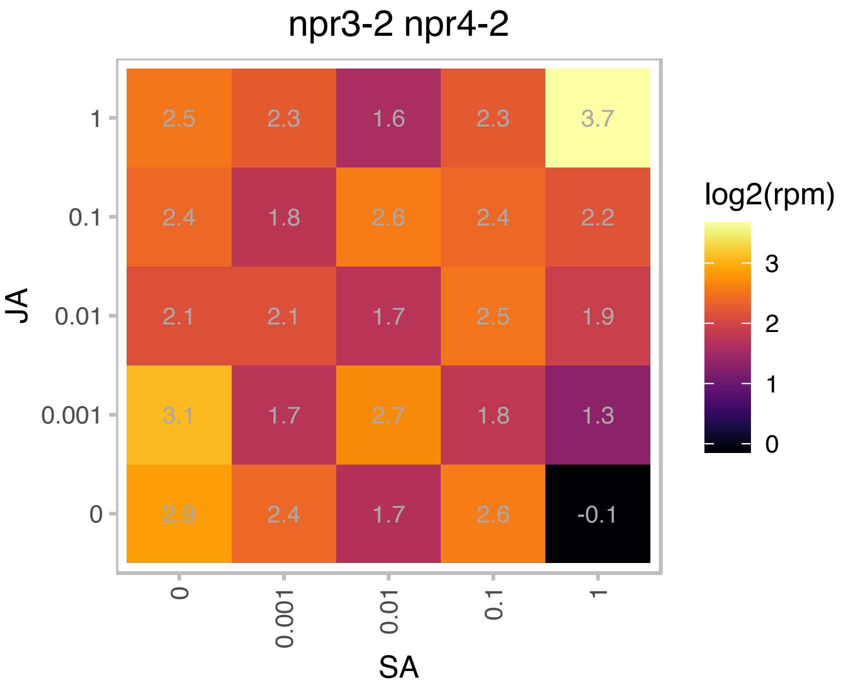


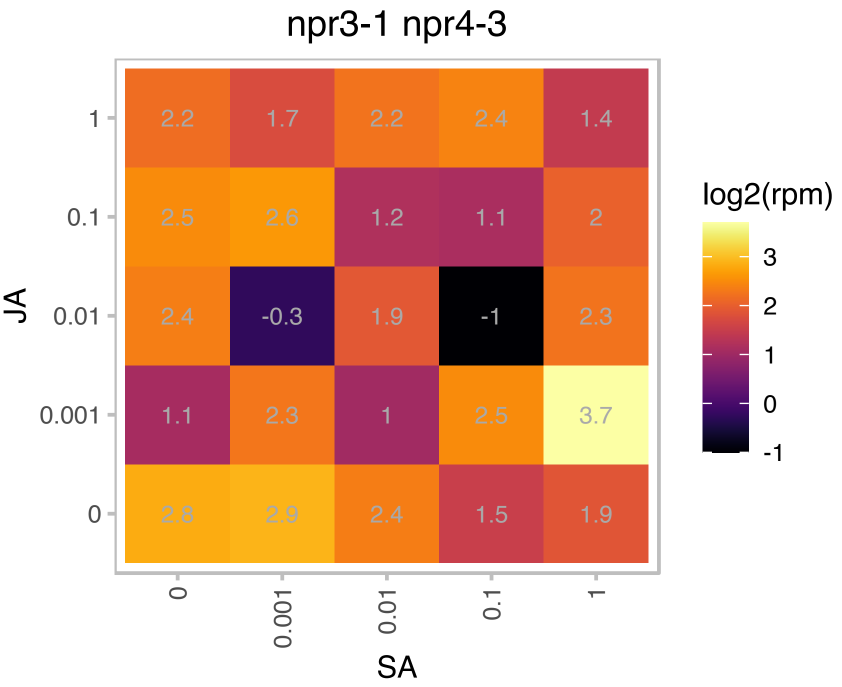

# AT3G06840.1 : NA

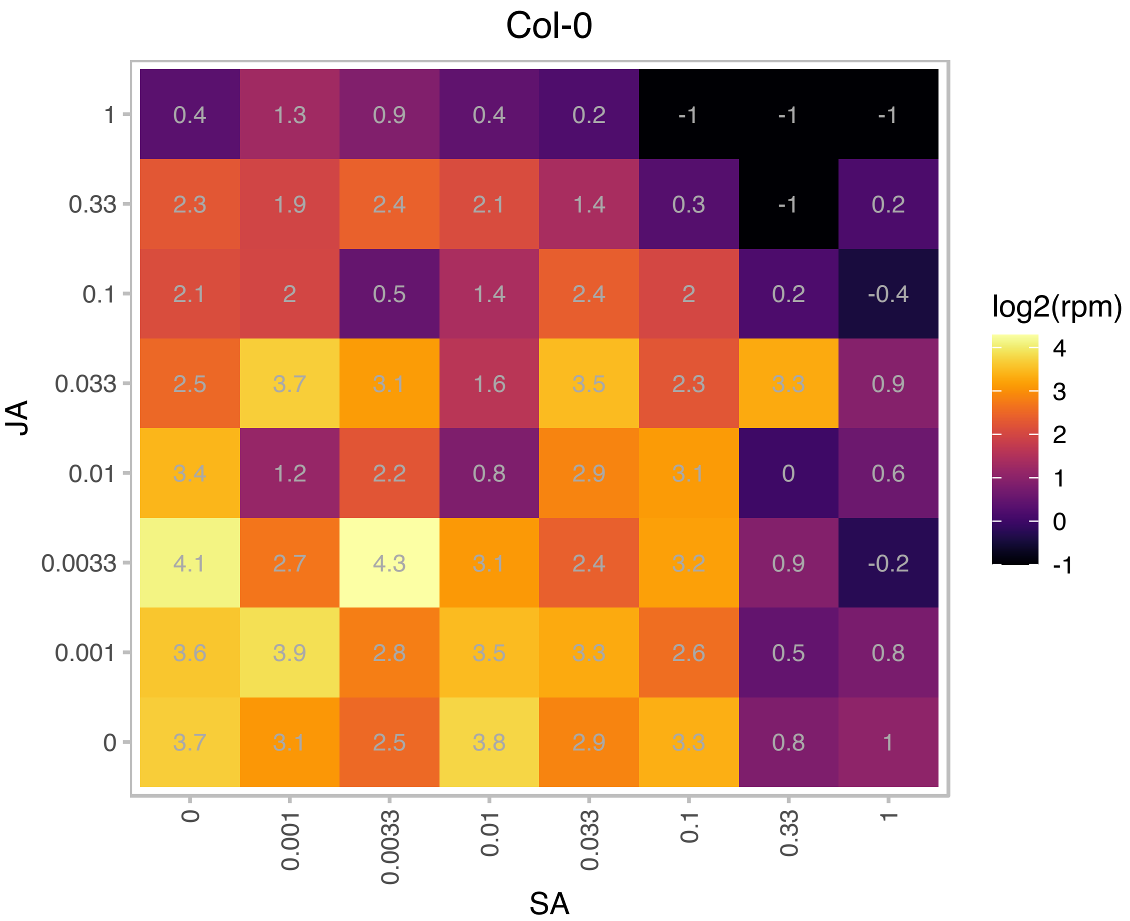


| | |
|---|---|
| Gene | AT3G06840 |
| is.expressed_genes | TRUE |
| is.responsive_genes | TRUE |
| is.apcluster | F-11 |

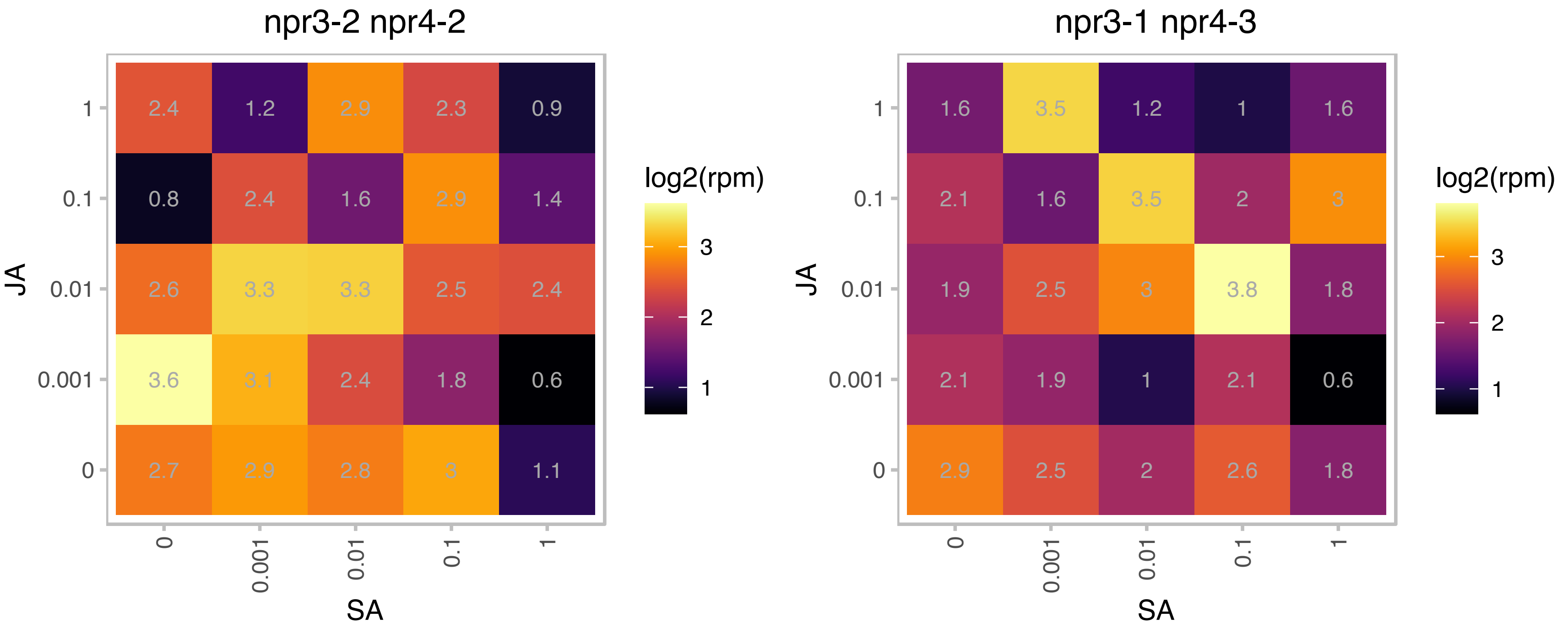

# AT3G06980.1 : NA

DEA(D/H)-box RNA helicase family protein

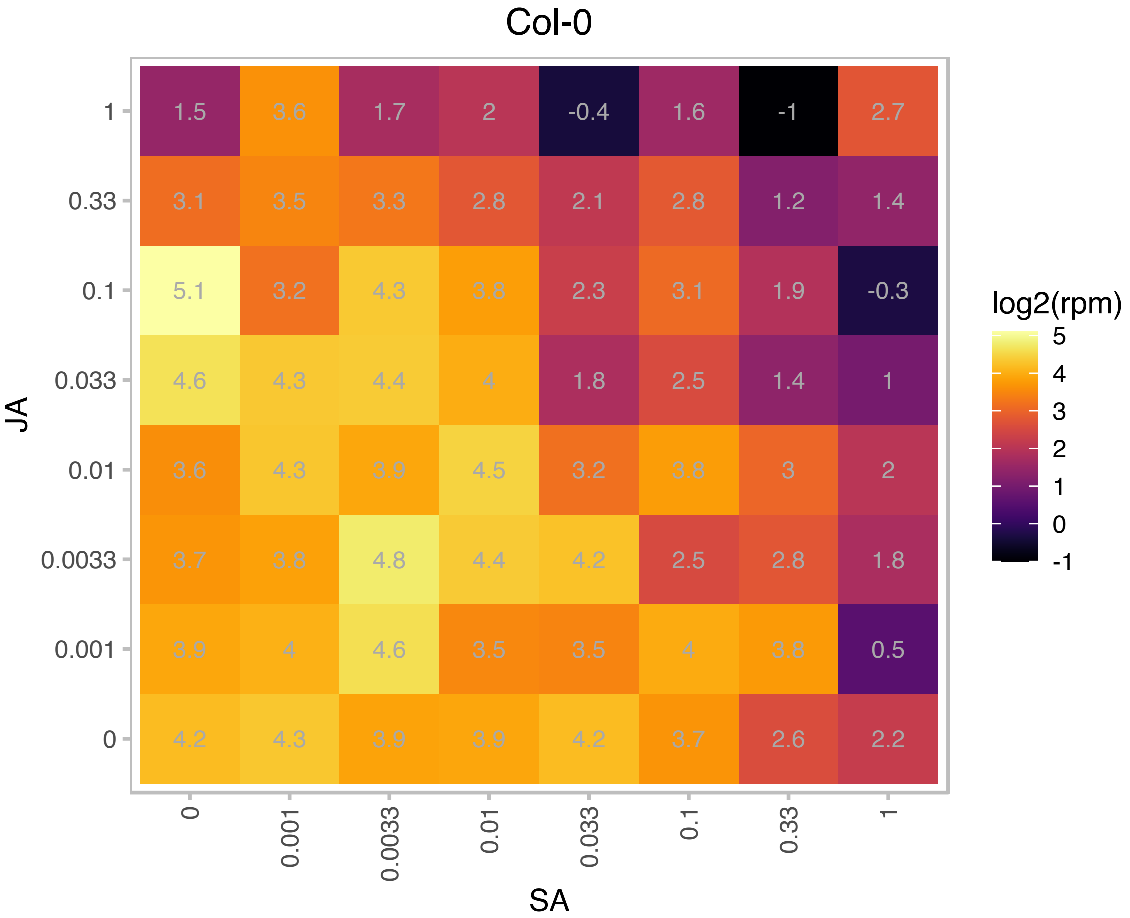


| | |
|---|---|
| Gene | AT3G06980 |
| is.expressed_genes | TRUE |
| is.responsive_genes | TRUE |
| is.apcluster | F-11 |

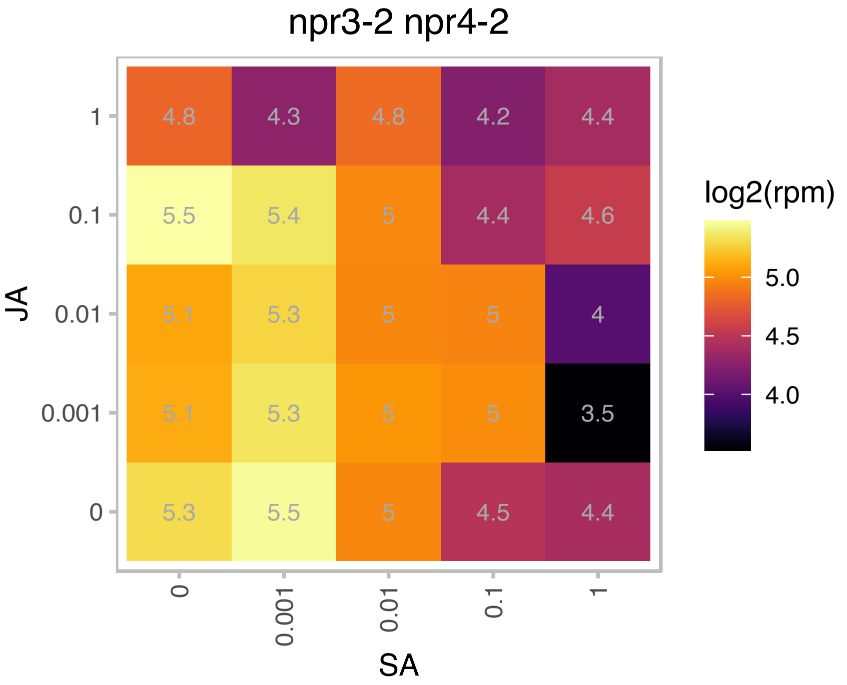


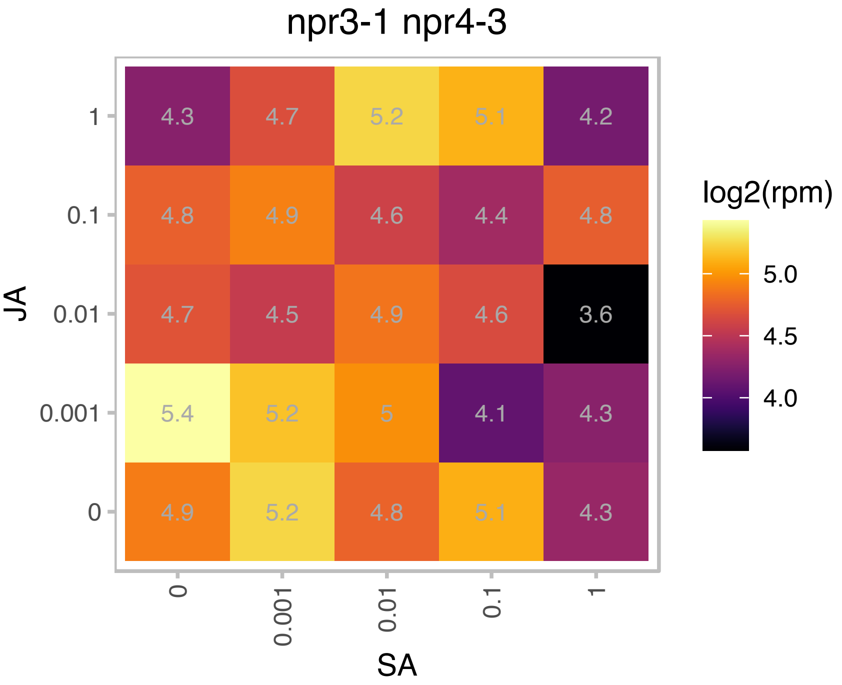

### AT3G08770.1 : AtLtpI-6, LTP6

lipid transfer protein 6

Predicted to encode a PR (pathogenesis-related) protein. Belongs to the lipid transfer protein (PR-14) family with the following members: At2g38540/LTP1, At2g38530/LTP2, At5g59320/LTP3, At5g59310/LTP4, At3g51600/LTP5, At3g08770/LTP6, At2g15050/LTP7, At2g18370/LTP8, At2g15325/LTP9, At5g01870/LTP10, At4g33355/LTP11, At3g51590/LTP12, At5g44265/LTP13, At5g62065/LTP14, At4g08530/LTP15.

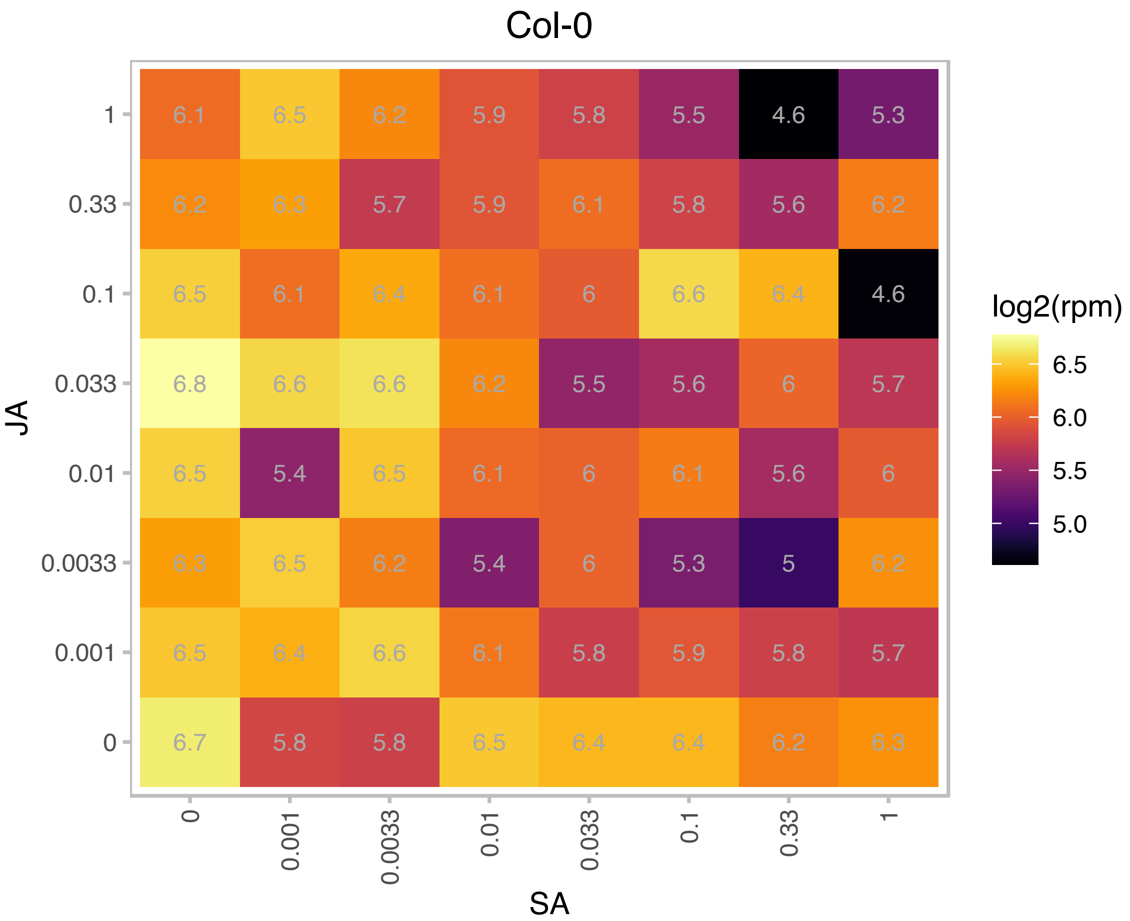


| | |
|---|---|
| Gene | AT3G08770 |
| is.expressed_genes | TRUE |
| is.responsive_genes | TRUE |
| is.apcluster | F-11 |

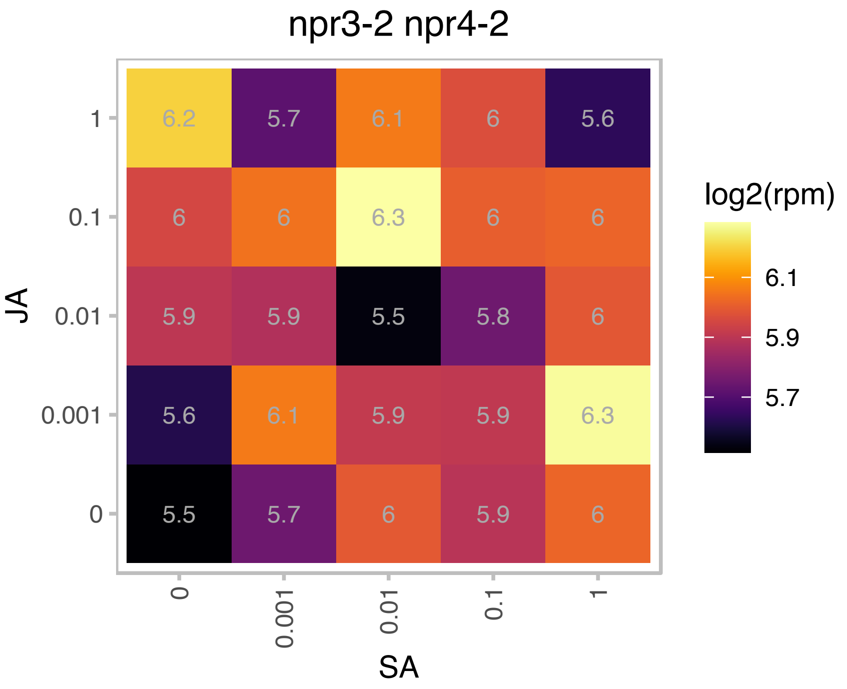


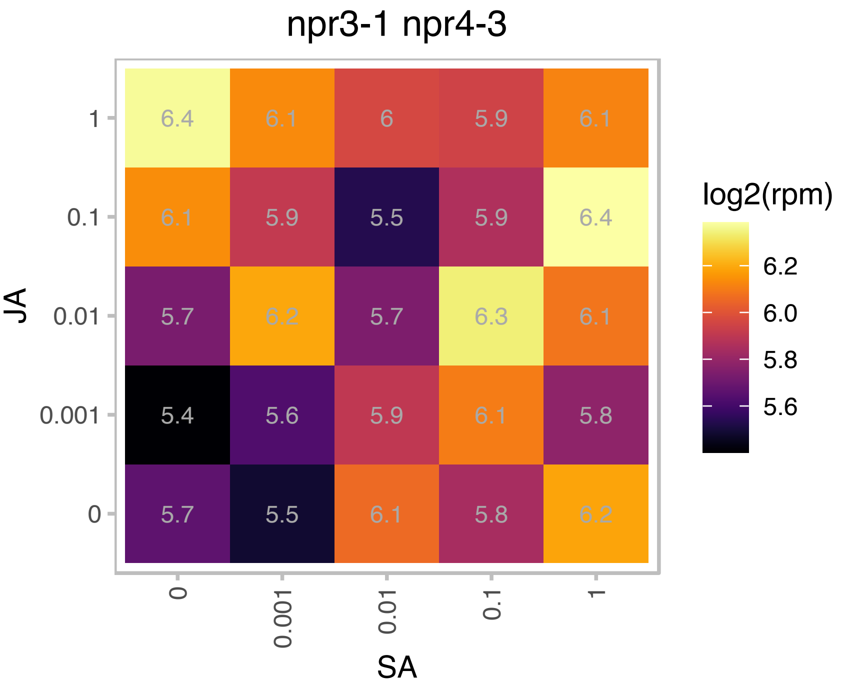

# AT3G09050.1 : NA

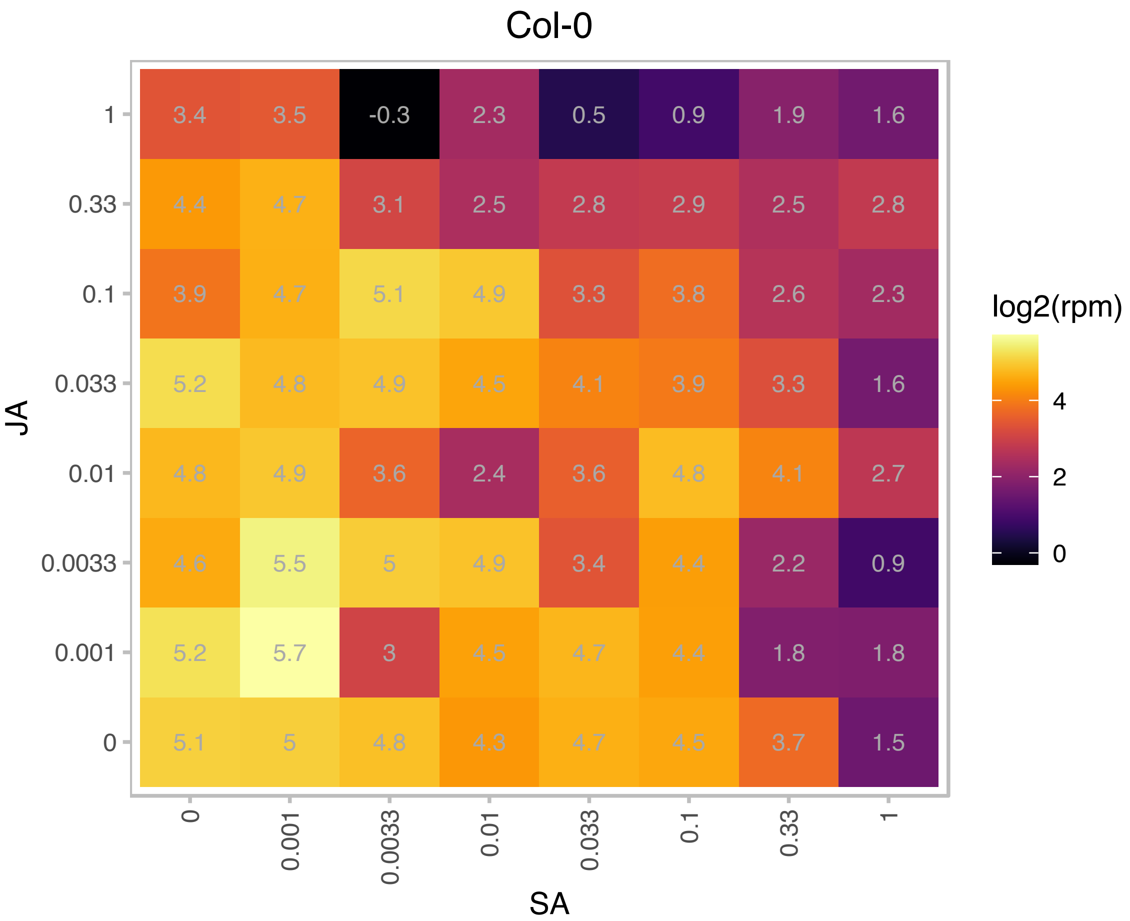


| | |
|---|---|
| Gene | AT3G09050 |
| is.expressed_genes | TRUE |
| is.responsive_genes | TRUE |
| is.apcluster | F-11 |

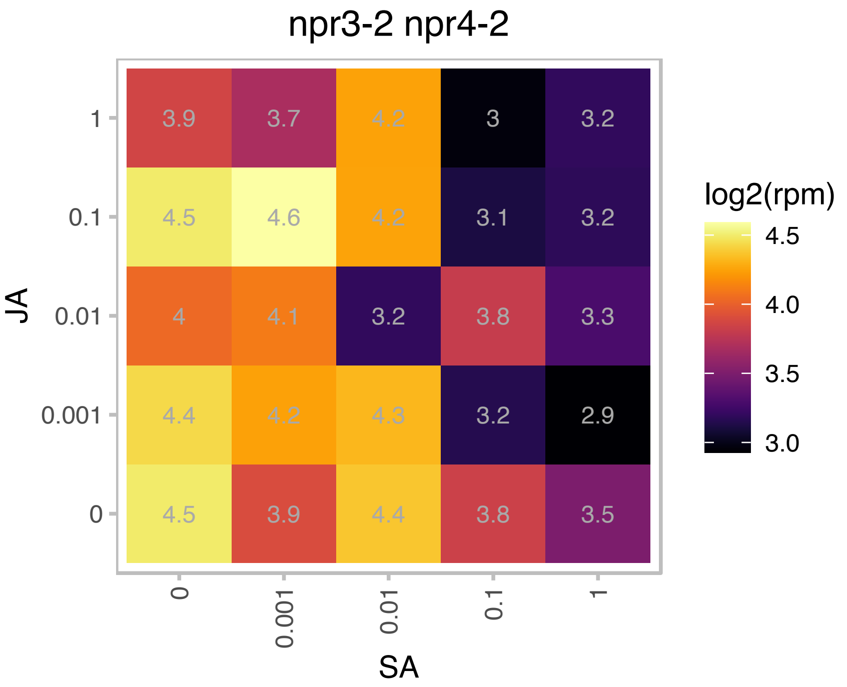


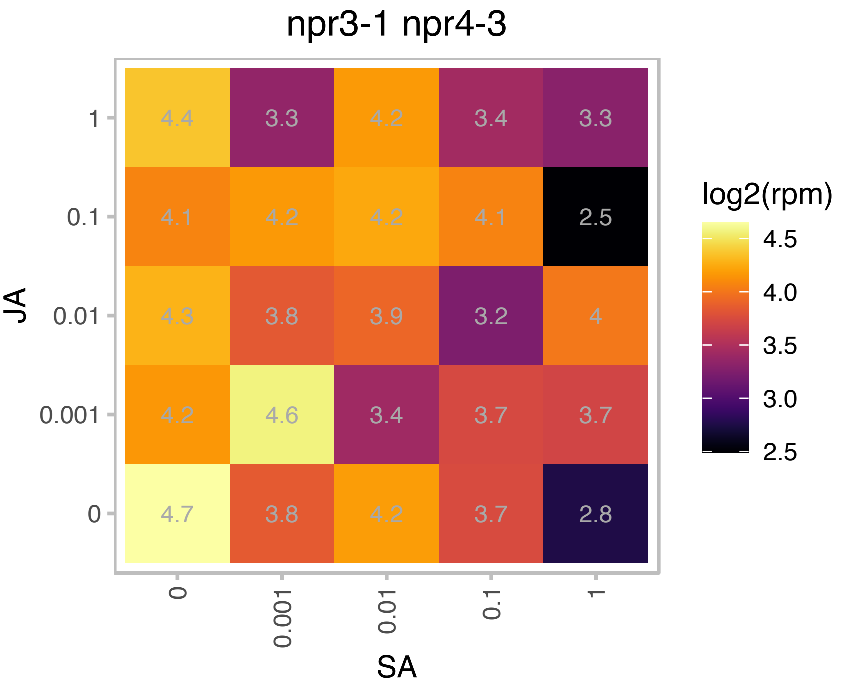

# AT3G09860.1 : NA

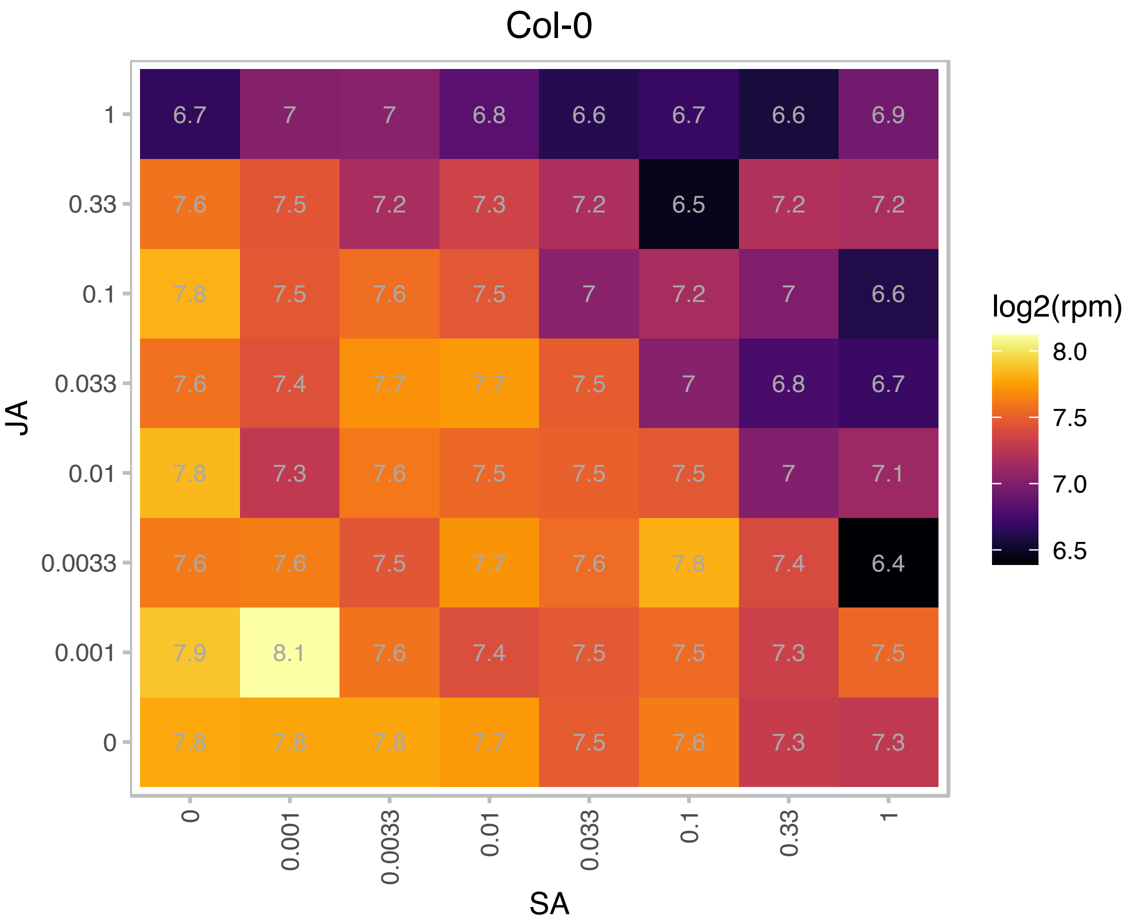


| | |
|---|---|
| Gene | AT3G09860 |
| is.expressed_genes | TRUE |
| is.responsive_genes | TRUE |
| is.apcluster | F-11 |

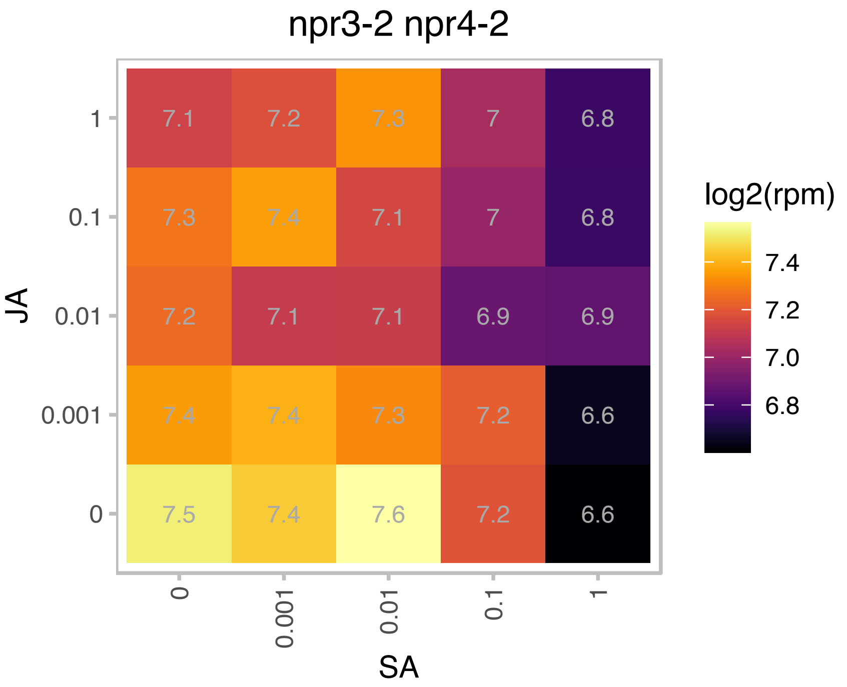


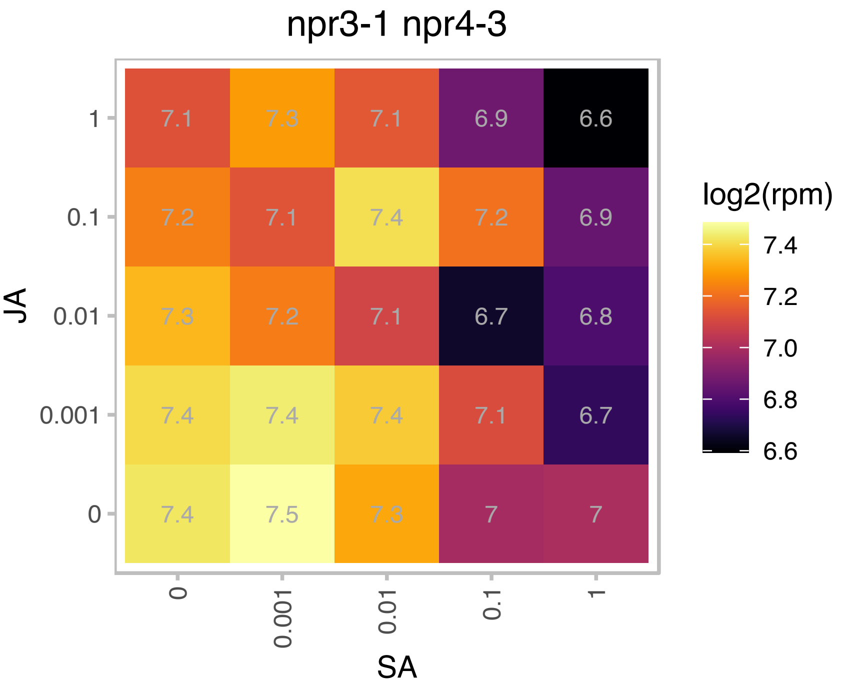

### AT3G10230.1 : AtLCY, LYC, SZL1

lycopene cyclase
Encodes a protein with lycopene β-cyclase activity. This enzyme uses the linear, symmetrical lycopene as substrate. However, unlike the ε-cyclase which adds only one ring, the β-cyclase introduces a ring at both ends of lycopene to form the bicyclic β-carotene.

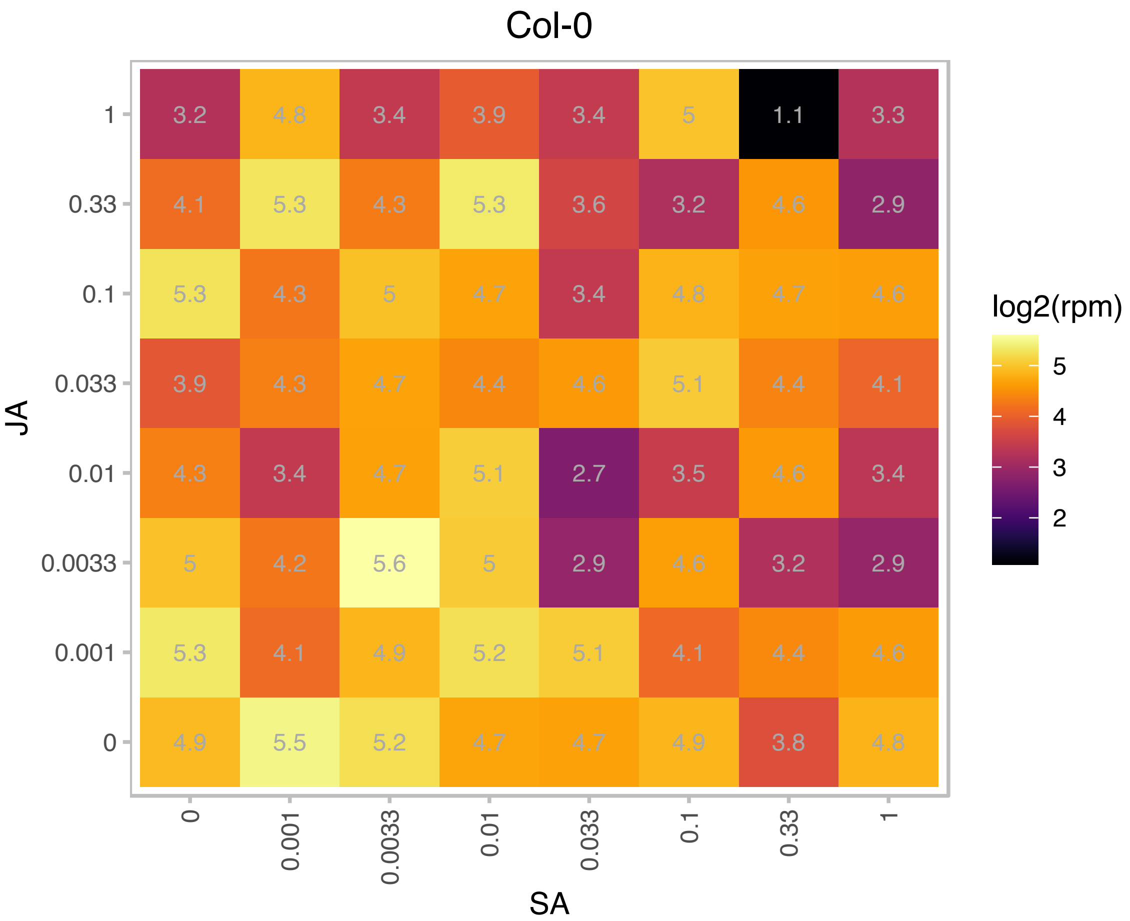


| | |
|---|---|
| Gene | AT3G10230 |
| is.expressed_genes | TRUE |
| is.responsive_genes | TRUE |
| is.apcluster | F-11 |

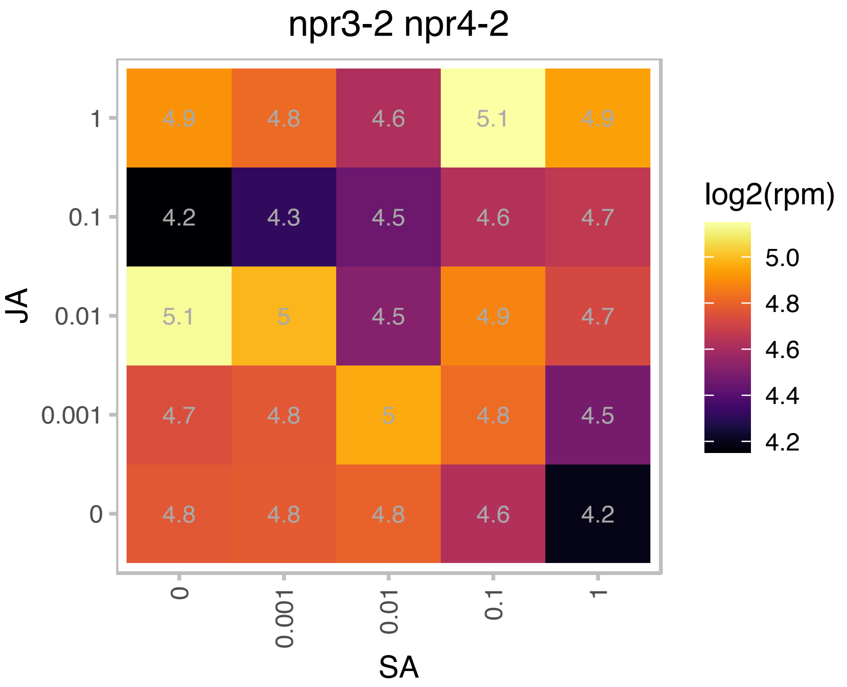


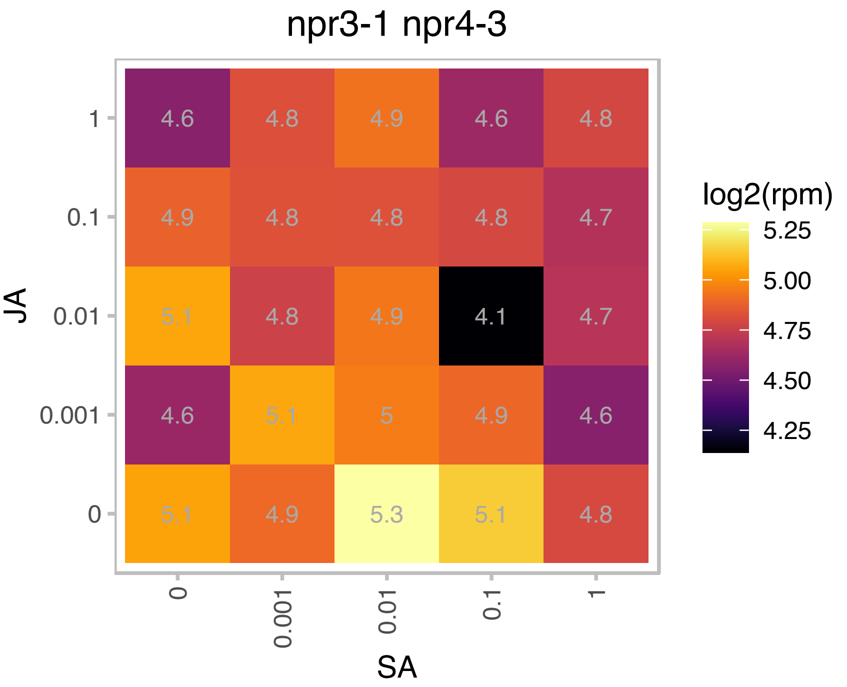

# AT3G12685.1 : NA

Acid phosphatase/vanadium-dependent haloperoxidase-related protein

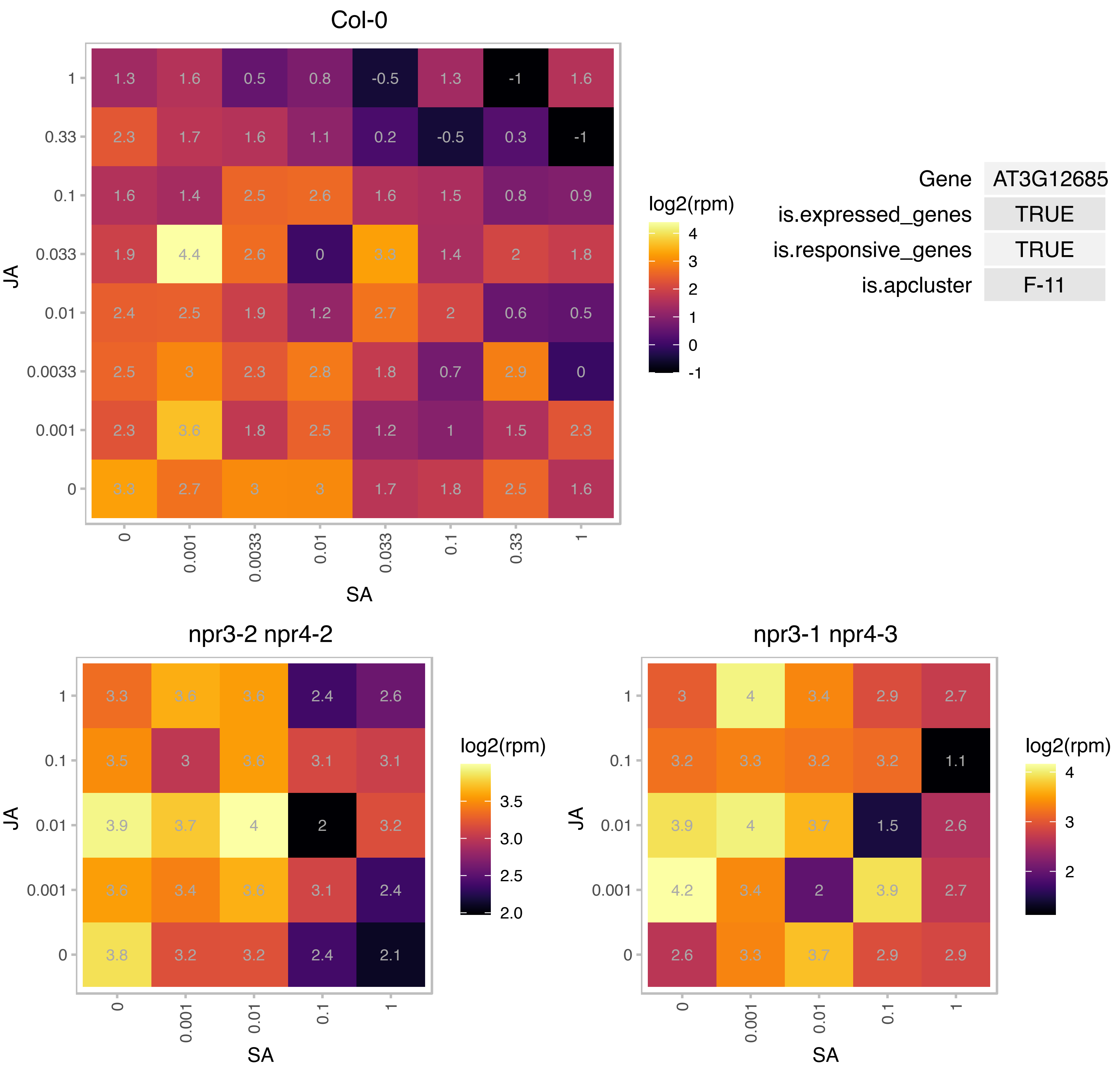


| | |
|---|---|
| Gene | AT3G12685 |
| is.expressed_genes | TRUE |
| is.responsive_genes | TRUE |
| is.apcluster | F-11 |

### AT3G16910.1 : AAE7, ACN1

acyl-activating enzyme 7

Encodes a peroxisomal protein with acetyl-CoA synthetase activity that is responsible for the activation of acetate for entry into the glyoxylate cycle.

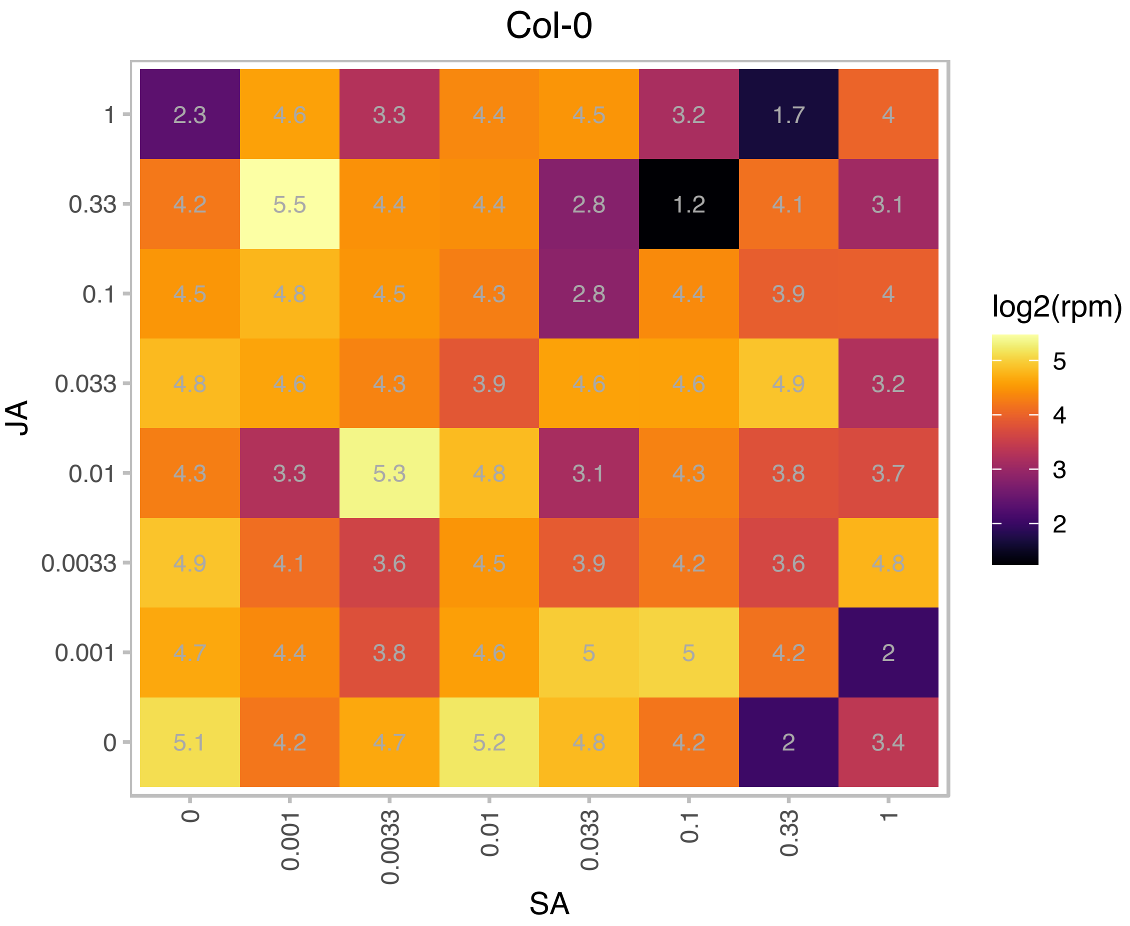


| | |
|---|---|
| Gene | AT3G16910 |
| is.expressed_genes | TRUE |
| is.responsive_genes | TRUE |
| is.apcluster | F-11 |

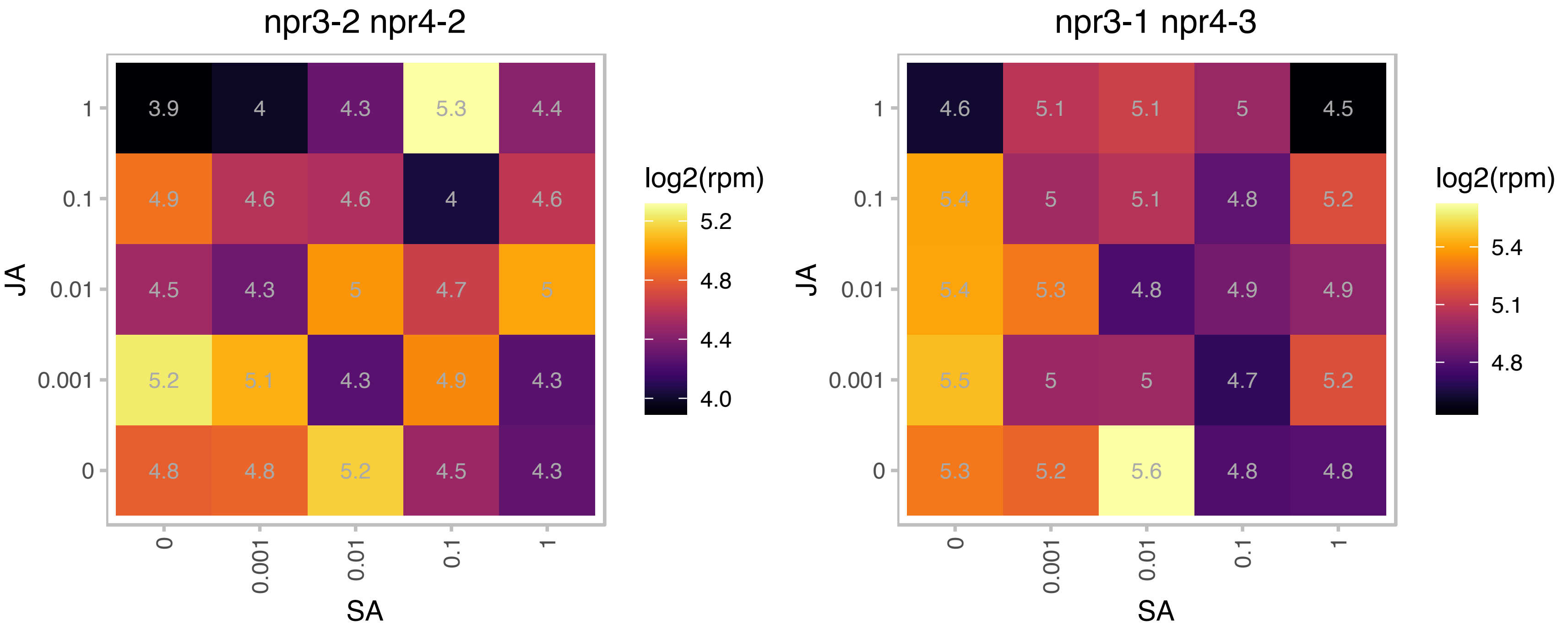

### AT3G17930.1 : DAC

Encodes a thylakoid membrane protein involved in the accumulation of the cytochrome b6/f complex.

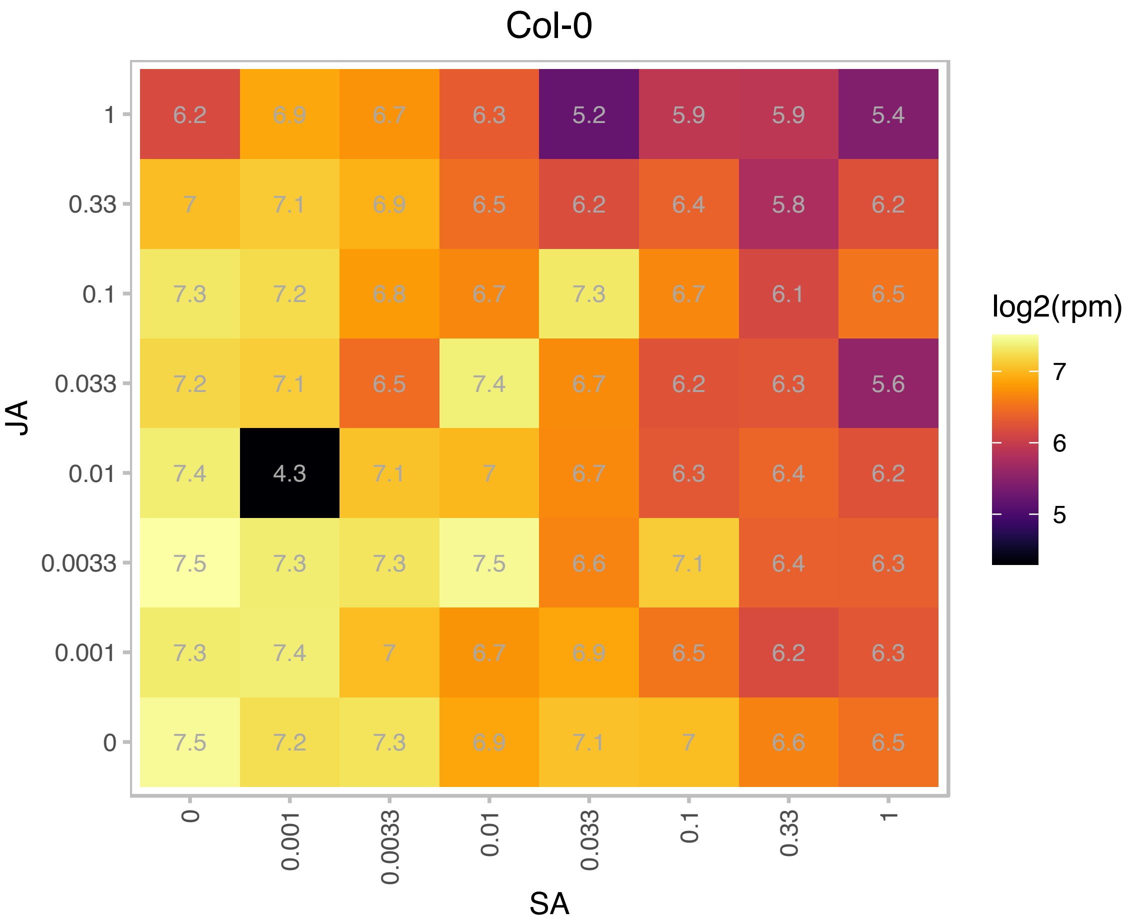


| Gene | AT3G17930 |
|---|---|
| is.expressed_genes | TRUE |
| is.responsive_genes | TRUE |
| is.apcluster | F-11 |

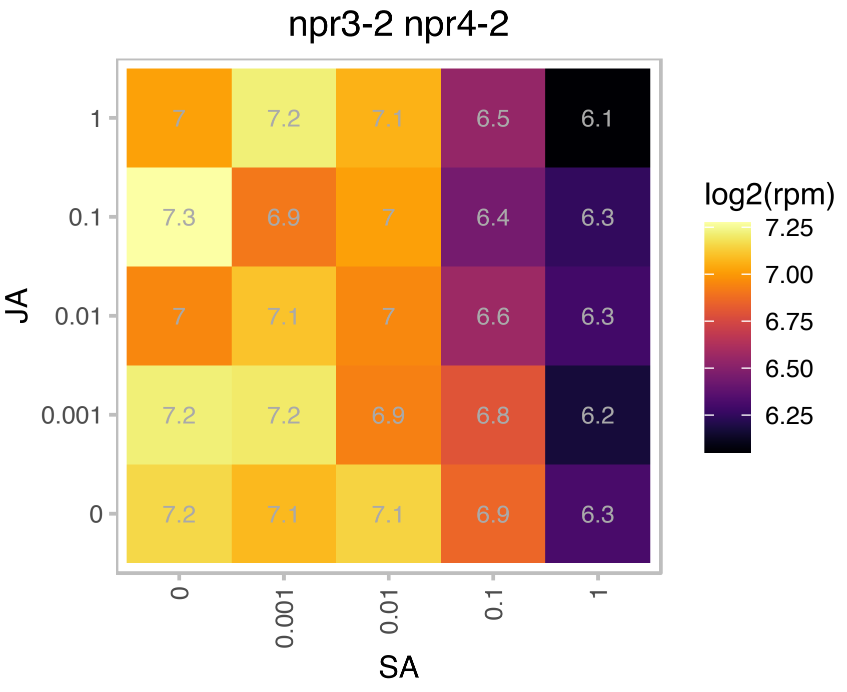


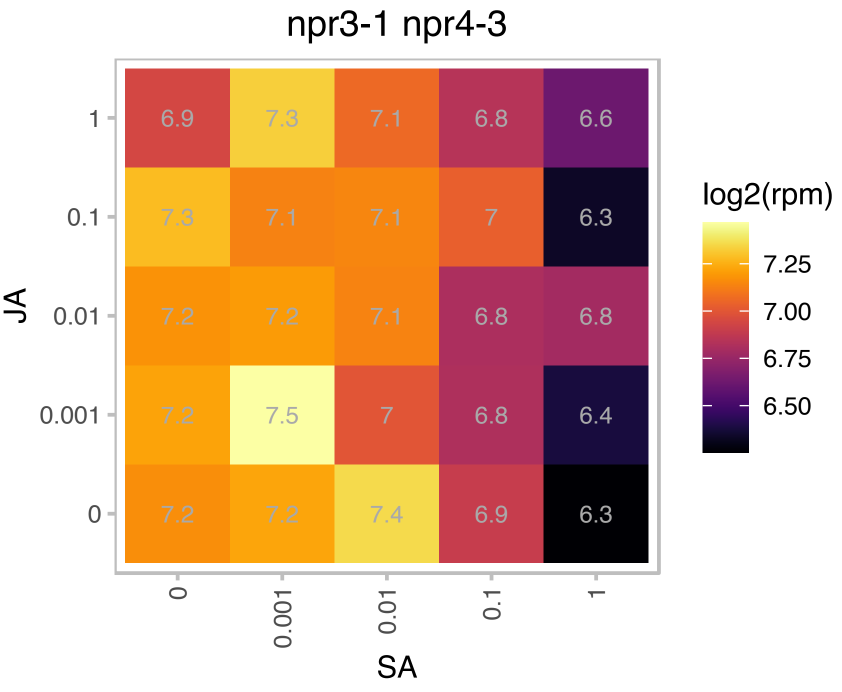

### AT3G23170.1 : PRP

PRP is a proline/serine rich protein of unknown function. It interacts with defense related MAP kinase MPK6 and others. It's expression is induced by PAMP elicitors. May play a role in response to pathogens.

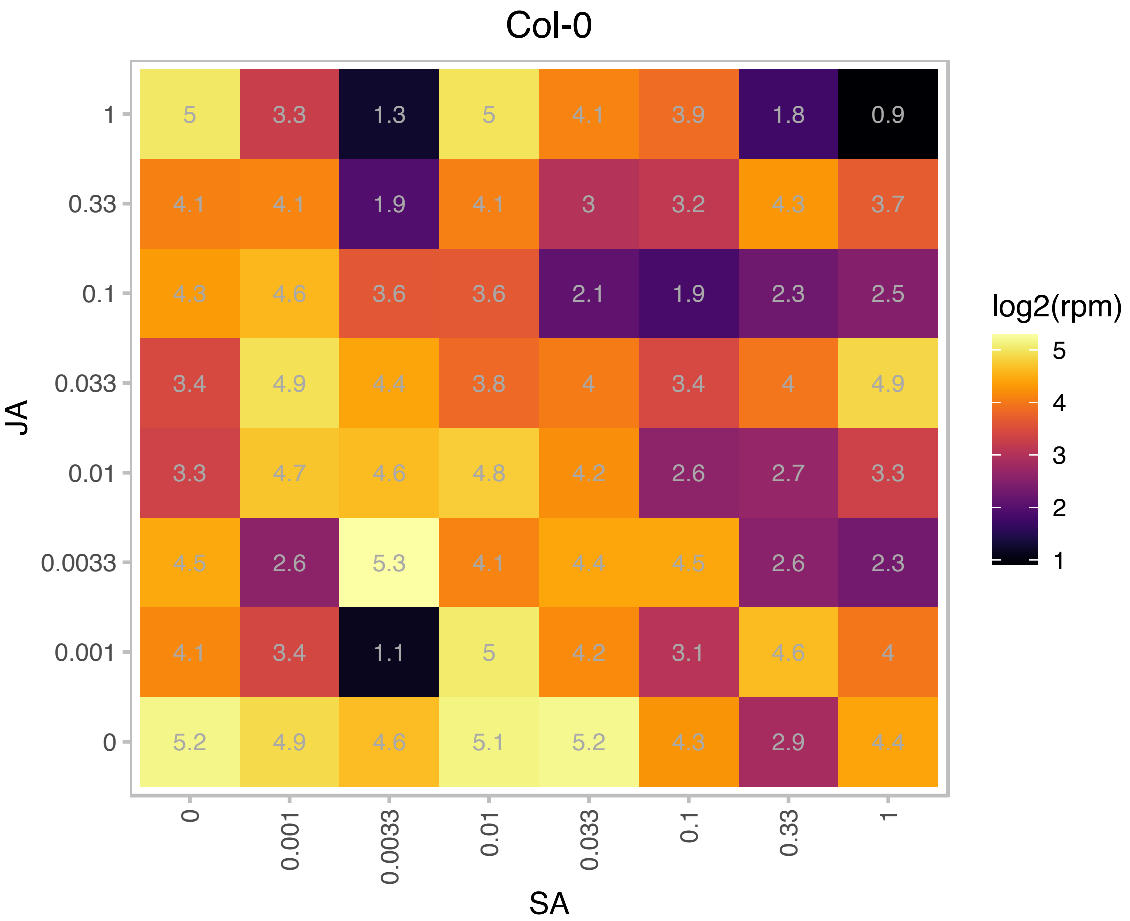


| | |
|---|---|
| Gene | AT3G23170 |
| is.expressed_genes | TRUE |
| is.responsive_genes | TRUE |
| is.apcluster | F-11 |

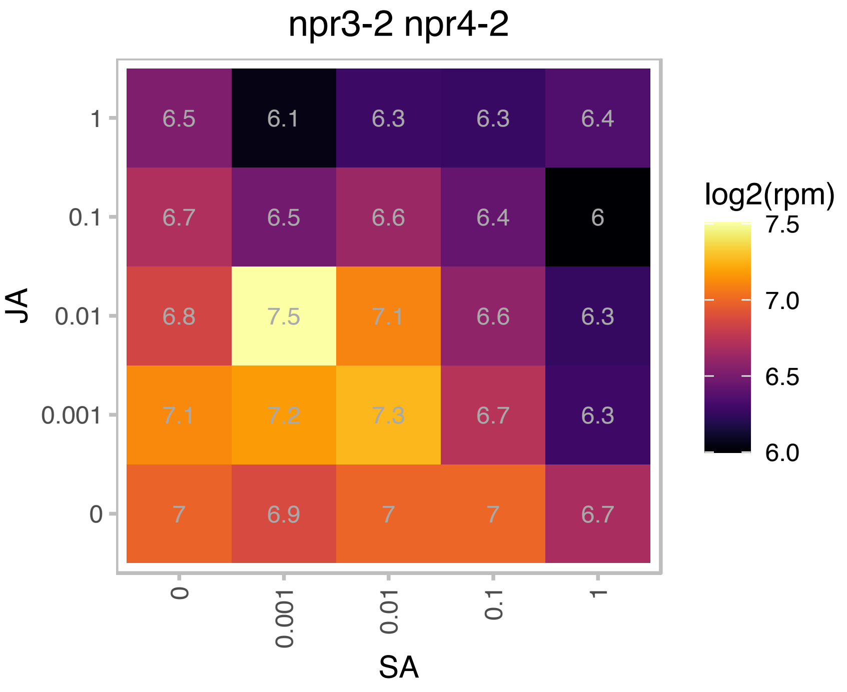


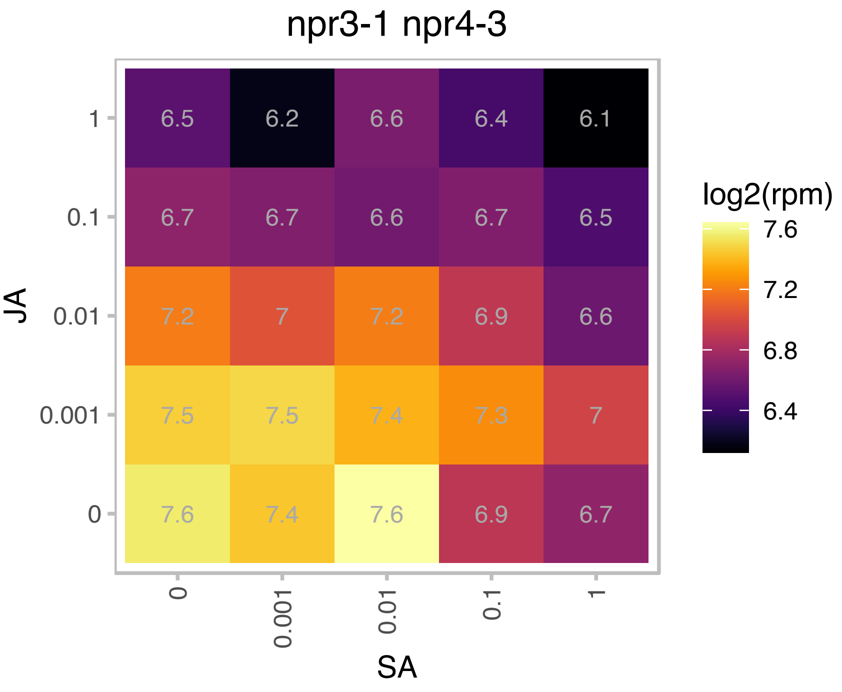

### AT3G23805.1 : RALFL24

ralf-like 24
Member of a diversely expressed predicted peptide family showing sequence similarity to tobacco Rapid Alkalinization Factor (RALF), and is believed to play an essential role in the physiology of Arabidopsis. Consists of a single exon and is characterized by a conserved C-terminal motif and N-terminal signal peptide.

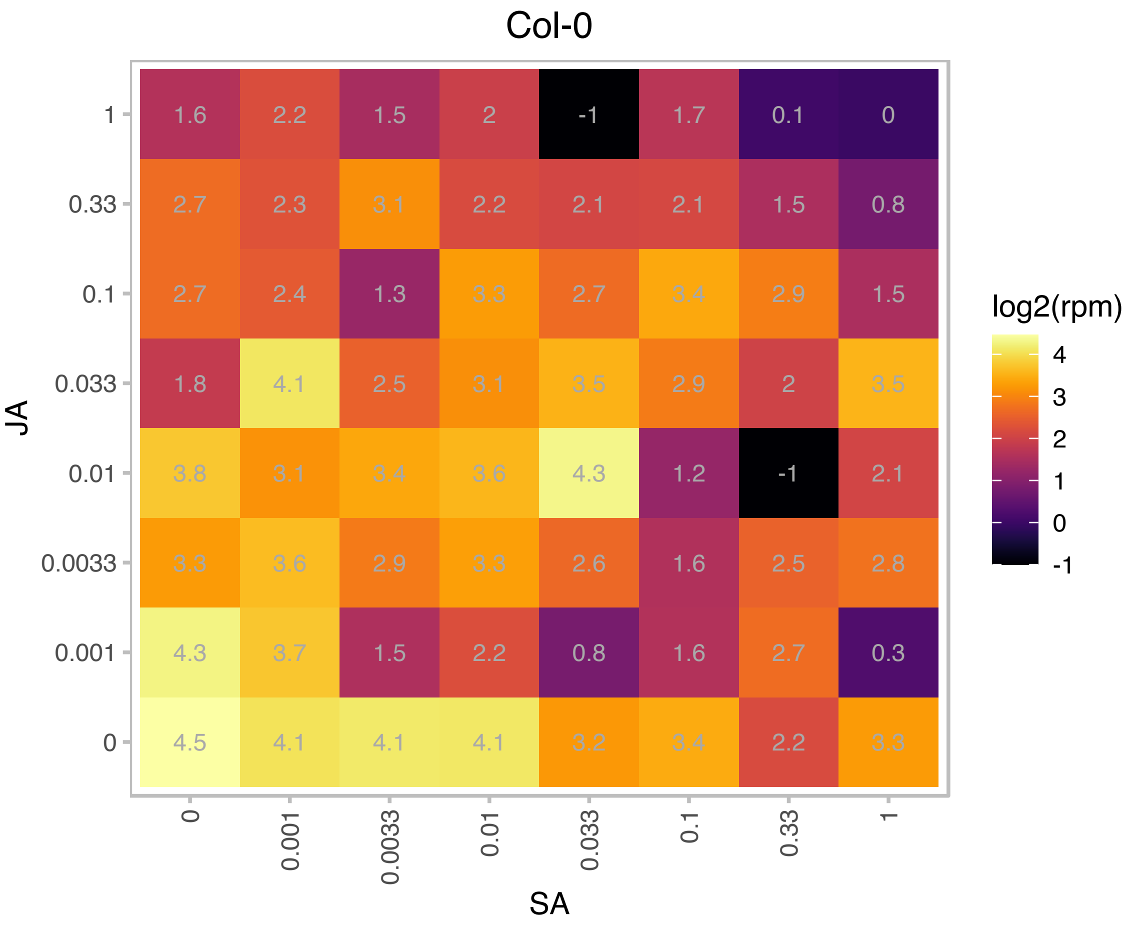


| | |
|---|---|
| Gene | AT3G23805 |
| is.expressed_genes | TRUE |
| is.responsive_genes | TRUE |
| is.apcluster | F-11 |

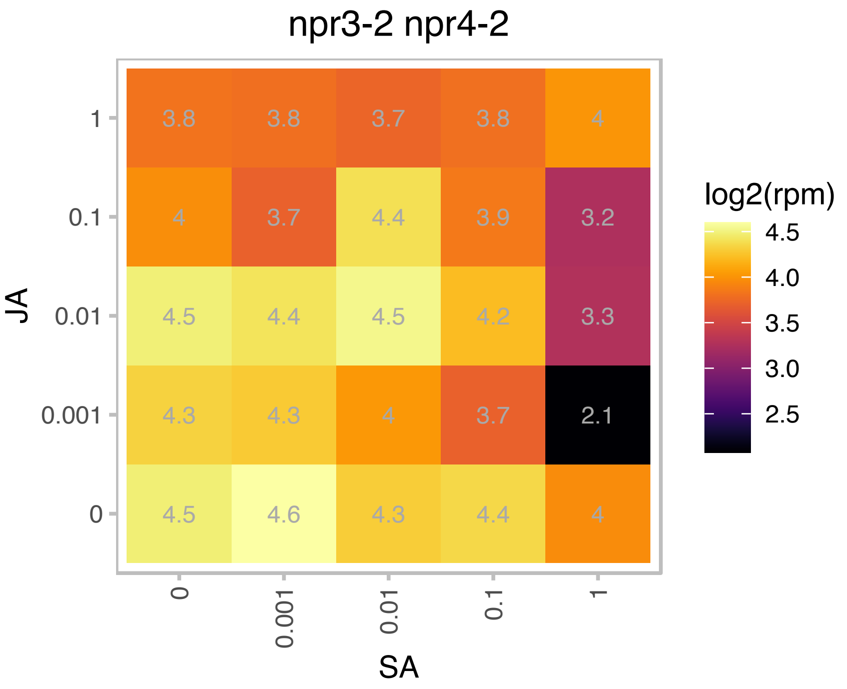


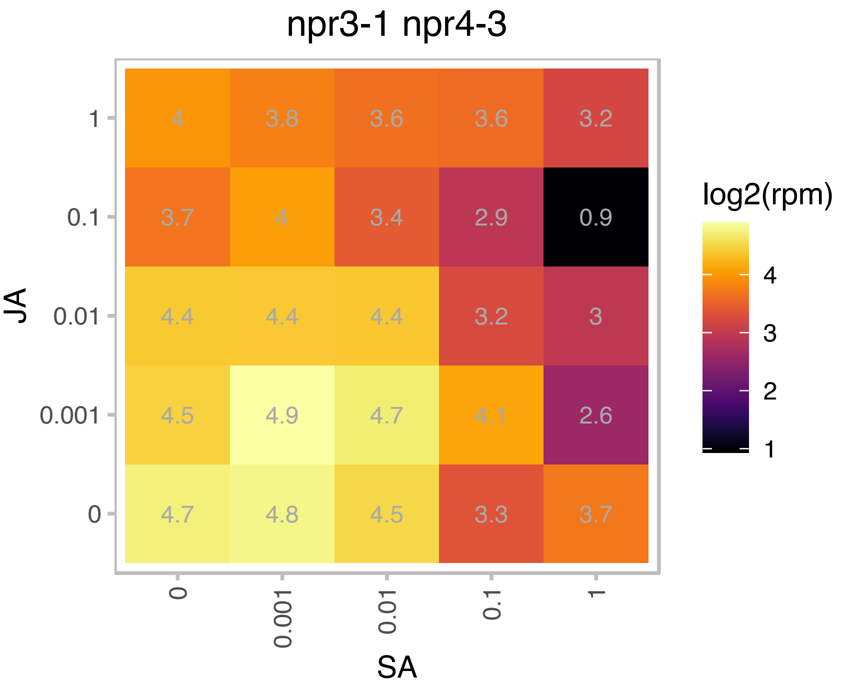

# AT3G26070.1 : NA

Plastid-lipid associated protein PAP / fibrillin family protein

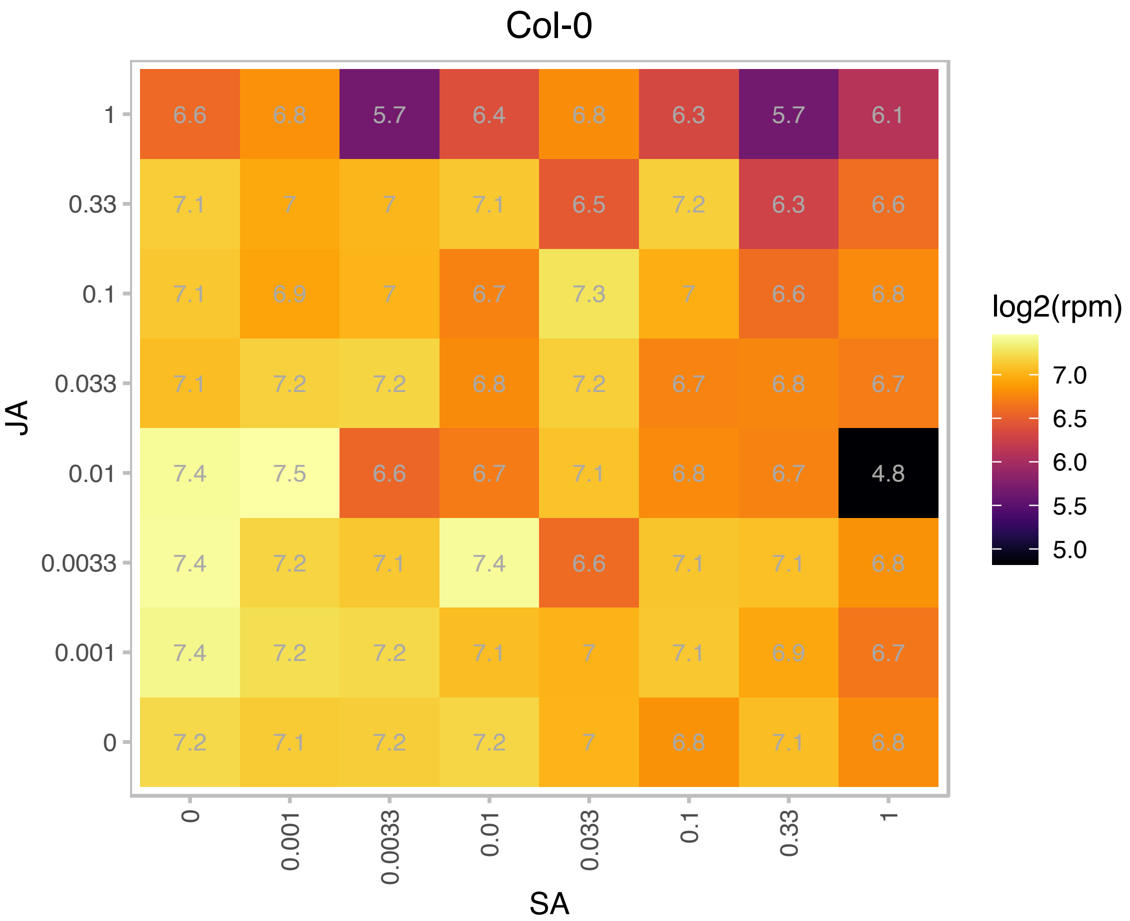


| | |
|---|---|
| Gene | AT3G26070 |
| is.expressed_genes | TRUE |
| is.responsive_genes | TRUE |
| is.apcluster | F-11 |

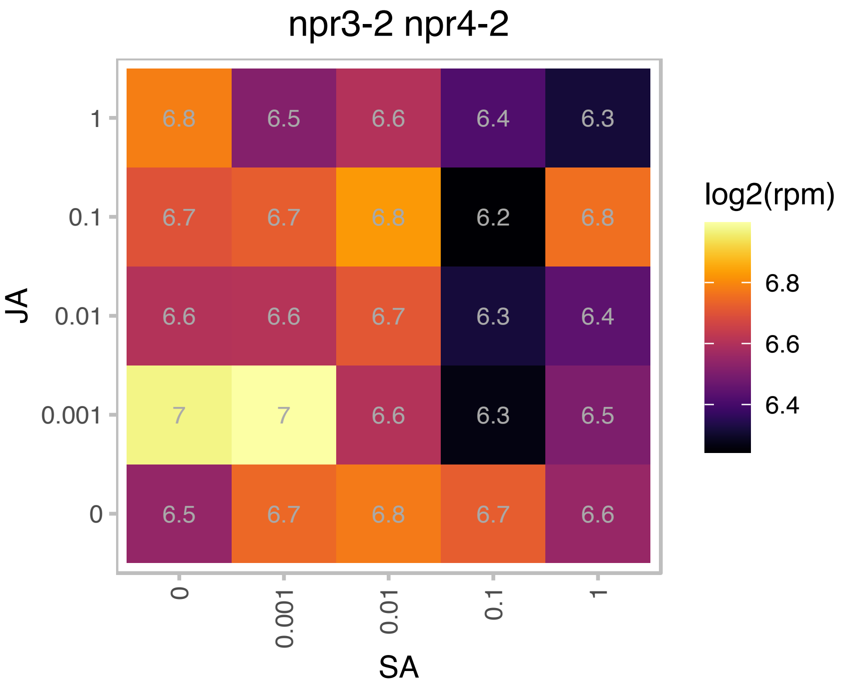


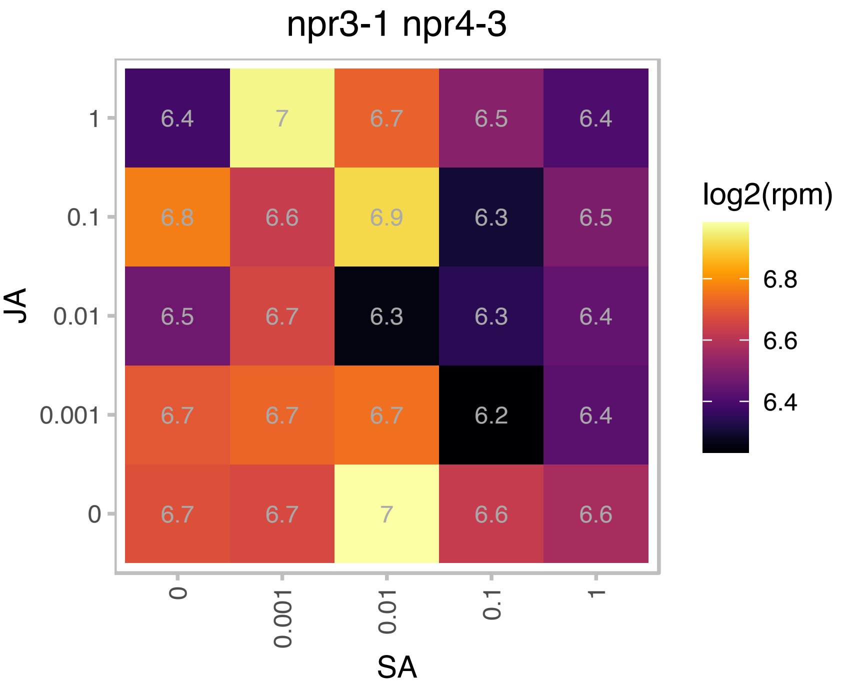

# AT3G44020.1 : NA

thylakoid lumenal P17.1 protein

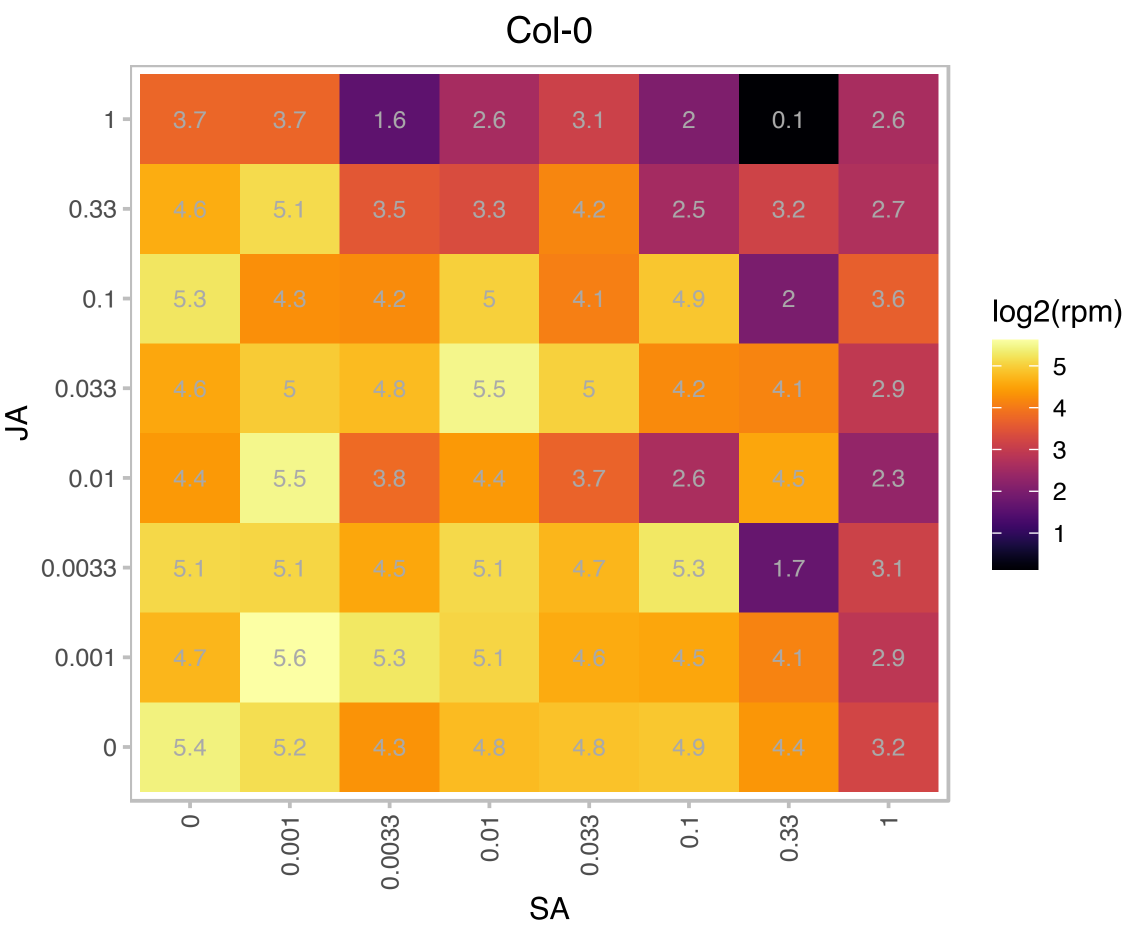


| | |
|---|---|
| Gene | AT3G44020 |
| is.expressed_genes | TRUE |
| is.responsive_genes | TRUE |
| is.apcluster | F-11 |

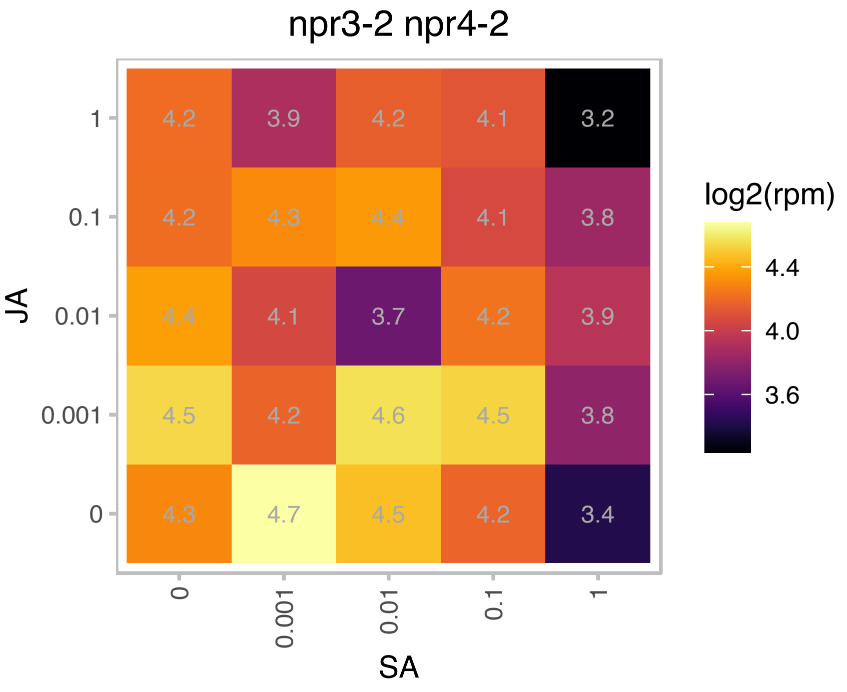


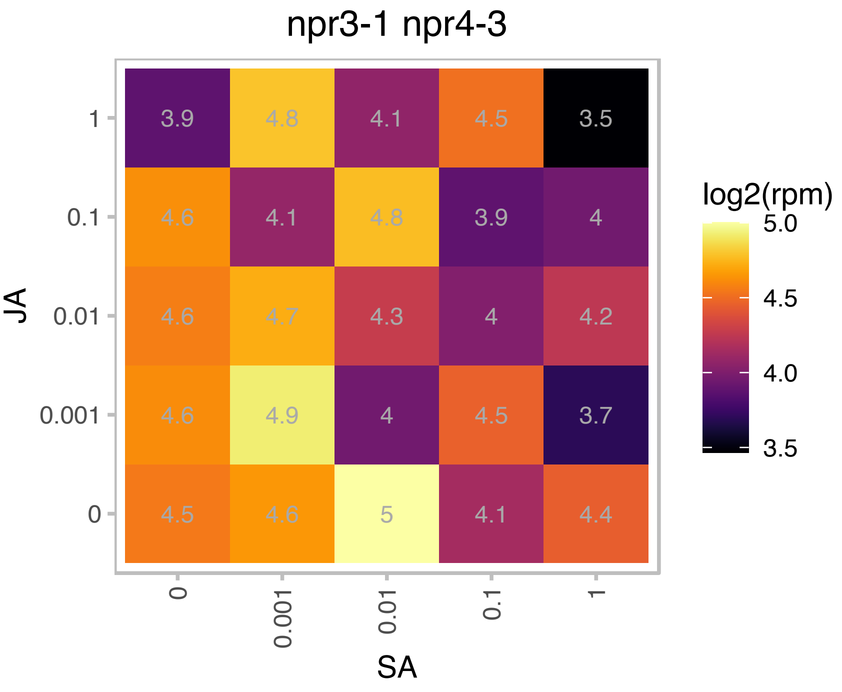

# AT3G51720.1 : NA

Plant protein of unknown function (DUF827)

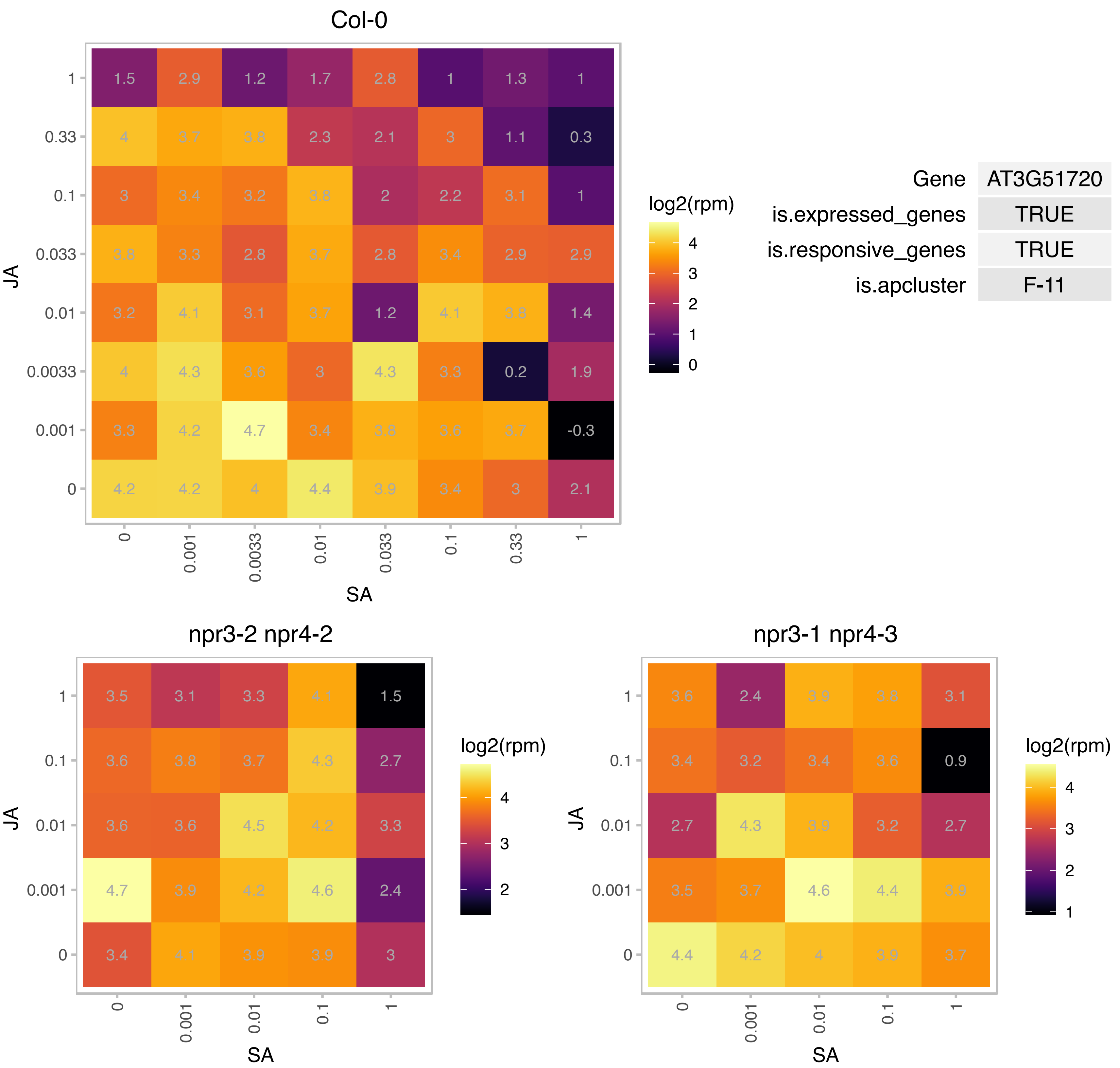


| Col-0: JA \ SA | 0 | 0.001 | 0.0033 | 0.01 | 0.033 | 0.1 | 0.33 | 1 |
|---|---|---|---|---|---|---|---|---|
| 1 | 1.5 | 2.9 | 1.2 | 1.7 | 2.8 | 1 | 1.3 | 1 |
| 0.33 | 4 | 3.7 | 3.8 | 2.3 | 2.1 | 3 | 1.1 | 0.3 |
| 0.1 | 3 | 3.4 | 3.2 | 3.8 | 2 | 2.2 | 3.1 | 1 |
| 0.033 | 3.8 | 3.3 | 2.8 | 3.7 | 2.8 | 3.4 | 2.9 | 2.9 |
| 0.01 | 3.2 | 4.1 | 3.1 | 3.7 | 1.2 | 4.1 | 3.8 | 1.4 |
| 0.0033 | 4 | 4.3 | 3.6 | 3 | 4.3 | 3.3 | 0.2 | 1.9 |
| 0.001 | 3.3 | 4.2 | 4.7 | 3.4 | 3.8 | 3.6 | 3.7 | -0.3 |
| 0 | 4.2 | 4.2 | 4 | 4.4 | 3.9 | 3.4 | 3 | 2.1 |

| npr3-2 npr4-2: JA \ SA | 0 | 0.001 | 0.01 | 0.1 | 1 |
|---|---|---|---|---|---|
| 1 | 3.5 | 3.1 | 3.3 | 4.1 | 1.5 |
| 0.1 | 3.6 | 3.8 | 3.7 | 4.3 | 2.7 |
| 0.01 | 3.6 | 3.6 | 4.5 | 4.2 | 3.3 |
| 0.001 | 4.7 | 3.9 | 4.2 | 4.6 | 2.4 |
| 0 | 3.4 | 4.1 | 3.9 | 3.9 | 3 |

| npr3-1 npr4-3: JA \ SA | 0 | 0.001 | 0.01 | 0.1 | 1 |
|---|---|---|---|---|---|
| 1 | 3.6 | 2.4 | 3.9 | 3.8 | 3.1 |
| 0.1 | 3.4 | 3.2 | 3.4 | 3.6 | 0.9 |
| 0.01 | 2.7 | 4.3 | 3.9 | 3.2 | 2.7 |
| 0.001 | 3.5 | 3.7 | 4.6 | 4.4 | 3.9 |
| 0 | 4.4 | 4.2 | 4 | 3.9 | 3.7 |

# AT3G52072.1 : NA

other RNA

Potential natural antisense gene, locus overlaps with AT3G52070

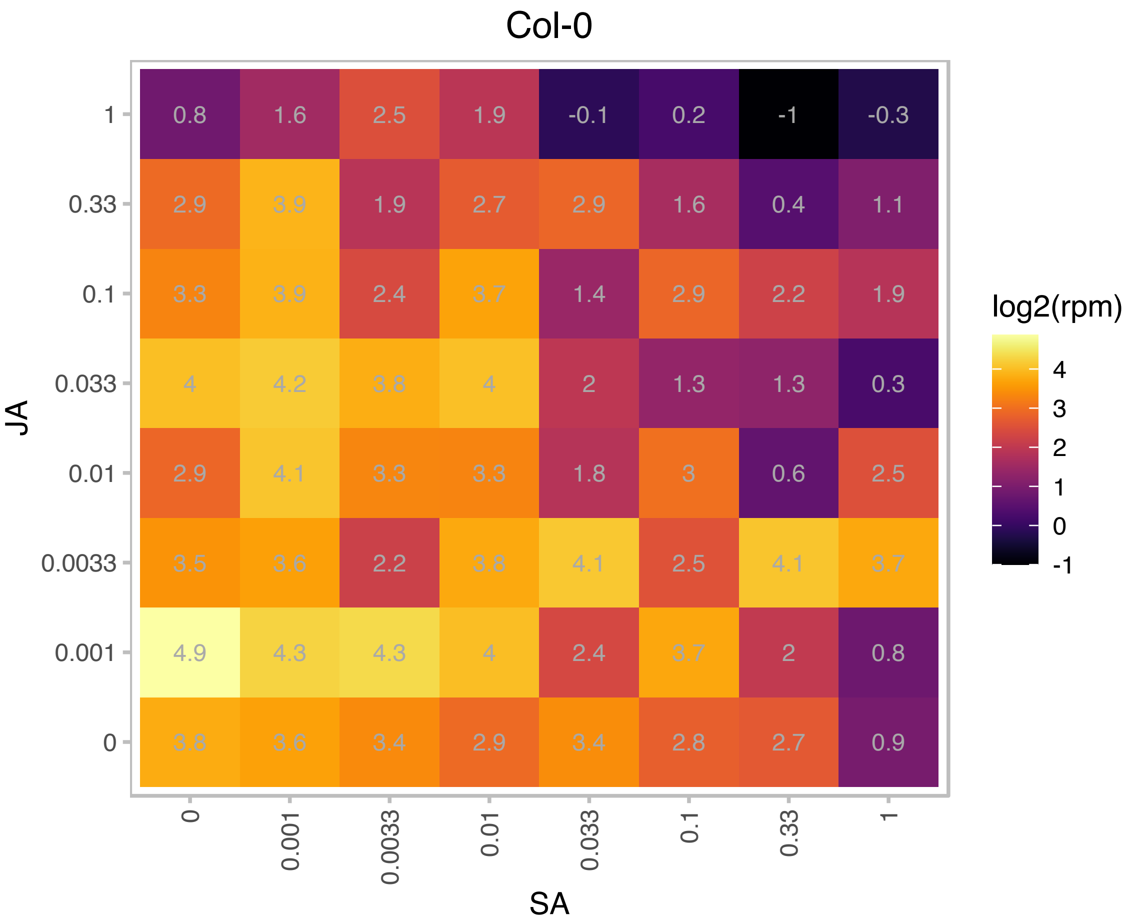


| | |
|---|---|
| Gene | AT3G52072 |
| is.expressed_genes | TRUE |
| is.responsive_genes | TRUE |
| is.apcluster | F-11 |

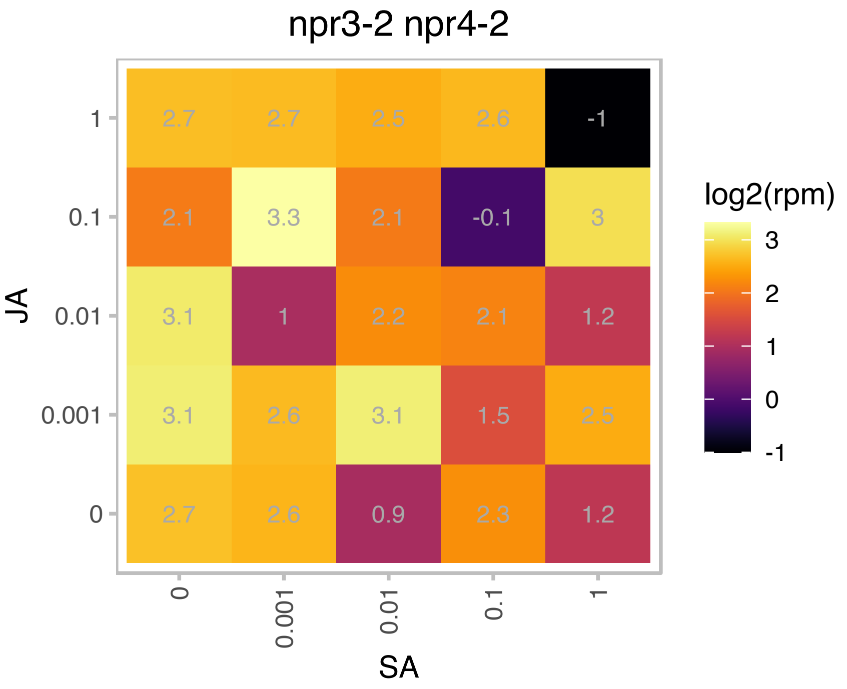


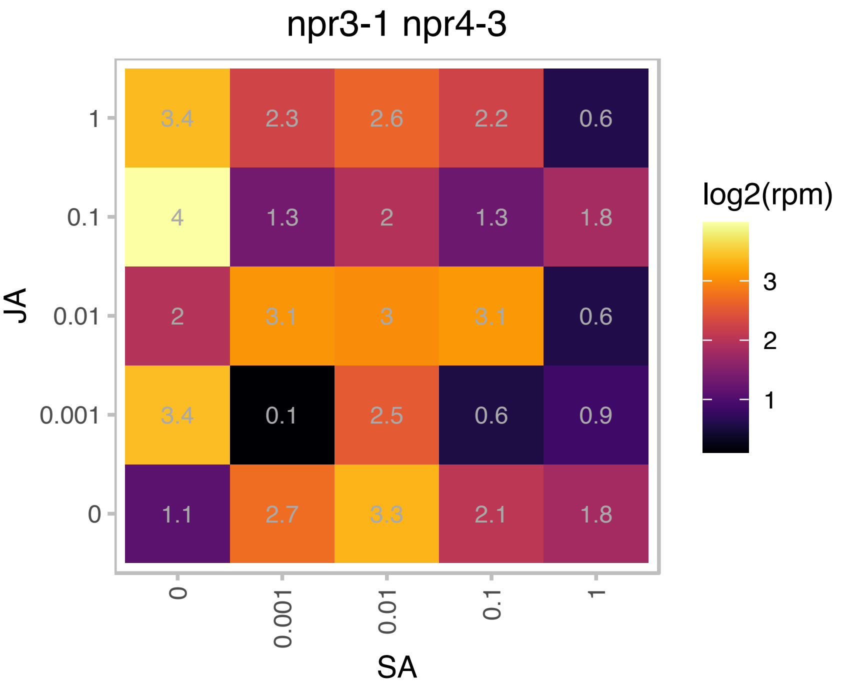

# AT3G62550.1 : NA

Adenine nucleotide alpha hydrolases-like superfamily protein

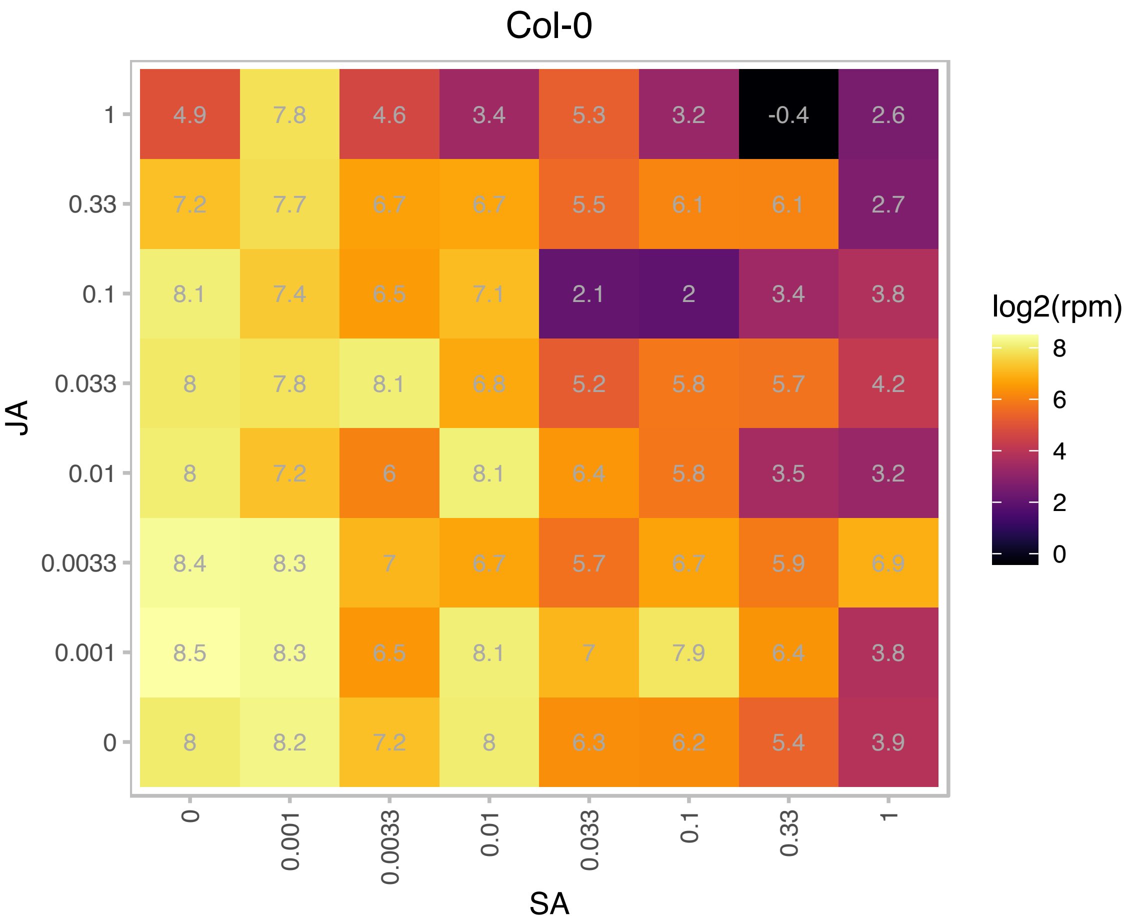


| | |
|---|---|
| Gene | AT3G62550 |
| is.expressed_genes | TRUE |
| is.responsive_genes | TRUE |
| is.apcluster | F-11 |

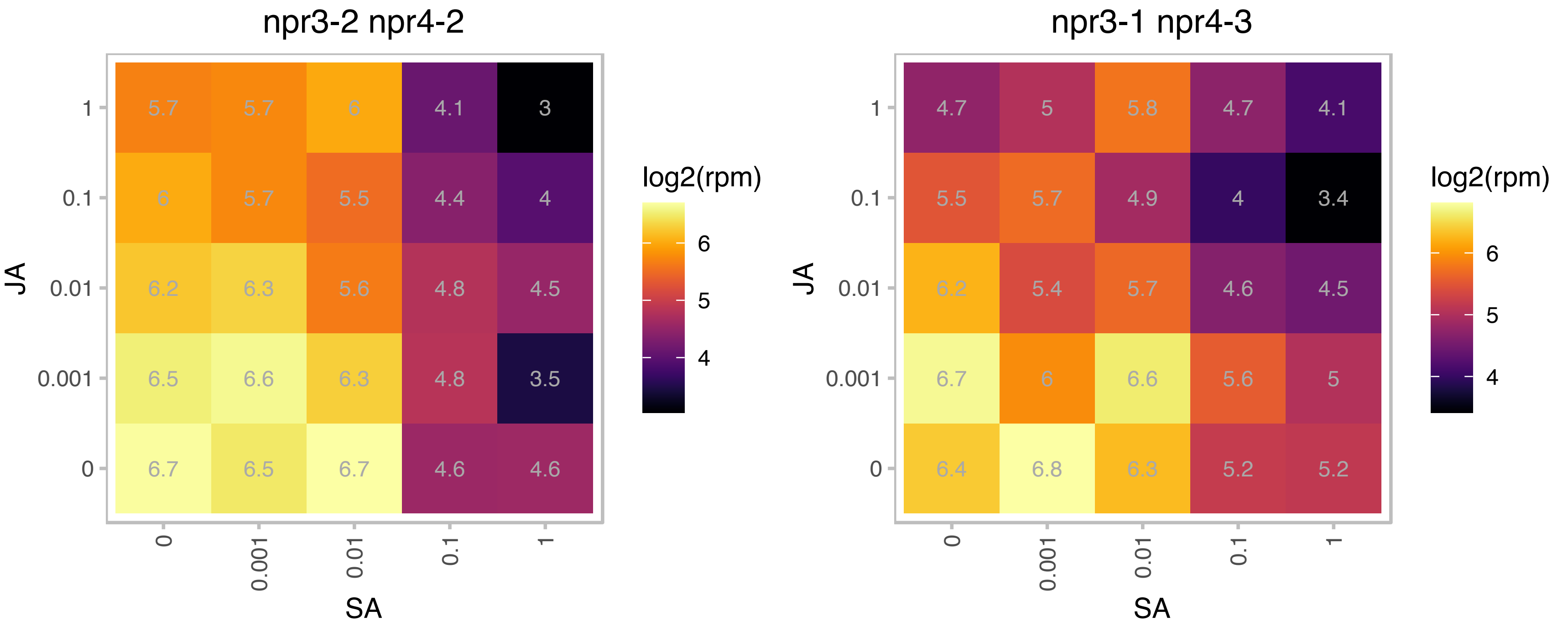

# AT4G00026.1 : SD3

Encodes SD3 (Segregation Distortion 3), a protein with high similarity to yeast translocase on the inner mitochondrial membrane 21 (TIM21). sd3 mutants show seedling-lethal phenotype in light-grown seedlings and shorter hypocotyls in dark-grown seedlings. SD3 overexpression plants show increase in cell number and cell size, as well as elevated ATP level.

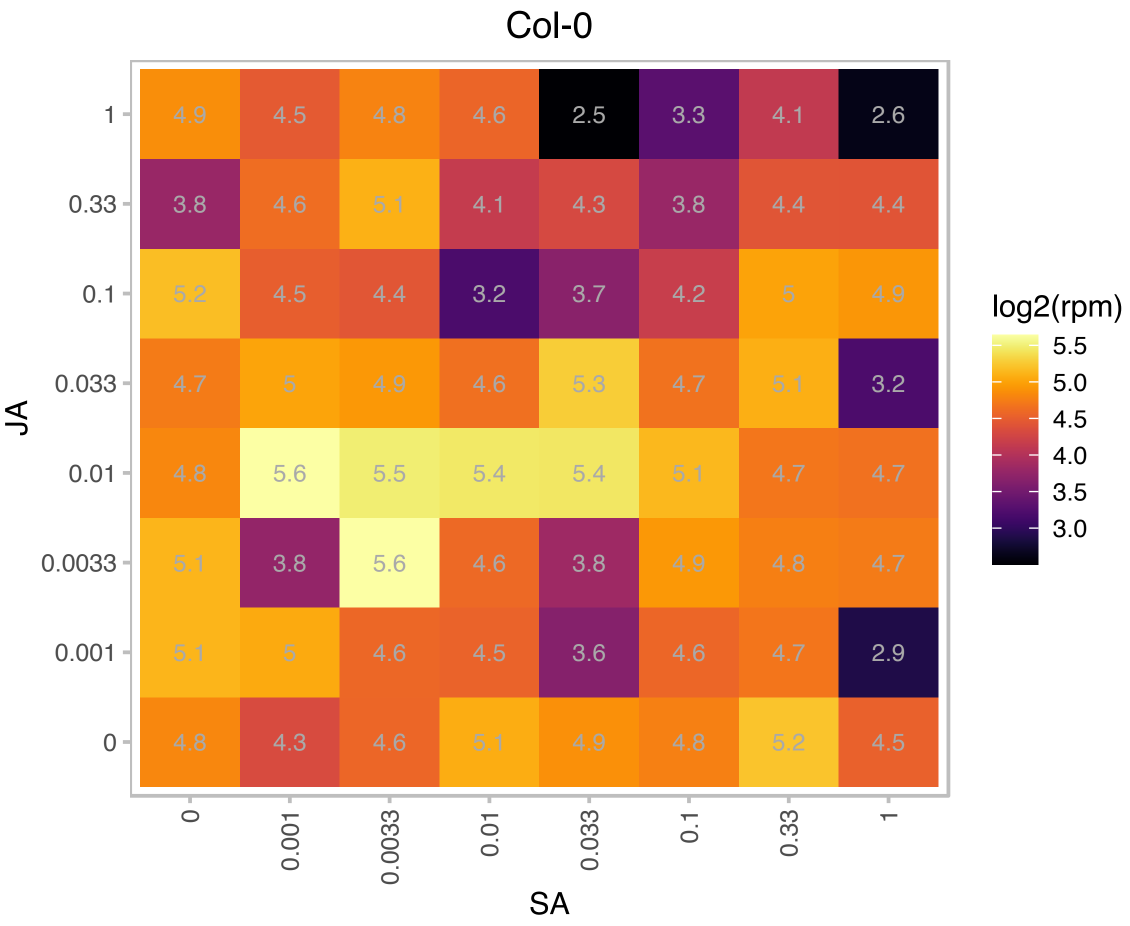


| | |
|---|---|
| Gene | AT4G00026 |
| is.expressed_genes | TRUE |
| is.responsive_genes | TRUE |
| is.apcluster | F-11 |

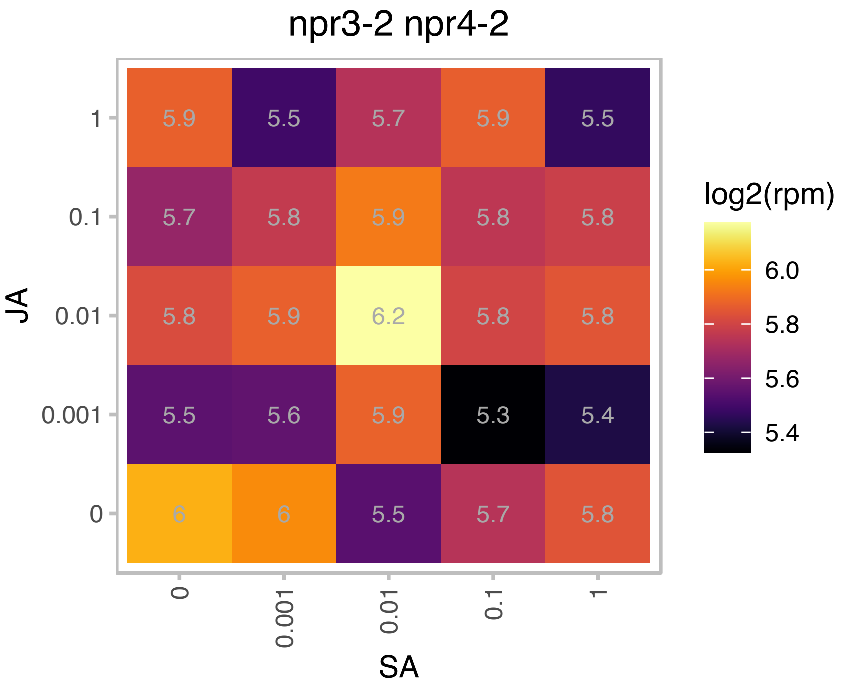


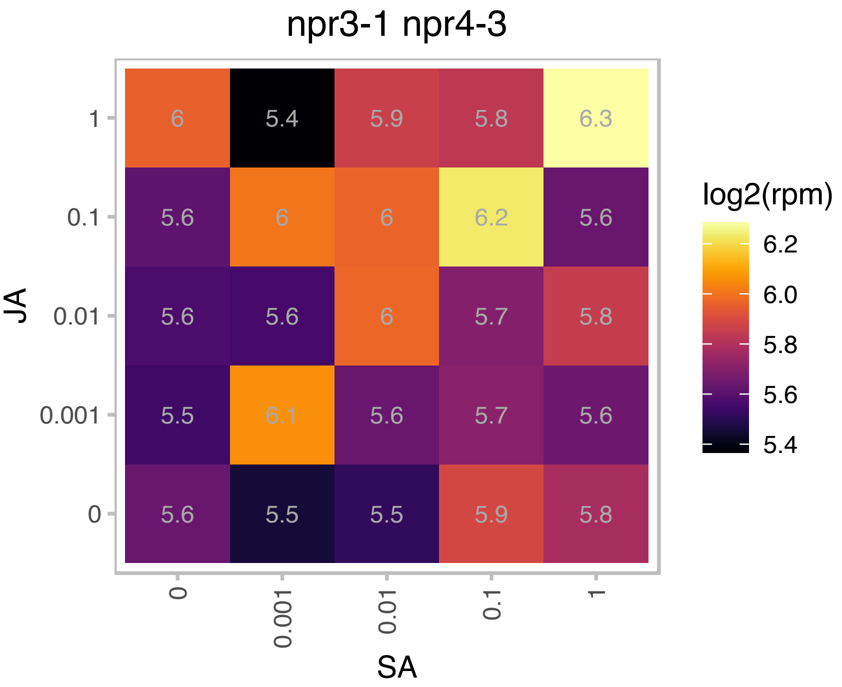

### AT4G00180.1 : YAB3

Plant-specific transcription factor YABBY family protein

YABBY gene family member, likely has transcription factor activity, involved in specifying abaxial cell fate. Along with FIL, involved in patterning of the fruit. GUS reporter gene expression in seedlings is observed in the young leaves and as the leaf matures, expression is restricted to the abaxial tissues of leaves, expression is also observed on either side of the leaf margin in the younger tissues of leaf blades.

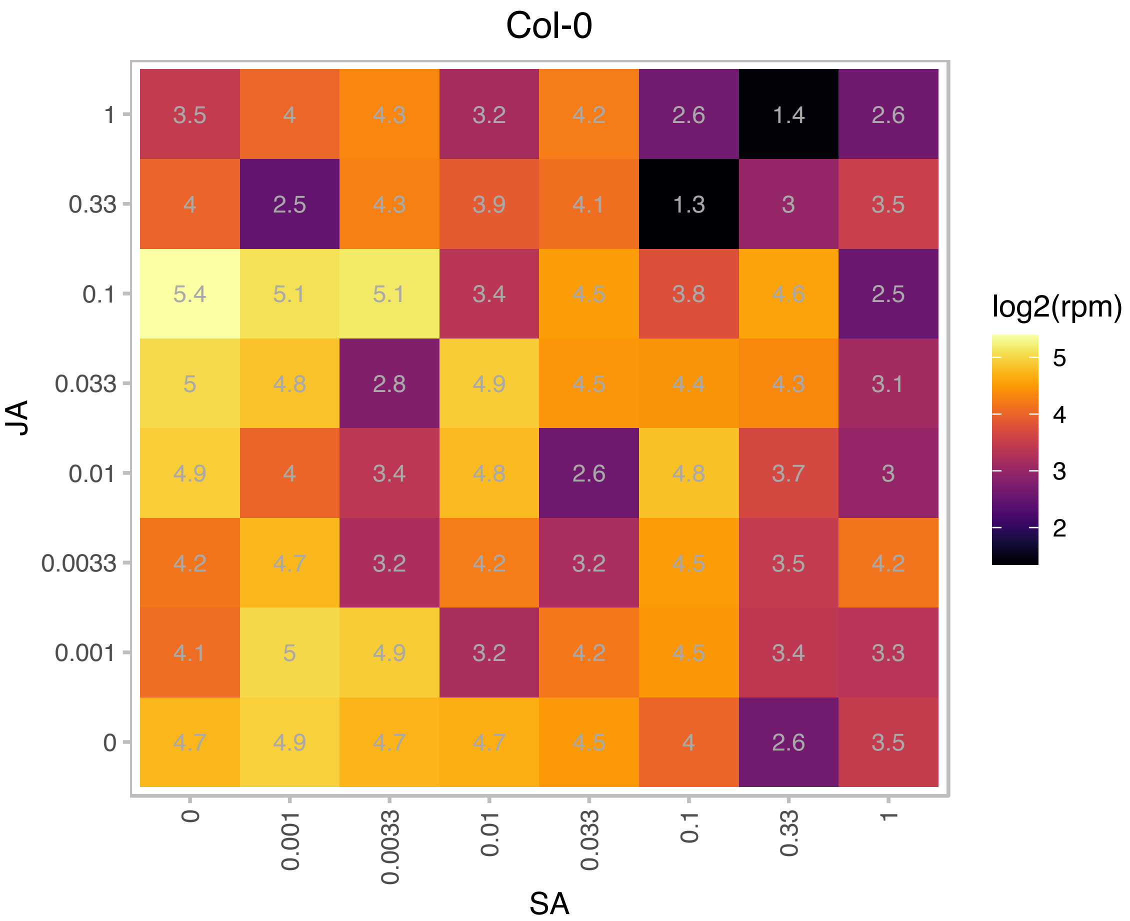


| | |
|---|---|
| Gene | AT4G00180 |
| is.expressed_genes | TRUE |
| is.responsive_genes | TRUE |
| is.apcluster | F-11 |

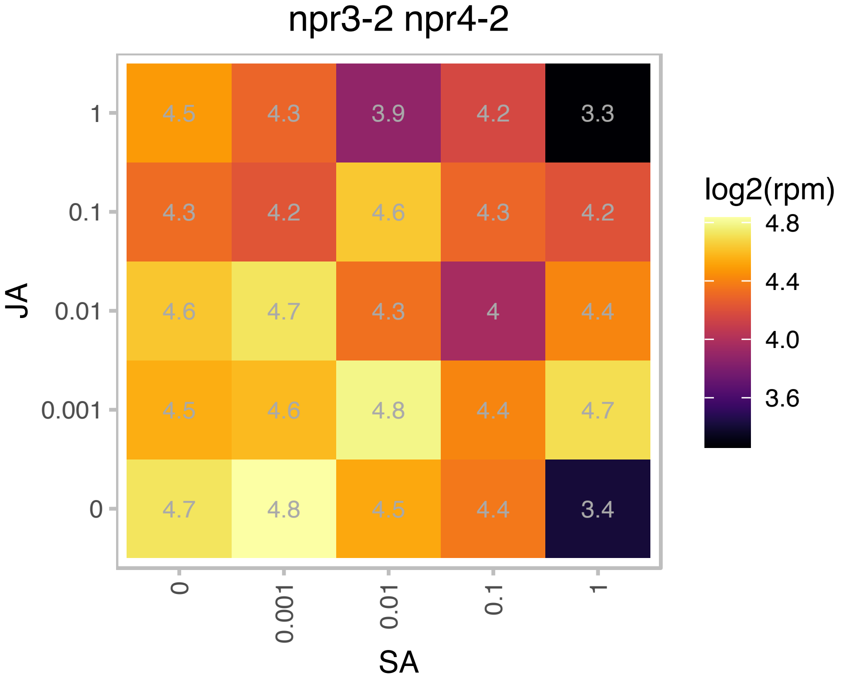


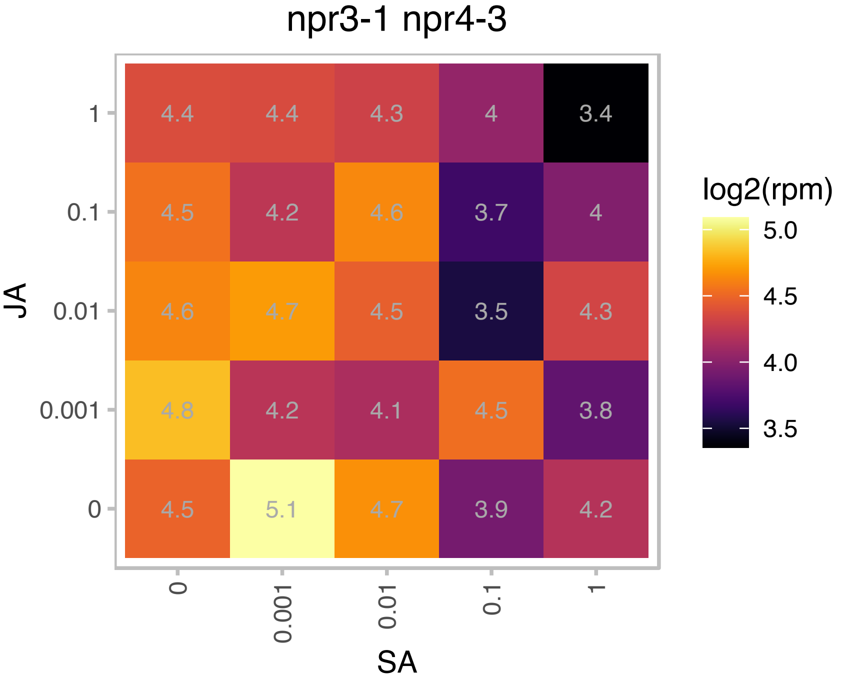

### AT4G00680.1 : ADF8

actin depolymerizing factor 8

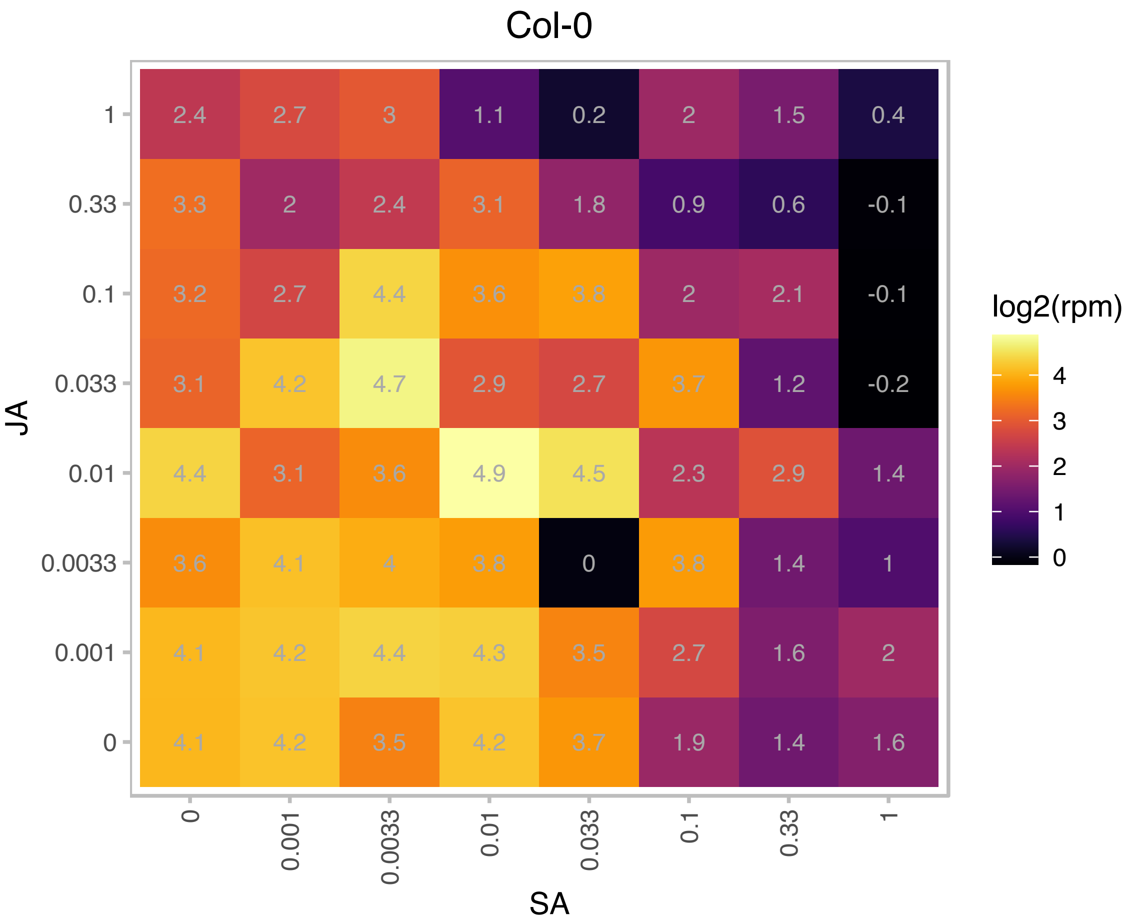


| | |
|---|---|
| Gene | AT4G00680 |
| is.expressed_genes | TRUE |
| is.responsive_genes | TRUE |
| is.apcluster | F-11 |

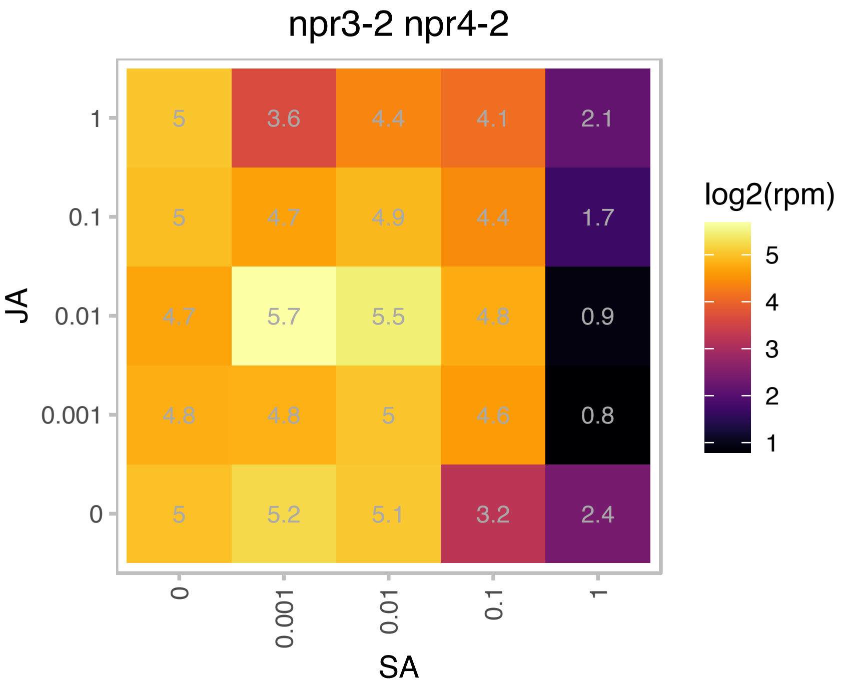


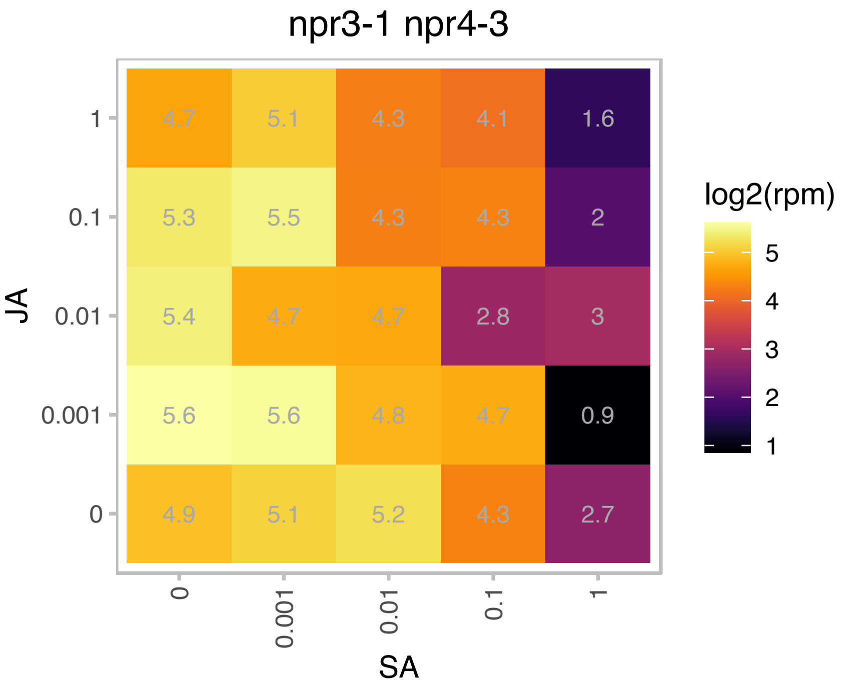

# AT4G00895.1 : NA

ATPase, F1 complex, OSCP/delta subunit protein

| | |
|---|---|
| Gene | AT4G00895 |
| is.expressed_genes | TRUE |
| is.responsive_genes | TRUE |
| is.apcluster | F-11 |

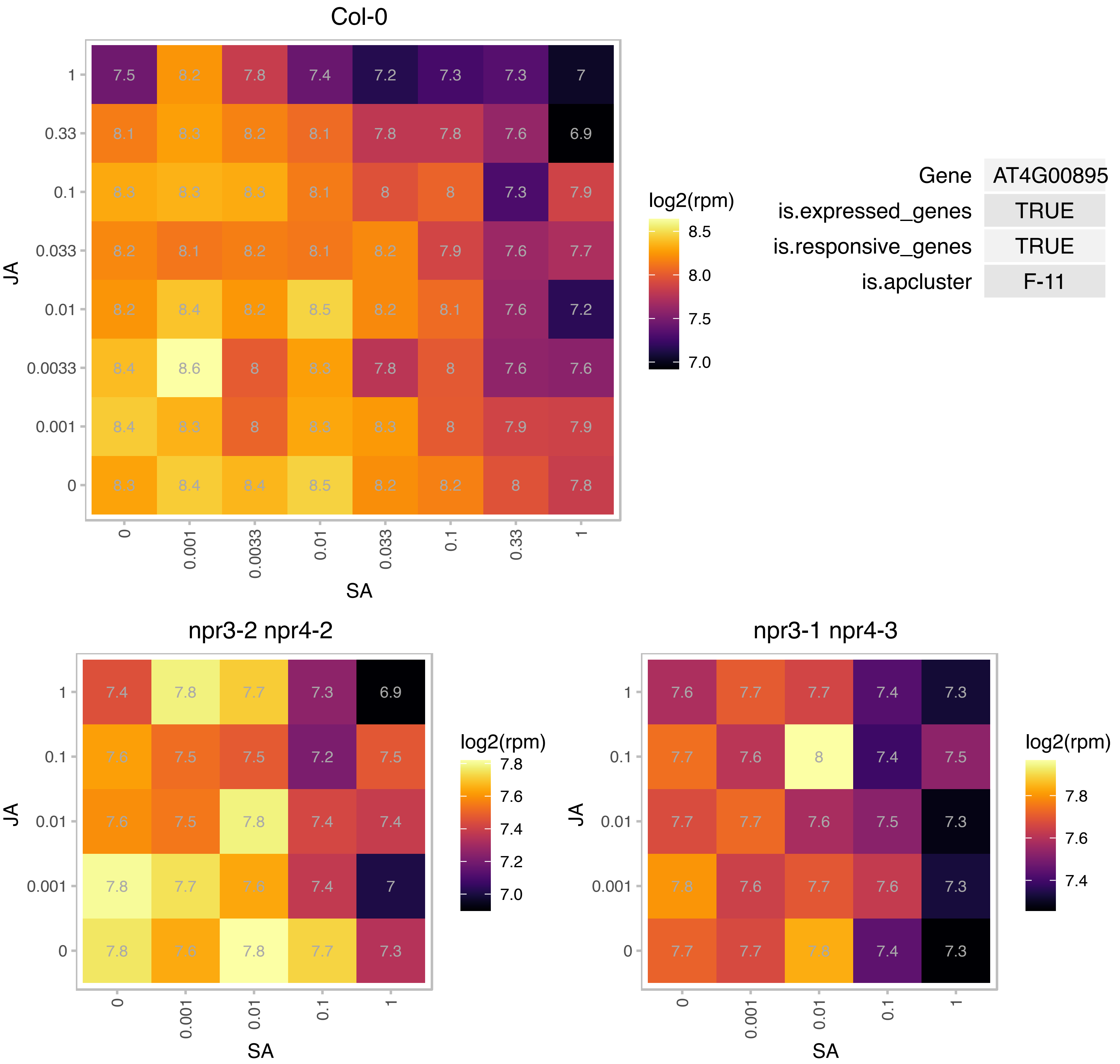

### AT4G02800.1 : MadA1

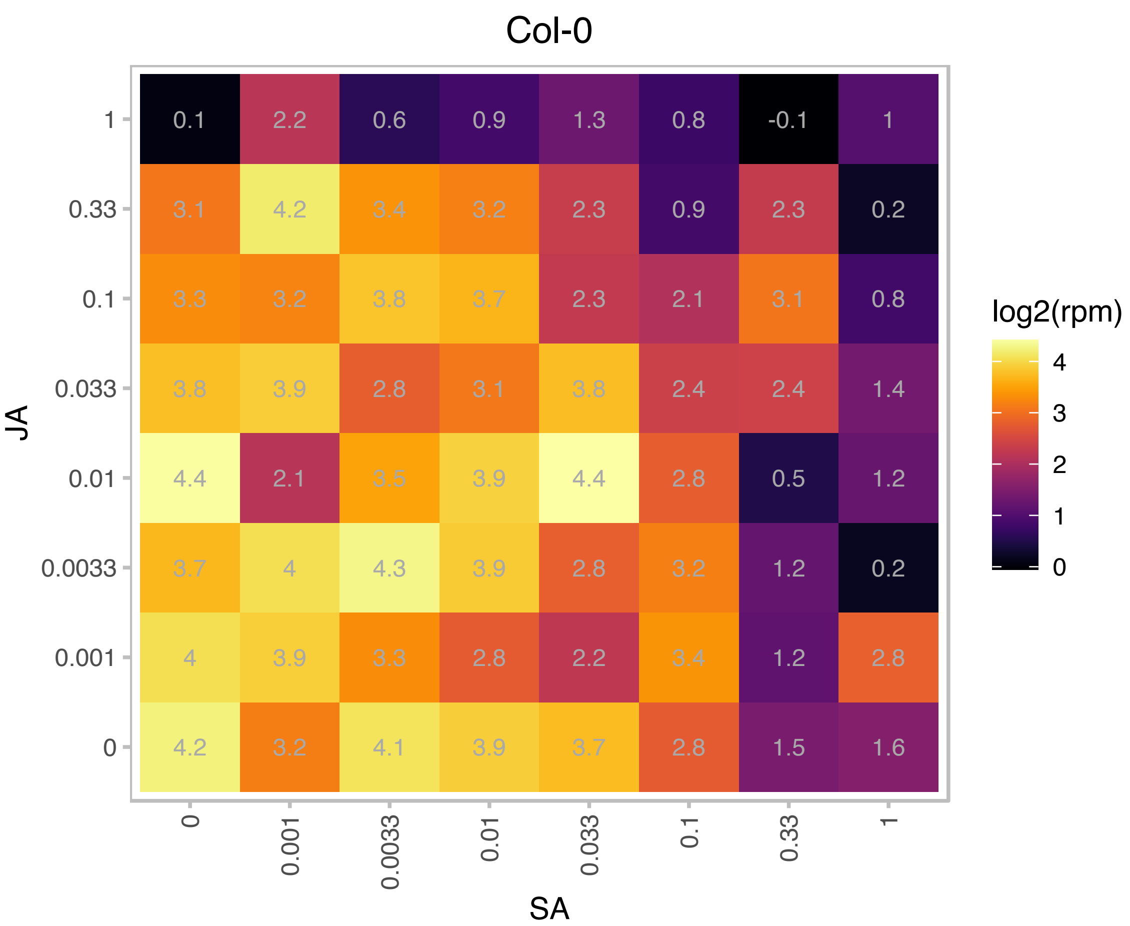


| | |
|---|---|
| Gene | AT4G02800 |
| is.expressed_genes | TRUE |
| is.responsive_genes | TRUE |
| is.apcluster | F-11 |

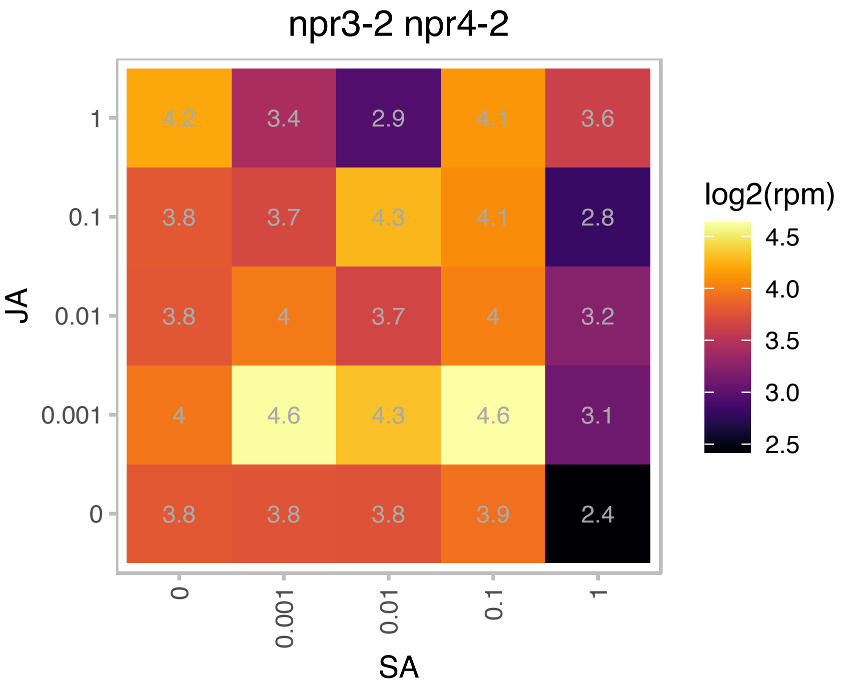


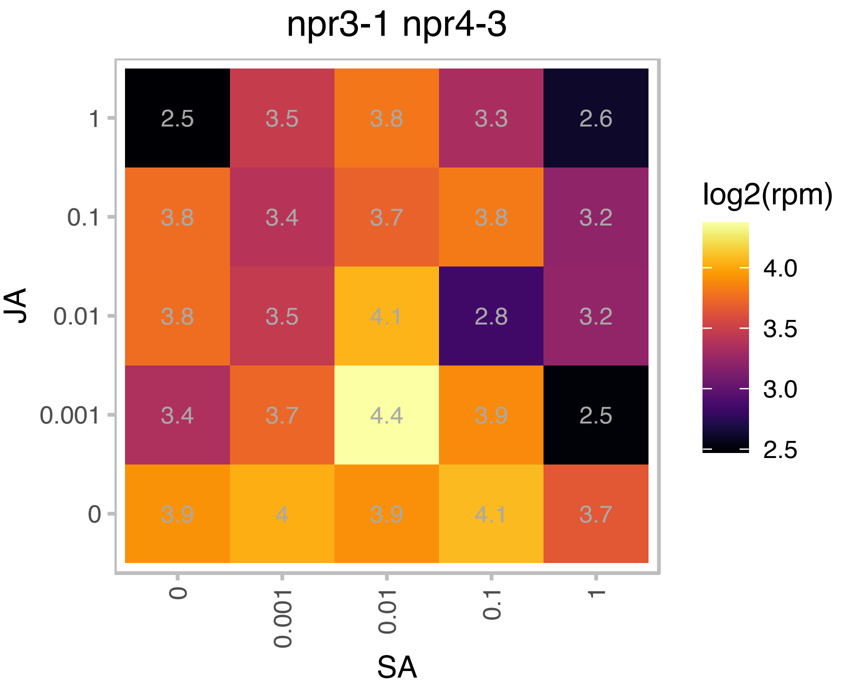

### AT4G03210.1 : XTH9

xyloglucan endotransglucosylase/hydrolase 9
encodes a member of xyloglucan endotransglucosylase/hydrolases (XTHs) that catalyze the cleavage and molecular grafting of xyloglucan chains function in loosening and rearrangement of the cell wall. Gene is expressed in shoot apex region, flower buds, flower stalks and internodes bearing flowers.

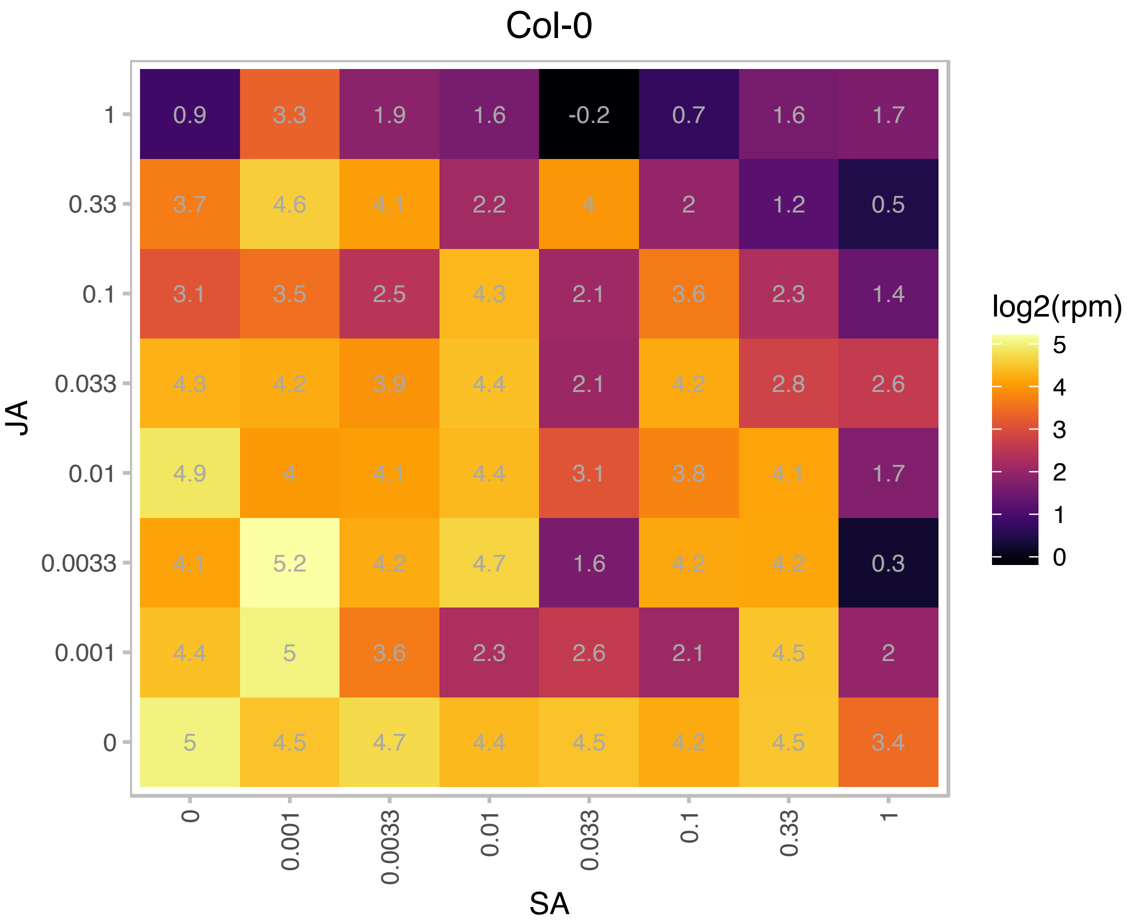


| | |
|---|---|
| Gene | AT4G03210 |
| is.expressed_genes | TRUE |
| is.responsive_genes | TRUE |
| is.apcluster | F-11 |

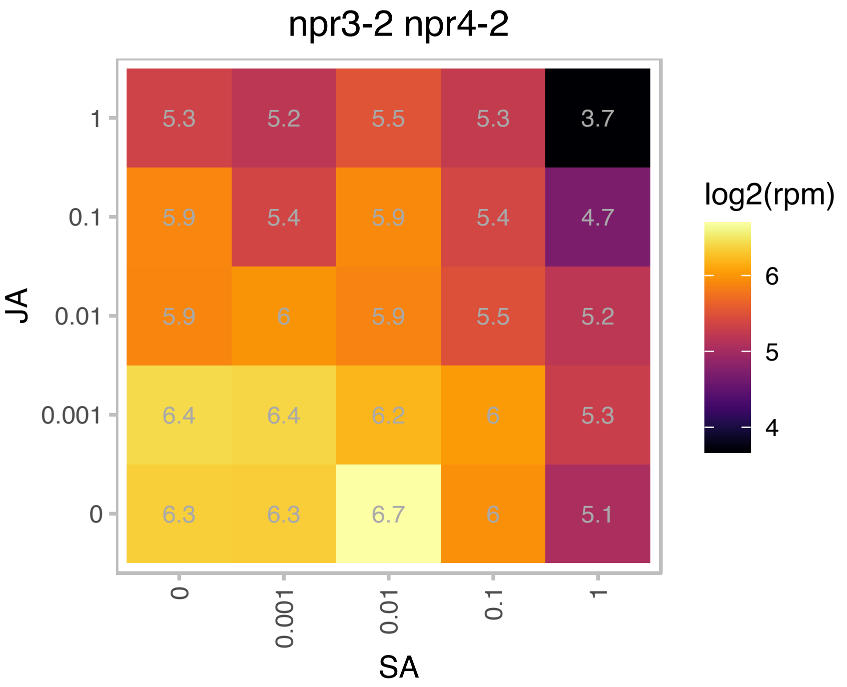


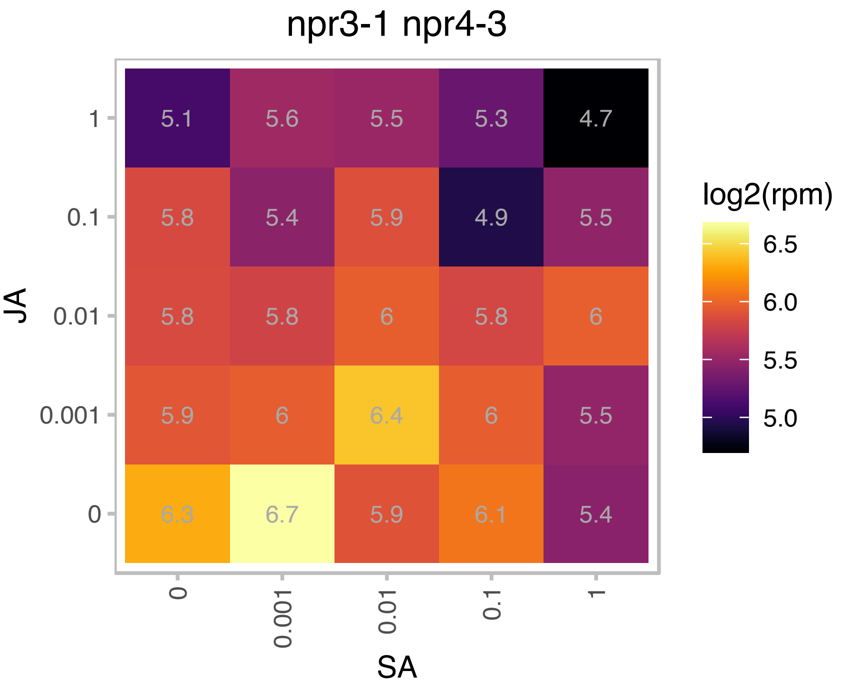

# AT4G04595.1 : NA

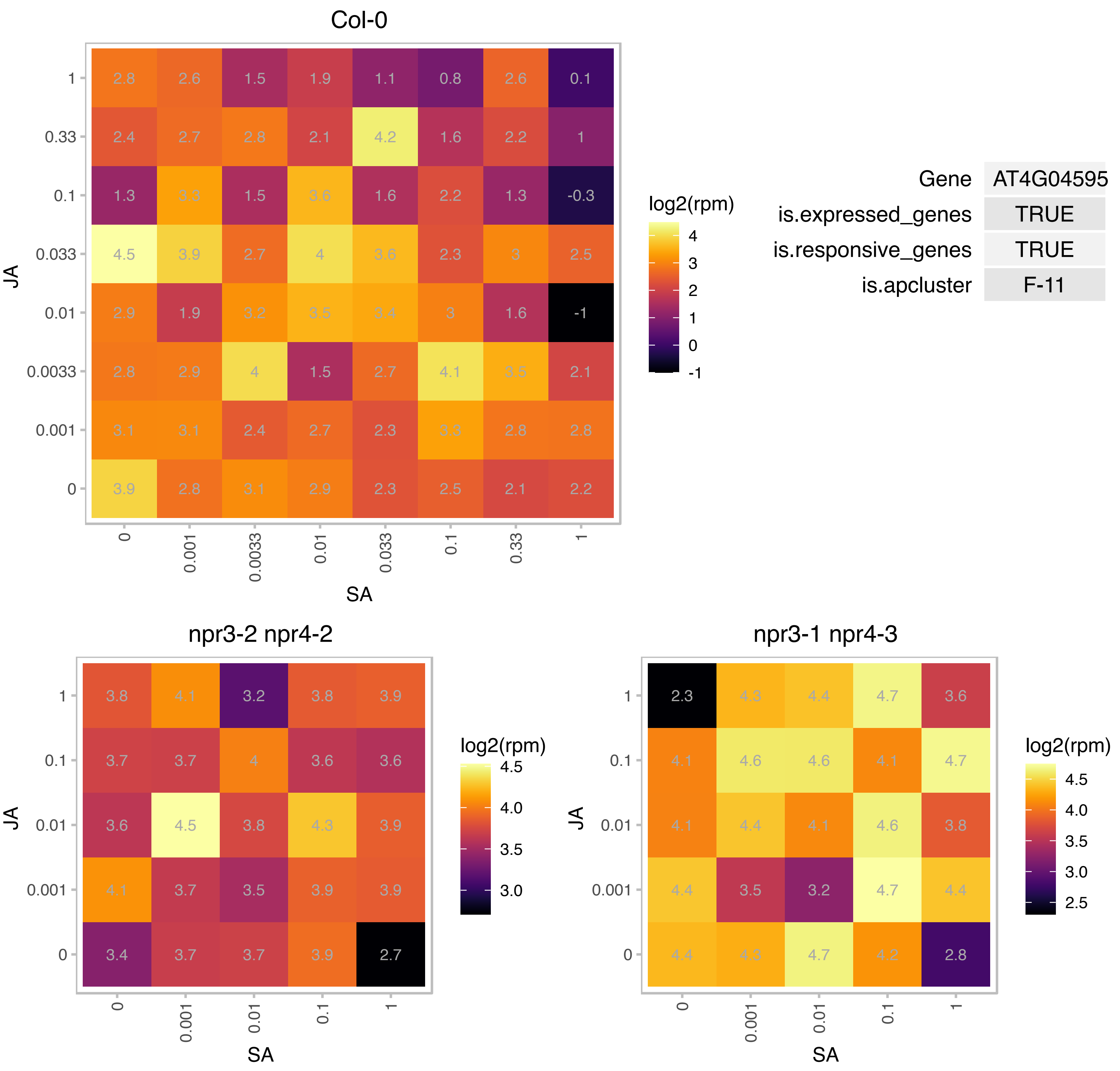

### AT4G08950.1 : EXO

Phosphate-responsive 1 family protein

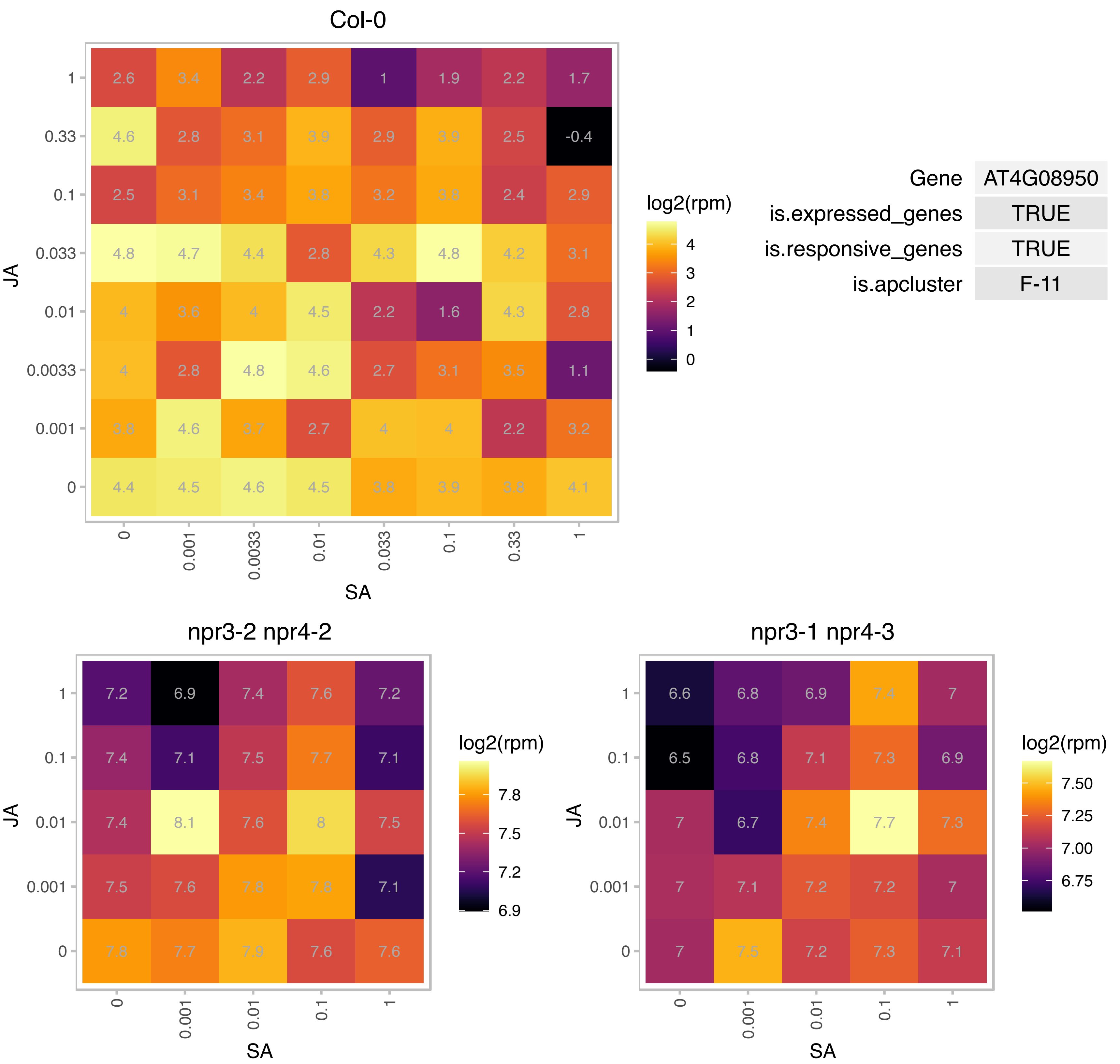


| | |
|---|---|
| Gene | AT4G08950 |
| is.expressed_genes | TRUE |
| is.responsive_genes | TRUE |
| is.apcluster | F-11 |

### AT4G12420.1 : SKU5

Cupredoxin superfamily protein.

Encodes a protein of unknown function involved in directed root tip growth. It is a member of 19-member gene family and is distantly related structurally to the multiple-copper oxidases ascorbate oxidase and laccase, though it lacks the copper-binding domains. The protein is glycosylated and GPI-anchored. It is localized to the plasma membrane and the cell wall. The gene is expressed most strongly in expanding tissues.

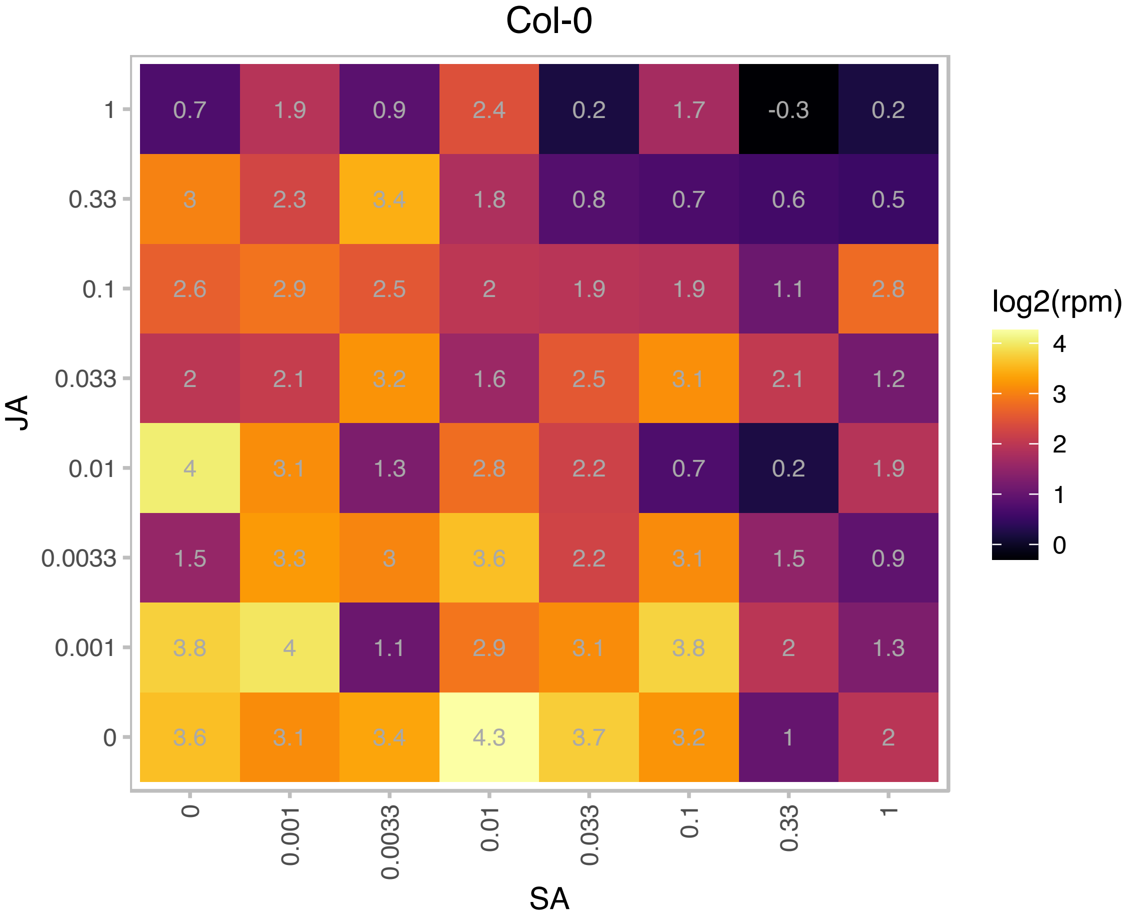


| | |
|---|---|
| Gene | AT4G12420 |
| is.expressed_genes | TRUE |
| is.responsive_genes | TRUE |
| is.apcluster | F-11 |

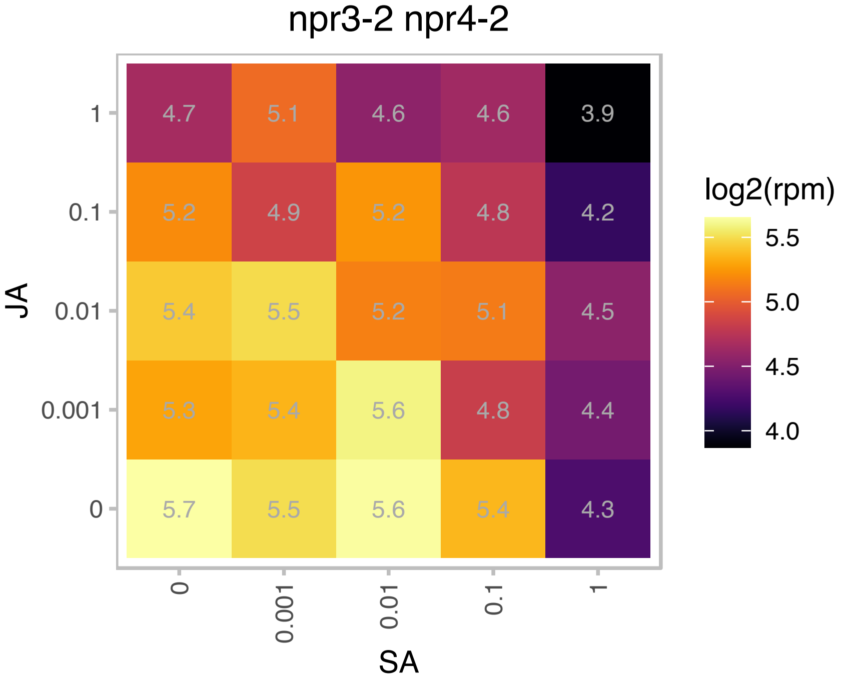


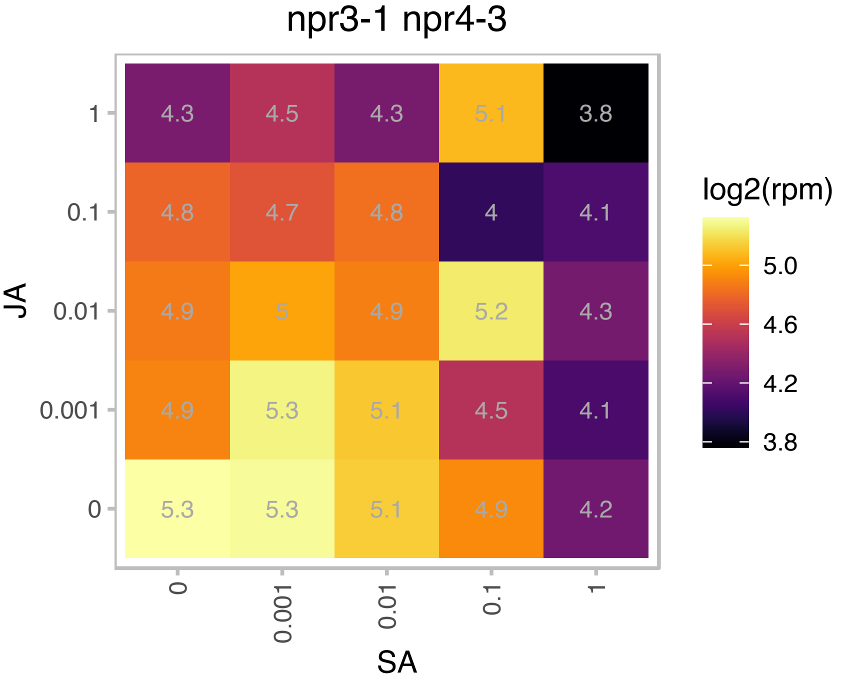

### AT4G12510.1 : AZI5

Bifunctional inhibitor/lipid-transfer protein/seed storage 2S albumin superfamily protein

Encodes a member of the AZI family of lipid transfer proteins.

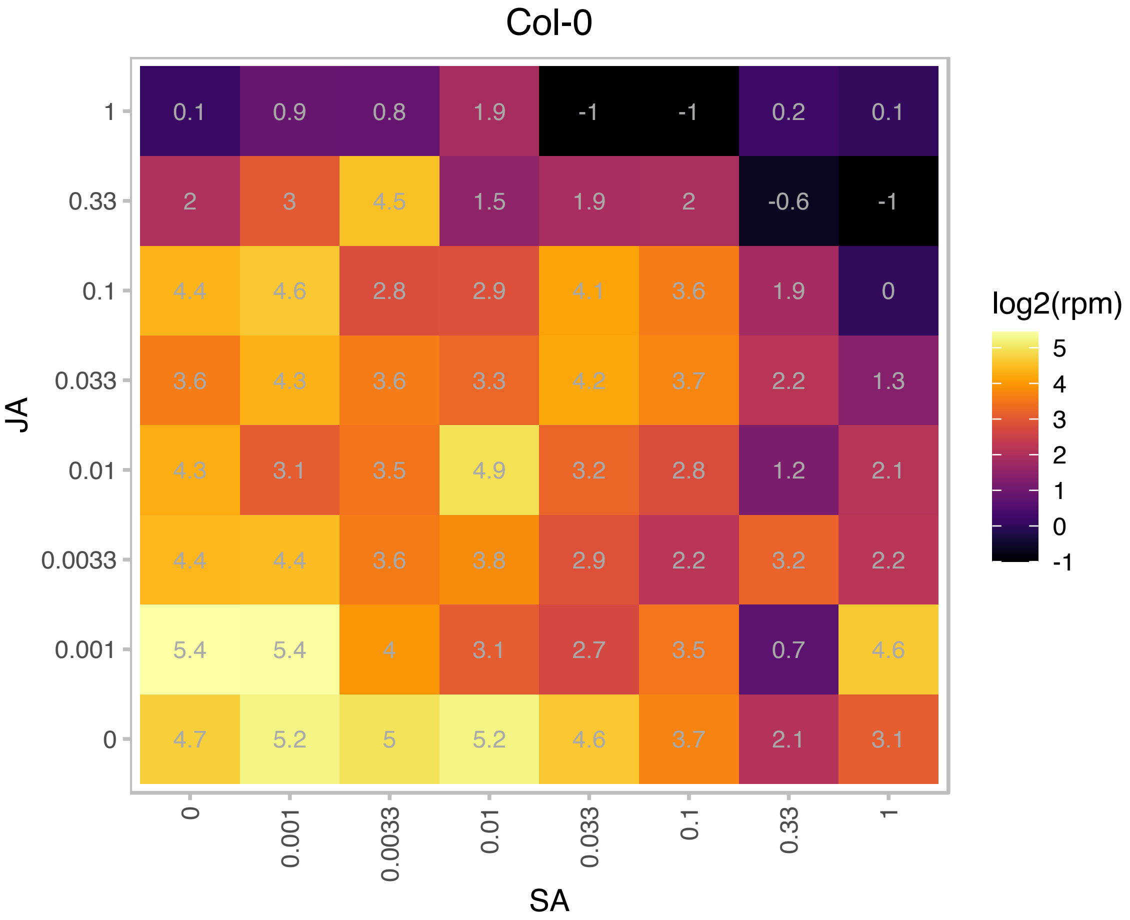


| | |
|---|---|
| Gene | AT4G12510 |
| is.expressed_genes | TRUE |
| is.responsive_genes | TRUE |
| is.apcluster | F-11 |

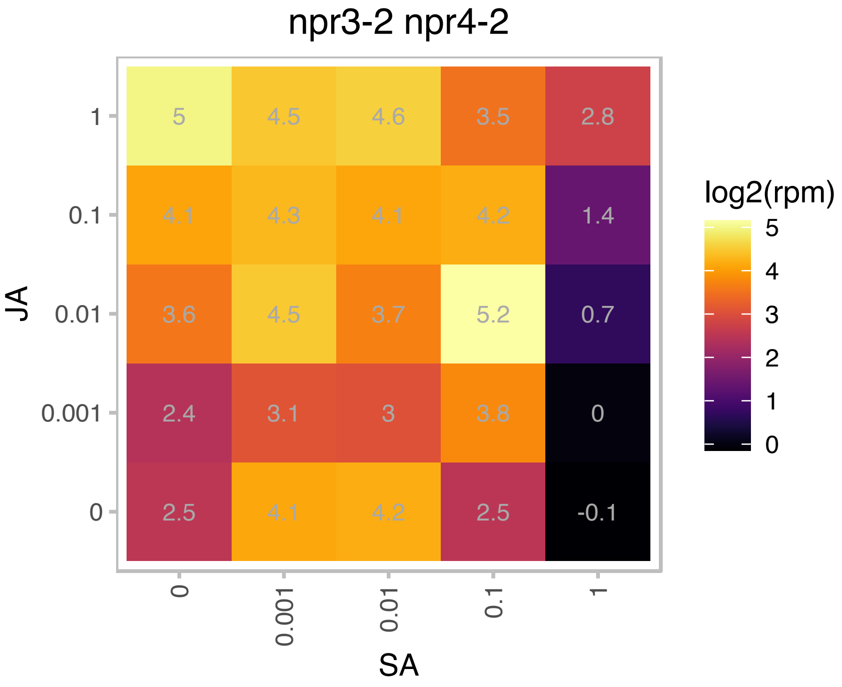


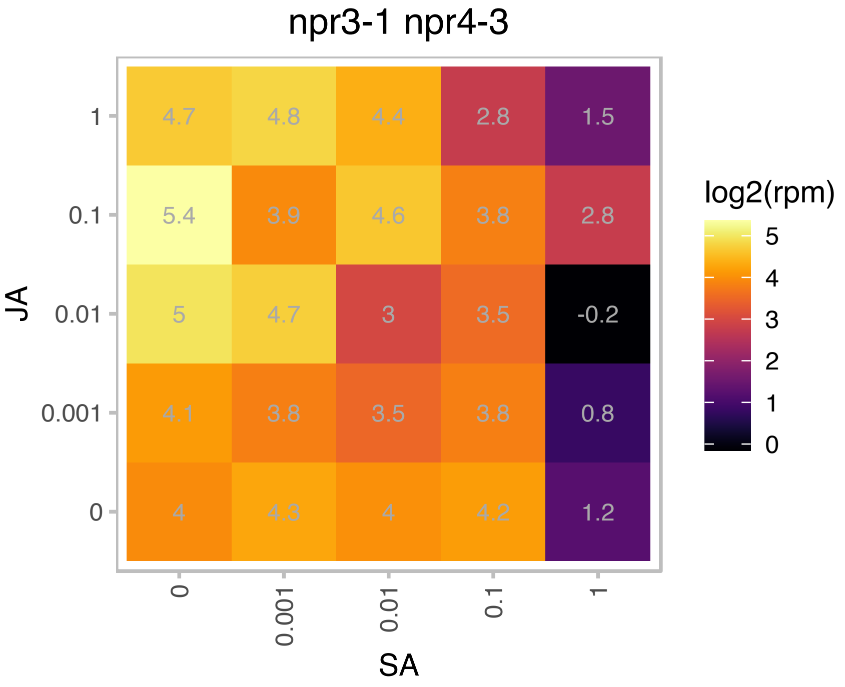

### AT4G12880.1 : AtENODL19, ENODL19

early nodulin-like protein 19

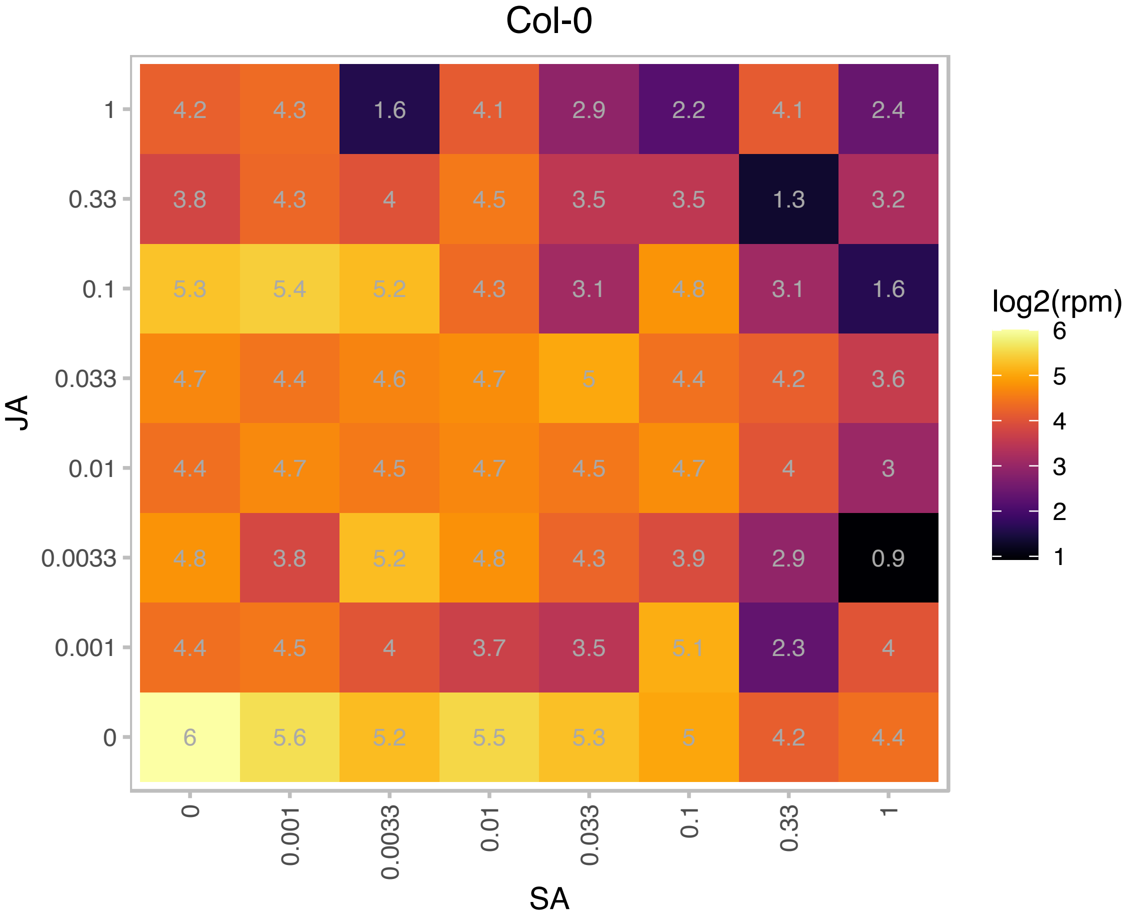


| | |
|---|---|
| Gene | AT4G12880 |
| is.expressed_genes | TRUE |
| is.responsive_genes | TRUE |
| is.apcluster | F-11 |

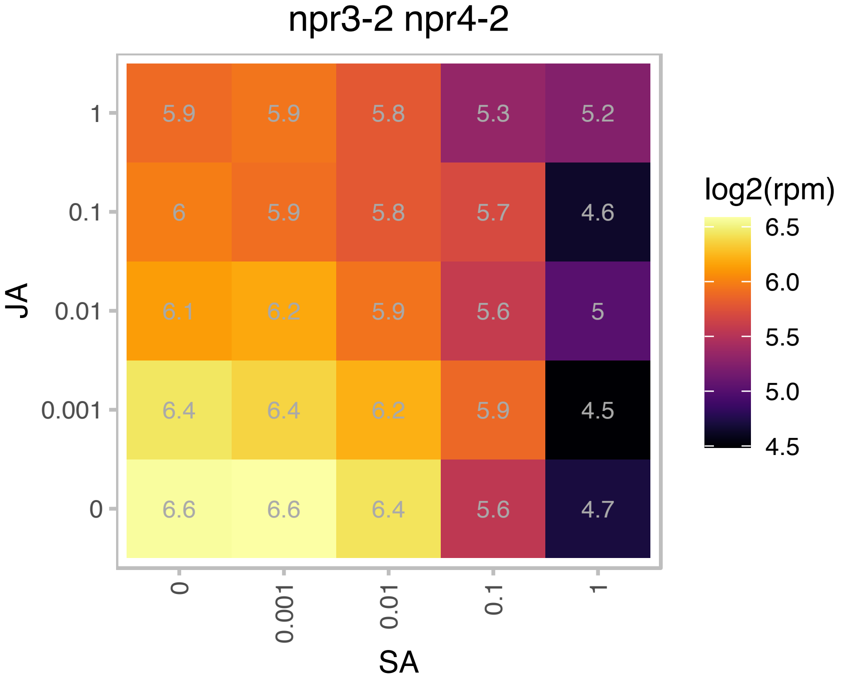


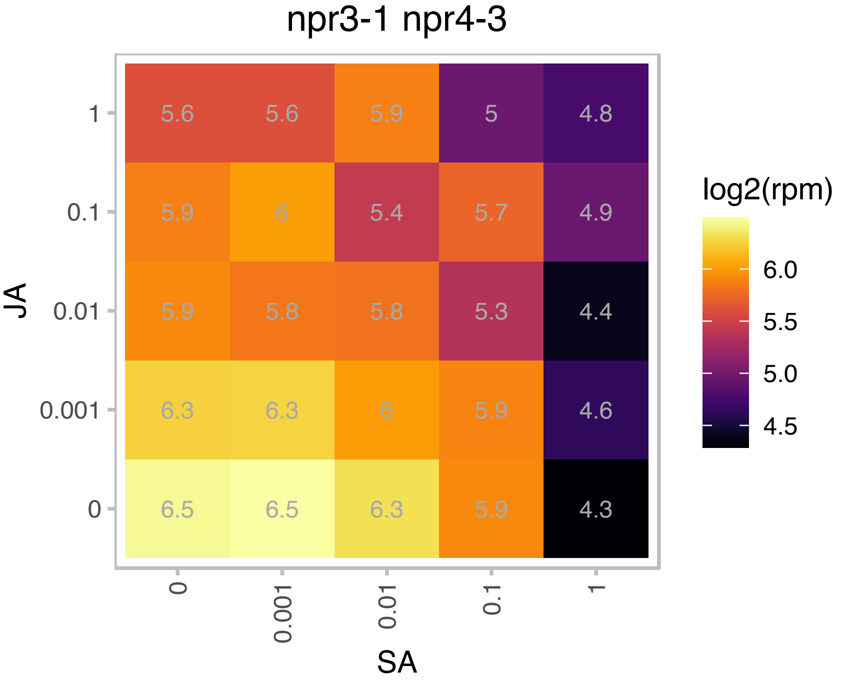

### AT4G12970.1 : AtEPFL9, EPFL9, STOMAGEN

Encodes a cysteine-rich peptide, a secretory factor that is produced in the mesophyll cells and acts on the
stomagen
epidermis to increase stomatal formation.  Its mature form is a 45-aa peptide with three intramolecular disulfide bonds.  It is proposed that STOMAGEN increases stomatal number by competing with two negative regulators of stomatal density, EPF1 and EPF2. STOMAGEN has been shown to compete with EPF2 for binding to the ER and TMM receptor kinases.Binding of STOMAGEN to ER prevents induction of the EPF2-ER MAPK cascade
It's transcript levels change after inducing MUTE expression in a mute background.

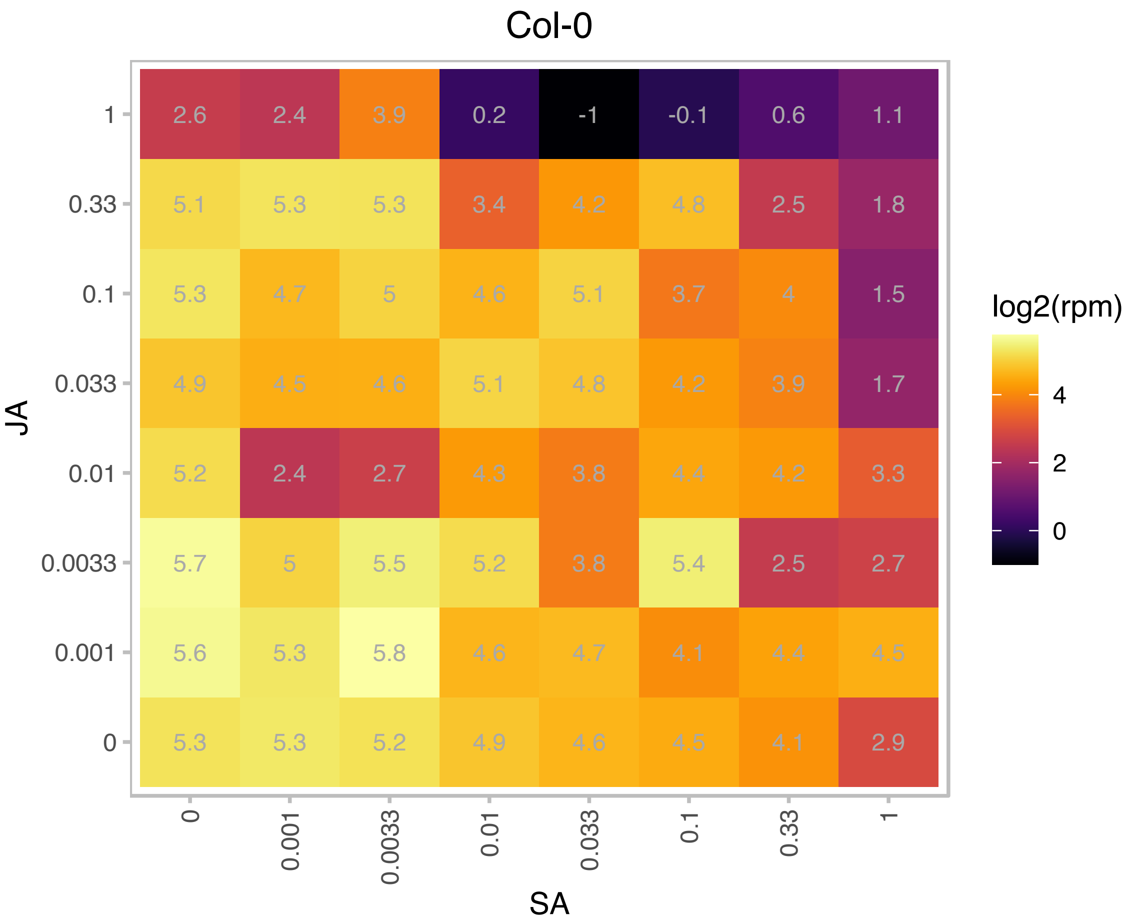


| | |
|---|---|
| Gene | AT4G12970 |
| is.expressed_genes | TRUE |
| is.responsive_genes | TRUE |
| is.apcluster | F-11 |

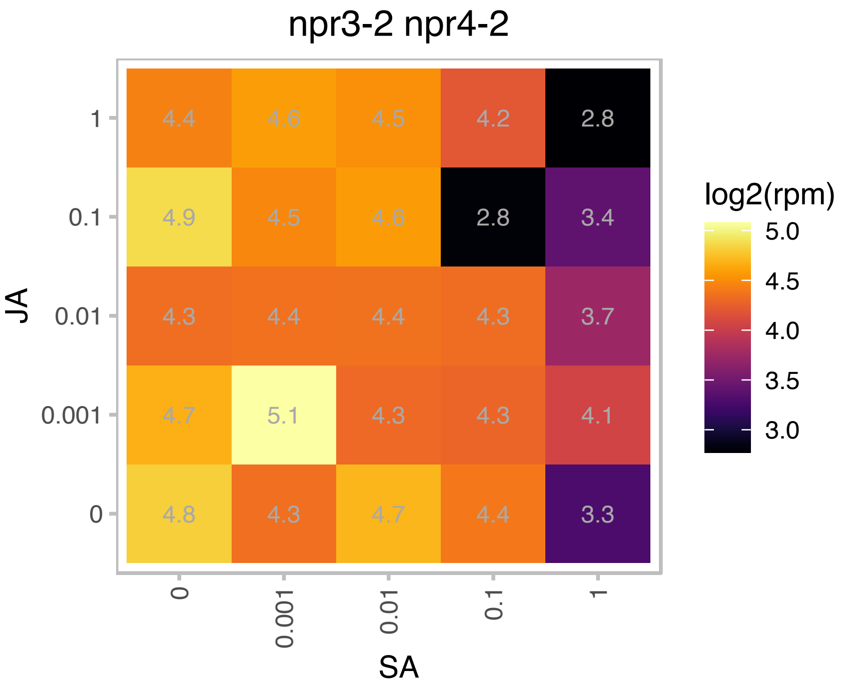


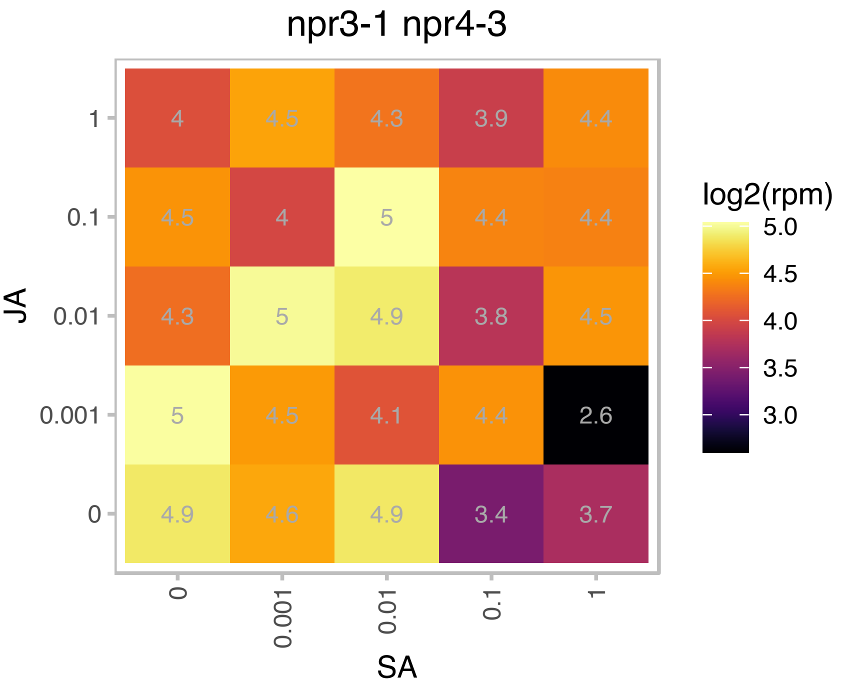

# AT4G13100.1 : NA

RING/U-box superfamily protein

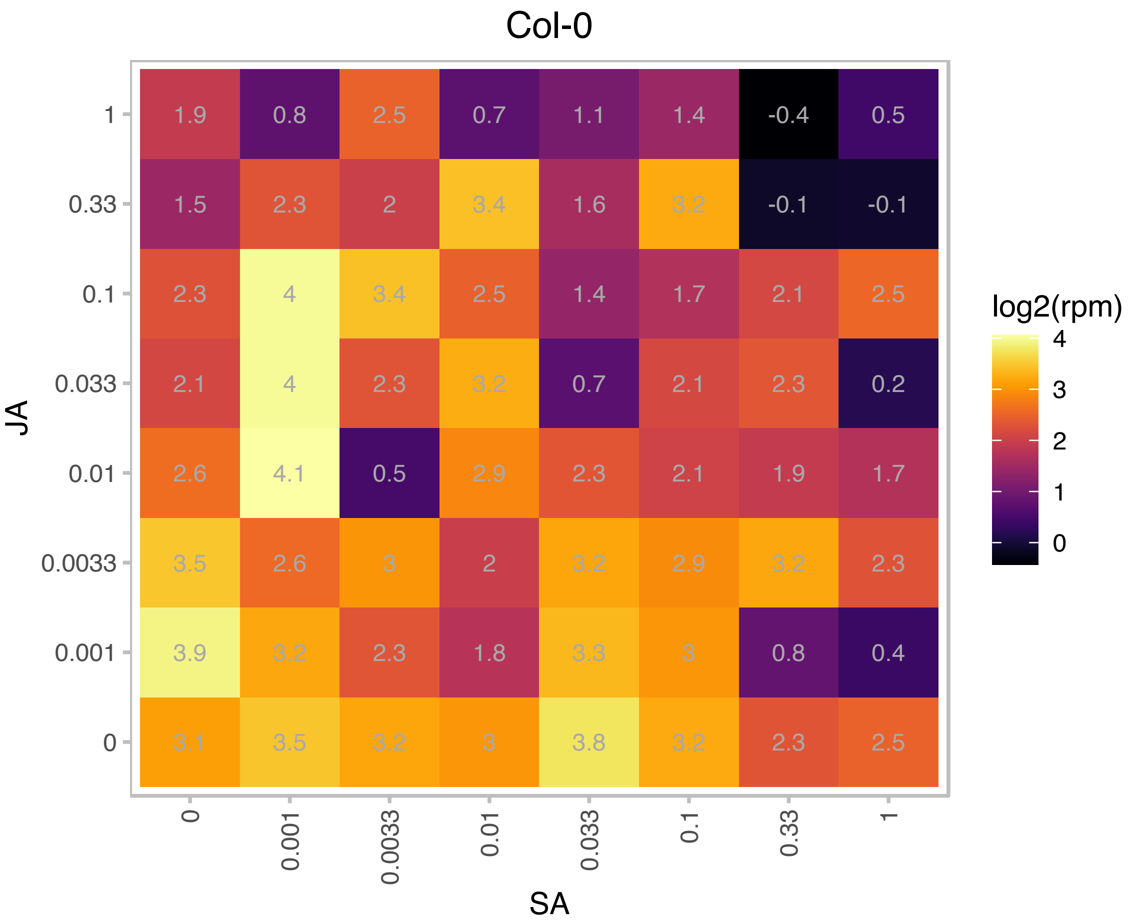


| | |
|---|---|
| Gene | AT4G13100 |
| is.expressed_genes | TRUE |
| is.responsive_genes | TRUE |
| is.apcluster | F-11 |

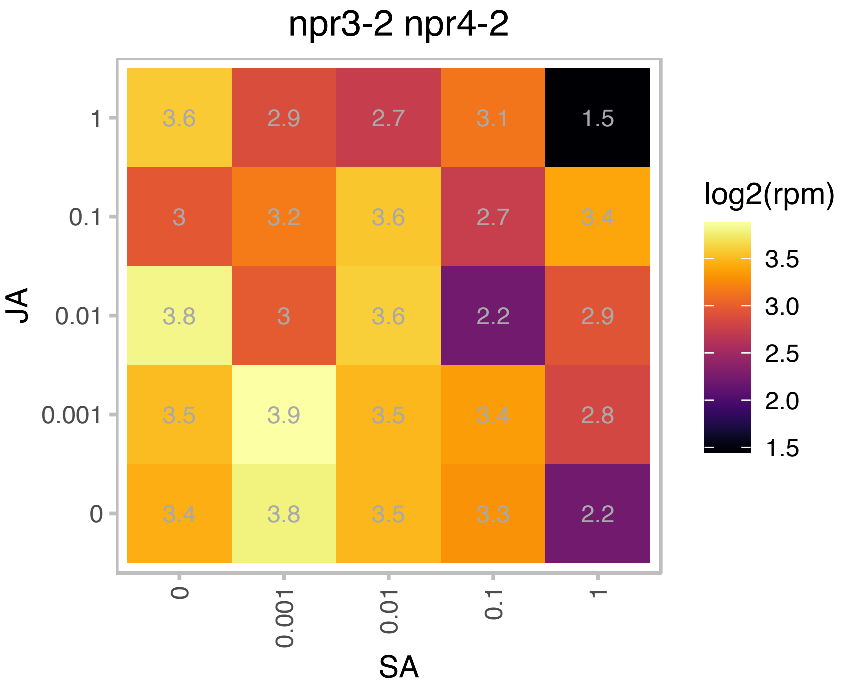


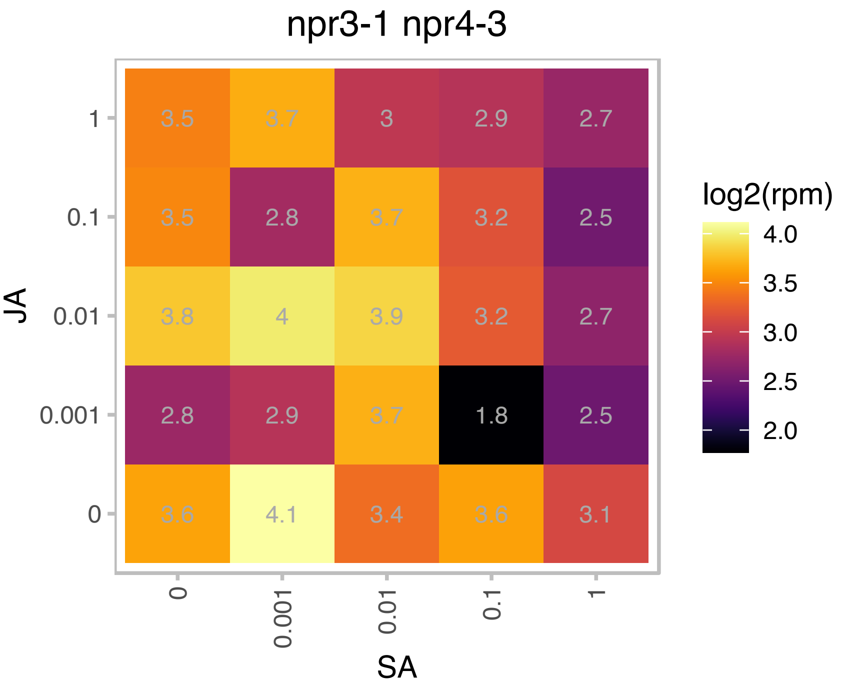

# AT4G14540.1 : NF-YB3

nuclear factor Y, subunit B3

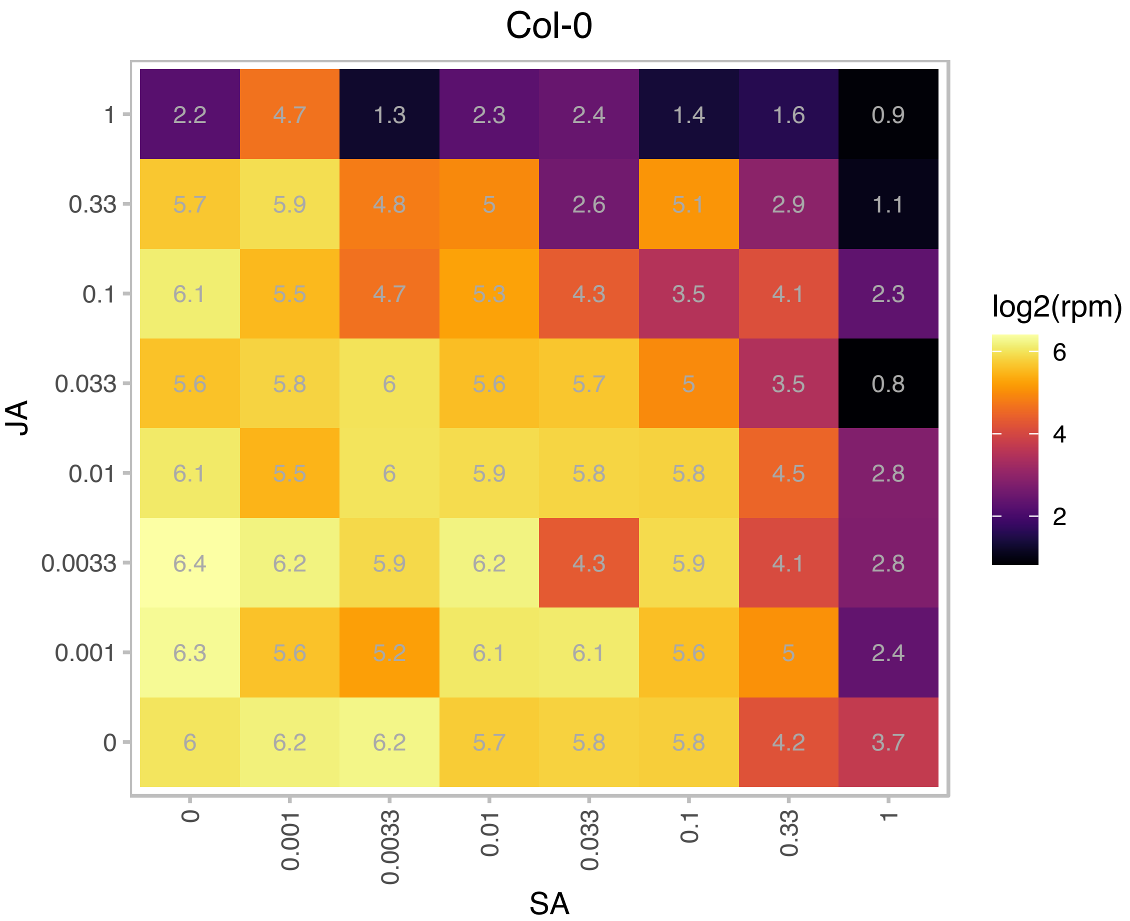


| | |
|---|---|
| Gene | AT4G14540 |
| is.expressed_genes | TRUE |
| is.responsive_genes | TRUE |
| is.apcluster | F-11 |

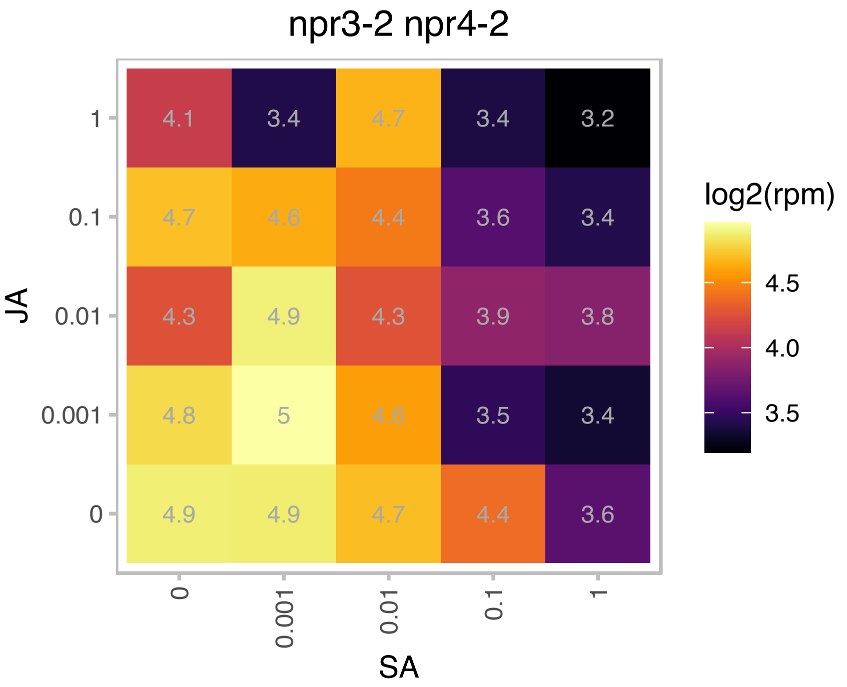


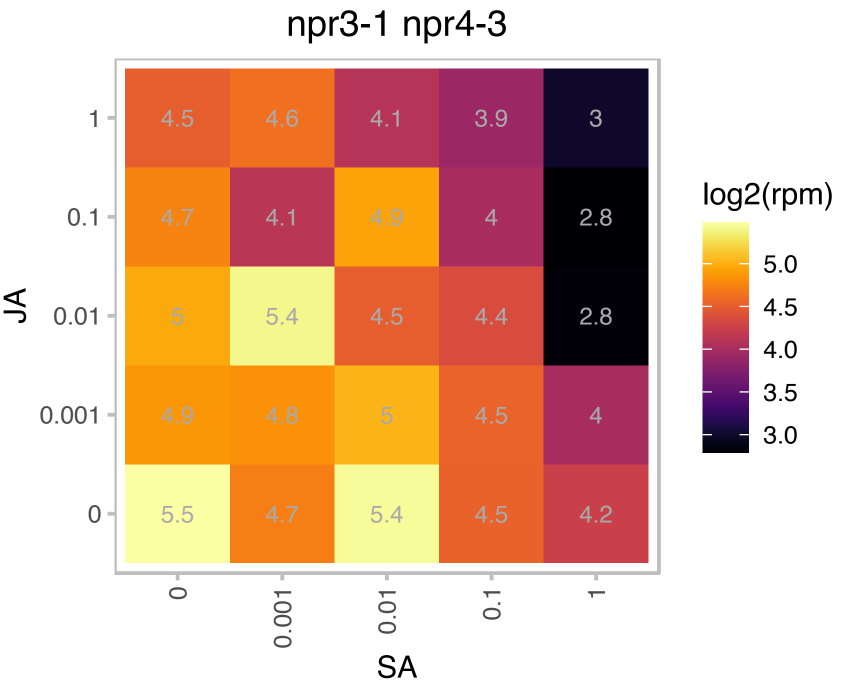

# AT4G15790.1 : NA

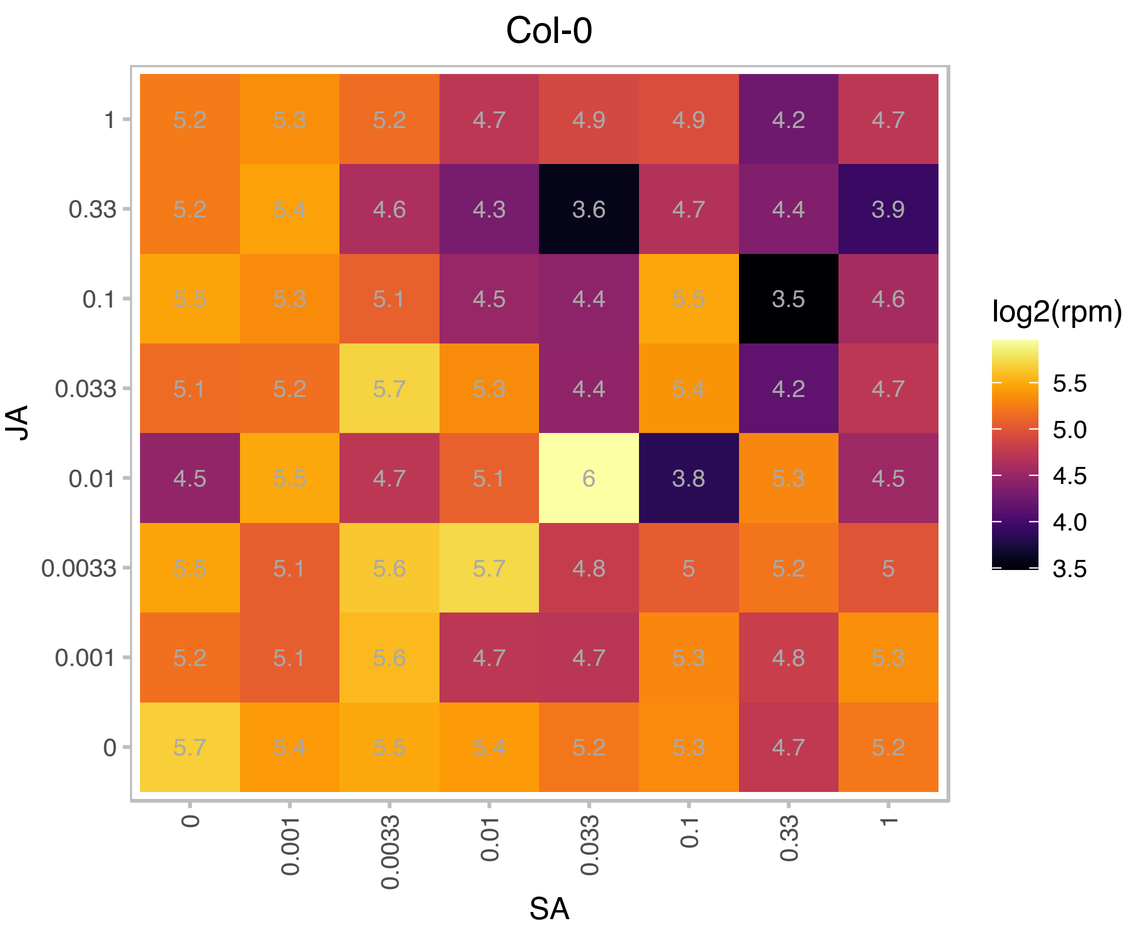


| | |
|---|---|
| Gene | AT4G15790 |
| is.expressed_genes | TRUE |
| is.responsive_genes | TRUE |
| is.apcluster | F-11 |

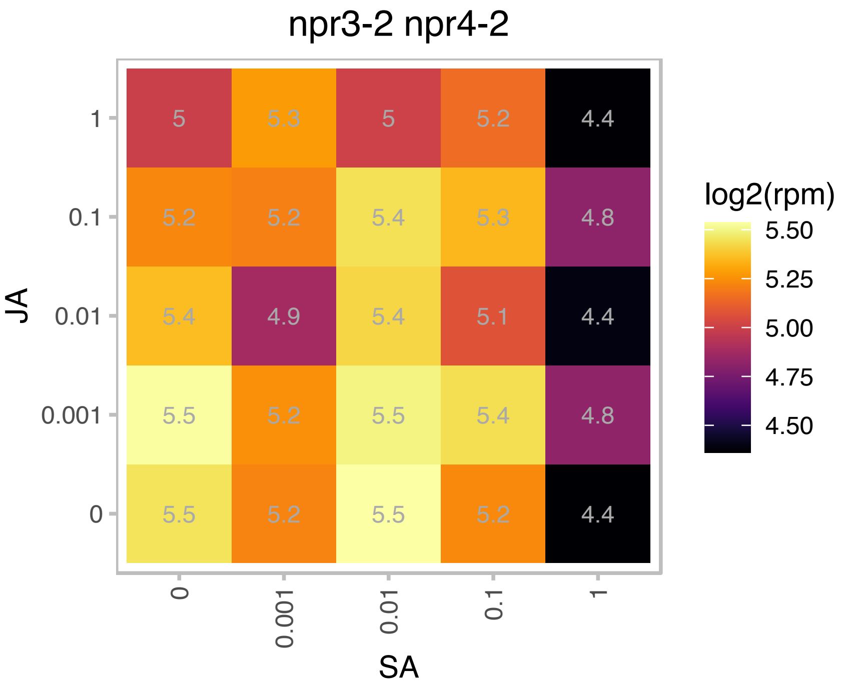


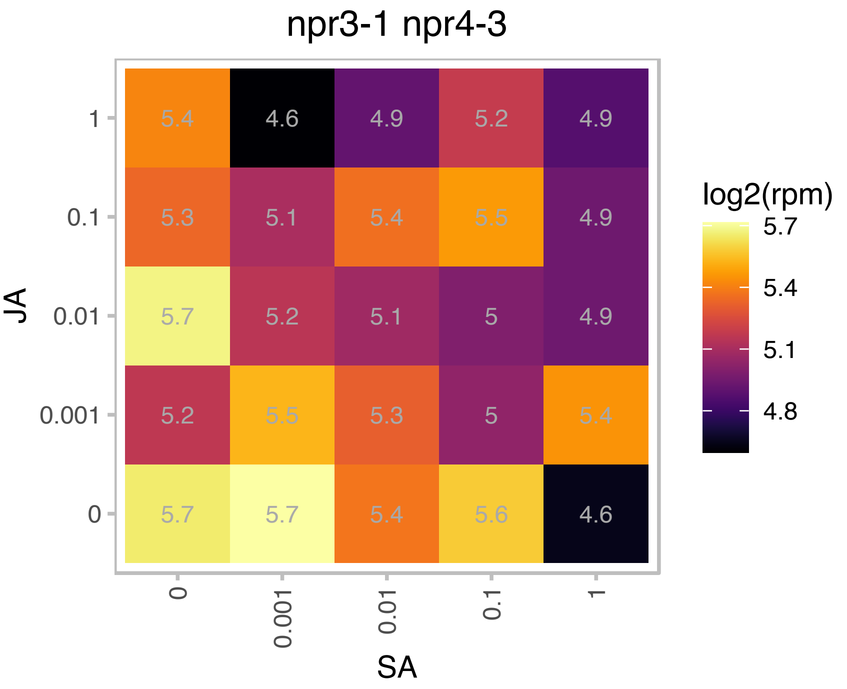

# AT4G16410.1 : NA

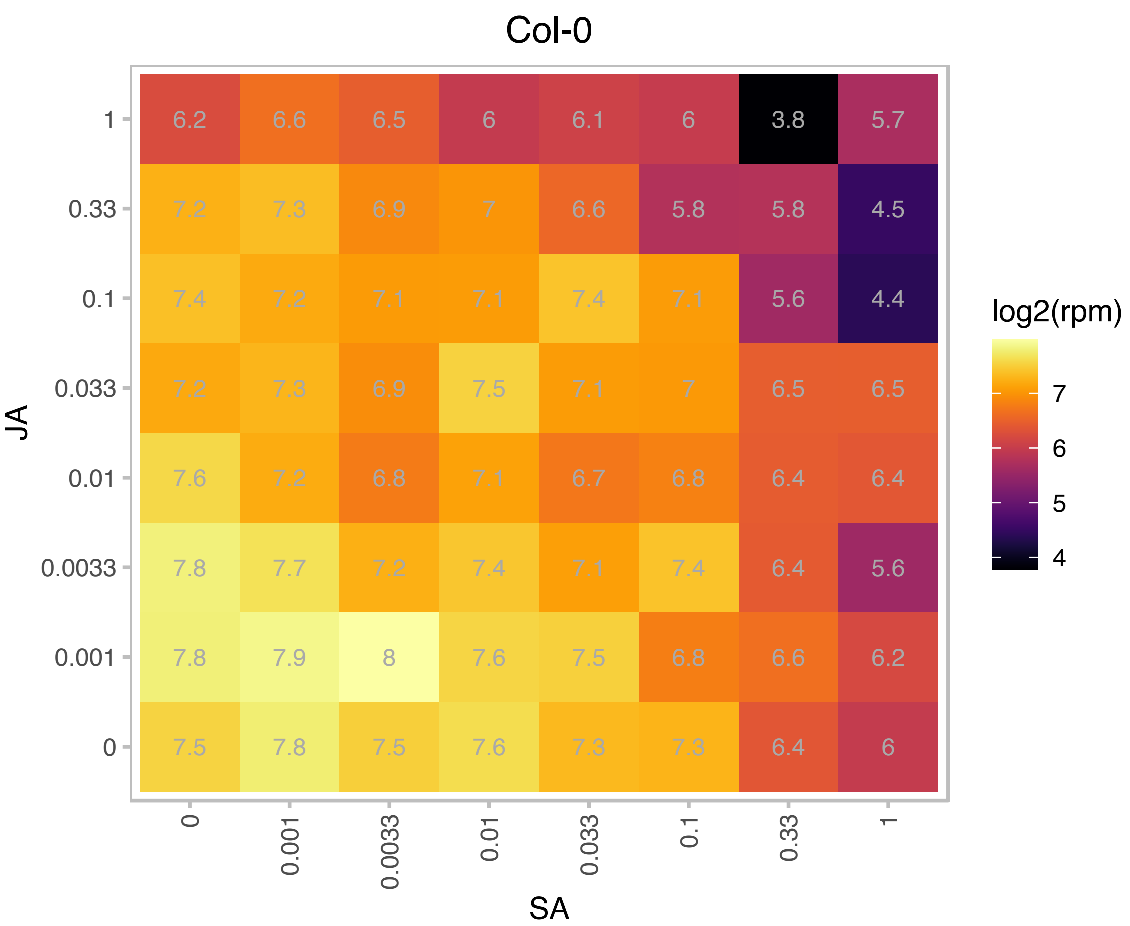


| | |
|---|---|
| Gene | AT4G16410 |
| is.expressed_genes | TRUE |
| is.responsive_genes | TRUE |
| is.apcluster | F-11 |

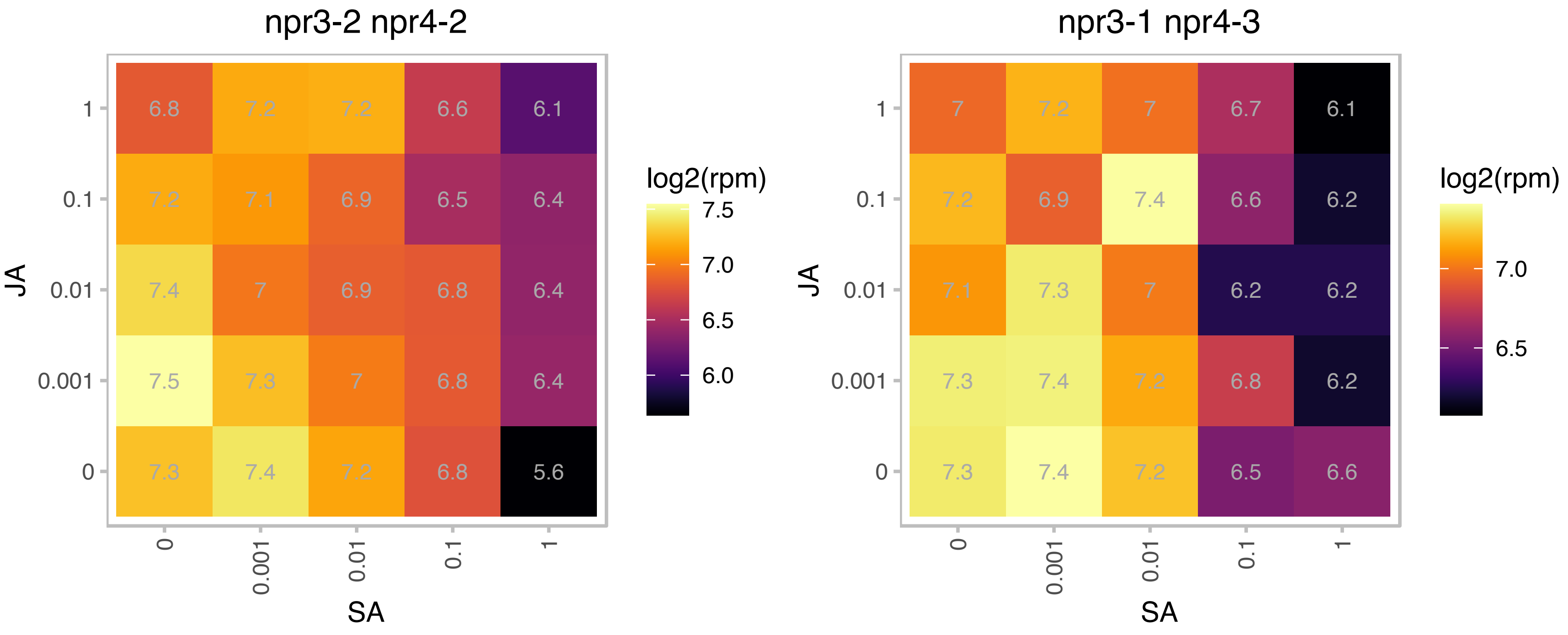

### AT4G17600.1 : LIL3:1

Chlorophyll A-B binding family protein
Encodes Lil3:1 (light-harvesting-like) protein. Belongs to the Lhc super-gene family encodes the light-harvesting chlorophyll a/b-binding (LHC) proteins that constitute the antenna system of the photosynthetic apparatus. A generic LHC motif is present in Lil3:1. The mRNA is cell-to-cell mobile.

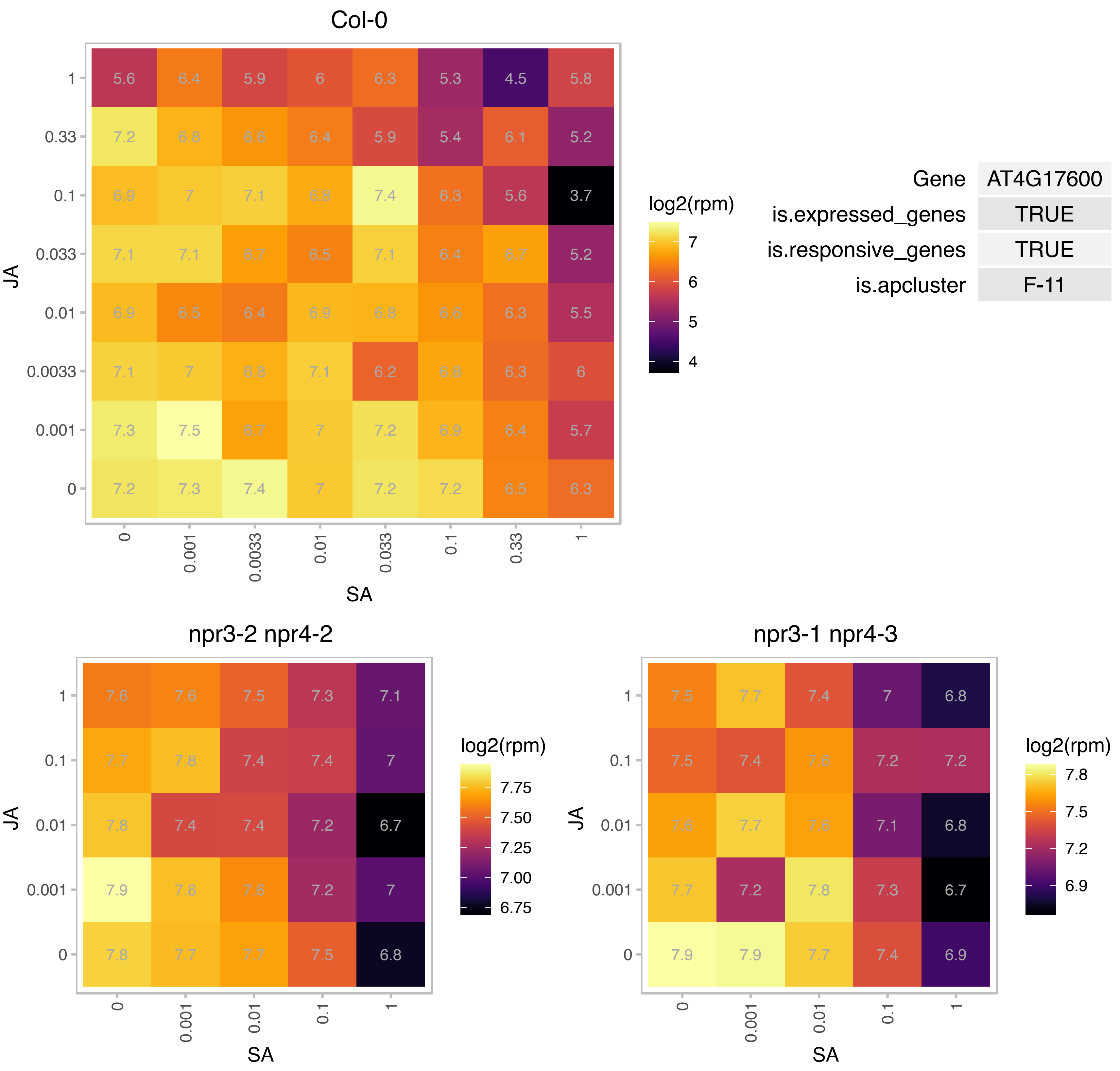


| | |
|---|---|
| Gene | AT4G17600 |
| is.expressed_genes | TRUE |
| is.responsive_genes | TRUE |
| is.apcluster | F-11 |

# AT4G17810.1 : NA

C2H2 and C2HC zinc fingers superfamily protein

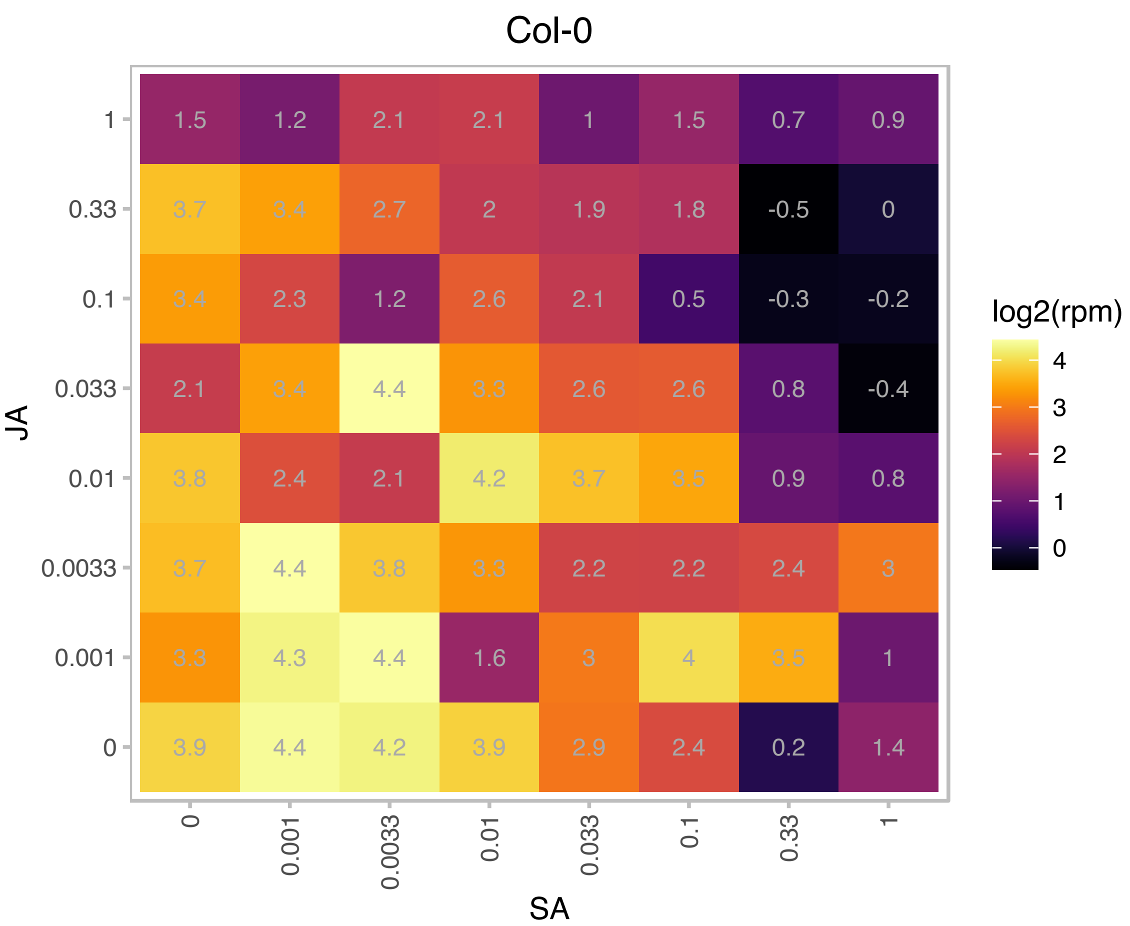


| | |
|---|---|
| Gene | AT4G17810 |
| is.expressed_genes | TRUE |
| is.responsive_genes | TRUE |
| is.apcluster | F-11 |

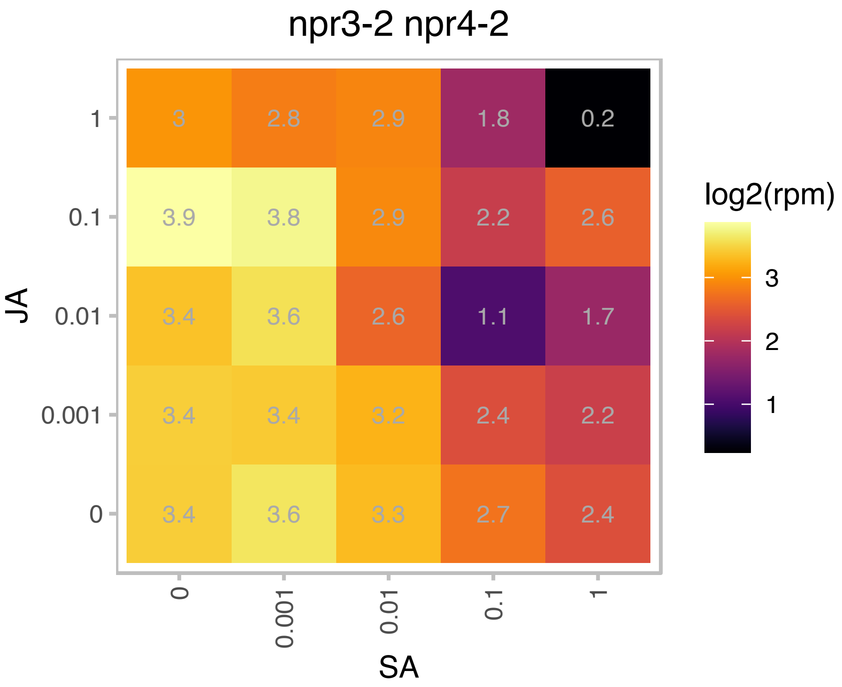


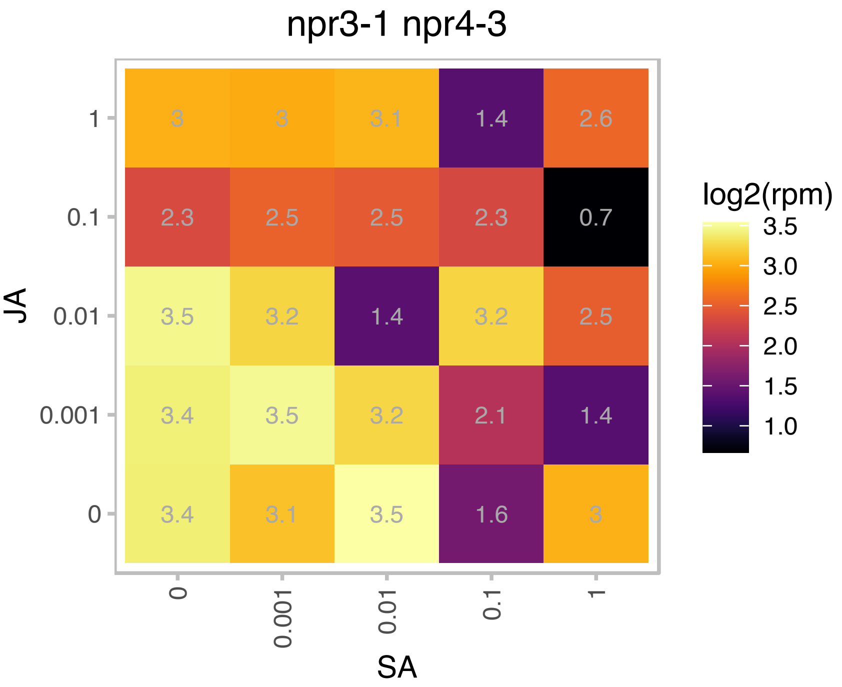

### AT4G21960.1 : PRXR1

Peroxidase superfamily protein

Encodes AT4g21960 (AT4g21960/T8O5_170). The mRNA is cell-to-cell mobile.

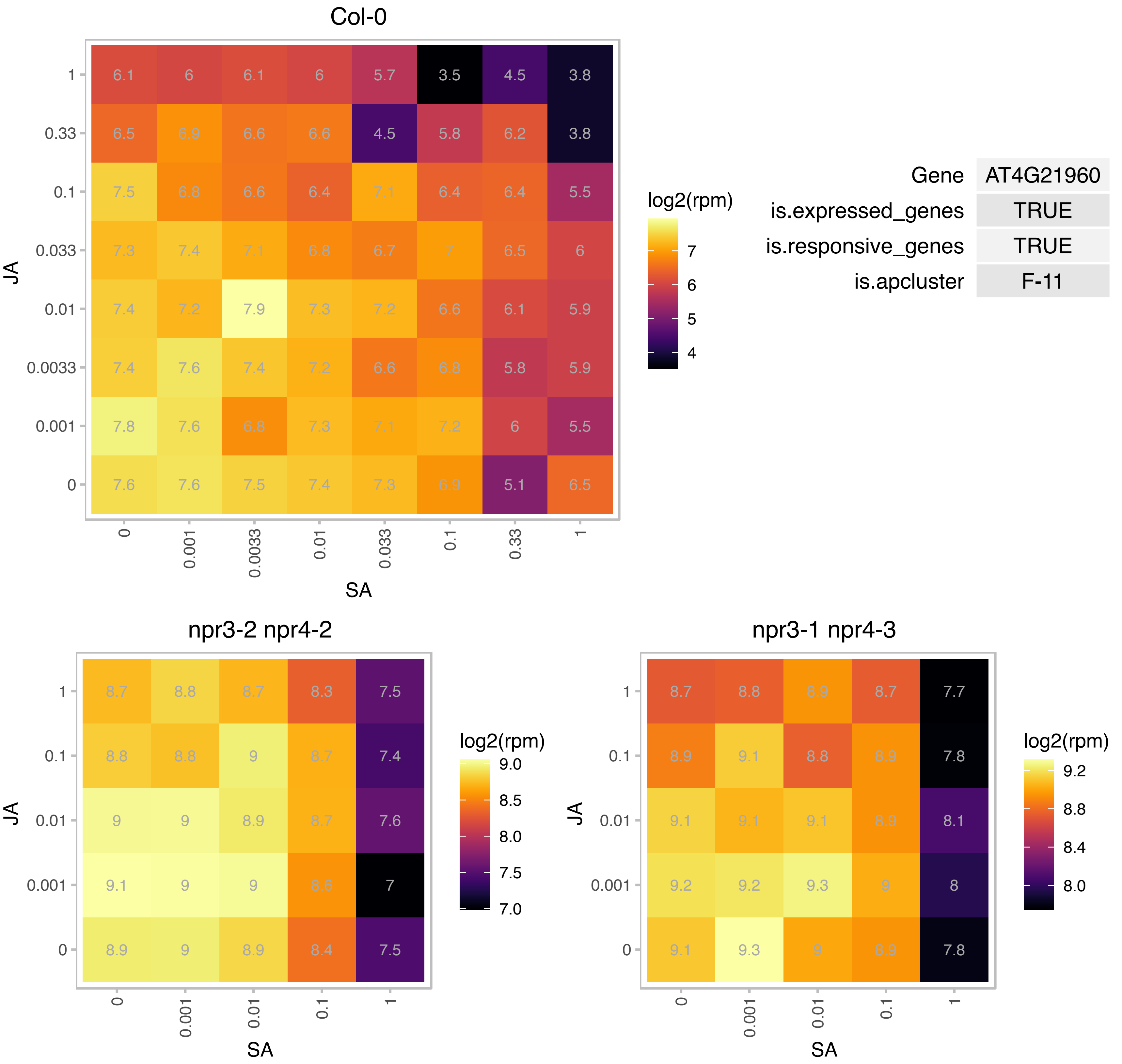


| | |
|---|---|
| Gene | AT4G21960 |
| is.expressed_genes | TRUE |
| is.responsive_genes | TRUE |
| is.apcluster | F-11 |

### AT4G23800.1 : 3xHMG-box2

HMG (high mobility group) box protein
Encodes a protein containing three copies of the HMG (high mobility group)-box domain. The two Arabidopsis 3xHMG-box proteins are: AT4G11080 (3xHMG-box1) and AT4G23800 (3xHMG-box2). Interacts with mitotic and meiotic chromosomes.

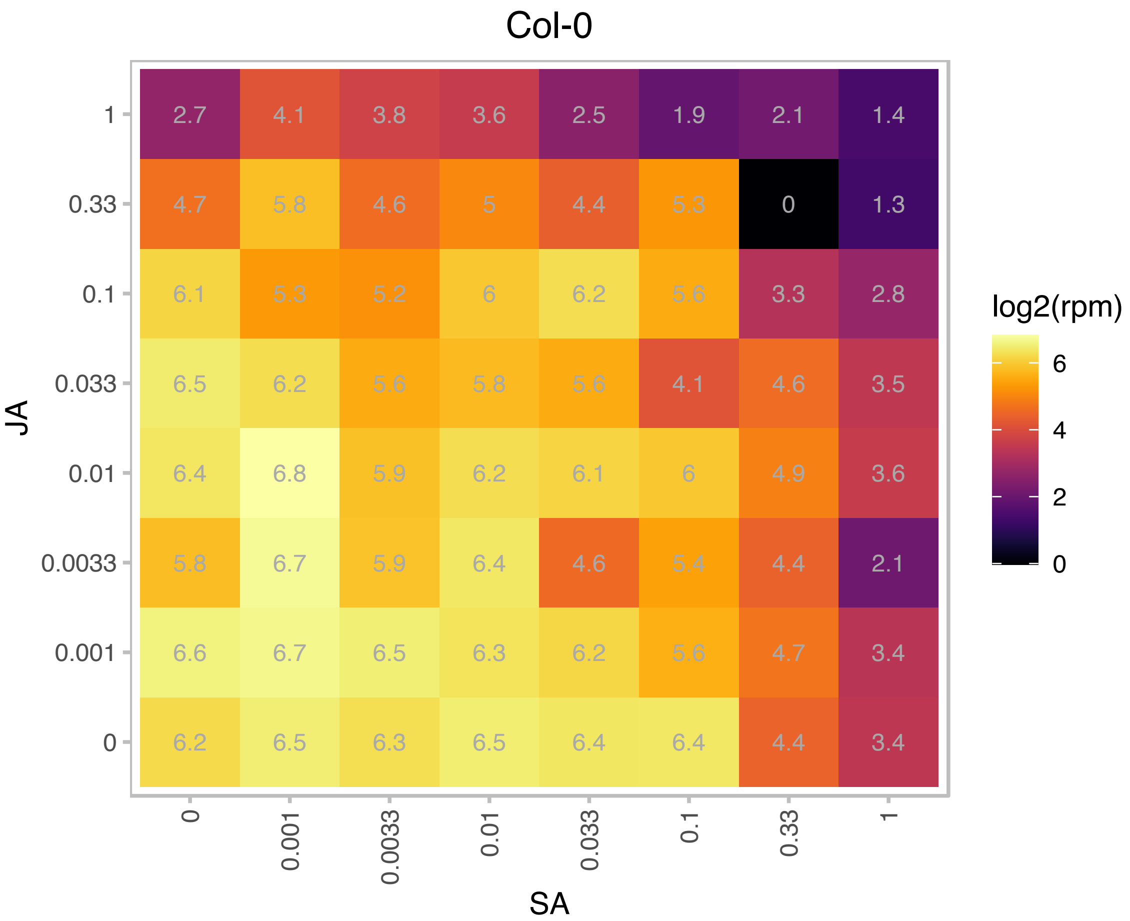


| JA \ SA | 0 | 0.001 | 0.0033 | 0.01 | 0.033 | 0.1 | 0.33 | 1 |
|---|---|---|---|---|---|---|---|---|
| 1 | 2.7 | 4.1 | 3.8 | 3.6 | 2.5 | 1.9 | 2.1 | 1.4 |
| 0.33 | 4.7 | 5.8 | 4.6 | 5 | 4.4 | 5.3 | 0 | 1.3 |
| 0.1 | 6.1 | 5.3 | 5.2 | 6 | 6.2 | 5.6 | 3.3 | 2.8 |
| 0.033 | 6.5 | 6.2 | 5.6 | 5.8 | 5.6 | 4.1 | 4.6 | 3.5 |
| 0.01 | 6.4 | 6.8 | 5.9 | 6.2 | 6.1 | 6 | 4.9 | 3.6 |
| 0.0033 | 5.8 | 6.7 | 5.9 | 6.4 | 4.6 | 5.4 | 4.4 | 2.1 |
| 0.001 | 6.6 | 6.7 | 6.5 | 6.3 | 6.2 | 5.6 | 4.7 | 3.4 |
| 0 | 6.2 | 6.5 | 6.3 | 6.5 | 6.4 | 6.4 | 4.4 | 3.4 |

| | |
|---|---|
| Gene | AT4G23800 |
| is.expressed_genes | TRUE |
| is.responsive_genes | TRUE |
| is.apcluster | F-11 |

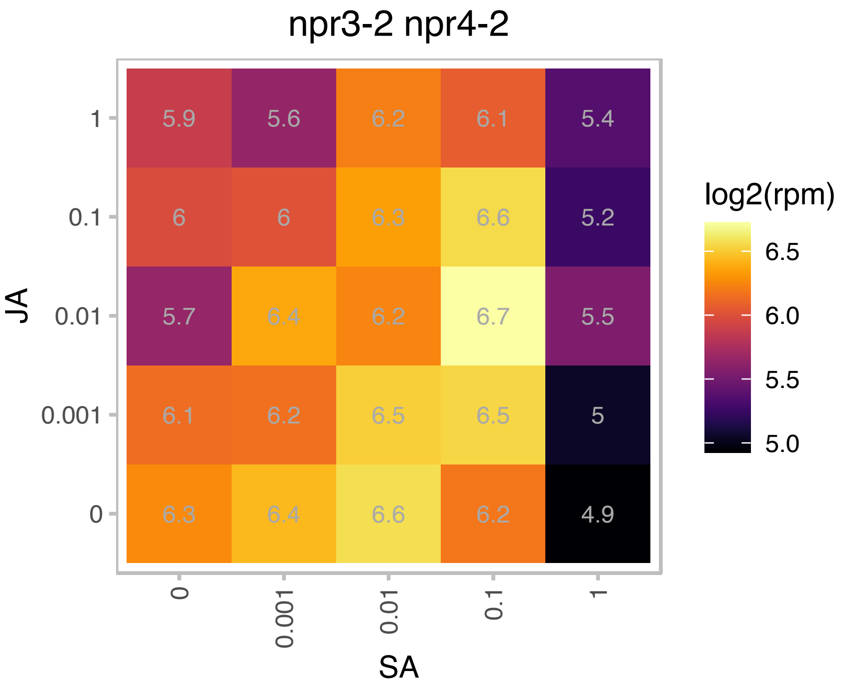


| JA \ SA | 0 | 0.001 | 0.01 | 0.1 | 1 |
|---|---|---|---|---|---|
| 1 | 5.9 | 5.6 | 6.2 | 6.1 | 5.4 |
| 0.1 | 6 | 6 | 6.3 | 6.6 | 5.2 |
| 0.01 | 5.7 | 6.4 | 6.2 | 6.7 | 5.5 |
| 0.001 | 6.1 | 6.2 | 6.5 | 6.5 | 5 |
| 0 | 6.3 | 6.4 | 6.6 | 6.2 | 4.9 |

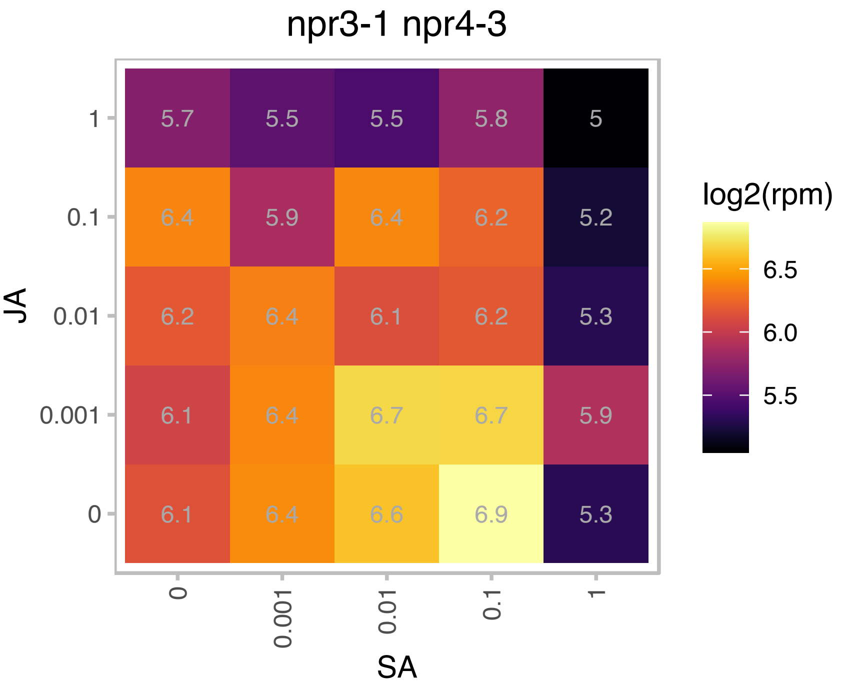


| JA \ SA | 0 | 0.001 | 0.01 | 0.1 | 1 |
|---|---|---|---|---|---|
| 1 | 5.7 | 5.5 | 5.5 | 5.8 | 5 |
| 0.1 | 6.4 | 5.9 | 6.4 | 6.2 | 5.2 |
| 0.01 | 6.2 | 6.4 | 6.1 | 6.2 | 5.3 |
| 0.001 | 6.1 | 6.4 | 6.7 | 6.7 | 5.9 |
| 0 | 6.1 | 6.4 | 6.6 | 6.9 | 5.3 |

# AT4G24090.1 : NA

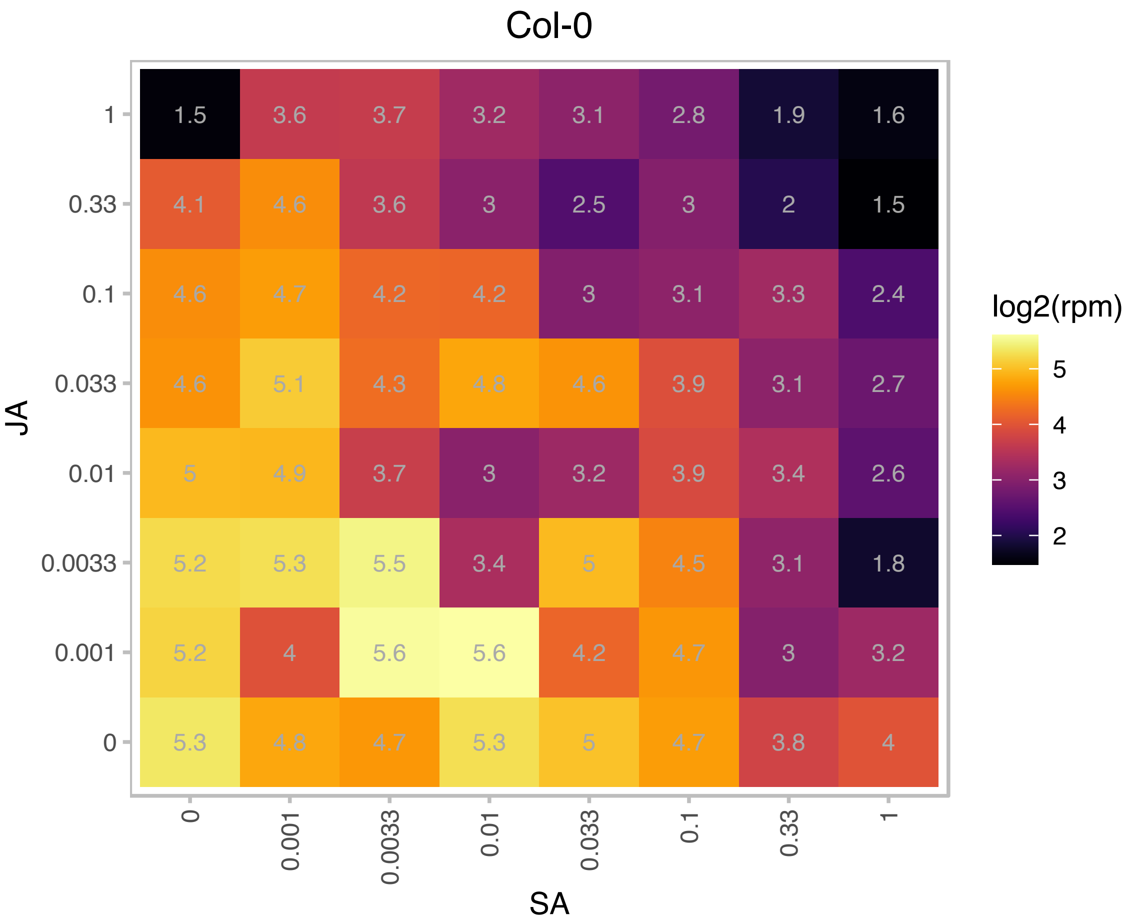


| | |
|---|---|
| Gene | AT4G24090 |
| is.expressed_genes | TRUE |
| is.responsive_genes | TRUE |
| is.apcluster | F-11 |

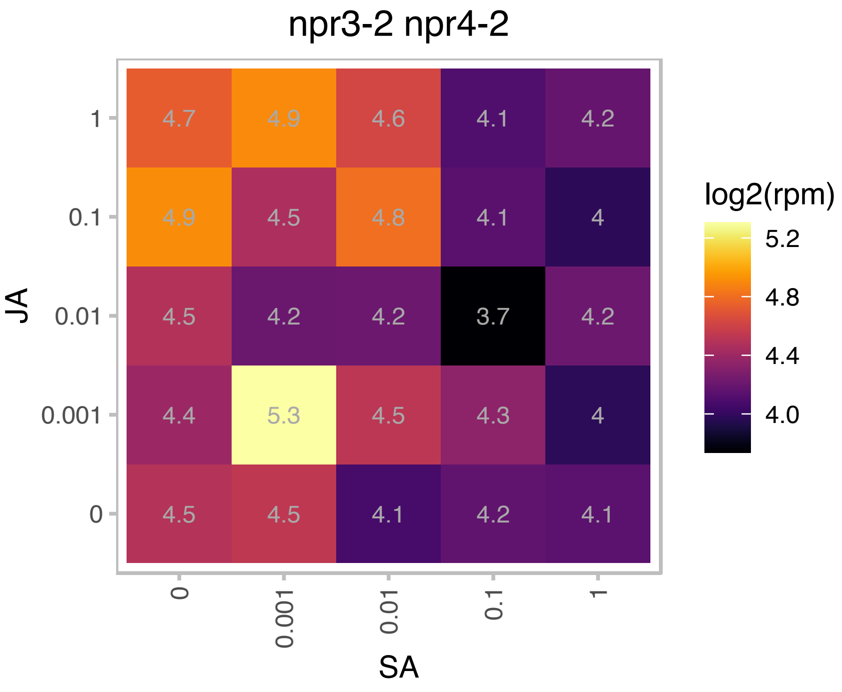


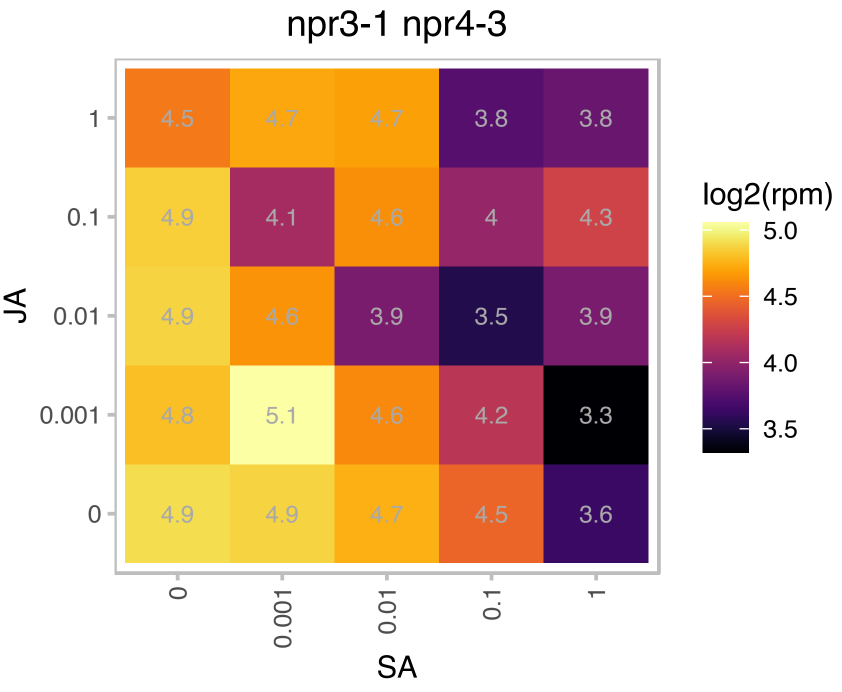

### AT4G25910.1 : ATCNFU3, NFU3

NFU domain protein 3
Encodes a protein containing the NFU domain that may be involved in iron-sulfur cluster assembly. Part of a five member gene family, more closely related to NFU1 and 2 than to NFU4 and 5. Targeted to the chloroplast. The mRNA is cell-to-cell mobile.

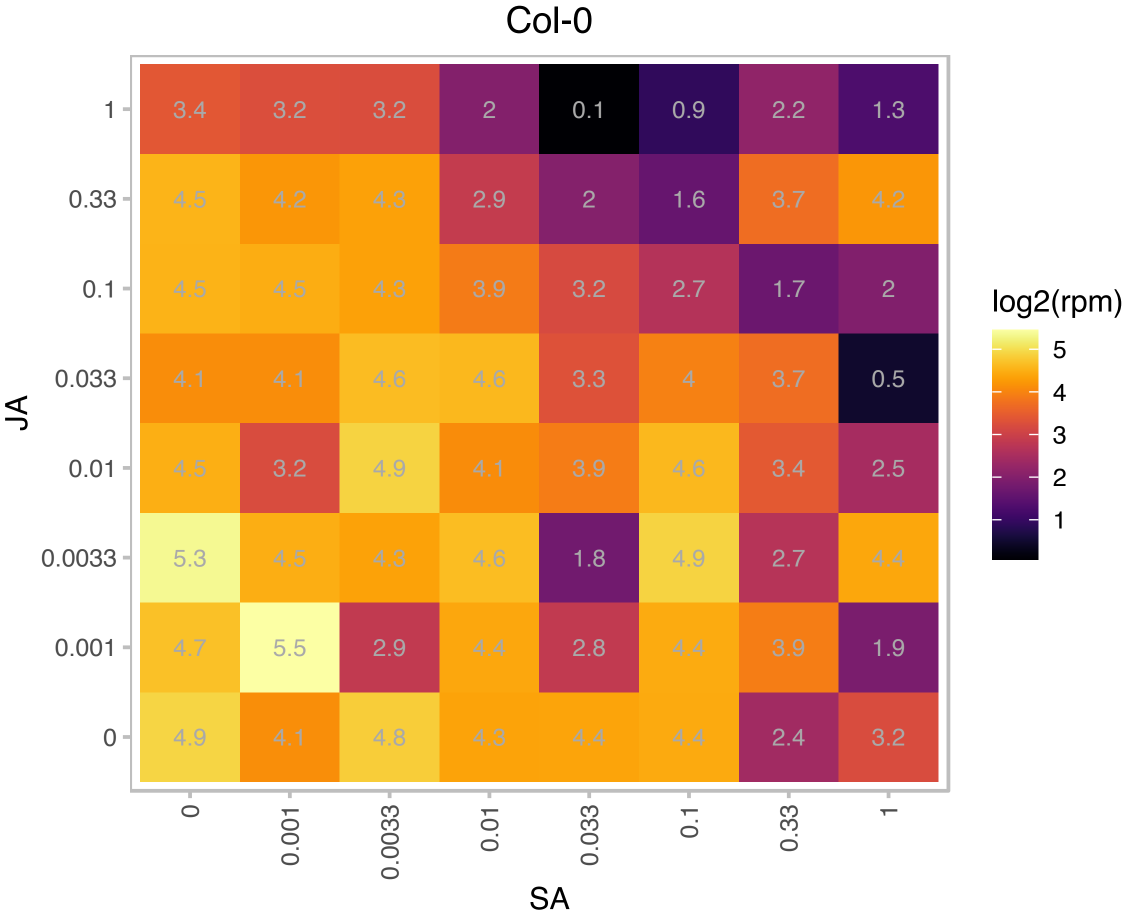


| | |
|---|---|
| Gene | AT4G25910 |
| is.expressed_genes | TRUE |
| is.responsive_genes | TRUE |
| is.apcluster | F-11 |

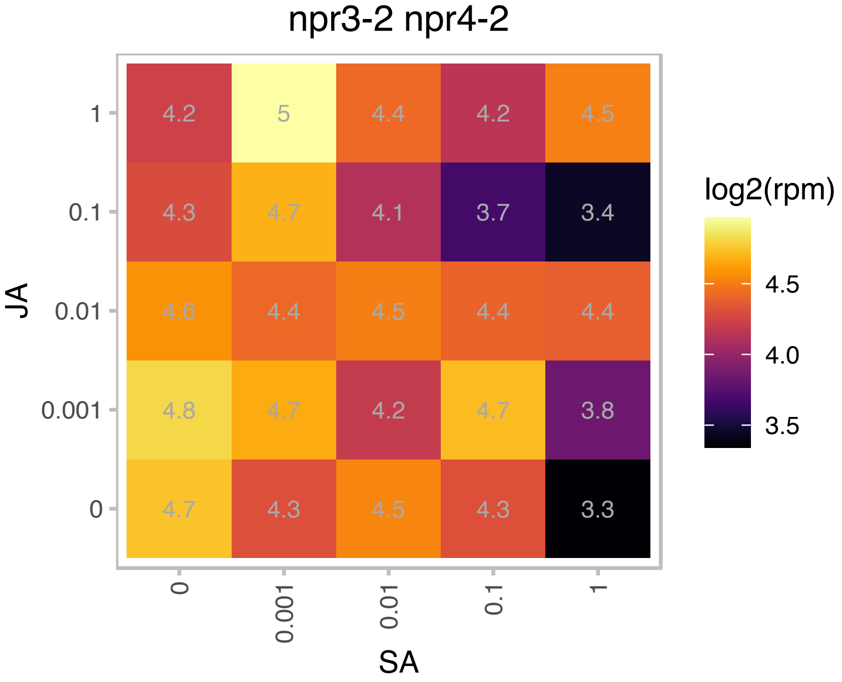


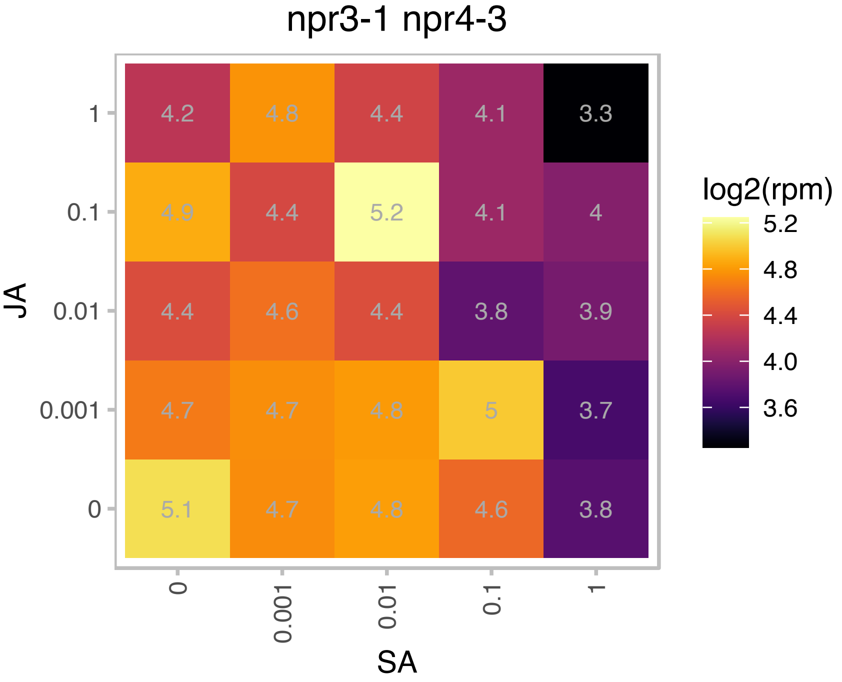

# AT4G26010.1 : NA

Peroxidase superfamily protein

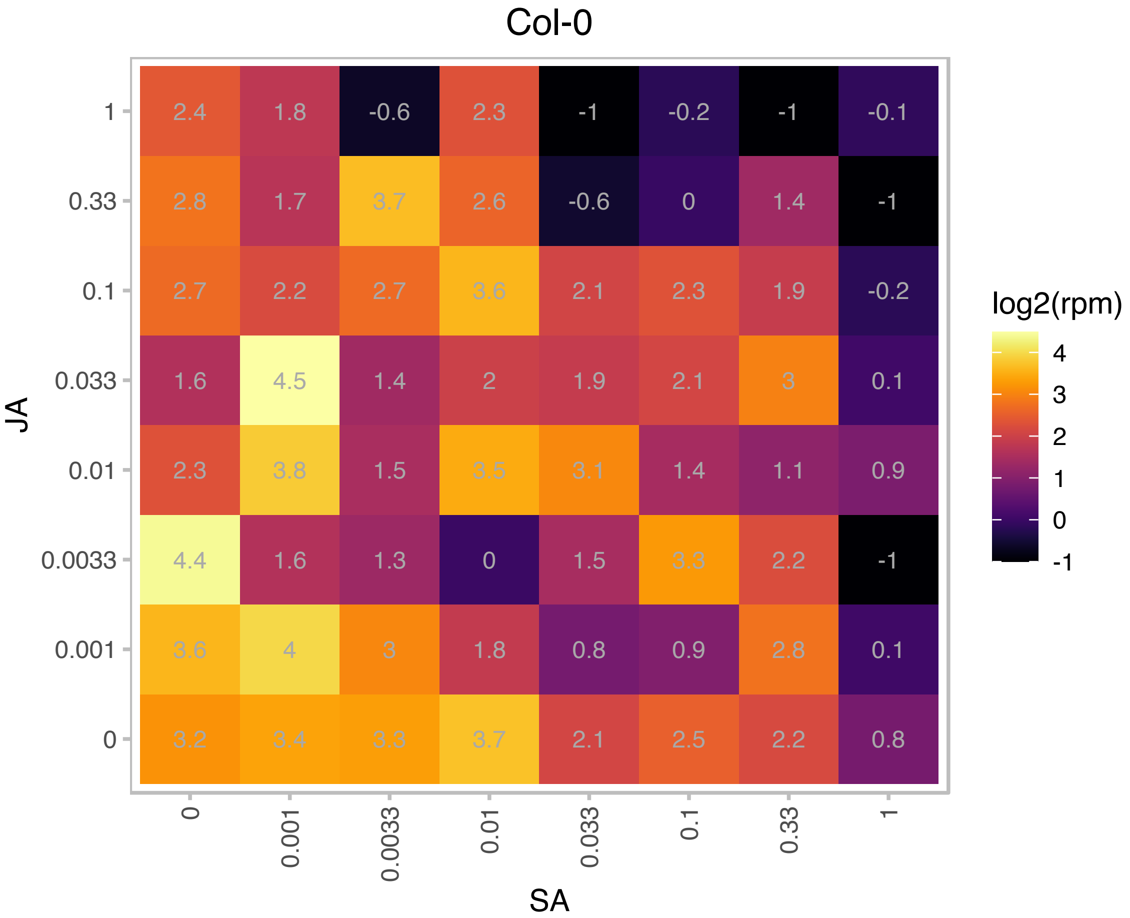


| | |
|---|---|
| Gene | AT4G26010 |
| is.expressed_genes | TRUE |
| is.responsive_genes | TRUE |
| is.apcluster | F-11 |

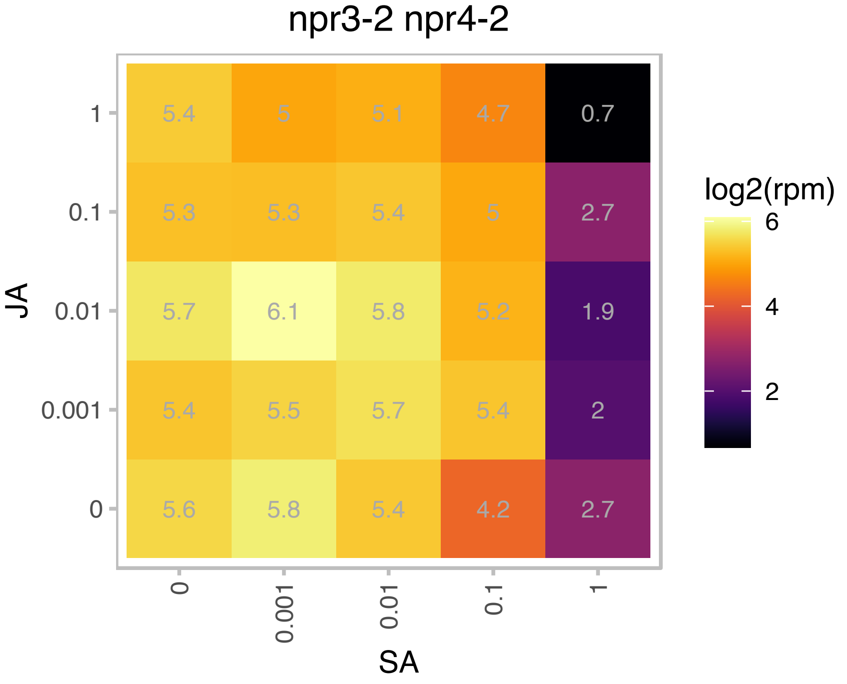


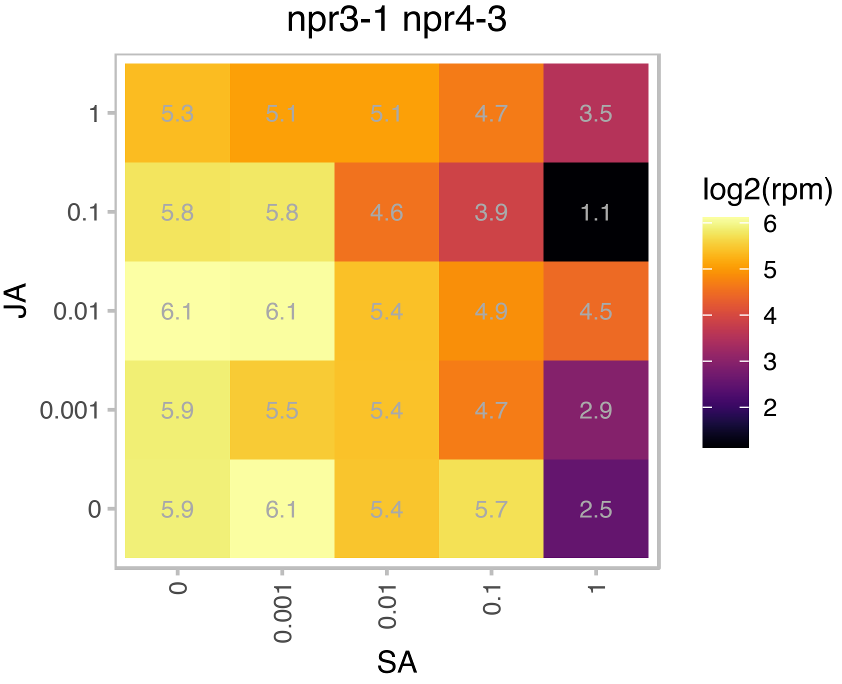

# AT4G26670.1 : HP20

Mitochondrial import inner membrane translocase subunit Tim17/Tim22/Tim23 family protein

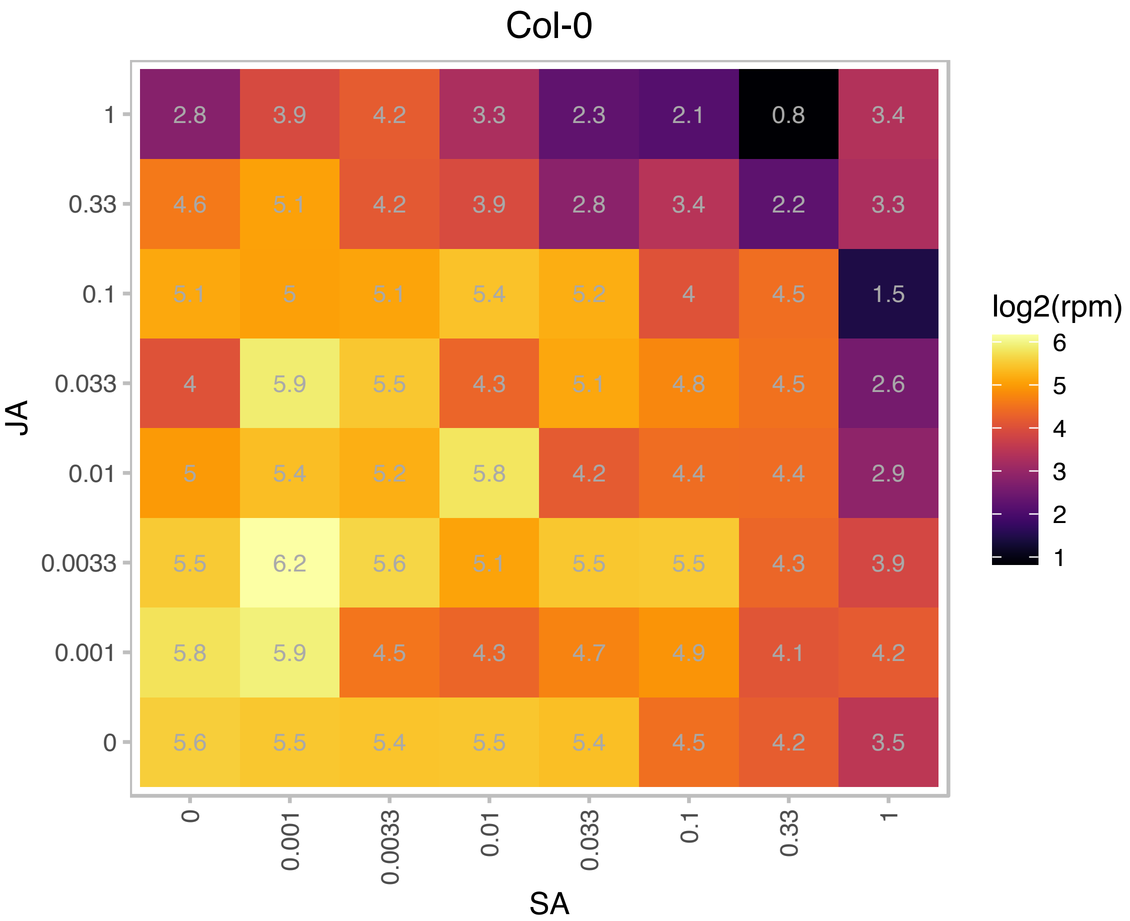


| | |
|---|---|
| Gene | AT4G26670 |
| is.expressed_genes | TRUE |
| is.responsive_genes | TRUE |
| is.apcluster | F-11 |

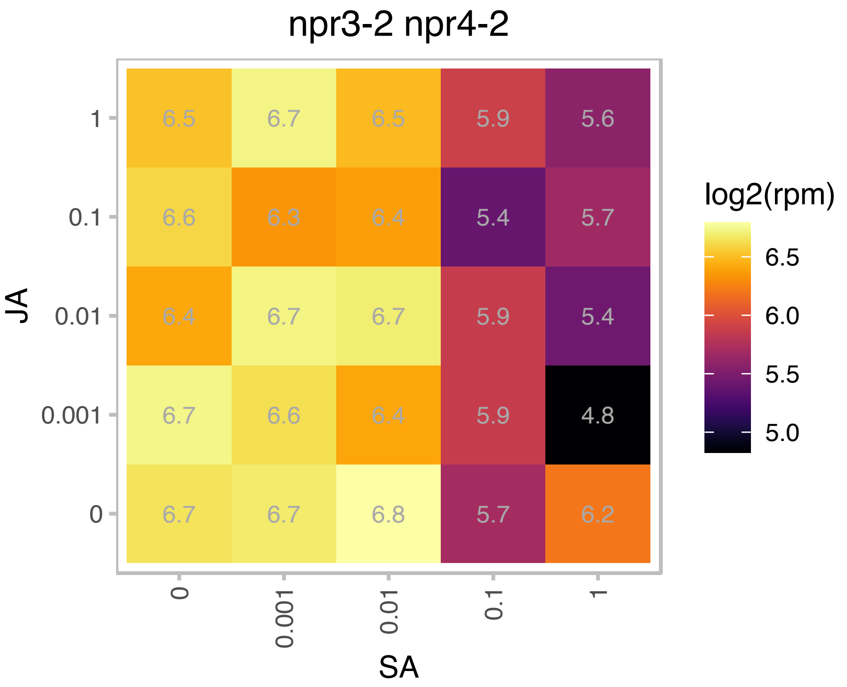


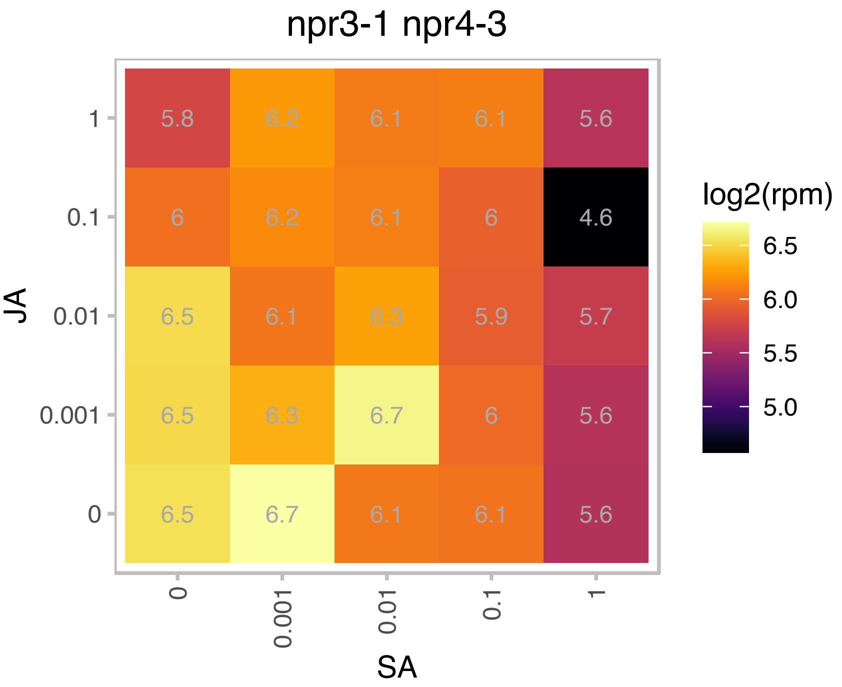

### AT4G26760.1 : MAP65-2

microtubule-associated protein 65-2

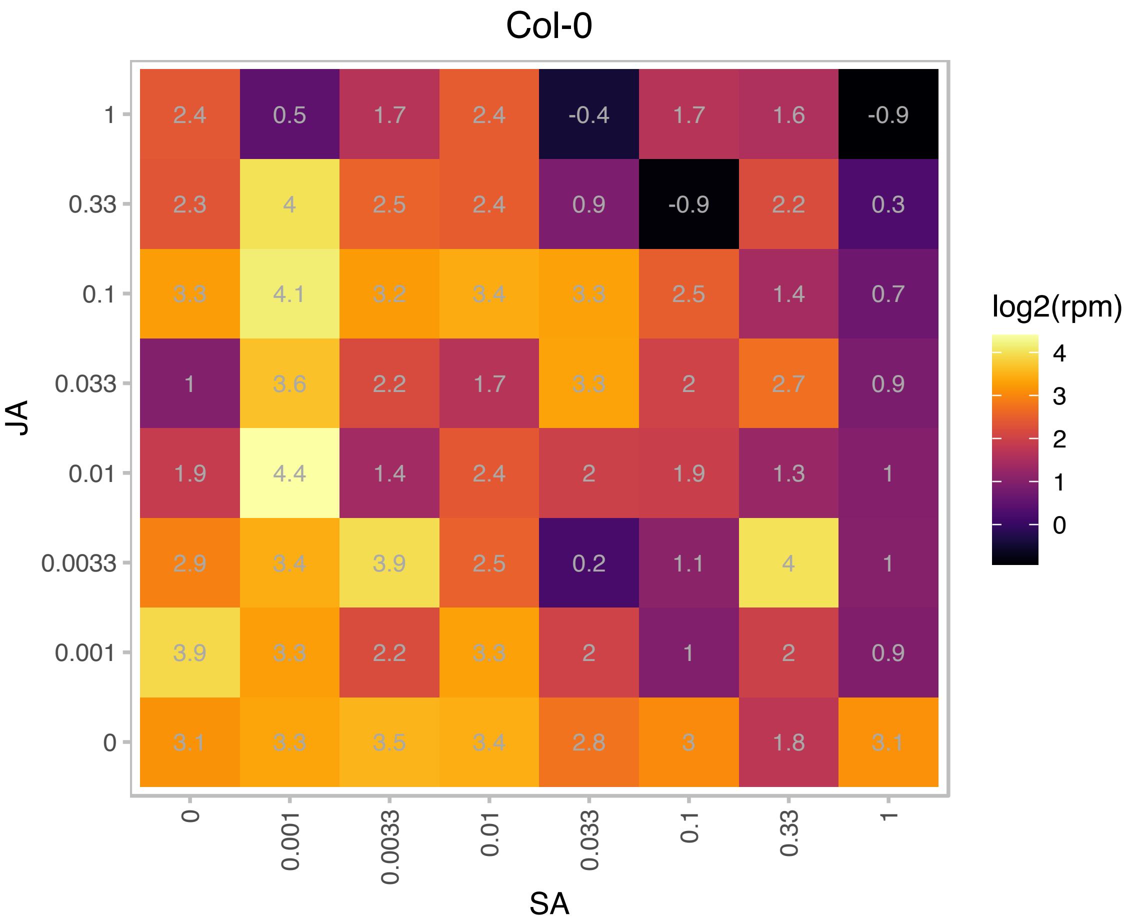


| | |
|---|---|
| Gene | AT4G26760 |
| is.expressed_genes | TRUE |
| is.responsive_genes | TRUE |
| is.apcluster | F-11 |

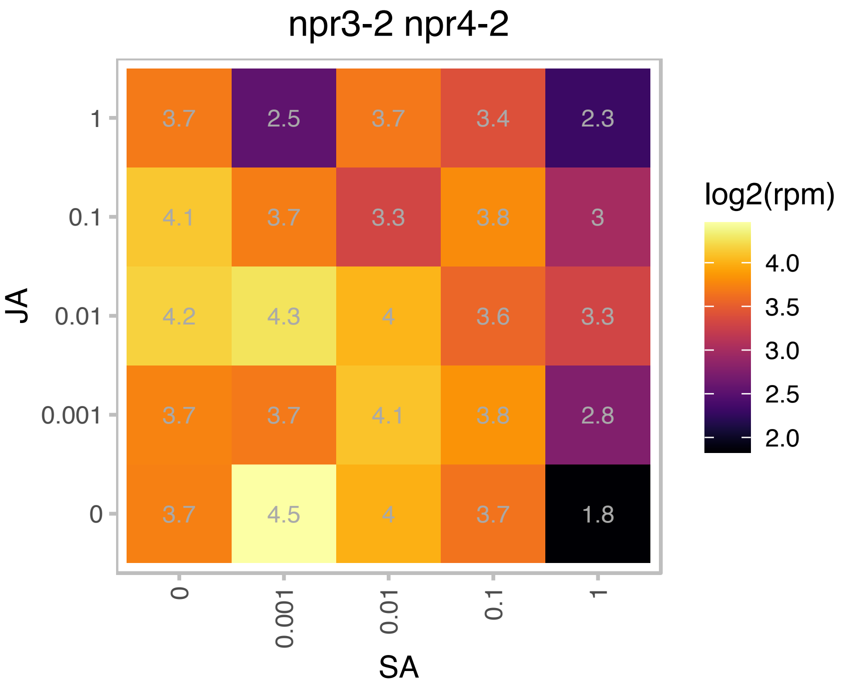


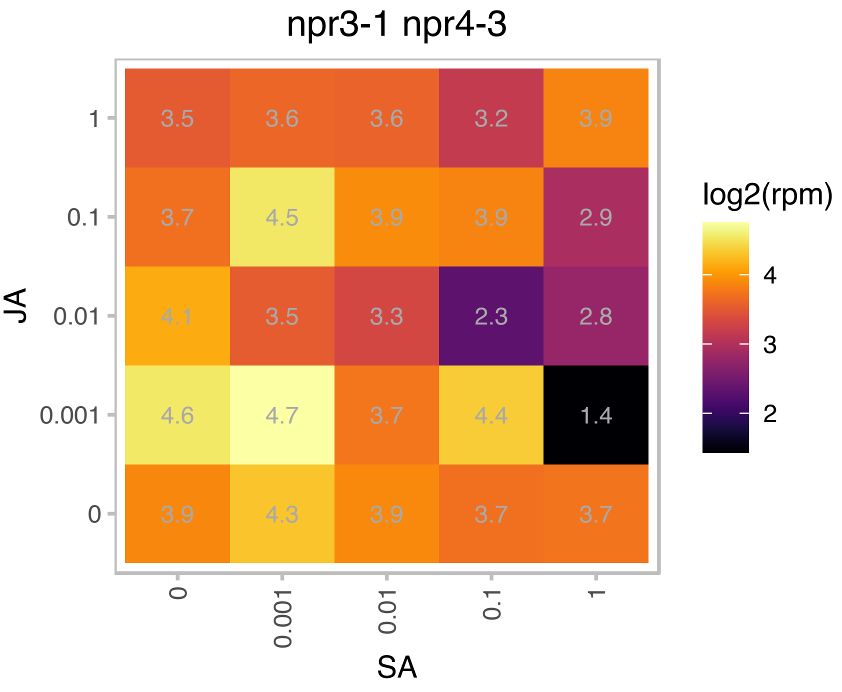

### AT4G27440.1 : PORB

protochlorophyllide oxidoreductase B

light-dependent NADPH:protochlorophyllide oxidoreductase B The mRNA is cell-to-cell mobile.

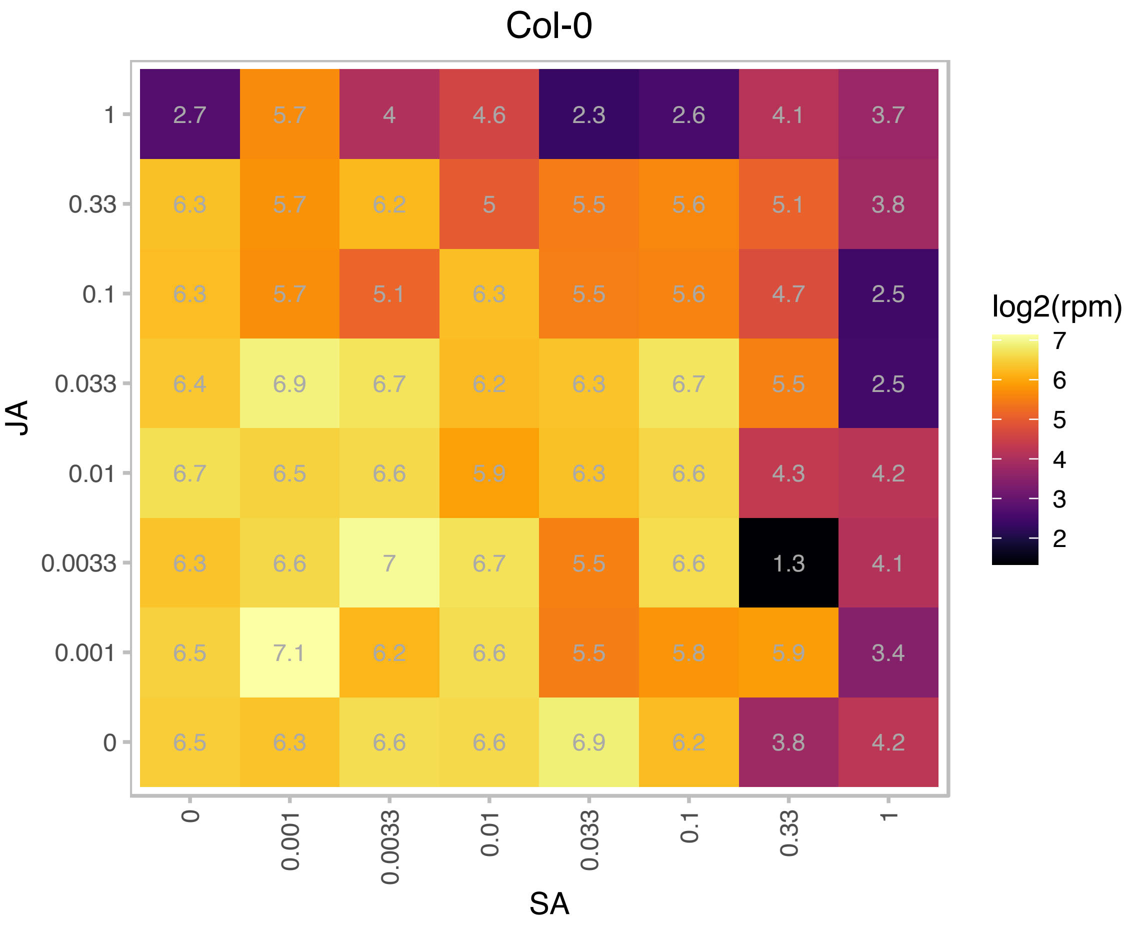


| | |
|---|---|
| Gene | AT4G27440 |
| is.expressed_genes | TRUE |
| is.responsive_genes | TRUE |
| is.apcluster | F-11 |

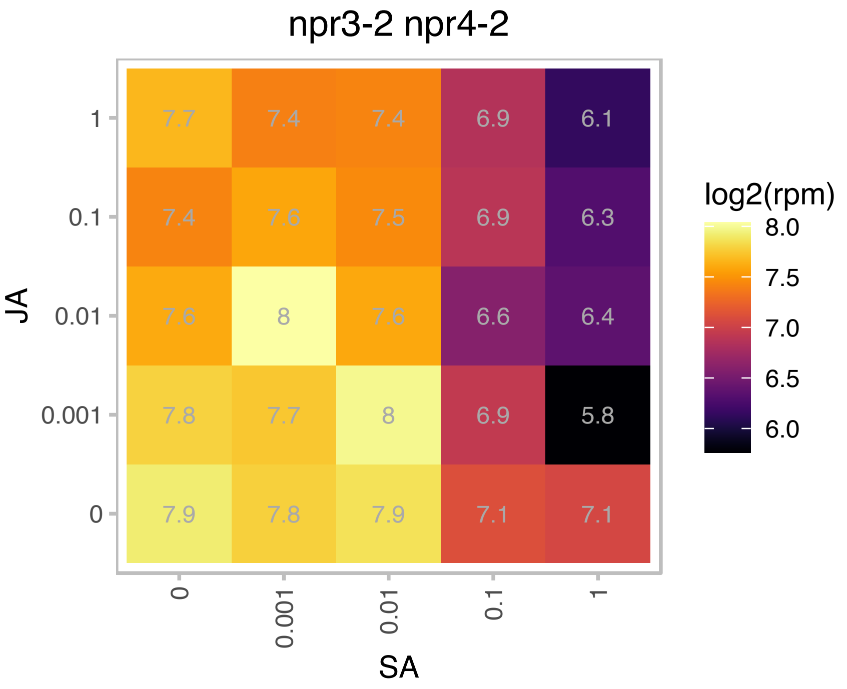


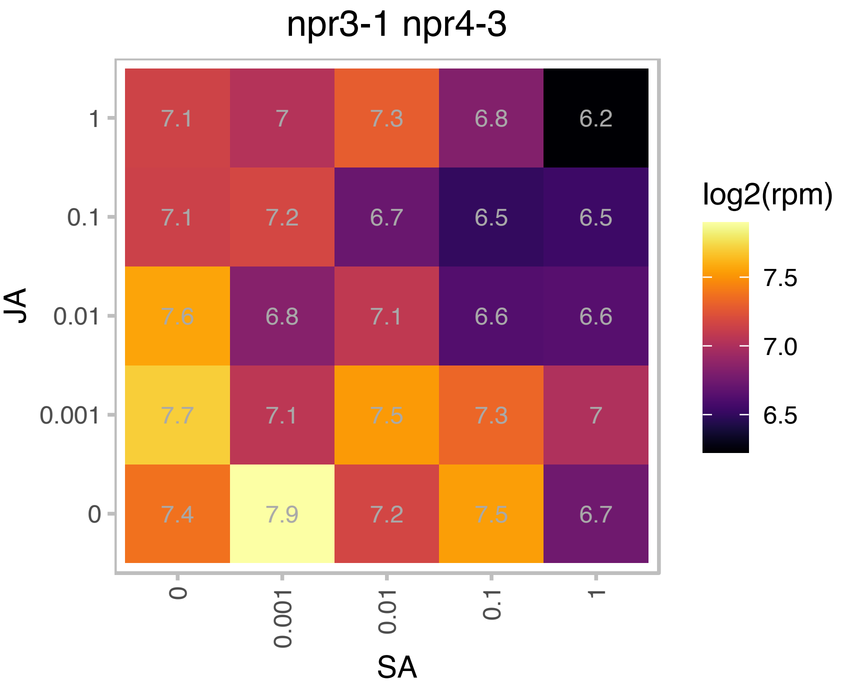

# AT4G27700.1 : NA

Rhodanese/Cell cycle control phosphatase superfamily protein

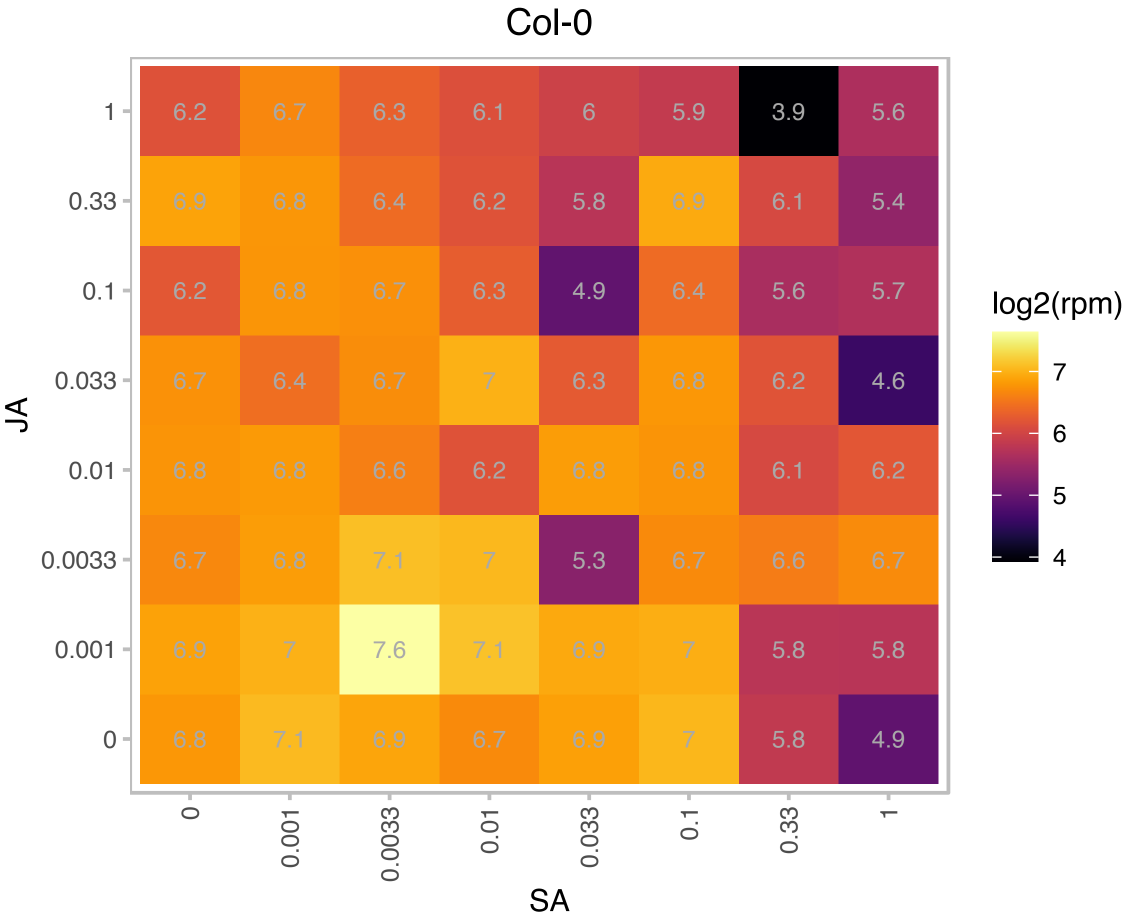


| | |
|---|---|
| Gene | AT4G27700 |
| is.expressed_genes | TRUE |
| is.responsive_genes | TRUE |
| is.apcluster | F-11 |

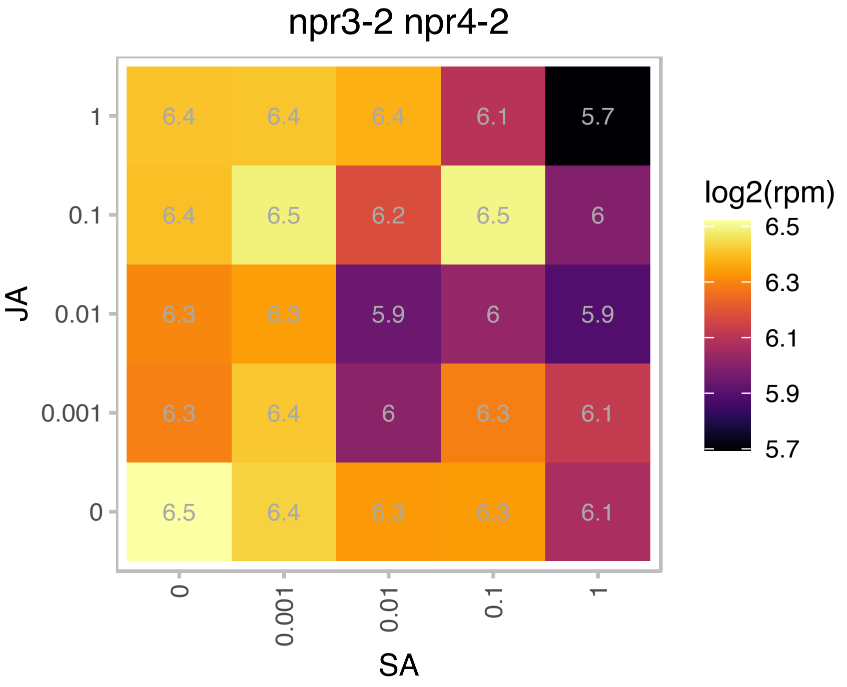


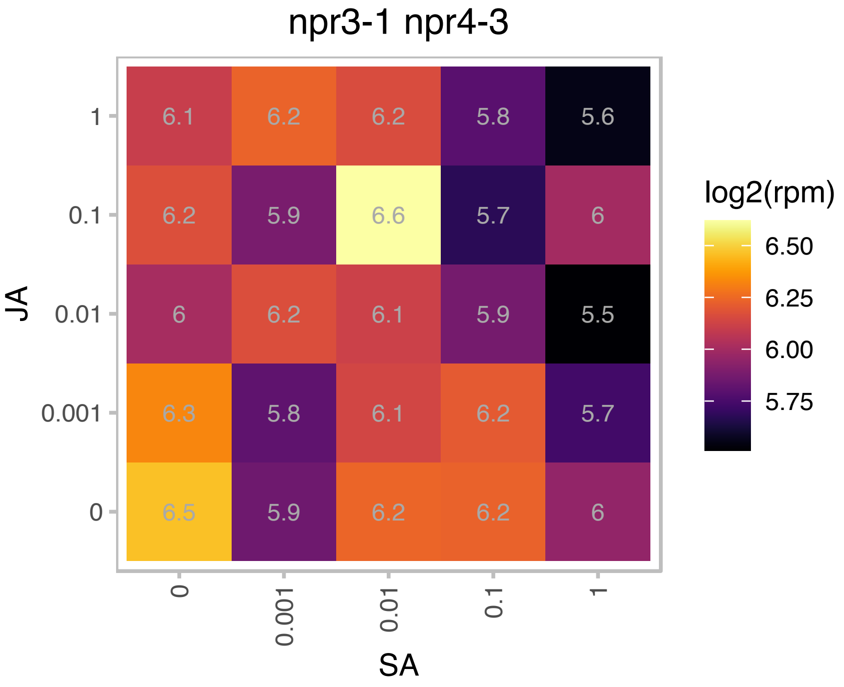

# AT4G28025.1 : NA

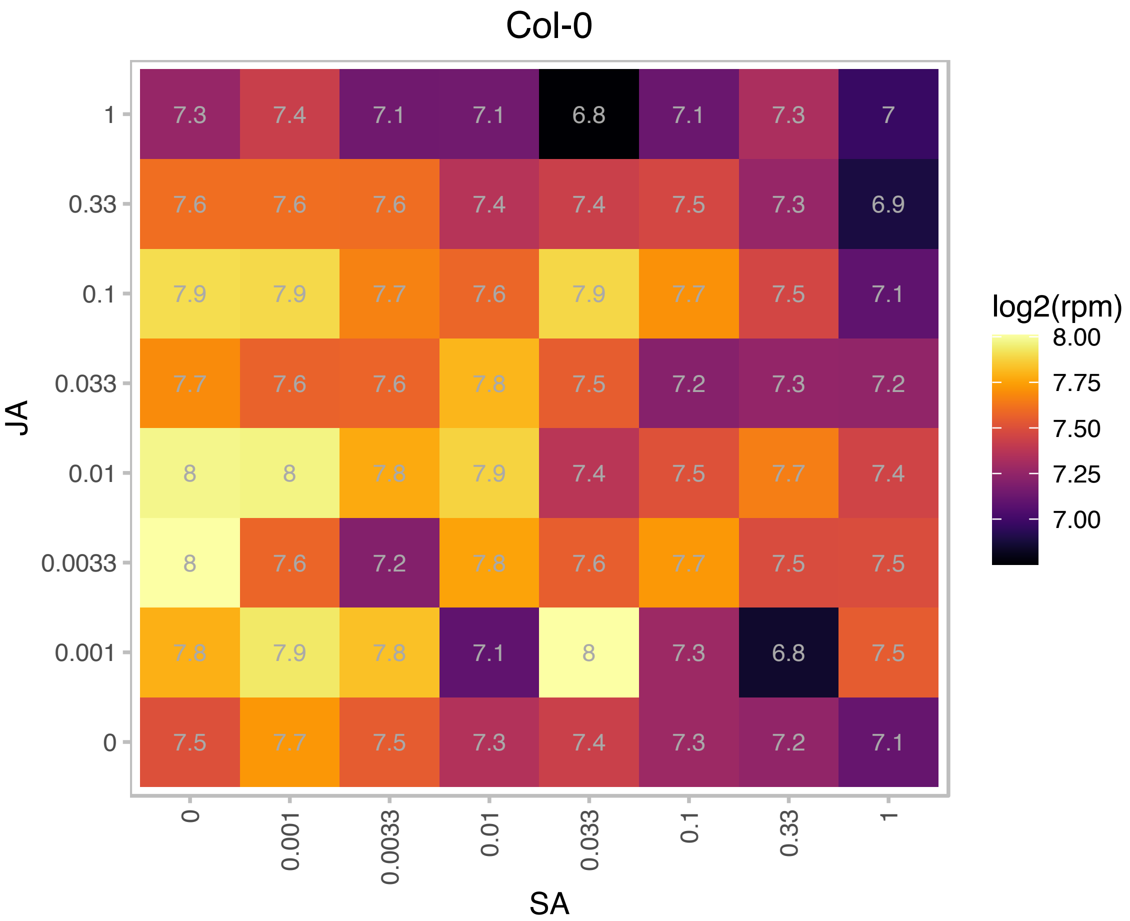


| | |
|---|---|
| Gene | AT4G28025 |
| is.expressed_genes | TRUE |
| is.responsive_genes | TRUE |
| is.apcluster | F-11 |

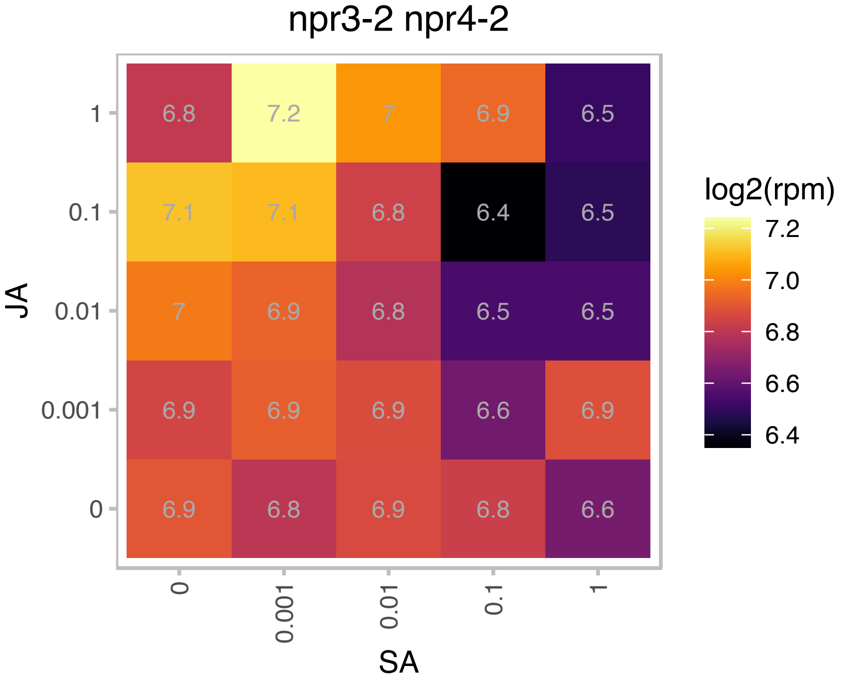


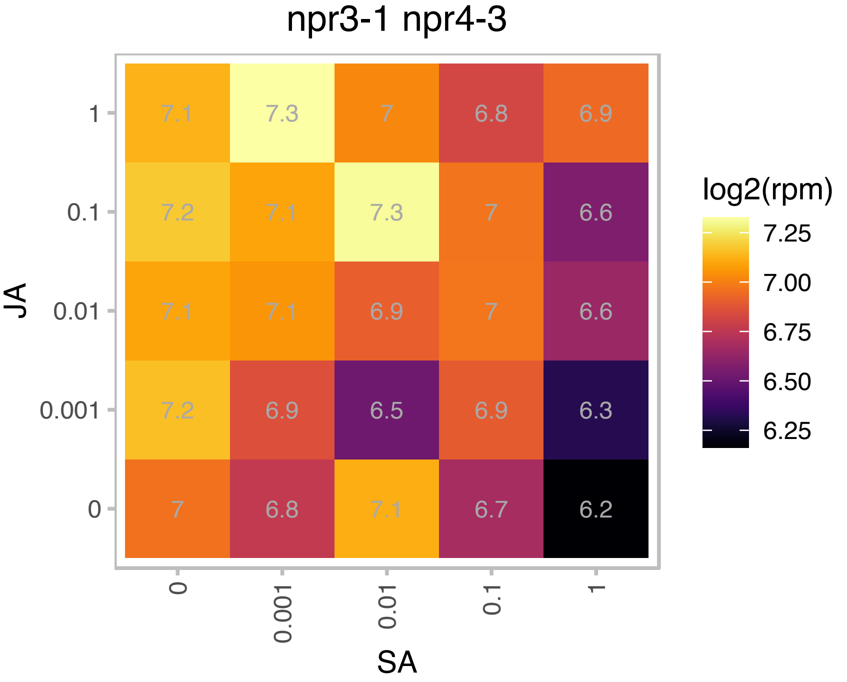

# AT4G30660.1 : NA

Low temperature and salt responsive protein family

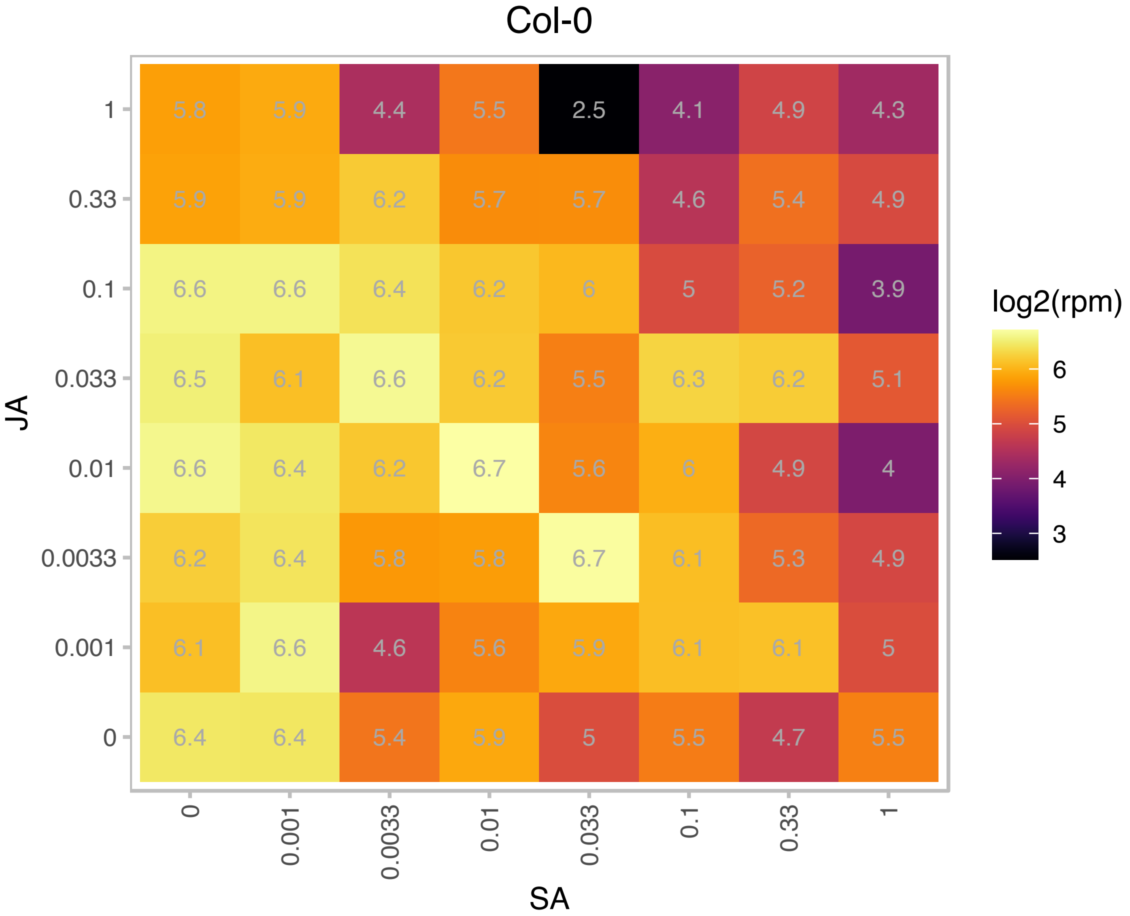


| | |
|---|---|
| Gene | AT4G30660 |
| is.expressed_genes | TRUE |
| is.responsive_genes | TRUE |
| is.apcluster | F-11 |

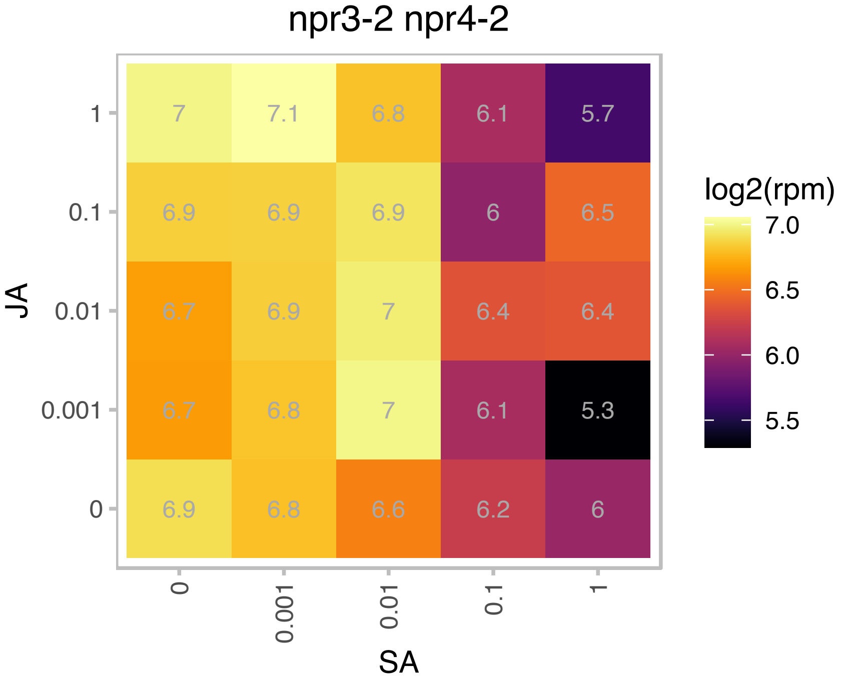


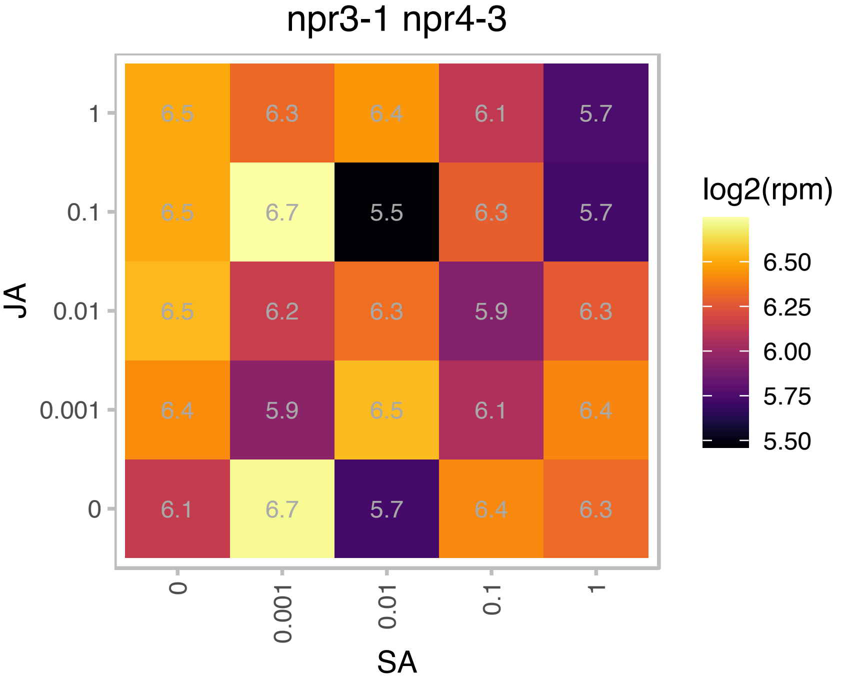

# AT4G30845.1 : NA

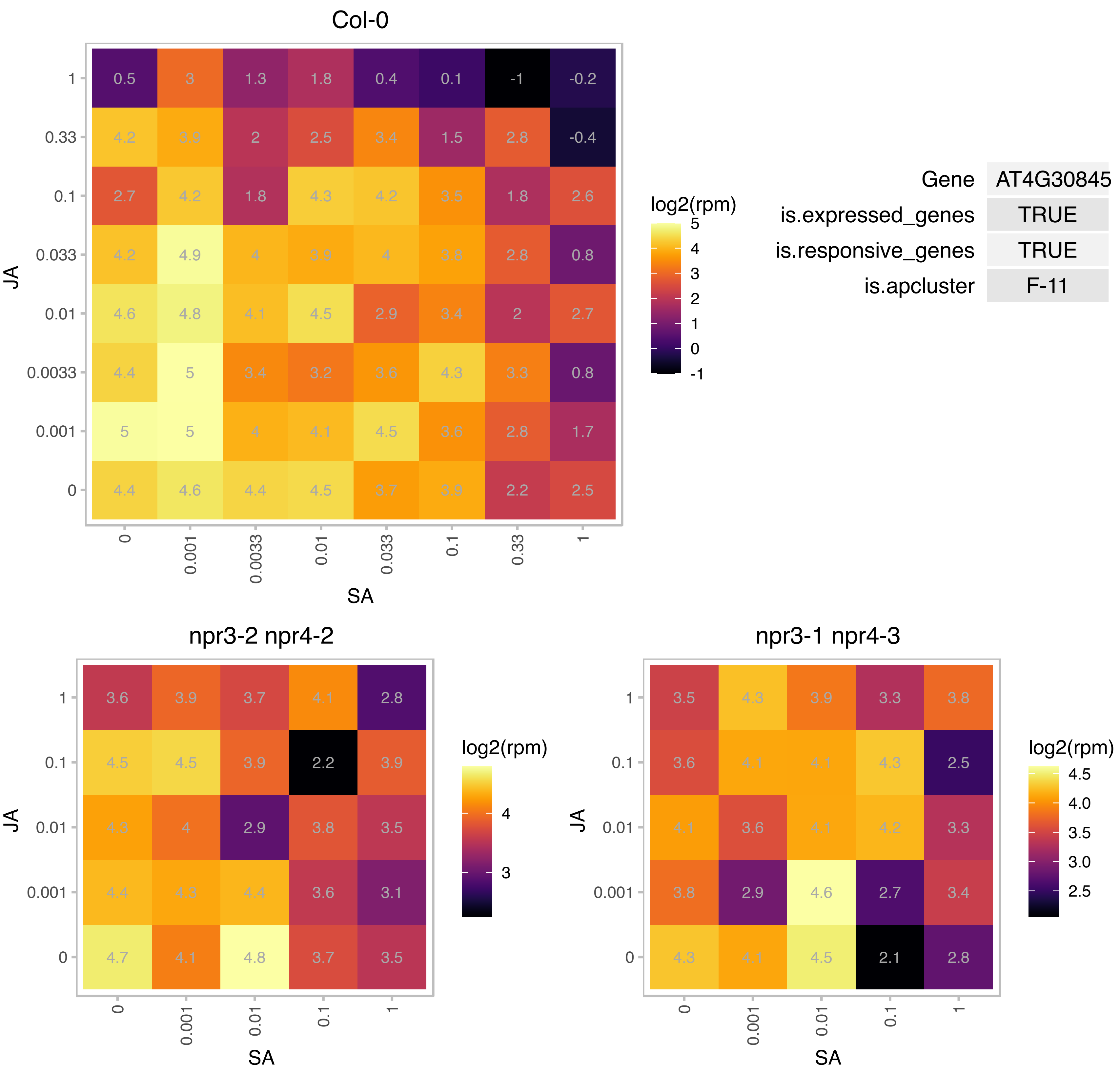

# AT4G31040.1 : NA

CemA-like proton extrusion protein-related

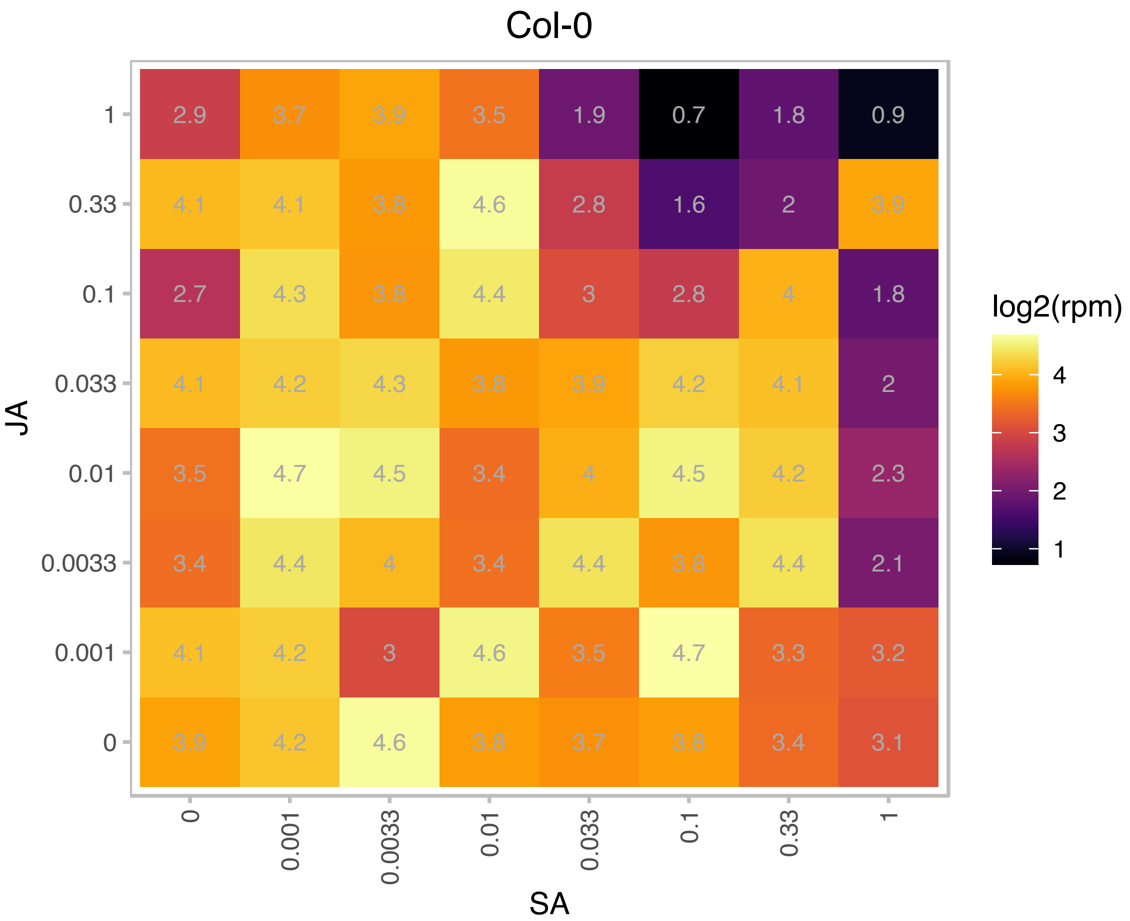


| | |
|---|---|
| Gene | AT4G31040 |
| is.expressed_genes | TRUE |
| is.responsive_genes | TRUE |
| is.apcluster | F-11 |

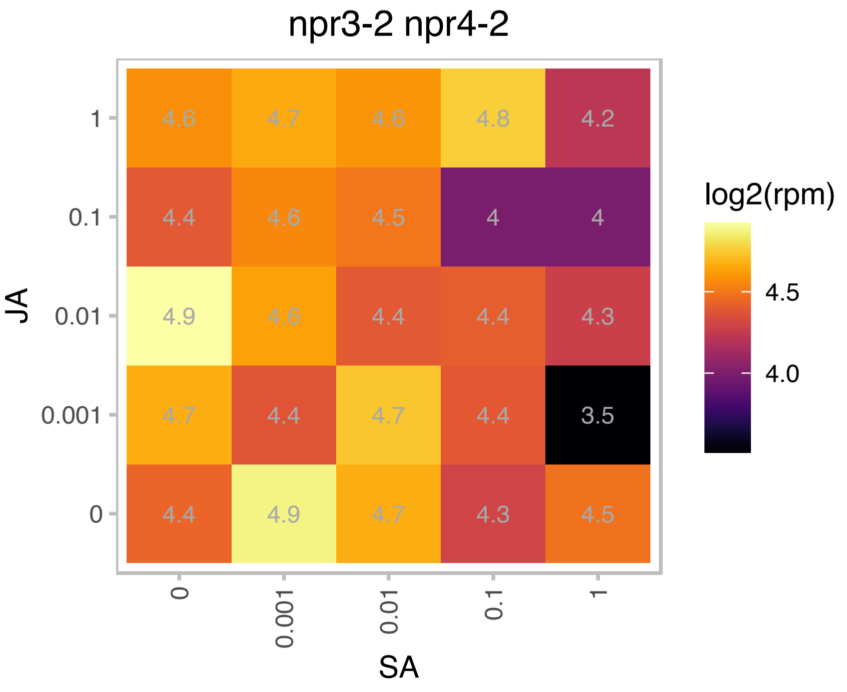


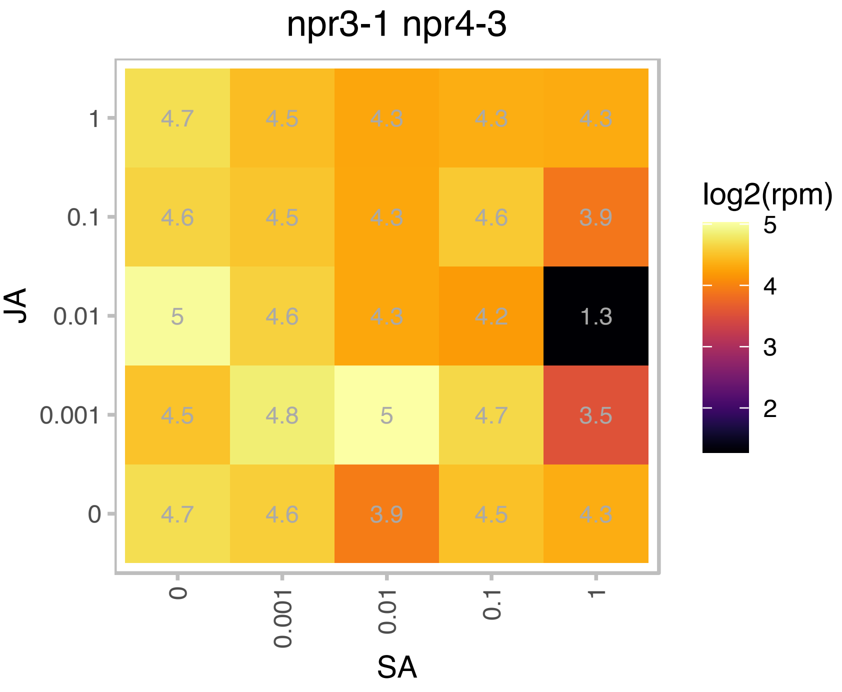

### AT4G31840.1 : AtENODL15, ENODL15

early nodulin-like protein 15

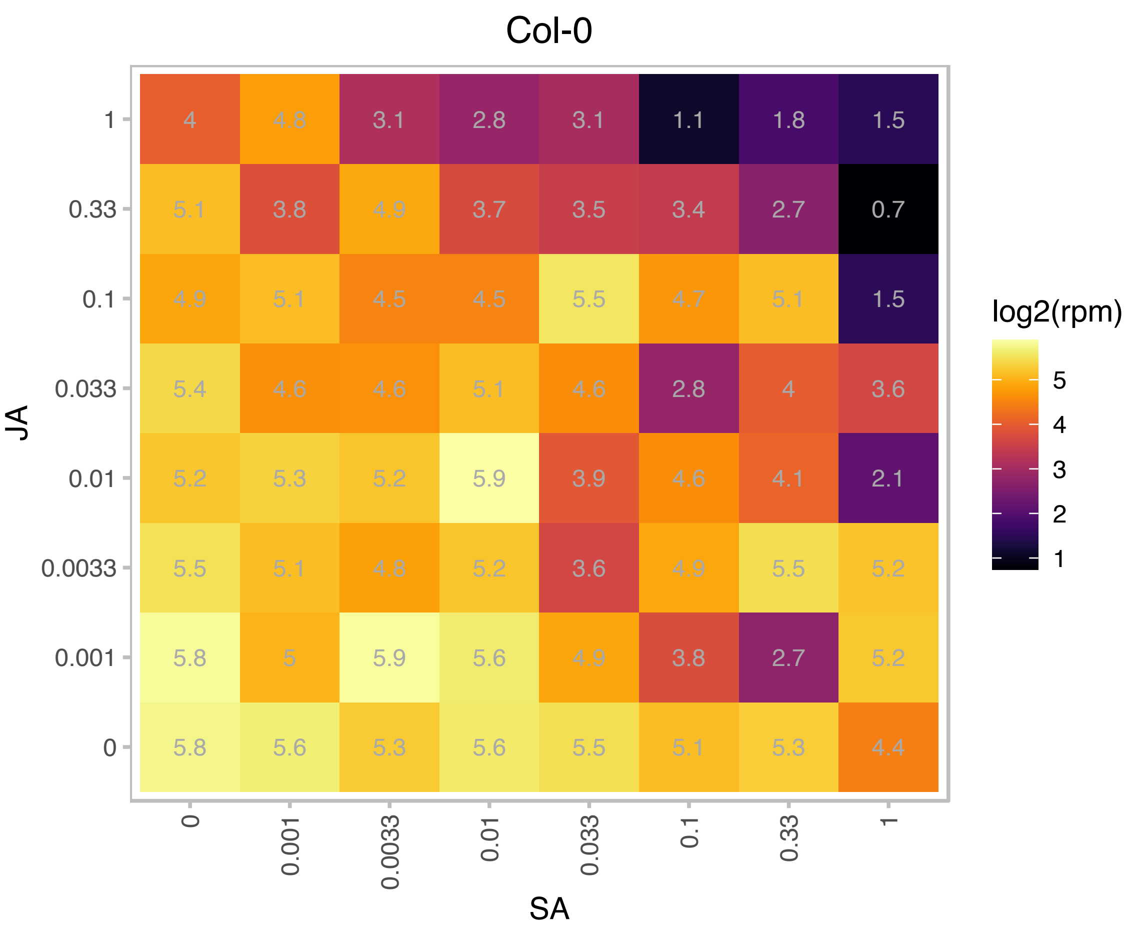


| | |
|---|---|
| Gene | AT4G31840 |
| is.expressed_genes | TRUE |
| is.responsive_genes | TRUE |
| is.apcluster | F-11 |

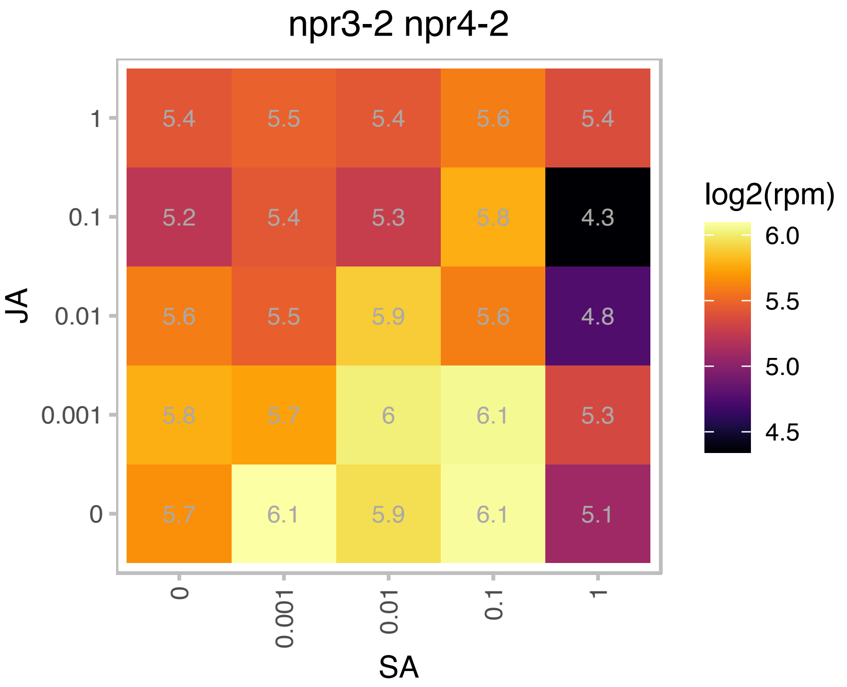


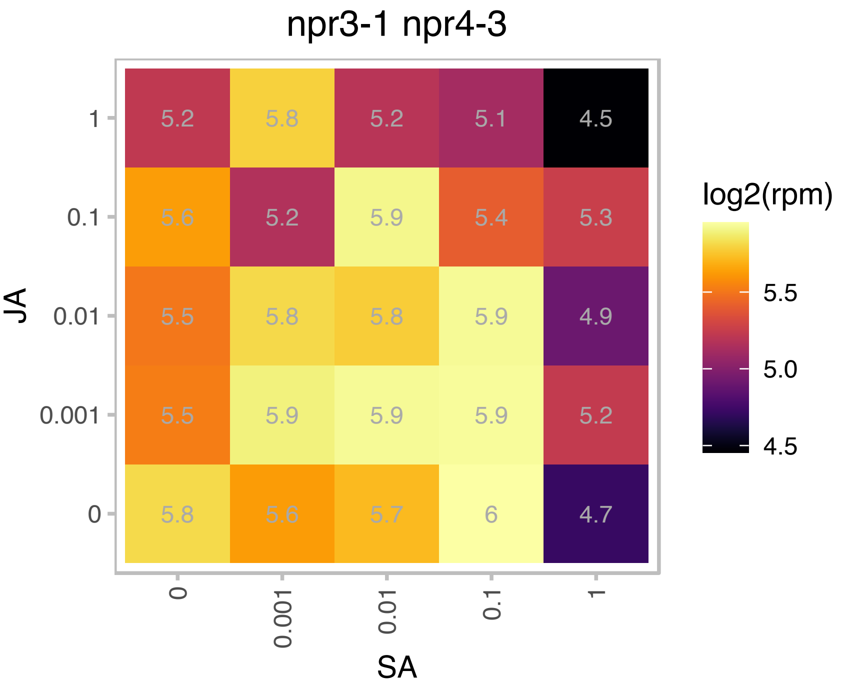

### AT4G33660.1 : AthCYSTM11

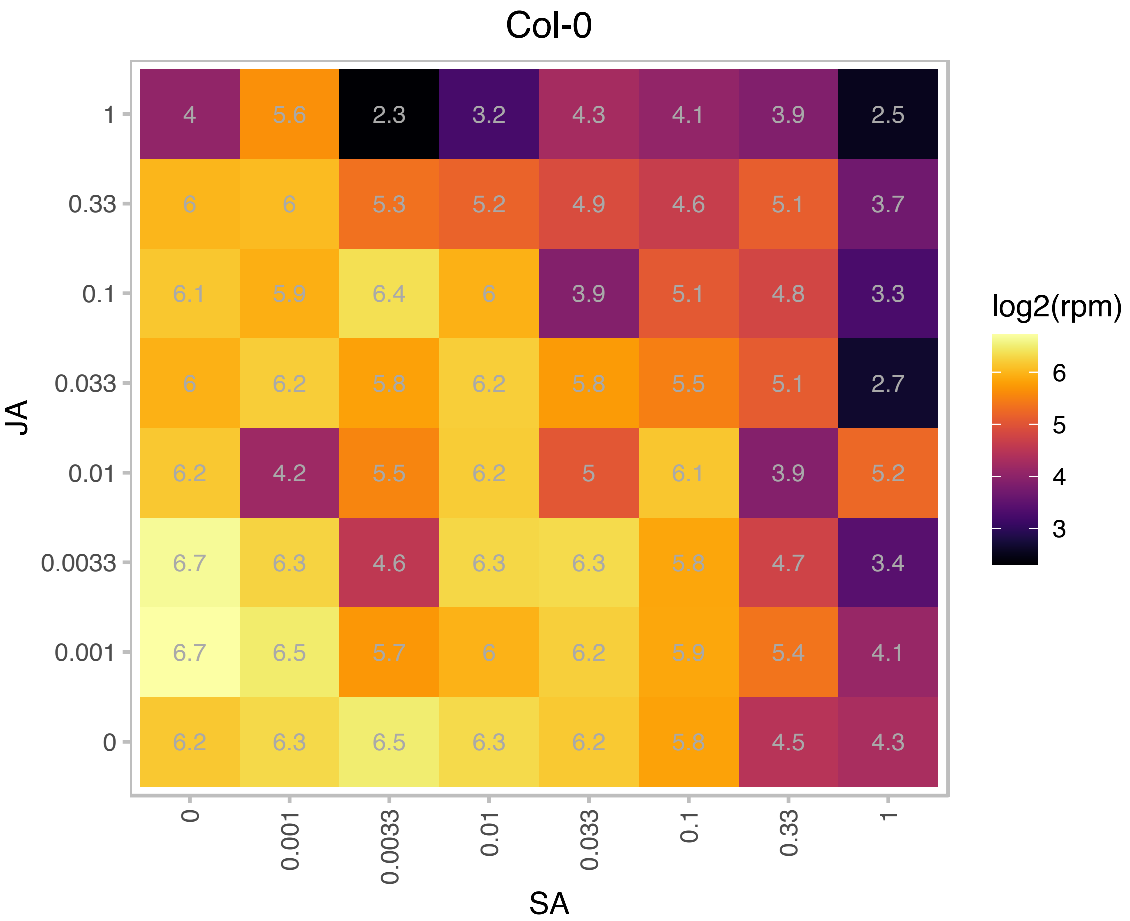


| | |
|---|---|
| Gene | AT4G33660 |
| is.expressed_genes | TRUE |
| is.responsive_genes | TRUE |
| is.apcluster | F-11 |

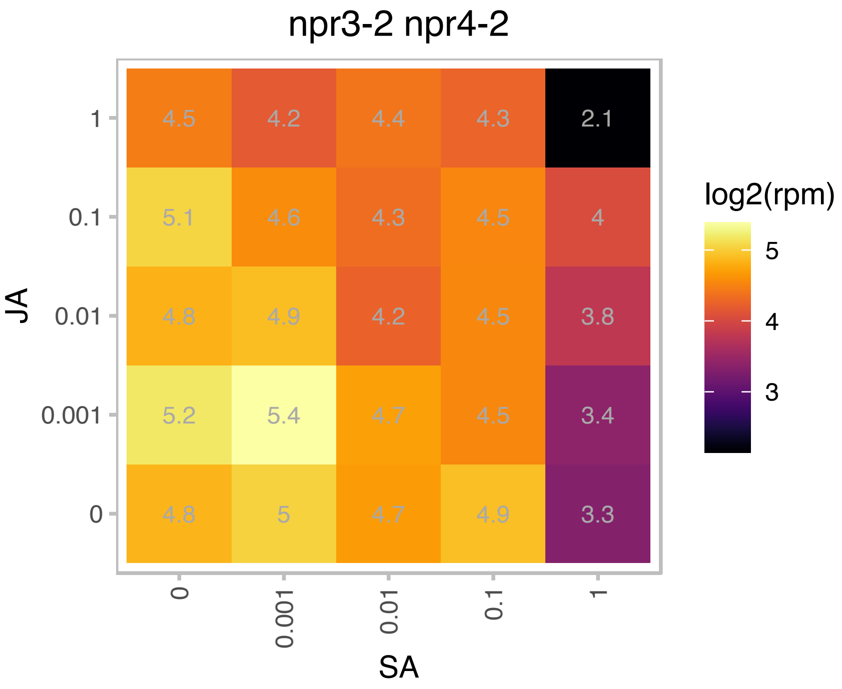


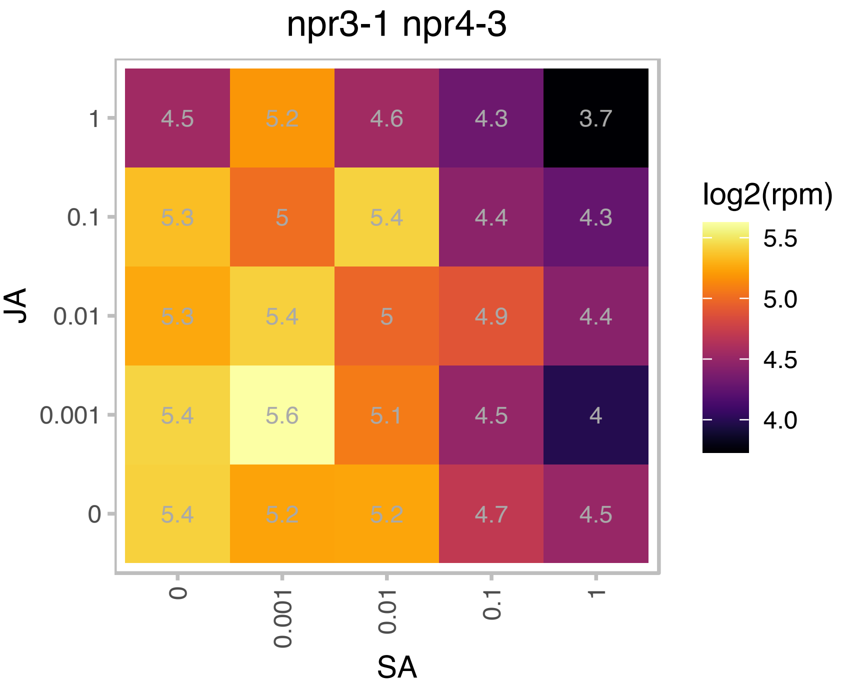

# AT4G34555.1 : NA

Ribosomal protein S25 family protein

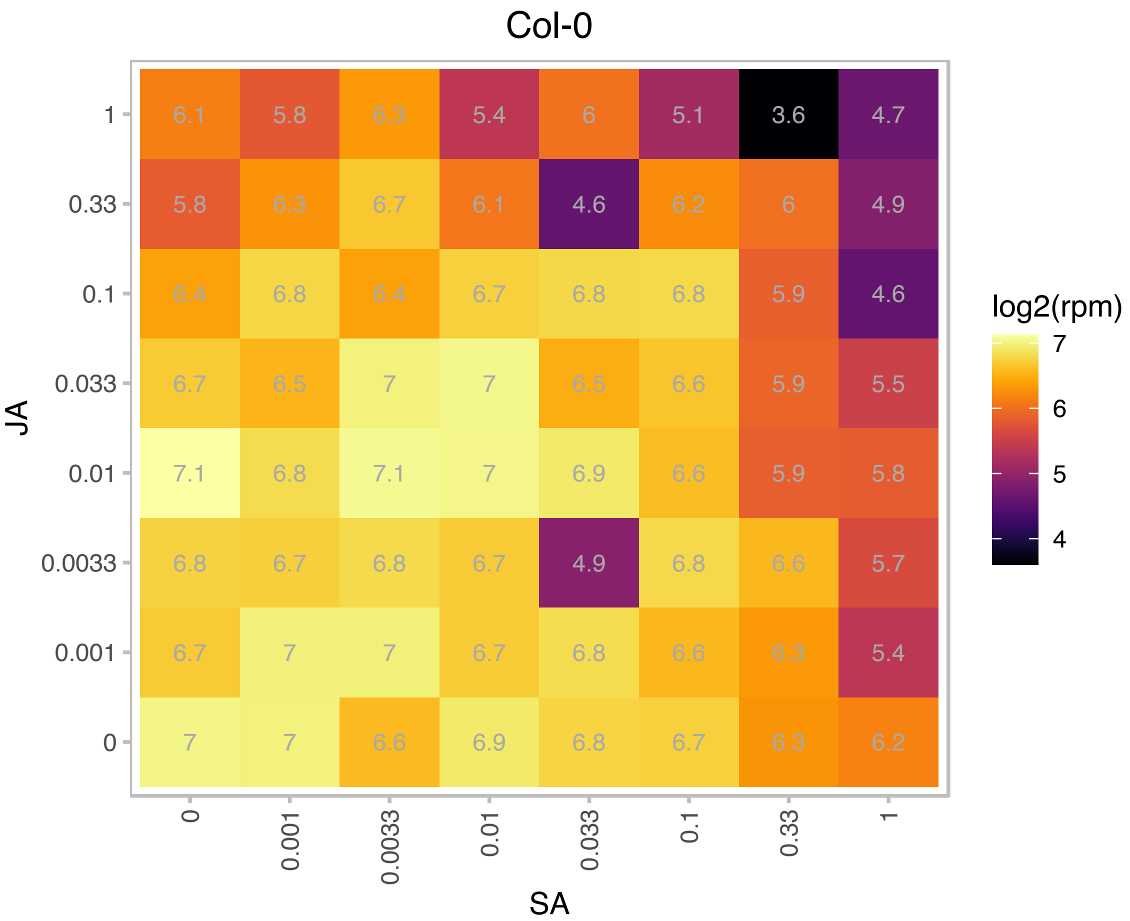


| | |
|---|---|
| Gene | AT4G34555 |
| is.expressed_genes | TRUE |
| is.responsive_genes | TRUE |
| is.apcluster | F-11 |

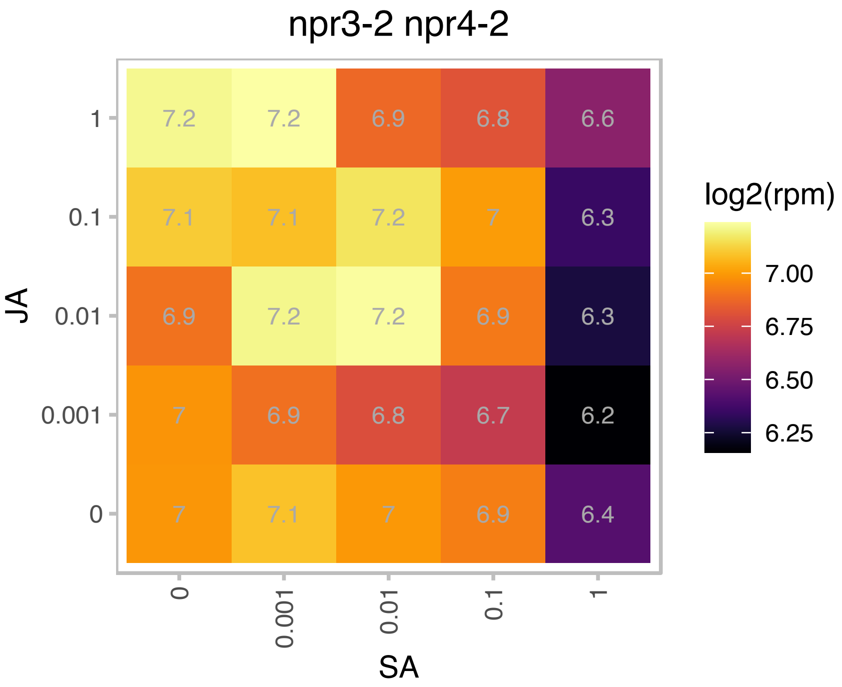


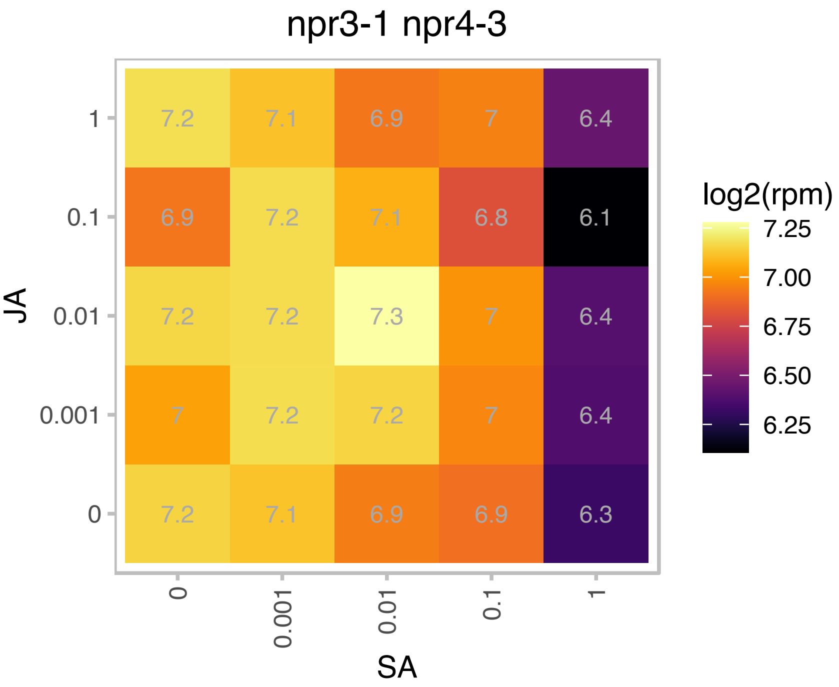

### AT4G35950.1 : ARAC6, ATRAC6, ATROP5, RAC2, RAC6,

RAC-like 6

A member of ROP GTPases gene family-like. GTP binding protein Arac6.

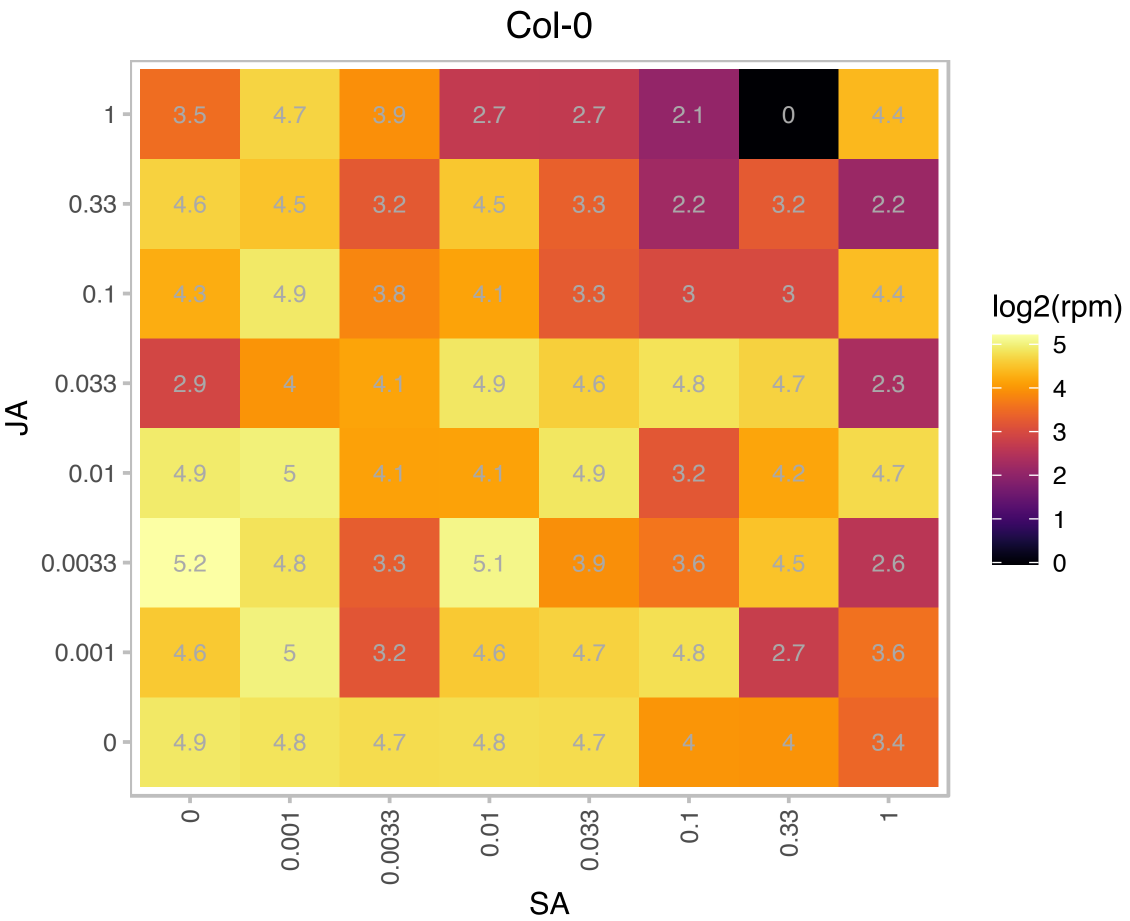


| | |
|---|---|
| Gene | AT4G35950 |
| is.expressed_genes | TRUE |
| is.responsive_genes | TRUE |
| is.apcluster | F-11 |

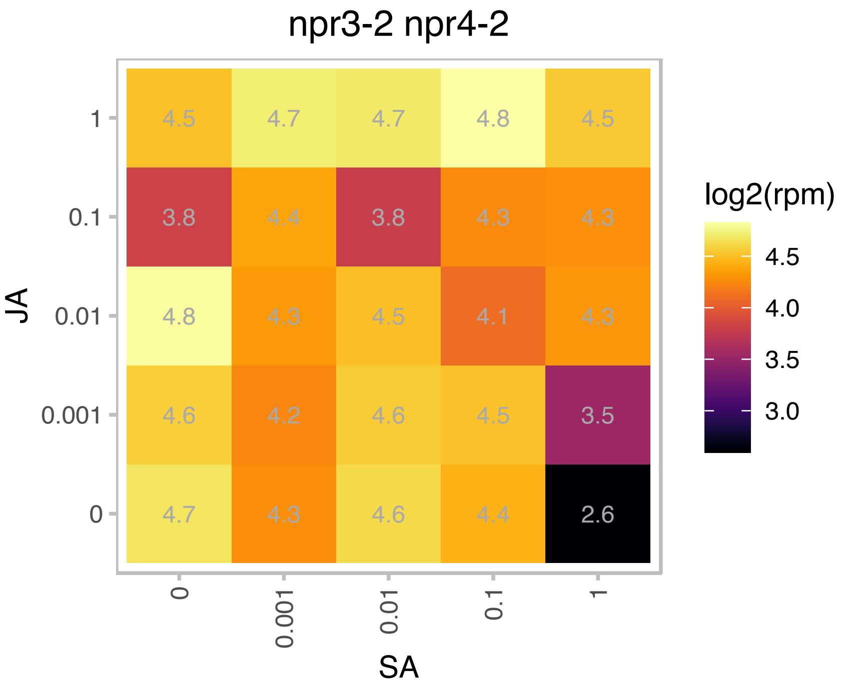


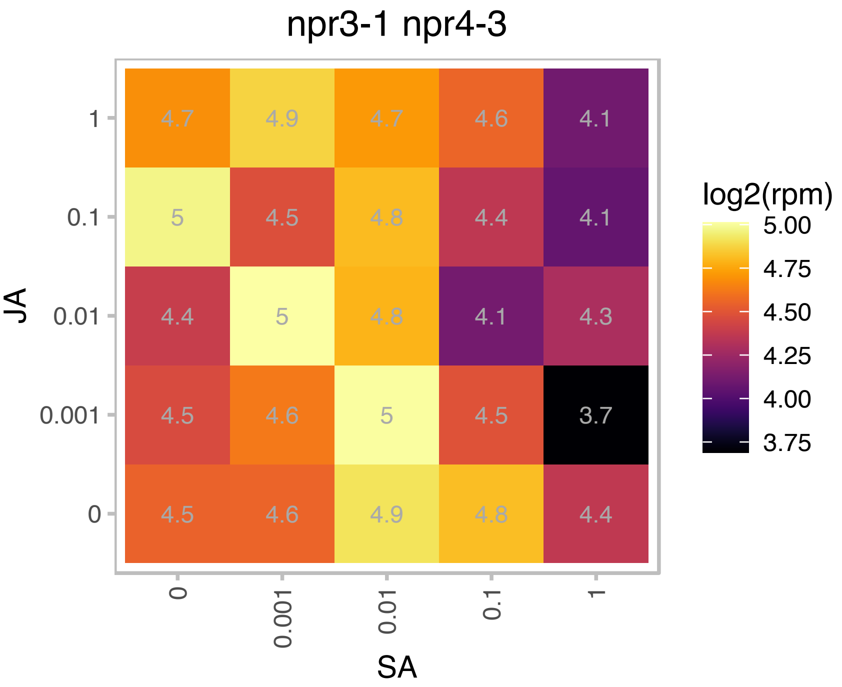

# AT4G36970.1 : NA

Remorin family protein

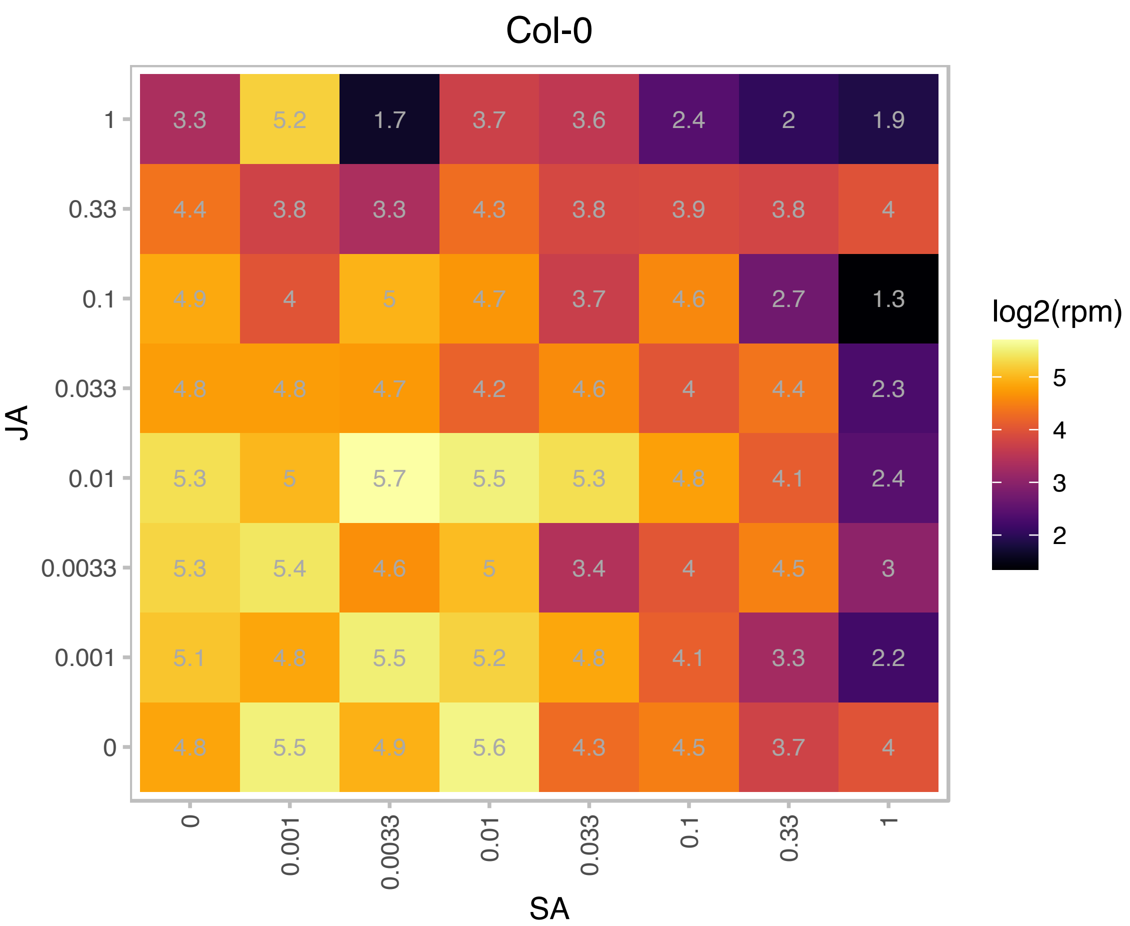


| | |
|---|---|
| Gene | AT4G36970 |
| is.expressed_genes | TRUE |
| is.responsive_genes | TRUE |
| is.apcluster | F-11 |

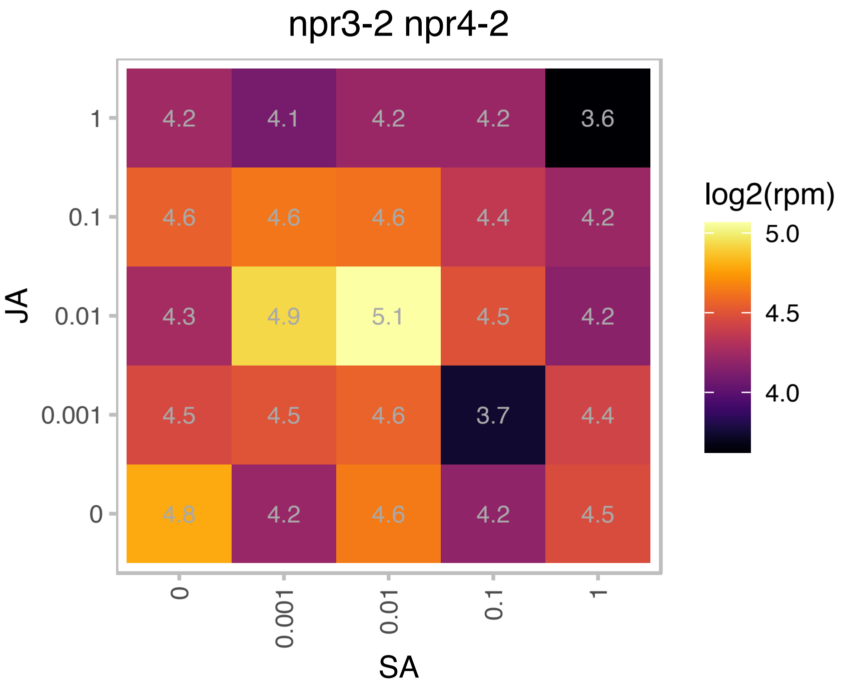


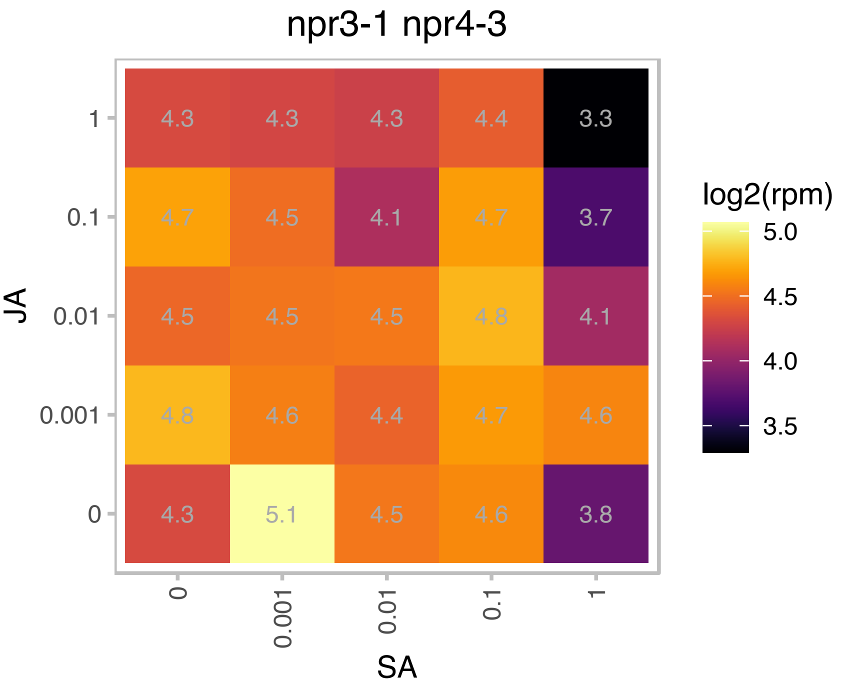

### AT4G37925.1 : NDH-M, NdhM

subunit NDH-M of NAD(P)H:plastoquinone dehydrogenase complex

Encodes subunit NDH-M of NAD(P)H:plastoquinone dehydrogenase complex (Ndh complex) present in the thylakoi
membrane of chloroplasts. This subunit is thought to be required for Ndh complex assembly.

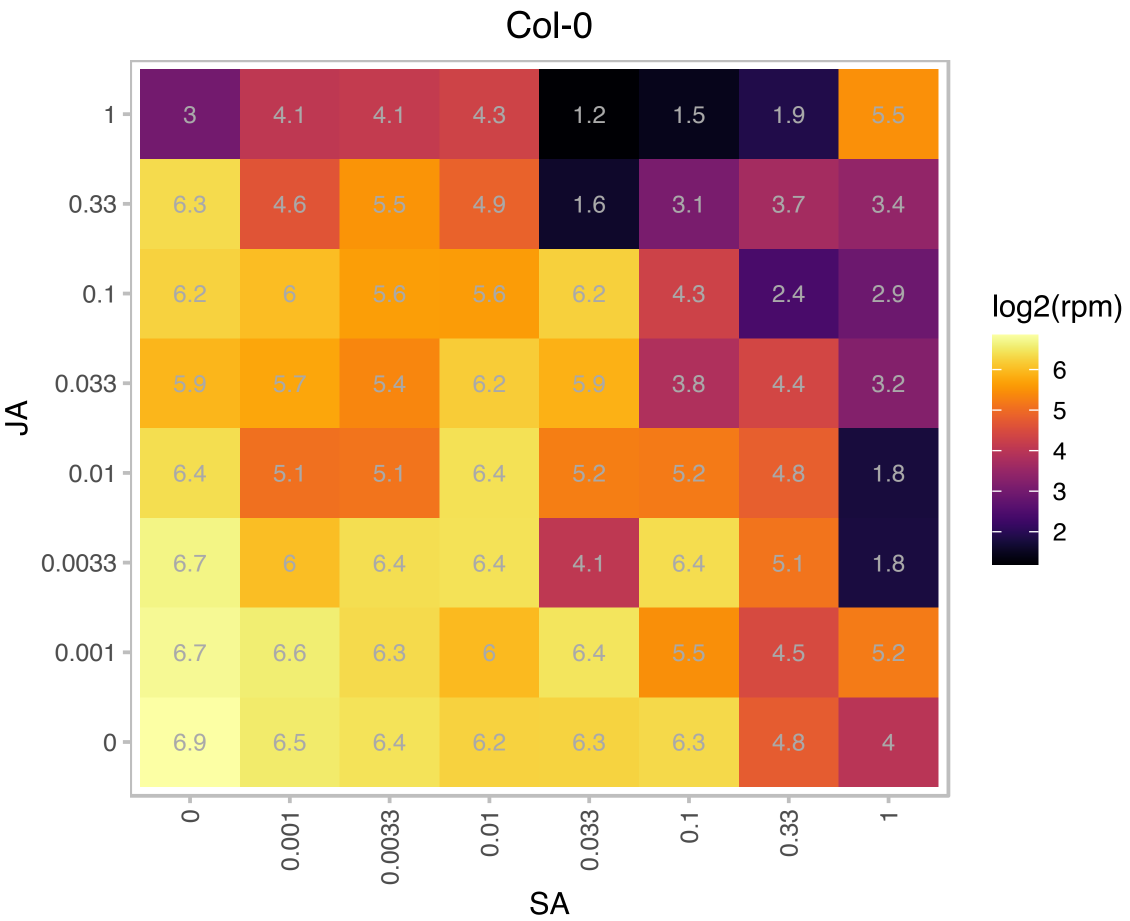


| | |
|---|---|
| Gene | AT4G37925 |
| is.expressed_genes | TRUE |
| is.responsive_genes | TRUE |
| is.apcluster | F-11 |

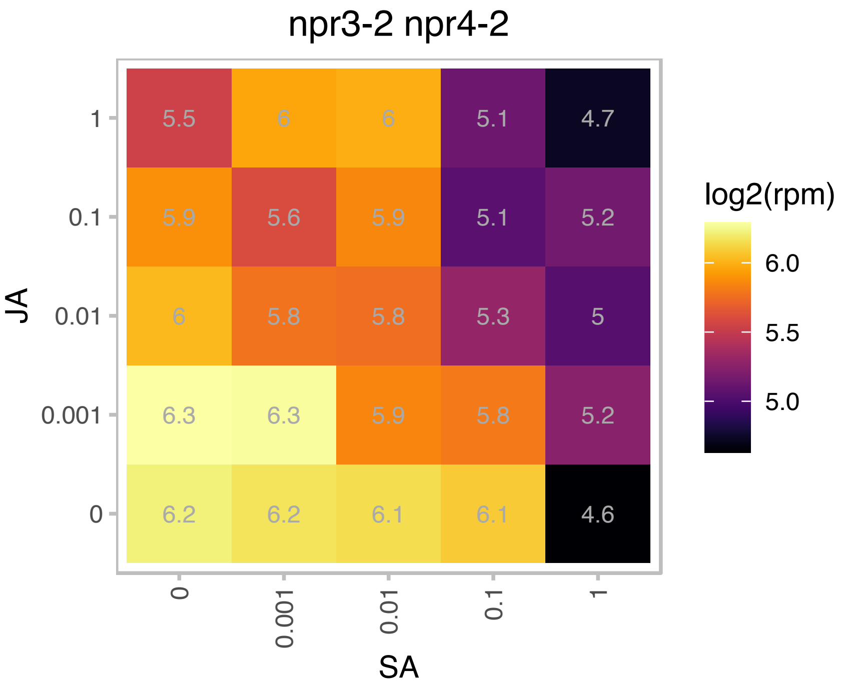


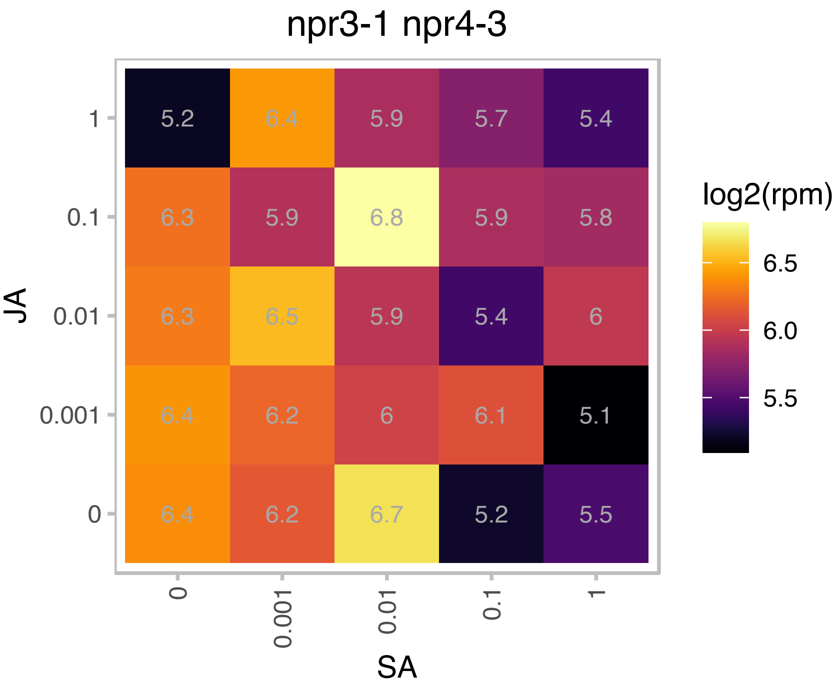

### AT5G02120.1 : OHP, OHP1, PDE335

one helix protein
Encodes a one helix protein homologous to cyanobacterial high-light inducible proteins.  The protein is localized to the thylakoid membrane and its transcript is transiently induced by exposure to high light conditions. The mRNA is cell-to-cell mobile.

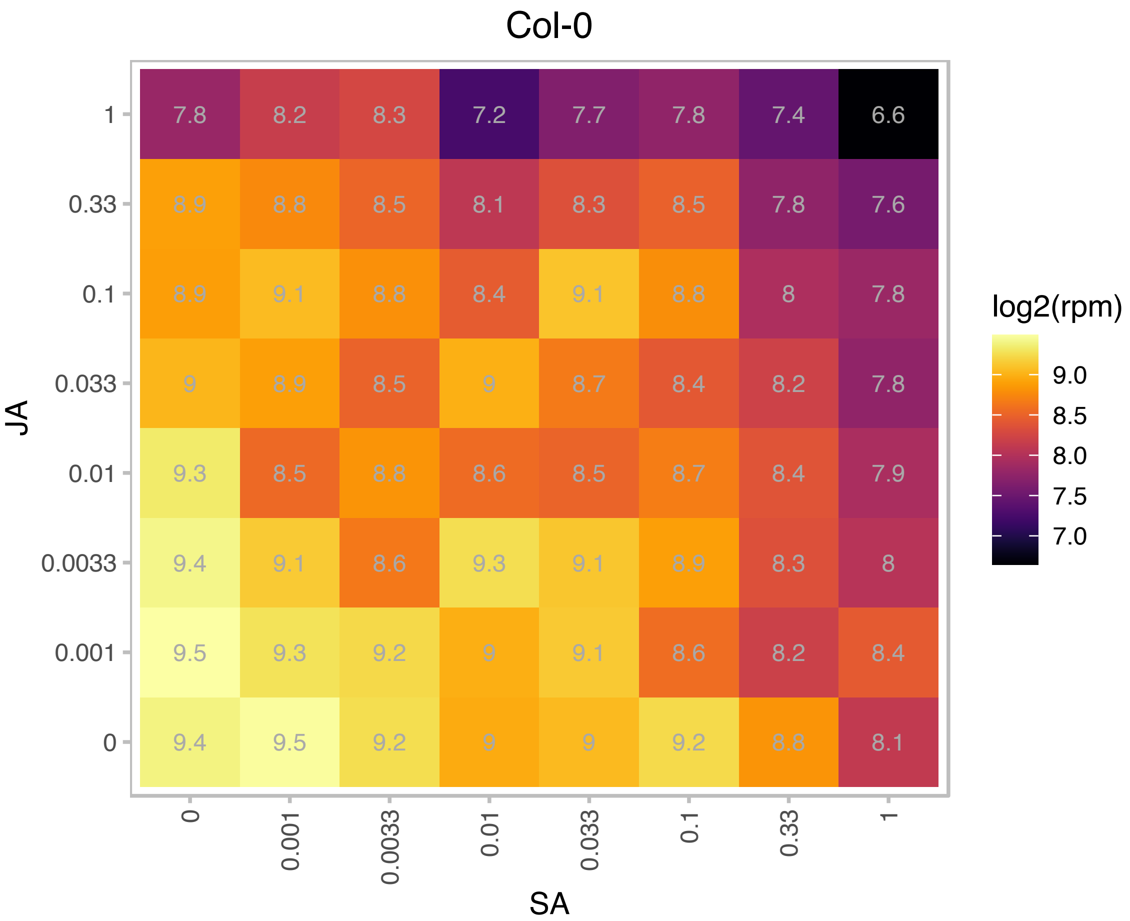


| | |
|---|---|
| Gene | AT5G02120 |
| is.expressed_genes | TRUE |
| is.responsive_genes | TRUE |
| is.apcluster | F-11 |

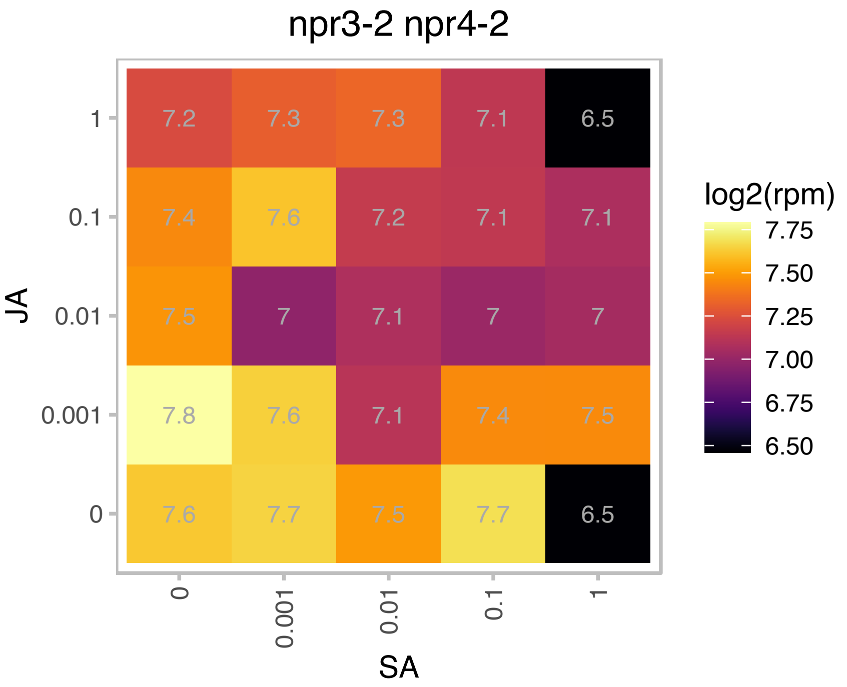


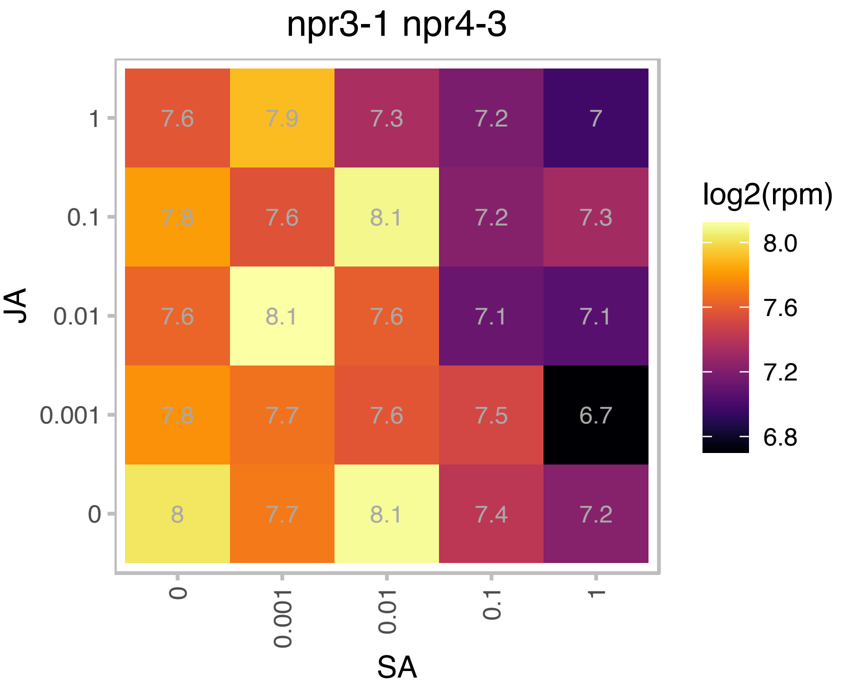

### AT5G08000.1 : E13L3, PDCB2

glucan endo-1,3-beta-glucosidase-like protein 3

Encodes a member of the X8-GPI family of proteins. It localizes to the plasmodesmata and binds callose.

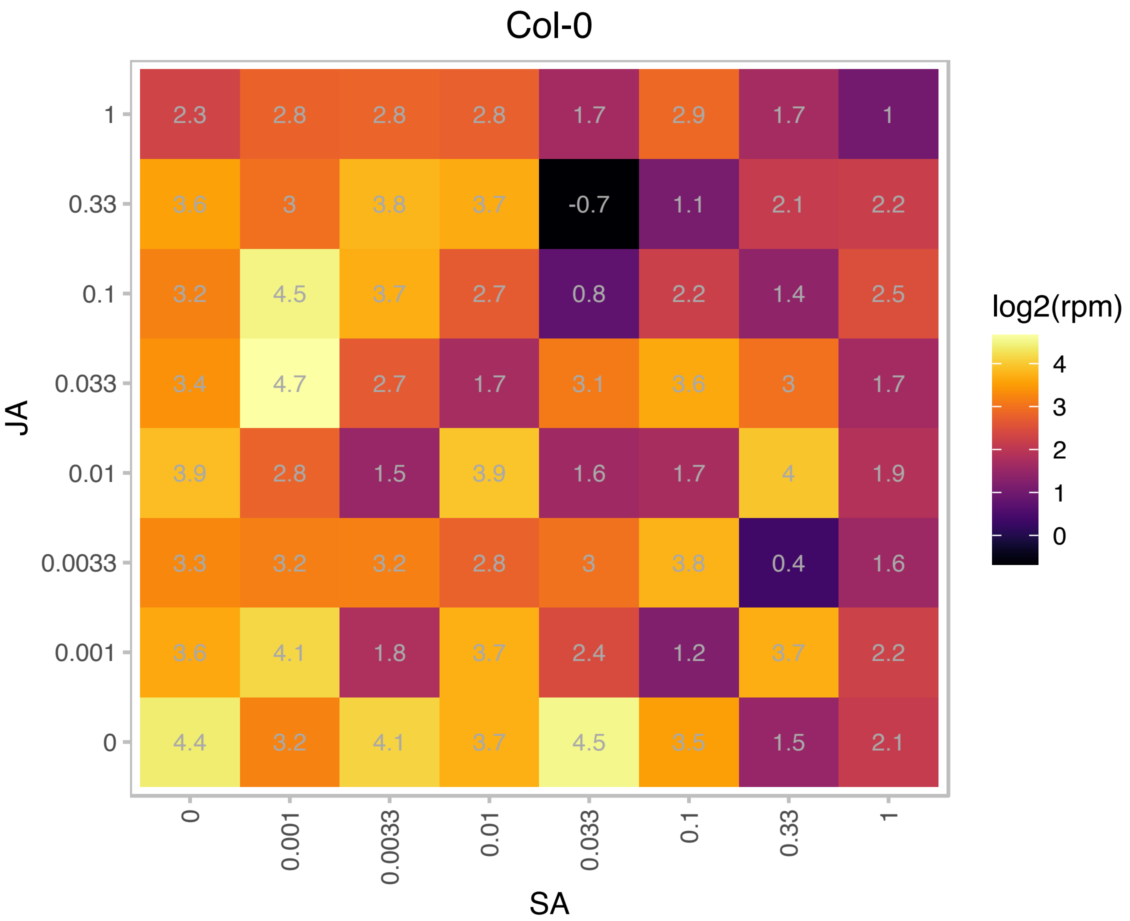


| Gene | AT5G08000 |
|---|---|
| is.expressed_genes | TRUE |
| is.responsive_genes | TRUE |
| is.apcluster | F-11 |

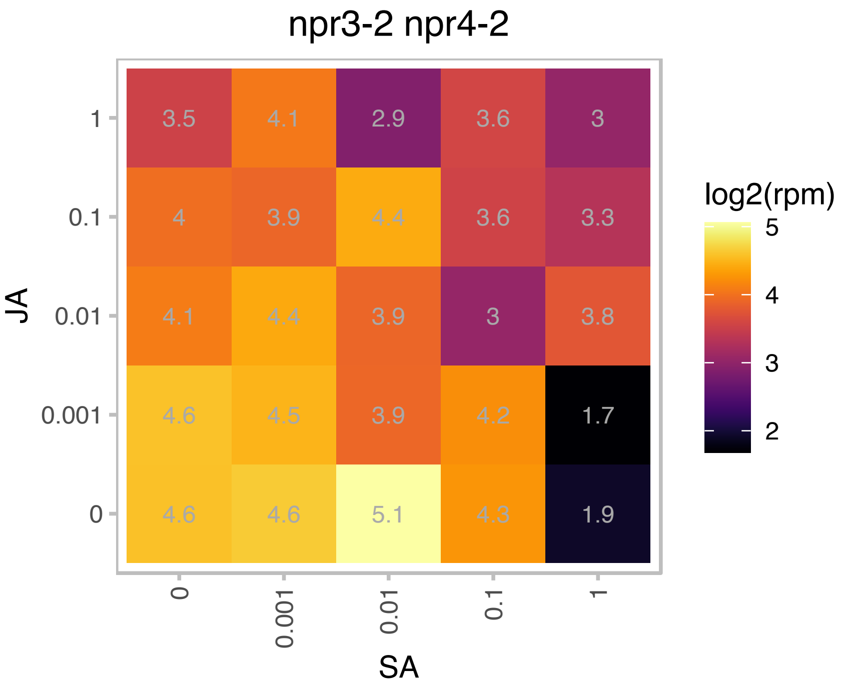


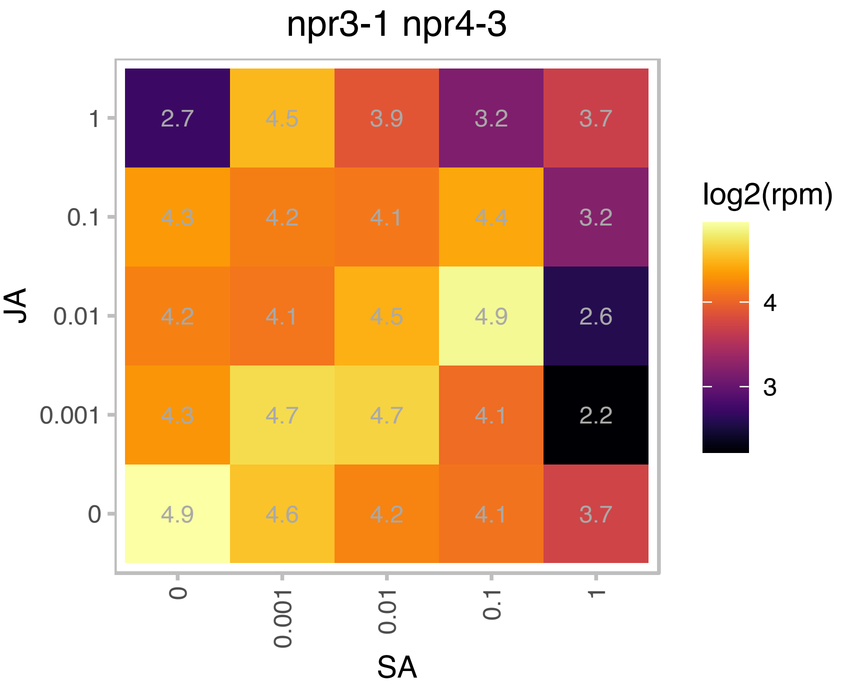

# AT5G09960.1 : NA

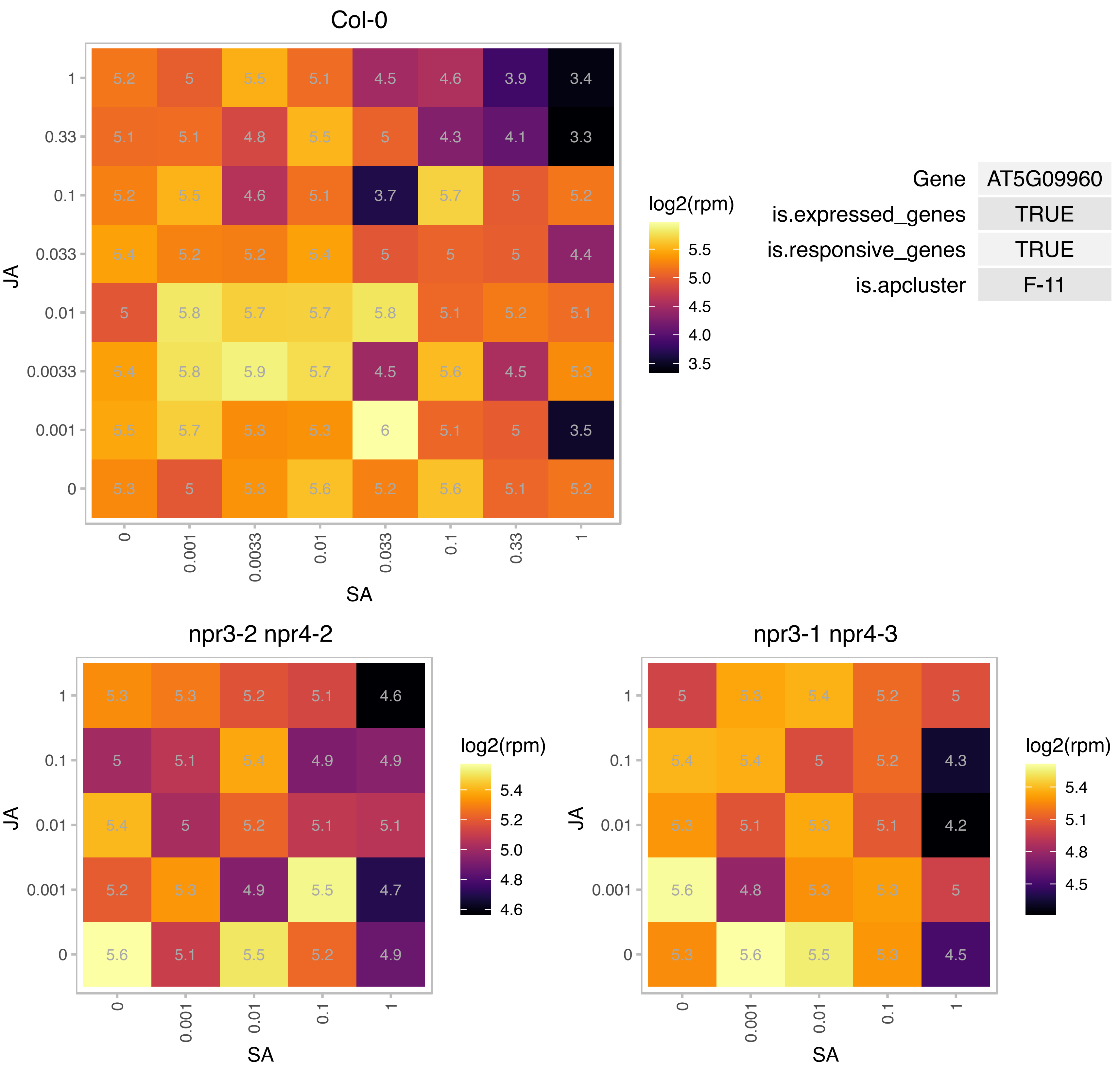

### AT5G11270.1 : OCP3

overexpressor of cationic peroxidase 3
Encodes a homeodomain transcription factor involved in mediating resistance to infection by necrotrophic pathogens dependent on perception of jasmonic acid through COI1. Expressed in the nucleus. Downregulated upor fungal infection. Also involved in drought tolerance.

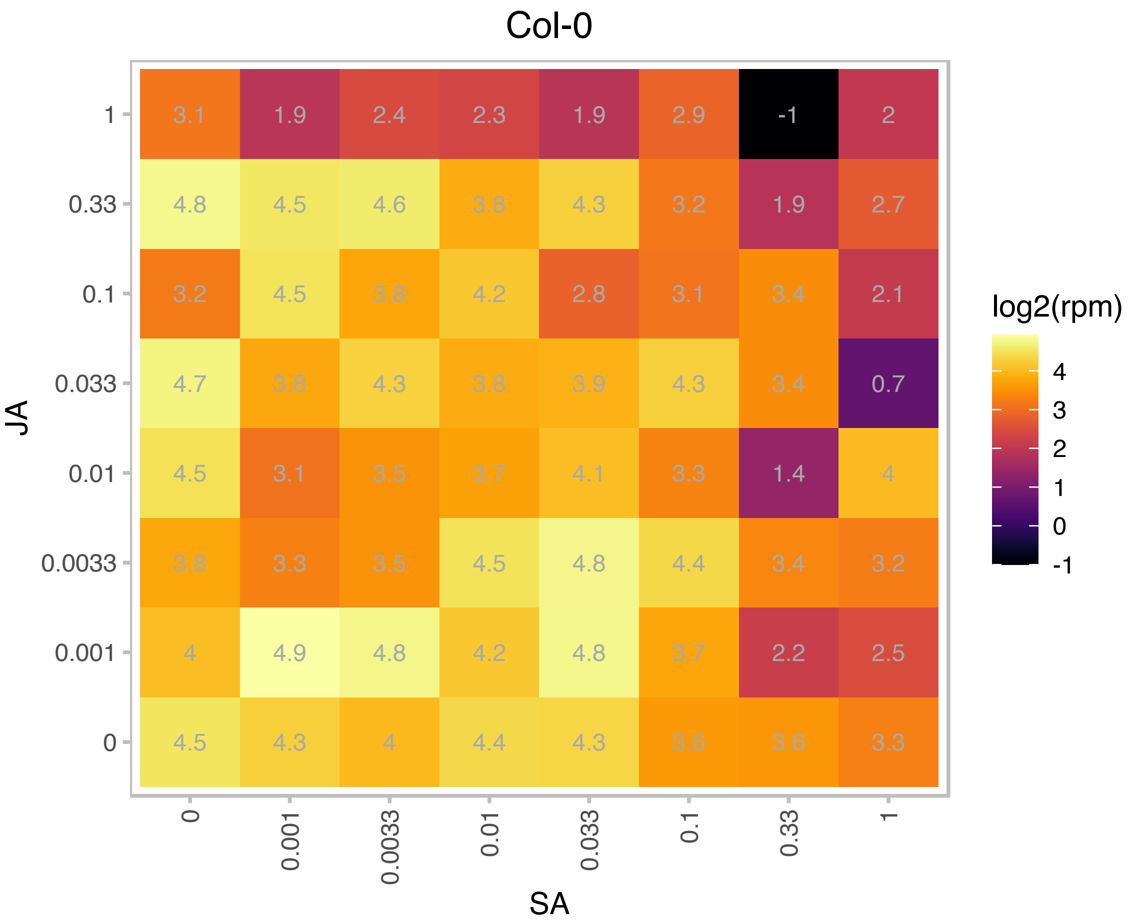


| | |
|---|---|
| Gene | AT5G11270 |
| is.expressed_genes | TRUE |
| is.responsive_genes | TRUE |
| is.apcluster | F-11 |

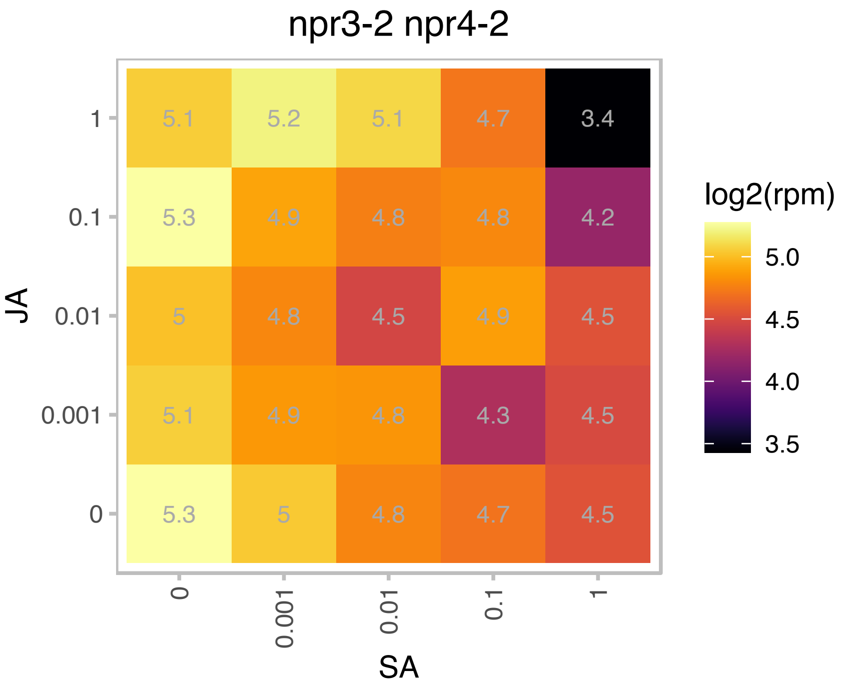


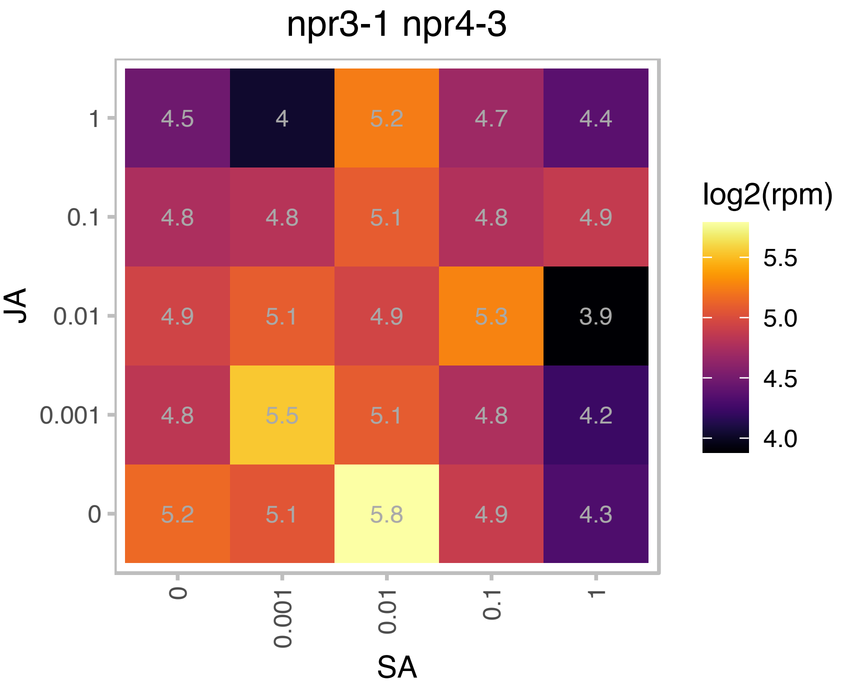

# AT5G11750.1 : NA

Ribosomal protein L19 family protein

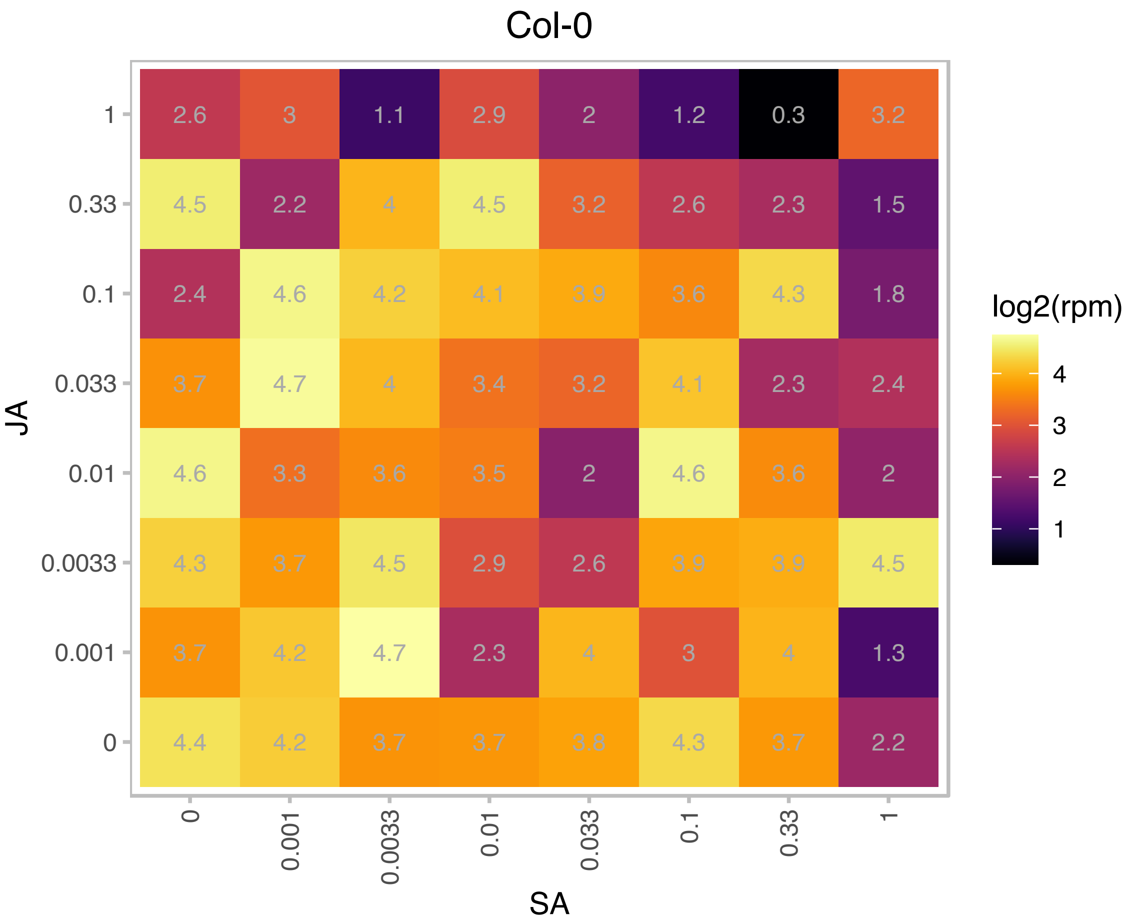


| | |
|---|---|
| Gene | AT5G11750 |
| is.expressed_genes | TRUE |
| is.responsive_genes | TRUE |
| is.apcluster | F-11 |

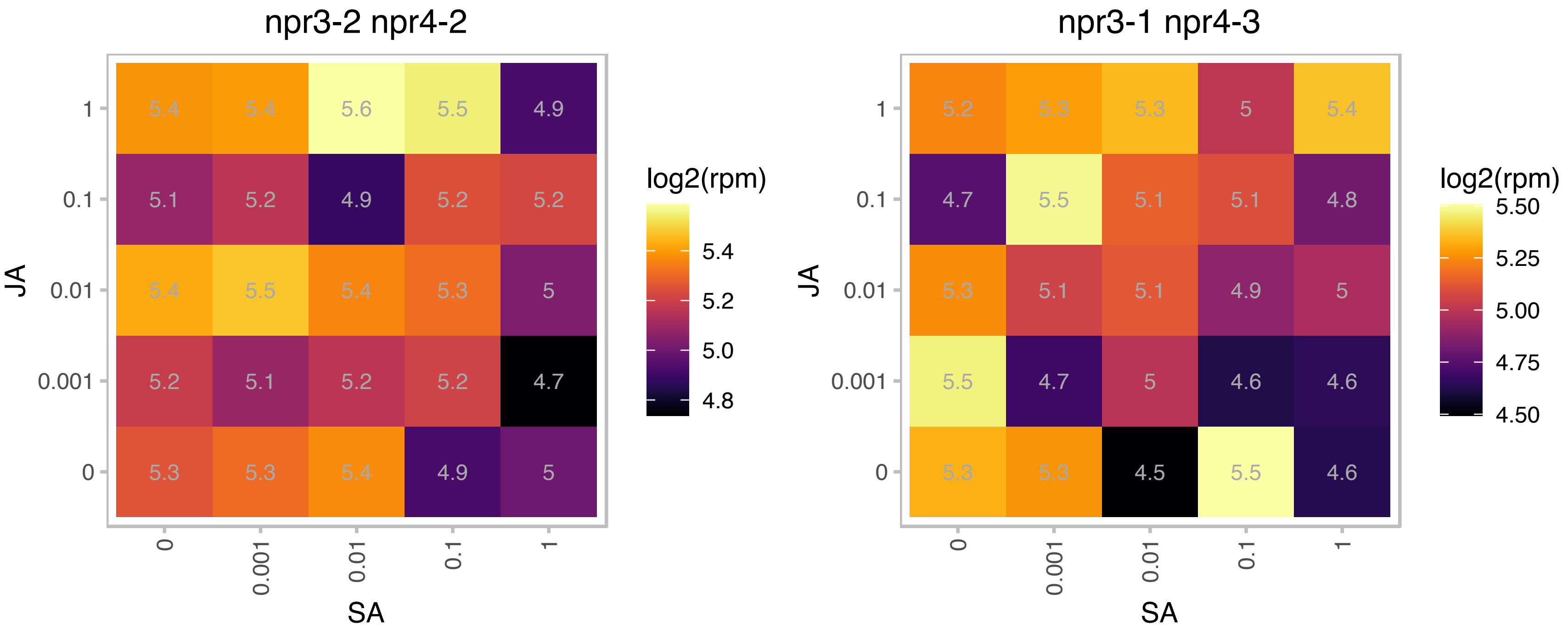

### AT5G13730.1 : SIG4, SIGD

sigma factor 4

Encodes sigma 4 factor, involved in regulating the activity of the plastid-encoded RNA polymerase PEP. Regulates the overall quantity of NDH complexes and thus influences NDH activity.

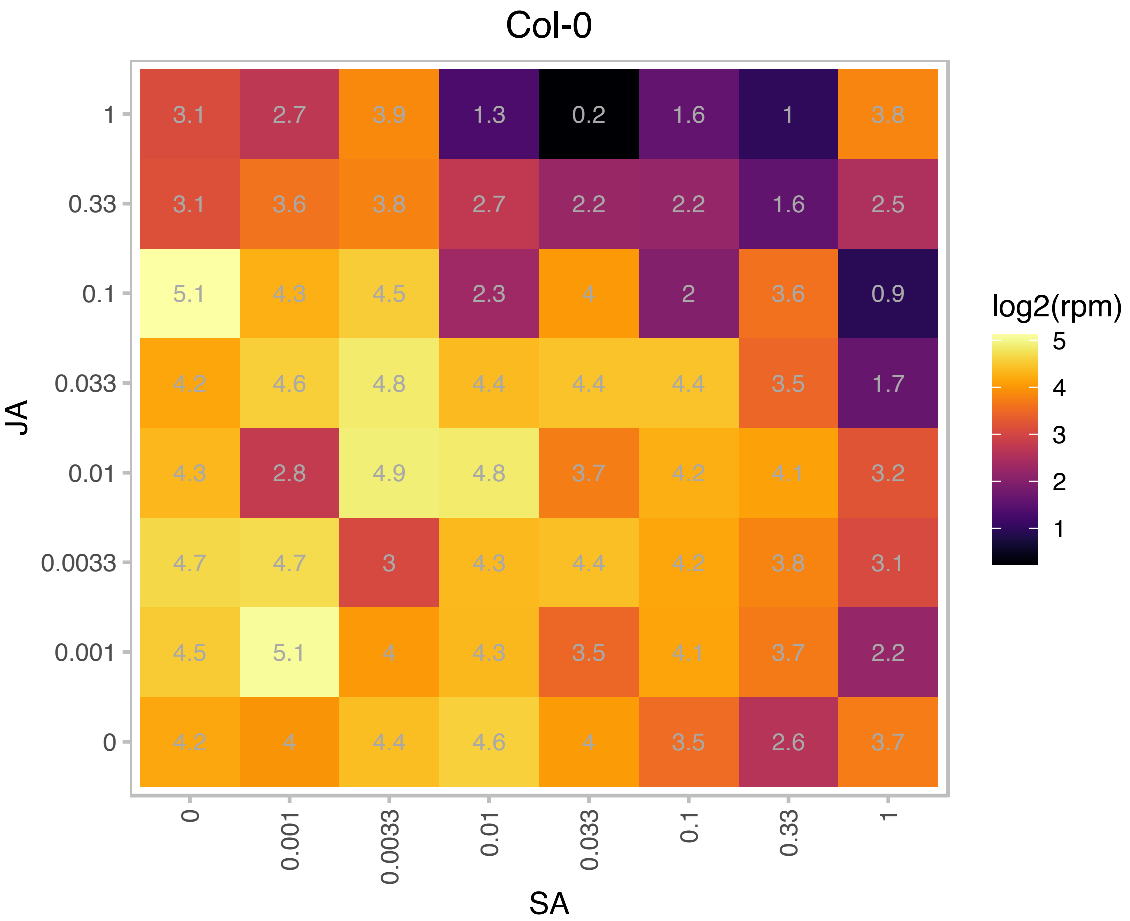


| | |
|---|---|
| Gene | AT5G13730 |
| is.expressed_genes | TRUE |
| is.responsive_genes | TRUE |
| is.apcluster | F-11 |

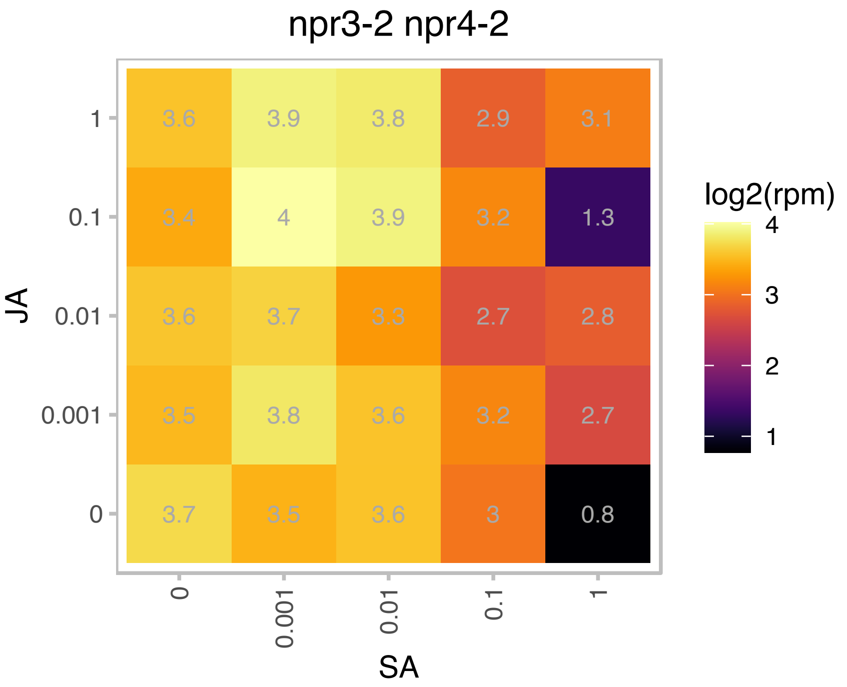


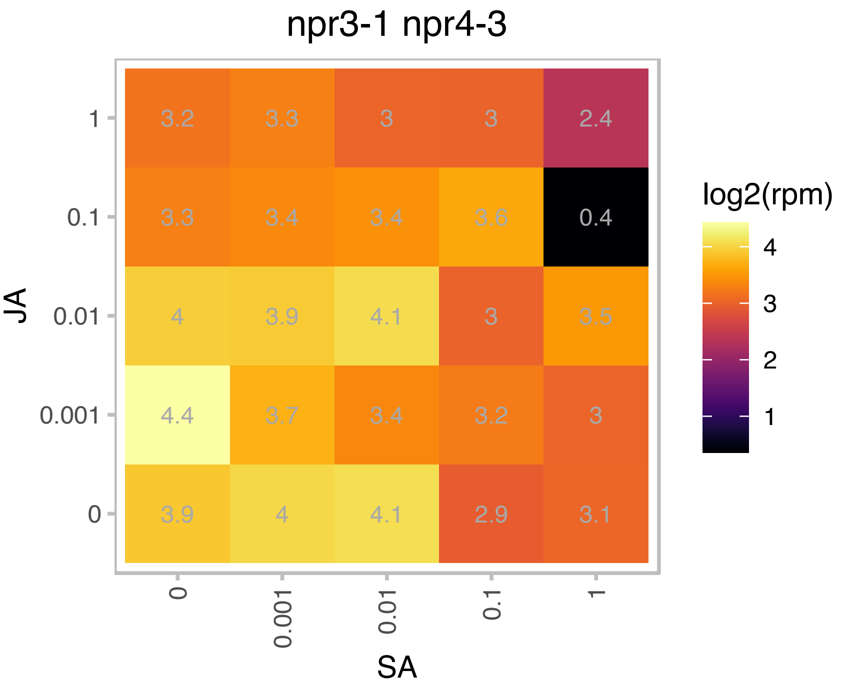

### AT5G15760.1 : PSRP3/2

Ribosomal protein PSRP-3/Ycf65

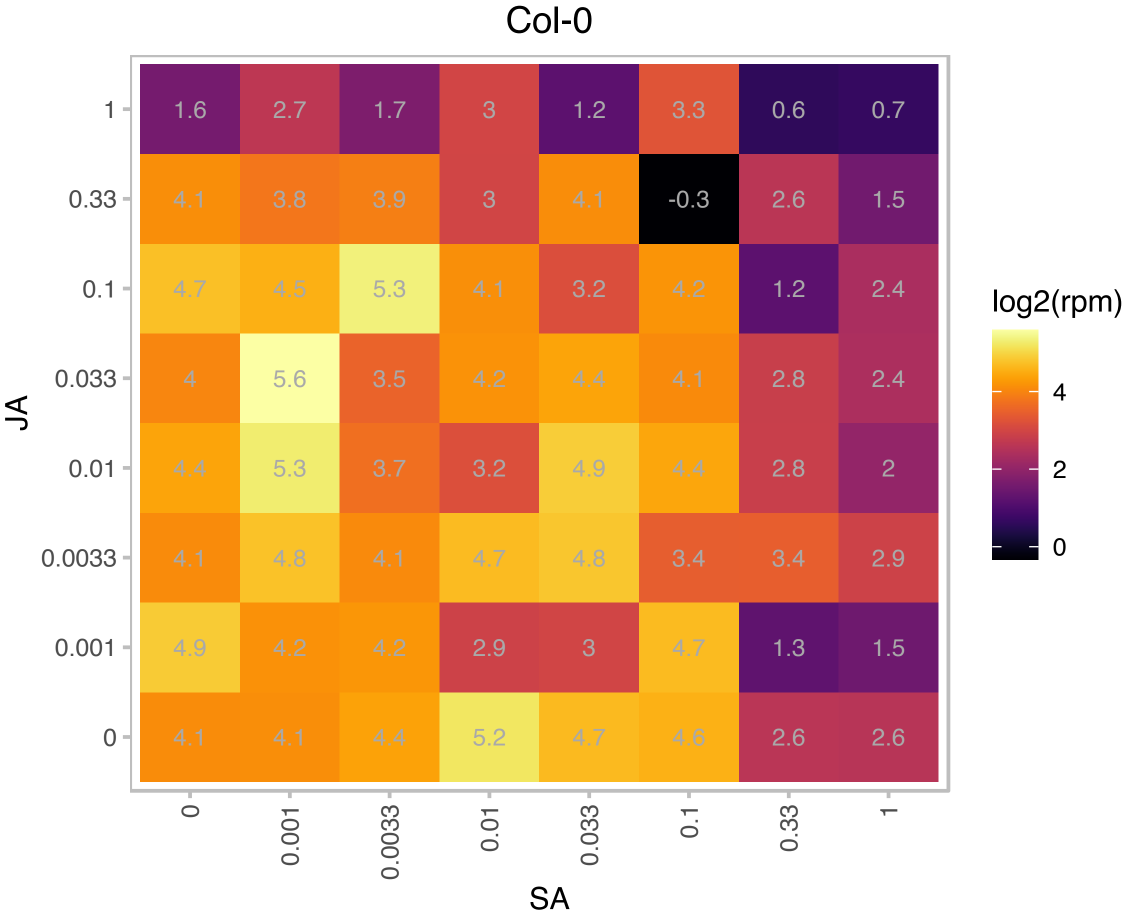


| | |
|---|---|
| Gene | AT5G15760 |
| is.expressed_genes | TRUE |
| is.responsive_genes | TRUE |
| is.apcluster | F-11 |

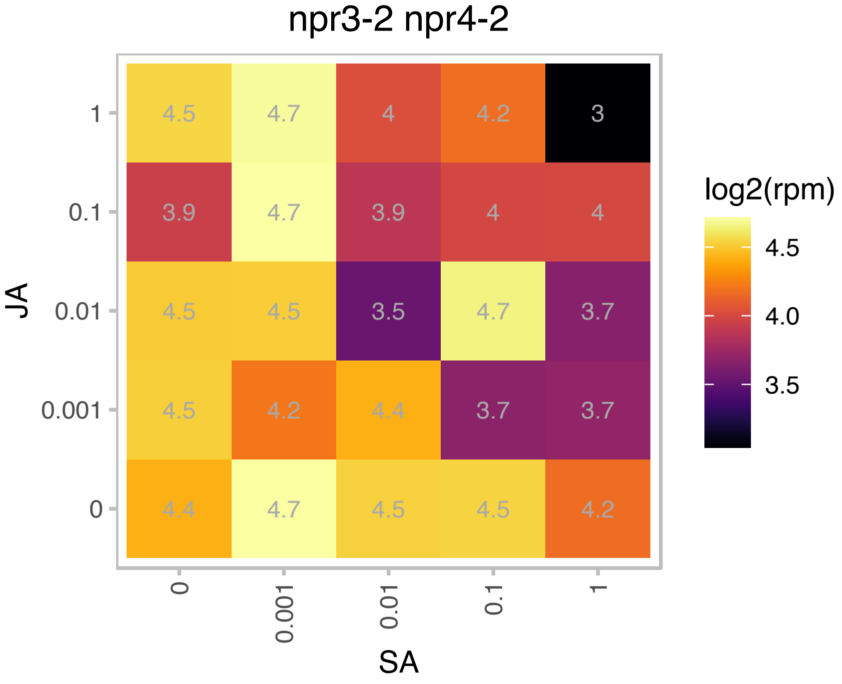


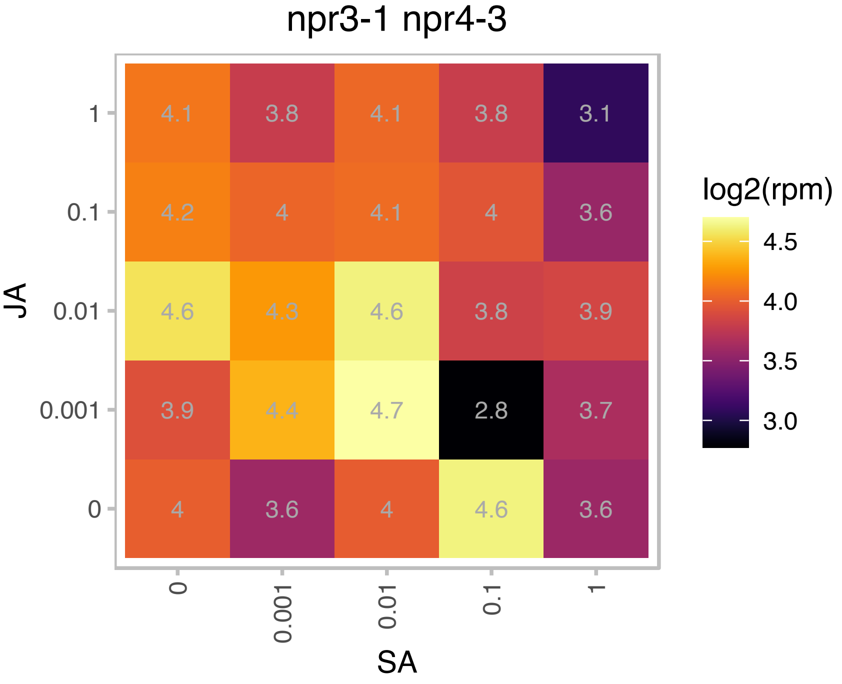

### AT5G18100.1 : CSD3

copper/zinc superoxide dismutase 3

A putative peroxisomal CuZnSOD inducible by a high-light pulse.

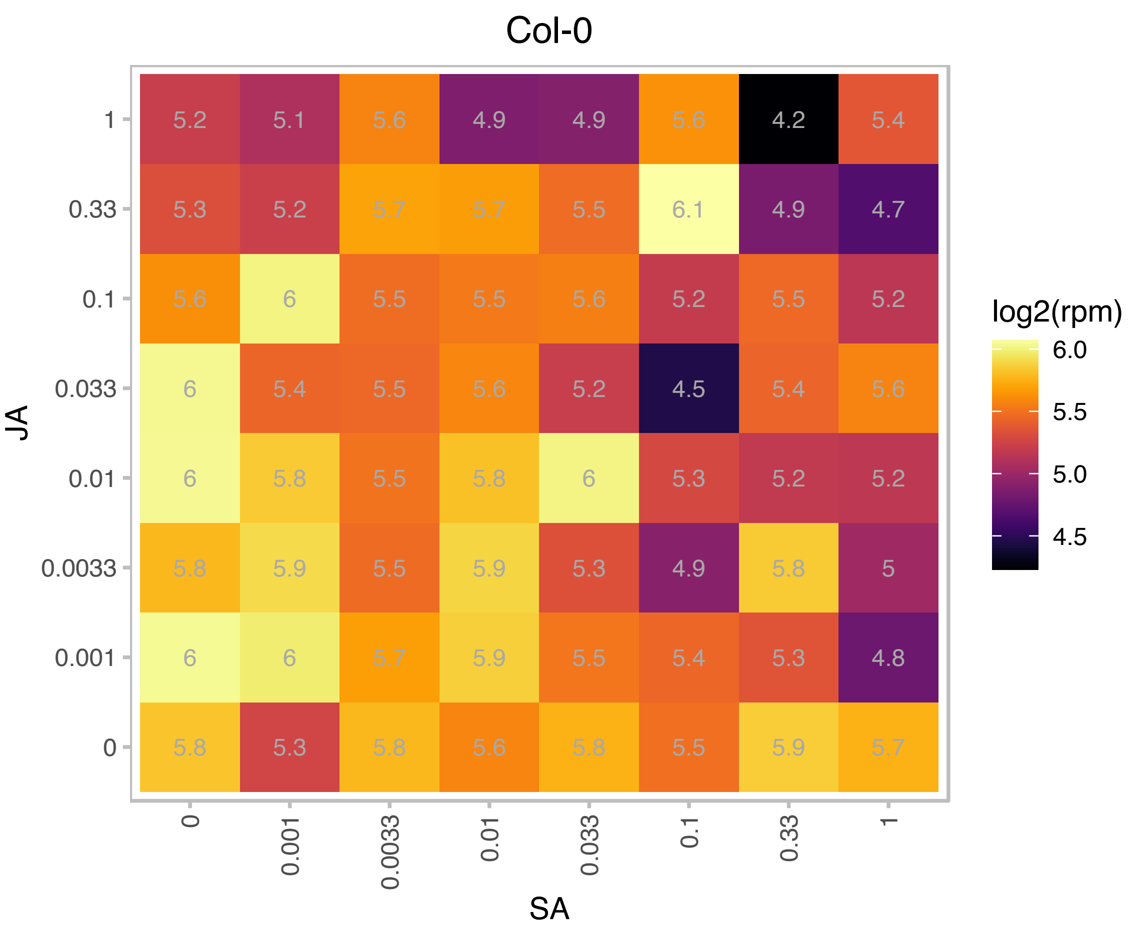


| | |
|---|---|
| Gene | AT5G18100 |
| is.expressed_genes | TRUE |
| is.responsive_genes | TRUE |
| is.apcluster | F-11 |

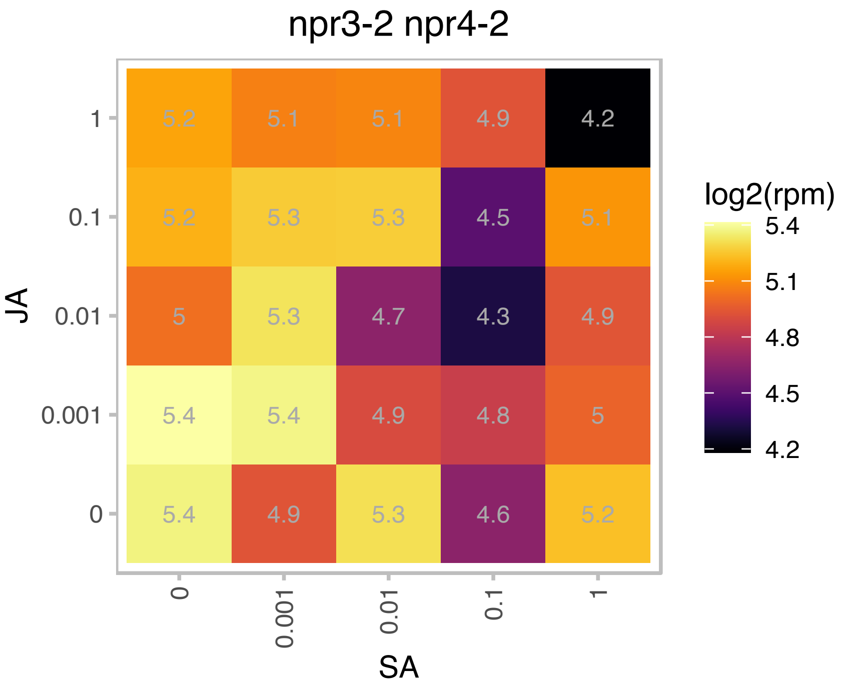


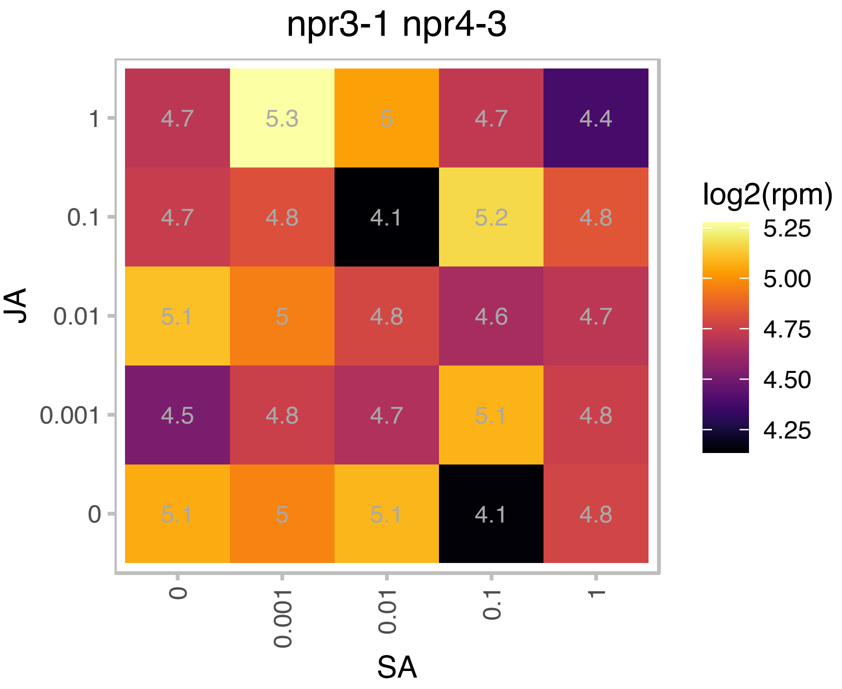

# AT5G19340.1 : NA

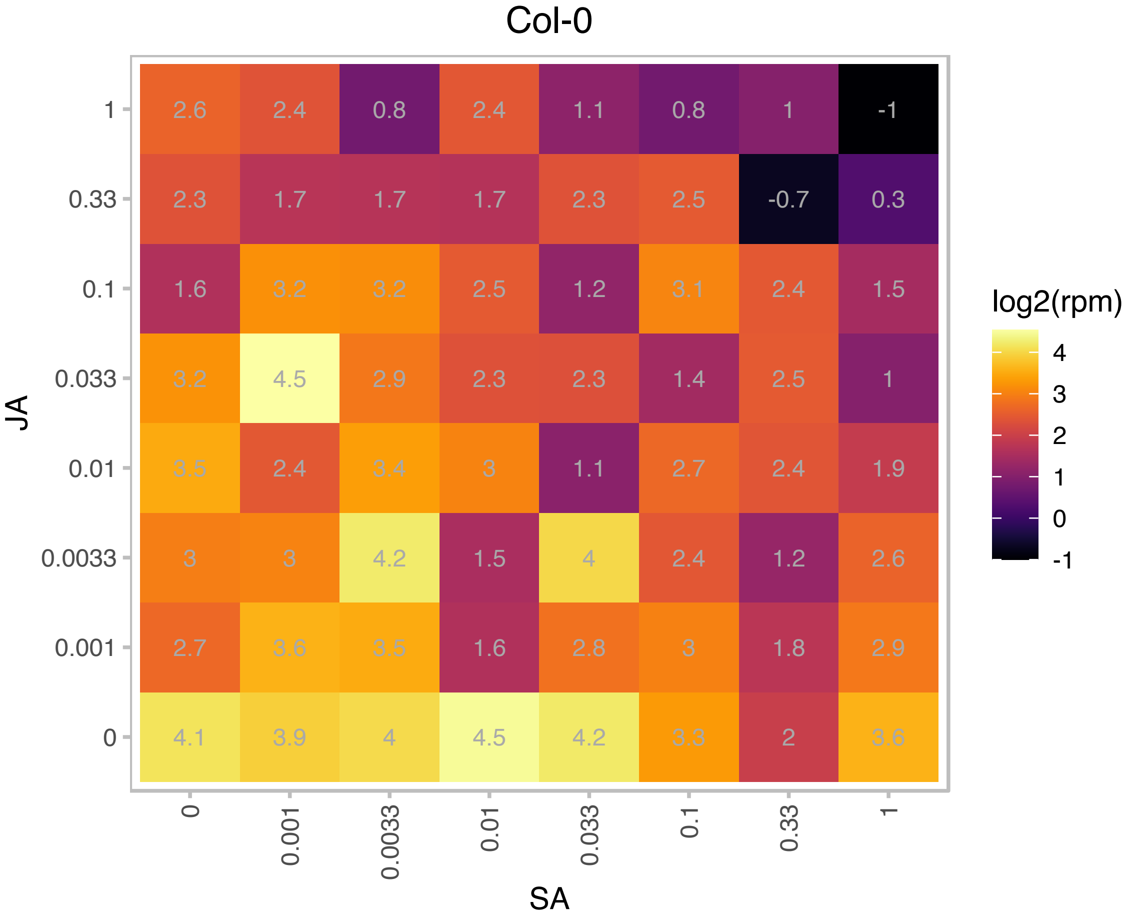


| | |
|---|---|
| Gene | AT5G19340 |
| is.expressed_genes | TRUE |
| is.responsive_genes | TRUE |
| is.apcluster | F-11 |

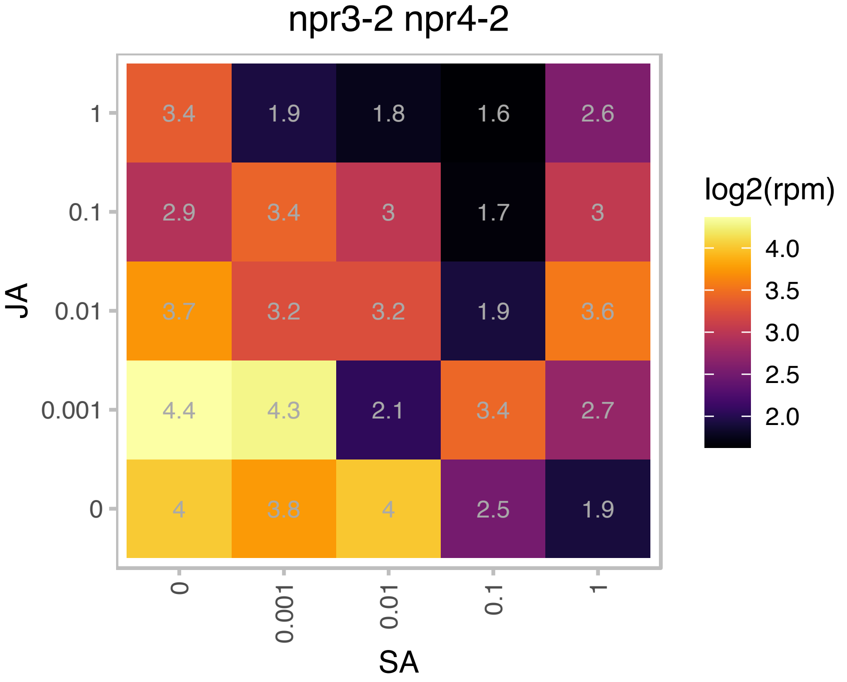


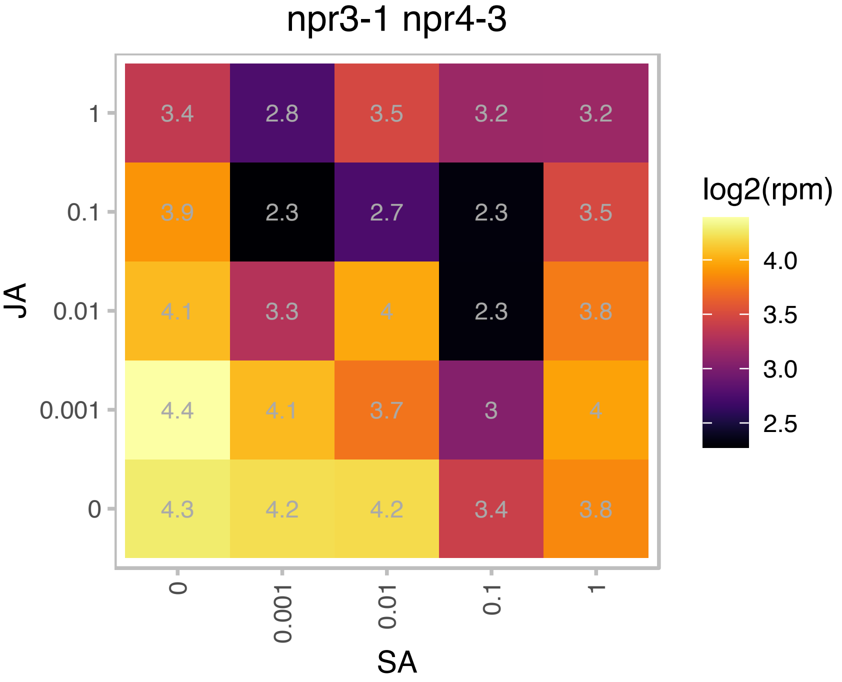

# AT5G19940.1 : NA

Plastid-lipid associated protein PAP / fibrillin family protein

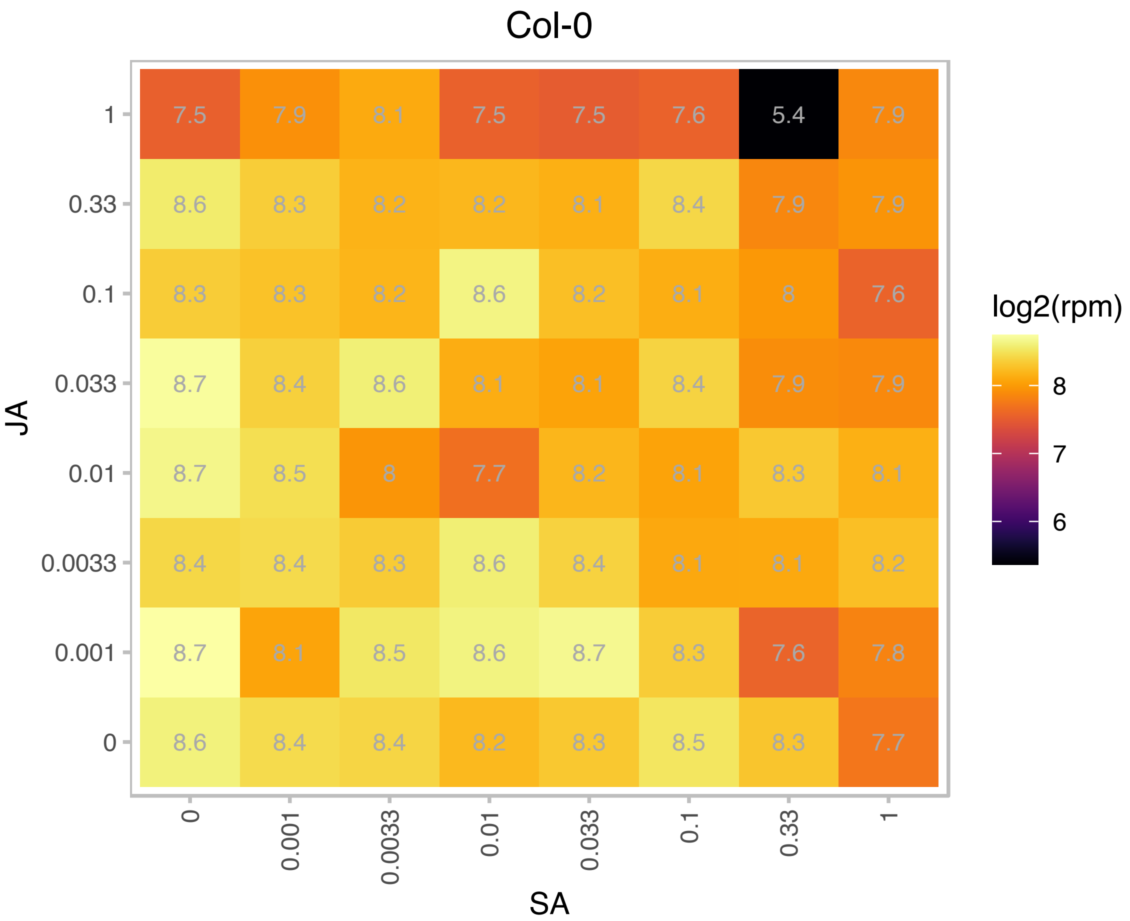


| | |
|---|---|
| Gene | AT5G19940 |
| is.expressed_genes | TRUE |
| is.responsive_genes | TRUE |
| is.apcluster | F-11 |

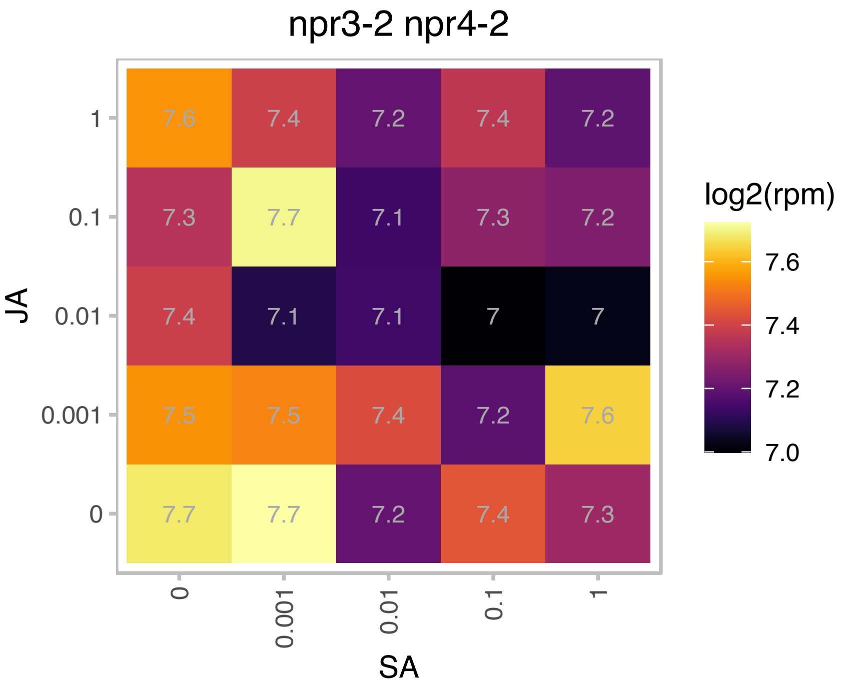


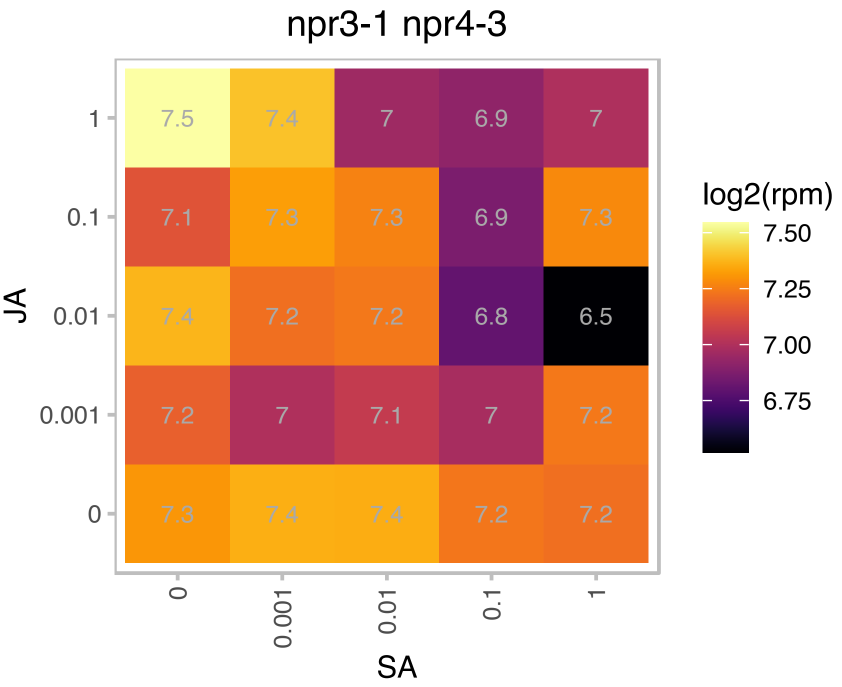

### AT5G20630.1 : ATGER3, GER3, GLP3, GLP3A, GLP3B

germin 3

Encodes a germin-like protein. Its transcripts are more abundant in RNA from leaves collected in the evening, suggesting some kind of circadian regulation.

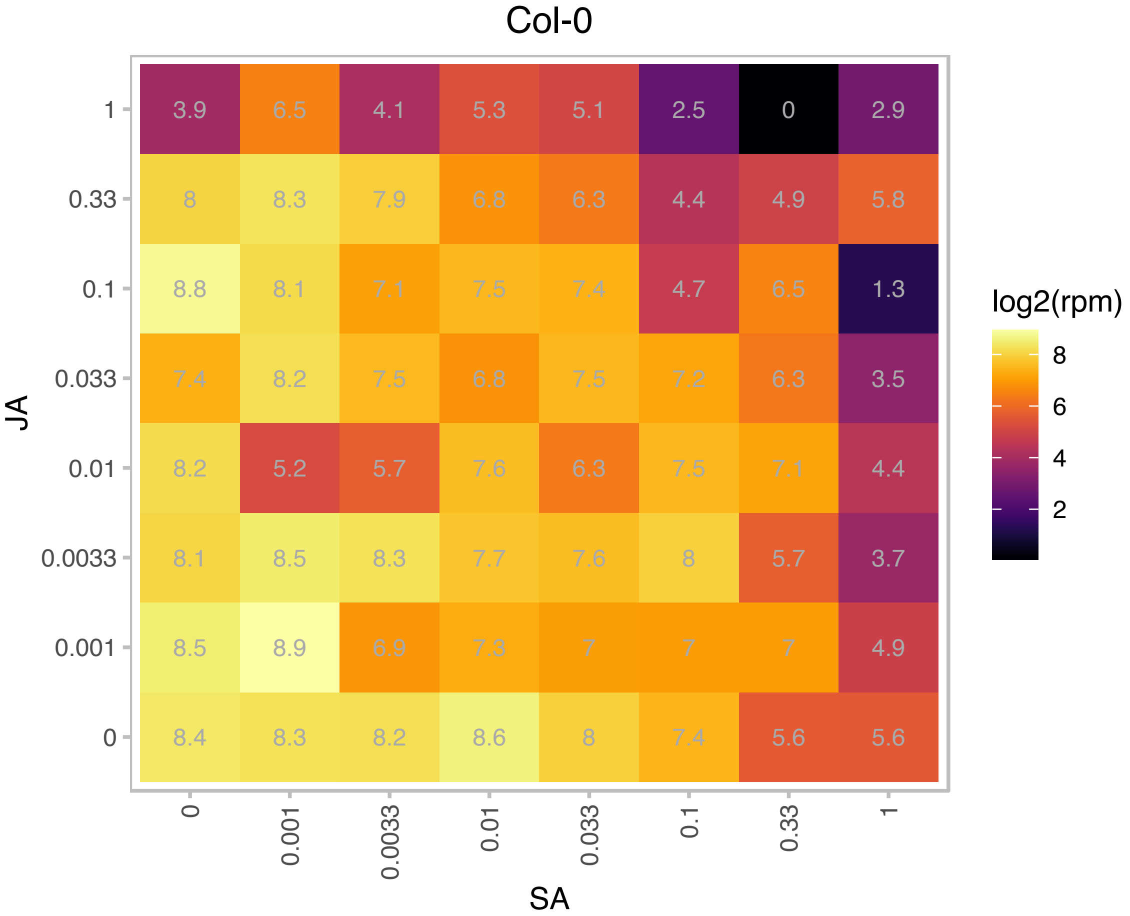


| | |
|---|---|
| Gene | AT5G20630 |
| is.expressed_genes | TRUE |
| is.responsive_genes | TRUE |
| is.apcluster | F-11 |

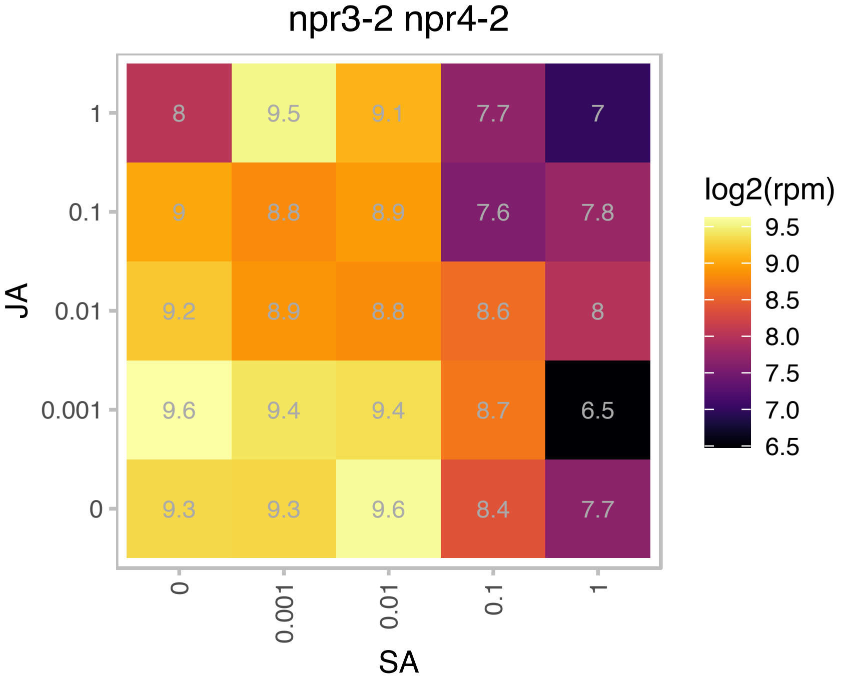


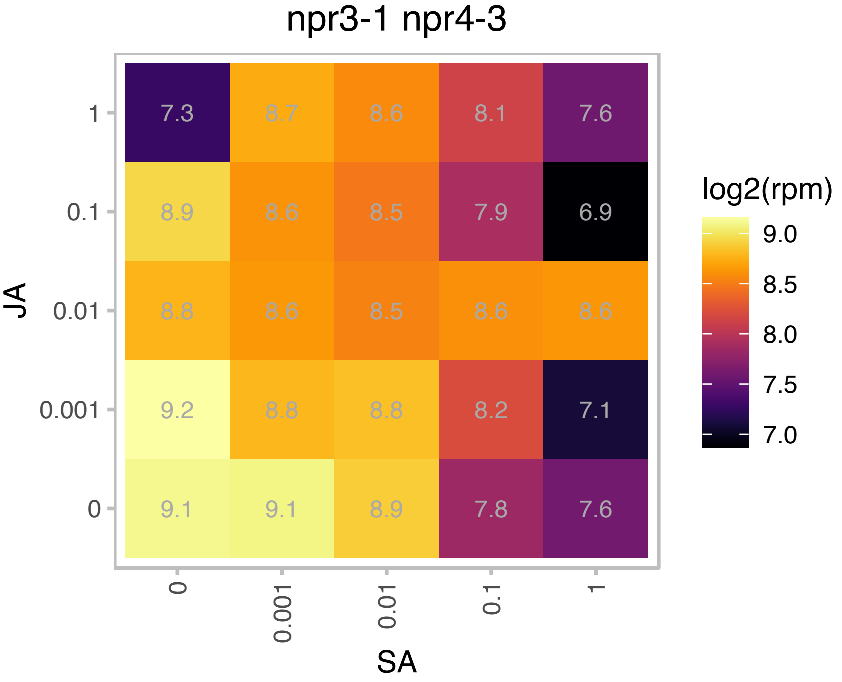

# AT5G22210.1 : NA

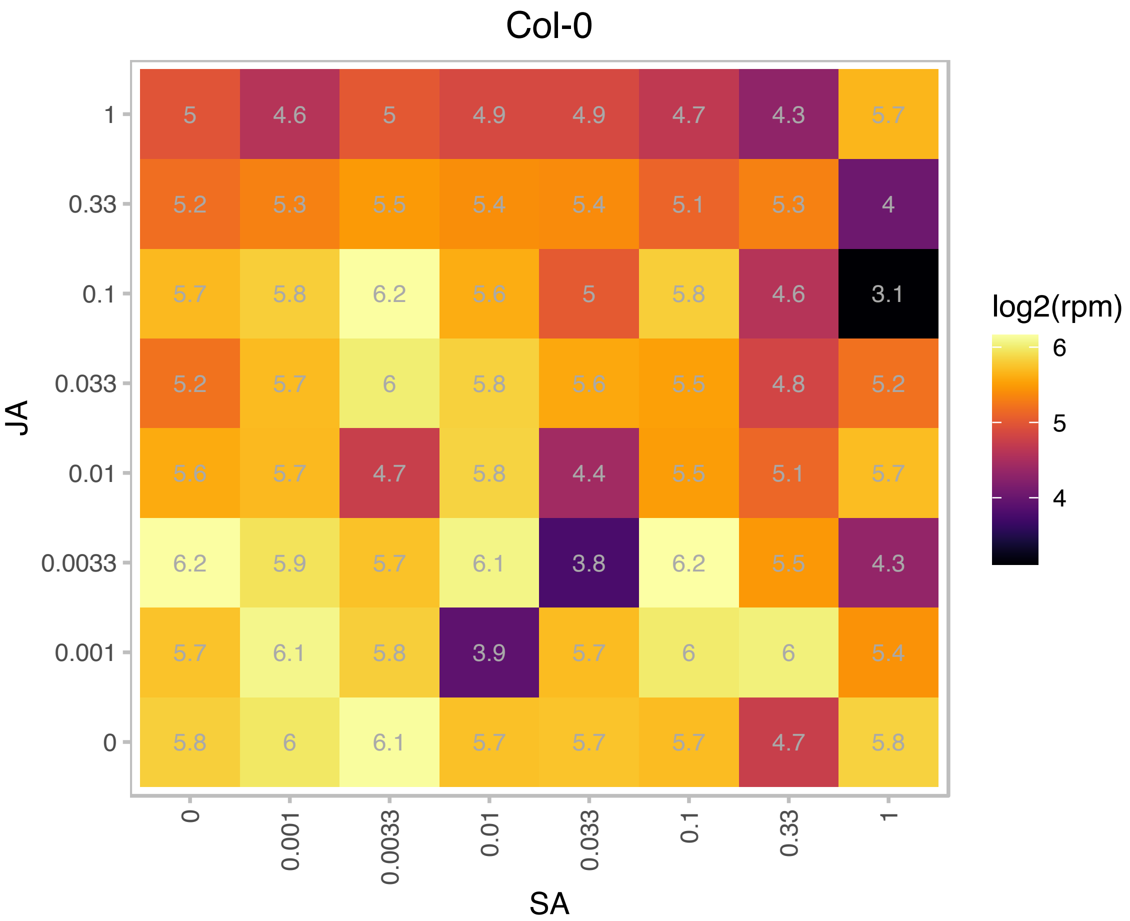


| Gene | AT5G22210 |
|---|---|
| is.expressed_genes | TRUE |
| is.responsive_genes | TRUE |
| is.apcluster | F-11 |

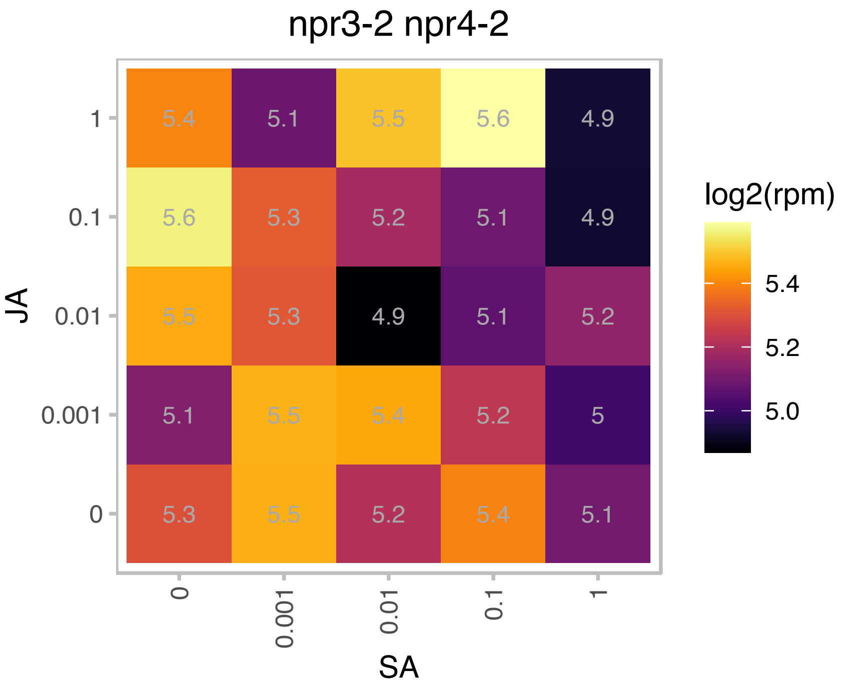


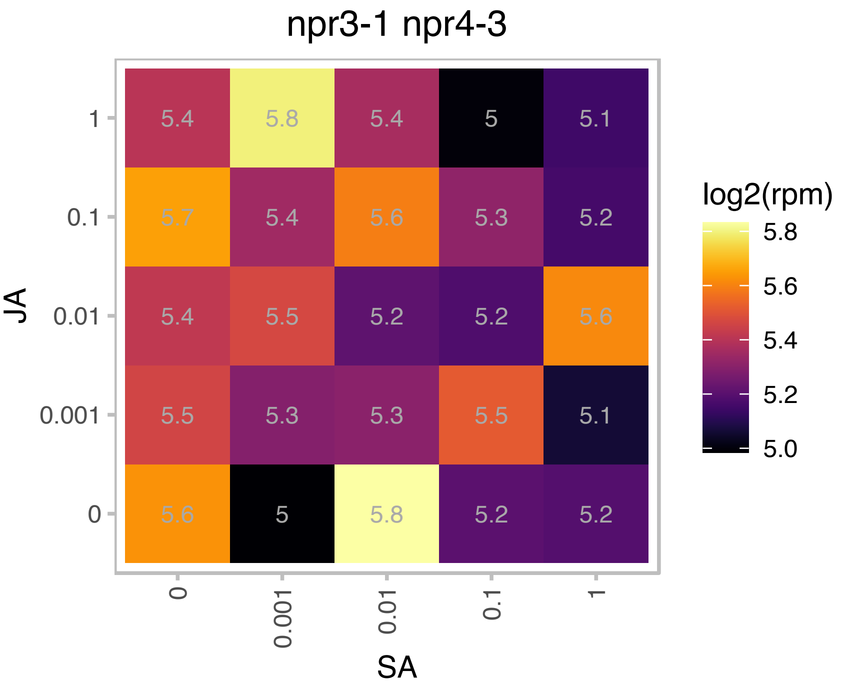

# AT5G24165.1 : NA

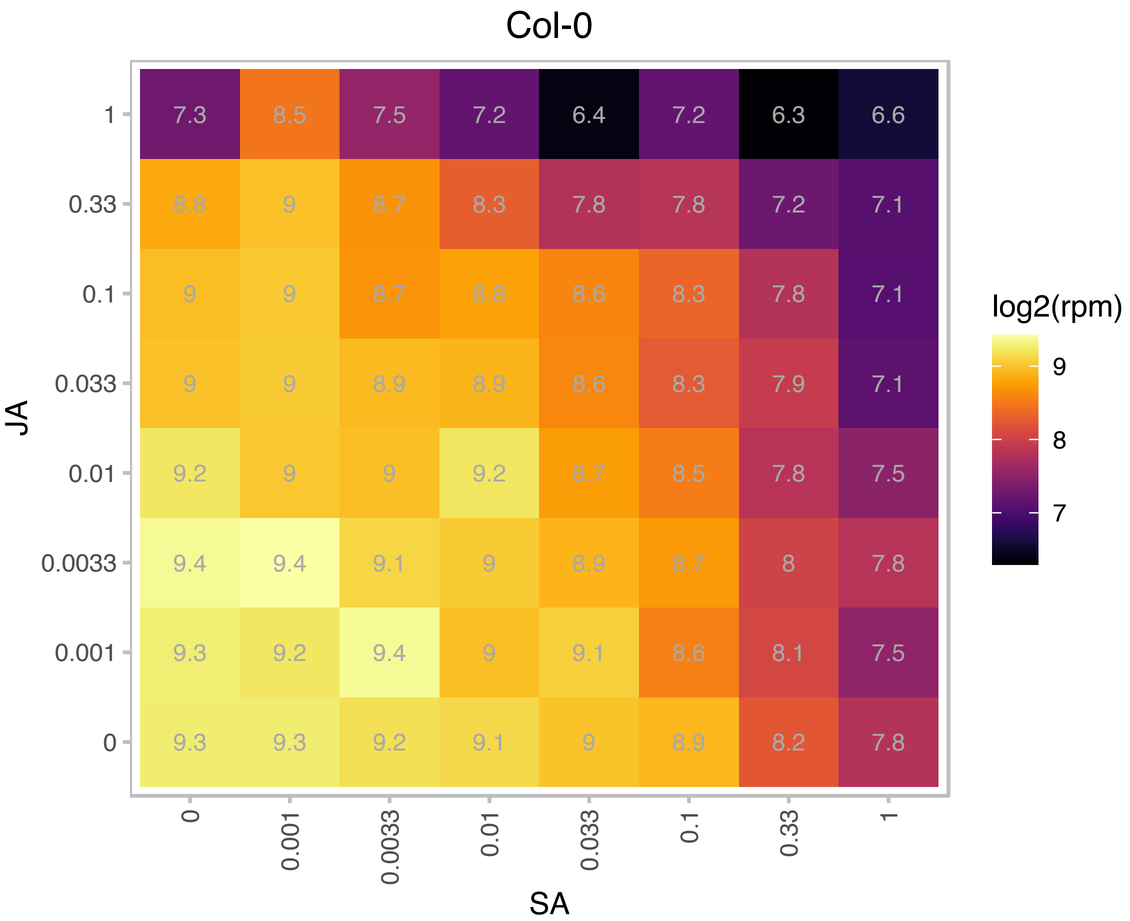


| | |
|---|---|
| Gene | AT5G24165 |
| is.expressed_genes | TRUE |
| is.responsive_genes | TRUE |
| is.apcluster | F-11 |

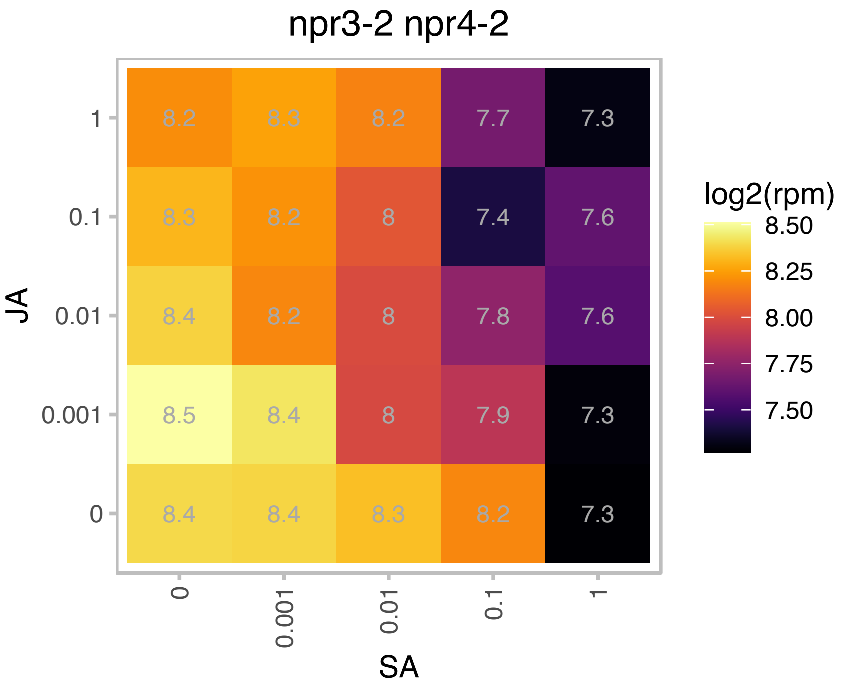


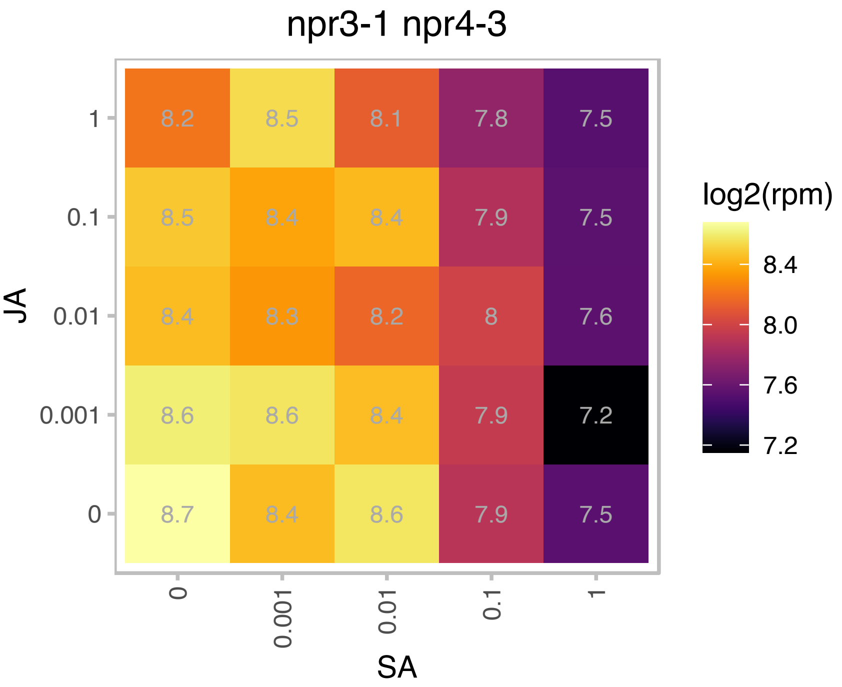

# AT5G25757.1 : NA

RNA polymerase I-associated factor PAF67

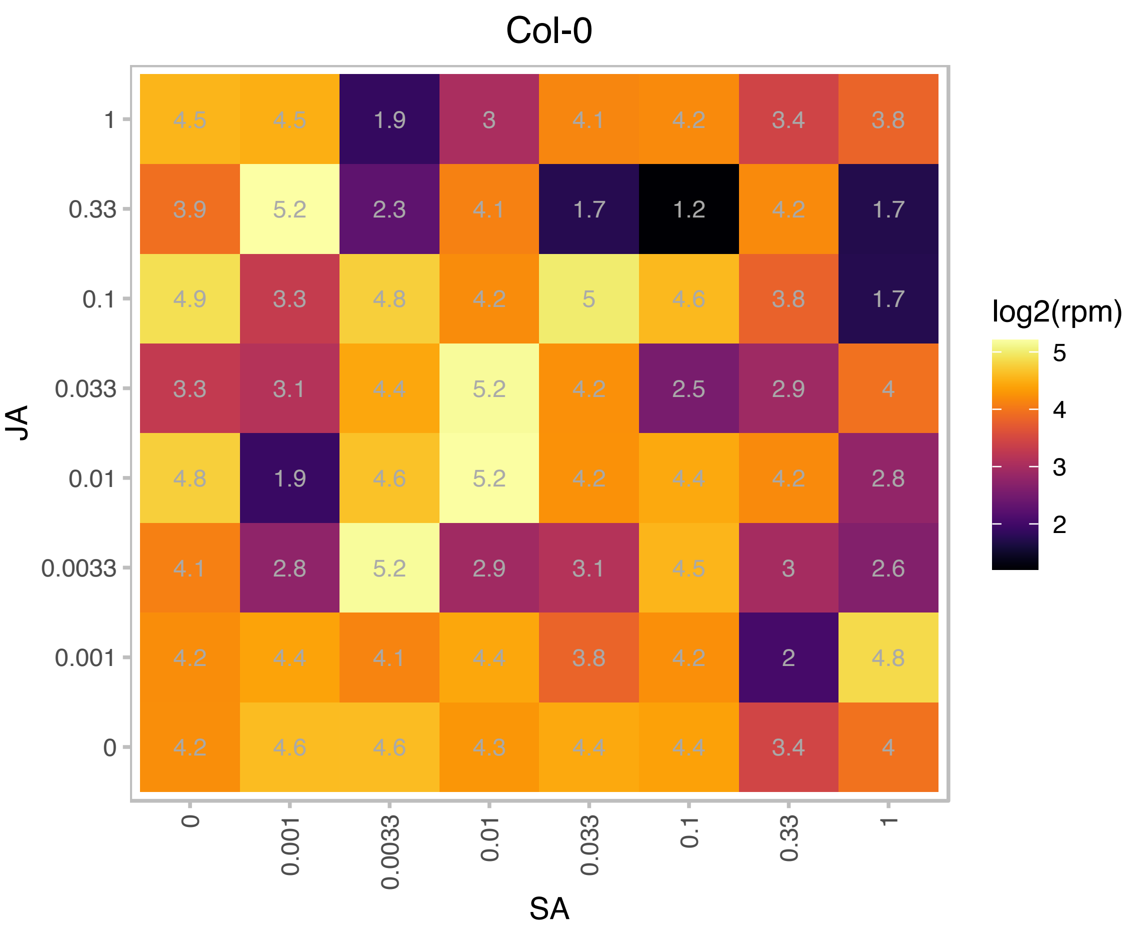


| | |
|---|---|
| Gene | AT5G25757 |
| is.expressed_genes | TRUE |
| is.responsive_genes | TRUE |
| is.apcluster | F-11 |

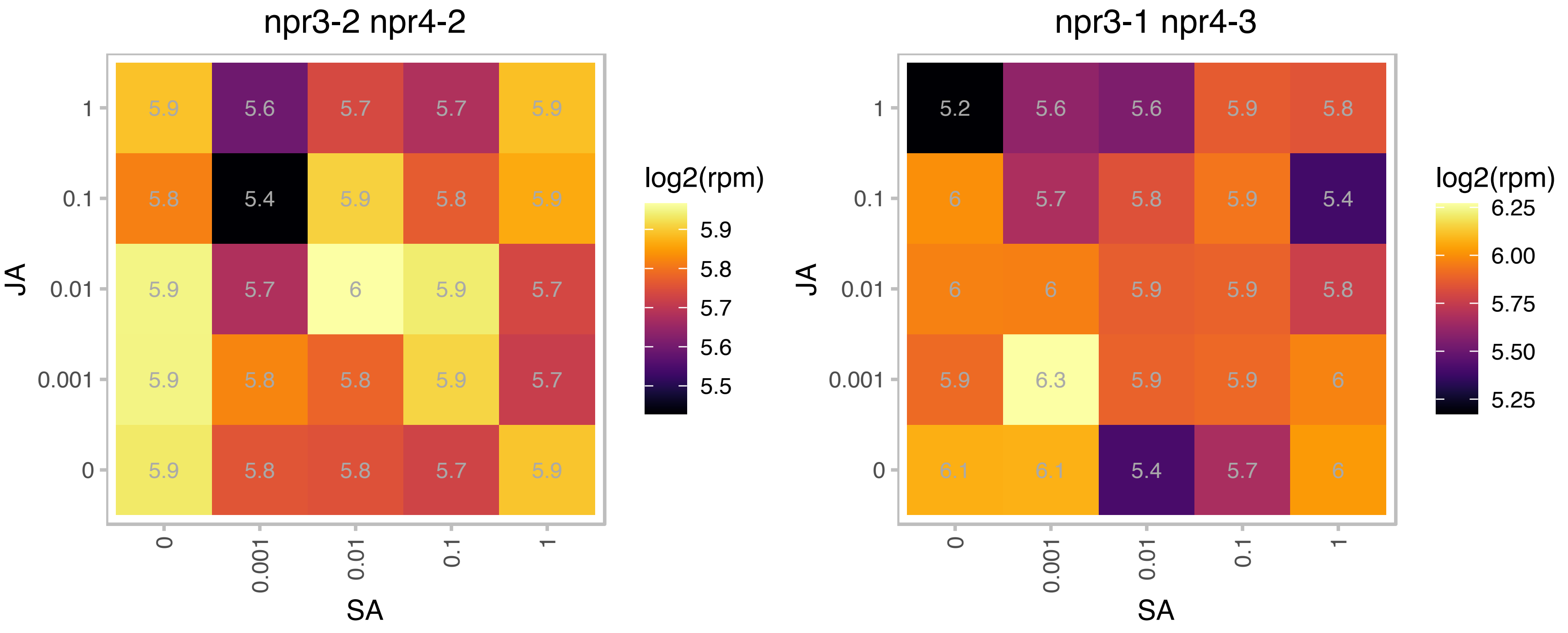

# AT5G35790.1 : G6PD1

glucose-6-phosphate dehydrogenase 1

Encodes a plastidic glucose-6-phosphate dehydrogenase that is sensitive to reduction by DTT and whose mRNA is more prevalent in developing organs but absent in the root.

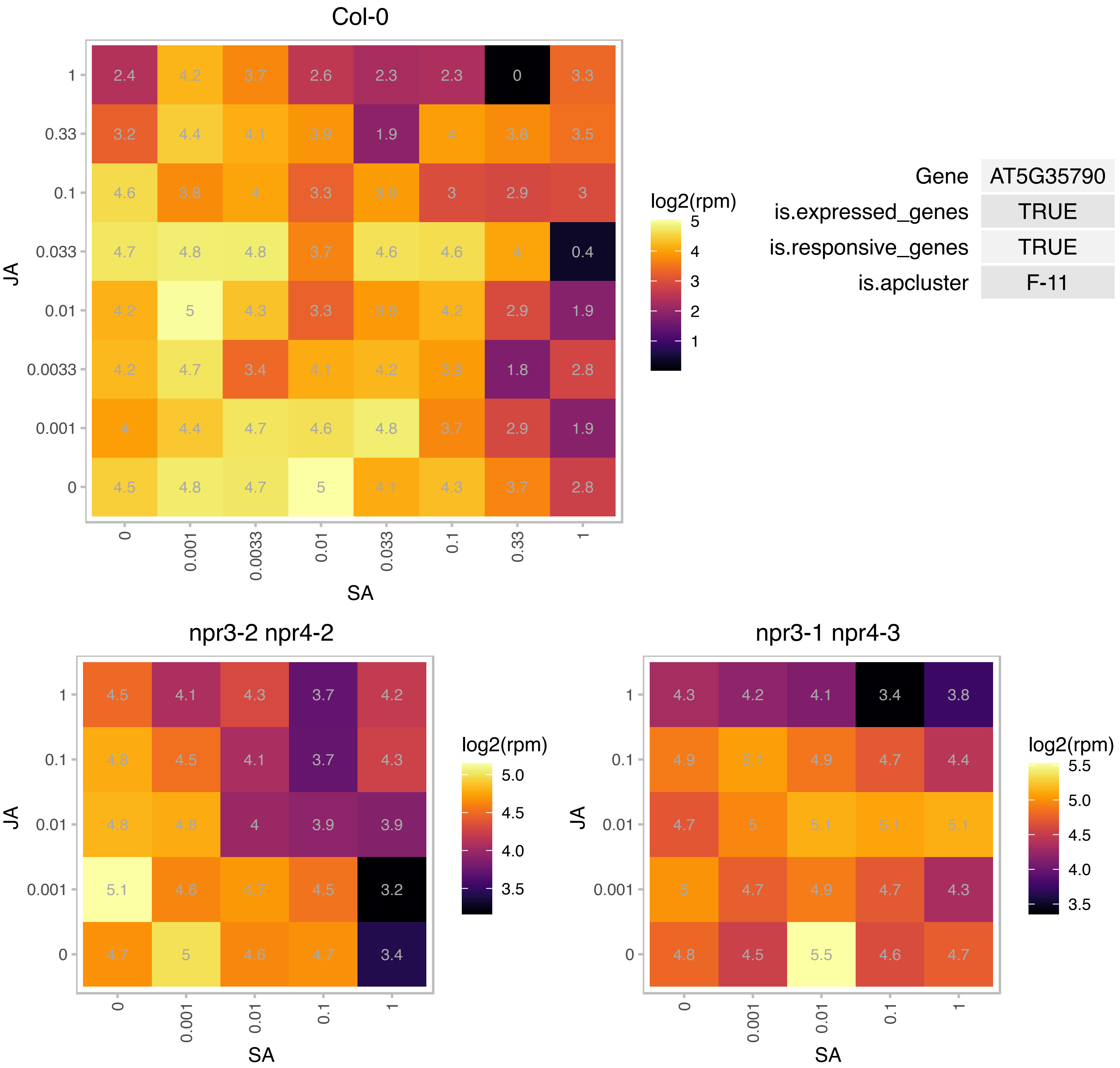


| | |
|---|---|
| Gene | AT5G35790 |
| is.expressed_genes | TRUE |
| is.responsive_genes | TRUE |
| is.apcluster | F-11 |

### AT5G39210.1 : CRR7

chlororespiratory reduction 7
Encodes a protein of the chloroplastic NAD(P)H dehydrogenase complex (NDH Complex) involved in respiration, photosystem I (PSI) cyclic electron transport and CO2 uptake. The product of this gene appears to be essential for the stable formation of the NDH Complex. The mRNA is cell-to-cell mobile.

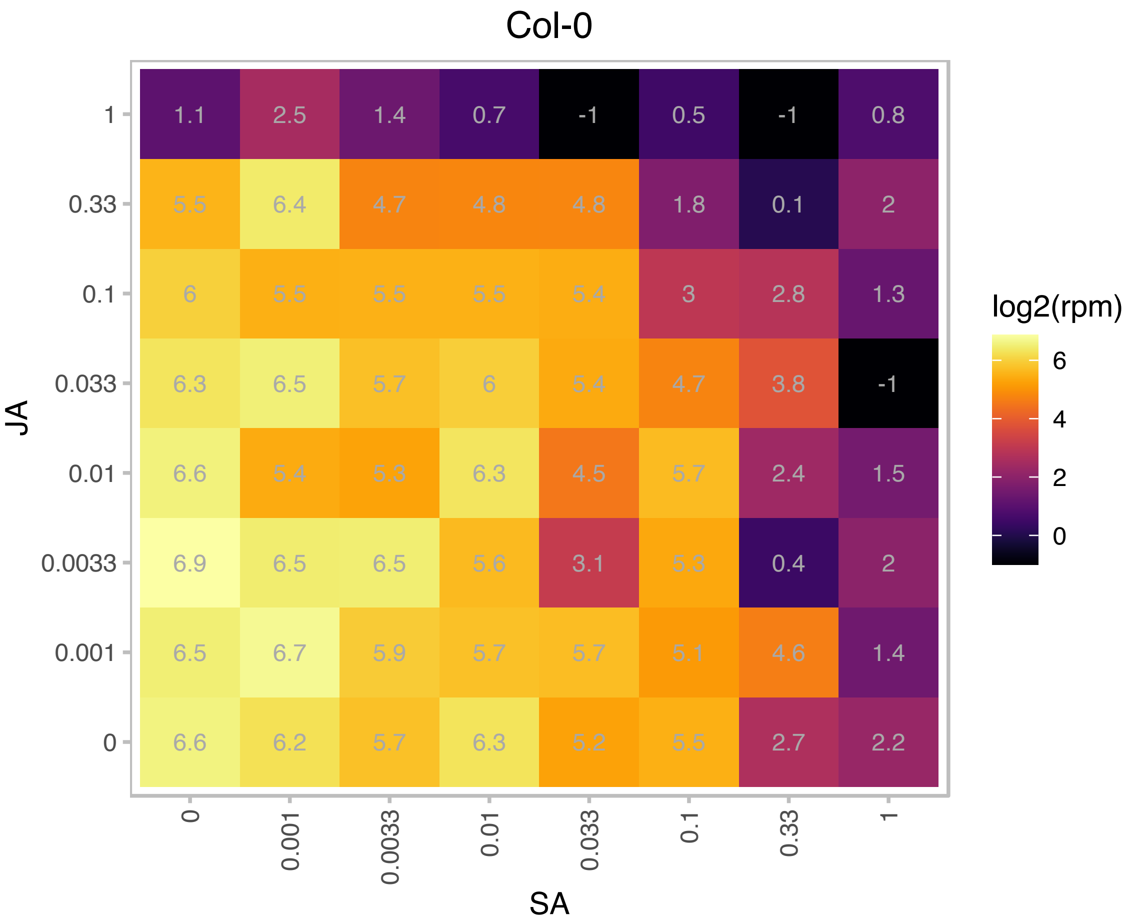


| | |
|---|---|
| Gene | AT5G39210 |
| is.expressed_genes | TRUE |
| is.responsive_genes | TRUE |
| is.apcluster | F-11 |

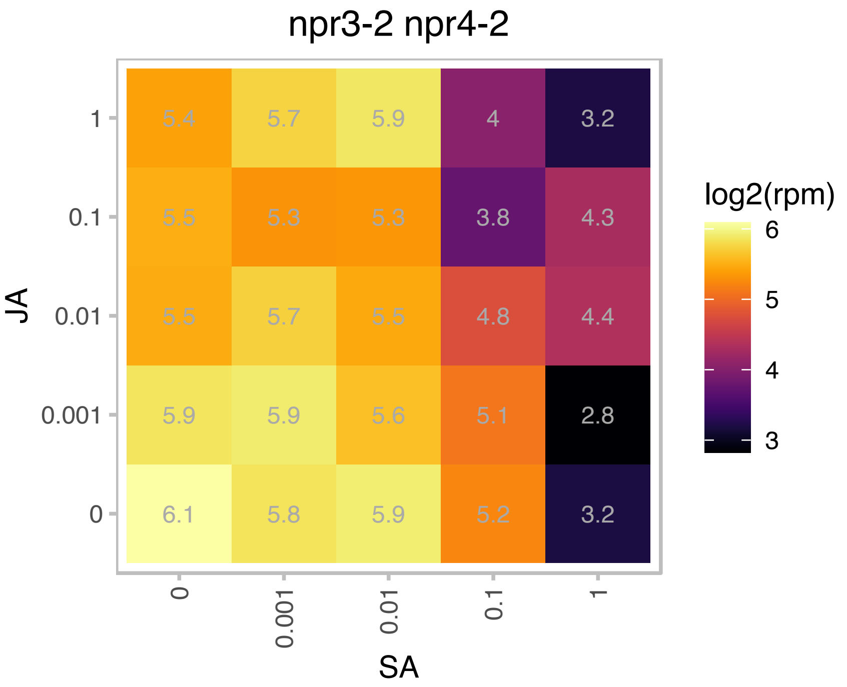


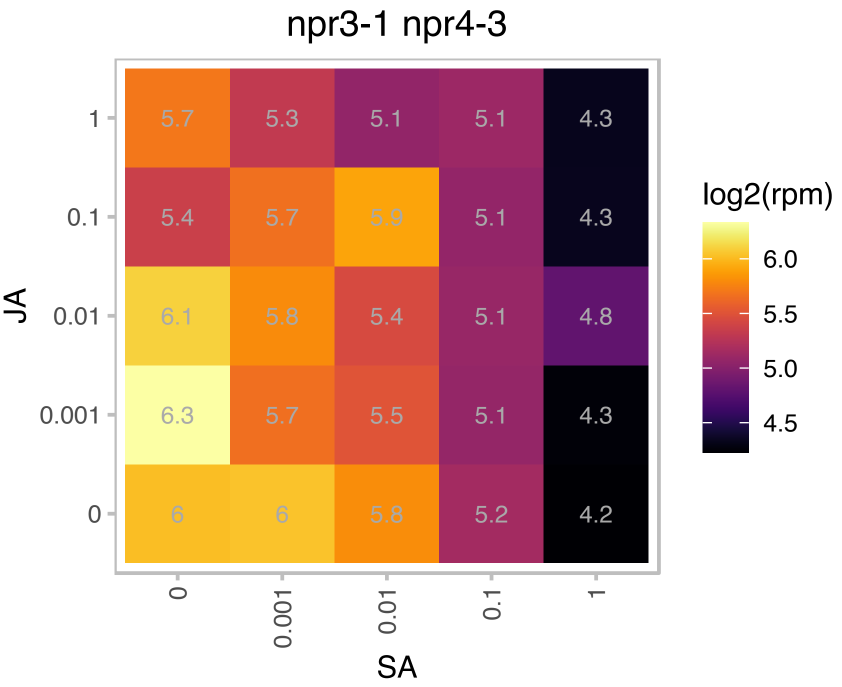

### AT5G40610.1 : GPDHp

NAD-dependent glycerol-3-phosphate dehydrogenase family protein

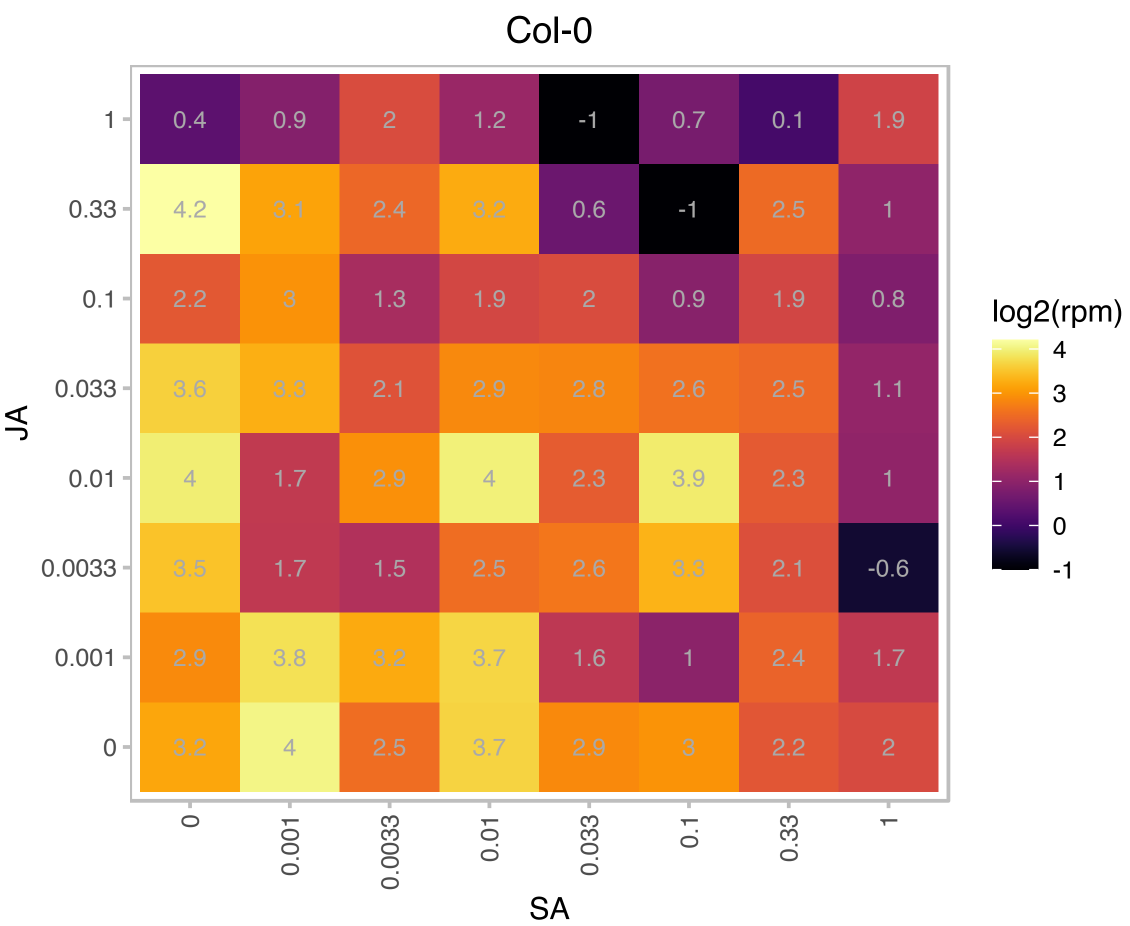


| | |
|---|---|
| Gene | AT5G40610 |
| is.expressed_genes | TRUE |
| is.responsive_genes | TRUE |
| is.apcluster | F-11 |

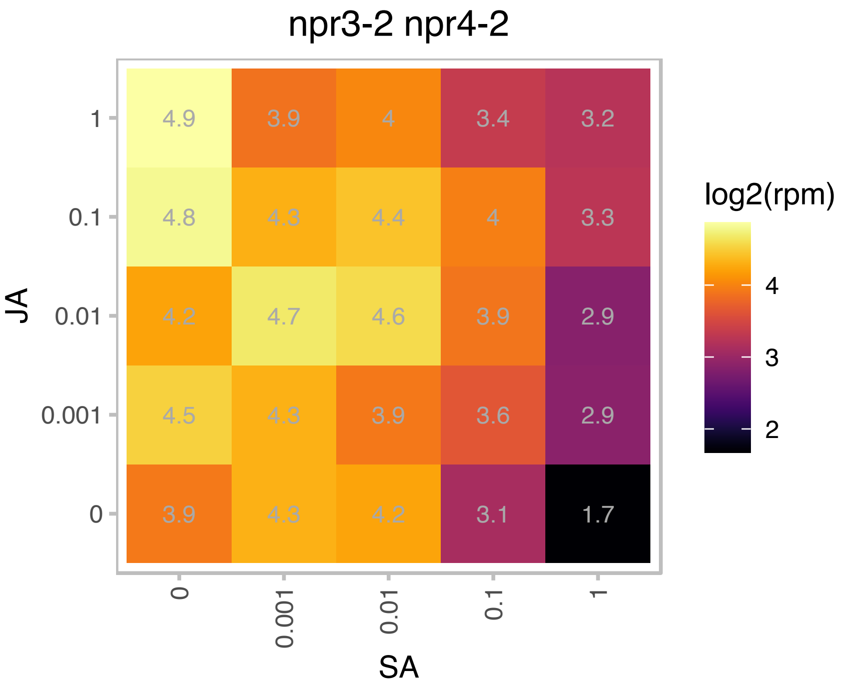


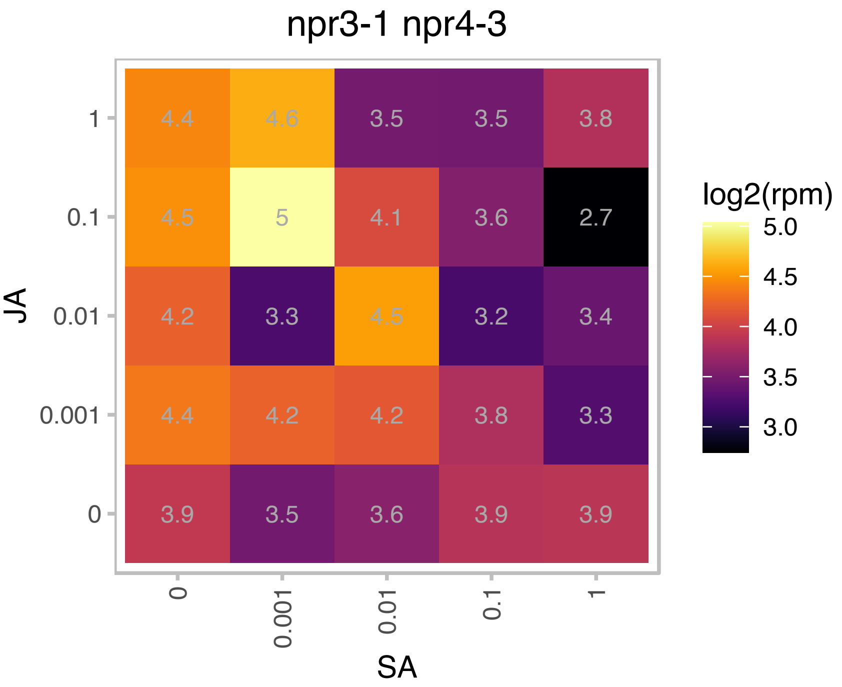

### AT5G42030.1 : ABIL4

ABL interactor-like protein 4

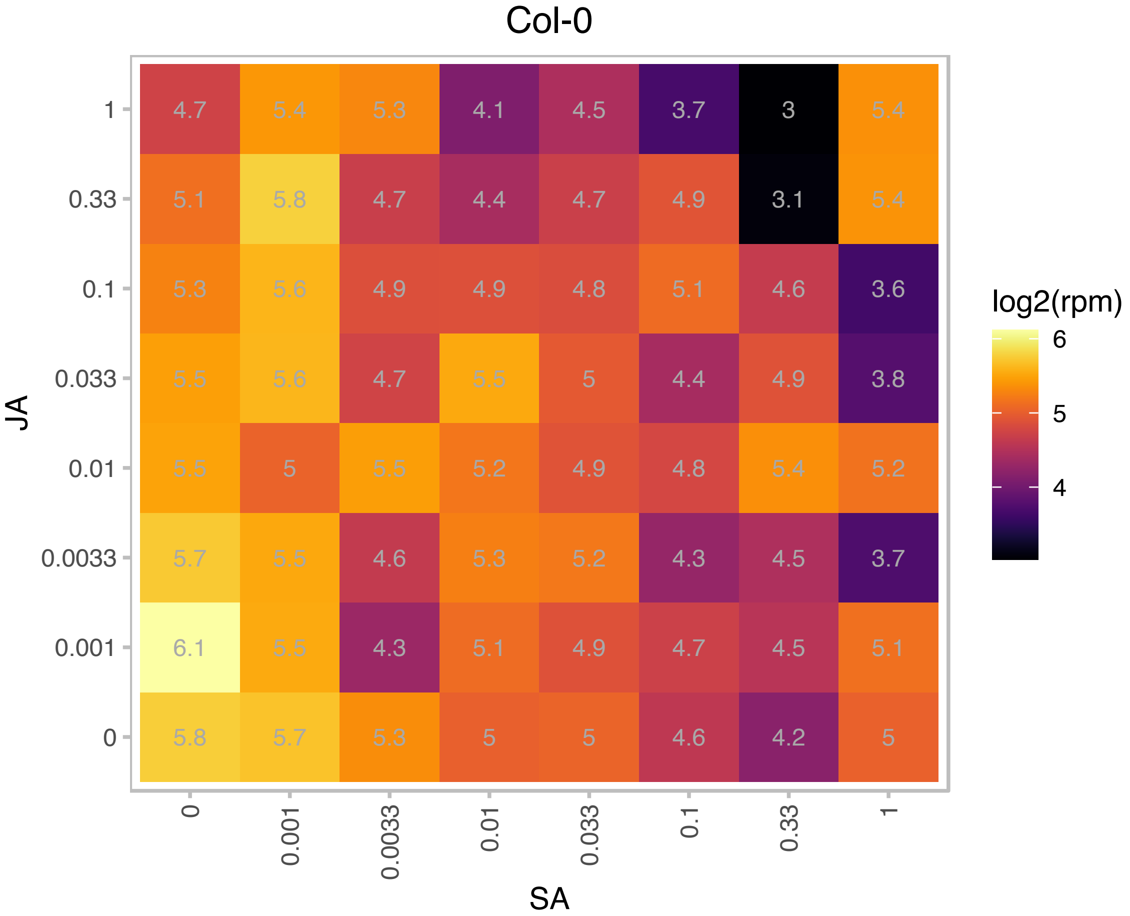


| | |
|---|---|
| Gene | AT5G42030 |
| is.expressed_genes | TRUE |
| is.responsive_genes | TRUE |
| is.apcluster | F-11 |

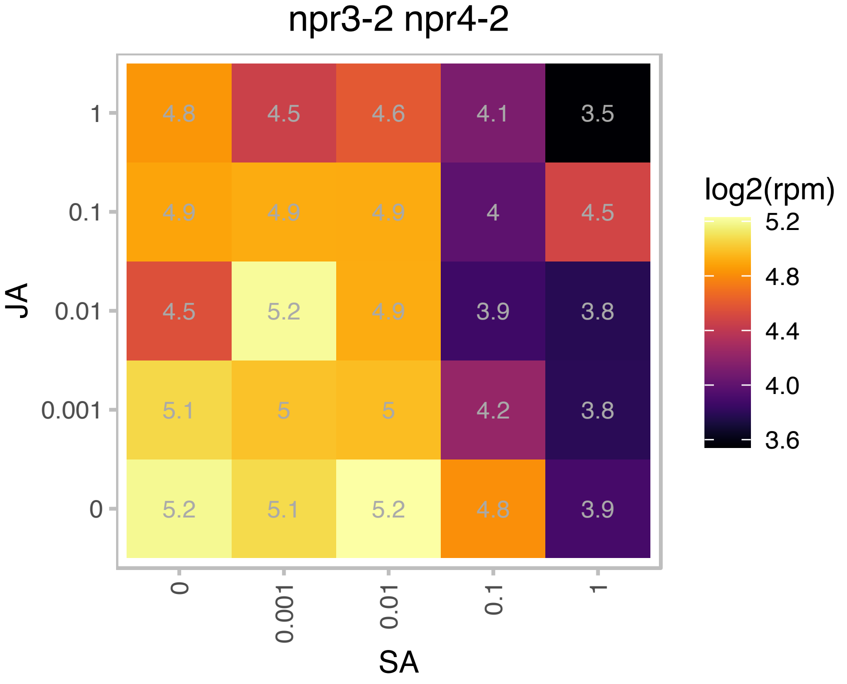


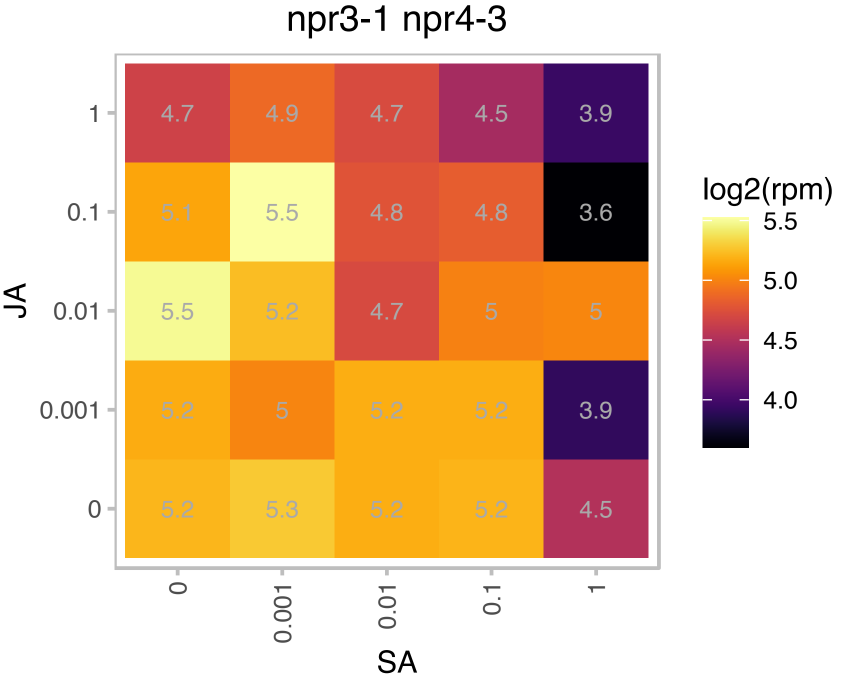

# AT5G42330.1 : NA

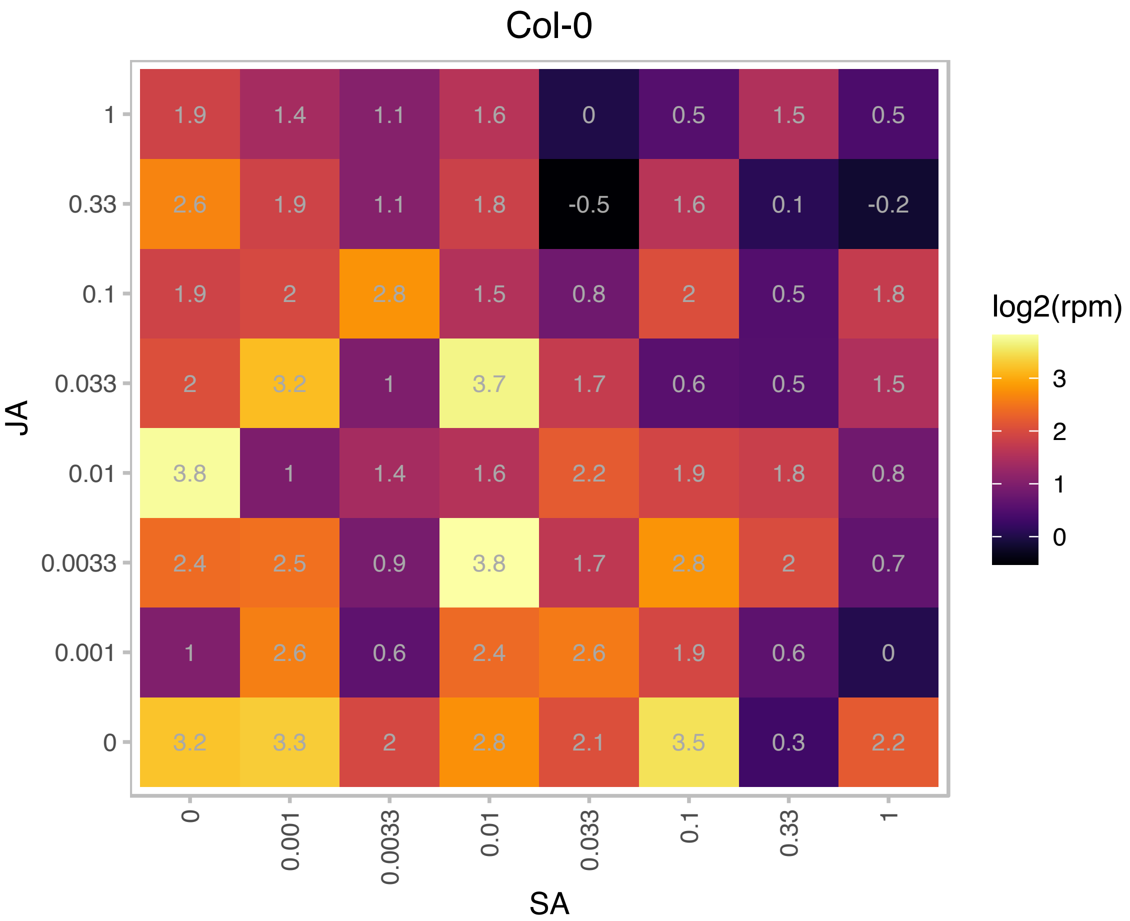


| | |
|---|---|
| Gene | AT5G42330 |
| is.expressed_genes | TRUE |
| is.responsive_genes | TRUE |
| is.apcluster | F-11 |

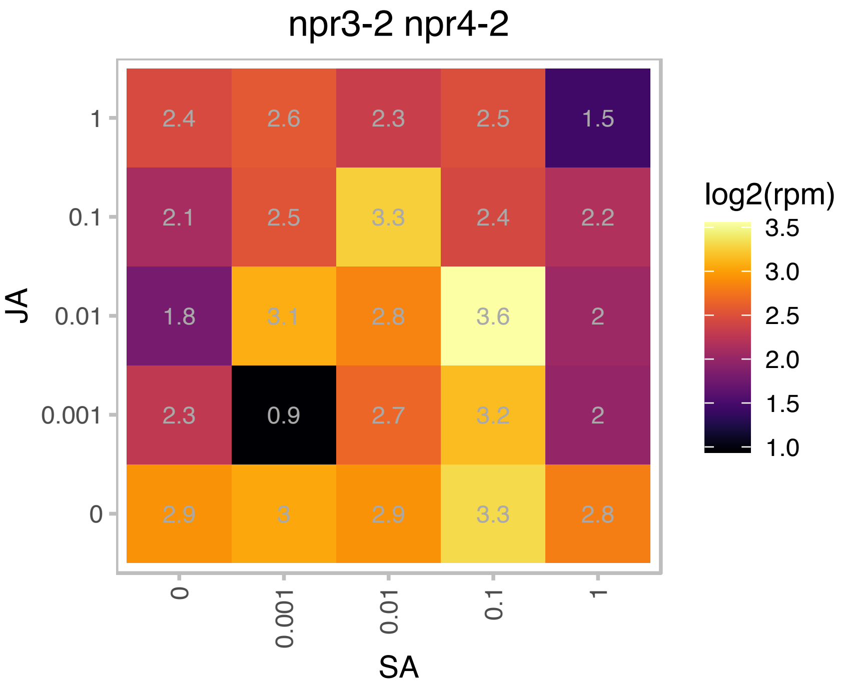


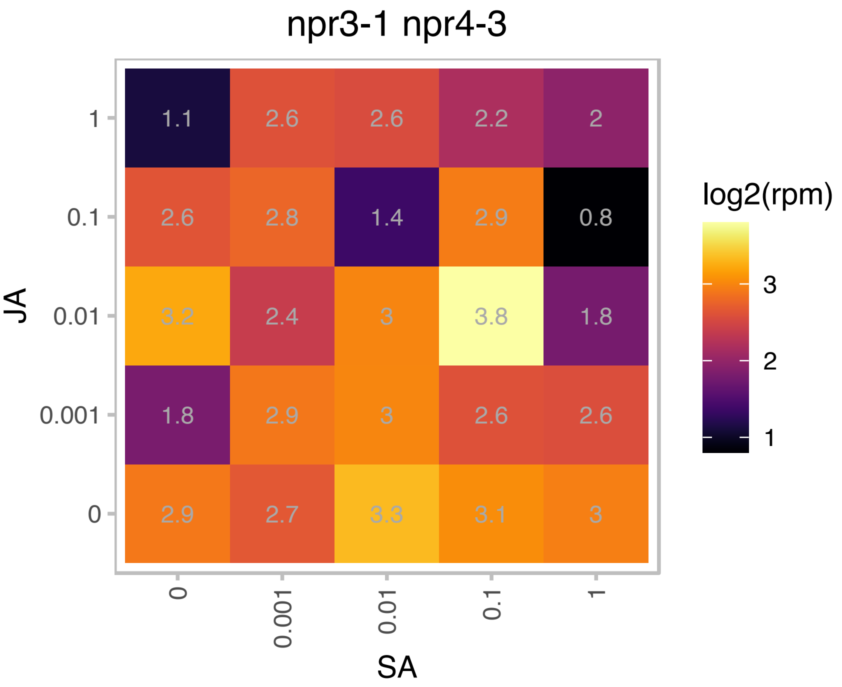

# AT5G42770.1 : NA

Maf-like protein

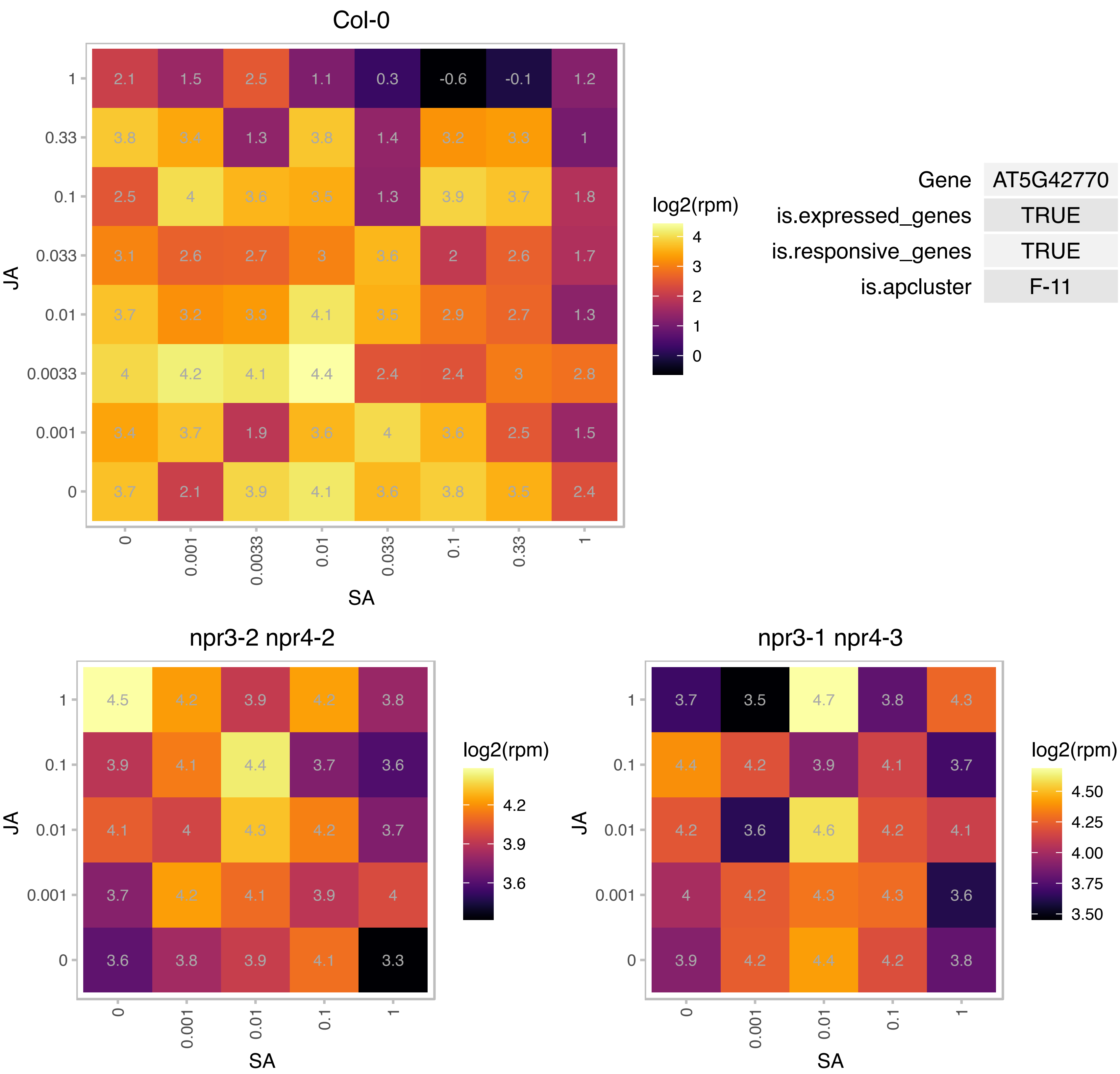


| | |
|---|---|
| Gene | AT5G42770 |
| is.expressed_genes | TRUE |
| is.responsive_genes | TRUE |
| is.apcluster | F-11 |

### AT5G43700.1 : ATAUX2-11, IAA4

AUX/IAA transcriptional regulator family protein

Auxin inducible protein similar to transcription factors.

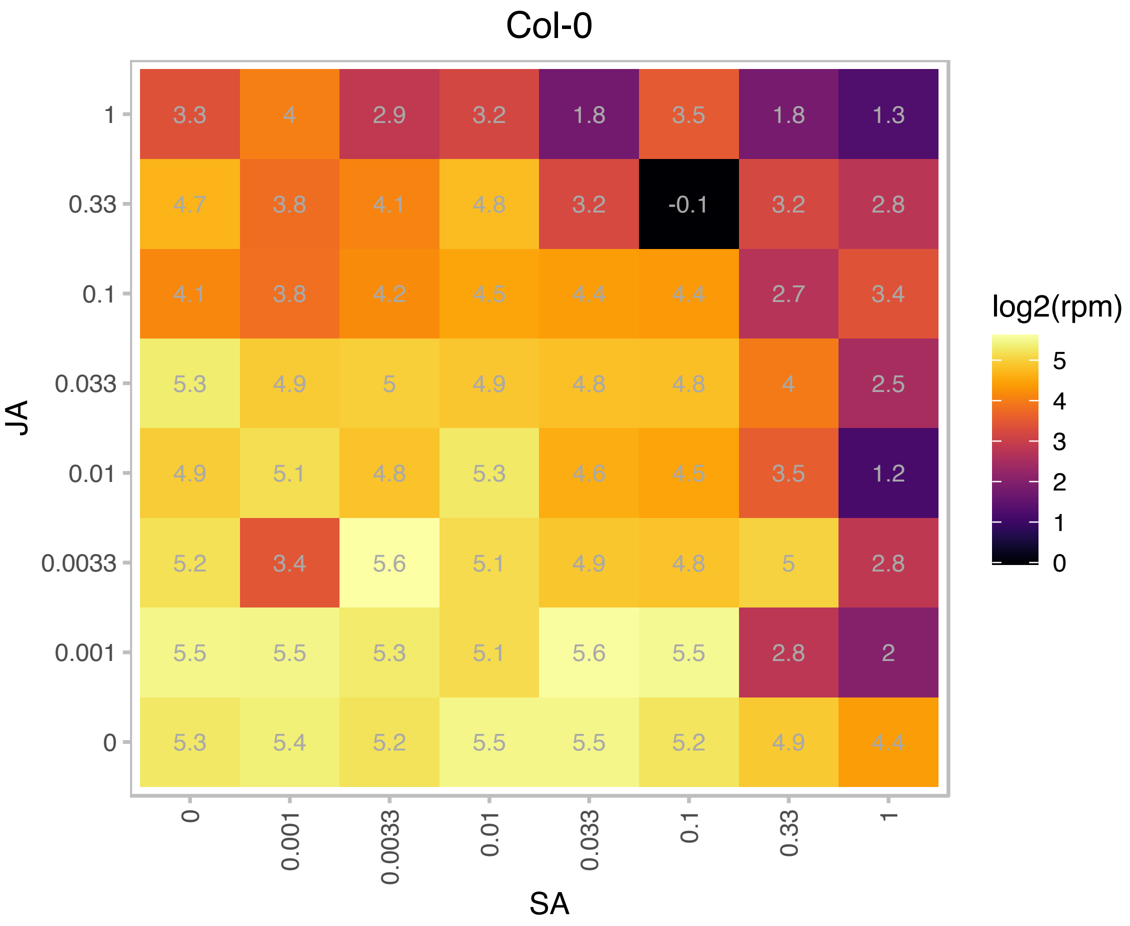


| | |
|---|---|
| Gene | AT5G43700 |
| is.expressed_genes | TRUE |
| is.responsive_genes | TRUE |
| is.apcluster | F-11 |

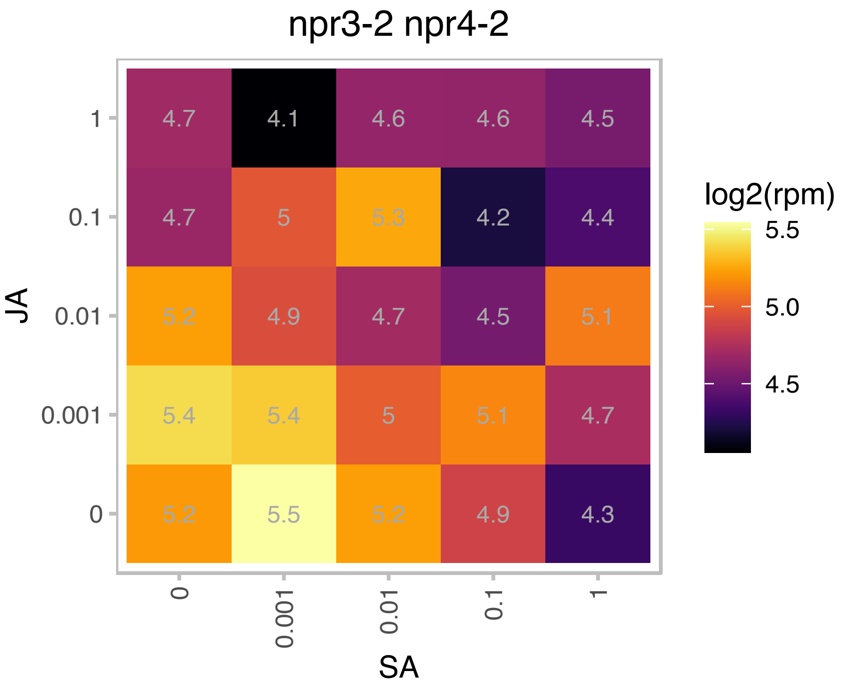


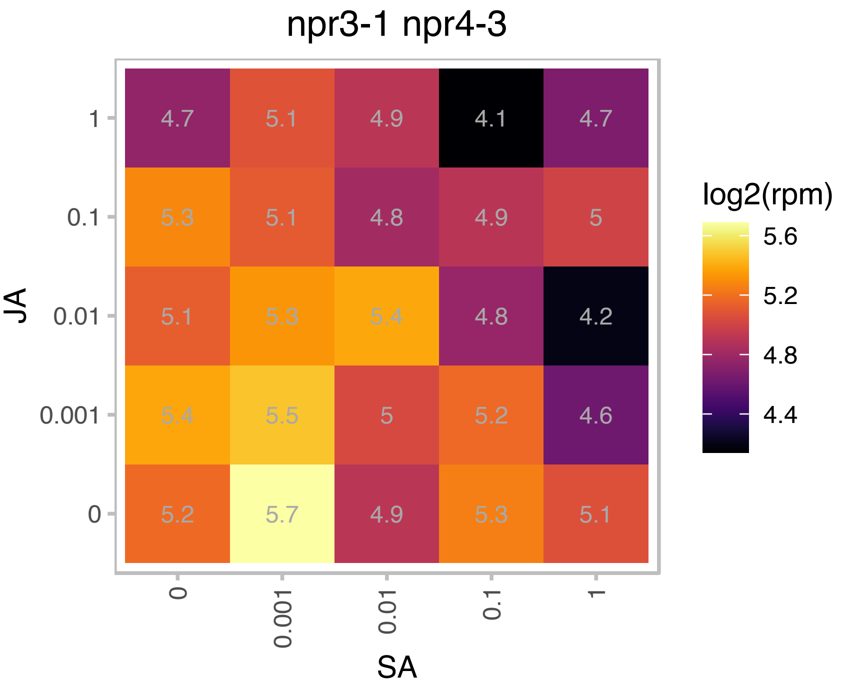

### AT5G44130.1 : FLA13

FASCICLIN-like arabinogalactan protein 13 precursor

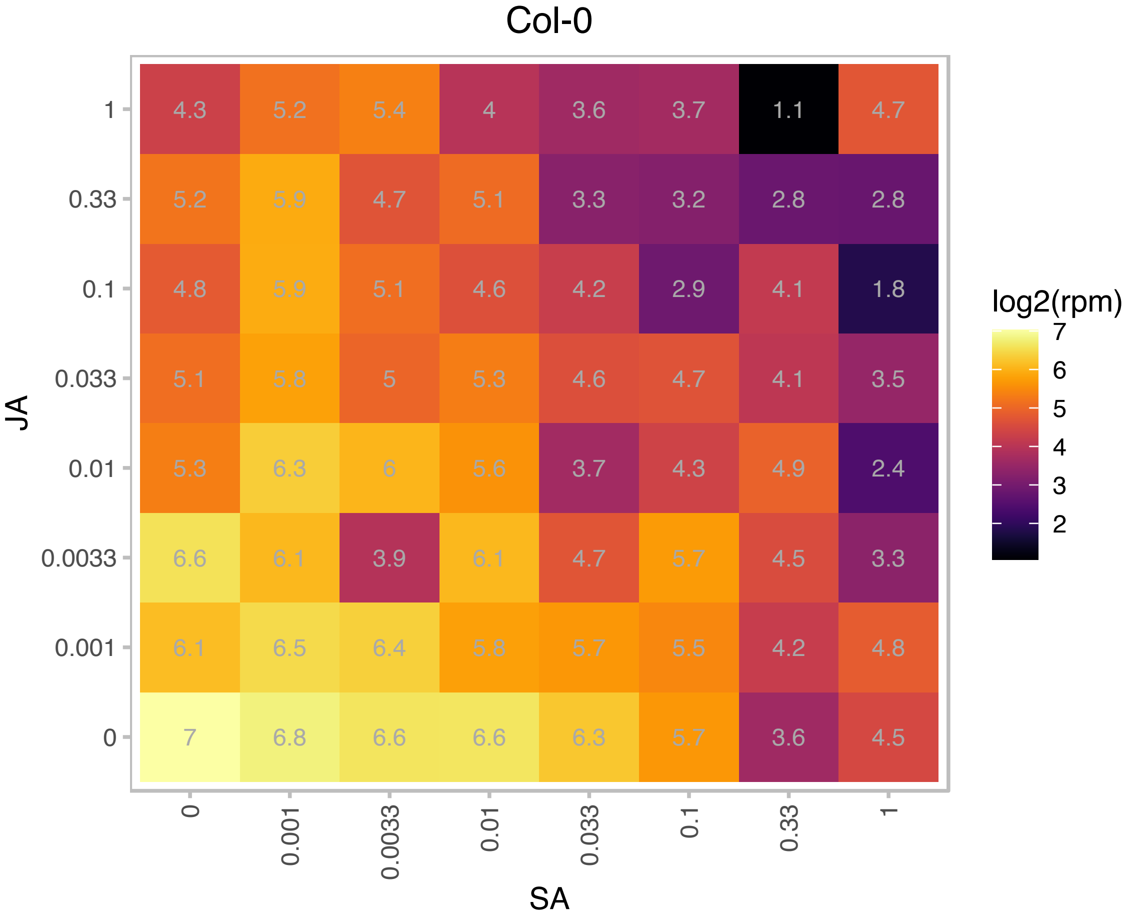


| | |
|---|---|
| Gene | AT5G44130 |
| is.expressed_genes | TRUE |
| is.responsive_genes | TRUE |
| is.apcluster | F-11 |

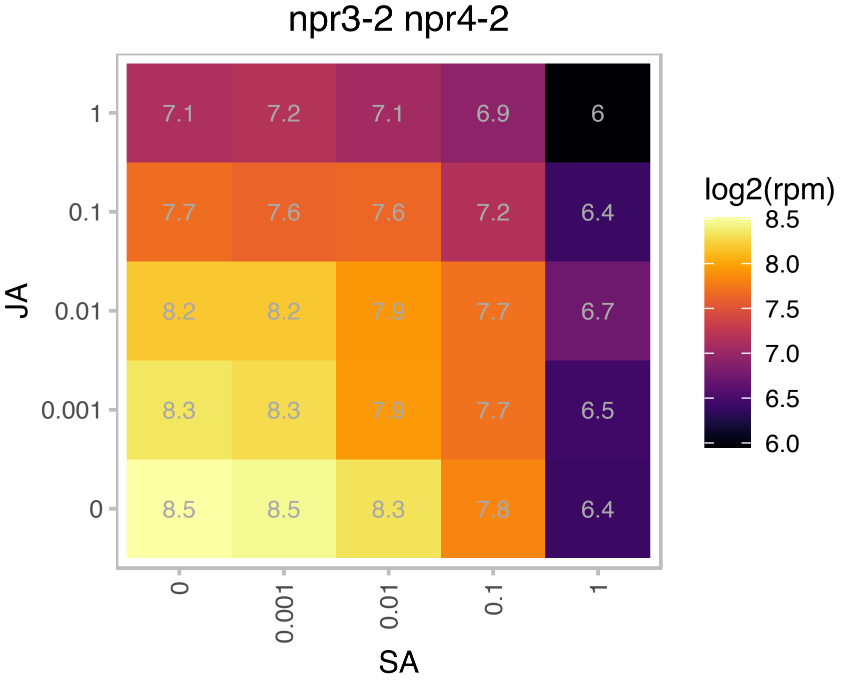


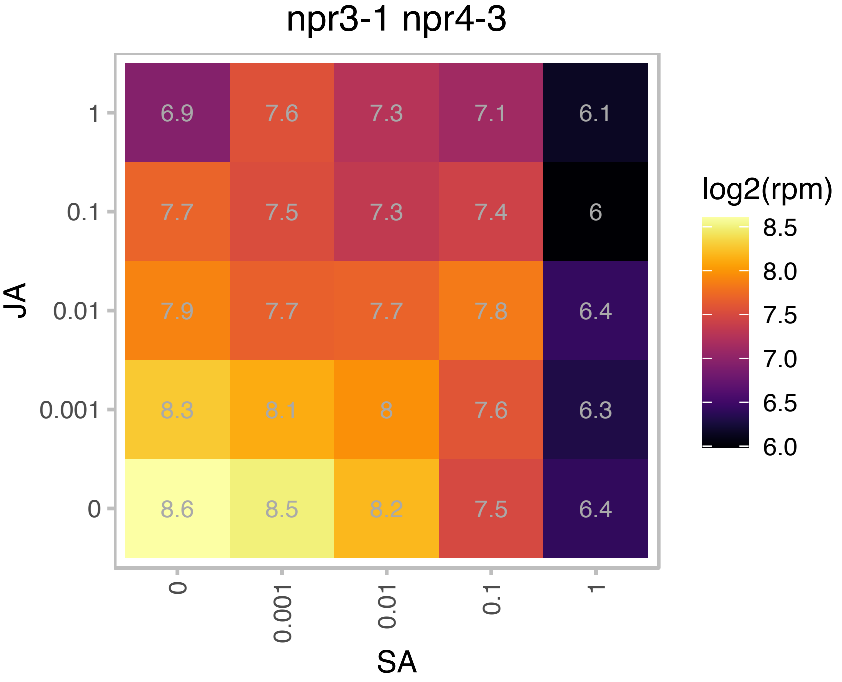

### AT5G48545.1 : HINT3

histidine triad nucleotide-binding 3

Encodes a protein that has adenylylsulfate sulfohydrolase activity (E.C. 3.6.2.1) in vitro.

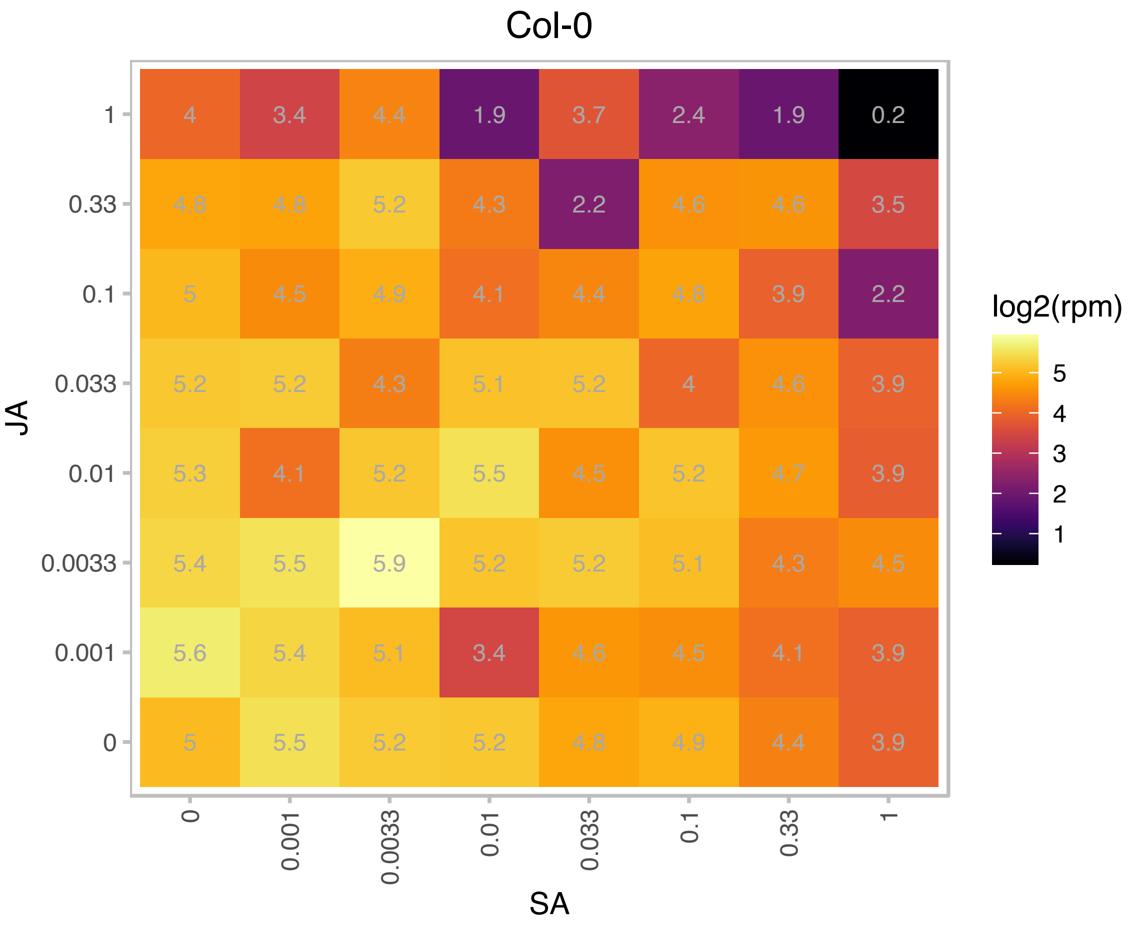


| | |
|---|---|
| Gene | AT5G48545 |
| is.expressed_genes | TRUE |
| is.responsive_genes | TRUE |
| is.apcluster | F-11 |

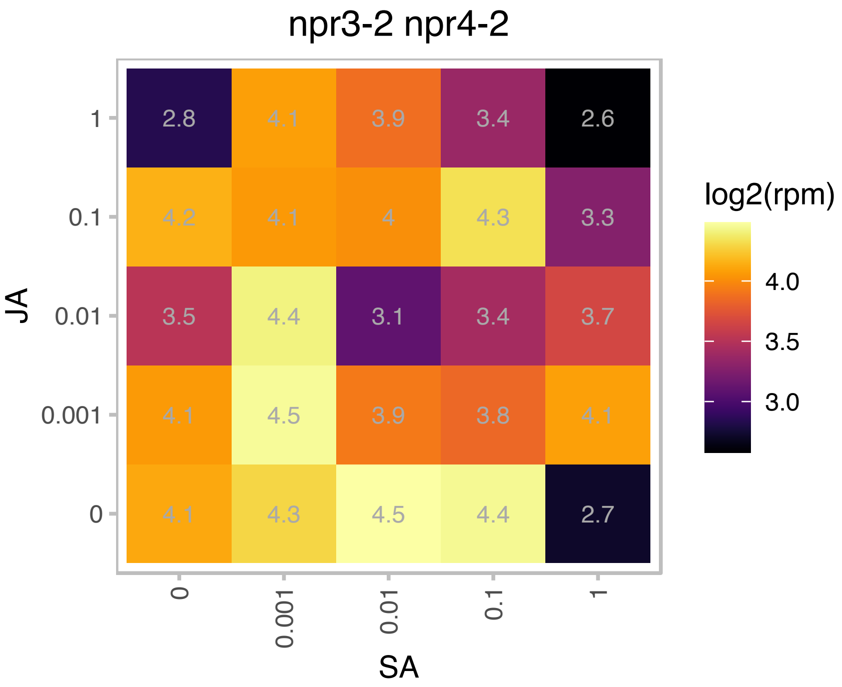


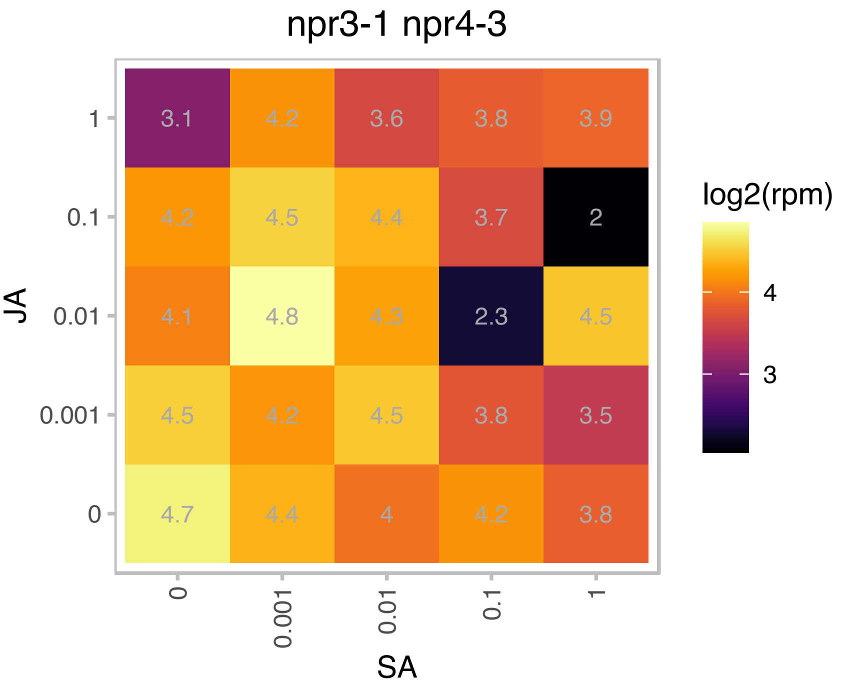

# AT5G52190.1 : NA

Sugar isomerase (SIS) family protein

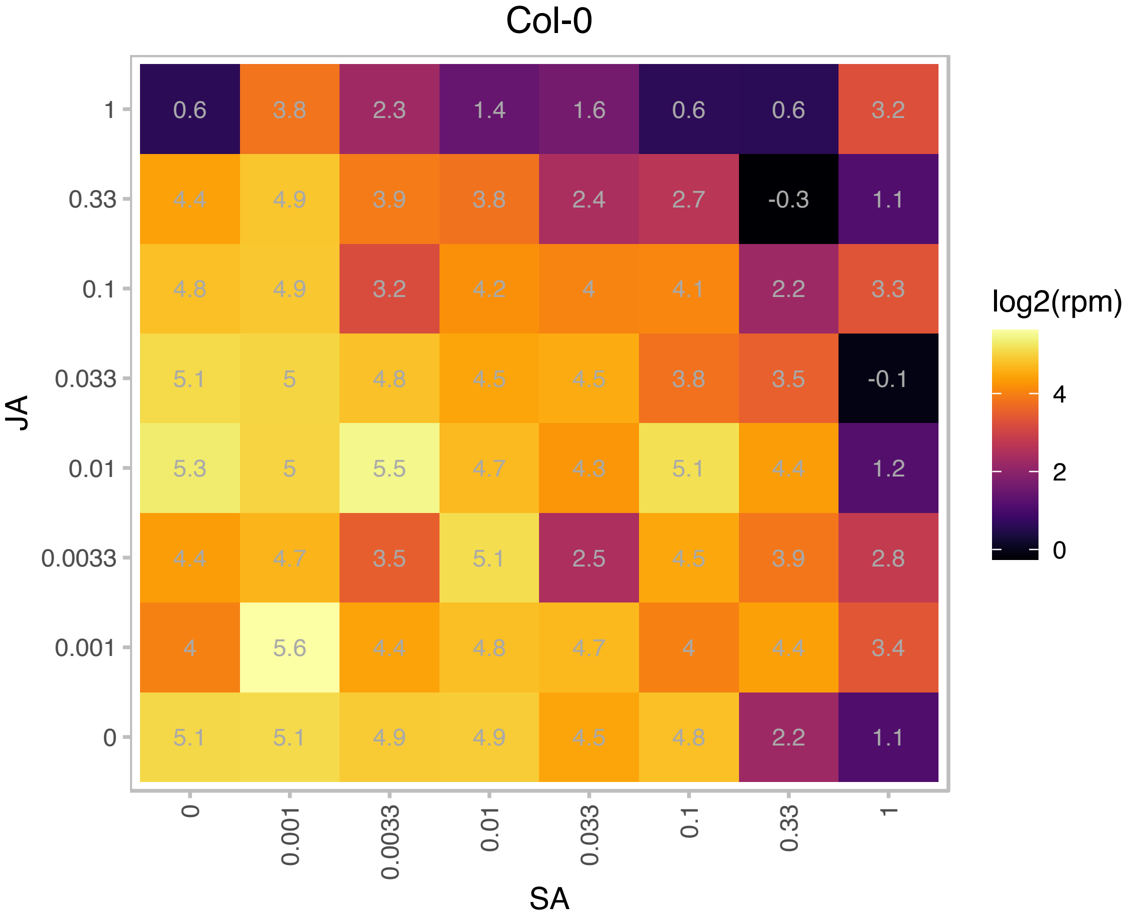


| | |
|---|---|
| Gene | AT5G52190 |
| is.expressed_genes | TRUE |
| is.responsive_genes | TRUE |
| is.apcluster | F-11 |

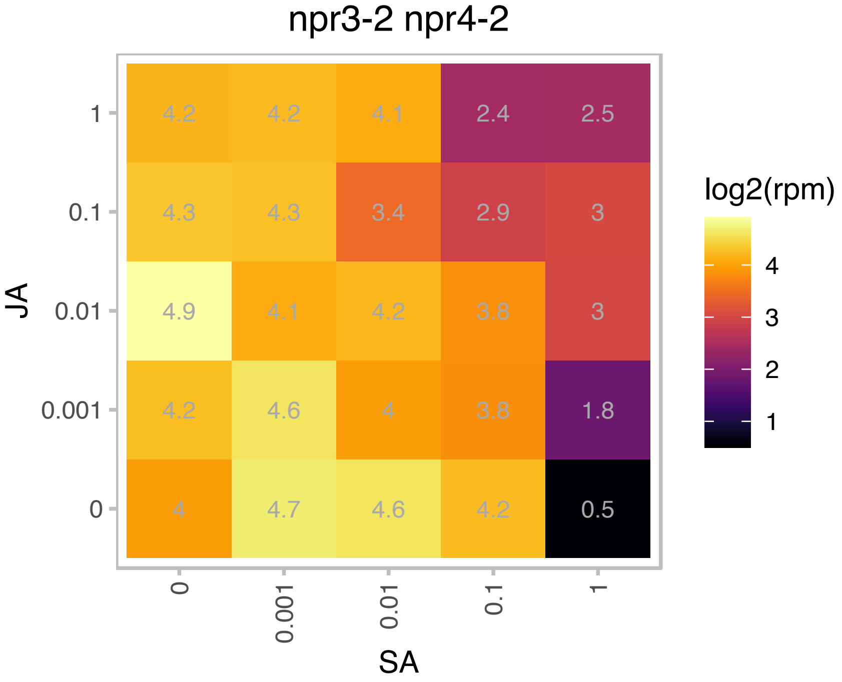


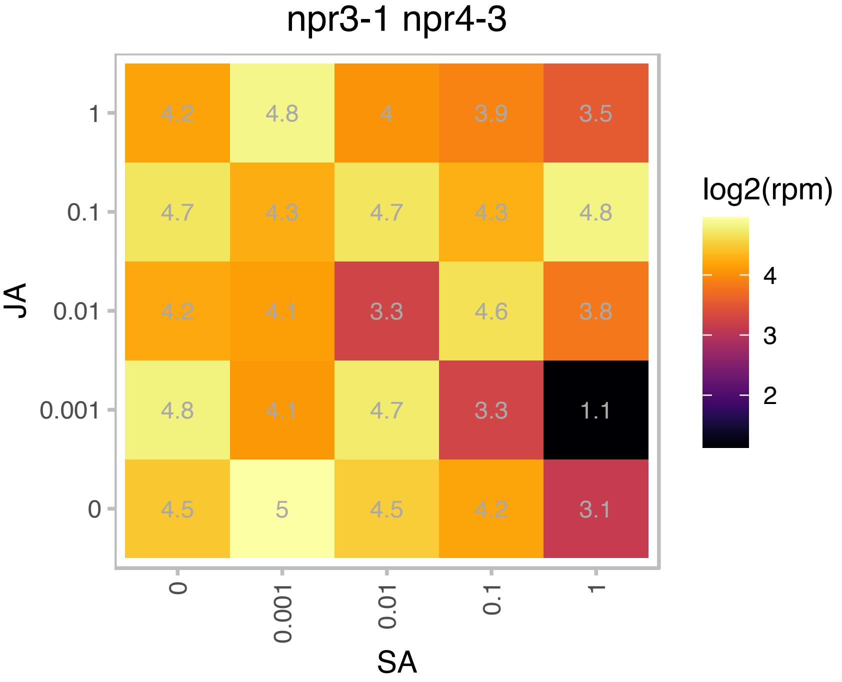

# AT5G53880.1 : NA

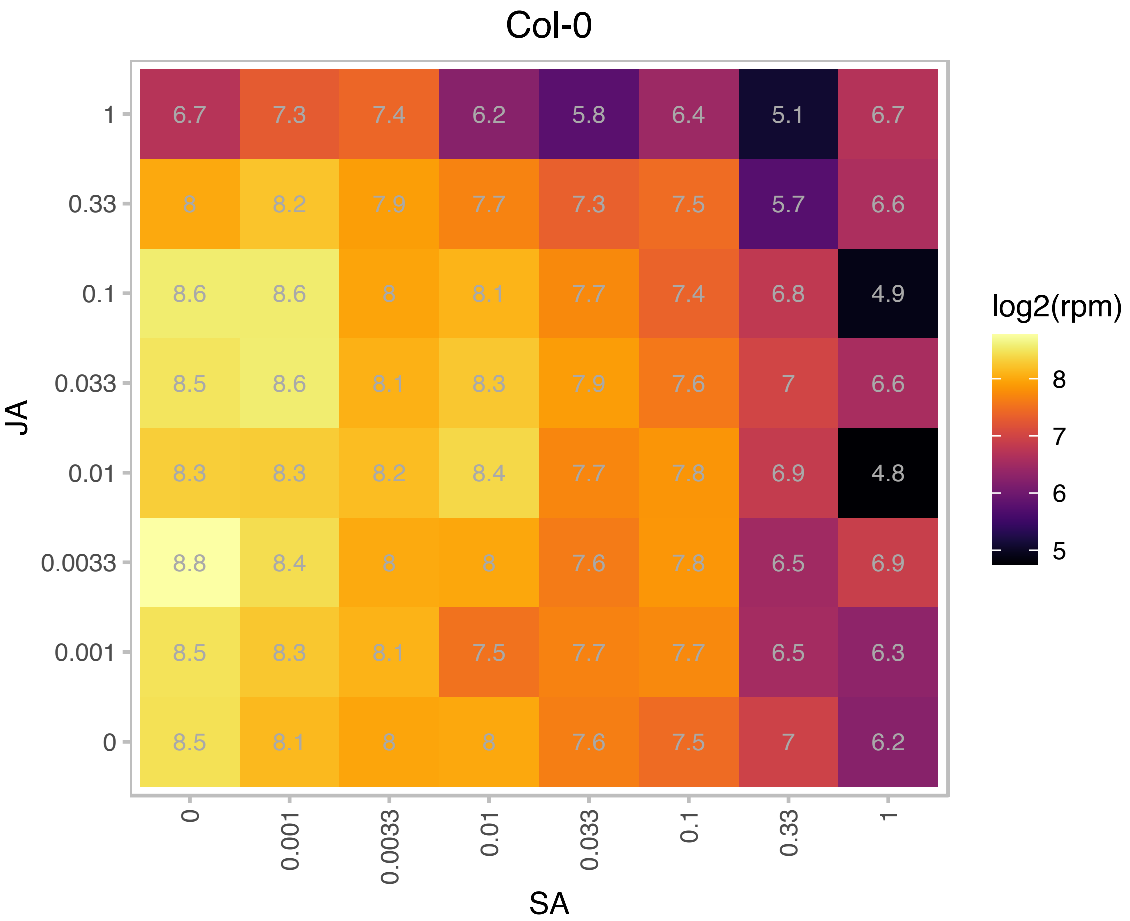


| | |
|---|---|
| Gene | AT5G53880 |
| is.expressed_genes | TRUE |
| is.responsive_genes | TRUE |
| is.apcluster | F-11 |

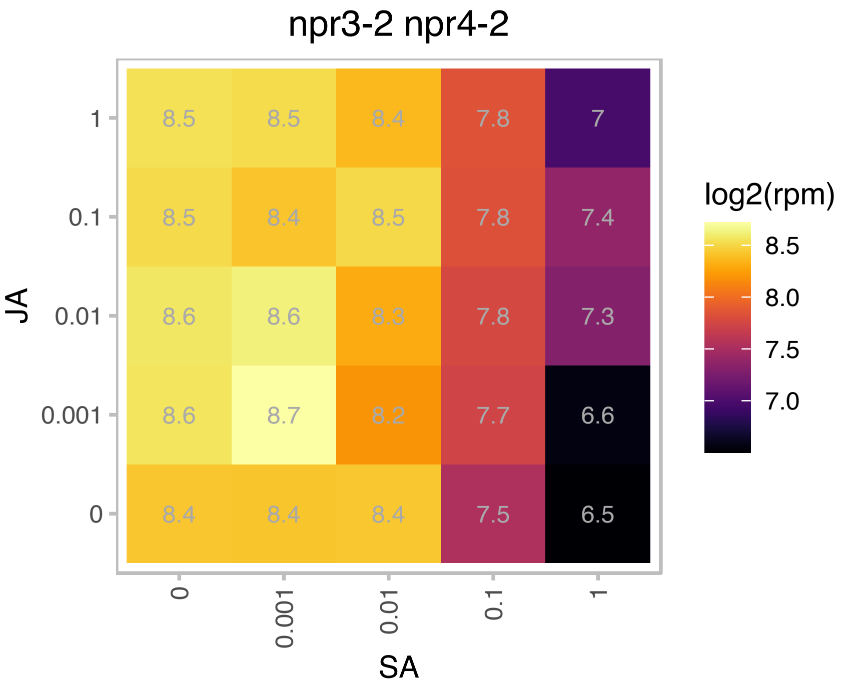


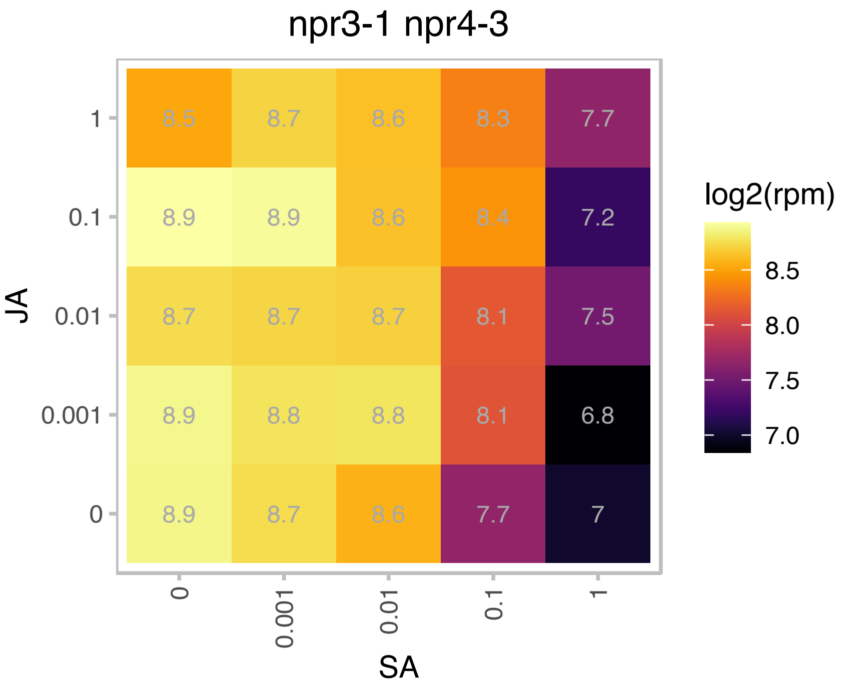

### AT5G55710.1 : AtTic20-V, Tic20-V

Encodes a component of the TIC (translocon at the inner envelope membrane of chloroplasts) protein translocation machinery mediating the protein translocation across the inner envelope of plastids. The Arabidopsis genome encodes four Tic20 homologous proteins, AT1G04940(Tic20-I), AT2G47840(Tic20-II), AT4G03320(Tic20-IV) and AT5G55710(Tic20-V).

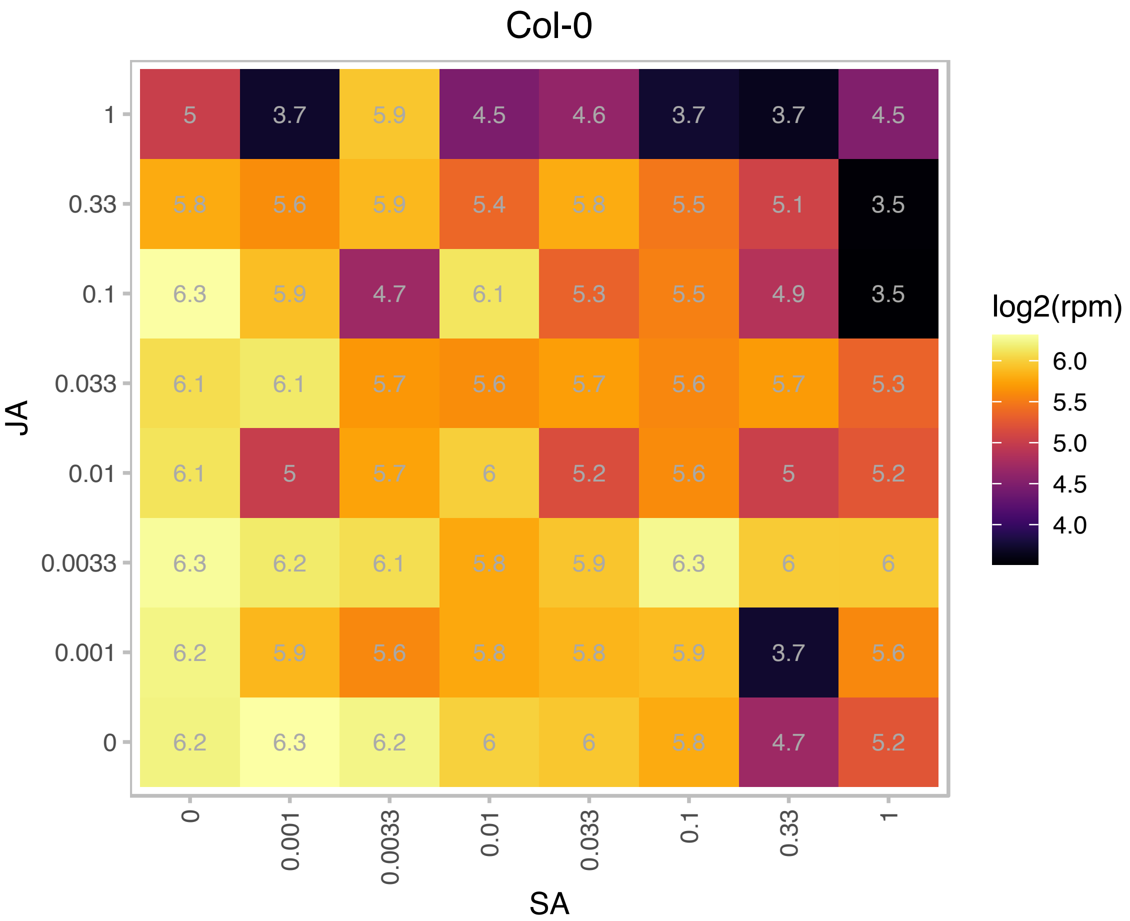


| | |
|---|---|
| Gene | AT5G55710 |
| is.expressed_genes | TRUE |
| is.responsive_genes | TRUE |
| is.apcluster | F-11 |

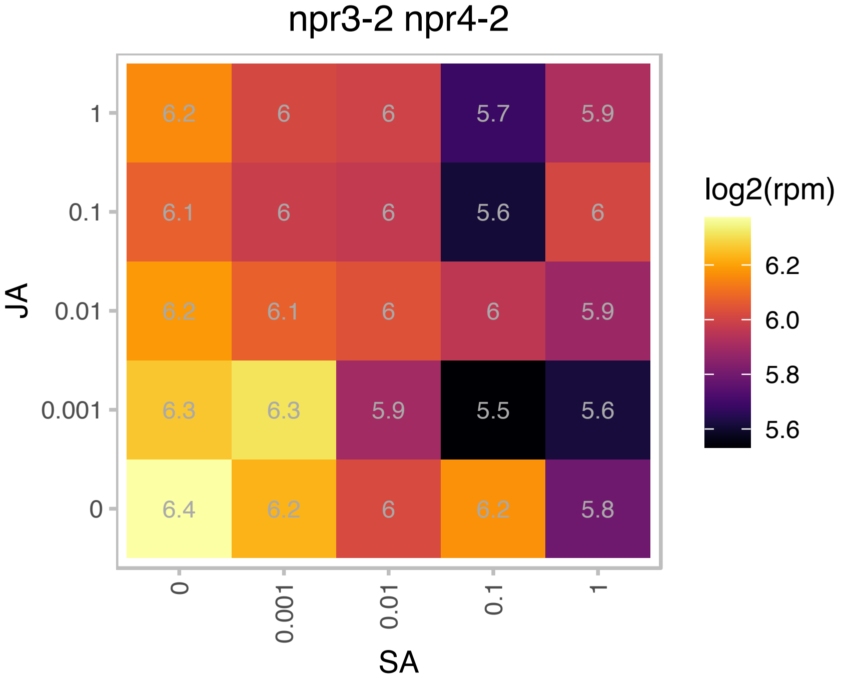


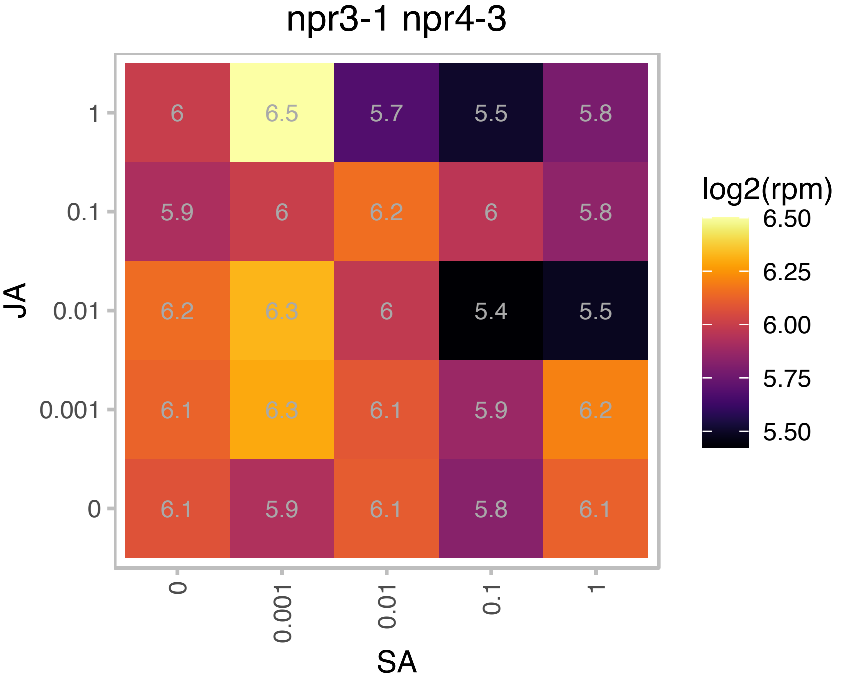

# AT5G55790.1 : NA

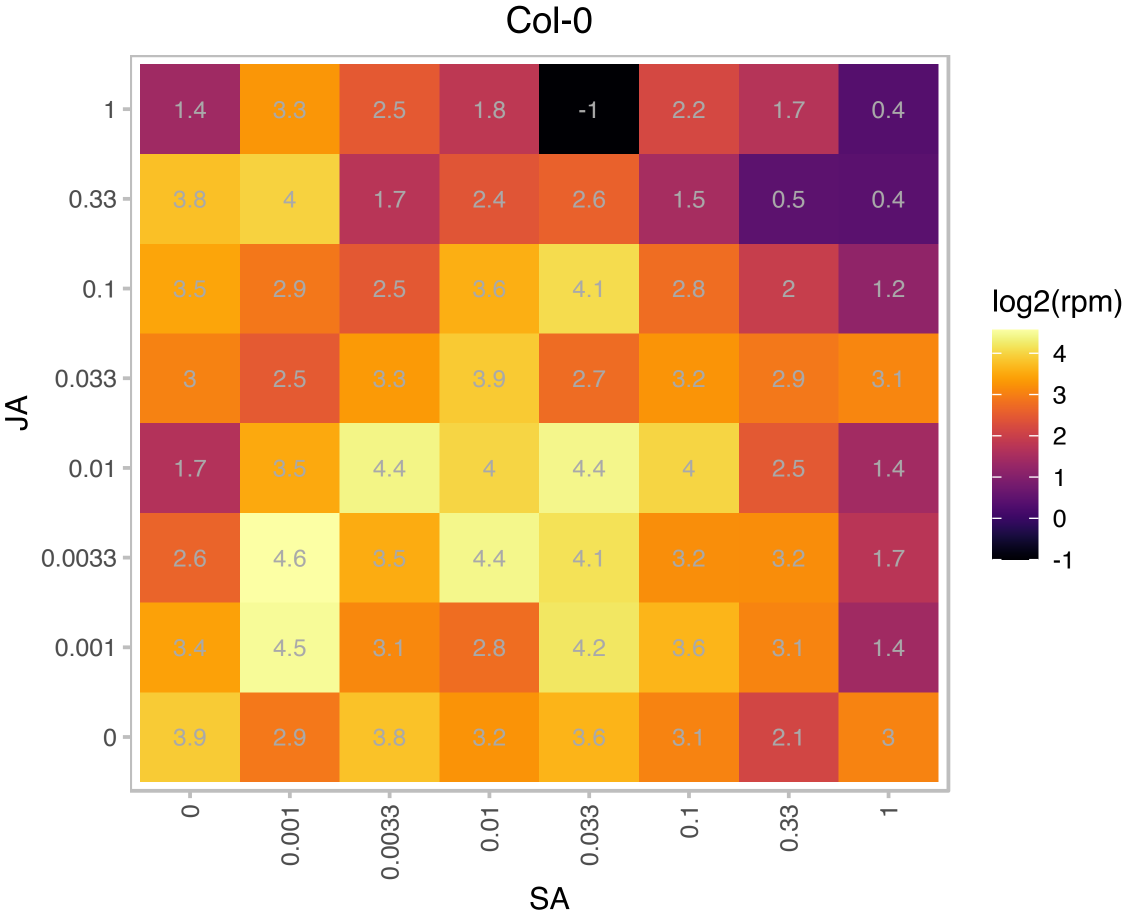


| | |
|---|---|
| Gene | AT5G55790 |
| is.expressed_genes | TRUE |
| is.responsive_genes | TRUE |
| is.apcluster | F-11 |

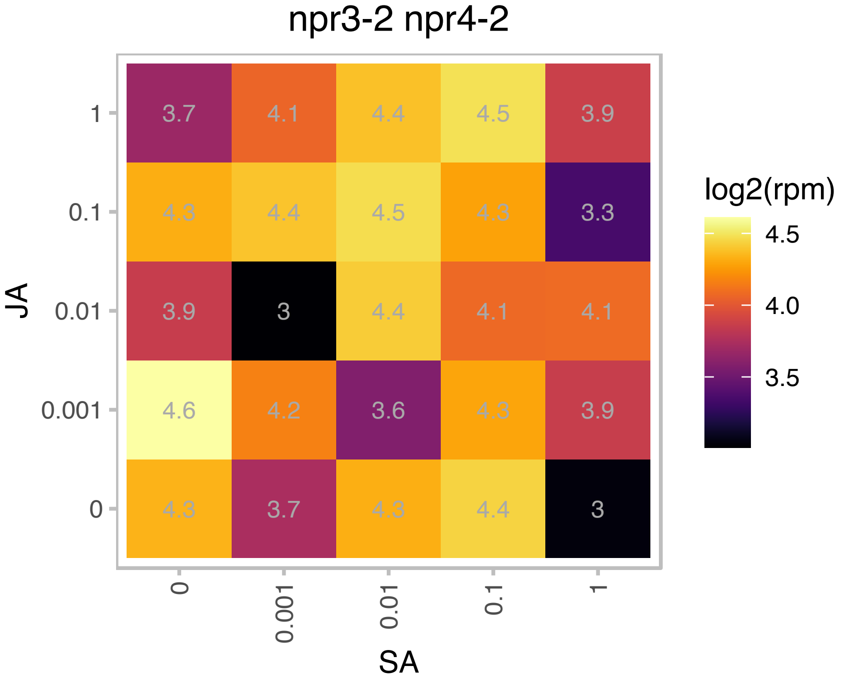


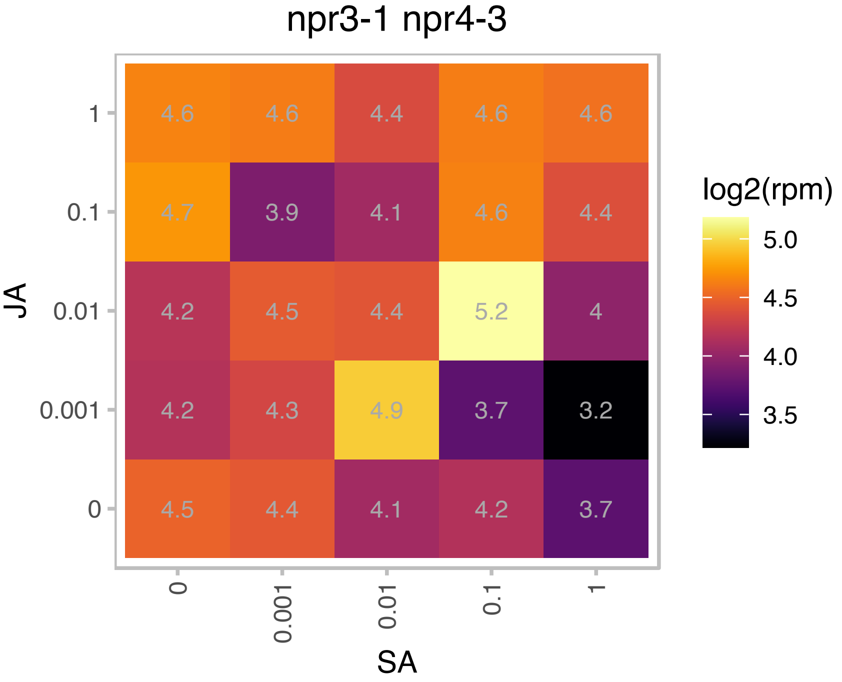

# AT5G62140.1 : NA

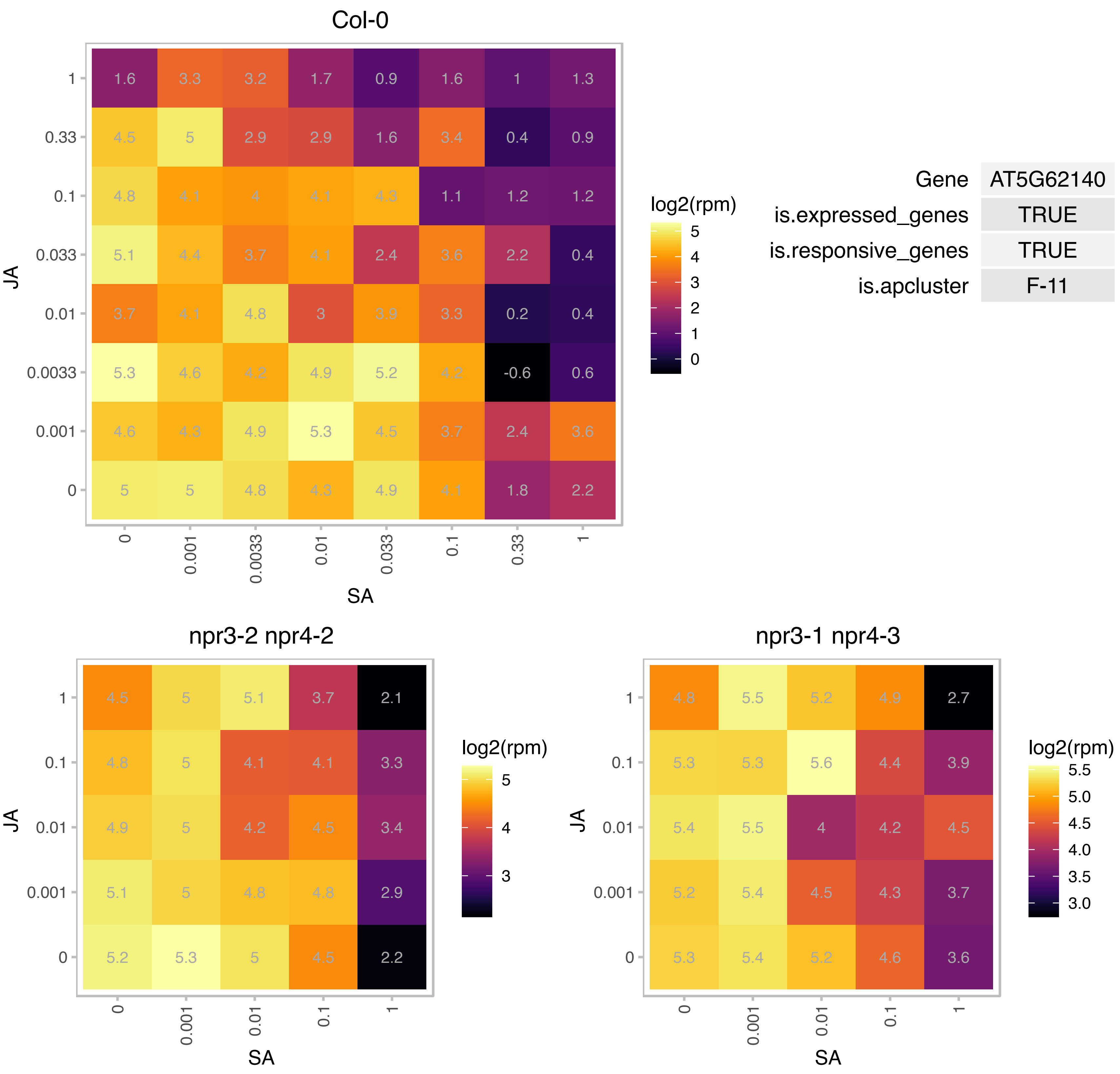

# AT5G63500.1 : NA

Protein of unknown function (DUF 3339)

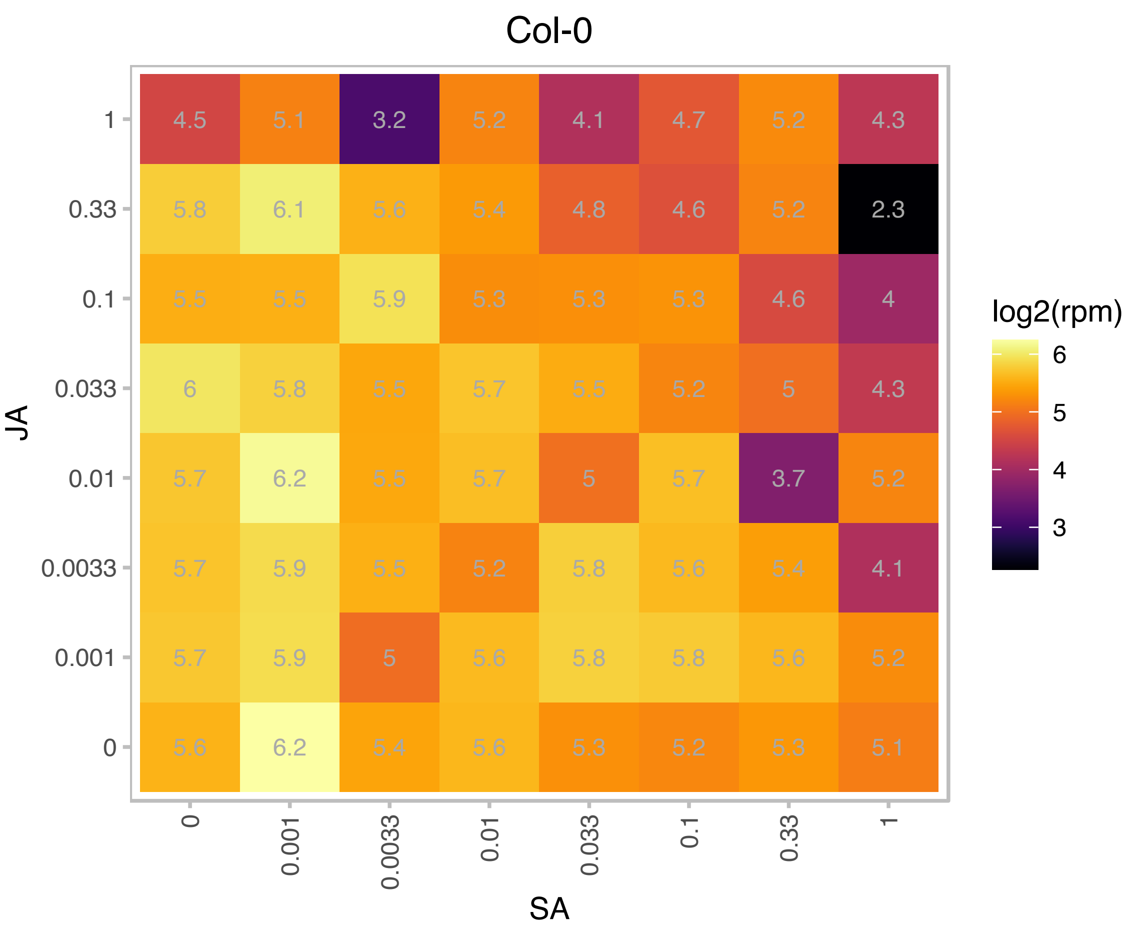


| | |
|---|---|
| Gene | AT5G63500 |
| is.expressed_genes | TRUE |
| is.responsive_genes | TRUE |
| is.apcluster | F-11 |

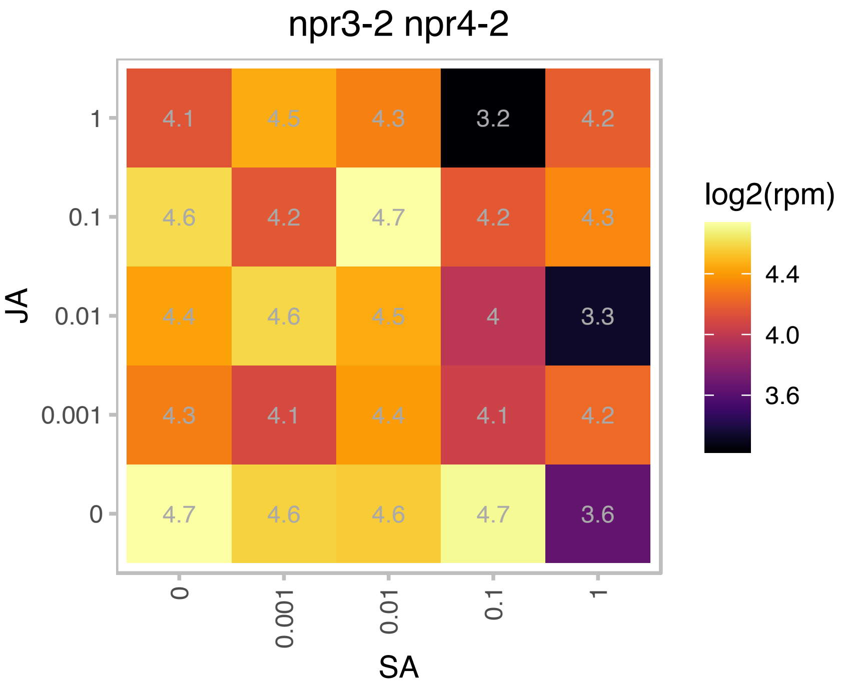


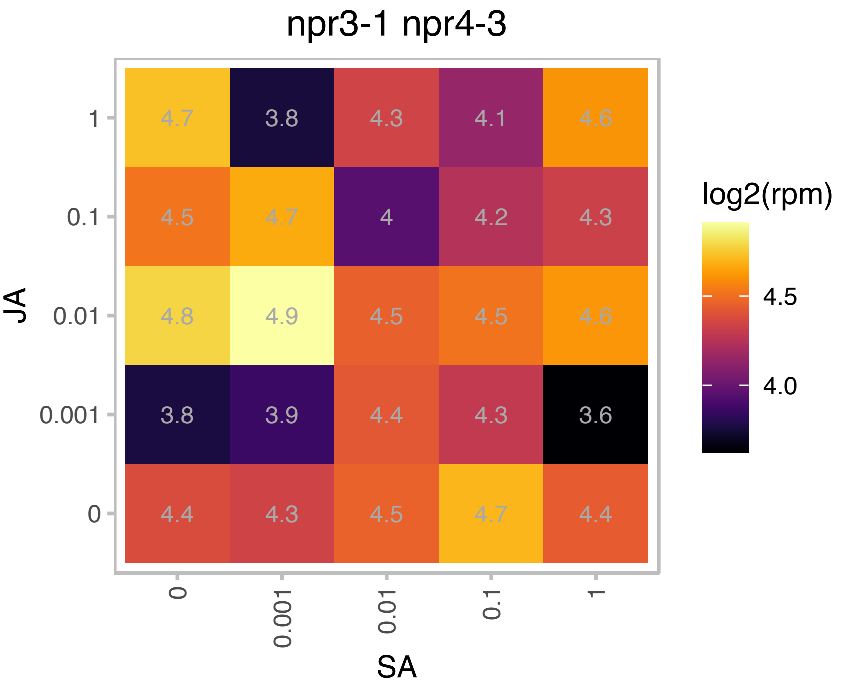

# AT5G64480.1 : NA

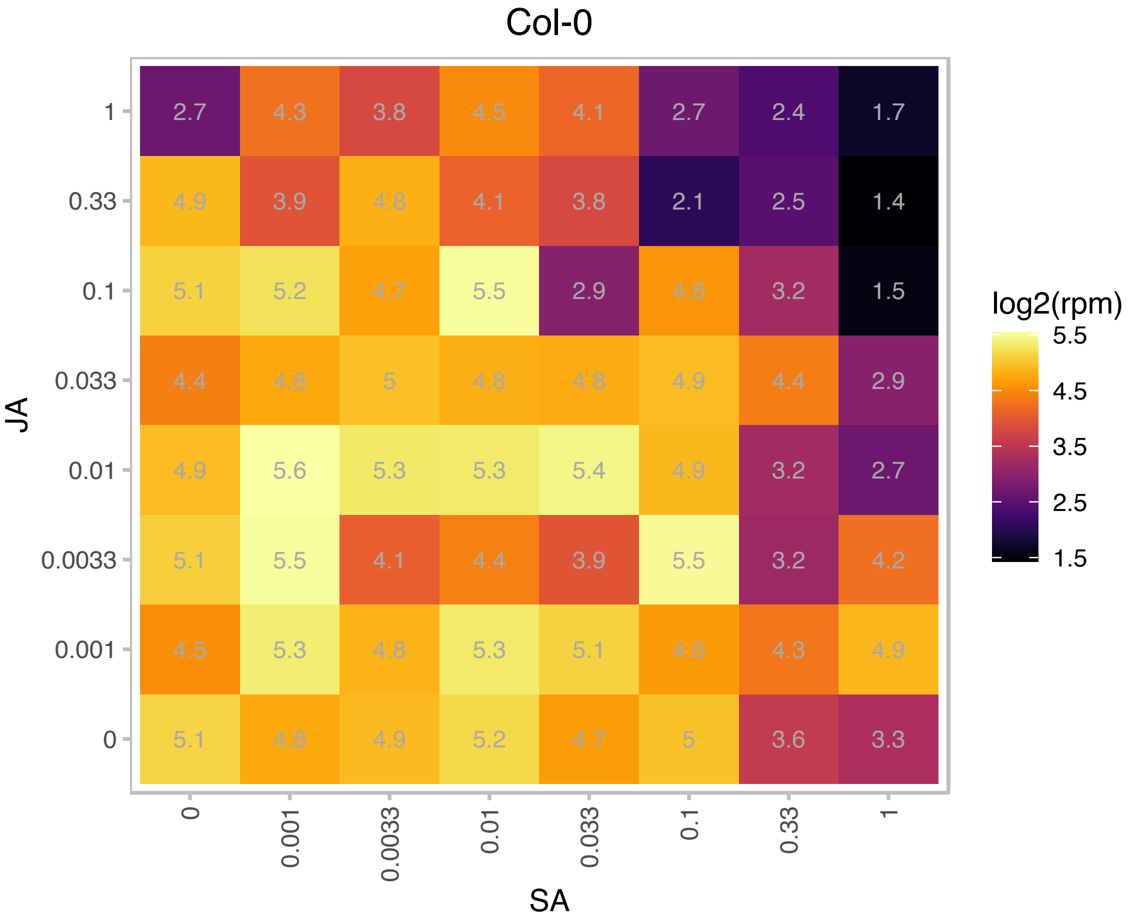


| | |
|---|---|
| Gene | AT5G64480 |
| is.expressed_genes | TRUE |
| is.responsive_genes | TRUE |
| is.apcluster | F-11 |

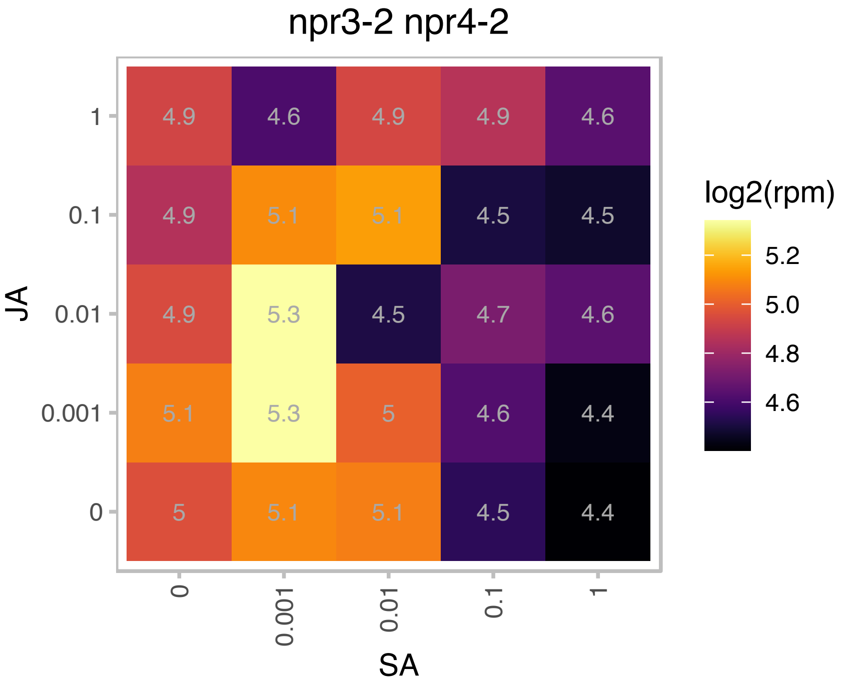


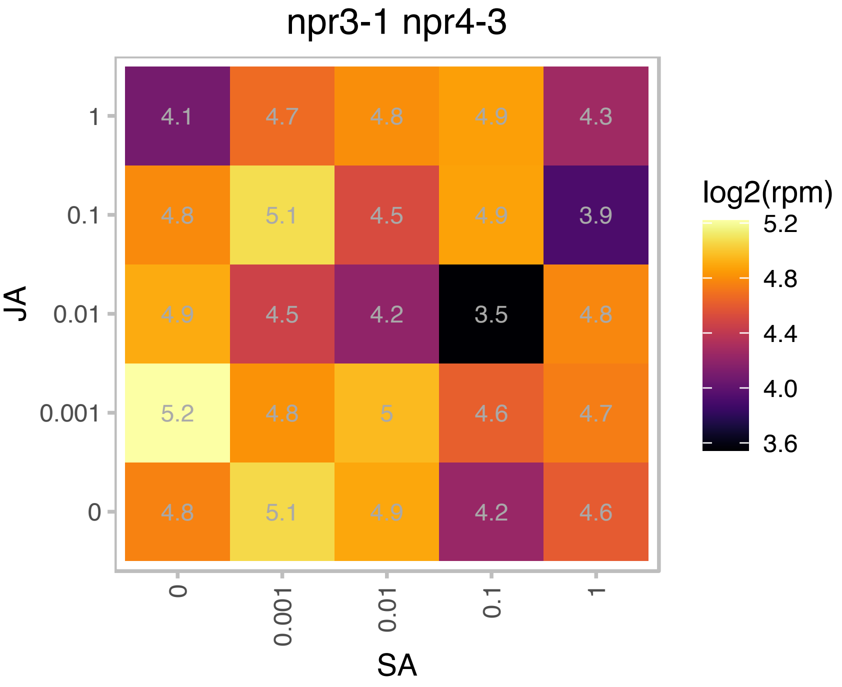

# AT1G23710.1 : NA

Protein of unknown function (DUF1645)

| | |
|---|---|
| Gene | AT1G23710 |
| is.expressed_genes | TRUE |
| is.responsive_genes | TRUE |
| is.apcluster | G-01 |

Col-0

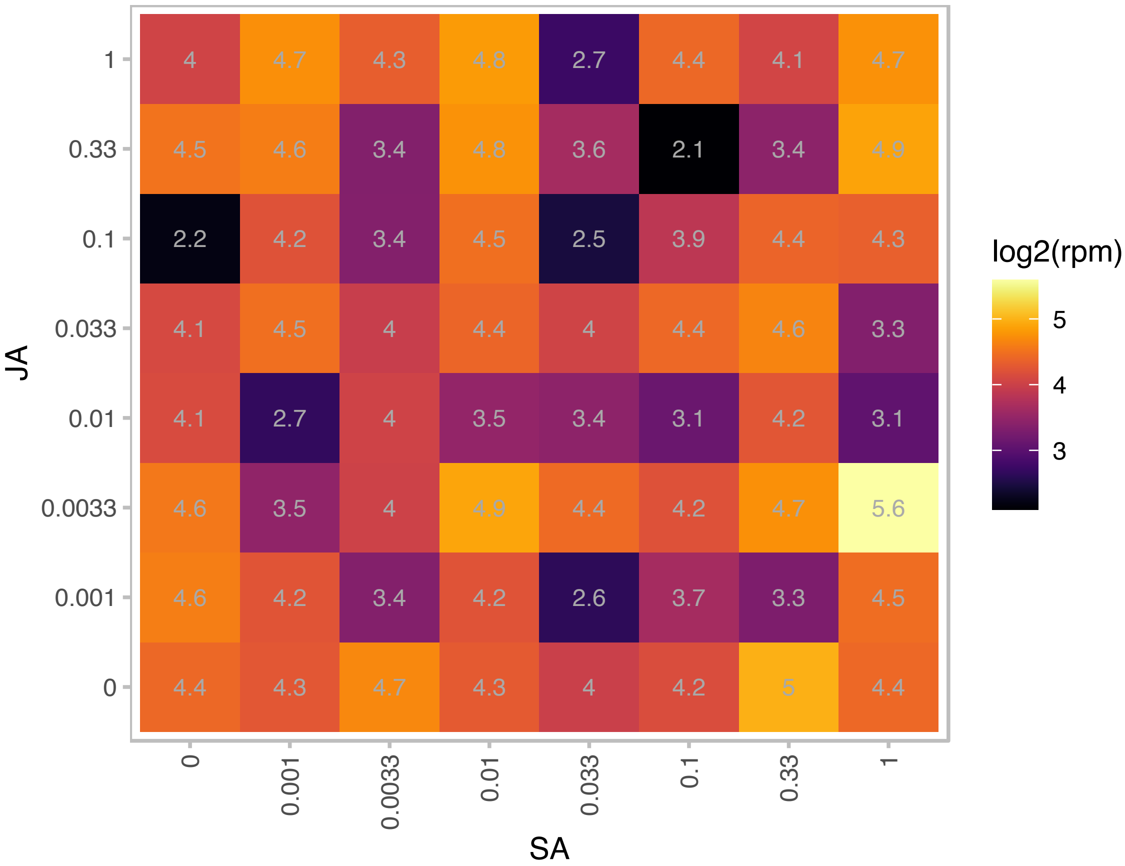


npr3-2 npr4-2

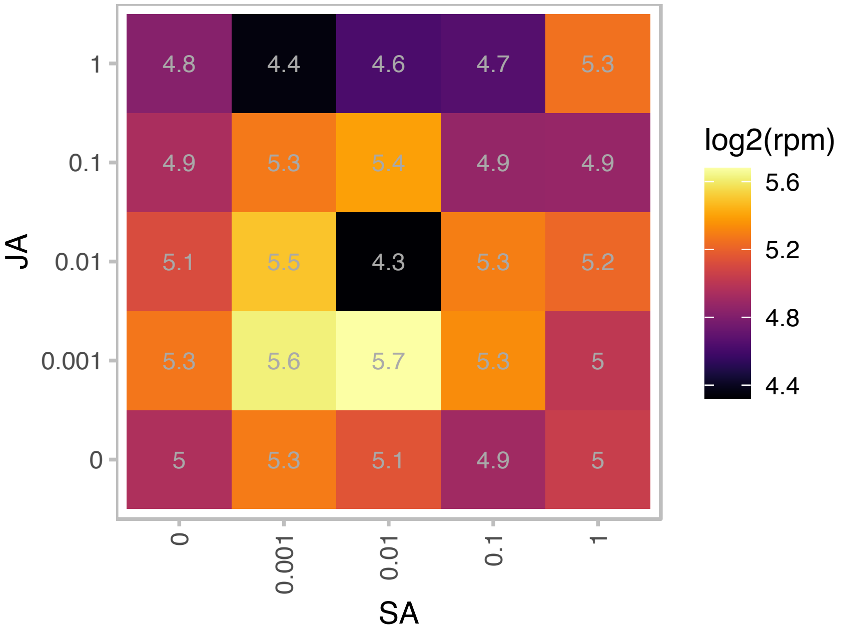


npr3-1 npr4-3

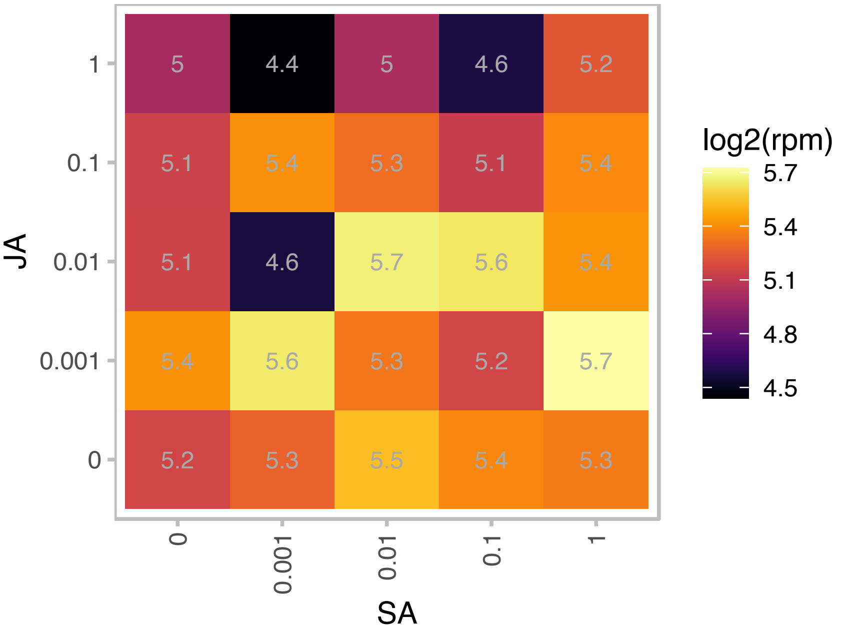

# AT1G42960.1 : NA

expressed protein localized to the inner membrane of the chloroplast.

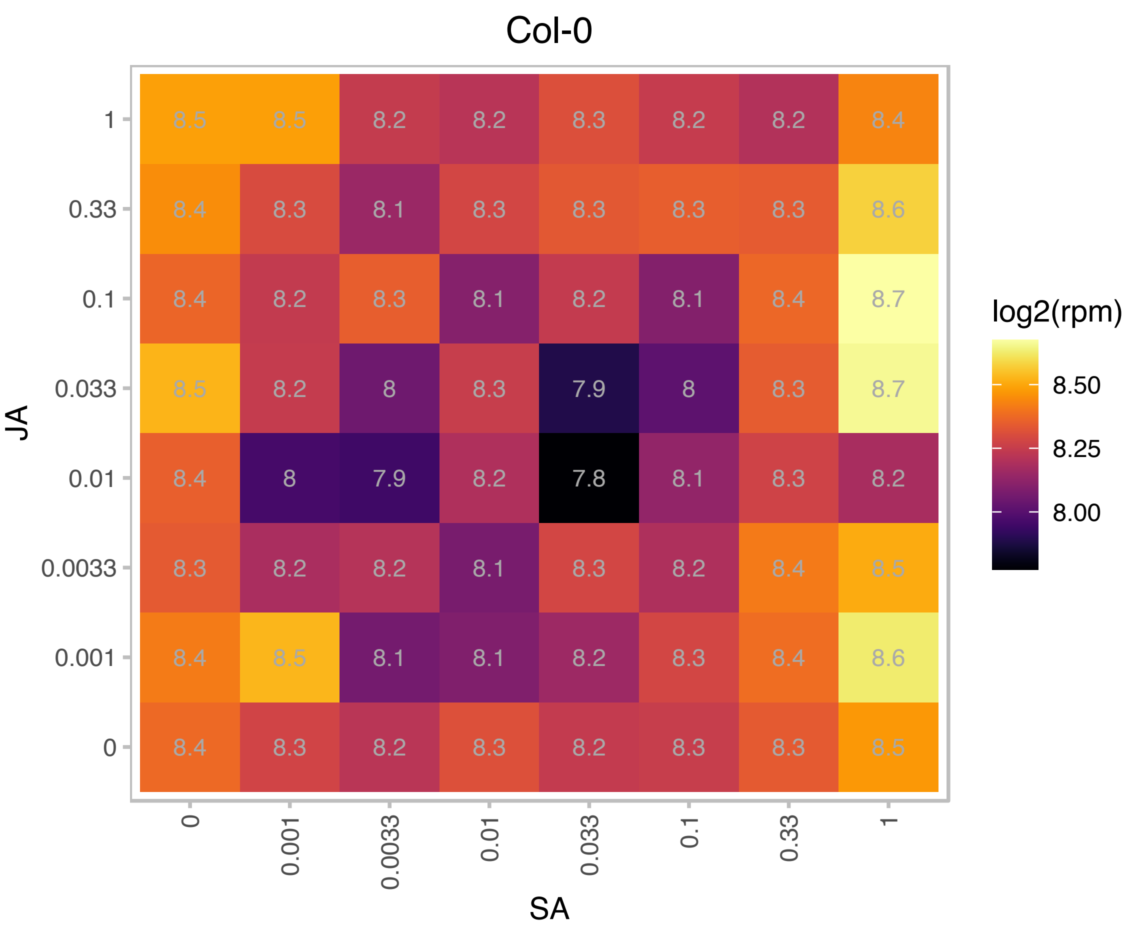


| | |
|---|---|
| Gene | AT1G42960 |
| is.expressed_genes | TRUE |
| is.responsive_genes | TRUE |
| is.apcluster | G-01 |

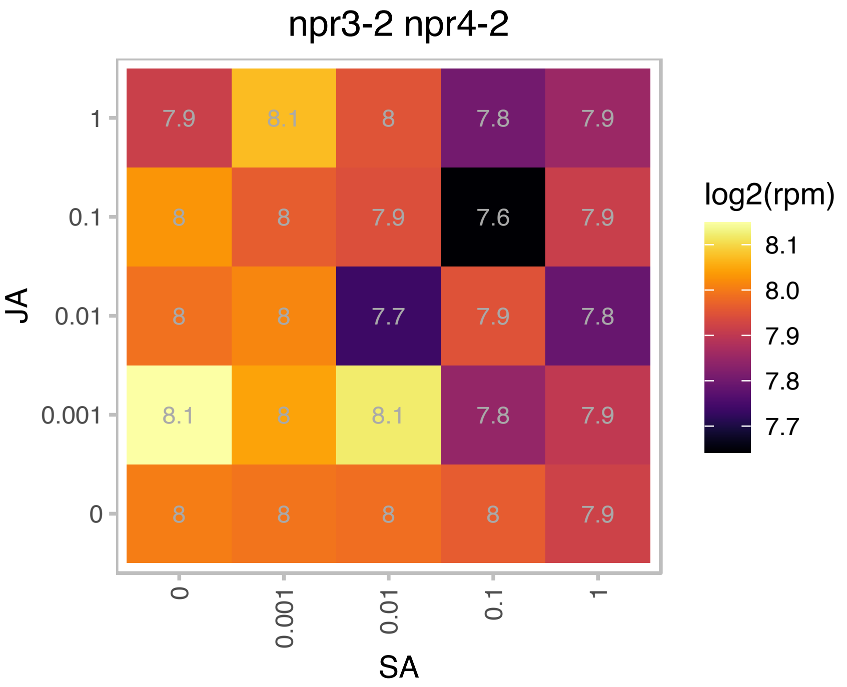


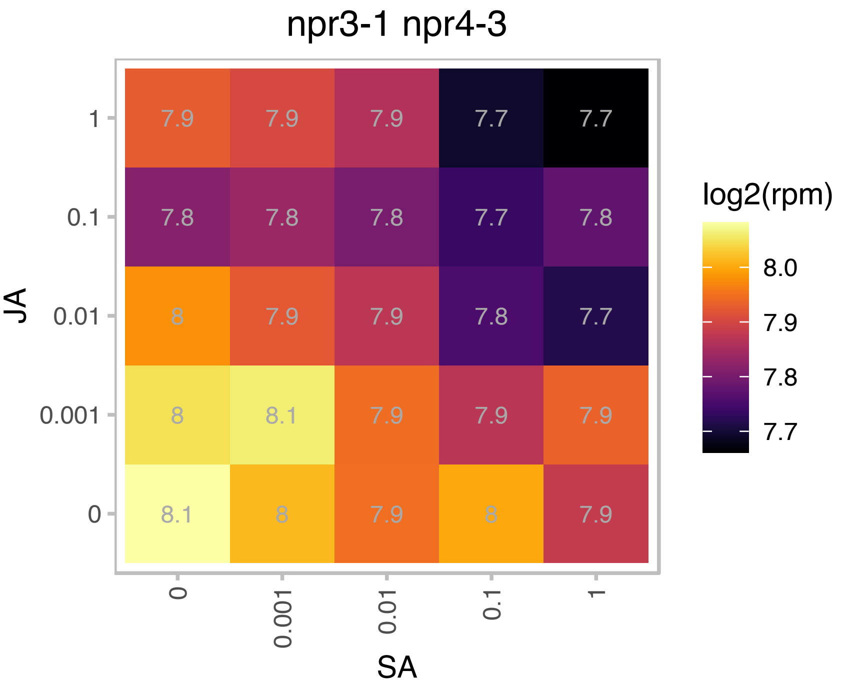

### AT1G65960.1 : GAD2

glutamate decarboxylase 2

glutamate decarboxylase (GAD2) The mRNA is cell-to-cell mobile.

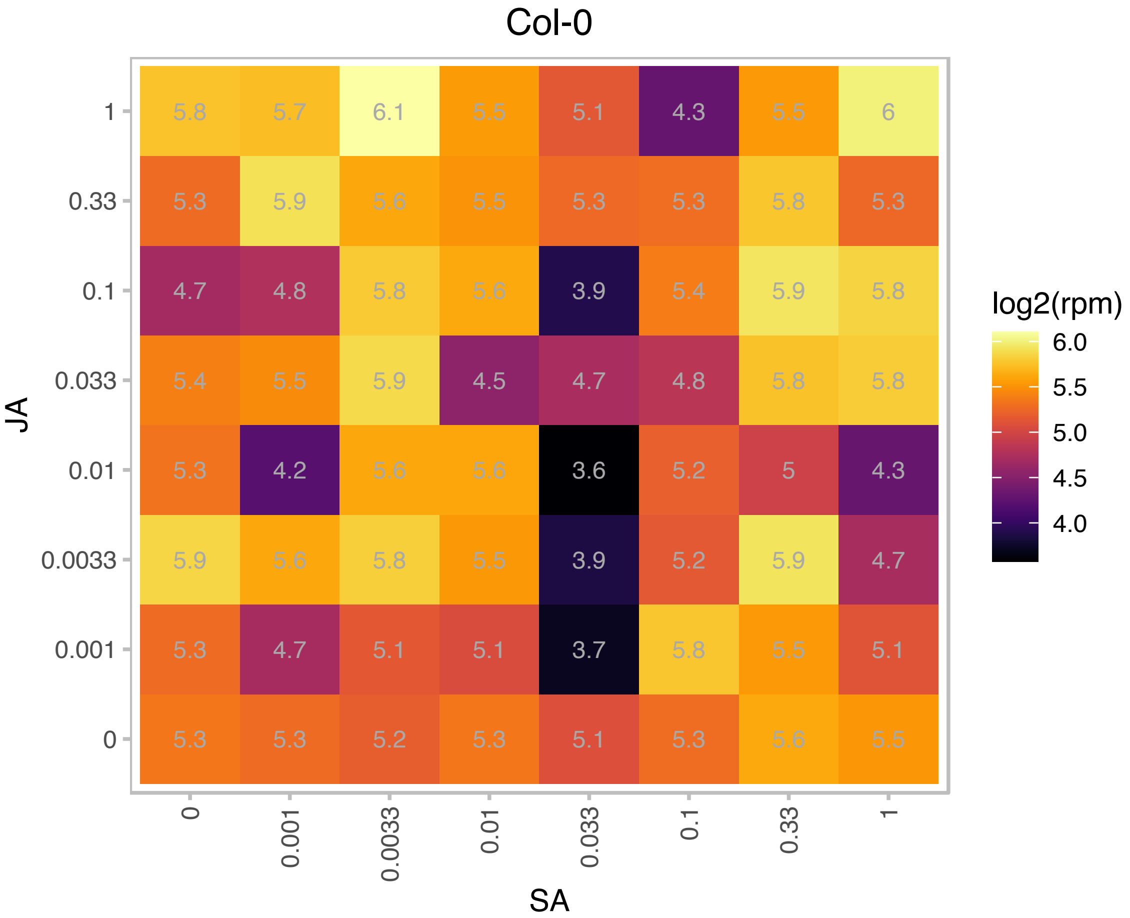


| | |
|---|---|
| Gene | AT1G65960 |
| is.expressed_genes | TRUE |
| is.responsive_genes | TRUE |
| is.apcluster | G-01 |

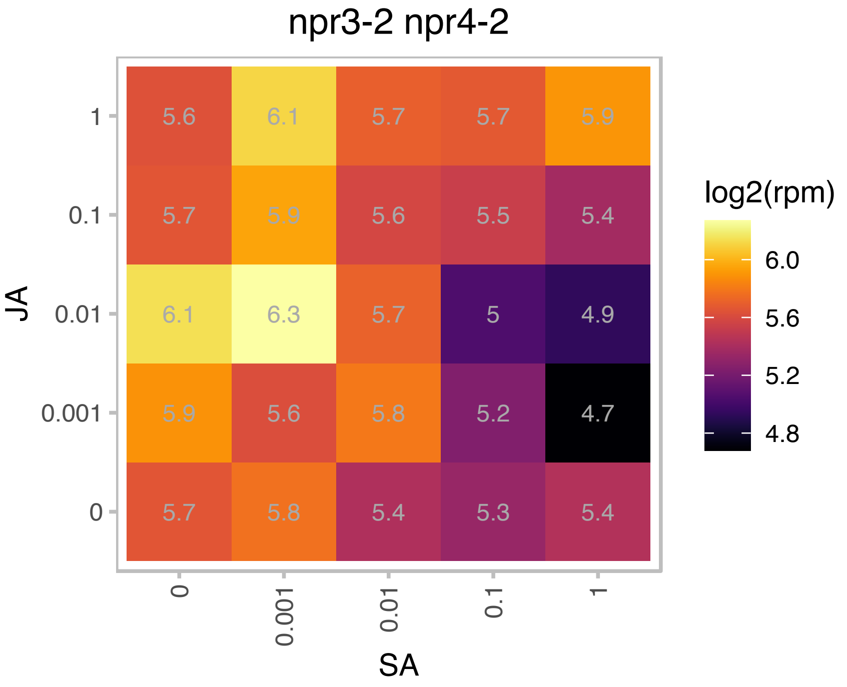


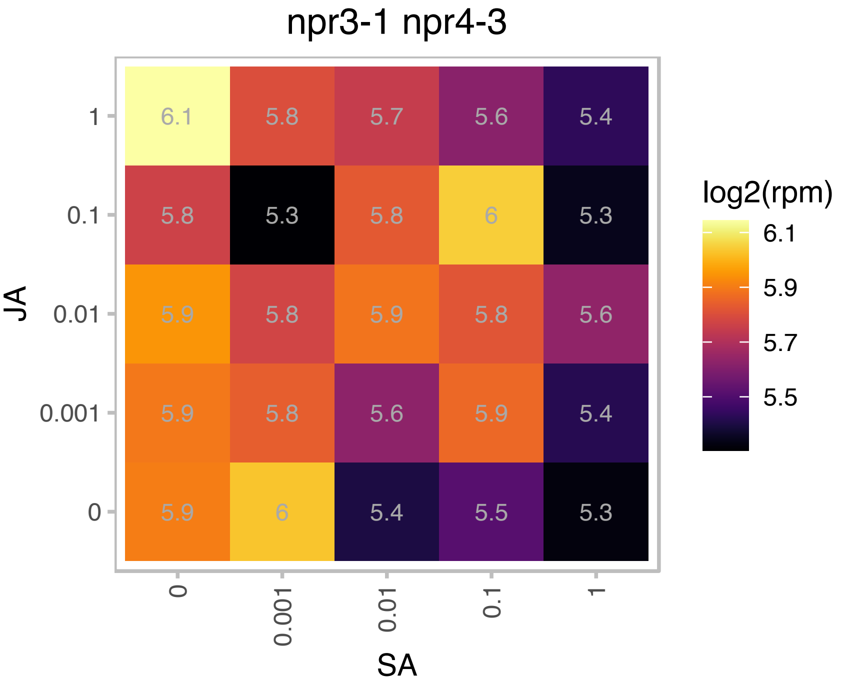

# AT2G02060.1 : NA

Homeodomain-like superfamily protein

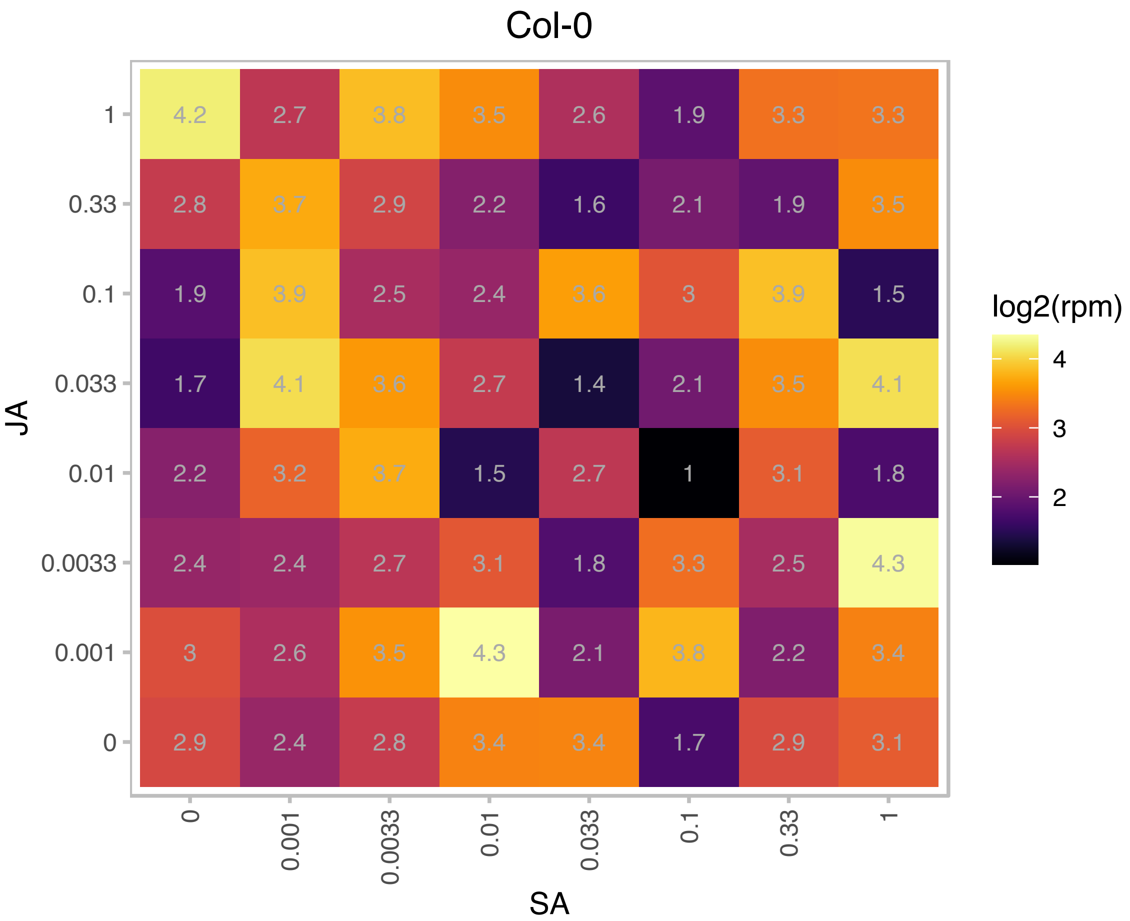


| | |
|---|---|
| Gene | AT2G02060 |
| is.expressed_genes | TRUE |
| is.responsive_genes | TRUE |
| is.apcluster | G-01 |

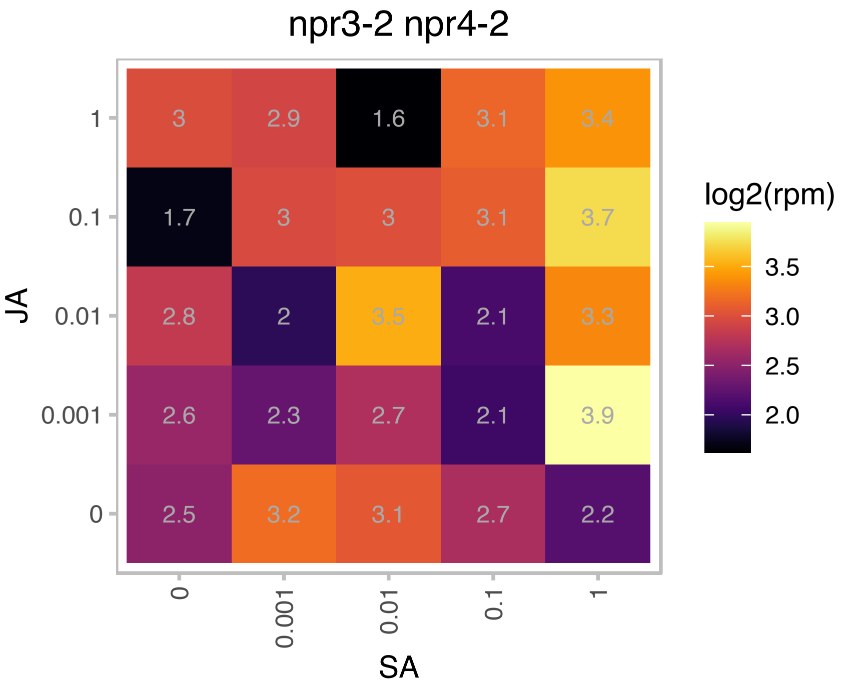


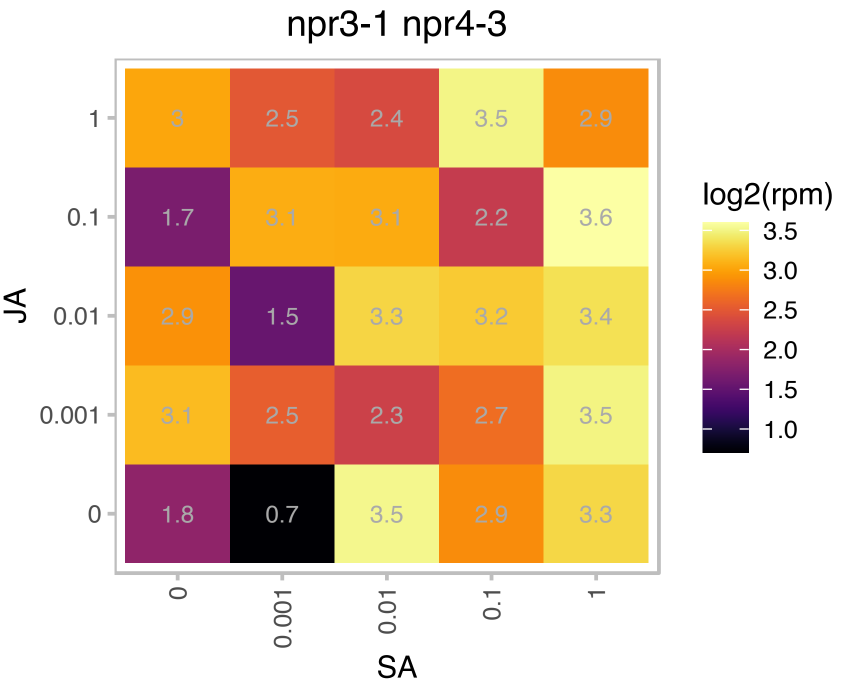

# AT2G20724.1 : NA

Annotated as pseudogene of unknown protein.Possibly not a pseudogene based on evidence for transcription (RNA-seq) and translation (Ribo-seq) described in PMID:27791167 .

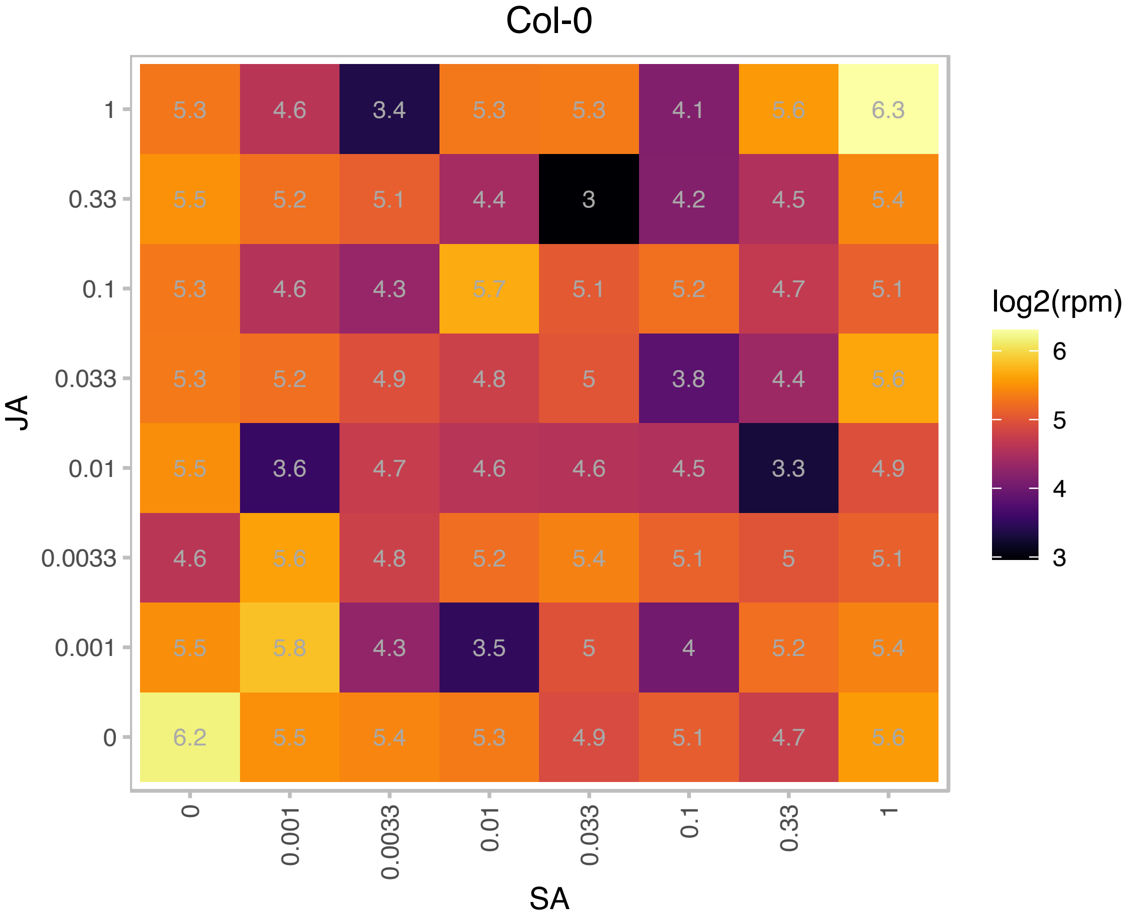


| | |
|---|---|
| Gene | AT2G20724 |
| is.expressed_genes | TRUE |
| is.responsive_genes | TRUE |
| is.apcluster | G-01 |

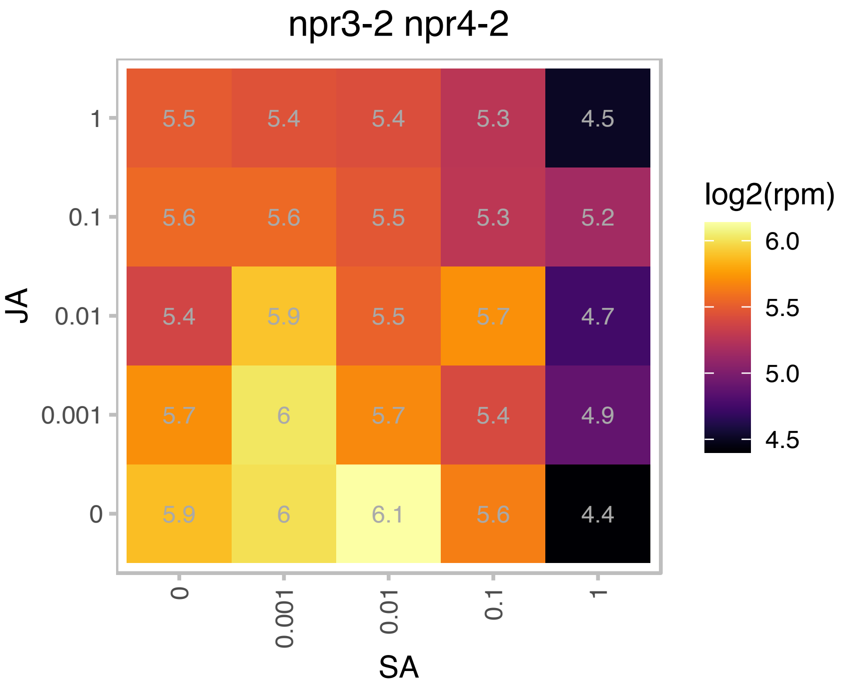


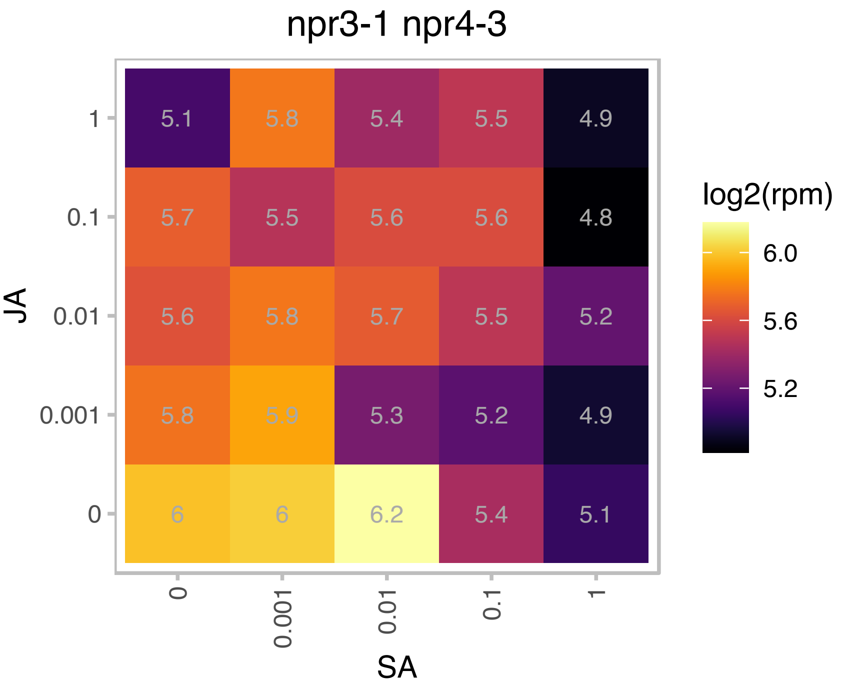

### AT2G25840.1 : OVA4

Nucleotidylyl transferase superfamily protein

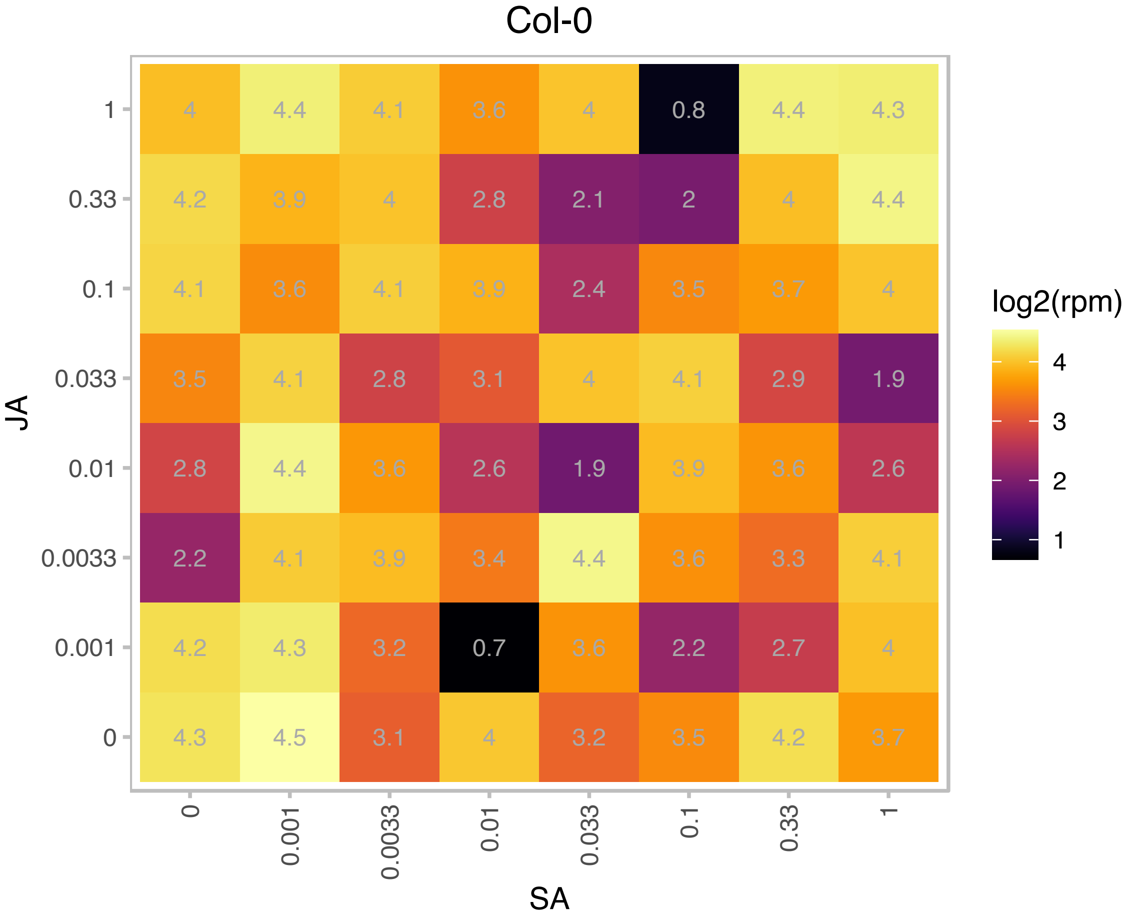


| | |
|---|---|
| Gene | AT2G25840 |
| is.expressed_genes | TRUE |
| is.responsive_genes | TRUE |
| is.apcluster | G-01 |

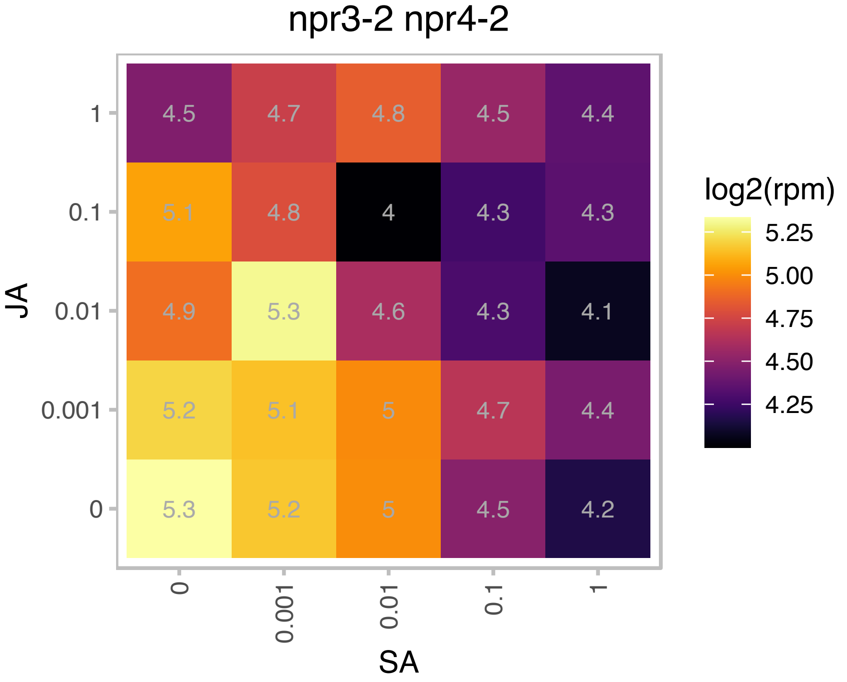


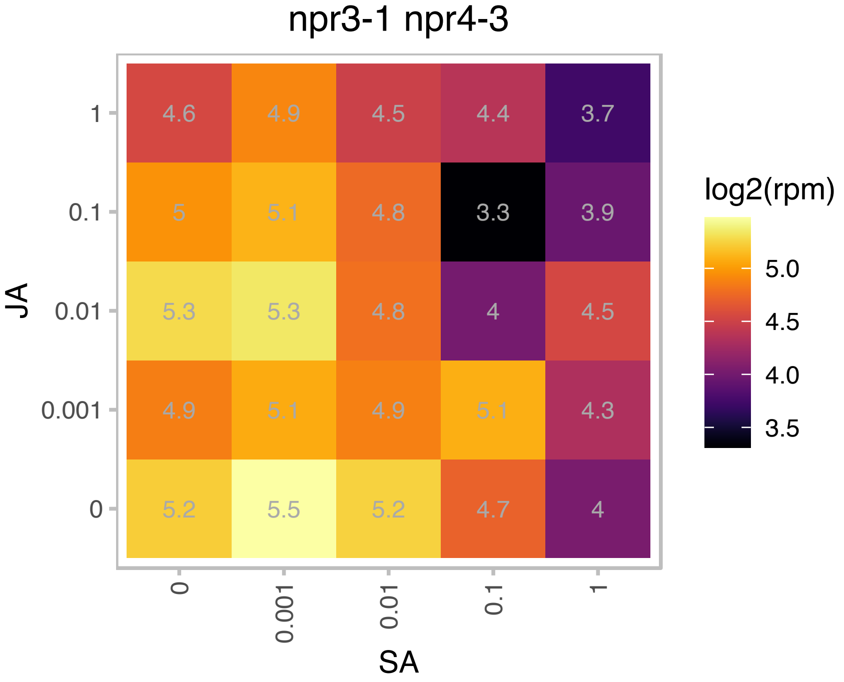

### AT2G28900.1 : ATOEP16-1, ATOEP16-L, OEP16, OEP16-1

outer plastid envelope protein 16-1

Encodes AtOEP16, a 16-KDa plastid outer membrane protein involved in plastid import of protochlorophyllide oxidoreductase A.  Predominantly expressed in leaves and is also inducible by cold treatment.

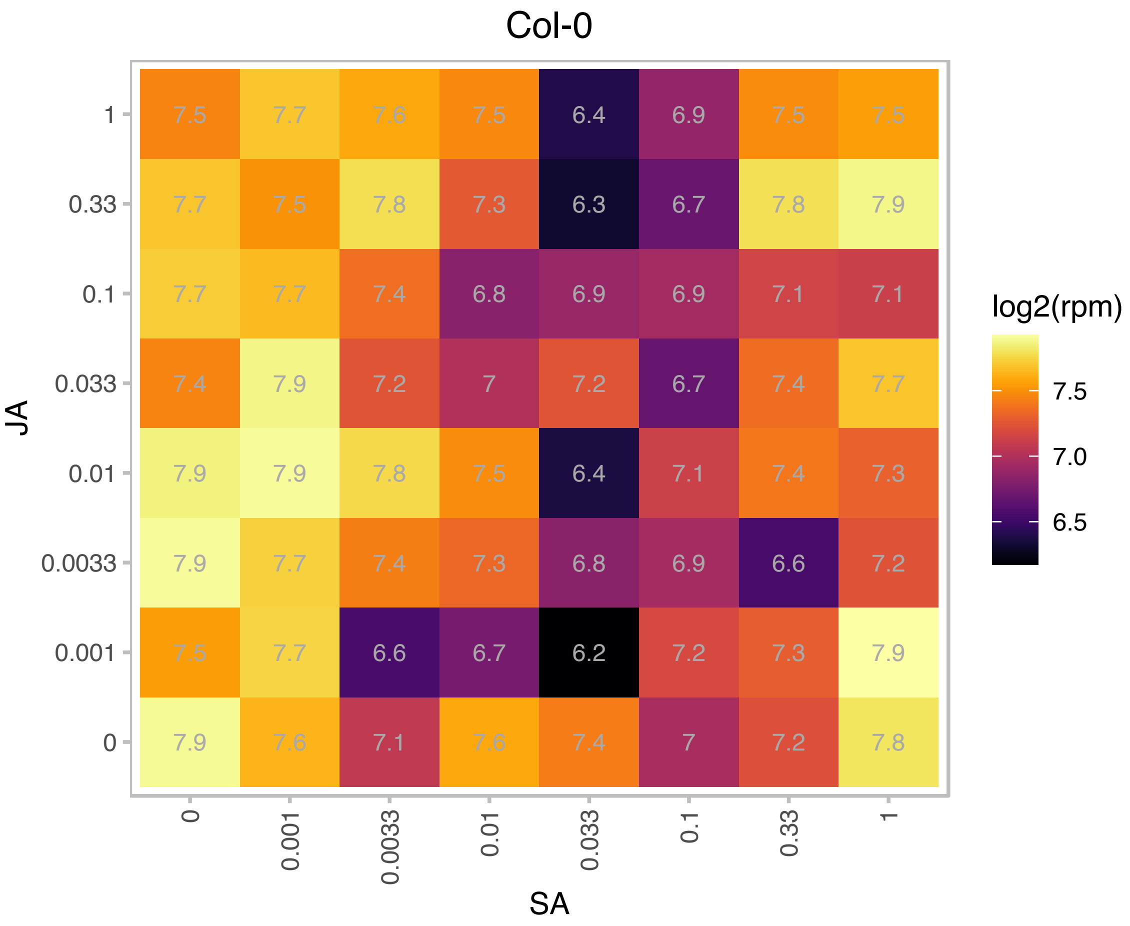


| | |
|---|---|
| Gene | AT2G28900 |
| is.expressed_genes | TRUE |
| is.responsive_genes | TRUE |
| is.apcluster | G-01 |

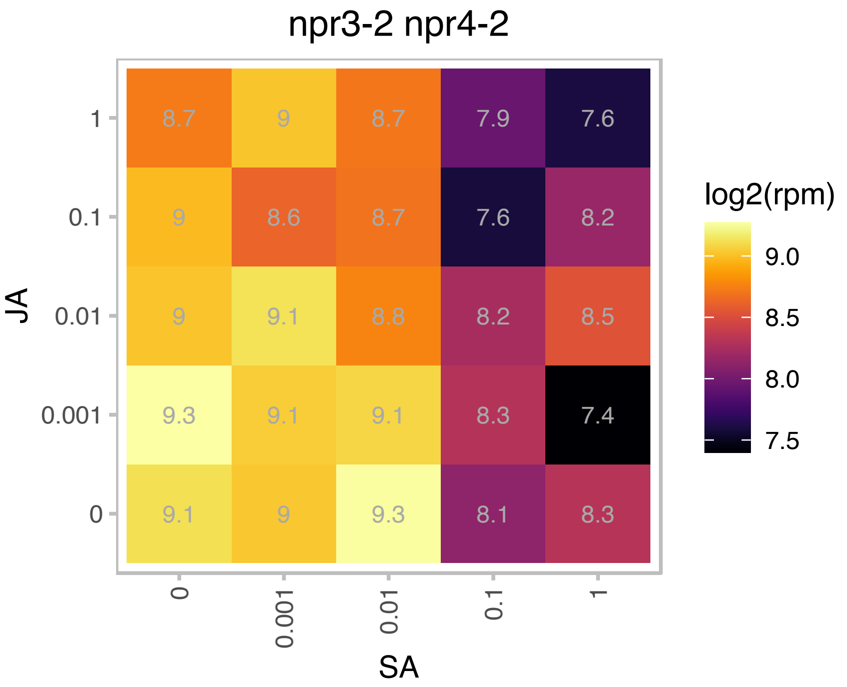


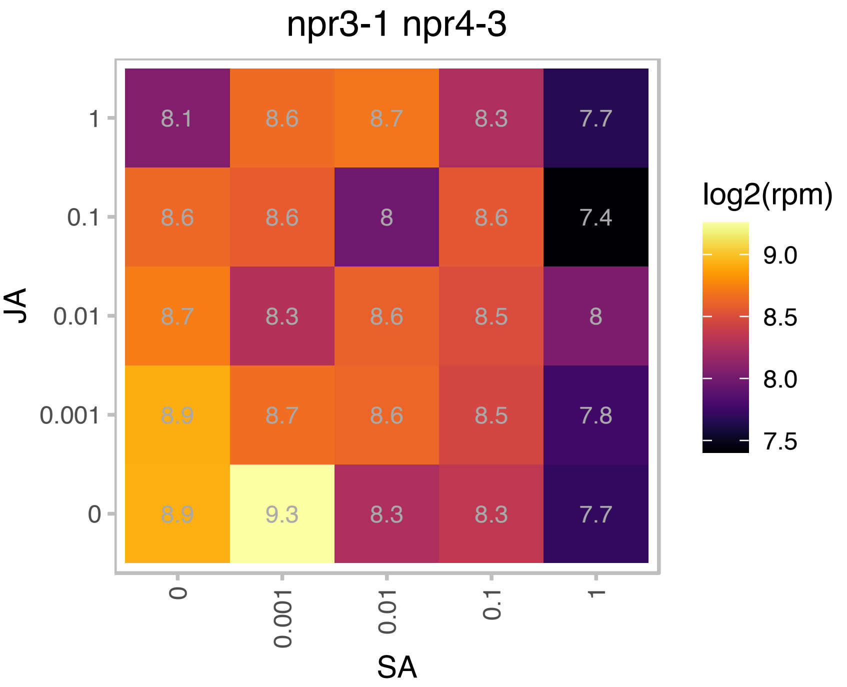

### AT2G33800.1 : EMB3113, PRPS5, RPS5, SCA1

Ribosomal protein S5 family protein

Encodes SCABRA1 (SCA1), a nuclear gene encoding a plastid-type ribosomal protein that functions  as a structural component of the 70S plastid ribosome. The sca1-rps5 allele exhibits defects in plastid 16SrRNA processing and a resulting decrease in accumulation of photosynthetic proteins. Loss-of-function mutations enhance the polarity defects of the as2 mutants.

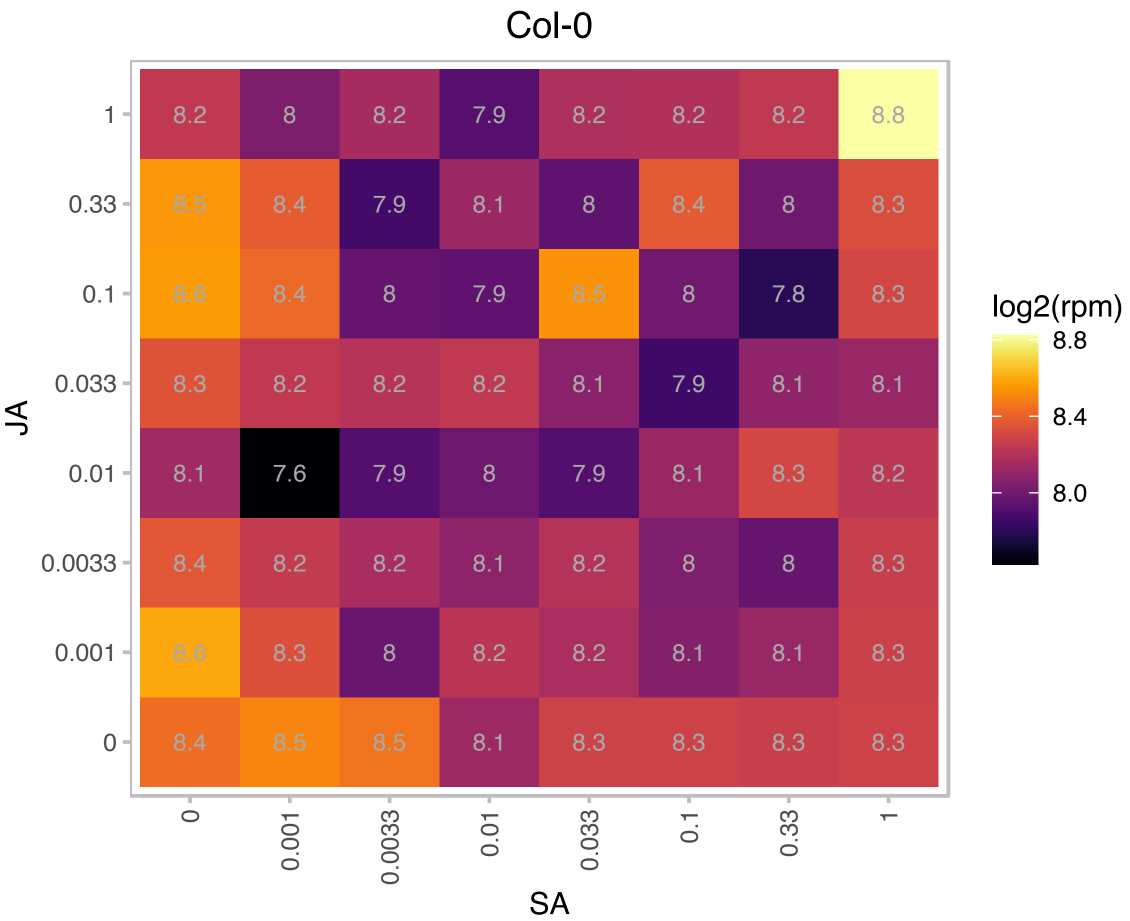


| | |
|---|---|
| Gene | AT2G33800 |
| is.expressed_genes | TRUE |
| is.responsive_genes | TRUE |
| is.apcluster | G-01 |

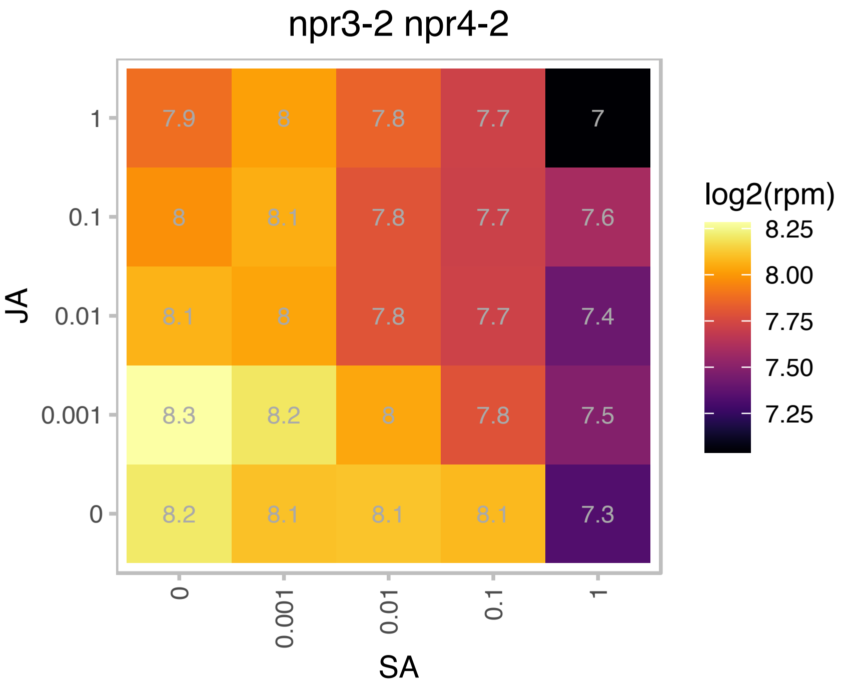


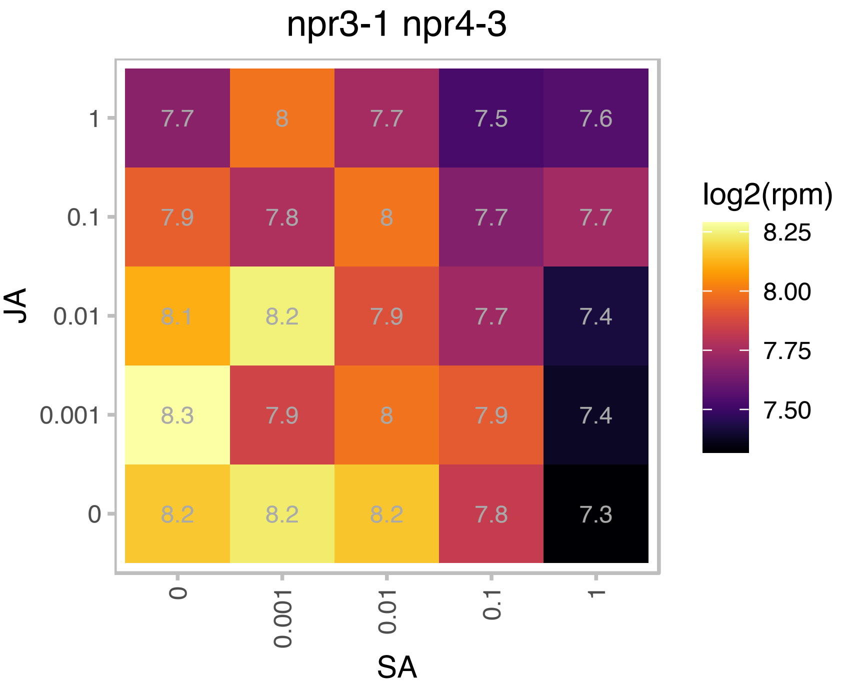

# AT3G07470.1 : NA

Protein of unknown function, DUF538

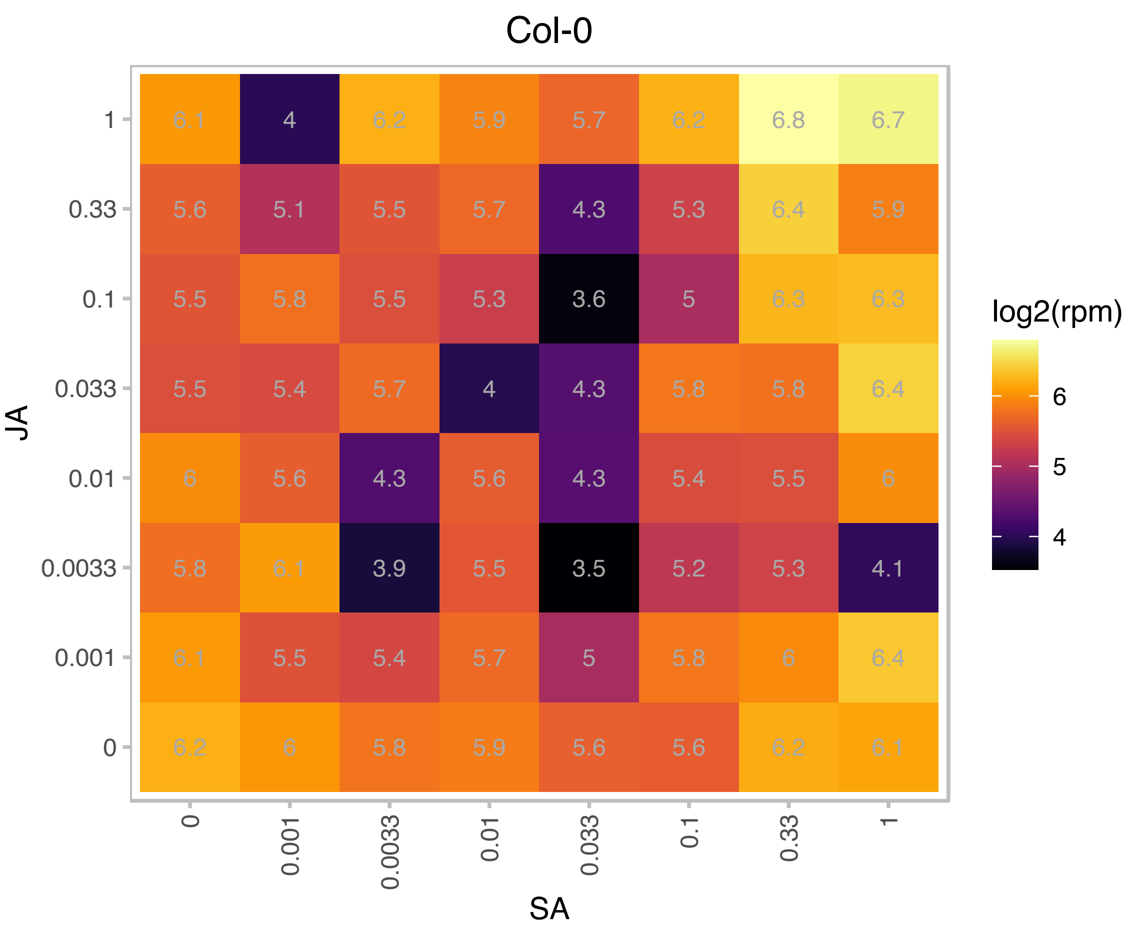


| | |
|---|---|
| Gene | AT3G07470 |
| is.expressed_genes | TRUE |
| is.responsive_genes | TRUE |
| is.apcluster | G-01 |

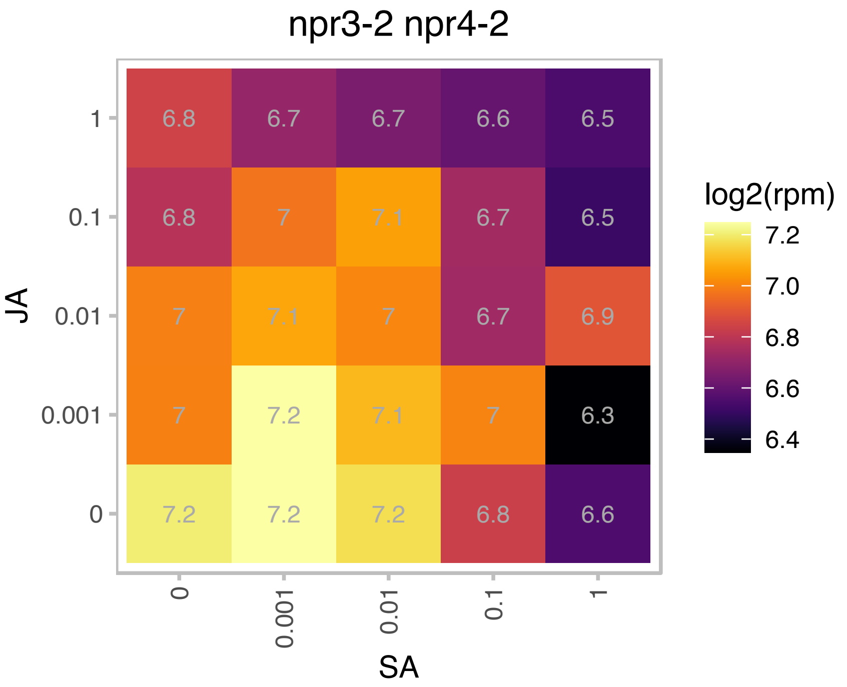


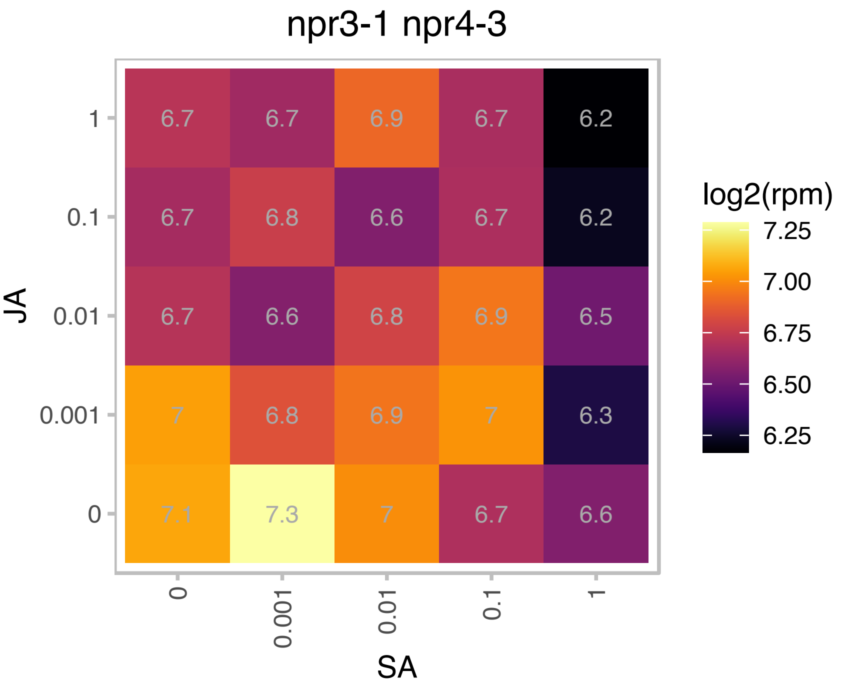

# AT3G11560.1 : NA

LETM1-like protein

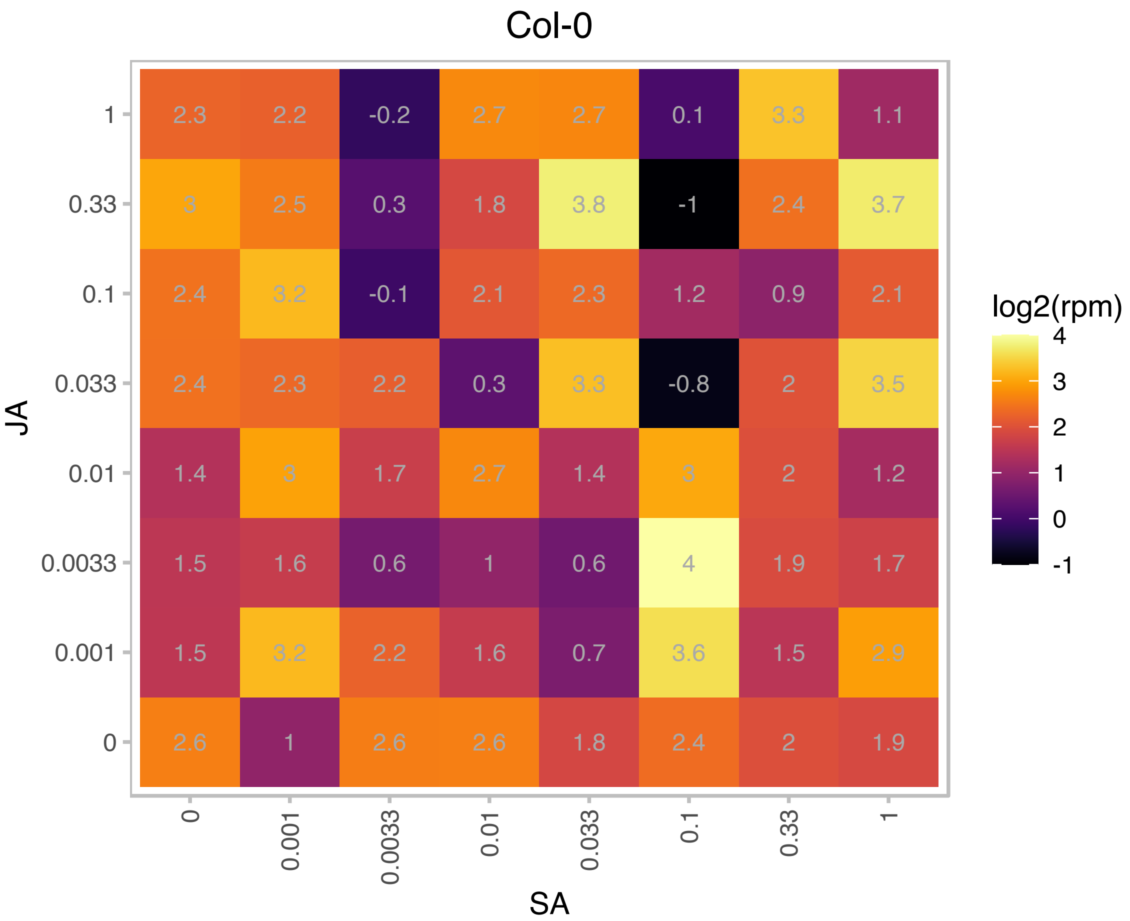


| | |
|---|---|
| Gene | AT3G11560 |
| is.expressed_genes | TRUE |
| is.responsive_genes | TRUE |
| is.apcluster | G-01 |

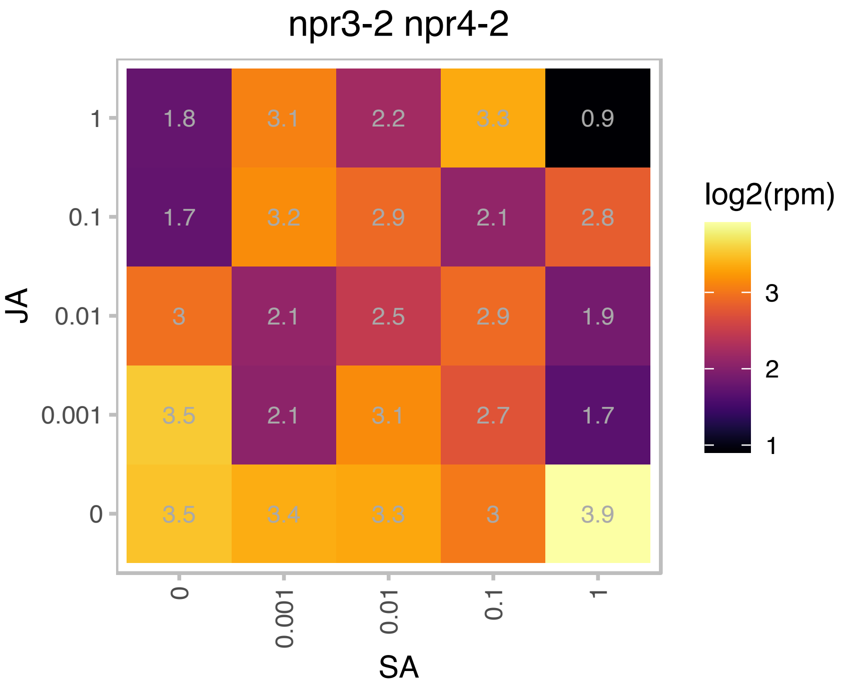


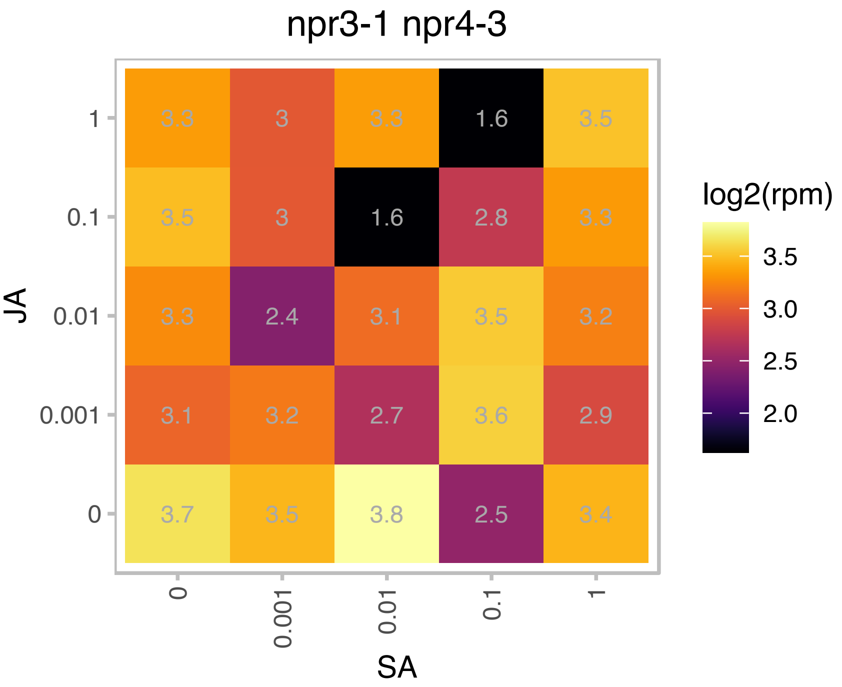

### AT3G25920.1 : RPL15

ribosomal protein L15

encodes a plastid ribosomal protein CL15, a constituent of the large subunit of the ribosomal complex

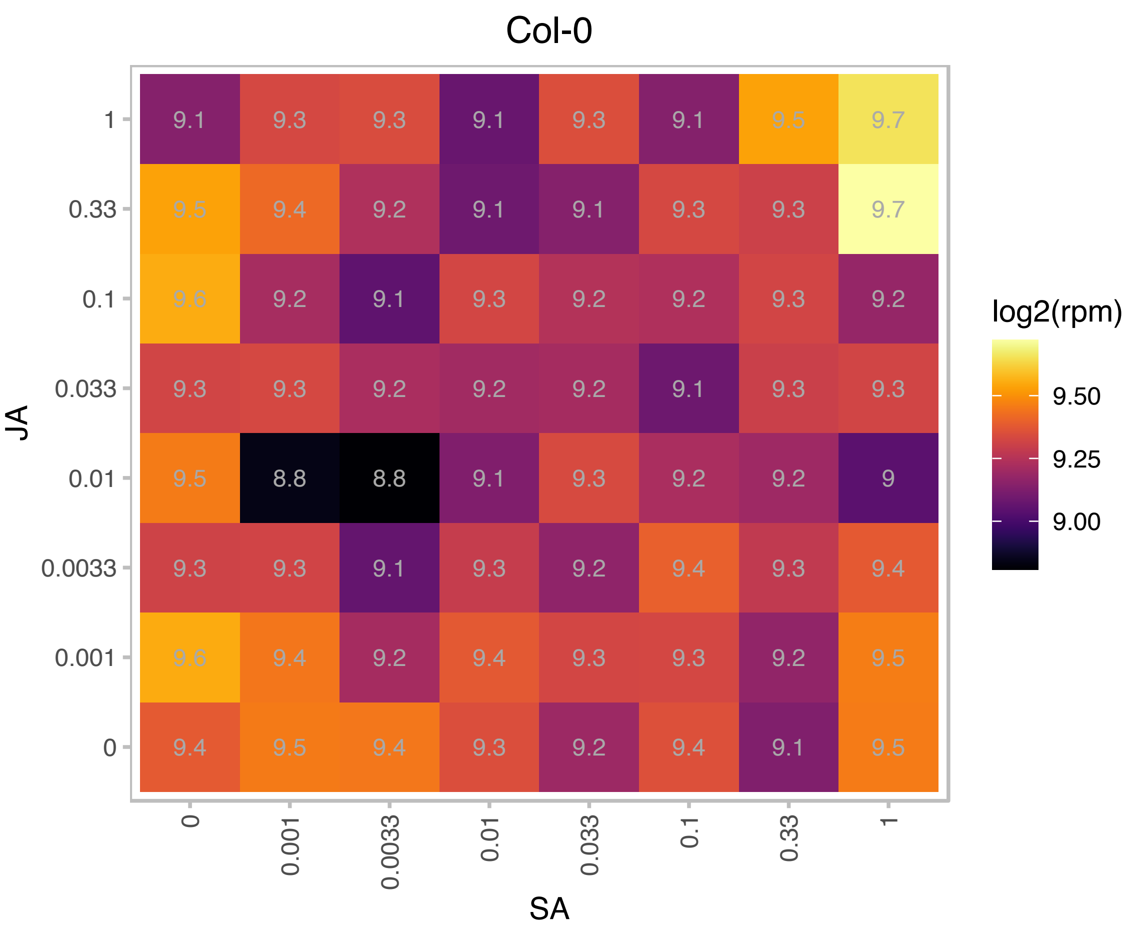


| | |
|---|---|
| Gene | AT3G25920 |
| is.expressed_genes | TRUE |
| is.responsive_genes | TRUE |
| is.apcluster | G-01 |

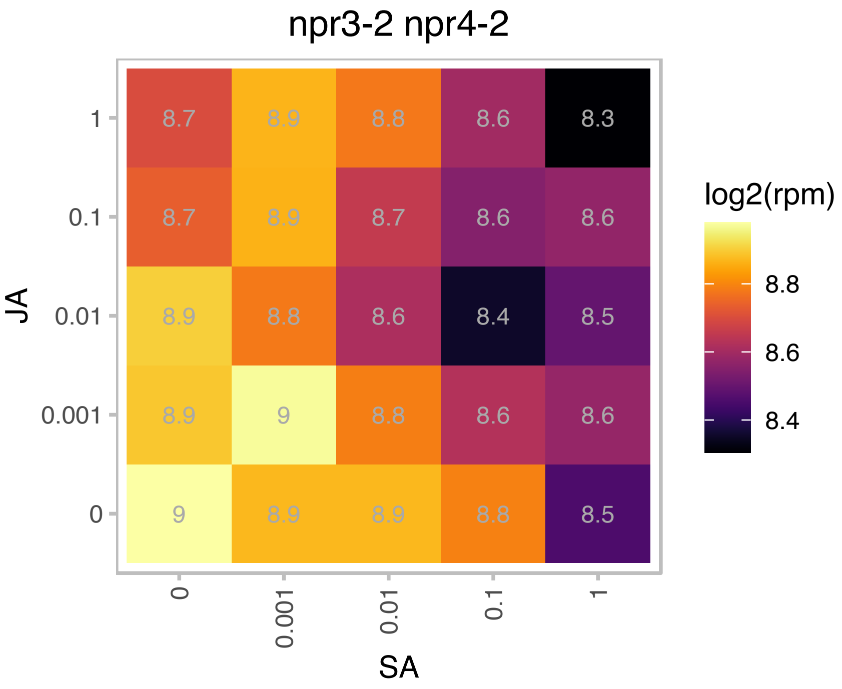


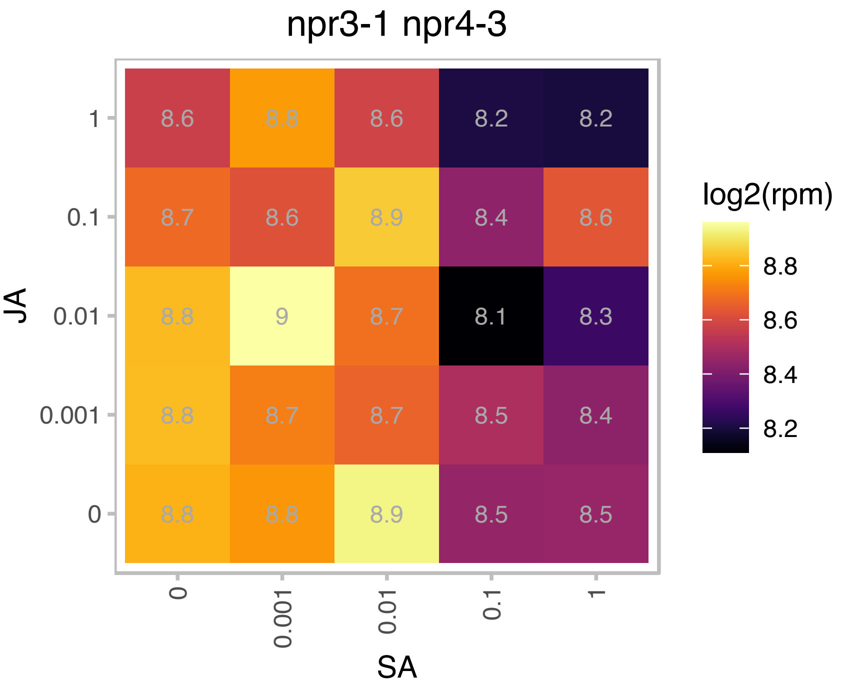

### AT3G50820.1 : OEC33, PSBO-2, PSBO2

photosystem II subunit O-2

Encodes a protein which is an extrinsic subunit of photosystem II and which has been proposed to play a central role in stabilization of the catalytic manganese cluster. In <i>Arabidopsis thaliana</i> the PsbO proteins are encoded by two genes: <i>psbO1</i> and <i>psbO2</i>. PsbO2 is the minor isoform in the wild-type. Mutants defective in this gene have been shown to be affected in the dephosphorylation of the D1 protein of PSII.

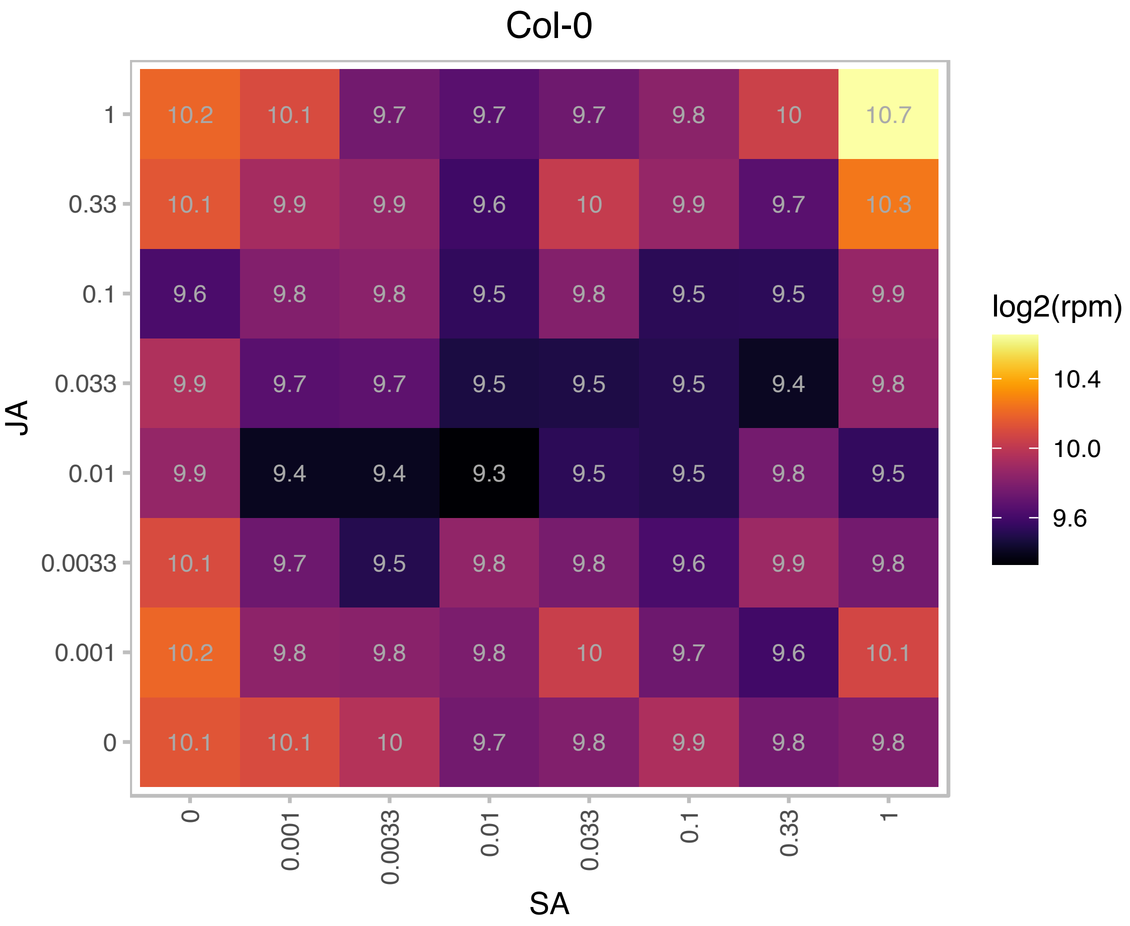


| | |
|---|---|
| Gene | AT3G50820 |
| is.expressed_genes | TRUE |
| is.responsive_genes | TRUE |
| is.apcluster | G-01 |

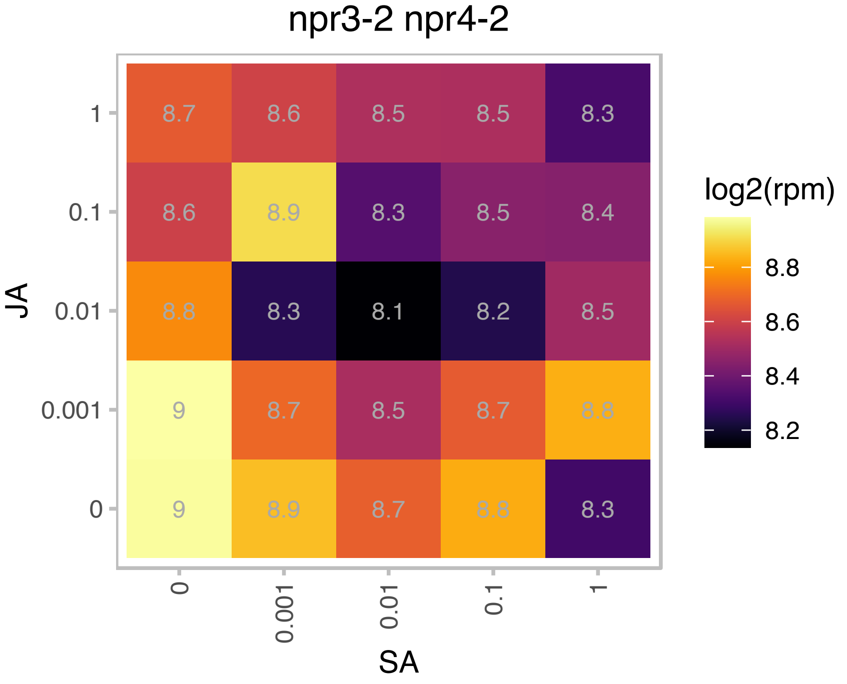


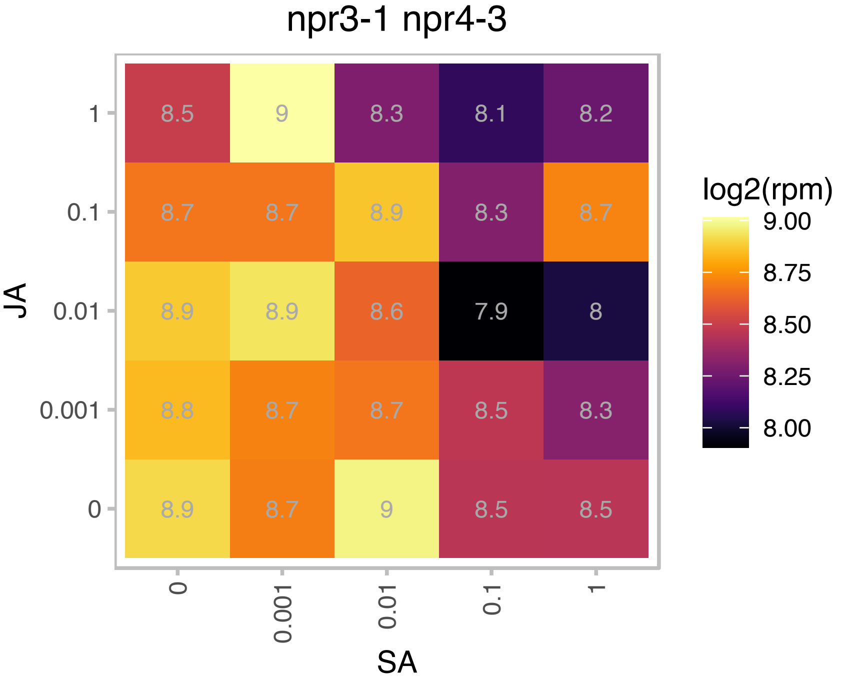

### AT3G52180.1 : ATPTPKIS1, ATSEX4, DSP4, SEX4

dual specificity protein phosphatase (DsPTP1) family protein

Encodes a plant-specific glucan phosphatase that contains a noncatalytic carbohydrate-binding module as well as a dual specificity protein phosphatase domain. SEX4 can dephosphorylate C6- and C3-glucosyl residues on native starch grains and related maltodextrin compounds in vitro. This protein interacts with the plant SnRK AKIN11, binds starch, and is localized in the chloroplast. sex4 mutants have elevated levels of starch.

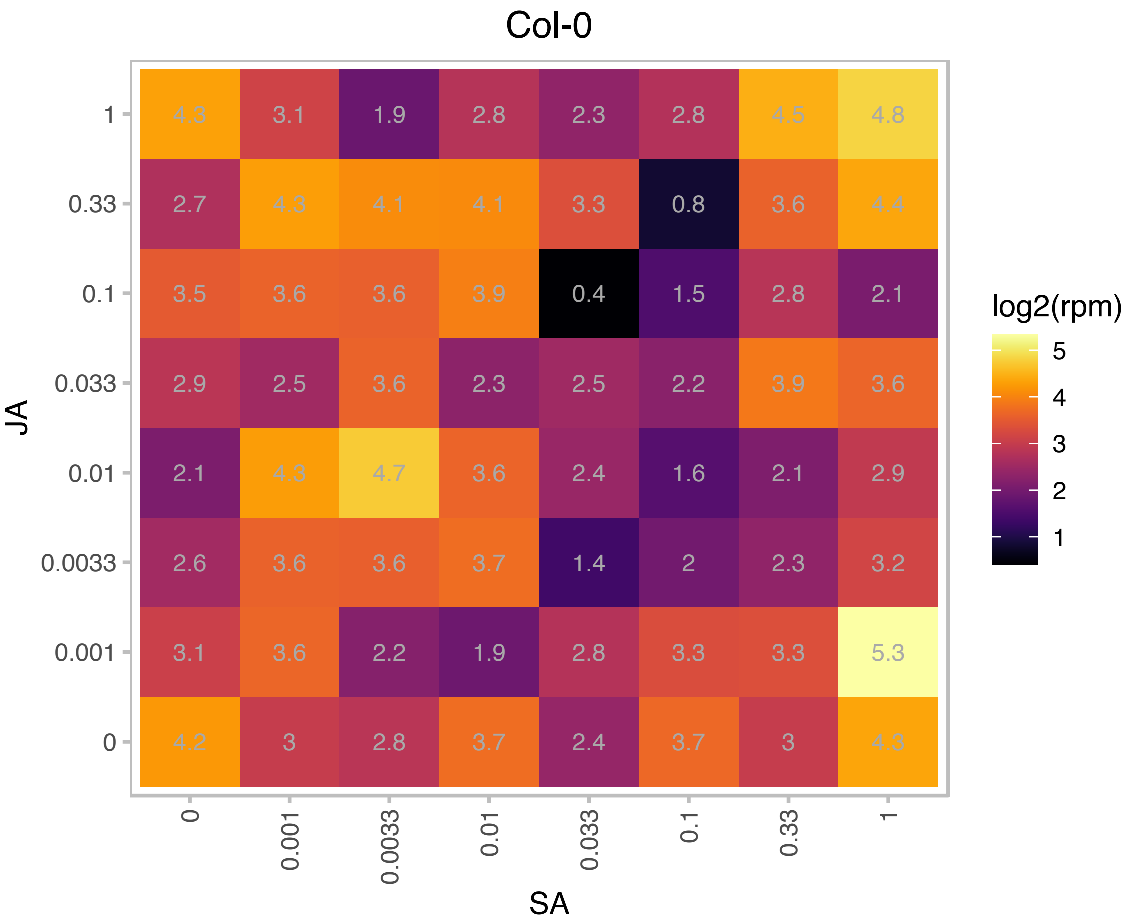


| | |
|---|---|
| Gene | AT3G52180 |
| is.expressed_genes | TRUE |
| is.responsive_genes | TRUE |
| is.apcluster | G-01 |

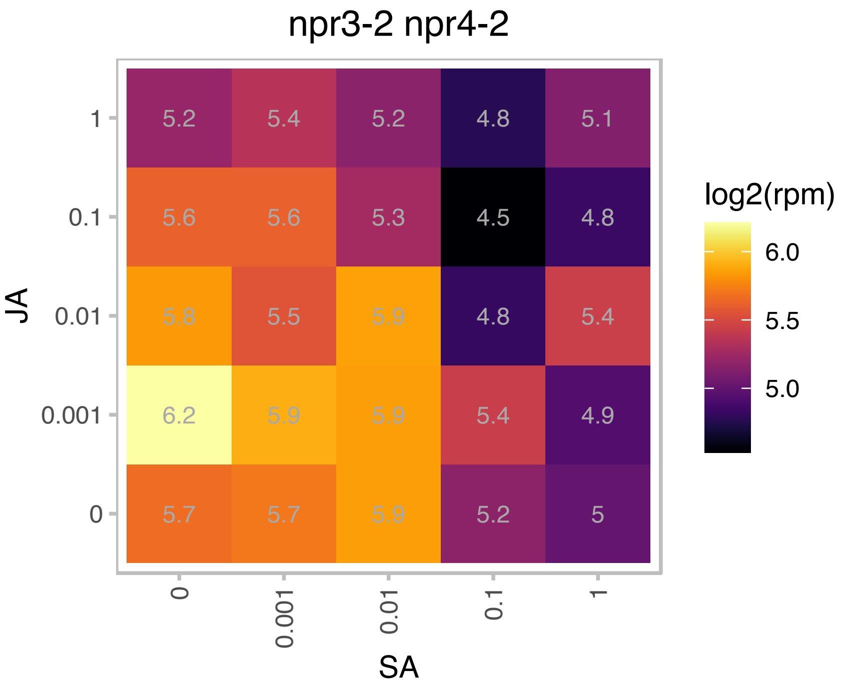


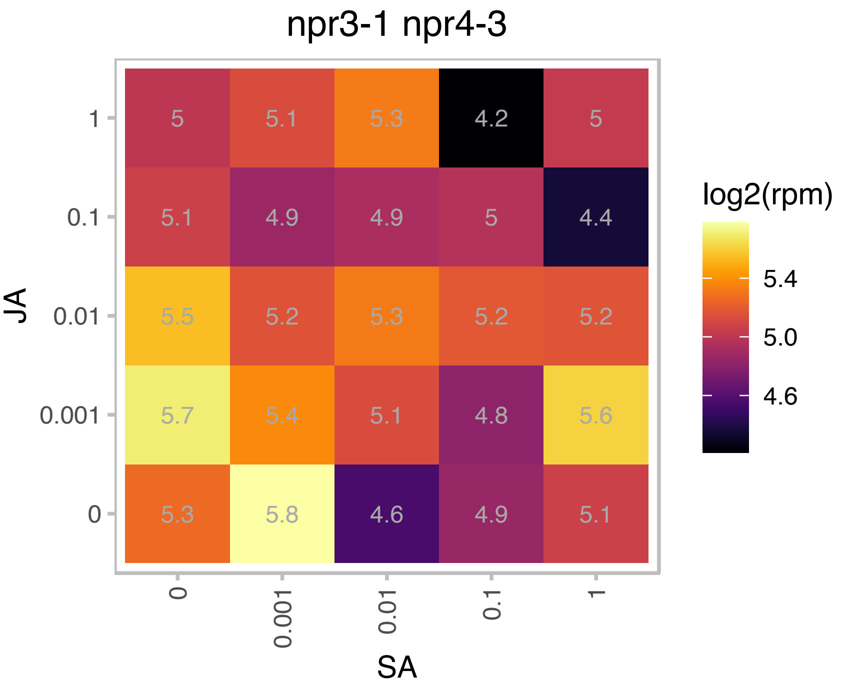

# AT4G24930.1 : NA

thylakoid lumenal 17.9 kDa protein, chloroplast

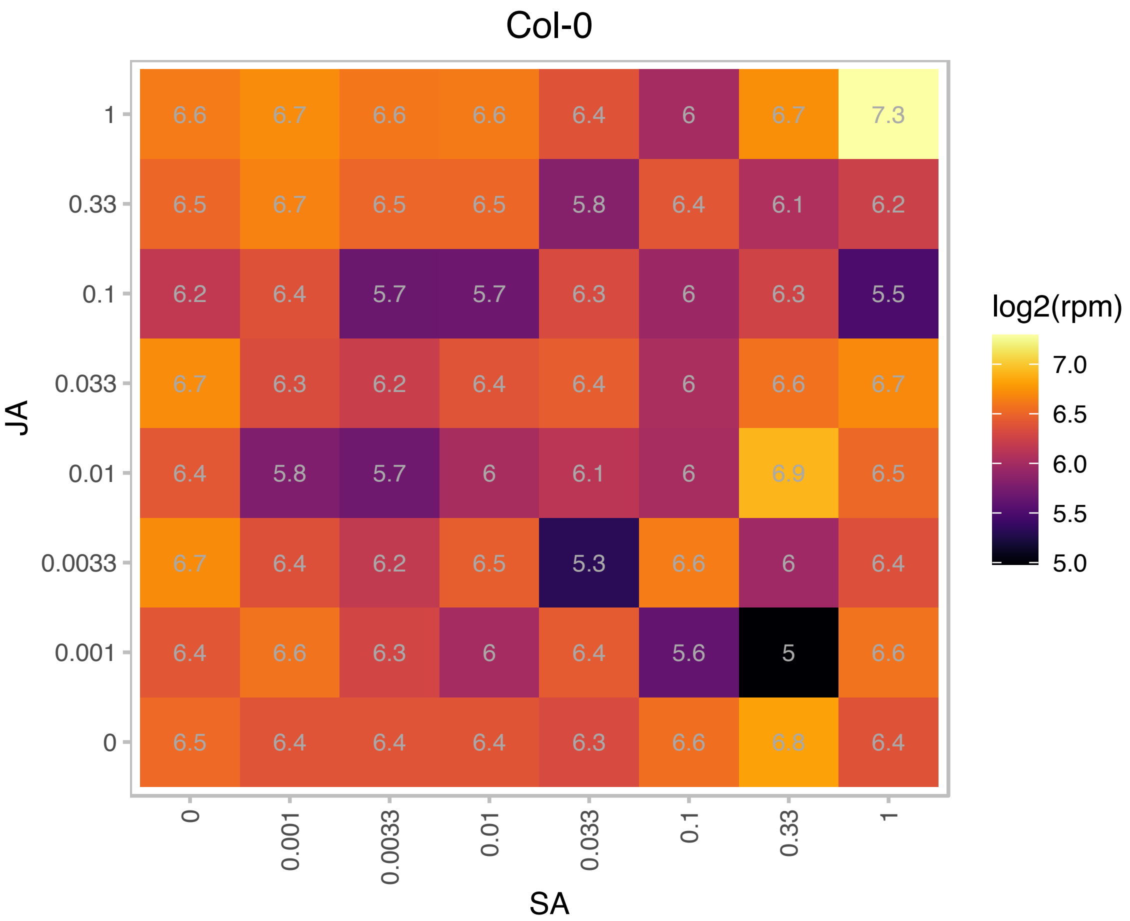


| | |
|---|---|
| Gene | AT4G24930 |
| is.expressed_genes | TRUE |
| is.responsive_genes | TRUE |
| is.apcluster | G-01 |

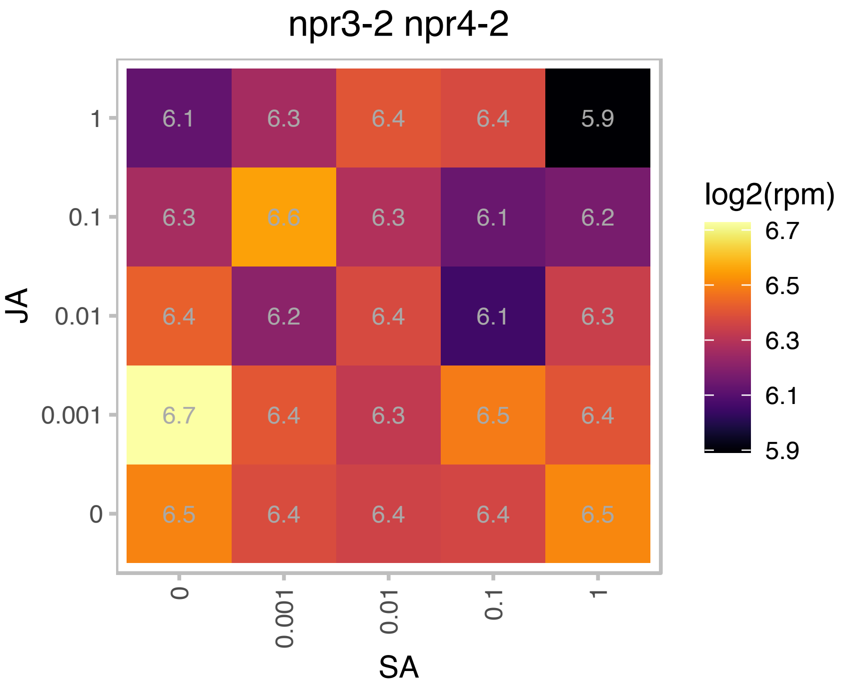


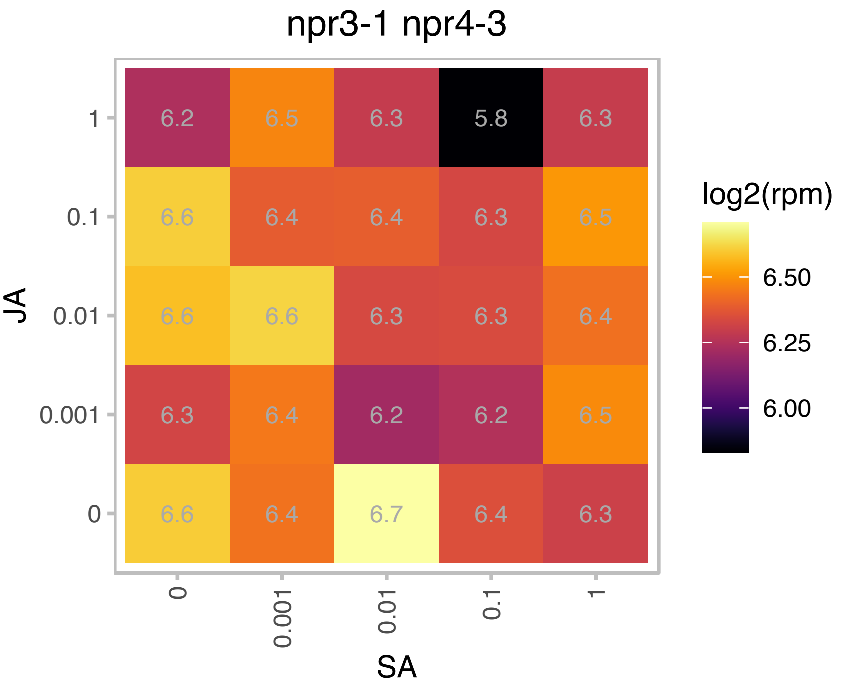

### AT4G29670.1 : ACHT2

atypical CYS HIS rich thioredoxin 2

Encodes a member of the thioredoxin family protein.  Located in the chloroplast.  Shows high activity towards the chloroplast 2-Cys peroxiredoxin A, and poor activity towards the chloroplast NADP-malate dehydrogenase.

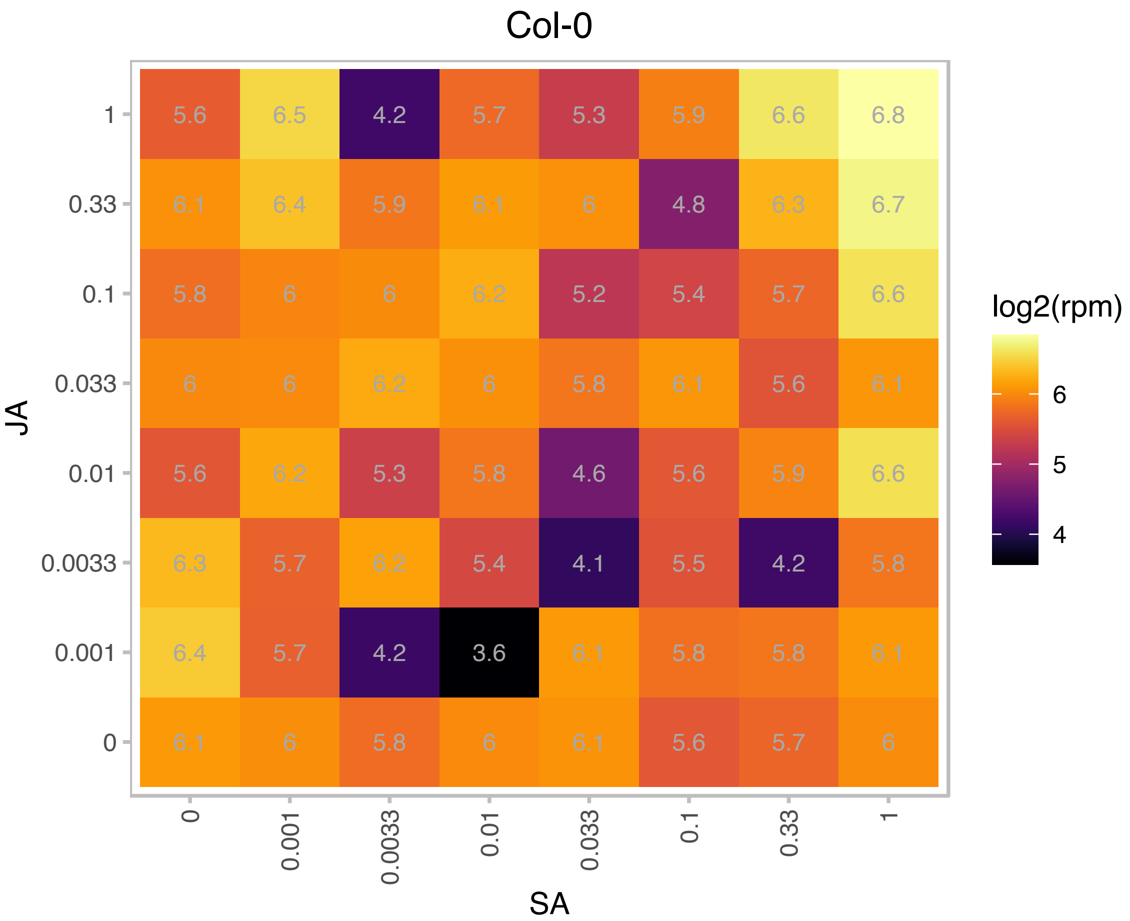


| | |
|---|---|
| Gene | AT4G29670 |
| is.expressed_genes | TRUE |
| is.responsive_genes | TRUE |
| is.apcluster | G-01 |

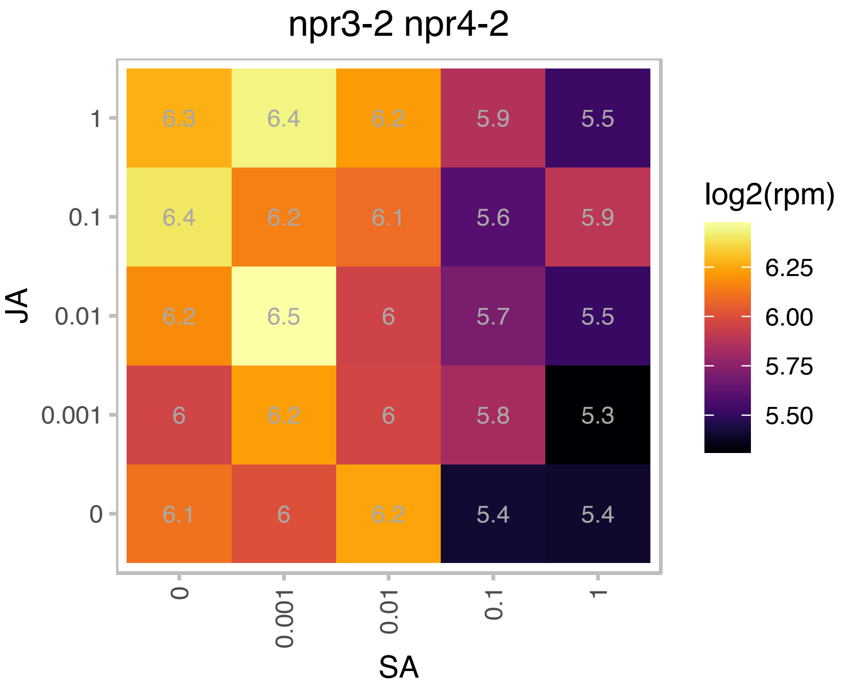


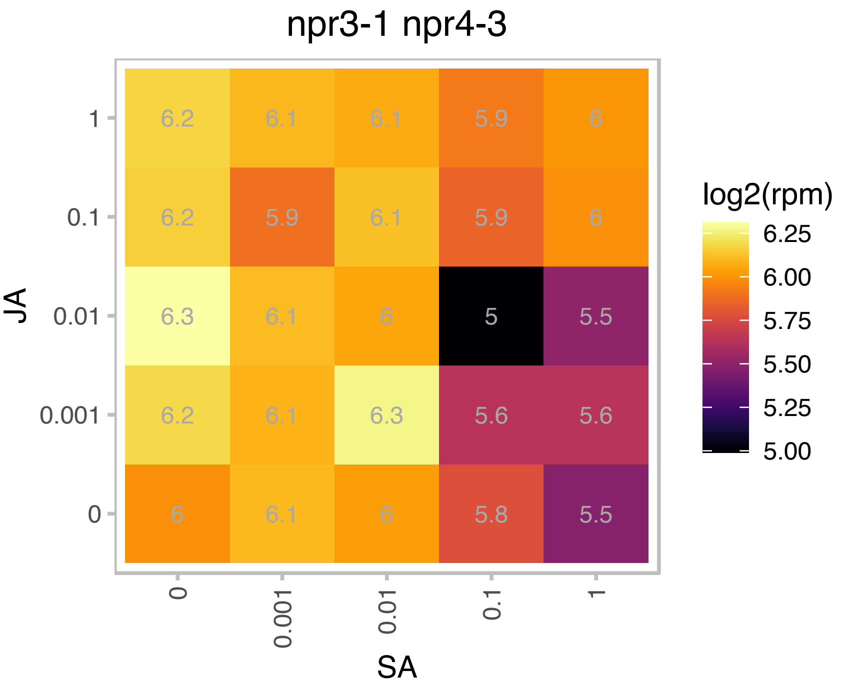

### AT5G08410.1 : FTRA2

ferredoxin/thioredoxin reductase subunit A (variable subunit) 2

| | |
|---|---|
| Gene | AT5G08410 |
| is.expressed_genes | TRUE |
| is.responsive_genes | TRUE |
| is.apcluster | G-01 |

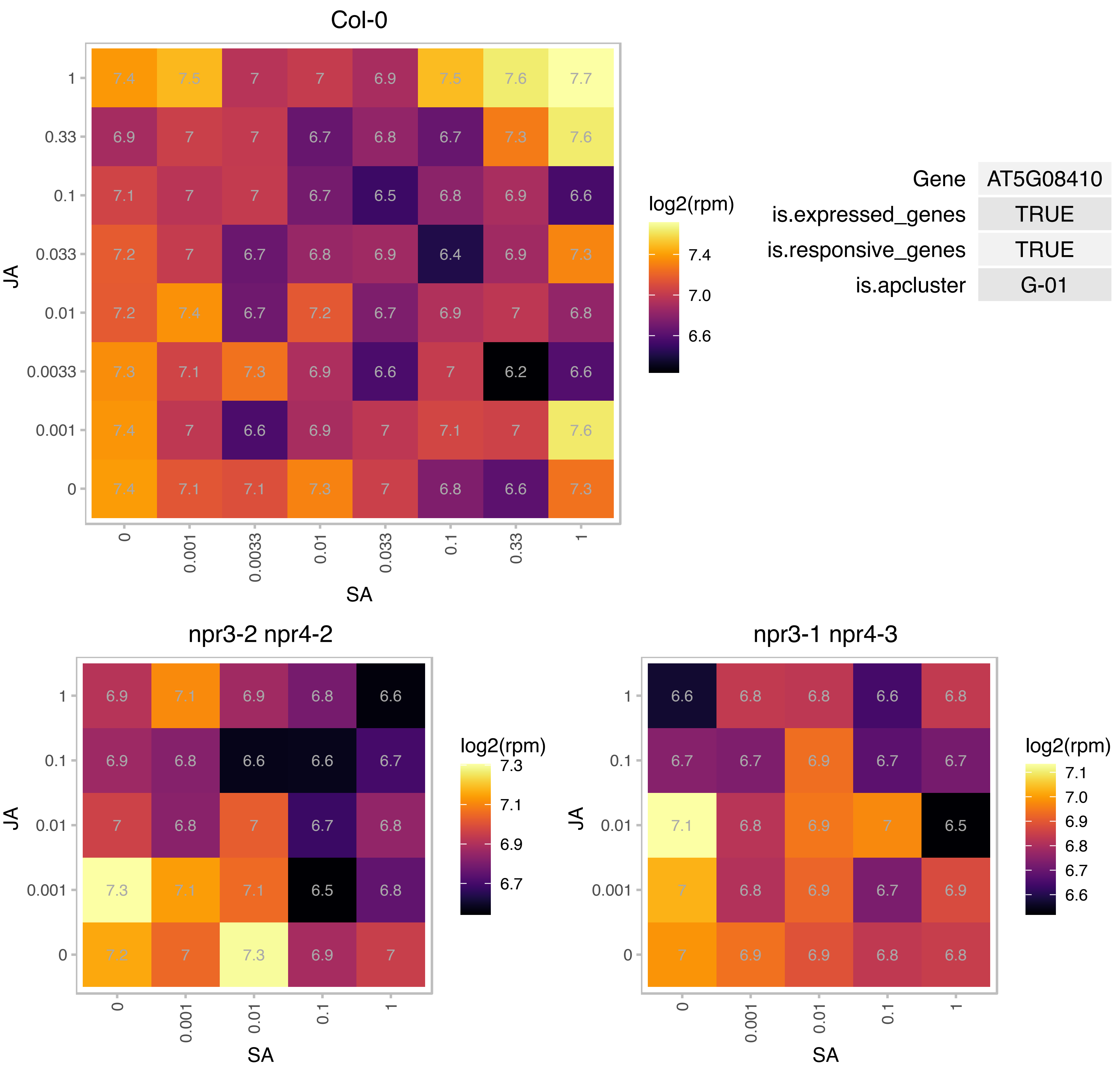

# AT5G11900.1 : NA

Translation initiation factor SUI1 family protein

| | |
|---|---|
| Gene | AT5G11900 |
| is.expressed_genes | TRUE |
| is.responsive_genes | TRUE |
| is.apcluster | G-01 |

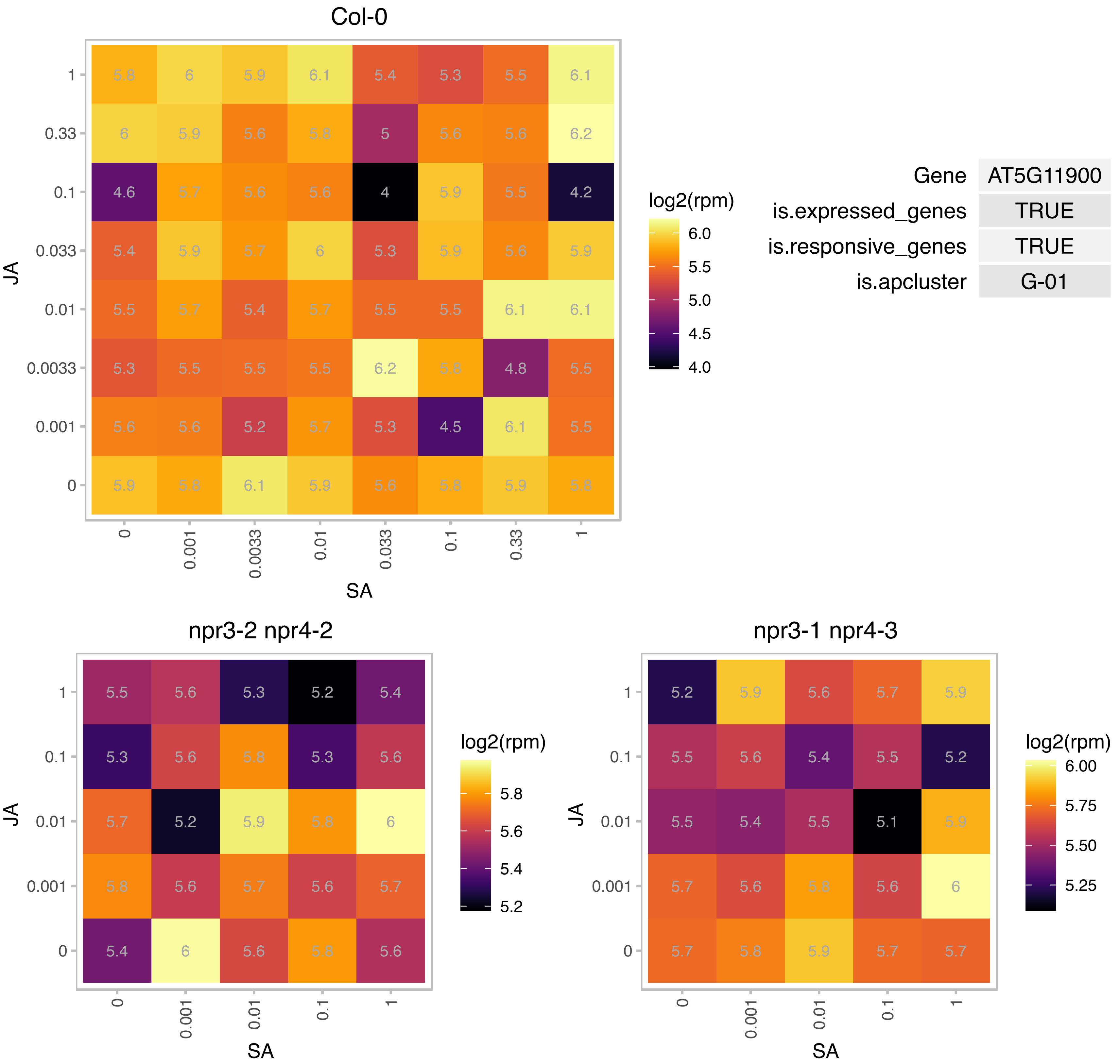

### AT5G12130.1 : ATTERC, PDE149

integral membrane TerC family protein

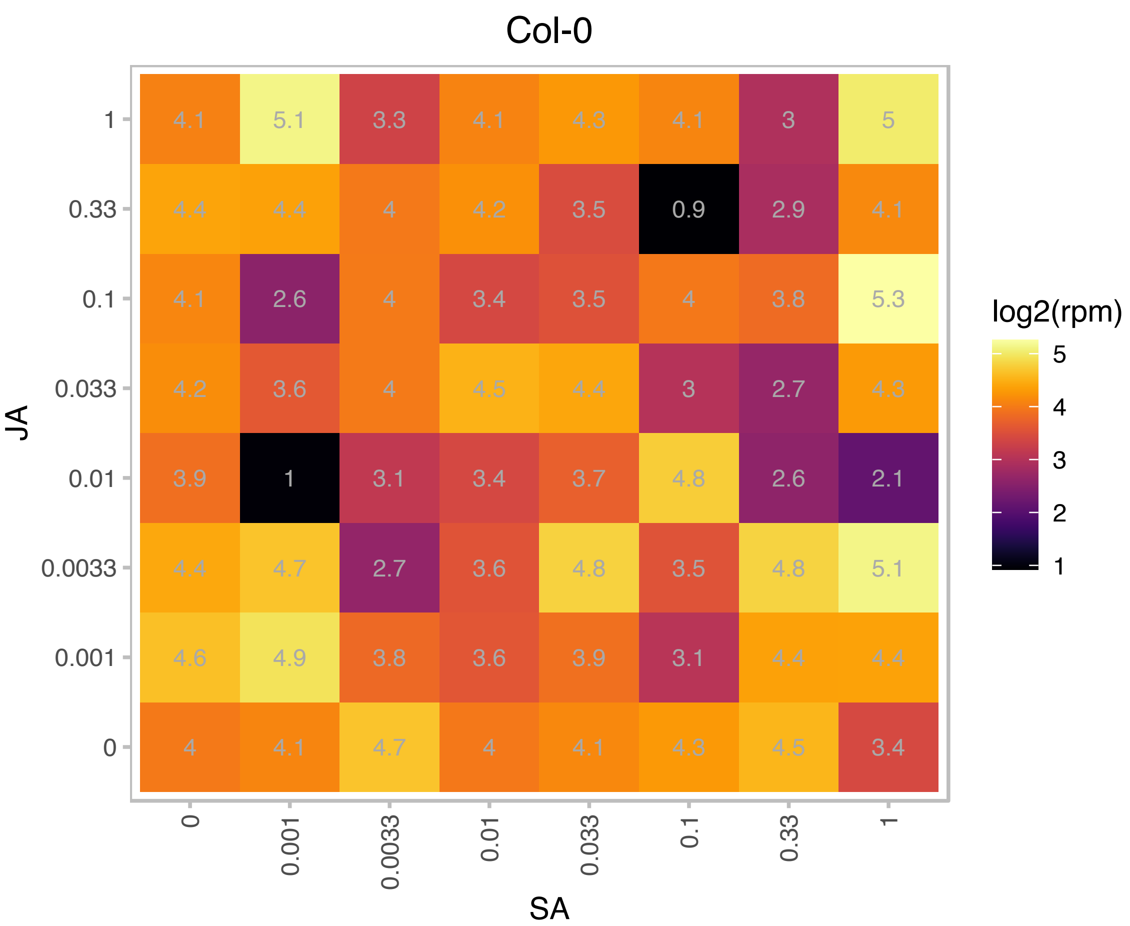


| | |
|---|---|
| Gene | AT5G12130 |
| is.expressed_genes | TRUE |
| is.responsive_genes | TRUE |
| is.apcluster | G-01 |

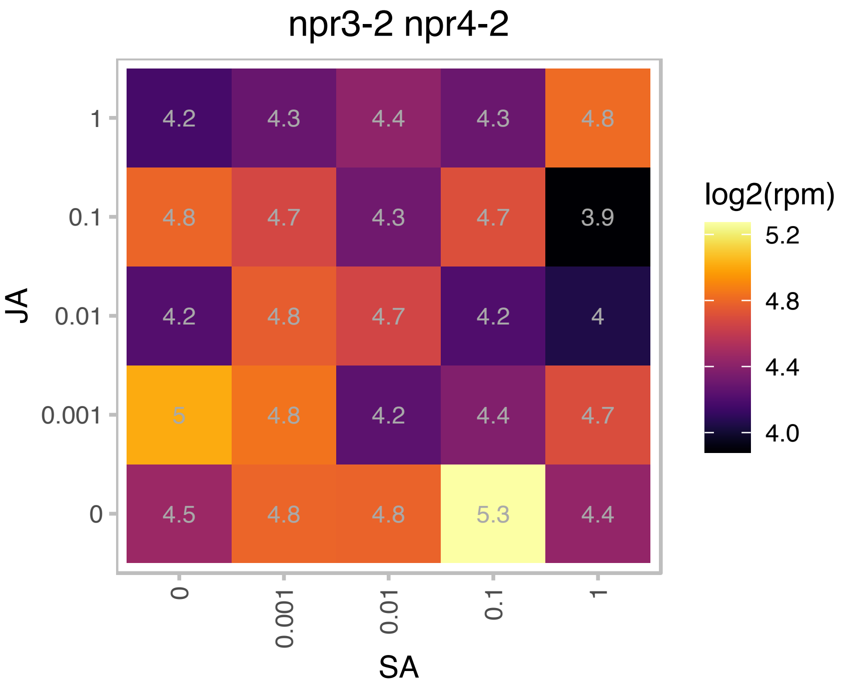


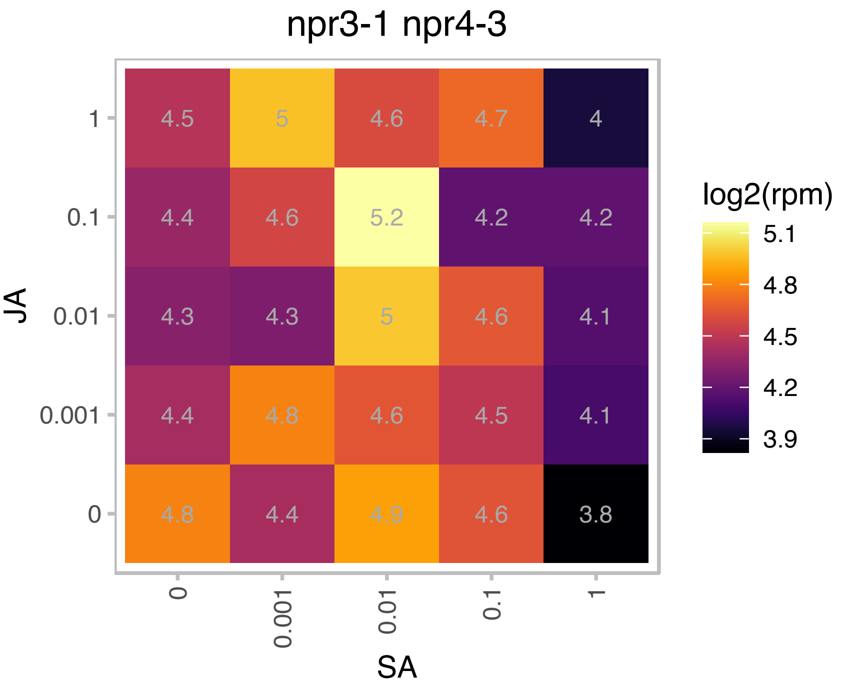

### AT5G21920.1 : ATYLMG2, YLMG2

YGGT family protein

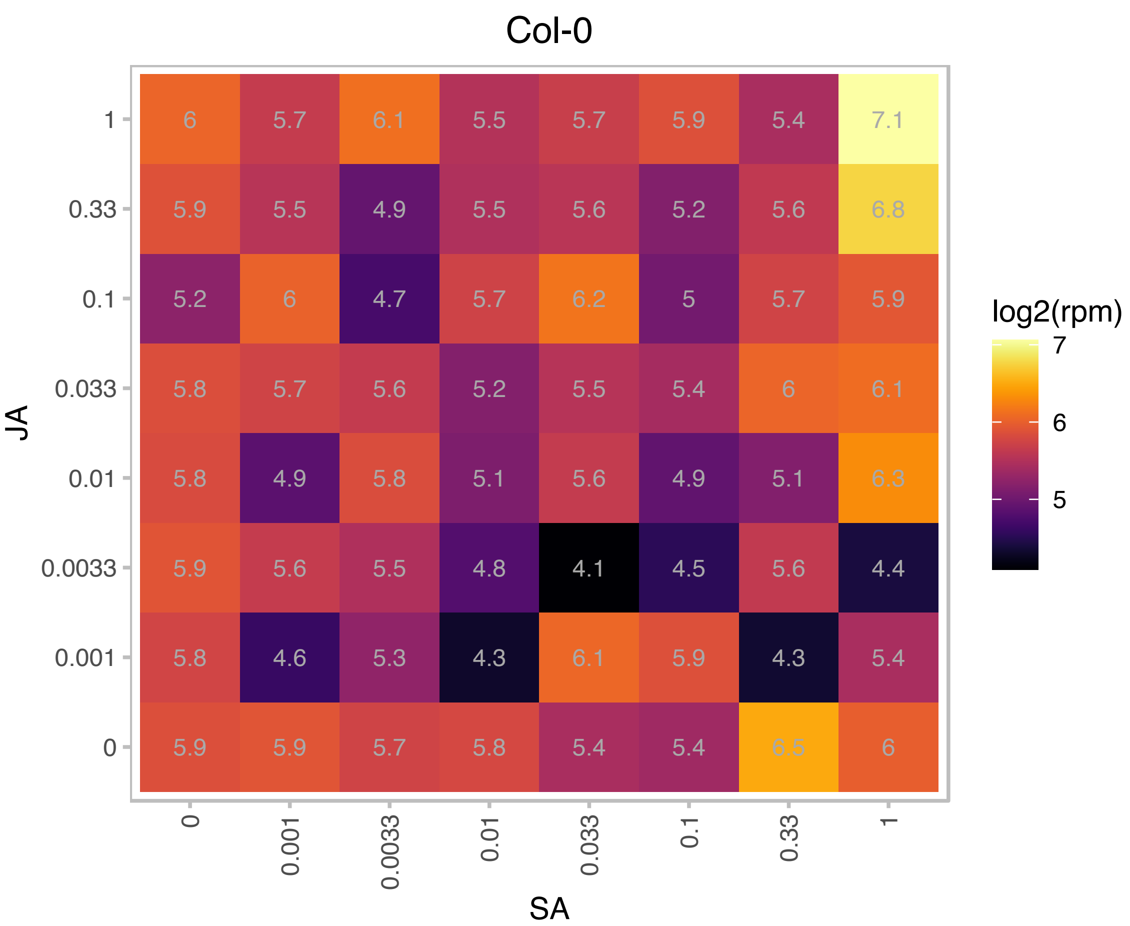


| | |
|---|---|
| Gene | AT5G21920 |
| is.expressed_genes | TRUE |
| is.responsive_genes | TRUE |
| is.apcluster | G-01 |

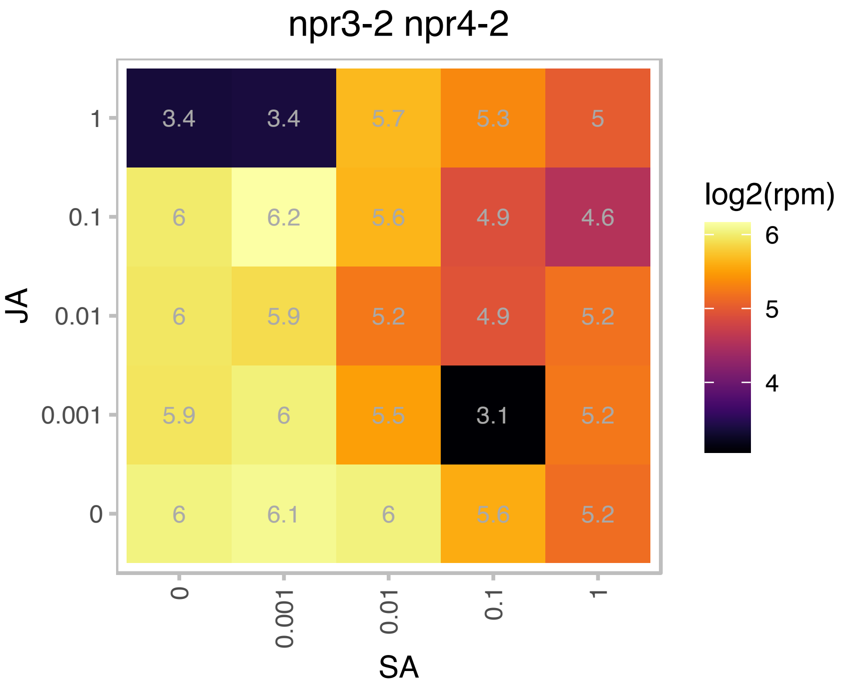


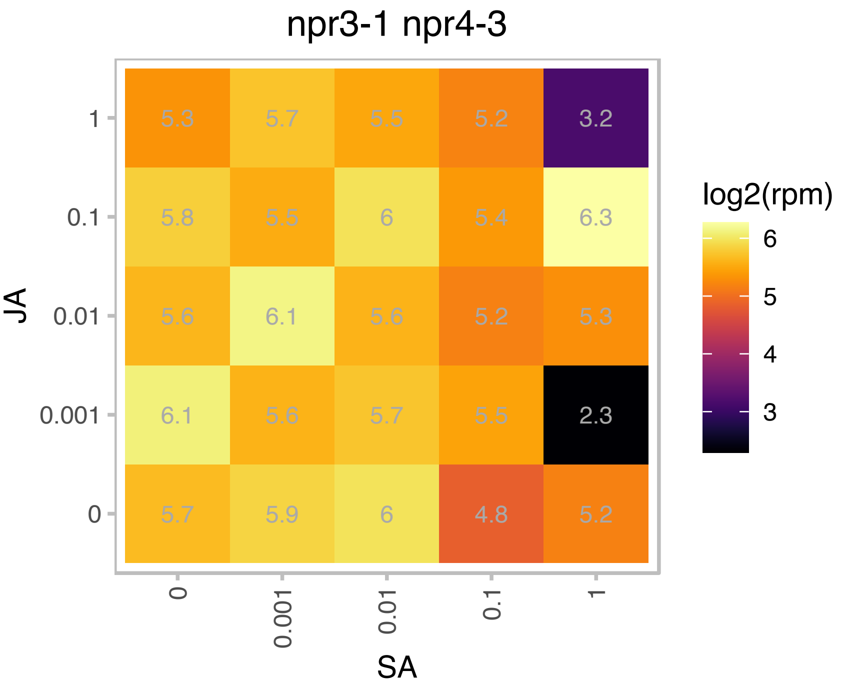

### AT5G42900.1 : COR27

cold regulated gene 27

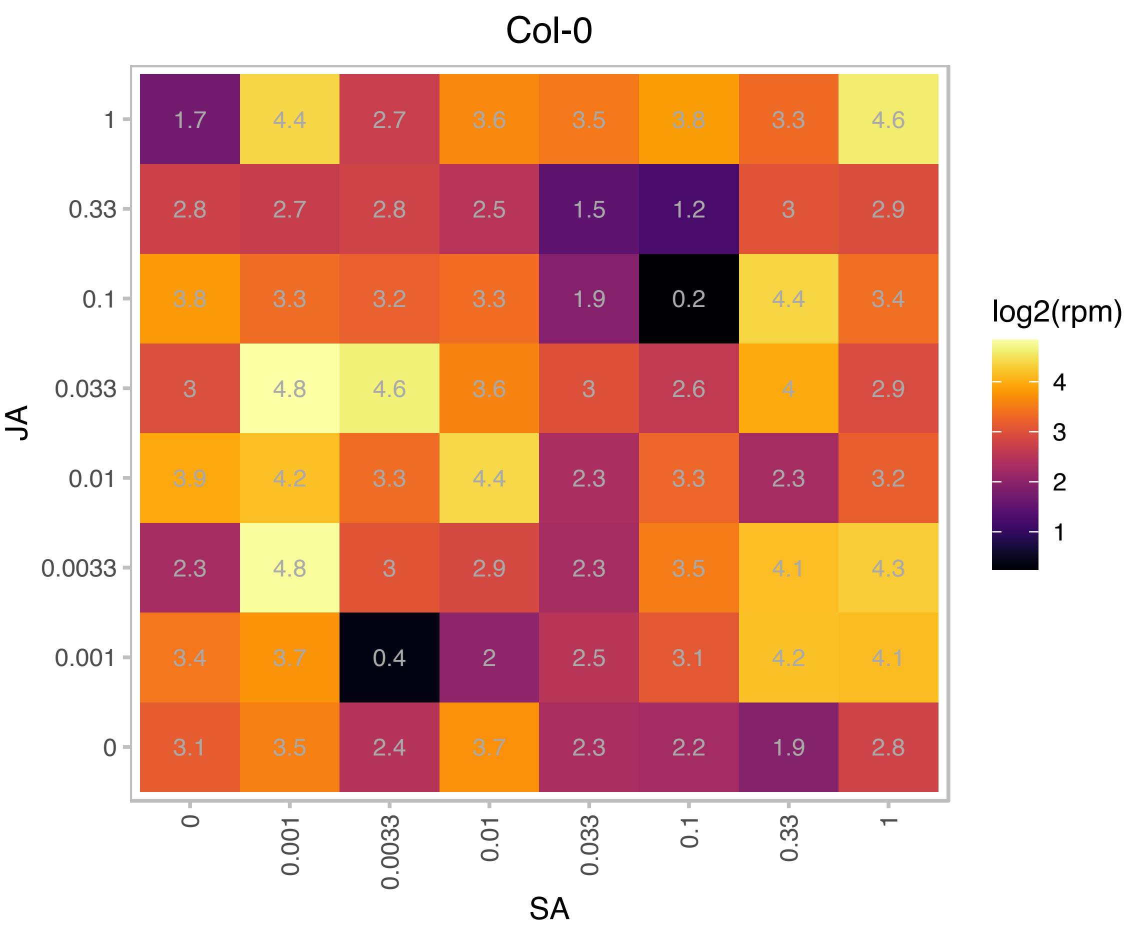


| | |
|---|---|
| Gene | AT5G42900 |
| is.expressed_genes | TRUE |
| is.responsive_genes | TRUE |
| is.apcluster | G-01 |

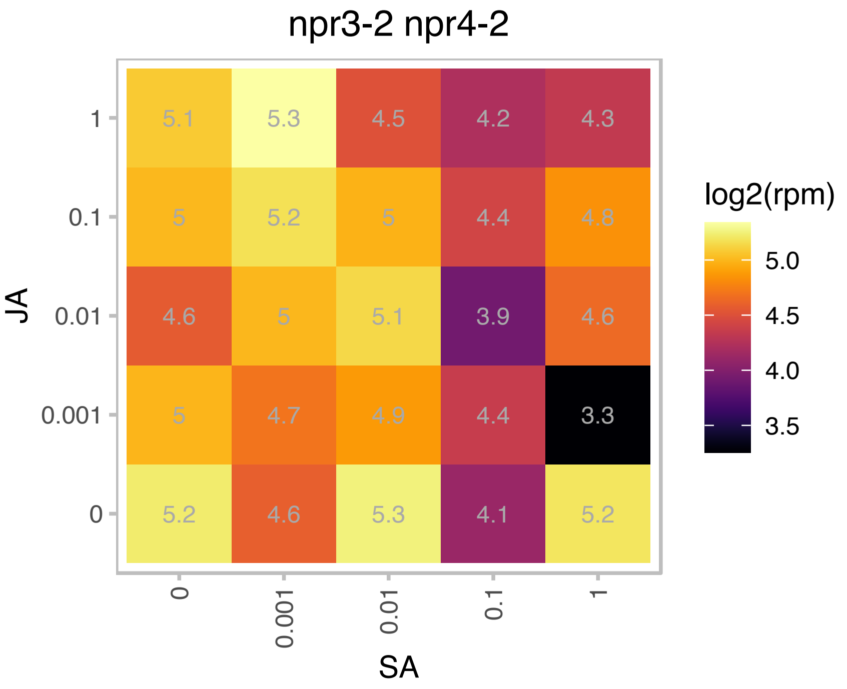


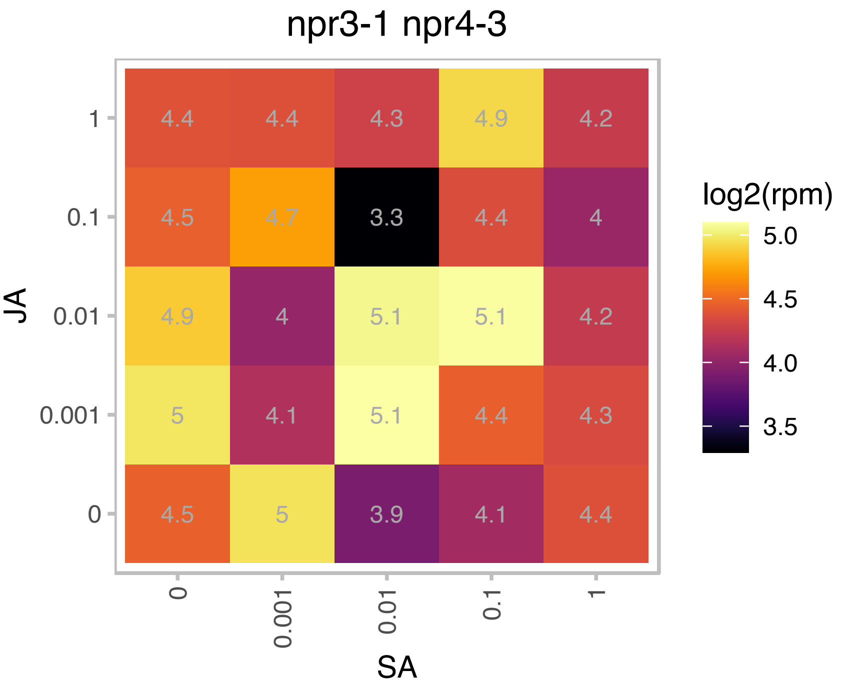

### AT5G46110.1 : APE2, TPT

Glucose-6-phosphate/phosphate translocator-related
Encodes a chloroplast triose phosphate / 3-phosphoglycerate translocator that transports triose phosphates derived from the Calvin cycle in the stroma to the cytosol for use in sucrose synthesis and other biosynthetic processes. A tpt mutant has altered acclimation responses. The mRNA is cell-to-cell mobile.

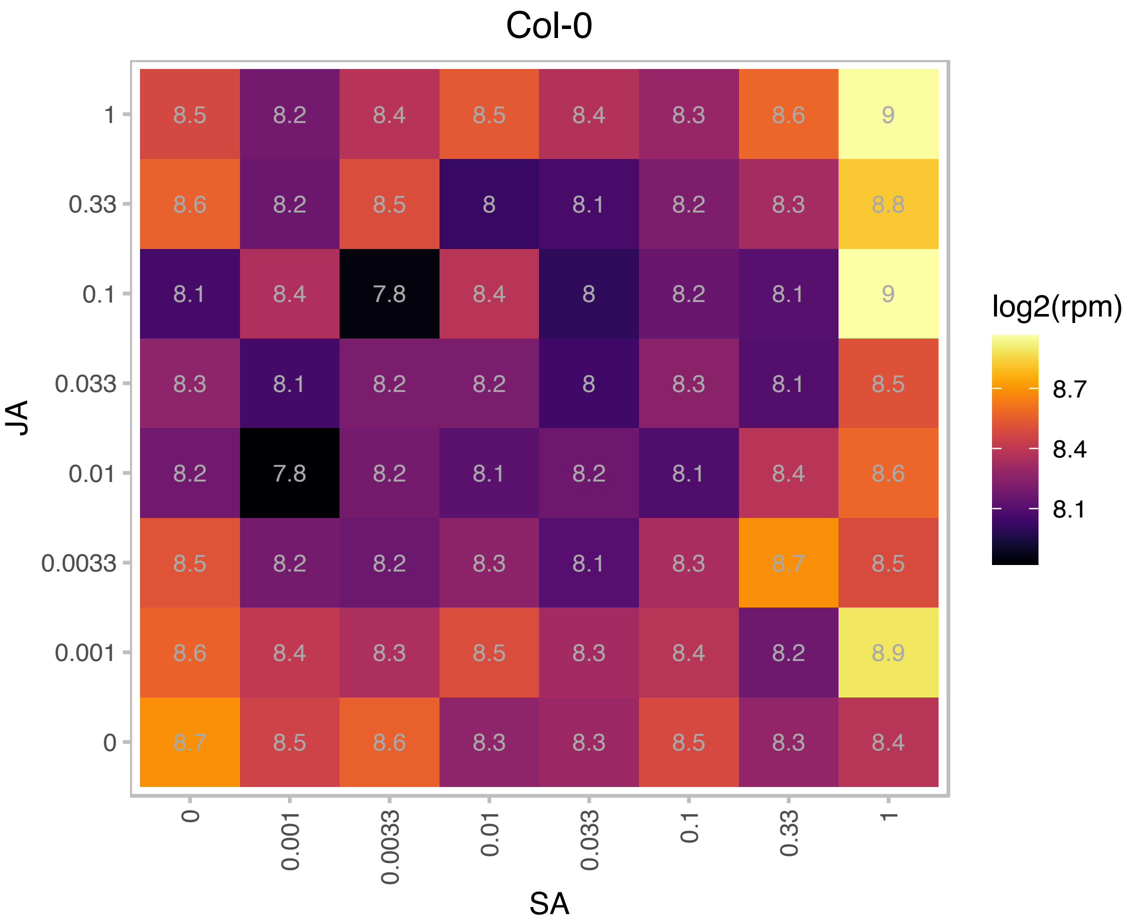


| | |
|---|---|
| Gene | AT5G46110 |
| is.expressed_genes | TRUE |
| is.responsive_genes | TRUE |
| is.apcluster | G-01 |

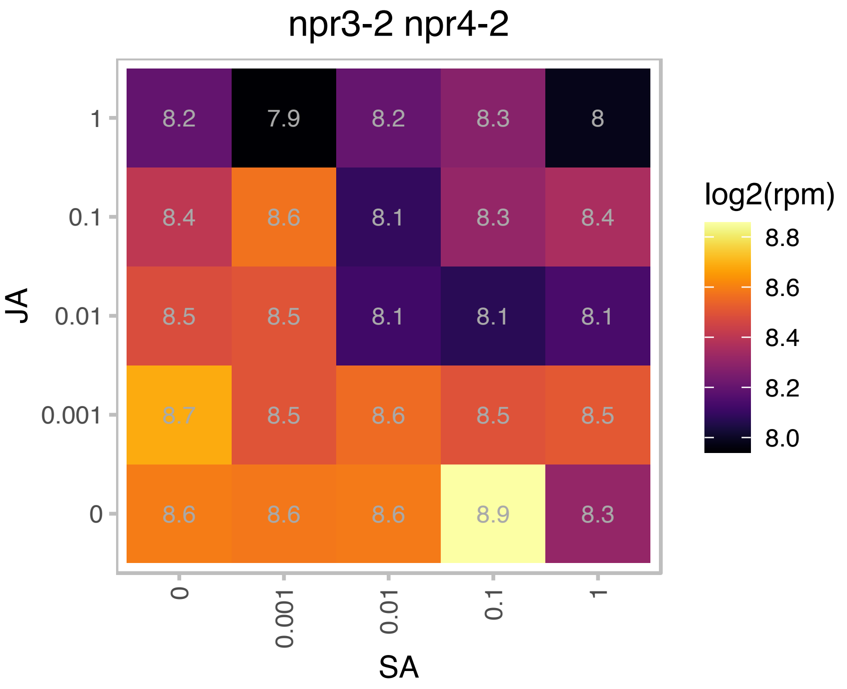


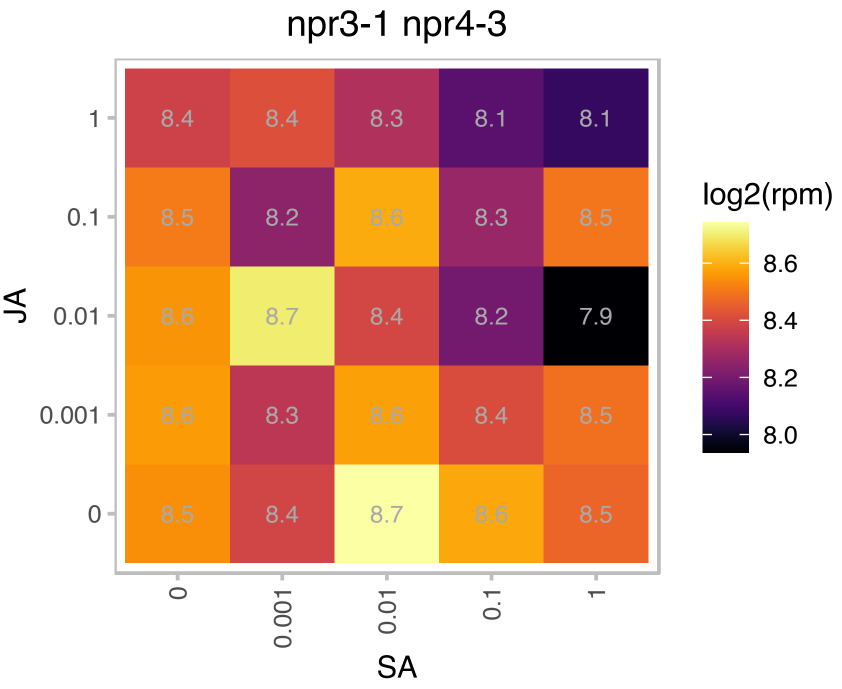

### AT5G48470.1 : PRDA1

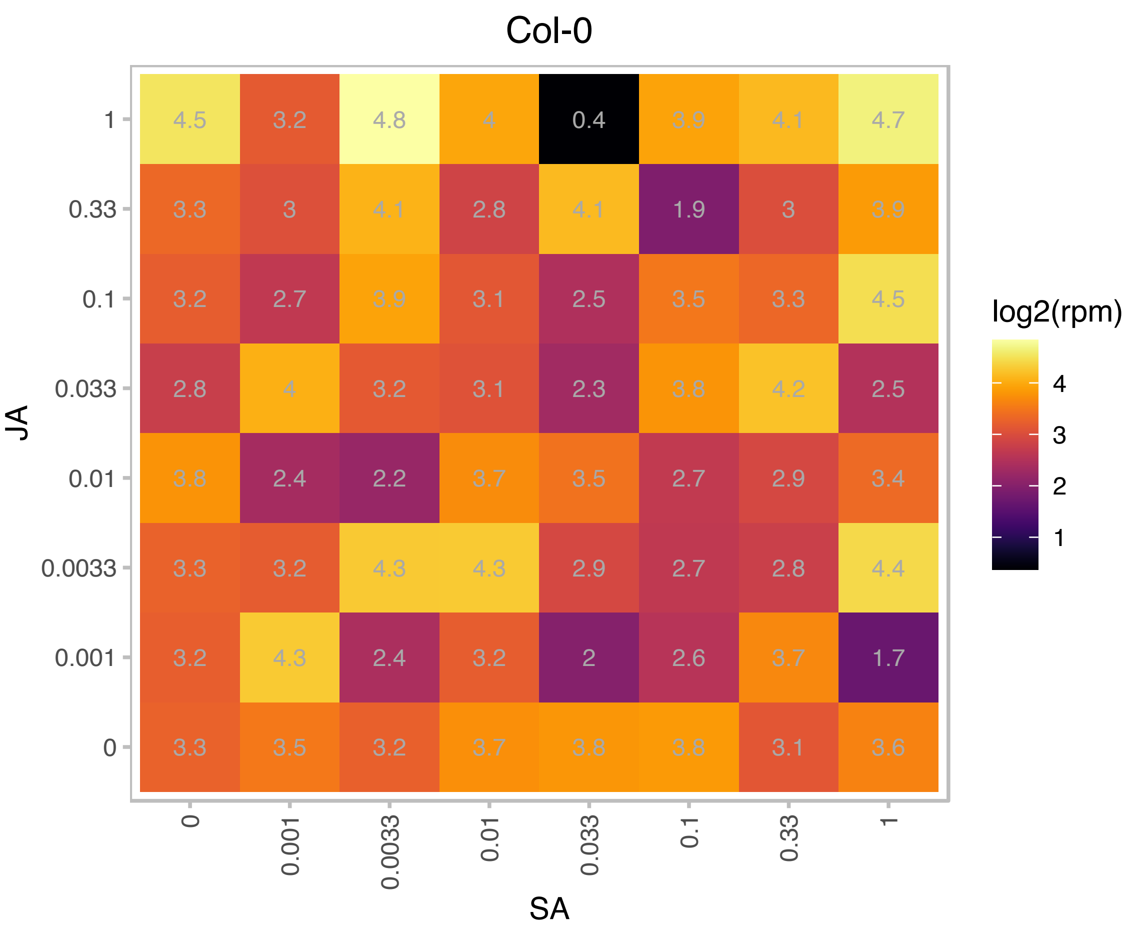


| | |
|---|---|
| Gene | AT5G48470 |
| is.expressed_genes | TRUE |
| is.responsive_genes | TRUE |
| is.apcluster | G-01 |

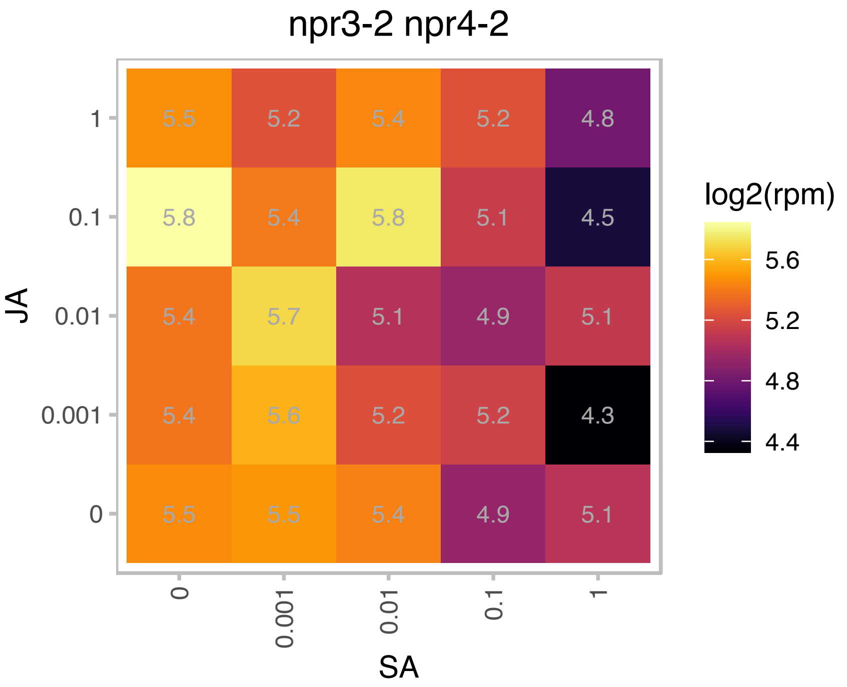


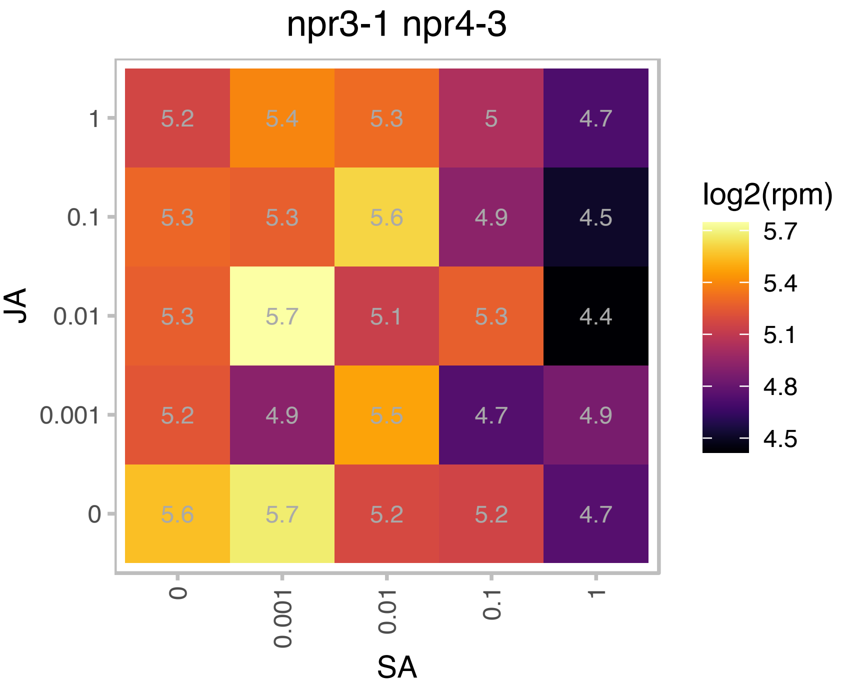

### AT5G54290.1 : CcdA

cytochrome c biogenesis protein family

Encodes CcdA, a thylakoid membrane protein required for the transfer of reducing equivalents from stroma to thylakoid lumen.

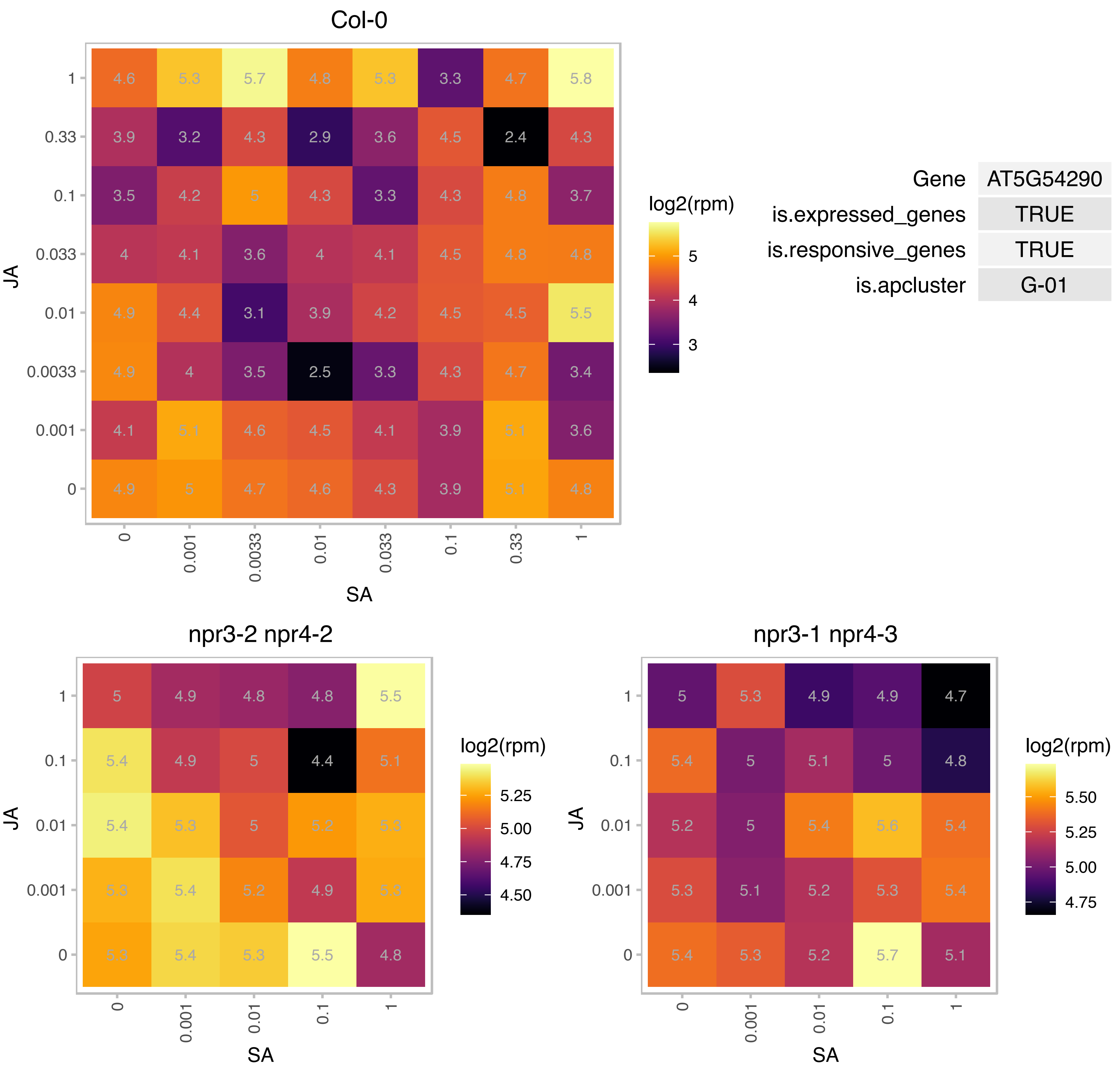


| | |
|---|---|
| Gene | AT5G54290 |
| is.expressed_genes | TRUE |
| is.responsive_genes | TRUE |
| is.apcluster | G-01 |

### AT5G54770.1 : THI1, THI4, TZ

thiazole biosynthetic enzyme, chloroplast (ARA6) (THI1) (THI4)

Encodes a thiamine biosynthetic gene that has a dual function in thiamine biosynthesis and mitochondrial DNA damage tolerance. It appears to be involved in producing the thiazole portion of thiamine (vitamin B1). A crystal structure of the protein reveals that it forms a 2-ring homo-octamer. The mRNA is cell-to-cell mobile.

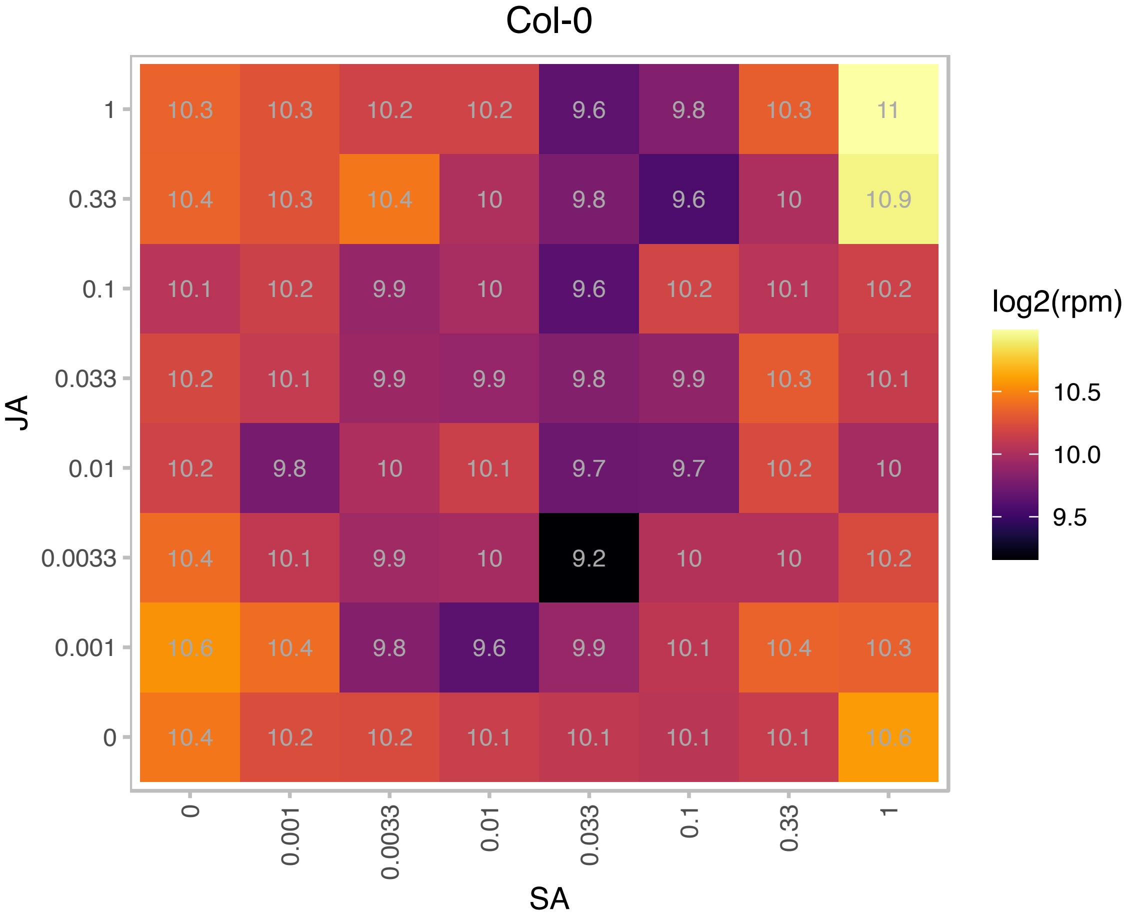


| | |
|---|---|
| Gene | AT5G54770 |
| is.expressed_genes | TRUE |
| is.responsive_genes | TRUE |
| is.apcluster | G-01 |

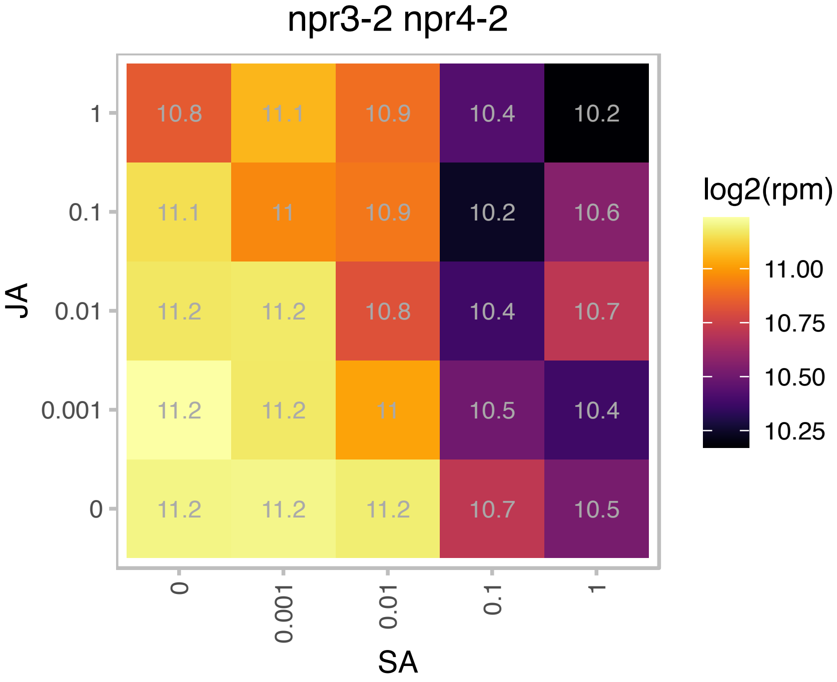


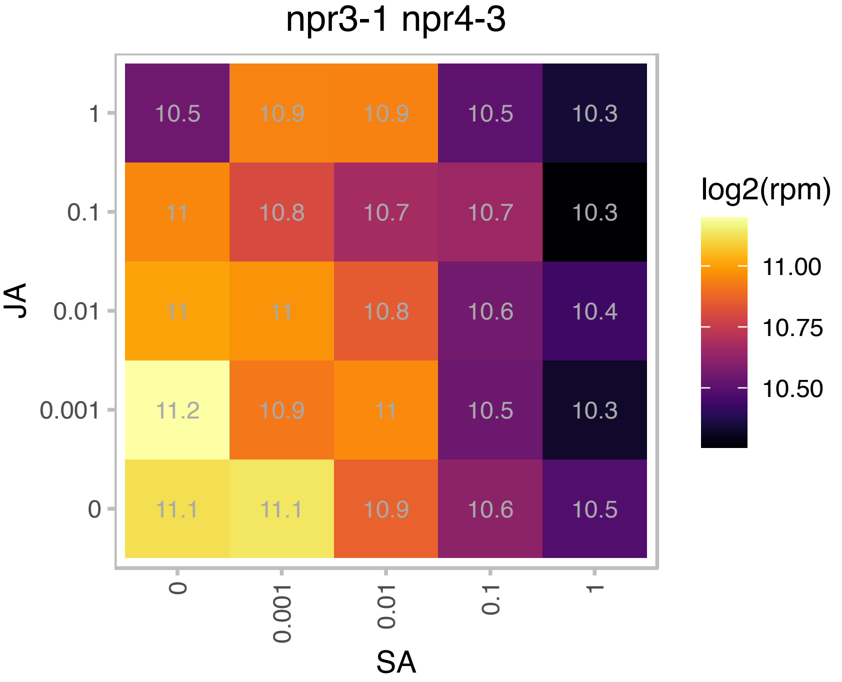

### AT5G56600.1 : PFN3, PRF3

profilin 3
Encodes profilin3, a low-molecular weight, actin monomer-binding protein that regulates the organization of actin cytoskeleton. Originally known as profilin5, and later named profilin3. Expressed in vegetative organs. Mutants have slightly elongated petioles.

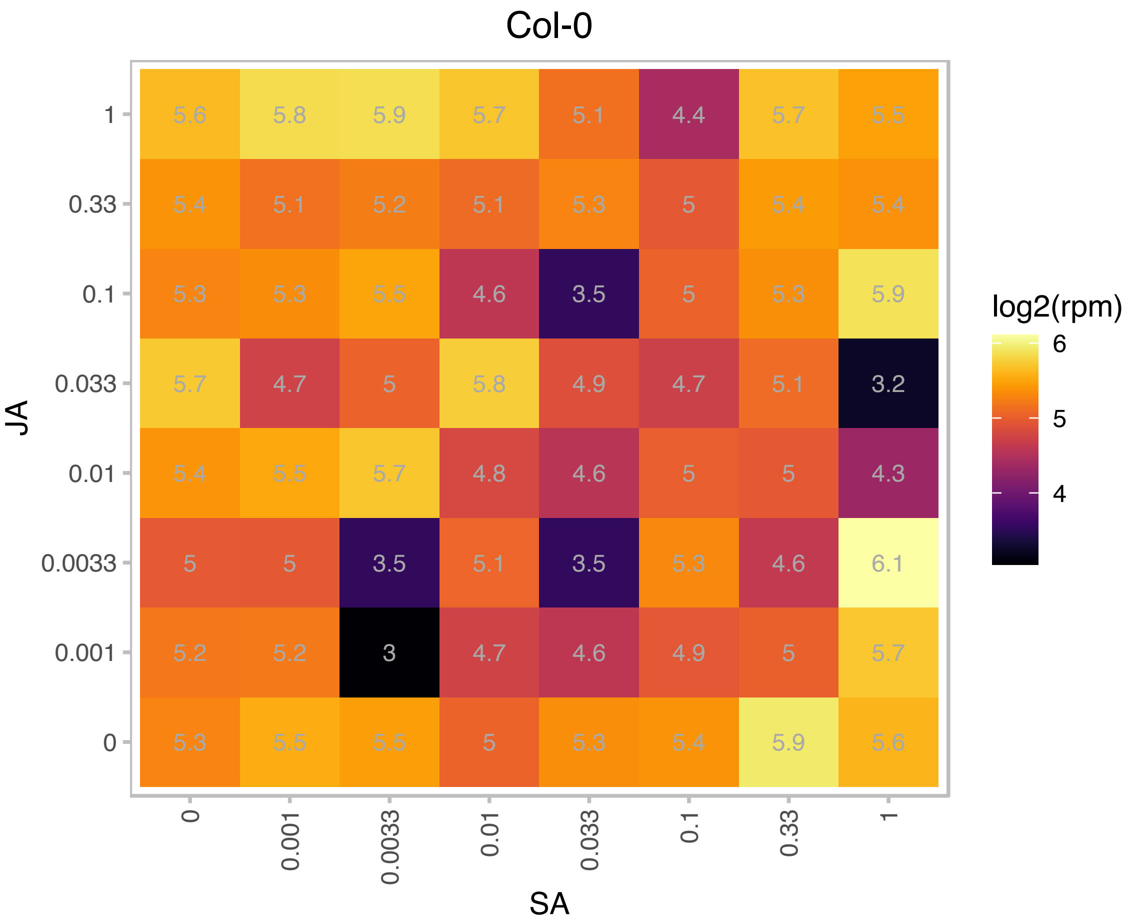


| | |
|---|---|
| Gene | AT5G56600 |
| is.expressed_genes | TRUE |
| is.responsive_genes | TRUE |
| is.apcluster | G-01 |

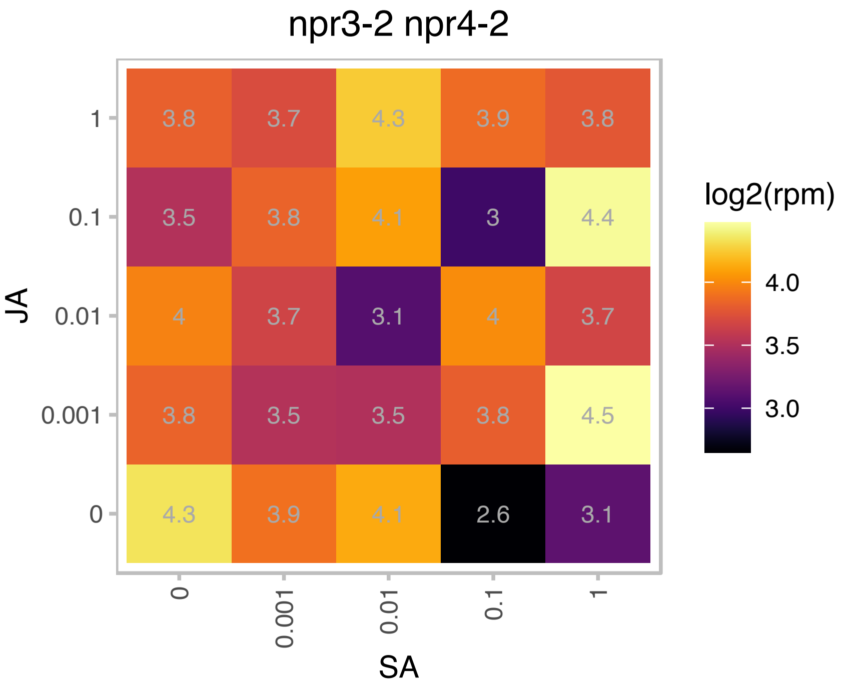


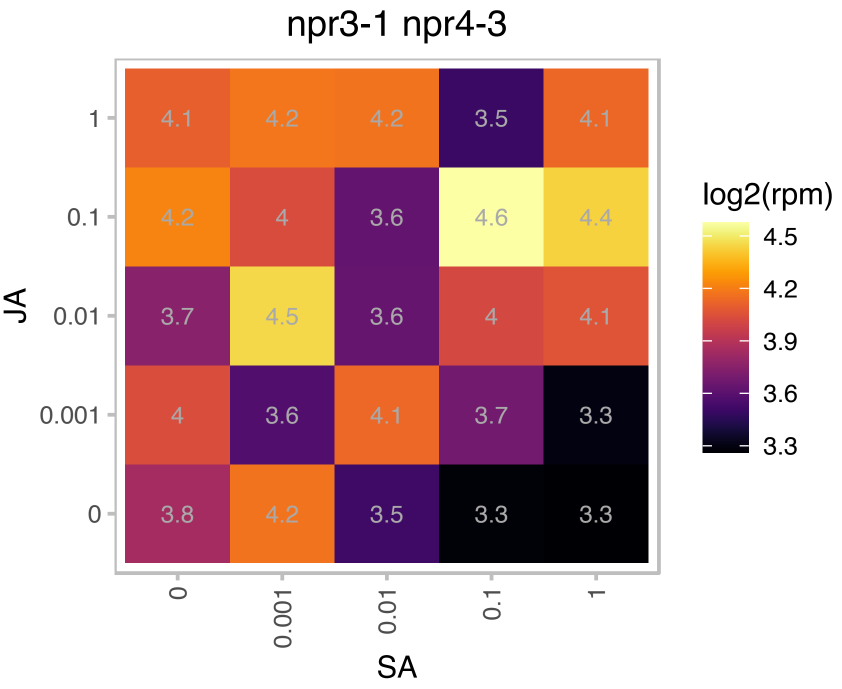

### ATCG01110.1 : NDHH

NAD(P)H dehydrogenase subunit H

Encodes the 49KDa plastid NAD(P)H dehydrogenase subunit H protein.  Its transcription is regulated by an ndhF-specific plastid sigma factor, SIG4.

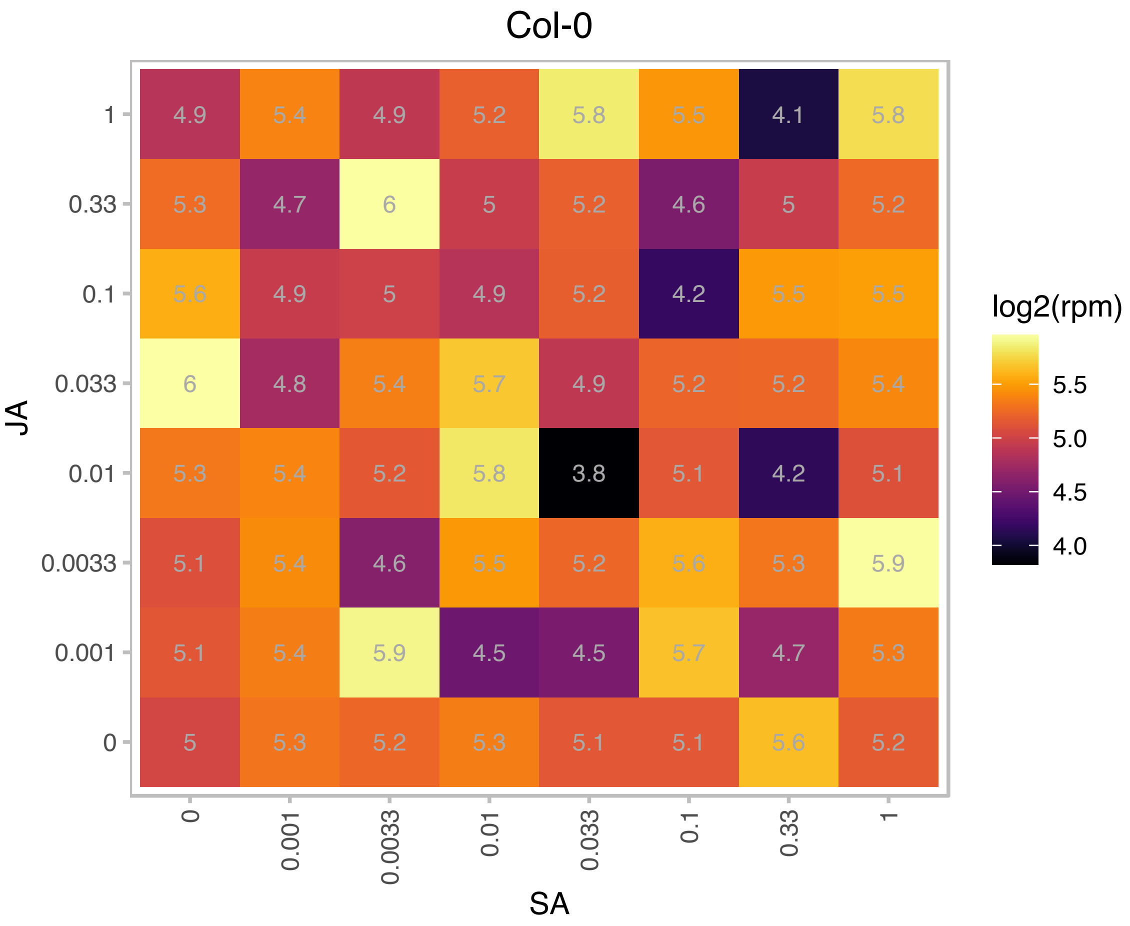


| | |
|---|---|
| Gene | ATCG01110 |
| is.expressed_genes | TRUE |
| is.responsive_genes | TRUE |
| is.apcluster | G-01 |

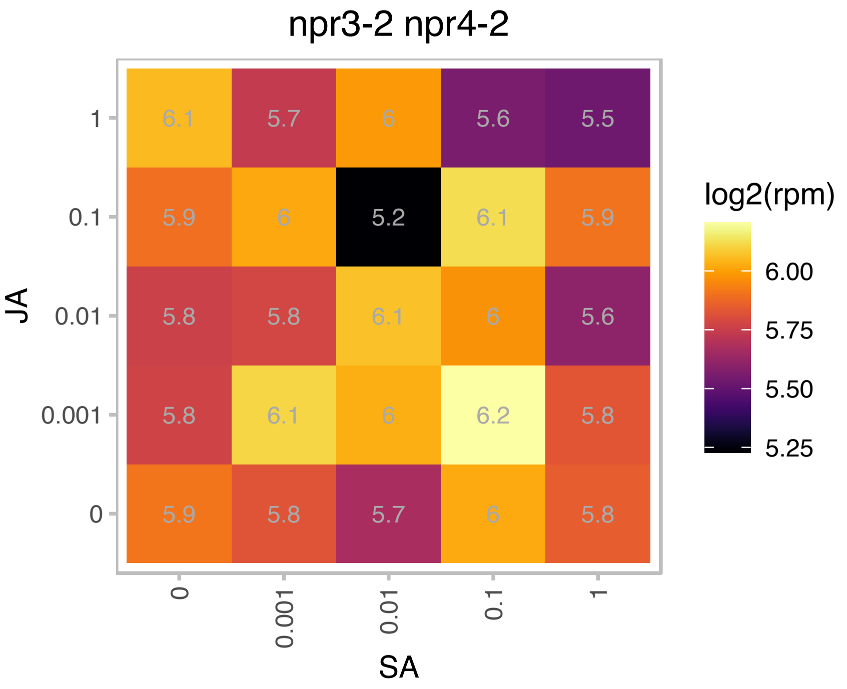


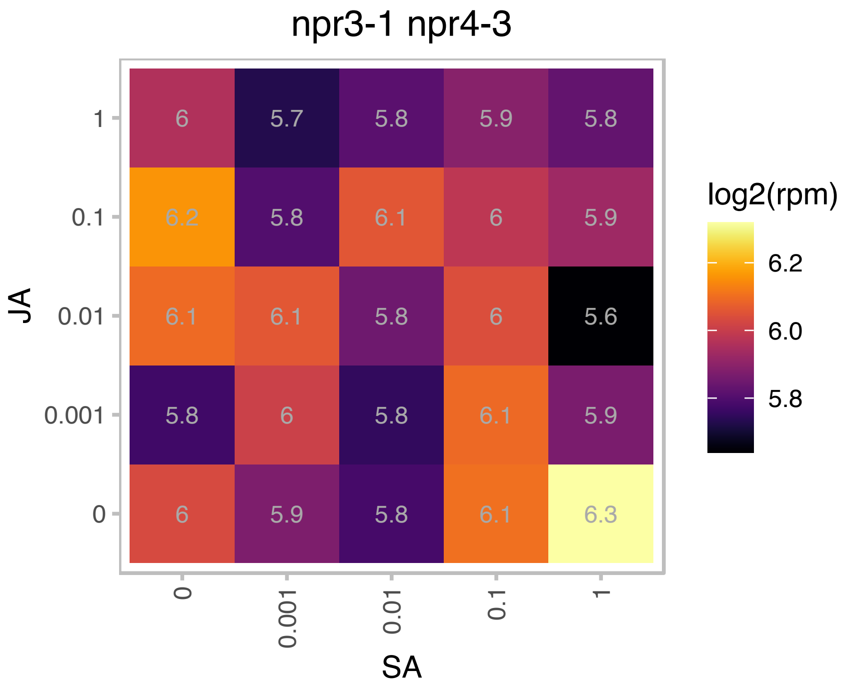

### AT1G06830.1 : CEPD1, ROXY6

Glutaredoxin family protein

Encodes a member of the CC-type glutaredoxin (ROXY) family that has been shown to interact with the transcription factor TGA2. CEPD1 is a non secreted polypeptide that is highly similar to CEPD2 which is another member of a novel family related to glutaredoxins. It is induced by nitrogen starvation. It acts downstream of the CEP1 peptide in systemic N-demand signalling .The RNA is expressed in the phloem of cotelydon and leaf vasculature but the peptide is graft transmissible, traveling from the shoot to the root.

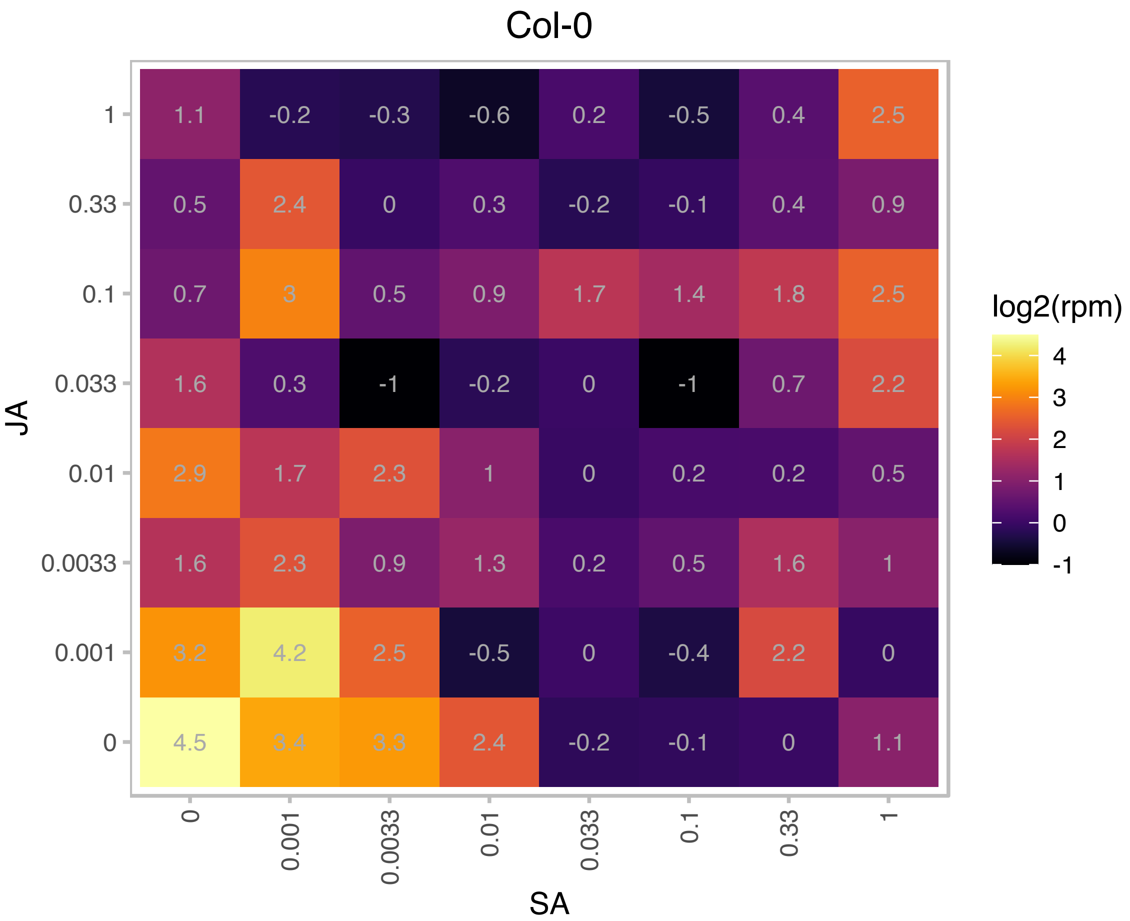


| | |
|---|---|
| Gene | AT1G06830 |
| is.expressed_genes | TRUE |
| is.responsive_genes | TRUE |
| is.apcluster | G-02 |

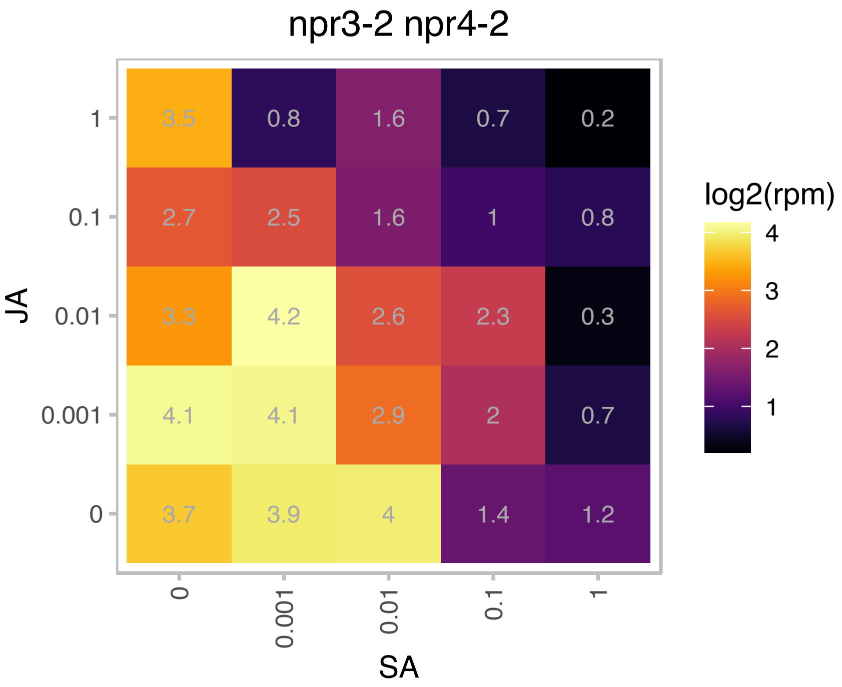


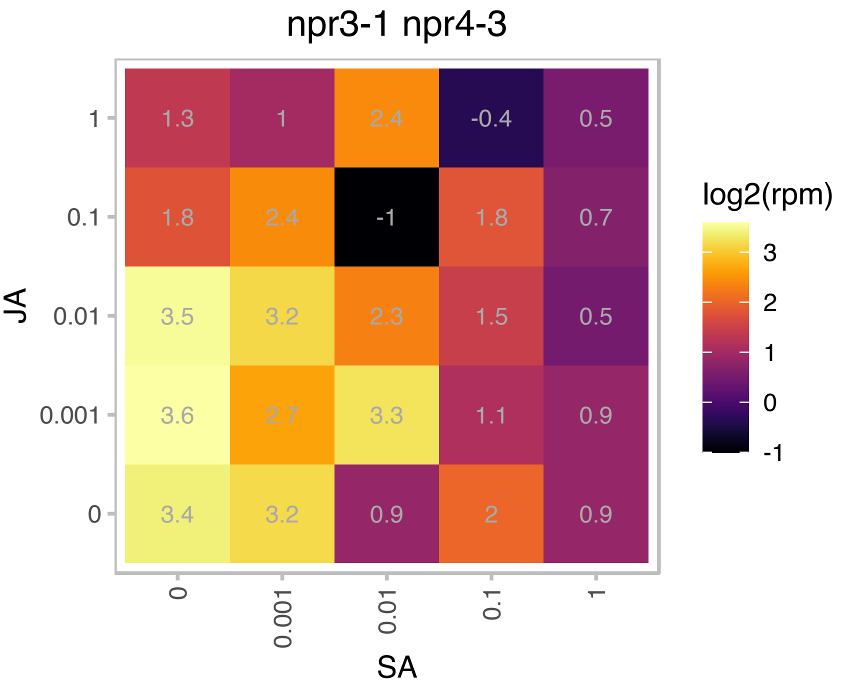

### AT1G16460.1 : ATMST2, ATRDH2, MST2, RDH2, ST2, STI

rhodanese homologue 2

encodes a cytoplasmic thiosulfate:cyanide sulfurtransferase, activity of which increased the rhodanese activity of transgenic yeast.  Can also act as a mercaptopyruvate sulfurtransferase.

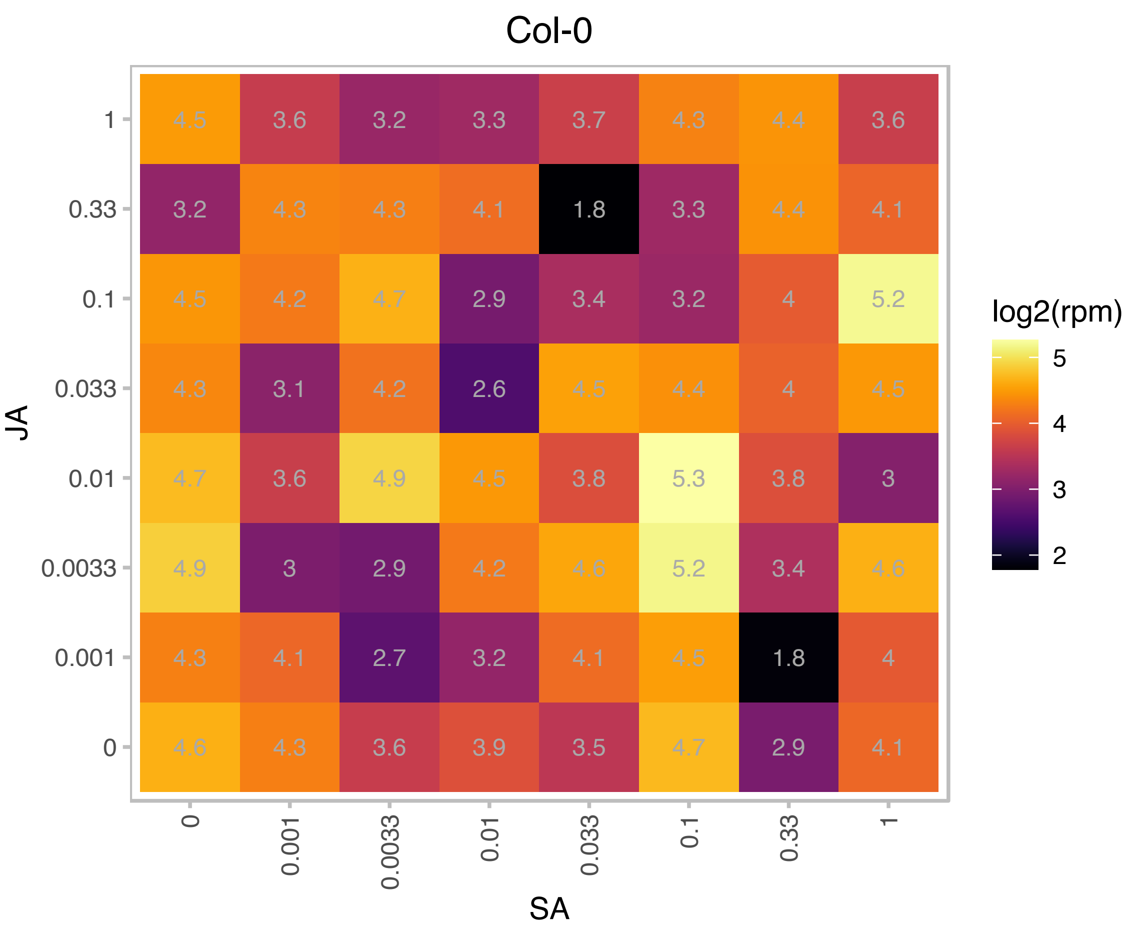


| | |
|---|---|
| Gene | AT1G16460 |
| is.expressed_genes | TRUE |
| is.responsive_genes | TRUE |
| is.apcluster | G-02 |

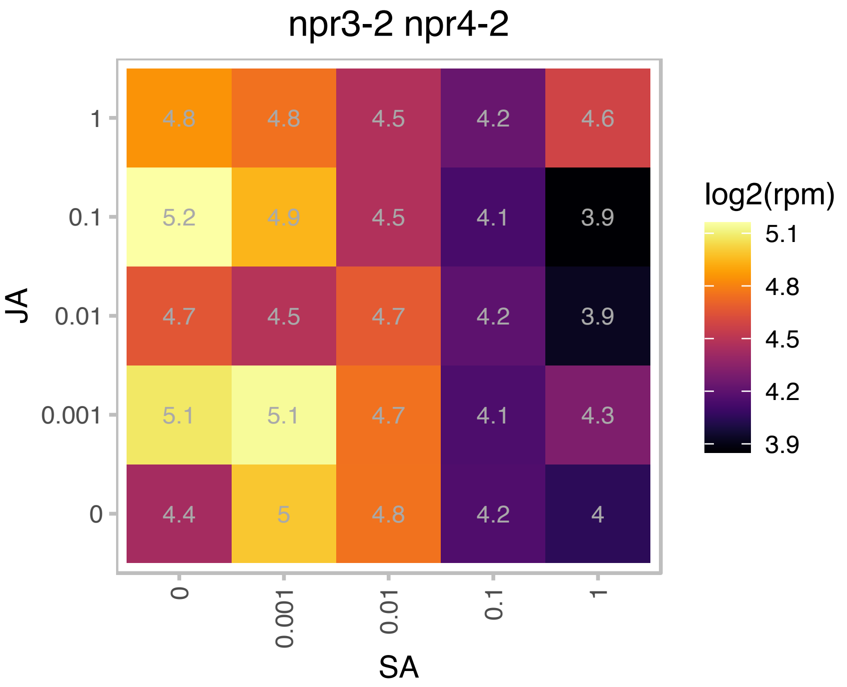


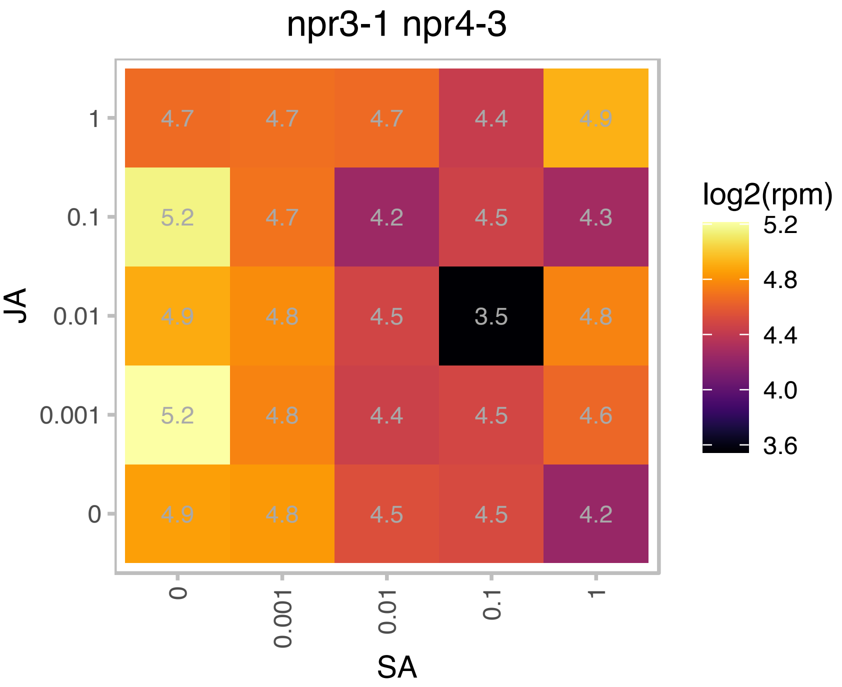

### AT1G32550.1 : FdC2

2Fe-2S ferredoxin-like superfamily protein

Encodes FdC2, a ferredoxin protein capable of alternative electron partitioning.  FdC1 level increases in conditions of acceptor limitation at PSI.

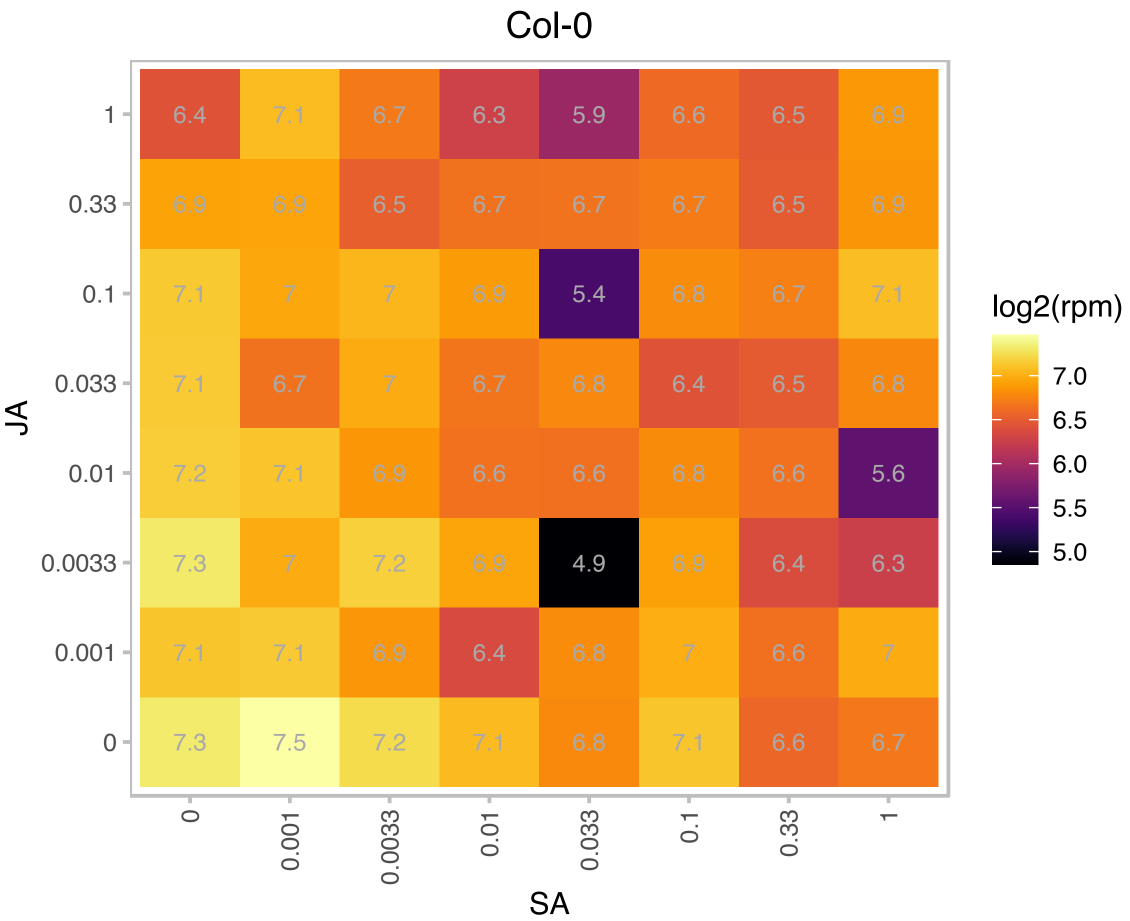


| | |
|---|---|
| Gene | AT1G32550 |
| is.expressed_genes | TRUE |
| is.responsive_genes | TRUE |
| is.apcluster | G-02 |

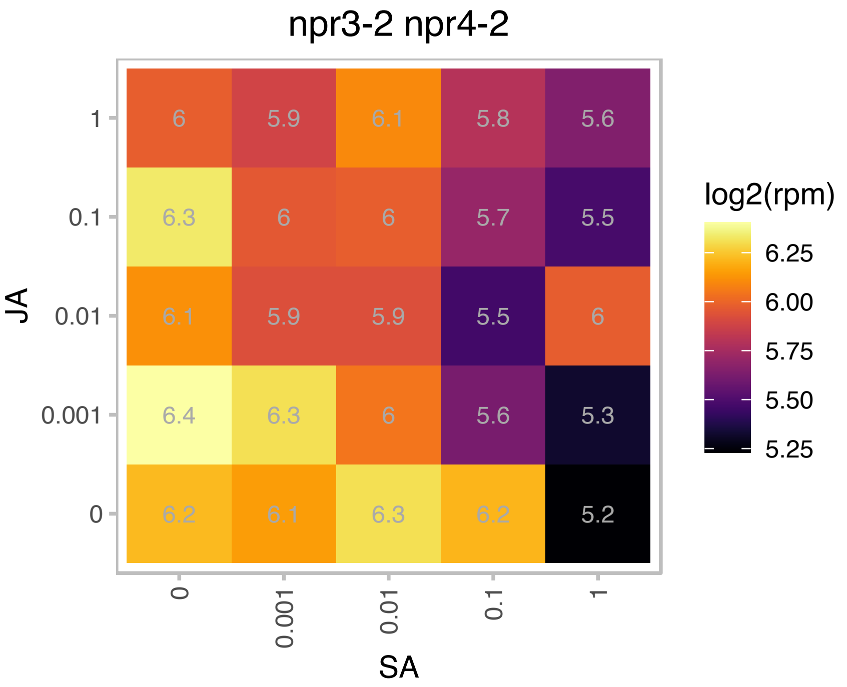


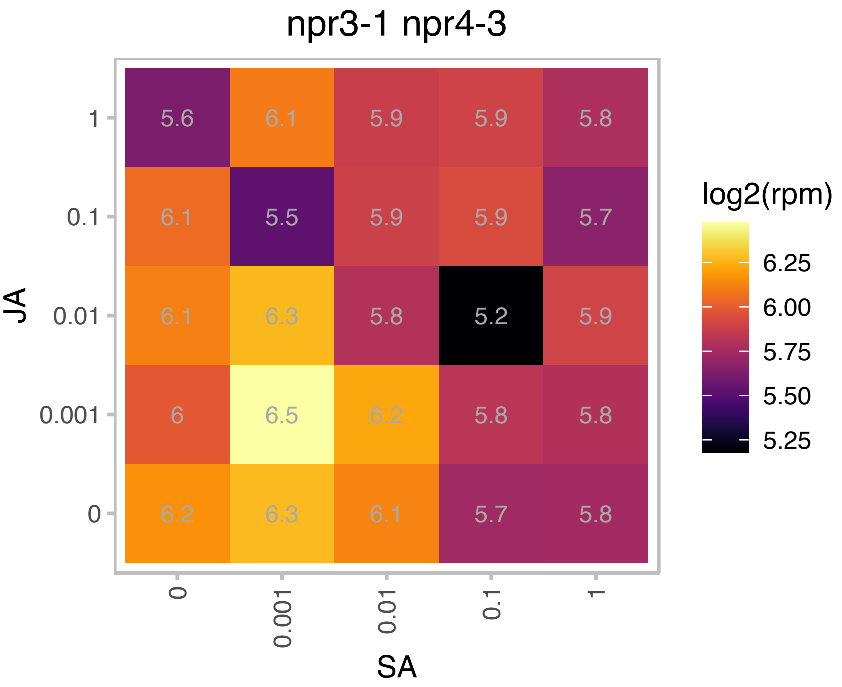

### AT1G43560.1 : Aty2, ty2

thioredoxin Y2

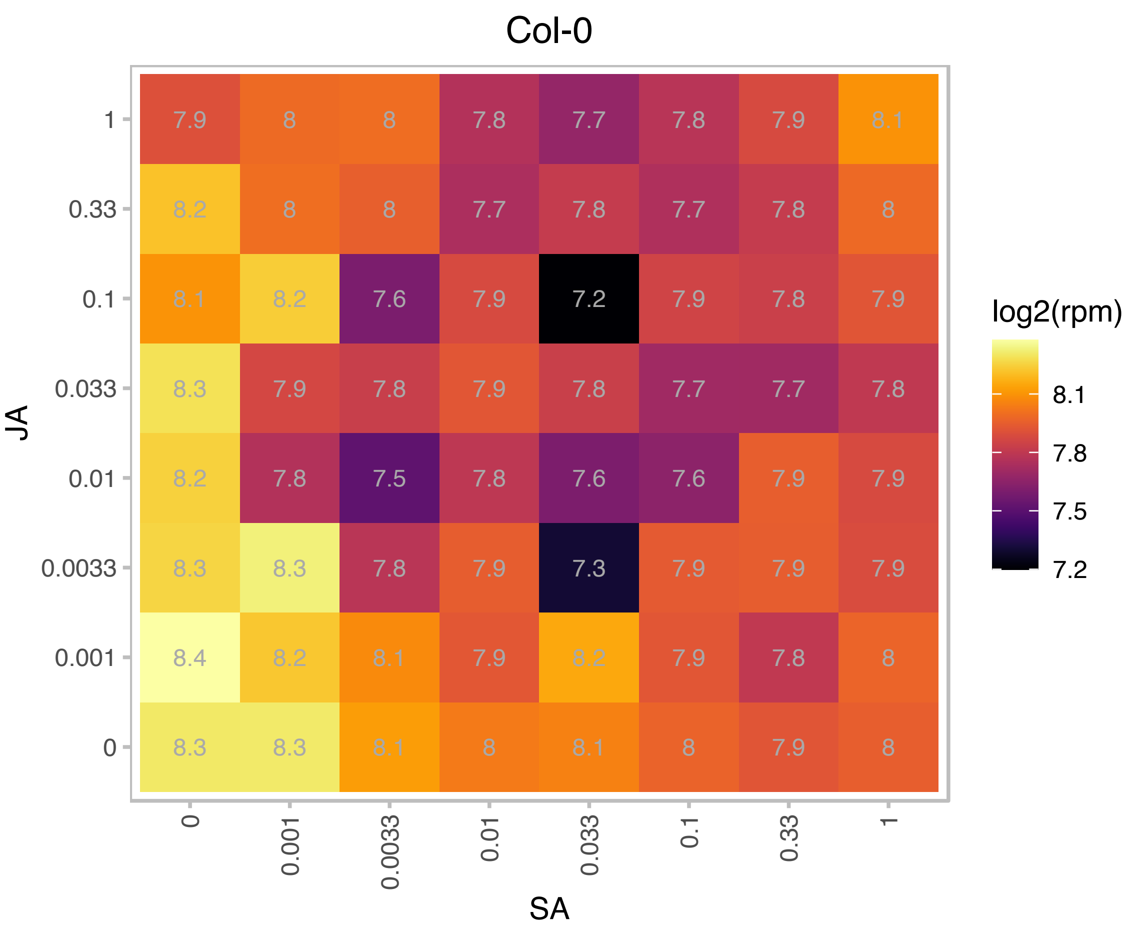


| | |
|---|---|
| Gene | AT1G43560 |
| is.expressed_genes | TRUE |
| is.responsive_genes | TRUE |
| is.apcluster | G-02 |

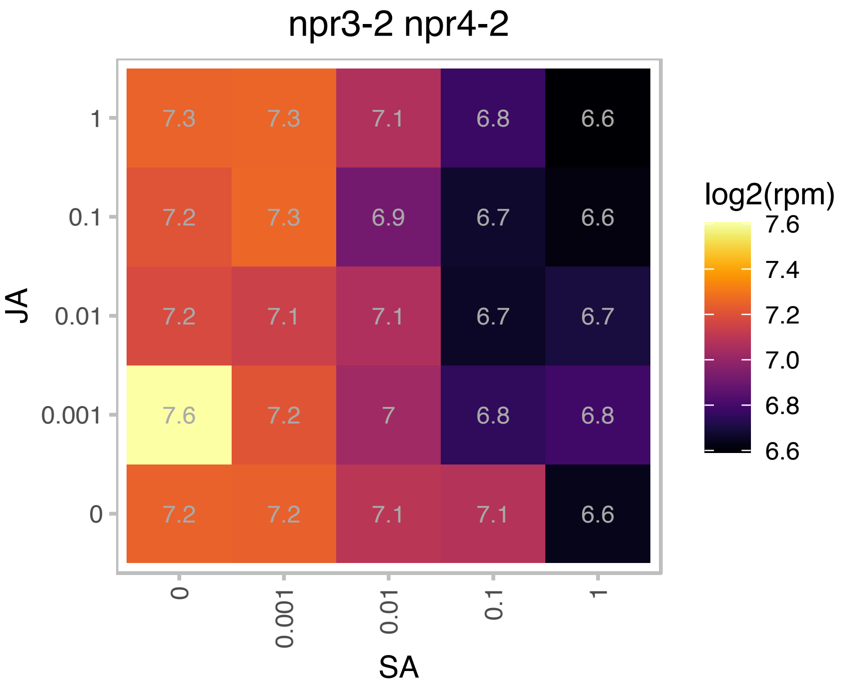


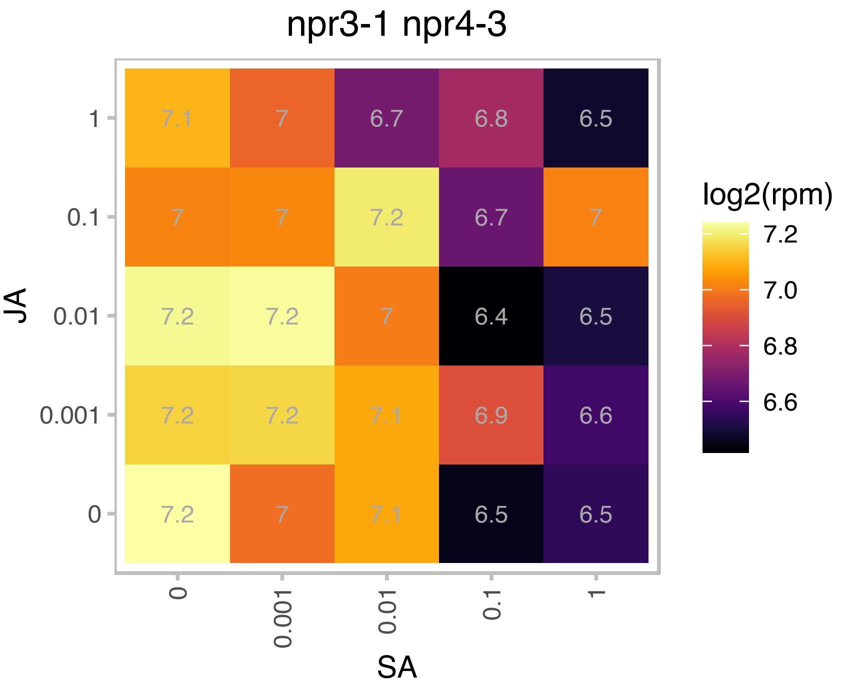

# AT1G75800.1 : NA

Pathogenesis-related thaumatin superfamily protein

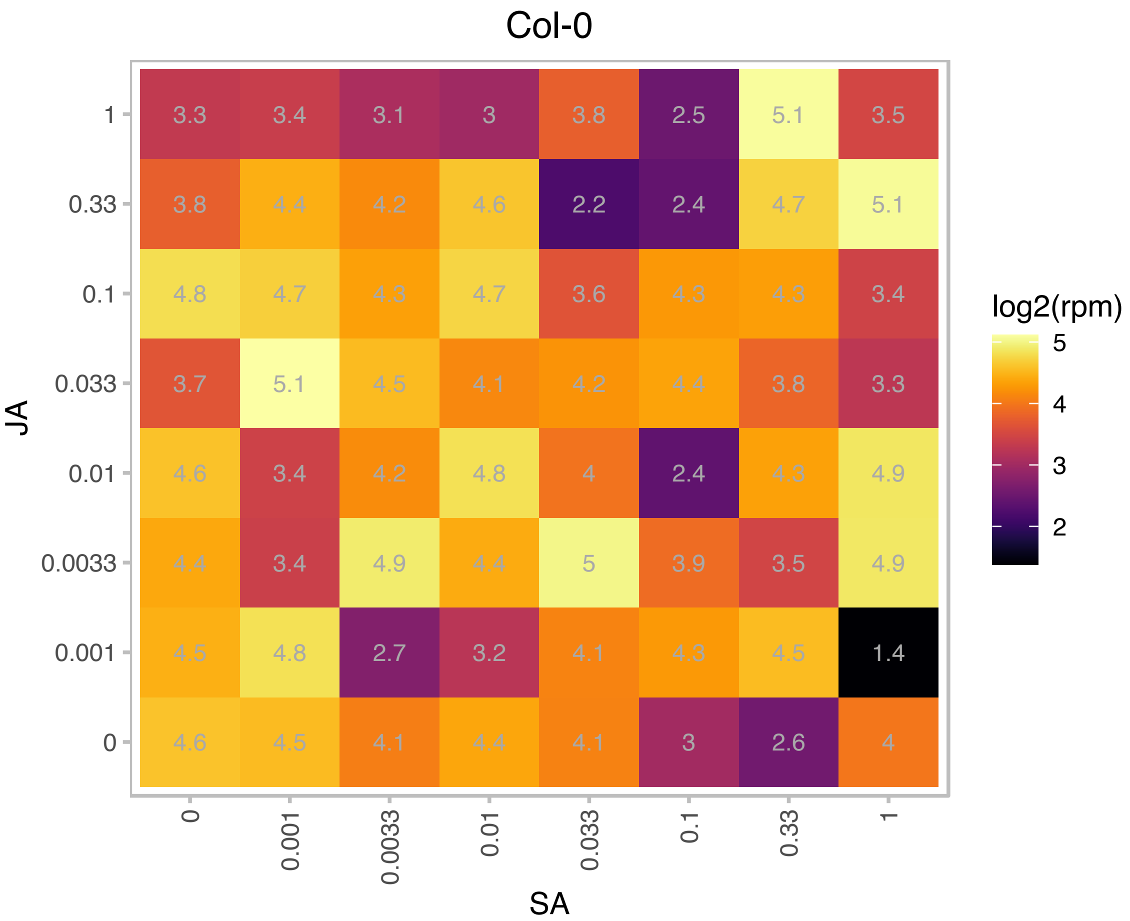


| | |
|---|---|
| Gene | AT1G75800 |
| is.expressed_genes | TRUE |
| is.responsive_genes | TRUE |
| is.apcluster | G-02 |

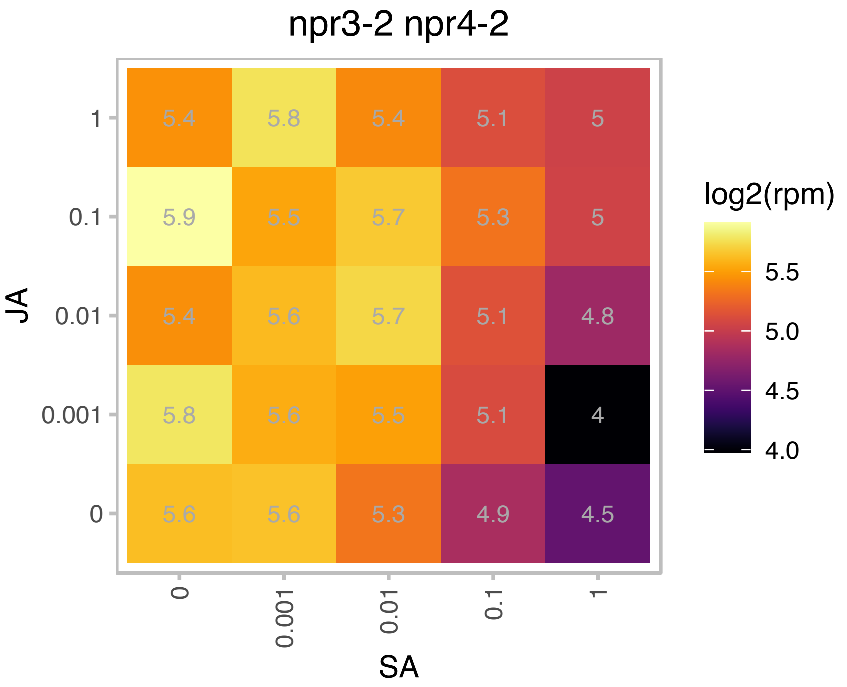


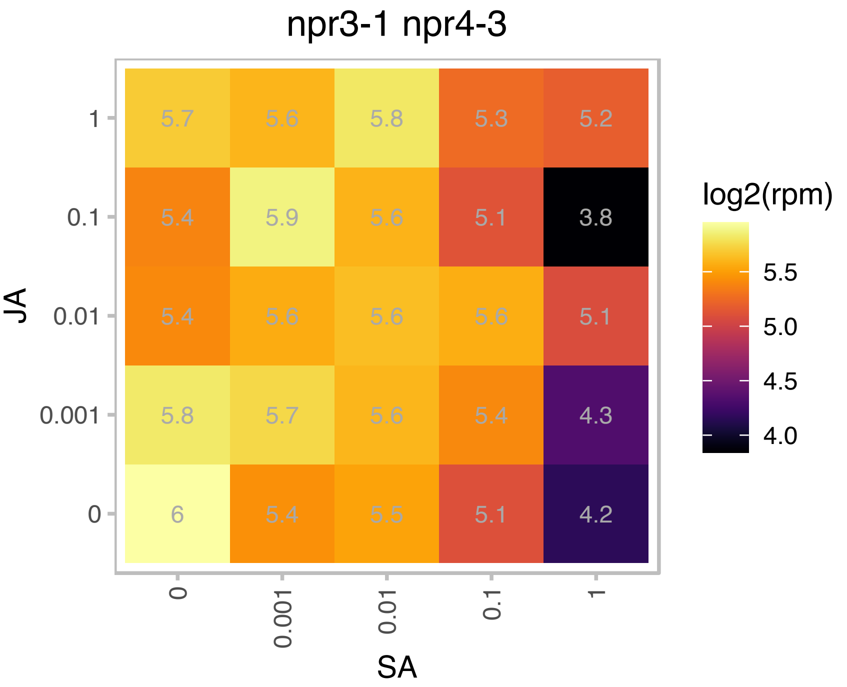

# AT2G21250.1 : NA

NAD(P)-linked oxidoreductase superfamily protein

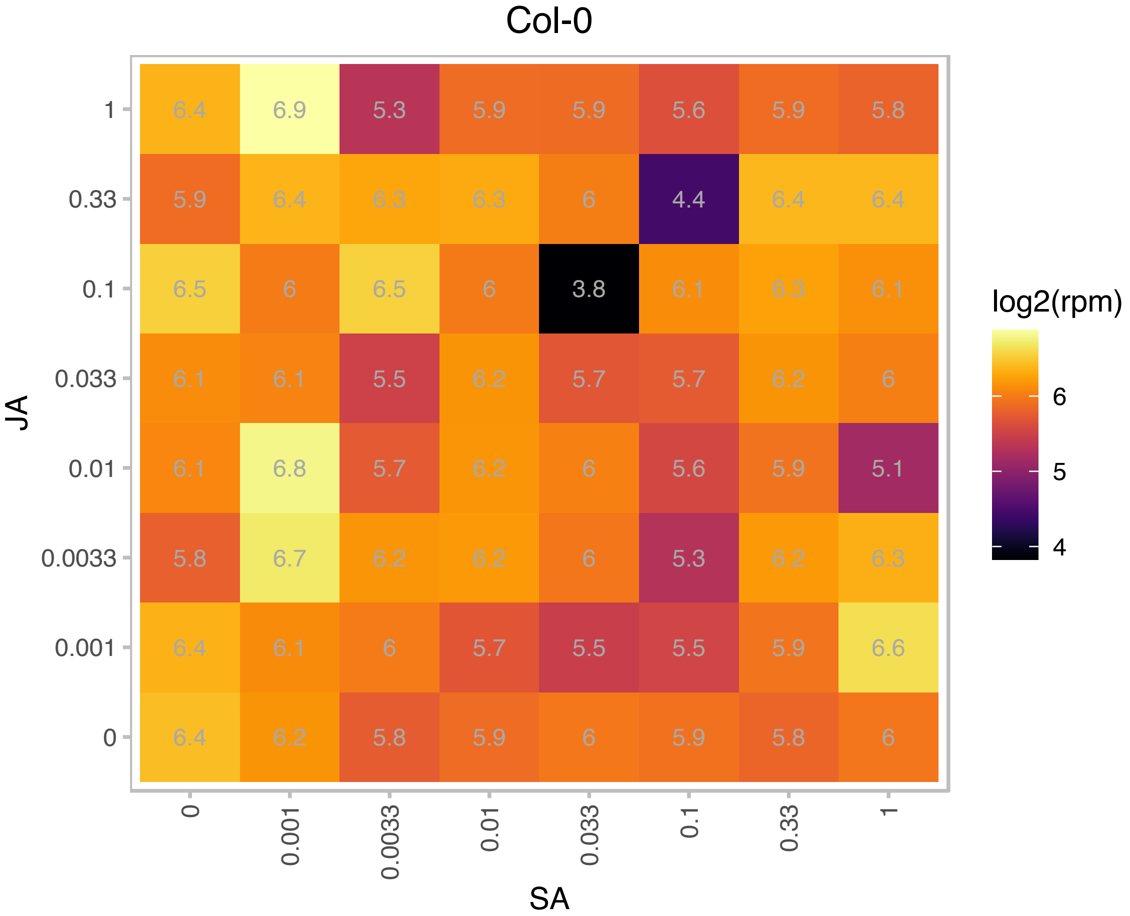


| | |
|---|---|
| Gene | AT2G21250 |
| is.expressed_genes | TRUE |
| is.responsive_genes | TRUE |
| is.apcluster | G-02 |

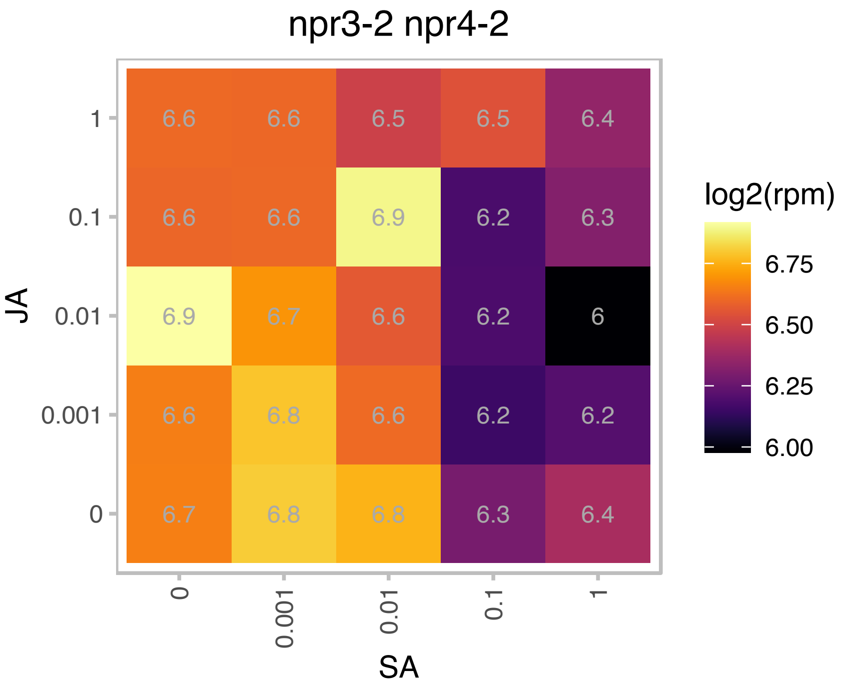


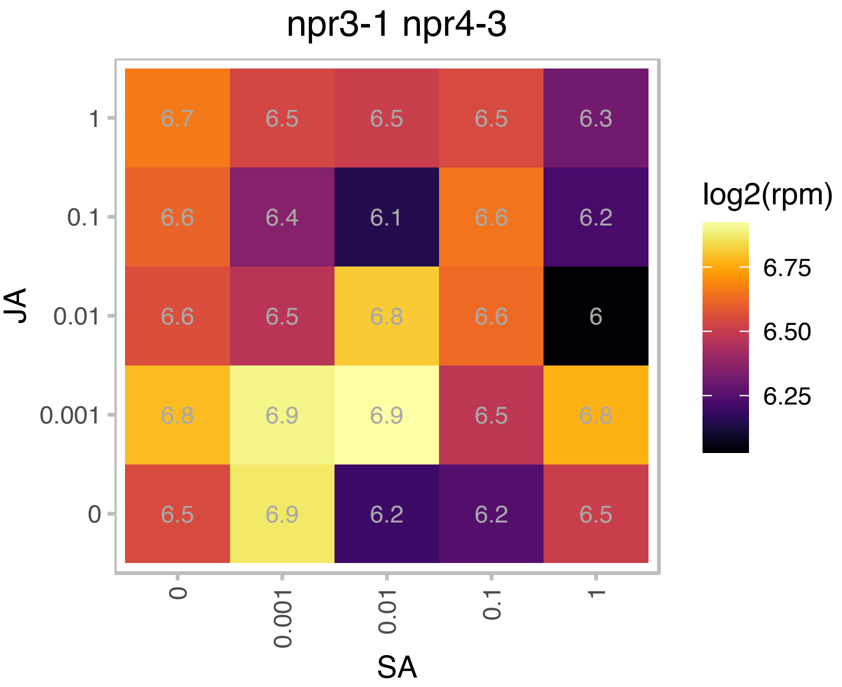

### AT2G35370.1 : GDCH

glycine decarboxylase complex H

Encodes glycine decarboxylase complex H protein. Involved in photorespiration. The mRNA is cell-to-cell mobile.

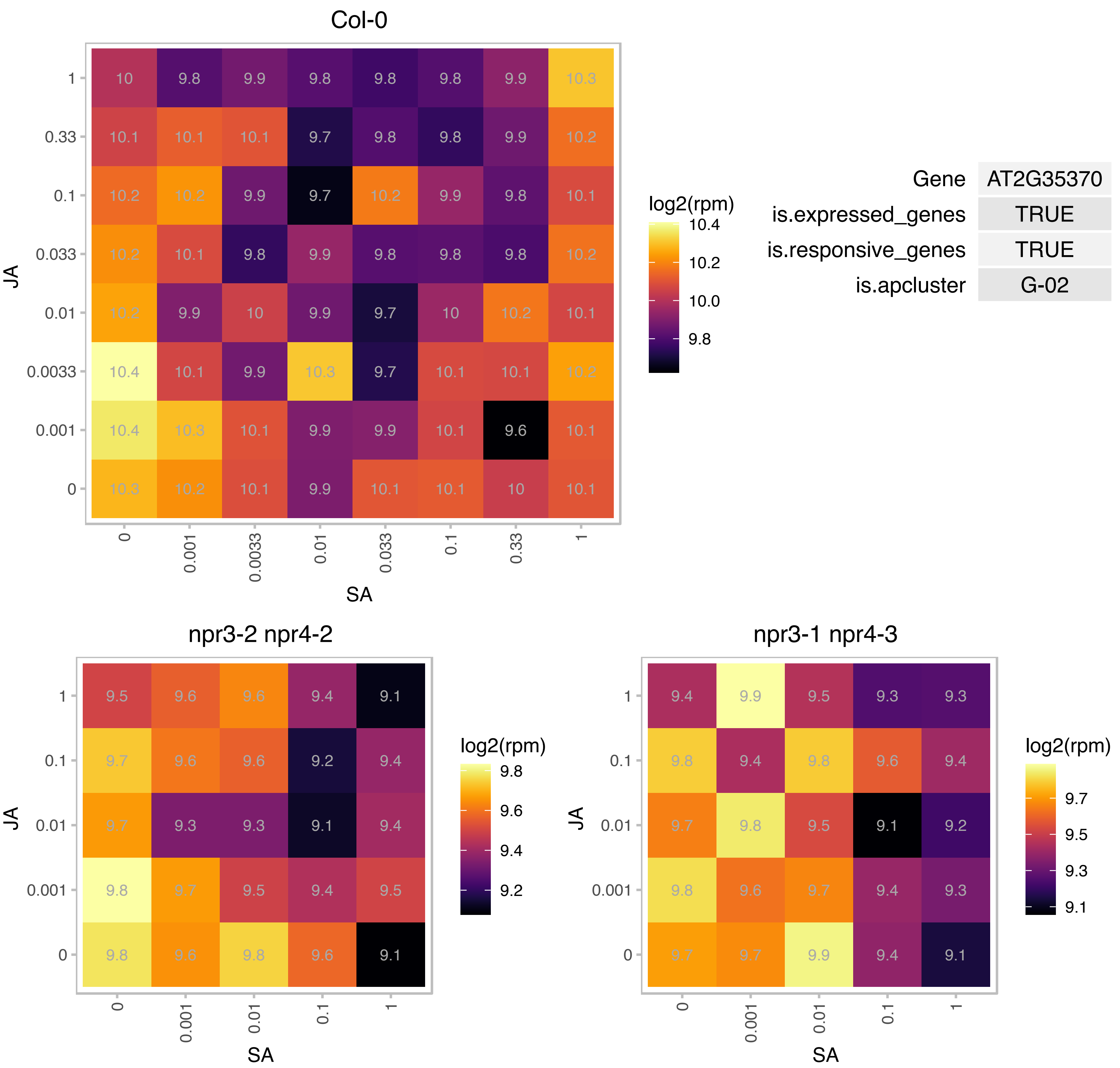


| | |
|---|---|
| Gene | AT2G35370 |
| is.expressed_genes | TRUE |
| is.responsive_genes | TRUE |
| is.apcluster | G-02 |

### AT2G45820.1 : Rem1.3

Remorin family protein

| | |
|---|---|
| Gene | AT2G45820 |
| is.expressed_genes | TRUE |
| is.responsive_genes | TRUE |
| is.apcluster | G-02 |

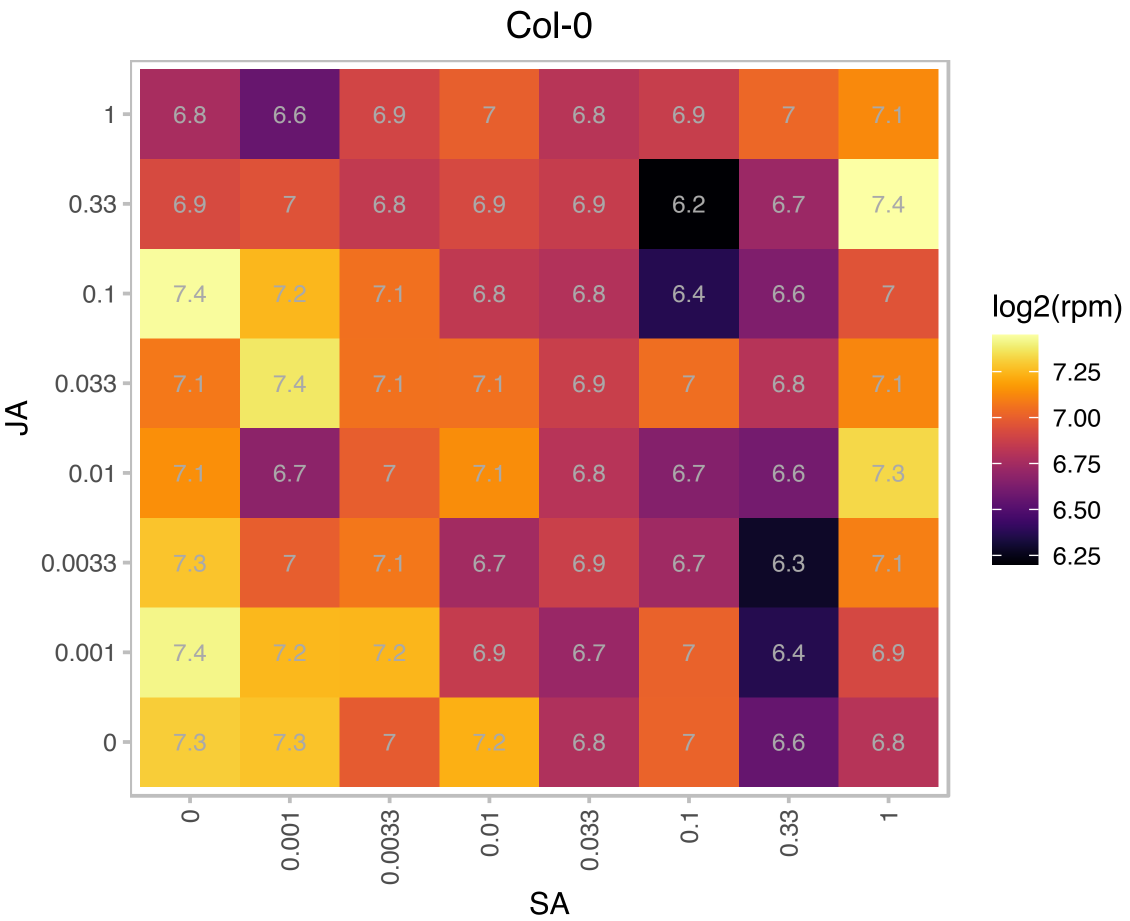


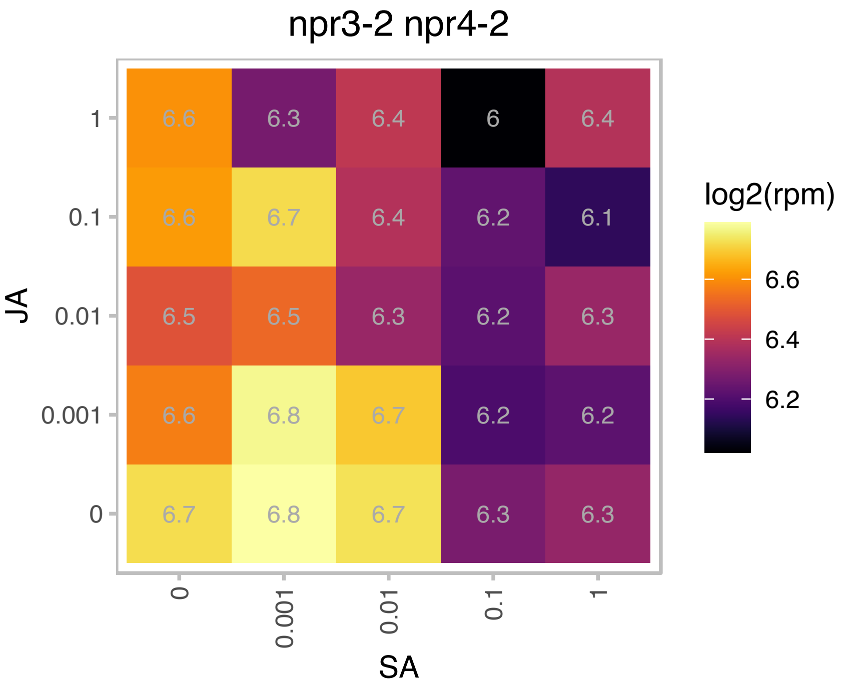


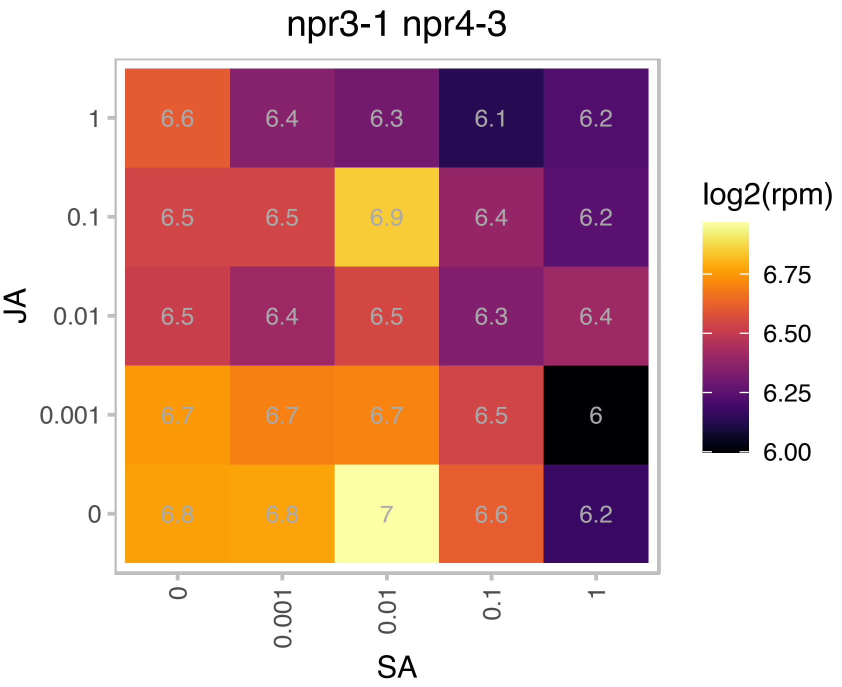

# AT3G07460.1 : NA

Protein of unknown function, DUF538

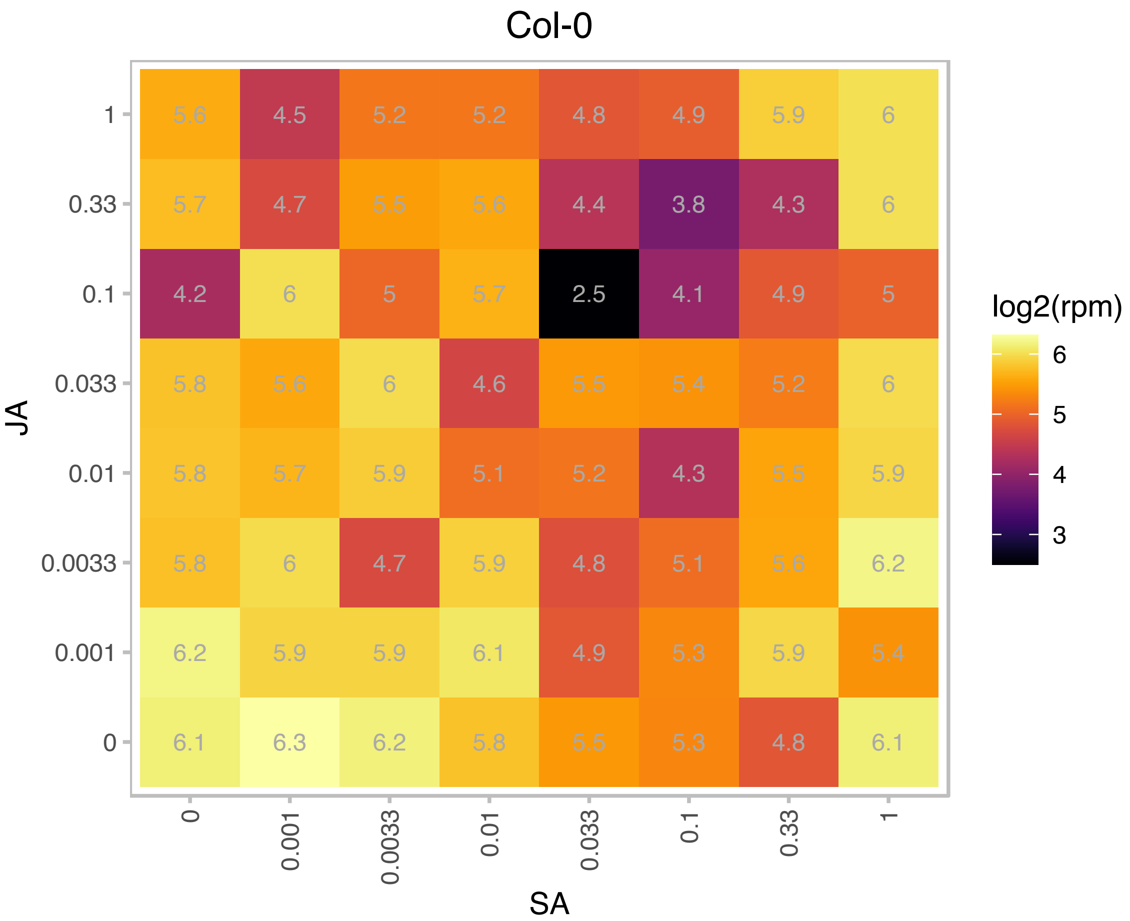


| | |
|---|---|
| Gene | AT3G07460 |
| is.expressed_genes | TRUE |
| is.responsive_genes | TRUE |
| is.apcluster | G-02 |

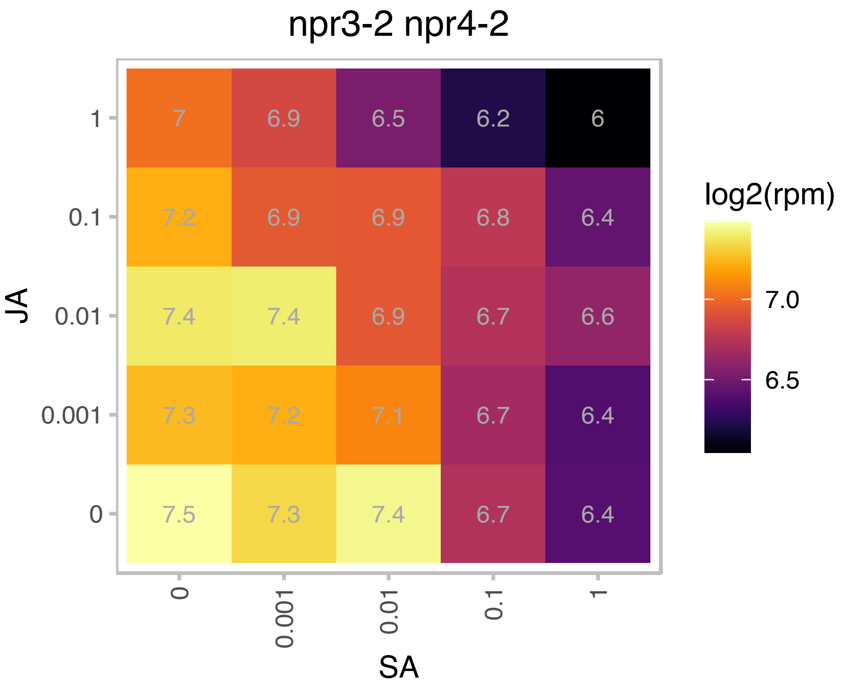


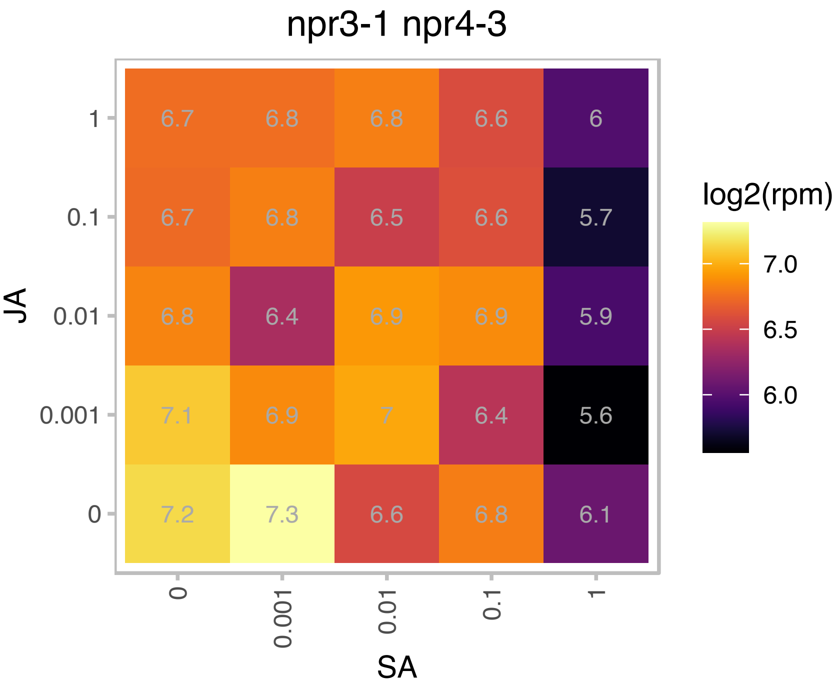

### AT3G16240.1 : AQP1, ATTIP2;1, DELTA-TIP, DELTA-TIP1,

delta tonoplast integral protein
Delta tonoplast intrinsic protein, functions as a water channel and ammonium (NH3) transporter. Highly expressed in flower, shoot, and stem. Expression shows diurnal regulation and is induced by ammonium (NH3). Protein localized to vacuolar membrane. The mRNA is cell-to-cell mobile.

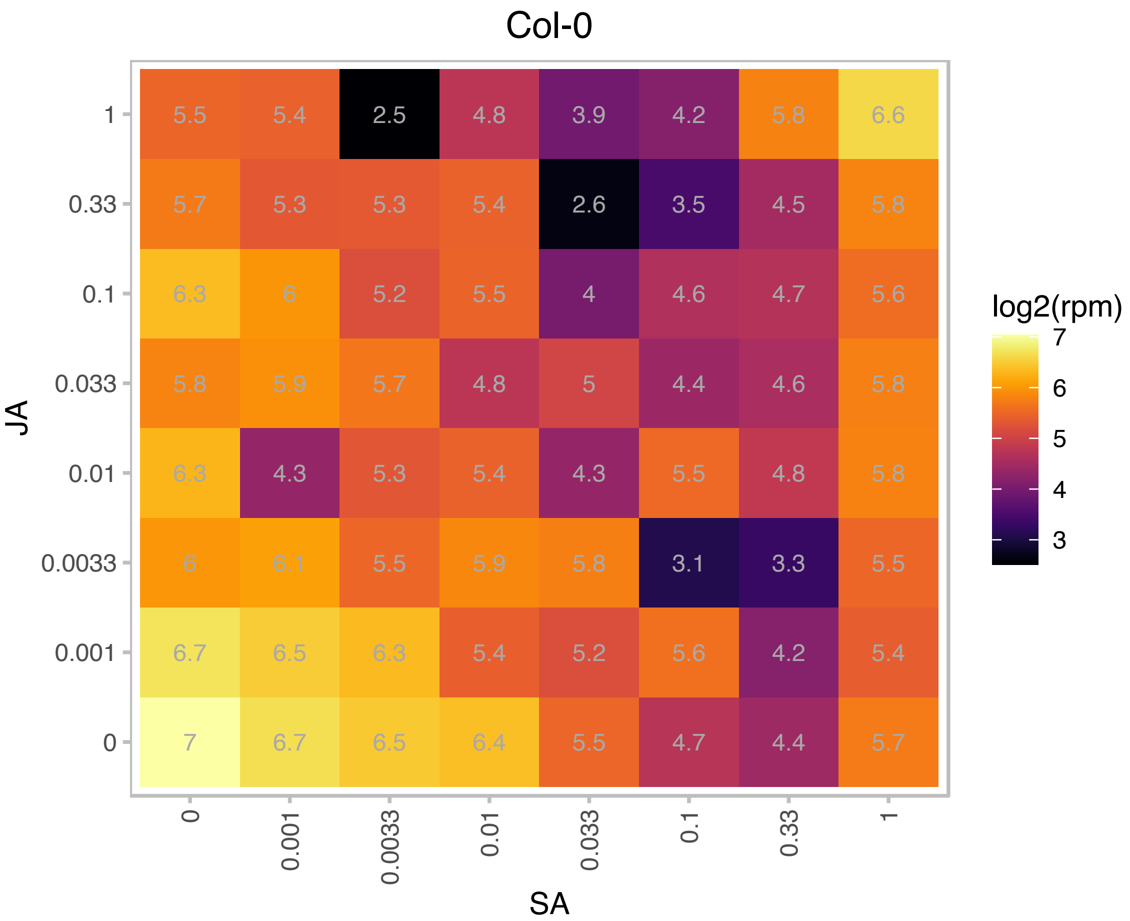


| | |
|---|---|
| Gene | AT3G16240 |
| is.expressed_genes | TRUE |
| is.responsive_genes | TRUE |
| is.apcluster | G-02 |

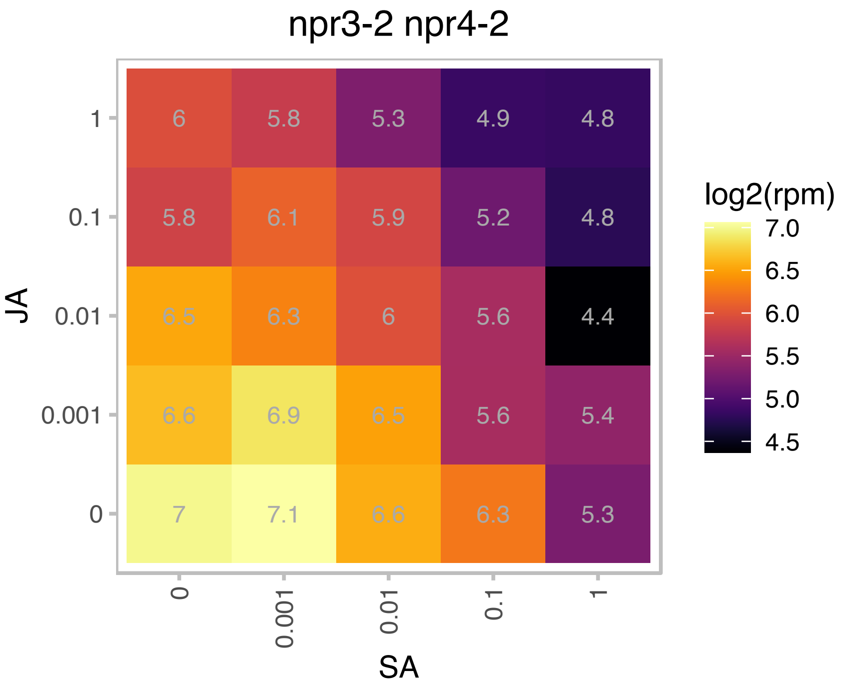


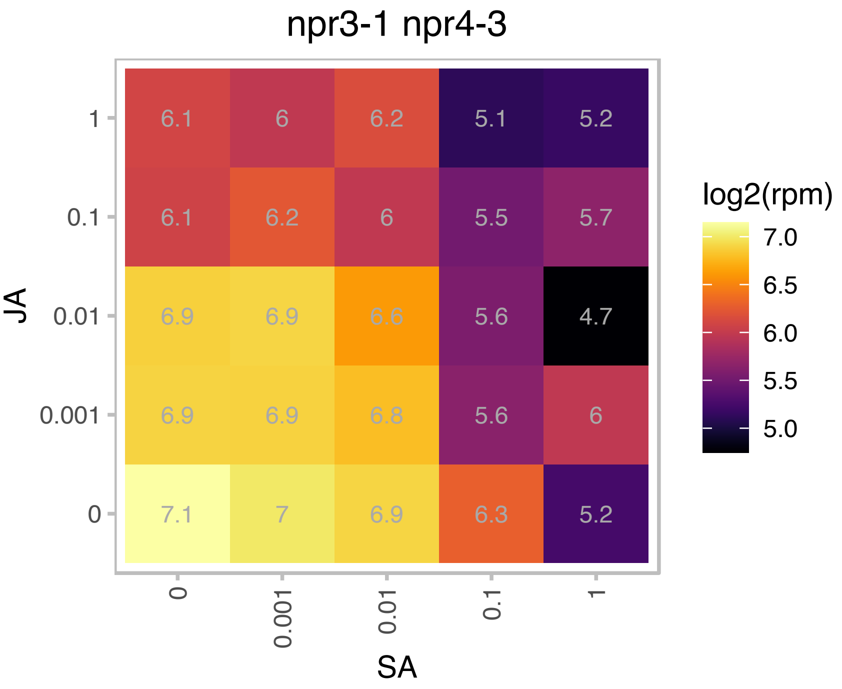

### AT3G24430.1 : HCF101

ATP binding
Encodes chloroplast protein HCF101 (high chlorophyll fluorescence 101).  Serves as a chloroplast scaffold protein that specifically assembles [4Fe-4S] clusters and transfers them to the chloroplast membrane and soluble target proteins.

Col-0

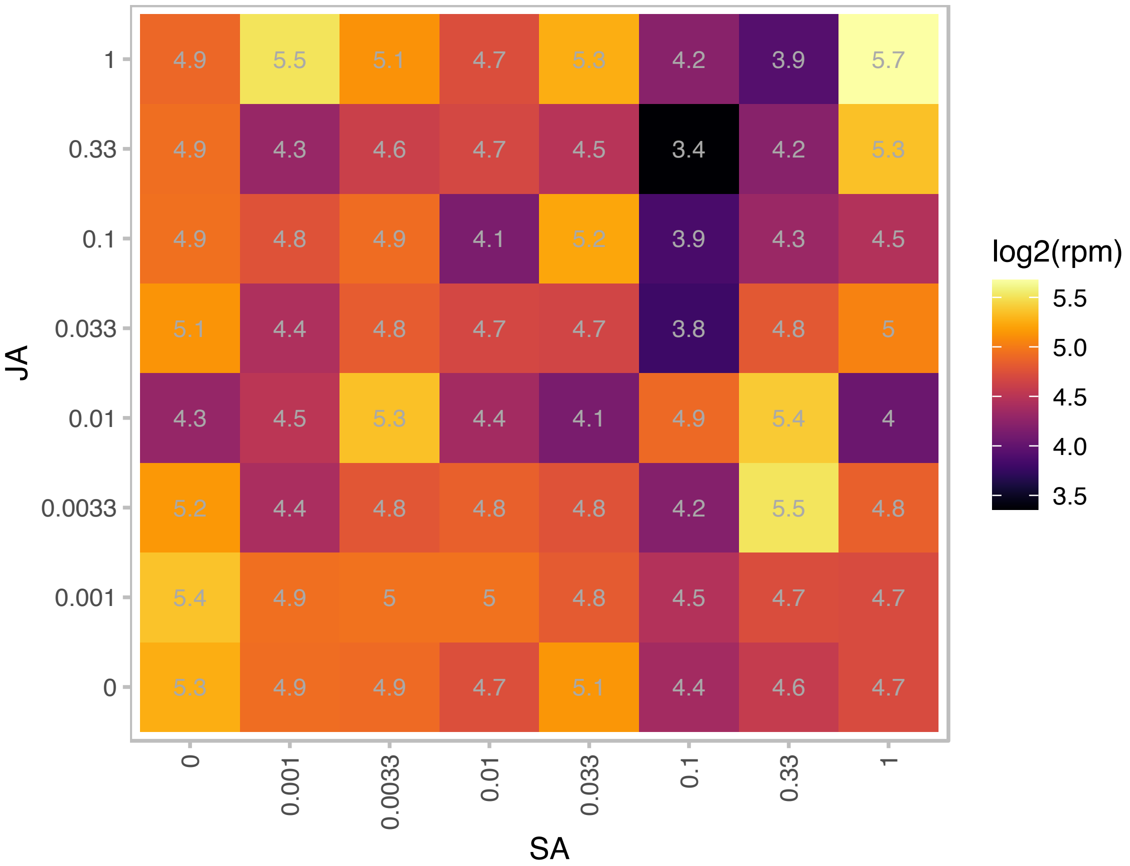


| Gene | AT3G24430 |
|---|---|
| is.expressed_genes | TRUE |
| is.responsive_genes | TRUE |
| is.apcluster | G-02 |

npr3-2 npr4-2

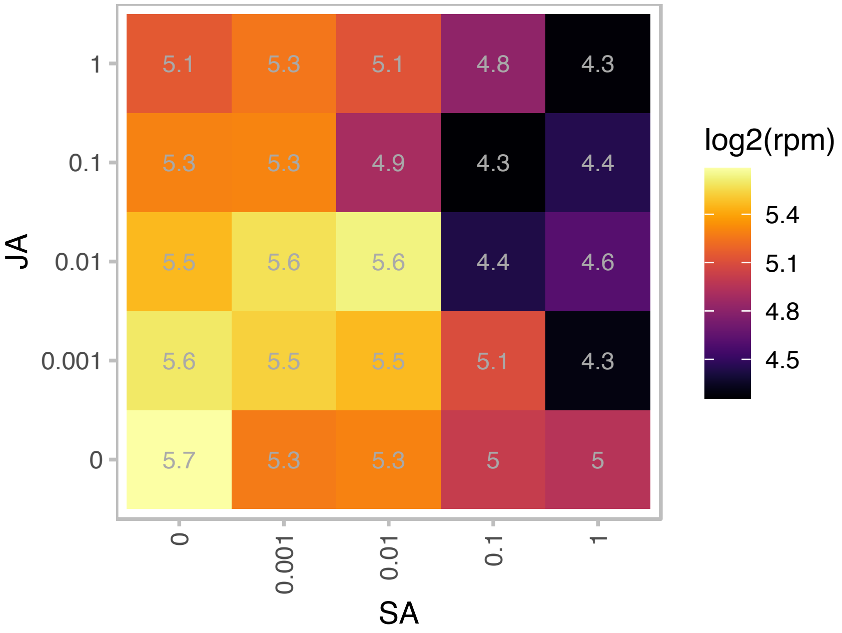


npr3-1 npr4-3

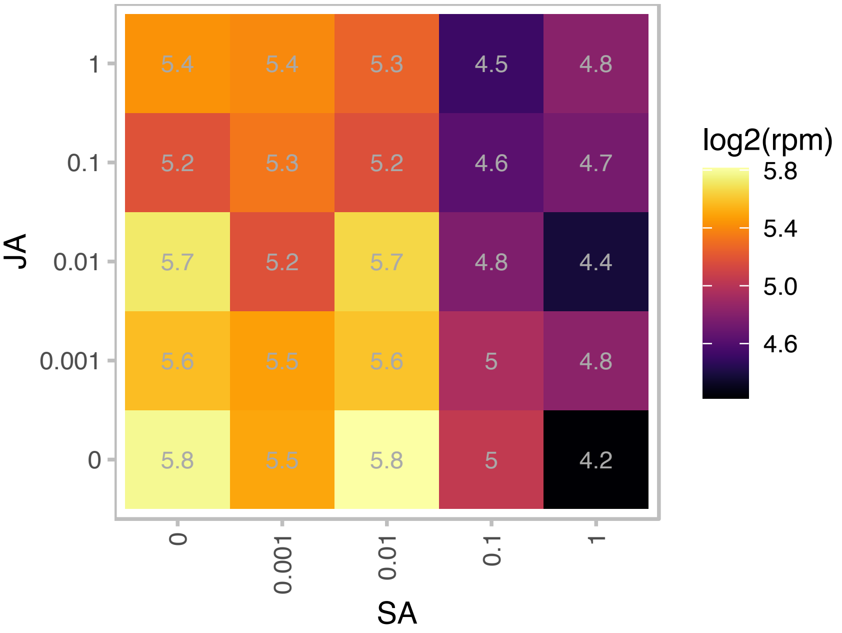

# AT3G25805.1 : NA

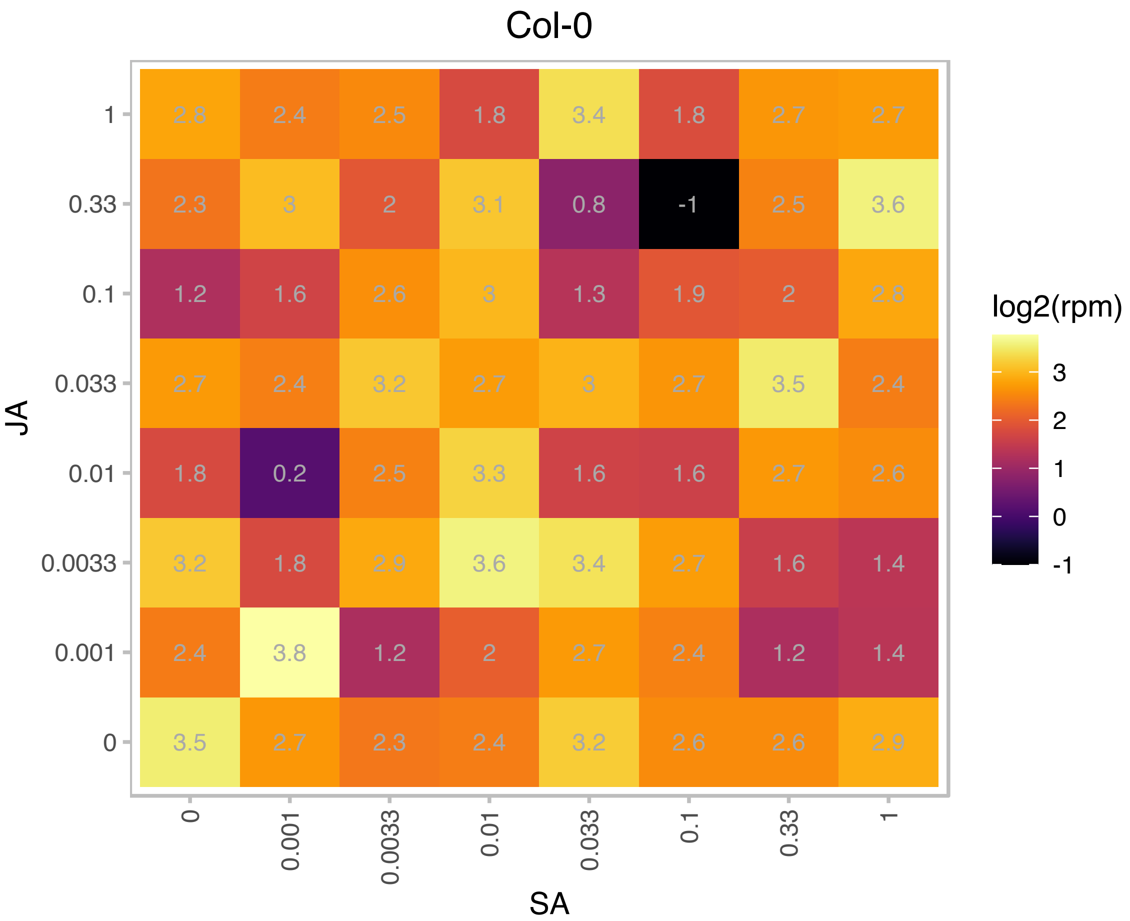


| | |
|---|---|
| Gene | AT3G25805 |
| is.expressed_genes | TRUE |
| is.responsive_genes | TRUE |
| is.apcluster | G-02 |

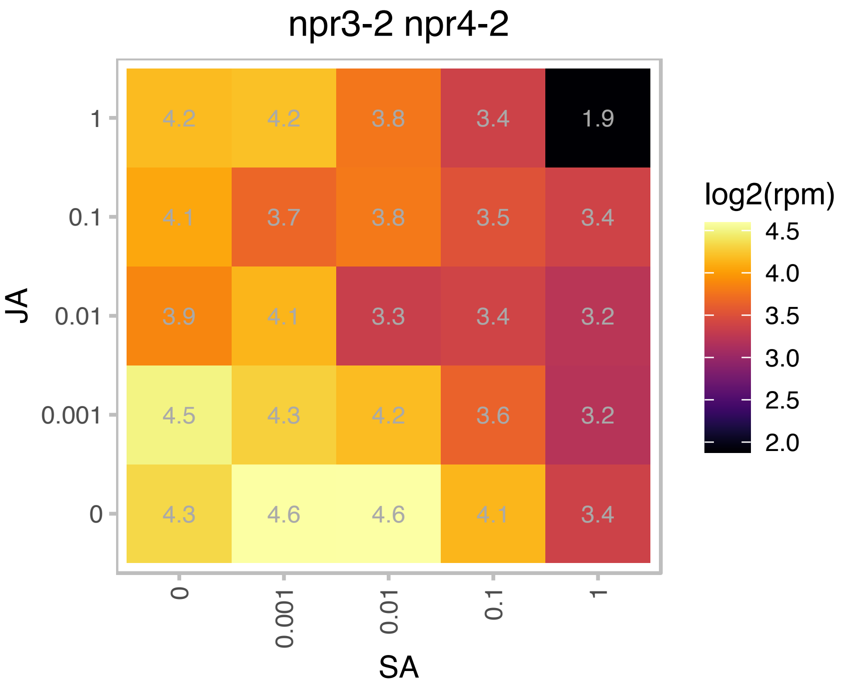


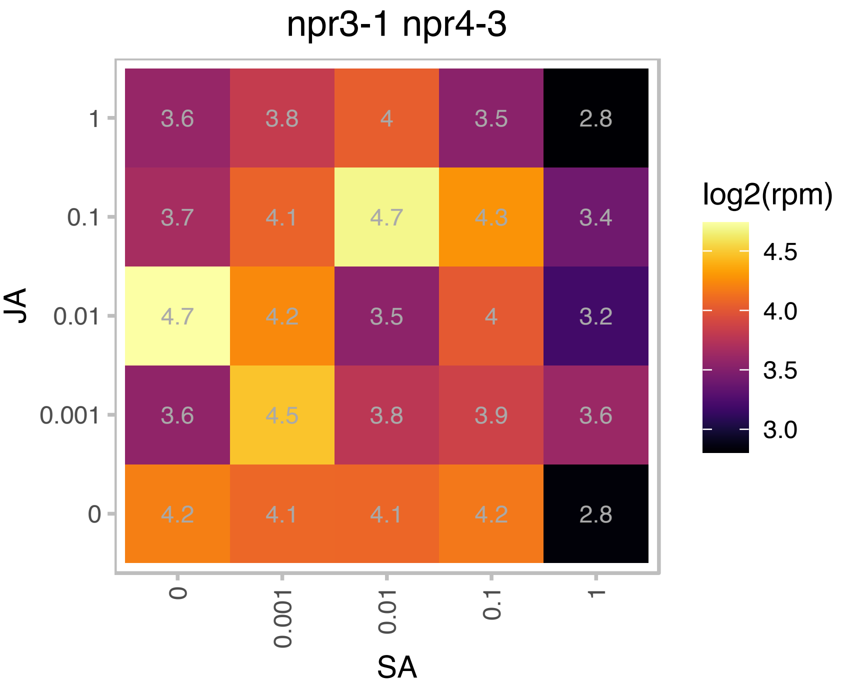

### AT3G46970.1 : ATPHS2, PHS2

alpha-glucan phosphorylase 2

Encodes a cytosolic alpha-glucan phosphorylase. In vitro, the enzyme has a preference for branched polysaccharides, such as glycogen.

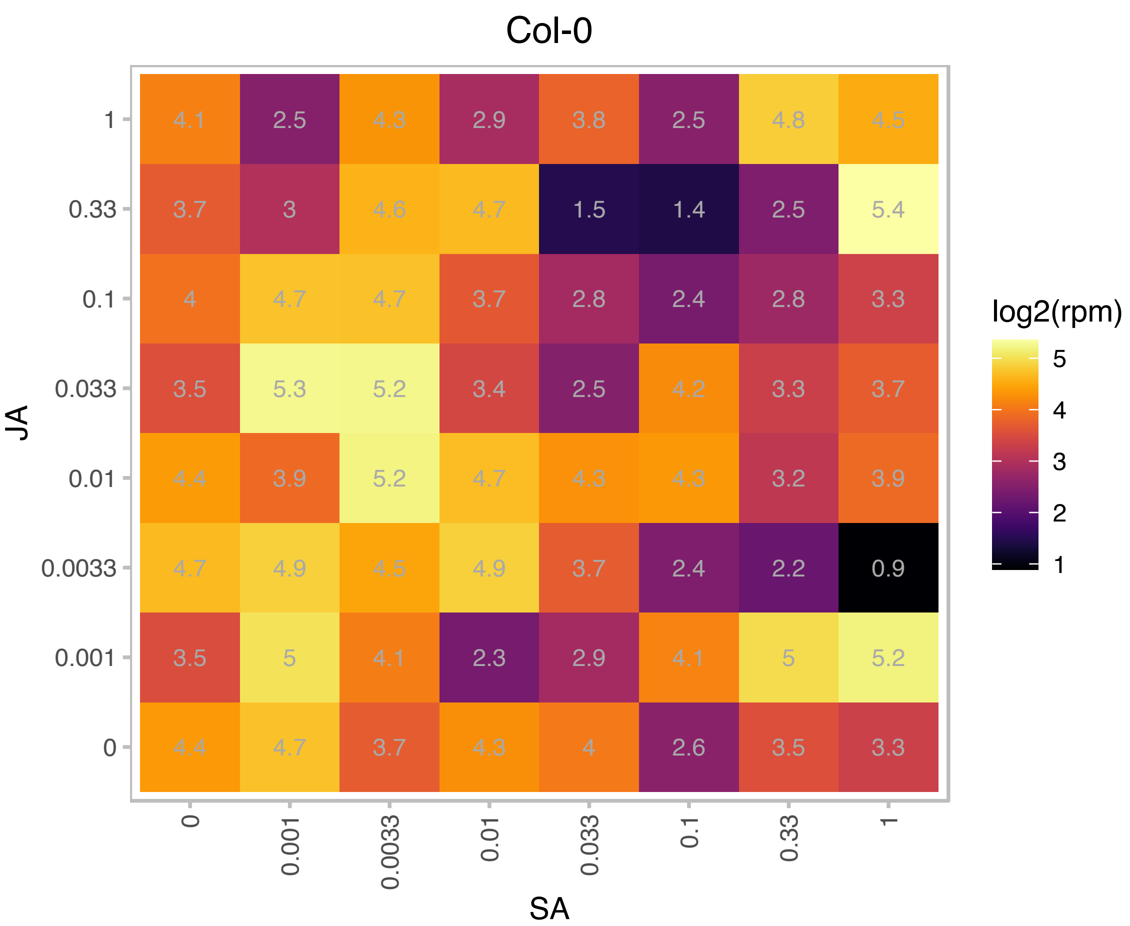


| | |
|---|---|
| Gene | AT3G46970 |
| is.expressed_genes | TRUE |
| is.responsive_genes | TRUE |
| is.apcluster | G-02 |

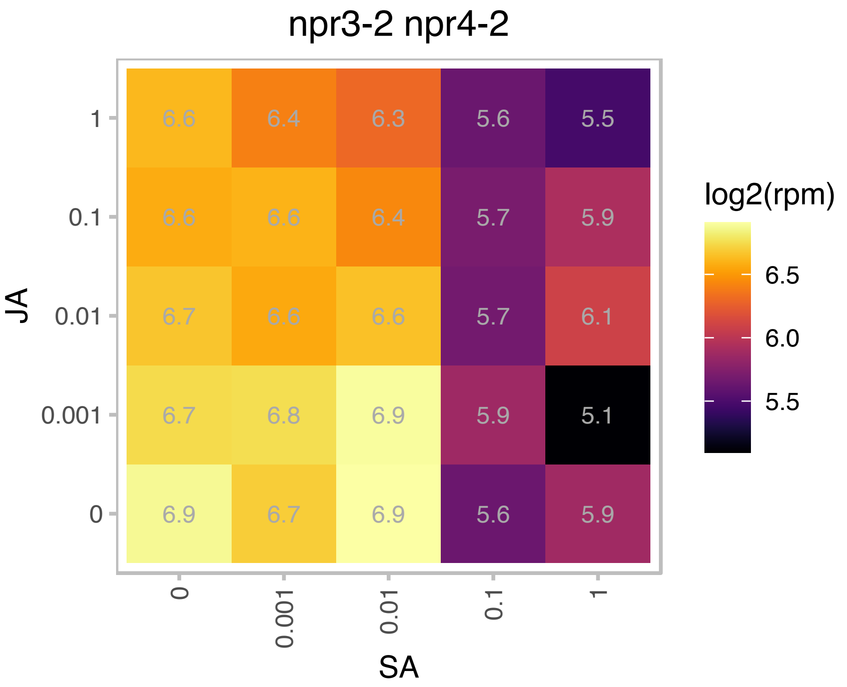


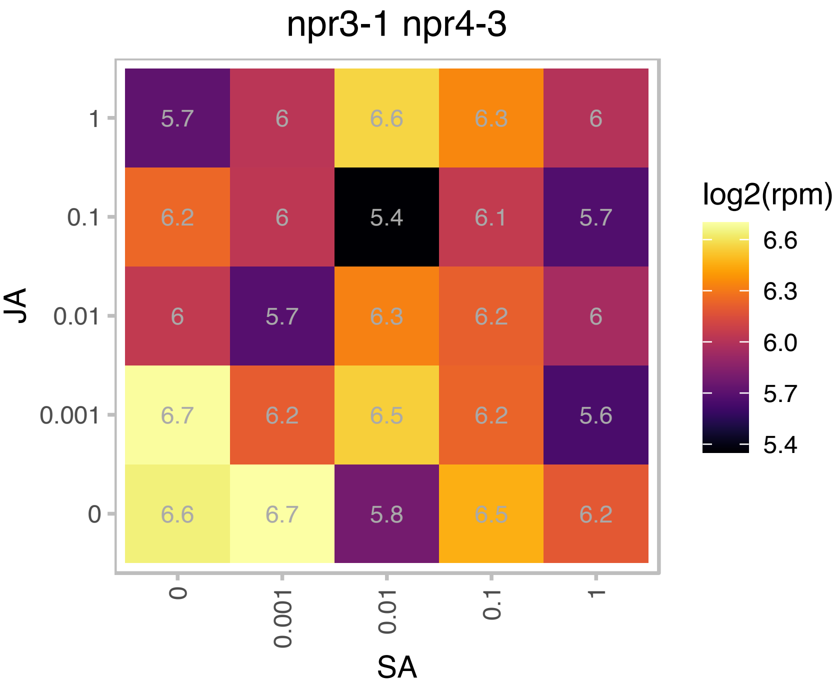

# AT3G47590.1 : NA

alpha/beta-Hydrolases superfamily protein

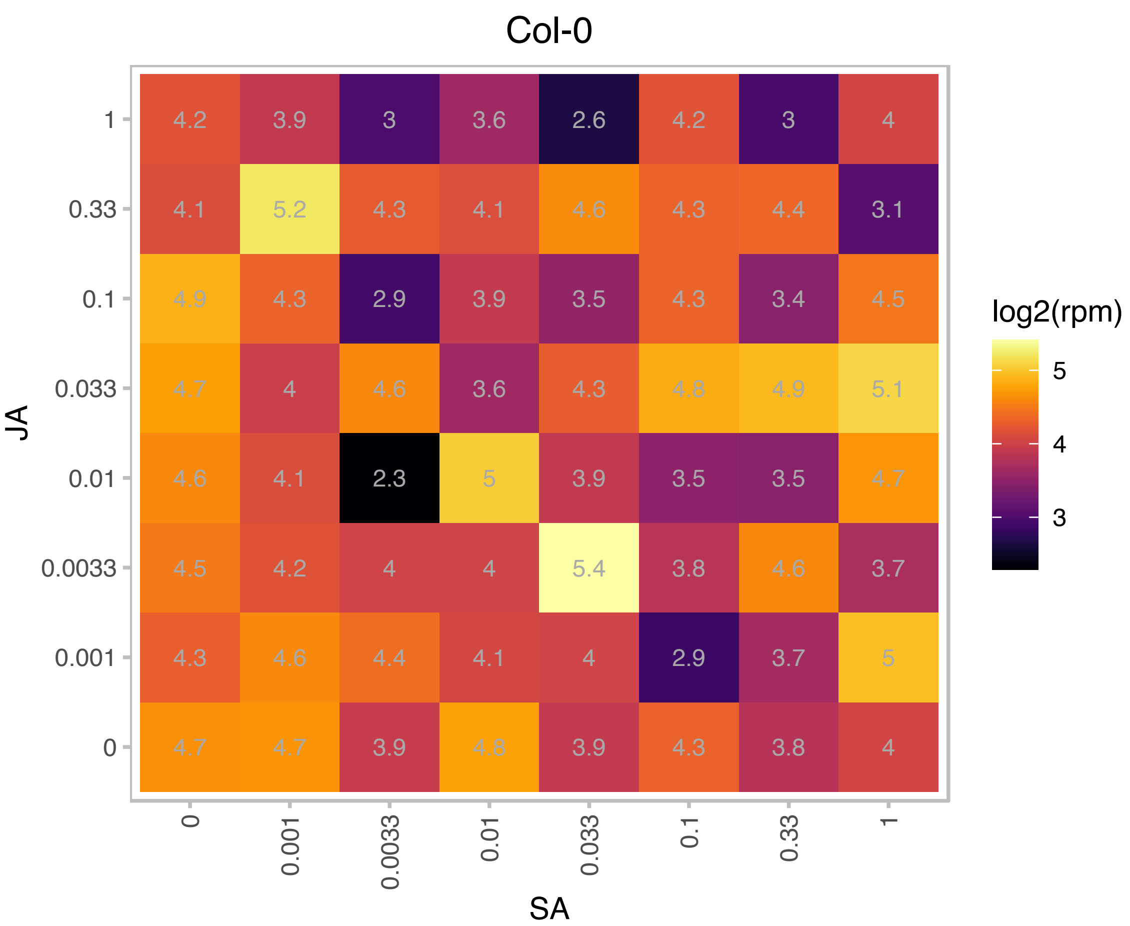


| | |
|---|---|
| Gene | AT3G47590 |
| is.expressed_genes | TRUE |
| is.responsive_genes | TRUE |
| is.apcluster | G-02 |

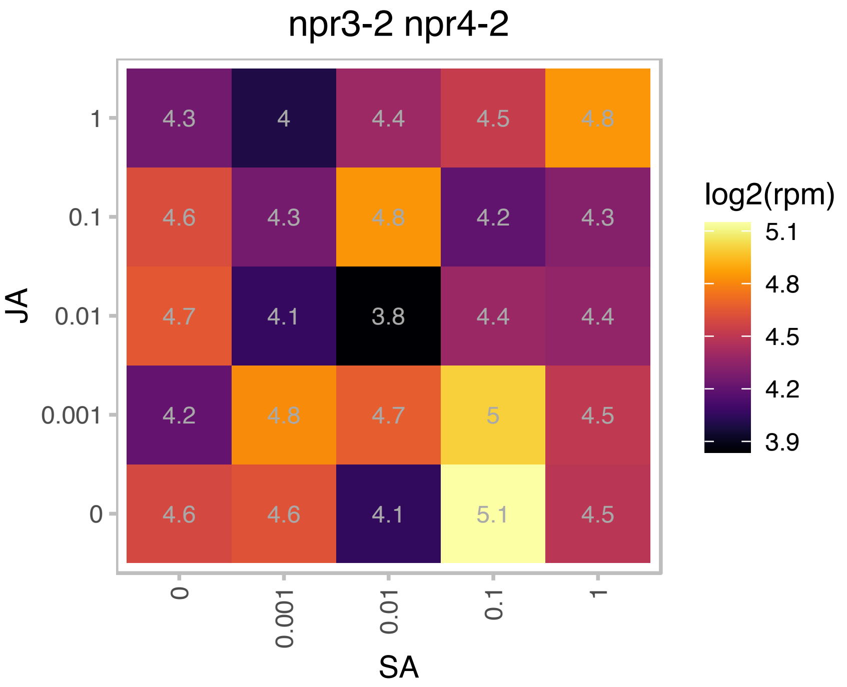


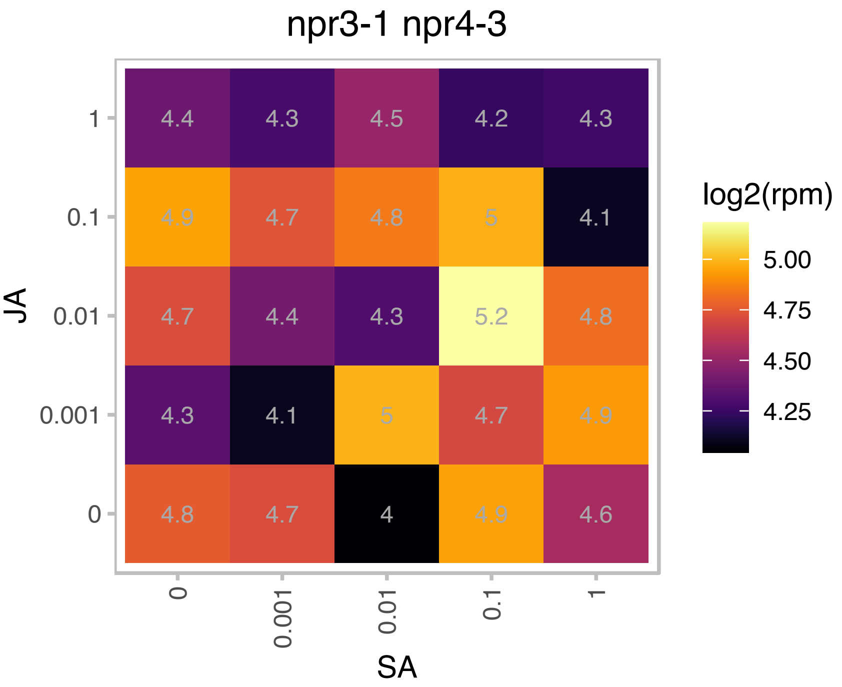

### AT4G04330.1 : AtRbcX1, RbcX1

Chaperonin-like RbcX protein

Encodes a chloroplast thylakoid localized RbcX protein that acts as a chaperone in the folding of Rubisco.

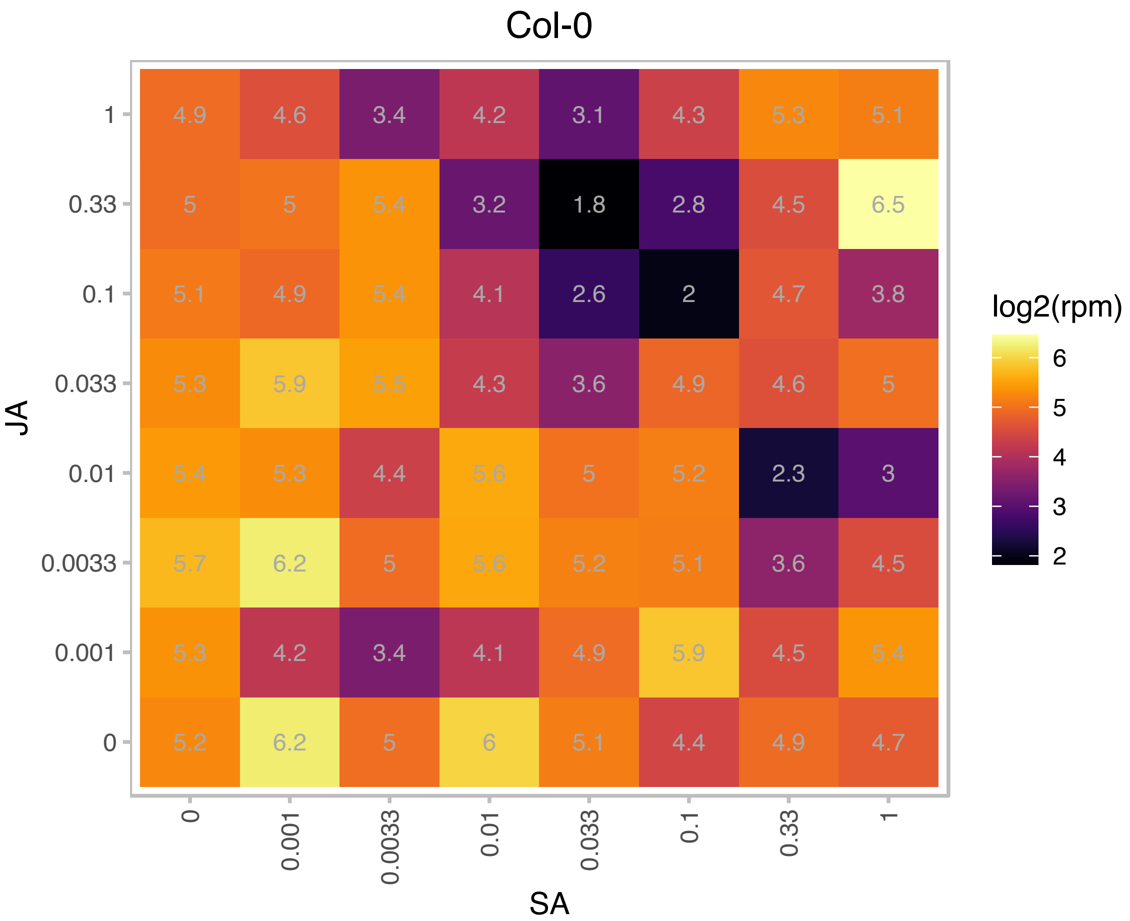


| | |
|---|---|
| Gene | AT4G04330 |
| is.expressed_genes | TRUE |
| is.responsive_genes | TRUE |
| is.apcluster | G-02 |

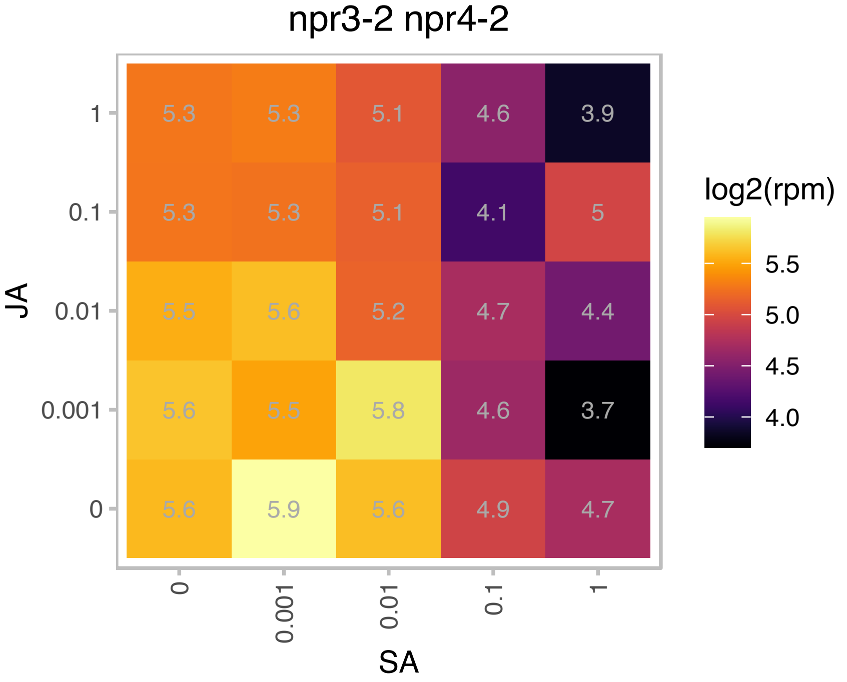


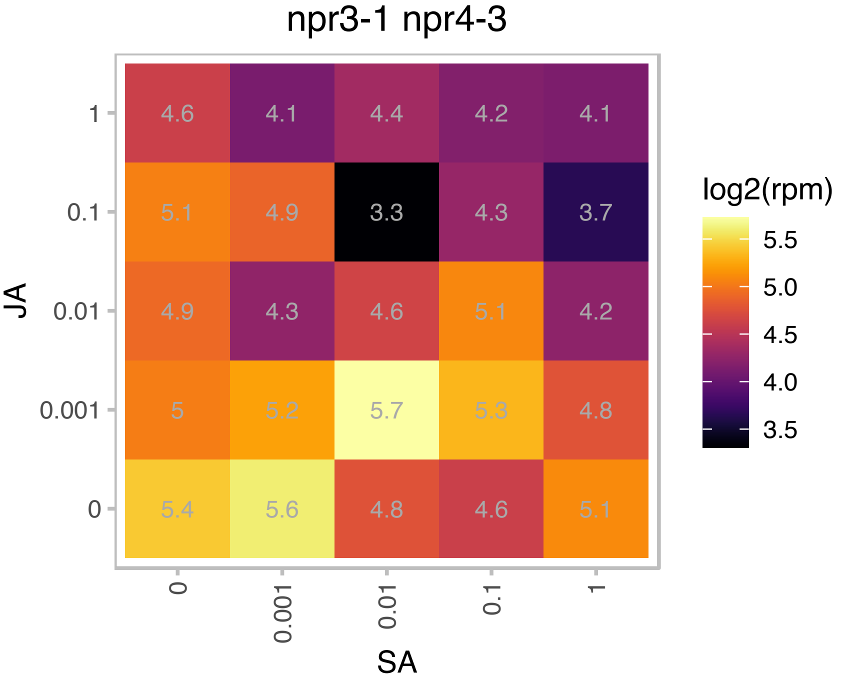

### AT4G15110.1 : CYP97B3

cytochrome P450, family 97, subfamily B, polypeptide 3

member of CYP97B

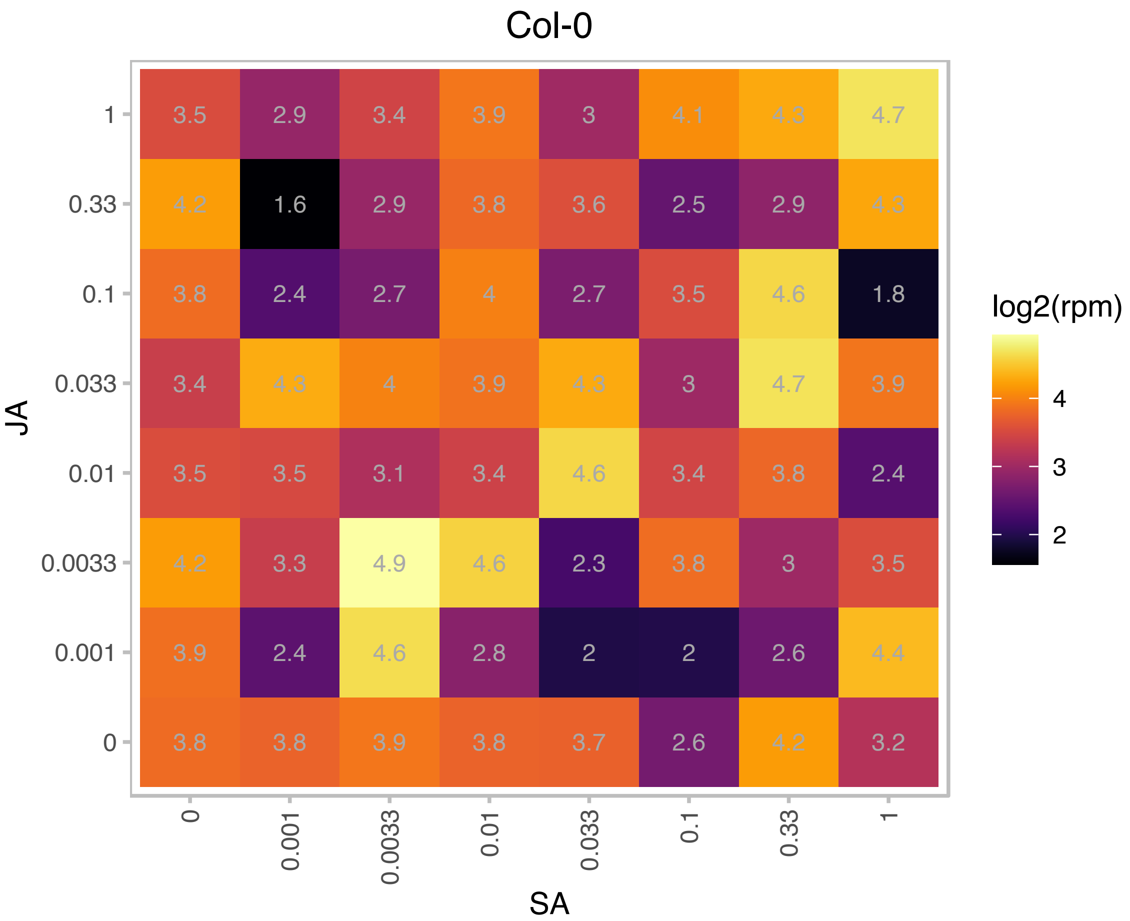


| | |
|---|---|
| Gene | AT4G15110 |
| is.expressed_genes | TRUE |
| is.responsive_genes | TRUE |
| is.apcluster | G-02 |

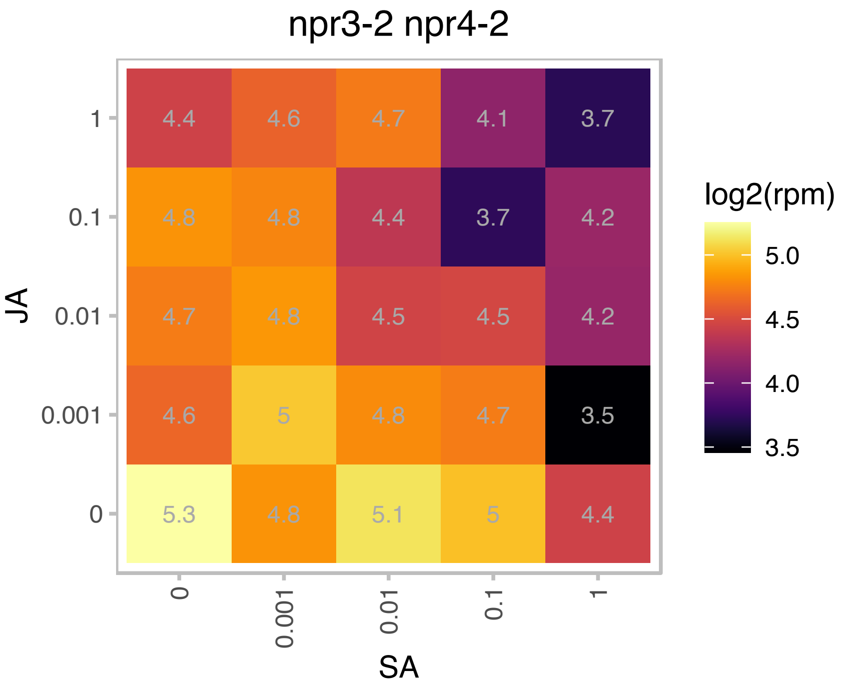


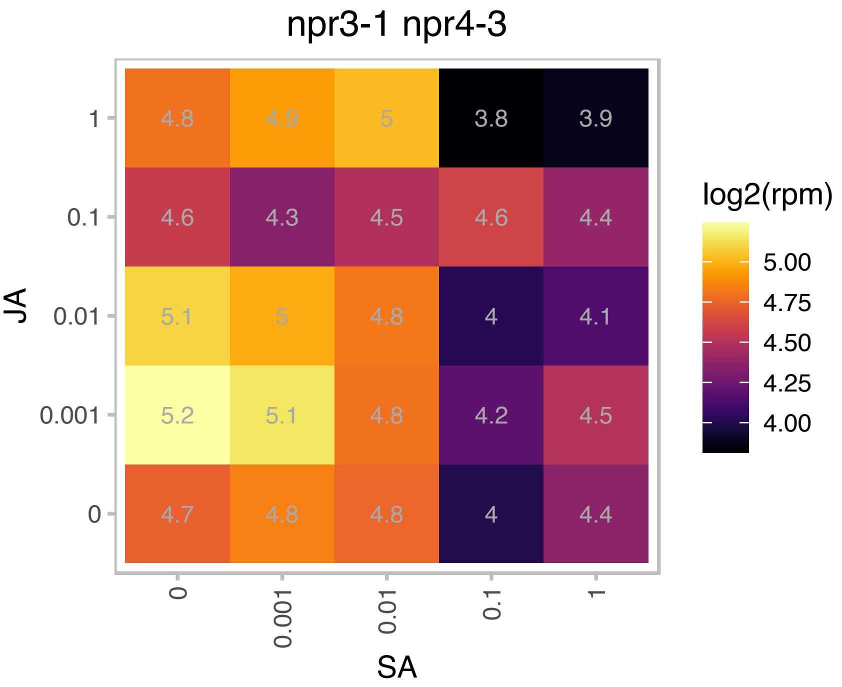

### AT4G28590.1 : AtECB1, ECB1, MRL7, SVR4

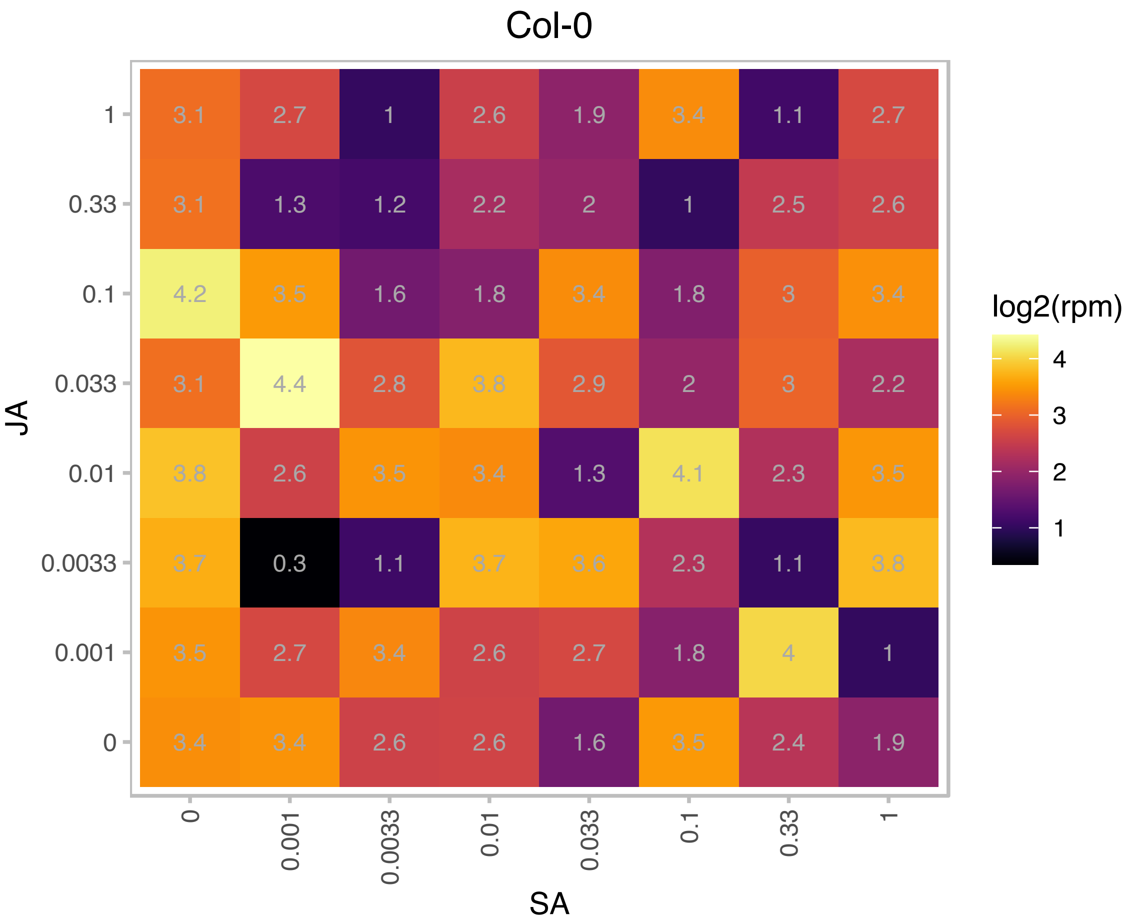


| | |
|---|---|
| Gene | AT4G28590 |
| is.expressed_genes | TRUE |
| is.responsive_genes | TRUE |
| is.apcluster | G-02 |

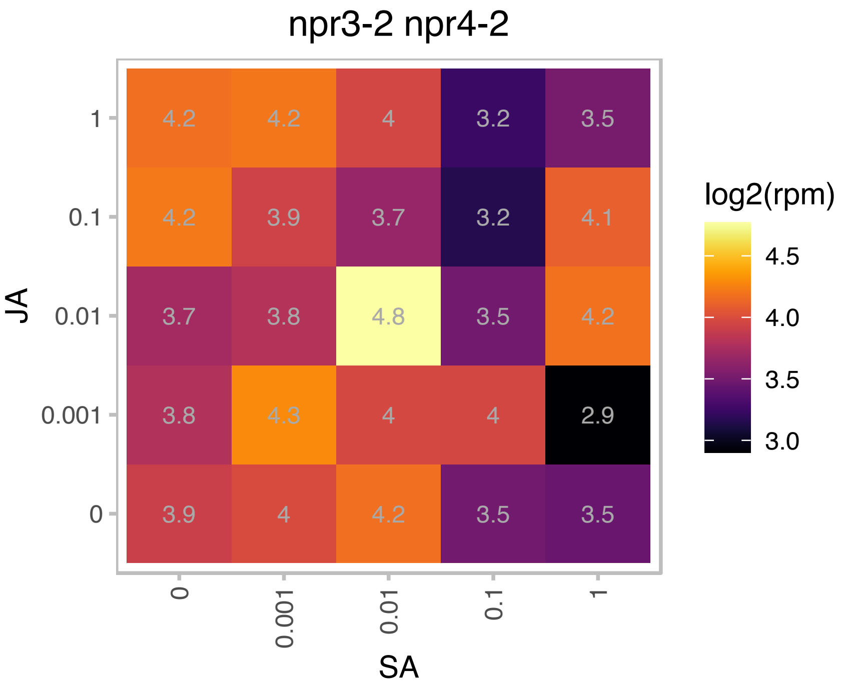


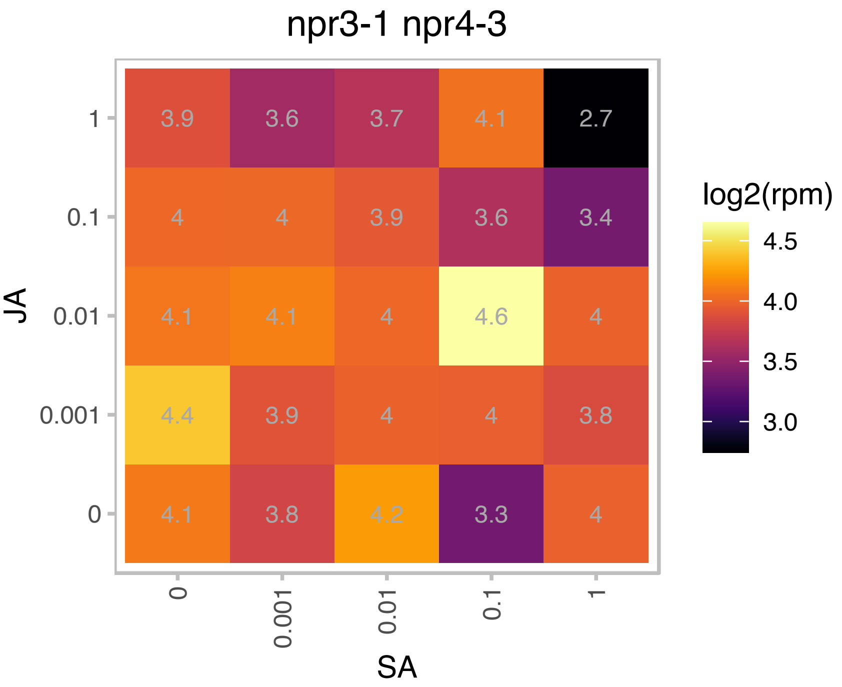

### AT4G35450.1 : AFT, AKR2, AKR2A

ankyrin repeat-containing protein 2
Involved in targeting of chloroplast outer membrane proteins to the chloroplast. Double mutants of AKR2A and the highly homologous AKR2B have yellow leaves, significantly reduced chloroplast proteins, and no thylakoid membranes.

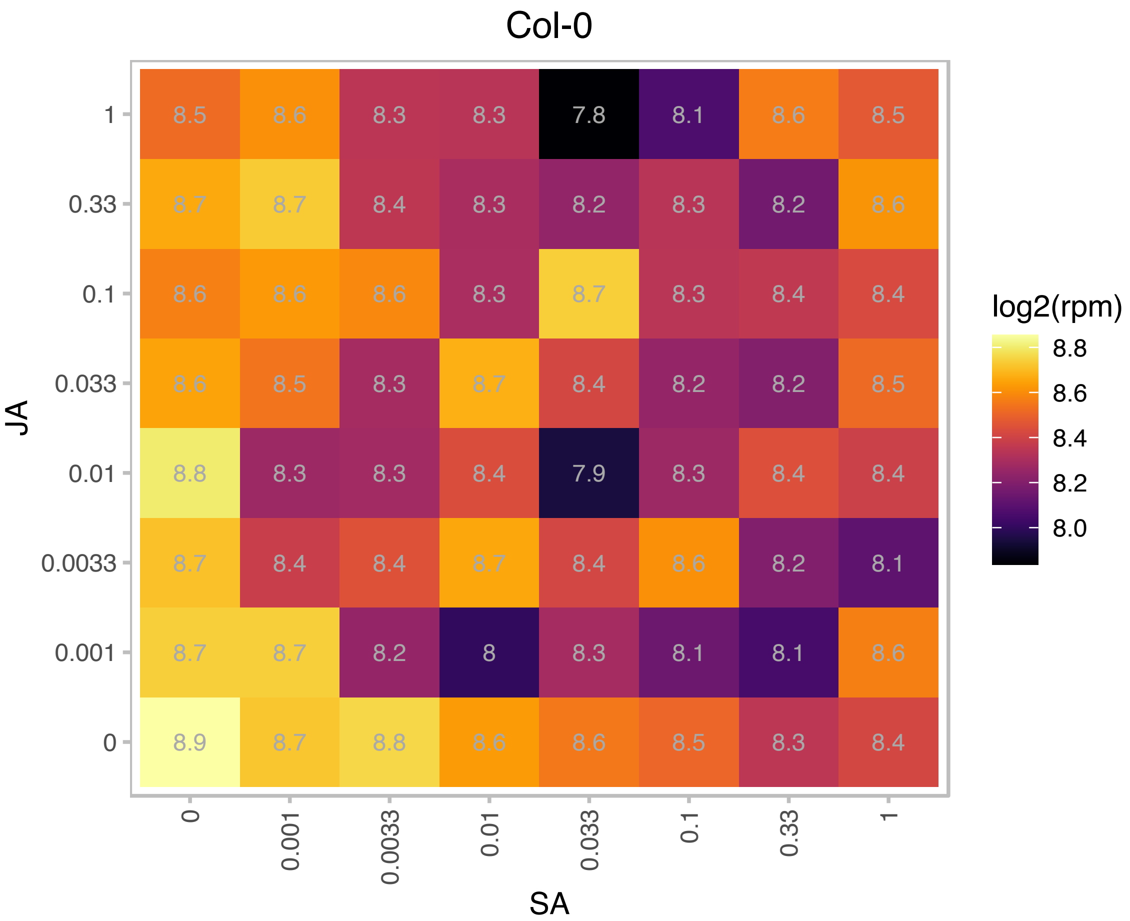


| Gene | AT4G35450 |
|---|---|
| is.expressed_genes | TRUE |
| is.responsive_genes | TRUE |
| is.apcluster | G-02 |

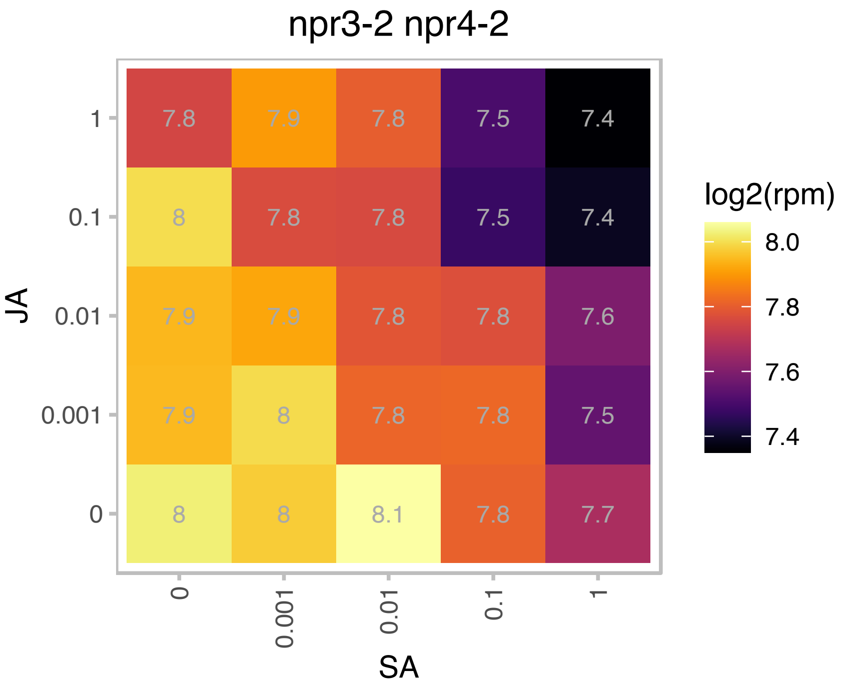


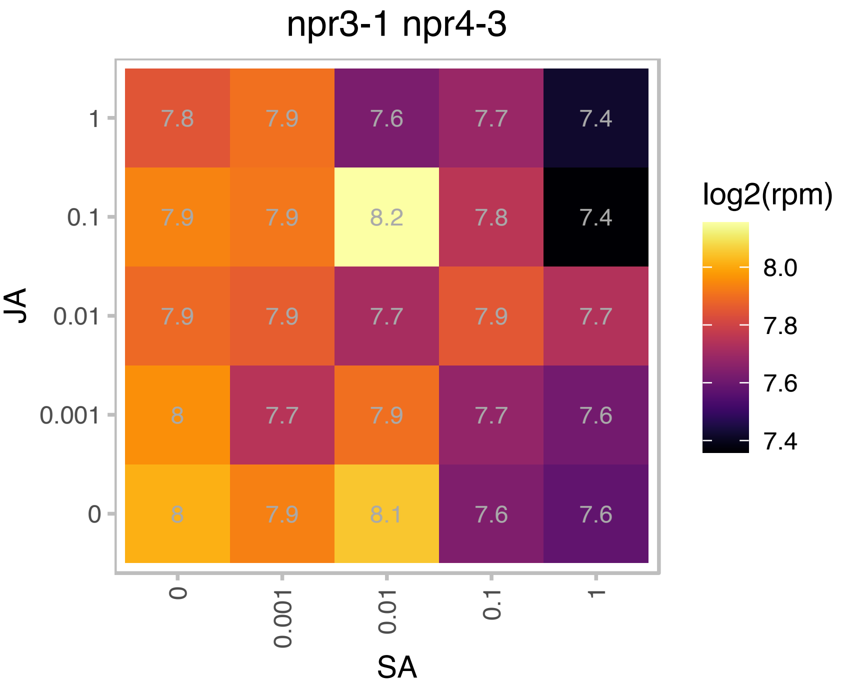

# AT4G38932.1 : NA

Potential natural antisense gene, locus overlaps with AT4G38930

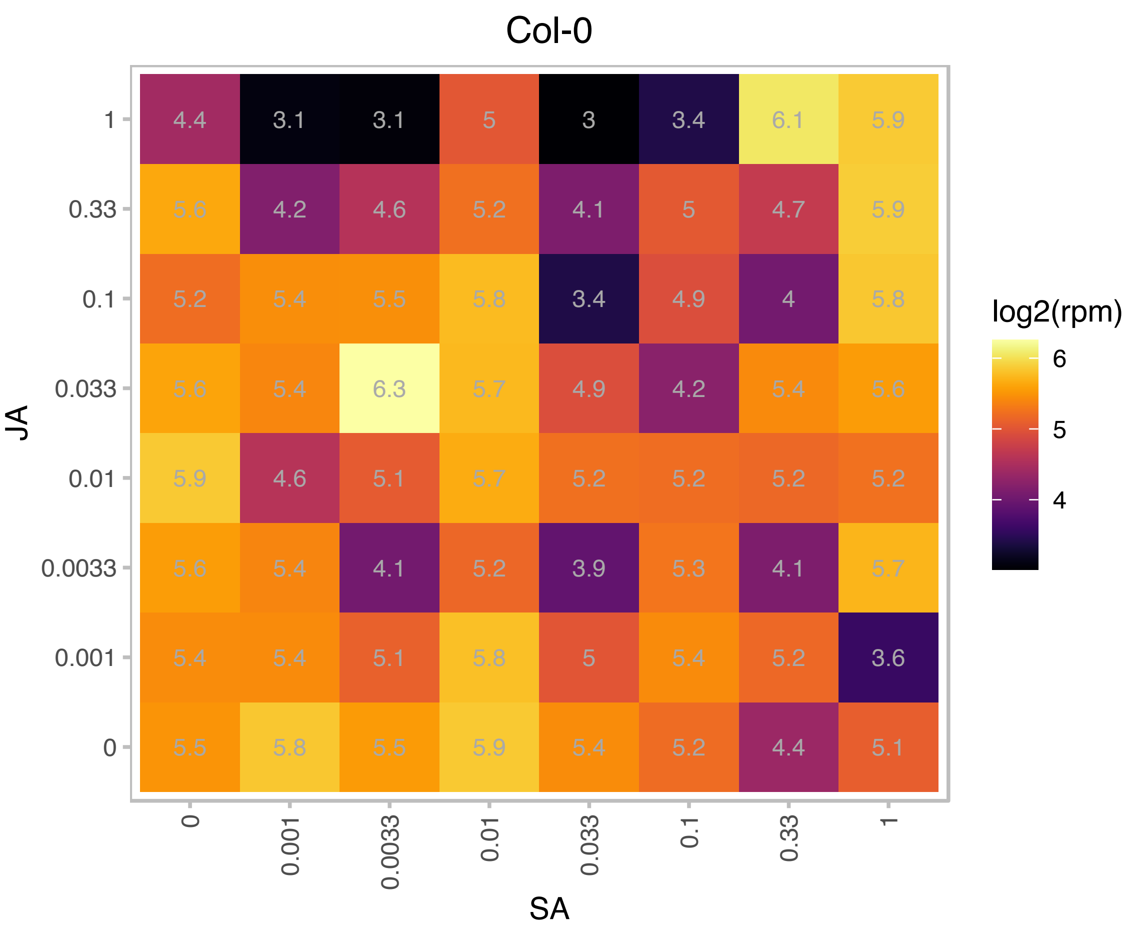


| | |
|---|---|
| Gene | AT4G38932 |
| is.expressed_genes | TRUE |
| is.responsive_genes | TRUE |
| is.apcluster | G-02 |

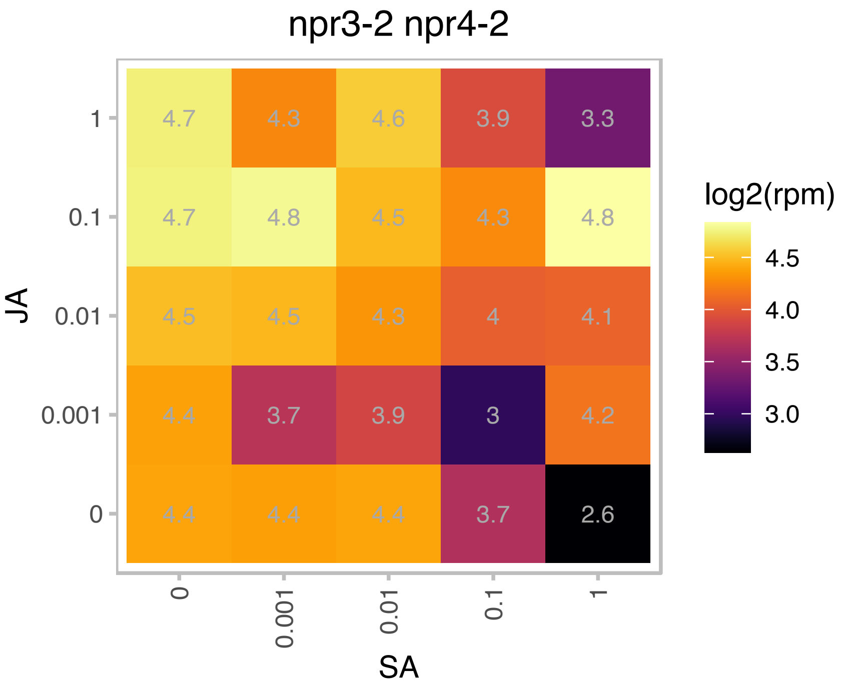


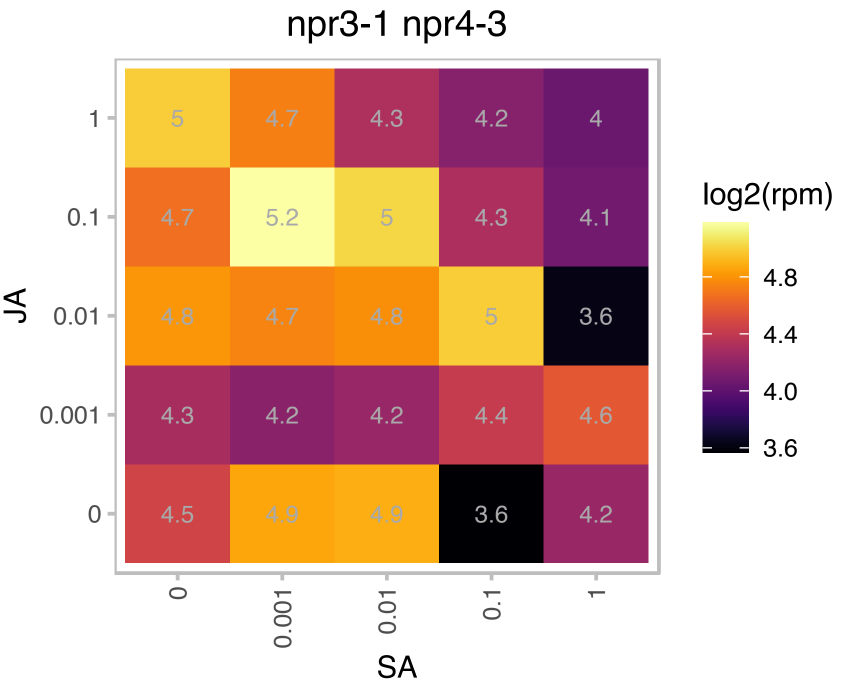

### AT5G51970.1 : ATSDH

GroES-like zinc-binding alcohol dehydrogenase family protein

Encodes a putative sorbitol dehydrogenase that can be thiolated in vitro.

Col-0

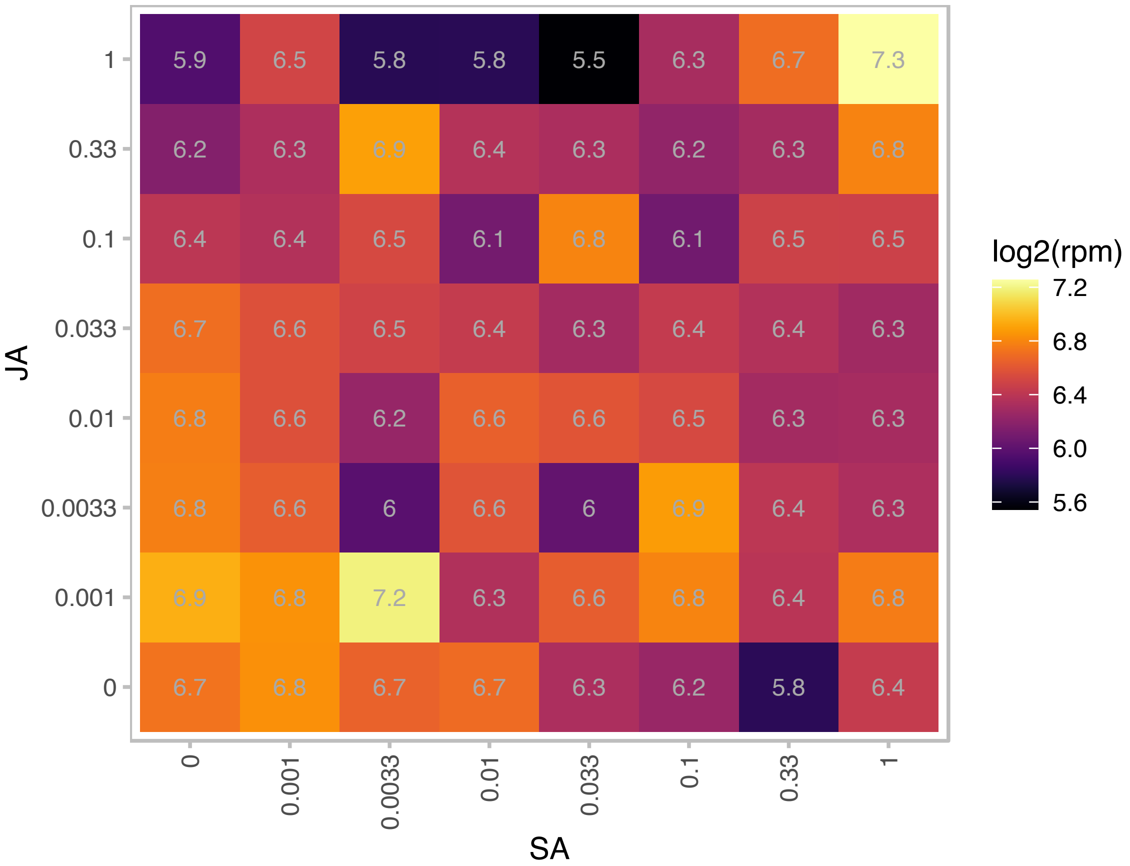


| | |
|---|---|
| Gene | AT5G51970 |
| is.expressed_genes | TRUE |
| is.responsive_genes | TRUE |
| is.apcluster | G-02 |

npr3-2 npr4-2

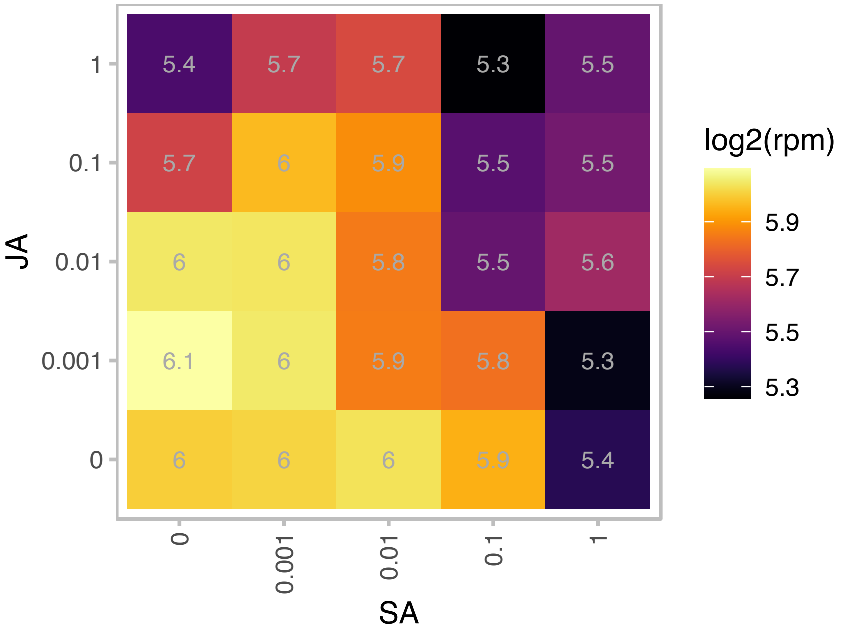


npr3-1 npr4-3

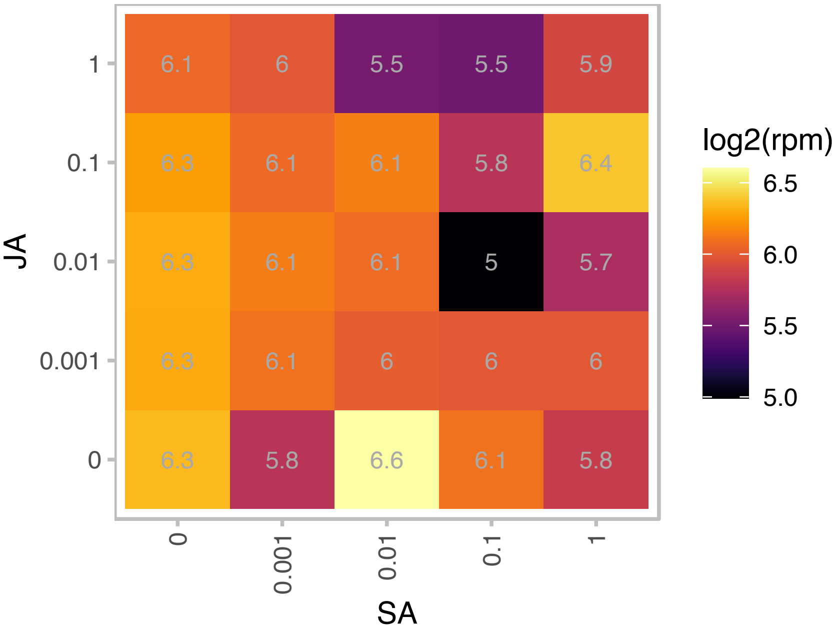

# AT5G53310.1 : NA

myosin heavy chain-related

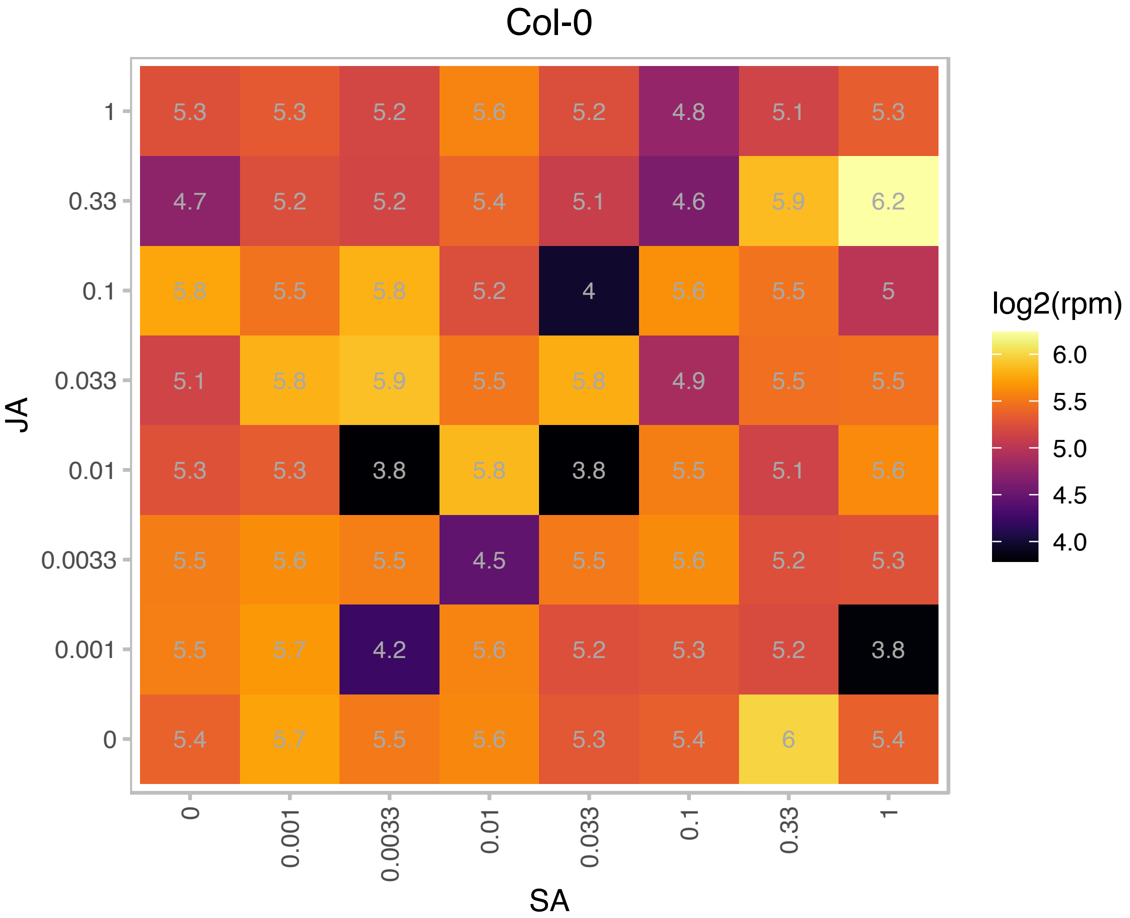


| | |
|---|---|
| Gene | AT5G53310 |
| is.expressed_genes | TRUE |
| is.responsive_genes | TRUE |
| is.apcluster | G-02 |

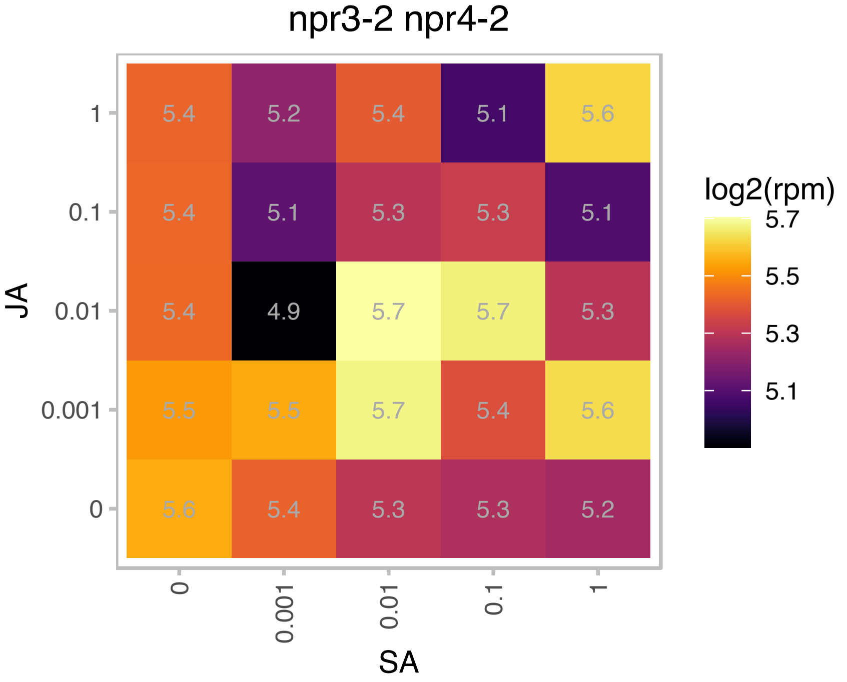


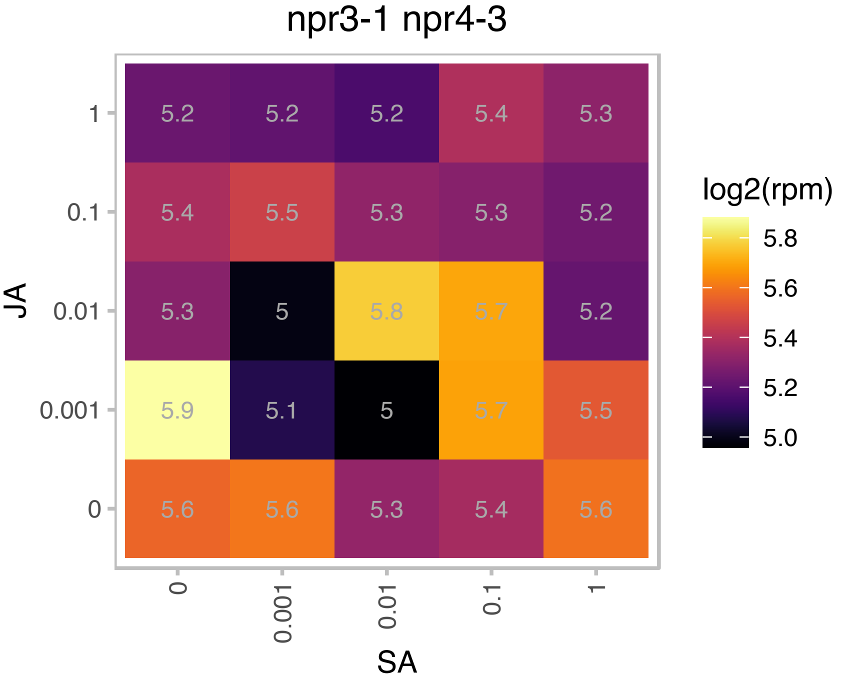

# AT5G64100.1 : NA

Peroxidase superfamily protein

| | |
|---|---|
| Gene | AT5G64100 |
| is.expressed_genes | TRUE |
| is.responsive_genes | TRUE |
| is.apcluster | G-02 |

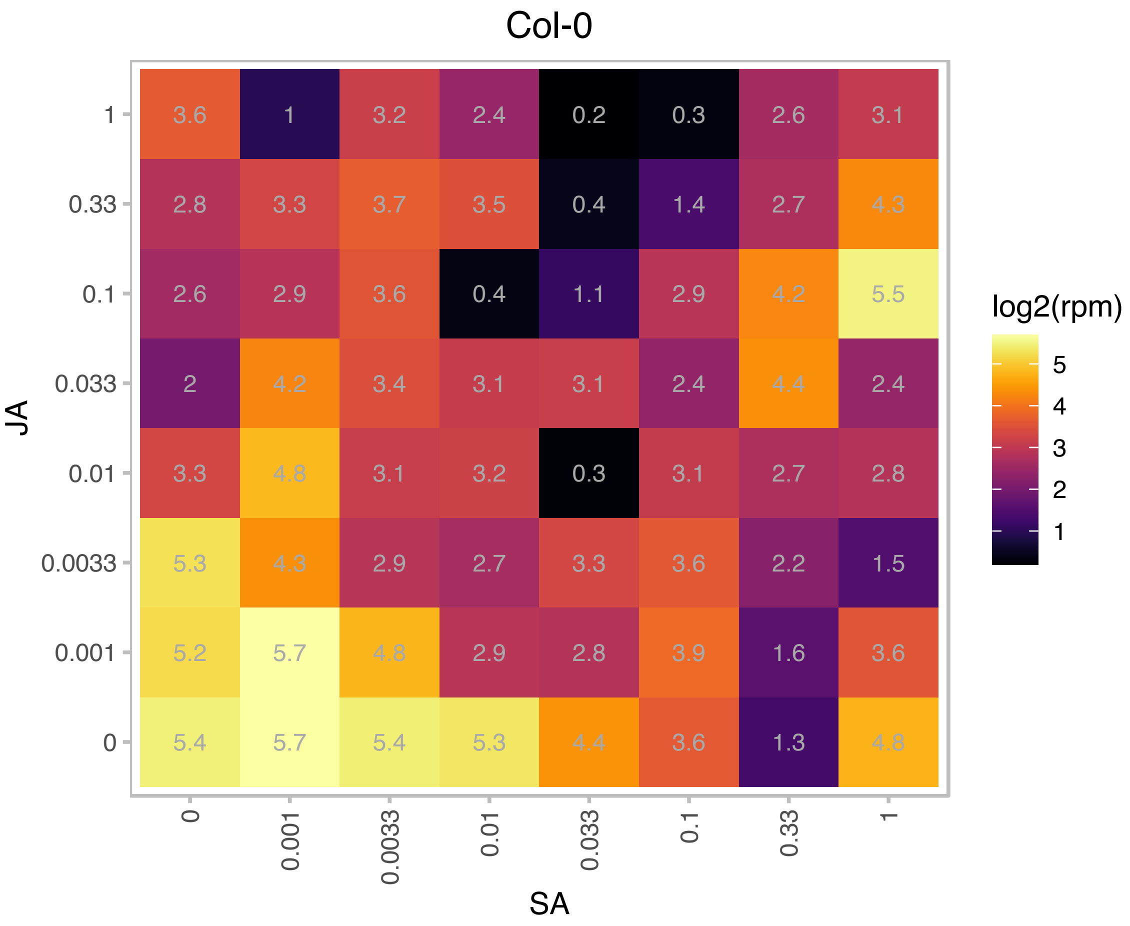


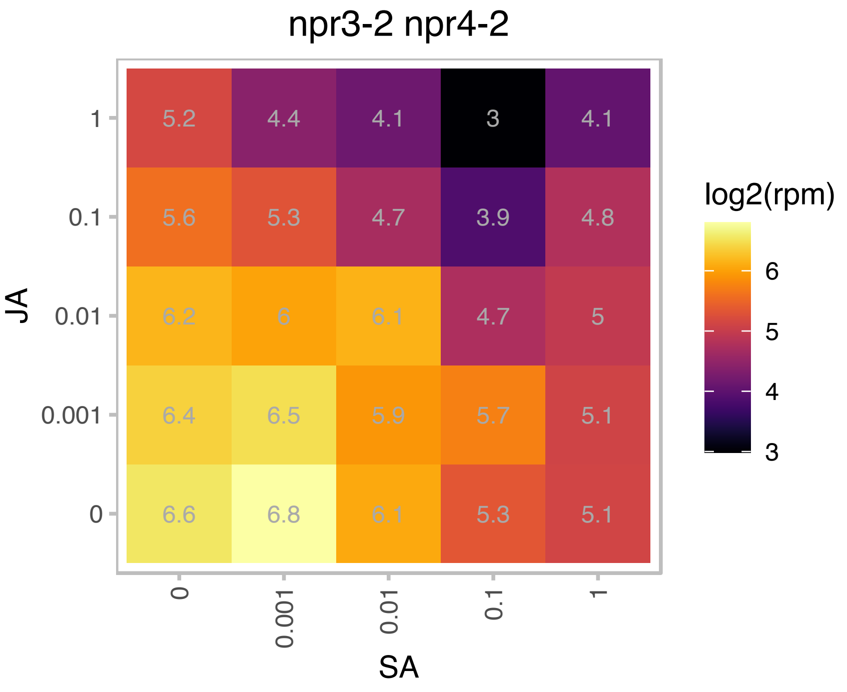


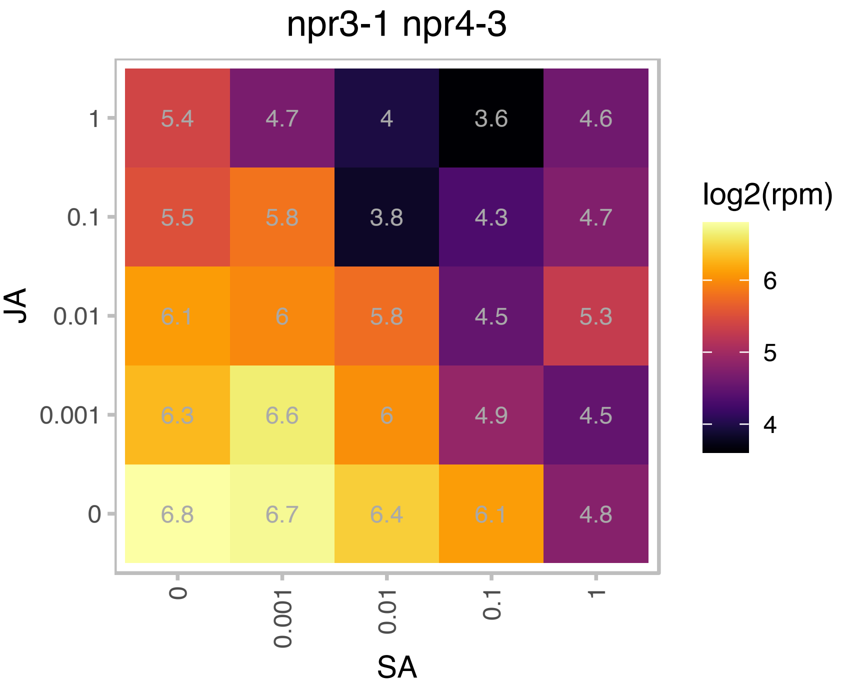

### ATCG00680.1 : PSBB

photosystem II reaction center protein B

encodes for CP47, subunit of the photosystem II reaction center.

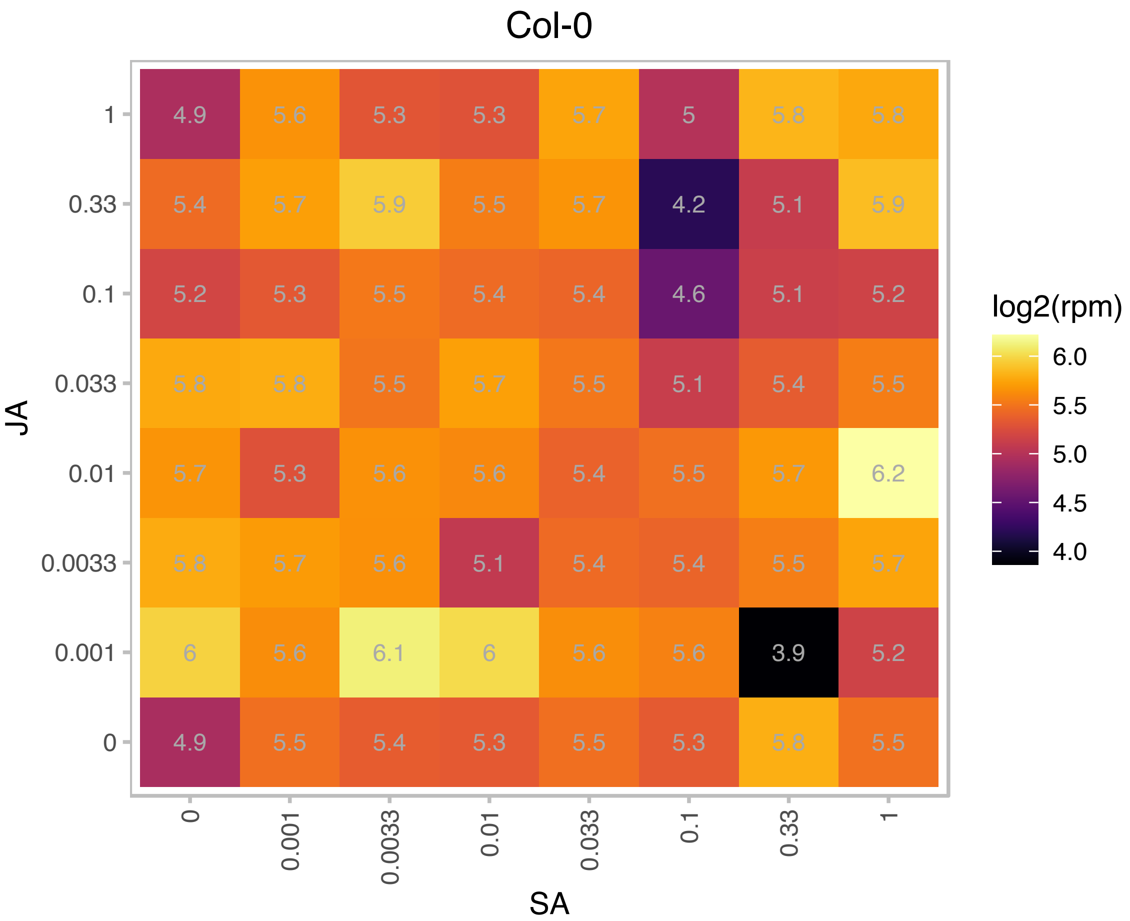


| | |
|---|---|
| Gene | ATCG00680 |
| is.expressed_genes | TRUE |
| is.responsive_genes | TRUE |
| is.apcluster | G-02 |

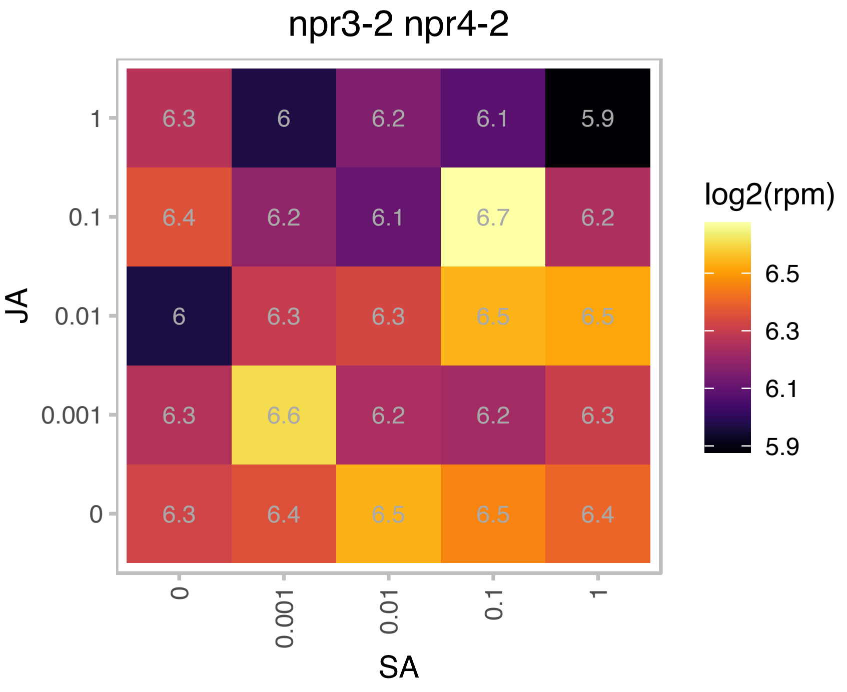


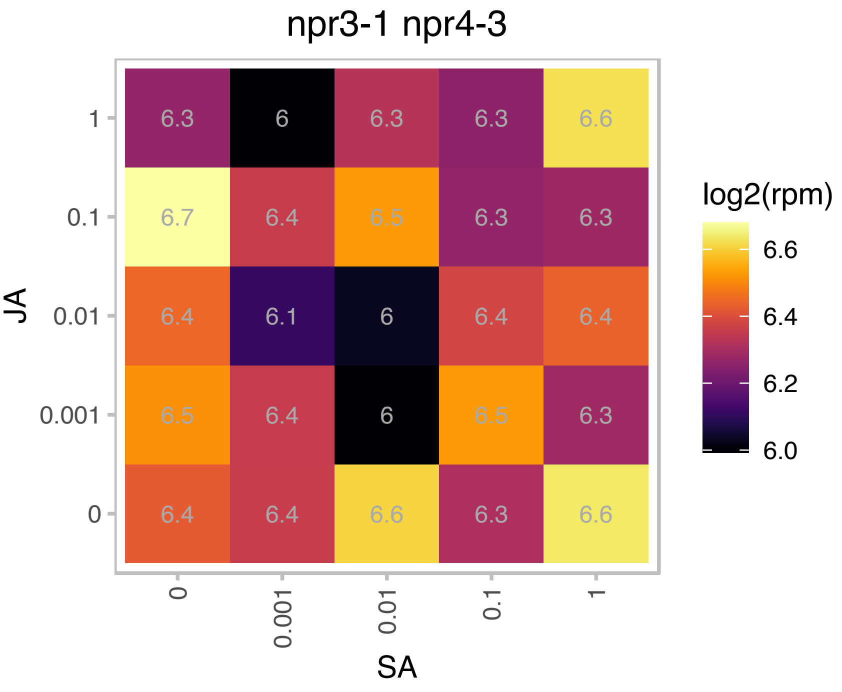

### AT1G08570.1 : ACHT4

atypical CYS HIS rich thioredoxin 4
Encodes a member of the thioredoxin family protein. Located in the chloroplast. Shows high activity towards the chloroplast 2-Cys peroxiredoxin A, and poor activity towards the chloroplast NADP-malate dehydrogenase. The mRNA is cell-to-cell mobile.

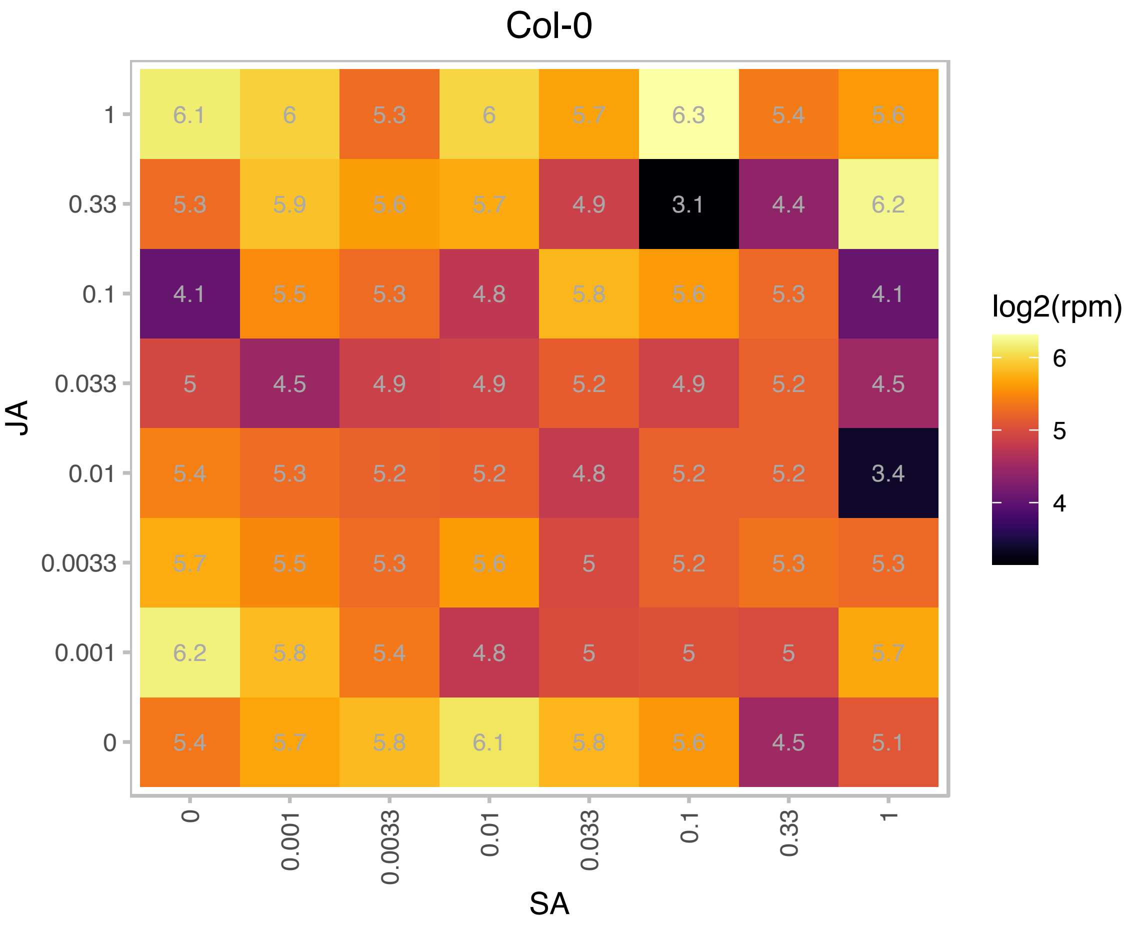


| | |
|---|---|
| Gene | AT1G08570 |
| is.expressed_genes | TRUE |
| is.responsive_genes | TRUE |
| is.apcluster | G-03 |

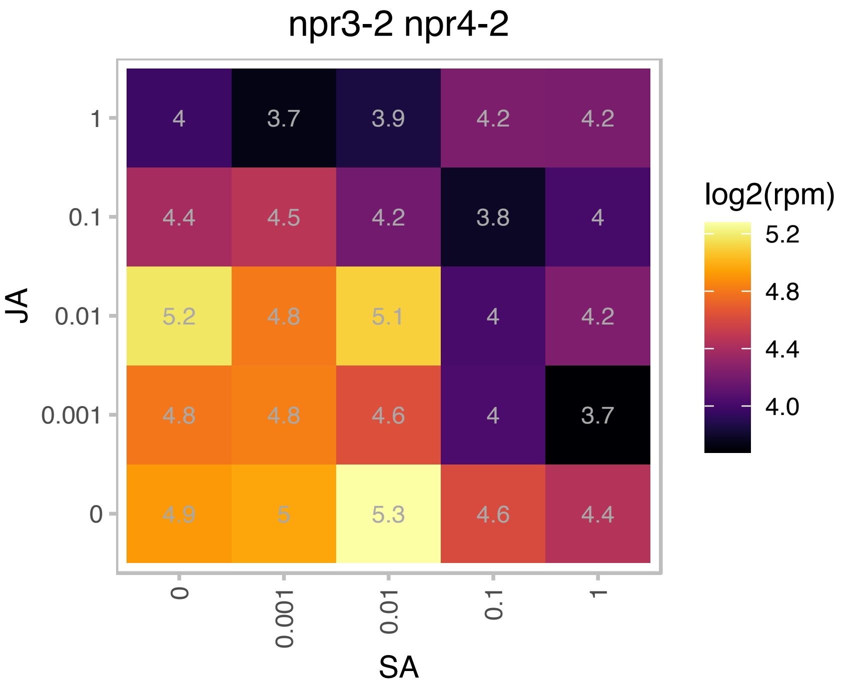


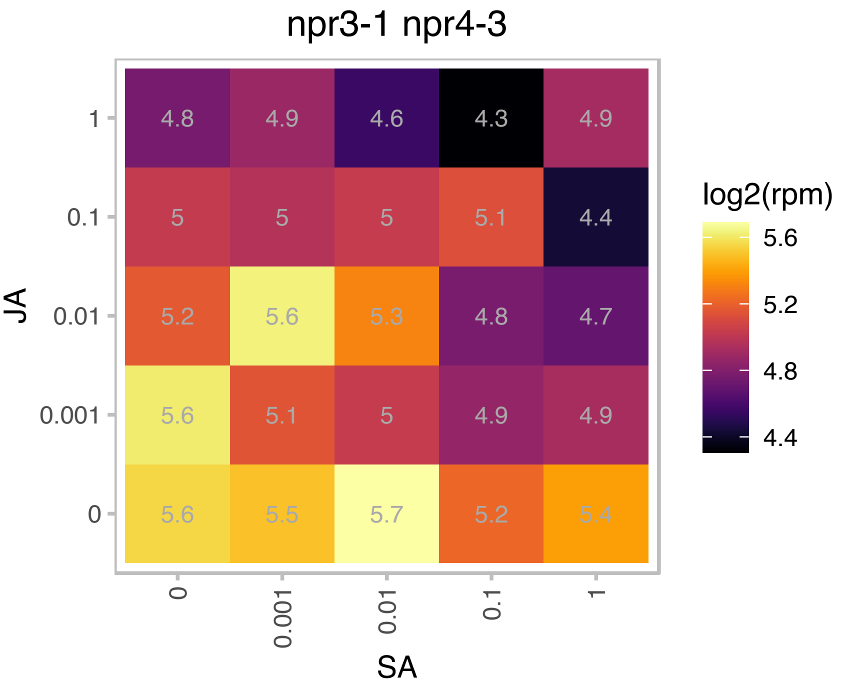

### AT1G17470.1 : ATDRG, ATDRG1, DRG1, Drg1-2

developmentally regulated G-protein 1

Encodes a member of the DRG (developmentally regulated G-protein) family expressed throughout the plant, with highest expression in actively growing tissues. Has GTPase activity.

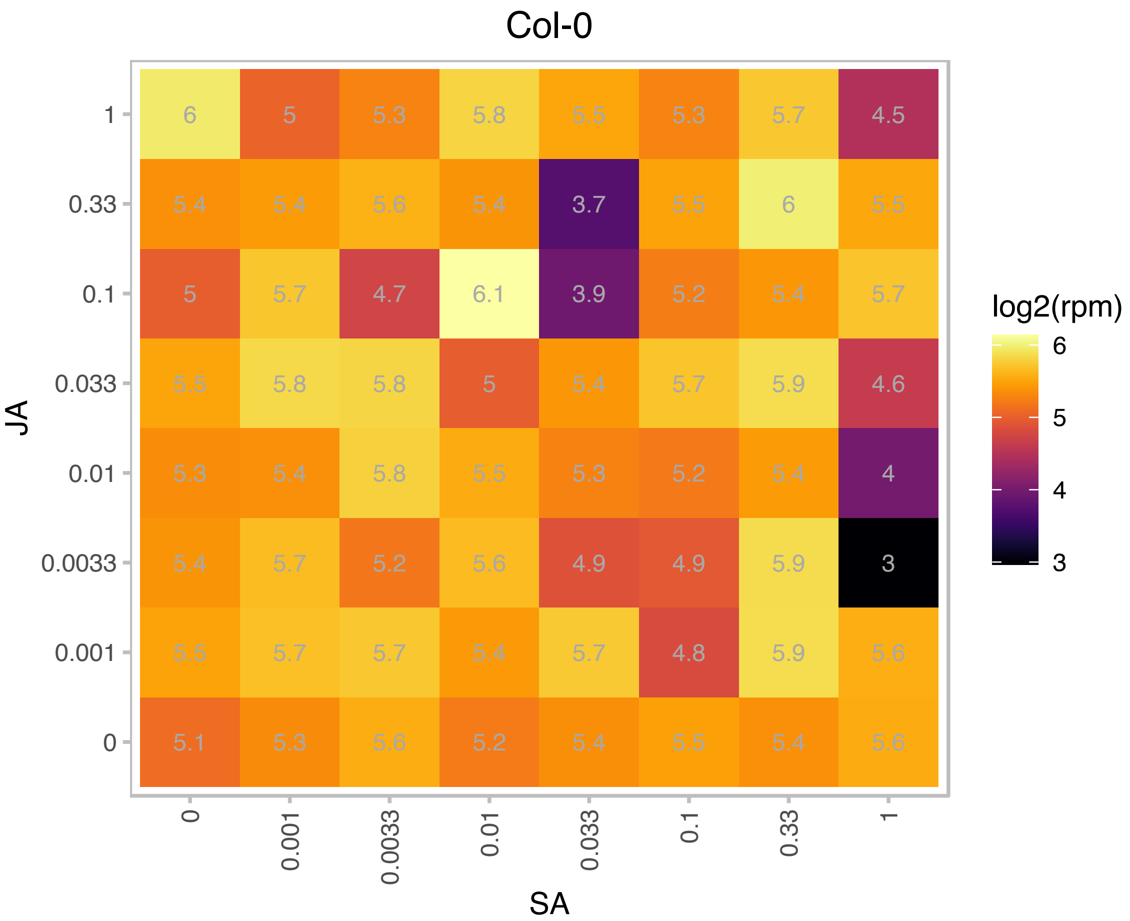


| | |
|---|---|
| Gene | AT1G17470 |
| is.expressed_genes | TRUE |
| is.responsive_genes | TRUE |
| is.apcluster | G-03 |

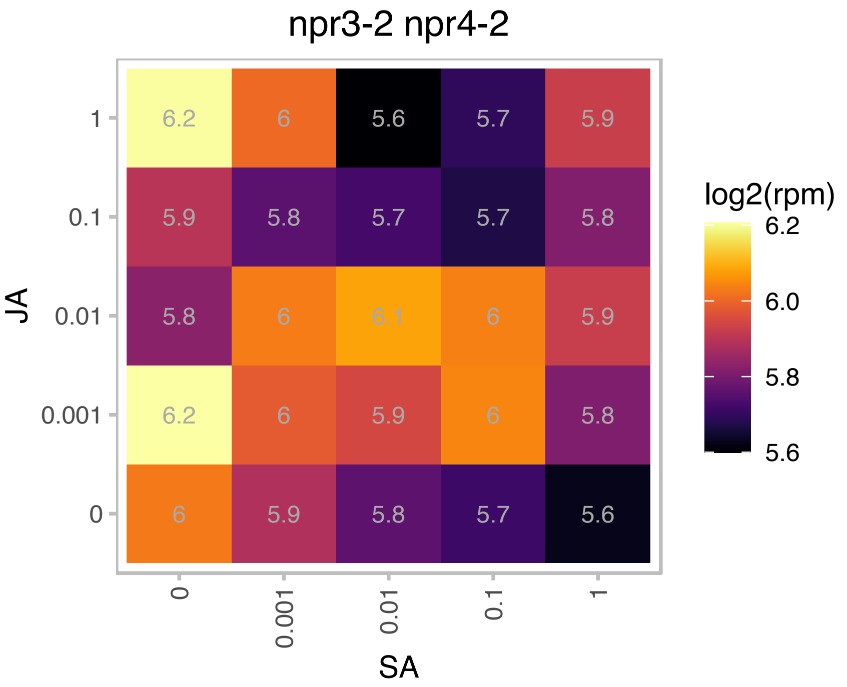


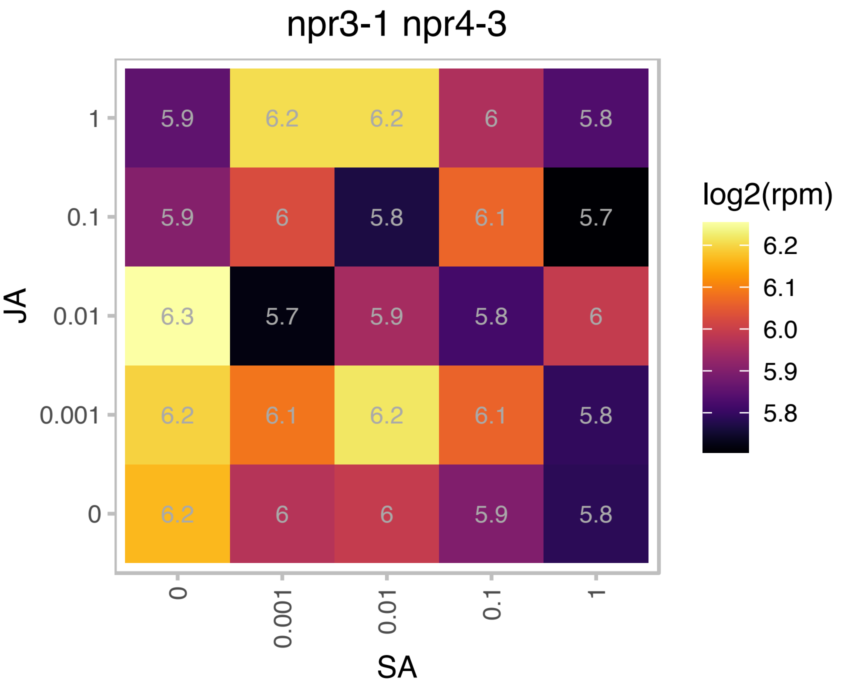

### AT1G54780.1 : AtTLP18.3, TLP18.3

thylakoid lumen 18.3 kDa protein

Encodes a thylakoid lumen protein regulating photosystem II repair cycle. Has acid phosphatase activity. The mRNA is cell-to-cell mobile.

| | |
|---|---|
| Gene | AT1G54780 |
| is.expressed_genes | TRUE |
| is.responsive_genes | TRUE |
| is.apcluster | G-03 |

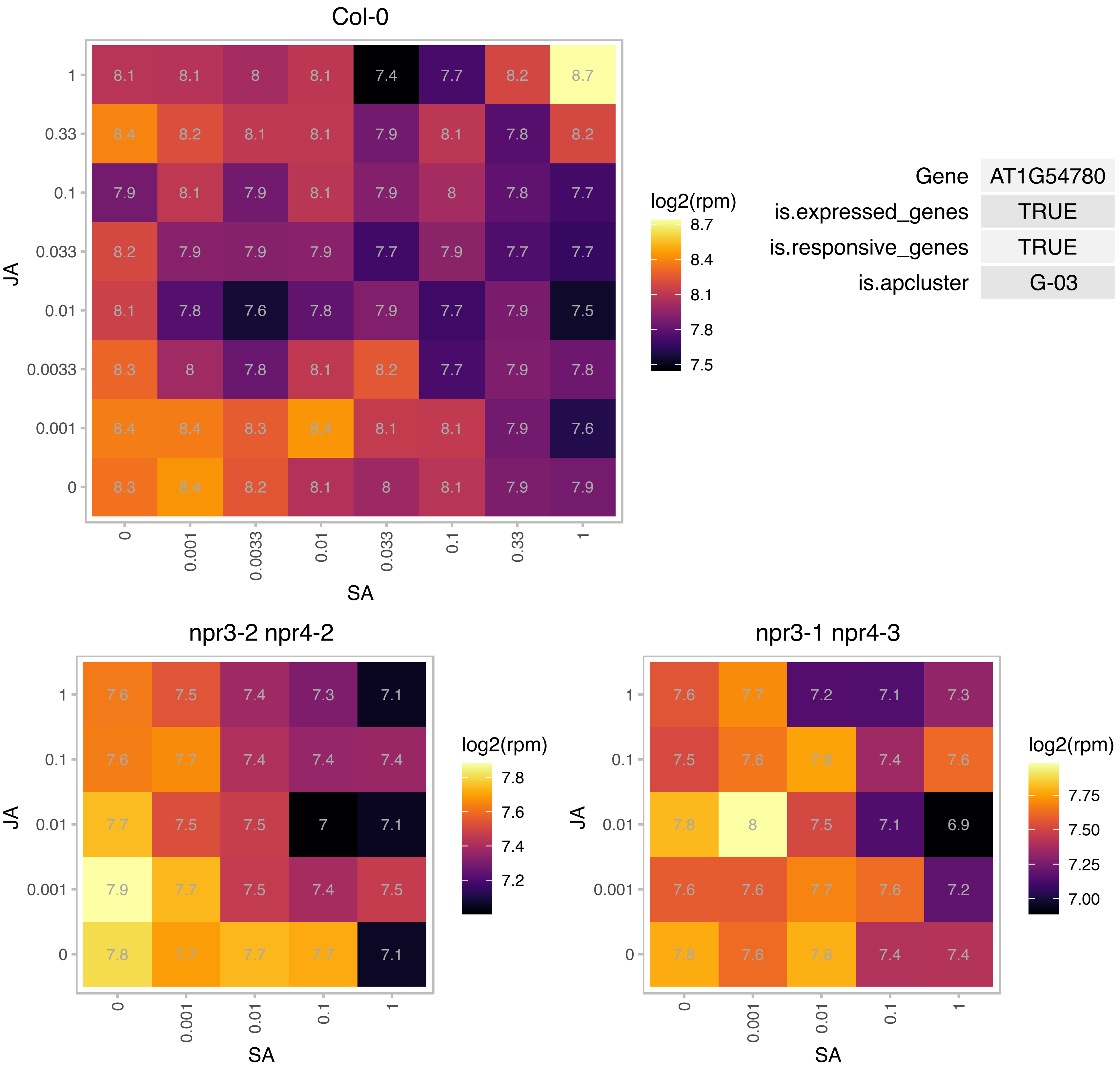

# AT2G03925.1 : NA

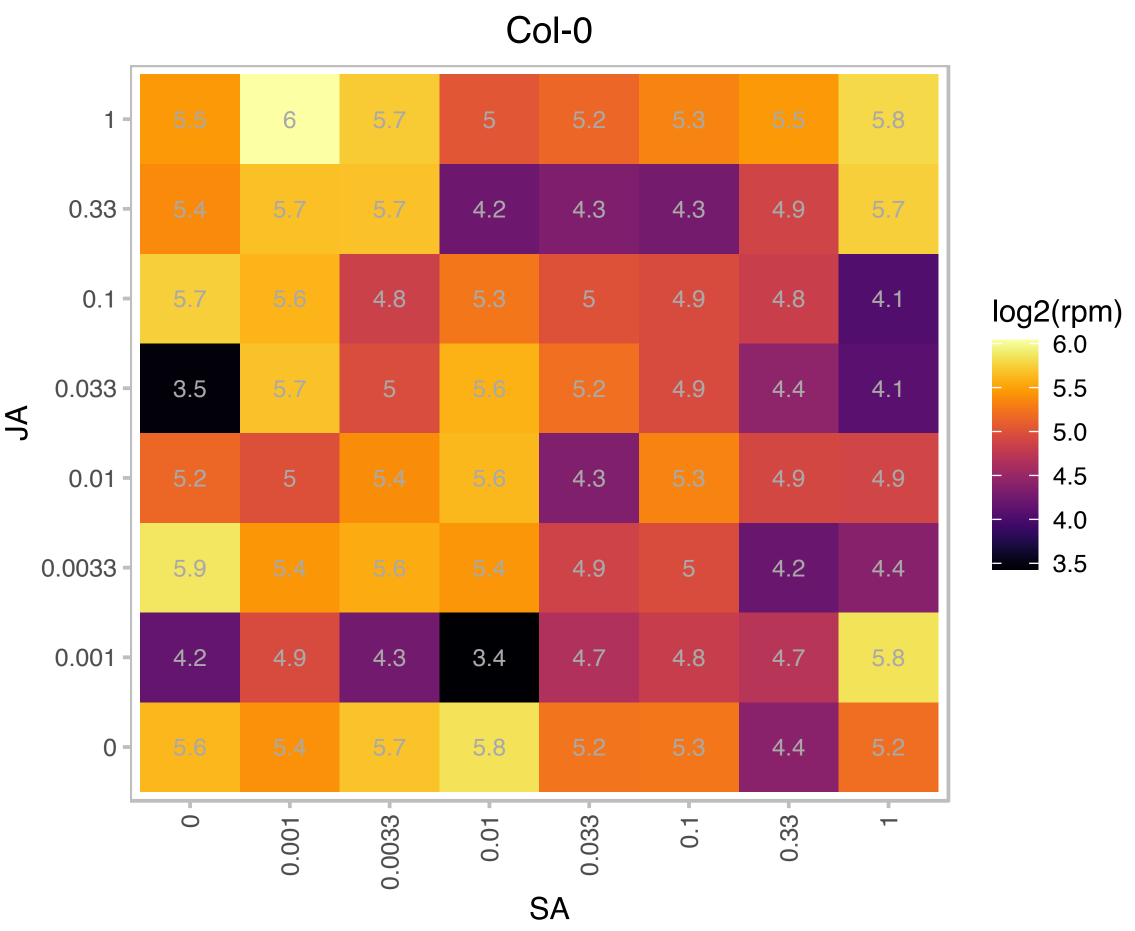


| | |
|---|---|
| Gene | AT2G03925 |
| is.expressed_genes | TRUE |
| is.responsive_genes | TRUE |
| is.apcluster | G-03 |

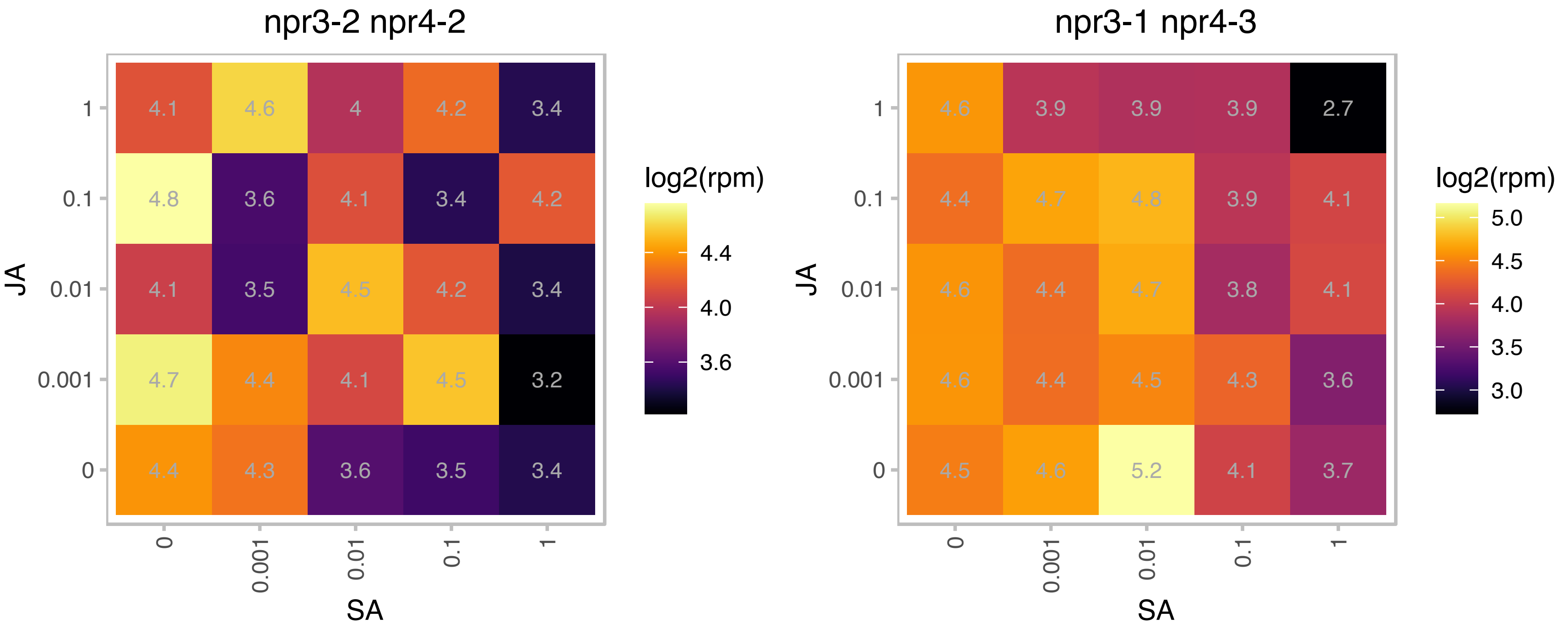

### AT2G06850.1 : EXGT-A1, EXT, XTH4

xyloglucan endotransglucosylase/hydrolase 4

endoxyloglucan transferase (EXGT-A1) gene

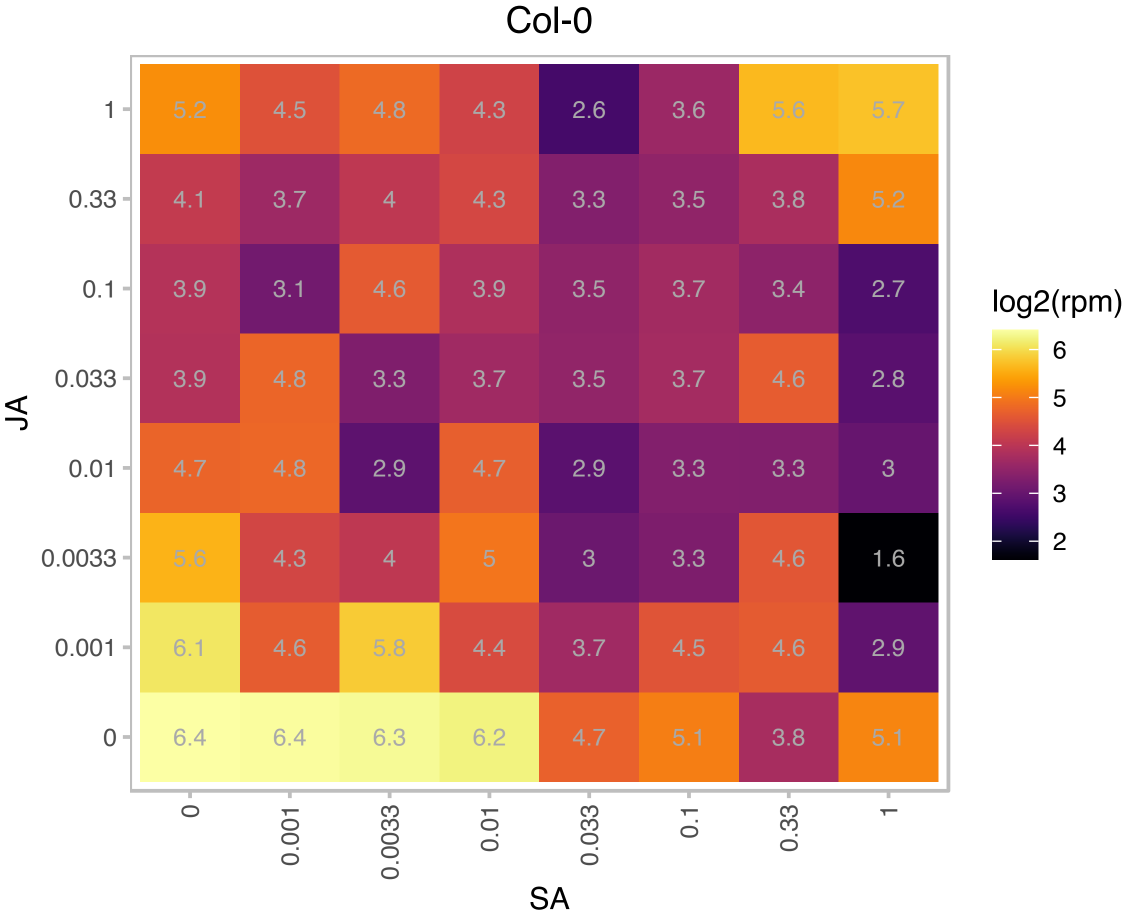


| | |
|---|---|
| Gene | AT2G06850 |
| is.expressed_genes | TRUE |
| is.responsive_genes | TRUE |
| is.apcluster | G-03 |

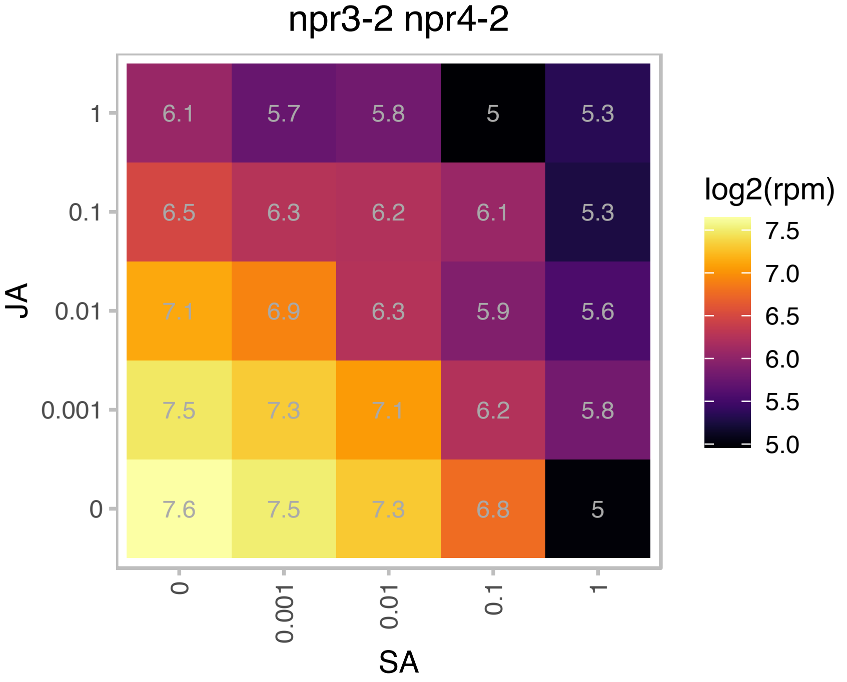


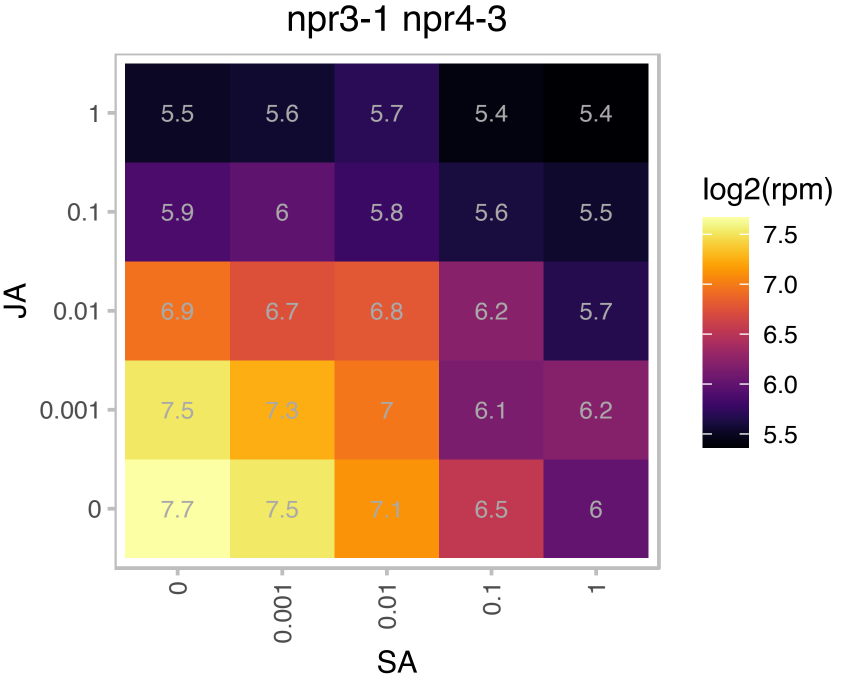

### AT2G29630.1 : PY, THIC

Encodes a protein involved in thiamine biosynthesis. The protein is an iron-sulfur cluster protein predicted to catalyze the conversion of 5-aminoimidazole ribonucleotide (AIR) to hydroxymethylpyrimidine phosphate thiaminC
(HMP-P). A severe reduction of THIC levels in plants decreases vitamin B1 (thiamin diphosphate (TPP)) levels and also leads to changes in the levels of numerous other metabolites since so many primary metabolic enzymes require a TPP co-factor. thiC mutants are chlorotic and arrest in their development at the cotyledon stage. A N-terminal targeting sequence directs the THIC protein to the chloroplast stroma. The THIC transcript is regulated by a TPP riboswitch in the 3' UTR. THIC transcripts are observed in seedlings 5 or more days after

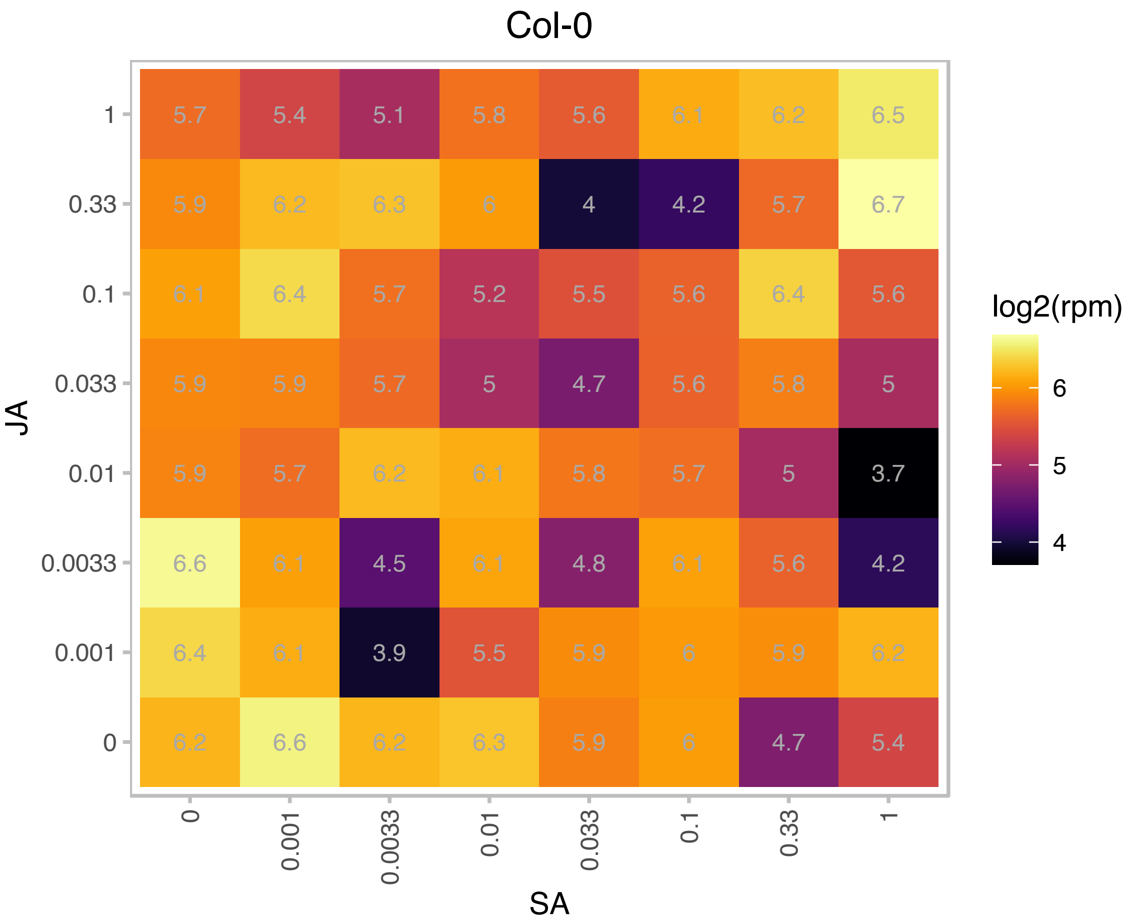


| | |
|---|---|
| Gene | AT2G29630 |
| is.expressed_genes | TRUE |
| is.responsive_genes | TRUE |
| is.apcluster | G-03 |

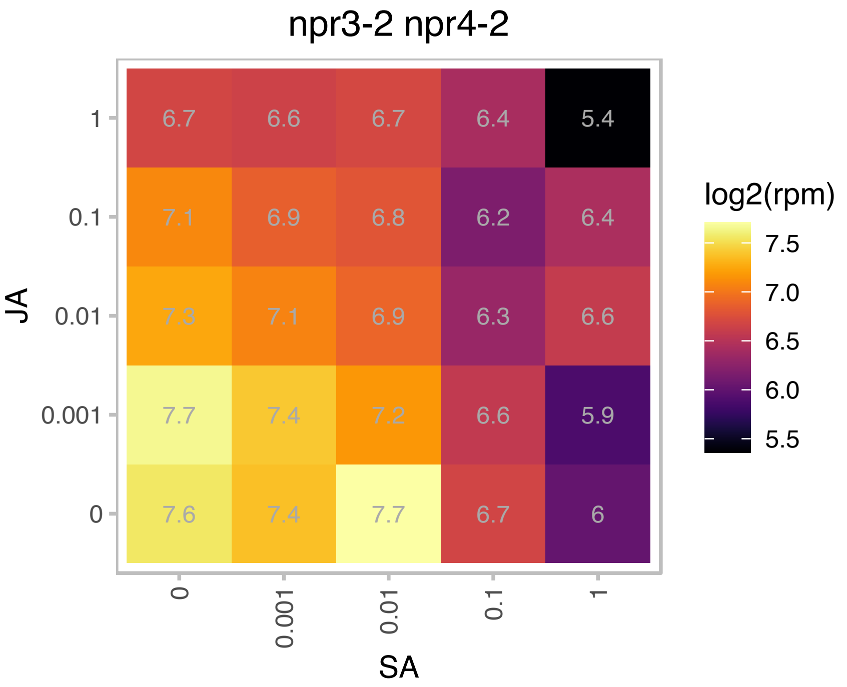


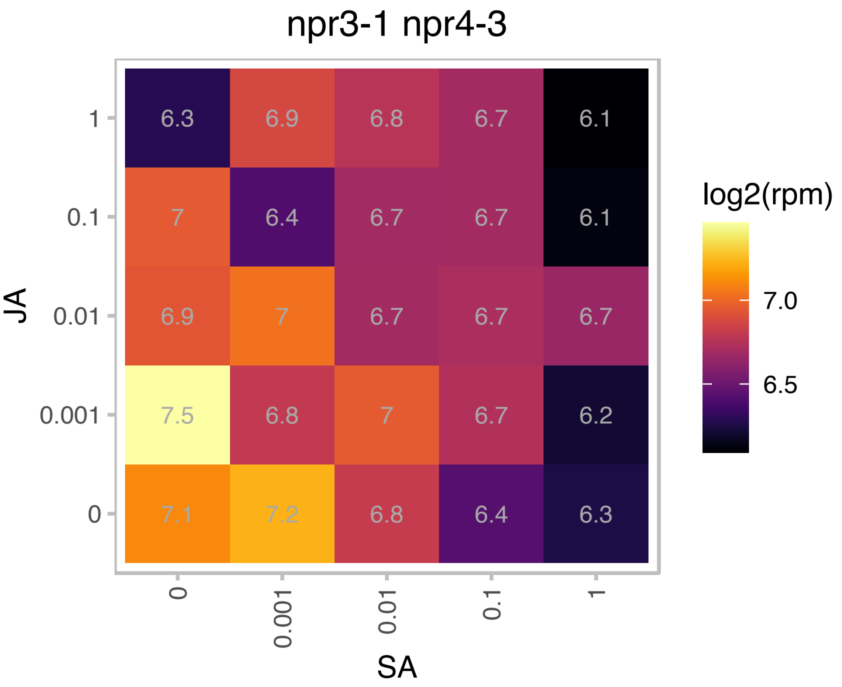

### AT2G38230.1 : ATPDX1.1, PDX1.1

pyridoxine biosynthesis 1.1

Encodes a protein predicted to function in tandem with PDX2 to form glutamine amidotransferase complex with involved in vitamin B6 biosynthesis. The mRNA is cell-to-cell mobile.

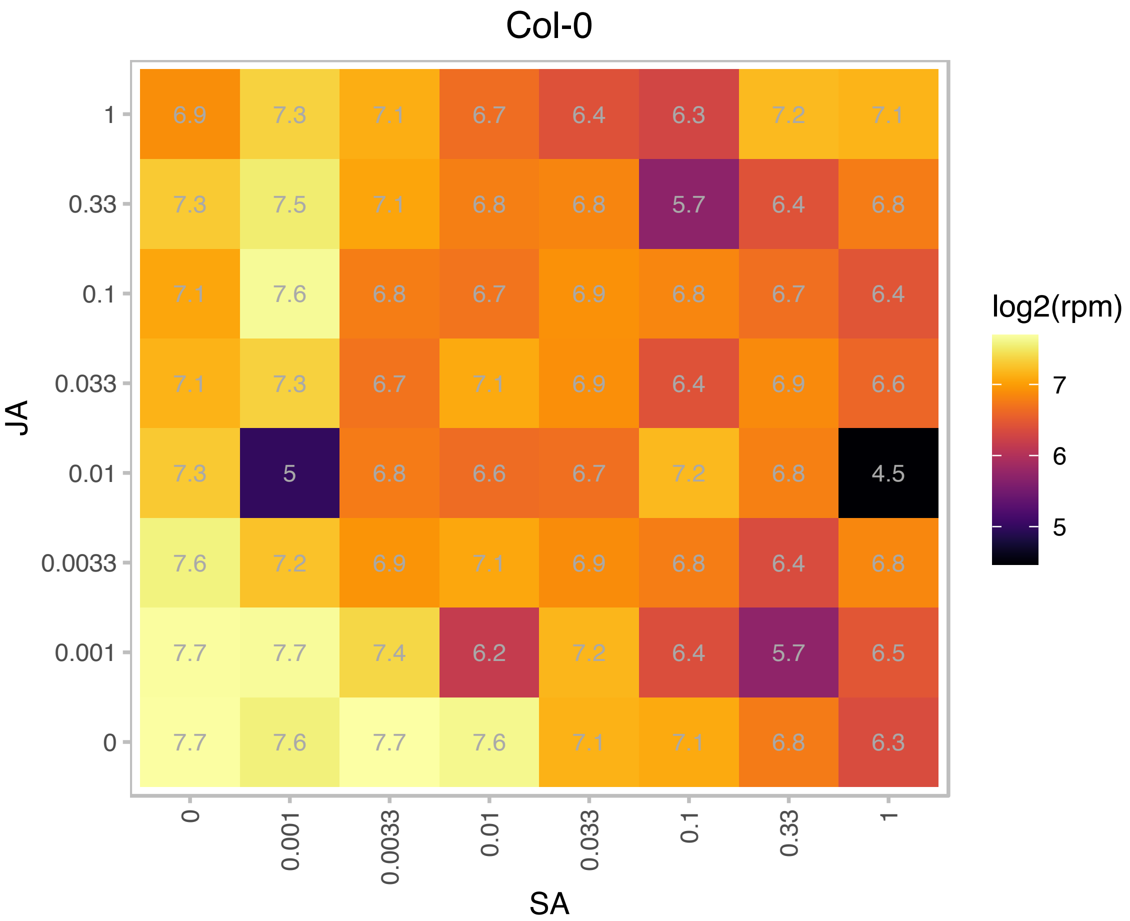


| | |
|---|---|
| Gene | AT2G38230 |
| is.expressed_genes | TRUE |
| is.responsive_genes | TRUE |
| is.apcluster | G-03 |

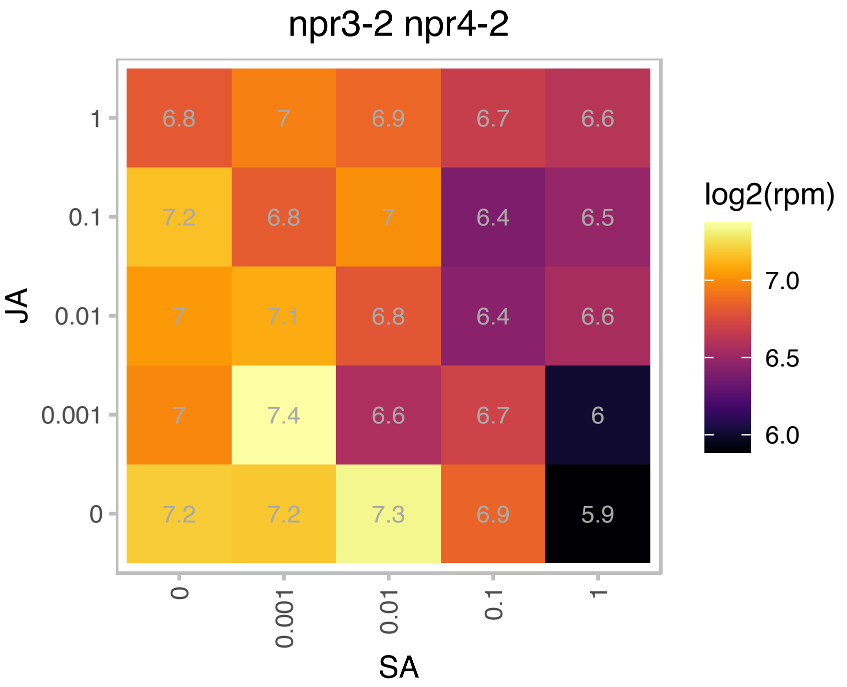


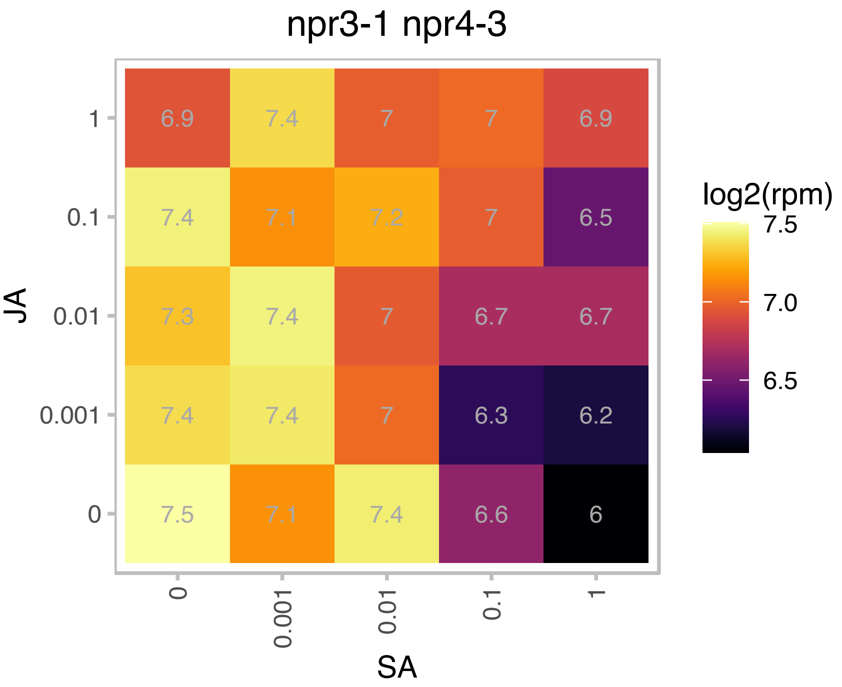

### AT2G42600.1 : ATPPC2, PPC2

phosphoenolpyruvate carboxylase 2

Encodes one of four Arabidopsis phosphoenolpyruvate carboxylase proteins.PPC1 and PPC2 are crucial for balancing carbon and nitrogen metabolism.

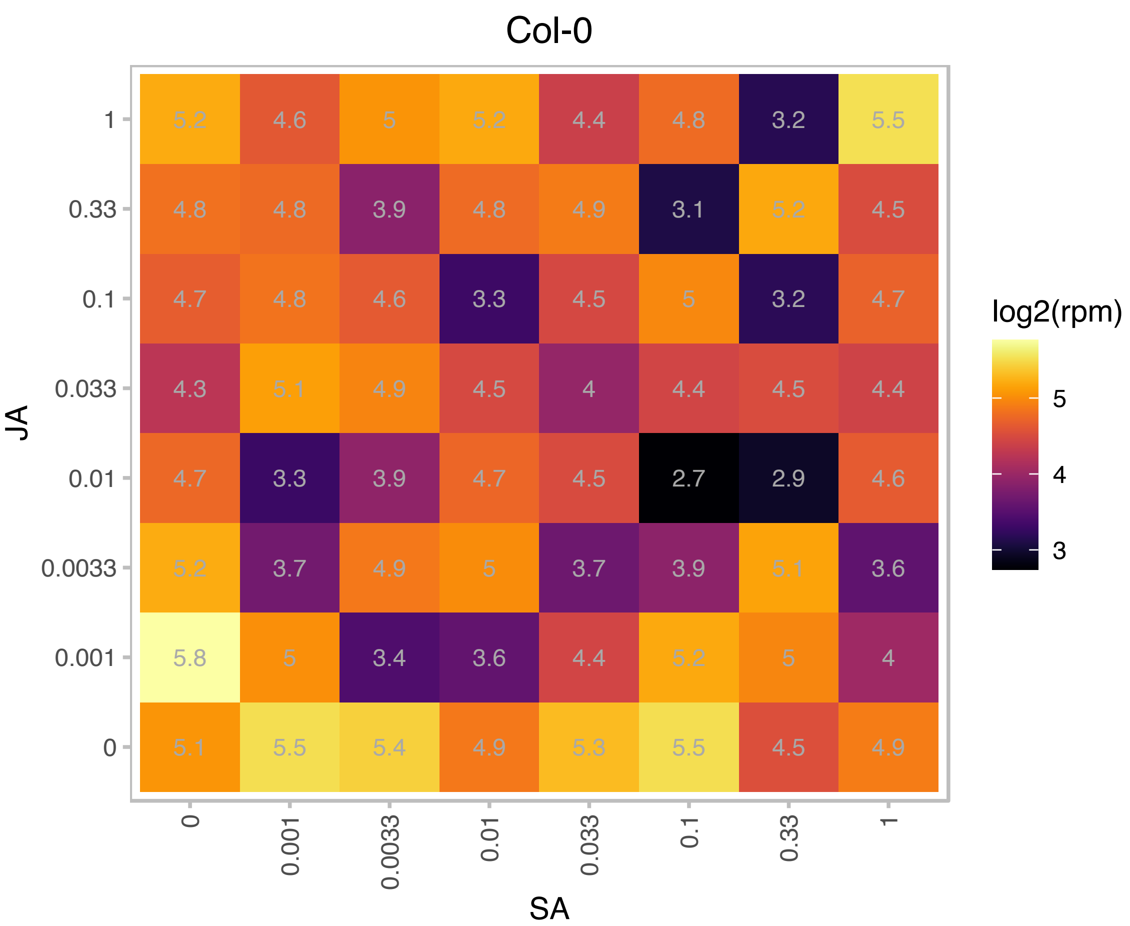


| | |
|---|---|
| Gene | AT2G42600 |
| is.expressed_genes | TRUE |
| is.responsive_genes | TRUE |
| is.apcluster | G-03 |

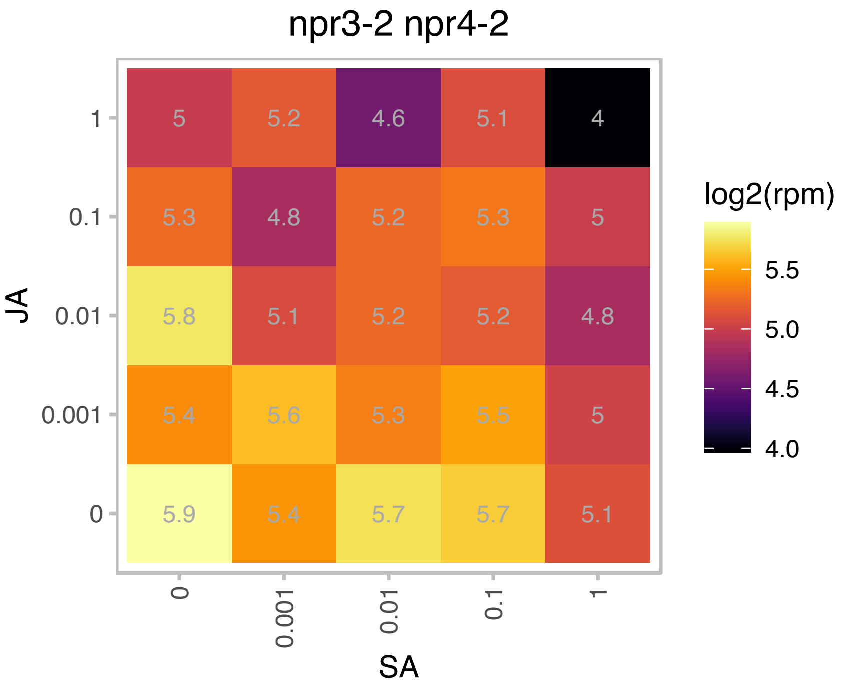


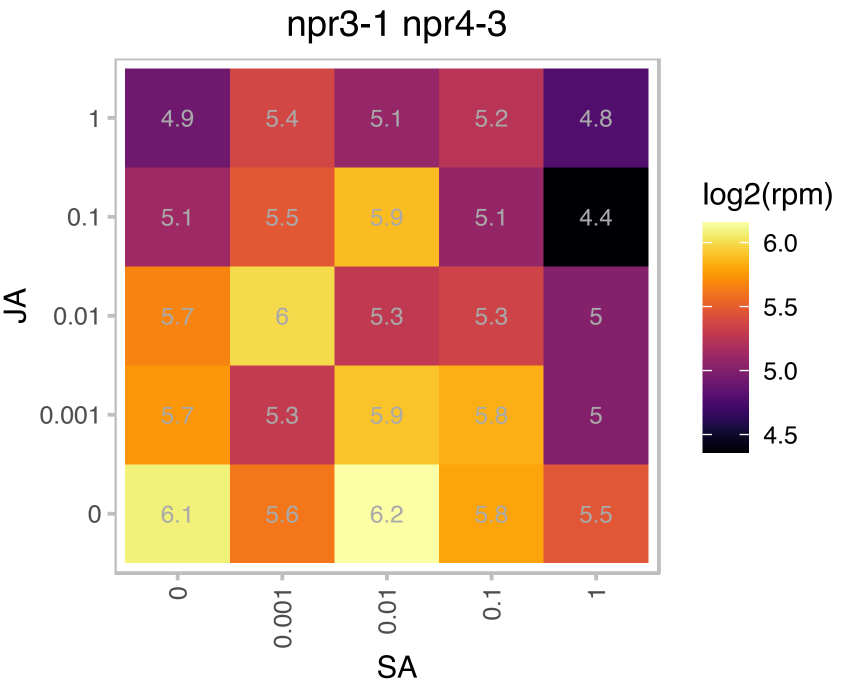

### AT2G42880.1 : ATMPK20, MPK20

MAP kinase 20

member of MAP Kinase

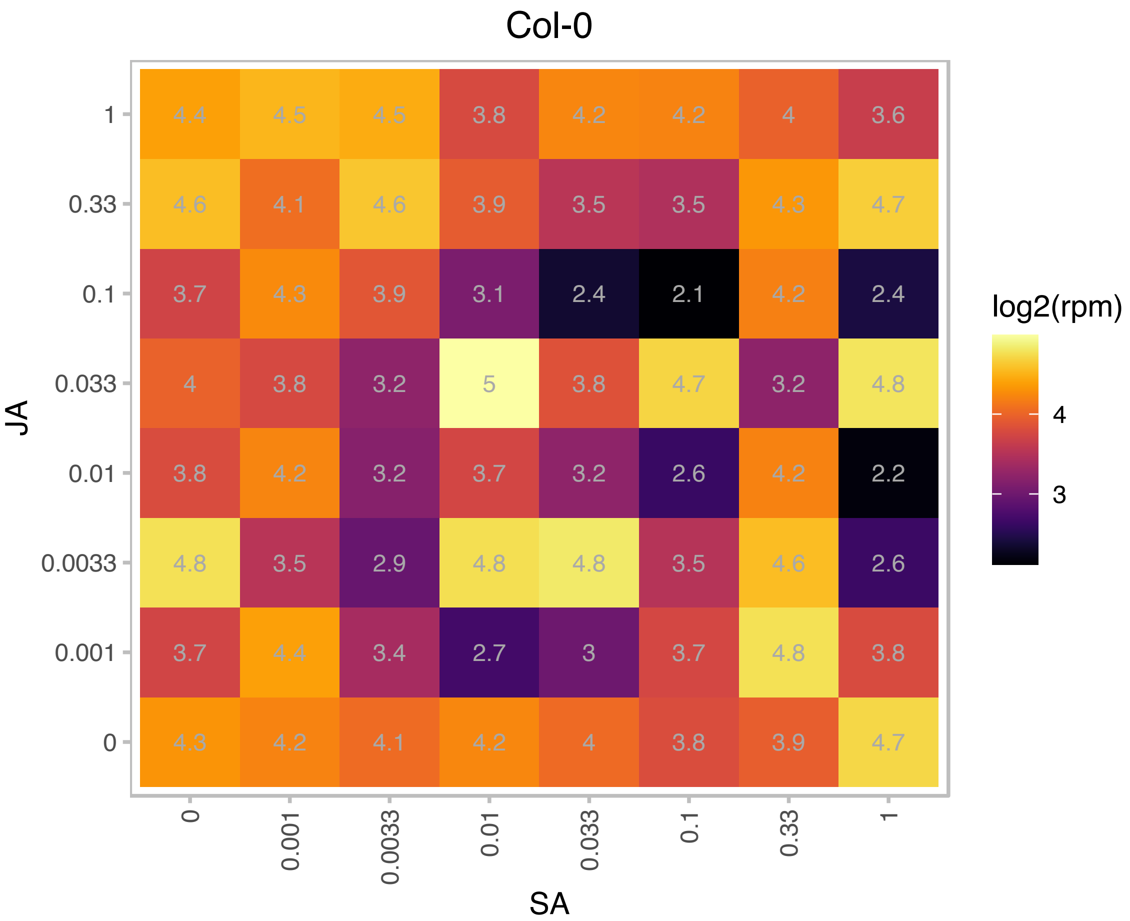


| | |
|---|---|
| Gene | AT2G42880 |
| is.expressed_genes | TRUE |
| is.responsive_genes | TRUE |
| is.apcluster | G-03 |

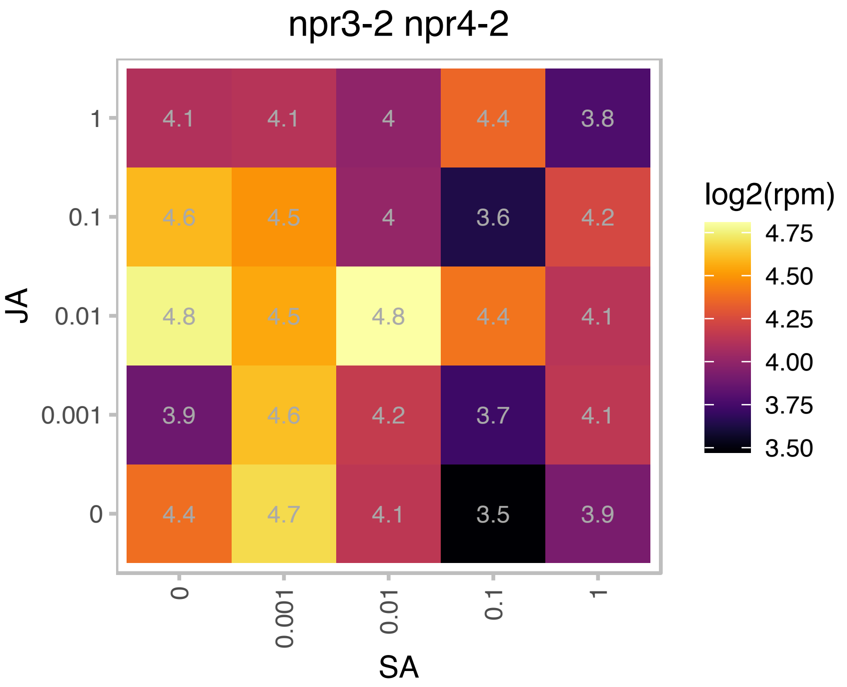


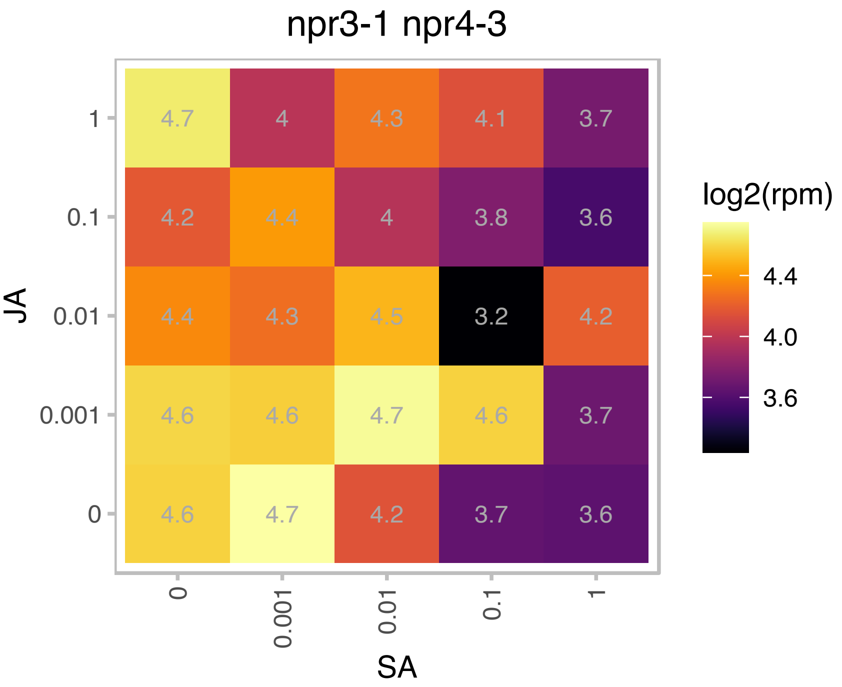

# AT3G06660.1 : NA

PAPA-1-like family protein / zinc finger (HIT type) family protein

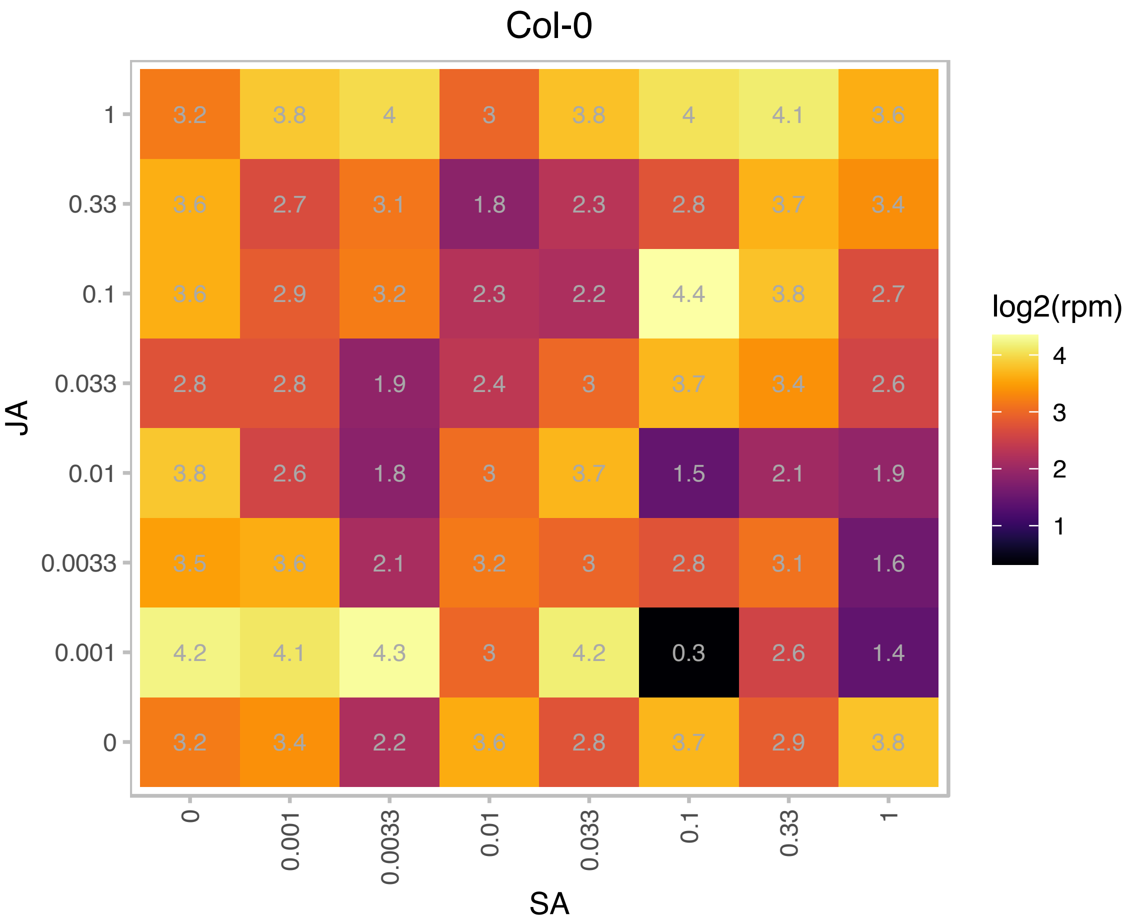


| | |
|---|---|
| Gene | AT3G06660 |
| is.expressed_genes | TRUE |
| is.responsive_genes | TRUE |
| is.apcluster | G-03 |

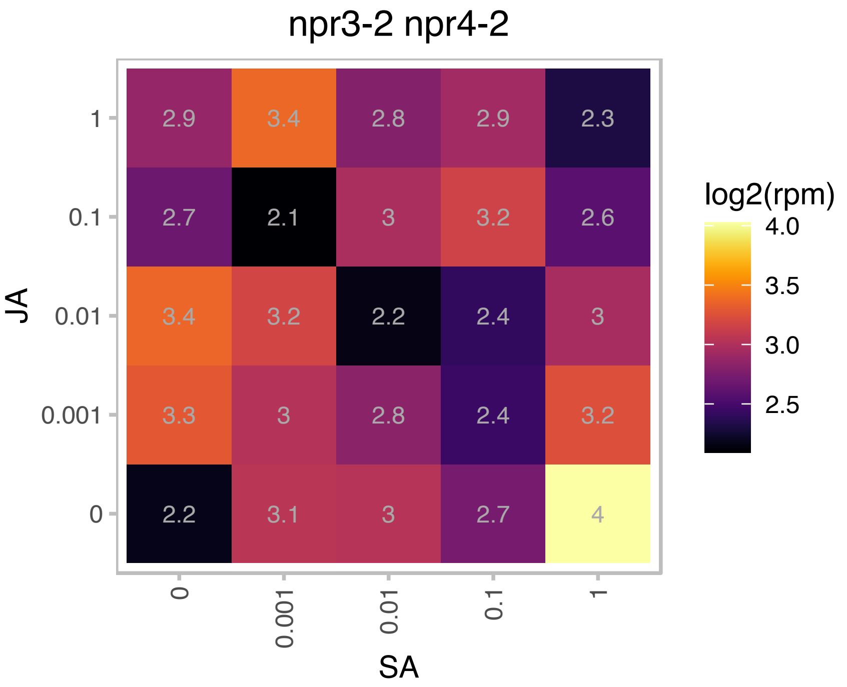


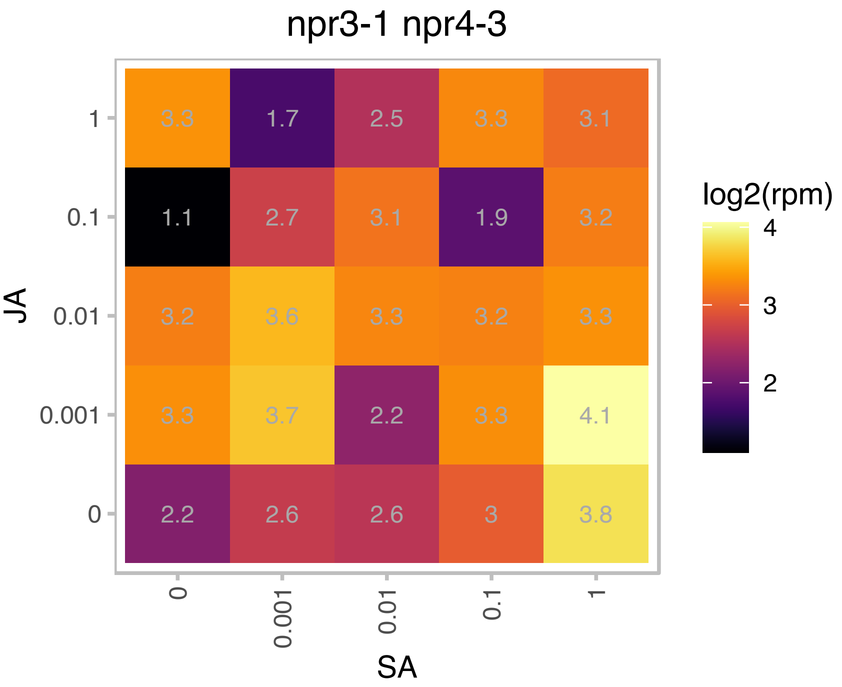

### AT3G13672.1 : SINA2, SINAT6

TRAF-like superfamily protein

SINAT homolog with truncated RING finger and zinc finger domains.

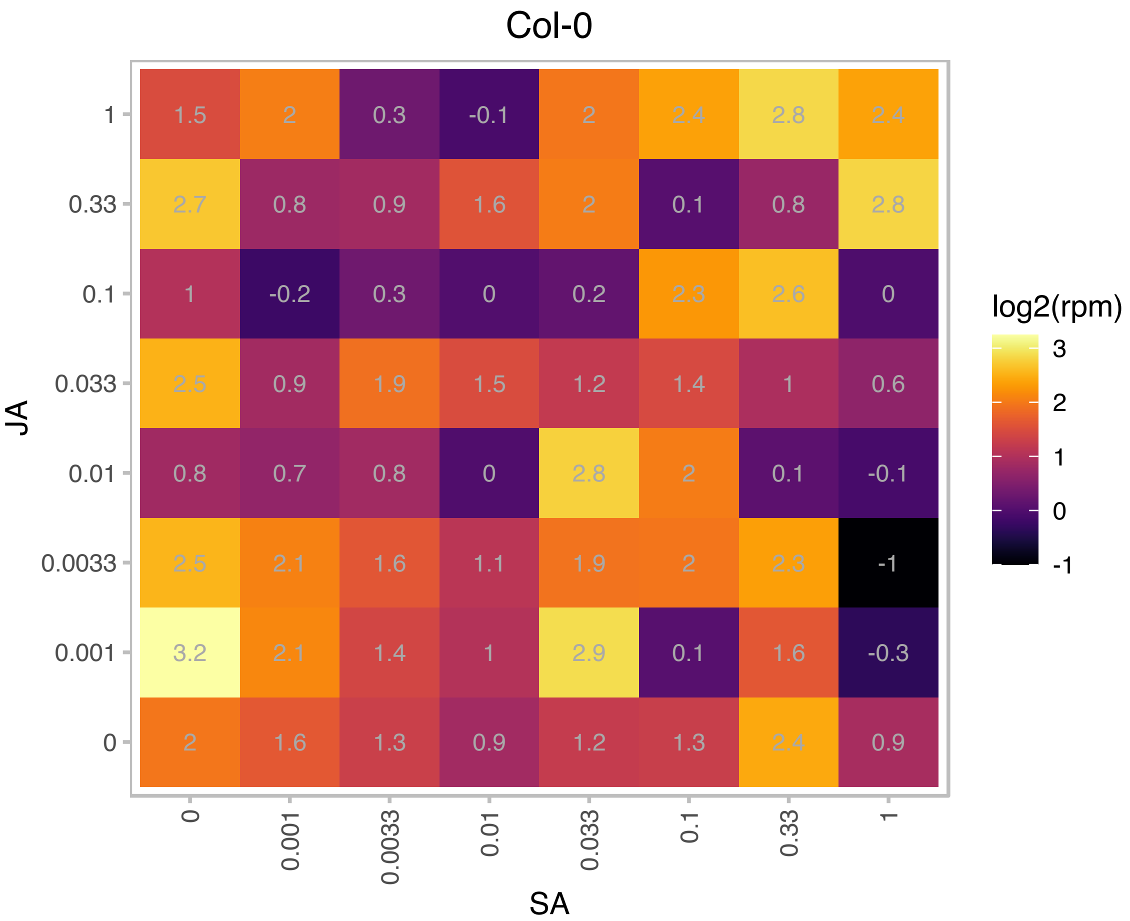


| | |
|---|---|
| Gene | AT3G13672 |
| is.expressed_genes | TRUE |
| is.responsive_genes | TRUE |
| is.apcluster | G-03 |

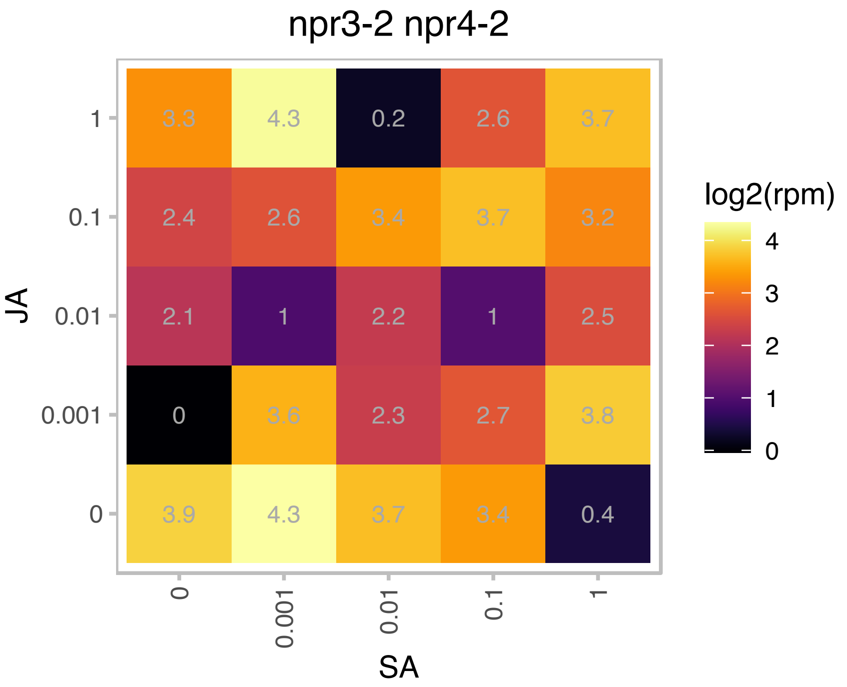


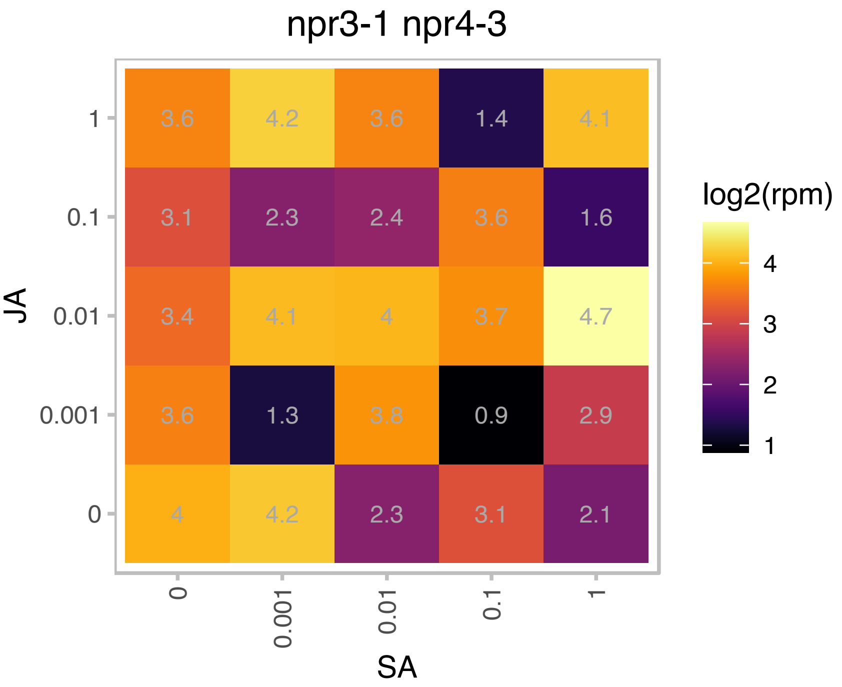

### AT3G14770.1 : AtSWEET2, SWEET2

Nodulin MtN3 family protein

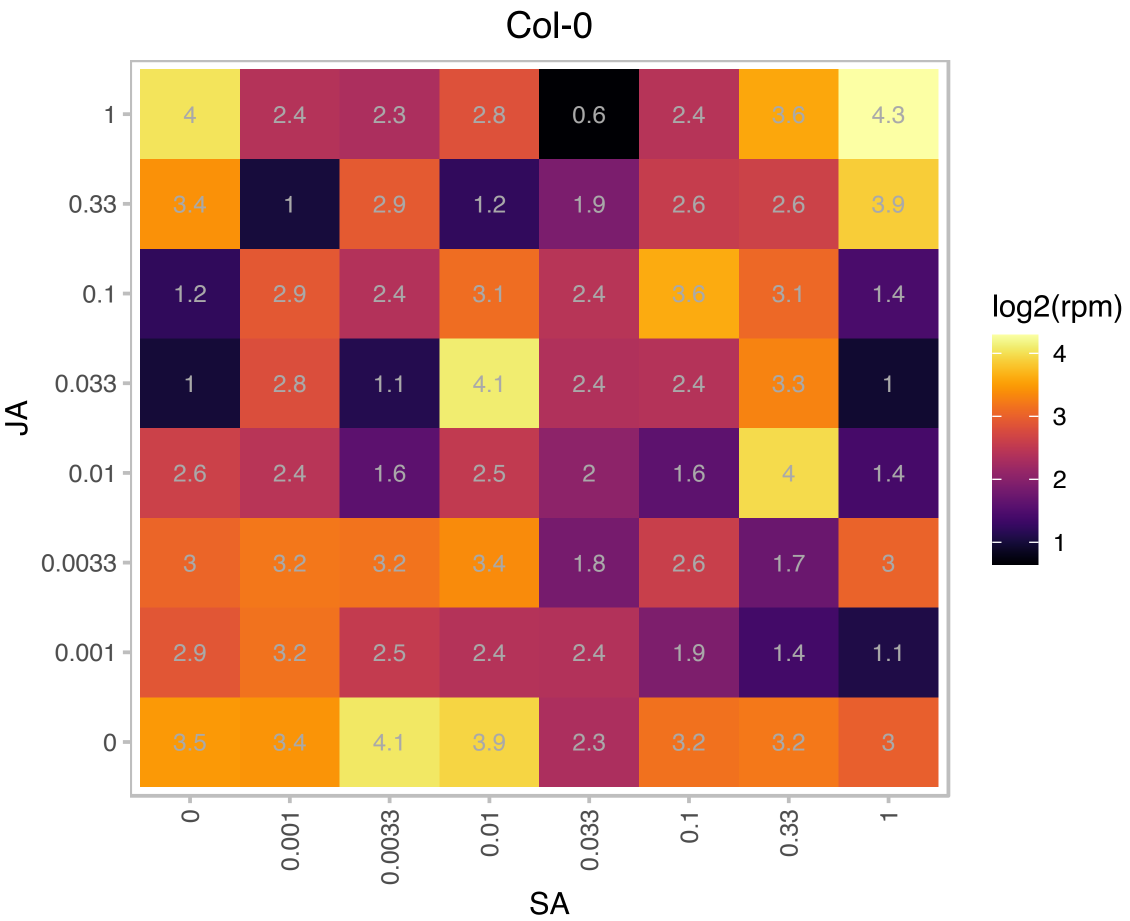


| | |
|---|---|
| Gene | AT3G14770 |
| is.expressed_genes | TRUE |
| is.responsive_genes | TRUE |
| is.apcluster | G-03 |

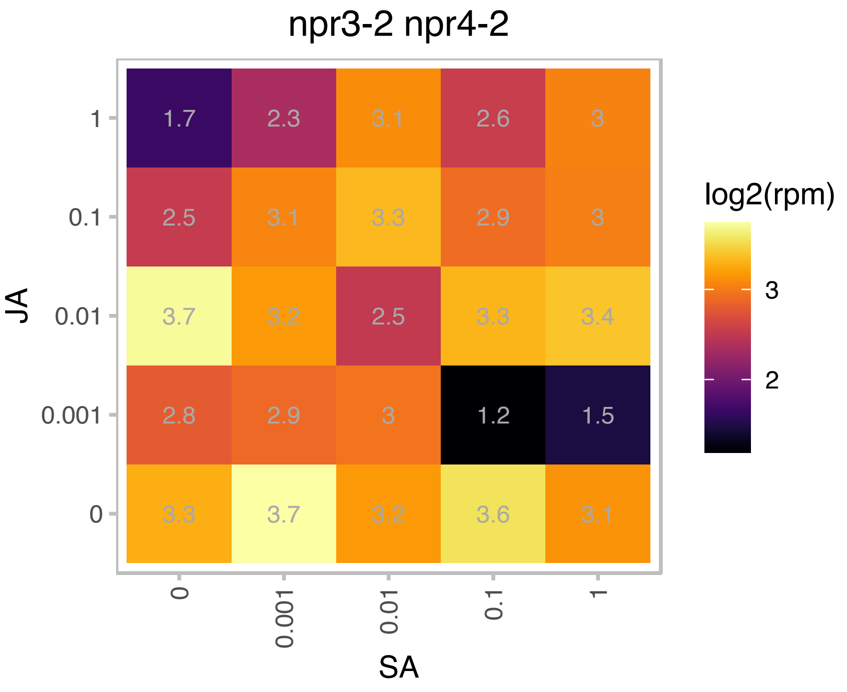


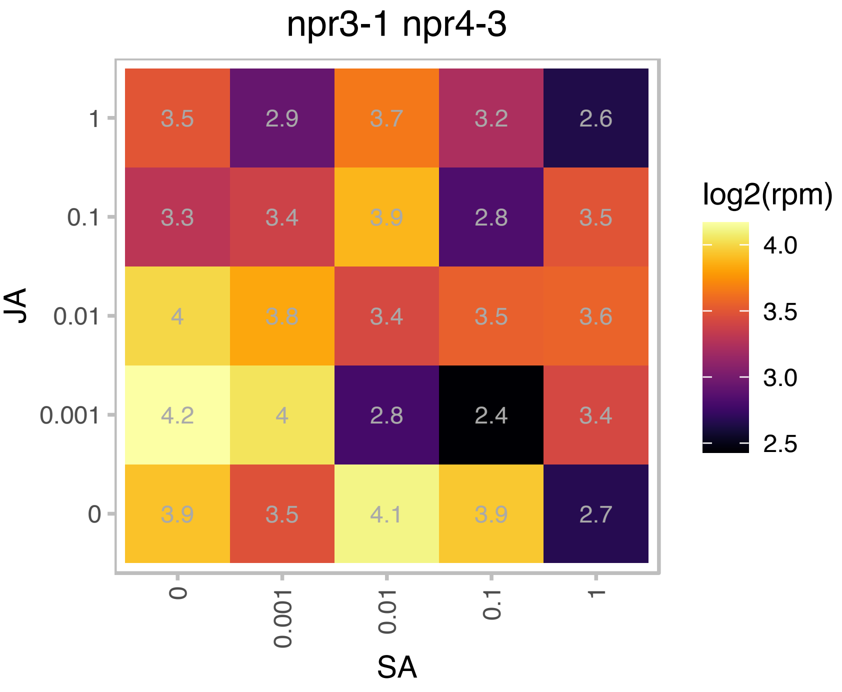

# AT3G16060.1 : NA

ATP binding microtubule motor family protein

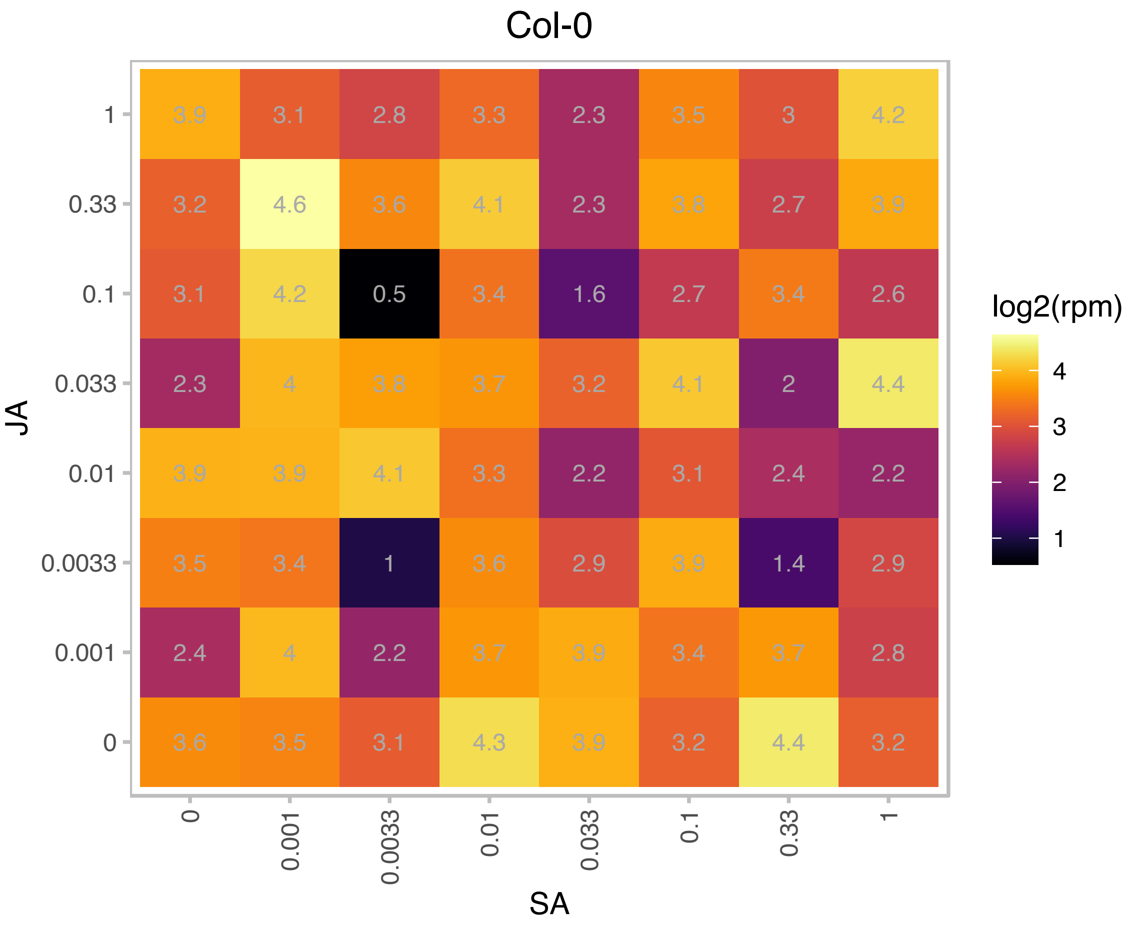


| | |
|---|---|
| Gene | AT3G16060 |
| is.expressed_genes | TRUE |
| is.responsive_genes | TRUE |
| is.apcluster | G-03 |

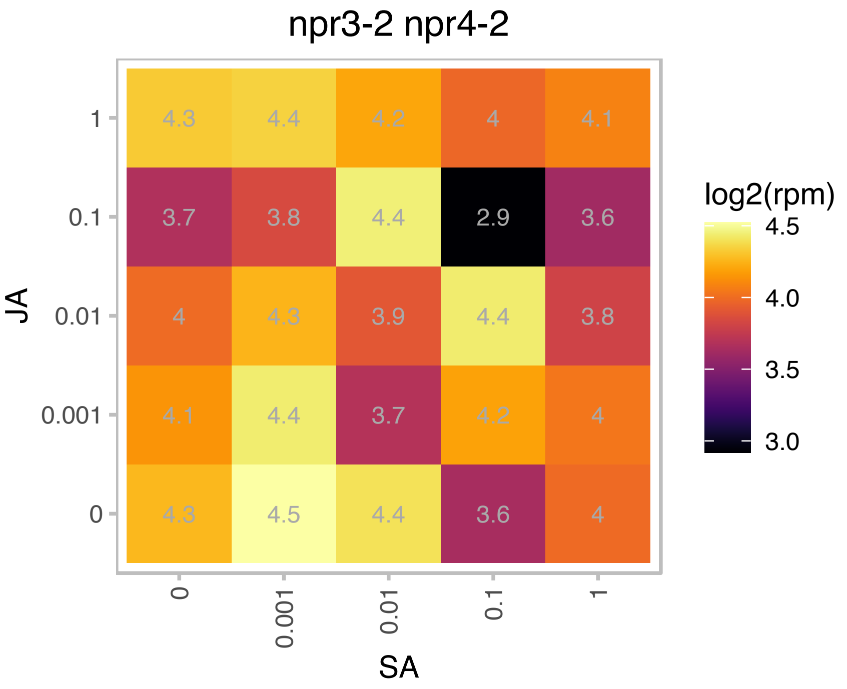


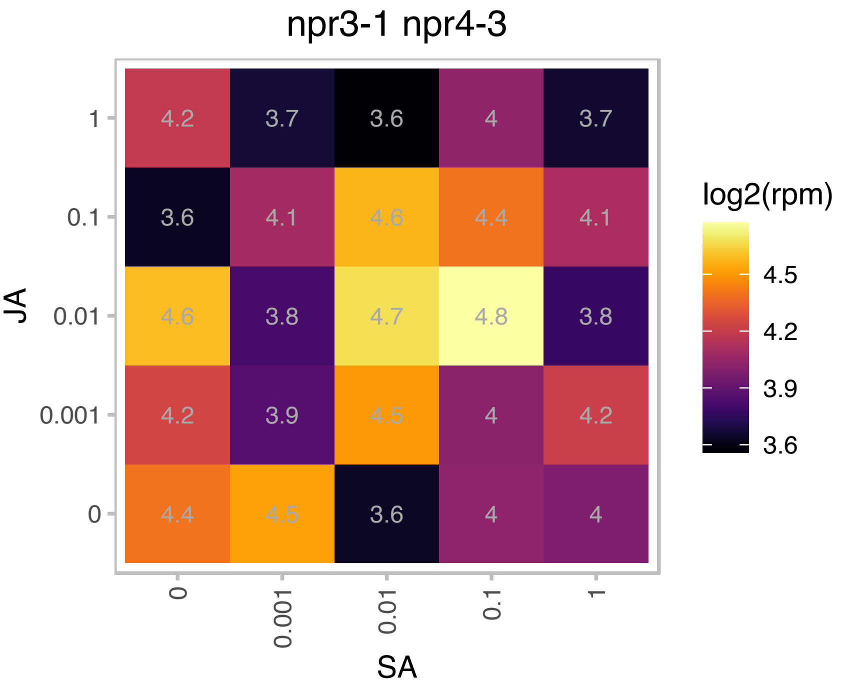

# AT3G27320.1 : NA

alpha/beta-Hydrolases superfamily protein

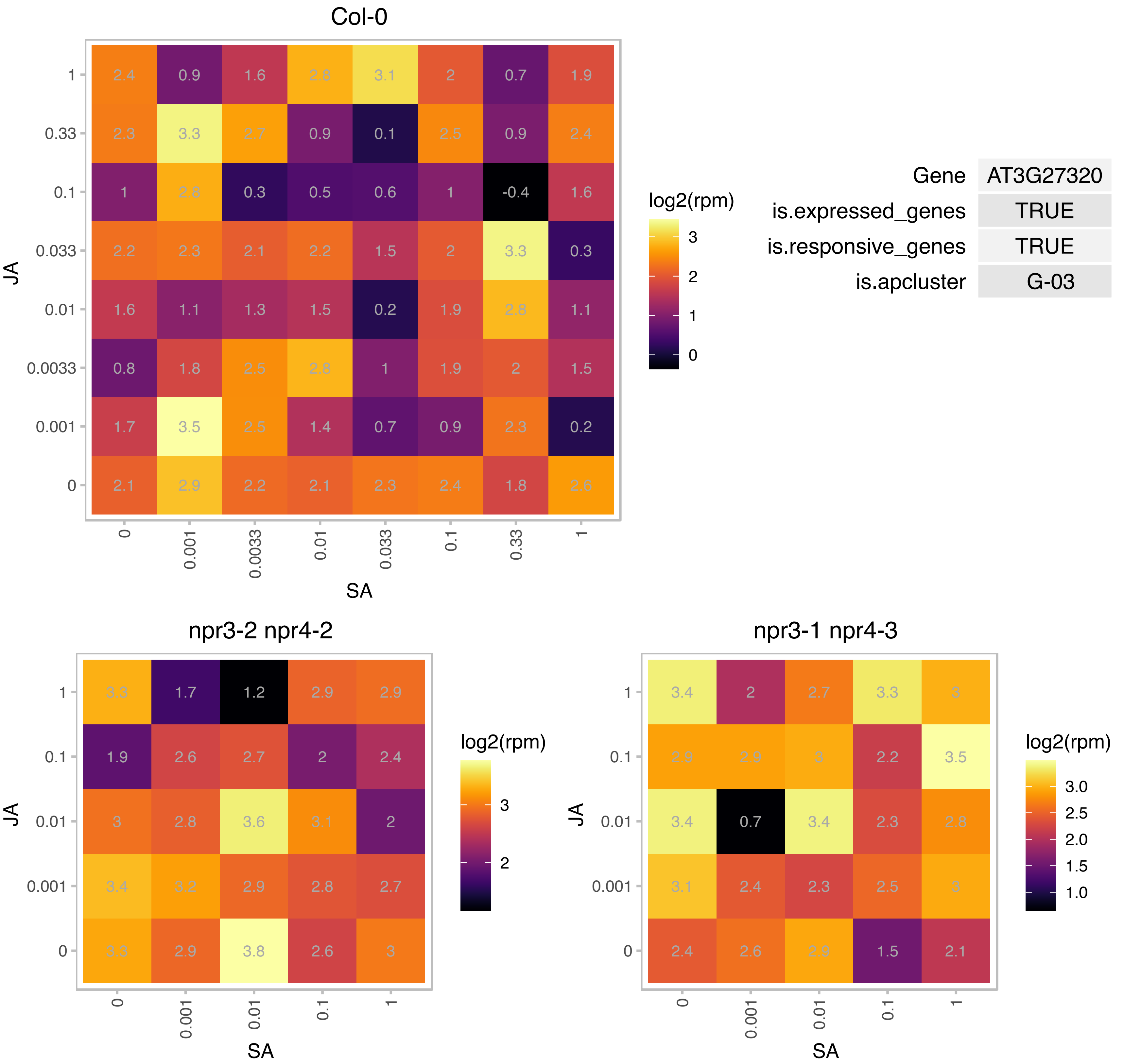


| | |
|---|---|
| Gene | AT3G27320 |
| is.expressed_genes | TRUE |
| is.responsive_genes | TRUE |
| is.apcluster | G-03 |

### AT3G62910.1 : APG3, AtcpRF1, cpRF1

Peptide chain release factor 1

Encodes a plastid-localized ribosome release factor 1 that is essential in chloroplast development. Pale green, albino mutant seedlings arrest early in seedling development.

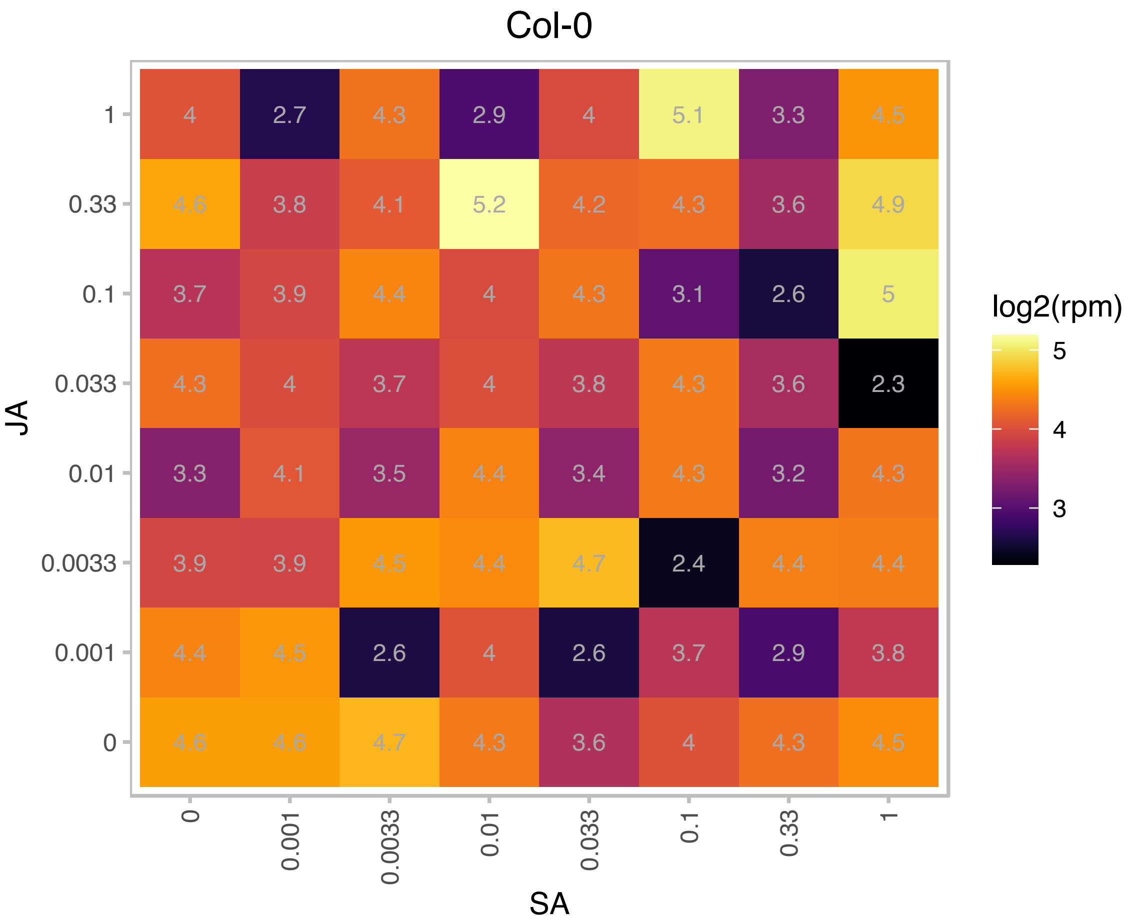


| | |
|---|---|
| Gene | AT3G62910 |
| is.expressed_genes | TRUE |
| is.responsive_genes | TRUE |
| is.apcluster | G-03 |

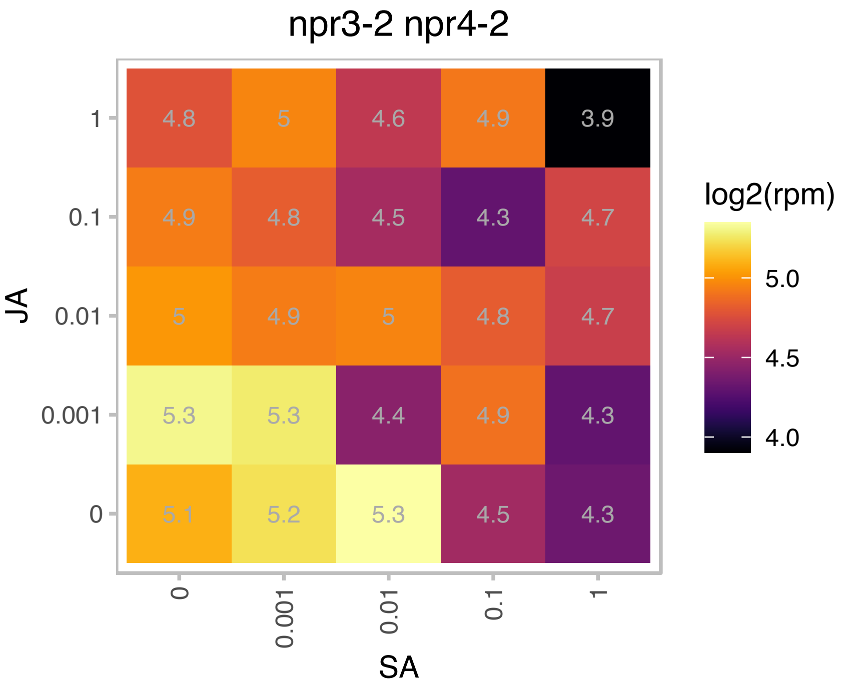


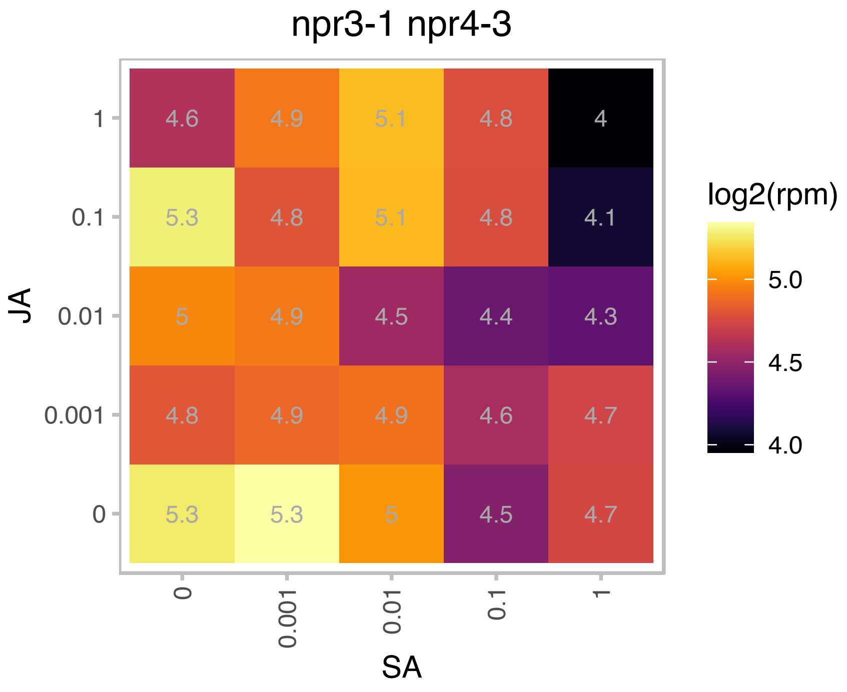

### AT4G26510.1 : UKL4

uridine kinase-like 4

One of the homologous genes predicted to encode proteins with UPRT domains (Uracil phosphoribosyltransferase). Five of these genes (At5g40870, At3g27190, At1g55810, At4g26510 and At3g27440) show a high level of identity, and are annotated as also containing a N-terminal uracil kinase (UK) domain. These genes are referred to as UKL1 (UK-like 1), UKL2, UKL3, UKL4 and UKL5, respectively.

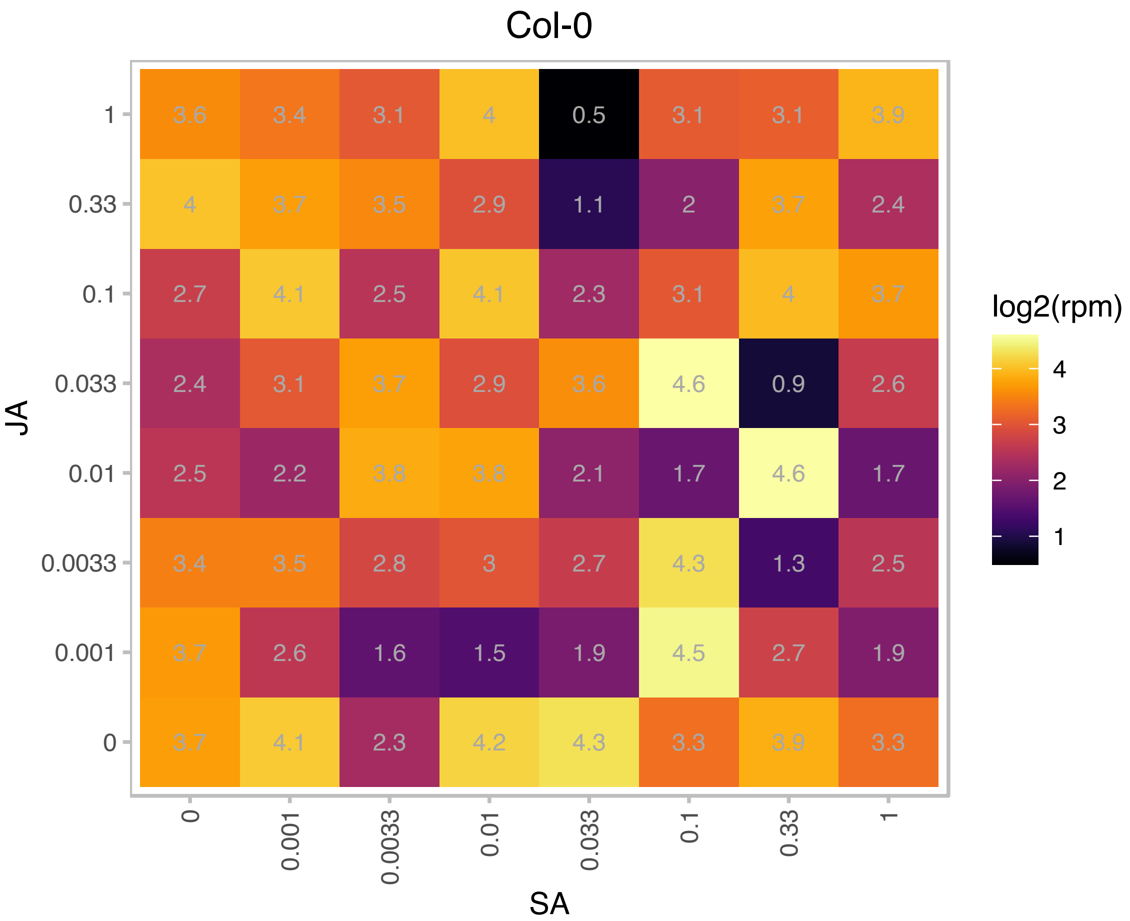


| | |
|---|---|
| Gene | AT4G26510 |
| is.expressed_genes | TRUE |
| is.responsive_genes | TRUE |
| is.apcluster | G-03 |

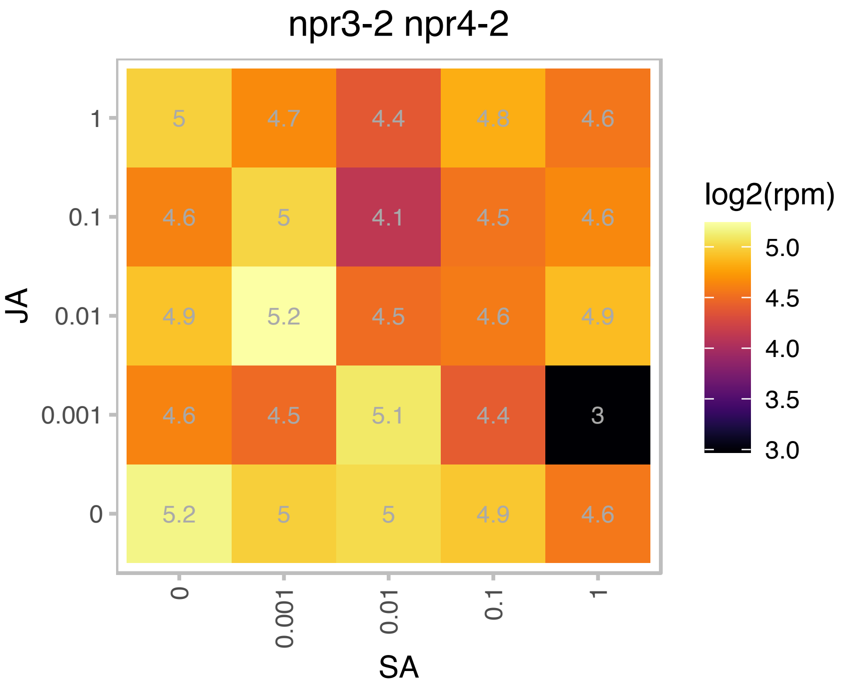


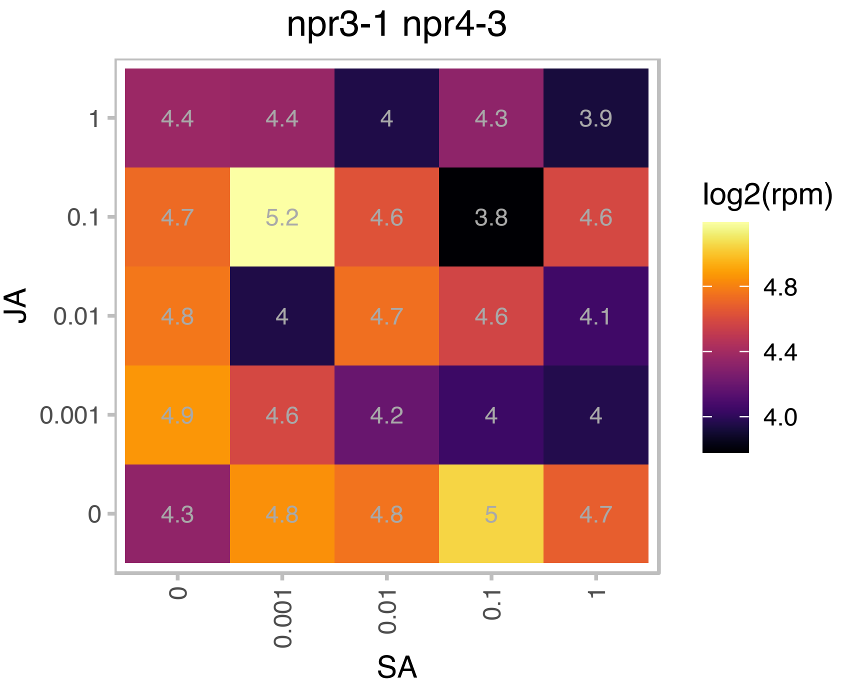

### AT5G13650.1 : SVR3

elongation factor family protein
Encodes SVR3, a putative chloroplast TypA translation elongation GTPase. Loss of SVR3 suppresses variegation mediated by var2. SVR3 is essential for plants' ability to develop functional chloroplasts under chilling stress (8C), but not at normal temperature (22C).

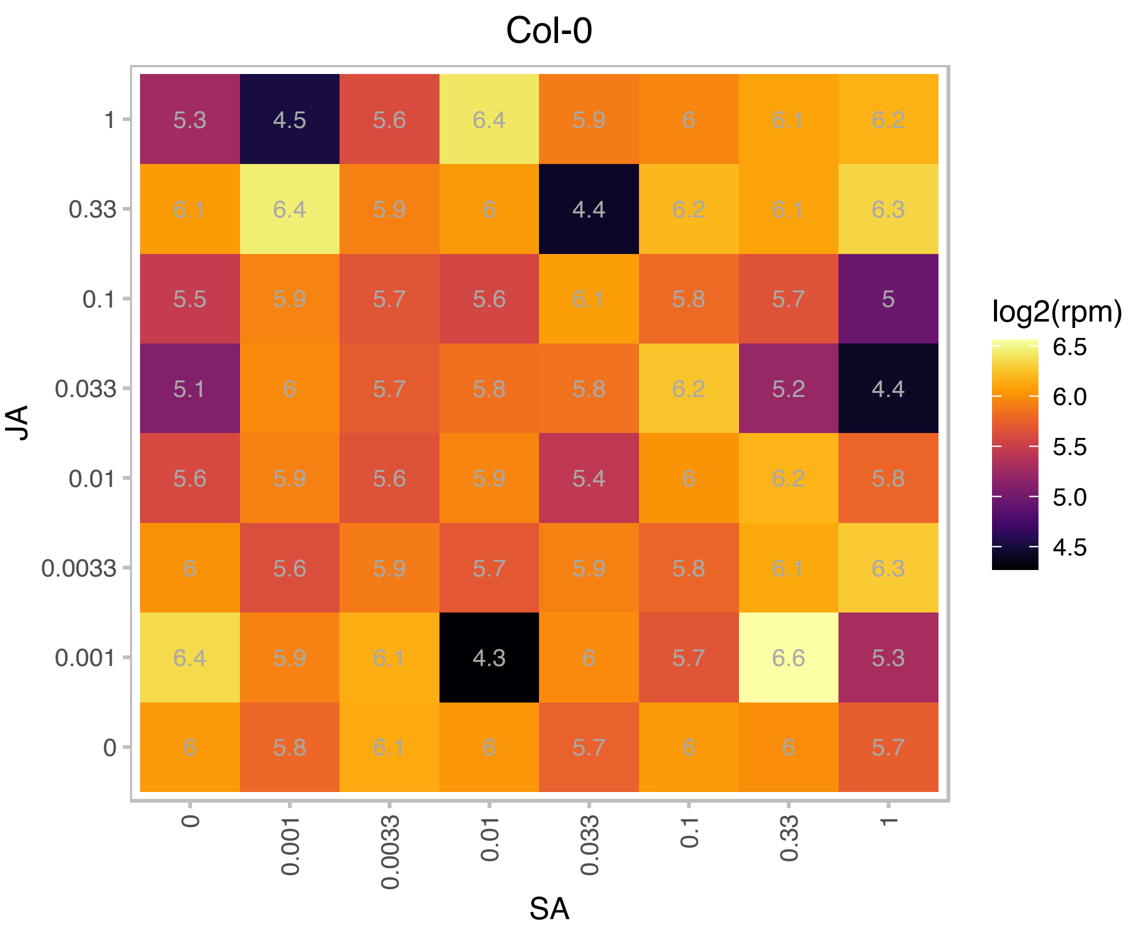


| | |
|---|---|
| Gene | AT5G13650 |
| is.expressed_genes | TRUE |
| is.responsive_genes | TRUE |
| is.apcluster | G-03 |

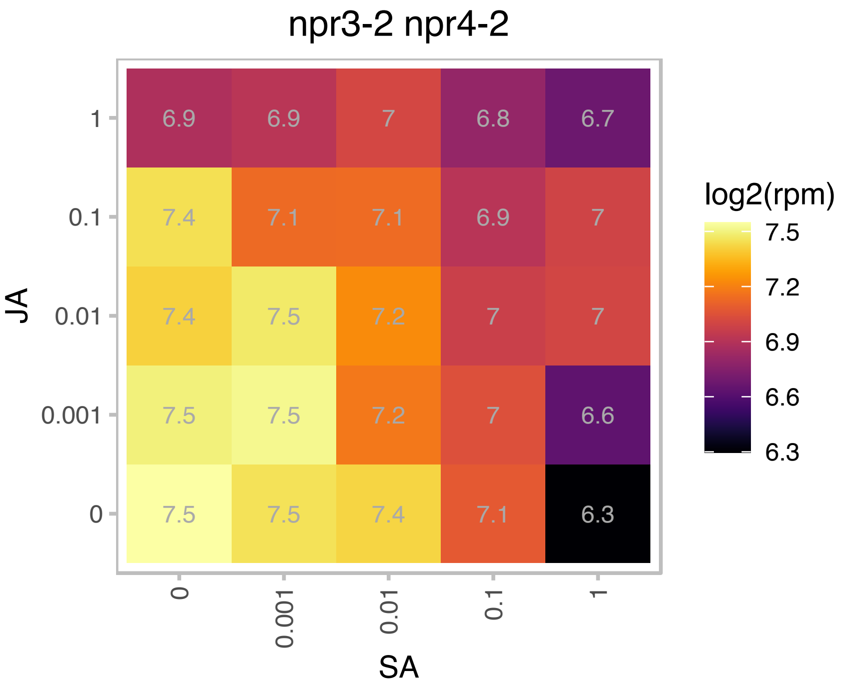


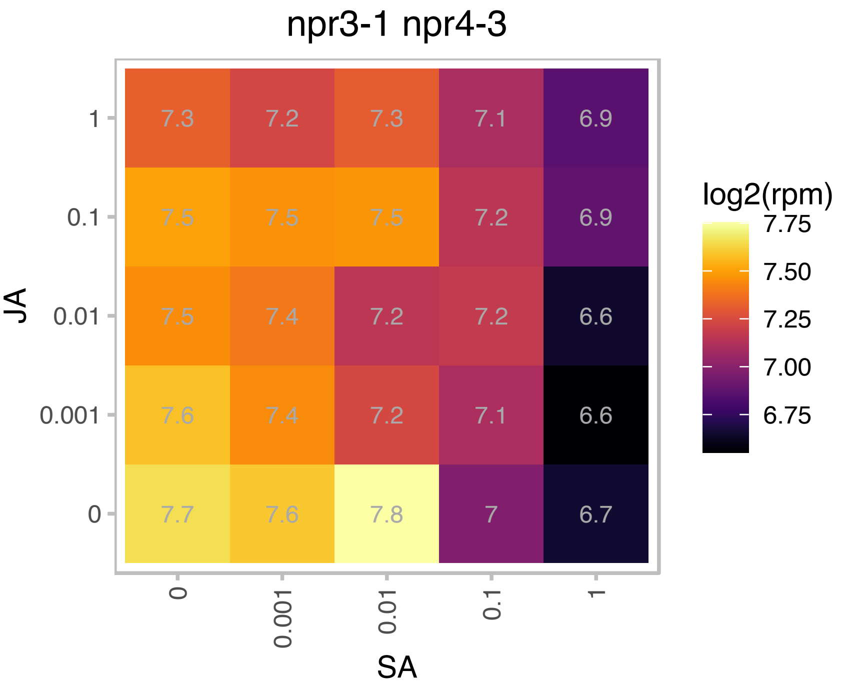

### AT5G14740.1 : BETA CA2, CA18, CA2

carbonic anhydrase 2

Encodes a beta carbonic anhydrase likely to be localized in the cytoplasm.  Expression of its mRNA is seen in etiolated seedlings and points to a possible nonphotosynthetic role for this isoform.

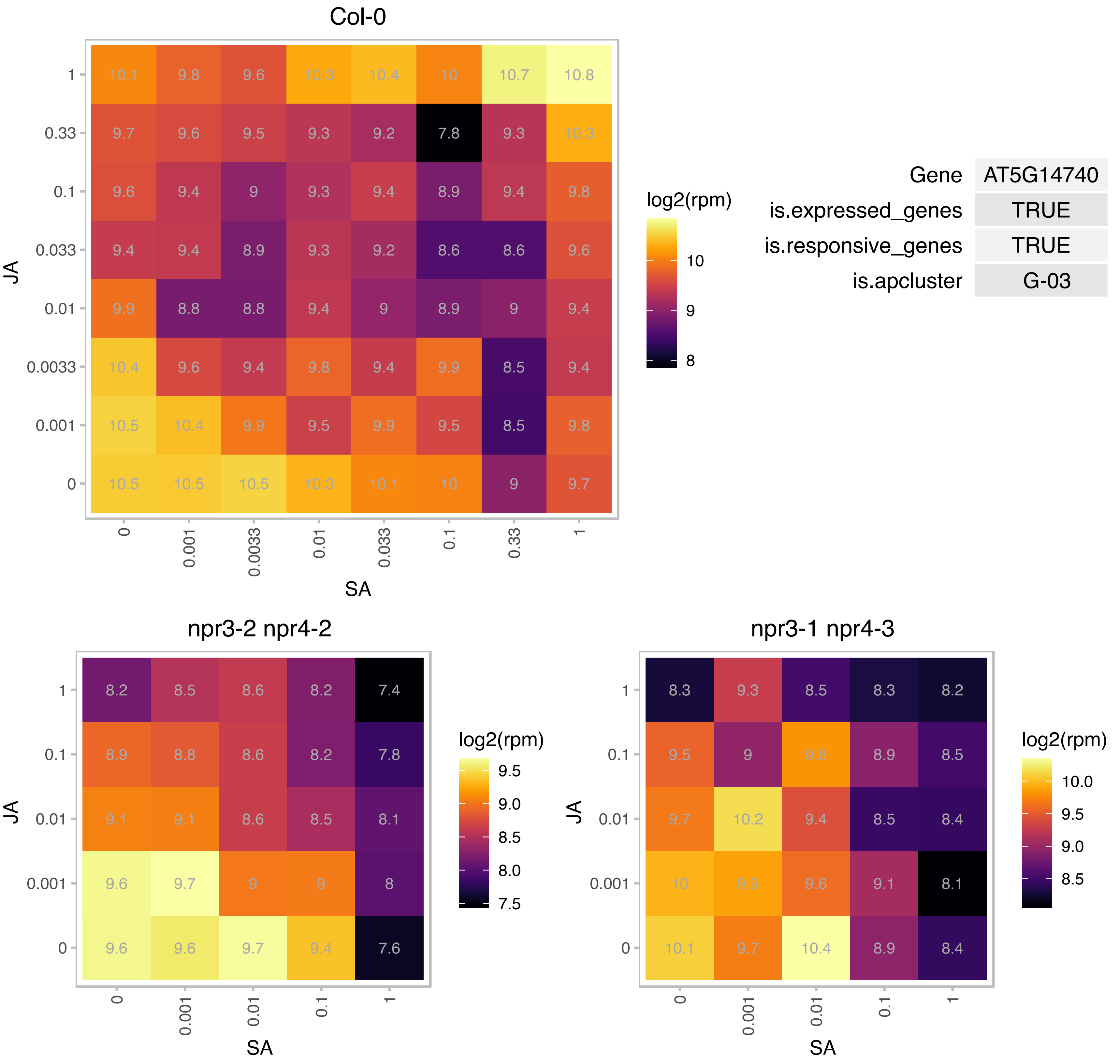


| | |
|---|---|
| Gene | AT5G14740 |
| is.expressed_genes | TRUE |
| is.responsive_genes | TRUE |
| is.apcluster | G-03 |

# AT5G14970.1 : NA

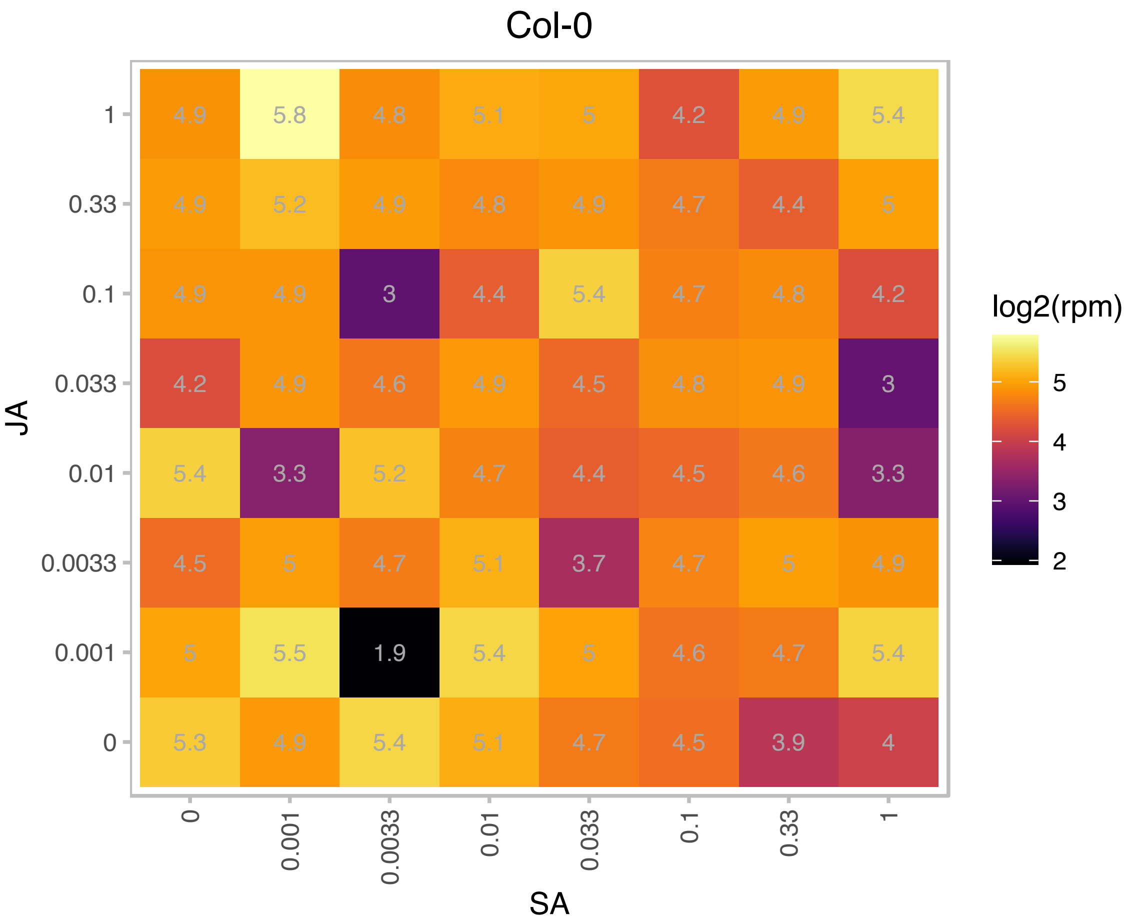


| | |
|---|---|
| Gene | AT5G14970 |
| is.expressed_genes | TRUE |
| is.responsive_genes | TRUE |
| is.apcluster | G-03 |

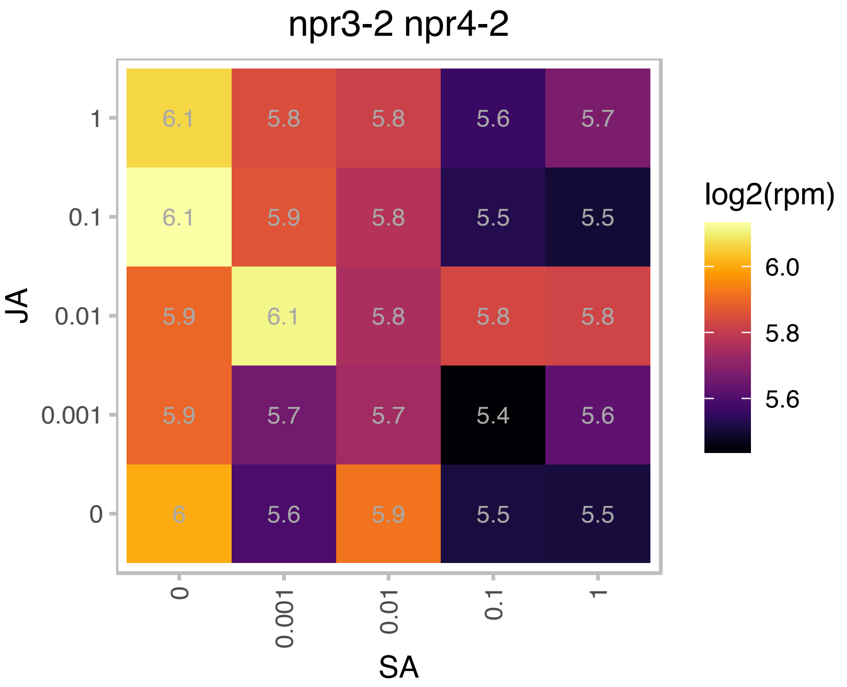


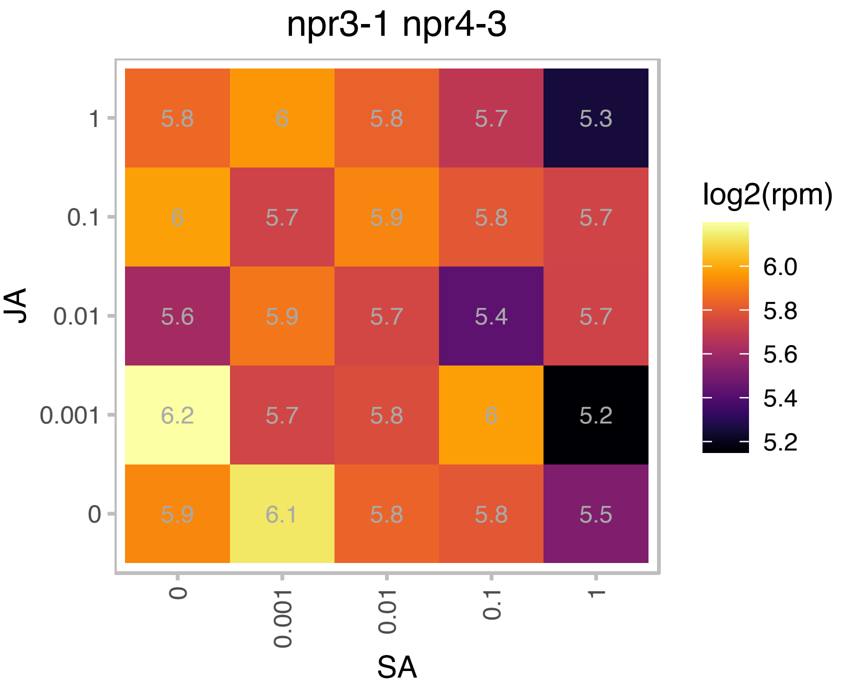

### AT5G16715.1 : EMB2247

ATP binding;valine-tRNA ligases;aminoacyl-tRNA ligases;nucleotide binding;ATP binding;aminoacyl-tRNA ligases

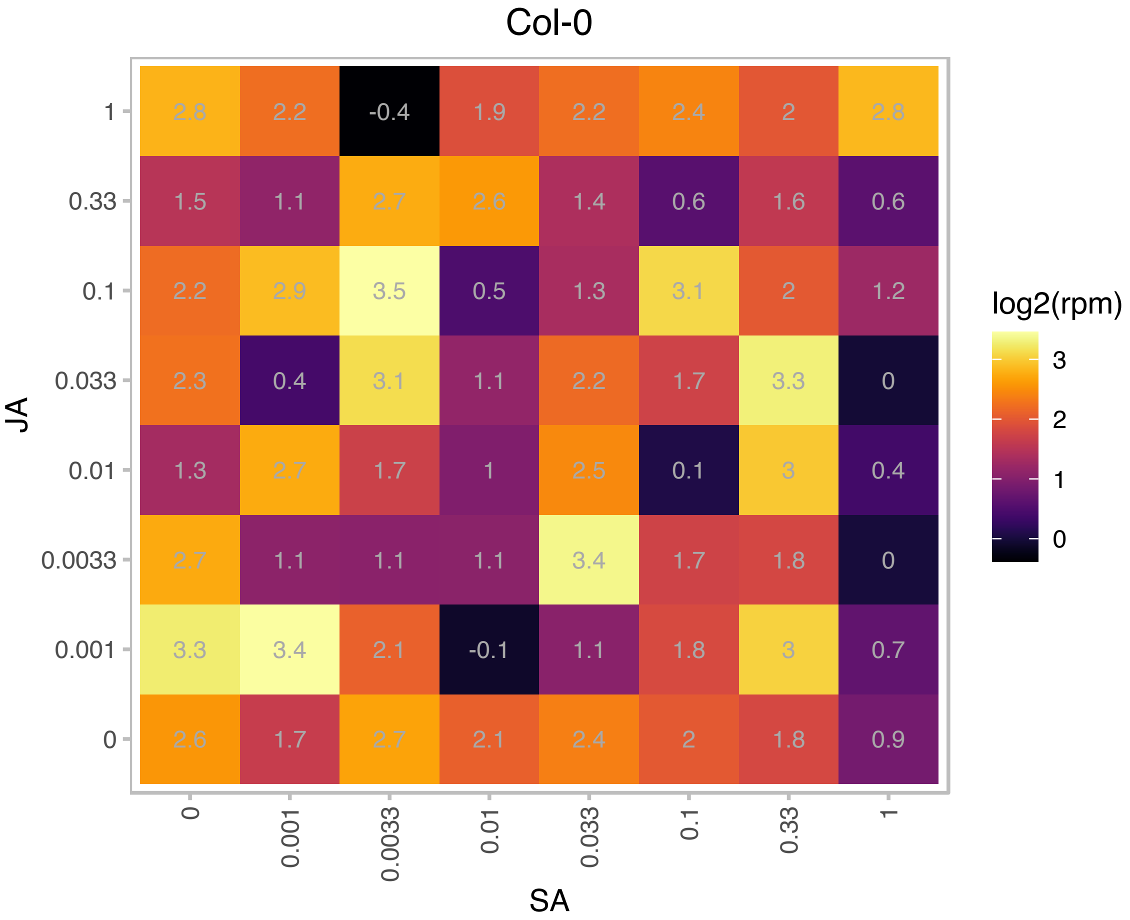


| | |
|---|---|
| Gene | AT5G16715 |
| is.expressed_genes | TRUE |
| is.responsive_genes | TRUE |
| is.apcluster | G-03 |

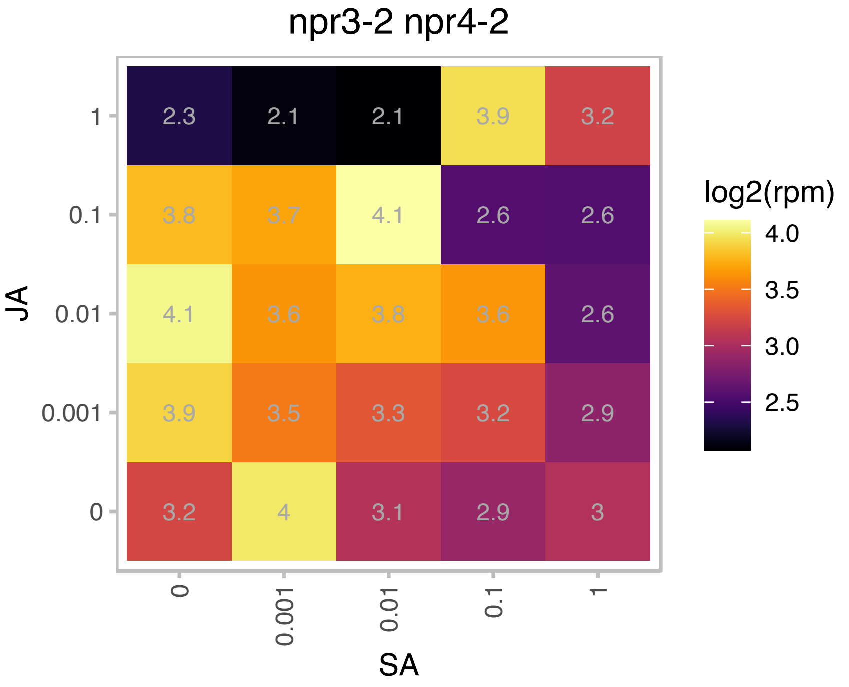


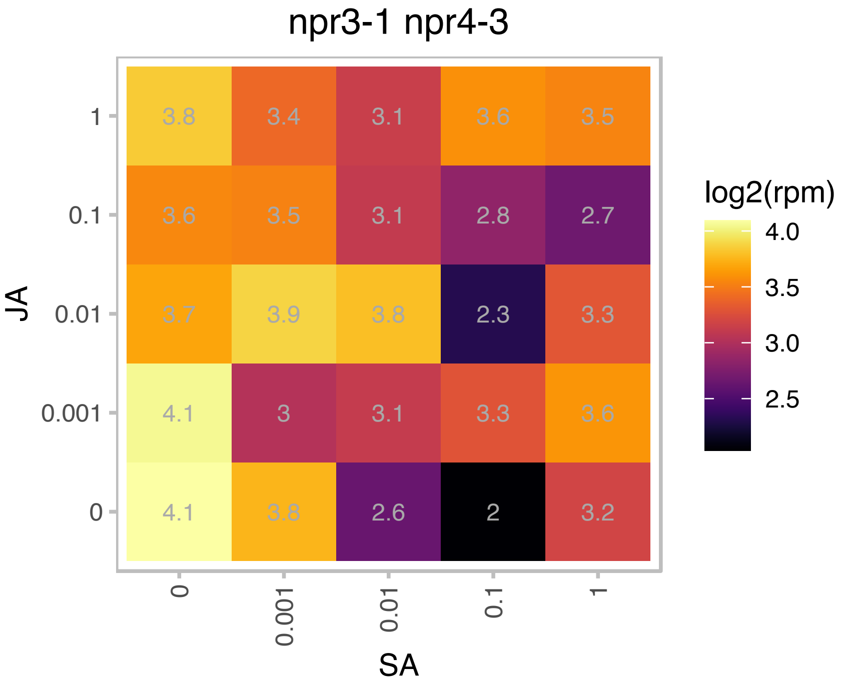

### AT5G21430.1 : CRRL, NdhU

Chaperone DnaJ-domain superfamily protein

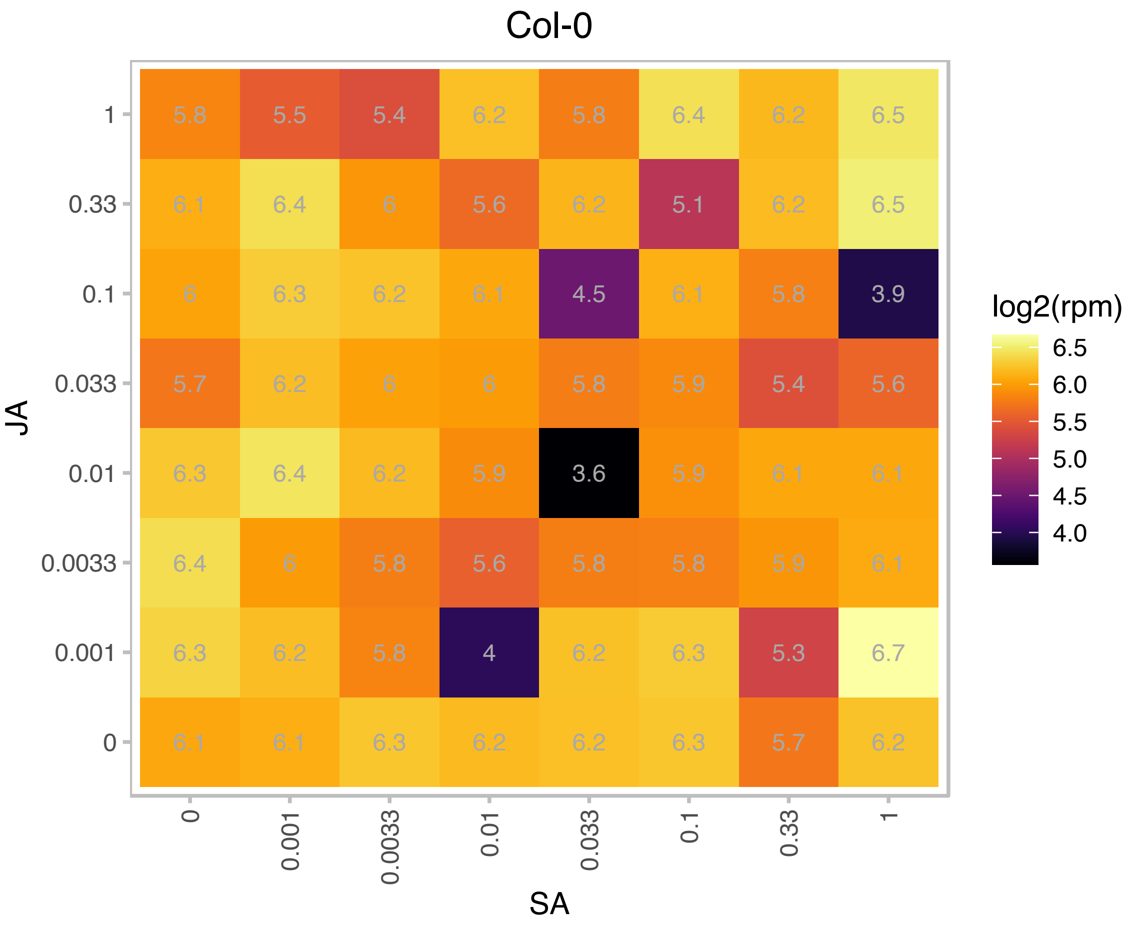


| | |
|---|---|
| Gene | AT5G21430 |
| is.expressed_genes | TRUE |
| is.responsive_genes | TRUE |
| is.apcluster | G-03 |

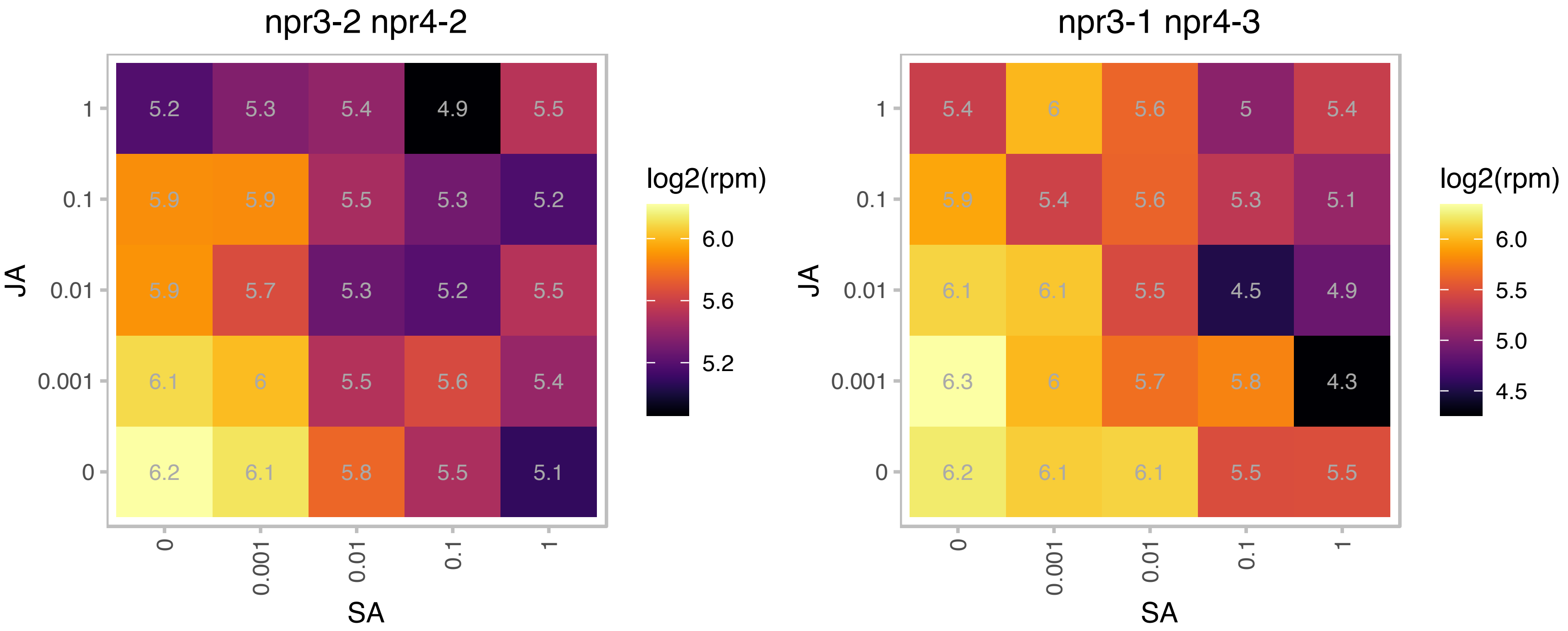

# AT5G22340.1 : NA

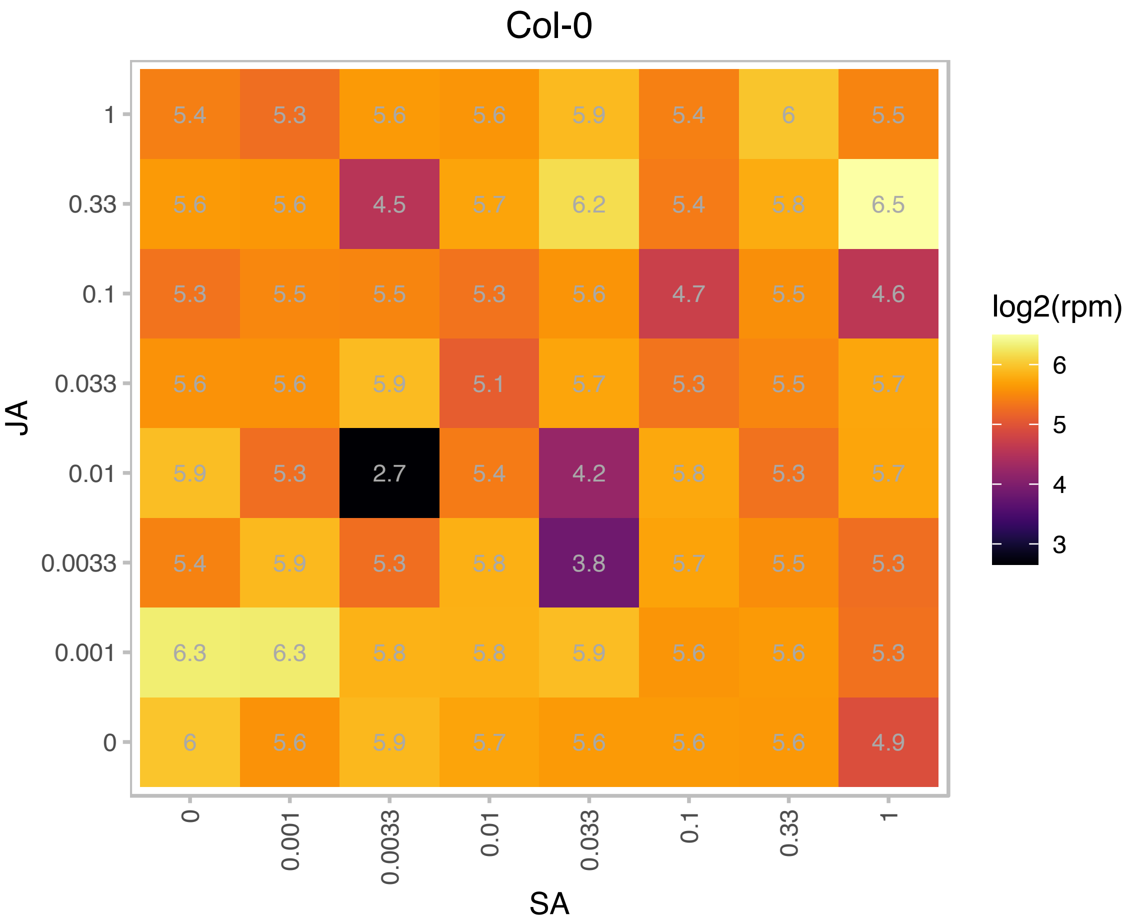


| | |
|---|---|
| Gene | AT5G22340 |
| is.expressed_genes | TRUE |
| is.responsive_genes | TRUE |
| is.apcluster | G-03 |

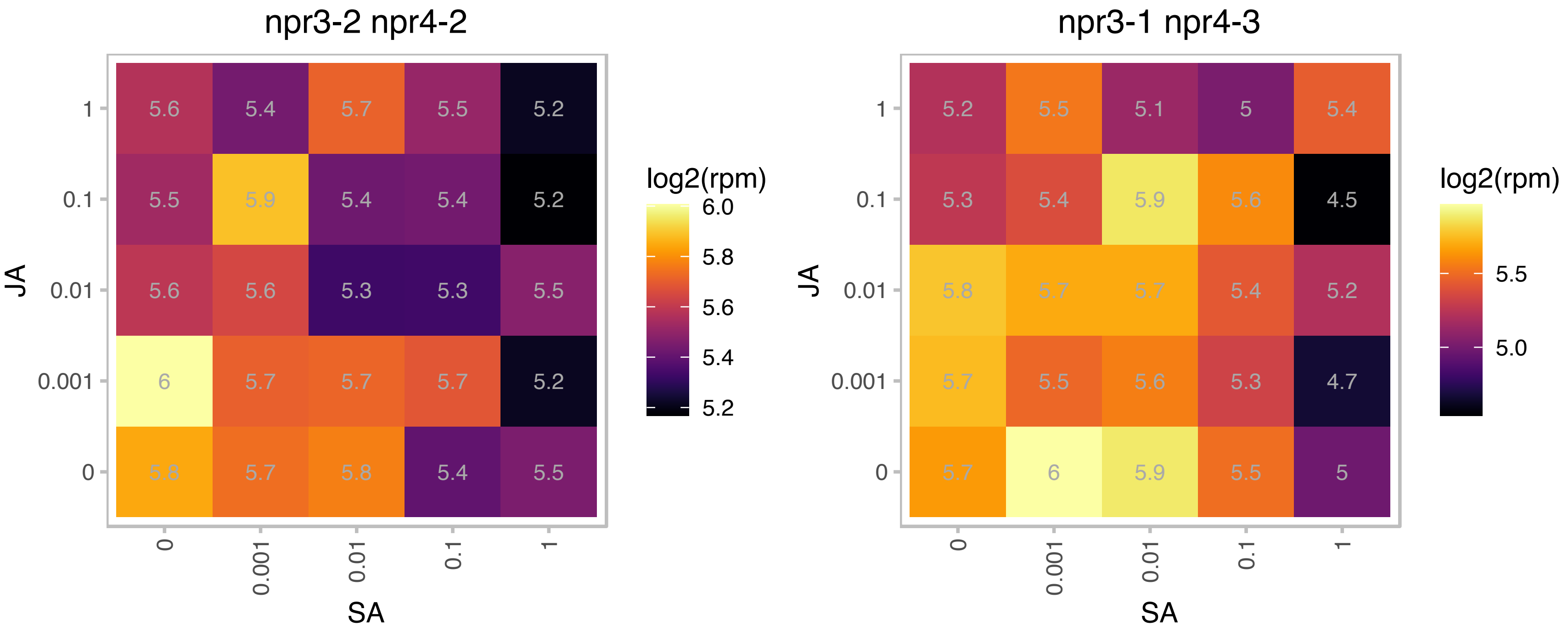

# AT5G50250.1 : CP31B

chloroplast RNA-binding protein 31B

Encodes a RNA binding protein. A substrate of the type III effector HopU1 (mono-ADP-ribosyltransferase). Supports editing of specific CP31A-dependent sites.

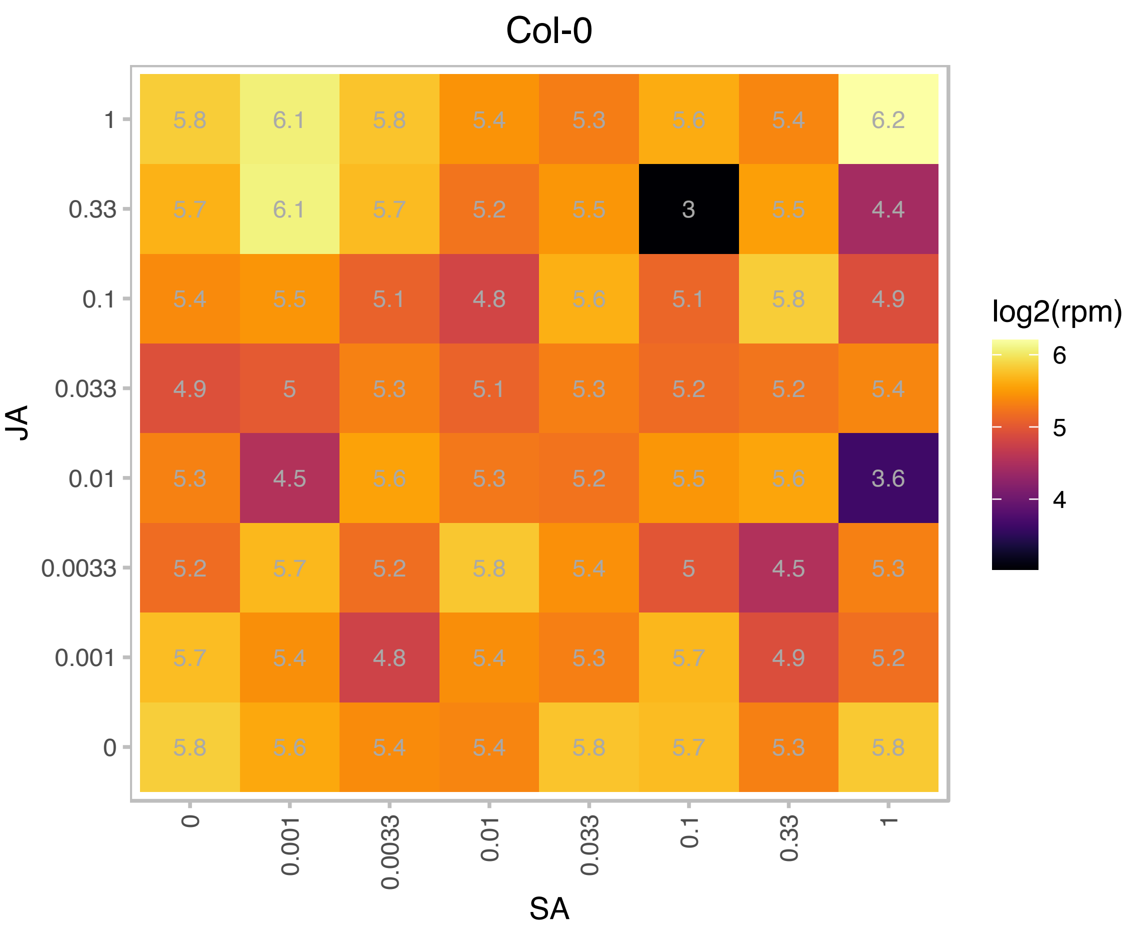


| | |
|---|---|
| Gene | AT5G50250 |
| is.expressed_genes | TRUE |
| is.responsive_genes | TRUE |
| is.apcluster | G-03 |

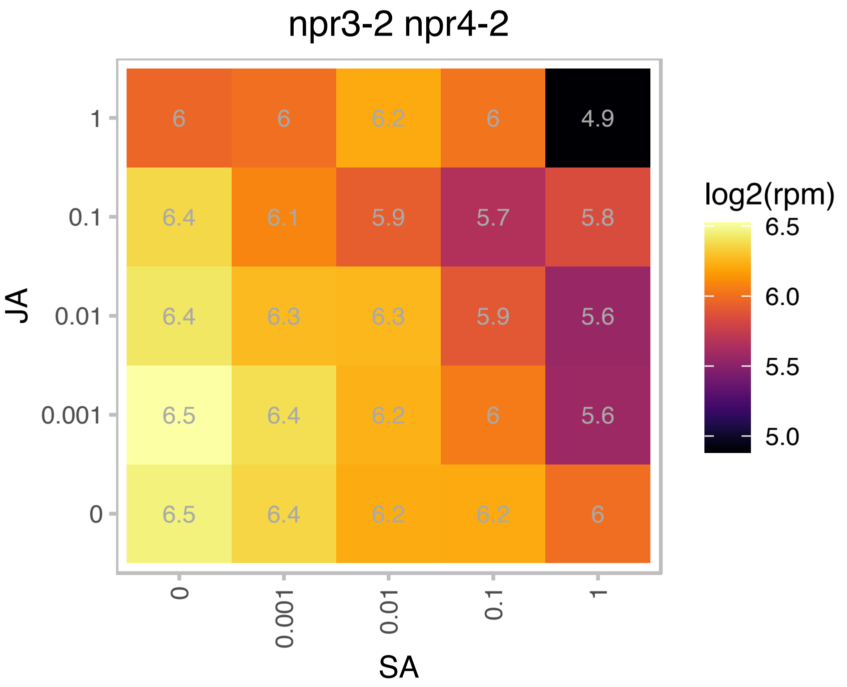


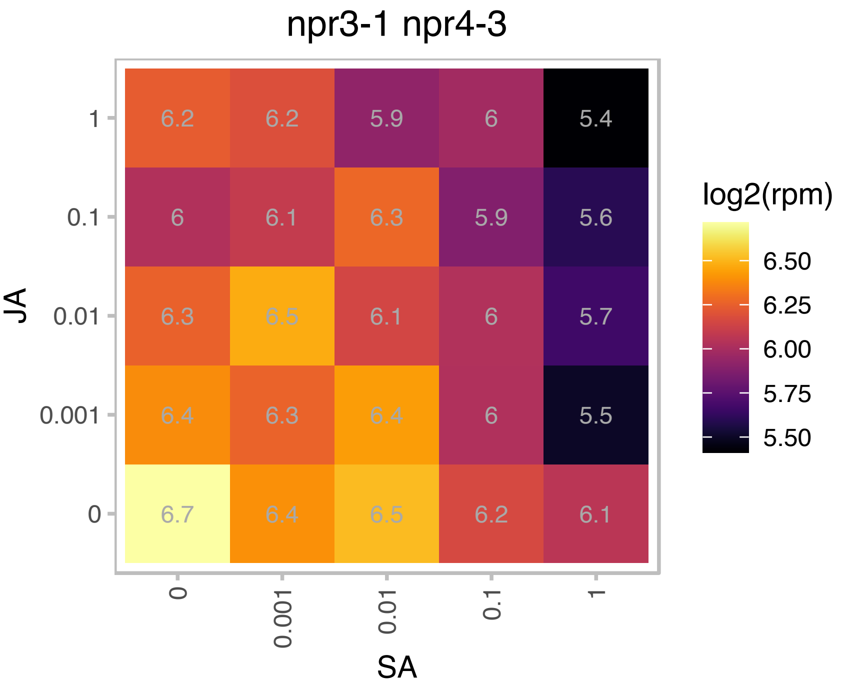

# AT5G52880.1 : NA

F-box family protein

| | |
|---|---|
| Gene | AT5G52880 |
| is.expressed_genes | TRUE |
| is.responsive_genes | TRUE |
| is.apcluster | G-03 |

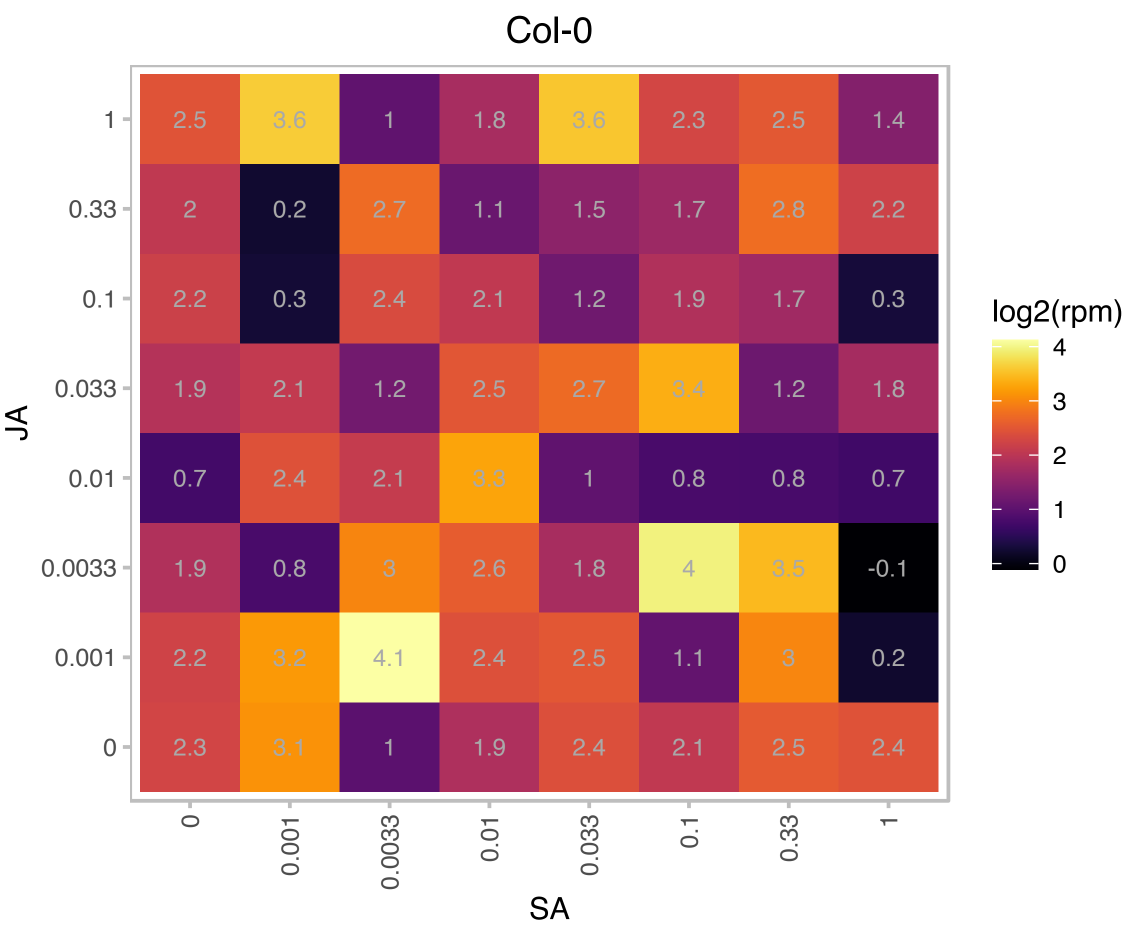


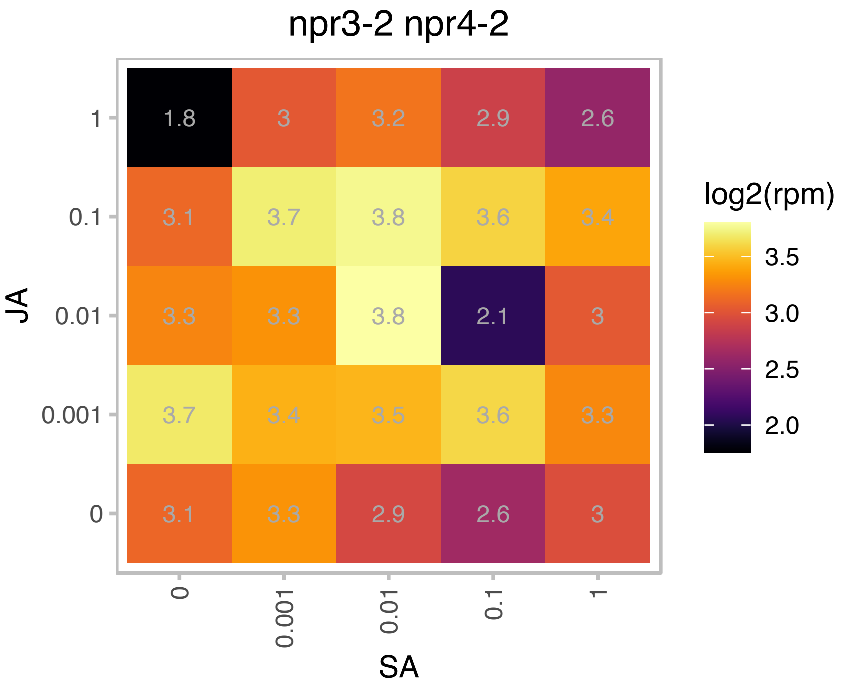


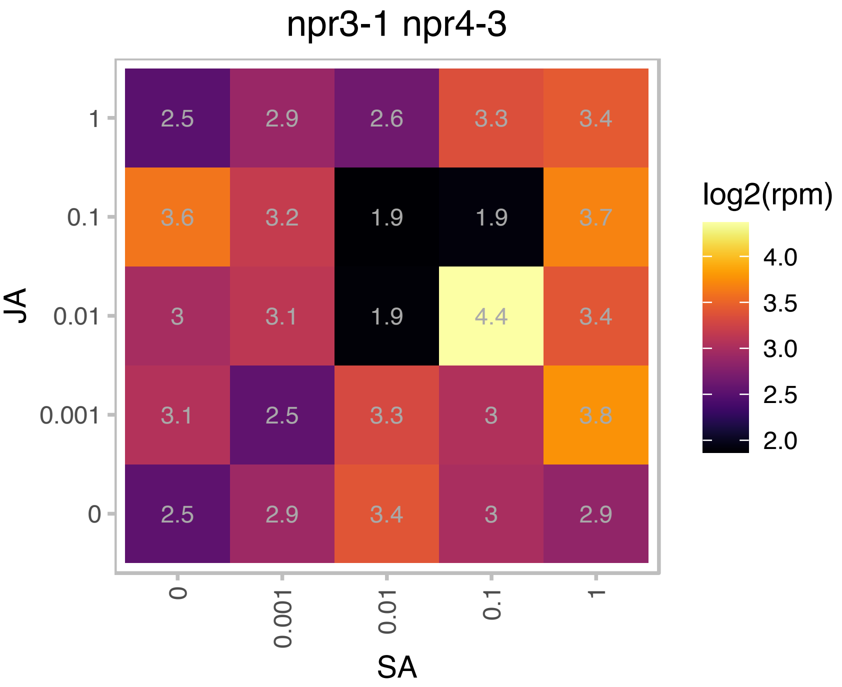

# AT5G53490.1 : TL17

Tetratricopeptide repeat (TPR)-like superfamily protein

thylakoid lumenal 17.4 kDa protein, chloroplast, identical to SP:P81760 Thylakoid lumenal 17.4 kDa protein, chloroplast precursor (P17.4) {Arabidopsis thaliana}. Putative pentapeptide protein.

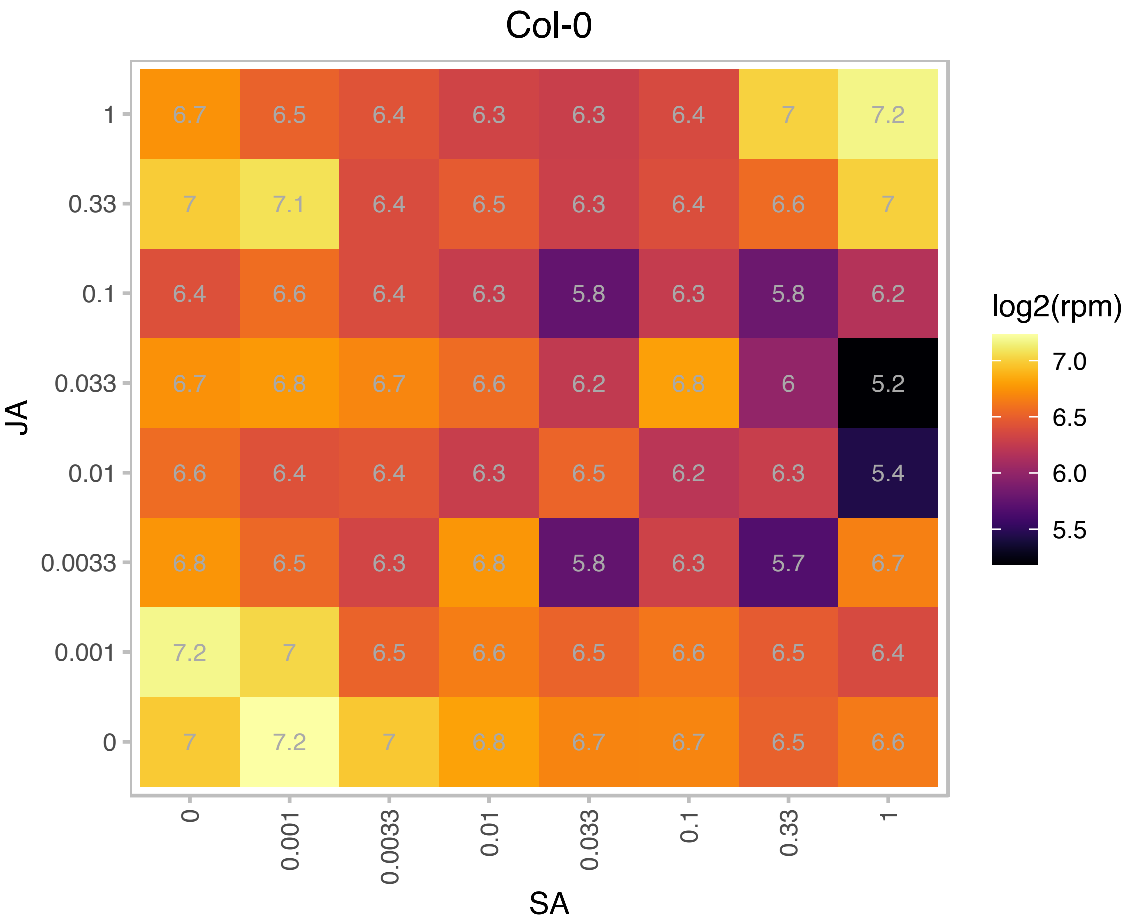


| | |
|---|---|
| Gene | AT5G53490 |
| is.expressed_genes | TRUE |
| is.responsive_genes | TRUE |
| is.apcluster | G-03 |

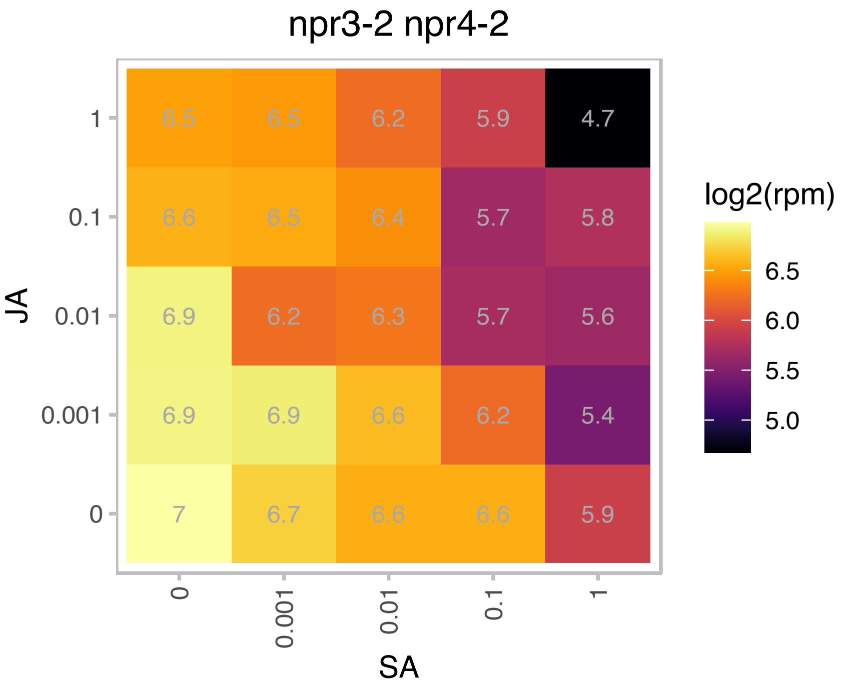


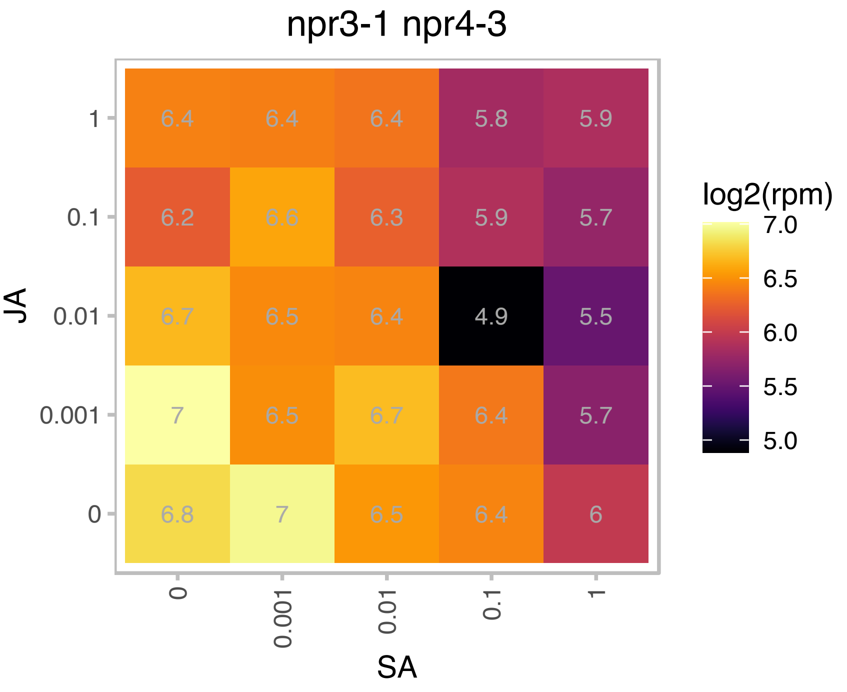

# AT1G05070.1 : NA

Protein of unknown function (DUF1068)

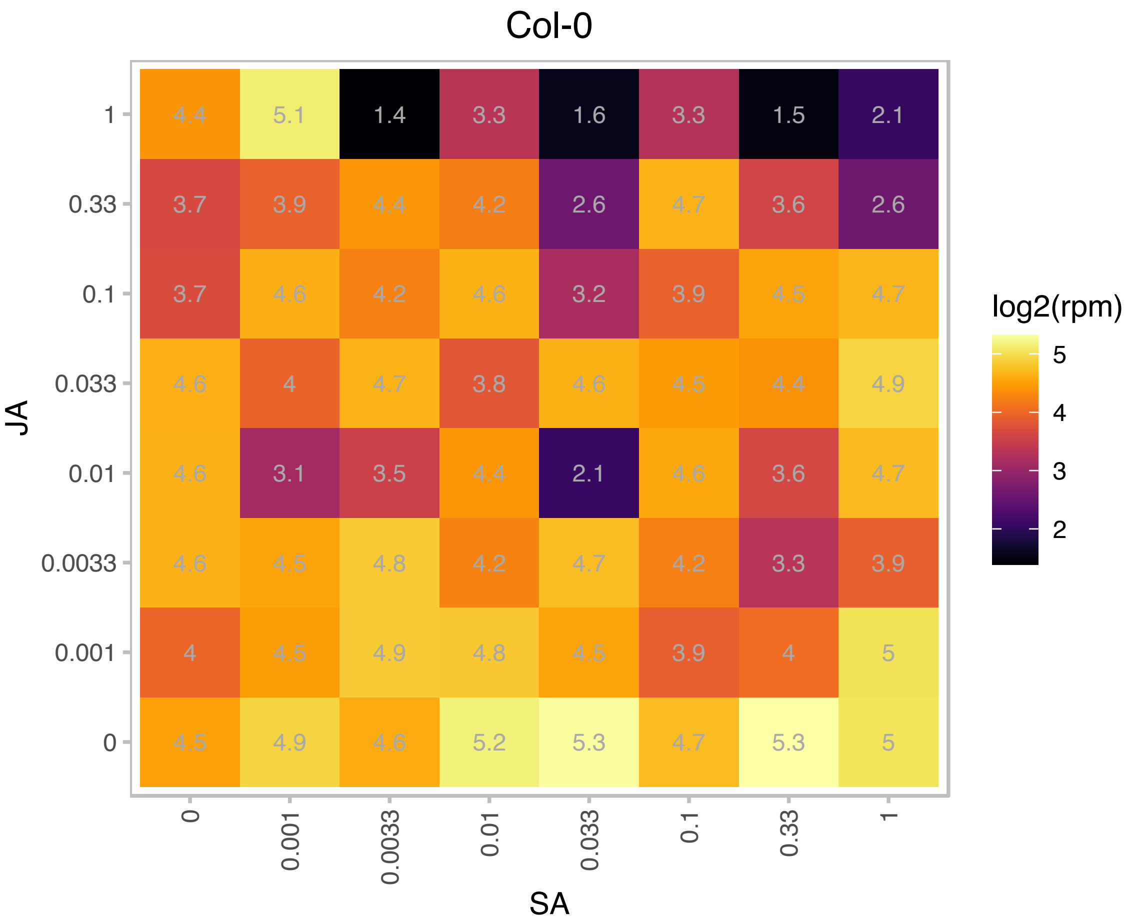


| | |
|---|---|
| Gene | AT1G05070 |
| is.expressed_genes | TRUE |
| is.responsive_genes | TRUE |
| is.apcluster | H-01 |

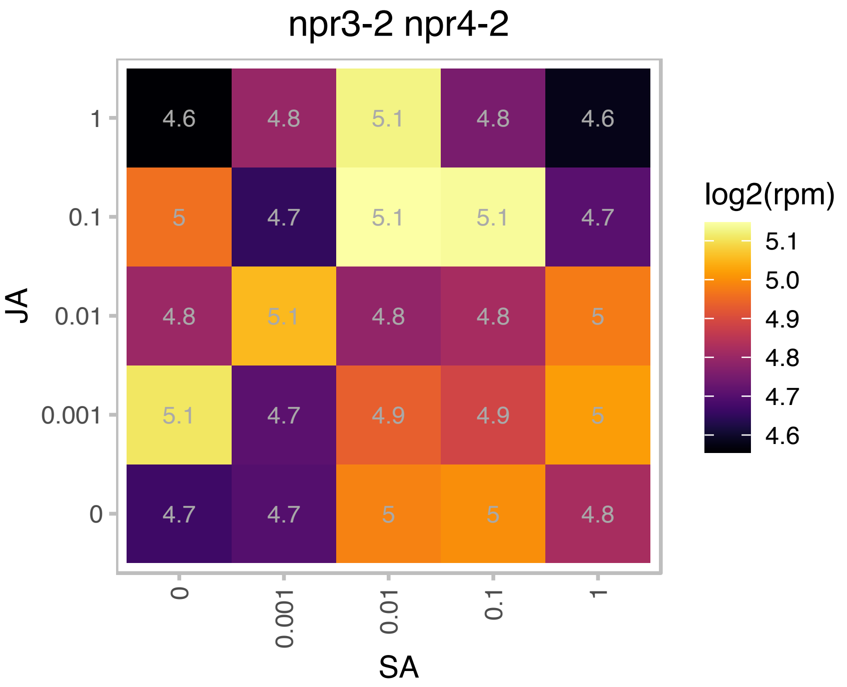


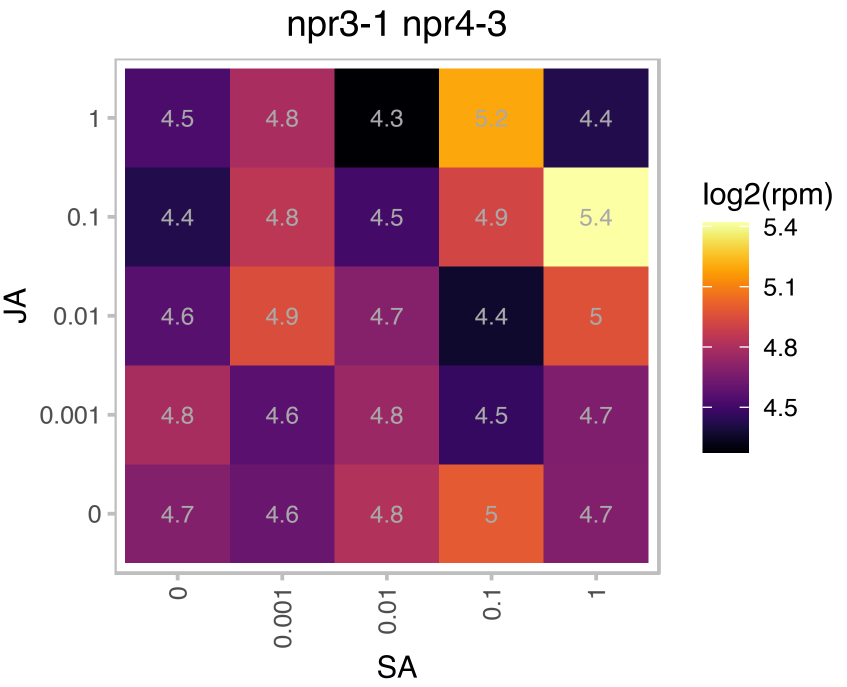

# AT1G07280.1 : NA

Tetratricopeptide repeat (TPR)-like superfamily protein

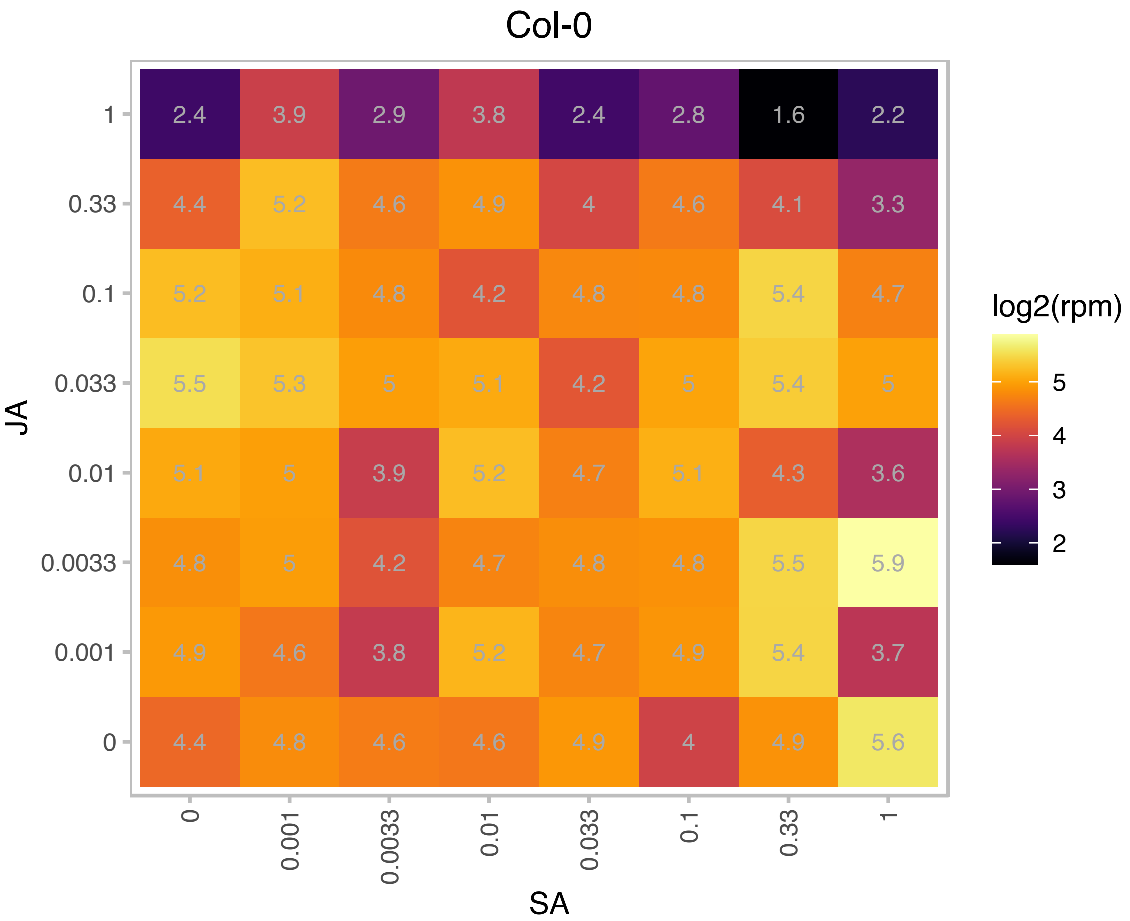


| | |
|---|---|
| Gene | AT1G07280 |
| is.expressed_genes | TRUE |
| is.responsive_genes | TRUE |
| is.apcluster | H-01 |

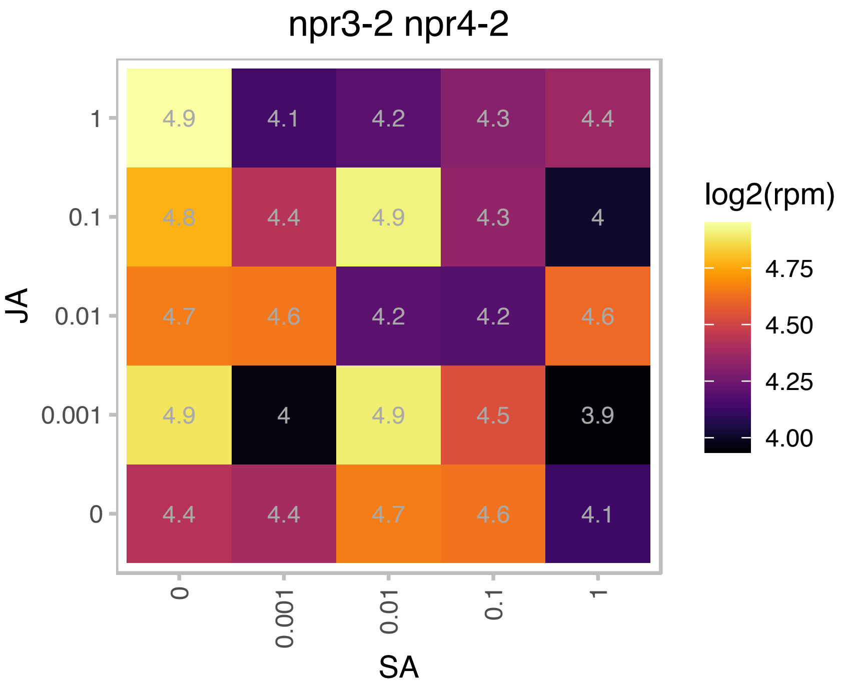


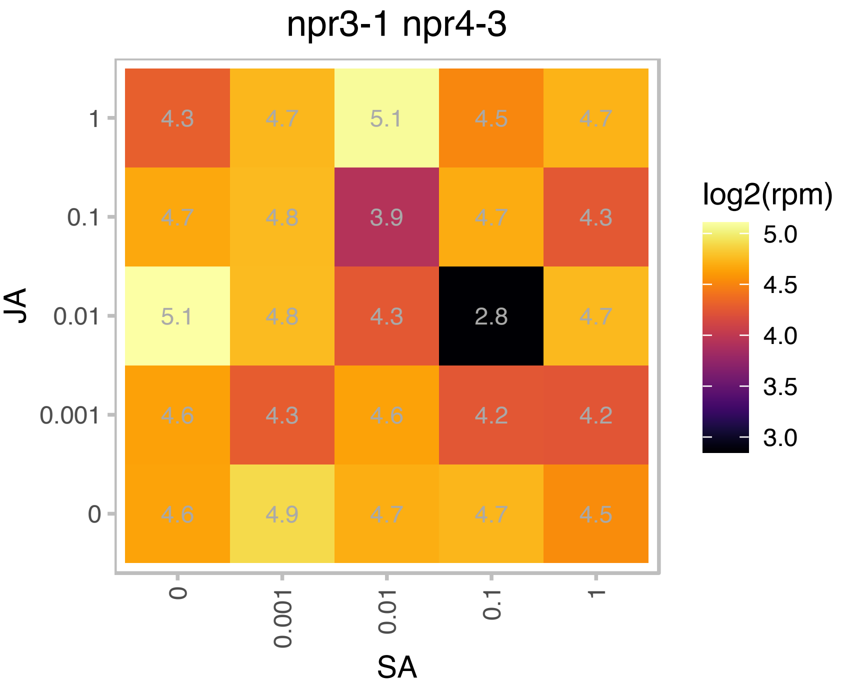

# AT1G07830.1 : NA

ribosomal protein L29 family protein

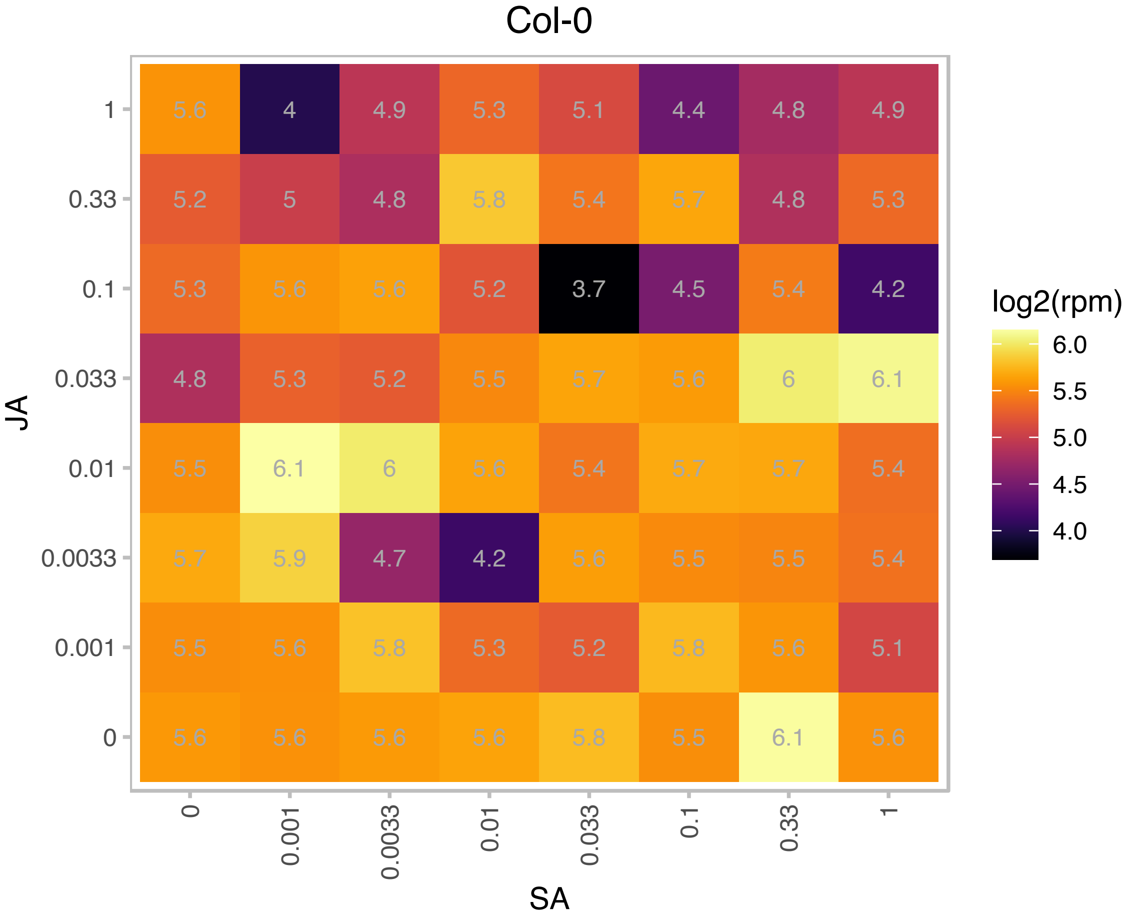


| | |
|---|---|
| Gene | AT1G07830 |
| is.expressed_genes | TRUE |
| is.responsive_genes | TRUE |
| is.apcluster | H-01 |

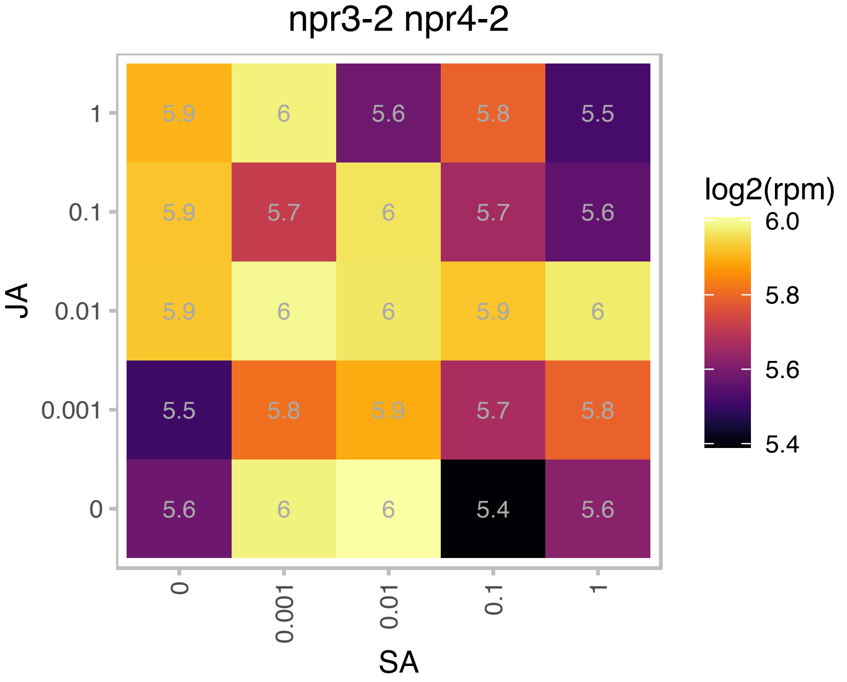


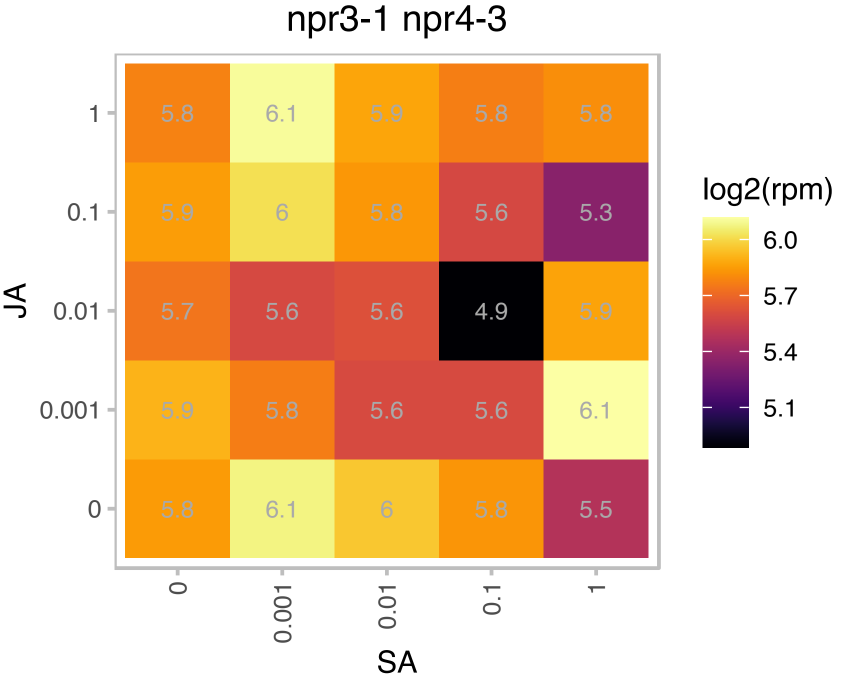

# AT1G11820.1 : NA

O-Glycosyl hydrolases family 17 protein

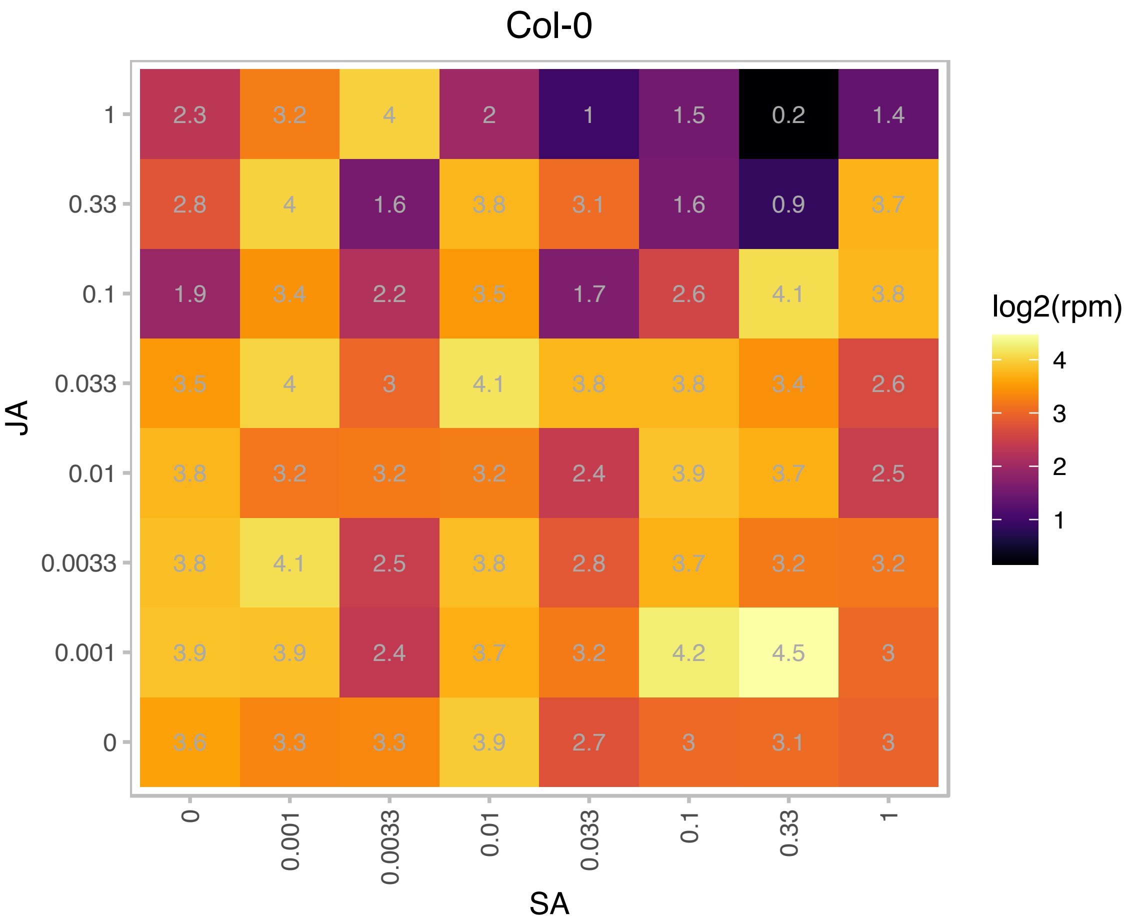


| | |
|---|---|
| Gene | AT1G11820 |
| is.expressed_genes | TRUE |
| is.responsive_genes | TRUE |
| is.apcluster | H-01 |

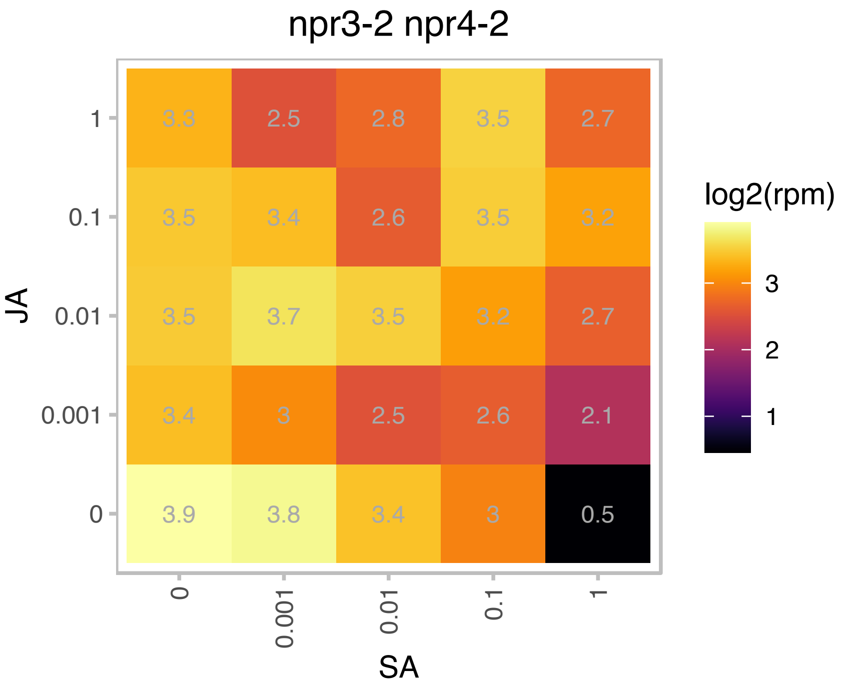


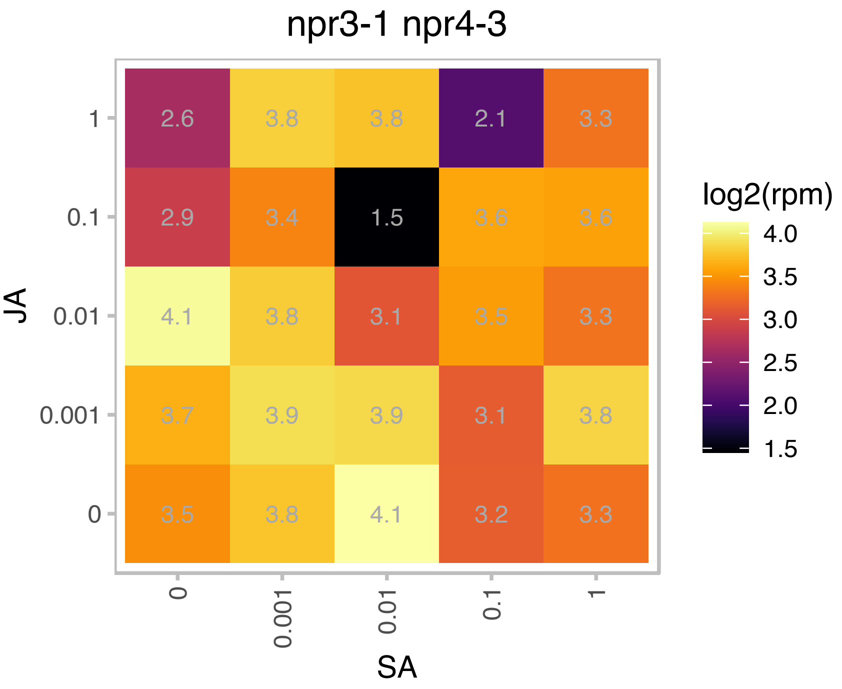

### AT1G20050.1 : HYD1, MAD4

C-8,7 sterol isomerase

C-8 sterol isomerase that also plays a role in miRNA function.

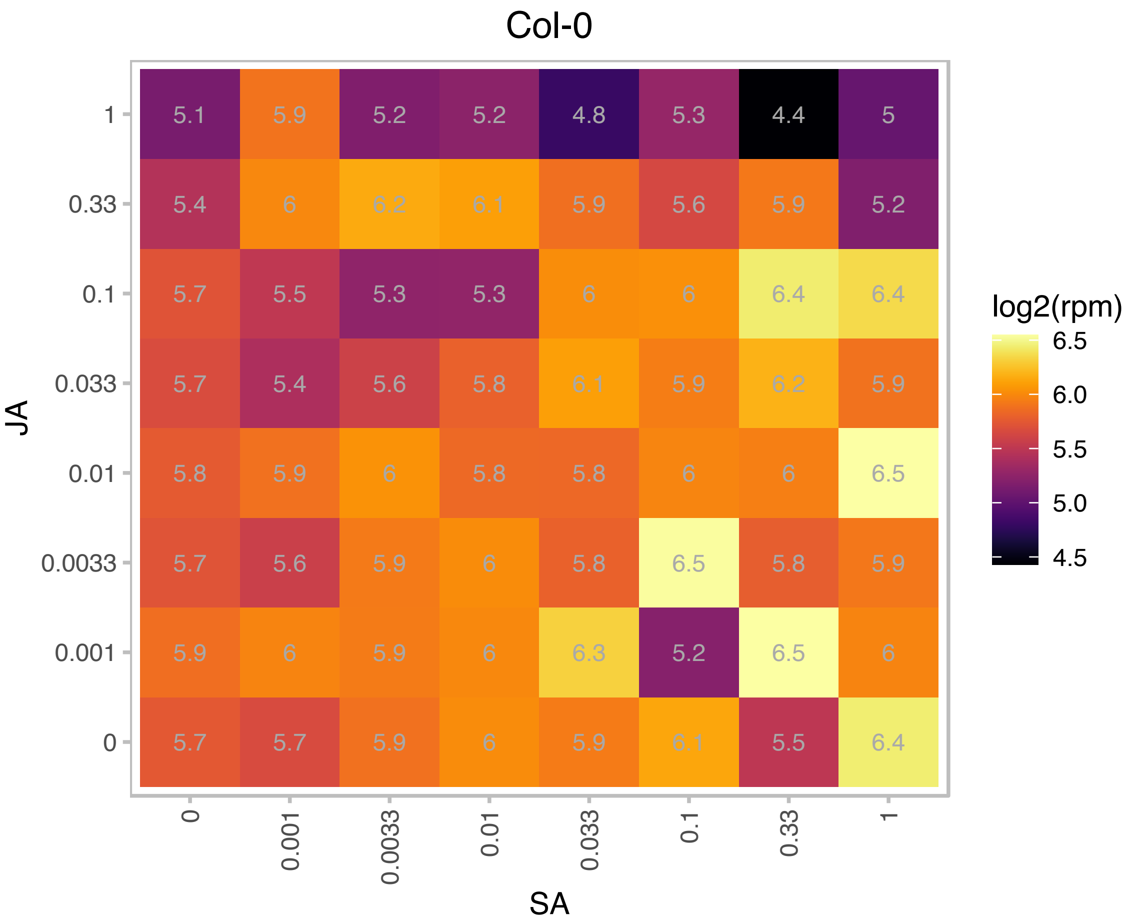


| | |
|---|---|
| Gene | AT1G20050 |
| is.expressed_genes | TRUE |
| is.responsive_genes | TRUE |
| is.apcluster | H-01 |

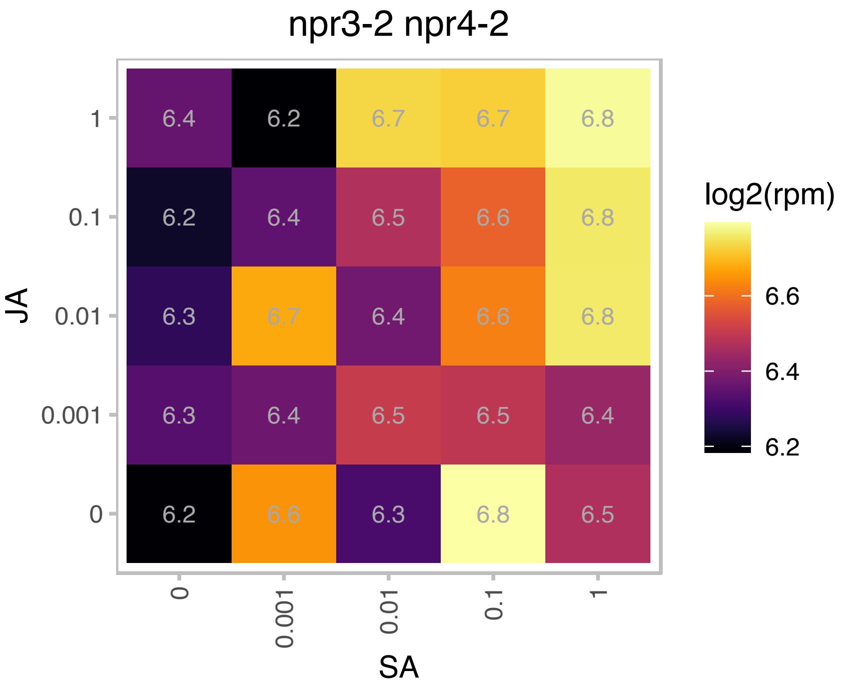


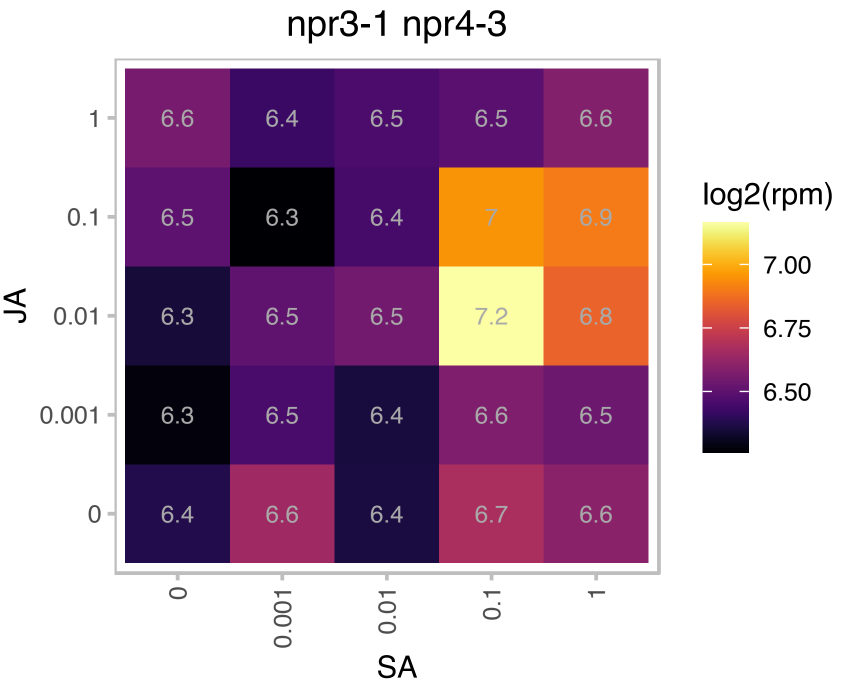

# AT1G23100.1 : NA

GroES-like family protein

| | |
|---|---|
| Gene | AT1G23100 |
| is.expressed_genes | TRUE |
| is.responsive_genes | TRUE |
| is.apcluster | H-01 |

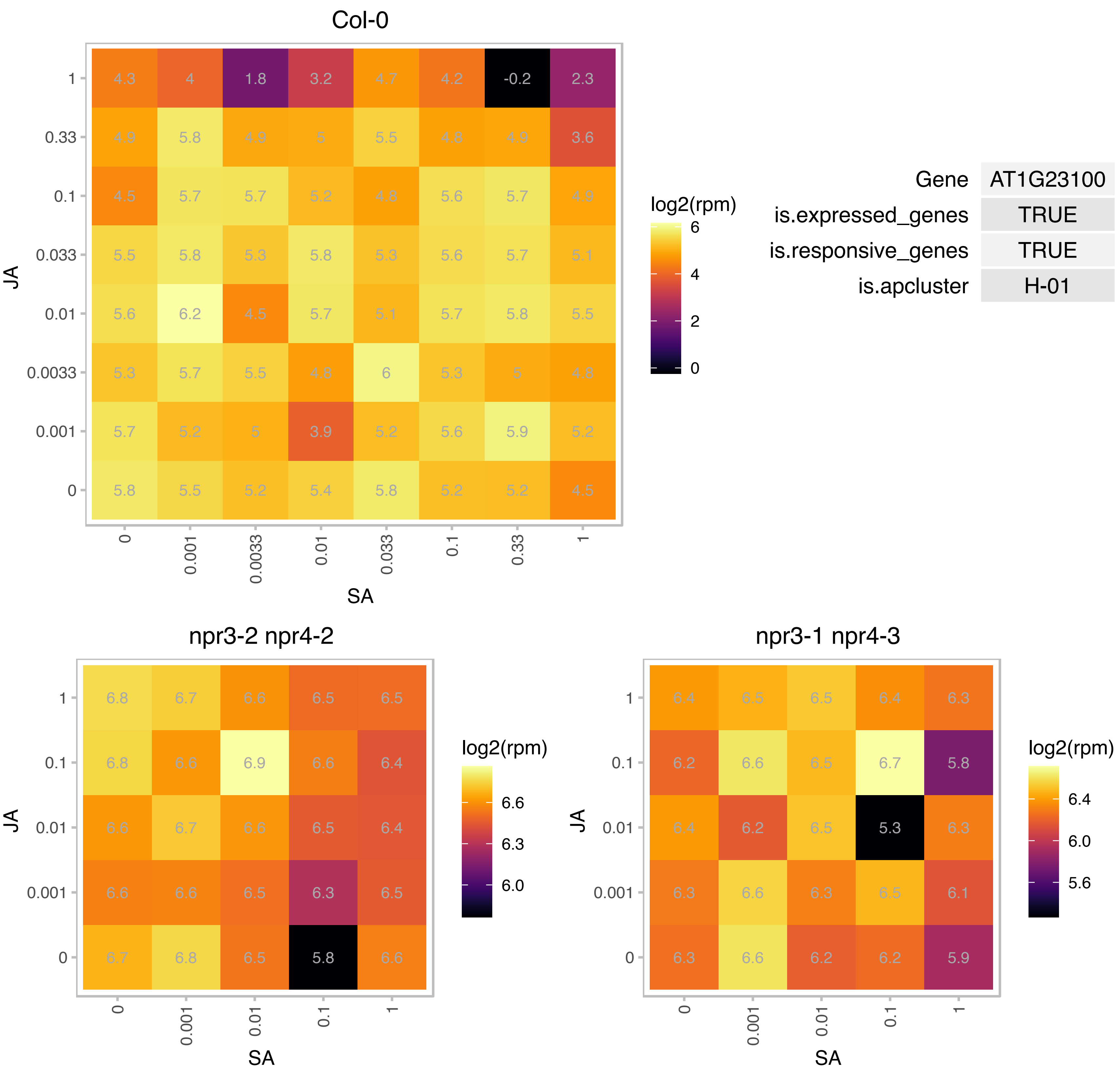

# AT1G28395.1 : NA

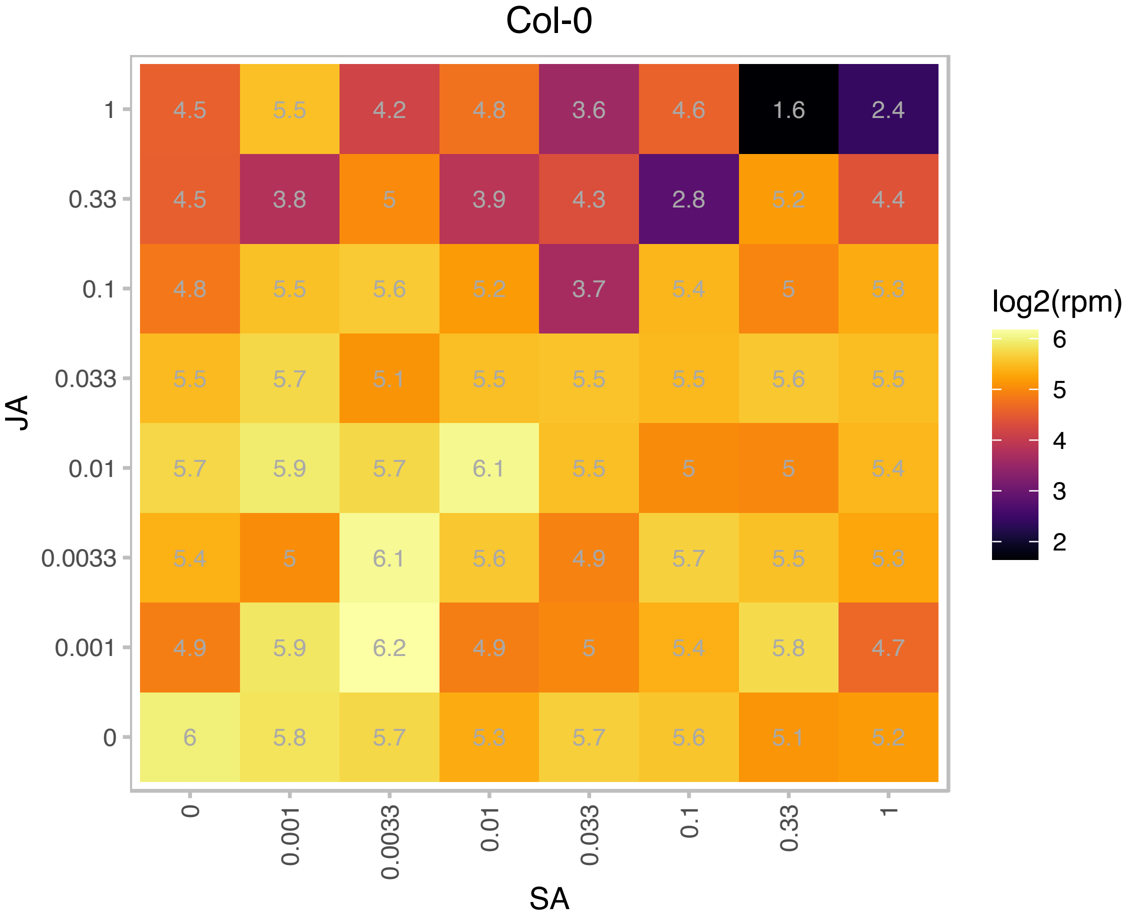


| | |
|---|---|
| Gene | AT1G28395 |
| is.expressed_genes | TRUE |
| is.responsive_genes | TRUE |
| is.apcluster | H-01 |

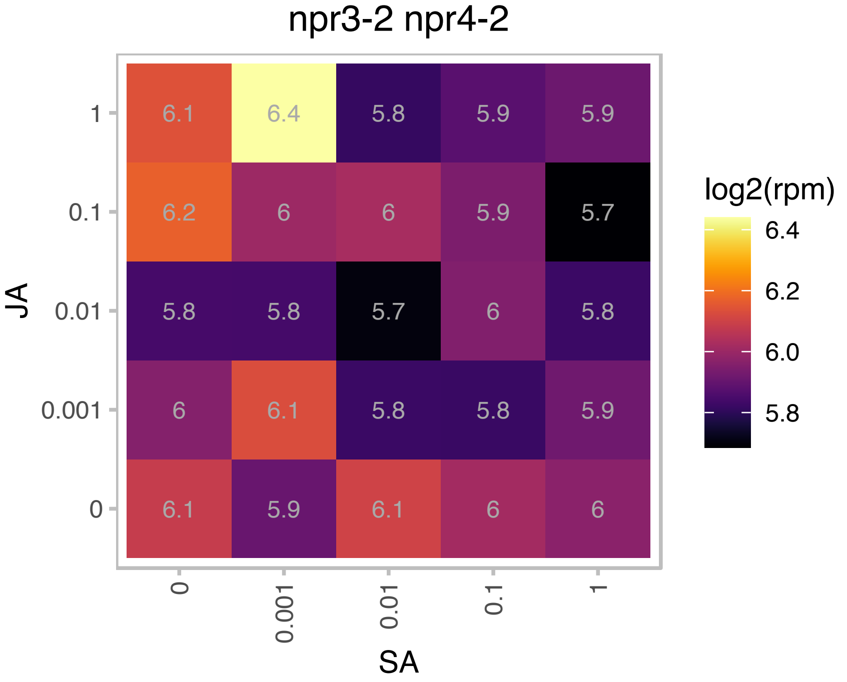


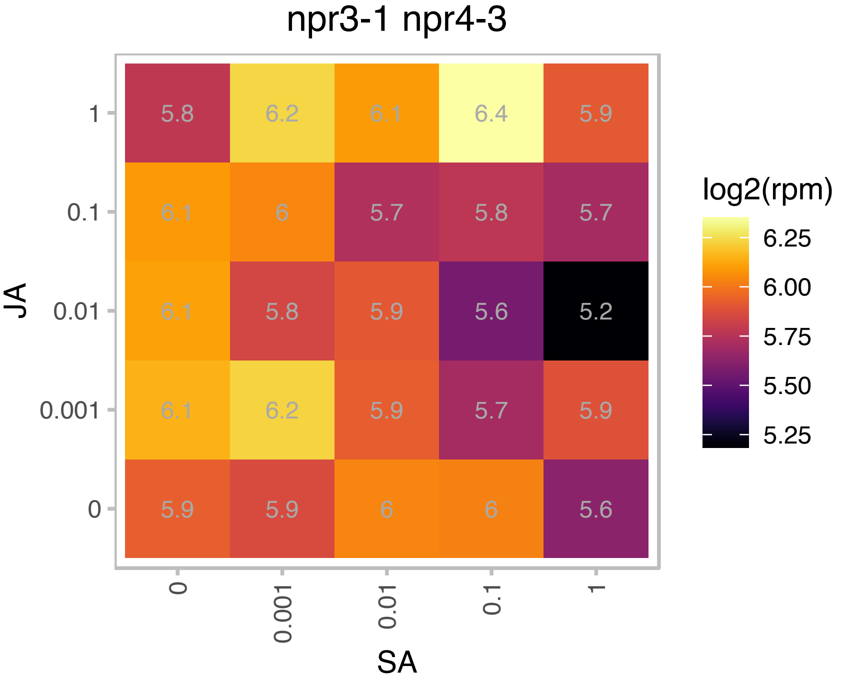

# AT1G29320.1 : NA

Transducin/WD40 repeat-like superfamily protein

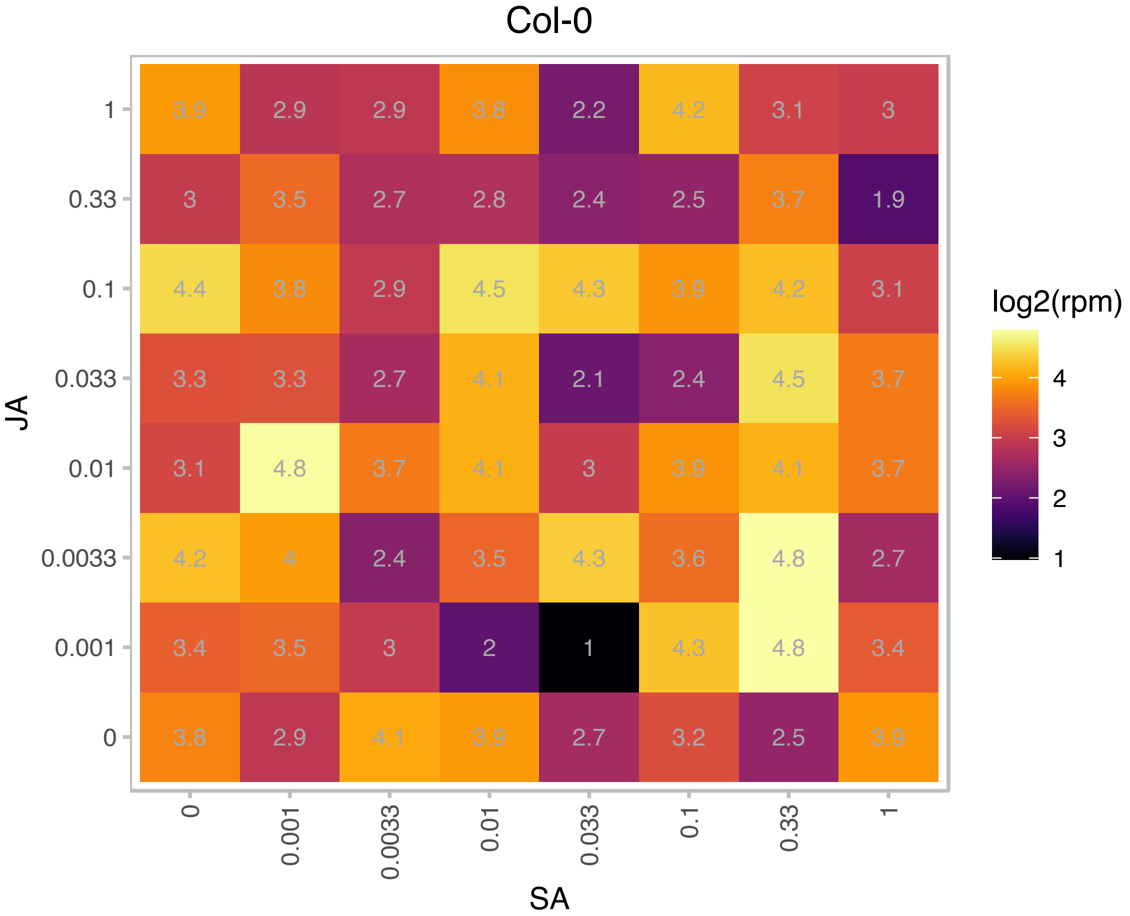


| | |
|---|---|
| Gene | AT1G29320 |
| is.expressed_genes | TRUE |
| is.responsive_genes | TRUE |
| is.apcluster | H-01 |

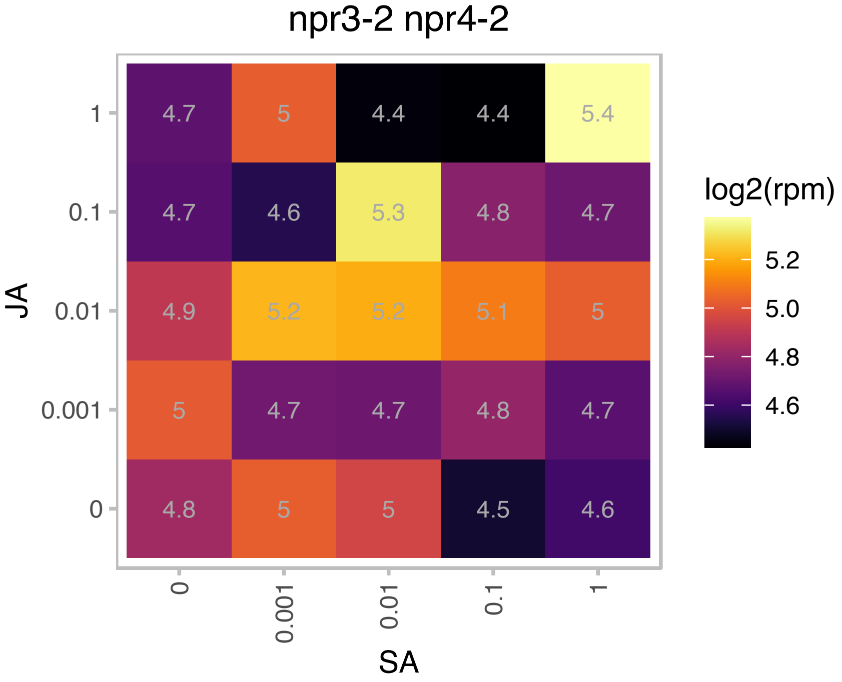


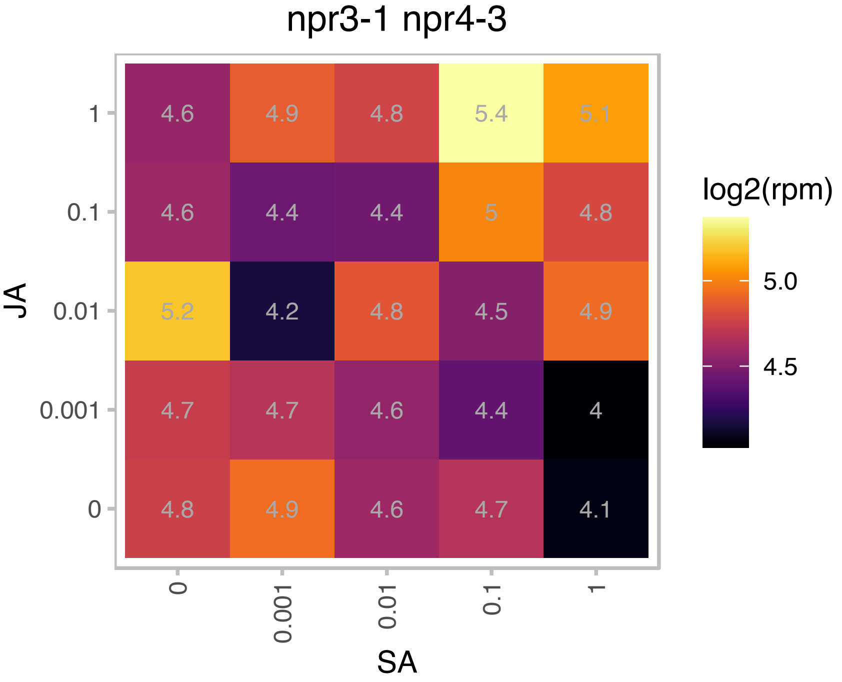

### AT1G57620.1 : CYB

emp24/gp25L/p24 family/GOLD family protein

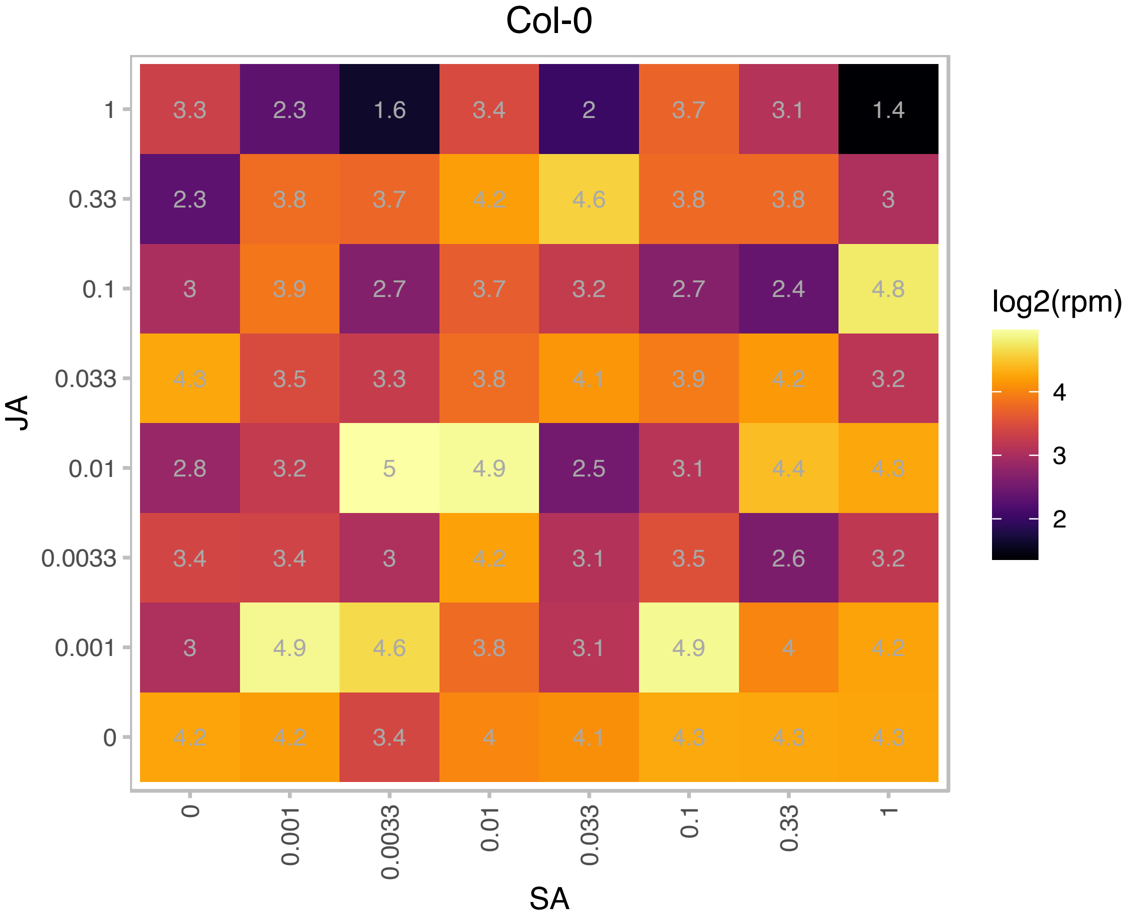


| | |
|---|---|
| Gene | AT1G57620 |
| is.expressed_genes | TRUE |
| is.responsive_genes | TRUE |
| is.apcluster | H-01 |

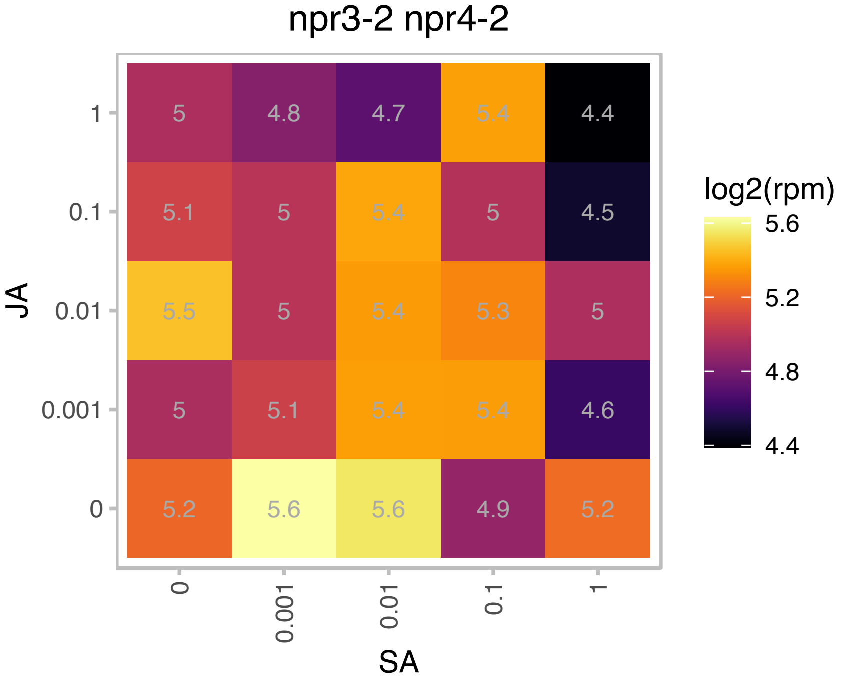


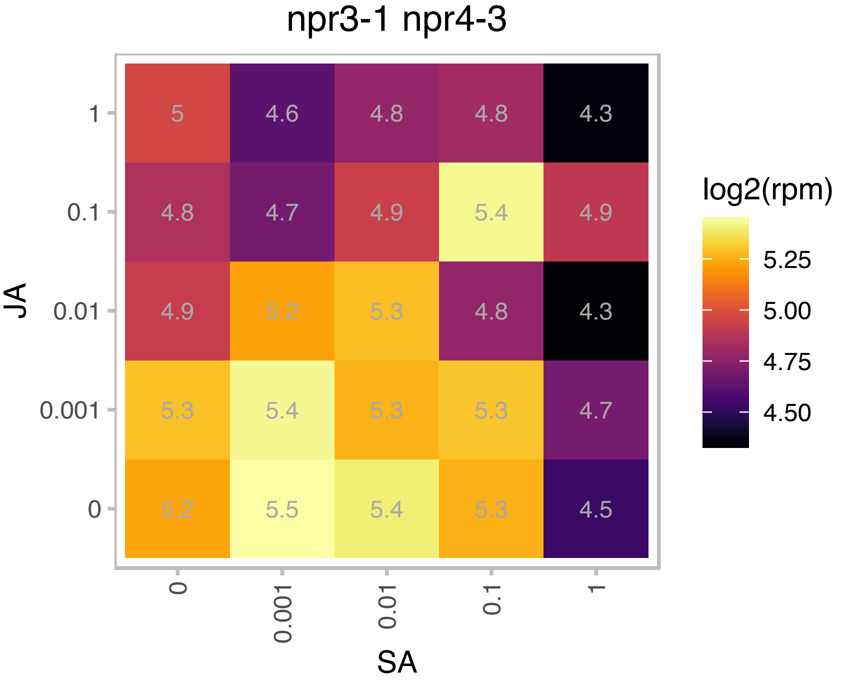

# AT1G68680.1 : NA

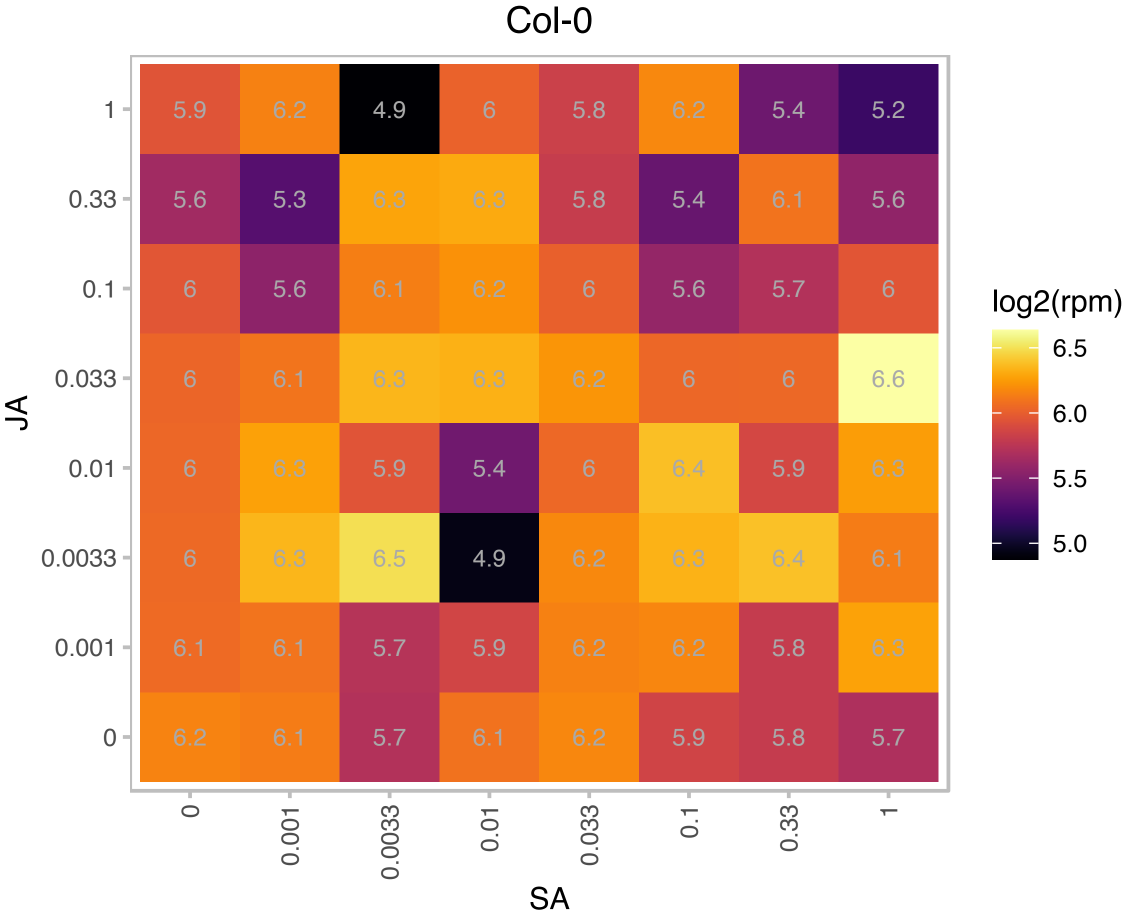


| | |
|---|---|
| Gene | AT1G68680 |
| is.expressed_genes | TRUE |
| is.responsive_genes | TRUE |
| is.apcluster | H-01 |

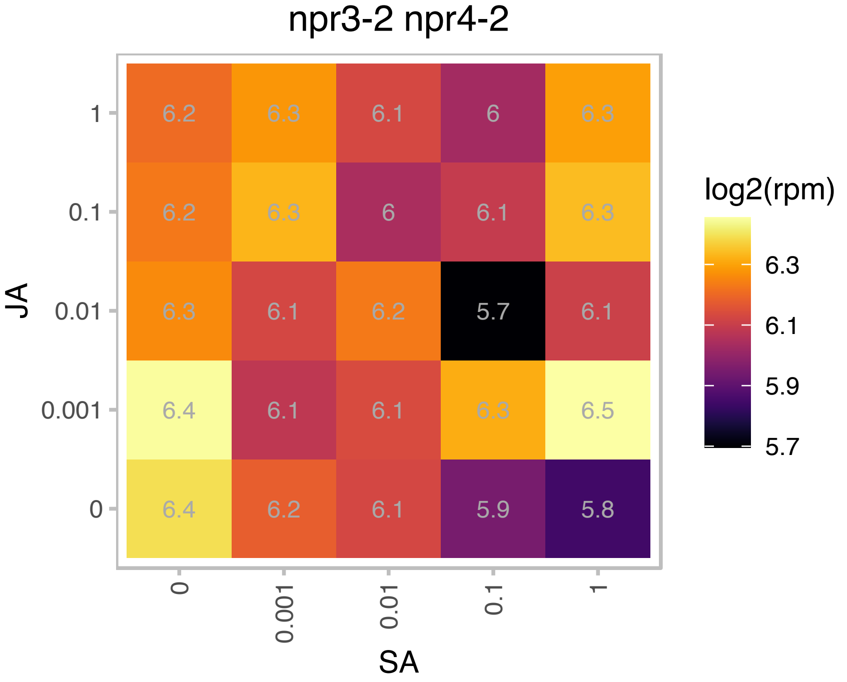


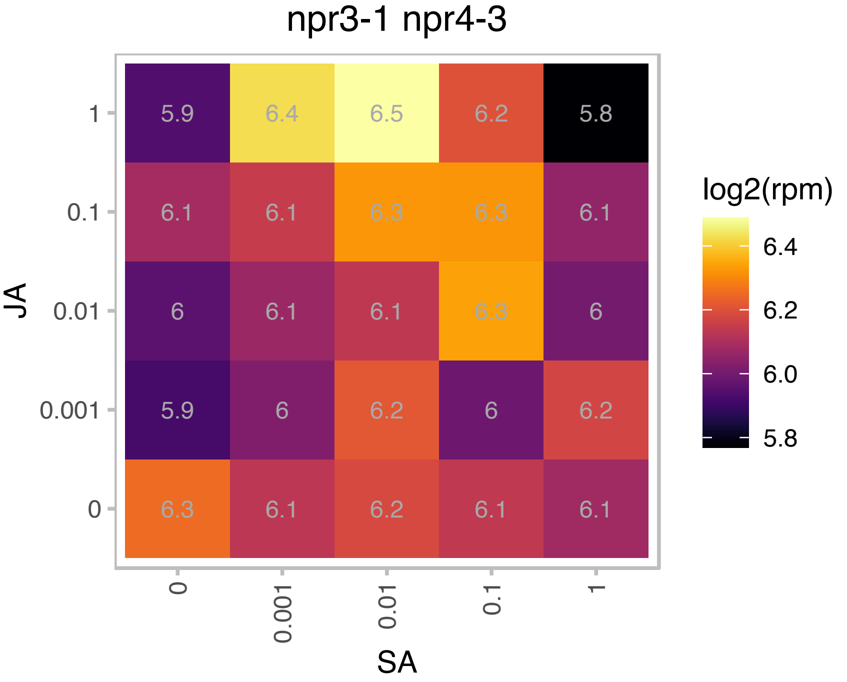

### AT1G70890.1 : MLP43, MLP43

MLP-like protein 43

| | |
|---|---|
| Gene | AT1G70890 |
| is.expressed_genes | TRUE |
| is.responsive_genes | TRUE |
| is.apcluster | H-01 |

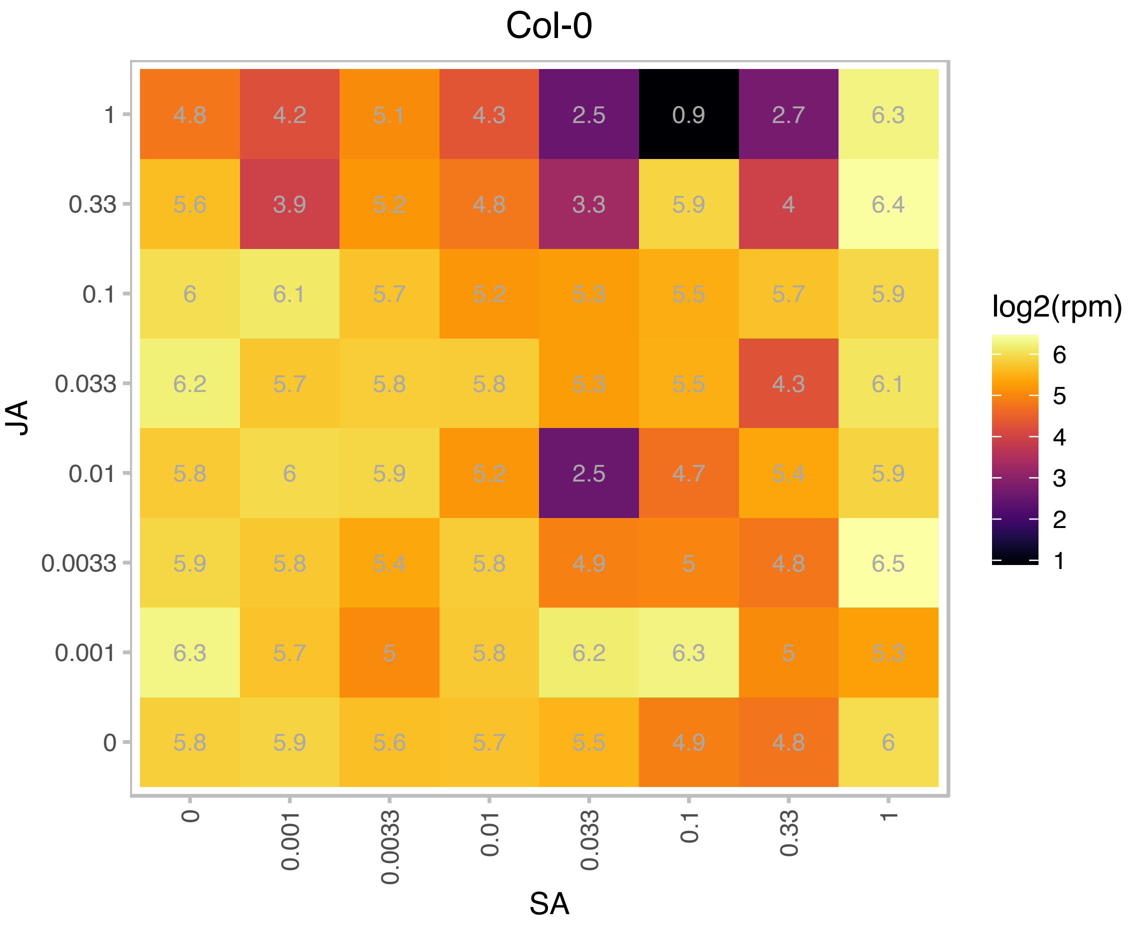


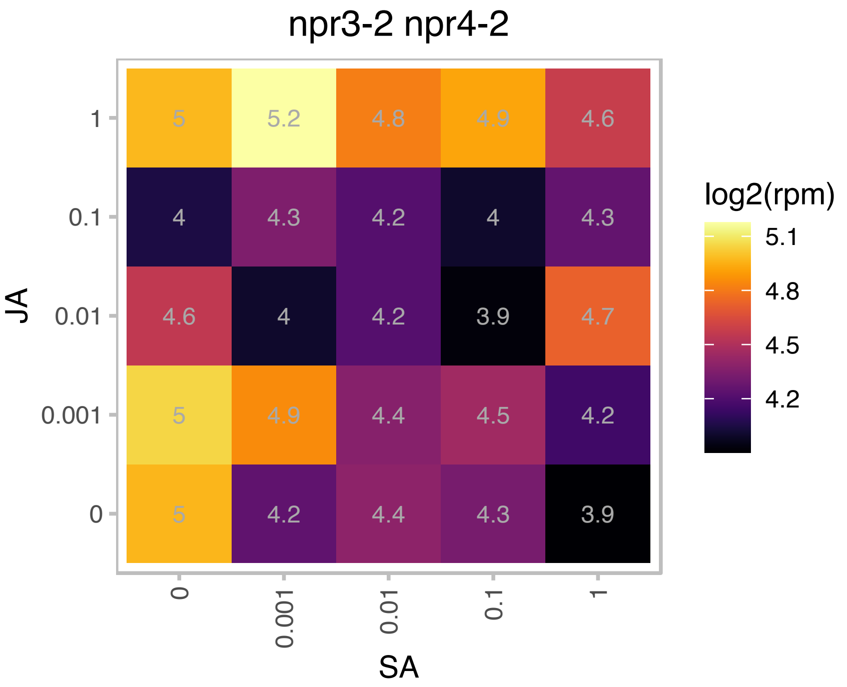


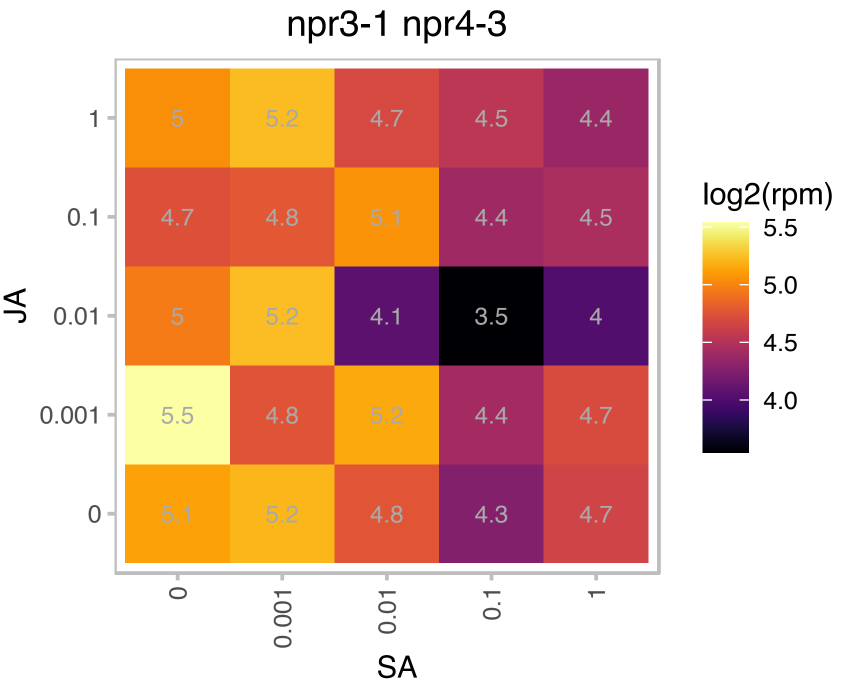

### AT1G71790.1 : AtCPB, CPB

Subunits of heterodimeric actin filament capping protein Capz superfamily

Encodes a heterodimeric actin binding protein composed of an alpha and a beta sumunit. Stabilizes actin filament cytoskeleton by capping.

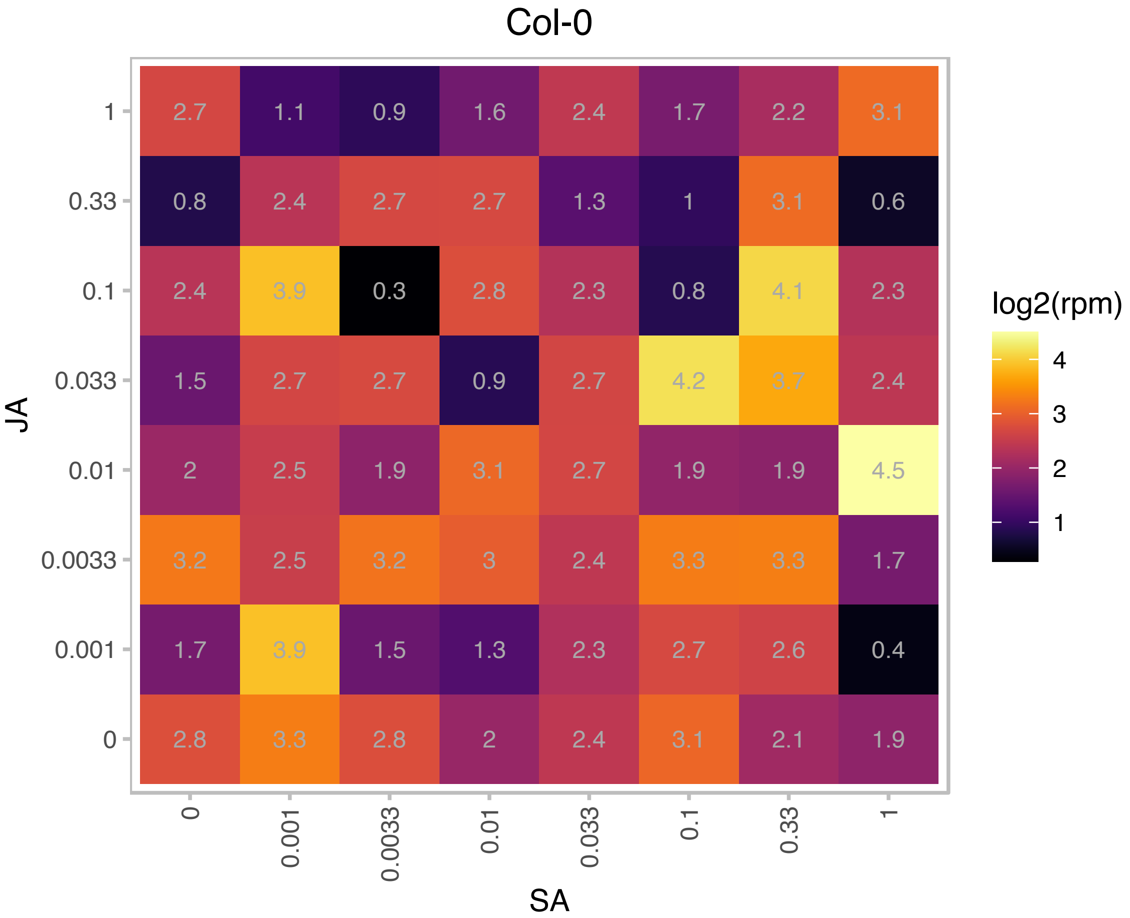


| | |
|---|---|
| Gene | AT1G71790 |
| is.expressed_genes | TRUE |
| is.responsive_genes | TRUE |
| is.apcluster | H-01 |

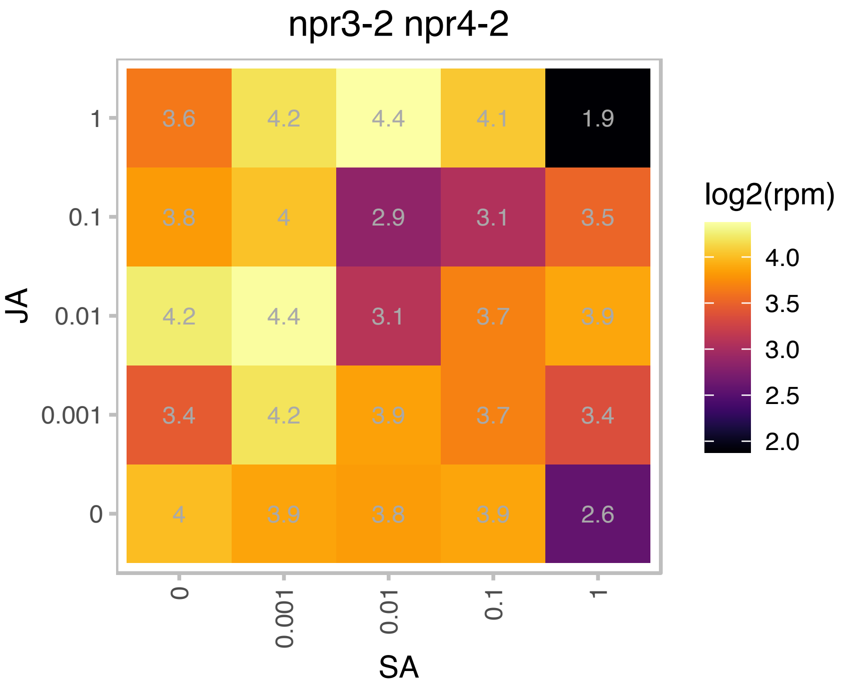


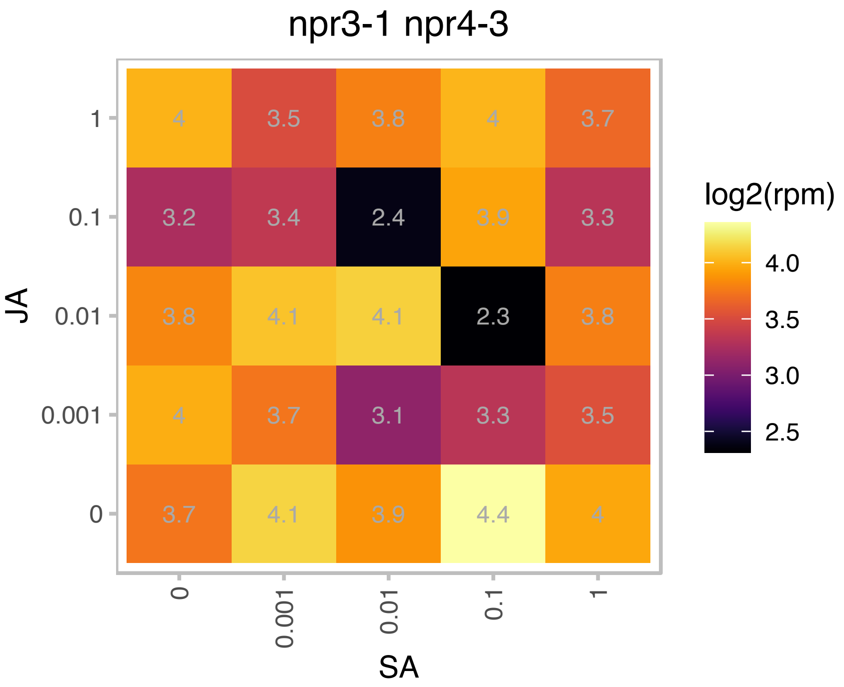

### AT1G80560.1 : ATIMD2, IMD2

isopropylmalate dehydrogenase 2
The AtIMD2 is one out of 3 genes encoding the enzyme 3-isopropylmalate dehydrogenase involved in leucine biosynthesis in Arabidopsis. Its subcellular location has been targeted to plastids. The mRNA is cell-to-cell mobile.

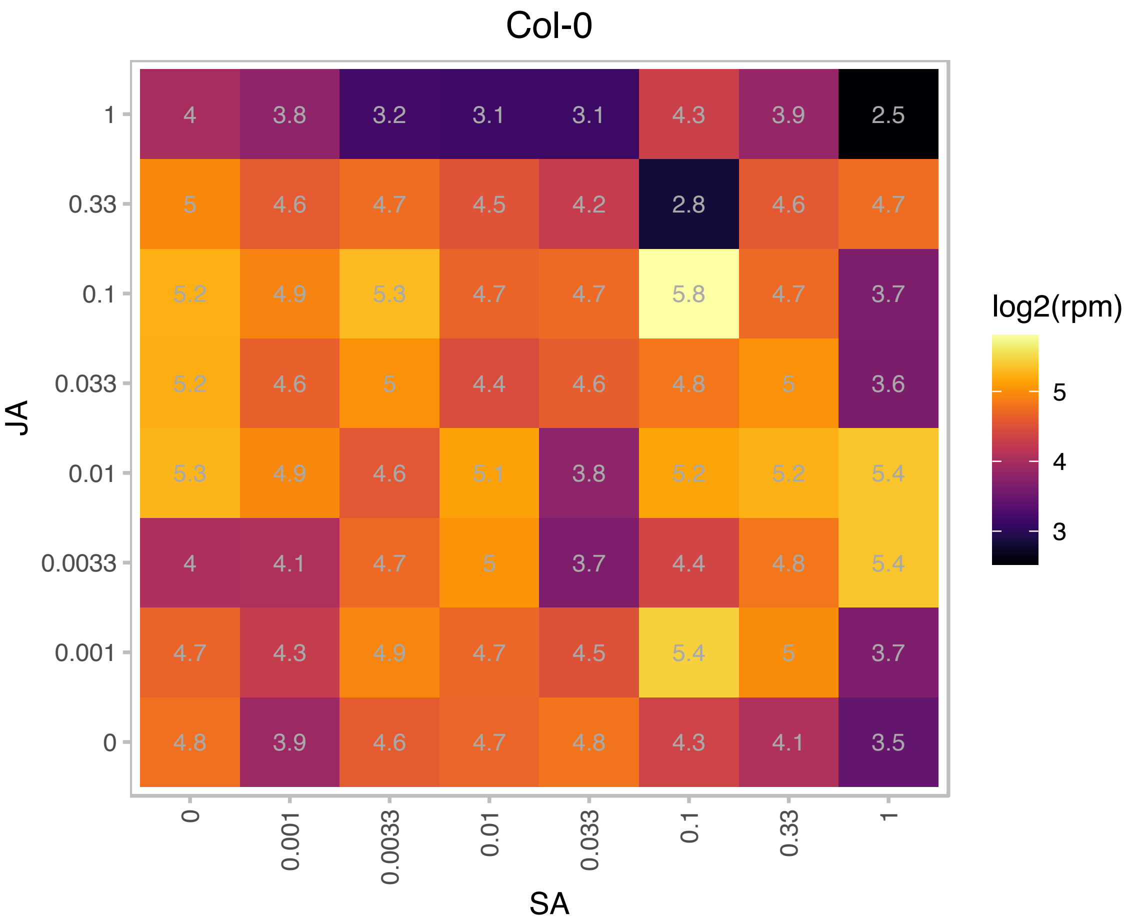


| | |
|---|---|
| Gene | AT1G80560 |
| is.expressed_genes | TRUE |
| is.responsive_genes | TRUE |
| is.apcluster | H-01 |

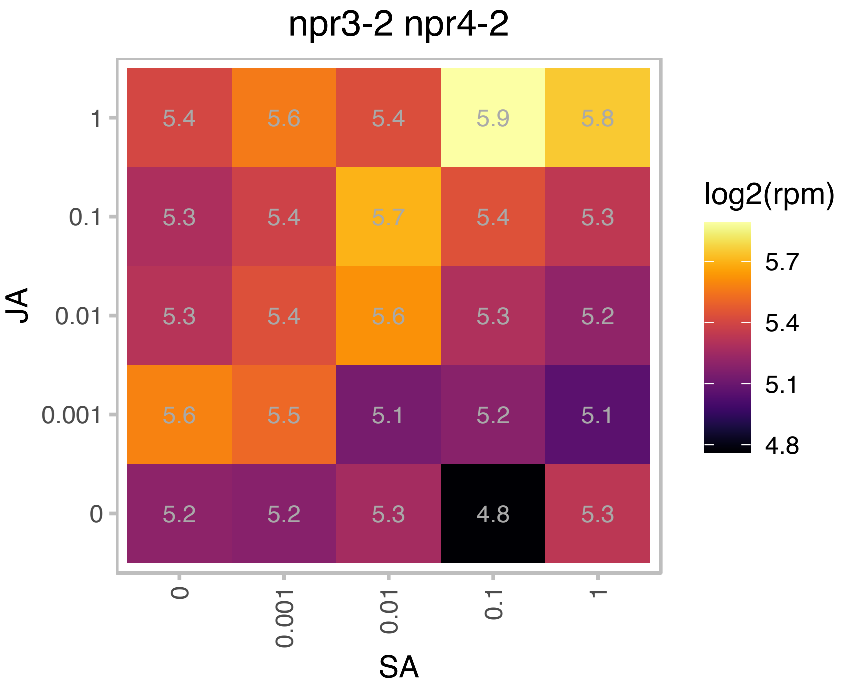


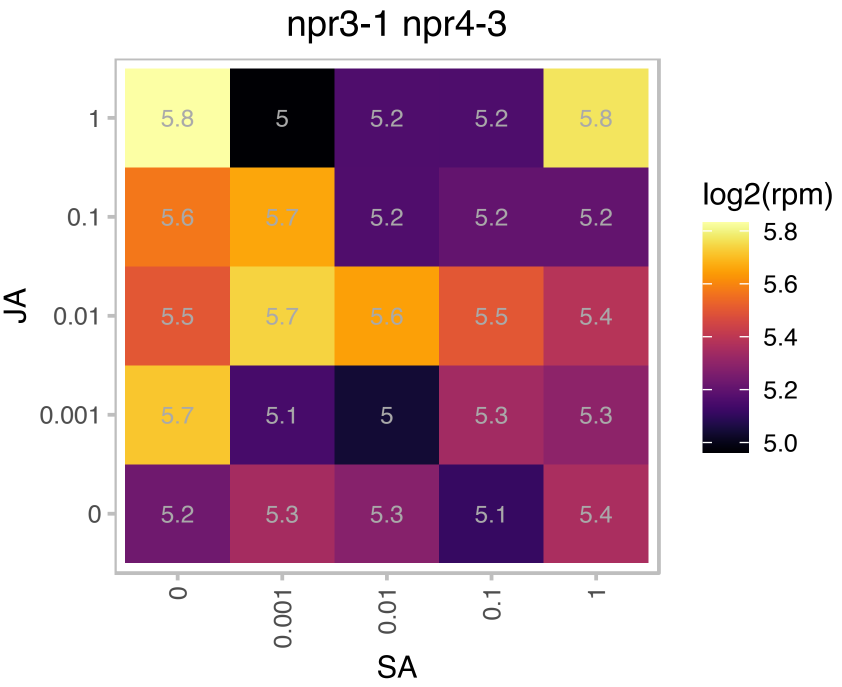

# AT2G01400.1 : NA

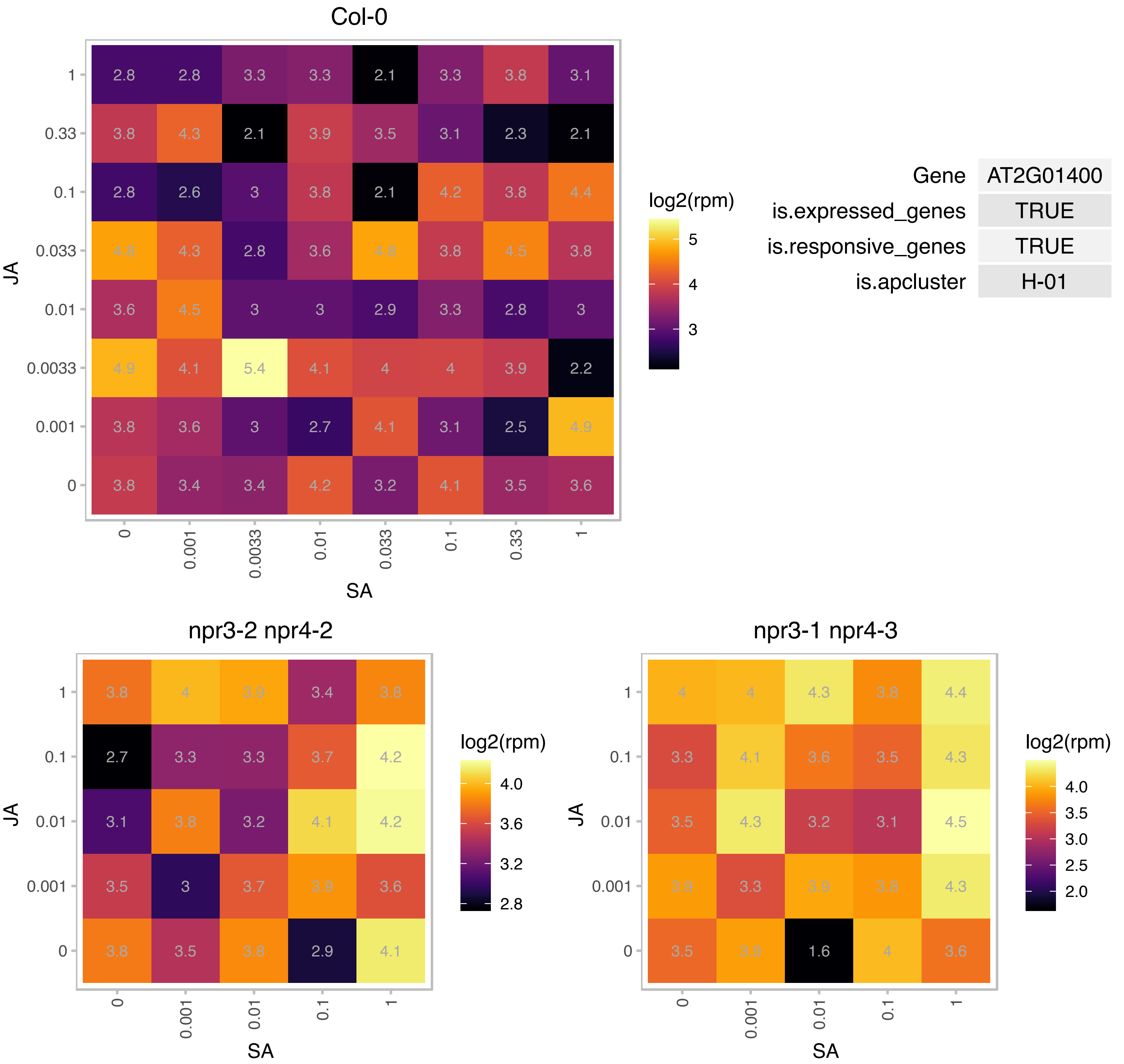


| Gene | AT2G01400 |
|---|---|
| is.expressed_genes | TRUE |
| is.responsive_genes | TRUE |
| is.apcluster | H-01 |

### AT2G02740.1 : ATWHY3, PTAC11, WHY3

ssDNA-binding transcriptional regulator

Encodes a homolog of the potato p24 protein.  It shares the conserved KGKAAL domain, a putative DNA-binding domain, with potato p24 and is localized to the plastid and not the nucleus.

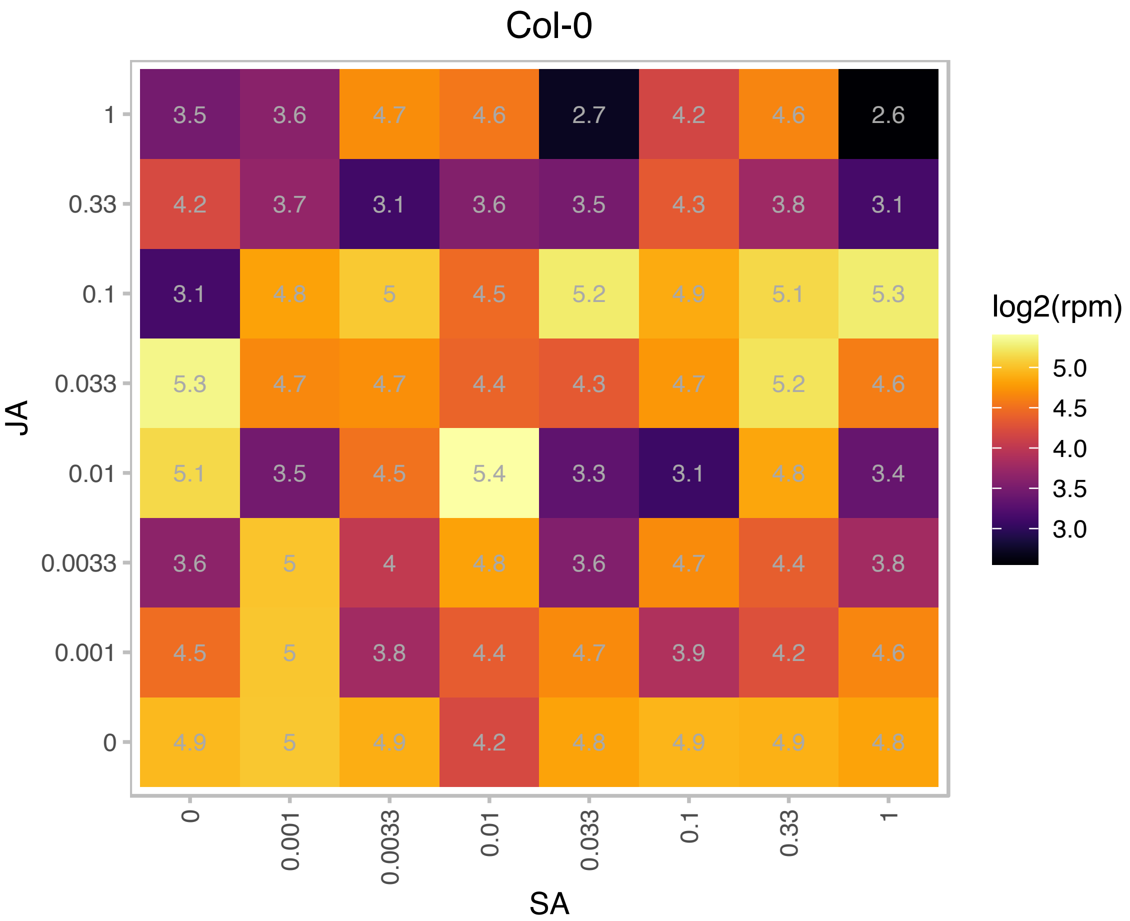


| Gene | AT2G02740 |
|---|---|
| is.expressed_genes | TRUE |
| is.responsive_genes | TRUE |
| is.apcluster | H-01 |

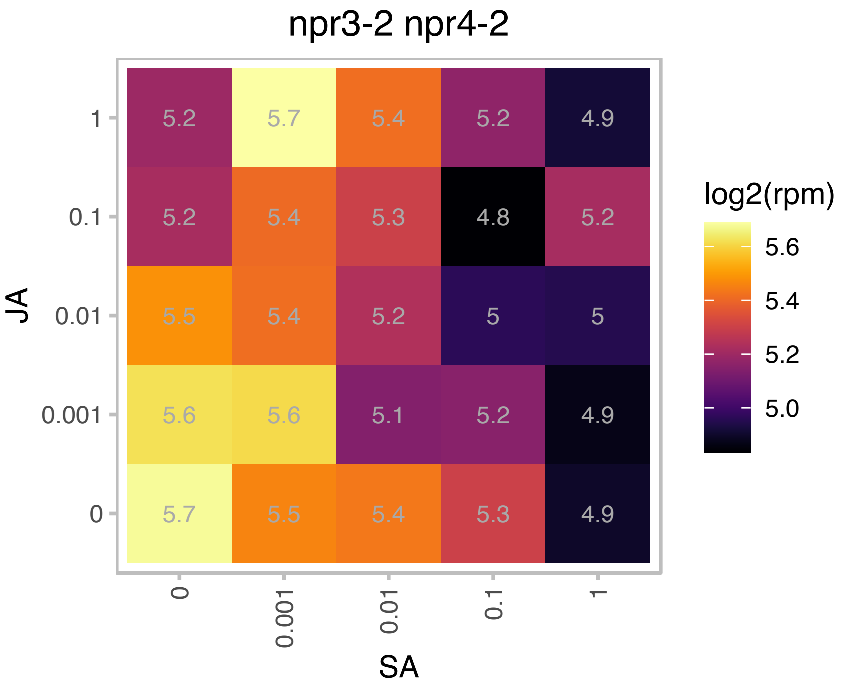


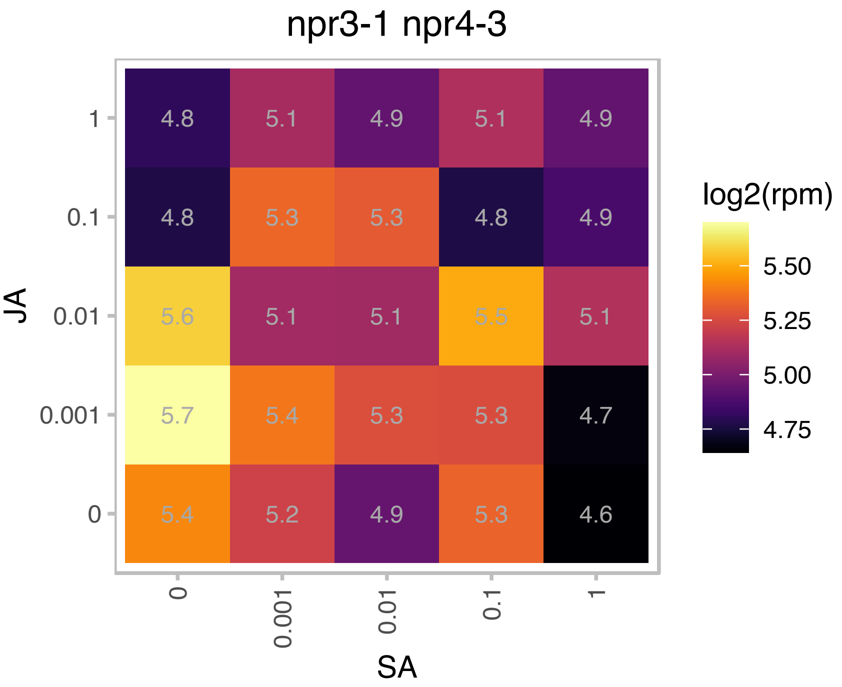

### AT2G06925.1 : ATSPLA2-ALPHA, PLA2-ALPHA

Phospholipase A2 family protein
Encodes a secretory phospholipase A2 enzyme, which specifically hydrolyzes the sn-2 position of phospholipids. The enzyme has a preference towards linoleoyl acyl chain over palmitoyl acyl chain. It also has a slight preference for phosphatidylcholine over phosphatidylethanolamine.

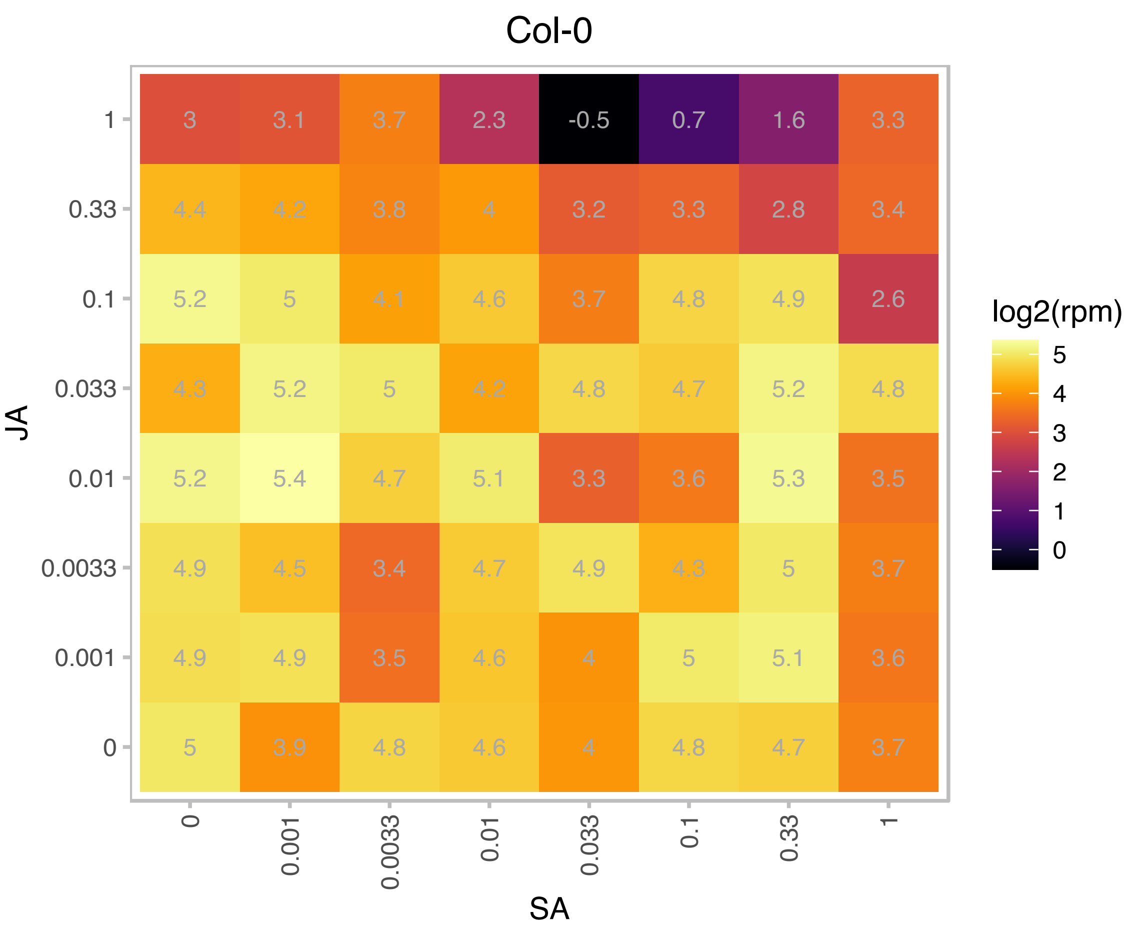


| | |
|---|---|
| Gene | AT2G06925 |
| is.expressed_genes | TRUE |
| is.responsive_genes | TRUE |
| is.apcluster | H-01 |

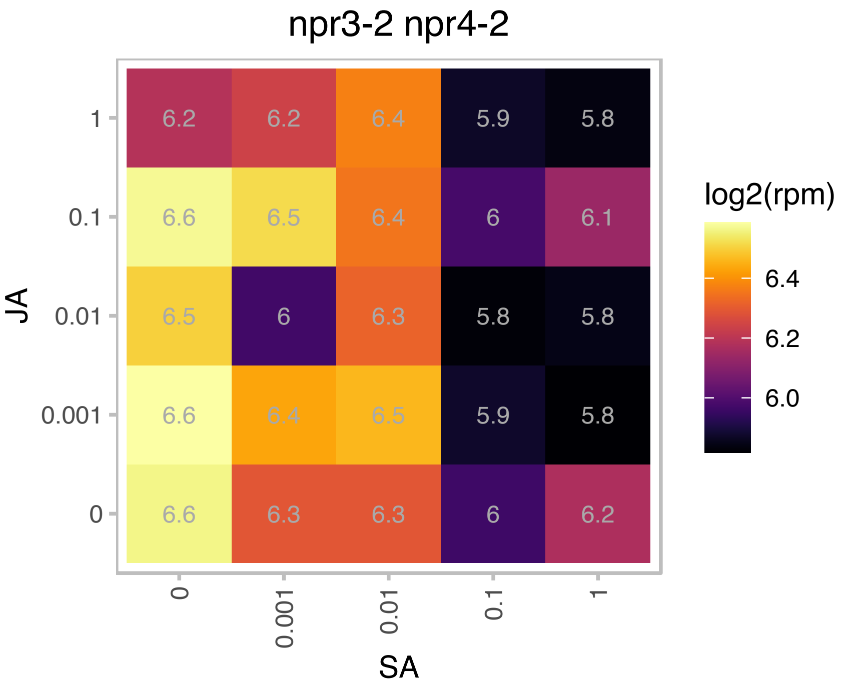


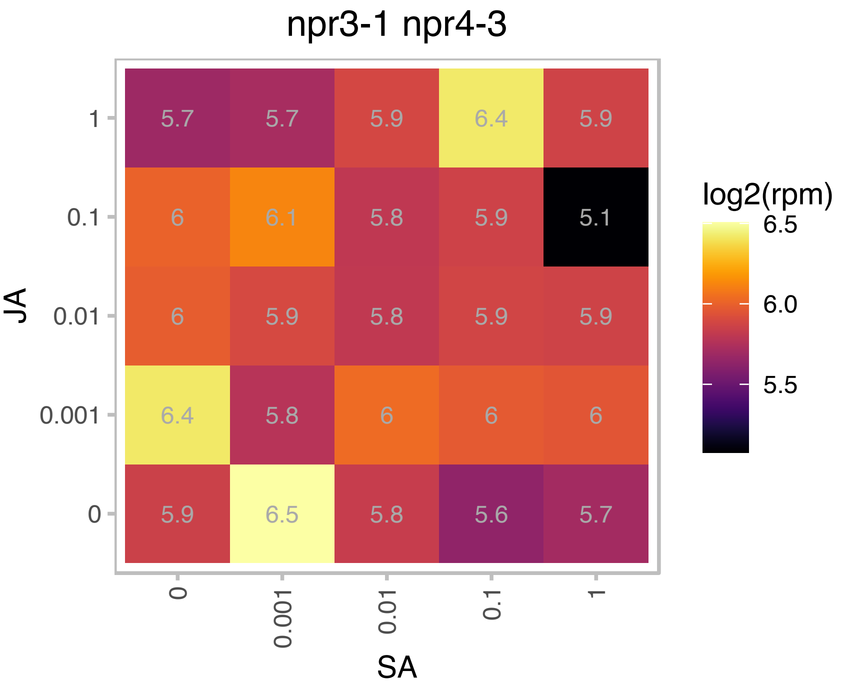

# AT2G06950.1 : NA

transposable element gene

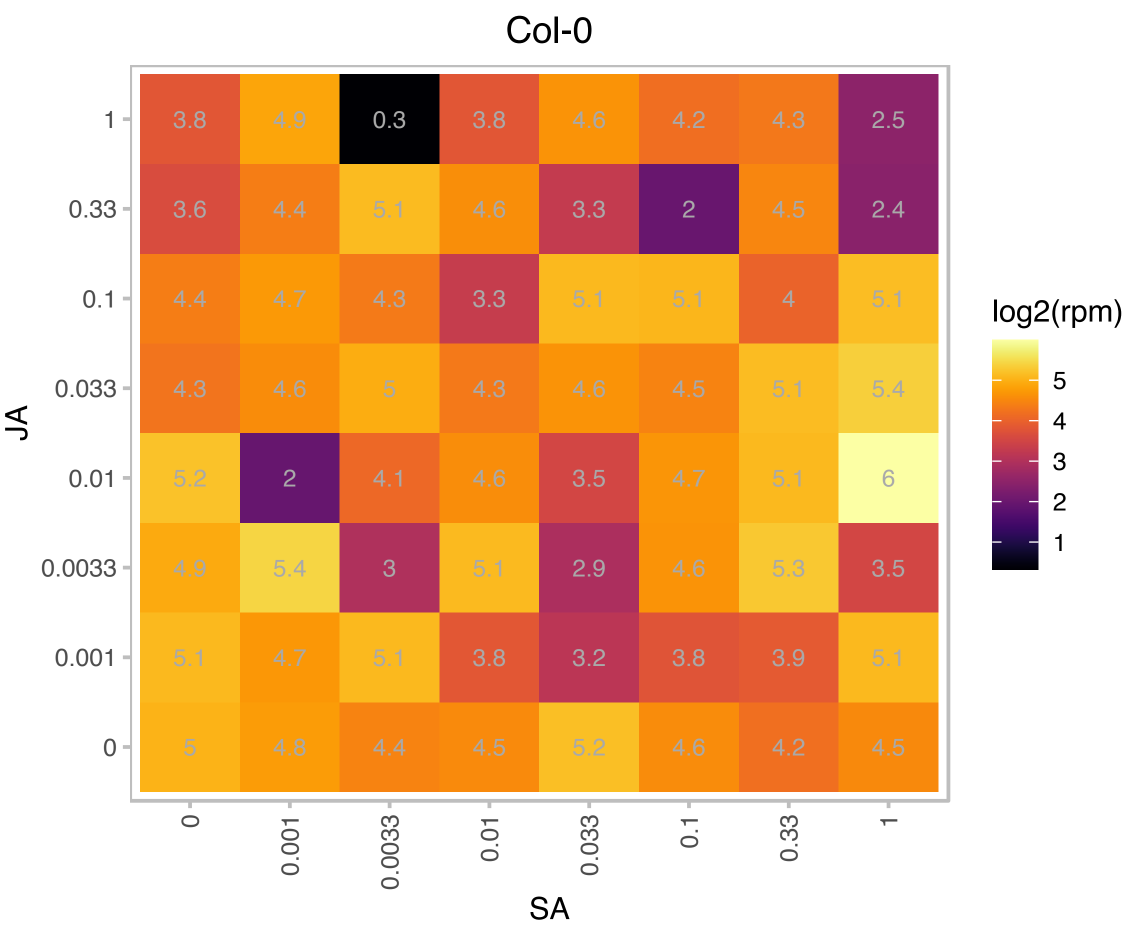


| | |
|---|---|
| Gene | AT2G06950 |
| is.expressed_genes | TRUE |
| is.responsive_genes | TRUE |
| is.apcluster | H-01 |

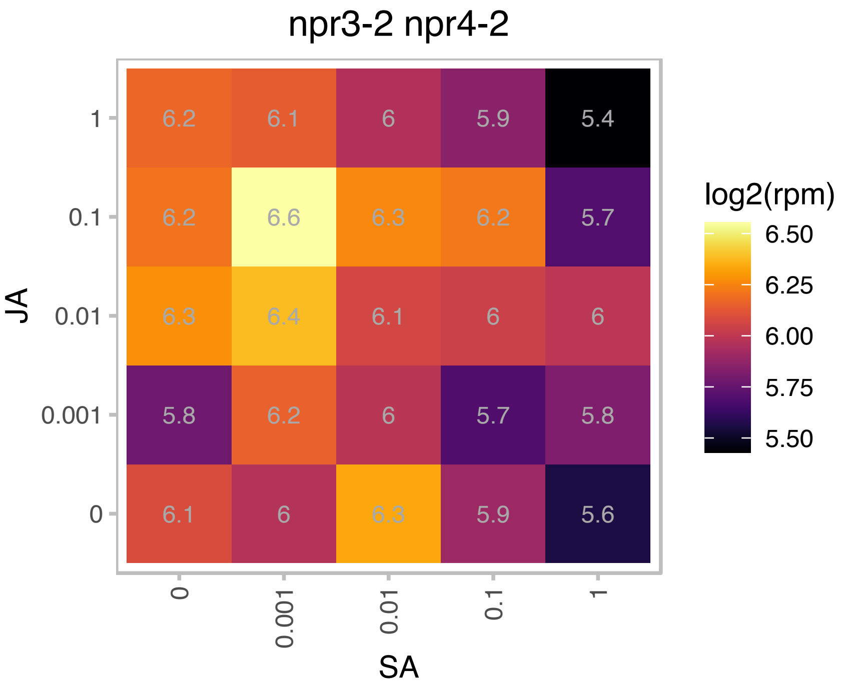


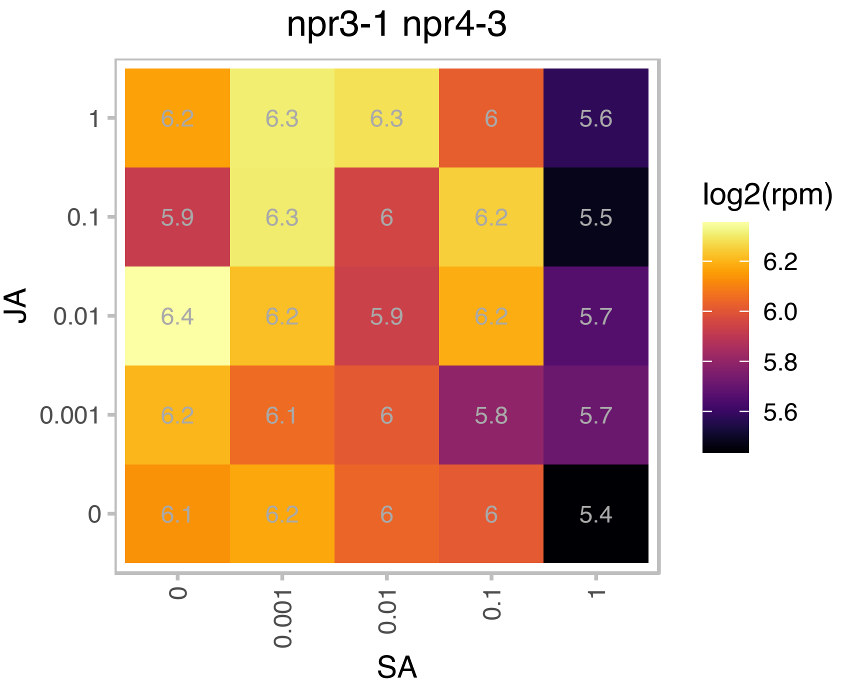

# AT2G07739.1 : NA

Ycf1 protein

| | |
|---|---|
| Gene | AT2G07739 |
| is.expressed_genes | TRUE |
| is.responsive_genes | TRUE |
| is.apcluster | H-01 |

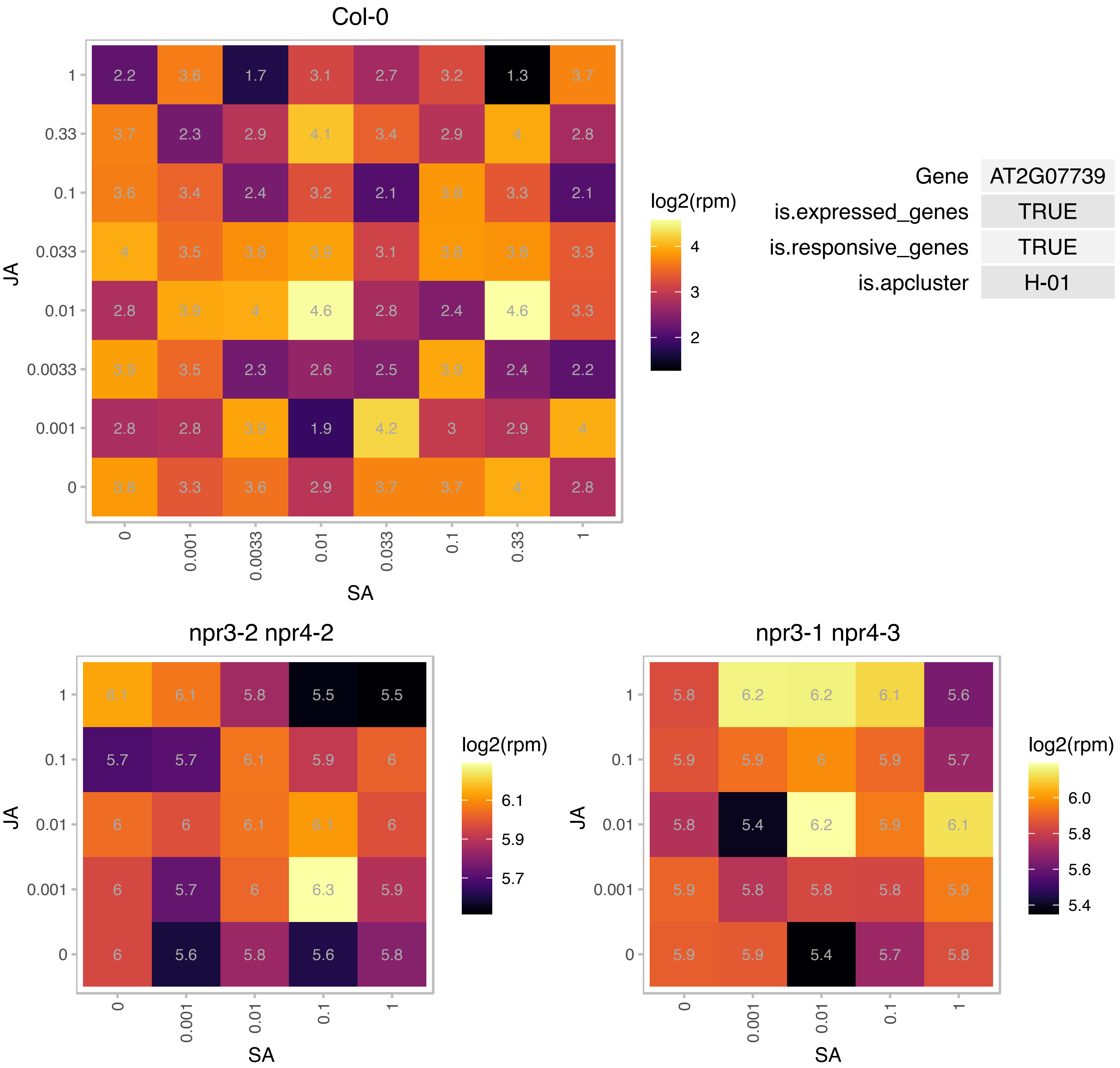

### AT2G26900.1 : BASS2

Sodium Bile acid symporter family

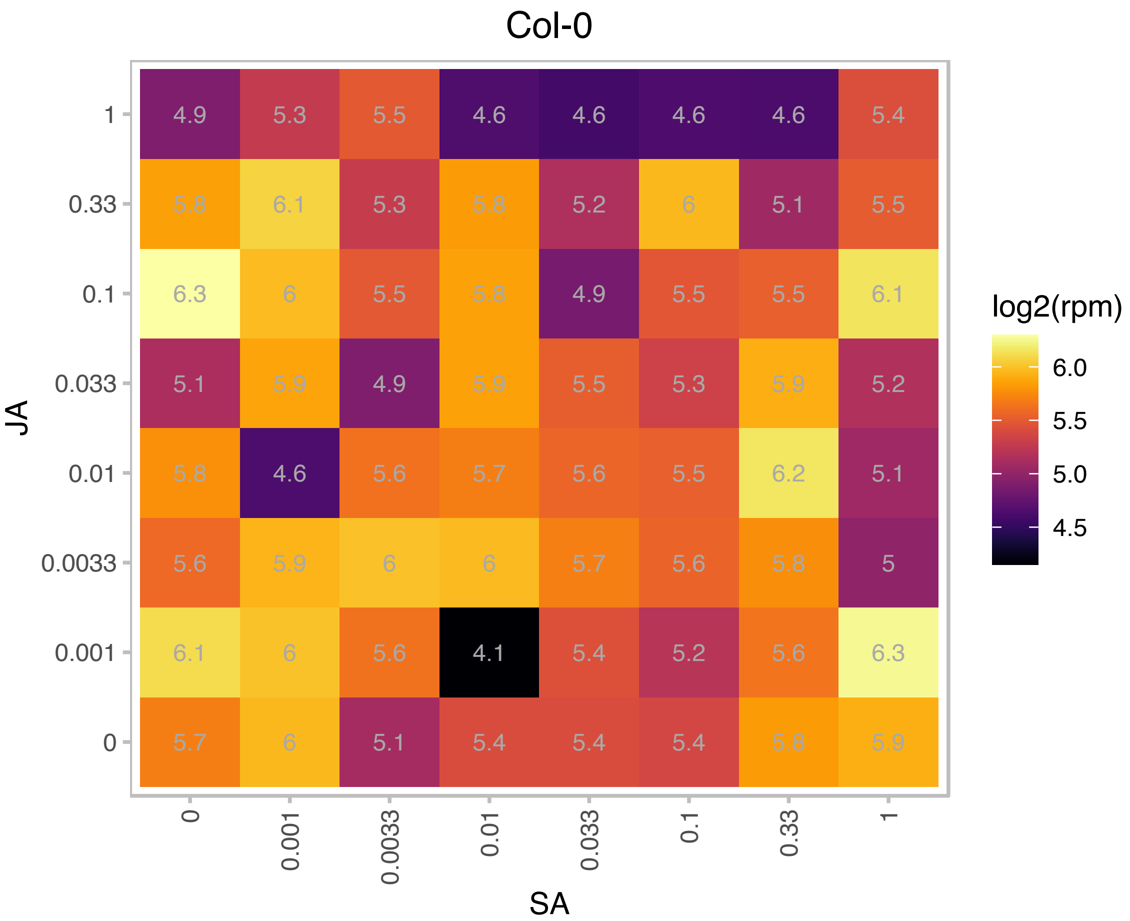


| | |
|---|---|
| Gene | AT2G26900 |
| is.expressed_genes | TRUE |
| is.responsive_genes | TRUE |
| is.apcluster | H-01 |

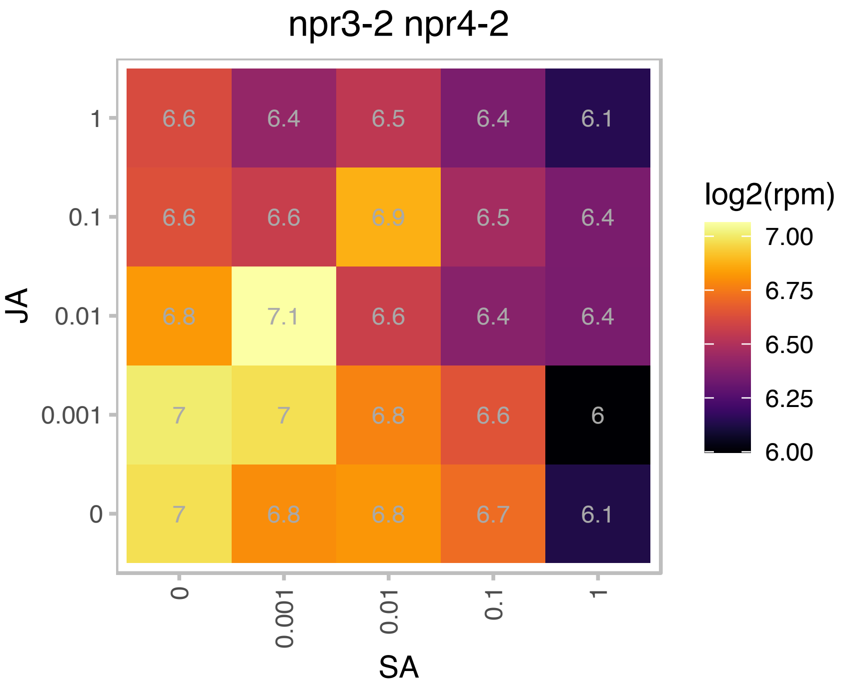


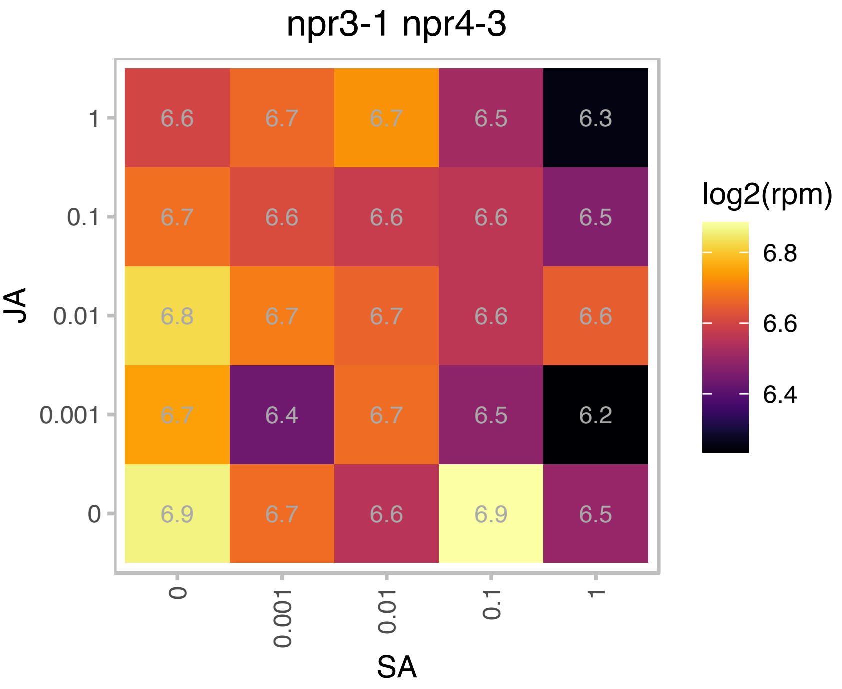

### AT2G30942.1 : ssSPTb

Protein of unknown function (DUF3317)
Encodes a 56-amino acid polypeptide with low but significant similarity to human small subunit of serine palmitoyltransferase that localizes to the ER and physically interacts with and greatly stimulates the activity of LCB1/LCB2 heterodimer ser palmitoyltransferase complex.

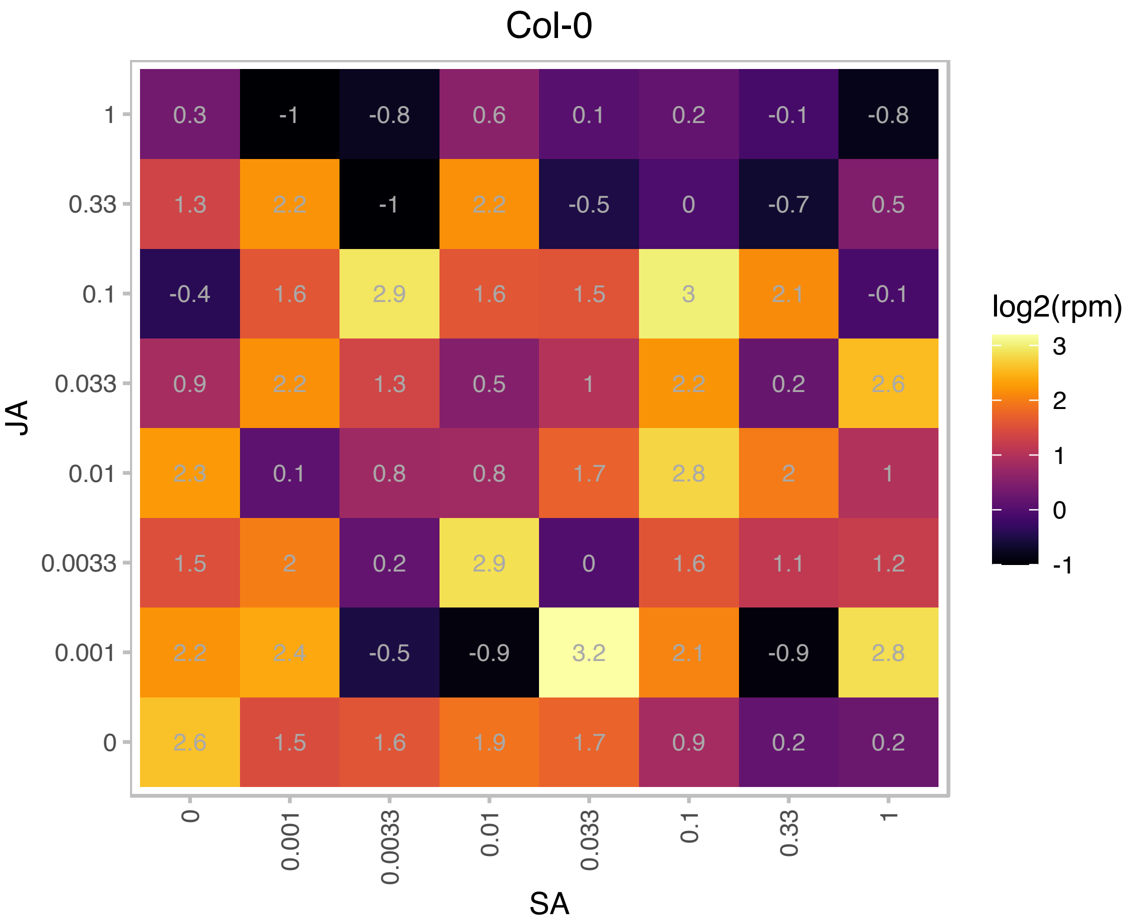


| | |
|---|---|
| Gene | AT2G30942 |
| is.expressed_genes | TRUE |
| is.responsive_genes | TRUE |
| is.apcluster | H-01 |

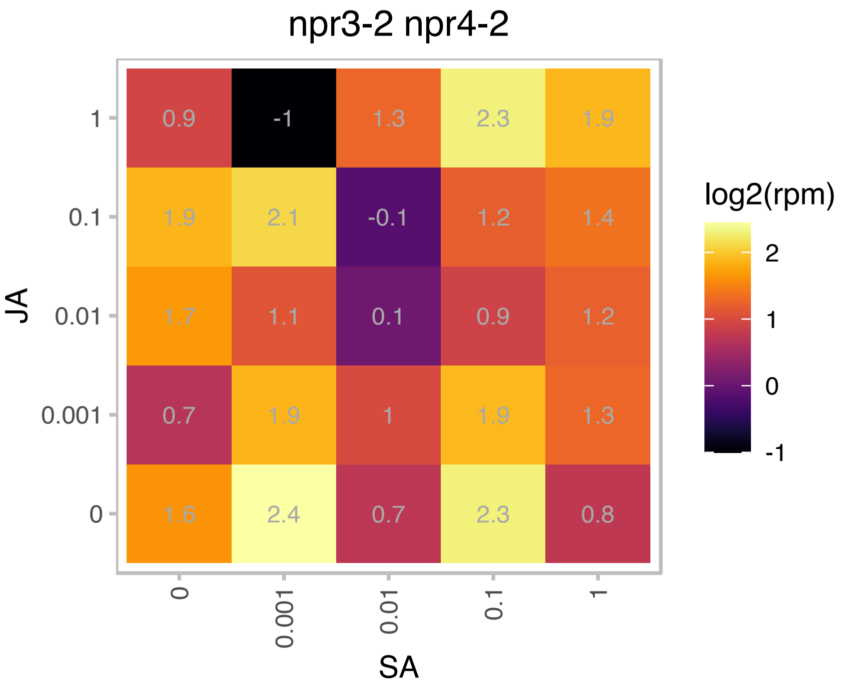


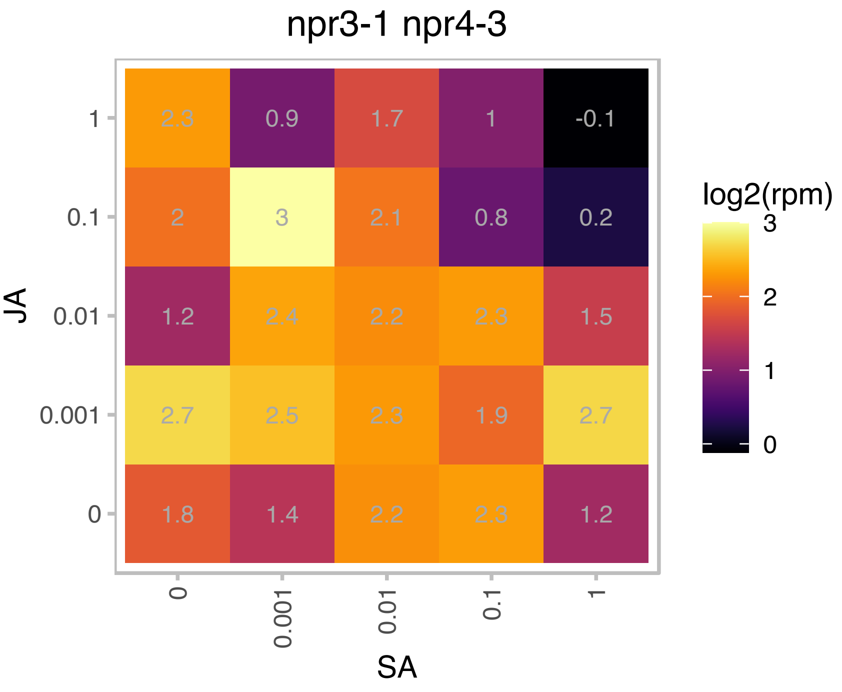

# AT2G34585.1 : NA

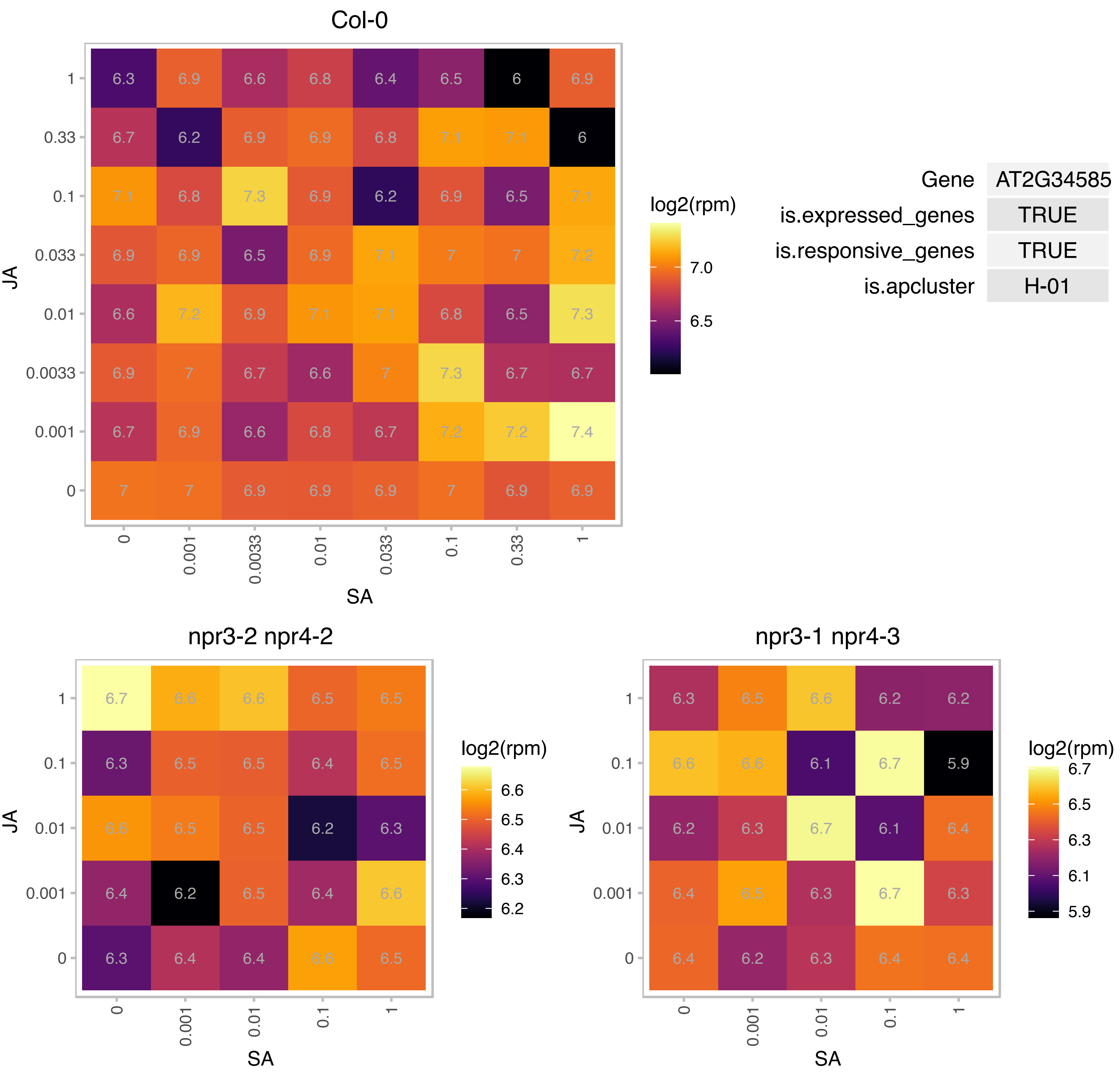

# AT2G37510.1 : NA

RNA-binding (RRM/RBD/RNP motifs) family protein

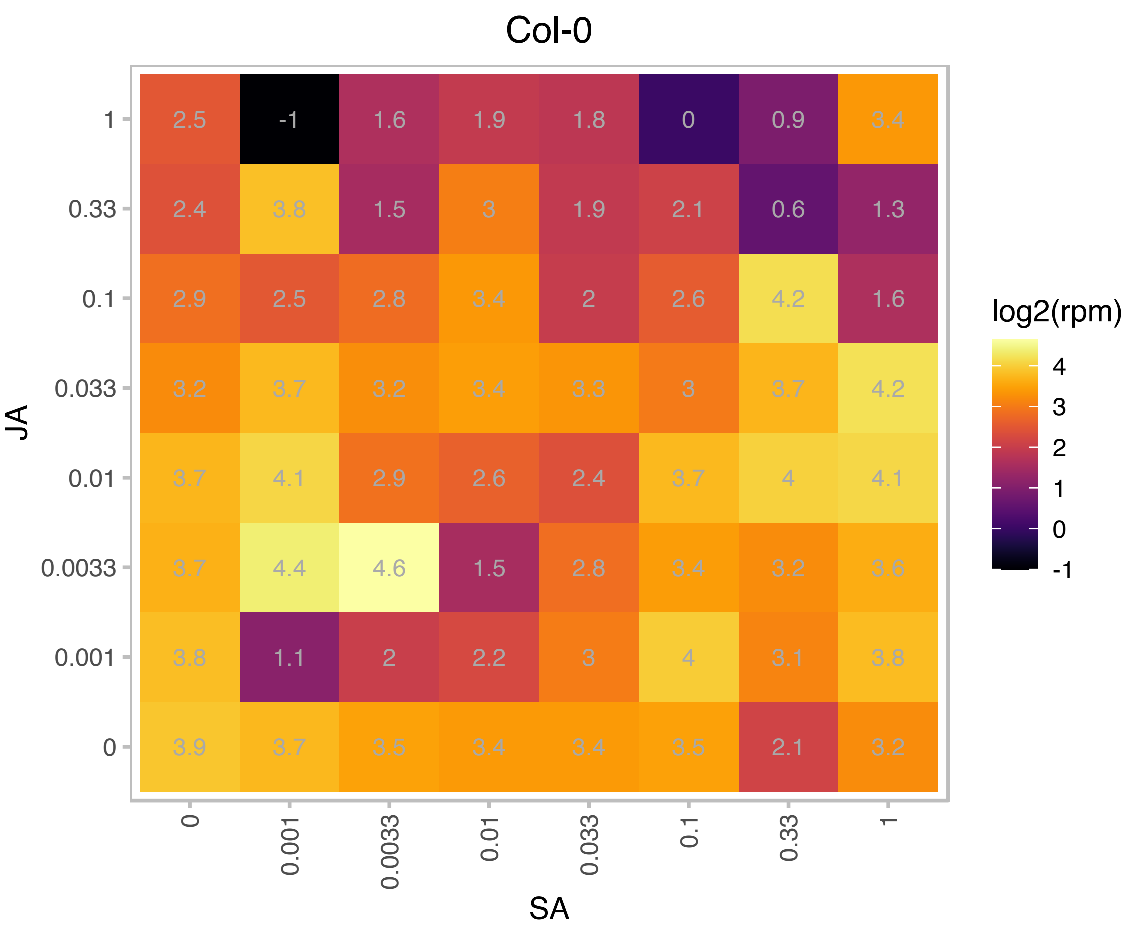


| | |
|---|---|
| Gene | AT2G37510 |
| is.expressed_genes | TRUE |
| is.responsive_genes | TRUE |
| is.apcluster | H-01 |

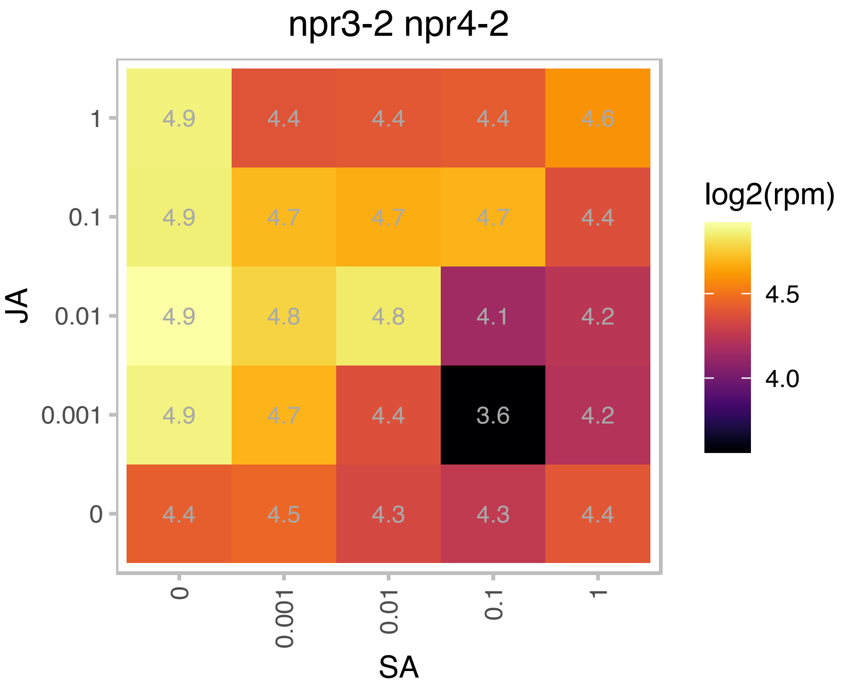


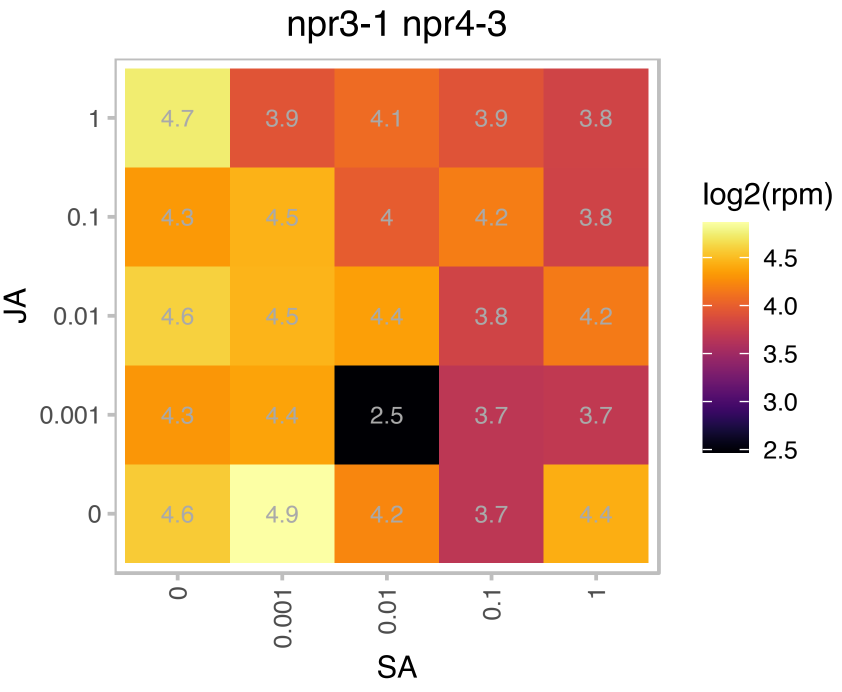

# AT3G02910.1 : NA

AIG2-like (avirulence induced gene) family protein

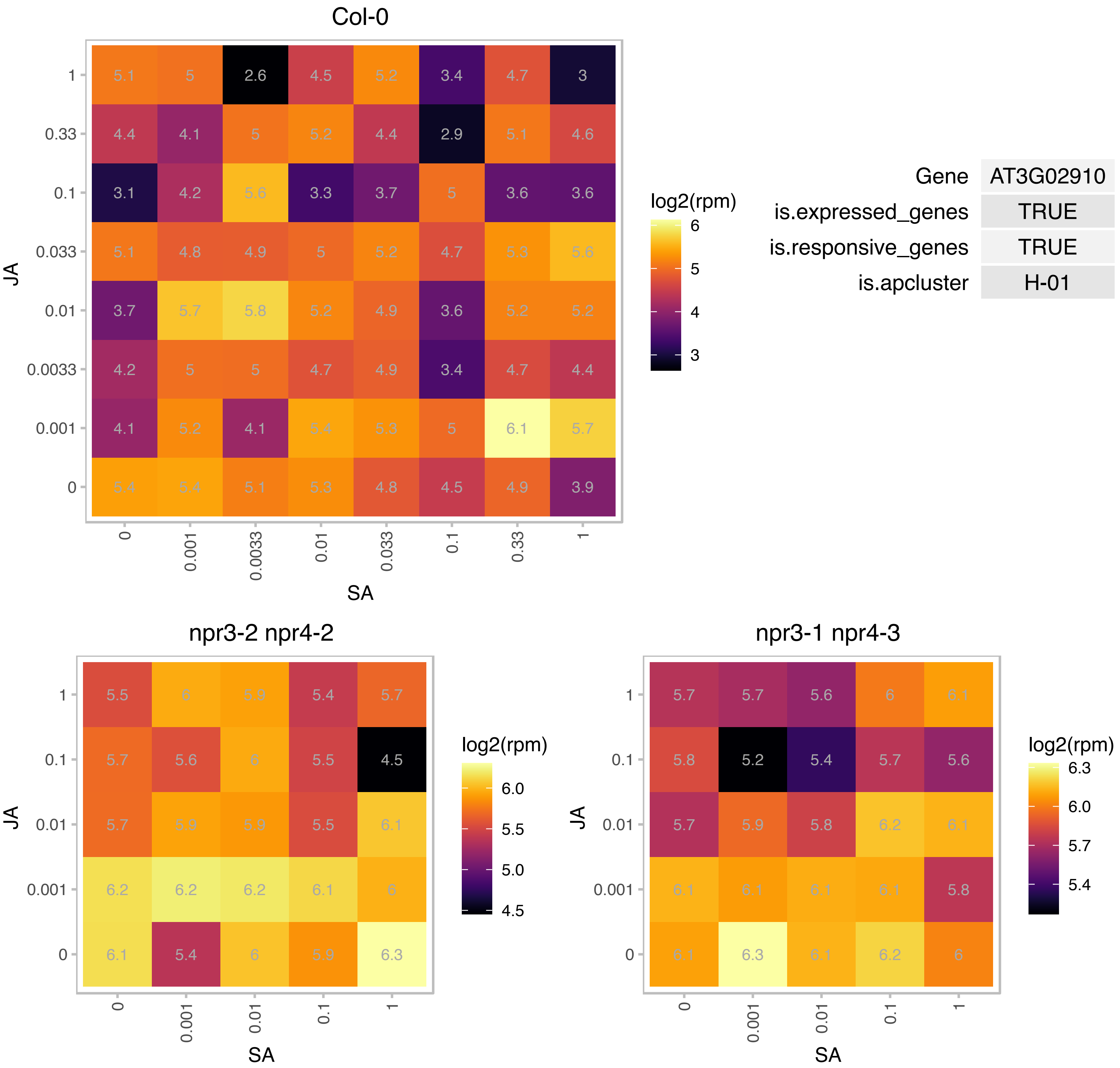


| | |
|---|---|
| Gene | AT3G02910 |
| is.expressed_genes | TRUE |
| is.responsive_genes | TRUE |
| is.apcluster | H-01 |

# AT3G05727.1 : NA

S locus-related glycoprotein 1 (SLR1) binding pollen coat protein family

Encodes a defensin-like (DEFL) family protein.

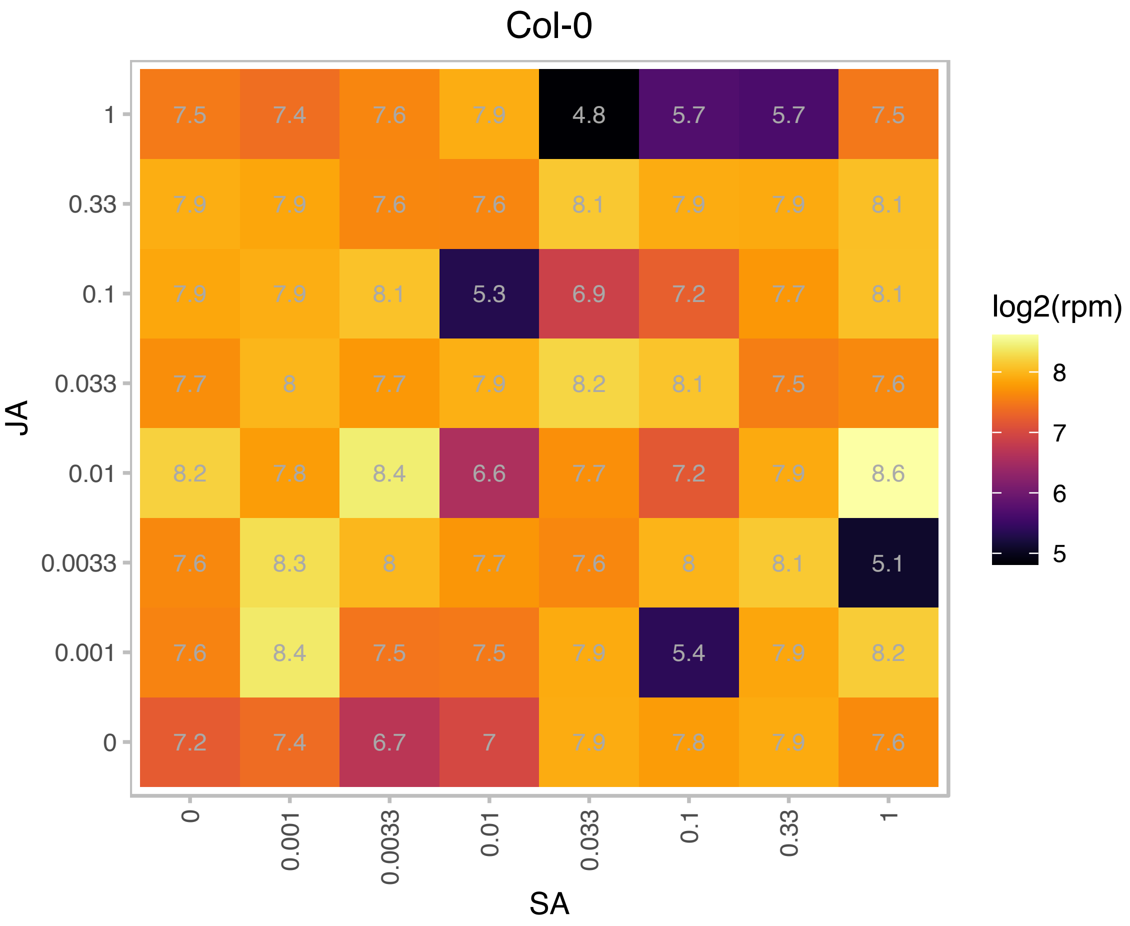


| | |
|---|---|
| Gene | AT3G05727 |
| is.expressed_genes | TRUE |
| is.responsive_genes | TRUE |
| is.apcluster | H-01 |

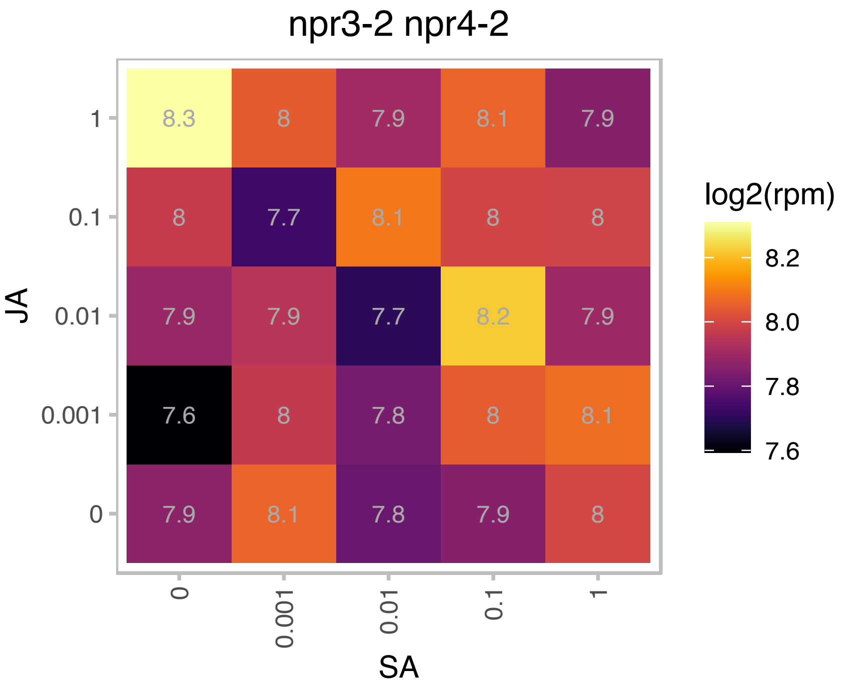


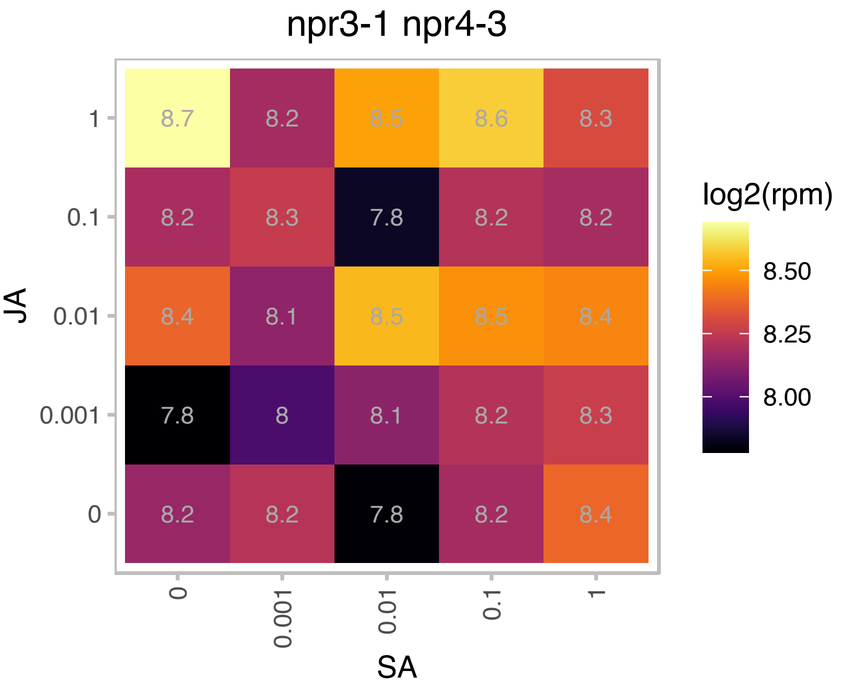

# AT3G06320.1 : NA

Ribosomal protein L33 family protein

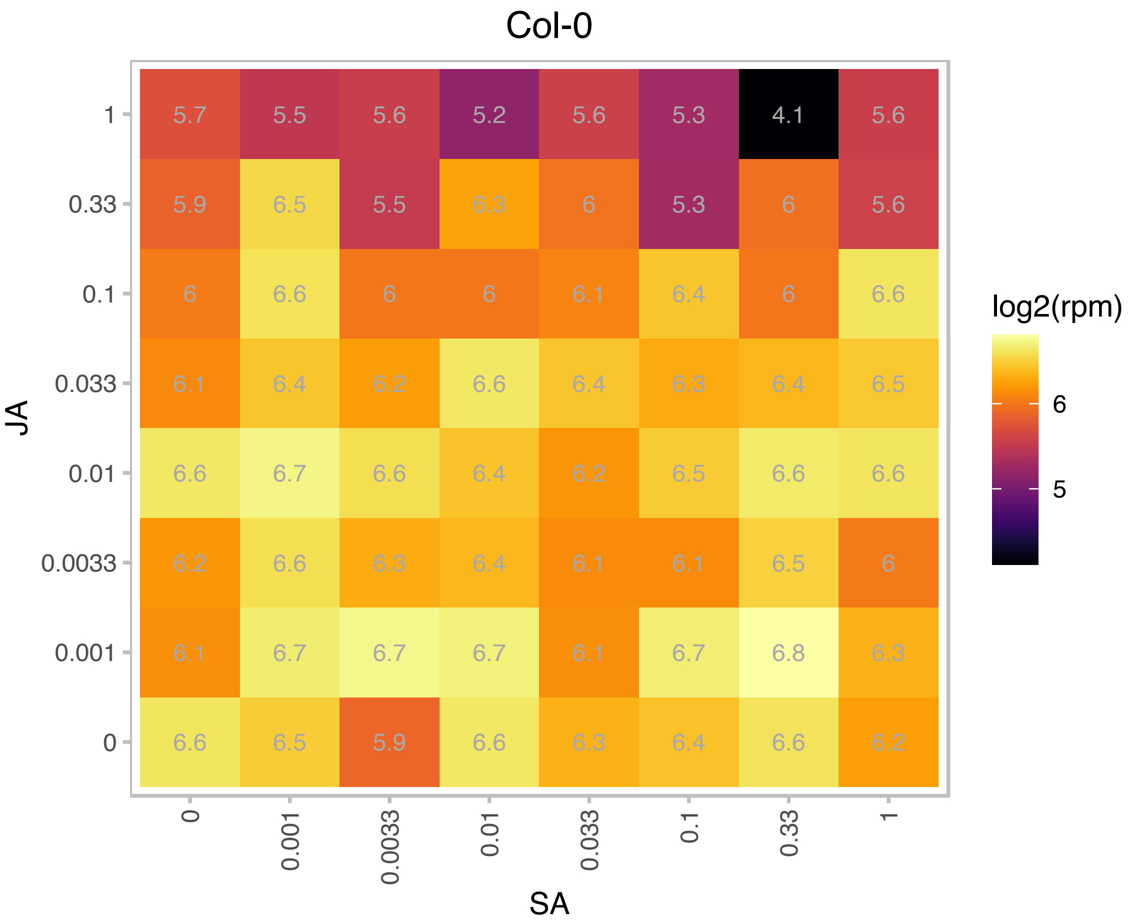


| | |
|---|---|
| Gene | AT3G06320 |
| is.expressed_genes | TRUE |
| is.responsive_genes | TRUE |
| is.apcluster | H-01 |

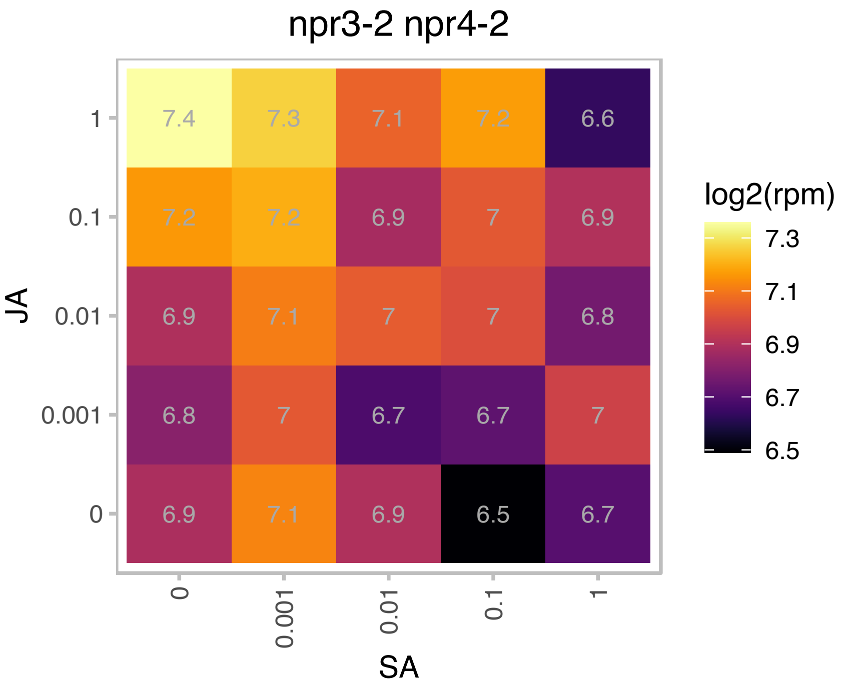


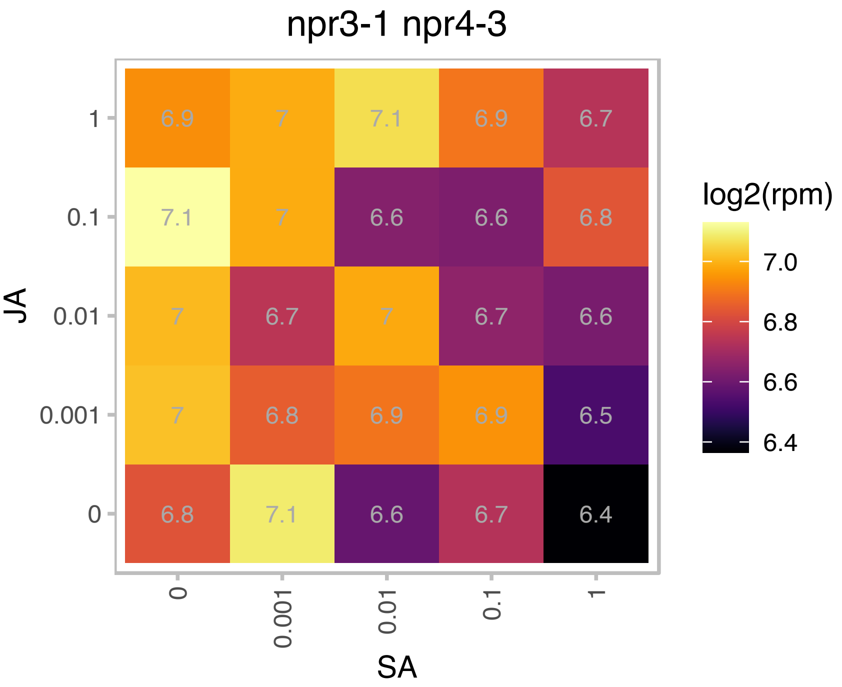

# AT3G15710.1 : NA

Peptidase S24/S26A/S26B/S26C family protein

| | |
|---|---|
| Gene | AT3G15710 |
| is.expressed_genes | TRUE |
| is.responsive_genes | TRUE |
| is.apcluster | H-01 |

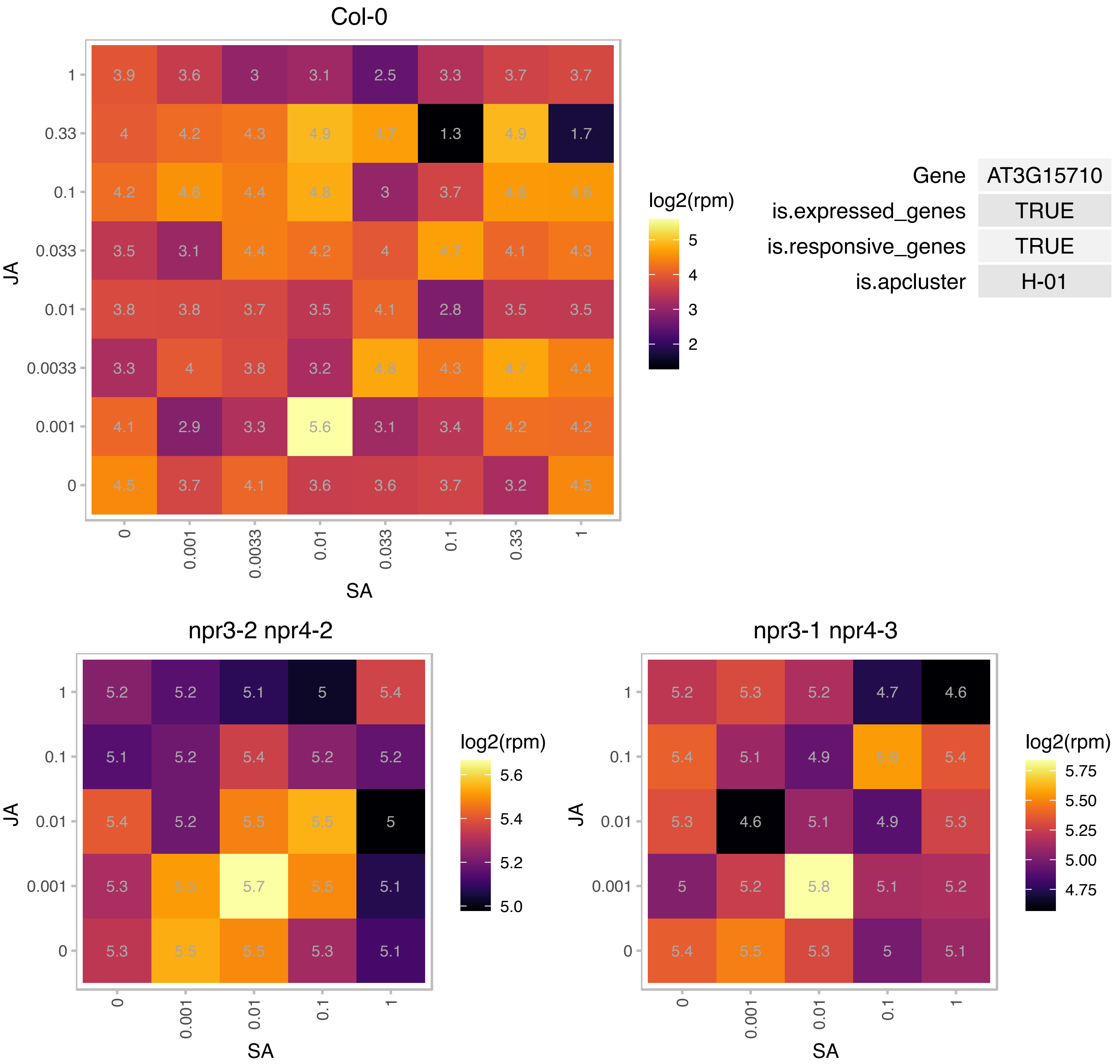

### AT3G55005.1 : TON1B

tonneau 1b (TON1b)

Encodes a protein of unknown function that is involved in cortical microtubule organization. Mutants exhibit abnormal cell growth and patterns of division. TON1B appears to be redundant with TON1A. Encodes a novel protein that is similar to human FOP and OFD1 centrosomal proteins. Localizes to the preprophase band, cytoplasm and cell cortex where it is probably associated with the cortical cytoskeleton. TON1B associates with plant centrin CEN1.

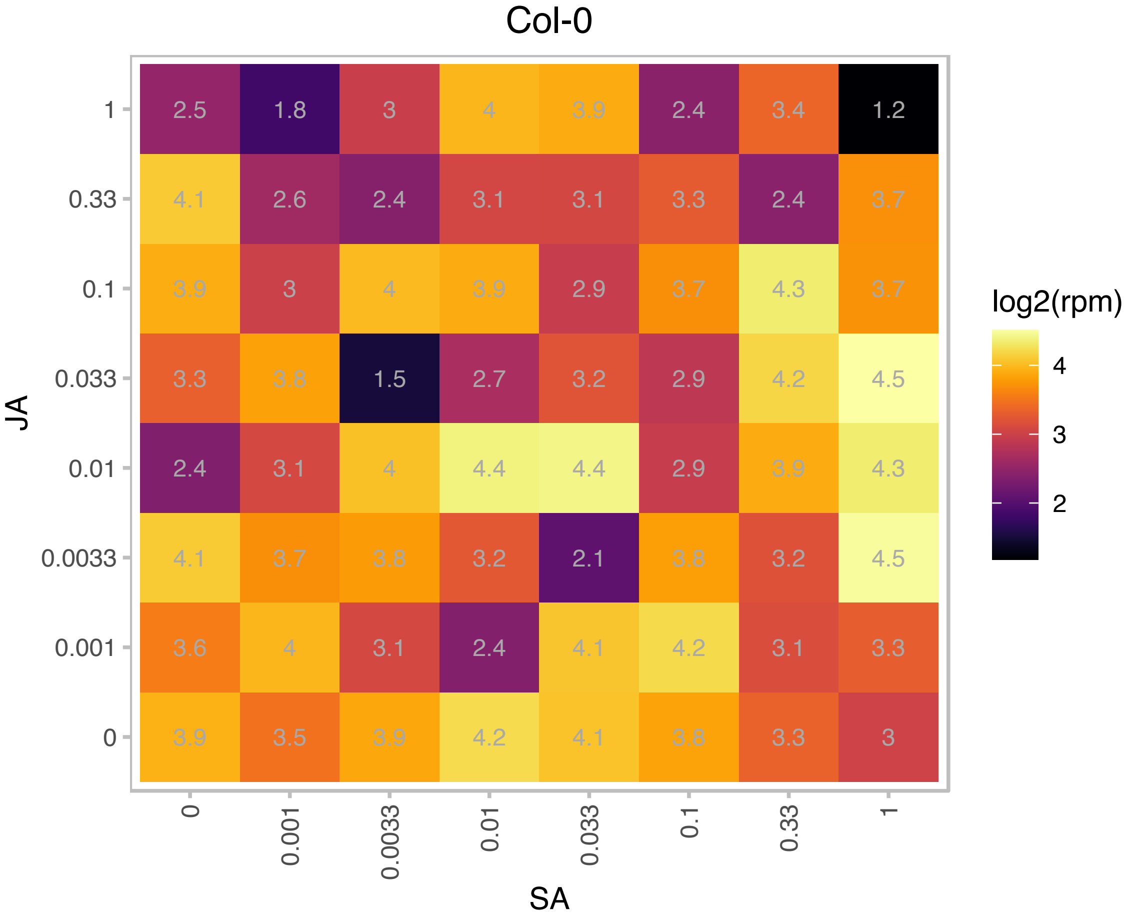


| | |
|---|---|
| Gene | AT3G55005 |
| is.expressed_genes | TRUE |
| is.responsive_genes | TRUE |
| is.apcluster | H-01 |

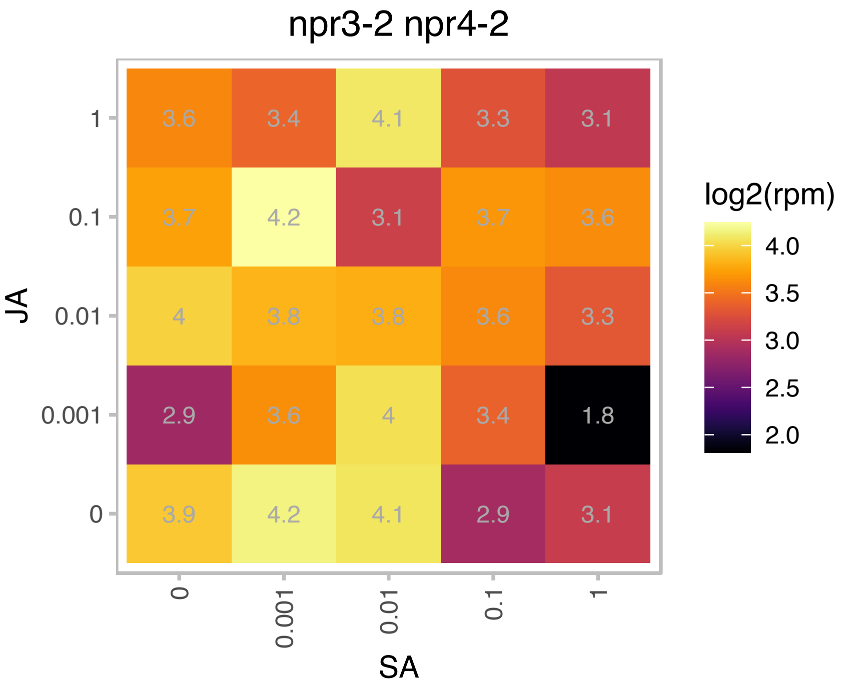


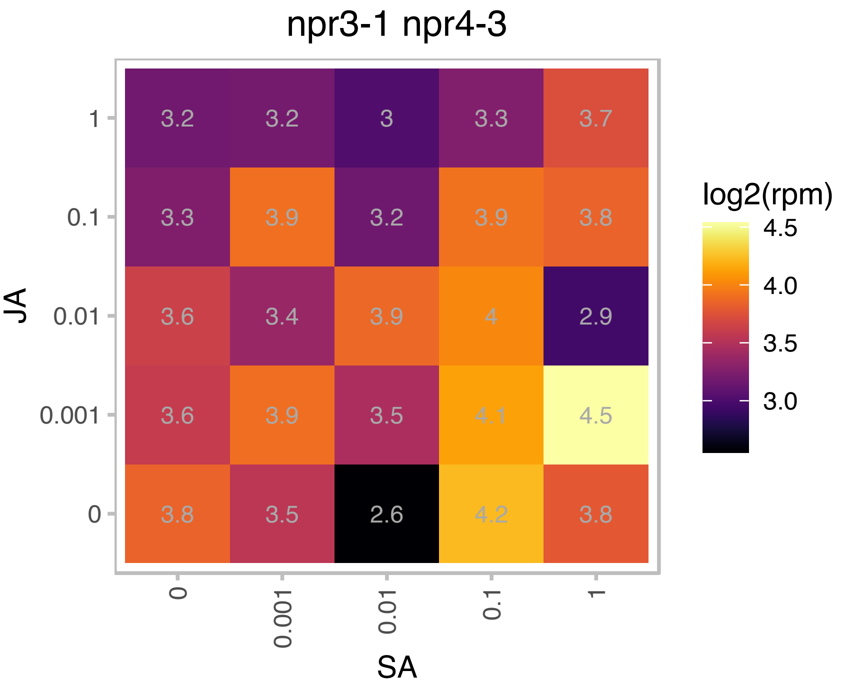

# AT3G61100.1 : NA

Putative endonuclease or glycosyl hydrolase

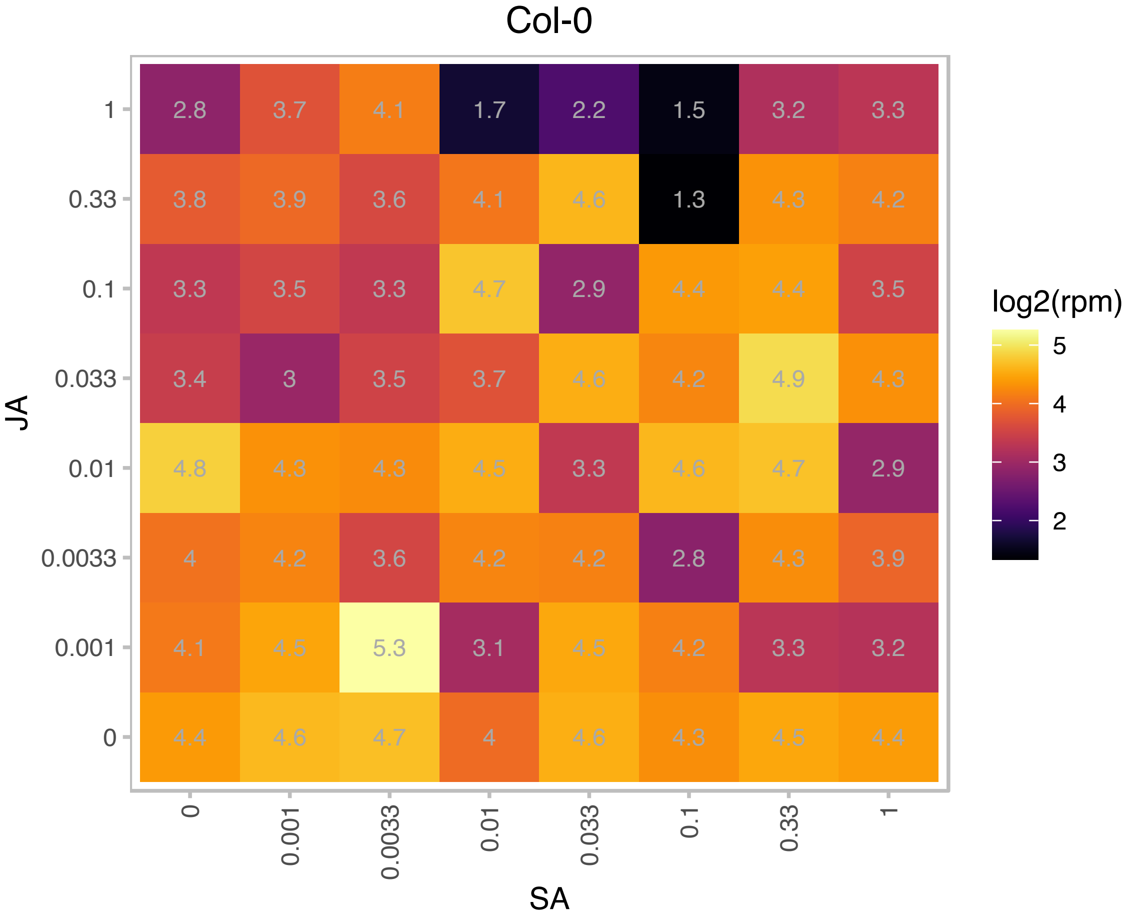


| | |
|---|---|
| Gene | AT3G61100 |
| is.expressed_genes | TRUE |
| is.responsive_genes | TRUE |
| is.apcluster | H-01 |

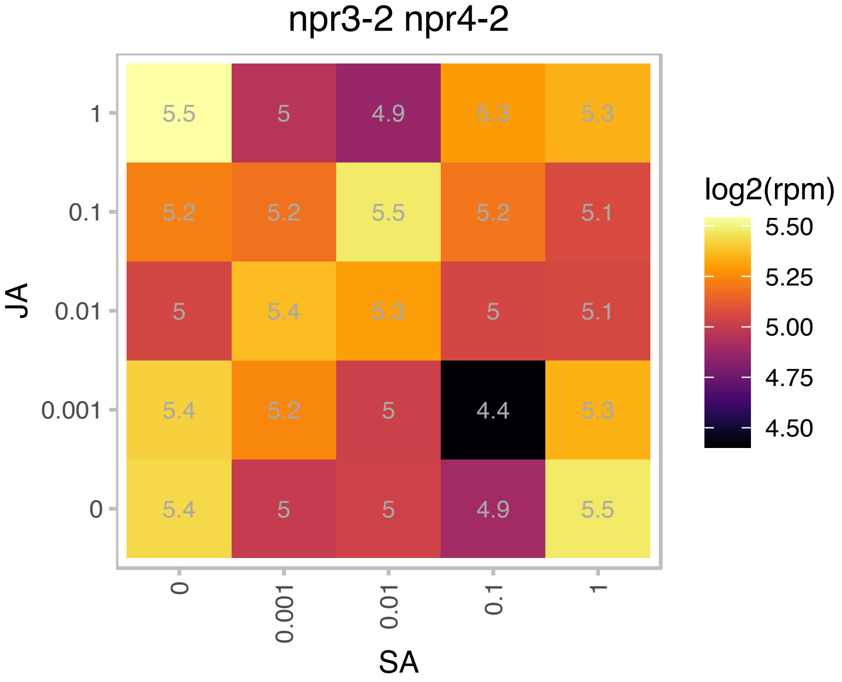


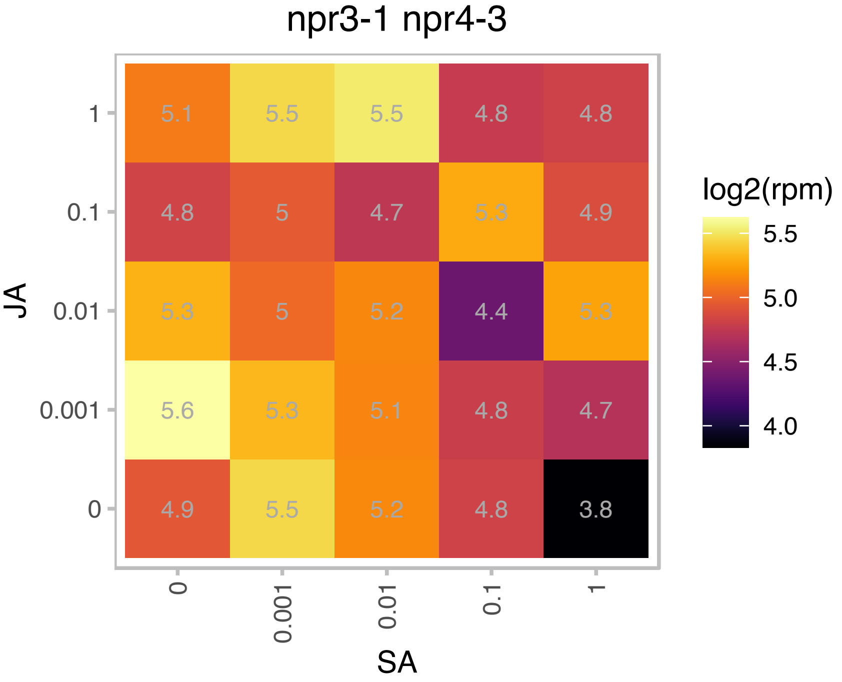

# AT4G18460.1 : NA

D-Tyr-tRNA(Tyr) deacylase family protein

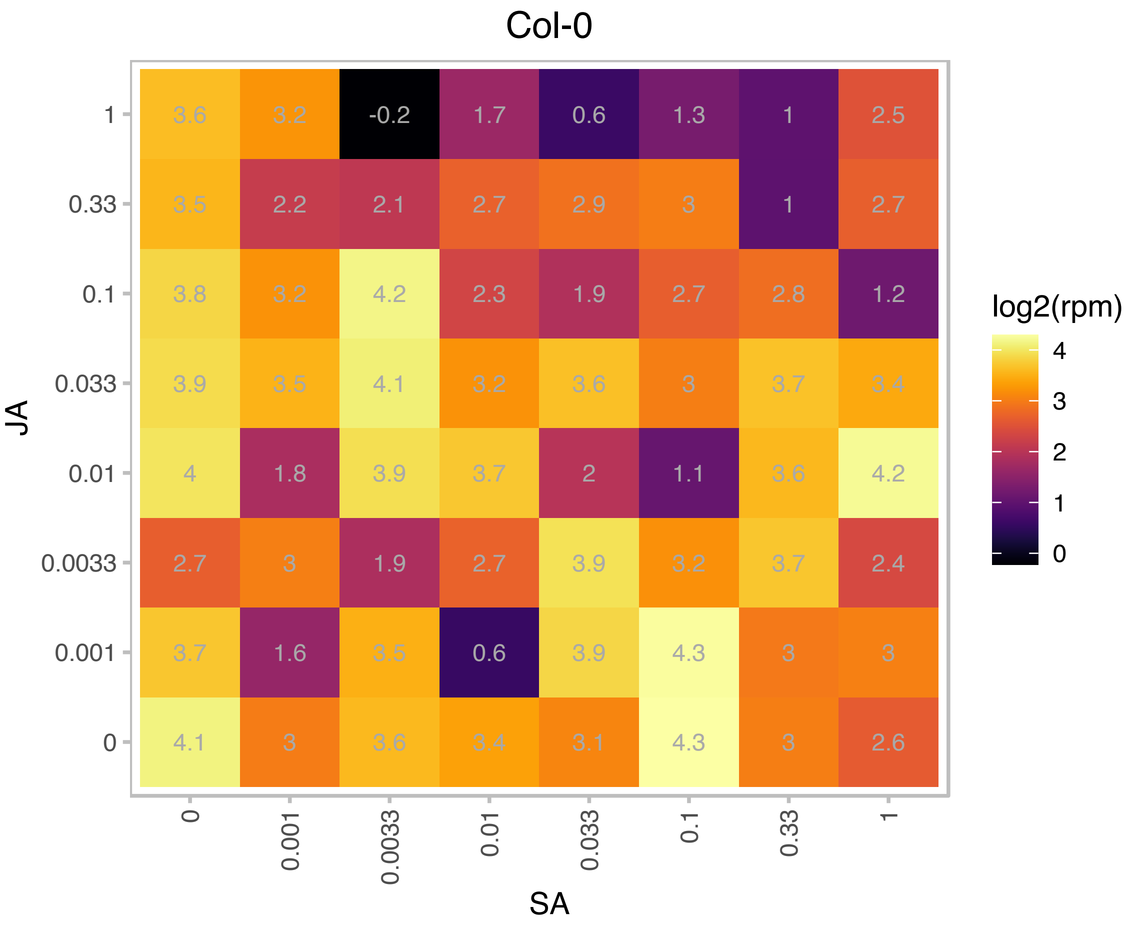


| | |
|---|---|
| Gene | AT4G18460 |
| is.expressed_genes | TRUE |
| is.responsive_genes | TRUE |
| is.apcluster | H-01 |

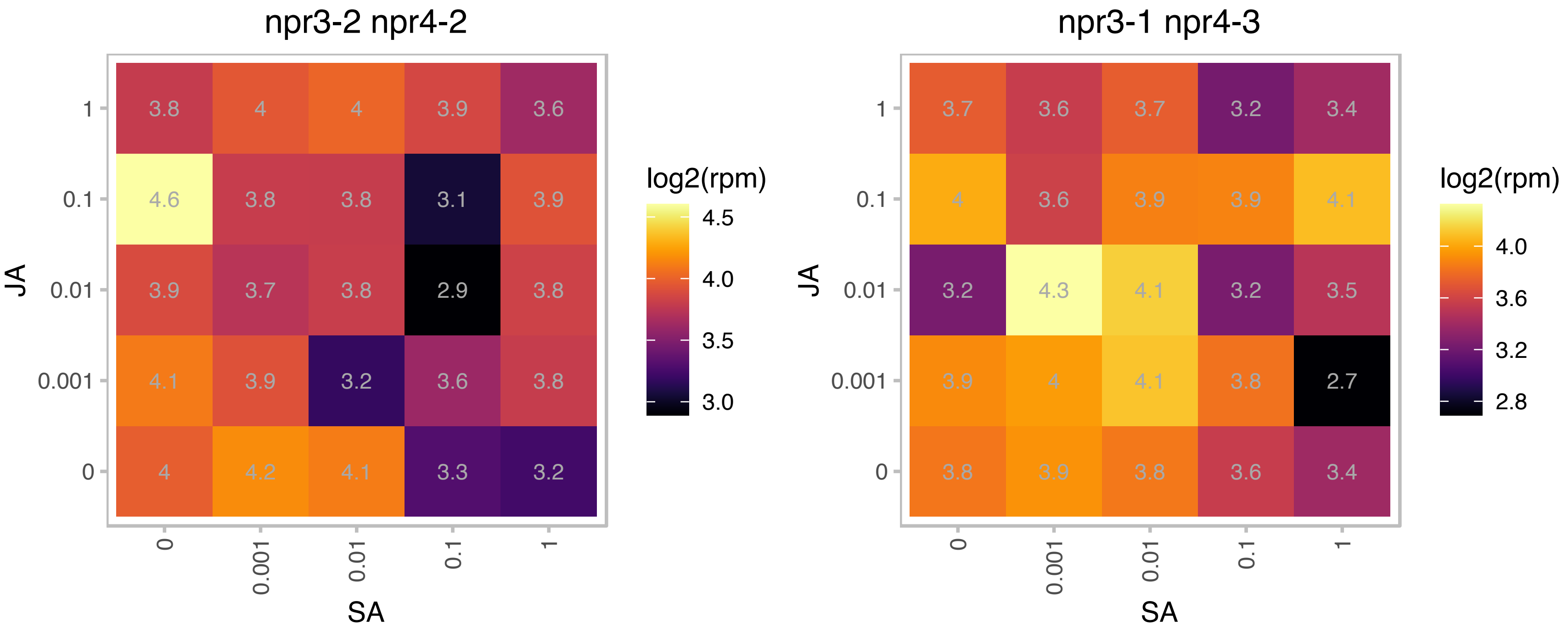

### AT4G19003.1 : VPS25

E2F/DP family winged-helix DNA-binding domain

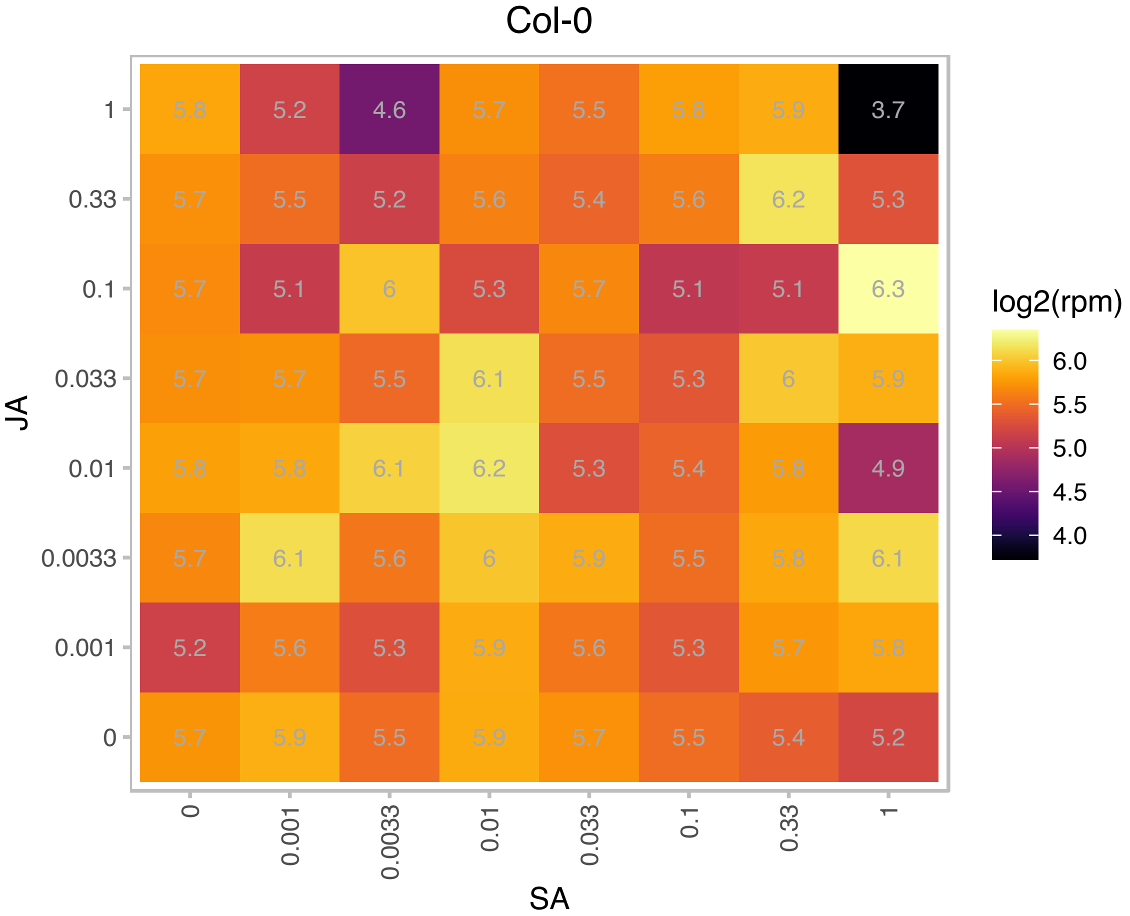


| | |
|---|---|
| Gene | AT4G19003 |
| is.expressed_genes | TRUE |
| is.responsive_genes | TRUE |
| is.apcluster | H-01 |

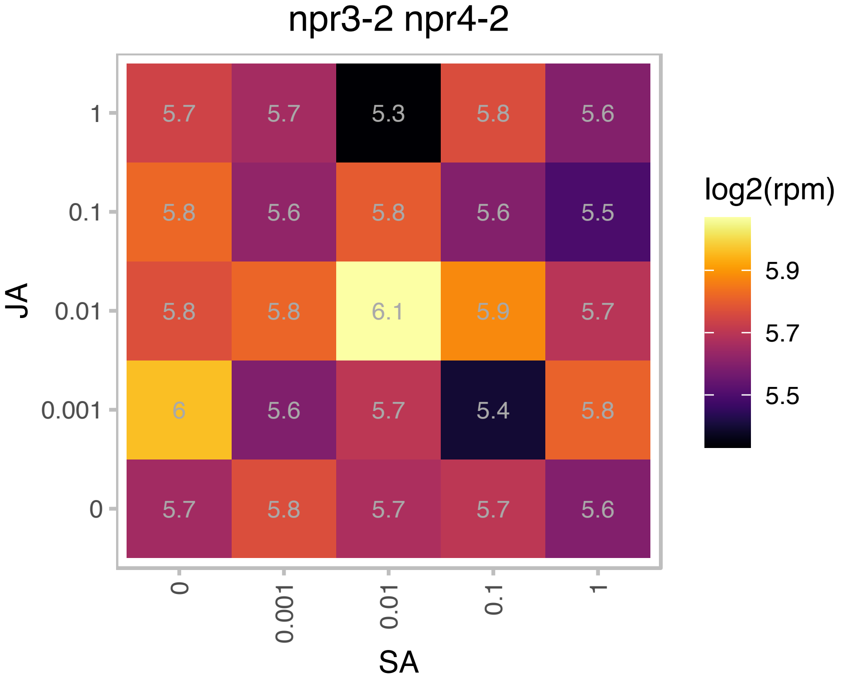


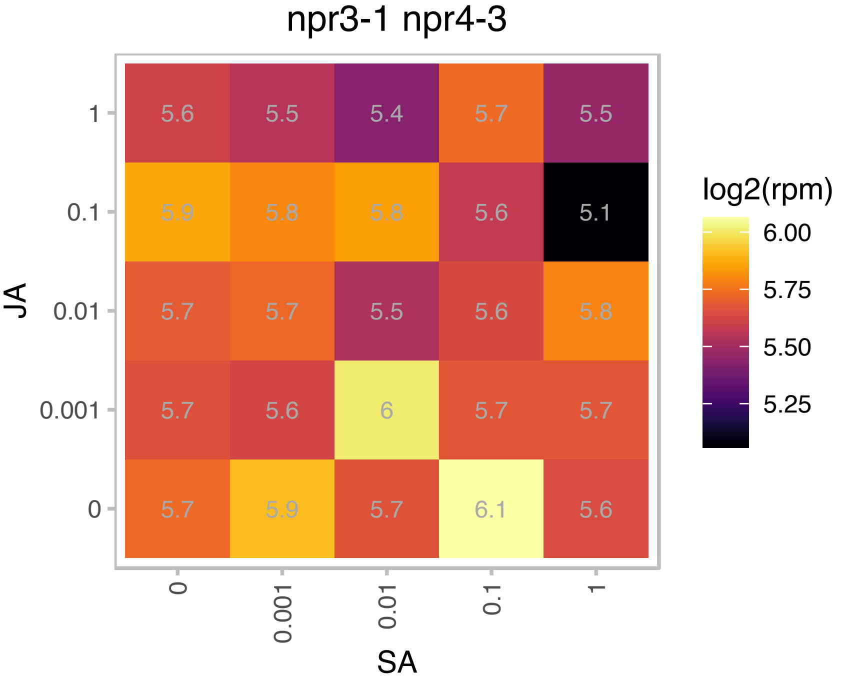

### AT4G26220.1 : CCoAOMT7

S-adenosyl-L-methionine-dependent methyltransferases superfamily protein
Encodes a caffeoyl-coenzyme A O-methyltransferase (CCoAOMT)-like protein with a strong preference for methylating the para position of flavanones and dihydroflavonols, whereas flavones and flavonols are methylated in the meta-position.

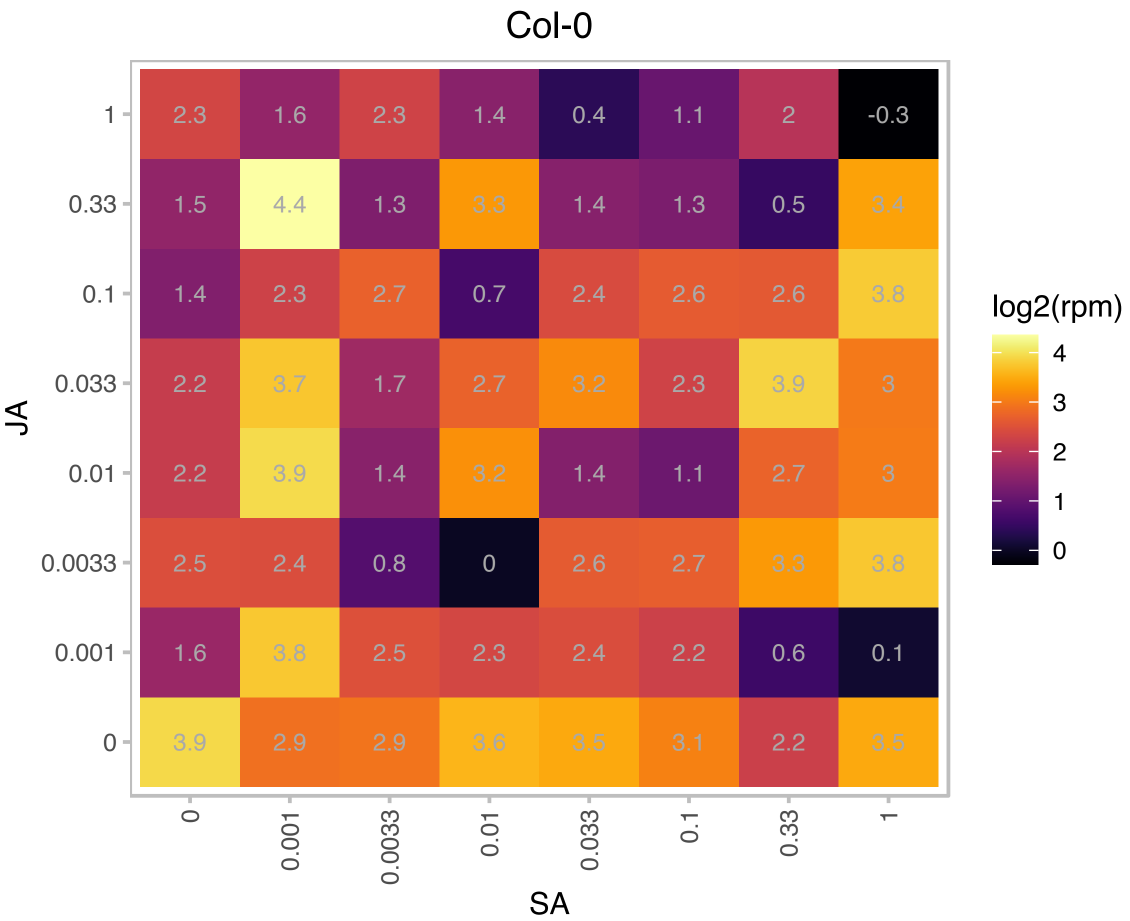


| | |
|---|---|
| Gene | AT4G26220 |
| is.expressed_genes | TRUE |
| is.responsive_genes | TRUE |
| is.apcluster | H-01 |

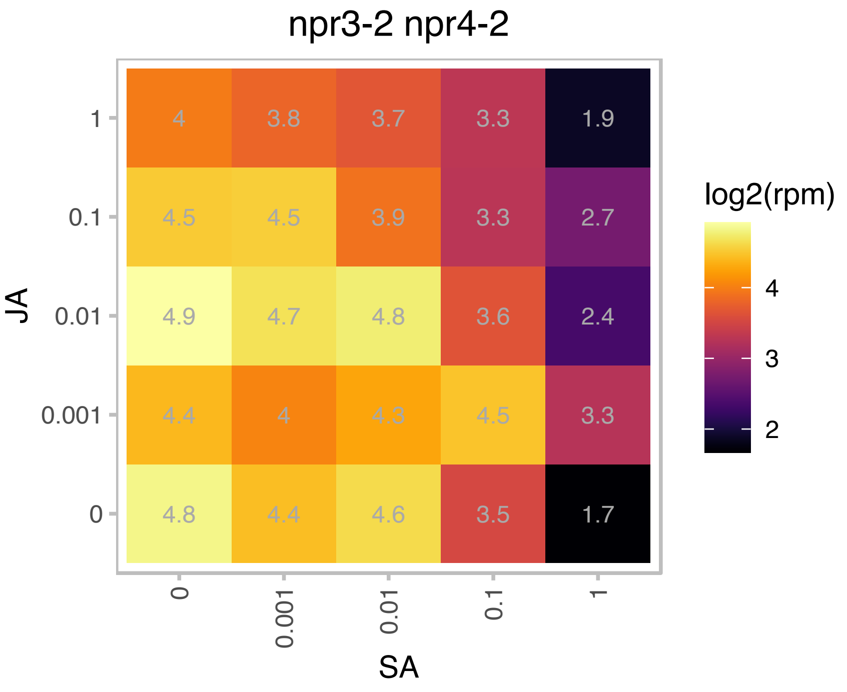


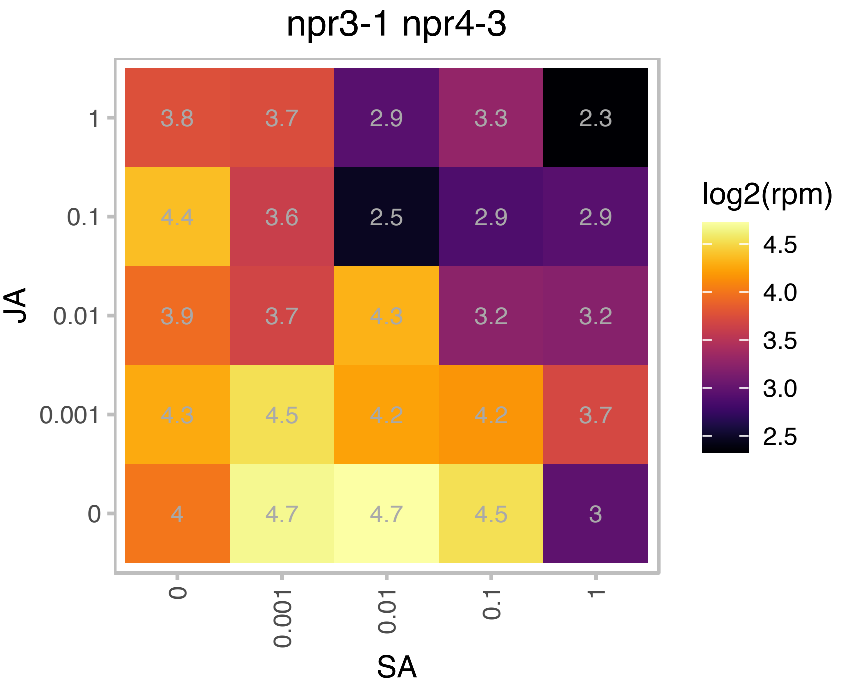

### AT4G27490.1 : RRP41L

3'-5'-exoribonuclease family protein

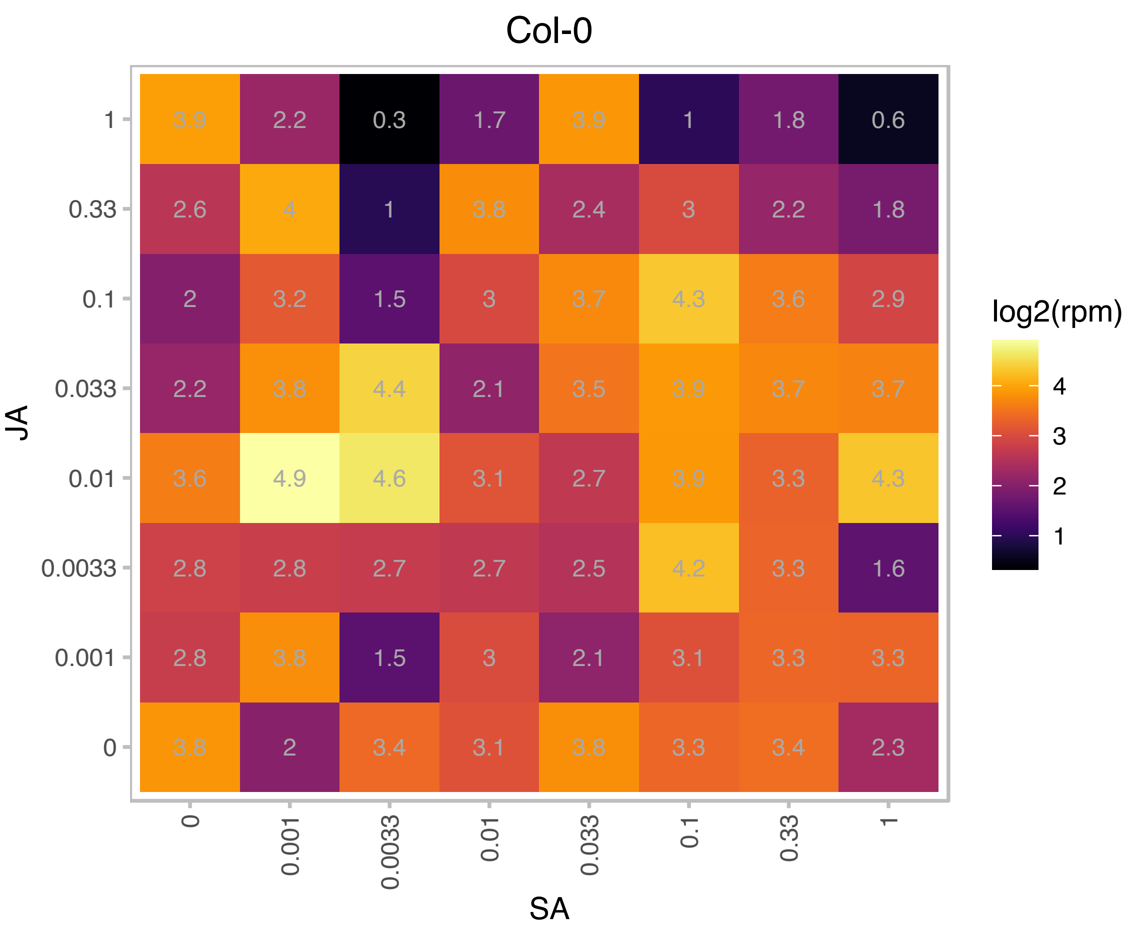


| | |
|---|---|
| Gene | AT4G27490 |
| is.expressed_genes | TRUE |
| is.responsive_genes | TRUE |
| is.apcluster | H-01 |

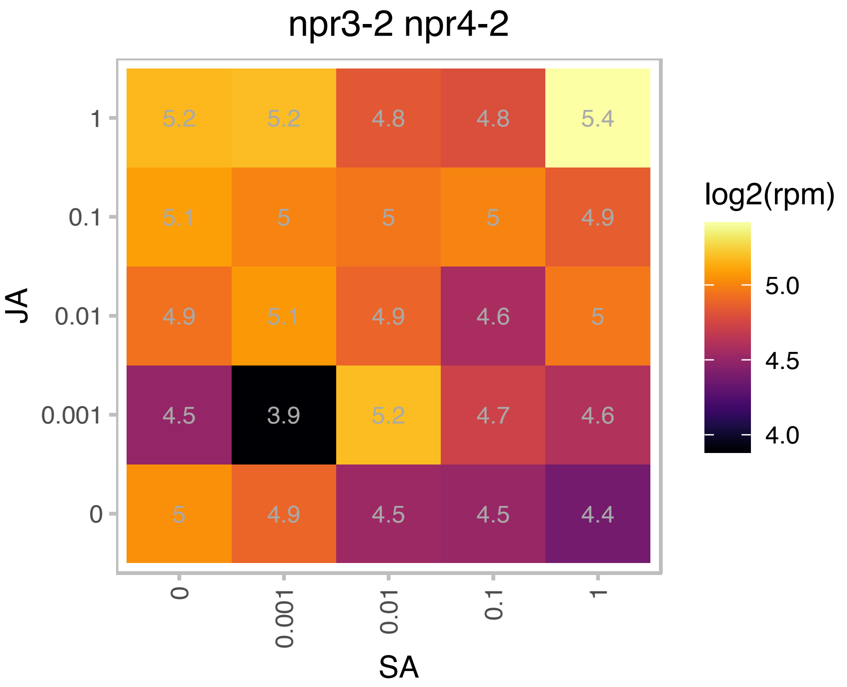


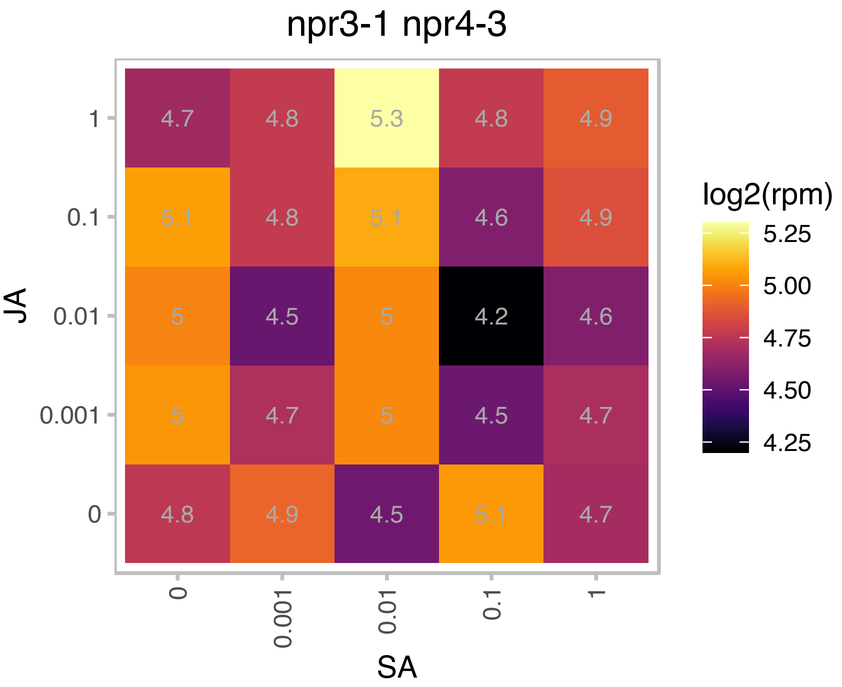

# AT4G31310.1 : NA

AIG2-like (avirulence induced gene) family protein

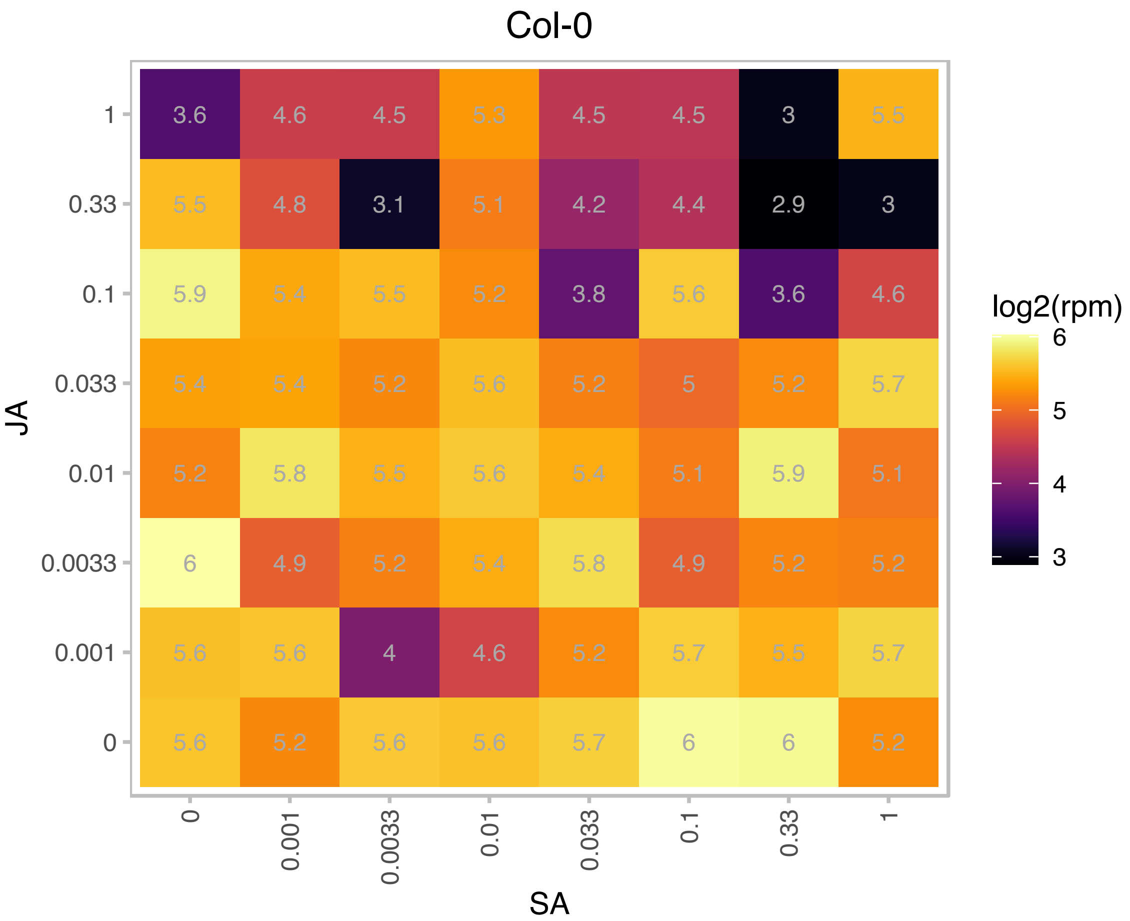


| | |
|---|---|
| Gene | AT4G31310 |
| is.expressed_genes | TRUE |
| is.responsive_genes | TRUE |
| is.apcluster | H-01 |

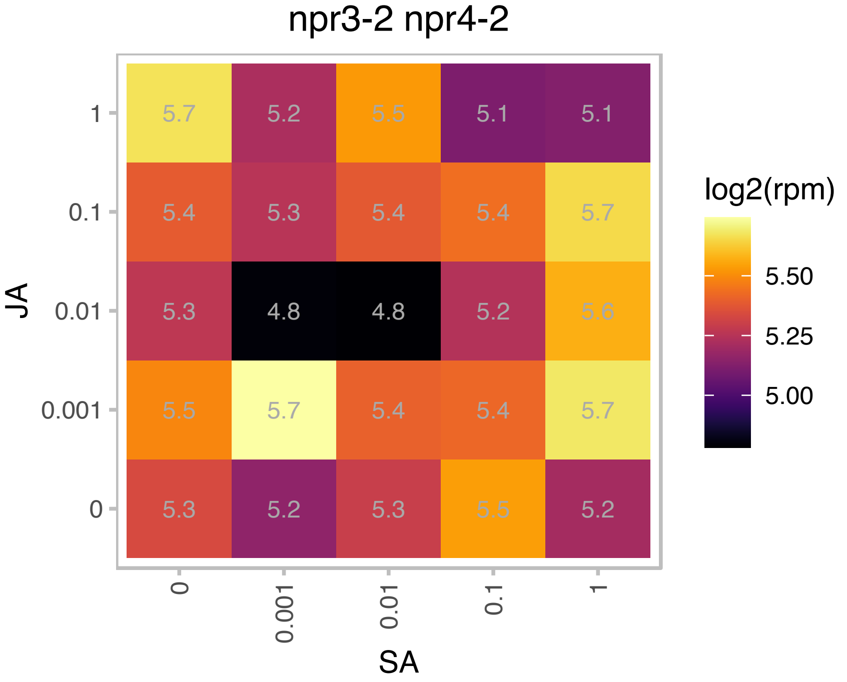


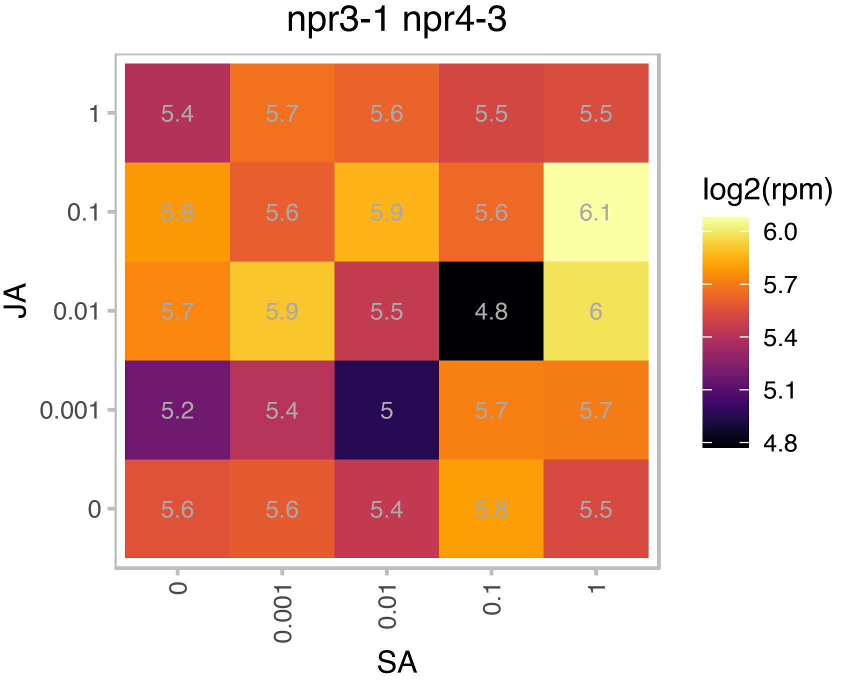

# AT4G34070.1 : NA

Calcium-binding EF-hand family protein

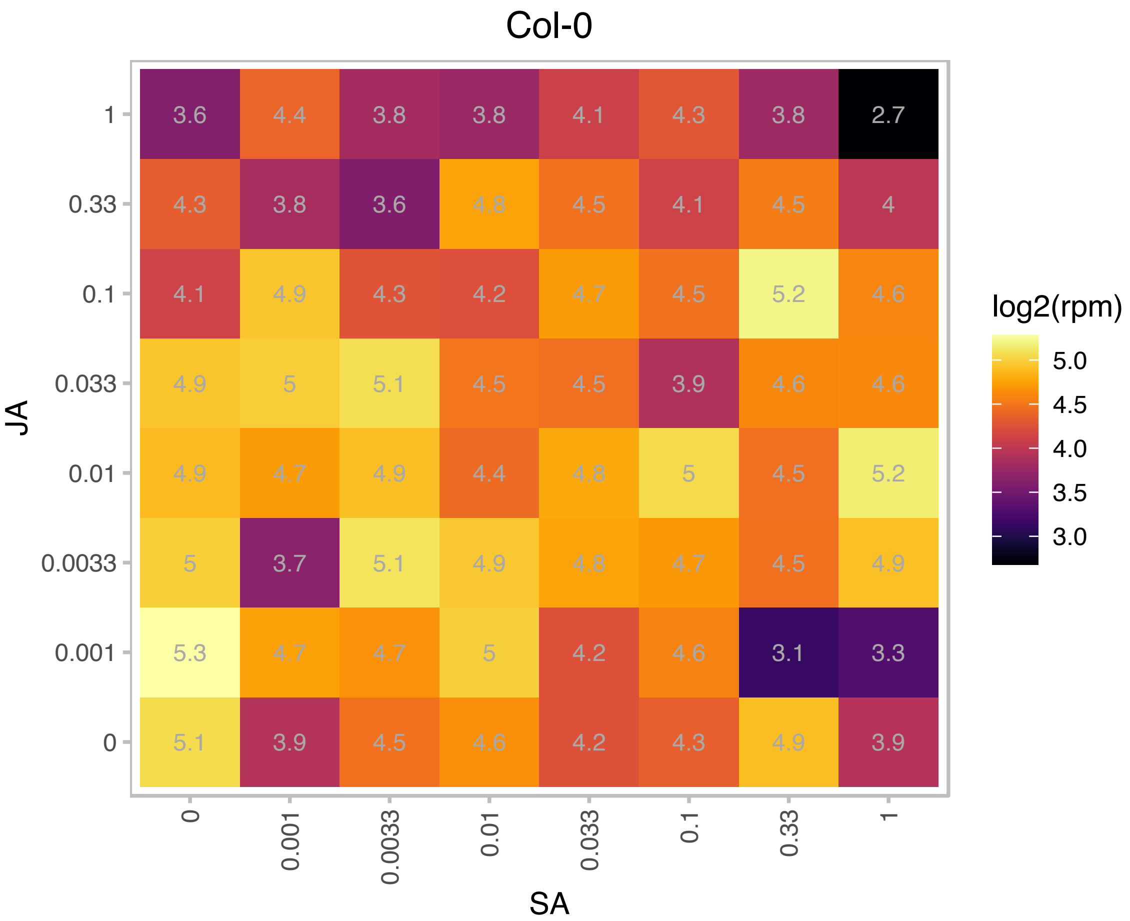


| | |
|---|---|
| Gene | AT4G34070 |
| is.expressed_genes | TRUE |
| is.responsive_genes | TRUE |
| is.apcluster | H-01 |

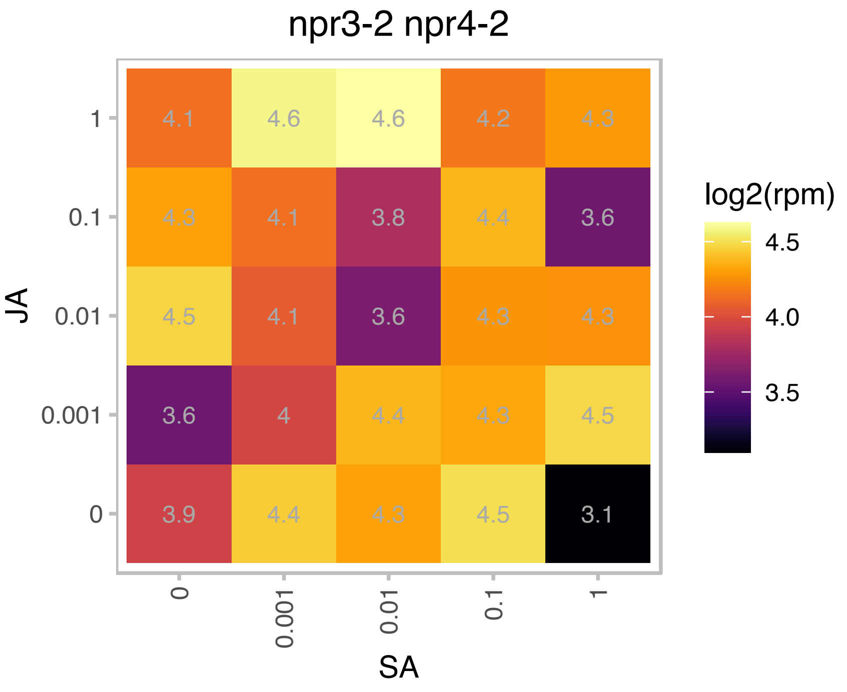


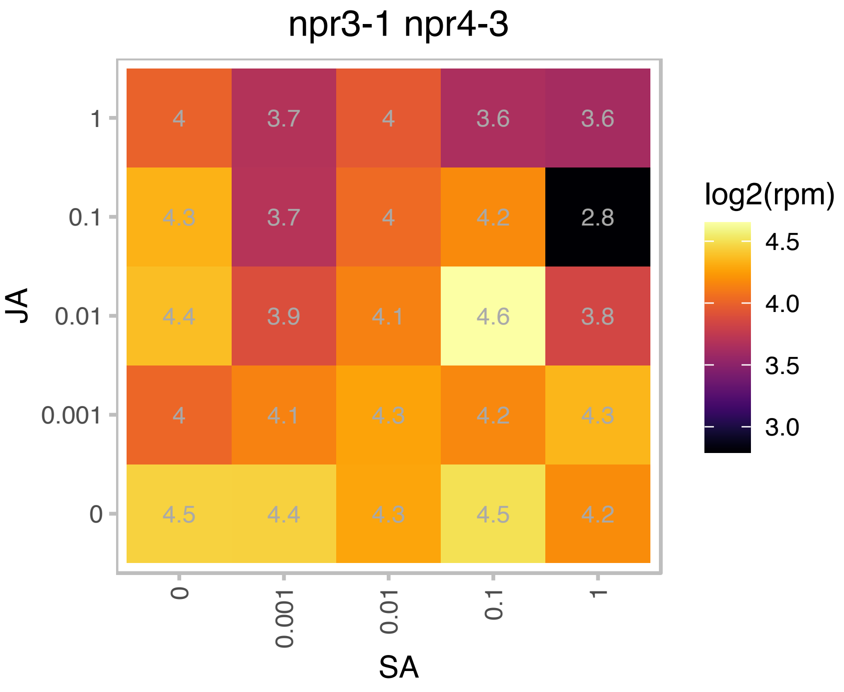

# AT4G35785.1 : NA

RNA-binding (RRM/RBD/RNP motifs) family protein

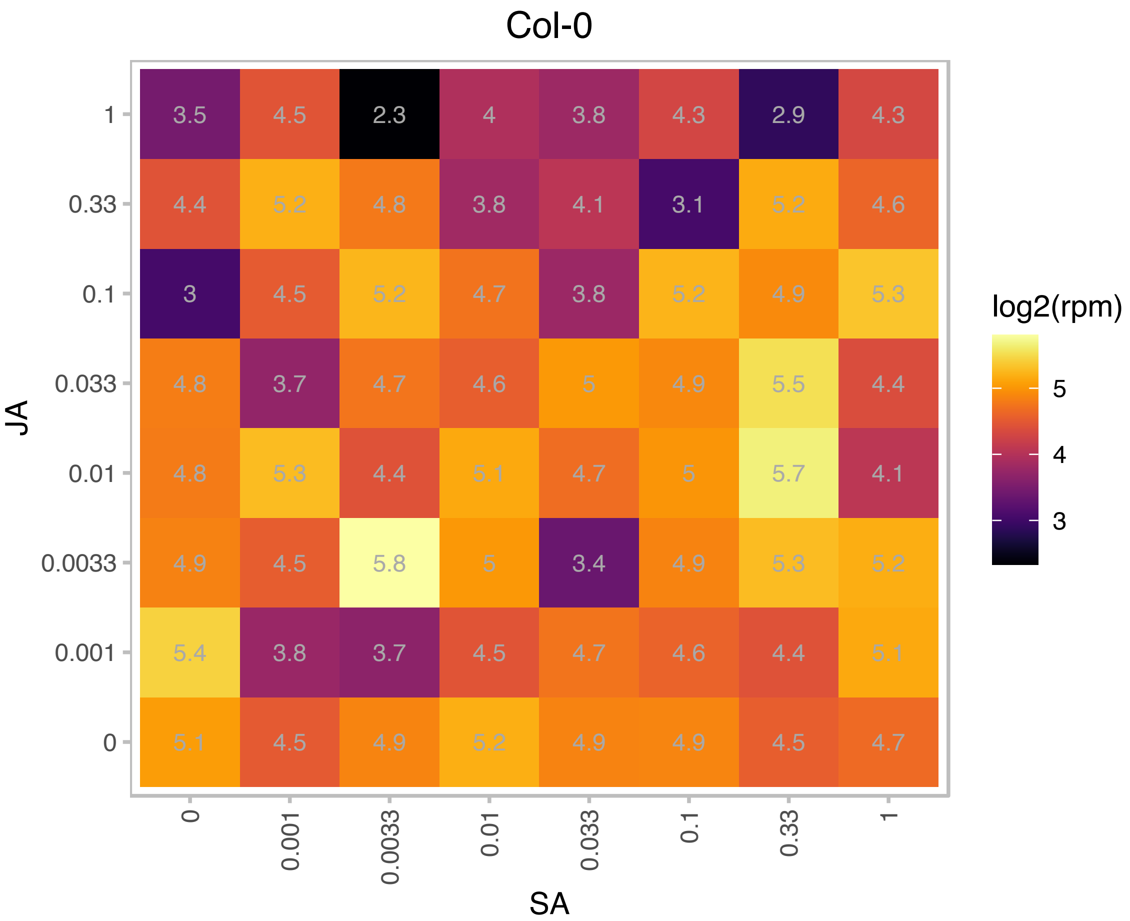


| | |
|---|---|
| Gene | AT4G35785 |
| is.expressed_genes | TRUE |
| is.responsive_genes | TRUE |
| is.apcluster | H-01 |

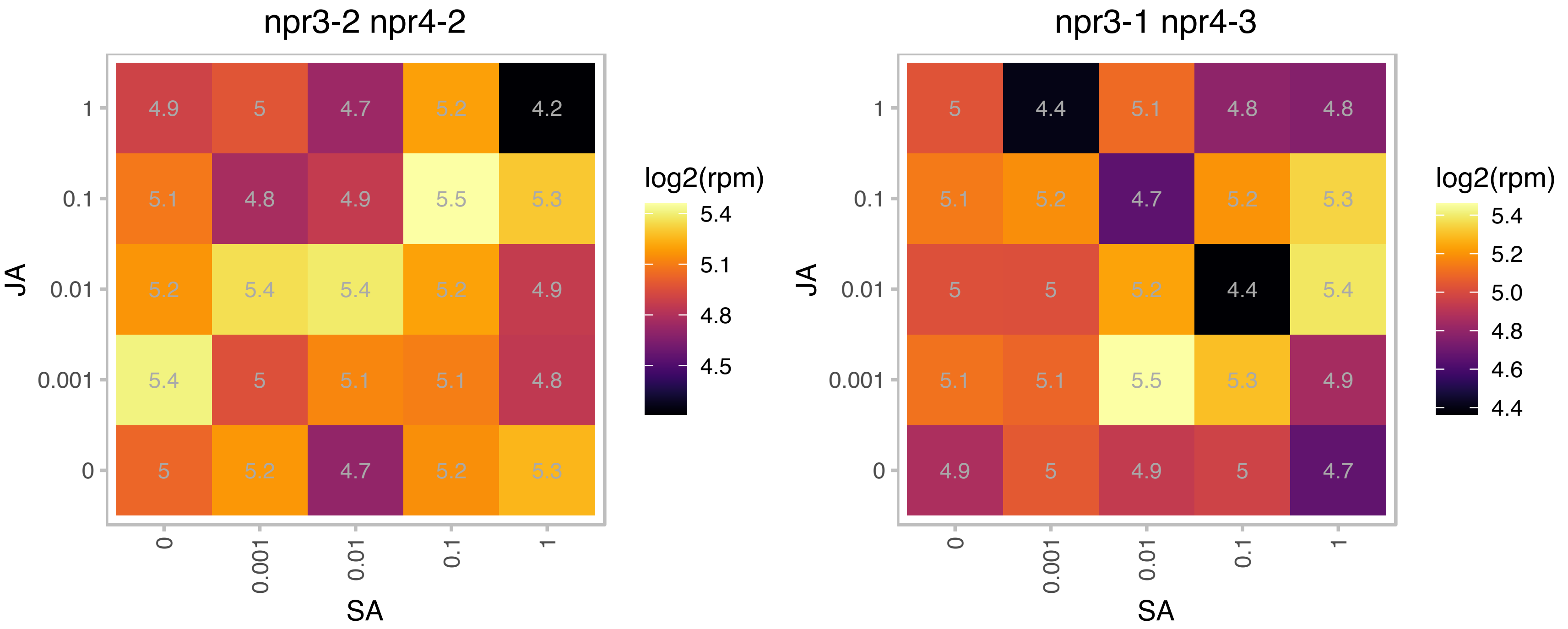

### AT4G39330.1 : ATCAD9, CAD9

cinnamyl alcohol dehydrogenase 9

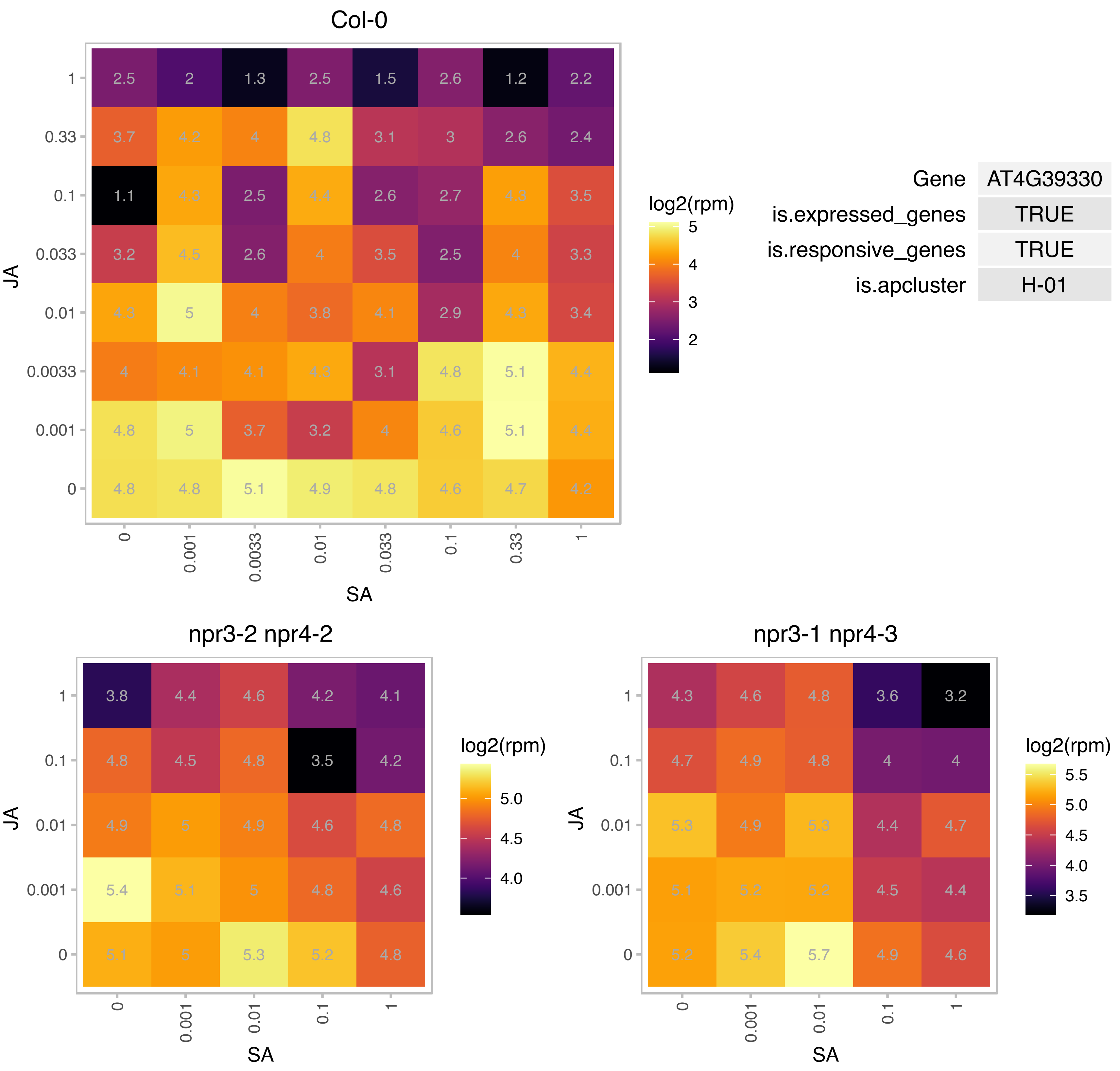

# AT5G02050.1 : NA

Mitochondrial glycoprotein family protein

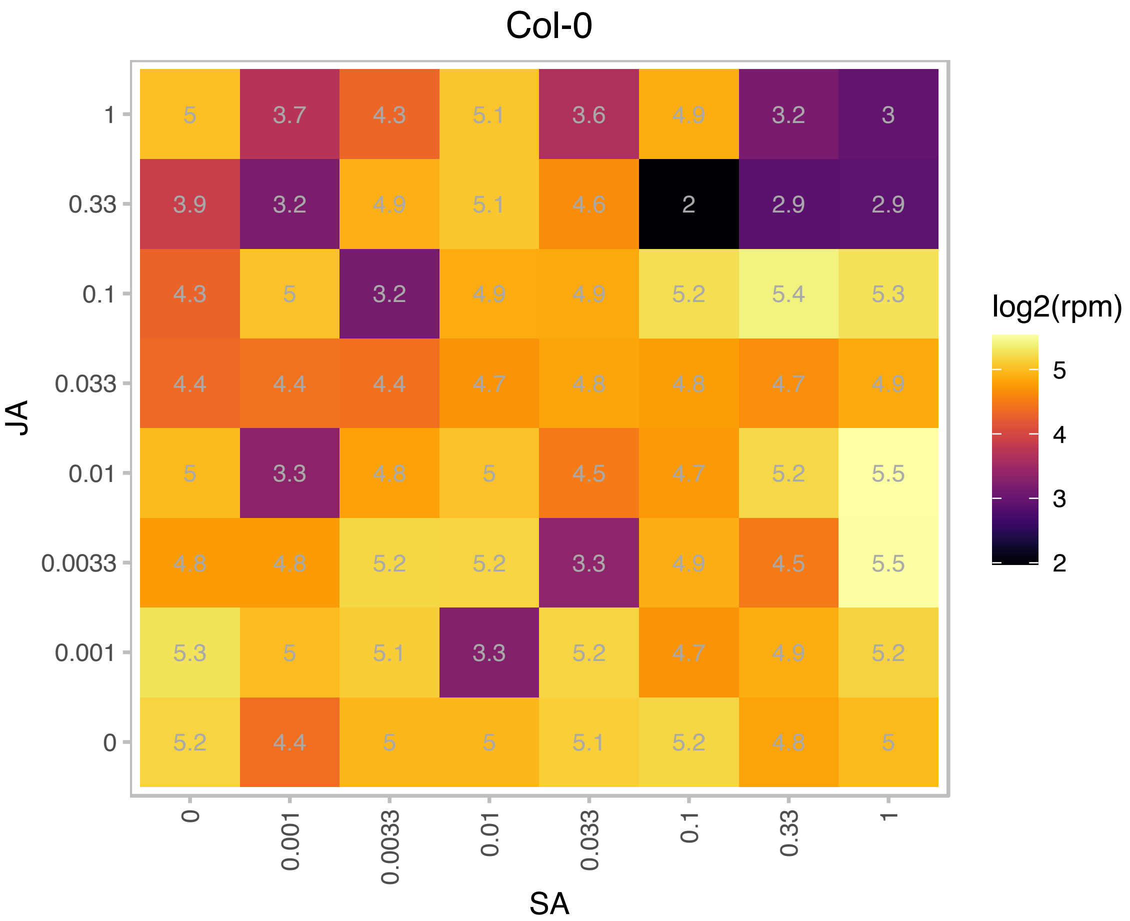


| | |
|---|---|
| Gene | AT5G02050 |
| is.expressed_genes | TRUE |
| is.responsive_genes | TRUE |
| is.apcluster | H-01 |

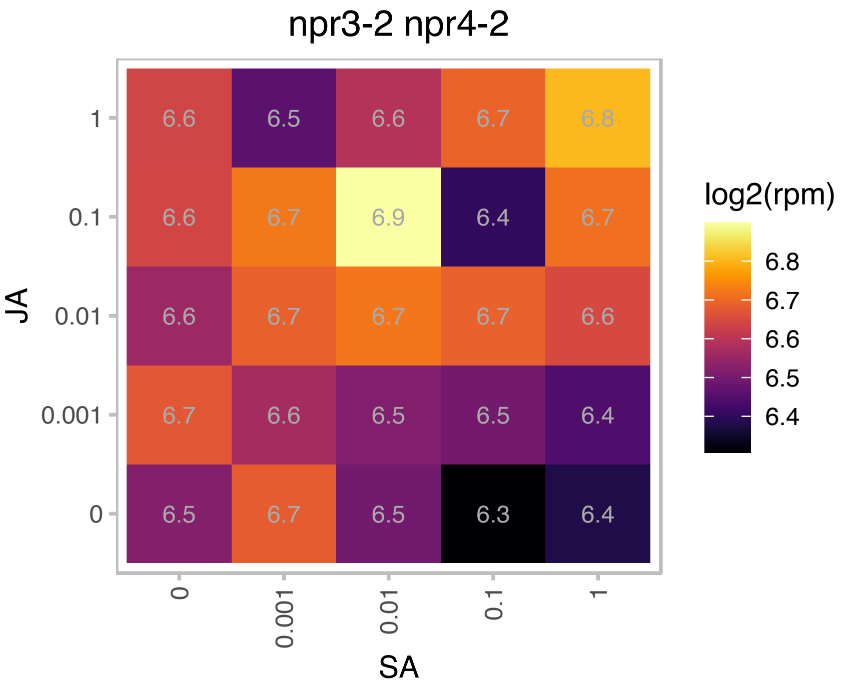


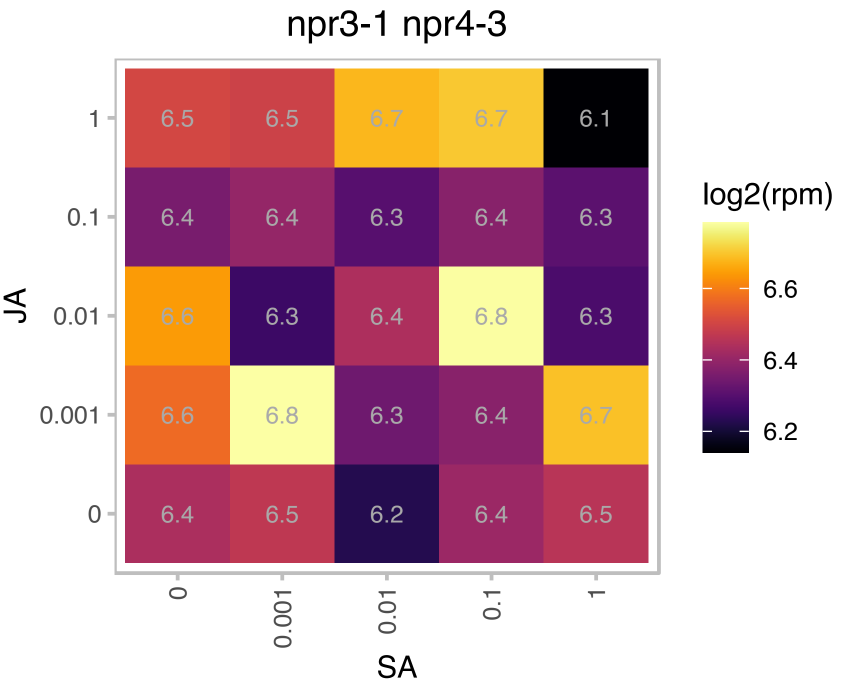

# AT5G26800.1 : NA

| | |
|---|---|
| Gene | AT5G26800 |
| is.expressed_genes | TRUE |
| is.responsive_genes | TRUE |
| is.apcluster | H-01 |

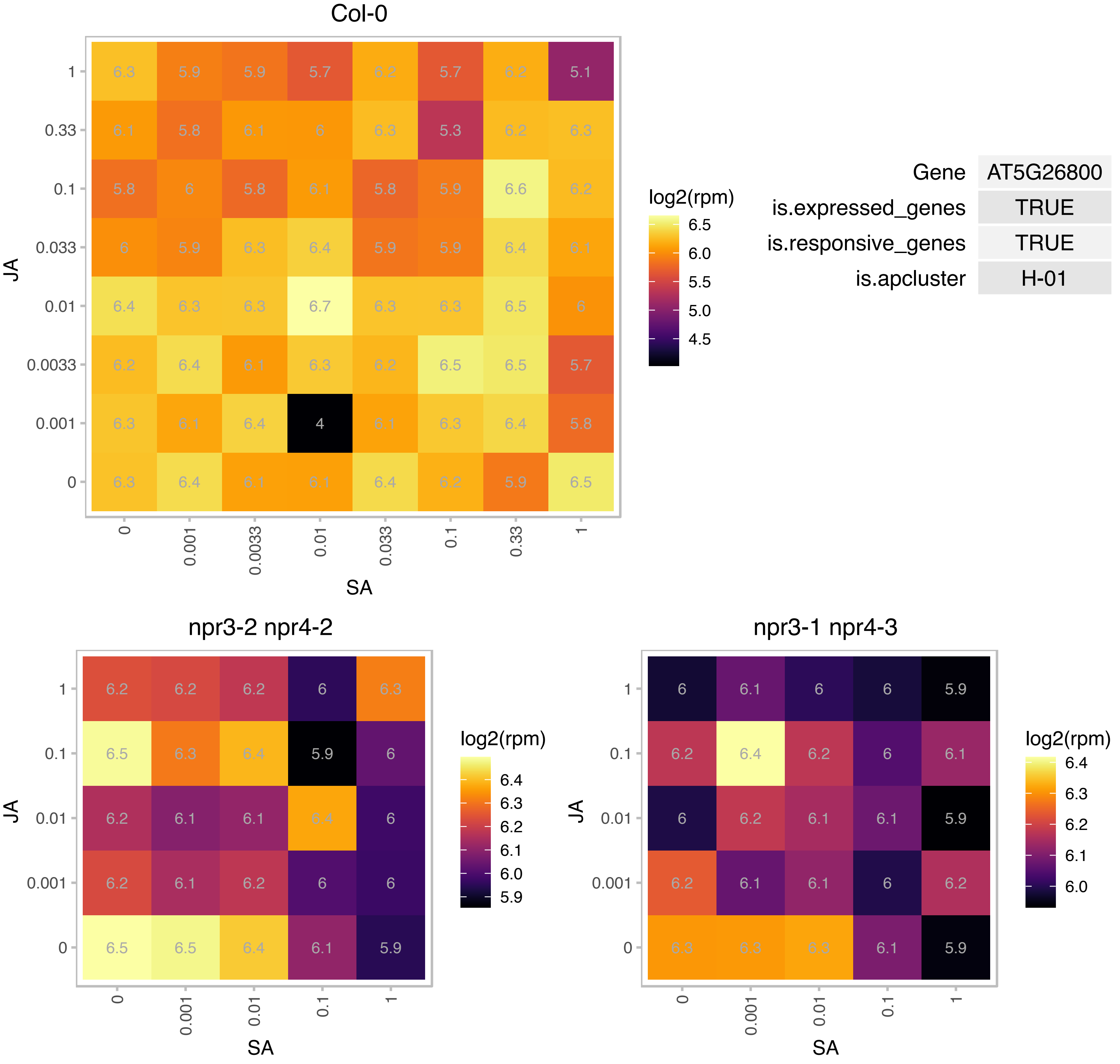

### AT5G28770.1 : AtbZIP63, BZO2H3

bZIP transcription factor family protein

bZIP protein BZO2H3 mRNA, partial cds

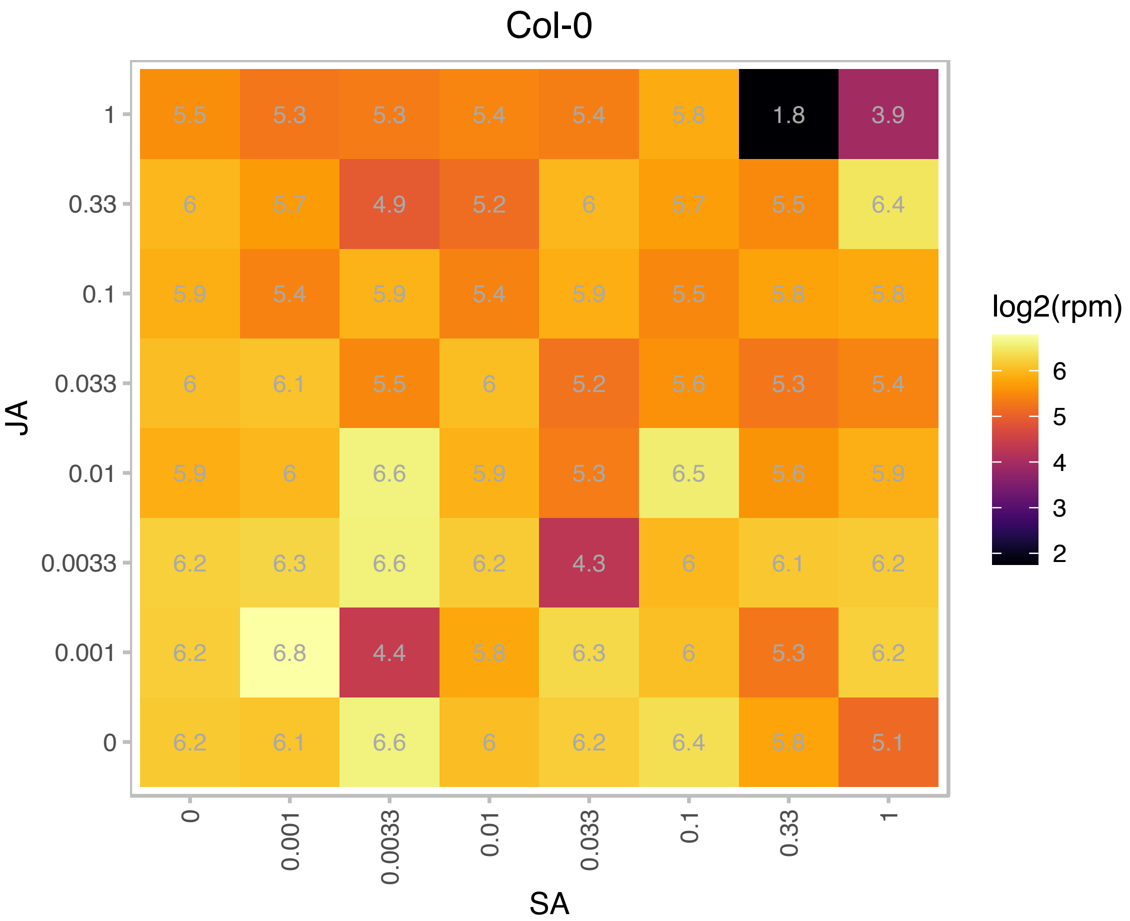


| | |
|---|---|
| Gene | AT5G28770 |
| is.expressed_genes | TRUE |
| is.responsive_genes | TRUE |
| is.apcluster | H-01 |

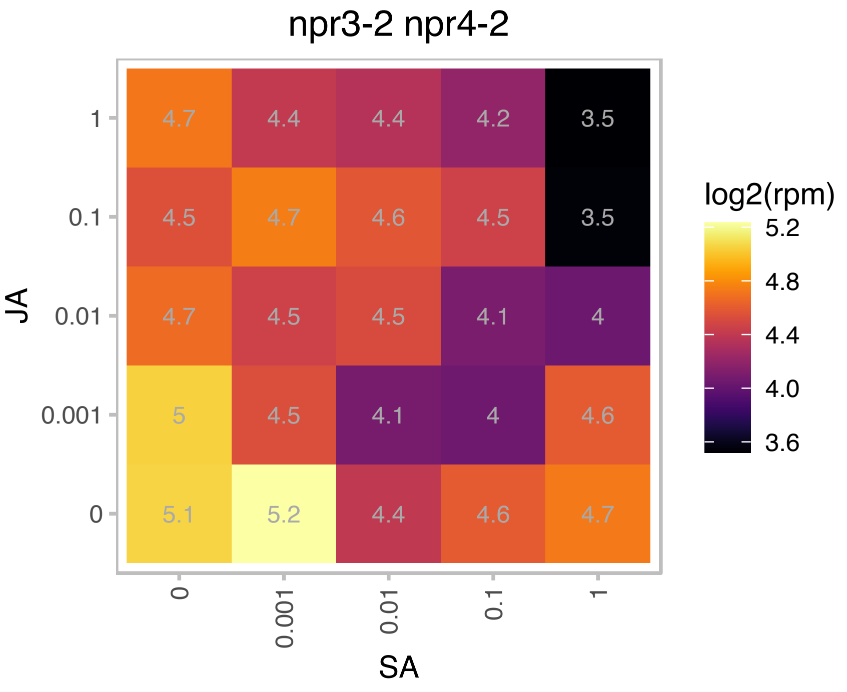


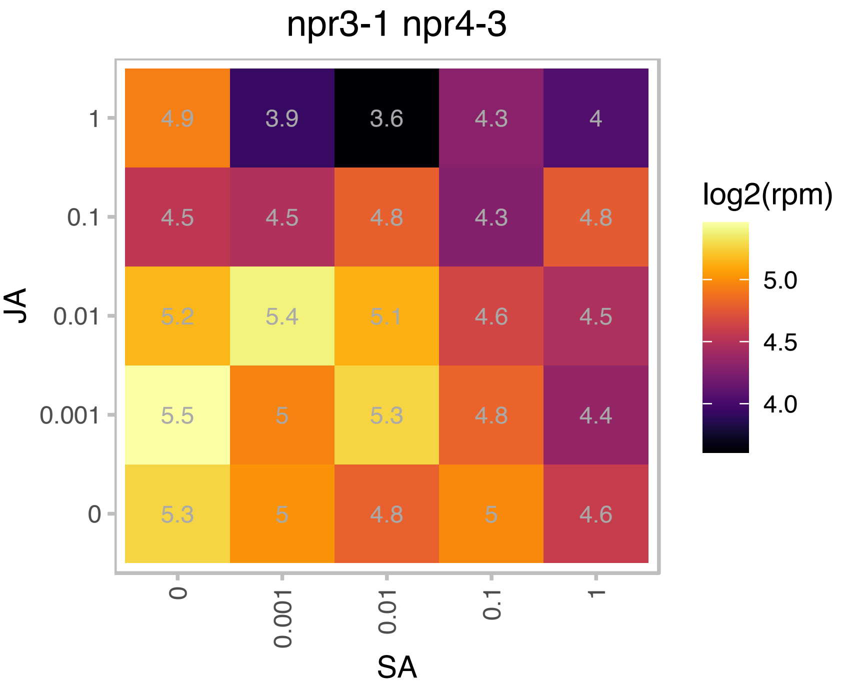

### AT5G54270.1 : LHCB3, LHCB3*1

light-harvesting chlorophyll B-binding protein 3

Lhcb3 protein is a component of the main light harvesting chlorophyll a/b-protein complex of Photosystem II (LHC II).

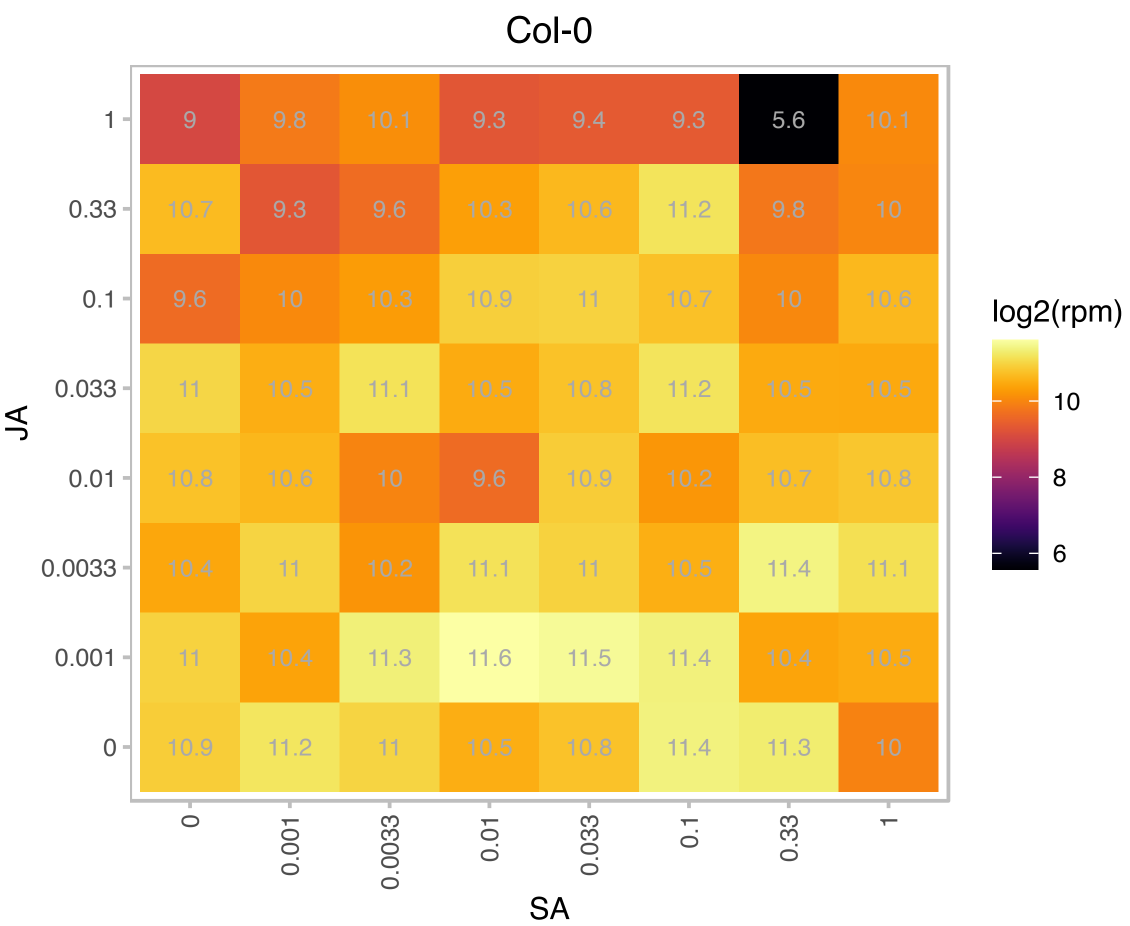


| | |
|---|---|
| Gene | AT5G54270 |
| is.expressed_genes | TRUE |
| is.responsive_genes | TRUE |
| is.apcluster | H-01 |

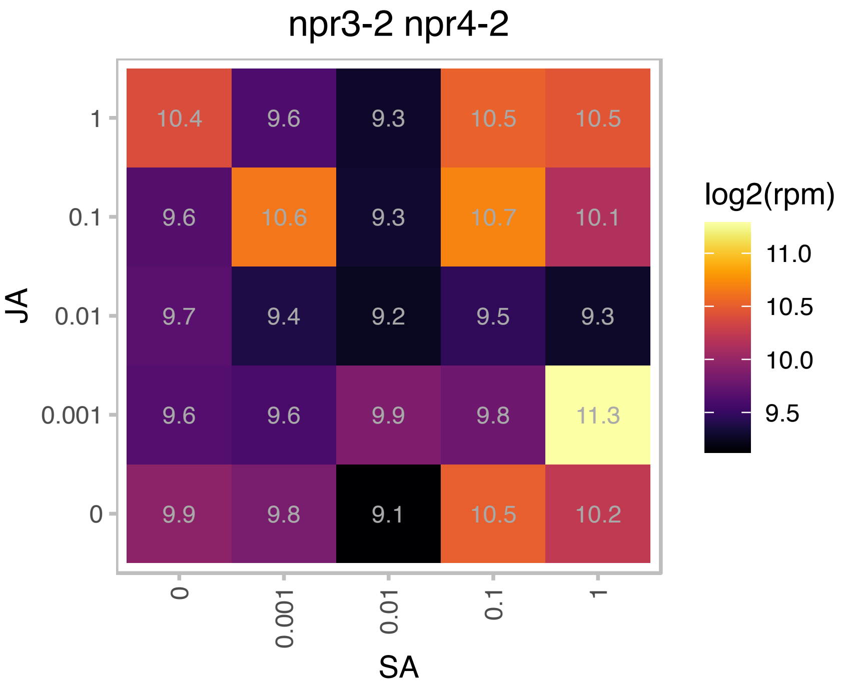


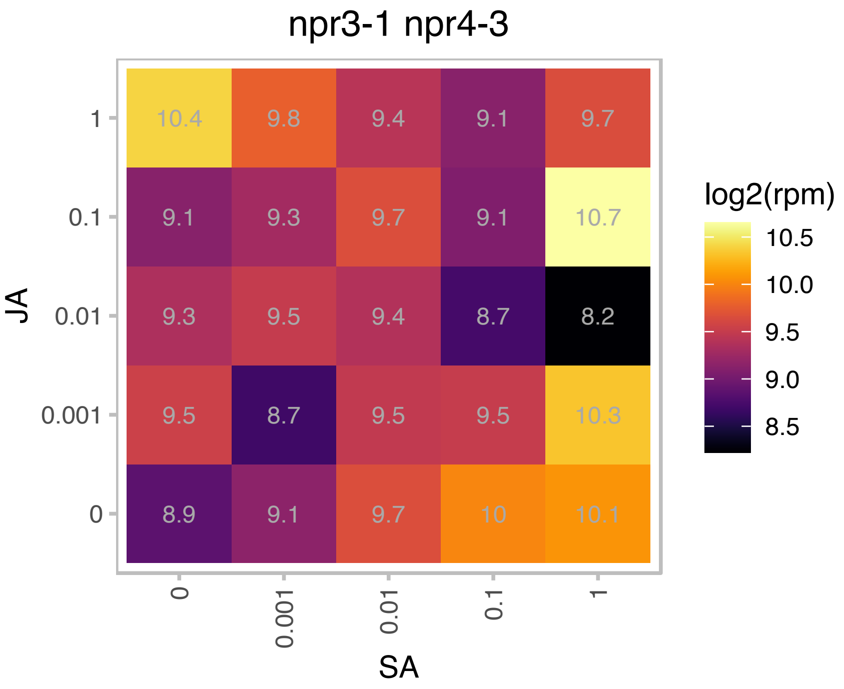

### AT5G56540.1 : AGP14, ATAGP14

arabinogalactan protein 14

Encodes arabinogalactan protein (AGP14). Mutants exhibit longer root hairs. The mRNA is cell-to-cell mobile.

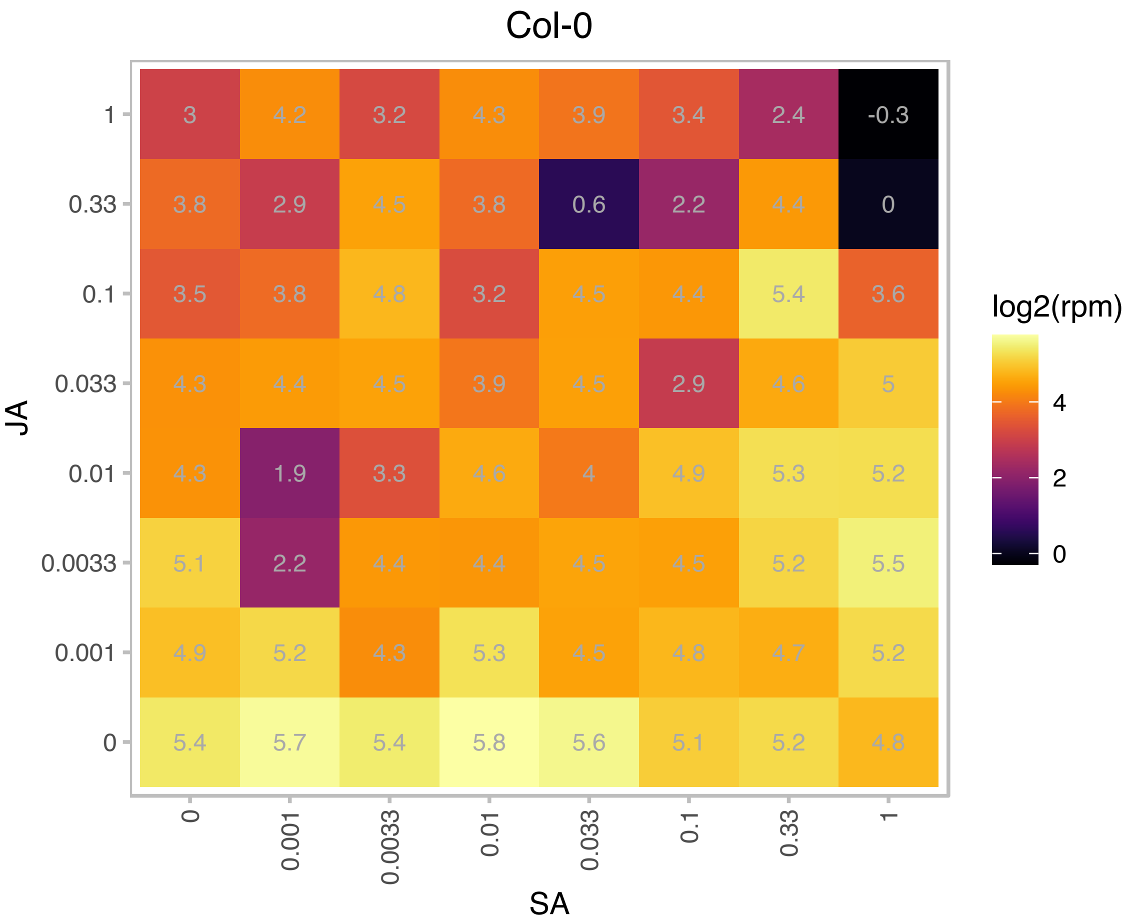


| | |
|---|---|
| Gene | AT5G56540 |
| is.expressed_genes | TRUE |
| is.responsive_genes | TRUE |
| is.apcluster | H-01 |

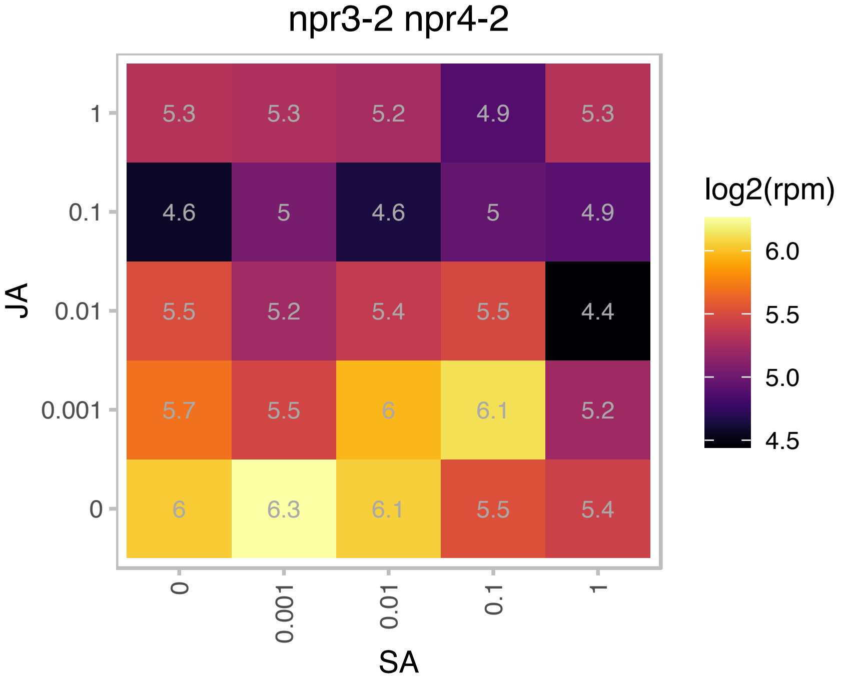


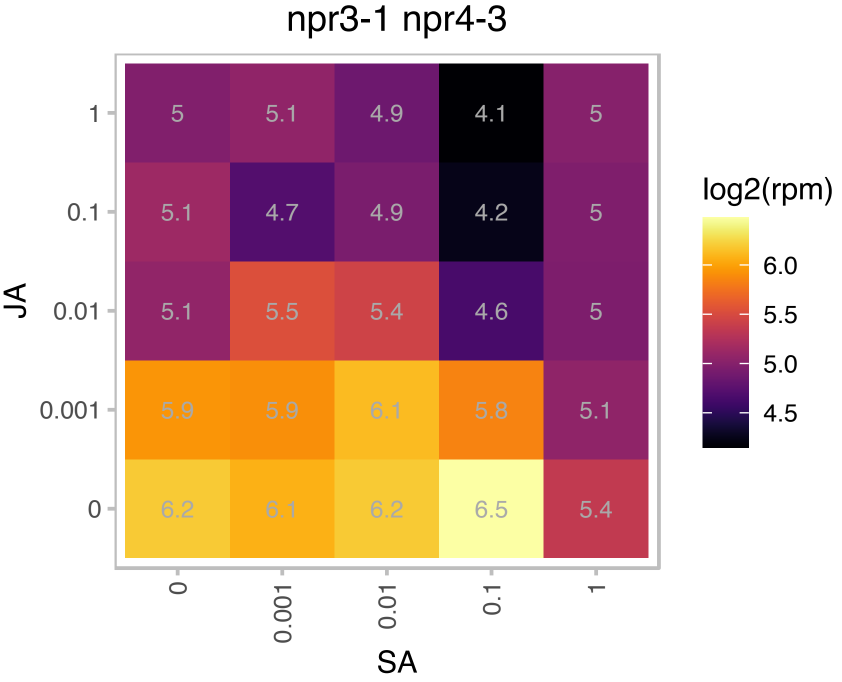

### AT5G60970.1 : TCP5

TEOSINTE BRANCHED 1, cycloidea and PCF transcription factor 5

TCP gene involved in heterochronic control of leaf differentiation.

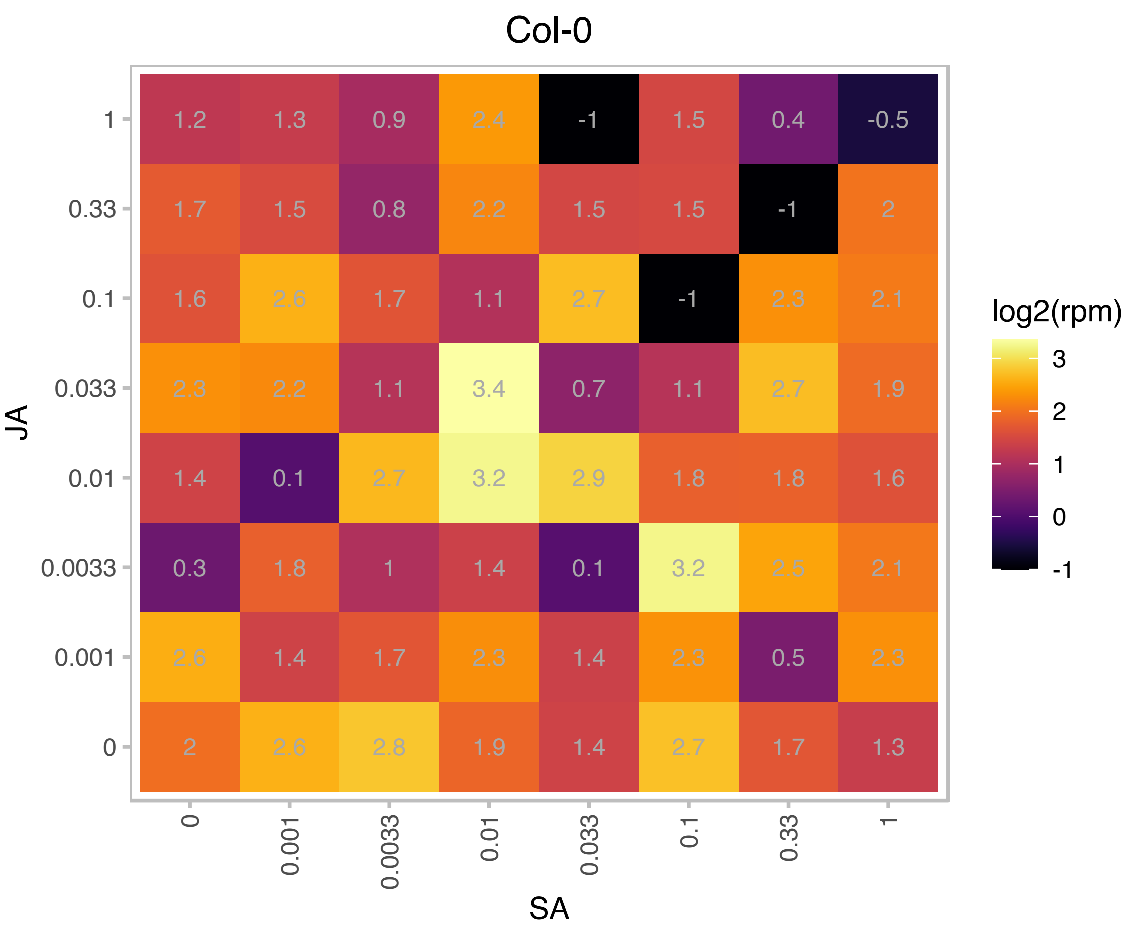


| | |
|---|---|
| Gene | AT5G60970 |
| is.expressed_genes | TRUE |
| is.responsive_genes | TRUE |
| is.apcluster | H-01 |

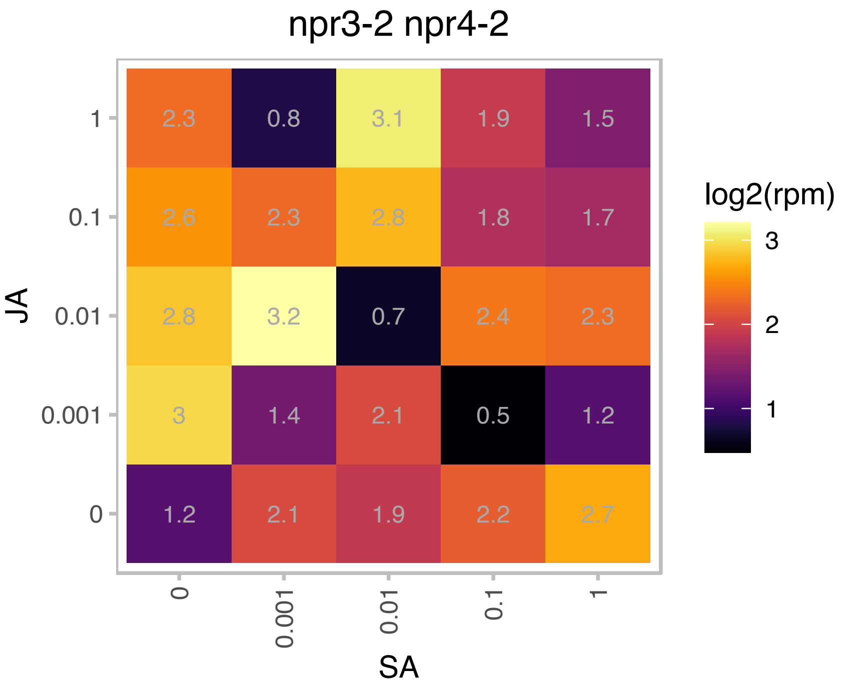


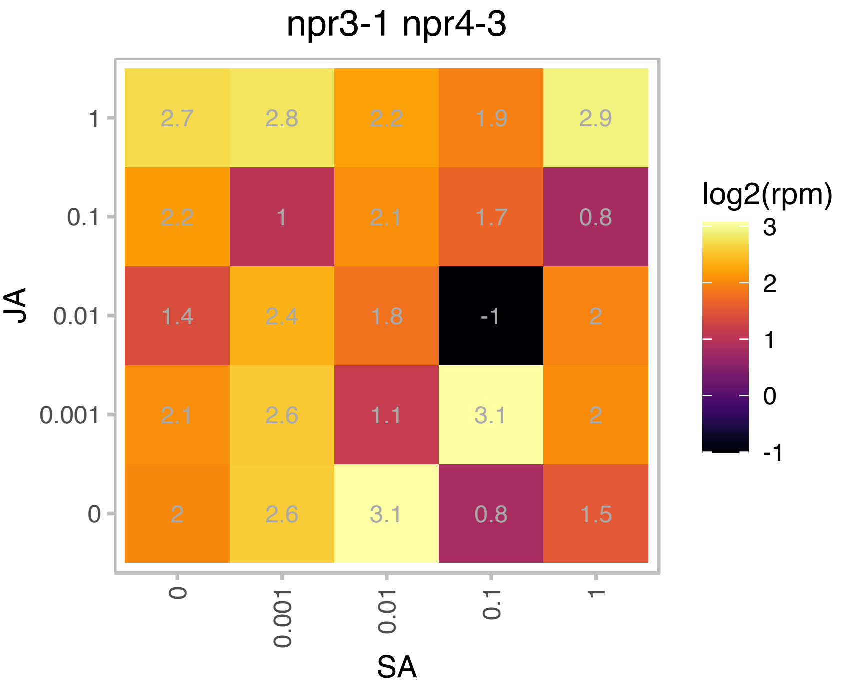

### AT5G61230.1 : ANK6, PIA2

Ankyrin repeat family protein

Encodes ANK6, a mitochondrial ankyrin repeat protein required for male-female gamete recognition.

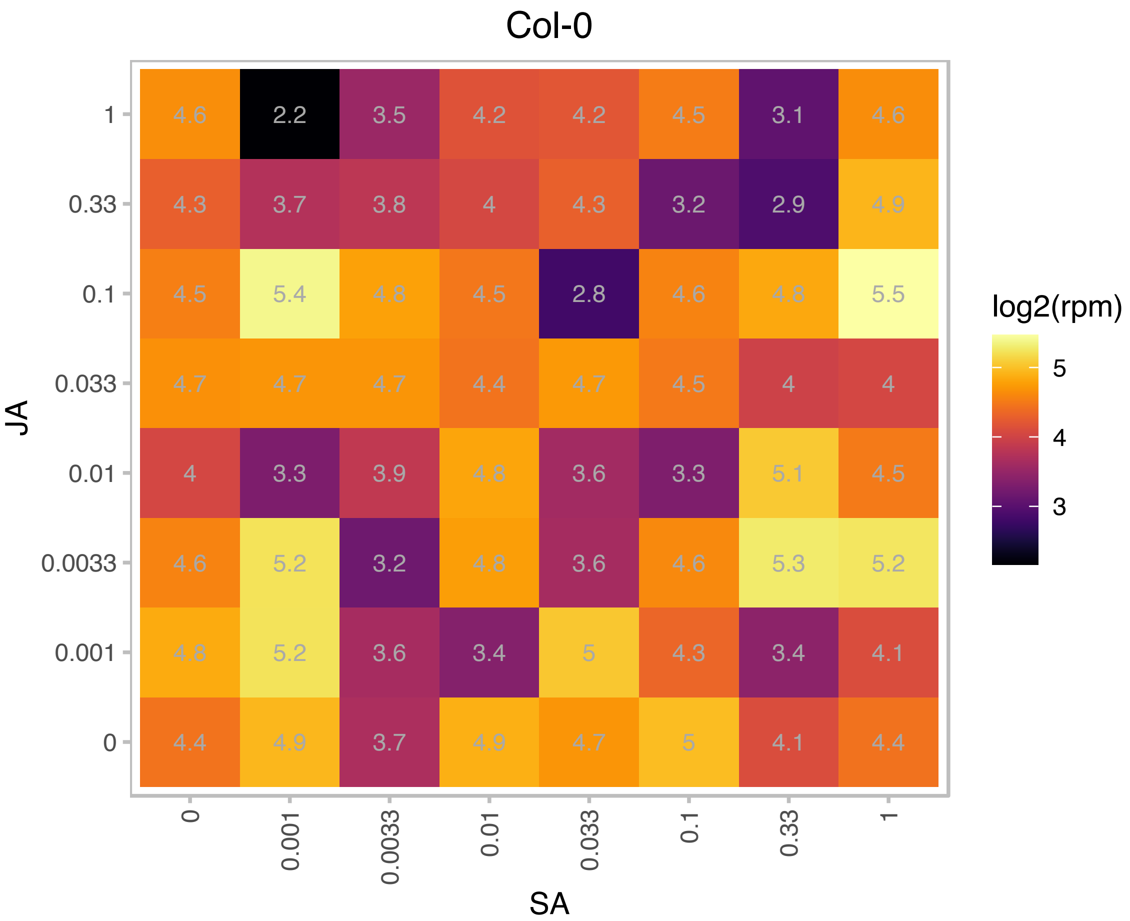


| | |
|---|---|
| Gene | AT5G61230 |
| is.expressed_genes | TRUE |
| is.responsive_genes | TRUE |
| is.apcluster | H-01 |

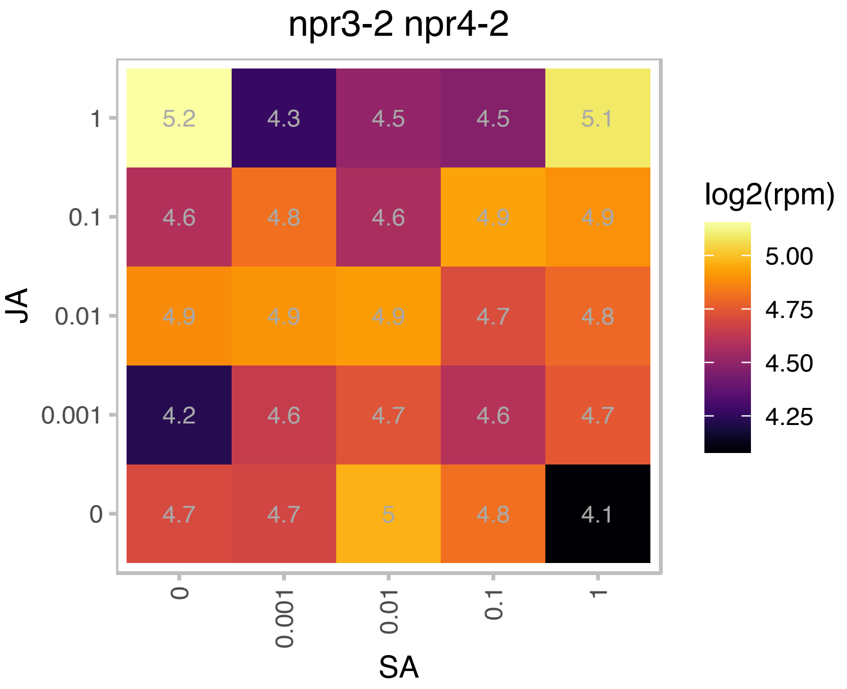


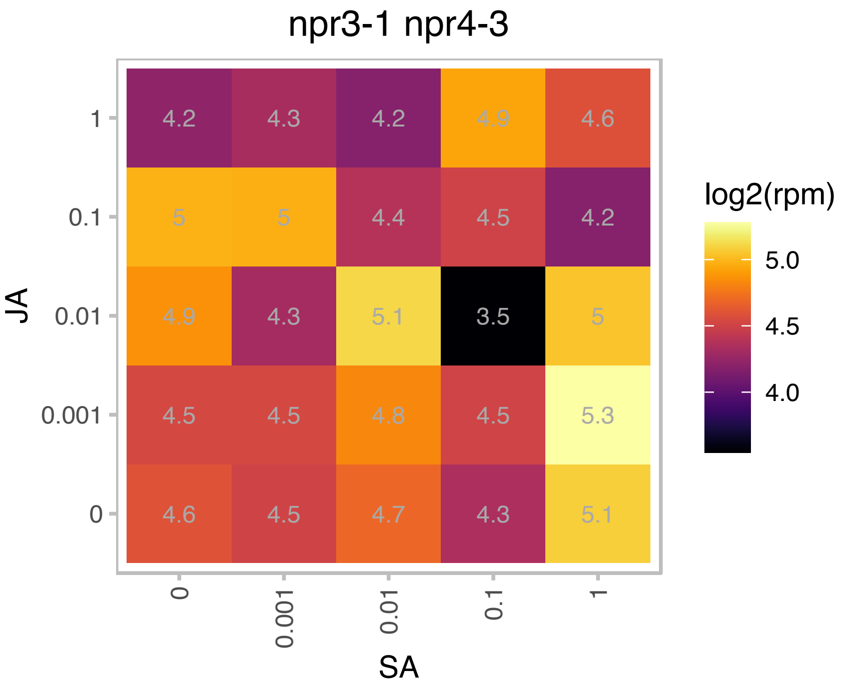

### AT5G64770.1 : CLEL 9, GLV2, RGF9

Encodes a root meristem growth factor (RGF). Belongs to a family of functionally redundant homologous peptides that are secreted, tyrosine-sulfated, and expressed mainly in the stem cell area and the innermost layer of central columella cells. RGFs are required for maintenance of the root stem cell niche and transit amplifying cell proliferation. Members of this family include: At5g60810 (RGF1), At1g13620 (RGF2), At2g04025 (RGF3), At3g30350 (RGF4), At5g51451 (RGF5), At4g16515 (RGF6), At3g02240 (RGF7), At2g03830 (RGF8) and At5g64770 (RGF9).

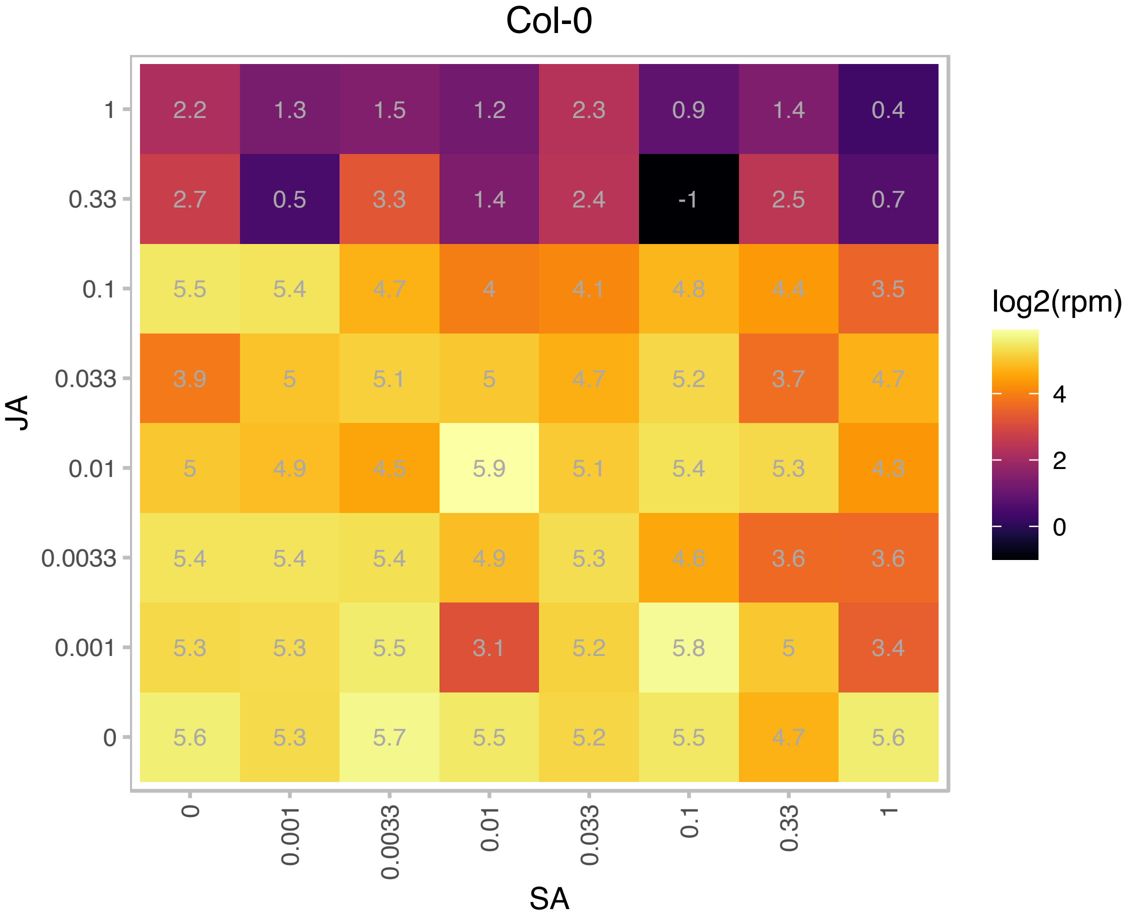


| | |
|---|---|
| Gene | AT5G64770 |
| is.expressed_genes | TRUE |
| is.responsive_genes | TRUE |
| is.apcluster | H-01 |

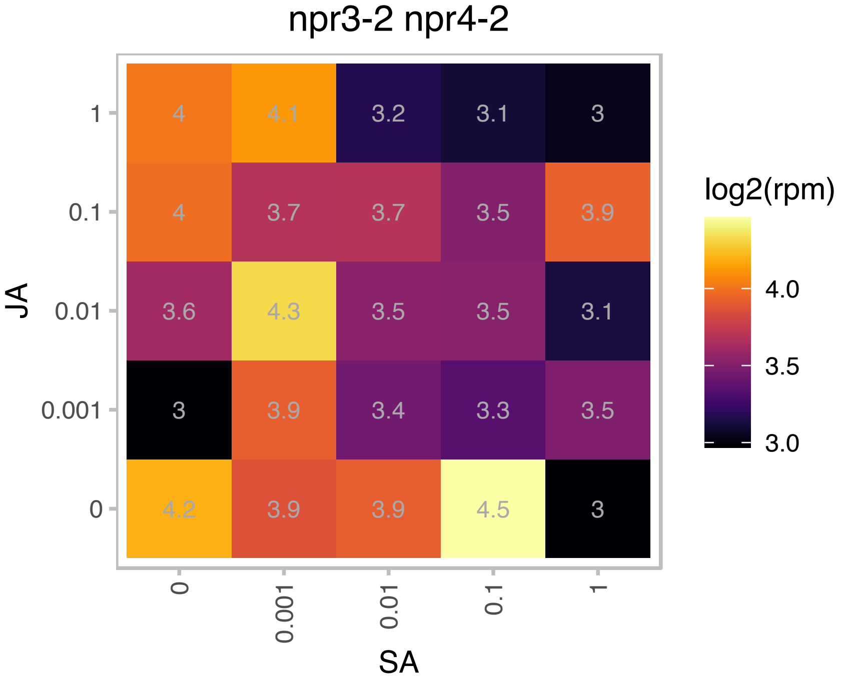


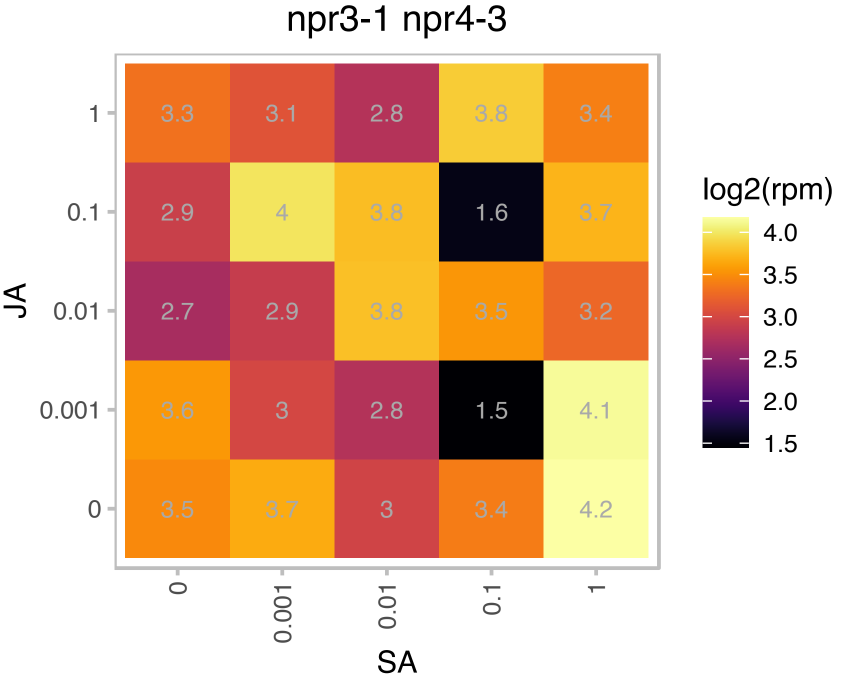

### ATCG00470.1 : ATPE

ATP synthase epsilon chain

ATPase epsilon subunit

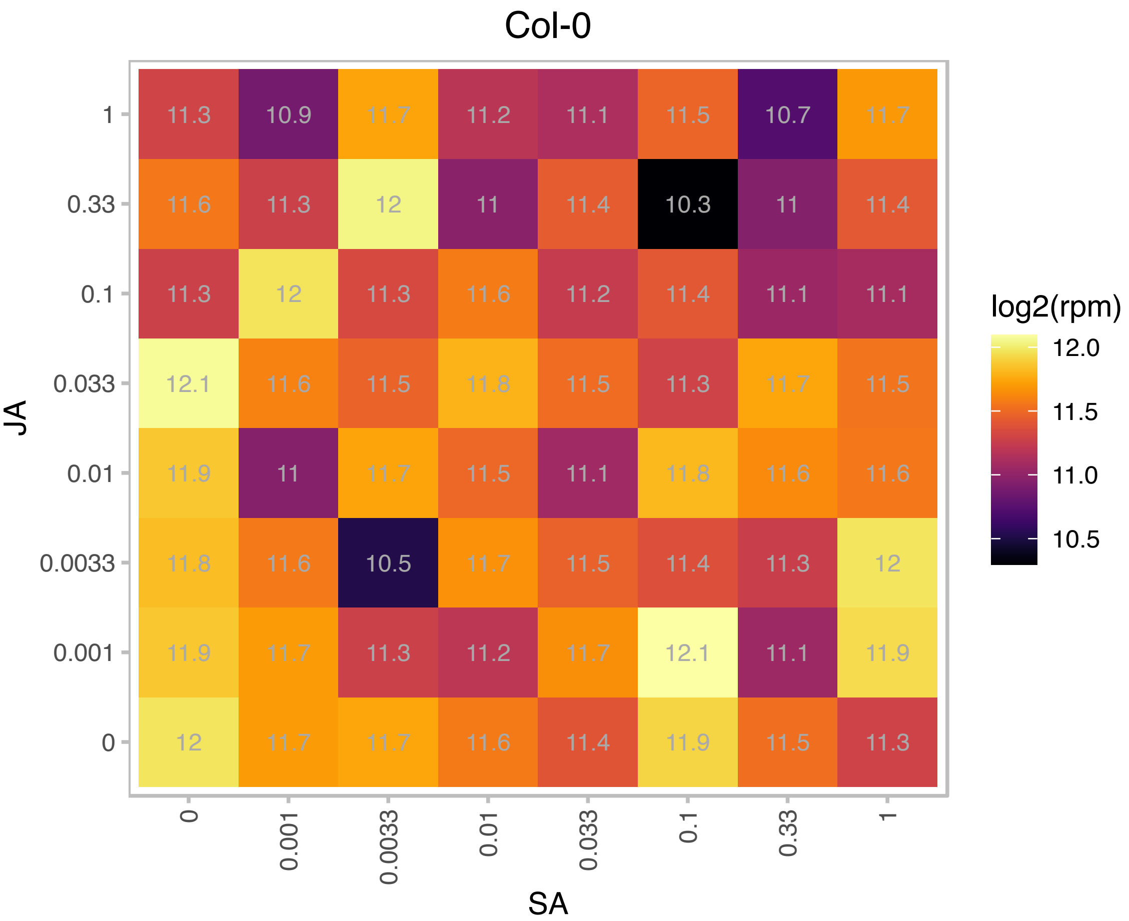


| | |
|---|---|
| Gene | ATCG00470 |
| is.expressed_genes | TRUE |
| is.responsive_genes | TRUE |
| is.apcluster | H-01 |

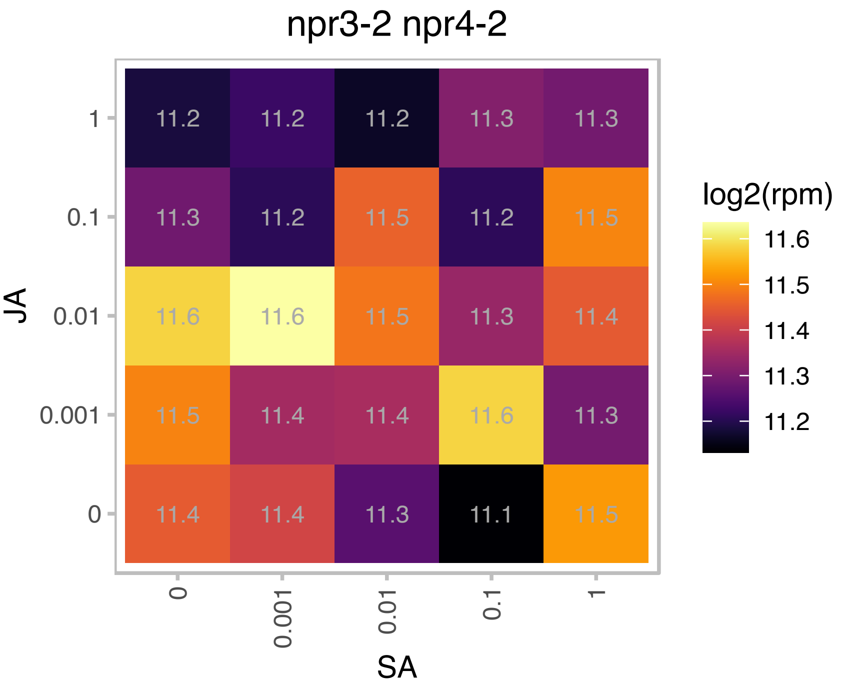


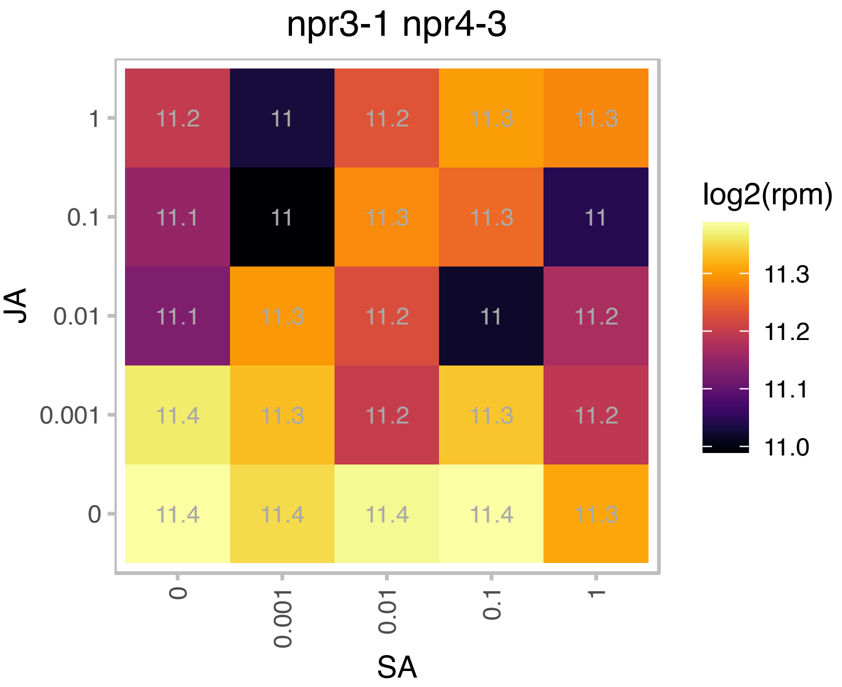

### ATCG00480.1 : AthCF1beta, ATPB, CF1beta, PB

ATP synthase subunit beta

chloroplast-encoded gene for beta subunit of ATP synthase

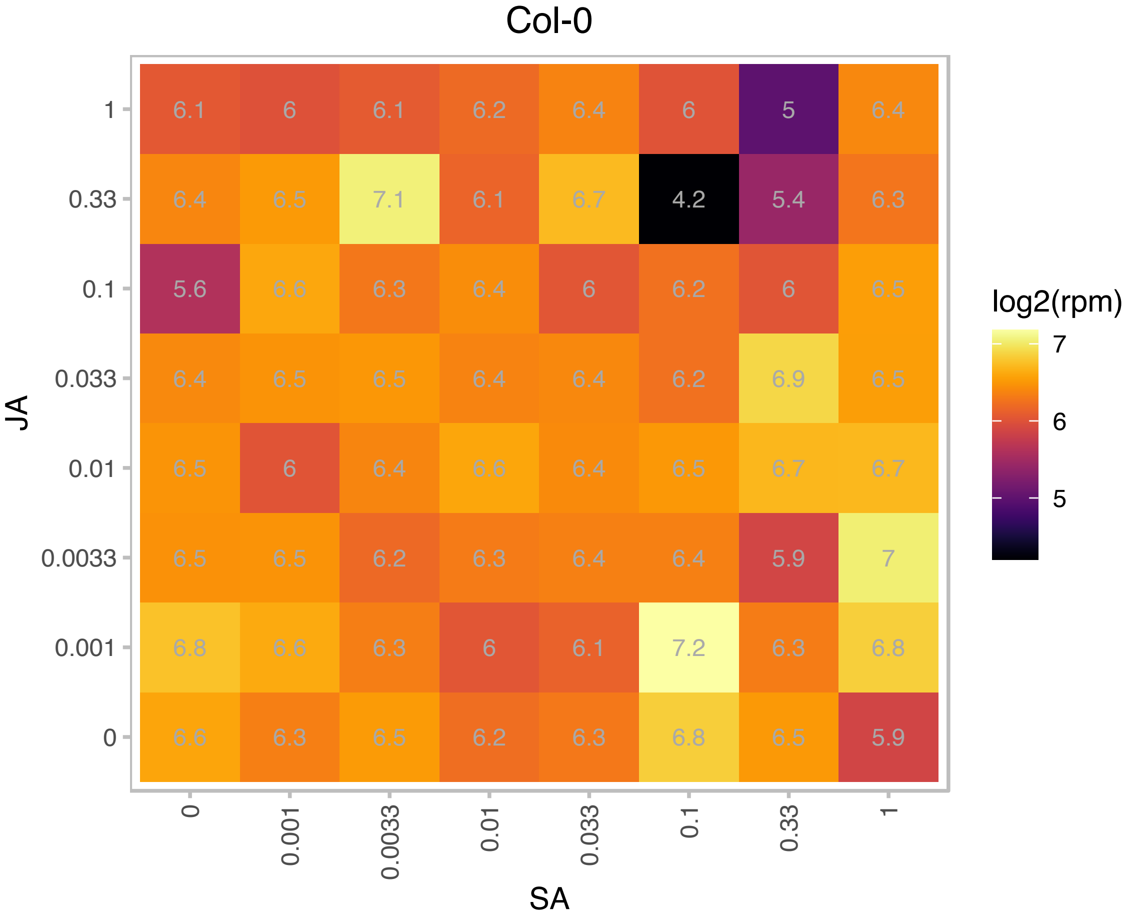


| | |
|---|---|
| Gene | ATCG00480 |
| is.expressed_genes | TRUE |
| is.responsive_genes | TRUE |
| is.apcluster | H-01 |

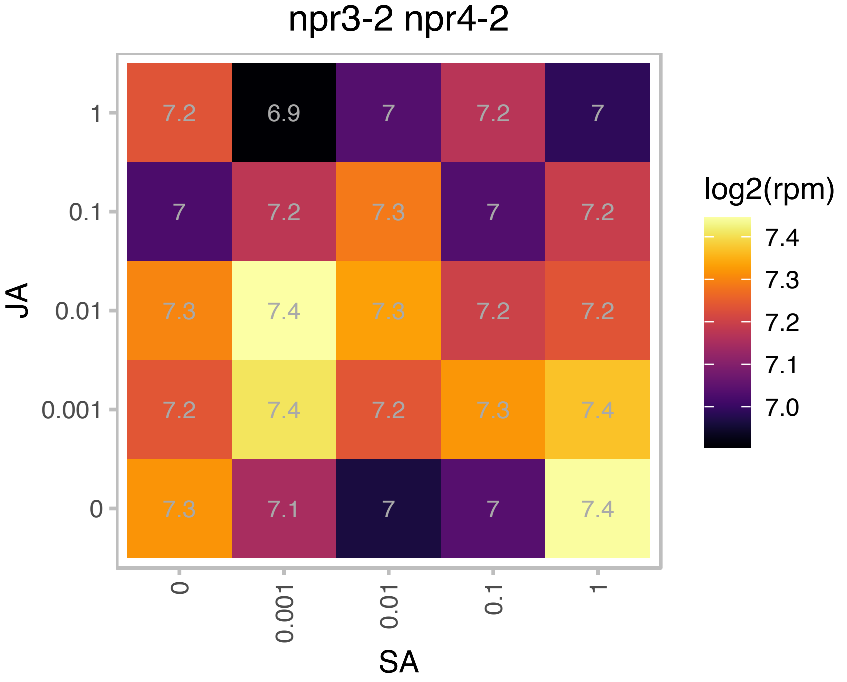


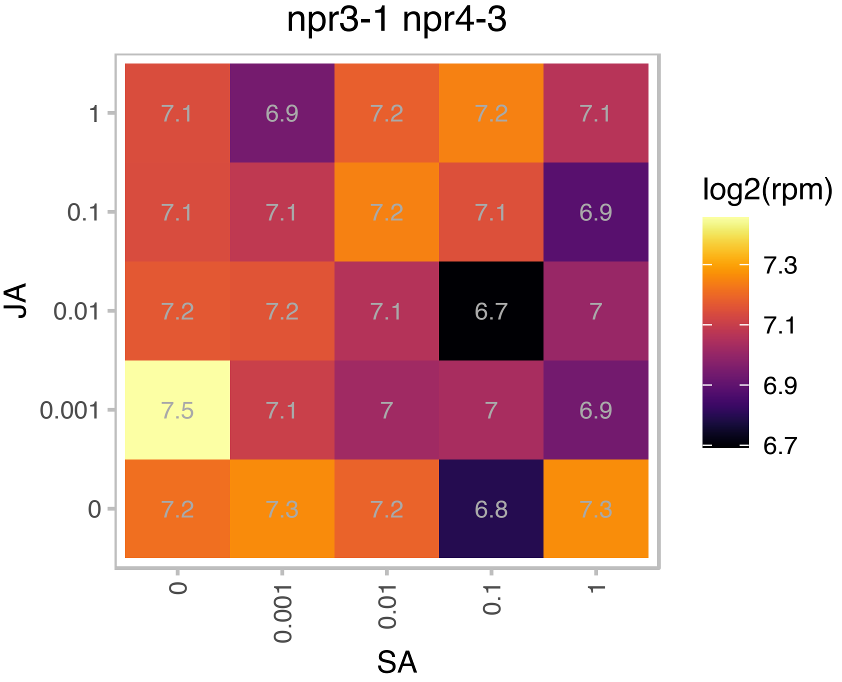

### ATCG00750.1 : RPS11

ribosomal protein S11

30S chloroplast ribosomal protein S11

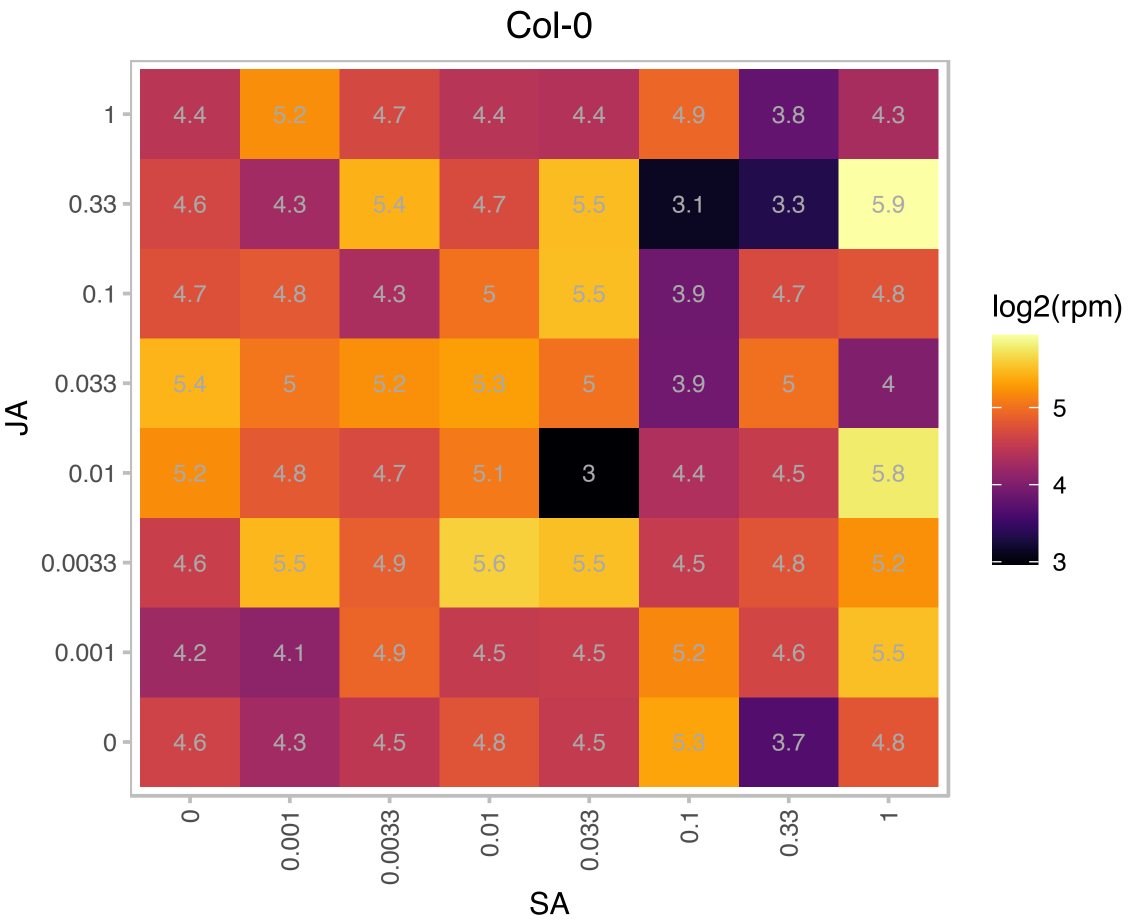


| | |
|---|---|
| Gene | ATCG00750 |
| is.expressed_genes | TRUE |
| is.responsive_genes | TRUE |
| is.apcluster | H-01 |

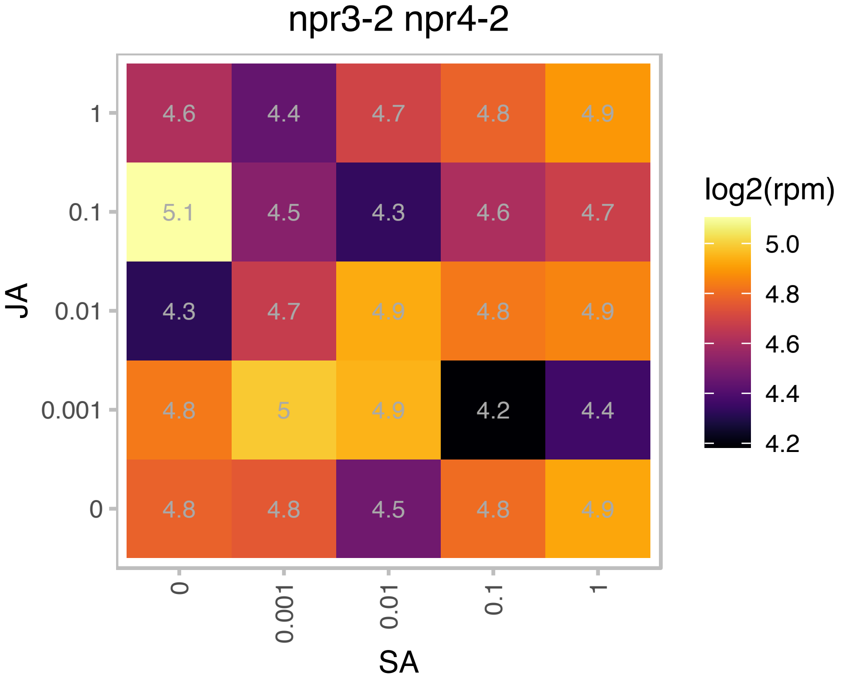


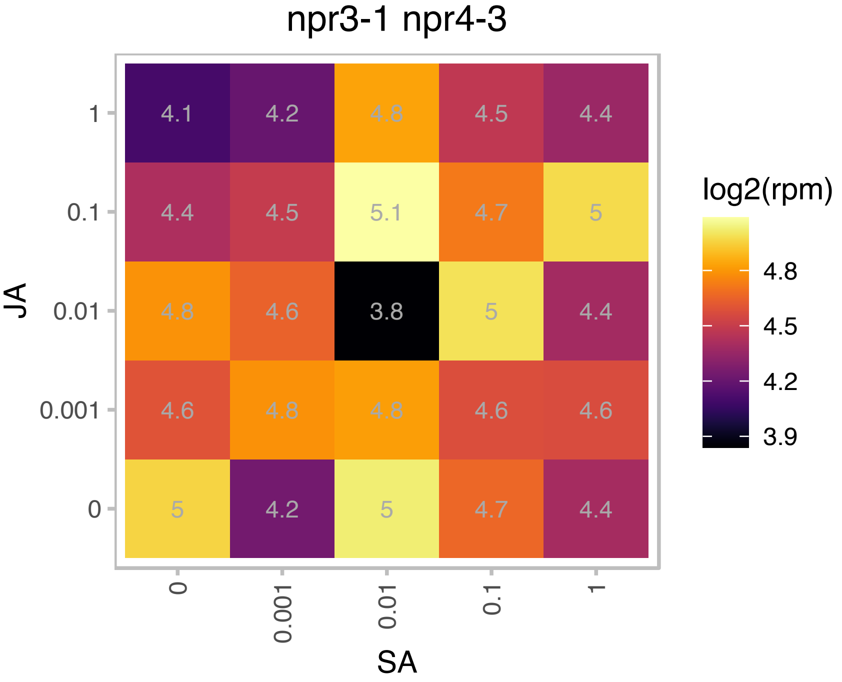

# AT1G05063.1 : NA

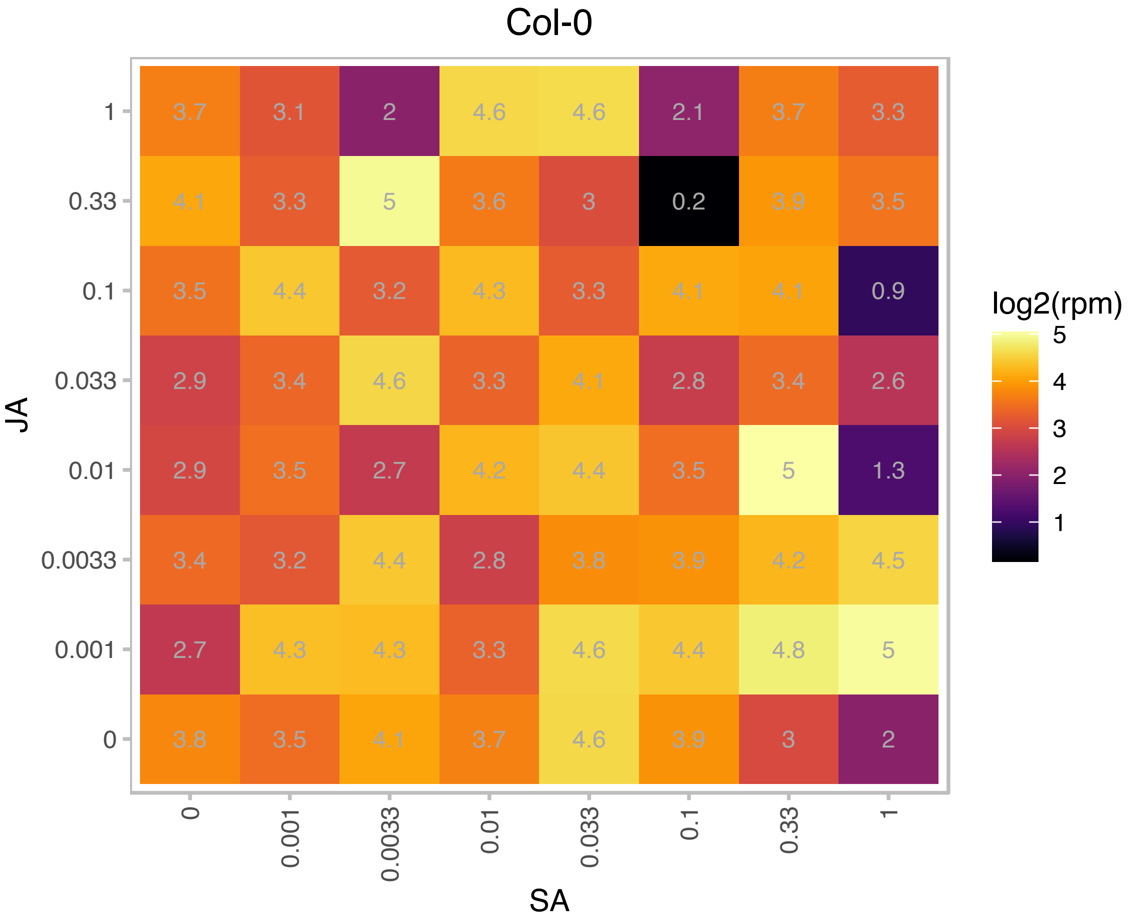


| | |
|---|---|
| Gene | AT1G05063 |
| is.expressed_genes | TRUE |
| is.responsive_genes | TRUE |
| is.apcluster | H-02 |

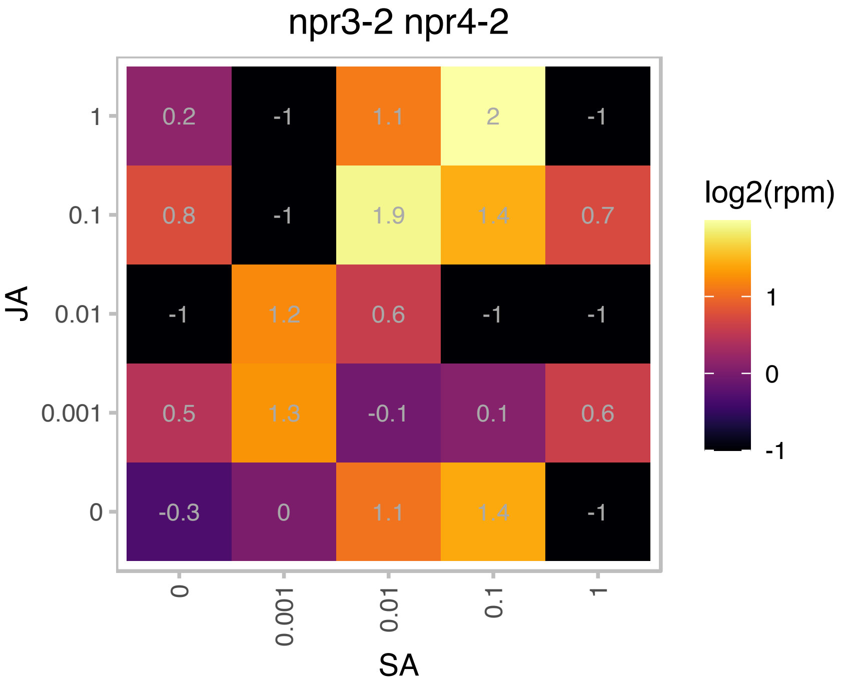


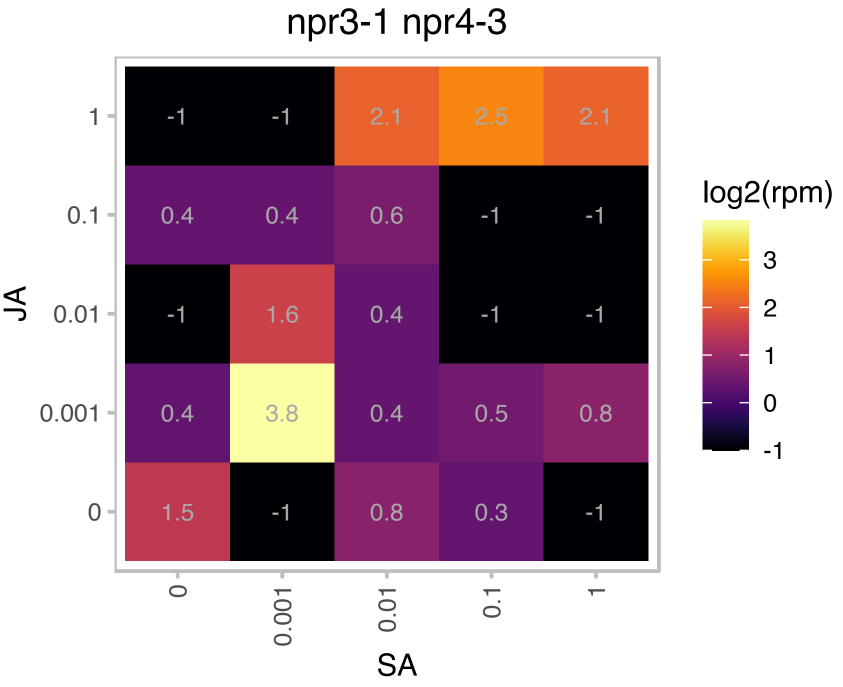

### AT1G05805.1 : AKS2

basic helix-loop-helix (bHLH) DNA-binding superfamily protein

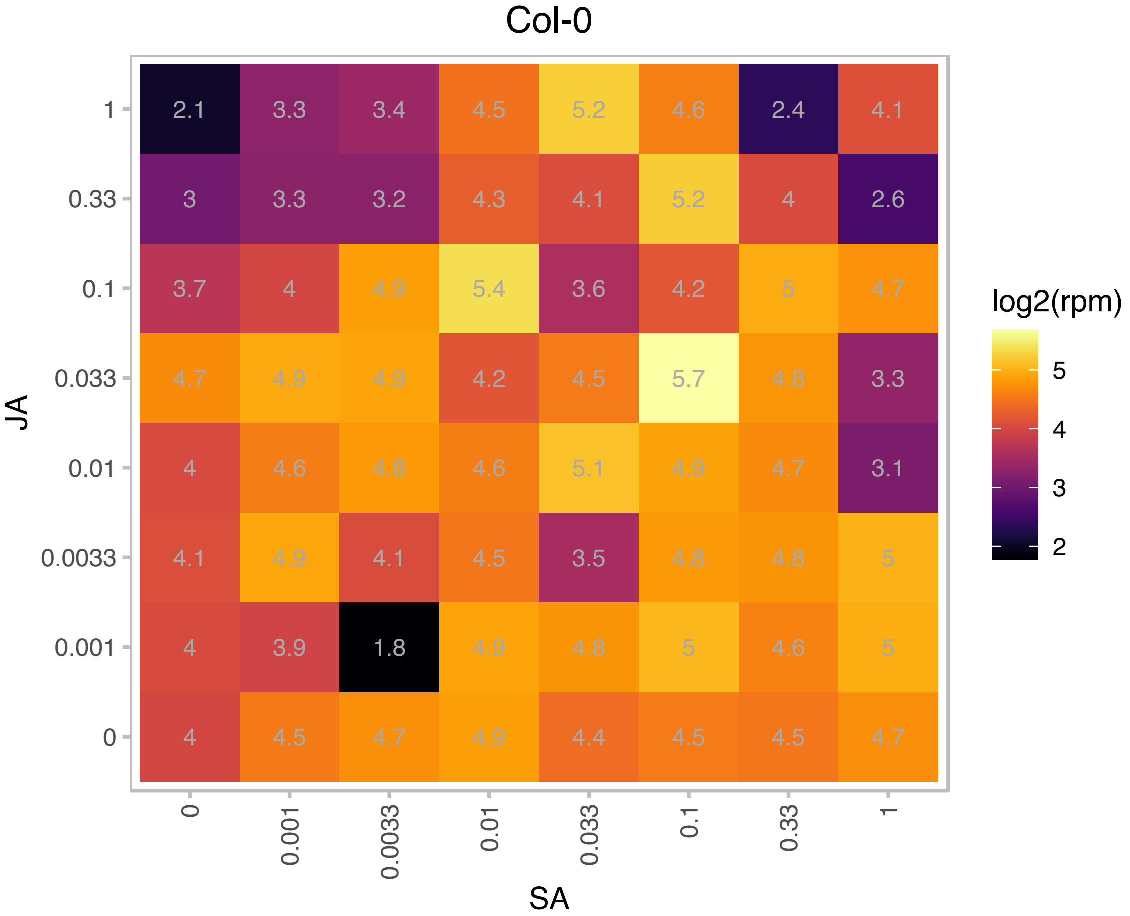


| | |
|---|---|
| Gene | AT1G05805 |
| is.expressed_genes | TRUE |
| is.responsive_genes | TRUE |
| is.apcluster | H-02 |

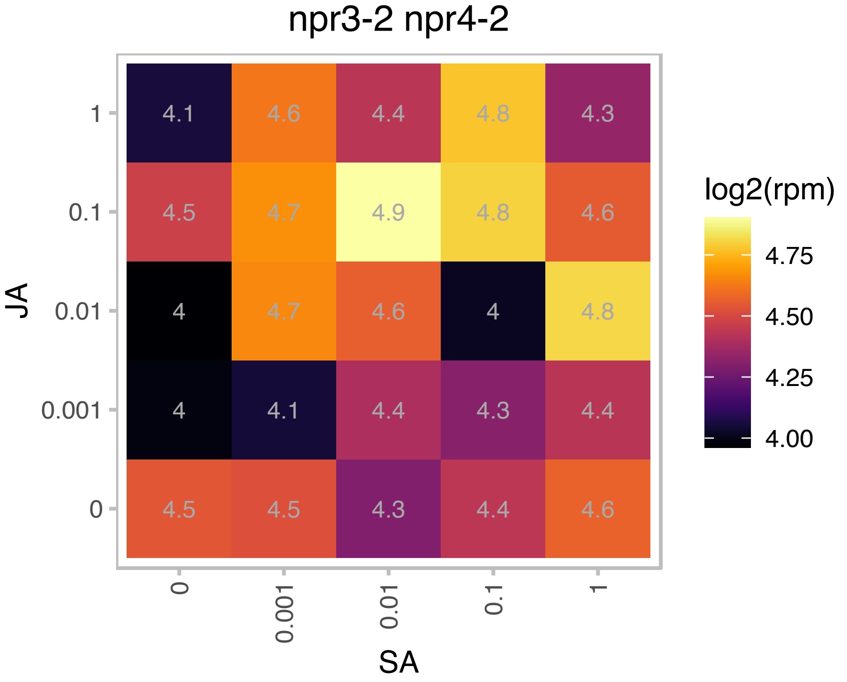


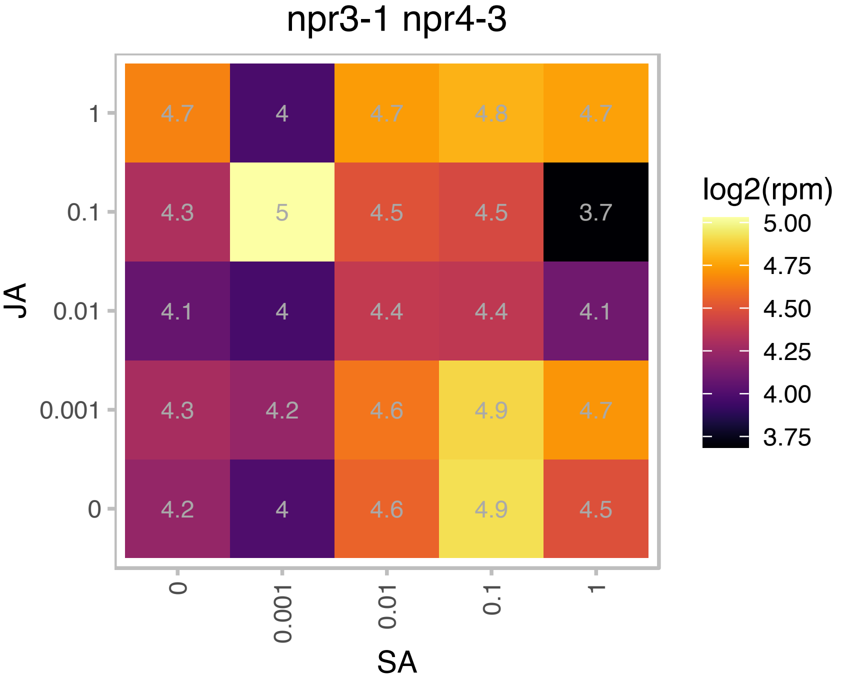

### AT1G09630.1 : ATRAB-A2A, ATRAB11C, ATRABA2A, RAB

RAB GTPase 11C

Encodes a putative GTP-binding protein. Associates with organelles on a pathway from the Golgi to the plasma membrane in interphase. In dividing cells acts at the cell plate.

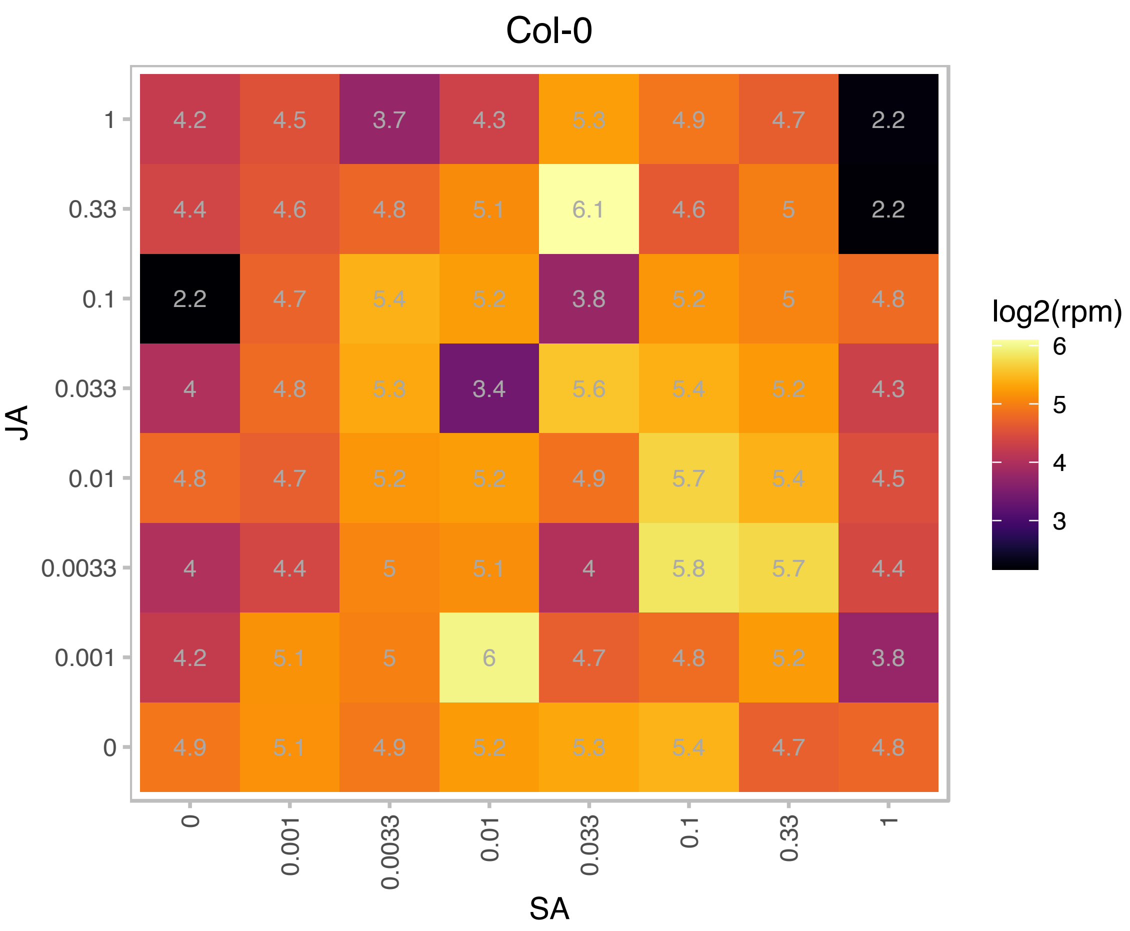


| | |
|---|---|
| Gene | AT1G09630 |
| is.expressed_genes | TRUE |
| is.responsive_genes | TRUE |
| is.apcluster | H-02 |

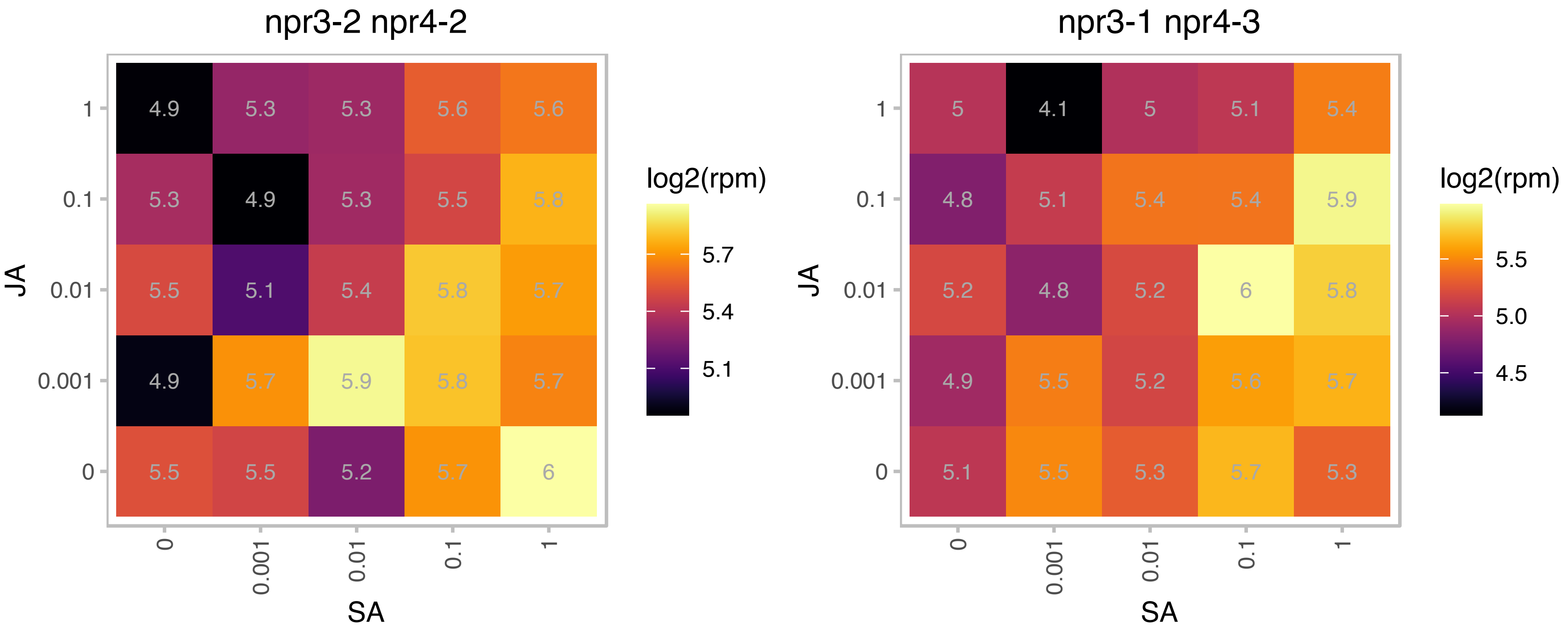

### AT1G13750.1 : ATPAP1, PAP1

Purple acid phosphatases superfamily protein

Encodes a purple acid phosphatase whose expression is responsive to both phosphate (Pi) and phosphite (Phi) in roots.

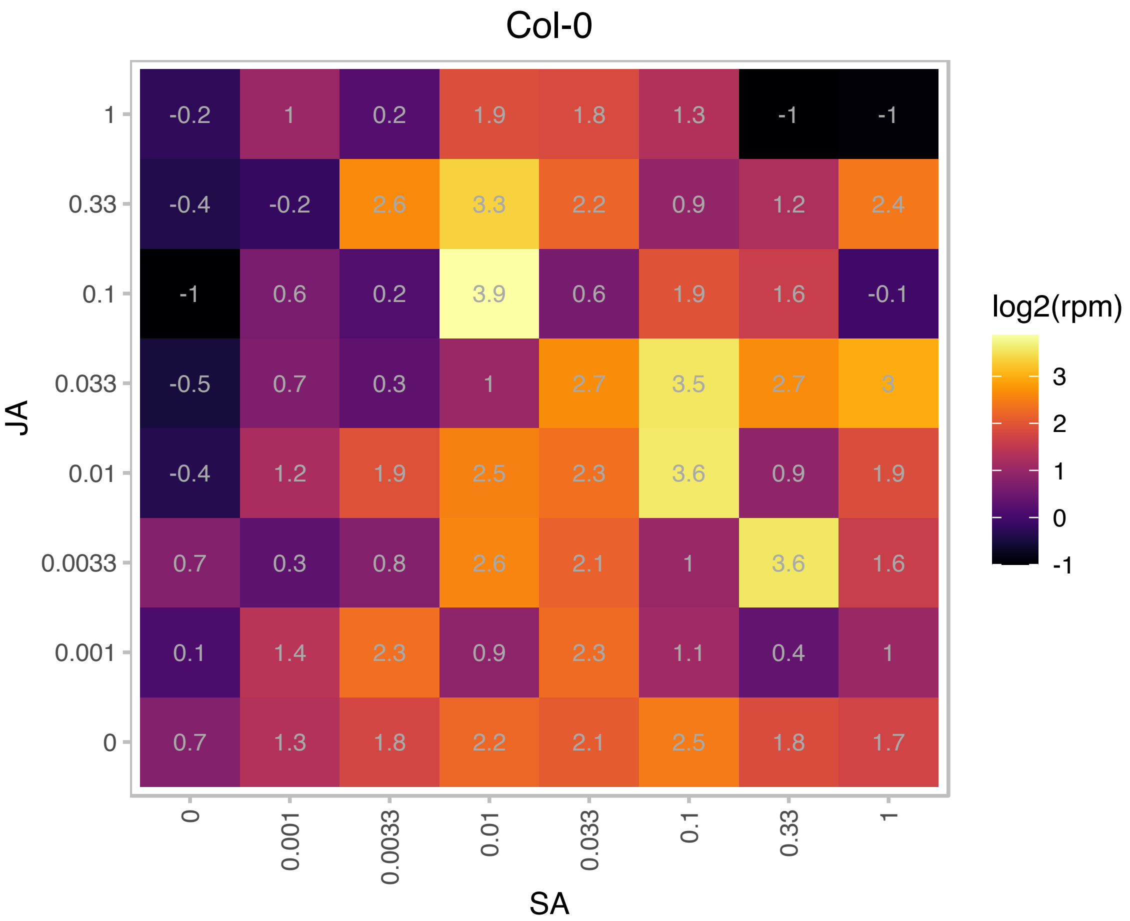


| | |
|---|---|
| Gene | AT1G13750 |
| is.expressed_genes | TRUE |
| is.responsive_genes | TRUE |
| is.apcluster | H-02 |

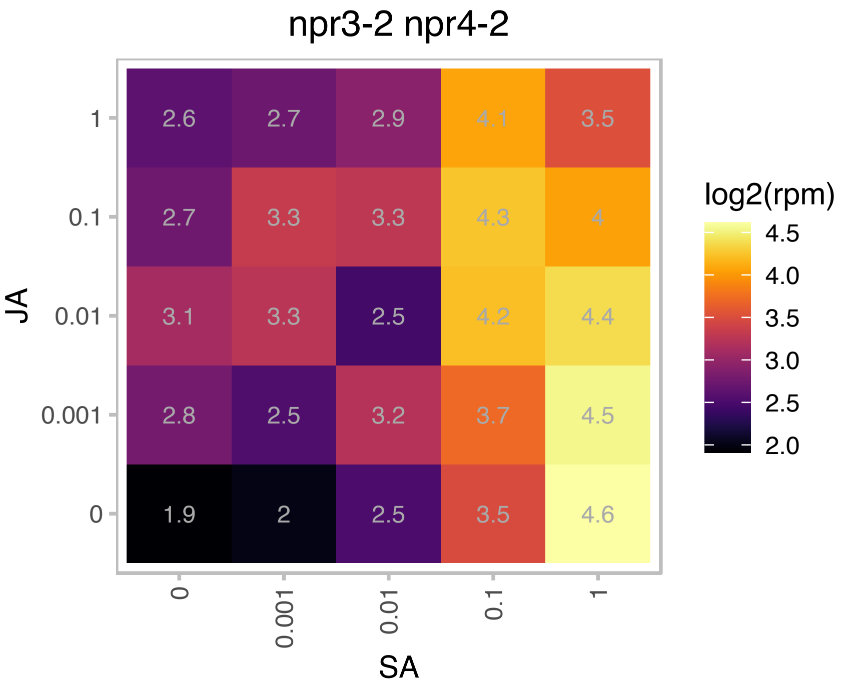


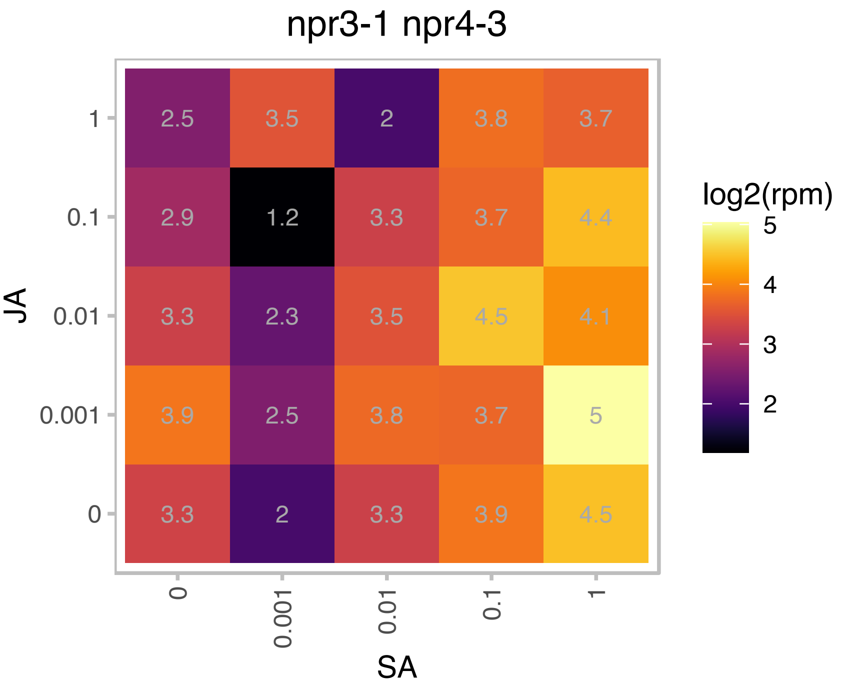

### AT1G14380.1 : IQD28

IQ-domain 28

Encodes a microtubule-associated protein.

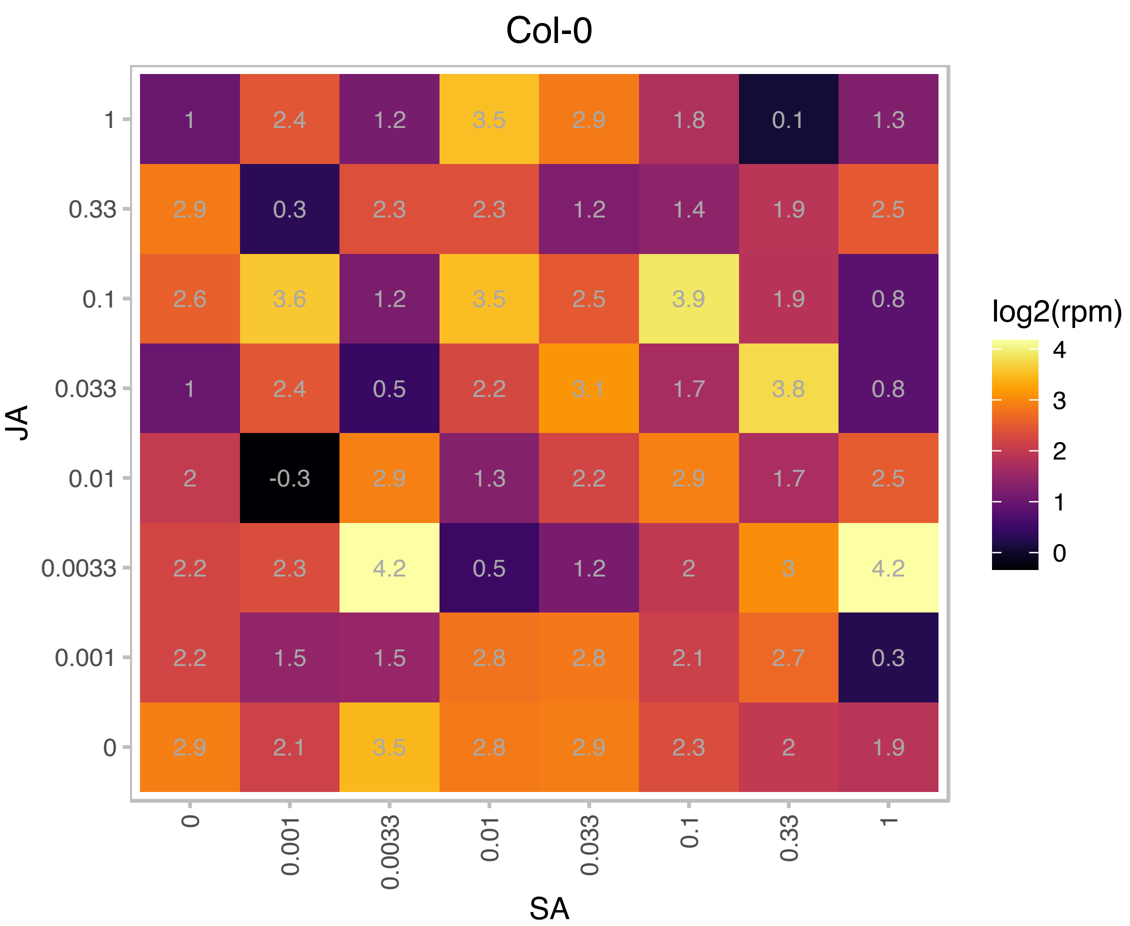


| | |
|---|---|
| Gene | AT1G14380 |
| is.expressed_genes | TRUE |
| is.responsive_genes | TRUE |
| is.apcluster | H-02 |

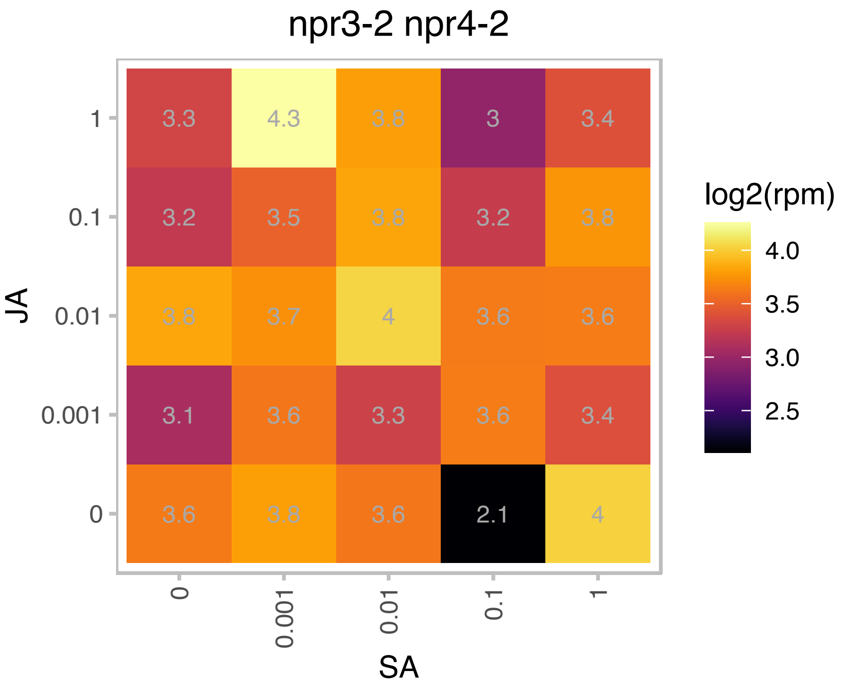


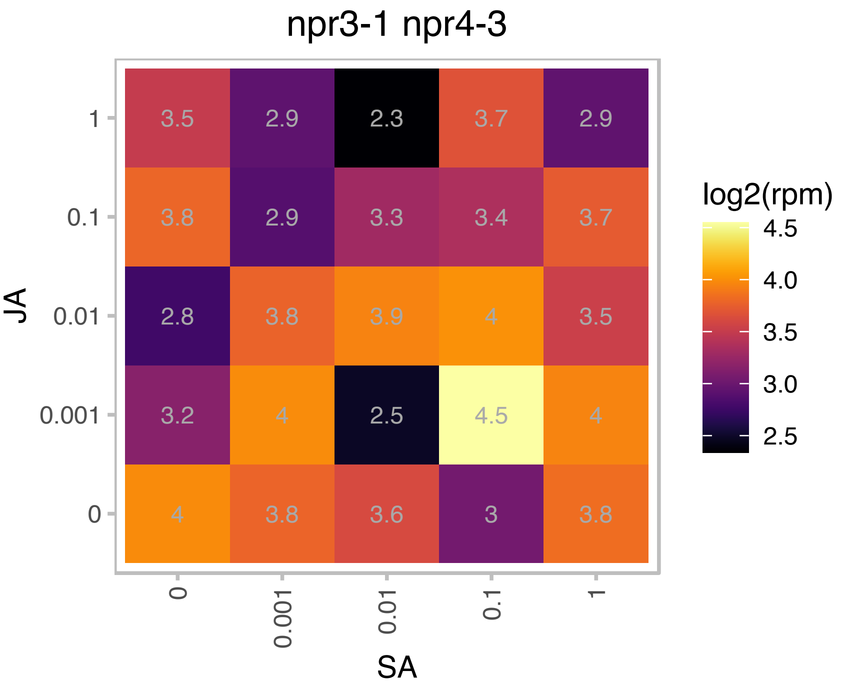

# AT1G19960.1 : NA

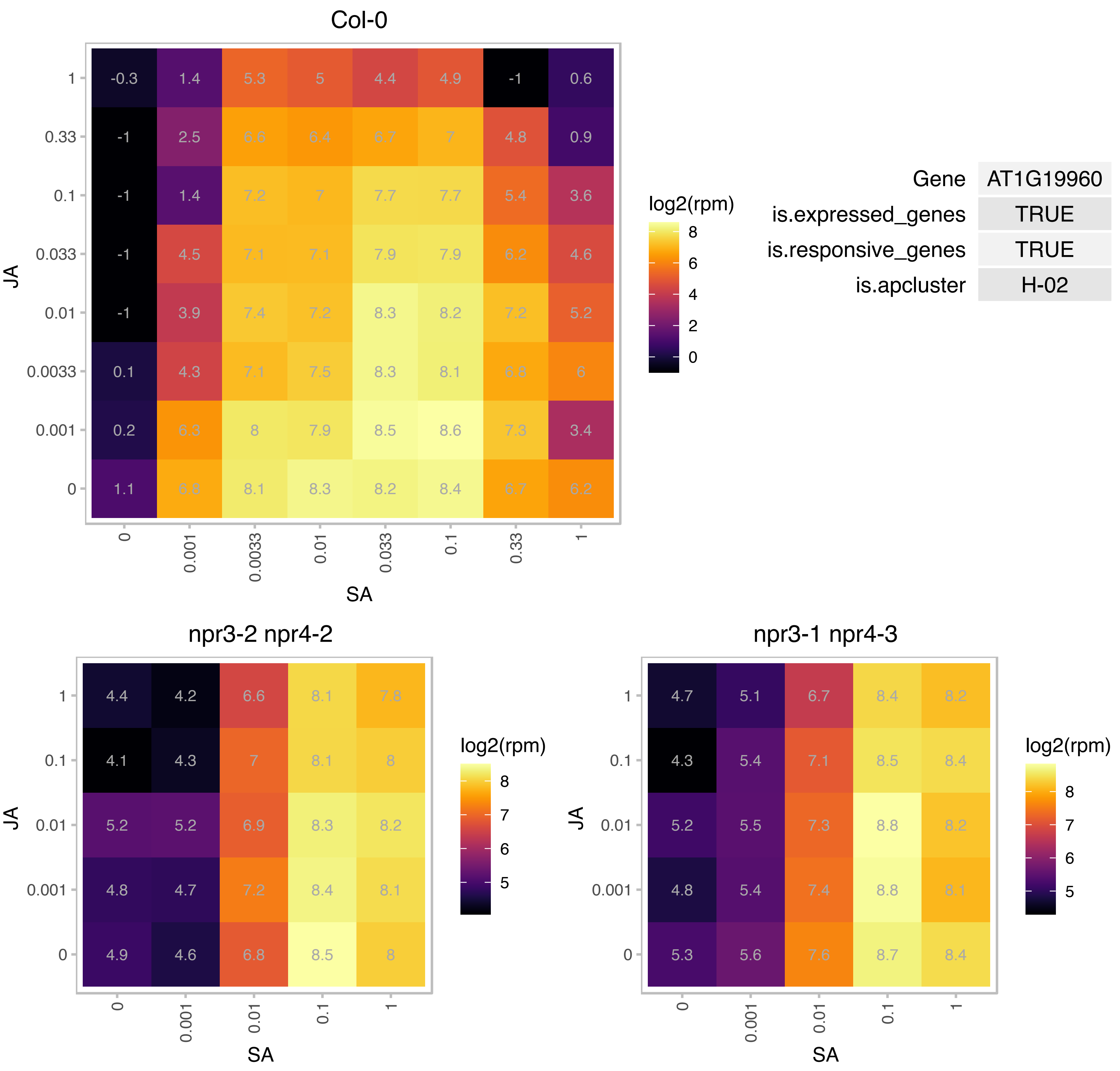


| | |
|---|---|
| Gene | AT1G19960 |
| is.expressed_genes | TRUE |
| is.responsive_genes | TRUE |
| is.apcluster | H-02 |

### AT1G21250.1 : AtWAK1, PRO25, WAK1

cell wall-associated kinase

Encodes a cell wall-associated kinase that interacts with AtGRP3 and may function as a signaling receptor of extracellular matrix component such as oligogalacturonides. The mRNA is cell-to-cell mobile.

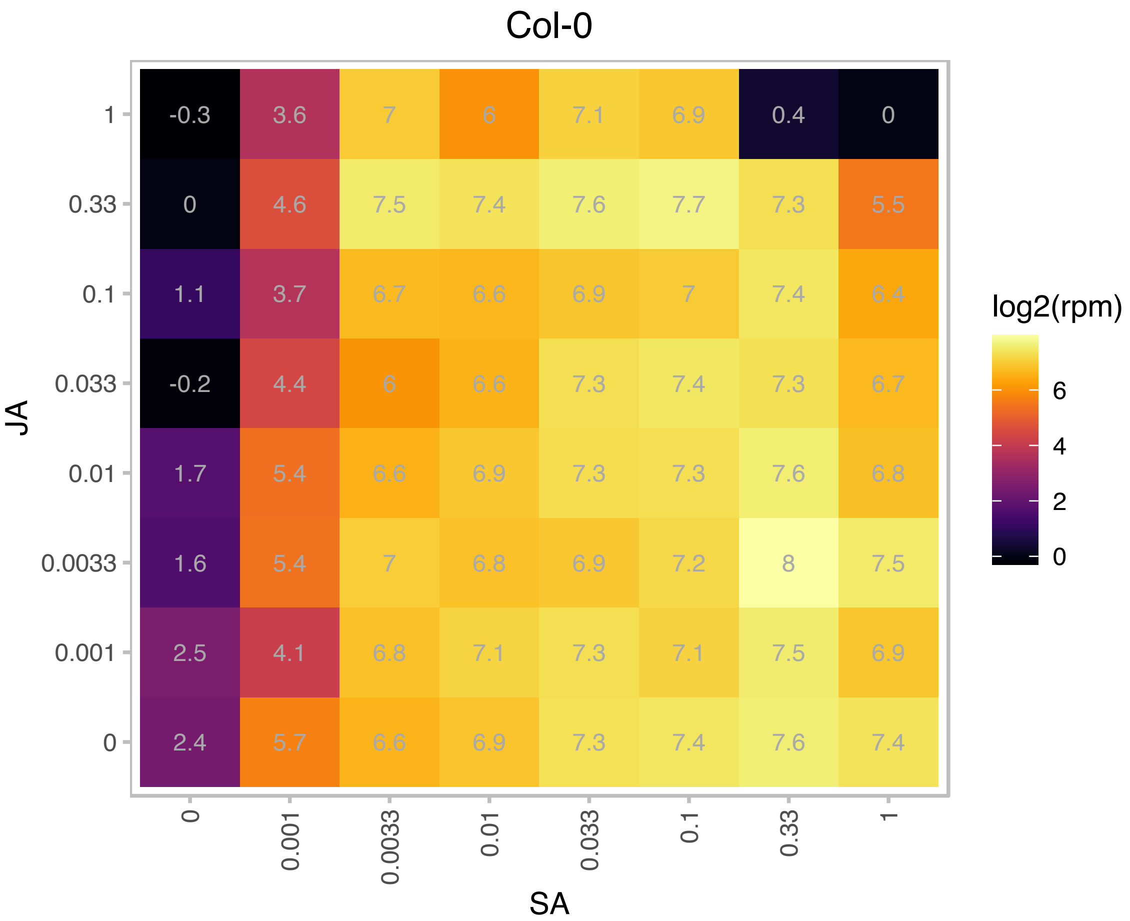


| | |
|---|---|
| Gene | AT1G21250 |
| is.expressed_genes | TRUE |
| is.responsive_genes | TRUE |
| is.apcluster | H-02 |

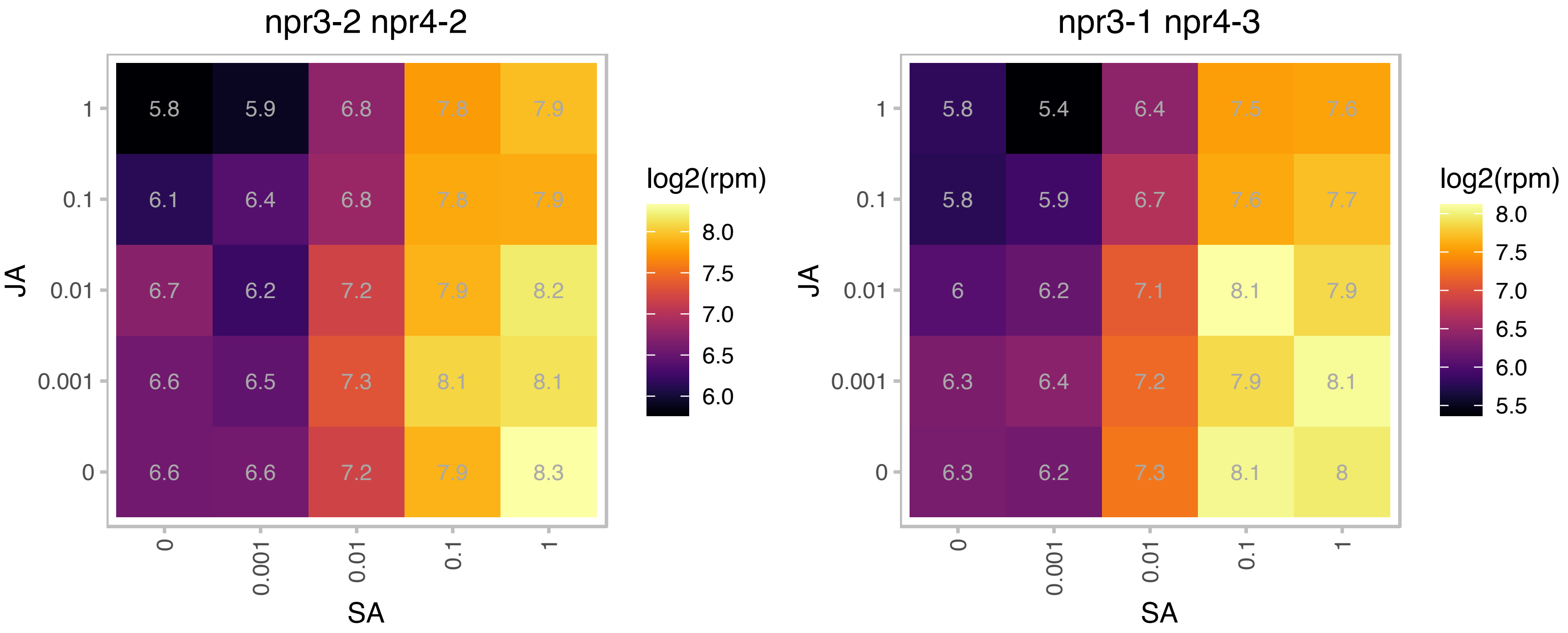

# AT1G24147.1 : NA

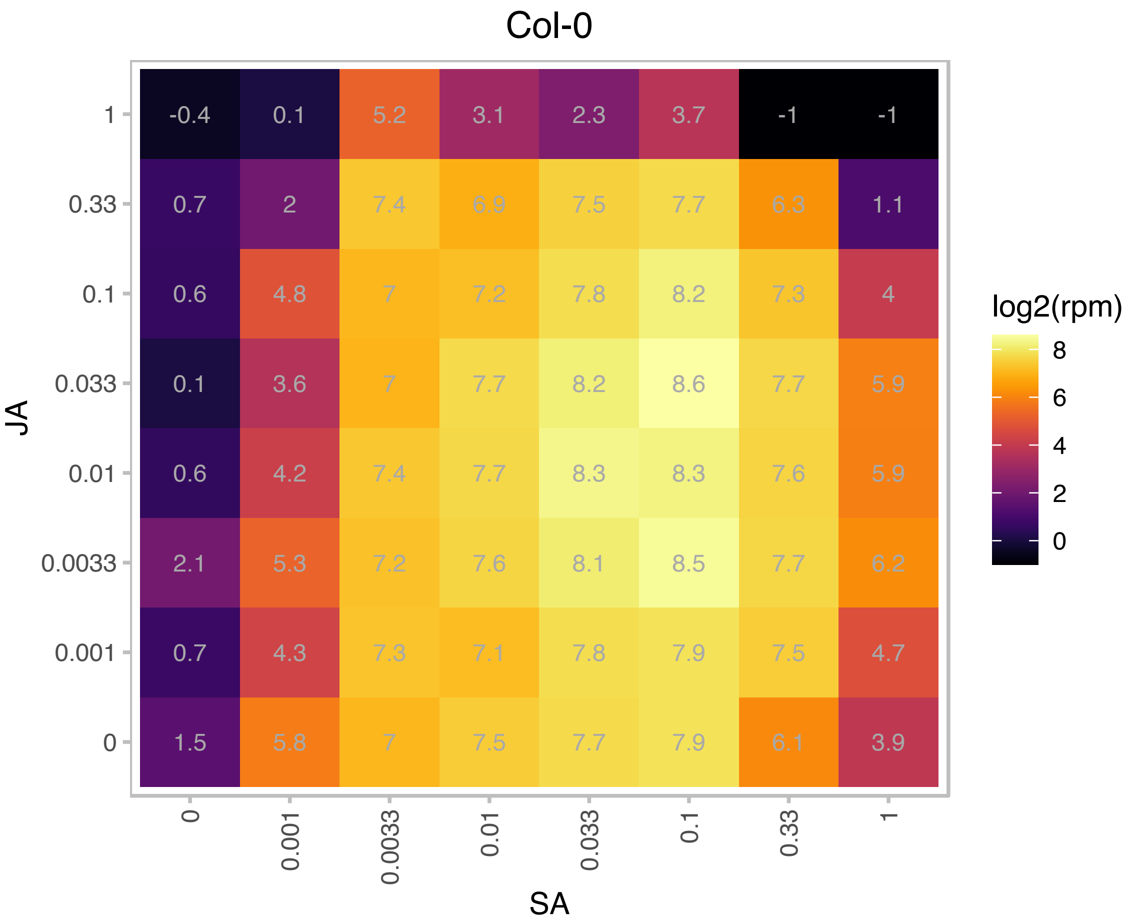


| | |
|---|---|
| Gene | AT1G24147 |
| is.expressed_genes | TRUE |
| is.responsive_genes | TRUE |
| is.apcluster | H-02 |

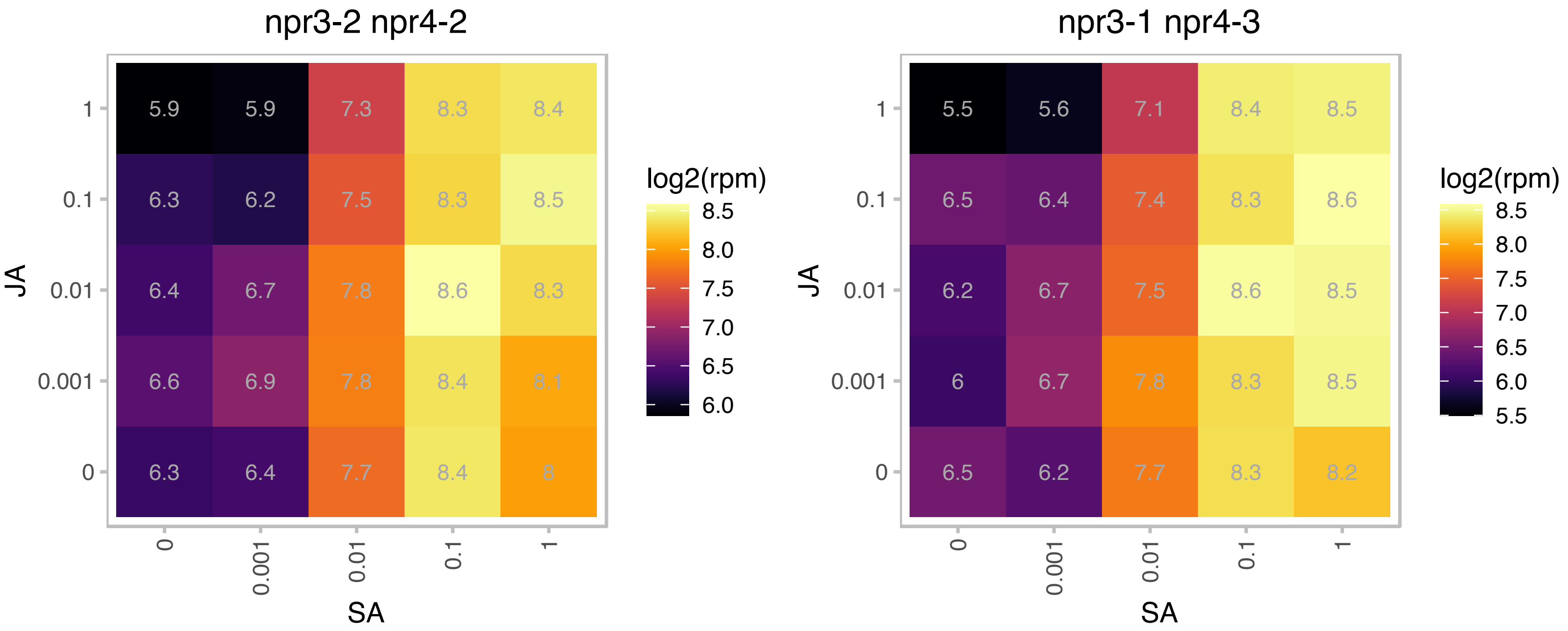

# AT1G35710.1 : NA

Protein kinase family protein with leucine-rich repeat domain

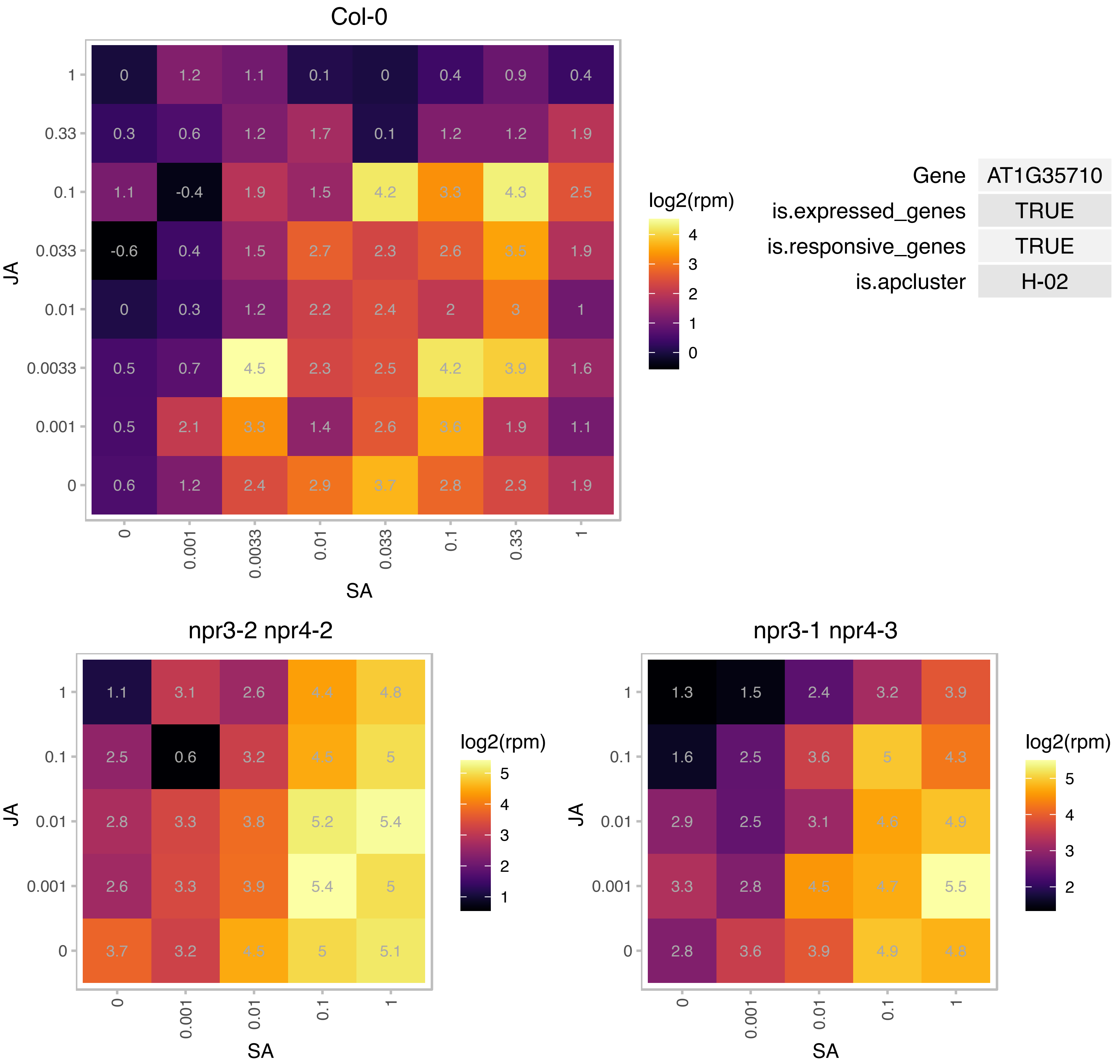


| | |
|---|---|
| Gene | AT1G35710 |
| is.expressed_genes | TRUE |
| is.responsive_genes | TRUE |
| is.apcluster | H-02 |

# AT1G49750.1 : NA

Leucine-rich repeat (LRR) family protein

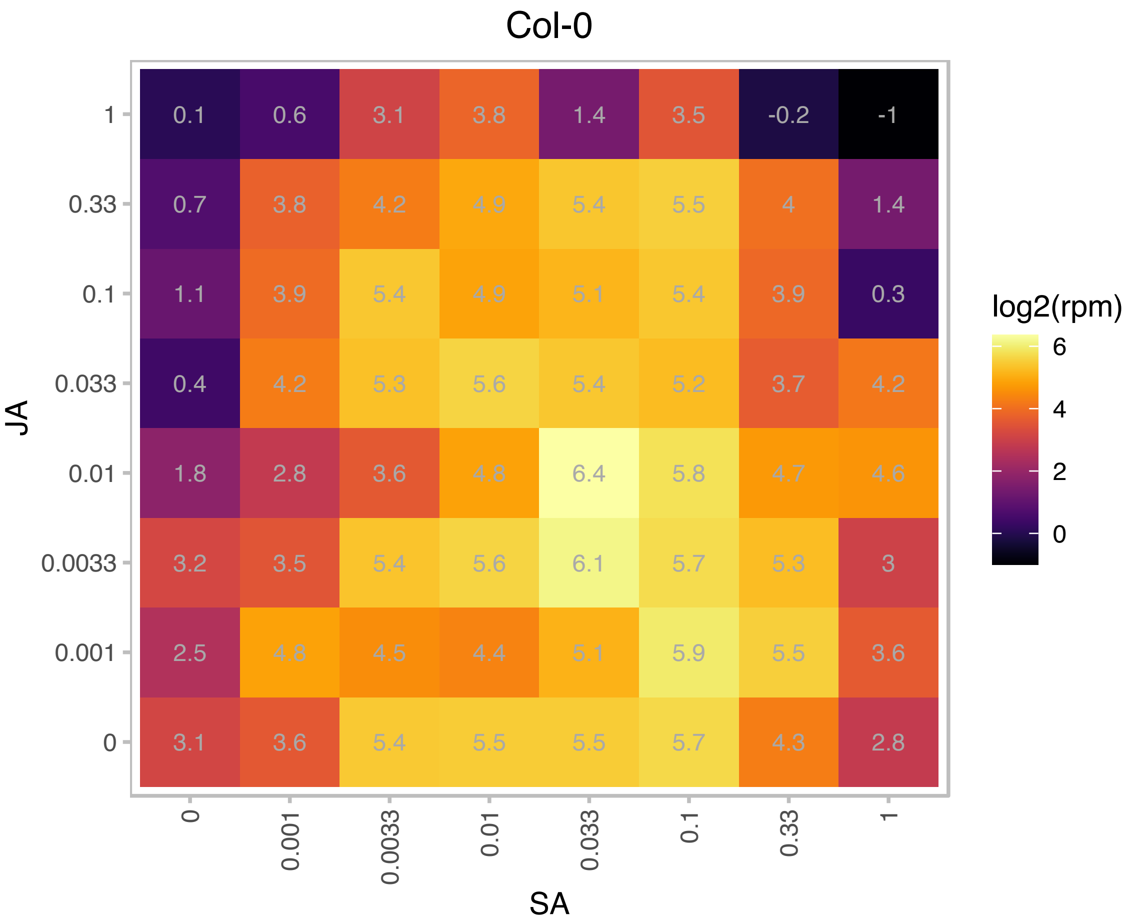


| | |
|---|---|
| Gene | AT1G49750 |
| is.expressed_genes | TRUE |
| is.responsive_genes | TRUE |
| is.apcluster | H-02 |

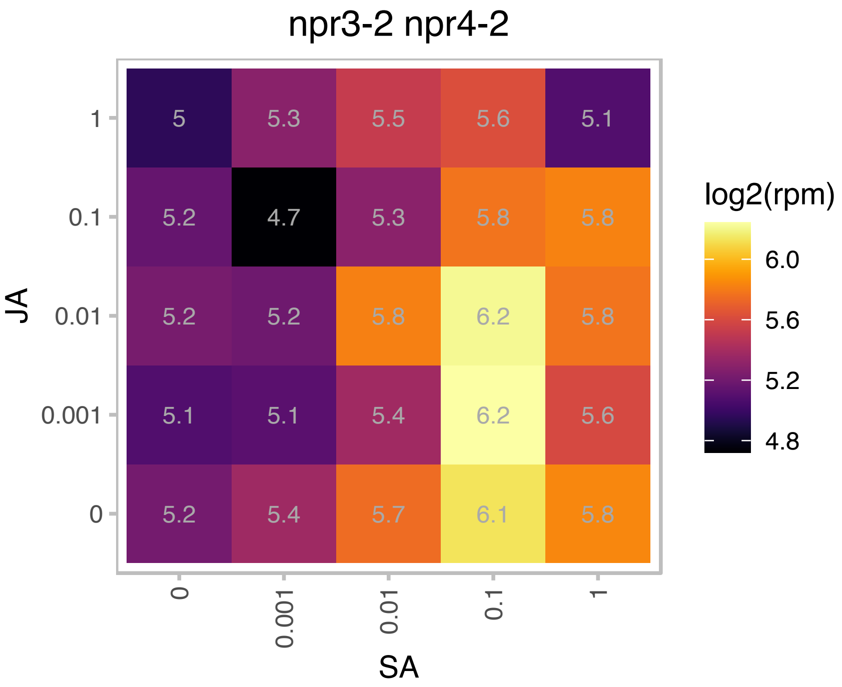


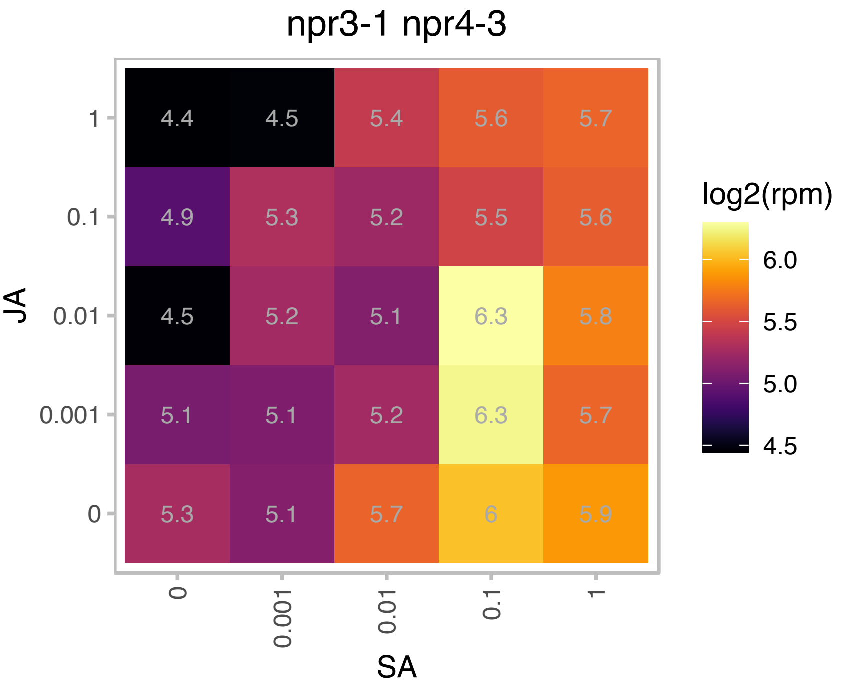

# AT1G61170.1 : NA

| | |
|---|---|
| Gene | AT1G61170 |
| is.expressed_genes | TRUE |
| is.responsive_genes | TRUE |
| is.apcluster | H-02 |

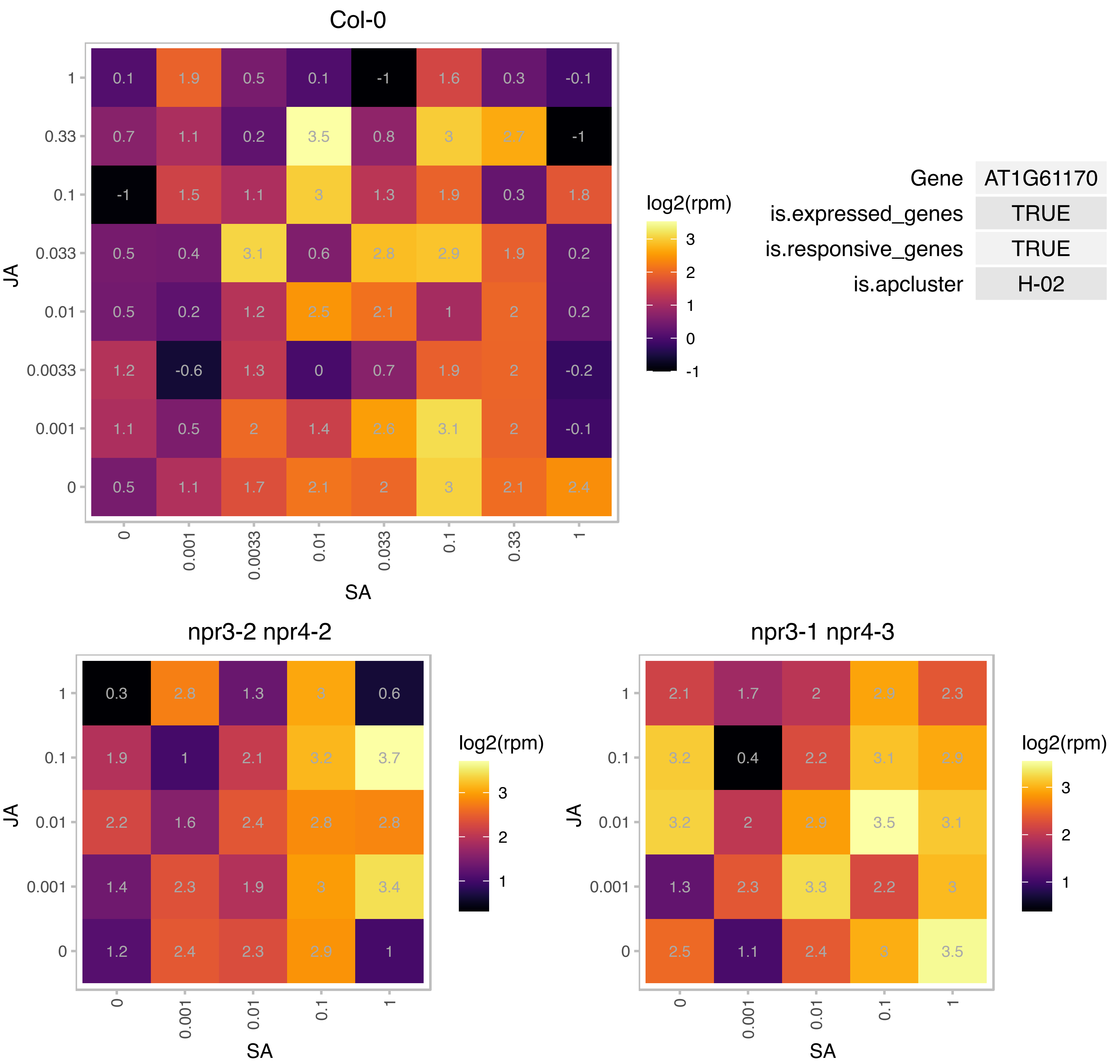

### AT1G65800.1 : ARK2, AtARK2, RK2

receptor kinase 2

Encodes a putative receptor-like serine/threonine protein kinases that is similar to brassica self-incompatibility (S) locus. expressed in specifically in cotyledons, leaves, and sepals, in correlation with the maturation of these structures. Together with AtPUB9, it is required for auxin-mediated lateral root development under phosphate-starved conditions. The mRNA is cell-to-cell mobile.

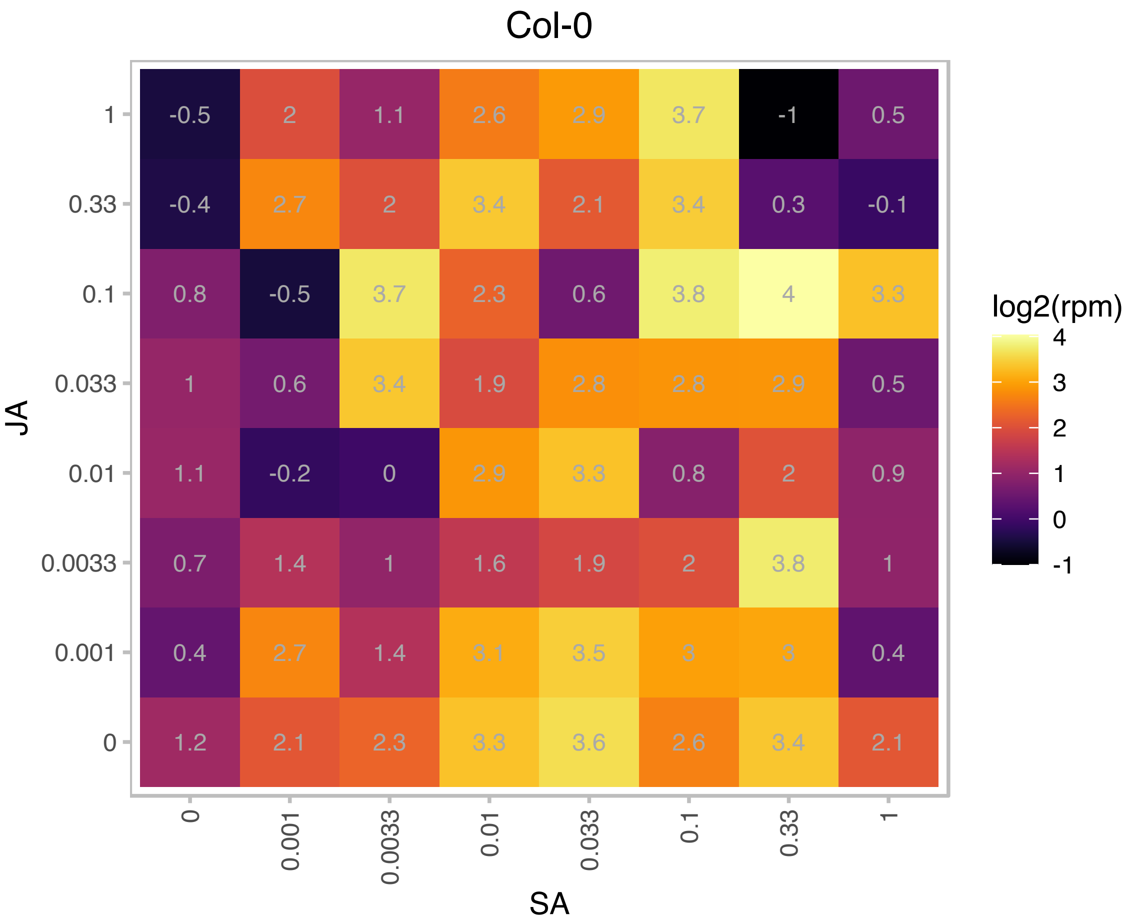


| | |
|---|---|
| Gene | AT1G65800 |
| is.expressed_genes | TRUE |
| is.responsive_genes | TRUE |
| is.apcluster | H-02 |

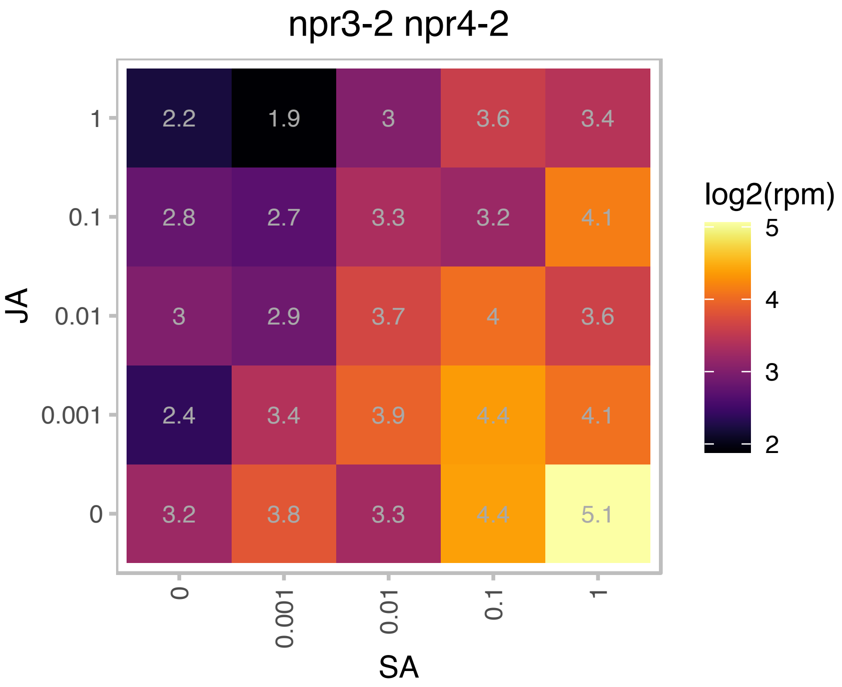


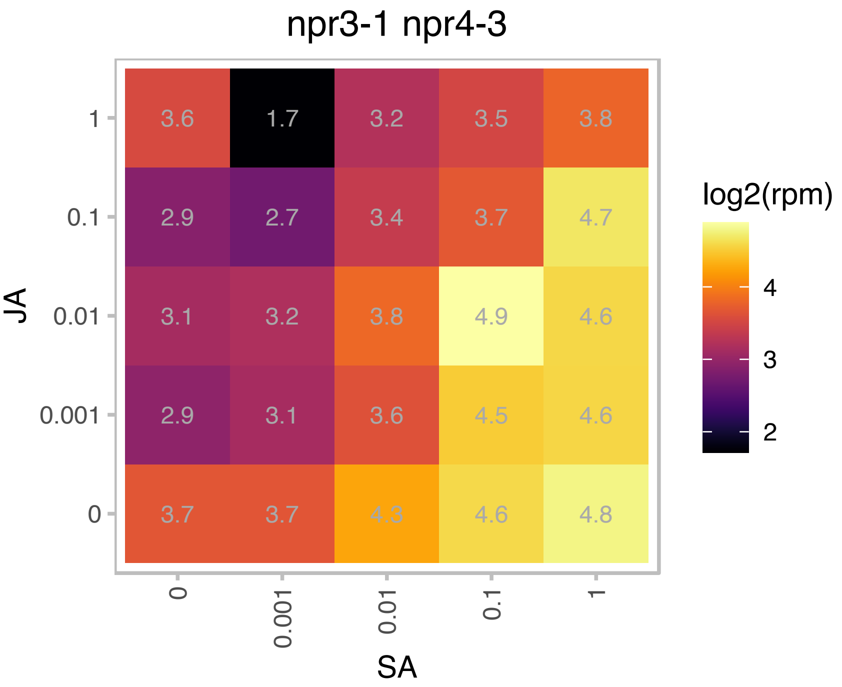

### AT1G66970.1 : GDPDL1, SVL2

SHV3-like 2

Encodes a member of the glycerophosphodiester phosphodiesterase like (GDPD-like) family.

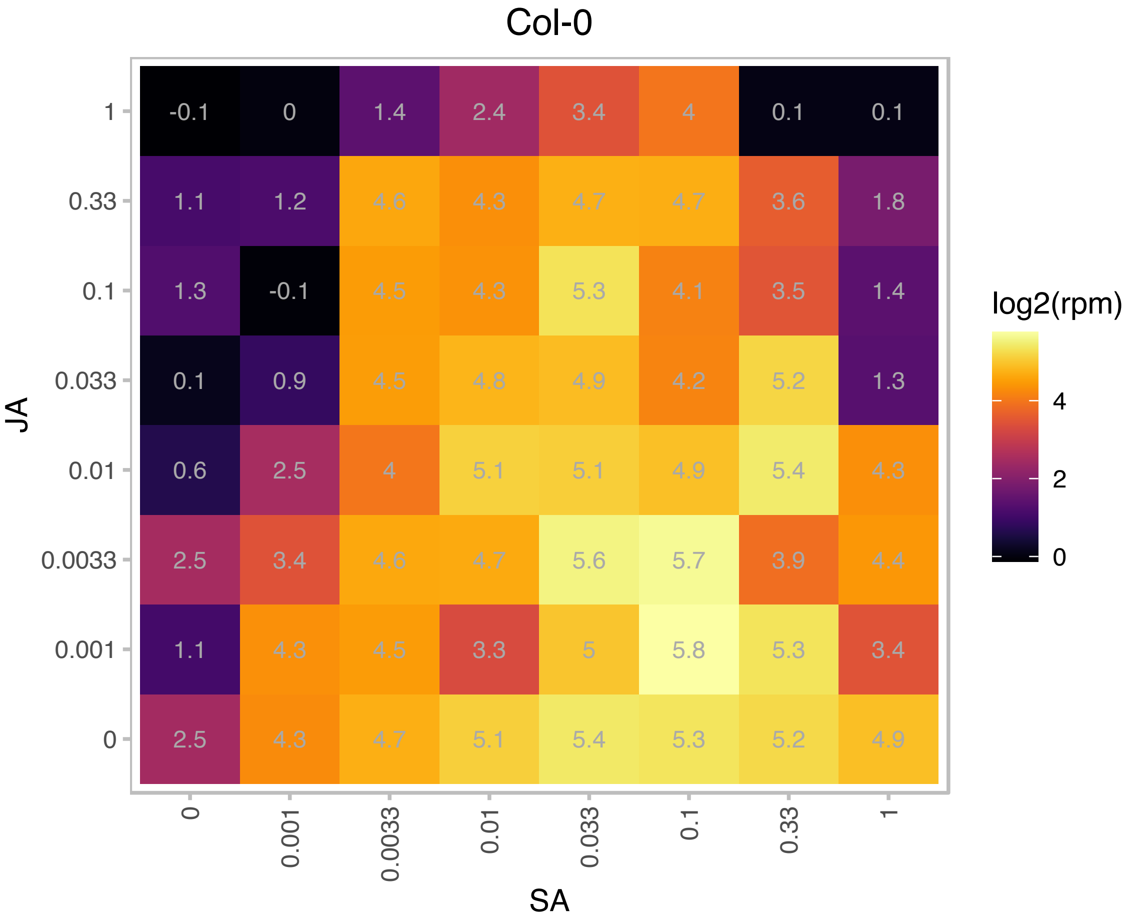


| | |
|---|---|
| Gene | AT1G66970 |
| is.expressed_genes | TRUE |
| is.responsive_genes | TRUE |
| is.apcluster | H-02 |

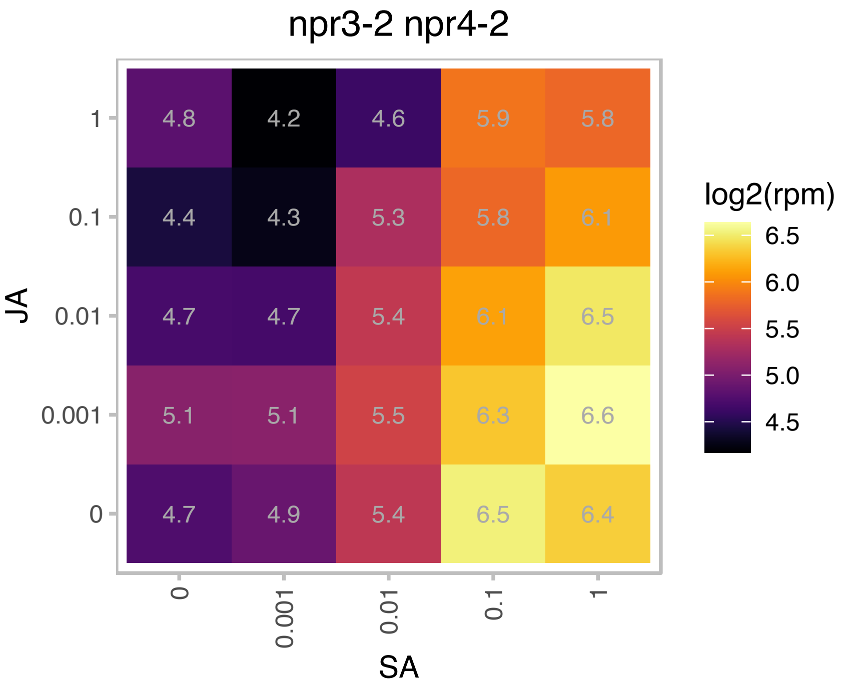


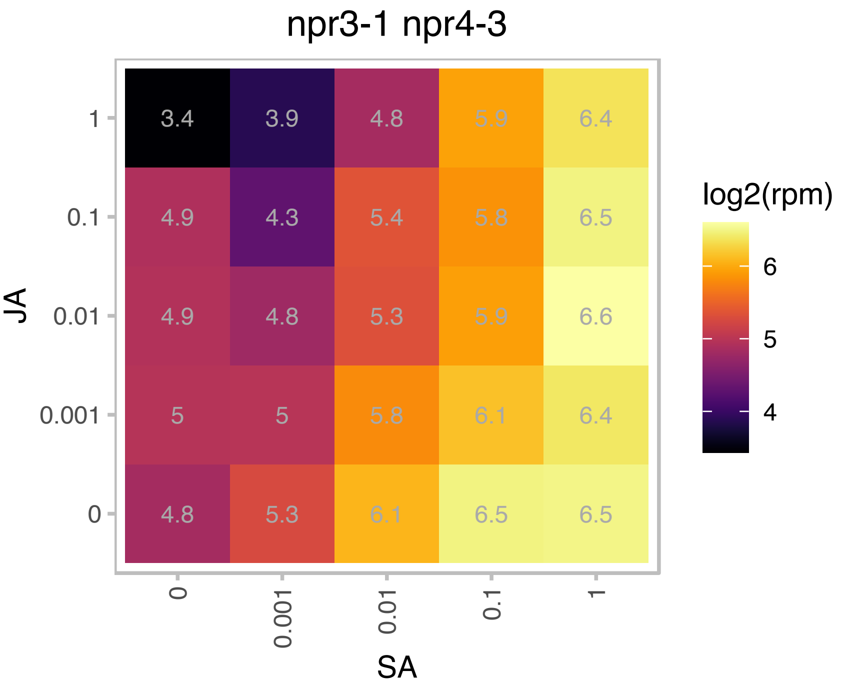

# AT1G67148.1 : NA

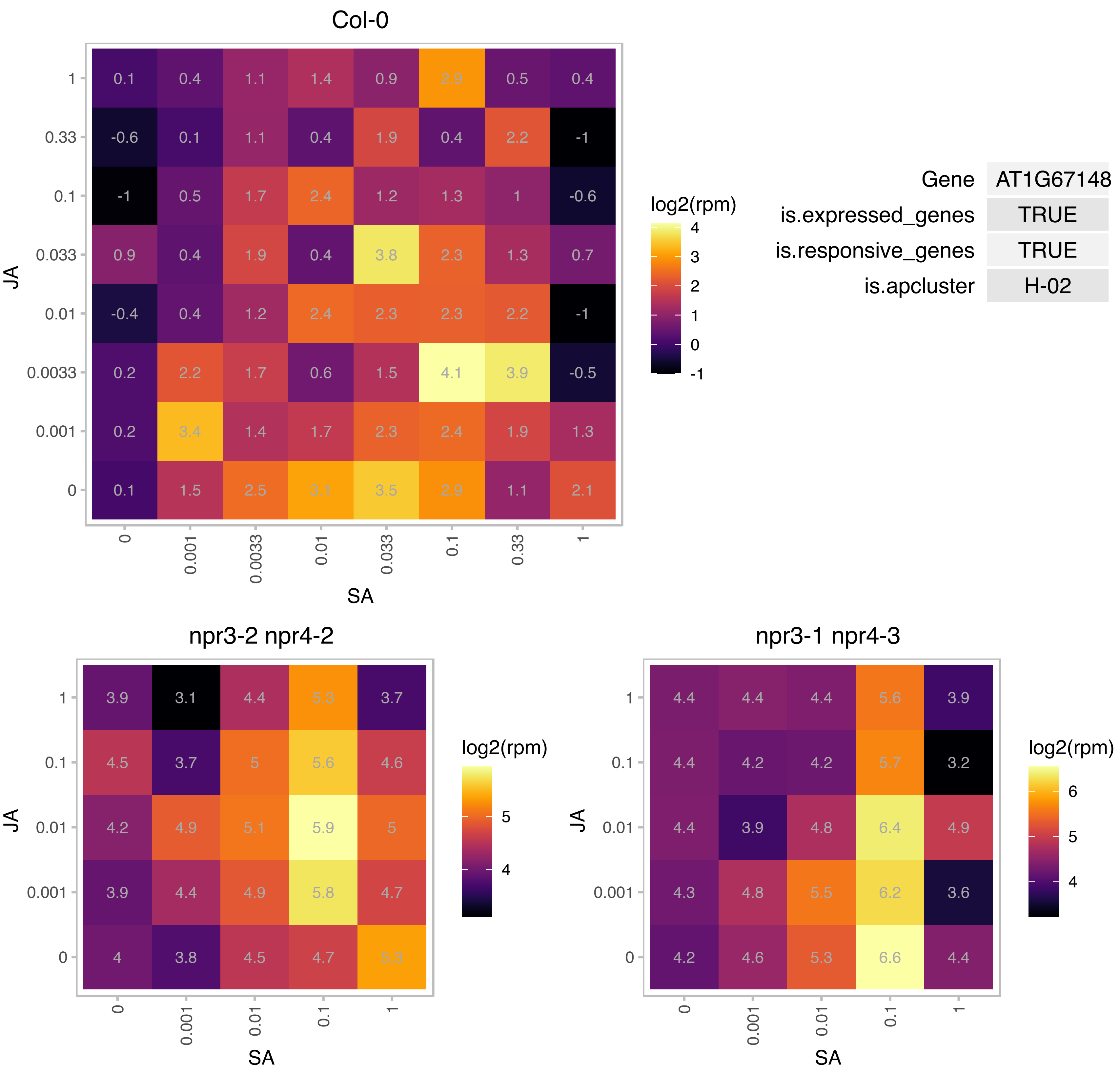

### AT1G79380.1 : RGLG4

Ca(2)-dependent phospholipid-binding protein (Copine) family

Encodes a ubiquitin ligase that is an essential upstream modulator of JA signaling in response to various stimuli.

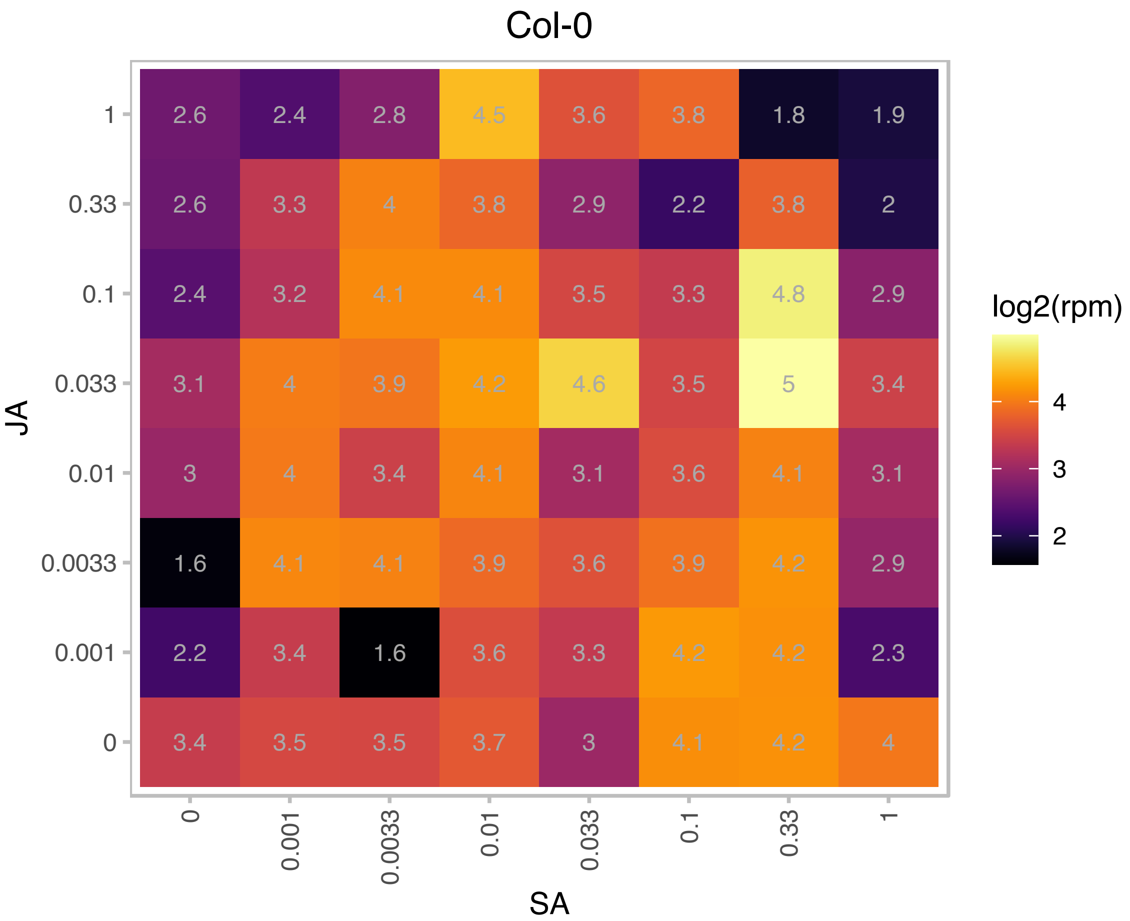


| | |
|---|---|
| Gene | AT1G79380 |
| is.expressed_genes | TRUE |
| is.responsive_genes | TRUE |
| is.apcluster | H-02 |

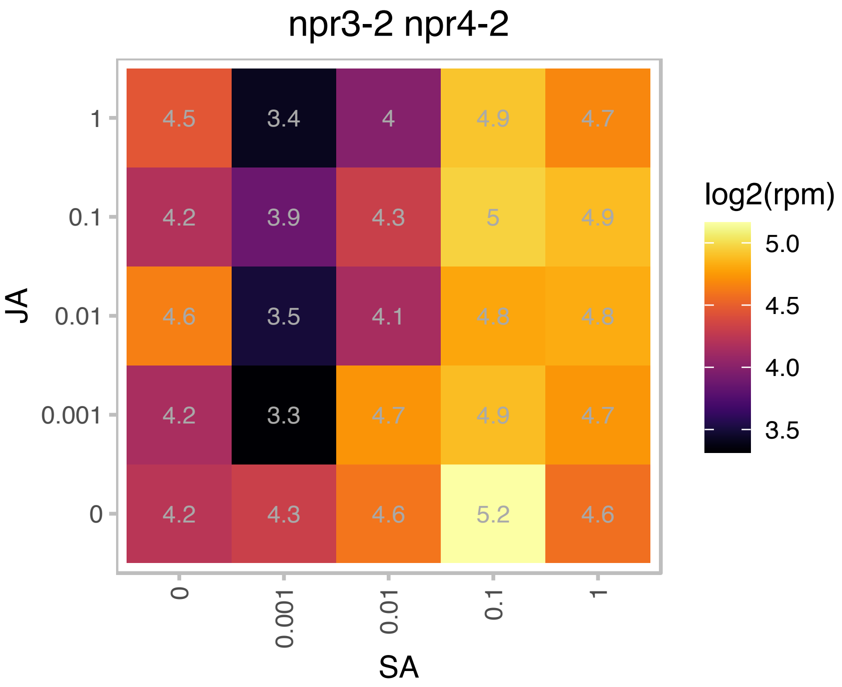


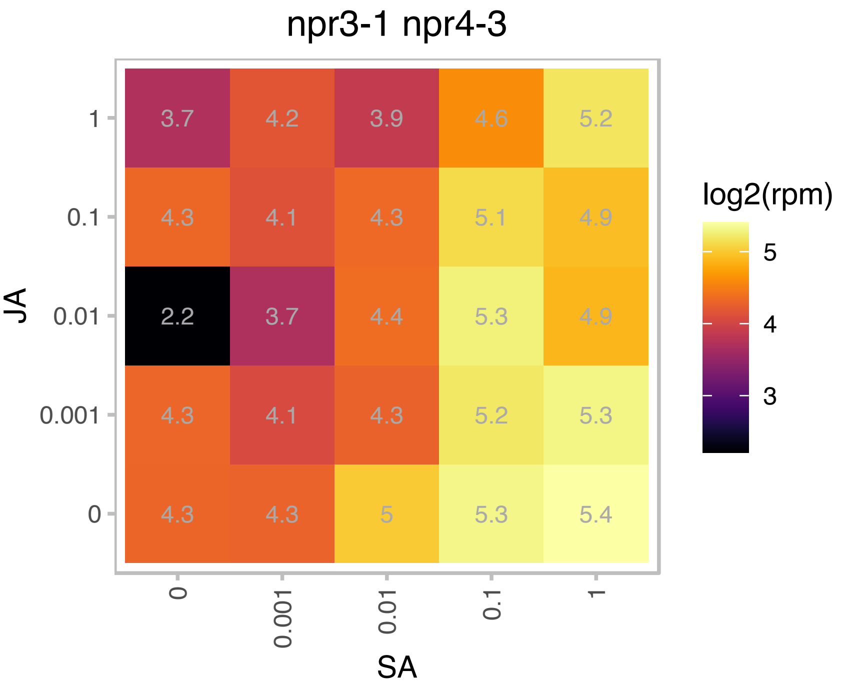

### AT2G03680.1 : SKU6, SPR1

spiral1

The SPR1 gene encodes a plant-specific 12-kD protein which has a repeated motif at both ends, separated by a predicted rod-like domain, suggesting that it may act as an intermolecular linker. Ubiquitously expressed and belongs to a six-member gene family in Arabidopsis; expressed in transgenic seedlings localized to microtubules within the cortical array, preprophase band, phragmoplast, and mitotic spindle.

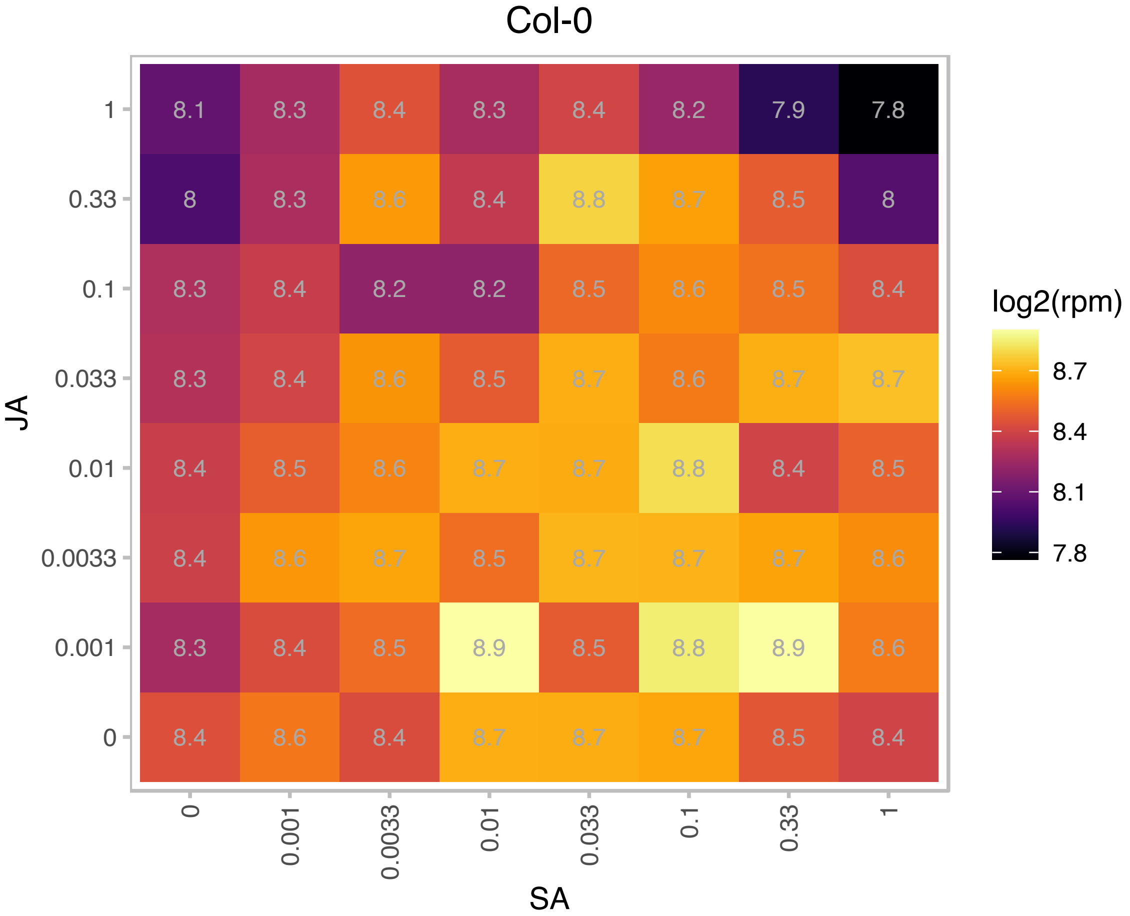


| | |
|---|---|
| Gene | AT2G03680 |
| is.expressed_genes | TRUE |
| is.responsive_genes | TRUE |
| is.apcluster | H-02 |

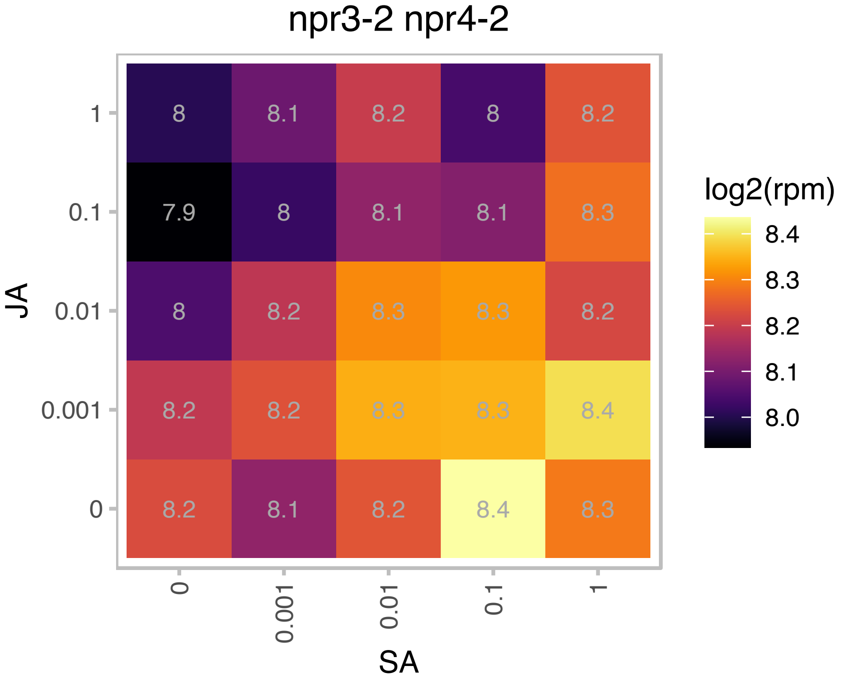


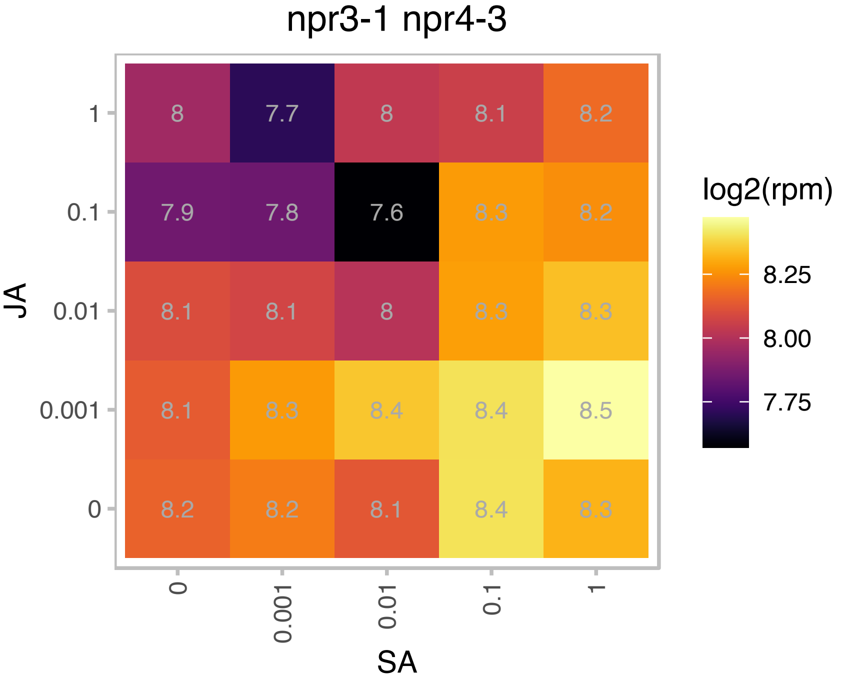

# AT2G04900.1 : NA

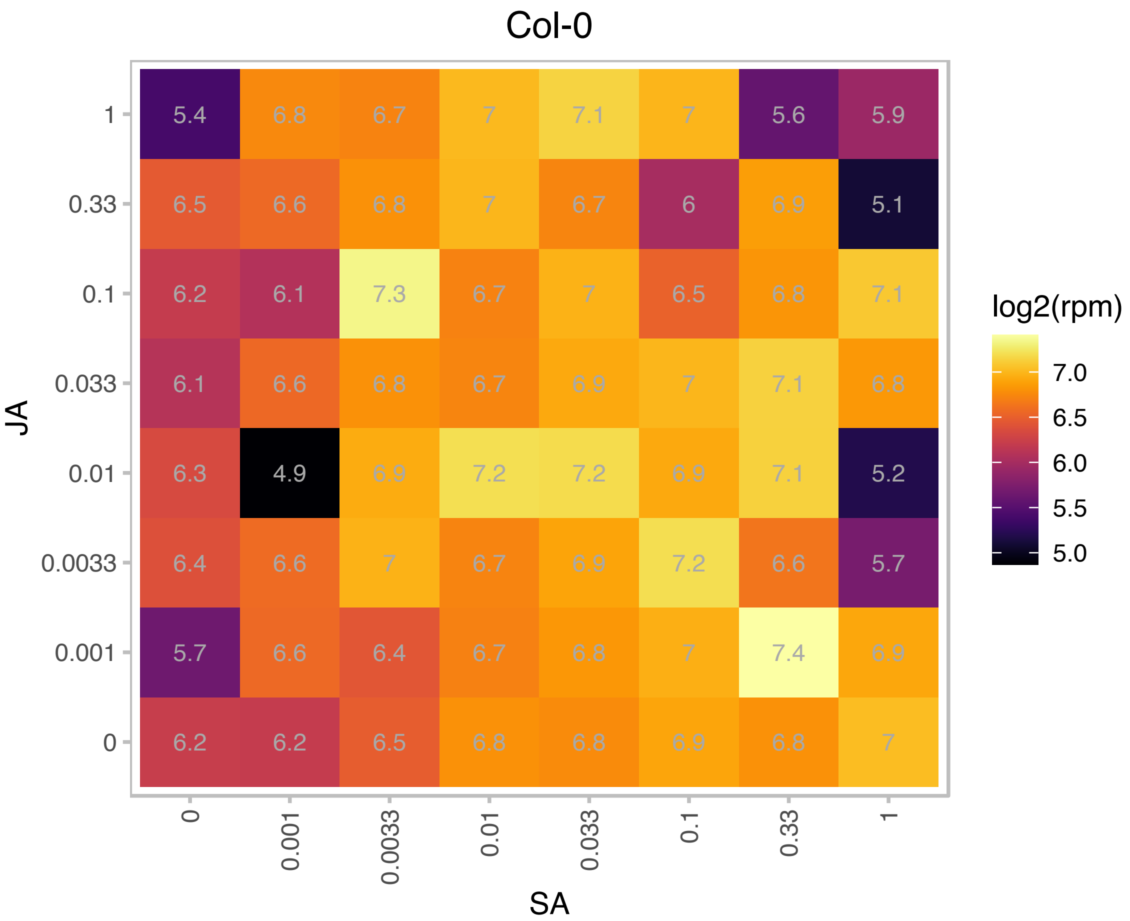


| | |
|---|---|
| Gene | AT2G04900 |
| is.expressed_genes | TRUE |
| is.responsive_genes | TRUE |
| is.apcluster | H-02 |

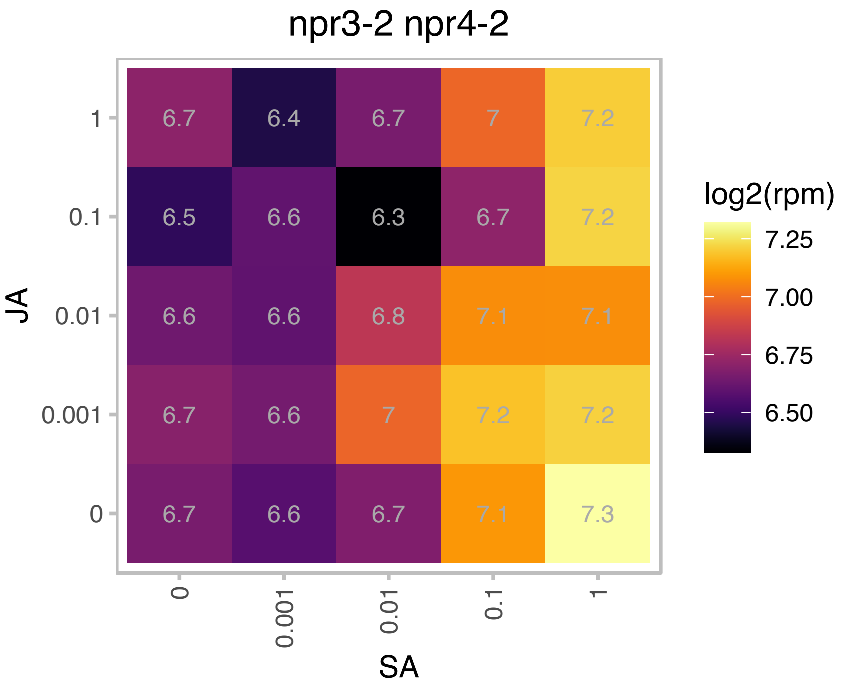


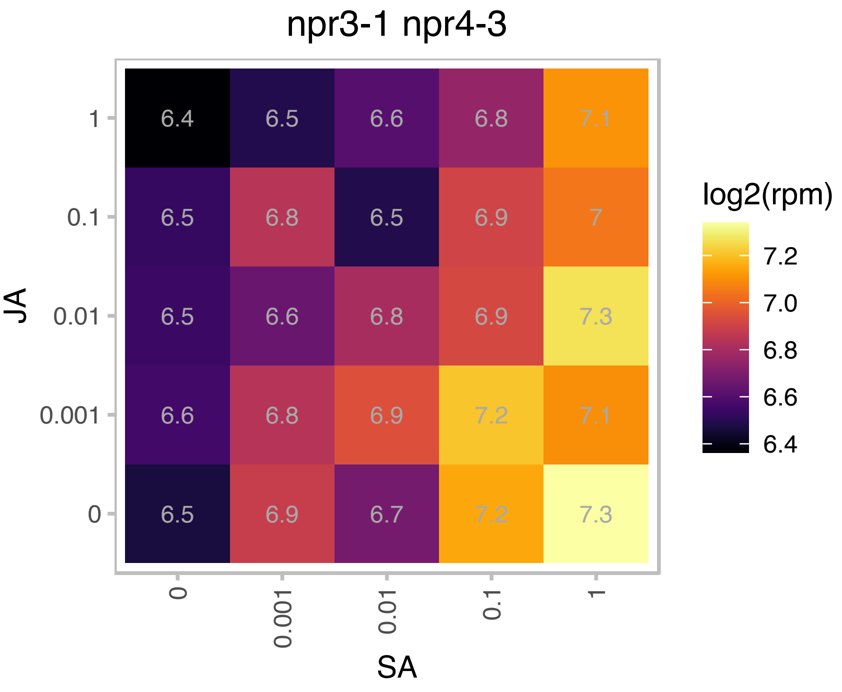

### AT2G16430.1 : ATPAP10, PAP10

purple acid phosphatase 10

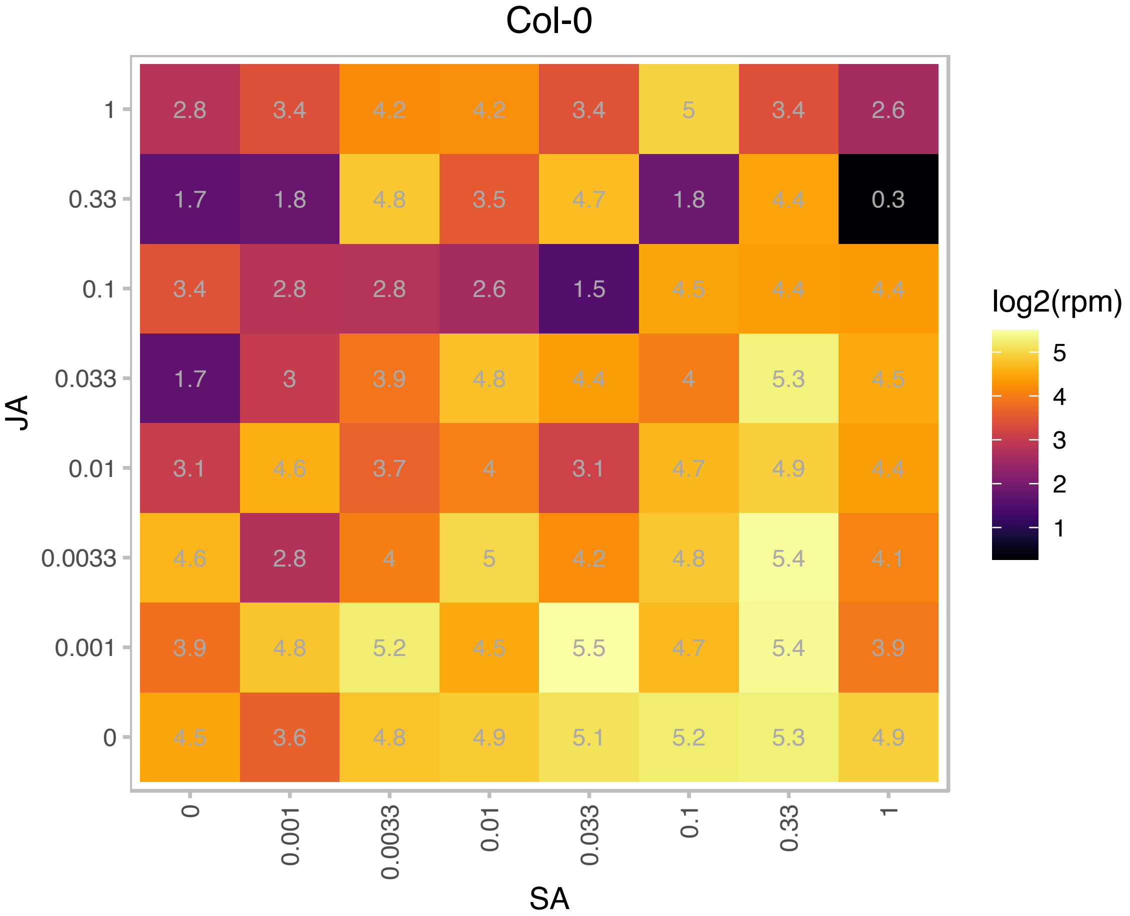


| Gene | AT2G16430 |
|---|---|
| is.expressed_genes | TRUE |
| is.responsive_genes | TRUE |
| is.apcluster | H-02 |

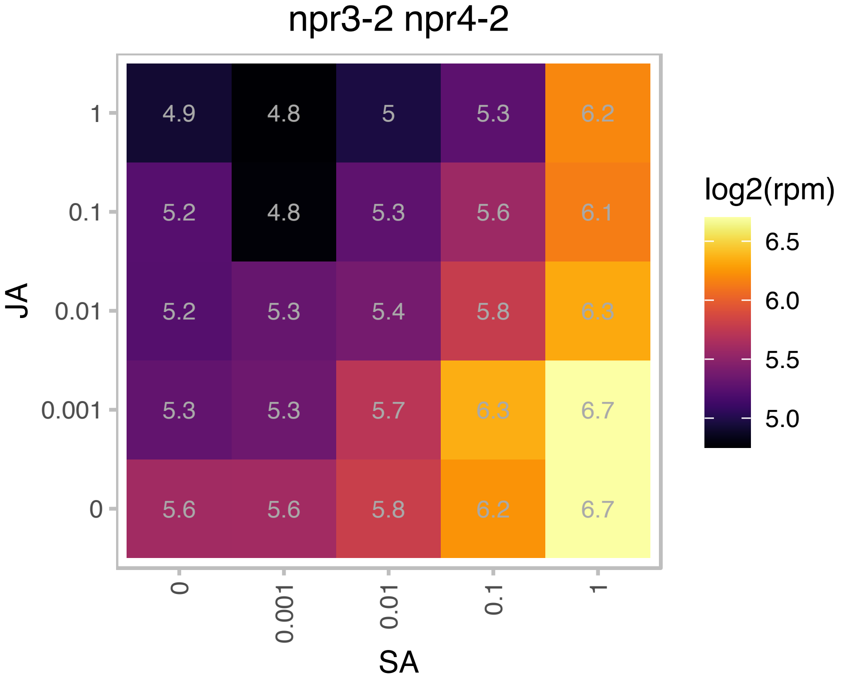


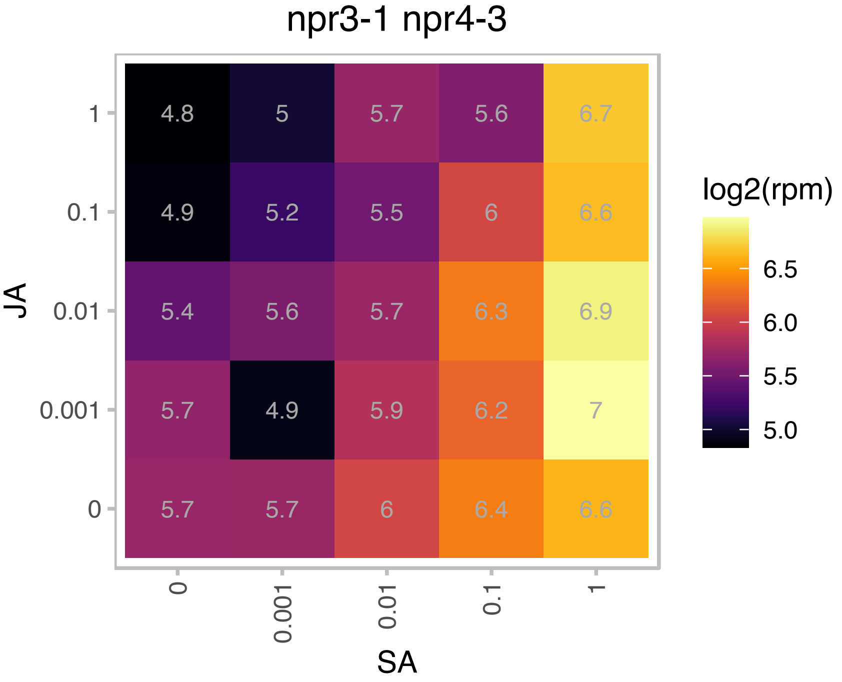

### AT2G16600.1 : AtCYP19-1, CYP19, ROC3

rotamase CYP 3

Encodes cytosolic cyclophilin ROC3. The mRNA is cell-to-cell mobile.

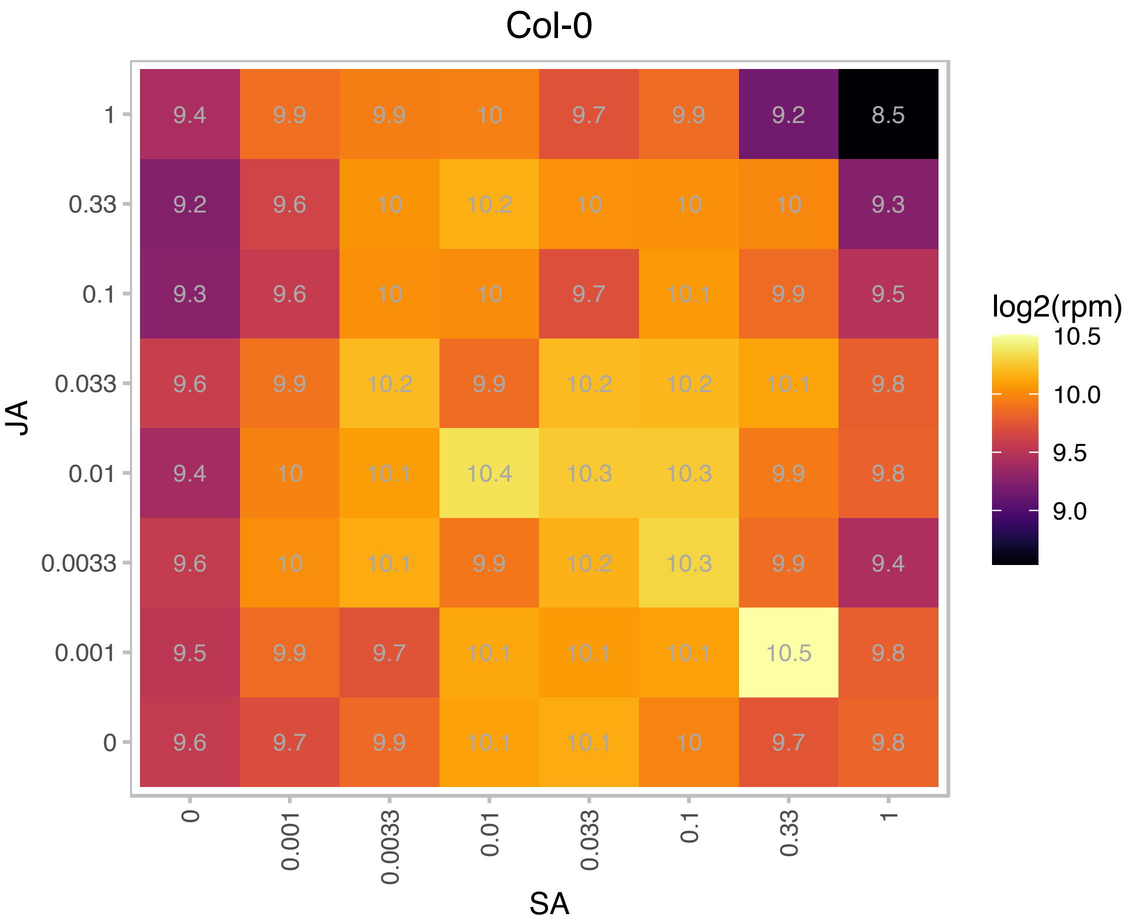


| | |
|---|---|
| Gene | AT2G16600 |
| is.expressed_genes | TRUE |
| is.responsive_genes | TRUE |
| is.apcluster | H-02 |

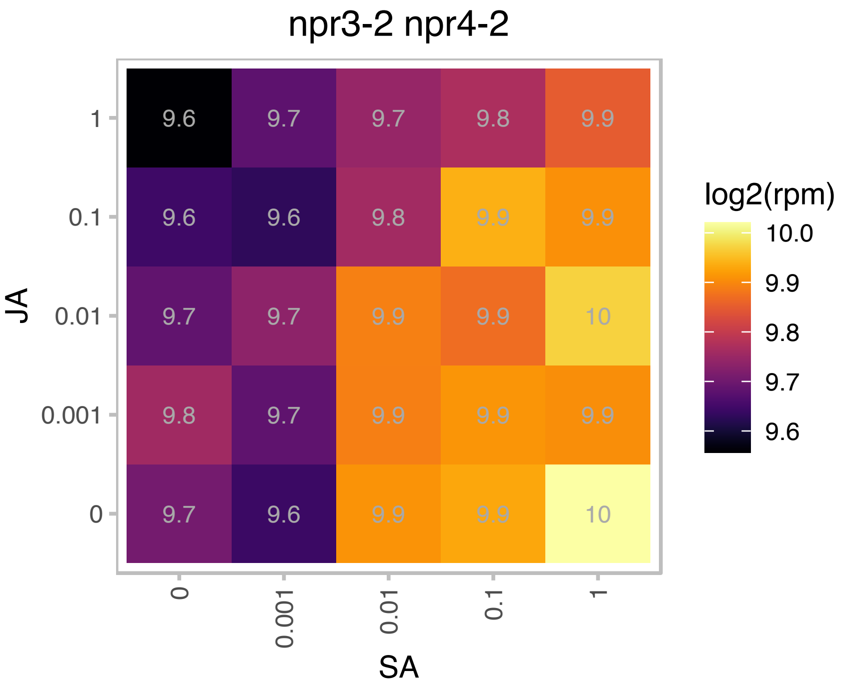


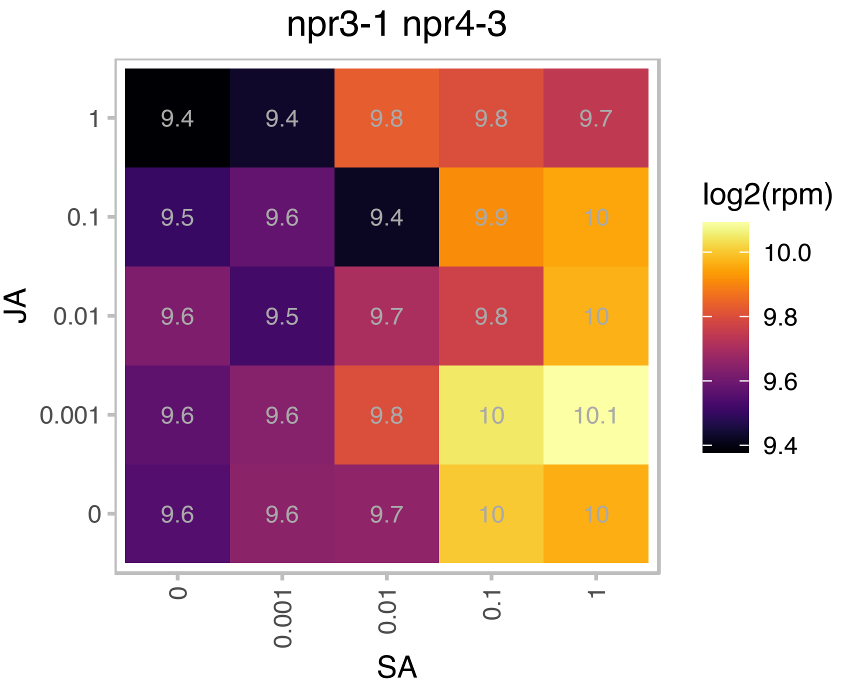

### AT2G25000.1 : ATWRKY60, WRKY60

WRKY DNA-binding protein 60

Pathogen-induced transcription factor. Forms protein complexes with itself and with WRKY40. Coexpression with WRKY18 or WRKY40 made plants more susceptible to both P. syringae and B. cinerea. WRKY18, WRKY40, and W have partially redundant roles in response to the hemibiotrophic bacterial pathogen Pseudomonas syringae and the necrotrophic fungal pathogen Botrytis cinerea, with WRKY18 playing a more important role than the other two.

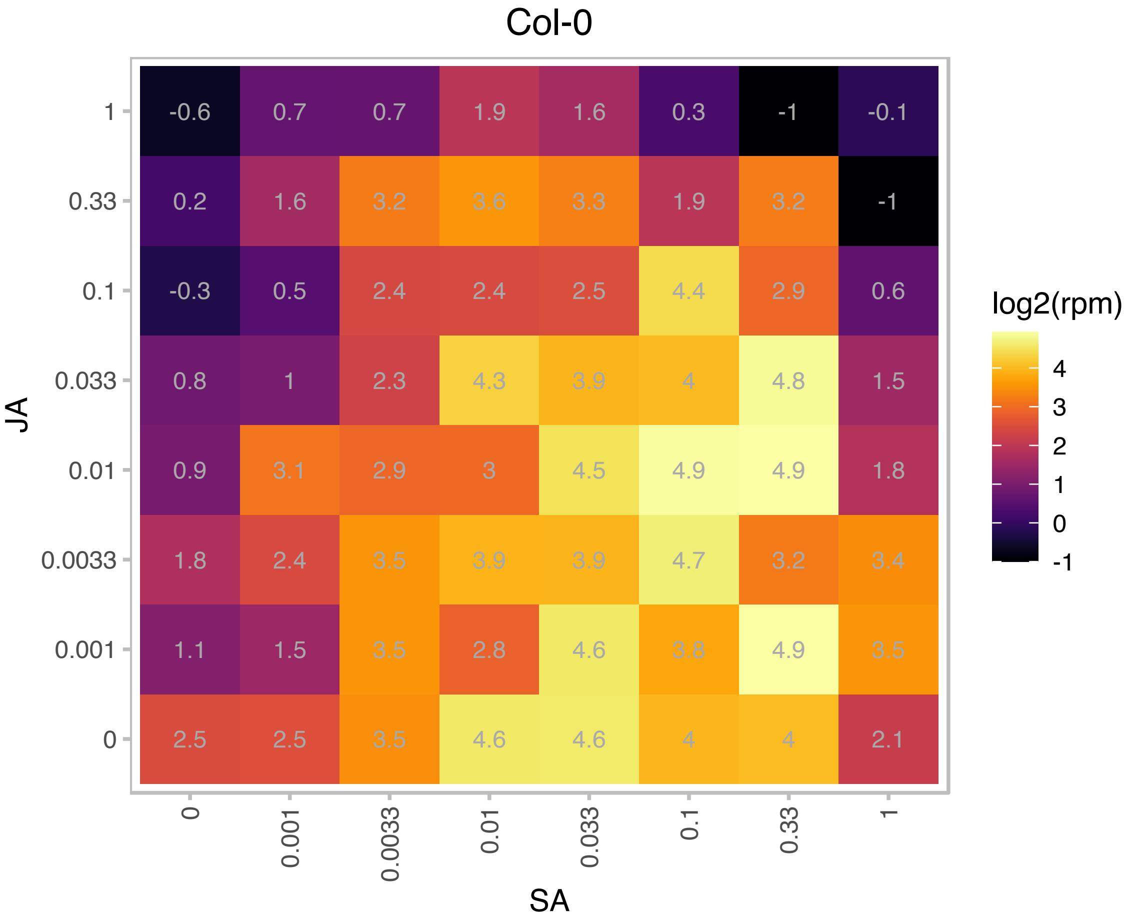


| | |
|---|---|
| Gene | AT2G25000 |
| is.expressed_genes | TRUE |
| is.responsive_genes | TRUE |
| is.apcluster | H-02 |

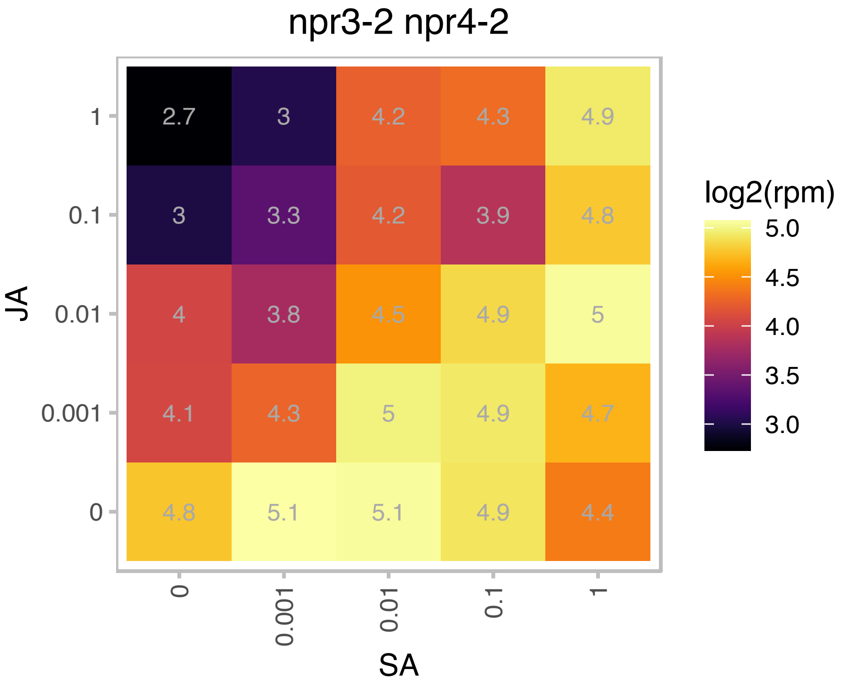


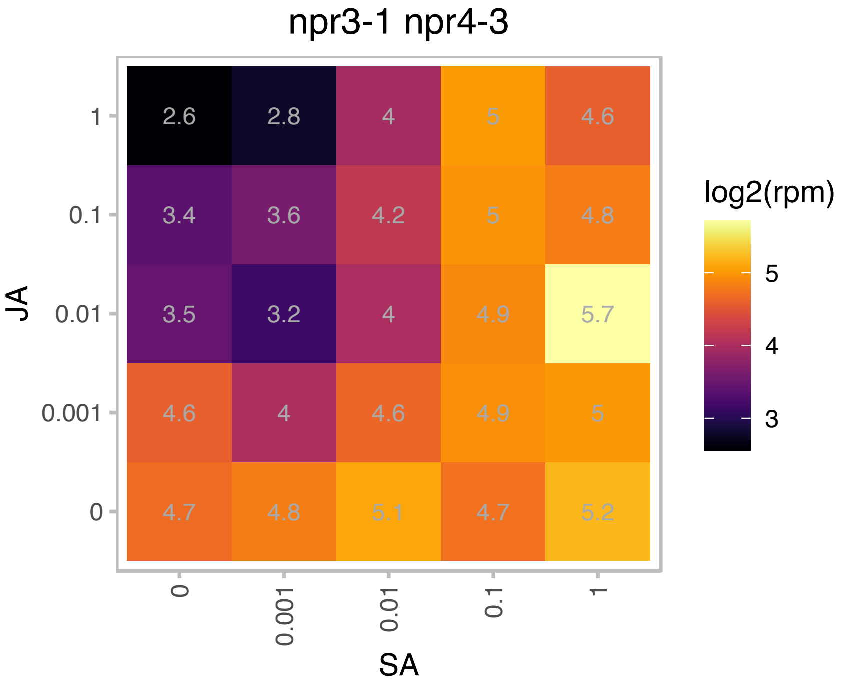

### AT2G26440.1 : PME12

Plant invertase/pectin methylesterase inhibitor superfamily

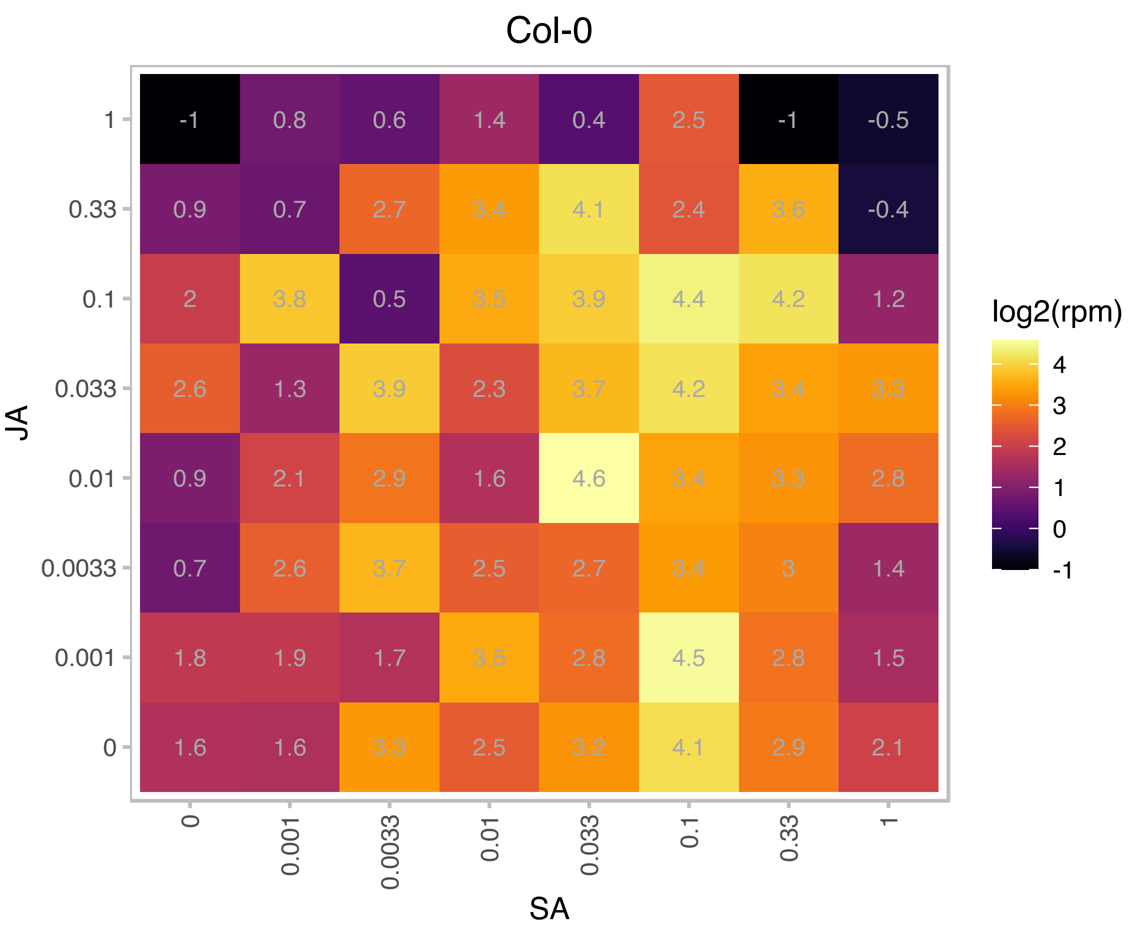


| | |
|---|---|
| Gene | AT2G26440 |
| is.expressed_genes | TRUE |
| is.responsive_genes | TRUE |
| is.apcluster | H-02 |

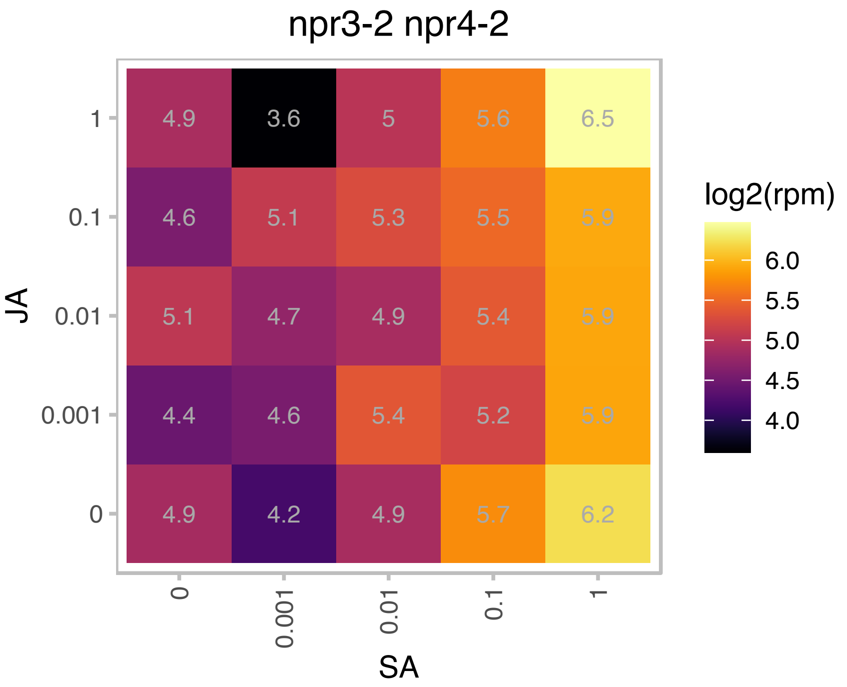


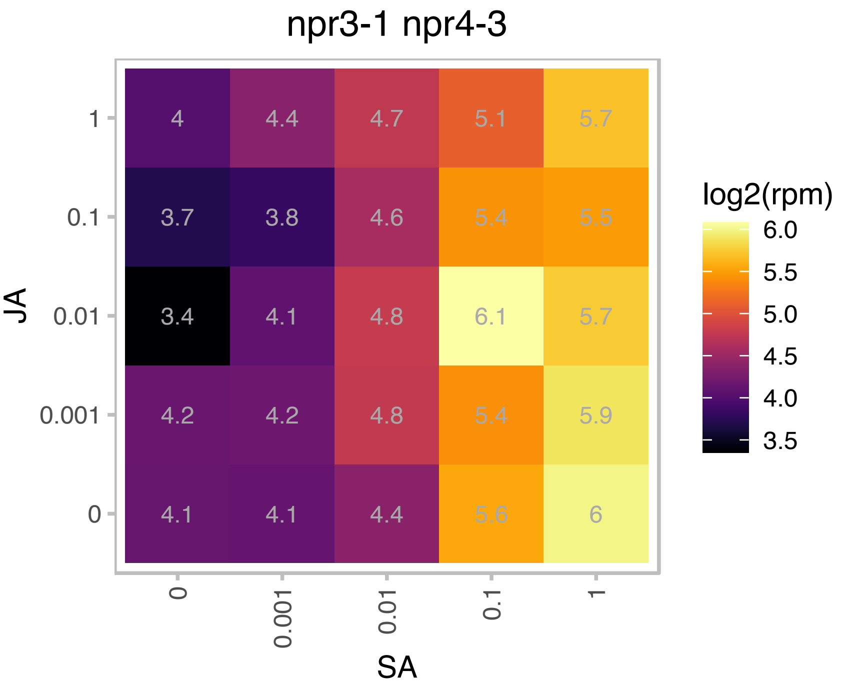

### AT2G30500.1 : NET4B

Kinase interacting (KIP1-like) family protein

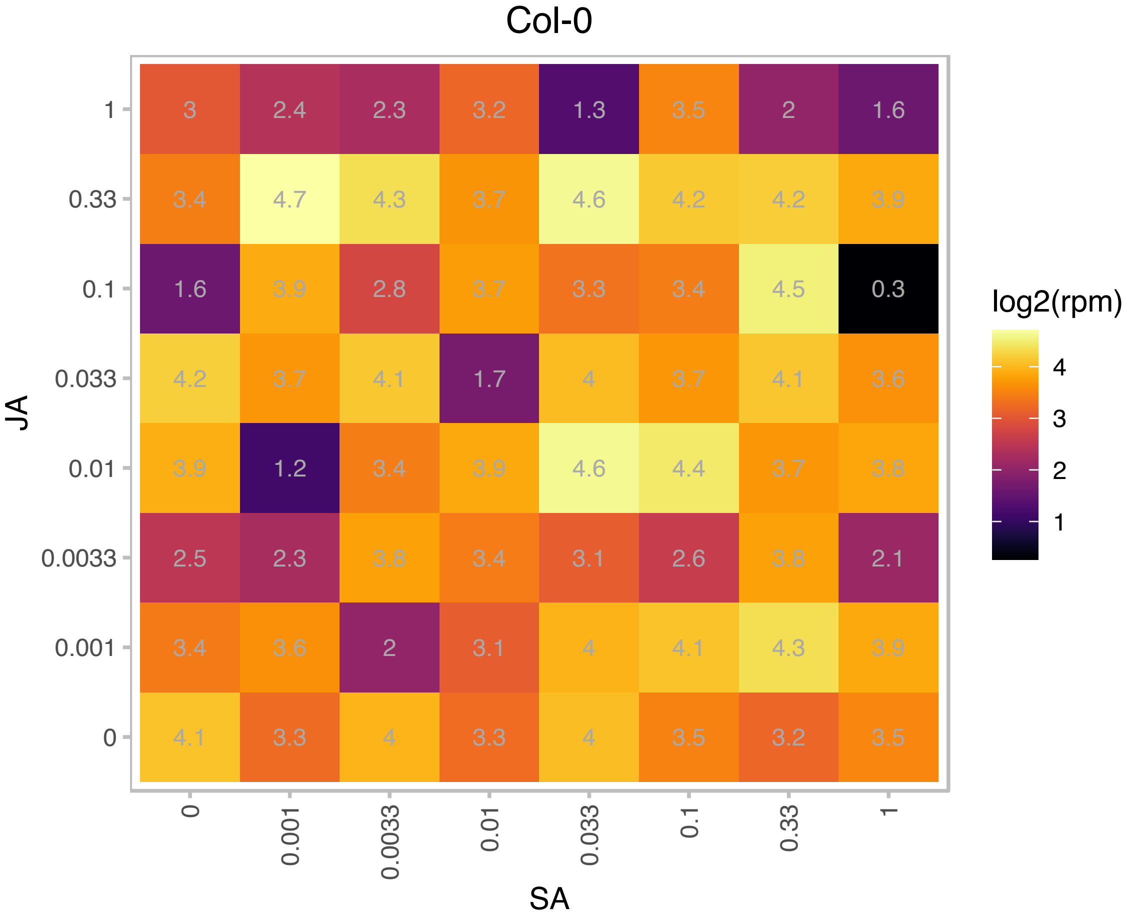


| | |
|---|---|
| Gene | AT2G30500 |
| is.expressed_genes | TRUE |
| is.responsive_genes | TRUE |
| is.apcluster | H-02 |

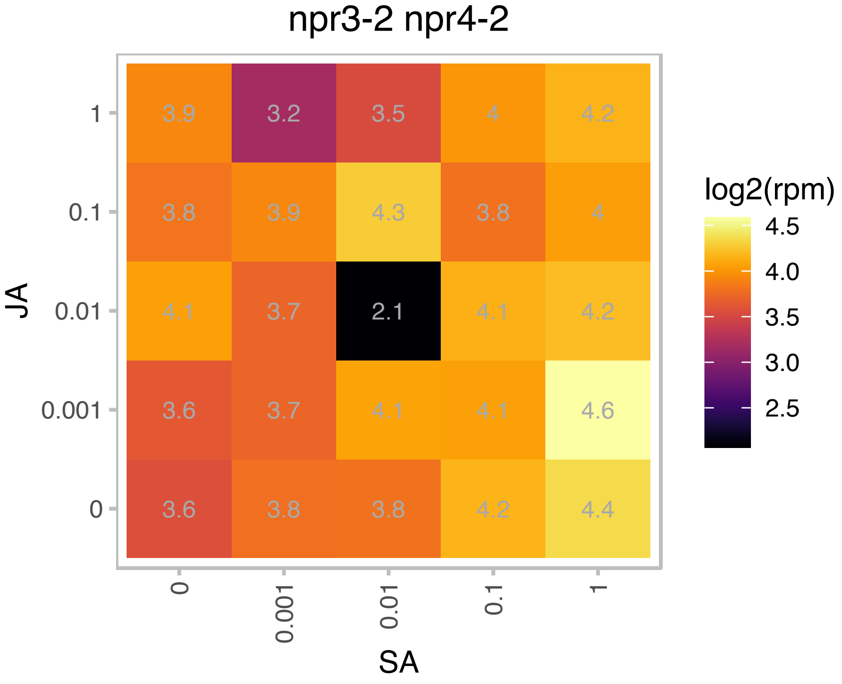


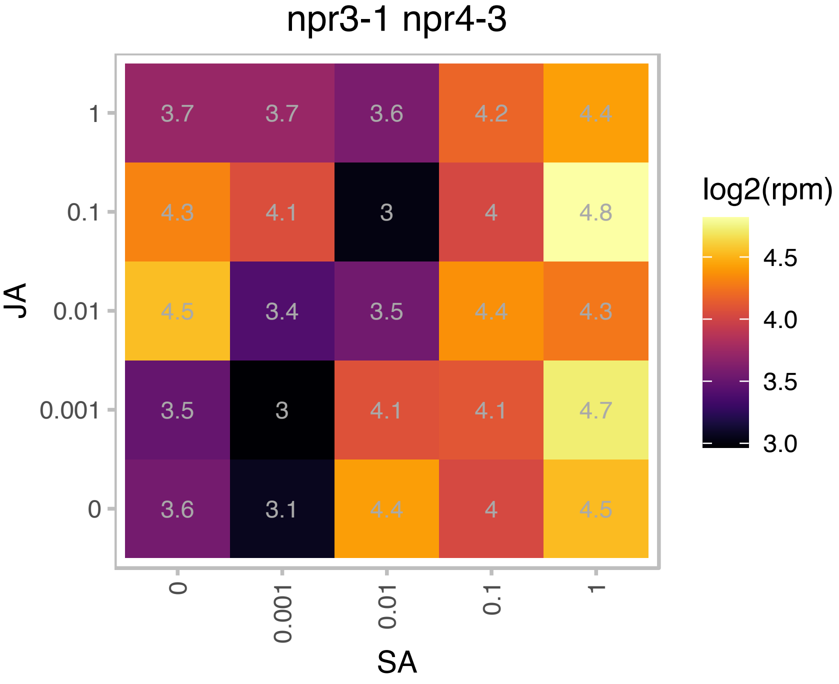

# AT2G33845.1 : NA

Nucleic acid-binding, OB-fold-like protein

| Gene | AT2G33845 |
|---|---|
| is.expressed_genes | TRUE |
| is.responsive_genes | TRUE |
| is.apcluster | H-02 |

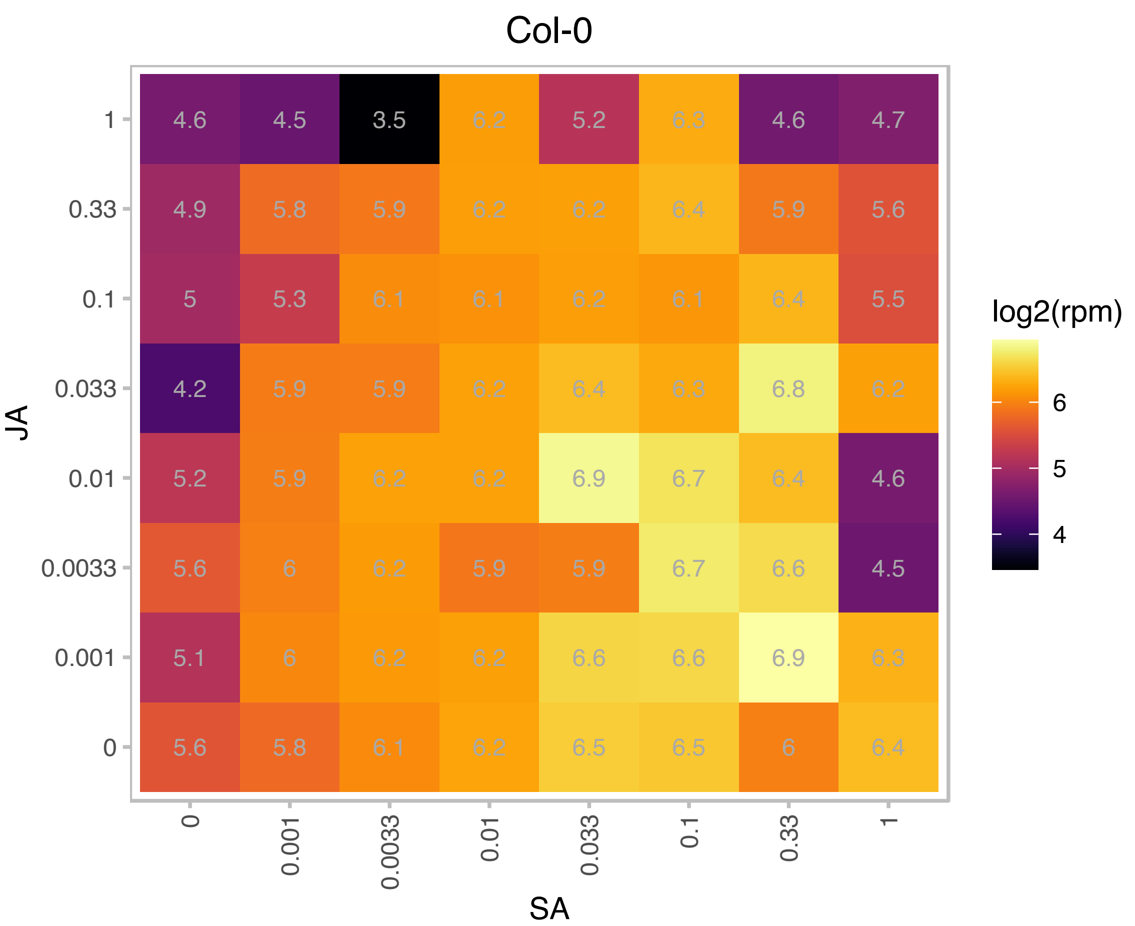


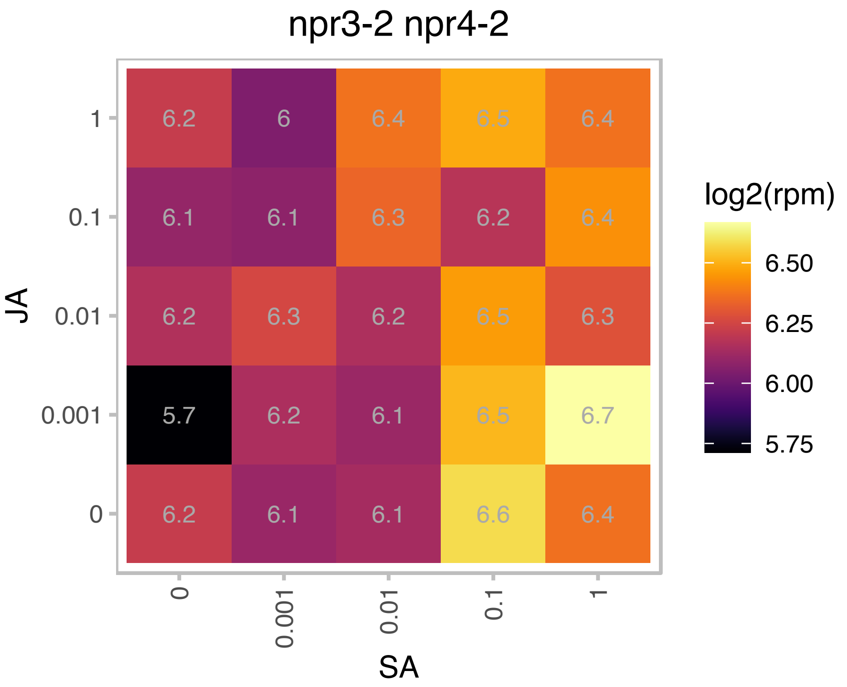


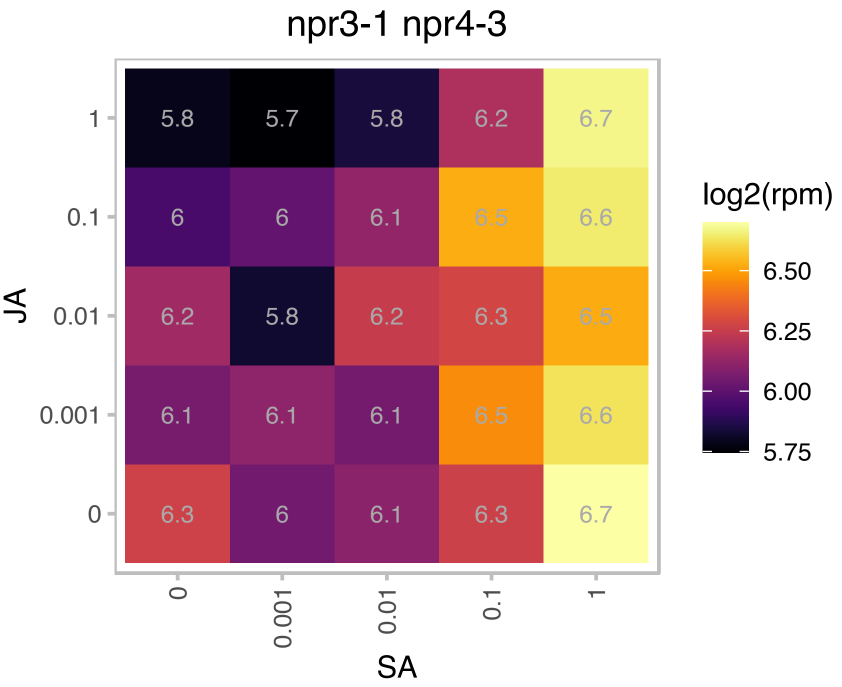

# AT2G39210.1 : NA

Major facilitator superfamily protein

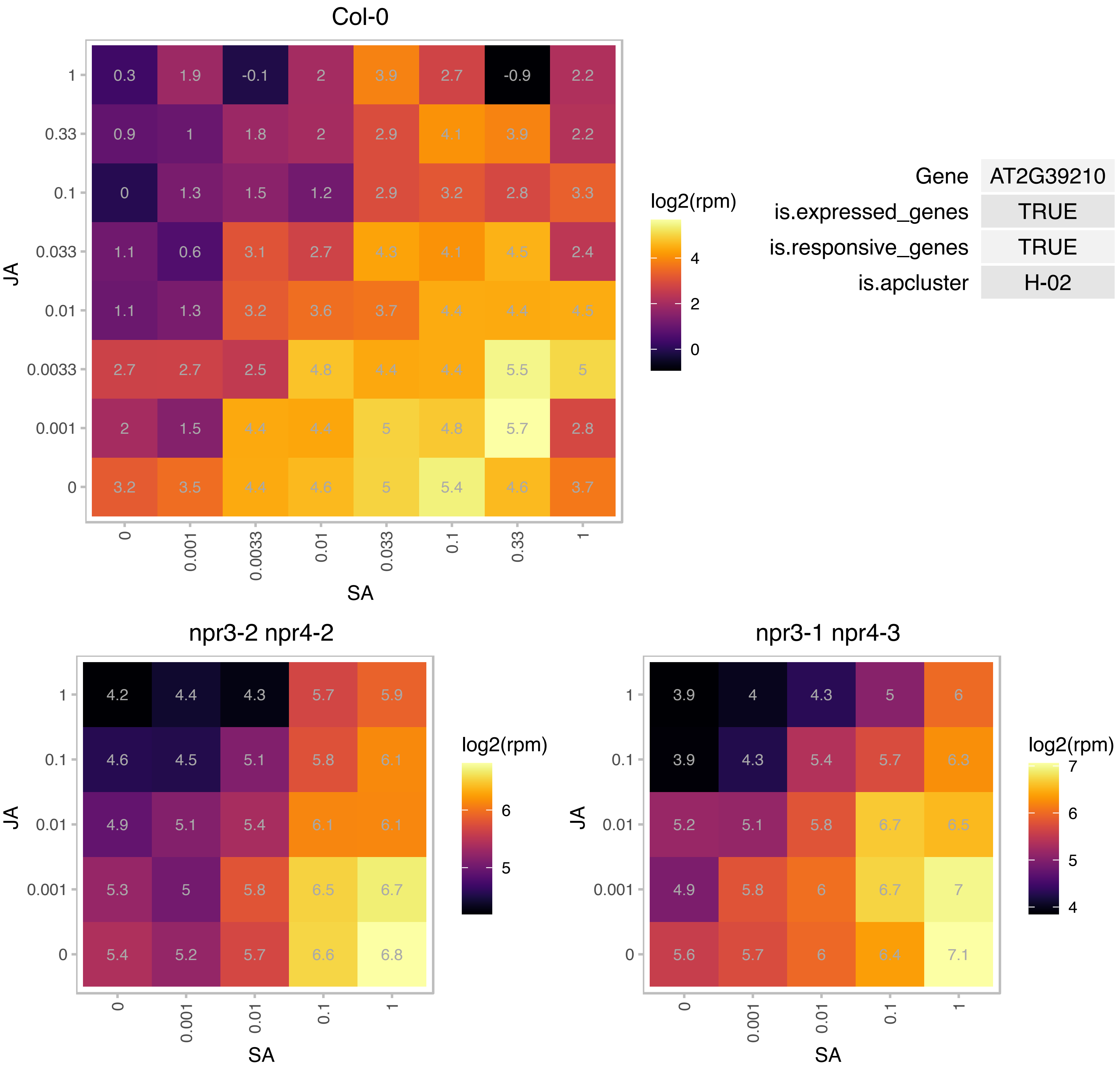


| | |
|---|---|
| Gene | AT2G39210 |
| is.expressed_genes | TRUE |
| is.responsive_genes | TRUE |
| is.apcluster | H-02 |

# AT2G40600.1 : NA

appr-1-p processing enzyme family protein

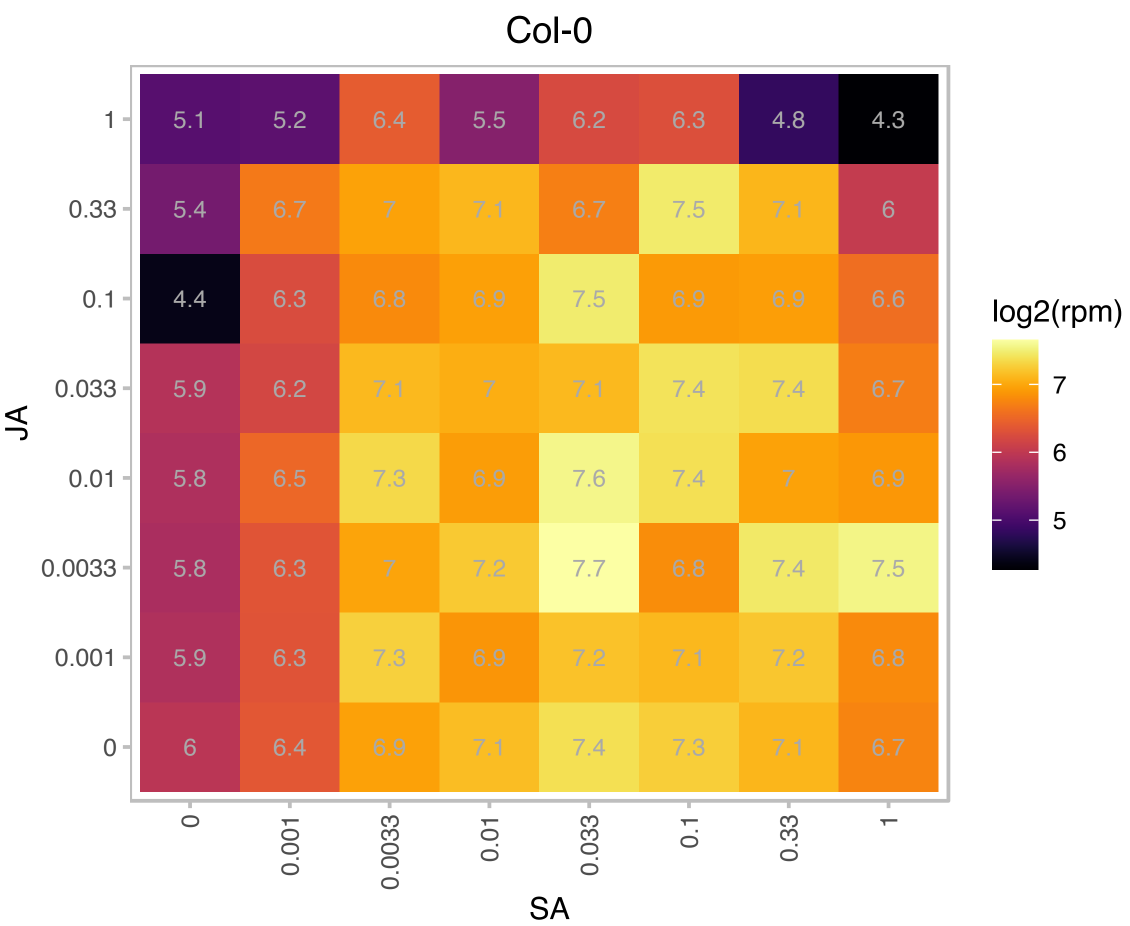


| | |
|---|---|
| Gene | AT2G40600 |
| is.expressed_genes | TRUE |
| is.responsive_genes | TRUE |
| is.apcluster | H-02 |

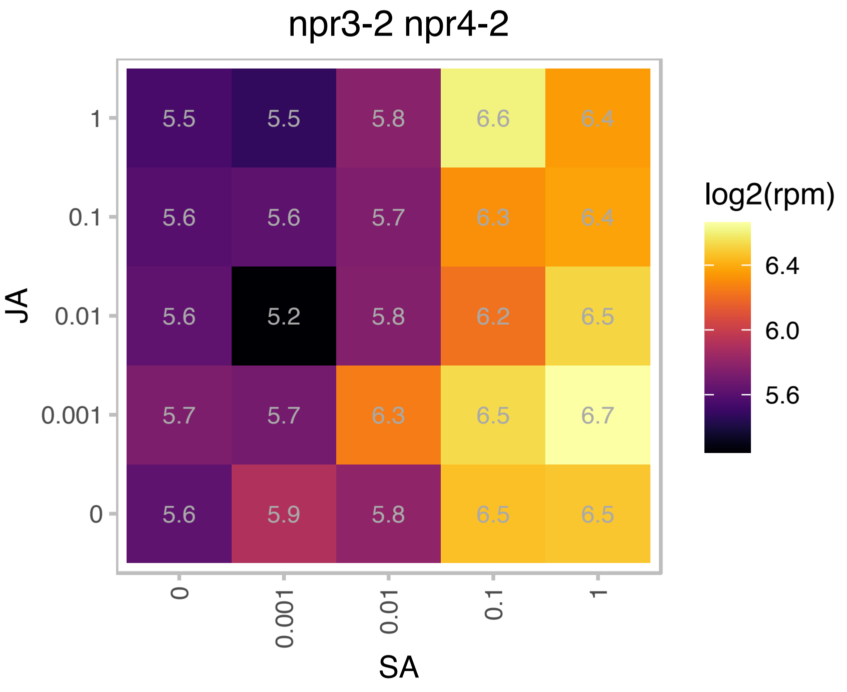


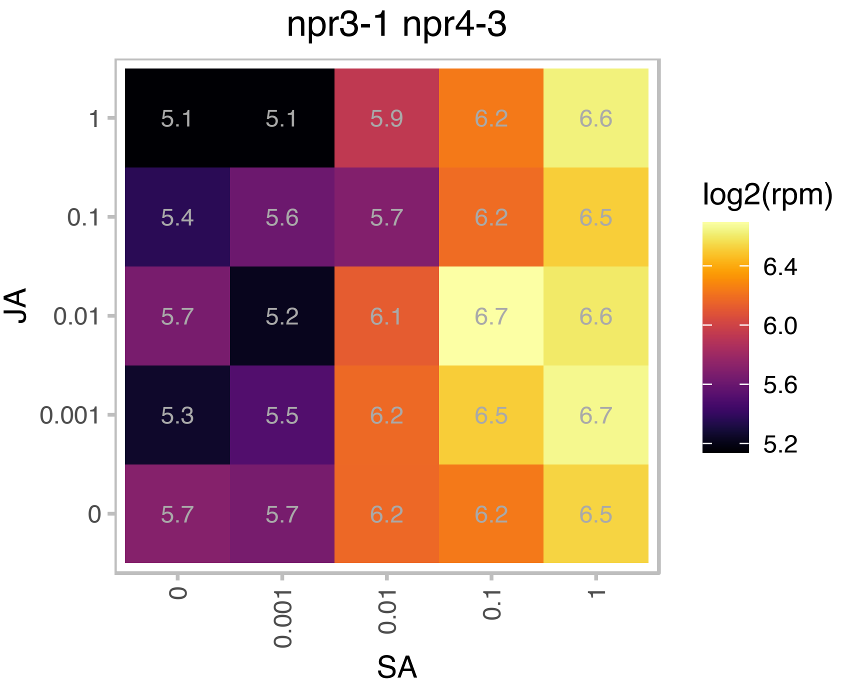

# AT2G44210.1 : NA

Protein of Unknown Function (DUF239)

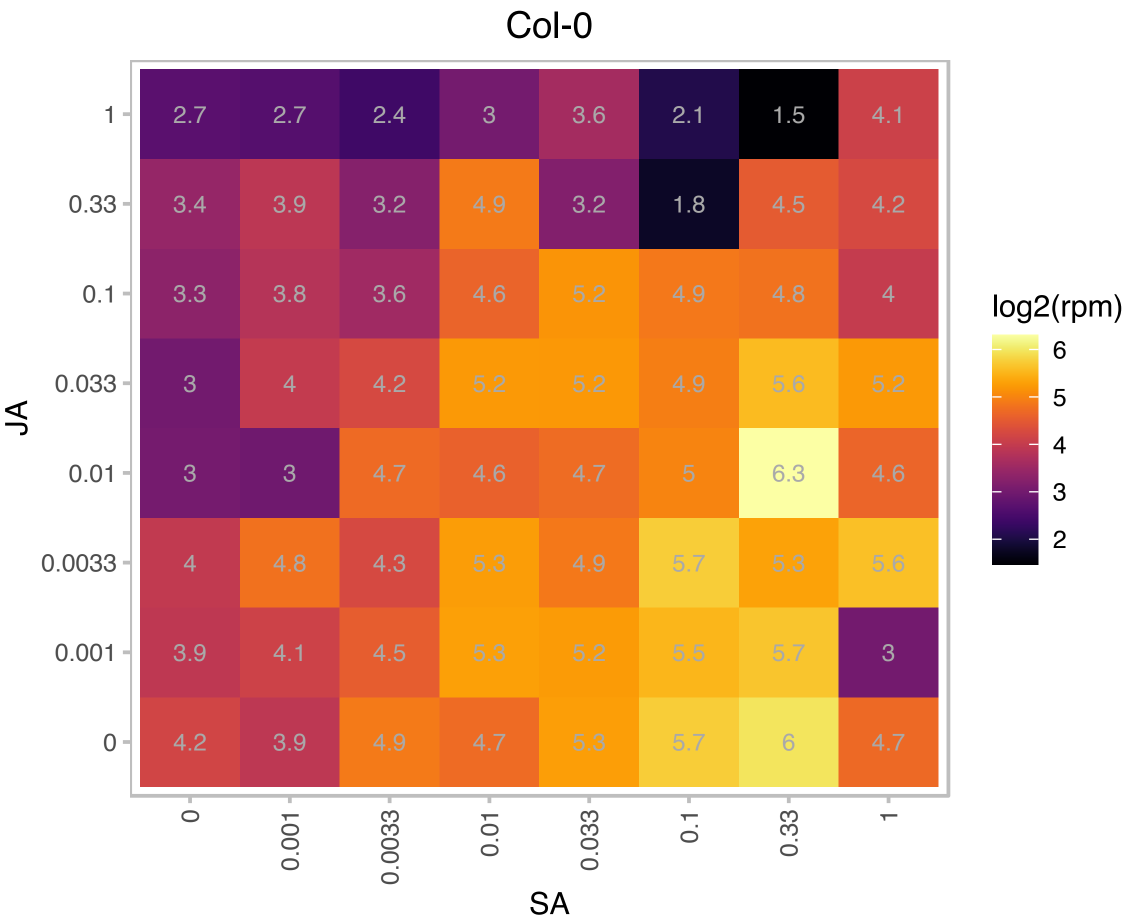


| | |
|---|---|
| Gene | AT2G44210 |
| is.expressed_genes | TRUE |
| is.responsive_genes | TRUE |
| is.apcluster | H-02 |

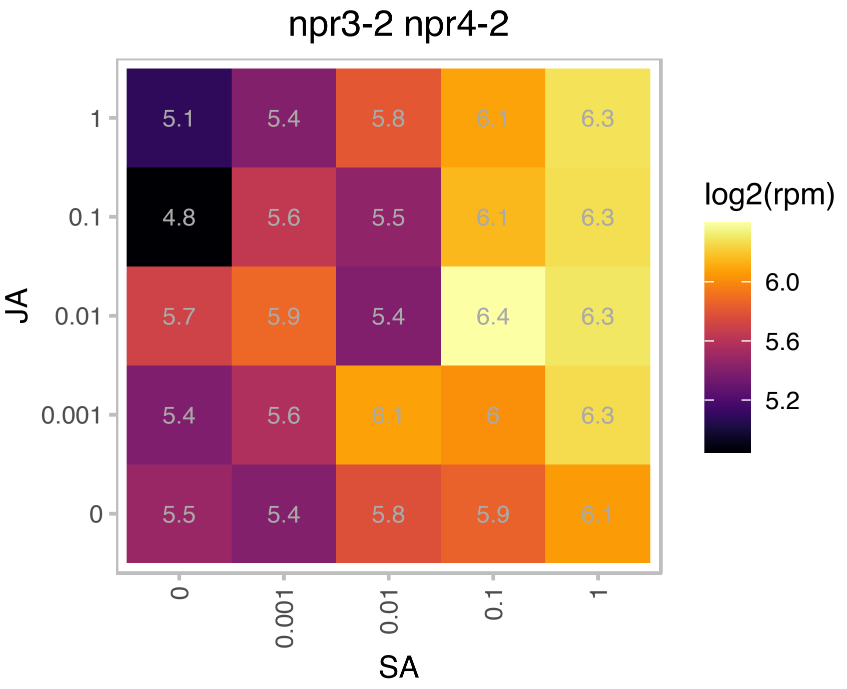


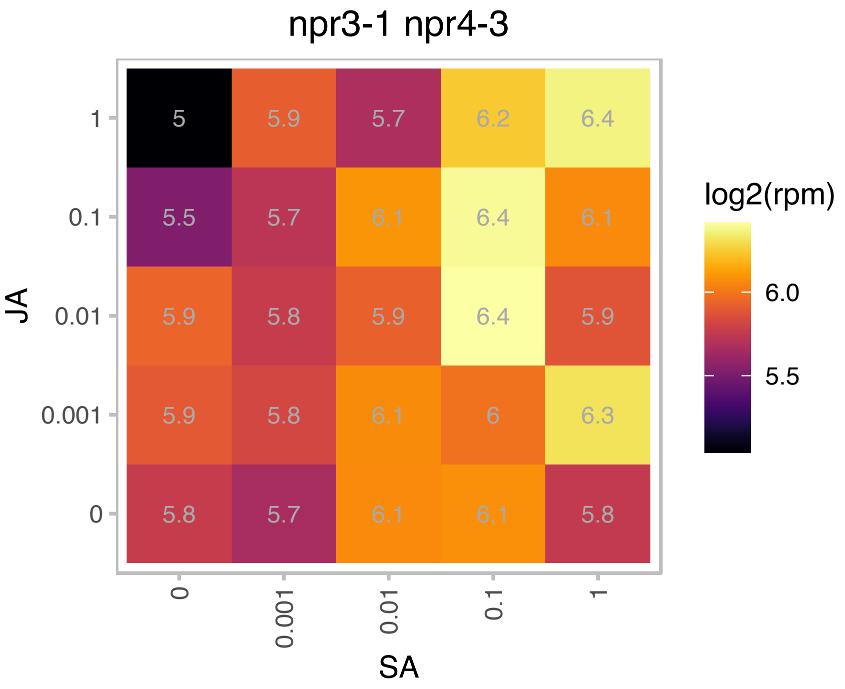

### AT2G46270.1 : GBF3

G-box binding factor 3
encodes a bZIP G-box binding protein whose expression is induced by ABA. It has been shown to bind to Adh that contains the G-box and is induced by cold and water deprivation. GBF3 has been shown to be expressed mostly in the root and dark-grown leaves. GBF3 can act as homodimers and as heterodimers with GFB1, GBF2 and GBF4. In addition, GBF3□ □ □ □ s DNA binding activity is enhanced by GIP1, GPRI1 and GPRI2.

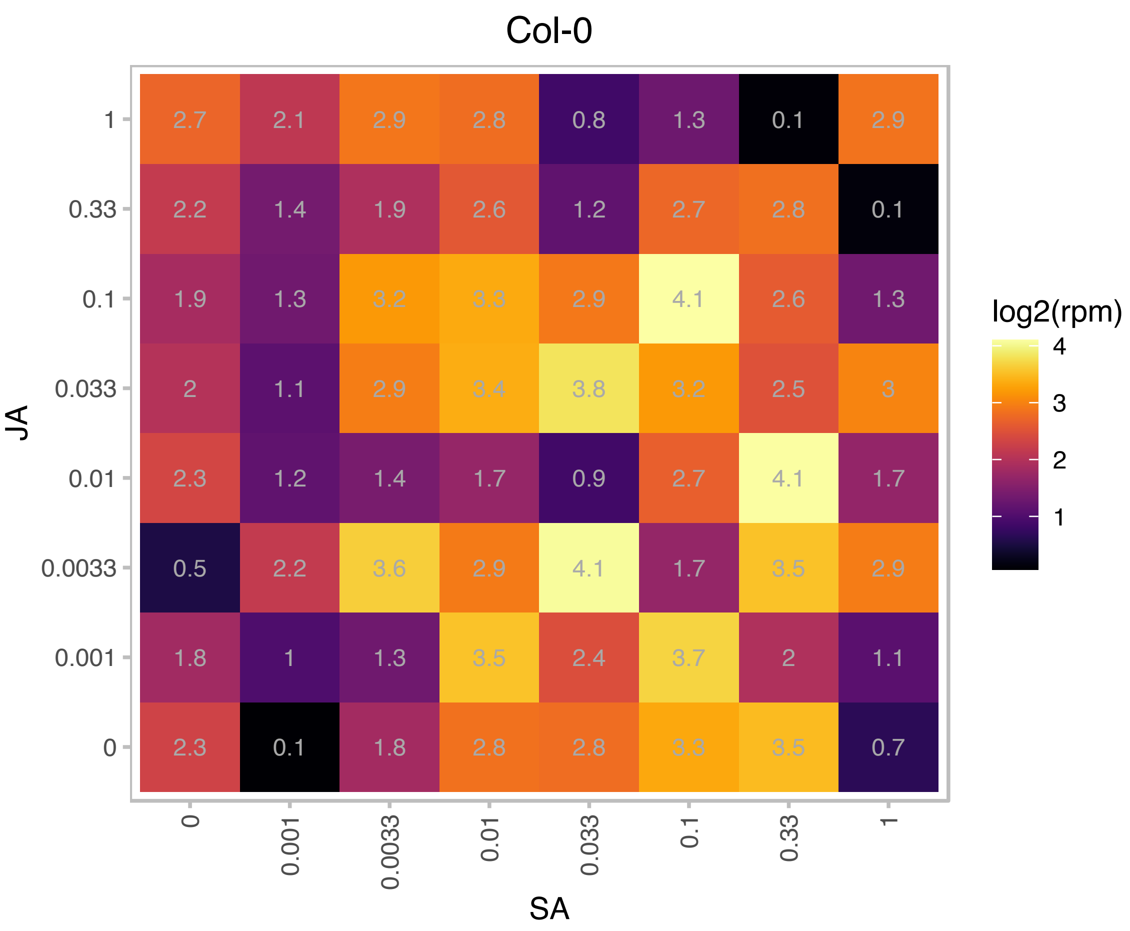


| | |
|---|---|
| Gene | AT2G46270 |
| is.expressed_genes | TRUE |
| is.responsive_genes | TRUE |
| is.apcluster | H-02 |

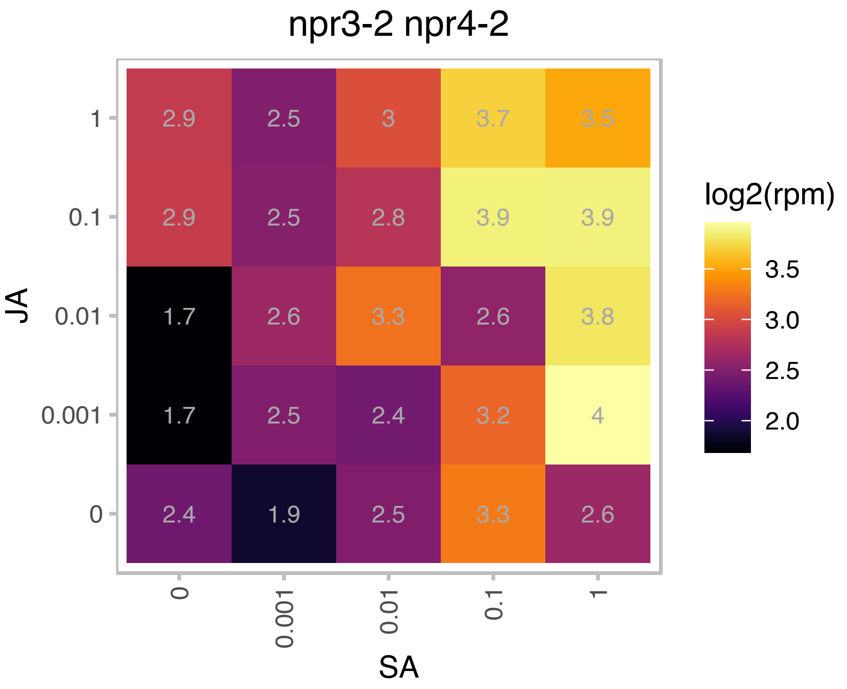


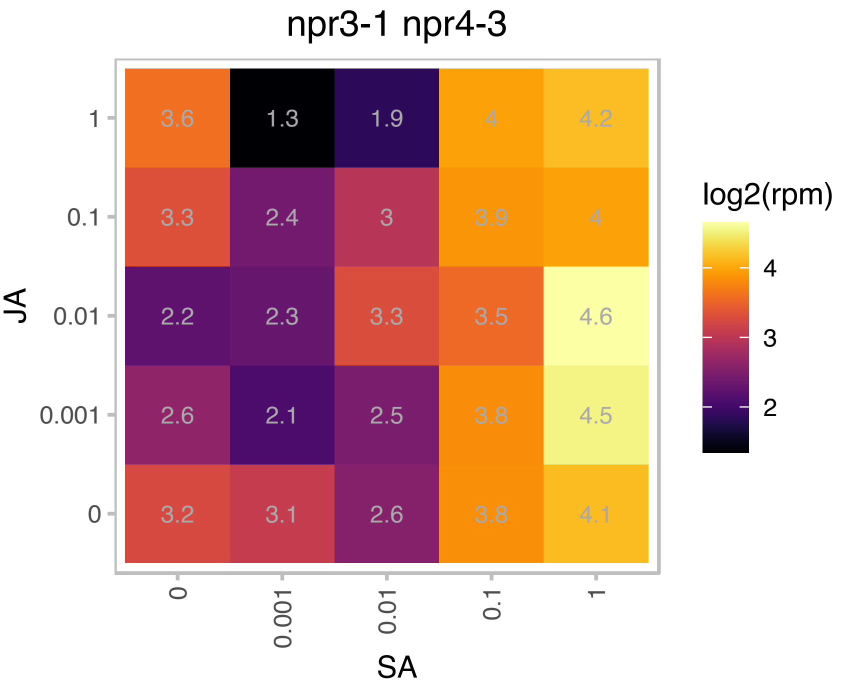

# AT2G46420.1 : NA

Plant protein 1589 of unknown function

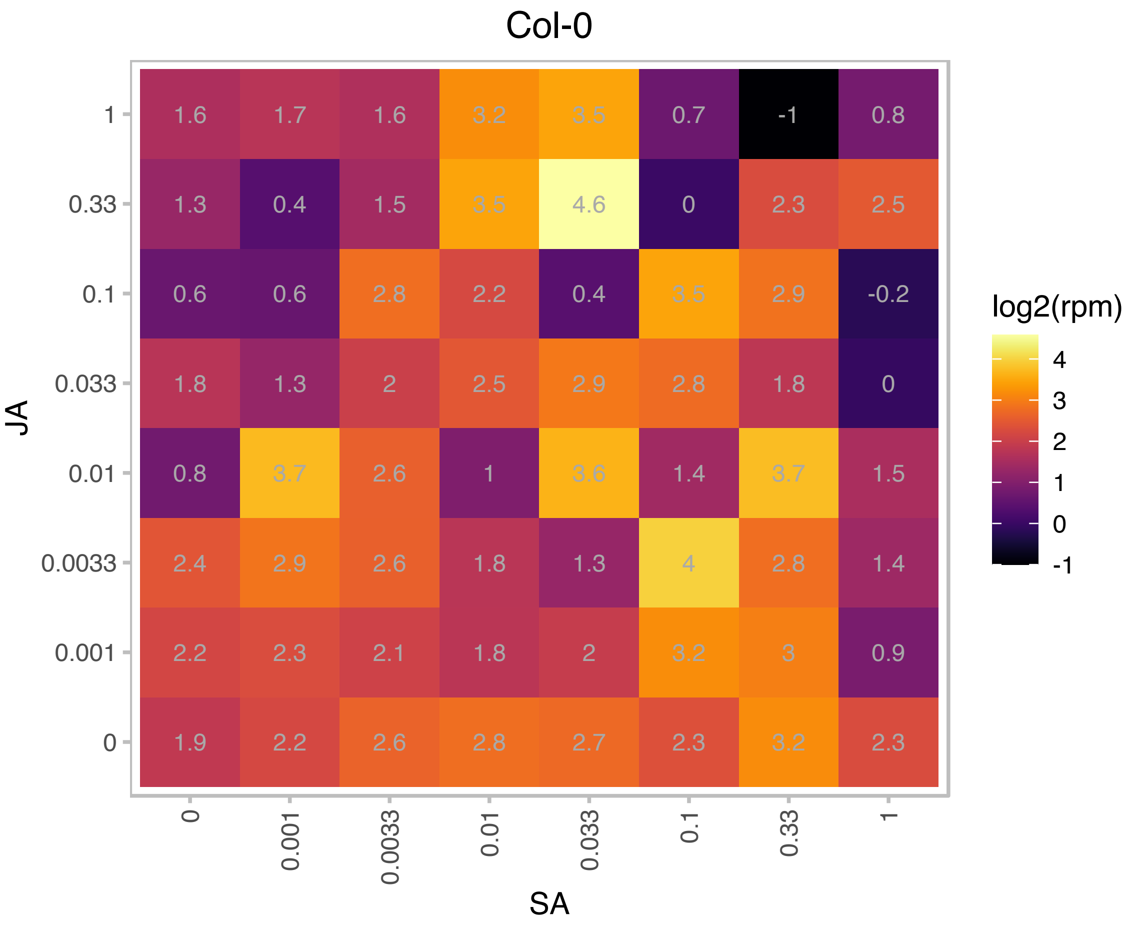


| | |
|---|---|
| Gene | AT2G46420 |
| is.expressed_genes | TRUE |
| is.responsive_genes | TRUE |
| is.apcluster | H-02 |

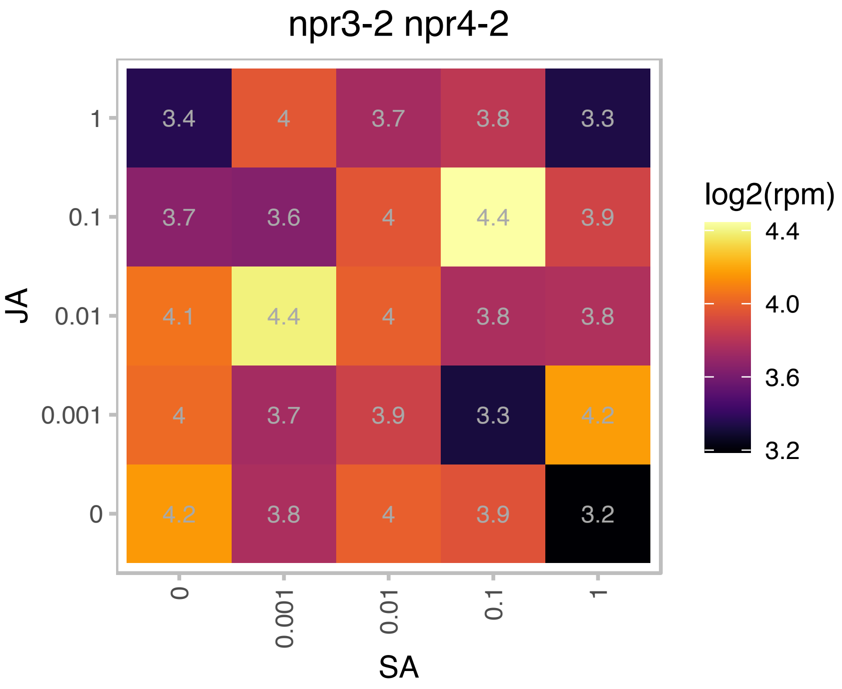


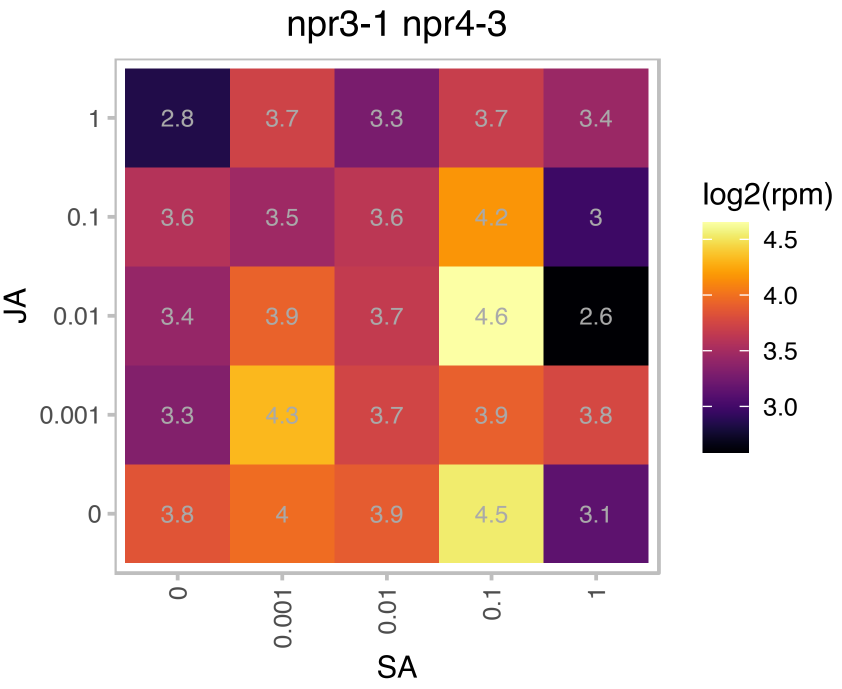

### AT2G47130.1 : AtSDR3, SDR3

NAD(P)-binding Rossmann-fold superfamily protein

Encodes a short-chain dehydrogenase/reductase that is not involved in ABA biosynthesis but plays an important role in plant defense response to bacteria.

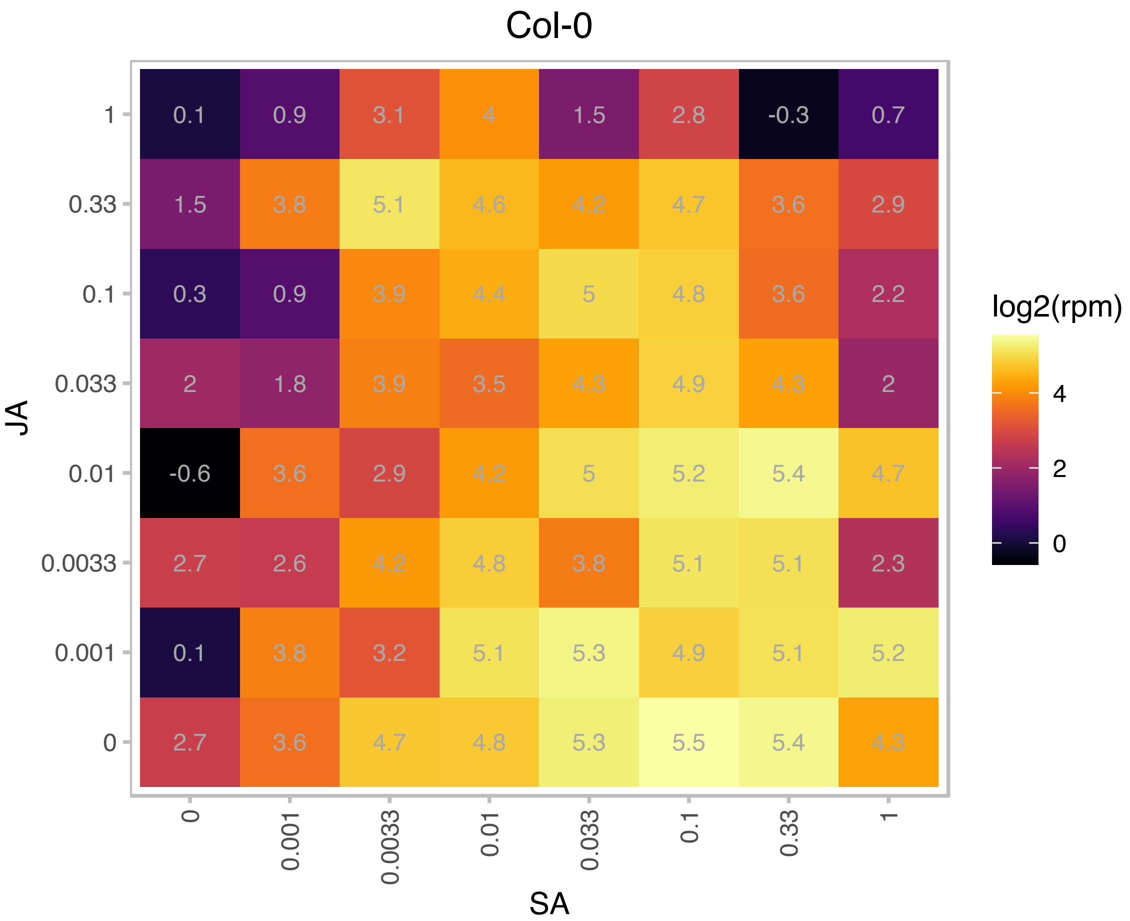


| | |
|---|---|
| Gene | AT2G47130 |
| is.expressed_genes | TRUE |
| is.responsive_genes | TRUE |
| is.apcluster | H-02 |

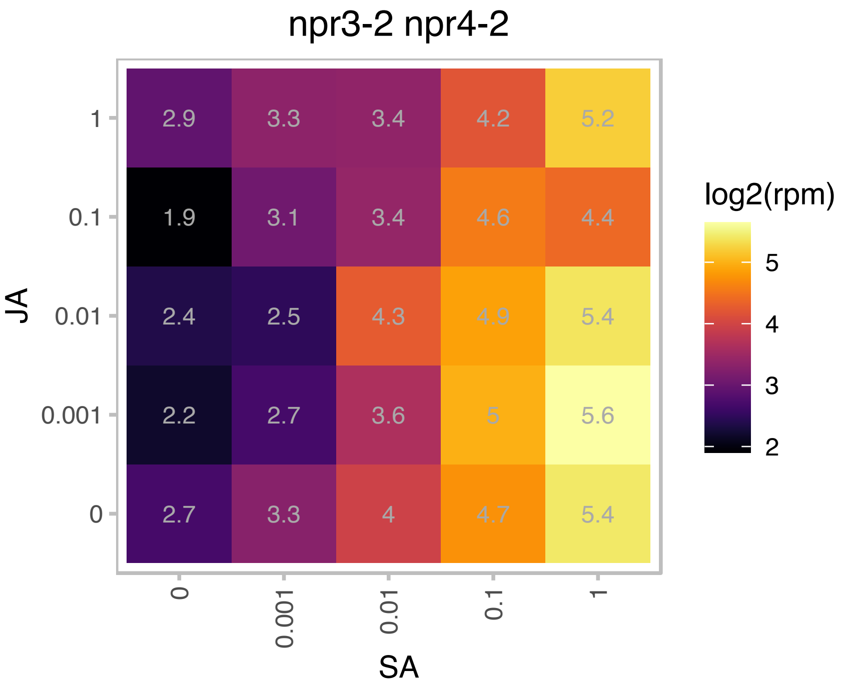


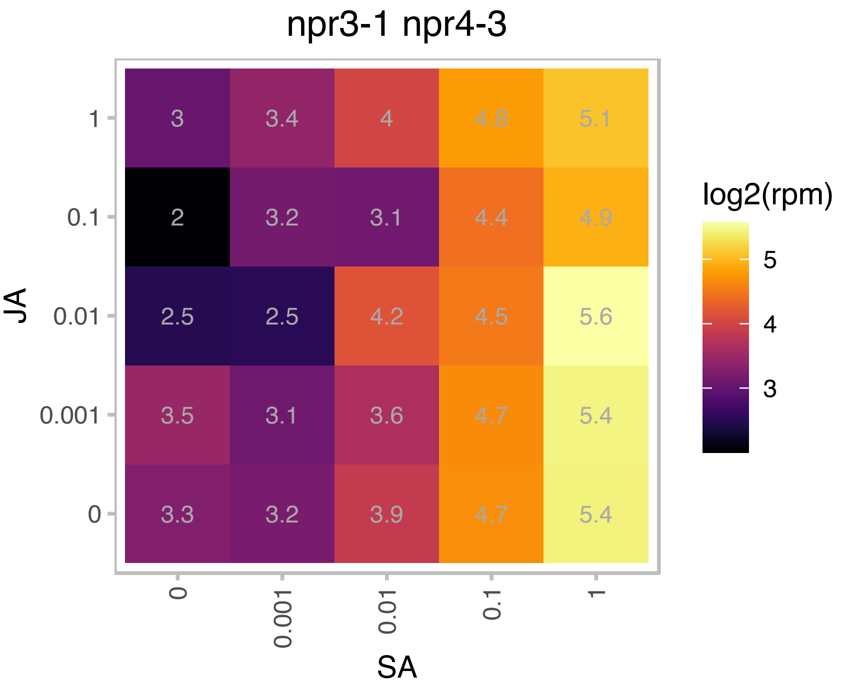

# AT3G07195.1 : NA

RPM1-interacting protein 4 (RIN4) family protein

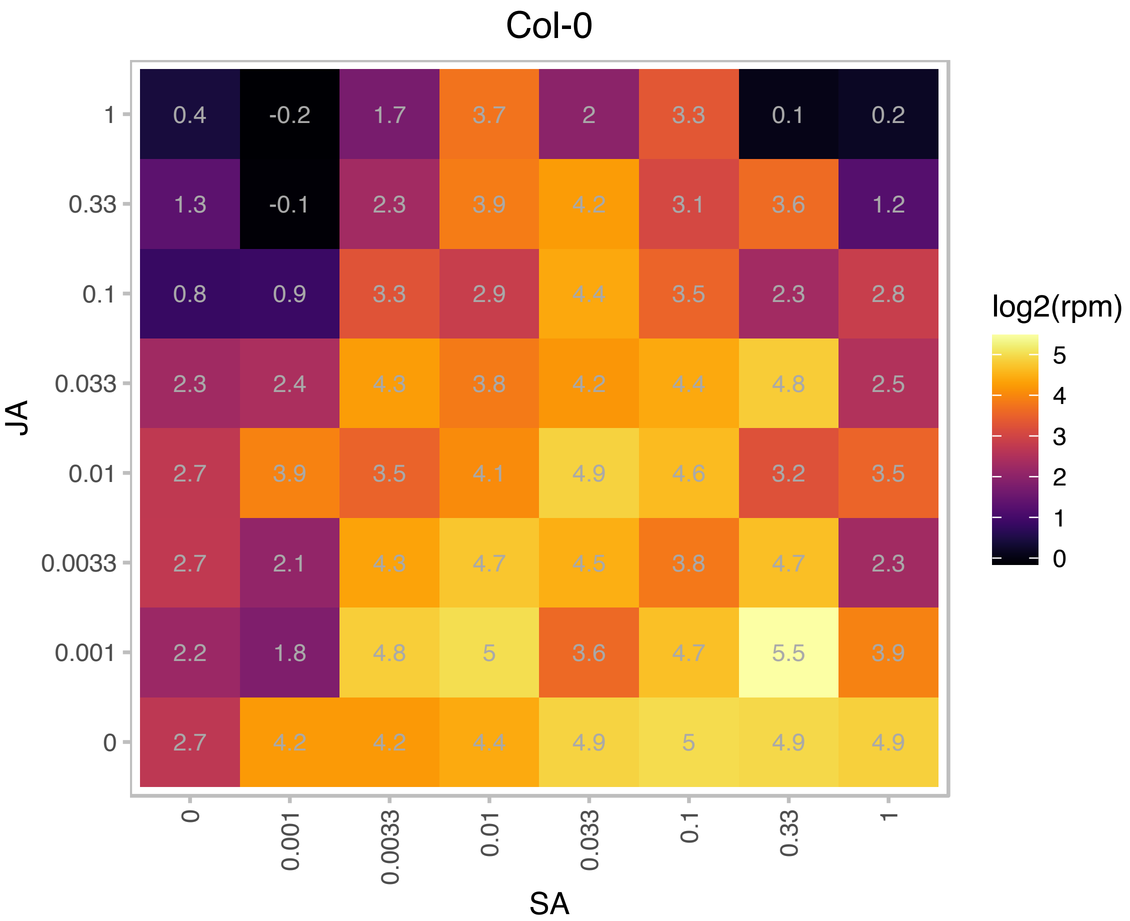


| | |
|---|---|
| Gene | AT3G07195 |
| is.expressed_genes | TRUE |
| is.responsive_genes | TRUE |
| is.apcluster | H-02 |

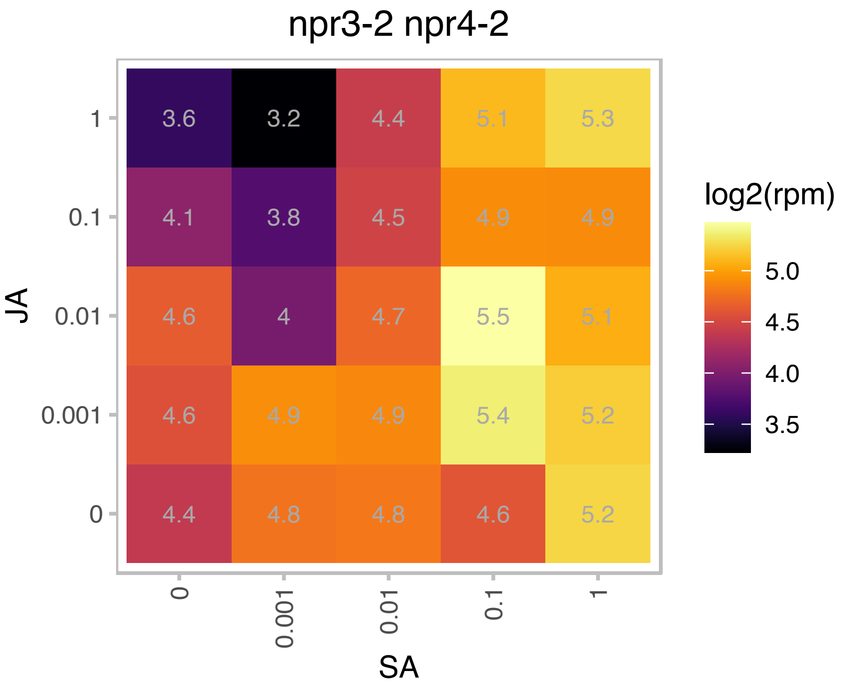


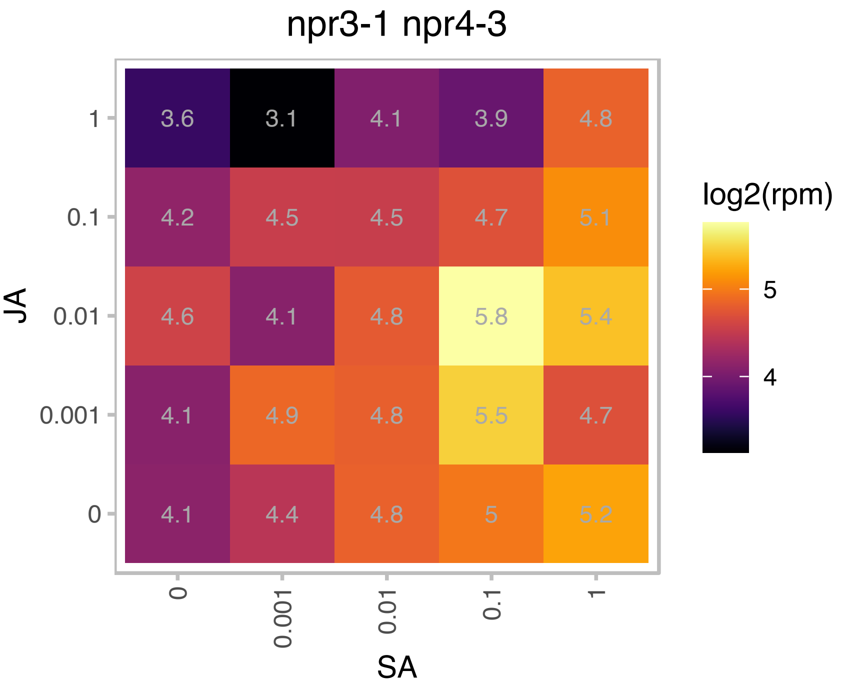

# AT3G07600.1 : NA

Heavy metal transport/detoxification superfamily protein

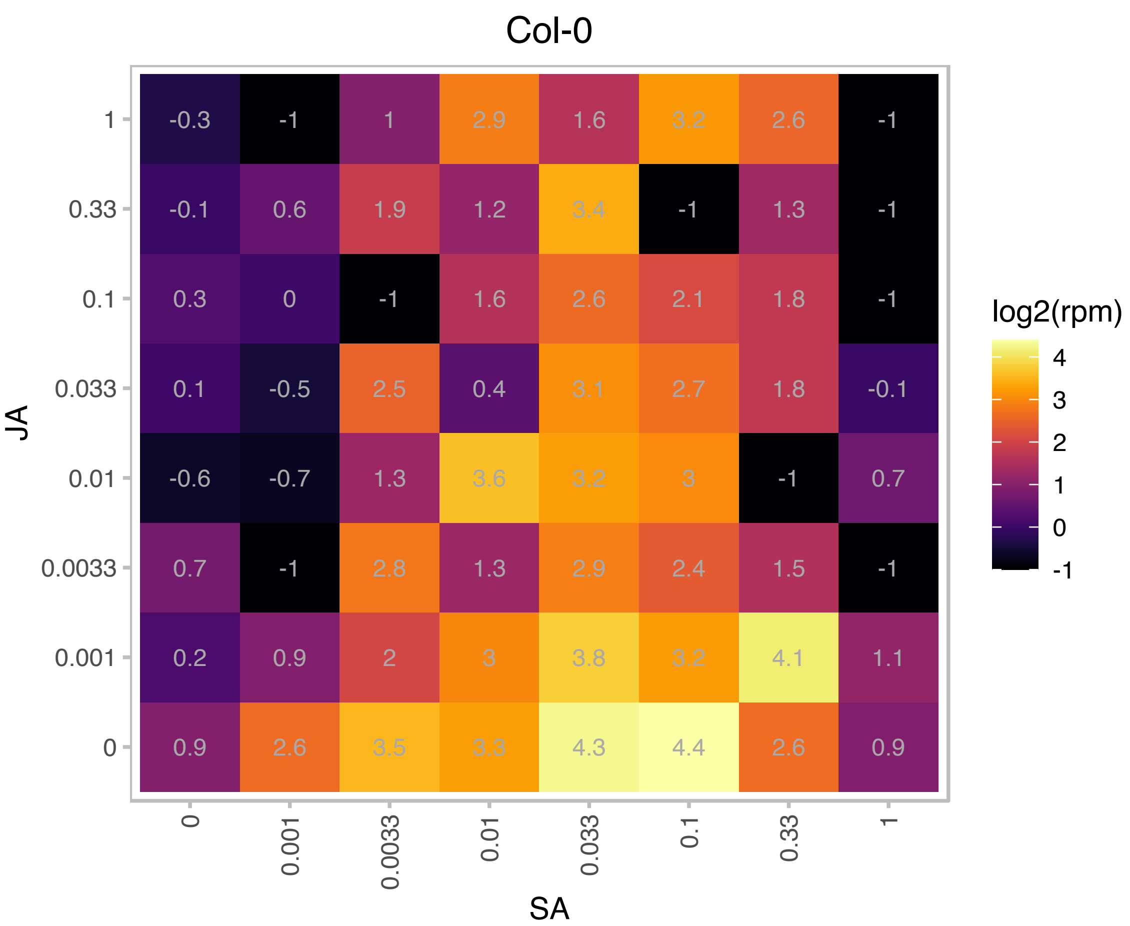


| | |
|---|---|
| Gene | AT3G07600 |
| is.expressed_genes | TRUE |
| is.responsive_genes | TRUE |
| is.apcluster | H-02 |

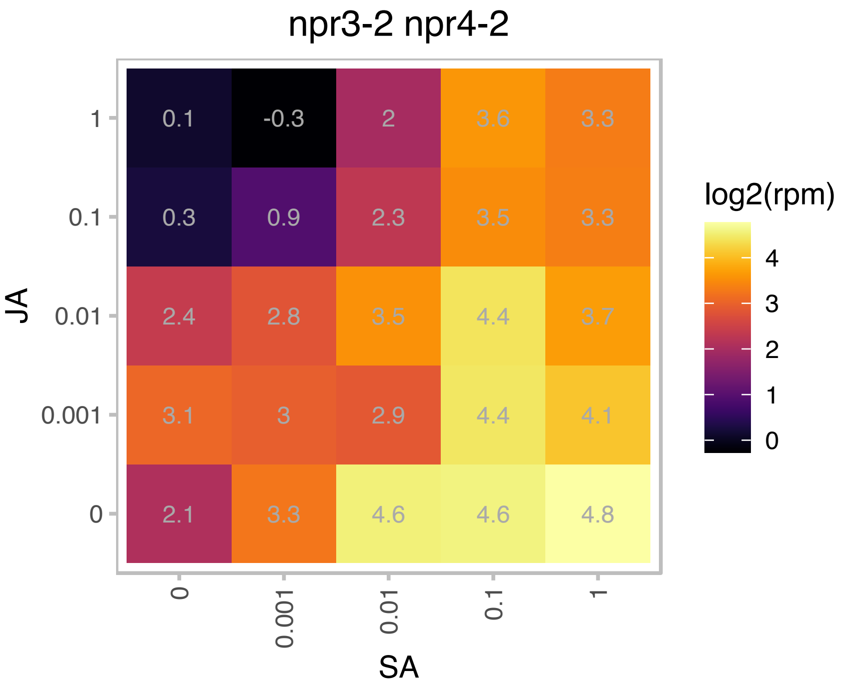


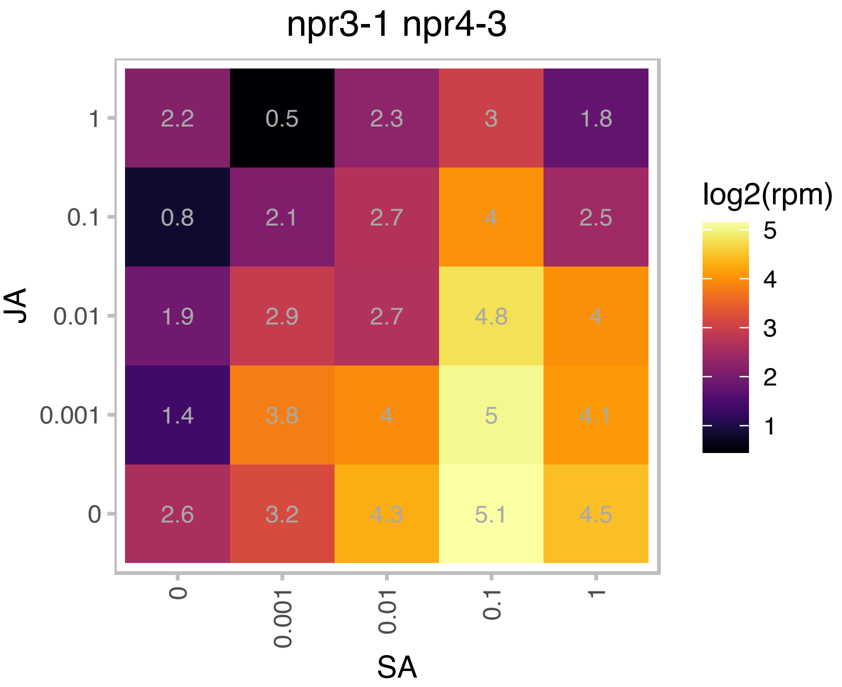

# AT3G29010.1 : NA

Biotin/lipoate A/B protein ligase family

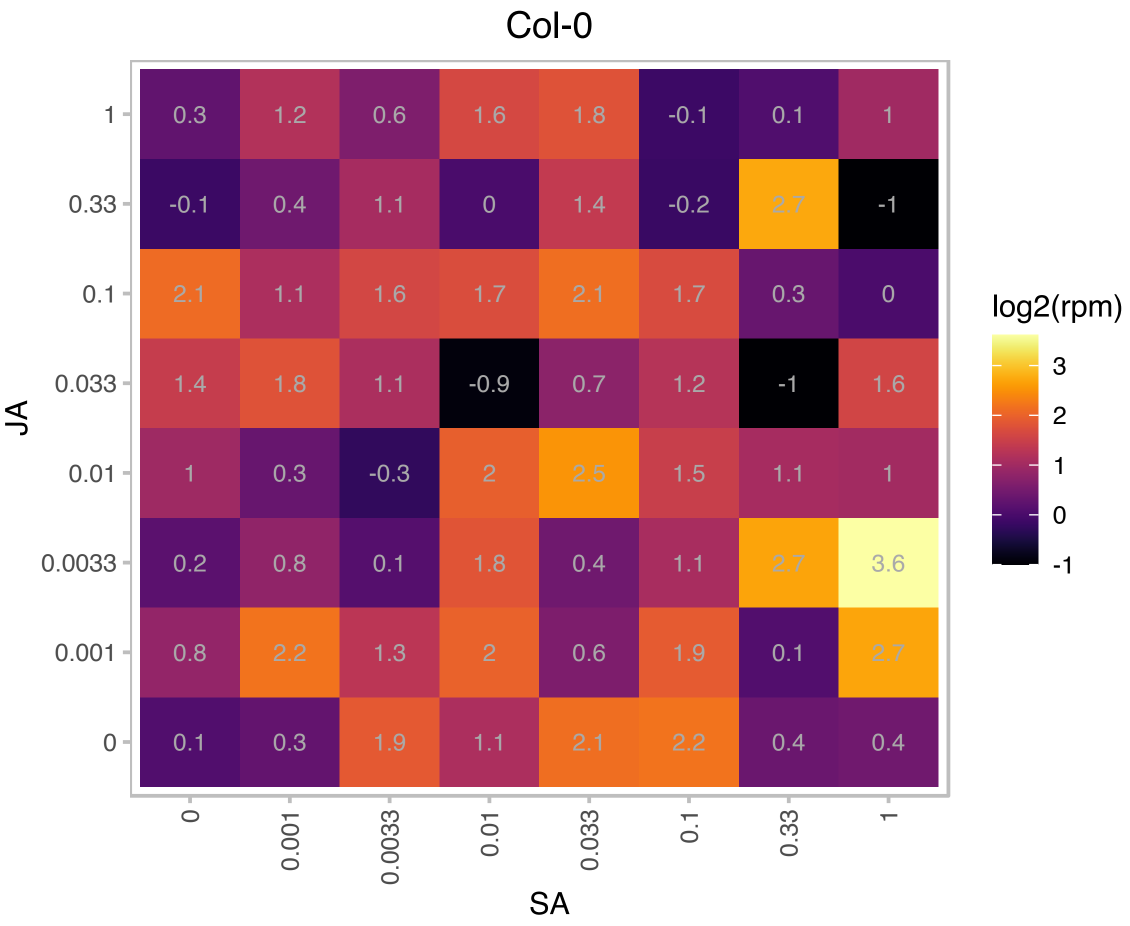


| | |
|---|---|
| Gene | AT3G29010 |
| is.expressed_genes | TRUE |
| is.responsive_genes | TRUE |
| is.apcluster | H-02 |

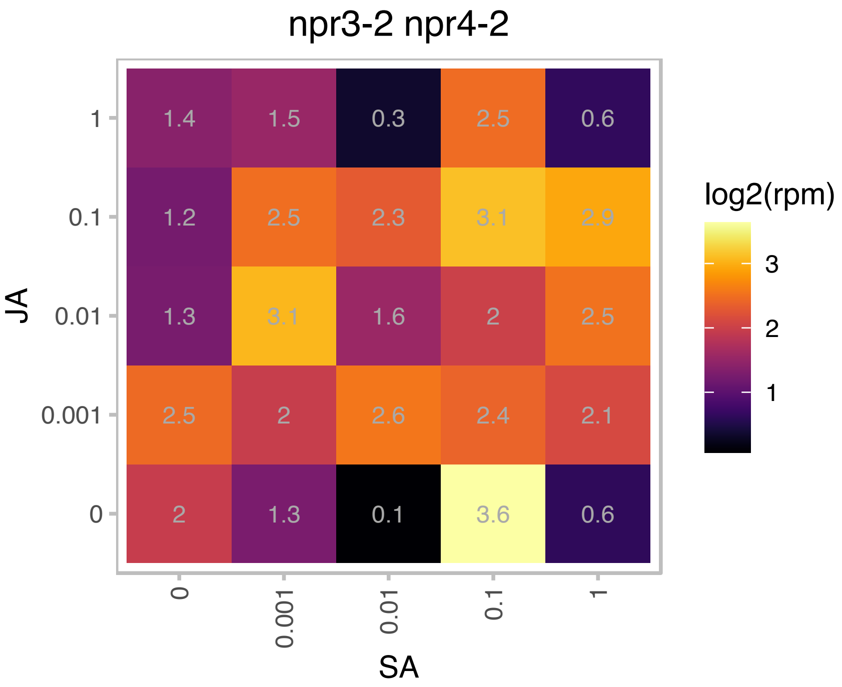


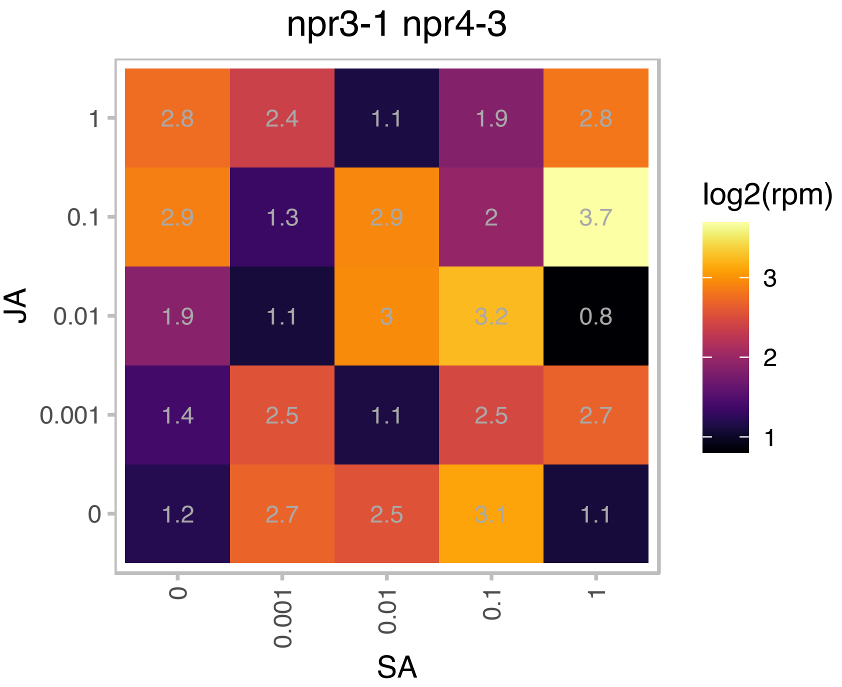

### AT3G52090.1 : ATRPB13.6, NRPB11, NRPD11, NRPE11

DNA-directed RNA polymerase, RBP11-like

Non-catalytic subunit common to nuclear DNA-dependent RNA polymerases II, IV and V; homologous to budding yeast RPB11 and the E. oli RNA polymerase alpha subunit.

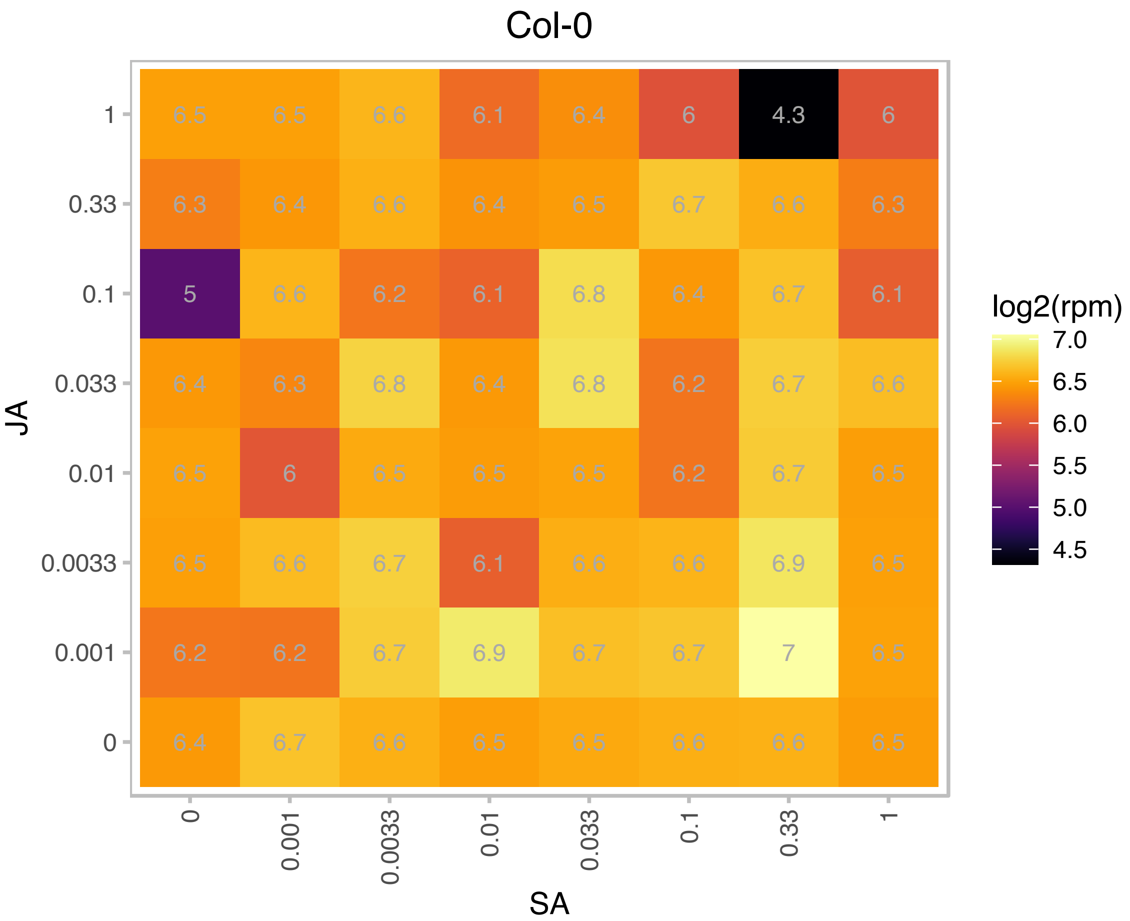


| | |
|---|---|
| Gene | AT3G52090 |
| is.expressed_genes | TRUE |
| is.responsive_genes | TRUE |
| is.apcluster | H-02 |

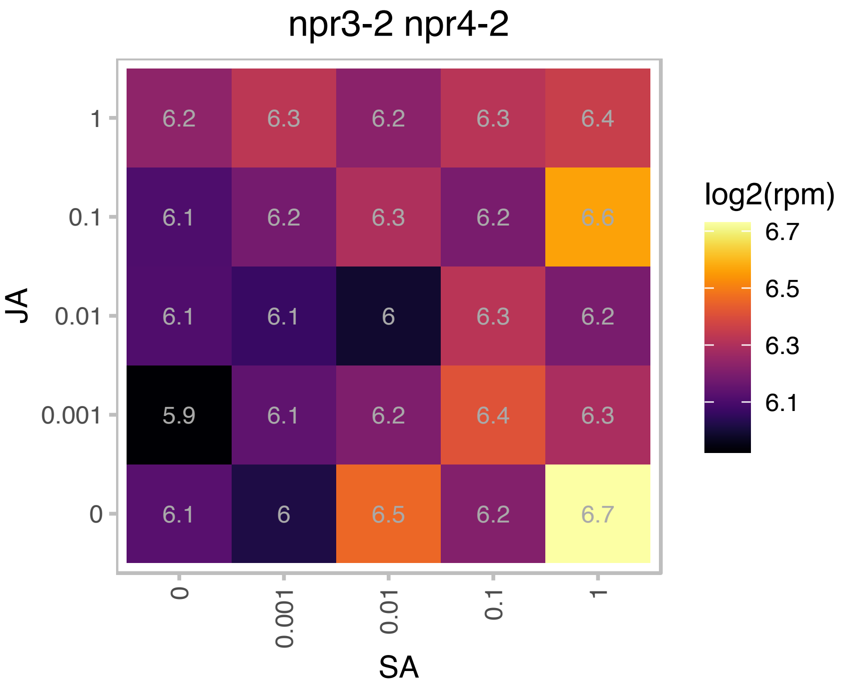


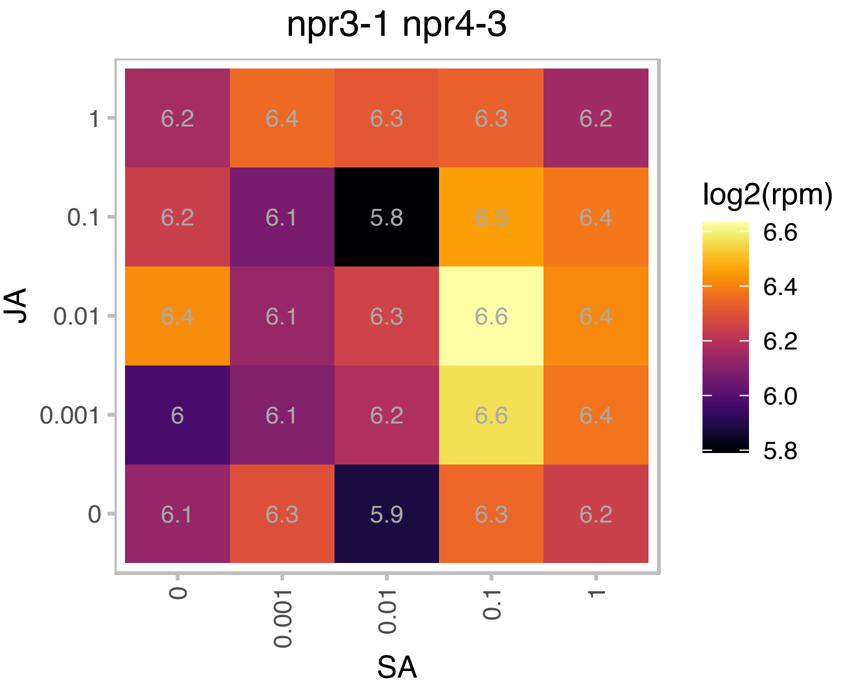

### AT3G52860.1 : MED28

Encodes a mediator subunit,, important for development and senescence.

Col-0

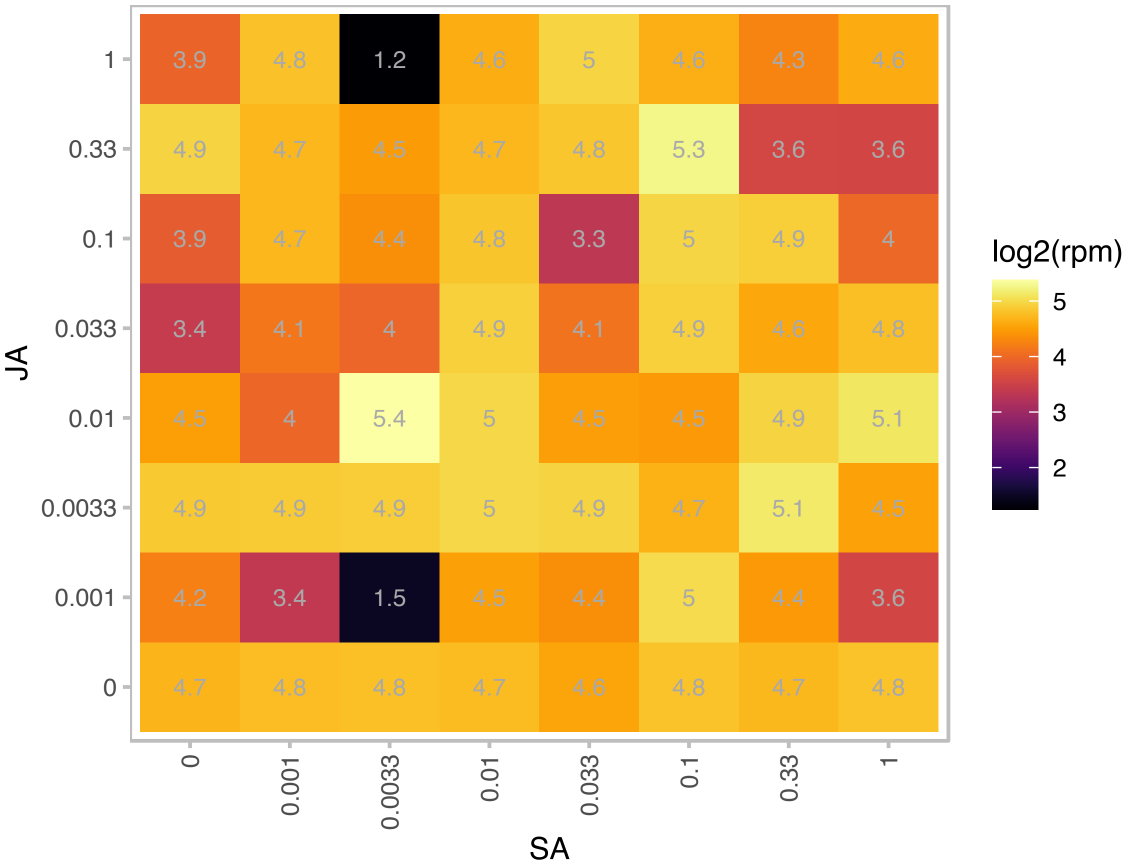


| | |
|---|---|
| Gene | AT3G52860 |
| is.expressed_genes | TRUE |
| is.responsive_genes | TRUE |
| is.apcluster | H-02 |

npr3-2 npr4-2

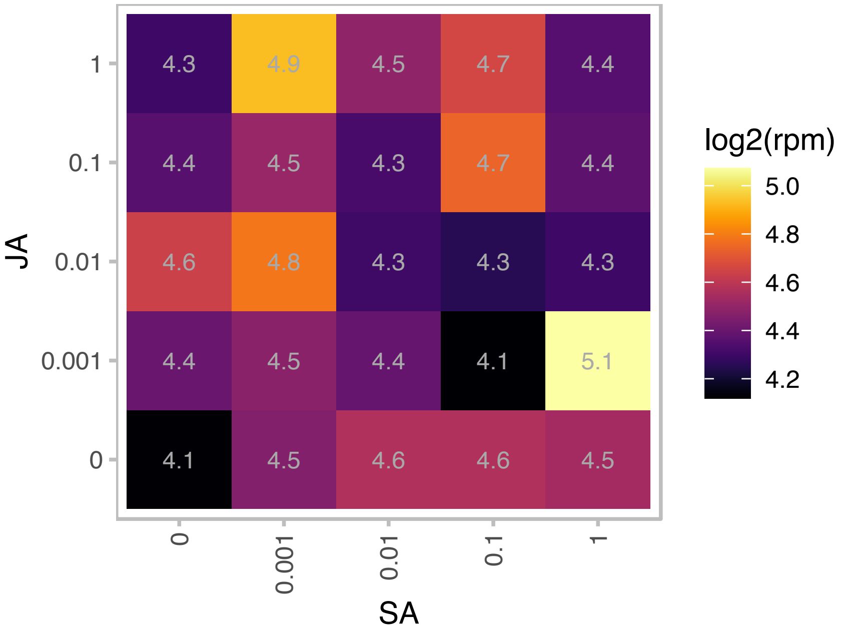


npr3-1 npr4-3

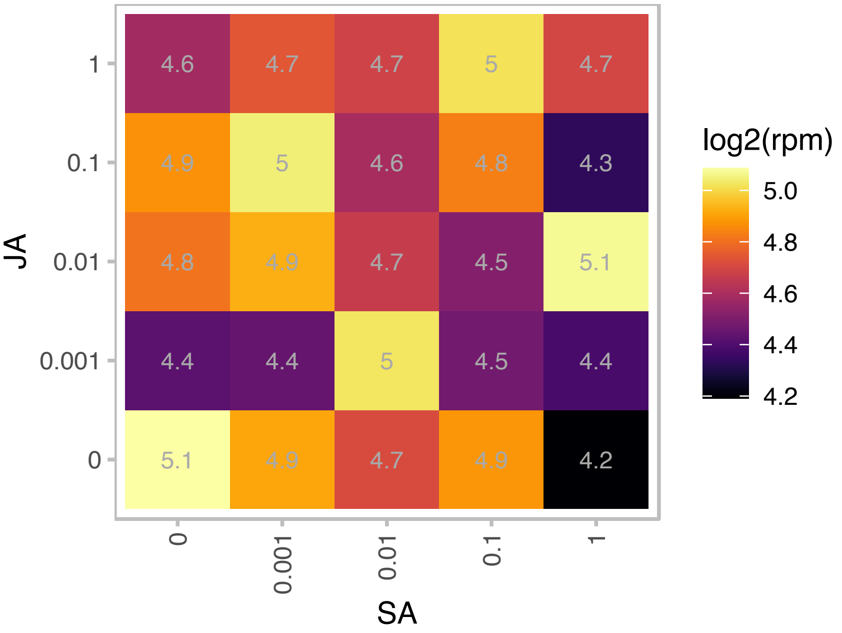

# AT3G61700.1 : NA

Plant protein 1589 of unknown function

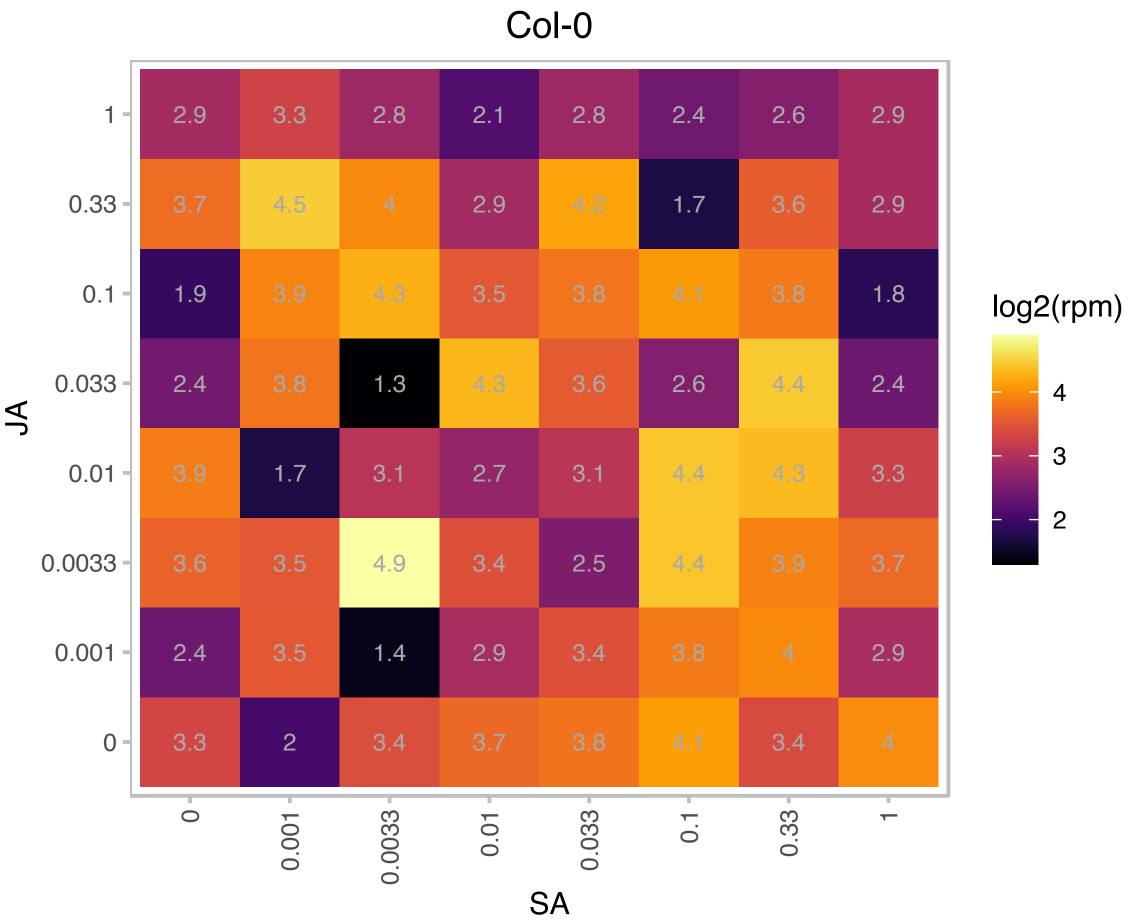


| | |
|---|---|
| Gene | AT3G61700 |
| is.expressed_genes | TRUE |
| is.responsive_genes | TRUE |
| is.apcluster | H-02 |

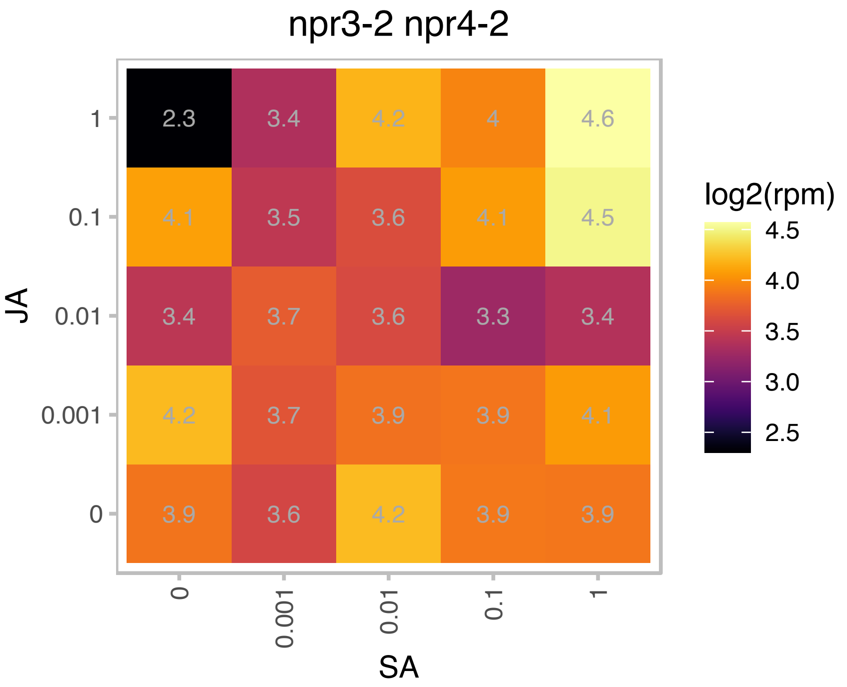


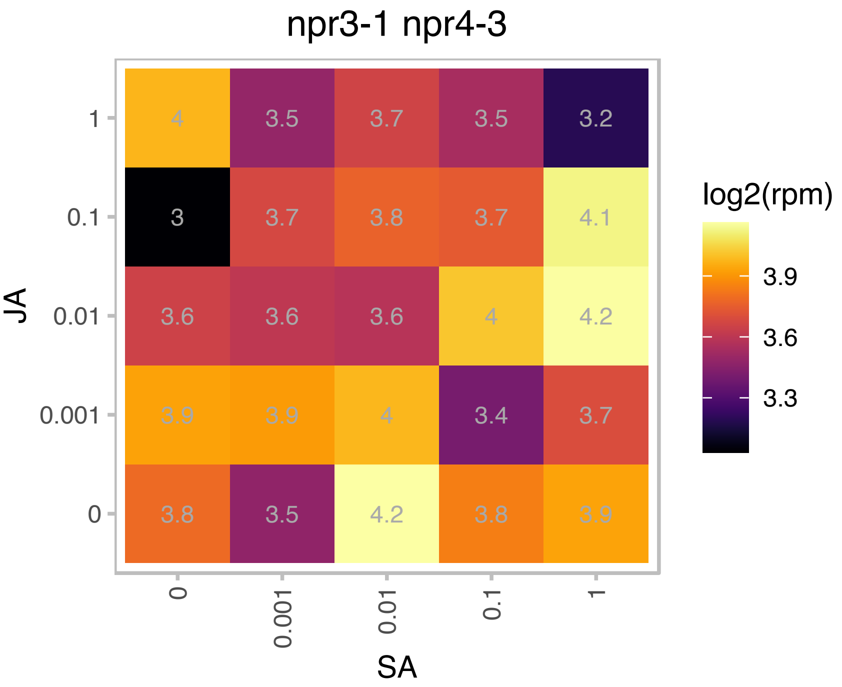

### AT3G63030.1 : MBD4

methyl-CPG-binding domain 4

Protein containing methyl-CpG-binding domain.Has sequence similarity to human MBD proteins.

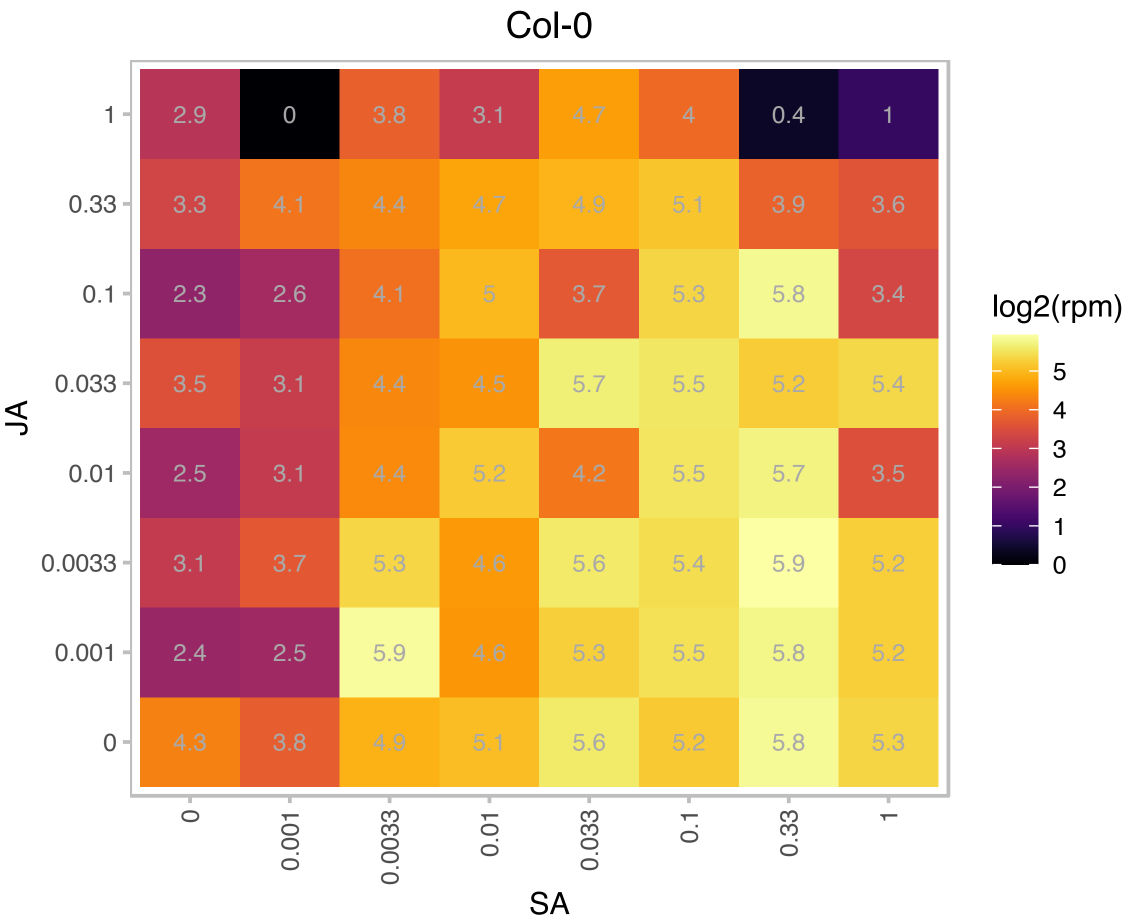


| | |
|---|---|
| Gene | AT3G63030 |
| is.expressed_genes | TRUE |
| is.responsive_genes | TRUE |
| is.apcluster | H-02 |

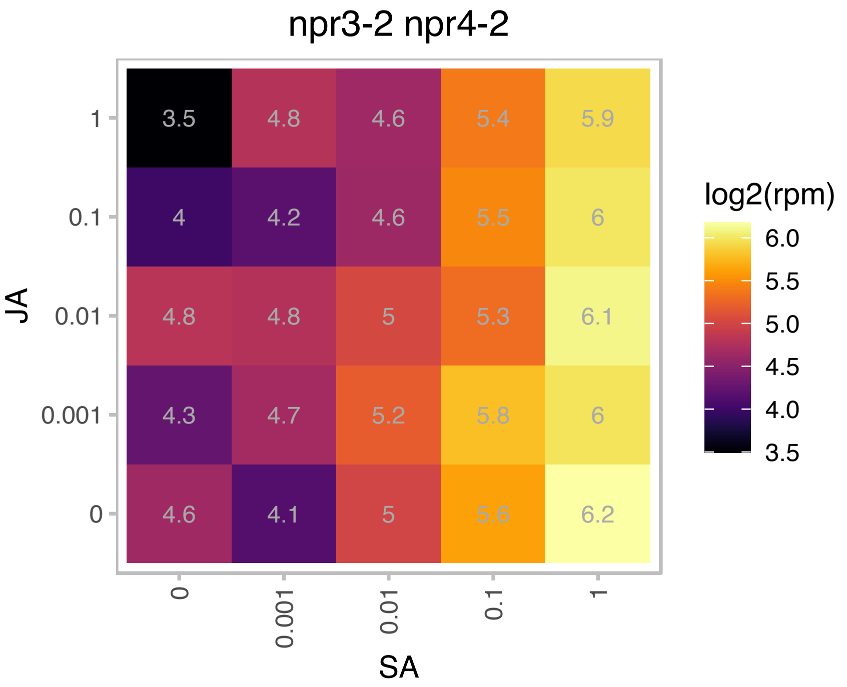


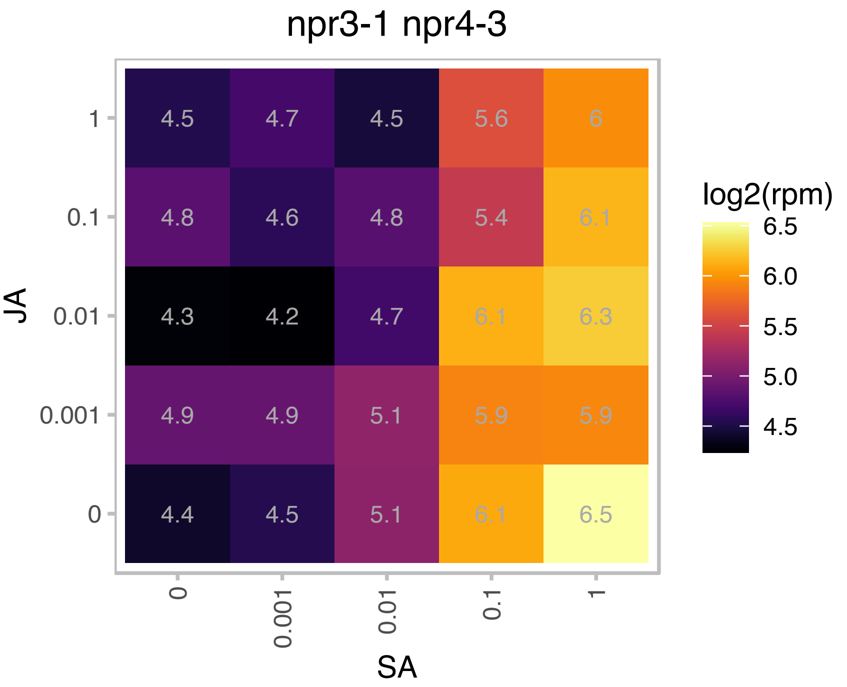

### AT3G63420.1 : AGG1, ATAGG1, GG1

Ggamma-subunit 1

heterotrimeric G protein gamma-subunit (AGG1) mRNA,

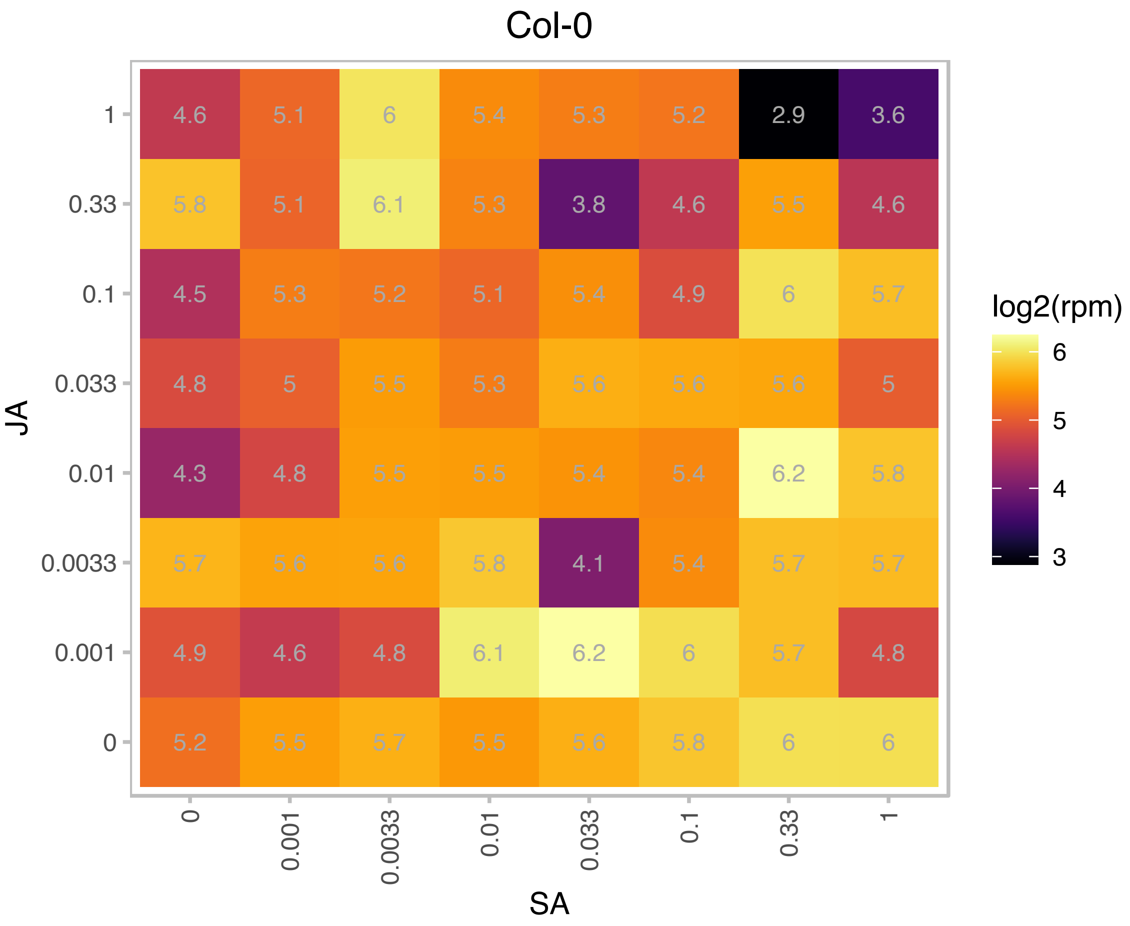


| | |
|---|---|
| Gene | AT3G63420 |
| is.expressed_genes | TRUE |
| is.responsive_genes | TRUE |
| is.apcluster | H-02 |

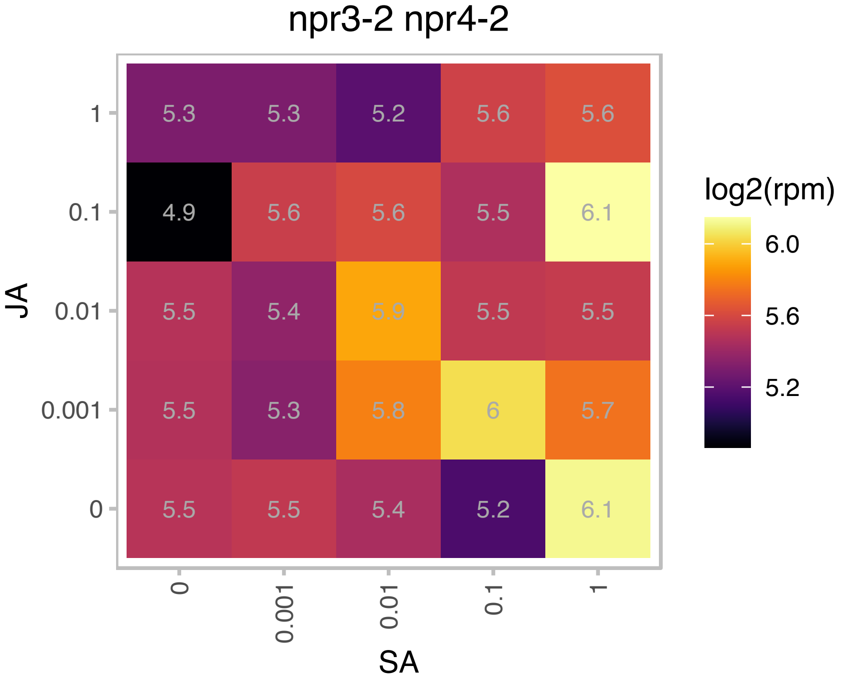


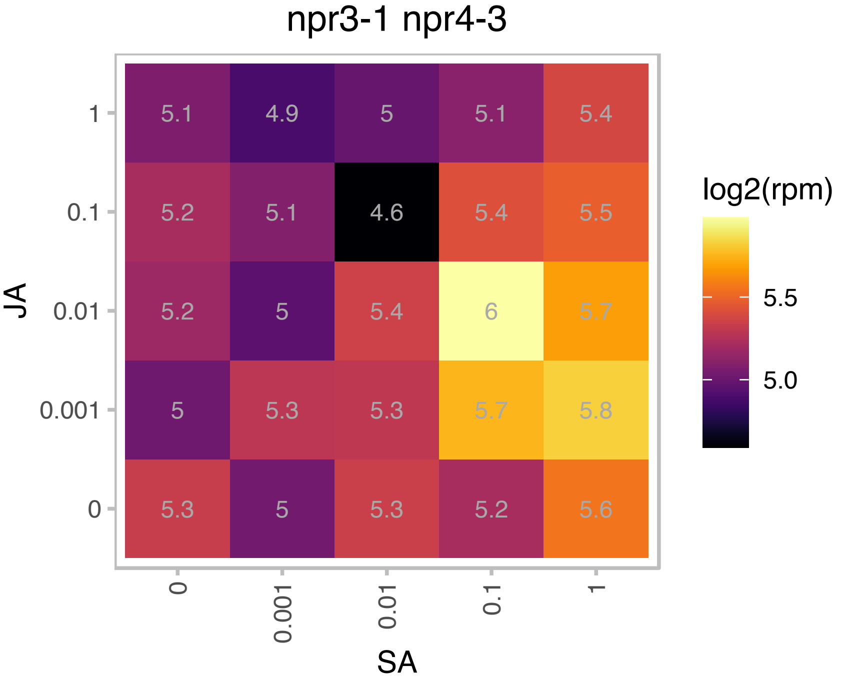

# AT4G02370.1 : NA

Protein of unknown function, DUF538

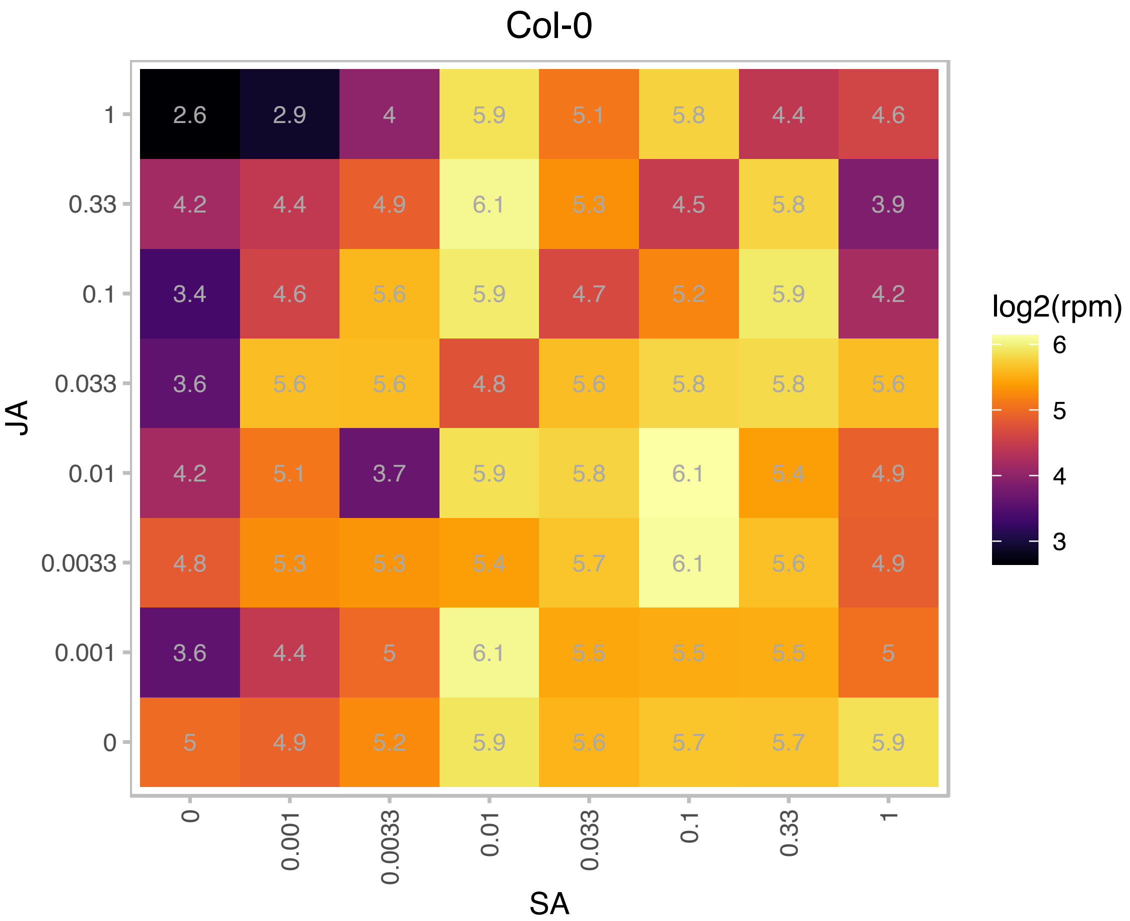


| | |
|---|---|
| Gene | AT4G02370 |
| is.expressed_genes | TRUE |
| is.responsive_genes | TRUE |
| is.apcluster | H-02 |

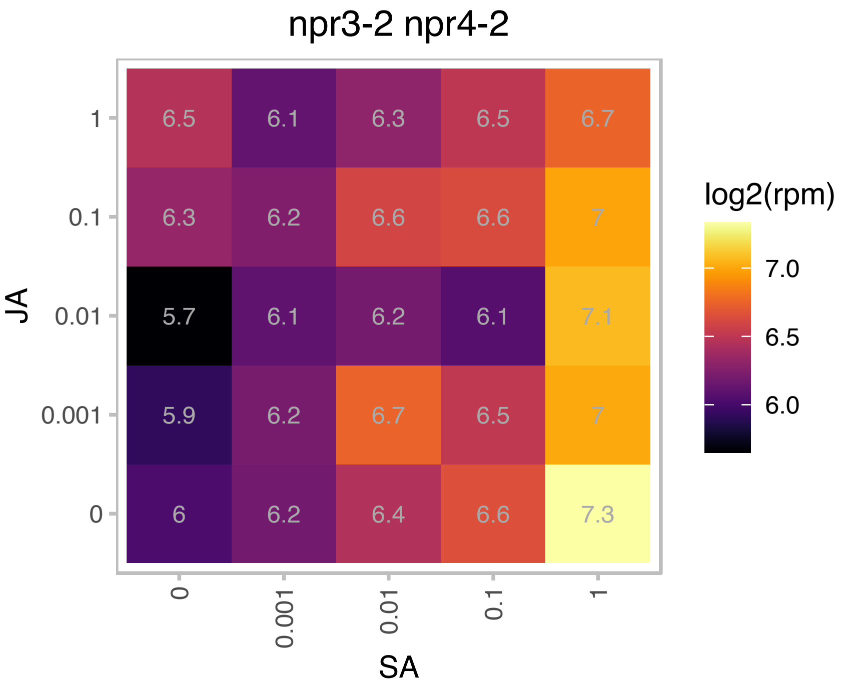


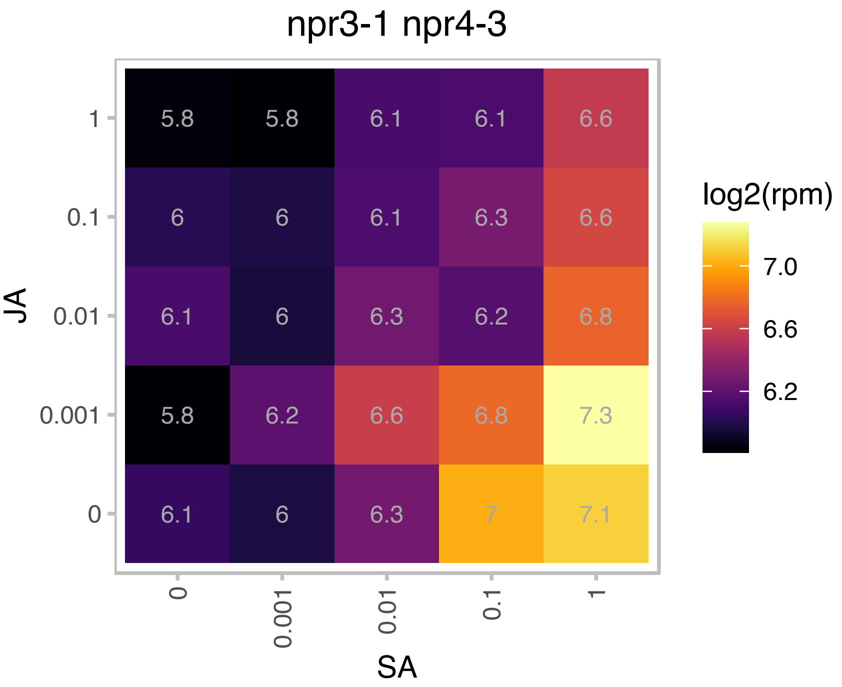

### AT4G11890.1 : ARCK1, CRK45

Protein kinase superfamily protein

Encodes a receptor-like cytosolic kinase ARCK1. Negatively controls abscisic acid and osmotic stress signal transduction.

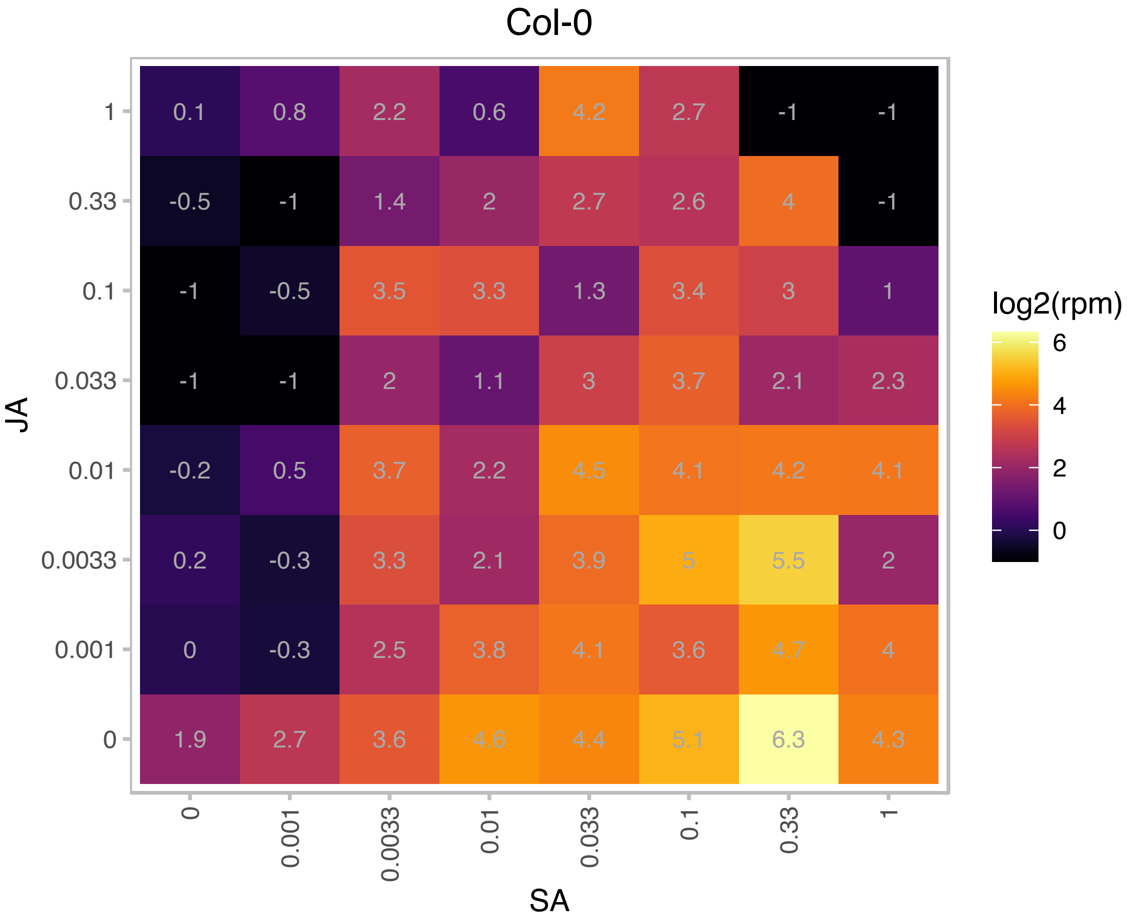


| | |
|---|---|
| Gene | AT4G11890 |
| is.expressed_genes | TRUE |
| is.responsive_genes | TRUE |
| is.apcluster | H-02 |

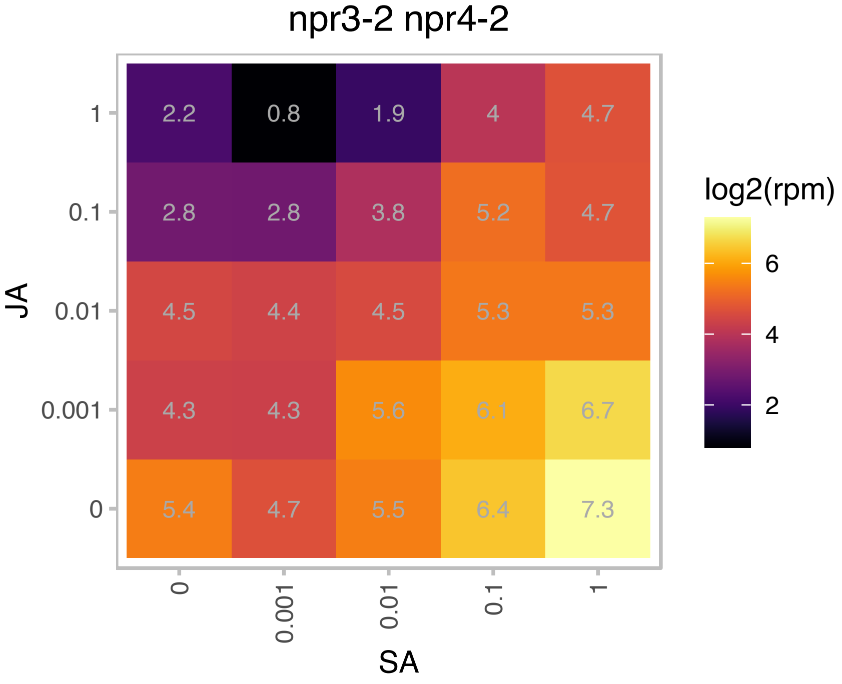


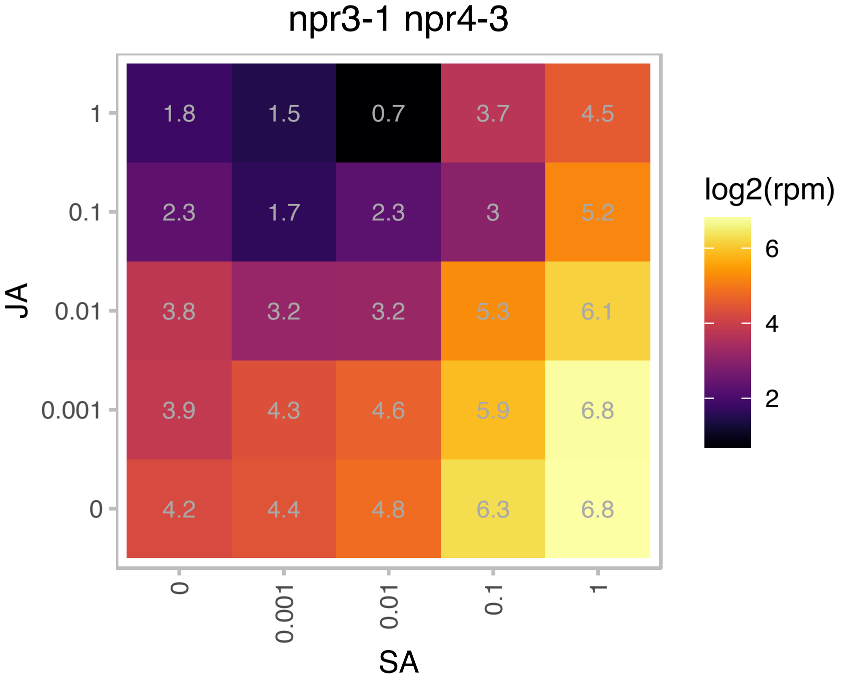

### AT4G14400.1 : ACD6

ankyrin repeat family protein

encodes a novel protein with putative ankyrin and transmembrane regions. It is a member of one of the largest uncharacterized gene families in higher plants. The gene is involved in resistance to Pseudomonas syringae.

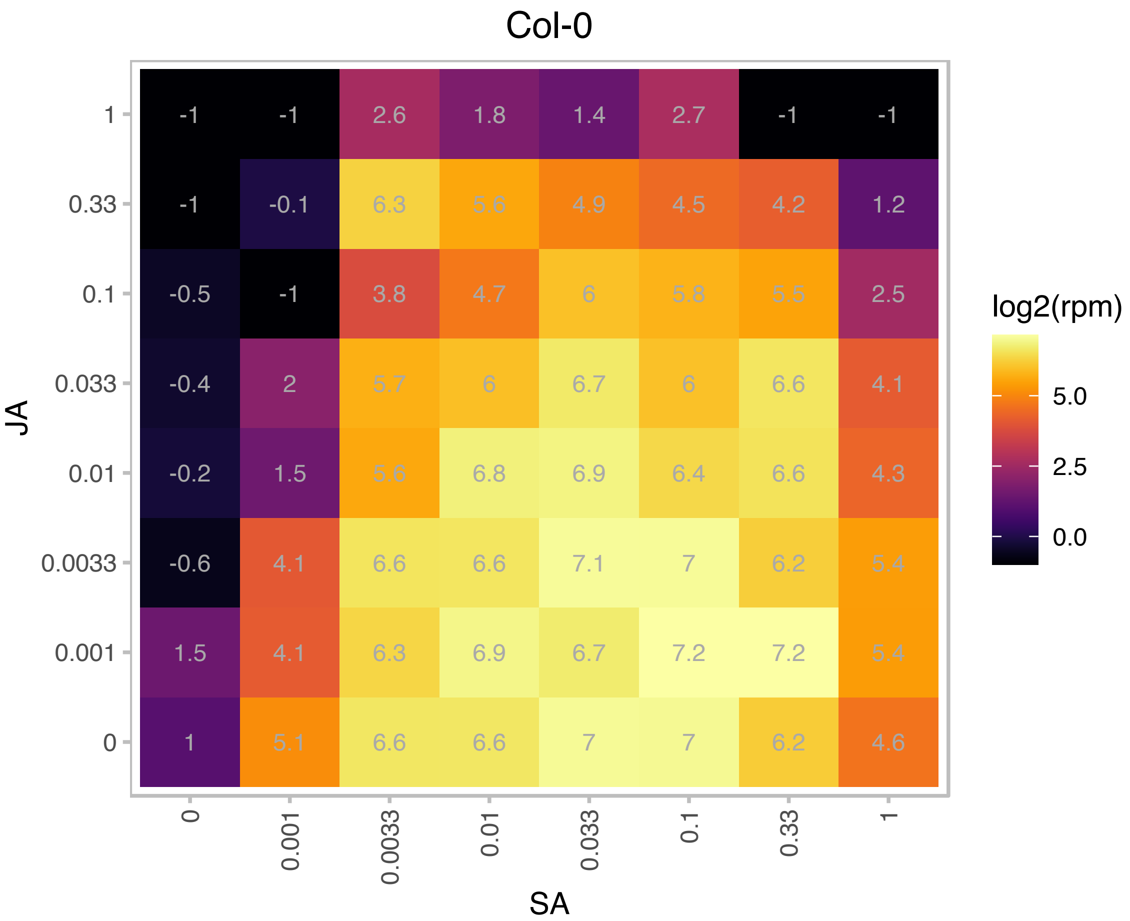


| | |
|---|---|
| Gene | AT4G14400 |
| is.expressed_genes | TRUE |
| is.responsive_genes | TRUE |
| is.apcluster | H-02 |

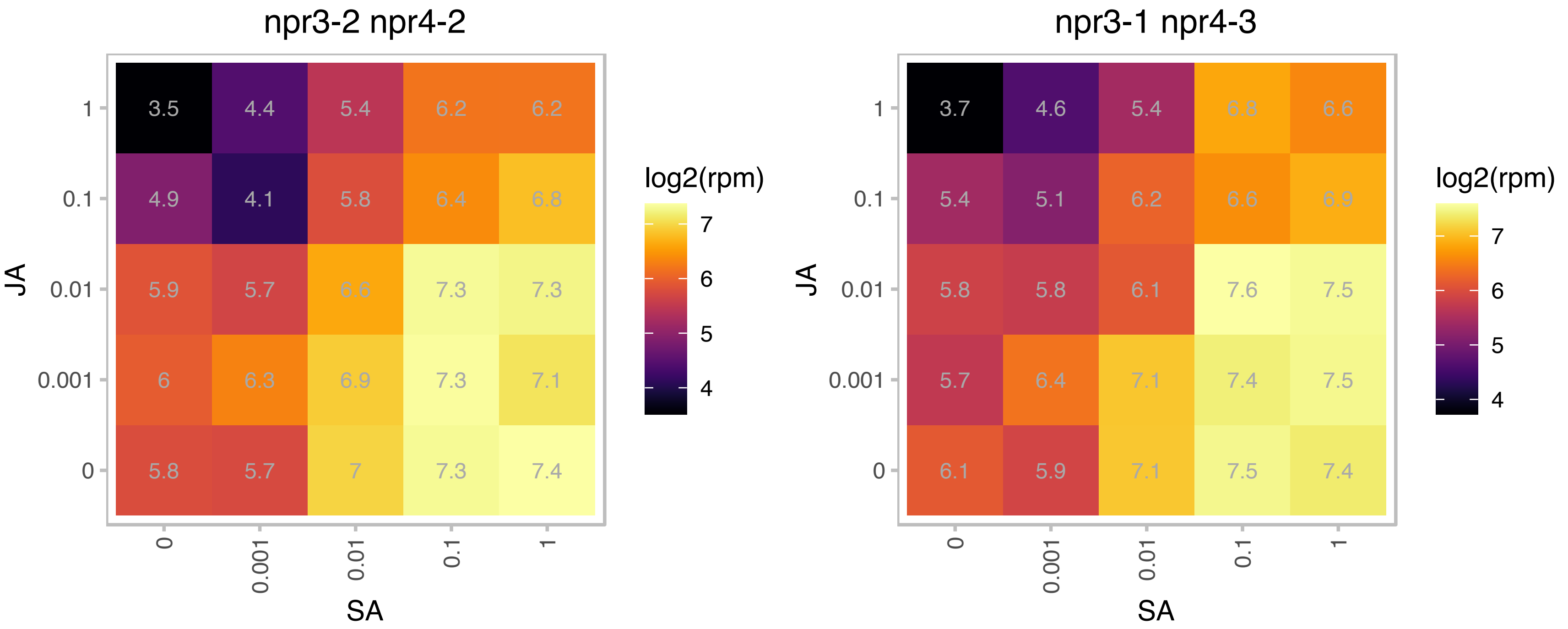

### AT4G16990.1 : RLM3

disease resistance protein (TIR-NBS class), putative

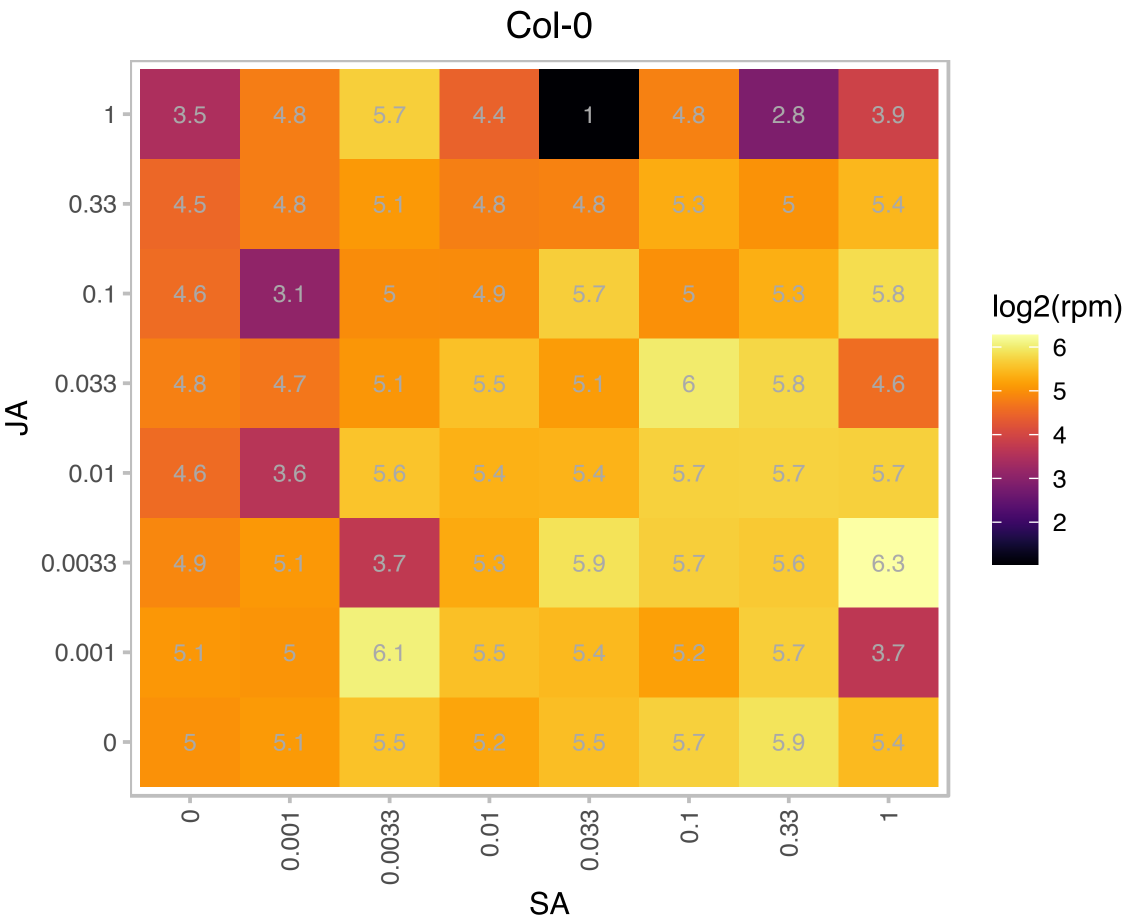


| | |
|---|---|
| Gene | AT4G16990 |
| is.expressed_genes | TRUE |
| is.responsive_genes | TRUE |
| is.apcluster | H-02 |

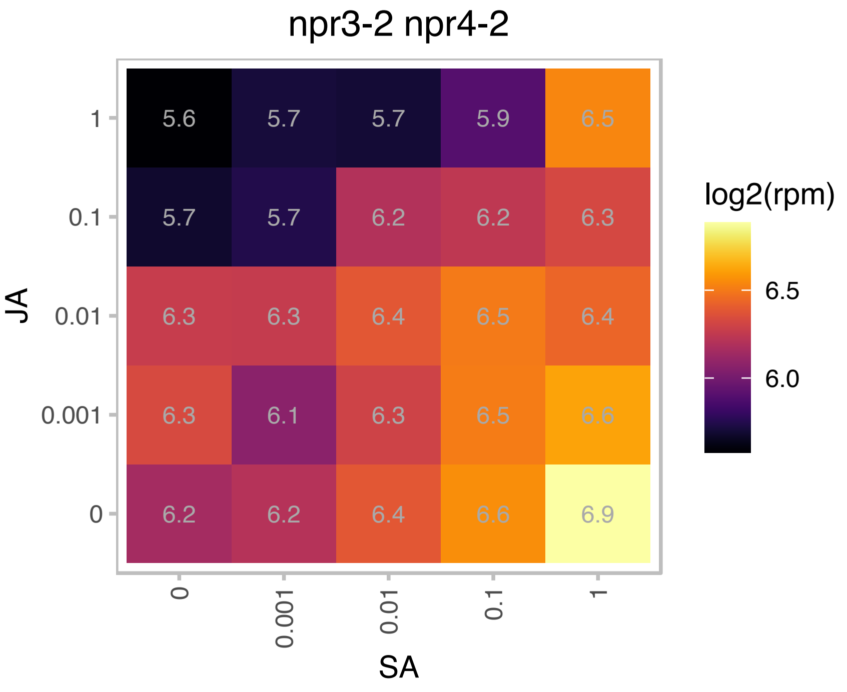


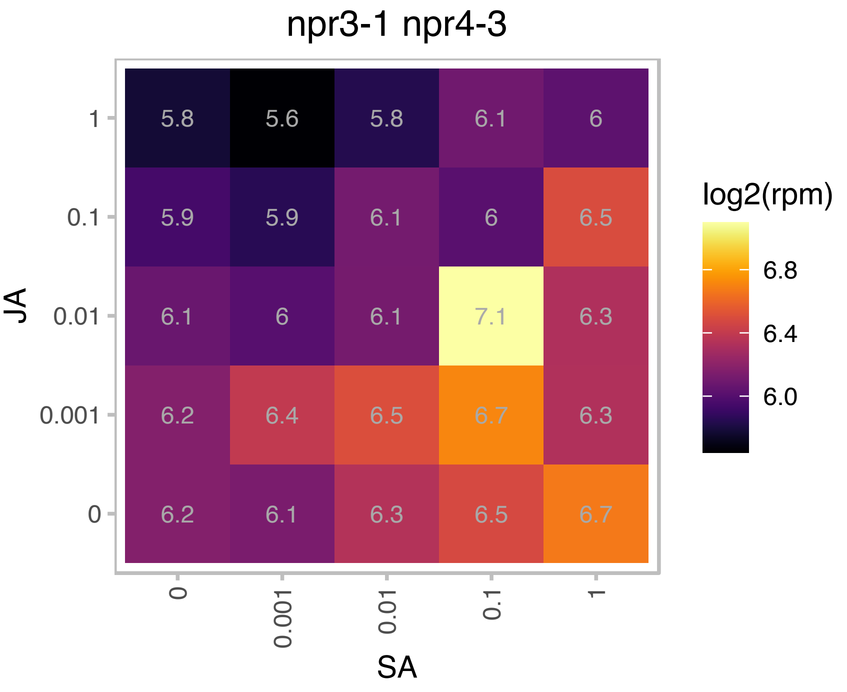

### AT4G19350.1 : EMB3006

embryo defective 3006

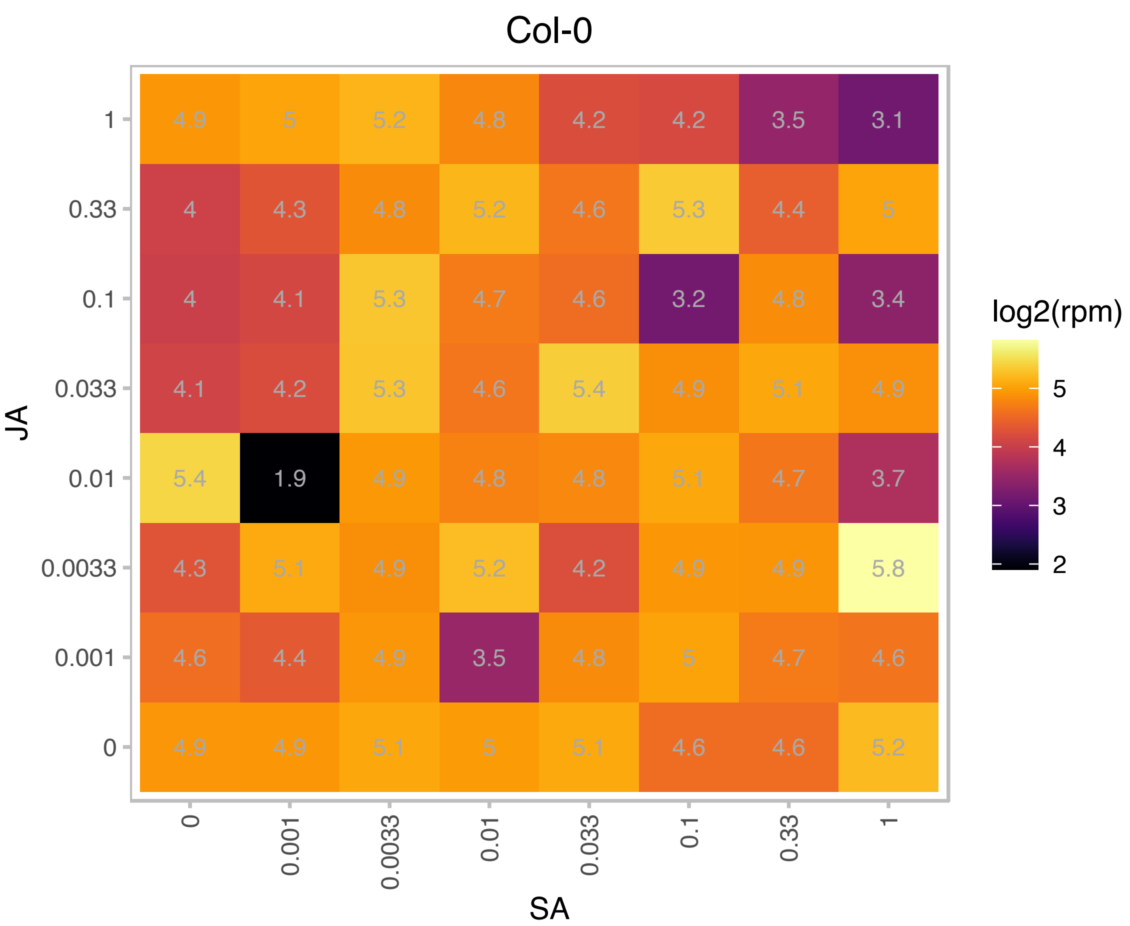


| | |
|---|---|
| Gene | AT4G19350 |
| is.expressed_genes | TRUE |
| is.responsive_genes | TRUE |
| is.apcluster | H-02 |

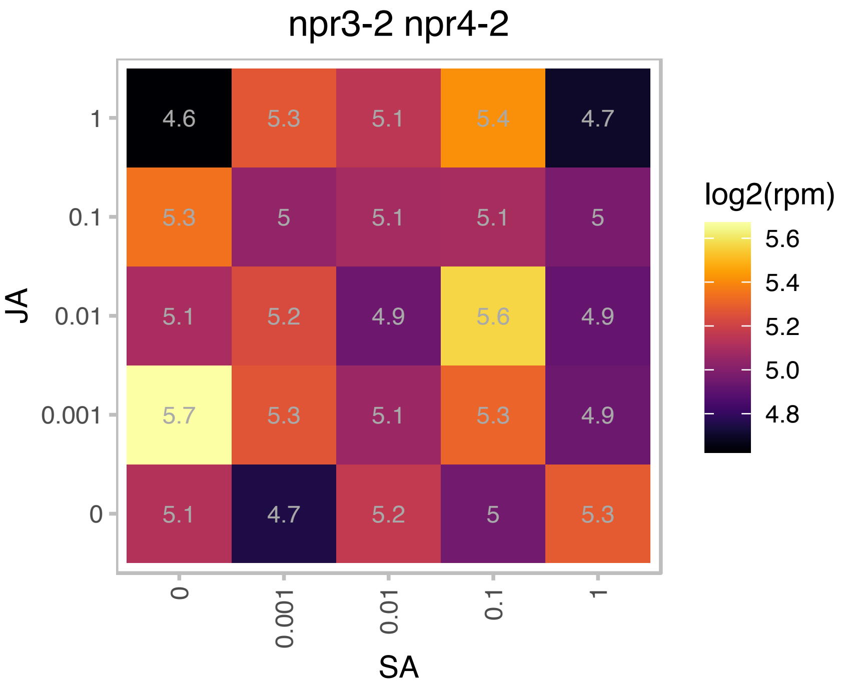


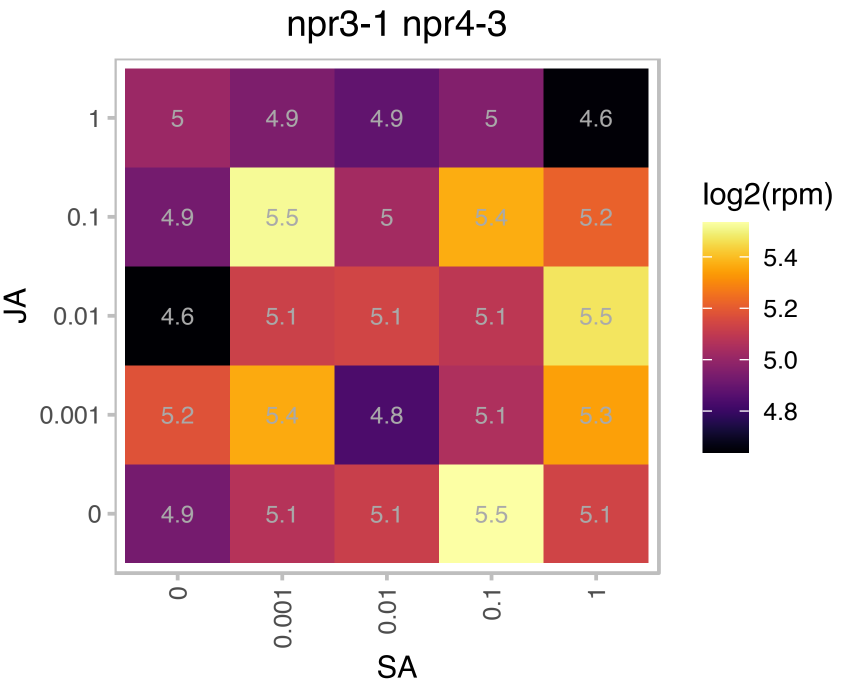

### AT4G23170.1 : CRK9, EP1

receptor-like protein kinase-related family protein
Induced in response to Salicylic acid.Similar to receptor-like kinase 4 and 5. NPR1, a known positive regulator of the SA signaling pathway is responsible for the SA-dependent induction and constitutive repression of EP1 gene's basal expression. The mRNA is cell-to-cell mobile.

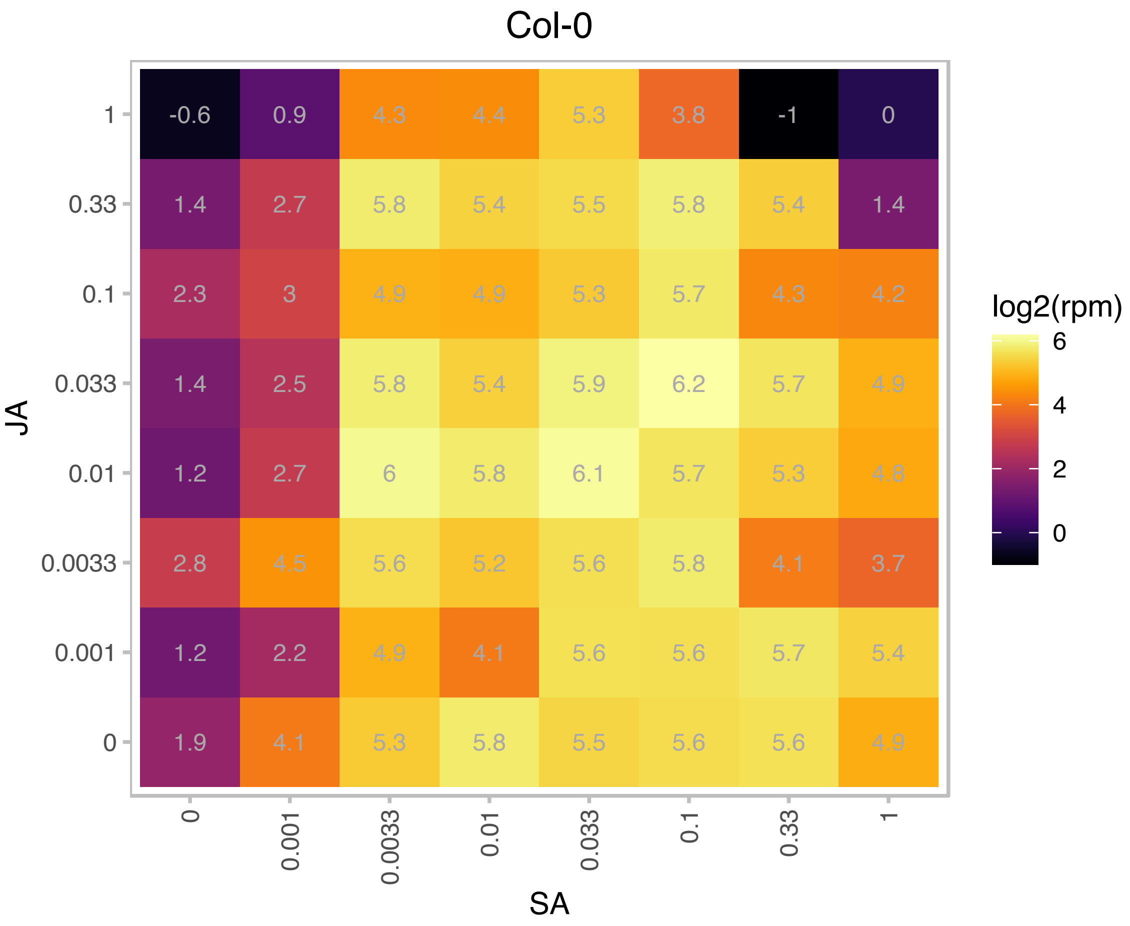


| | |
|---|---|
| Gene | AT4G23170 |
| is.expressed_genes | TRUE |
| is.responsive_genes | TRUE |
| is.apcluster | H-02 |

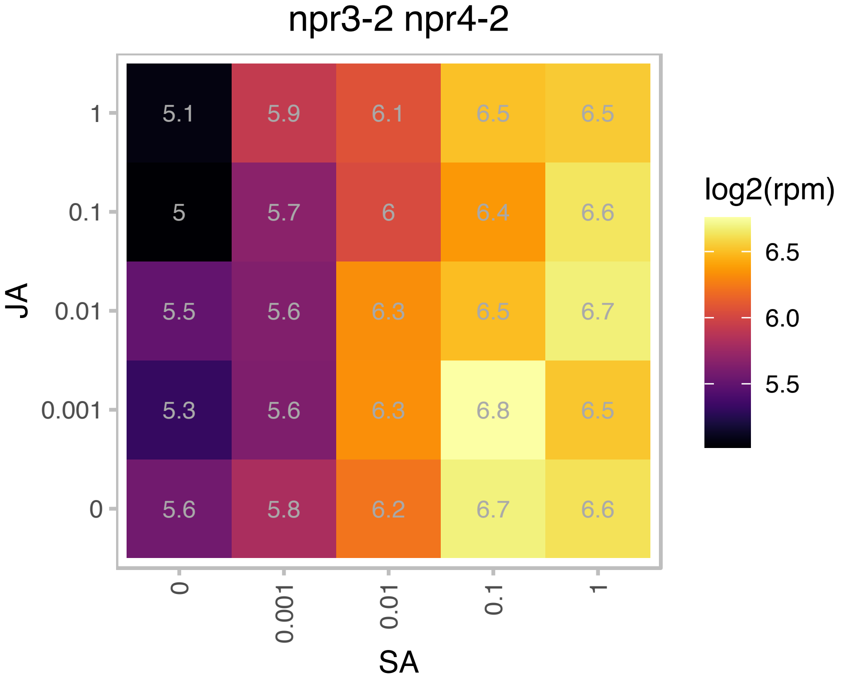


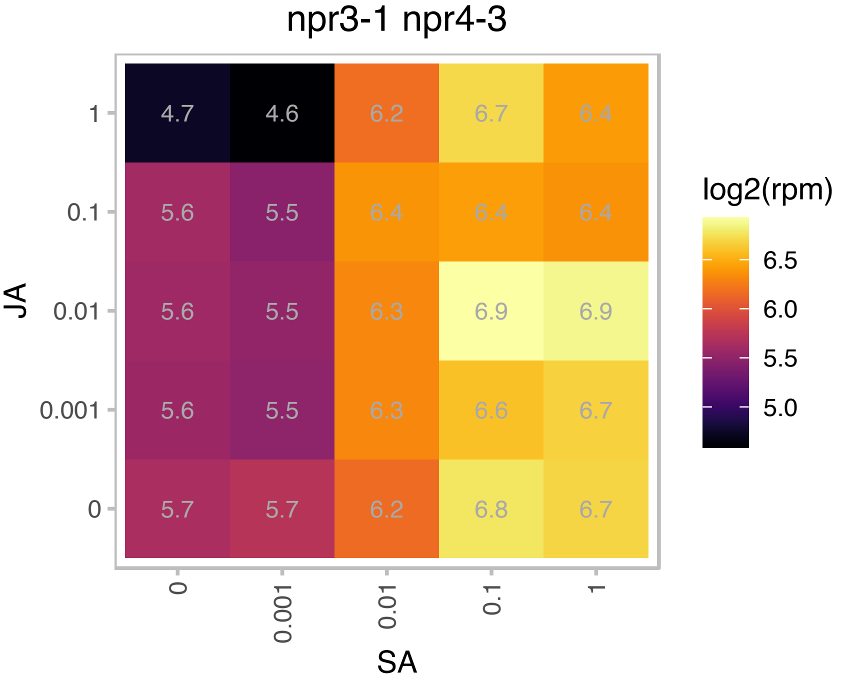

### AT4G23260.1 : CRK18

cysteine-rich RLK (RECEPTOR-like protein kinase) 18

Encodes a cysteine-rich receptor-like protein kinase.

| | |
|---|---|
| Gene | AT4G23260 |
| is.expressed_genes | TRUE |
| is.responsive_genes | TRUE |
| is.apcluster | H-02 |

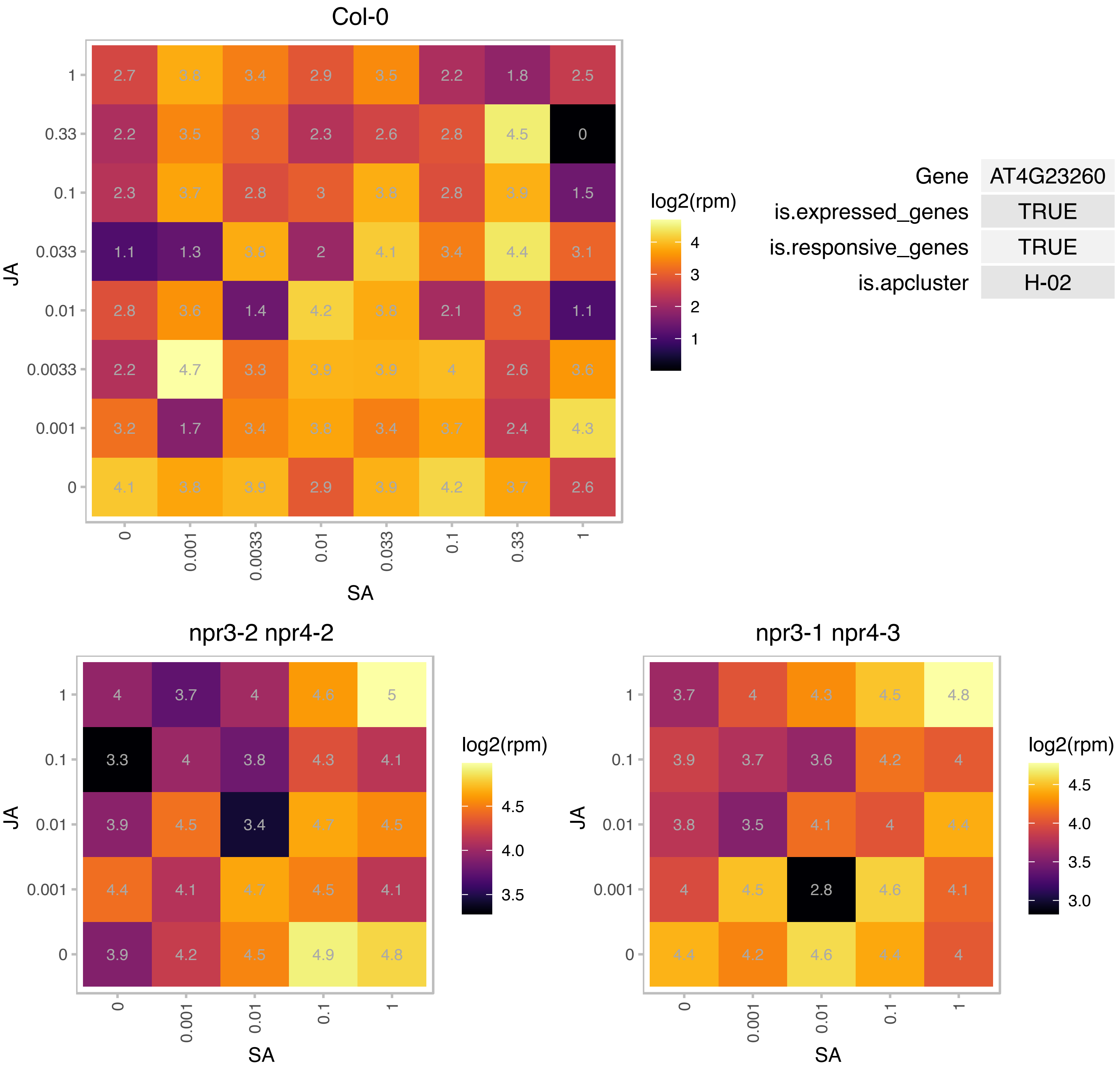

### AT4G25110.1 : AtMC2, AtMCP1c, MC2, MCP1c

metacaspase 2

Encodes a type I metacaspase. Two Arabidopsis metacaspases, AT1G02170 (MC1) and AT4G25110 (MC2) antagonistically control programmed cell death in Arabidopsis. MC1 is a positive regulator of cell death and requires conserved caspase-like putative catalytic residues for its function. MC2 negatively regulates cell death. This function is independent of the putative catalytic residues.  A third type I Arabidopsis metacaspase is MC3 (AT5g64240).

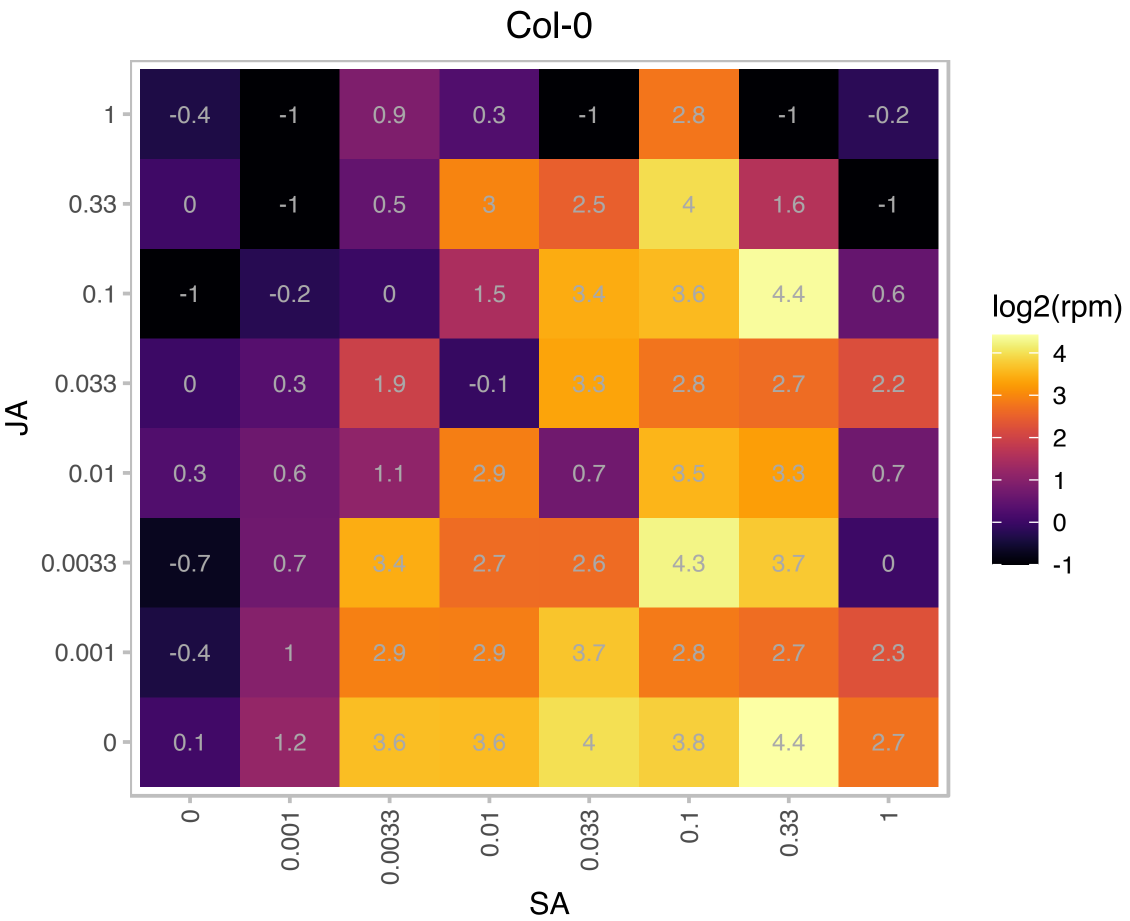


| | |
|---|---|
| Gene | AT4G25110 |
| is.expressed_genes | TRUE |
| is.responsive_genes | TRUE |
| is.apcluster | H-02 |

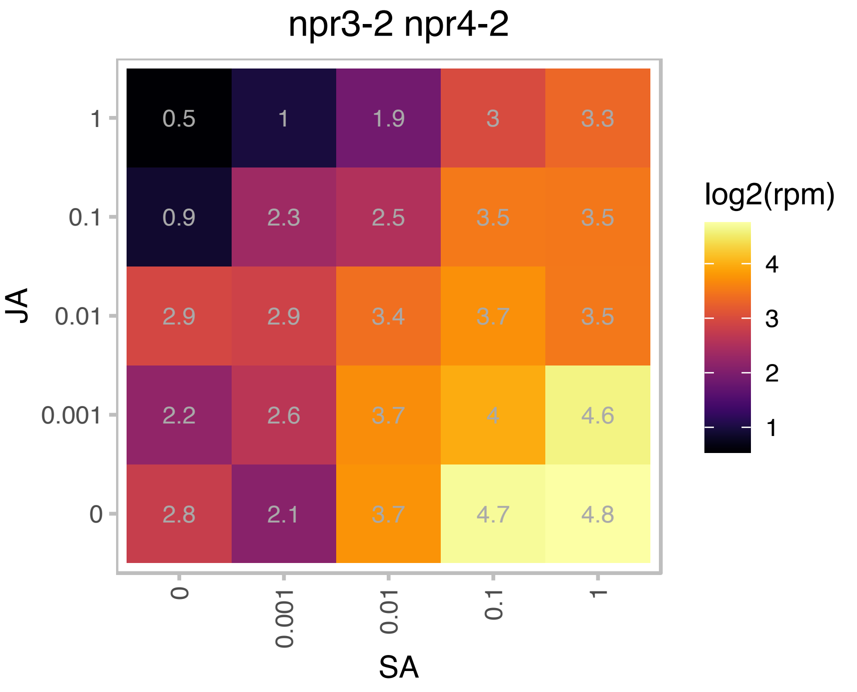


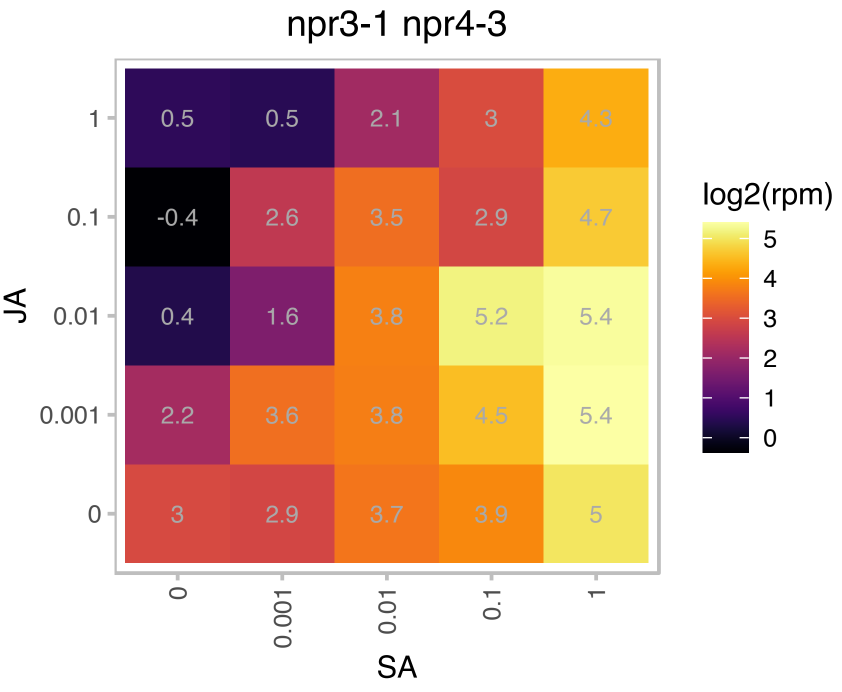

### AT4G31600.1 : UTr7

UDP-N-acetylglucosamine (UAA) transporter family

Encodes a Golgi-localized UDP□ □ lucose/UDP□ □ alactose transporter that affects lateral root emergence.

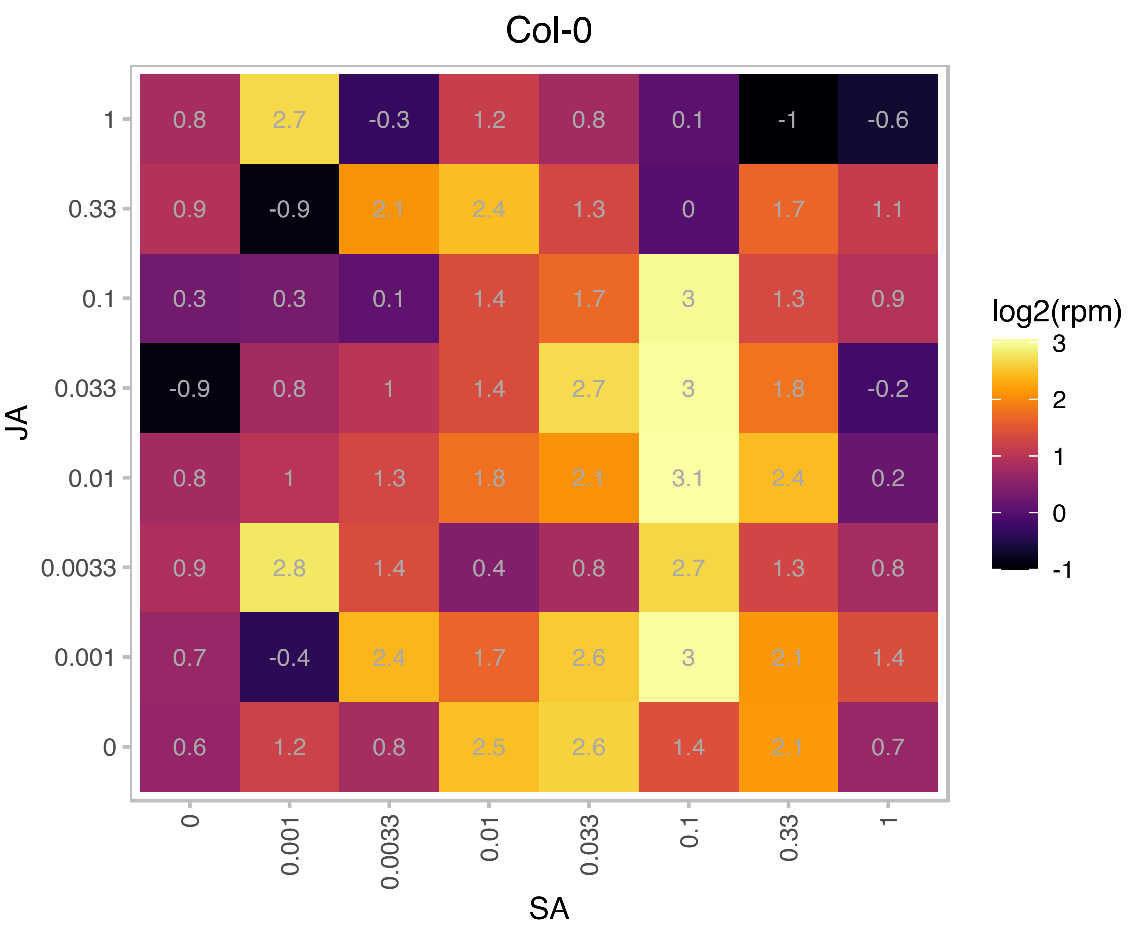


| | |
|---|---|
| Gene | AT4G31600 |
| is.expressed_genes | TRUE |
| is.responsive_genes | TRUE |
| is.apcluster | H-02 |

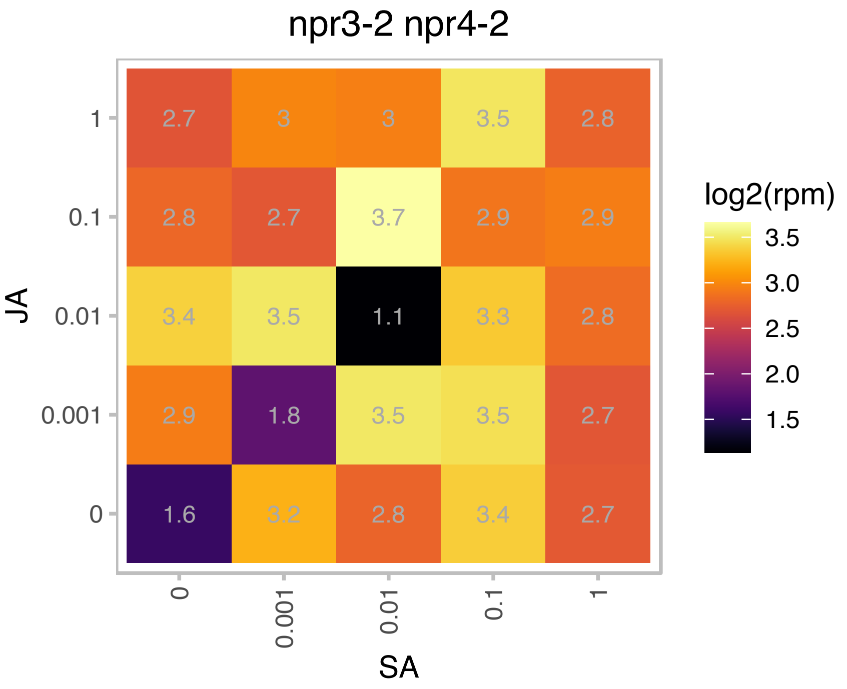


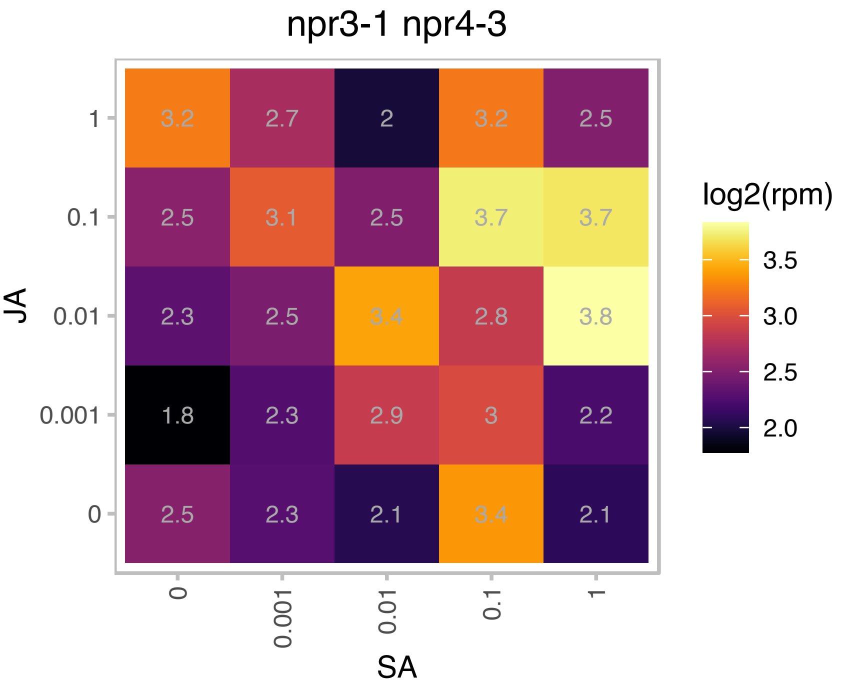

### AT4G35020.1 : ARAC3, ATROP6, RAC3, RHO1PS, ROP6

RAC-like 3

A member of ROP GTPase gene family; Encodes a Rho-like GTP binding protein.

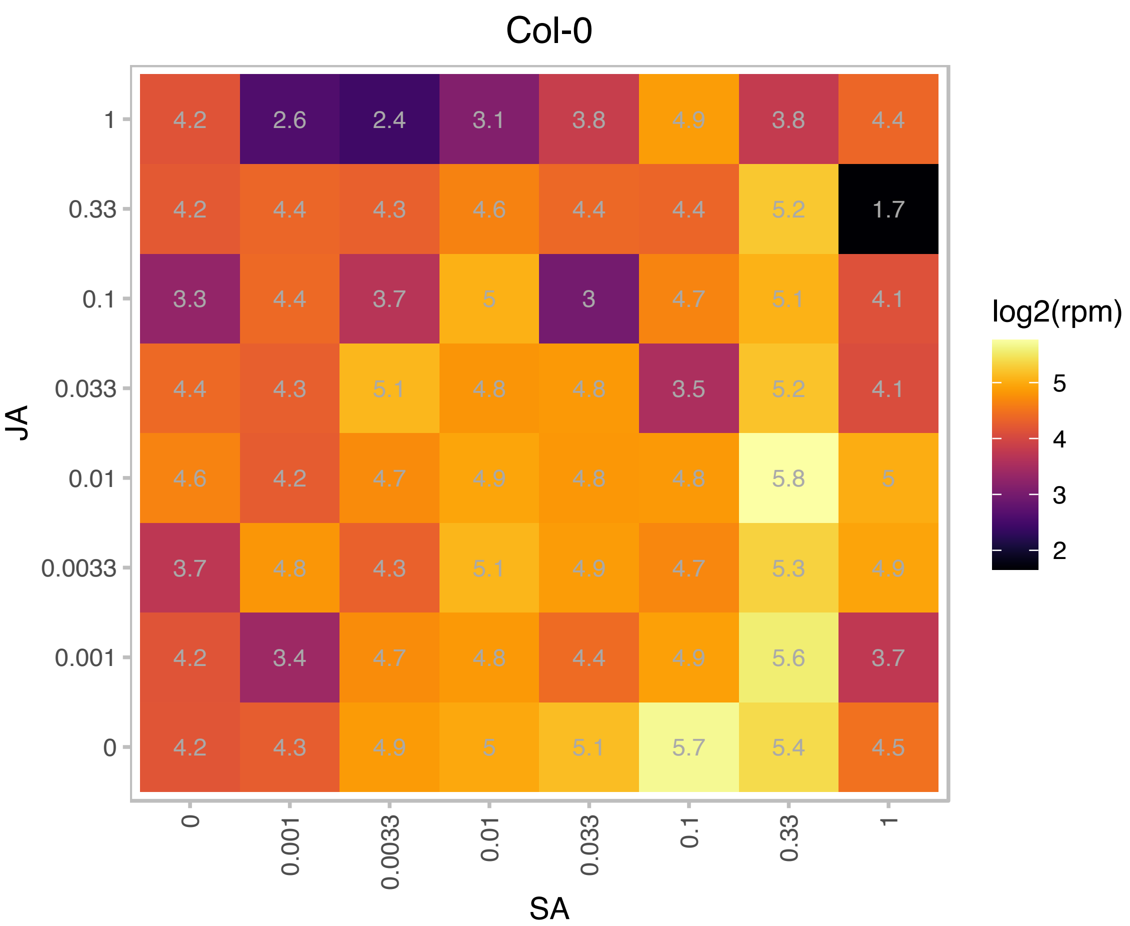


| | |
|---|---|
| Gene | AT4G35020 |
| is.expressed_genes | TRUE |
| is.responsive_genes | TRUE |
| is.apcluster | H-02 |

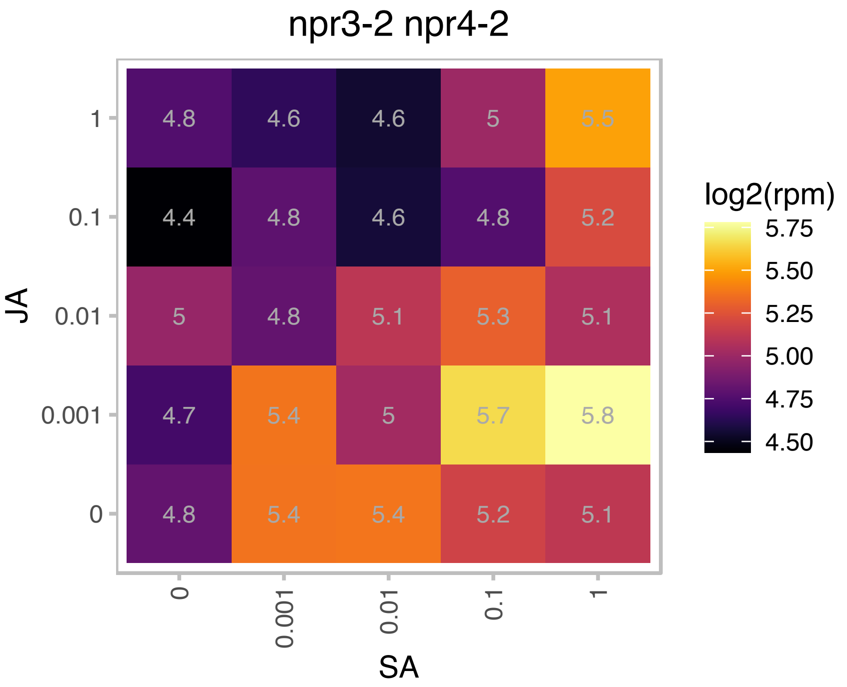


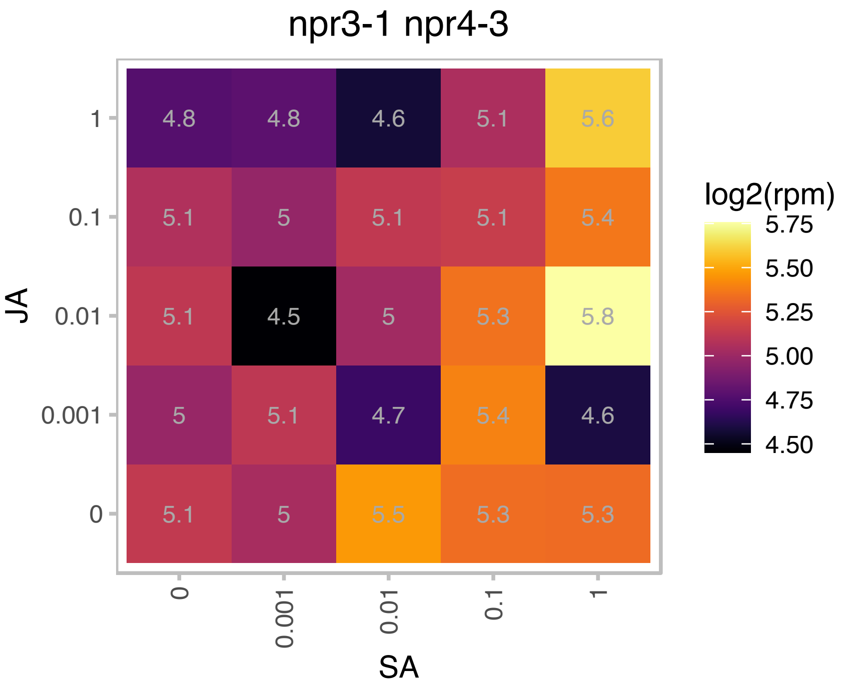

# AT5G03440.1 : NA

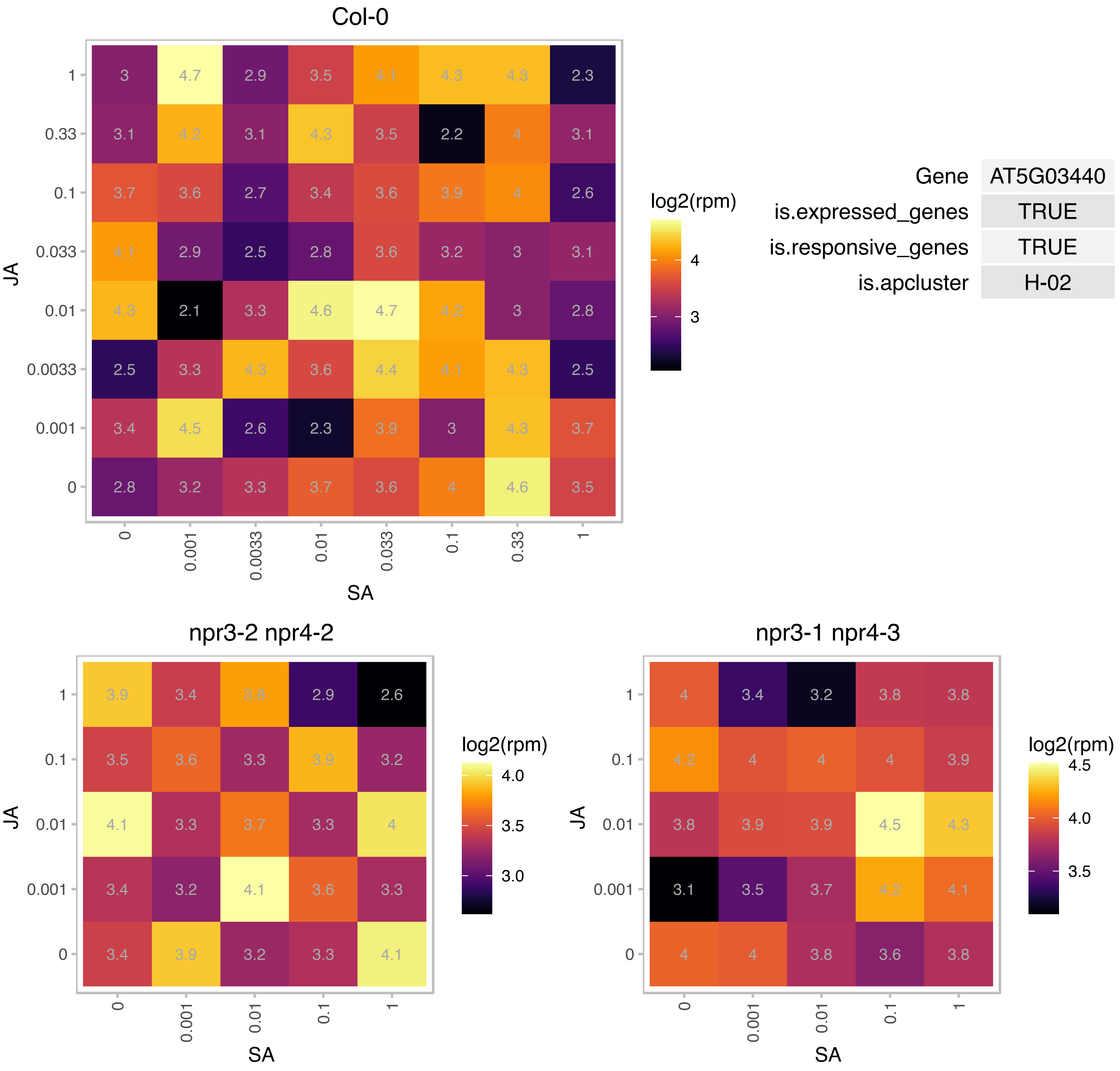

### AT5G08120.1 : MPB2C

movement protein binding protein 2C
Microtubule-associated and viral movement protein binding protein. Negatively regulates KN1 association to plasmodesmata and, consequently, cell-to-cell transport. Involved in the alignment of cortical microtubules, the patterning of stomata and in restricting tobamoviral infections.

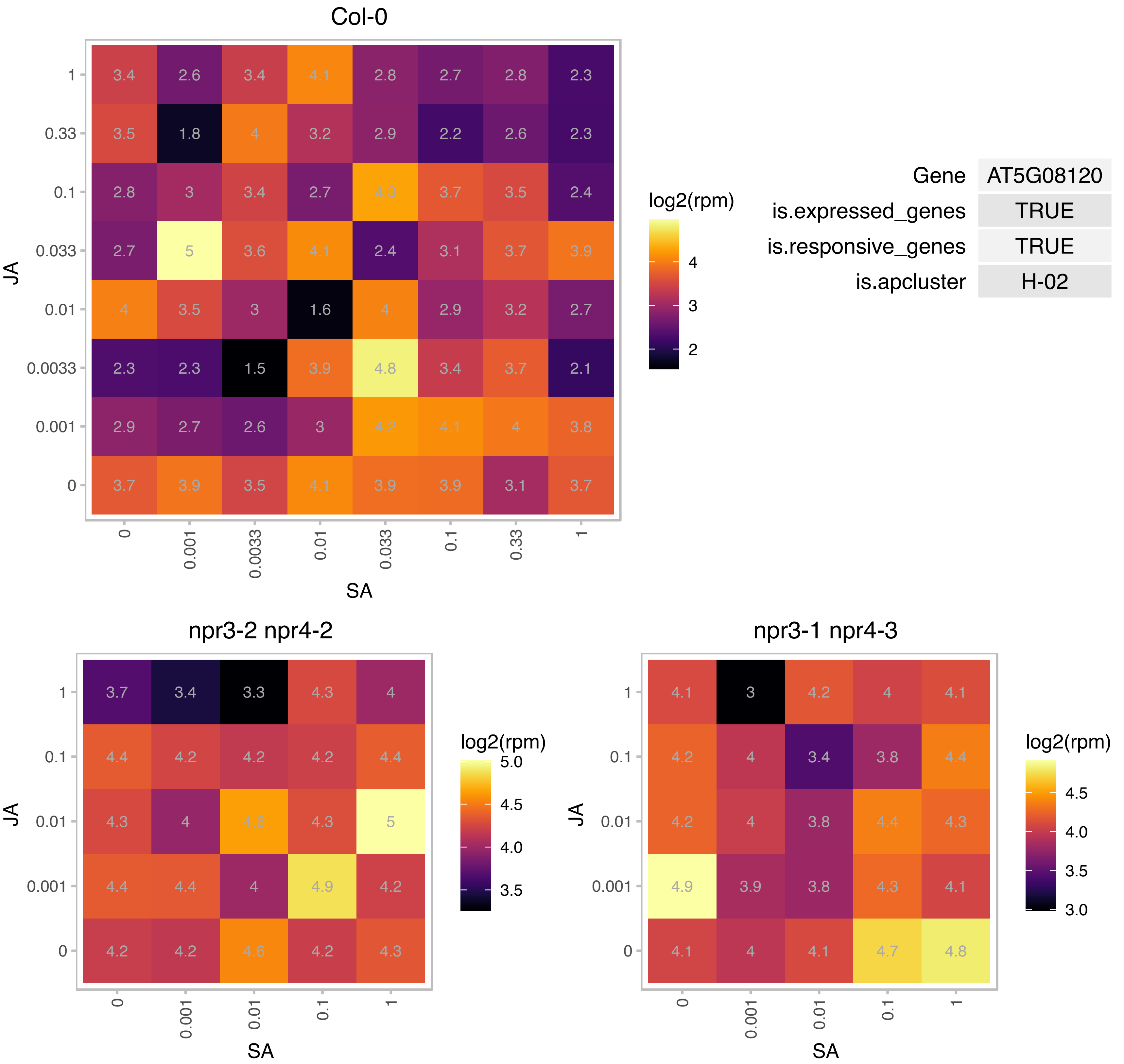


| | |
|---|---|
| Gene | AT5G08120 |
| is.expressed_genes | TRUE |
| is.responsive_genes | TRUE |
| is.apcluster | H-02 |

# AT5G08760.1 : NA

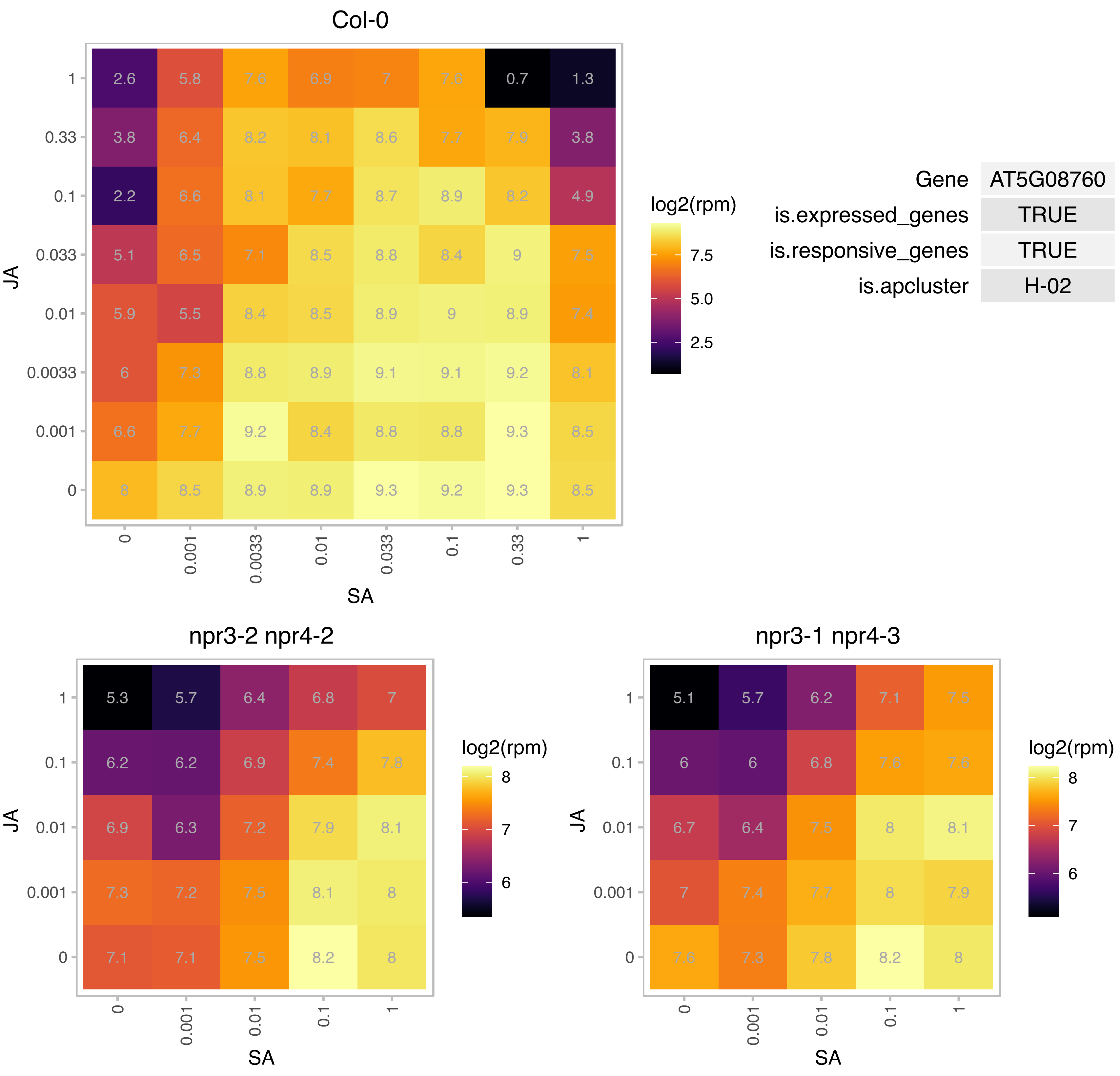

# AT5G09290.1 : NA

Inositol monophosphatase family protein

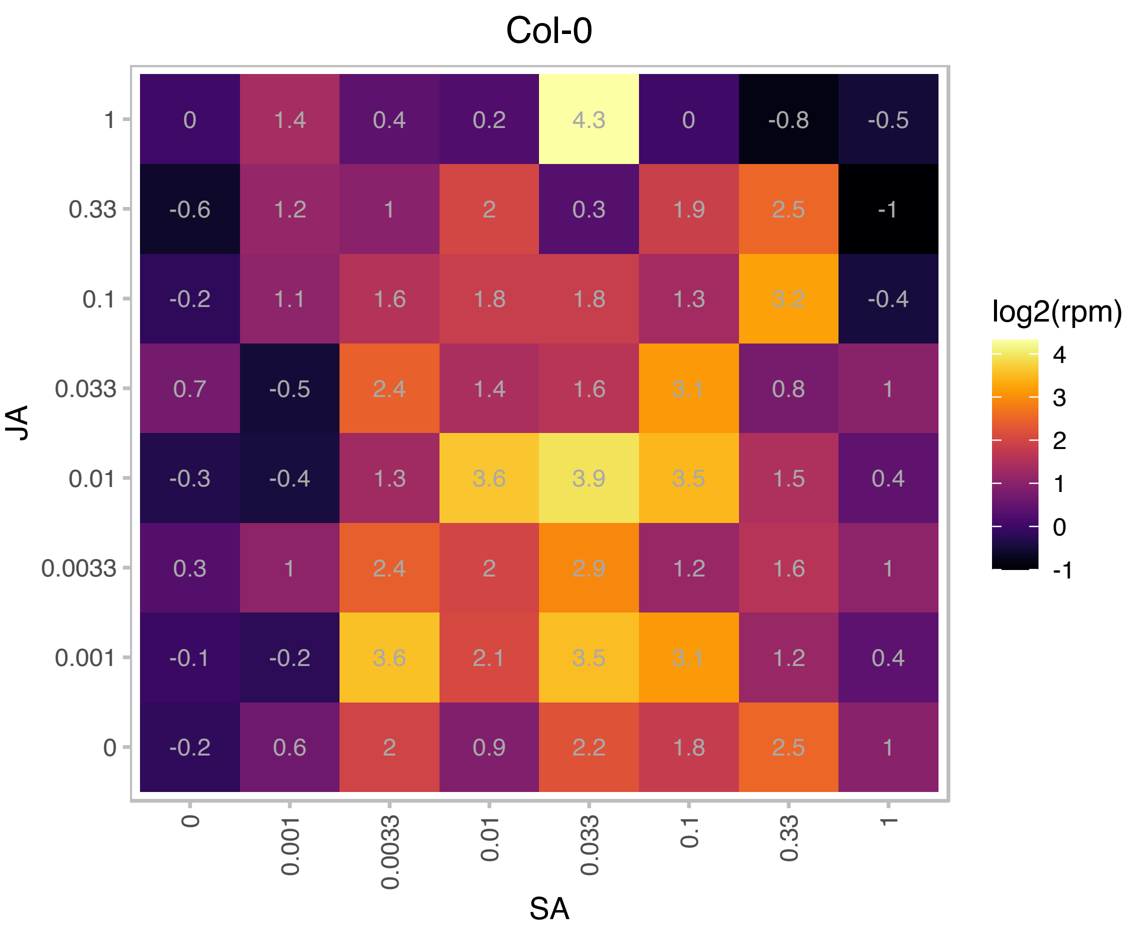


| | |
|---|---|
| Gene | AT5G09290 |
| is.expressed_genes | TRUE |
| is.responsive_genes | TRUE |
| is.apcluster | H-02 |

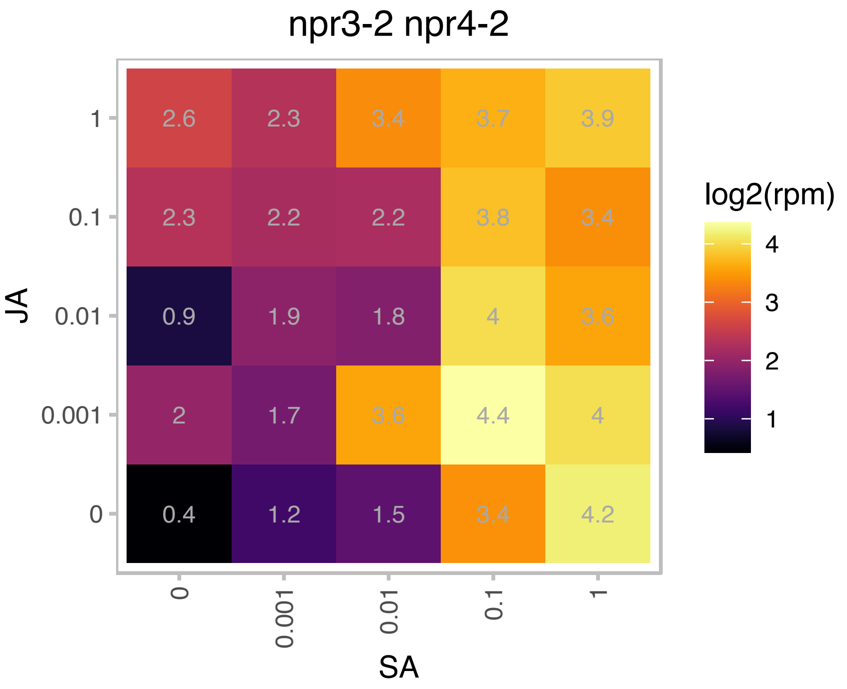


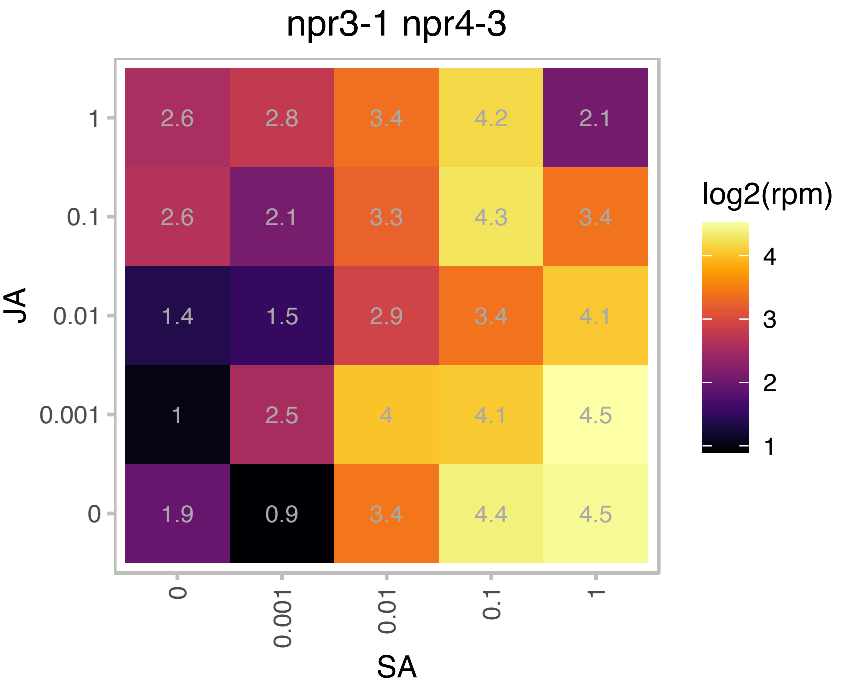

### AT5G10810.1 : ATER, ER

enhancer of rudimentary protein, putative

enhancer of rudimentary homolog ATER

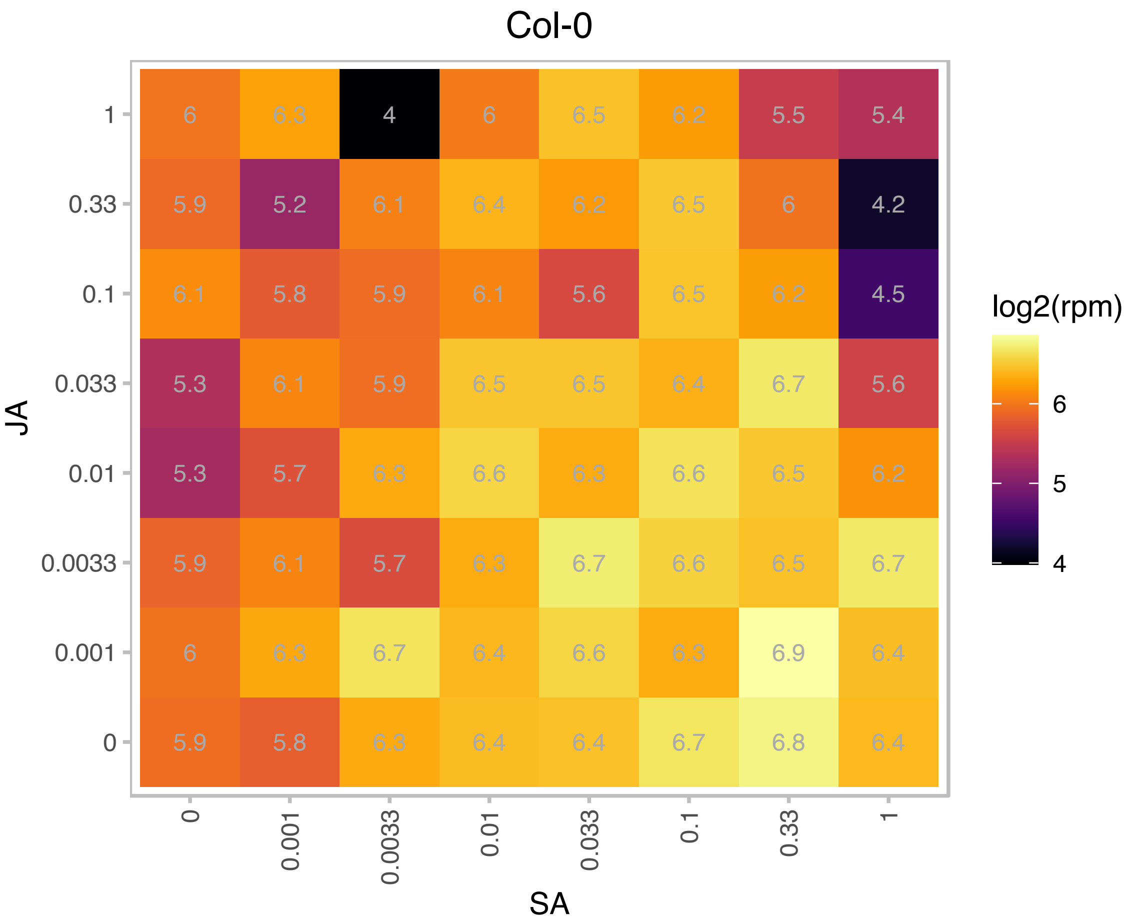


| | |
|---|---|
| Gene | AT5G10810 |
| is.expressed_genes | TRUE |
| is.responsive_genes | TRUE |
| is.apcluster | H-02 |

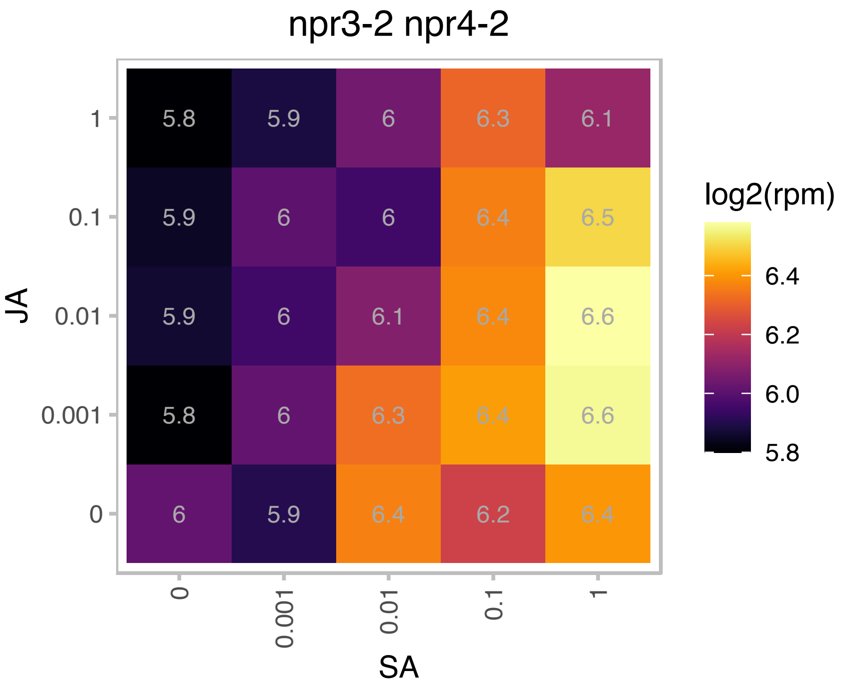


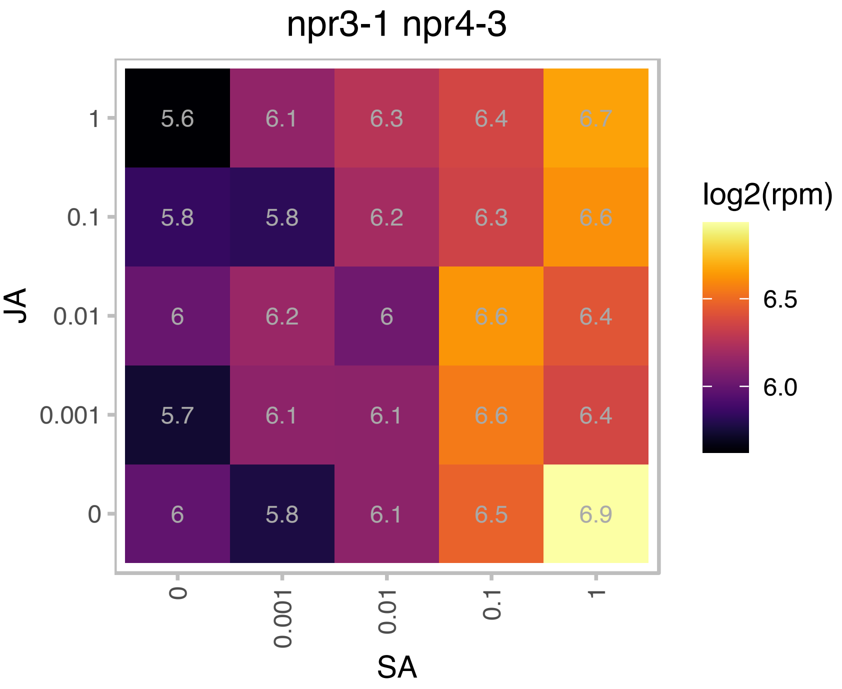

# AT5G10960.1 : NA

Polynucleotidyl transferase, ribonuclease H-like superfamily protein

| | |
|---|---|
| Gene | AT5G10960 |
| is.expressed_genes | TRUE |
| is.responsive_genes | TRUE |
| is.apcluster | H-02 |

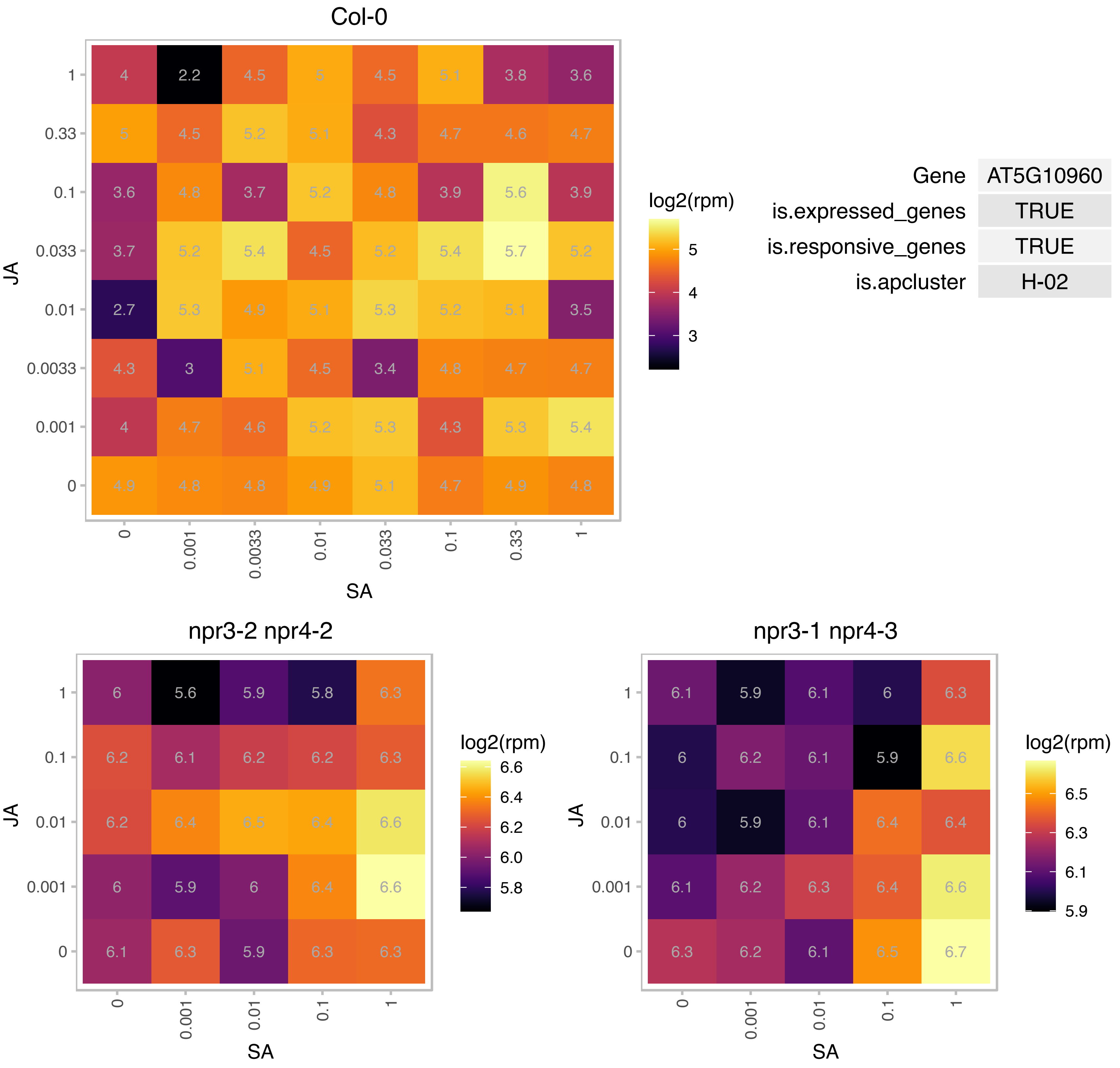

# AT5G19240.1 : NA

Glycoprotein membrane precursor GPI-anchored

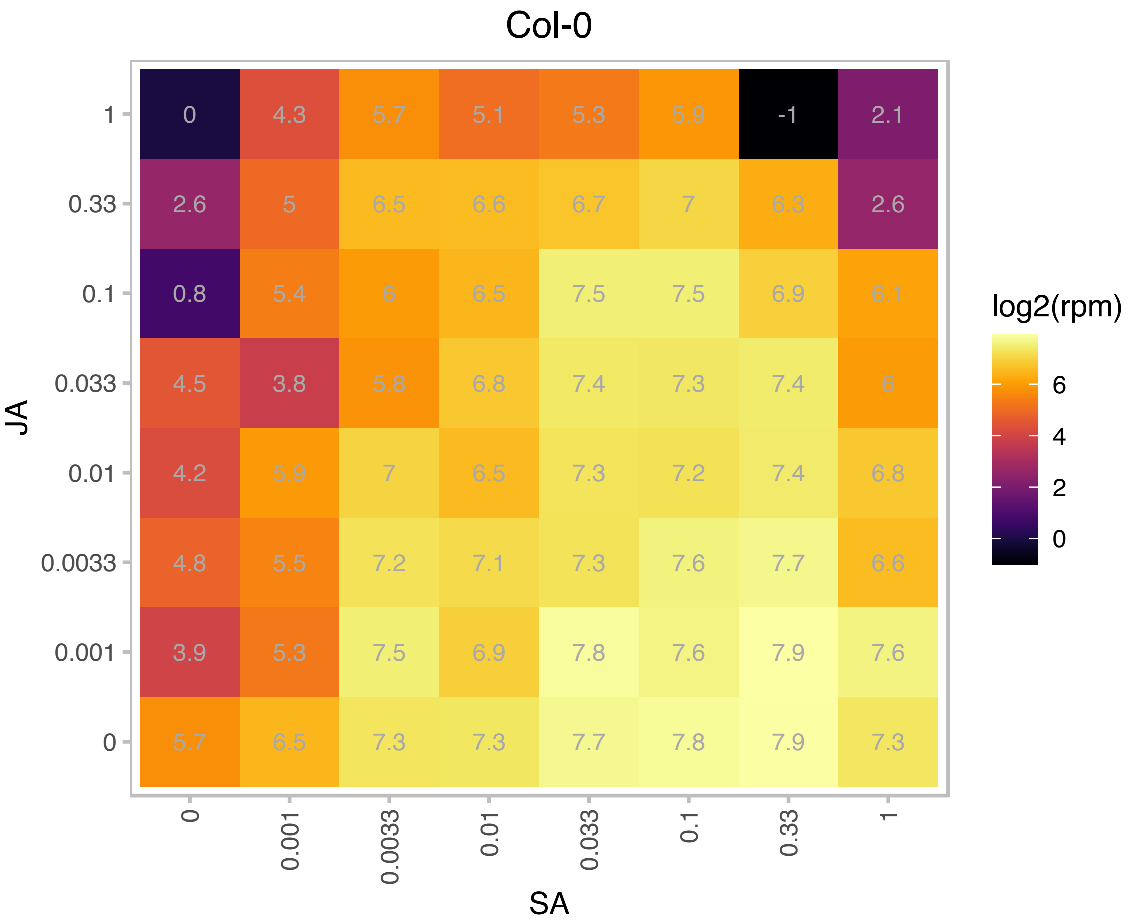


| | |
|---|---|
| Gene | AT5G19240 |
| is.expressed_genes | TRUE |
| is.responsive_genes | TRUE |
| is.apcluster | H-02 |

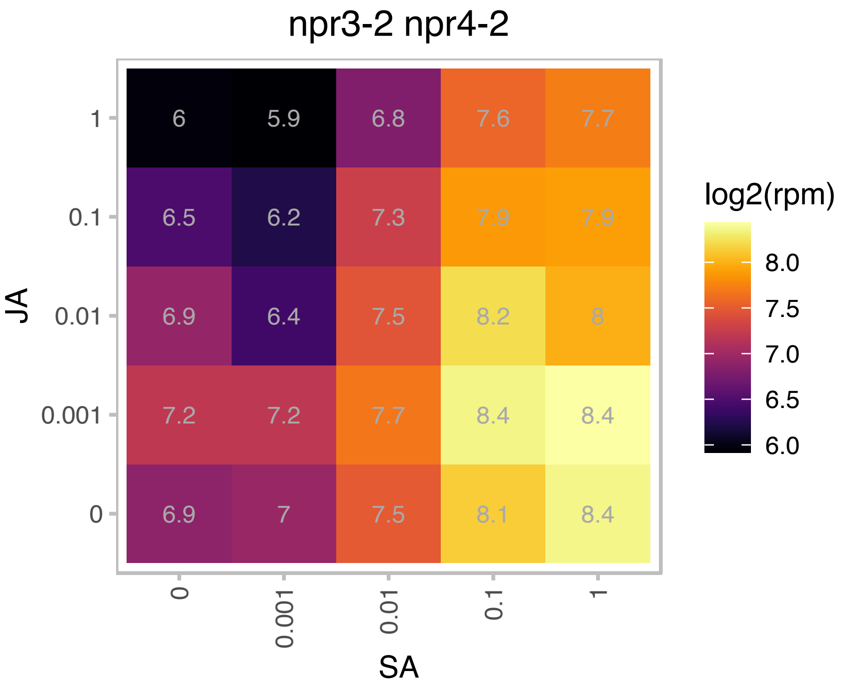


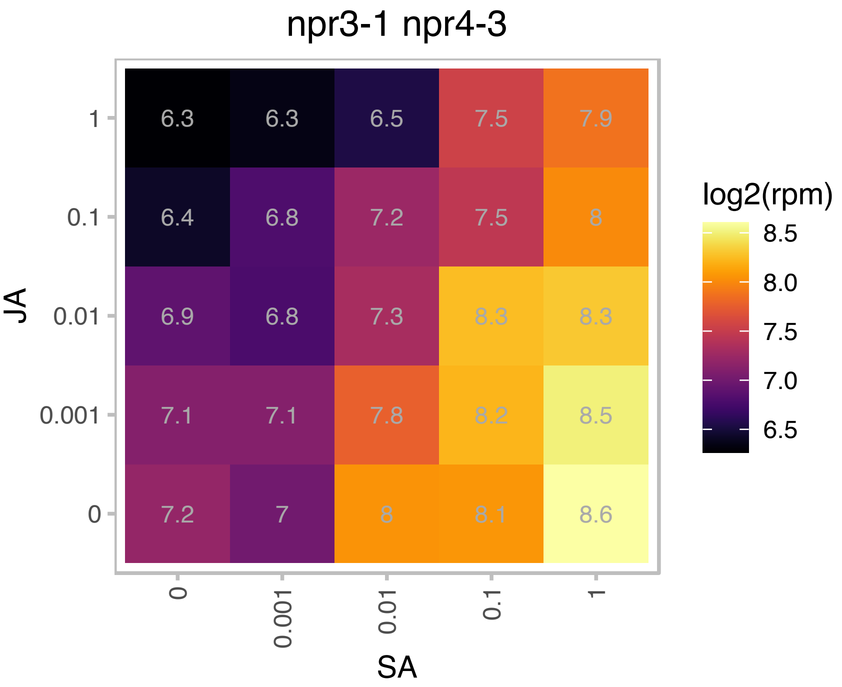

# AT5G38990.1 : NA

Malectin/receptor-like protein kinase family protein

| | |
|---|---|
| Gene | AT5G38990 |
| is.expressed_genes | TRUE |
| is.responsive_genes | TRUE |
| is.apcluster | H-02 |

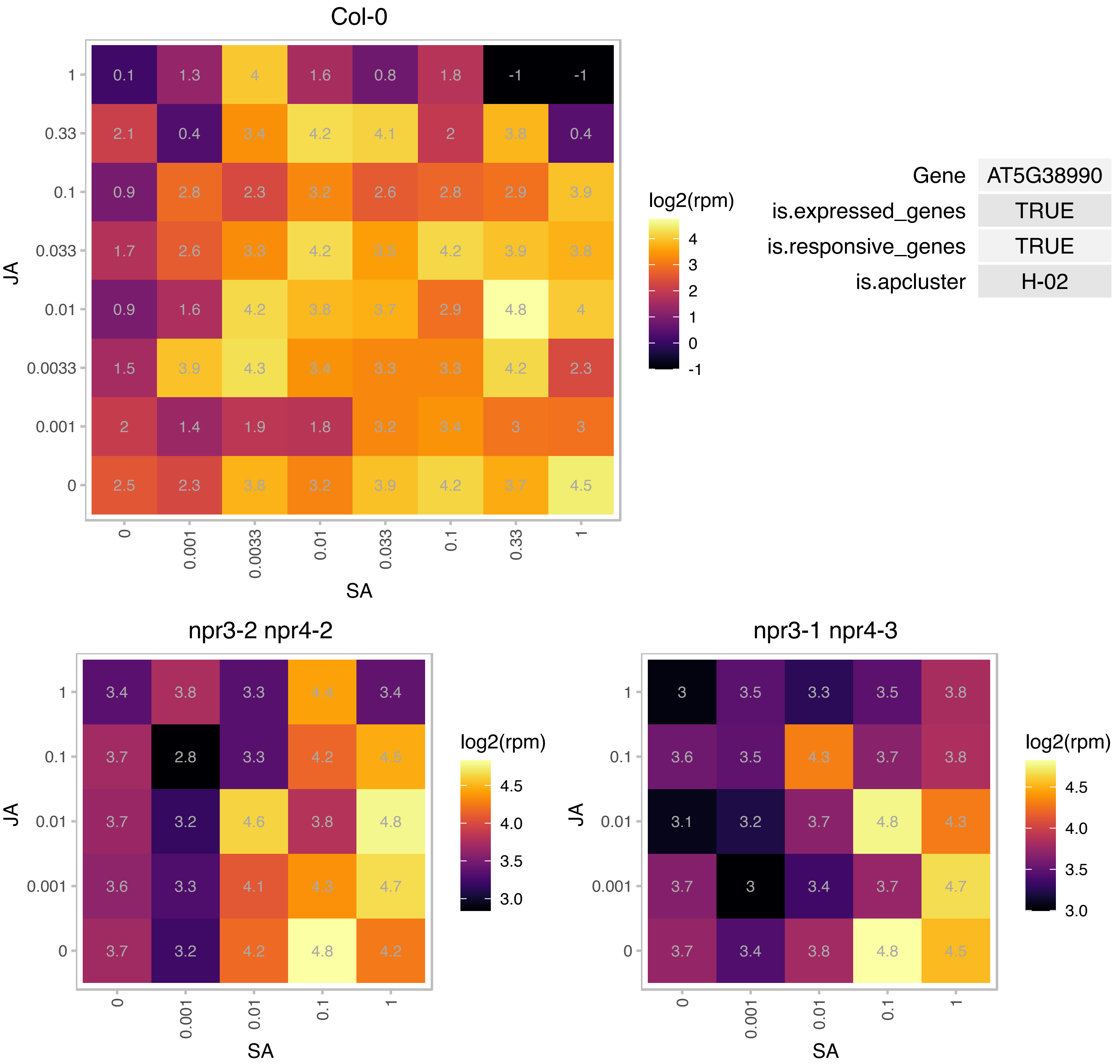

# AT5G39670.1 : NA

Calcium-binding EF-hand family protein

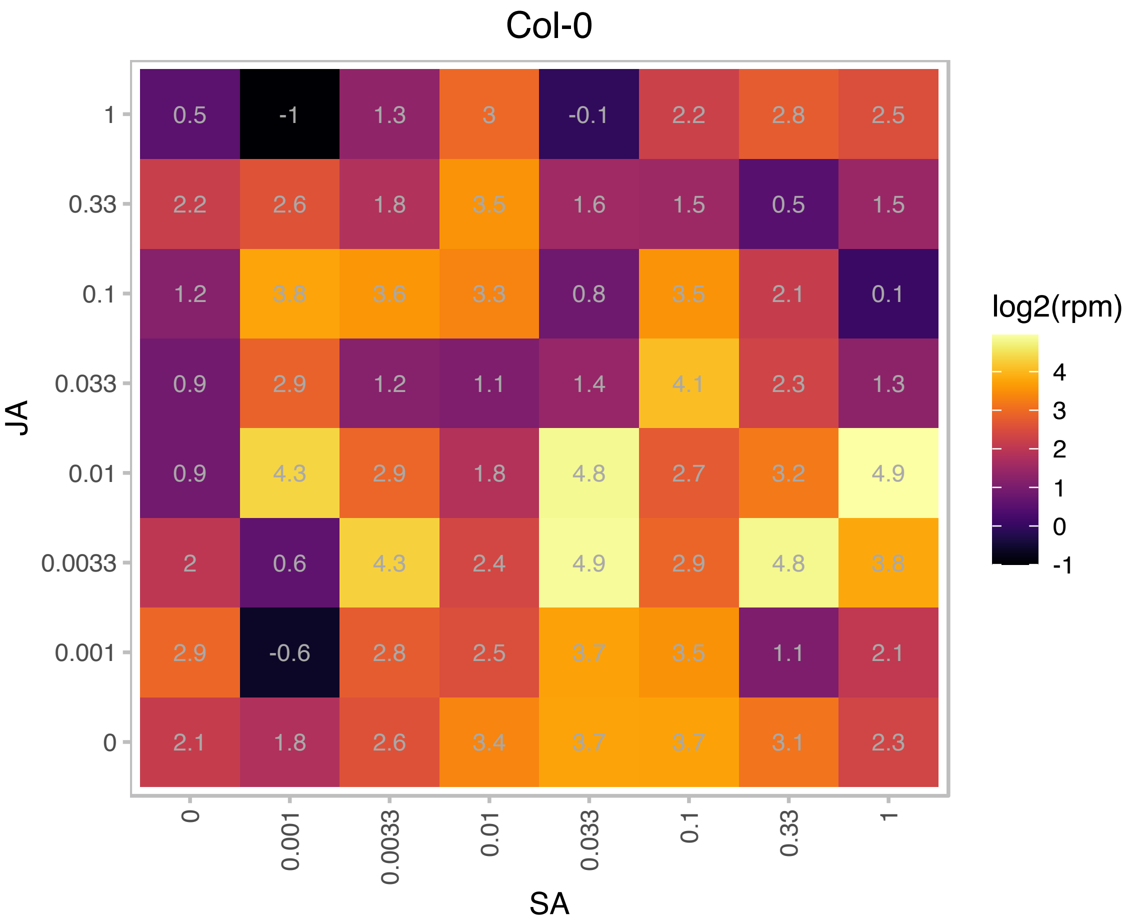


| | |
|---|---|
| Gene | AT5G39670 |
| is.expressed_genes | TRUE |
| is.responsive_genes | TRUE |
| is.apcluster | H-02 |

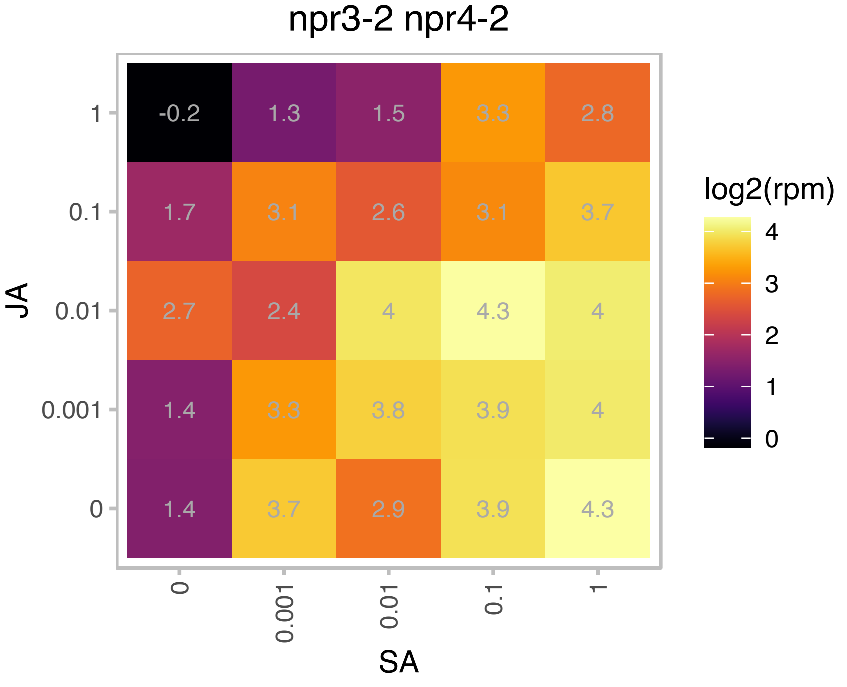


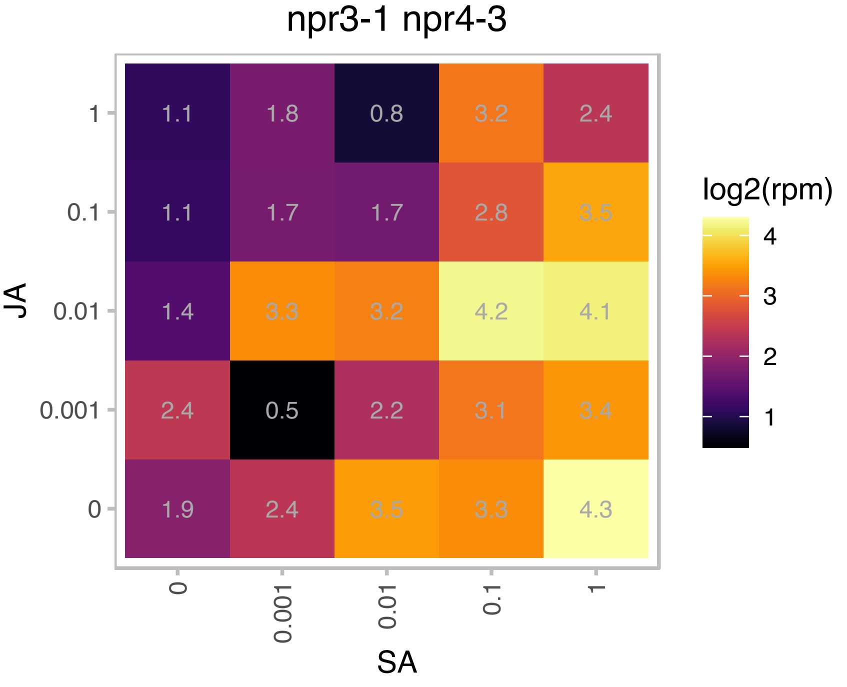

# AT5G42530.1 : NA

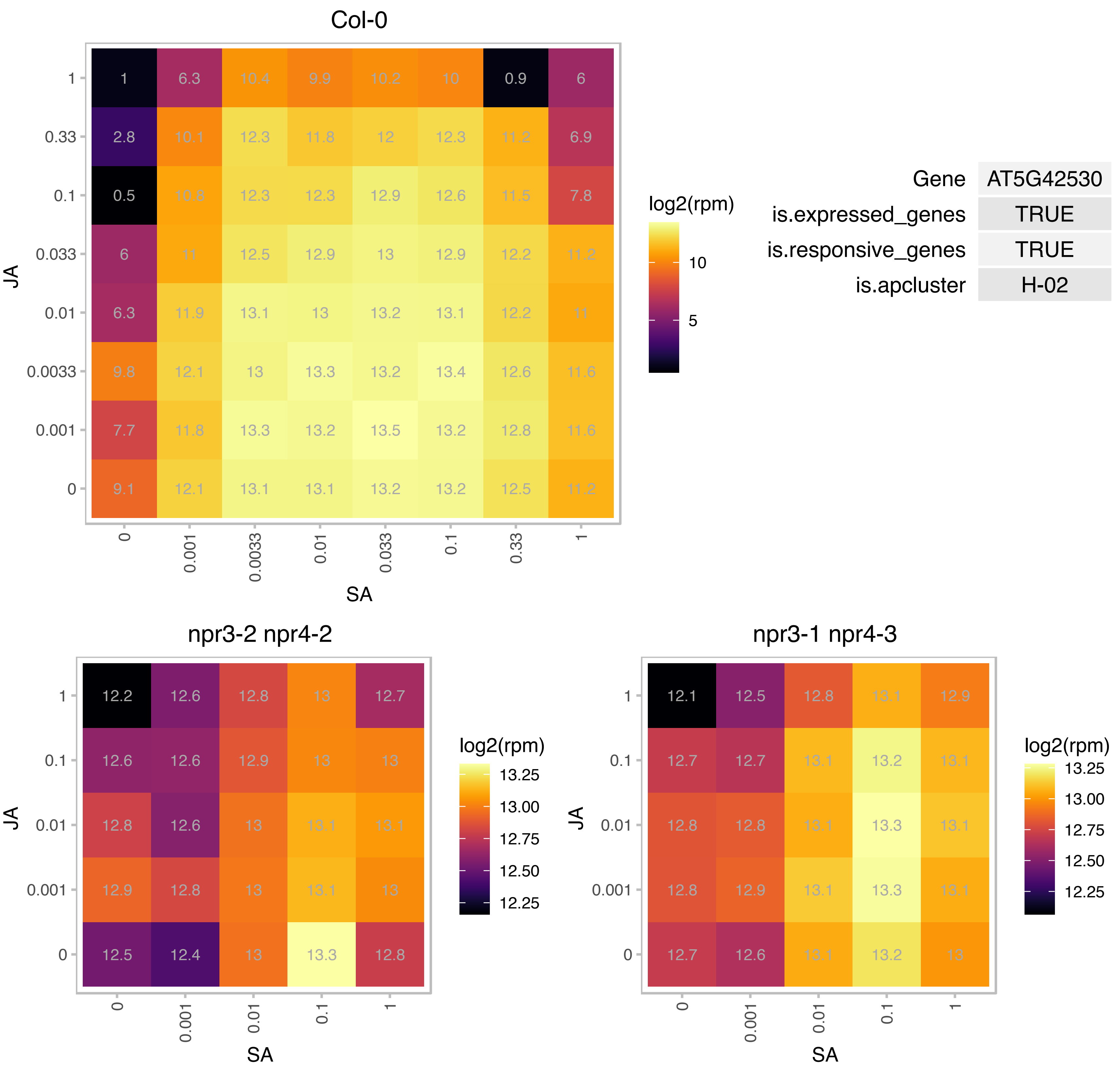


| | |
|---|---|
| Gene | AT5G42530 |
| is.expressed_genes | TRUE |
| is.responsive_genes | TRUE |
| is.apcluster | H-02 |

# AT5G44568.1 : NA

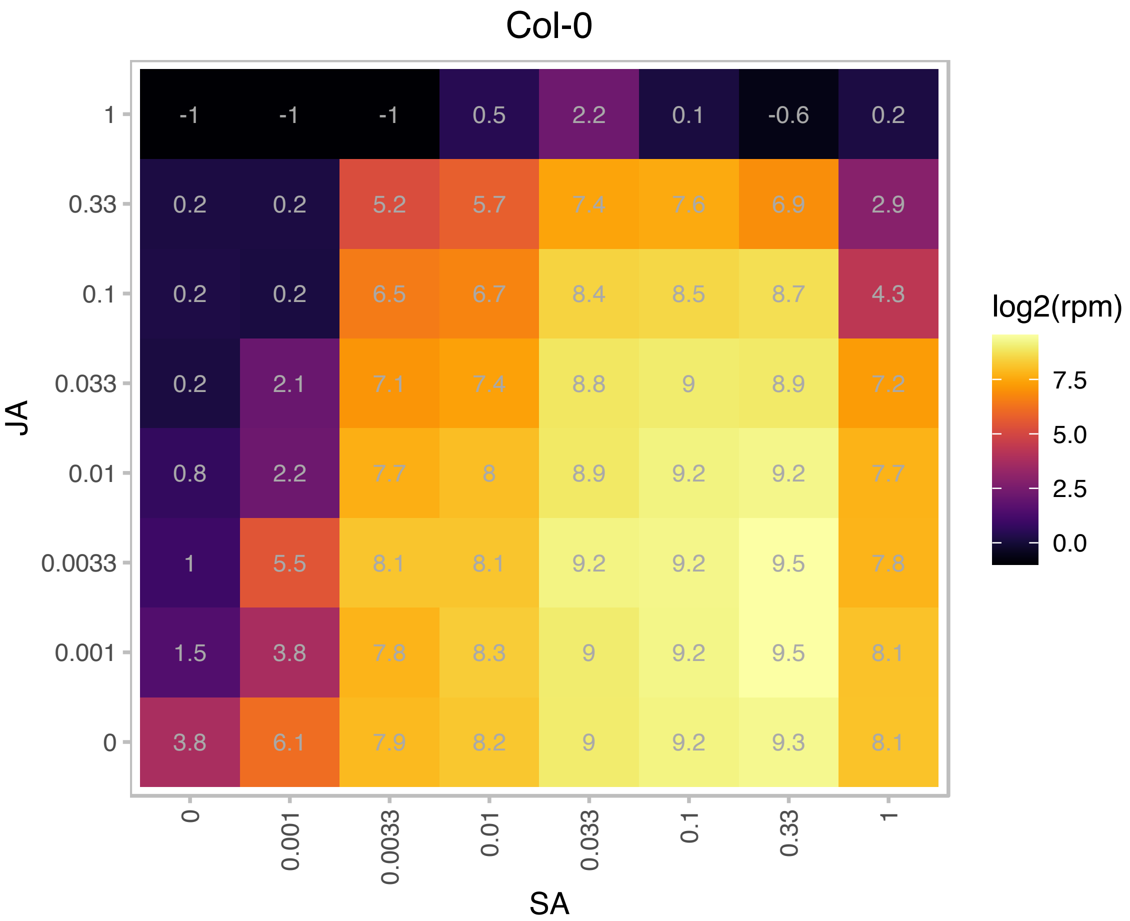


| | |
|---|---|
| Gene | AT5G44568 |
| is.expressed_genes | TRUE |
| is.responsive_genes | TRUE |
| is.apcluster | H-02 |

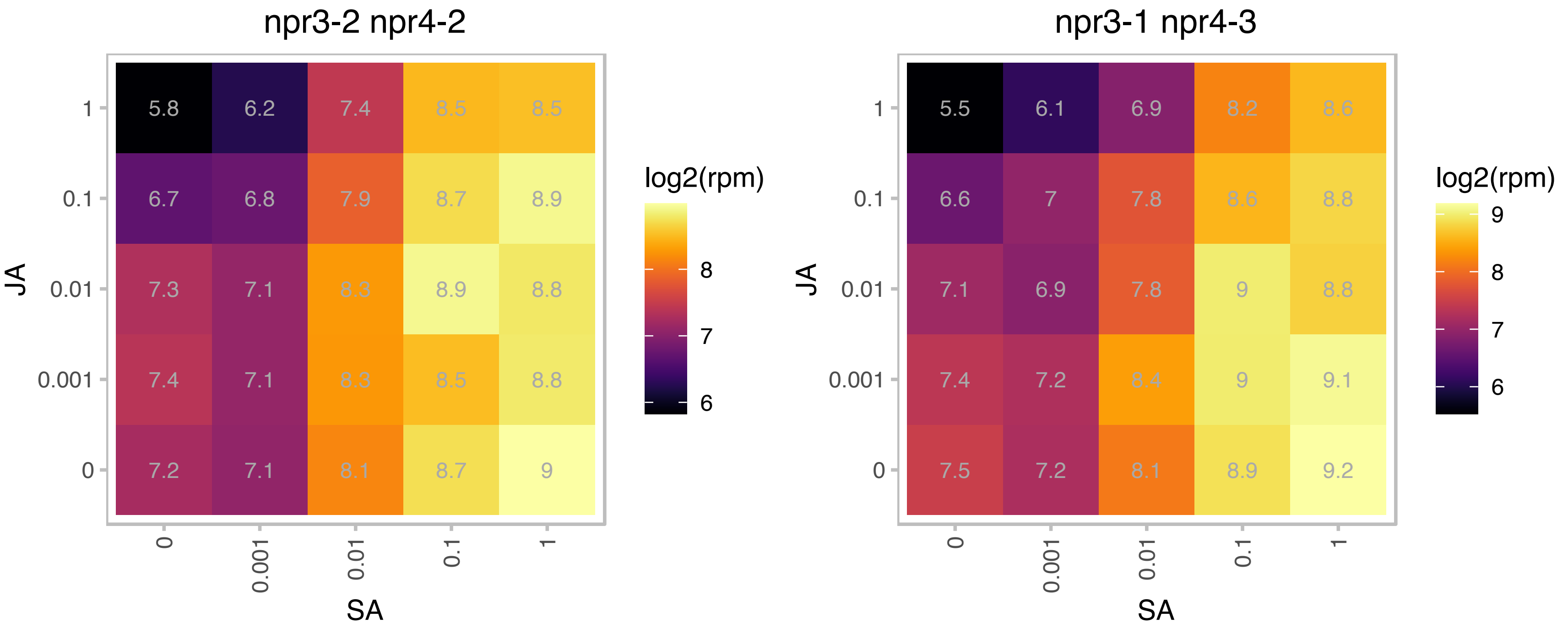

# AT5G50460.1 : NA

secE/sec61-gamma protein transport protein

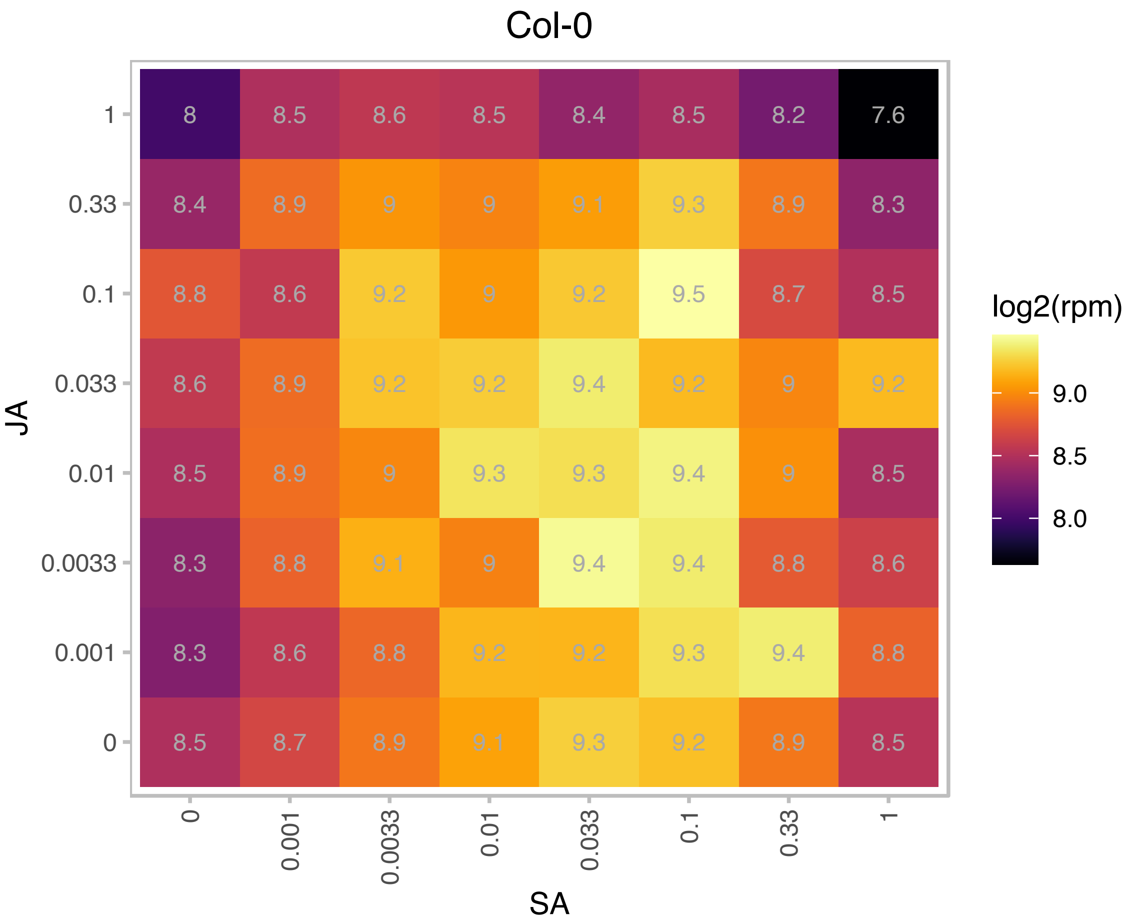


| | |
|---|---|
| Gene | AT5G50460 |
| is.expressed_genes | TRUE |
| is.responsive_genes | TRUE |
| is.apcluster | H-02 |

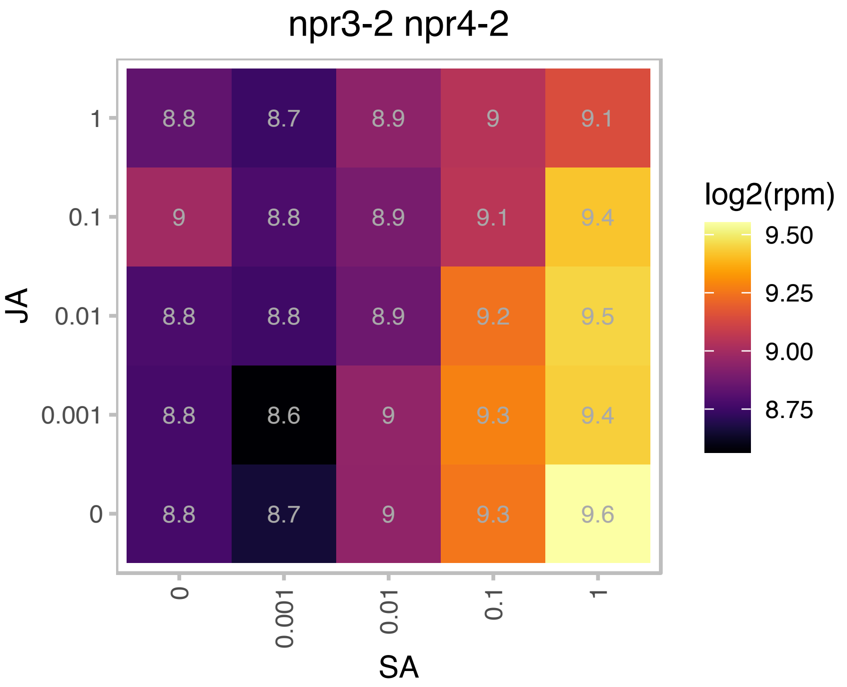


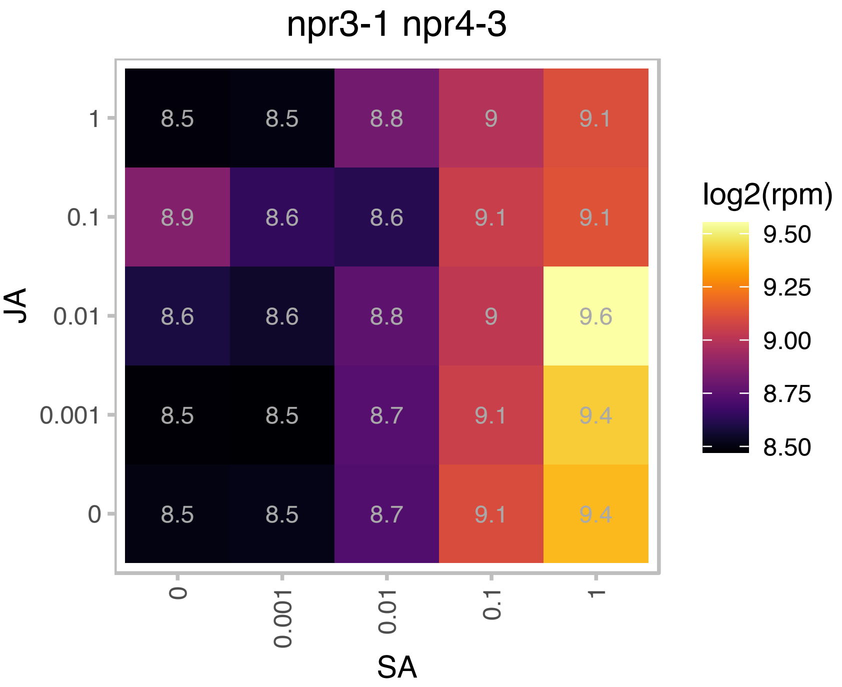

# AT5G52760.1 : NA

Copper transport protein family

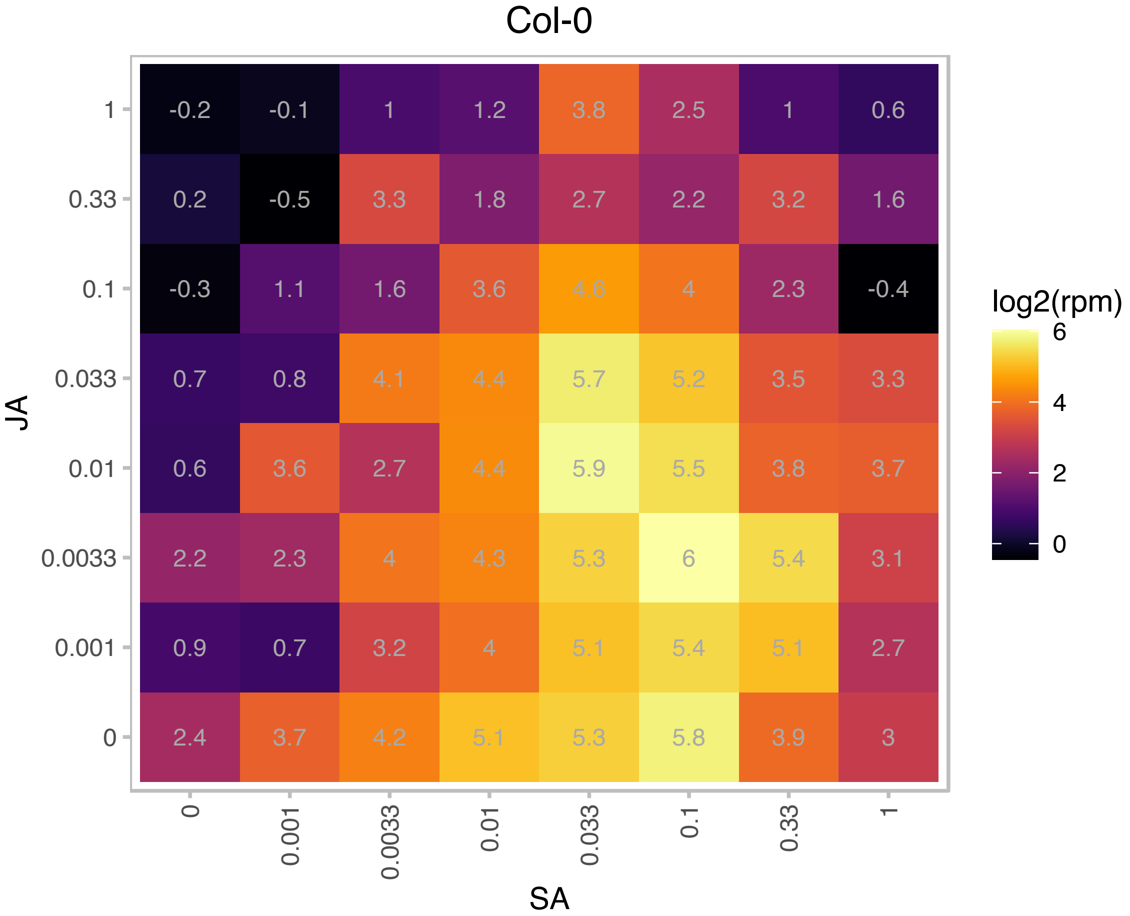


| | |
|---|---|
| Gene | AT5G52760 |
| is.expressed_genes | TRUE |
| is.responsive_genes | TRUE |
| is.apcluster | H-02 |

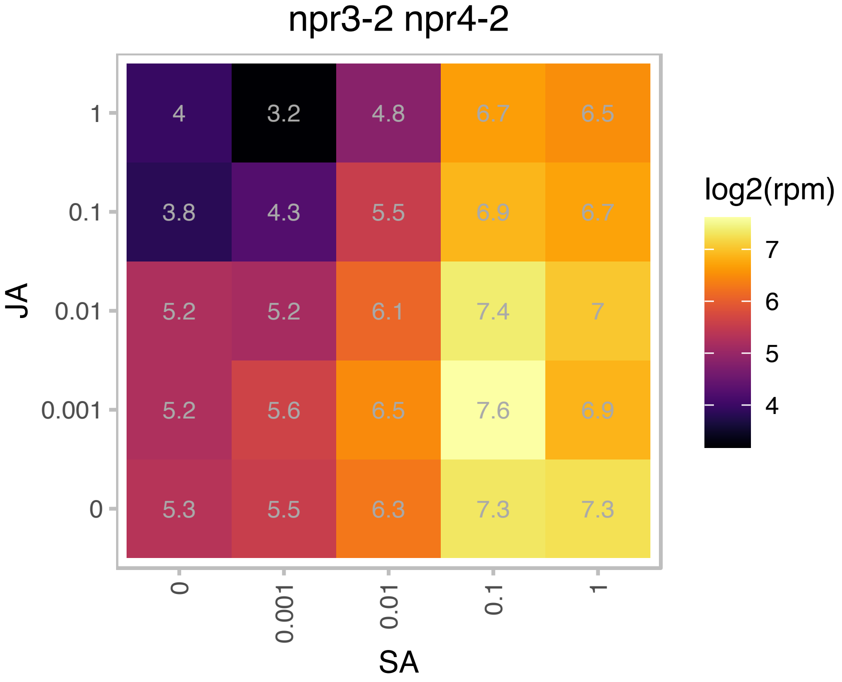


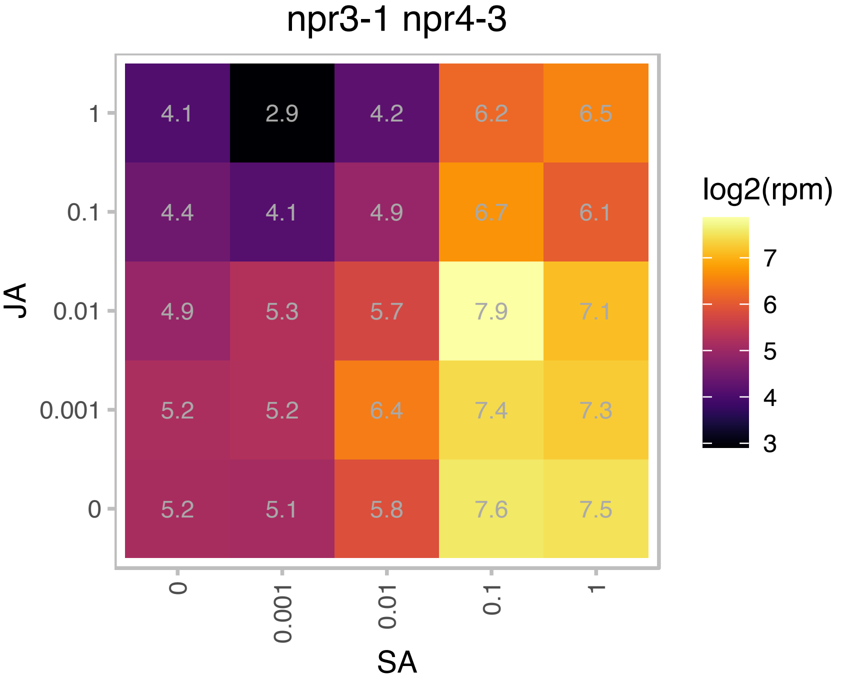

### AT5G53550.1 : ATYSL3, YSL3

YELLOW STRIPE like 3

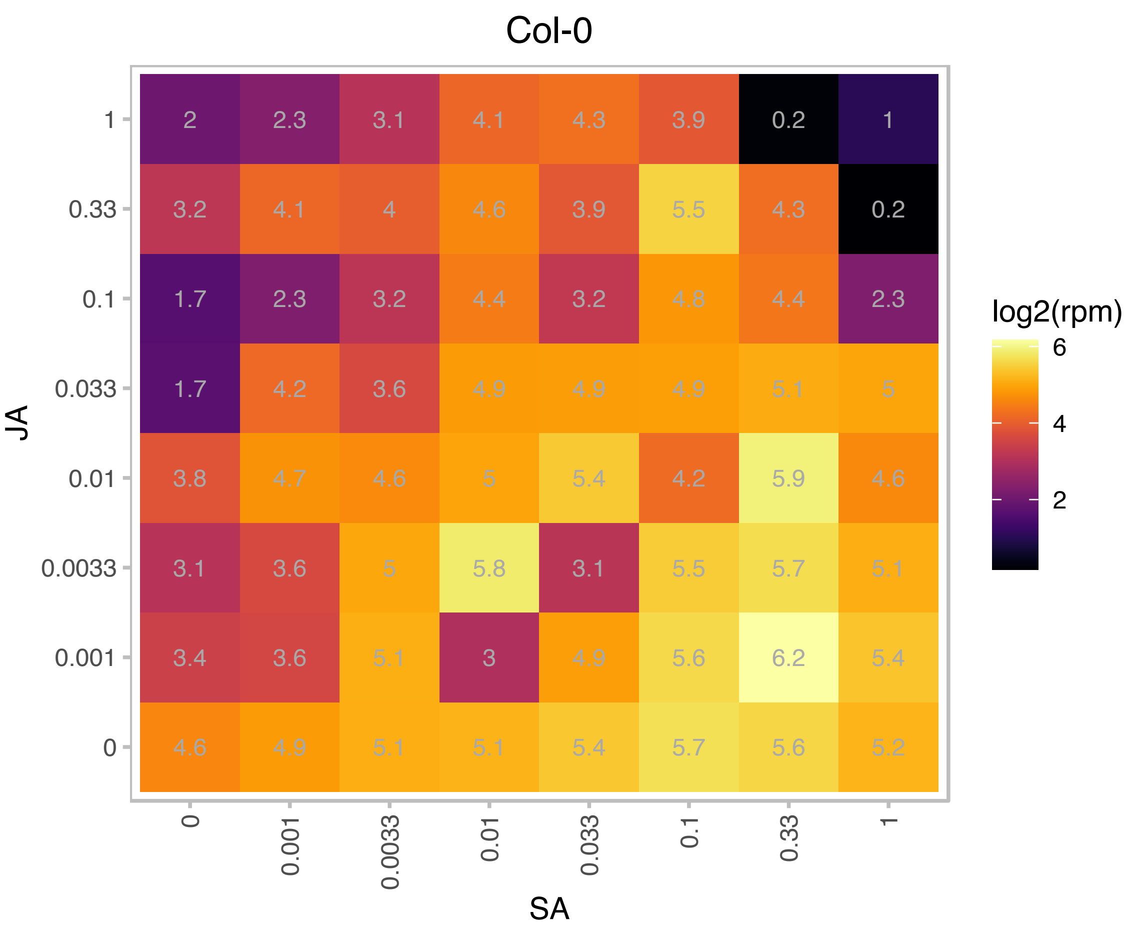


| | |
|---|---|
| Gene | AT5G53550 |
| is.expressed_genes | TRUE |
| is.responsive_genes | TRUE |
| is.apcluster | H-02 |

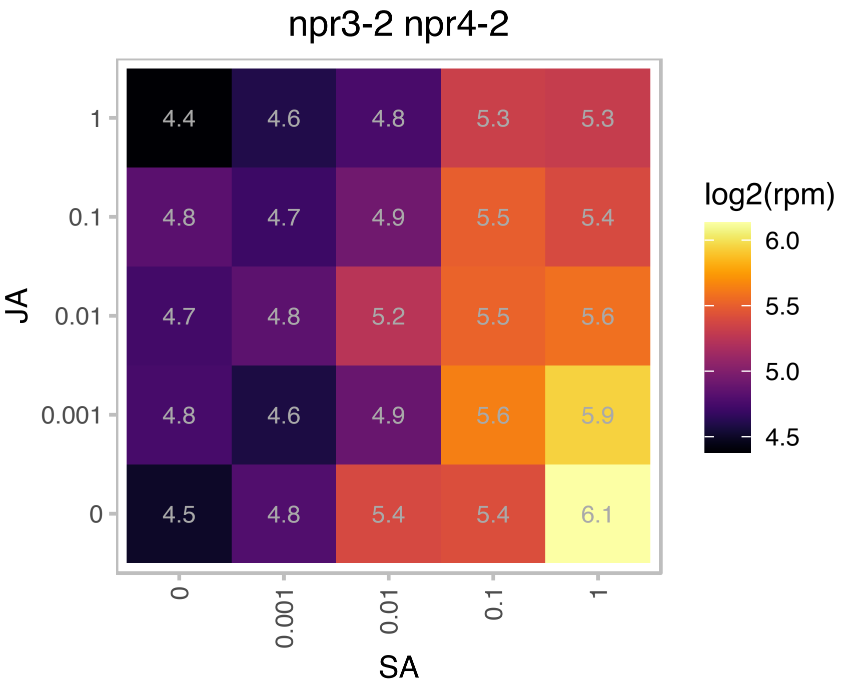


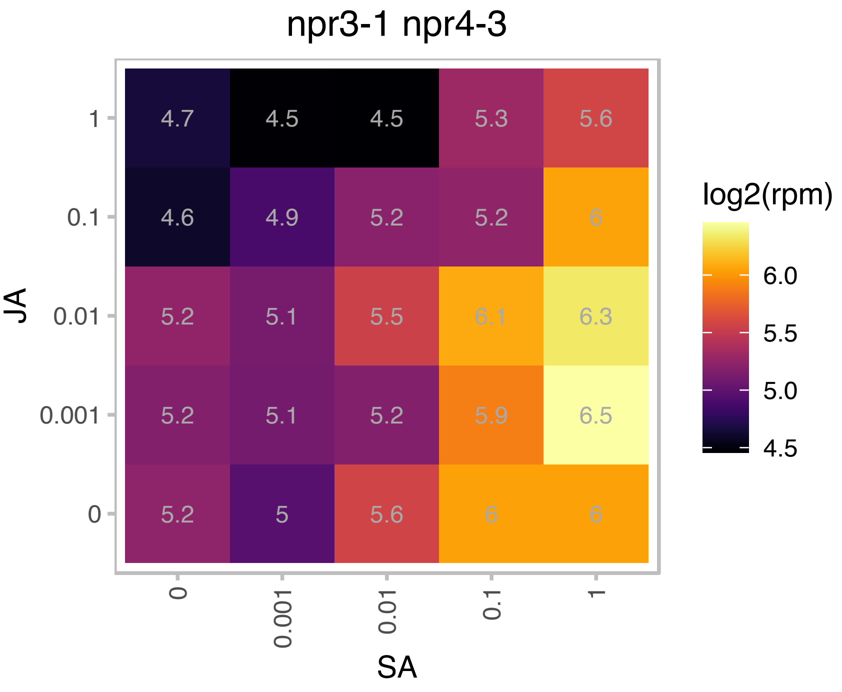

### AT5G55170.1 : ATSUMO3, SUM3, SUMO 3, SUMO3

small ubiquitin-like modifier 3
Encodes a small ubiquitin-like modifier (SUMO) polypeptide that becomes covalently attached to various intracellular protein targets, much like ubiquitination, leading to post-translational modification of those targets.

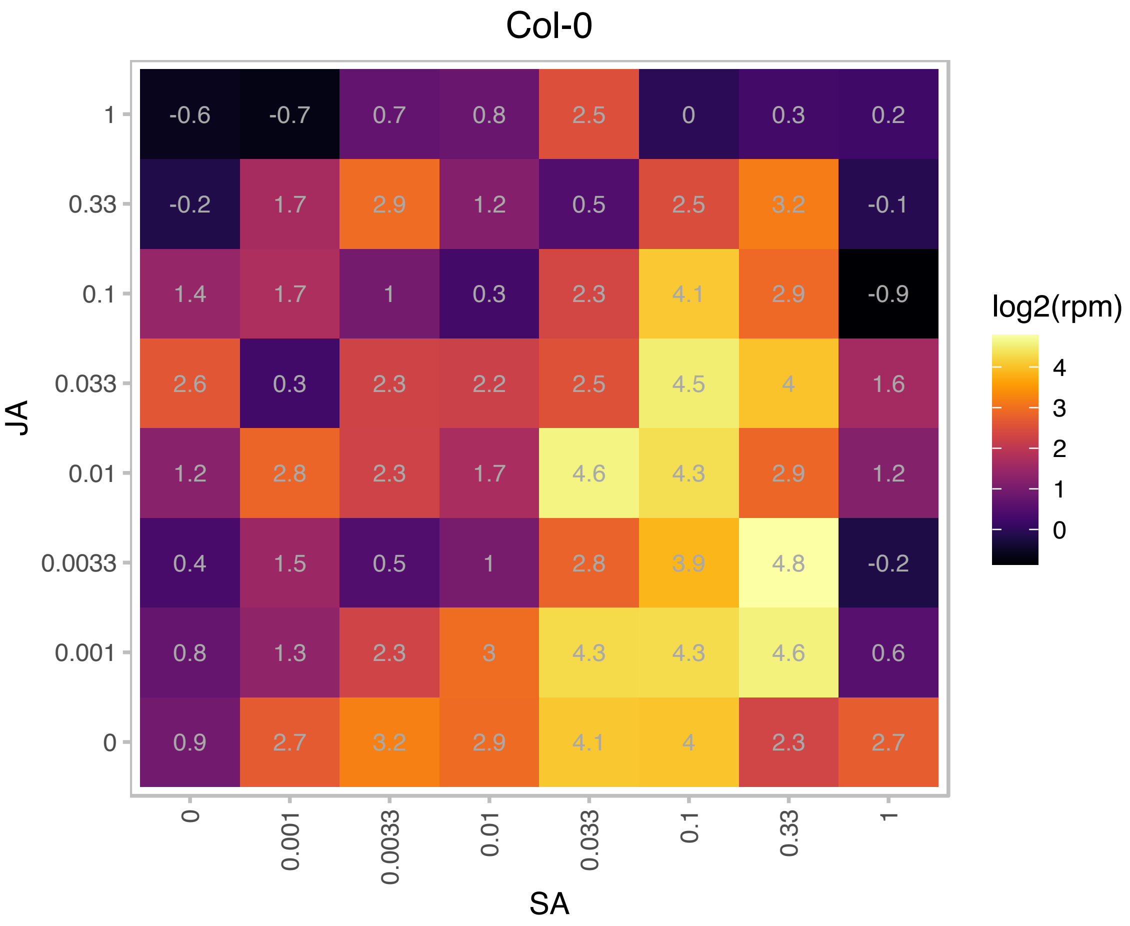


| | |
|---|---|
| Gene | AT5G55170 |
| is.expressed_genes | TRUE |
| is.responsive_genes | TRUE |
| is.apcluster | H-02 |

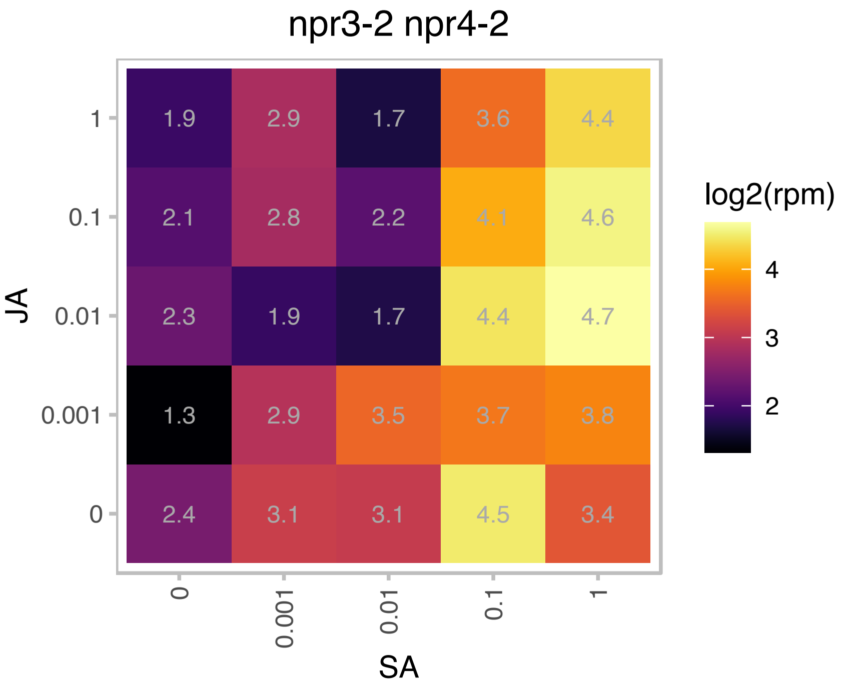


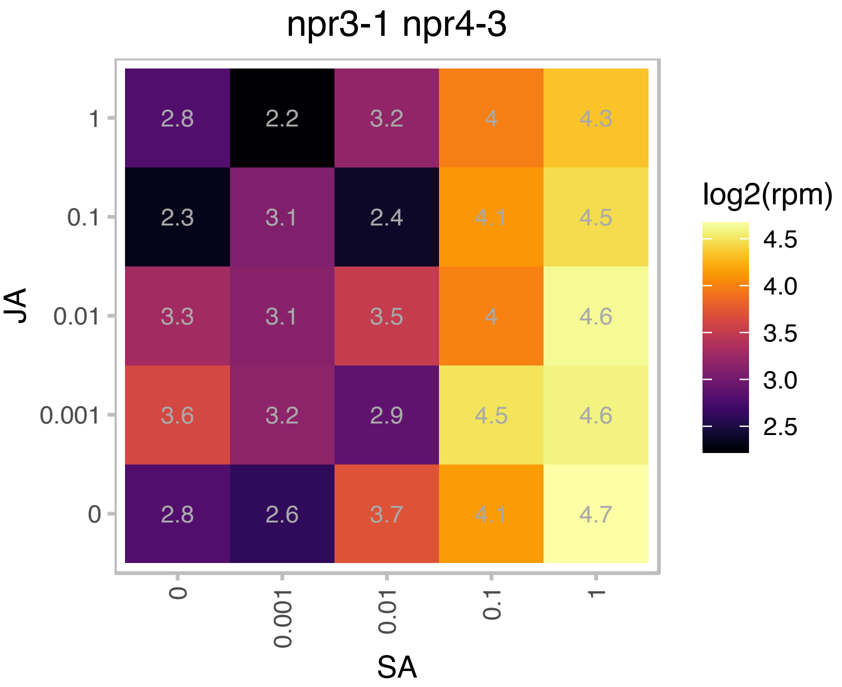

# AT5G58640.1 : NA

Selenoprotein, Rdx type

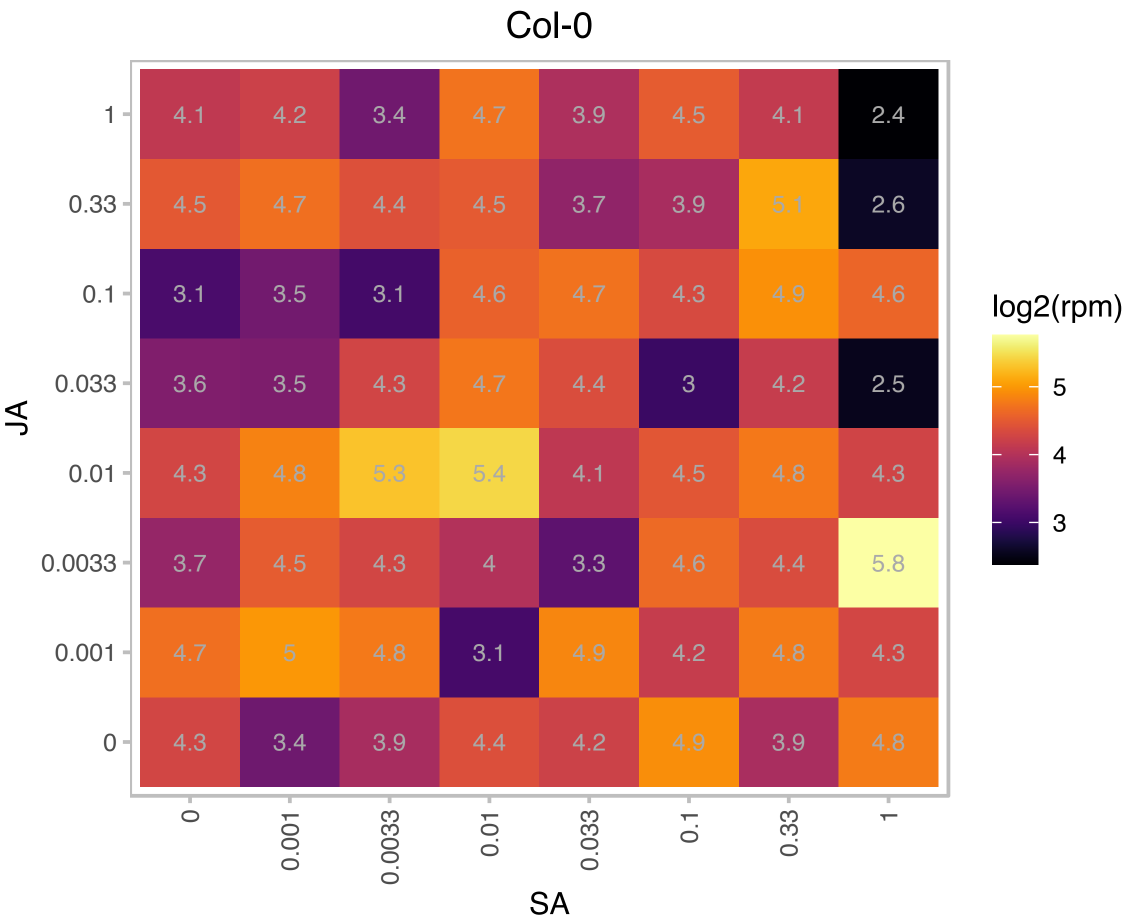


| | |
|---|---|
| Gene | AT5G58640 |
| is.expressed_genes | TRUE |
| is.responsive_genes | TRUE |
| is.apcluster | H-02 |

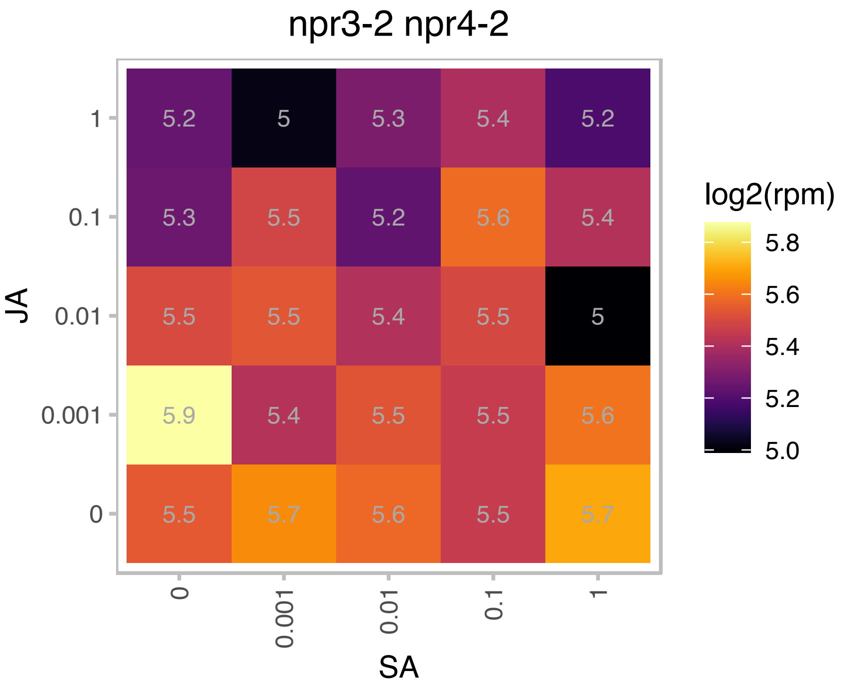


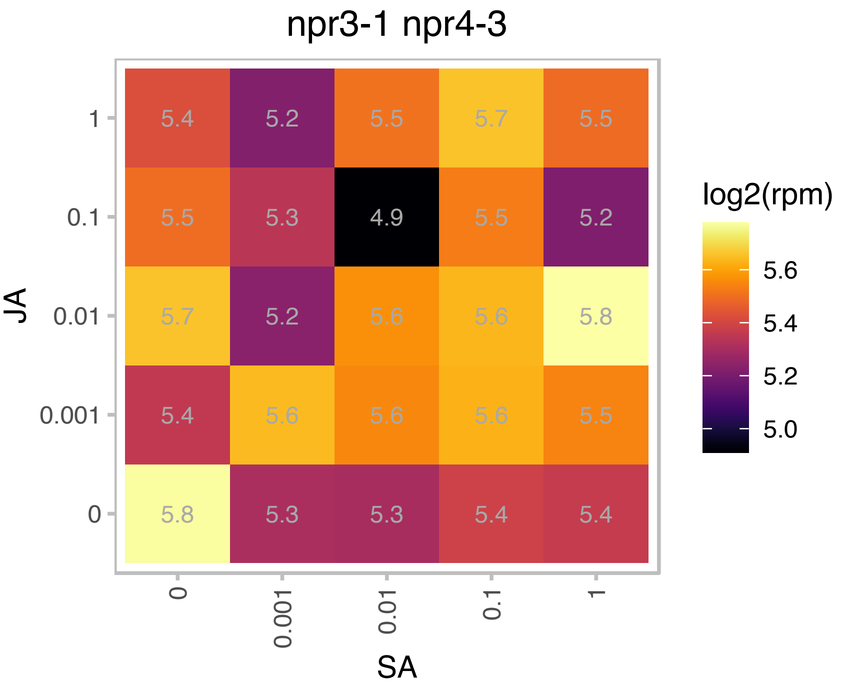

### AT5G62070.1 : IQD23

IQ-domain 23

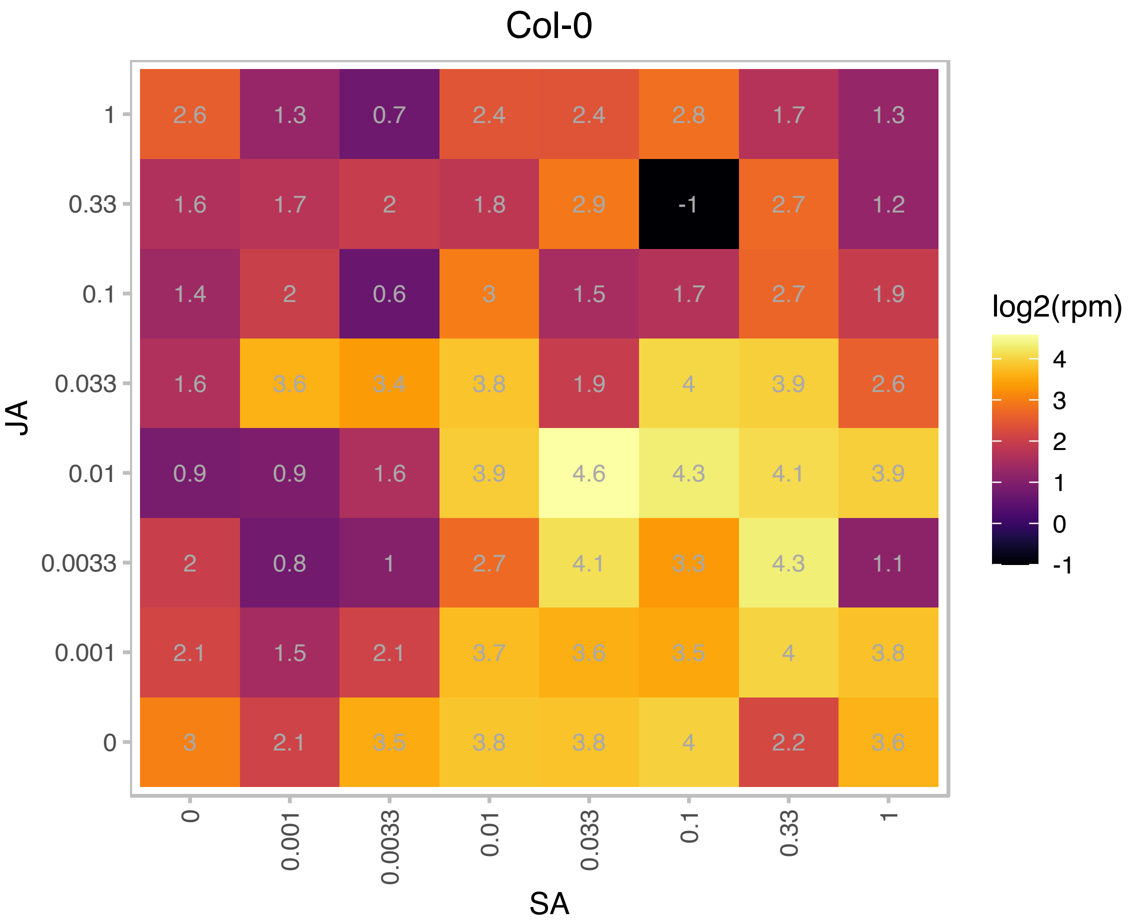


| | |
|---|---|
| Gene | AT5G62070 |
| is.expressed_genes | TRUE |
| is.responsive_genes | TRUE |
| is.apcluster | H-02 |

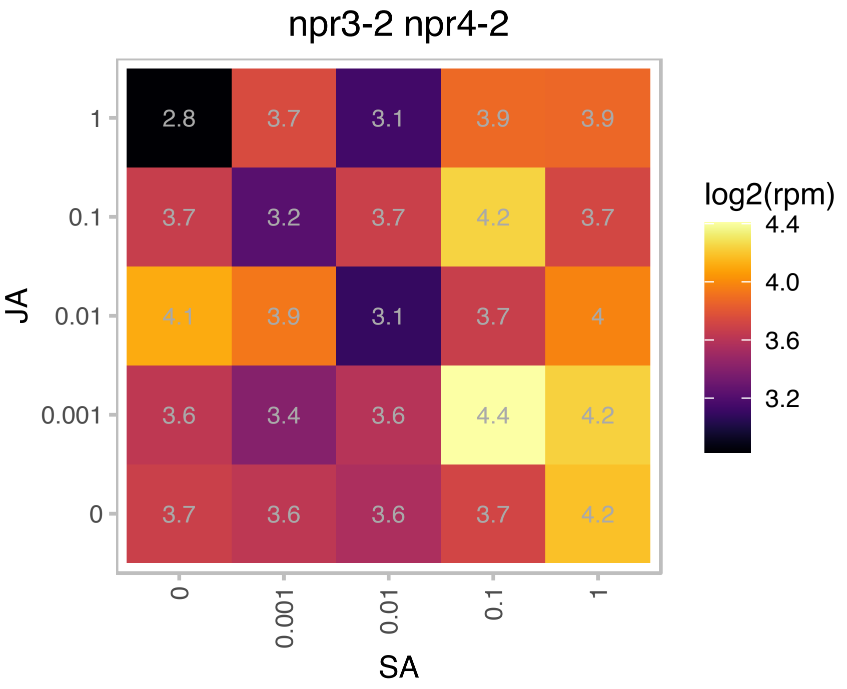


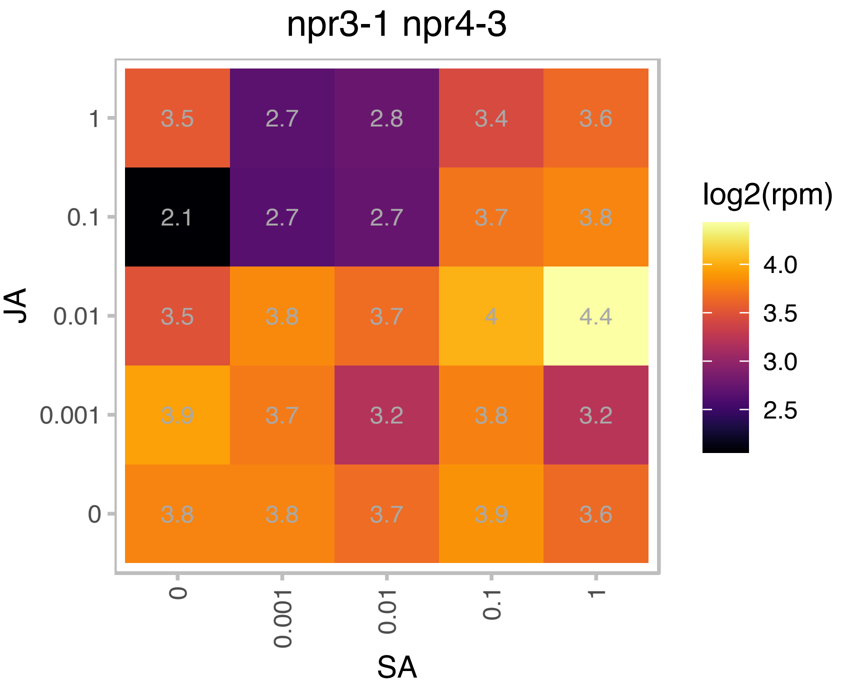

### AT1G14290.1 : AtSBH2, SBH2

sphingoid base hydroxylase 2
Encodes one of the two redundant sphingoid base hydroxylases (SBH). Involved in sphingolipid trihydroxy long-chain base (4-hydroxysphinganine) biosynthesis. Double mutants of SBHs were dwarfed and not able to progress from vegetative to reproductive growth.

Col-0

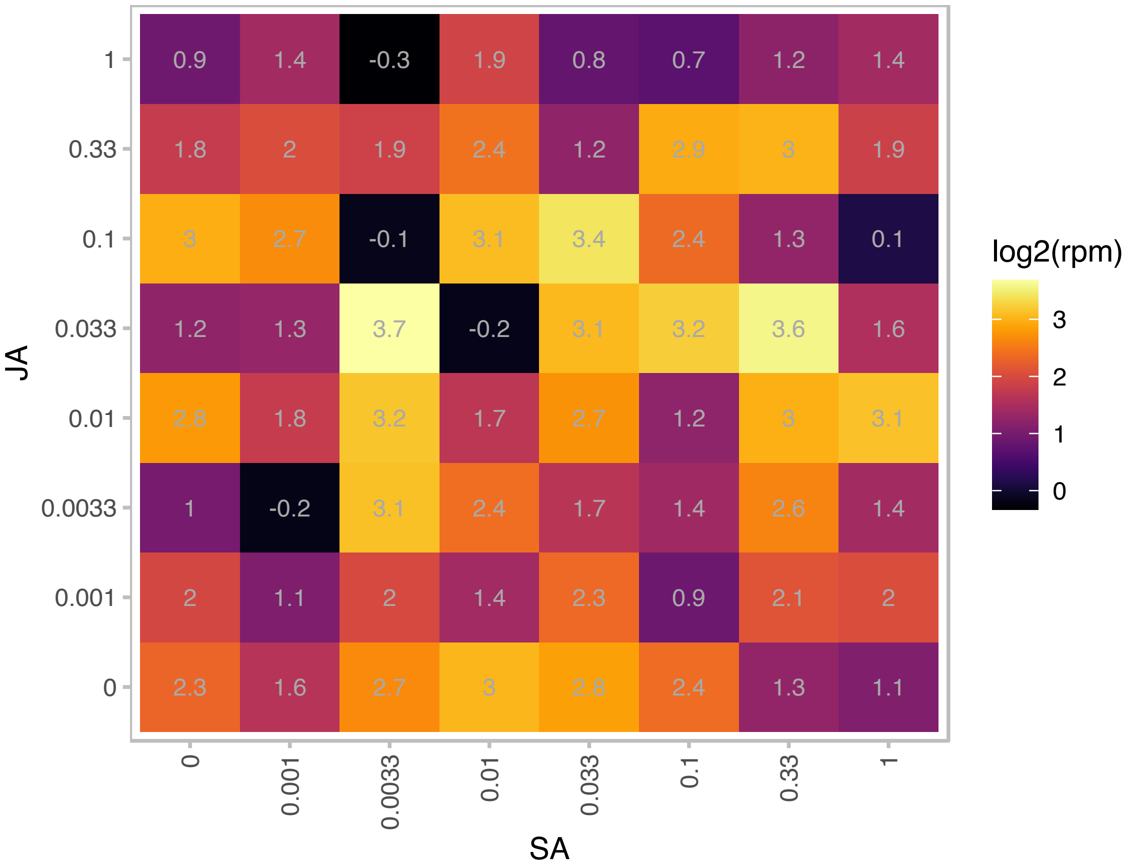


| | |
|---|---|
| Gene | AT1G14290 |
| is.expressed_genes | TRUE |
| is.responsive_genes | TRUE |
| is.apcluster | H-03 |

npr3-2 npr4-2

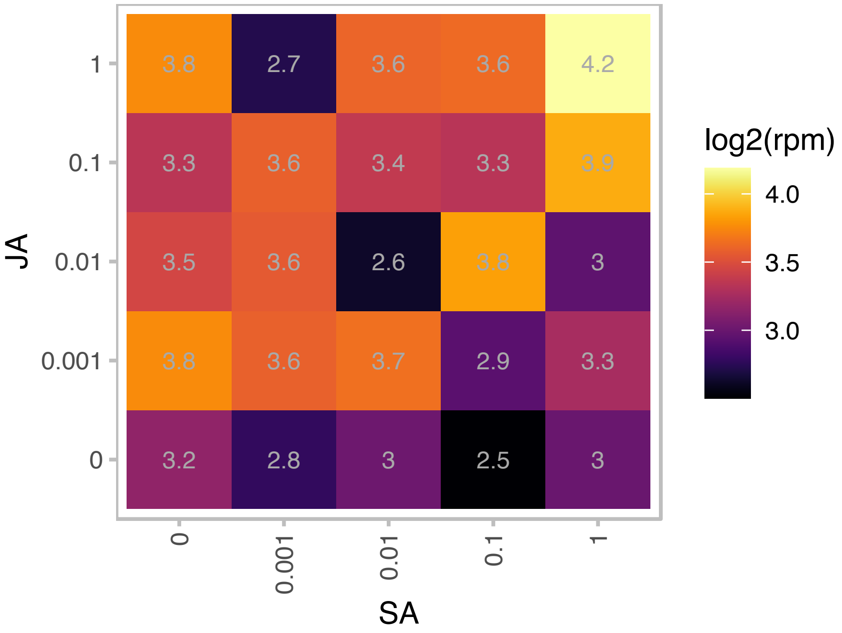


npr3-1 npr4-3

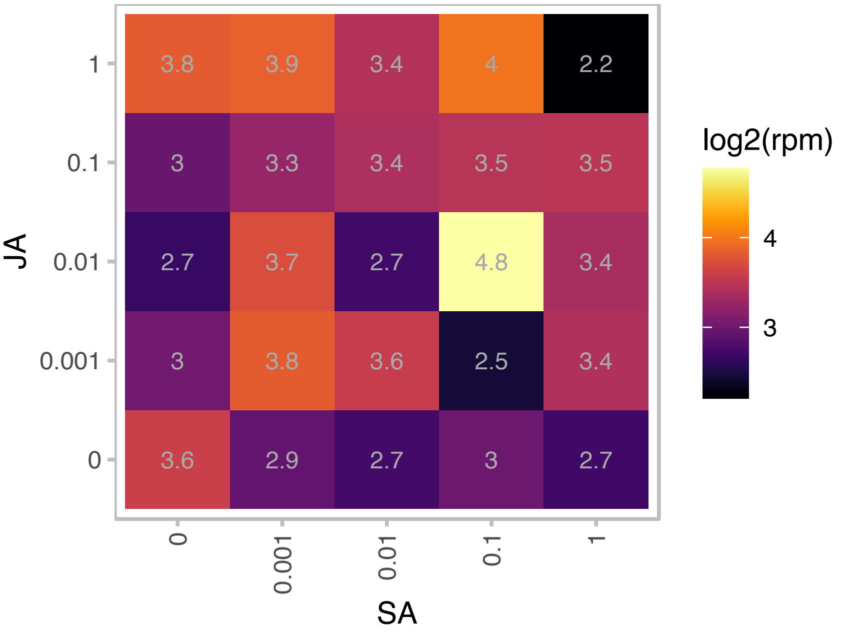

# AT1G14310.1 : NA

Haloacid dehalogenase-like hydrolase (HAD) superfamily protein

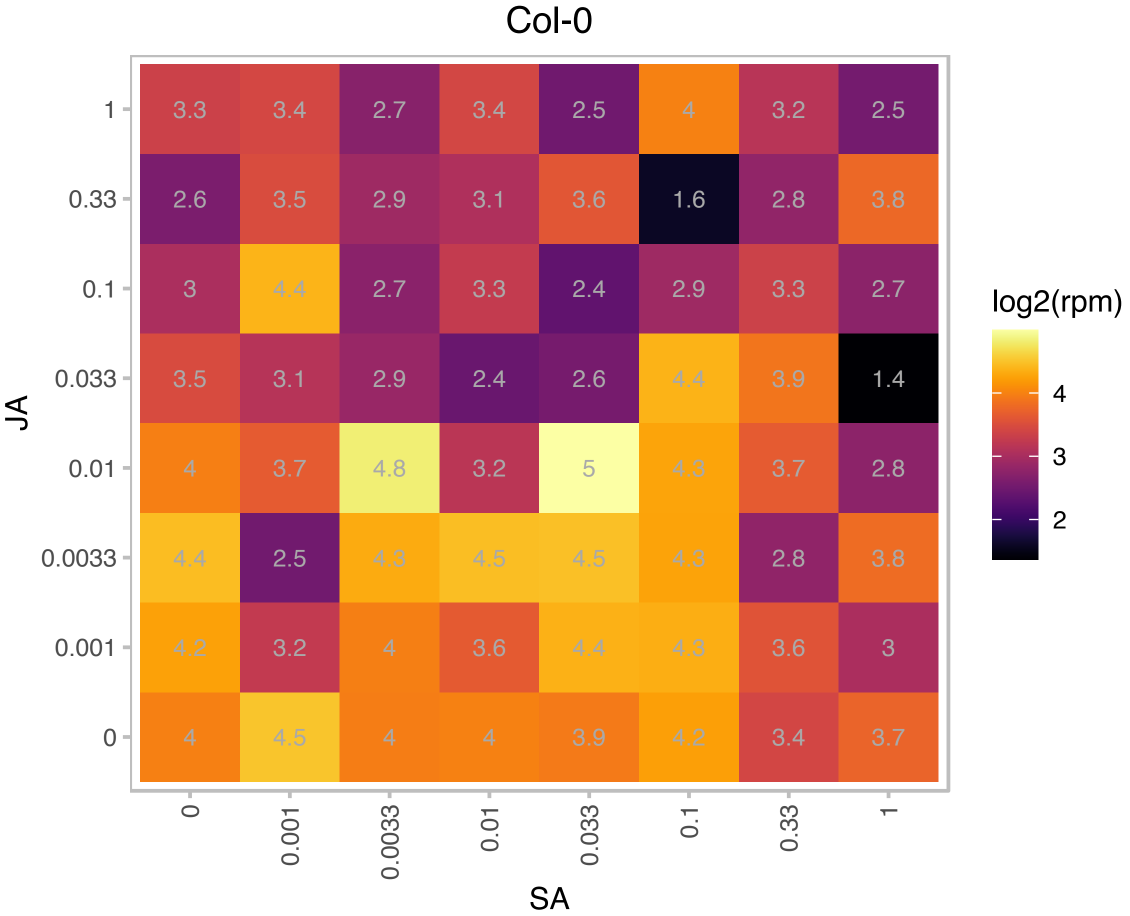


| | |
|---|---|
| Gene | AT1G14310 |
| is.expressed_genes | TRUE |
| is.responsive_genes | TRUE |
| is.apcluster | H-03 |

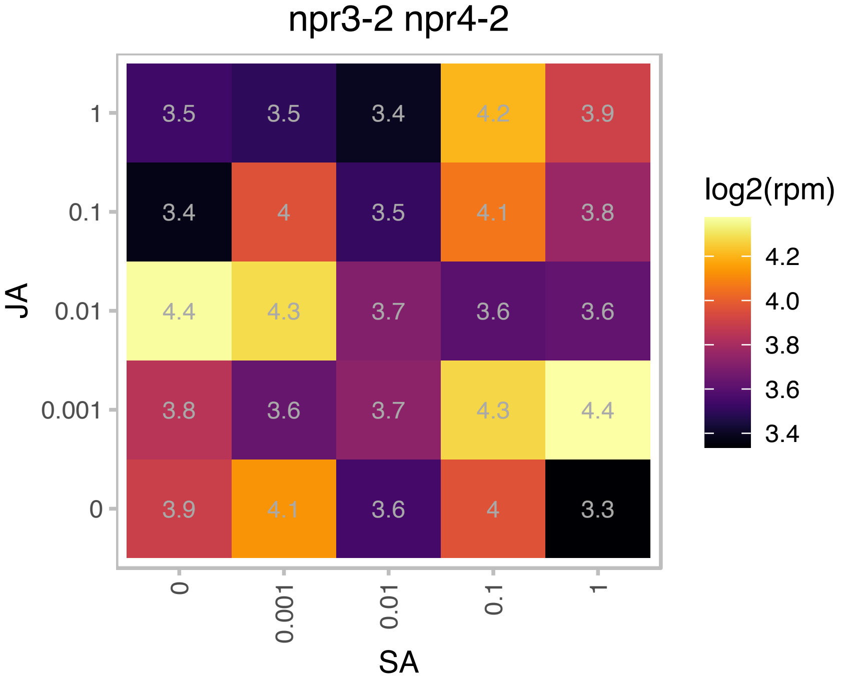


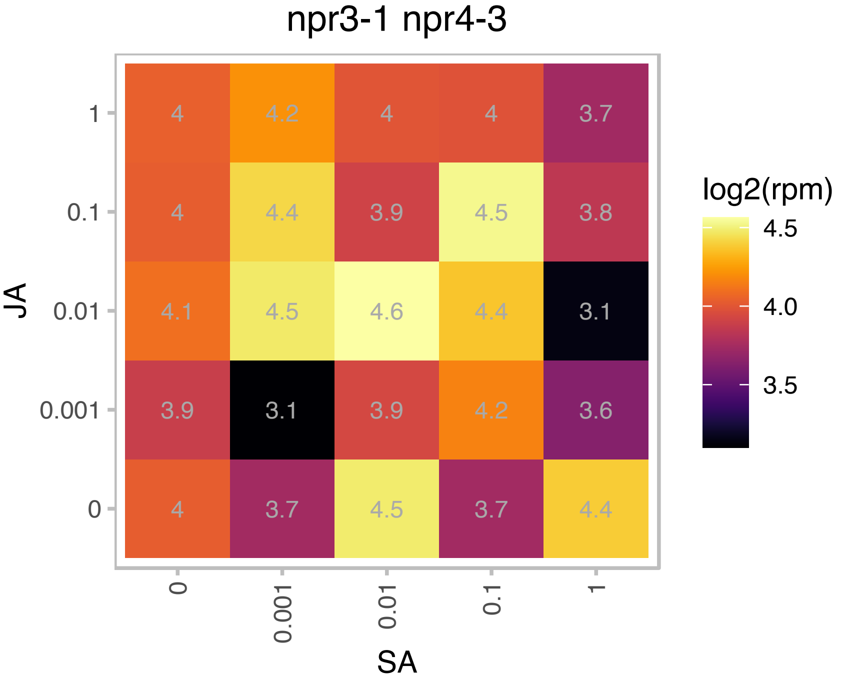

### AT1G14620.1 : DECOY

decoy

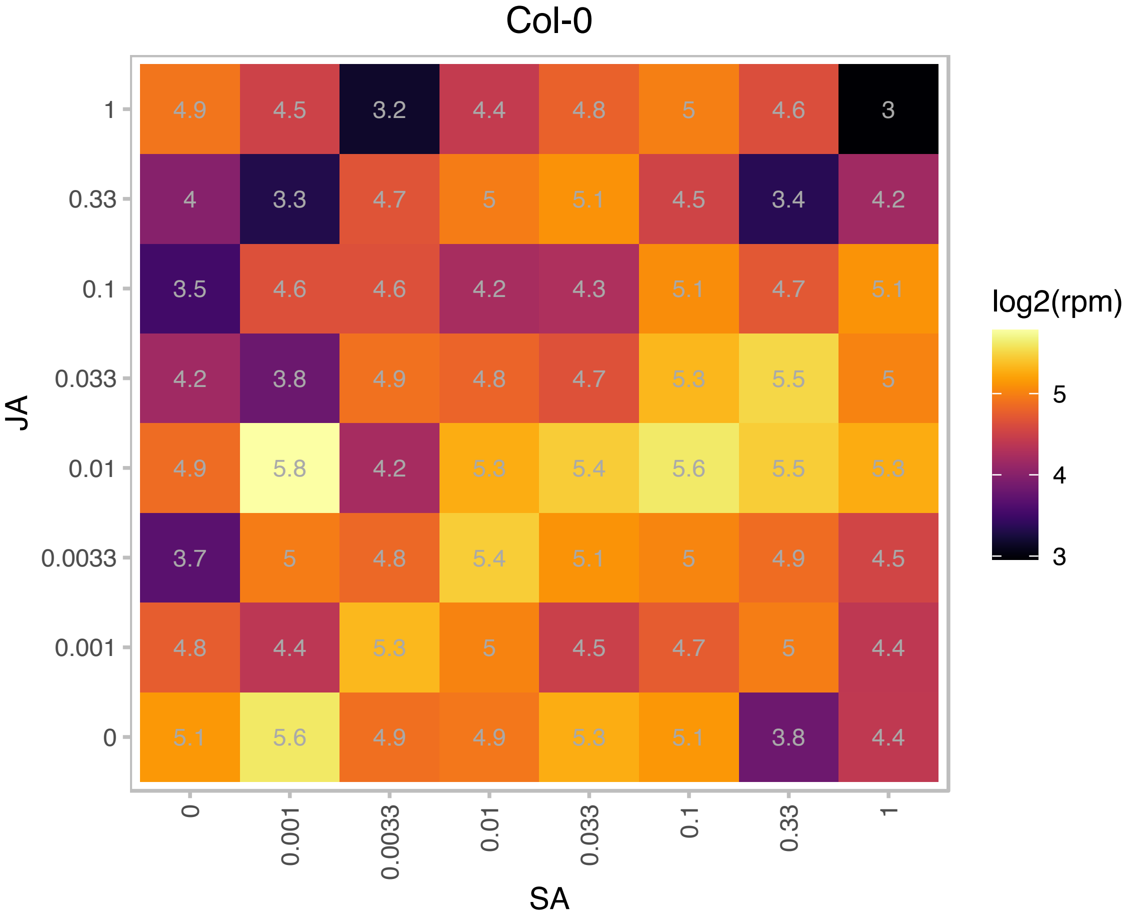


| | |
|---|---|
| Gene | AT1G14620 |
| is.expressed_genes | TRUE |
| is.responsive_genes | TRUE |
| is.apcluster | H-03 |

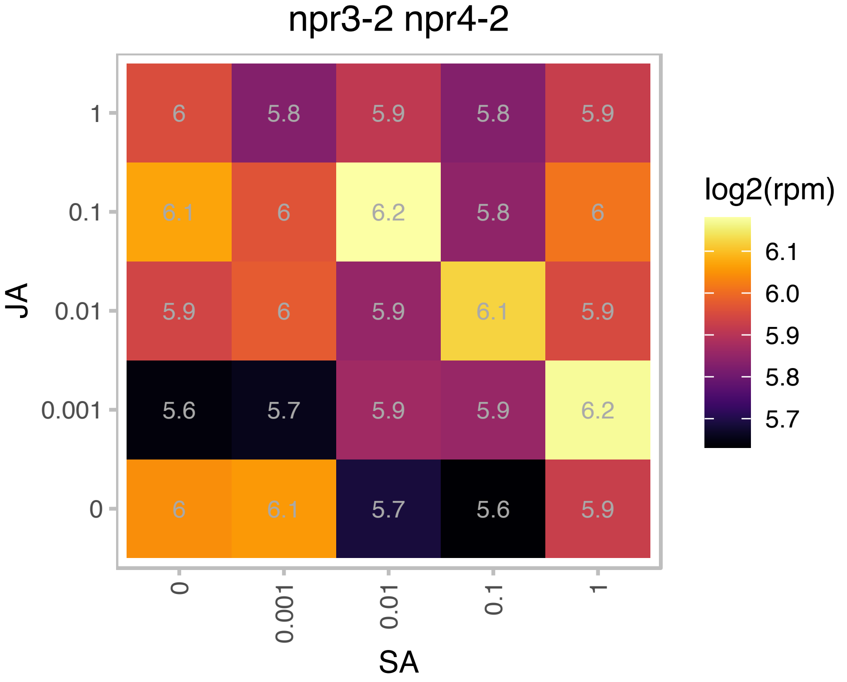


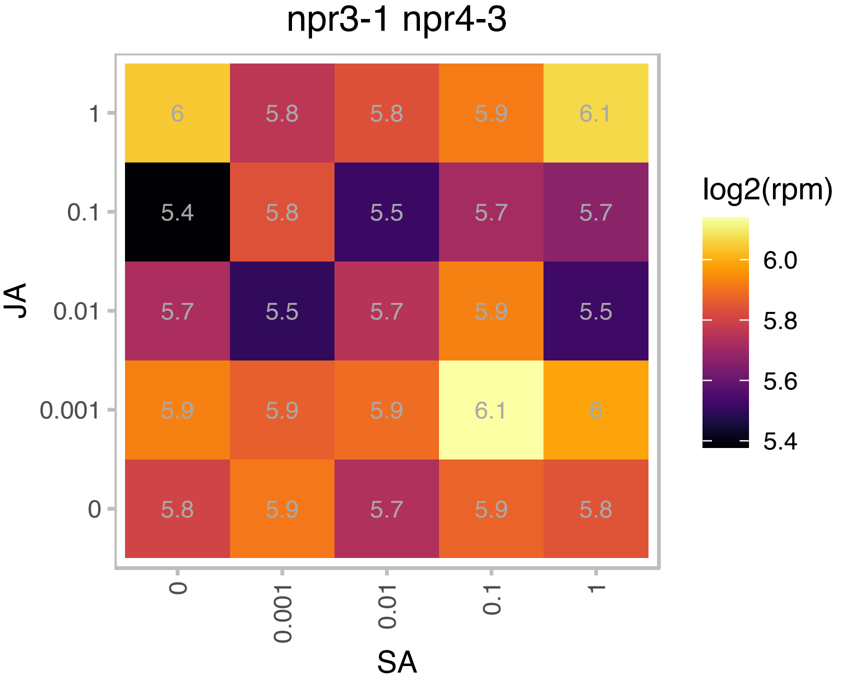

### AT1G23390.1 : KFB

Kelch repeat-containing F-box family protein
A kelch domain-containing F-box protein. Its N terminus contains a typical F-box motif but its C-terminal domain only consists of one predicted kelch motif. Predicted to be stu Interacts with chalcone synthase CHS to mediate CHS ubiquitination and degradation.

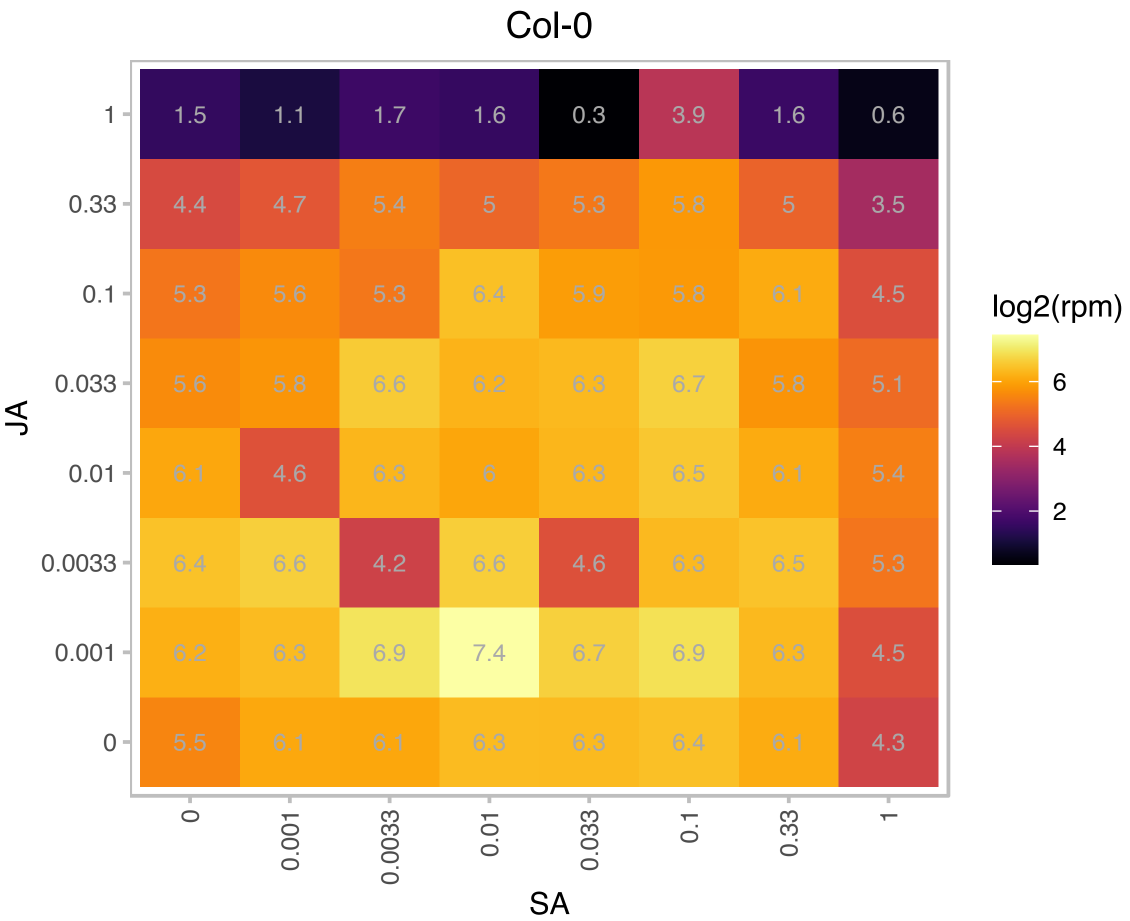


| | |
|---|---|
| Gene | AT1G23390 |
| is.expressed_genes | TRUE |
| is.responsive_genes | TRUE |
| is.apcluster | H-03 |

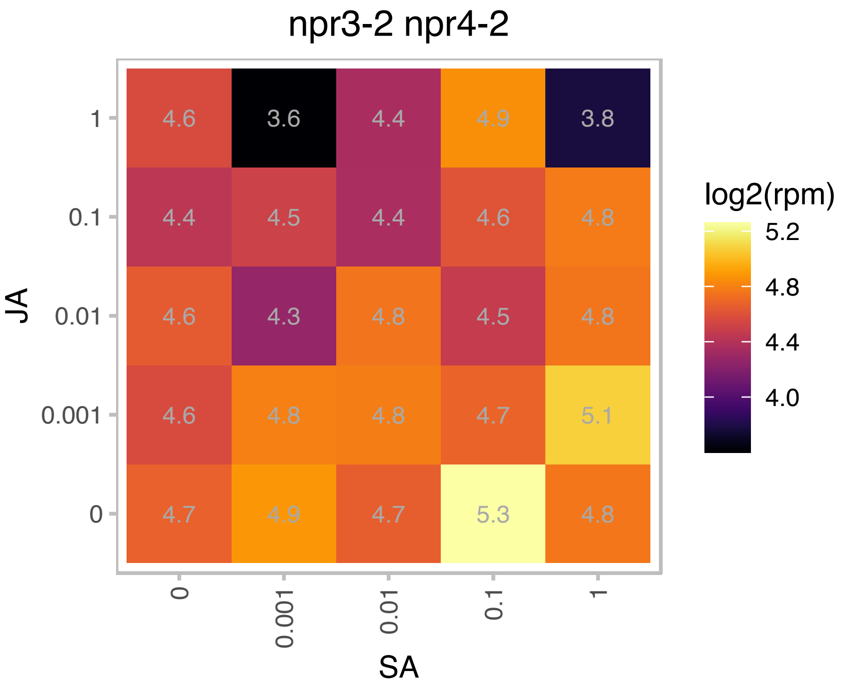


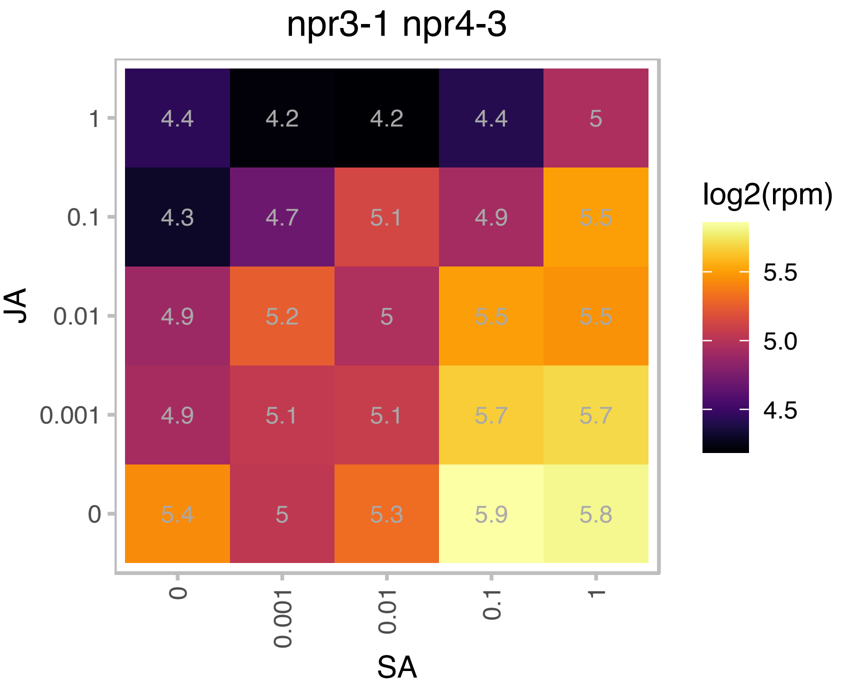

### AT1G29920.1 : AB165, CAB2, LHCB1.1

chlorophyll A/B-binding protein 2

Encodes lhcb1.1 a component of the LHCIIb light harvesting complex associated with photosystem II.

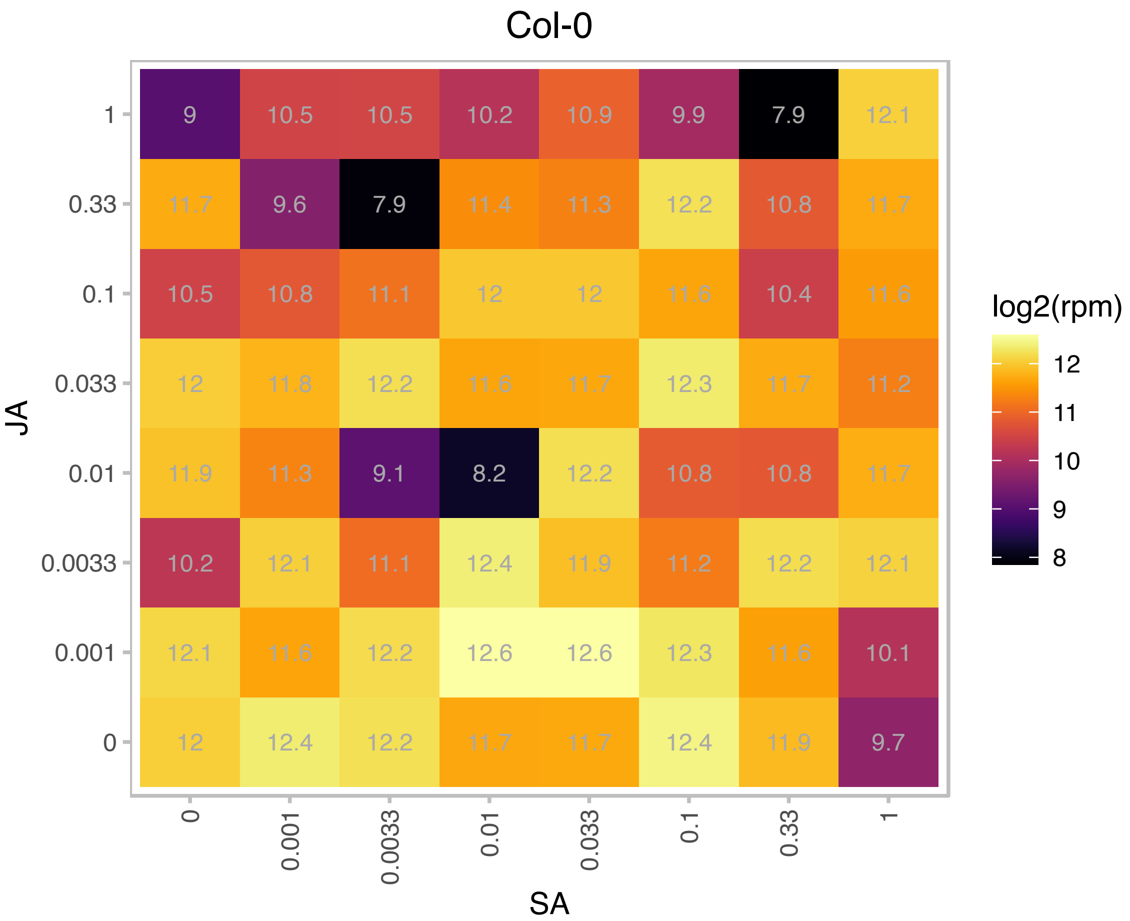


| | |
|---|---|
| Gene | AT1G29920 |
| is.expressed_genes | TRUE |
| is.responsive_genes | TRUE |
| is.apcluster | H-03 |

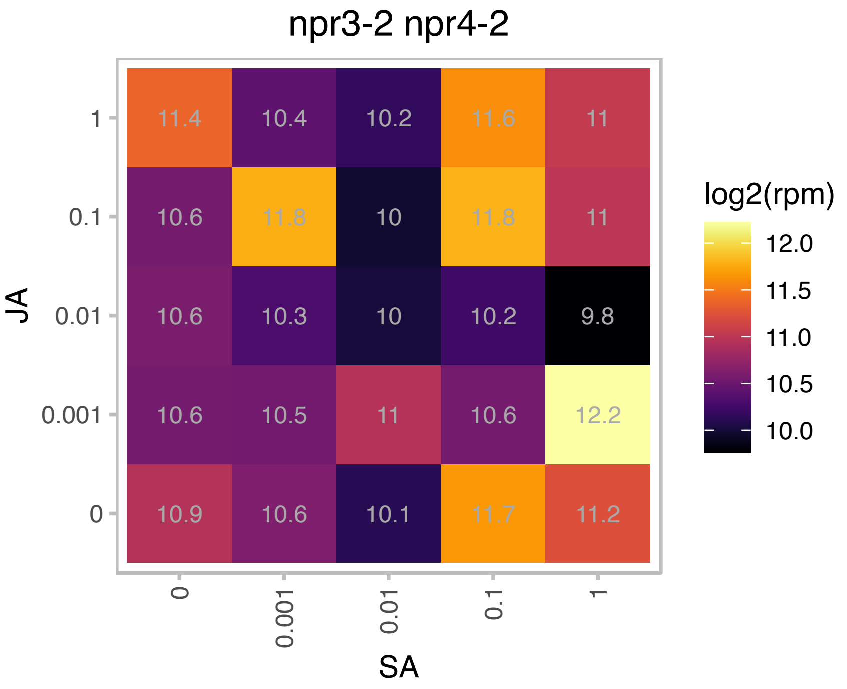


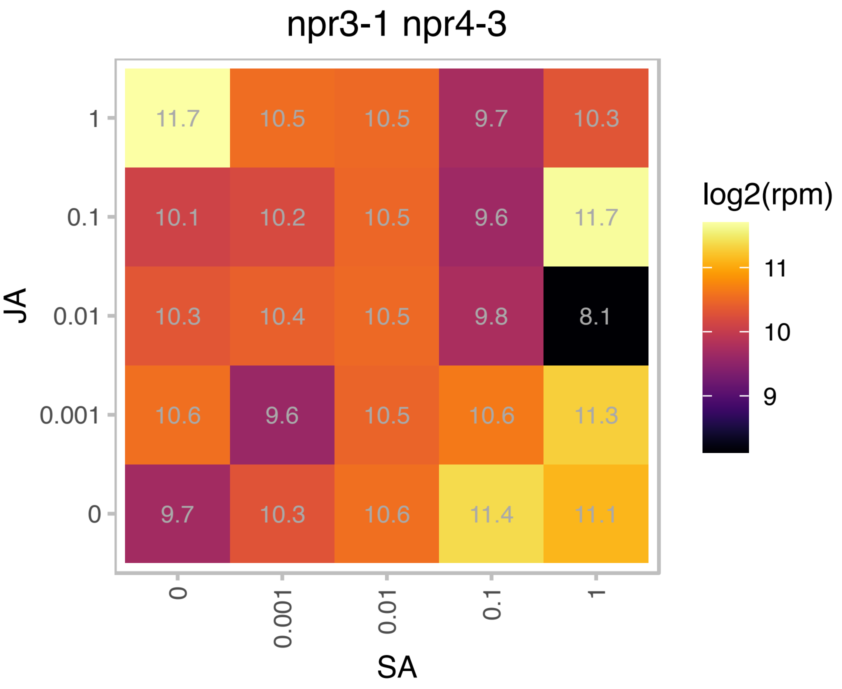

### AT1G29930.1 : AB140, CAB1, CAB140, LHCB1.3

chlorophyll A/B binding protein 1

Subunit of light-harvesting complex II (LHCII),which absorbs light and transfers energy to the photosynthetic reaction center. The mRNA is cell-to-cell mobile.

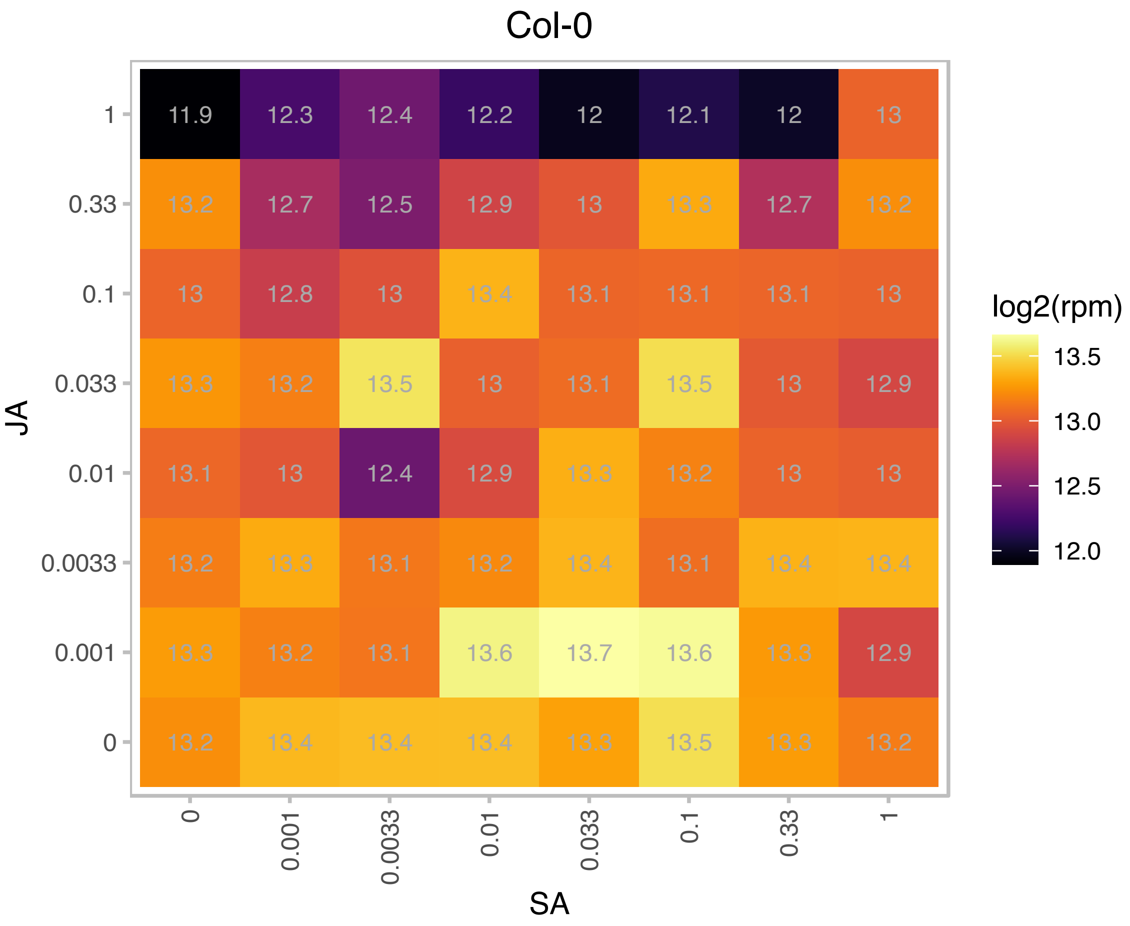


| | |
|---|---|
| Gene | AT1G29930 |
| is.expressed_genes | TRUE |
| is.responsive_genes | TRUE |
| is.apcluster | H-03 |

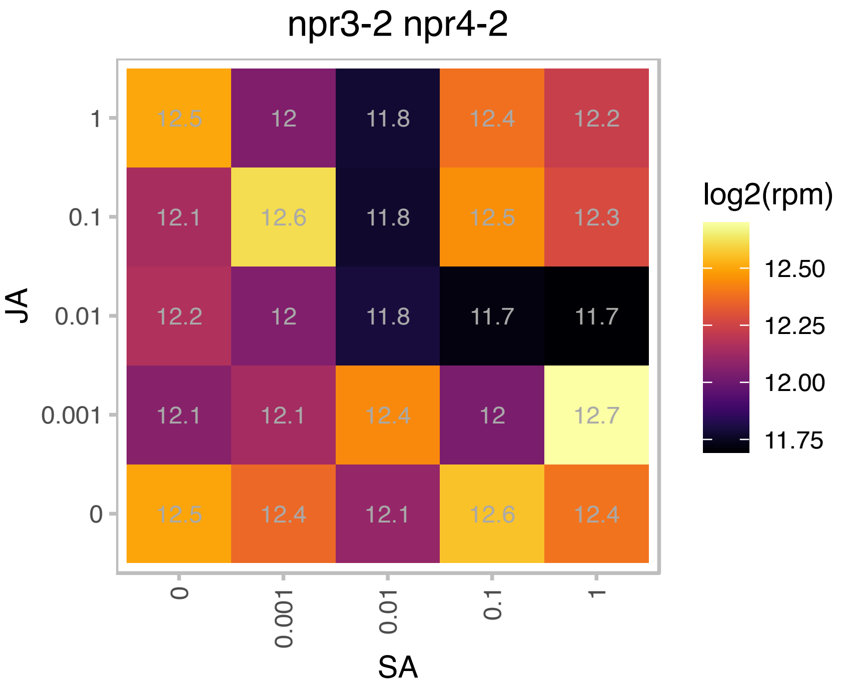


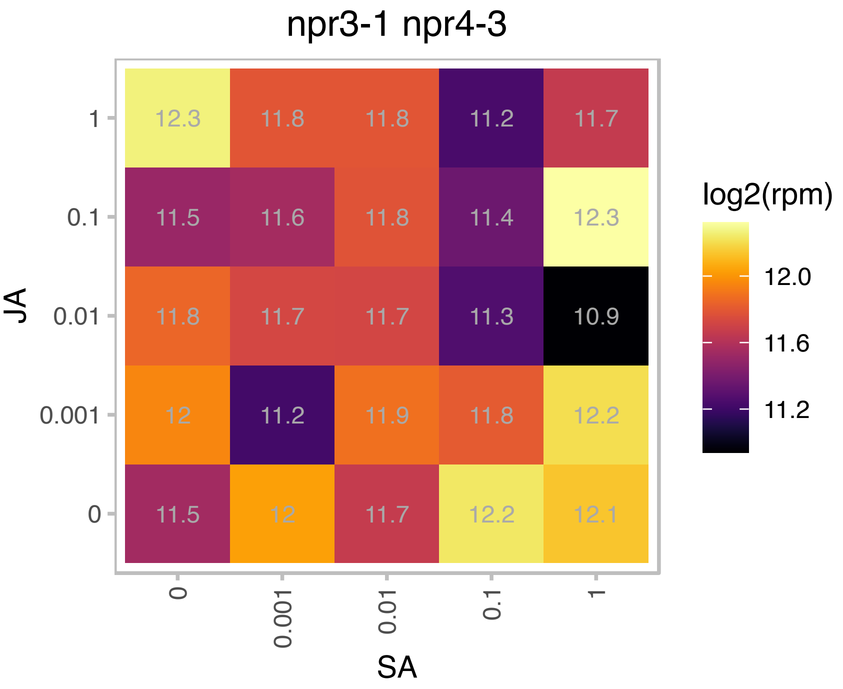

### AT1G33960.1 : AIG1

P-loop containing nucleoside triphosphate hydrolases superfamily protein

Identified as a gene that is induced by avirulence gene avrRpt2 and RPS2 after infection with Pseudomonas syringae pv maculicola strain ES4326 carrying avrRpt2

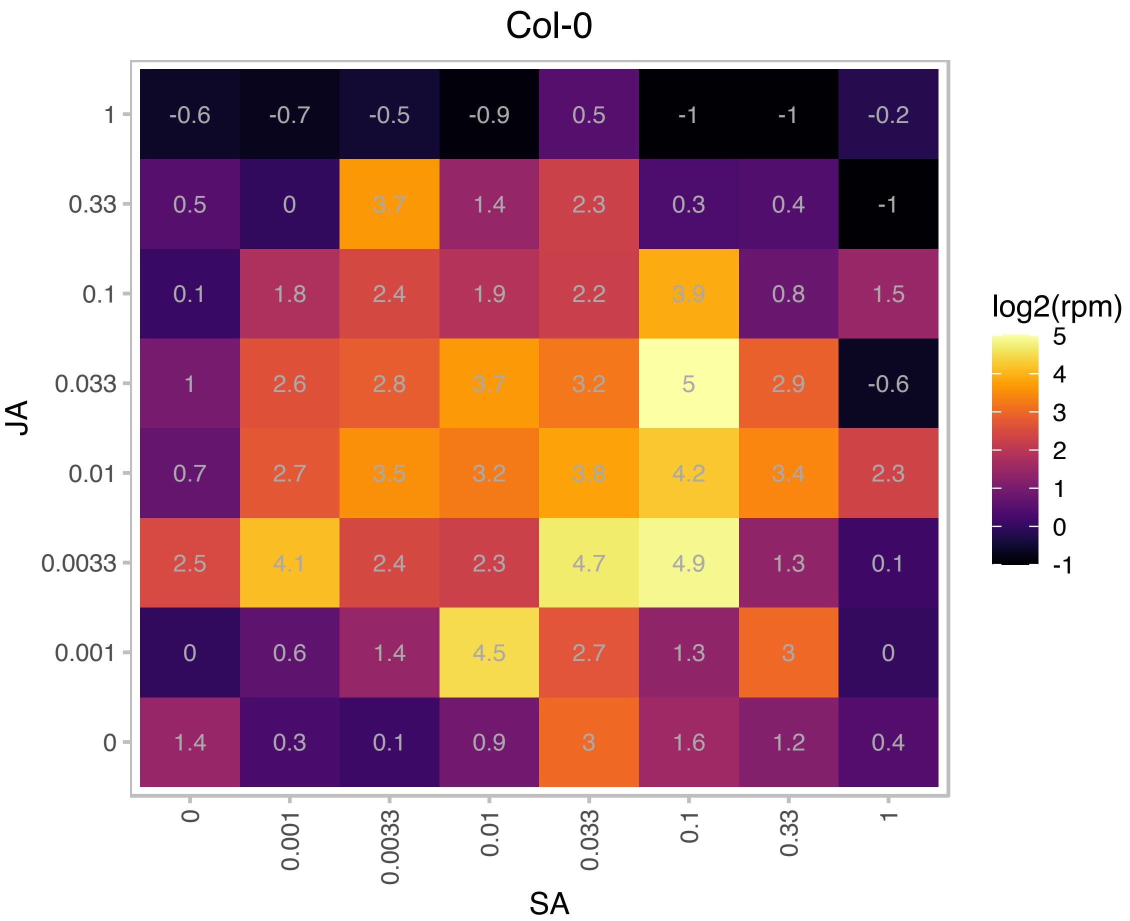


| | |
|---|---|
| Gene | AT1G33960 |
| is.expressed_genes | TRUE |
| is.responsive_genes | TRUE |
| is.apcluster | H-03 |

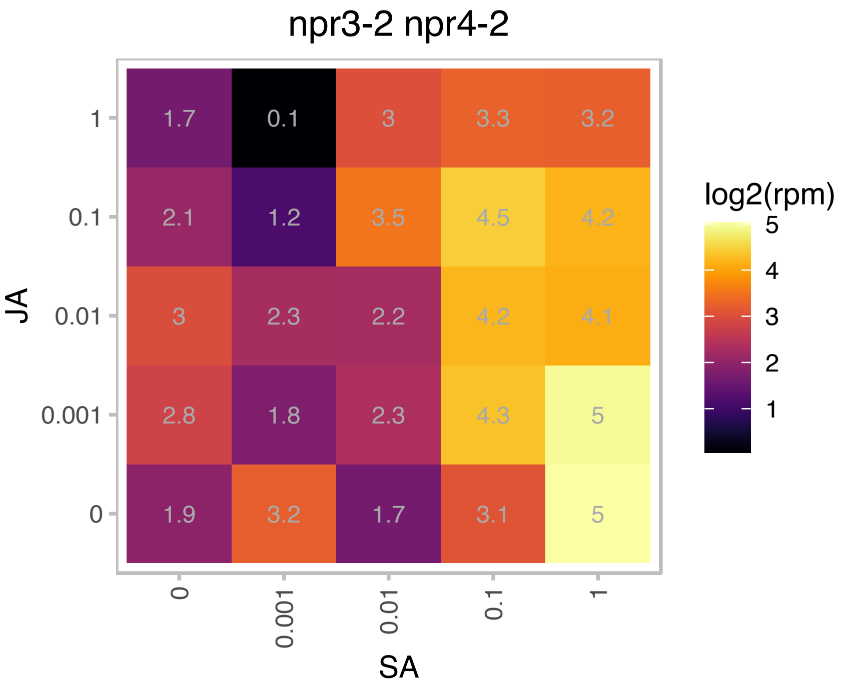


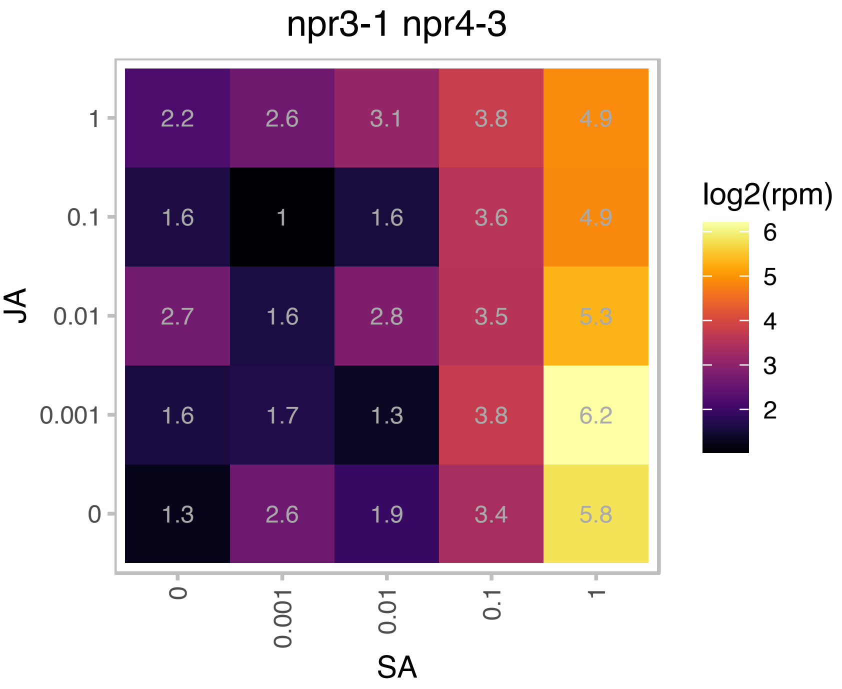

# AT1G55450.1 : NA

S-adenosyl-L-methionine-dependent methyltransferases superfamily protein

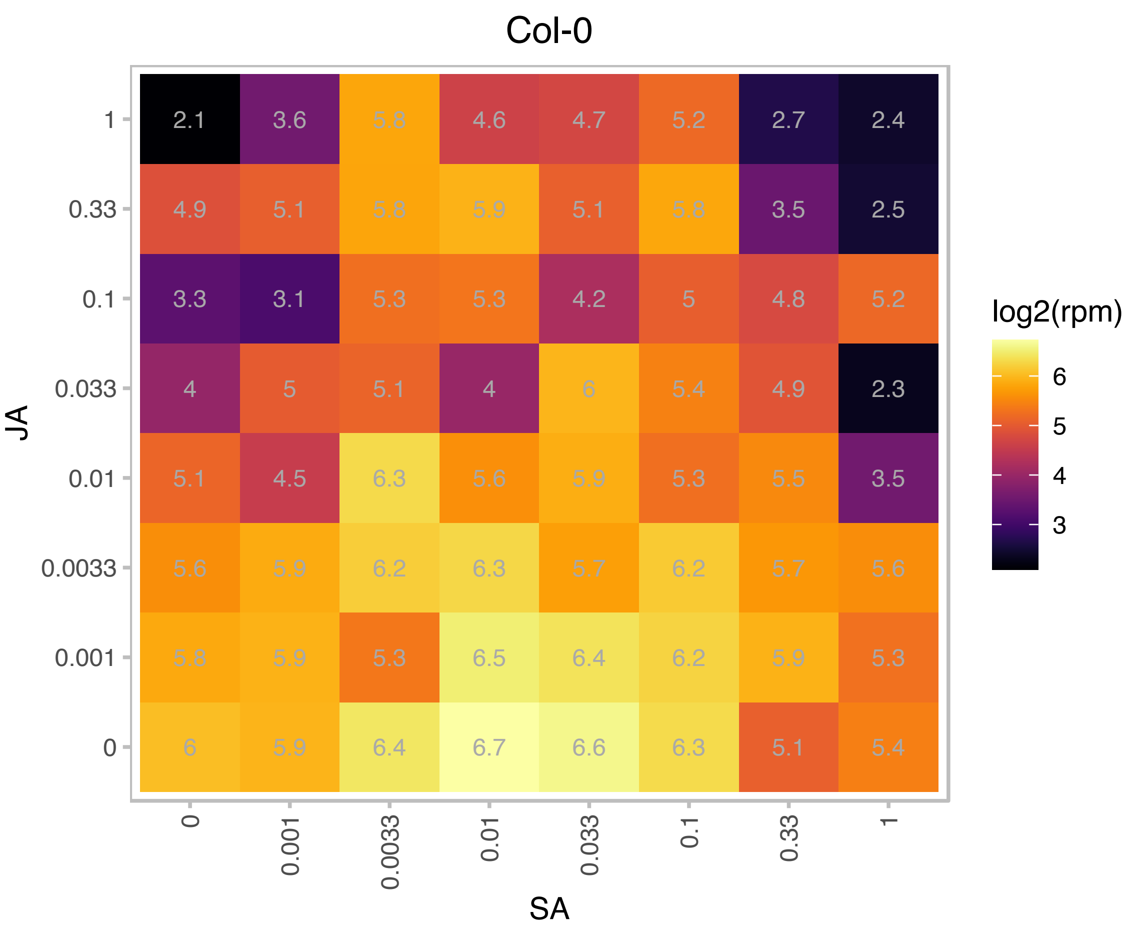


| | |
|---|---|
| Gene | AT1G55450 |
| is.expressed_genes | TRUE |
| is.responsive_genes | TRUE |
| is.apcluster | H-03 |

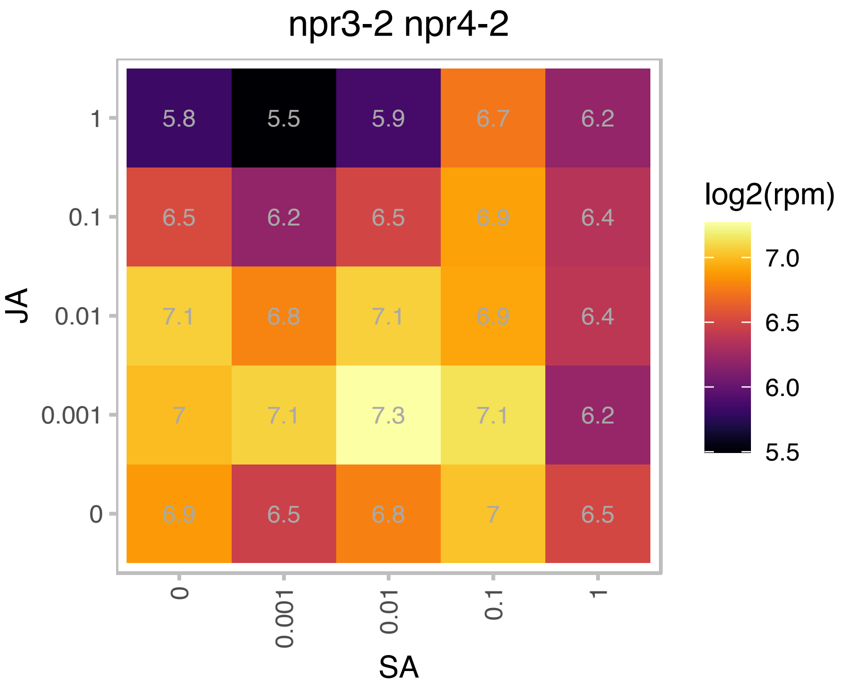


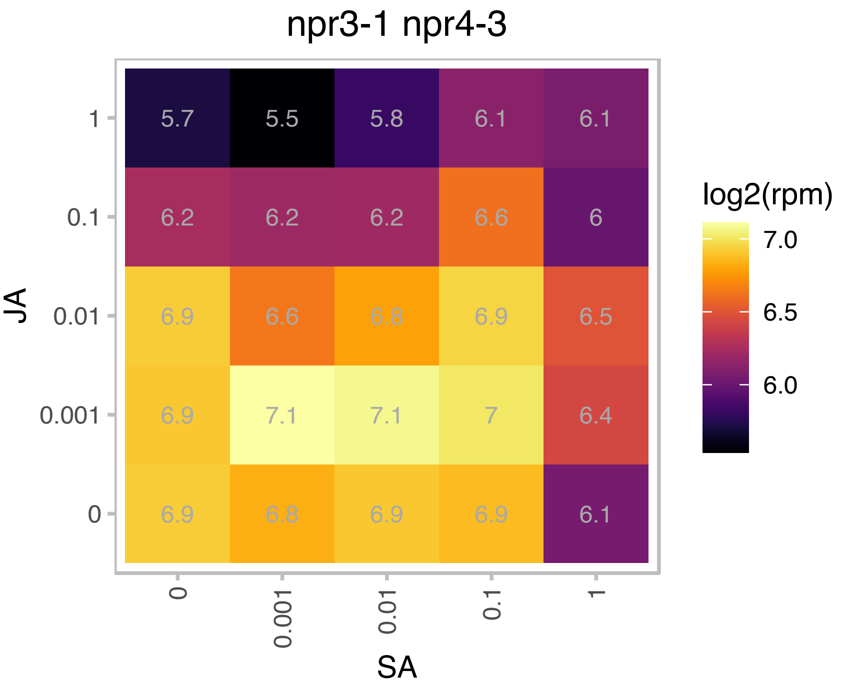

# AT1G65486.1 : NA

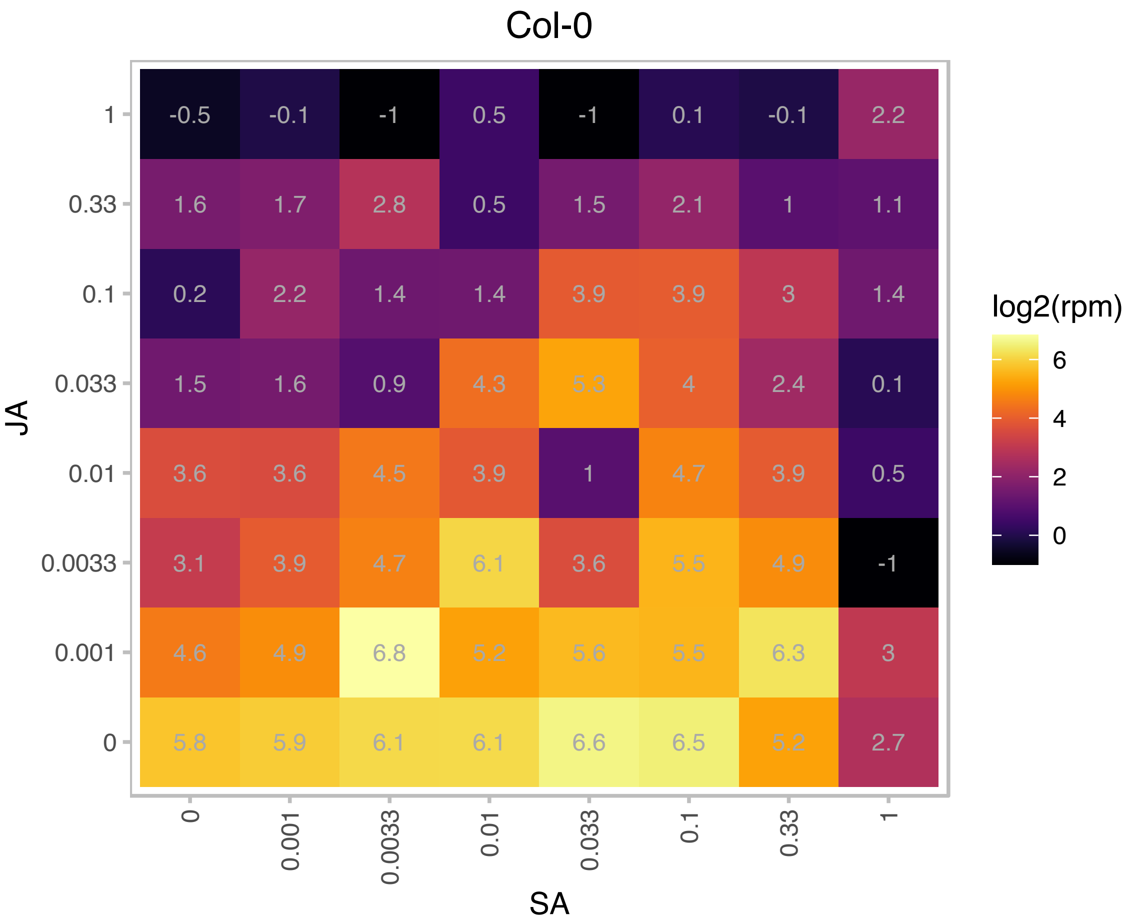


| | |
|---|---|
| Gene | AT1G65486 |
| is.expressed_genes | TRUE |
| is.responsive_genes | TRUE |
| is.apcluster | H-03 |

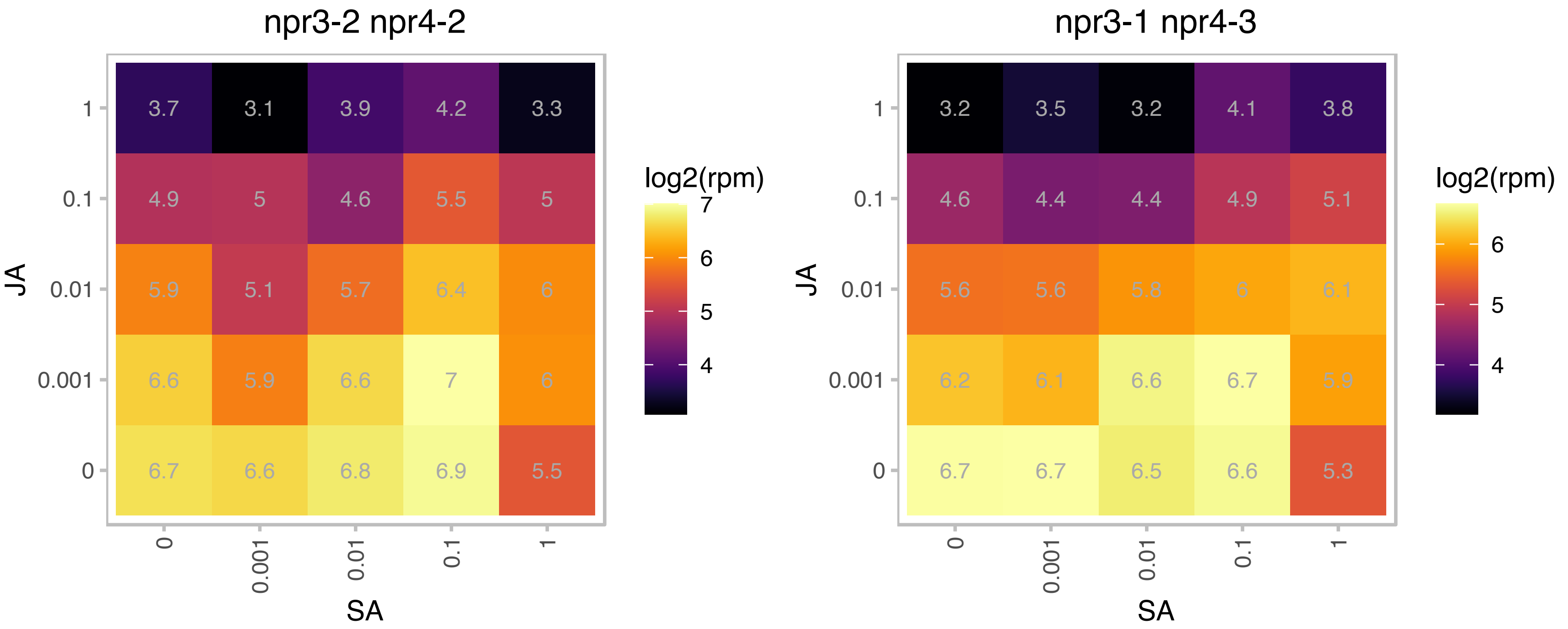

# AT1G65845.1 : NA

| | |
|---|---|
| Gene | AT1G65845 |
| is.expressed_genes | TRUE |
| is.responsive_genes | TRUE |
| is.apcluster | H-03 |

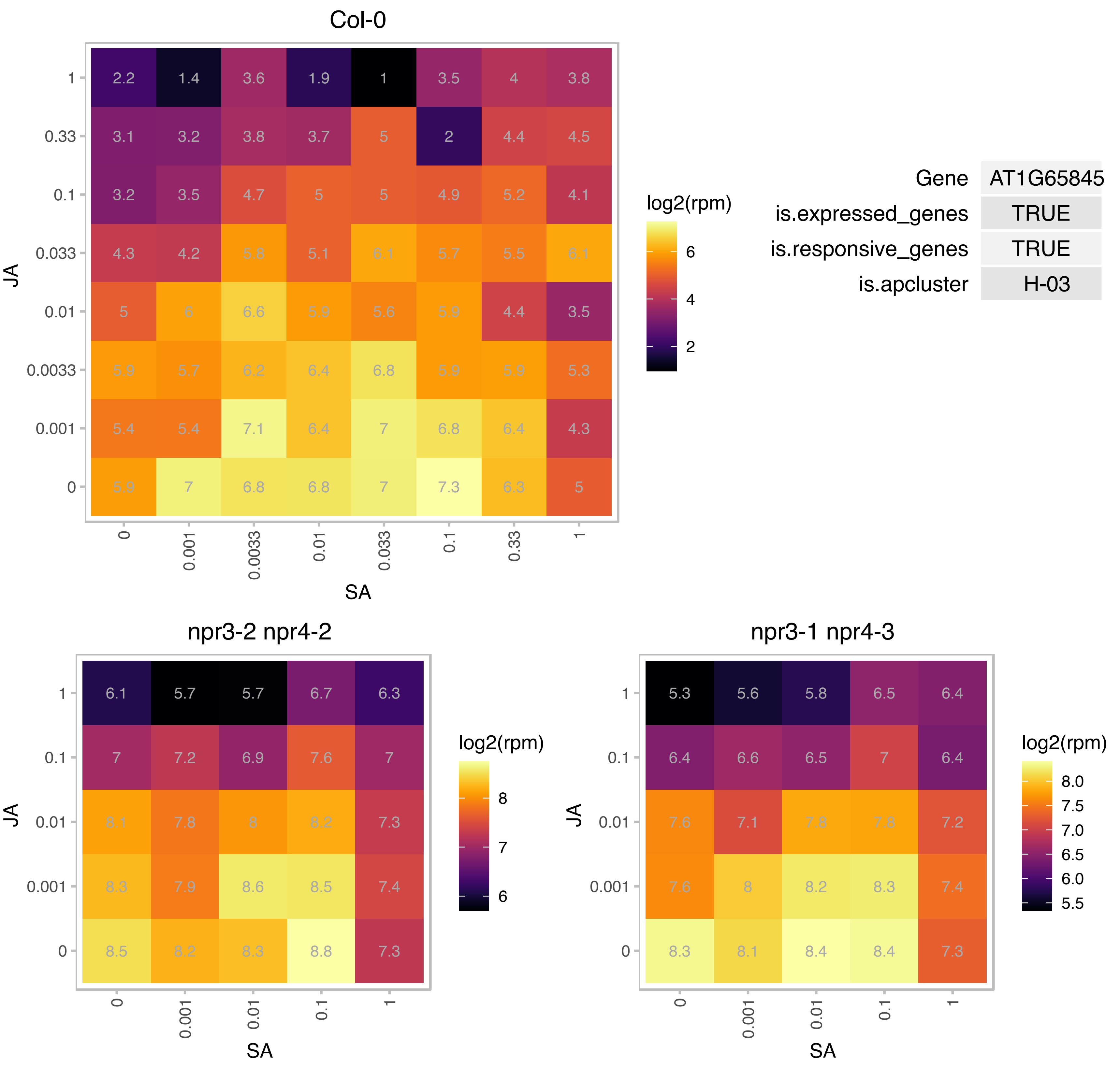

# AT1G68585.1 : NA

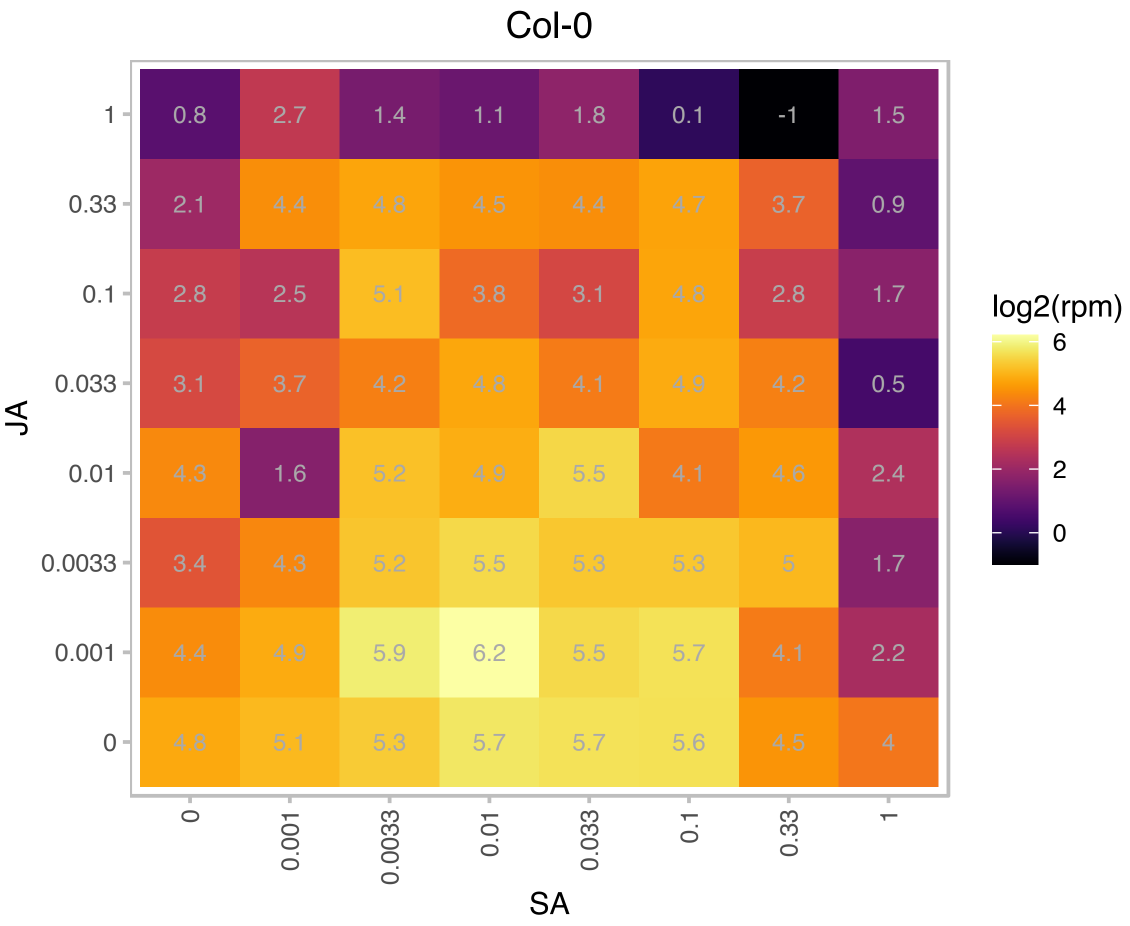


| | |
|---|---|
| Gene | AT1G68585 |
| is.expressed_genes | TRUE |
| is.responsive_genes | TRUE |
| is.apcluster | H-03 |

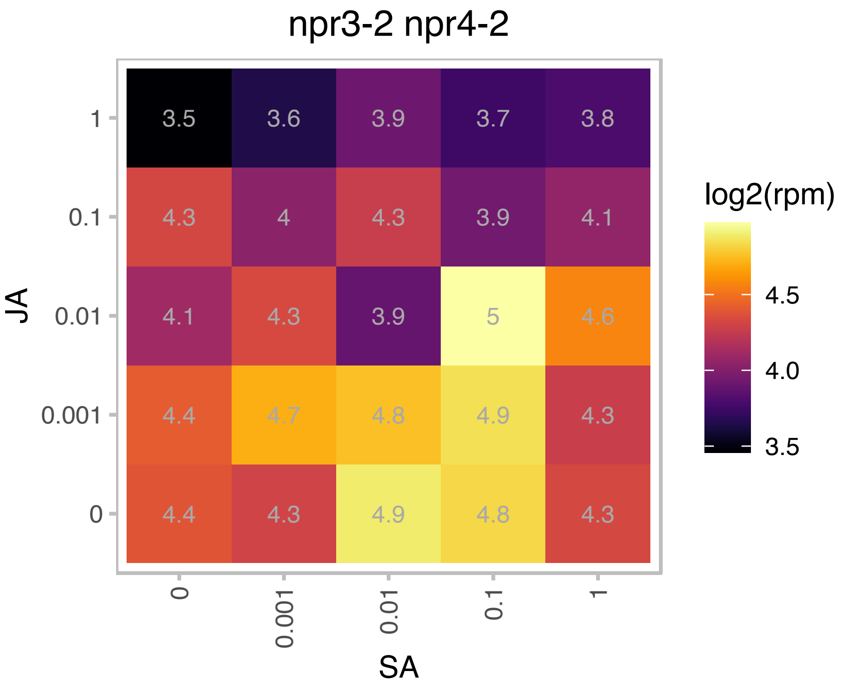


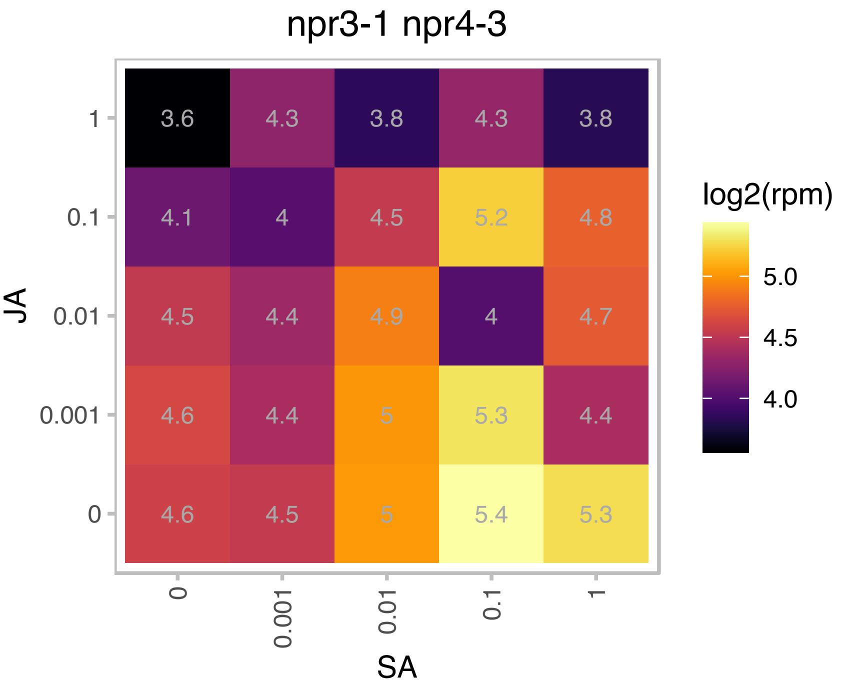

### AT1G72010.1 : AtTCP22, TCP22

TCP family transcription factor

Col-0

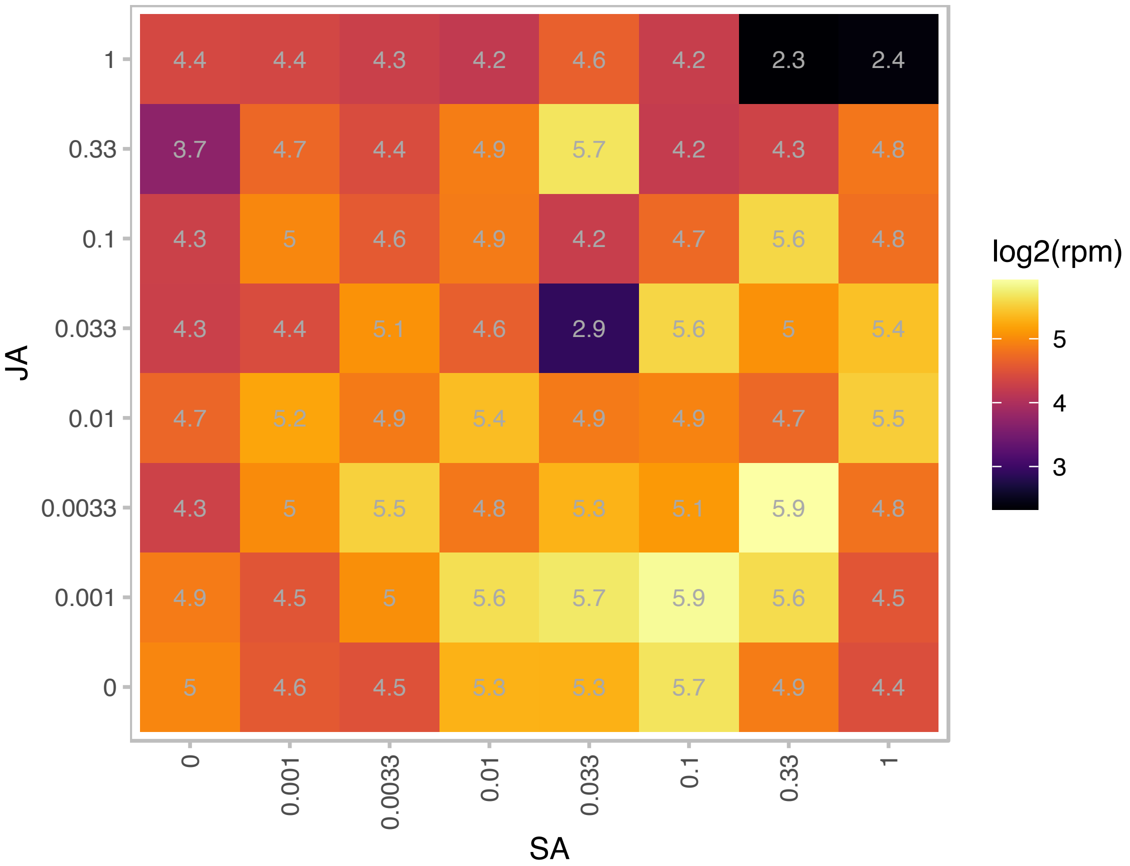


| | |
|---|---|
| Gene | AT1G72010 |
| is.expressed_genes | TRUE |
| is.responsive_genes | TRUE |
| is.apcluster | H-03 |

npr3-2 npr4-2

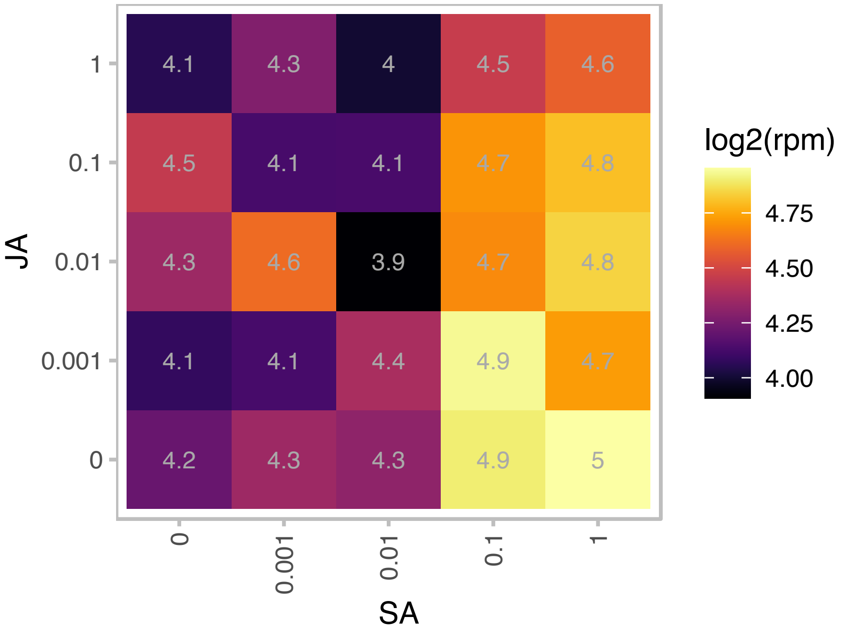


npr3-1 npr4-3

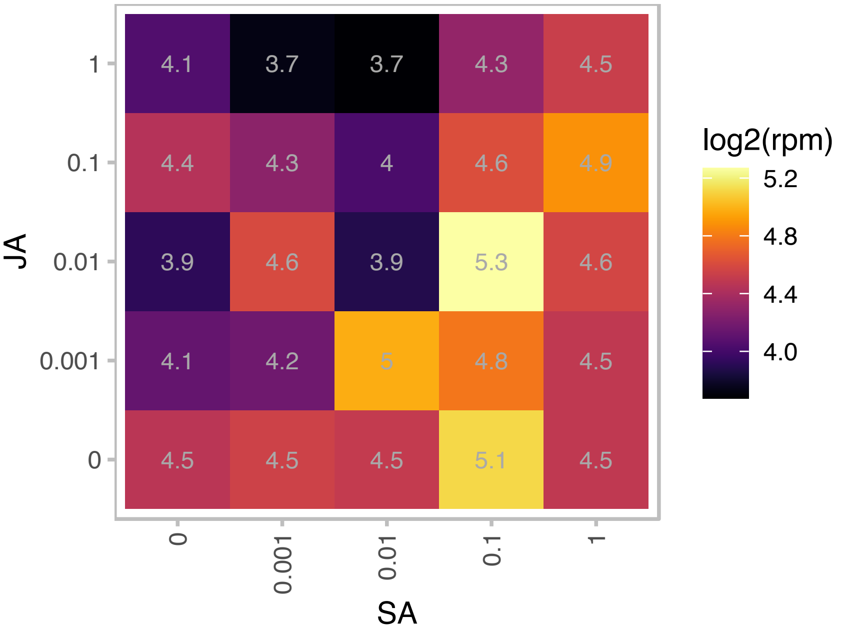

# AT1G72060.1 : NA

serine-type endopeptidase inhibitors

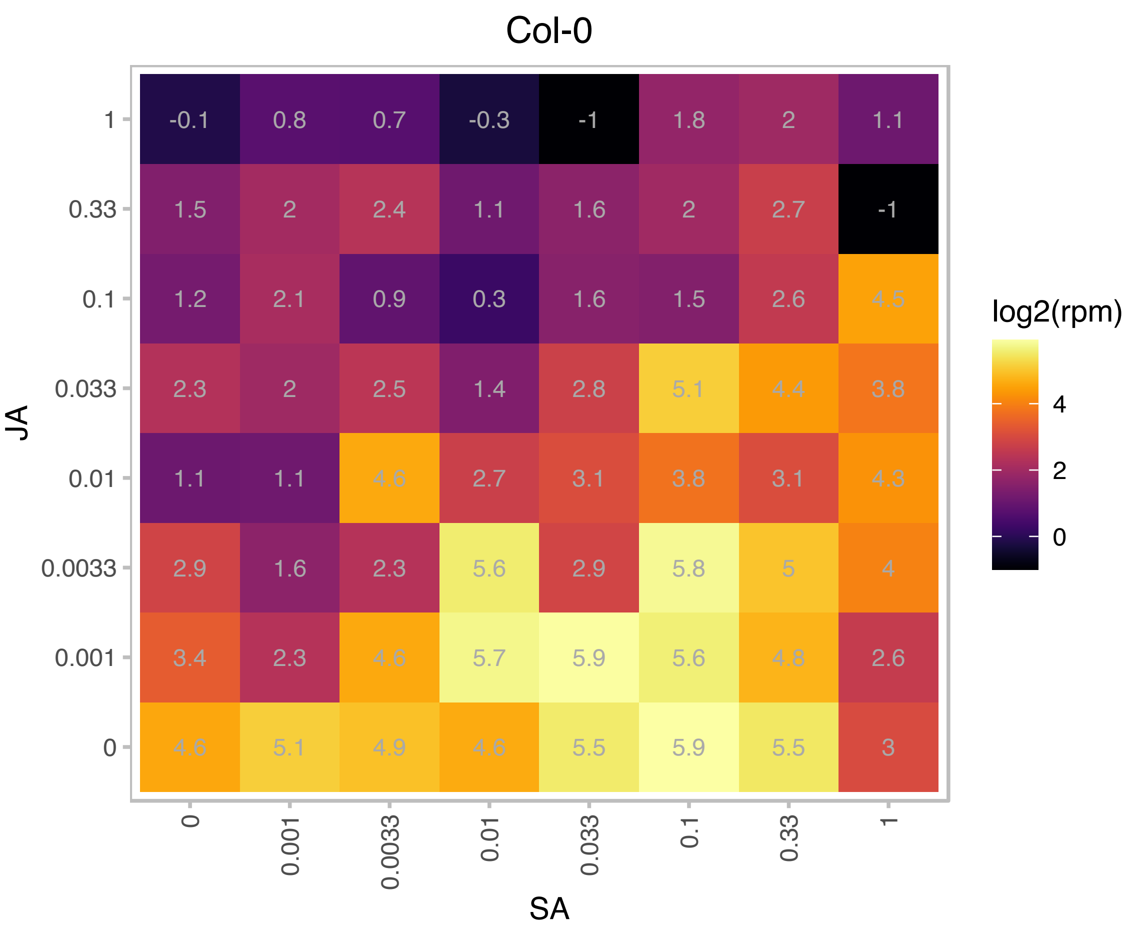


| | |
|---|---|
| Gene | AT1G72060 |
| is.expressed_genes | TRUE |
| is.responsive_genes | TRUE |
| is.apcluster | H-03 |

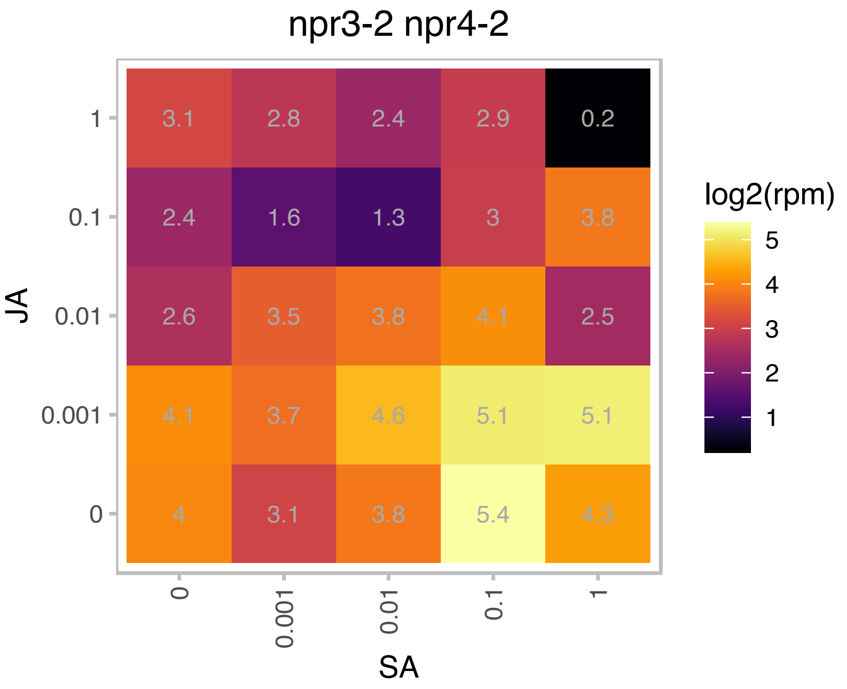


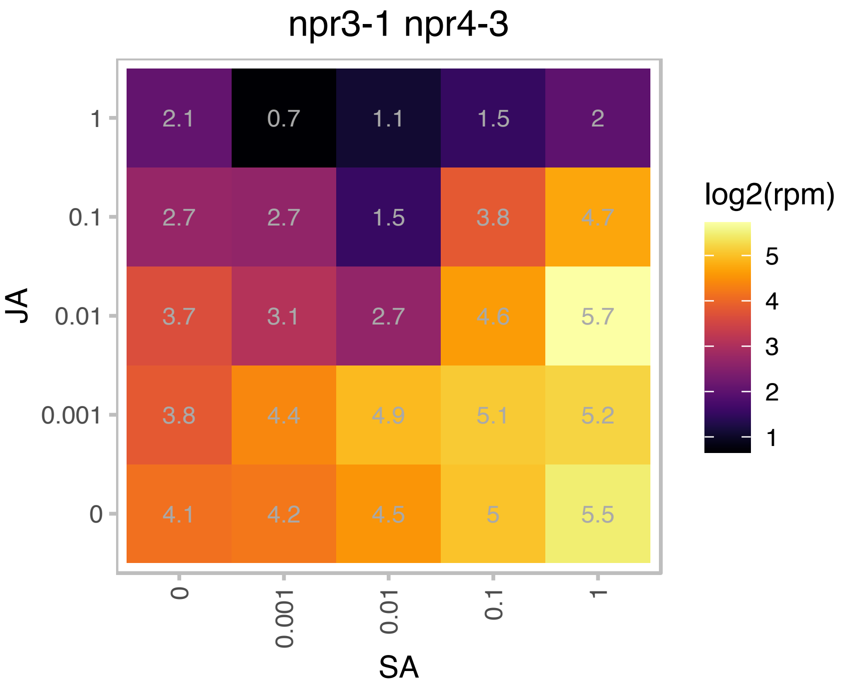

### AT1G76930.1 : ATEXT1, ATEXT4, EXT1, EXT4, ORG5

extensin 4
Encodes an Arabidopsis extensin gene that belongs to cell-wall hydroxyproline-rich glycoproteins. The cross-link of extensins enforces cell wall strength. Transgenic plants overexpressing this gene show an increase in stem thickness.

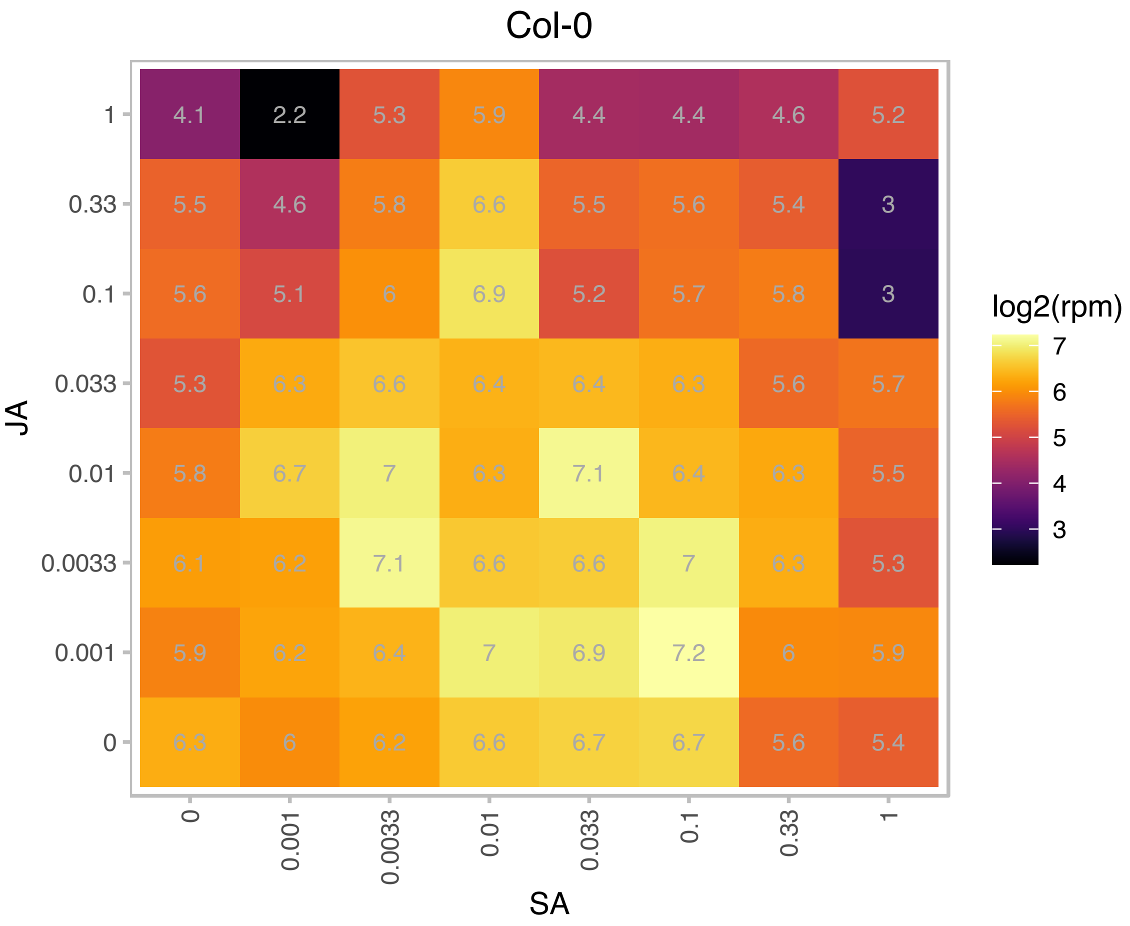


| | |
|---|---|
| Gene | AT1G76930 |
| is.expressed_genes | TRUE |
| is.responsive_genes | TRUE |
| is.apcluster | H-03 |

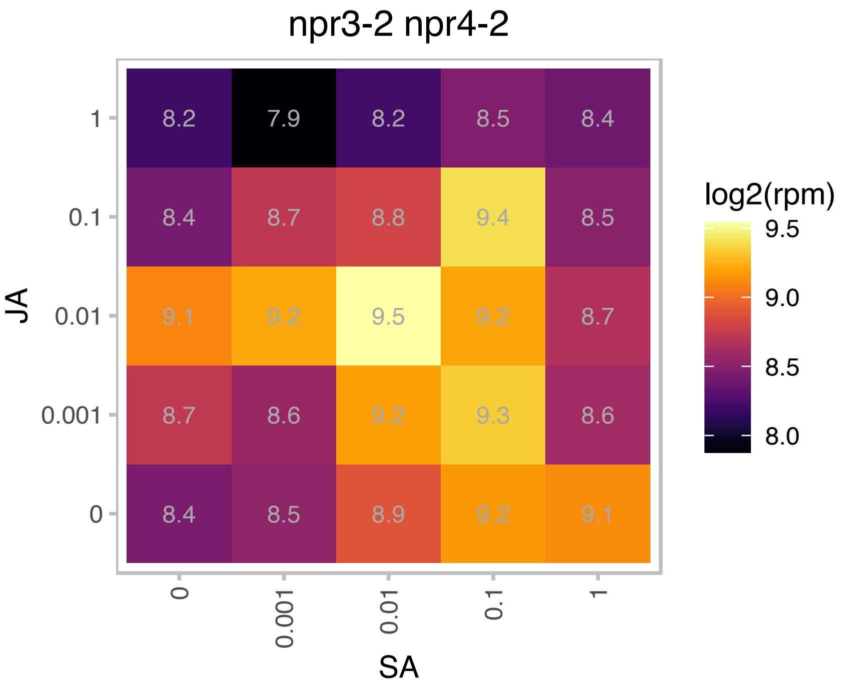


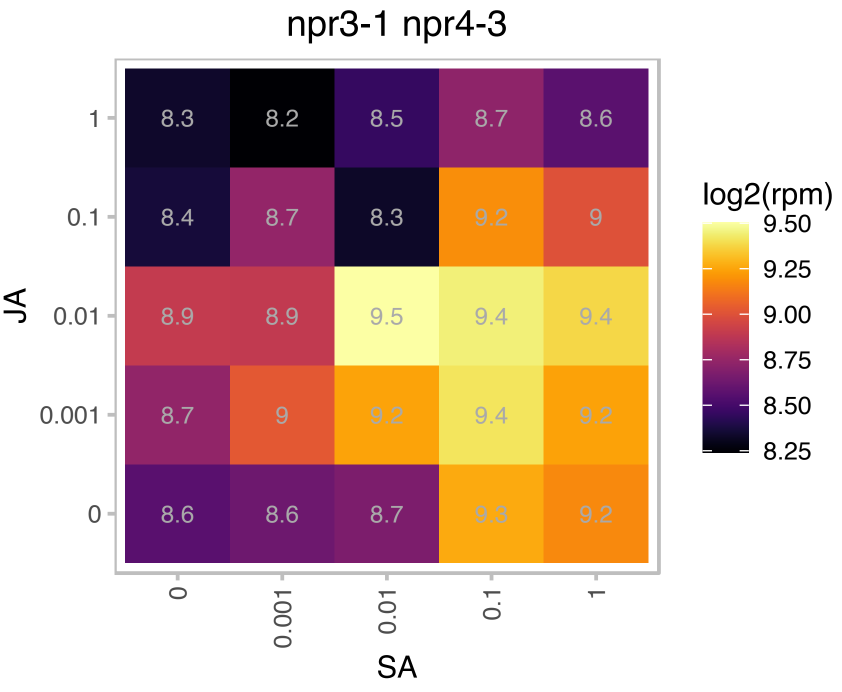

# AT1G76960.1 : NA

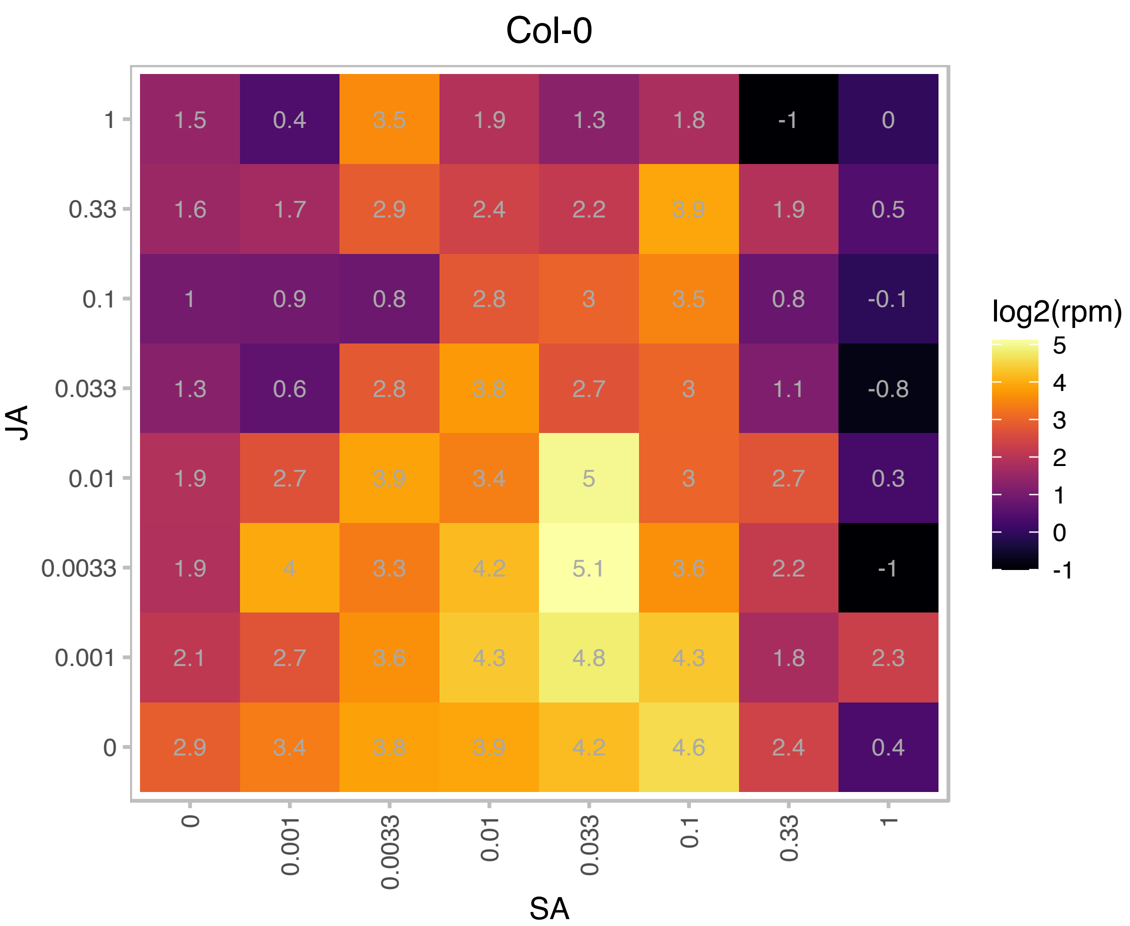


| | |
|---|---|
| Gene | AT1G76960 |
| is.expressed_genes | TRUE |
| is.responsive_genes | TRUE |
| is.apcluster | H-03 |

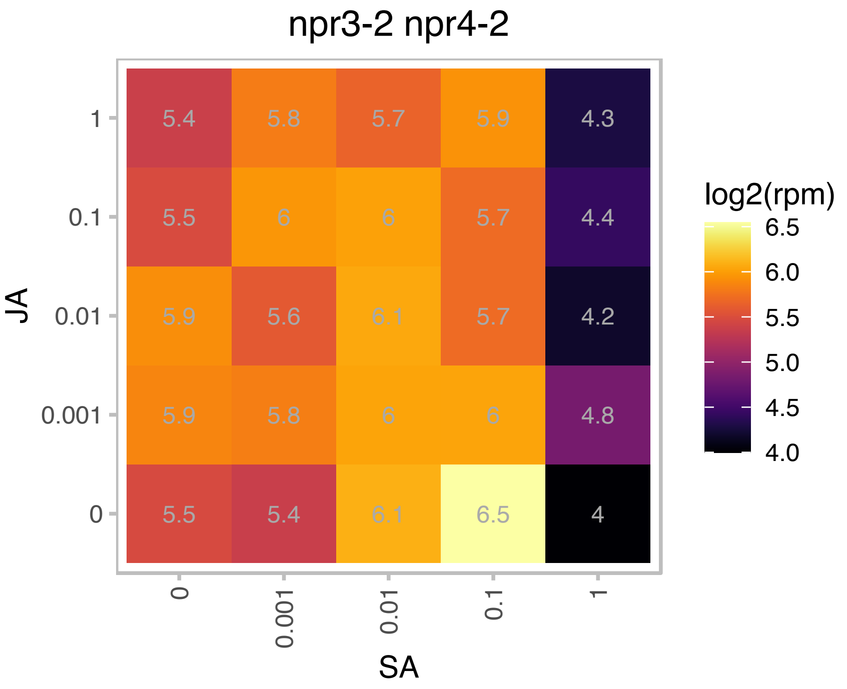


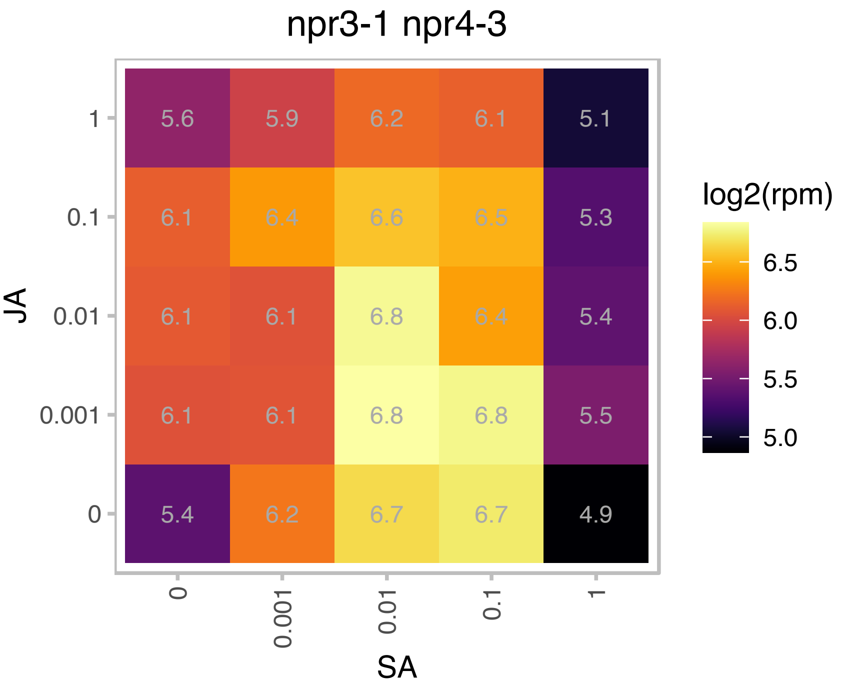

### AT2G04450.1 : ATNUDT6, ATNUDX6, NUDT6, NUDX6

nudix hydrolase homolog 6

Encodes a protein with NADH pyrophosphatase activity. Although this protein was also shown to have ADP-ribose diphosphatase activity in vitro, mutant analyses suggest that NUDX6 is involved in NADH metabolism in vivo.

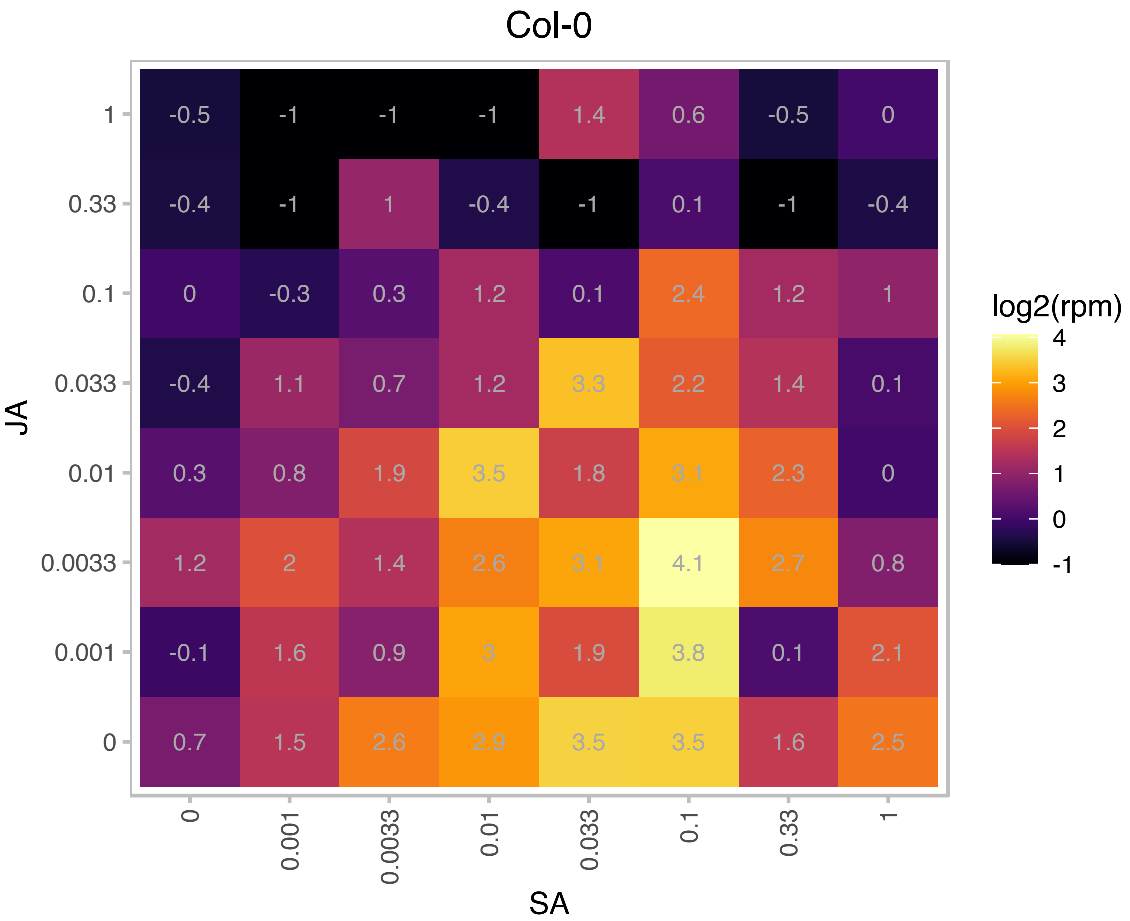


| | |
|---|---|
| Gene | AT2G04450 |
| is.expressed_genes | TRUE |
| is.responsive_genes | TRUE |
| is.apcluster | H-03 |

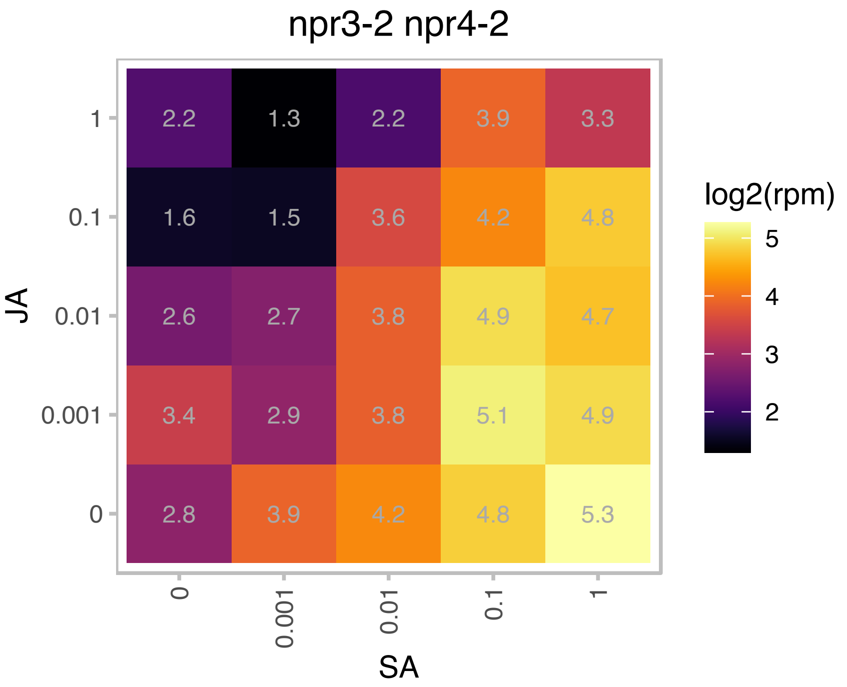


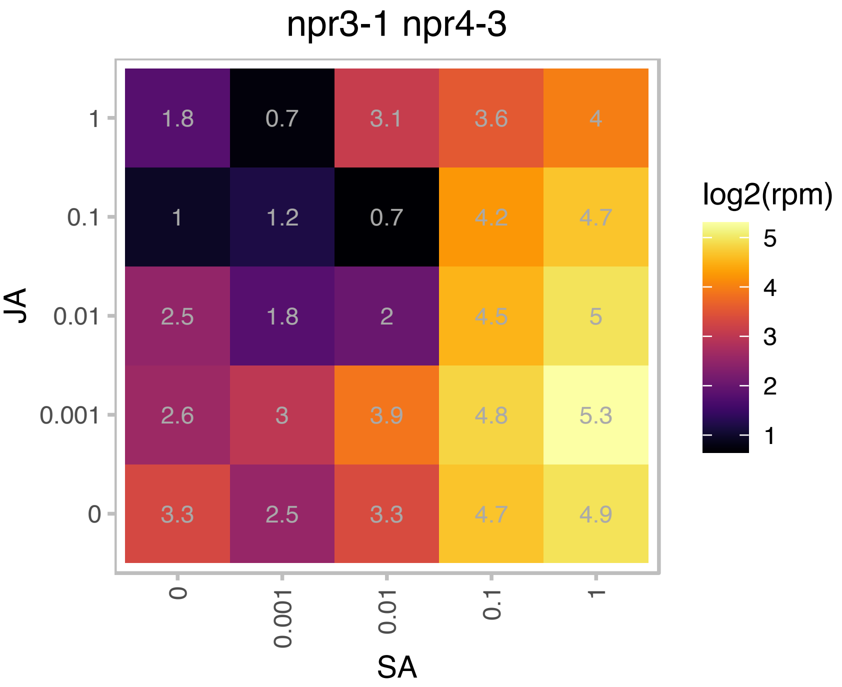

### AT2G05100.1 : LHCB2, LHCB2.1

photosystem II light harvesting complex gene 2.1
Lhcb2.1 protein encoding a subunit of the light harvesting complex II. Member of a gene family with high degree of sequence similarity. Initially LHCB2.3 was considered as a separate gene but appears to be an allele of LHCB2.1.

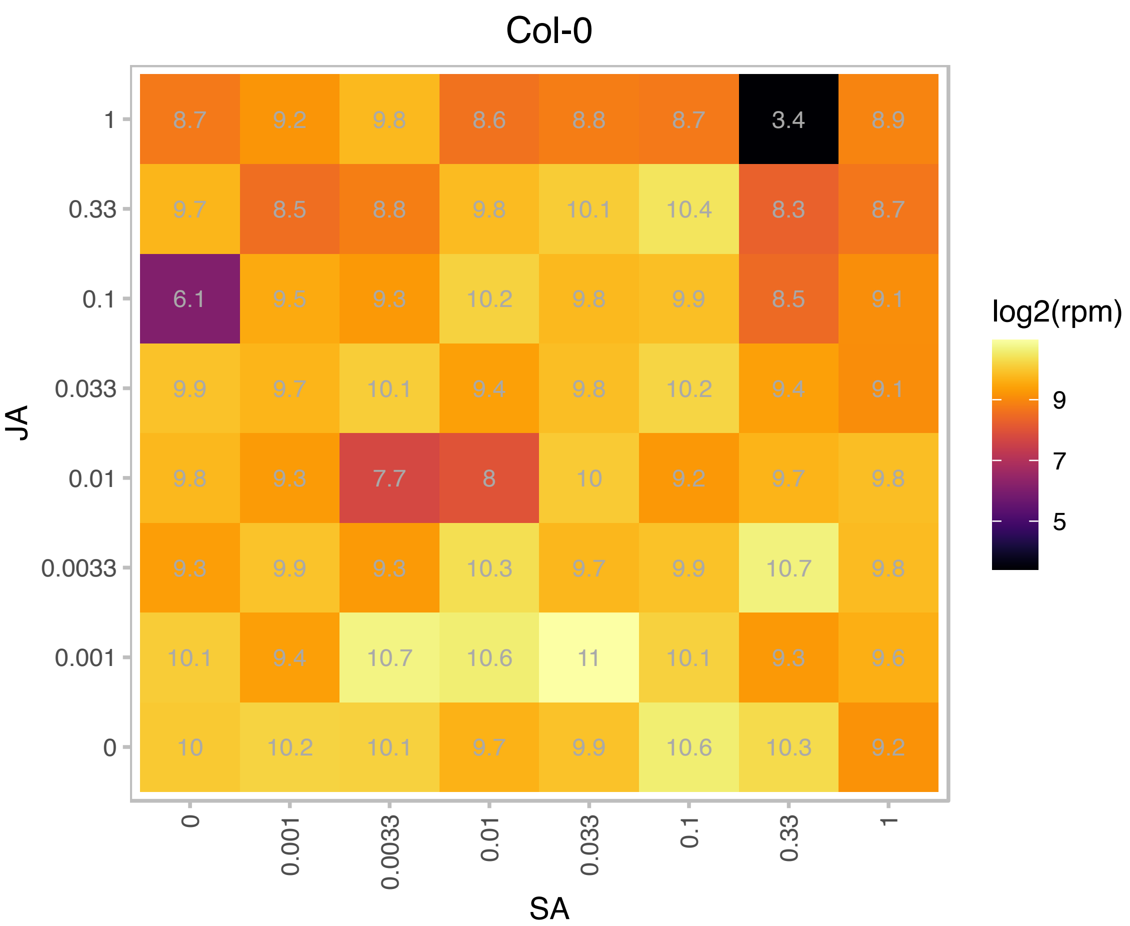


| | |
|---|---|
| Gene | AT2G05100 |
| is.expressed_genes | TRUE |
| is.responsive_genes | TRUE |
| is.apcluster | H-03 |

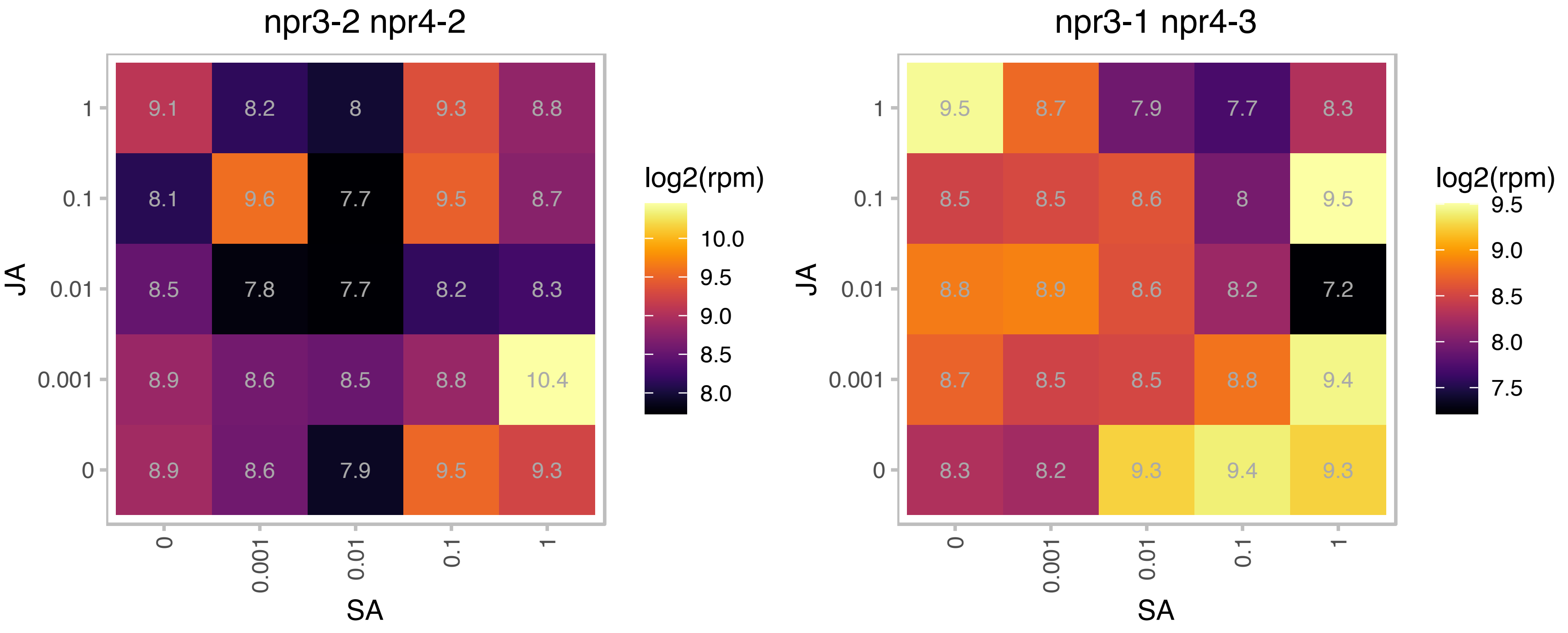

### AT2G16060.1 : AHB1, ARATH GLB1, ATGLB1, GLB1, HB1

hemoglobin 1
Encodes a class 1 nonsymbiotic hemoglobin induced by low oxygen levels with very high oxygen affinity. It is not likely to be a hemoglobin transporter because of its extremely high affinity for oxygen. Overexpression impairs cold stress-induced nitric oxide (NO) production.

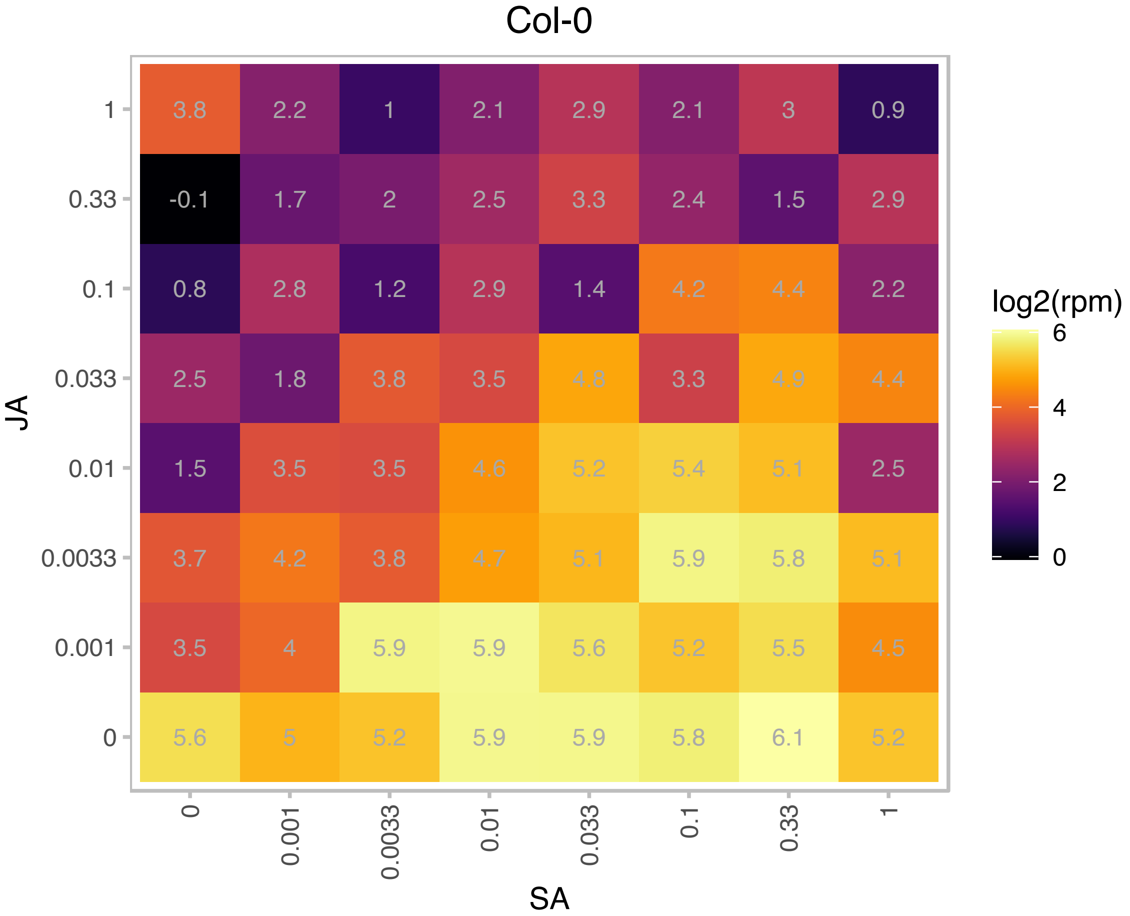


| | |
|---|---|
| Gene | AT2G16060 |
| is.expressed_genes | TRUE |
| is.responsive_genes | TRUE |
| is.apcluster | H-03 |

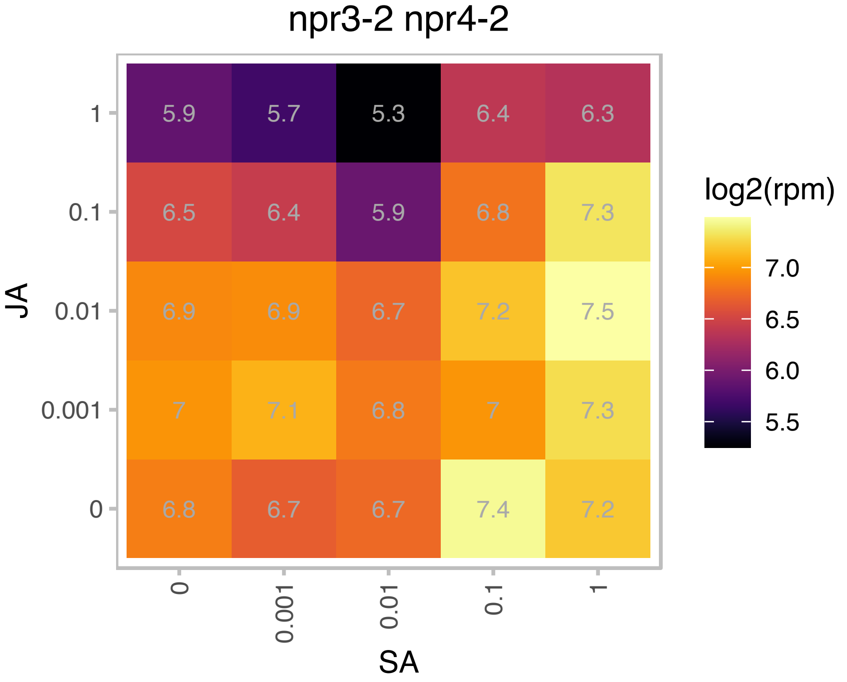


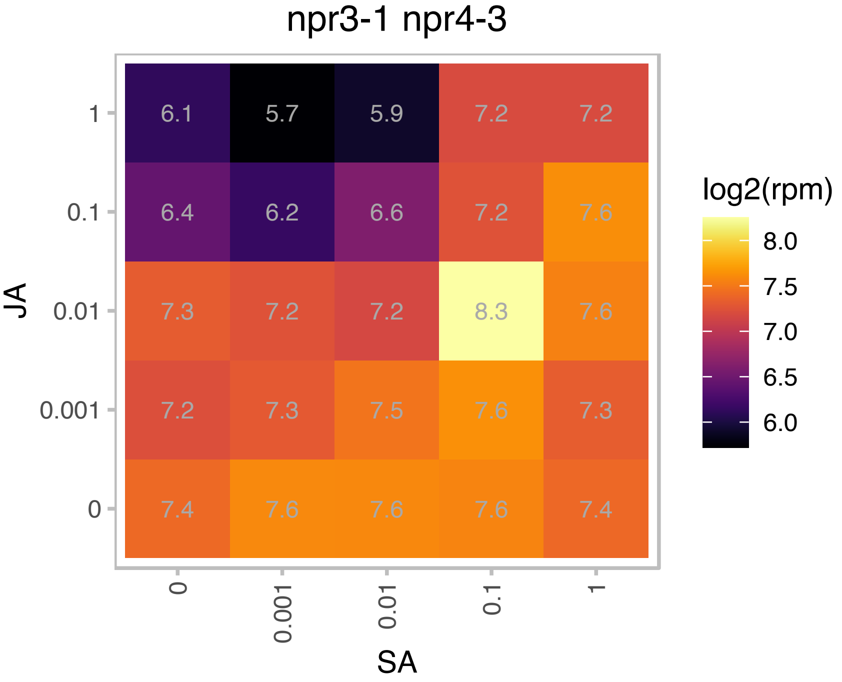

# AT2G20725.1 : NA

CAAX amino terminal protease family protein

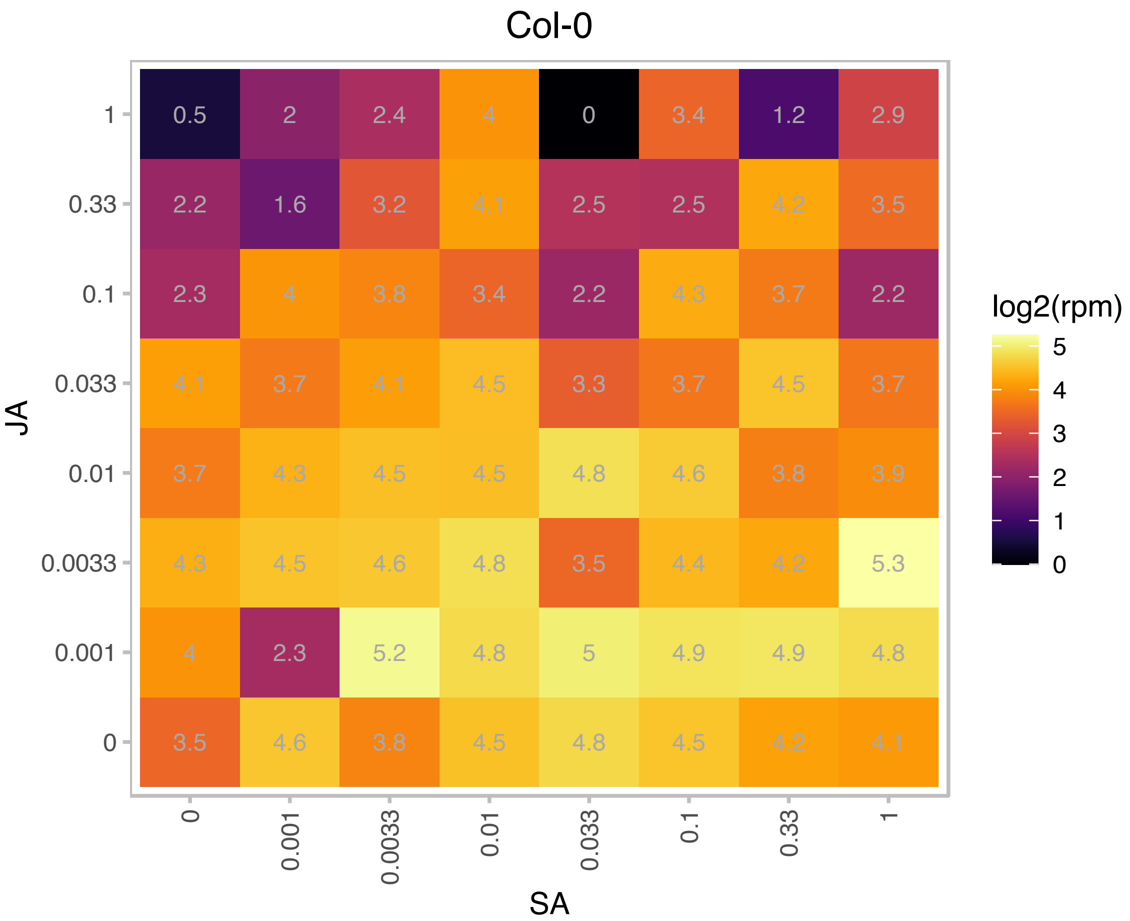


| Gene | AT2G20725 |
|---|---|
| is.expressed_genes | TRUE |
| is.responsive_genes | TRUE |
| is.apcluster | H-03 |

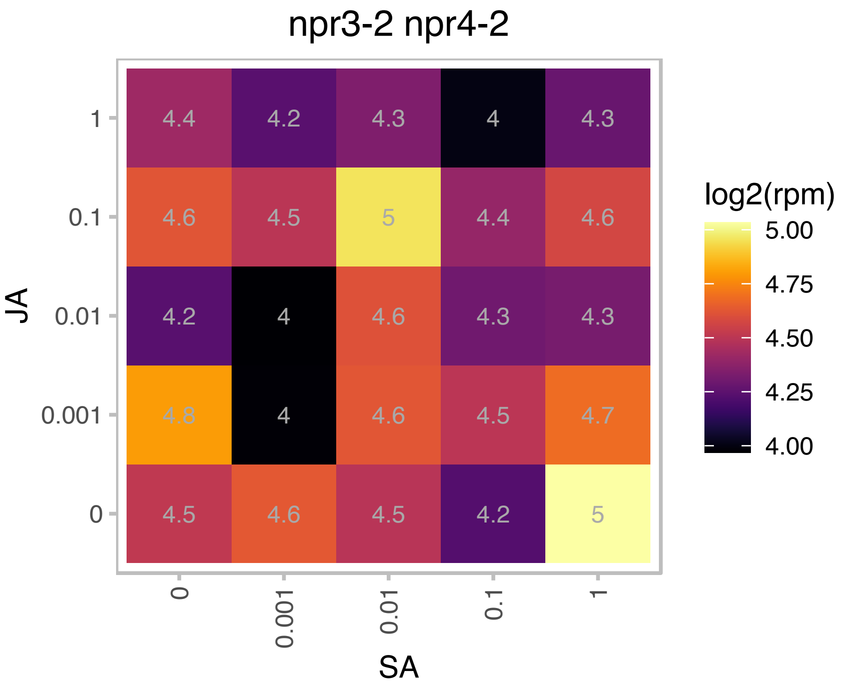


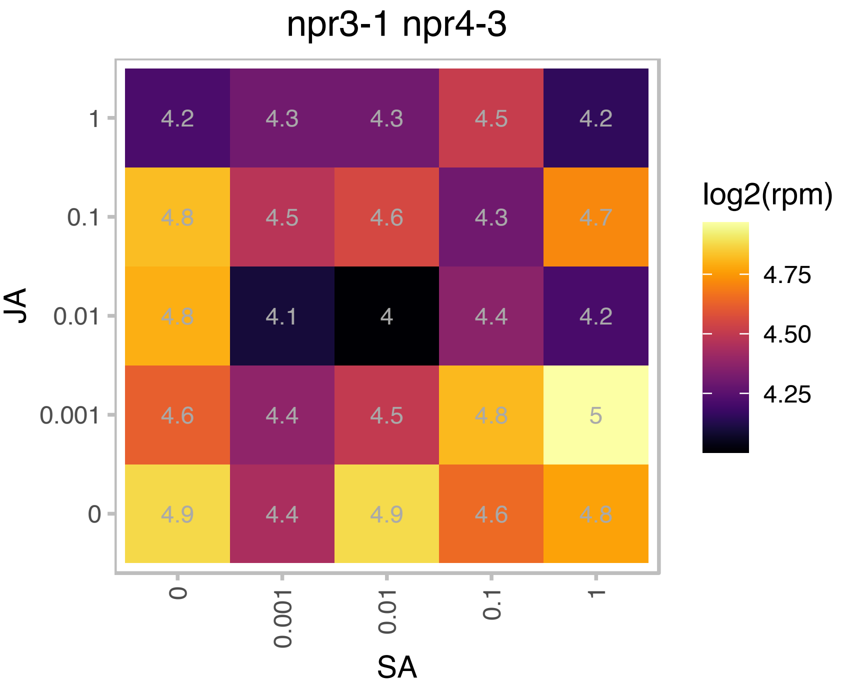

# AT2G24390.1 : NA

AIG2-like (avirulence induced gene) family protein

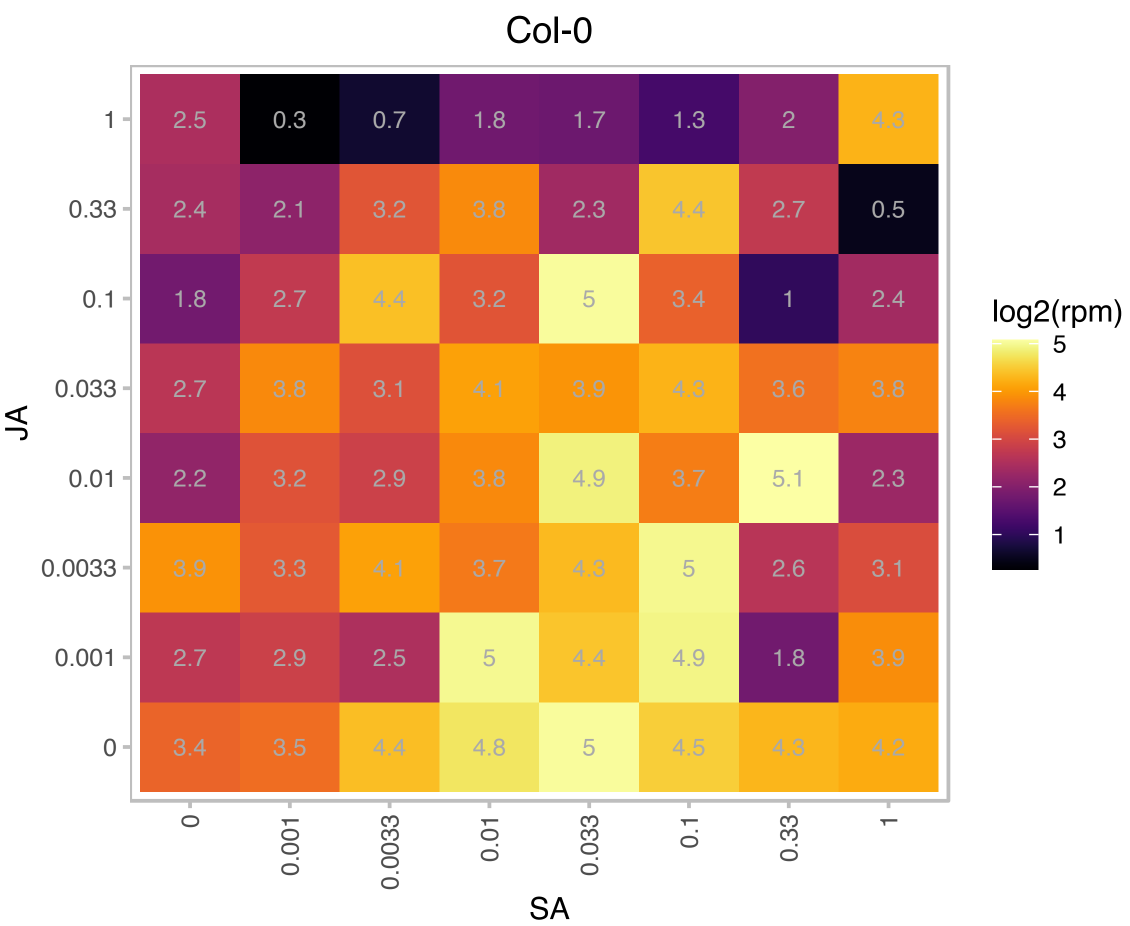


| | |
|---|---|
| Gene | AT2G24390 |
| is.expressed_genes | TRUE |
| is.responsive_genes | TRUE |
| is.apcluster | H-03 |

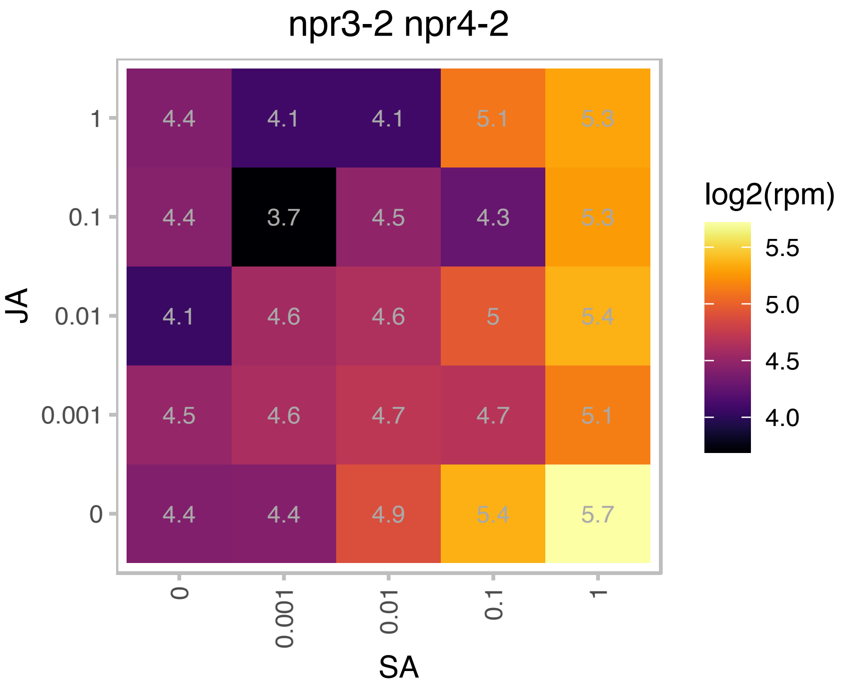


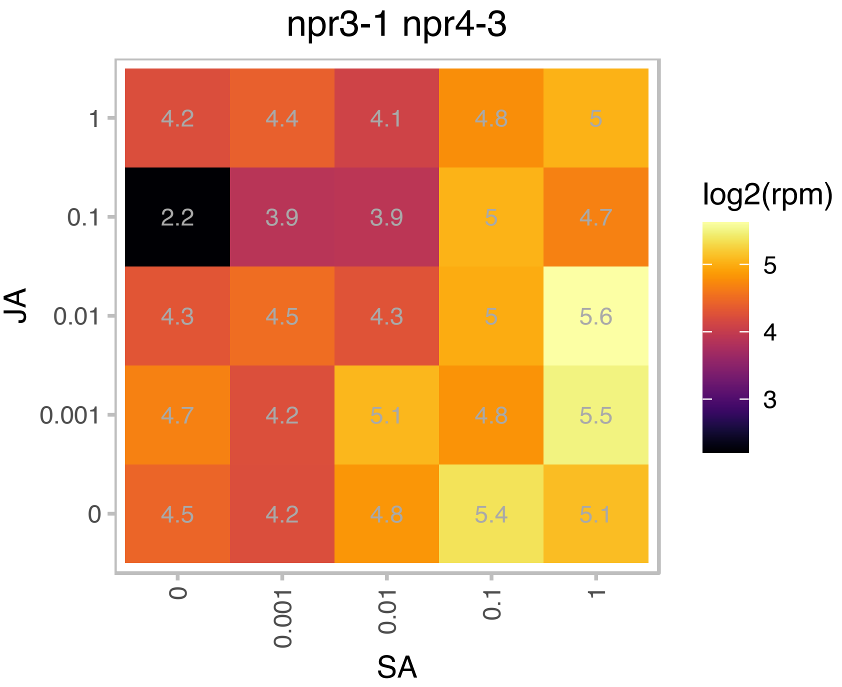

# AT2G24970.1 : NA

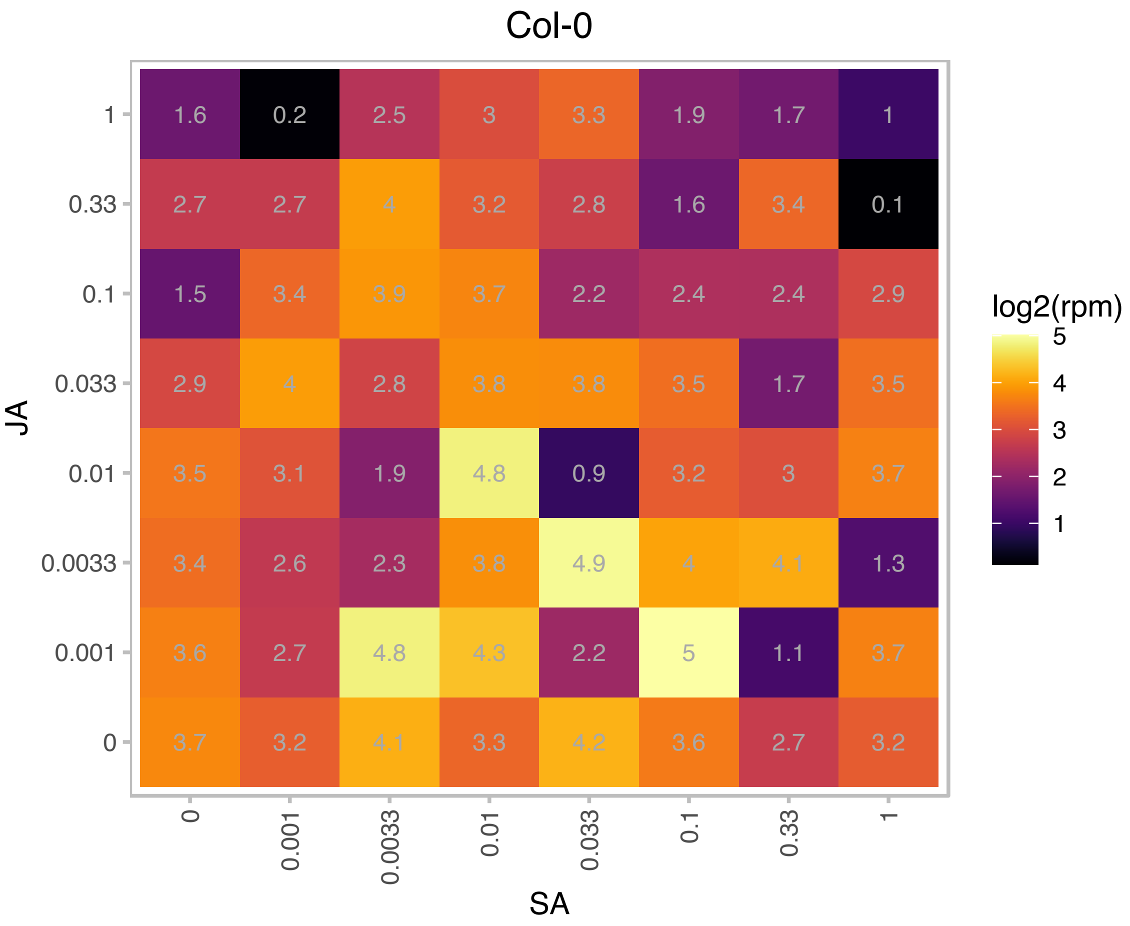


| | |
|---|---|
| Gene | AT2G24970 |
| is.expressed_genes | TRUE |
| is.responsive_genes | TRUE |
| is.apcluster | H-03 |

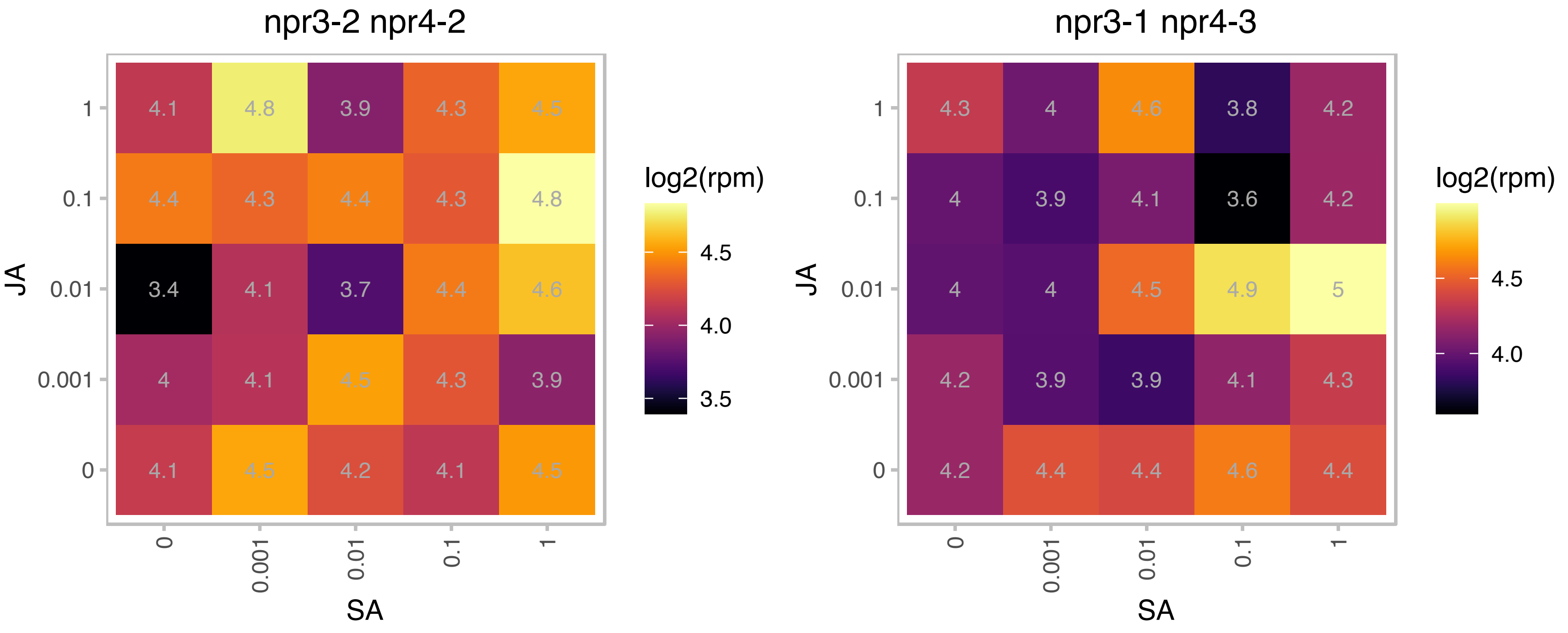

# AT2G26240.1 : NA

Transmembrane proteins 14C

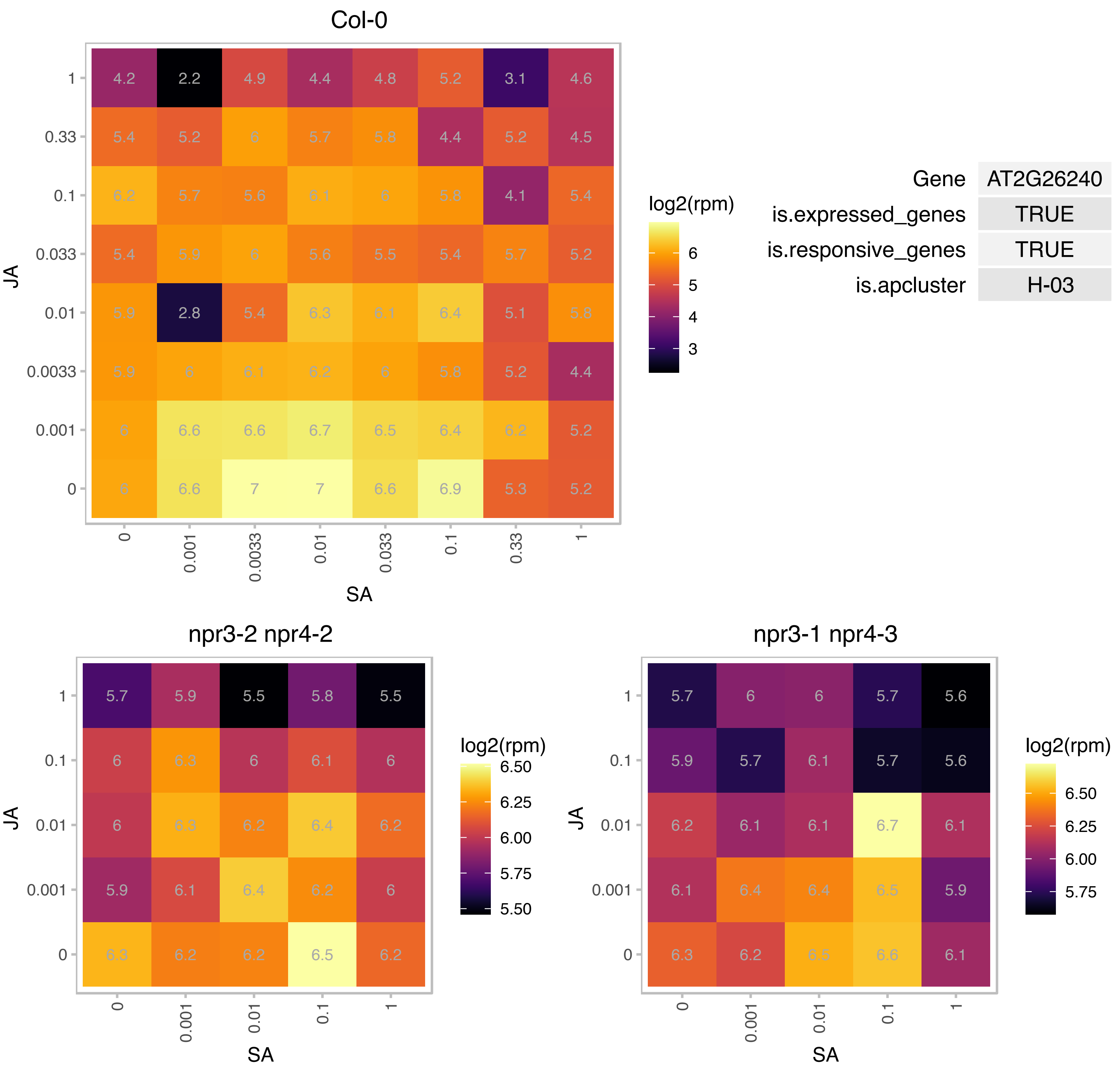


| | |
|---|---|
| Gene | AT2G26240 |
| is.expressed_genes | TRUE |
| is.responsive_genes | TRUE |
| is.apcluster | H-03 |

# AT2G27590.1 : NA

S-adenosyl-L-methionine-dependent methyltransferases superfamily protein

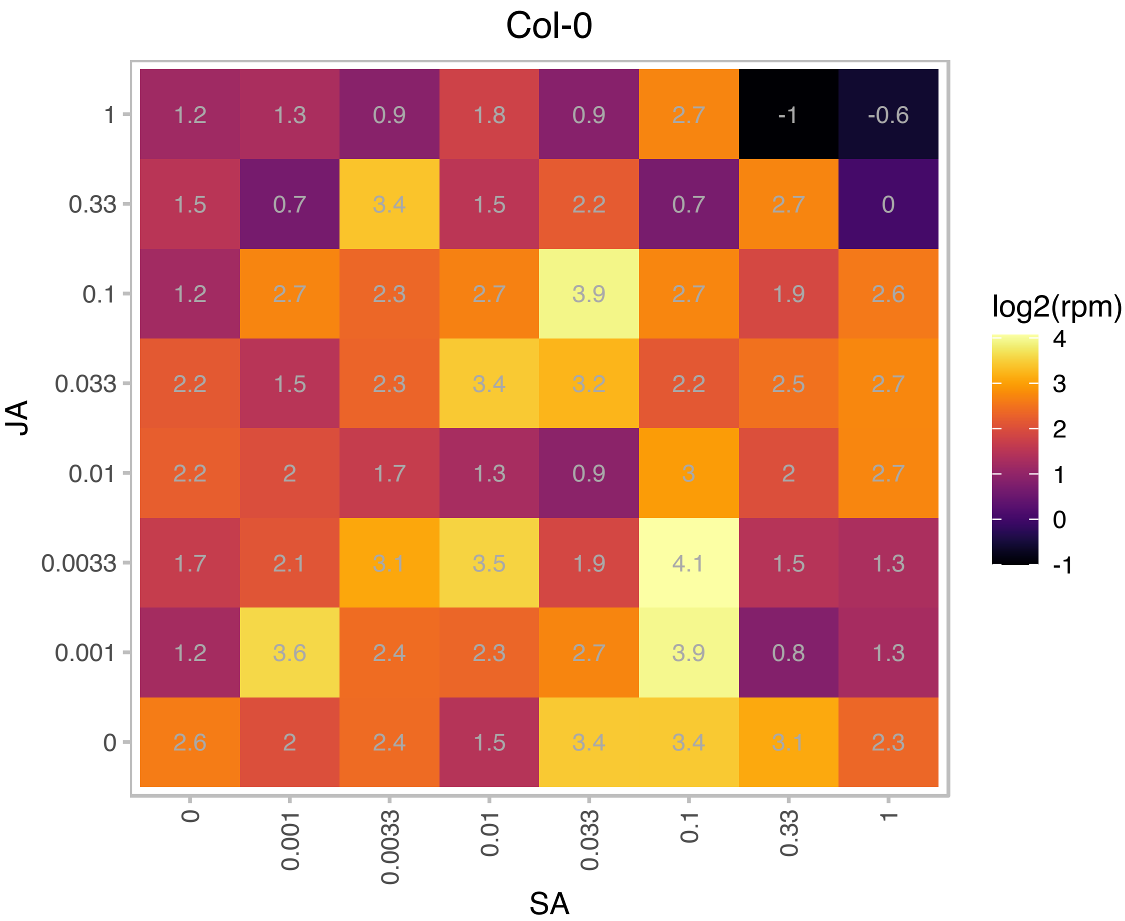


| | |
|---|---|
| Gene | AT2G27590 |
| is.expressed_genes | TRUE |
| is.responsive_genes | TRUE |
| is.apcluster | H-03 |

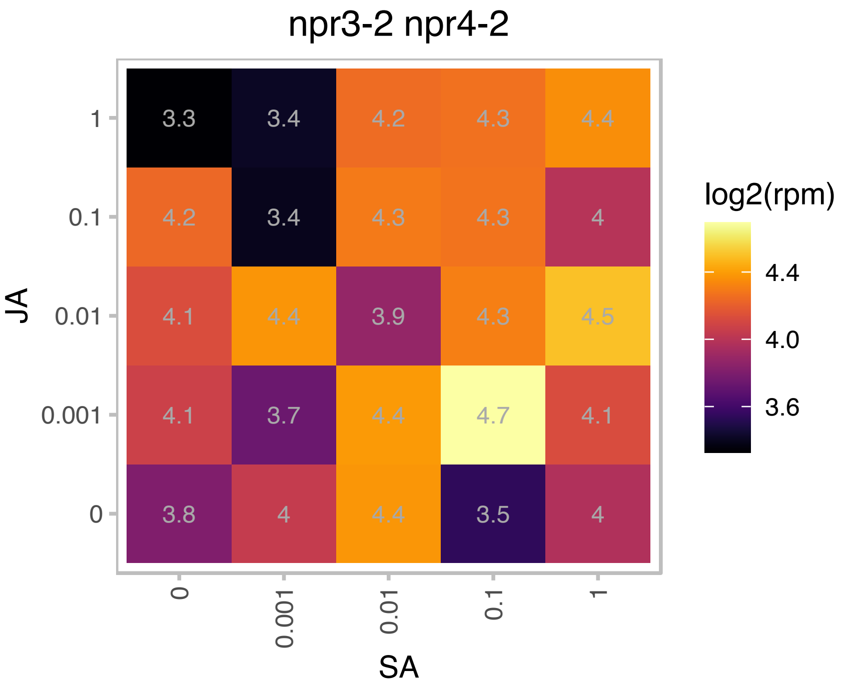


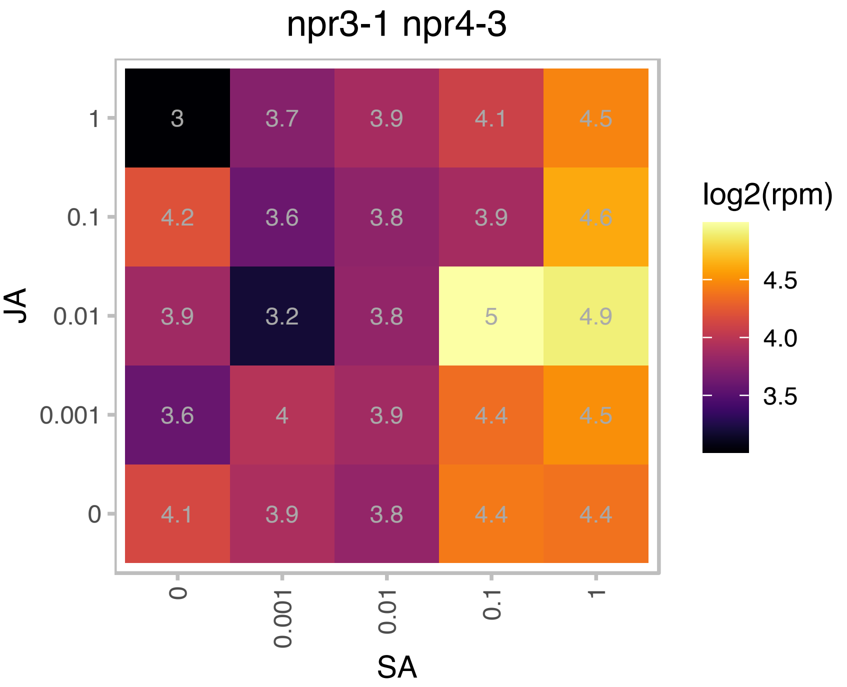

### AT2G34430.1 : LHB1B1, LHCB1.4

light-harvesting chlorophyll-protein complex II subunit B1

Photosystem II type I chlorophyll a/b-binding protein The mRNA is cell-to-cell mobile.

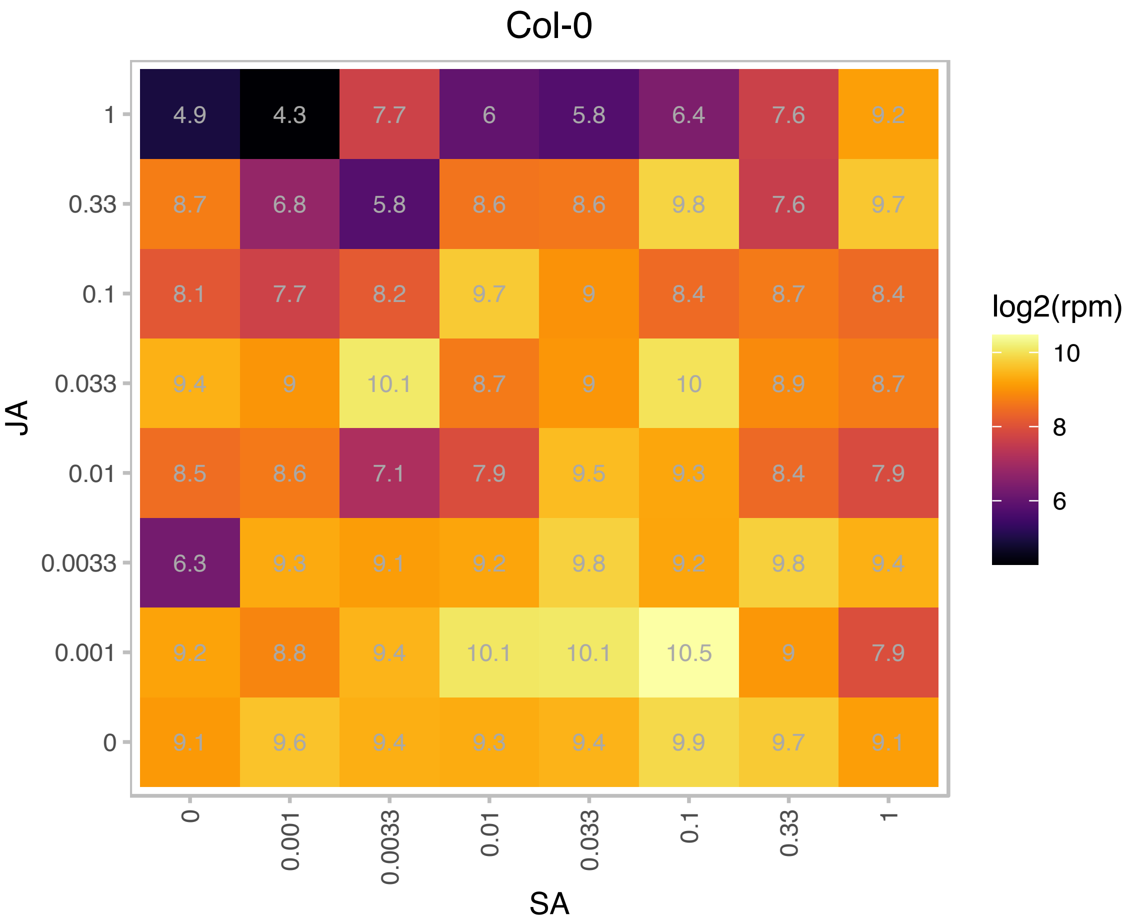


| | |
|---|---|
| Gene | AT2G34430 |
| is.expressed_genes | TRUE |
| is.responsive_genes | TRUE |
| is.apcluster | H-03 |

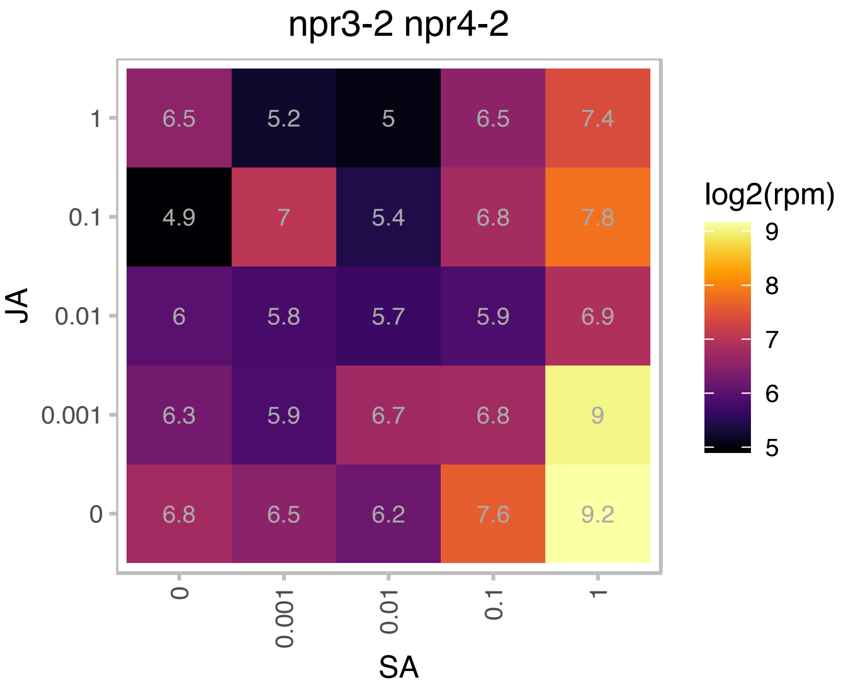


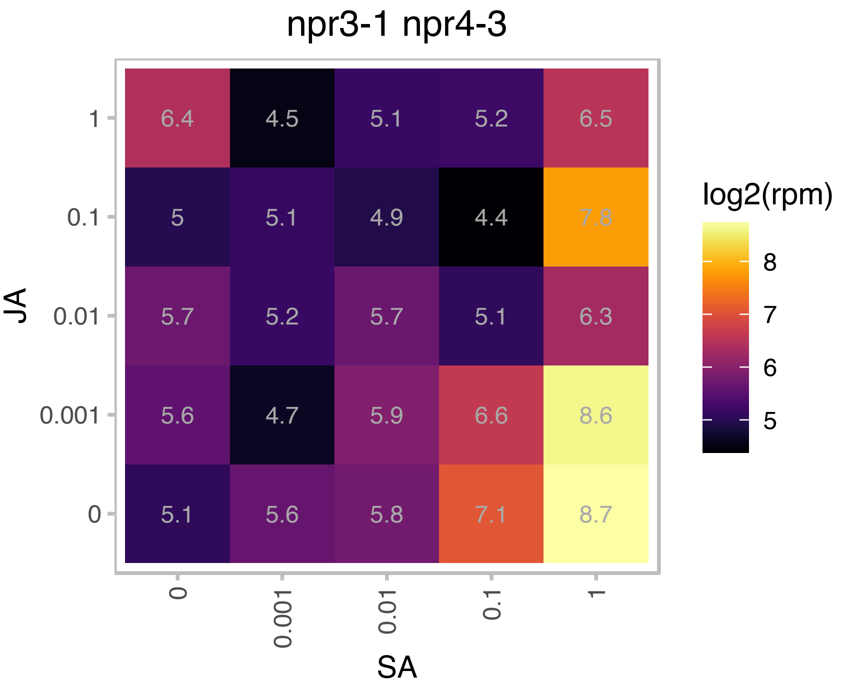

# AT2G38260.1 : NA

Probably not a pseudogene based on evidence for transcription (RNA-seq) and translation (Ribo-seq) described in PMID:27791167

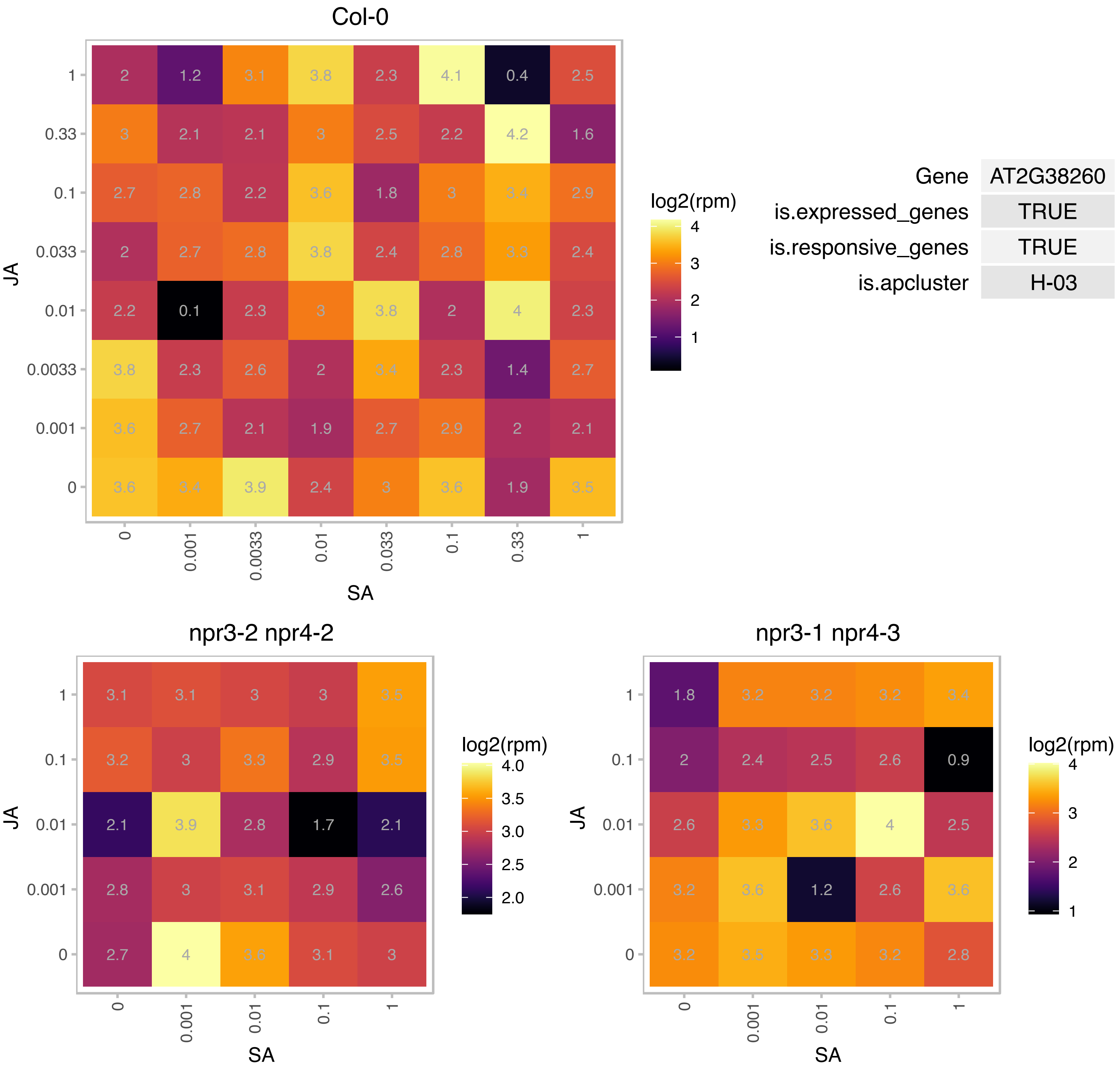


| | |
|---|---|
| Gene | AT2G38260 |
| is.expressed_genes | TRUE |
| is.responsive_genes | TRUE |
| is.apcluster | H-03 |

### AT2G39400.1 : MAGL6

alpha/beta-Hydrolases superfamily protein

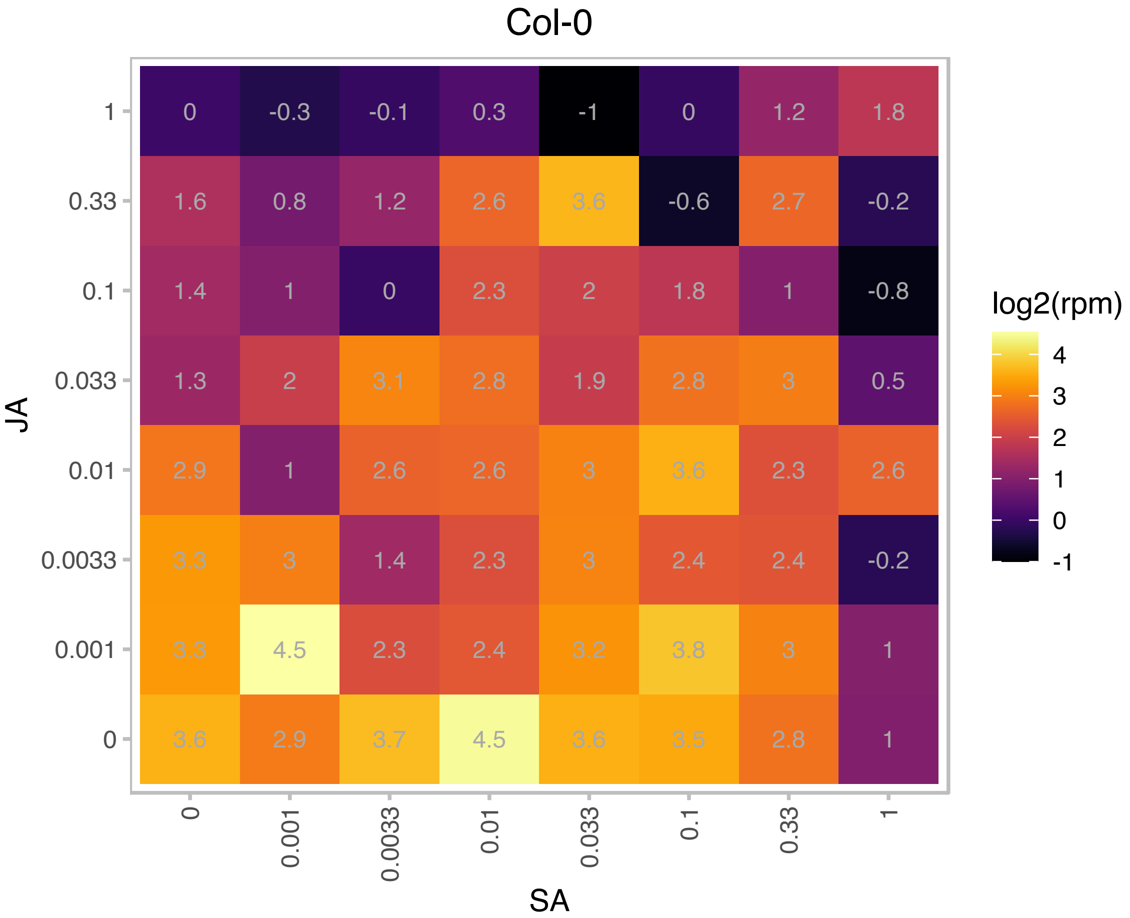


| | |
|---|---|
| Gene | AT2G39400 |
| is.expressed_genes | TRUE |
| is.responsive_genes | TRUE |
| is.apcluster | H-03 |

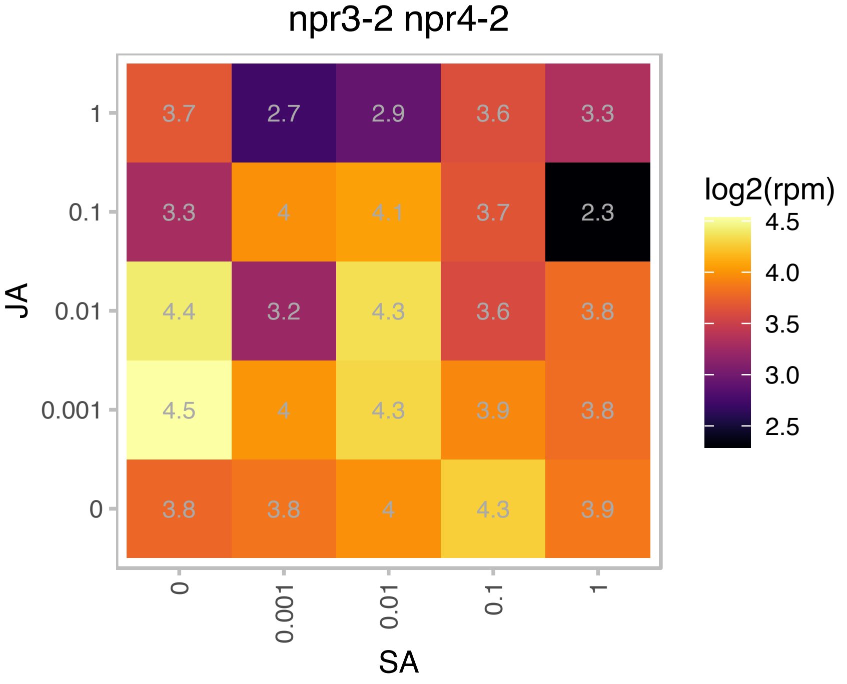


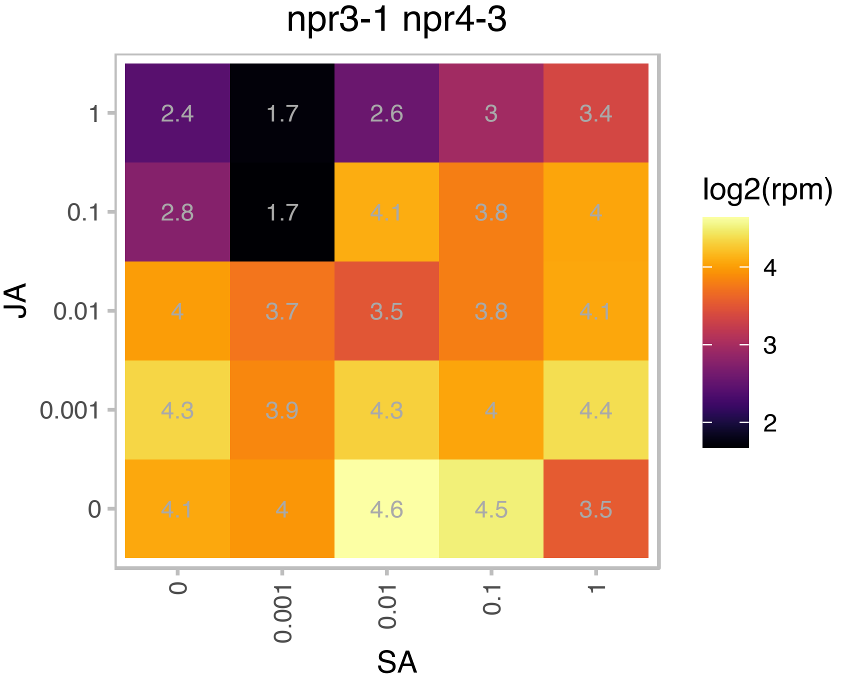

# AT2G39795.1 : NA

Mitochondrial glycoprotein family protein

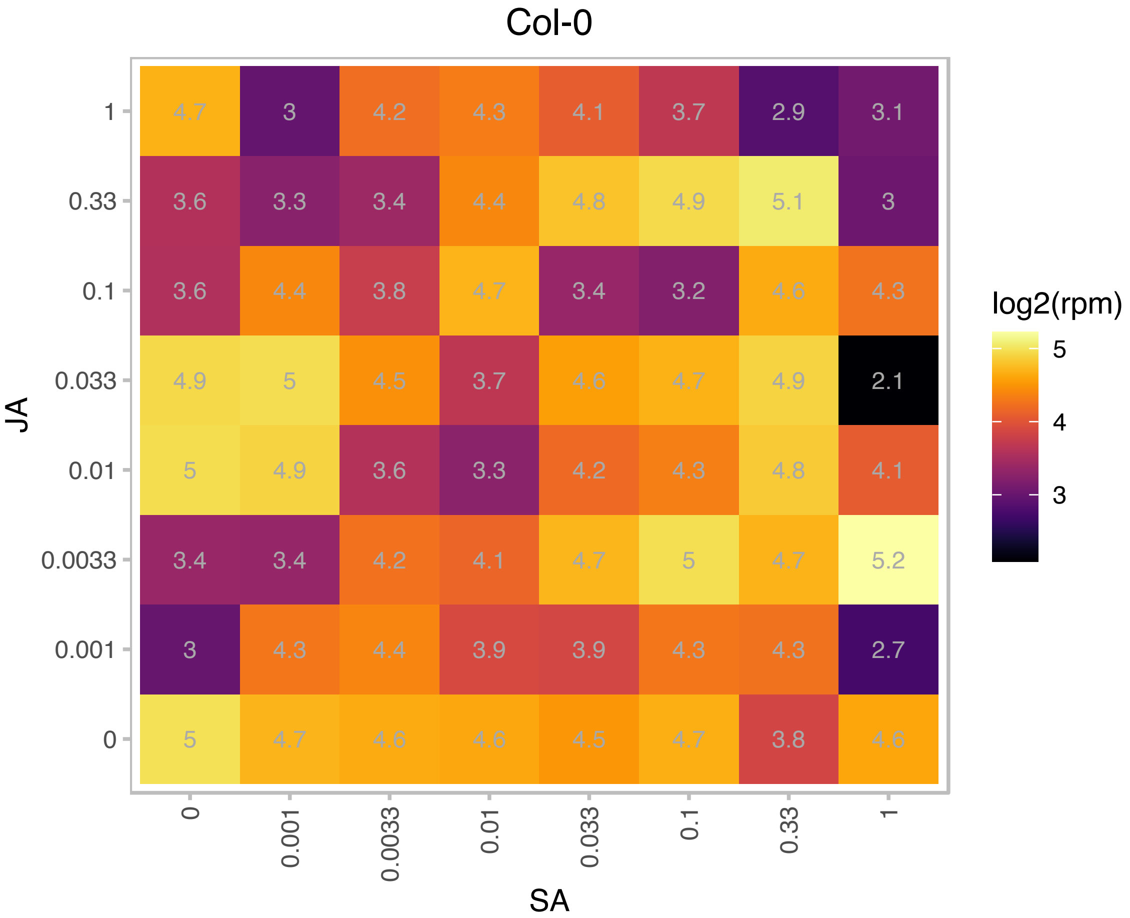


| | |
|---|---|
| Gene | AT2G39795 |
| is.expressed_genes | TRUE |
| is.responsive_genes | TRUE |
| is.apcluster | H-03 |

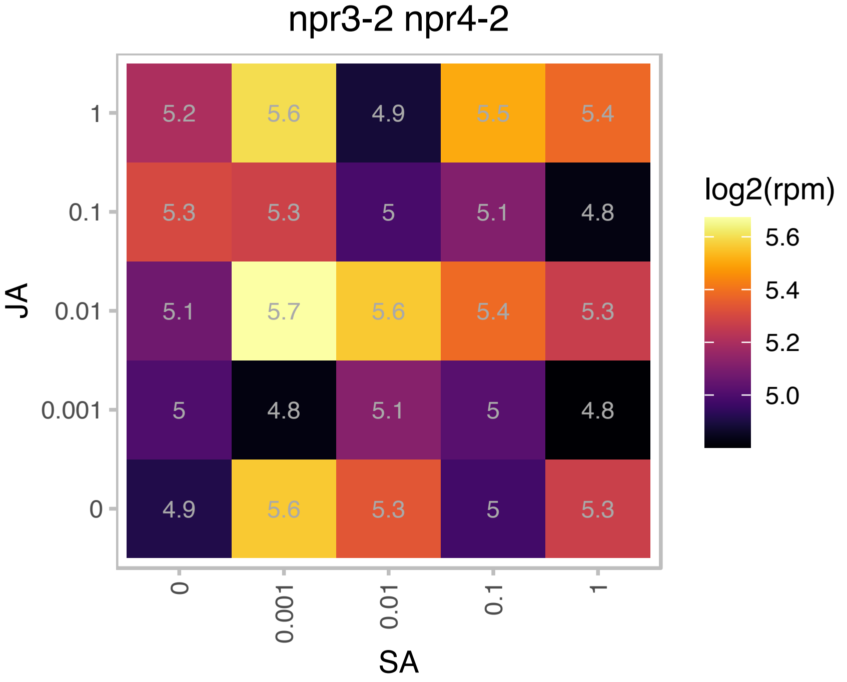


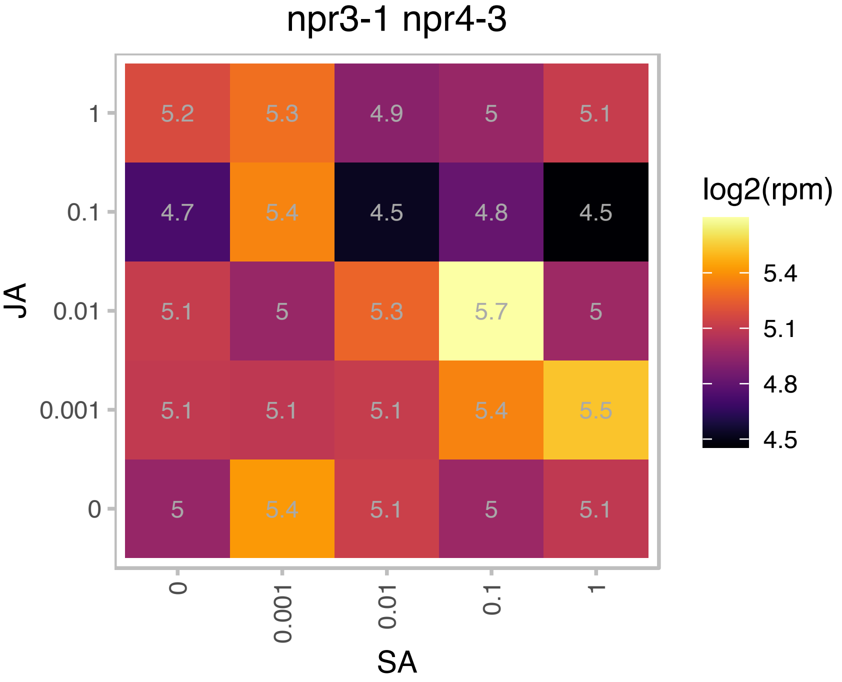

# AT2G40316.1 : NA

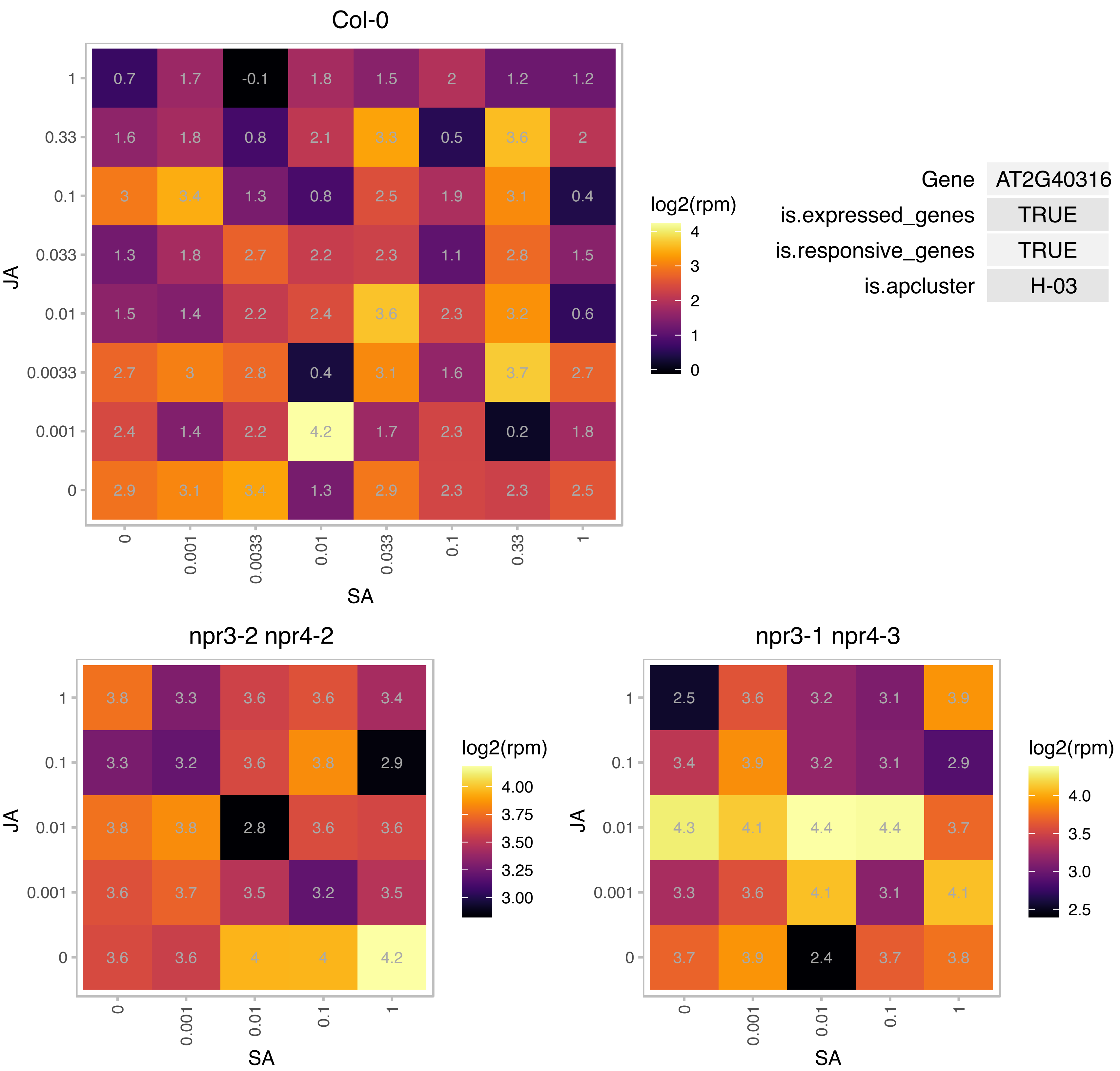

### AT2G43910.1 : ATHOL1, HOL1

HARMLESS TO OZONE LAYER 1

Col-0

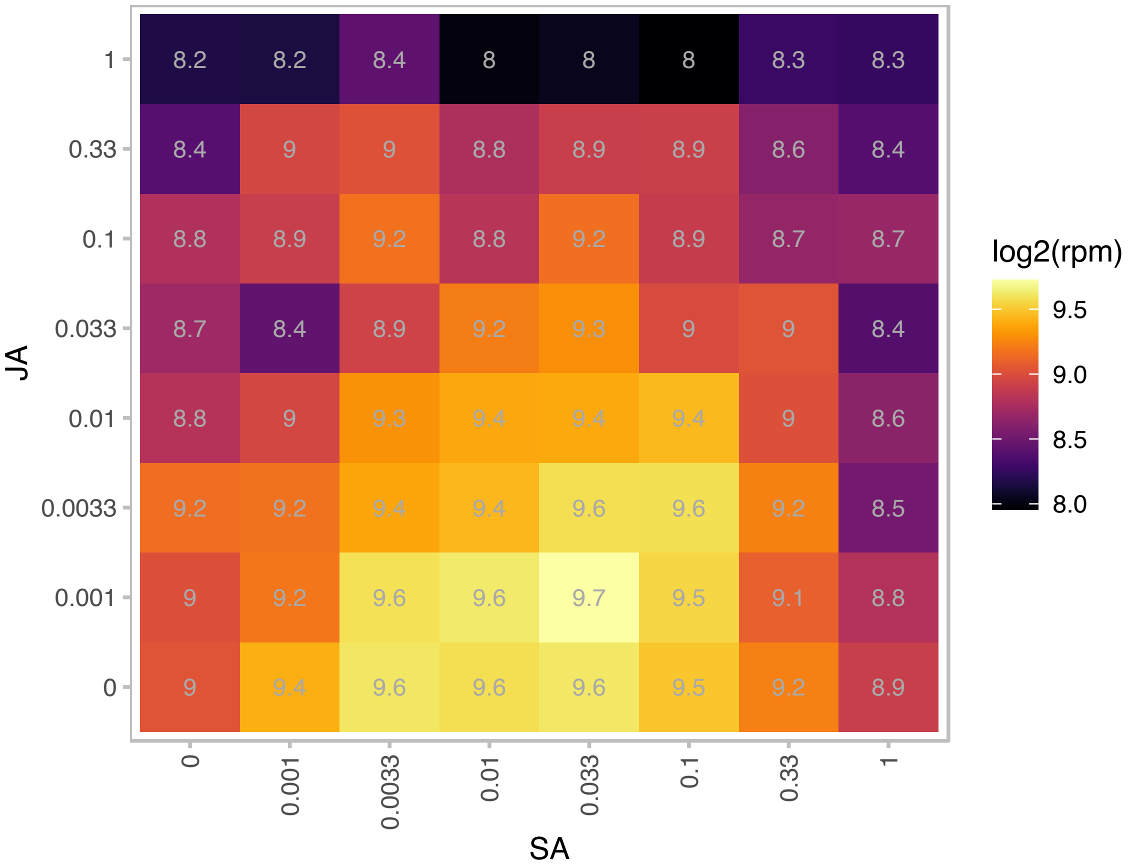


| | |
|---|---|
| Gene | AT2G43910 |
| is.expressed_genes | TRUE |
| is.responsive_genes | TRUE |
| is.apcluster | H-03 |

npr3-2 npr4-2

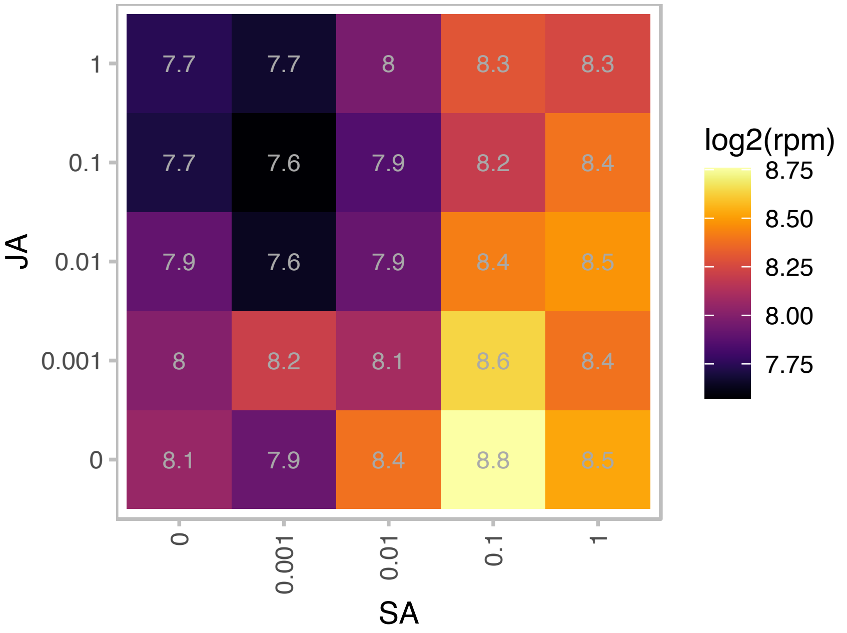


npr3-1 npr4-3

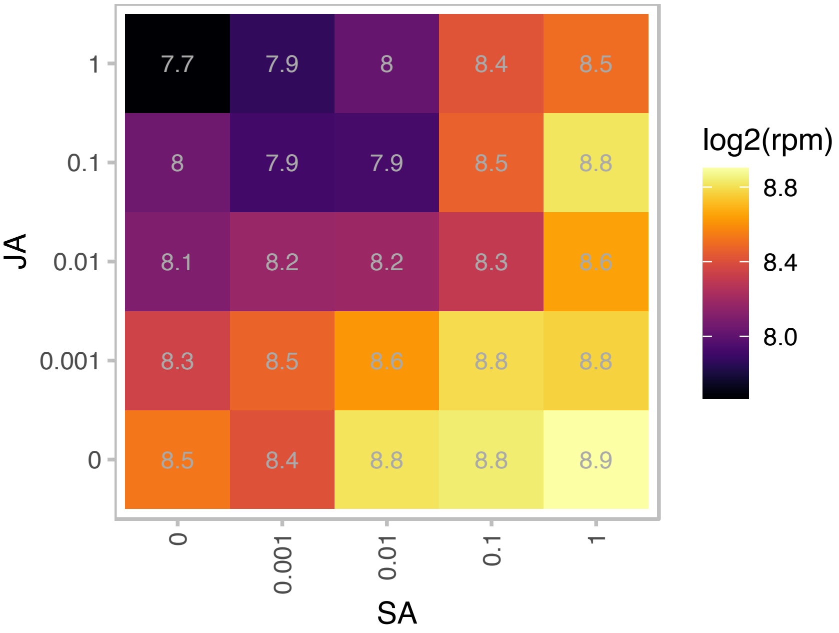

# AT2G44380.1 : NA

Cysteine/Histidine-rich C1 domain family protein

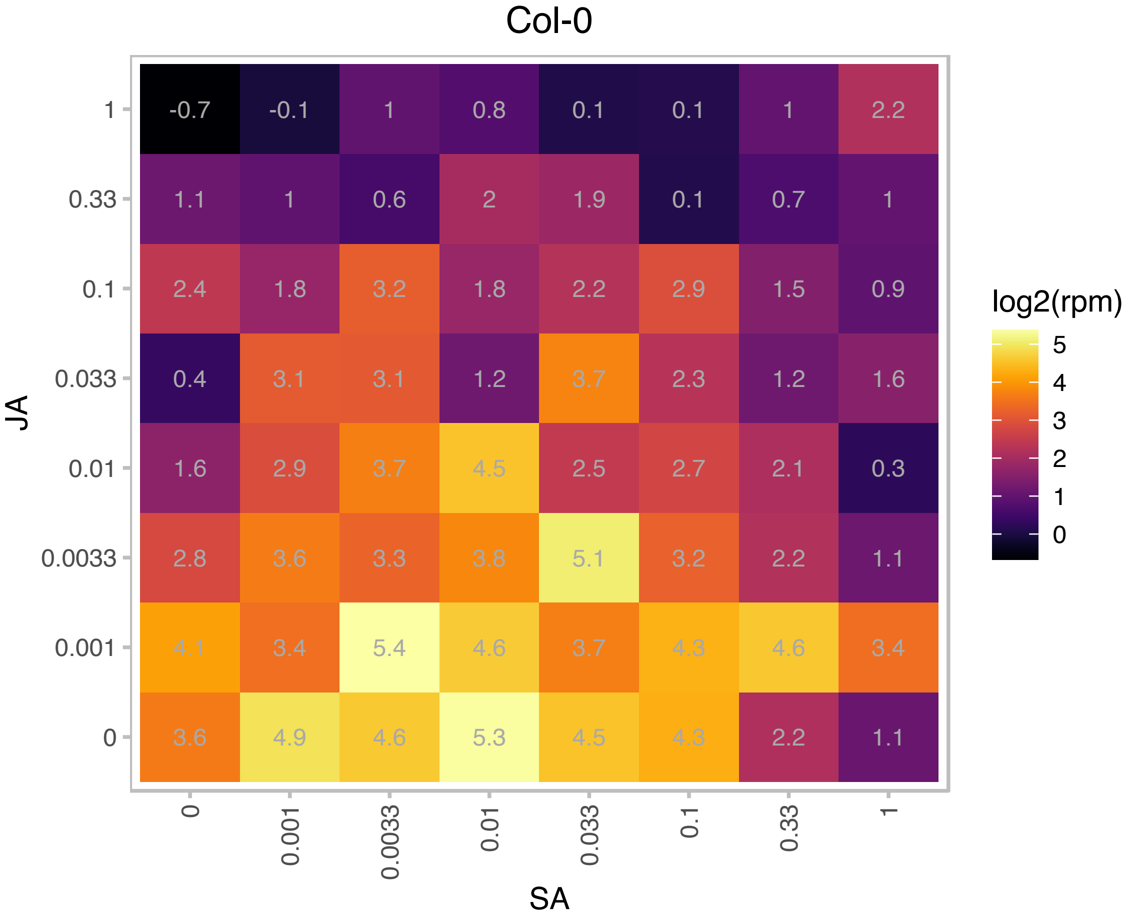


| | |
|---|---|
| Gene | AT2G44380 |
| is.expressed_genes | TRUE |
| is.responsive_genes | TRUE |
| is.apcluster | H-03 |

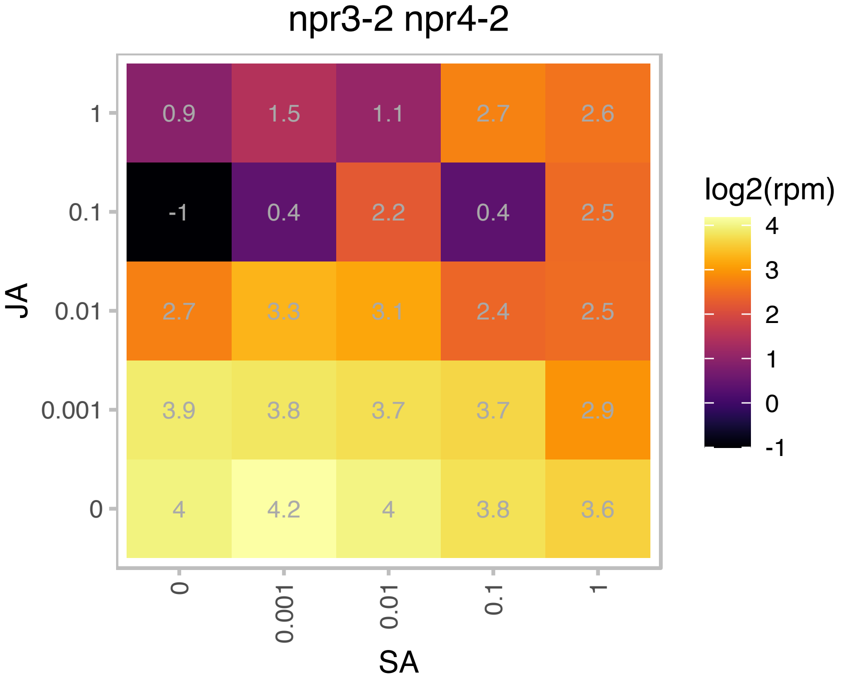


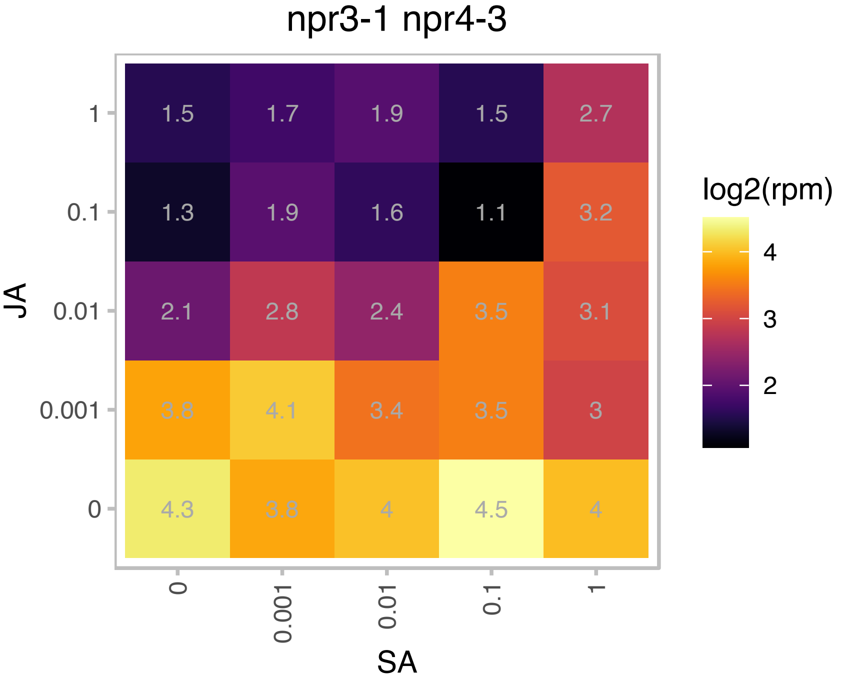

# AT3G03630.1 : CS26

cysteine synthase 26

Encodes a protein that possesses S-sulfocysteine synthase activity and lacks O-acetylserien(thiol)lyase activity.

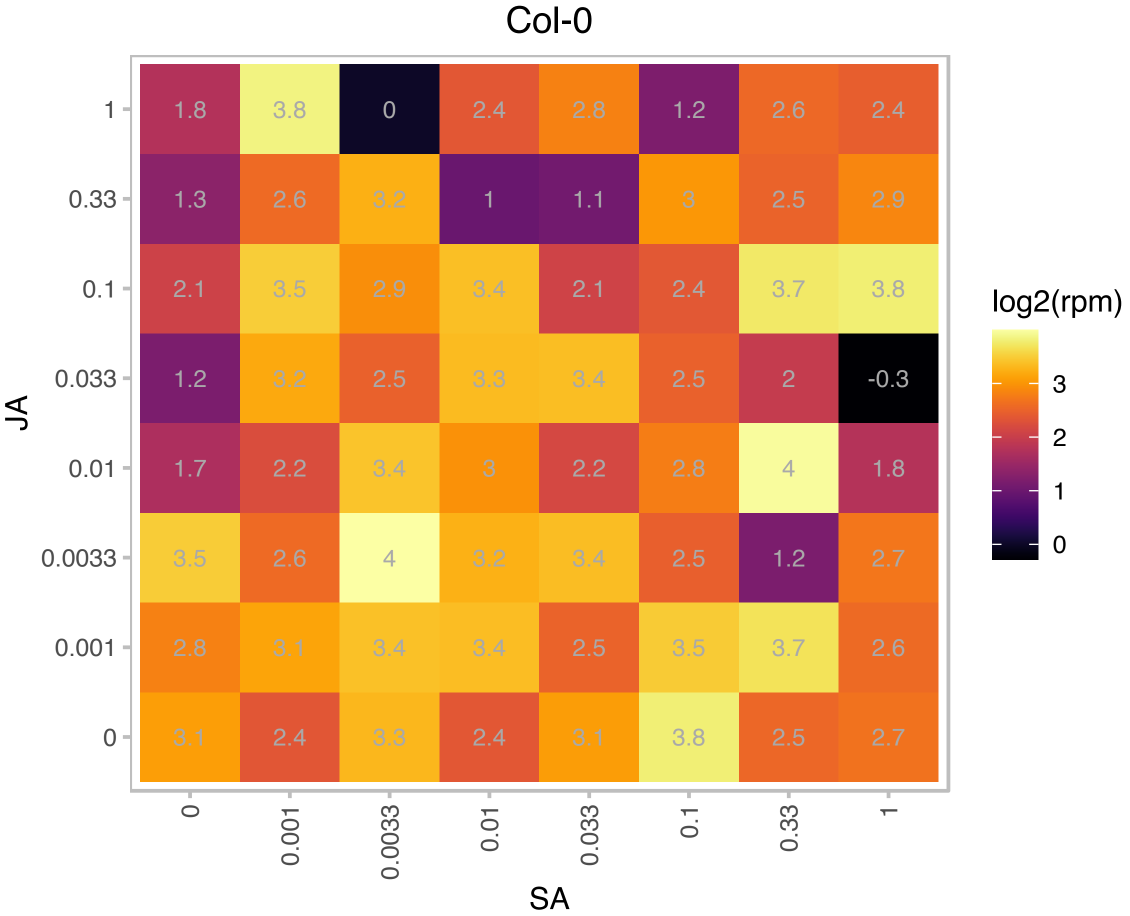


| | |
|---|---|
| Gene | AT3G03630 |
| is.expressed_genes | TRUE |
| is.responsive_genes | TRUE |
| is.apcluster | H-03 |

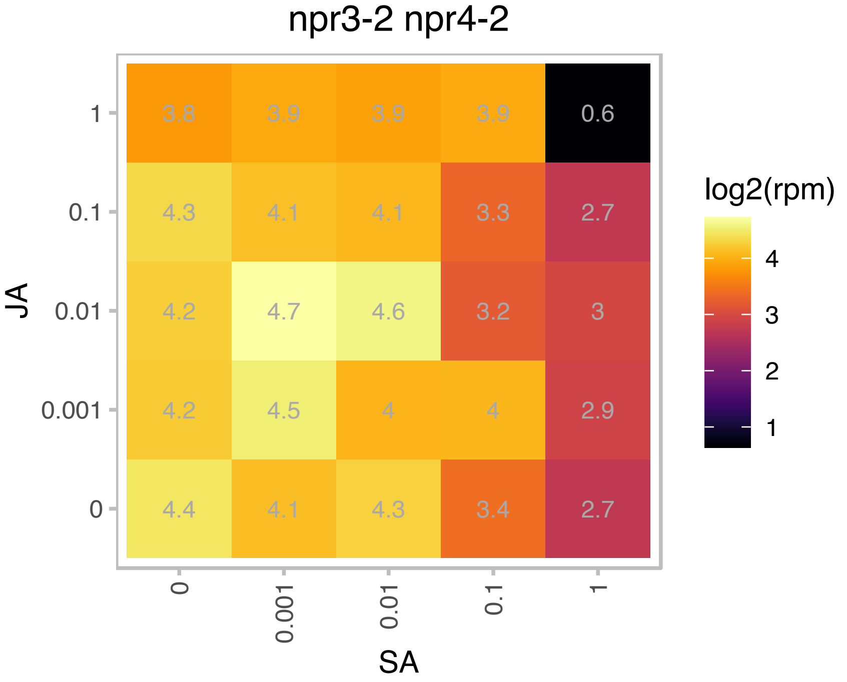


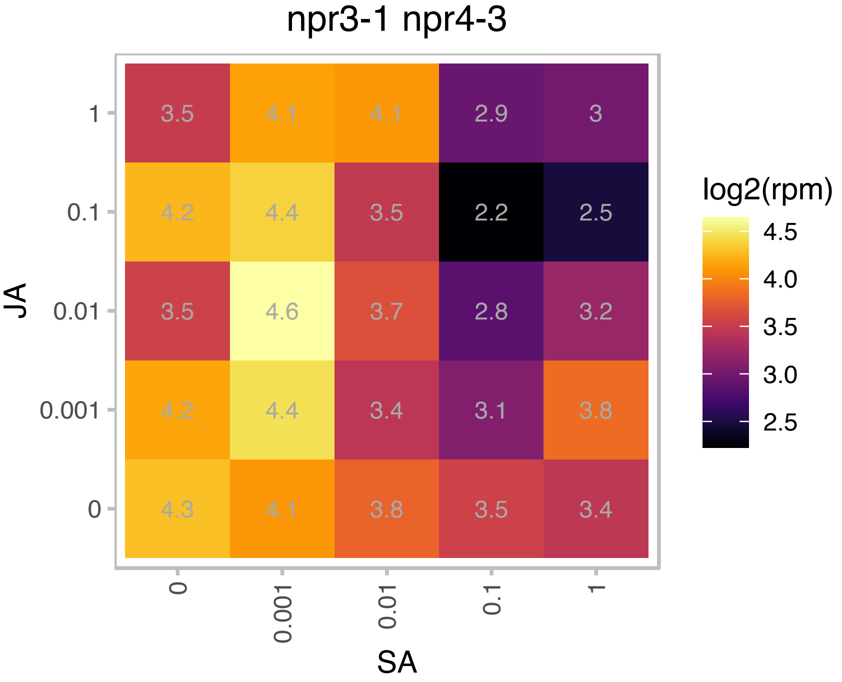

# AT3G06070.1 : NA

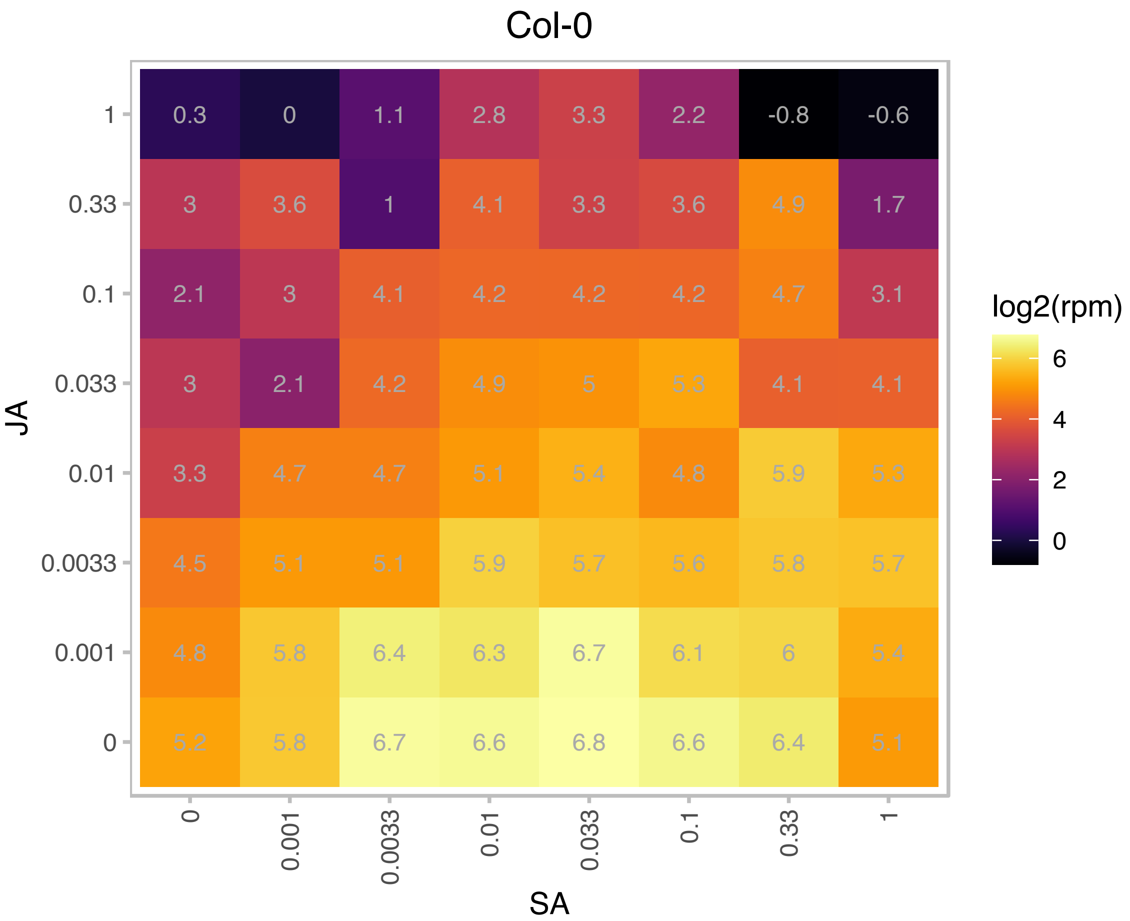


| | |
|---|---|
| Gene | AT3G06070 |
| is.expressed_genes | TRUE |
| is.responsive_genes | TRUE |
| is.apcluster | H-03 |

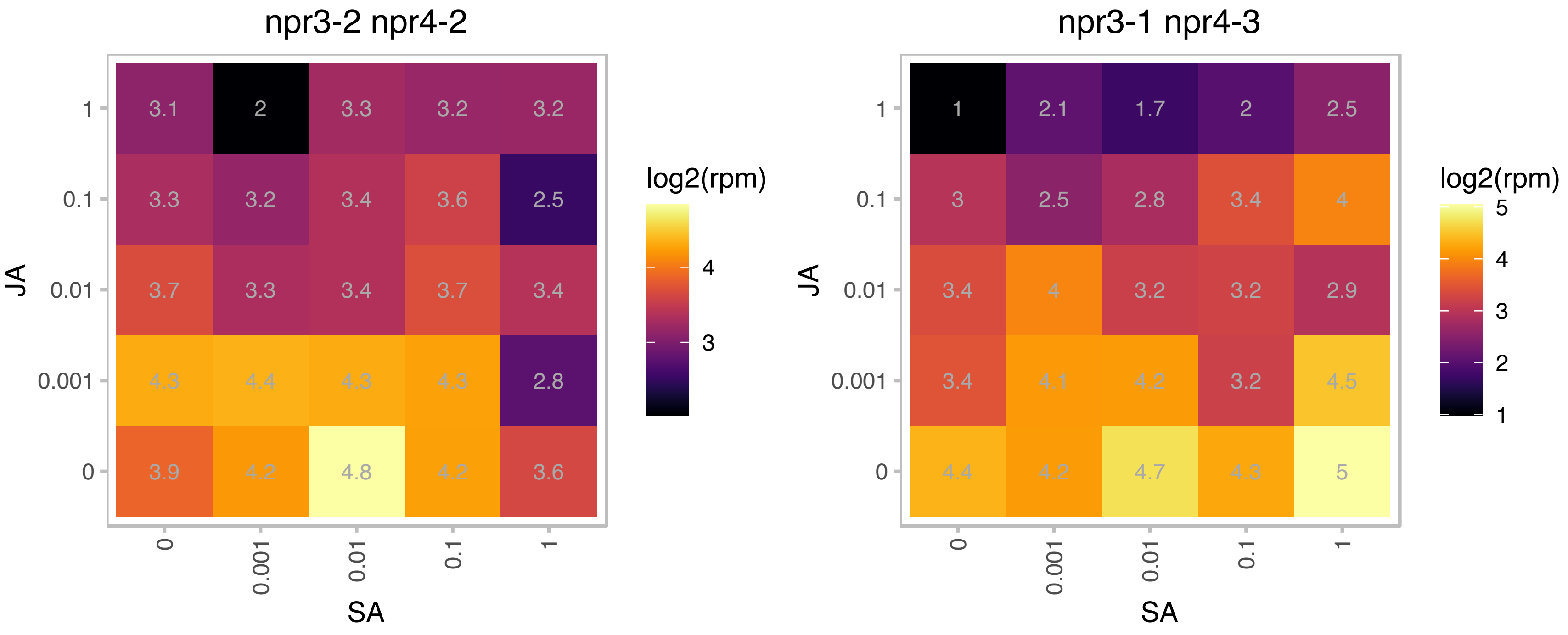

### AT3G11090.1 : LBD21

LOB domain-containing protein 21

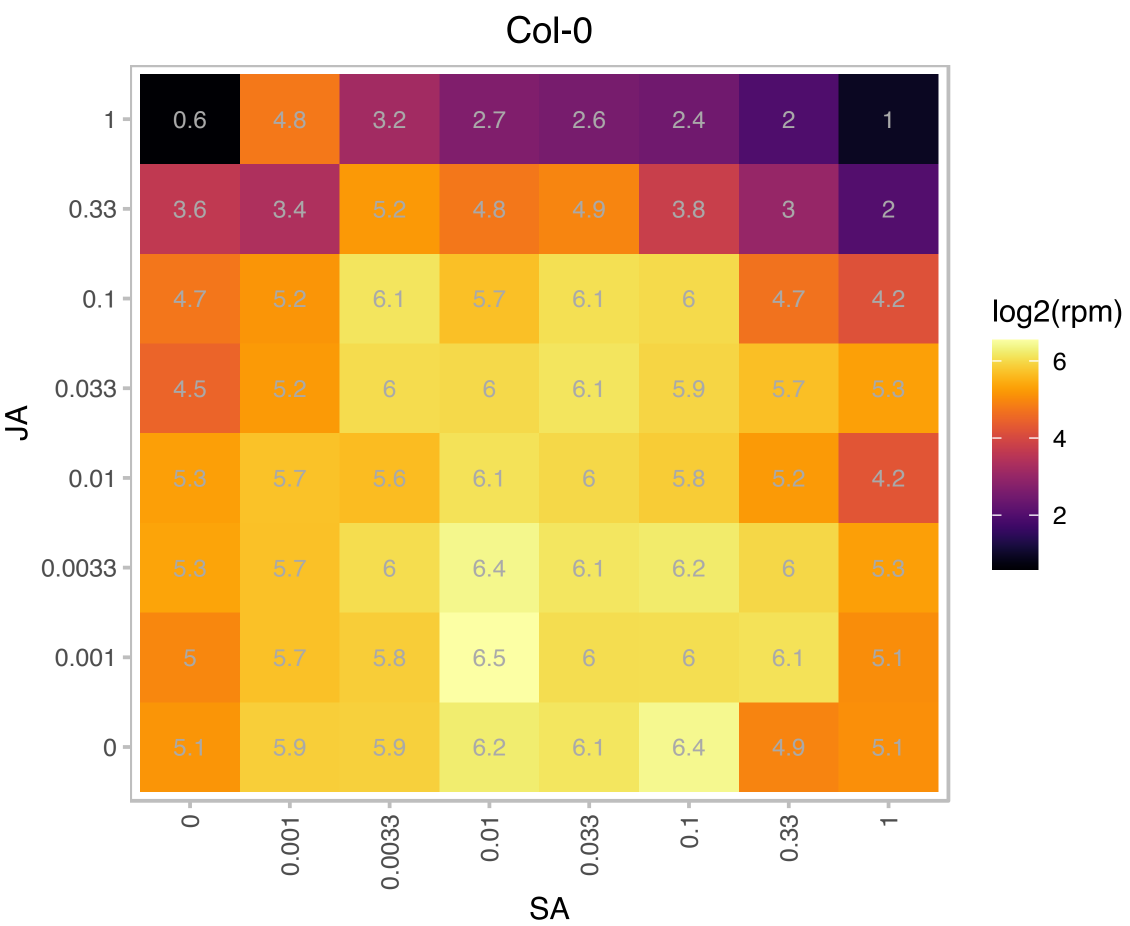


| | |
|---|---|
| Gene | AT3G11090 |
| is.expressed_genes | TRUE |
| is.responsive_genes | TRUE |
| is.apcluster | H-03 |

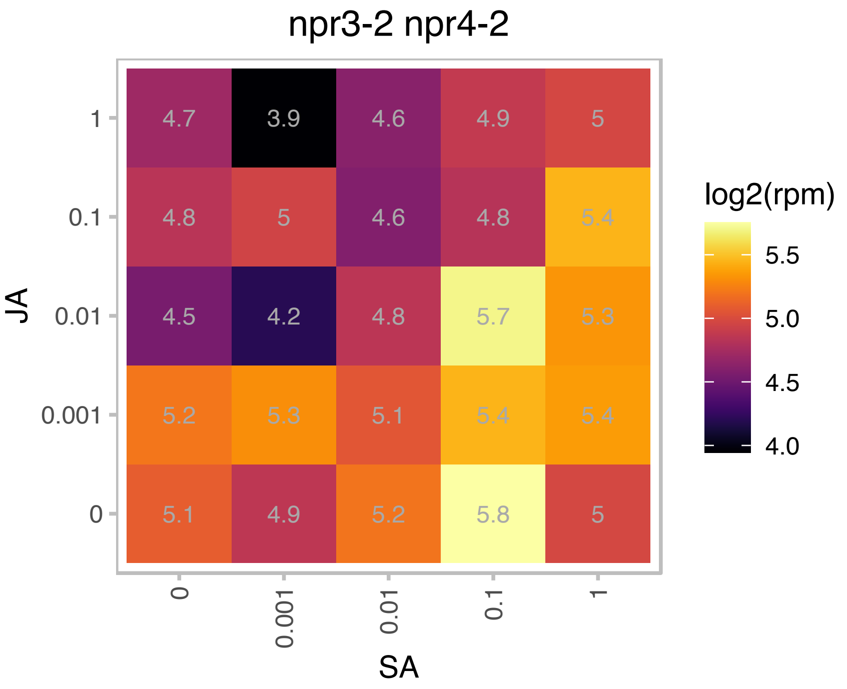


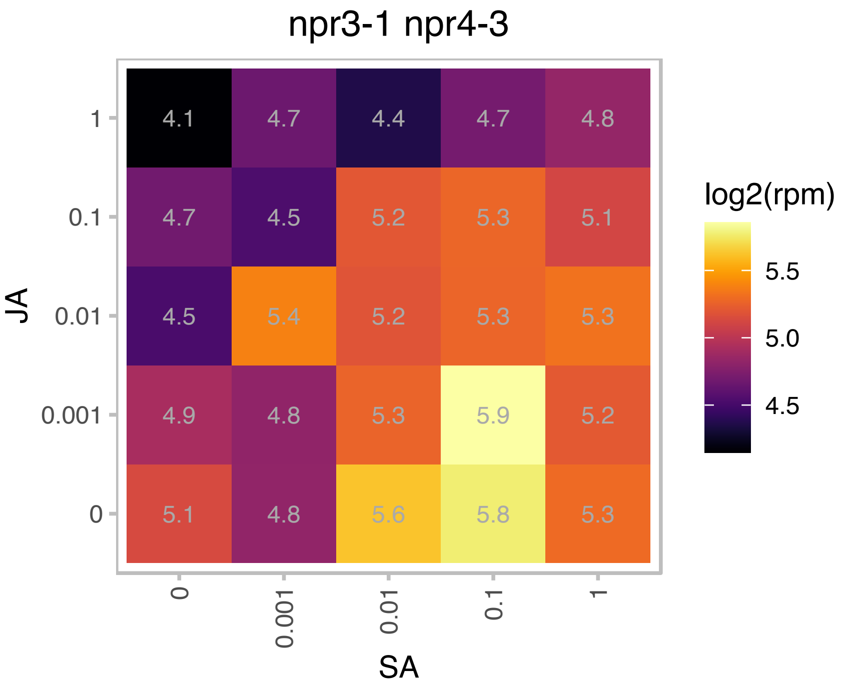

# AT3G11280.1 : NA

Duplicated homeodomain-like superfamily protein

Putative transcription factors interacting with the gene product of VHA-B1 (vacuolar ATPase subunit B1; as shown through yeast two-hybrid assay).

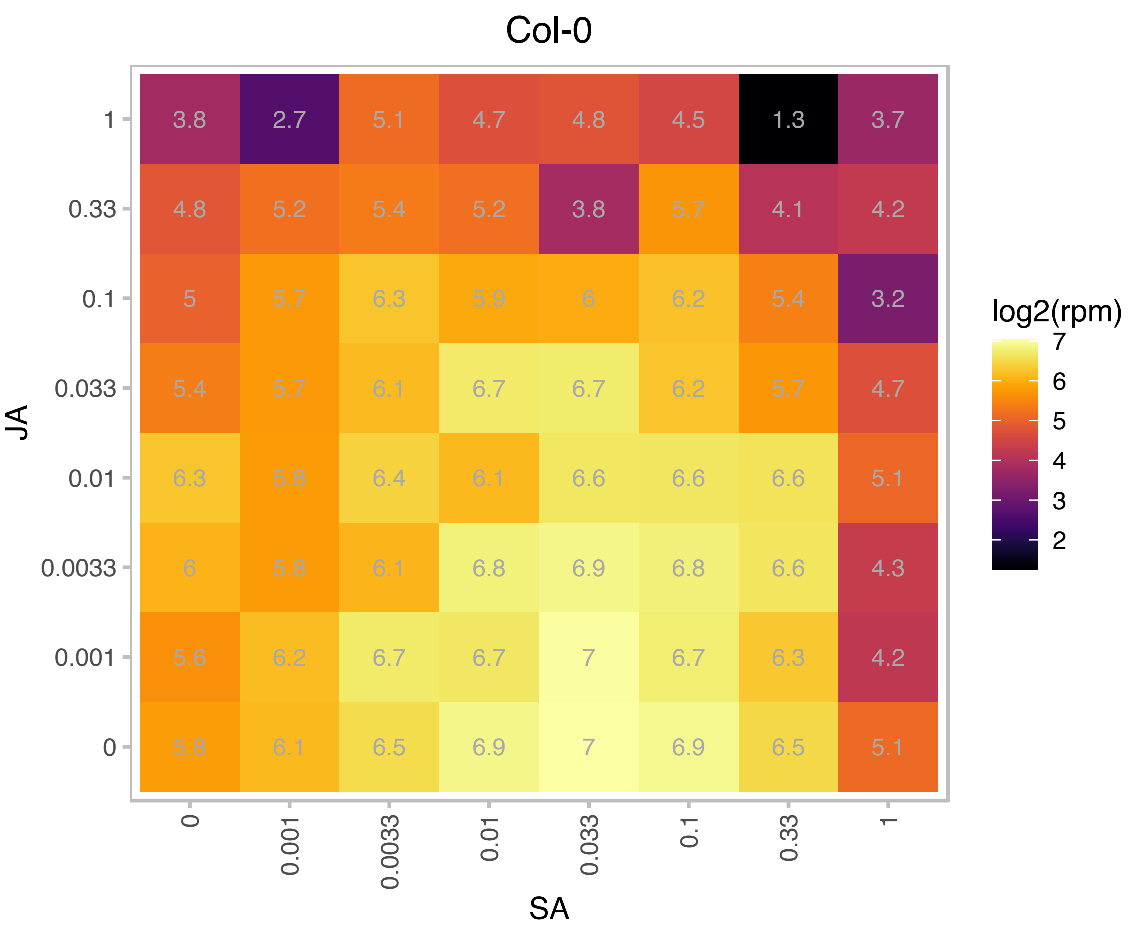


| | |
|---|---|
| Gene | AT3G11280 |
| is.expressed_genes | TRUE |
| is.responsive_genes | TRUE |
| is.apcluster | H-03 |

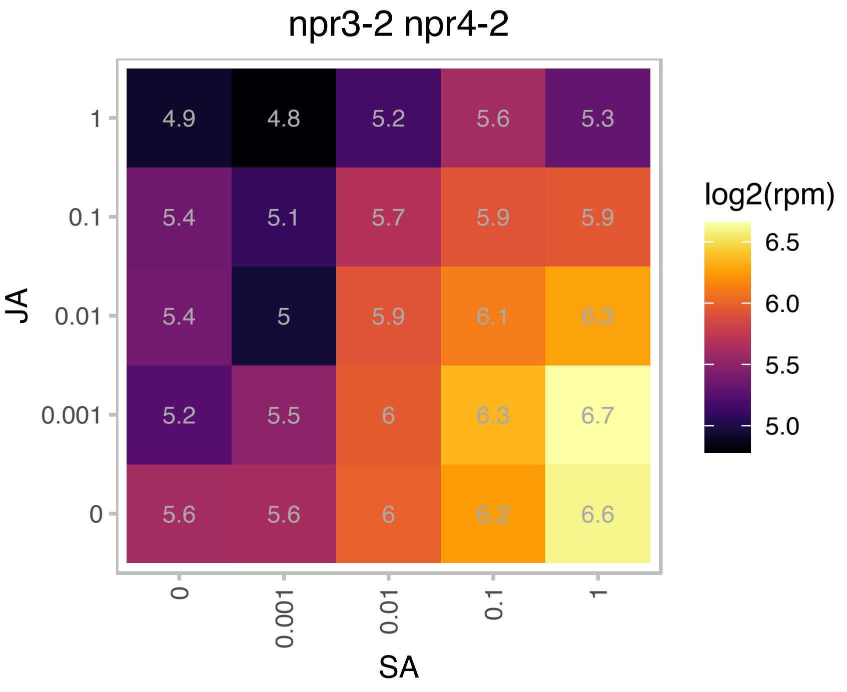


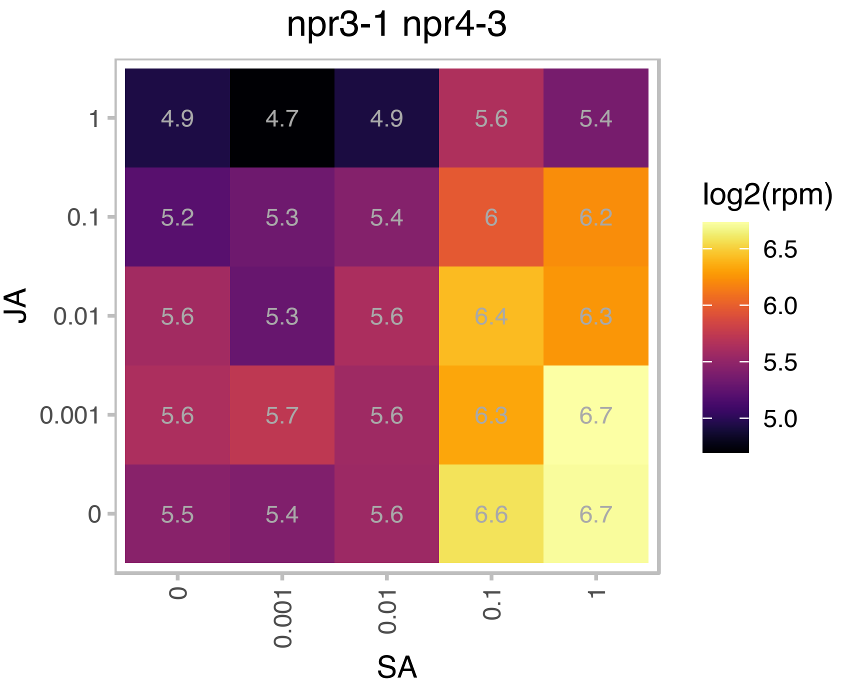

### AT3G22320.1 : ATRPABC24.3, NRPB5, NRPD5, RPB5A

Eukaryotic rpb5 RNA polymerase subunit family protein

Non-catalytic subunit common to DNA-dependent RNA polymerases I, II, III and IV; homologous to budding yeast RPB5.

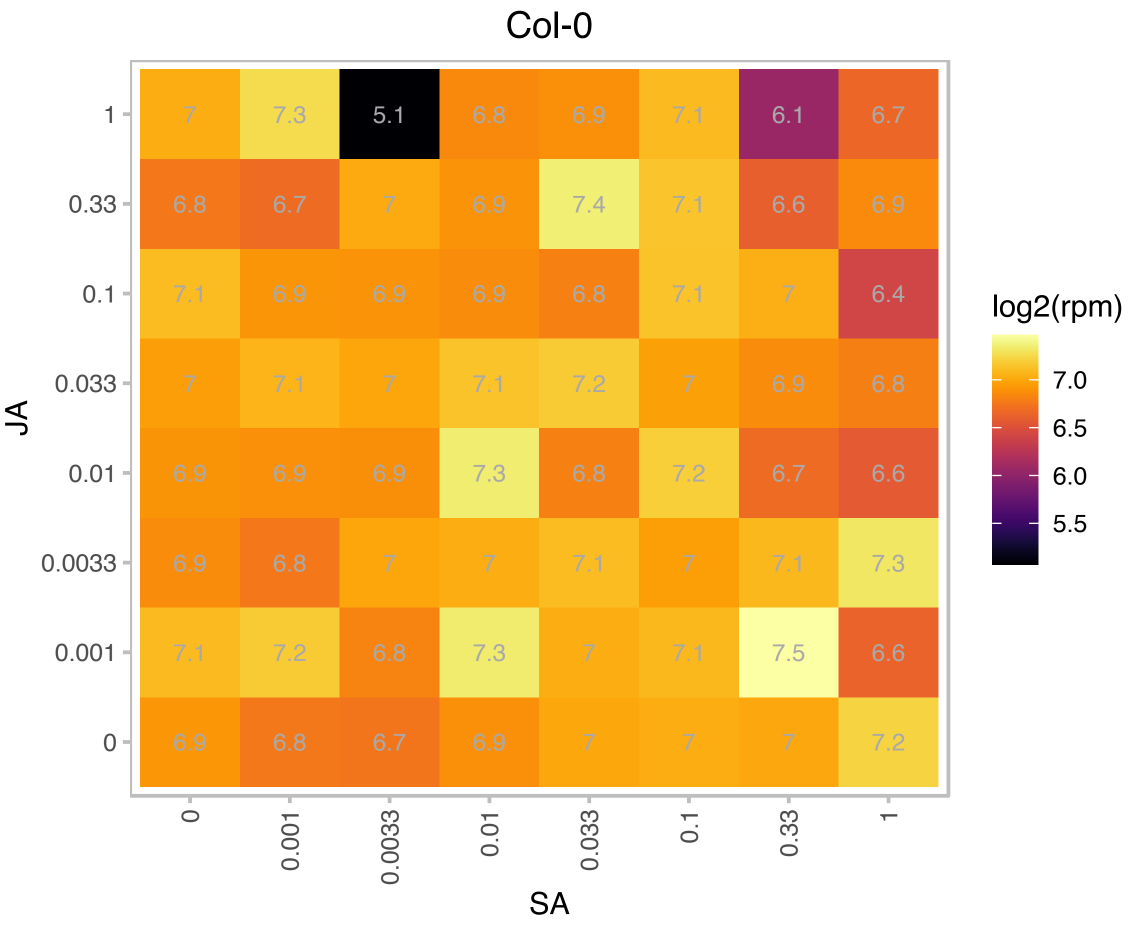


| | |
|---|---|
| Gene | AT3G22320 |
| is.expressed_genes | TRUE |
| is.responsive_genes | TRUE |
| is.apcluster | H-03 |

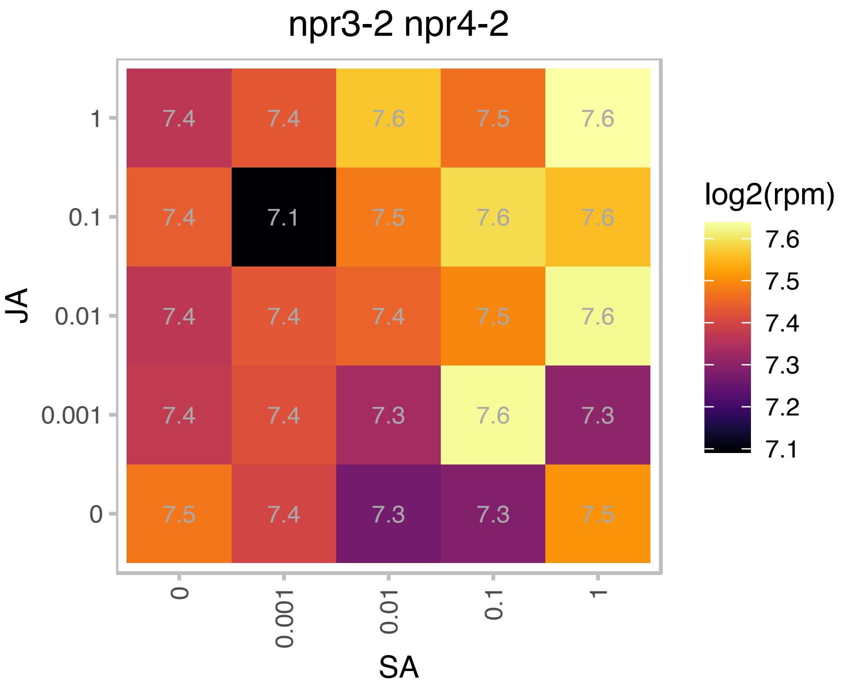


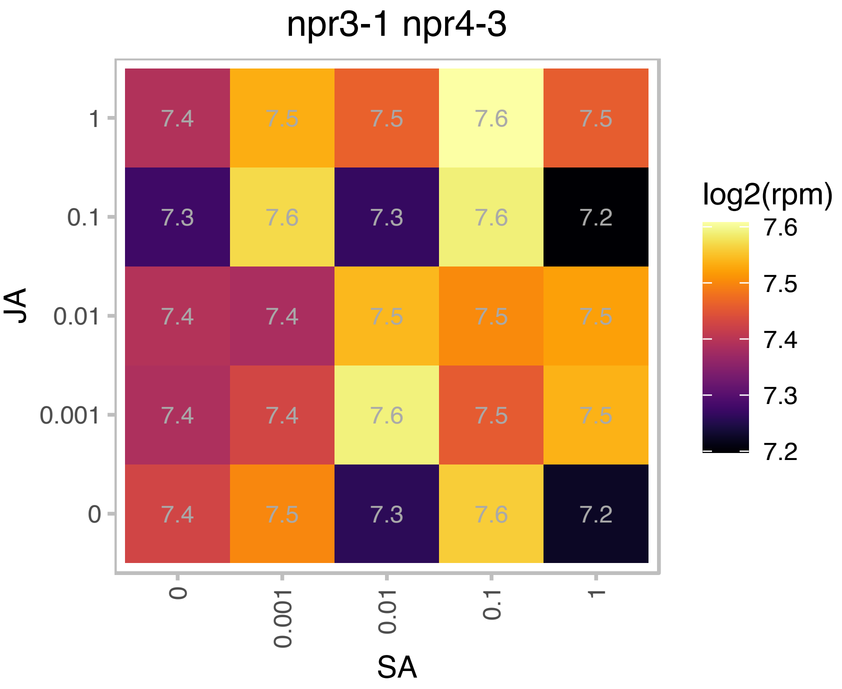

# AT3G25400.1 : NA

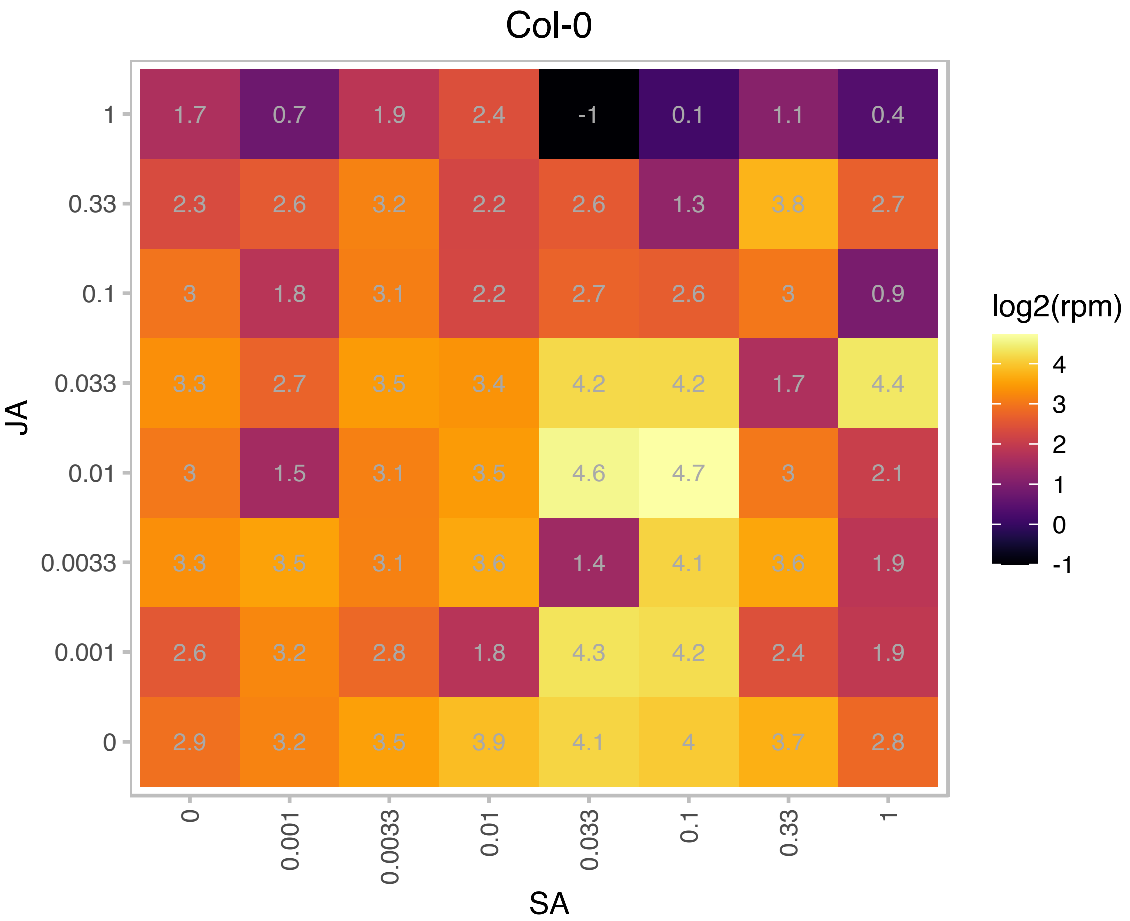


| | |
|---|---|
| Gene | AT3G25400 |
| is.expressed_genes | TRUE |
| is.responsive_genes | TRUE |
| is.apcluster | H-03 |

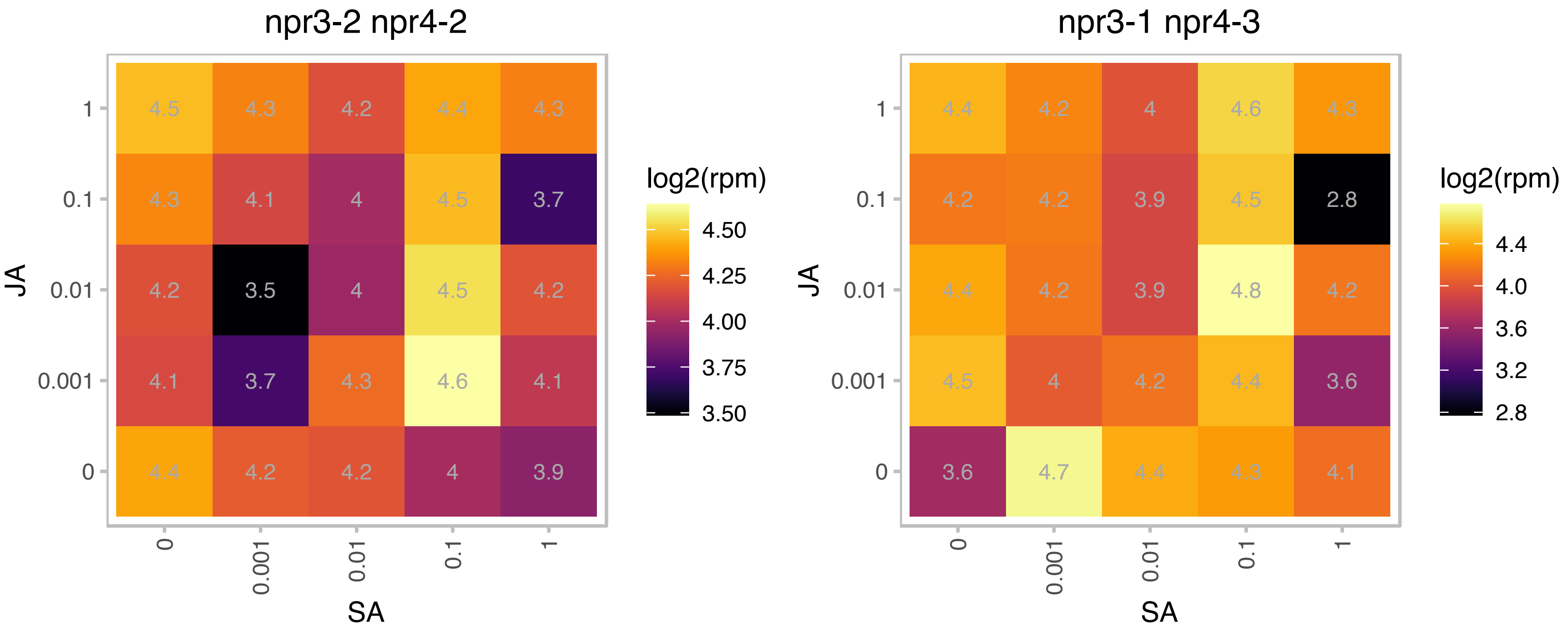

### AT3G27100.1 : ENY2

ENY2 is a component of the deubiquitination module of the SAGA complex and is involved in the deubiquitination of histone 2B .

Col-0

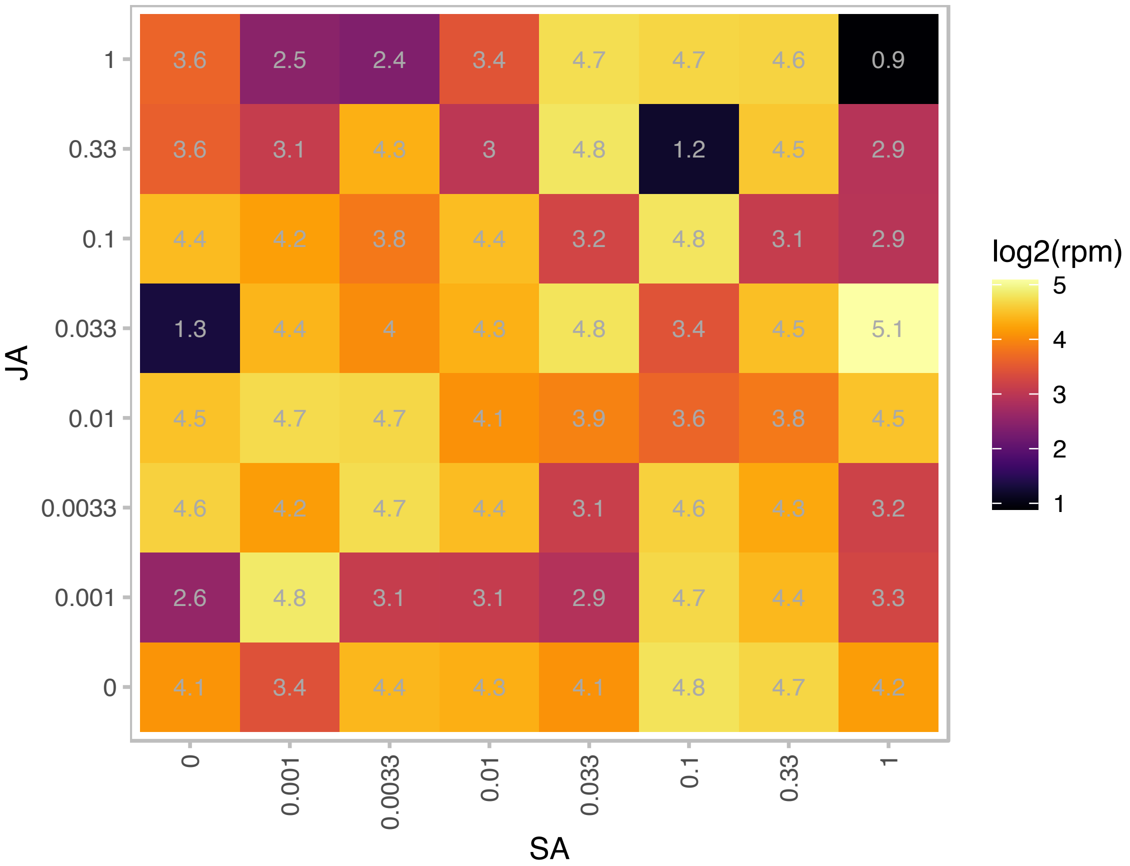


| | |
|---|---|
| Gene | AT3G27100 |
| is.expressed_genes | TRUE |
| is.responsive_genes | TRUE |
| is.apcluster | H-03 |

npr3-2 npr4-2

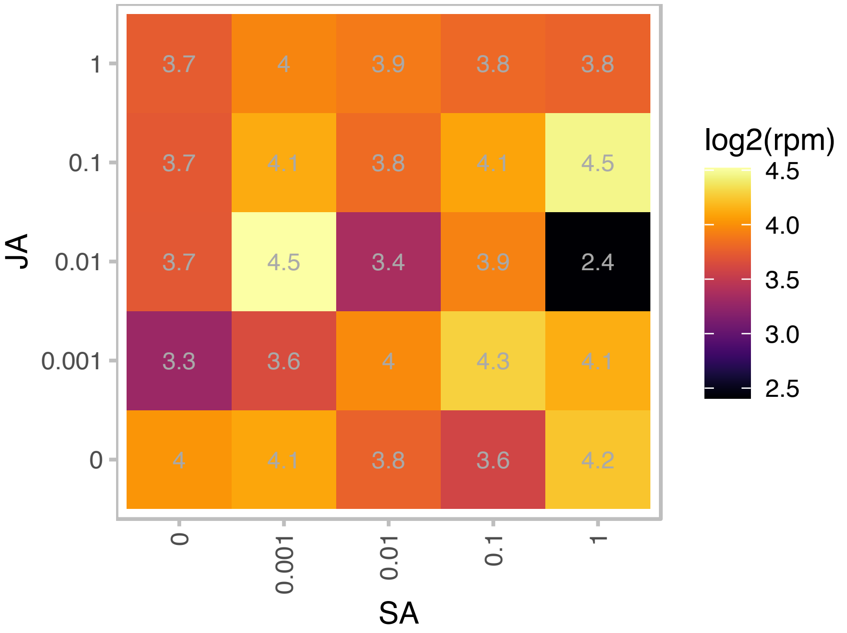


npr3-1 npr4-3

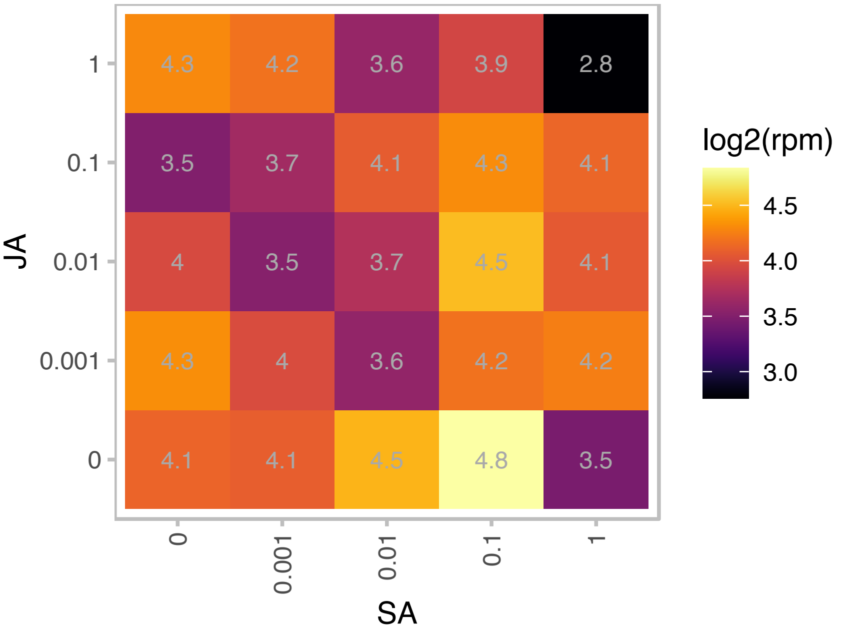

# AT3G46540.1 : NA

ENTH/VHS family protein

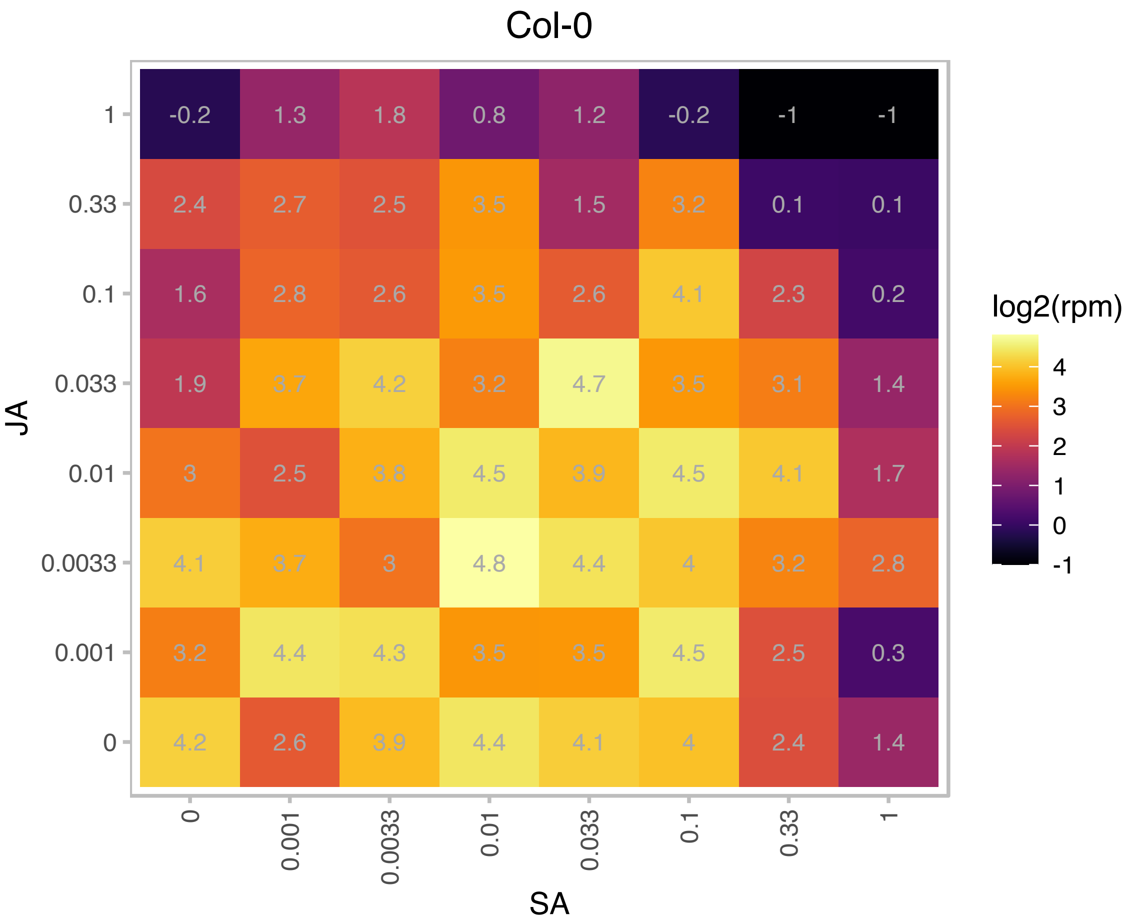


| | |
|---|---|
| Gene | AT3G46540 |
| is.expressed_genes | TRUE |
| is.responsive_genes | TRUE |
| is.apcluster | H-03 |

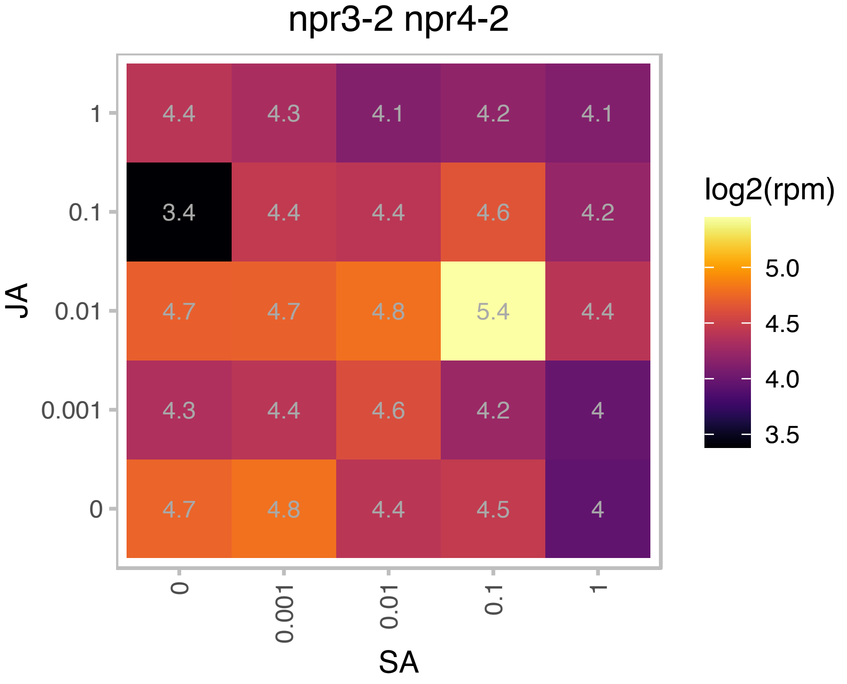


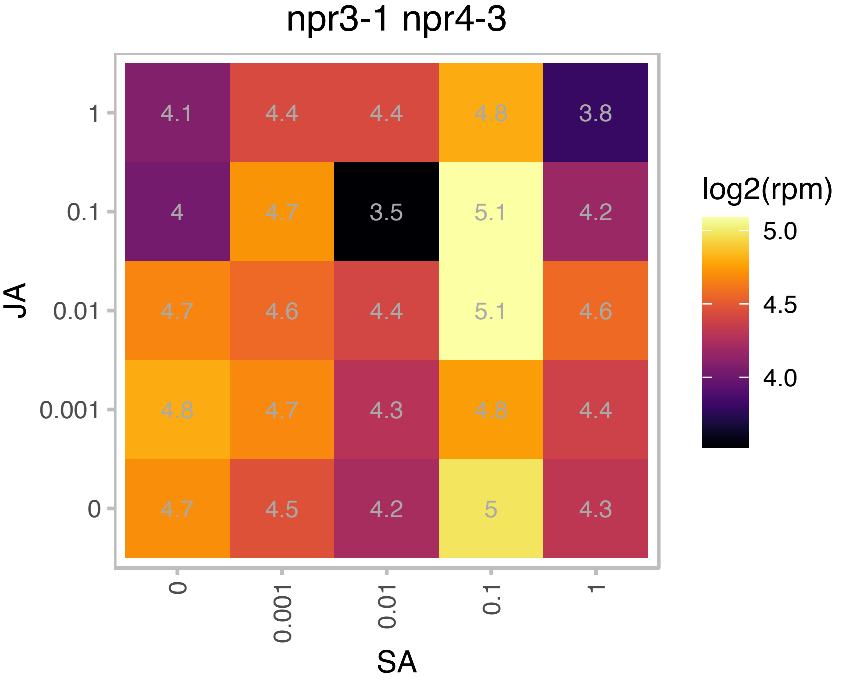

### AT3G52430.1 : ATPAD4, PAD4

alpha/beta-Hydrolases superfamily protein

Encodes a lipase-like gene that is important for salicylic acid signaling and function in resistance (R) gene-mediated and basal plant disease resistance. PAD4 can interact directly with EDS1, another disease resistance signaling protein. Expressed at elevated level in response to green peach aphid (GPA) feeding, and modulates the GPA feeding-induced leaf senescence through a mechanism that doesn't require camalexin synthesis and salicylic acid (SA) signaling. Required for the ssi2-dependent heightened resistance to GPA. The mRNA is cell-to-cell mobile.

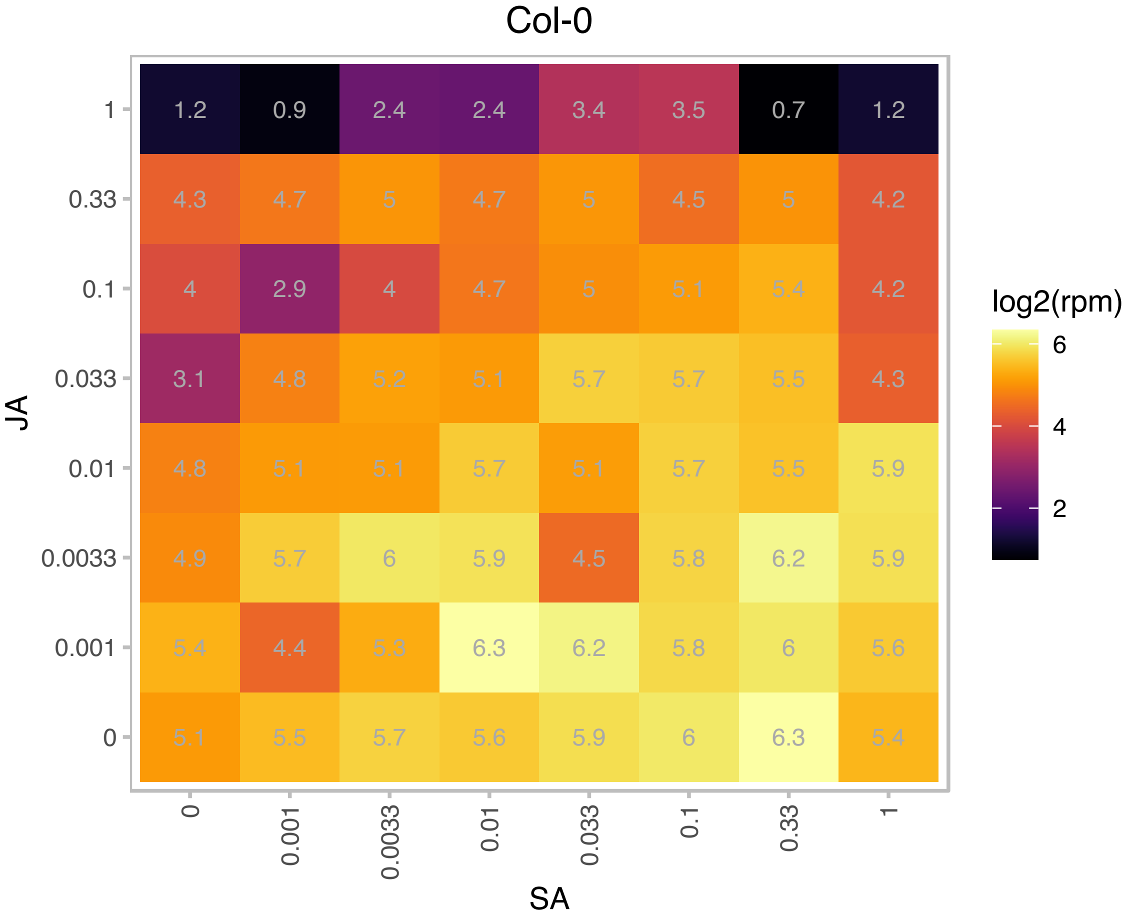


| | |
|---|---|
| Gene | AT3G52430 |
| is.expressed_genes | TRUE |
| is.responsive_genes | TRUE |
| is.apcluster | H-03 |

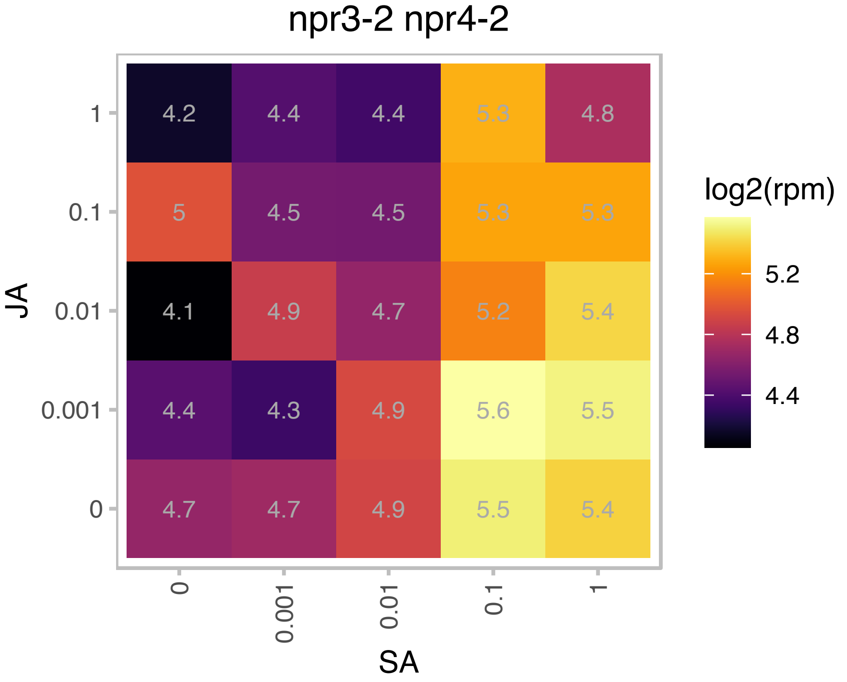


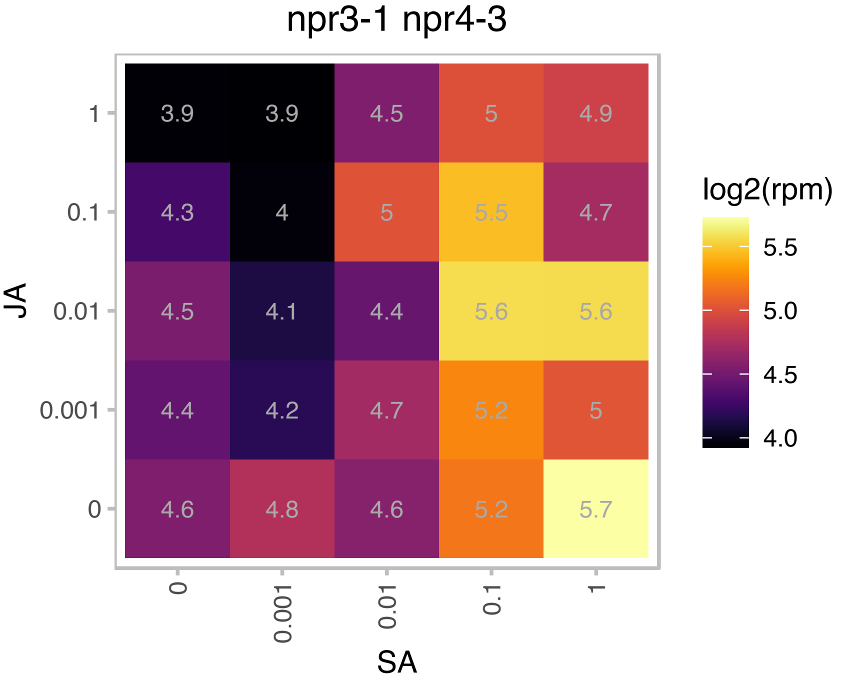

### AT4G03110.1 : AtBRN1, AtRBP-DR1, BRN1, RBP-DR1

RNA-binding protein-defense related 1

Encodes a putative RNA-binding protein that is located in the cytoplasm and is involved in the hypersensitive response and positively regulates salicylic acid-mediated immunity.

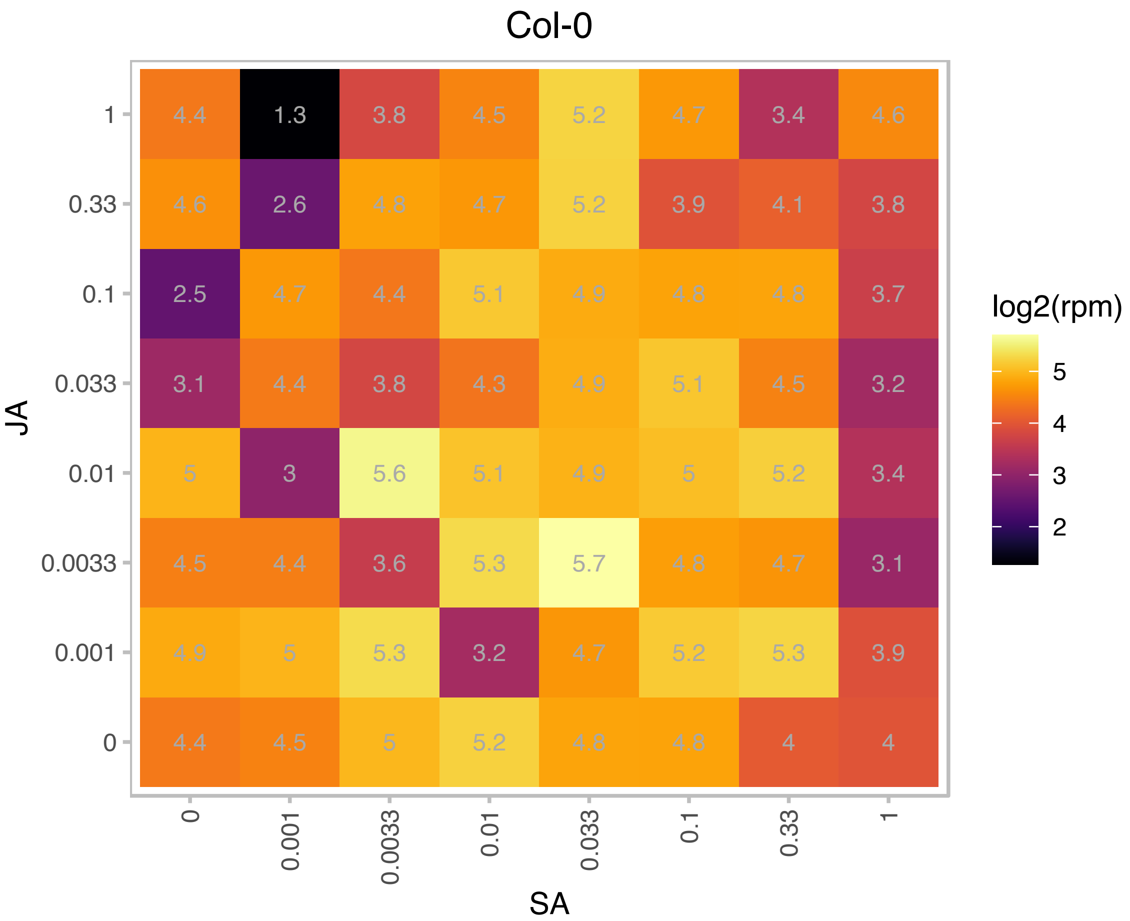


| | |
|---|---|
| Gene | AT4G03110 |
| is.expressed_genes | TRUE |
| is.responsive_genes | TRUE |
| is.apcluster | H-03 |

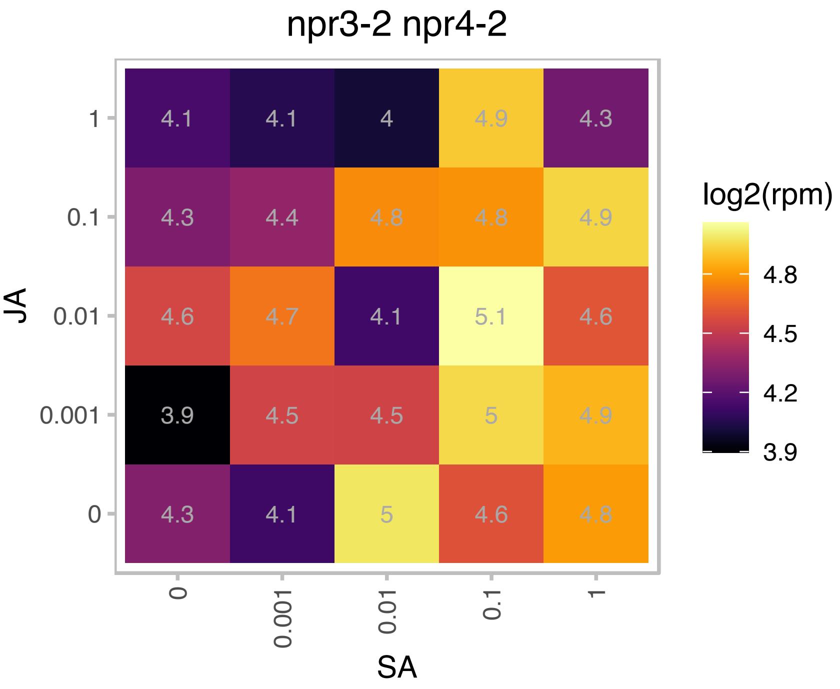


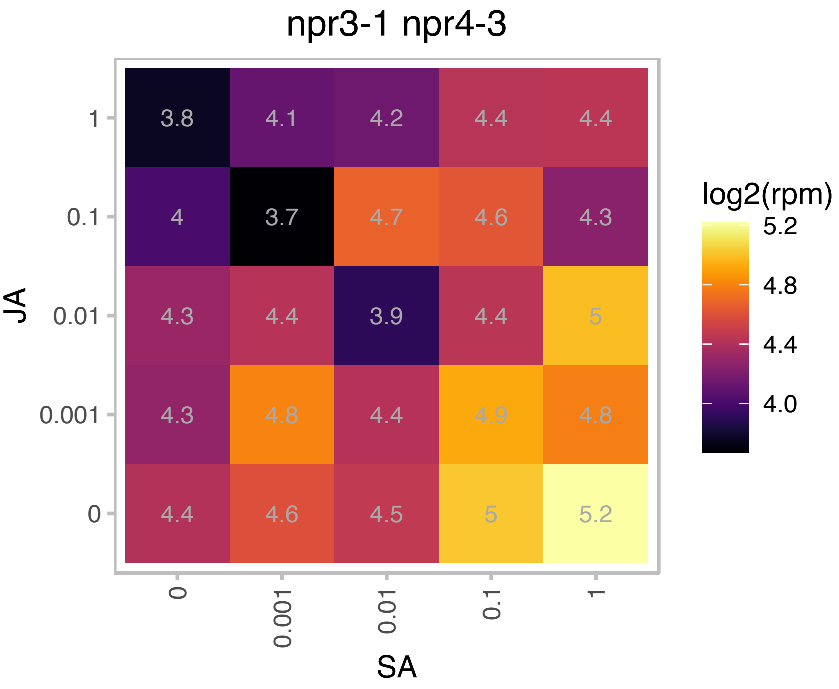

### AT4G12480.1 : EARLI1, pEARLI 1

Bifunctional inhibitor/lipid-transfer protein/seed storage 2S albumin superfamily protein

Encodes a putative lipid transfer protein, vernalization-responsive and cold-induced. It is involved in priming the SAR and ISR responses, specifically in propagating the cell-to-cell mobile signal.

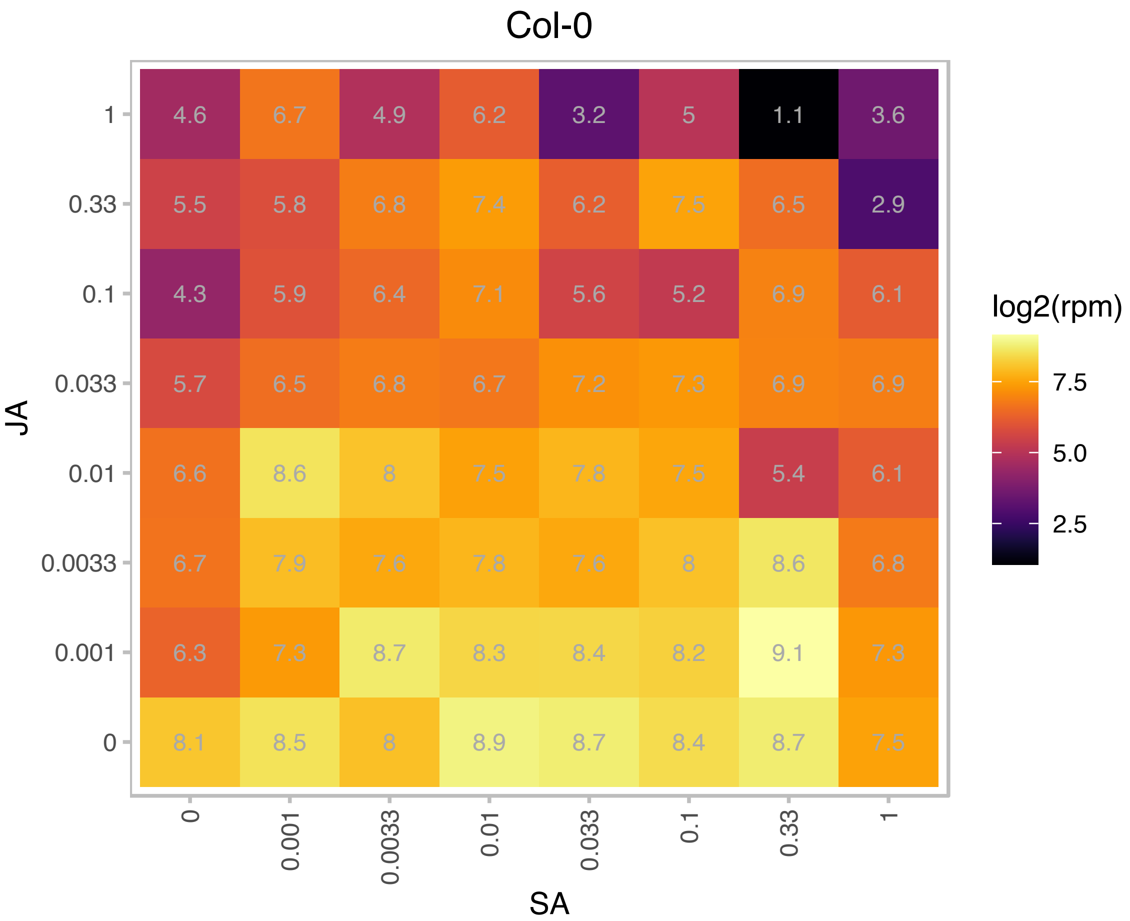


| | |
|---|---|
| Gene | AT4G12480 |
| is.expressed_genes | TRUE |
| is.responsive_genes | TRUE |
| is.apcluster | H-03 |

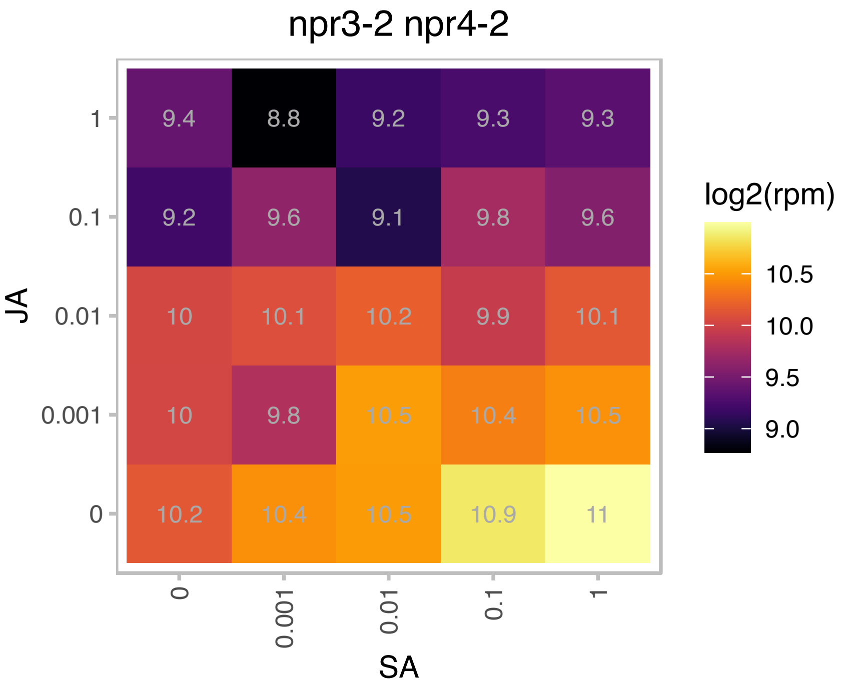


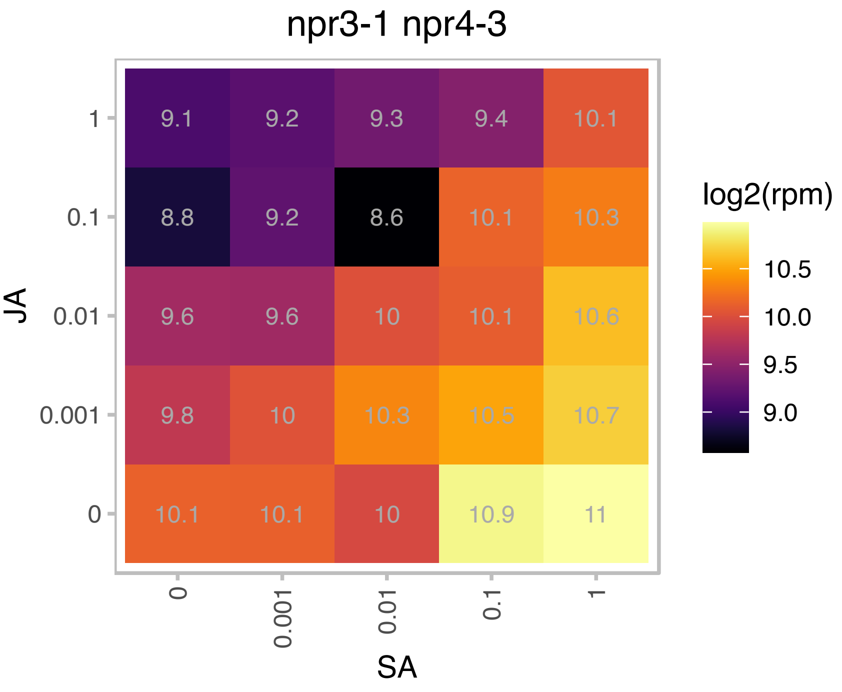

### AT4G16265.1 : DMS13, NRPB9B, NRPD9B, NRPE9B

RNA polymerases M/15 Kd subunit

One of two highly similar, non-catalytic subunits common to nuclear DNA-directed RNA polymerases II, IV and V; homologous to budding yeast RPB9. Appears to be redundant with At3g16980

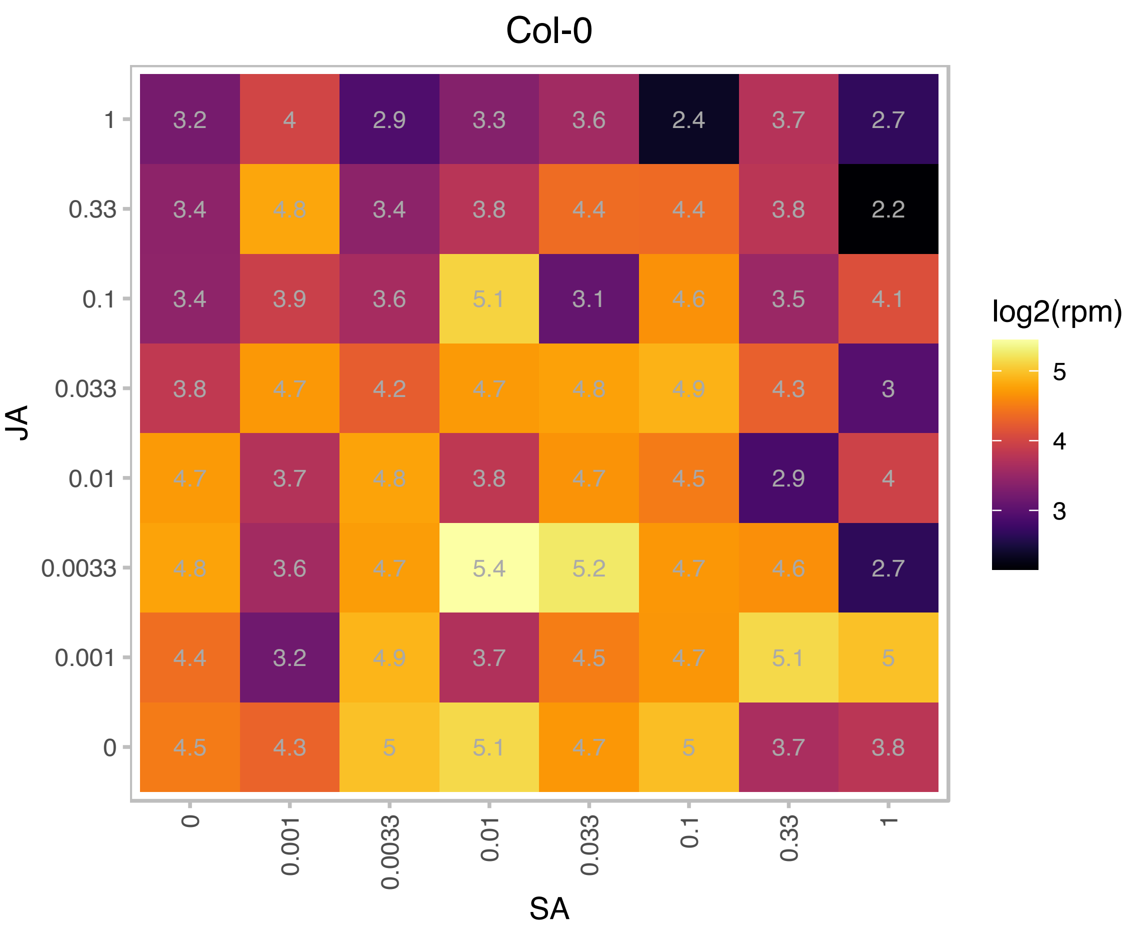


| | |
|---|---|
| Gene | AT4G16265 |
| is.expressed_genes | TRUE |
| is.responsive_genes | TRUE |
| is.apcluster | H-03 |

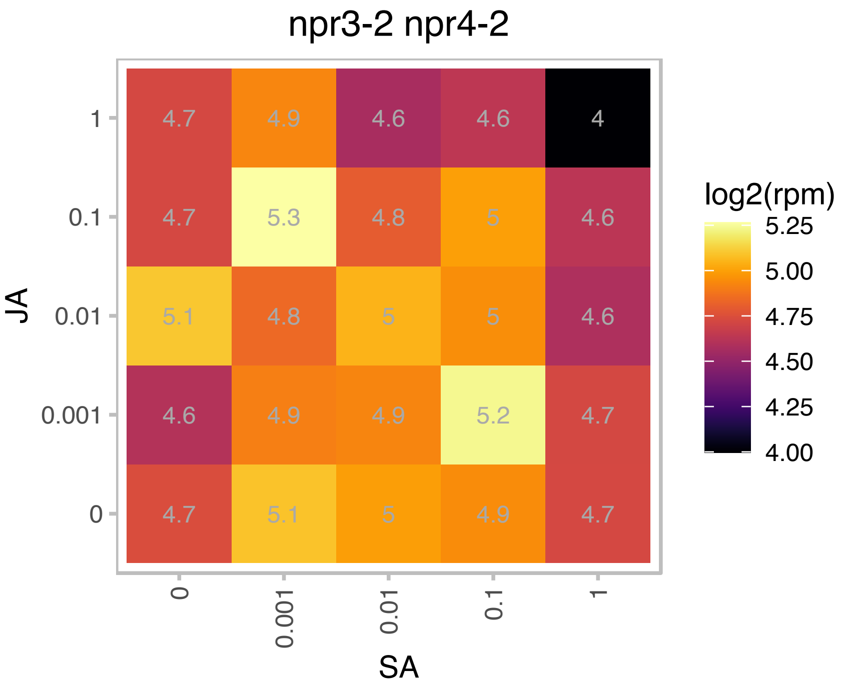


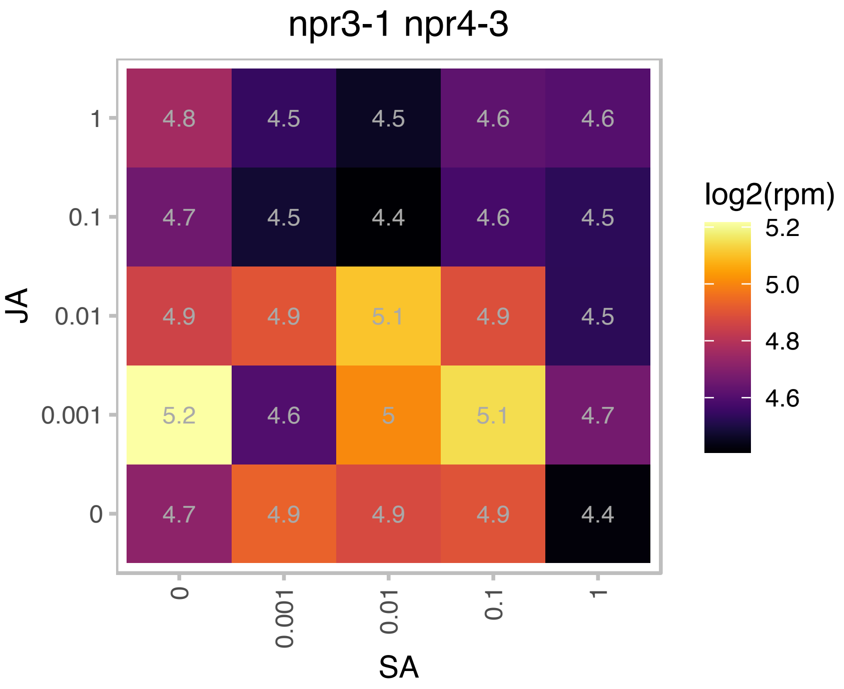

### AT4G28270.1 : ATRMA2, RMA2

RING membrane-anchor 2

Encodes a RING finger E3 ubiquitin ligase. Binds and ubiquitinates ABP1 in vivo and in vitro.

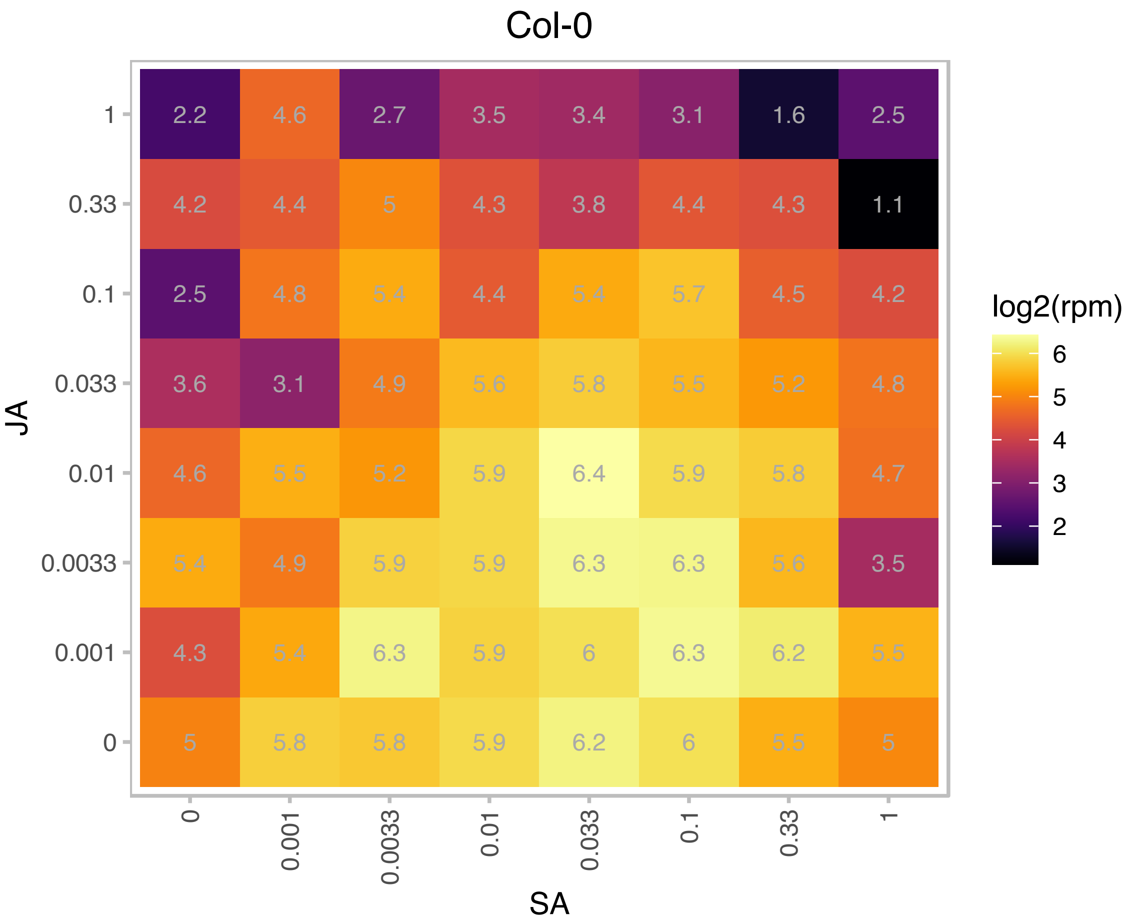


| | |
|---|---|
| Gene | AT4G28270 |
| is.expressed_genes | TRUE |
| is.responsive_genes | TRUE |
| is.apcluster | H-03 |

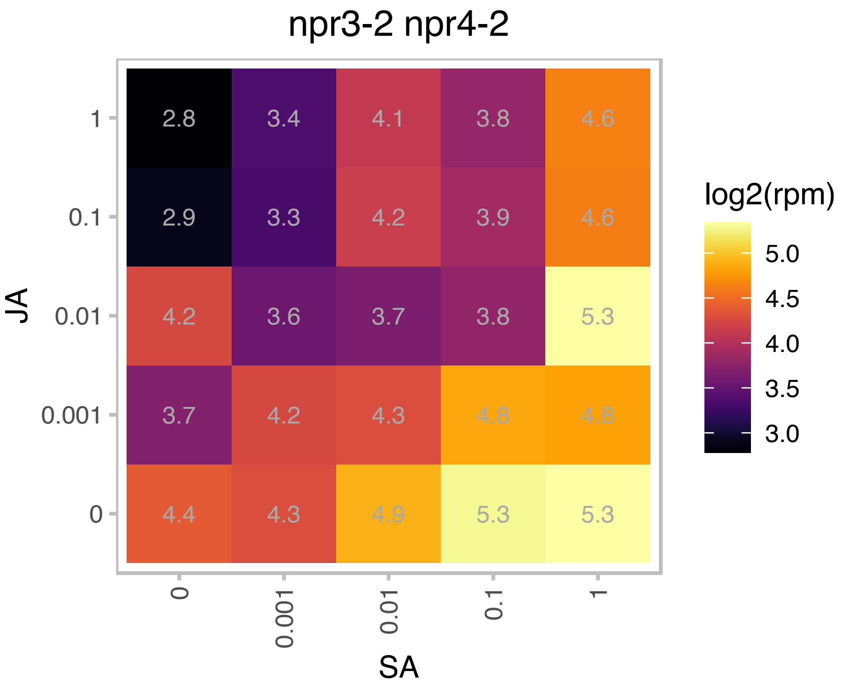


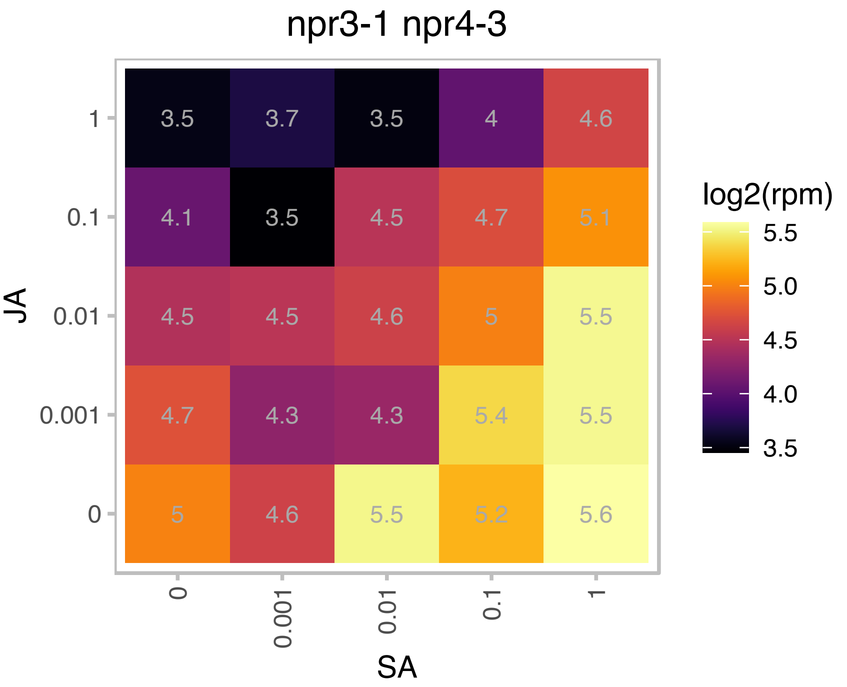

### AT5G14060.1 : CARAB-AK-LYS

Aspartate kinase family protein

lysine-sensitive aspartate kinase

Col-0

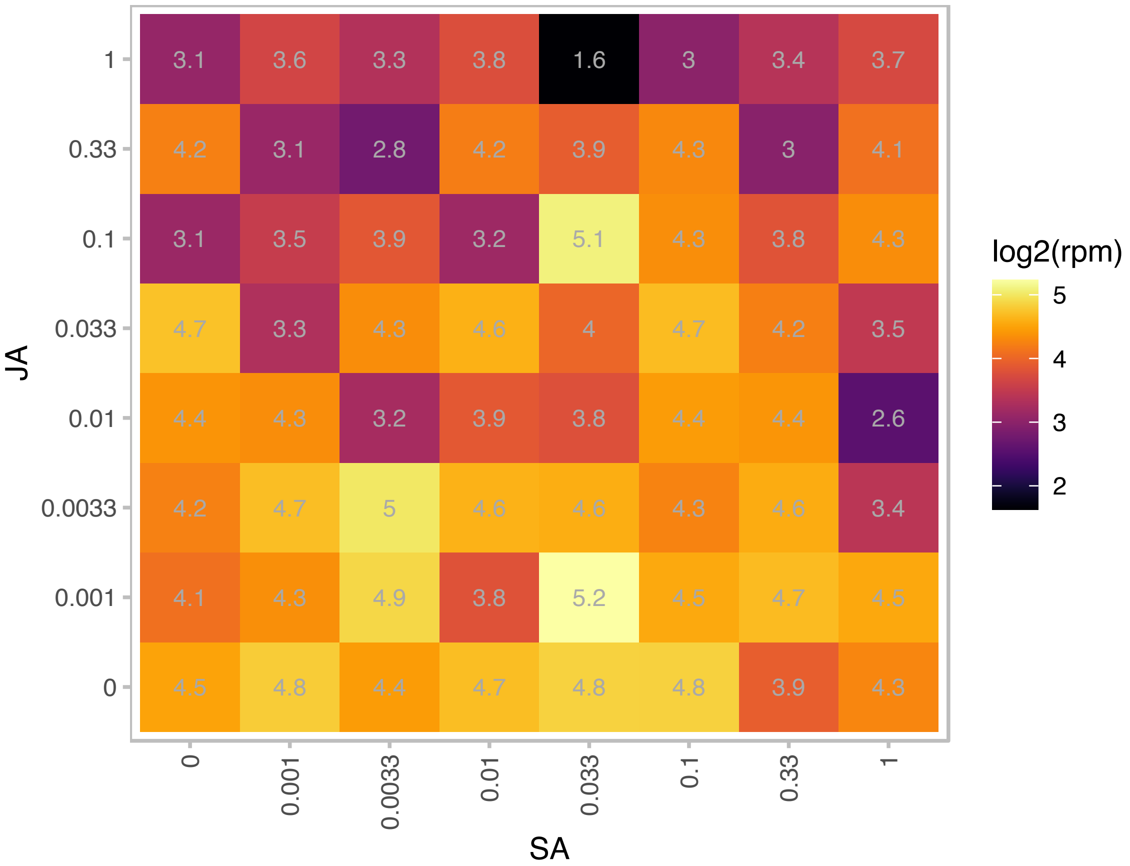


| | |
|---|---|
| Gene | AT5G14060 |
| is.expressed_genes | TRUE |
| is.responsive_genes | TRUE |
| is.apcluster | H-03 |

npr3-2 npr4-2

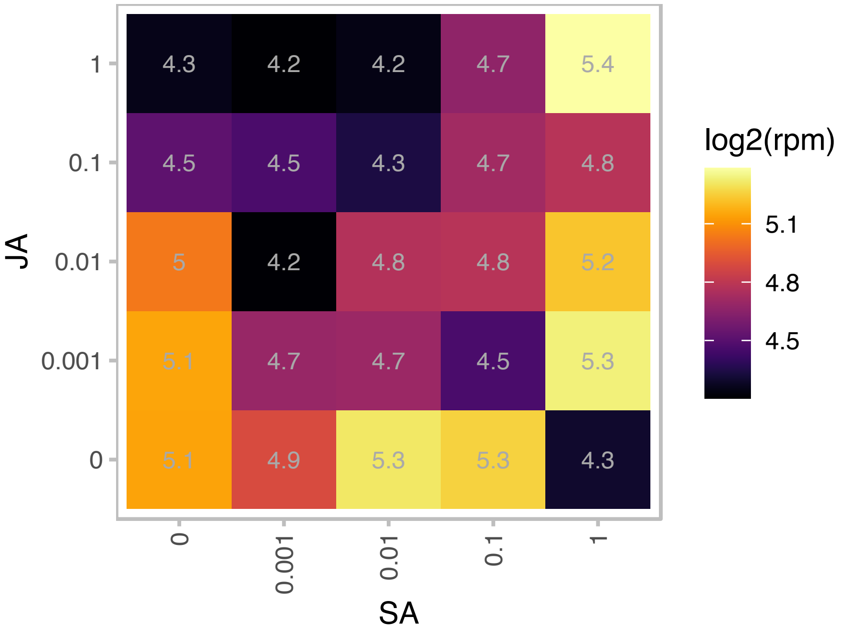


npr3-1 npr4-3

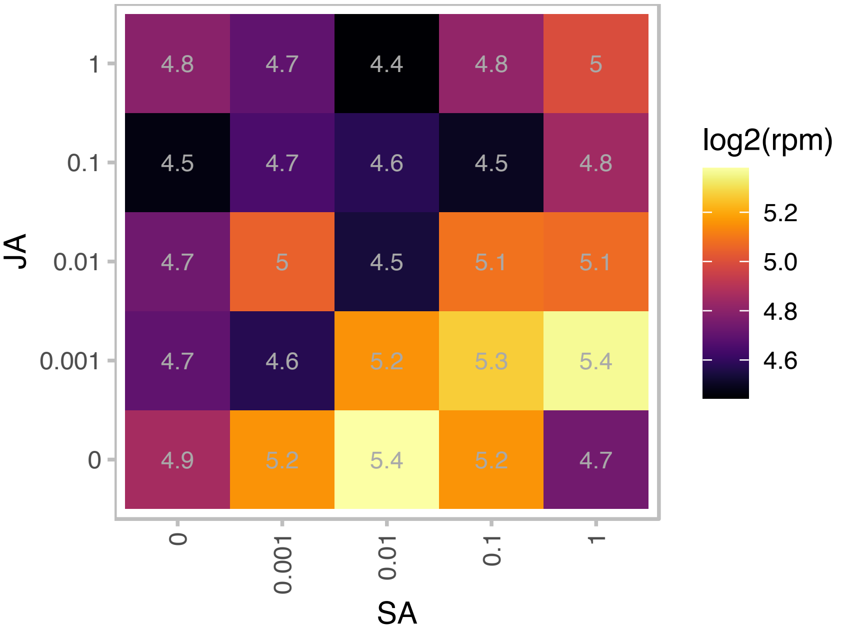

# AT5G19190.1 : NA

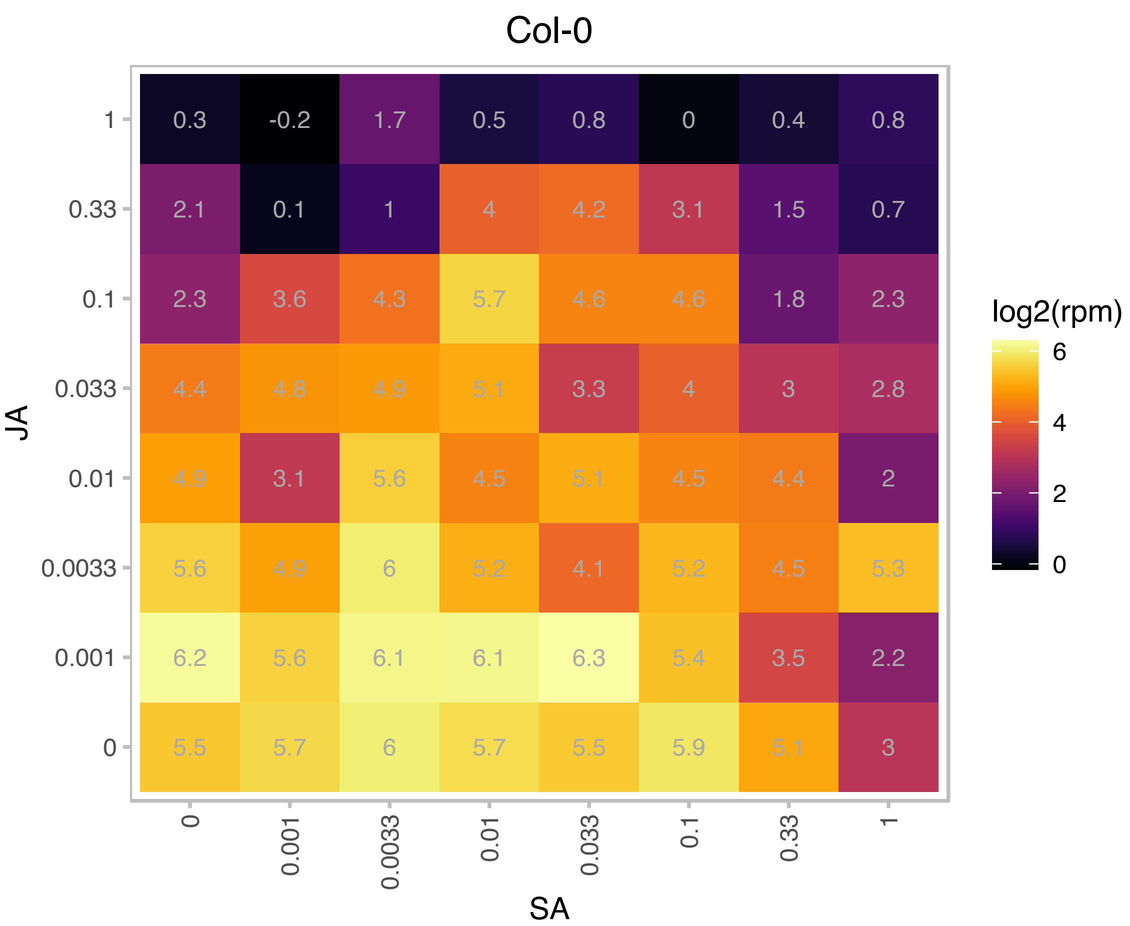


| | |
|---|---|
| Gene | AT5G19190 |
| is.expressed_genes | TRUE |
| is.responsive_genes | TRUE |
| is.apcluster | H-03 |

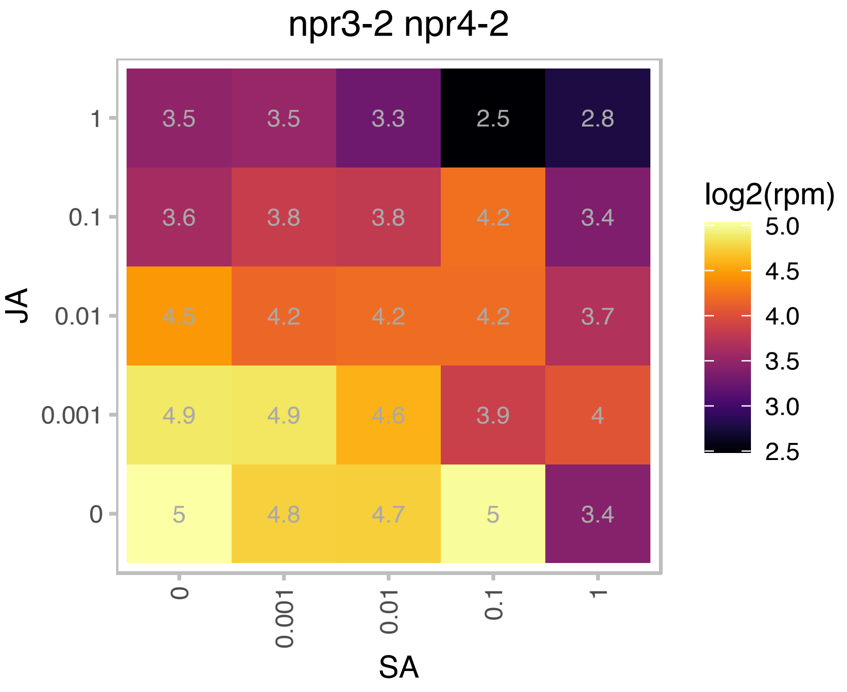


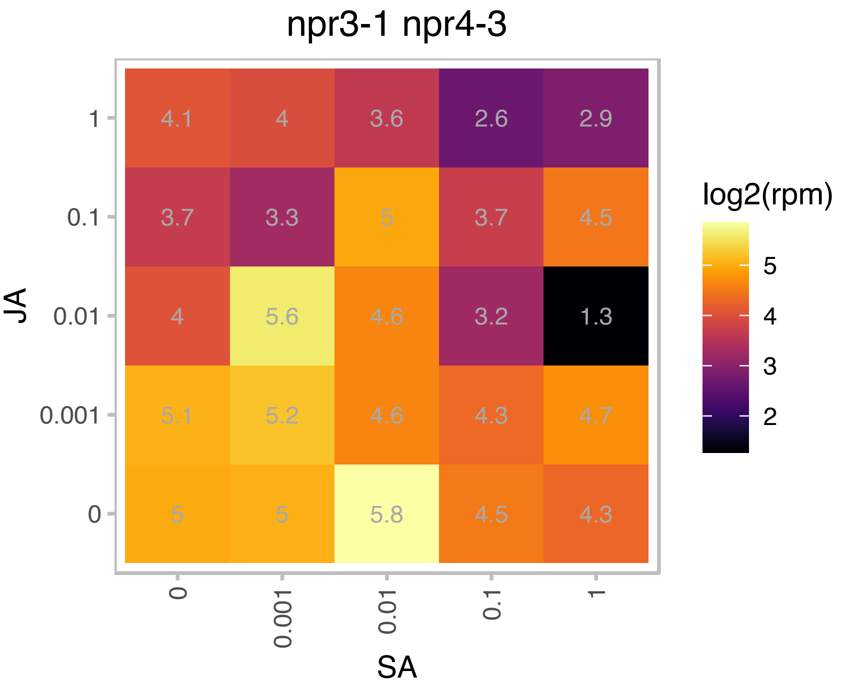

# AT5G19750.1 : NA

Peroxisomal membrane 22 kDa (Mpv17/PMP22) family protein

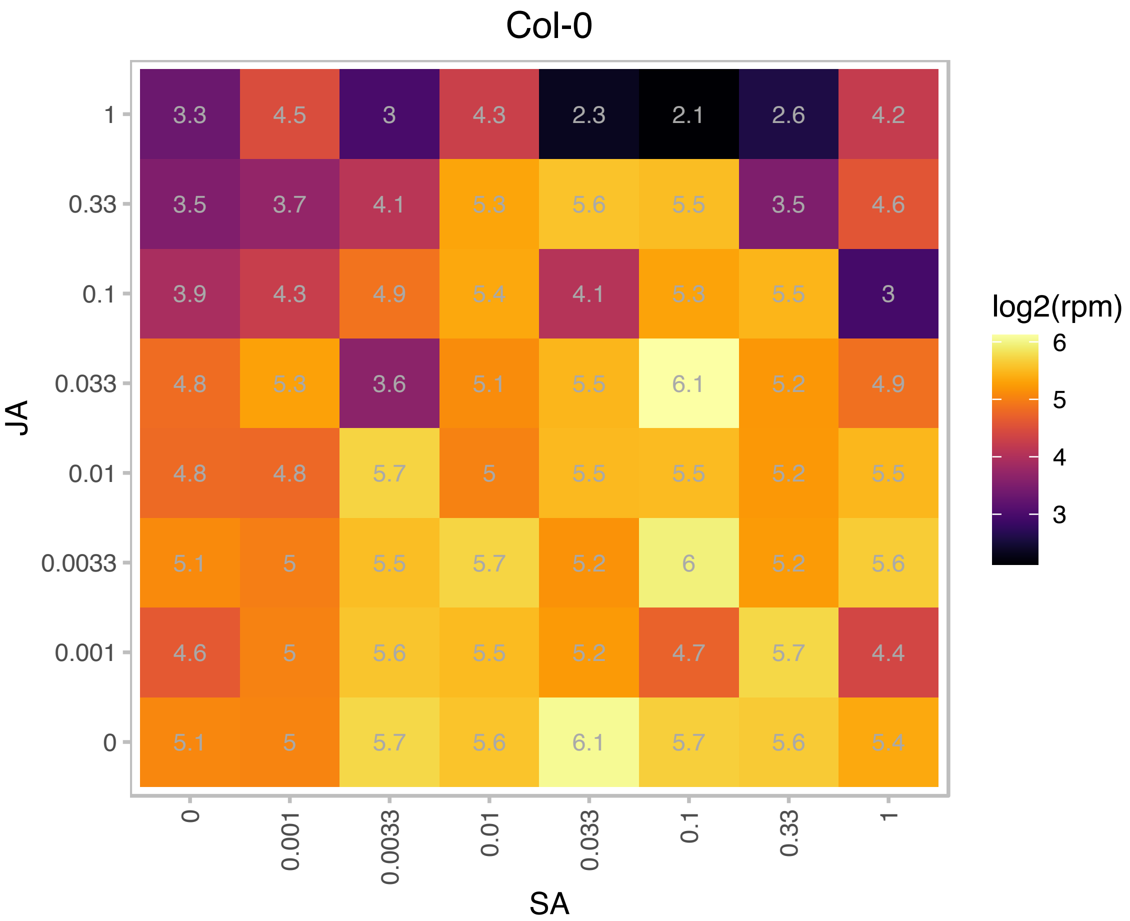


| | |
|---|---|
| Gene | AT5G19750 |
| is.expressed_genes | TRUE |
| is.responsive_genes | TRUE |
| is.apcluster | H-03 |

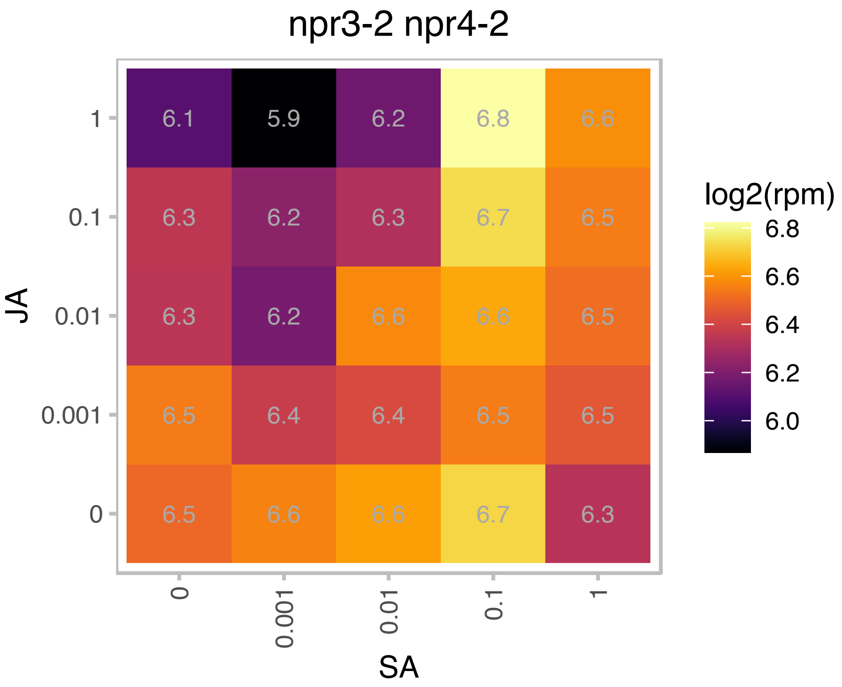


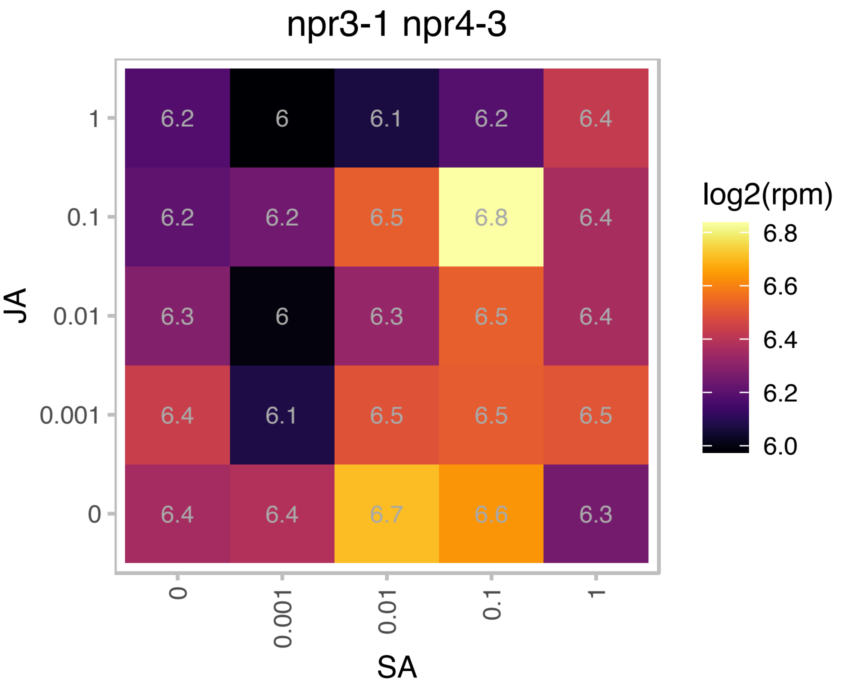

# AT5G22390.1 : NA

Protein of unknown function (DUF3049)

| | |
|---|---|
| Gene | AT5G22390 |
| is.expressed_genes | TRUE |
| is.responsive_genes | TRUE |
| is.apcluster | H-03 |

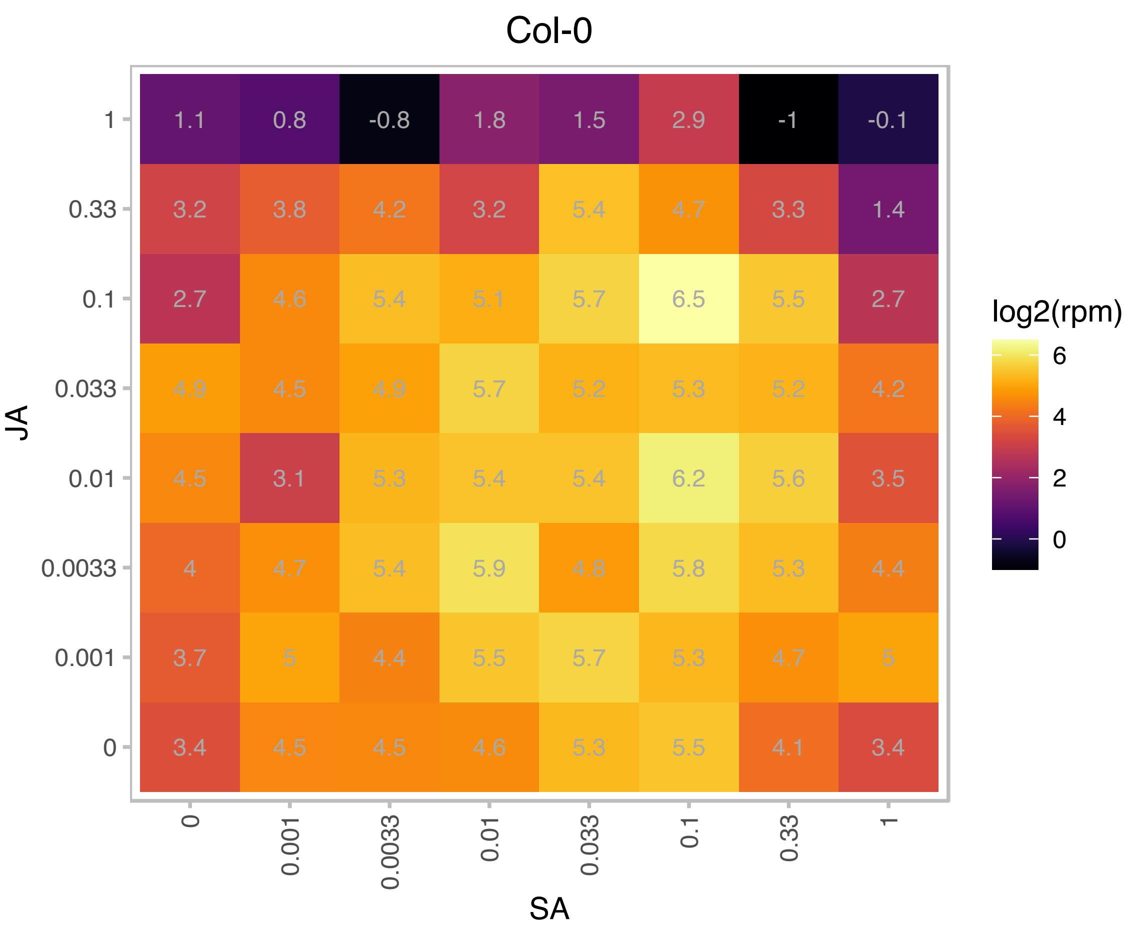


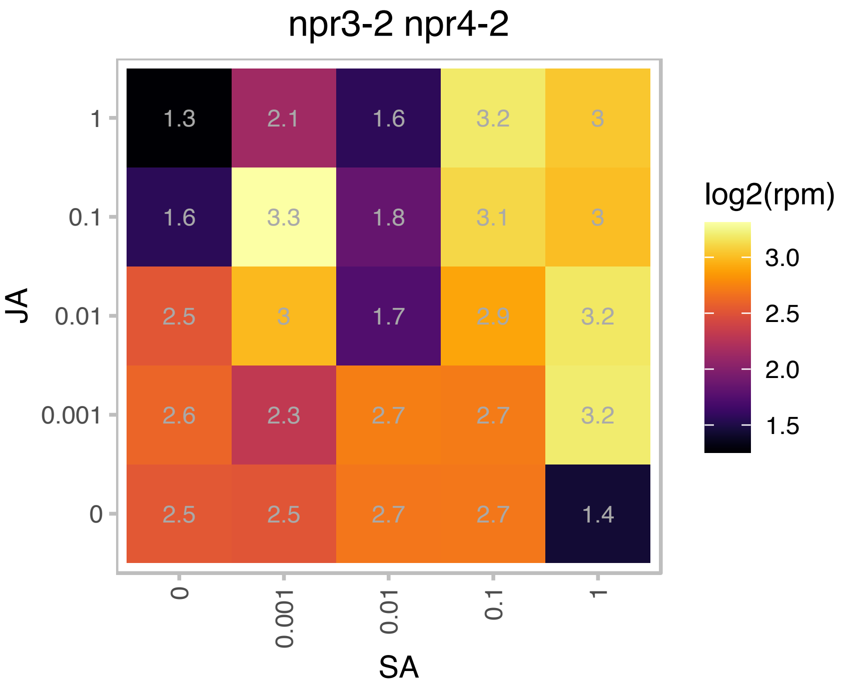


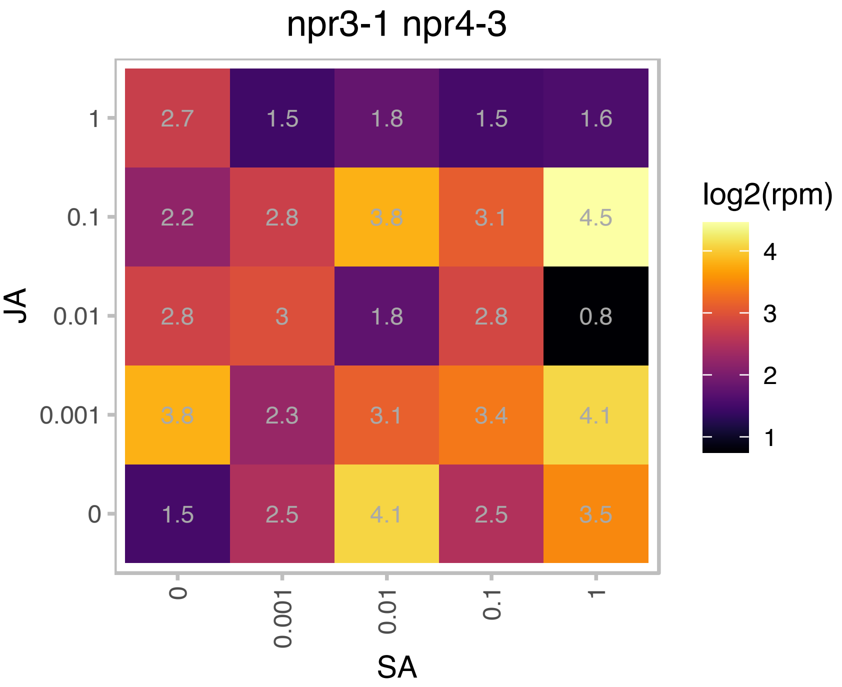

# AT5G22580.1 : NA

Stress responsive A/B Barrel Domain

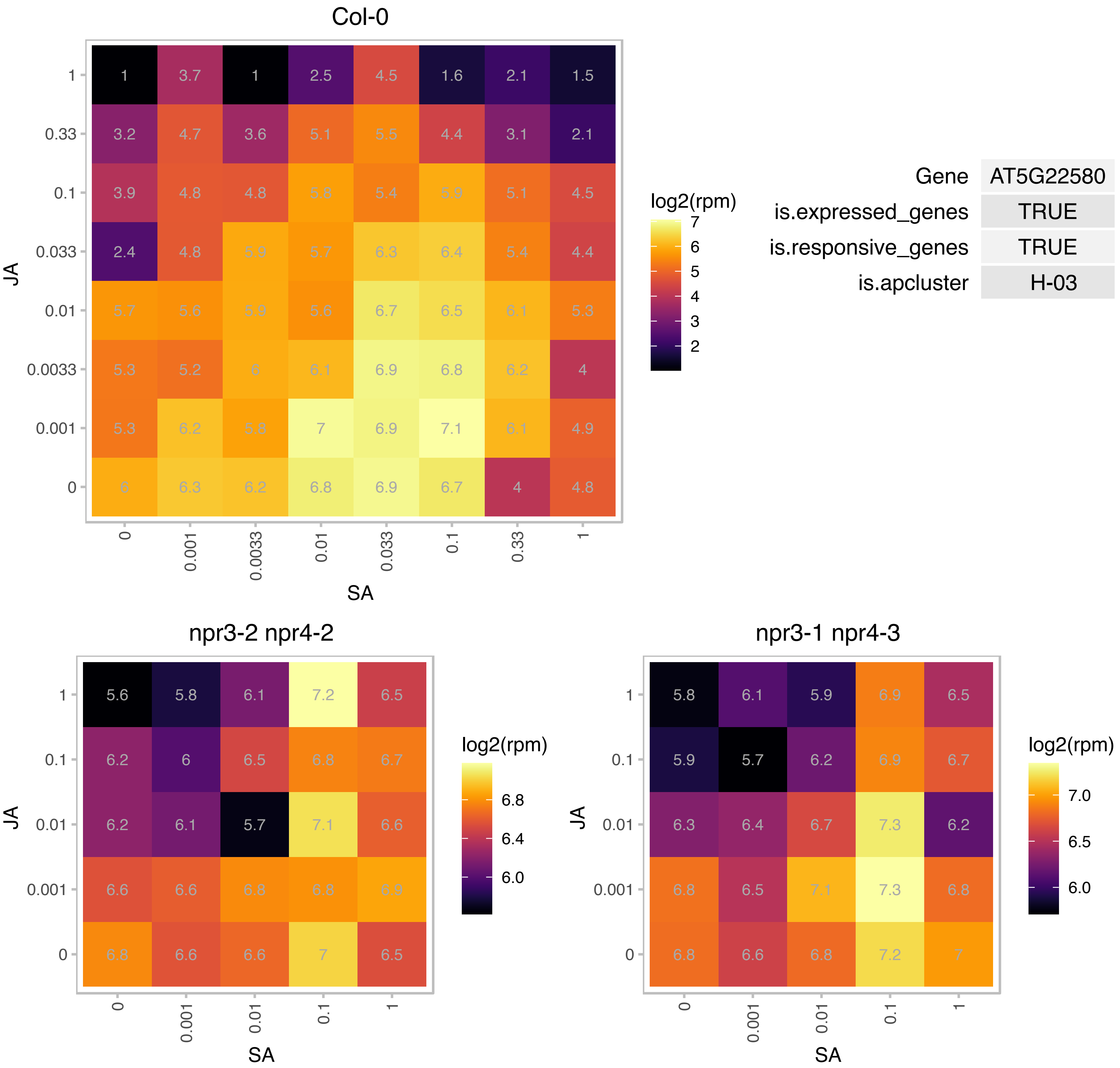


| | |
|---|---|
| Gene | AT5G22580 |
| is.expressed_genes | TRUE |
| is.responsive_genes | TRUE |
| is.apcluster | H-03 |

# AT5G23405.1 : NA

HMG-box (high mobility group) DNA-binding family protein

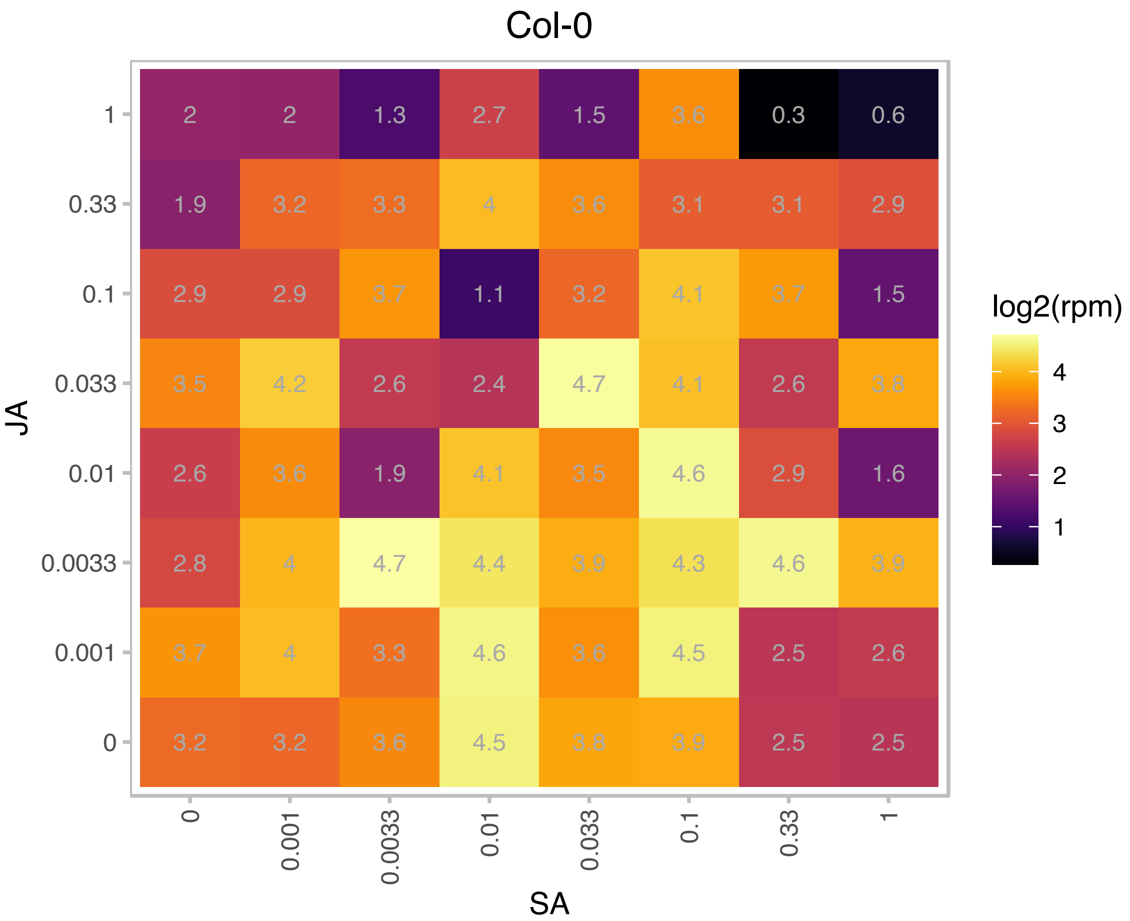


| | |
|---|---|
| Gene | AT5G23405 |
| is.expressed_genes | TRUE |
| is.responsive_genes | TRUE |
| is.apcluster | H-03 |

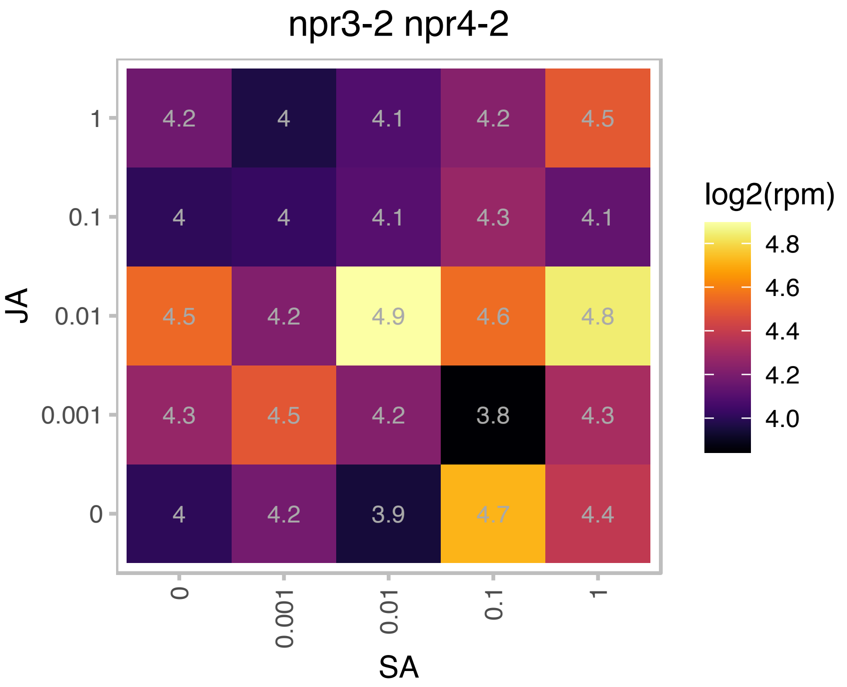


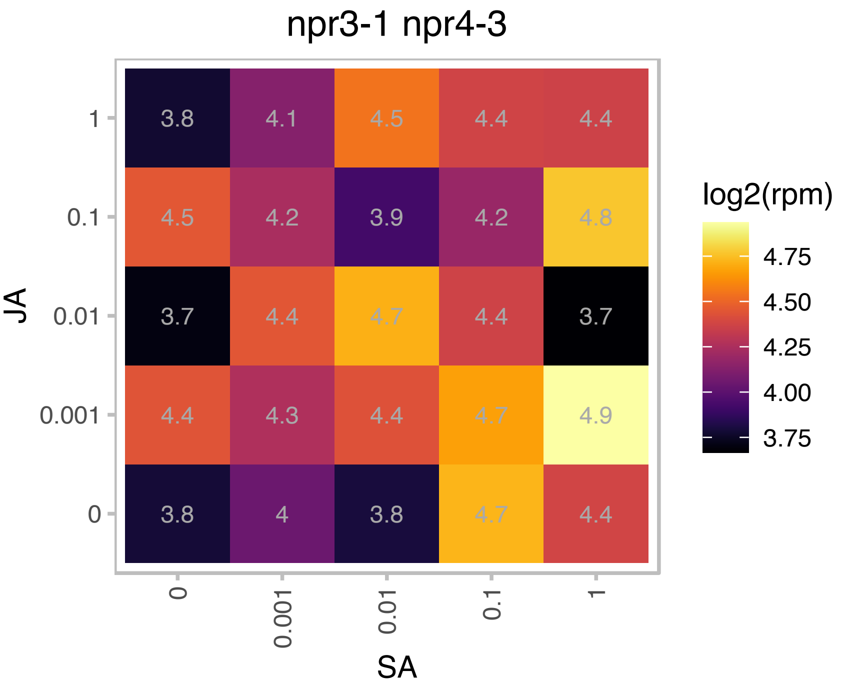

### AT5G25460.1 : DGR2

Protein of unknown function, DUF642

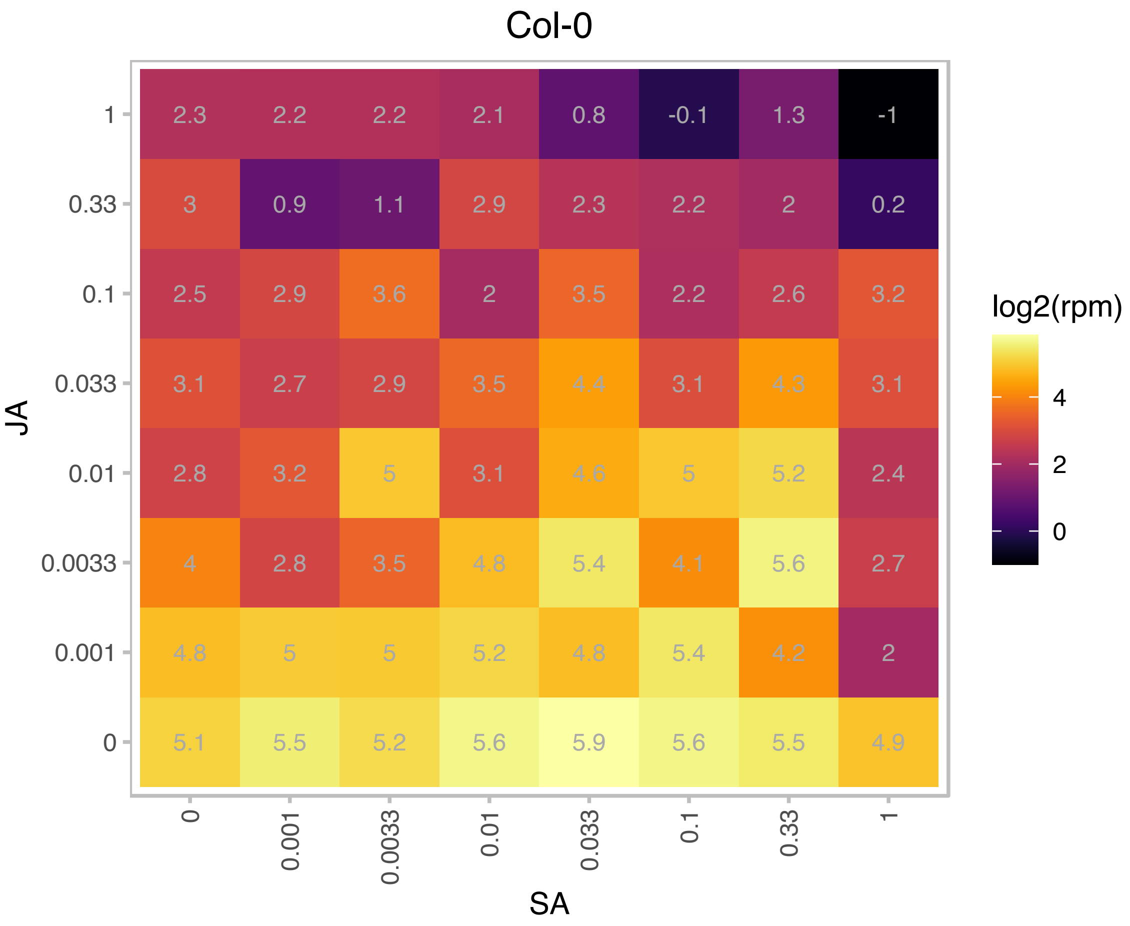


| | |
|---|---|
| Gene | AT5G25460 |
| is.expressed_genes | TRUE |
| is.responsive_genes | TRUE |
| is.apcluster | H-03 |

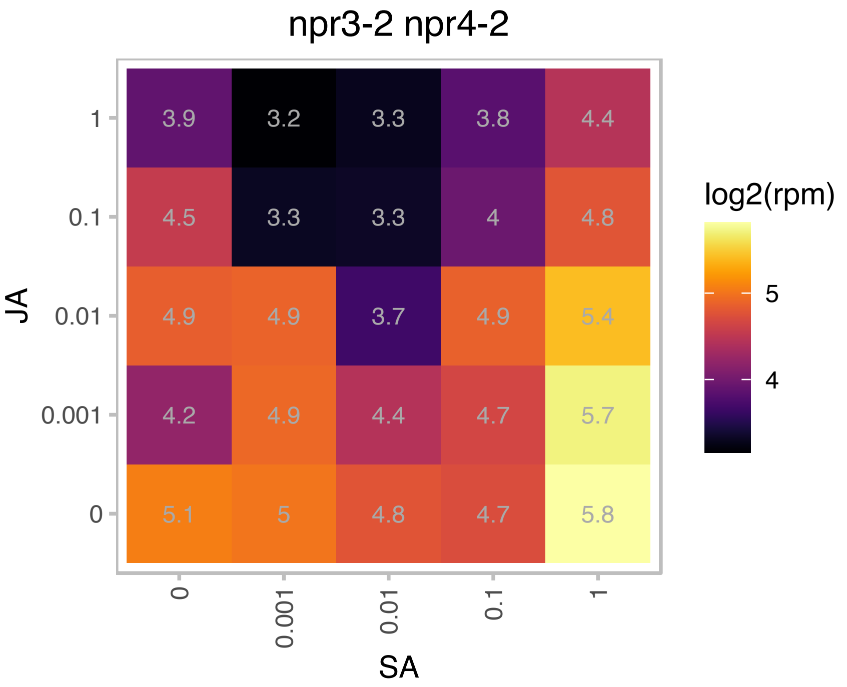


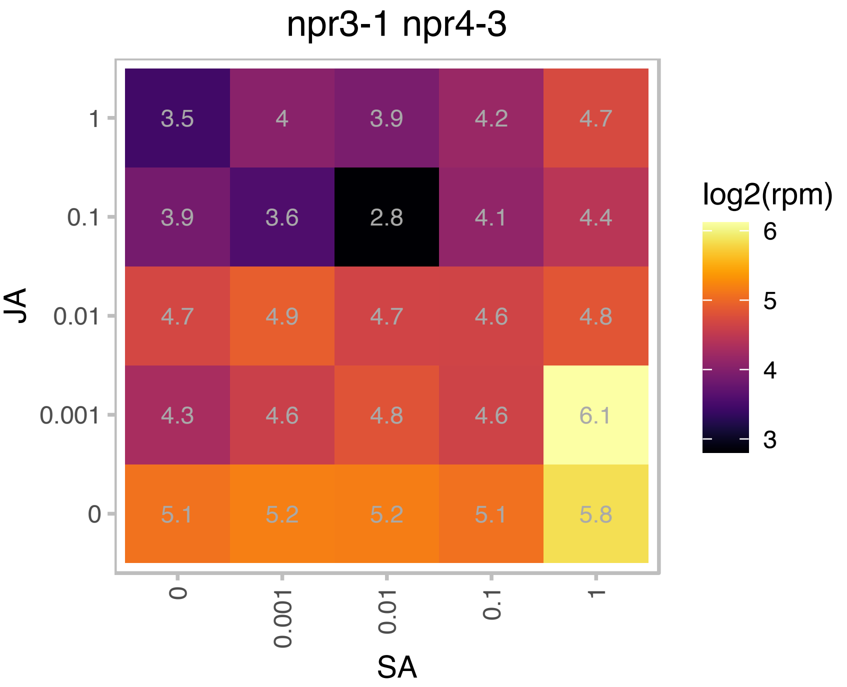

# AT5G44572.1 : NA

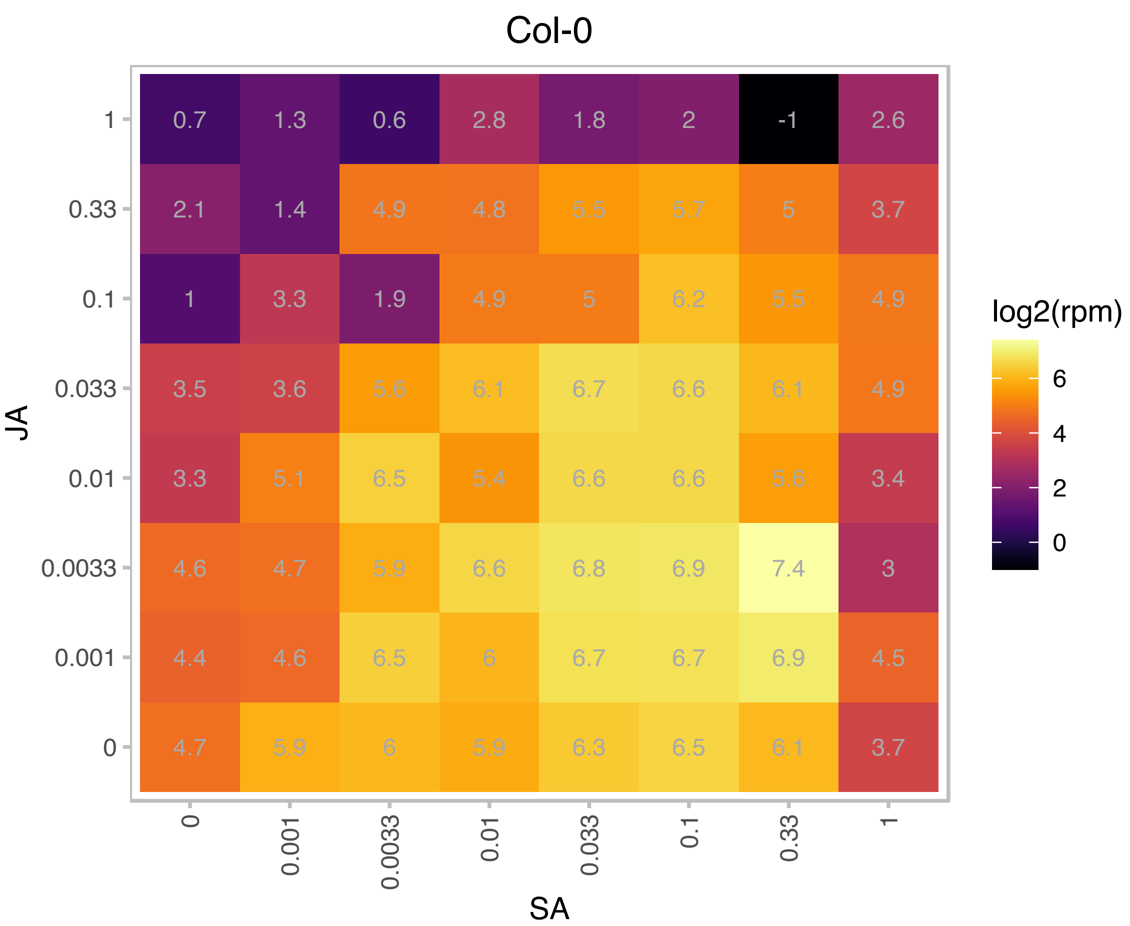


| | |
|---|---|
| Gene | AT5G44572 |
| is.expressed_genes | TRUE |
| is.responsive_genes | TRUE |
| is.apcluster | H-03 |

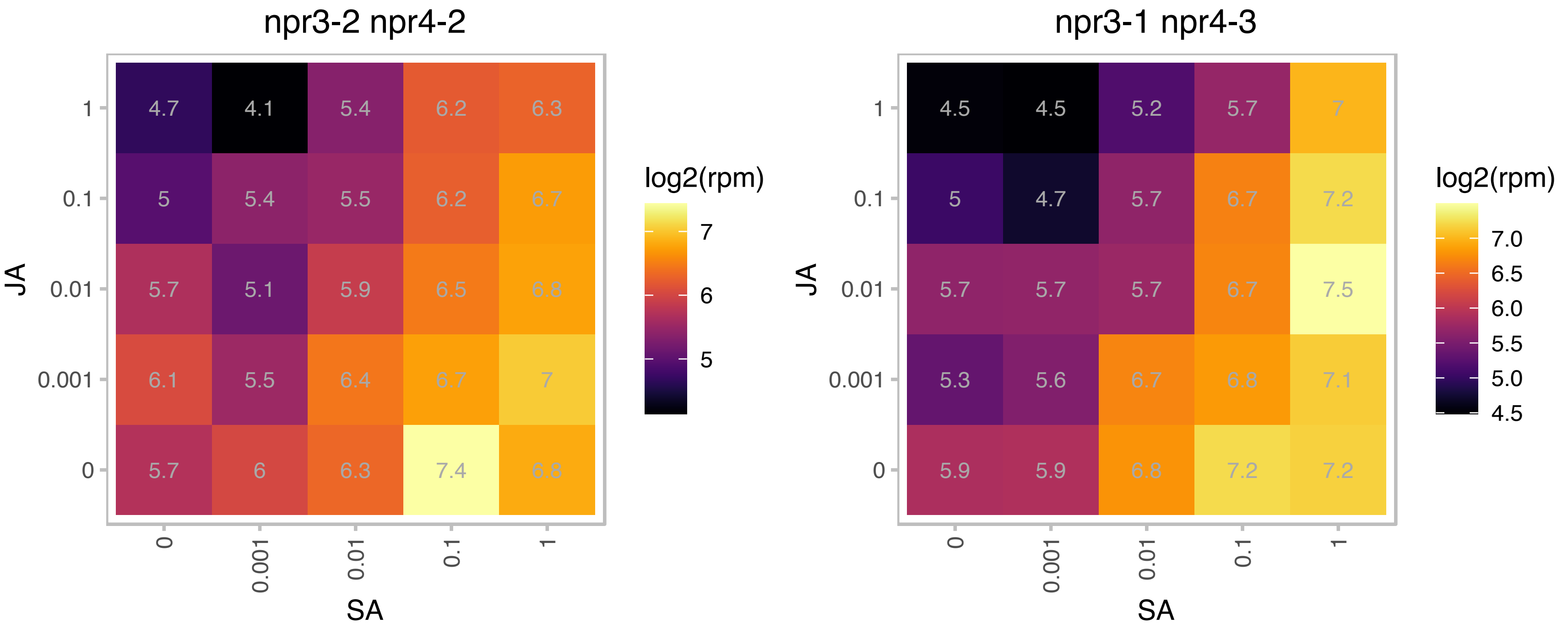

# AT5G44574.1 : NA

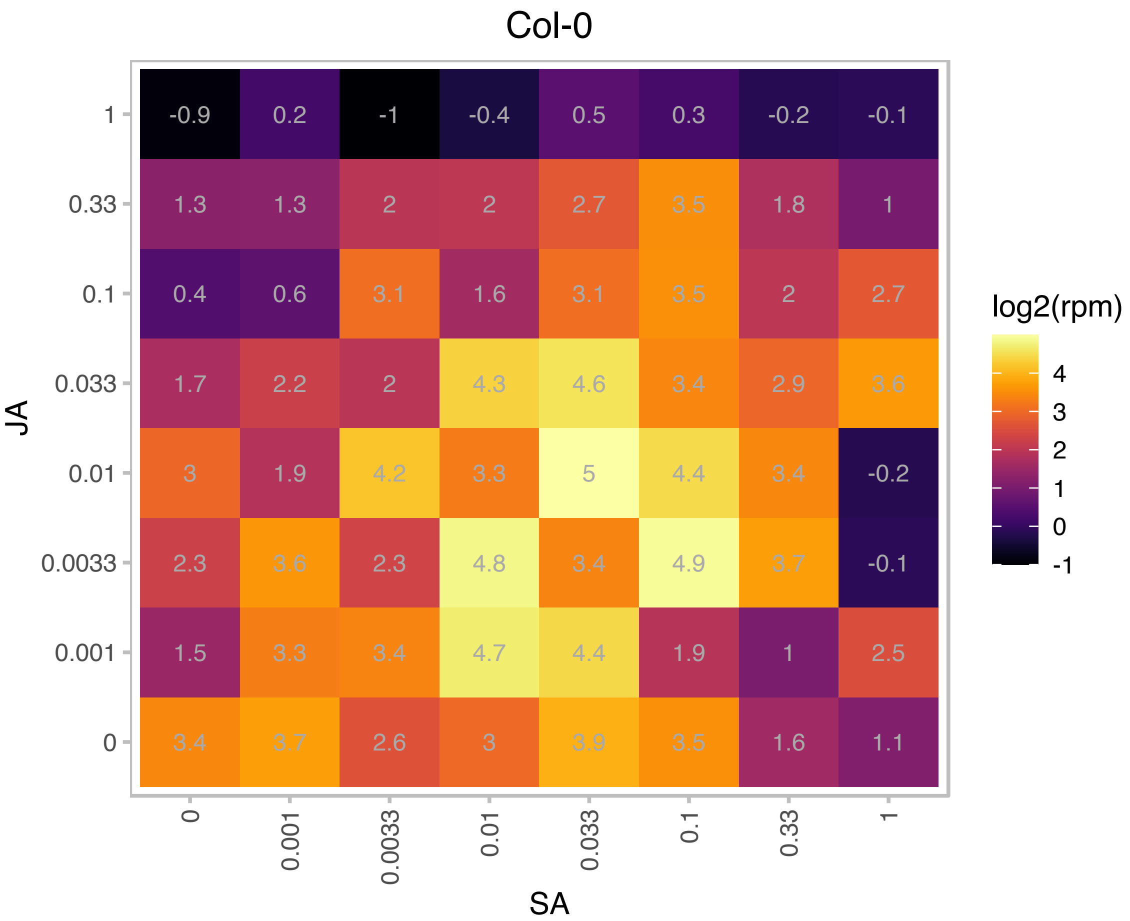


| | |
|---|---|
| Gene | AT5G44574 |
| is.expressed_genes | TRUE |
| is.responsive_genes | TRUE |
| is.apcluster | H-03 |

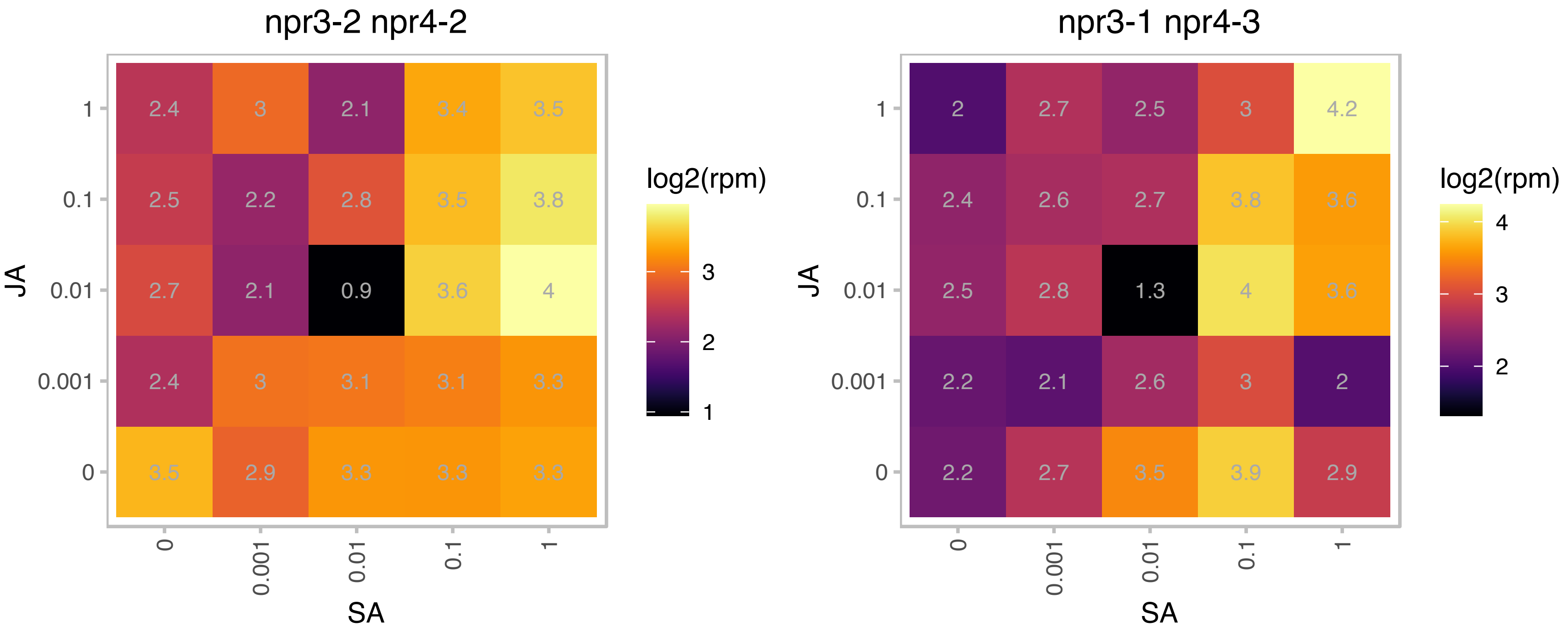

# AT5G44575.1 : NA

| | |
|---|---|
| Gene | AT5G44575 |
| is.expressed_genes | TRUE |
| is.responsive_genes | TRUE |
| is.apcluster | H-03 |

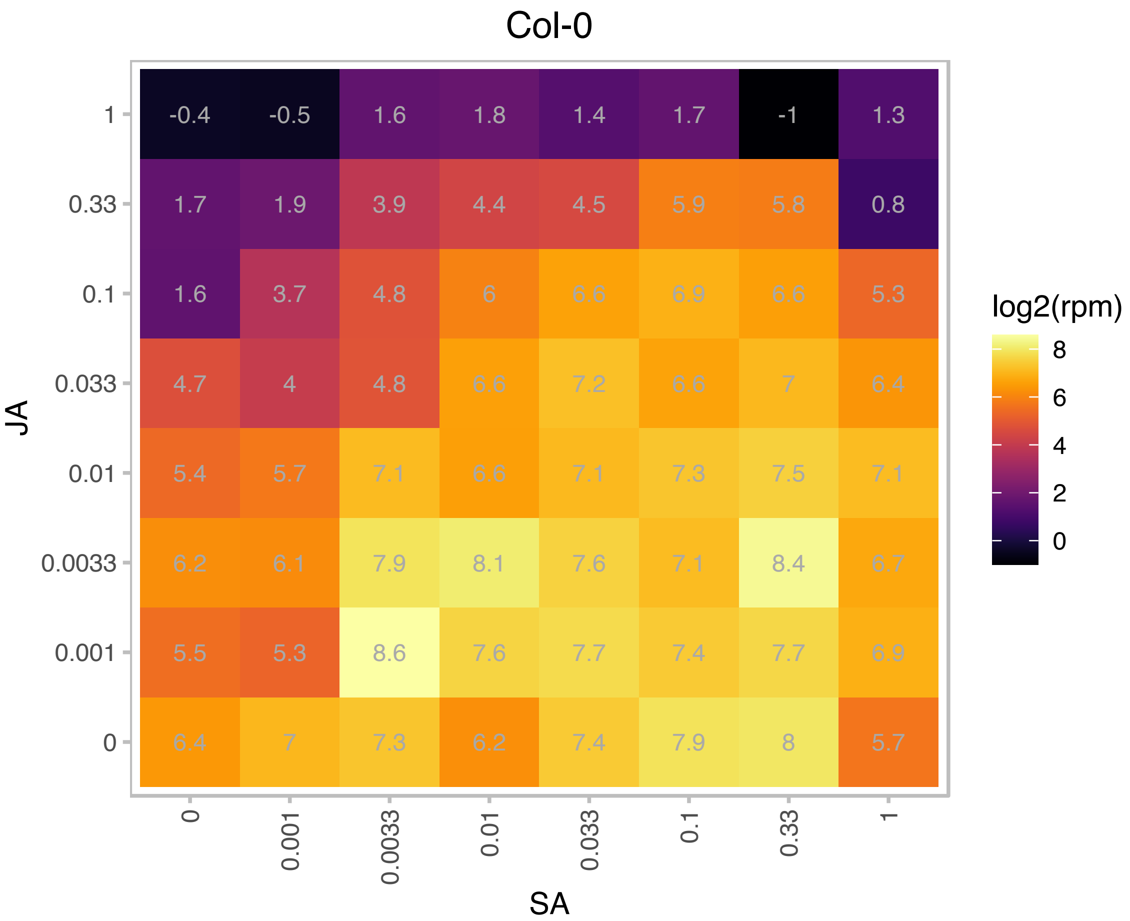


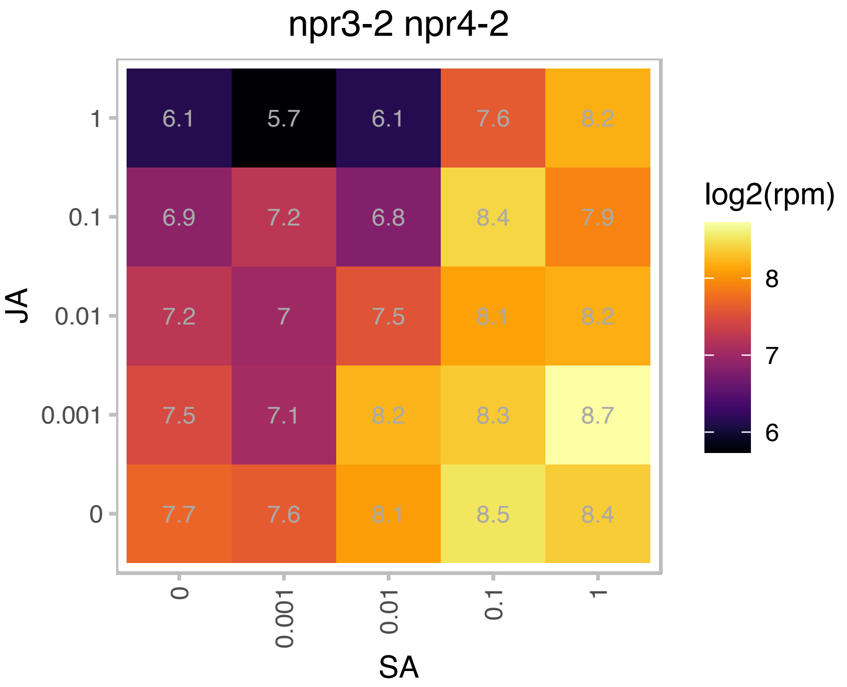


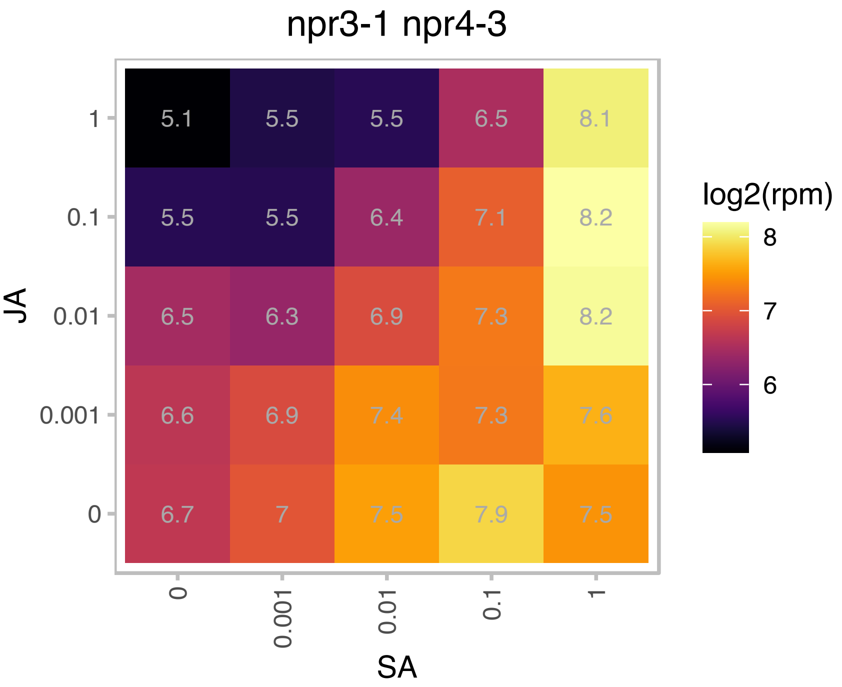

### AT5G44610.1 : MAP18, PCAP2

Encodes a protein with seven repeated VEEKK motifs. RNAi and overexpression experiments suggest that the gene is not involved in cell division but might be consequential for cell shape of epidermal and cortical cells.
microtubule-associated protein 18
The protein encoded by this gene binds to cortical microtubules and inhibits tubulin polymerization. Associates to the plasma membrane and interacts with calmodulin and phosphatidylinositol phosphates, indicating an involvement in cellular signal transduction. Expression is enhanced by abiotic and hormonal factors. Induced during senescence.Interacts with Ca2+/calmodulin complex, phosphatidylinositol phosphates, [illegible]

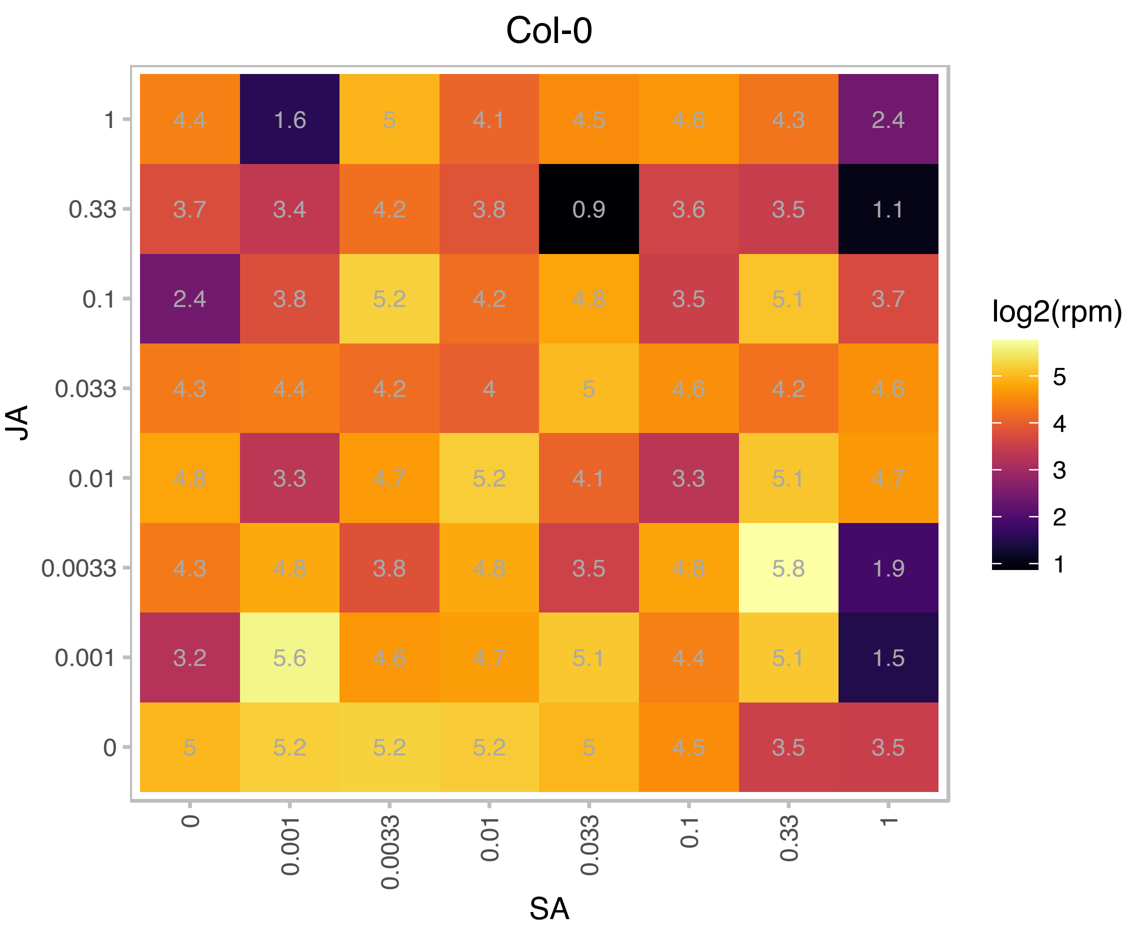


| | |
|---|---|
| Gene | AT5G44610 |
| is.expressed_genes | TRUE |
| is.responsive_genes | TRUE |
| is.apcluster | H-03 |

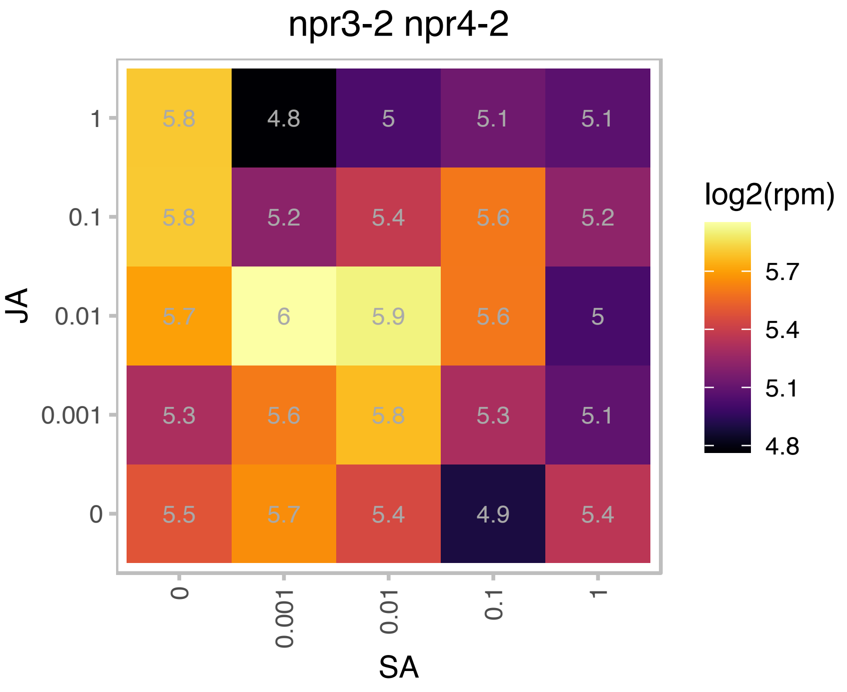


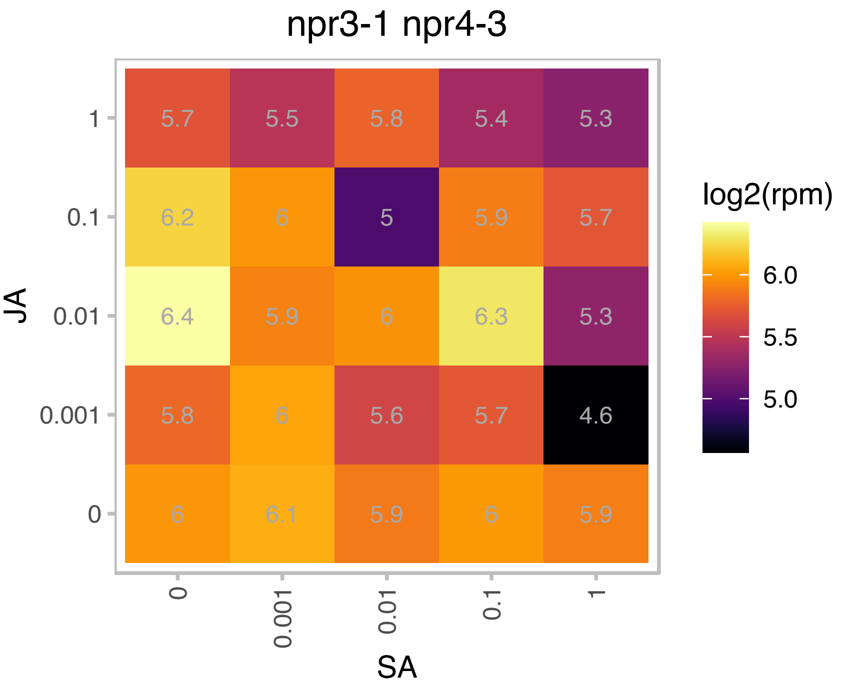

# AT5G48590.1 : NA

Protein of unknown function (DUF760)

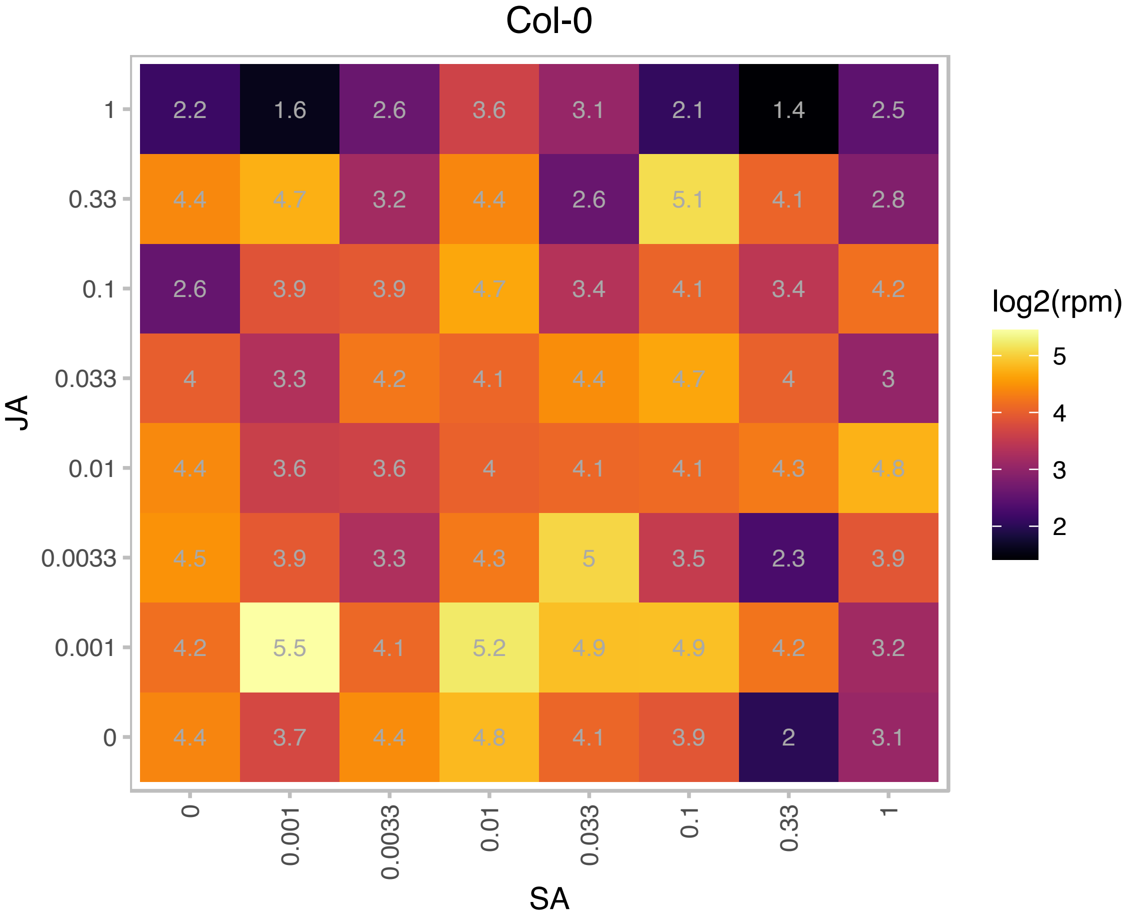


| | |
|---|---|
| Gene | AT5G48590 |
| is.expressed_genes | TRUE |
| is.responsive_genes | TRUE |
| is.apcluster | H-03 |

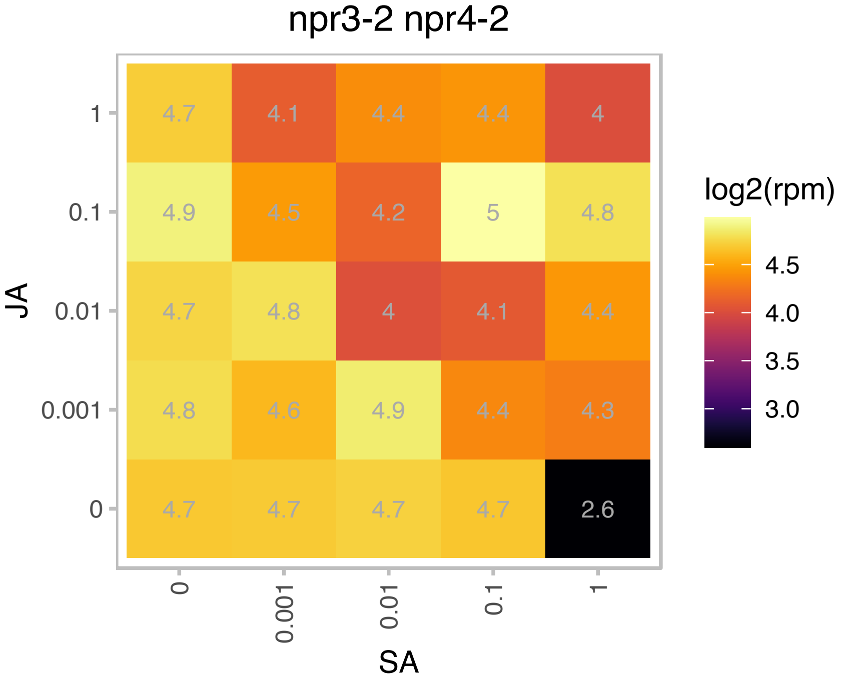


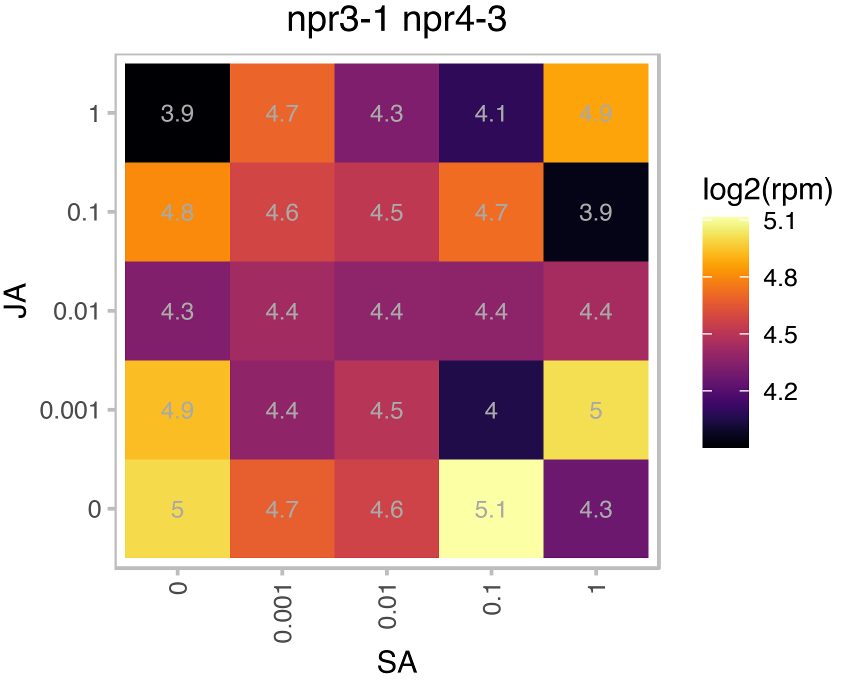

### AT5G55450.1 : AtLTP4.4

Bifunctional inhibitor/lipid-transfer protein/seed storage 2S albumin superfamily protein

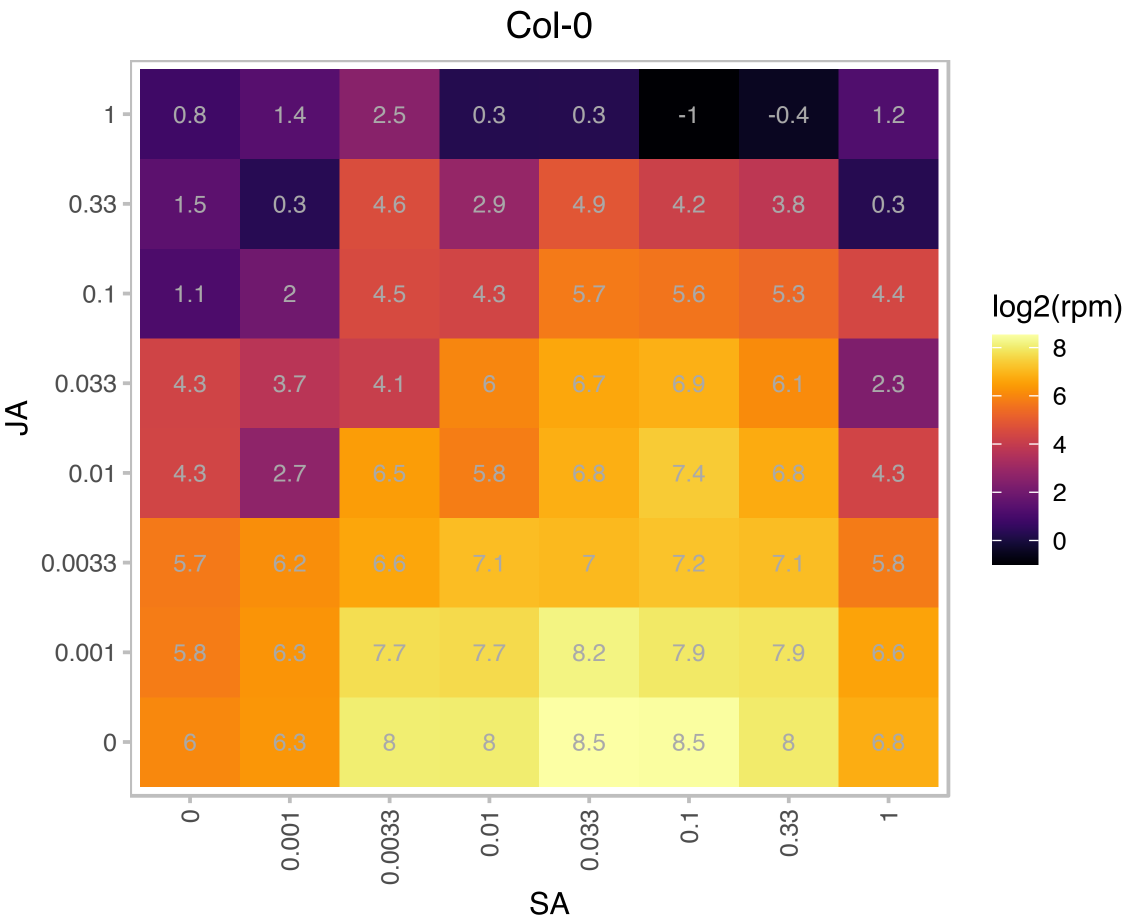


| | |
|---|---|
| Gene | AT5G55450 |
| is.expressed_genes | TRUE |
| is.responsive_genes | TRUE |
| is.apcluster | H-03 |

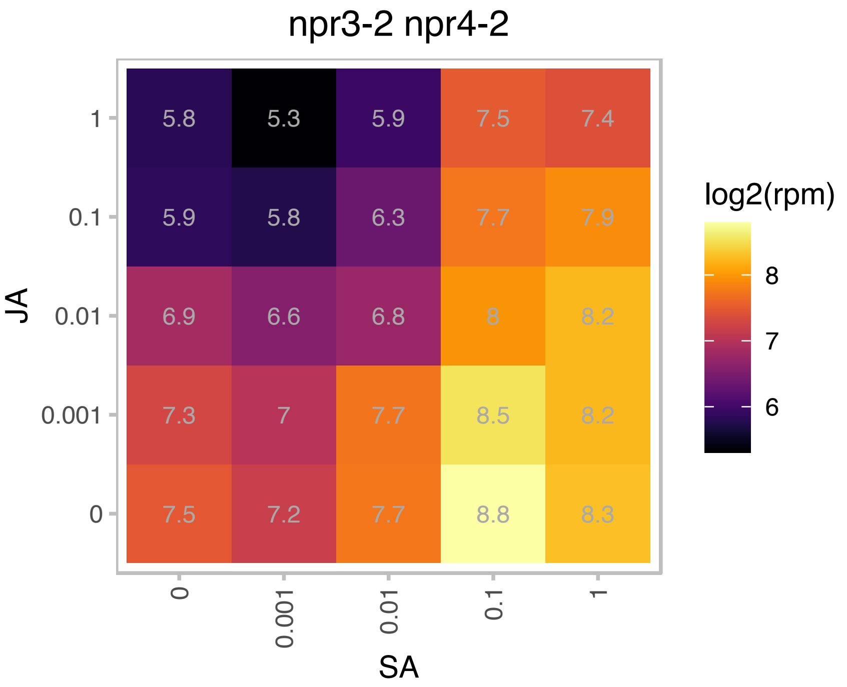


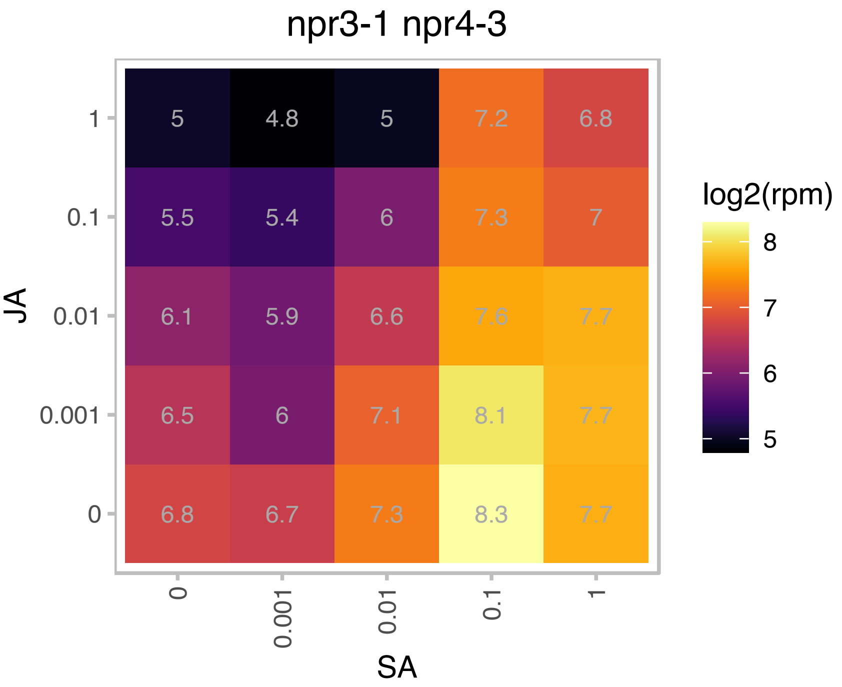

### AT5G56870.1 : BGAL4

beta-galactosidase 4

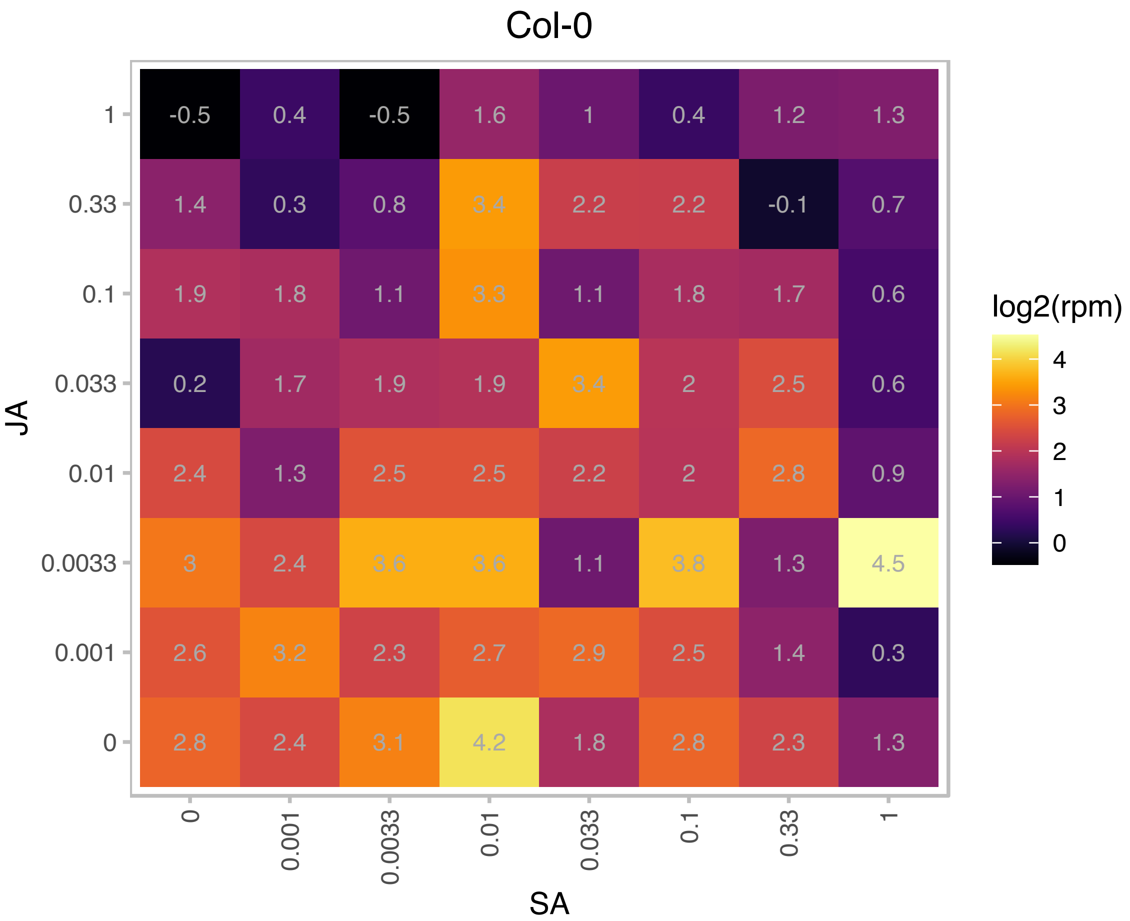


| | |
|---|---|
| Gene | AT5G56870 |
| is.expressed_genes | TRUE |
| is.responsive_genes | TRUE |
| is.apcluster | H-03 |

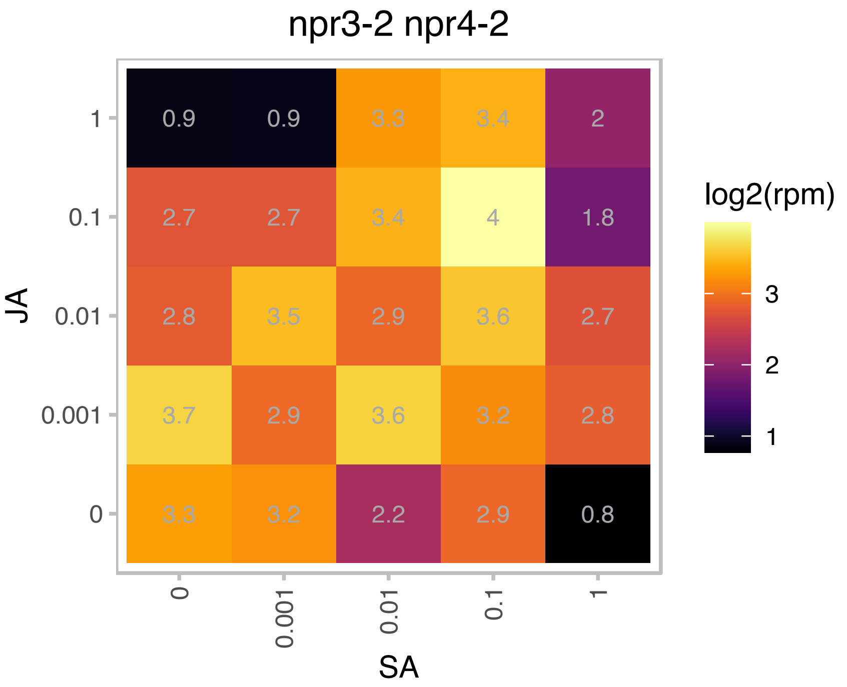


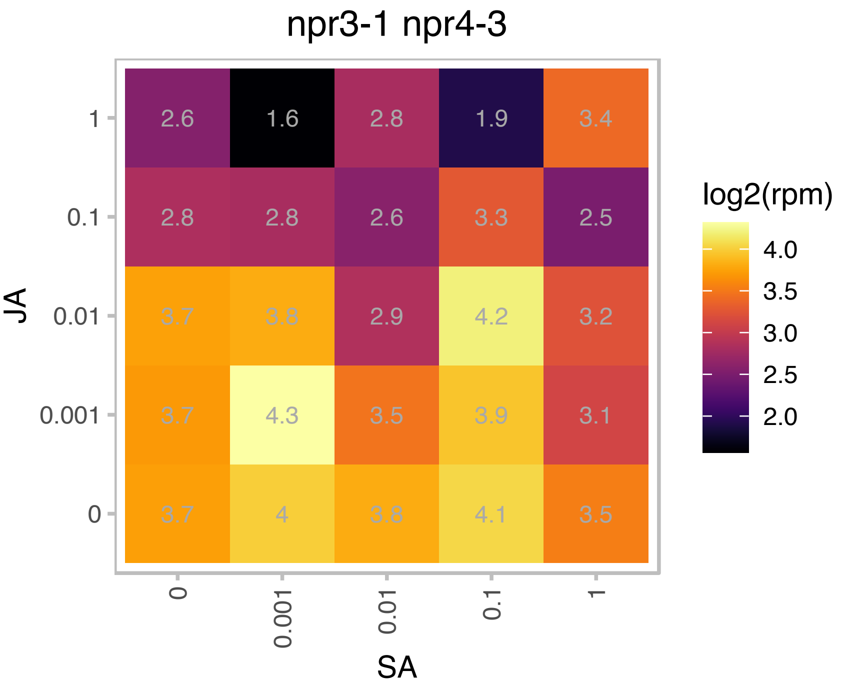

# AT5G57785.1 : NA

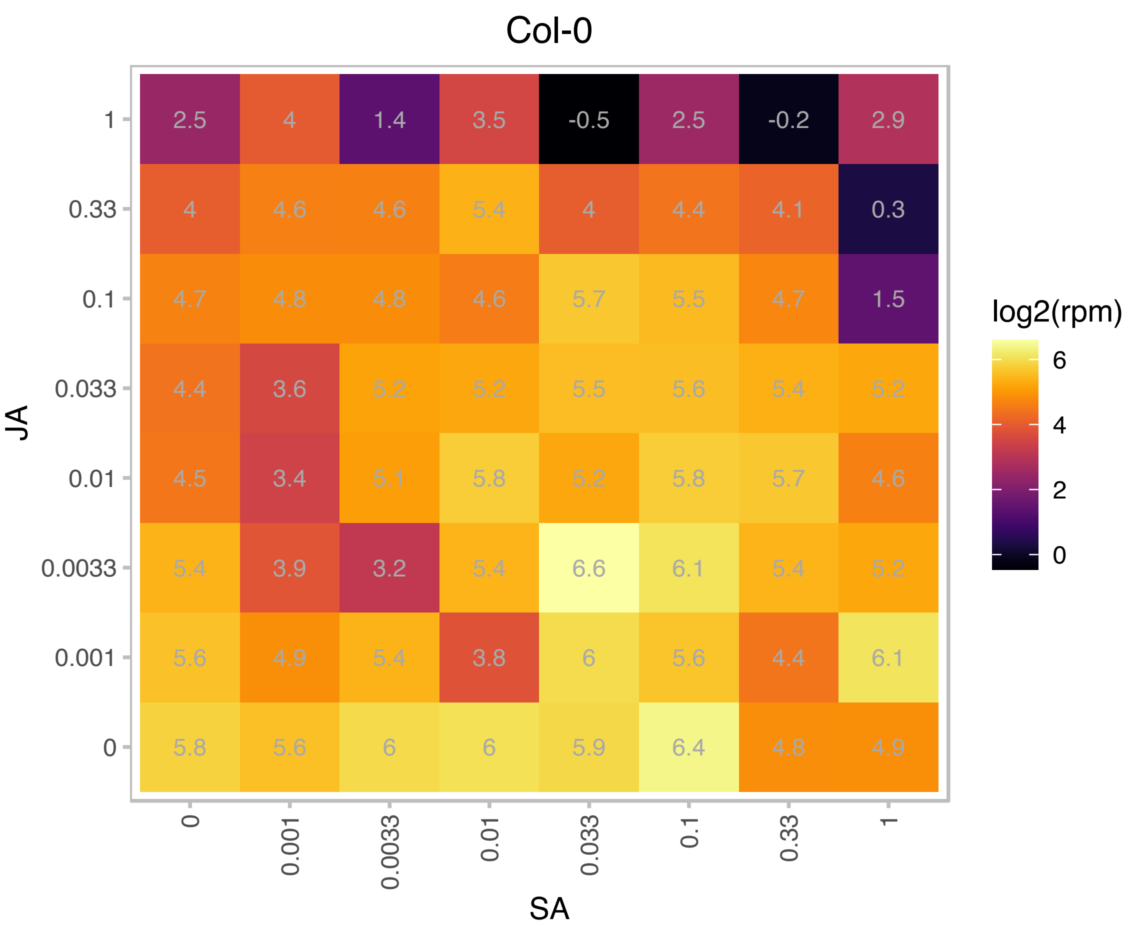


| | |
|---|---|
| Gene | AT5G57785 |
| is.expressed_genes | TRUE |
| is.responsive_genes | TRUE |
| is.apcluster | H-03 |

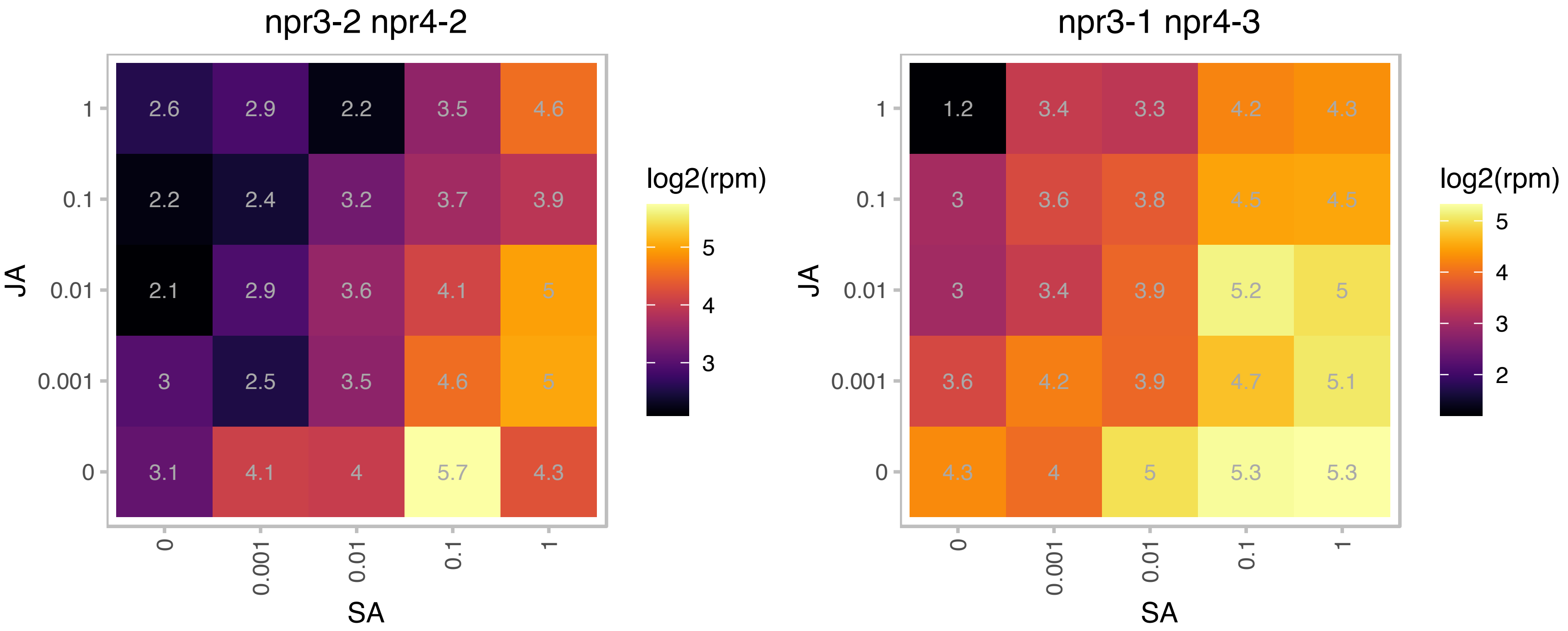

# AT5G62340.1 : NA

Plant invertase/pectin methylesterase inhibitor superfamily protein

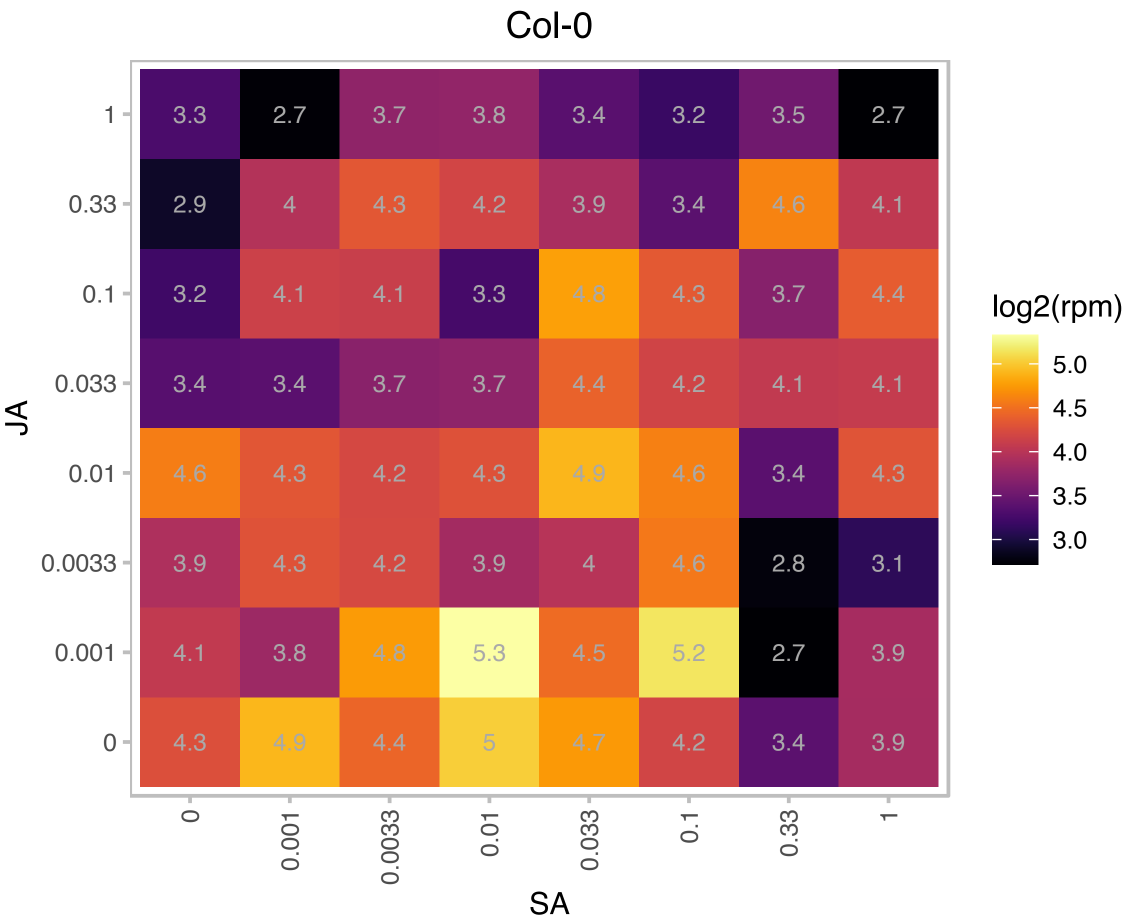


| | |
|---|---|
| Gene | AT5G62340 |
| is.expressed_genes | TRUE |
| is.responsive_genes | TRUE |
| is.apcluster | H-03 |

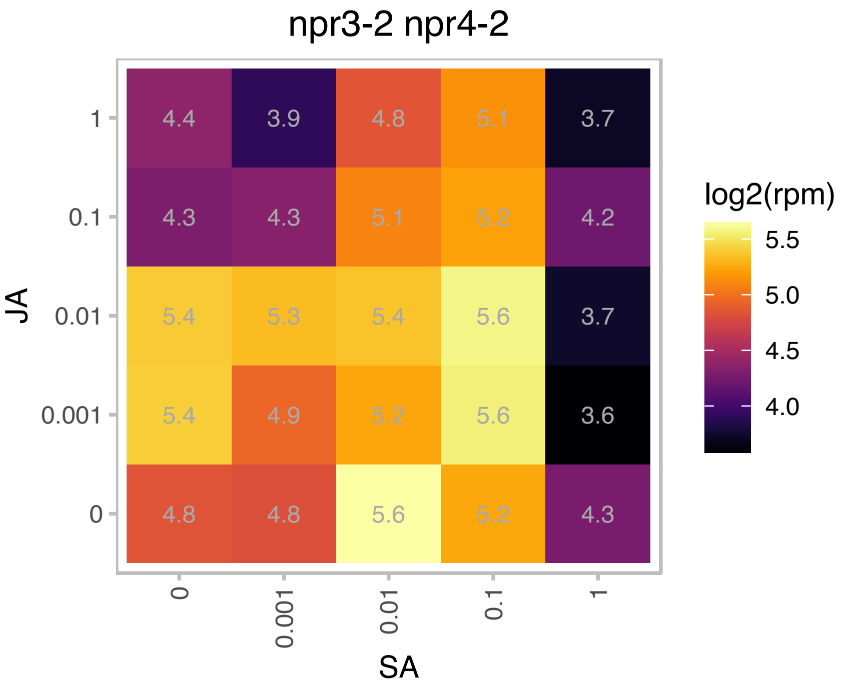


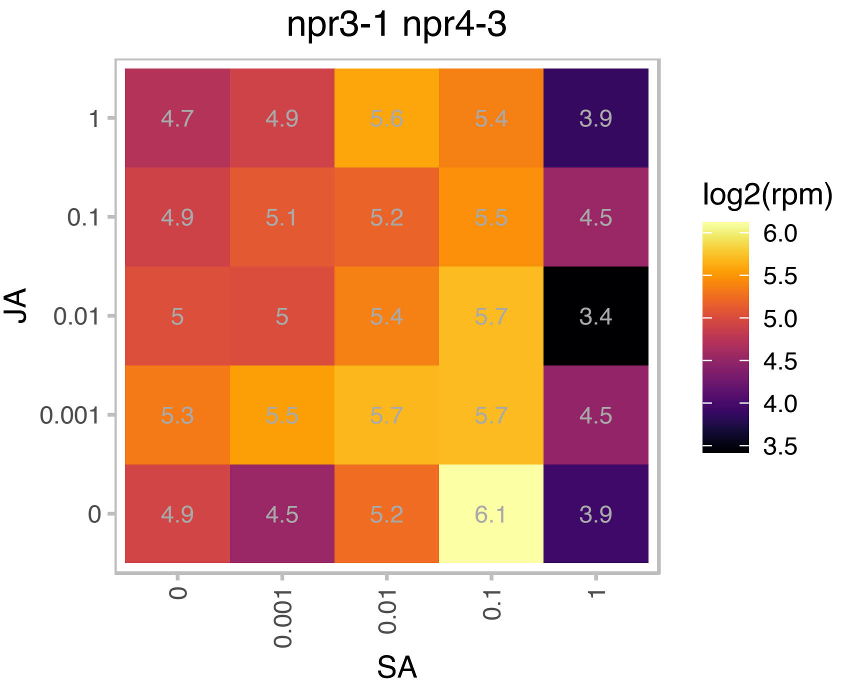

### AT5G63570.1 : GSA1

glutamate-1-semialdehyde-2,1-aminomutase
Encodes a protein with homology to glutamate-1-semialdehyde 2,1-aminomutase catalyzing the conversion of glutamate-1-semialdehyde (GSA) into 5-amino levulinate. The expression of this gene was demonstrated to be light-induced. The mRNA is cell-to-cell mobile.

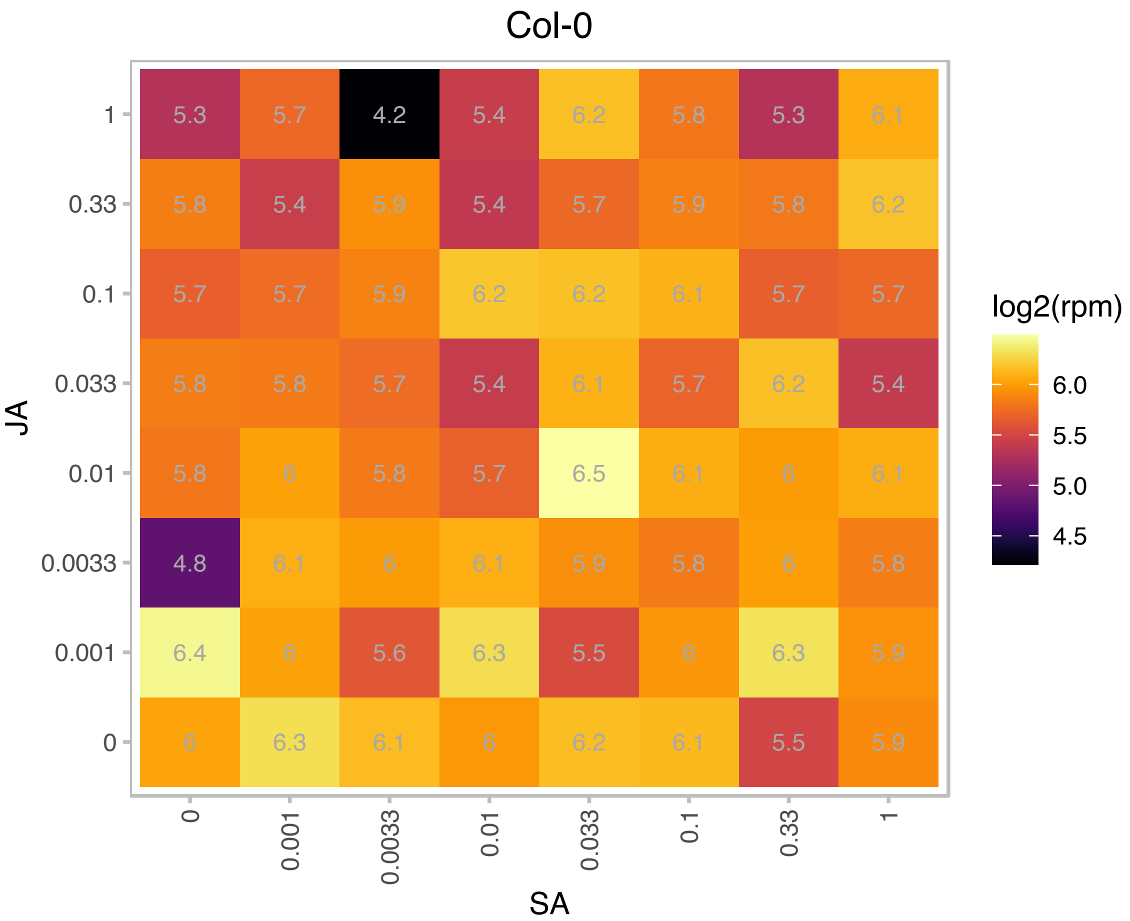


| | |
|---|---|
| Gene | AT5G63570 |
| is.expressed_genes | TRUE |
| is.responsive_genes | TRUE |
| is.apcluster | H-03 |

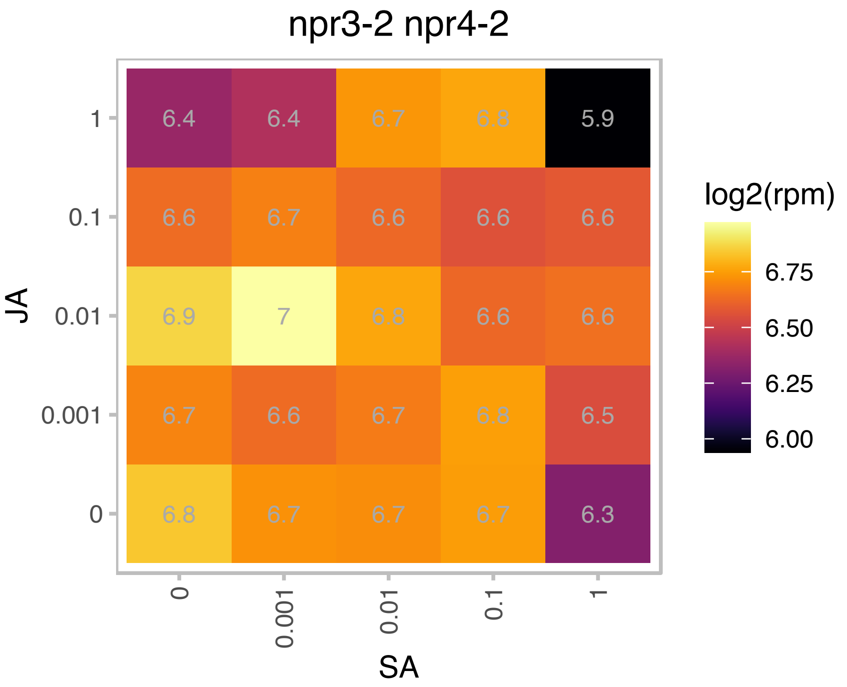


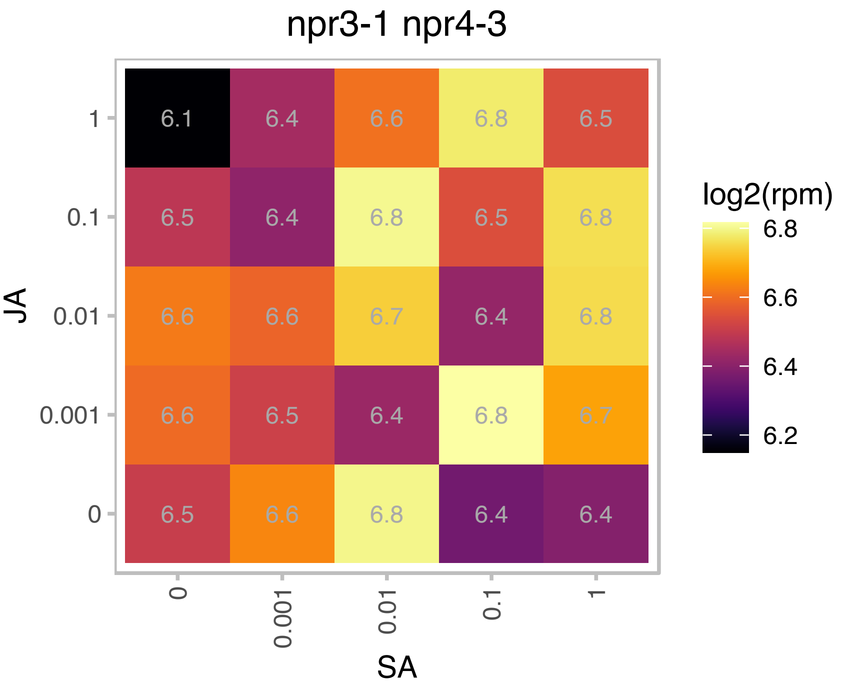

### AT1G01470.1 : AtLEA14, LEA1, LEA14, LSR3

Late embryogenesis abundant protein

Encodes late-embryogenesis abundant protein whose mRNA levels are induced in response to wounding and light stress. Might be involved in protection against desiccation.

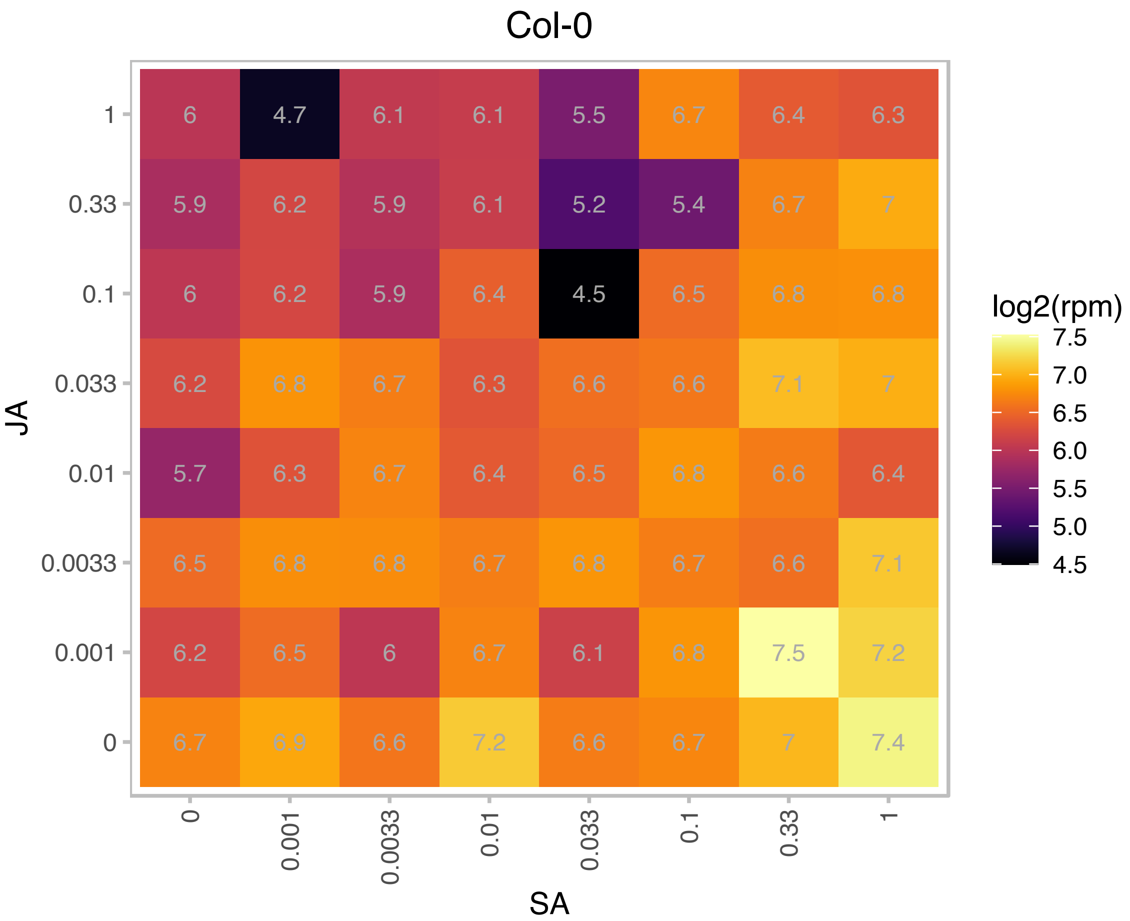


| | |
|---|---|
| Gene | AT1G01470 |
| is.expressed_genes | TRUE |
| is.responsive_genes | TRUE |
| is.apcluster | H-04 |

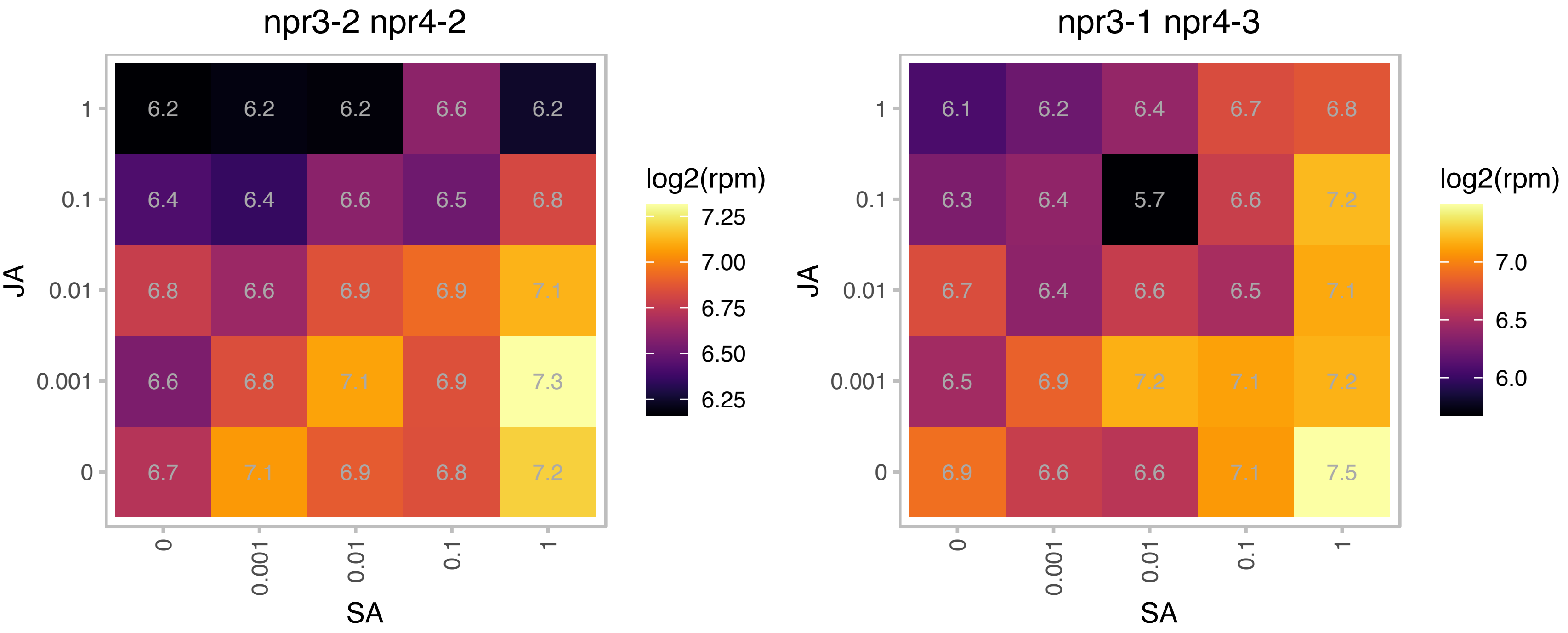

# AT1G09927.1 : NA

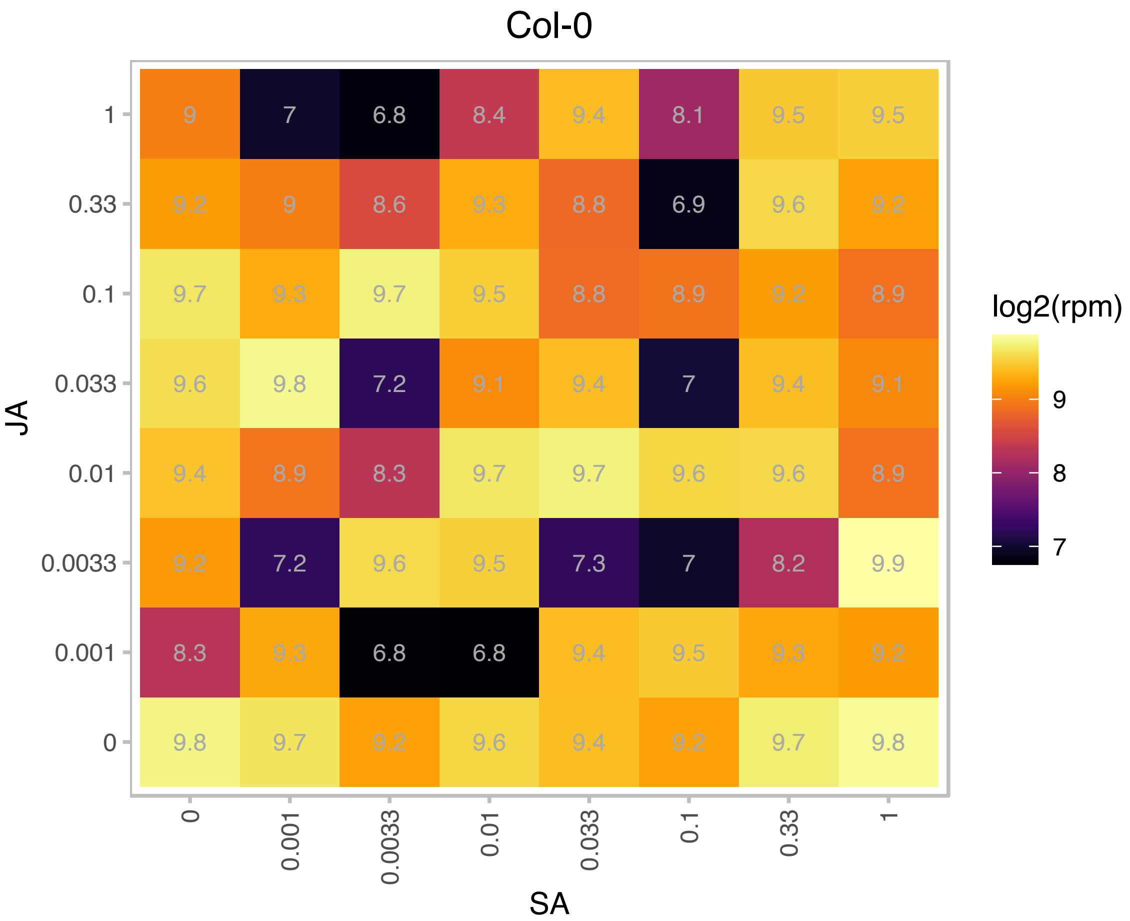


| | |
|---|---|
| Gene | AT1G09927 |
| is.expressed_genes | TRUE |
| is.responsive_genes | TRUE |
| is.apcluster | H-04 |

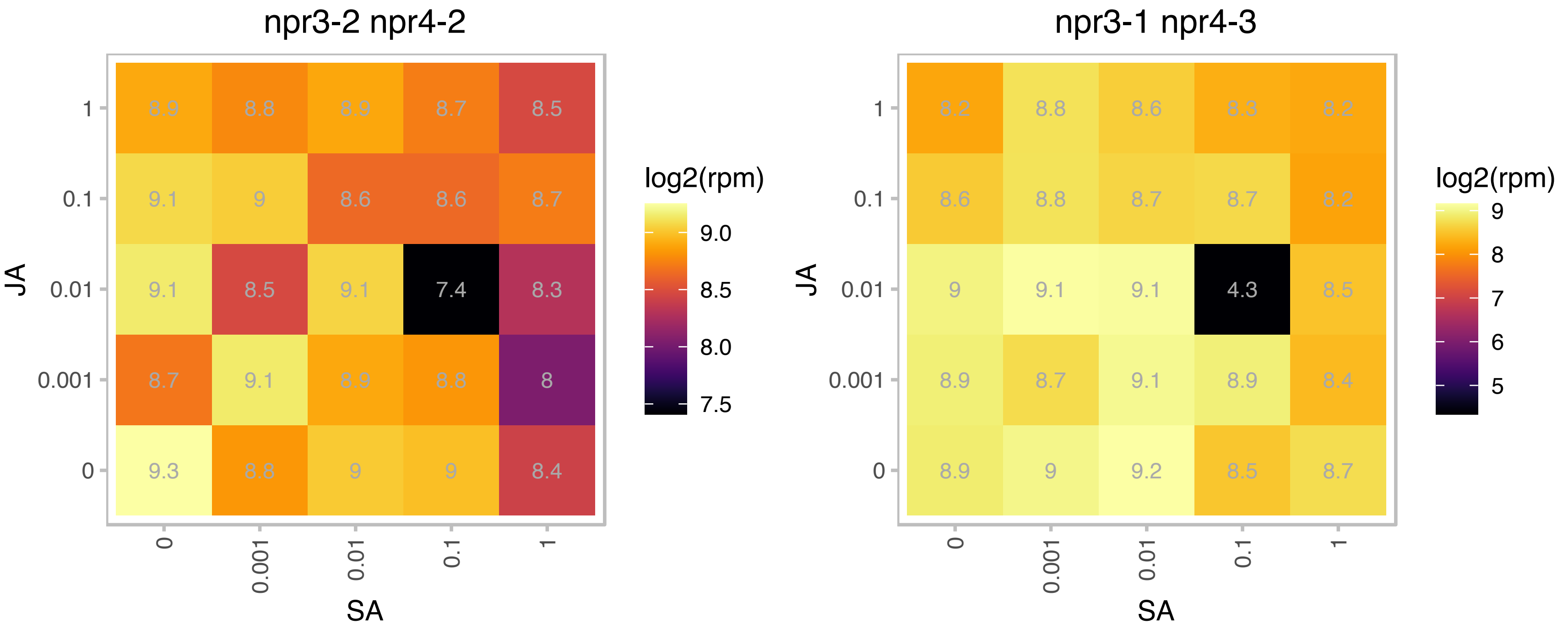

# AT1G24996.1 : NA

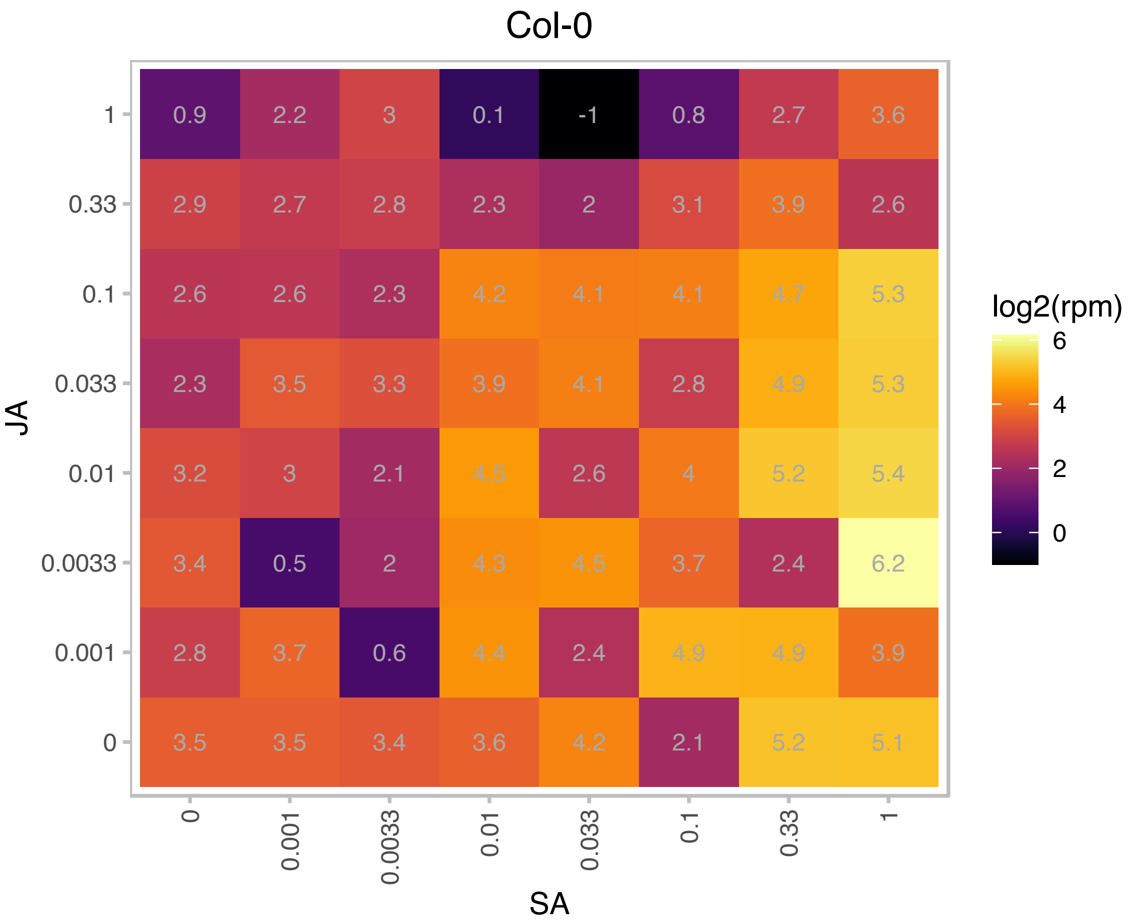


| | |
|---|---|
| Gene | AT1G24996 |
| is.expressed_genes | TRUE |
| is.responsive_genes | TRUE |
| is.apcluster | H-04 |

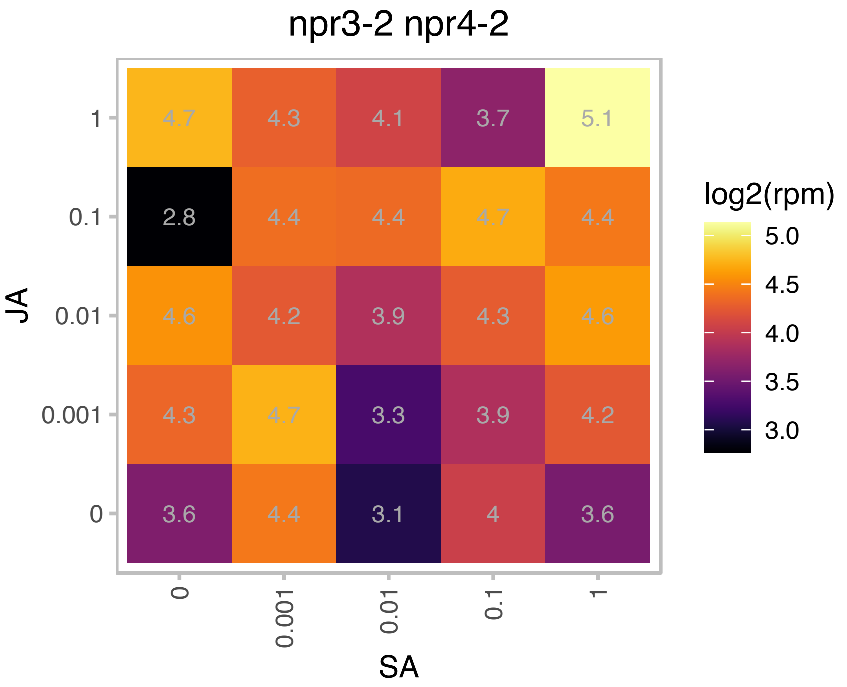


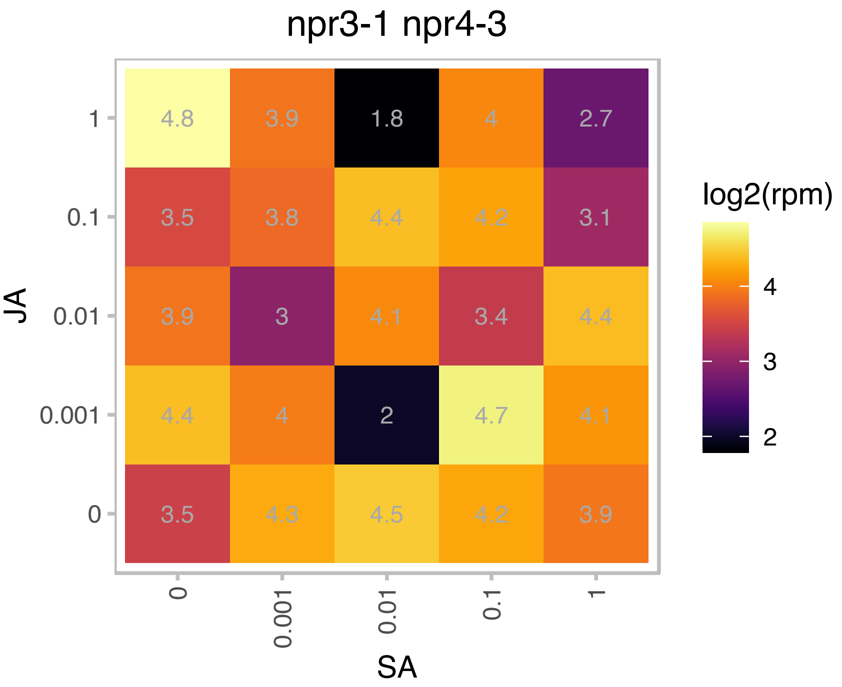

# AT1G53633.1 : NA

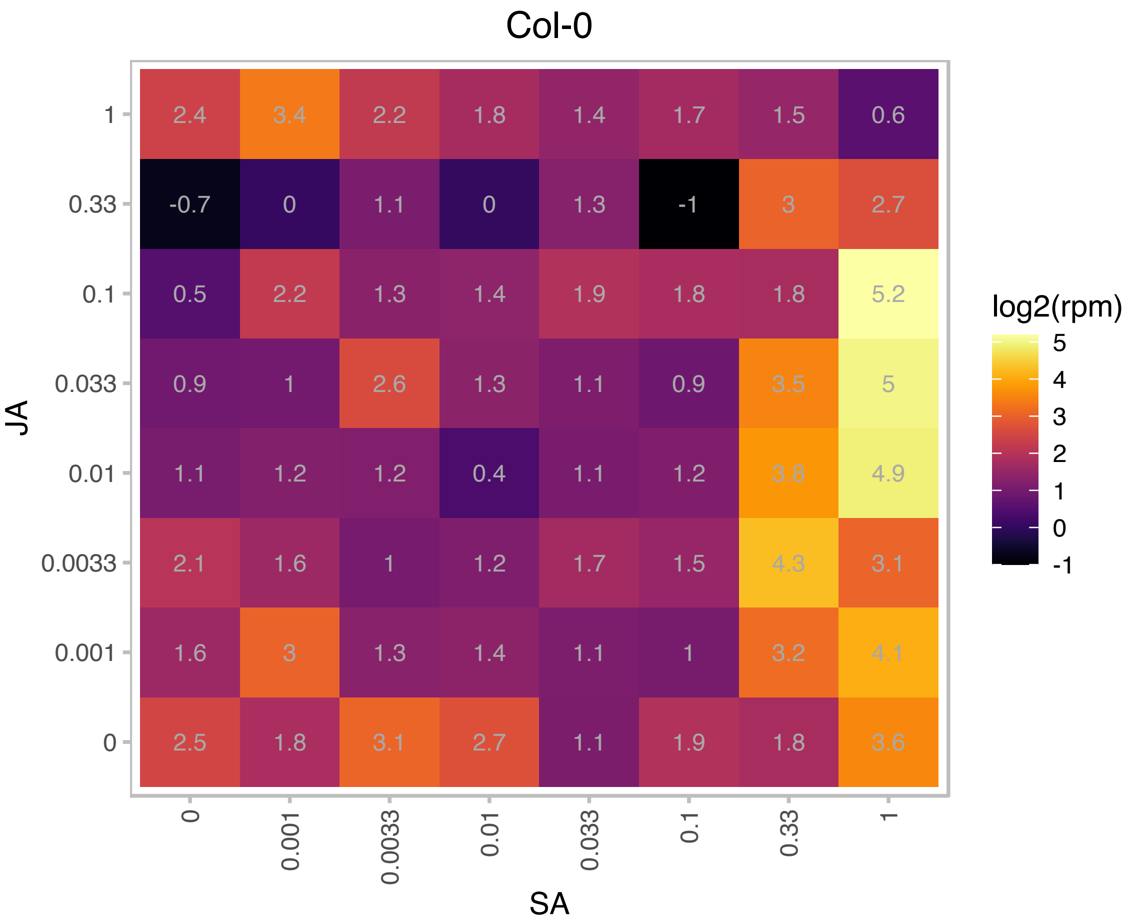


| | |
|---|---|
| Gene | AT1G53633 |
| is.expressed_genes | TRUE |
| is.responsive_genes | TRUE |
| is.apcluster | H-04 |

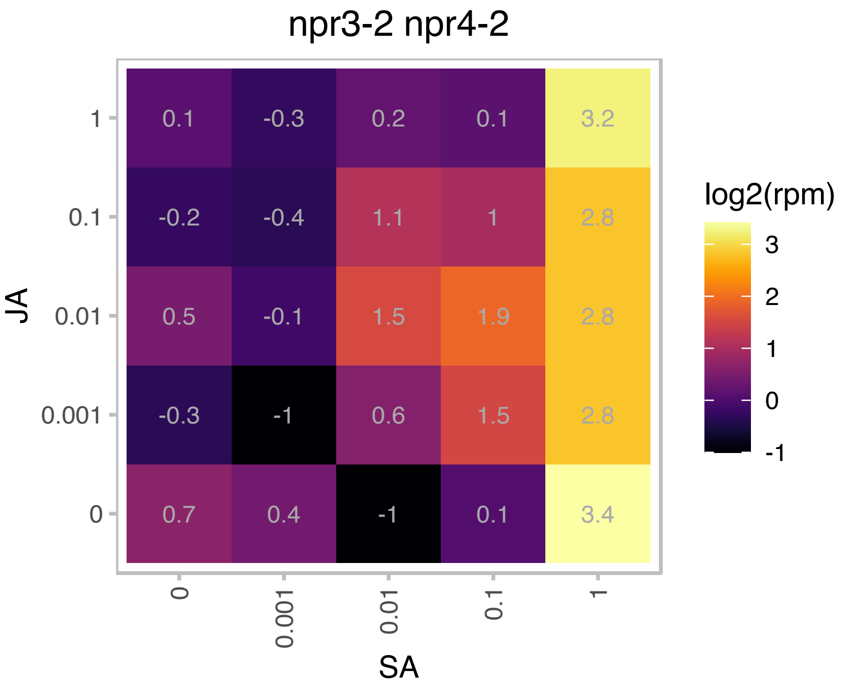


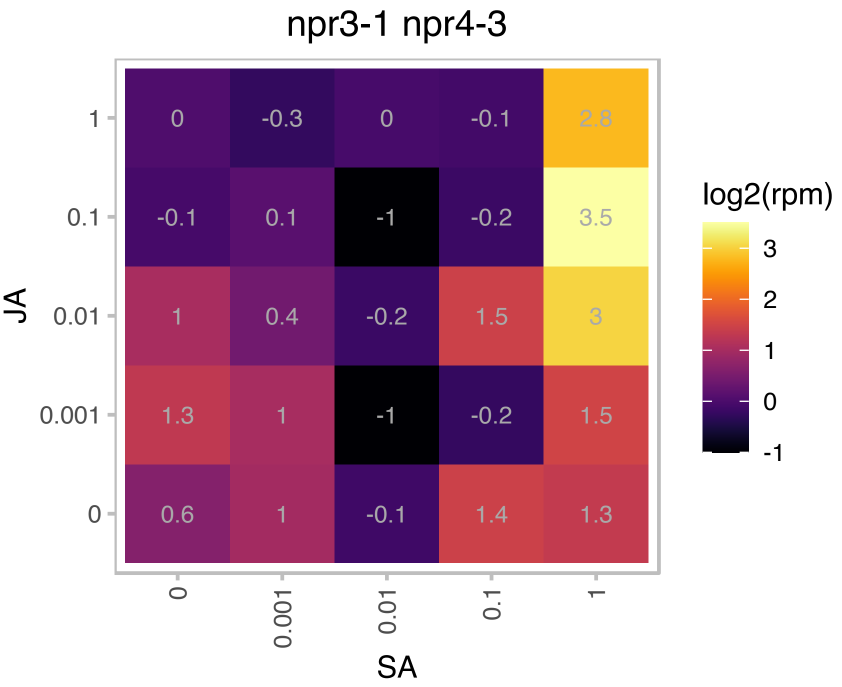

# AT1G55675.1 : NA

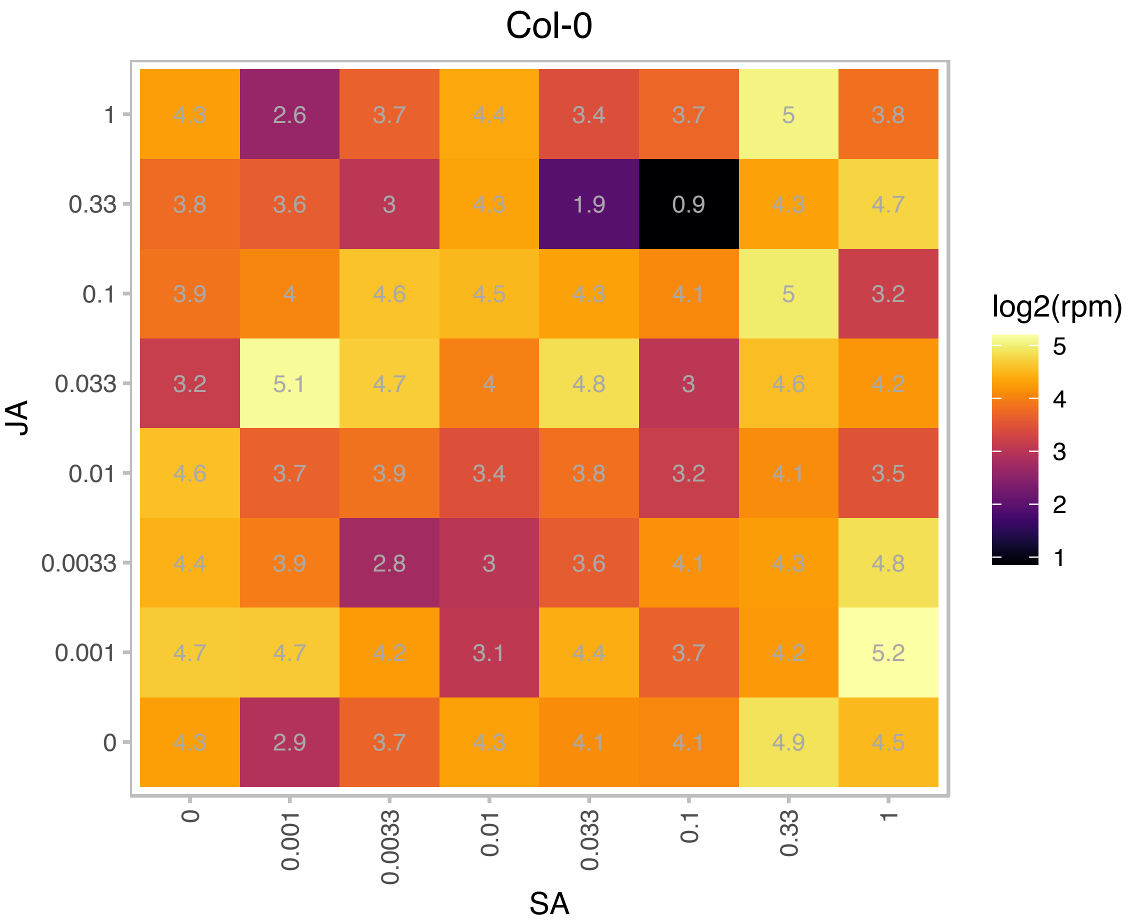


| | |
|---|---|
| Gene | AT1G55675 |
| is.expressed_genes | TRUE |
| is.responsive_genes | TRUE |
| is.apcluster | H-04 |

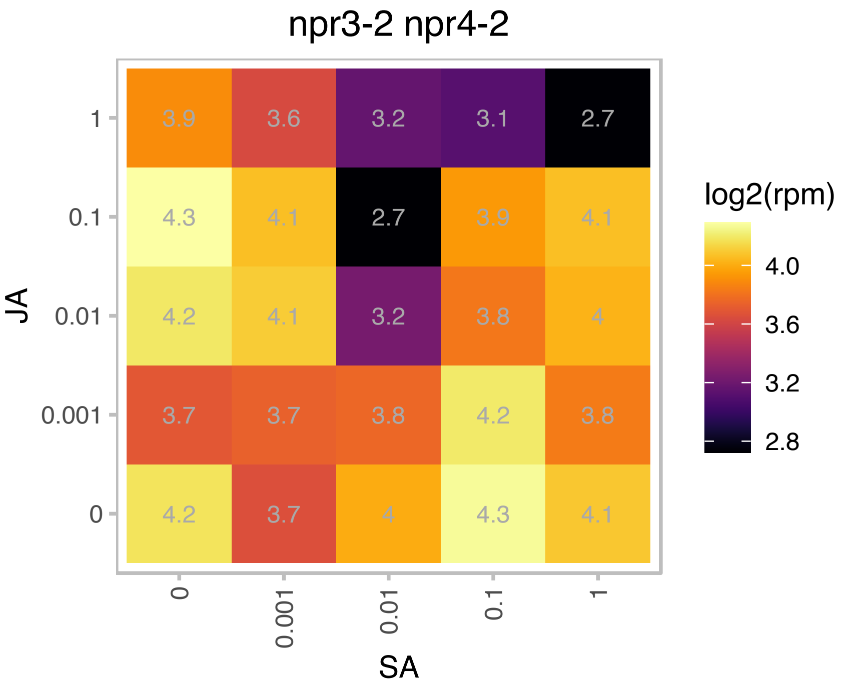


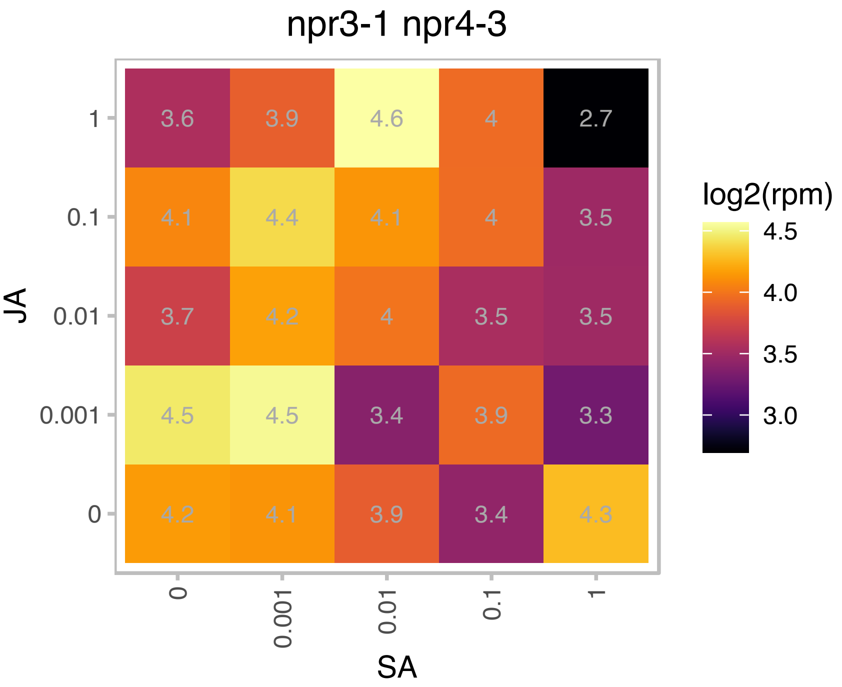

# AT1G67060.1 : NA

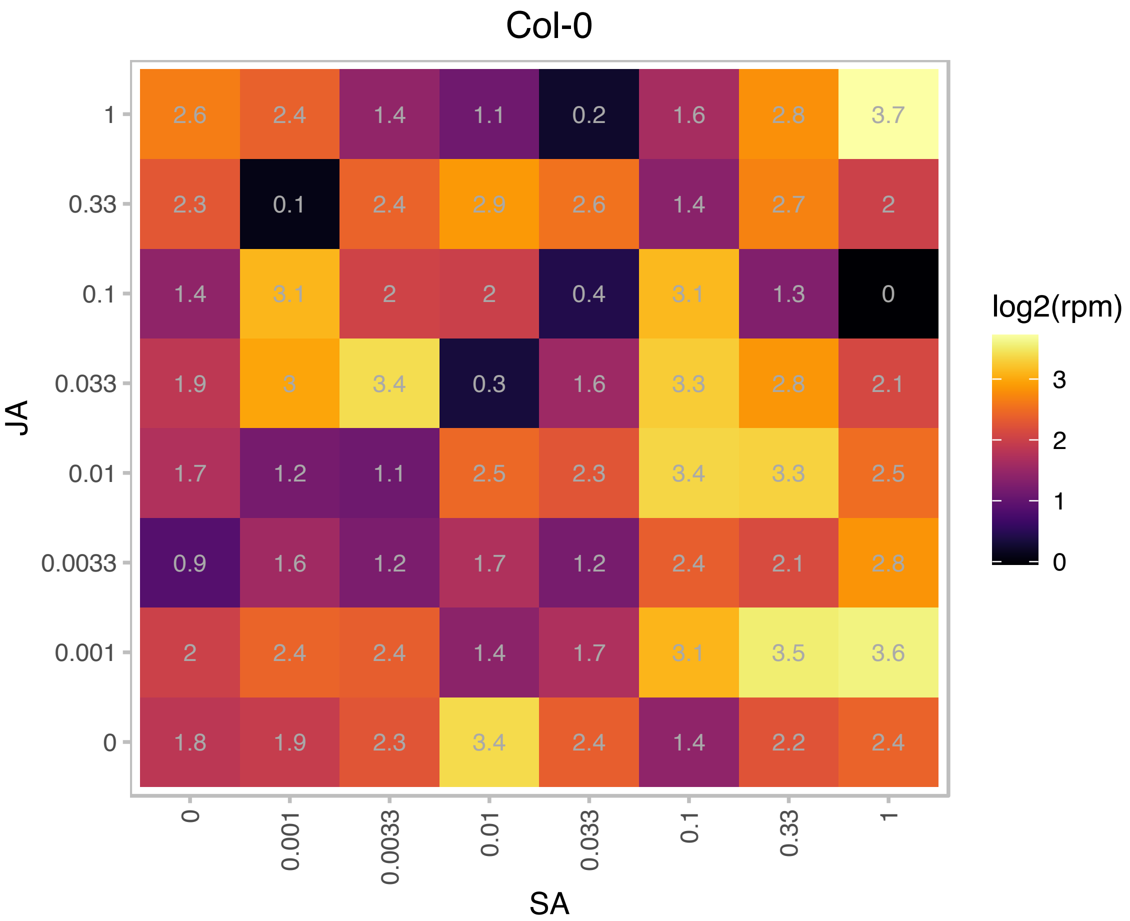


| | |
|---|---|
| Gene | AT1G67060 |
| is.expressed_genes | TRUE |
| is.responsive_genes | TRUE |
| is.apcluster | H-04 |

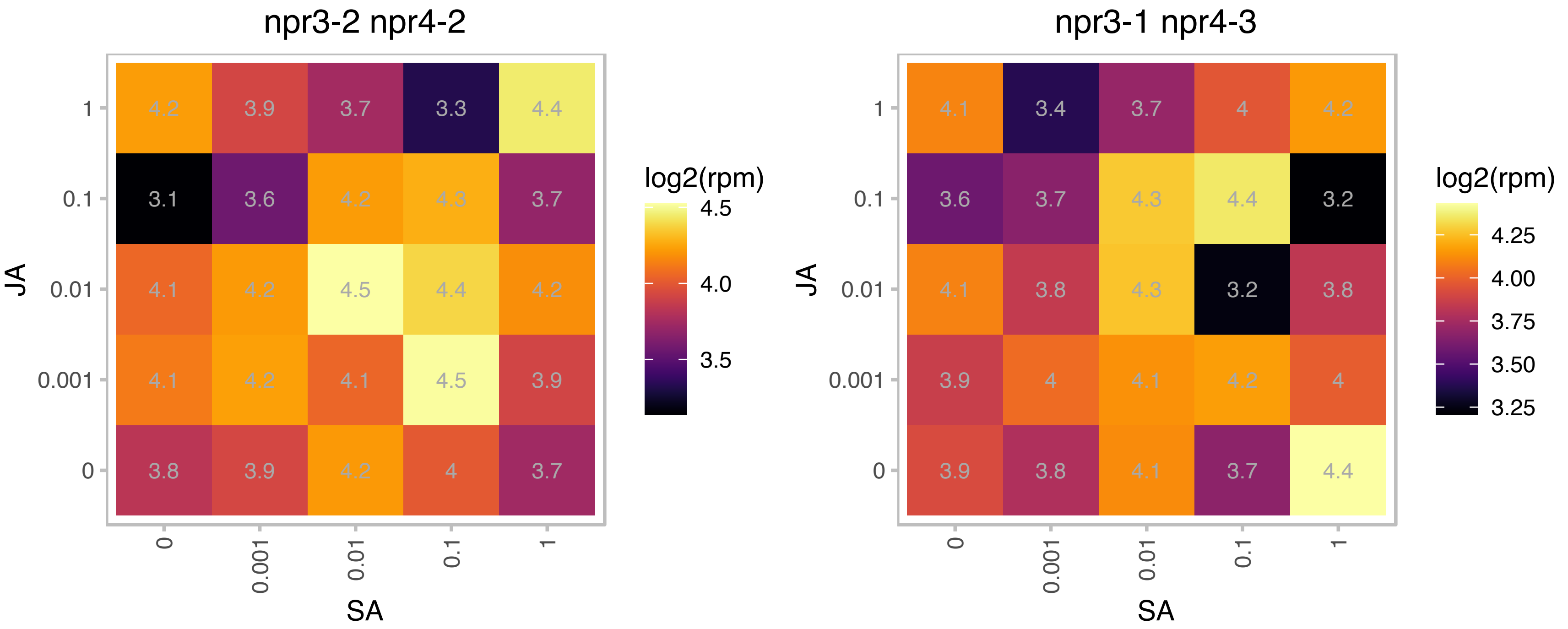

# AT1G80780.1 : NA

Polynucleotidyl transferase, ribonuclease H-like superfamily protein

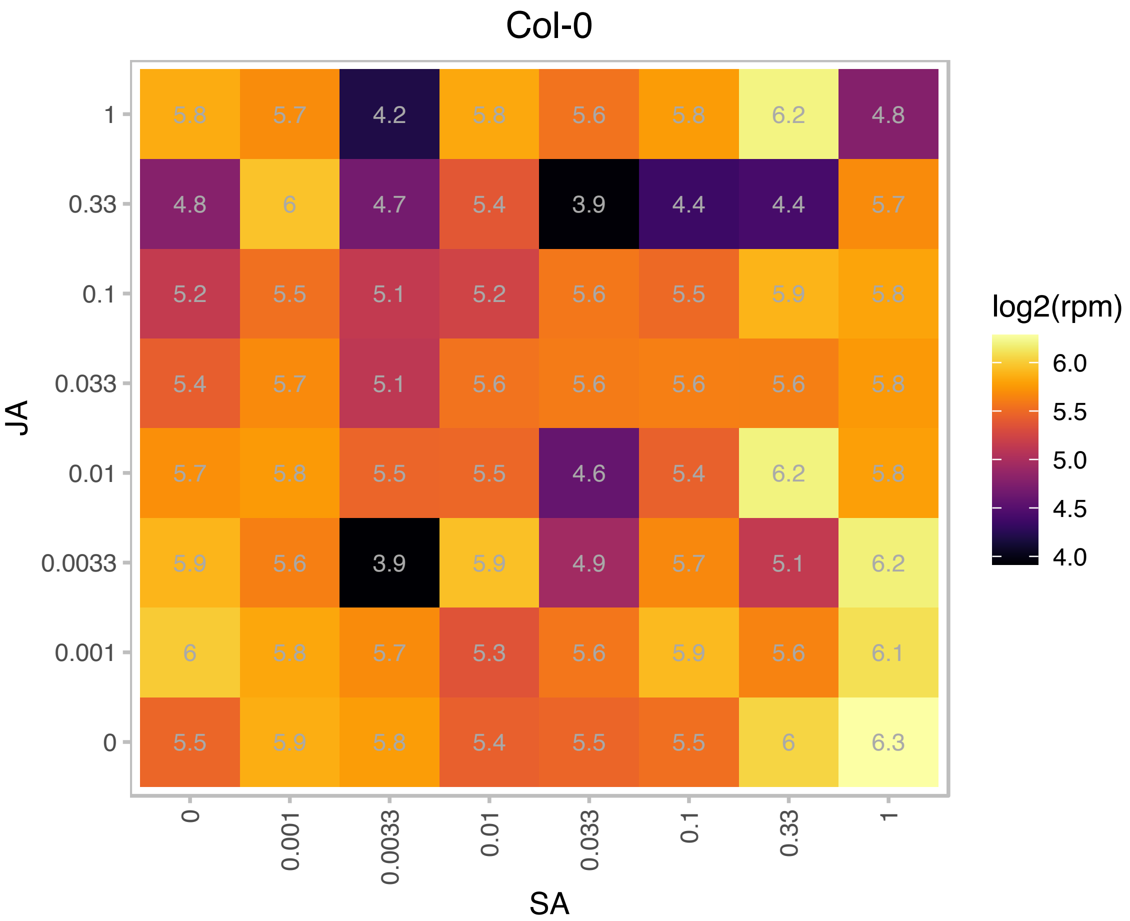


| | |
|---|---|
| Gene | AT1G80780 |
| is.expressed_genes | TRUE |
| is.responsive_genes | TRUE |
| is.apcluster | H-04 |

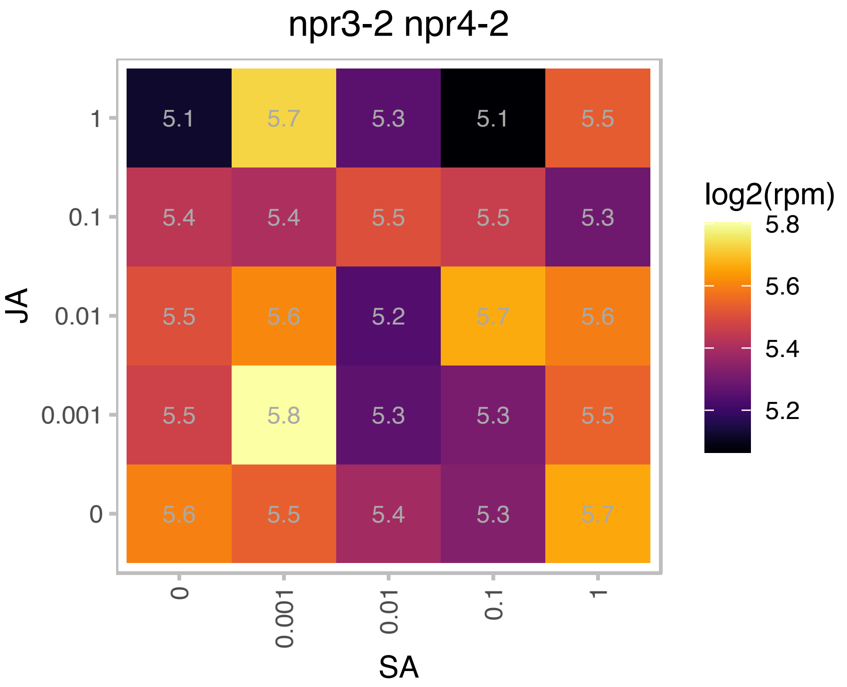


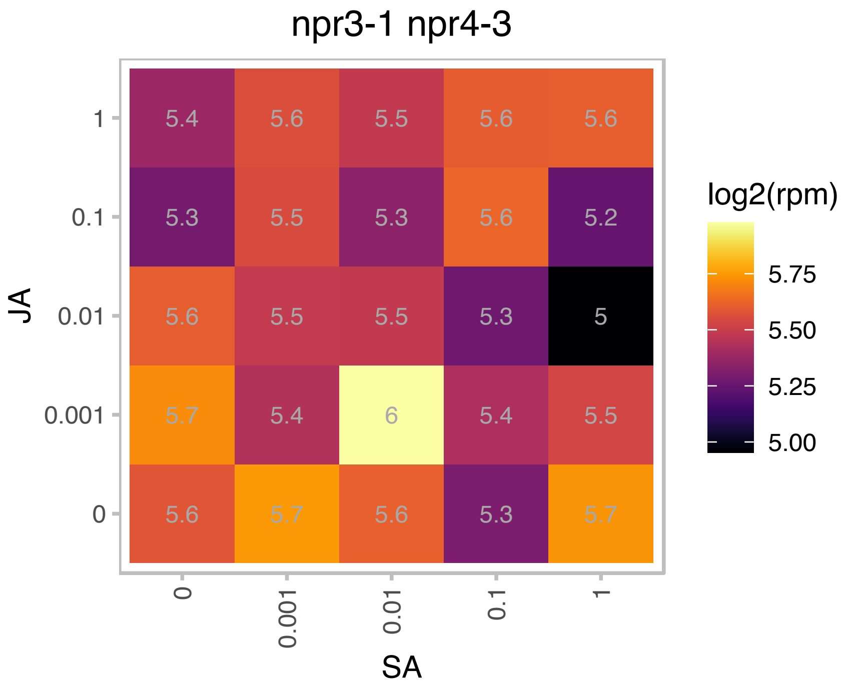

# AT2G05325.1 : NA

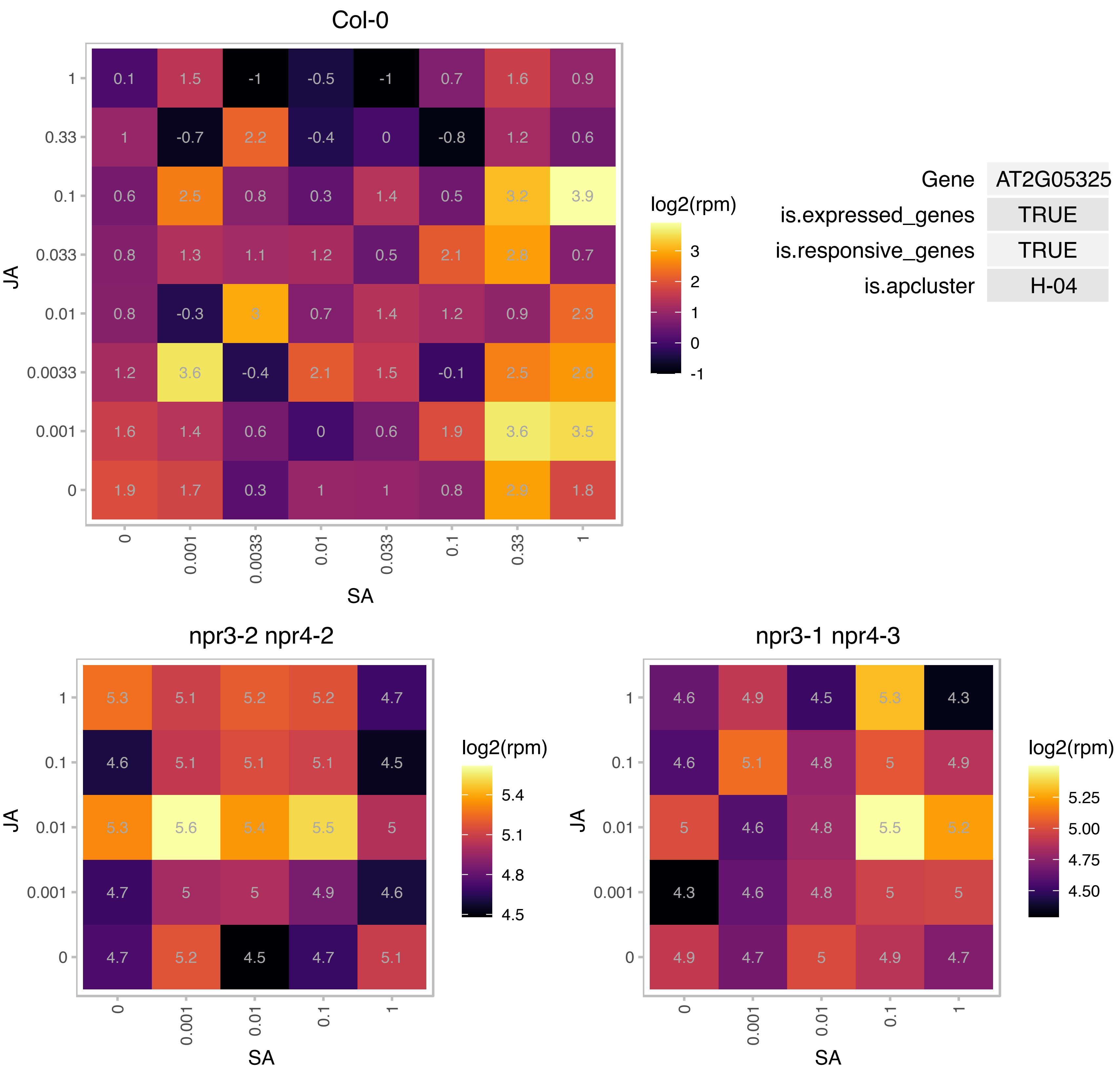

# AT2G05375.1 : NA

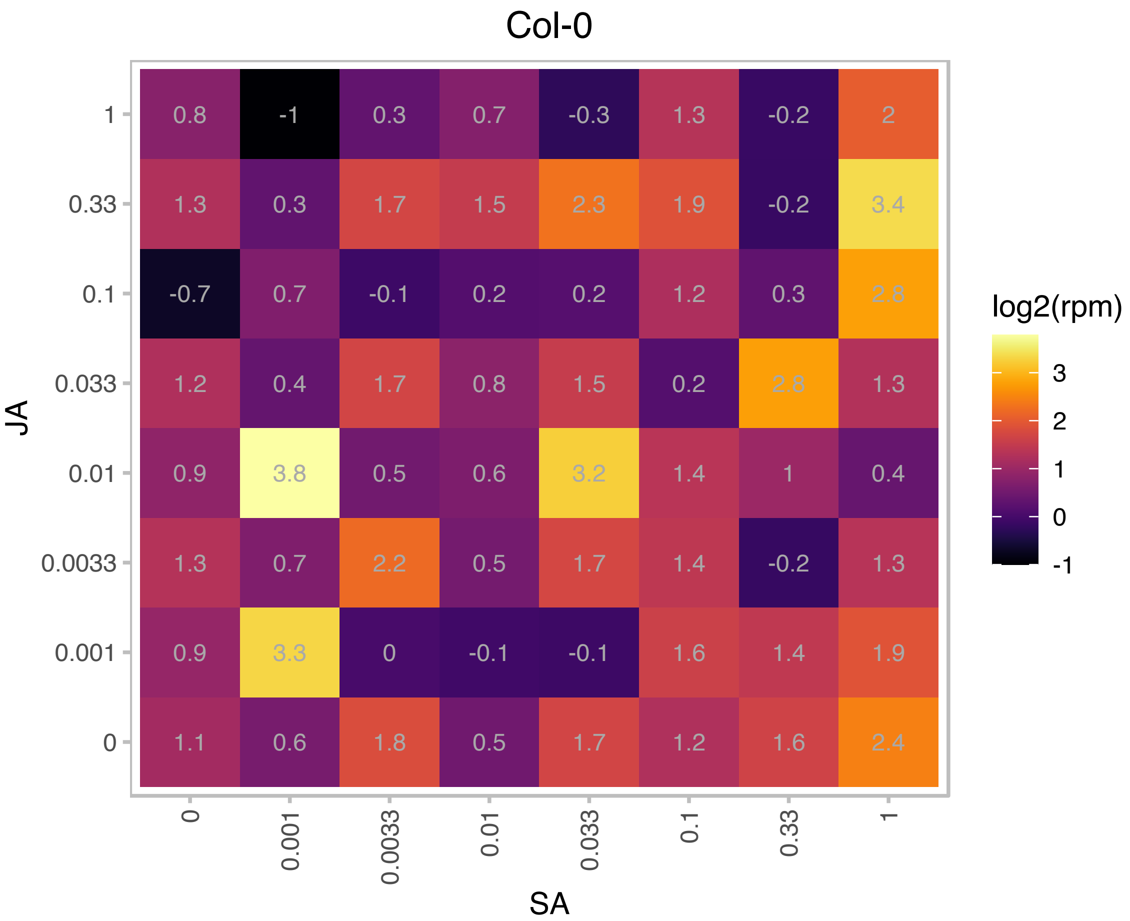


| | |
|---|---|
| Gene | AT2G05375 |
| is.expressed_genes | TRUE |
| is.responsive_genes | TRUE |
| is.apcluster | H-04 |

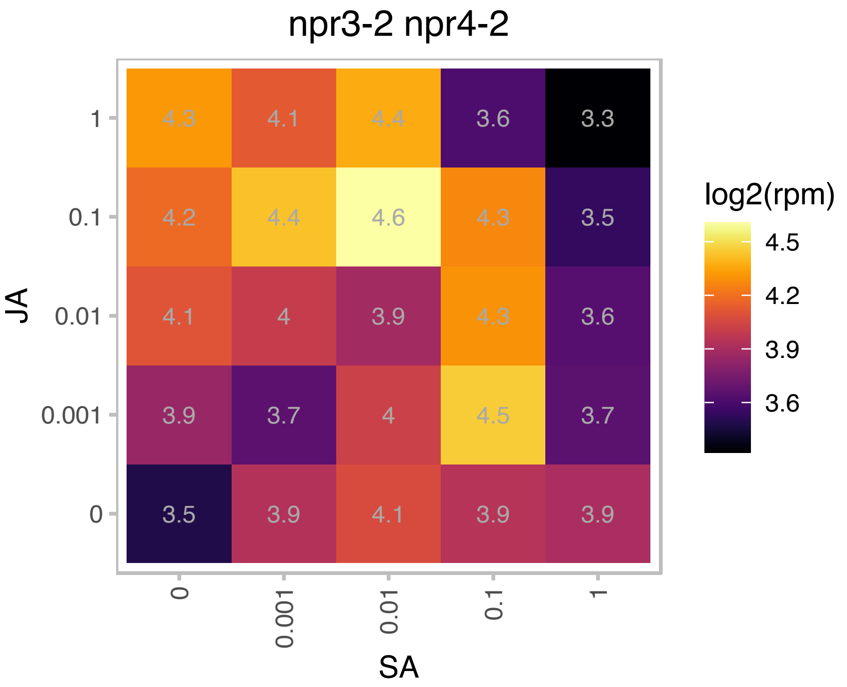


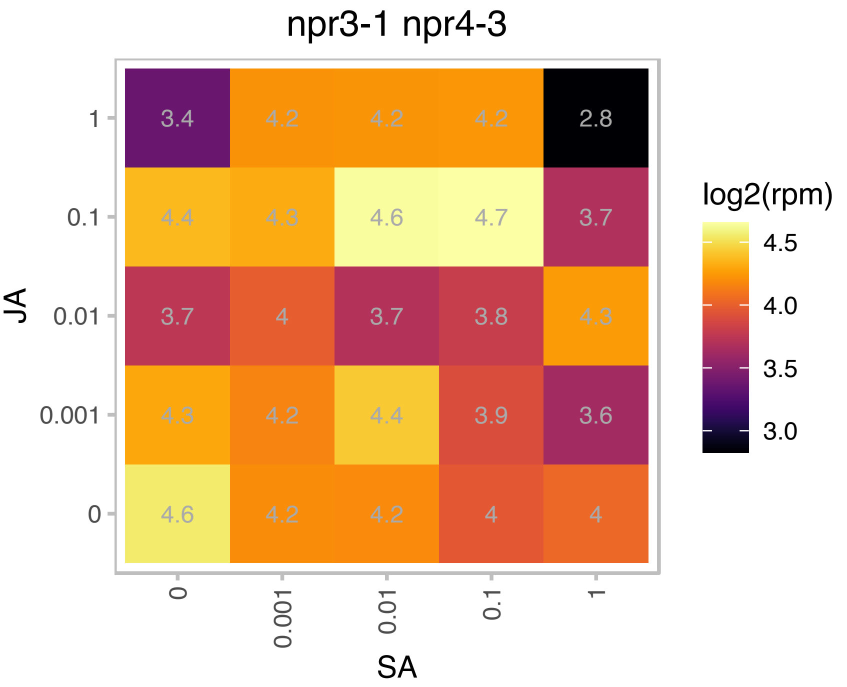

# AT2G16586.1 : NA

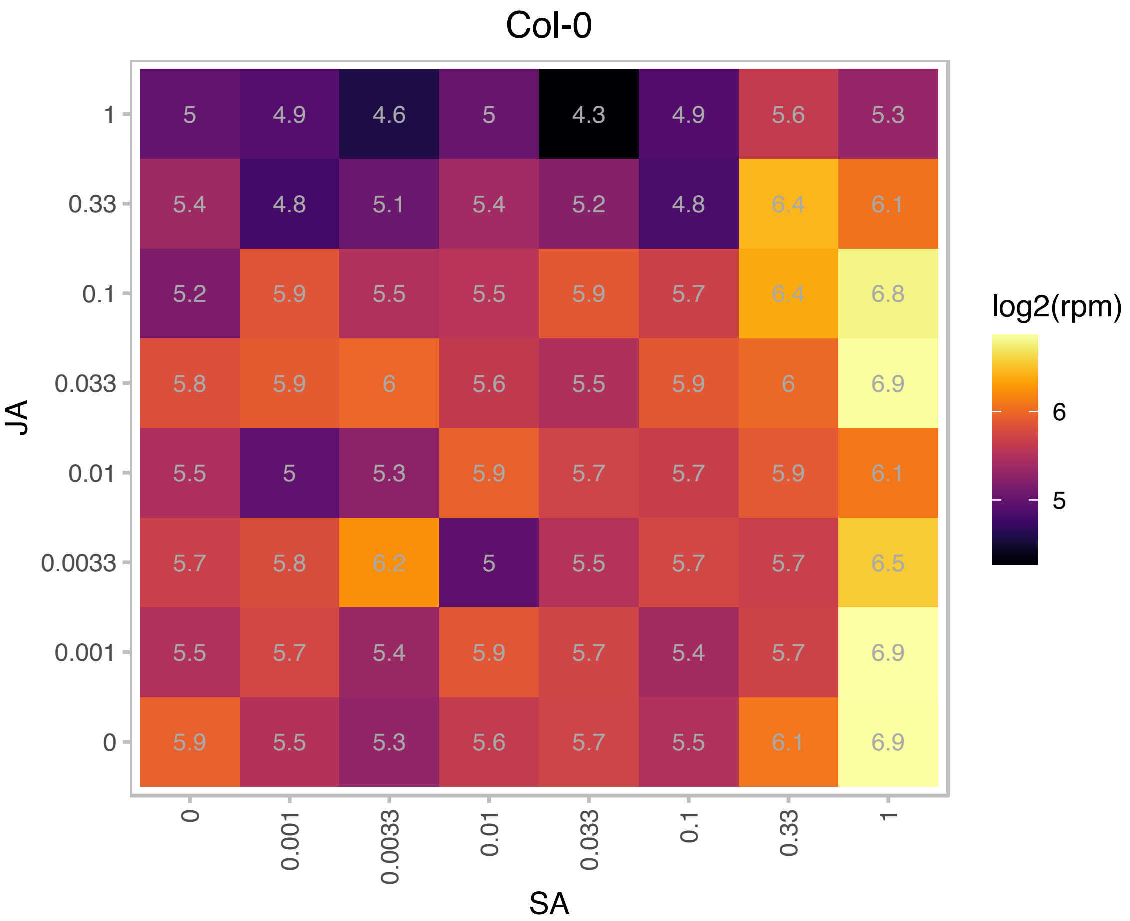


| | |
|---|---|
| Gene | AT2G16586 |
| is.expressed_genes | TRUE |
| is.responsive_genes | TRUE |
| is.apcluster | H-04 |

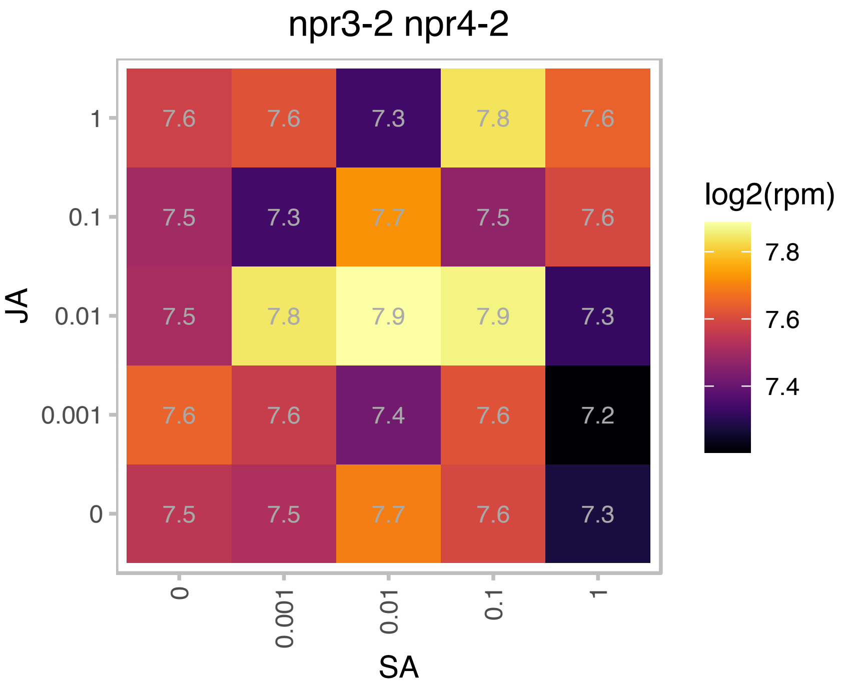


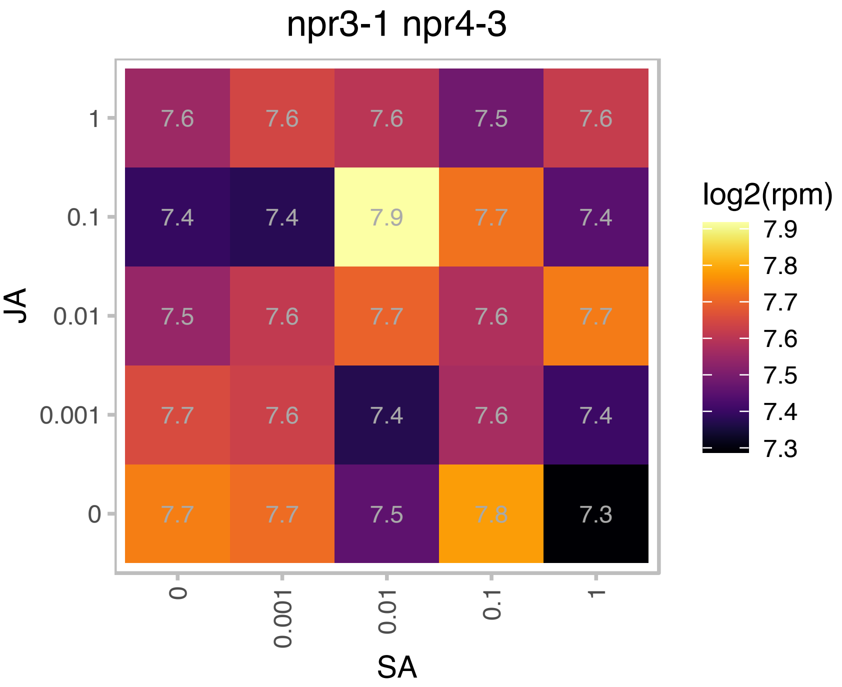

# AT2G35450.1 : NA

catalytics;hydrolases

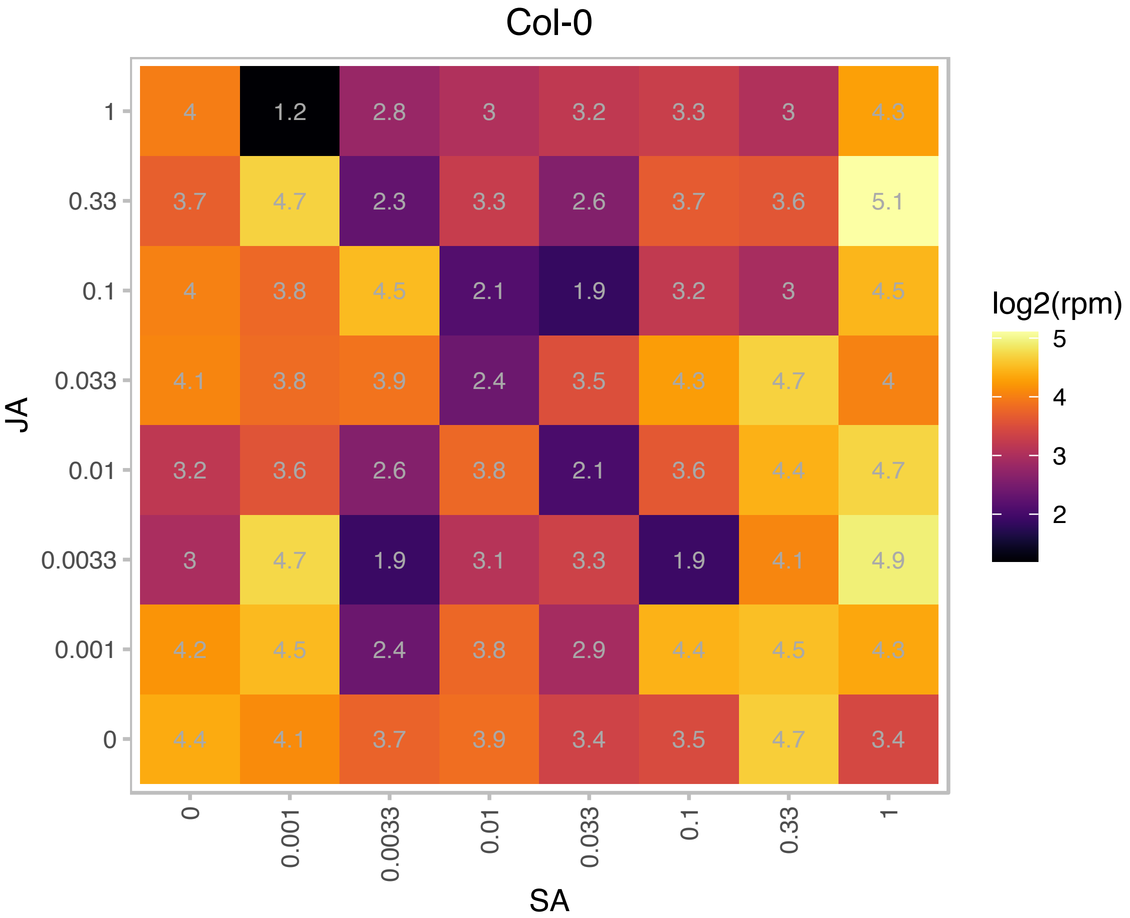


| | |
|---|---|
| Gene | AT2G35450 |
| is.expressed_genes | TRUE |
| is.responsive_genes | TRUE |
| is.apcluster | H-04 |

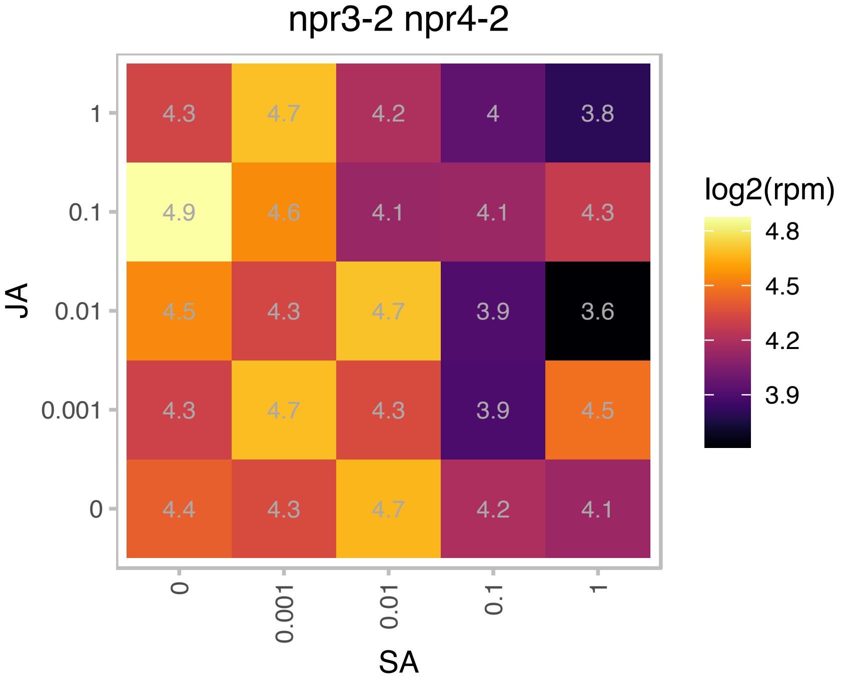


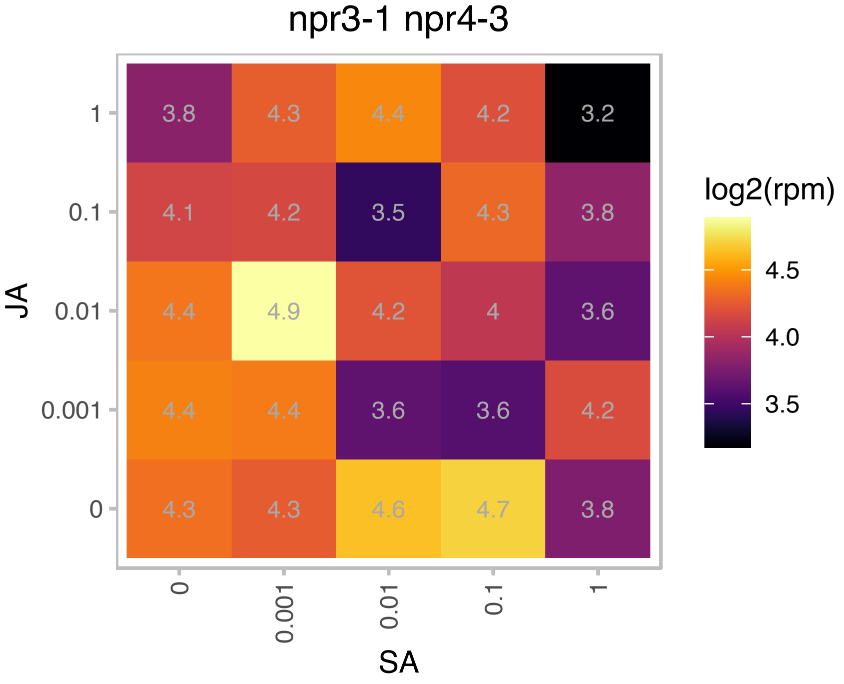

# AT2G39670.1 : NA

Radical SAM superfamily protein

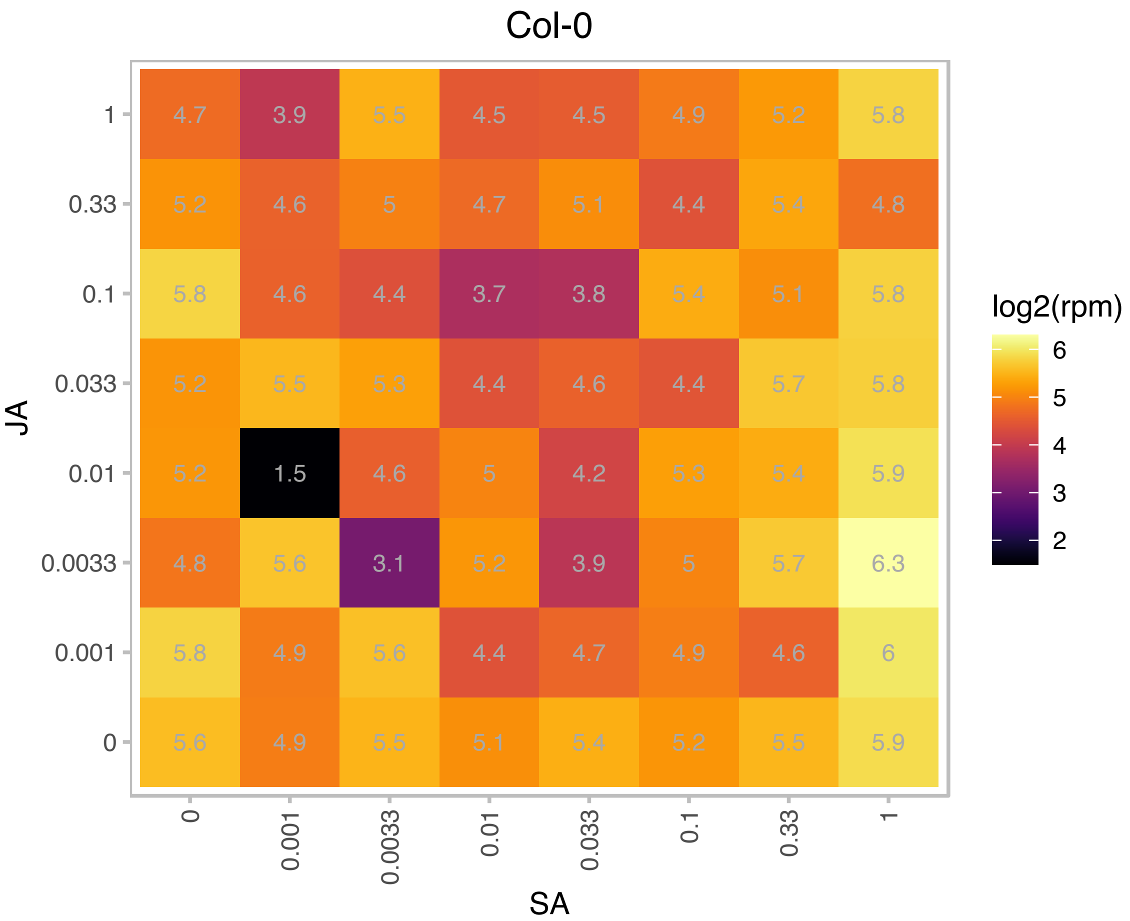


| | |
|---|---|
| Gene | AT2G39670 |
| is.expressed_genes | TRUE |
| is.responsive_genes | TRUE |
| is.apcluster | H-04 |

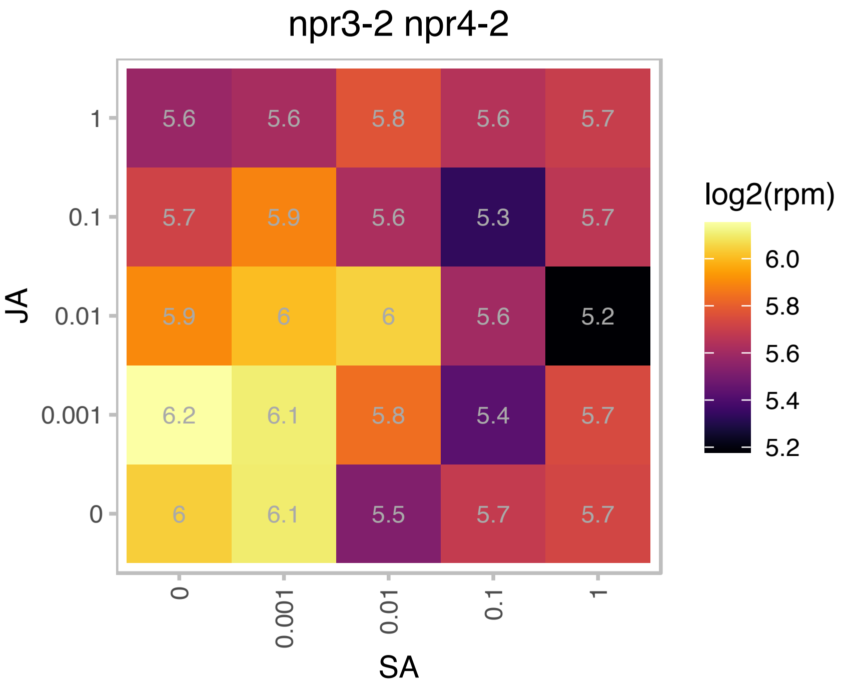


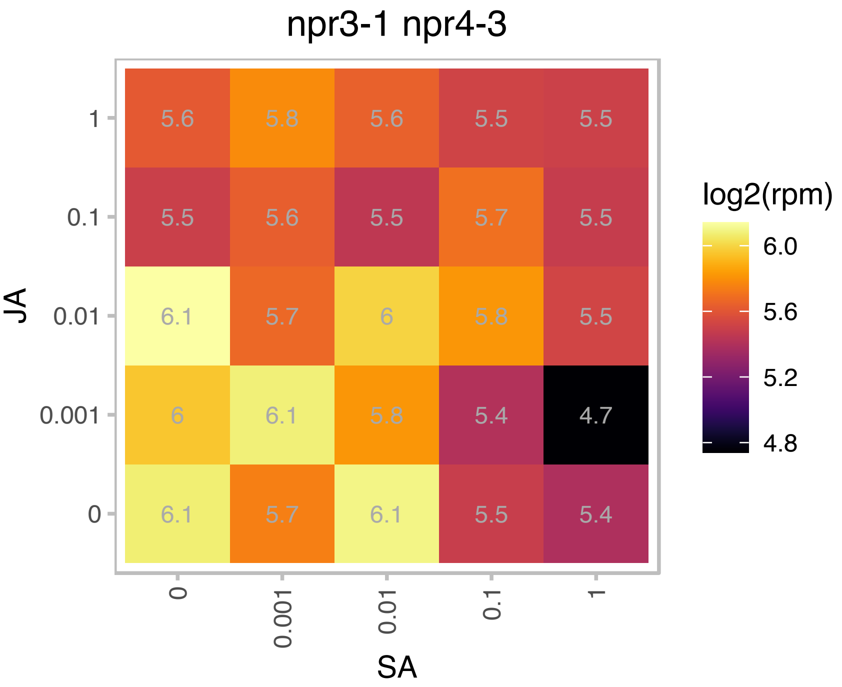

### AT3G51520.1 : AtDGAT2, DGAT2

diacylglycerol acyltransferase family
Encodes a functional acyl-CoA:diacylglycerol acyltransferase with different acyl-CoA substrate preferences and shows higher DAG to TAG conversion rate than AtDGAT1. It increases both C18:2 and C18:3 polyunsaturated fatty acids at the expense of C16:0.

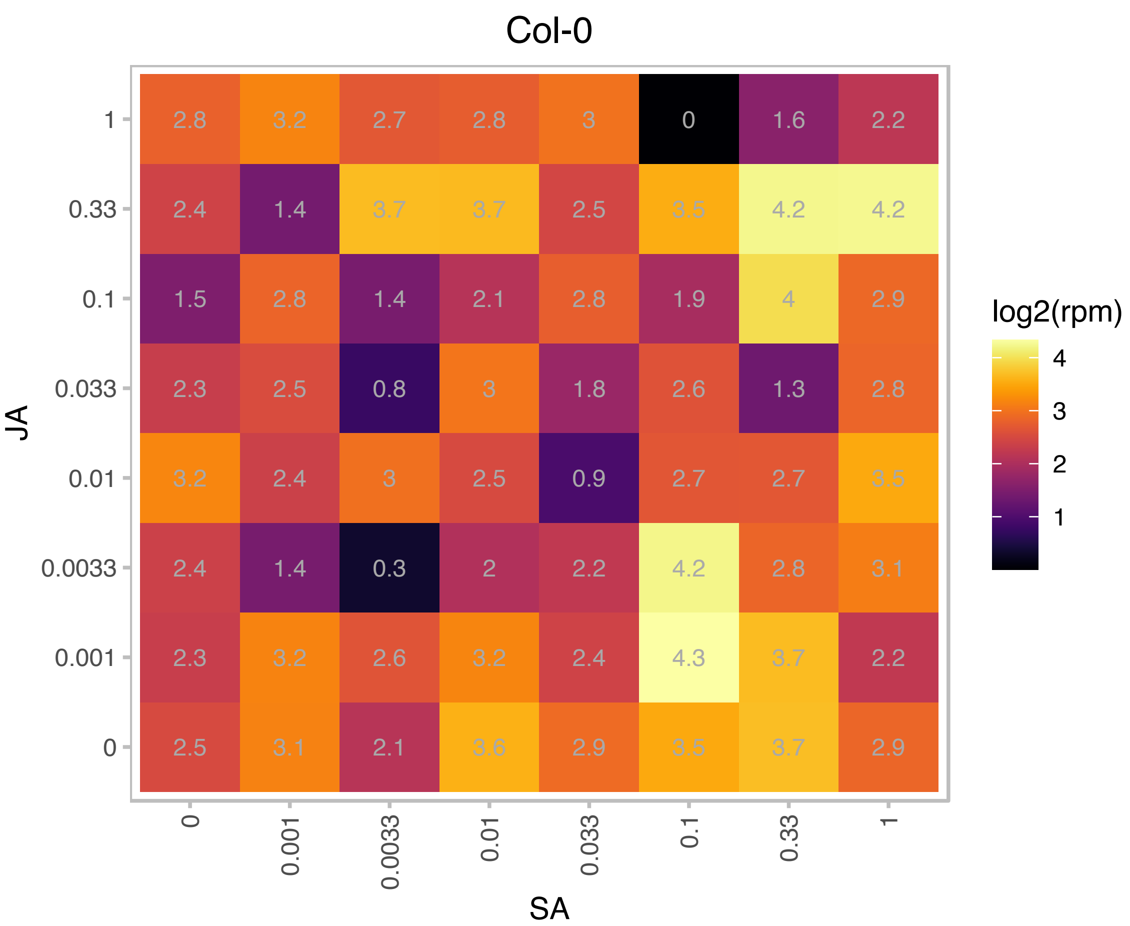


| | |
|---|---|
| Gene | AT3G51520 |
| is.expressed_genes | TRUE |
| is.responsive_genes | TRUE |
| is.apcluster | H-04 |

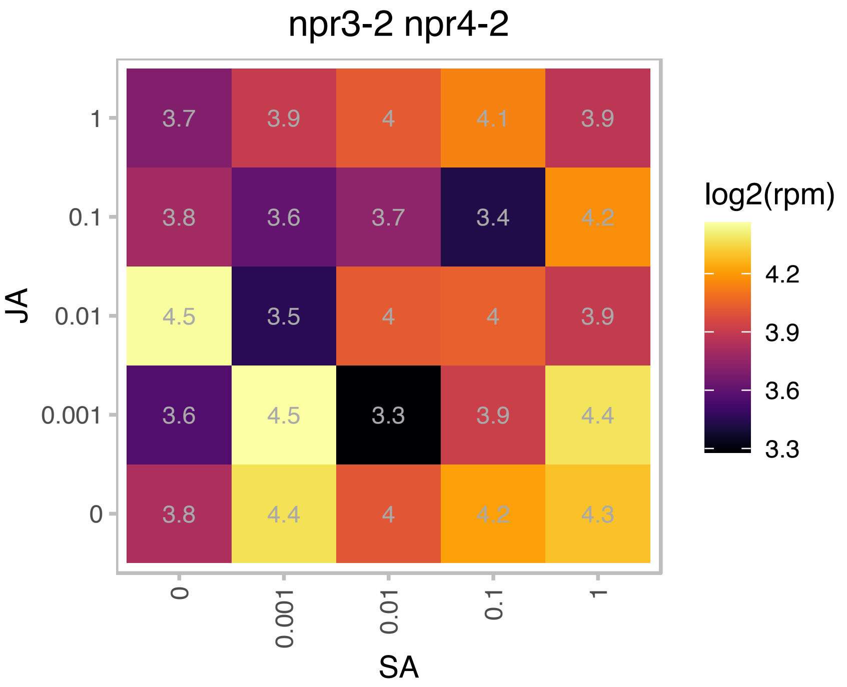


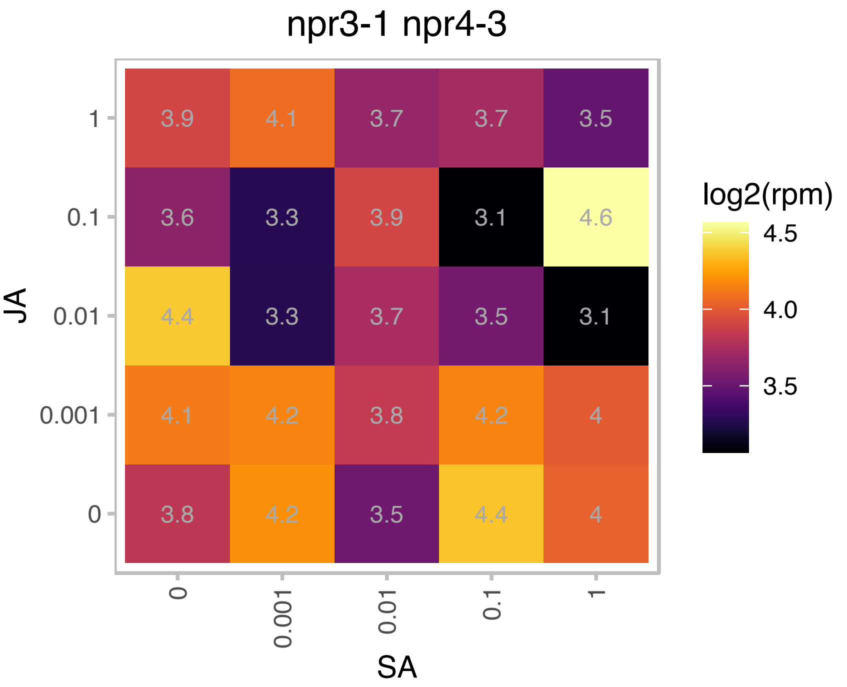

### AT3G51820.1 : ATG4, CHLG, G4, PDE325

UbiA prenyltransferase family protein

Encodes a protein with chlorophyll synthase activity. This enzyme has been shown to perform the esterification of chlorophyllide (a and b), the last step of chlorophyll biosynthesis. Although it can use either geranylgeranyl pyrophosphate (GGPP) or phytyl pyrophosphate (PhyPP) as substrates, the esterification reaction was faster with GGPP than with PhyPP.

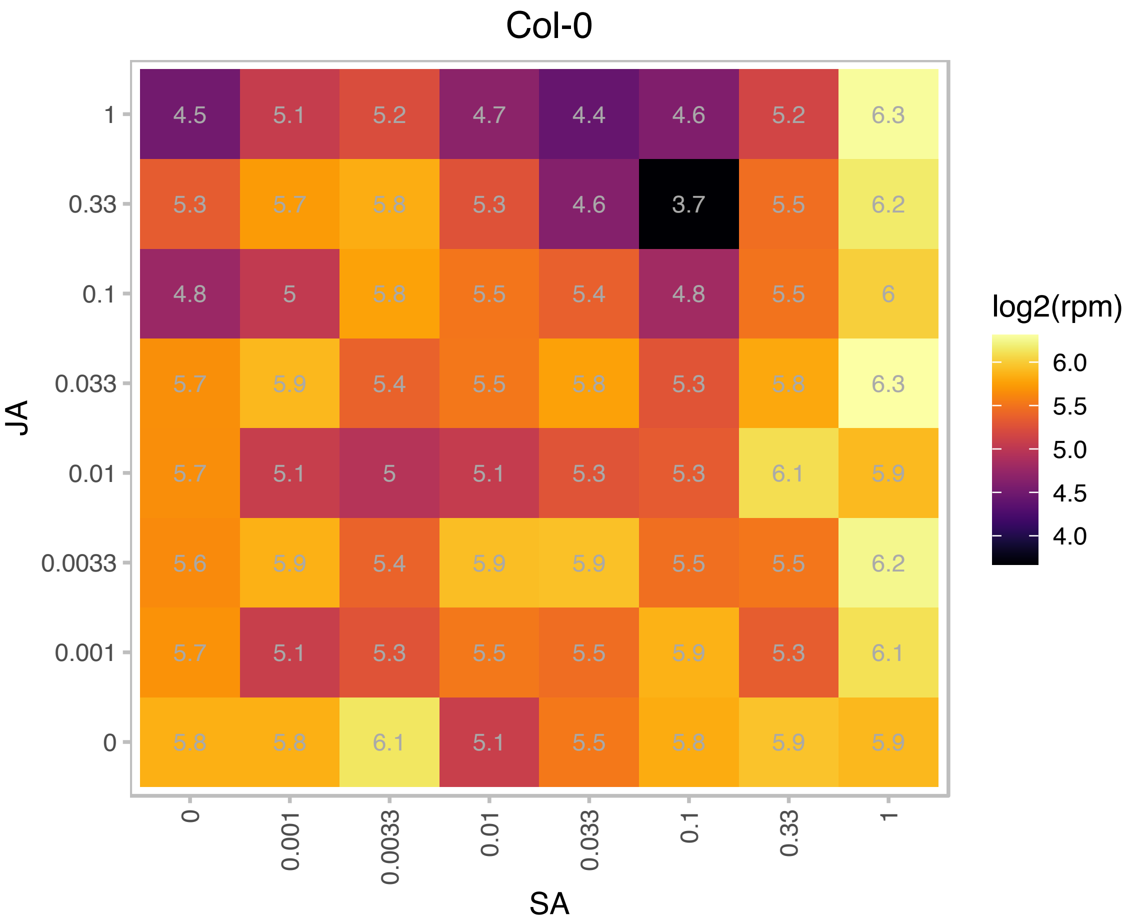


| | |
|---|---|
| Gene | AT3G51820 |
| is.expressed_genes | TRUE |
| is.responsive_genes | TRUE |
| is.apcluster | H-04 |

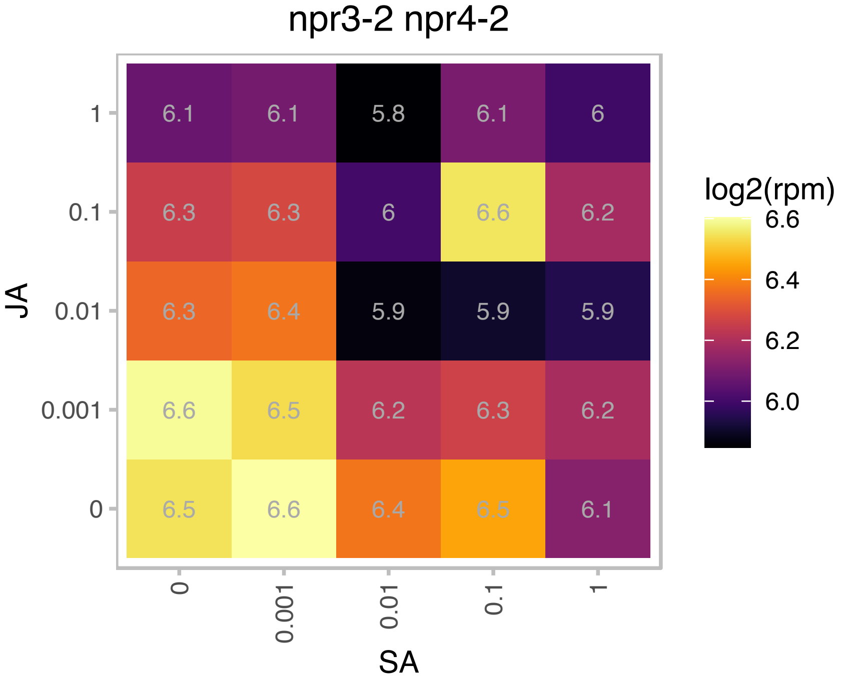


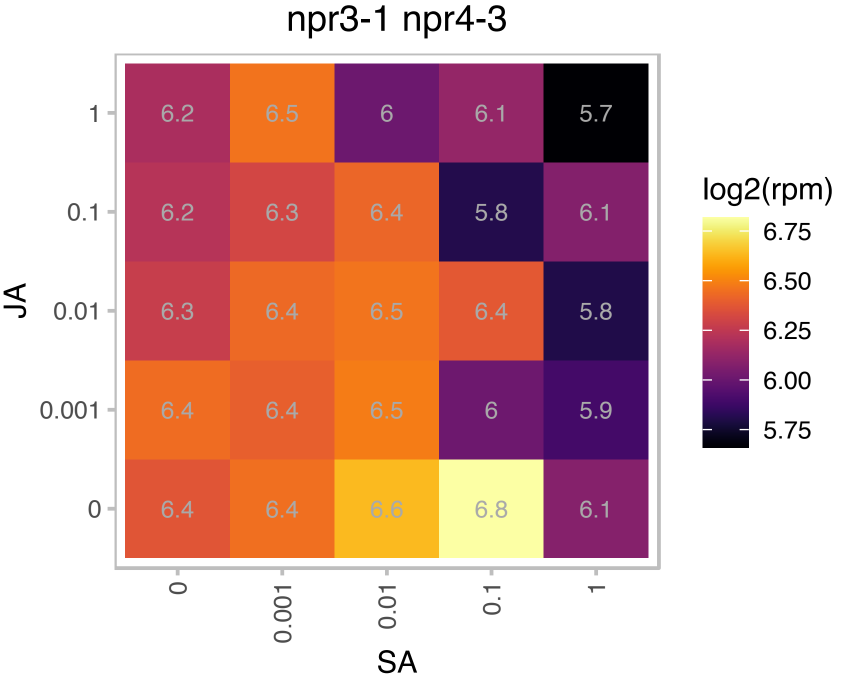

# AT3G53560.1 : NA

Tetratricopeptide repeat (TPR)-like superfamily protein

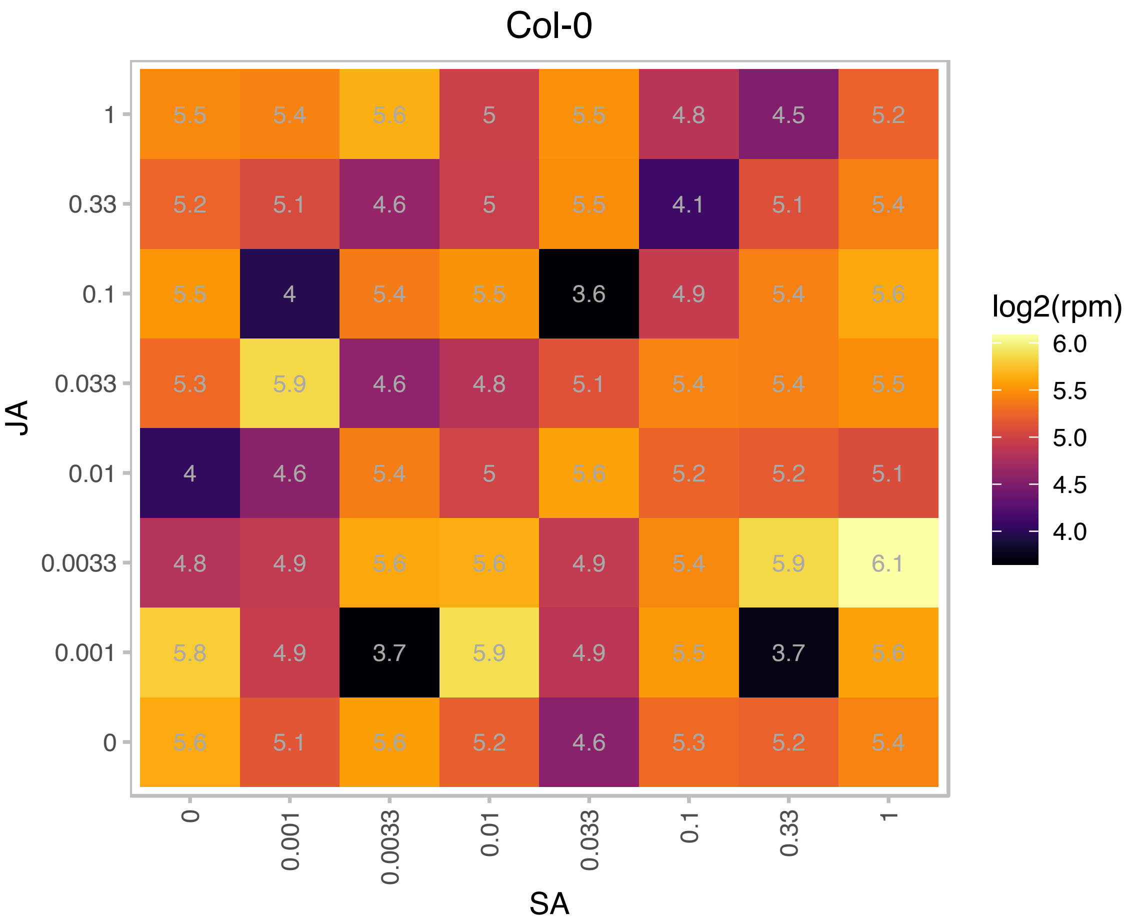


| | |
|---|---|
| Gene | AT3G53560 |
| is.expressed_genes | TRUE |
| is.responsive_genes | TRUE |
| is.apcluster | H-04 |

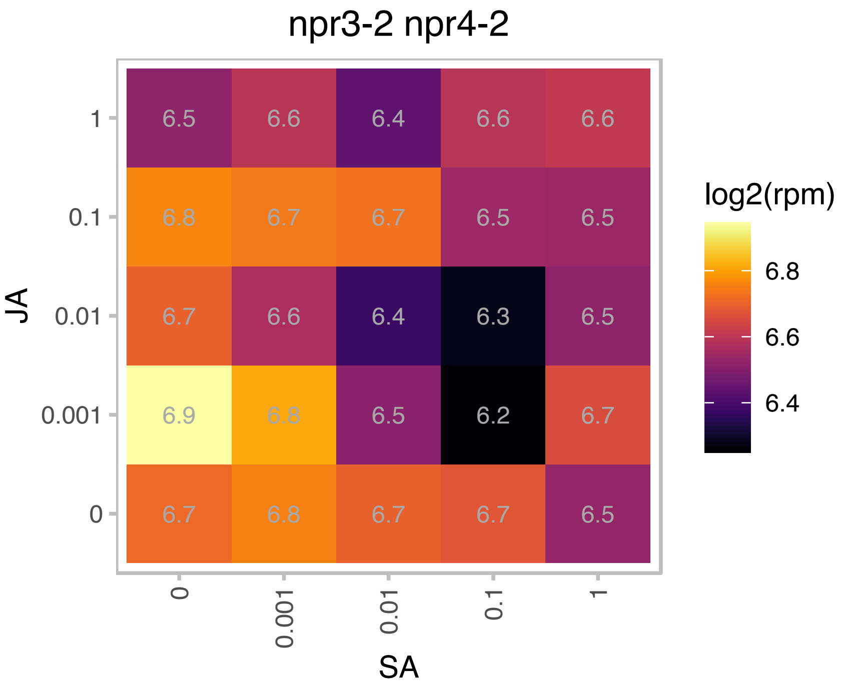


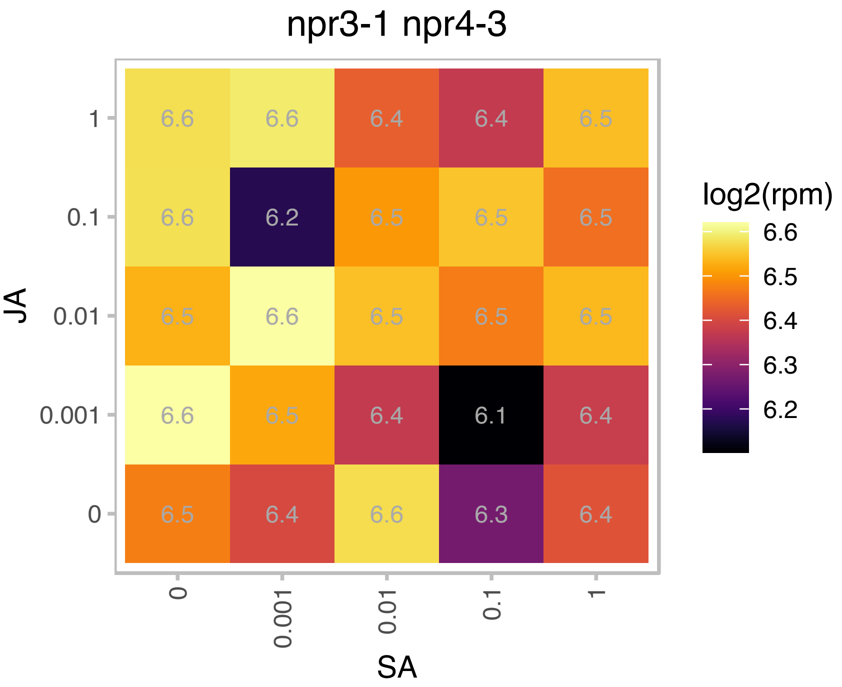

# AT4G17250.1 : NA

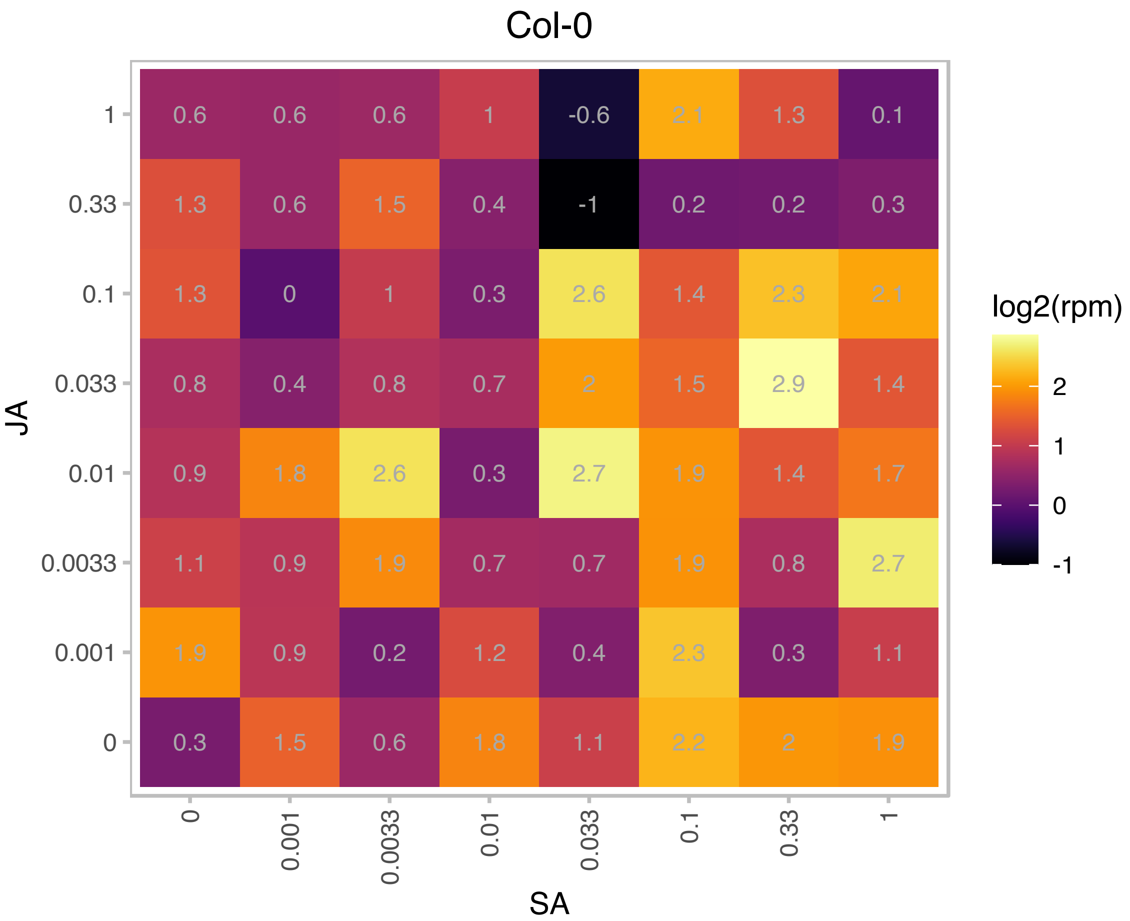


| | |
|---|---|
| Gene | AT4G17250 |
| is.expressed_genes | TRUE |
| is.responsive_genes | TRUE |
| is.apcluster | H-04 |

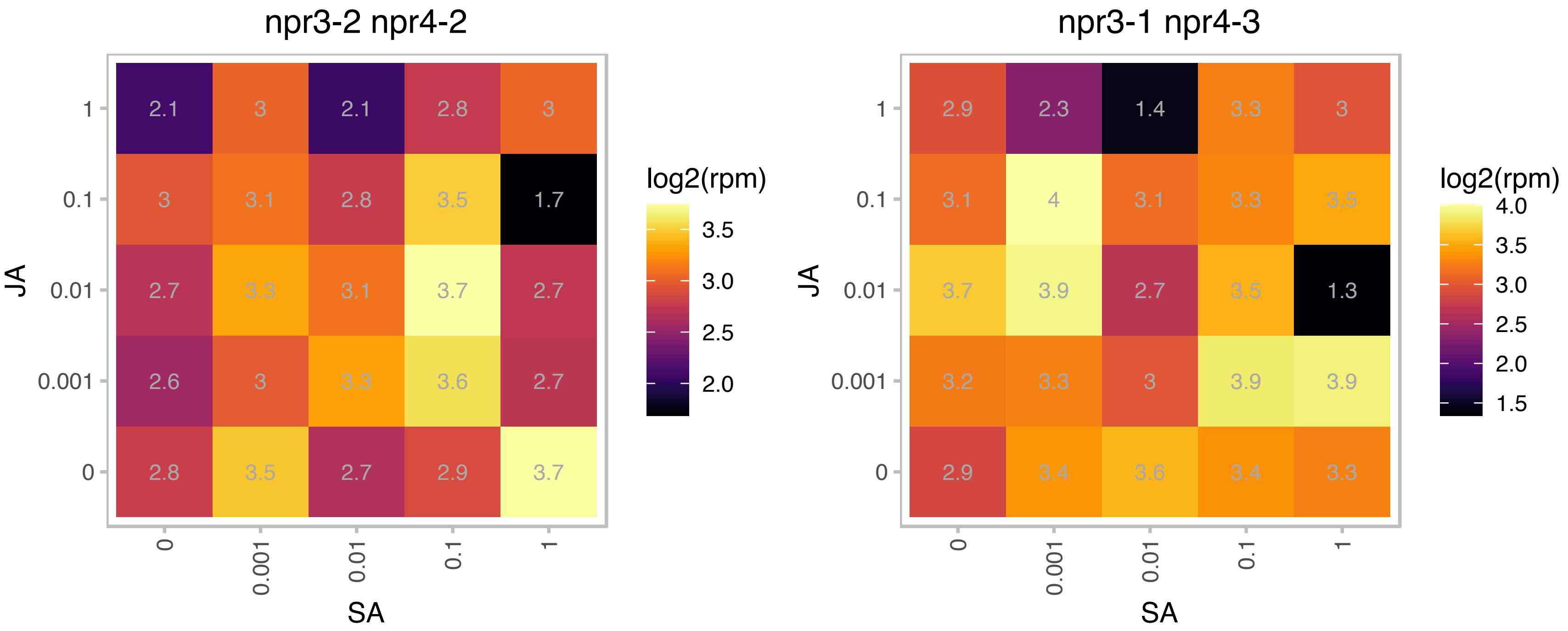

### AT4G17460.1 : HAT1, JAB

Homeobox-leucine zipper protein 4 (HB-4) / HD-ZIP protein

Encodes a class II HD-ZIP protein that regulates meristematic activity in different tissues, and that it is necessary for the correct formation of the gynoecium.

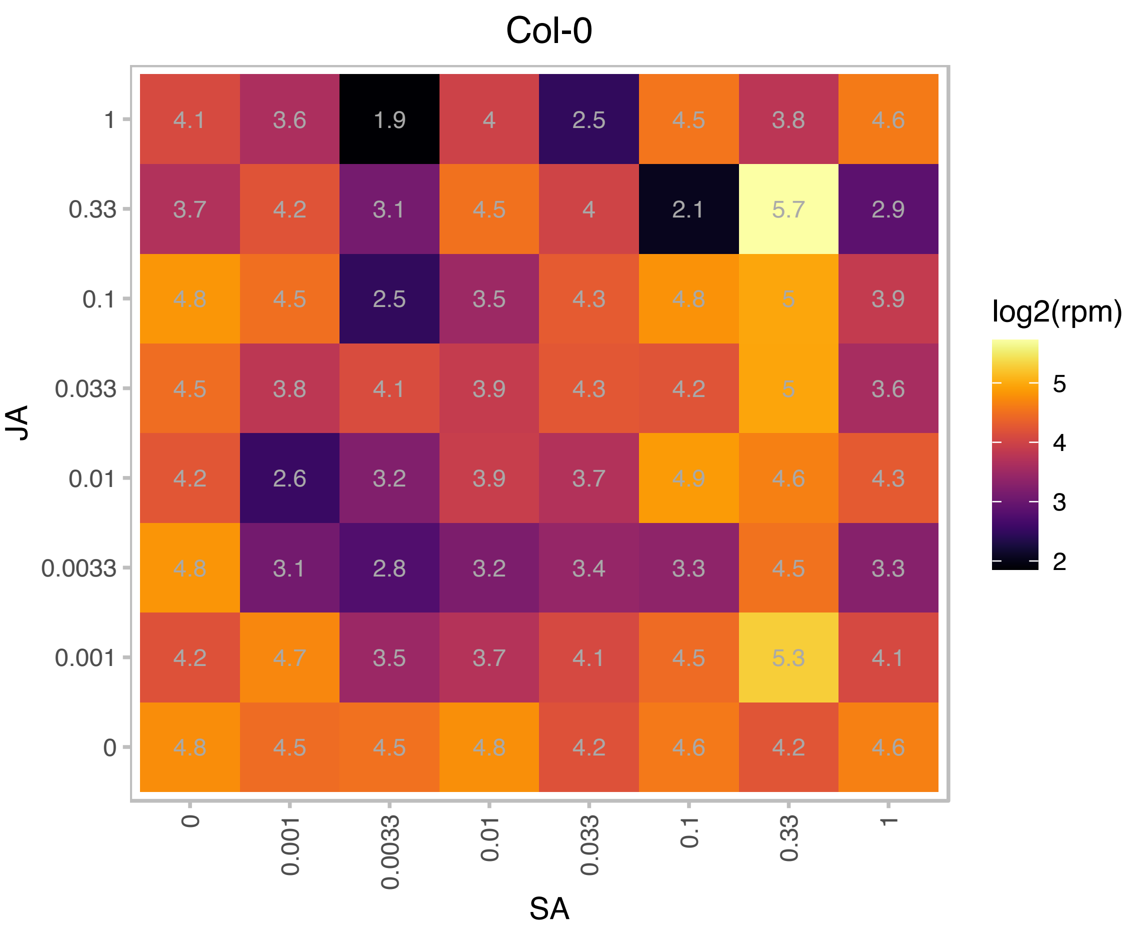


| | |
|---|---|
| Gene | AT4G17460 |
| is.expressed_genes | TRUE |
| is.responsive_genes | TRUE |
| is.apcluster | H-04 |

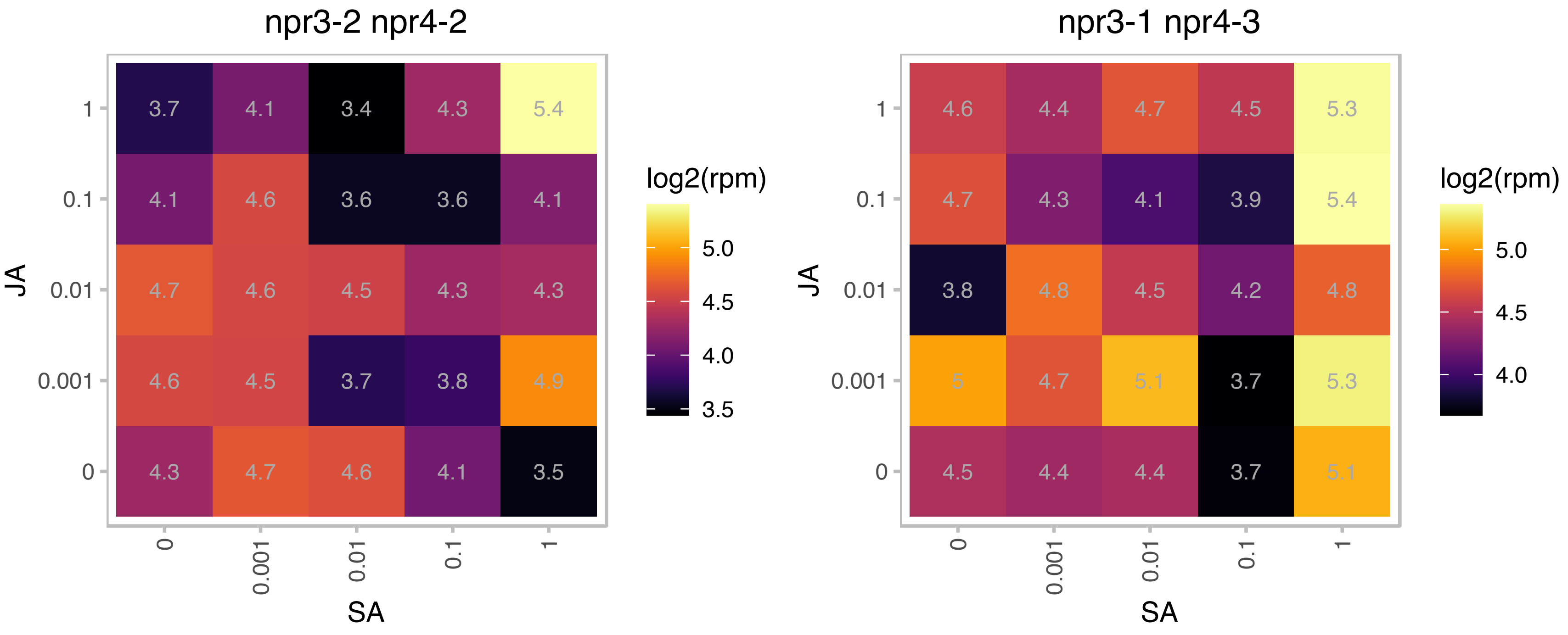

### AT4G23400.1 : PIP1;5, PIP1D

plasma membrane intrinsic protein 1;5

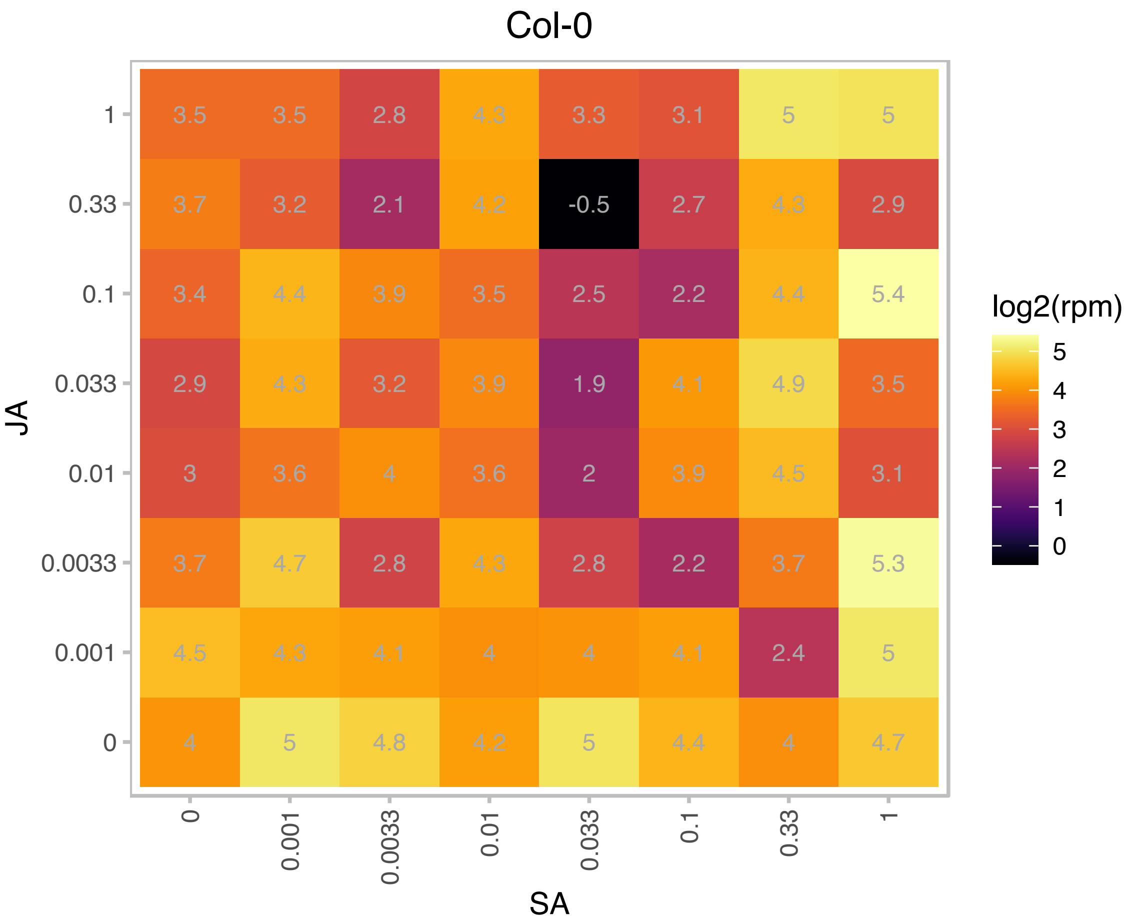


| | |
|---|---|
| Gene | AT4G23400 |
| is.expressed_genes | TRUE |
| is.responsive_genes | TRUE |
| is.apcluster | H-04 |

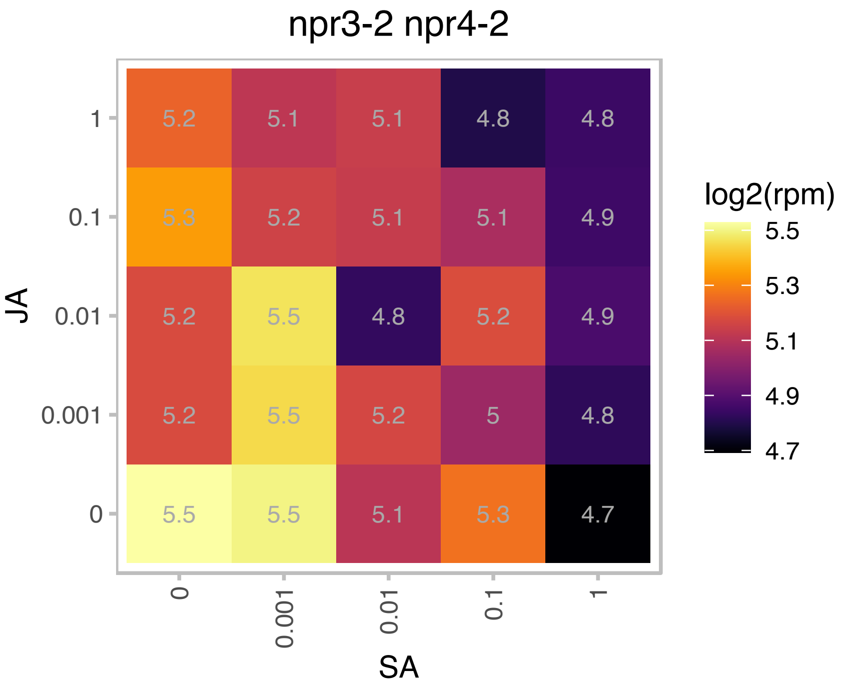


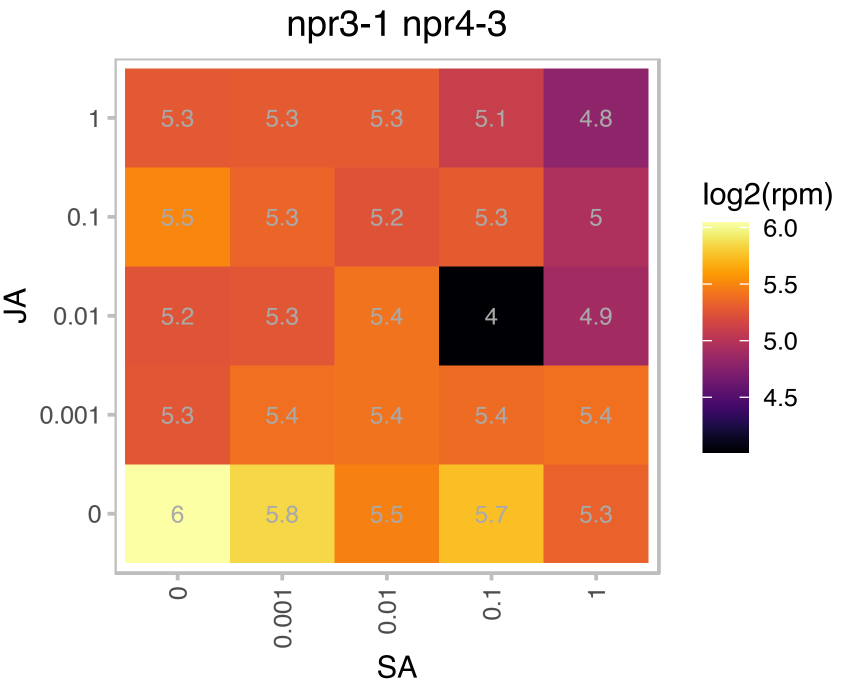

# AT4G32930.1 : NA

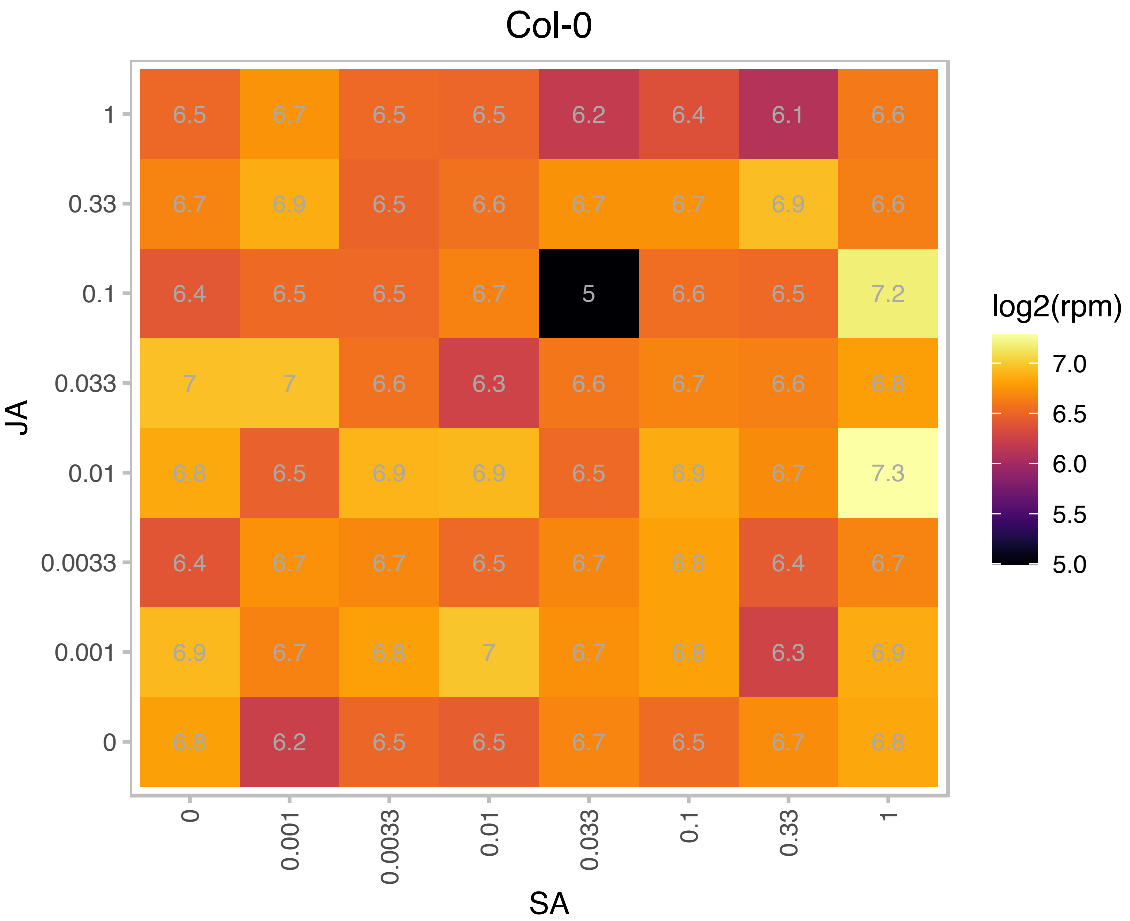


| | |
|---|---|
| Gene | AT4G32930 |
| is.expressed_genes | TRUE |
| is.responsive_genes | TRUE |
| is.apcluster | H-04 |

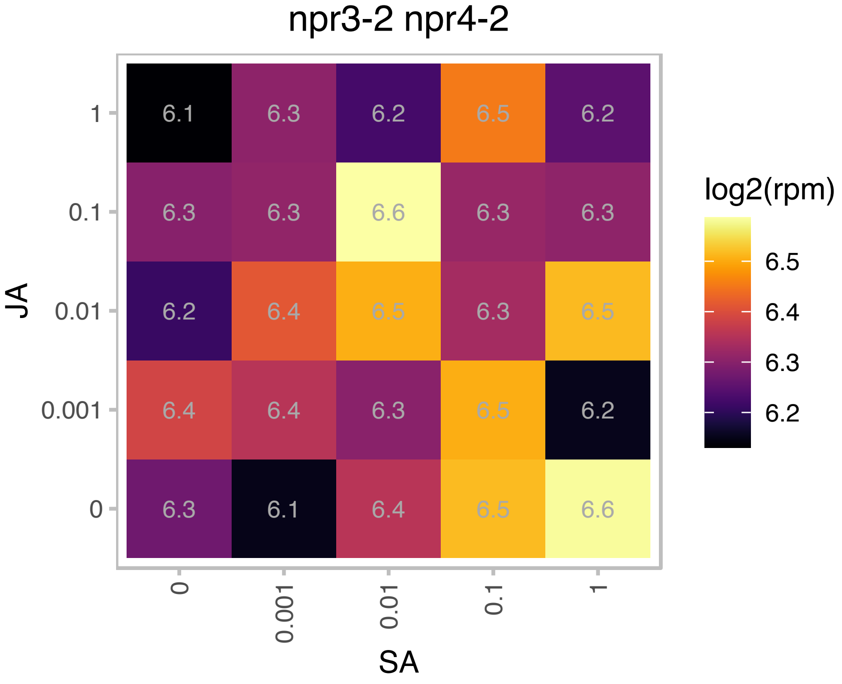


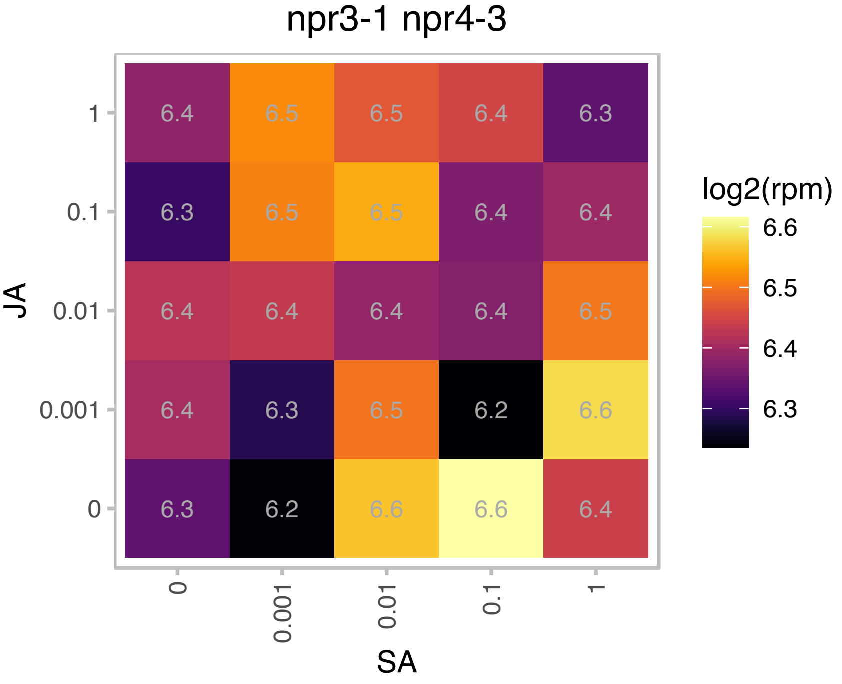

# AT4G34360.1 : NA

S-adenosyl-L-methionine-dependent methyltransferases superfamily protein

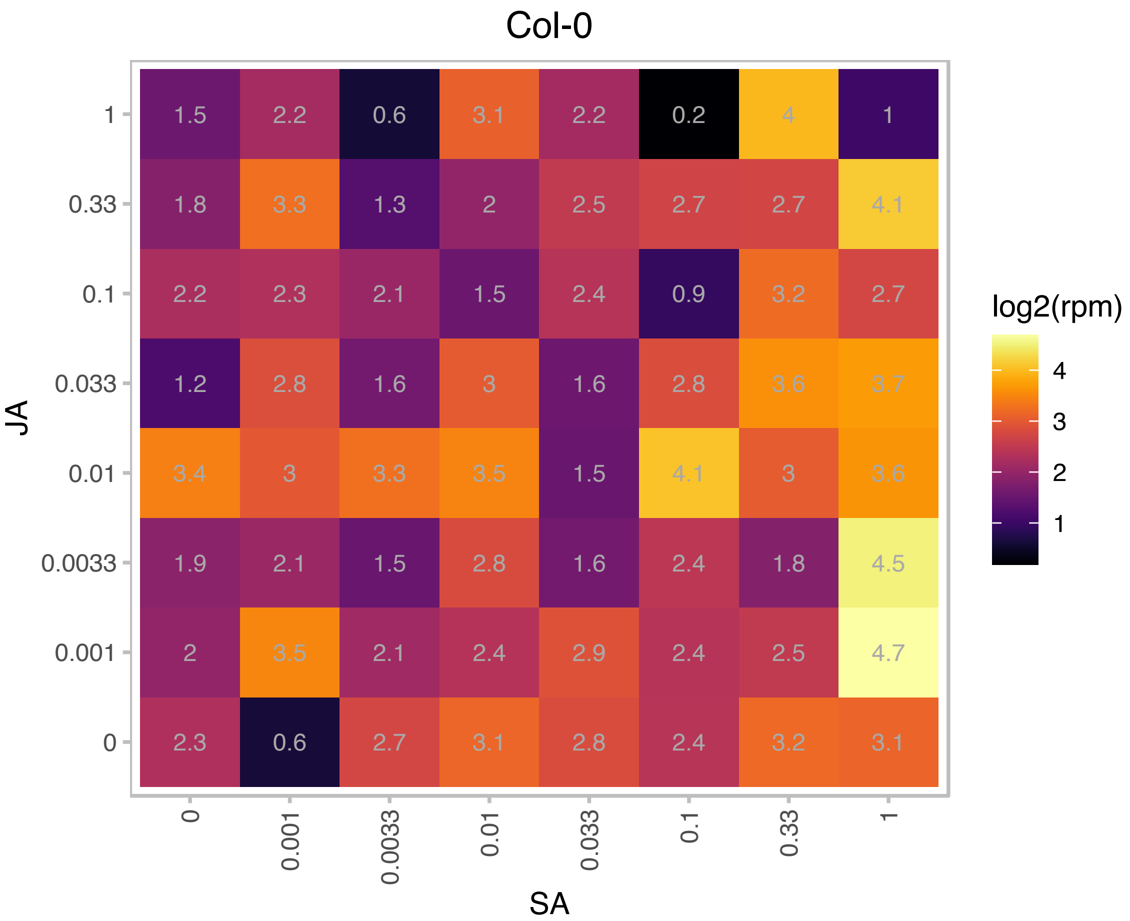


| | |
|---|---|
| Gene | AT4G34360 |
| is.expressed_genes | TRUE |
| is.responsive_genes | TRUE |
| is.apcluster | H-04 |

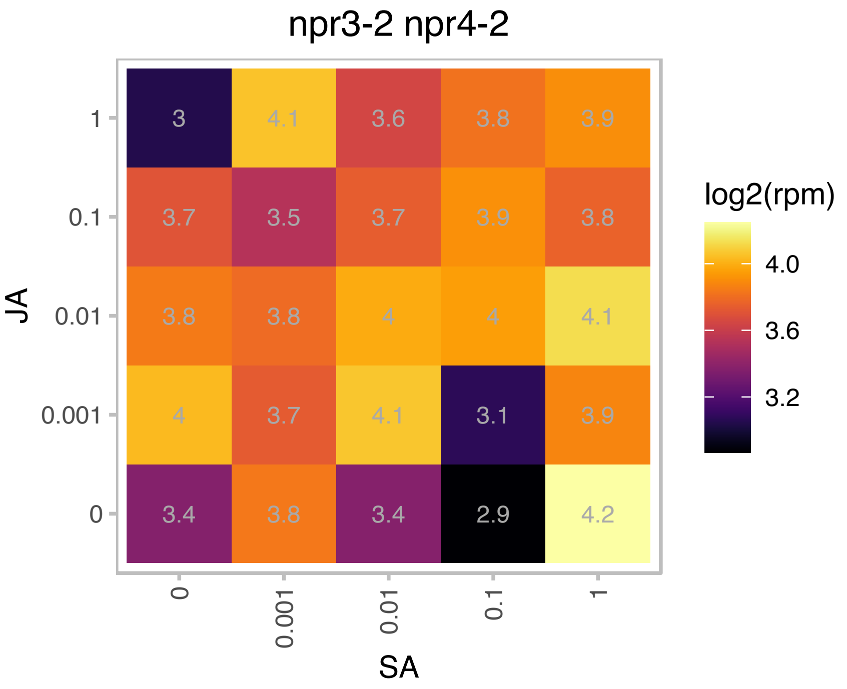


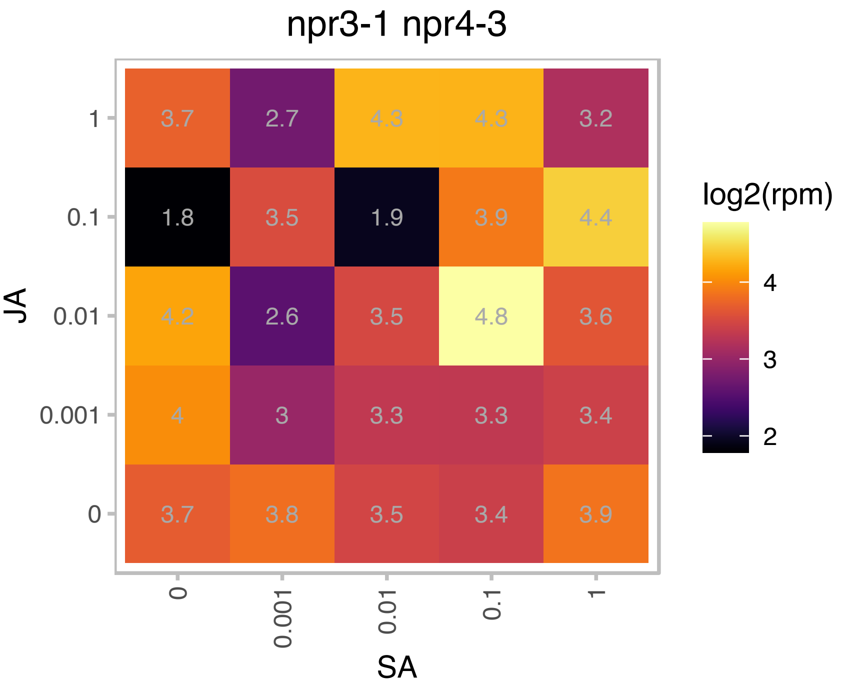

# AT4G39235.1 : NA

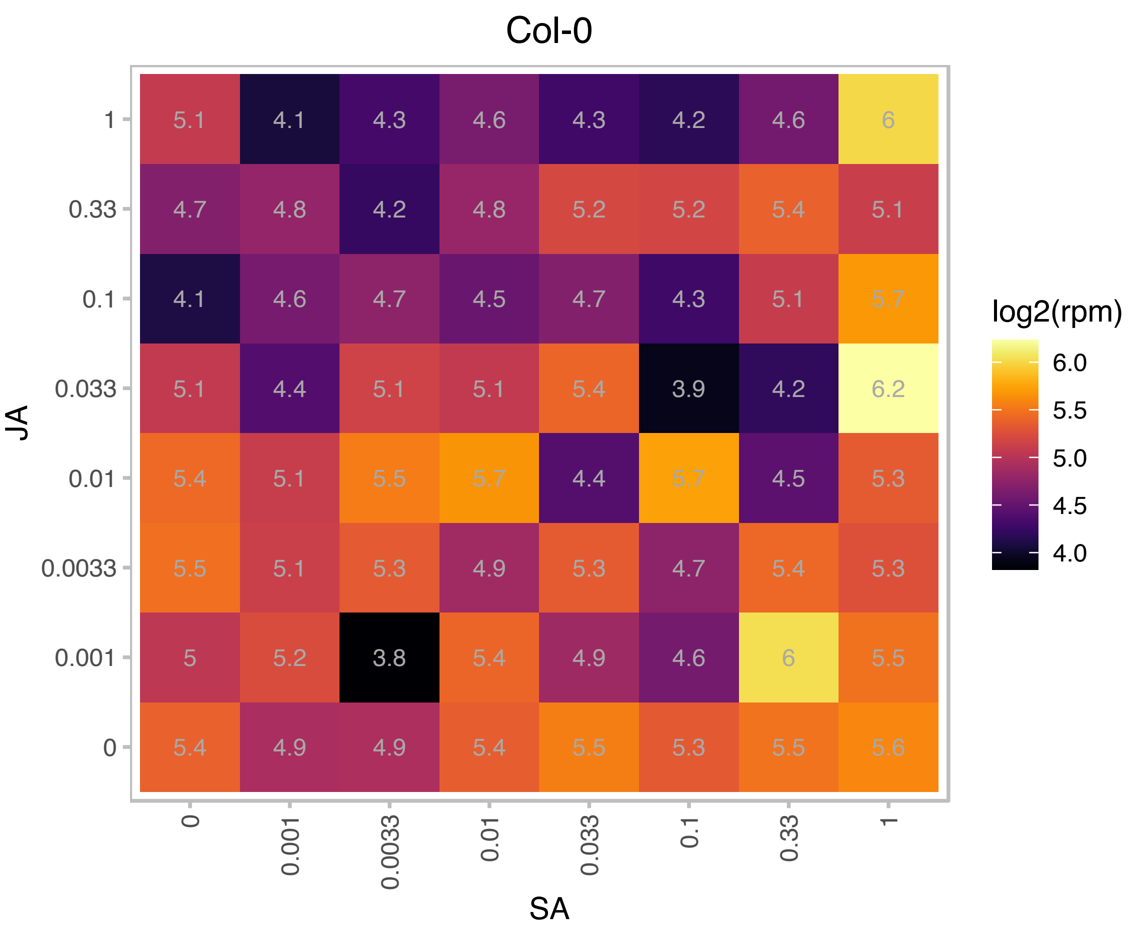


| | |
|---|---|
| Gene | AT4G39235 |
| is.expressed_genes | TRUE |
| is.responsive_genes | TRUE |
| is.apcluster | H-04 |

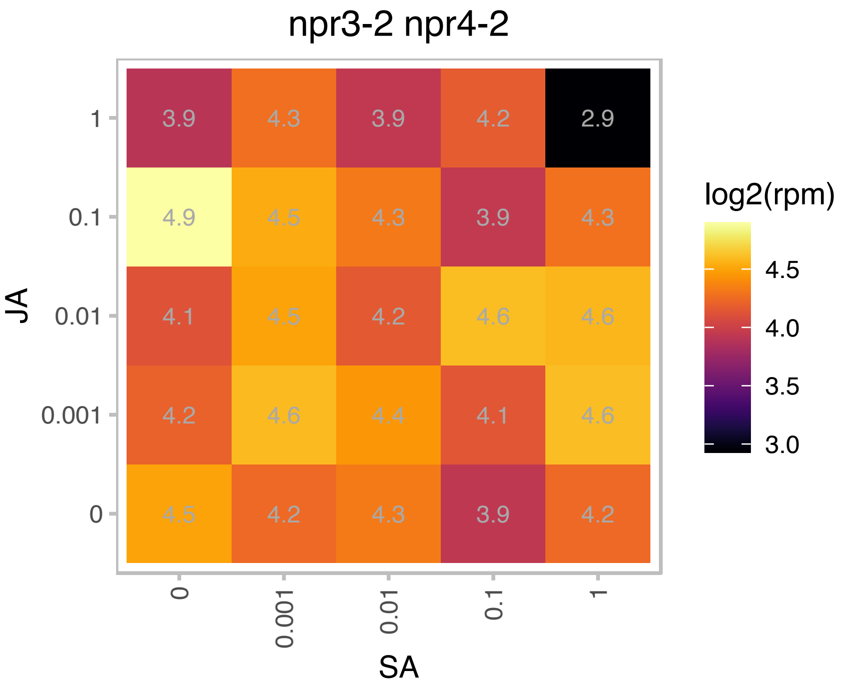


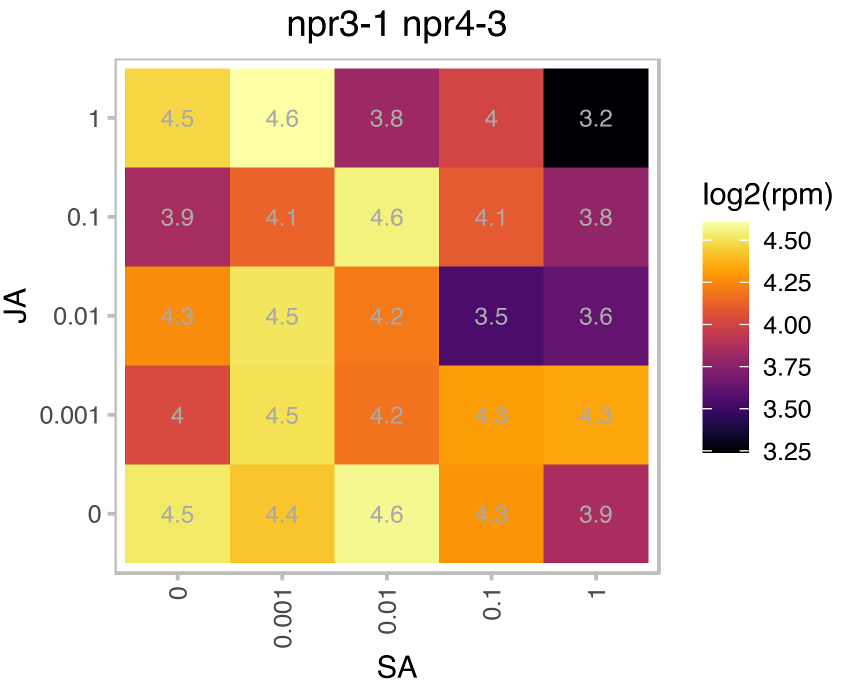

# AT5G01750.1 : NA

Protein of unknown function (DUF567)

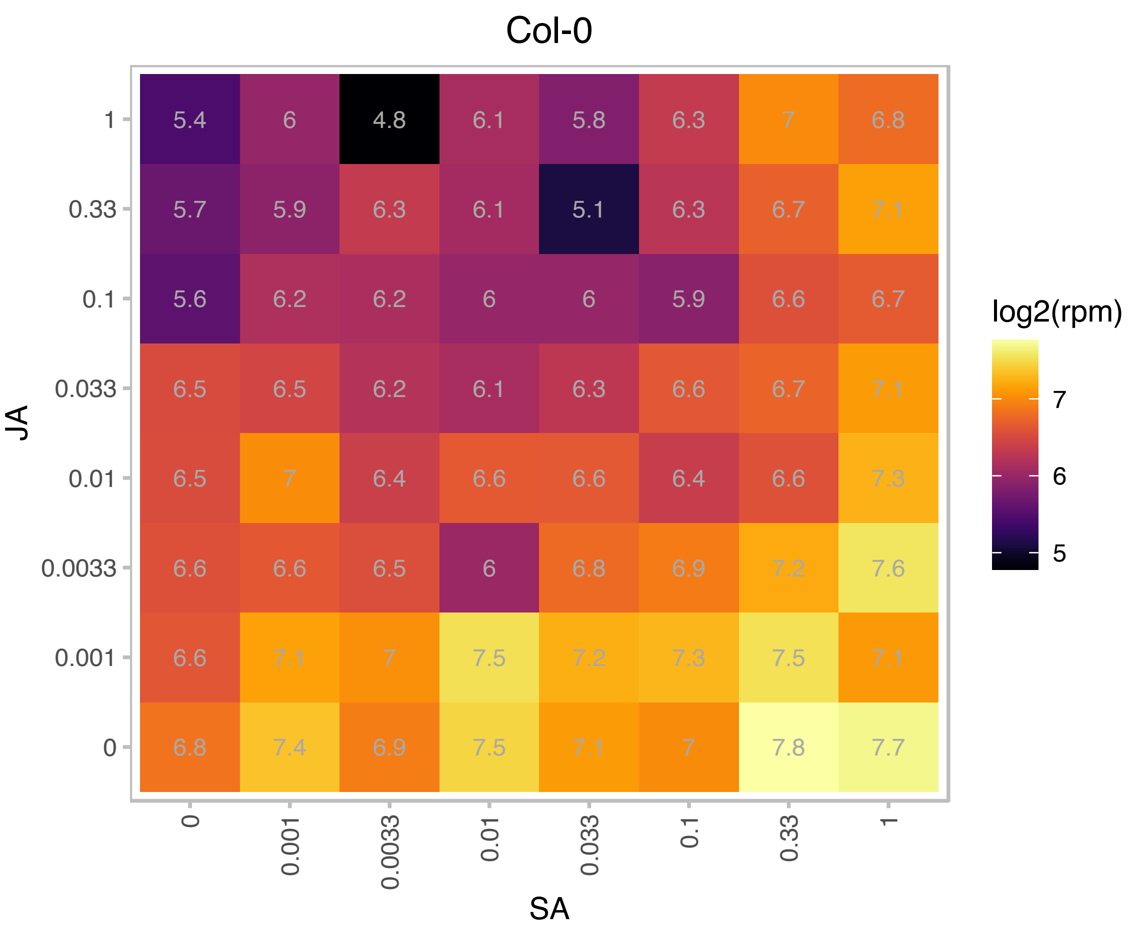


| | |
|---|---|
| Gene | AT5G01750 |
| is.expressed_genes | TRUE |
| is.responsive_genes | TRUE |
| is.apcluster | H-04 |

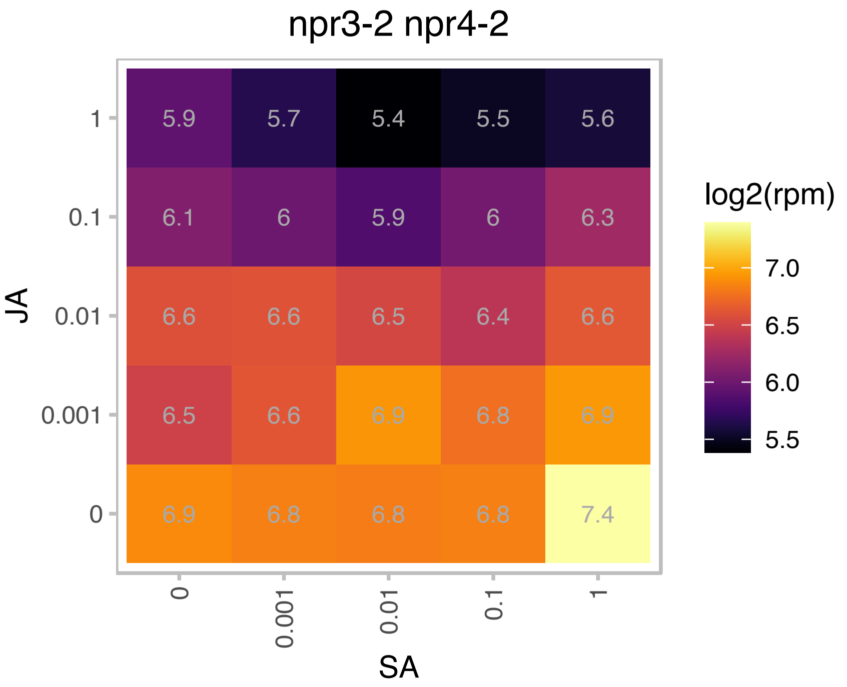


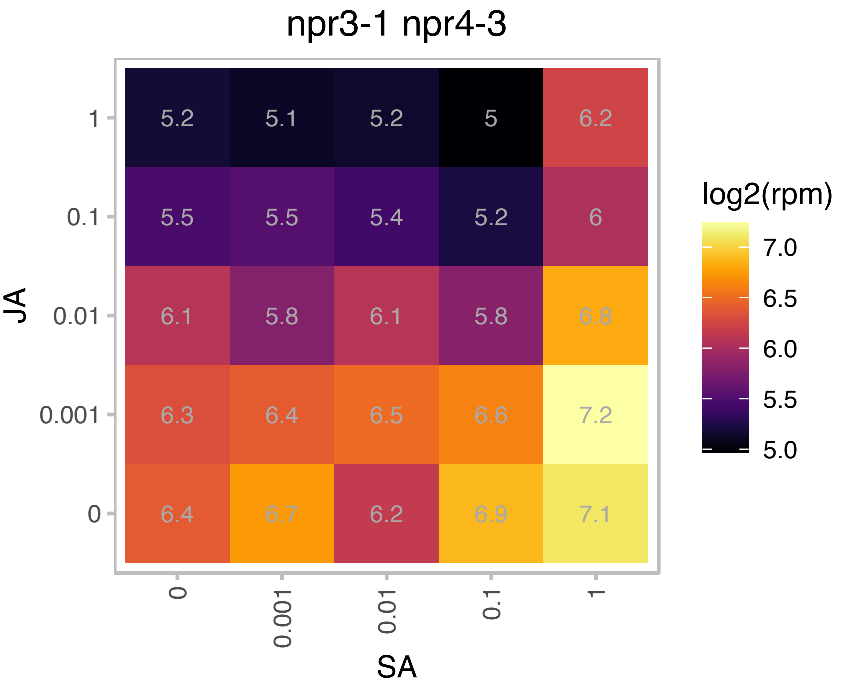

### AT5G09220.1 : AAP2

amino acid permease 2

member of AAAP family The mRNA is cell-to-cell mobile.

Col-0

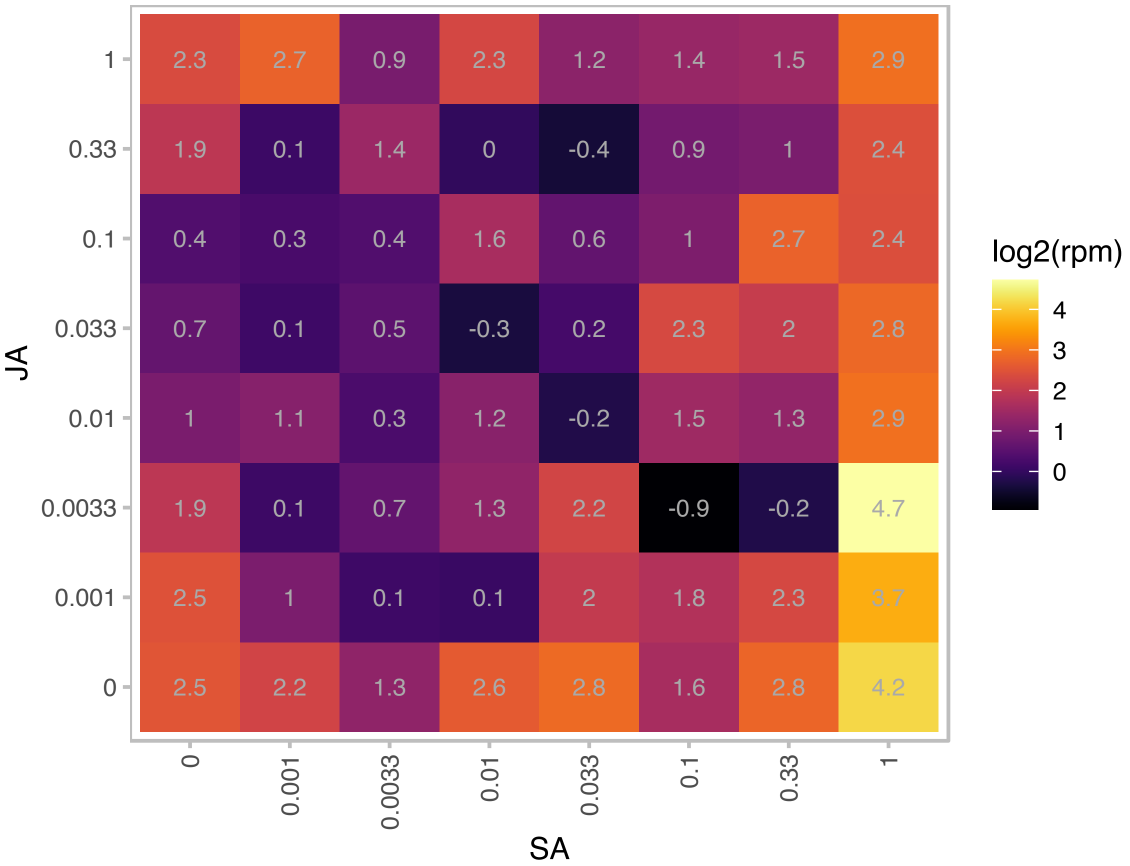


| | |
|---|---|
| Gene | AT5G09220 |
| is.expressed_genes | TRUE |
| is.responsive_genes | TRUE |
| is.apcluster | H-04 |

npr3-2 npr4-2

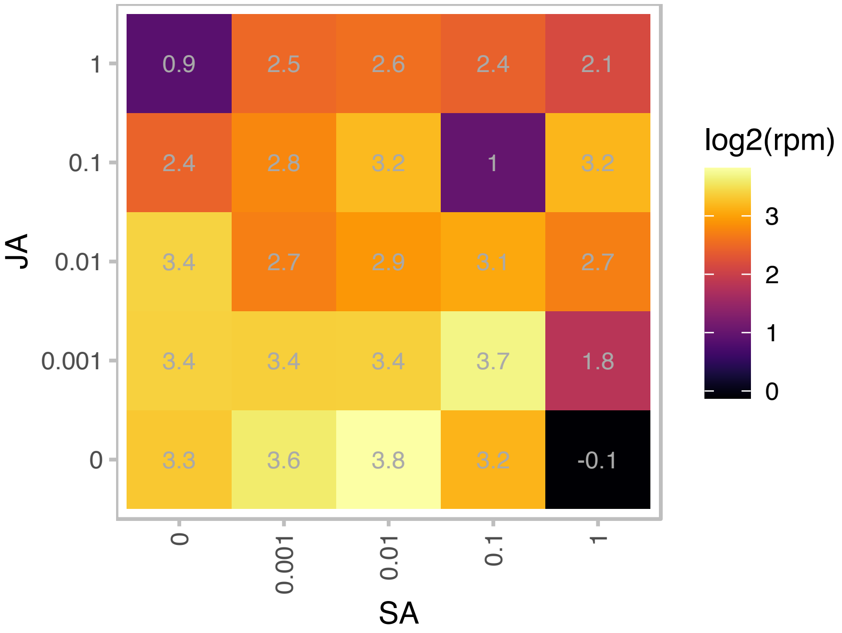


npr3-1 npr4-3

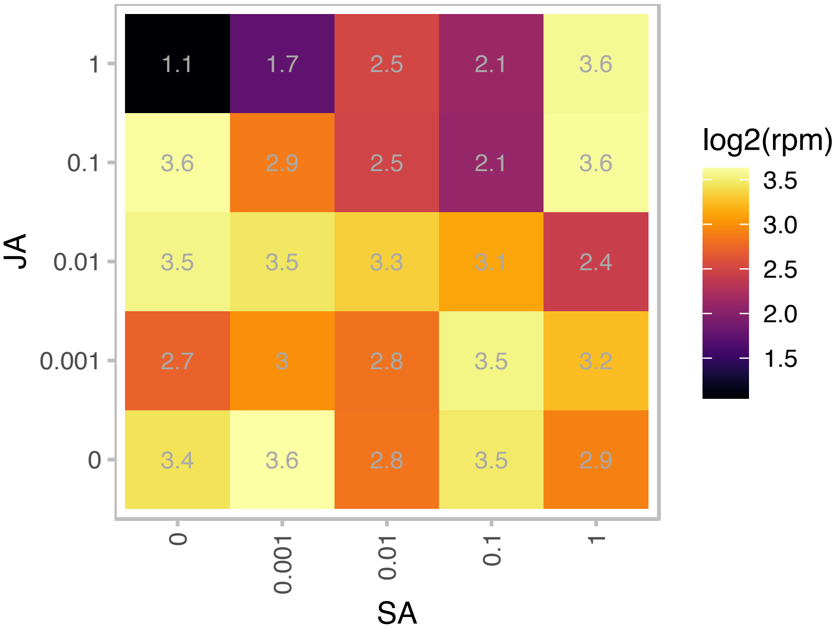

# AT5G22100.1 : NA

RNA cyclase family protein

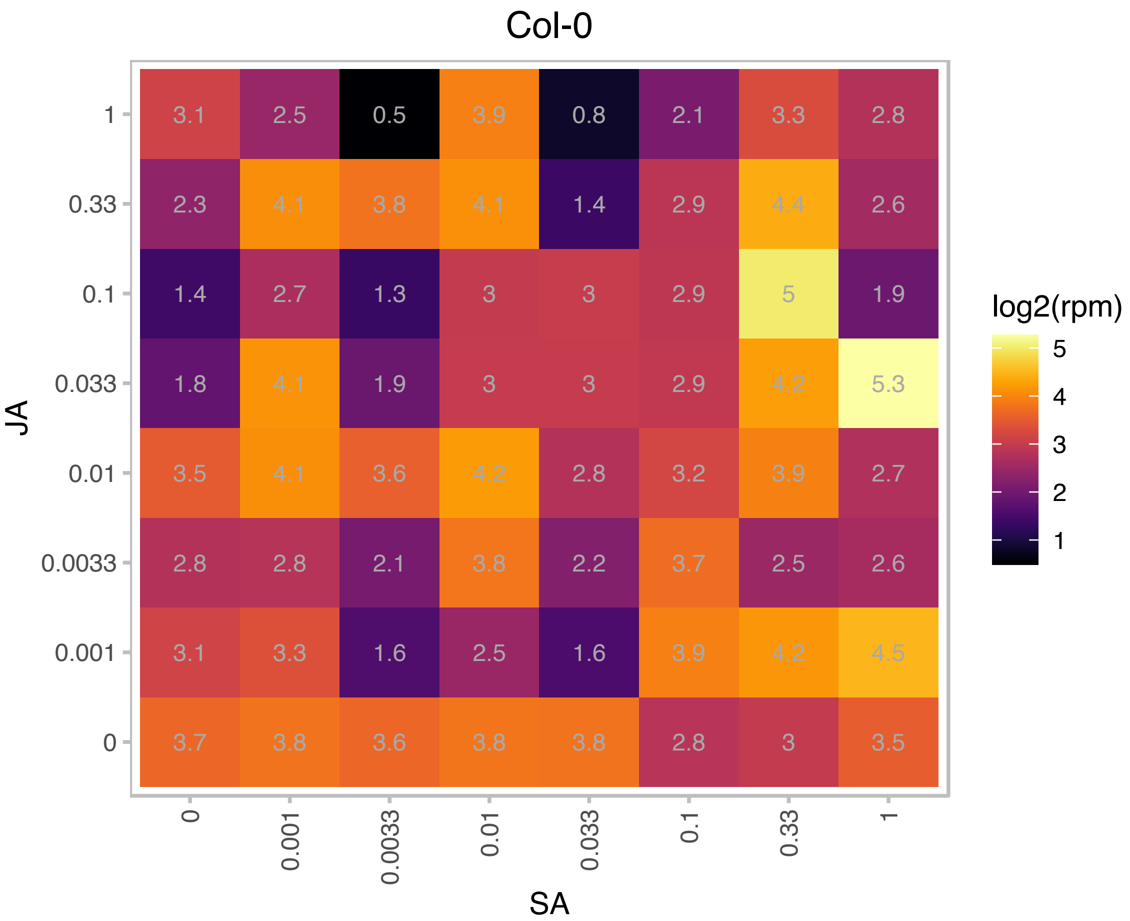


| | |
|---|---|
| Gene | AT5G22100 |
| is.expressed_genes | TRUE |
| is.responsive_genes | TRUE |
| is.apcluster | H-04 |

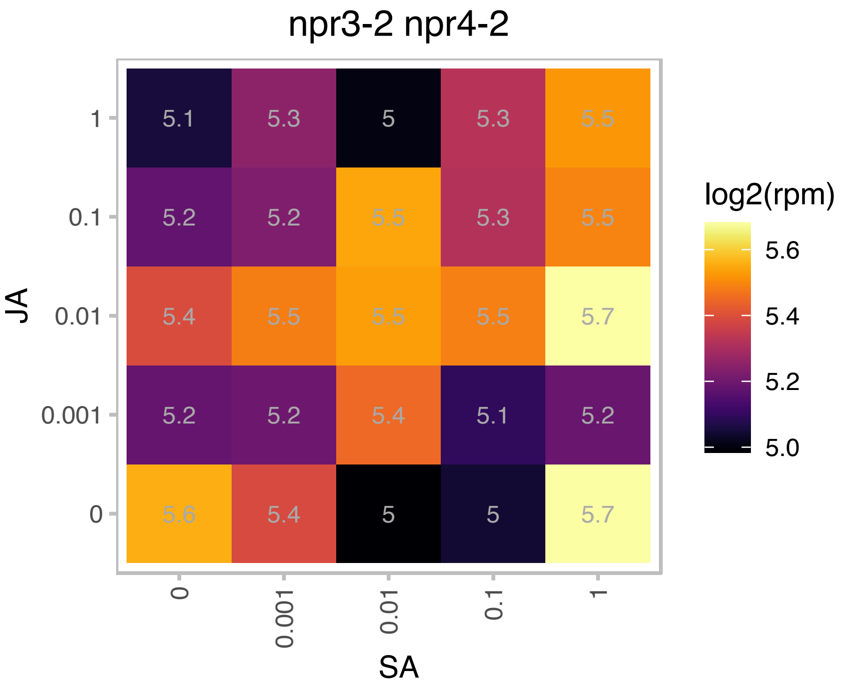


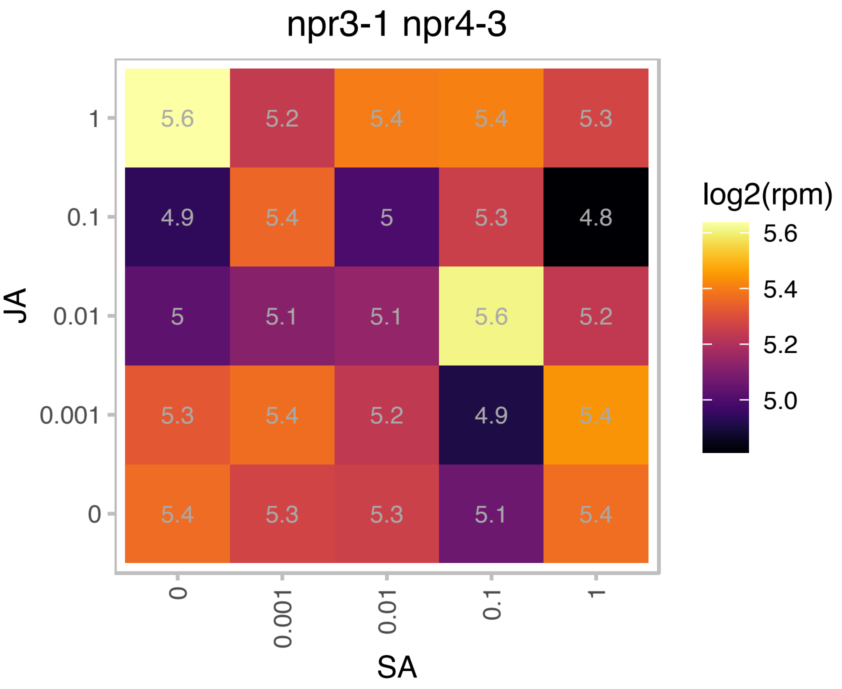

### AT5G38520.1 : CLD1

alpha/beta-Hydrolases superfamily protein
CLD1 is involved in steady-state chlorophyll turnover; CLD1 dephytylates chlorophyll a, chlorophyll b, and pheophytin a in vitro; CLD1 and CHLG form a salvage cycle in recycling chlorophyll. Suppression of CLD1 expression results in reduced tolerance to moderately high temperature.

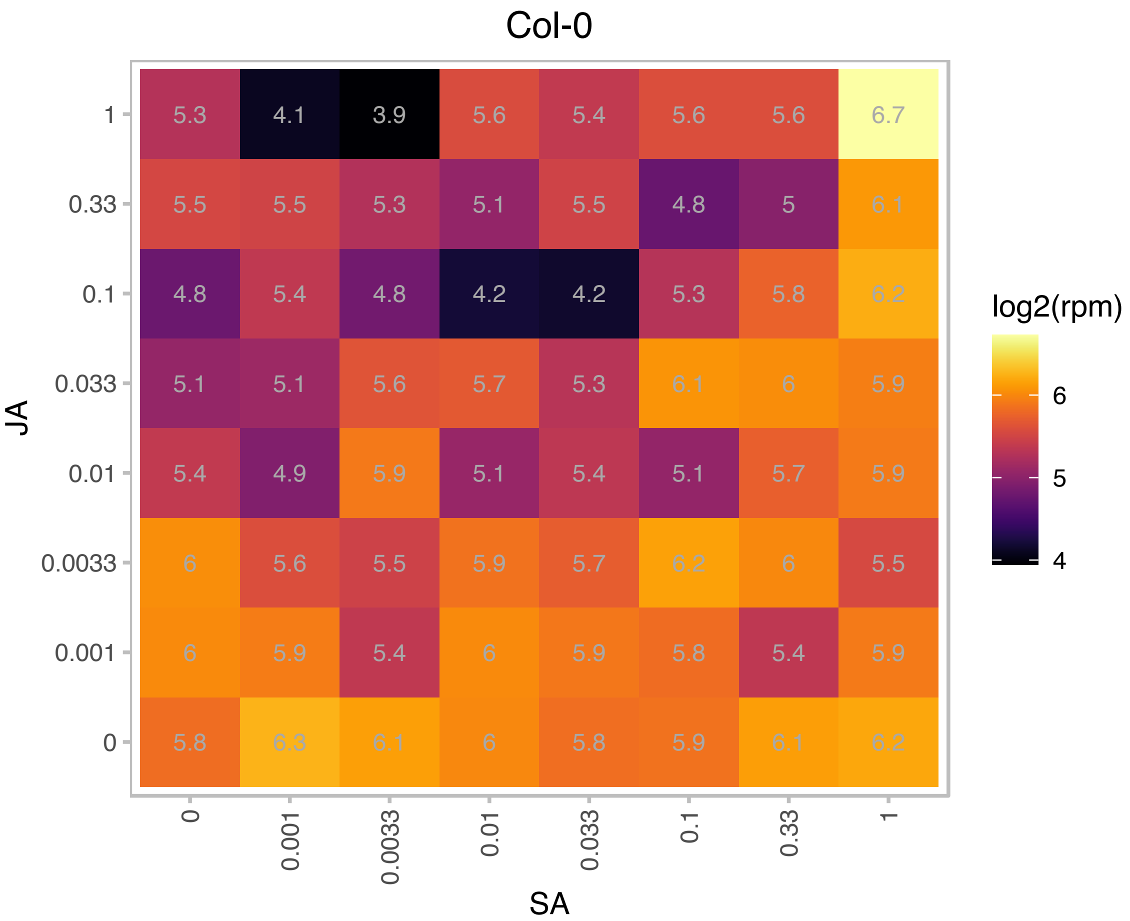


| | |
|---|---|
| Gene | AT5G38520 |
| is.expressed_genes | TRUE |
| is.responsive_genes | TRUE |
| is.apcluster | H-04 |

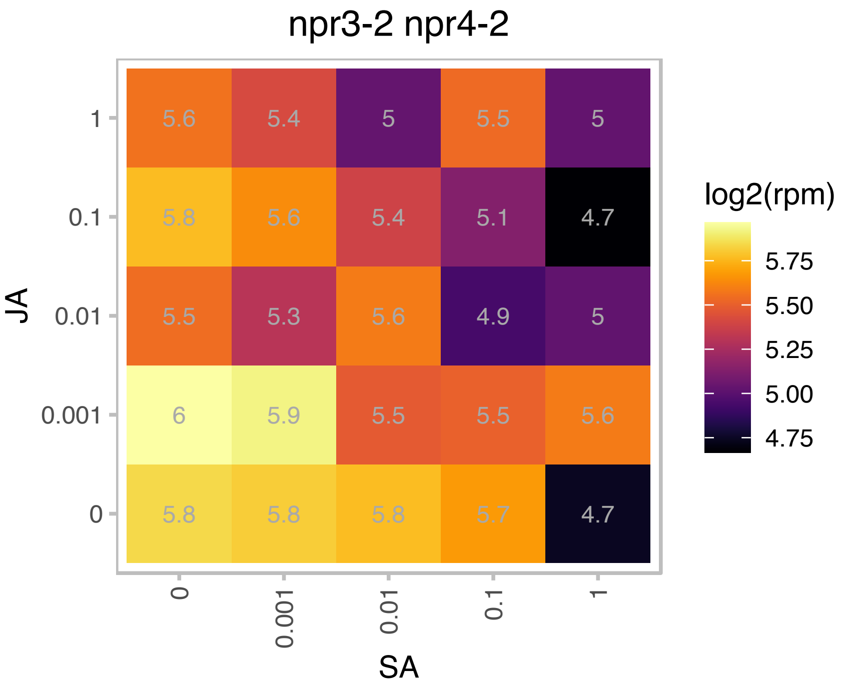


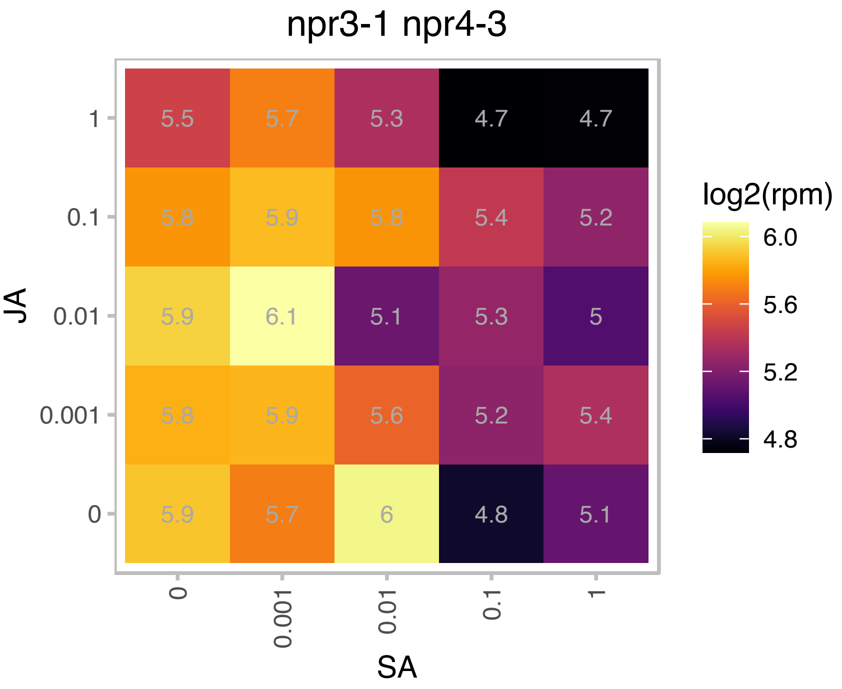

# AT5G42825.1 : NA

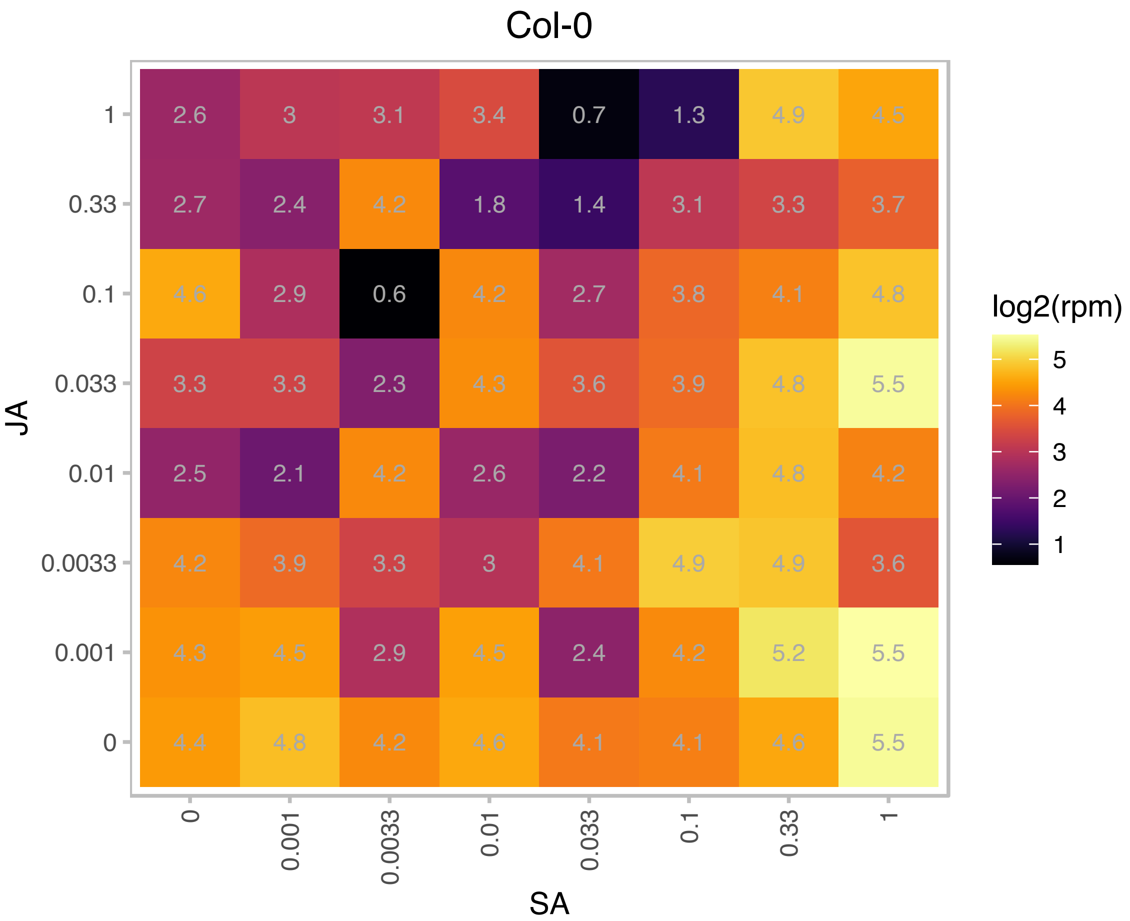


| | |
|---|---|
| Gene | AT5G42825 |
| is.expressed_genes | TRUE |
| is.responsive_genes | TRUE |
| is.apcluster | H-04 |

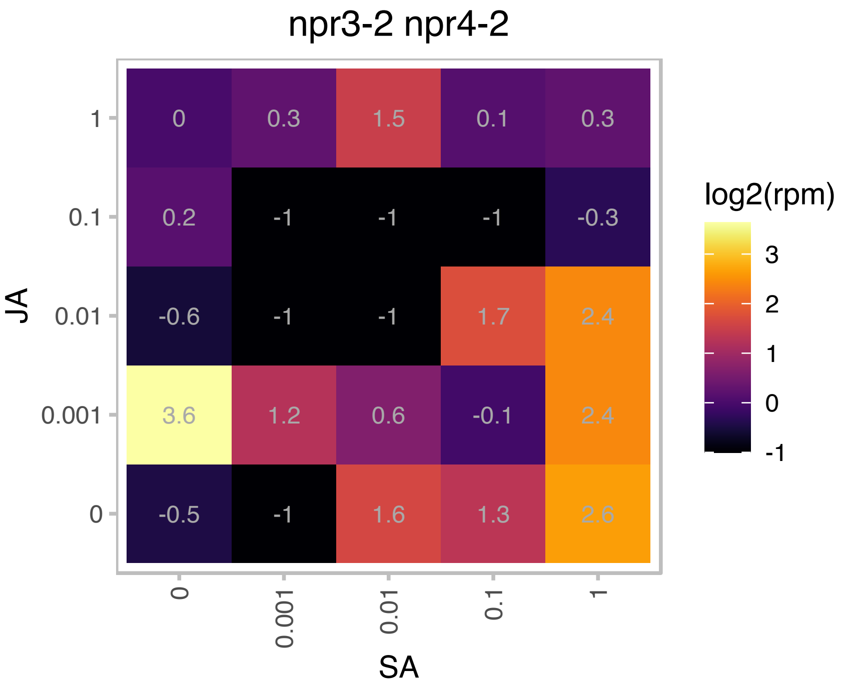


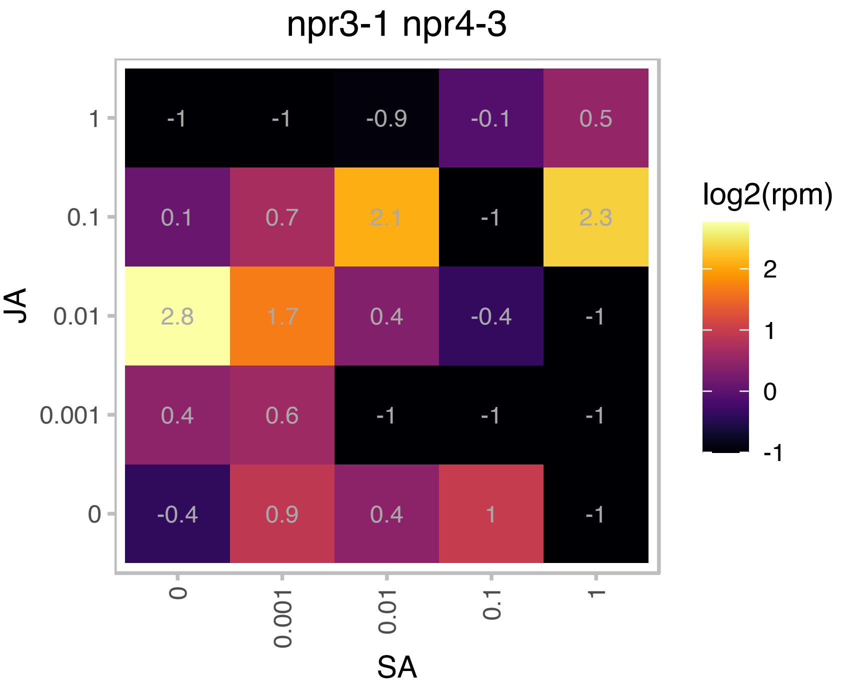

### AT5G65940.1 : CHY1

beta-hydroxyisobutyryl-CoA hydrolase 1

hydrolyzes beta-hydroxyisobutyryl-CoA

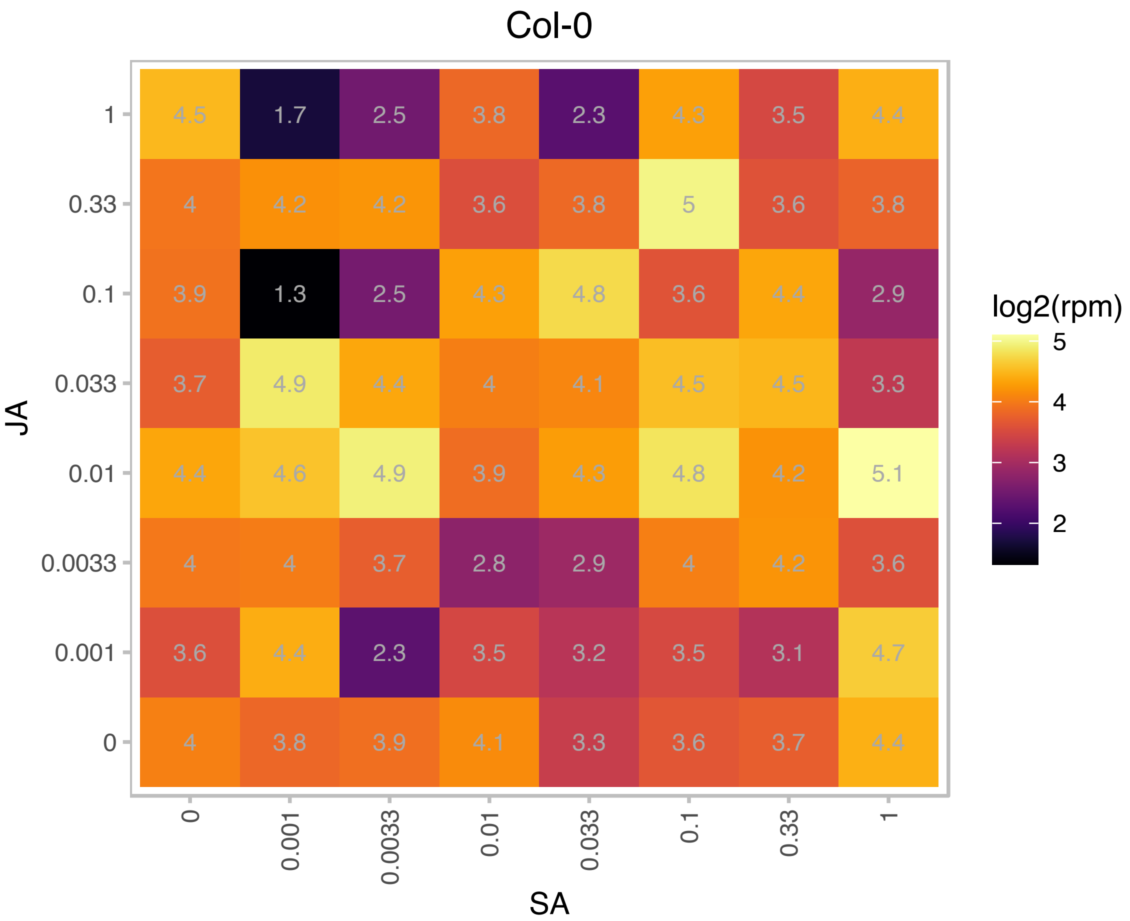


| | |
|---|---|
| Gene | AT5G65940 |
| is.expressed_genes | TRUE |
| is.responsive_genes | TRUE |
| is.apcluster | H-04 |

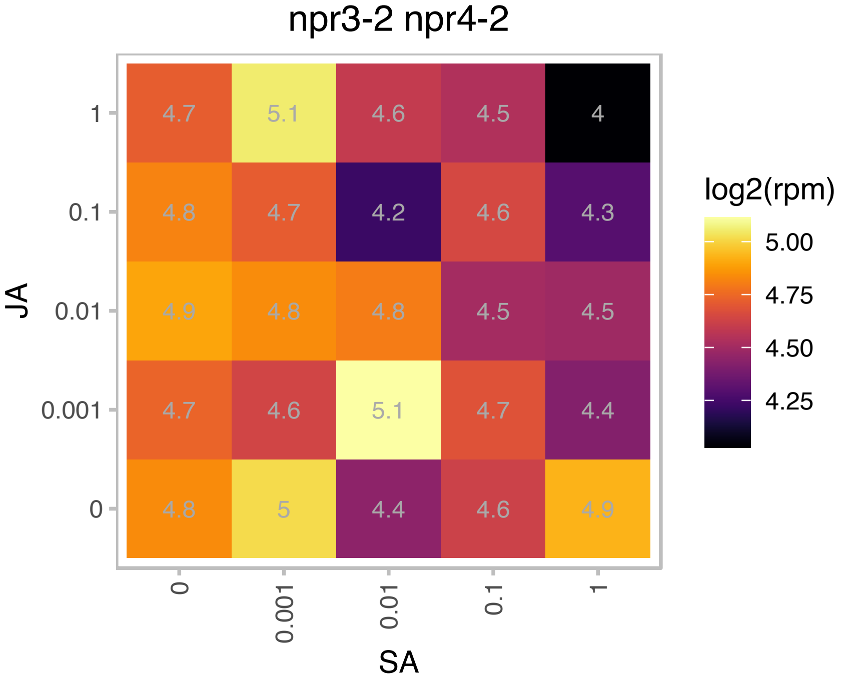


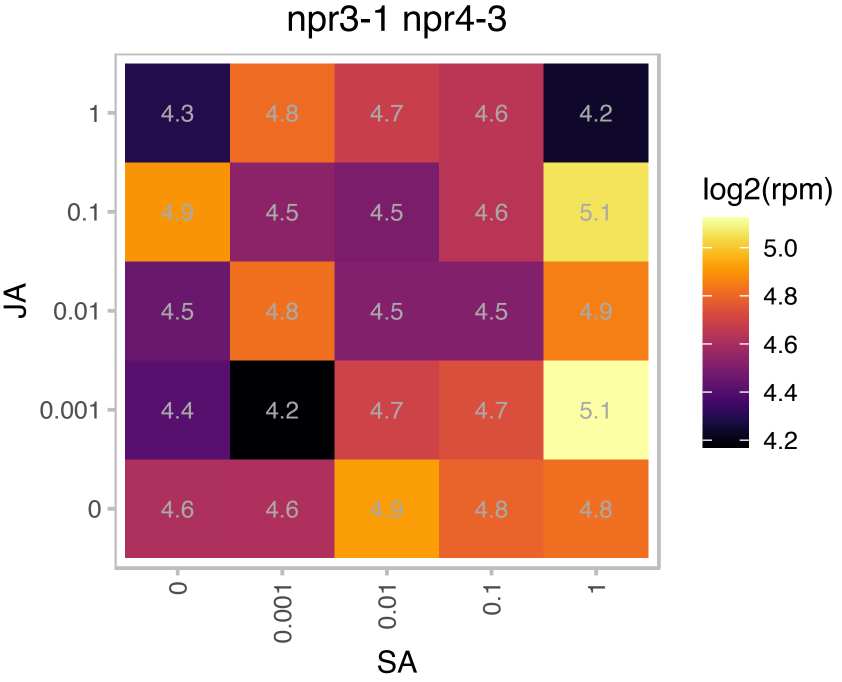

# AT1G09310.1 : NA

Protein of unknown function, DUF538

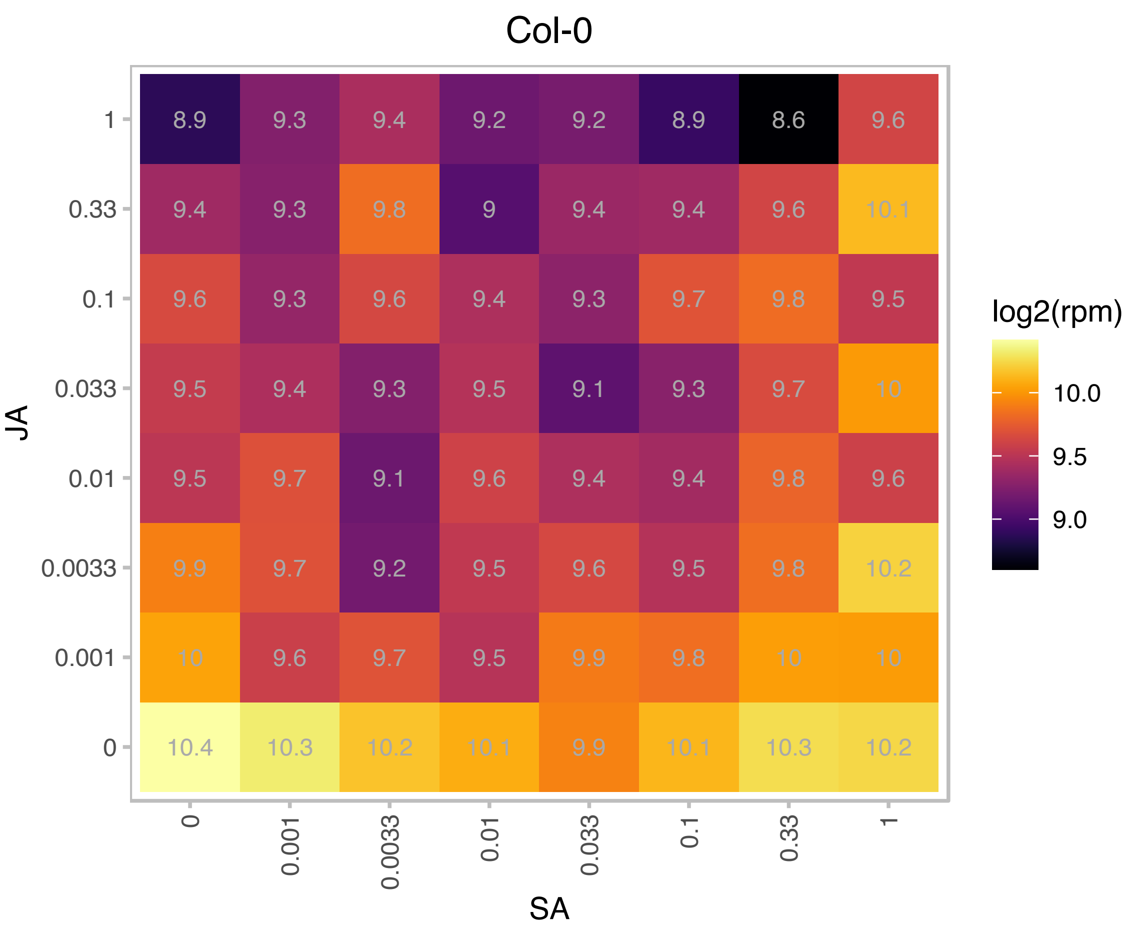


| | |
|---|---|
| Gene | AT1G09310 |
| is.expressed_genes | TRUE |
| is.responsive_genes | TRUE |
| is.apcluster | H-05 |

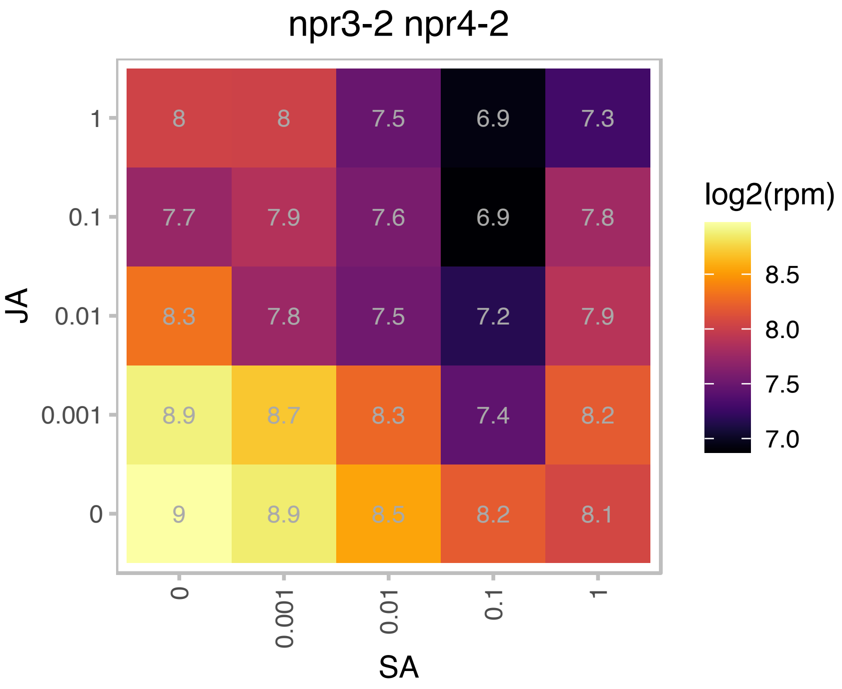


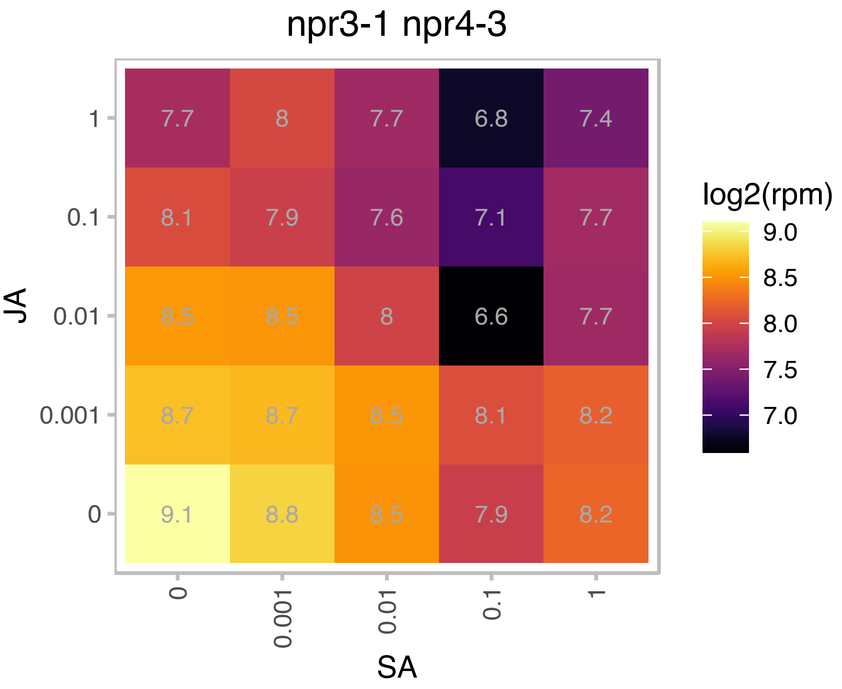

### AT1G11545.1 : XTH8

xyloglucan endotransglucosylase/hydrolase 8

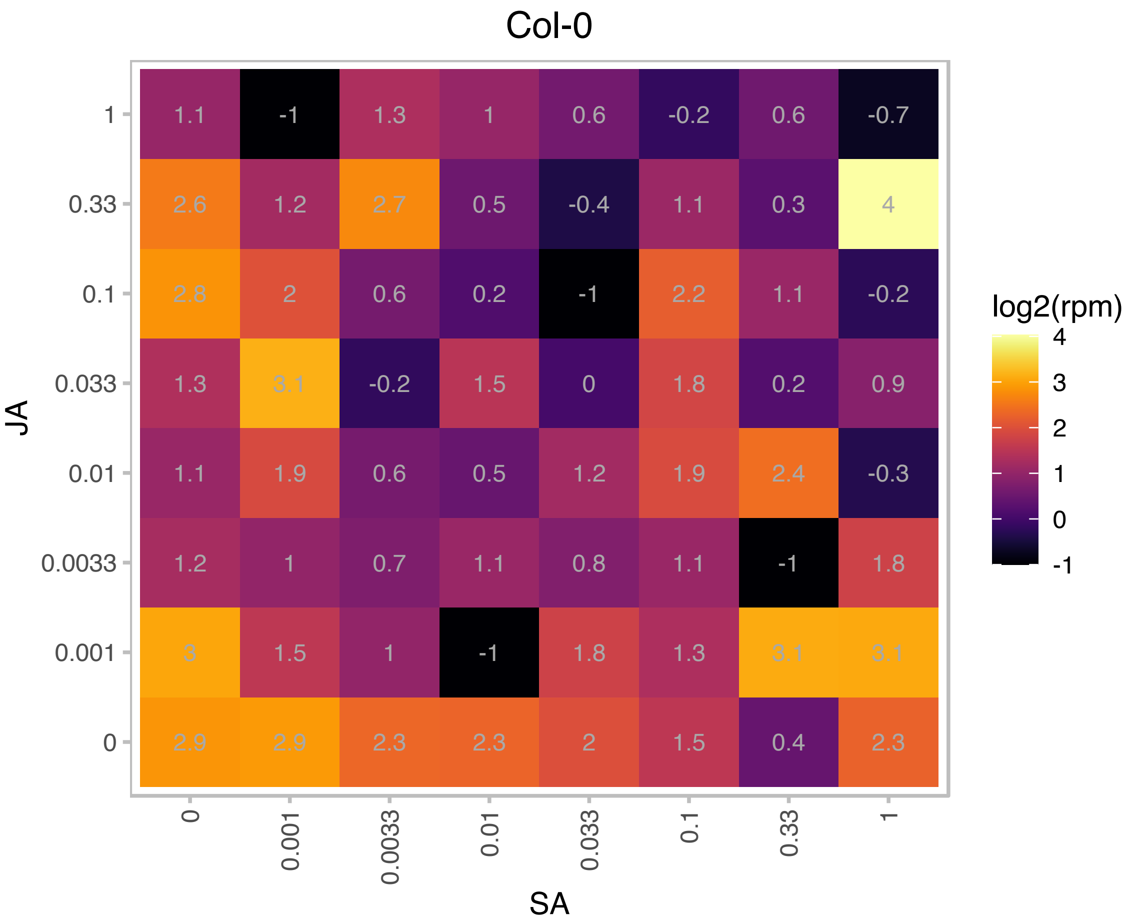


| | |
|---|---|
| Gene | AT1G11545 |
| is.expressed_genes | TRUE |
| is.responsive_genes | TRUE |
| is.apcluster | H-05 |

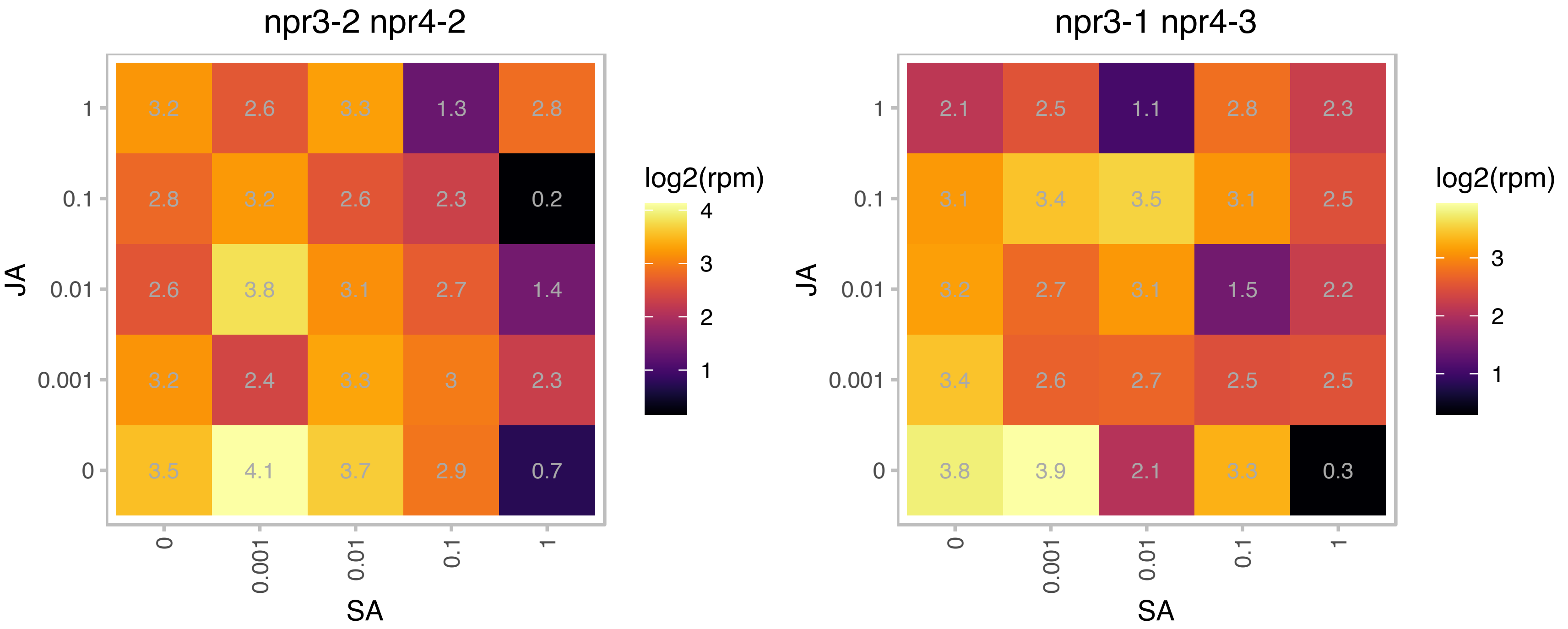

### AT1G20340.1 : DRT112, PETE2

recombination and DNA-damage resistance protein (DRT112) One of two Arabidopsis plastocyanin genes. Predominant form, expressed 10x higher than PETE1. PETE2 is thought to be post-transcriptionally regulated via copper accumulation and is involved in copper homeostasis. Mutation of this gene does not have obvious effect on photosynthesis. In plsp1-1 mutant plastids, the nonmature form of the protein localizes in the membrane.

Cupredoxin superfamily protein

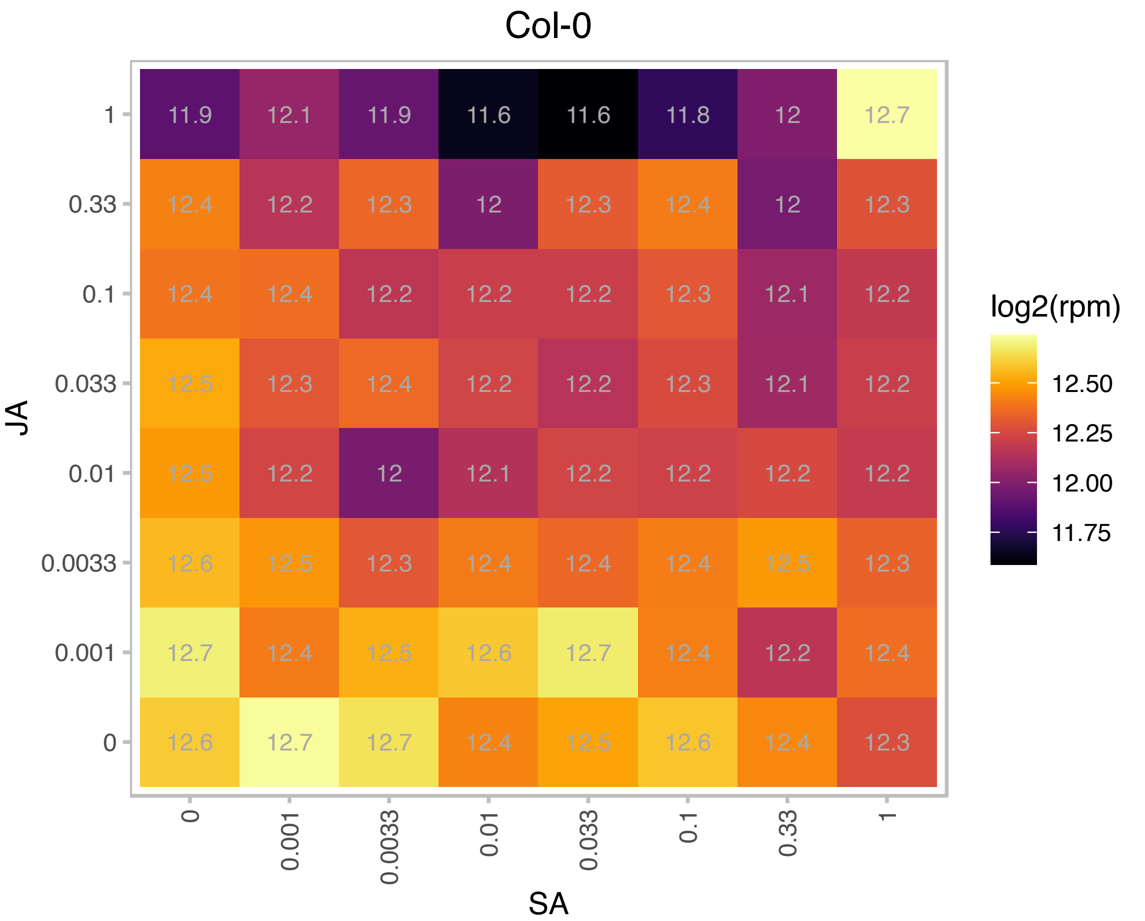


| | |
|---|---|
| Gene | AT1G20340 |
| is.expressed_genes | TRUE |
| is.responsive_genes | TRUE |
| is.apcluster | H-05 |

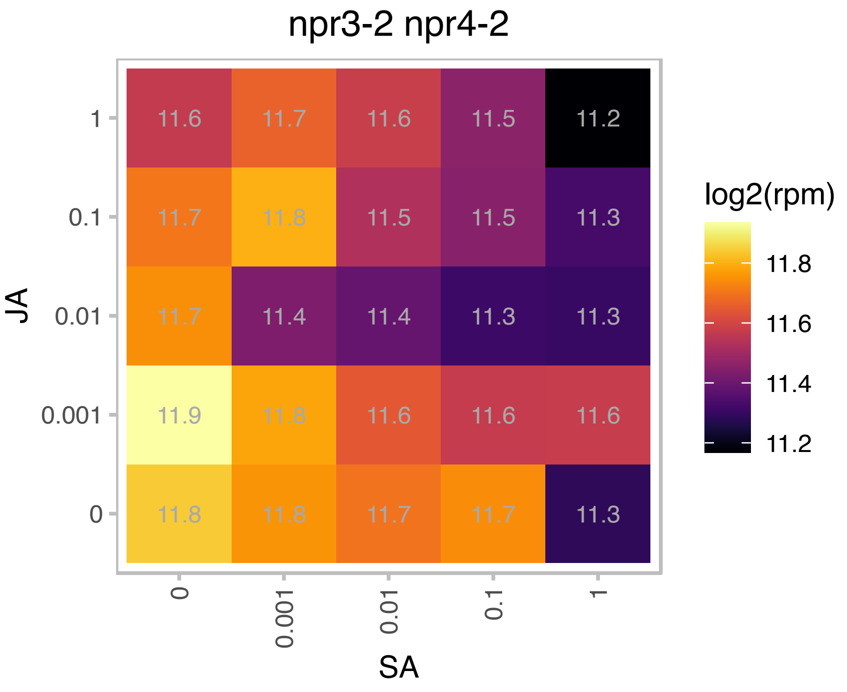


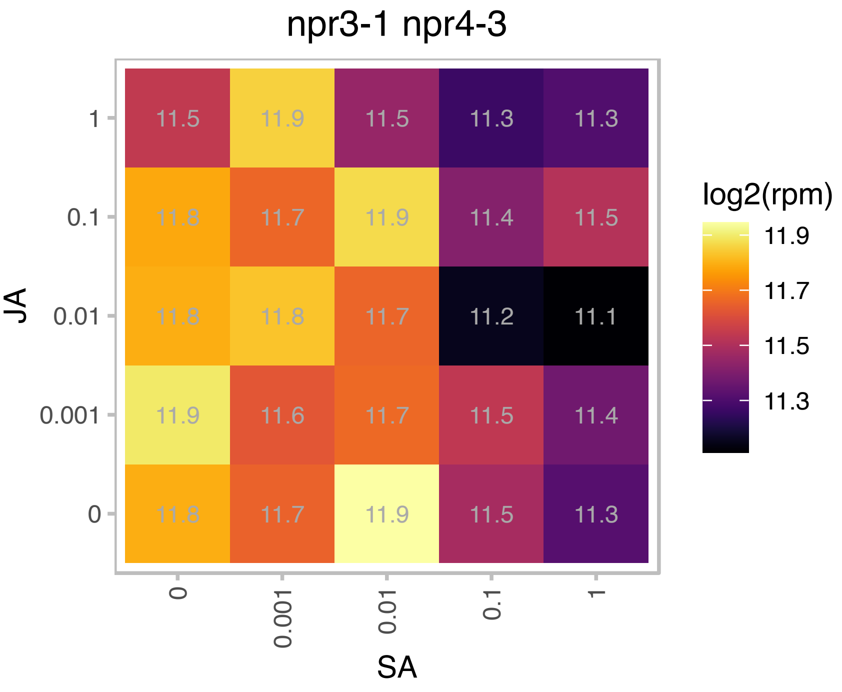

# AT1G28150.1 : NA

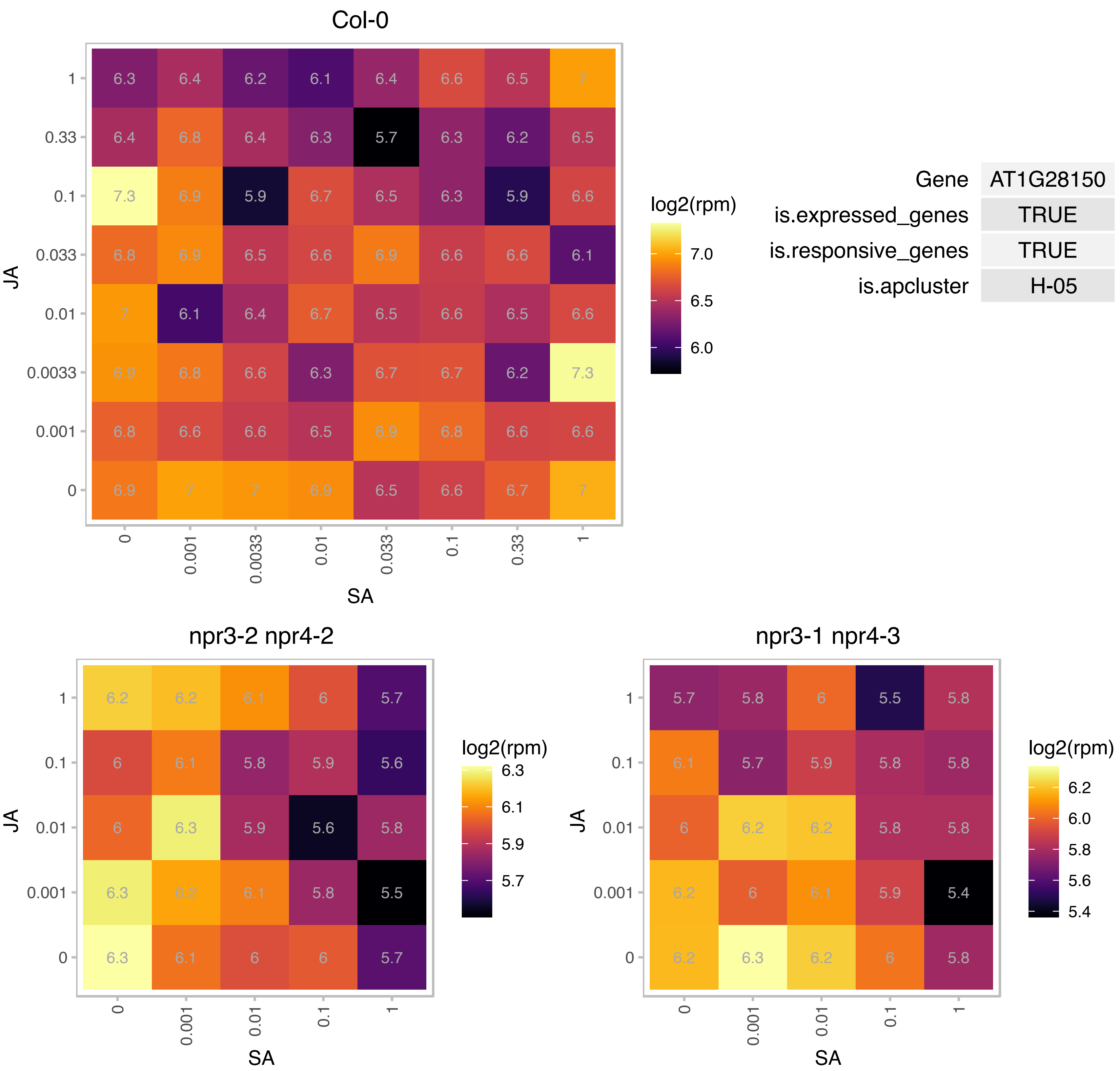

# AT1G48355.1 : NA

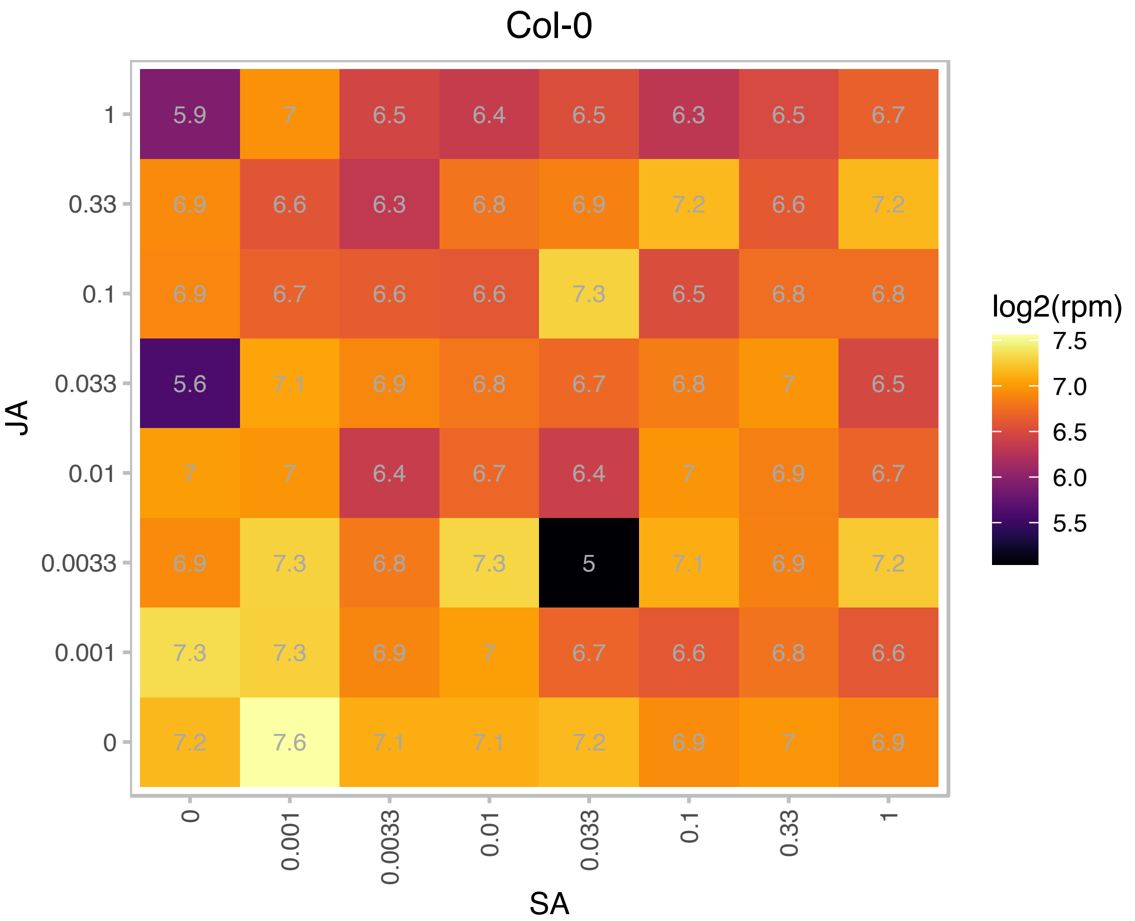


| | |
|---|---|
| Gene | AT1G48355 |
| is.expressed_genes | TRUE |
| is.responsive_genes | TRUE |
| is.apcluster | H-05 |

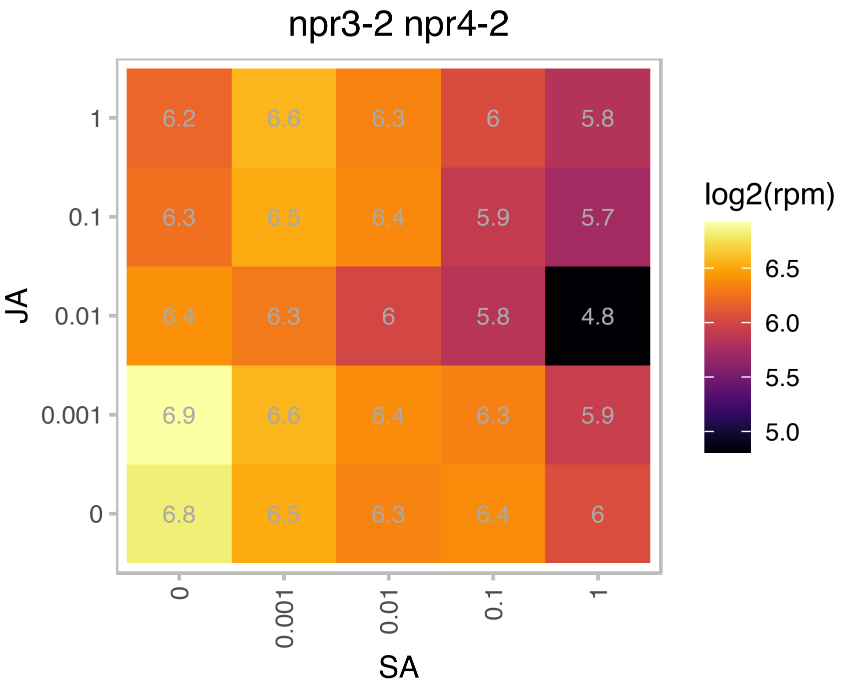


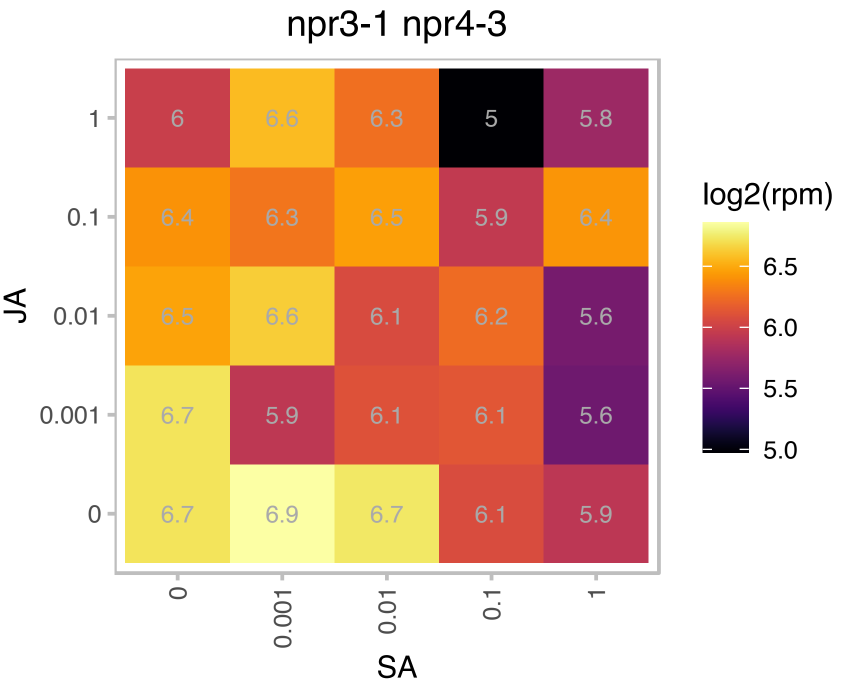

# AT1G68220.1 : NA

Protein of unknown function (DUF1218)

| | |
|---|---|
| Gene | AT1G68220 |
| is.expressed_genes | TRUE |
| is.responsive_genes | TRUE |
| is.apcluster | H-05 |

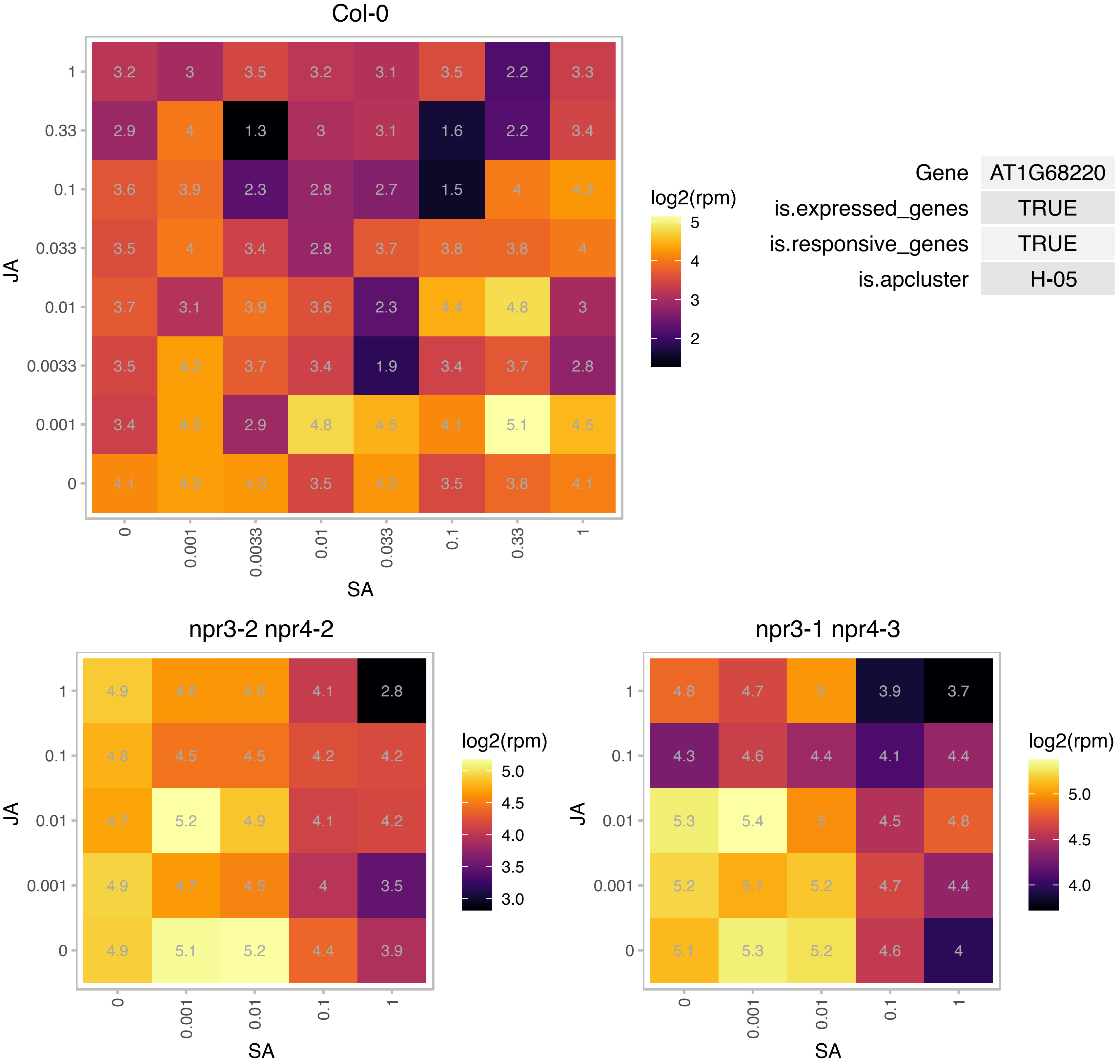

### AT1G73600.1 : AtPMT3, NMT, NMT3

S-adenosyl-L-methionine-dependent methyltransferases superfamily protein

Encodes a S-adenosyl-L-methionine-dependent phosphoethanolamine N-methyltransferase whose expression is responsive to both phosphate (Pi) and phosphite (Phi) in roots.

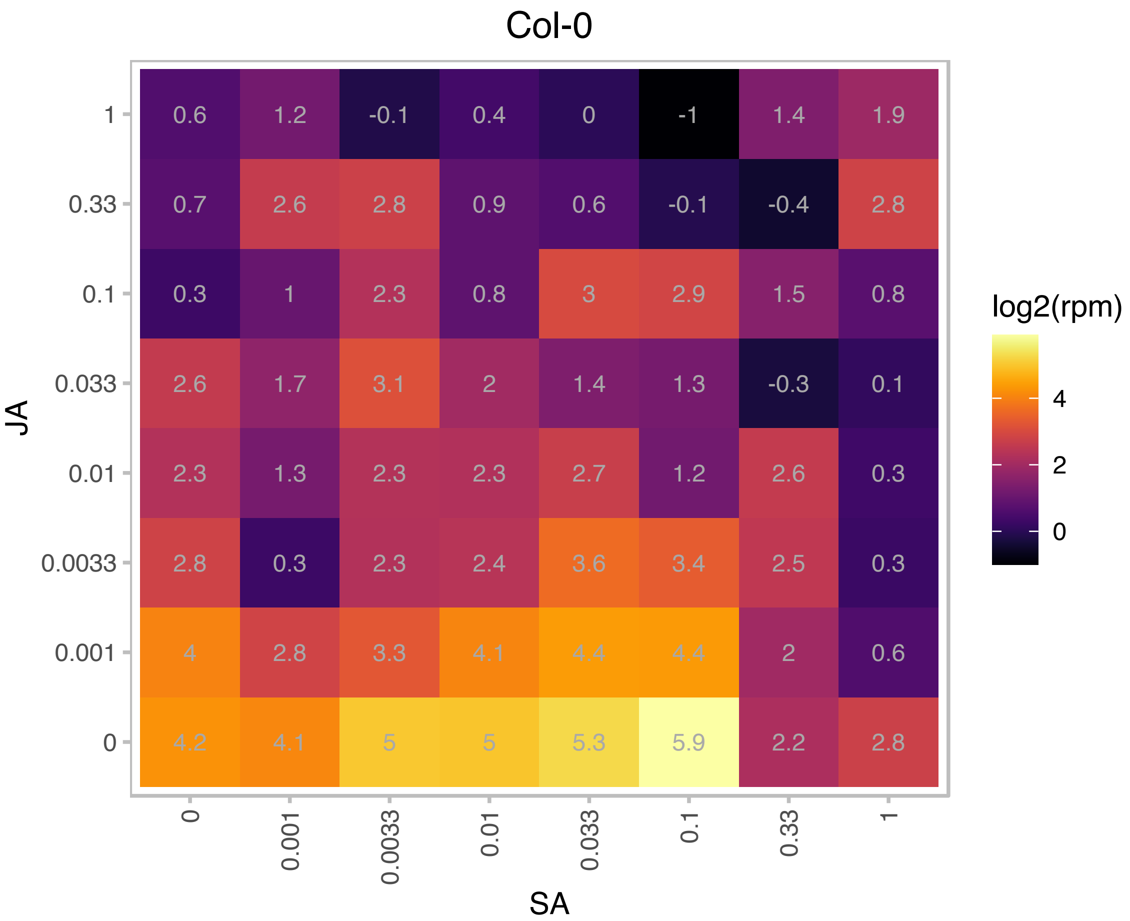


| Gene | AT1G73600 |
|---|---|
| is.expressed_genes | TRUE |
| is.responsive_genes | TRUE |
| is.apcluster | H-05 |

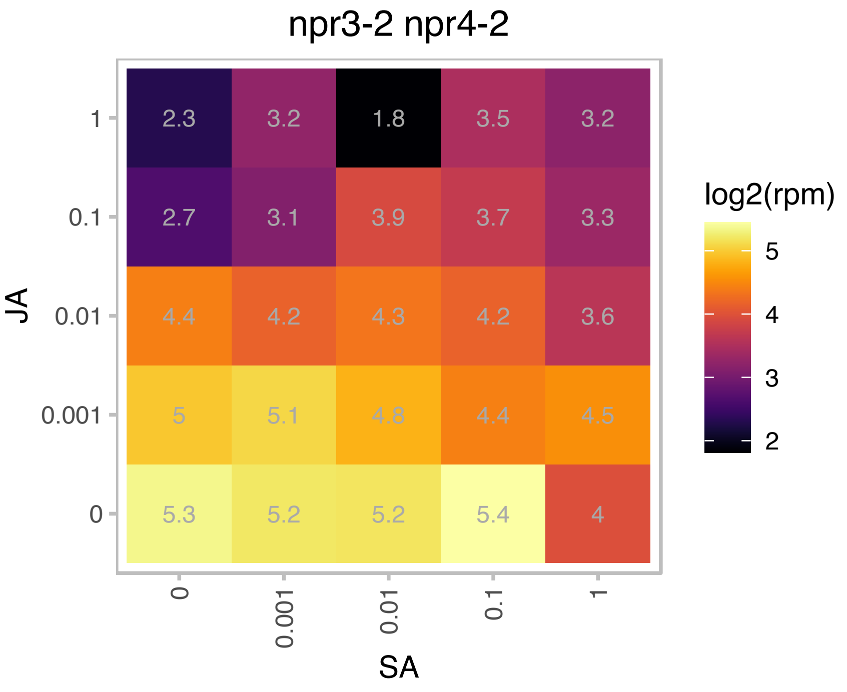


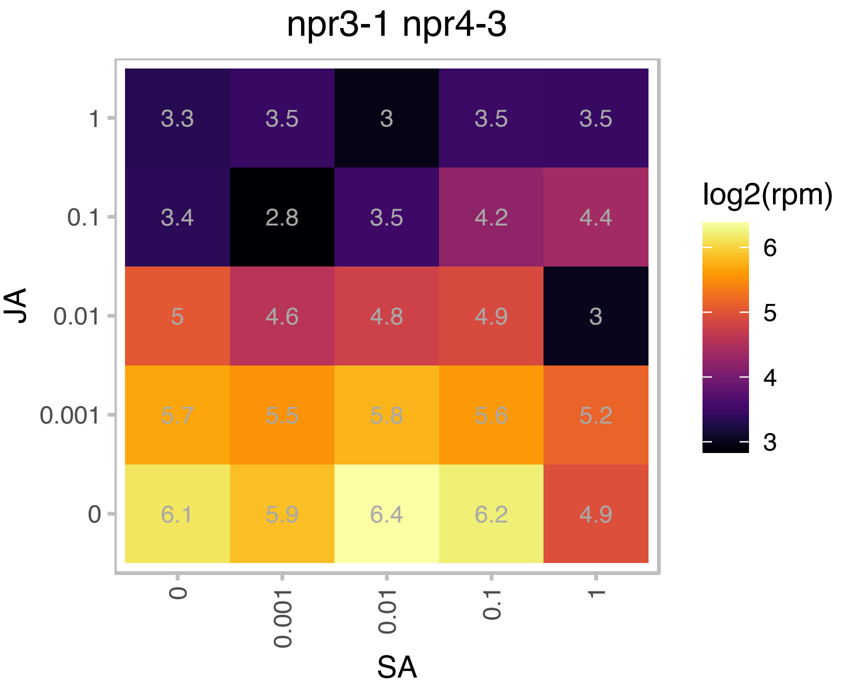

### AT1G80460.1 : GLI1, NHO1

Actin-like ATPase superfamily protein

Encodes a protein similar to glycerol kinase, which converts glycerol to glycerol 3-phosphate and performs a rate-limiting step in glycerol metabolism. This gene is required for both general and specific resistance against bacteria and fungi. Arabidopsis thaliana glycerol kinase (GLR1) mRNA.Involved in flagellin-induced non-host resistance to Pseudomonas. Coronatine partially suppresses flagellin-induced expression of NHO1.

Col-0

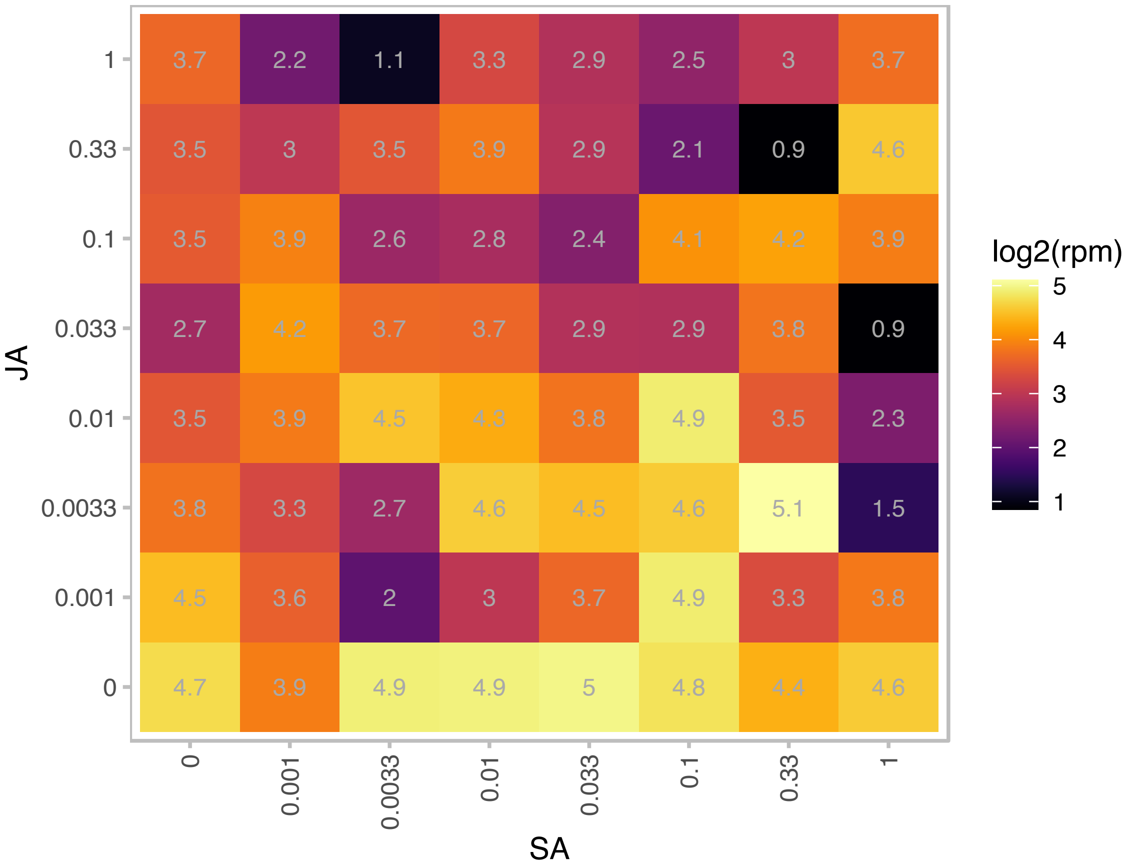


| | |
|---|---|
| Gene | AT1G80460 |
| is.expressed_genes | TRUE |
| is.responsive_genes | TRUE |
| is.apcluster | H-05 |

npr3-2 npr4-2

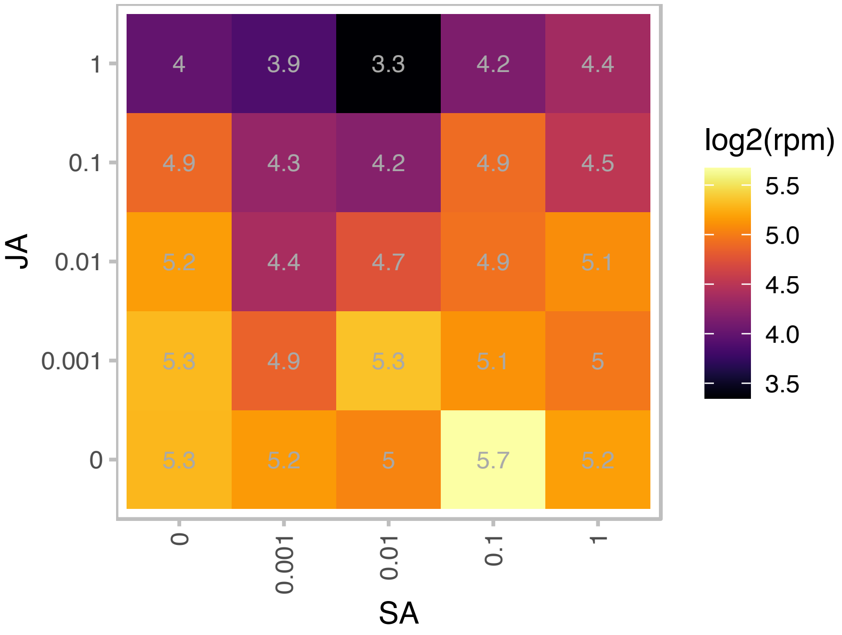


npr3-1 npr4-3

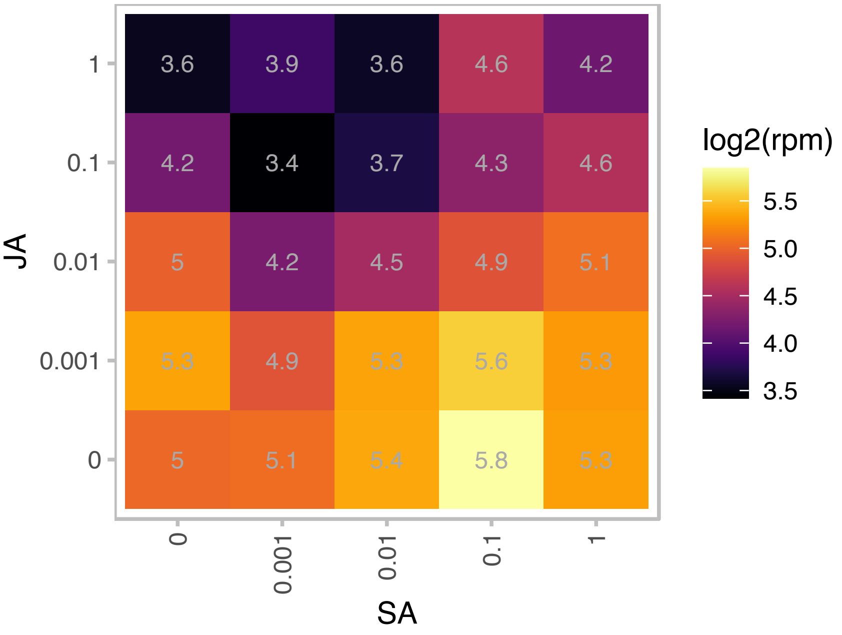

### AT2G02950.1 : PKS1

phytochrome kinase substrate 1
Encodes a basic soluble protein which can independently bind to either PHYA or PHYB, regardless of whether the phytochromes are in the Pr or Pfr state. PKS1 can be phosphorylated by oat phyA in vitro in a light regulated manner. It is postulated to be a negative regulator of phyB signalling.

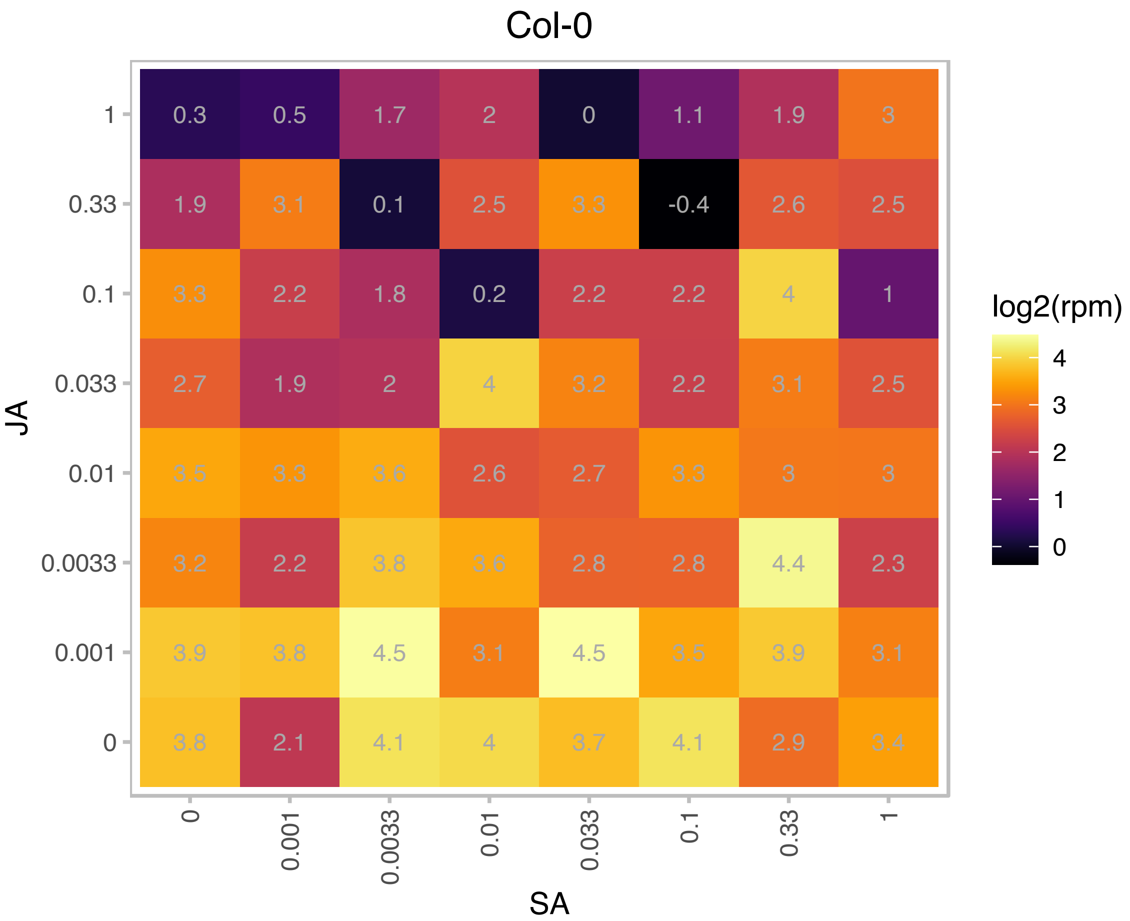


| | |
|---|---|
| Gene | AT2G02950 |
| is.expressed_genes | TRUE |
| is.responsive_genes | TRUE |
| is.apcluster | H-05 |

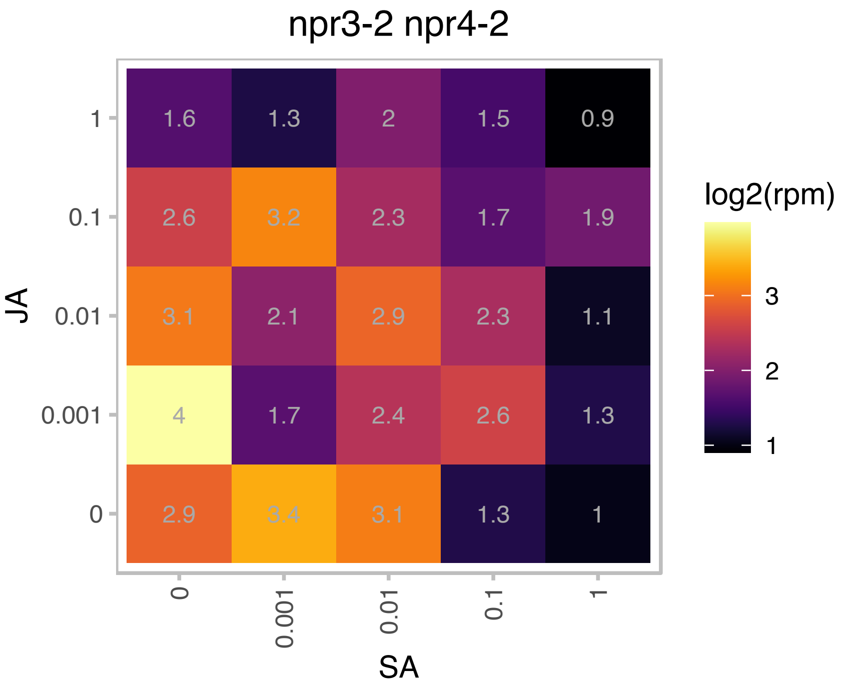


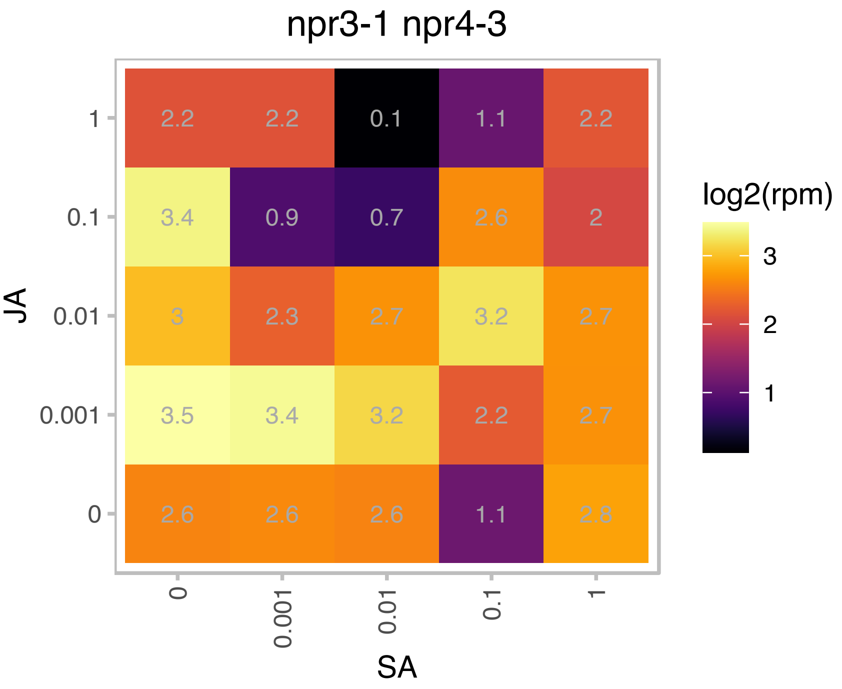

### AT2G05620.1 : AtPGR5, PGR5

proton gradient regulation 5

Involved in electron flow in Photosystem I. Essential for photoprotection.

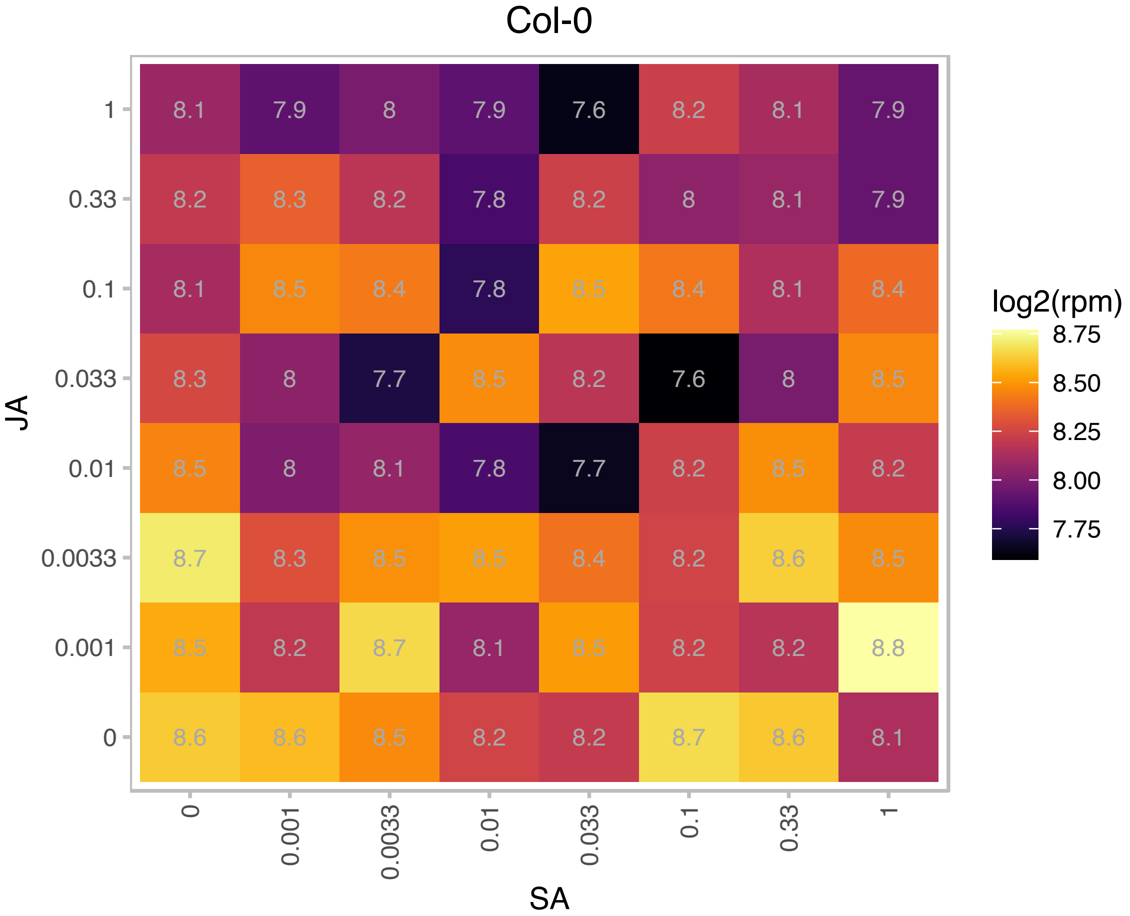


| | |
|---|---|
| Gene | AT2G05620 |
| is.expressed_genes | TRUE |
| is.responsive_genes | TRUE |
| is.apcluster | H-05 |

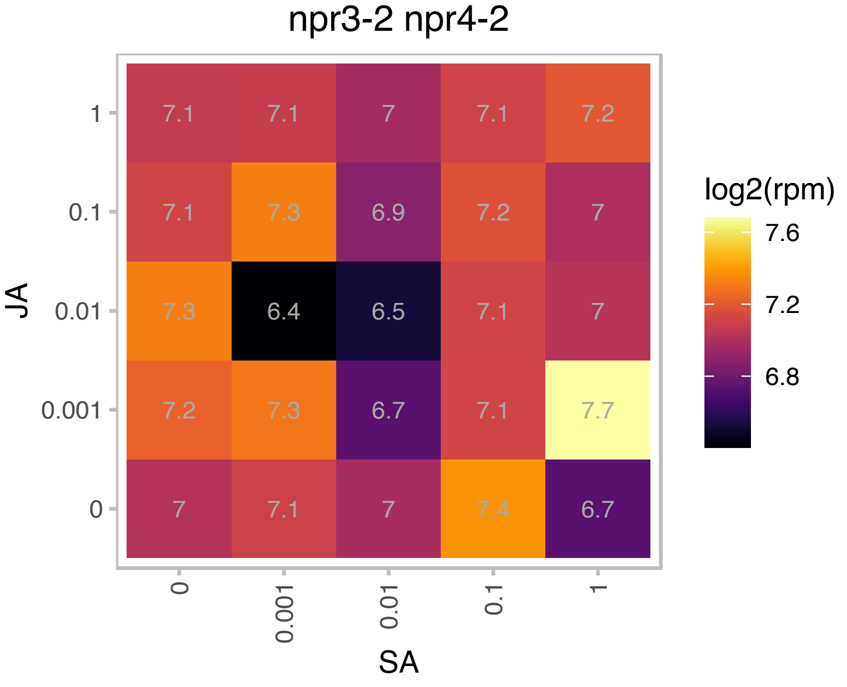


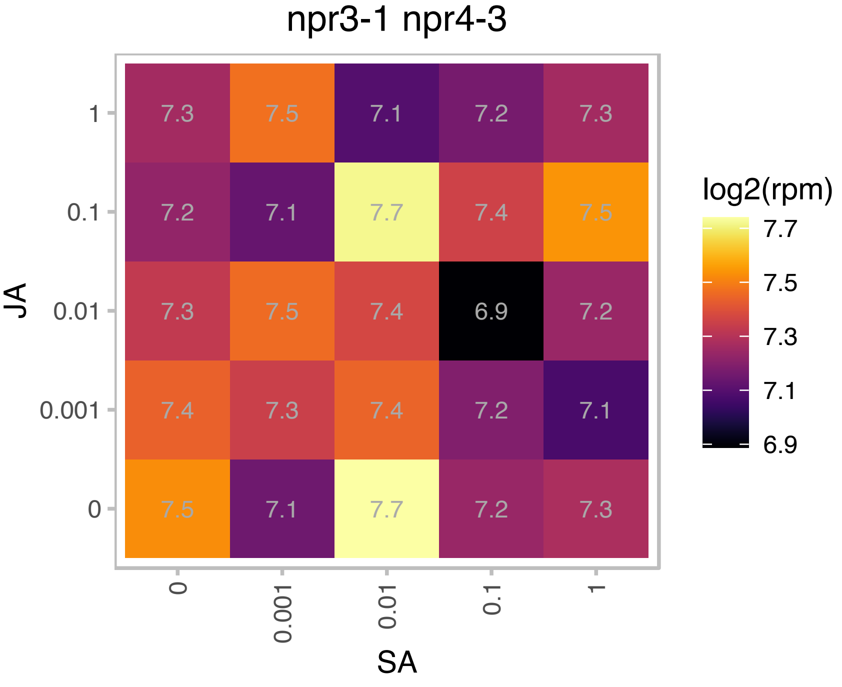

### AT2G18390.1 : ARL2, ATARLC1, HAL, TTN5

ADP-ribosylation factor family protein

Encodes a member of ARF-like GTPase family. A thaliana has 21 members, in two subfamilies, ARF and ARF-like (ARL) GTPases.  Mutant has abnormal mitosis and cell cycle control during seed development.

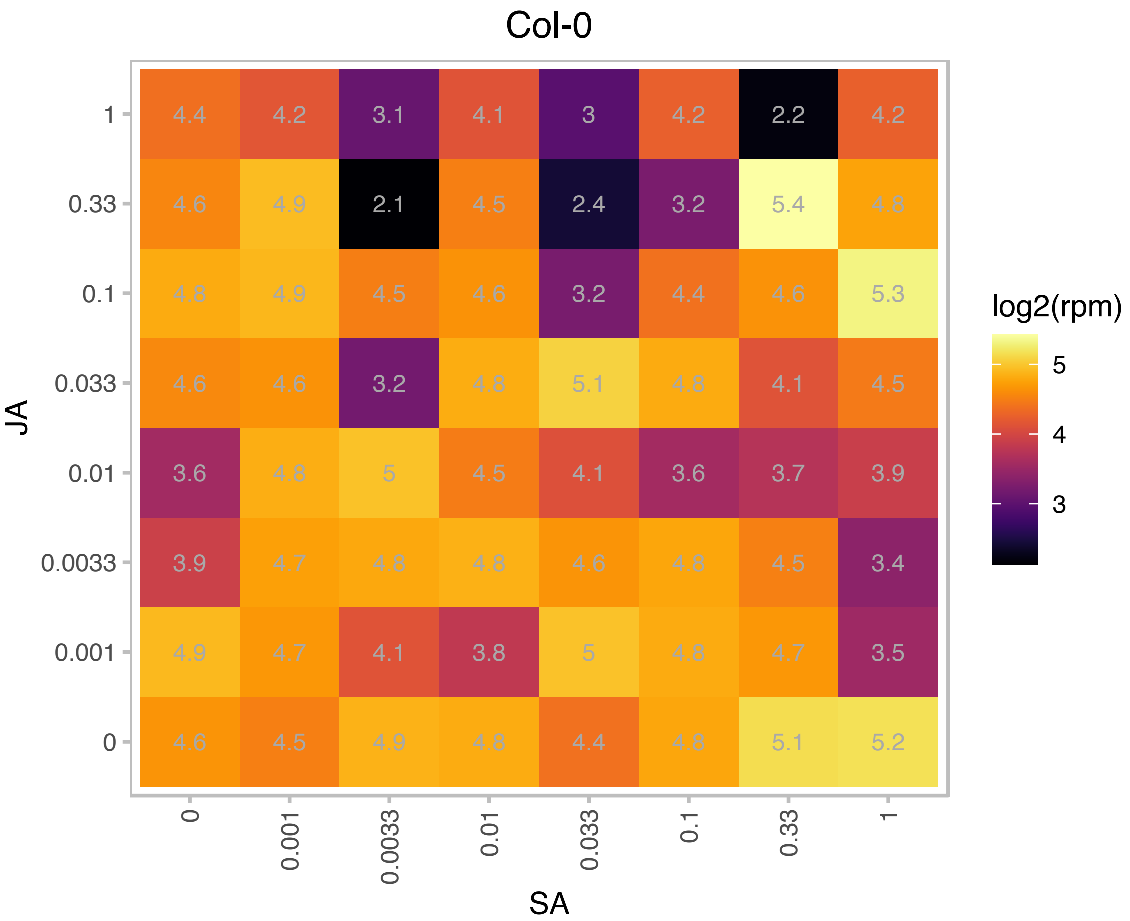


| | |
|---|---|
| Gene | AT2G18390 |
| is.expressed_genes | TRUE |
| is.responsive_genes | TRUE |
| is.apcluster | H-05 |

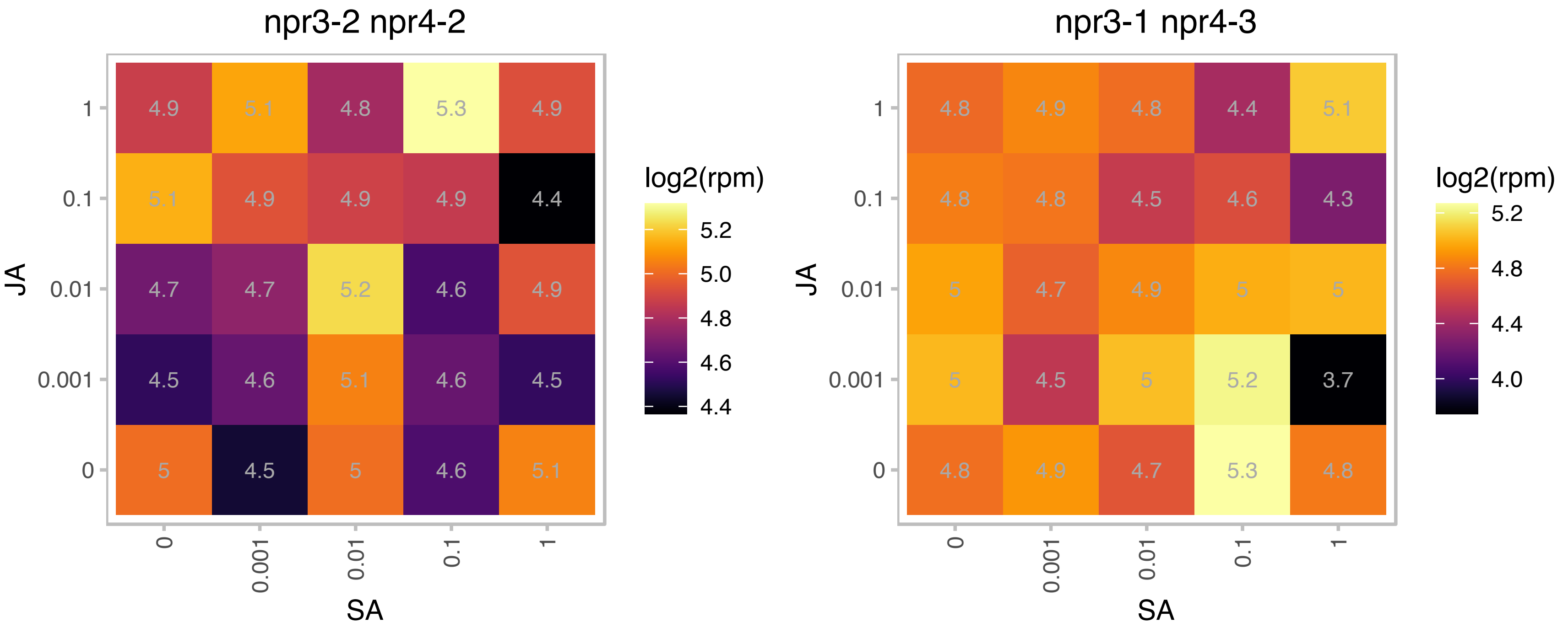

# AT2G36145.1 : NA

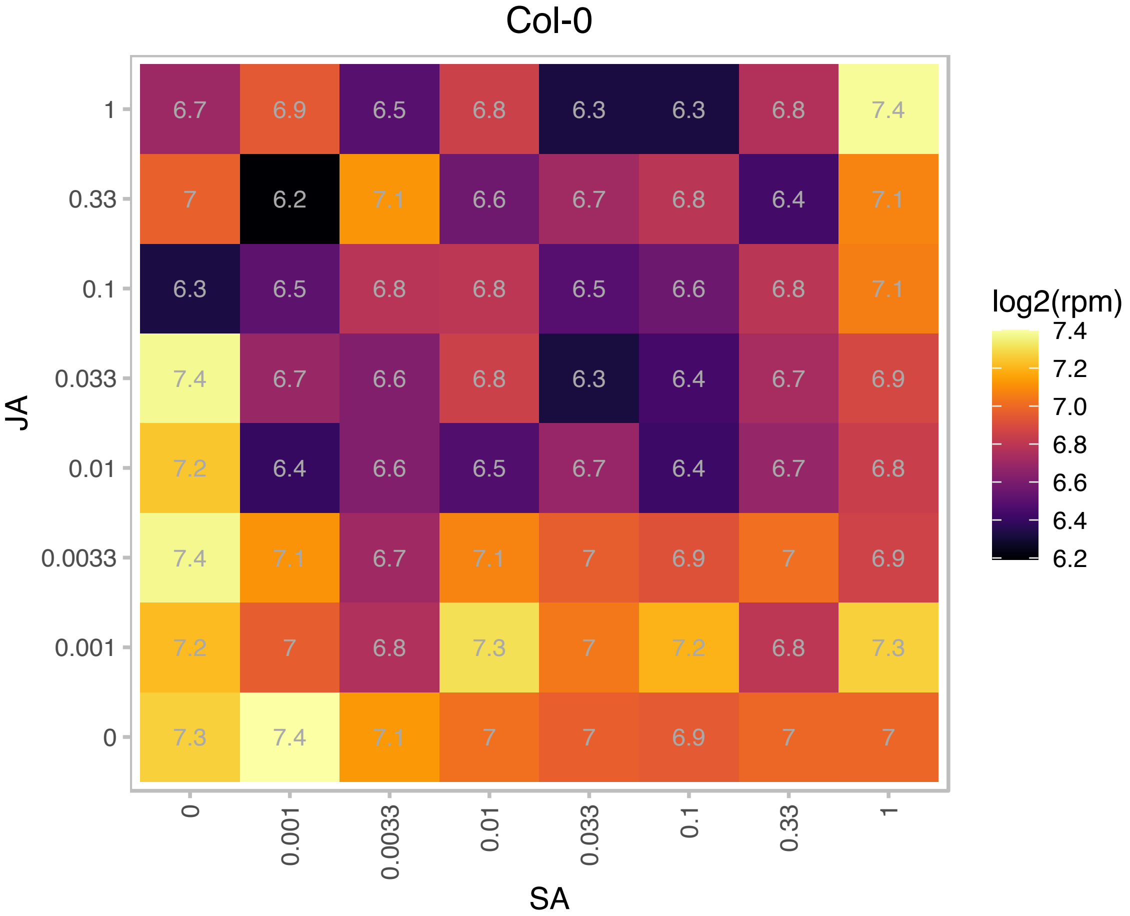


| | |
|---|---|
| Gene | AT2G36145 |
| is.expressed_genes | TRUE |
| is.responsive_genes | TRUE |
| is.apcluster | H-05 |

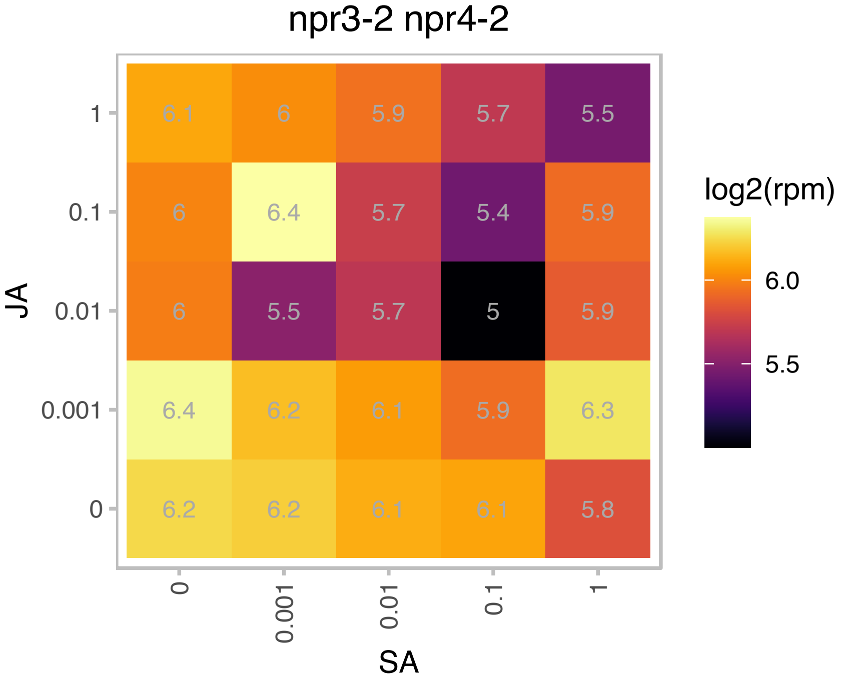


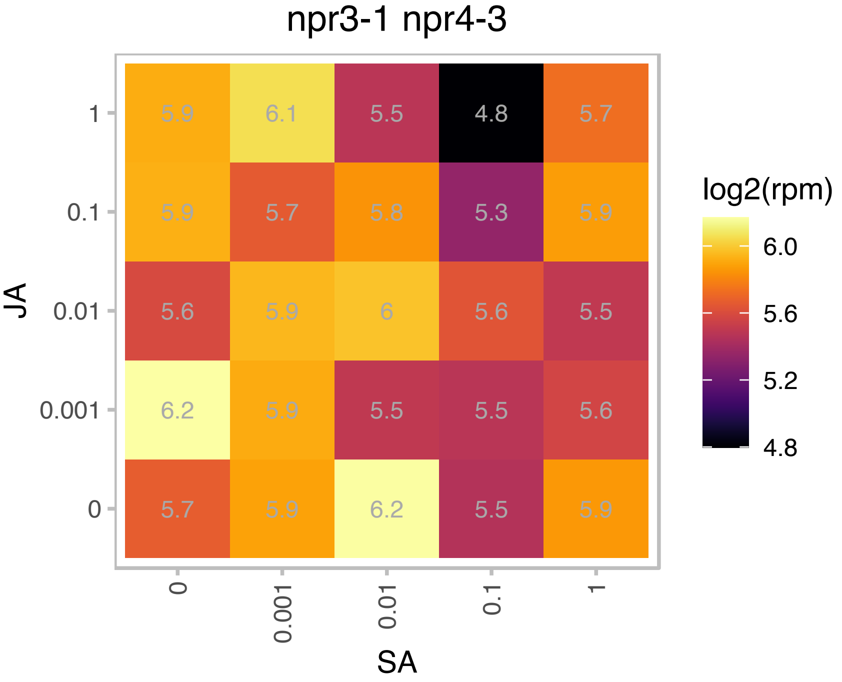

### AT2G43030.1 : PRPL3

Ribosomal protein L3 family protein

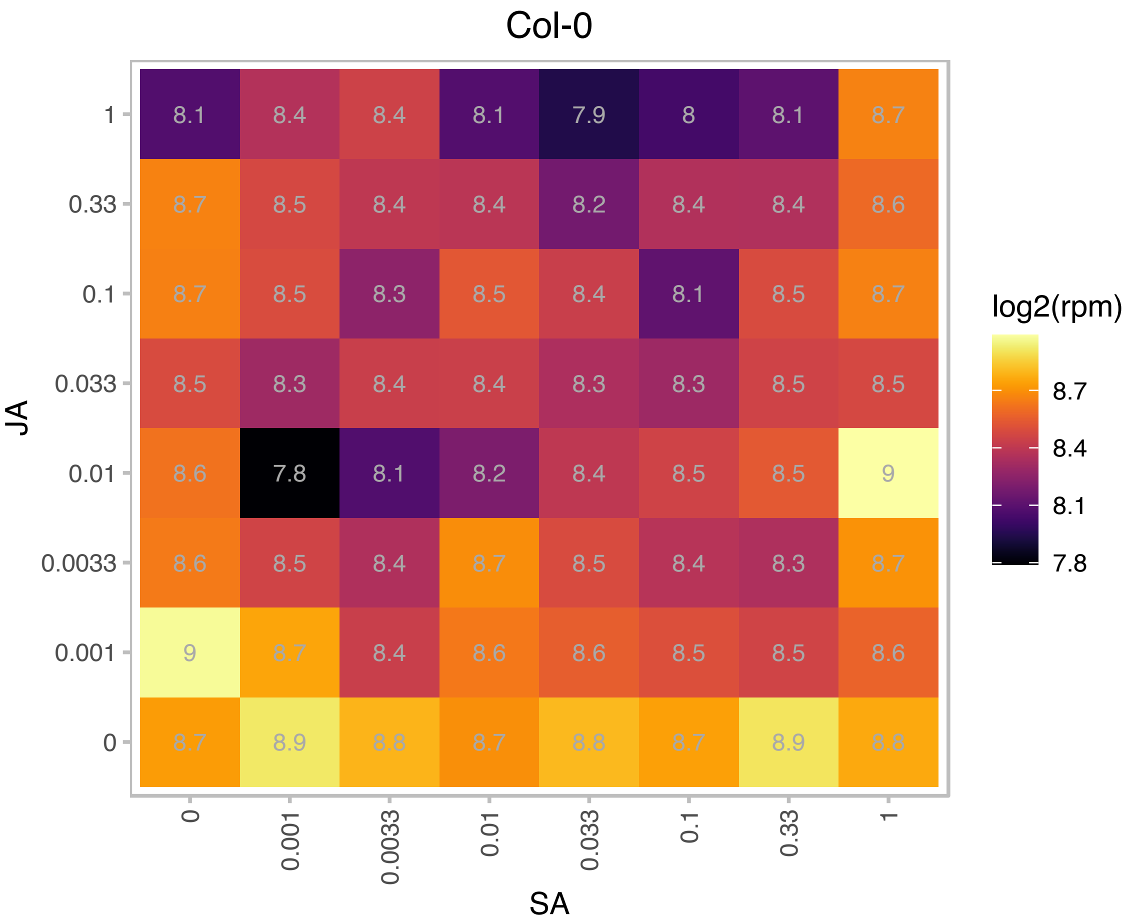


| | |
|---|---|
| Gene | AT2G43030 |
| is.expressed_genes | TRUE |
| is.responsive_genes | TRUE |
| is.apcluster | H-05 |

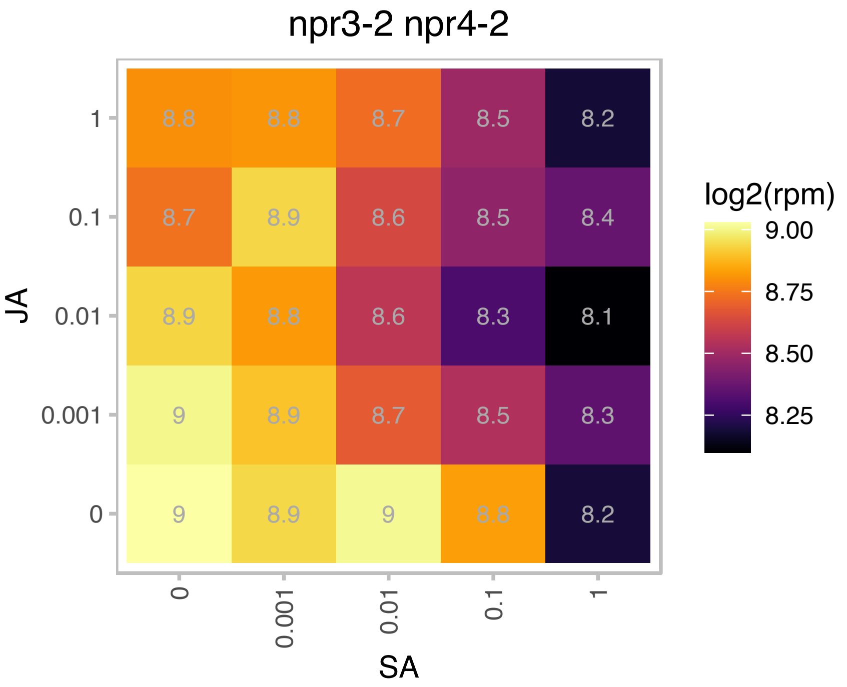


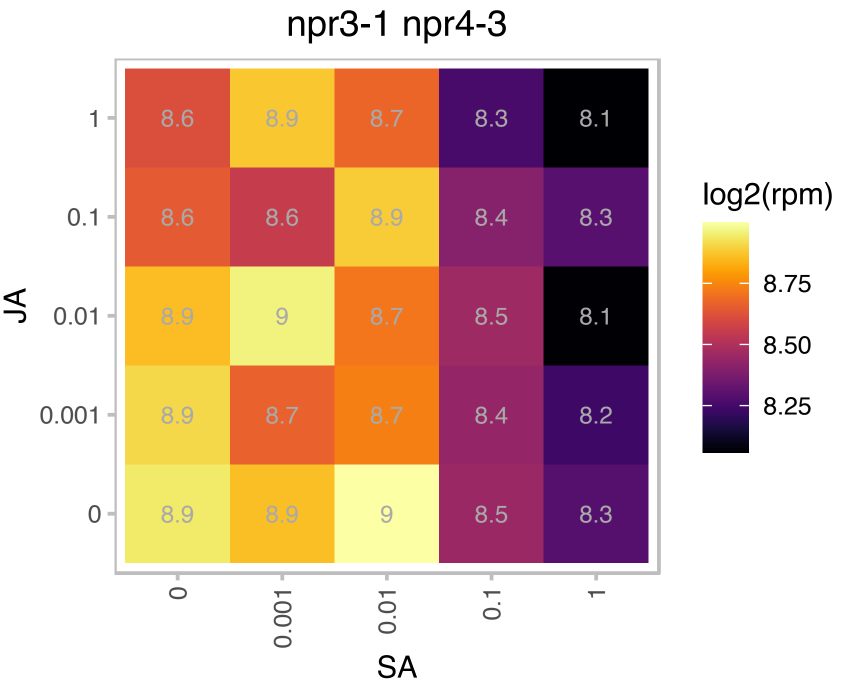

### AT3G02730.1 : ATF1, TRXF1

thioredoxin F-type 1

Encodes a type-f thioredoxin.  Has a role in the short-term activation of carbon metabolism. Loss affects growth under short-day conditions.

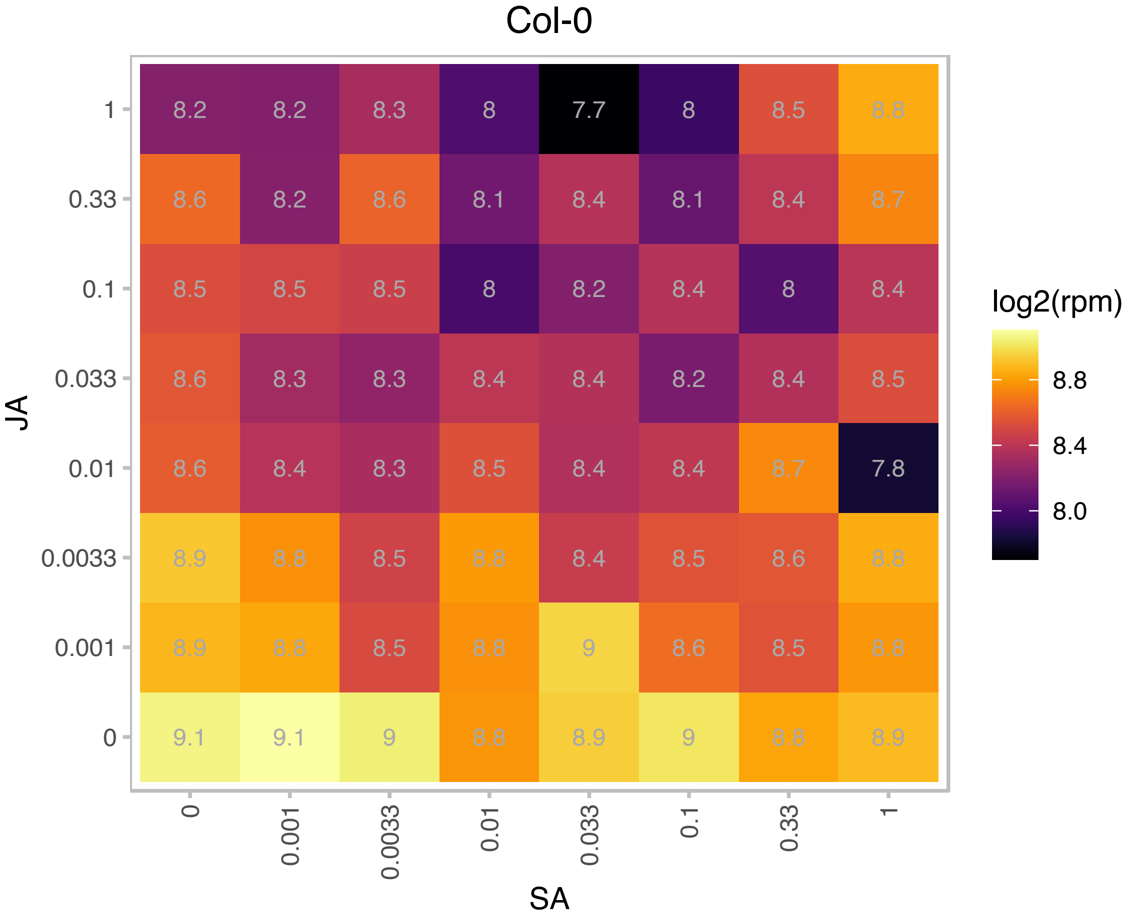


| | |
|---|---|
| Gene | AT3G02730 |
| is.expressed_genes | TRUE |
| is.responsive_genes | TRUE |
| is.apcluster | H-05 |

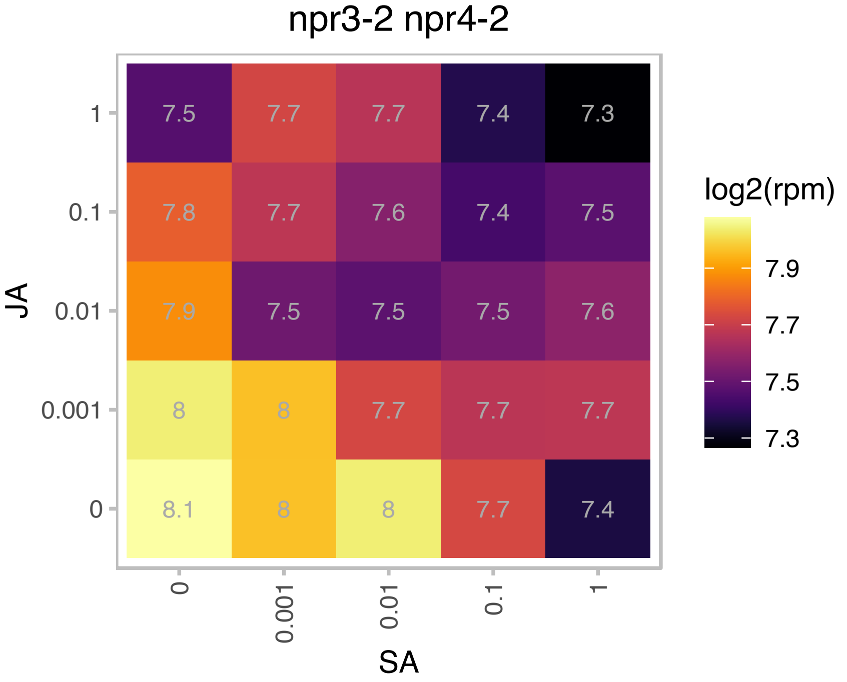


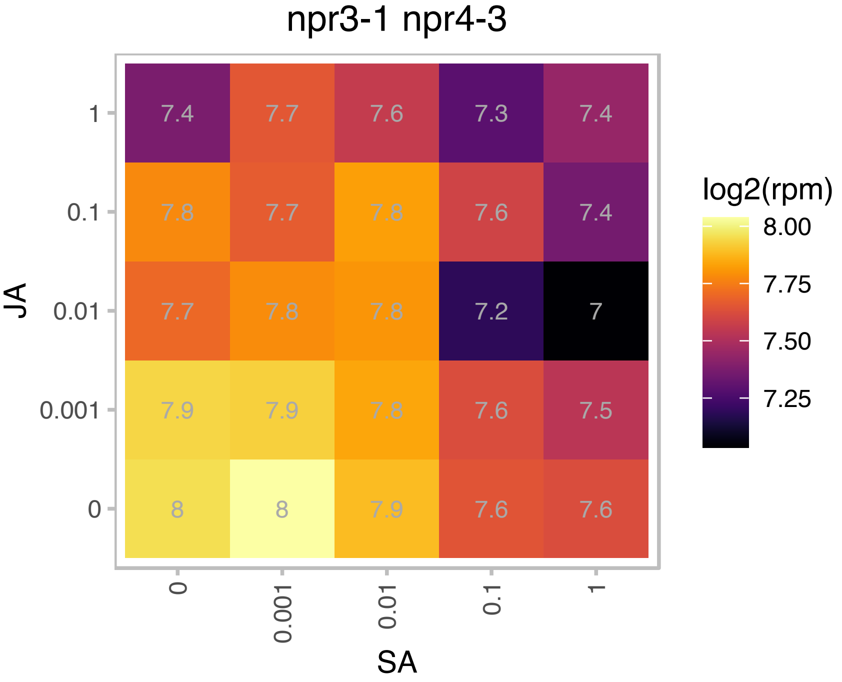

### AT3G04790.1 : EMB3119

Ribose 5-phosphate isomerase, type A protein

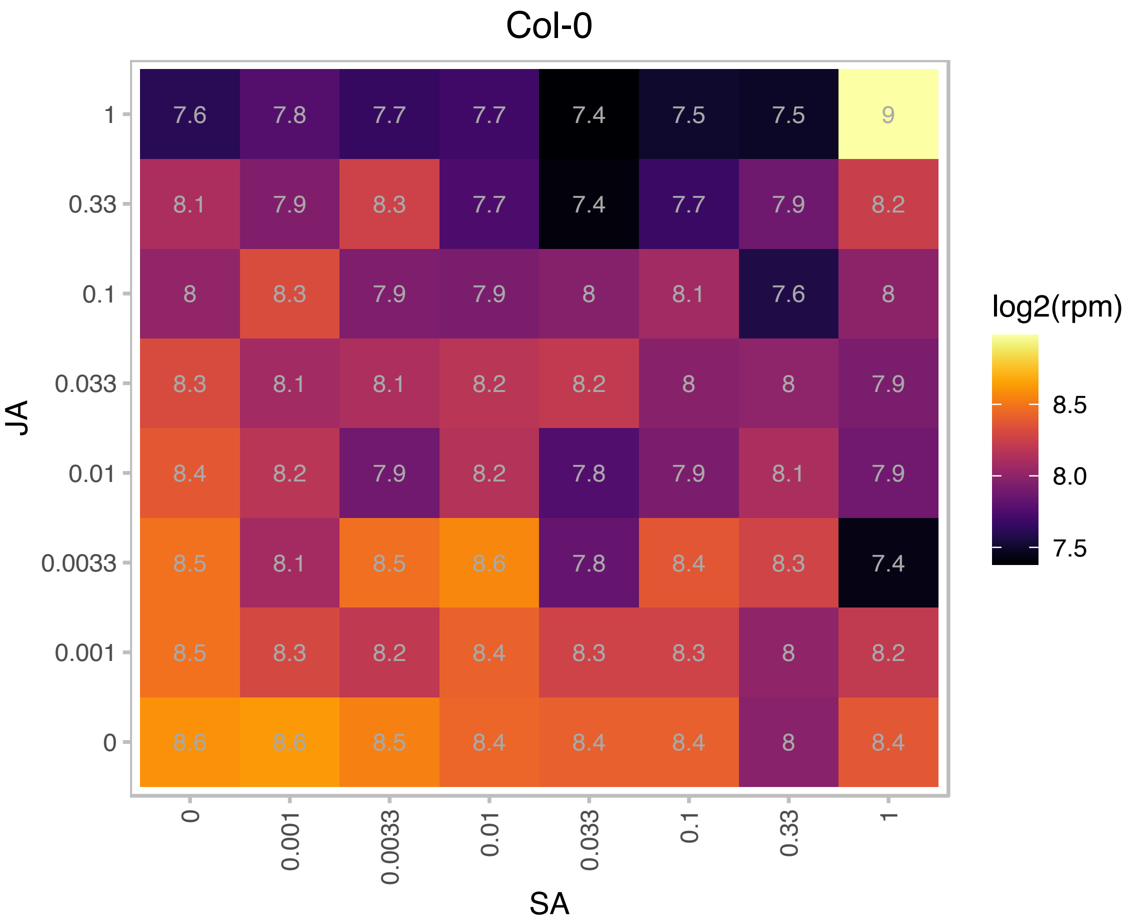


| | |
|---|---|
| Gene | AT3G04790 |
| is.expressed_genes | TRUE |
| is.responsive_genes | TRUE |
| is.apcluster | H-05 |

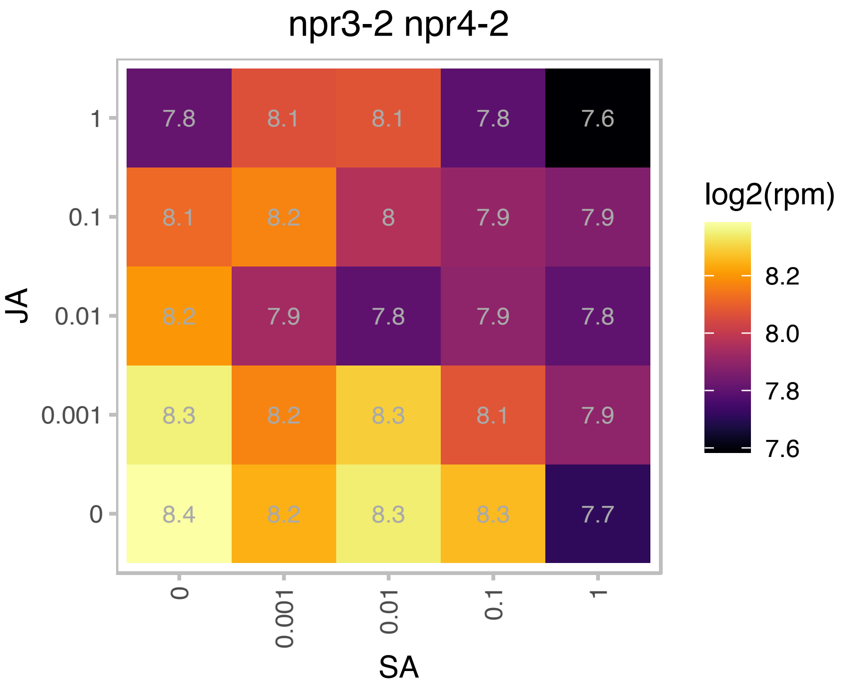


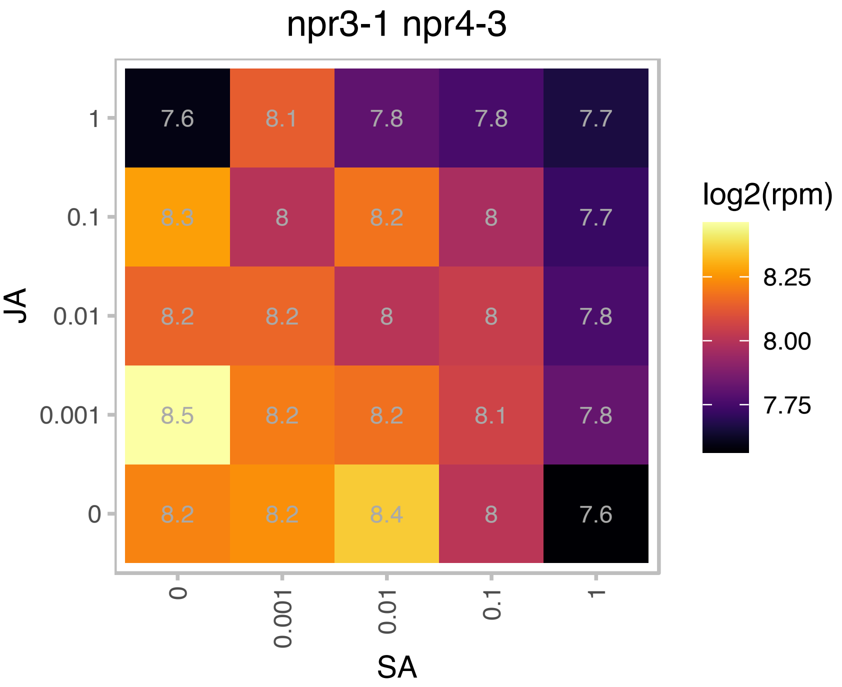

### AT3G07880.1 : SCN1

Immunoglobulin E-set superfamily protein

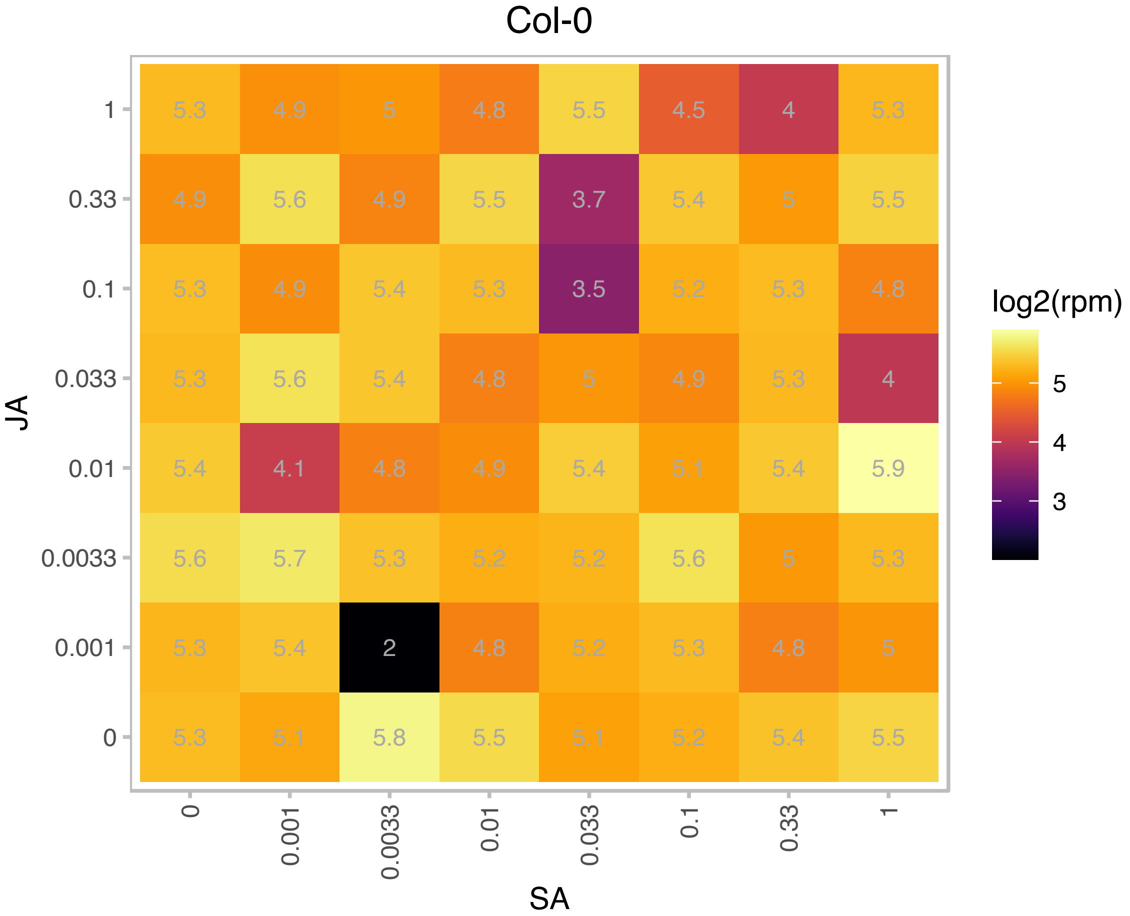


| | |
|---|---|
| Gene | AT3G07880 |
| is.expressed_genes | TRUE |
| is.responsive_genes | TRUE |
| is.apcluster | H-05 |

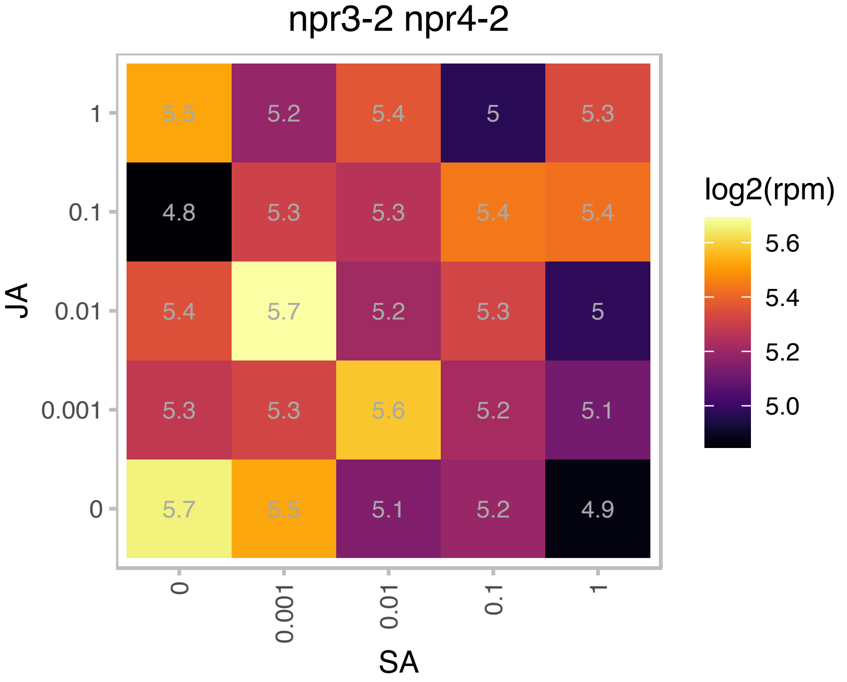


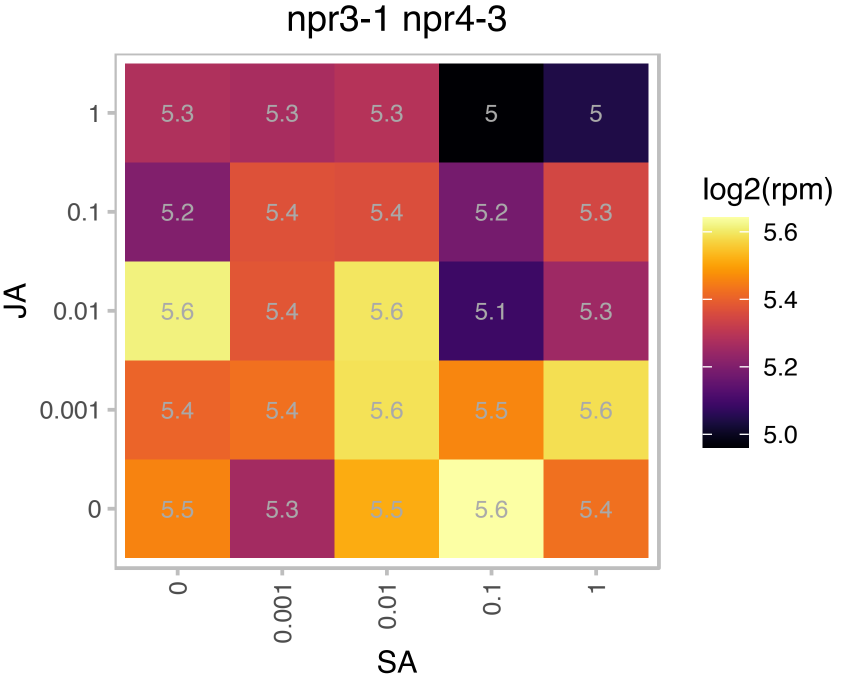

# AT3G07910.1 : NA

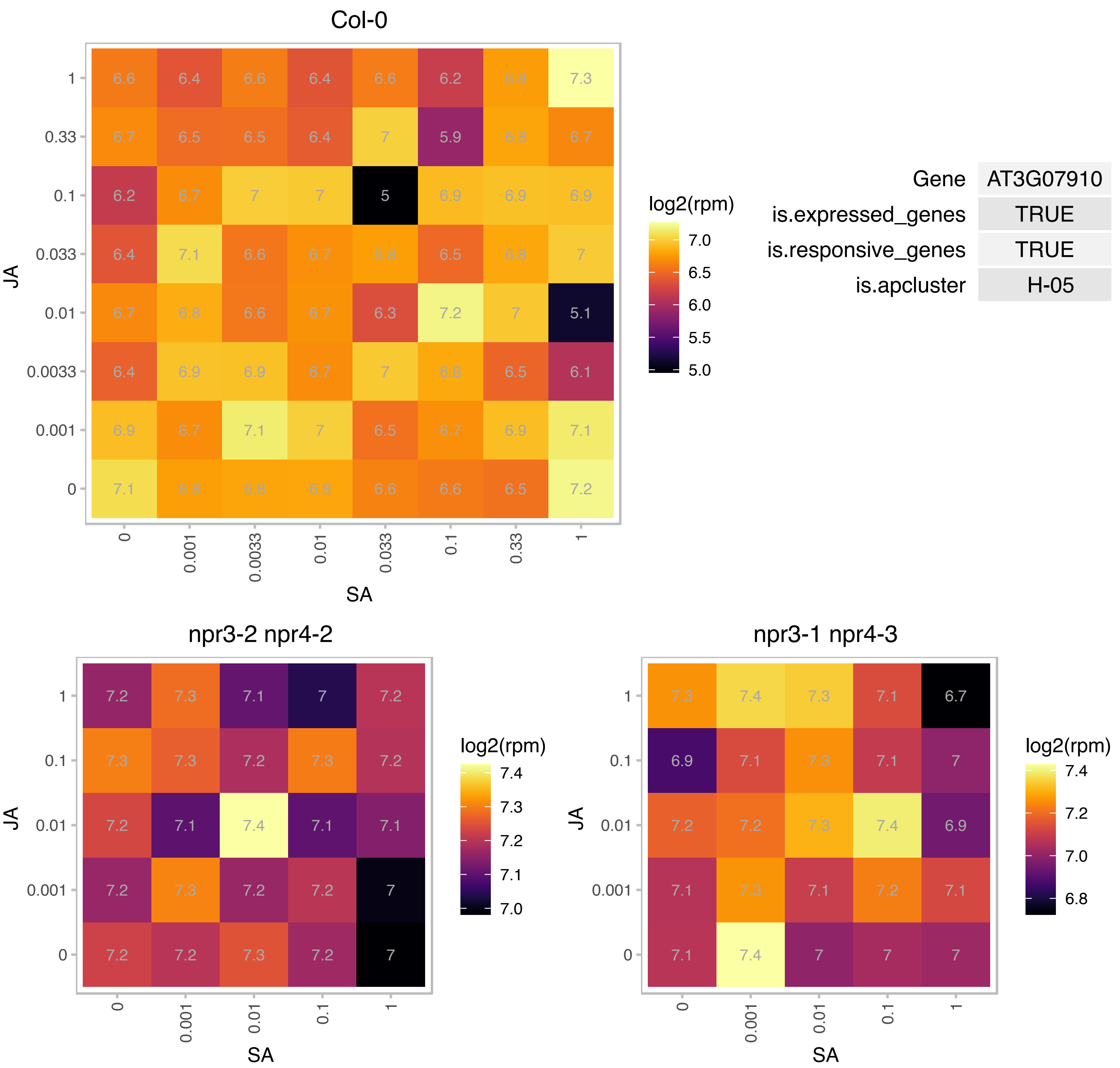

# AT3G08740.1 : NA

elongation factor P (EF-P) family protein

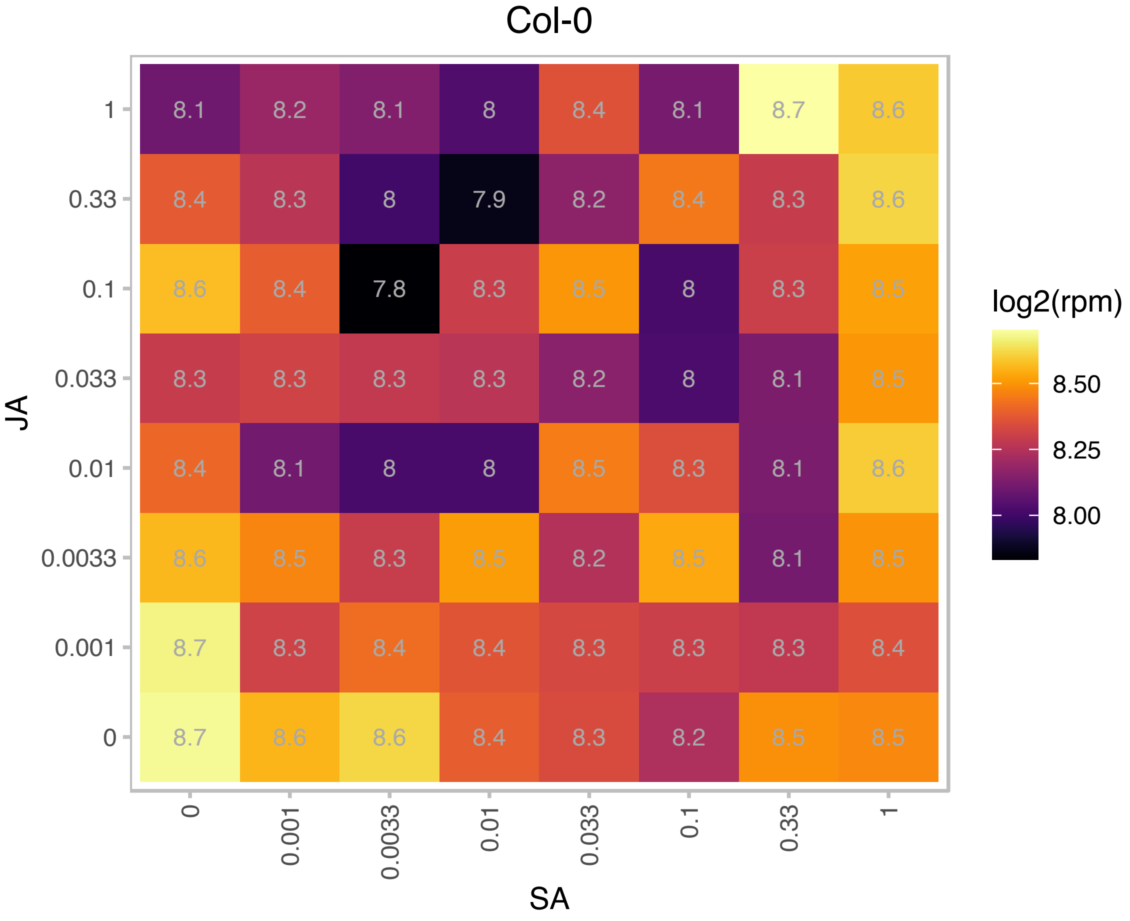


| | |
|---|---|
| Gene | AT3G08740 |
| is.expressed_genes | TRUE |
| is.responsive_genes | TRUE |
| is.apcluster | H-05 |

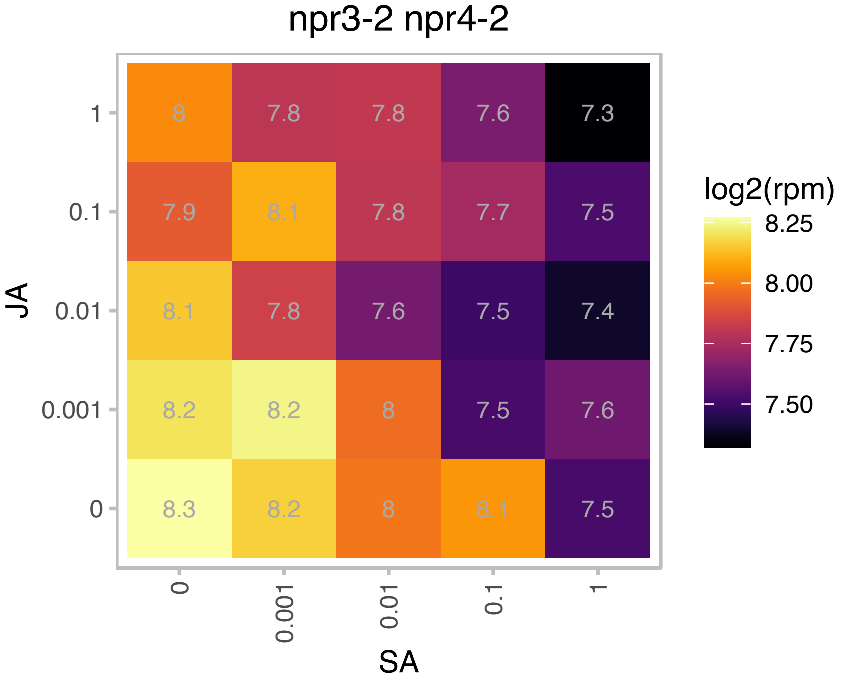


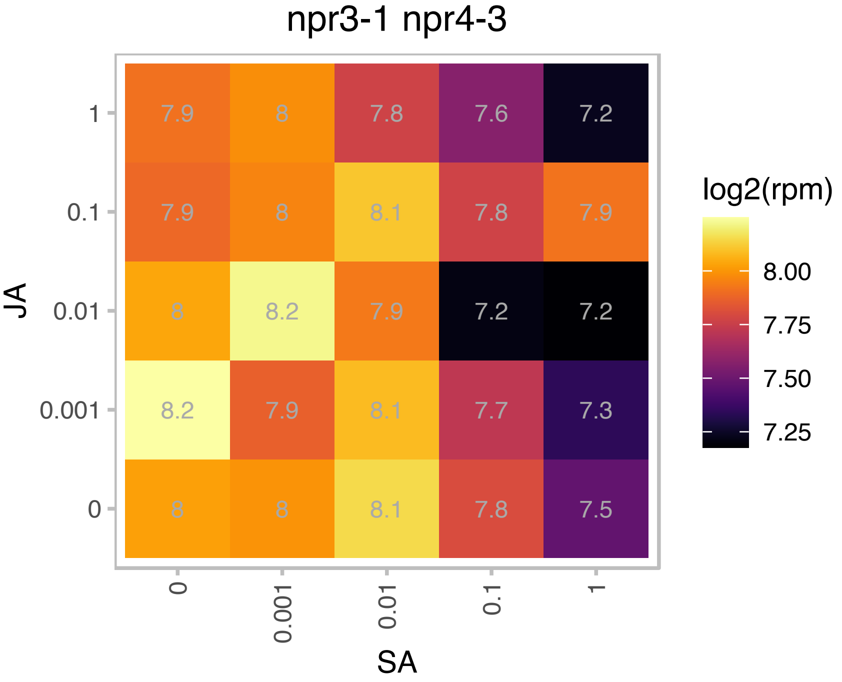

### AT3G10520.1 : AHB2, ARATH GLB2, ATGLB2, GLB2, HB2

haemoglobin 2
Encodes a class 2 non-symbiotic hemoglobin.  Over-expression of AHb2 in seeds led to a 40% increase in the total fatty acid content of developing and mature seeds in three subsequent generations. This was mainly due to an increase in the poly-unsaturated C18:2 (omega-6) linoleic and C18:3 (omega-3) alpha-linolenic acids.

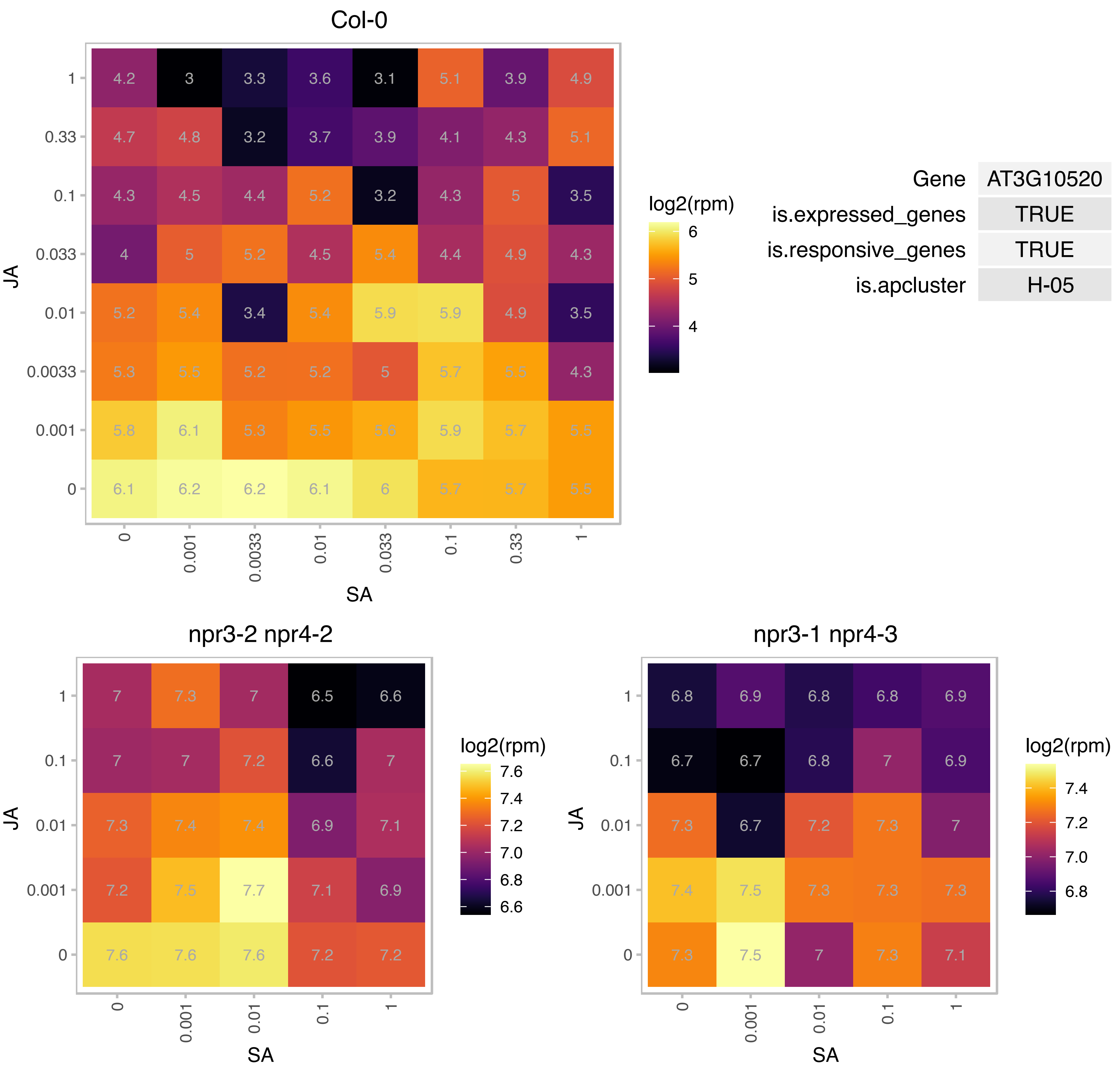


| | |
|---|---|
| Gene | AT3G10520 |
| is.expressed_genes | TRUE |
| is.responsive_genes | TRUE |
| is.apcluster | H-05 |

# AT3G13435.1 : NA

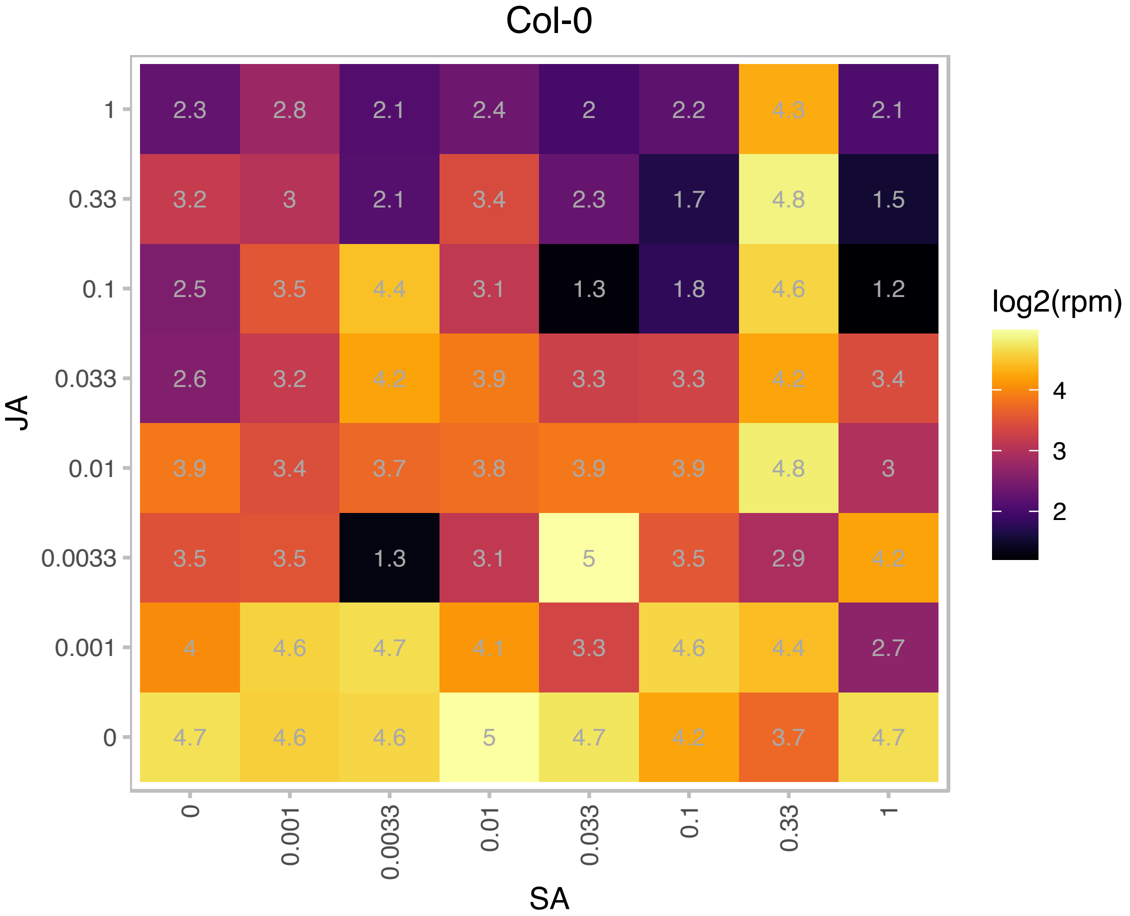


| | |
|---|---|
| Gene | AT3G13435 |
| is.expressed_genes | TRUE |
| is.responsive_genes | TRUE |
| is.apcluster | H-05 |

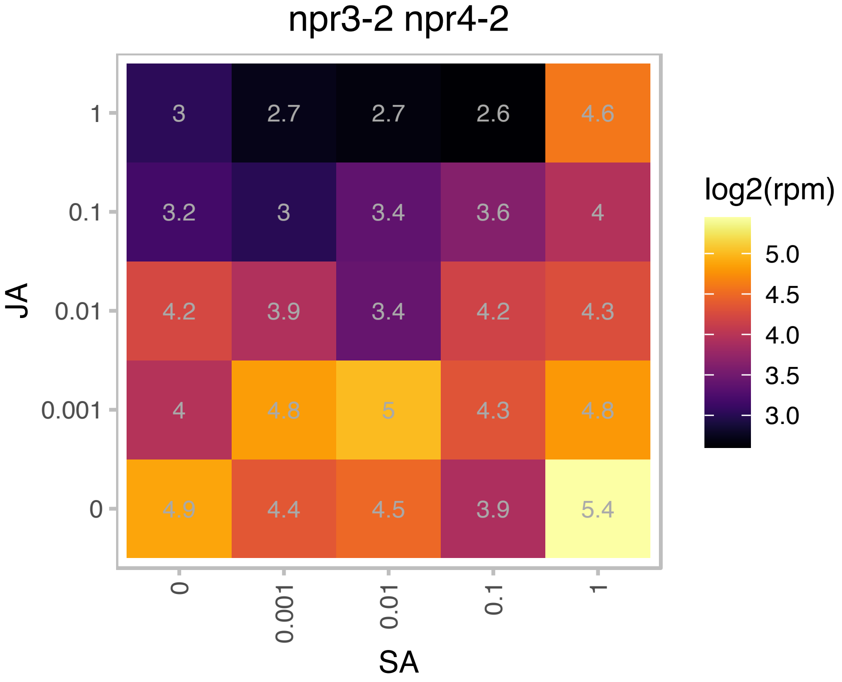


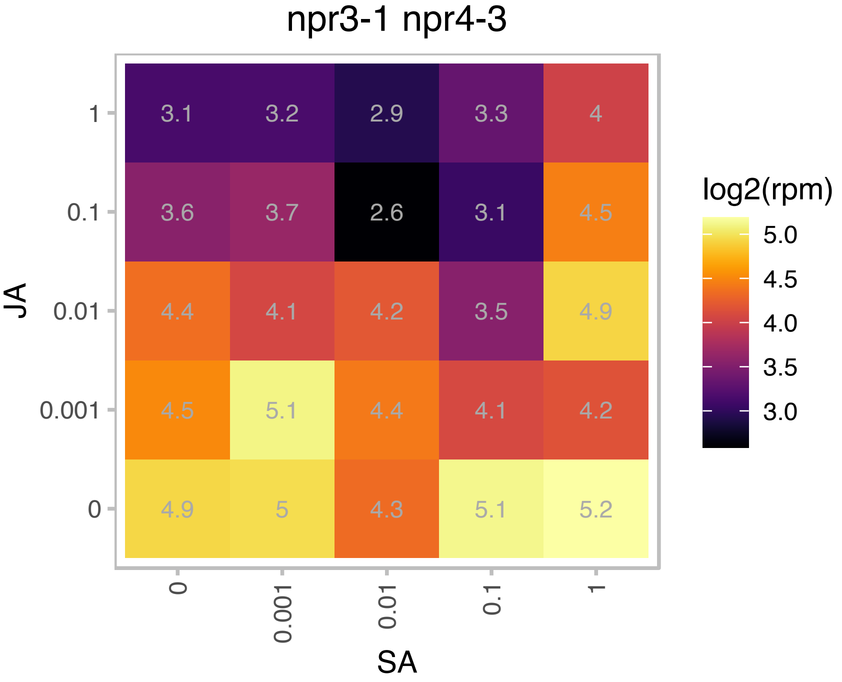

# AT3G15480.1 : NA

Protein of unknown function (DUF1218)

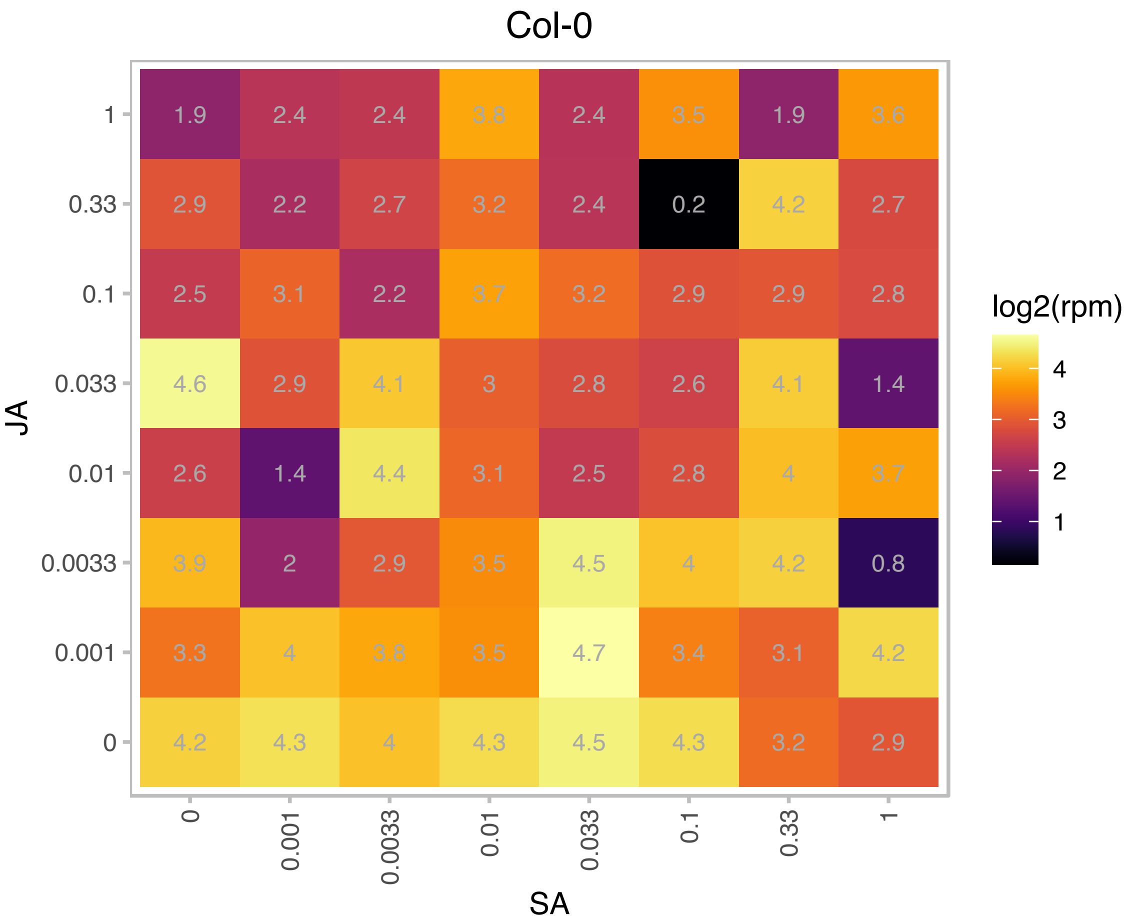


| | |
|---|---|
| Gene | AT3G15480 |
| is.expressed_genes | TRUE |
| is.responsive_genes | TRUE |
| is.apcluster | H-05 |

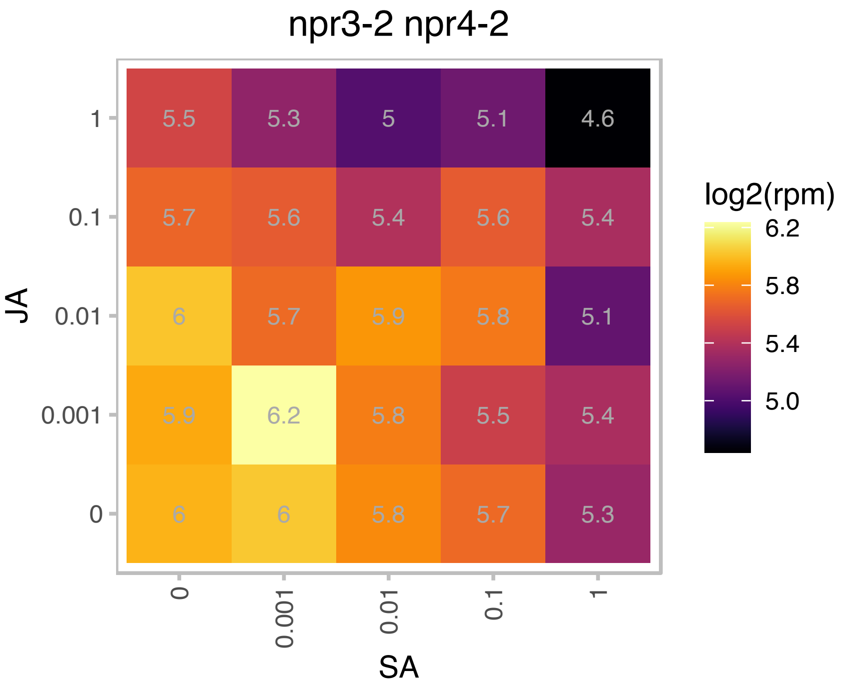


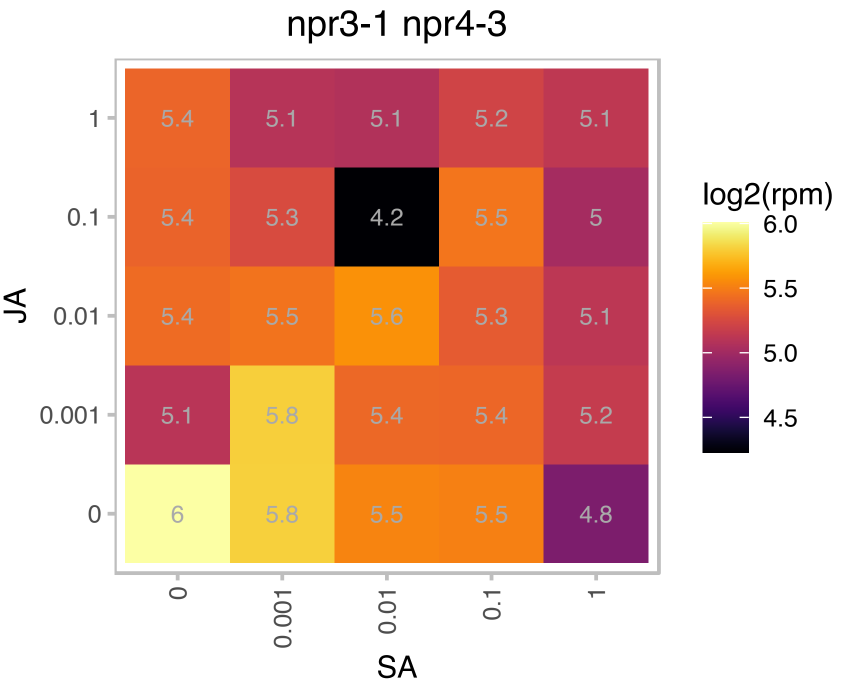

### AT3G23000.1 : ATSR2, ATSRPK1, CIPK7, PKS7, SnRK3.1

CBL-interacting protein kinase 7

Encodes a serine/threonine protein kinase with similarities to CBL-interacting protein kinases, SNF1 and SOS2. The mRNA is cell-to-cell mobile.

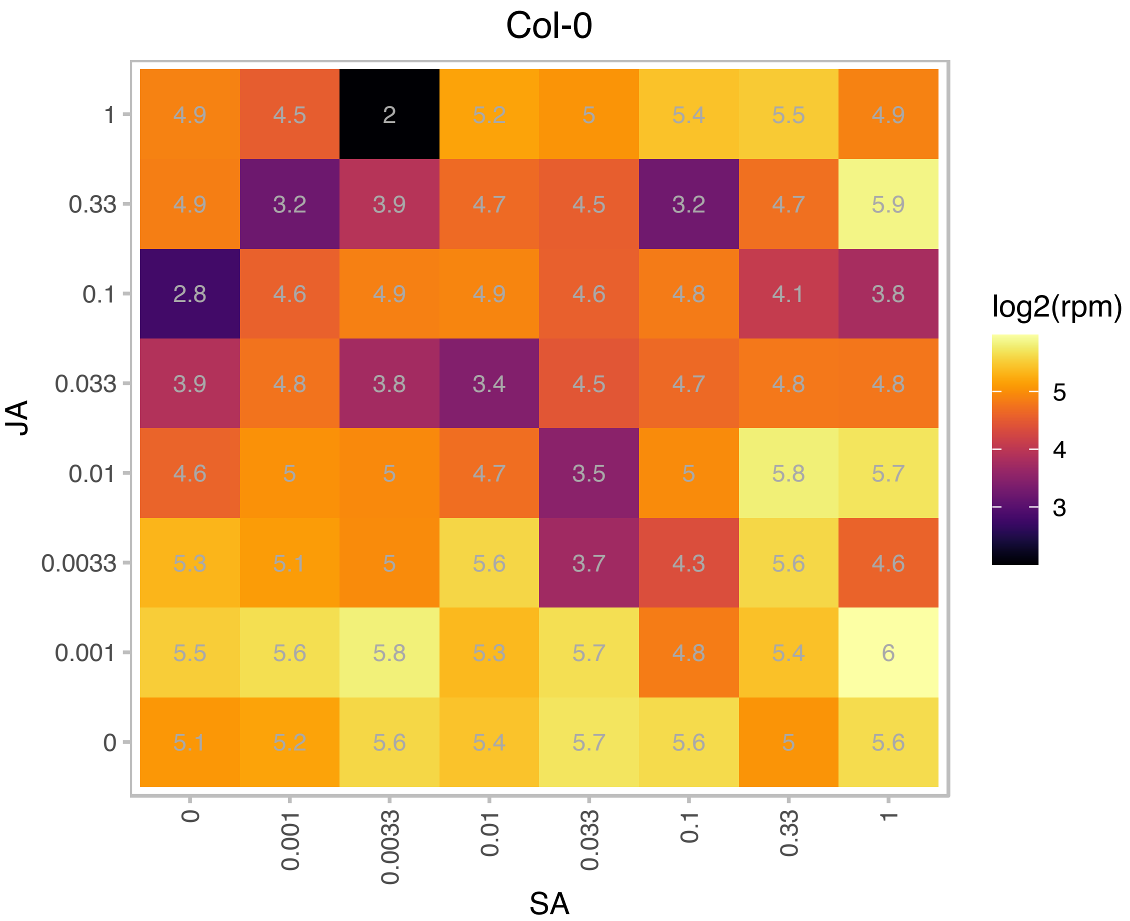


| | |
|---|---|
| Gene | AT3G23000 |
| is.expressed_genes | TRUE |
| is.responsive_genes | TRUE |
| is.apcluster | H-05 |

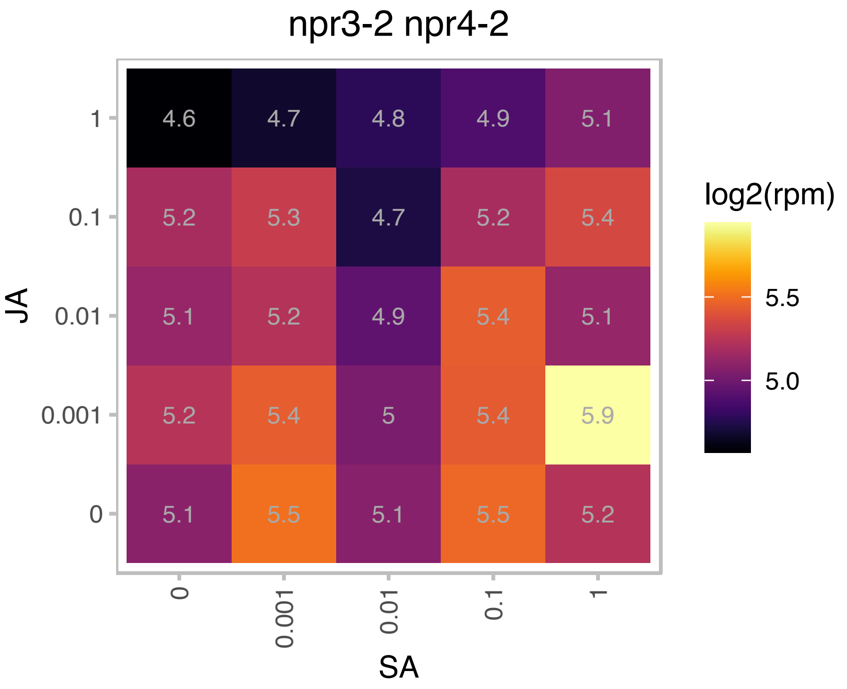


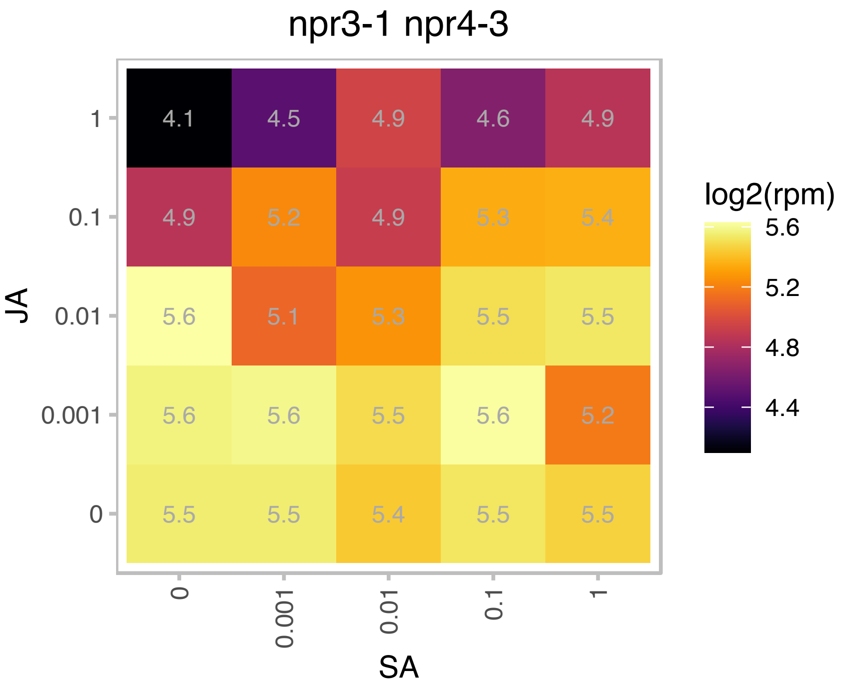

### AT3G26710.1 : CCB1

cofactor assembly of complex C

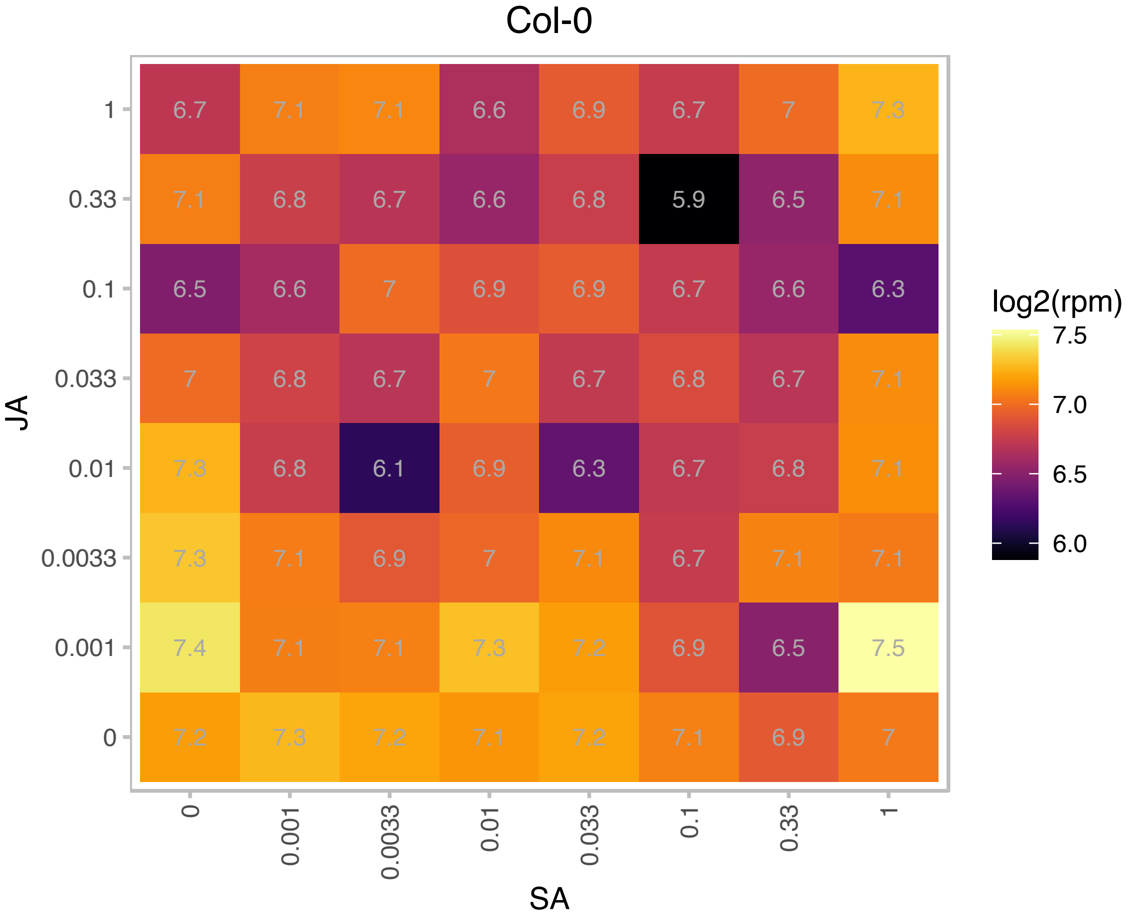


| | |
|---|---|
| Gene | AT3G26710 |
| is.expressed_genes | TRUE |
| is.responsive_genes | TRUE |
| is.apcluster | H-05 |

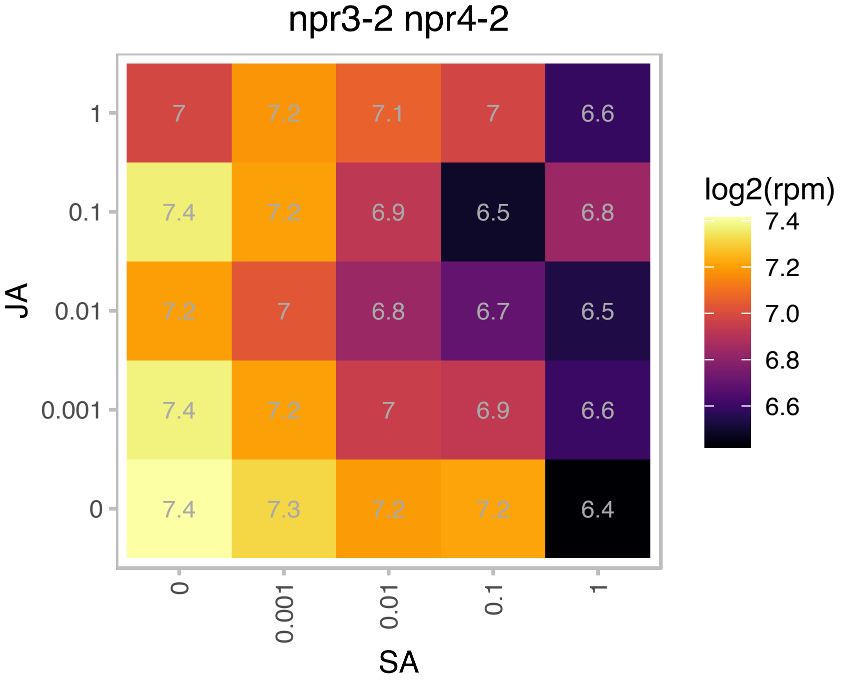


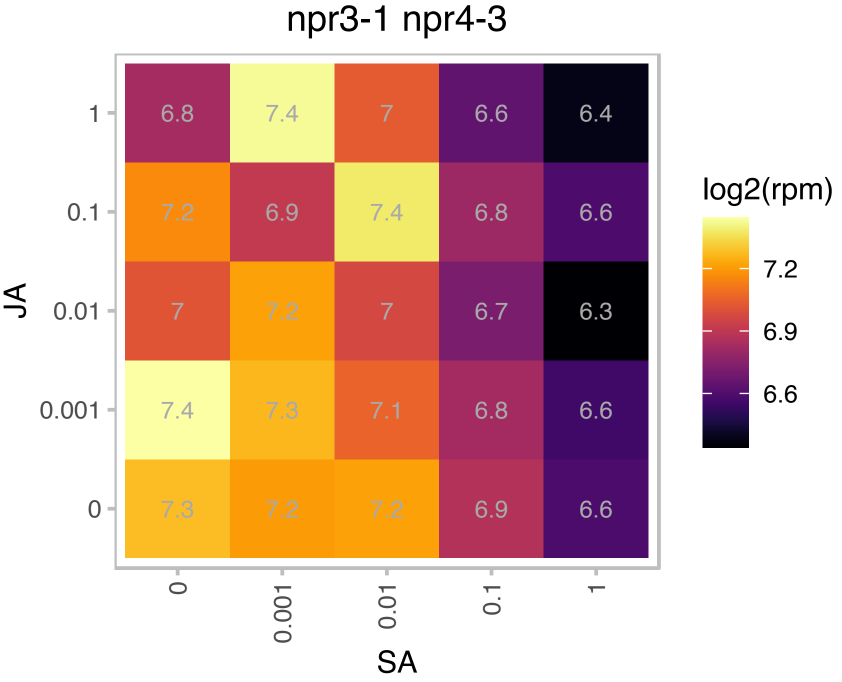

### AT3G26980.1 : MUB4

membrane-anchored ubiquitin-fold protein 4 precursor

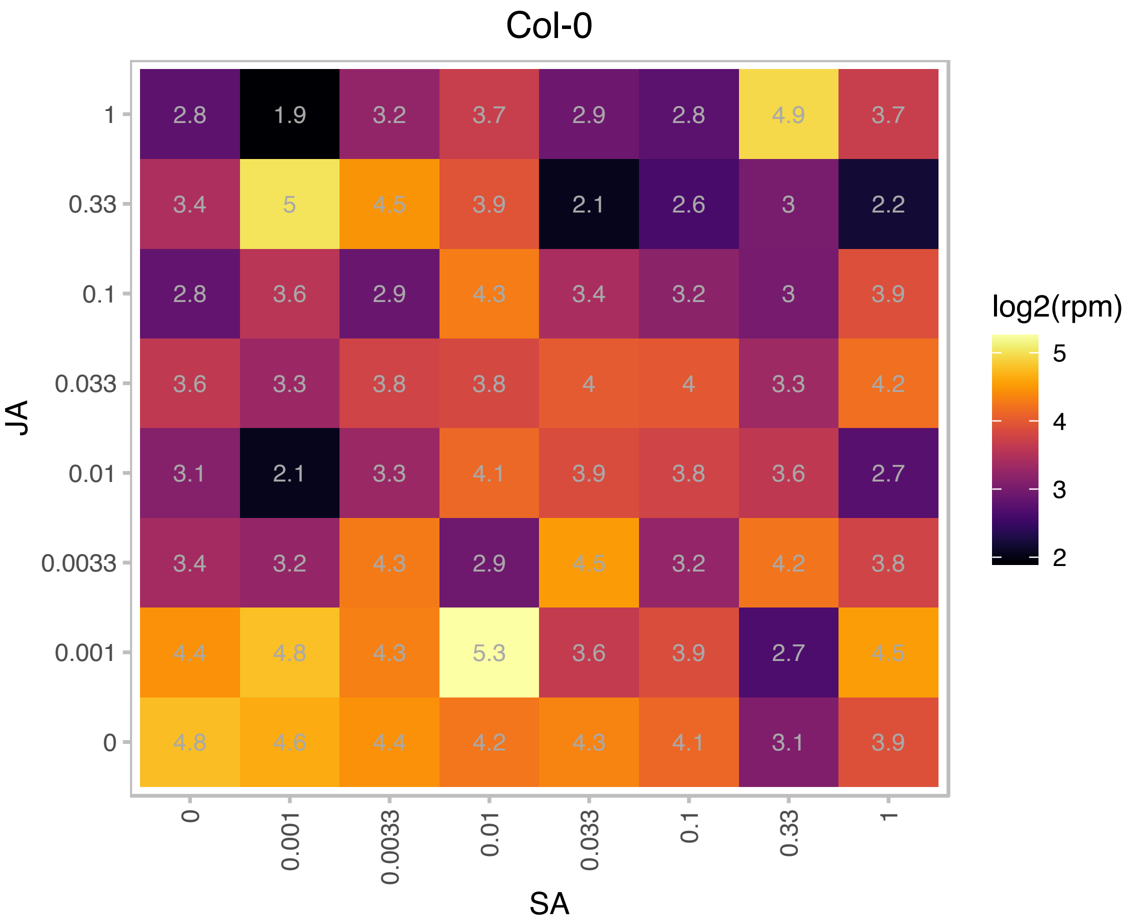


| | |
|---|---|
| Gene | AT3G26980 |
| is.expressed_genes | TRUE |
| is.responsive_genes | TRUE |
| is.apcluster | H-05 |

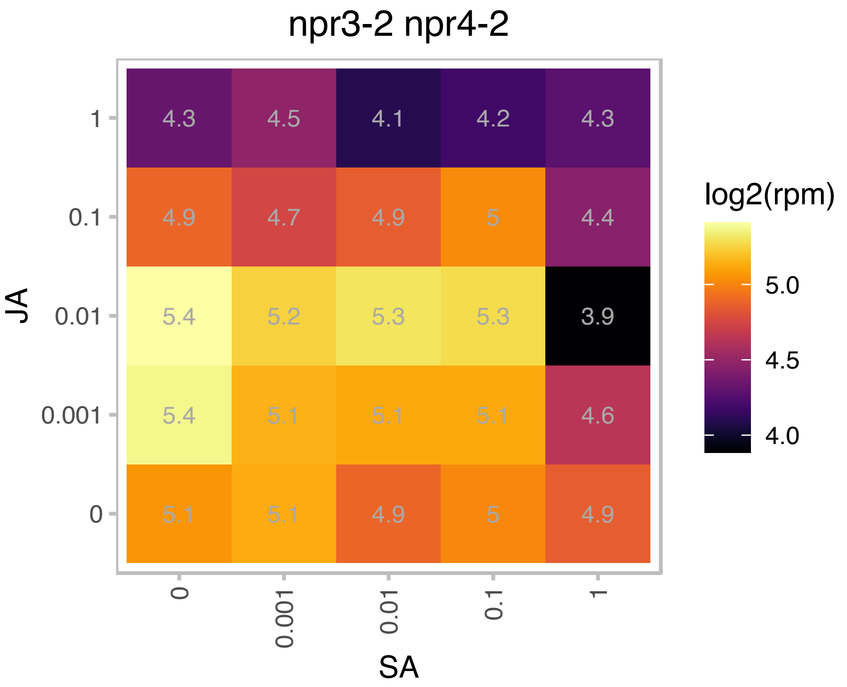


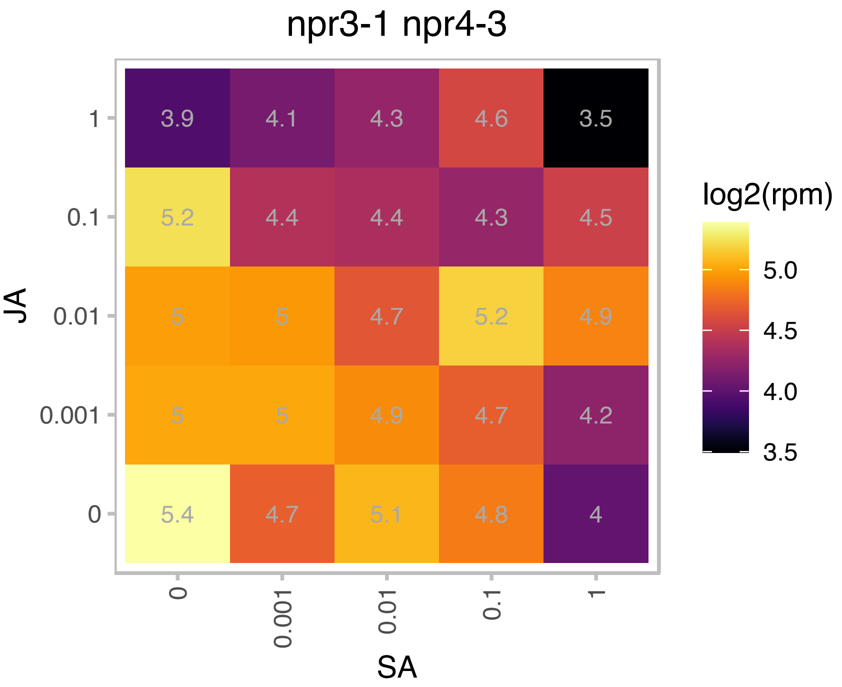

### AT3G27830.1 : RPL12, RPL12-A

ribosomal protein L12-A

50S ribosomal protein L12-A The mRNA is cell-to-cell mobile.

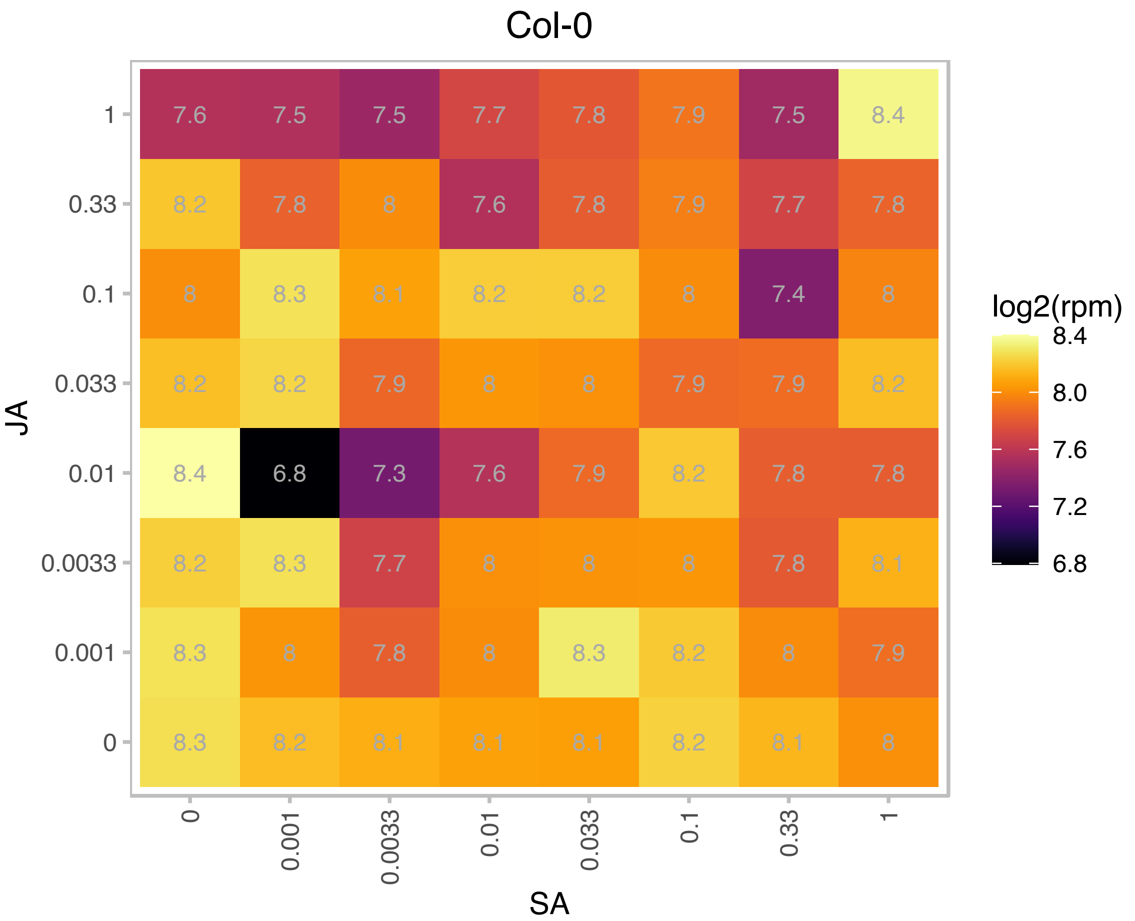


| | |
|---|---|
| Gene | AT3G27830 |
| is.expressed_genes | TRUE |
| is.responsive_genes | TRUE |
| is.apcluster | H-05 |

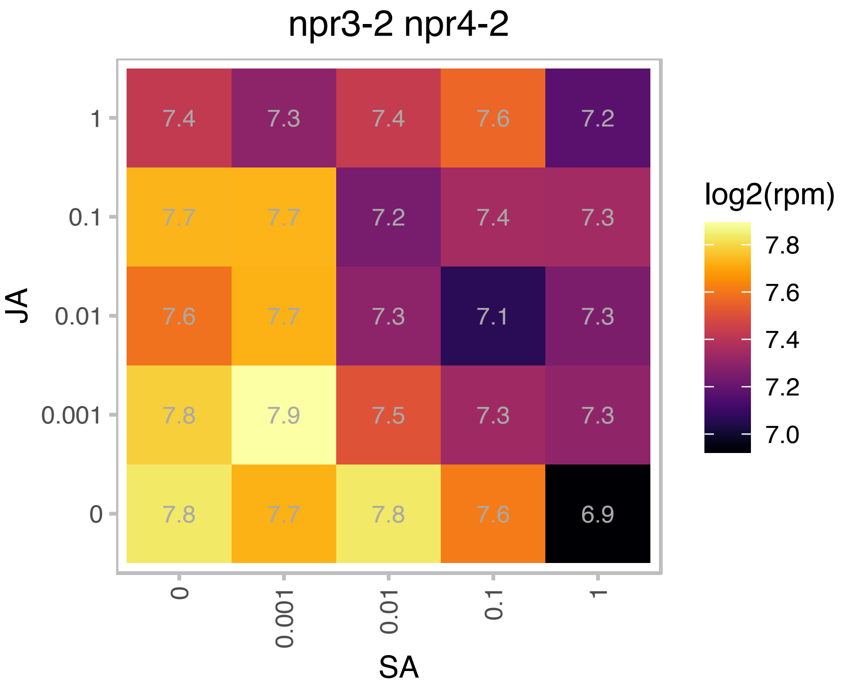


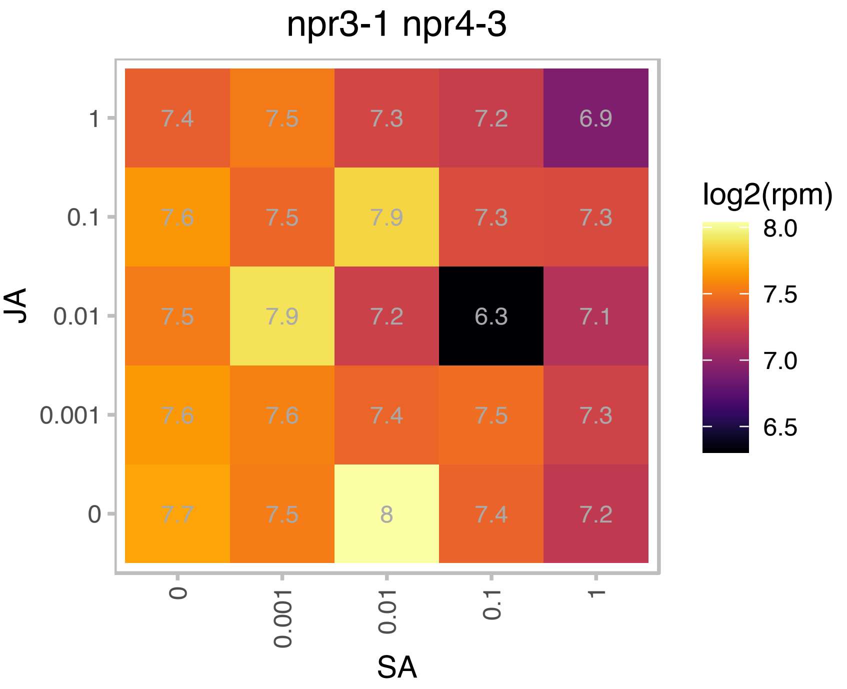

# AT3G45160.1 : NA

Putative membrane lipoprotein

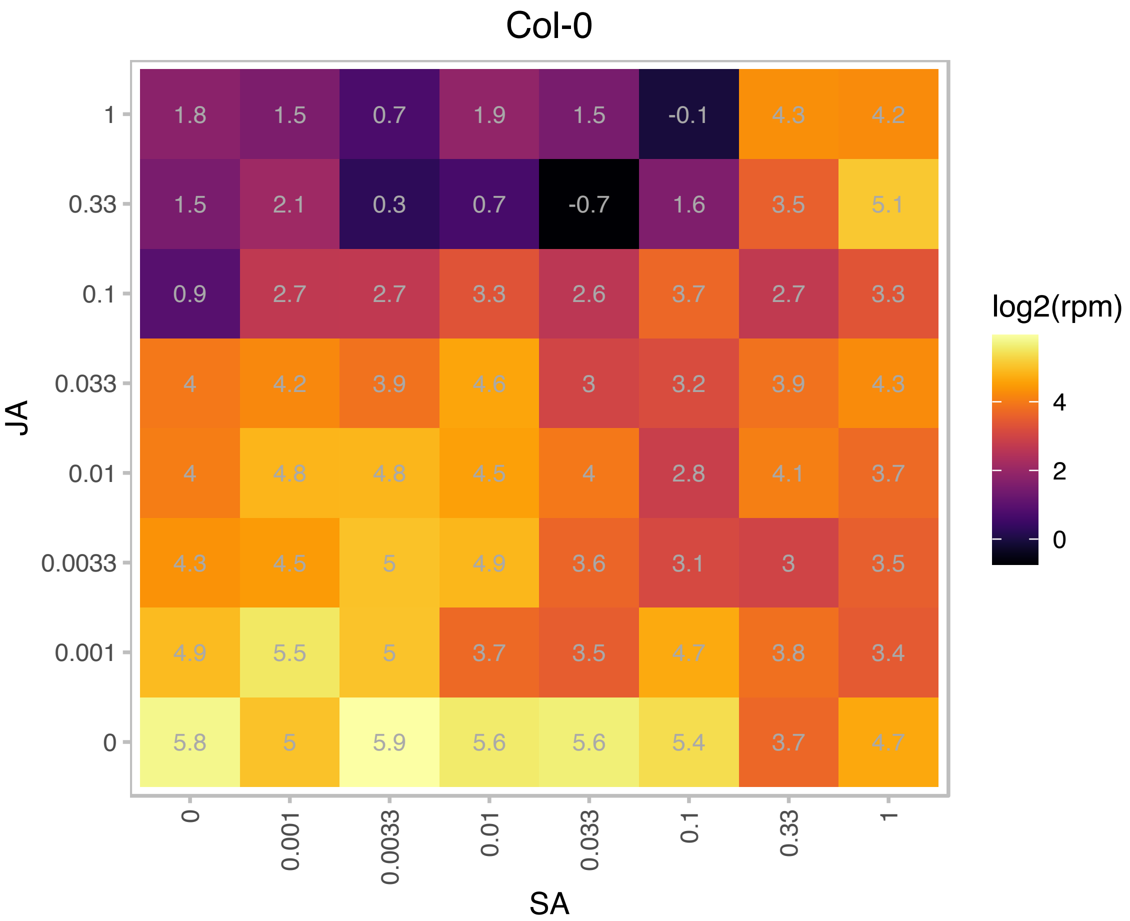


| | |
|---|---|
| Gene | AT3G45160 |
| is.expressed_genes | TRUE |
| is.responsive_genes | TRUE |
| is.apcluster | H-05 |

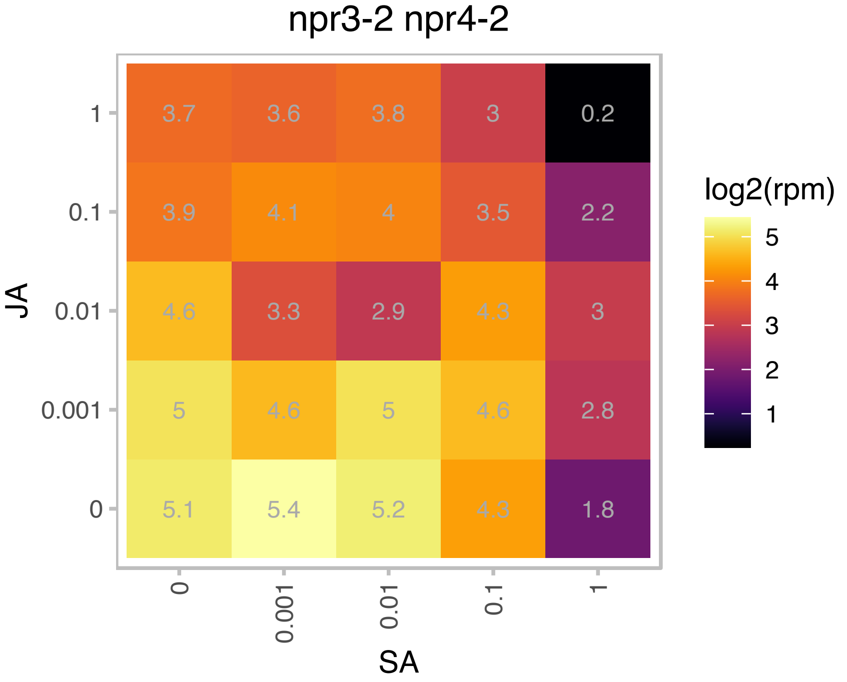


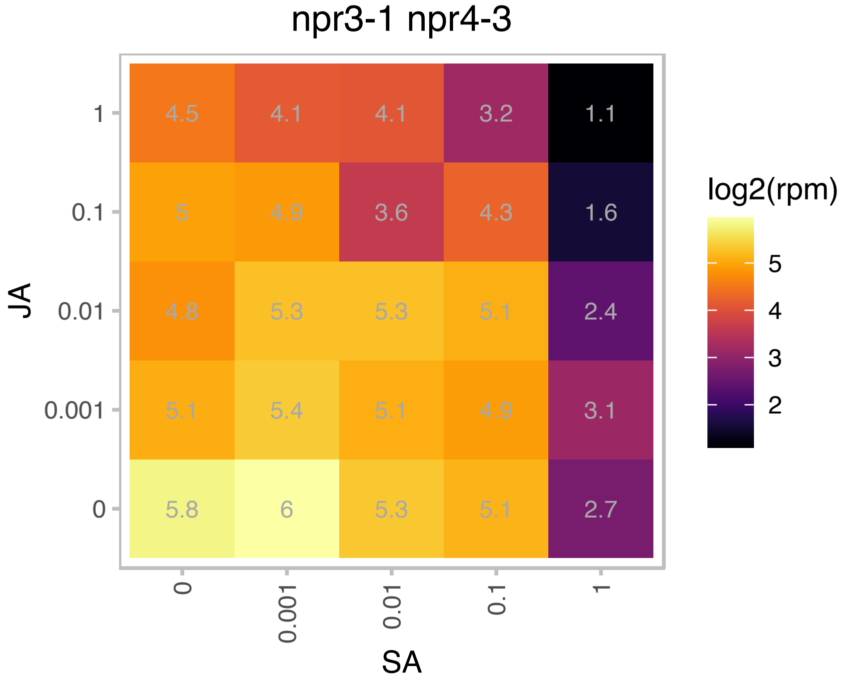

### AT3G55250.1 : PDE329, PSA3

Encodes a nucleus-encoded protein, Photosystem I Assembly 3 (PSA3), that is required for PSI accumulation.

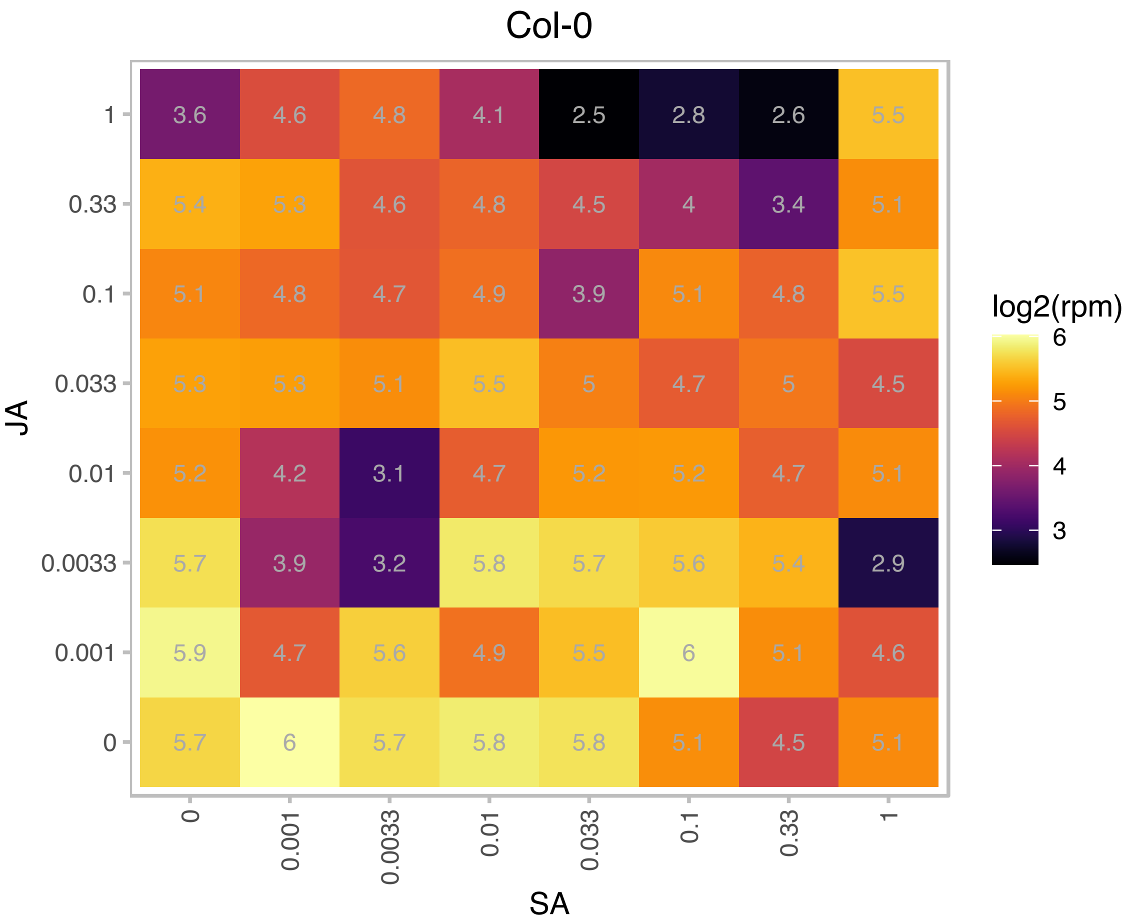


| | |
|---|---|
| Gene | AT3G55250 |
| is.expressed_genes | TRUE |
| is.responsive_genes | TRUE |
| is.apcluster | H-05 |

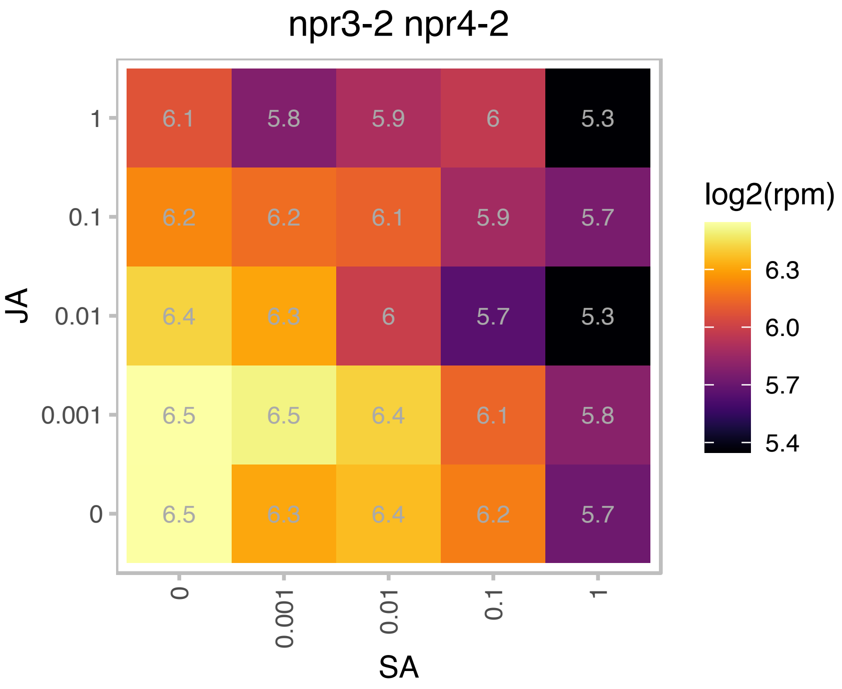


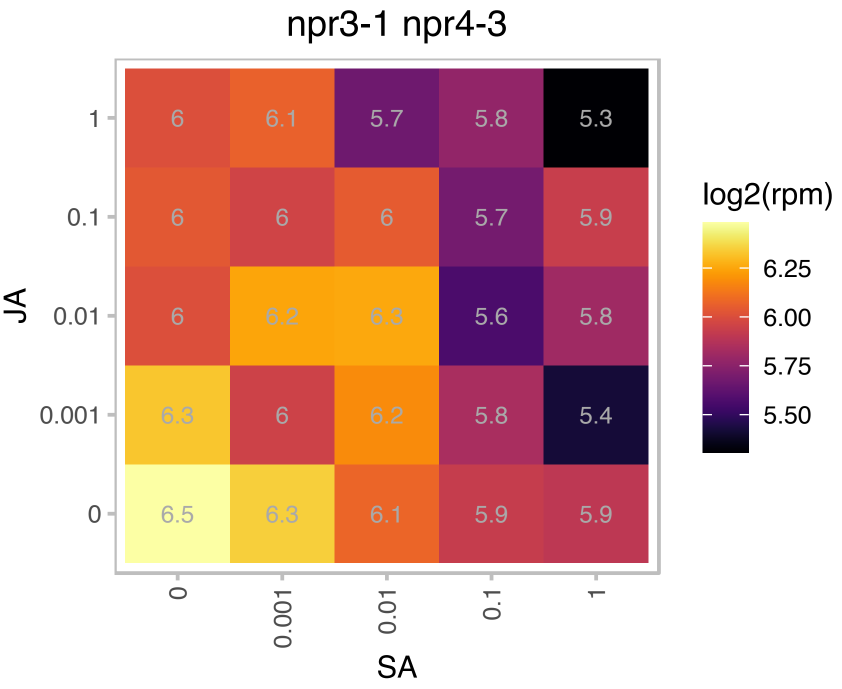

# AT4G02725.1 : NA

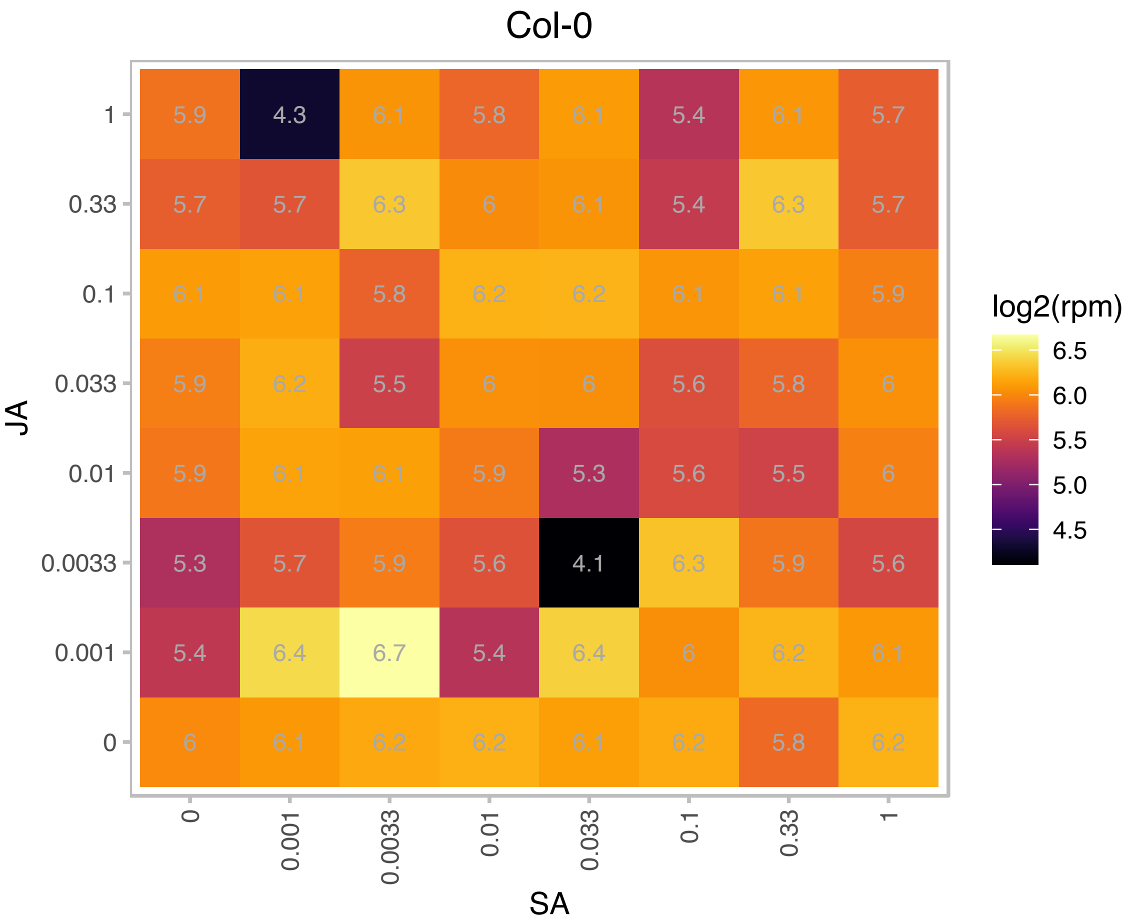


| | |
|---|---|
| Gene | AT4G02725 |
| is.expressed_genes | TRUE |
| is.responsive_genes | TRUE |
| is.apcluster | H-05 |

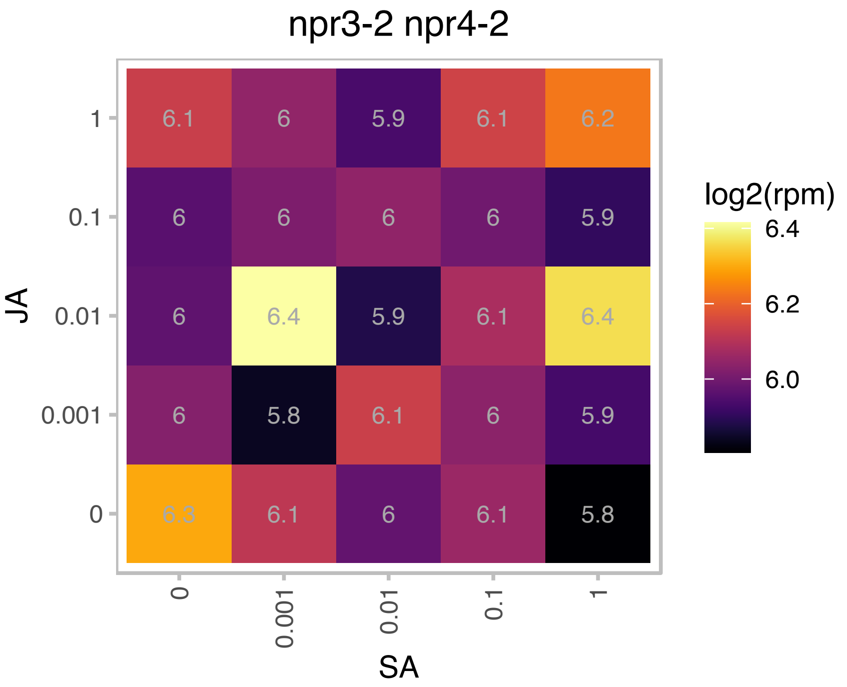


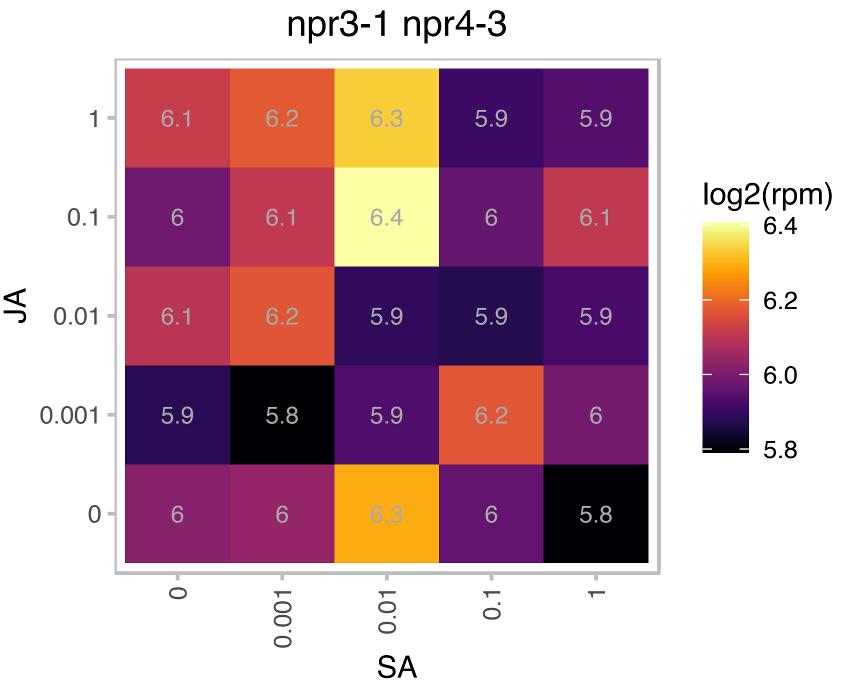

# AT4G02920.1 : NA

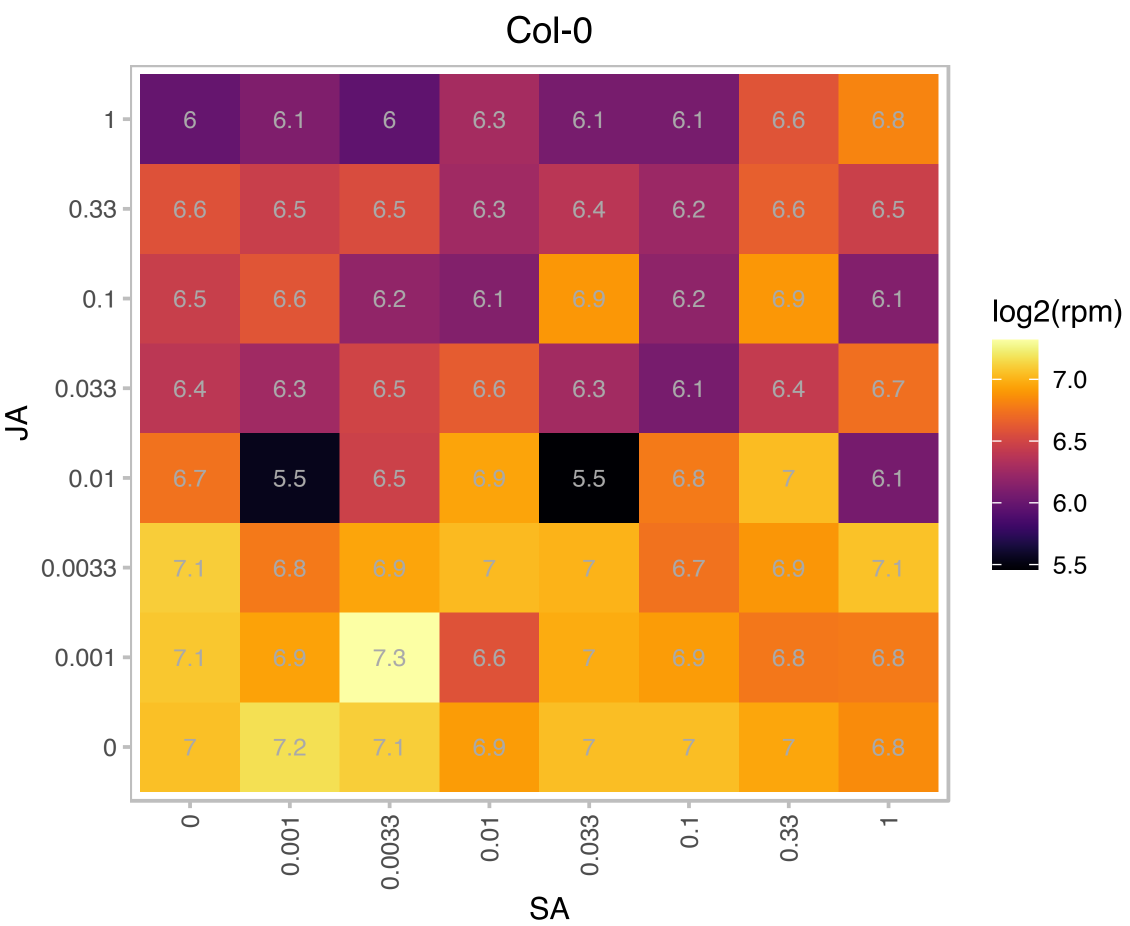


| | |
|---|---|
| Gene | AT4G02920 |
| is.expressed_genes | TRUE |
| is.responsive_genes | TRUE |
| is.apcluster | H-05 |

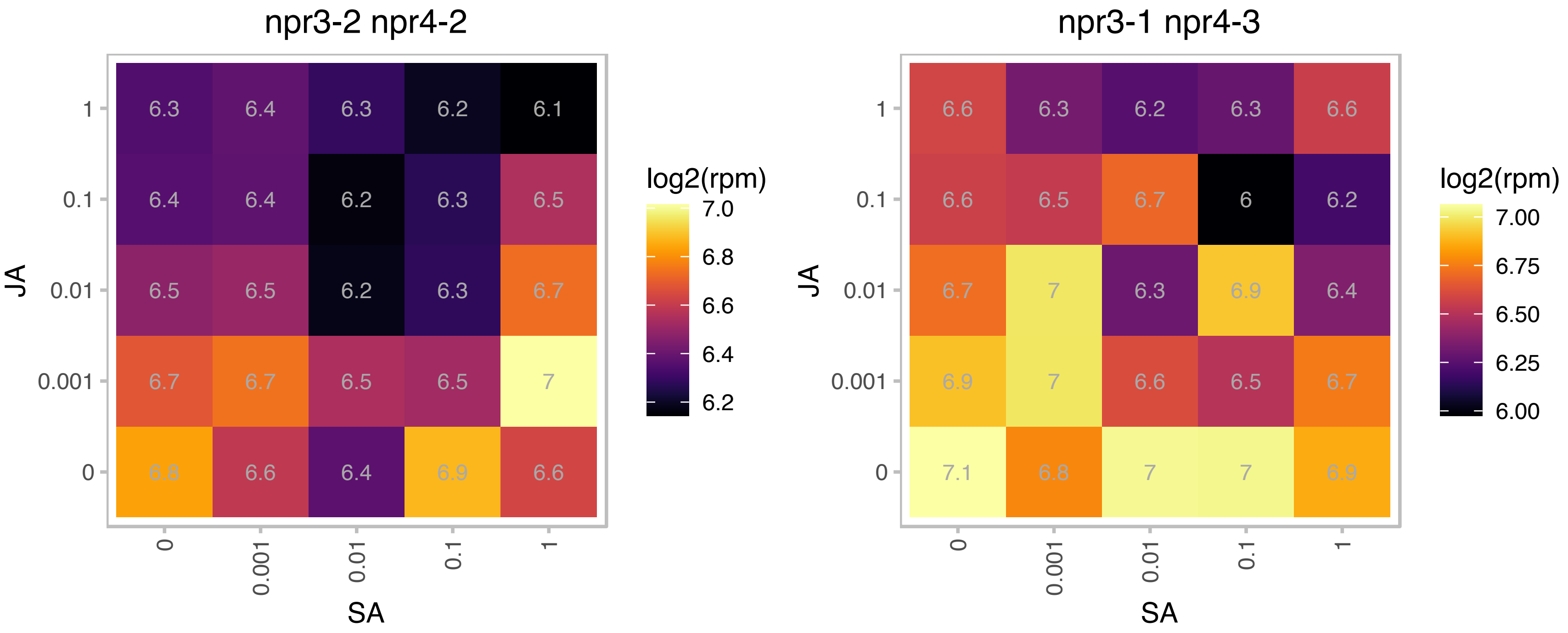

# AT4G09040.1 : NA

RNA-binding (RRM/RBD/RNP motifs) family protein

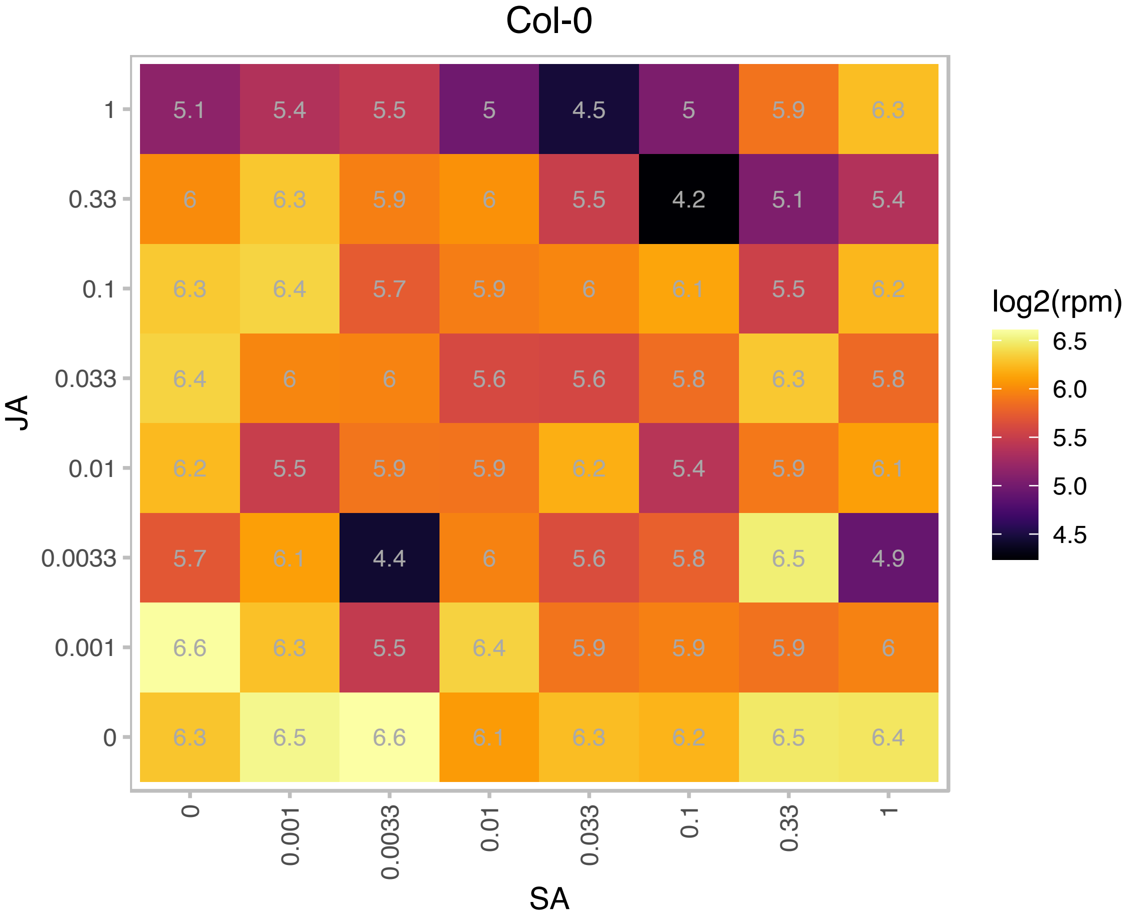


| | |
|---|---|
| Gene | AT4G09040 |
| is.expressed_genes | TRUE |
| is.responsive_genes | TRUE |
| is.apcluster | H-05 |

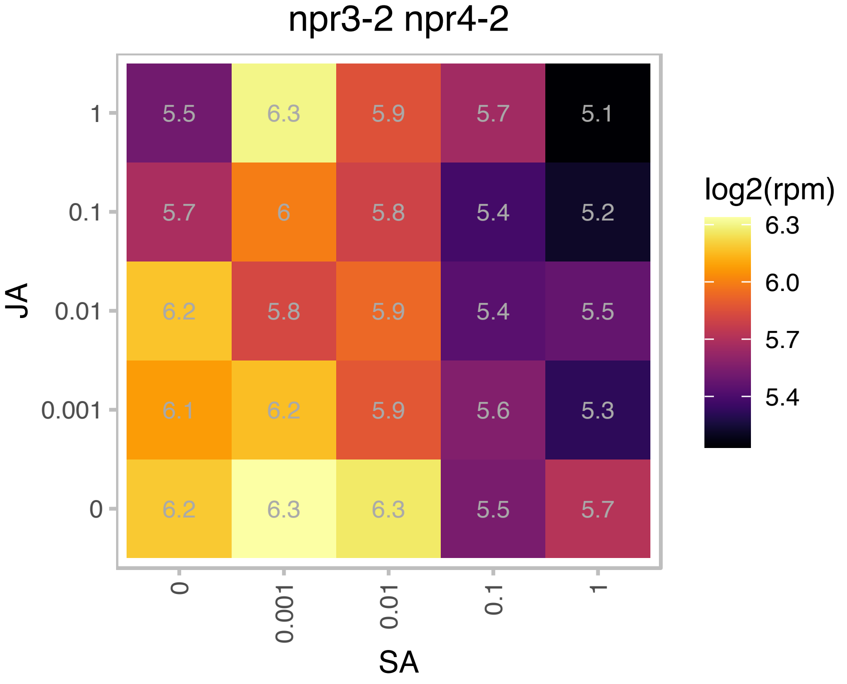


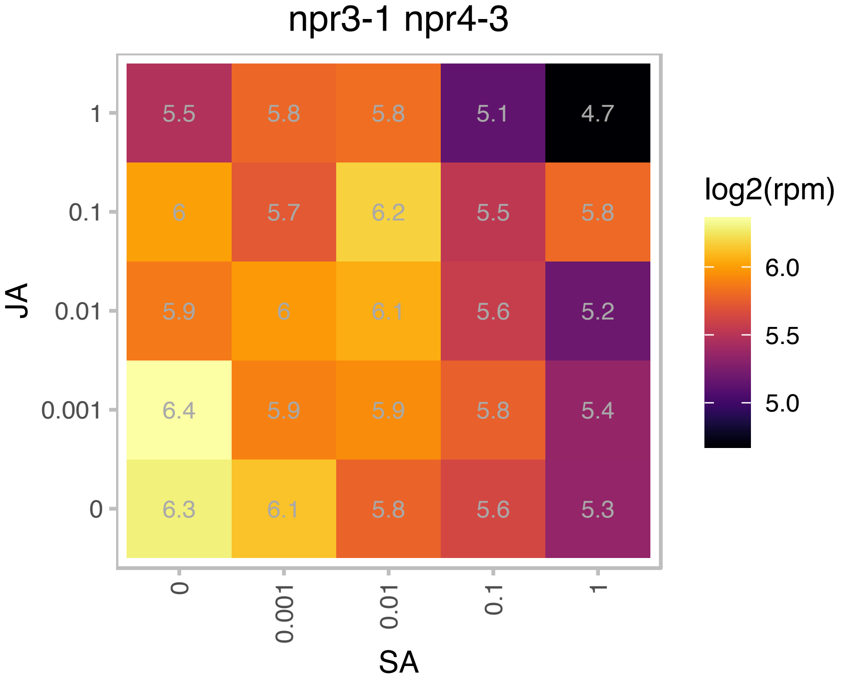

# AT4G29905.1 : NA

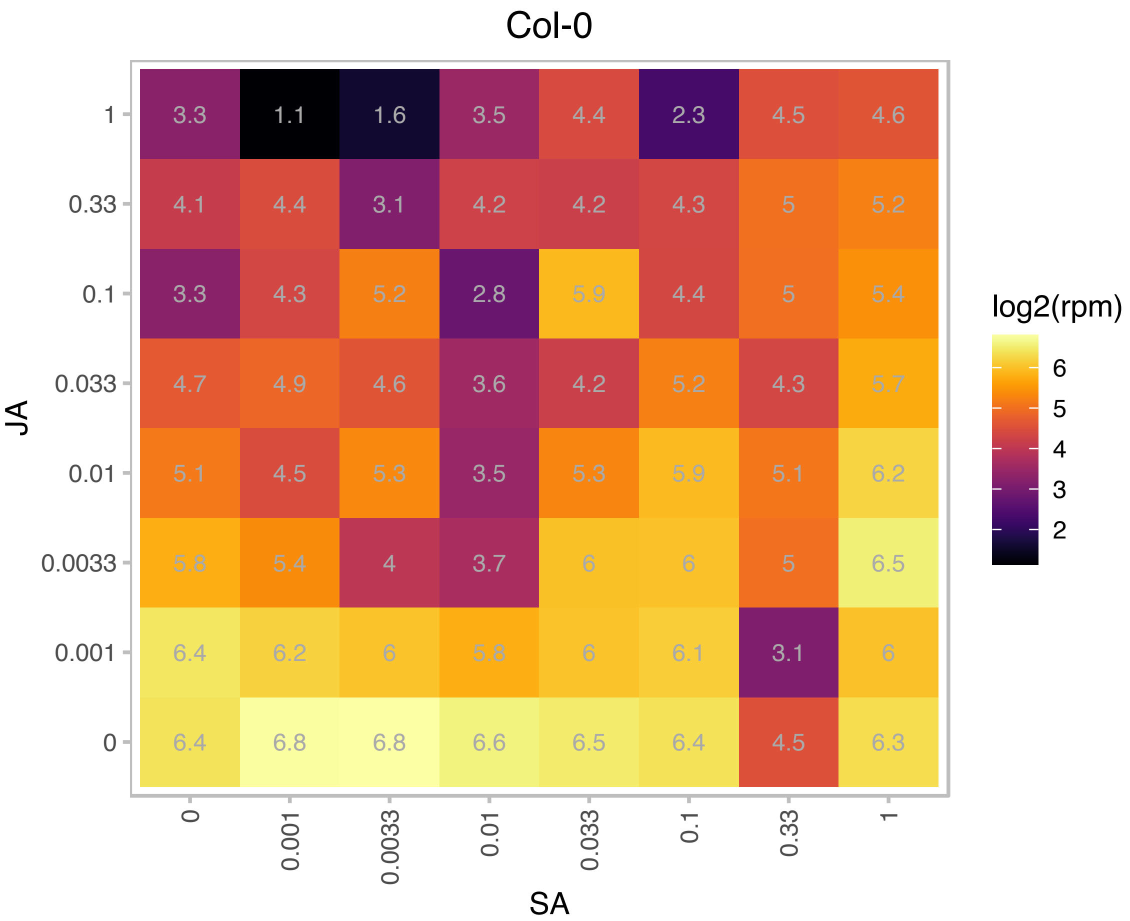


| Gene | AT4G29905 |
|---|---|
| is.expressed_genes | TRUE |
| is.responsive_genes | TRUE |
| is.apcluster | H-05 |

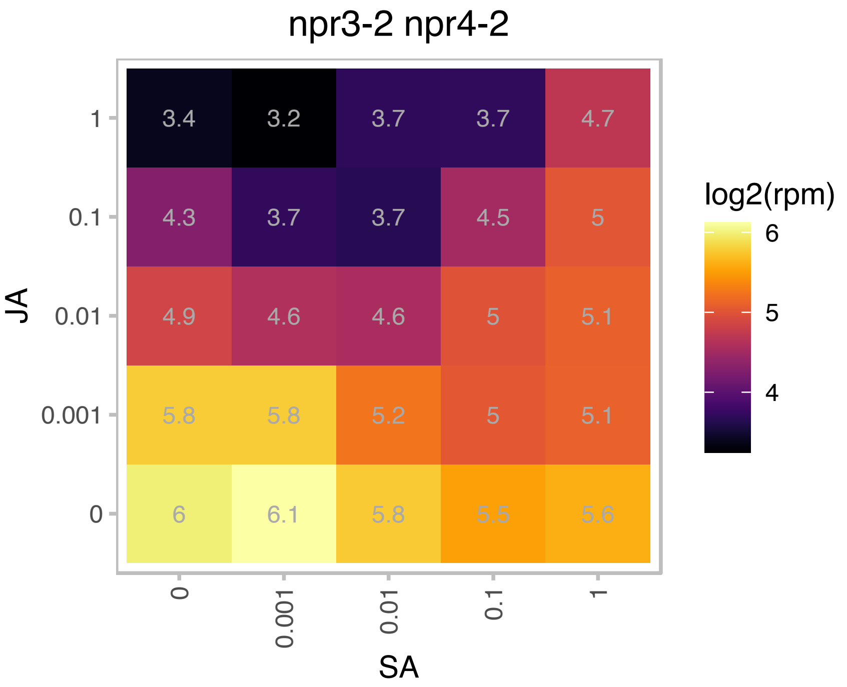


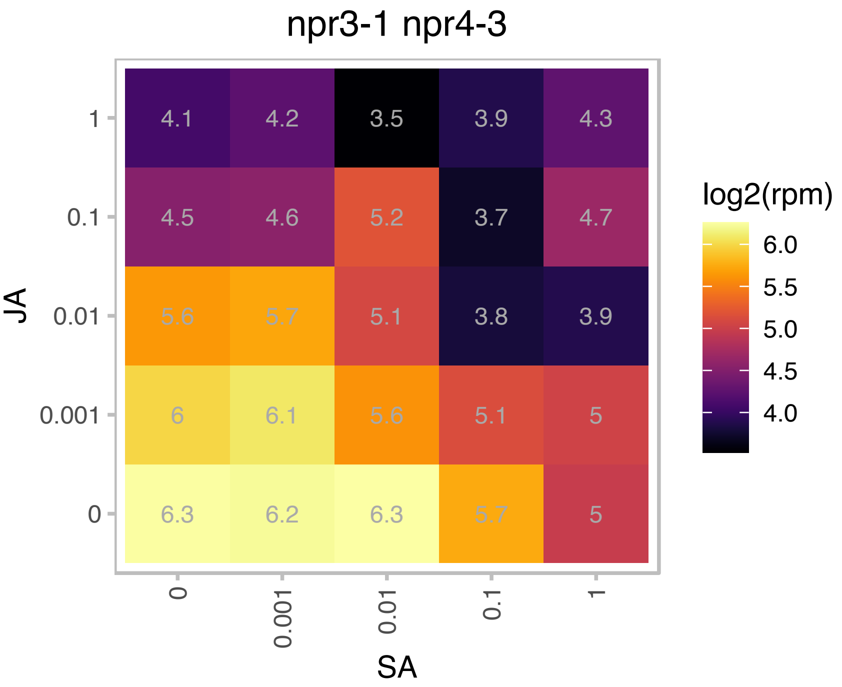

### AT4G33030.1 : SQD1

sulfoquinovosyldiacylglycerol 1

involved in sulfolipid biosynthesis The mRNA is cell-to-cell mobile.

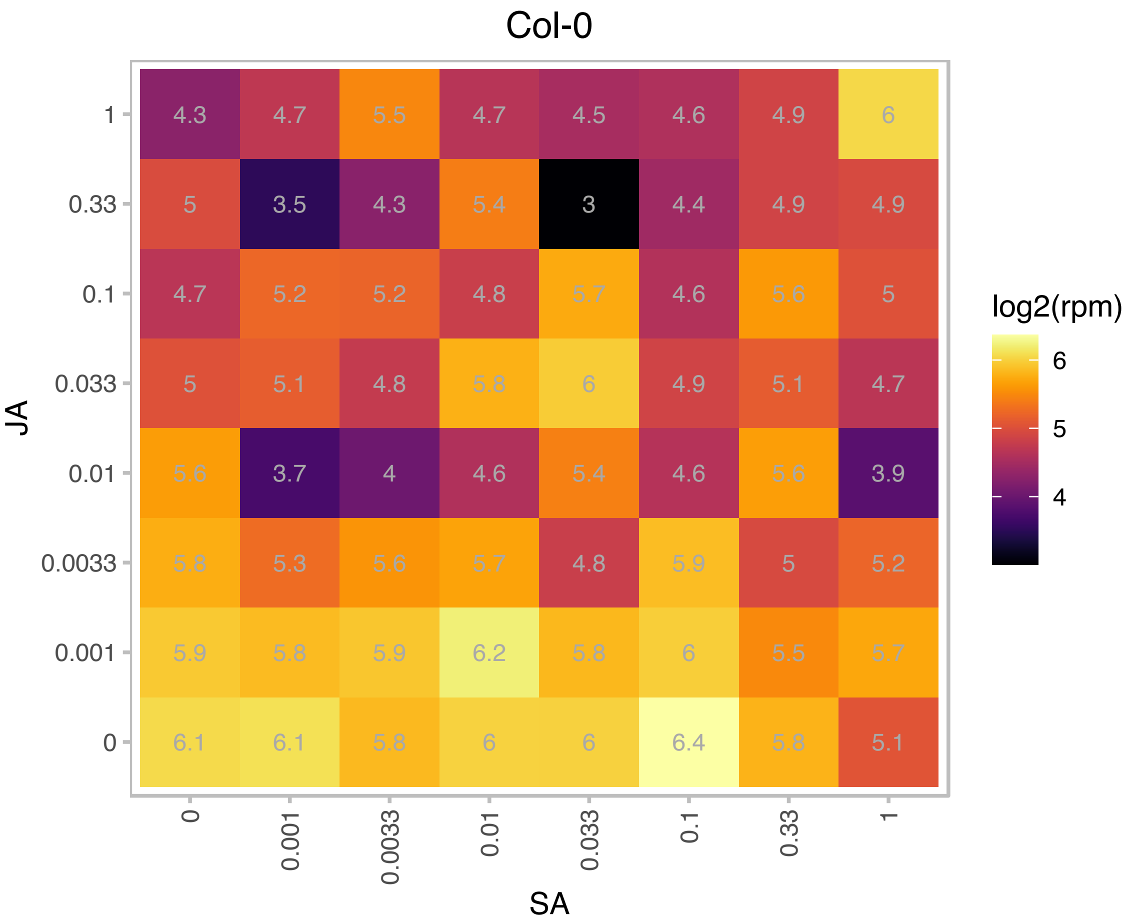


| | |
|---|---|
| Gene | AT4G33030 |
| is.expressed_genes | TRUE |
| is.responsive_genes | TRUE |
| is.apcluster | H-05 |

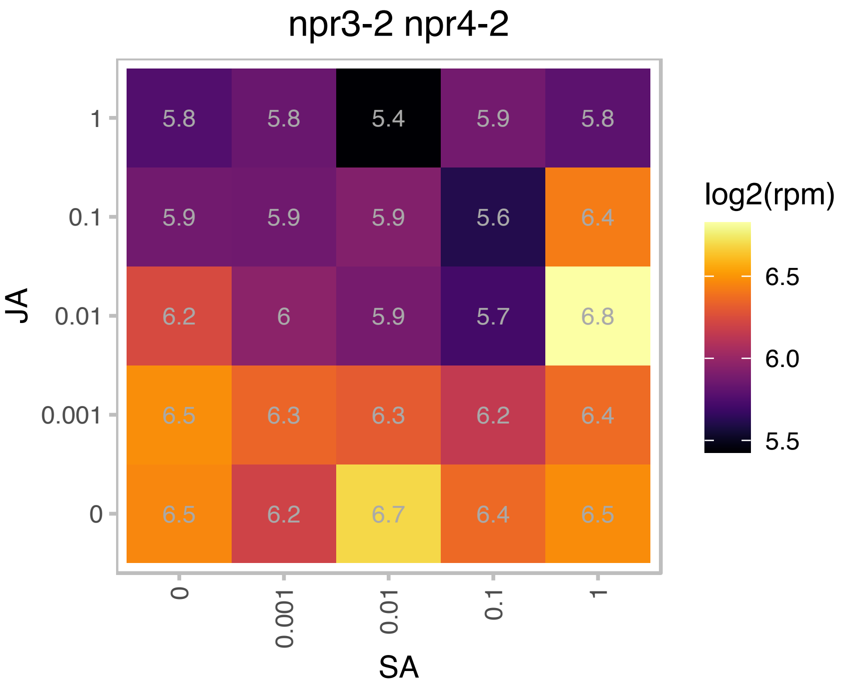


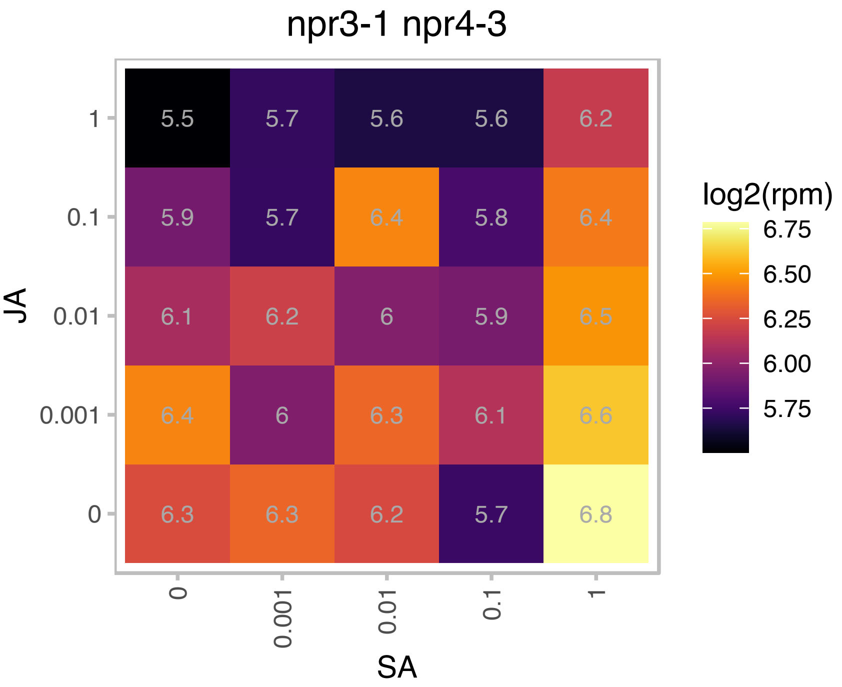

### AT4G34730.1 : RBF1

ribosome-binding factor A family protein

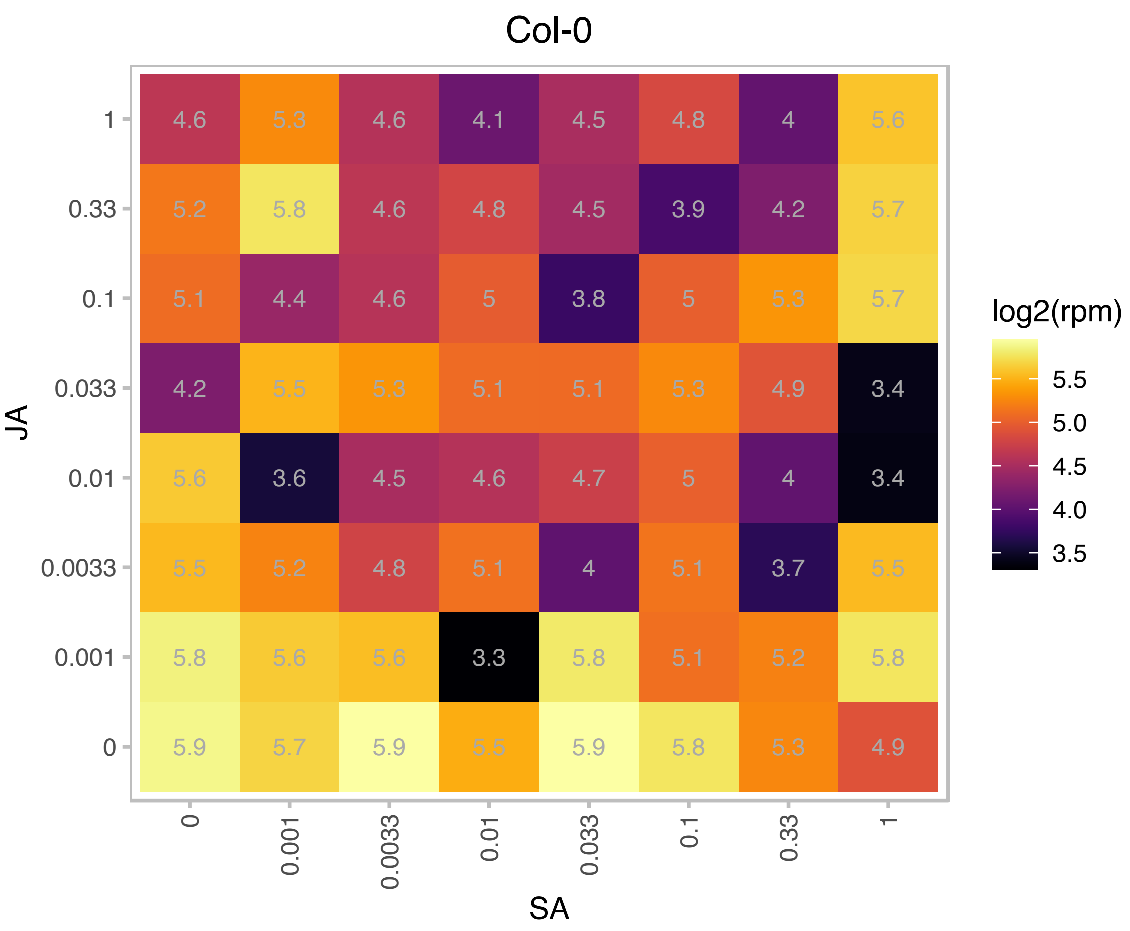


| | |
|---|---|
| Gene | AT4G34730 |
| is.expressed_genes | TRUE |
| is.responsive_genes | TRUE |
| is.apcluster | H-05 |

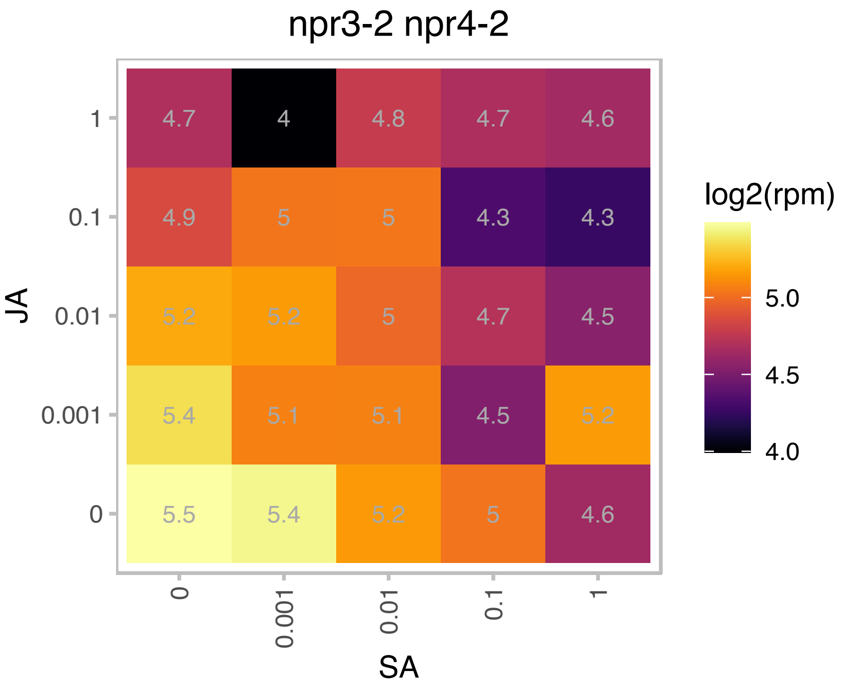


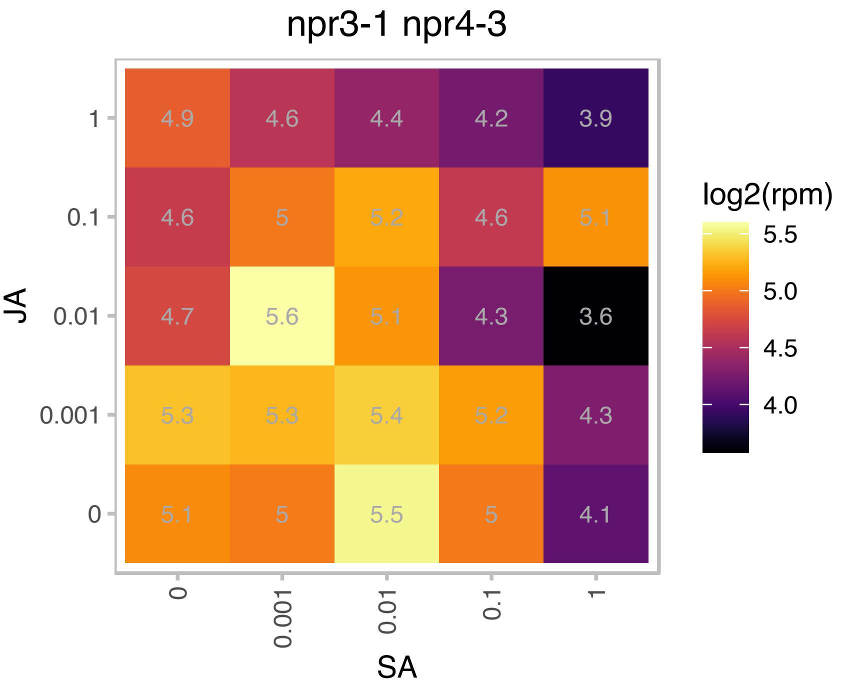

### AT5G17400.1 : ER-ANT1

This gene is predicted to encode an ER-localized adenine nucleotide transporter with six putative
endoplasmic reticulum-adenine nucleotide transporter 1
transmembrane helices. It appears to act as a ATP:ADP antiporter when expressed in E.coli plasma membranes. Transcript levels for several ER-localized chaperones (e.g. BIP1/2) and other ATP-requiring ER proteins (e.g. CPK2) are reduced in er-ant1 knock-out lines, suggesting a lack of adequate ATP transport into the ER in these mutants. They also have reduced seed oil and seed protein levels.

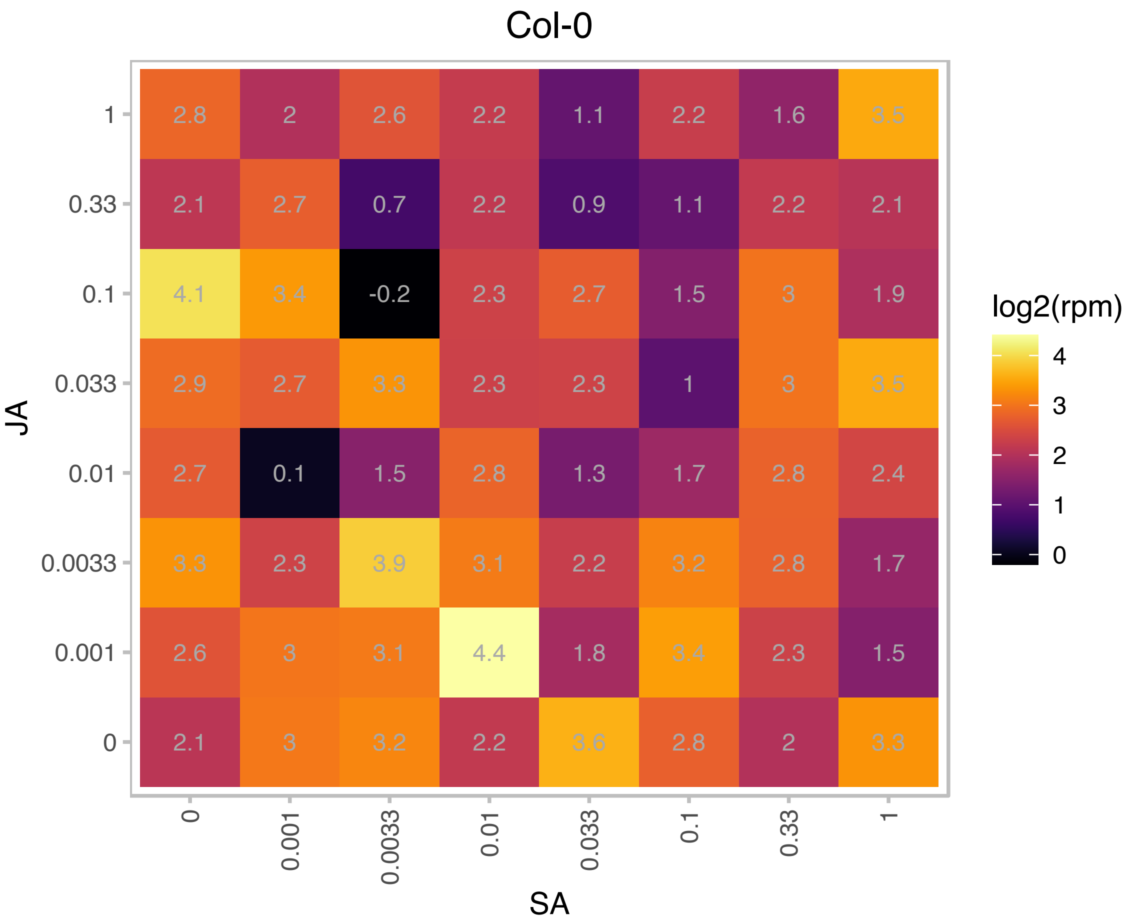


| | |
|---|---|
| Gene | AT5G17400 |
| is.expressed_genes | TRUE |
| is.responsive_genes | TRUE |
| is.apcluster | H-05 |

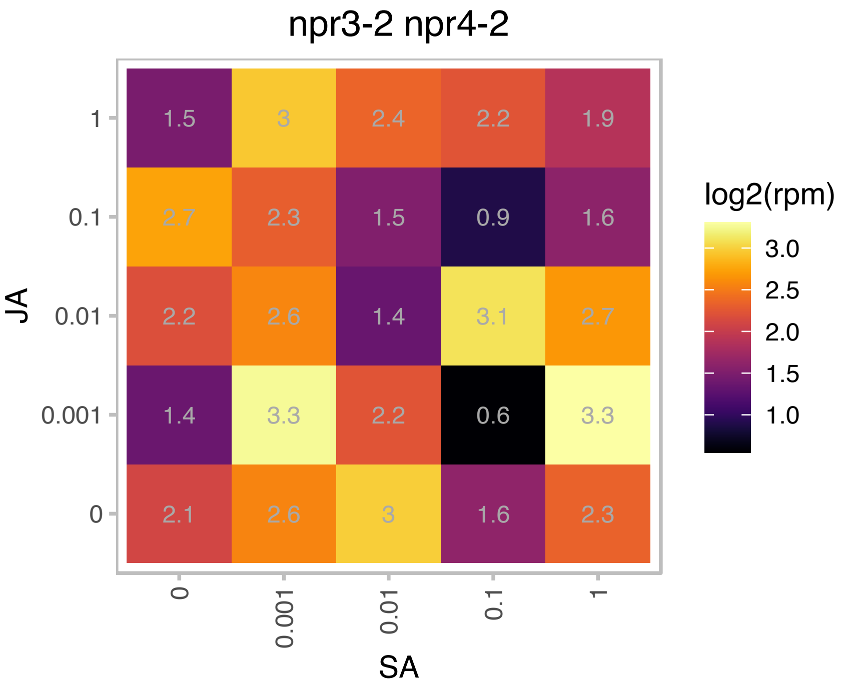


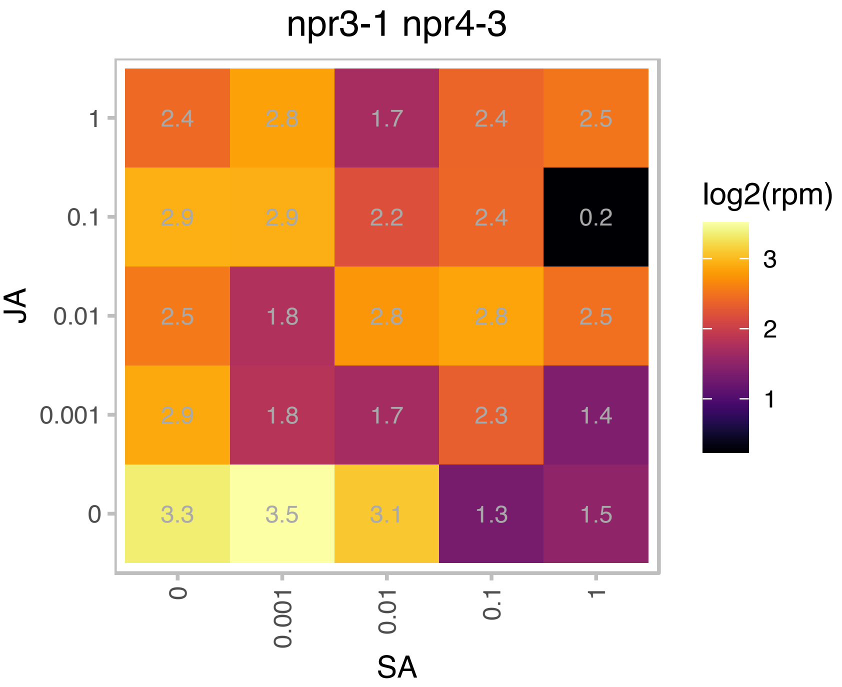

### AT5G18660.1 : DVR, PCB2

NAD(P)-binding Rossmann-fold superfamily protein

Encodes a protein with 3,8-divinyl protochlorophyllide a 8-vinyl reductase activity. Mutants accumulate divinyl chlorophyll rather than monovinyl chlorophyll.

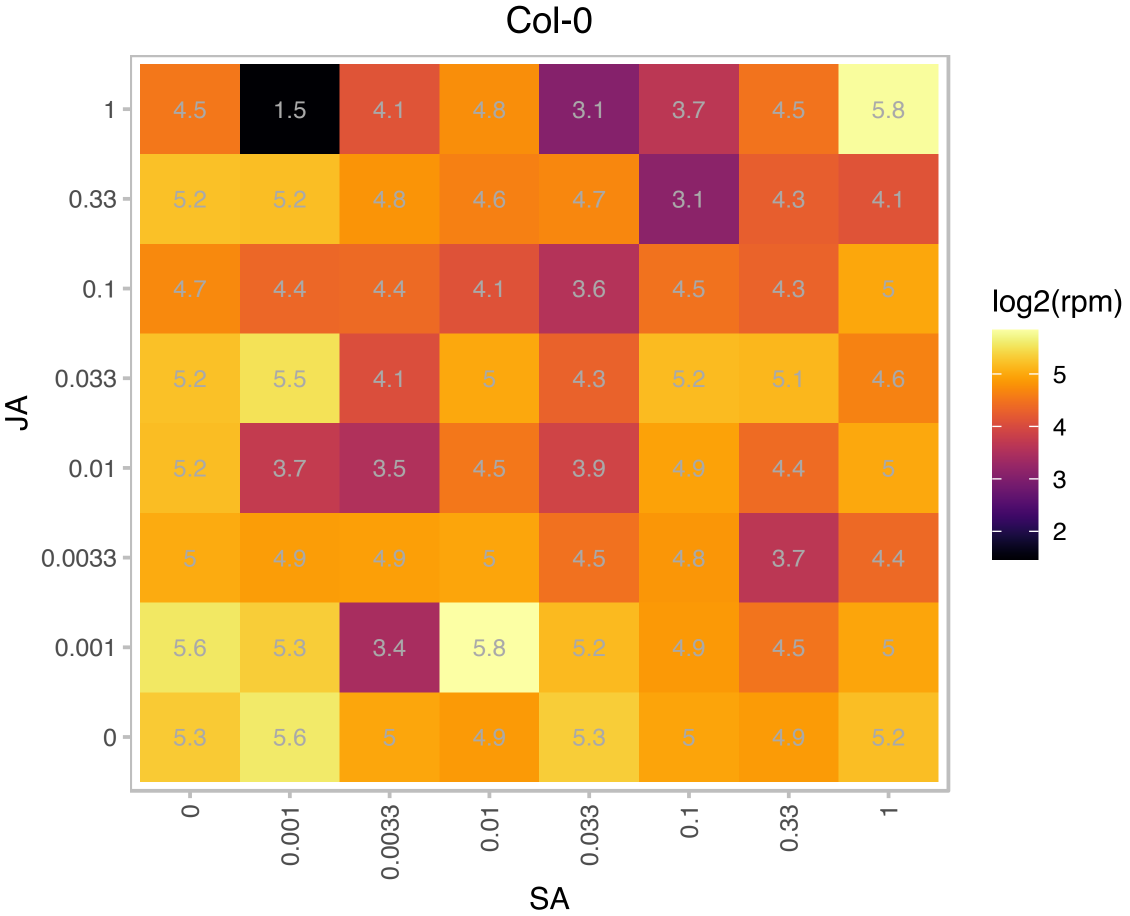


| | |
|---|---|
| Gene | AT5G18660 |
| is.expressed_genes | TRUE |
| is.responsive_genes | TRUE |
| is.apcluster | H-05 |

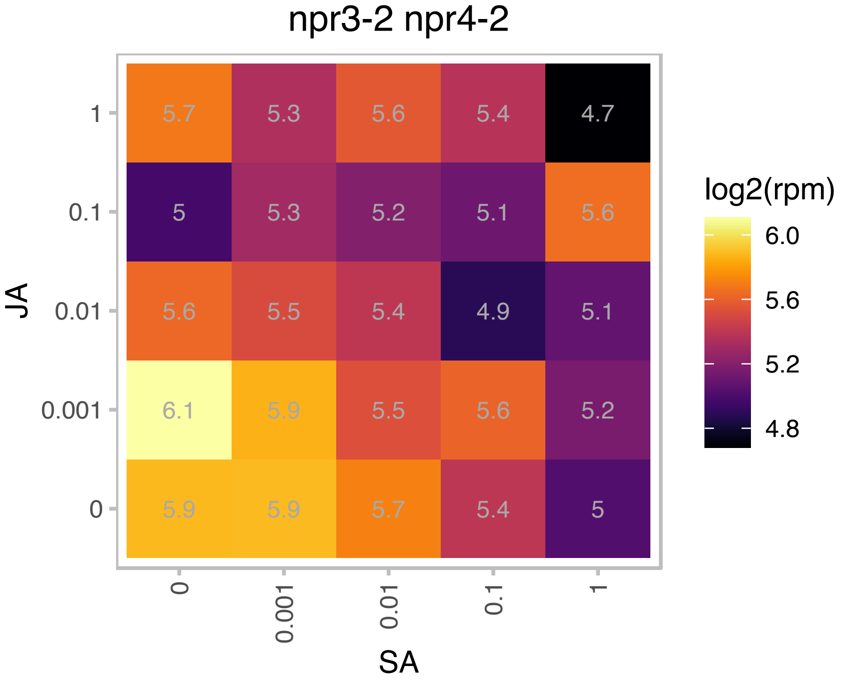


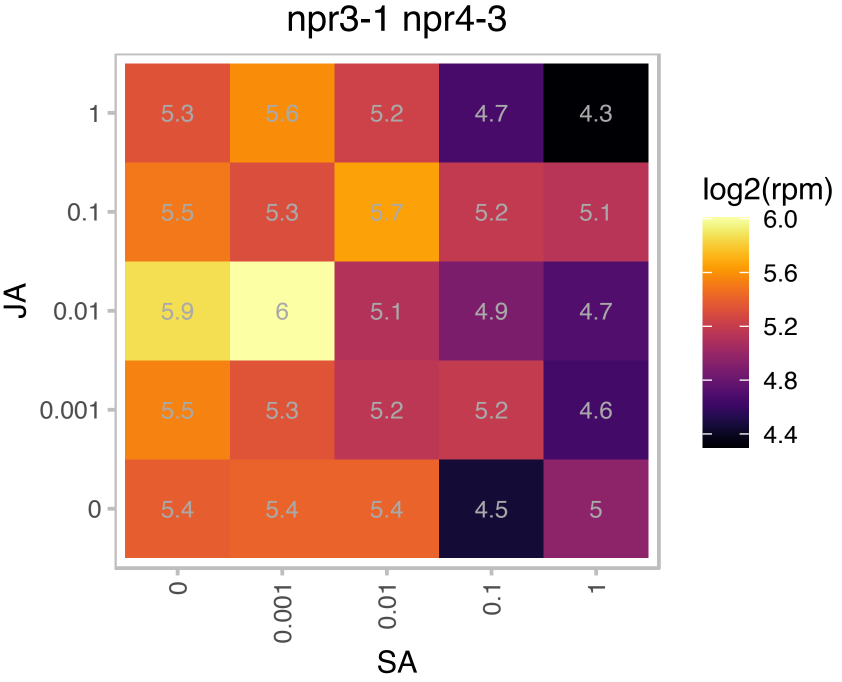

# AT5G27290.1 : NA

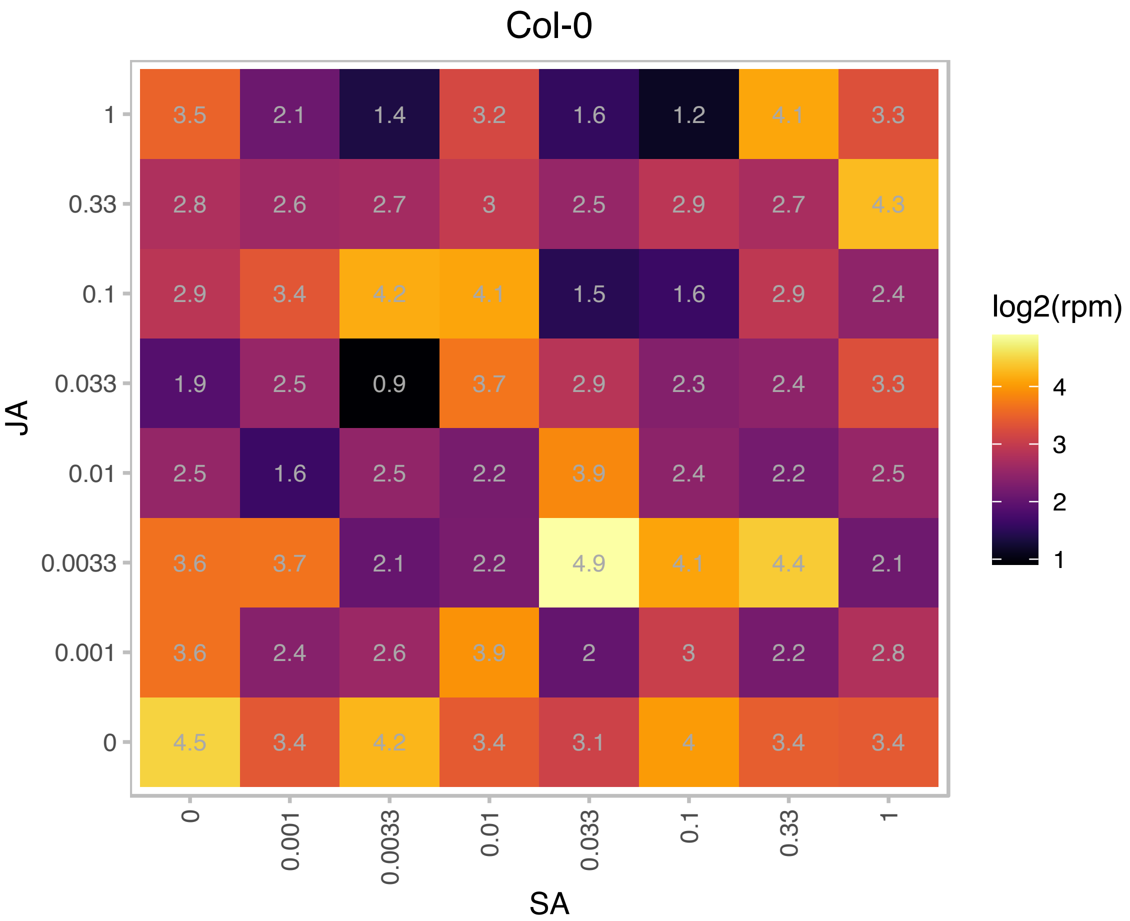


| | |
|---|---|
| Gene | AT5G27290 |
| is.expressed_genes | TRUE |
| is.responsive_genes | TRUE |
| is.apcluster | H-05 |

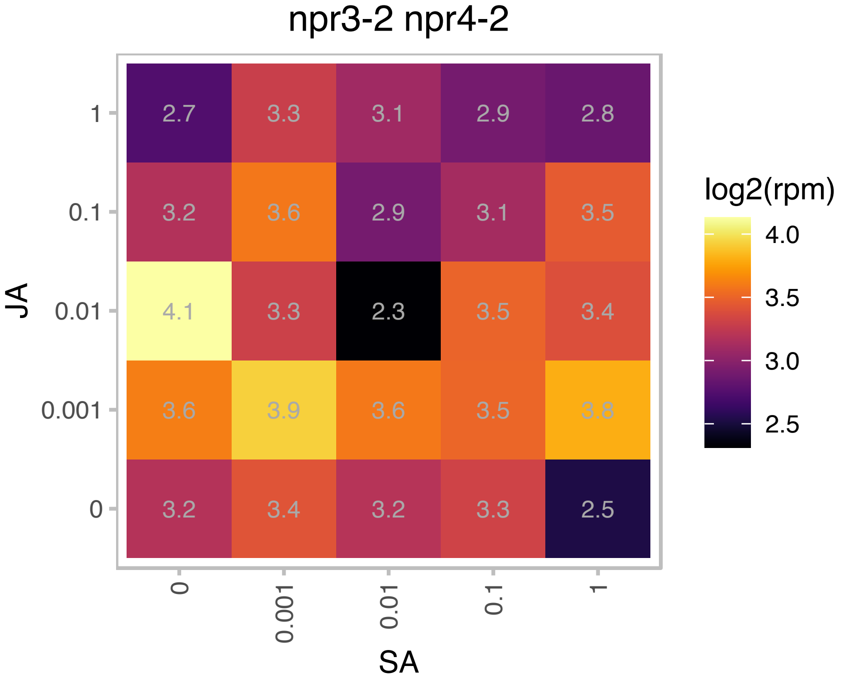


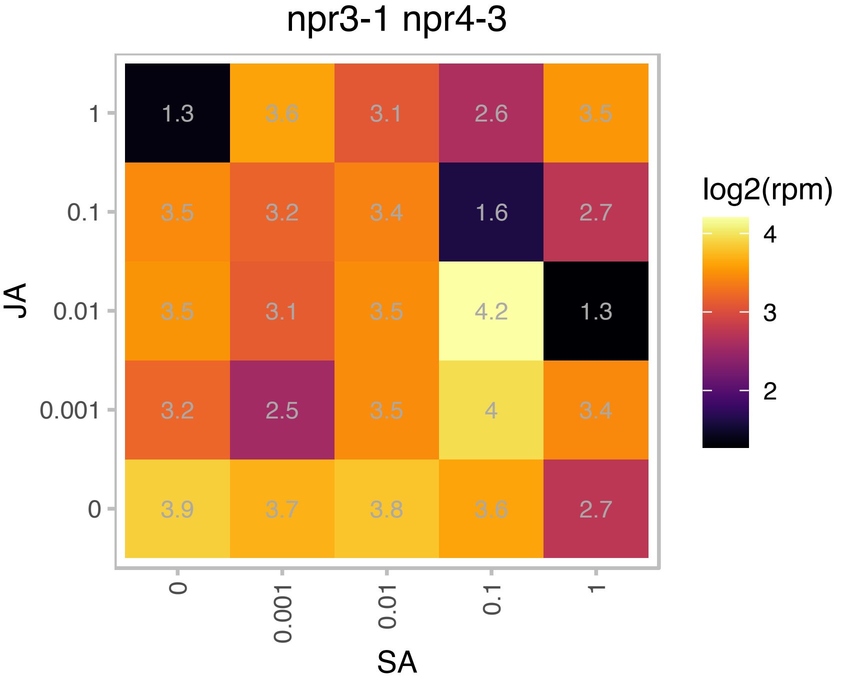

# AT5G28450.1 : NA

Chlorophyll A-B binding family protein

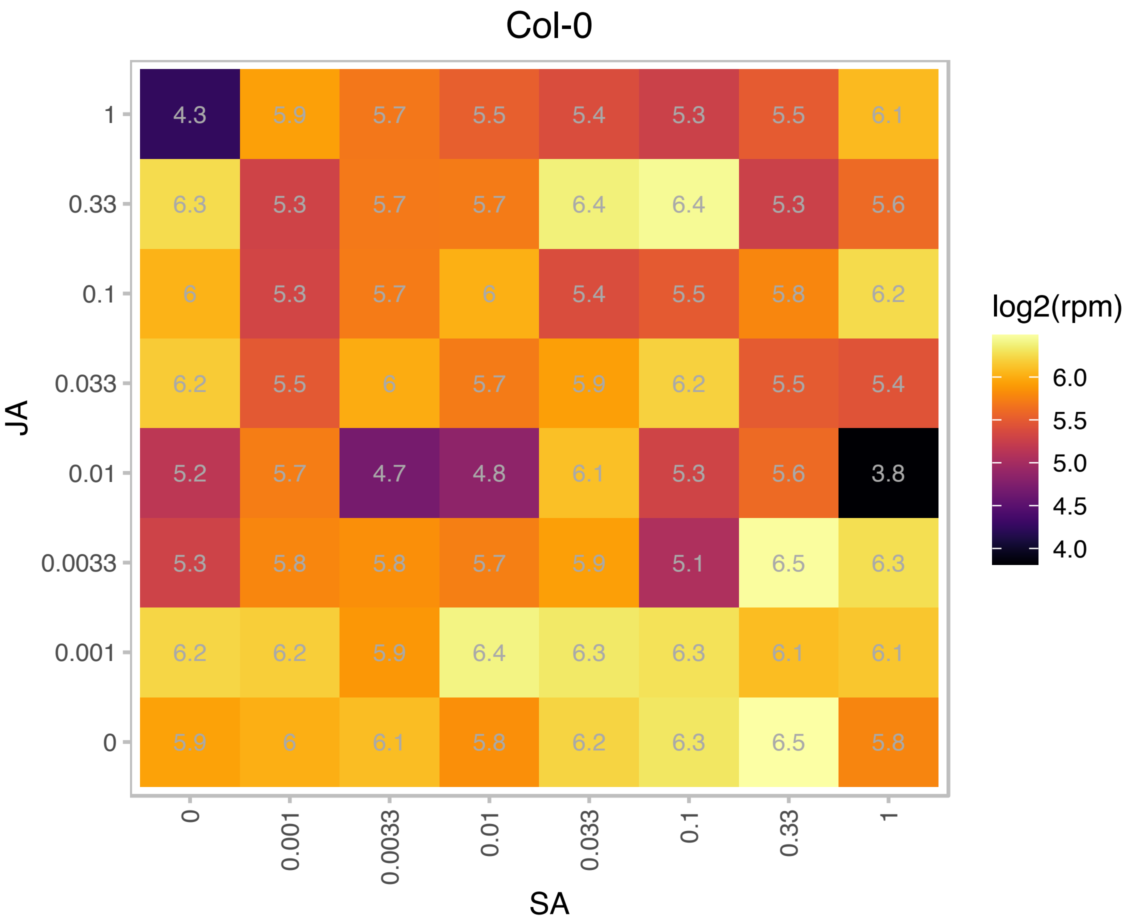


| | |
|---|---|
| Gene | AT5G28450 |
| is.expressed_genes | TRUE |
| is.responsive_genes | TRUE |
| is.apcluster | H-05 |

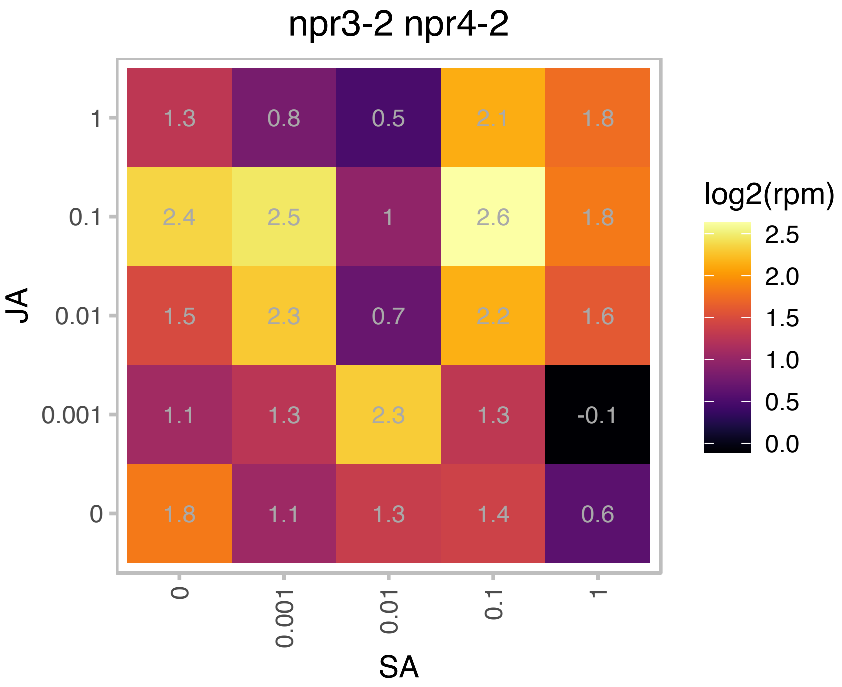


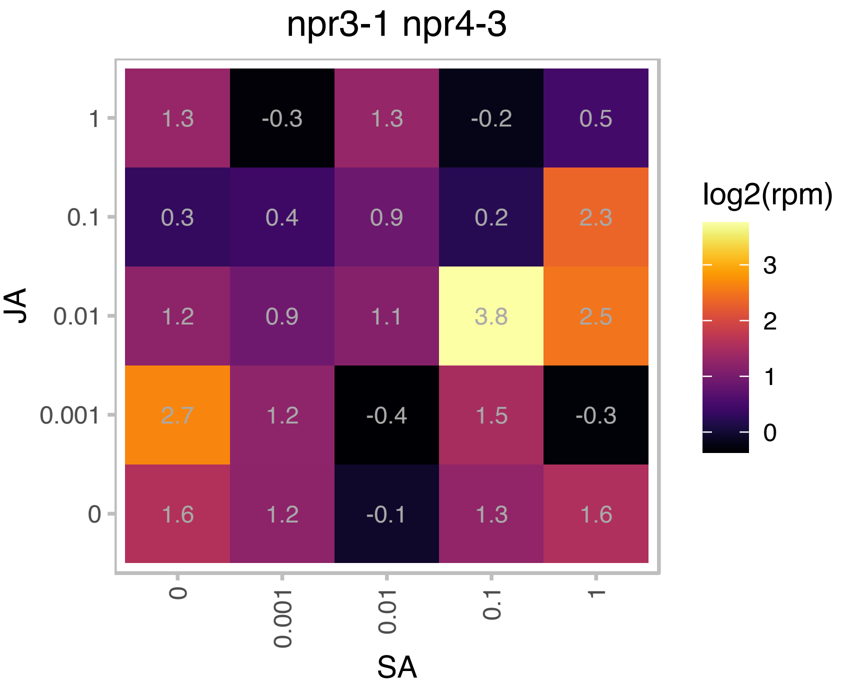

# AT5G46900.1 : NA

Bifunctional inhibitor/lipid-transfer protein/seed storage 2S albumin superfamily protein

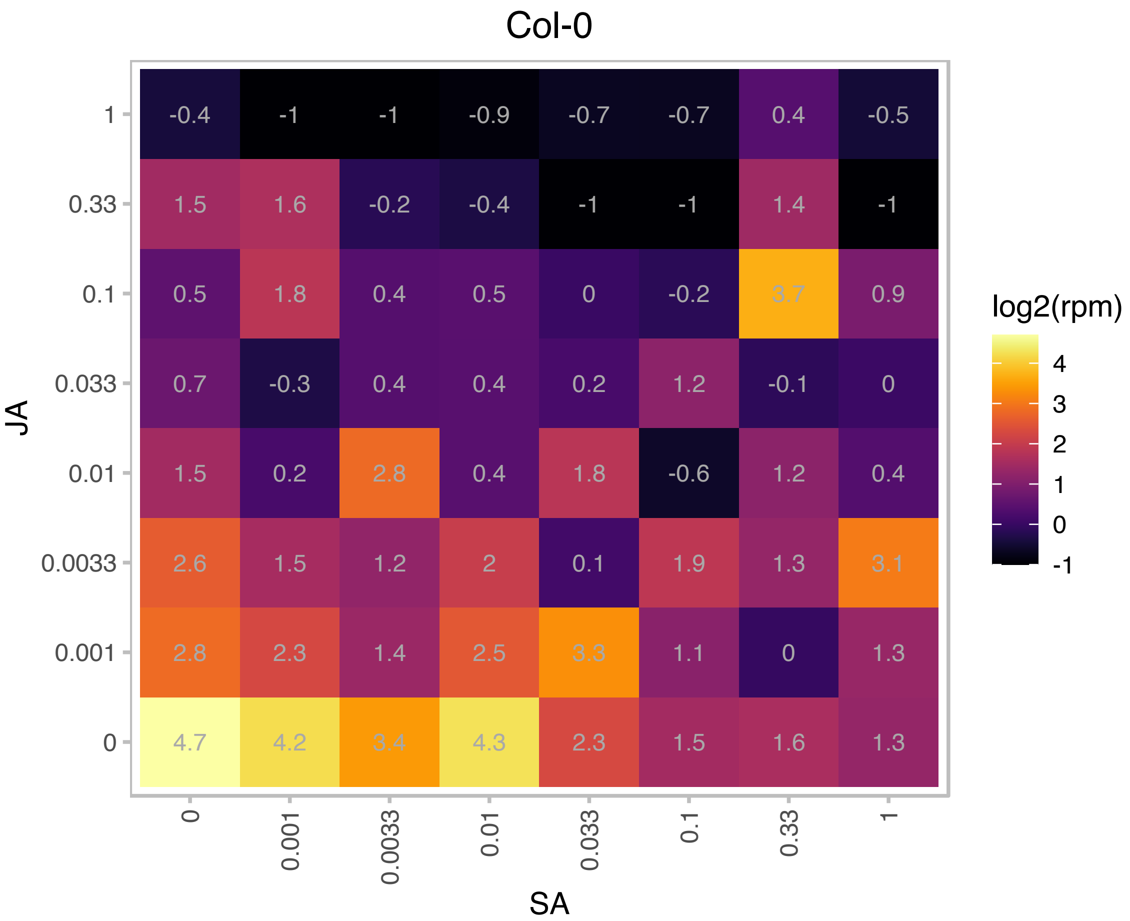


| | |
|---|---|
| Gene | AT5G46900 |
| is.expressed_genes | TRUE |
| is.responsive_genes | TRUE |
| is.apcluster | H-05 |

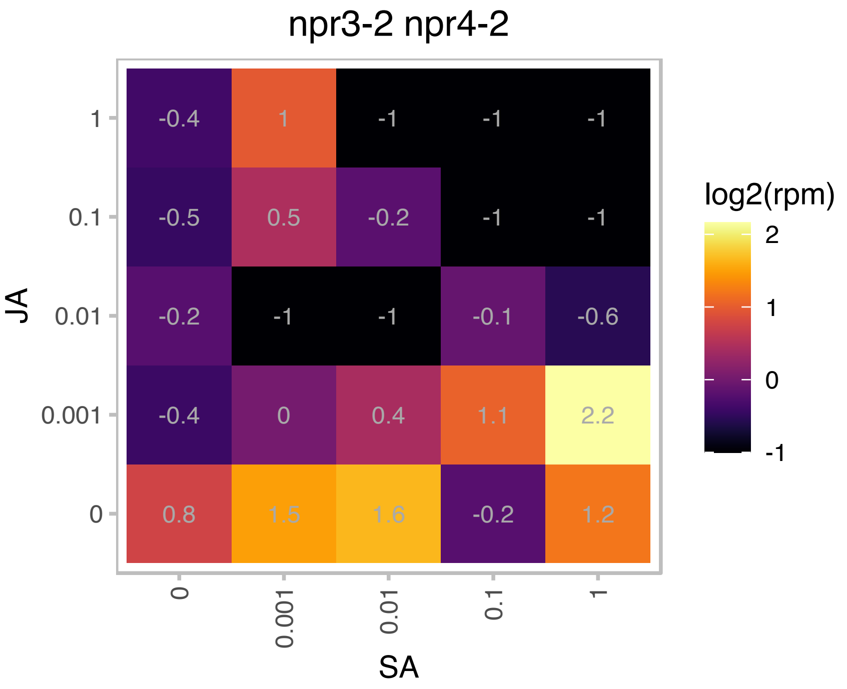


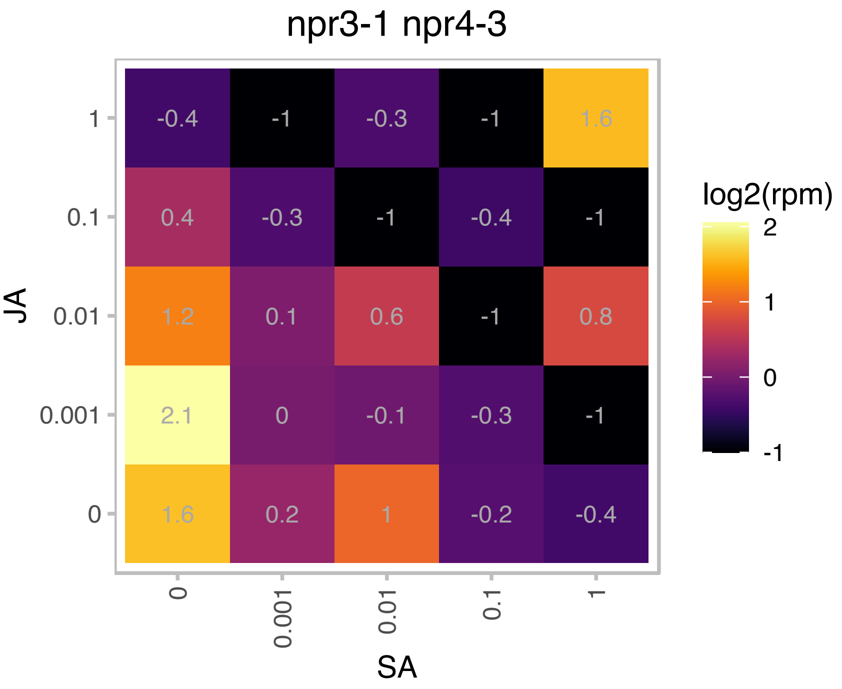

# AT5G52970.1 : NA

thylakoid lumen 15.0 kDa protein

| | |
|---|---|
| Gene | AT5G52970 |
| is.expressed_genes | TRUE |
| is.responsive_genes | TRUE |
| is.apcluster | H-05 |

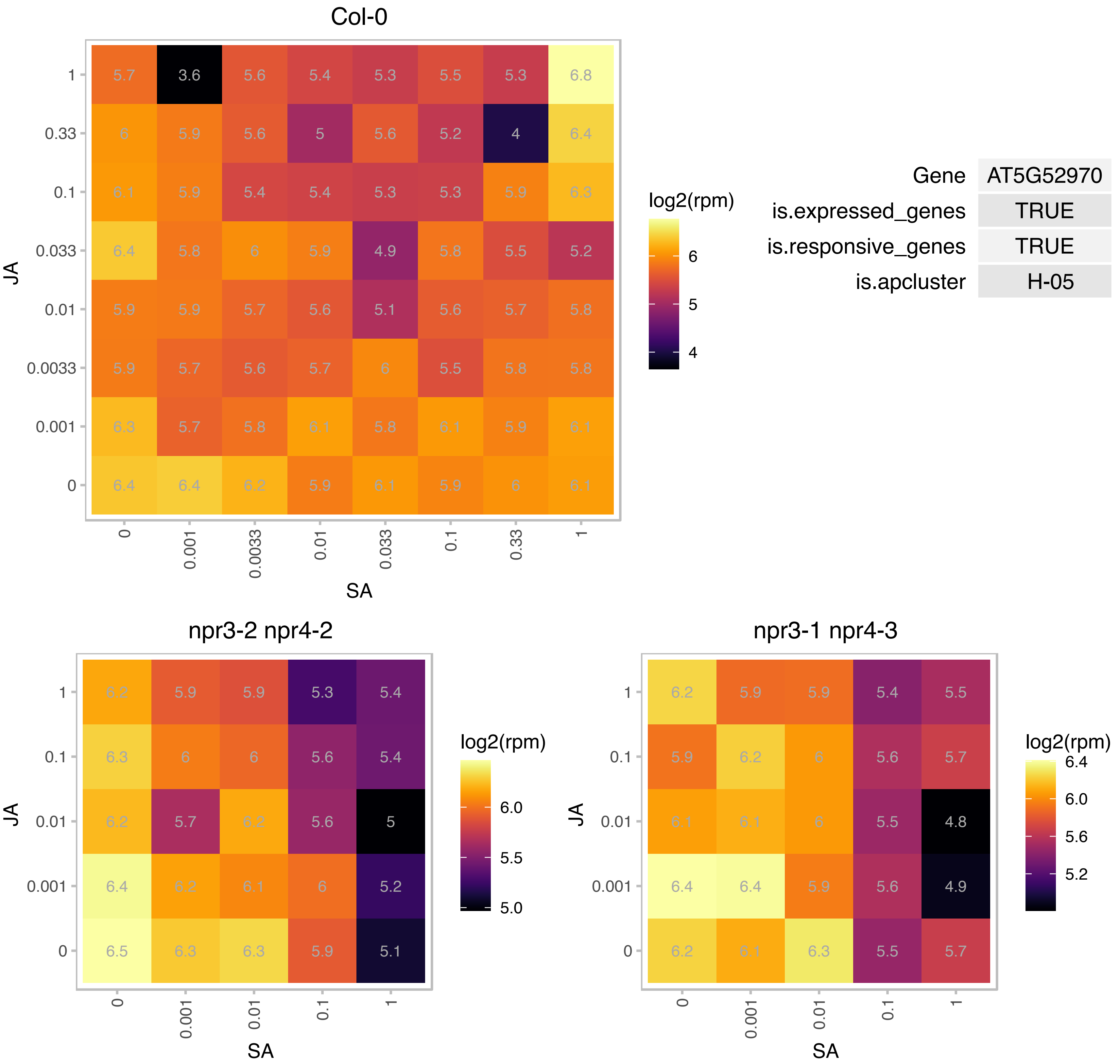

# AT5G55620.1 : NA

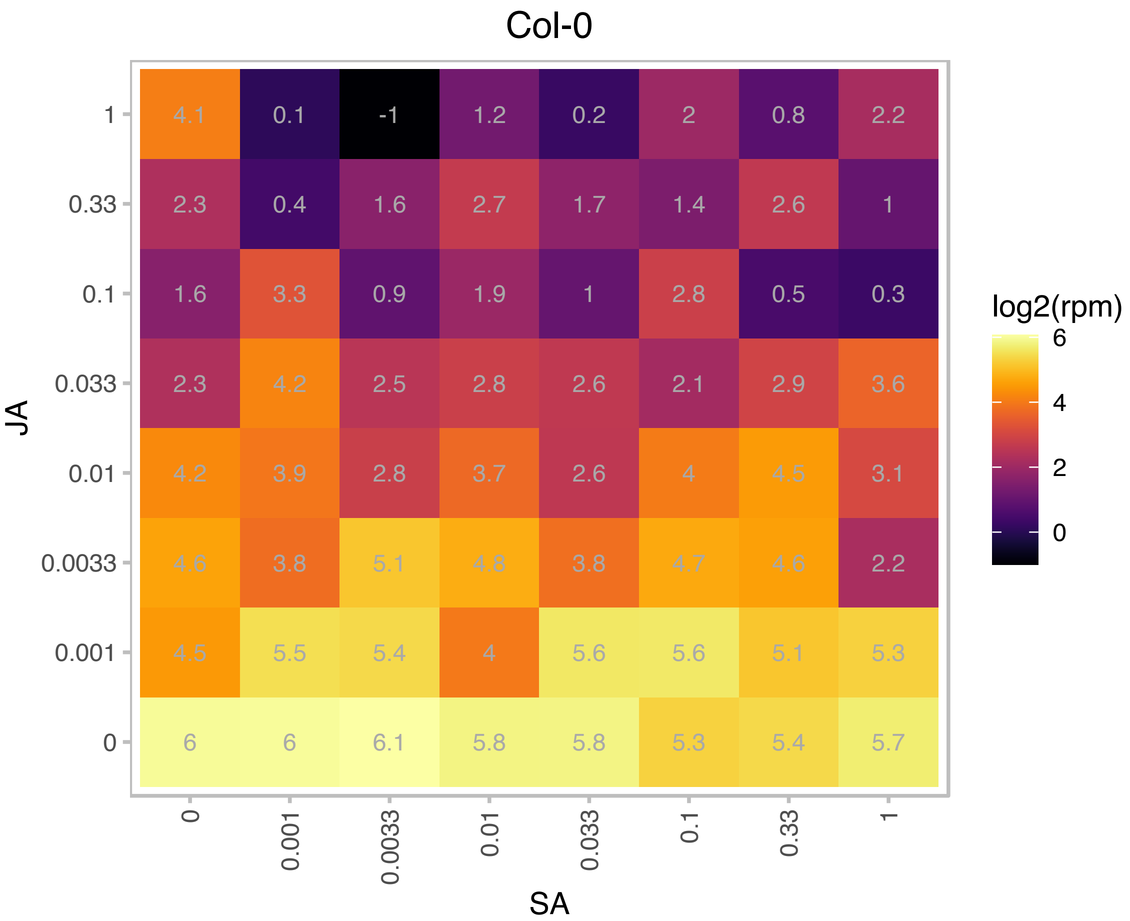


| | |
|---|---|
| Gene | AT5G55620 |
| is.expressed_genes | TRUE |
| is.responsive_genes | TRUE |
| is.apcluster | H-05 |

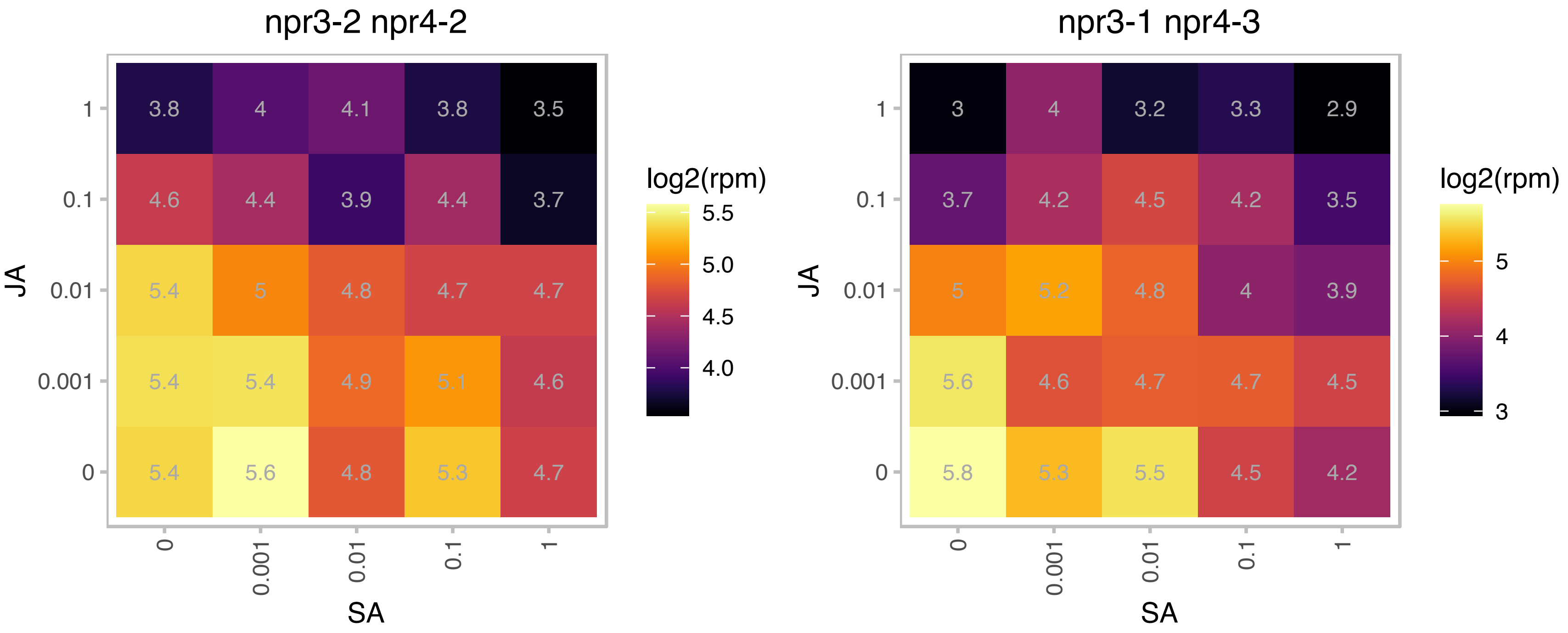

### AT5G57440.1 : GPP2, GS1

Haloacid dehalogenase-like hydrolase (HAD) superfamily protein

a member of haloacid dehalogenase-like hydrolase family

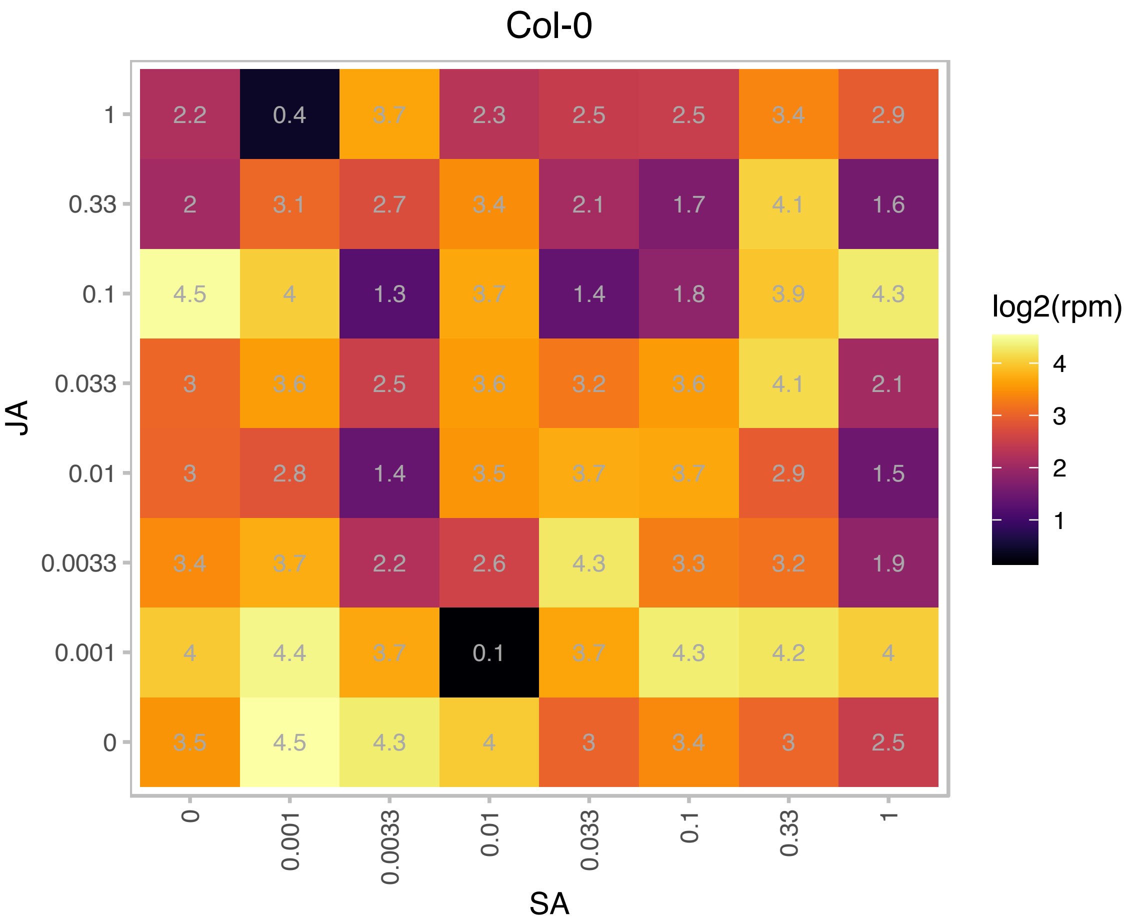


| | |
|---|---|
| Gene | AT5G57440 |
| is.expressed_genes | TRUE |
| is.responsive_genes | TRUE |
| is.apcluster | H-05 |

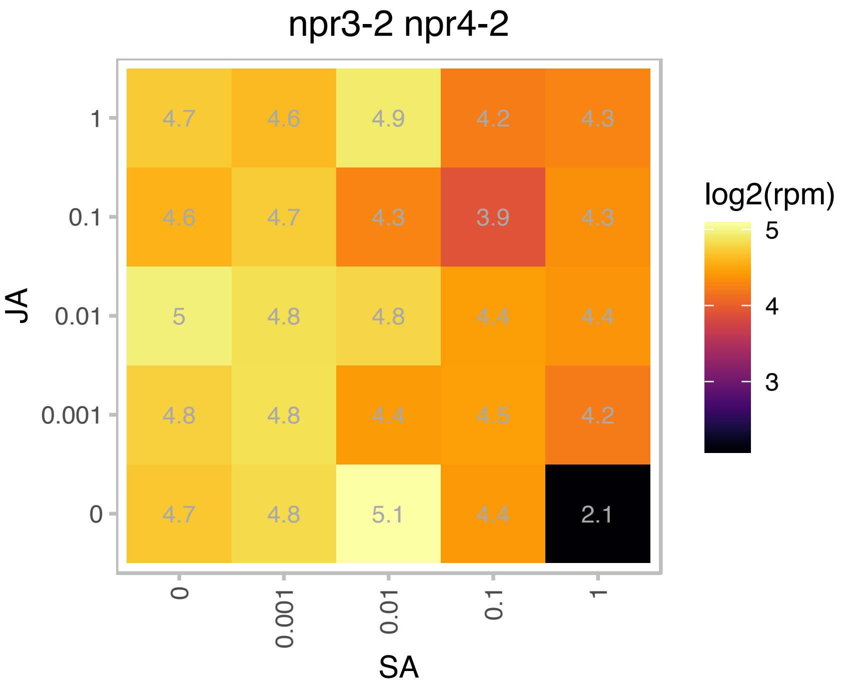


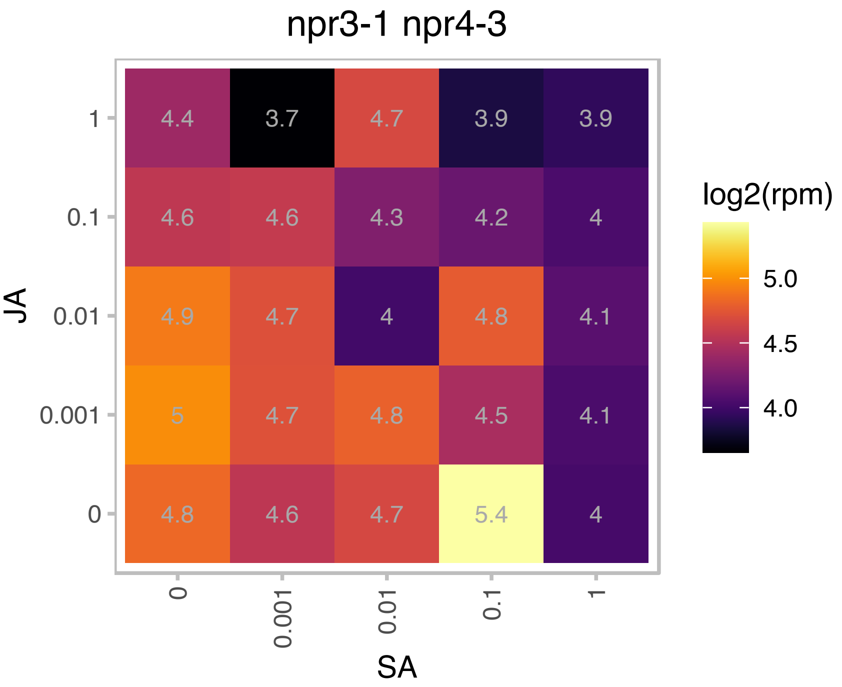

### AT5G64120.1 : AtPRX71, PRX71

Peroxidase superfamily protein

Encodes a cell wall bound peroxidase that is induced by hypo-osmolarity and is involved in the lignification of cell walls.

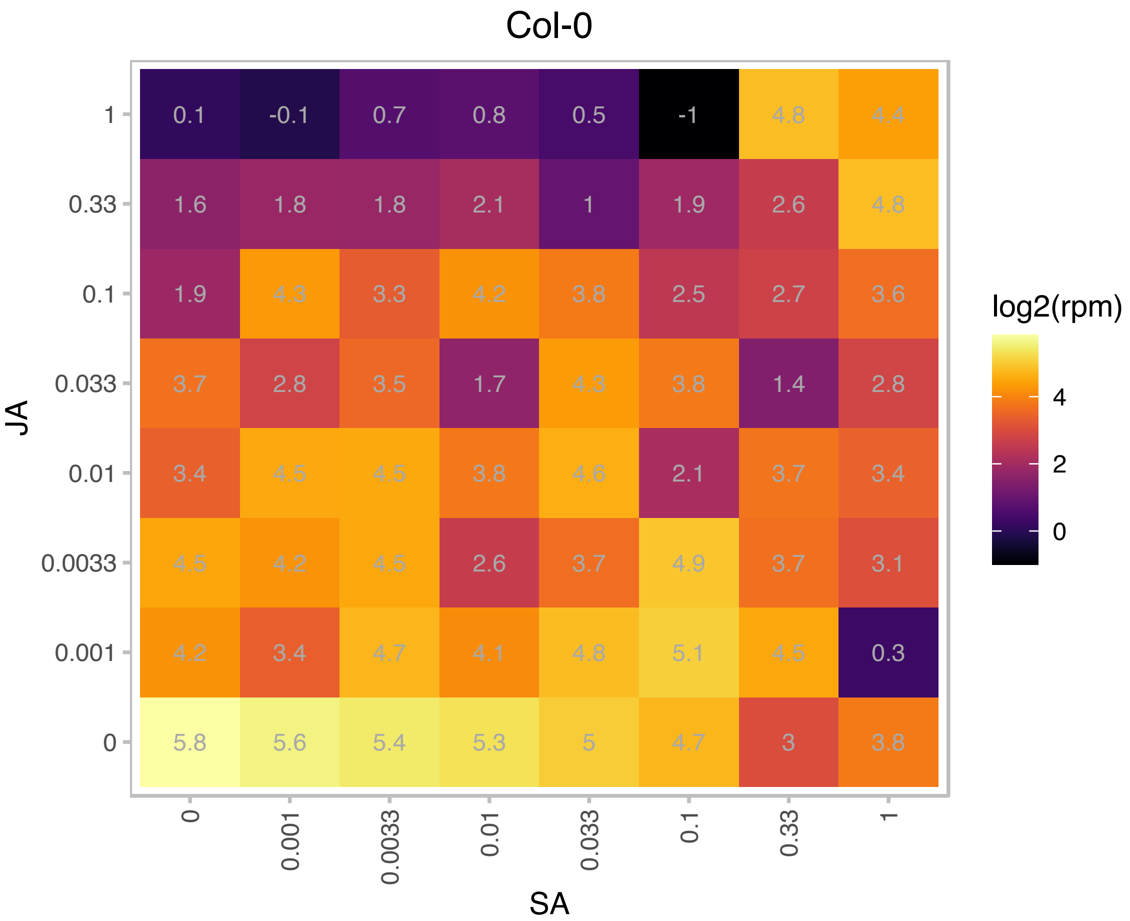


| | |
|---|---|
| Gene | AT5G64120 |
| is.expressed_genes | TRUE |
| is.responsive_genes | TRUE |
| is.apcluster | H-05 |

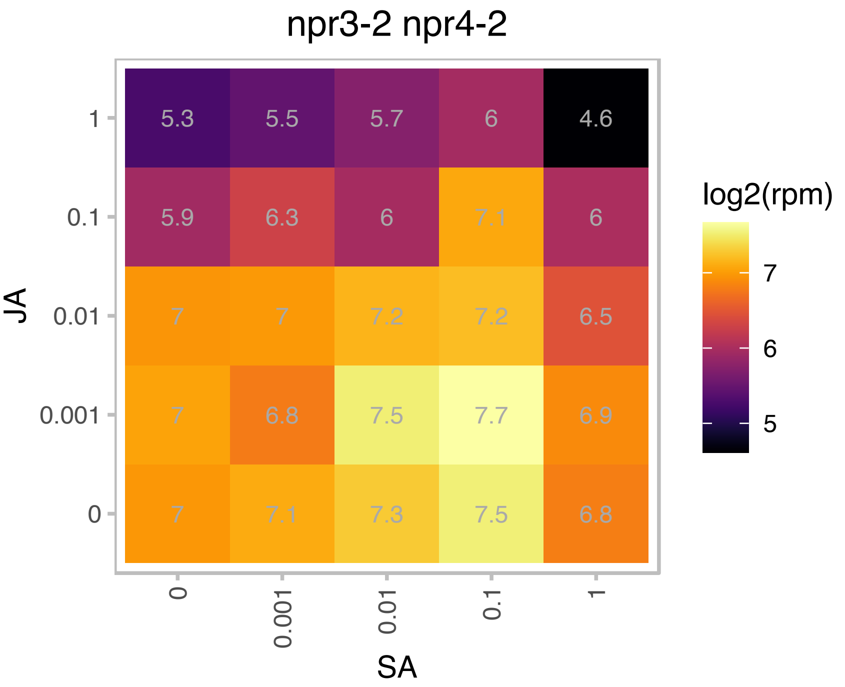


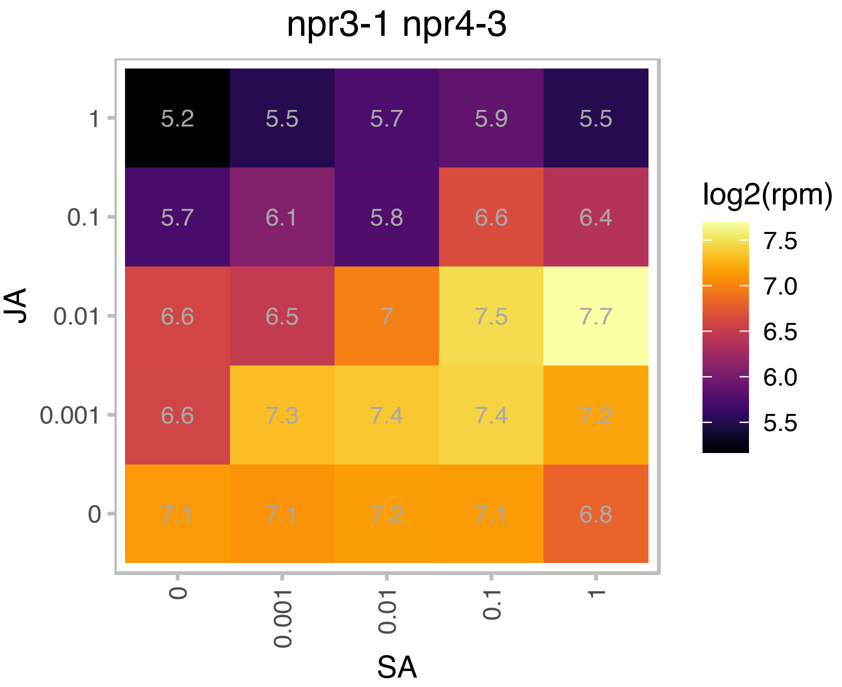

### AT5G66055.1 : AKRP, EMB16, EMB2036

ankyrin repeat protein

A locus involved in embryogenesis. Mutations in this locus result in embryo lethality.

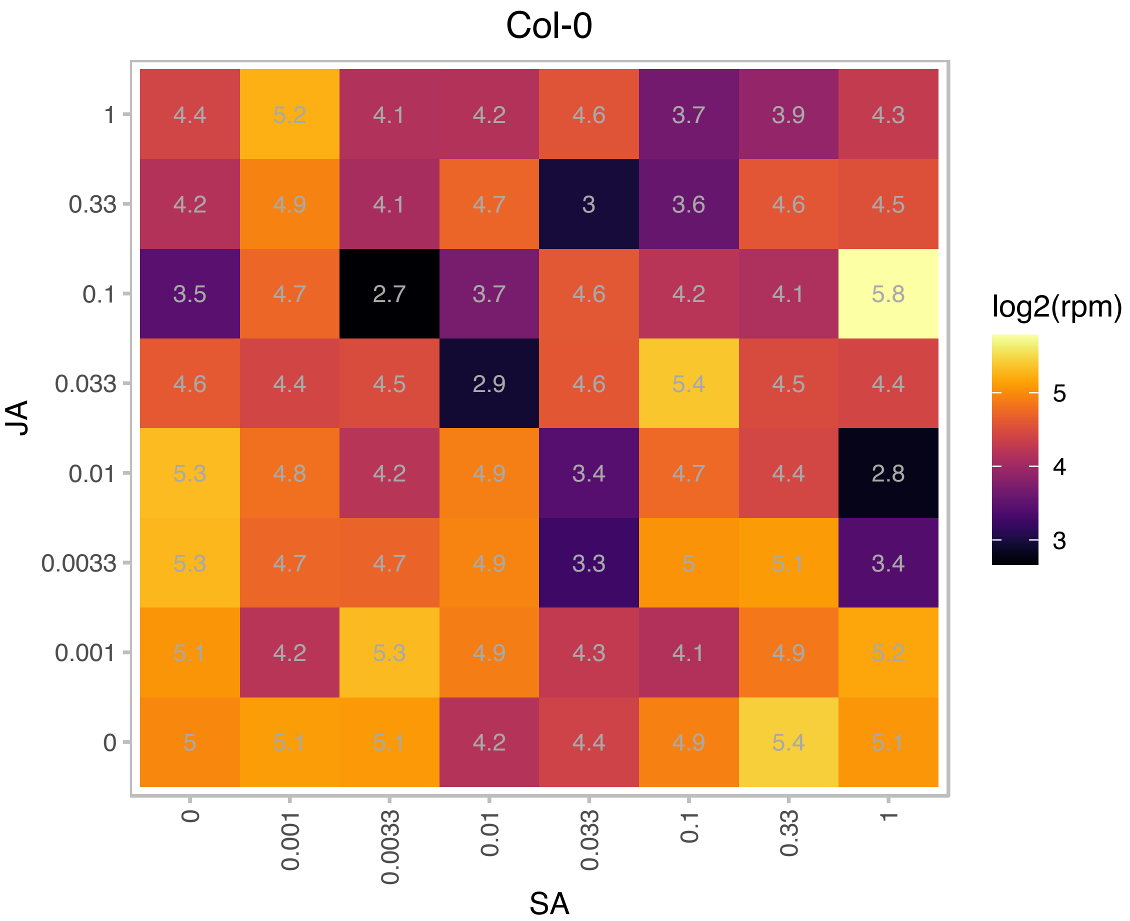


| | |
|---|---|
| Gene | AT5G66055 |
| is.expressed_genes | TRUE |
| is.responsive_genes | TRUE |
| is.apcluster | H-05 |

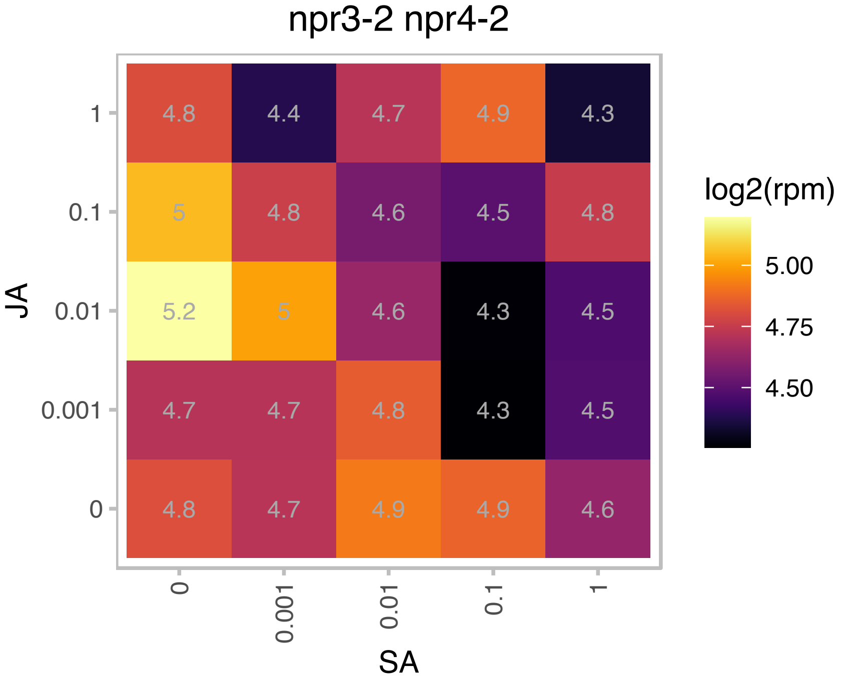


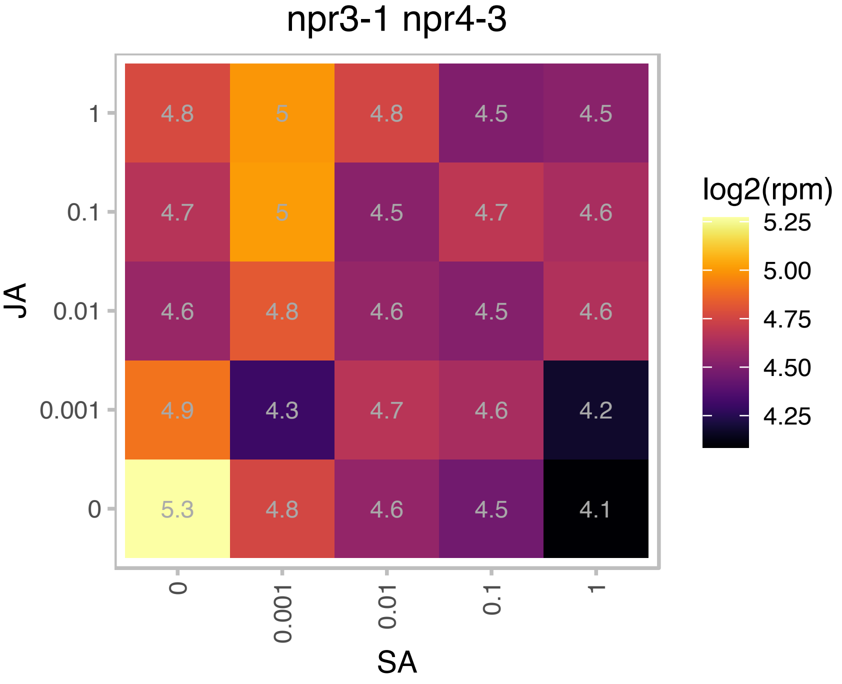

### AT1G14140.1 : UCP3

Mitochondrial substrate carrier family protein

| | |
|---|---|
| Gene | AT1G14140 |
| is.expressed_genes | TRUE |
| is.responsive_genes | TRUE |
| is.apcluster | H-06 |

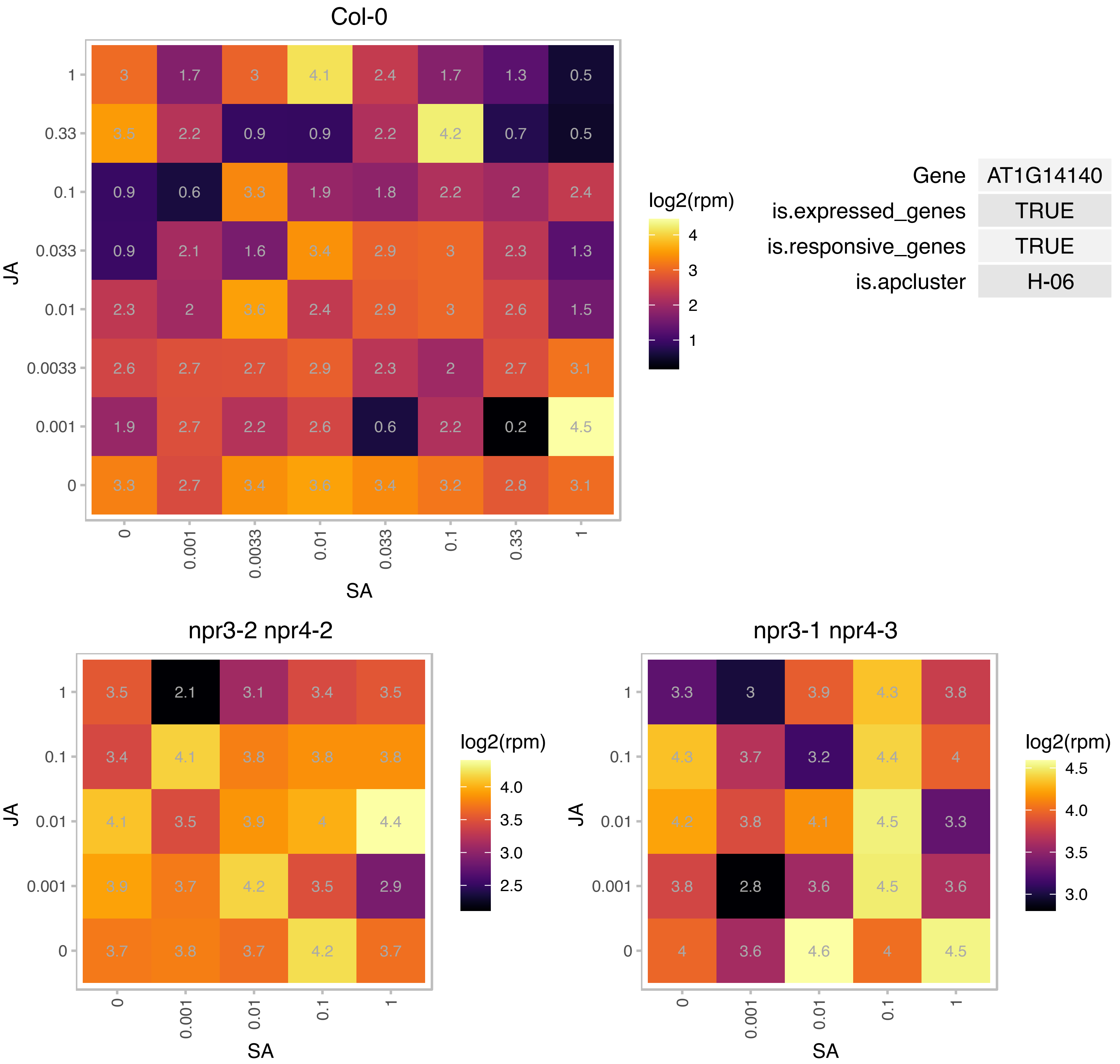

### AT1G23140.1 : CAR8

Calcium-dependent lipid-binding (CaLB domain) family protein

Expression is upregulated in the shoot of cax1/cax3 mutant.

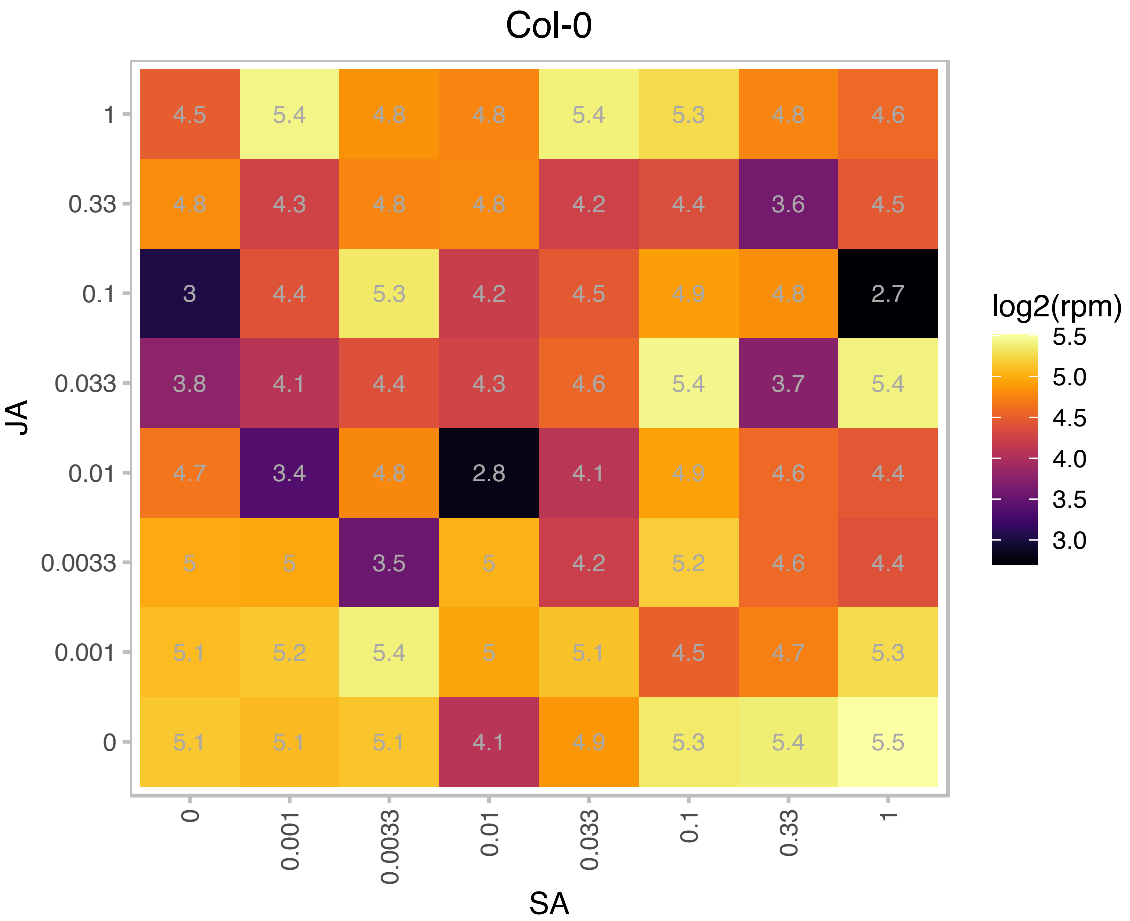

| | |
|---|---|
| Gene | AT1G23140 |
| is.expressed_genes | TRUE |
| is.responsive_genes | TRUE |
| is.apcluster | H-06 |

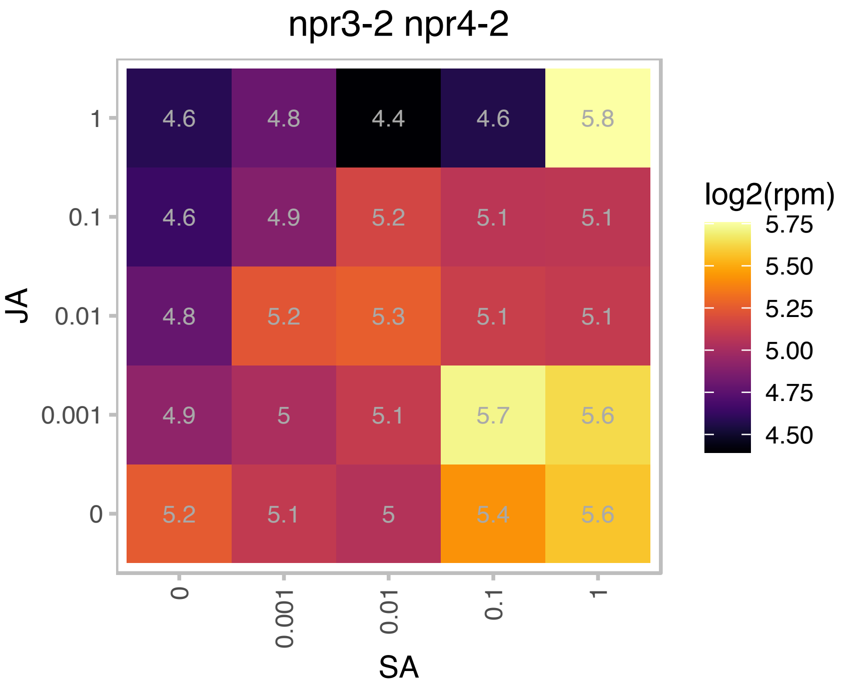

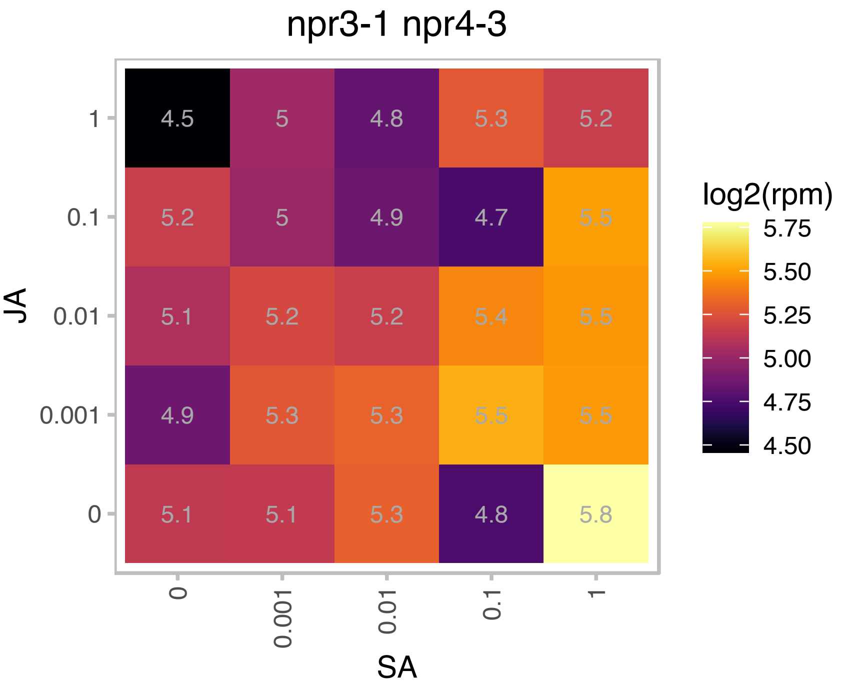

### AT1G27950.1 : LTPG1

glycosylphosphatidylinositol-anchored lipid protein transfer 1

Encodes LTPG1, a lipid transfer protein with a predicted GPI (glycosylphosphatidylinositol)-anchor domain. Localized in the plasma membrane. Disruption of the LTPG1 gene causes alterations of cuticular lipid composition, but no significant changes on total wax and cutin monomer loads are seen. The mRNA is cell-to-cell mobile.

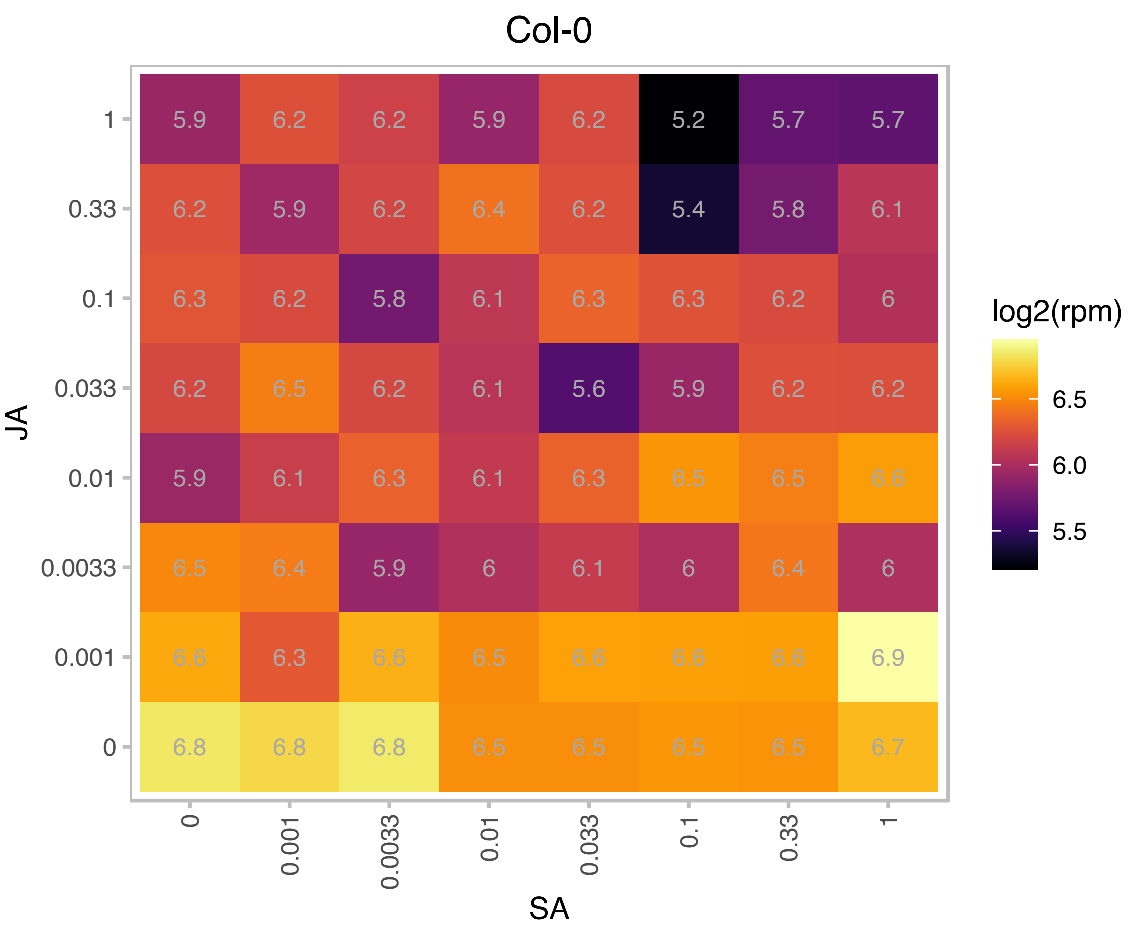


| | |
|---|---|
| Gene | AT1G27950 |
| is.expressed_genes | TRUE |
| is.responsive_genes | TRUE |
| is.apcluster | H-06 |

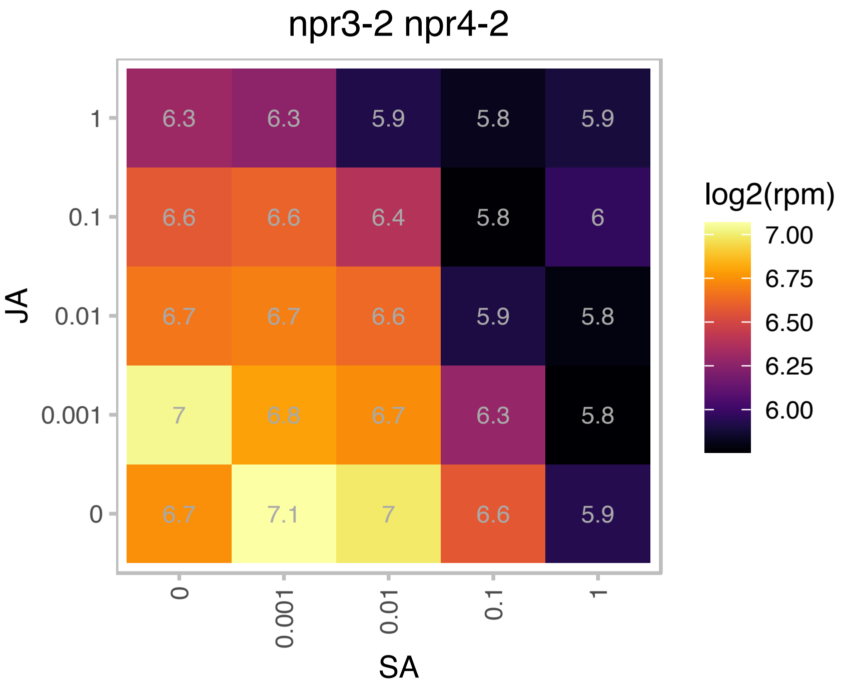


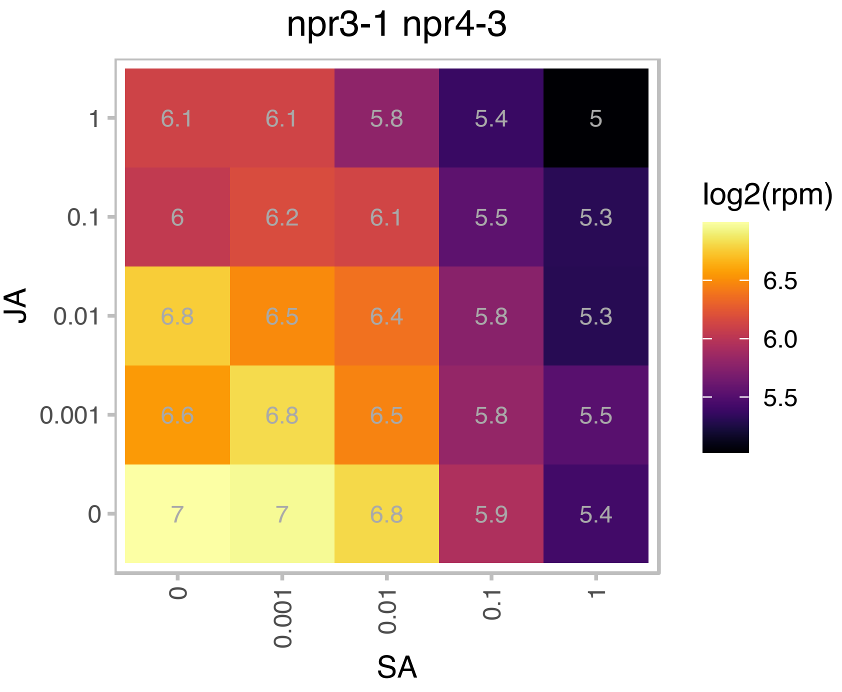

### AT1G47200.1 : WPP2

WPP domain protein 2
WPP family members contains an NE targeting domain. This domain, called the WPP domain after a highly conserved Trp-Pro-Pro motif, is necessary for NE targeting of WPP1. RNAi suppression of WPP2 resulted in reduced mitotic activity.

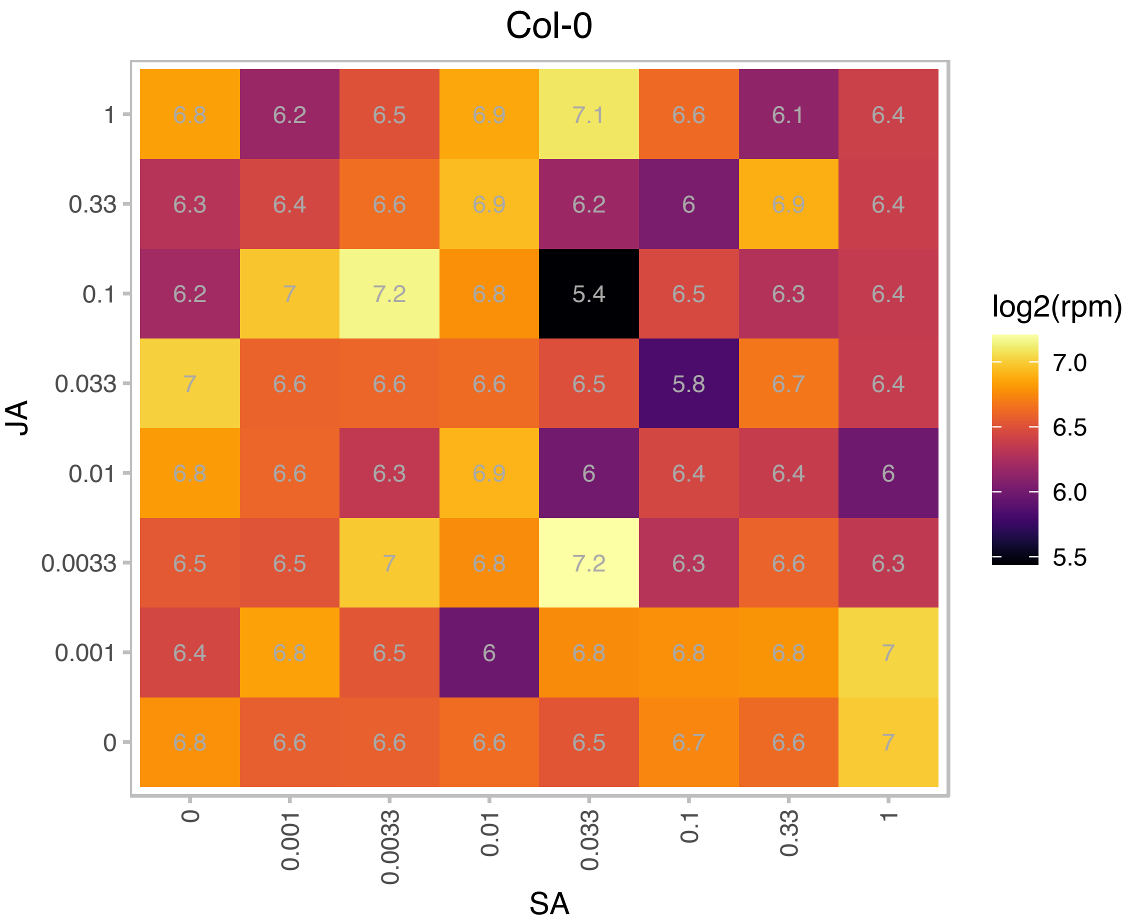


| | |
|---|---|
| Gene | AT1G47200 |
| is.expressed_genes | TRUE |
| is.responsive_genes | TRUE |
| is.apcluster | H-06 |

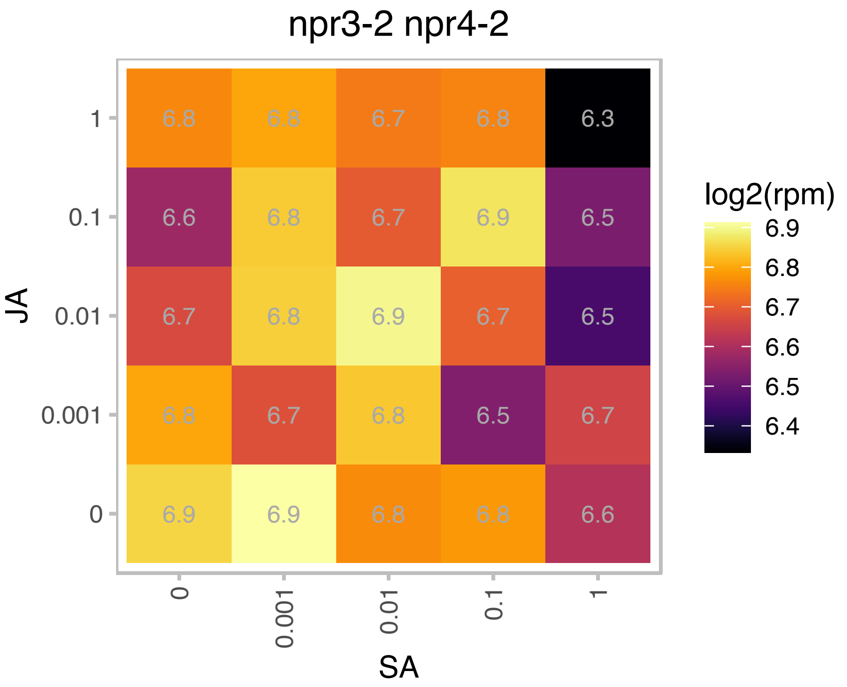


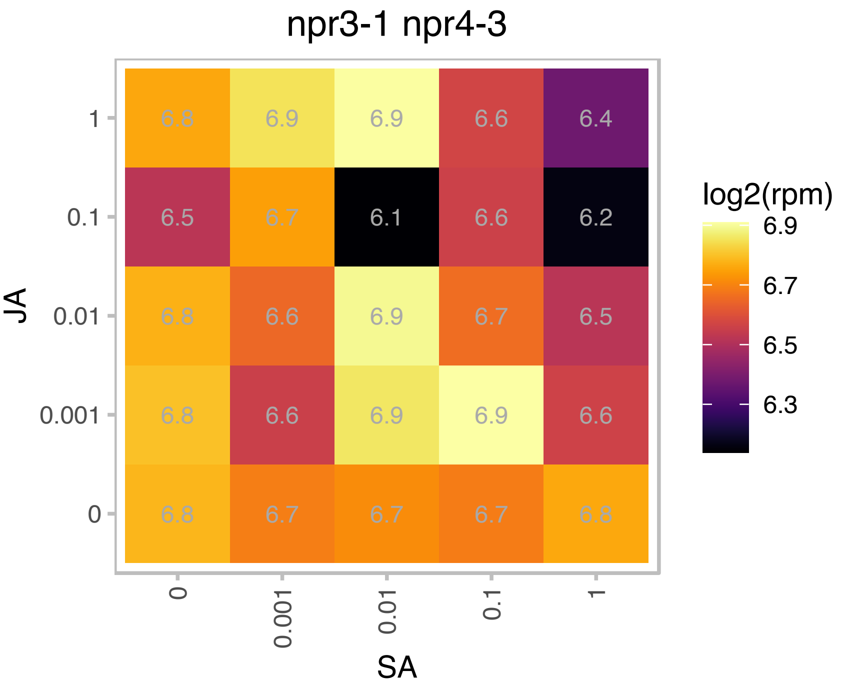

### AT1G75630.1 : AVA-P4

vacuolar H+-pumping ATPase 16 kDa proteolipid subunit 4

vacuolar H+-pumping ATPase 16 kD proteolipid (ava-p) mRNA, The mRNA is cell-to-cell mobile.

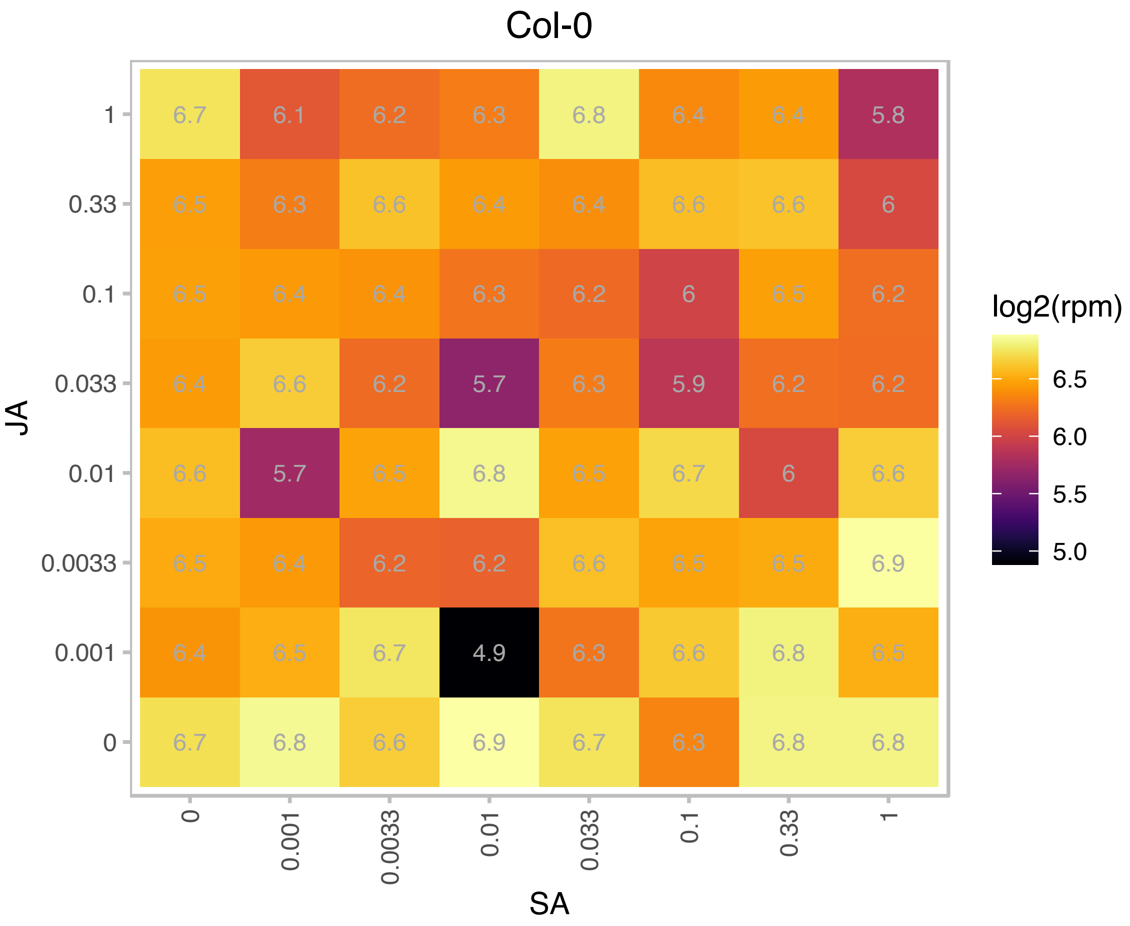


| | |
|---|---|
| Gene | AT1G75630 |
| is.expressed_genes | TRUE |
| is.responsive_genes | TRUE |
| is.apcluster | H-06 |

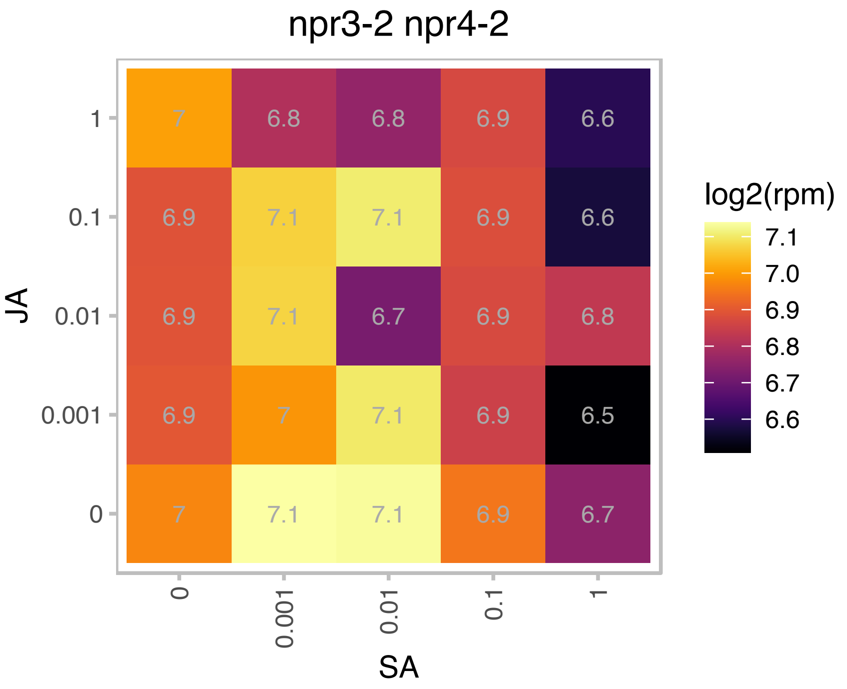


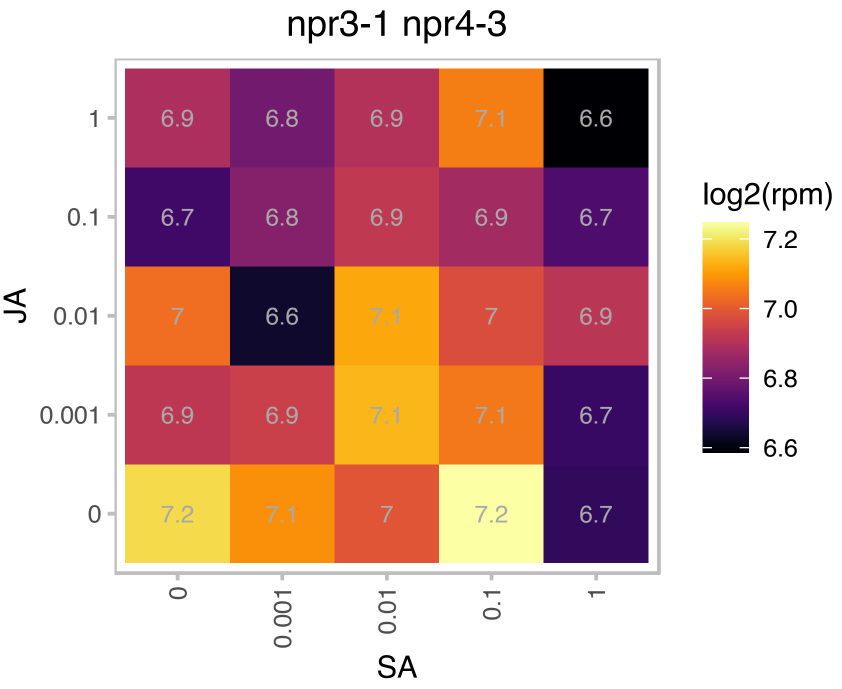

### AT2G18328.1 : ATRL4, RL4

RAD-like 4

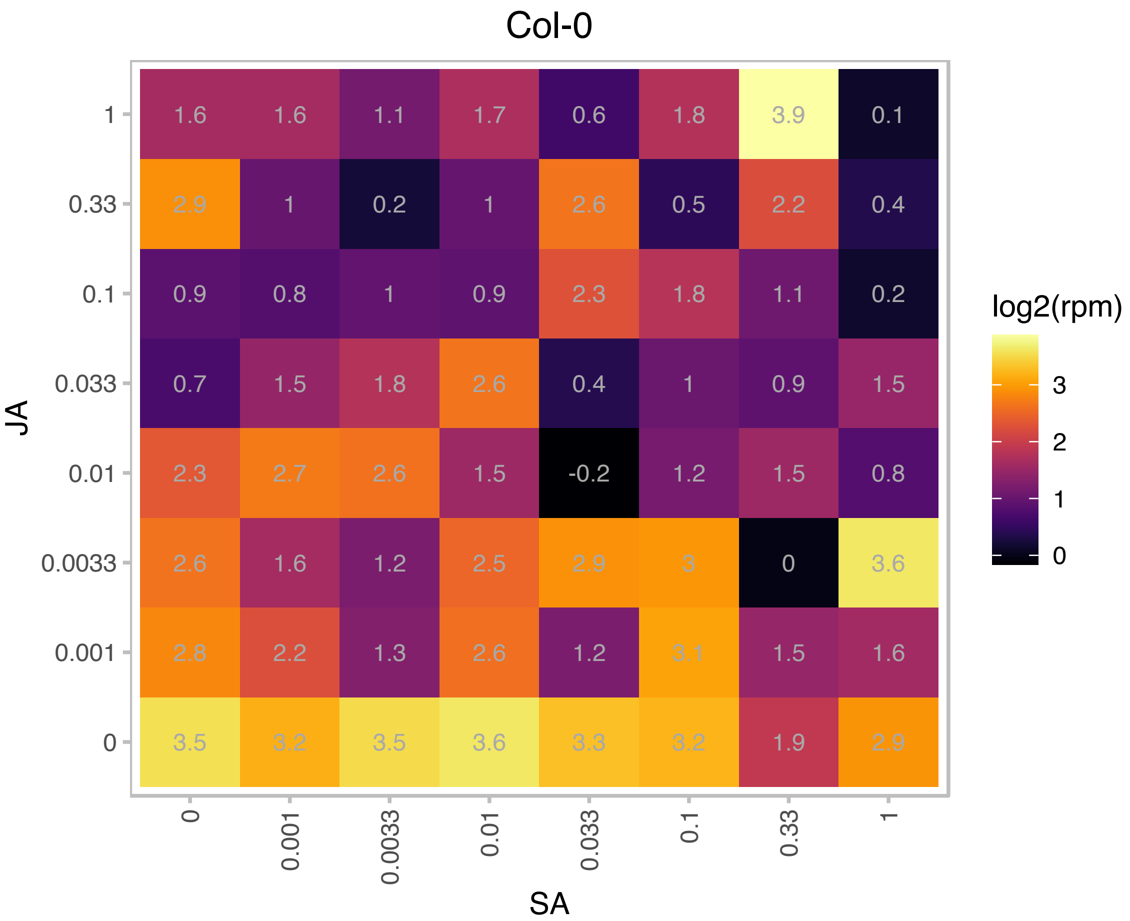


| | |
|---|---|
| Gene | AT2G18328 |
| is.expressed_genes | TRUE |
| is.responsive_genes | TRUE |
| is.apcluster | H-06 |

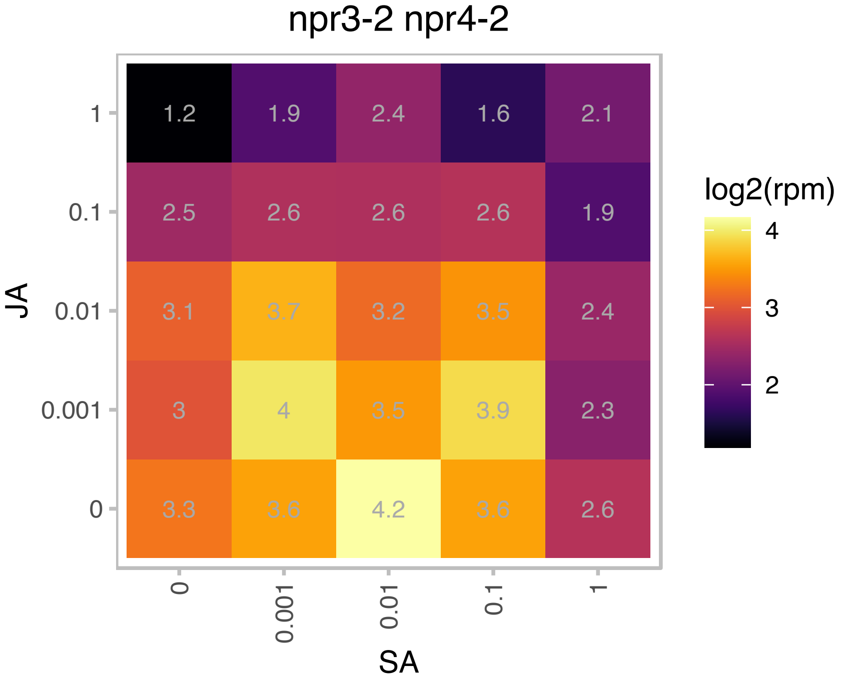


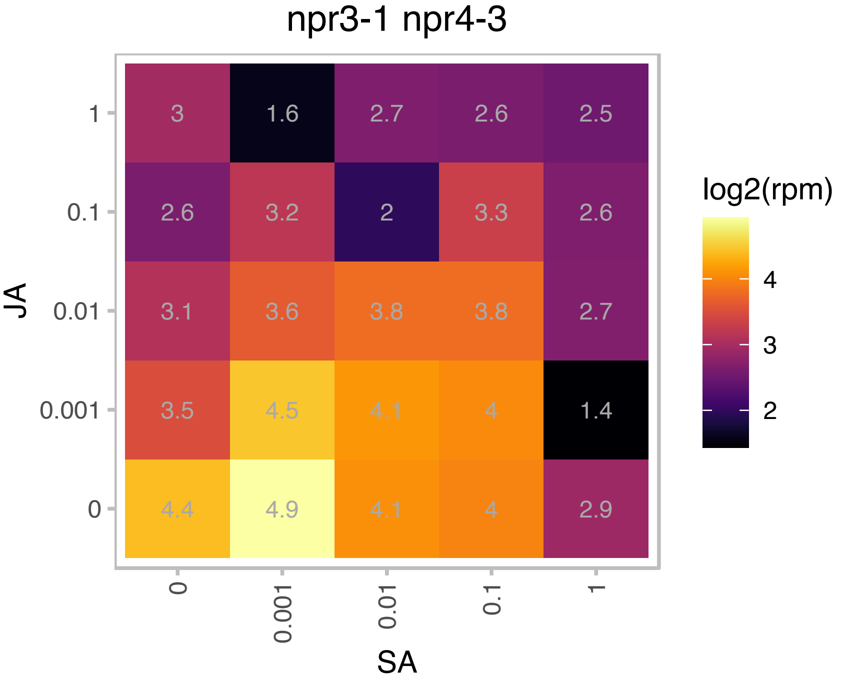

### AT2G26070.1 : AtRTE1, RTE1

Protein of unknown function (DUF778).

Encodes a predicted membrane protein. Similar sequences are widely distributed and conserved in plants, animals and protists but absent in fungi and prokaryotes. The sequence has no known motifs and no biological function has been assigned in any species. In Arabidopsis, it appears to be involved in the negative regulation of the response to ethylene, is localized to the Golgi and is a positive regulator of ETR1.

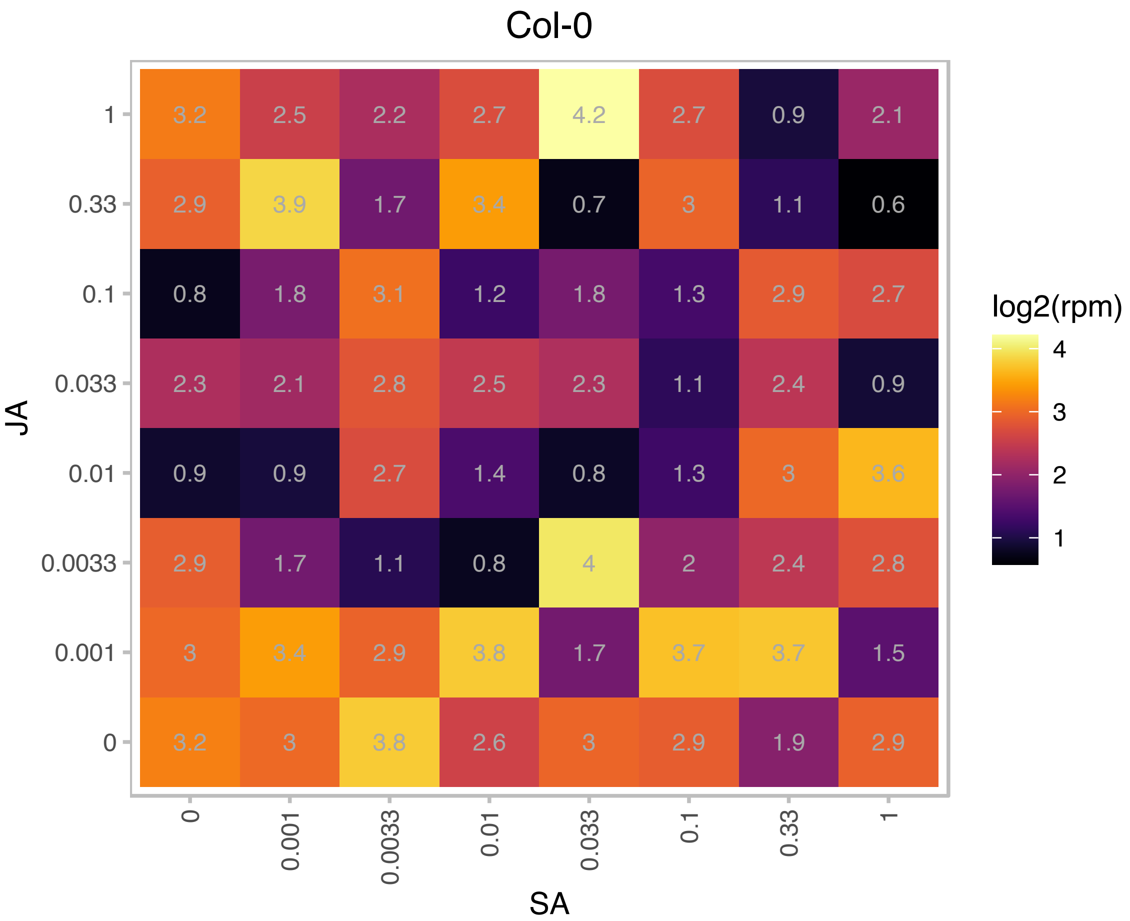


| | |
|---|---|
| Gene | AT2G26070 |
| is.expressed_genes | TRUE |
| is.responsive_genes | TRUE |
| is.apcluster | H-06 |

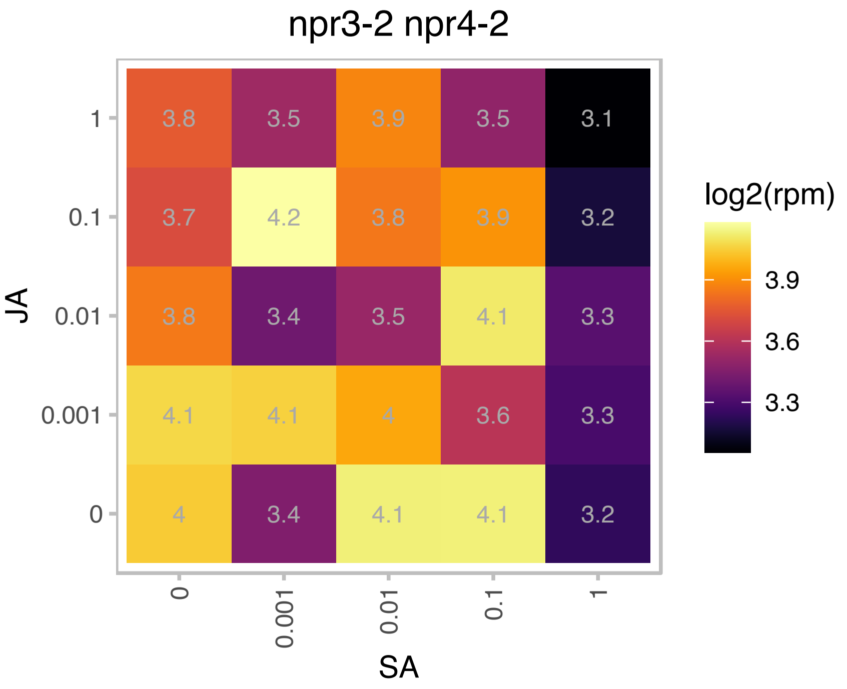


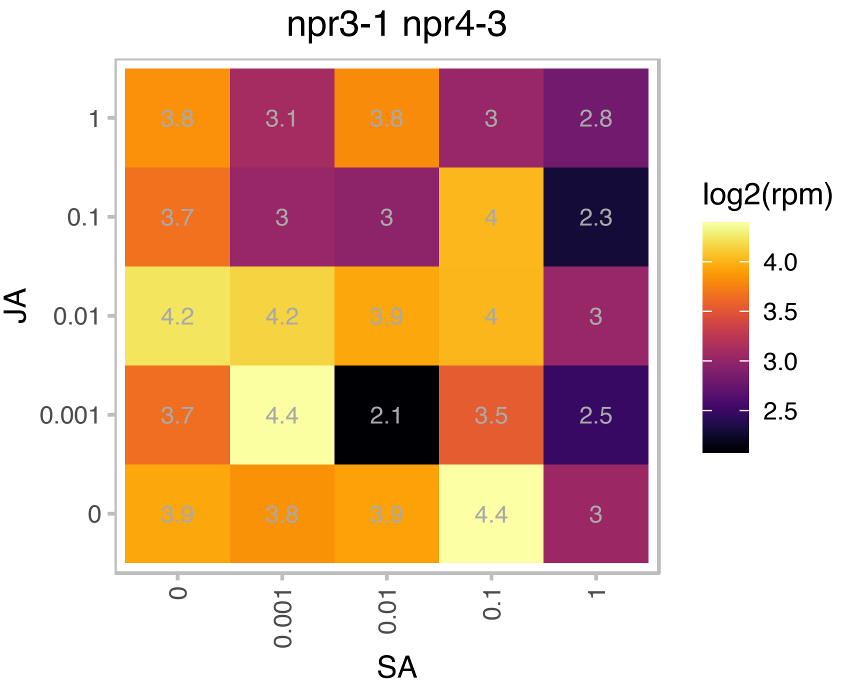

### AT2G27050.1 : AtEIL1, EIL1

ETHYLENE-INSENSITIVE3-like 1

ethylene-insensitive3-like1 (EIL1) The mRNA is cell-to-cell mobile.

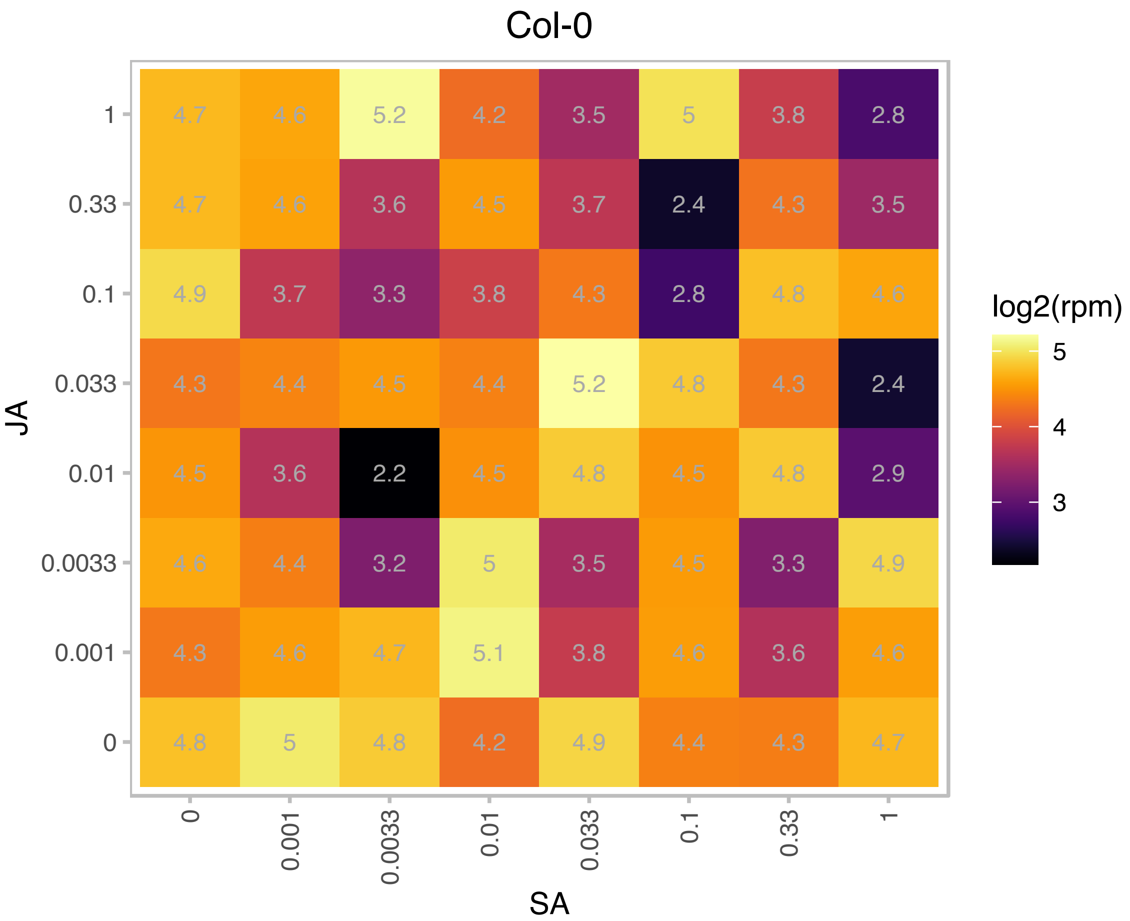


| | |
|---|---|
| Gene | AT2G27050 |
| is.expressed_genes | TRUE |
| is.responsive_genes | TRUE |
| is.apcluster | H-06 |

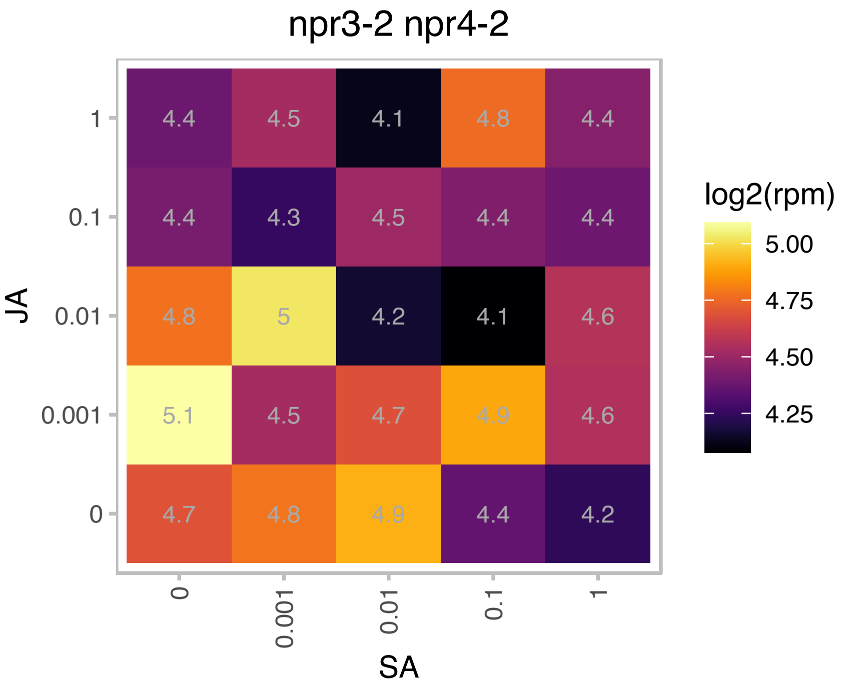


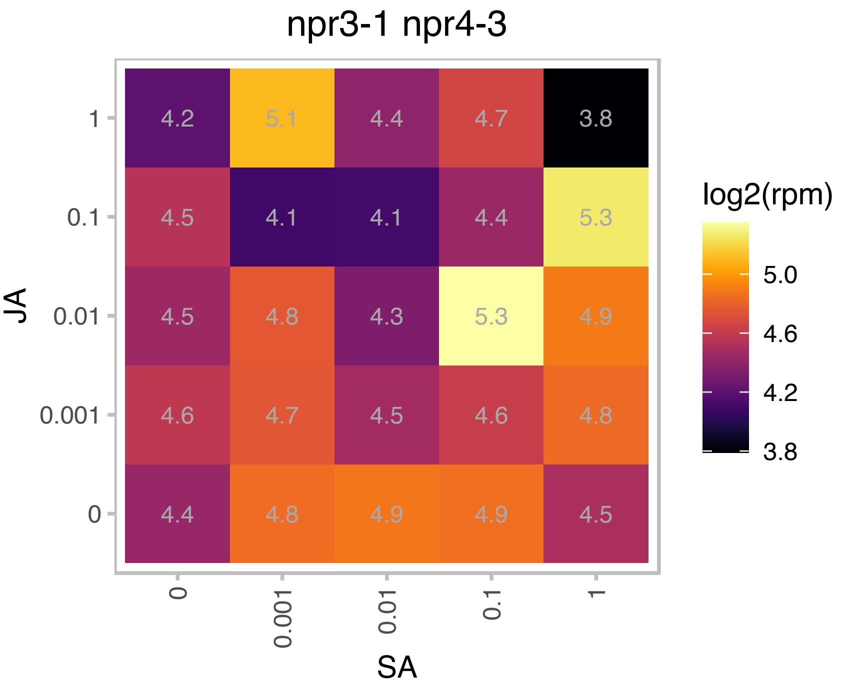

# AT2G31730.1 : NA

basic helix-loop-helix (bHLH) DNA-binding superfamily protein

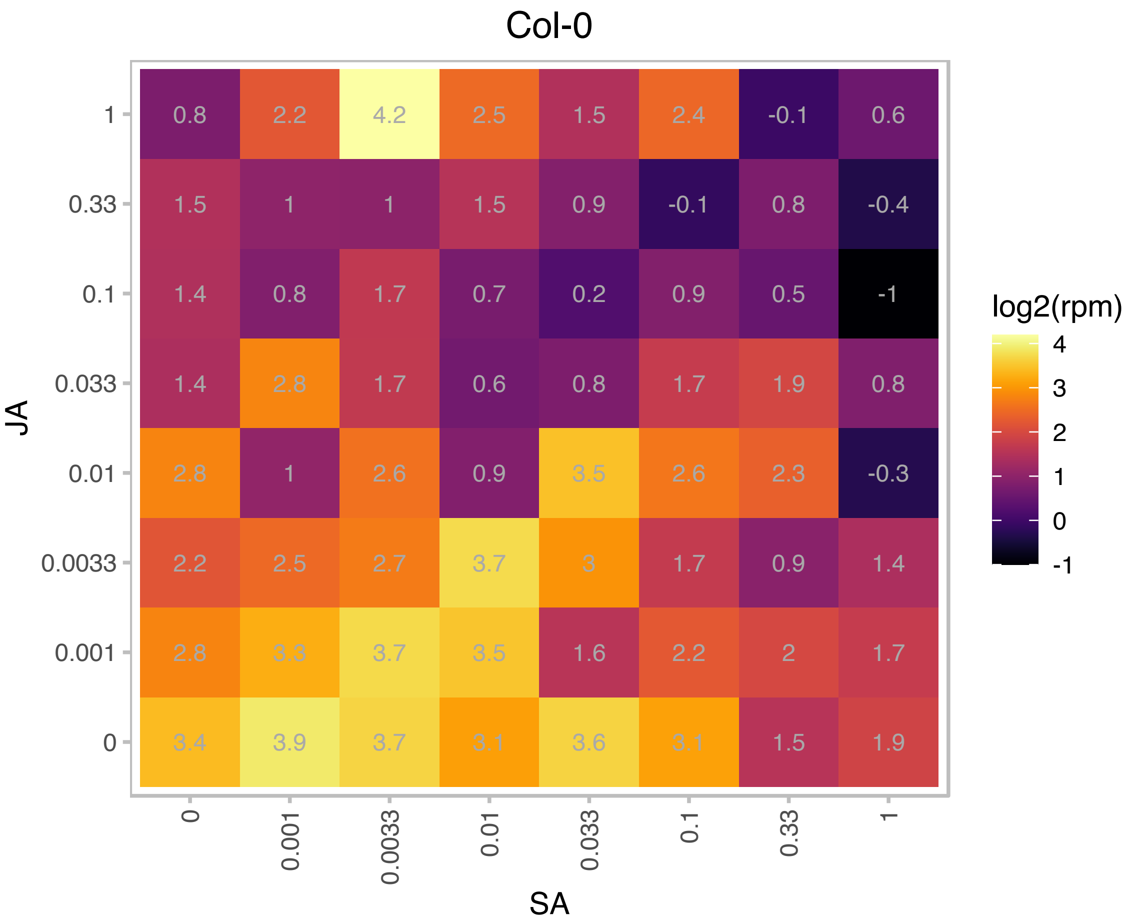


| | |
|---|---|
| Gene | AT2G31730 |
| is.expressed_genes | TRUE |
| is.responsive_genes | TRUE |
| is.apcluster | H-06 |

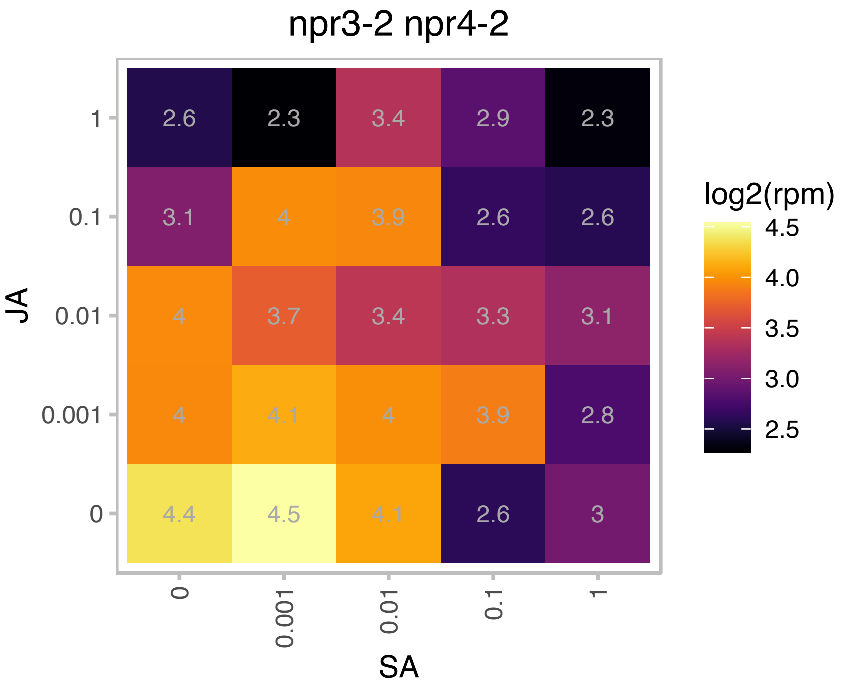


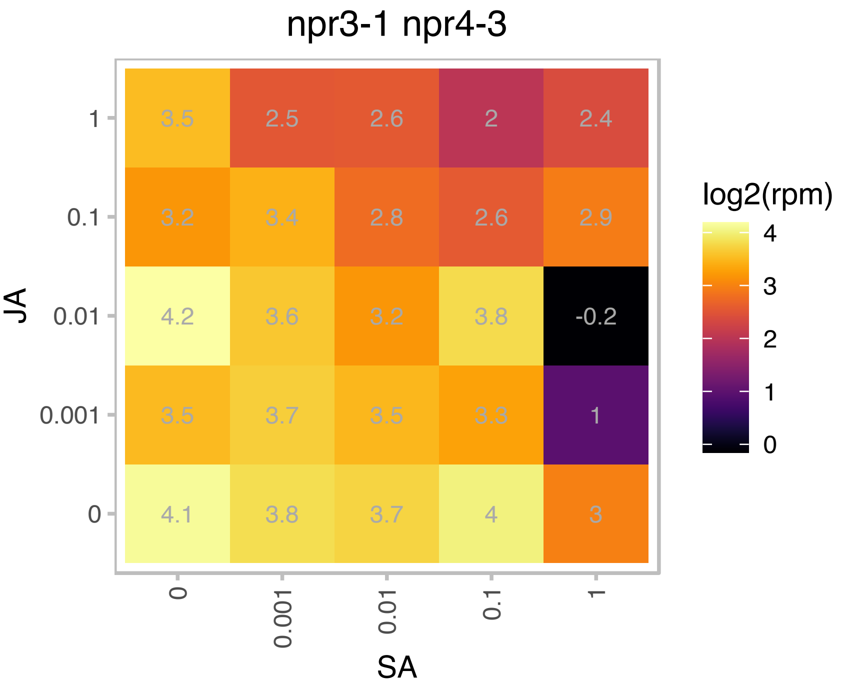

# AT2G33250.1 : NA

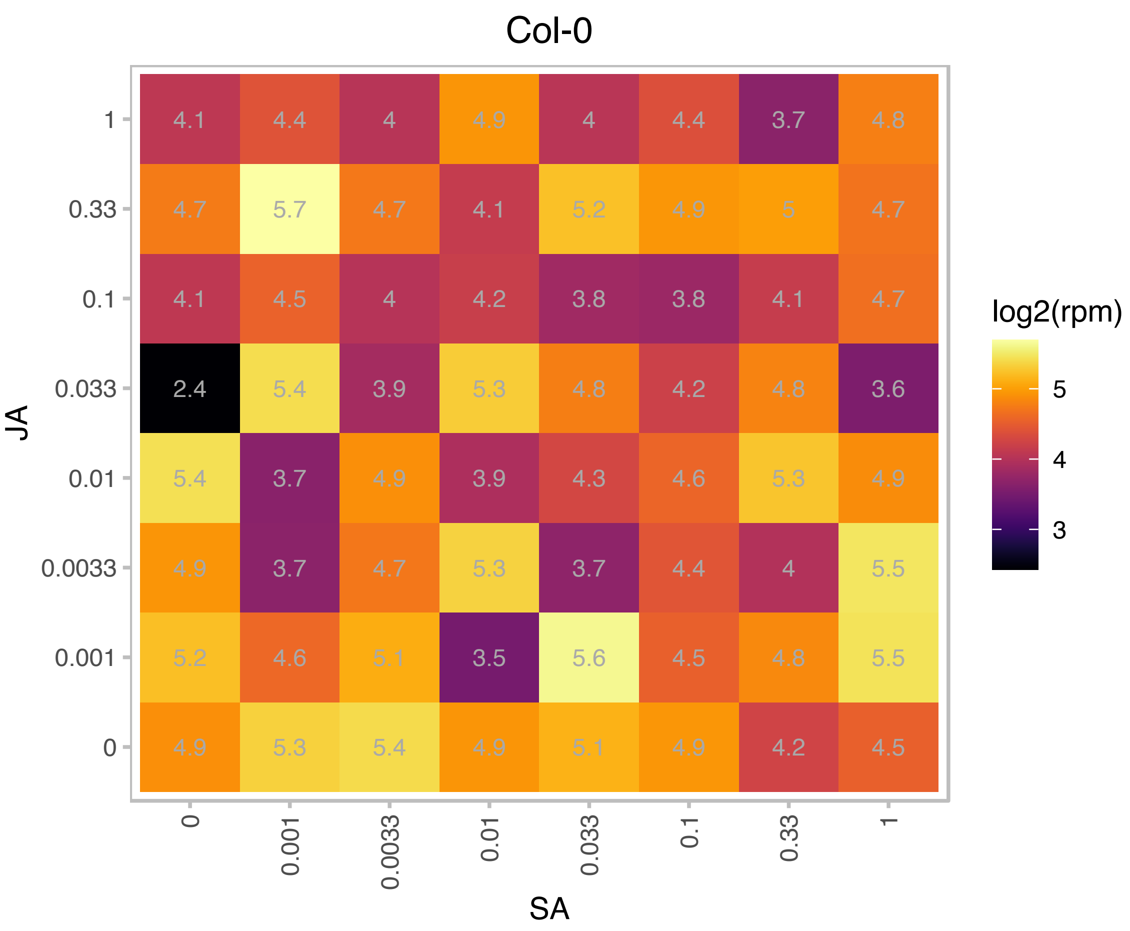


| | |
|---|---|
| Gene | AT2G33250 |
| is.expressed_genes | TRUE |
| is.responsive_genes | TRUE |
| is.apcluster | H-06 |

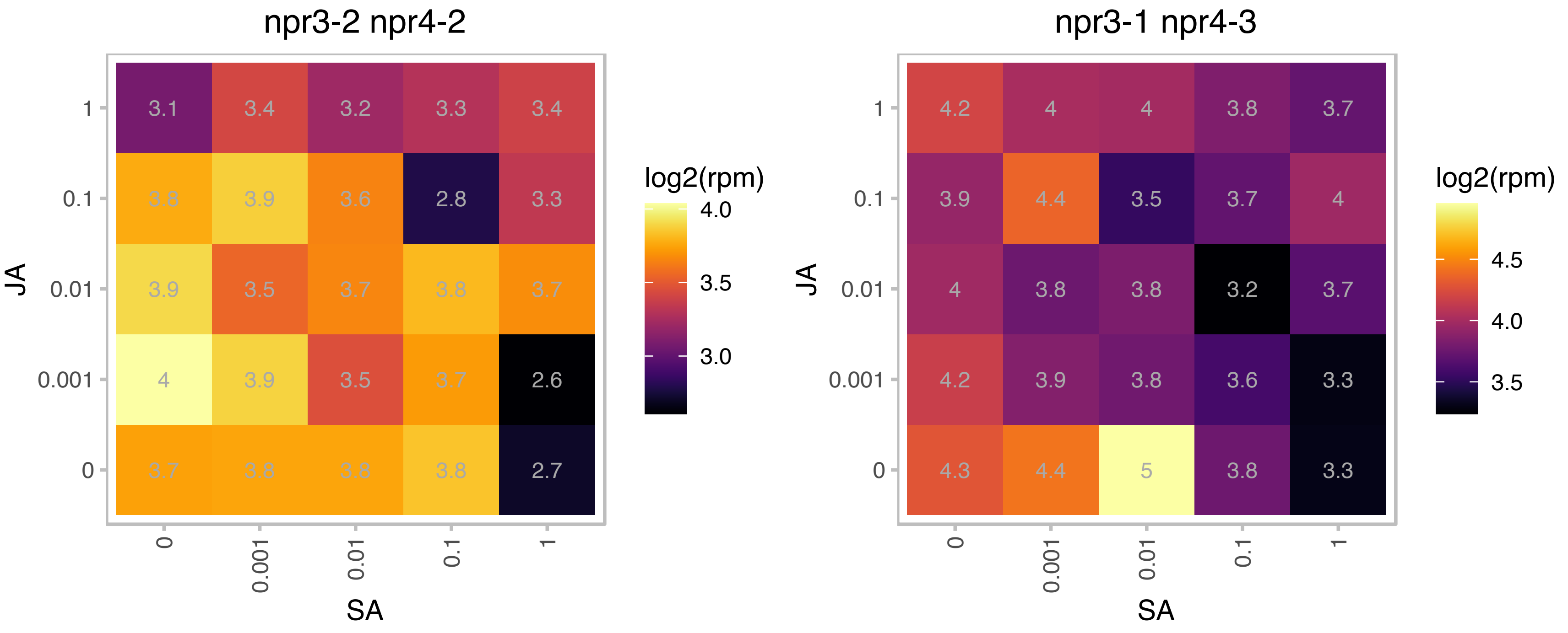

### AT2G41230.1 : ARL2, OSR1

Encodes an ER-localized plant hormone-responsive gene and appears to act redundantly with ARGOS and ARL during organ growth. Over-expression modifies plant sensitivity to ethylene, leading to improved drought tolerance.

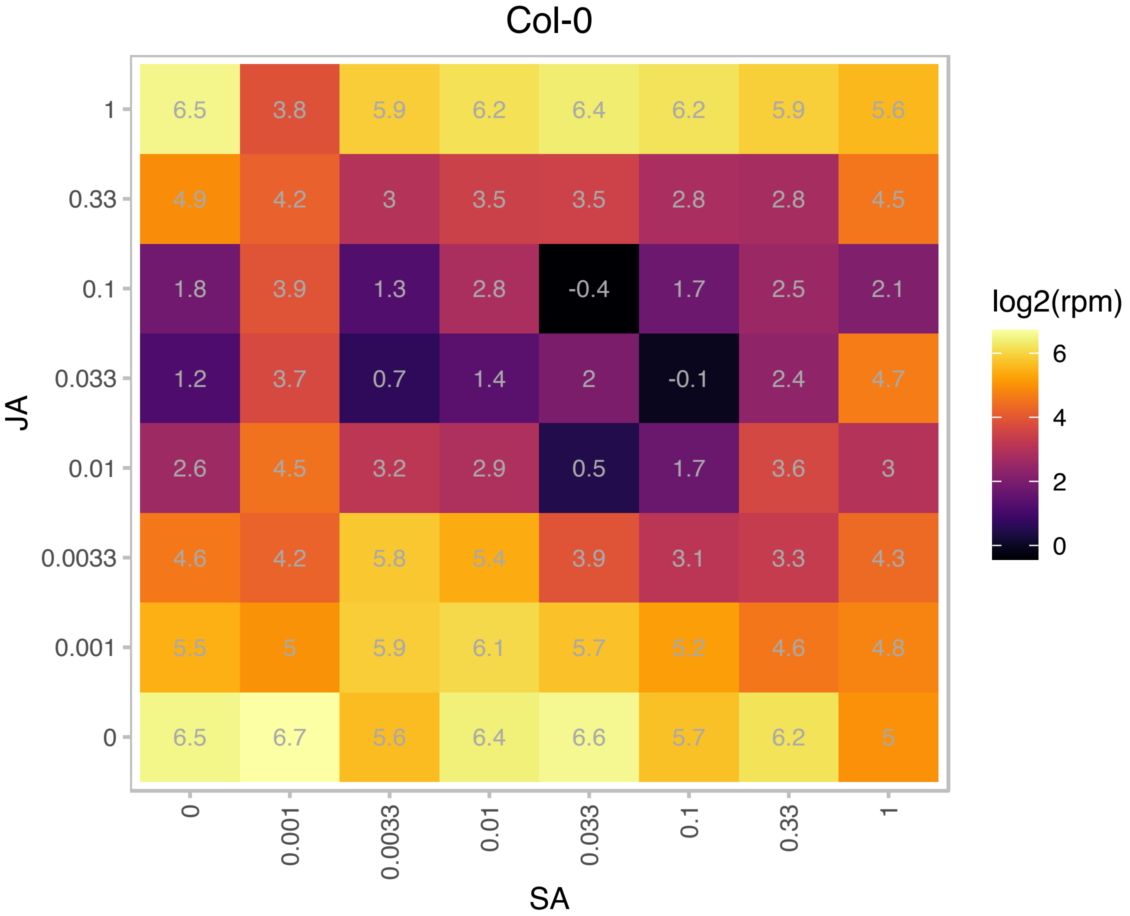


| | |
|---|---|
| Gene | AT2G41230 |
| is.expressed_genes | TRUE |
| is.responsive_genes | TRUE |
| is.apcluster | H-06 |

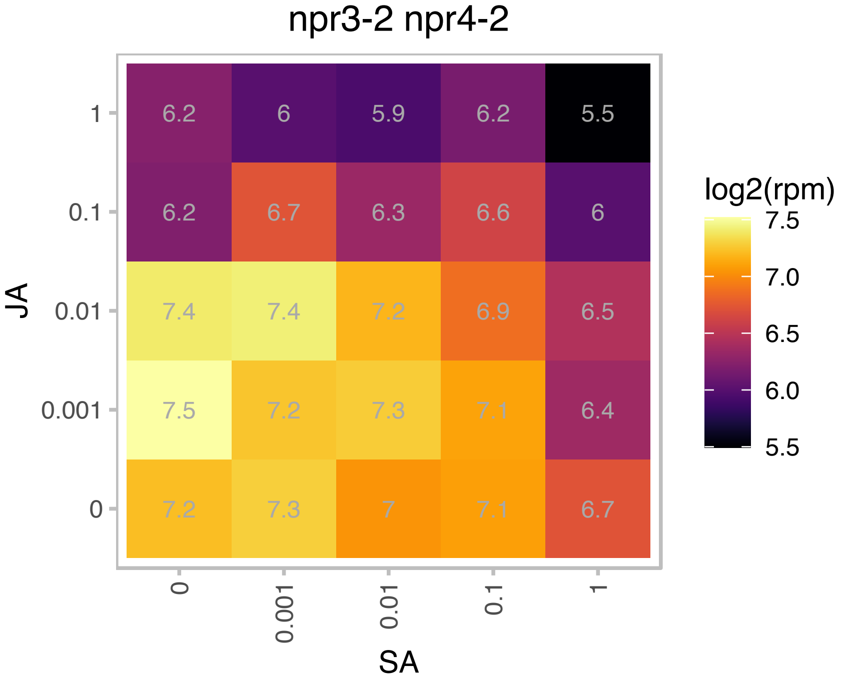


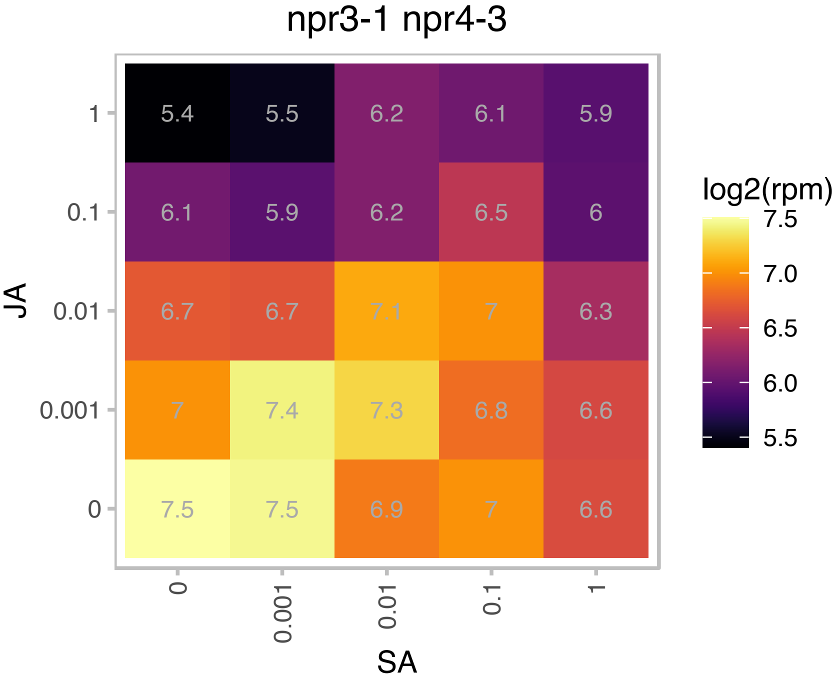

# AT2G42975.1 : NA

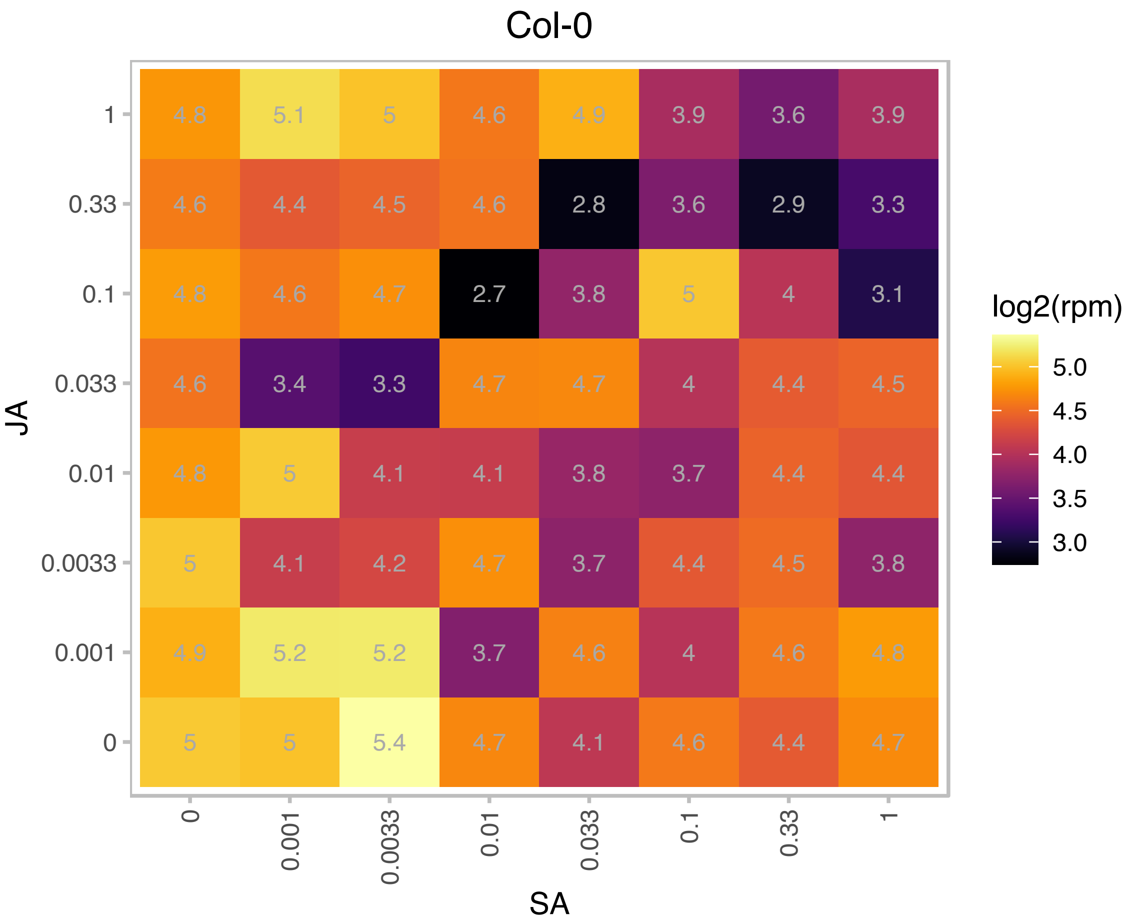


| | |
|---|---|
| Gene | AT2G42975 |
| is.expressed_genes | TRUE |
| is.responsive_genes | TRUE |
| is.apcluster | H-06 |

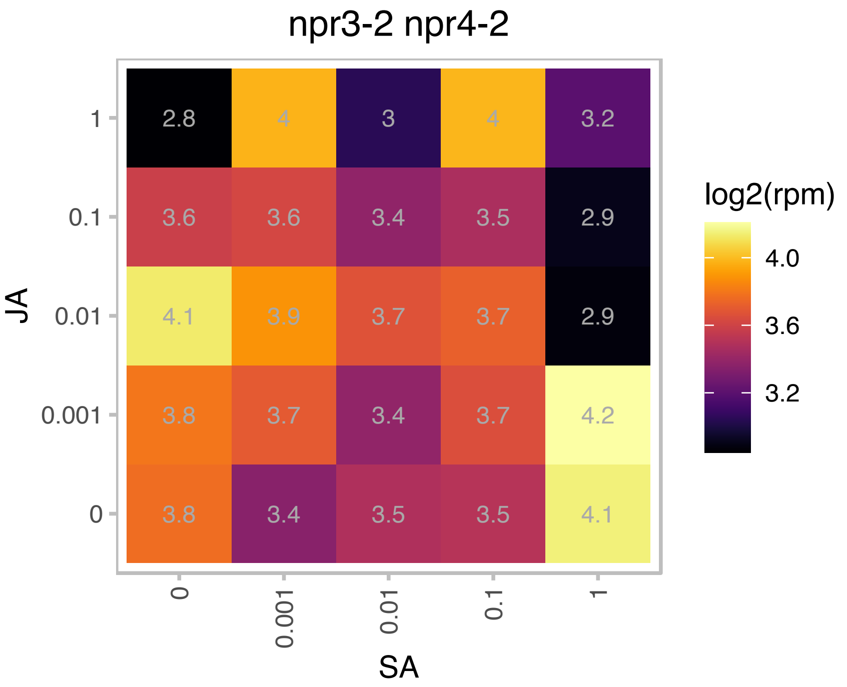


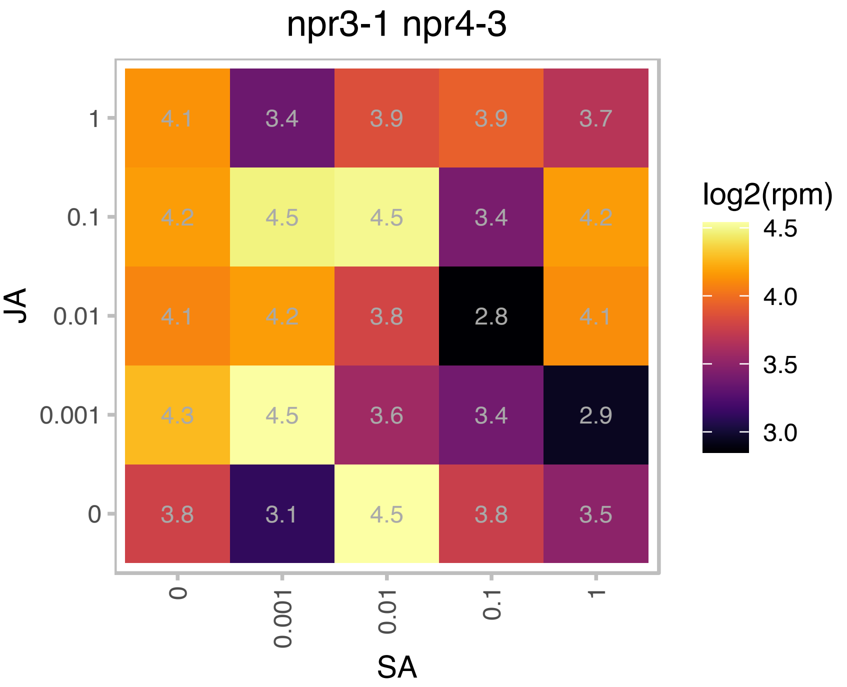

### AT2G44080.1 : ARL

ARGOS-like

Encodes ARL, a gene similar to ARGOS involved in cell expansion-dependent organ growth. Upregulated by brassinosteroid. Acts downstream of BRI1. The mRNA is cell-to-cell mobile.

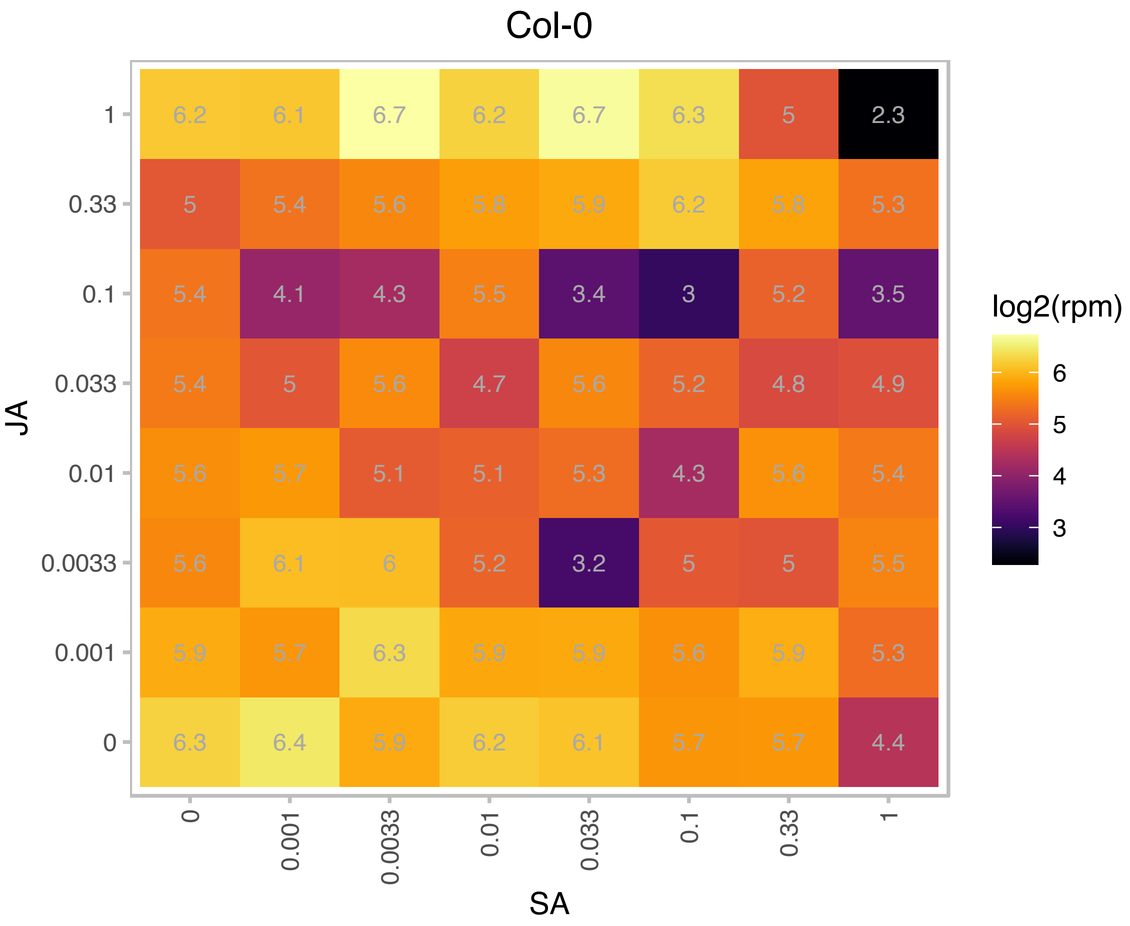


| | |
|---|---|
| Gene | AT2G44080 |
| is.expressed_genes | TRUE |
| is.responsive_genes | TRUE |
| is.apcluster | H-06 |

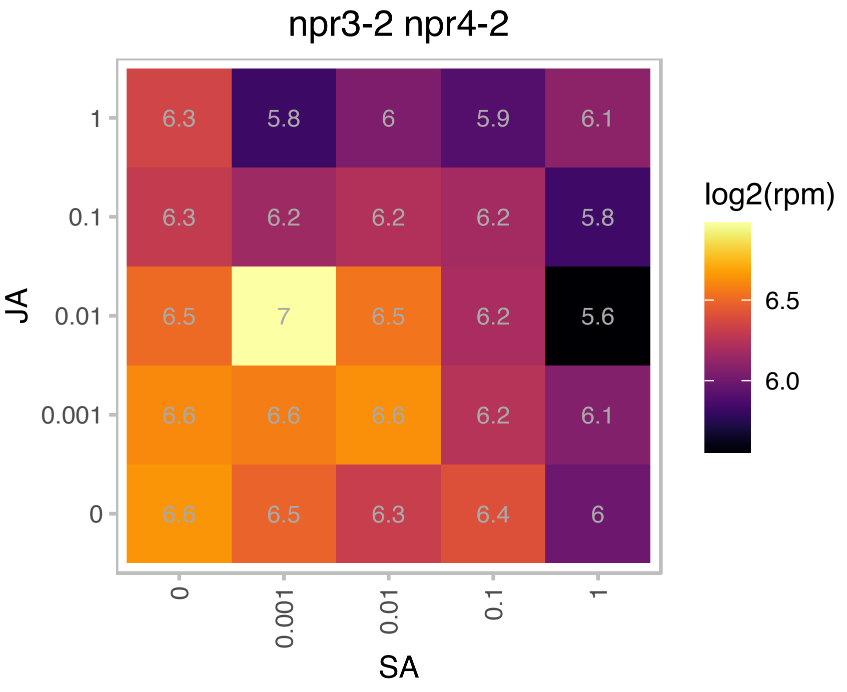


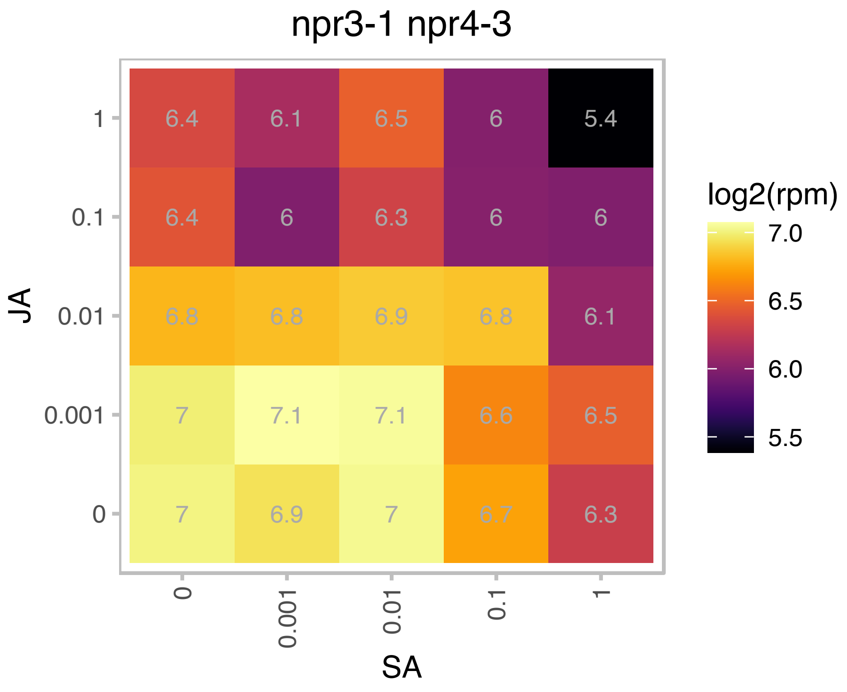

# AT2G46000.1 : NA

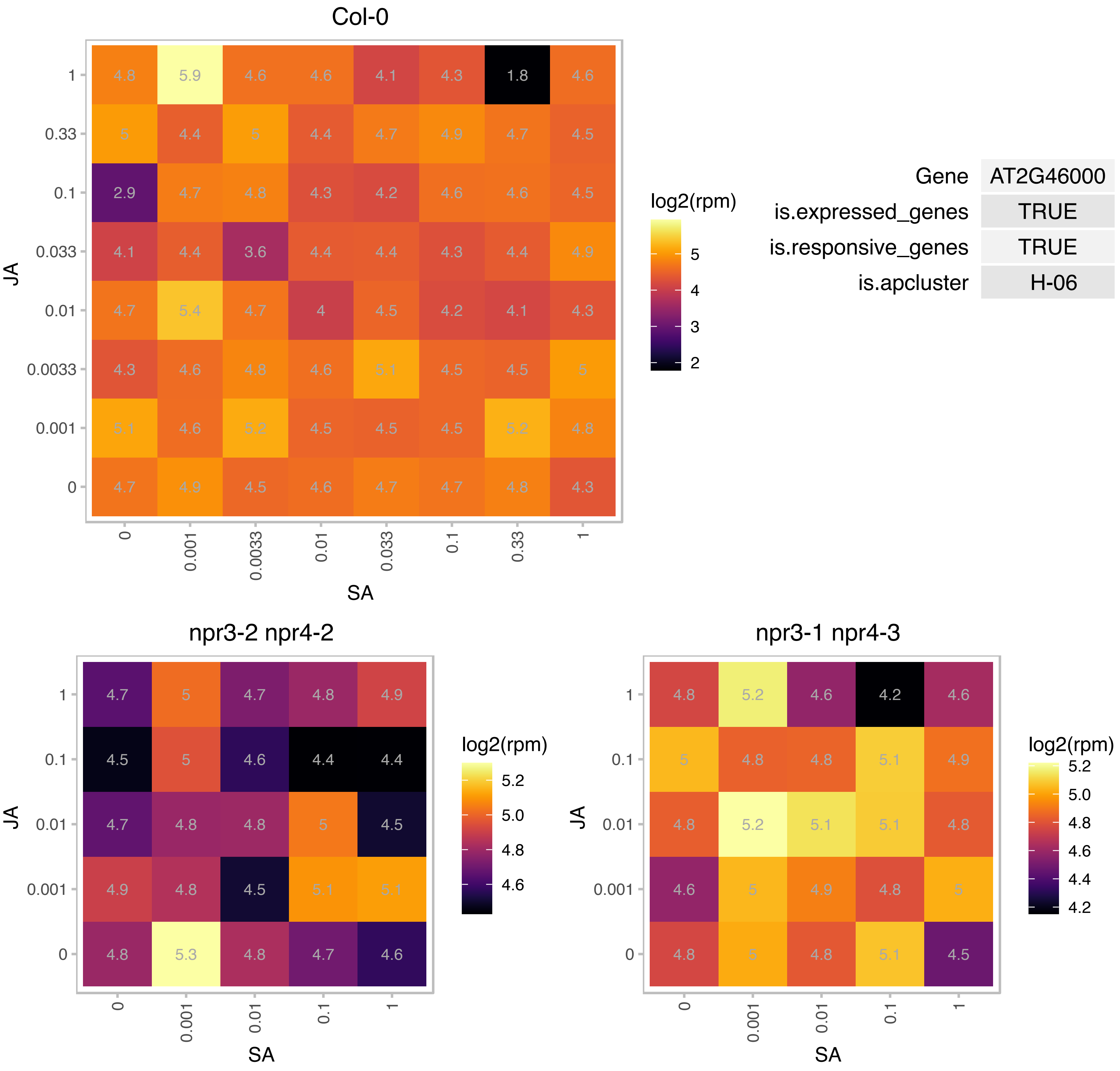

### AT3G06590.1 : AIF2, RITF1

basic helix-loop-helix (bHLH) DNA-binding superfamily protein

Encodes RITF1, a bHLH transcription factor that regulates the transcription of several genes involved in the detoxification of reactive oxygen species generated by salt stress.

Col-0

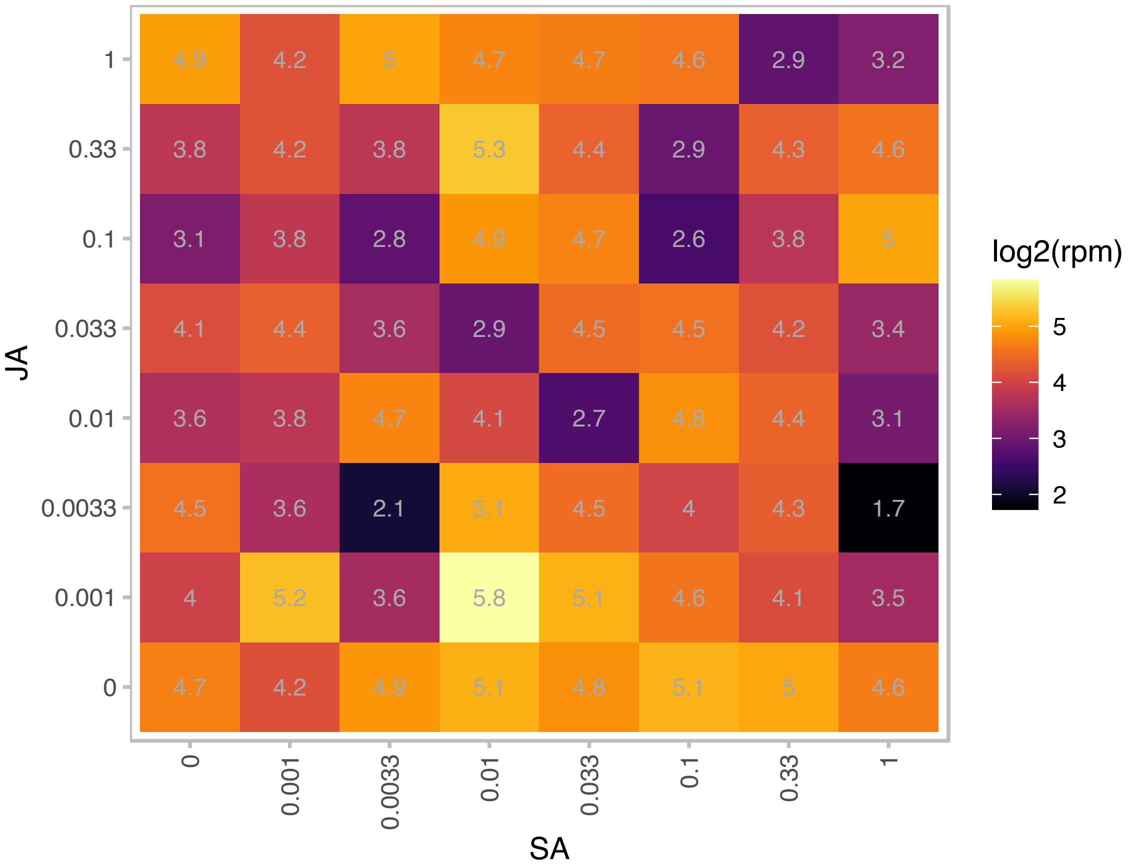


| | |
|---|---|
| Gene | AT3G06590 |
| is.expressed_genes | TRUE |
| is.responsive_genes | TRUE |
| is.apcluster | H-06 |

npr3-2 npr4-2

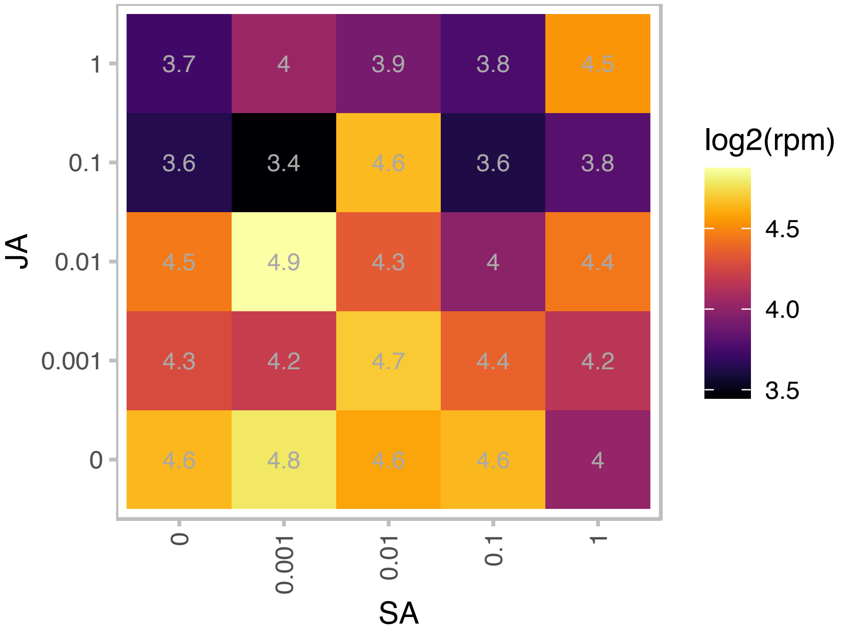


npr3-1 npr4-3

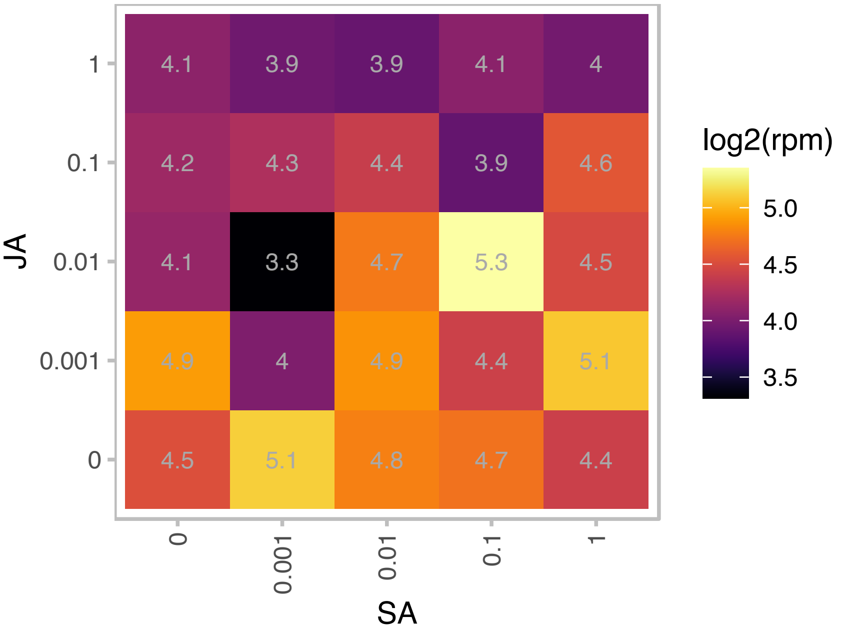

# AT3G10720.1 : NA

Plant invertase/pectin methylesterase inhibitor superfamily

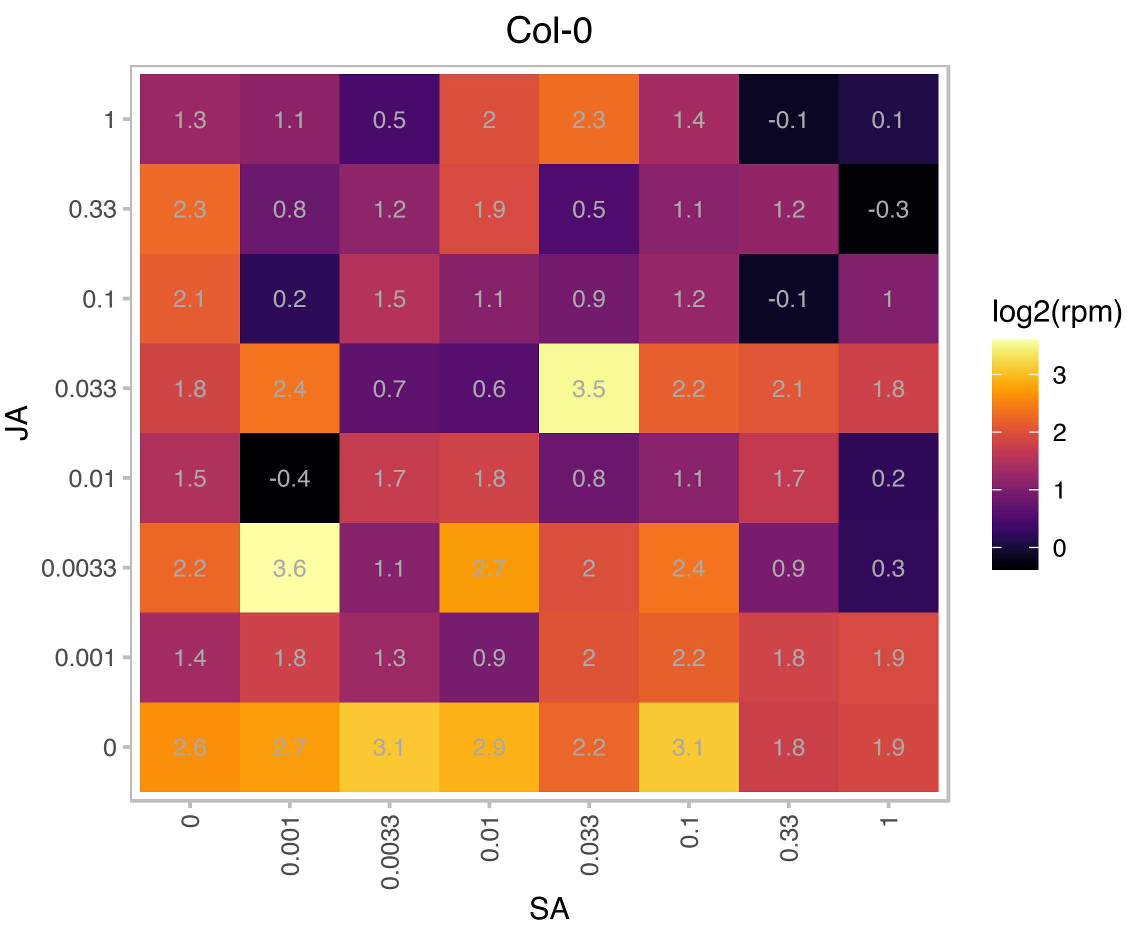


| | |
|---|---|
| Gene | AT3G10720 |
| is.expressed_genes | TRUE |
| is.responsive_genes | TRUE |
| is.apcluster | H-06 |

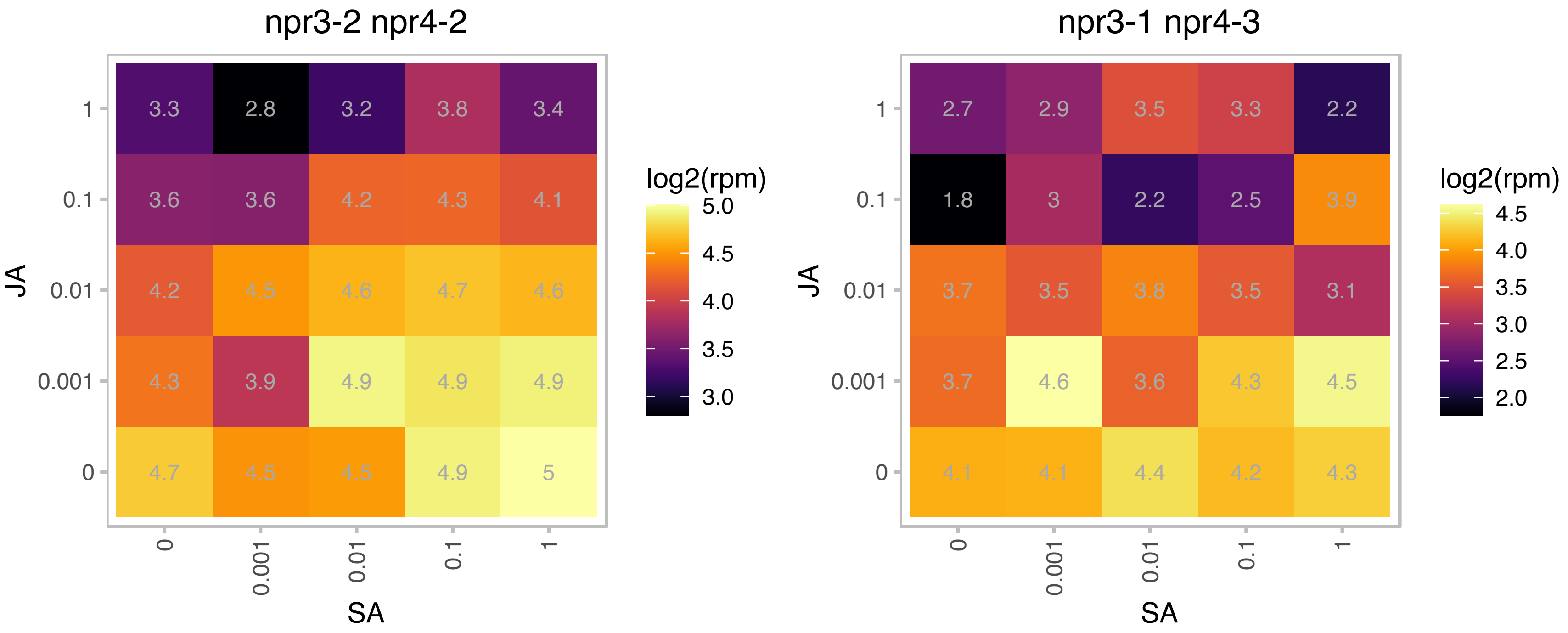

# AT3G16370.1 : NA

GDSL-like Lipase/Acylhydrolase superfamily protein

GDSL-motif esterase/acyltransferase/lipase. Enzyme group with broad substrate specificity that may catalyze acyltransfer or hydrolase reactions with lipid and non-lipid substrates. The mRNA is cell-to-cell mobile.

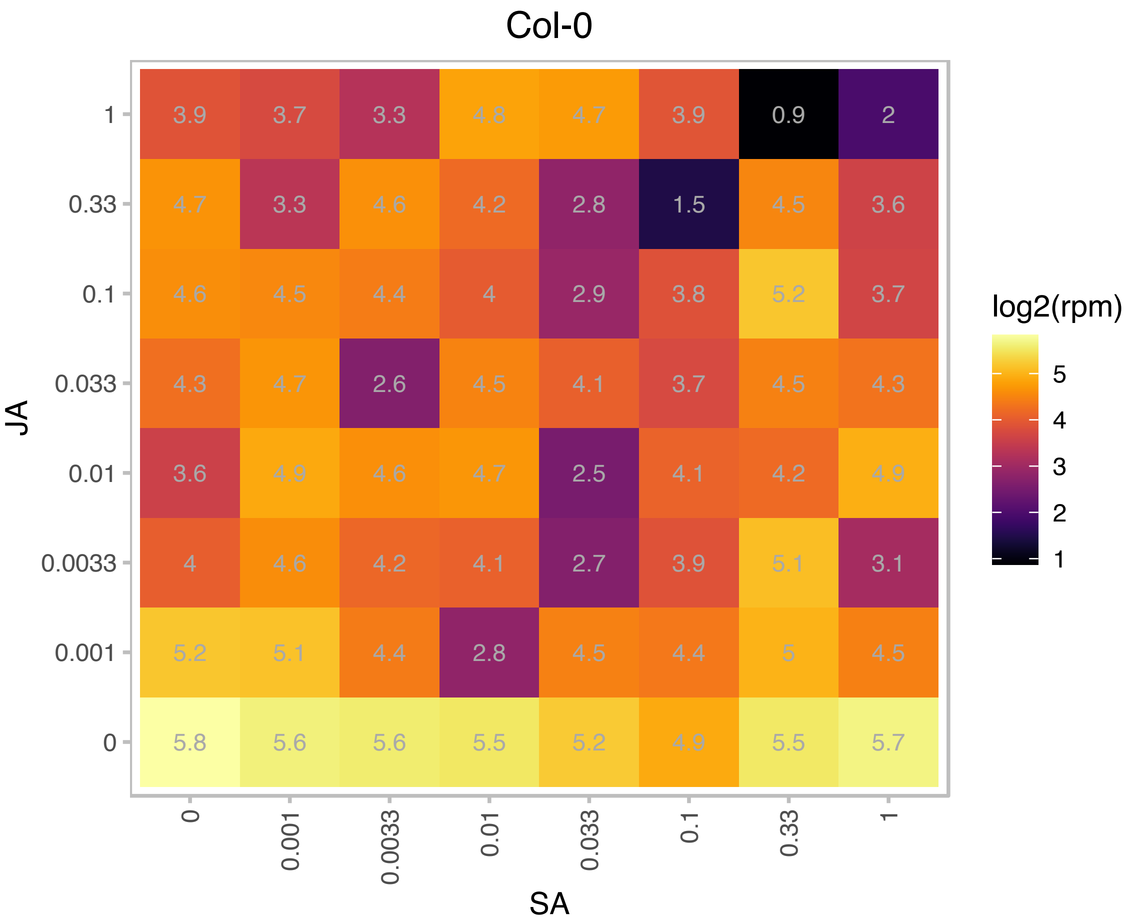


| | |
|---|---|
| Gene | AT3G16370 |
| is.expressed_genes | TRUE |
| is.responsive_genes | TRUE |
| is.apcluster | H-06 |

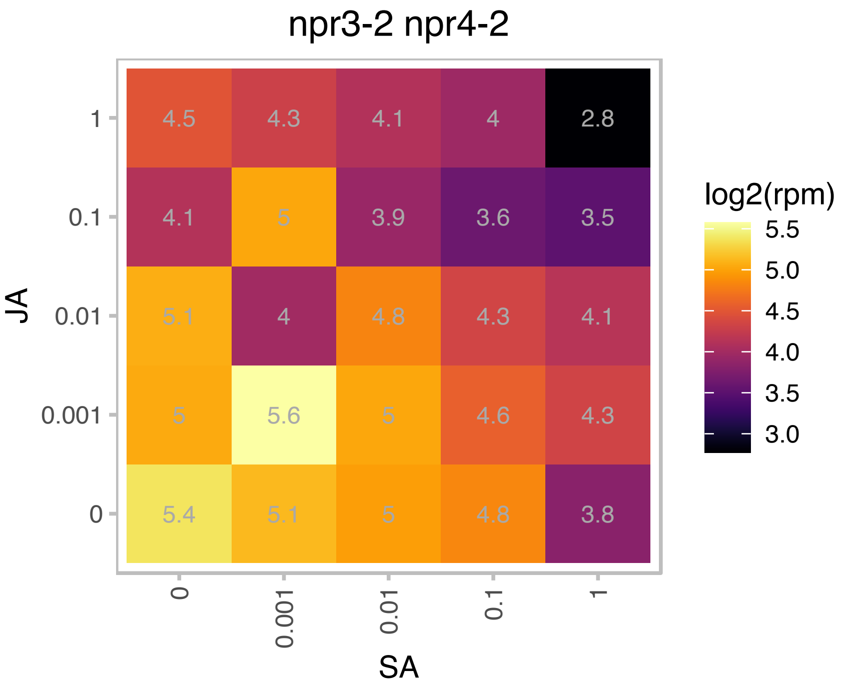


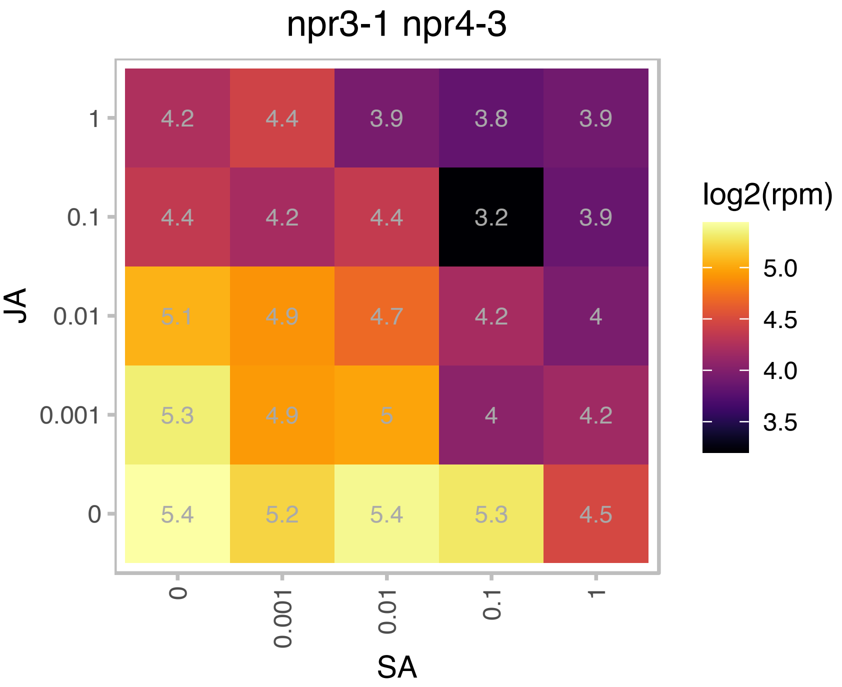

### AT3G24770.1 : CLE41

CLAVATA3/ESR-RELATED 41

Belongs to a large gene family, called CLE for CLAVATA3/ESR-related, encoding small peptides with conserved carboxyl termini. The C-terminal 12 amino acid sequence of CLE41 is identical to that of a dodeca peptide (TDIF, tracheary element differentiation inhibitory factor) isolated from Arabidopsis and functions as a suppressor of plant stem cell differentiation. TDIF sequence is also identical to the C-terminal 12 amino acids of CLE44 (At4g13195). The protein is expressed in the vascular system and is involved in axillary bud formation. The mRNA is cell-to-cell mobile.

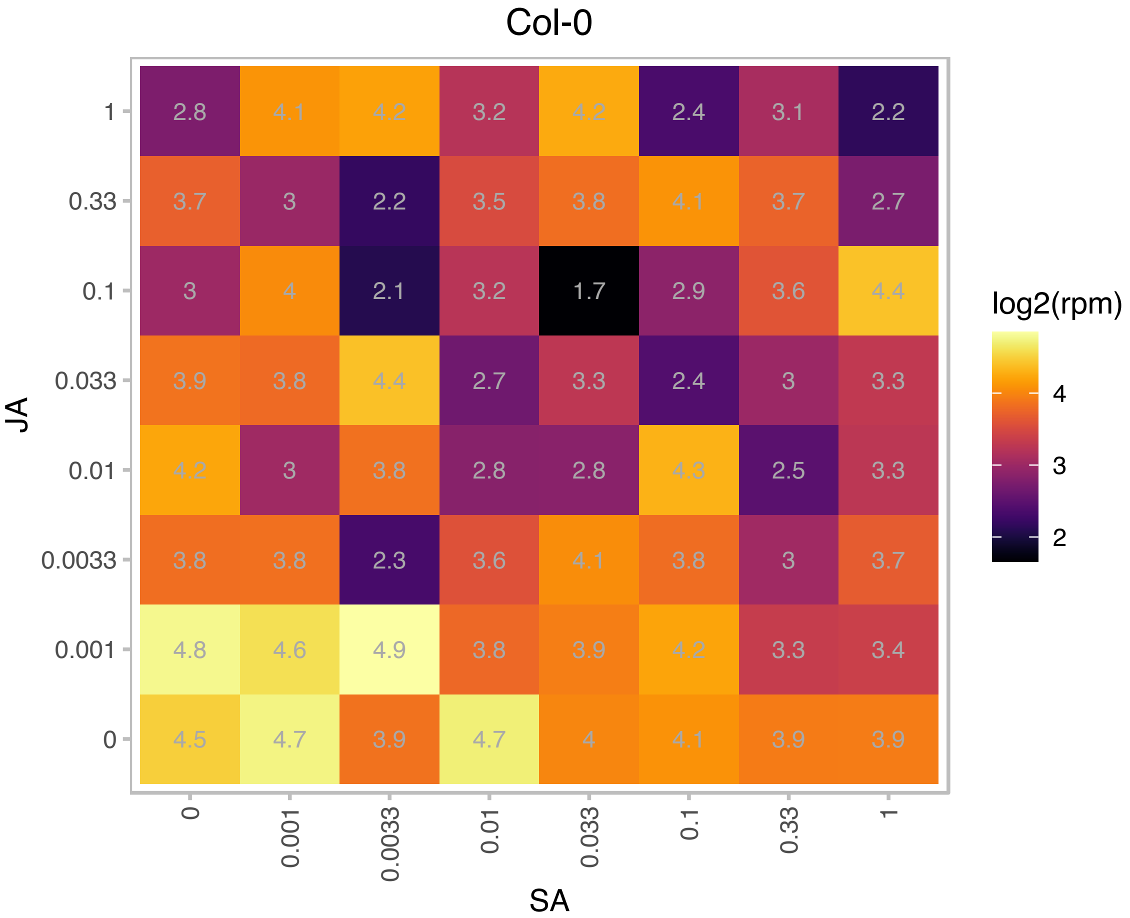


| | |
|---|---|
| Gene | AT3G24770 |
| is.expressed_genes | TRUE |
| is.responsive_genes | TRUE |
| is.apcluster | H-06 |

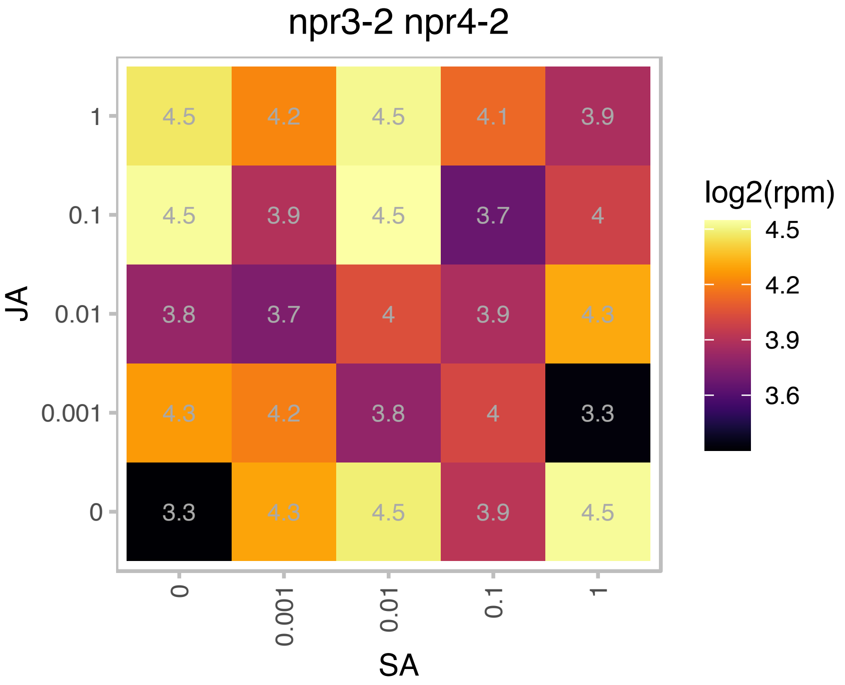


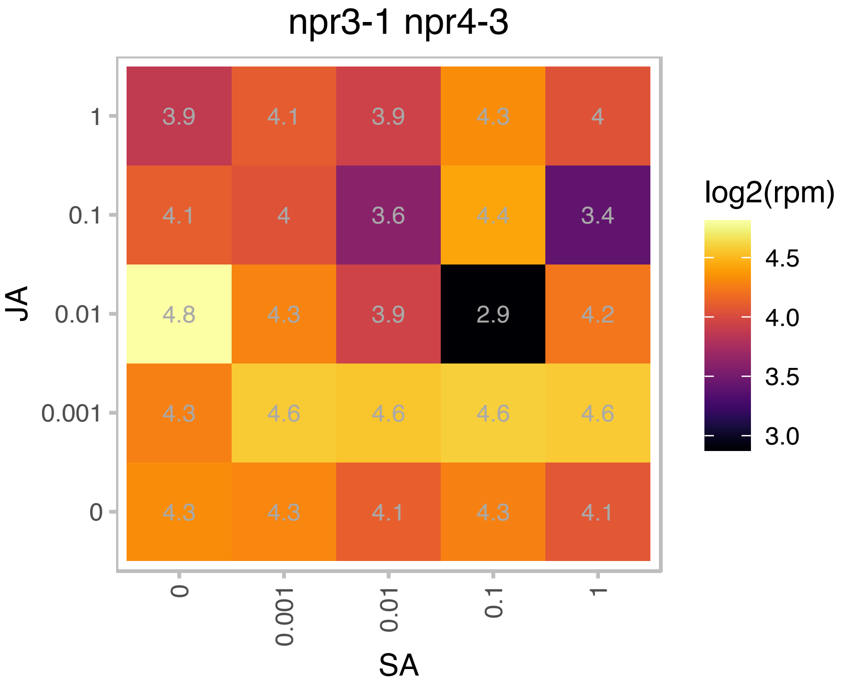

### AT3G44735.1 : ATPSK3, PSK1, PSK3

PHYTOSULFOKINE 3 PRECURSOR

Phytosulfokine 3 precursor, coding for a unique plant peptide growth factor.

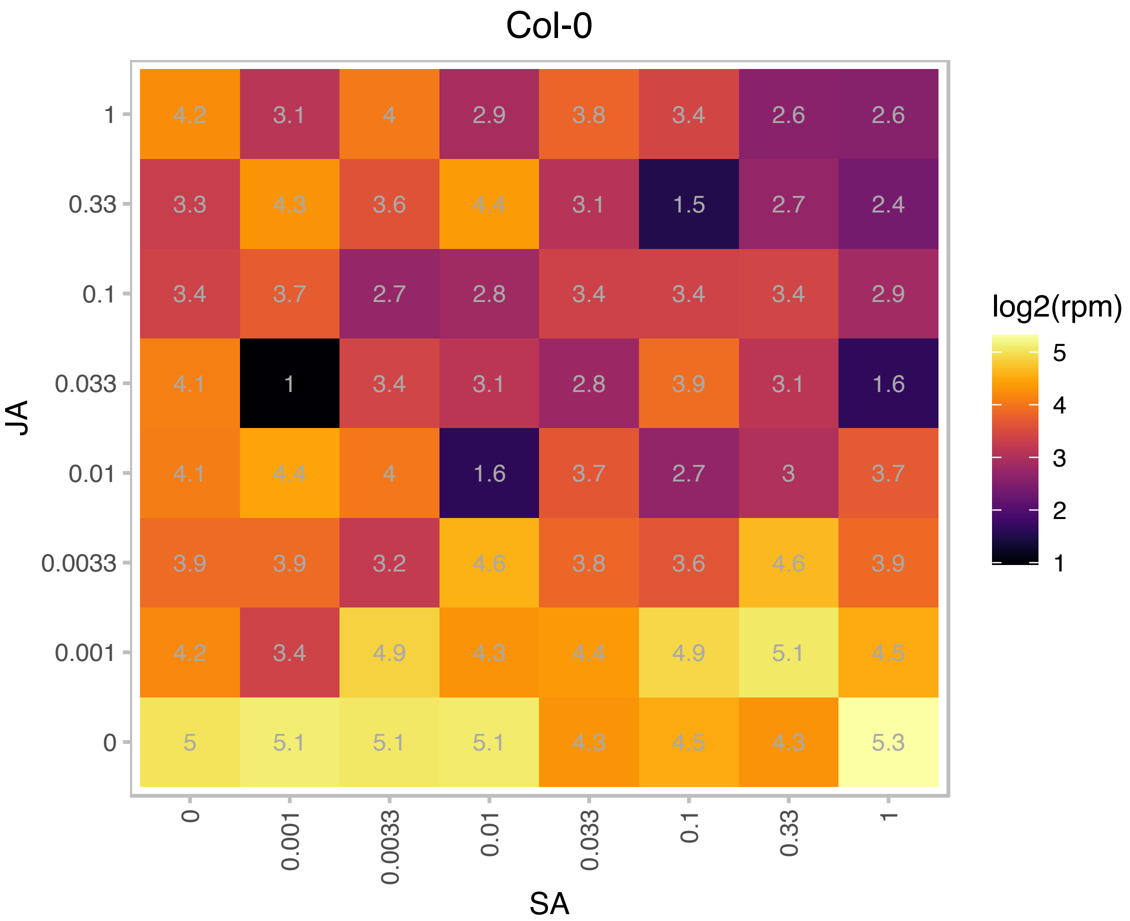


| | |
|---|---|
| Gene | AT3G44735 |
| is.expressed_genes | TRUE |
| is.responsive_genes | TRUE |
| is.apcluster | H-06 |

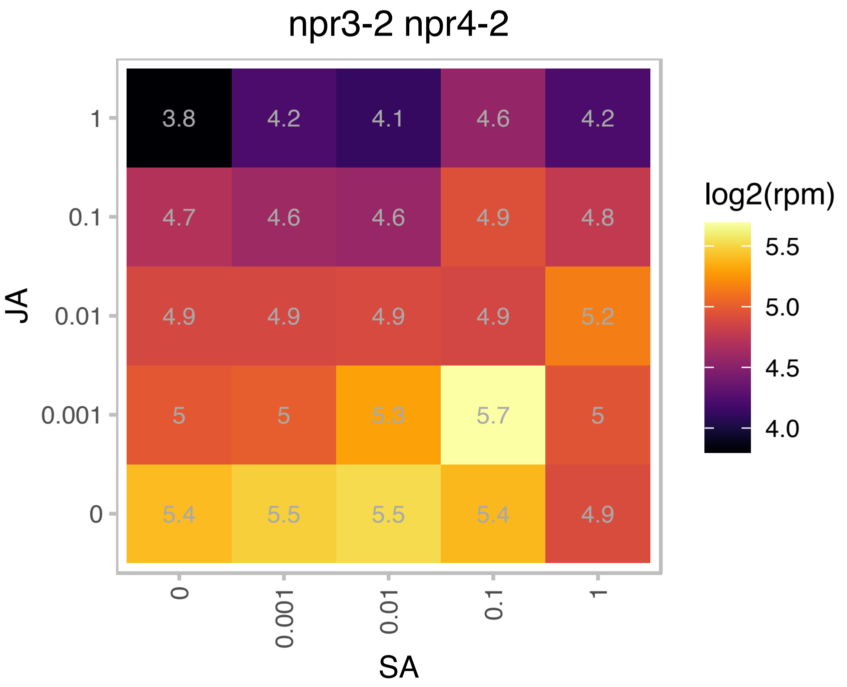


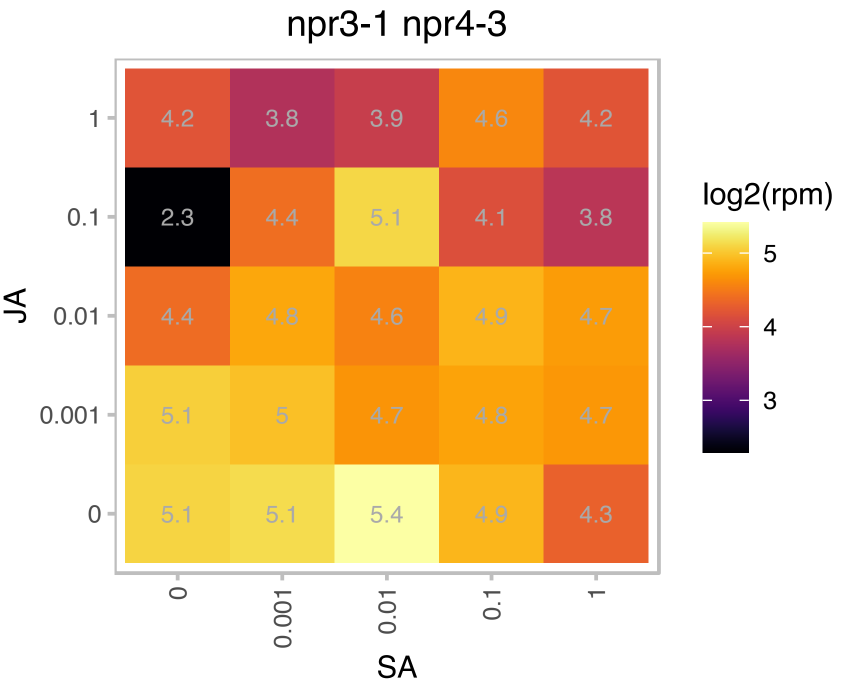

### AT3G51920.1 : ATCML9, CAM9, CML9

calmodulin 9

encodes a divergent member of calmodulin, which is an EF-hand family of Ca2+-binding proteins. This gene is expressed in leaves, flowers and siliques. The gene functionally complements yeast calmodulin 1 (CAM1) but only when selected against the plasmid harboring wild-type yeast sequences. Also the protein does not form formed a complex with a basic amphiphilic helical peptide in the presence of Ca2+ in vitro. Authors suggest that this gene may represent a Ca2+-binding sensor protein that interacts with a more limited set of target proteins than do more conventional CaM isoforms. Mutations in this gene alter plant responses to abiotic [illegible]

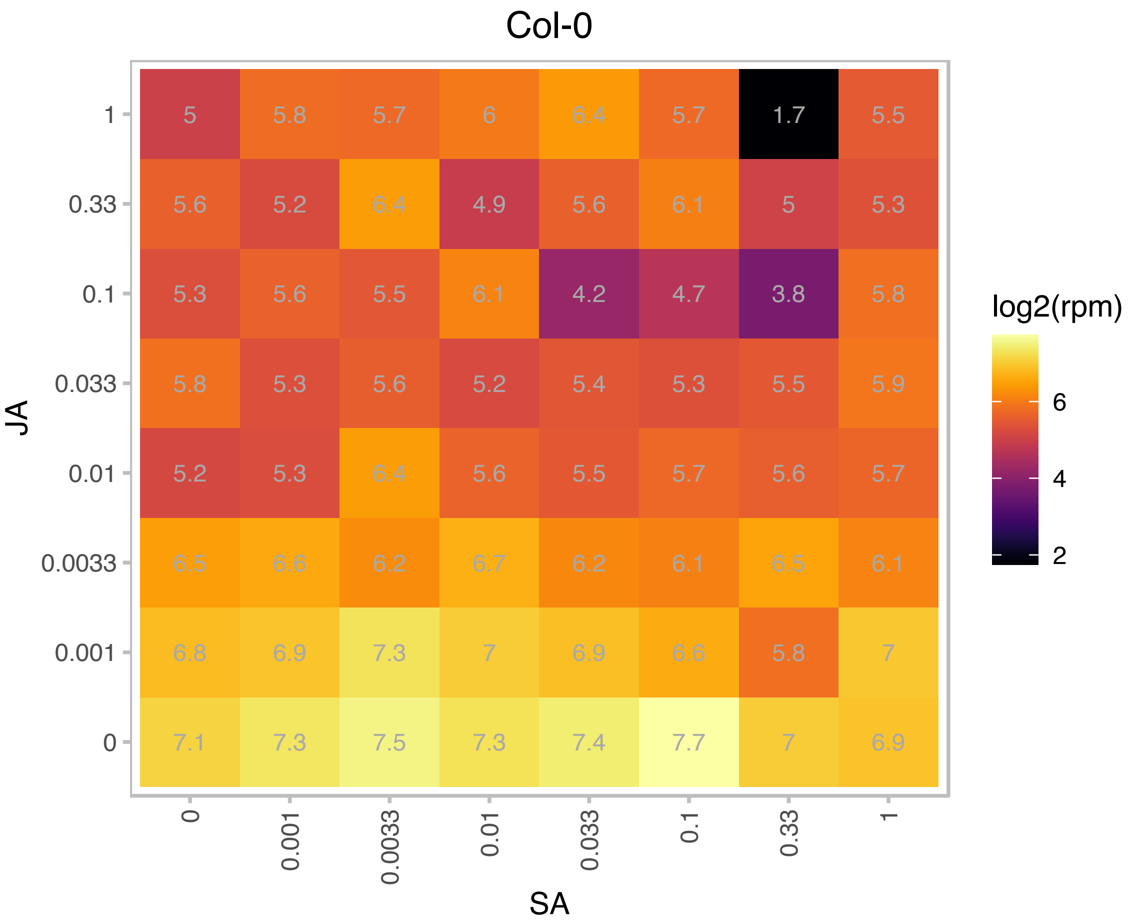


| | |
|---|---|
| Gene | AT3G51920 |
| is.expressed_genes | TRUE |
| is.responsive_genes | TRUE |
| is.apcluster | H-06 |

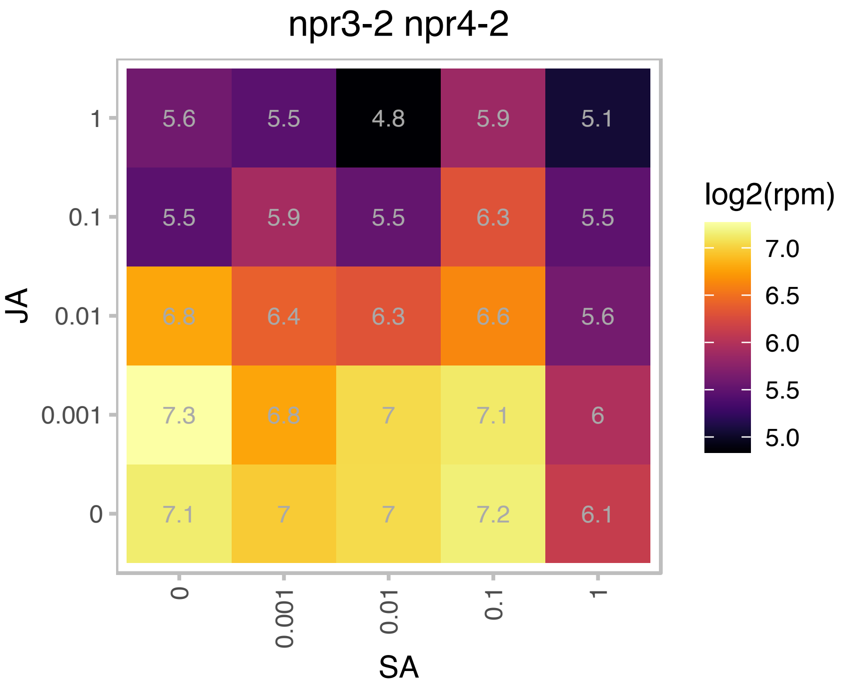


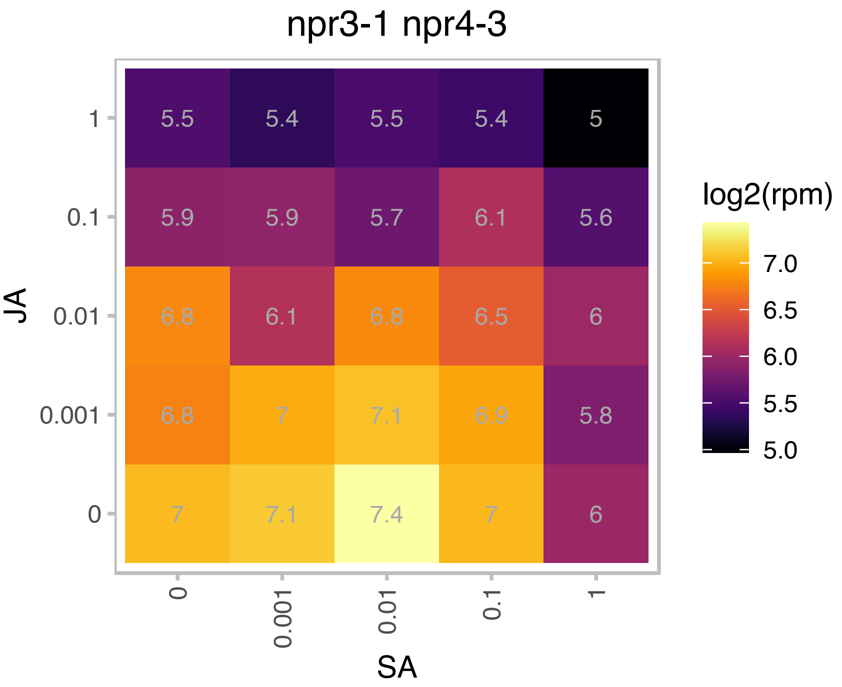

# AT3G56290.1 : NA

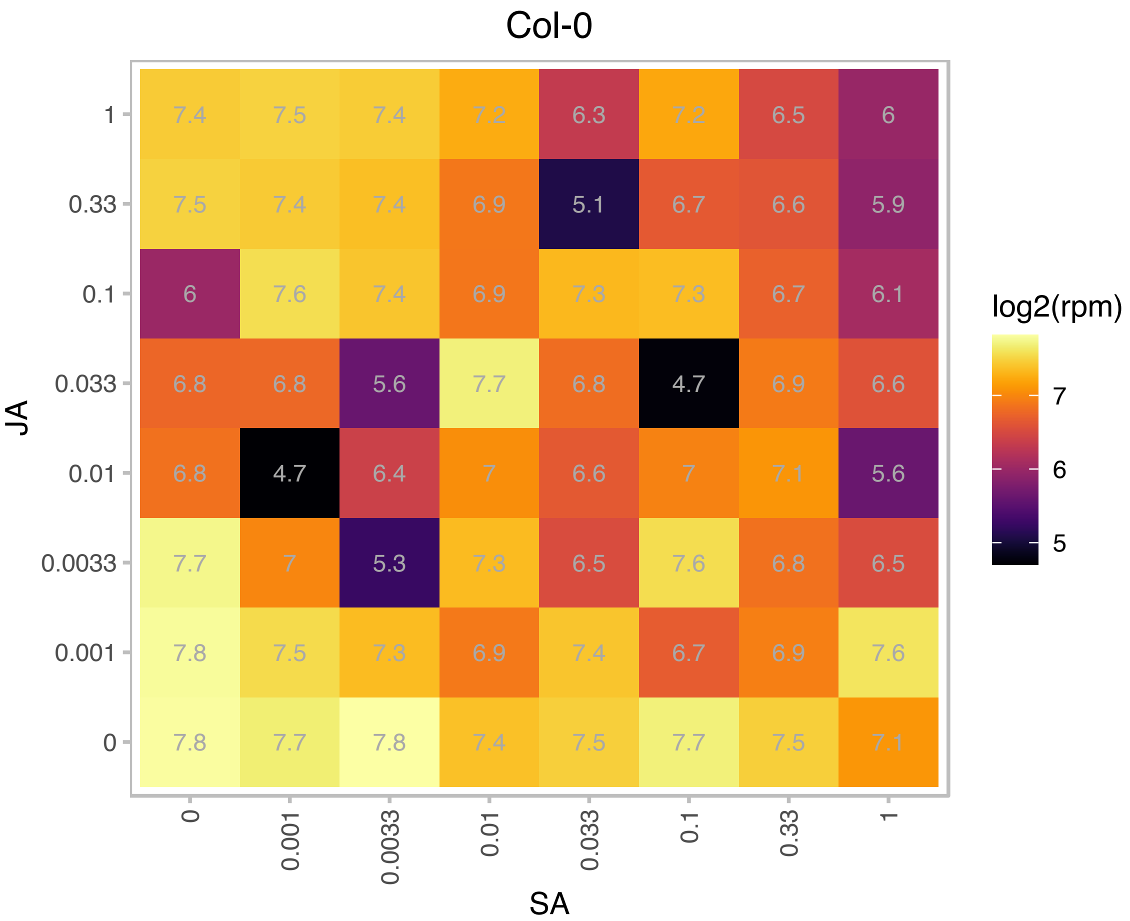


| | |
|---|---|
| Gene | AT3G56290 |
| is.expressed_genes | TRUE |
| is.responsive_genes | TRUE |
| is.apcluster | H-06 |

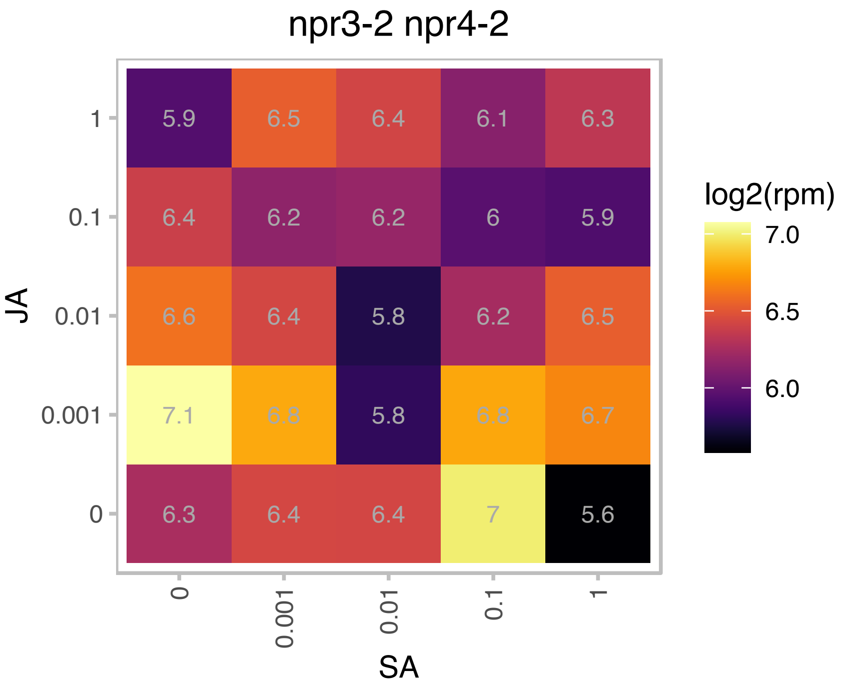


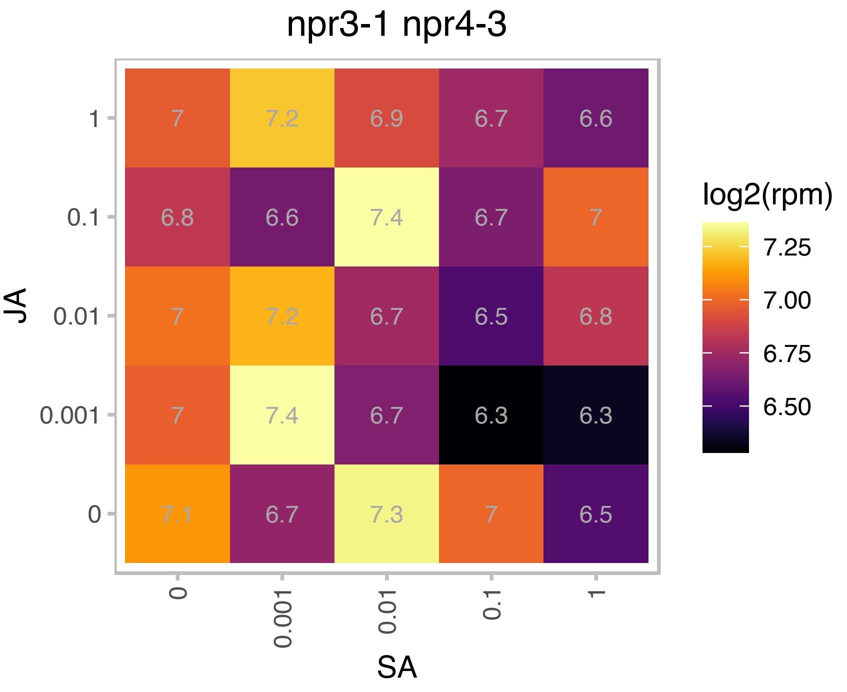

# AT3G56880.1 : NA

VQ motif-containing protein

Col-0

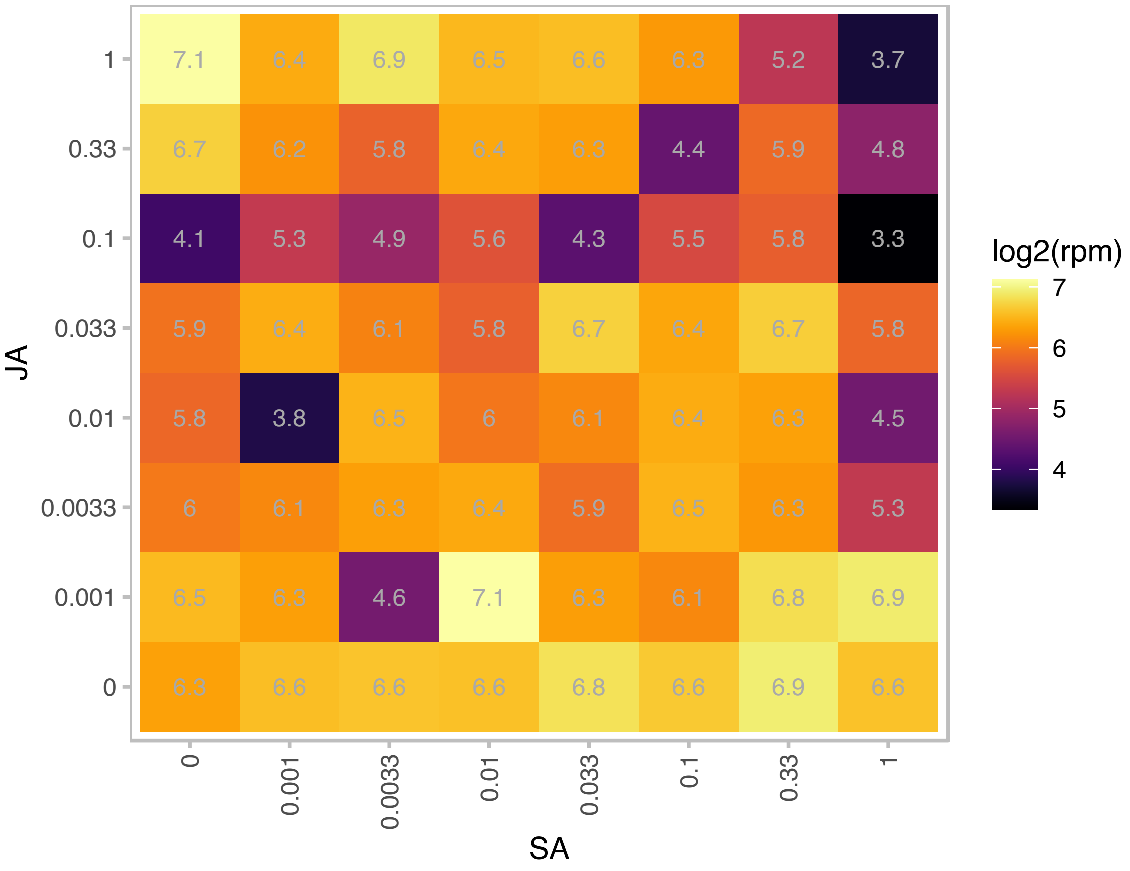


| | |
|---|---|
| Gene | AT3G56880 |
| is.expressed_genes | TRUE |
| is.responsive_genes | TRUE |
| is.apcluster | H-06 |

npr3-2 npr4-2

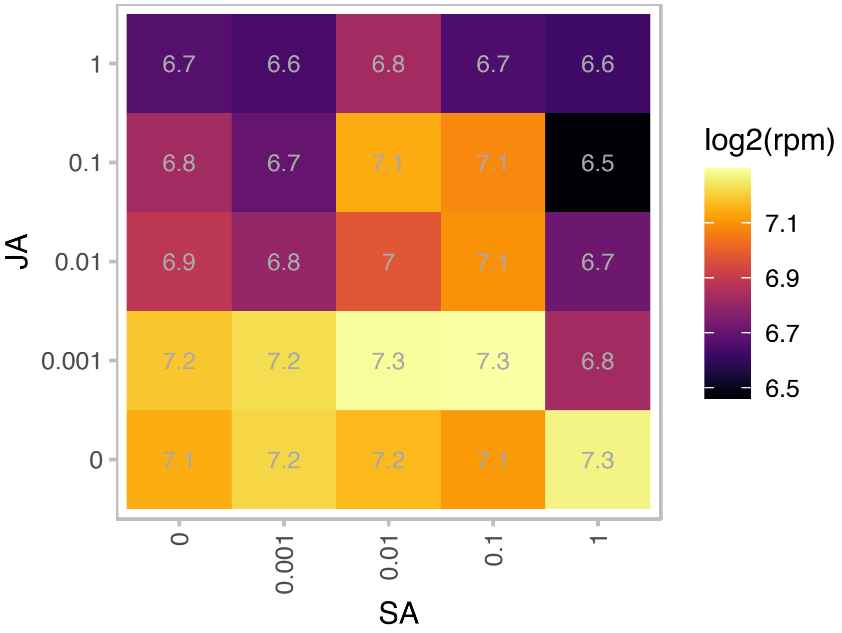


npr3-1 npr4-3

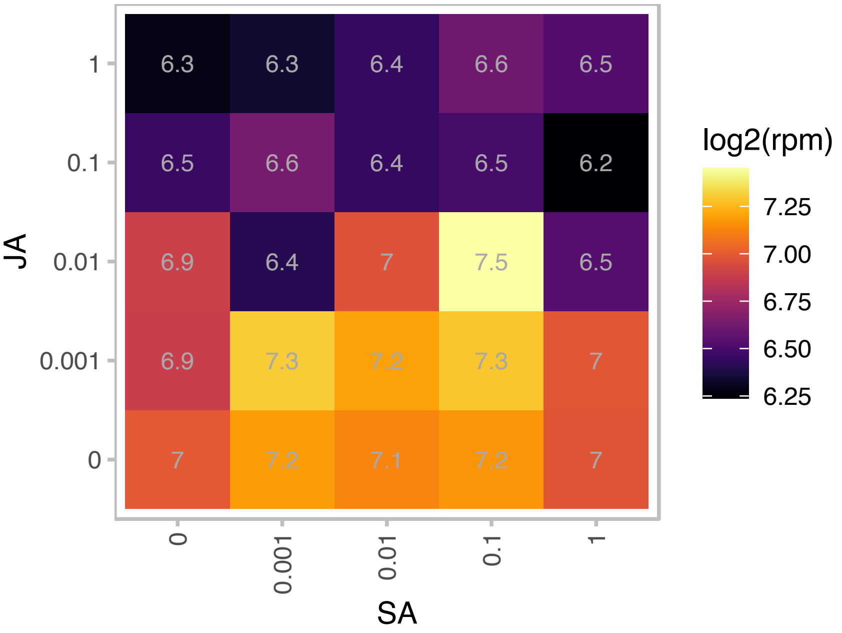

### AT4G14440.1 : ATECI3, ECI3, HCD1

3-hydroxyacyl-CoA dehydratase 1

encodes a cytosolic delta3, delta2-enoyl CoA isomerase, involved in unsaturated fatty acid degradation

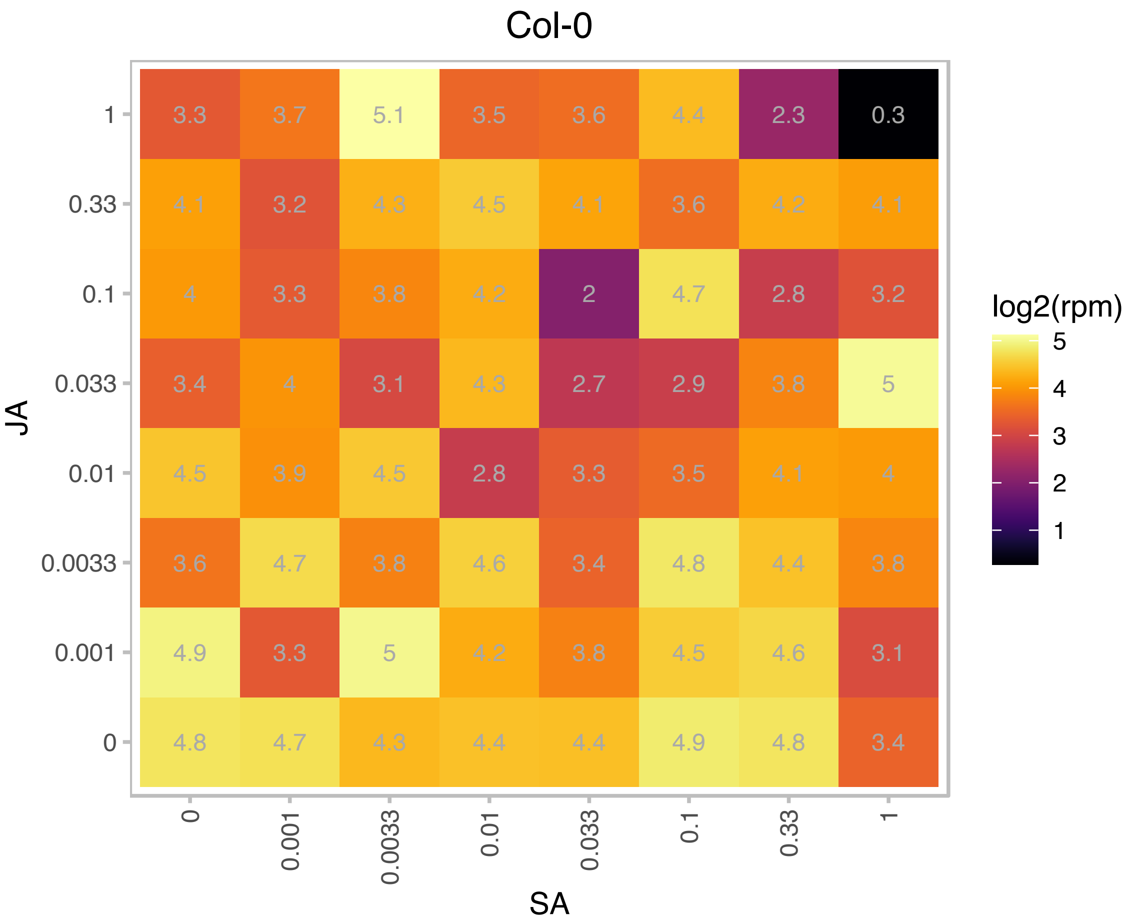


| | |
|---|---|
| Gene | AT4G14440 |
| is.expressed_genes | TRUE |
| is.responsive_genes | TRUE |
| is.apcluster | H-06 |

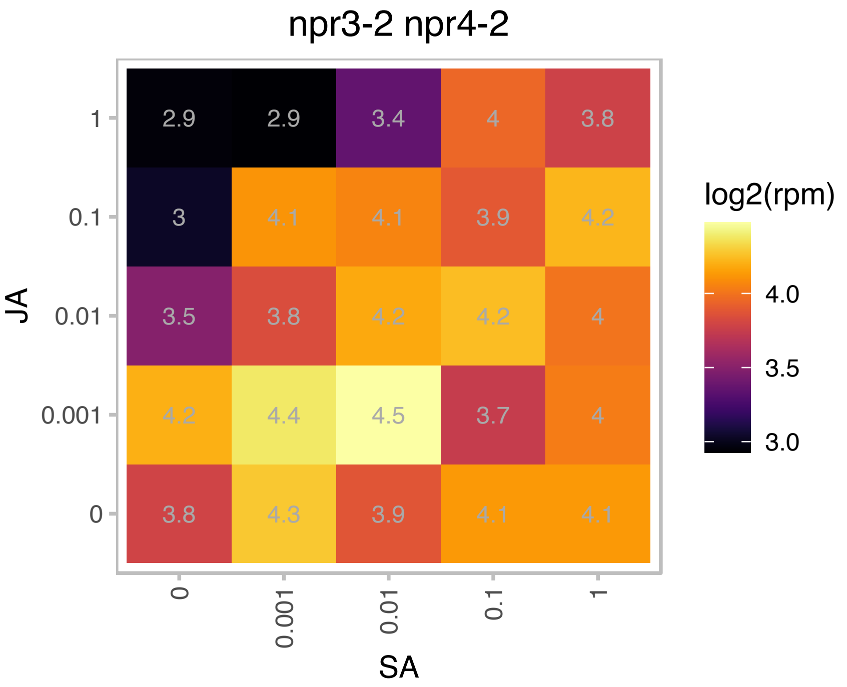


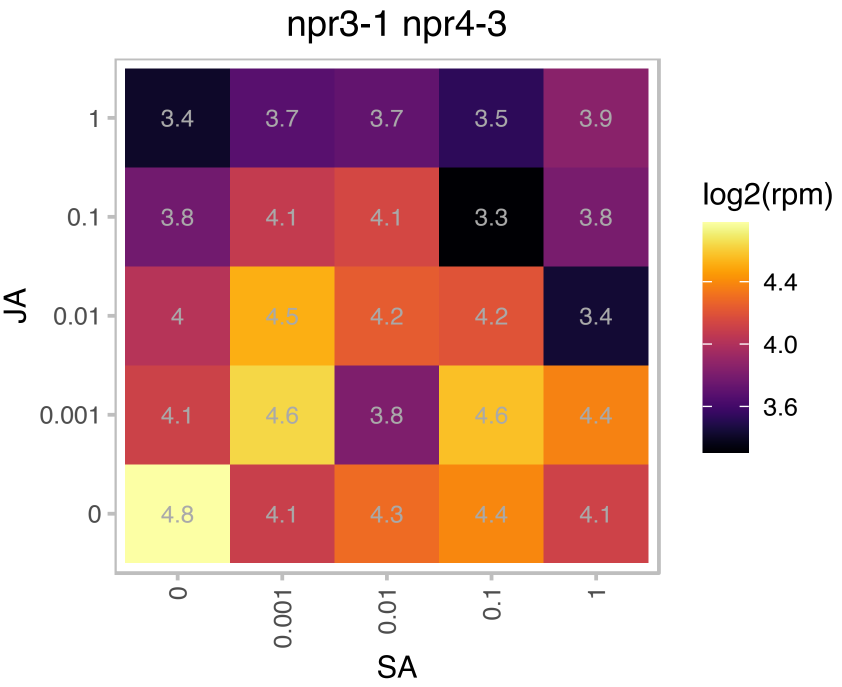

# AT4G24730.1 : NA

Calcineurin-like metallo-phosphoesterase superfamily protein

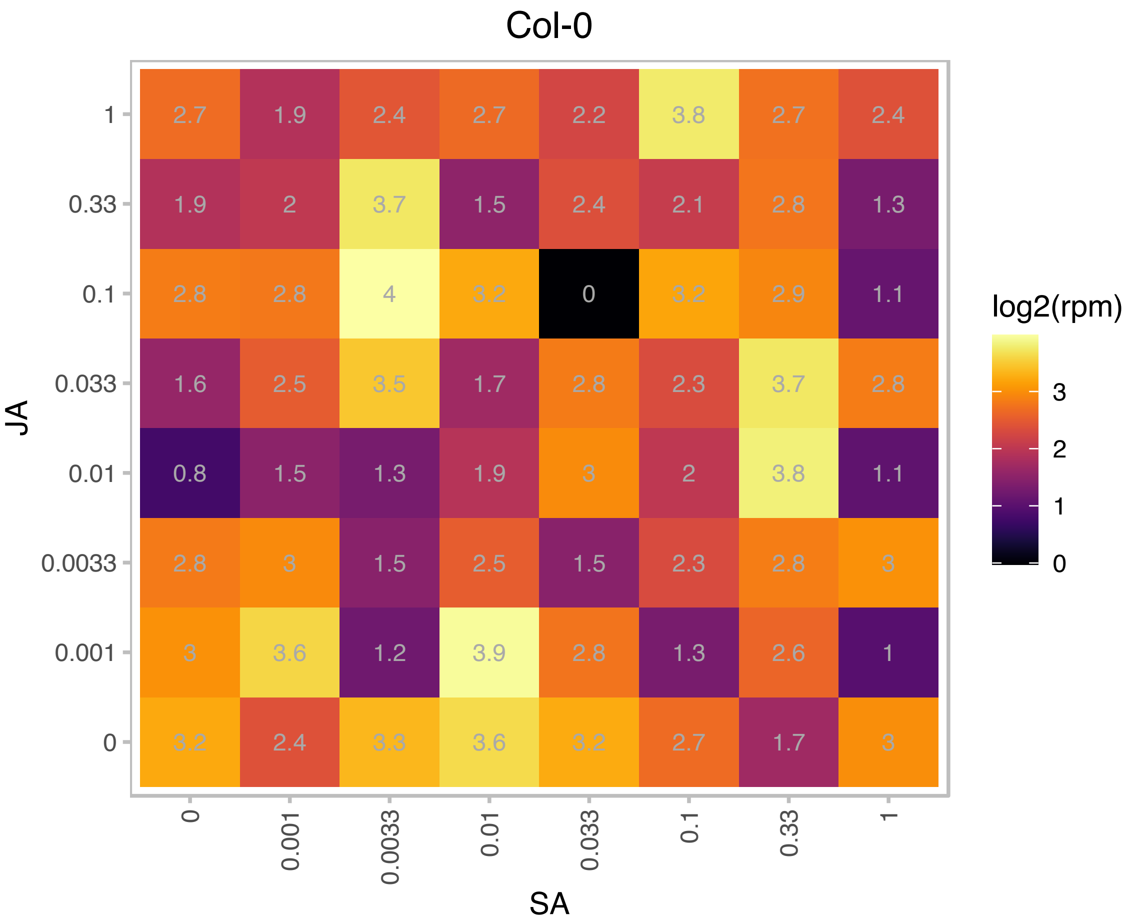


| | |
|---|---|
| Gene | AT4G24730 |
| is.expressed_genes | TRUE |
| is.responsive_genes | TRUE |
| is.apcluster | H-06 |

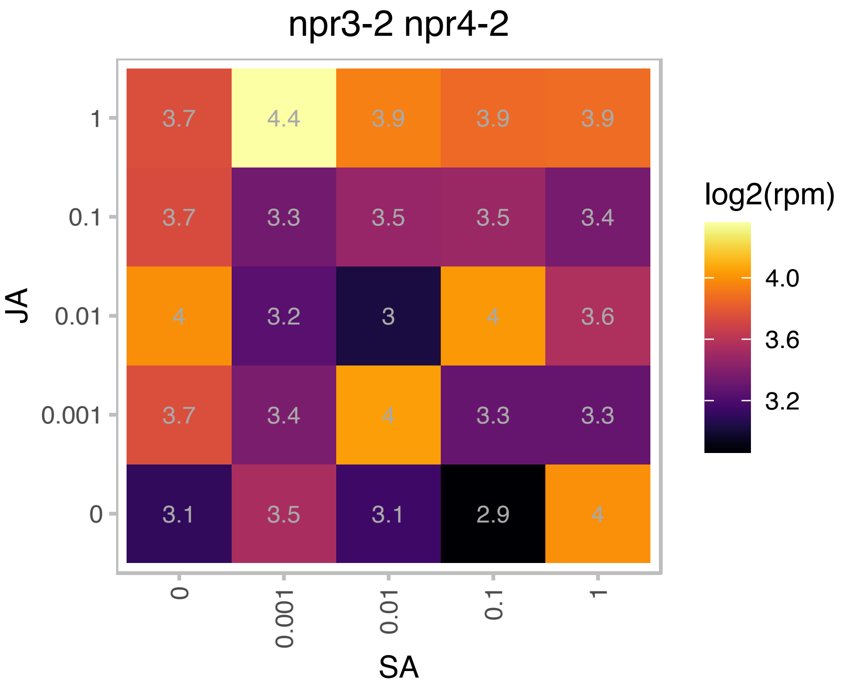


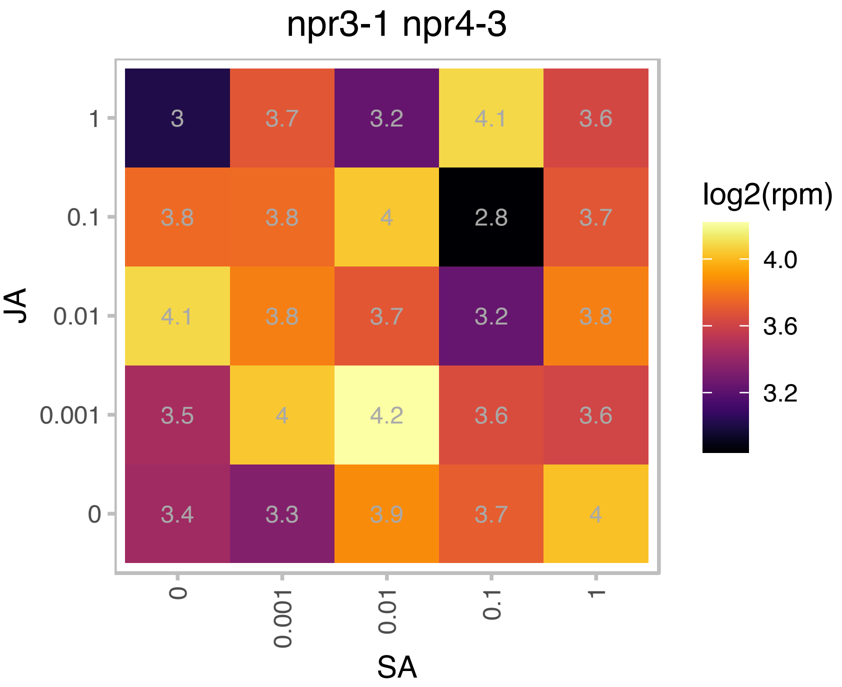

### AT4G34190.1 : SEP1

stress enhanced protein 1

Encodes a stress enhanced protein that localizes to the thylakoid membrane and whose mRNA is upregulated in response to high light intensity.  It may be involved in chlorophyll binding.

Col-0

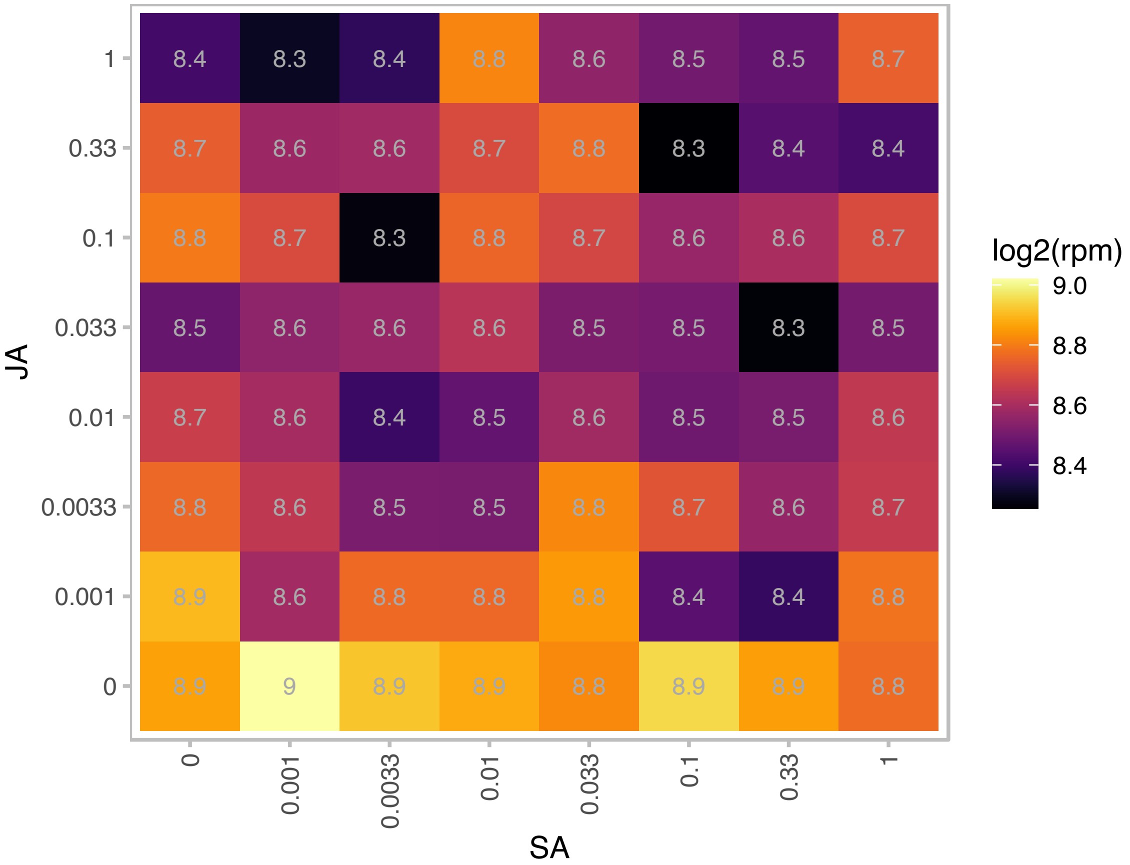


| | |
|---|---|
| Gene | AT4G34190 |
| is.expressed_genes | TRUE |
| is.responsive_genes | TRUE |
| is.apcluster | H-06 |

npr3-2 npr4-2

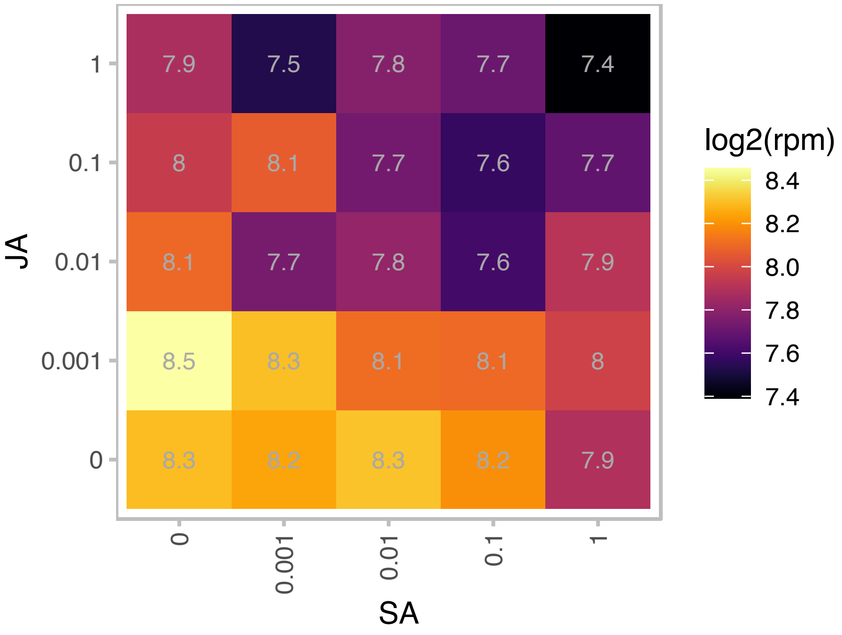


npr3-1 npr4-3

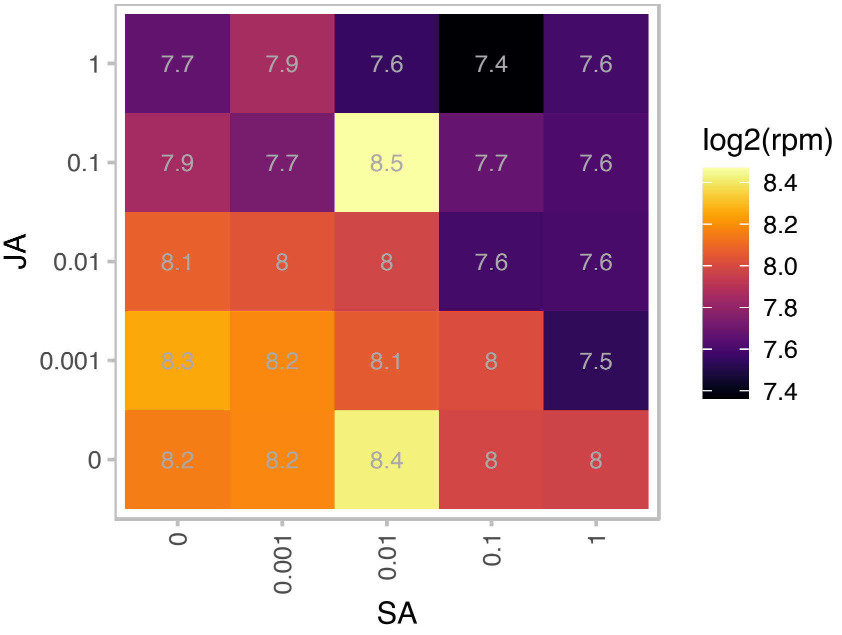

### AT4G37470.1 : KAI2

alpha/beta-Hydrolases superfamily protein

HTL belonging to the alpha/beta fold hydrolase superfamily. Mutant and over-expression studies indicates its involvement in seedling de-etiolation process.

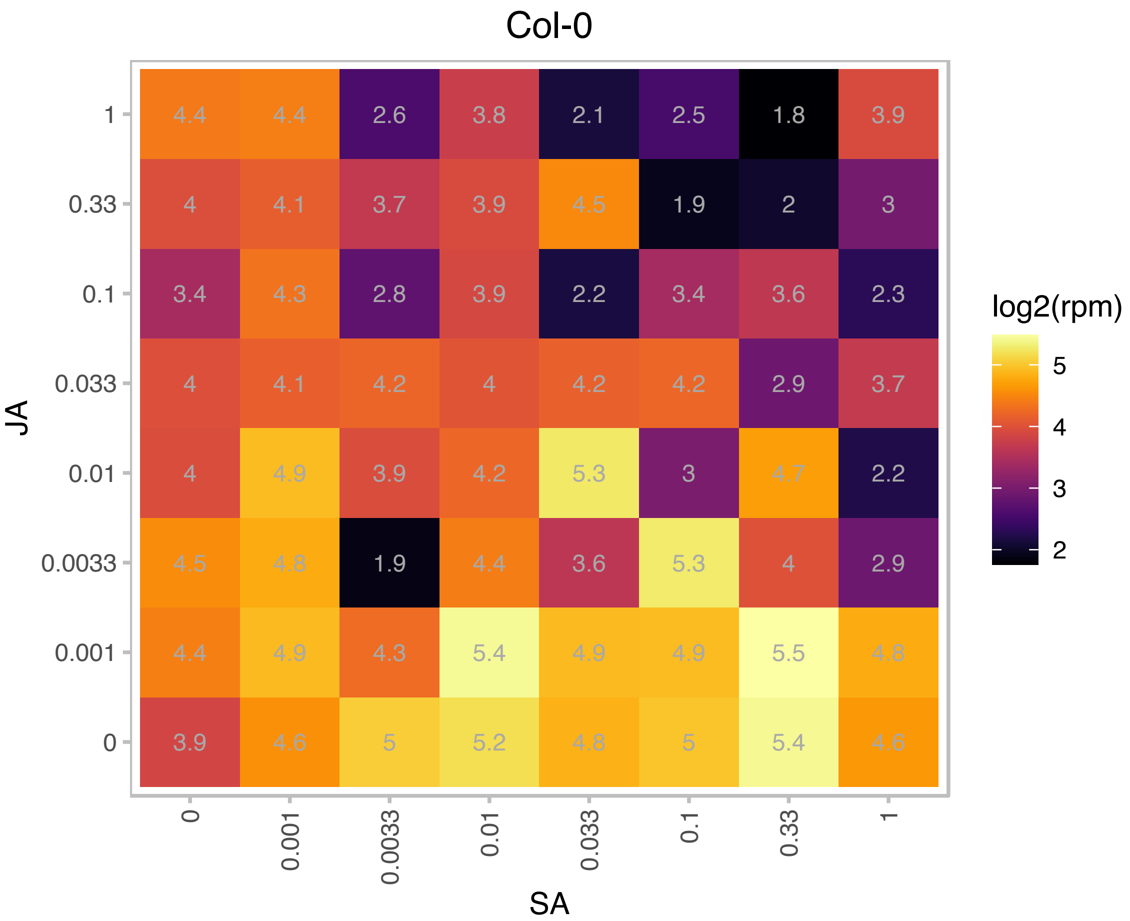


| | |
|---|---|
| Gene | AT4G37470 |
| is.expressed_genes | TRUE |
| is.responsive_genes | TRUE |
| is.apcluster | H-06 |

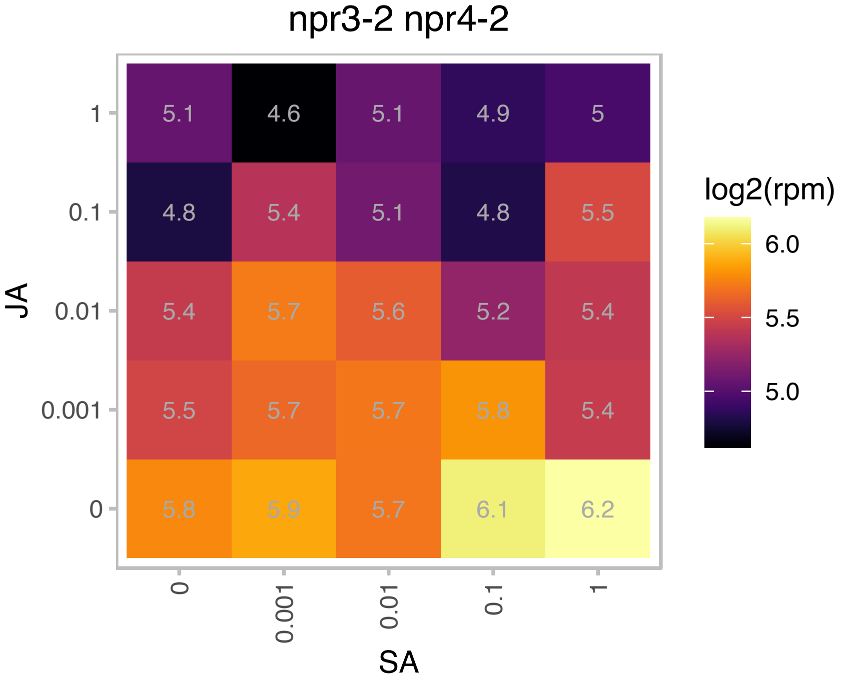


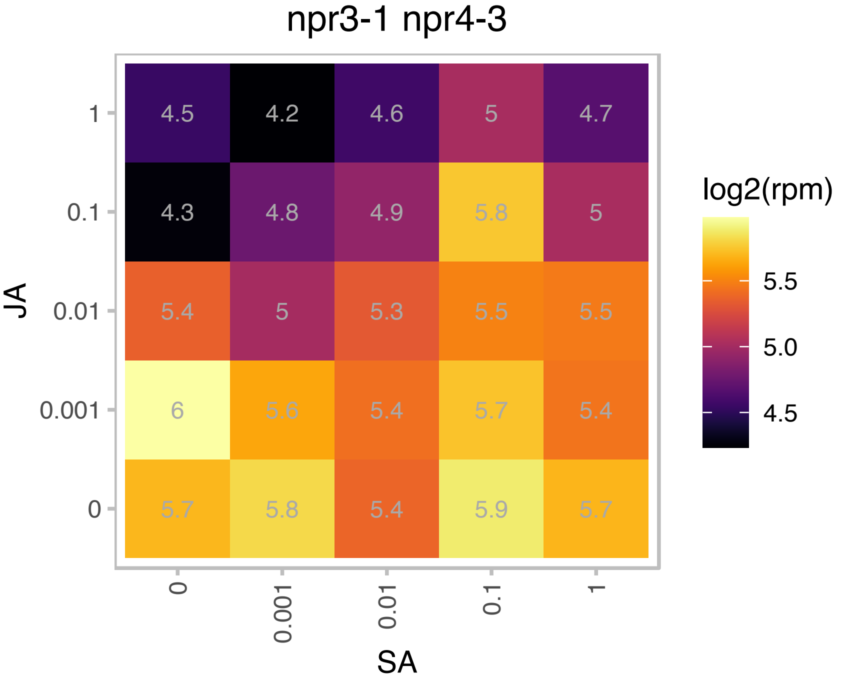

### AT5G03650.1 : SBE2.2

starch branching enzyme 2.2

Encodes starch branching enzyme (E.C.2.4.1.18) similar to SBE2 from maize and rice. Expressed throughout the plant and highest in seedlings and cauline leaves.

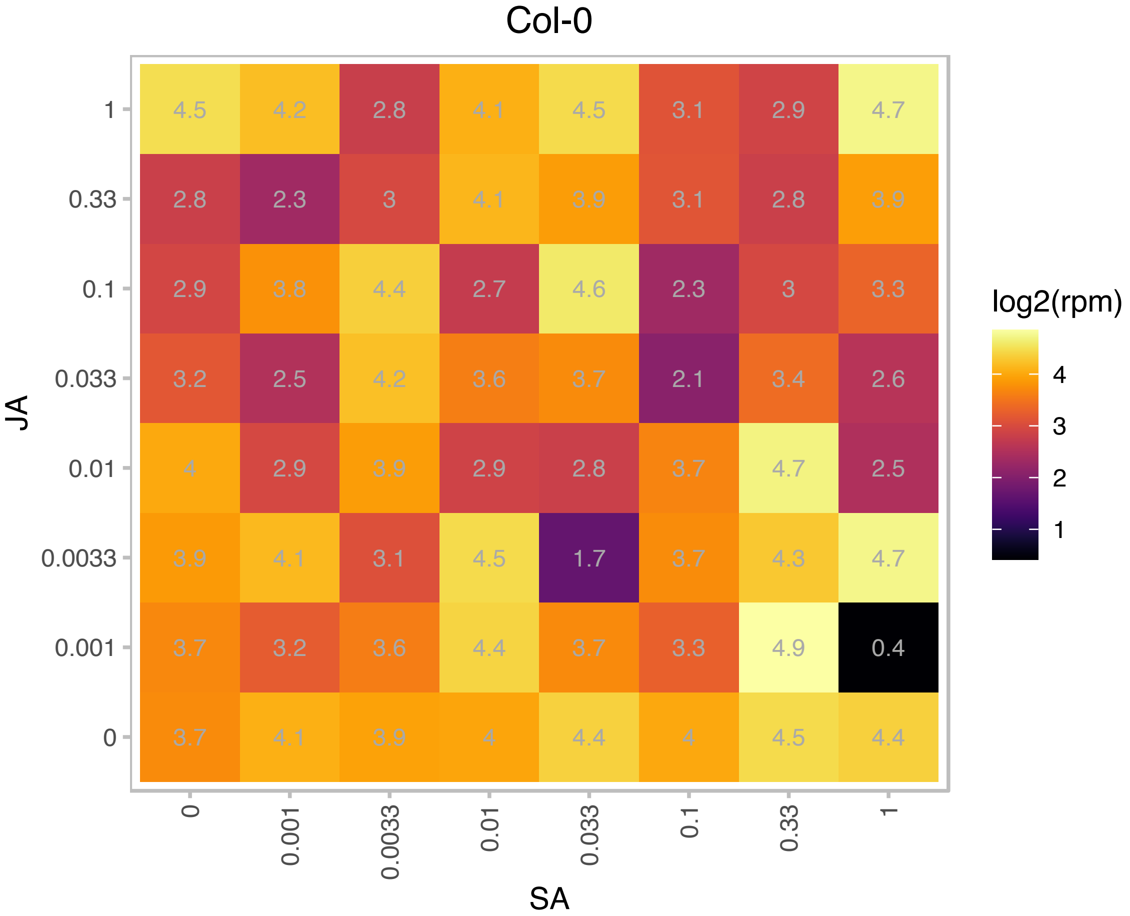


| | |
|---|---|
| Gene | AT5G03650 |
| is.expressed_genes | TRUE |
| is.responsive_genes | TRUE |
| is.apcluster | H-06 |

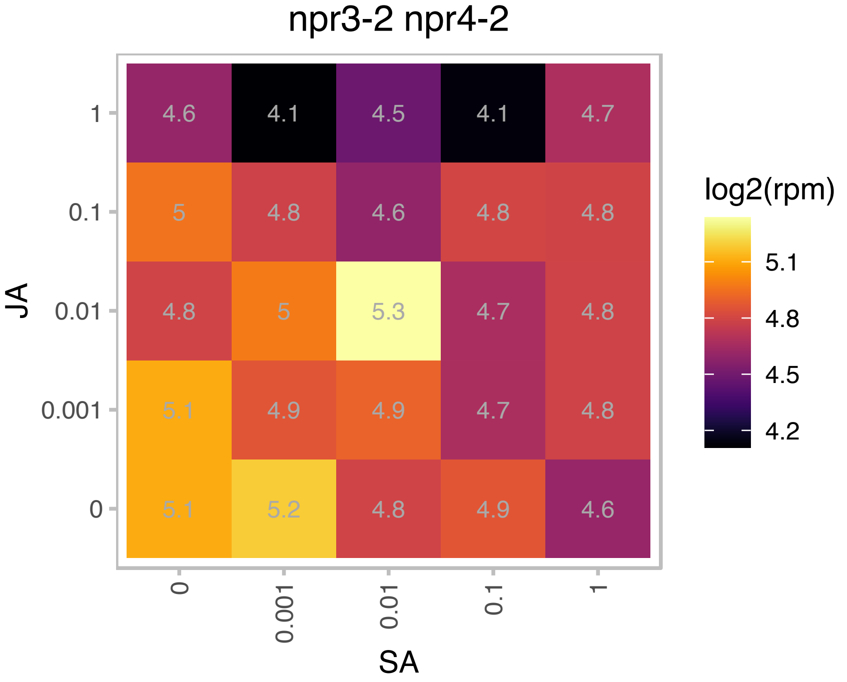


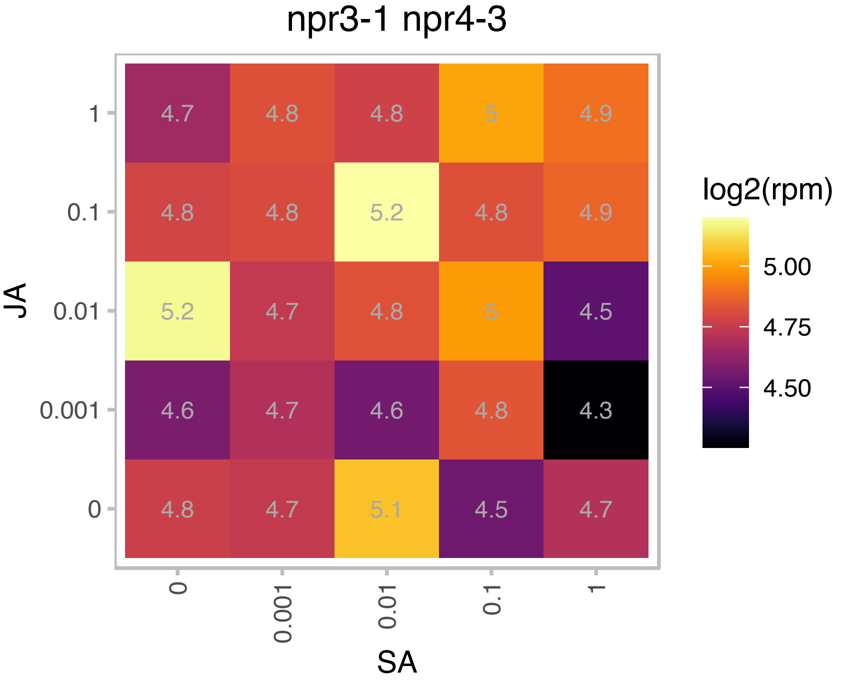

# AT5G08540.1 : NA

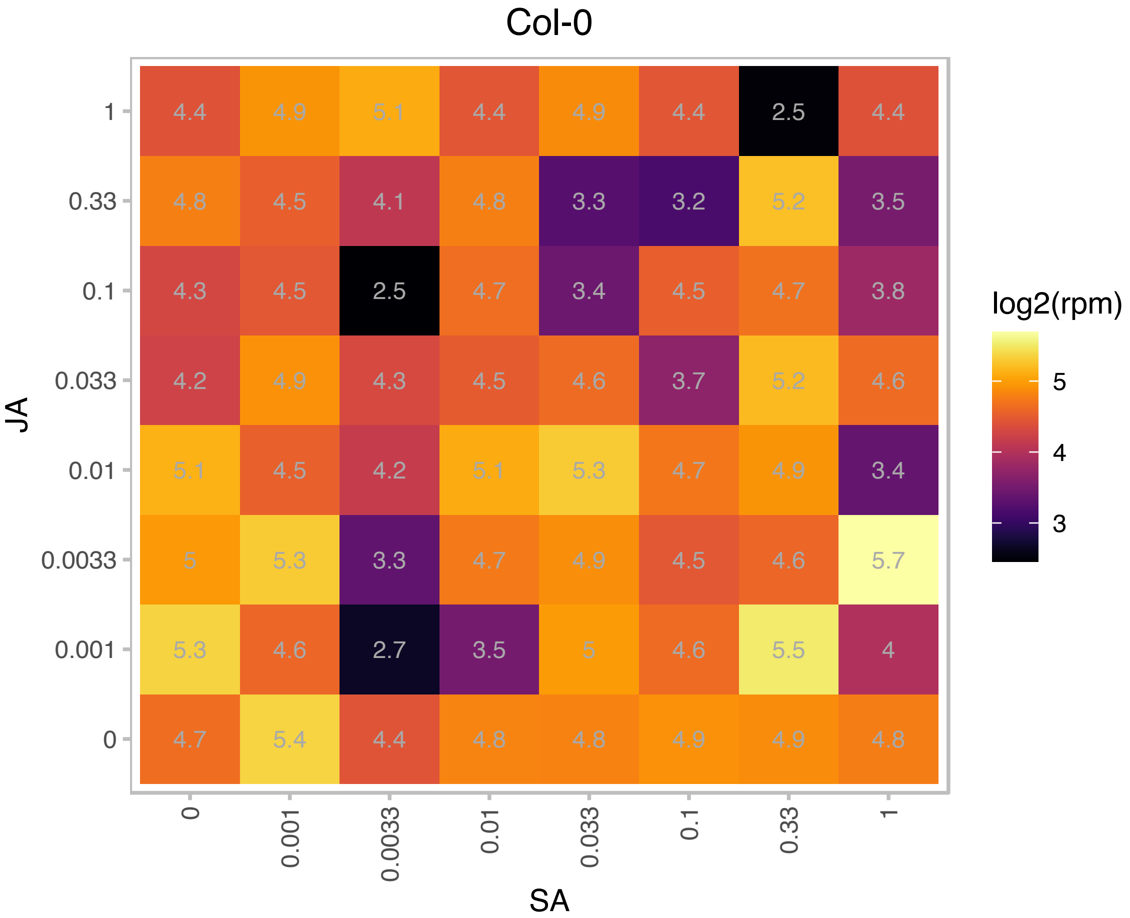


| | |
|---|---|
| Gene | AT5G08540 |
| is.expressed_genes | TRUE |
| is.responsive_genes | TRUE |
| is.apcluster | H-06 |

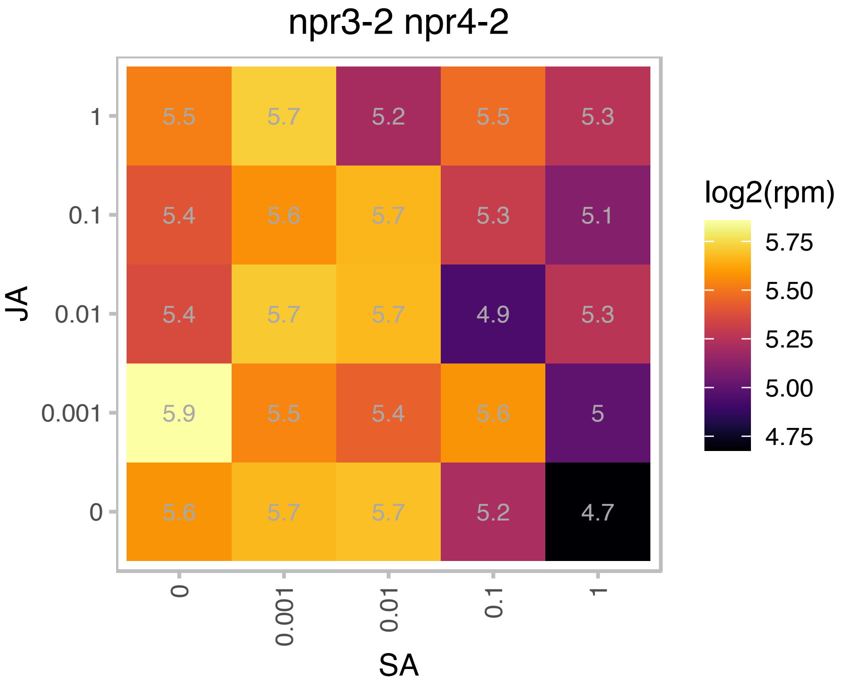


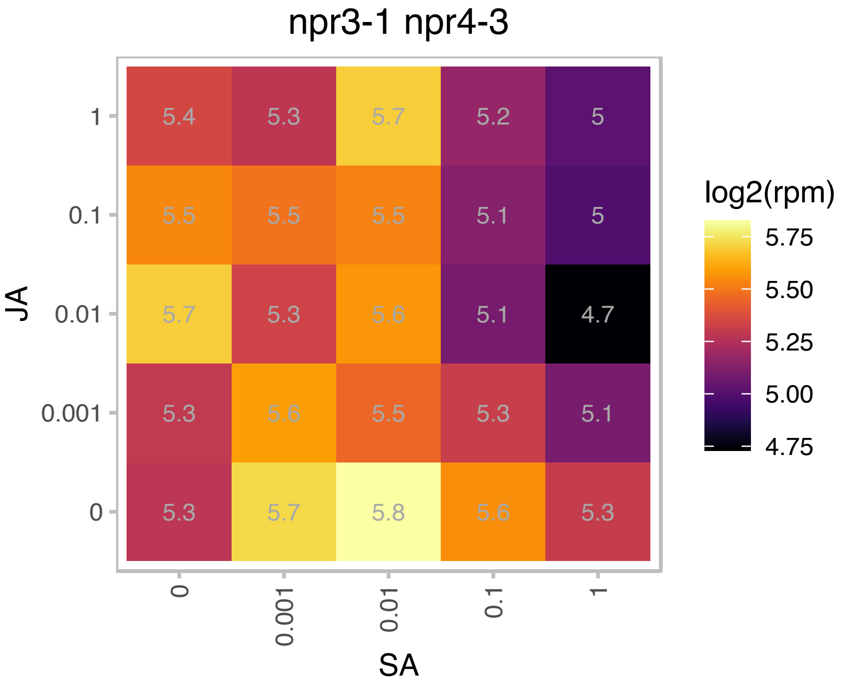

### AT5G16400.1 : ATF2, TRXF2

thioredoxin F2

Encodes an f-type thioredoxin (Trx-f2) localized in chloroplast stroma.

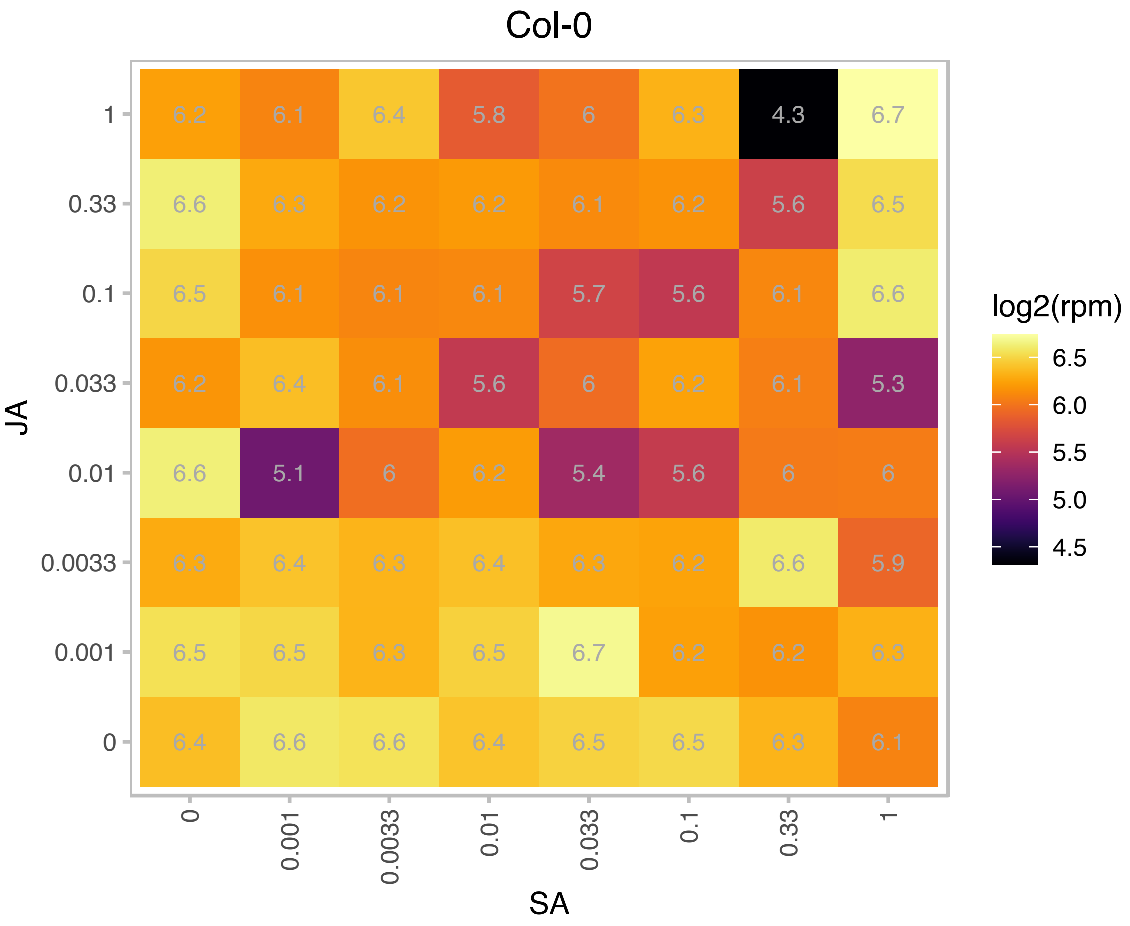


| | |
|---|---|
| Gene | AT5G16400 |
| is.expressed_genes | TRUE |
| is.responsive_genes | TRUE |
| is.apcluster | H-06 |

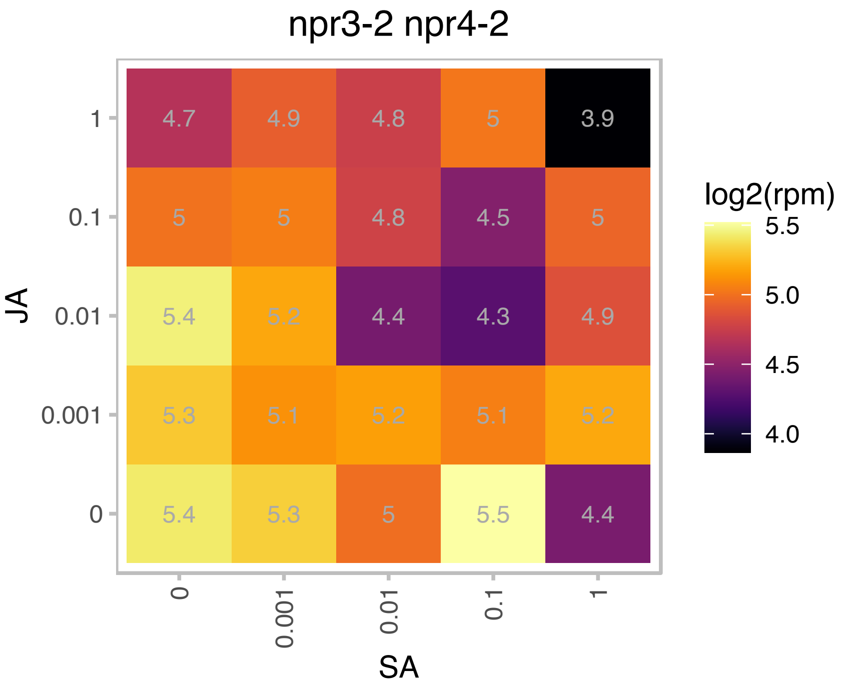


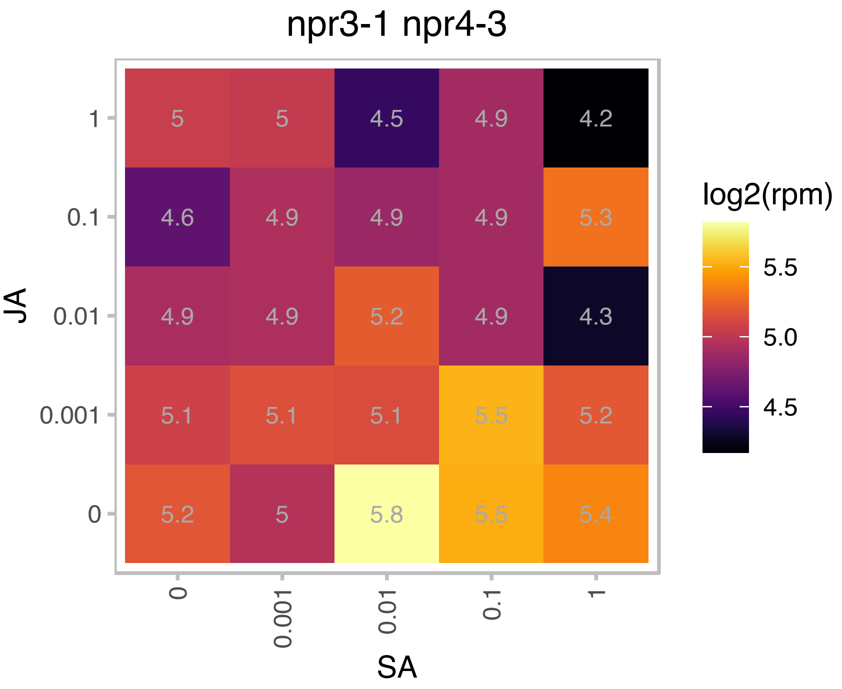

# AT5G43440.1 : NA

2-oxoglutarate (2OG) and Fe(II)-dependent oxygenase superfamily protein

encodes a protein whose sequence is similar to ACC oxidase

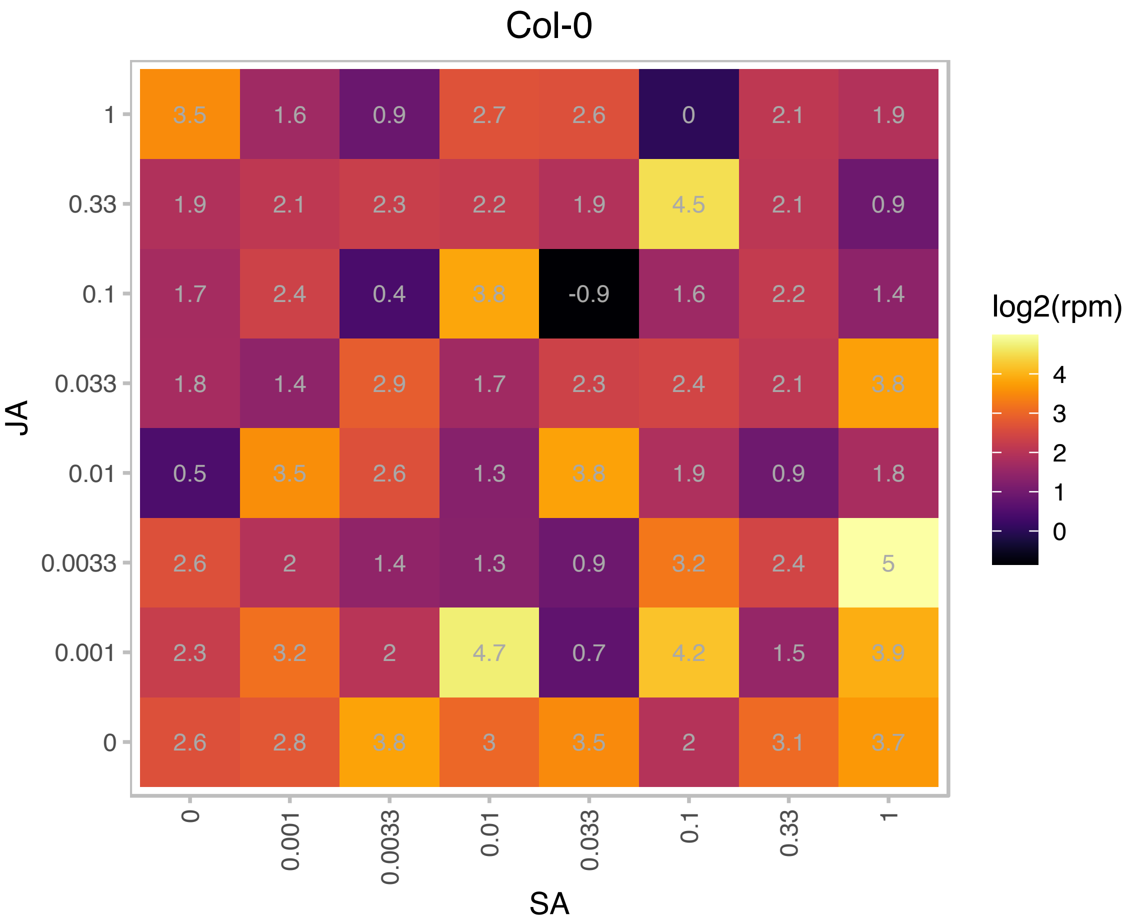


| | |
|---|---|
| Gene | AT5G43440 |
| is.expressed_genes | TRUE |
| is.responsive_genes | TRUE |
| is.apcluster | H-06 |

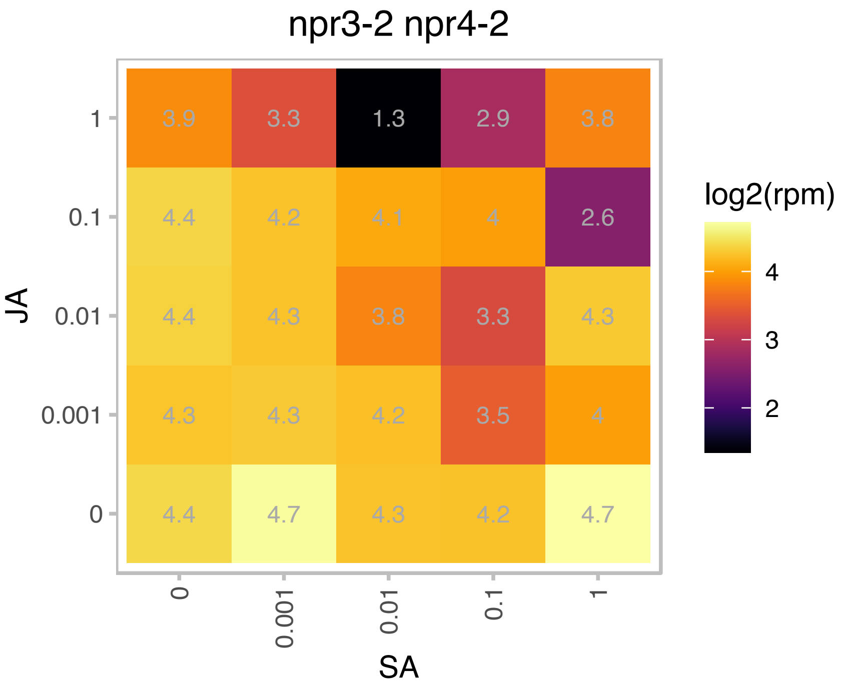


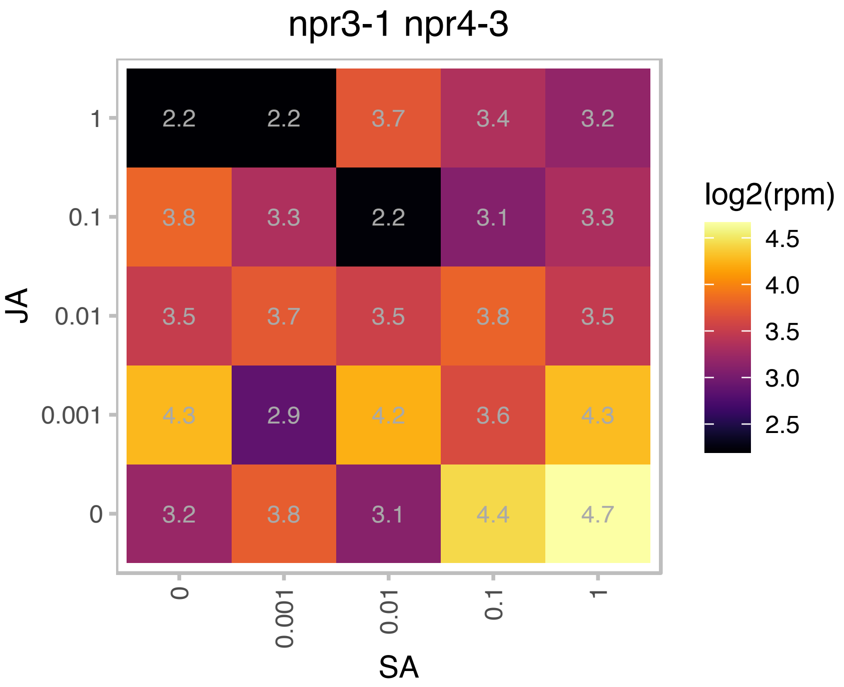

### AT5G64940.1 : abc1k8, ATATH13, ATH13, ATOSA1, OSA1

ABC2 homolog 13
ABC1K8 is a member of an atypical protein kinase family that is induced by heavy metals. Loss of function mutations affect the metabolic profile of chloroplast lipids. It appears to function along with ABC1K7 in mediating lipid membrane changes in response to stress. The mRNA is cell-to-cell mobile.

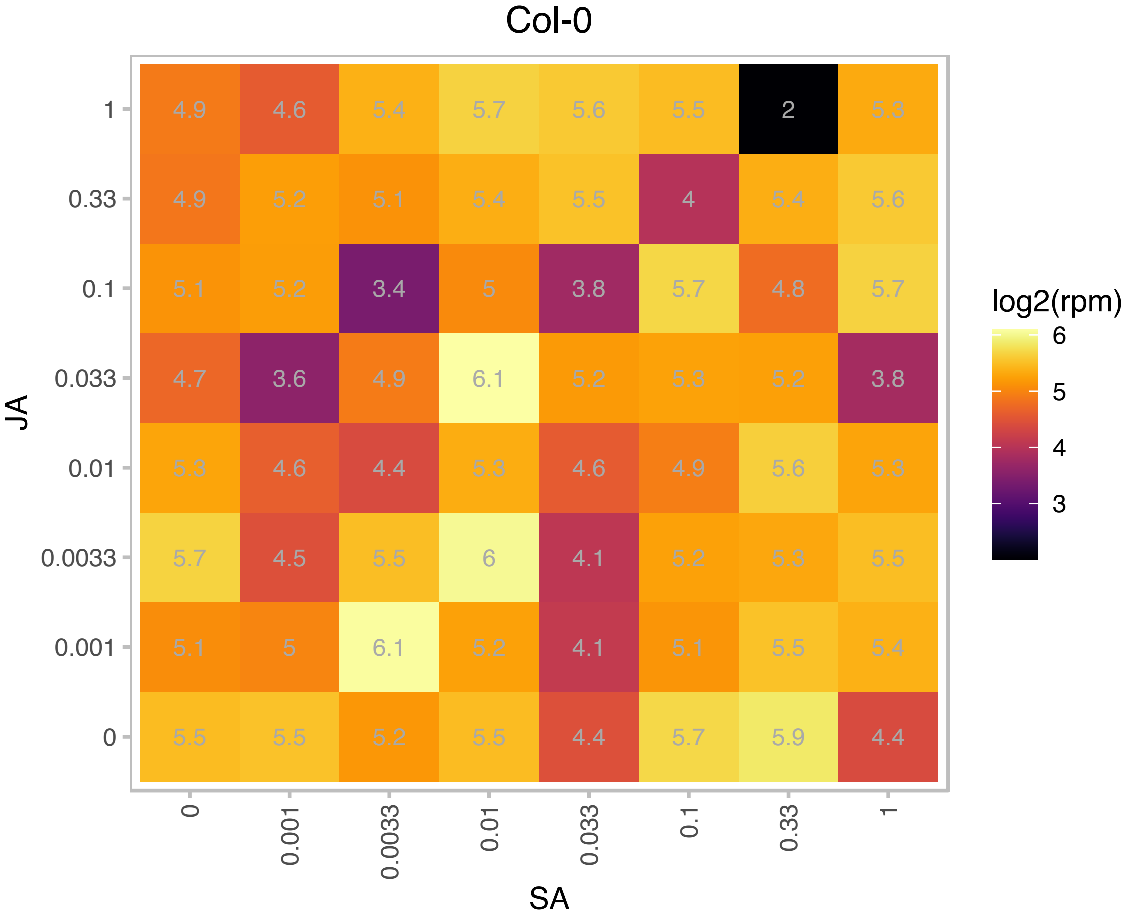


| | |
|---|---|
| Gene | AT5G64940 |
| is.expressed_genes | TRUE |
| is.responsive_genes | TRUE |
| is.apcluster | H-06 |

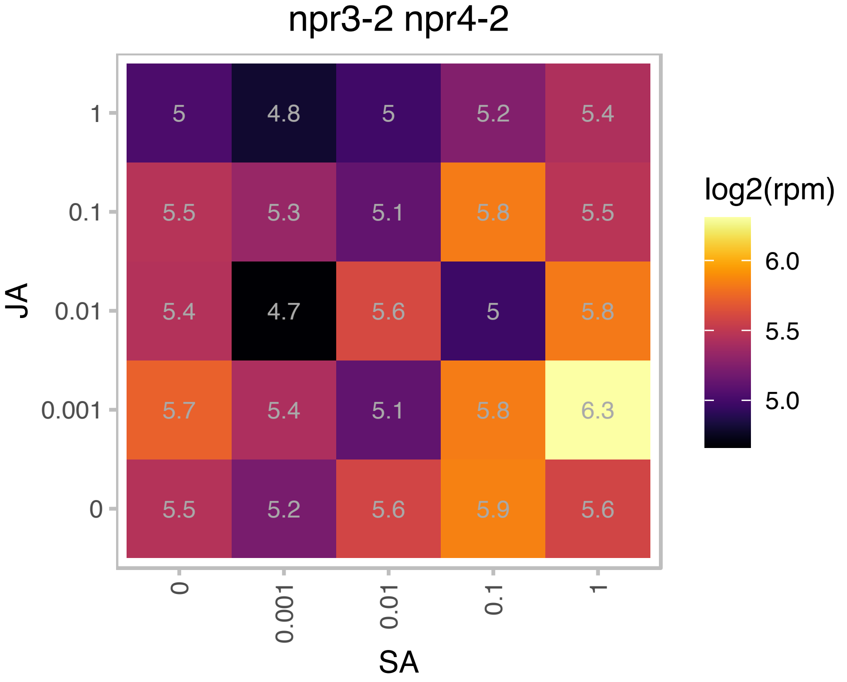


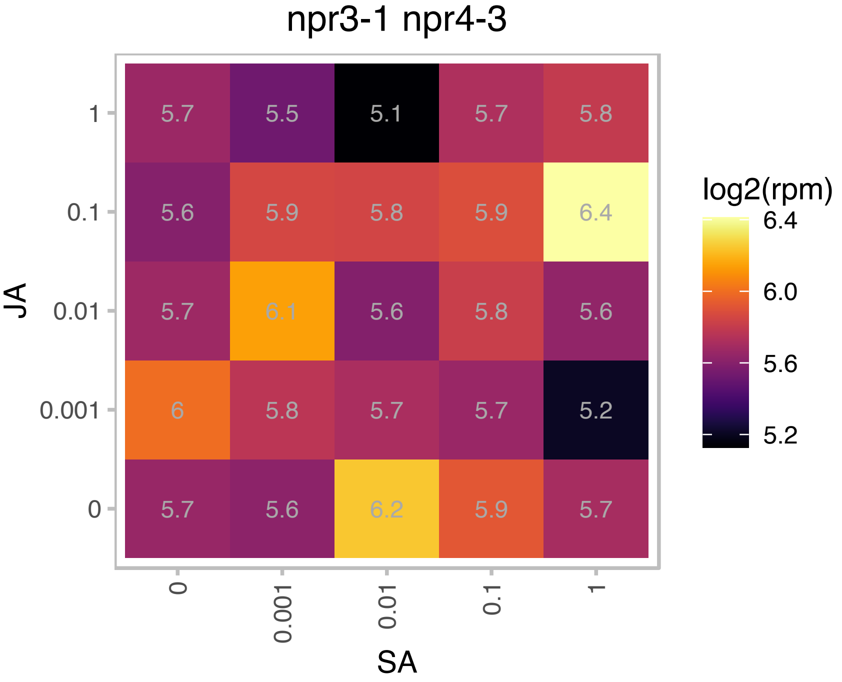

### AT5G67370.1 : CGLD27

Protein of unknown function (DUF1230)

Col-0

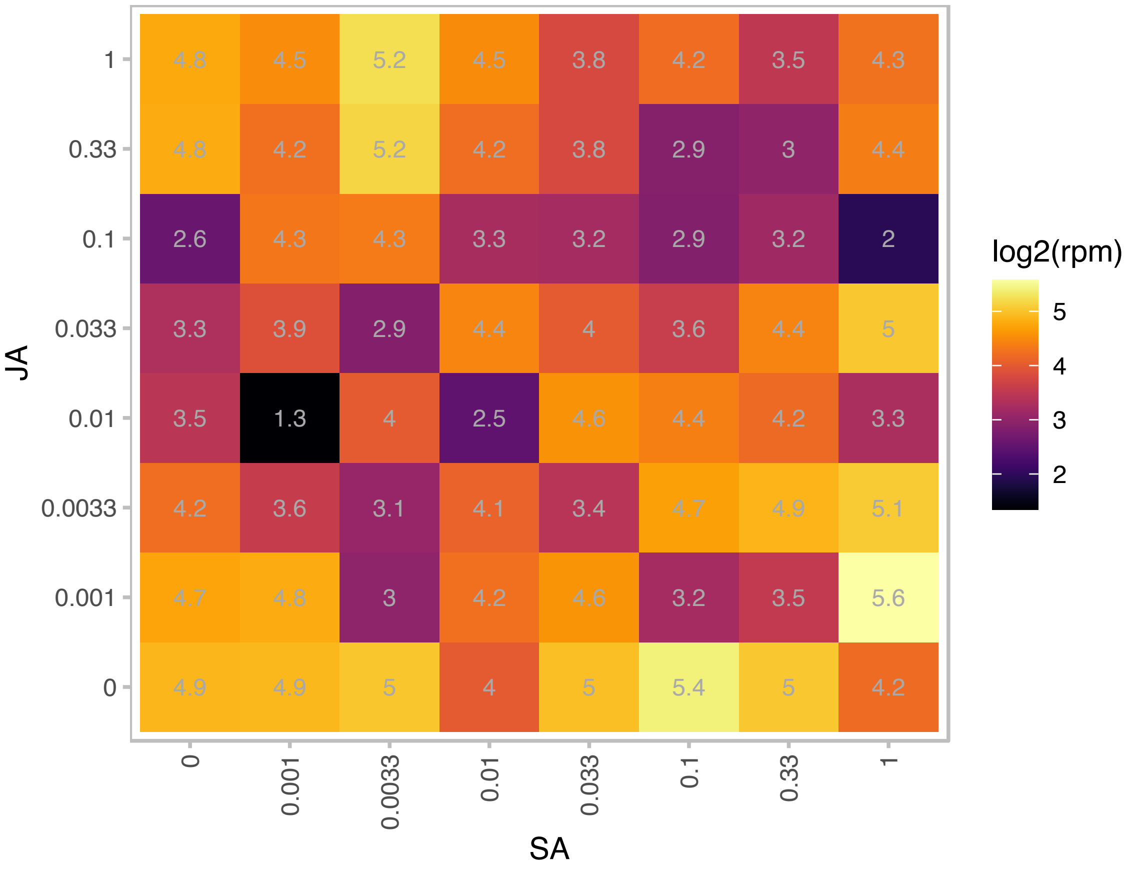


| | |
|---|---|
| Gene | AT5G67370 |
| is.expressed_genes | TRUE |
| is.responsive_genes | TRUE |
| is.apcluster | H-06 |

npr3-2 npr4-2

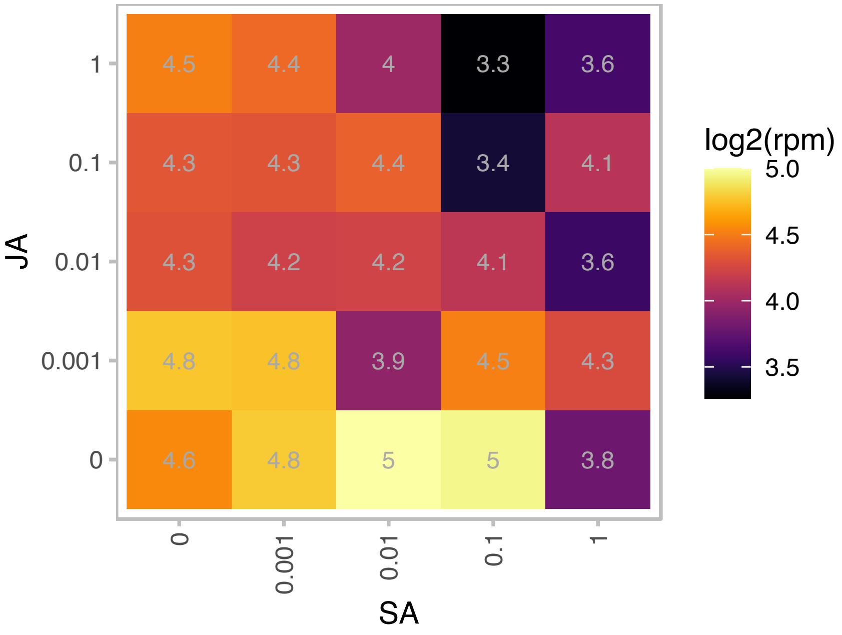


npr3-1 npr4-3

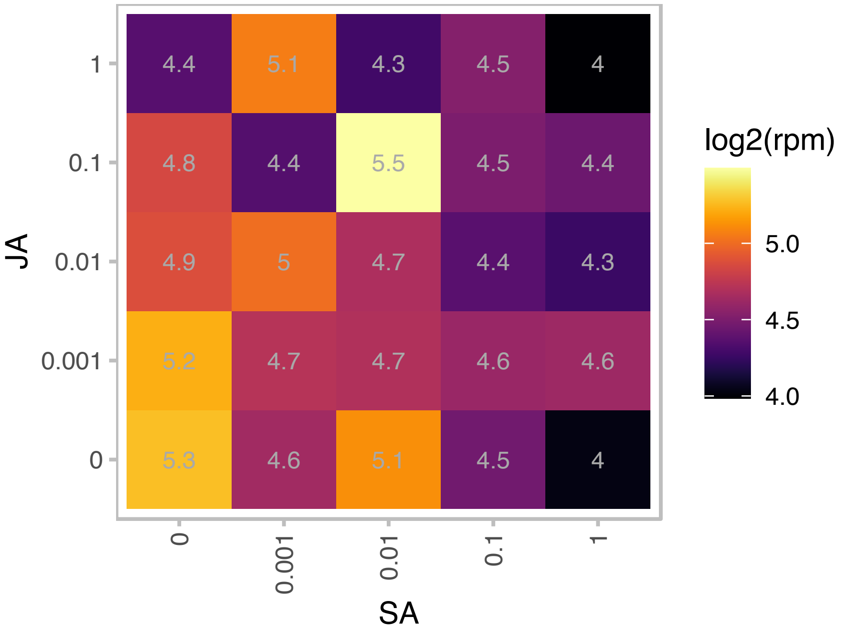
